# The Everything Bagel Feature Finder: Ultra-fast automated feature finding for untargeted metabolomics

**DOI:** 10.64898/2026.08.17.744735

**Authors:** Yourae Shin, Yasin El Abiead, Alan K. Jarmusch, Michael Strobel, Paul E. Abraham, Sarah Thurmon, Deepa D. Acharya, Allegra Aron, Aivett Bilbao, Benjamin P. Bowen, Corey D. Broeckling, Chris J. Brown, Vincent Charron-Lamoureux, Xingyuan Chen, Tito Damiani, Alex Doty, Xiuxia Du, Neha Garg, Stilianos Papadopoulos Lambidis, Laura-Isobel McCall, Kaylie I. Kirkwood-Donelson, Trent Northen, Jessica Prenni, Emma E. Rennie, Oliver B. Vining, Crystal X. Wang, Quanbo Xiong, Haoqi Nina Zhao, Pieter C Dorrestein, Daniel Petras, Vanessa V Phelan, Mingxun Wang

**Author notes:** Correspondence should be addressed to M.W.

## Abstract

Metabolomics studies are increasingly being applied with hundreds to thousands, even tens of thousands of samples that demand rapid, automated data processing while maintaining analytical sensitivity or quantitative accuracy. A major computational bottleneck is feature finding, which is the transformation of LC-MS and LC-MS/MS data into a set of analyte signals aligned and quantified across samples. Feature finding can be computationally intensive and often requires manual iterative parameter optimization. To accelerate this process, we present the Everything Bagel (EB) feature finder, an ultra-fast automated feature finding tool that integrates feature detection, retention-time alignment, and gap filling designed for run-time and memory efficiency. We benchmarked EB against two automated feature finding methods on eight benchmarking datasets. Specifically, we evaluated these three feature finding methods by measuring spike-in standard detection coverage, dilution series quantification accuracy, and yeast ^12^C/^13^C credentialed features. In this evaluation, the EB feature finder achieved performance comparable to, and often exceeding, existing methods while requiring up to 150-fold lower CPU hours and up to 113-fold lower wall time. We further demonstrated the bioanalytical validity of EB by reanalyzing published datasets used for biomarker discovery and reproduced biologically significant features that matched the published findings using manually tuned feature finding settings. Taken along with the speed improvements, we anticipate EB will enhance the ability to automatically analyze datasets with thousands to tens of thousands of samples for the community.

## Introduction

Untargeted mass spectrometry-based metabolomics is a key analytical tool to delineate the small molecule composition of biological and environmental systems^1–3^. Liquid chromatography (LC) coupled to mass spectrometry (MS) separates small molecules into an orthogonal space of chromatographic retention time (RT) and mass-to-charge ratio (*m/z*) for quantification. Tandem mass spectrometry (MS/MS), when performed, generates spectra that provide structural information by isolating and fragmenting individual precursor ions (defined by their *m/z*)^4^. LC-MS experiments can capture thousands to tens of thousands of these signals within a study^5^. Converting raw data into quantitative measurements suitable for statistical and biological analysis requires feature finding (FF), the computational pre-processing step that transforms the raw signal into an integrated peak area used for quantification^6^. In this FF process, a feature is defined as a set of *m/z*, retention time, and peak intensities that represent an analyte for small molecules within a dataset.

When tandem mass spectrum is acquired, FF links MS/MS spectra to the corresponding precursor features to support molecular annotation. Feature-finding workflows commonly include feature detection, retention-time correction and alignment across multiple LC-MS/MS runs in a dataset, gap-filling across runs (to fill in missing peak areas), adduct assignment, and in-source fragment assignment. The quality of the resulting feature table depends strongly on parameters governing mass accuracy, intensity thresholds, chromatographic peak shape and width, retention-time tolerances, and minimum detection frequency across samples. Inappropriate parameter settings can result in missed peaks, artificial peak splitting or merging, inaccurate cross-sample alignment, and ultimately distorted quantitative measurements. Numerous FF tools are available, for example XCMS^7^, MS-DIAL^8^, MZmine^9^, SLAW^10^, asari^11^, and MassCube^12^. However, existing methods have limited throughput^13^, and many workflows require iterative manual parameter optimization to achieve acceptable performance across different instruments and chromatographic methods^14^.

Here, we present the Everything Bagel feature finder (EB), an ultra-fast automated FF workflow. Our method used up to 150 times less CPU hours and up to 113 times less wall time compared to other methods, enabling FF of hundreds of files in minutes using 16 CPU cores and < 30 GB of memory on benchmark datasets. We benchmarked EB against XCMS with Isotopologue Parameter Optimization (XCMS-IPO) and MassCube (MC) using eight datasets from the PR002464 metabolomics benchmark. EB performance was comparable or improved on the detection of spike-in standards, yeast ^12^C/^13^C credentialed features ^15^, and quantitative performance across different ionization modes, chromatography techniques, and instruments, demonstrating its robustness and applicability across various experimental settings.

## Results

### Software architecture

We developed the Everything Bagel feature finder (EB) to be automated based on Rust^16^. To reduce complexity, we integrated multiple computational steps for feature finding into one tool that comprises modules to handle reading input spectra, parameter optimization, feature detection, alignment, gap-filling, tandem mass spectra assignment, and feature grouping (**Figure 1**). The overall ethos was that EB would accelerate the task of feature finding in two ways: automated parameter optimization removes the burden of iterative manual parameter optimization, and highly optimized algorithms provide ultra-fast data processing.

**Figure 1.**
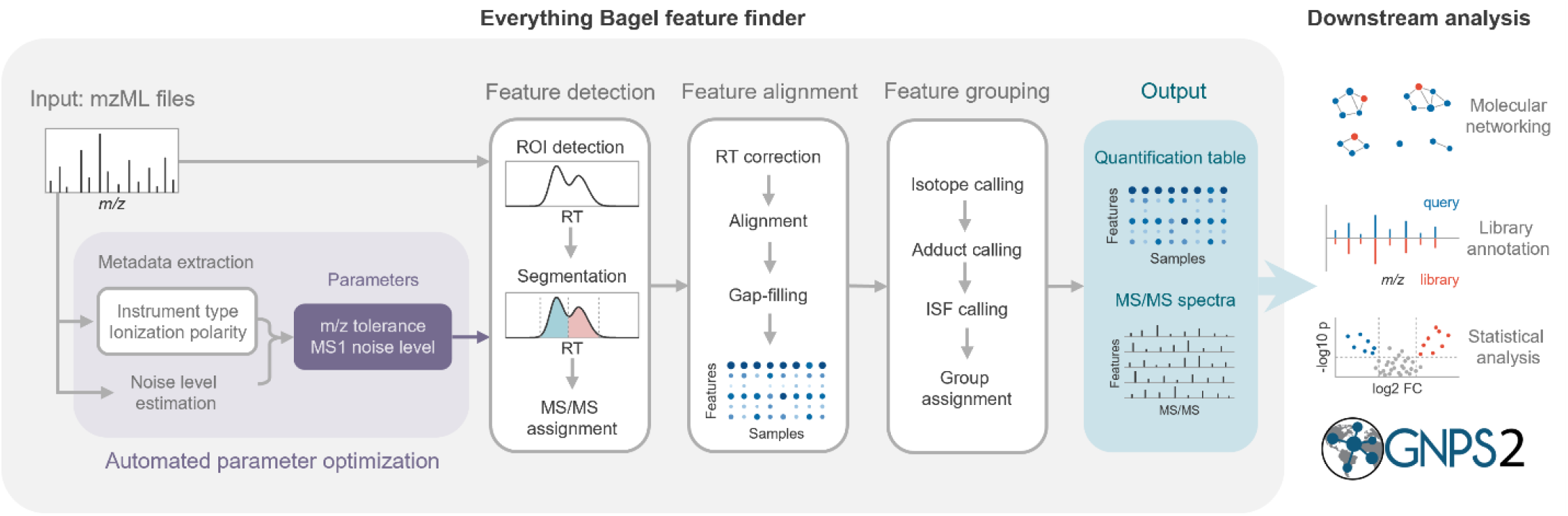
A schematic diagram of the Everything Bagel feature finder.

EB is accessible through an open-source software, a desktop application, a web application, and a Nextflow^28^ workflow built in the Global Natural Products Social Molecular Networking 2 (GNPS2)^26^ infrastructure (See **Software/Code Availability**). We made EB accessible through a variety of channels to meet diverse computational demands among metabolomics researchers, with varying degrees of resource availability and understanding of system architecture.

Although EB does not directly support the annotation of features, EB is integrated into a workflow that encompasses feature finding, library annotation, and feature-based molecular networking^29^ in the GNPS2 infrastructure. The workflow offers streamlined processing of experimental data with compatibility to associated workflows on the GNPS2 ecosystem, as well as effortless visualization and accessibility through the dashboard.

### Feature finding evaluation on metabolomics benchmark

The performance of feature finding was evaluated over a constructed metabolomics benchmark (PR002464)^24^ composed of 8 sub-datasets (**Table 1**). Across these datasets we benchmarked three variants of EB: standard mode (retaining peaks that appear in at least 10% of samples, identical to Masscube’s defaults), reproducible mode (retaining peaks that appear in at least 3 samples), rare mode (retaining peaks that appear in at least one sample). We selected two widely used algorithms, XCMS-IPO (XCMS)^7,14^ and MassCube (MC)^12^ to benchmark against EB for CPU time, wall time and memory usage (**Figure 2**). All jobs were allocated with 16 CPU cores (AMD EPYC 7713 Zen 3) and 256GB of memory.

**Table 1.** Benchmarking datasets and evaluation criteria. The comparison of XCMS, MassCube, and EB was performed over the PR002464 metabolomics benchmark. The performance evaluation of EB on biological samples was performed on *Trypanosoma cruzi* infection biomarker datasets. RP: reverse-phase. HILIC: hydrophilic interaction liquid chromatography.

| Dataset | Instrument | Column | Ion mode | Description | Measurements |  |  |
| --- | --- | --- | --- | --- | --- | --- | --- |
|  |  |  |  |  | Computational cost | Sensitivity | Quantitative accuracy |
| PR002464 metabolomics benchmark |  |  |  |  |  |  |  |
| ST004500 | Orbitrap | RP | negative | LC-MS; a hybrid dataset with MS1-only samples paired with pooled MS/MS acquisition samples | CPU hours, wall time, and memory usage | Spike-in standard detection coverage | Spike-in standard dilution linearity and yeast <sup>12</sup>C/<sup>13</sup>C pair detection coverage with dilution linearity test |
| ST004501 | Orbitrap | HILIC | positive |  |  |  |  |
| ST004502 | Orbitrap | HILIC | negative |  |  |  |  |
| ST004503 | Q-ToF | RP | positive |  |  |  |  |
| ST004504 | Orbitrap | RP | positive |  |  |  |  |
| ST004626 | Q-ToF | RP | negative |  |  |  |  |
| ST004650 | Q-ToF | HILIC | positive |  |  |  |  |
| ST004651 | Q-ToF | HILIC | negative |  |  |  |  |
| Trypanosoma cruzi infection biomarker discovery and validation |  |  |  |  |  |  |  |
| MSV000089112 | Orbitrap | RP | positive | LC-MS/MS samples | - | Biomarker detection at reported m/z and RT | Biomarker fold-change evaluation |

**Figure 2.**
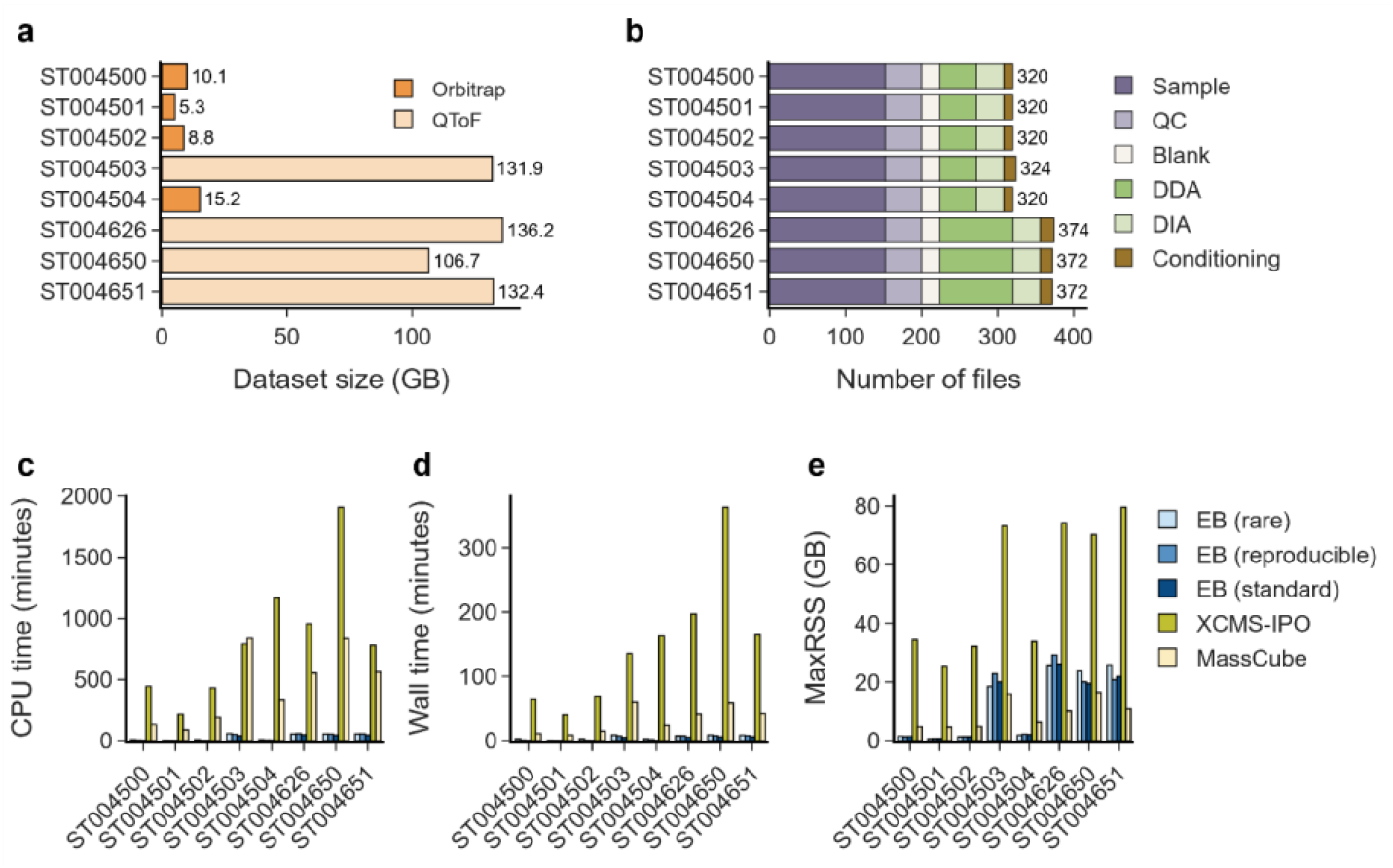
Benchmarking results. **(a)** The size and **(b)** The number of files in each dataset. Orbitrap datasets were 5∼15 GB, and Q-ToF datasets were 105∼136 GB. In each dataset, data-independent acquisition (DIA) MS/MS and conditioning runs were excluded from the analysis. **(c)** The CPU time, **(d)** the wall time, and **(e)** maximum resident set size (MaxRSS) of EB, XCMS, and MassCube in each dataset. EB was tested in three reproducibility modes: rare, reproducible, and standard. EB CPU time ranged from 2.05 minutes to 9.87 minutes in Orbitrap datasets, and from 40.87 minutes to 59.52 minutes in Q-ToF datasets. EB wall time (on 16 CPU Cores) ranged from 0.35 minutes to 3.19 minutes in Orbitrap datasets, and from 5.25 minutes to 9.21 minutes in Q-ToF datasets. The peak memory usage (MaxRSS) of EB, XCMS, and MassCube in each dataset. The memory usage of EB ranged from 0.69 GB to 2.3 GB in Orbitrap datasets, and from 18.51 GB to 29.31 GB in Q-ToF datasets.

EB completed the analysis faster in terms of both CPU time and wall time across all datasets (**Figure 2**). EB standard mode required between 16-192 times (16-40 times in Q-ToF datasets, 85-192 times in Orbitrap datasets) fewer CPU - hours compared to XCMS, and 11-55 times (11-20 times in Q-ToF datasets, 25-55 times in Orbitrap datasets) fewer CPU-hours compared to MassCube. In wall time, EB standard modeachieved 26-177 fold improvement compared to XCMS and 6-26 fold improvement compared to MassCube. In Orbitrap datasets, the memory usage was 2.7-30 times lower compared to both XCMS and MassCube. In the Q-ToF datasets, memory usage was 2.5-3.2 times lower compared to XCMS, but 1.4-2.9 times higher than MassCube. We concluded that the minimum resident size of the MS1 peaks was larger in EB because the adaptive noise thresholds EB used for these datasets, ranging from 141 to 190, were much lower than MassCube. MassCube uses a fixed MS1 intensity tolerance of 1,000 in Q-ToF datasets, which caused the removal of > 98% of the MS1 peaks in these datasets, significantly reducing the size of working peaks kept in memory. Overall, in these benchmark datasets, EB demonstrated faster data processing compared to other methods with < 30 GB of memory usage, enabling untargeted feature finding on hundreds of files at the seconds to minutes scale.

We assessed the sensitivity of three feature finding tools using spike-in standards in PR002464 datasets. We used MS/MS based library search for spike-in standard annotation as it provides higher-quality and confidence identifications than simple *m/z* match on precursor ions. Out of 96 spike-in standards, we found MS/MS spectra matching 88 of these structures in NIST20 and GNPS2 spectral libraries (**SI Table 5 and Methods - Public data acquisition**). First, we searched all MS/MS spectra acquired in each dataset against library MS/MS spectra of these 88 structures and found spectral matches to 66 structures. This set of 66 structures constitutes the best-case scenario for feature finding results. Then, we annotated consensus features in each tool result using MS/MS spectra linked to consensus features. We found that the EB rare mode, when searching the features against the same library, found 63 structure matches (**Figure 3a**), followed by the EB reproducible mode that showed 62 matches. Notably, EB rare mode recovered a structure, Lariciresinol (theoretical [M-H] ^1-^. *m/z*=359.1500), that was not matched in the original full LC-MS/MS search because EB corrected the precursor *m/z* based upon the MS1 feature *m/z*. The original isolation window target *m/z* of the MS/MS scan was 359.1140, while EB assigned the MS/MS scan to the feature *m/z*=359.1486, which closely approximates the library *m/z*=359.1500. The four missing standards that EB rare mode did not detect (1,4-cinole, S-allyl-L-cysteine, (-)-hydroxycitric acid lactone, and 2-phenylethanol) were not detected in any other method (See XICs of these missed standards in **SI Figure 1**). EB standard mode, the most conservative detection mode for EB, showed a total number of 60 library annotated standards, outperforming the other methods by eight more annotated standard matches (**Figure 3a**).

**Figure 3.**
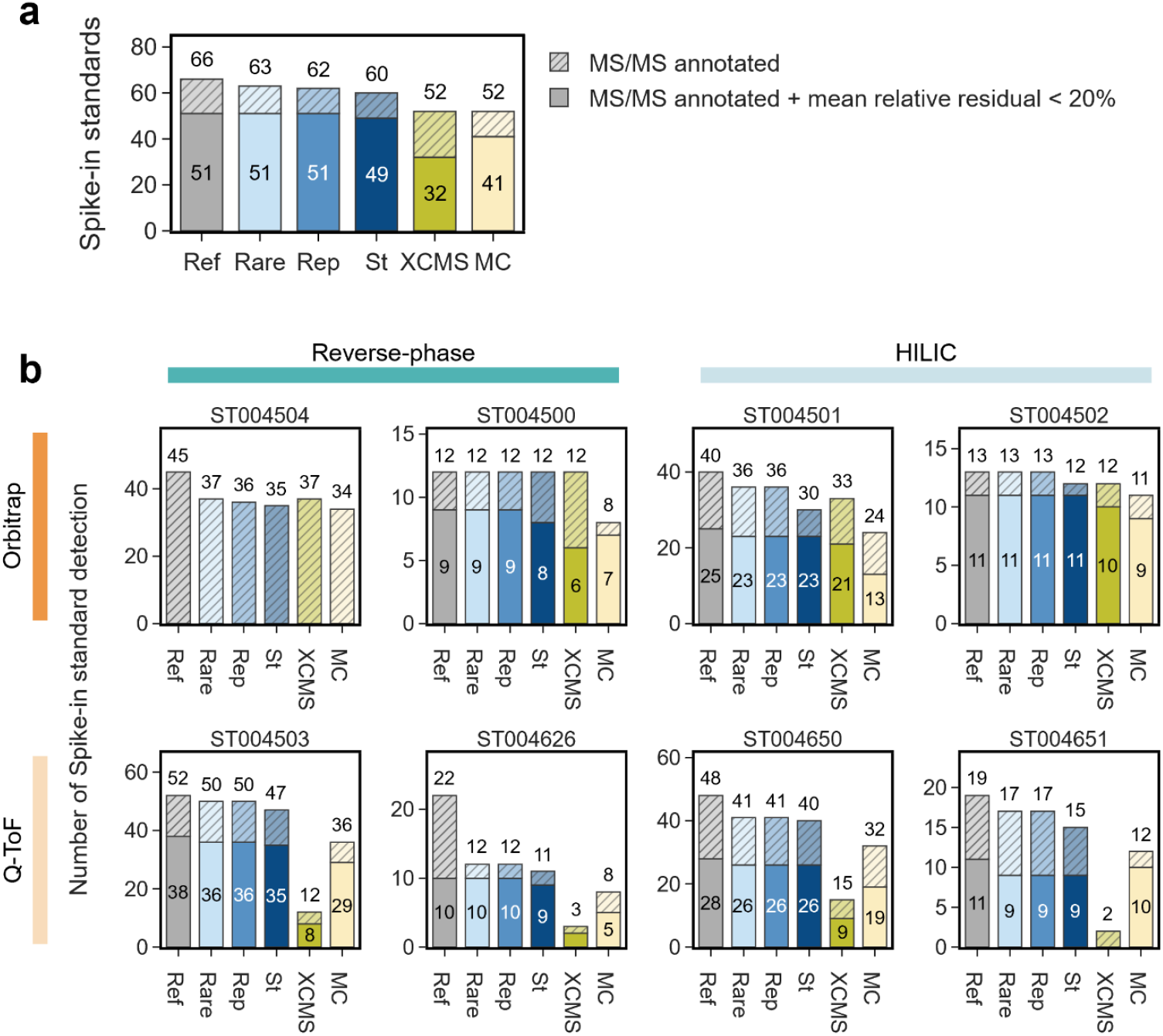
Number of spike-in standards detected in feature finding results using EB, XCMS, and MassCube. **(a)** The number of unique structures across all datasets. **(b)** The number of unique structures in each dataset. All MS/MS spectra in each dataset were searched against the library of 78 structures to measure the upper limit of detection (Ref). MS/MS annotated (hatched) groups show the number of spike-in standards that were annotated in consensus feature MS/MS spectra. MS/MS annotated + mean relative residual < 20% (solid) groups show the number of spike-in standards that were annotated through library search and followed expected abundance series across 5-level dilution with mean relative residual < 20%. Ref: all MS/MS spectra. St: EB with standard mode. Rep: EB with reproducible mode. Rare: EB with rare mode. MC: MassCube.

Next, we evaluated the quantification of consensus features matching spike-in standards in the PR002464 datasets. Out of 66 standards found in library search against all MS/MS spectra, 51 standards were detected and showed dilution linearity (defined as having mean relative residual < 20%) in at least one of the feature finding tools (**Figure 3a**). EB rare mode and EB reproducible mode detected all the 51 standards with dilution linearity, while EB standard mode detected 49 standards with dilution linearity. EB standard mode showed better quantitative performance compared to XCMS and MassCube in each dataset, except for ST004651 where MassCube detected 10 standards with dilution linearity while EB detected 9 standards with dilution linearity (**Figure 3b**). The spike-in standard features that passed dilution linearity test in each tool are listed in **SI Table 1** . Detailed spike-in standard quantification results on the dilution series are shown in **SI Data Files 2-9**.

We attempted to investigate the precision of feature finding tools at a fixed sensitivity of the spike-in standards, but the number of true positive features was not known for each dataset. Instead, we decided to compare the total number of features detected in each tool to the number of spike-in standards that passed dilution linearity test as a proxy for the true positive features. We selected the number of spike-in standards that passed dilution linearity test to represent both precise detection and quantification of feature finding tools. We hypothesized that the feature finding tool that found more spike-in standards with dilution linearity and a smaller total number of features is preferable, visualized as the upper left quadrant in **Figure 4** . Across the 8 datasets of the metabolomics benchmark where this evaluation was possible, EB standard mode detected a total number of features similar to XCMS and MassCube while recovering more of the spike-in standards (**Figure 4**). While EB reproducible and EB rare exhibited slightly higher detection of the spike- in standards, this caused the total number of features to rise. At least in this benchmark set, where spike-in standards are repeatedly measured within a dataset, EB standard, with its 10% detection rate floor, seemed to be an optimal trade off. Overall, EB demonstrated high sensitivity across all datasets compared to other tools without excessively inflating the total number of features.

**Figure 4.**
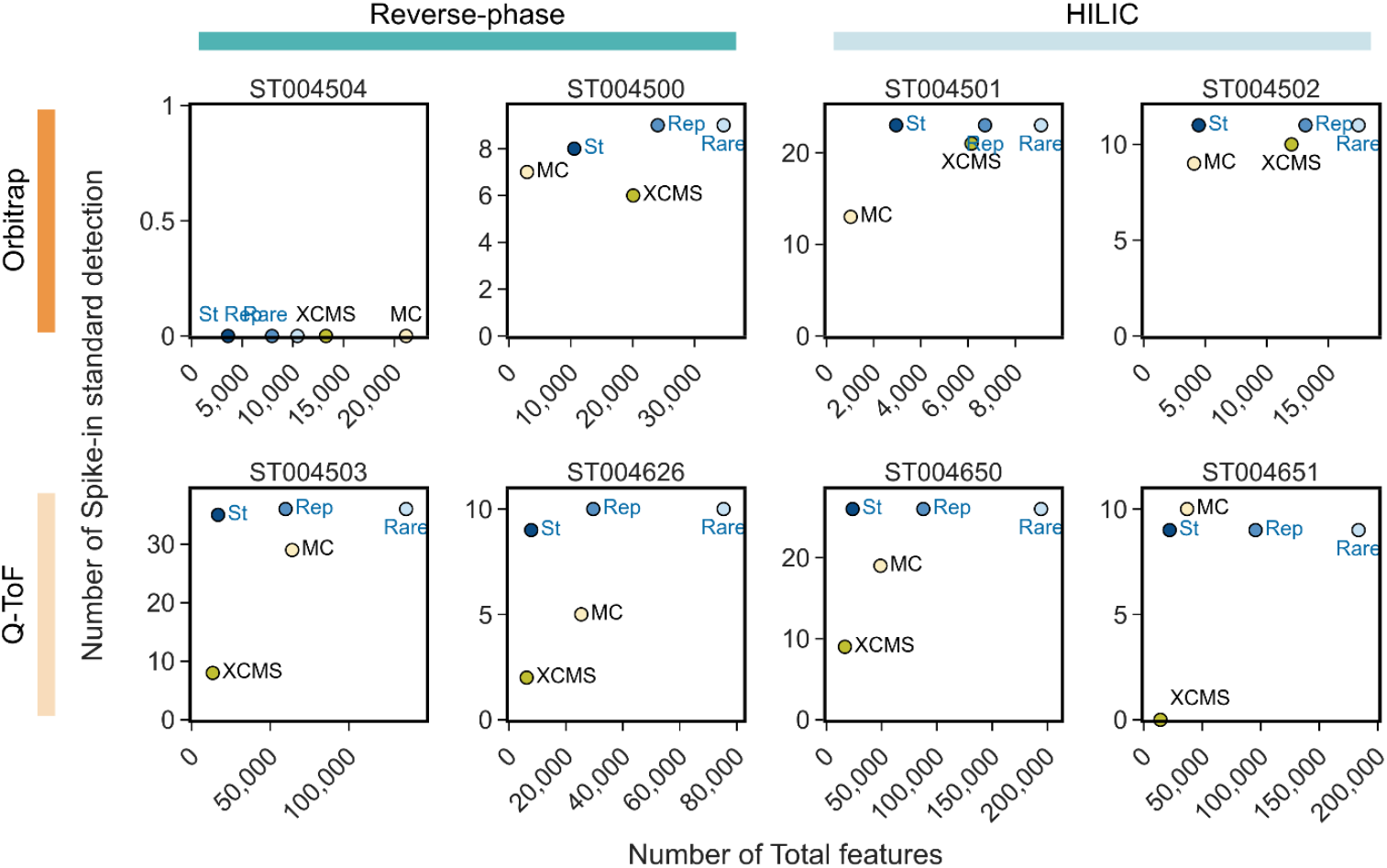
Total features and spike-in standard detections across tools in each dataset. Each data point represents the number of spike-in standards with dilution linearity (mean relative residual < 20%) annotated through library search and quantitative evaluation compared to the total feature detections in each tool. A data point towards the upper left direction indicates more spike-in standard detection with a lower number of features.

While the number of features and recovery of spike-in standards provided insights on EB performance, we found it difficult to comprehensively evaluate the quality of the total set of features that each tool detected, as feature quality assessment often relies on manual investigation by experts. Instead, we compared the features identified in each tool and extracted the XICs of features that are uniquely detected in EB standard mode, XCMS, and MassCube to allow manual interpretations of detection gaps (See **SI Data Files 10-33**). We used EB standard mode to assess the discrepancy in core algorithms, rather than the alignment detection rate threshold. We matched features across different tool results using instrument-specific *m/z* tolerance (10 ppm for Orbitrap, 25 ppm for Q-ToF) and RT tolerance of 0.2 minutes expanded to the RT boundaries detected in EB standard mode. A tool - unique feature was defined as a feature from one tool that is not matched to any feature in other tool results within the tolerance.

We found that none of the tools could find and quantify any spike-in standards in the Orbitrap reverse-phase (RP) dataset in the positive ion-mode (ST004504) with < 20% residual error in the abundance correlation (**Figure 3**). In comparing ST004504 (positive mode) versus its counterpart in negative mode (ST004500), we found that most features in ST004504 were very abundant in the blanks (**SI Figure 2**). In contrast, ST004500 exhibited significantly higher abundance in sample and quality control (QC) runs compared to the blank runs for most features. Moreover, the number of features detected in ST004504 was ∼3 fold fewer than in ST004500. In contrast, for the pos-neg pairs of Q-ToF RP datasets, the total number of features detected in the positiveion mode was comparable or greater (**Figure 4**). We concluded that dominant signals found in ST004504 are background or matrix signals, and thus target analytes were not identified in any feature finding method.

The metabolomics benchmark (PR002464) datasets include the commercial yeast extract samples to provide biological analytes as a part of the benchmark. To assess the performance of each feature finding method in samples with a biological matrix, we further evaluated the quantitative performance of feature finding methods using the set of yeast extract dilution series samples in the benchmark datasets.

We extracted the reference yeast analytes (see **Methods Commercial yeast extract dilution series evaluation**) in each dataset, a set of yeast ^12^C native ion features that showed dilution linearity in at least one of the tool results. We used the standard mode of EB to compare with the other tools. Out of 1,024 reference yeast analytes detected across all datasets, EB standard mode detected 689 analytes, followed by XCMS (674 analytes) and MassCube (595 analytes). The number of overlapping reference yeast analytes detected in each method is shown in **Figure 5** .

**Figure 5.**
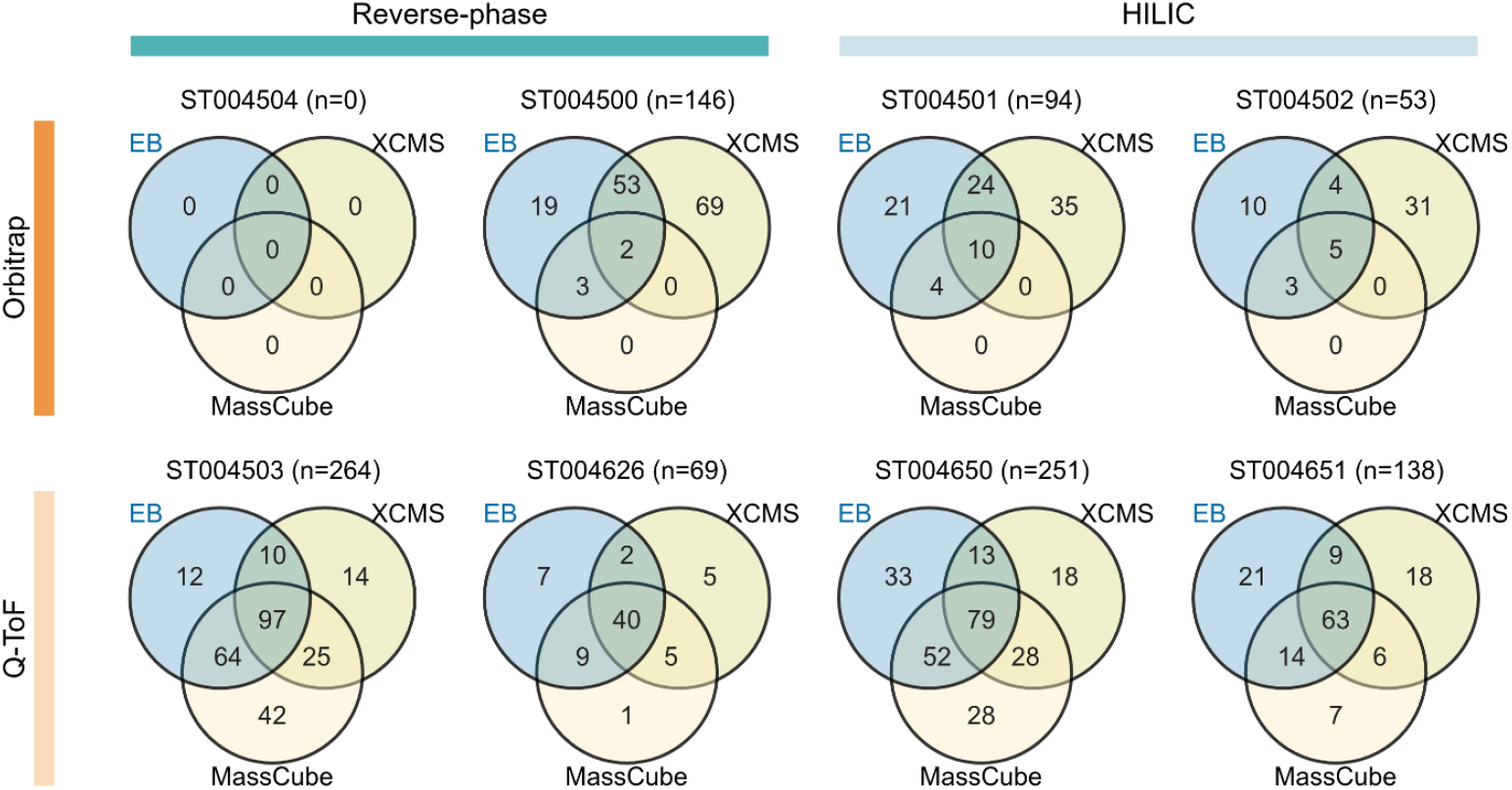
The reference yeast analytes detected in XCMS, MassCube, and EB standard mode. In Orbitrap datasets, EB and XCMS showed the highest overlap between two tools. In Q-ToF datasets, EB and MassCube showed the highest overlap between two tools, except for ST004651. The XICs of features matched with reference yeast analytes that are uniquely detected in each tool are shown in **SI Data Files 34-36**.

We then evaluated the quantitative performance of each tool on the 4-level dilution series of reference yeast analytes. The mean abundance across replicates was computed at each dilution level, and the top-level abundance was used as an anchor to compute the expected abundance at each lower dilution level. If two or more values are missing for the top dilution level, the analyte is considered to be not detected. The mean relative residual value was computed on all dilution levels below the top level. We compared the mean relative residual of the reference yeast analytes that were detected and had no more than one missing value at the top level in all three tools for each dataset (**Figure 6**). We performed a Wilcoxon signed-rank test on mean relative residual distributions of the shared reference yeast analytes in each dataset and confirmed that EB standard mode showed either comparable (*p*-value ≥ 0.05 or |median difference| ≤ 5%) or exceeding (*p*-value < 0.05 and |median difference| > 5%) quantitative performance to XCMS and MassCube. The number of reference yeast analytes that passed dilution linearity test compared to the total number of features is shown in **SI Figure 3** .

**Figure 6.**
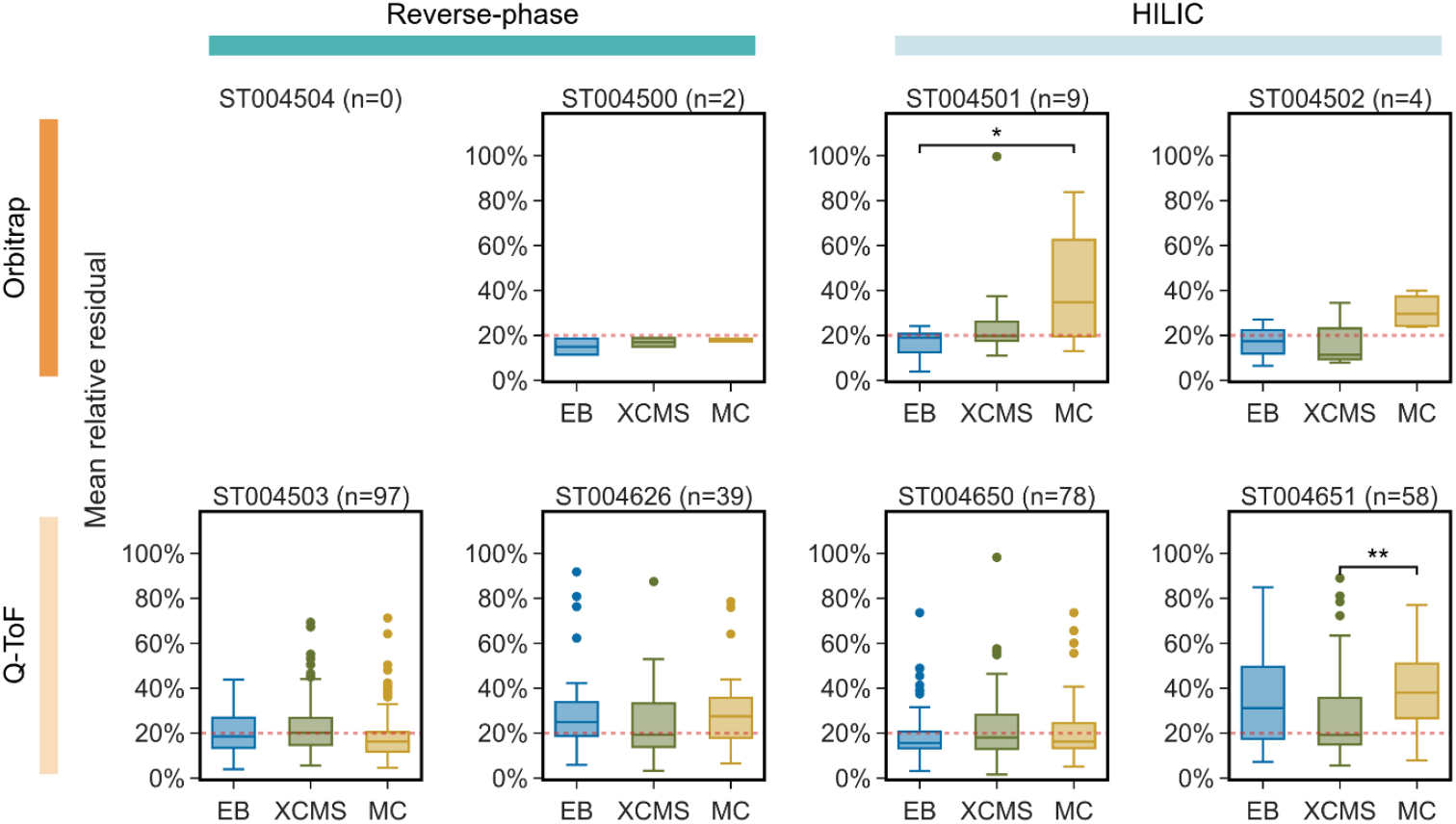
The mean relative residual values of yeast ^12^C native ion features matched the yeast analyte reference set in XCMS, MassCube, and EB standard mode. The mean relative residual value is evaluated on expected abundance along the 4-level dilution series. The mean relative residual is zero when the mean feature abundance across replicates perfectly follows the expected abundance. The number of reference yeast analytes detected in all three tools is shown in each dataset subtitle. The y-axis is capped to [0, 100%] range for visibility. ST004504 is not shown in the figure due to the lack of yeast ^12^C native ion features that passed blank removal. *: *p*-value < 0.05 and |median difference| > 5%. **: *p*-value < 0.01 and |median difference| > 5%.

### Reproducing biological signatures in a public dataset

We performed reanalysis on publicly available untargeted dataset (MSV000089112) to validate that EB reproduces biological signatures reported in the published study^27^. In the reanalysis using EB reproducible mode, 22 out of 23 biomarkers reported as indicators of *T*rypanosoma cruzi infection and Chagas disease was determined to be significant, replicating the targeted Skyline^30^ analysis in the original paper. Out of those 22 biomarkers, 18 biomarkers showed a clean one-to-one match to the validated biomarkers within *m/z* tolerance of 10 ppm and RT tolerance of 0.2 minute (**SI Table 3**).

Among five biomarkers that did not show a one-to-one match, one biomarker (*m/z* 272.1488, RT=3.77) was matched to two features in the reanalysis within the *m/z* and RT tolerance. The feature, found at RT=3.78, showed an increasing pattern in week 2 samples which is opposite to the literature reported fold-change direction, but the isomer at RT=3.85 reproduced the decreasing pattern in both female and male week 2 samples (**Figure 7**). In the original study, Skyline did not distinguish between these two isomers, and no significant time point was reported in female samples. EB reanalysis reported the isomers into two separate features and revealed the decreasing pattern of the isomer at RT=3.85 in both female and male week 2 samples. Two biomarkers (*m/z* 343.222, RT=3.64 and *m/z* 497.1334, RT=3.1) lost the increasing pattern reported in the original study. One of these biomarkers (*m/z* 343.222, RT=3.64) showed an increasing pattern in week 5 female samples, concurring with original findings (*p*-value=0.001), but the significance did not pass Benjamini-Hochberg (BH) correction (**SI Figure 4**). In the other biomarker (*m/z* 497.1334, RT=3.1), an increasing trend was detected in week 3 female samples (concurring with original manuscript findings, *p*-value=0.011) as well, but it did not survive BH correction. One biomarker (*m/z* 134.06, RT=1.83) was reported with mixed directions in male samples, increasing in week 1 and decreasing in week 10. The EB reanalysis showed decreasing patterns in both week 1 (*p*-value=0.026) and week 10 (*p*-value=0.031) male samples, although both were not significant after BH correction. We examined the XICs of the feature matched to the biomarker and confirmed that the majority of week 1 infected male samples exhibited a decreasing pattern in the untargete d dataset, opposed to the original targeted analysis (**SI Figure 5a**). One biomarker (*m/z* 411.1543, RT=3.53) was reported with mixed direction in female samples by the original work, increasing in week 5 and decreasing in week 10 and17 samples. In the EB reanalysis of the untargeted data, the feature was only detected in three out of 16 weeks 5 infected samples, with no signal in the MS1 for any other untargeted sample leading to the flat fold-change (**SI Figure 5b**).

**Figure 7.**
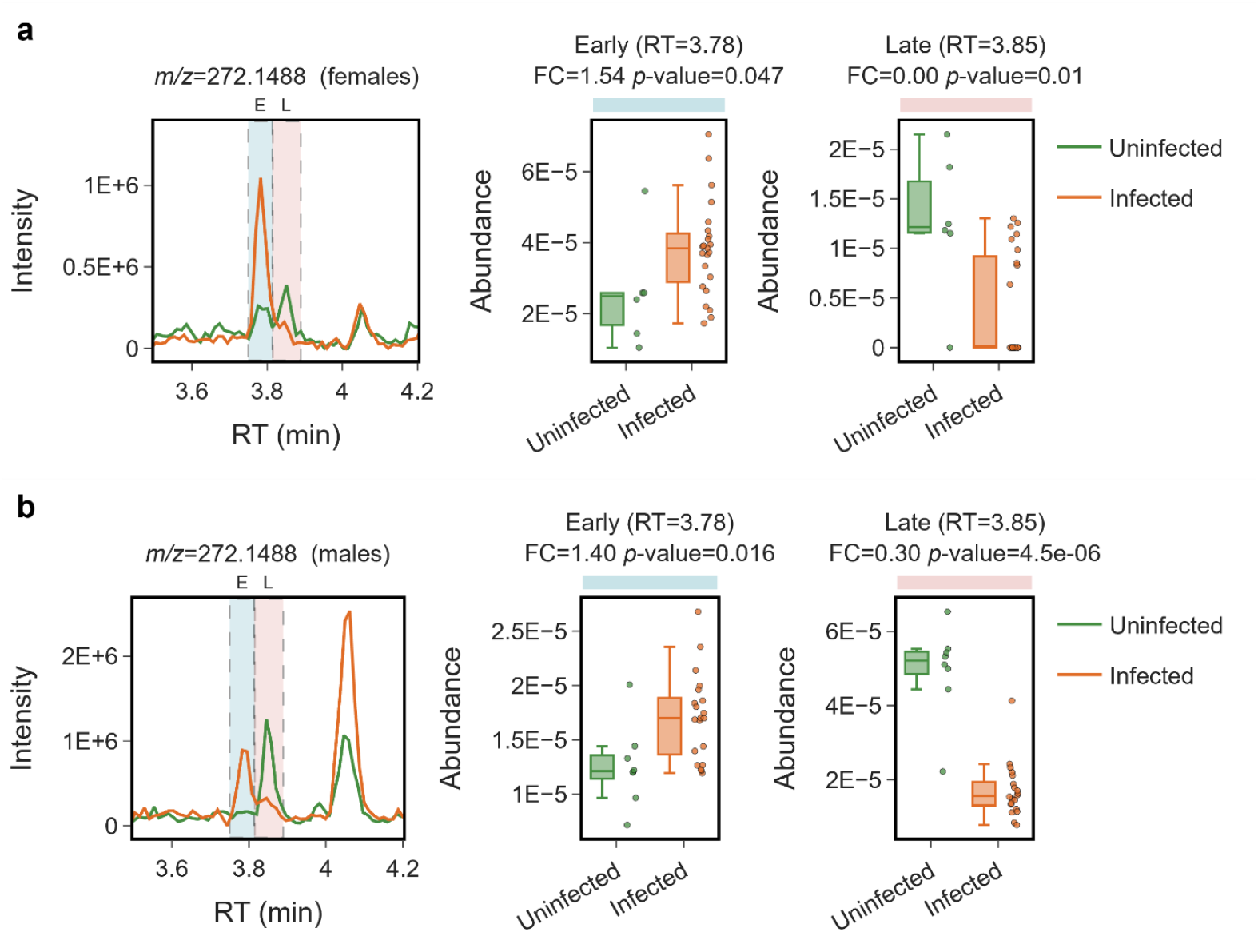
A biomarker (*m/z* 272.1488, RT=3.77) split into two features in the reanalysis, **(a)** week 2 females, **(b)** week 2 males. The XICs show an early peak detected at RT=3.78 (shown in the blue shaded area) and a late isomer detected at RT=3.85 (shown in the pink shaded area). The fold-change direction of the late isomer conforms with the original study. FC: fold-change. E: early. L: late.

We hypothesized that EB was fundamentally limited by the sensitivity of untargeted metabolomics data. MS1 signals that would have indicated the presence of the analyte were absent in most samples from the untargeted metabolomics data and were only observed in a higher sensitivity targeted method. The main gap between the published work and our EB reanalysis is that the original study performed the validation through the targeted analysis on a diagnostic fragment, rather than MS1 peak area of the precursor over the untargeted dataset. In the vast majority of the biomarkers reported, EB was able to resolve the isomers in the untargeted LC-MS/MS data, EB was able to recover the biomarker pattern in the original publication (**Figure 7b**).

### Scalability on collaboration and public datasets

While we demonstrate analysis and benchmarking on a modest sample size (hundreds of samples), in collaboration with industry partners, we have scaled EB Feature Finding to over 17K LC-MS samples in an industrial setting. EB was able to complete the analysis within 10 CPU-Hours and ∼100GB of memory for a diverse natural product collection in Corteva Agriscience. Additionally, on open data, we have re-analyzed a 2900 sample dataset (MSV000094173) that originally required over a day of parameter tweaking and runtime with MZmine^9^, that now completes in 12 minutes with EB. We anticipate the fully automated nature of EB combined with the scalable algorithms presented in this manuscript will enable feature finding analysis from a few samples at instrument computers all the way up to industrial sized datasets for the community.

## Discussion

This work demonstrates that feature finding with EB can be substantially accelerated. In the benchmarking datasets in this manuscript, we demonstrated significant speed improvements on datasets in the hundreds of samples.

Additionally, while manual parameter tuning is often recommended in other feature finding methods, EB aims to be a fully automated feature finder that determines the parameters solely based on the input dataset. Taken together, these features of EB alleviate the time-intensive manual effort of parameter optimization and the computational overhead of feature finding in untargeted metabolomics studies. During the development of EB, we found that the early and substantive feedback from the community significantly enhanced the performance and robustness of the tool. This dovetailed with rapid algorithmic iterations to improve the tool and keep the community testers engaged. At the end, this feedback combined with algorithmic improvements led to significantly accelerating the speed for EB feature finding.

We evaluated feature finding methods on their spike-in standard detection coverage (sensitivity), dilution-series quantification accuracy, and yeast ^12^C/^13^C credentialed feature detection (sensitivity on biological matrix). EB showed high sensitivity and quantification accuracy in spike-in standards detection and high sensitivity in yeast ^12^C/^13^C credentialed analytes across all datasets in the benchmark, demonstrating its robustness across these experimental settings. However, we acknowledge that these datasets might not encompass the full spectrum of sample types and experimental conditions tha t the untargeted metabolomics community generates. To enhance robustness, we collaborated and tested EB across a wide cross-section of the community. This included different instrument manufacturers and models, sample matrices, and chromatography/instrument methods. As noted above, we found this feedback invaluable to improve EB.

One of the limitations of the EB FF is that it currently only supports centroided LC-MS and data dependent LC - MS/MS datasets and does not support GC-MS data. Further, more complicated data acquisition strategies such as polarity switching, MS^n^ ^31^, mass spectrometry coupled with ion mobility spectrometry, and acquisition using different columns or varying LC gradient, and data-independent MS/MS acquisition are not supported. However, these are exciting directions that could build upon the modular framework of the EB FF. Finally, we want to acknowledge that feature finding is only one step of the data analysis pipeline for untargeted metabolomics. To ensure compatibility with downstream computational tools for annotation, visualization, molecular networking, and statistical analysis we have ensured that the data file formats are in line with existing feature finding tools in the field.

## Software/Code Availability

The Everything Bagel feature finder is provided as an open - source software - DOI in Zenodo upon publication. The software is command-line executable in Linux and intended for computationally demanding data processing on HPC clusters.

A graphical user interface coupled for EB is also accessible through a desktop application for offline environments - DOI in Zenodo upon publication.

A web application that is ∼30x computationally slower than the Linux or desktop application but does not require local installation or data transfer is available at featurefinding.gnps2.org

A Nextflow workflow built in the GNPS2 infrastructure (https://gnps2.org/workflowinput?workflowname=everything_bagel_workflow) is publicly available to GNPS2 users.

The GNPS2 workflow provides easier access to computational resources and streamlined downstream analysis for large scale feature finding in the tens of thousands of sample sized datasets.

## Acknowledgements

We acknowledge Kambria Philips, Tal Luzatto, and Madeleine Ernst for her early testing of our software. This paper was typeset with the bioRxiv word template by @Chrelli: www.github.com/chrelli/bioRxiv-word-template.

## Author contributions

MW conceptualized the project, supervised, and wrote the manuscript. YS developed computational approaches and wrote the manuscript. YE supervised, benchmarked, evaluated, and edited the manuscript. BPB, PEA, AKJ, PCD: Validation (benchmarking and evaluation of the software). TRN, LIM, CXW, VCL, HNZ, PEA, AHD, CDB, XD, NG, ATA, AKJ, XC: software evaluation and manuscript editing. VVP conceptualization, software evaluation, and manuscript editing. MS evaluation, software development, and manuscript editing. EER provided data and edited the manuscript.

## Materials and Methods

### Software Architecture and supported inputs

The Everything Bagel feature finder (EB) is a Rust^16^-based software for automated preprocessing of LC-MS and LC-MS/MS based untargeted metabolomics dataset that integrates parameter optimization, feature detection, alignment, and feature grouping. EB supports centroid MS dataset in mzML^17^ or mzXML^18^ formats and handles datasets acquired using a wide range of instrumentation and methods. Within a single dataset, the same instrumentation and methodology (e.g. ionization polarity, chromatographic method, column, acquisition settings, etc.) must be used to generate a valid result. EB supports datasets consisting of MS1-only samples paired with MS/MS acquisition on pooled quality controls^19^.

### Automated parameter optimization

Parameter optimization in EB is based on instrumentation description in the mzML metadata. The instrument type is extracted from the analyzer and instrument configuration section of the first file in the input dataset.

#### *m/z* tolerance

The *m/z* tolerance is selected based on the extracted instrument type: 10 ppm for Orbitrap instruments, 25 ppm for Q-ToF instruments, 0.2 Da for ion-trap instruments, and 25 ppm default when the type of instrument cannot be determined. An *m/z* tolerance of 0.5 Da is used in alignment and feature grouping for ion-trap instruments. For high resolution data (Orbitrap and Q-ToF), the *m/z* tolerance is clamped to a minimum absolute value, 0.002 Da in region-of-interest (ROI) extraction and 0.005 Da in alignment and feature grouping.

#### MS1 noise intensity threshold

The MS1 noise intensity threshold is estimated using MS1 scans sampled from a maximum of 30 files. Files are sorted lexicographically by filename and sampled at approximately even intervals across the dataset. Within each selected file, the first MS1 scan from every 10 consecutive scans is retained. The first and final 50 MS1 scans are ignored to mitigate the influence of column-void and end-of-run signals. For Orbitrap data, the noise threshold is computed as the 75% quantile intensity across all MS1 peaks in the sampled scans. For Q-ToF data, all MS1 peaks in the chosen scans are log10 transformed, and any peak with intensity values under 0.1% and over 99.9% quantile is removed from the distribution. The intensities are histogrammed into 256 bins, and the resulting distribution is smoothed by gaussian filtering with σ=4. The nearest local maximum from the median value is selected as the apex. From this local maximum, the higher of the half-maximum intensities of this apex peak is set as the noise threshold. Any MS1 peak intensity below the noise threshold is considered to be noise in the data and is not considered as part of feature finding.

### Feature detection

Feature detection begins with a region of interest (ROI) from MS1 signals within each file. An ROI is a sequence of MS1 peaks *p* = (*m*_1_, *p*_1_), …, (*m*_*n*_, *p*_*n*_), where *m*_*i*_ is the m/z of the peak and *p*_*i*_ is the intensity of the peak, with maximum intensity of *p*_k_. Each MS1 scan is processed in the acquisition order and peaks within each scan are processed in descending order of intensity. Each peak joins an ROI at the closest *m/z* within the *m/z* tolerance and assignment occurs in a one-to-one manner. If a peak from the same scan already occupies the ROI, the next closest ROI is considered. A new ROI is seeded when a peak is not matched to any available ROIs. An ROI remains active until it reaches > 10 consecutive MS1 scans with no matching peak. After extension is complete, an ROI is removed from the analysis if *p*_k_ is smaller than 5 times the noise intensity threshold. When the noise intensity threshold is set to zero, an ROI is dropped if either *p*_1_ > 0.2 × *p*_k_ or *p*_*n*_ > 0.2 × *p*_k_. A noise score is calculated by the relative sum of total variance across the ROI compared to the maximum intensity.

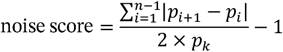

An ROI is not segmented if (1) the ROI includes less than 5 consecutive scans, (2) the ROI is shorter than or equal to 60 seconds and its noise score is greater than 3.0, (3) the ROI is longer than 60 seconds and either its median intensity is greater than 40% of its maximum intensity or its noise score is greater than 20.0, (4) the mean relative residual of the raw ROI compared to its gaussian-filtered signal (σ=6) is less than 0.15, or (5) the mean relative residual of the raw ROI compared to its σ=0.6 ga ussian-filtered signal is greater than 0.15. Otherwise, the ROI is processed through iterative segmentation to produce gaussian-shaped peaks. The desired prominence of the ROI is 0.4 × *p*_k_ in the first iteration. In the second iteration, the desired prominence is tuned to 0.03 × *p*_k_ if noise score of the ROI is below 0.1, otherwise 0.15 × *p*_k_. The gaussian-filtered peak intensities with σ= 0.6 are computed for the ROI, *p*′ = *p*′_1_, …, *p*′_*n*_. A local maximum *p*′_*t*_ is collected if *p*′_*t*_ *> p*′_*t−*1_, *p*′_*t*_ *> p*′_*t+*1_ and *p*′_*t*_ *>* 0.2 × *p*_k_. The prominence of a local maximum *p*′_*t*_ is *prom* = 1 *−* (*p*′_*v*_/*p*′_*t*_) where *p*′_*v*_ is the valley on either the left or the right side of the local maximum. The left valley is defined as the minimum value between *p*′_*t*_ and its adjacent local maximum, *p*′_*l*_, where l < t. If there is no such *p*′_*l*_, *p*′_*v*_ = *min*(*p*′_1_ … *p*′_*t*_). The right valley is defined as the minimum value between *p*′_*t*_ and its adjacent local maximum, *p*′_*r*_, where t < r. If there is no such *p*′_*r*_, *p*′_*v*_ = *min*(*p*′_*t*_ … *p*′_*n*_). The overall prominence of a local maximum *p*′_*t*_ is *min*(*prom*_*l*_, *prom*_*r*_). The set of local maxima detected in an ROI is pruned by removing local maxima satisfying any of the following conditions: (1) the overall prominence of the local maximum is below the desired prominence, or (2) the overall prominence of the local maximum is below 5 × the maximum overall prominence of surrounding local maxima within 10 scans window.

A surviving local maximum is extracted as a feature, and its gaussian similarity is computed as a Pearson correlation coefficient between raw intensity *p* and a bi-gaussian fitted intensity 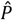.

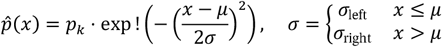

The full width at half maximum (FWHM) is computed by walking away from the apex until it reaches either an adjacent valley or the end of the ROI. The FWHM is obtained separately on the left and the right side to handle asymmetric peak shape. The signal-to-noise ratio (SNR) is computed using the following formula, where *p*_*off−pea*k_ represents peak intensities outside the inclusive FWHM interval, [*apex − FWHM*_*left*_, *apex + FWHM*_*right*_].

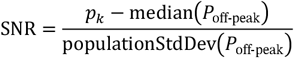

All features are tested against the following gates and removed if any of the gate is not satisfied: gaussian similarity ≥ 0.7, noise score ≤ 2, number of scans ≥ 6, mean relative residual of the raw ROI compared to its σ=0.6 gaussian filtered signal ≤ 0.15, SNR ≥ 3.0, and number of scans equal to or above half max-intensity > 1.

### MS/MS assignment

Each MS/MS scan is processed sequentially and assigned to a detected feature using precursor *m/z* and retention time. EB first identifies candidate features whose *m/z* values match the reported precursor *m/z* of the MS/MS scan within the instrument-specific mass tolerance. If no selected ion *m/z* is provided in the MS/MS scan, the *m/z* search is expanded to the precursor isolation window of the MS/MS scan. Among these candidate features, the MS/MS scan is assigned to the highest intensity feature whose chromatographic boundaries contain the scan acquisition time. If none of the candidate features contain the scan time, the MS/MS scan is assigned to the highest intensity candidate feature within 0.1 minutes RT tolerance.

### Linear RT correction

EB corrects retention time variation across files automatically by identifying high-confidence features as alignment anchors and mapping each file into a common retention time space. Candidate anchors are selected within each file based on four criteria: a low noise score (noise score < 0.1), a relative high intensity (intensity > 75% quantile of all features detected in the file), a narrow shape (FWHM < the median FWHM of all features in the file), and an isolated *m/z* value that has no confounding features within the instrument-specific *m/z* tolerance. The anchor coverage of each file is computed as the percentage of 0.5-minute RT bins that have at least one anchor. The file with the highest anchor coverage is selected as the RT correction reference.

For each non-reference file, a feature is matched to a reference anchor within instrument-specific *m/z* tolerance and an RT tolerance of 0.5 minutes. A match is retained only when the hit is unique to the anchor. The RT gap between anchors and matched features is computed, and outlier anchor-feature matches are dropped from the correction (median absolute deviation k>3.5). The pairs of adjacent control points are iteratively scanned until all non-monotonic corrections are resolved. A pair of points with an inversion of less than 1.8 seconds are averaged into one control point. A pair of points with a larger inversion are both dropped. The resolved control points are smoothed using a density-adaptive k=5 nearest median. The control points that escape the original RT range are removed, and identity anchors are inserted at (0, 0) and (max rt, max rt). The RT correction model is computed as linear transformation between control points.

### Alignment

Following file-specific retention time correction, all detected features are mapped into a consensus RT space using the file-specific RT correction model and pooled together into a descending intensity order. The features are sequentially aligned into consensus clusters, where each cluster represents a set of features, *c* = *f*_1_, …, *f*_k_. Each feature searches for clusters within *m/z* tolerance and RT tolerance of 0.2 minutes. The cluster *m/z* and RT are computed as the weighted sum of its members, and the distance is computed as 2 × (*Δm*/*z*) / (*m*/*z* tolerance*) +* (*Δ*RT) / (RT *tolerance)* . The feature joins the matched cluster if (1) no member of the cluster is from the same file of the feature, (2) its raw RT is in between the raw RT of the nearest monotonicity guards with 3 seconds tolerance, and (3) its consensus RT bounds overlap with the cluster’s RT bounds. When multiple clusters satisfy the conditions, the cluster with the smallest distance is selected. The monotonicity guards are pairs of per-file raw RTs and consensus RTs, *g* = (*raw*_1_, *con*_1_), …, (*raw*_*m*_, *con*_*m*_). A feature with a narrow (FWHM < 3 seconds) and clean gaussian shape (gaussian similarity > 0.9) is collected as a monotonicity guard when it merges with a consensus feature cluster. During clustering, each feature’s raw RT, *raw*_*f*_, is tested against its adjacent monotonicity guards, (*raw*_*l*_, *con*_*l*_) and (*raw*_*r*_, *con*_*r*_), where *raw*_*l*_ < *raw*_*f*_ and *raw*_*r*_ *> raw*_*f*_, to ensure that the candidate cluster consensus RT, *con*_*f*_, satisfies *con*_*l*_ *−* (3 seconds) < *con*_*f*_ < *con*_*r*_ + (3 seconds). If not, the merge is rejected.

A feature that matches no existing cluster forms its own cluster. Once all pooled features are processed, clusters lying within the instrument-specific *m/z* tolerance and an RT tolerance of 0.2 minutes are grouped into a family *C* = *c*_1_ … *c*_*k*_ . Each pair of clusters (*c*_*a*_, *c*_*b*_) in the family, with its member features *c*_*a*_ = *f*_*a*1_, … and *c*_*b*_ = *f*_*b*1_, …, is evaluated to determine whether the pair should remain segmented or be merged at the alignment level. The pair is merged if (1) the members of *c*_*a*_ and *c*_*b*_ are jointly detected in fewer than two files, or (2) the number of files in which a member of both *c*_*a*_ and *c*_*b*_ is detected is below 0.05 × the number of files in which either is detected. When the pair passes both criteria, relative prominence is collected from each file in which the two clusters are both detected and a member of *c*_*a*_ and *c*_*b*_ is adjacent to each other without any other feature detected in between. The relative prominence is computed as a maximum of the right prominence of the early peak and the left prominence of the late peak. The segmentation between *c*_*a*_ and *c*_*b*_ is accepted when the median of relative prominence across collected files > 0.2.

In the pooled set of clusters, a cluster is considered isolated when its consensus RT bounds do not overlap with any other clusters’ RT bounds or when its members are not detected in any other clusters and its consensus FWHM does not overlap with any other clusters’ FWHM in the set. The consensus FWHM was considered to complement the strict no overlap in RT bounds criterion. In the pooled set of clusters, all features that are not assigned to a segmented or isolated cluster are merged into its closest segmented or isolated cluster based o n the consensus RT distance. After reclustering, all features assigned to a segmented or isolated cluster are reassigned to its closest isolated or segmented cluster for further refinement. Refined clusters detected in the number of files larger than the detection threshold are extracted into consensus features. Each consensus feature *m/z* and retention time is assigned from its maximum intensity member. The RT bounds are computed as the median of consensus RT bounds of its members.

A consensus feature is retained based on the minimum detection rate or minimum detection count threshold across runs. The default threshold is a minimum detection rate of 10%. A consensus feature is retained if its detection rate is equal to or greater than the threshold. Otherwise, a feature is removed from the analysis.

### Gap-filling

After the alignment step, features detected in each file are grouped into consensus features across runs. Gap-filling is performed to estimate the peak area of consensus features in the files where no corresponding per-file feature was linked to a given consensus feature. For each consensus feature, intensity values for all files are checked for missingness and filled if a value is missing. The value is computed from the extracted ion chromatogram (XIC) of the file in which the value is missing at consensus *m/z* and RT bounds. The consensus RT bounds are mapped into the original RT coordinates of the file using the file-specific correction model. Signals below the estimated MS1 noise threshold are excluded, and the remaining chromatographic signals are used to es timate the peak area.

### Feature grouping

Consensus features are grouped into feature groups through isotope, adduct, and in-source fragmentation (ISF) calling. The per-file coelution graph that connects consensus features detected within a 3 second RT tolerance is built for each file. Each featur e in a connected component is enumerated to search for possible isotope chain matches within the connected component using z=1…3 and isotope count=1…5. The isotope chain match is accepted if (1) both features are detected in at least two files, (2) the pai r is connected and have intensity ratio within [0.002, 1.5 ×(*m/z*/13)×0.011×isotope count]^20,21^ in at least 75% of the files in which both are detected. A set of disjoint isotope chains that maximizes total explained intensity is selected via branch -and-bound with greedy upper bound estimation.

Each consensus feature assigned as a native ion is tested on the set of canonical primary adducts ([M+H] ^1+^, [M+Na] ^1+^, [M+K] ^1+^, [M+NH3+H] ^1+^, [M+2H] ^2+^, [M+3H] ^3+^ for positive ion mode, [M-H] ^1-^, [M+Cl] ^1-^, [M+HCOO] ^1-^, [M-2H] ^2-^, [M-3H] ^3-^ for negative ion mode)^22,23^ and enumerated to search for consensus features matched within the known *m/z* difference between the candidate primary adduct and other adduct forms within *m/z* tolerance. The adduct match is accepted when (1) both features are detected in at least two files, (2) the pair has the XIC correlation score > 0.7 and is connected in at least 75% of files in which both are detected. The XIC correlation score was computed using Pearson correlation coefficient on a pair of XICs, *p*_*a*_ = *p*_*a*1_, …, *p*_*an*_ and *p*_*b*_ = *p*_*b*1_, … *p*_*bm*_, trimmed on the shared scan range of both features. A set of disjoint adduct chains that maximize the sum of feature intensity weighted by the canonical adduct classes (**SI Table 4**) is selected via greedy disjoint selection.

ISF is evaluated among consensus features assigned as native ions.Each consensus feature is paired with another consensus feature from a different adduct group within 0.1 minutes of the consensus RT and an *m/z* difference > 0.1 Da. The lower *m/z* feature is considered as the candidate fragment, and the higher *m/z* is considered as the candidate source. The ISF is identified if (1) the product ion *m/z* (i.e. fragment) appears in the precursor ion MS/MS spectrum within *m/z* tolerance, (2) both features are detected in at least two files, and (3) a pair coelutes within a 1.5 second RT tolerance in at least 75% of files in which both were detected.

### Tool comparisons

Untargeted feature finding was performed using XCMS-IPO, MassCube, and EB to compare computational cost and feature-detection performance across the benchmarking datasets. We used XCMS (v4.10.0; Bioconductor 3.23, R 4.6) in a containerized R environment with the xcmsSet interface. For each dataset, centWave peak detection parameters were tuned by Isotopologue Parameter Optimization (IPO v1.38.0). We optimized over ppm and peak width on three representative QC files, with ppm range set by instrument (3-12 ppm for Orbitrap, 5-40 ppm for Q-ToF) and the remaining parameters fixed (snthresh=10, noise=10 00, prefiler=3/1000, mzdiff=-0.001). For grouping, we used parameters equivalent to EB (minfrac=0.1, mzwid=0.01). The remaining parameters were fixed to the package default. We used the untargeted metabolomics workflow in MassCube (v1.2.20), with tool defa ults for noise threshold (MS1 absolute threshold=50,000, MS2=10,000 for Orbitrap; MS1/MS2=1,000/500 for Q - ToF). The remaining parameters were fixed to the tool default (MS1 *m/z* tolerance=0.01 Da, alignment *m/z* tolerance=0.01 Da and RT tolerance=0.2 min, minimum detection rate=0.1). We performed feature finding with EB with the package default parameters and three reproducibility modes: rare (minimum detection count=1), reproducible (minimum detection count=3), and standard (minimum detection rate=0.1) to compare the results across different settings. The EB standard mode has the most similar detection rate filtration compared to the MassCube.

### Metabolomics Workbench datasets

Untargeted metabolomics benchmark datasets (PR002464) were obtained from the Metabolomics Workbench^24^. The benchmark contains datasets acquired using multiple chromatographic methods, ionization modes, and mass-spectrometer platforms. The GC-MS/MS dataset (ST003934) was excluded because it was acquired using an experimental configuration not supported by EB. We used the remaining eight LC-MS/MS datasets for feature finding evaluation. Performance was evaluated based on spike-in standard detection coverage (sensitivity), dilution-series quantification accuracy (quantitative performance), and recovery of yeast ^12^C/^13^C credentialed features (sensitivity in biological analytes). In each dataset, pooled QC runs with data-independent acquisition (DIA) MS/MS spectra and conditioning runs were excluded from the analysis.

### Spike-in standard spectral library

We created a dataset-specific MS/MS library of compounds spiked into PR002464. For the 96 structures spiked in, we searched NIST20 for identical planar InChIKeys^25^, with exception of 2’-epitaxol and paclitaxel, which used full InChiKeys to distinguish stereochemistry. If the structures were not found, we searched for the Global Natural Products Social Molecular Networking 2 (GNPS2)^26^ public MS/MS libraries using InChIKey as well. Out of 96 standards, a total number of 78 standards were covered in the NIST20 library. From the GNPS2 library, 10 standards were uniquely covered in addition to the NIST20 library, yielding MS/MS library for a total 88 structures of the 96 structures.

### Spike-in standard annotation and dilution series evaluation

A set of MS/MS spectra assigned to features in each tool result was annotated through the dataset-specific MS/MS library of spike-in compounds. The library search was performed on MS/MS spectra assigned to consensus features in each tool output using top 10 peaks in 50 Da window, fragment mass tolerance of 0.01 Da, minimum matched peak count of four, and cosine similarity threshold=0.7. The precursor mass tolerance of 0.02 Da was used for Orbitrap datasets, and 0.5 Da was used for Q-ToF datasets. To determine the upper limit of the spike-in standard detection, we performed library search on all MS/MS spectra in each dataset with the same parameters.

We further measured spike-in standards dilution-series accuracy across spike-in 1000 µL samples with different spike A solution and spike B solution ratios, denoted as spike A 1000 µL, spike A 750 µL:spike B 250 µL, spike A 500 µL:spike B 500 µL, spike A 250 µL:spike B 750 µL, and spike B 1000 µL. The mean intensity series of each feature across 5 mixture groups, *S* = [*s*_1_, *s*_2_, *s*_3_, *s*_4_, *s*_5_], was computed from the median value over replicates in each group. For each spike-in standard, we computed the expected abundance in each mixture using the abundance in a dominantly loaded mixture, either *s*_1_ or *s*_5_, and expected relative ratio based on the design table specifications (**SI Table 2**). Then, we computed the relative residual of measured abundance to the expected abundance in each mixture across all levels except for the dominant level, either *s*_1_ or *s*_5_. The spike-in standard features were tested for mean relative residual (∑_*i*_|*s*_*i*_ *− s*′_*i*_| / ∑_*i*_ *s*_*i*_) < 20% to determine the dilution linearity of the quantification.

### Commercial yeast extract dilution series evaluation

We utilized Cambridge Isotope Labs yeast extract (^12^C and^13^C) dilution samples for the quantitative evaluation of feature finding results. We extracted features that were detected in yeast ^12^C 1000 µL samples and showed mean yeast ^12^C 1000 µL sample intensity > 3 × mean process and solvent blank intensity, and mean yeast ^12^C 1000 µL sample intensity > mean plasma 1000 µL sample intensity. We extracted features using yeast^13^C 1000 µL samples with the same yeast sample dominance filters. Each retained yeast ^12^C feature was sorted in the intensity descending order and paired with yeast^13^C features at *m/z* difference of 1.003355 × number of carbons (between 3-60 and ≤ *m/z* / 12) within instrument-specific *m/z* tolerance (10 ppm for Orbitrap, 25 ppm for Q-ToF) and 0.1 minute RT tolerance^15^. The match is rejected if *mean*(*native*/*heavy*) ≤ 0.75 or *mean*(*native*/ *heavy*) ≥ 5. If multiple yeast^13^C features are matched, the closest feature to the estimated *m/z* and RT was selected. The raw intensity reported from the tool was used for evaluation without normalization.

We calculated the quantification accuracy of the 4-level yeast dilution series on yeast ^12^C/^13^C balanced samples. In each pair, the mean intensities of the yeast ^12^C native ion feature were calculated across replicates at each dilution level of yeast ^12^C/^13^C balanced dilution samples. The mean intensity series, *S* = [*s*_1_, *s*_2_, *s*_3_, *s*_4_], was compared to the expected intensity series, *S*′ = [*0*.1 × *s*_4_, 0.2 × *s*_4_, *0*.5 × *s*_4_, *s*_4_], and was evaluated upon the mean relative residual across all levels below the top level (*s*_4_). The mean relative residual was not computed if two or more samples did not have intensity measured in the top level. We collected yeast ^12^C native ion features that passed mean relative residual < 20% in XCMS, MassCube, and EB standard mode. The collected features were deduplicated using instrument-specific *m/z* tolerance (10 ppm for Orbitrap, 25 ppm for Q-ToF) and RT tolerance of 0.2 minutes. All matches within the tolerance were merged to handle duplicate and split detections. The deduplicated set of features was considered as the reference yeast analytes, which represents a set of yeast ^12^C native ion features that exhibited dilution linearity in at least one of the tool results.

### Reanalysis of a published biological dataset

We obtained an untargeted biomarker validation dataset from published work^27^ to validate that EB reproduces the validated biological signatures. We used EB reproducible mode with package default parameters for feature finding and followed the 3-fold blank removal and TIC normalization steps described in the original publication for statistical analysis. We performed differential abundance analysis between uninfected and infected samples for each week in which the samples were collected. We compared feature abundance with a two-sided Mann-Whitney test (minimum 3 samples required for each group) and converted *p*-values into Benjamini-Hochberg false discovery rate (*q*-values) across all features under analysis at the time point. A feature was called significant if a *q*-value < 0.05 in at least one time point was identified. The fold change direction was assigned accordingly to the median fold change at the time point.

## Supplementary Information

**SI Data File 1**. Detailed report on metabolomics benchmark results. This is an aggregate with an index to explore all unique comparisons.

**SI Data File 2**. Spike-in standards dilution series (ST004500).

**SI Data File 3**. Spike-in standards dilution series (ST004501).

**SI Data File 4**. Spike-in standards dilution series (ST004502).

**SI Data File 5**. Spike-in standards dilution series (ST004503).

**SI Data File 6**. Spike-in standards dilution series (ST004504).

**SI Data File 7**. Spike-in standards dilution series (ST004626).

**SI Data File 8**. Spike-in standards dilution series (ST004650).

**SI Data File 9**. Spike-in standards dilution series (ST004651).

**SI Data File 10**. The representative XICs of EB-unique features (ST004500).

**SI Data File 11**. The representative XICs of XCMS-unique features (ST004500).

**SI Data File 12**. The representative XICs of MassCube-unique features (ST004500).

**SI Data File 13**. The representative XICs of EB-unique features (ST004501).

**SI Data File 14**. The representative XICs of XCMS-unique features (ST004501).

**SI Data File 15**. The representative XICs of MassCube-unique features (ST004501).

**SI Data File 16**. The representative XICs of EB-unique features (ST004502).

**SI Data File 17**. The representative XICs of XCMS-unique features (ST004502).

**SI Data File 18**. The representative XICs of MassCube-unique features (ST004502).

**SI Data File 19**. The representative XICs of EB-unique features (ST004503).

**SI Data File 20**. The representative XICs of XCMS-unique features (ST004503).

**SI Data File 21**. The representative XICs of MassCube-unique features (ST004503).

**SI Data File 22**. The representative XICs of EB-unique features (ST004504).

**SI Data File 23**. The representative XICs of XCMS-unique features (ST004504).

**SI Data File 24**. The representative XICs of MassCube-unique features (ST004504).

**SI Data File 25**. The representative XICs of EB-unique features (ST004626).

**SI Data File 26**. The representative XICs of XCMS-unique features (ST004626).

**SI Data File 27**. The representative XICs of MassCube-unique features (ST004626).

**SI Data File 28**. The representative XICs of EB-unique features (ST004650).

**SI Data File 29**. The representative XICs of XCMS-unique features (ST004650).

**SI Data File 30**. The representative XICs of MassCube-unique features (ST004650).

**SI Data File 31**. The representative XICs of EB-unique features (ST004651).

**SI Data File 32**. The representative XICs of XCMS-unique features (ST004651).

**SI Data File 33**. The representative XICs of MassCube-unique features (ST004651).

**SI Data File 34**. The representative XICs of EB-unique reference yeast analyte features.

**SI Data File 35**. The representative XICs of XCMS-unique reference yeast analyte features.

**SI Data File 36**. The representative XICs of MassCube-unique reference yeast analyte features.

**SI Table 1**. 96 spike-in detection *m/z*, RT, and library annotation.

**SI Table 2**. Spike-in standards dilution series design.

**SI Table 3**. Validated biomarkers reproduction. The EB reanalysis results that matched the reported results in the original publication are bolded. N/S: not significant, *: isomer at RT=3.85.

**SI Table 4**. The canonical adduct class weights assigned for the weighted intensity sum calculation.

**SI Table 5**. Spike-in standard library extracted from NIST20 and GNPS2 public libraries.

**SI Figure 1**. The XICs of spike-in standards that were not detected in EB.

**SI Figure 2**. The consensus features abundance across runs in the Orbitrap RP datasets.

**SI Figure 3**. Total features and yeast reference analytes with dilution linearity across tools in each dataset.

**SI Figure 4**. A validated biomarker (*m/z* 343.222, RT=3.64) reproduced in the EB reanalysis.

**SI Figure 5**. The XICs of two biomarkers with different fold change patterns compared to the original study.

