## Supplementary material for "The Everything Bagel Feature Finder: Ultra-fast automated feature finding for untargeted metabolomics": SI Figure

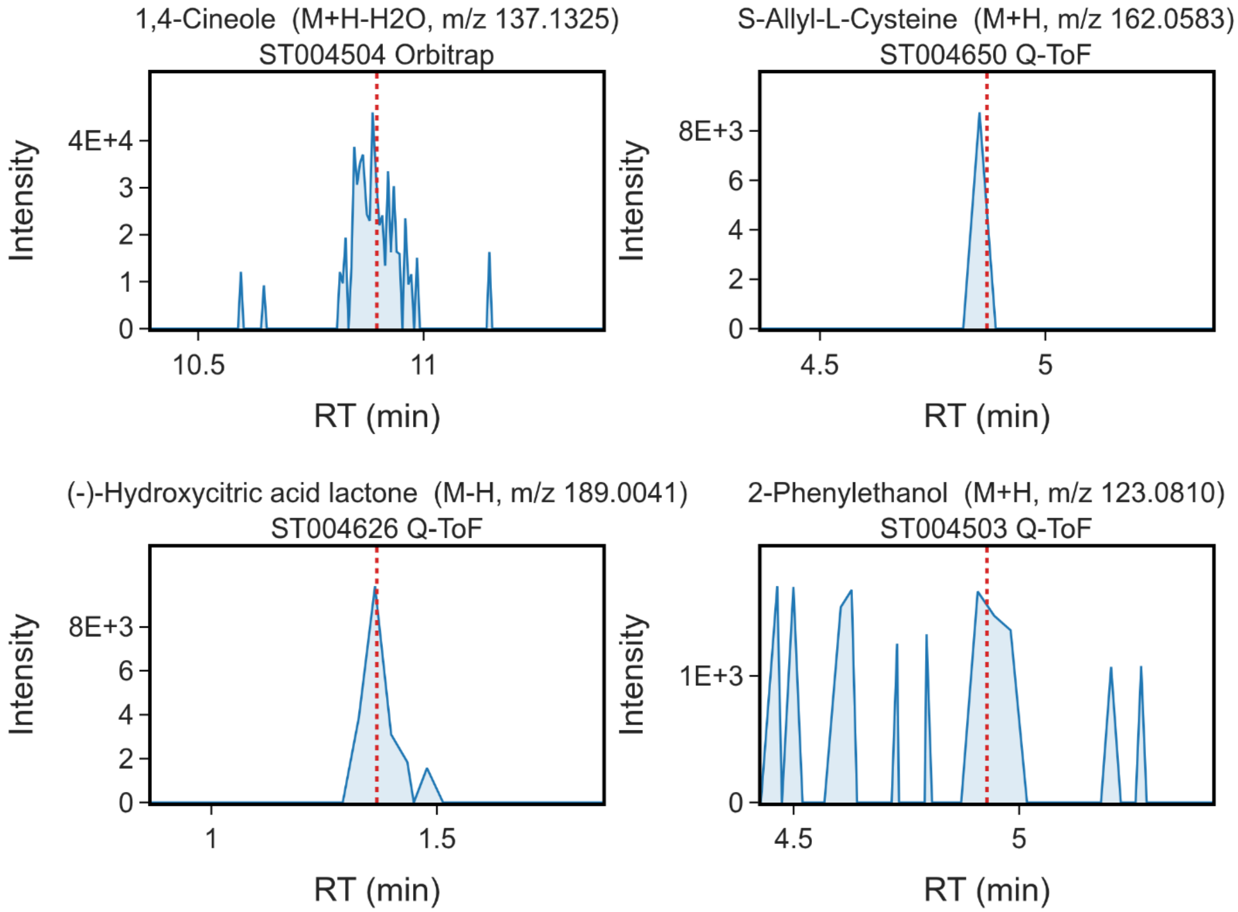


**SI Figure 1**. The XICs of spike-in standards that were not detected in EB. The red dash line indicates the RT of MS/MS acquisition. The XICs are extracted from the source file of library annotated MS/MS scans on library *m/z* value with instrument-specific *m/z* tolerance (10 ppm for Orbitrap, 25 ppm for Q-ToF) with the minimum threshold of 0.002 Da.

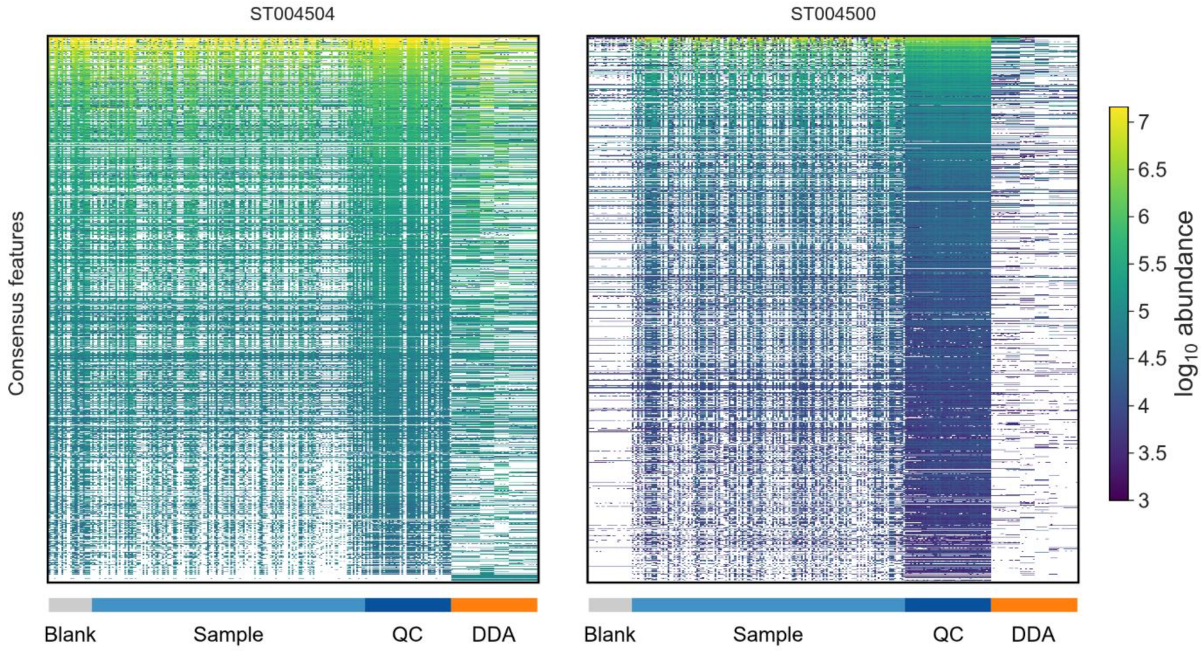


**SI Figure 2**. The consensus features abundance across runs in the Orbitrap RP datasets. Each row represents a consensus feature found in each dataset using EB standard mode (3,594 features in ST004504 and 10,555 features in ST004500). Each column represents an LC-MS/MS run in the dataset, grouped by their sample type (Blank: blank samples, Sample: MS1 acquisition samples, QC: pooled quality control samples, DDA: data-dependent MS/MS acquisition samples).

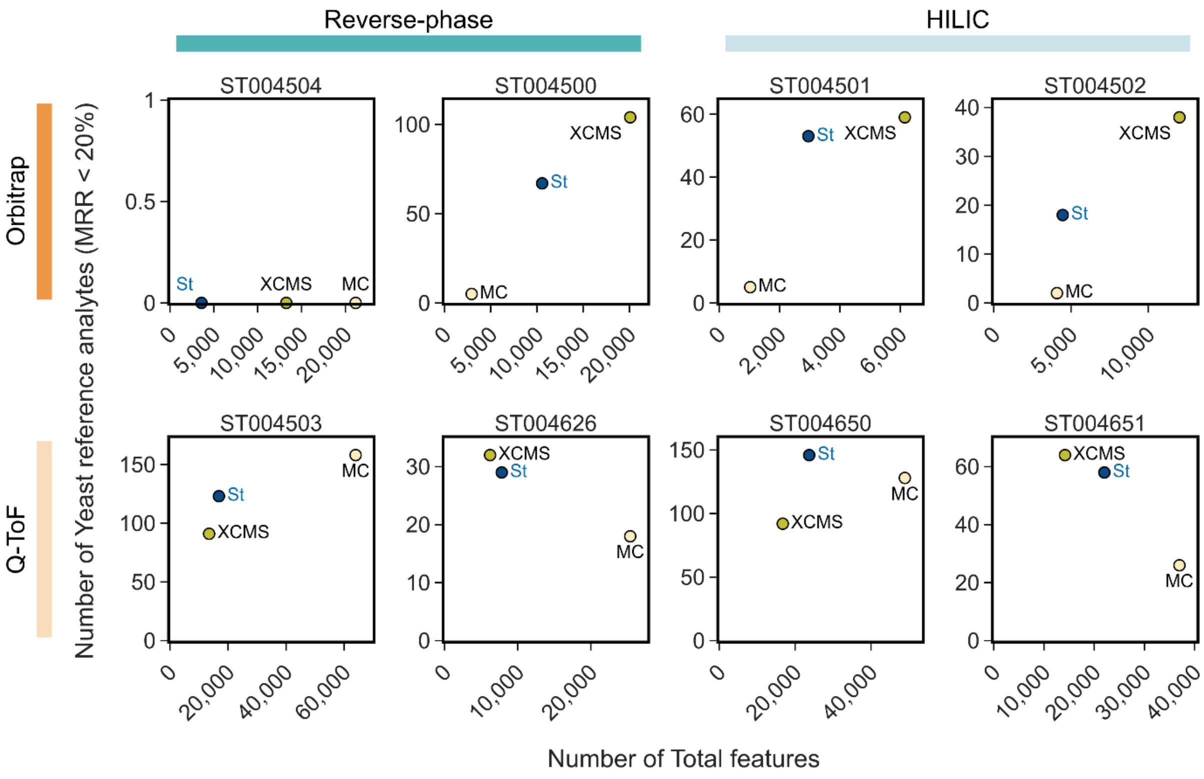


**SI Figure 3**. Total features and yeast reference analytes with dilution linearity across tools in each dataset. Each data point represents the number of yeast reference analytes with dilution linearity (mean relative residual < 20%) compared to the total feature detections in each tool. A data point towards the upper left direction indicates more yeast reference analyte detection with less total number of features. MRR: mean relative residual.

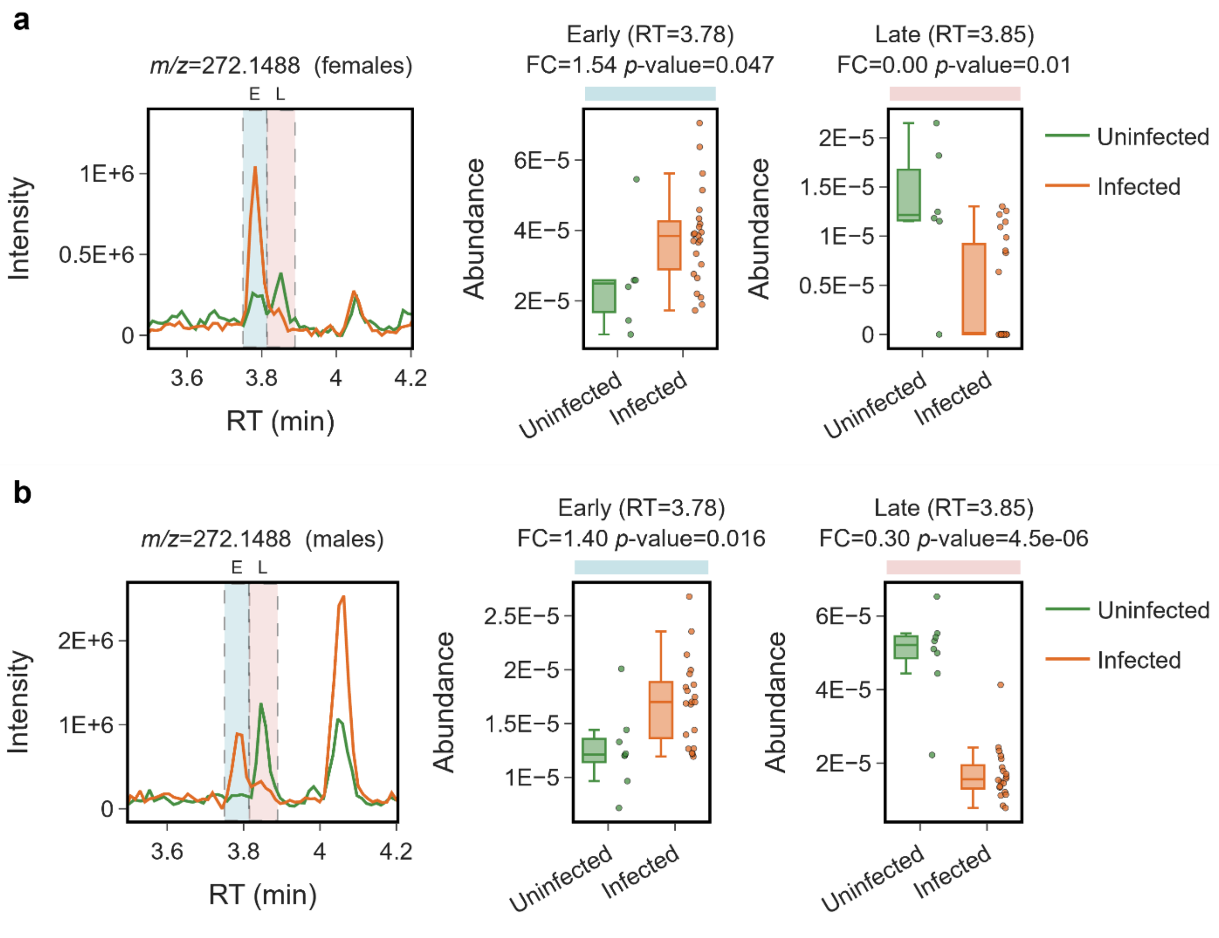


**SI Figure 4**. A validated biomarker (*m/z* 343.222, RT=3.64) reproduced in the EB reanalysis. The biomarker showed an increasing pattern in week 5 female infected samples and week 4, 5 male samples in the original study (*q*-value < 0.05). In the reanalysis, none of the groups showed *q*-value < 0.05 (*p*-value=0.001 in week 5 females, *p*-value=0.171 in week 4 males, and *p*-value=0.150 in week 5 males).

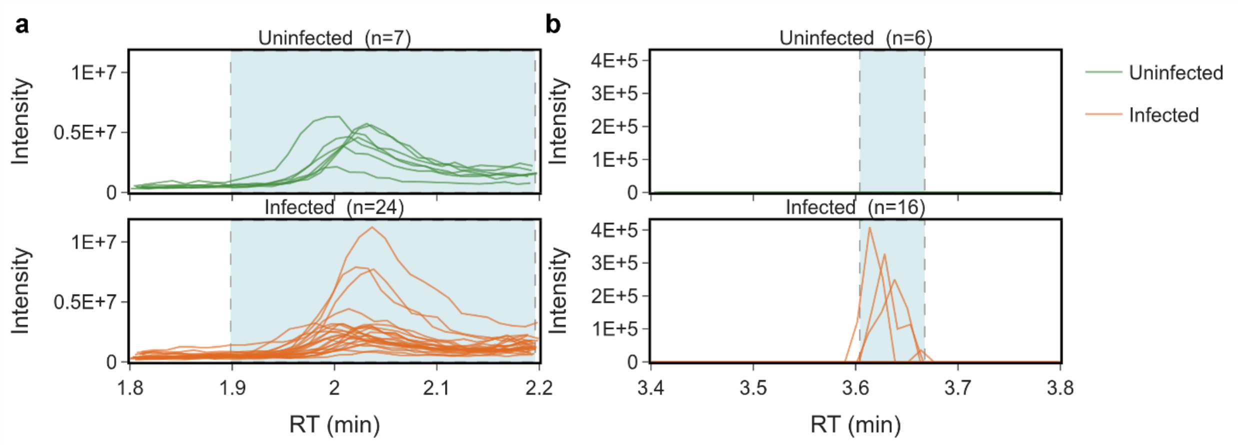


**SI Figure 5**. The XICs of two biomarkers with different fold change patterns compared to the original study. The XICs are extracted with instrument-specific *m/z* tolerance (10 ppm for Orbitrap, 25 ppm for Q-ToF) with the minimum threshold of 0.002 Da. The yellow shade represents the RT boundaries of each feature. **(a)** The biomarker (*m/z* 134.06, RT=1.83) showed a decreasing pattern in week 1 male samples, opposite of the original work. **(b)** The biomarker (*m/z* 411.1543, RT=3.53) was only detected in three infected samples, resulting in a flat fold-change in the reanalysis.
