## Supplementary material for "The Everything Bagel Feature Finder: Ultra-fast automated feature finding for untargeted metabolomics": SI Data File: index.html

Supplementary Data

### Supplementary Data

67 evidence pages comparing EB, XCMS and MassCube across PR002464 metabolomics benchmark.

Spike-in standards dilution series Tool-unique features Tool-unique native yeast analytes

Every page is a self-contained HTML file with inline SVG — no
network access needed. Some XIC reports are tens of MB and take a moment to render.

#### Spike-in standards dilution series

spikein\_boxplot\_report\_<dataset>.html

The abundance of features annotated as spike-in standards across sample groups in EB reproducible mode, XCMS, and MassCube. One representative feature for each spike-in standard was selected, either the one that has the lowest mean relative residual, or the one with the highest cosine similarity in library search if none of the annotated features passed mean relative residual < 20%.

| Dataset | Report |
| --- | --- |
| ST004500Orbitrap · RP · neg · 320 files | Supplementary Data 2408 KB |
| ST004501Orbitrap · HILIC · pos · 320 files | Supplementary Data 31.2 MB |
| ST004502Orbitrap · HILIC · neg · 320 files | Supplementary Data 4457 KB |
| ST004503Q-ToF · RP · pos · 324 files | Supplementary Data 51.2 MB |
| ST004504Orbitrap · RP · pos · 320 files | Supplementary Data 61.3 MB |
| ST004626Q-ToF · RP · neg · 374 files | Supplementary Data 7297 KB |
| ST004650Q-ToF · HILIC · pos · 372 files | Supplementary Data 81.1 MB |
| ST004651Q-ToF · HILIC · neg · 372 files | Supplementary Data 9397 KB |

#### Tool-unique features

tool\_unique\_xic\_<tool>\_<dataset>.html

The XICs of features detected by only one of the three tools across EB standard mode (min\_detection\_rate = 0.1), XCMS, and MassCube. Matching uses instrument-tuned m/z tolerance and an RT tolerance of 0.2 minute. Each XIC is drawn at the feature's m/z (± instrument ppm, floored at 0.002 Da for extraction only) from the raw mzML of that tool's highest-intensity detection.

| Dataset | EB | XCMS | MassCube |
| --- | --- | --- | --- |
| ST004500Orbitrap · RP · neg · 320 files | Supplementary Data 104.2 MB | Supplementary Data 1142.2 MB | Supplementary Data 121.3 MB |
| ST004501Orbitrap · HILIC · pos · 320 files | Supplementary Data 131.4 MB | Supplementary Data 1416.9 MB | Supplementary Data 15731 KB |
| ST004502Orbitrap · HILIC · neg · 320 files | Supplementary Data 162.5 MB | Supplementary Data 1738.2 MB | Supplementary Data 184.5 MB |
| ST004503Q-ToF · RP · pos · 324 files | Supplementary Data 1911.9 MB | Supplementary Data 202.0 MB | Supplementary Data 21145.6 MB |
| ST004504Orbitrap · RP · pos · 320 files | Supplementary Data 22407 KB | Supplementary Data 2341.8 MB | Supplementary Data 2447.7 MB |
| ST004626Q-ToF · RP · neg · 374 files | Supplementary Data 2510.4 MB | Supplementary Data 261.7 MB | Supplementary Data 2750.4 MB |
| ST004650Q-ToF · HILIC · pos · 372 files | Supplementary Data 2824.7 MB | Supplementary Data 293.5 MB | Supplementary Data 30108.8 MB |
| ST004651Q-ToF · HILIC · neg · 372 files | Supplementary Data 3123.1 MB | Supplementary Data 322.3 MB | Supplementary Data 3373.1 MB |

#### Tool-unique native yeast analytes

yeast\_native\_unique\_xic\_<tool>.html

The XICs of features matched to reference yeast analytes found in one tool that was not detected in the other tools. The feature with the highest abundance in yeast 12C 1000 µL samples is selected as the representative feature for each analyte. The XICs are extracted from the yeast 12C 1000 µL sample with the highest abundance.

| EB | XCMS | MassCube |
| --- | --- | --- |
| Supplementary Data 34640 KB | Supplementary Data 35833 KB | Supplementary Data 36465 KB |
