## Supplementary material for "The Everything Bagel Feature Finder: Ultra-fast automated feature finding for untargeted metabolomics": SI Data File: spikein_boxplot_report_ST004500.html

Spike-in standards dilution series - ST004500

### Spike-in standards dilution series - ST004500

12 standards · 32 tool detections

A1000A750:B250A500:B500A250:B750B1000|process.blanksolvent.blankplasma onlyyeast onlyplasma+yeast|expected, top levelexpected at each level

The y axis is linear and autoscaled per cell. **resid** = that feature's mean relative residual over the 5-level dilution series (pass = resid < 0.20). The **×** beside each box is the expectated abudance at the level, in the corresponding own colour.

| Standard | EB (reproducible mode) | XCMS | MassCube |
| --- | --- | --- | --- |
| #1 Acesulfame potassium  mix 12 · A:B 8:1 neutral mass 162.9939288  YGCFIWIQZPHFLU-UHFFFAOYSA-N | pass05e41e5 m/z 161.9867 · RT 3.35 min · cos 0.952 · id 287 · **resid 0.192** | fail05e61e7 m/z 161.9867 · RT 2.96 min · cos 0.951 · id 684 · **resid 267.597** | fail01e62e63e6 m/z 161.9866 · RT 2.98 min · cos 0.951 · id 332 · **resid 0.800** |
| #5 ALITAME HYDRATE  mix 14 · A:B 1:4 neutral mass 331.1565775  IVBOUFAWPCPFTQ-UHFFFAOYSA-N | pass02e64e6 m/z 330.1490 · RT 6.34 min · cos 0.899 · id 78 · **resid 0.071** | pass01e72e7 m/z 330.1490 · RT 6.34 min · cos 0.899 · id 3972 · **resid 0.161** | pass02e64e6 m/z 330.1490 · RT 6.32 min · cos 0.887 · id 95 · **resid 0.071** |
| #14 Baohuoside I  mix 5 · A:B 4:1 neutral mass 514.1838972  NGMYNFJANBHLKA-LVKFHIPRSA-N | pass02.5e65e67.5e6 m/z 513.1765 · RT 9.19 min · cos 0.907 · id 119 · **resid 0.106** | pass02e74e7 m/z 513.1764 · RT 9.19 min · cos 0.907 · id 8749 · **resid 0.116** | pass02.5e65e67.5e6 m/z 513.1764 · RT 9.17 min · cos 0.922 · id 145 · **resid 0.106** |
| #53 Lecanoric acid  mix 4 · A:B 8:1 neutral mass 318.0739528  HEMSJKZDHNSSEW-UHFFFAOYSA-N | pass02e64e66e6 m/z 317.0663 · RT 8.48 min · cos 0.949 · id 147 · **resid 0.102** | pass05e61e7 m/z 167.0350 · RT 8.90 min · cos 0.946 · id 769 · **resid 0.121** | pass02e64e66e6 m/z 317.0664 · RT 8.48 min · cos 0.936 · id 185 · **resid 0.102** |
| #54 Leonurine  mix 10 · A:B 1:2 neutral mass 311.1481208  WNGSUWLDMZFYNZ-UHFFFAOYSA-N | pass02e64e66e6 m/z 310.1406 · RT 5.34 min · cos 0.913 · id 83 · **resid 0.043** | pass01e72e73e7 m/z 310.1405 · RT 5.34 min · cos 0.913 · id 3550 · **resid 0.073** | pass02e64e66e6 m/z 310.1405 · RT 5.34 min · cos 0.888 · id 102 · **resid 0.043** |
| #62 N-Acetyl-DL-methionine  mix 6 · A:B 1:4 neutral mass 191.0616145  XUYPXLNMDZIRQH-UHFFFAOYSA-N | pass01e62e63e6 m/z 190.0544 · RT 4.89 min · cos 0.865 · id 153 · **resid 0.113** | fail02e64e66e6 m/z 190.0546 · RT 4.91 min · cos 0.865 · id 1168 · **resid 1.198** | pass01e62e63e6 m/z 190.0545 · RT 4.91 min · cos 0.885 · id 192 · **resid 0.113** |
| #63 NEOTAME MONOHYDRATE  mix 22 · A:B 1:4 neutral mass 378.2154721  HLIAVLHNDJUHFG-HOTGVXAUSA-N | pass05e61e7 m/z 377.2080 · RT 8.01 min · cos 0.887 · id 34 · **resid 0.111** | pass02e74e76e7 m/z 377.2079 · RT 8.02 min · cos 0.887 · id 5068 · **resid 0.098** | pass05e61e7 m/z 377.2080 · RT 8.00 min · cos 0.857 · id 36 · **resid 0.111** |
| #69 Oroxindin  mix 23 · A:B 1:16 neutral mass 460.1005615  LNOHXHDWGCMVCO-NTKSAMNMSA-N | fail01e52e5 m/z 459.0926 · RT 7.17 min · cos 0.724 · id 1895 · **resid 0.304** | fail05e51e61.5e6 m/z 459.0928 · RT 7.19 min · cos 0.724 · id 7278 · **resid 0.248** | not detected |
| #70 [(1R,2S,3R,4S,7R,9S,10S,12R,15R)-4,12-Diacetyloxy-15-[(2R,3S)-3-benzamido-2-hydroxy-3-phenylpropanoyl]oxy-1,9-dihydroxy-10,14,17,17-tetramethyl-11-oxo-6-oxatetracyclo[11.3.1.03,10.04,7]heptadec-13-en-2-yl] benzoate  mix 11 · A:B 1:8 neutral mass 853.3309553  RCINICONZNJXQF-WTEIXIHQSA-N | fail02e64e66e6 m/z 898.3292 · RT 9.11 min · cos 0.901 · id 55 · **resid 0.294** | fail01e72e73e7 m/z 898.3291 · RT 9.10 min · cos 0.901 · id 17833 · **resid 0.416** | not detected |
| #76 4-HYDROXYBENZOIC ACID PROPYLESTER  mix 6 · A:B 1:4 neutral mass 180.0786442  QELSKZZBTMNZEB-UHFFFAOYSA-N | pass02.5e55e57.5e5 m/z 179.0716 · RT 8.76 min · cos 0.724 · id 500 · **resid 0.179** | fail02e64e6 m/z 179.0714 · RT 8.76 min · cos 0.724 · id 974 · **resid 0.297** | pass02.5e55e57.5e5 m/z 179.0714 · RT 8.74 min · cos 0.980 · id 654 · **resid 0.179** |
| #81 Salidroside  mix 13 · A:B 4:1 neutral mass 300.120903  ILRCGYURZSFMEG-RKQHYHRCSA-N | pass01e52e53e5 m/z 299.1133 · RT 4.77 min · cos 0.921 · id 2021 · **resid 0.132** | pass05e51e6 m/z 299.1132 · RT 4.79 min · cos 0.921 · id 3335 · **resid 0.123** | not detected |
| #88 L(+)-Tartaric acid  mix 12 · A:B 8:1 neutral mass 150.0164379  FEWJPZIEWOKRBE-JCYAYHJZSA-N | fail02e54e5 m/z 149.0278 · RT 1.41 min · cos 0.943 · id 1881 · **resid 1.093** | fail05e51e61.5e6 m/z 149.0276 · RT 1.40 min · cos 0.943 · id 514 · **resid 1.483** | not detected |
