## Supplementary material for "The Everything Bagel Feature Finder: Ultra-fast automated feature finding for untargeted metabolomics": SI Data File: spikein_boxplot_report_ST004501.html

Spike-in standards dilution series - ST004501

### Spike-in standards dilution series - ST004501

36 standards · 93 tool detections

A1000A750:B250A500:B500A250:B750B1000|process.blanksolvent.blankplasma onlyyeast onlyplasma+yeast|expected, top levelexpected at each level

The y axis is linear and autoscaled per cell. **resid** = that feature's mean relative residual over the 5-level dilution series (pass = resid < 0.20). The **×** beside each box is the expectated abudance at the level, in the corresponding own colour.

| Standard | EB (reproducible mode) | XCMS | MassCube |
| --- | --- | --- | --- |
| #5 ALITAME HYDRATE  mix 14 · A:B 1:4 neutral mass 331.1565775  IVBOUFAWPCPFTQ-UHFFFAOYSA-N | pass02.5e55e57.5e5 m/z 332.1641 · RT 4.68 min · cos 0.923 · id 349 · **resid 0.161** | pass01e62e6 m/z 332.1640 · RT 4.70 min · cos 0.923 · id 4188 · **resid 0.032** | pass02.5e55e57.5e5 m/z 332.1640 · RT 4.72 min · cos 0.910 · id 447 · **resid 0.098** |
| #9 Anemone camphor  mix 7 · A:B 1:16 neutral mass 192.0422587  JLUQTCXCAFSSLD-NXEZZACHSA-N | fail01e42e43e4 m/z 193.0495 · RT 1.22 min · cos 0.703 · id 4743 · **resid 0.503** | not detected | not detected |
| #10 Apocarotenal  mix 19 · A:B 1:8 neutral mass 416.3079159  DFMMVLFMMAQXHZ-DOKBYWHISA-N | fail01e62e63e6 m/z 417.3153 · RT 0.80 min · cos 0.832 · id 135 · **resid 0.586** | fail02e64e6 m/z 417.3155 · RT 0.82 min · cos 0.832 · id 5153 · **resid 0.485** | fail01e62e63e6 m/z 417.3155 · RT 0.81 min · cos 0.840 · id 166 · **resid 0.586** |
| #11 Artemetin  mix 17 · A:B 2:1 neutral mass 388.1158176  RIGYMJVFEJNCKD-UHFFFAOYSA-N | pass05e61e71.5e7 m/z 389.1234 · RT 0.87 min · cos 0.929 · id 32 · **resid 0.164** | pass01e72e73e7 m/z 389.1234 · RT 0.87 min · cos 0.929 · id 4935 · **resid 0.197** | pass05e61e71.5e7 m/z 389.1234 · RT 0.88 min · cos 0.946 · id 44 · **resid 0.164** |
| #12 Aspartame  mix 12 · A:B 8:1 neutral mass 294.1215717  IAOZJIPTCAWIRG-QWRGUYRKSA-N | pass01e62e6 m/z 295.1290 · RT 4.43 min · cos 0.927 · id 166 · **resid 0.183** | pass02.5e65e67.5e6 m/z 295.1290 · RT 4.44 min · cos 0.927 · id 3532 · **resid 0.135** | pass01e62e6 m/z 295.1290 · RT 4.42 min · cos 0.931 · id 202 · **resid 0.183** |
| #14 Baohuoside I  mix 5 · A:B 4:1 neutral mass 514.1838972  NGMYNFJANBHLKA-LVKFHIPRSA-N | pass05e51e6 m/z 369.1335 · RT 1.43 min · cos 0.915 · id 303 · **resid 0.121** | pass01e62e63e6 m/z 369.1336 · RT 1.38 min · cos 0.915 · id 4707 · **resid 0.058** | not detected |
| #16 beta-Maltose  mix 13 · A:B 4:1 neutral mass 342.1162115  GUBGYTABKSRVRQ-QUYVBRFLSA-N | fail01e42e4 m/z 325.1129 · RT 5.03 min · cos 0.830 · id 4118 · **resid 7.762** | fail02e44e4 m/z 325.1131 · RT 5.02 min · cos 0.830 · id 4084 · **resid 6.997** | not detected |
| #18 Byakangelicol  mix 20 · A:B 8:1 neutral mass 316.0946882  ORBITTMJKIGFNH-LLVKDONJSA-N | pass01e72e73e7 m/z 317.1021 · RT 0.90 min · cos 0.938 · id 15 · **resid 0.148** | pass02.5e75e77.5e7 m/z 317.1021 · RT 0.91 min · cos 0.938 · id 3949 · **resid 0.117** | pass01e72e73e7 m/z 317.1020 · RT 0.91 min · cos 0.942 · id 22 · **resid 0.148** |
| #20 Cannabidivarin  mix 1 · A:B 2:1 neutral mass 286.1932801  REOZWEGFPHTFEI-JKSUJKDBSA-N | pass05e41e51.5e5 m/z 287.2007 · RT 0.85 min · cos 0.865 · id 497 · **resid 0.182** | fail01e52e53e5 m/z 287.2006 · RT 0.86 min · cos 0.865 · id 3403 · **resid 0.216** | fail05e41e51.5e5 m/z 287.2006 · RT 0.85 min · cos 0.835 · id 646 · **resid 0.380** |
| #21 Cantharidin  mix 2 · A:B 1:2 neutral mass 196.0735589  DHZBEENLJMYSHQ-XCVPVQRUSA-N | pass02e54e56e5 m/z 197.0809 · RT 0.94 min · cos 0.931 · id 584 · **resid 0.137** | pass01e62e63e6 m/z 197.0808 · RT 1.20 min · cos 0.923 · id 1805 · **resid 0.072** | pass05e51e6 m/z 197.0809 · RT 1.21 min · cos 0.912 · id 386 · **resid 0.099** |
| #25 Creatinolfosfate  mix 16 · A:B 16:1 neutral mass 197.0565429  FOIPWTMKYXWFGC-UHFFFAOYSA-N | fail02e44e46e4 m/z 198.0638 · RT 7.46 min · cos 0.855 · id 4162 · **resid 0.708** | fail02.5e55e57.5e5 m/z 198.0638 · RT 7.40 min · cos 0.855 · id 1826 · **resid 0.645** | not detected |
| #28 1,2,3,4,5,6-Hexahydro-1,5-methano-pyrido[1,2-a][1,5]diazocin-8-one  mix 15 · A:B 1:16 neutral mass 190.1106131  ANJTVLIZGCUXLD-BDAKNGLRSA-N | pass05e61e7 m/z 191.1180 · RT 5.30 min · cos 0.906 · id 48 · **resid 0.188** | fail02e64e66e6 m/z 191.1133 · RT 5.31 min · cos 0.906 · id 1711 · **resid 29.091** | not detected |
| #29 1-Deoxynojirimycin  mix 14 · A:B 1:4 neutral mass 163.0844579  LXBIFEVIBLOUGU-JGWLITMVSA-N | pass02e54e5 m/z 164.0916 · RT 6.73 min · cos 0.994 · id 512 · **resid 0.052** | pass05e51e61.5e6 m/z 164.0918 · RT 6.75 min · cos 0.994 · id 1201 · **resid 0.042** | not detected |
| #30 Digoxigenin  mix 11 · A:B 1:8 neutral mass 390.2406242  SHIBSTMRCDJXLN-KCZCNTNESA-N | pass01e52e53e5 m/z 355.2272 · RT 1.24 min · cos 0.899 · id 921 · **resid 0.093** | pass02e54e56e5 m/z 355.2270 · RT 1.25 min · cos 0.899 · id 4535 · **resid 0.060** | pass05e51e61.5e6 m/z 391.2483 · RT 1.27 min · cos 0.896 · id 329 · **resid 0.196** |
| #31 Dihydrocapsaicin  mix 9 · A:B 2:1 neutral mass 307.2147438  XJQPQKLURWNAAH-UHFFFAOYSA-N | fail01e52e53e5 m/z 615.4370 · RT 0.88 min · cos 0.957 · id 1192 · **resid 0.764** | fail05e61e7 m/z 308.2221 · RT 0.90 min · cos 0.961 · id 3781 · **resid 0.320** | fail02e64e6 m/z 137.0597 · RT 0.91 min · cos 0.954 · id 123 · **resid 0.358** |
| #34 2'-epitaxol  mix 9 · A:B 2:1 neutral mass 853.3309553  RCINICONZNJXQF-MEUUVHMJSA-N | fail01e42e4 m/z 286.1074 · RT 1.18 min · cos 0.848 · id 4732 · **resid 0.695** | not detected | not detected |
| #36 Evodiamine  mix 20 · A:B 8:1 neutral mass 303.1371622  TXDUTHBFYKGSAH-SFHVURJKSA-N | pass02.5e45e47.5e4 m/z 134.0599 · RT 0.92 min · cos 0.828 · id 1460 · **resid 0.186** | pass02.5e55e57.5e5 m/z 134.0600 · RT 0.82 min · cos 0.726 · id 711 · **resid 0.176** | fail05e51e61.5e6 m/z 304.1445 · RT 0.92 min · cos 0.900 · id 248 · **resid 0.332** |
| #38 (1'R,2'S,3S,6'S,8'R)-2'-Ethenyl-4'-methylspiro[1H-indole-3,7'-9-oxa-4-azatetracyclo[6.3.1.02,6.05,11]dodecane]-2-one  mix 3 · A:B 1:8 neutral mass 322.1681279  NFYYATWFXNPTRM-WKRGLROPSA-N | pass05e71e81.5e8 m/z 323.1756 · RT 3.83 min · cos 0.979 · id 3 · **resid 0.041** | pass02e84e8 m/z 323.1756 · RT 3.83 min · cos 0.925 · id 4051 · **resid 0.028** | pass05e71e81.5e8 m/z 323.1756 · RT 3.86 min · cos 0.889 · id 4 · **resid 0.041** |
| #45 (+)-Hydrastine  mix 18 · A:B 1:2 neutral mass 383.1368874  JZUTXVTYJDCMDU-RBUKOAKNSA-N | fail05e41e51.5e5 m/z 406.1270 · RT 0.98 min · cos 0.913 · id 1667 · **resid 0.724** | pass01e72e73e7 m/z 384.1444 · RT 0.99 min · cos 0.889 · id 4888 · **resid 0.116** | fail05e61e71.5e7 m/z 384.1444 · RT 1.00 min · cos 0.892 · id 38 · **resid 0.214** |
| #49 Isopimpinellin  mix 4 · A:B 8:1 neutral mass 246.0528234  DFMAXQKDIGCMTL-UHFFFAOYSA-N | pass05e61e71.5e7 m/z 247.0602 · RT 0.88 min · cos 0.914 · id 38 · **resid 0.114** | pass01e72e7 m/z 247.0602 · RT 0.88 min · cos 0.914 · id 2679 · **resid 0.081** | fail01e62e63e6 m/z 232.0322 · RT 0.91 min · cos 0.721 · id 160 · **resid 0.202** |
| #51 Kirenol  mix 8 · A:B 16:1 neutral mass 338.2457096  NRYNTARIOIRWAB-JPDRSCFKSA-N | fail05e31e41.5e4 m/z 321.2424 · RT 1.62 min · cos 0.763 · id 4702 · **resid 0.307** | fail02e44e46e4 m/z 321.2426 · RT 1.60 min · cos 0.763 · id 4027 · **resid 0.227** | not detected |
| #54 Leonurine  mix 10 · A:B 1:2 neutral mass 311.1481208  WNGSUWLDMZFYNZ-UHFFFAOYSA-N | pass02.5e75e77.5e7 m/z 312.1556 · RT 4.01 min · cos 0.939 · id 6 · **resid 0.076** | pass01e82e8 m/z 312.1556 · RT 4.02 min · cos 0.939 · id 3848 · **resid 0.045** | pass02.5e75e77.5e7 m/z 312.1458 · RT 4.02 min · cos 0.953 · id 9 · **resid 0.153** |
| #57 Loganin  mix 15 · A:B 1:16 neutral mass 390.152597  AMBQHHVBBHTQBF-UOUCRYGSSA-N | pass05e41e51.5e5 m/z 408.1868 · RT 2.93 min · cos 0.906 · id 1413 · **resid 0.192** | pass01e52e53e5 m/z 179.0703 · RT 2.90 min · cos 0.890 · id 1464 · **resid 0.090** | fail01e52e53e5 m/z 229.1071 · RT 2.98 min · cos 0.732 · id 851 · **resid 0.497** |
| #58 2'-Methoxyflavone  mix 20 · A:B 8:1 neutral mass 252.0786442  YEHDMSUNJUONMW-UHFFFAOYSA-N | pass05e51e6 m/z 238.0620 · RT 0.88 min · cos 0.887 · id 312 · **resid 0.154** | fail05e51e61.5e6 m/z 238.0624 · RT 0.90 min · cos 0.889 · id 2517 · **resid 2.074** | fail05e51e6 m/z 238.0624 · RT 0.93 min · cos 0.891 · id 487 · **resid 1.315** |
| #61 N-Acetyl-D-Glucosamine  mix 22 · A:B 1:4 neutral mass 221.0899372  OVRNDRQMDRJTHS-RTRLPJTCSA-N | pass02.5e45e47.5e4 m/z 138.0550 · RT 4.68 min · cos 0.962 · id 2078 · **resid 0.063** | pass01e52e53e5 m/z 186.0761 · RT 4.68 min · cos 0.959 · id 1601 · **resid 0.104** | pass01e52e53e5 m/z 204.0866 · RT 4.70 min · cos 0.918 · id 833 · **resid 0.078** |
| #62 N-Acetyl-DL-methionine  mix 6 · A:B 1:4 neutral mass 191.0616145  XUYPXLNMDZIRQH-UHFFFAOYSA-N | fail01e52e5 m/z 192.0689 · RT 1.41 min · cos 0.837 · id 1079 · **resid 0.284** | fail02e54e5 m/z 192.0689 · RT 1.41 min · cos 0.837 · id 1723 · **resid 0.345** | not detected |
| #63 NEOTAME MONOHYDRATE  mix 22 · A:B 1:4 neutral mass 378.2154721  HLIAVLHNDJUHFG-HOTGVXAUSA-N | pass05e41e5 m/z 401.2051 · RT 2.45 min · cos 0.777 · id 2144 · **resid 0.107** | pass02e74e7 m/z 379.2231 · RT 2.45 min · cos 0.943 · id 4834 · **resid 0.198** | fail01e52e53e5 m/z 379.2231 · RT 1.90 min · cos 0.939 · id 717 · **resid 0.417** |
| #65 (4As)-4,4a-dimethyl-6-prop-1-en-2-yl-3,4,5,6,7,8-hexahydronaphthalen-2-one  mix 3 · A:B 1:8 neutral mass 218.1670653  WTOYNNBCKUYIKC-QOZQQMKHSA-N | fail05e41e5 m/z 219.1745 · RT 0.85 min · cos 0.887 · id 1652 · **resid 0.972** | fail01e52e5 m/z 219.1743 · RT 0.86 min · cos 0.887 · id 2193 · **resid 0.507** | not detected |
| #70 [(1R,2S,3R,4S,7R,9S,10S,12R,15R)-4,12-Diacetyloxy-15-[(2R,3S)-3-benzamido-2-hydroxy-3-phenylpropanoyl]oxy-1,9-dihydroxy-10,14,17,17-tetramethyl-11-oxo-6-oxatetracyclo[11.3.1.03,10.04,7]heptadec-13-en-2-yl] benzoate  mix 11 · A:B 1:8 neutral mass 853.3309553  RCINICONZNJXQF-WTEIXIHQSA-N | fail05e41e51.5e5 m/z 286.1072 · RT 0.93 min · cos 0.862 · id 1354 · **resid 0.550** | not detected | not detected |
| #71 Palatinose Hydrate  mix 14 · A:B 1:4 neutral mass 342.1162115  PVXPPJIGRGXGCY-TZLCEDOOSA-N | not tested02e44e4 m/z 360.1501 · RT 7.02 min · cos 0.888 · id 3058 · **resid —** | fail05e41e51.5e5 m/z 360.1503 · RT 6.98 min · cos 0.888 · id 4601 · **resid 0.248** | not detected |
| #85 (2E)-2-[(2S,3R,4S,6R)-2-Hydroxy-3,4-dimethoxy-6-[(2R,3R,4S,5S,6R)-3,4,5-trihydroxy-6-(hydroxymethyl)oxan-2-yl]oxycyclohexylidene]acetonitrile  mix 16 · A:B 16:1 neutral mass 375.1529314  KURSRHBVYUACKS-LUIBMDKYSA-N | pass02e54e56e5 m/z 393.1870 · RT 3.33 min · cos 0.953 · id 418 · **resid 0.071** | pass01e62e63e6 m/z 393.1872 · RT 3.34 min · cos 0.953 · id 4974 · **resid 0.060** | fail02e54e56e5 m/z 393.1871 · RT 3.36 min · cos 0.955 · id 534 · **resid 0.306** |
| #86 Sweroside  mix 22 · A:B 1:4 neutral mass 358.1263823  VSJGJMKGNMDJCI-ZASXJUAOSA-N | pass02e54e56e5 m/z 197.0809 · RT 2.51 min · cos 0.938 · id 510 · **resid 0.053** | pass01e62e63e6 m/z 197.0809 · RT 2.52 min · cos 0.938 · id 1806 · **resid 0.051** | pass02e54e56e5 m/z 197.0809 · RT 2.53 min · cos 0.942 · id 654 · **resid 0.053** |
| #87 l-Tetrahydrocoptisine  mix 21 · A:B 4:1 neutral mass 323.115758  UXYJCYXWJGAKQY-OAHLLOKOSA-N | pass05e71e81.5e8 m/z 324.1232 · RT 0.99 min · cos 0.980 · id 2 · **resid 0.021** | pass01e82e83e8 m/z 324.1232 · RT 0.99 min · cos 0.933 · id 4063 · **resid 0.033** | pass05e71e81.5e8 m/z 324.1232 · RT 1.00 min · cos 0.933 · id 2 · **resid 0.021** |
| #94 Verarine  mix 2 · A:B 1:2 neutral mass 393.3031649  FSMHPYDCTDZJLV-CGARSJMXSA-N | pass02e74e76e7 m/z 394.3108 · RT 3.00 min · cos 0.864 · id 7 · **resid 0.089** | fail05e71e8 m/z 394.3108 · RT 2.52 min · cos 0.864 · id 4991 · **resid 5.111** | pass02e74e76e7 m/z 394.3107 · RT 3.03 min · cos 0.826 · id 11 · **resid 0.089** |
| #95 Vinpocetine  mix 24 · A:B 16:1 neutral mass 350.1994281  DDNCQMVWWZOMLN-IRLDBZIGSA-N | pass01e72e73e7 m/z 351.2068 · RT 1.97 min · cos 0.873 · id 13 · **resid 0.189** | pass05e71e8 m/z 351.2069 · RT 1.40 min · cos 0.871 · id 4475 · **resid 0.161** | pass01e72e73e7 m/z 351.2069 · RT 1.99 min · cos 0.882 · id 21 · **resid 0.189** |
| #96 alpha-Tocopherol acetate  mix 24 · A:B 16:1 neutral mass 472.3916455  ZAKOWWREFLAJOT-CEFNRUSXSA-N | fail01e52e53e5 m/z 473.3995 · RT 0.80 min · cos 0.878 · id 433 · **resid 0.247** | pass02e54e56e5 m/z 473.3993 · RT 0.80 min · cos 0.878 · id 5443 · **resid 0.125** | fail01e52e53e5 m/z 473.3994 · RT 0.81 min · cos 0.882 · id 558 · **resid 0.393** |
