## Supplementary material for "The Everything Bagel Feature Finder: Ultra-fast automated feature finding for untargeted metabolomics": SI Data File: spikein_boxplot_report_ST004502.html

Spike-in standards dilution series - ST004502

### Spike-in standards dilution series - ST004502

14 standards · 36 tool detections

A1000A750:B250A500:B500A250:B750B1000|process.blanksolvent.blankplasma onlyyeast onlyplasma+yeast|expected, top levelexpected at each level

The y axis is linear and autoscaled per cell. **resid** = that feature's mean relative residual over the 5-level dilution series (pass = resid < 0.20). The **×** beside each box is the expectated abudance at the level, in the corresponding own colour.

| Standard | EB (reproducible mode) | XCMS | MassCube |
| --- | --- | --- | --- |
| #1 Acesulfame potassium  mix 12 · A:B 8:1 neutral mass 162.9939288  YGCFIWIQZPHFLU-UHFFFAOYSA-N | fail02e74e7 m/z 161.9866 · RT 1.23 min · cos 0.961 · id 15 · **resid 0.233** | fail02.5e75e77.5e7 m/z 161.9866 · RT 1.23 min · cos 0.961 · id 1600 · **resid 73.279** | not detected |
| #5 ALITAME HYDRATE  mix 14 · A:B 1:4 neutral mass 331.1565775  IVBOUFAWPCPFTQ-UHFFFAOYSA-N | pass01e62e6 m/z 312.1386 · RT 4.60 min · cos 0.937 · id 243 · **resid 0.041** | pass01e52e53e5 m/z 312.1436 · RT 4.03 min · cos 0.937 · id 6326 · **resid 0.030** | pass01e62e6 m/z 312.1387 · RT 4.59 min · cos 0.938 · id 605 · **resid 0.041** |
| #14 Baohuoside I  mix 5 · A:B 4:1 neutral mass 514.1838972  NGMYNFJANBHLKA-LVKFHIPRSA-N | pass02e64e6 m/z 513.1766 · RT 1.44 min · cos 0.935 · id 170 · **resid 0.089** | pass05e61e7 m/z 513.1771 · RT 1.42 min · cos 0.935 · id 10310 · **resid 0.067** | pass02e64e6 m/z 513.1768 · RT 1.43 min · cos 0.943 · id 427 · **resid 0.089** |
| #16 beta-Maltose  mix 13 · A:B 4:1 neutral mass 342.1162115  GUBGYTABKSRVRQ-QUYVBRFLSA-N | pass02e44e46e4 m/z 341.1087 · RT 6.61 min · cos 0.892 · id 5436 · **resid 0.200** | pass05e61e71.5e7 m/z 377.0855 · RT 6.41 min · cos 0.877 · id 8060 · **resid 0.079** | pass02e54e56e5 m/z 377.0854 · RT 6.47 min · cos 0.889 · id 1918 · **resid 0.085** |
| #40 Ginsenoside-Rg5  mix 1 · A:B 2:1 neutral mass 766.4867277  NJUXRKMKOFXMRX-UHFFFAOYSA-N | not tested01e0 m/z 811.4845 · RT 4.06 min · cos 0.921 · id 3688 · **resid —** | not detected | not detected |
| #53 Lecanoric acid  mix 4 · A:B 8:1 neutral mass 318.0739528  HEMSJKZDHNSSEW-UHFFFAOYSA-N | pass01e72e7 m/z 317.0665 · RT 1.26 min · cos 0.951 · id 27 · **resid 0.118** | pass02e74e76e7 m/z 317.0665 · RT 1.26 min · cos 0.951 · id 6475 · **resid 0.052** | pass05e61e7 m/z 317.0665 · RT 1.32 min · cos 0.955 · id 310 · **resid 0.041** |
| #54 Leonurine  mix 10 · A:B 1:2 neutral mass 311.1481208  WNGSUWLDMZFYNZ-UHFFFAOYSA-N | pass02.5e65e67.5e6 m/z 310.1407 · RT 3.84 min · cos 0.929 · id 82 · **resid 0.073** | pass01e72e73e7 m/z 310.1407 · RT 3.85 min · cos 0.929 · id 6277 · **resid 0.094** | pass02.5e65e67.5e6 m/z 310.1407 · RT 3.88 min · cos 0.935 · id 225 · **resid 0.073** |
| #57 Loganin  mix 15 · A:B 1:16 neutral mass 390.152597  AMBQHHVBBHTQBF-UOUCRYGSSA-N | pass02e64e6 m/z 435.1507 · RT 2.89 min · cos 0.733 · id 140 · **resid 0.014** | pass01e72e73e7 m/z 435.1508 · RT 2.87 min · cos 0.733 · id 9184 · **resid 0.089** | pass02e64e6 m/z 435.1506 · RT 2.90 min · cos 0.741 · id 360 · **resid 0.014** |
| #62 N-Acetyl-DL-methionine  mix 6 · A:B 1:4 neutral mass 191.0616145  XUYPXLNMDZIRQH-UHFFFAOYSA-N | pass01e72e7 m/z 190.0543 · RT 1.41 min · cos 0.987 · id 24 · **resid 0.096** | pass02.5e75e77.5e7 m/z 190.0542 · RT 1.40 min · cos 0.987 · id 2417 · **resid 0.146** | pass01e72e7 m/z 190.0544 · RT 1.40 min · cos 0.974 · id 86 · **resid 0.096** |
| #63 NEOTAME MONOHYDRATE  mix 22 · A:B 1:4 neutral mass 378.2154721  HLIAVLHNDJUHFG-HOTGVXAUSA-N | pass02e64e66e6 m/z 377.2083 · RT 2.35 min · cos 0.931 · id 132 · **resid 0.061** | pass02e54e5 m/z 345.1819 · RT 2.33 min · cos 0.761 · id 7272 · **resid 0.169** | pass02e64e66e6 m/z 377.2082 · RT 2.33 min · cos 0.932 · id 346 · **resid 0.061** |
| #69 Oroxindin  mix 23 · A:B 1:16 neutral mass 460.1005615  LNOHXHDWGCMVCO-NTKSAMNMSA-N | pass02e54e5 m/z 459.0932 · RT 3.64 min · cos 0.915 · id 1893 · **resid 0.073** | pass01e62e6 m/z 459.0940 · RT 3.66 min · cos 0.915 · id 9595 · **resid 0.111** | fail02e54e5 m/z 459.0934 · RT 3.64 min · cos 0.907 · id 2912 · **resid 0.384** |
| #71 Palatinose Hydrate  mix 14 · A:B 1:4 neutral mass 342.1162115  PVXPPJIGRGXGCY-TZLCEDOOSA-N | not detected | not detected | not tested02e54e56e5 m/z 377.0854 · RT 6.81 min · cos 0.834 · id 2484 · **resid —** |
| #76 4-HYDROXYBENZOIC ACID PROPYLESTER  mix 6 · A:B 1:4 neutral mass 180.0786442  QELSKZZBTMNZEB-UHFFFAOYSA-N | pass01e52e5 m/z 179.0713 · RT 0.91 min · cos 0.910 · id 1579 · **resid 0.151** | fail02e54e5 m/z 179.0712 · RT 1.11 min · cos 0.910 · id 2086 · **resid 13.593** | not detected |
| #81 Salidroside  mix 13 · A:B 4:1 neutral mass 300.120903  ILRCGYURZSFMEG-RKQHYHRCSA-N | pass02e54e56e5 m/z 299.1136 · RT 3.03 min · cos 0.957 · id 1453 · **resid 0.087** | pass01e62e63e6 m/z 299.1136 · RT 2.99 min · cos 0.957 · id 5961 · **resid 0.133** | pass02e54e56e5 m/z 299.1135 · RT 3.00 min · cos 0.985 · id 2569 · **resid 0.025** |
