## Supplementary material for "The Everything Bagel Feature Finder: Ultra-fast automated feature finding for untargeted metabolomics": SI Data File: spikein_boxplot_report_ST004503.html

Spike-in standards dilution series - ST004503

### Spike-in standards dilution series - ST004503

50 standards · 98 tool detections

A1000A750:B250A500:B500A250:B750B1000|process.blanksolvent.blankplasma onlyyeast onlyplasma+yeast|expected, top levelexpected at each level

The y axis is linear and autoscaled per cell. **resid** = that feature's mean relative residual over the 5-level dilution series (pass = resid < 0.20). The **×** beside each box is the expectated abudance at the level, in the corresponding own colour.

| Standard | EB (reproducible mode) | XCMS | MassCube |
| --- | --- | --- | --- |
| #2 1-(3-Hydroxy-4-methoxyphenyl)ethanone  mix 5 · A:B 4:1 neutral mass 166.0629942  YLTGFGDODHXMFB-UHFFFAOYSA-N | pass01e62e63e6 m/z 167.0708 · RT 5.60 min · cos 0.944 · id 332 · **resid 0.104** | pass02e64e66e6 m/z 167.0708 · RT 5.61 min · cos 0.944 · id 640 · **resid 0.092** | pass01e62e63e6 m/z 167.0709 · RT 5.65 min · cos 0.952 · id 479 · **resid 0.104** |
| #5 ALITAME HYDRATE  mix 14 · A:B 1:4 neutral mass 331.1565775  IVBOUFAWPCPFTQ-UHFFFAOYSA-N | pass01e52e53e5 m/z 332.1635 · RT 5.91 min · cos 0.910 · id 1713 · **resid 0.113** | not detected | pass01e52e53e5 m/z 332.1642 · RT 5.92 min · cos 0.890 · id 2805 · **resid 0.113** |
| #8 (3Z)-3-[2-[(1R,4As,6R,8aS)-6-hydroxy-5-(hydroxymethyl)-5,8a-dimethyl-2-methylidene-3,4,4a,6,7,8-hexahydro-1H-naphthalen-1-yl]ethylidene]-4-hydroxyoxolan-2-one  mix 8 · A:B 16:1 neutral mass 350.2093241  BOJKULTULYSRAS-RYAUHTGOSA-N | pass05e51e6 m/z 297.1854 · RT 6.76 min · cos 0.915 · id 687 · **resid 0.061** | not detected | pass05e51e6 m/z 297.1853 · RT 6.81 min · cos 0.919 · id 1019 · **resid 0.061** |
| #10 Apocarotenal  mix 19 · A:B 1:8 neutral mass 416.3079159  DFMMVLFMMAQXHZ-DOKBYWHISA-N | pass01e62e63e6 m/z 417.3163 · RT 13.76 min · cos 0.770 · id 317 · **resid 0.107** | not detected | pass01e62e63e6 m/z 417.3152 · RT 13.78 min · cos 0.782 · id 461 · **resid 0.107** |
| #11 Artemetin  mix 17 · A:B 2:1 neutral mass 388.1158176  RIGYMJVFEJNCKD-UHFFFAOYSA-N | fail02e64e66e6 m/z 389.1237 · RT 7.69 min · cos 0.911 · id 155 · **resid 0.323** | not detected | not detected |
| #12 Aspartame  mix 12 · A:B 8:1 neutral mass 294.1215717  IAOZJIPTCAWIRG-QWRGUYRKSA-N | pass05e51e61.5e6 m/z 295.1292 · RT 5.18 min · cos 0.847 · id 544 · **resid 0.197** | not detected | pass05e51e61.5e6 m/z 295.1293 · RT 5.21 min · cos 0.863 · id 800 · **resid 0.197** |
| #14 Baohuoside I  mix 5 · A:B 4:1 neutral mass 514.1838972  NGMYNFJANBHLKA-LVKFHIPRSA-N | pass02.5e55e57.5e5 m/z 369.1337 · RT 8.49 min · cos 0.906 · id 873 · **resid 0.056** | not detected | not detected |
| #15 Benzyl alcohol  mix 6 · A:B 1:4 neutral mass 108.0575149  WVDDGKGOMKODPV-UHFFFAOYSA-N | fail05e51e6 m/z 91.0551 · RT 4.77 min · cos 0.844 · id 697 · **resid 14.035** | not detected | not detected |
| #16 beta-Maltose  mix 13 · A:B 4:1 neutral mass 342.1162115  GUBGYTABKSRVRQ-QUYVBRFLSA-N | fail02e54e56e5 m/z 365.1055 · RT 1.34 min · cos 0.773 · id 1275 · **resid 0.545** | not detected | not detected |
| #18 Byakangelicol  mix 20 · A:B 8:1 neutral mass 316.0946882  ORBITTMJKIGFNH-LLVKDONJSA-N | pass05e51e61.5e6 m/z 299.0916 · RT 6.43 min · cos 0.938 · id 524 · **resid 0.068** | not detected | pass02e64e66e6 m/z 317.1024 · RT 7.12 min · cos 0.920 · id 400 · **resid 0.162** |
| #19 Cannabidiol  mix 7 · A:B 1:16 neutral mass 314.2245802  QHMBSVQNZZTUGM-ZWKOTPCHSA-N | fail05e31e4 m/z 193.1237 · RT 10.96 min · cos 0.723 · id 21747 · **resid 0.718** | not detected | pass05e51e6 m/z 315.2323 · RT 10.92 min · cos 0.930 · id 1371 · **resid 0.189** |
| #20 Cannabidivarin  mix 1 · A:B 2:1 neutral mass 286.1932801  REOZWEGFPHTFEI-JKSUJKDBSA-N | pass05e51e61.5e6 m/z 287.2007 · RT 10.33 min · cos 0.882 · id 543 · **resid 0.172** | not detected | pass05e51e6 m/z 287.2009 · RT 10.48 min · cos 0.885 · id 1020 · **resid 0.103** |
| #21 Cantharidin  mix 2 · A:B 1:2 neutral mass 196.0735589  DHZBEENLJMYSHQ-XCVPVQRUSA-N | fail02e44e4 m/z 179.0704 · RT 4.88 min · cos 0.777 · id 5308 · **resid 0.582** | pass02e44e46e4 m/z 197.0917 · RT 4.50 min · cos 0.901 · id 994 · **resid 0.179** | pass01e52e5 m/z 197.0812 · RT 5.75 min · cos 0.851 · id 3493 · **resid 0.142** |
| #25 Creatinolfosfate  mix 16 · A:B 16:1 neutral mass 197.0565429  FOIPWTMKYXWFGC-UHFFFAOYSA-N | fail02e54e5 m/z 198.0638 · RT 1.34 min · cos 0.943 · id 1651 · **resid 0.474** | not detected | fail02e54e5 m/z 198.0649 · RT 1.36 min · cos 0.933 · id 2697 · **resid 0.488** |
| #28 1,2,3,4,5,6-Hexahydro-1,5-methano-pyrido[1,2-a][1,5]diazocin-8-one  mix 15 · A:B 1:16 neutral mass 190.1106131  ANJTVLIZGCUXLD-BDAKNGLRSA-N | pass01e52e53e5 m/z 148.0762 · RT 1.43 min · cos 0.857 · id 1891 · **resid 0.106** | pass05e51e6 m/z 148.0763 · RT 1.44 min · cos 0.851 · id 434 · **resid 0.069** | pass01e52e53e5 m/z 148.0764 · RT 1.50 min · cos 0.864 · id 3098 · **resid 0.106** |
| #29 1-Deoxynojirimycin  mix 14 · A:B 1:4 neutral mass 163.0844579  LXBIFEVIBLOUGU-JGWLITMVSA-N | fail02e54e56e5 m/z 164.0921 · RT 1.19 min · cos 0.895 · id 1059 · **resid 0.252** | fail05e51e61.5e6 m/z 164.0923 · RT 1.21 min · cos 0.895 · id 607 · **resid 0.223** | fail02e54e56e5 m/z 164.0928 · RT 1.20 min · cos 0.893 · id 1618 · **resid 0.252** |
| #30 Digoxigenin  mix 11 · A:B 1:8 neutral mass 390.2406242  SHIBSTMRCDJXLN-KCZCNTNESA-N | pass05e51e61.5e6 m/z 355.2266 · RT 5.92 min · cos 0.907 · id 555 · **resid 0.018** | not detected | pass05e51e61.5e6 m/z 355.2270 · RT 5.96 min · cos 0.906 · id 822 · **resid 0.018** |
| #31 Dihydrocapsaicin  mix 9 · A:B 2:1 neutral mass 307.2147438  XJQPQKLURWNAAH-UHFFFAOYSA-N | pass05e51e61.5e6 m/z 308.2224 · RT 9.18 min · cos 0.969 · id 468 · **resid 0.090** | pass01e72e7 m/z 137.0602 · RT 9.15 min · cos 0.972 · id 326 · **resid 0.117** | pass05e51e61.5e6 m/z 308.2226 · RT 9.19 min · cos 0.967 · id 1517 · **resid 0.093** |
| #32 Diosgenin  mix 18 · A:B 1:2 neutral mass 414.3133952  WQLVFSAGQJTQCK-VKROHFNGSA-N | pass02e54e5 m/z 415.3208 · RT 11.80 min · cos 0.910 · id 1470 · **resid 0.199** | not detected | fail02e54e5 m/z 415.3208 · RT 11.94 min · cos 0.880 · id 2246 · **resid 0.228** |
| #33 Echinacoside  mix 23 · A:B 1:16 neutral mass 786.2582439  FSBUXLDOLNLABB-ISAKITKMSA-N | pass05e41e5 m/z 804.2917 · RT 4.67 min · cos 0.915 · id 5671 · **resid 0.101** | not detected | not detected |
| #34 2'-epitaxol  mix 9 · A:B 2:1 neutral mass 853.3309553  RCINICONZNJXQF-MEUUVHMJSA-N | fail02.5e45e47.5e4 m/z 286.1072 · RT 8.36 min · cos 0.919 · id 6242 · **resid 1.548** | not detected | not detected |
| #36 Evodiamine  mix 20 · A:B 8:1 neutral mass 303.1371622  TXDUTHBFYKGSAH-SFHVURJKSA-N | pass01e62e6 m/z 304.1442 · RT 8.71 min · cos 0.931 · id 446 · **resid 0.036** | pass05e41e51.5e5 m/z 134.0606 · RT 8.72 min · cos 0.865 · id 307 · **resid 0.118** | pass01e62e6 m/z 304.1448 · RT 8.75 min · cos 0.916 · id 665 · **resid 0.036** |
| #38 (1'R,2'S,3S,6'S,8'R)-2'-Ethenyl-4'-methylspiro[1H-indole-3,7'-9-oxa-4-azatetracyclo[6.3.1.02,6.05,11]dodecane]-2-one  mix 3 · A:B 1:8 neutral mass 322.1681279  NFYYATWFXNPTRM-WKRGLROPSA-N | pass01e72e73e7 m/z 323.1767 · RT 4.51 min · cos 0.851 · id 28 · **resid 0.106** | not detected | pass01e52e53e5 m/z 236.1075 · RT 4.49 min · cos 0.780 · id 3708 · **resid 0.109** |
| #45 (+)-Hydrastine  mix 18 · A:B 1:2 neutral mass 383.1368874  JZUTXVTYJDCMDU-RBUKOAKNSA-N | pass05e61e7 m/z 384.1448 · RT 4.93 min · cos 0.934 · id 97 · **resid 0.059** | not detected | pass05e61e7 m/z 384.1445 · RT 4.91 min · cos 0.919 · id 134 · **resid 0.059** |
| #48 3-Hydroxy-6-methoxyflavone  mix 17 · A:B 2:1 neutral mass 268.0735589  OGURJSOPVFCIOO-UHFFFAOYSA-N | pass05e51e6 m/z 269.0811 · RT 9.29 min · cos 0.836 · id 727 · **resid 0.093** | not detected | not detected |
| #49 Isopimpinellin  mix 4 · A:B 8:1 neutral mass 246.0528234  DFMAXQKDIGCMTL-UHFFFAOYSA-N | pass02e74e7 m/z 247.0605 · RT 7.10 min · cos 0.963 · id 20 · **resid 0.125** | not detected | not detected |
| #51 Kirenol  mix 8 · A:B 16:1 neutral mass 338.2457096  NRYNTARIOIRWAB-JPDRSCFKSA-N | pass01e52e5 m/z 321.2426 · RT 7.38 min · cos 0.783 · id 3144 · **resid 0.149** | not detected | pass01e52e5 m/z 321.2428 · RT 7.41 min · cos 0.819 · id 5418 · **resid 0.149** |
| #54 Leonurine  mix 10 · A:B 1:2 neutral mass 311.1481208  WNGSUWLDMZFYNZ-UHFFFAOYSA-N | pass01e72e7 m/z 312.1558 · RT 4.94 min · cos 0.917 · id 56 · **resid 0.037** | pass05e51e6 m/z 181.0505 · RT 4.96 min · cos 0.722 · id 805 · **resid 0.057** | pass01e72e7 m/z 312.1559 · RT 4.91 min · cos 0.927 · id 72 · **resid 0.037** |
| #55 Limonin  mix 19 · A:B 1:8 neutral mass 470.1940679  KBDSLGBFQAGHBE-MSGMIQHVSA-N | pass02.5e55e57.5e5 m/z 471.2020 · RT 6.74 min · cos 0.860 · id 1013 · **resid 0.067** | not detected | pass02.5e55e57.5e5 m/z 471.2018 · RT 6.78 min · cos 0.851 · id 1542 · **resid 0.067** |
| #56 Linustatin  mix 14 · A:B 1:4 neutral mass 409.1584107  FERSMFQBWVBKQK-CXTTVELOSA-N | fail05e51e61.5e6 m/z 432.1489 · RT 4.41 min · cos 0.867 · id 456 · **resid 0.231** | not detected | not detected |
| #57 Loganin  mix 15 · A:B 1:16 neutral mass 390.152597  AMBQHHVBBHTQBF-UOUCRYGSSA-N | pass05e41e51.5e5 m/z 229.1071 · RT 4.89 min · cos 0.925 · id 3970 · **resid 0.099** | not detected | pass05e41e51.5e5 m/z 229.1074 · RT 4.89 min · cos 0.903 · id 7040 · **resid 0.121** |
| #58 2'-Methoxyflavone  mix 20 · A:B 8:1 neutral mass 252.0786442  YEHDMSUNJUONMW-UHFFFAOYSA-N | pass02.5e55e57.5e5 m/z 238.0625 · RT 8.52 min · cos 0.927 · id 891 · **resid 0.048** | not detected | pass02.5e55e57.5e5 m/z 238.0626 · RT 8.56 min · cos 0.940 · id 1323 · **resid 0.048** |
| #59 5-Methoxyflavone  mix 7 · A:B 1:16 neutral mass 252.0786442  XRQSPUXANRGDAV-UHFFFAOYSA-N | pass01e72e7 m/z 253.0862 · RT 8.03 min · cos 0.770 · id 50 · **resid 0.062** | not detected | not detected |
| #60 7-Methoxyflavonol  mix 18 · A:B 1:2 neutral mass 268.0735589  IPRIGHIBTRMTDP-UHFFFAOYSA-N | pass05e51e6 m/z 269.0811 · RT 9.10 min · cos 0.764 · id 809 · **resid 0.031** | not detected | not detected |
| #61 N-Acetyl-D-Glucosamine  mix 22 · A:B 1:4 neutral mass 221.0899372  OVRNDRQMDRJTHS-RTRLPJTCSA-N | fail05e41e51.5e5 m/z 138.0551 · RT 1.36 min · cos 0.856 · id 4326 · **resid 0.304** | fail02e54e56e5 m/z 138.0555 · RT 1.31 min · cos 0.856 · id 339 · **resid 2.658** | fail02e44e46e4 m/z 204.0876 · RT 1.41 min · cos 0.894 · id 16837 · **resid 0.251** |
| #62 N-Acetyl-DL-methionine  mix 6 · A:B 1:4 neutral mass 191.0616145  XUYPXLNMDZIRQH-UHFFFAOYSA-N | pass02e44e46e4 m/z 192.0691 · RT 4.62 min · cos 0.949 · id 6968 · **resid 0.126** | not detected | pass02e44e46e4 m/z 192.0695 · RT 4.64 min · cos 0.955 · id 13038 · **resid 0.126** |
| #63 NEOTAME MONOHYDRATE  mix 22 · A:B 1:4 neutral mass 378.2154721  HLIAVLHNDJUHFG-HOTGVXAUSA-N | pass02e64e66e6 m/z 379.2231 · RT 7.44 min · cos 0.942 · id 131 · **resid 0.143** | not detected | pass02e64e66e6 m/z 379.2233 · RT 7.47 min · cos 0.942 · id 183 · **resid 0.143** |
| #65 (4As)-4,4a-dimethyl-6-prop-1-en-2-yl-3,4,5,6,7,8-hexahydronaphthalen-2-one  mix 3 · A:B 1:8 neutral mass 218.1670653  WTOYNNBCKUYIKC-QOZQQMKHSA-N | pass05e31e4 m/z 145.1024 · RT 13.76 min · cos 0.827 · id 23343 · **resid 0.147** | pass02.5e45e47.5e4 m/z 145.1016 · RT 13.76 min · cos 0.827 · id 415 · **resid 0.193** | fail01e62e63e6 m/z 219.1752 · RT 9.89 min · cos 0.856 · id 456 · **resid 0.384** |
| #70 [(1R,2S,3R,4S,7R,9S,10S,12R,15R)-4,12-Diacetyloxy-15-[(2R,3S)-3-benzamido-2-hydroxy-3-phenylpropanoyl]oxy-1,9-dihydroxy-10,14,17,17-tetramethyl-11-oxo-6-oxatetracyclo[11.3.1.03,10.04,7]heptadec-13-en-2-yl] benzoate  mix 11 · A:B 1:8 neutral mass 853.3309553  RCINICONZNJXQF-WTEIXIHQSA-N | fail05e41e51.5e5 m/z 794.3165 · RT 8.35 min · cos 0.742 · id 4469 · **resid 2.366** | not detected | not detected |
| #71 Palatinose Hydrate  mix 14 · A:B 1:4 neutral mass 342.1162115  PVXPPJIGRGXGCY-TZLCEDOOSA-N | fail01e42e4 m/z 360.1503 · RT 1.30 min · cos 0.742 · id 10805 · **resid 0.458** | not detected | fail02e44e46e4 m/z 360.1503 · RT 1.37 min · cos 0.865 · id 16989 · **resid 0.419** |
| #82 Salvinorin A  mix 5 · A:B 4:1 neutral mass 432.1784179  OBSYBRPAKCASQB-AGQYDFLVSA-N | pass01e62e63e6 m/z 373.1647 · RT 7.68 min · cos 0.859 · id 330 · **resid 0.149** | fail02e64e66e6 m/z 373.2377 · RT 7.78 min · cos 0.792 · id 3679 · **resid 2.616** | pass01e62e63e6 m/z 373.1648 · RT 7.71 min · cos 0.755 · id 475 · **resid 0.149** |
| #83 Sclareolide  mix 19 · A:B 1:8 neutral mass 250.1932801  IMKJGXCIJJXALX-SHUKQUCYSA-N | pass05e41e51.5e5 m/z 215.1797 · RT 10.21 min · cos 0.913 · id 4403 · **resid 0.190** | not detected | pass05e51e6 m/z 191.1800 · RT 10.28 min · cos 0.908 · id 1093 · **resid 0.183** |
| #85 (2E)-2-[(2S,3R,4S,6R)-2-Hydroxy-3,4-dimethoxy-6-[(2R,3R,4S,5S,6R)-3,4,5-trihydroxy-6-(hydroxymethyl)oxan-2-yl]oxycyclohexylidene]acetonitrile  mix 16 · A:B 16:1 neutral mass 375.1529314  KURSRHBVYUACKS-LUIBMDKYSA-N | pass02e54e5 m/z 393.1873 · RT 4.39 min · cos 0.918 · id 1502 · **resid 0.094** | not detected | pass02e54e56e5 m/z 214.1078 · RT 4.42 min · cos 0.930 · id 2184 · **resid 0.070** |
| #86 Sweroside  mix 22 · A:B 1:4 neutral mass 358.1263823  VSJGJMKGNMDJCI-ZASXJUAOSA-N | pass05e41e51.5e5 m/z 359.1343 · RT 4.96 min · cos 0.946 · id 3394 · **resid 0.032** | not detected | pass05e41e51.5e5 m/z 359.1343 · RT 5.00 min · cos 0.951 · id 5908 · **resid 0.032** |
| #87 l-Tetrahydrocoptisine  mix 21 · A:B 4:1 neutral mass 323.115758  UXYJCYXWJGAKQY-OAHLLOKOSA-N | pass02e74e7 m/z 324.1234 · RT 5.59 min · cos 0.941 · id 17 · **resid 0.089** | pass02e64e6 m/z 176.0711 · RT 5.62 min · cos 0.746 · id 743 · **resid 0.124** | pass02e54e5 m/z 176.0706 · RT 5.61 min · cos 0.969 · id 3154 · **resid 0.042** |
| #89 [4,12-Diacetyloxy-1,9-dihydroxy-15-[2-hydroxy-3-phenyl-3-[(2-phenylacetyl)amino]propanoyl]oxy-10,14,17,17-tetramethyl-11-oxo-6-oxatetracyclo[11.3.1.03,10.04,7]heptadec-13-en-2-yl] benzoate  mix 2 · A:B 1:2 neutral mass 867.3466054  ZNHNNRQMRNHUFB-UHFFFAOYSA-N | fail02e44e4 m/z 300.1230 · RT 8.52 min · cos 0.882 · id 8917 · **resid 0.253** | not detected | not detected |
| #93 URSOLIC ACID  mix 9 · A:B 2:1 neutral mass 456.3603454  WCGUUGGRBIKTOS-YKKLBAJLSA-N | fail05e31e41.5e4 m/z 439.3731 · RT 11.10 min · cos 0.828 · id 8978 · **resid 0.201** | not detected | fail05e41e51.5e5 m/z 439.3578 · RT 11.22 min · cos 0.713 · id 5655 · **resid 0.246** |
| #94 Verarine  mix 2 · A:B 1:2 neutral mass 393.3031649  FSMHPYDCTDZJLV-CGARSJMXSA-N | pass02e74e7 m/z 394.3109 · RT 7.18 min · cos 0.771 · id 16 · **resid 0.075** | not detected | not detected |
| #95 Vinpocetine  mix 24 · A:B 16:1 neutral mass 350.1994281  DDNCQMVWWZOMLN-IRLDBZIGSA-N | pass02e74e7 m/z 351.2074 · RT 6.76 min · cos 0.927 · id 18 · **resid 0.106** | not detected | pass02e74e7 m/z 351.2073 · RT 6.76 min · cos 0.861 · id 22 · **resid 0.106** |
| #96 alpha-Tocopherol acetate  mix 24 · A:B 16:1 neutral mass 472.3916455  ZAKOWWREFLAJOT-CEFNRUSXSA-N | pass01e72e7 m/z 473.3988 · RT 13.93 min · cos 0.890 · id 53 · **resid 0.193** | fail01e52e5 m/z 473.5059 · RT 13.94 min · cos 0.894 · id 5154 · **resid 0.475** | pass01e72e7 m/z 473.3992 · RT 13.95 min · cos 0.910 · id 65 · **resid 0.193** |
