## Supplementary material for "The Everything Bagel Feature Finder: Ultra-fast automated feature finding for untargeted metabolomics": SI Data File: spikein_boxplot_report_ST004504.html

Spike-in standards dilution series - ST004504

### Spike-in standards dilution series - ST004504

41 standards · 107 tool detections

A1000A750:B250A500:B500A250:B750B1000|process.blanksolvent.blankplasma onlyyeast onlyplasma+yeast|expected, top levelexpected at each level

The y axis is linear and autoscaled per cell. **resid** = that feature's mean relative residual over the 5-level dilution series (pass = resid < 0.20). The **×** beside each box is the expectated abudance at the level, in the corresponding own colour.

| Standard | EB (reproducible mode) | XCMS | MassCube |
| --- | --- | --- | --- |
| #5 ALITAME HYDRATE  mix 14 · A:B 1:4 neutral mass 331.1565775  IVBOUFAWPCPFTQ-UHFFFAOYSA-N | fail02.5e55e57.5e5 m/z 332.1639 · RT 6.15 min · cos 0.926 · id 1766 · **resid 1.094** | fail02e64e6 m/z 332.1640 · RT 6.16 min · cos 0.926 · id 5967 · **resid 0.869** | fail02.5e55e57.5e5 m/z 332.1640 · RT 6.18 min · cos 0.916 · id 14715 · **resid 1.348** |
| #8 (3Z)-3-[2-[(1R,4As,6R,8aS)-6-hydroxy-5-(hydroxymethyl)-5,8a-dimethyl-2-methylidene-3,4,4a,6,7,8-hexahydro-1H-naphthalen-1-yl]ethylidene]-4-hydroxyoxolan-2-one  mix 8 · A:B 16:1 neutral mass 350.2093241  BOJKULTULYSRAS-RYAUHTGOSA-N | fail02e54e56e5 m/z 333.2062 · RT 7.05 min · cos 0.935 · id 2236 · **resid 24.801** | fail01e62e63e6 m/z 333.2062 · RT 7.07 min · cos 0.935 · id 5996 · **resid 23.848** | not tested02e54e56e5 m/z 333.2062 · RT 7.09 min · cos 0.946 · id 18635 · **resid —** |
| #10 Apocarotenal  mix 19 · A:B 1:8 neutral mass 416.3079159  DFMMVLFMMAQXHZ-DOKBYWHISA-N | fail01e72e73e7 m/z 417.3154 · RT 14.47 min · cos 0.848 · id 79 · **resid 2.705** | fail05e71e81.5e8 m/z 417.3154 · RT 14.48 min · cos 0.855 · id 7690 · **resid 2.463** | fail01e72e73e7 m/z 417.3153 · RT 14.60 min · cos 0.854 · id 824 · **resid 3.487** |
| #11 Artemetin  mix 17 · A:B 2:1 neutral mass 388.1158176  RIGYMJVFEJNCKD-UHFFFAOYSA-N | fail01e62e63e6 m/z 389.1232 · RT 8.10 min · cos 0.830 · id 601 · **resid 1.357** | fail05e61e71.5e7 m/z 389.1232 · RT 8.12 min · cos 0.830 · id 7140 · **resid 1.412** | fail01e62e63e6 m/z 389.1232 · RT 8.11 min · cos 0.942 · id 5142 · **resid 1.357** |
| #12 Aspartame  mix 12 · A:B 8:1 neutral mass 294.1215717  IAOZJIPTCAWIRG-QWRGUYRKSA-N | fail05e51e61.5e6 m/z 295.1288 · RT 5.40 min · cos 0.917 · id 977 · **resid 6.423** | fail02e64e66e6 m/z 295.1289 · RT 5.42 min · cos 0.917 · id 5126 · **resid 6.339** | fail05e51e61.5e6 m/z 295.1288 · RT 5.45 min · cos 0.921 · id 8321 · **resid 6.436** |
| #14 Baohuoside I  mix 5 · A:B 4:1 neutral mass 514.1838972  NGMYNFJANBHLKA-LVKFHIPRSA-N | fail05e51e6 m/z 369.1333 · RT 8.95 min · cos 0.797 · id 1357 · **resid 5.087** | fail02e64e66e6 m/z 369.1334 · RT 8.97 min · cos 0.797 · id 6730 · **resid 4.099** | not tested05e51e61.5e6 m/z 515.1910 · RT 8.85 min · cos 0.758 · id 8665 · **resid —** |
| #18 Byakangelicol  mix 20 · A:B 8:1 neutral mass 316.0946882  ORBITTMJKIGFNH-LLVKDONJSA-N | fail01e62e6 m/z 299.0915 · RT 7.51 min · cos 0.940 · id 956 · **resid 7.945** | fail05e61e7 m/z 299.0915 · RT 7.53 min · cos 0.940 · id 5212 · **resid 8.074** | fail02e64e6 m/z 317.1020 · RT 8.41 min · cos 0.940 · id 4511 · **resid 19.970** |
| #19 Cannabidiol  mix 7 · A:B 1:16 neutral mass 314.2245802  QHMBSVQNZZTUGM-ZWKOTPCHSA-N | not detected | fail02e74e76e7 m/z 315.2320 · RT 11.83 min · cos 0.917 · id 5590 · **resid 15.921** | fail02.5e65e67.5e6 m/z 315.2320 · RT 11.89 min · cos 0.917 · id 2188 · **resid 3.954** |
| #20 Cannabidivarin  mix 1 · A:B 2:1 neutral mass 286.1932801  REOZWEGFPHTFEI-JKSUJKDBSA-N | fail05e61e7 m/z 287.2006 · RT 11.12 min · cos 0.914 · id 260 · **resid 2.336** | fail01e72e73e7 m/z 287.2006 · RT 11.13 min · cos 0.899 · id 4949 · **resid 2.359** | fail02e64e66e6 m/z 287.2006 · RT 11.83 min · cos 0.934 · id 3523 · **resid 0.535** |
| #21 Cantharidin  mix 2 · A:B 1:2 neutral mass 196.0735589  DHZBEENLJMYSHQ-XCVPVQRUSA-N | fail01e52e53e5 m/z 197.0807 · RT 5.98 min · cos 0.857 · id 3033 · **resid 0.929** | not detected | fail01e52e53e5 m/z 197.0808 · RT 6.04 min · cos 0.849 · id 19612 · **resid 1.000** |
| #25 Creatinolfosfate  mix 16 · A:B 16:1 neutral mass 197.0565429  FOIPWTMKYXWFGC-UHFFFAOYSA-N | fail02e54e56e5 m/z 198.0638 · RT 1.24 min · cos 0.878 · id 2764 · **resid 13.517** | fail01e72e73e7 m/z 198.0638 · RT 1.26 min · cos 0.878 · id 2514 · **resid 16.507** | fail02.5e55e57.5e5 m/z 198.0639 · RT 1.24 min · cos 0.886 · id 14427 · **resid 10.148** |
| #28 1,2,3,4,5,6-Hexahydro-1,5-methano-pyrido[1,2-a][1,5]diazocin-8-one  mix 15 · A:B 1:16 neutral mass 190.1106131  ANJTVLIZGCUXLD-BDAKNGLRSA-N | not tested01e0 m/z 191.1178 · RT 2.93 min · cos 0.944 · id 545 · **resid —** | fail02e74e76e7 m/z 191.1178 · RT 1.66 min · cos 0.919 · id 2338 · **resid 1.169** | not detected |
| #30 Digoxigenin  mix 11 · A:B 1:8 neutral mass 390.2406242  SHIBSTMRCDJXLN-KCZCNTNESA-N | fail05e51e61.5e6 m/z 355.2269 · RT 6.20 min · cos 0.934 · id 996 · **resid 2.845** | fail02e64e66e6 m/z 373.2375 · RT 6.21 min · cos 0.867 · id 6831 · **resid 2.463** | fail05e51e6 m/z 373.2375 · RT 6.24 min · cos 0.867 · id 11059 · **resid 2.700** |
| #31 Dihydrocapsaicin  mix 9 · A:B 2:1 neutral mass 307.2147438  XJQPQKLURWNAAH-UHFFFAOYSA-N | fail05e61e71.5e7 m/z 308.2221 · RT 9.74 min · cos 0.954 · id 179 · **resid 1.414** | fail02.5e75e77.5e7 m/z 308.2220 · RT 9.76 min · cos 0.954 · id 5433 · **resid 0.838** | fail05e61e71.5e7 m/z 308.2220 · RT 9.74 min · cos 0.967 · id 1479 · **resid 1.414** |
| #32 Diosgenin  mix 18 · A:B 1:2 neutral mass 414.3133952  WQLVFSAGQJTQCK-VKROHFNGSA-N | not detected | not detected | fail02e54e56e5 m/z 415.3209 · RT 12.79 min · cos 0.866 · id 17163 · **resid 0.602** |
| #33 Echinacoside  mix 23 · A:B 1:16 neutral mass 786.2582439  FSBUXLDOLNLABB-ISAKITKMSA-N | not detected | fail01e52e5 m/z 804.2927 · RT 4.96 min · cos 0.915 · id 12181 · **resid 8.462** | not detected |
| #34 2'-epitaxol  mix 9 · A:B 2:1 neutral mass 853.3309553  RCINICONZNJXQF-MEUUVHMJSA-N | not detected | fail02e54e56e5 m/z 794.3174 · RT 8.85 min · cos 0.741 · id 12035 · **resid 7.588** | not detected |
| #36 Evodiamine  mix 20 · A:B 8:1 neutral mass 303.1371622  TXDUTHBFYKGSAH-SFHVURJKSA-N | fail02e64e66e6 m/z 304.1444 · RT 9.16 min · cos 0.891 · id 356 · **resid 6.267** | fail01e72e73e7 m/z 304.1444 · RT 9.65 min · cos 0.874 · id 5331 · **resid 2.582** | not tested02e64e66e6 m/z 304.1445 · RT 9.04 min · cos 0.877 · id 3459 · **resid —** |
| #38 (1'R,2'S,3S,6'S,8'R)-2'-Ethenyl-4'-methylspiro[1H-indole-3,7'-9-oxa-4-azatetracyclo[6.3.1.02,6.05,11]dodecane]-2-one  mix 3 · A:B 1:8 neutral mass 322.1681279  NFYYATWFXNPTRM-WKRGLROPSA-N | fail05e71e81.5e8 m/z 323.1753 · RT 4.70 min · cos 0.975 · id 15 · **resid 5.594** | fail02e84e8 m/z 323.1754 · RT 4.72 min · cos 0.951 · id 5777 · **resid 4.743** | fail05e71e81.5e8 m/z 323.1753 · RT 4.75 min · cos 0.891 · id 64 · **resid 5.594** |
| #45 (+)-Hydrastine  mix 18 · A:B 1:2 neutral mass 383.1368874  JZUTXVTYJDCMDU-RBUKOAKNSA-N | fail01e52e53e5 m/z 406.1261 · RT 5.17 min · cos 0.914 · id 3620 · **resid 0.574** | fail02.5e75e77.5e7 m/z 384.1442 · RT 5.19 min · cos 0.926 · id 7056 · **resid 0.512** | fail01e72e7 m/z 384.1442 · RT 5.21 min · cos 0.927 · id 798 · **resid 0.555** |
| #48 3-Hydroxy-6-methoxyflavone  mix 17 · A:B 2:1 neutral mass 268.0735589  OGURJSOPVFCIOO-UHFFFAOYSA-N | fail01e62e63e6 m/z 269.0809 · RT 9.84 min · cos 0.701 · id 727 · **resid 1.377** | fail01e72e73e7 m/z 269.0809 · RT 9.82 min · cos 0.746 · id 4487 · **resid 1.002** | not detected |
| #49 Isopimpinellin  mix 4 · A:B 8:1 neutral mass 246.0528234  DFMAXQKDIGCMTL-UHFFFAOYSA-N | fail05e71e8 m/z 247.0600 · RT 7.42 min · cos 0.964 · id 17 · **resid 7.427** | fail01e82e83e8 m/z 247.0670 · RT 7.43 min · cos 0.948 · id 3892 · **resid 10.935** | not detected |
| #54 Leonurine  mix 10 · A:B 1:2 neutral mass 311.1481208  WNGSUWLDMZFYNZ-UHFFFAOYSA-N | fail02e74e7 m/z 312.1553 · RT 5.17 min · cos 0.990 · id 40 · **resid 0.584** | fail01e82e8 m/z 312.1553 · RT 5.19 min · cos 0.958 · id 5513 · **resid 0.534** | fail02e74e7 m/z 312.1553 · RT 5.22 min · cos 0.955 · id 342 · **resid 0.584** |
| #55 Limonin  mix 19 · A:B 1:8 neutral mass 470.1940679  KBDSLGBFQAGHBE-MSGMIQHVSA-N | fail02e54e5 m/z 471.2012 · RT 7.05 min · cos 0.881 · id 2752 · **resid 3.600** | fail01e62e6 m/z 471.2013 · RT 7.07 min · cos 0.881 · id 8490 · **resid 3.779** | not tested02e54e5 m/z 471.2012 · RT 7.10 min · cos 0.845 · id 19381 · **resid —** |
| #57 Loganin  mix 15 · A:B 1:16 neutral mass 390.152597  AMBQHHVBBHTQBF-UOUCRYGSSA-N | fail02e54e56e5 m/z 229.1071 · RT 5.15 min · cos 0.930 · id 1933 · **resid 12.390** | fail01e62e63e6 m/z 229.1071 · RT 5.17 min · cos 0.930 · id 3400 · **resid 2.497** | not tested02e54e56e5 m/z 229.1071 · RT 5.20 min · cos 0.933 · id 16091 · **resid —** |
| #58 2'-Methoxyflavone  mix 20 · A:B 8:1 neutral mass 252.0786442  YEHDMSUNJUONMW-UHFFFAOYSA-N | fail05e71e81.5e8 m/z 253.0859 · RT 8.94 min · cos 0.888 · id 8 · **resid 7.333** | fail02e84e86e8 m/z 253.0859 · RT 8.94 min · cos 0.879 · id 4069 · **resid 6.771** | fail05e71e8 m/z 253.0859 · RT 9.03 min · cos 0.922 · id 33 · **resid 0.931** |
| #62 N-Acetyl-DL-methionine  mix 6 · A:B 1:4 neutral mass 191.0616145  XUYPXLNMDZIRQH-UHFFFAOYSA-N | fail01e62e6 m/z 192.0689 · RT 4.73 min · cos 0.722 · id 863 · **resid 1.502** | not detected | not detected |
| #63 NEOTAME MONOHYDRATE  mix 22 · A:B 1:4 neutral mass 378.2154721  HLIAVLHNDJUHFG-HOTGVXAUSA-N | fail02e64e66e6 m/z 379.2229 · RT 7.79 min · cos 0.919 · id 381 · **resid 1.329** | fail01e72e73e7 m/z 379.2229 · RT 7.81 min · cos 0.919 · id 6956 · **resid 1.283** | fail02e64e66e6 m/z 379.2229 · RT 7.84 min · cos 0.922 · id 3242 · **resid 1.329** |
| #65 (4As)-4,4a-dimethyl-6-prop-1-en-2-yl-3,4,5,6,7,8-hexahydronaphthalen-2-one  mix 3 · A:B 1:8 neutral mass 218.1670653  WTOYNNBCKUYIKC-QOZQQMKHSA-N | fail05e61e7 m/z 219.1743 · RT 10.42 min · cos 0.922 · id 204 · **resid 2.321** | fail02e74e76e7 m/z 219.1744 · RT 10.43 min · cos 0.922 · id 3138 · **resid 2.917** | fail05e61e7 m/z 219.1744 · RT 10.48 min · cos 0.922 · id 1808 · **resid 1.927** |
| #70 [(1R,2S,3R,4S,7R,9S,10S,12R,15R)-4,12-Diacetyloxy-15-[(2R,3S)-3-benzamido-2-hydroxy-3-phenylpropanoyl]oxy-1,9-dihydroxy-10,14,17,17-tetramethyl-11-oxo-6-oxatetracyclo[11.3.1.03,10.04,7]heptadec-13-en-2-yl] benzoate  mix 11 · A:B 1:8 neutral mass 853.3309553  RCINICONZNJXQF-WTEIXIHQSA-N | fail05e41e51.5e5 m/z 286.1075 · RT 8.88 min · cos 0.710 · id 5960 · **resid 1.448** | fail02.5e55e57.5e5 m/z 286.1074 · RT 8.88 min · cos 0.710 · id 4912 · **resid 1.100** | not detected |
| #71 Palatinose Hydrate  mix 14 · A:B 1:4 neutral mass 342.1162115  PVXPPJIGRGXGCY-TZLCEDOOSA-N | fail02e54e5 m/z 360.1502 · RT 1.26 min · cos 0.940 · id 2232 · **resid 1.286** | not detected | fail02e54e5 m/z 360.1502 · RT 1.26 min · cos 0.949 · id 19249 · **resid 1.768** |
| #79 Resorcinol  mix 1 · A:B 2:1 neutral mass 110.0367794  GHMLBKRAJCXXBS-UHFFFAOYSA-N | not detected | fail05e61e7 m/z 111.0440 · RT 13.29 min · cos 0.811 · id 682 · **resid 0.411** | fail05e51e61.5e6 m/z 111.0440 · RT 13.32 min · cos 0.850 · id 7229 · **resid 0.395** |
| #82 Salvinorin A  mix 5 · A:B 4:1 neutral mass 432.1784179  OBSYBRPAKCASQB-AGQYDFLVSA-N | fail02e64e6 m/z 373.1646 · RT 8.07 min · cos 0.843 · id 506 · **resid 2.825** | fail01e72e7 m/z 373.1647 · RT 8.09 min · cos 0.843 · id 6825 · **resid 3.145** | fail02e64e6 m/z 373.1646 · RT 8.26 min · cos 0.846 · id 4420 · **resid 5.360** |
| #83 Sclareolide  mix 19 · A:B 1:8 neutral mass 250.1932801  IMKJGXCIJJXALX-SHUKQUCYSA-N | fail05e51e6 m/z 233.1900 · RT 10.86 min · cos 0.933 · id 1397 · **resid 1.725** | fail02e64e66e6 m/z 233.1900 · RT 10.85 min · cos 0.933 · id 3527 · **resid 2.984** | fail05e51e6 m/z 233.1900 · RT 10.93 min · cos 0.933 · id 11657 · **resid 2.756** |
| #85 (2E)-2-[(2S,3R,4S,6R)-2-Hydroxy-3,4-dimethoxy-6-[(2R,3R,4S,5S,6R)-3,4,5-trihydroxy-6-(hydroxymethyl)oxan-2-yl]oxycyclohexylidene]acetonitrile  mix 16 · A:B 16:1 neutral mass 375.1529314  KURSRHBVYUACKS-LUIBMDKYSA-N | fail02e54e56e5 m/z 376.1603 · RT 4.54 min · cos 0.935 · id 2397 · **resid 16.302** | fail05e51e61.5e6 m/z 376.1604 · RT 4.55 min · cos 0.935 · id 6885 · **resid 22.294** | fail02e54e5 m/z 214.1074 · RT 4.58 min · cos 0.928 · id 19394 · **resid 15.501** |
| #86 Sweroside  mix 22 · A:B 1:4 neutral mass 358.1263823  VSJGJMKGNMDJCI-ZASXJUAOSA-N | fail05e51e61.5e6 m/z 197.0808 · RT 5.19 min · cos 0.883 · id 1153 · **resid 1.531** | fail02e64e66e6 m/z 197.0808 · RT 5.22 min · cos 0.883 · id 2495 · **resid 1.418** | fail05e51e61.5e6 m/z 359.1338 · RT 5.23 min · cos 0.931 · id 9590 · **resid 1.562** |
| #87 l-Tetrahydrocoptisine  mix 21 · A:B 4:1 neutral mass 323.115758  UXYJCYXWJGAKQY-OAHLLOKOSA-N | fail02.5e75e77.5e7 m/z 324.1230 · RT 5.85 min · cos 0.941 · id 23 · **resid 3.956** | fail01e82e83e8 m/z 324.1230 · RT 5.86 min · cos 0.924 · id 5793 · **resid 3.460** | fail02.5e75e77.5e7 m/z 324.1230 · RT 5.89 min · cos 0.928 · id 144 · **resid 3.956** |
| #89 [4,12-Diacetyloxy-1,9-dihydroxy-15-[2-hydroxy-3-phenyl-3-[(2-phenylacetyl)amino]propanoyl]oxy-10,14,17,17-tetramethyl-11-oxo-6-oxatetracyclo[11.3.1.03,10.04,7]heptadec-13-en-2-yl] benzoate  mix 2 · A:B 1:2 neutral mass 867.3466054  ZNHNNRQMRNHUFB-UHFFFAOYSA-N | fail02e54e56e5 m/z 868.3539 · RT 9.05 min · cos 0.842 · id 2304 · **resid 0.977** | fail01e62e63e6 m/z 868.3540 · RT 9.07 min · cos 0.842 · id 12708 · **resid 0.624** | fail02e54e5 m/z 868.3538 · RT 9.12 min · cos 0.846 · id 19100 · **resid 1.037** |
| #94 Verarine  mix 2 · A:B 1:2 neutral mass 393.3031649  FSMHPYDCTDZJLV-CGARSJMXSA-N | fail02e74e76e7 m/z 394.3103 · RT 7.47 min · cos 0.826 · id 32 · **resid 0.490** | fail01e82e83e8 m/z 394.3103 · RT 7.48 min · cos 0.826 · id 7254 · **resid 0.453** | fail02e74e76e7 m/z 394.3103 · RT 7.51 min · cos 0.825 · id 214 · **resid 0.490** |
| #95 Vinpocetine  mix 24 · A:B 16:1 neutral mass 350.1994281  DDNCQMVWWZOMLN-IRLDBZIGSA-N | fail02e74e7 m/z 351.2067 · RT 7.03 min · cos 0.879 · id 55 · **resid 12.167** | fail01e82e8 m/z 351.2068 · RT 7.05 min · cos 0.878 · id 6373 · **resid 13.137** | fail02e74e7 m/z 351.2068 · RT 7.08 min · cos 0.872 · id 513 · **resid 11.777** |
| #96 alpha-Tocopherol acetate  mix 24 · A:B 16:1 neutral mass 472.3916455  ZAKOWWREFLAJOT-CEFNRUSXSA-N | fail02e74e7 m/z 473.3988 · RT 14.68 min · cos 0.898 · id 51 · **resid 10.874** | fail01e82e8 m/z 473.3990 · RT 14.70 min · cos 0.898 · id 8528 · **resid 11.786** | fail02e74e7 m/z 473.3990 · RT 14.58 min · cos 0.912 · id 494 · **resid 59.139** |
