## Supplementary material for "The Everything Bagel Feature Finder: Ultra-fast automated feature finding for untargeted metabolomics": SI Data File: spikein_boxplot_report_ST004626.html

Spike-in standards dilution series - ST004626

### Spike-in standards dilution series - ST004626

13 standards · 23 tool detections

A1000A750:B250A500:B500A250:B750B1000|process.blanksolvent.blankplasma onlyyeast onlyplasma+yeast|expected, top levelexpected at each level

The y axis is linear and autoscaled per cell. **resid** = that feature's mean relative residual over the 5-level dilution series (pass = resid < 0.20). The **×** beside each box is the expectated abudance at the level, in the corresponding own colour.

| Standard | EB (reproducible mode) | XCMS | MassCube |
| --- | --- | --- | --- |
| #5 ALITAME HYDRATE  mix 14 · A:B 1:4 neutral mass 331.1565775  IVBOUFAWPCPFTQ-UHFFFAOYSA-N | pass02e54e5 m/z 330.1486 · RT 5.93 min · cos 0.830 · id 265 · **resid 0.172** | not detected | pass02e54e5 m/z 330.1486 · RT 5.90 min · cos 0.920 · id 658 · **resid 0.172** |
| #14 Baohuoside I  mix 5 · A:B 4:1 neutral mass 514.1838972  NGMYNFJANBHLKA-LVKFHIPRSA-N | pass02.5e55e57.5e5 m/z 513.1763 · RT 8.51 min · cos 0.825 · id 120 · **resid 0.150** | pass02e64e66e6 m/z 513.1759 · RT 8.51 min · cos 0.825 · id 2097 · **resid 0.025** | not detected |
| #33 Echinacoside  mix 23 · A:B 1:16 neutral mass 786.2582439  FSBUXLDOLNLABB-ISAKITKMSA-N | pass01e42e43e4 m/z 392.1201 · RT 4.68 min · cos 0.782 · id 3173 · **resid 0.123** | not detected | not detected |
| #40 Ginsenoside-Rg5  mix 1 · A:B 2:1 neutral mass 766.4867277  NJUXRKMKOFXMRX-UHFFFAOYSA-N | pass02.5e45e47.5e4 m/z 811.4854 · RT 10.39 min · cos 0.859 · id 984 · **resid 0.078** | not detected | not detected |
| #44 Honokiol  mix 5 · A:B 4:1 neutral mass 266.1306798  FVYXIJYOAGAUQK-UHFFFAOYSA-N | pass01e52e53e5 m/z 265.1236 · RT 9.39 min · cos 0.756 · id 277 · **resid 0.160** | pass01e62e63e6 m/z 265.1249 · RT 9.39 min · cos 0.711 · id 761 · **resid 0.136** | not detected |
| #53 Lecanoric acid  mix 4 · A:B 8:1 neutral mass 318.0739528  HEMSJKZDHNSSEW-UHFFFAOYSA-N | pass05e51e6 m/z 317.0663 · RT 7.89 min · cos 0.949 · id 74 · **resid 0.190** | not detected | pass05e51e61.5e6 m/z 167.0344 · RT 7.80 min · cos 0.952 · id 113 · **resid 0.176** |
| #54 Leonurine  mix 10 · A:B 1:2 neutral mass 311.1481208  WNGSUWLDMZFYNZ-UHFFFAOYSA-N | pass01e52e53e5 m/z 310.1405 · RT 4.95 min · cos 0.859 · id 393 · **resid 0.059** | not detected | pass01e52e53e5 m/z 310.1402 · RT 4.98 min · cos 0.873 · id 1027 · **resid 0.059** |
| #61 N-Acetyl-D-Glucosamine  mix 22 · A:B 1:4 neutral mass 221.0899372  OVRNDRQMDRJTHS-RTRLPJTCSA-N | pass02.5e35e37.5e3 m/z 266.0826 · RT 1.37 min · cos 0.765 · id 7645 · **resid 0.152** | not detected | not detected |
| #62 N-Acetyl-DL-methionine  mix 6 · A:B 1:4 neutral mass 191.0616145  XUYPXLNMDZIRQH-UHFFFAOYSA-N | not detected | not detected | fail02.5e55e57.5e5 m/z 190.0537 · RT 4.55 min · cos 0.977 · id 305 · **resid 0.254** |
| #63 NEOTAME MONOHYDRATE  mix 22 · A:B 1:4 neutral mass 378.2154721  HLIAVLHNDJUHFG-HOTGVXAUSA-N | pass02.5e55e57.5e5 m/z 377.2080 · RT 7.47 min · cos 0.882 · id 123 · **resid 0.101** | not detected | pass02.5e55e57.5e5 m/z 377.2074 · RT 7.41 min · cos 0.900 · id 284 · **resid 0.101** |
| #70 [(1R,2S,3R,4S,7R,9S,10S,12R,15R)-4,12-Diacetyloxy-15-[(2R,3S)-3-benzamido-2-hydroxy-3-phenylpropanoyl]oxy-1,9-dihydroxy-10,14,17,17-tetramethyl-11-oxo-6-oxatetracyclo[11.3.1.03,10.04,7]heptadec-13-en-2-yl] benzoate  mix 11 · A:B 1:8 neutral mass 853.3309553  RCINICONZNJXQF-WTEIXIHQSA-N | fail01e52e53e5 m/z 898.3297 · RT 8.38 min · cos 0.811 · id 297 · **resid 0.460** | fail01e62e63e6 m/z 898.3282 · RT 8.38 min · cos 0.751 · id 5129 · **resid 0.761** | fail01e52e53e5 m/z 898.3282 · RT 8.35 min · cos 0.771 · id 738 · **resid 0.610** |
| #76 4-HYDROXYBENZOIC ACID PROPYLESTER  mix 6 · A:B 1:4 neutral mass 180.0786442  QELSKZZBTMNZEB-UHFFFAOYSA-N | pass05e51e6 m/z 179.0709 · RT 8.10 min · cos 0.951 · id 66 · **resid 0.177** | not detected | pass05e51e6 m/z 179.0708 · RT 8.11 min · cos 0.982 · id 141 · **resid 0.180** |
| #81 Salidroside  mix 13 · A:B 4:1 neutral mass 300.120903  ILRCGYURZSFMEG-RKQHYHRCSA-N | fail02.5e45e47.5e4 m/z 345.1187 · RT 4.46 min · cos 0.754 · id 1227 · **resid 0.214** | not detected | fail02.5e45e47.5e4 m/z 345.1184 · RT 4.45 min · cos 0.852 · id 3360 · **resid 0.214** |
