## Supplementary material for "The Everything Bagel Feature Finder: Ultra-fast automated feature finding for untargeted metabolomics": SI Data File: spikein_boxplot_report_ST004650.html

Spike-in standards dilution series - ST004650

### Spike-in standards dilution series - ST004650

43 standards · 88 tool detections

A1000A750:B250A500:B500A250:B750B1000|process.blanksolvent.blankplasma onlyyeast onlyplasma+yeast|expected, top levelexpected at each level

The y axis is linear and autoscaled per cell. **resid** = that feature's mean relative residual over the 5-level dilution series (pass = resid < 0.20). The **×** beside each box is the expectated abudance at the level, in the corresponding own colour.

| Standard | EB (reproducible mode) | XCMS | MassCube |
| --- | --- | --- | --- |
| #5 ALITAME HYDRATE  mix 14 · A:B 1:4 neutral mass 331.1565775  IVBOUFAWPCPFTQ-UHFFFAOYSA-N | pass02e54e56e5 m/z 354.1459 · RT 4.54 min · cos 0.925 · id 1024 · **resid 0.100** | not detected | fail02.5e55e57.5e5 m/z 332.1638 · RT 4.52 min · cos 0.917 · id 1457 · **resid 0.243** |
| #9 Anemone camphor  mix 7 · A:B 1:16 neutral mass 192.0422587  JLUQTCXCAFSSLD-NXEZZACHSA-N | pass02e44e46e4 m/z 193.0505 · RT 1.28 min · cos 0.832 · id 5140 · **resid 0.080** | not detected | pass02e44e46e4 m/z 193.0498 · RT 1.25 min · cos 0.824 · id 5255 · **resid 0.080** |
| #10 Apocarotenal  mix 19 · A:B 1:8 neutral mass 416.3079159  DFMMVLFMMAQXHZ-DOKBYWHISA-N | fail01e62e63e6 m/z 417.3156 · RT 0.84 min · cos 0.821 · id 234 · **resid 0.224** | fail02e64e66e6 m/z 417.3137 · RT 0.84 min · cos 0.814 · id 6401 · **resid 0.241** | fail01e62e63e6 m/z 417.3139 · RT 0.82 min · cos 0.822 · id 325 · **resid 0.224** |
| #11 Artemetin  mix 17 · A:B 2:1 neutral mass 388.1158176  RIGYMJVFEJNCKD-UHFFFAOYSA-N | fail02e44e4 m/z 389.1835 · RT 0.98 min · cos 0.958 · id 10734 · **resid 0.329** | pass01e72e7 m/z 389.1236 · RT 0.90 min · cos 0.964 · id 5755 · **resid 0.149** | not detected |
| #12 Aspartame  mix 12 · A:B 8:1 neutral mass 294.1215717  IAOZJIPTCAWIRG-QWRGUYRKSA-N | pass01e62e6 m/z 295.1290 · RT 4.33 min · cos 0.896 · id 126 · **resid 0.064** | not detected | pass01e62e6 m/z 295.1290 · RT 4.29 min · cos 0.893 · id 149 · **resid 0.064** |
| #14 Baohuoside I  mix 5 · A:B 4:1 neutral mass 514.1838972  NGMYNFJANBHLKA-LVKFHIPRSA-N | pass01e32e33e3 m/z 369.0804 · RT 1.29 min · cos 0.716 · id 49334 · **resid 0.136** | pass02e64e66e6 m/z 515.1913 · RT 1.58 min · cos 0.878 · id 8158 · **resid 0.081** | not detected |
| #15 Benzyl alcohol  mix 6 · A:B 1:4 neutral mass 108.0575149  WVDDGKGOMKODPV-UHFFFAOYSA-N | fail01e62e63e6 m/z 91.0553 · RT 3.58 min · cos 0.842 · id 275 · **resid 9.048** | not detected | not detected |
| #16 beta-Maltose  mix 13 · A:B 4:1 neutral mass 342.1162115  GUBGYTABKSRVRQ-QUYVBRFLSA-N | pass05e51e6 m/z 365.1056 · RT 6.45 min · cos 0.829 · id 739 · **resid 0.196** | fail01e62e63e6 m/z 365.1057 · RT 6.77 min · cos 0.761 · id 5220 · **resid 0.410** | not detected |
| #18 Byakangelicol  mix 20 · A:B 8:1 neutral mass 316.0946882  ORBITTMJKIGFNH-LLVKDONJSA-N | pass02.5e65e67.5e6 m/z 233.0449 · RT 0.94 min · cos 0.729 · id 124 · **resid 0.173** | not detected | pass02.5e65e67.5e6 m/z 317.1018 · RT 0.97 min · cos 0.926 · id 121 · **resid 0.047** |
| #19 Cannabidiol  mix 7 · A:B 1:16 neutral mass 314.2245802  QHMBSVQNZZTUGM-ZWKOTPCHSA-N | pass02e54e56e5 m/z 315.2327 · RT 0.89 min · cos 0.929 · id 984 · **resid 0.095** | not detected | pass02e54e56e5 m/z 315.2323 · RT 0.89 min · cos 0.936 · id 1564 · **resid 0.173** |
| #20 Cannabidivarin  mix 1 · A:B 2:1 neutral mass 286.1932801  REOZWEGFPHTFEI-JKSUJKDBSA-N | pass02e54e56e5 m/z 287.2009 · RT 0.89 min · cos 0.870 · id 1296 · **resid 0.090** | not detected | pass02e54e56e5 m/z 287.2004 · RT 0.88 min · cos 0.871 · id 2044 · **resid 0.090** |
| #21 Cantharidin  mix 2 · A:B 1:2 neutral mass 196.0735589  DHZBEENLJMYSHQ-XCVPVQRUSA-N | pass02e54e56e5 m/z 197.0818 · RT 1.25 min · cos 0.942 · id 1085 · **resid 0.120** | not detected | pass02e54e56e5 m/z 197.0814 · RT 1.25 min · cos 0.943 · id 1733 · **resid 0.120** |
| #25 Creatinolfosfate  mix 16 · A:B 16:1 neutral mass 197.0565429  FOIPWTMKYXWFGC-UHFFFAOYSA-N | fail02e44e4 m/z 198.0643 · RT 6.67 min · cos 0.927 · id 2003 · **resid 0.552** | not detected | not detected |
| #28 1,2,3,4,5,6-Hexahydro-1,5-methano-pyrido[1,2-a][1,5]diazocin-8-one  mix 15 · A:B 1:16 neutral mass 190.1106131  ANJTVLIZGCUXLD-BDAKNGLRSA-N | pass02.5e65e67.5e6 m/z 191.1185 · RT 5.12 min · cos 0.923 · id 106 · **resid 0.110** | not detected | pass02e54e5 m/z 148.0764 · RT 5.09 min · cos 0.862 · id 2771 · **resid 0.137** |
| #29 1-Deoxynojirimycin  mix 14 · A:B 1:4 neutral mass 163.0844579  LXBIFEVIBLOUGU-JGWLITMVSA-N | pass01e52e53e5 m/z 164.0925 · RT 6.50 min · cos 0.915 · id 1105 · **resid 0.126** | not detected | pass01e52e53e5 m/z 164.0922 · RT 6.46 min · cos 0.882 · id 1764 · **resid 0.126** |
| #30 Digoxigenin  mix 11 · A:B 1:8 neutral mass 390.2406242  SHIBSTMRCDJXLN-KCZCNTNESA-N | pass05e51e6 m/z 373.2385 · RT 1.32 min · cos 0.899 · id 738 · **resid 0.051** | pass05e51e61.5e6 m/z 373.2376 · RT 1.10 min · cos 0.899 · id 5422 · **resid 0.133** | pass05e51e6 m/z 373.2374 · RT 1.31 min · cos 0.901 · id 1102 · **resid 0.051** |
| #31 Dihydrocapsaicin  mix 9 · A:B 2:1 neutral mass 307.2147438  XJQPQKLURWNAAH-UHFFFAOYSA-N | fail02e64e6 m/z 137.0606 · RT 0.94 min · cos 0.978 · id 206 · **resid 1.003** | fail02e64e66e6 m/z 137.0604 · RT 1.18 min · cos 0.978 · id 551 · **resid 0.739** | fail02e64e6 m/z 137.0603 · RT 0.93 min · cos 0.961 · id 276 · **resid 1.003** |
| #32 Diosgenin  mix 18 · A:B 1:2 neutral mass 414.3133952  WQLVFSAGQJTQCK-VKROHFNGSA-N | fail01e52e5 m/z 415.2995 · RT 0.84 min · cos 0.757 · id 2887 · **resid 0.380** | not detected | fail05e41e5 m/z 415.3174 · RT 0.91 min · cos 0.746 · id 9365 · **resid 0.516** |
| #33 Echinacoside  mix 23 · A:B 1:16 neutral mass 786.2582439  FSBUXLDOLNLABB-ISAKITKMSA-N | pass01e42e43e4 m/z 804.2933 · RT 5.36 min · cos 0.739 · id 13622 · **resid 0.154** | not detected | pass01e42e43e4 m/z 804.2948 · RT 5.35 min · cos 0.886 · id 22376 · **resid 0.188** |
| #34 2'-epitaxol  mix 9 · A:B 2:1 neutral mass 853.3309553  RCINICONZNJXQF-MEUUVHMJSA-N | fail01e32e33e3 m/z 286.2145 · RT 1.41 min · cos 0.846 · id 54142 · **resid 0.890** | not detected | not detected |
| #36 Evodiamine  mix 20 · A:B 8:1 neutral mass 303.1371622  TXDUTHBFYKGSAH-SFHVURJKSA-N | pass01e52e5 m/z 171.0906 · RT 0.96 min · cos 0.885 · id 3325 · **resid 0.200** | not detected | pass01e52e5 m/z 171.0916 · RT 0.94 min · cos 0.923 · id 5264 · **resid 0.200** |
| #38 (1'R,2'S,3S,6'S,8'R)-2'-Ethenyl-4'-methylspiro[1H-indole-3,7'-9-oxa-4-azatetracyclo[6.3.1.02,6.05,11]dodecane]-2-one  mix 3 · A:B 1:8 neutral mass 322.1681279  NFYYATWFXNPTRM-WKRGLROPSA-N | fail02e74e76e7 m/z 323.1761 · RT 3.71 min · cos 0.863 · id 2 · **resid 0.326** | pass01e82e83e8 m/z 323.1756 · RT 3.67 min · cos 0.856 · id 4162 · **resid 0.107** | fail02e54e5 m/z 236.1073 · RT 3.61 min · cos 0.781 · id 2453 · **resid 0.313** |
| #45 (+)-Hydrastine  mix 18 · A:B 1:2 neutral mass 383.1368874  JZUTXVTYJDCMDU-RBUKOAKNSA-N | pass02.5e65e67.5e6 m/z 384.1425 · RT 1.45 min · cos 0.943 · id 117 · **resid 0.090** | fail02e74e7 m/z 384.1442 · RT 1.04 min · cos 0.936 · id 5660 · **resid 0.321** | pass02e54e56e5 m/z 323.0916 · RT 1.38 min · cos 0.934 · id 2064 · **resid 0.059** |
| #49 Isopimpinellin  mix 4 · A:B 8:1 neutral mass 246.0528234  DFMAXQKDIGCMTL-UHFFFAOYSA-N | fail01e32e33e3 m/z 246.9808 · RT 0.89 min · cos 0.946 · id 14654 · **resid 0.204** | not detected | not detected |
| #54 Leonurine  mix 10 · A:B 1:2 neutral mass 311.1481208  WNGSUWLDMZFYNZ-UHFFFAOYSA-N | pass05e31e41.5e4 m/z 312.0772 · RT 3.81 min · cos 0.921 · id 22704 · **resid 0.101** | not detected | pass02e74e7 m/z 312.1560 · RT 3.76 min · cos 0.917 · id 10 · **resid 0.061** |
| #56 Linustatin  mix 14 · A:B 1:4 neutral mass 409.1584107  FERSMFQBWVBKQK-CXTTVELOSA-N | fail01e62e6 m/z 432.1484 · RT 4.97 min · cos 0.895 · id 366 · **resid 0.452** | not detected | not detected |
| #57 Loganin  mix 15 · A:B 1:16 neutral mass 390.152597  AMBQHHVBBHTQBF-UOUCRYGSSA-N | pass01e52e5 m/z 413.1428 · RT 3.27 min · cos 0.737 · id 3020 · **resid 0.075** | not detected | fail02e44e46e4 m/z 408.1870 · RT 3.29 min · cos 0.964 · id 14404 · **resid 0.360** |
| #58 2'-Methoxyflavone  mix 20 · A:B 8:1 neutral mass 252.0786442  YEHDMSUNJUONMW-UHFFFAOYSA-N | fail05e51e61.5e6 m/z 238.0809 · RT 0.82 min · cos 0.933 · id 591 · **resid 4.165** | not detected | fail02e74e76e7 m/z 253.0862 · RT 0.96 min · cos 0.947 · id 6 · **resid 14.453** |
| #59 5-Methoxyflavone  mix 7 · A:B 1:16 neutral mass 252.0786442  XRQSPUXANRGDAV-UHFFFAOYSA-N | fail02e44e4 m/z 238.0036 · RT 0.82 min · cos 0.919 · id 10407 · **resid 5.259** | not detected | not detected |
| #60 7-Methoxyflavonol  mix 18 · A:B 1:2 neutral mass 268.0735589  IPRIGHIBTRMTDP-UHFFFAOYSA-N | fail01e52e5 m/z 268.9828 · RT 0.82 min · cos 0.920 · id 2509 · **resid 0.372** | not detected | not detected |
| #61 N-Acetyl-D-Glucosamine  mix 22 · A:B 1:4 neutral mass 221.0899372  OVRNDRQMDRJTHS-RTRLPJTCSA-N | pass02.5e45e47.5e4 m/z 186.0769 · RT 4.73 min · cos 0.932 · id 2999 · **resid 0.166** | not detected | fail01e52e53e5 m/z 204.0870 · RT 4.68 min · cos 0.943 · id 3608 · **resid 0.204** |
| #62 N-Acetyl-DL-methionine  mix 6 · A:B 1:4 neutral mass 191.0616145  XUYPXLNMDZIRQH-UHFFFAOYSA-N | pass05e41e51.5e5 m/z 192.0686 · RT 1.53 min · cos 0.944 · id 4292 · **resid 0.195** | not detected | pass05e41e51.5e5 m/z 192.0698 · RT 1.52 min · cos 0.953 · id 7075 · **resid 0.195** |
| #63 NEOTAME MONOHYDRATE  mix 22 · A:B 1:4 neutral mass 378.2154721  HLIAVLHNDJUHFG-HOTGVXAUSA-N | pass05e61e71.5e7 m/z 379.2229 · RT 3.01 min · cos 0.921 · id 57 · **resid 0.123** | pass02.5e75e77.5e7 m/z 379.2228 · RT 2.95 min · cos 0.939 · id 5546 · **resid 0.174** | pass02.5e55e57.5e5 m/z 379.2229 · RT 2.72 min · cos 0.944 · id 1637 · **resid 0.121** |
| #65 (4As)-4,4a-dimethyl-6-prop-1-en-2-yl-3,4,5,6,7,8-hexahydronaphthalen-2-one  mix 3 · A:B 1:8 neutral mass 218.1670653  WTOYNNBCKUYIKC-QOZQQMKHSA-N | fail05e51e6 m/z 219.1732 · RT 0.88 min · cos 0.883 · id 655 · **resid 0.534** | not detected | fail05e51e6 m/z 219.1744 · RT 0.88 min · cos 0.885 · id 978 · **resid 0.534** |
| #70 [(1R,2S,3R,4S,7R,9S,10S,12R,15R)-4,12-Diacetyloxy-15-[(2R,3S)-3-benzamido-2-hydroxy-3-phenylpropanoyl]oxy-1,9-dihydroxy-10,14,17,17-tetramethyl-11-oxo-6-oxatetracyclo[11.3.1.03,10.04,7]heptadec-13-en-2-yl] benzoate  mix 11 · A:B 1:8 neutral mass 853.3309553  RCINICONZNJXQF-WTEIXIHQSA-N | not detected | not detected | fail02e44e46e4 m/z 286.1077 · RT 1.29 min · cos 0.913 · id 6987 · **resid 4.125** |
| #71 Palatinose Hydrate  mix 14 · A:B 1:4 neutral mass 342.1162115  PVXPPJIGRGXGCY-TZLCEDOOSA-N | pass01e42e4 m/z 360.1509 · RT 6.30 min · cos 0.707 · id 13057 · **resid 0.166** | fail02e44e46e4 m/z 360.1501 · RT 6.57 min · cos 0.748 · id 5133 · **resid 0.292** | not tested01e42e4 m/z 360.1499 · RT 6.86 min · cos 0.797 · id 26824 · **resid —** |
| #85 (2E)-2-[(2S,3R,4S,6R)-2-Hydroxy-3,4-dimethoxy-6-[(2R,3R,4S,5S,6R)-3,4,5-trihydroxy-6-(hydroxymethyl)oxan-2-yl]oxycyclohexylidene]acetonitrile  mix 16 · A:B 16:1 neutral mass 375.1529314  KURSRHBVYUACKS-LUIBMDKYSA-N | fail05e51e6 m/z 376.1606 · RT 3.60 min · cos 0.937 · id 653 · **resid 0.368** | not detected | fail05e51e6 m/z 376.1601 · RT 3.54 min · cos 0.935 · id 974 · **resid 0.368** |
| #86 Sweroside  mix 22 · A:B 1:4 neutral mass 358.1263823  VSJGJMKGNMDJCI-ZASXJUAOSA-N | pass02e54e5 m/z 359.1334 · RT 2.95 min · cos 0.953 · id 1579 · **resid 0.044** | pass05e51e61.5e6 m/z 359.1337 · RT 2.96 min · cos 0.953 · id 5109 · **resid 0.064** | pass02e54e5 m/z 359.1337 · RT 2.94 min · cos 0.962 · id 2490 · **resid 0.044** |
| #87 l-Tetrahydrocoptisine  mix 21 · A:B 4:1 neutral mass 323.115758  UXYJCYXWJGAKQY-OAHLLOKOSA-N | pass02e34e36e3 m/z 324.1787 · RT 1.53 min · cos 0.952 · id 37843 · **resid 0.175** | pass05e71e81.5e8 m/z 324.1230 · RT 1.07 min · cos 0.933 · id 4187 · **resid 0.089** | pass01e72e7 m/z 324.1232 · RT 1.42 min · cos 0.958 · id 33 · **resid 0.179** |
| #89 [4,12-Diacetyloxy-1,9-dihydroxy-15-[2-hydroxy-3-phenyl-3-[(2-phenylacetyl)amino]propanoyl]oxy-10,14,17,17-tetramethyl-11-oxo-6-oxatetracyclo[11.3.1.03,10.04,7]heptadec-13-en-2-yl] benzoate  mix 2 · A:B 1:2 neutral mass 867.3466054  ZNHNNRQMRNHUFB-UHFFFAOYSA-N | not detected | not detected | fail02e44e4 m/z 300.1235 · RT 1.28 min · cos 0.893 · id 10619 · **resid 1.801** |
| #94 Verarine  mix 2 · A:B 1:2 neutral mass 393.3031649  FSMHPYDCTDZJLV-CGARSJMXSA-N | pass02e74e76e7 m/z 394.3113 · RT 3.05 min · cos 0.796 · id 4 · **resid 0.083** | pass01e82e8 m/z 394.3107 · RT 2.96 min · cos 0.775 · id 5917 · **resid 0.114** | not detected |
| #95 Vinpocetine  mix 24 · A:B 16:1 neutral mass 350.1994281  DDNCQMVWWZOMLN-IRLDBZIGSA-N | pass01e72e7 m/z 351.2075 · RT 2.16 min · cos 0.933 · id 30 · **resid 0.114** | fail02.5e75e77.5e7 m/z 351.2070 · RT 1.55 min · cos 0.933 · id 4890 · **resid 0.401** | pass02e64e6 m/z 351.2068 · RT 1.42 min · cos 0.863 · id 243 · **resid 0.083** |
| #96 alpha-Tocopherol acetate  mix 24 · A:B 16:1 neutral mass 472.3916455  ZAKOWWREFLAJOT-CEFNRUSXSA-N | pass02e64e66e6 m/z 473.3998 · RT 0.83 min · cos 0.902 · id 141 · **resid 0.104** | pass05e61e71.5e7 m/z 473.3978 · RT 0.83 min · cos 0.914 · id 7419 · **resid 0.056** | pass02e64e66e6 m/z 473.3986 · RT 0.81 min · cos 0.914 · id 166 · **resid 0.104** |
