## Supplementary material for "The Everything Bagel Feature Finder: Ultra-fast automated feature finding for untargeted metabolomics": SI Data File: spikein_boxplot_report_ST004651.html

Spike-in standards dilution series - ST004651

### Spike-in standards dilution series - ST004651

18 standards · 31 tool detections

A1000A750:B250A500:B500A250:B750B1000|process.blanksolvent.blankplasma onlyyeast onlyplasma+yeast|expected, top levelexpected at each level

The y axis is linear and autoscaled per cell. **resid** = that feature's mean relative residual over the 5-level dilution series (pass = resid < 0.20). The **×** beside each box is the expectated abudance at the level, in the corresponding own colour.

| Standard | EB (reproducible mode) | XCMS | MassCube |
| --- | --- | --- | --- |
| #1 Acesulfame potassium  mix 12 · A:B 8:1 neutral mass 162.9939288  YGCFIWIQZPHFLU-UHFFFAOYSA-N | fail05e31e4 m/z 162.0245 · RT 0.99 min · cos 0.978 · id 29013 · **resid 10.001** | not detected | not detected |
| #5 ALITAME HYDRATE  mix 14 · A:B 1:4 neutral mass 331.1565775  IVBOUFAWPCPFTQ-UHFFFAOYSA-N | pass02e54e56e5 m/z 312.1386 · RT 4.54 min · cos 0.890 · id 629 · **resid 0.149** | not detected | pass01e62e6 m/z 330.1485 · RT 4.55 min · cos 0.909 · id 115 · **resid 0.175** |
| #7 Aloe-emodin  mix 19 · A:B 1:8 neutral mass 270.0528234  YDQWDHRMZQUTBA-UHFFFAOYSA-N | fail01e52e53e5 m/z 269.0447 · RT 0.97 min · cos 0.959 · id 1170 · **resid 0.262** | not detected | fail01e52e53e5 m/z 269.0448 · RT 0.98 min · cos 0.958 · id 1346 · **resid 0.262** |
| #14 Baohuoside I  mix 5 · A:B 4:1 neutral mass 514.1838972  NGMYNFJANBHLKA-LVKFHIPRSA-N | fail02e64e66e6 m/z 513.1755 · RT 1.58 min · cos 0.962 · id 27 · **resid 0.303** | fail01e62e63e6 m/z 513.1759 · RT 1.34 min · cos 0.952 · id 6027 · **resid 0.228** | not detected |
| #16 beta-Maltose  mix 13 · A:B 4:1 neutral mass 342.1162115  GUBGYTABKSRVRQ-QUYVBRFLSA-N | pass01e42e43e4 m/z 387.1137 · RT 6.43 min · cos 0.807 · id 12594 · **resid 0.128** | not detected | pass01e42e43e4 m/z 387.1140 · RT 6.86 min · cos 0.925 · id 16465 · **resid 0.079** |
| #40 Ginsenoside-Rg5  mix 1 · A:B 2:1 neutral mass 766.4867277  NJUXRKMKOFXMRX-UHFFFAOYSA-N | pass05e51e61.5e6 m/z 811.4839 · RT 4.00 min · cos 0.916 · id 256 · **resid 0.047** | not detected | not detected |
| #44 Honokiol  mix 5 · A:B 4:1 neutral mass 266.1306798  FVYXIJYOAGAUQK-UHFFFAOYSA-N | fail02e34e36e3 m/z 265.0756 · RT 1.00 min · cos 0.893 · id 79973 · **resid 1.191** | not detected | not detected |
| #52 (-)-Lariciresinol  mix 3 · A:B 1:8 neutral mass 360.1572885  MHXCIKYXNYCMHY-SXGZJXTBSA-N | fail05e41e5 m/z 359.1486 · RT 1.12 min · cos 0.770 · id 3837 · **resid 0.290** | not detected | not detected |
| #53 Lecanoric acid  mix 4 · A:B 8:1 neutral mass 318.0739528  HEMSJKZDHNSSEW-UHFFFAOYSA-N | pass01e62e63e6 m/z 167.0348 · RT 0.98 min · cos 0.950 · id 117 · **resid 0.071** | not detected | pass01e62e63e6 m/z 167.0343 · RT 0.99 min · cos 0.958 · id 133 · **resid 0.071** |
| #54 Leonurine  mix 10 · A:B 1:2 neutral mass 311.1481208  WNGSUWLDMZFYNZ-UHFFFAOYSA-N | pass05e41e51.5e5 m/z 295.1163 · RT 3.82 min · cos 0.817 · id 3109 · **resid 0.077** | not detected | pass01e62e6 m/z 310.1402 · RT 3.90 min · cos 0.906 · id 150 · **resid 0.084** |
| #57 Loganin  mix 15 · A:B 1:16 neutral mass 390.152597  AMBQHHVBBHTQBF-UOUCRYGSSA-N | not detected | not detected | pass01e62e6 m/z 435.1500 · RT 3.27 min · cos 0.999 · id 168 · **resid 0.090** |
| #62 N-Acetyl-DL-methionine  mix 6 · A:B 1:4 neutral mass 191.0616145  XUYPXLNMDZIRQH-UHFFFAOYSA-N | fail05e41e5 m/z 381.1155 · RT 1.51 min · cos 0.905 · id 4406 · **resid 0.319** | not detected | pass02e64e66e6 m/z 190.0537 · RT 1.50 min · cos 0.974 · id 26 · **resid 0.121** |
| #63 NEOTAME MONOHYDRATE  mix 22 · A:B 1:4 neutral mass 378.2154721  HLIAVLHNDJUHFG-HOTGVXAUSA-N | pass01e62e63e6 m/z 377.2078 · RT 3.03 min · cos 0.860 · id 48 · **resid 0.035** | not detected | pass02e64e6 m/z 377.2077 · RT 3.06 min · cos 0.897 · id 52 · **resid 0.145** |
| #69 Oroxindin  mix 23 · A:B 1:16 neutral mass 460.1005615  LNOHXHDWGCMVCO-NTKSAMNMSA-N | pass02e54e5 m/z 459.0926 · RT 3.63 min · cos 0.824 · id 1015 · **resid 0.102** | not detected | pass02e54e5 m/z 459.0924 · RT 3.62 min · cos 0.863 · id 1118 · **resid 0.100** |
| #70 [(1R,2S,3R,4S,7R,9S,10S,12R,15R)-4,12-Diacetyloxy-15-[(2R,3S)-3-benzamido-2-hydroxy-3-phenylpropanoyl]oxy-1,9-dihydroxy-10,14,17,17-tetramethyl-11-oxo-6-oxatetracyclo[11.3.1.03,10.04,7]heptadec-13-en-2-yl] benzoate  mix 11 · A:B 1:8 neutral mass 853.3309553  RCINICONZNJXQF-WTEIXIHQSA-N | fail05e41e51.5e5 m/z 888.2991 · RT 1.22 min · cos 0.749 · id 2633 · **resid 0.326** | fail05e51e61.5e6 m/z 898.3288 · RT 1.02 min · cos 0.787 · id 11902 · **resid 0.513** | fail05e51e6 m/z 898.3284 · RT 0.99 min · cos 0.774 · id 434 · **resid 0.551** |
| #76 4-HYDROXYBENZOIC ACID PROPYLESTER  mix 6 · A:B 1:4 neutral mass 180.0786442  QELSKZZBTMNZEB-UHFFFAOYSA-N | pass02e64e66e6 m/z 179.0707 · RT 0.93 min · cos 0.976 · id 30 · **resid 0.099** | not detected | pass02e64e66e6 m/z 179.0707 · RT 0.94 min · cos 0.981 · id 35 · **resid 0.099** |
| #81 Salidroside  mix 13 · A:B 4:1 neutral mass 300.120903  ILRCGYURZSFMEG-RKQHYHRCSA-N | pass02e54e5 m/z 345.1184 · RT 3.33 min · cos 0.922 · id 972 · **resid 0.131** | not detected | pass02e54e56e5 m/z 345.1184 · RT 3.32 min · cos 0.936 · id 607 · **resid 0.053** |
| #84 Scutellarin  mix 11 · A:B 1:8 neutral mass 462.079826  DJSISFGPUUYILV-ZFORQUDYSA-N | not tested02e24e2 m/z 461.0709 · RT 4.79 min · cos 0.724 · id 37313 · **resid —** | not detected | not detected |
