## Supplementary material for "The Everything Bagel Feature Finder: Ultra-fast automated feature finding for untargeted metabolomics": SI Data File: tool_unique_xic_st_EB_ST004500.html

EB-unique features (not detected by the other two tools) - ST004500

### EB-unique features (not detected by the other two tools) - ST004500

1158 features · orbitrap · XIC window m/z ± 10 ppm (≥ 0.002 Da) · ± 0.5 min

Every panel is a feature **EB** detected that neither of the other two tools detected — three-way unique across EB (standard mode (min\_detection\_rate=0.1)), Masscube and XCMS, under the standard matching rule (m/z ± 10 ppm; apex within the reference RT window ± 0.2 min). It shows the MS1 extracted-ion chromatogram at the feature's m/z, drawn from the raw mzML named in `alignment_reference_file`, zoomed to ± 0.5 min around the feature's reported apex RT. **Gray solid line** = the feature's reported apex RT; **red dashed line** = EB's assigned MS/MS — `MS2_scan_id` (native scan number) in `MS2_source_file`; the panel widens to keep an off-apex MS/MS on-screen. Features with no assigned MS/MS show only the gray line. y is per-cell autoscaled. Rows keep the table order (m/z ascending).

m/z 102.9567

01E+41111.5RT (min)

lcorbi\_t3\_neg\_2\_005\_01

m/z 114.0197

01234E+411.5RT (min)

lcorbi\_t3\_neg\_3\_005\_23

m/z 116.0255

0246E+41.52RT (min)

lcorbi\_t3\_neg\_3\_002\_24

m/z 116.0255

01E+51.52RT (min)

lcorbi\_t3\_neg\_4\_011\_24

m/z 116.9209

01E+415.516RT (min)

lcorbi\_t3\_neg\_2\_043\_33

m/z 119.0170

02468E+422.5RT (min)

lcorbi\_t3\_neg\_4\_096\_60

m/z 119.0351

0123E+500.5RT (min)

lcorbi\_t3\_neg\_1\_039\_38

m/z 119.9075

0123E+400.5RT (min)

lcorbi\_t3\_neg\_4\_066\_57

m/z 125.8732

01234E+418.519RT (min)

lcorbi\_t3\_neg\_1\_044\_23

m/z 125.8733

012E+500.5RT (min)

lcorbi\_t3\_neg\_2\_029\_12

m/z 125.8733

012E+500.5RT (min)

lcorbi\_t3\_neg\_3\_039\_29

m/z 125.8734

01234E+411.5RT (min)

lcorbi\_t3\_neg\_2\_006\_10

m/z 126.8810

024E+400.5RT (min)

lcorbi\_t3\_neg\_4\_043\_11

m/z 127.8703

01E+500.5RT (min)

lcorbi\_t3\_neg\_2\_031\_25

m/z 127.8704

01E+500.5RT (min)

lcorbi\_t3\_neg\_3\_039\_29

m/z 128.8780

0123E+400.5RT (min)

lcorbi\_t3\_neg\_4\_043\_11

m/z 128.8781

012E+411.5RT (min)

lcorbi\_t3\_neg\_1\_011\_48

m/z 129.0385

01234E+41.52RT (min)

lcorbi\_t3\_neg\_2\_053\_23

m/z 130.0872

01234E+42.53RT (min)

lcorbi\_t3\_neg\_4\_034\_01

m/z 130.0872

01E+400.5RT (min)

lcorbi\_t3\_neg\_1\_017\_51

m/z 132.8680

012E+511.5RT (min)

lcorbi\_t3\_neg\_4\_021\_25

m/z 133.0507

02468E+400.5RT (min)

lcorbi\_t3\_neg\_1\_022\_34

m/z 134.8655

012E+511.5RT (min)

lcorbi\_t3\_neg\_4\_021\_25

m/z 135.0437

01234E+410.511RT (min)

lcorbi\_t3\_neg\_4\_034\_01

m/z 135.8948

01E+415.516RT (min)

lcorbi\_t3\_neg\_2\_039\_31

m/z 135.8948

012E+41515.5RT (min)

lcorbi\_t3\_neg\_1\_054\_37

m/z 136.0242

01E+511.5RT (min)

lcorbi\_t3\_neg\_2\_022\_24

m/z 136.8917

02468E+41515.5RT (min)

lcorbi\_t3\_neg\_1\_054\_37

m/z 136.8917

02468E+41515.5RT (min)

lcorbi\_t3\_neg\_1\_050\_45

m/z 136.8918

0246E+415.516RT (min)

lcorbi\_t3\_neg\_1\_053\_32

m/z 136.9099

0123E+400.5RT (min)

lcorbi\_t3\_neg\_1\_015\_27

m/z 136.9175

01234E+411.5RT (min)

lcorbi\_t3\_neg\_2\_052\_32

m/z 139.0070

01234E+41313.5RT (min)

lcorbi\_t3\_neg\_4\_034\_01

m/z 139.0070

0246E+41414.5RT (min)

lcorbi\_t3\_neg\_4\_048\_03

m/z 139.0070

01234E+41313.5RT (min)

lcorbi\_t3\_neg\_4\_007\_13

m/z 139.0070

0123E+41313.5RT (min)

lcorbi\_t3\_neg\_4\_081\_58

m/z 147.0301

012E+511.5RT (min)

lcorbi\_t3\_neg\_4\_054\_36

m/z 147.0301

0123E+52.53RT (min)

lcorbi\_t3\_neg\_4\_054\_36

m/z 151.0072

0246E+411.5RT (min)

lcorbi\_t3\_neg\_2\_048\_27

m/z 152.0353

0246E+488.5RT (min)

lcorbi\_t3\_neg\_4\_029\_31

m/z 152.0466

01E+52.53RT (min)

lcorbi\_t3\_neg\_4\_096\_60

m/z 152.0467

012E+511.5RT (min)

lcorbi\_t3\_neg\_4\_039\_32

m/z 152.0467

012E+52.53RT (min)

lcorbi\_t3\_neg\_4\_039\_32

m/z 152.9785

01E+400.5RT (min)

lcorbi\_t3\_neg\_1\_046\_11

m/z 152.9786

012E+411.5RT (min)

lcorbi\_t3\_neg\_4\_007\_13

m/z 155.0589

0246E+411.5RT (min)

lcorbi\_t3\_neg\_4\_011\_24

m/z 158.8472

01E+500.5RT (min)

lcorbi\_t3\_neg\_4\_035\_33

m/z 159.9756

0123E+41515.5RT (min)

lcorbi\_t3\_neg\_1\_049\_55

m/z 159.9756

0123E+414.515RT (min)

lcorbi\_t3\_neg\_2\_041\_54

m/z 159.9756

0123E+414.515RT (min)

lcorbi\_t3\_neg\_1\_050\_45

m/z 159.9757

012E+415.516RT (min)

lcorbi\_t3\_neg\_2\_036\_35

m/z 160.0438

012E+52.53RT (min)

lcorbi\_t3\_neg\_2\_027\_13

m/z 160.9355

0123E+400.5RT (min)

lcorbi\_t3\_neg\_4\_083\_59

m/z 161.8502

0246E+400.5RT (min)

lcorbi\_t3\_neg\_3\_020\_31

m/z 161.9866

01234E+44.55RT (min)

lcorbi\_t3\_neg\_3\_045\_34

m/z 162.9895

02468E+42.53RT (min)

lcorbi\_t3\_neg\_3\_045\_34

m/z 163.0343

0246E+42.53RT (min)

lcorbi\_t3\_neg\_2\_050\_17

m/z 163.0343

0246E+42.53RT (min)

lcorbi\_t3\_neg\_4\_036\_16

m/z 163.0344

01E+42.53RT (min)

lcorbi\_t3\_neg\_3\_019\_12

m/z 163.8395

01E+511.5RT (min)

lcorbi\_t3\_neg\_1\_032\_13

m/z 163.8396

01E+511.5RT (min)

lcorbi\_t3\_neg\_2\_027\_13

m/z 163.8471

02468E+400.5RT (min)

lcorbi\_t3\_neg\_3\_020\_31

m/z 163.8974

0123E+400.5RT (min)

lcorbi\_t3\_neg\_4\_083\_59

m/z 163.9823

01E+52.53RT (min)

lcorbi\_t3\_neg\_3\_045\_34

m/z 164.0716

0246E+433.5RT (min)

lcorbi\_t3\_neg\_4\_034\_01

m/z 164.8361

01234E+418.519RT (min)

lcorbi\_t3\_neg\_1\_021\_31

m/z 164.8362

0123E+418.519RT (min)

lcorbi\_t3\_neg\_1\_028\_10

m/z 164.8364

0246E+500.5RT (min)

lcorbi\_t3\_neg\_4\_035\_33

m/z 165.0228

01E+411.5RT (min)

lcorbi\_t3\_neg\_4\_029\_31

m/z 166.0176

024E+411.5RT (min)

lcorbi\_t3\_neg\_3\_052\_32

m/z 166.8335

024E+400.5RT (min)

lcorbi\_t3\_neg\_4\_035\_33

m/z 167.0656

0123E+52.53RT (min)

lcorbi\_t3\_neg\_4\_039\_32

m/z 168.0243

0246E+41.52RT (min)

lcorbi\_t3\_neg\_2\_048\_27

m/z 168.0278

012E+511.5RT (min)

lcorbi\_t3\_neg\_4\_046\_37

m/z 170.0919

0246E+433.5RT (min)

lcorbi\_t3\_neg\_3\_019\_12

m/z 170.8709

0246E+400.5RT (min)

lcorbi\_t3\_neg\_4\_013\_09

m/z 170.9443

01E+41111.5RT (min)

lcorbi\_t3\_neg\_2\_007\_14

m/z 170.9443

01E+511.5RT (min)

lcorbi\_t3\_neg\_2\_005\_01

m/z 172.8306

024E+411.5RT (min)

lcorbi\_t3\_neg\_3\_010\_15

m/z 172.8307

0246E+411.5RT (min)

lcorbi\_t3\_neg\_1\_051\_15

m/z 172.8679

01234E+400.5RT (min)

lcorbi\_t3\_neg\_4\_043\_11

m/z 173.0092

02468E+41.52RT (min)

lcorbi\_t3\_neg\_3\_053\_14

m/z 175.0400

0123E+488.5RT (min)

lcorbi\_t3\_neg\_4\_040\_19

m/z 177.0921

024E+499.5RT (min)

lcorbi\_t3\_neg\_3\_019\_12

m/z 179.0292

012E+511.5RT (min)

lcorbi\_t3\_neg\_4\_039\_32

m/z 180.0594

02468E+411.5RT (min)

lcorbi\_t3\_neg\_2\_048\_27

m/z 180.0610

024E+411.5RT (min)

lcorbi\_t3\_neg\_4\_042\_14

m/z 180.0666

0123E+42.53RT (min)

lcorbi\_t3\_neg\_4\_007\_13

m/z 180.0667

02468E+422.5RT (min)

lcorbi\_t3\_neg\_2\_048\_27

m/z 180.8996

012E+411.5RT (min)

lcorbi\_t3\_neg\_1\_017\_51

m/z 182.9949

0246E+444.5RT (min)

lcorbi\_t3\_neg\_4\_005\_27

m/z 184.8484

0123E+411.5RT (min)

lcorbi\_t3\_neg\_4\_072\_57

m/z 184.8768

02468E+411.5RT (min)

lcorbi\_t3\_neg\_3\_019\_12

m/z 186.0554

01234E+411.5RT (min)

lcorbi\_t3\_neg\_4\_039\_32

m/z 186.0867

024E+422.5RT (min)

lcorbi\_t3\_neg\_3\_019\_12

m/z 186.0868

0123E+411.5RT (min)

lcorbi\_t3\_neg\_4\_051\_40

m/z 187.0249

012E+511.5RT (min)

lcorbi\_t3\_neg\_3\_005\_23

m/z 187.8613

024E+400.5RT (min)

lcorbi\_t3\_neg\_1\_014\_28

m/z 188.0565

01E+522.5RT (min)

lcorbi\_t3\_neg\_4\_054\_36

m/z 188.0565

01E+522.5RT (min)

lcorbi\_t3\_neg\_4\_044\_38

m/z 188.9863

01E+544.5RT (min)

lcorbi\_t3\_neg\_2\_048\_27

m/z 188.9863

0246E+43.54RT (min)

lcorbi\_t3\_neg\_4\_034\_01

m/z 189.8581

024E+411.5RT (min)

lcorbi\_t3\_neg\_4\_102\_61

m/z 190.1292

02468E+411.5RT (min)

lcorbi\_t3\_neg\_2\_022\_24

m/z 191.0150

0123E+411.5RT (min)

lcorbi\_t3\_neg\_4\_017\_51

m/z 191.0250

012E+41.52RT (min)

lcorbi\_t3\_neg\_4\_044\_38

m/z 193.0236

02468E+41.52RT (min)

lcorbi\_t3\_neg\_2\_048\_27

m/z 194.0295

0246E+41.52RT (min)

lcorbi\_t3\_neg\_4\_050\_02

m/z 194.0300

0123E+41.52RT (min)

lcorbi\_t3\_neg\_1\_055\_05

m/z 194.0483

01E+522.5RT (min)

lcorbi\_t3\_neg\_2\_022\_24

m/z 195.0604

02468E+42.53RT (min)

lcorbi\_t3\_neg\_4\_039\_32

m/z 195.0797

01E+522.5RT (min)

lcorbi\_t3\_neg\_4\_039\_32

m/z 195.8109

012E+418.519RT (min)

lcorbi\_t3\_neg\_1\_046\_11

m/z 195.9761

01E+411.5RT (min)

lcorbi\_t3\_neg\_2\_002\_26

m/z 196.0365

024E+41.52RT (min)

lcorbi\_t3\_neg\_4\_011\_24

m/z 196.0365

02468E+41.52RT (min)

lcorbi\_t3\_neg\_4\_011\_24

m/z 196.0555

012E+411.5RT (min)

lcorbi\_t3\_neg\_2\_052\_32

m/z 196.9470

0246E+411.5RT (min)

lcorbi\_t3\_neg\_2\_022\_24

m/z 197.8080

0123E+418.519RT (min)

lcorbi\_t3\_neg\_1\_046\_11

m/z 197.9023

01E+400.5RT (min)

lcorbi\_t3\_neg\_3\_032\_17

m/z 199.8049

0246E+500.5RT (min)

lcorbi\_t3\_neg\_1\_022\_34

m/z 199.8055

012E+600.5RT (min)

lcorbi\_t3\_neg\_2\_049\_55

m/z 201.1420

012E+41414.5RT (min)

lcorbi\_t3\_neg\_4\_005\_27

m/z 201.8019

01E+500.5RT (min)

lcorbi\_t3\_neg\_1\_022\_34

m/z 202.0249

012E+41818.5RT (min)

lcorbi\_t3\_neg\_1\_022\_34

m/z 202.0383

02468E+411.5RT (min)

lcorbi\_t3\_neg\_2\_048\_27

m/z 203.0019

01E+54.55RT (min)

lcorbi\_t3\_neg\_2\_048\_27

m/z 203.0156

01E+41818.5RT (min)

lcorbi\_t3\_neg\_1\_013\_35

m/z 203.8445

01234E+400.5RT (min)

lcorbi\_t3\_neg\_3\_020\_31

m/z 205.0353

012E+522.5RT (min)

lcorbi\_t3\_neg\_4\_054\_36

m/z 205.0353

012E+522.5RT (min)

lcorbi\_t3\_neg\_4\_083\_59

m/z 205.8395

02468E+413.514RT (min)

lcorbi\_t3\_neg\_4\_040\_19

m/z 205.8397

0246E+418.519RT (min)

lcorbi\_t3\_neg\_1\_044\_23

m/z 207.8366

01234E+413.514RT (min)

lcorbi\_t3\_neg\_2\_037\_09

m/z 207.8367

0246E+418.519RT (min)

lcorbi\_t3\_neg\_1\_044\_23

m/z 209.8335

012E+500.5RT (min)

lcorbi\_t3\_neg\_2\_029\_12

m/z 209.8340

012E+500.5RT (min)

lcorbi\_t3\_neg\_3\_051\_46

m/z 210.0652

012E+411.5RT (min)

lcorbi\_t3\_neg\_3\_028\_35

m/z 210.8839

012E+400.5RT (min)

lcorbi\_t3\_neg\_1\_043\_14

m/z 212.0586

0123E+522.5RT (min)

lcorbi\_t3\_neg\_4\_039\_32

m/z 212.0588

01234E+522.5RT (min)

lcorbi\_t3\_neg\_4\_095\_60

m/z 214.0768

0123E+44.55RT (min)

lcorbi\_t3\_neg\_4\_087\_59

m/z 214.8101

0123E+411.5RT (min)

lcorbi\_t3\_neg\_4\_088\_59

m/z 214.9338

012E+411.5RT (min)

lcorbi\_t3\_neg\_3\_021\_38

m/z 215.0462

012E+411.5RT (min)

lcorbi\_t3\_neg\_2\_045\_30

m/z 215.0728

01E+52.53RT (min)

lcorbi\_t3\_neg\_2\_050\_17

m/z 215.8113

01234E+411.5RT (min)

lcorbi\_t3\_neg\_4\_088\_59

m/z 215.8684

024E+413.514RT (min)

lcorbi\_t3\_neg\_4\_040\_19

m/z 215.8685

012E+500.5RT (min)

lcorbi\_t3\_neg\_2\_031\_25

m/z 216.0181

01E+511.5RT (min)

lcorbi\_t3\_neg\_2\_022\_24

m/z 216.0335

01234E+41414.5RT (min)

lcorbi\_t3\_neg\_4\_019\_08

m/z 216.8094

0246E+411.5RT (min)

lcorbi\_t3\_neg\_4\_088\_59

m/z 217.0230

01234E+411.5RT (min)

lcorbi\_t3\_neg\_4\_005\_27

m/z 217.0299

012E+51414.5RT (min)

lcorbi\_t3\_neg\_4\_019\_08

m/z 217.0477

01234E+411.5RT (min)

lcorbi\_t3\_neg\_2\_002\_26

m/z 217.0525

0246E+41.52RT (min)

lcorbi\_t3\_neg\_4\_075\_58

m/z 217.8098

012E+411.5RT (min)

lcorbi\_t3\_neg\_4\_089\_59

m/z 217.8655

0123E+413.514RT (min)

lcorbi\_t3\_neg\_4\_040\_19

m/z 217.8655

01E+500.5RT (min)

lcorbi\_t3\_neg\_2\_031\_25

m/z 218.0263

01234E+51414.5RT (min)

lcorbi\_t3\_neg\_4\_019\_08

m/z 218.8091

0246E+411.5RT (min)

lcorbi\_t3\_neg\_4\_089\_59

m/z 219.0115

0246E+411.5RT (min)

lcorbi\_t3\_neg\_2\_048\_27

m/z 219.0274

024E+41212.5RT (min)

lcorbi\_t3\_neg\_2\_054\_15

m/z 219.0496

0246E+411.5RT (min)

lcorbi\_t3\_neg\_2\_052\_32

m/z 219.8084

0123E+411.5RT (min)

lcorbi\_t3\_neg\_4\_087\_59

m/z 219.8916

02468E+413.514RT (min)

lcorbi\_t3\_neg\_4\_040\_19

m/z 220.8079

0246E+411.5RT (min)

lcorbi\_t3\_neg\_4\_089\_59

m/z 221.0256

01E+41818.5RT (min)

lcorbi\_t3\_neg\_1\_055\_05

m/z 221.0601

012E+511.5RT (min)

lcorbi\_t3\_neg\_4\_032\_23

m/z 221.8424

01E+522.5RT (min)

lcorbi\_t3\_neg\_4\_036\_16

m/z 221.8425

0246E+400.5RT (min)

lcorbi\_t3\_neg\_4\_066\_57

m/z 221.8887

02468E+413.514RT (min)

lcorbi\_t3\_neg\_4\_040\_19

m/z 222.8064

012E+411.5RT (min)

lcorbi\_t3\_neg\_4\_088\_59

m/z 223.0282

01234E+41212.5RT (min)

lcorbi\_t3\_neg\_4\_034\_01

m/z 223.0282

0123E+51313.5RT (min)

lcorbi\_t3\_neg\_4\_045\_35

m/z 223.0975

01E+48.59RT (min)

lcorbi\_t3\_neg\_2\_048\_27

m/z 223.8393

01E+42.53RT (min)

lcorbi\_t3\_neg\_3\_039\_29

m/z 223.8395

012E+41.52RT (min)

lcorbi\_t3\_neg\_1\_047\_02

m/z 225.0615

02468E+44.55RT (min)

lcorbi\_t3\_neg\_4\_023\_18

m/z 225.8972

0246E+400.5RT (min)

lcorbi\_t3\_neg\_3\_029\_02

m/z 226.9783

012E+413.514RT (min)

lcorbi\_t3\_neg\_1\_014\_28

m/z 226.9784

012E+413.514RT (min)

lcorbi\_t3\_neg\_1\_010\_17

m/z 227.8942

01E+400.5RT (min)

lcorbi\_t3\_neg\_3\_004\_30

m/z 228.9633

0123E+411.5RT (min)

lcorbi\_t3\_neg\_4\_042\_14

m/z 229.8823

01E+511.5RT (min)

lcorbi\_t3\_neg\_3\_019\_12

m/z 229.9062

0123E+411.5RT (min)

lcorbi\_t3\_neg\_4\_012\_17

m/z 230.0125

0246E+42.53RT (min)

lcorbi\_t3\_neg\_4\_034\_01

m/z 231.0269

02468E+411.5RT (min)

lcorbi\_t3\_neg\_4\_032\_23

m/z 232.7637

024E+41.52RT (min)

lcorbi\_t3\_neg\_2\_021\_41

m/z 232.9023

01234E+511.5RT (min)

lcorbi\_t3\_neg\_1\_055\_05

m/z 233.0099

01E+411.5RT (min)

lcorbi\_t3\_neg\_4\_095\_60

m/z 234.1579

01E+510.511RT (min)

lcorbi\_t3\_neg\_4\_042\_14

m/z 238.8581

0123E+411.5RT (min)

lcorbi\_t3\_neg\_1\_026\_19

m/z 238.8627

012E+411.5RT (min)

lcorbi\_t3\_neg\_4\_039\_32

m/z 238.9317

012E+41111.5RT (min)

lcorbi\_t3\_neg\_4\_074\_58

m/z 238.9372

0123E+411.5RT (min)

lcorbi\_t3\_neg\_3\_018\_13

m/z 239.0595

01E+511.512RT (min)

lcorbi\_t3\_neg\_4\_007\_13

m/z 239.0595

0246E+51313.5RT (min)

lcorbi\_t3\_neg\_4\_045\_35

m/z 239.0595

024E+41212.5RT (min)

lcorbi\_t3\_neg\_4\_007\_13

m/z 239.0801

0246E+411.5RT (min)

lcorbi\_t3\_neg\_4\_039\_32

m/z 241.9320

012E+411.5RT (min)

lcorbi\_t3\_neg\_4\_005\_27

m/z 242.9433

01E+410.511RT (min)

lcorbi\_t3\_neg\_2\_014\_29

m/z 243.0620

012E+422.5RT (min)

lcorbi\_t3\_neg\_4\_036\_16

m/z 243.0774

0123E+45.56RT (min)

lcorbi\_t3\_neg\_4\_023\_18

m/z 243.0805

0246E+455.5RT (min)

lcorbi\_t3\_neg\_1\_055\_05

m/z 244.0493

0246E+44.55RT (min)

lcorbi\_t3\_neg\_2\_035\_46

m/z 244.0493

0246E+44.55RT (min)

lcorbi\_t3\_neg\_4\_026\_46

m/z 245.0473

024E+411.5RT (min)

lcorbi\_t3\_neg\_4\_054\_36

m/z 245.0473

024E+41.52RT (min)

lcorbi\_t3\_neg\_4\_054\_36

m/z 247.0279

012E+41818.5RT (min)

lcorbi\_t3\_neg\_1\_020\_36

m/z 247.0592

0246E+44.55RT (min)

lcorbi\_t3\_neg\_2\_022\_24

m/z 247.0595

02468E+44.55RT (min)

lcorbi\_t3\_neg\_4\_097\_60

m/z 248.9020

024E+47.58RT (min)

lcorbi\_t3\_neg\_4\_034\_01

m/z 248.9182

0246E+411.5RT (min)

lcorbi\_t3\_neg\_1\_055\_05

m/z 249.0565

0123E+44.55RT (min)

lcorbi\_t3\_neg\_4\_097\_60

m/z 253.0569

0123E+511.5RT (min)

lcorbi\_t3\_neg\_3\_002\_24

m/z 253.0748

01234E+411.5RT (min)

lcorbi\_t3\_neg\_4\_039\_32

m/z 253.0925

0123E+410.511RT (min)

lcorbi\_t3\_neg\_4\_019\_08

m/z 253.1443

01E+599.5RT (min)

lcorbi\_t3\_neg\_2\_048\_27

m/z 255.8219

01234E+422.5RT (min)

lcorbi\_t3\_neg\_3\_031\_45

m/z 255.8222

01E+522.5RT (min)

lcorbi\_t3\_neg\_4\_036\_16

m/z 257.0657

012E+511.5RT (min)

lcorbi\_t3\_neg\_4\_032\_23

m/z 257.1140

01E+51.52RT (min)

lcorbi\_t3\_neg\_2\_053\_23

m/z 257.1141

02468E+422.5RT (min)

lcorbi\_t3\_neg\_4\_032\_23

m/z 257.8188

01234E+422.5RT (min)

lcorbi\_t3\_neg\_3\_031\_45

m/z 257.8191

01E+522.5RT (min)

lcorbi\_t3\_neg\_4\_036\_16

m/z 259.0122

01234E+411.5RT (min)

lcorbi\_t3\_neg\_2\_035\_46

m/z 259.0127

01E+511.5RT (min)

lcorbi\_t3\_neg\_4\_007\_13

m/z 259.0160

01E+511.5RT (min)

lcorbi\_t3\_neg\_3\_005\_23

m/z 259.0754

01E+52.53RT (min)

lcorbi\_t3\_neg\_1\_055\_05

m/z 259.0815

01234E+41313.5RT (min)

lcorbi\_t3\_neg\_3\_018\_13

m/z 259.8158

012E+42.53RT (min)

lcorbi\_t3\_neg\_4\_034\_01

m/z 259.8159

01234E+422.5RT (min)

lcorbi\_t3\_neg\_4\_036\_16

m/z 260.0779

01E+513.514RT (min)

lcorbi\_t3\_neg\_4\_040\_19

m/z 260.8922

0246E+411.5RT (min)

lcorbi\_t3\_neg\_4\_039\_32

m/z 261.0414

01E+41111.5RT (min)

lcorbi\_t3\_neg\_2\_048\_27

m/z 261.0742

02468E+413.514RT (min)

lcorbi\_t3\_neg\_4\_040\_19

m/z 261.0878

0123E+46.57RT (min)

lcorbi\_t3\_neg\_4\_044\_38

m/z 261.1368

01E+566.5RT (min)

lcorbi\_t3\_neg\_4\_039\_32

m/z 263.0341

012E+41111.5RT (min)

lcorbi\_t3\_neg\_2\_048\_27

m/z 263.1434

02468E+466.5RT (min)

lcorbi\_t3\_neg\_4\_039\_32

m/z 263.8522

01E+42.53RT (min)

lcorbi\_t3\_neg\_3\_022\_27

m/z 265.1231

0246E+410.511RT (min)

lcorbi\_t3\_neg\_4\_004\_34

m/z 265.1476

02468E+41313.5RT (min)

lcorbi\_t3\_neg\_4\_034\_01

m/z 265.1477

02468E+41313.5RT (min)

lcorbi\_t3\_neg\_4\_034\_01

m/z 266.0393

01E+511.5RT (min)

lcorbi\_t3\_neg\_3\_002\_24

m/z 266.0825

01E+411.5RT (min)

lcorbi\_t3\_neg\_4\_017\_51

m/z 266.1265

012E+410.511RT (min)

lcorbi\_t3\_neg\_4\_023\_18

m/z 266.9324

012E+411.5RT (min)

lcorbi\_t3\_neg\_4\_033\_53

m/z 269.0932

01E+511.5RT (min)

lcorbi\_t3\_neg\_4\_035\_33

m/z 272.0887

024E+41.52RT (min)

lcorbi\_t3\_neg\_4\_029\_31

m/z 273.8267

012E+44.55RT (min)

lcorbi\_t3\_neg\_1\_054\_37

m/z 273.8271

012E+400.5RT (min)

lcorbi\_t3\_neg\_4\_066\_57

m/z 274.1043

01E+51.52RT (min)

lcorbi\_t3\_neg\_3\_051\_46

m/z 275.1686

01E+51010.5RT (min)

lcorbi\_t3\_neg\_4\_102\_61

m/z 275.8238

012E+44.55RT (min)

lcorbi\_t3\_neg\_2\_004\_28

m/z 275.8242

01E+400.5RT (min)

lcorbi\_t3\_neg\_4\_066\_57

m/z 277.1208

0246E+410.511RT (min)

lcorbi\_t3\_neg\_2\_007\_14

m/z 278.1577

0246E+410.511RT (min)

lcorbi\_t3\_neg\_4\_042\_14

m/z 279.8587

01234E+411.5RT (min)

lcorbi\_t3\_neg\_4\_048\_03

m/z 279.9079

0123E+411.5RT (min)

lcorbi\_t3\_neg\_1\_051\_15

m/z 281.1303

0123E+51010.5RT (min)

lcorbi\_t3\_neg\_4\_023\_18

m/z 281.8557

01234E+411.5RT (min)

lcorbi\_t3\_neg\_4\_048\_03

m/z 281.9391

0246E+42.53RT (min)

lcorbi\_t3\_neg\_2\_039\_31

m/z 284.9156

01234E+411.5RT (min)

lcorbi\_t3\_neg\_2\_052\_32

m/z 288.8766

01234E+41.52RT (min)

lcorbi\_t3\_neg\_3\_005\_23

m/z 288.8768

01E+522.5RT (min)

lcorbi\_t3\_neg\_3\_037\_36

m/z 289.0327

012E+511.5RT (min)

lcorbi\_t3\_neg\_3\_005\_23

m/z 289.0373

0123E+51.52RT (min)

lcorbi\_t3\_neg\_4\_079\_58

m/z 289.1233

0123E+411.5RT (min)

lcorbi\_t3\_neg\_3\_005\_23

m/z 289.8618

012E+411.5RT (min)

lcorbi\_t3\_neg\_2\_048\_27

m/z 289.9378

01E+511.5RT (min)

lcorbi\_t3\_neg\_2\_048\_27

m/z 289.9441

01234E+433.5RT (min)

lcorbi\_t3\_neg\_1\_032\_13

m/z 290.8735

01234E+41.52RT (min)

lcorbi\_t3\_neg\_3\_005\_23

m/z 290.8737

01E+522.5RT (min)

lcorbi\_t3\_neg\_3\_037\_36

m/z 290.9337

01E+511.5RT (min)

lcorbi\_t3\_neg\_2\_048\_27

m/z 291.0659

024E+488.5RT (min)

lcorbi\_t3\_neg\_4\_056\_56

m/z 291.1964

01234E+510.511RT (min)

lcorbi\_t3\_neg\_1\_055\_05

m/z 291.9411

012E+433.5RT (min)

lcorbi\_t3\_neg\_1\_032\_13

m/z 292.0860

01E+511.5RT (min)

lcorbi\_t3\_neg\_4\_039\_32

m/z 293.1239

012E+45.56RT (min)

lcorbi\_t3\_neg\_4\_023\_18

m/z 293.1392

0246E+488.5RT (min)

lcorbi\_t3\_neg\_2\_048\_27

m/z 293.1691

02468E+59.510RT (min)

lcorbi\_t3\_neg\_4\_019\_08

m/z 293.1791

01E+41313.5RT (min)

lcorbi\_t3\_neg\_4\_034\_01

m/z 293.8932

0246E+41.52RT (min)

lcorbi\_t3\_neg\_3\_031\_45

m/z 294.9530

01E+51818.5RT (min)

lcorbi\_t3\_neg\_1\_001\_49

m/z 294.9928

02468E+411.5RT (min)

lcorbi\_t3\_neg\_3\_053\_14

m/z 295.1744

0123E+410.511RT (min)

lcorbi\_t3\_neg\_4\_046\_37

m/z 295.8901

0246E+41.52RT (min)

lcorbi\_t3\_neg\_3\_031\_45

m/z 295.9642

01234E+433.5RT (min)

lcorbi\_t3\_neg\_3\_019\_12

m/z 297.0816

012E+511.5RT (min)

lcorbi\_t3\_neg\_1\_055\_05

m/z 297.1004

01234E+411.5RT (min)

lcorbi\_t3\_neg\_4\_035\_33

m/z 297.1527

0123E+41313.5RT (min)

lcorbi\_t3\_neg\_4\_044\_38

m/z 297.8935

024E+411.5RT (min)

lcorbi\_t3\_neg\_4\_007\_13

m/z 297.9610

012E+433.5RT (min)

lcorbi\_t3\_neg\_4\_013\_09

m/z 298.0694

024E+42.53RT (min)

lcorbi\_t3\_neg\_3\_005\_23

m/z 298.9355

01E+57.58RT (min)

lcorbi\_t3\_neg\_4\_023\_18

m/z 299.0502

0123E+411.5RT (min)

lcorbi\_t3\_neg\_3\_045\_34

m/z 299.0831

01E+511.5RT (min)

lcorbi\_t3\_neg\_4\_011\_24

m/z 299.1564

012E+522.5RT (min)

lcorbi\_t3\_neg\_4\_011\_24

m/z 299.1565

012E+522.5RT (min)

lcorbi\_t3\_neg\_2\_022\_24

m/z 299.2588

01E+511.512RT (min)

lcorbi\_t3\_neg\_3\_005\_23

m/z 299.2592

0246E+41212.5RT (min)

lcorbi\_t3\_neg\_4\_097\_60

m/z 300.0086

0246E+411.5RT (min)

lcorbi\_t3\_neg\_2\_048\_27

m/z 300.2621

01E+411.512RT (min)

lcorbi\_t3\_neg\_3\_005\_23

m/z 305.0542

0246E+42.53RT (min)

lcorbi\_t3\_neg\_2\_053\_23

m/z 305.0639

01E+511.5RT (min)

lcorbi\_t3\_neg\_3\_005\_23

m/z 305.2482

012E+41313.5RT (min)

lcorbi\_t3\_neg\_4\_034\_01

m/z 306.1023

024E+411.5RT (min)

lcorbi\_t3\_neg\_4\_038\_15

m/z 306.9188

012E+41111.5RT (min)

lcorbi\_t3\_neg\_2\_010\_02

m/z 307.0795

02468E+41.52RT (min)

lcorbi\_t3\_neg\_4\_029\_31

m/z 307.0994

02468E+42.53RT (min)

lcorbi\_t3\_neg\_4\_011\_24

m/z 308.9063

01234E+411.5RT (min)

lcorbi\_t3\_neg\_2\_048\_27

m/z 308.9237

012E+411.5RT (min)

lcorbi\_t3\_neg\_4\_037\_29

m/z 309.1703

024E+49.510RT (min)

lcorbi\_t3\_neg\_4\_017\_51

m/z 309.1703

0246E+49.510RT (min)

lcorbi\_t3\_neg\_2\_048\_27

m/z 309.1704

01234E+499.5RT (min)

lcorbi\_t3\_neg\_4\_017\_51

m/z 310.0665

012E+511.5RT (min)

lcorbi\_t3\_neg\_2\_051\_34

m/z 310.0950

02468E+411.5RT (min)

lcorbi\_t3\_neg\_2\_052\_32

m/z 310.1738

01234E+41111.5RT (min)

lcorbi\_t3\_neg\_4\_023\_18

m/z 310.7773

012E+411.5RT (min)

lcorbi\_t3\_neg\_4\_042\_14

m/z 313.0781

012E+51313.5RT (min)

lcorbi\_t3\_neg\_4\_045\_35

m/z 313.2878

01234E+410.511RT (min)

lcorbi\_t3\_neg\_1\_055\_05

m/z 315.0872

02468E+42.53RT (min)

lcorbi\_t3\_neg\_3\_002\_24

m/z 315.2537

0246E+41111.5RT (min)

lcorbi\_t3\_neg\_4\_029\_31

m/z 316.0445

01E+511.5RT (min)

lcorbi\_t3\_neg\_1\_023\_03

m/z 316.0879

01E+511.5RT (min)

lcorbi\_t3\_neg\_3\_002\_24

m/z 316.1096

024E+51.52RT (min)

lcorbi\_t3\_neg\_4\_039\_32

m/z 316.8634

024E+411.5RT (min)

lcorbi\_t3\_neg\_1\_055\_05

m/z 316.9078

02468E+42.53RT (min)

lcorbi\_t3\_neg\_3\_045\_34

m/z 316.9475

0246E+41111.5RT (min)

lcorbi\_t3\_neg\_3\_053\_14

m/z 317.0865

0123E+411.5RT (min)

lcorbi\_t3\_neg\_2\_039\_31

m/z 317.2118

012E+411.512RT (min)

lcorbi\_t3\_neg\_4\_042\_14

m/z 317.3191

01E+511.512RT (min)

lcorbi\_t3\_neg\_4\_011\_24

m/z 318.0761

01E+511.5RT (min)

lcorbi\_t3\_neg\_4\_029\_31

m/z 318.1365

024E+411.5RT (min)

lcorbi\_t3\_neg\_1\_055\_05

m/z 318.9047

02468E+42.53RT (min)

lcorbi\_t3\_neg\_3\_045\_34

m/z 320.8269

012E+411.5RT (min)

lcorbi\_t3\_neg\_2\_052\_32

m/z 322.9038

01E+41818.5RT (min)

lcorbi\_t3\_neg\_1\_055\_05

m/z 326.0801

0246E+411.5RT (min)

lcorbi\_t3\_neg\_1\_055\_05

m/z 326.0877

02468E+42.53RT (min)

lcorbi\_t3\_neg\_1\_032\_13

m/z 327.1349

012E+411.5RT (min)

lcorbi\_t3\_neg\_2\_035\_46

m/z 327.2900

012E+41212.5RT (min)

lcorbi\_t3\_neg\_3\_005\_23

m/z 329.2329

01E+58.59RT (min)

lcorbi\_t3\_neg\_4\_042\_14

m/z 330.2363

0123E+499.5RT (min)

lcorbi\_t3\_neg\_4\_029\_31

m/z 330.2364

0246E+499.5RT (min)

lcorbi\_t3\_neg\_2\_048\_27

m/z 333.2068

012E+499.5RT (min)

lcorbi\_t3\_neg\_3\_048\_08

m/z 333.9250

0123E+411.5RT (min)

lcorbi\_t3\_neg\_3\_006\_48

m/z 334.0621

0246E+47.58RT (min)

lcorbi\_t3\_neg\_4\_005\_27

m/z 334.9196

0123E+411.5RT (min)

lcorbi\_t3\_neg\_3\_009\_50

m/z 335.8666

02468E+411.5RT (min)

lcorbi\_t3\_neg\_3\_034\_21

m/z 336.0504

01E+511.5RT (min)

lcorbi\_t3\_neg\_2\_048\_27

m/z 337.0624

01E+511.5RT (min)

lcorbi\_t3\_neg\_4\_029\_31

m/z 337.8635

02468E+411.5RT (min)

lcorbi\_t3\_neg\_3\_034\_21

m/z 338.0905

012E+44.55RT (min)

lcorbi\_t3\_neg\_4\_023\_18

m/z 338.8665

012E+411.5RT (min)

lcorbi\_t3\_neg\_3\_053\_14

m/z 339.2147

012E+41212.5RT (min)

lcorbi\_t3\_neg\_4\_029\_31

m/z 339.2206

01E+51313.5RT (min)

lcorbi\_t3\_neg\_4\_007\_13

m/z 340.8293

012E+411.5RT (min)

lcorbi\_t3\_neg\_2\_052\_32

m/z 342.2468

0123E+41212.5RT (min)

lcorbi\_t3\_neg\_4\_029\_31

m/z 343.9951

024E+400.5RT (min)

lcorbi\_t3\_neg\_4\_035\_33

m/z 348.8954

01234E+411.5RT (min)

lcorbi\_t3\_neg\_3\_051\_46

m/z 349.2413

01234E+41212.5RT (min)

lcorbi\_t3\_neg\_2\_056\_56

m/z 350.1299

024E+41010.5RT (min)

lcorbi\_t3\_neg\_4\_023\_18

m/z 351.0208

01E+511.5RT (min)

lcorbi\_t3\_neg\_2\_048\_27

m/z 351.1373

01E+410.511RT (min)

lcorbi\_t3\_neg\_4\_046\_37

m/z 353.0001

012E+411.5RT (min)

lcorbi\_t3\_neg\_1\_053\_32

m/z 353.1237

024E+45.56RT (min)

lcorbi\_t3\_neg\_4\_001\_49

m/z 354.1498

01E+412.513RT (min)

lcorbi\_t3\_neg\_4\_007\_13

m/z 354.9970

012E+411.5RT (min)

lcorbi\_t3\_neg\_1\_053\_32

m/z 355.1046

0123E+411.5RT (min)

lcorbi\_t3\_neg\_3\_037\_36

m/z 355.3214

01E+412.513RT (min)

lcorbi\_t3\_neg\_3\_005\_23

m/z 355.7385

012E+411.5RT (min)

lcorbi\_t3\_neg\_2\_008\_16

m/z 356.1654

012E+41313.5RT (min)

lcorbi\_t3\_neg\_4\_050\_02

m/z 357.1621

012E+41313.5RT (min)

lcorbi\_t3\_neg\_2\_001\_49

m/z 359.0443

01E+511.5RT (min)

lcorbi\_t3\_neg\_3\_028\_35

m/z 362.0122

01234E+49.510RT (min)

lcorbi\_t3\_neg\_4\_017\_51

m/z 362.1087

0246E+44.55RT (min)

lcorbi\_t3\_neg\_4\_023\_18

m/z 363.0289

012E+411.5RT (min)

lcorbi\_t3\_neg\_2\_052\_32

m/z 364.8085

012E+411.5RT (min)

lcorbi\_t3\_neg\_4\_046\_37

m/z 365.2362

0123E+411.512RT (min)

lcorbi\_t3\_neg\_4\_032\_23

m/z 365.2696

01234E+41212.5RT (min)

lcorbi\_t3\_neg\_4\_034\_01

m/z 369.1736

012E+577.5RT (min)

lcorbi\_t3\_neg\_4\_034\_01

m/z 370.1768

01234E+477.5RT (min)

lcorbi\_t3\_neg\_4\_034\_01

m/z 374.9060

01E+41111.5RT (min)

lcorbi\_t3\_neg\_4\_050\_02

m/z 376.0222

02468E+411.5RT (min)

lcorbi\_t3\_neg\_4\_039\_32

m/z 376.8959

01234E+411.5RT (min)

lcorbi\_t3\_neg\_2\_048\_27

m/z 377.1967

0123E+477.5RT (min)

lcorbi\_t3\_neg\_4\_023\_18

m/z 379.9713

012E+411.5RT (min)

lcorbi\_t3\_neg\_1\_043\_14

m/z 382.1815

01E+413.514RT (min)

lcorbi\_t3\_neg\_4\_053\_43

m/z 383.2201

0246E+410.511RT (min)

lcorbi\_t3\_neg\_2\_053\_23

m/z 384.9347

0123E+41111.5RT (min)

lcorbi\_t3\_neg\_3\_053\_14

m/z 384.9349

01234E+41111.5RT (min)

lcorbi\_t3\_neg\_4\_050\_02

m/z 385.2171

01E+410.511RT (min)

lcorbi\_t3\_neg\_3\_020\_31

m/z 388.8144

01234E+411.5RT (min)

lcorbi\_t3\_neg\_3\_052\_32

m/z 394.0490

012E+410.511RT (min)

lcorbi\_t3\_neg\_4\_034\_01

m/z 394.8917

01234E+411.5RT (min)

lcorbi\_t3\_neg\_2\_007\_14

m/z 396.8669

0246E+411.5RT (min)

lcorbi\_t3\_neg\_4\_039\_32

m/z 397.2124

01E+577.5RT (min)

lcorbi\_t3\_neg\_4\_023\_18

m/z 397.2203

012E+49.510RT (min)

lcorbi\_t3\_neg\_2\_048\_27

m/z 397.2206

012E+499.5RT (min)

lcorbi\_t3\_neg\_2\_048\_27

m/z 397.3683

01234E+41414.5RT (min)

lcorbi\_t3\_neg\_2\_020\_05

m/z 399.2206

0123E+48.59RT (min)

lcorbi\_t3\_neg\_4\_034\_01

m/z 399.3223

01E+511.512RT (min)

lcorbi\_t3\_neg\_4\_007\_13

m/z 400.2340

02468E+41313.5RT (min)

lcorbi\_t3\_neg\_1\_055\_05

m/z 401.1500

01234E+41313.5RT (min)

lcorbi\_t3\_neg\_4\_038\_15

m/z 402.9165

01E+511.5RT (min)

lcorbi\_t3\_neg\_2\_048\_27

m/z 403.0950

0246E+41111.5RT (min)

lcorbi\_t3\_neg\_2\_035\_46

m/z 404.0156

012E+411.5RT (min)

lcorbi\_t3\_neg\_3\_032\_17

m/z 404.1604

012E+41313.5RT (min)

lcorbi\_t3\_neg\_2\_007\_14

m/z 406.8543

01234E+411.5RT (min)

lcorbi\_t3\_neg\_3\_053\_14

m/z 407.0112

012E+410.511RT (min)

lcorbi\_t3\_neg\_1\_043\_14

m/z 407.1067

012E+55.56RT (min)

lcorbi\_t3\_neg\_4\_023\_18

m/z 407.1990

02468E+48.59RT (min)

lcorbi\_t3\_neg\_4\_017\_51

m/z 408.1506

012E+52.53RT (min)

lcorbi\_t3\_neg\_4\_040\_19

m/z 408.7694

0246E+411.5RT (min)

lcorbi\_t3\_neg\_4\_007\_13

m/z 411.0972

0123E+41010.5RT (min)

lcorbi\_t3\_neg\_4\_023\_18

m/z 411.3837

0123E+413.514RT (min)

lcorbi\_t3\_neg\_2\_007\_14

m/z 411.3839

0123E+413.514RT (min)

lcorbi\_t3\_neg\_3\_053\_14

m/z 413.2721

012E+41313.5RT (min)

lcorbi\_t3\_neg\_4\_034\_01

m/z 416.8830

0246E+411.5RT (min)

lcorbi\_t3\_neg\_3\_051\_46

m/z 417.0027

024E+411.5RT (min)

lcorbi\_t3\_neg\_3\_028\_35

m/z 417.2278

0123E+47.58RT (min)

lcorbi\_t3\_neg\_4\_017\_51

m/z 418.8940

0123E+511.5RT (min)

lcorbi\_t3\_neg\_2\_048\_27

m/z 419.0478

012E+49.510RT (min)

lcorbi\_t3\_neg\_1\_039\_38

m/z 421.1112

01E+54.55RT (min)

lcorbi\_t3\_neg\_4\_039\_32

m/z 421.1494

0123E+410.511RT (min)

lcorbi\_t3\_neg\_2\_048\_27

m/z 421.1497

01234E+410.511RT (min)

lcorbi\_t3\_neg\_2\_007\_14

m/z 424.7441

012E+411.5RT (min)

lcorbi\_t3\_neg\_4\_042\_14

m/z 426.3134

0123E+41313.5RT (min)

lcorbi\_t3\_neg\_4\_040\_19

m/z 426.6944

012E+411.5RT (min)

lcorbi\_t3\_neg\_1\_043\_14

m/z 427.8105

012E+411.5RT (min)

lcorbi\_t3\_neg\_4\_052\_26

m/z 429.1536

0123E+410.511RT (min)

lcorbi\_t3\_neg\_4\_054\_36

m/z 429.1537

02468E+44.55RT (min)

lcorbi\_t3\_neg\_2\_048\_27

m/z 429.8073

012E+411.5RT (min)

lcorbi\_t3\_neg\_3\_013\_28

m/z 430.7684

01E+411.5RT (min)

lcorbi\_t3\_neg\_4\_005\_27

m/z 430.9280

02468E+411.5RT (min)

lcorbi\_t3\_neg\_2\_007\_14

m/z 433.1319

01E+599.5RT (min)

lcorbi\_t3\_neg\_2\_048\_27

m/z 433.3252

024E+41313.5RT (min)

lcorbi\_t3\_neg\_1\_055\_05

m/z 433.8682

024E+411.5RT (min)

lcorbi\_t3\_neg\_3\_018\_13

m/z 434.7472

01234E+411.5RT (min)

lcorbi\_t3\_neg\_4\_039\_32

m/z 434.8714

012E+511.5RT (min)

lcorbi\_t3\_neg\_2\_048\_27

m/z 436.8590

01E+511.5RT (min)

lcorbi\_t3\_neg\_3\_018\_13

m/z 437.1610

0246E+47.58RT (min)

lcorbi\_t3\_neg\_4\_034\_01

m/z 437.3548

01E+51212.5RT (min)

lcorbi\_t3\_neg\_2\_022\_24

m/z 437.9904

01E+41010.5RT (min)

lcorbi\_t3\_neg\_3\_037\_36

m/z 438.1602

01E+41010.5RT (min)

lcorbi\_t3\_neg\_4\_023\_18

m/z 438.7256

0123E+411.5RT (min)

lcorbi\_t3\_neg\_1\_055\_05

m/z 438.9890

0246E+410.511RT (min)

lcorbi\_t3\_neg\_1\_043\_14

m/z 440.1903

024E+44.55RT (min)

lcorbi\_t3\_neg\_4\_034\_01

m/z 440.3127

01234E+51212.5RT (min)

lcorbi\_t3\_neg\_1\_055\_05

m/z 440.7230

0123E+411.5RT (min)

lcorbi\_t3\_neg\_3\_002\_24

m/z 440.9861

0246E+410.511RT (min)

lcorbi\_t3\_neg\_1\_043\_14

m/z 441.0451

0246E+411.5RT (min)

lcorbi\_t3\_neg\_2\_052\_32

m/z 441.2872

02468E+413.514RT (min)

lcorbi\_t3\_neg\_4\_034\_01

m/z 441.9311

02468E+488.5RT (min)

lcorbi\_t3\_neg\_1\_054\_37

m/z 441.9799

01E+510.511RT (min)

lcorbi\_t3\_neg\_1\_043\_14

m/z 442.8937

01E+41111.5RT (min)

lcorbi\_t3\_neg\_2\_010\_02

m/z 443.2795

012E+41414.5RT (min)

lcorbi\_t3\_neg\_3\_023\_07

m/z 443.3162

0246E+41212.5RT (min)

lcorbi\_t3\_neg\_4\_032\_23

m/z 443.8973

02468E+411.5RT (min)

lcorbi\_t3\_neg\_4\_007\_13

m/z 444.7758

024E+411.5RT (min)

lcorbi\_t3\_neg\_4\_039\_32

m/z 445.1287

024E+41010.5RT (min)

lcorbi\_t3\_neg\_1\_007\_26

m/z 446.7733

0246E+411.5RT (min)

lcorbi\_t3\_neg\_3\_052\_32

m/z 446.9055

0123E+511.5RT (min)

lcorbi\_t3\_neg\_2\_048\_27

m/z 447.3325

0123E+41313.5RT (min)

lcorbi\_t3\_neg\_4\_017\_51

m/z 448.3064

01E+58.59RT (min)

lcorbi\_t3\_neg\_2\_048\_27

m/z 449.3098

01E+59.510RT (min)

lcorbi\_t3\_neg\_4\_023\_18

m/z 449.3098

0123E+48.59RT (min)

lcorbi\_t3\_neg\_2\_048\_27

m/z 450.8569

02468E+411.5RT (min)

lcorbi\_t3\_neg\_4\_044\_38

m/z 451.3367

01E+412.513RT (min)

lcorbi\_t3\_neg\_4\_017\_51

m/z 452.2779

0123E+410.511RT (min)

lcorbi\_t3\_neg\_4\_007\_13

m/z 452.9224

012E+41111.5RT (min)

lcorbi\_t3\_neg\_2\_048\_27

m/z 452.9224

0123E+41111.5RT (min)

lcorbi\_t3\_neg\_4\_050\_02

m/z 454.0445

02468E+488.5RT (min)

lcorbi\_t3\_neg\_4\_052\_26

m/z 454.8049

02468E+411.5RT (min)

lcorbi\_t3\_neg\_4\_039\_32

m/z 454.9268

02468E+411.5RT (min)

lcorbi\_t3\_neg\_2\_027\_13

m/z 455.1593

01E+52.53RT (min)

lcorbi\_t3\_neg\_4\_040\_19

m/z 455.2734

01E+49.510RT (min)

lcorbi\_t3\_neg\_4\_054\_36

m/z 455.3044

024E+412.513RT (min)

lcorbi\_t3\_neg\_3\_024\_26

m/z 456.2992

012E+413.514RT (min)

lcorbi\_t3\_neg\_4\_034\_01

m/z 456.3633

02468E+411.512RT (min)

lcorbi\_t3\_neg\_4\_007\_13

m/z 456.7811

01234E+411.5RT (min)

lcorbi\_t3\_neg\_4\_042\_14

m/z 456.8020

01234E+411.5RT (min)

lcorbi\_t3\_neg\_4\_039\_32

m/z 459.3244

01E+51313.5RT (min)

lcorbi\_t3\_neg\_1\_055\_05

m/z 459.3268

0123E+41414.5RT (min)

lcorbi\_t3\_neg\_1\_056\_56

m/z 464.8130

024E+411.5RT (min)

lcorbi\_t3\_neg\_4\_042\_14

m/z 465.2133

012E+41010.5RT (min)

lcorbi\_t3\_neg\_4\_044\_38

m/z 465.3580

01234E+41313.5RT (min)

lcorbi\_t3\_neg\_4\_042\_14

m/z 466.2782

0123E+41212.5RT (min)

lcorbi\_t3\_neg\_4\_032\_23

m/z 466.3301

0123E+411.512RT (min)

lcorbi\_t3\_neg\_4\_034\_01

m/z 466.8114

024E+411.5RT (min)

lcorbi\_t3\_neg\_1\_055\_05

m/z 466.8310

01234E+411.5RT (min)

lcorbi\_t3\_neg\_1\_053\_32

m/z 469.1860

0246E+477.5RT (min)

lcorbi\_t3\_neg\_2\_048\_27

m/z 469.8997

024E+411.5RT (min)

lcorbi\_t3\_neg\_3\_017\_51

m/z 470.1479

0246E+488.5RT (min)

lcorbi\_t3\_neg\_4\_023\_18

m/z 471.3412

0123E+412.513RT (min)

lcorbi\_t3\_neg\_3\_007\_16

m/z 472.3453

012E+511.512RT (min)

lcorbi\_t3\_neg\_1\_055\_05

m/z 473.2822

012E+41313.5RT (min)

lcorbi\_t3\_neg\_2\_020\_05

m/z 474.8417

0246E+411.5RT (min)

lcorbi\_t3\_neg\_3\_053\_14

m/z 474.8625

012E+511.5RT (min)

lcorbi\_t3\_neg\_2\_052\_32

m/z 476.7319

0123E+411.5RT (min)

lcorbi\_t3\_neg\_4\_044\_38

m/z 476.9584

012E+411.5RT (min)

lcorbi\_t3\_neg\_3\_037\_36

m/z 478.2935

01234E+410.511RT (min)

lcorbi\_t3\_neg\_1\_055\_05

m/z 480.7261

01234E+411.5RT (min)

lcorbi\_t3\_neg\_4\_054\_36

m/z 480.8703

01E+511.5RT (min)

lcorbi\_t3\_neg\_4\_035\_33

m/z 482.3152

012E+41212.5RT (min)

lcorbi\_t3\_neg\_4\_034\_01

m/z 482.9751

012E+511.5RT (min)

lcorbi\_t3\_neg\_2\_048\_27

m/z 484.3258

024E+41212.5RT (min)

lcorbi\_t3\_neg\_4\_010\_28

m/z 484.8705

01E+511.5RT (min)

lcorbi\_t3\_neg\_3\_049\_55

m/z 485.2728

01E+51414.5RT (min)

lcorbi\_t3\_neg\_2\_048\_27

m/z 485.6665

01234E+411.5RT (min)

lcorbi\_t3\_neg\_2\_039\_31

m/z 486.7603

024E+411.5RT (min)

lcorbi\_t3\_neg\_4\_039\_32

m/z 486.7860

0246E+411.5RT (min)

lcorbi\_t3\_neg\_4\_007\_13

m/z 488.7580

0246E+411.5RT (min)

lcorbi\_t3\_neg\_4\_029\_31

m/z 489.3585

0123E+411.512RT (min)

lcorbi\_t3\_neg\_4\_005\_27

m/z 492.6548

012E+411.5RT (min)

lcorbi\_t3\_neg\_2\_024\_37

m/z 492.7316

024E+411.5RT (min)

lcorbi\_t3\_neg\_4\_042\_14

m/z 494.5932

0123E+411.5RT (min)

lcorbi\_t3\_neg\_2\_007\_14

m/z 494.7029

01234E+411.5RT (min)

lcorbi\_t3\_neg\_4\_039\_32

m/z 495.3091

01E+41313.5RT (min)

lcorbi\_t3\_neg\_4\_029\_31

m/z 495.6503

0123E+411.5RT (min)

lcorbi\_t3\_neg\_3\_020\_31

m/z 496.7896

02468E+411.5RT (min)

lcorbi\_t3\_neg\_3\_020\_31

m/z 498.2625

024E+411.512RT (min)

lcorbi\_t3\_neg\_4\_047\_30

m/z 498.2892

012E+58.59RT (min)

lcorbi\_t3\_neg\_4\_023\_18

m/z 500.3064

012E+41111.5RT (min)

lcorbi\_t3\_neg\_4\_034\_01

m/z 501.7636

012E+411.5RT (min)

lcorbi\_t3\_neg\_3\_020\_31

m/z 501.8560

02468E+411.5RT (min)

lcorbi\_t3\_neg\_1\_032\_13

m/z 502.2940

01E+510.511RT (min)

lcorbi\_t3\_neg\_1\_055\_05

m/z 503.3863

01E+511.512RT (min)

lcorbi\_t3\_neg\_1\_055\_05

m/z 503.6674

01E+411.5RT (min)

lcorbi\_t3\_neg\_2\_005\_01

m/z 504.7110

0246E+411.5RT (min)

lcorbi\_t3\_neg\_3\_053\_14

m/z 504.8466

01E+511.5RT (min)

lcorbi\_t3\_neg\_3\_018\_13

m/z 504.8641

0246E+411.5RT (min)

lcorbi\_t3\_neg\_4\_042\_14

m/z 505.6657

012E+411.5RT (min)

lcorbi\_t3\_neg\_3\_037\_36

m/z 507.9775

01E+54.55RT (min)

lcorbi\_t3\_neg\_2\_048\_27

m/z 508.1416

012E+411.5RT (min)

lcorbi\_t3\_neg\_4\_012\_17

m/z 508.8156

02468E+411.5RT (min)

lcorbi\_t3\_neg\_3\_028\_35

m/z 510.7921

02468E+411.5RT (min)

lcorbi\_t3\_neg\_4\_042\_14

m/z 510.8811

01E+41111.5RT (min)

lcorbi\_t3\_neg\_3\_029\_02

m/z 511.2910

01234E+411.512RT (min)

lcorbi\_t3\_neg\_4\_005\_27

m/z 511.8848

01E+511.5RT (min)

lcorbi\_t3\_neg\_3\_018\_13

m/z 512.7427

01234E+411.5RT (min)

lcorbi\_t3\_neg\_1\_043\_14

m/z 512.7634

01234E+411.5RT (min)

lcorbi\_t3\_neg\_4\_039\_32

m/z 513.3115

0123E+41212.5RT (min)

lcorbi\_t3\_neg\_4\_023\_18

m/z 514.6461

01234E+411.5RT (min)

lcorbi\_t3\_neg\_4\_046\_37

m/z 514.7397

0246E+411.5RT (min)

lcorbi\_t3\_neg\_4\_042\_14

m/z 514.8929

01234E+511.5RT (min)

lcorbi\_t3\_neg\_3\_010\_15

m/z 515.1920

012E+47.58RT (min)

lcorbi\_t3\_neg\_4\_037\_29

m/z 515.3334

0246E+412.513RT (min)

lcorbi\_t3\_neg\_4\_007\_13

m/z 515.3709

0246E+412.513RT (min)

lcorbi\_t3\_neg\_4\_011\_24

m/z 516.2939

01234E+49.510RT (min)

lcorbi\_t3\_neg\_4\_023\_18

m/z 516.6434

0123E+411.5RT (min)

lcorbi\_t3\_neg\_4\_046\_37

m/z 517.3896

0123E+41212.5RT (min)

lcorbi\_t3\_neg\_2\_007\_14

m/z 517.6431

012E+411.5RT (min)

lcorbi\_t3\_neg\_4\_054\_36

m/z 518.2441

01E+511.512RT (min)

lcorbi\_t3\_neg\_4\_007\_13

m/z 518.2520

01E+51010.5RT (min)

lcorbi\_t3\_neg\_1\_055\_05

m/z 518.2521

01234E+41010.5RT (min)

lcorbi\_t3\_neg\_4\_029\_31

m/z 518.8444

0246E+411.5RT (min)

lcorbi\_t3\_neg\_4\_044\_38

m/z 519.0616

01E+511.5RT (min)

lcorbi\_t3\_neg\_2\_052\_32

m/z 519.2475

0123E+411.512RT (min)

lcorbi\_t3\_neg\_4\_007\_13

m/z 519.9912

0123E+45.56RT (min)

lcorbi\_t3\_neg\_3\_014\_19

m/z 520.9100

012E+41111.5RT (min)

lcorbi\_t3\_neg\_3\_029\_02

m/z 521.6343

01234E+411.5RT (min)

lcorbi\_t3\_neg\_4\_029\_31

m/z 522.7718

0246E+411.5RT (min)

lcorbi\_t3\_neg\_4\_042\_14

m/z 522.7925

0246E+411.5RT (min)

lcorbi\_t3\_neg\_4\_039\_32

m/z 522.9146

01E+511.5RT (min)

lcorbi\_t3\_neg\_3\_018\_13

m/z 524.7691

0246E+411.5RT (min)

lcorbi\_t3\_neg\_1\_043\_14

m/z 524.7894

024E+411.5RT (min)

lcorbi\_t3\_neg\_4\_017\_51

m/z 529.3047

02468E+41010.5RT (min)

lcorbi\_t3\_neg\_4\_032\_23

m/z 529.3127

0246E+411.512RT (min)

lcorbi\_t3\_neg\_4\_038\_15

m/z 529.3127

01E+511.512RT (min)

lcorbi\_t3\_neg\_4\_034\_01

m/z 529.3127

02468E+41212.5RT (min)

lcorbi\_t3\_neg\_4\_007\_13

m/z 530.3082

012E+41010.5RT (min)

lcorbi\_t3\_neg\_4\_032\_23

m/z 530.6199

0123E+411.5RT (min)

lcorbi\_t3\_neg\_4\_017\_51

m/z 530.8668

02468E+411.5RT (min)

lcorbi\_t3\_neg\_2\_007\_14

m/z 531.6183

0123E+411.5RT (min)

lcorbi\_t3\_neg\_2\_035\_46

m/z 532.6168

012E+411.5RT (min)

lcorbi\_t3\_neg\_4\_039\_32

m/z 532.8005

02468E+411.5RT (min)

lcorbi\_t3\_neg\_3\_053\_14

m/z 532.8213

02468E+411.5RT (min)

lcorbi\_t3\_neg\_4\_039\_32

m/z 534.2947

01E+512.513RT (min)

lcorbi\_t3\_neg\_3\_007\_16

m/z 534.3024

01234E+413.514RT (min)

lcorbi\_t3\_neg\_1\_023\_03

m/z 534.6459

012E+411.5RT (min)

lcorbi\_t3\_neg\_2\_052\_32

m/z 534.6899

01E+411.5RT (min)

lcorbi\_t3\_neg\_2\_043\_33

m/z 536.6876

01234E+411.5RT (min)

lcorbi\_t3\_neg\_4\_039\_32

m/z 537.4158

01E+411.512RT (min)

lcorbi\_t3\_neg\_4\_034\_01

m/z 537.8873

02468E+411.5RT (min)

lcorbi\_t3\_neg\_3\_051\_46

m/z 538.6843

0123E+411.5RT (min)

lcorbi\_t3\_neg\_4\_046\_37

m/z 538.8293

01E+511.5RT (min)

lcorbi\_t3\_neg\_4\_044\_38

m/z 540.0542

01E+511.5RT (min)

lcorbi\_t3\_neg\_4\_046\_37

m/z 540.3619

01E+51111.5RT (min)

lcorbi\_t3\_neg\_4\_034\_01

m/z 540.3623

01E+511.512RT (min)

lcorbi\_t3\_neg\_4\_005\_27

m/z 540.3685

0246E+41111.5RT (min)

lcorbi\_t3\_neg\_2\_007\_14

m/z 540.3828

0123E+411.512RT (min)

lcorbi\_t3\_neg\_4\_042\_14

m/z 542.2498

02468E+410.511RT (min)

lcorbi\_t3\_neg\_4\_021\_25

m/z 542.3325

01E+51010.5RT (min)

lcorbi\_t3\_neg\_1\_055\_05

m/z 542.6121

0123E+411.5RT (min)

lcorbi\_t3\_neg\_1\_043\_14

m/z 542.6583

012E+411.5RT (min)

lcorbi\_t3\_neg\_4\_005\_27

m/z 542.8294

01E+511.5RT (min)

lcorbi\_t3\_neg\_3\_053\_14

m/z 542.8500

01E+511.5RT (min)

lcorbi\_t3\_neg\_2\_052\_32

m/z 543.6250

0123E+411.5RT (min)

lcorbi\_t3\_neg\_3\_045\_34

m/z 544.6093

01234E+411.5RT (min)

lcorbi\_t3\_neg\_4\_038\_15

m/z 545.6221

01234E+411.5RT (min)

lcorbi\_t3\_neg\_2\_012\_36

m/z 548.5574

0123E+411.5RT (min)

lcorbi\_t3\_neg\_4\_042\_14

m/z 548.8518

0246E+411.5RT (min)

lcorbi\_t3\_neg\_2\_048\_27

m/z 548.8580

012E+511.5RT (min)

lcorbi\_t3\_neg\_4\_035\_33

m/z 550.3513

012E+411.512RT (min)

lcorbi\_t3\_neg\_4\_005\_27

m/z 550.6142

01234E+411.5RT (min)

lcorbi\_t3\_neg\_3\_047\_37

m/z 550.6901

0246E+411.5RT (min)

lcorbi\_t3\_neg\_4\_038\_15

m/z 552.3668

01234E+51212.5RT (min)

lcorbi\_t3\_neg\_4\_005\_27

m/z 552.6411

024E+411.5RT (min)

lcorbi\_t3\_neg\_4\_042\_14

m/z 552.7765

0123E+511.5RT (min)

lcorbi\_t3\_neg\_4\_007\_13

m/z 552.8580

01E+511.5RT (min)

lcorbi\_t3\_neg\_3\_051\_46

m/z 553.3759

0246E+411.512RT (min)

lcorbi\_t3\_neg\_4\_029\_31

m/z 554.6073

01234E+411.5RT (min)

lcorbi\_t3\_neg\_4\_045\_35

m/z 555.3020

0123E+41010.5RT (min)

lcorbi\_t3\_neg\_4\_032\_23

m/z 556.2996

024E+41111.5RT (min)

lcorbi\_t3\_neg\_4\_007\_13

m/z 556.2997

024E+41111.5RT (min)

lcorbi\_t3\_neg\_4\_007\_13

m/z 557.3029

01E+510.511RT (min)

lcorbi\_t3\_neg\_4\_042\_14

m/z 557.3368

0123E+41111.5RT (min)

lcorbi\_t3\_neg\_4\_046\_37

m/z 559.3427

012E+41313.5RT (min)

lcorbi\_t3\_neg\_4\_017\_51

m/z 559.5992

01234E+411.5RT (min)

lcorbi\_t3\_neg\_4\_001\_49

m/z 560.7194

024E+411.5RT (min)

lcorbi\_t3\_neg\_4\_005\_27

m/z 560.8269

012E+411.5RT (min)

lcorbi\_t3\_neg\_2\_051\_34

m/z 561.3784

01234E+41212.5RT (min)

lcorbi\_t3\_neg\_2\_002\_26

m/z 561.5966

01234E+411.5RT (min)

lcorbi\_t3\_neg\_2\_043\_33

m/z 562.6700

0246E+411.5RT (min)

lcorbi\_t3\_neg\_4\_042\_14

m/z 562.8055

012E+511.5RT (min)

lcorbi\_t3\_neg\_3\_018\_13

m/z 564.2969

012E+41111.5RT (min)

lcorbi\_t3\_neg\_4\_044\_38

m/z 564.2971

0123E+41111.5RT (min)

lcorbi\_t3\_neg\_4\_034\_01

m/z 564.3718

01E+41111.5RT (min)

lcorbi\_t3\_neg\_4\_005\_27

m/z 564.8322

0246E+411.5RT (min)

lcorbi\_t3\_neg\_2\_052\_32

m/z 565.6224

012E+411.5RT (min)

lcorbi\_t3\_neg\_4\_046\_37

m/z 566.3301

01E+510.511RT (min)

lcorbi\_t3\_neg\_1\_055\_05

m/z 566.3303

024E+410.511RT (min)

lcorbi\_t3\_neg\_3\_052\_32

m/z 566.5156

01E+51414.5RT (min)

lcorbi\_t3\_neg\_4\_032\_23

m/z 566.5156

0246E+41414.5RT (min)

lcorbi\_t3\_neg\_2\_039\_31

m/z 568.3210

0123E+41212.5RT (min)

lcorbi\_t3\_neg\_4\_034\_01

m/z 568.3461

02468E+41111.5RT (min)

lcorbi\_t3\_neg\_3\_002\_24

m/z 568.7509

01E+511.5RT (min)

lcorbi\_t3\_neg\_4\_038\_15

m/z 570.2759

01E+51010.5RT (min)

lcorbi\_t3\_neg\_4\_023\_18

m/z 571.1450

01234E+533.5RT (min)

lcorbi\_t3\_neg\_2\_048\_27

m/z 572.6054

0123E+411.5RT (min)

lcorbi\_t3\_neg\_4\_044\_38

m/z 572.6993

0246E+411.5RT (min)

lcorbi\_t3\_neg\_4\_042\_14

m/z 573.4525

02468E+412.513RT (min)

lcorbi\_t3\_neg\_2\_007\_14

m/z 574.4555

0123E+412.513RT (min)

lcorbi\_t3\_neg\_4\_042\_14

m/z 574.6017

01234E+411.5RT (min)

lcorbi\_t3\_neg\_4\_044\_38

m/z 577.4839

0246E+41414.5RT (min)

lcorbi\_t3\_neg\_4\_034\_01

m/z 578.3781

012E+41313.5RT (min)

lcorbi\_t3\_neg\_4\_034\_01

m/z 578.3820

012E+41212.5RT (min)

lcorbi\_t3\_neg\_2\_007\_14

m/z 578.3821

012E+41212.5RT (min)

lcorbi\_t3\_neg\_4\_034\_01

m/z 578.7798

02468E+411.5RT (min)

lcorbi\_t3\_neg\_1\_043\_14

m/z 579.5930

01234E+411.5RT (min)

lcorbi\_t3\_neg\_4\_046\_37

m/z 579.8724

01E+511.5RT (min)

lcorbi\_t3\_neg\_1\_032\_13

m/z 580.8239

0123E+411.5RT (min)

lcorbi\_t3\_neg\_1\_053\_32

m/z 581.8698

01234E+411.5RT (min)

lcorbi\_t3\_neg\_2\_002\_26

m/z 582.3413

012E+413.514RT (min)

lcorbi\_t3\_neg\_4\_023\_18

m/z 582.7482

0123E+411.5RT (min)

lcorbi\_t3\_neg\_2\_052\_32

m/z 582.8212

0123E+411.5RT (min)

lcorbi\_t3\_neg\_3\_052\_32

m/z 582.8808

0123E+511.5RT (min)

lcorbi\_t3\_neg\_4\_034\_01

m/z 583.3002

0246E+411.512RT (min)

lcorbi\_t3\_neg\_4\_007\_13

m/z 584.3121

01E+51212.5RT (min)

lcorbi\_t3\_neg\_4\_007\_13

m/z 584.8350

01E+511.5RT (min)

lcorbi\_t3\_neg\_4\_039\_32

m/z 586.3093

01234E+41212.5RT (min)

lcorbi\_t3\_neg\_4\_007\_13

m/z 586.3154

0123E+41111.5RT (min)

lcorbi\_t3\_neg\_4\_012\_17

m/z 587.3762

01234E+413.514RT (min)

lcorbi\_t3\_neg\_4\_023\_18

m/z 588.3586

01E+41111.5RT (min)

lcorbi\_t3\_neg\_4\_017\_51

m/z 588.3797

01E+413.514RT (min)

lcorbi\_t3\_neg\_3\_024\_26

m/z 588.8978

01E+41111.5RT (min)

lcorbi\_t3\_neg\_3\_029\_02

m/z 589.5780

01234E+411.5RT (min)

lcorbi\_t3\_neg\_3\_030\_33

m/z 589.9012

0123E+511.5RT (min)

lcorbi\_t3\_neg\_4\_007\_13

m/z 591.2728

012E+49.510RT (min)

lcorbi\_t3\_neg\_4\_017\_51

m/z 592.6290

024E+411.5RT (min)

lcorbi\_t3\_neg\_4\_012\_17

m/z 593.3920

01E+414.515RT (min)

lcorbi\_t3\_neg\_1\_005\_07

m/z 593.3974

01234E+41414.5RT (min)

lcorbi\_t3\_neg\_4\_019\_08

m/z 593.4104

0246E+41414.5RT (min)

lcorbi\_t3\_neg\_3\_023\_07

m/z 594.4125

012E+41414.5RT (min)

lcorbi\_t3\_neg\_4\_040\_19

m/z 594.5850

0246E+411.5RT (min)

lcorbi\_t3\_neg\_3\_018\_13

m/z 594.6716

0246E+411.5RT (min)

lcorbi\_t3\_neg\_3\_007\_16

m/z 594.8639

01E+511.5RT (min)

lcorbi\_t3\_neg\_4\_044\_38

m/z 596.3257

024E+413.514RT (min)

lcorbi\_t3\_neg\_4\_040\_19

m/z 596.4902

02468E+41414.5RT (min)

lcorbi\_t3\_neg\_4\_032\_23

m/z 596.8400

012E+511.5RT (min)

lcorbi\_t3\_neg\_4\_038\_15

m/z 597.4031

01E+51111.5RT (min)

lcorbi\_t3\_neg\_2\_039\_31

m/z 599.3145

01234E+410.511RT (min)

lcorbi\_t3\_neg\_4\_017\_51

m/z 600.6174

024E+411.5RT (min)

lcorbi\_t3\_neg\_2\_048\_27

m/z 602.6575

0246E+411.5RT (min)

lcorbi\_t3\_neg\_4\_038\_15

m/z 604.5673

024E+411.5RT (min)

lcorbi\_t3\_neg\_2\_048\_27

m/z 604.7001

0123E+411.5RT (min)

lcorbi\_t3\_neg\_4\_005\_27

m/z 606.0742

01E+511.5RT (min)

lcorbi\_t3\_neg\_4\_046\_37

m/z 606.2969

01E+511.512RT (min)

lcorbi\_t3\_neg\_4\_007\_13

m/z 606.8170

01E+511.5RT (min)

lcorbi\_t3\_neg\_4\_039\_32

m/z 607.3058

012E+410.511RT (min)

lcorbi\_t3\_neg\_4\_007\_13

m/z 609.5709

024E+411.5RT (min)

lcorbi\_t3\_neg\_2\_028\_38

m/z 610.4404

0246E+41414.5RT (min)

lcorbi\_t3\_neg\_1\_035\_25

m/z 610.5696

0246E+411.5RT (min)

lcorbi\_t3\_neg\_4\_045\_35

m/z 610.5999

024E+411.5RT (min)

lcorbi\_t3\_neg\_4\_042\_14

m/z 610.6462

0123E+411.5RT (min)

lcorbi\_t3\_neg\_2\_048\_27

m/z 610.8174

01E+511.5RT (min)

lcorbi\_t3\_neg\_3\_053\_14

m/z 611.5344

01234E+411.5RT (min)

lcorbi\_t3\_neg\_1\_043\_14

m/z 612.1479

0246E+411.5RT (min)

lcorbi\_t3\_neg\_4\_029\_31

m/z 612.5660

024E+411.5RT (min)

lcorbi\_t3\_neg\_3\_028\_35

m/z 612.7324

02468E+411.5RT (min)

lcorbi\_t3\_neg\_4\_007\_13

m/z 613.1406

01E+411.5RT (min)

lcorbi\_t3\_neg\_4\_029\_31

m/z 616.3749

0123E+411.512RT (min)

lcorbi\_t3\_neg\_4\_029\_31

m/z 616.8458

012E+511.5RT (min)

lcorbi\_t3\_neg\_4\_044\_38

m/z 618.6778

0246E+411.5RT (min)

lcorbi\_t3\_neg\_1\_043\_14

m/z 620.7641

012E+511.5RT (min)

lcorbi\_t3\_neg\_3\_018\_13

m/z 620.8458

01E+511.5RT (min)

lcorbi\_t3\_neg\_3\_053\_14

m/z 621.4437

012E+41414.5RT (min)

lcorbi\_t3\_neg\_3\_023\_07

m/z 622.6270

0246E+411.5RT (min)

lcorbi\_t3\_neg\_4\_042\_14

m/z 622.7151

01E+511.5RT (min)

lcorbi\_t3\_neg\_1\_043\_14

m/z 628.7059

024E+411.5RT (min)

lcorbi\_t3\_neg\_3\_004\_30

m/z 630.2083

012E+511.5RT (min)

lcorbi\_t3\_neg\_1\_055\_05

m/z 630.6581

02468E+411.5RT (min)

lcorbi\_t3\_neg\_4\_042\_14

m/z 630.7933

012E+511.5RT (min)

lcorbi\_t3\_neg\_1\_032\_13

m/z 632.7439

01E+511.5RT (min)

lcorbi\_t3\_neg\_2\_004\_28

m/z 633.2072

01E+511.5RT (min)

lcorbi\_t3\_neg\_1\_055\_05

m/z 634.5696

024E+411.5RT (min)

lcorbi\_t3\_neg\_2\_048\_27

m/z 634.6140

024E+411.5RT (min)

lcorbi\_t3\_neg\_4\_007\_13

m/z 635.2169

01E+455.5RT (min)

lcorbi\_t3\_neg\_4\_017\_51

m/z 636.7385

01E+511.5RT (min)

lcorbi\_t3\_neg\_2\_004\_28

m/z 638.5502

01234E+411.5RT (min)

lcorbi\_t3\_neg\_4\_046\_37

m/z 638.7166

02468E+411.5RT (min)

lcorbi\_t3\_neg\_4\_007\_13

m/z 638.8249

0246E+511.5RT (min)

lcorbi\_t3\_neg\_2\_027\_13

m/z 639.5490

0246E+411.5RT (min)

lcorbi\_t3\_neg\_3\_037\_36

m/z 640.5471

01234E+411.5RT (min)

lcorbi\_t3\_neg\_3\_020\_31

m/z 641.5457

01234E+411.5RT (min)

lcorbi\_t3\_neg\_2\_039\_31

m/z 642.6019

0246E+411.5RT (min)

lcorbi\_t3\_neg\_4\_007\_13

m/z 643.1172

0246E+42.53RT (min)

lcorbi\_t3\_neg\_3\_005\_23

m/z 645.2275

01234E+42.53RT (min)

lcorbi\_t3\_neg\_3\_001\_49

m/z 646.2692

02468E+410.511RT (min)

lcorbi\_t3\_neg\_2\_007\_14

m/z 646.2696

02468E+410.511RT (min)

lcorbi\_t3\_neg\_4\_042\_14

m/z 648.8540

01234E+511.5RT (min)

lcorbi\_t3\_neg\_3\_018\_13

m/z 650.5540

01234E+41414.5RT (min)

lcorbi\_t3\_neg\_3\_005\_23

m/z 650.5636

0123E+411.5RT (min)

lcorbi\_t3\_neg\_4\_044\_38

m/z 650.6132

01E+51414.5RT (min)

lcorbi\_t3\_neg\_2\_052\_32

m/z 650.6133

01234E+51414.5RT (min)

lcorbi\_t3\_neg\_2\_022\_24

m/z 652.6299

0246E+411.5RT (min)

lcorbi\_t3\_neg\_4\_007\_13

m/z 652.8017

0123E+511.5RT (min)

lcorbi\_t3\_neg\_3\_053\_14

m/z 654.5406

01234E+411.5RT (min)

lcorbi\_t3\_neg\_4\_007\_13

m/z 654.6709

01E+511.5RT (min)

lcorbi\_t3\_neg\_4\_007\_13

m/z 654.7989

012E+511.5RT (min)

lcorbi\_t3\_neg\_3\_053\_14

m/z 655.3121

01E+41111.5RT (min)

lcorbi\_t3\_neg\_4\_041\_54

m/z 656.5786

0246E+411.5RT (min)

lcorbi\_t3\_neg\_2\_045\_30

m/z 657.8885

012E+511.5RT (min)

lcorbi\_t3\_neg\_3\_018\_13

m/z 659.5139

012E+41414.5RT (min)

lcorbi\_t3\_neg\_4\_007\_13

m/z 660.5294

024E+411.5RT (min)

lcorbi\_t3\_neg\_2\_048\_27

m/z 662.1022

012E+511.5RT (min)

lcorbi\_t3\_neg\_4\_029\_31

m/z 662.5272

01234E+411.5RT (min)

lcorbi\_t3\_neg\_4\_034\_01

m/z 662.8303

01234E+511.5RT (min)

lcorbi\_t3\_neg\_2\_045\_30

m/z 663.1052

024E+411.5RT (min)

lcorbi\_t3\_neg\_4\_029\_31

m/z 663.1829

01E+52.53RT (min)

lcorbi\_t3\_neg\_3\_002\_24

m/z 664.7000

01E+511.5RT (min)

lcorbi\_t3\_neg\_4\_007\_13

m/z 664.8282

01E+511.5RT (min)

lcorbi\_t3\_neg\_3\_053\_14

m/z 666.3937

01234E+41111.5RT (min)

lcorbi\_t3\_neg\_4\_029\_31

m/z 666.5488

012E+414.515RT (min)

lcorbi\_t3\_neg\_3\_005\_23

m/z 666.6076

02468E+411.5RT (min)

lcorbi\_t3\_neg\_4\_038\_15

m/z 668.3182

0123E+513.514RT (min)

lcorbi\_t3\_neg\_3\_005\_23

m/z 669.4943

012E+411.5RT (min)

lcorbi\_t3\_neg\_1\_051\_15

m/z 670.5562

01234E+411.5RT (min)

lcorbi\_t3\_neg\_1\_043\_14

m/z 670.6451

0246E+411.5RT (min)

lcorbi\_t3\_neg\_3\_032\_17

m/z 670.6912

01E+511.5RT (min)

lcorbi\_t3\_neg\_3\_018\_13

m/z 672.8587

0246E+511.5RT (min)

lcorbi\_t3\_neg\_3\_053\_14

m/z 674.5239

0123E+411.5RT (min)

lcorbi\_t3\_neg\_4\_039\_32

m/z 674.7285

012E+511.5RT (min)

lcorbi\_t3\_neg\_4\_007\_13

m/z 675.4849

01234E+411.5RT (min)

lcorbi\_t3\_neg\_1\_043\_14

m/z 676.6366

02468E+411.5RT (min)

lcorbi\_t3\_neg\_4\_042\_14

m/z 678.5524

01234E+413.514RT (min)

lcorbi\_t3\_neg\_2\_022\_24

m/z 678.7226

0123E+511.5RT (min)

lcorbi\_t3\_neg\_4\_007\_13

m/z 678.8051

01E+511.5RT (min)

lcorbi\_t3\_neg\_4\_042\_14

m/z 680.5494

02468E+413.514RT (min)

lcorbi\_t3\_neg\_4\_040\_19

m/z 680.5855

024E+411.5RT (min)

lcorbi\_t3\_neg\_4\_042\_14

m/z 682.5823

01234E+411.5RT (min)

lcorbi\_t3\_neg\_4\_042\_14

m/z 682.6709

01E+511.5RT (min)

lcorbi\_t3\_neg\_2\_007\_14

m/z 683.1714

012E+511.5RT (min)

lcorbi\_t3\_neg\_1\_053\_32

m/z 684.6682

01E+511.5RT (min)

lcorbi\_t3\_neg\_4\_012\_17

m/z 685.5340

0123E+414.515RT (min)

lcorbi\_t3\_neg\_4\_044\_38

m/z 686.6651

0246E+411.5RT (min)

lcorbi\_t3\_neg\_4\_005\_27

m/z 688.2675

012E+41313.5RT (min)

lcorbi\_t3\_neg\_1\_055\_05

m/z 688.2679

0123E+413.514RT (min)

lcorbi\_t3\_neg\_4\_007\_13

m/z 688.7518

012E+511.5RT (min)

lcorbi\_t3\_neg\_3\_018\_13

m/z 690.7023

01E+511.5RT (min)

lcorbi\_t3\_neg\_2\_004\_28

m/z 692.5088

0246E+41414.5RT (min)

lcorbi\_t3\_neg\_4\_054\_36

m/z 692.6997

01E+511.5RT (min)

lcorbi\_t3\_neg\_4\_038\_15

m/z 693.4590

01E+51414.5RT (min)

lcorbi\_t3\_neg\_2\_043\_33

m/z 694.2884

01E+41111.5RT (min)

lcorbi\_t3\_neg\_3\_032\_17

m/z 694.5687

01E+511.5RT (min)

lcorbi\_t3\_neg\_4\_007\_13

m/z 694.6969

01E+511.5RT (min)

lcorbi\_t3\_neg\_3\_022\_27

m/z 694.7864

0246E+511.5RT (min)

lcorbi\_t3\_neg\_1\_032\_13

m/z 695.5280

01E+514.515RT (min)

lcorbi\_t3\_neg\_4\_054\_36

m/z 695.5833

012E+414.515RT (min)

lcorbi\_t3\_neg\_3\_005\_23

m/z 696.3044

012E+411.512RT (min)

lcorbi\_t3\_neg\_4\_034\_01

m/z 696.5319

01234E+414.515RT (min)

lcorbi\_t3\_neg\_3\_020\_31

m/z 696.7833

0246E+511.5RT (min)

lcorbi\_t3\_neg\_3\_018\_13

m/z 697.2733

01234E+41313.5RT (min)

lcorbi\_t3\_neg\_4\_039\_32

m/z 698.5134

01E+51414.5RT (min)

lcorbi\_t3\_neg\_4\_039\_32

m/z 700.5144

01234E+411.5RT (min)

lcorbi\_t3\_neg\_2\_048\_27

m/z 700.5602

0246E+411.5RT (min)

lcorbi\_t3\_neg\_4\_007\_13

m/z 702.4930

01E+51414.5RT (min)

lcorbi\_t3\_neg\_4\_019\_08

m/z 702.5002

0246E+414.515RT (min)

lcorbi\_t3\_neg\_4\_046\_37

m/z 702.7286

012E+511.5RT (min)

lcorbi\_t3\_neg\_3\_053\_14

m/z 704.4896

0246E+411.5RT (min)

lcorbi\_t3\_neg\_4\_007\_13

m/z 704.5151

02468E+41414.5RT (min)

lcorbi\_t3\_neg\_4\_032\_23

m/z 704.5977

01E+511.5RT (min)

lcorbi\_t3\_neg\_4\_007\_13

m/z 704.8152

02468E+511.5RT (min)

lcorbi\_t3\_neg\_4\_007\_13

m/z 706.5052

024E+411.5RT (min)

lcorbi\_t3\_neg\_2\_048\_27

m/z 706.5244

01E+514.515RT (min)

lcorbi\_t3\_neg\_4\_044\_38

m/z 706.5264

0123E+411.5RT (min)

lcorbi\_t3\_neg\_3\_028\_35

m/z 706.8123

024E+511.5RT (min)

lcorbi\_t3\_neg\_4\_007\_13

m/z 708.5919

01E+511.5RT (min)

lcorbi\_t3\_neg\_4\_007\_13

m/z 710.5430

01234E+411.5RT (min)

lcorbi\_t3\_neg\_2\_048\_27

m/z 710.7600

012E+511.5RT (min)

lcorbi\_t3\_neg\_2\_054\_15

m/z 711.4788

0123E+411.5RT (min)

lcorbi\_t3\_neg\_4\_007\_13

m/z 712.5294

01E+414.515RT (min)

lcorbi\_t3\_neg\_2\_052\_32

m/z 712.5406

02468E+411.5RT (min)

lcorbi\_t3\_neg\_2\_048\_27

m/z 712.7571

012E+511.5RT (min)

lcorbi\_t3\_neg\_2\_054\_15

m/z 713.5586

012E+41414.5RT (min)

lcorbi\_t3\_neg\_4\_007\_13

m/z 714.5078

0123E+413.514RT (min)

lcorbi\_t3\_neg\_2\_051\_34

m/z 714.5081

012E+713.514RT (min)

lcorbi\_t3\_neg\_2\_053\_23

m/z 714.5372

024E+411.5RT (min)

lcorbi\_t3\_neg\_4\_038\_15

m/z 714.8439

01E+611.5RT (min)

lcorbi\_t3\_neg\_3\_018\_13

m/z 716.5243

0246E+61414.5RT (min)

lcorbi\_t3\_neg\_2\_036\_35

m/z 718.4669

012E+411.5RT (min)

lcorbi\_t3\_neg\_4\_034\_01

m/z 718.5143

01E+51414.5RT (min)

lcorbi\_t3\_neg\_4\_019\_08

m/z 718.6205

01E+511.5RT (min)

lcorbi\_t3\_neg\_4\_007\_13

m/z 720.2940

01E+51313.5RT (min)

lcorbi\_t3\_neg\_1\_023\_03

m/z 720.4644

0123E+411.5RT (min)

lcorbi\_t3\_neg\_2\_048\_27

m/z 720.5355

024E+41414.5RT (min)

lcorbi\_t3\_neg\_3\_005\_23

m/z 720.5717

0246E+411.5RT (min)

lcorbi\_t3\_neg\_2\_048\_27

m/z 720.7890

0123E+511.5RT (min)

lcorbi\_t3\_neg\_3\_053\_14

m/z 721.2975

0123E+41313.5RT (min)

lcorbi\_t3\_neg\_1\_023\_03

m/z 721.4645

01234E+413.514RT (min)

lcorbi\_t3\_neg\_4\_044\_38

m/z 721.5241

012E+414.515RT (min)

lcorbi\_t3\_neg\_4\_032\_23

m/z 722.4711

012E+41515.5RT (min)

lcorbi\_t3\_neg\_4\_046\_37

m/z 722.5688

02468E+411.5RT (min)

lcorbi\_t3\_neg\_2\_048\_27

m/z 723.5217

0123E+413.514RT (min)

lcorbi\_t3\_neg\_4\_007\_13

m/z 724.4870

0123E+414.515RT (min)

lcorbi\_t3\_neg\_4\_007\_13

m/z 724.5254

01E+413.514RT (min)

lcorbi\_t3\_neg\_4\_007\_13

m/z 724.5660

0246E+411.5RT (min)

lcorbi\_t3\_neg\_4\_012\_17

m/z 724.6550

012E+511.5RT (min)

lcorbi\_t3\_neg\_4\_007\_13

m/z 724.8726

01E+611.5RT (min)

lcorbi\_t3\_neg\_3\_018\_13

m/z 726.2599

0123E+41414.5RT (min)

lcorbi\_t3\_neg\_4\_030\_04

m/z 726.5345

01E+414.515RT (min)

lcorbi\_t3\_neg\_2\_039\_31

m/z 726.6525

012E+511.5RT (min)

lcorbi\_t3\_neg\_4\_007\_13

m/z 728.6497

01E+511.5RT (min)

lcorbi\_t3\_neg\_3\_018\_13

m/z 729.5618

012E+413.514RT (min)

lcorbi\_t3\_neg\_4\_039\_32

m/z 729.6420

024E+414.515RT (min)

lcorbi\_t3\_neg\_4\_039\_32

m/z 730.5833

012E+41414.5RT (min)

lcorbi\_t3\_neg\_4\_011\_24

m/z 730.6006

02468E+411.5RT (min)

lcorbi\_t3\_neg\_3\_032\_17

m/z 730.8176

01234E+511.5RT (min)

lcorbi\_t3\_neg\_4\_038\_15

m/z 731.5343

01234E+413.514RT (min)

lcorbi\_t3\_neg\_4\_007\_13

m/z 732.5973

01E+511.5RT (min)

lcorbi\_t3\_neg\_3\_032\_17

m/z 733.5037

01E+513.514RT (min)

lcorbi\_t3\_neg\_4\_029\_31

m/z 734.6844

012E+511.5RT (min)

lcorbi\_t3\_neg\_4\_007\_13

m/z 736.5255

01E+513.514RT (min)

lcorbi\_t3\_neg\_4\_007\_13

m/z 736.5294

02468E+414.515RT (min)

lcorbi\_t3\_neg\_4\_039\_32

m/z 736.6809

012E+511.5RT (min)

lcorbi\_t3\_neg\_4\_007\_13

m/z 737.2919

0246E+41313.5RT (min)

lcorbi\_t3\_neg\_4\_030\_04

m/z 737.5328

01234E+41414.5RT (min)

lcorbi\_t3\_neg\_4\_039\_32

m/z 737.6722

01234E+414.515RT (min)

lcorbi\_t3\_neg\_1\_053\_32

m/z 738.5079

024E+413.514RT (min)

lcorbi\_t3\_neg\_4\_017\_51

m/z 738.5088

01E+713.514RT (min)

lcorbi\_t3\_neg\_3\_047\_37

m/z 738.5450

0123E+414.515RT (min)

lcorbi\_t3\_neg\_4\_007\_13

m/z 739.4598

02468E+411.5RT (min)

lcorbi\_t3\_neg\_4\_007\_13

m/z 739.5113

01E+413.514RT (min)

lcorbi\_t3\_neg\_4\_034\_01

m/z 739.5125

024E+613.514RT (min)

lcorbi\_t3\_neg\_4\_045\_35

m/z 740.2635

01234E+41313.5RT (min)

lcorbi\_t3\_neg\_1\_023\_03

m/z 740.8463

024E+511.5RT (min)

lcorbi\_t3\_neg\_3\_029\_02

m/z 741.4691

0123E+41414.5RT (min)

lcorbi\_t3\_neg\_1\_044\_23

m/z 741.5709

01E+41515.5RT (min)

lcorbi\_t3\_neg\_2\_050\_17

m/z 742.6263

01E+511.5RT (min)

lcorbi\_t3\_neg\_2\_048\_27

m/z 742.7158

01E+511.5RT (min)

lcorbi\_t3\_neg\_2\_027\_13

m/z 743.5345

012E+413.514RT (min)

lcorbi\_t3\_neg\_4\_007\_13

m/z 744.4958

01E+511.5RT (min)

lcorbi\_t3\_neg\_4\_007\_13

m/z 744.6236

024E+411.5RT (min)

lcorbi\_t3\_neg\_3\_032\_17

m/z 744.7131

0123E+511.5RT (min)

lcorbi\_t3\_neg\_2\_027\_13

m/z 745.5499

02468E+61313.5RT (min)

lcorbi\_t3\_neg\_4\_007\_13

m/z 745.6396

012E+414.515RT (min)

lcorbi\_t3\_neg\_3\_020\_31

m/z 746.4931

01E+511.5RT (min)

lcorbi\_t3\_neg\_4\_007\_13

m/z 746.5515

024E+41414.5RT (min)

lcorbi\_t3\_neg\_3\_005\_23

m/z 746.7925

01E+511.5RT (min)

lcorbi\_t3\_neg\_3\_053\_14

m/z 747.5640

01234E+414.515RT (min)

lcorbi\_t3\_neg\_2\_053\_23

m/z 747.5652

01E+713.514RT (min)

lcorbi\_t3\_neg\_4\_007\_13

m/z 748.5685

0246E+613.514RT (min)

lcorbi\_t3\_neg\_4\_007\_13

m/z 752.5159

01234E+413.514RT (min)

lcorbi\_t3\_neg\_4\_046\_37

m/z 754.4646

012E+513.514RT (min)

lcorbi\_t3\_neg\_3\_005\_23

m/z 754.5754

012E+414.515RT (min)

lcorbi\_t3\_neg\_4\_035\_33

m/z 754.7420

0123E+511.5RT (min)

lcorbi\_t3\_neg\_1\_032\_13

m/z 755.6598

012E+414.515RT (min)

lcorbi\_t3\_neg\_2\_039\_31

m/z 756.3196

01E+414.515RT (min)

lcorbi\_t3\_neg\_4\_012\_17

m/z 756.5216

02468E+411.5RT (min)

lcorbi\_t3\_neg\_4\_007\_13

m/z 758.5612

02468E+413.514RT (min)

lcorbi\_t3\_neg\_4\_044\_38

m/z 759.4906

01E+41313.5RT (min)

lcorbi\_t3\_neg\_3\_007\_16

m/z 760.4906

0123E+411.5RT (min)

lcorbi\_t3\_neg\_4\_046\_37

m/z 760.6047

01234E+413.514RT (min)

lcorbi\_t3\_neg\_2\_052\_32

m/z 761.4995

012E+414.515RT (min)

lcorbi\_t3\_neg\_4\_029\_31

m/z 762.2759

01E+41111.5RT (min)

lcorbi\_t3\_neg\_2\_007\_14

m/z 762.4218

0123E+411.5RT (min)

lcorbi\_t3\_neg\_3\_053\_14

m/z 762.5284

0246E+413.514RT (min)

lcorbi\_t3\_neg\_4\_007\_13

m/z 762.5382

01234E+41414.5RT (min)

lcorbi\_t3\_neg\_3\_005\_23

m/z 762.5443

012E+41414.5RT (min)

lcorbi\_t3\_neg\_4\_034\_01

m/z 762.5486

01E+413.514RT (min)

lcorbi\_t3\_neg\_3\_028\_35

m/z 762.7734

01234E+511.5RT (min)

lcorbi\_t3\_neg\_2\_027\_13

m/z 763.5107

01E+513.514RT (min)

lcorbi\_t3\_neg\_2\_039\_31

m/z 763.5607

024E+41313.5RT (min)

lcorbi\_t3\_neg\_4\_007\_13

m/z 764.4442

0246E+411.5RT (min)

lcorbi\_t3\_neg\_4\_007\_13

m/z 764.5230

0123E+413.514RT (min)

lcorbi\_t3\_neg\_4\_034\_01

m/z 765.5756

012E+413.514RT (min)

lcorbi\_t3\_neg\_4\_007\_13

m/z 766.5808

01234E+41414.5RT (min)

lcorbi\_t3\_neg\_4\_007\_13

m/z 767.4409

0246E+411.5RT (min)

lcorbi\_t3\_neg\_4\_007\_13

m/z 767.4869

01234E+413.514RT (min)

lcorbi\_t3\_neg\_4\_054\_36

m/z 767.4908

012E+414.515RT (min)

lcorbi\_t3\_neg\_4\_034\_01

m/z 768.3564

012E+413.514RT (min)

lcorbi\_t3\_neg\_4\_017\_51

m/z 769.5501

01234E+413.514RT (min)

lcorbi\_t3\_neg\_4\_007\_13

m/z 769.5577

0246E+414.515RT (min)

lcorbi\_t3\_neg\_3\_007\_16

m/z 771.4328

0123E+411.5RT (min)

lcorbi\_t3\_neg\_2\_048\_27

m/z 771.5643

01E+51414.5RT (min)

lcorbi\_t3\_neg\_1\_054\_37

m/z 772.5767

0123E+51414.5RT (min)

lcorbi\_t3\_neg\_3\_005\_23

m/z 772.6347

024E+413.514RT (min)

lcorbi\_t3\_neg\_2\_052\_32

m/z 772.6401

012E+41515.5RT (min)

lcorbi\_t3\_neg\_2\_012\_36

m/z 772.8024

0246E+511.5RT (min)

lcorbi\_t3\_neg\_4\_007\_13

m/z 773.5798

0246E+41414.5RT (min)

lcorbi\_t3\_neg\_4\_032\_23

m/z 773.5814

012E+613.514RT (min)

lcorbi\_t3\_neg\_4\_007\_13

m/z 773.5836

012E+41515.5RT (min)

lcorbi\_t3\_neg\_4\_034\_01

m/z 774.5197

0123E+513.514RT (min)

lcorbi\_t3\_neg\_2\_036\_35

m/z 774.7996

0123E+511.5RT (min)

lcorbi\_t3\_neg\_2\_027\_13

m/z 776.4629

0123E+411.5RT (min)

lcorbi\_t3\_neg\_3\_045\_34

m/z 776.6348

01234E+614.515RT (min)

lcorbi\_t3\_neg\_1\_052\_24

m/z 776.6693

01234E+41414.5RT (min)

lcorbi\_t3\_neg\_2\_052\_32

m/z 778.5400

0246E+41414.5RT (min)

lcorbi\_t3\_neg\_4\_005\_27

m/z 778.6436

0123E+613.514RT (min)

lcorbi\_t3\_neg\_2\_022\_24

m/z 779.5428

0123E+41414.5RT (min)

lcorbi\_t3\_neg\_4\_042\_14

m/z 780.4187

01234E+411.5RT (min)

lcorbi\_t3\_neg\_2\_048\_27

m/z 780.5494

01E+41515.5RT (min)

lcorbi\_t3\_neg\_4\_029\_31

m/z 781.4832

01E+51414.5RT (min)

lcorbi\_t3\_neg\_3\_055\_06

m/z 781.5585

01E+51414.5RT (min)

lcorbi\_t3\_neg\_4\_042\_14

m/z 782.4788

012E+513.514RT (min)

lcorbi\_t3\_neg\_4\_019\_08

m/z 782.7163

0246E+41414.5RT (min)

lcorbi\_t3\_neg\_3\_002\_24

m/z 782.8312

02468E+511.5RT (min)

lcorbi\_t3\_neg\_3\_018\_13

m/z 783.4141

0123E+411.5RT (min)

lcorbi\_t3\_neg\_4\_012\_17

m/z 783.4990

02468E+41414.5RT (min)

lcorbi\_t3\_neg\_3\_048\_08

m/z 783.6309

02468E+413.514RT (min)

lcorbi\_t3\_neg\_2\_052\_32

m/z 785.5145

02468E+51414.5RT (min)

lcorbi\_t3\_neg\_3\_005\_23

m/z 786.6082

0246E+411.5RT (min)

lcorbi\_t3\_neg\_1\_032\_13

m/z 788.2811

0123E+413.514RT (min)

lcorbi\_t3\_neg\_4\_007\_13

m/z 788.5245

02468E+41414.5RT (min)

lcorbi\_t3\_neg\_3\_018\_13

m/z 788.5245

01E+51414.5RT (min)

lcorbi\_t3\_neg\_4\_007\_13

m/z 789.5279

01234E+413.514RT (min)

lcorbi\_t3\_neg\_4\_007\_13

m/z 790.5782

01E+413.514RT (min)

lcorbi\_t3\_neg\_4\_007\_13

m/z 792.5289

01E+414.515RT (min)

lcorbi\_t3\_neg\_4\_029\_31

m/z 792.5570

02468E+41515.5RT (min)

lcorbi\_t3\_neg\_4\_029\_31

m/z 792.8599

01E+611.5RT (min)

lcorbi\_t3\_neg\_3\_018\_13

m/z 793.5431

0123E+413.514RT (min)

lcorbi\_t3\_neg\_4\_044\_38

m/z 793.6069

01234E+413.514RT (min)

lcorbi\_t3\_neg\_2\_052\_32

m/z 794.4960

01E+513.514RT (min)

lcorbi\_t3\_neg\_1\_044\_23

m/z 794.5094

01E+41313.5RT (min)

lcorbi\_t3\_neg\_4\_034\_01

m/z 796.5112

02468E+41414.5RT (min)

lcorbi\_t3\_neg\_4\_032\_23

m/z 797.5148

0246E+41414.5RT (min)

lcorbi\_t3\_neg\_1\_029\_21

m/z 797.6320

01E+514.515RT (min)

lcorbi\_t3\_neg\_2\_039\_31

m/z 798.6351

0123E+414.515RT (min)

lcorbi\_t3\_neg\_2\_039\_31

m/z 798.6883

01E+513.514RT (min)

lcorbi\_t3\_neg\_1\_053\_32

m/z 799.9320

0123E+411.5RT (min)

lcorbi\_t3\_neg\_2\_002\_26

m/z 800.4570

02468E+411.5RT (min)

lcorbi\_t3\_neg\_4\_007\_13

m/z 800.6830

024E+413.514RT (min)

lcorbi\_t3\_neg\_4\_035\_33

m/z 800.9304

0123E+411.5RT (min)

lcorbi\_t3\_neg\_4\_005\_27

m/z 802.4641

01E+513.514RT (min)

lcorbi\_t3\_neg\_1\_013\_35

m/z 802.5135

0123E+414.515RT (min)

lcorbi\_t3\_neg\_2\_053\_23

m/z 802.5822

01234E+411.5RT (min)

lcorbi\_t3\_neg\_3\_024\_26

m/z 803.4674

024E+413.514RT (min)

lcorbi\_t3\_neg\_3\_005\_23

m/z 803.5116

0123E+414.515RT (min)

lcorbi\_t3\_neg\_2\_051\_34

m/z 803.5987

02468E+41414.5RT (min)

lcorbi\_t3\_neg\_2\_053\_23

m/z 804.2761

0123E+41313.5RT (min)

lcorbi\_t3\_neg\_4\_048\_03

m/z 804.5295

024E+414.515RT (min)

lcorbi\_t3\_neg\_2\_024\_37

m/z 804.6482

0246E+413.514RT (min)

lcorbi\_t3\_neg\_2\_052\_32

m/z 805.4833

01E+513.514RT (min)

lcorbi\_t3\_neg\_3\_047\_37

m/z 805.5067

012E+413.514RT (min)

lcorbi\_t3\_neg\_4\_019\_08

m/z 805.5270

012E+414.515RT (min)

lcorbi\_t3\_neg\_4\_046\_37

m/z 806.4781

01E+513.514RT (min)

lcorbi\_t3\_neg\_4\_019\_08

m/z 806.5711

012E+514.515RT (min)

lcorbi\_t3\_neg\_4\_005\_27

m/z 807.4008

0123E+411.5RT (min)

lcorbi\_t3\_neg\_4\_034\_01

m/z 807.4990

01E+513.514RT (min)

lcorbi\_t3\_neg\_4\_045\_35

m/z 807.4991

01E+51414.5RT (min)

lcorbi\_t3\_neg\_1\_044\_23

m/z 807.5223

012E+413.514RT (min)

lcorbi\_t3\_neg\_2\_008\_16

m/z 807.5397

02468E+413.514RT (min)

lcorbi\_t3\_neg\_4\_029\_31

m/z 807.5744

0123E+414.515RT (min)

lcorbi\_t3\_neg\_4\_005\_27

m/z 808.4455

02468E+411.5RT (min)

lcorbi\_t3\_neg\_4\_007\_13

m/z 808.5444

01234E+41414.5RT (min)

lcorbi\_t3\_neg\_2\_024\_37

m/z 808.9166

0123E+411.5RT (min)

lcorbi\_t3\_neg\_2\_048\_27

m/z 809.5144

024E+51414.5RT (min)

lcorbi\_t3\_neg\_3\_045\_34

m/z 809.5148

01E+51414.5RT (min)

lcorbi\_t3\_neg\_3\_005\_23

m/z 809.5905

012E+414.515RT (min)

lcorbi\_t3\_neg\_3\_053\_14

m/z 811.5302

012E+51414.5RT (min)

lcorbi\_t3\_neg\_3\_005\_23

m/z 811.6301

01E+514.515RT (min)

lcorbi\_t3\_neg\_2\_052\_32

m/z 812.4830

01E+511.5RT (min)

lcorbi\_t3\_neg\_4\_007\_13

m/z 813.4290

012E+411.5RT (min)

lcorbi\_t3\_neg\_2\_054\_15

m/z 813.5456

01E+514.515RT (min)

lcorbi\_t3\_neg\_2\_053\_23

m/z 813.5458

02468E+414.515RT (min)

lcorbi\_t3\_neg\_4\_032\_23

m/z 814.4800

01E+511.5RT (min)

lcorbi\_t3\_neg\_4\_007\_13

m/z 815.6368

012E+414.515RT (min)

lcorbi\_t3\_neg\_4\_011\_24

m/z 816.4318

01234E+411.5RT (min)

lcorbi\_t3\_neg\_3\_032\_17

m/z 816.4592

0123E+411.5RT (min)

lcorbi\_t3\_neg\_4\_042\_14

m/z 816.5101

012E+414.515RT (min)

lcorbi\_t3\_neg\_4\_029\_31

m/z 816.5148

01E+513.514RT (min)

lcorbi\_t3\_neg\_1\_020\_36

m/z 816.5824

0123E+41313.5RT (min)

lcorbi\_t3\_neg\_4\_039\_32

m/z 816.6343

0123E+414.515RT (min)

lcorbi\_t3\_neg\_4\_011\_24

m/z 817.5801

012E+41515.5RT (min)

lcorbi\_t3\_neg\_4\_029\_31

m/z 817.5947

01E+513.514RT (min)

lcorbi\_t3\_neg\_2\_052\_32

m/z 818.5164

0123E+413.514RT (min)

lcorbi\_t3\_neg\_2\_027\_13

m/z 818.5308

02468E+41414.5RT (min)

lcorbi\_t3\_neg\_1\_013\_35

m/z 819.3828

01234E+411.5RT (min)

lcorbi\_t3\_neg\_4\_042\_14

m/z 819.5204

01E+413.514RT (min)

lcorbi\_t3\_neg\_2\_027\_13

m/z 819.5358

012E+414.515RT (min)

lcorbi\_t3\_neg\_3\_030\_33

m/z 819.5586

01E+51313.5RT (min)

lcorbi\_t3\_neg\_2\_007\_14

m/z 819.6111

012E+51414.5RT (min)

lcorbi\_t3\_neg\_4\_011\_24

m/z 820.4474

0123E+411.5RT (min)

lcorbi\_t3\_neg\_2\_039\_31

m/z 820.5144

01E+511.5RT (min)

lcorbi\_t3\_neg\_4\_007\_13

m/z 820.5461

0123E+414.515RT (min)

lcorbi\_t3\_neg\_3\_021\_38

m/z 820.6069

012E+51313.5RT (min)

lcorbi\_t3\_neg\_2\_043\_33

m/z 820.6190

012E+414.515RT (min)

lcorbi\_t3\_neg\_1\_007\_26

m/z 821.6287

01234E+413.514RT (min)

lcorbi\_t3\_neg\_2\_022\_24

m/z 822.5118

01E+511.5RT (min)

lcorbi\_t3\_neg\_4\_007\_13

m/z 822.5756

01E+513.514RT (min)

lcorbi\_t3\_neg\_4\_029\_31

m/z 822.5771

01234E+413.514RT (min)

lcorbi\_t3\_neg\_4\_007\_13

m/z 822.5886

024E+413.514RT (min)

lcorbi\_t3\_neg\_1\_039\_38

m/z 822.6346

0123E+414.515RT (min)

lcorbi\_t3\_neg\_2\_007\_14

m/z 822.6346

0123E+414.515RT (min)

lcorbi\_t3\_neg\_4\_036\_16

m/z 823.5790

0246E+413.514RT (min)

lcorbi\_t3\_neg\_4\_029\_31

m/z 823.6447

01E+513.514RT (min)

lcorbi\_t3\_neg\_2\_052\_32

m/z 824.3999

02468E+411.5RT (min)

lcorbi\_t3\_neg\_4\_007\_13

m/z 824.4196

024E+411.5RT (min)

lcorbi\_t3\_neg\_4\_005\_27

m/z 825.6220

02468E+413.514RT (min)

lcorbi\_t3\_neg\_2\_022\_24

m/z 826.5062

01E+511.5RT (min)

lcorbi\_t3\_neg\_4\_007\_13

m/z 828.4574

01E+511.5RT (min)

lcorbi\_t3\_neg\_2\_048\_27

m/z 828.6255

024E+413.514RT (min)

lcorbi\_t3\_neg\_2\_052\_32

m/z 829.3926

01234E+411.5RT (min)

lcorbi\_t3\_neg\_4\_007\_13

m/z 830.4543

01E+511.5RT (min)

lcorbi\_t3\_neg\_2\_048\_27

m/z 830.5268

012E+413.514RT (min)

lcorbi\_t3\_neg\_2\_039\_31

m/z 830.6416

02468E+413.514RT (min)

lcorbi\_t3\_neg\_1\_052\_24

m/z 830.6417

02468E+413.514RT (min)

lcorbi\_t3\_neg\_2\_052\_32

m/z 831.6539

01234E+41414.5RT (min)

lcorbi\_t3\_neg\_1\_053\_32

m/z 832.2886

01234E+413.514RT (min)

lcorbi\_t3\_neg\_1\_023\_03

m/z 832.4516

0246E+411.5RT (min)

lcorbi\_t3\_neg\_2\_048\_27

m/z 832.6391

0123E+413.514RT (min)

lcorbi\_t3\_neg\_2\_022\_24

m/z 832.6401

012E+413.514RT (min)

lcorbi\_t3\_neg\_3\_052\_32

m/z 832.6572

02468E+41414.5RT (min)

lcorbi\_t3\_neg\_3\_002\_24

m/z 833.5644

012E+414.515RT (min)

lcorbi\_t3\_neg\_4\_034\_01

m/z 833.6696

0123E+41414.5RT (min)

lcorbi\_t3\_neg\_3\_002\_24

m/z 834.4235

0123E+411.5RT (min)

lcorbi\_t3\_neg\_4\_046\_37

m/z 834.5652

01234E+41414.5RT (min)

lcorbi\_t3\_neg\_4\_039\_32

m/z 834.6541

01234E+41414.5RT (min)

lcorbi\_t3\_neg\_4\_019\_08

m/z 834.6732

01E+51414.5RT (min)

lcorbi\_t3\_neg\_3\_002\_24

m/z 835.5300

01E+41414.5RT (min)

lcorbi\_t3\_neg\_3\_053\_14

m/z 835.6350

01E+51313.5RT (min)

lcorbi\_t3\_neg\_4\_039\_32

m/z 836.6455

024E+41414.5RT (min)

lcorbi\_t3\_neg\_2\_022\_24

m/z 836.6475

01E+513.514RT (min)

lcorbi\_t3\_neg\_2\_052\_32

m/z 836.6702

0123E+41414.5RT (min)

lcorbi\_t3\_neg\_3\_002\_24

m/z 838.4168

0123E+411.5RT (min)

lcorbi\_t3\_neg\_4\_046\_37

m/z 838.4860

02468E+411.5RT (min)

lcorbi\_t3\_neg\_2\_048\_27

m/z 840.4832

01E+511.5RT (min)

lcorbi\_t3\_neg\_2\_048\_27

m/z 841.6436

01234E+414.515RT (min)

lcorbi\_t3\_neg\_3\_052\_32

m/z 842.5552

0246E+413.514RT (min)

lcorbi\_t3\_neg\_4\_007\_13

m/z 843.5586

0123E+413.514RT (min)

lcorbi\_t3\_neg\_4\_007\_13

m/z 843.6013

0246E+413.514RT (min)

lcorbi\_t3\_neg\_4\_039\_32

m/z 843.6013

024E+41313.514RT (min)

lcorbi\_t3\_neg\_4\_039\_32

m/z 844.6044

01E+51313.5RT (min)

lcorbi\_t3\_neg\_2\_052\_32

m/z 844.6135

01E+513.514RT (min)

lcorbi\_t3\_neg\_2\_022\_24

m/z 844.6534

0246E+41414.5RT (min)

lcorbi\_t3\_neg\_4\_039\_32

m/z 845.4896

01234E+477.5RT (min)

lcorbi\_t3\_neg\_2\_048\_27

m/z 845.6174

012E+513.514RT (min)

lcorbi\_t3\_neg\_2\_052\_32

m/z 845.6211

01E+51313.5RT (min)

lcorbi\_t3\_neg\_4\_039\_32

m/z 845.6488

02468E+413.514RT (min)

lcorbi\_t3\_neg\_4\_039\_32

m/z 846.6346

01234E+414.515RT (min)

lcorbi\_t3\_neg\_2\_014\_29

m/z 846.6620

012E+513.514RT (min)

lcorbi\_t3\_neg\_4\_039\_32

m/z 847.6369

01E+513.514RT (min)

lcorbi\_t3\_neg\_4\_039\_32

m/z 848.3354

01234E+411.5RT (min)

lcorbi\_t3\_neg\_3\_053\_14

m/z 848.5924

0123E+413.514RT (min)

lcorbi\_t3\_neg\_2\_007\_14

m/z 848.6360

01234E+513.514RT (min)

lcorbi\_t3\_neg\_1\_052\_24

m/z 848.6451

02468E+41414.5RT (min)

lcorbi\_t3\_neg\_4\_011\_24

m/z 849.5396

024E+41313.5RT (min)

lcorbi\_t3\_neg\_4\_029\_31

m/z 849.6500

02468E+413.514RT (min)

lcorbi\_t3\_neg\_4\_039\_32

m/z 850.6516

012E+51414.5RT (min)

lcorbi\_t3\_neg\_3\_002\_24

m/z 850.6607

012E+51414.5RT (min)

lcorbi\_t3\_neg\_3\_002\_24

m/z 854.5281

01234E+41414.5RT (min)

lcorbi\_t3\_neg\_4\_005\_27

m/z 855.6378

024E+414.515RT (min)

lcorbi\_t3\_neg\_2\_052\_32

m/z 855.6454

0246E+413.514RT (min)

lcorbi\_t3\_neg\_3\_052\_32

m/z 857.6615

01E+513.514RT (min)

lcorbi\_t3\_neg\_2\_052\_32

m/z 859.3696

01234E+411.5RT (min)

lcorbi\_t3\_neg\_2\_048\_27

m/z 859.6766

0246E+41414.5RT (min)

lcorbi\_t3\_neg\_4\_039\_32

m/z 860.6404

0246E+413.514RT (min)

lcorbi\_t3\_neg\_4\_039\_32

m/z 861.6437

012E+513.514RT (min)

lcorbi\_t3\_neg\_4\_039\_32

m/z 861.6751

012E+41414.5RT (min)

lcorbi\_t3\_neg\_1\_052\_24

m/z 861.6914

01234E+41414.5RT (min)

lcorbi\_t3\_neg\_3\_002\_24

m/z 862.6581

0246E+413.514RT (min)

lcorbi\_t3\_neg\_4\_011\_24

m/z 862.6779

02468E+41414.5RT (min)

lcorbi\_t3\_neg\_1\_053\_32

m/z 862.6953

01E+51414.5RT (min)

lcorbi\_t3\_neg\_1\_052\_24

m/z 863.6002

012E+413.514RT (min)

lcorbi\_t3\_neg\_3\_002\_24

m/z 864.6156

01E+414.515RT (min)

lcorbi\_t3\_neg\_3\_005\_23

m/z 864.6928

01234E+41414.5RT (min)

lcorbi\_t3\_neg\_3\_002\_24

m/z 865.6200

02468E+413.514RT (min)

lcorbi\_t3\_neg\_3\_002\_24

m/z 866.7535

0123E+413.514RT (min)

lcorbi\_t3\_neg\_1\_053\_32

m/z 867.6392

02468E+413.514RT (min)

lcorbi\_t3\_neg\_4\_039\_32

m/z 868.5770

01E+51414.5RT (min)

lcorbi\_t3\_neg\_1\_044\_23

m/z 869.6475

01234E+41414.5RT (min)

lcorbi\_t3\_neg\_2\_022\_24

m/z 870.3340

0246E+48.59RT (min)

lcorbi\_t3\_neg\_2\_030\_47

m/z 870.6572

012E+51414.5RT (min)

lcorbi\_t3\_neg\_4\_039\_32

m/z 871.3375

0123E+48.59RT (min)

lcorbi\_t3\_neg\_4\_006\_47

m/z 871.3503

01234E+411.5RT (min)

lcorbi\_t3\_neg\_2\_048\_27

m/z 872.6000

01E+513.514RT (min)

lcorbi\_t3\_neg\_4\_042\_14

m/z 872.6405

01E+513.514RT (min)

lcorbi\_t3\_neg\_4\_039\_32

m/z 873.5118

012E+413.514RT (min)

lcorbi\_t3\_neg\_3\_053\_14

m/z 873.5121

012E+413.514RT (min)

lcorbi\_t3\_neg\_1\_047\_02

m/z 875.6676

01E+414.515RT (min)

lcorbi\_t3\_neg\_4\_005\_27

m/z 875.6678

01E+414.515RT (min)

lcorbi\_t3\_neg\_4\_036\_16

m/z 878.5847

0123E+413.514RT (min)

lcorbi\_t3\_neg\_4\_007\_13

m/z 882.3325

0123E+411.5RT (min)

lcorbi\_t3\_neg\_1\_043\_14

m/z 885.6818

012E+413.514RT (min)

lcorbi\_t3\_neg\_1\_053\_32

m/z 888.6625

0246E+413.514RT (min)

lcorbi\_t3\_neg\_4\_039\_32

m/z 891.3439

0123E+411.5RT (min)

lcorbi\_t3\_neg\_2\_048\_27

m/z 896.7020

0123E+41414.5RT (min)

lcorbi\_t3\_neg\_3\_002\_24

m/z 897.6754

0246E+41414.5RT (min)

lcorbi\_t3\_neg\_1\_055\_05

m/z 898.5568

0246E+413.514RT (min)

lcorbi\_t3\_neg\_4\_011\_24

m/z 904.6074

0123E+41414.5RT (min)

lcorbi\_t3\_neg\_3\_007\_16

m/z 906.3109

01234E+48.59RT (min)

lcorbi\_t3\_neg\_2\_030\_47

m/z 906.6230

024E+414.515RT (min)

lcorbi\_t3\_neg\_4\_007\_13

m/z 907.3146

012E+48.59RT (min)

lcorbi\_t3\_neg\_2\_030\_47

m/z 908.3426

024E+411.5RT (min)

lcorbi\_t3\_neg\_4\_007\_13

m/z 911.5543

0123E+413.514RT (min)

lcorbi\_t3\_neg\_2\_008\_16

m/z 912.5923

024E+41313.5RT (min)

lcorbi\_t3\_neg\_2\_052\_32

m/z 917.3430

0246E+48.59RT (min)

lcorbi\_t3\_neg\_2\_030\_47

m/z 918.3456

012E+48.59RT (min)

lcorbi\_t3\_neg\_2\_030\_47

m/z 918.6422

012E+413.514RT (min)

lcorbi\_t3\_neg\_2\_052\_32

m/z 919.6450

01E+513.514RT (min)

lcorbi\_t3\_neg\_2\_052\_32

m/z 920.6698

01E+513.514RT (min)

lcorbi\_t3\_neg\_3\_002\_24

m/z 922.6857

0123E+413.514RT (min)

lcorbi\_t3\_neg\_1\_052\_24

m/z 926.6613

02468E+41414.5RT (min)

lcorbi\_t3\_neg\_3\_005\_23

m/z 929.5587

012E+413.514RT (min)

lcorbi\_t3\_neg\_4\_038\_15

m/z 930.3556

024E+48.59RT (min)

lcorbi\_t3\_neg\_2\_030\_47

m/z 930.3557

024E+48.59RT (min)

lcorbi\_t3\_neg\_2\_030\_47

m/z 931.3585

01E+48.59RT (min)

lcorbi\_t3\_neg\_4\_033\_53

m/z 931.3588

012E+48.59RT (min)

lcorbi\_t3\_neg\_2\_030\_47

m/z 932.3045

0123E+411.5RT (min)

lcorbi\_t3\_neg\_3\_004\_30

m/z 932.6056

012E+413.514RT (min)

lcorbi\_t3\_neg\_1\_053\_32

m/z 934.6210

0123E+413.514RT (min)

lcorbi\_t3\_neg\_1\_052\_24

m/z 938.6830

0123E+413.514RT (min)

lcorbi\_t3\_neg\_2\_052\_32

m/z 939.6738

01E+51414.5RT (min)

lcorbi\_t3\_neg\_2\_053\_23

m/z 941.6309

024E+413.514RT (min)

lcorbi\_t3\_neg\_3\_052\_32

m/z 956.6311

01E+41414.5RT (min)

lcorbi\_t3\_neg\_2\_021\_41

m/z 961.5153

012E+413.514RT (min)

lcorbi\_t3\_neg\_4\_029\_31

m/z 963.5392

012E+41414.5RT (min)

lcorbi\_t3\_neg\_2\_011\_04

m/z 976.5875

01E+513.514RT (min)

lcorbi\_t3\_neg\_1\_052\_24

m/z 978.2839

0123E+411.5RT (min)

lcorbi\_t3\_neg\_2\_048\_27

m/z 984.3265

0123E+48.59RT (min)

lcorbi\_t3\_neg\_2\_030\_47

m/z 985.3303

0123E+48.59RT (min)

lcorbi\_t3\_neg\_4\_040\_19

m/z 989.4933

01234E+41414.5RT (min)

lcorbi\_t3\_neg\_1\_043\_14

m/z 991.6320

01E+513.514RT (min)

lcorbi\_t3\_neg\_4\_005\_27

m/z 1064.1965

012E+411.5RT (min)

lcorbi\_t3\_neg\_2\_007\_14

m/z 1135.8878

01E+515.516RT (min)

lcorbi\_t3\_neg\_4\_005\_27
