## Supplementary material for "The Everything Bagel Feature Finder: Ultra-fast automated feature finding for untargeted metabolomics": SI Data File: tool_unique_xic_st_EB_ST004501.html

EB-unique features (not detected by the other two tools) - ST004501

### EB-unique features (not detected by the other two tools) - ST004501

343 features · orbitrap · XIC window m/z ± 10 ppm (≥ 0.002 Da) · ± 0.5 min

Every panel is a feature **EB** detected that neither of the other two tools detected — three-way unique across EB (standard mode (min\_detection\_rate=0.1)), Masscube and XCMS, under the standard matching rule (m/z ± 10 ppm; apex within the reference RT window ± 0.2 min). It shows the MS1 extracted-ion chromatogram at the feature's m/z, drawn from the raw mzML named in `alignment_reference_file`, zoomed to ± 0.5 min around the feature's reported apex RT. **Gray solid line** = the feature's reported apex RT; **red dashed line** = EB's assigned MS/MS — `MS2_scan_id` (native scan number) in `MS2_source_file`; the panel widens to keep an off-apex MS/MS on-screen. Features with no assigned MS/MS show only the gray line. y is per-cell autoscaled. Rows keep the table order (m/z ascending).

m/z 58.0651

01E+53.54RT (min)

lcorbi\_hilic\_pos\_4\_077\_58

m/z 59.0604

0123E+44.55RT (min)

lcorbi\_hilic\_pos\_1\_020\_13

m/z 60.0798

012E+43.54RT (min)

lcorbi\_hilic\_pos\_2\_012\_37

m/z 70.0641

024E+43.54RT (min)

lcorbi\_hilic\_pos\_1\_014\_27

m/z 70.0651

01E+555.5RT (min)

lcorbi\_hilic\_pos\_2\_007\_28

m/z 70.0651

02468E+44.55RT (min)

lcorbi\_hilic\_pos\_4\_020\_37

m/z 74.0236

0123E+41.52RT (min)

lcorbi\_hilic\_pos\_4\_067\_57

m/z 74.0601

02468E+444.5RT (min)

lcorbi\_hilic\_pos\_1\_054\_23

m/z 74.0786

01234E+55.56RT (min)

lcorbi\_hilic\_pos\_1\_003\_24

m/z 80.9478

0246E+400.5RT (min)

lcorbi\_hilic\_pos\_3\_013\_30

m/z 80.9478

024E+455.5RT (min)

lcorbi\_hilic\_pos\_4\_068\_57

m/z 80.9478

024E+44.55RT (min)

lcorbi\_hilic\_pos\_4\_082\_59

m/z 80.9478

024E+44.55RT (min)

lcorbi\_hilic\_pos\_4\_066\_57

m/z 80.9478

0123E+466.5RT (min)

lcorbi\_hilic\_pos\_1\_046\_30

m/z 81.0306

012E+511.5RT (min)

lcorbi\_hilic\_pos\_2\_016\_18

m/z 81.0445

0246E+43.54RT (min)

lcorbi\_hilic\_pos\_4\_004\_38

m/z 83.0603

0123E+46.57RT (min)

lcorbi\_hilic\_pos\_1\_016\_33

m/z 84.0444

0123E+444.5RT (min)

lcorbi\_hilic\_pos\_4\_088\_59

m/z 85.0283

01E+455.5RT (min)

lcorbi\_hilic\_pos\_2\_051\_15

m/z 85.0283

0123E+46.57RT (min)

lcorbi\_hilic\_pos\_4\_083\_59

m/z 85.0663

0123E+511.5RT (min)

lcorbi\_hilic\_pos\_1\_036\_32

m/z 86.0601

0123E+411.5RT (min)

lcorbi\_hilic\_pos\_1\_019\_01

m/z 86.0964

01234E+43.54RT (min)

lcorbi\_hilic\_pos\_4\_070\_57

m/z 87.8015

01E+411.5RT (min)

lcorbi\_hilic\_pos\_4\_099\_61

m/z 88.0393

01E+56.57RT (min)

lcorbi\_hilic\_pos\_2\_013\_36

m/z 88.0578

02468E+433.5RT (min)

lcorbi\_hilic\_pos\_1\_036\_32

m/z 95.0855

0123E+411.5RT (min)

lcorbi\_hilic\_pos\_4\_107\_62

m/z 96.0444

0246E+41.52RT (min)

lcorbi\_hilic\_pos\_1\_020\_13

m/z 98.9842

02468E+477.5RT (min)

lcorbi\_hilic\_pos\_1\_003\_24

m/z 101.0612

01234E+411.5RT (min)

lcorbi\_hilic\_pos\_4\_093\_60

m/z 102.0550

01E+511.5RT (min)

lcorbi\_hilic\_pos\_3\_054\_16

m/z 103.0542

012E+455.5RT (min)

lcorbi\_hilic\_pos\_1\_034\_37

m/z 103.0542

012E+455.5RT (min)

lcorbi\_hilic\_pos\_4\_077\_58

m/z 103.0753

02468E+40.51RT (min)

lcorbi\_hilic\_pos\_4\_107\_62

m/z 104.0706

012E+45.56RT (min)

lcorbi\_hilic\_pos\_1\_008\_38

m/z 106.0683

012E+51.52RT (min)

lcorbi\_hilic\_pos\_4\_096\_60

m/z 106.0863

012E+44.55RT (min)

lcorbi\_hilic\_pos\_1\_044\_17

m/z 106.0863

01234E+40.51RT (min)

lcorbi\_hilic\_pos\_3\_002\_04

m/z 107.0873

012E+44.55RT (min)

lcorbi\_hilic\_pos\_3\_020\_31

m/z 107.0893

01E+544.5RT (min)

lcorbi\_hilic\_pos\_4\_110\_62

m/z 107.0895

0123E+455.5RT (min)

lcorbi\_hilic\_pos\_2\_007\_28

m/z 109.0760

0246E+43.54RT (min)

lcorbi\_hilic\_pos\_3\_048\_38

m/z 112.0504

0123E+43.54RT (min)

lcorbi\_hilic\_pos\_4\_082\_59

m/z 112.0504

0246E+444.5RT (min)

lcorbi\_hilic\_pos\_4\_089\_59

m/z 113.1073

012E+41.52RT (min)

lcorbi\_hilic\_pos\_4\_018\_31

m/z 114.0549

01234E+444.5RT (min)

lcorbi\_hilic\_pos\_4\_001\_49

m/z 115.0880

0246E+47.58RT (min)

lcorbi\_hilic\_pos\_3\_010\_32

m/z 116.0704

0123E+43.54RT (min)

lcorbi\_hilic\_pos\_4\_080\_58

m/z 116.0706

0246E+40.51RT (min)

lcorbi\_hilic\_pos\_1\_045\_12

m/z 116.1070

012E+42.53RT (min)

lcorbi\_hilic\_pos\_4\_075\_58

m/z 117.0546

0123E+444.5RT (min)

lcorbi\_hilic\_pos\_1\_054\_23

m/z 117.0573

01E+455.5RT (min)

lcorbi\_hilic\_pos\_4\_014\_27

m/z 117.0912

02468E+43.54RT (min)

lcorbi\_hilic\_pos\_4\_087\_59

m/z 118.0651

024E+444.5RT (min)

lcorbi\_hilic\_pos\_4\_067\_57

m/z 118.0651

024E+444.5RT (min)

lcorbi\_hilic\_pos\_4\_066\_57

m/z 118.0820

01E+53.54RT (min)

lcorbi\_hilic\_pos\_4\_107\_62

m/z 118.0862

0246E+411.5RT (min)

lcorbi\_hilic\_pos\_1\_044\_17

m/z 118.0920

02468E+455.5RT (min)

lcorbi\_hilic\_pos\_4\_080\_58

m/z 119.0716

0246E+444.5RT (min)

lcorbi\_hilic\_pos\_4\_001\_49

m/z 119.0718

01E+55.56RT (min)

lcorbi\_hilic\_pos\_4\_091\_60

m/z 119.0832

024E+455.5RT (min)

lcorbi\_hilic\_pos\_4\_072\_57

m/z 119.0891

0123E+43.54RT (min)

lcorbi\_hilic\_pos\_4\_107\_62

m/z 120.0808

02468E+444.5RT (min)

lcorbi\_hilic\_pos\_4\_099\_61

m/z 120.0809

01E+544.5RT (min)

lcorbi\_hilic\_pos\_1\_037\_18

m/z 120.0903

024E+455.5RT (min)

lcorbi\_hilic\_pos\_4\_069\_57

m/z 120.1034

01E+555.5RT (min)

lcorbi\_hilic\_pos\_4\_092\_60

m/z 122.0362

01E+50.51RT (min)

lcorbi\_hilic\_pos\_1\_055\_20

m/z 122.0811

0123E+444.5RT (min)

lcorbi\_hilic\_pos\_1\_008\_38

m/z 122.0811

01234E+444.5RT (min)

lcorbi\_hilic\_pos\_4\_084\_59

m/z 122.0965

012E+41.52RT (min)

lcorbi\_hilic\_pos\_1\_044\_17

m/z 125.0903

012E+46.57RT (min)

lcorbi\_hilic\_pos\_2\_008\_25

m/z 128.0705

01234E+422.5RT (min)

lcorbi\_hilic\_pos\_4\_087\_59

m/z 128.0707

01E+544.5RT (min)

lcorbi\_hilic\_pos\_1\_008\_38

m/z 128.0722

012E+511.5RT (min)

lcorbi\_hilic\_pos\_3\_010\_32

m/z 128.0723

012E+511.5RT (min)

lcorbi\_hilic\_pos\_1\_036\_32

m/z 130.0499

0246E+422.5RT (min)

lcorbi\_hilic\_pos\_4\_070\_57

m/z 130.0652

0123E+444.5RT (min)

lcorbi\_hilic\_pos\_1\_037\_18

m/z 130.0863

01234E+45.56RT (min)

lcorbi\_hilic\_pos\_4\_083\_59

m/z 130.0863

0123E+41.52RT (min)

lcorbi\_hilic\_pos\_4\_073\_57

m/z 130.0863

0123E+45.56RT (min)

lcorbi\_hilic\_pos\_1\_031\_36

m/z 130.0863

02468E+455.5RT (min)

lcorbi\_hilic\_pos\_4\_077\_58

m/z 131.0896

01234E+43.54RT (min)

lcorbi\_hilic\_pos\_2\_001\_49

m/z 131.0910

02468E+46.57RT (min)

lcorbi\_hilic\_pos\_2\_003\_33

m/z 132.0655

01E+55.56RT (min)

lcorbi\_hilic\_pos\_4\_083\_59

m/z 133.0760

02468E+411.5RT (min)

lcorbi\_hilic\_pos\_4\_094\_60

m/z 133.0760

02468E+411.5RT (min)

lcorbi\_hilic\_pos\_4\_094\_60

m/z 133.0858

01E+50.51RT (min)

lcorbi\_hilic\_pos\_4\_015\_20

m/z 133.0860

0123E+50.51RT (min)

lcorbi\_hilic\_pos\_4\_086\_59

m/z 134.0600

012E+511.5RT (min)

lcorbi\_hilic\_pos\_3\_004\_27

m/z 137.0652

01E+533.5RT (min)

lcorbi\_hilic\_pos\_1\_054\_23

m/z 137.0710

012E+54.55RT (min)

lcorbi\_hilic\_pos\_4\_020\_37

m/z 138.0550

02468E+44.55RT (min)

lcorbi\_hilic\_pos\_4\_108\_62

m/z 139.0503

012E+511.5RT (min)

lcorbi\_hilic\_pos\_3\_035\_22

m/z 140.0705

01234E+41.52RT (min)

lcorbi\_hilic\_pos\_2\_012\_37

m/z 141.9917

0123E+488.5RT (min)

lcorbi\_hilic\_pos\_1\_042\_40

m/z 142.0863

0246E+43.54RT (min)

lcorbi\_hilic\_pos\_1\_020\_13

m/z 143.0814

012E+455.5RT (min)

lcorbi\_hilic\_pos\_1\_031\_36

m/z 144.0809

024E+455.5RT (min)

lcorbi\_hilic\_pos\_1\_023\_28

m/z 144.1018

01E+64.55RT (min)

lcorbi\_hilic\_pos\_4\_020\_37

m/z 144.1019

0246E+55.56RT (min)

lcorbi\_hilic\_pos\_4\_081\_58

m/z 146.0600

0123E+455.5RT (min)

lcorbi\_hilic\_pos\_4\_014\_27

m/z 147.1144

02468E+46.57RT (min)

lcorbi\_hilic\_pos\_2\_003\_33

m/z 148.0798

01E+56.57RT (min)

lcorbi\_hilic\_pos\_1\_052\_35

m/z 148.1161

01E+47.58RT (min)

lcorbi\_hilic\_pos\_4\_020\_37

m/z 149.0822

01E+54.55RT (min)

lcorbi\_hilic\_pos\_2\_001\_49

m/z 149.0823

01E+544.5RT (min)

lcorbi\_hilic\_pos\_2\_001\_49

m/z 149.1187

012E+533.5RT (min)

lcorbi\_hilic\_pos\_2\_037\_24

m/z 150.0646

0123E+46.57RT (min)

lcorbi\_hilic\_pos\_1\_043\_25

m/z 150.1146

02468E+444.5RT (min)

lcorbi\_hilic\_pos\_4\_107\_62

m/z 151.0614

0123E+42.53RT (min)

lcorbi\_hilic\_pos\_2\_023\_34

m/z 152.0567

0246E+444.5RT (min)

lcorbi\_hilic\_pos\_4\_089\_59

m/z 152.0697

012E+433.5RT (min)

lcorbi\_hilic\_pos\_4\_056\_56

m/z 152.0739

01E+56.57RT (min)

lcorbi\_hilic\_pos\_1\_016\_33

m/z 152.0999

01E+50.51RT (min)

lcorbi\_hilic\_pos\_4\_088\_59

m/z 155.0814

012E+411.5RT (min)

lcorbi\_hilic\_pos\_2\_012\_37

m/z 155.0815

01234E+422.5RT (min)

lcorbi\_hilic\_pos\_4\_078\_58

m/z 155.0815

024E+40.51RT (min)

lcorbi\_hilic\_pos\_4\_073\_57

m/z 156.0265

02468E+46.57RT (min)

lcorbi\_hilic\_pos\_1\_032\_31

m/z 158.0812

01E+51.52RT (min)

lcorbi\_hilic\_pos\_2\_020\_31

m/z 158.0925

0246E+466.5RT (min)

lcorbi\_hilic\_pos\_1\_021\_46

m/z 158.0941

0123E+43.54RT (min)

lcorbi\_hilic\_pos\_4\_102\_61

m/z 160.0369

01E+555.5RT (min)

lcorbi\_hilic\_pos\_4\_071\_57

m/z 160.0400

01E+56.57RT (min)

lcorbi\_hilic\_pos\_1\_036\_32

m/z 160.0968

024E+455.5RT (min)

lcorbi\_hilic\_pos\_4\_070\_57

m/z 160.0970

01E+544.5RT (min)

lcorbi\_hilic\_pos\_2\_009\_50

m/z 162.0761

01E+544.5RT (min)

lcorbi\_hilic\_pos\_1\_054\_23

m/z 162.0761

012E+566.5RT (min)

lcorbi\_hilic\_pos\_1\_054\_23

m/z 162.0914

0123E+455.5RT (min)

lcorbi\_hilic\_pos\_4\_112\_62

m/z 164.0738

01234E+41.52RT (min)

lcorbi\_hilic\_pos\_4\_069\_57

m/z 164.0785

02468E+40.51RT (min)

lcorbi\_hilic\_pos\_4\_087\_59

m/z 164.1170

01E+555.5RT (min)

lcorbi\_hilic\_pos\_3\_004\_27

m/z 164.1171

01E+555.5RT (min)

lcorbi\_hilic\_pos\_1\_031\_36

m/z 165.0513

012E+55.56RT (min)

lcorbi\_hilic\_pos\_1\_036\_32

m/z 167.0895

012E+455.5RT (min)

lcorbi\_hilic\_pos\_4\_020\_37

m/z 167.0896

0123E+455.5RT (min)

lcorbi\_hilic\_pos\_1\_034\_37

m/z 167.0927

02468E+43.54RT (min)

lcorbi\_hilic\_pos\_4\_020\_37

m/z 167.0928

024E+45.56RT (min)

lcorbi\_hilic\_pos\_4\_001\_49

m/z 167.0928

01E+555.5RT (min)

lcorbi\_hilic\_pos\_1\_030\_48

m/z 167.0929

0246E+444.5RT (min)

lcorbi\_hilic\_pos\_2\_001\_49

m/z 168.0963

02468E+466.5RT (min)

lcorbi\_hilic\_pos\_1\_003\_24

m/z 169.0972

0246E+41.52RT (min)

lcorbi\_hilic\_pos\_4\_071\_57

m/z 170.0768

02468E+43.54RT (min)

lcorbi\_hilic\_pos\_4\_092\_60

m/z 170.0923

024E+477.5RT (min)

lcorbi\_hilic\_pos\_1\_036\_32

m/z 172.1413

024E+40.51RT (min)

lcorbi\_hilic\_pos\_3\_003\_37

m/z 174.1489

0123E+42.53RT (min)

lcorbi\_hilic\_pos\_4\_066\_57

m/z 174.1853

024E+40.51RT (min)

lcorbi\_hilic\_pos\_2\_014\_20

m/z 176.0295

01E+55.56RT (min)

lcorbi\_hilic\_pos\_4\_086\_59

m/z 176.0608

0123E+444.5RT (min)

lcorbi\_hilic\_pos\_2\_005\_23

m/z 176.0705

01234E+511.5RT (min)

lcorbi\_hilic\_pos\_4\_099\_61

m/z 176.0918

0246E+422.5RT (min)

lcorbi\_hilic\_pos\_4\_001\_49

m/z 176.1282

012E+53.54RT (min)

lcorbi\_hilic\_pos\_4\_072\_57

m/z 179.0814

0123E+41.52RT (min)

lcorbi\_hilic\_pos\_3\_011\_34

m/z 179.0855

01234E+50.51RT (min)

lcorbi\_hilic\_pos\_4\_089\_59

m/z 179.0856

0246E+50.51RT (min)

lcorbi\_hilic\_pos\_4\_029\_41

m/z 182.0578

02468E+455.5RT (min)

lcorbi\_hilic\_pos\_4\_020\_37

m/z 182.0779

0123E+47.58RT (min)

lcorbi\_hilic\_pos\_1\_040\_06

m/z 182.0787

01234E+455.5RT (min)

lcorbi\_hilic\_pos\_4\_071\_57

m/z 182.0813

01E+55.56RT (min)

lcorbi\_hilic\_pos\_4\_020\_37

m/z 184.0732

0123E+46.57RT (min)

lcorbi\_hilic\_pos\_4\_020\_37

m/z 186.1125

01E+544.5RT (min)

lcorbi\_hilic\_pos\_1\_014\_27

m/z 187.0785

01E+55.56RT (min)

lcorbi\_hilic\_pos\_1\_003\_24

m/z 187.1077

01E+55.56RT (min)

lcorbi\_hilic\_pos\_4\_083\_59

m/z 188.0707

01E+544.5RT (min)

lcorbi\_hilic\_pos\_4\_069\_57

m/z 188.2009

01E+52.53RT (min)

lcorbi\_hilic\_pos\_4\_054\_45

m/z 191.1277

0246E+411.5RT (min)

lcorbi\_hilic\_pos\_4\_103\_61

m/z 191.1286

0246E+455.5RT (min)

lcorbi\_hilic\_pos\_4\_113\_62

m/z 196.0653

01234E+41.52RT (min)

lcorbi\_hilic\_pos\_1\_036\_32

m/z 197.0809

02468E+50.51RT (min)

lcorbi\_hilic\_pos\_1\_012\_19

m/z 199.1886

0123E+41.52RT (min)

lcorbi\_hilic\_pos\_1\_051\_26

m/z 200.2009

0123E+52.53RT (min)

lcorbi\_hilic\_pos\_4\_014\_27

m/z 201.0546

0123E+40.51RT (min)

lcorbi\_hilic\_pos\_4\_031\_17

m/z 202.1803

0123E+433.5RT (min)

lcorbi\_hilic\_pos\_3\_014\_08

m/z 203.0525

0123E+54.55RT (min)

lcorbi\_hilic\_pos\_4\_078\_58

m/z 203.1501

0246E+466.5RT (min)

lcorbi\_hilic\_pos\_4\_089\_59

m/z 204.1175

01E+54.55RT (min)

lcorbi\_hilic\_pos\_4\_020\_37

m/z 205.0971

0123E+444.5RT (min)

lcorbi\_hilic\_pos\_4\_072\_57

m/z 206.0399

01E+566.5RT (min)

lcorbi\_hilic\_pos\_4\_087\_59

m/z 206.1752

01E+53.54RT (min)

lcorbi\_hilic\_pos\_4\_050\_16

m/z 210.1487

01234E+41.52RT (min)

lcorbi\_hilic\_pos\_4\_084\_59

m/z 214.0508

0246E+411.5RT (min)

lcorbi\_hilic\_pos\_2\_029\_19

m/z 216.2323

0246E+433.5RT (min)

lcorbi\_hilic\_pos\_1\_034\_37

m/z 216.9226

024E+455.5RT (min)

lcorbi\_hilic\_pos\_4\_070\_57

m/z 216.9227

01234E+466.5RT (min)

lcorbi\_hilic\_pos\_4\_070\_57

m/z 217.1986

01234E+433.5RT (min)

lcorbi\_hilic\_pos\_3\_047\_28

m/z 217.1991

01E+522.5RT (min)

lcorbi\_hilic\_pos\_4\_013\_23

m/z 217.1992

0123E+433.5RT (min)

lcorbi\_hilic\_pos\_4\_012\_29

m/z 218.1953

012E+533.5RT (min)

lcorbi\_hilic\_pos\_4\_099\_61

m/z 218.2179

0246E+533.5RT (min)

lcorbi\_hilic\_pos\_3\_026\_47

m/z 218.2190

02468E+533.5RT (min)

lcorbi\_hilic\_pos\_1\_008\_38

m/z 218.2310

012E+533.5RT (min)

lcorbi\_hilic\_pos\_4\_091\_60

m/z 218.2377

02468E+433.5RT (min)

lcorbi\_hilic\_pos\_2\_056\_56

m/z 219.0231

0246E+46.57RT (min)

lcorbi\_hilic\_pos\_4\_094\_60

m/z 220.1907

012E+43.54RT (min)

lcorbi\_hilic\_pos\_4\_050\_16

m/z 223.0924

0123E+455.5RT (min)

lcorbi\_hilic\_pos\_2\_020\_31

m/z 223.0925

0123E+455.5RT (min)

lcorbi\_hilic\_pos\_2\_020\_31

m/z 226.0869

01E+56.57RT (min)

lcorbi\_hilic\_pos\_4\_092\_60

m/z 226.1802

0123E+50.51RT (min)

lcorbi\_hilic\_pos\_3\_019\_43

m/z 226.9464

02468E+45.56RT (min)

lcorbi\_hilic\_pos\_4\_066\_57

m/z 229.1071

012E+50.51RT (min)

lcorbi\_hilic\_pos\_1\_012\_19

m/z 231.1257

01E+511.5RT (min)

lcorbi\_hilic\_pos\_4\_091\_60

m/z 232.0275

0246E+411.5RT (min)

lcorbi\_hilic\_pos\_4\_014\_27

m/z 232.1330

0246E+40.51RT (min)

lcorbi\_hilic\_pos\_2\_053\_27

m/z 236.1111

01234E+433.5RT (min)

lcorbi\_hilic\_pos\_4\_041\_54

m/z 240.2686

01234E+444.5RT (min)

lcorbi\_hilic\_pos\_2\_046\_46

m/z 241.2717

01E+511.5RT (min)

lcorbi\_hilic\_pos\_2\_020\_31

m/z 242.2843

01E+52.53RT (min)

lcorbi\_hilic\_pos\_1\_054\_23

m/z 244.2634

0246E+40.51RT (min)

lcorbi\_hilic\_pos\_4\_026\_11

m/z 245.1412

01E+511.5RT (min)

lcorbi\_hilic\_pos\_4\_092\_60

m/z 245.1496

012E+422.5RT (min)

lcorbi\_hilic\_pos\_4\_018\_31

m/z 245.1860

02468E+44.55RT (min)

lcorbi\_hilic\_pos\_4\_086\_59

m/z 245.2305

012E+41.52RT (min)

lcorbi\_hilic\_pos\_1\_034\_37

m/z 245.2305

01234E+42.53RT (min)

lcorbi\_hilic\_pos\_1\_034\_37

m/z 245.2668

0123E+422.5RT (min)

lcorbi\_hilic\_pos\_4\_097\_60

m/z 246.0775

01E+55.56RT (min)

lcorbi\_hilic\_pos\_4\_073\_57

m/z 246.0777

01E+566.5RT (min)

lcorbi\_hilic\_pos\_4\_068\_57

m/z 246.1337

012E+50.51RT (min)

lcorbi\_hilic\_pos\_4\_111\_62

m/z 246.2291

01E+533.5RT (min)

lcorbi\_hilic\_pos\_4\_105\_61

m/z 246.2376

01234E+52.53RT (min)

lcorbi\_hilic\_pos\_1\_054\_23

m/z 246.2507

01E+52.53RT (min)

lcorbi\_hilic\_pos\_1\_020\_13

m/z 246.2509

01E+533.5RT (min)

lcorbi\_hilic\_pos\_2\_019\_16

m/z 247.0787

01234E+43.54RT (min)

lcorbi\_hilic\_pos\_2\_001\_49

m/z 247.0787

01E+455.5RT (min)

lcorbi\_hilic\_pos\_4\_020\_37

m/z 247.1481

024E+411.5RT (min)

lcorbi\_hilic\_pos\_1\_036\_32

m/z 247.1732

012E+43.54RT (min)

lcorbi\_hilic\_pos\_1\_031\_36

m/z 250.1759

0123E+40.51RT (min)

lcorbi\_hilic\_pos\_3\_019\_43

m/z 253.0551

01E+50.51RT (min)

lcorbi\_hilic\_pos\_3\_047\_28

m/z 253.1645

02468E+40.51RT (min)

lcorbi\_hilic\_pos\_4\_099\_61

m/z 254.2842

0123E+411.5RT (min)

lcorbi\_hilic\_pos\_2\_012\_37

m/z 258.2788

02468E+40.51RT (min)

lcorbi\_hilic\_pos\_4\_109\_62

m/z 258.2792

012E+52.53RT (min)

lcorbi\_hilic\_pos\_1\_054\_23

m/z 259.1289

024E+46.57RT (min)

lcorbi\_hilic\_pos\_4\_084\_59

m/z 261.0485

0123E+45.56RT (min)

lcorbi\_hilic\_pos\_4\_076\_58

m/z 264.1090

0246E+433.5RT (min)

lcorbi\_hilic\_pos\_1\_050\_05

m/z 266.1598

01E+55.56RT (min)

lcorbi\_hilic\_pos\_1\_048\_02

m/z 266.2842

01E+511.5RT (min)

lcorbi\_hilic\_pos\_2\_012\_37

m/z 267.1720

024E+40.51RT (min)

lcorbi\_hilic\_pos\_2\_014\_20

m/z 267.2875

012E+411.5RT (min)

lcorbi\_hilic\_pos\_2\_012\_37

m/z 268.2998

012E+511.5RT (min)

lcorbi\_hilic\_pos\_2\_012\_37

m/z 269.3032

01234E+411.5RT (min)

lcorbi\_hilic\_pos\_3\_024\_35

m/z 272.2583

01E+50.51RT (min)

lcorbi\_hilic\_pos\_1\_024\_03

m/z 272.2586

01E+511.5RT (min)

lcorbi\_hilic\_pos\_1\_020\_13

m/z 273.1420

012E+555.5RT (min)

lcorbi\_hilic\_pos\_1\_044\_17

m/z 273.2615

0123E+40.51RT (min)

lcorbi\_hilic\_pos\_4\_014\_27

m/z 273.2619

01234E+42.53RT (min)

lcorbi\_hilic\_pos\_1\_034\_37

m/z 274.1650

01234E+50.51RT (min)

lcorbi\_hilic\_pos\_4\_107\_62

m/z 274.2581

024E+42.53RT (min)

lcorbi\_hilic\_pos\_4\_105\_61

m/z 276.0689

0123E+444.5RT (min)

lcorbi\_hilic\_pos\_4\_086\_59

m/z 276.1806

012E+44.55RT (min)

lcorbi\_hilic\_pos\_2\_049\_55

m/z 279.2320

01E+511.5RT (min)

lcorbi\_hilic\_pos\_4\_080\_58

m/z 281.2585

0123E+40.51RT (min)

lcorbi\_hilic\_pos\_2\_021\_17

m/z 282.2791

012E+544.5RT (min)

lcorbi\_hilic\_pos\_4\_013\_23

m/z 283.2742

0246E+511.5RT (min)

lcorbi\_hilic\_pos\_3\_034\_26

m/z 284.2776

01E+511.5RT (min)

lcorbi\_hilic\_pos\_1\_008\_38

m/z 284.2949

0123E+43.54RT (min)

lcorbi\_hilic\_pos\_4\_032\_35

m/z 286.0911

02468E+433.5RT (min)

lcorbi\_hilic\_pos\_2\_008\_25

m/z 286.1074

0123E+411.5RT (min)

lcorbi\_hilic\_pos\_3\_026\_47

m/z 286.2741

0123E+411.5RT (min)

lcorbi\_hilic\_pos\_1\_029\_14

m/z 286.2741

01234E+411.5RT (min)

lcorbi\_hilic\_pos\_1\_054\_23

m/z 286.3102

01234E+433.5RT (min)

lcorbi\_hilic\_pos\_4\_085\_59

m/z 288.2363

012E+444.5RT (min)

lcorbi\_hilic\_pos\_4\_082\_59

m/z 288.2897

012E+50.51RT (min)

lcorbi\_hilic\_pos\_1\_042\_40

m/z 289.2929

012E+52.53RT (min)

lcorbi\_hilic\_pos\_1\_054\_23

m/z 290.1598

012E+444.5RT (min)

lcorbi\_hilic\_pos\_4\_021\_13

m/z 291.1067

012E+544.5RT (min)

lcorbi\_hilic\_pos\_2\_030\_05

m/z 291.1068

012E+544.5RT (min)

lcorbi\_hilic\_pos\_1\_050\_05

m/z 293.0981

012E+57.58RT (min)

lcorbi\_hilic\_pos\_1\_036\_32

m/z 294.1912

01E+544.5RT (min)

lcorbi\_hilic\_pos\_1\_048\_02

m/z 295.0899

01234E+433.5RT (min)

lcorbi\_hilic\_pos\_4\_013\_23

m/z 295.2267

0246E+411.5RT (min)

lcorbi\_hilic\_pos\_4\_077\_58

m/z 295.2268

0246E+411.5RT (min)

lcorbi\_hilic\_pos\_4\_077\_58

m/z 296.1217

01E+53.54RT (min)

lcorbi\_hilic\_pos\_4\_066\_57

m/z 296.1494

0246E+411.5RT (min)

lcorbi\_hilic\_pos\_3\_054\_16

m/z 297.2902

01234E+411.5RT (min)

lcorbi\_hilic\_pos\_1\_034\_37

m/z 299.9961

0123E+45.56RT (min)

lcorbi\_hilic\_pos\_4\_084\_59

m/z 300.0927

02468E+422.5RT (min)

lcorbi\_hilic\_pos\_2\_022\_35

m/z 300.2896

01E+533.5RT (min)

lcorbi\_hilic\_pos\_4\_087\_59

m/z 300.2896

01E+50.51RT (min)

lcorbi\_hilic\_pos\_4\_018\_31

m/z 300.2899

01E+511.5RT (min)

lcorbi\_hilic\_pos\_4\_085\_59

m/z 301.2933

012E+433.5RT (min)

lcorbi\_hilic\_pos\_4\_089\_59

m/z 303.2319

0123E+411.5RT (min)

lcorbi\_hilic\_pos\_4\_102\_61

m/z 303.3086

0246E+43.54RT (min)

lcorbi\_hilic\_pos\_4\_004\_38

m/z 304.2928

02468E+40.51RT (min)

lcorbi\_hilic\_pos\_1\_032\_31

m/z 304.3000

01E+51.52RT (min)

lcorbi\_hilic\_pos\_4\_020\_37

m/z 305.3033

01E+50.51RT (min)

lcorbi\_hilic\_pos\_4\_110\_62

m/z 309.2902

012E+511.5RT (min)

lcorbi\_hilic\_pos\_3\_003\_37

m/z 310.2306

02468E+40.51RT (min)

lcorbi\_hilic\_pos\_4\_018\_31

m/z 310.2937

024E+411.5RT (min)

lcorbi\_hilic\_pos\_4\_032\_35

m/z 311.3059

0123E+511.5RT (min)

lcorbi\_hilic\_pos\_1\_008\_38

m/z 312.1374

0246E+544.5RT (min)

lcorbi\_hilic\_pos\_4\_109\_62

m/z 312.3094

0246E+411.5RT (min)

lcorbi\_hilic\_pos\_2\_012\_37

m/z 316.2118

02468E+41.52RT (min)

lcorbi\_hilic\_pos\_4\_090\_60

m/z 316.2121

01E+52.53RT (min)

lcorbi\_hilic\_pos\_2\_024\_32

m/z 316.3210

01E+52.53RT (min)

lcorbi\_hilic\_pos\_1\_054\_23

m/z 317.2156

012E+42.53RT (min)

lcorbi\_hilic\_pos\_2\_024\_32

m/z 318.1471

0123E+444.5RT (min)

lcorbi\_hilic\_pos\_3\_005\_48

m/z 318.3500

02468E+444.5RT (min)

lcorbi\_hilic\_pos\_3\_010\_32

m/z 322.1078

012E+511.5RT (min)

lcorbi\_hilic\_pos\_4\_099\_61

m/z 323.0916

0246E+40.51RT (min)

lcorbi\_hilic\_pos\_4\_008\_19

m/z 323.1636

02468E+444.5RT (min)

lcorbi\_hilic\_pos\_3\_010\_32

m/z 324.1120

02468E+511.5RT (min)

lcorbi\_hilic\_pos\_4\_030\_18

m/z 324.1355

0246E+511.5RT (min)

lcorbi\_hilic\_pos\_2\_016\_18

m/z 324.1672

01E+53.54RT (min)

lcorbi\_hilic\_pos\_3\_004\_27

m/z 325.1914

0246E+43.54RT (min)

lcorbi\_hilic\_pos\_4\_113\_62

m/z 326.1291

012E+611.5RT (min)

lcorbi\_hilic\_pos\_4\_101\_61

m/z 327.2014

012E+50.51RT (min)

lcorbi\_hilic\_pos\_3\_019\_43

m/z 328.2592

01E+50.51RT (min)

lcorbi\_hilic\_pos\_1\_052\_35

m/z 331.3036

012E+511.5RT (min)

lcorbi\_hilic\_pos\_1\_031\_36

m/z 332.3315

01234E+400.5RT (min)

lcorbi\_hilic\_pos\_1\_013\_42

m/z 332.3315

012E+51.52RT (min)

lcorbi\_hilic\_pos\_4\_020\_37

m/z 333.3345

01E+50.51RT (min)

lcorbi\_hilic\_pos\_4\_106\_62

m/z 337.1107

01E+566.5RT (min)

lcorbi\_hilic\_pos\_2\_046\_46

m/z 337.1108

024E+45.56RT (min)

lcorbi\_hilic\_pos\_1\_049\_55

m/z 339.3371

01234E+40.51RT (min)

lcorbi\_hilic\_pos\_4\_068\_57

m/z 340.1545

01E+51.52RT (min)

lcorbi\_hilic\_pos\_4\_099\_61

m/z 342.1274

01234E+566.5RT (min)

lcorbi\_hilic\_pos\_1\_021\_46

m/z 344.1492

01234E+444.5RT (min)

lcorbi\_hilic\_pos\_2\_003\_33

m/z 344.2280

012E+50.51RT (min)

lcorbi\_hilic\_pos\_3\_019\_43

m/z 345.3192

01E+51.52RT (min)

lcorbi\_hilic\_pos\_4\_082\_59

m/z 350.0208

01E+57.58RT (min)

lcorbi\_hilic\_pos\_1\_029\_14

m/z 351.1979

0246E+411.5RT (min)

lcorbi\_hilic\_pos\_3\_034\_26

m/z 351.2199

024E+411.5RT (min)

lcorbi\_hilic\_pos\_1\_028\_34

m/z 355.2269

012E+50.51RT (min)

lcorbi\_hilic\_pos\_4\_008\_19

m/z 358.3317

012E+41.52RT (min)

lcorbi\_hilic\_pos\_4\_082\_59

m/z 360.3624

02468E+40.51RT (min)

lcorbi\_hilic\_pos\_4\_111\_62

m/z 362.3764

01E+51.52RT (min)

lcorbi\_hilic\_pos\_4\_090\_60

m/z 363.0336

0123E+46.57RT (min)

lcorbi\_hilic\_pos\_1\_044\_17

m/z 370.1365

01E+511.5RT (min)

lcorbi\_hilic\_pos\_3\_023\_18

m/z 372.1589

024E+41.52RT (min)

lcorbi\_hilic\_pos\_4\_066\_57

m/z 372.3479

01234E+411.5RT (min)

lcorbi\_hilic\_pos\_4\_086\_59

m/z 373.2376

01E+50.51RT (min)

lcorbi\_hilic\_pos\_4\_112\_62

m/z 386.1505

01234E+511.5RT (min)

lcorbi\_hilic\_pos\_4\_110\_62

m/z 392.2516

01E+50.51RT (min)

lcorbi\_hilic\_pos\_2\_029\_19

m/z 392.3374

024E+40.51RT (min)

lcorbi\_hilic\_pos\_2\_020\_31

m/z 393.3841

012E+40.51RT (min)

lcorbi\_hilic\_pos\_4\_031\_17

m/z 405.4149

02468E+40.51RT (min)

lcorbi\_hilic\_pos\_1\_036\_32

m/z 406.1270

01E+50.51RT (min)

lcorbi\_hilic\_pos\_4\_100\_61

m/z 415.3000

01E+50.51RT (min)

lcorbi\_hilic\_pos\_4\_008\_19

m/z 436.3061

0123E+50.51RT (min)

lcorbi\_hilic\_pos\_4\_106\_62

m/z 453.3437

01234E+41.52RT (min)

lcorbi\_hilic\_pos\_3\_029\_23

m/z 499.3548

0246E+444.5RT (min)

lcorbi\_hilic\_pos\_2\_030\_05

m/z 515.1921

01234E+511.5RT (min)

lcorbi\_hilic\_pos\_4\_099\_61

m/z 584.2639

0123E+40.51RT (min)

lcorbi\_hilic\_pos\_3\_004\_27

m/z 758.5695

024E+41.52RT (min)

lcorbi\_hilic\_pos\_4\_018\_31
