## Supplementary material for "The Everything Bagel Feature Finder: Ultra-fast automated feature finding for untargeted metabolomics": SI Data File: tool_unique_xic_st_EB_ST004502.html

EB-unique features (not detected by the other two tools) - ST004502

### EB-unique features (not detected by the other two tools) - ST004502

645 features · orbitrap · XIC window m/z ± 10 ppm (≥ 0.002 Da) · ± 0.5 min

Every panel is a feature **EB** detected that neither of the other two tools detected — three-way unique across EB (standard mode (min\_detection\_rate=0.1)), Masscube and XCMS, under the standard matching rule (m/z ± 10 ppm; apex within the reference RT window ± 0.2 min). It shows the MS1 extracted-ion chromatogram at the feature's m/z, drawn from the raw mzML named in `alignment_reference_file`, zoomed to ± 0.5 min around the feature's reported apex RT. **Gray solid line** = the feature's reported apex RT; **red dashed line** = EB's assigned MS/MS — `MS2_scan_id` (native scan number) in `MS2_source_file`; the panel widens to keep an off-apex MS/MS on-screen. Features with no assigned MS/MS show only the gray line. y is per-cell autoscaled. Rows keep the table order (m/z ascending).

m/z 59.0138

02468E+444.5RT (min)

lcorbi\_hilic\_neg\_4\_085\_59

m/z 61.9883

02468E+40.51RT (min)

lcorbi\_hilic\_neg\_2\_011\_29

m/z 61.9883

01E+511.5RT (min)

lcorbi\_hilic\_neg\_1\_052\_41

m/z 61.9885

0123E+44.55RT (min)

lcorbi\_hilic\_neg\_2\_007\_08

m/z 68.9958

01234E+533.5RT (min)

lcorbi\_hilic\_neg\_1\_005\_36

m/z 70.2853

024E+40.51RT (min)

lcorbi\_hilic\_neg\_2\_013\_18

m/z 73.0296

01234E+444.5RT (min)

lcorbi\_hilic\_neg\_4\_087\_59

m/z 74.0239

0123E+511.5RT (min)

lcorbi\_hilic\_neg\_3\_011\_24

m/z 74.0239

01234E+411.5RT (min)

lcorbi\_hilic\_neg\_2\_001\_49

m/z 76.0315

01E+566.5RT (min)

lcorbi\_hilic\_neg\_1\_012\_32

m/z 79.9573

0123E+40.51RT (min)

lcorbi\_hilic\_neg\_4\_078\_58

m/z 79.9573

0123E+477.5RT (min)

lcorbi\_hilic\_neg\_1\_001\_49

m/z 82.9541

024E+444.5RT (min)

lcorbi\_hilic\_neg\_4\_047\_07

m/z 82.9542

024E+477.5RT (min)

lcorbi\_hilic\_neg\_4\_048\_37

m/z 82.9542

024E+477.5RT (min)

lcorbi\_hilic\_neg\_4\_048\_37

m/z 82.9542

01234E+511.5RT (min)

lcorbi\_hilic\_neg\_3\_024\_27

m/z 82.9543

01E+54.55RT (min)

lcorbi\_hilic\_neg\_3\_051\_46

m/z 82.9544

01E+566.5RT (min)

lcorbi\_hilic\_neg\_3\_048\_33

m/z 84.9512

01E+511.5RT (min)

lcorbi\_hilic\_neg\_3\_024\_27

m/z 85.0295

0246E+444.5RT (min)

lcorbi\_hilic\_neg\_4\_082\_59

m/z 85.0295

01E+511.5RT (min)

lcorbi\_hilic\_neg\_2\_014\_31

m/z 89.0429

01E+51.52RT (min)

lcorbi\_hilic\_neg\_4\_005\_32

m/z 90.0188

01E+51.52RT (min)

lcorbi\_hilic\_neg\_4\_095\_60

m/z 90.0471

02468E+40.51RT (min)

lcorbi\_hilic\_neg\_3\_037\_12

m/z 91.0504

01E+566.5RT (min)

lcorbi\_hilic\_neg\_4\_015\_24

m/z 92.2882

012E+433.5RT (min)

lcorbi\_hilic\_neg\_4\_069\_57

m/z 92.9260

01E+55.56RT (min)

lcorbi\_hilic\_neg\_1\_051\_33

m/z 96.9580

0123E+54.55RT (min)

lcorbi\_hilic\_neg\_1\_022\_48

m/z 98.9067

0246E+444.5RT (min)

lcorbi\_hilic\_neg\_3\_011\_24

m/z 100.9351

012E+611.5RT (min)

lcorbi\_hilic\_neg\_2\_039\_07

m/z 102.9567

024E+411.5RT (min)

lcorbi\_hilic\_neg\_4\_030\_05

m/z 102.9567

01234E+411.5RT (min)

lcorbi\_hilic\_neg\_4\_046\_15

m/z 102.9567

012E+54.55RT (min)

lcorbi\_hilic\_neg\_2\_021\_05

m/z 103.0221

012E+51.52RT (min)

lcorbi\_hilic\_neg\_4\_092\_60

m/z 104.9538

024E+46.57RT (min)

lcorbi\_hilic\_neg\_4\_084\_59

m/z 104.9539

01234E+444.5RT (min)

lcorbi\_hilic\_neg\_4\_038\_27

m/z 108.8997

01234E+54.55RT (min)

lcorbi\_hilic\_neg\_2\_025\_52

m/z 108.9020

0123E+43.54RT (min)

lcorbi\_hilic\_neg\_1\_019\_06

m/z 108.9020

0123E+43.54RT (min)

lcorbi\_hilic\_neg\_4\_079\_58

m/z 109.0407

01E+56.57RT (min)

lcorbi\_hilic\_neg\_1\_016\_37

m/z 110.8992

0246E+411.5RT (min)

lcorbi\_hilic\_neg\_1\_006\_08

m/z 111.0090

01E+522.5RT (min)

lcorbi\_hilic\_neg\_1\_004\_17

m/z 111.0199

012E+411.5RT (min)

lcorbi\_hilic\_neg\_3\_036\_17

m/z 111.0200

01234E+43.54RT (min)

lcorbi\_hilic\_neg\_4\_004\_13

m/z 111.9774

01E+511.5RT (min)

lcorbi\_hilic\_neg\_4\_040\_17

m/z 112.9716

024E+511.5RT (min)

lcorbi\_hilic\_neg\_4\_040\_17

m/z 112.9800

0246E+411.5RT (min)

lcorbi\_hilic\_neg\_4\_040\_17

m/z 112.9935

0246E+455.5RT (min)

lcorbi\_hilic\_neg\_4\_074\_58

m/z 112.9991

02468E+411.5RT (min)

lcorbi\_hilic\_neg\_4\_071\_57

m/z 113.0243

02468E+444.5RT (min)

lcorbi\_hilic\_neg\_4\_031\_14

m/z 113.0244

01E+544.5RT (min)

lcorbi\_hilic\_neg\_4\_088\_59

m/z 114.0572

0123E+55.56RT (min)

lcorbi\_hilic\_neg\_1\_055\_24

m/z 114.9340

01234E+44.55RT (min)

lcorbi\_hilic\_neg\_4\_070\_57

m/z 115.0037

01234E+511.5RT (min)

lcorbi\_hilic\_neg\_2\_022\_37

m/z 115.0333

01234E+41.52RT (min)

lcorbi\_hilic\_neg\_4\_094\_60

m/z 115.0333

02468E+455.5RT (min)

lcorbi\_hilic\_neg\_1\_012\_32

m/z 115.0333

01E+55.56RT (min)

lcorbi\_hilic\_neg\_3\_011\_24

m/z 115.0401

02468E+41.52RT (min)

lcorbi\_hilic\_neg\_4\_036\_36

m/z 115.9207

01E+56.57RT (min)

lcorbi\_hilic\_neg\_4\_055\_38

m/z 116.0255

01E+522.5RT (min)

lcorbi\_hilic\_neg\_3\_044\_32

m/z 117.9287

02468E+44.555.5RT (min)

lcorbi\_hilic\_neg\_2\_011\_29

m/z 118.9266

0246E+411.5RT (min)

lcorbi\_hilic\_neg\_4\_107\_62

m/z 119.0171

012E+511.5RT (min)

lcorbi\_hilic\_neg\_2\_055\_32

m/z 119.0349

024E+45.56RT (min)

lcorbi\_hilic\_neg\_3\_040\_19

m/z 119.0502

01E+50.51RT (min)

lcorbi\_hilic\_neg\_2\_020\_27

m/z 119.0502

0246E+433.5RT (min)

lcorbi\_hilic\_neg\_2\_019\_21

m/z 119.9443

01234E+46.57RT (min)

lcorbi\_hilic\_neg\_4\_006\_06

m/z 119.9469

0246E+45.56RT (min)

lcorbi\_hilic\_neg\_4\_078\_58

m/z 119.9469

01234E+44.55RT (min)

lcorbi\_hilic\_neg\_4\_085\_59

m/z 119.9469

0246E+44.55RT (min)

lcorbi\_hilic\_neg\_4\_067\_57

m/z 121.9131

01234E+411.5RT (min)

lcorbi\_hilic\_neg\_3\_039\_31

m/z 121.9439

024E+411.5RT (min)

lcorbi\_hilic\_neg\_4\_018\_12

m/z 121.9440

0246E+41.52RT (min)

lcorbi\_hilic\_neg\_1\_027\_18

m/z 122.0328

01E+50.51RT (min)

lcorbi\_hilic\_neg\_3\_010\_18

m/z 123.0450

02468E+50.51RT (min)

lcorbi\_hilic\_neg\_2\_019\_21

m/z 123.0451

0246E+40.51RT (min)

lcorbi\_hilic\_neg\_2\_020\_27

m/z 124.0151

012E+45.56RT (min)

lcorbi\_hilic\_neg\_4\_068\_57

m/z 124.9840

012E+51.52RT (min)

lcorbi\_hilic\_neg\_4\_004\_13

m/z 128.9567

02468E+644.5RT (min)

lcorbi\_hilic\_neg\_2\_047\_48

m/z 129.0134

01E+52.53RT (min)

lcorbi\_hilic\_neg\_2\_019\_21

m/z 129.0193

0246E+511.5RT (min)

lcorbi\_hilic\_neg\_2\_022\_37

m/z 129.0323

024E+433.5RT (min)

lcorbi\_hilic\_neg\_4\_089\_59

m/z 129.0491

01234E+411.5RT (min)

lcorbi\_hilic\_neg\_1\_001\_49

m/z 129.0491

02468E+42.53RT (min)

lcorbi\_hilic\_neg\_1\_020\_13

m/z 129.9335

0123E+488.5RT (min)

lcorbi\_hilic\_neg\_4\_005\_32

m/z 130.0226

01234E+41.52RT (min)

lcorbi\_hilic\_neg\_4\_031\_14

m/z 130.9630

01E+54.55RT (min)

lcorbi\_hilic\_neg\_4\_074\_58

m/z 131.0445

012E+51.52RT (min)

lcorbi\_hilic\_neg\_4\_015\_24

m/z 132.0217

02468E+411.5RT (min)

lcorbi\_hilic\_neg\_2\_020\_27

m/z 132.8997

0246E+43.54RT (min)

lcorbi\_hilic\_neg\_4\_006\_06

m/z 132.9001

0123E+422.5RT (min)

lcorbi\_hilic\_neg\_4\_028\_03

m/z 132.9002

0246E+466.5RT (min)

lcorbi\_hilic\_neg\_4\_023\_34

m/z 132.9002

01E+53.54RT (min)

lcorbi\_hilic\_neg\_4\_074\_58

m/z 132.9234

02468E+46.57RT (min)

lcorbi\_hilic\_neg\_4\_036\_36

m/z 133.0296

0246E+50.51RT (min)

lcorbi\_hilic\_neg\_2\_015\_19

m/z 133.0495

0246E+577.5RT (min)

lcorbi\_hilic\_neg\_2\_005\_24

m/z 134.0306

0123E+422.5RT (min)

lcorbi\_hilic\_neg\_2\_054\_13

m/z 134.0307

01234E+422.5RT (min)

lcorbi\_hilic\_neg\_4\_008\_35

m/z 134.0473

02468E+422.5RT (min)

lcorbi\_hilic\_neg\_1\_005\_36

m/z 134.8625

024E+55.56RT (min)

lcorbi\_hilic\_neg\_2\_049\_55

m/z 135.0408

01234E+577.5RT (min)

lcorbi\_hilic\_neg\_1\_012\_32

m/z 135.8947

01E+566.5RT (min)

lcorbi\_hilic\_neg\_4\_023\_34

m/z 135.9208

0246E+411.5RT (min)

lcorbi\_hilic\_neg\_4\_107\_62

m/z 135.9208

0246E+41.52RT (min)

lcorbi\_hilic\_neg\_4\_098\_61

m/z 135.9208

024E+411.5RT (min)

lcorbi\_hilic\_neg\_4\_103\_61

m/z 135.9709

024E+411.5RT (min)

lcorbi\_hilic\_neg\_1\_006\_08

m/z 136.0484

0246E+50.51RT (min)

lcorbi\_hilic\_neg\_1\_010\_22

m/z 136.0517

024E+511.5RT (min)

lcorbi\_hilic\_neg\_4\_093\_60

m/z 136.8978

024E+44.55RT (min)

lcorbi\_hilic\_neg\_4\_108\_62

m/z 137.0248

02468E+666.5RT (min)

lcorbi\_hilic\_neg\_2\_005\_24

m/z 137.8917

024E+466.5RT (min)

lcorbi\_hilic\_neg\_4\_038\_27

m/z 137.8917

0246E+43.54RT (min)

lcorbi\_hilic\_neg\_4\_074\_58

m/z 137.8917

01E+566.5RT (min)

lcorbi\_hilic\_neg\_4\_082\_59

m/z 138.0475

01E+56.57RT (min)

lcorbi\_hilic\_neg\_1\_012\_32

m/z 138.8600

0123E+50.51RT (min)

lcorbi\_hilic\_neg\_1\_046\_19

m/z 138.9848

02468E+466.5RT (min)

lcorbi\_hilic\_neg\_4\_082\_59

m/z 139.0638

01E+522.5RT (min)

lcorbi\_hilic\_neg\_3\_011\_24

m/z 140.9940

01E+522.5RT (min)

lcorbi\_hilic\_neg\_4\_071\_57

m/z 142.9290

024E+411.5RT (min)

lcorbi\_hilic\_neg\_4\_019\_09

m/z 143.0349

02468E+444.5RT (min)

lcorbi\_hilic\_neg\_4\_031\_14

m/z 143.0461

012E+444.5RT (min)

lcorbi\_hilic\_neg\_4\_085\_59

m/z 143.0462

0123E+45.56RT (min)

lcorbi\_hilic\_neg\_4\_073\_57

m/z 143.9758

01E+577.5RT (min)

lcorbi\_hilic\_neg\_4\_089\_59

m/z 145.0684

01E+56.57RT (min)

lcorbi\_hilic\_neg\_4\_071\_57

m/z 145.9825

01E+577.5RT (min)

lcorbi\_hilic\_neg\_4\_094\_60

m/z 146.0578

02468E+46.57RT (min)

lcorbi\_hilic\_neg\_1\_029\_31

m/z 146.9587

024E+411.5RT (min)

lcorbi\_hilic\_neg\_3\_032\_42

m/z 146.9669

01234E+44.55RT (min)

lcorbi\_hilic\_neg\_4\_072\_57

m/z 147.0298

0123E+411.5RT (min)

lcorbi\_hilic\_neg\_2\_001\_49

m/z 148.0436

024E+455.5RT (min)

lcorbi\_hilic\_neg\_1\_004\_17

m/z 149.0255

0123E+566.5RT (min)

lcorbi\_hilic\_neg\_3\_053\_13

m/z 149.0272

01E+566.5RT (min)

lcorbi\_hilic\_neg\_4\_089\_59

m/z 149.0455

01E+55.56RT (min)

lcorbi\_hilic\_neg\_3\_018\_38

m/z 151.0432

02468E+411.5RT (min)

lcorbi\_hilic\_neg\_2\_001\_49

m/z 151.0697

012E+566.5RT (min)

lcorbi\_hilic\_neg\_4\_093\_60

m/z 153.8653

012E+577.5RT (min)

lcorbi\_hilic\_neg\_3\_001\_49

m/z 153.8654

012E+57.58RT (min)

lcorbi\_hilic\_neg\_1\_025\_52

m/z 154.0274

01234E+46.57RT (min)

lcorbi\_hilic\_neg\_3\_055\_26

m/z 154.9282

0123E+477.5RT (min)

lcorbi\_hilic\_neg\_1\_006\_08

m/z 154.9282

0123E+47.58RT (min)

lcorbi\_hilic\_neg\_1\_019\_06

m/z 154.9473

01234E+46.57RT (min)

lcorbi\_hilic\_neg\_2\_001\_49

m/z 154.9474

0246E+46.57RT (min)

lcorbi\_hilic\_neg\_1\_037\_46

m/z 155.0556

01E+64.55RT (min)

lcorbi\_hilic\_neg\_2\_005\_24

m/z 155.8657

01E+56.57RT (min)

lcorbi\_hilic\_neg\_1\_039\_38

m/z 156.0570

01234E+444.5RT (min)

lcorbi\_hilic\_neg\_4\_097\_60

m/z 156.0853

01E+644.5RT (min)

lcorbi\_hilic\_neg\_4\_093\_60

m/z 156.0924

01E+544.5RT (min)

lcorbi\_hilic\_neg\_4\_097\_60

m/z 156.9233

01234E+47.58RT (min)

lcorbi\_hilic\_neg\_1\_022\_48

m/z 158.9389

01234E+46.57RT (min)

lcorbi\_hilic\_neg\_4\_097\_60

m/z 158.9391

0246E+44.55RT (min)

lcorbi\_hilic\_neg\_4\_076\_58

m/z 159.0074

01234E+40.51RT (min)

lcorbi\_hilic\_neg\_4\_109\_62

m/z 160.0615

024E+41.52RT (min)

lcorbi\_hilic\_neg\_1\_023\_23

m/z 160.8387

0246E+46.57RT (min)

lcorbi\_hilic\_neg\_3\_001\_49

m/z 161.0244

0123E+50.51RT (min)

lcorbi\_hilic\_neg\_3\_040\_19

m/z 161.0455

01E+53.54RT (min)

lcorbi\_hilic\_neg\_4\_004\_13

m/z 161.0458

024E+444.5RT (min)

lcorbi\_hilic\_neg\_4\_031\_14

m/z 162.8357

0246E+46.57RT (min)

lcorbi\_hilic\_neg\_3\_001\_49

m/z 162.8410

01E+533.5RT (min)

lcorbi\_hilic\_neg\_1\_020\_13

m/z 162.8420

024E+588.5RT (min)

lcorbi\_hilic\_neg\_1\_018\_21

m/z 162.9398

01E+511.5RT (min)

lcorbi\_hilic\_neg\_4\_024\_39

m/z 162.9896

01E+60.51RT (min)

lcorbi\_hilic\_neg\_4\_014\_18

m/z 163.0343

012E+511.5RT (min)

lcorbi\_hilic\_neg\_1\_004\_17

m/z 163.9904

01E+50.51RT (min)

lcorbi\_hilic\_neg\_4\_014\_18

m/z 163.9910

01234E+46.57RT (min)

lcorbi\_hilic\_neg\_4\_072\_57

m/z 165.0039

024E+46.57RT (min)

lcorbi\_hilic\_neg\_4\_093\_60

m/z 165.0223

02468E+411.5RT (min)

lcorbi\_hilic\_neg\_1\_004\_17

m/z 165.0226

02468E+466.5RT (min)

lcorbi\_hilic\_neg\_1\_029\_31

m/z 166.0226

0246E+50.51RT (min)

lcorbi\_hilic\_neg\_4\_075\_58

m/z 166.8356

01E+57.58RT (min)

lcorbi\_hilic\_neg\_3\_050\_03

m/z 166.8357

01E+588.5RT (min)

lcorbi\_hilic\_neg\_1\_024\_11

m/z 167.0210

01234E+655.5RT (min)

lcorbi\_hilic\_neg\_2\_020\_27

m/z 167.0229

02468E+46.57RT (min)

lcorbi\_hilic\_neg\_1\_023\_23

m/z 167.8312

024E+58.59RT (min)

lcorbi\_hilic\_neg\_4\_035\_48

m/z 167.8331

02468E+544.5RT (min)

lcorbi\_hilic\_neg\_1\_025\_52

m/z 167.8367

02468E+57.58RT (min)

lcorbi\_hilic\_neg\_3\_004\_41

m/z 167.8368

01E+566.5RT (min)

lcorbi\_hilic\_neg\_1\_002\_26

m/z 168.0243

012E+55.56RT (min)

lcorbi\_hilic\_neg\_4\_068\_57

m/z 168.0666

01E+51.52RT (min)

lcorbi\_hilic\_neg\_2\_020\_27

m/z 169.0554

0123E+566.5RT (min)

lcorbi\_hilic\_neg\_4\_095\_60

m/z 169.8304

01E+644.5RT (min)

lcorbi\_hilic\_neg\_1\_025\_52

m/z 169.8342

01E+566.5RT (min)

lcorbi\_hilic\_neg\_1\_002\_26

m/z 170.8332

01E+56.57RT (min)

lcorbi\_hilic\_neg\_4\_085\_59

m/z 170.8333

0123E+466.5RT (min)

lcorbi\_hilic\_neg\_1\_005\_36

m/z 171.0299

024E+411.5RT (min)

lcorbi\_hilic\_neg\_4\_087\_59

m/z 171.0411

01E+52.53RT (min)

lcorbi\_hilic\_neg\_4\_071\_57

m/z 171.8315

01E+566.5RT (min)

lcorbi\_hilic\_neg\_4\_076\_58

m/z 172.9580

0246E+44.55RT (min)

lcorbi\_hilic\_neg\_1\_003\_14

m/z 173.0389

02468E+41.52RT (min)

lcorbi\_hilic\_neg\_4\_067\_57

m/z 173.0455

01E+511.5RT (min)

lcorbi\_hilic\_neg\_4\_037\_31

m/z 173.9947

01234E+40.51RT (min)

lcorbi\_hilic\_neg\_2\_020\_27

m/z 174.0240

01234E+422.5RT (min)

lcorbi\_hilic\_neg\_4\_085\_59

m/z 174.0323

01E+50.51RT (min)

lcorbi\_hilic\_neg\_1\_036\_09

m/z 174.0409

01E+533.5RT (min)

lcorbi\_hilic\_neg\_1\_002\_26

m/z 175.0249

01E+52.53RT (min)

lcorbi\_hilic\_neg\_1\_026\_34

m/z 175.0805

01E+55.56RT (min)

lcorbi\_hilic\_neg\_3\_028\_23

m/z 175.0918

01E+56.57RT (min)

lcorbi\_hilic\_neg\_1\_029\_31

m/z 176.0226

0246E+422.5RT (min)

lcorbi\_hilic\_neg\_4\_100\_61

m/z 176.0281

01E+51.52RT (min)

lcorbi\_hilic\_neg\_4\_102\_61

m/z 176.0433

0246E+40.51RT (min)

lcorbi\_hilic\_neg\_4\_112\_62

m/z 176.9979

01E+50.51RT (min)

lcorbi\_hilic\_neg\_4\_043\_11

m/z 177.0404

01E+511.5RT (min)

lcorbi\_hilic\_neg\_2\_054\_13

m/z 177.0404

01E+511.5RT (min)

lcorbi\_hilic\_neg\_3\_016\_16

m/z 177.0404

0246E+42.53RT (min)

lcorbi\_hilic\_neg\_1\_035\_29

m/z 177.0557

02468E+50.51RT (min)

lcorbi\_hilic\_neg\_2\_015\_19

m/z 177.8993

01234E+56.57RT (min)

lcorbi\_hilic\_neg\_3\_001\_49

m/z 178.9030

0246E+46.57RT (min)

lcorbi\_hilic\_neg\_4\_031\_14

m/z 178.9031

02468E+466.5RT (min)

lcorbi\_hilic\_neg\_3\_002\_15

m/z 178.9031

01234E+477.5RT (min)

lcorbi\_hilic\_neg\_1\_005\_36

m/z 178.9838

0246E+466.5RT (min)

lcorbi\_hilic\_neg\_1\_005\_36

m/z 179.0341

02468E+422.5RT (min)

lcorbi\_hilic\_neg\_2\_011\_29

m/z 179.0342

01234E+422.5RT (min)

lcorbi\_hilic\_neg\_2\_028\_33

m/z 179.0344

01234E+422.5RT (min)

lcorbi\_hilic\_neg\_4\_031\_14

m/z 179.0561

0246E+466.5RT (min)

lcorbi\_hilic\_neg\_3\_018\_38

m/z 179.0561

0246E+43.54RT (min)

lcorbi\_hilic\_neg\_2\_020\_27

m/z 180.0192

02468E+466.5RT (min)

lcorbi\_hilic\_neg\_2\_005\_24

m/z 180.0665

024E+40.51RT (min)

lcorbi\_hilic\_neg\_3\_039\_31

m/z 180.0666

01E+55.56RT (min)

lcorbi\_hilic\_neg\_2\_020\_27

m/z 180.1166

01E+566.5RT (min)

lcorbi\_hilic\_neg\_4\_095\_60

m/z 180.9730

01234E+46.57RT (min)

lcorbi\_hilic\_neg\_4\_084\_59

m/z 181.0450

0123E+566.5RT (min)

lcorbi\_hilic\_neg\_3\_044\_32

m/z 181.0510

01E+51.52RT (min)

lcorbi\_hilic\_neg\_1\_003\_14

m/z 181.0618

01E+51.52RT (min)

lcorbi\_hilic\_neg\_2\_001\_49

m/z 181.1005

012E+55.56RT (min)

lcorbi\_hilic\_neg\_4\_096\_60

m/z 181.9376

0246E+41.52RT (min)

lcorbi\_hilic\_neg\_2\_008\_39

m/z 181.9861

012E+511.5RT (min)

lcorbi\_hilic\_neg\_1\_039\_38

m/z 182.0439

012E+566.5RT (min)

lcorbi\_hilic\_neg\_1\_023\_23

m/z 182.0797

01E+566.5RT (min)

lcorbi\_hilic\_neg\_1\_012\_32

m/z 182.0846

01234E+56.57RT (min)

lcorbi\_hilic\_neg\_3\_011\_24

m/z 182.8616

012E+44.55RT (min)

lcorbi\_hilic\_neg\_4\_075\_58

m/z 182.9946

0123E+611.5RT (min)

lcorbi\_hilic\_neg\_2\_020\_27

m/z 182.9949

01E+611.5RT (min)

lcorbi\_hilic\_neg\_4\_106\_62

m/z 182.9971

012E+522.5RT (min)

lcorbi\_hilic\_neg\_4\_004\_13

m/z 183.0065

012E+51.52RT (min)

lcorbi\_hilic\_neg\_3\_028\_23

m/z 183.0159

0246E+444.5RT (min)

lcorbi\_hilic\_neg\_4\_078\_58

m/z 183.9955

01E+511.5RT (min)

lcorbi\_hilic\_neg\_2\_015\_19

m/z 185.0067

0123E+41.52RT (min)

lcorbi\_hilic\_neg\_4\_089\_59

m/z 185.0125

0246E+422.5RT (min)

lcorbi\_hilic\_neg\_4\_097\_60

m/z 186.0414

02468E+44.55RT (min)

lcorbi\_hilic\_neg\_4\_053\_23

m/z 187.0612

0246E+411.5RT (min)

lcorbi\_hilic\_neg\_4\_087\_59

m/z 187.9419

0246E+400.5RT (min)

lcorbi\_hilic\_neg\_3\_006\_28

m/z 188.0564

01234E+42.53RT (min)

lcorbi\_hilic\_neg\_4\_089\_59

m/z 188.0565

012E+522.5RT (min)

lcorbi\_hilic\_neg\_1\_012\_32

m/z 188.0612

01E+51.52RT (min)

lcorbi\_hilic\_neg\_4\_078\_58

m/z 188.0636

02468E+466.5RT (min)

lcorbi\_hilic\_neg\_1\_055\_24

m/z 189.0029

02468E+45.56RT (min)

lcorbi\_hilic\_neg\_4\_073\_57

m/z 189.0193

01E+511.5RT (min)

lcorbi\_hilic\_neg\_2\_020\_27

m/z 189.0403

01E+52.53RT (min)

lcorbi\_hilic\_neg\_2\_020\_27

m/z 190.0390

01E+522.5RT (min)

lcorbi\_hilic\_neg\_3\_011\_24

m/z 190.0478

01E+566.5RT (min)

lcorbi\_hilic\_neg\_2\_005\_24

m/z 190.0491

01E+511.5RT (min)

lcorbi\_hilic\_neg\_2\_020\_27

m/z 191.0197

012E+56.57RT (min)

lcorbi\_hilic\_neg\_1\_016\_37

m/z 191.0199

0246E+466.5RT (min)

lcorbi\_hilic\_neg\_4\_045\_25

m/z 191.0494

0246E+455.5RT (min)

lcorbi\_hilic\_neg\_1\_004\_17

m/z 191.0495

01234E+43.54RT (min)

lcorbi\_hilic\_neg\_4\_071\_57

m/z 191.0732

01E+51.52RT (min)

lcorbi\_hilic\_neg\_4\_107\_62

m/z 191.1077

0246E+411.5RT (min)

lcorbi\_hilic\_neg\_4\_004\_13

m/z 191.1078

024E+400.5RT (min)

lcorbi\_hilic\_neg\_1\_017\_51

m/z 192.0495

01E+544.5RT (min)

lcorbi\_hilic\_neg\_4\_030\_05

m/z 192.0544

012E+55.56RT (min)

lcorbi\_hilic\_neg\_1\_012\_32

m/z 192.0577

0246E+411.5RT (min)

lcorbi\_hilic\_neg\_2\_020\_27

m/z 192.9457

024E+411.5RT (min)

lcorbi\_hilic\_neg\_4\_039\_42

m/z 192.9496

01234E+422.5RT (min)

lcorbi\_hilic\_neg\_4\_095\_60

m/z 193.0285

01234E+54.55RT (min)

lcorbi\_hilic\_neg\_2\_005\_24

m/z 193.0385

0123E+55.56RT (min)

lcorbi\_hilic\_neg\_1\_023\_23

m/z 193.0451

01E+51.52RT (min)

lcorbi\_hilic\_neg\_4\_095\_60

m/z 193.0532

024E+411.5RT (min)

lcorbi\_hilic\_neg\_2\_020\_27

m/z 193.0644

012E+54.55RT (min)

lcorbi\_hilic\_neg\_1\_055\_24

m/z 193.0869

01234E+411.5RT (min)

lcorbi\_hilic\_neg\_1\_039\_38

m/z 193.8157

02468E+400.5RT (min)

lcorbi\_hilic\_neg\_4\_080\_58

m/z 194.0513

02468E+466.5RT (min)

lcorbi\_hilic\_neg\_2\_055\_32

m/z 194.0585

024E+544.5RT (min)

lcorbi\_hilic\_neg\_4\_091\_60

m/z 194.0767

01E+533.5RT (min)

lcorbi\_hilic\_neg\_4\_096\_60

m/z 194.9016

024E+56.57RT (min)

lcorbi\_hilic\_neg\_3\_022\_06

m/z 194.9016

01234E+577.5RT (min)

lcorbi\_hilic\_neg\_1\_025\_52

m/z 194.9179

0123E+47.58RT (min)

lcorbi\_hilic\_neg\_4\_096\_60

m/z 194.9285

02468E+477.5RT (min)

lcorbi\_hilic\_neg\_1\_001\_49

m/z 194.9420

024E+677.5RT (min)

lcorbi\_hilic\_neg\_4\_005\_32

m/z 194.9522

0246E+477.5RT (min)

lcorbi\_hilic\_neg\_3\_042\_05

m/z 194.9524

0246E+477.5RT (min)

lcorbi\_hilic\_neg\_1\_047\_05

m/z 195.1042

024E+42.53RT (min)

lcorbi\_hilic\_neg\_1\_022\_48

m/z 196.0557

01E+544.5RT (min)

lcorbi\_hilic\_neg\_4\_097\_60

m/z 196.0557

01E+544.5RT (min)

lcorbi\_hilic\_neg\_4\_095\_60

m/z 196.9324

0123E+477.5RT (min)

lcorbi\_hilic\_neg\_1\_023\_23

m/z 197.0222

01E+51.52RT (min)

lcorbi\_hilic\_neg\_4\_067\_57

m/z 197.0680

0246E+42.53RT (min)

lcorbi\_hilic\_neg\_4\_004\_13

m/z 198.0495

01E+55.56RT (min)

lcorbi\_hilic\_neg\_4\_077\_58

m/z 198.8853

024E+488.5RT (min)

lcorbi\_hilic\_neg\_1\_046\_19

m/z 198.9178

024E+477.5RT (min)

lcorbi\_hilic\_neg\_4\_086\_59

m/z 199.0070

0246E+411.5RT (min)

lcorbi\_hilic\_neg\_4\_097\_60

m/z 199.0070

0123E+50.51RT (min)

lcorbi\_hilic\_neg\_2\_020\_27

m/z 199.0281

0123E+43.54RT (min)

lcorbi\_hilic\_neg\_4\_076\_58

m/z 199.0474

01234E+43.54RT (min)

lcorbi\_hilic\_neg\_4\_081\_58

m/z 199.0583

0123E+55.56RT (min)

lcorbi\_hilic\_neg\_1\_055\_24

m/z 201.0396

01E+533.5RT (min)

lcorbi\_hilic\_neg\_4\_094\_60

m/z 201.9061

0123E+41.52RT (min)

lcorbi\_hilic\_neg\_4\_074\_58

m/z 202.0192

02468E+422.5RT (min)

lcorbi\_hilic\_neg\_4\_082\_59

m/z 202.0608

01E+522.5RT (min)

lcorbi\_hilic\_neg\_4\_074\_58

m/z 202.0610

01E+522.5RT (min)

lcorbi\_hilic\_neg\_4\_055\_38

m/z 202.0714

01E+544.5RT (min)

lcorbi\_hilic\_neg\_3\_044\_32

m/z 202.0833

01234E+744.5RT (min)

lcorbi\_hilic\_neg\_3\_044\_32

m/z 202.0833

01234E+744.5RT (min)

lcorbi\_hilic\_neg\_1\_055\_24

m/z 202.0946

0246E+544.5RT (min)

lcorbi\_hilic\_neg\_4\_095\_60

m/z 202.9289

01E+54.55RT (min)

lcorbi\_hilic\_neg\_4\_075\_58

m/z 202.9289

0246E+46.57RT (min)

lcorbi\_hilic\_neg\_4\_097\_60

m/z 203.0469

01E+56.57RT (min)

lcorbi\_hilic\_neg\_4\_091\_60

m/z 203.0673

01234E+46.57RT (min)

lcorbi\_hilic\_neg\_1\_016\_37

m/z 204.9464

0246E+40.51RT (min)

lcorbi\_hilic\_neg\_1\_002\_26

m/z 204.9963

01234E+40.51RT (min)

lcorbi\_hilic\_neg\_4\_001\_49

m/z 205.0106

012E+56.57RT (min)

lcorbi\_hilic\_neg\_4\_097\_60

m/z 205.9947

0123E+566.5RT (min)

lcorbi\_hilic\_neg\_4\_090\_60

m/z 207.0120

0123E+511.5RT (min)

lcorbi\_hilic\_neg\_4\_026\_33

m/z 207.0541

01E+51.52RT (min)

lcorbi\_hilic\_neg\_4\_073\_57

m/z 208.9346

0246E+40.51RT (min)

lcorbi\_hilic\_neg\_1\_006\_08

m/z 209.0042

0246E+40.51RT (min)

lcorbi\_hilic\_neg\_1\_030\_04

m/z 209.0609

0123E+43.54RT (min)

lcorbi\_hilic\_neg\_3\_031\_14

m/z 210.9315

012E+555.5RT (min)

lcorbi\_hilic\_neg\_1\_050\_07

m/z 211.0012

01E+55.56RT (min)

lcorbi\_hilic\_neg\_3\_012\_35

m/z 211.0724

0246E+422.5RT (min)

lcorbi\_hilic\_neg\_4\_089\_59

m/z 211.0747

01E+54.55RT (min)

lcorbi\_hilic\_neg\_4\_096\_60

m/z 212.0049

01E+51.52RT (min)

lcorbi\_hilic\_neg\_1\_018\_21

m/z 213.0055

012E+511.5RT (min)

lcorbi\_hilic\_neg\_1\_051\_33

m/z 213.9270

0123E+411.5RT (min)

lcorbi\_hilic\_neg\_4\_017\_51

m/z 214.0019

024E+41.52RT (min)

lcorbi\_hilic\_neg\_4\_103\_61

m/z 214.0260

0123E+40.51RT (min)

lcorbi\_hilic\_neg\_4\_080\_58

m/z 214.9190

01E+54.55RT (min)

lcorbi\_hilic\_neg\_4\_078\_58

m/z 214.9190

01E+54.55RT (min)

lcorbi\_hilic\_neg\_4\_074\_58

m/z 215.0019

0246E+40.51RT (min)

lcorbi\_hilic\_neg\_2\_020\_27

m/z 215.0260

01E+555.5RT (min)

lcorbi\_hilic\_neg\_1\_015\_30

m/z 215.8767

0123E+46.57RT (min)

lcorbi\_hilic\_neg\_1\_048\_35

m/z 215.8768

0246E+466.5RT (min)

lcorbi\_hilic\_neg\_2\_037\_15

m/z 215.8768

01234E+466.5RT (min)

lcorbi\_hilic\_neg\_4\_040\_17

m/z 215.8770

0123E+46.57RT (min)

lcorbi\_hilic\_neg\_1\_005\_36

m/z 215.9251

0123E+411.5RT (min)

lcorbi\_hilic\_neg\_4\_017\_51

m/z 216.0182

01E+577.5RT (min)

lcorbi\_hilic\_neg\_1\_055\_24

m/z 216.0361

024E+466.5RT (min)

lcorbi\_hilic\_neg\_4\_067\_57

m/z 216.0383

0246E+422.5RT (min)

lcorbi\_hilic\_neg\_4\_083\_59

m/z 216.8537

01E+577.5RT (min)

lcorbi\_hilic\_neg\_4\_082\_59

m/z 216.8537

0246E+477.5RT (min)

lcorbi\_hilic\_neg\_4\_047\_07

m/z 217.0297

012E+566.5RT (min)

lcorbi\_hilic\_neg\_4\_080\_58

m/z 217.0297

024E+65.56RT (min)

lcorbi\_hilic\_neg\_1\_021\_28

m/z 217.0298

012E+566.5RT (min)

lcorbi\_hilic\_neg\_4\_067\_57

m/z 217.0298

02468E+46.57RT (min)

lcorbi\_hilic\_neg\_4\_095\_60

m/z 218.0200

01234E+466.5RT (min)

lcorbi\_hilic\_neg\_4\_085\_59

m/z 218.0331

0246E+444.5RT (min)

lcorbi\_hilic\_neg\_4\_076\_58

m/z 218.0539

0246E+4123RT (min)

lcorbi\_hilic\_neg\_1\_016\_37

m/z 218.0659

0123E+56.57RT (min)

lcorbi\_hilic\_neg\_4\_015\_24

m/z 218.0821

01E+56.57RT (min)

lcorbi\_hilic\_neg\_4\_095\_60

m/z 218.9631

01E+566.5RT (min)

lcorbi\_hilic\_neg\_1\_012\_32

m/z 218.9633

01234E+44.55RT (min)

lcorbi\_hilic\_neg\_1\_019\_06

m/z 219.0459

01E+54.55RT (min)

lcorbi\_hilic\_neg\_1\_023\_23

m/z 219.0726

01E+50.51RT (min)

lcorbi\_hilic\_neg\_2\_015\_19

m/z 219.0733

01E+544.5RT (min)

lcorbi\_hilic\_neg\_3\_044\_32

m/z 219.0734

01E+511.5RT (min)

lcorbi\_hilic\_neg\_2\_020\_27

m/z 219.9453

0246E+40.51RT (min)

lcorbi\_hilic\_neg\_4\_104\_61

m/z 221.0257

02468E+422.5RT (min)

lcorbi\_hilic\_neg\_4\_083\_59

m/z 221.0528

02468E+43.54RT (min)

lcorbi\_hilic\_neg\_1\_012\_32

m/z 221.0666

0123E+41.52RT (min)

lcorbi\_hilic\_neg\_4\_085\_59

m/z 221.0720

0246E+40.51RT (min)

lcorbi\_hilic\_neg\_4\_109\_62

m/z 221.8424

0246E+43.54RT (min)

lcorbi\_hilic\_neg\_4\_082\_59

m/z 221.8424

0246E+43.54RT (min)

lcorbi\_hilic\_neg\_3\_022\_06

m/z 222.9834

0123E+411.5RT (min)

lcorbi\_hilic\_neg\_1\_026\_34

m/z 223.0197

012E+411.5RT (min)

lcorbi\_hilic\_neg\_2\_048\_38

m/z 223.0636

0246E+555.5RT (min)

lcorbi\_hilic\_neg\_2\_005\_24

m/z 223.0638

0246E+54.55RT (min)

lcorbi\_hilic\_neg\_2\_005\_24

m/z 223.0823

01234E+43.54RT (min)

lcorbi\_hilic\_neg\_4\_001\_49

m/z 223.1339

012E+511.5RT (min)

lcorbi\_hilic\_neg\_4\_004\_13

m/z 224.0617

01E+544.5RT (min)

lcorbi\_hilic\_neg\_2\_038\_36

m/z 225.0607

012E+54.55RT (min)

lcorbi\_hilic\_neg\_2\_005\_24

m/z 225.0615

012E+566.5RT (min)

lcorbi\_hilic\_neg\_4\_004\_13

m/z 225.0615

024E+444.5RT (min)

lcorbi\_hilic\_neg\_4\_055\_38

m/z 225.0672

0123E+455.5RT (min)

lcorbi\_hilic\_neg\_4\_054\_30

m/z 227.0383

0246E+40.51RT (min)

lcorbi\_hilic\_neg\_4\_074\_58

m/z 228.0642

0246E+466.5RT (min)

lcorbi\_hilic\_neg\_4\_089\_59

m/z 228.0786

024E+57.58RT (min)

lcorbi\_hilic\_neg\_2\_005\_24

m/z 228.9506

01E+511.5RT (min)

lcorbi\_hilic\_neg\_1\_025\_52

m/z 228.9926

0246E+40.51RT (min)

lcorbi\_hilic\_neg\_1\_002\_26

m/z 229.0934

012E+51.52RT (min)

lcorbi\_hilic\_neg\_4\_096\_60

m/z 230.0072

0123E+577.5RT (min)

lcorbi\_hilic\_neg\_4\_084\_59

m/z 230.9485

01E+511.5RT (min)

lcorbi\_hilic\_neg\_1\_025\_52

m/z 231.0630

01234E+477.5RT (min)

lcorbi\_hilic\_neg\_4\_070\_57

m/z 233.0819

01E+50.51RT (min)

lcorbi\_hilic\_neg\_4\_113\_62

m/z 233.0819

01234E+511.5RT (min)

lcorbi\_hilic\_neg\_1\_046\_19

m/z 233.0930

012E+42.53RT (min)

lcorbi\_hilic\_neg\_2\_001\_49

m/z 233.9244

012E+44.55RT (min)

lcorbi\_hilic\_neg\_1\_009\_50

m/z 235.9688

012E+577.5RT (min)

lcorbi\_hilic\_neg\_4\_097\_60

m/z 236.0095

024E+411.5RT (min)

lcorbi\_hilic\_neg\_4\_070\_57

m/z 236.0296

012E+655.5RT (min)

lcorbi\_hilic\_neg\_2\_005\_24

m/z 237.1165

012E+50.51RT (min)

lcorbi\_hilic\_neg\_2\_038\_36

m/z 238.0293

01E+544.5RT (min)

lcorbi\_hilic\_neg\_2\_005\_24

m/z 238.0396

0246E+411.5RT (min)

lcorbi\_hilic\_neg\_2\_005\_24

m/z 238.1085

02468E+411.5RT (min)

lcorbi\_hilic\_neg\_2\_020\_27

m/z 238.9315

0123E+46.57RT (min)

lcorbi\_hilic\_neg\_3\_005\_08

m/z 239.0923

012E+50.51RT (min)

lcorbi\_hilic\_neg\_3\_039\_31

m/z 241.1446

024E+411.5RT (min)

lcorbi\_hilic\_neg\_4\_077\_58

m/z 241.9762

024E+433.5RT (min)

lcorbi\_hilic\_neg\_4\_085\_59

m/z 242.0646

01E+55.56RT (min)

lcorbi\_hilic\_neg\_4\_083\_59

m/z 242.0758

02468E+46.57RT (min)

lcorbi\_hilic\_neg\_4\_088\_59

m/z 242.0799

01E+566.5RT (min)

lcorbi\_hilic\_neg\_4\_089\_59

m/z 243.0310

0123E+422.5RT (min)

lcorbi\_hilic\_neg\_4\_102\_61

m/z 243.0622

0123E+533.5RT (min)

lcorbi\_hilic\_neg\_4\_080\_58

m/z 243.0774

01234E+444.5RT (min)

lcorbi\_hilic\_neg\_1\_002\_26

m/z 245.0488

0123E+50.51RT (min)

lcorbi\_hilic\_neg\_3\_024\_27

m/z 245.0488

024E+511.5RT (min)

lcorbi\_hilic\_neg\_2\_012\_14

m/z 245.0489

01234E+50.51RT (min)

lcorbi\_hilic\_neg\_4\_078\_58

m/z 246.0259

01234E+422.5RT (min)

lcorbi\_hilic\_neg\_4\_083\_59

m/z 246.0468

0246E+466.5RT (min)

lcorbi\_hilic\_neg\_1\_016\_37

m/z 247.0403

01E+55.56RT (min)

lcorbi\_hilic\_neg\_1\_046\_19

m/z 247.1065

01E+577.5RT (min)

lcorbi\_hilic\_neg\_4\_005\_32

m/z 248.0904

01E+56.57RT (min)

lcorbi\_hilic\_neg\_2\_005\_24

m/z 248.9551

02468E+655.5RT (min)

lcorbi\_hilic\_neg\_1\_051\_33

m/z 248.9605

024E+46.57RT (min)

lcorbi\_hilic\_neg\_4\_088\_59

m/z 248.9732

0123E+400.5RT (min)

lcorbi\_hilic\_neg\_4\_070\_57

m/z 248.9795

01E+54.55RT (min)

lcorbi\_hilic\_neg\_3\_023\_36

m/z 249.0981

012E+566.5RT (min)

lcorbi\_hilic\_neg\_2\_005\_24

m/z 250.9764

02468E+44.55RT (min)

lcorbi\_hilic\_neg\_4\_077\_58

m/z 252.0665

01E+555.5RT (min)

lcorbi\_hilic\_neg\_2\_055\_32

m/z 252.1004

01E+511.5RT (min)

lcorbi\_hilic\_neg\_2\_005\_24

m/z 253.0235

02468E+53.54RT (min)

lcorbi\_hilic\_neg\_3\_055\_26

m/z 253.1445

0123E+411.5RT (min)

lcorbi\_hilic\_neg\_4\_076\_58

m/z 254.0686

02468E+45.56RT (min)

lcorbi\_hilic\_neg\_2\_032\_25

m/z 254.0782

01234E+46.57RT (min)

lcorbi\_hilic\_neg\_1\_029\_31

m/z 254.0800

01E+51.52RT (min)

lcorbi\_hilic\_neg\_4\_089\_59

m/z 255.8222

024E+43.54RT (min)

lcorbi\_hilic\_neg\_4\_089\_59

m/z 256.0539

01E+64.55RT (min)

lcorbi\_hilic\_neg\_3\_027\_45

m/z 257.1143

012E+57.58RT (min)

lcorbi\_hilic\_neg\_1\_023\_23

m/z 257.8189

01234E+43.54RT (min)

lcorbi\_hilic\_neg\_3\_051\_46

m/z 257.8192

024E+43.54RT (min)

lcorbi\_hilic\_neg\_4\_088\_59

m/z 258.0577

0123E+555.5RT (min)

lcorbi\_hilic\_neg\_2\_040\_35

m/z 258.0578

0246E+455.5RT (min)

lcorbi\_hilic\_neg\_1\_029\_31

m/z 258.9952

012E+444.5RT (min)

lcorbi\_hilic\_neg\_2\_044\_46

m/z 259.0128

012E+566.5RT (min)

lcorbi\_hilic\_neg\_4\_073\_57

m/z 259.0128

01234E+54.55RT (min)

lcorbi\_hilic\_neg\_4\_066\_57

m/z 259.0129

012E+566.5RT (min)

lcorbi\_hilic\_neg\_4\_004\_13

m/z 259.0515

01234E+555.5RT (min)

lcorbi\_hilic\_neg\_1\_029\_31

m/z 259.0515

012E+555.5RT (min)

lcorbi\_hilic\_neg\_2\_040\_35

m/z 261.0560

01E+54.55RT (min)

lcorbi\_hilic\_neg\_1\_009\_50

m/z 261.0561

02468E+43.54RT (min)

lcorbi\_hilic\_neg\_2\_001\_49

m/z 262.0914

02468E+444.5RT (min)

lcorbi\_hilic\_neg\_1\_027\_18

m/z 262.0952

012E+533.5RT (min)

lcorbi\_hilic\_neg\_2\_032\_25

m/z 262.9244

02468E+488.5RT (min)

lcorbi\_hilic\_neg\_2\_009\_50

m/z 263.0725

0246E+42.53RT (min)

lcorbi\_hilic\_neg\_1\_037\_46

m/z 263.0860

012E+566.5RT (min)

lcorbi\_hilic\_neg\_4\_009\_50

m/z 263.0924

012E+511.5RT (min)

lcorbi\_hilic\_neg\_1\_001\_49

m/z 263.1102

01E+51.52RT (min)

lcorbi\_hilic\_neg\_4\_093\_60

m/z 263.1287

0246E+40.51RT (min)

lcorbi\_hilic\_neg\_1\_003\_14

m/z 263.1345

01E+51.52RT (min)

lcorbi\_hilic\_neg\_4\_084\_59

m/z 264.0802

024E+54.55RT (min)

lcorbi\_hilic\_neg\_2\_005\_24

m/z 264.9342

01234E+44.55RT (min)

lcorbi\_hilic\_neg\_4\_003\_08

m/z 265.0367

01E+57.58RT (min)

lcorbi\_hilic\_neg\_2\_005\_24

m/z 265.0887

01E+566.5RT (min)

lcorbi\_hilic\_neg\_3\_042\_05

m/z 265.1321

02468E+40.51RT (min)

lcorbi\_hilic\_neg\_4\_089\_59

m/z 266.0830

012E+54.55RT (min)

lcorbi\_hilic\_neg\_4\_097\_60

m/z 266.1511

01E+611.5RT (min)

lcorbi\_hilic\_neg\_1\_023\_23

m/z 267.0777

01E+65.56RT (min)

lcorbi\_hilic\_neg\_4\_009\_50

m/z 267.1238

01234E+50.51RT (min)

lcorbi\_hilic\_neg\_4\_075\_58

m/z 268.0868

01234E+455.5RT (min)

lcorbi\_hilic\_neg\_3\_039\_31

m/z 268.1511

01E+56.57RT (min)

lcorbi\_hilic\_neg\_4\_094\_60

m/z 269.0545

01E+53.54RT (min)

lcorbi\_hilic\_neg\_3\_028\_23

m/z 269.0779

0123E+511.5RT (min)

lcorbi\_hilic\_neg\_1\_023\_23

m/z 269.0876

0246E+45.56RT (min)

lcorbi\_hilic\_neg\_4\_072\_57

m/z 269.0876

0246E+45.56RT (min)

lcorbi\_hilic\_neg\_4\_068\_57

m/z 269.0877

012E+54.55RT (min)

lcorbi\_hilic\_neg\_3\_009\_50

m/z 269.0879

01E+53.54RT (min)

lcorbi\_hilic\_neg\_2\_001\_49

m/z 269.1212

01234E+566.5RT (min)

lcorbi\_hilic\_neg\_4\_005\_32

m/z 270.0705

012E+544.5RT (min)

lcorbi\_hilic\_neg\_1\_055\_24

m/z 270.7903

01234E+44.55RT (min)

lcorbi\_hilic\_neg\_4\_078\_58

m/z 271.0595

01E+522.5RT (min)

lcorbi\_hilic\_neg\_4\_004\_13

m/z 271.0939

0246E+433.5RT (min)

lcorbi\_hilic\_neg\_3\_028\_23

m/z 271.0953

012E+51.52RT (min)

lcorbi\_hilic\_neg\_4\_015\_24

m/z 272.8831

02468E+46.57RT (min)

lcorbi\_hilic\_neg\_4\_083\_59

m/z 273.0636

0246E+422.5RT (min)

lcorbi\_hilic\_neg\_4\_075\_58

m/z 275.1077

02468E+42.53RT (min)

lcorbi\_hilic\_neg\_1\_037\_46

m/z 277.0812

01E+53.54RT (min)

lcorbi\_hilic\_neg\_4\_090\_60

m/z 277.0953

01234E+577.5RT (min)

lcorbi\_hilic\_neg\_2\_005\_24

m/z 277.1525

024E+511.5RT (min)

lcorbi\_hilic\_neg\_2\_050\_17

m/z 279.1237

0246E+40.51RT (min)

lcorbi\_hilic\_neg\_2\_019\_21

m/z 280.1038

012E+56.57RT (min)

lcorbi\_hilic\_neg\_4\_087\_59

m/z 280.1147

0123E+50.51RT (min)

lcorbi\_hilic\_neg\_4\_091\_60

m/z 280.9023

02468E+44.55RT (min)

lcorbi\_hilic\_neg\_3\_022\_06

m/z 281.0455

0123E+50.51RT (min)

lcorbi\_hilic\_neg\_4\_111\_62

m/z 281.0576

01E+53.54RT (min)

lcorbi\_hilic\_neg\_4\_089\_59

m/z 281.1301

01E+50.51RT (min)

lcorbi\_hilic\_neg\_2\_019\_21

m/z 281.1426

0246E+40.51RT (min)

lcorbi\_hilic\_neg\_4\_089\_59

m/z 281.1485

01E+50.51RT (min)

lcorbi\_hilic\_neg\_2\_013\_18

m/z 281.1485

024E+511.5RT (min)

lcorbi\_hilic\_neg\_1\_002\_26

m/z 283.0610

0246E+411.5RT (min)

lcorbi\_hilic\_neg\_4\_103\_61

m/z 283.0678

0123E+40.51RT (min)

lcorbi\_hilic\_neg\_3\_006\_28

m/z 283.1446

01234E+50.51RT (min)

lcorbi\_hilic\_neg\_1\_027\_18

m/z 285.1259

01E+50.51RT (min)

lcorbi\_hilic\_neg\_4\_078\_58

m/z 286.1556

012E+57.58RT (min)

lcorbi\_hilic\_neg\_2\_005\_24

m/z 286.9118

01E+555.5RT (min)

lcorbi\_hilic\_neg\_1\_027\_18

m/z 287.0503

01234E+577.5RT (min)

lcorbi\_hilic\_neg\_1\_005\_36

m/z 288.0660

01234E+46.57RT (min)

lcorbi\_hilic\_neg\_1\_029\_31

m/z 288.8446

012E+54.55RT (min)

lcorbi\_hilic\_neg\_1\_051\_33

m/z 289.0696

01E+54.55RT (min)

lcorbi\_hilic\_neg\_4\_073\_57

m/z 289.1072

01E+55.56RT (min)

lcorbi\_hilic\_neg\_1\_047\_05

m/z 291.0833

01E+57.58RT (min)

lcorbi\_hilic\_neg\_2\_055\_32

m/z 291.0946

01E+52.53RT (min)

lcorbi\_hilic\_neg\_2\_044\_46

m/z 291.1027

01E+566.5RT (min)

lcorbi\_hilic\_neg\_4\_005\_32

m/z 293.0489

01E+65.56RT (min)

lcorbi\_hilic\_neg\_1\_003\_14

m/z 293.1078

02468E+544.5RT (min)

lcorbi\_hilic\_neg\_1\_028\_45

m/z 293.1693

0123E+40.51RT (min)

lcorbi\_hilic\_neg\_1\_023\_23

m/z 293.1884

0123E+40.51RT (min)

lcorbi\_hilic\_neg\_3\_028\_23

m/z 295.1121

01E+54.55RT (min)

lcorbi\_hilic\_neg\_2\_020\_27

m/z 295.1816

01E+511.5RT (min)

lcorbi\_hilic\_neg\_4\_107\_62

m/z 295.1819

01E+50.51RT (min)

lcorbi\_hilic\_neg\_4\_113\_62

m/z 295.2277

012E+511.5RT (min)

lcorbi\_hilic\_neg\_4\_075\_58

m/z 296.0499

02468E+57.58RT (min)

lcorbi\_hilic\_neg\_4\_005\_32

m/z 297.0789

01234E+55.56RT (min)

lcorbi\_hilic\_neg\_2\_005\_24

m/z 297.2434

0246E+411.5RT (min)

lcorbi\_hilic\_neg\_4\_075\_58

m/z 298.8735

012E+54.55RT (min)

lcorbi\_hilic\_neg\_4\_005\_32

m/z 299.1504

01E+66.57RT (min)

lcorbi\_hilic\_neg\_4\_005\_32

m/z 300.1169

024E+433.5RT (min)

lcorbi\_hilic\_neg\_4\_099\_61

m/z 302.9957

02468E+45.56RT (min)

lcorbi\_hilic\_neg\_4\_066\_57

m/z 303.0487

024E+46.57RT (min)

lcorbi\_hilic\_neg\_4\_089\_59

m/z 304.0522

01234E+422.5RT (min)

lcorbi\_hilic\_neg\_4\_083\_59

m/z 305.0641

012E+56.57RT (min)

lcorbi\_hilic\_neg\_4\_089\_59

m/z 305.0833

01E+53.54RT (min)

lcorbi\_hilic\_neg\_2\_021\_05

m/z 306.0699

01E+66.57RT (min)

lcorbi\_hilic\_neg\_1\_051\_33

m/z 306.9191

02468E+466.5RT (min)

lcorbi\_hilic\_neg\_4\_067\_57

m/z 307.1583

02468E+40.51RT (min)

lcorbi\_hilic\_neg\_2\_021\_05

m/z 309.1635

01234E+40.51RT (min)

lcorbi\_hilic\_neg\_2\_040\_35

m/z 309.1705

024E+411.5RT (min)

lcorbi\_hilic\_neg\_2\_048\_38

m/z 309.1740

0246E+611.5RT (min)

lcorbi\_hilic\_neg\_1\_023\_23

m/z 309.1840

024E+40.51RT (min)

lcorbi\_hilic\_neg\_2\_022\_37

m/z 309.2073

012E+50.51RT (min)

lcorbi\_hilic\_neg\_4\_045\_25

m/z 310.1302

01234E+43.54RT (min)

lcorbi\_hilic\_neg\_3\_040\_19

m/z 310.1505

02468E+455.5RT (min)

lcorbi\_hilic\_neg\_1\_021\_28

m/z 313.0843

012E+56.57RT (min)

lcorbi\_hilic\_neg\_2\_005\_24

m/z 313.1139

012E+55.56RT (min)

lcorbi\_hilic\_neg\_4\_001\_49

m/z 313.1643

01234E+50.51RT (min)

lcorbi\_hilic\_neg\_4\_112\_62

m/z 314.8986

01E+577.5RT (min)

lcorbi\_hilic\_neg\_3\_029\_25

m/z 315.1067

01E+56.57RT (min)

lcorbi\_hilic\_neg\_1\_012\_32

m/z 316.0811

01234E+566.5RT (min)

lcorbi\_hilic\_neg\_4\_005\_32

m/z 316.0875

01234E+566.5RT (min)

lcorbi\_hilic\_neg\_2\_055\_32

m/z 316.9408

024E+655.5RT (min)

lcorbi\_hilic\_neg\_1\_051\_33

m/z 317.0839

01E+522.5RT (min)

lcorbi\_hilic\_neg\_4\_055\_38

m/z 317.1158

012E+50.51RT (min)

lcorbi\_hilic\_neg\_2\_013\_18

m/z 318.0699

012E+60.51RT (min)

lcorbi\_hilic\_neg\_2\_013\_18

m/z 318.1036

01E+55.56RT (min)

lcorbi\_hilic\_neg\_4\_027\_47

m/z 318.1306

012E+52.53RT (min)

lcorbi\_hilic\_neg\_3\_051\_46

m/z 319.0838

01E+56.57RT (min)

lcorbi\_hilic\_neg\_1\_029\_31

m/z 319.1338

012E+566.5RT (min)

lcorbi\_hilic\_neg\_1\_037\_46

m/z 323.1394

02468E+56.57RT (min)

lcorbi\_hilic\_neg\_2\_005\_24

m/z 325.1251

0246E+45.56RT (min)

lcorbi\_hilic\_neg\_4\_068\_57

m/z 327.1797

0123E+511.5RT (min)

lcorbi\_hilic\_neg\_4\_111\_62

m/z 327.2904

02468E+411.5RT (min)

lcorbi\_hilic\_neg\_4\_072\_57

m/z 330.1418

01E+64.55RT (min)

lcorbi\_hilic\_neg\_1\_028\_45

m/z 331.0450

0123E+422.5RT (min)

lcorbi\_hilic\_neg\_4\_098\_61

m/z 332.0514

0246E+56.57RT (min)

lcorbi\_hilic\_neg\_2\_005\_24

m/z 332.1949

0246E+40.51RT (min)

lcorbi\_hilic\_neg\_4\_105\_61

m/z 333.0592

012E+50.51RT (min)

lcorbi\_hilic\_neg\_1\_010\_22

m/z 333.0827

01E+52.53RT (min)

lcorbi\_hilic\_neg\_4\_107\_62

m/z 334.0559

0123E+422.5RT (min)

lcorbi\_hilic\_neg\_4\_089\_59

m/z 335.0963

024E+422.5RT (min)

lcorbi\_hilic\_neg\_4\_051\_16

m/z 335.2224

01234E+411.5RT (min)

lcorbi\_hilic\_neg\_2\_012\_14

m/z 337.2497

01E+50.51RT (min)

lcorbi\_hilic\_neg\_4\_080\_58

m/z 338.0846

012E+56.57RT (min)

lcorbi\_hilic\_neg\_4\_005\_32

m/z 339.2080

012E+50.51RT (min)

lcorbi\_hilic\_neg\_4\_070\_57

m/z 341.0955

0246E+46.57RT (min)

lcorbi\_hilic\_neg\_1\_019\_06

m/z 341.1088

01E+56.57RT (min)

lcorbi\_hilic\_neg\_4\_068\_57

m/z 341.1088

0123E+577.5RT (min)

lcorbi\_hilic\_neg\_4\_088\_59

m/z 341.1966

0246E+411.5RT (min)

lcorbi\_hilic\_neg\_2\_032\_25

m/z 342.0819

012E+57.58RT (min)

lcorbi\_hilic\_neg\_2\_005\_24

m/z 343.9310

01E+50.51RT (min)

lcorbi\_hilic\_neg\_3\_040\_19

m/z 343.9870

01E+666.57RT (min)

lcorbi\_hilic\_neg\_1\_028\_45

m/z 345.1383

02468E+411.5RT (min)

lcorbi\_hilic\_neg\_4\_014\_18

m/z 346.9931

012E+566.5RT (min)

lcorbi\_hilic\_neg\_1\_003\_14

m/z 346.9957

01234E+67.58RT (min)

lcorbi\_hilic\_neg\_1\_051\_33

m/z 347.9934

0123E+67.58RT (min)

lcorbi\_hilic\_neg\_1\_051\_33

m/z 349.0951

01E+50.51RT (min)

lcorbi\_hilic\_neg\_3\_037\_12

m/z 350.0941

01E+566.5RT (min)

lcorbi\_hilic\_neg\_1\_037\_46

m/z 351.0875

01E+55.56RT (min)

lcorbi\_hilic\_neg\_4\_001\_49

m/z 353.1244

012E+522.5RT (min)

lcorbi\_hilic\_neg\_3\_011\_24

m/z 353.1490

01234E+577.5RT (min)

lcorbi\_hilic\_neg\_4\_092\_60

m/z 353.2001

0123E+611.5RT (min)

lcorbi\_hilic\_neg\_1\_023\_23

m/z 355.1104

012E+566.5RT (min)

lcorbi\_hilic\_neg\_1\_037\_46

m/z 355.1244

01E+55.56RT (min)

lcorbi\_hilic\_neg\_3\_043\_02

m/z 359.1190

0123E+566.5RT (min)

lcorbi\_hilic\_neg\_1\_037\_46

m/z 359.1193

012E+455.5RT (min)

lcorbi\_hilic\_neg\_1\_021\_28

m/z 360.1198

01E+666.5RT (min)

lcorbi\_hilic\_neg\_2\_005\_24

m/z 362.1090

01E+533.5RT (min)

lcorbi\_hilic\_neg\_4\_103\_61

m/z 362.1167

02468E+466.5RT (min)

lcorbi\_hilic\_neg\_4\_005\_32

m/z 364.8694

01E+54.55RT (min)

lcorbi\_hilic\_neg\_4\_090\_60

m/z 368.1617

0246E+50.51RT (min)

lcorbi\_hilic\_neg\_4\_080\_58

m/z 369.1538

0246E+40.51RT (min)

lcorbi\_hilic\_neg\_4\_080\_58

m/z 370.1771

012E+50.51RT (min)

lcorbi\_hilic\_neg\_4\_068\_57

m/z 370.8066

01E+54.55RT (min)

lcorbi\_hilic\_neg\_4\_005\_32

m/z 371.0772

01E+50.51RT (min)

lcorbi\_hilic\_neg\_3\_046\_39

m/z 377.0856

0123E+577.5RT (min)

lcorbi\_hilic\_neg\_1\_005\_36

m/z 378.0525

02468E+466.5RT (min)

lcorbi\_hilic\_neg\_4\_089\_59

m/z 380.8350

0246E+54.55RT (min)

lcorbi\_hilic\_neg\_4\_005\_32

m/z 381.1059

012E+50.51RT (min)

lcorbi\_hilic\_neg\_3\_037\_12

m/z 382.8332

01E+54.55RT (min)

lcorbi\_hilic\_neg\_4\_005\_32

m/z 383.1145

0246E+46.57RT (min)

lcorbi\_hilic\_neg\_4\_089\_59

m/z 383.1531

0123E+511.5RT (min)

lcorbi\_hilic\_neg\_4\_031\_14

m/z 384.9267

0123E+655.5RT (min)

lcorbi\_hilic\_neg\_4\_005\_32

m/z 385.1691

0246E+41.52RT (min)

lcorbi\_hilic\_neg\_4\_078\_58

m/z 385.1692

0246E+41.52RT (min)

lcorbi\_hilic\_neg\_4\_077\_58

m/z 385.1730

012E+511.5RT (min)

lcorbi\_hilic\_neg\_4\_075\_58

m/z 385.1731

012E+50.51RT (min)

lcorbi\_hilic\_neg\_4\_074\_58

m/z 387.1141

0123E+566.5RT (min)

lcorbi\_hilic\_neg\_2\_013\_18

m/z 387.1144

012E+577.5RT (min)

lcorbi\_hilic\_neg\_4\_088\_59

m/z 389.1172

01E+577.5RT (min)

lcorbi\_hilic\_neg\_1\_028\_45

m/z 389.1257

01234E+577.5RT (min)

lcorbi\_hilic\_neg\_1\_012\_32

m/z 394.1000

012E+522.5RT (min)

lcorbi\_hilic\_neg\_4\_055\_38

m/z 395.1321

01E+60.51RT (min)

lcorbi\_hilic\_neg\_1\_027\_18

m/z 396.2109

012E+50.51RT (min)

lcorbi\_hilic\_neg\_2\_013\_18

m/z 396.3167

0246E+50.51RT (min)

lcorbi\_hilic\_neg\_3\_037\_12

m/z 397.2054

01234E+50.51RT (min)

lcorbi\_hilic\_neg\_4\_080\_58

m/z 397.2263

012E+611.5RT (min)

lcorbi\_hilic\_neg\_1\_026\_34

m/z 397.3201

01E+50.51RT (min)

lcorbi\_hilic\_neg\_3\_037\_12

m/z 398.1433

0123E+566.5RT (min)

lcorbi\_hilic\_neg\_4\_045\_25

m/z 399.1458

012E+56.57RT (min)

lcorbi\_hilic\_neg\_2\_005\_24

m/z 399.1459

012E+577.5RT (min)

lcorbi\_hilic\_neg\_2\_005\_24

m/z 399.1548

0123E+577.5RT (min)

lcorbi\_hilic\_neg\_4\_096\_60

m/z 399.2212

01E+511.5RT (min)

lcorbi\_hilic\_neg\_4\_080\_58

m/z 400.1458

01E+566.5RT (min)

lcorbi\_hilic\_neg\_4\_088\_59

m/z 401.1300

01E+55.56RT (min)

lcorbi\_hilic\_neg\_3\_043\_02

m/z 413.2003

0123E+511.5RT (min)

lcorbi\_hilic\_neg\_4\_031\_14

m/z 414.8376

02468E+477.5RT (min)

lcorbi\_hilic\_neg\_2\_005\_24

m/z 416.9919

01E+50.51RT (min)

lcorbi\_hilic\_neg\_4\_098\_61

m/z 432.0169

01E+51.52RT (min)

lcorbi\_hilic\_neg\_4\_104\_61

m/z 432.1241

0246E+45.56RT (min)

lcorbi\_hilic\_neg\_1\_001\_49

m/z 442.3223

01234E+50.51RT (min)

lcorbi\_hilic\_neg\_3\_037\_12

m/z 443.3256

01E+50.51RT (min)

lcorbi\_hilic\_neg\_3\_037\_12

m/z 449.3633

012E+511.5RT (min)

lcorbi\_hilic\_neg\_4\_077\_58

m/z 452.9130

0123E+655.5RT (min)

lcorbi\_hilic\_neg\_1\_051\_33

m/z 464.7947

012E+54.55RT (min)

lcorbi\_hilic\_neg\_4\_005\_32

m/z 465.3584

01E+511.5RT (min)

lcorbi\_hilic\_neg\_4\_078\_58

m/z 466.7932

0246E+44.55RT (min)

lcorbi\_hilic\_neg\_2\_005\_24

m/z 467.3742

0123E+511.5RT (min)

lcorbi\_hilic\_neg\_4\_077\_58

m/z 473.3636

012E+511.5RT (min)

lcorbi\_hilic\_neg\_4\_079\_58

m/z 489.3034

012E+52.53RT (min)

lcorbi\_hilic\_neg\_4\_089\_59

m/z 512.1969

012E+50.51RT (min)

lcorbi\_hilic\_neg\_4\_100\_61

m/z 514.1804

01E+50.51RT (min)

lcorbi\_hilic\_neg\_4\_112\_62

m/z 518.3306

024E+555.5RT (min)

lcorbi\_hilic\_neg\_2\_005\_24

m/z 537.4166

02468E+411.5RT (min)

lcorbi\_hilic\_neg\_4\_080\_58

m/z 548.7548

01E+54.55RT (min)

lcorbi\_hilic\_neg\_4\_005\_32

m/z 573.4531

0246E+511.5RT (min)

lcorbi\_hilic\_neg\_4\_031\_14

m/z 593.4792

0246E+511.5RT (min)

lcorbi\_hilic\_neg\_4\_080\_58

m/z 594.4824

01E+511.5RT (min)

lcorbi\_hilic\_neg\_4\_079\_58

m/z 595.4942

01234E+511.5RT (min)

lcorbi\_hilic\_neg\_4\_079\_58

m/z 772.5136

01E+544.5RT (min)

lcorbi\_hilic\_neg\_4\_084\_59

m/z 786.5284

01E+544.5RT (min)

lcorbi\_hilic\_neg\_4\_086\_59
