## Supplementary material for "The Everything Bagel Feature Finder: Ultra-fast automated feature finding for untargeted metabolomics": SI Data File: tool_unique_xic_st_EB_ST004503.html

EB-unique features (not detected by the other two tools) - ST004503

### EB-unique features (not detected by the other two tools) - ST004503

2133 features · qtof · XIC window m/z ± 25 ppm (≥ 0.002 Da) · ± 0.5 min

Every panel is a feature **EB** detected that neither of the other two tools detected — three-way unique across EB (standard mode (min\_detection\_rate=0.1)), Masscube and XCMS, under the standard matching rule (m/z ± 25 ppm; apex within the reference RT window ± 0.2 min). It shows the MS1 extracted-ion chromatogram at the feature's m/z, drawn from the raw mzML named in `alignment_reference_file`, zoomed to ± 0.5 min around the feature's reported apex RT. **Gray solid line** = the feature's reported apex RT; **red dashed line** = EB's assigned MS/MS — `MS2_scan_id` (native scan number) in `MS2_source_file`; the panel widens to keep an off-apex MS/MS on-screen. Features with no assigned MS/MS show only the gray line. y is per-cell autoscaled. Rows keep the table order (m/z ascending).

m/z 50.6266

01234E+41515.5RT (min)

lcqtof\_t3\_pos\_3\_022\_03

m/z 59.0497

0123E+34.55RT (min)

lcqtof\_t3\_pos\_2\_010\_26

m/z 60.0813

012E+311.5RT (min)

lcqtof\_t3\_pos\_4\_074\_58

m/z 61.0593

024E+311.5RT (min)

lcqtof\_t3\_pos\_4\_014\_24

m/z 62.8446

0246E+414.515RT (min)

lcqtof\_t3\_pos\_4\_012\_12

m/z 65.0398

0123E+34.55RT (min)

lcqtof\_t3\_pos\_1\_047\_26

m/z 78.0362

01234E+311.5RT (min)

lcqtof\_t3\_pos\_4\_014\_24

m/z 81.0577

01234E+317.518RT (min)

lcqtof\_t3\_pos\_2\_007\_07

m/z 85.0487

0123E+344.5RT (min)

lcqtof\_t3\_pos\_4\_053\_26

m/z 86.9964

0246E+311.5RT (min)

lcqtof\_t3\_pos\_4\_049\_55

m/z 87.0552

0246E+311.5RT (min)

lcqtof\_t3\_pos\_4\_091\_60

m/z 88.1128

0123E+311.5RT (min)

lcqtof\_t3\_pos\_4\_021\_01

m/z 88.1148

0123E+311.5RT (min)

lcqtof\_t3\_pos\_4\_042\_18

m/z 89.0427

0246E+311.5RT (min)

lcqtof\_t3\_pos\_4\_092\_60

m/z 89.0984

024E+311.5RT (min)

lcqtof\_t3\_pos\_3\_013\_24

m/z 93.0695

024E+344.5RT (min)

lcqtof\_t3\_pos\_4\_021\_01

m/z 94.0660

012E+344.5RT (min)

lcqtof\_t3\_pos\_4\_021\_01

m/z 96.0468

012E+344.5RT (min)

lcqtof\_t3\_pos\_2\_008\_31

m/z 96.9629

01E+311.5RT (min)

lcqtof\_t3\_pos\_1\_011\_43

m/z 97.0686

01234E+344.5RT (min)

lcqtof\_t3\_pos\_4\_012\_12

m/z 98.0637

0123E+311.5RT (min)

lcqtof\_t3\_pos\_2\_022\_32

m/z 98.0969

0123E+311.5RT (min)

lcqtof\_t3\_pos\_1\_010\_27

m/z 99.0277

01234E+344.5RT (min)

lcqtof\_t3\_pos\_1\_018\_30

m/z 100.0875

01E+411.5RT (min)

lcqtof\_t3\_pos\_4\_042\_18

m/z 100.1127

01234E+344.5RT (min)

lcqtof\_t3\_pos\_4\_052\_06

m/z 101.0614

0123E+311.5RT (min)

lcqtof\_t3\_pos\_4\_097\_60

m/z 102.0491

02468E+344.5RT (min)

lcqtof\_t3\_pos\_4\_021\_01

m/z 102.1300

0123E+344.5RT (min)

lcqtof\_t3\_pos\_2\_026\_40

m/z 104.0502

024E+344.5RT (min)

lcqtof\_t3\_pos\_4\_042\_18

m/z 104.0516

01234E+311.5RT (min)

lcqtof\_t3\_pos\_2\_020\_27

m/z 105.0345

012E+36.57RT (min)

lcqtof\_t3\_pos\_4\_111\_62

m/z 106.0662

012E+34.55RT (min)

lcqtof\_t3\_pos\_2\_010\_26

m/z 106.1129

012E+311.5RT (min)

lcqtof\_t3\_pos\_4\_003\_05

m/z 106.9510

012E+317.518RT (min)

lcqtof\_t3\_pos\_2\_005\_17

m/z 108.0841

01234E+344.5RT (min)

lcqtof\_t3\_pos\_4\_049\_55

m/z 108.9493

012E+311.5RT (min)

lcqtof\_t3\_pos\_1\_014\_14

m/z 109.0754

0123E+311.5RT (min)

lcqtof\_t3\_pos\_4\_034\_23

m/z 109.0765

012E+311.5RT (min)

lcqtof\_t3\_pos\_2\_003\_16

m/z 110.0611

01234E+311.5RT (min)

lcqtof\_t3\_pos\_1\_029\_19

m/z 111.2002

01234E+317.518RT (min)

lcqtof\_t3\_pos\_2\_007\_07

m/z 112.0464

0246E+311.5RT (min)

lcqtof\_t3\_pos\_4\_082\_59

m/z 112.0872

024E+311.5RT (min)

lcqtof\_t3\_pos\_1\_020\_35

m/z 112.1010

0123E+317.518RT (min)

lcqtof\_t3\_pos\_4\_100\_61

m/z 114.0040

01234E+311.5RT (min)

lcqtof\_t3\_pos\_4\_079\_58

m/z 114.0926

012E+311.5RT (min)

lcqtof\_t3\_pos\_2\_012\_35

m/z 114.1034

024E+311.5RT (min)

lcqtof\_t3\_pos\_2\_012\_35

m/z 116.0643

0123E+36.57RT (min)

lcqtof\_t3\_pos\_4\_042\_18

m/z 117.0624

02468E+35.56RT (min)

lcqtof\_t3\_pos\_4\_049\_55

m/z 117.1282

02468E+314.515RT (min)

lcqtof\_t3\_pos\_4\_094\_60

m/z 118.0673

01E+544.5RT (min)

lcqtof\_t3\_pos\_4\_021\_01

m/z 118.3440

02468E+311.5RT (min)

lcqtof\_t3\_pos\_2\_015\_13

m/z 120.0037

012E+311.5RT (min)

lcqtof\_t3\_pos\_3\_015\_13

m/z 122.0591

01E+344.5RT (min)

lcqtof\_t3\_pos\_2\_012\_35

m/z 122.1008

0123E+444.5RT (min)

lcqtof\_t3\_pos\_4\_049\_55

m/z 124.0953

012E+317.518RT (min)

lcqtof\_t3\_pos\_2\_013\_29

m/z 124.1133

0246E+35.56RT (min)

lcqtof\_t3\_pos\_4\_021\_01

m/z 124.9393

012E+311.5RT (min)

lcqtof\_t3\_pos\_1\_008\_31

m/z 125.0214

012E+311.5RT (min)

lcqtof\_t3\_pos\_1\_032\_13

m/z 125.0656

0123E+344.5RT (min)

lcqtof\_t3\_pos\_2\_002\_37

m/z 126.0354

024E+388.5RT (min)

lcqtof\_t3\_pos\_1\_051\_38

m/z 126.0636

0246E+35.56RT (min)

lcqtof\_t3\_pos\_4\_105\_61

m/z 126.1041

01E+311.5RT (min)

lcqtof\_t3\_pos\_3\_002\_17

m/z 127.0080

0246E+311.5RT (min)

lcqtof\_t3\_pos\_2\_023\_05

m/z 127.0919

012E+444.5RT (min)

lcqtof\_t3\_pos\_4\_021\_01

m/z 127.0998

02468E+344.5RT (min)

lcqtof\_t3\_pos\_4\_021\_01

m/z 128.0464

01234E+34.55RT (min)

lcqtof\_t3\_pos\_4\_021\_01

m/z 128.0630

024E+34.55RT (min)

lcqtof\_t3\_pos\_3\_113\_62

m/z 128.0718

0246E+344.5RT (min)

lcqtof\_t3\_pos\_2\_015\_13

m/z 128.1443

0123E+344.5RT (min)

lcqtof\_t3\_pos\_4\_011\_10

m/z 128.9545

012E+317.518RT (min)

lcqtof\_t3\_pos\_1\_024\_40

m/z 129.0542

0123E+355.5RT (min)

lcqtof\_t3\_pos\_4\_003\_05

m/z 130.0415

0246E+311.5RT (min)

lcqtof\_t3\_pos\_4\_078\_58

m/z 132.0216

0123E+366.5RT (min)

lcqtof\_t3\_pos\_4\_042\_18

m/z 132.0446

0246E+34.55RT (min)

lcqtof\_t3\_pos\_4\_021\_01

m/z 132.0507

0123E+34.55RT (min)

lcqtof\_t3\_pos\_4\_049\_55

m/z 132.1019

01E+411.5RT (min)

lcqtof\_t3\_pos\_4\_012\_12

m/z 132.1133

0123E+34.55RT (min)

lcqtof\_t3\_pos\_4\_024\_20

m/z 133.0699

012E+311.5RT (min)

lcqtof\_t3\_pos\_1\_014\_14

m/z 133.0985

01234E+311.5RT (min)

lcqtof\_t3\_pos\_2\_039\_23

m/z 134.0197

024E+311.5RT (min)

lcqtof\_t3\_pos\_4\_029\_37

m/z 134.0611

024E+355.5RT (min)

lcqtof\_t3\_pos\_4\_054\_31

m/z 135.0669

01234E+344.5RT (min)

lcqtof\_t3\_pos\_4\_091\_60

m/z 135.0921

01E+317.518RT (min)

lcqtof\_t3\_pos\_2\_033\_53

m/z 135.0955

01E+411.5RT (min)

lcqtof\_t3\_pos\_4\_024\_20

m/z 136.0172

01234E+388.5RT (min)

lcqtof\_t3\_pos\_1\_051\_38

m/z 136.0397

0246E+34.55RT (min)

lcqtof\_t3\_pos\_4\_021\_01

m/z 136.0411

01234E+344.5RT (min)

lcqtof\_t3\_pos\_1\_027\_16

m/z 136.0679

024E+311.5RT (min)

lcqtof\_t3\_pos\_1\_026\_32

m/z 136.1174

0123E+444.5RT (min)

lcqtof\_t3\_pos\_4\_049\_55

m/z 136.1472

0123E+311.5RT (min)

lcqtof\_t3\_pos\_2\_022\_32

m/z 137.0521

012E+311.5RT (min)

lcqtof\_t3\_pos\_2\_022\_32

m/z 137.0782

012E+411.5RT (min)

lcqtof\_t3\_pos\_4\_097\_60

m/z 137.0827

024E+411.5RT (min)

lcqtof\_t3\_pos\_1\_048\_24

m/z 137.1010

0246E+344.5RT (min)

lcqtof\_t3\_pos\_1\_025\_52

m/z 137.9890

0123E+311.5RT (min)

lcqtof\_t3\_pos\_3\_005\_18

m/z 139.0946

0246E+344.5RT (min)

lcqtof\_t3\_pos\_4\_021\_01

m/z 139.1126

0123E+355.5RT (min)

lcqtof\_t3\_pos\_4\_021\_01

m/z 139.1288

01234E+344.5RT (min)

lcqtof\_t3\_pos\_2\_031\_01

m/z 139.9888

01234E+311.5RT (min)

lcqtof\_t3\_pos\_3\_096\_60

m/z 140.9043

0123E+311.5RT (min)

lcqtof\_t3\_pos\_4\_021\_01

m/z 141.1105

01E+444.5RT (min)

lcqtof\_t3\_pos\_4\_021\_01

m/z 141.1277

0123E+366.5RT (min)

lcqtof\_t3\_pos\_4\_001\_49

m/z 141.9601

0246E+311.5RT (min)

lcqtof\_t3\_pos\_1\_013\_20

m/z 143.0705

01E+411.5RT (min)

lcqtof\_t3\_pos\_1\_026\_32

m/z 143.0854

01234E+35.56RT (min)

lcqtof\_t3\_pos\_4\_029\_37

m/z 143.1196

012E+311.5RT (min)

lcqtof\_t3\_pos\_4\_021\_01

m/z 143.1574

01E+36.57RT (min)

lcqtof\_t3\_pos\_2\_043\_48

m/z 144.1071

01234E+511.5RT (min)

lcqtof\_t3\_pos\_4\_049\_55

m/z 144.1763

012E+34.55RT (min)

lcqtof\_t3\_pos\_2\_045\_10

m/z 145.0659

024E+35.56RT (min)

lcqtof\_t3\_pos\_4\_029\_37

m/z 145.0787

012E+444.5RT (min)

lcqtof\_t3\_pos\_4\_021\_01

m/z 145.1513

0246E+31414.5RT (min)

lcqtof\_t3\_pos\_4\_093\_60

m/z 146.0651

01E+45.56RT (min)

lcqtof\_t3\_pos\_4\_049\_55

m/z 147.0865

02468E+411.5RT (min)

lcqtof\_t3\_pos\_2\_034\_24

m/z 147.1176

01234E+35.56RT (min)

lcqtof\_t3\_pos\_4\_024\_20

m/z 148.0767

012E+35.56RT (min)

lcqtof\_t3\_pos\_4\_088\_59

m/z 148.9361

0123E+311.5RT (min)

lcqtof\_t3\_pos\_2\_031\_01

m/z 149.0711

024E+34.55RT (min)

lcqtof\_t3\_pos\_4\_029\_37

m/z 149.0800

01234E+33.54RT (min)

lcqtof\_t3\_pos\_2\_020\_27

m/z 149.1024

01E+411.5RT (min)

lcqtof\_t3\_pos\_2\_034\_24

m/z 150.0448

0123E+377.5RT (min)

lcqtof\_t3\_pos\_4\_024\_20

m/z 150.0750

0246E+311.5RT (min)

lcqtof\_t3\_pos\_2\_030\_33

m/z 151.0665

024E+344.5RT (min)

lcqtof\_t3\_pos\_2\_020\_27

m/z 151.1299

012E+344.5RT (min)

lcqtof\_t3\_pos\_2\_007\_07

m/z 152.0184

01234E+35.56RT (min)

lcqtof\_t3\_pos\_4\_024\_20

m/z 152.0619

01E+45.56RT (min)

lcqtof\_t3\_pos\_4\_029\_37

m/z 152.0748

0123E+44.55RT (min)

lcqtof\_t3\_pos\_4\_049\_55

m/z 152.1072

01234E+35.56RT (min)

lcqtof\_t3\_pos\_4\_073\_57

m/z 153.1081

01E+444.5RT (min)

lcqtof\_t3\_pos\_4\_021\_01

m/z 154.0817

01234E+344.5RT (min)

lcqtof\_t3\_pos\_1\_047\_26

m/z 155.0732

01E+45.56RT (min)

lcqtof\_t3\_pos\_4\_029\_37

m/z 155.0743

01E+444.5RT (min)

lcqtof\_t3\_pos\_4\_011\_10

m/z 155.0796

01234E+311.5RT (min)

lcqtof\_t3\_pos\_2\_012\_35

m/z 155.0865

0246E+377.5RT (min)

lcqtof\_t3\_pos\_2\_011\_19

m/z 155.0930

024E+311.5RT (min)

lcqtof\_t3\_pos\_1\_026\_32

m/z 155.1181

012E+344.5RT (min)

lcqtof\_t3\_pos\_1\_038\_15

m/z 156.0669

01234E+344.5RT (min)

lcqtof\_t3\_pos\_1\_043\_23

m/z 157.0610

012E+344.5RT (min)

lcqtof\_t3\_pos\_4\_101\_61

m/z 157.0800

01E+411.5RT (min)

lcqtof\_t3\_pos\_2\_028\_34

m/z 157.1013

0123E+35.56RT (min)

lcqtof\_t3\_pos\_4\_020\_19

m/z 157.1367

0123E+344.5RT (min)

lcqtof\_t3\_pos\_1\_014\_14

m/z 157.1705

0123E+317.518RT (min)

lcqtof\_t3\_pos\_4\_085\_59

m/z 158.1178

024E+34.55RT (min)

lcqtof\_t3\_pos\_4\_021\_01

m/z 158.1537

01E+344.5RT (min)

lcqtof\_t3\_pos\_1\_007\_07

m/z 159.0446

0246E+37.58RT (min)

lcqtof\_t3\_pos\_2\_010\_26

m/z 159.0519

0246E+311.5RT (min)

lcqtof\_t3\_pos\_4\_010\_36

m/z 159.0944

024E+311.5RT (min)

lcqtof\_t3\_pos\_2\_028\_34

m/z 159.1135

0123E+344.5RT (min)

lcqtof\_t3\_pos\_1\_038\_15

m/z 160.0405

0123E+411.5RT (min)

lcqtof\_t3\_pos\_1\_026\_32

m/z 160.9686

01234E+311.5RT (min)

lcqtof\_t3\_pos\_4\_072\_57

m/z 161.1285

012E+311.5RT (min)

lcqtof\_t3\_pos\_2\_003\_16

m/z 162.1087

012E+317.518RT (min)

lcqtof\_t3\_pos\_2\_001\_49

m/z 163.0017

01234E+377.5RT (min)

lcqtof\_t3\_pos\_4\_105\_61

m/z 163.0401

01234E+34.55RT (min)

lcqtof\_t3\_pos\_4\_042\_18

m/z 163.0554

02468E+34.55RT (min)

lcqtof\_t3\_pos\_3\_113\_62

m/z 163.0994

0246E+311.5RT (min)

lcqtof\_t3\_pos\_4\_090\_60

m/z 164.0617

01E+444.5RT (min)

lcqtof\_t3\_pos\_1\_018\_30

m/z 164.0836

02468E+344.5RT (min)

lcqtof\_t3\_pos\_4\_002\_46

m/z 164.1131

01E+388.5RT (min)

lcqtof\_t3\_pos\_2\_012\_35

m/z 164.1138

0123E+344.5RT (min)

lcqtof\_t3\_pos\_4\_021\_01

m/z 164.9204

01234E+411.5RT (min)

lcqtof\_t3\_pos\_4\_029\_37

m/z 166.0494

0246E+344.5RT (min)

lcqtof\_t3\_pos\_3\_089\_59

m/z 166.0768

01234E+35.56RT (min)

lcqtof\_t3\_pos\_4\_029\_37

m/z 166.0964

024E+311.5RT (min)

lcqtof\_t3\_pos\_2\_011\_19

m/z 166.1535

02468E+34.55RT (min)

lcqtof\_t3\_pos\_4\_014\_24

m/z 166.1596

02468E+377.5RT (min)

lcqtof\_t3\_pos\_4\_021\_01

m/z 167.0375

024E+311.5RT (min)

lcqtof\_t3\_pos\_2\_008\_31

m/z 167.0826

0246E+311.5RT (min)

lcqtof\_t3\_pos\_4\_082\_59

m/z 167.0879

01234E+311.5RT (min)

lcqtof\_t3\_pos\_4\_021\_01

m/z 168.0738

0123E+317.518RT (min)

lcqtof\_t3\_pos\_4\_018\_42

m/z 169.0609

01E+44.55RT (min)

lcqtof\_t3\_pos\_4\_047\_13

m/z 169.1254

012E+444.5RT (min)

lcqtof\_t3\_pos\_4\_021\_01

m/z 169.9862

012E+311.5RT (min)

lcqtof\_t3\_pos\_4\_015\_16

m/z 170.0601

0246E+311.5RT (min)

lcqtof\_t3\_pos\_2\_039\_23

m/z 170.1326

0123E+344.5RT (min)

lcqtof\_t3\_pos\_2\_026\_40

m/z 171.0290

02468E+311.5RT (min)

lcqtof\_t3\_pos\_1\_014\_14

m/z 172.0149

0123E+366.5RT (min)

lcqtof\_t3\_pos\_4\_042\_18

m/z 172.0765

02468E+355.5RT (min)

lcqtof\_t3\_pos\_4\_042\_18

m/z 173.1118

0246E+344.5RT (min)

lcqtof\_t3\_pos\_4\_042\_18

m/z 174.0308

024E+37.58RT (min)

lcqtof\_t3\_pos\_4\_053\_26

m/z 174.0614

01E+45.56RT (min)

lcqtof\_t3\_pos\_4\_049\_55

m/z 174.0840

0123E+355.5RT (min)

lcqtof\_t3\_pos\_4\_003\_05

m/z 175.1887

024E+34.55RT (min)

lcqtof\_t3\_pos\_4\_021\_01

m/z 175.1891

01E+414.515RT (min)

lcqtof\_t3\_pos\_4\_091\_60

m/z 176.0747

012E+344.5RT (min)

lcqtof\_t3\_pos\_4\_047\_13

m/z 177.0194

012E+366.5RT (min)

lcqtof\_t3\_pos\_4\_042\_18

m/z 177.1030

012E+411.5RT (min)

lcqtof\_t3\_pos\_4\_021\_01

m/z 177.9496

012E+311.5RT (min)

lcqtof\_t3\_pos\_1\_008\_31

m/z 178.0834

01E+45.56RT (min)

lcqtof\_t3\_pos\_4\_049\_55

m/z 178.0881

01E+411.5RT (min)

lcqtof\_t3\_pos\_4\_021\_01

m/z 179.0339

024E+366.5RT (min)

lcqtof\_t3\_pos\_4\_042\_18

m/z 179.1117

012E+36.57RT (min)

lcqtof\_t3\_pos\_2\_047\_02

m/z 179.1228

012E+317.518RT (min)

lcqtof\_t3\_pos\_2\_012\_35

m/z 181.0414

0246E+311.5RT (min)

lcqtof\_t3\_pos\_4\_037\_32

m/z 181.0543

0123E+311.5RT (min)

lcqtof\_t3\_pos\_2\_050\_14

m/z 181.0590

0246E+311.5RT (min)

lcqtof\_t3\_pos\_2\_012\_35

m/z 182.0576

0246E+31414.5RT (min)

lcqtof\_t3\_pos\_2\_039\_23

m/z 182.0583

024E+311.5RT (min)

lcqtof\_t3\_pos\_4\_021\_01

m/z 182.1163

01234E+355.5RT (min)

lcqtof\_t3\_pos\_4\_021\_01

m/z 182.1215

012E+344.5RT (min)

lcqtof\_t3\_pos\_1\_027\_16

m/z 182.1552

012E+344.5RT (min)

lcqtof\_t3\_pos\_1\_003\_33

m/z 183.0763

0123E+344.5RT (min)

lcqtof\_t3\_pos\_2\_012\_35

m/z 183.0798

012E+366.5RT (min)

lcqtof\_t3\_pos\_2\_026\_40

m/z 183.1952

012E+355.5RT (min)

lcqtof\_t3\_pos\_2\_022\_32

m/z 184.0765

01234E+34.55RT (min)

lcqtof\_t3\_pos\_3\_113\_62

m/z 184.0943

01E+411.5RT (min)

lcqtof\_t3\_pos\_1\_032\_13

m/z 184.0997

01E+444.5RT (min)

lcqtof\_t3\_pos\_4\_090\_60

m/z 185.0340

024E+311.5RT (min)

lcqtof\_t3\_pos\_1\_002\_36

m/z 185.0835

01E+444.5RT (min)

lcqtof\_t3\_pos\_4\_083\_59

m/z 186.1119

02468E+34.55RT (min)

lcqtof\_t3\_pos\_4\_021\_01

m/z 186.1506

0123E+355.5RT (min)

lcqtof\_t3\_pos\_4\_001\_49

m/z 186.6505

01234E+311.5RT (min)

lcqtof\_t3\_pos\_2\_034\_24

m/z 187.1493

012E+35.56RT (min)

lcqtof\_t3\_pos\_4\_024\_20

m/z 188.0463

0246E+366.5RT (min)

lcqtof\_t3\_pos\_4\_042\_18

m/z 188.0766

0123E+311.5RT (min)

lcqtof\_t3\_pos\_2\_040\_28

m/z 188.1119

0246E+311.5RT (min)

lcqtof\_t3\_pos\_2\_008\_31

m/z 188.1541

024E+36.57RT (min)

lcqtof\_t3\_pos\_4\_042\_18

m/z 189.1256

01234E+355.5RT (min)

lcqtof\_t3\_pos\_4\_096\_60

m/z 189.1632

01234E+35.56RT (min)

lcqtof\_t3\_pos\_4\_029\_37

m/z 190.0536

01234E+311.5RT (min)

lcqtof\_t3\_pos\_4\_042\_18

m/z 190.1629

01E+311.5RT (min)

lcqtof\_t3\_pos\_2\_012\_35

m/z 191.1888

01234E+411.5RT (min)

lcqtof\_t3\_pos\_4\_112\_62

m/z 191.3664

0246E+411.5RT (min)

lcqtof\_t3\_pos\_4\_112\_62

m/z 192.1041

01E+366.5RT (min)

lcqtof\_t3\_pos\_2\_001\_49

m/z 193.0364

0246E+311.5RT (min)

lcqtof\_t3\_pos\_4\_034\_23

m/z 193.0716

01E+44.55RT (min)

lcqtof\_t3\_pos\_4\_024\_20

m/z 193.0894

024E+34.55RT (min)

lcqtof\_t3\_pos\_4\_044\_38

m/z 193.1230

012E+34.55RT (min)

lcqtof\_t3\_pos\_1\_052\_06

m/z 193.1416

01E+344.5RT (min)

lcqtof\_t3\_pos\_2\_038\_21

m/z 193.1621

012E+311.5RT (min)

lcqtof\_t3\_pos\_4\_001\_49

m/z 194.1573

01E+344.5RT (min)

lcqtof\_t3\_pos\_2\_024\_45

m/z 194.1622

01234E+36.57RT (min)

lcqtof\_t3\_pos\_4\_006\_41

m/z 196.0583

012E+411.5RT (min)

lcqtof\_t3\_pos\_4\_037\_32

m/z 196.1180

012E+366.5RT (min)

lcqtof\_t3\_pos\_2\_030\_33

m/z 197.0827

0123E+344.5RT (min)

lcqtof\_t3\_pos\_2\_019\_08

m/z 197.9987

0123E+311.5RT (min)

lcqtof\_t3\_pos\_4\_037\_32

m/z 198.0404

0123E+311.5RT (min)

lcqtof\_t3\_pos\_2\_040\_28

m/z 198.0527

0123E+366.5RT (min)

lcqtof\_t3\_pos\_4\_021\_01

m/z 198.0756

01234E+344.5RT (min)

lcqtof\_t3\_pos\_4\_034\_23

m/z 198.1134

0123E+344.5RT (min)

lcqtof\_t3\_pos\_2\_035\_44

m/z 199.0069

0246E+355.5RT (min)

lcqtof\_t3\_pos\_4\_040\_28

m/z 199.0977

024E+344.5RT (min)

lcqtof\_t3\_pos\_2\_016\_39

m/z 200.0284

012E+311.5RT (min)

lcqtof\_t3\_pos\_2\_055\_18

m/z 200.0622

012E+377.5RT (min)

lcqtof\_t3\_pos\_2\_020\_27

m/z 200.0924

01234E+344.5RT (min)

lcqtof\_t3\_pos\_2\_027\_38

m/z 200.1546

01E+477.5RT (min)

lcqtof\_t3\_pos\_4\_049\_55

m/z 201.0088

012E+311.5RT (min)

lcqtof\_t3\_pos\_1\_028\_08

m/z 201.0920

024E+36.57RT (min)

lcqtof\_t3\_pos\_4\_042\_18

m/z 201.1324

01234E+311.5RT (min)

lcqtof\_t3\_pos\_2\_015\_13

m/z 202.0479

02468E+34.55RT (min)

lcqtof\_t3\_pos\_4\_021\_01

m/z 202.0762

01234E+34.55RT (min)

lcqtof\_t3\_pos\_3\_113\_62

m/z 202.0860

012E+34.55RT (min)

lcqtof\_t3\_pos\_2\_033\_53

m/z 202.1186

024E+311.5RT (min)

lcqtof\_t3\_pos\_2\_022\_32

m/z 202.8877

0246E+311.5RT (min)

lcqtof\_t3\_pos\_4\_001\_49

m/z 203.1419

024E+36.57RT (min)

lcqtof\_t3\_pos\_4\_021\_01

m/z 204.1141

0123E+344.5RT (min)

lcqtof\_t3\_pos\_2\_041\_54

m/z 204.1588

012E+344.5RT (min)

lcqtof\_t3\_pos\_1\_014\_14

m/z 204.1956

0246E+311.5RT (min)

lcqtof\_t3\_pos\_4\_021\_01

m/z 206.0238

0123E+366.5RT (min)

lcqtof\_t3\_pos\_4\_042\_18

m/z 206.1108

01E+41414.5RT (min)

lcqtof\_t3\_pos\_2\_022\_32

m/z 207.0275

01234E+366.5RT (min)

lcqtof\_t3\_pos\_4\_042\_18

m/z 207.1576

012E+344.5RT (min)

lcqtof\_t3\_pos\_2\_044\_25

m/z 207.1627

0246E+34.55RT (min)

lcqtof\_t3\_pos\_4\_014\_24

m/z 207.9982

0246E+311.5RT (min)

lcqtof\_t3\_pos\_4\_029\_37

m/z 208.2083

01234E+399.5RT (min)

lcqtof\_t3\_pos\_2\_029\_06

m/z 209.0302

024E+36.57RT (min)

lcqtof\_t3\_pos\_4\_021\_01

m/z 209.0550

0123E+311.5RT (min)

lcqtof\_t3\_pos\_4\_021\_01

m/z 209.0609

0123E+311.5RT (min)

lcqtof\_t3\_pos\_4\_029\_37

m/z 210.0624

0123E+311.5RT (min)

lcqtof\_t3\_pos\_4\_007\_14

m/z 210.1668

01234E+455.5RT (min)

lcqtof\_t3\_pos\_4\_029\_37

m/z 210.8896

024E+311.5RT (min)

lcqtof\_t3\_pos\_4\_021\_01

m/z 210.9792

01234E+311.5RT (min)

lcqtof\_t3\_pos\_2\_015\_13

m/z 210.9962

0246E+34.55RT (min)

lcqtof\_t3\_pos\_4\_021\_01

m/z 211.1098

02468E+344.5RT (min)

lcqtof\_t3\_pos\_2\_012\_35

m/z 211.1115

02468E+311.5RT (min)

lcqtof\_t3\_pos\_4\_082\_59

m/z 212.0916

01234E+344.5RT (min)

lcqtof\_t3\_pos\_2\_027\_38

m/z 212.1014

0123E+34.55RT (min)

lcqtof\_t3\_pos\_2\_027\_38

m/z 213.0757

012E+344.5RT (min)

lcqtof\_t3\_pos\_4\_053\_26

m/z 214.0641

0246E+36.57RT (min)

lcqtof\_t3\_pos\_4\_042\_18

m/z 214.0656

024E+34.55RT (min)

lcqtof\_t3\_pos\_2\_020\_27

m/z 214.0693

01234E+311.5RT (min)

lcqtof\_t3\_pos\_3\_079\_58

m/z 215.1015

024E+344.5RT (min)

lcqtof\_t3\_pos\_3\_018\_37

m/z 215.1408

012E+311.5RT (min)

lcqtof\_t3\_pos\_4\_078\_58

m/z 215.1802

024E+377.5RT (min)

lcqtof\_t3\_pos\_4\_001\_49

m/z 215.9685

0123E+311.5RT (min)

lcqtof\_t3\_pos\_4\_029\_37

m/z 216.0208

01E+411.5RT (min)

lcqtof\_t3\_pos\_4\_021\_01

m/z 217.0612

0123E+34.55RT (min)

lcqtof\_t3\_pos\_2\_039\_23

m/z 217.0701

0123E+34.55RT (min)

lcqtof\_t3\_pos\_2\_015\_13

m/z 217.9285

024E+311.5RT (min)

lcqtof\_t3\_pos\_2\_015\_13

m/z 218.9764

012E+311.5RT (min)

lcqtof\_t3\_pos\_2\_008\_31

m/z 219.0935

02468E+355.5RT (min)

lcqtof\_t3\_pos\_4\_014\_24

m/z 219.1346

012E+311.5RT (min)

lcqtof\_t3\_pos\_2\_012\_35

m/z 219.1414

0246E+311.5RT (min)

lcqtof\_t3\_pos\_4\_021\_01

m/z 220.0561

024E+311.5RT (min)

lcqtof\_t3\_pos\_2\_008\_31

m/z 220.0662

0123E+34.55RT (min)

lcqtof\_t3\_pos\_2\_010\_26

m/z 220.1644

0246E+344.5RT (min)

lcqtof\_t3\_pos\_2\_034\_24

m/z 221.0944

01E+411.5RT (min)

lcqtof\_t3\_pos\_4\_014\_24

m/z 221.1400

0246E+38.59RT (min)

lcqtof\_t3\_pos\_4\_049\_55

m/z 222.0808

0246E+344.5RT (min)

lcqtof\_t3\_pos\_4\_040\_28

m/z 222.0831

01234E+34.55RT (min)

lcqtof\_t3\_pos\_4\_029\_37

m/z 222.0923

0246E+34.55RT (min)

lcqtof\_t3\_pos\_3\_113\_62

m/z 224.1015

01E+411.5RT (min)

lcqtof\_t3\_pos\_2\_022\_32

m/z 224.1665

01E+344.5RT (min)

lcqtof\_t3\_pos\_1\_007\_07

m/z 225.0374

012E+311.5RT (min)

lcqtof\_t3\_pos\_1\_010\_27

m/z 225.0782

0246E+344.5RT (min)

lcqtof\_t3\_pos\_2\_011\_19

m/z 225.1683

024E+344.5RT (min)

lcqtof\_t3\_pos\_4\_014\_24

m/z 226.0679

024E+311.5RT (min)

lcqtof\_t3\_pos\_4\_014\_24

m/z 226.1900

0246E+344.5RT (min)

lcqtof\_t3\_pos\_4\_029\_37

m/z 227.2115

012E+366.5RT (min)

lcqtof\_t3\_pos\_4\_015\_16

m/z 227.9560

0123E+317.518RT (min)

lcqtof\_t3\_pos\_2\_005\_17

m/z 229.1702

0123E+377.5RT (min)

lcqtof\_t3\_pos\_4\_052\_06

m/z 230.0235

0246E+47.58RT (min)

lcqtof\_t3\_pos\_2\_010\_26

m/z 230.0634

01234E+34.55RT (min)

lcqtof\_t3\_pos\_2\_023\_05

m/z 230.0968

0123E+34.55RT (min)

lcqtof\_t3\_pos\_3\_113\_62

m/z 230.0991

01234E+36.57RT (min)

lcqtof\_t3\_pos\_3\_005\_18

m/z 230.1109

0246E+35.56RT (min)

lcqtof\_t3\_pos\_1\_029\_19

m/z 230.1844

0246E+34.55RT (min)

lcqtof\_t3\_pos\_4\_029\_37

m/z 231.0994

01234E+311.5RT (min)

lcqtof\_t3\_pos\_2\_039\_23

m/z 231.1469

012E+311.5RT (min)

lcqtof\_t3\_pos\_1\_008\_31

m/z 231.1829

01E+577.5RT (min)

lcqtof\_t3\_pos\_4\_042\_18

m/z 231.2126

0246E+314.515RT (min)

lcqtof\_t3\_pos\_4\_029\_37

m/z 232.0268

012E+344.5RT (min)

lcqtof\_t3\_pos\_4\_029\_37

m/z 232.0591

0246E+34.55RT (min)

lcqtof\_t3\_pos\_4\_047\_13

m/z 232.1065

024E+388.5RT (min)

lcqtof\_t3\_pos\_4\_021\_01

m/z 232.1751

012E+377.5RT (min)

lcqtof\_t3\_pos\_1\_042\_45

m/z 232.9805

01E+44.55RT (min)

lcqtof\_t3\_pos\_4\_021\_01

m/z 233.0451

024E+36.57RT (min)

lcqtof\_t3\_pos\_4\_008\_15

m/z 233.1164

012E+35.56RT (min)

lcqtof\_t3\_pos\_4\_052\_06

m/z 234.1655

0123E+355.5RT (min)

lcqtof\_t3\_pos\_4\_029\_37

m/z 234.2071

0246E+36.57RT (min)

lcqtof\_t3\_pos\_4\_015\_16

m/z 234.2096

0123E+355.5RT (min)

lcqtof\_t3\_pos\_4\_021\_01

m/z 234.2101

01E+377.5RT (min)

lcqtof\_t3\_pos\_2\_008\_31

m/z 235.0420

012E+34.55RT (min)

lcqtof\_t3\_pos\_4\_024\_20

m/z 236.0810

012E+34.55RT (min)

lcqtof\_t3\_pos\_4\_029\_37

m/z 237.0293

024E+311.5RT (min)

lcqtof\_t3\_pos\_2\_039\_23

m/z 237.0385

024E+34.55RT (min)

lcqtof\_t3\_pos\_4\_024\_20

m/z 237.0928

012E+34.55RT (min)

lcqtof\_t3\_pos\_4\_020\_19

m/z 237.0957

012E+311.5RT (min)

lcqtof\_t3\_pos\_2\_037\_46

m/z 238.0469

01234E+311.5RT (min)

lcqtof\_t3\_pos\_2\_002\_37

m/z 238.0589

012E+34.55RT (min)

lcqtof\_t3\_pos\_4\_029\_37

m/z 238.1202

01E+35.56RT (min)

lcqtof\_t3\_pos\_2\_027\_38

m/z 238.1486

0123E+44.55RT (min)

lcqtof\_t3\_pos\_4\_007\_14

m/z 238.1510

0123E+344.5RT (min)

lcqtof\_t3\_pos\_2\_010\_26

m/z 238.1810

012E+344.5RT (min)

lcqtof\_t3\_pos\_4\_052\_06

m/z 239.0695

0123E+34.55RT (min)

lcqtof\_t3\_pos\_4\_029\_37

m/z 239.0732

012E+34.55RT (min)

lcqtof\_t3\_pos\_2\_016\_39

m/z 239.0953

012E+344.5RT (min)

lcqtof\_t3\_pos\_2\_011\_19

m/z 239.1046

01234E+311.5RT (min)

lcqtof\_t3\_pos\_4\_082\_59

m/z 240.0731

0246E+311.5RT (min)

lcqtof\_t3\_pos\_4\_014\_24

m/z 242.2075

012E+35.56RT (min)

lcqtof\_t3\_pos\_1\_014\_14

m/z 242.6796

01234E+34.55RT (min)

lcqtof\_t3\_pos\_4\_029\_37

m/z 243.0547

02468E+344.5RT (min)

lcqtof\_t3\_pos\_4\_007\_14

m/z 243.0656

0123E+311.5RT (min)

lcqtof\_t3\_pos\_2\_022\_32

m/z 244.0704

01234E+34.55RT (min)

lcqtof\_t3\_pos\_4\_072\_57

m/z 244.0805

02468E+34.55RT (min)

lcqtof\_t3\_pos\_4\_029\_37

m/z 244.9274

01E+317.518RT (min)

lcqtof\_t3\_pos\_3\_005\_18

m/z 245.0996

02468E+311.5RT (min)

lcqtof\_t3\_pos\_2\_039\_23

m/z 245.1014

0246E+344.5RT (min)

lcqtof\_t3\_pos\_4\_029\_37

m/z 245.1093

024E+344.5RT (min)

lcqtof\_t3\_pos\_2\_009\_50

m/z 245.1306

01234E+35.56RT (min)

lcqtof\_t3\_pos\_4\_021\_01

m/z 245.1521

012E+311.5RT (min)

lcqtof\_t3\_pos\_4\_007\_14

m/z 246.0222

012E+311.5RT (min)

lcqtof\_t3\_pos\_2\_002\_37

m/z 246.0489

0123E+366.5RT (min)

lcqtof\_t3\_pos\_4\_042\_18

m/z 246.1938

01234E+344.5RT (min)

lcqtof\_t3\_pos\_4\_029\_37

m/z 247.0585

0246E+311.5RT (min)

lcqtof\_t3\_pos\_4\_034\_23

m/z 247.0606

0246E+344.5RT (min)

lcqtof\_t3\_pos\_4\_034\_23

m/z 247.0991

0123E+377.5RT (min)

lcqtof\_t3\_pos\_1\_036\_37

m/z 247.1315

01E+377.5RT (min)

lcqtof\_t3\_pos\_2\_012\_35

m/z 247.1360

012E+311.5RT (min)

lcqtof\_t3\_pos\_2\_012\_35

m/z 248.1000

0123E+311.5RT (min)

lcqtof\_t3\_pos\_2\_050\_14

m/z 248.1128

0246E+311.5RT (min)

lcqtof\_t3\_pos\_4\_042\_18

m/z 248.8822

012E+317.518RT (min)

lcqtof\_t3\_pos\_2\_011\_19

m/z 249.1568

012E+311.5RT (min)

lcqtof\_t3\_pos\_4\_047\_13

m/z 249.6870

0123E+355.5RT (min)

lcqtof\_t3\_pos\_4\_034\_23

m/z 250.0403

0123E+344.5RT (min)

lcqtof\_t3\_pos\_2\_010\_26

m/z 250.8694

02468E+311.5RT (min)

lcqtof\_t3\_pos\_4\_007\_14

m/z 252.1267

012E+311.5RT (min)

lcqtof\_t3\_pos\_2\_005\_17

m/z 252.1811

0123E+317.518RT (min)

lcqtof\_t3\_pos\_4\_074\_58

m/z 254.1194

0246E+311.5RT (min)

lcqtof\_t3\_pos\_4\_014\_24

m/z 255.1346

0246E+344.5RT (min)

lcqtof\_t3\_pos\_4\_029\_37

m/z 256.1448

0246E+36.57RT (min)

lcqtof\_t3\_pos\_4\_042\_18

m/z 257.0629

012E+344.5RT (min)

lcqtof\_t3\_pos\_3\_010\_01

m/z 257.1585

024E+311.5RT (min)

lcqtof\_t3\_pos\_2\_037\_46

m/z 257.9808

01234E+34.55RT (min)

lcqtof\_t3\_pos\_4\_007\_14

m/z 258.0536

012E+366.5RT (min)

lcqtof\_t3\_pos\_4\_046\_34

m/z 258.0949

01234E+344.5RT (min)

lcqtof\_t3\_pos\_2\_039\_23

m/z 258.9800

024E+311.5RT (min)

lcqtof\_t3\_pos\_4\_021\_01

m/z 259.0152

0246E+35.56RT (min)

lcqtof\_t3\_pos\_4\_021\_01

m/z 259.2024

012E+355.5RT (min)

lcqtof\_t3\_pos\_4\_029\_37

m/z 259.2849

0123E+39.510RT (min)

lcqtof\_t3\_pos\_4\_021\_01

m/z 259.9468

0246E+311.5RT (min)

lcqtof\_t3\_pos\_4\_029\_37

m/z 260.0294

012E+311.5RT (min)

lcqtof\_t3\_pos\_4\_046\_34

m/z 260.1192

0123E+34.55RT (min)

lcqtof\_t3\_pos\_2\_010\_26

m/z 260.1933

012E+311.5RT (min)

lcqtof\_t3\_pos\_4\_082\_59

m/z 260.8146

0246E+311.5RT (min)

lcqtof\_t3\_pos\_4\_021\_01

m/z 260.8445

0246E+311.5RT (min)

lcqtof\_t3\_pos\_4\_029\_37

m/z 261.0567

012E+34.55RT (min)

lcqtof\_t3\_pos\_4\_024\_20

m/z 261.1560

0246E+311.5RT (min)

lcqtof\_t3\_pos\_4\_089\_59

m/z 262.1498

0246E+35.56RT (min)

lcqtof\_t3\_pos\_4\_097\_60

m/z 262.1789

02468E+388.5RT (min)

lcqtof\_t3\_pos\_4\_004\_33

m/z 263.1238

01234E+311.5RT (min)

lcqtof\_t3\_pos\_2\_037\_46

m/z 263.1397

0123E+355.5RT (min)

lcqtof\_t3\_pos\_4\_035\_45

m/z 263.1516

024E+36.57RT (min)

lcqtof\_t3\_pos\_4\_052\_06

m/z 263.2429

0123E+37.58RT (min)

lcqtof\_t3\_pos\_4\_015\_16

m/z 264.1179

0246E+311.5RT (min)

lcqtof\_t3\_pos\_4\_015\_16

m/z 264.1983

012E+355.5RT (min)

lcqtof\_t3\_pos\_1\_032\_13

m/z 264.9908

0123E+311.5RT (min)

lcqtof\_t3\_pos\_4\_001\_49

m/z 265.0342

01E+344.5RT (min)

lcqtof\_t3\_pos\_4\_081\_58

m/z 265.1333

0123E+34.55RT (min)

lcqtof\_t3\_pos\_2\_039\_23

m/z 265.1567

0123E+311.5RT (min)

lcqtof\_t3\_pos\_2\_039\_23

m/z 265.1580

02468E+311.5RT (min)

lcqtof\_t3\_pos\_2\_022\_32

m/z 266.0557

012E+34.55RT (min)

lcqtof\_t3\_pos\_4\_042\_18

m/z 266.0782

024E+311.5RT (min)

lcqtof\_t3\_pos\_2\_027\_38

m/z 267.0811

024E+344.5RT (min)

lcqtof\_t3\_pos\_4\_077\_58

m/z 267.1666

0246E+311.5RT (min)

lcqtof\_t3\_pos\_4\_039\_02

m/z 268.0245

0123E+311.5RT (min)

lcqtof\_t3\_pos\_2\_003\_16

m/z 268.8472

01234E+311.5RT (min)

lcqtof\_t3\_pos\_4\_021\_01

m/z 269.1121

024E+344.5RT (min)

lcqtof\_t3\_pos\_4\_097\_60

m/z 269.1500

01234E+344.5RT (min)

lcqtof\_t3\_pos\_2\_027\_38

m/z 270.9786

01234E+317.518RT (min)

lcqtof\_t3\_pos\_2\_006\_04

m/z 271.0886

024E+344.5RT (min)

lcqtof\_t3\_pos\_4\_047\_13

m/z 272.1001

0246E+377.5RT (min)

lcqtof\_t3\_pos\_4\_042\_18

m/z 272.1883

01E+44.55RT (min)

lcqtof\_t3\_pos\_4\_001\_49

m/z 273.1459

02468E+344.5RT (min)

lcqtof\_t3\_pos\_4\_029\_37

m/z 273.9250

01234E+311.5RT (min)

lcqtof\_t3\_pos\_4\_046\_34

m/z 274.0605

0123E+311.5RT (min)

lcqtof\_t3\_pos\_2\_022\_32

m/z 274.0798

02468E+377.5RT (min)

lcqtof\_t3\_pos\_4\_042\_18

m/z 274.1701

01234E+34.55RT (min)

lcqtof\_t3\_pos\_4\_021\_01

m/z 275.0699

012E+34.55RT (min)

lcqtof\_t3\_pos\_4\_024\_20

m/z 276.2926

02468E+31414.5RT (min)

lcqtof\_t3\_pos\_4\_014\_24

m/z 277.0856

0123E+34.55RT (min)

lcqtof\_t3\_pos\_4\_029\_37

m/z 277.1241

0246E+37.58RT (min)

lcqtof\_t3\_pos\_4\_042\_18

m/z 277.1375

0123E+317.518RT (min)

lcqtof\_t3\_pos\_1\_001\_49

m/z 277.1758

012E+366.5RT (min)

lcqtof\_t3\_pos\_2\_032\_47

m/z 278.0426

012E+344.5RT (min)

lcqtof\_t3\_pos\_2\_018\_41

m/z 278.8817

024E+311.5RT (min)

lcqtof\_t3\_pos\_4\_003\_05

m/z 280.1625

01E+377.5RT (min)

lcqtof\_t3\_pos\_2\_015\_13

m/z 280.1948

012E+311.5RT (min)

lcqtof\_t3\_pos\_1\_002\_36

m/z 281.1562

01234E+344.5RT (min)

lcqtof\_t3\_pos\_2\_022\_32

m/z 282.0798

024E+344.5RT (min)

lcqtof\_t3\_pos\_4\_077\_58

m/z 282.1538

0246E+344.5RT (min)

lcqtof\_t3\_pos\_2\_012\_35

m/z 283.2269

01234E+34.55RT (min)

lcqtof\_t3\_pos\_4\_029\_37

m/z 284.0837

0123E+344.5RT (min)

lcqtof\_t3\_pos\_4\_083\_59

m/z 284.9108

0246E+311.5RT (min)

lcqtof\_t3\_pos\_4\_005\_30

m/z 285.1047

024E+344.5RT (min)

lcqtof\_t3\_pos\_4\_049\_55

m/z 285.1207

01234E+311.5RT (min)

lcqtof\_t3\_pos\_4\_089\_59

m/z 285.6971

01234E+355.5RT (min)

lcqtof\_t3\_pos\_4\_029\_37

m/z 286.0894

024E+311.5RT (min)

lcqtof\_t3\_pos\_4\_014\_24

m/z 286.0970

0246E+355.5RT (min)

lcqtof\_t3\_pos\_2\_010\_26

m/z 286.2145

0123E+34.55RT (min)

lcqtof\_t3\_pos\_4\_089\_59

m/z 286.9085

012E+311.5RT (min)

lcqtof\_t3\_pos\_4\_013\_29

m/z 287.0305

012E+34.55RT (min)

lcqtof\_t3\_pos\_1\_014\_14

m/z 287.1273

0123E+344.5RT (min)

lcqtof\_t3\_pos\_4\_014\_24

m/z 287.1985

0246E+311.5RT (min)

lcqtof\_t3\_pos\_4\_021\_01

m/z 288.0240

0123E+34.55RT (min)

lcqtof\_t3\_pos\_1\_014\_14

m/z 288.0845

02468E+311.5RT (min)

lcqtof\_t3\_pos\_4\_112\_62

m/z 288.2032

01234E+344.5RT (min)

lcqtof\_t3\_pos\_2\_027\_38

m/z 288.7721

01234E+311.5RT (min)

lcqtof\_t3\_pos\_4\_007\_14

m/z 289.0932

01E+355.5RT (min)

lcqtof\_t3\_pos\_2\_032\_47

m/z 290.0803

01234E+311.5RT (min)

lcqtof\_t3\_pos\_2\_039\_23

m/z 290.1481

0246E+311.5RT (min)

lcqtof\_t3\_pos\_2\_039\_23

m/z 290.2129

0123E+35.56RT (min)

lcqtof\_t3\_pos\_4\_003\_05

m/z 291.0269

01234E+366.5RT (min)

lcqtof\_t3\_pos\_4\_007\_14

m/z 292.2151

012E+35.56RT (min)

lcqtof\_t3\_pos\_4\_024\_20

m/z 292.8730

01234E+311.5RT (min)

lcqtof\_t3\_pos\_2\_028\_34

m/z 293.3247

02468E+314.515RT (min)

lcqtof\_t3\_pos\_4\_014\_24

m/z 294.1306

01234E+311.5RT (min)

lcqtof\_t3\_pos\_2\_015\_13

m/z 294.1578

0246E+34.55RT (min)

lcqtof\_t3\_pos\_4\_037\_32

m/z 294.1584

0246E+344.5RT (min)

lcqtof\_t3\_pos\_1\_026\_32

m/z 294.1697

0246E+344.5RT (min)

lcqtof\_t3\_pos\_4\_029\_37

m/z 295.9426

01E+311.5RT (min)

lcqtof\_t3\_pos\_1\_022\_28

m/z 296.1230

0123E+444.5RT (min)

lcqtof\_t3\_pos\_4\_001\_49

m/z 296.1496

0123E+444.5RT (min)

lcqtof\_t3\_pos\_4\_021\_01

m/z 296.1554

01E+46.57RT (min)

lcqtof\_t3\_pos\_4\_046\_34

m/z 297.0685

0123E+35.56RT (min)

lcqtof\_t3\_pos\_1\_001\_49

m/z 297.9687

01234E+311.5RT (min)

lcqtof\_t3\_pos\_4\_010\_36

m/z 298.1529

0246E+37.58RT (min)

lcqtof\_t3\_pos\_4\_042\_18

m/z 299.3071

012E+399.5RT (min)

lcqtof\_t3\_pos\_4\_083\_59

m/z 300.1658

0123E+311.5RT (min)

lcqtof\_t3\_pos\_1\_005\_46

m/z 300.7008

012E+344.5RT (min)

lcqtof\_t3\_pos\_1\_003\_33

m/z 300.7158

024E+355.5RT (min)

lcqtof\_t3\_pos\_4\_029\_37

m/z 300.8652

01234E+311.5RT (min)

lcqtof\_t3\_pos\_2\_022\_32

m/z 300.8952

012E+411.5RT (min)

lcqtof\_t3\_pos\_4\_029\_37

m/z 301.1124

01E+411.5RT (min)

lcqtof\_t3\_pos\_4\_069\_57

m/z 301.1154

0123E+311.5RT (min)

lcqtof\_t3\_pos\_2\_046\_30

m/z 301.2338

012E+36.57RT (min)

lcqtof\_t3\_pos\_4\_015\_16

m/z 302.2768

024E+311.5RT (min)

lcqtof\_t3\_pos\_2\_034\_24

m/z 302.6924

0123E+35.56RT (min)

lcqtof\_t3\_pos\_1\_002\_36

m/z 303.0470

024E+311.5RT (min)

lcqtof\_t3\_pos\_2\_012\_35

m/z 303.0873

0246E+366.5RT (min)

lcqtof\_t3\_pos\_4\_042\_18

m/z 303.1547

0246E+344.5RT (min)

lcqtof\_t3\_pos\_2\_027\_38

m/z 304.1494

02468E+344.5RT (min)

lcqtof\_t3\_pos\_4\_034\_23

m/z 304.1787

012E+37.58RT (min)

lcqtof\_t3\_pos\_2\_002\_37

m/z 306.1350

0246E+344.5RT (min)

lcqtof\_t3\_pos\_4\_021\_01

m/z 306.8573

02468E+311.5RT (min)

lcqtof\_t3\_pos\_2\_022\_32

m/z 308.0880

012E+344.5RT (min)

lcqtof\_t3\_pos\_3\_003\_14

m/z 308.1156

0123E+35.56RT (min)

lcqtof\_t3\_pos\_4\_003\_05

m/z 308.1808

0246E+344.5RT (min)

lcqtof\_t3\_pos\_2\_001\_49

m/z 308.5675

0123E+35.56RT (min)

lcqtof\_t3\_pos\_4\_010\_36

m/z 308.5768

01234E+355.5RT (min)

lcqtof\_t3\_pos\_4\_054\_31

m/z 308.6041

02468E+311.5RT (min)

lcqtof\_t3\_pos\_4\_014\_24

m/z 309.1498

0246E+37.58RT (min)

lcqtof\_t3\_pos\_4\_042\_18

m/z 310.1661

012E+34.55RT (min)

lcqtof\_t3\_pos\_2\_020\_27

m/z 310.9146

012E+317.518RT (min)

lcqtof\_t3\_pos\_2\_007\_07

m/z 311.2320

024E+35.56RT (min)

lcqtof\_t3\_pos\_4\_029\_37

m/z 311.6882

01234E+34.55RT (min)

lcqtof\_t3\_pos\_4\_021\_01

m/z 312.9009

01234E+355.5RT (min)

lcqtof\_t3\_pos\_4\_029\_37

m/z 313.5589

01234E+35.56RT (min)

lcqtof\_t3\_pos\_4\_029\_37

m/z 314.1340

0246E+311.5RT (min)

lcqtof\_t3\_pos\_4\_095\_60

m/z 314.1464

0246E+37.58RT (min)

lcqtof\_t3\_pos\_4\_042\_18

m/z 314.2460

01234E+355.5RT (min)

lcqtof\_t3\_pos\_4\_029\_37

m/z 314.3467

02468E+34.55RT (min)

lcqtof\_t3\_pos\_4\_029\_37

m/z 314.8547

0246E+311.5RT (min)

lcqtof\_t3\_pos\_4\_029\_37

m/z 315.0952

01234E+366.5RT (min)

lcqtof\_t3\_pos\_4\_006\_41

m/z 315.2274

01234E+366.5RT (min)

lcqtof\_t3\_pos\_4\_029\_37

m/z 316.1386

01234E+38.59RT (min)

lcqtof\_t3\_pos\_2\_011\_19

m/z 316.1795

02468E+344.5RT (min)

lcqtof\_t3\_pos\_2\_034\_24

m/z 317.2180

0123E+311.5RT (min)

lcqtof\_t3\_pos\_4\_034\_23

m/z 318.2157

012E+344.5RT (min)

lcqtof\_t3\_pos\_2\_022\_32

m/z 318.7773

0123E+311.5RT (min)

lcqtof\_t3\_pos\_4\_021\_01

m/z 318.8054

01234E+311.5RT (min)

lcqtof\_t3\_pos\_4\_001\_49

m/z 319.0593

012E+311.5RT (min)

lcqtof\_t3\_pos\_4\_045\_04

m/z 319.1289

012E+311.5RT (min)

lcqtof\_t3\_pos\_1\_032\_13

m/z 320.1718

012E+317.518RT (min)

lcqtof\_t3\_pos\_4\_027\_44

m/z 320.8038

012E+311.5RT (min)

lcqtof\_t3\_pos\_4\_037\_32

m/z 321.0746

024E+34.55RT (min)

lcqtof\_t3\_pos\_4\_024\_20

m/z 321.0949

02468E+311.5RT (min)

lcqtof\_t3\_pos\_4\_086\_59

m/z 321.2652

012E+355.5RT (min)

lcqtof\_t3\_pos\_4\_029\_37

m/z 322.1671

012E+34.55RT (min)

lcqtof\_t3\_pos\_2\_027\_38

m/z 322.1678

0246E+311.5RT (min)

lcqtof\_t3\_pos\_1\_048\_24

m/z 322.8142

0246E+311.5RT (min)

lcqtof\_t3\_pos\_4\_021\_01

m/z 322.8259

012E+317.518RT (min)

lcqtof\_t3\_pos\_1\_012\_42

m/z 322.9952

01234E+311.5RT (min)

lcqtof\_t3\_pos\_2\_039\_23

m/z 323.0134

0246E+311.5RT (min)

lcqtof\_t3\_pos\_4\_037\_32

m/z 323.0667

01234E+344.5RT (min)

lcqtof\_t3\_pos\_4\_034\_23

m/z 324.3209

012E+34.55RT (min)

lcqtof\_t3\_pos\_2\_040\_28

m/z 324.8084

02468E+311.5RT (min)

lcqtof\_t3\_pos\_4\_021\_01

m/z 325.0089

012E+311.5RT (min)

lcqtof\_t3\_pos\_2\_053\_15

m/z 325.5591

01E+55.56RT (min)

lcqtof\_t3\_pos\_4\_049\_55

m/z 326.1047

01234E+36.57RT (min)

lcqtof\_t3\_pos\_4\_013\_29

m/z 326.1407

0123E+311.5RT (min)

lcqtof\_t3\_pos\_1\_005\_46

m/z 326.1949

0246E+34.55RT (min)

lcqtof\_t3\_pos\_2\_027\_38

m/z 326.2104

012E+344.5RT (min)

lcqtof\_t3\_pos\_3\_007\_32

m/z 326.2412

024E+355.5RT (min)

lcqtof\_t3\_pos\_4\_029\_37

m/z 326.8070

01234E+311.5RT (min)

lcqtof\_t3\_pos\_4\_021\_01

m/z 327.1852

0123E+311.5RT (min)

lcqtof\_t3\_pos\_2\_041\_54

m/z 327.7437

01234E+36.57RT (min)

lcqtof\_t3\_pos\_4\_029\_37

m/z 328.0569

012E+311.5RT (min)

lcqtof\_t3\_pos\_4\_021\_01

m/z 328.0964

012E+355.5RT (min)

lcqtof\_t3\_pos\_3\_015\_13

m/z 328.1015

0123E+344.5RT (min)

lcqtof\_t3\_pos\_2\_008\_31

m/z 328.1607

01234E+344.5RT (min)

lcqtof\_t3\_pos\_4\_020\_19

m/z 328.1660

0246E+37.58RT (min)

lcqtof\_t3\_pos\_4\_042\_18

m/z 328.7599

012E+311.5RT (min)

lcqtof\_t3\_pos\_4\_021\_01

m/z 329.1797

0246E+344.5RT (min)

lcqtof\_t3\_pos\_4\_029\_37

m/z 329.1885

02468E+311.5RT (min)

lcqtof\_t3\_pos\_3\_011\_15

m/z 330.7553

0123E+311.5RT (min)

lcqtof\_t3\_pos\_4\_007\_14

m/z 332.1325

01234E+344.5RT (min)

lcqtof\_t3\_pos\_4\_023\_39

m/z 332.2248

0123E+355.5RT (min)

lcqtof\_t3\_pos\_4\_014\_24

m/z 332.2560

01234E+35.56RT (min)

lcqtof\_t3\_pos\_4\_003\_05

m/z 332.7509

012E+311.5RT (min)

lcqtof\_t3\_pos\_4\_021\_01

m/z 332.8400

0246E+311.5RT (min)

lcqtof\_t3\_pos\_4\_021\_01

m/z 333.2343

01234E+355.5RT (min)

lcqtof\_t3\_pos\_4\_029\_37

m/z 333.7024

01E+34.55RT (min)

lcqtof\_t3\_pos\_4\_044\_38

m/z 333.7061

024E+344.5RT (min)

lcqtof\_t3\_pos\_2\_050\_14

m/z 333.9381

01E+317.518RT (min)

lcqtof\_t3\_pos\_2\_004\_42

m/z 335.0967

01234E+311.5RT (min)

lcqtof\_t3\_pos\_4\_047\_13

m/z 336.0895

0123E+35.56RT (min)

lcqtof\_t3\_pos\_3\_029\_26

m/z 336.2414

0123E+377.5RT (min)

lcqtof\_t3\_pos\_4\_029\_37

m/z 336.2840

0123E+39.510RT (min)

lcqtof\_t3\_pos\_4\_082\_59

m/z 337.0815

024E+366.5RT (min)

lcqtof\_t3\_pos\_1\_052\_06

m/z 338.1818

0123E+344.5RT (min)

lcqtof\_t3\_pos\_4\_029\_37

m/z 338.1824

02468E+311.5RT (min)

lcqtof\_t3\_pos\_2\_047\_02

m/z 339.2764

01234E+344.5RT (min)

lcqtof\_t3\_pos\_4\_029\_37

m/z 339.7683

01234E+355.5RT (min)

lcqtof\_t3\_pos\_4\_029\_37

m/z 340.0415

0123E+311.5RT (min)

lcqtof\_t3\_pos\_2\_032\_47

m/z 340.1909

024E+311.5RT (min)

lcqtof\_t3\_pos\_3\_013\_24

m/z 340.2486

01E+466.5RT (min)

lcqtof\_t3\_pos\_4\_021\_01

m/z 340.3965

01234E+31111.5RT (min)

lcqtof\_t3\_pos\_2\_055\_18

m/z 340.9326

02468E+311.5RT (min)

lcqtof\_t3\_pos\_4\_021\_01

m/z 341.9301

01234E+311.5RT (min)

lcqtof\_t3\_pos\_4\_029\_37

m/z 342.0377

0246E+311.5RT (min)

lcqtof\_t3\_pos\_2\_022\_32

m/z 342.0906

012E+344.5RT (min)

lcqtof\_t3\_pos\_2\_010\_26

m/z 342.8001

0123E+311.5RT (min)

lcqtof\_t3\_pos\_2\_012\_35

m/z 343.0813

02468E+37.58RT (min)

lcqtof\_t3\_pos\_4\_042\_18

m/z 343.1970

0246E+344.5RT (min)

lcqtof\_t3\_pos\_4\_029\_37

m/z 343.2292

0123E+344.5RT (min)

lcqtof\_t3\_pos\_2\_034\_24

m/z 343.8824

01E+317.518RT (min)

lcqtof\_t3\_pos\_2\_024\_45

m/z 344.0859

024E+311.5RT (min)

lcqtof\_t3\_pos\_2\_008\_31

m/z 345.7367

01234E+36.57RT (min)

lcqtof\_t3\_pos\_4\_029\_37

m/z 346.1650

012E+34.55RT (min)

lcqtof\_t3\_pos\_2\_011\_19

m/z 346.1717

024E+37.58RT (min)

lcqtof\_t3\_pos\_4\_042\_18

m/z 346.1811

0246E+35.56RT (min)

lcqtof\_t3\_pos\_4\_029\_37

m/z 346.9549

01234E+311.5RT (min)

lcqtof\_t3\_pos\_2\_015\_13

m/z 347.2208

0246E+377.5RT (min)

lcqtof\_t3\_pos\_4\_021\_01

m/z 348.0191

012E+311.5RT (min)

lcqtof\_t3\_pos\_4\_080\_58

m/z 348.2581

024E+36.57RT (min)

lcqtof\_t3\_pos\_4\_029\_37

m/z 348.8320

0246E+311.5RT (min)

lcqtof\_t3\_pos\_2\_012\_35

m/z 349.2119

0246E+311.5RT (min)

lcqtof\_t3\_pos\_3\_013\_24

m/z 349.2129

0123E+311.5RT (min)

lcqtof\_t3\_pos\_3\_007\_32

m/z 349.2242

01234E+34.55RT (min)

lcqtof\_t3\_pos\_4\_021\_01

m/z 349.7380

0246E+35.56RT (min)

lcqtof\_t3\_pos\_4\_029\_37

m/z 350.0276

01E+317.518RT (min)

lcqtof\_t3\_pos\_2\_042\_12

m/z 350.1156

02468E+311.5RT (min)

lcqtof\_t3\_pos\_4\_047\_13

m/z 350.2403

024E+35.56RT (min)

lcqtof\_t3\_pos\_4\_003\_05

m/z 350.2424

0123E+37.58RT (min)

lcqtof\_t3\_pos\_4\_015\_16

m/z 351.0148

01E+411.5RT (min)

lcqtof\_t3\_pos\_4\_039\_02

m/z 351.1593

0123E+311.5RT (min)

lcqtof\_t3\_pos\_4\_021\_01

m/z 351.2407

0123E+366.5RT (min)

lcqtof\_t3\_pos\_4\_054\_31

m/z 351.5928

01234E+35.56RT (min)

lcqtof\_t3\_pos\_4\_029\_37

m/z 351.9911

012E+311.5RT (min)

lcqtof\_t3\_pos\_4\_021\_01

m/z 352.1072

0246E+311.5RT (min)

lcqtof\_t3\_pos\_2\_022\_32

m/z 352.1206

0123E+311.5RT (min)

lcqtof\_t3\_pos\_2\_017\_51

m/z 353.0040

01234E+311.5RT (min)

lcqtof\_t3\_pos\_2\_039\_23

m/z 353.0503

024E+39.510RT (min)

lcqtof\_t3\_pos\_4\_012\_12

m/z 353.1288

012E+311.5RT (min)

lcqtof\_t3\_pos\_4\_095\_60

m/z 353.2498

0123E+34.55RT (min)

lcqtof\_t3\_pos\_4\_029\_37

m/z 354.2292

0246E+344.5RT (min)

lcqtof\_t3\_pos\_2\_022\_32

m/z 354.7886

0246E+311.5RT (min)

lcqtof\_t3\_pos\_4\_003\_05

m/z 354.8396

012E+311.5RT (min)

lcqtof\_t3\_pos\_4\_037\_32

m/z 355.2660

024E+35.56RT (min)

lcqtof\_t3\_pos\_4\_029\_37

m/z 356.0621

024E+311.5RT (min)

lcqtof\_t3\_pos\_2\_012\_35

m/z 357.2281

01E+46.57RT (min)

lcqtof\_t3\_pos\_2\_010\_26

m/z 357.7382

012E+34.55RT (min)

lcqtof\_t3\_pos\_4\_008\_15

m/z 358.0477

02468E+311.5RT (min)

lcqtof\_t3\_pos\_2\_008\_31

m/z 358.1319

0123E+311.5RT (min)

lcqtof\_t3\_pos\_2\_003\_16

m/z 358.7106

0123E+344.5RT (min)

lcqtof\_t3\_pos\_4\_007\_14

m/z 358.7780

012E+311.5RT (min)

lcqtof\_t3\_pos\_2\_022\_32

m/z 359.1449

0246E+344.5RT (min)

lcqtof\_t3\_pos\_2\_022\_32

m/z 359.1494

024E+377.5RT (min)

lcqtof\_t3\_pos\_3\_005\_18

m/z 359.2049

01234E+36.57RT (min)

lcqtof\_t3\_pos\_4\_102\_61

m/z 359.2300

01234E+355.5RT (min)

lcqtof\_t3\_pos\_4\_029\_37

m/z 359.2345

0123E+35.56RT (min)

lcqtof\_t3\_pos\_2\_039\_23

m/z 359.2643

01234E+34.55RT (min)

lcqtof\_t3\_pos\_1\_004\_05

m/z 359.9745

012E+34.55RT (min)

lcqtof\_t3\_pos\_1\_029\_19

m/z 360.7413

01E+38.59RT (min)

lcqtof\_t3\_pos\_4\_103\_61

m/z 360.8464

0246E+311.5RT (min)

lcqtof\_t3\_pos\_2\_012\_35

m/z 361.0797

024E+411.5RT (min)

lcqtof\_t3\_pos\_2\_022\_32

m/z 361.1908

0246E+311.5RT (min)

lcqtof\_t3\_pos\_4\_087\_59

m/z 362.0952

024E+377.5RT (min)

lcqtof\_t3\_pos\_4\_046\_34

m/z 362.1133

012E+366.5RT (min)

lcqtof\_t3\_pos\_4\_042\_18

m/z 362.1544

0123E+311.5RT (min)

lcqtof\_t3\_pos\_1\_032\_13

m/z 362.2038

012E+36.57RT (min)

lcqtof\_t3\_pos\_4\_007\_14

m/z 362.2325

012E+366.5RT (min)

lcqtof\_t3\_pos\_4\_001\_49

m/z 363.9307

01E+317.518RT (min)

lcqtof\_t3\_pos\_2\_003\_16

m/z 363.9315

012E+317.518RT (min)

lcqtof\_t3\_pos\_2\_012\_35

m/z 364.2242

012E+34.55RT (min)

lcqtof\_t3\_pos\_4\_021\_01

m/z 364.3892

02468E+314.515RT (min)

lcqtof\_t3\_pos\_4\_014\_24

m/z 364.8279

02468E+311.5RT (min)

lcqtof\_t3\_pos\_4\_029\_37

m/z 366.1288

0123E+344.5RT (min)

lcqtof\_t3\_pos\_3\_009\_50

m/z 366.8217

0123E+311.5RT (min)

lcqtof\_t3\_pos\_4\_029\_37

m/z 368.1848

024E+344.5RT (min)

lcqtof\_t3\_pos\_4\_029\_37

m/z 369.1219

0123E+344.5RT (min)

lcqtof\_t3\_pos\_4\_007\_14

m/z 369.1436

02468E+311.5RT (min)

lcqtof\_t3\_pos\_2\_023\_05

m/z 369.2505

0123E+311.5RT (min)

lcqtof\_t3\_pos\_2\_012\_35

m/z 369.9260

02468E+311.5RT (min)

lcqtof\_t3\_pos\_4\_007\_14

m/z 370.1465

0246E+311.5RT (min)

lcqtof\_t3\_pos\_2\_041\_54

m/z 370.2979

0246E+377.5RT (min)

lcqtof\_t3\_pos\_4\_008\_15

m/z 370.7374

0123E+317.518RT (min)

lcqtof\_t3\_pos\_4\_082\_59

m/z 370.8692

01E+411.5RT (min)

lcqtof\_t3\_pos\_1\_014\_14

m/z 371.0333

01E+411.5RT (min)

lcqtof\_t3\_pos\_2\_002\_37

m/z 371.6791

02468E+34.55RT (min)

lcqtof\_t3\_pos\_4\_021\_01

m/z 372.0106

01234E+34.55RT (min)

lcqtof\_t3\_pos\_4\_113\_62

m/z 372.2686

01234E+344.5RT (min)

lcqtof\_t3\_pos\_4\_029\_37

m/z 373.1159

0246E+377.5RT (min)

lcqtof\_t3\_pos\_4\_042\_18

m/z 373.1518

024E+34.55RT (min)

lcqtof\_t3\_pos\_1\_010\_27

m/z 373.2041

01234E+344.5RT (min)

lcqtof\_t3\_pos\_4\_029\_37

m/z 373.2097

01234E+311.5RT (min)

lcqtof\_t3\_pos\_2\_037\_46

m/z 374.1054

024E+344.5RT (min)

lcqtof\_t3\_pos\_2\_020\_27

m/z 374.2552

0123E+34.55RT (min)

lcqtof\_t3\_pos\_4\_029\_37

m/z 374.2837

01E+46.57RT (min)

lcqtof\_t3\_pos\_4\_029\_37

m/z 374.6174

0246E+36.57RT (min)

lcqtof\_t3\_pos\_4\_029\_37

m/z 374.8366

012E+317.518RT (min)

lcqtof\_t3\_pos\_1\_012\_42

m/z 375.1345

01234E+344.5RT (min)

lcqtof\_t3\_pos\_2\_015\_13

m/z 375.1896

0123E+311.5RT (min)

lcqtof\_t3\_pos\_2\_050\_14

m/z 375.2254

02468E+344.5RT (min)

lcqtof\_t3\_pos\_2\_012\_35

m/z 375.2538

0123E+36.57RT (min)

lcqtof\_t3\_pos\_2\_010\_26

m/z 375.2744

0246E+311.5RT (min)

lcqtof\_t3\_pos\_2\_034\_24

m/z 376.0741

012E+366.5RT (min)

lcqtof\_t3\_pos\_2\_028\_34

m/z 377.0873

024E+36.57RT (min)

lcqtof\_t3\_pos\_4\_005\_30

m/z 378.7301

012E+311.5RT (min)

lcqtof\_t3\_pos\_4\_021\_01

m/z 378.9951

01234E+311.5RT (min)

lcqtof\_t3\_pos\_2\_014\_03

m/z 379.0760

012E+311.5RT (min)

lcqtof\_t3\_pos\_1\_002\_36

m/z 379.1006

0246E+34.55RT (min)

lcqtof\_t3\_pos\_4\_042\_18

m/z 379.2480

0123E+344.5RT (min)

lcqtof\_t3\_pos\_1\_008\_31

m/z 379.2496

01234E+35.56RT (min)

lcqtof\_t3\_pos\_4\_052\_06

m/z 379.3226

02468E+377.5RT (min)

lcqtof\_t3\_pos\_2\_011\_19

m/z 379.7520

01234E+34.55RT (min)

lcqtof\_t3\_pos\_4\_003\_05

m/z 379.9947

0123E+311.5RT (min)

lcqtof\_t3\_pos\_4\_019\_03

m/z 380.1685

0246E+37.58RT (min)

lcqtof\_t3\_pos\_1\_053\_18

m/z 381.0833

012E+317.518RT (min)

lcqtof\_t3\_pos\_2\_004\_42

m/z 381.1328

01234E+35.56RT (min)

lcqtof\_t3\_pos\_4\_036\_27

m/z 381.1777

012E+366.5RT (min)

lcqtof\_t3\_pos\_4\_051\_35

m/z 382.0835

0123E+311.5RT (min)

lcqtof\_t3\_pos\_3\_005\_18

m/z 382.1651

0123E+344.5RT (min)

lcqtof\_t3\_pos\_3\_001\_49

m/z 382.2093

01E+344.5RT (min)

lcqtof\_t3\_pos\_1\_007\_07

m/z 382.9975

0246E+377.5RT (min)

lcqtof\_t3\_pos\_4\_042\_18

m/z 383.0918

01234E+366.5RT (min)

lcqtof\_t3\_pos\_4\_042\_18

m/z 383.7921

0123E+35.56RT (min)

lcqtof\_t3\_pos\_4\_029\_37

m/z 384.1463

012E+317.518RT (min)

lcqtof\_t3\_pos\_2\_011\_19

m/z 384.6945

02468E+34.55RT (min)

lcqtof\_t3\_pos\_4\_024\_20

m/z 384.7524

0123E+344.5RT (min)

lcqtof\_t3\_pos\_2\_008\_31

m/z 384.8448

024E+311.5RT (min)

lcqtof\_t3\_pos\_2\_050\_14

m/z 384.9890

012E+311.5RT (min)

lcqtof\_t3\_pos\_4\_019\_03

m/z 385.0283

01234E+37.58RT (min)

lcqtof\_t3\_pos\_4\_104\_61

m/z 385.0812

0246E+344.5RT (min)

lcqtof\_t3\_pos\_2\_010\_26

m/z 385.0875

012E+355.5RT (min)

lcqtof\_t3\_pos\_3\_005\_18

m/z 385.2124

012E+344.5RT (min)

lcqtof\_t3\_pos\_4\_046\_34

m/z 385.8729

01E+317.518RT (min)

lcqtof\_t3\_pos\_2\_020\_27

m/z 385.9027

01E+411.5RT (min)

lcqtof\_t3\_pos\_4\_007\_14

m/z 386.0410

0246E+311.5RT (min)

lcqtof\_t3\_pos\_2\_022\_32

m/z 386.1745

0246E+36.57RT (min)

lcqtof\_t3\_pos\_4\_021\_01

m/z 386.2653

024E+344.5RT (min)

lcqtof\_t3\_pos\_4\_014\_24

m/z 386.2837

0246E+34.55RT (min)

lcqtof\_t3\_pos\_4\_029\_37

m/z 387.2672

012E+355.5RT (min)

lcqtof\_t3\_pos\_4\_029\_37

m/z 387.3101

0246E+311.5RT (min)

lcqtof\_t3\_pos\_4\_014\_24

m/z 388.0908

024E+377.5RT (min)

lcqtof\_t3\_pos\_4\_042\_18

m/z 388.2295

01234E+311.5RT (min)

lcqtof\_t3\_pos\_2\_008\_31

m/z 388.7990

024E+35.56RT (min)

lcqtof\_t3\_pos\_4\_029\_37

m/z 389.1001

0123E+311.5RT (min)

lcqtof\_t3\_pos\_2\_039\_23

m/z 389.2188

024E+311.5RT (min)

lcqtof\_t3\_pos\_2\_039\_23

m/z 389.2882

0123E+35.56RT (min)

lcqtof\_t3\_pos\_4\_089\_59

m/z 390.0563

012E+344.5RT (min)

lcqtof\_t3\_pos\_2\_019\_08

m/z 390.1030

012E+311.5RT (min)

lcqtof\_t3\_pos\_1\_018\_30

m/z 390.1629

02468E+355.5RT (min)

lcqtof\_t3\_pos\_2\_037\_46

m/z 391.2172

01234E+344.5RT (min)

lcqtof\_t3\_pos\_1\_014\_14

m/z 392.0716

01234E+344.5RT (min)

lcqtof\_t3\_pos\_4\_029\_37

m/z 392.1042

01234E+35.56RT (min)

lcqtof\_t3\_pos\_4\_020\_19

m/z 392.1816

0246E+36.57RT (min)

lcqtof\_t3\_pos\_4\_105\_61

m/z 392.2140

01234E+355.5RT (min)

lcqtof\_t3\_pos\_4\_042\_18

m/z 393.1364

0246E+344.5RT (min)

lcqtof\_t3\_pos\_2\_015\_13

m/z 393.2519

01234E+355.5RT (min)

lcqtof\_t3\_pos\_4\_008\_15

m/z 394.8753

012E+317.518RT (min)

lcqtof\_t3\_pos\_2\_007\_07

m/z 395.0147

024E+388.5RT (min)

lcqtof\_t3\_pos\_4\_088\_59

m/z 395.0386

0123E+311.5RT (min)

lcqtof\_t3\_pos\_2\_027\_38

m/z 395.0966

02468E+311.5RT (min)

lcqtof\_t3\_pos\_2\_047\_02

m/z 395.1334

0123E+34.55RT (min)

lcqtof\_t3\_pos\_3\_029\_26

m/z 396.1753

0123E+344.5RT (min)

lcqtof\_t3\_pos\_4\_097\_60

m/z 396.6809

02468E+377.5RT (min)

lcqtof\_t3\_pos\_2\_011\_19

m/z 396.8738

01234E+311.5RT (min)

lcqtof\_t3\_pos\_2\_050\_14

m/z 397.2729

024E+317.518RT (min)

lcqtof\_t3\_pos\_2\_007\_07

m/z 398.1168

0123E+35.56RT (min)

lcqtof\_t3\_pos\_2\_027\_38

m/z 398.1658

02468E+311.5RT (min)

lcqtof\_t3\_pos\_2\_034\_24

m/z 398.7430

02468E+311.5RT (min)

lcqtof\_t3\_pos\_4\_007\_14

m/z 399.2911

024E+34.55RT (min)

lcqtof\_t3\_pos\_4\_029\_37

m/z 399.9218

012E+355.5RT (min)

lcqtof\_t3\_pos\_4\_096\_60

m/z 400.0982

012E+34.55RT (min)

lcqtof\_t3\_pos\_2\_020\_27

m/z 400.1264

0246E+35.56RT (min)

lcqtof\_t3\_pos\_2\_011\_19

m/z 400.2425

012E+311.5RT (min)

lcqtof\_t3\_pos\_4\_014\_24

m/z 400.2564

02468E+314.515RT (min)

lcqtof\_t3\_pos\_4\_029\_37

m/z 401.1222

024E+35.56RT (min)

lcqtof\_t3\_pos\_2\_020\_27

m/z 401.2107

01234E+344.5RT (min)

lcqtof\_t3\_pos\_4\_029\_37

m/z 401.2590

01234E+317.518RT (min)

lcqtof\_t3\_pos\_4\_101\_61

m/z 401.2656

01234E+344.5RT (min)

lcqtof\_t3\_pos\_4\_054\_31

m/z 402.0700

012E+355.5RT (min)

lcqtof\_t3\_pos\_3\_005\_18

m/z 402.2466

024E+311.5RT (min)

lcqtof\_t3\_pos\_2\_039\_23

m/z 402.2804

024E+36.57RT (min)

lcqtof\_t3\_pos\_4\_029\_37

m/z 403.0287

012E+35.56RT (min)

lcqtof\_t3\_pos\_2\_011\_19

m/z 404.2457

01234E+377.5RT (min)

lcqtof\_t3\_pos\_4\_046\_34

m/z 404.6878

024E+311.5RT (min)

lcqtof\_t3\_pos\_4\_007\_14

m/z 405.1309

024E+311.5RT (min)

lcqtof\_t3\_pos\_2\_028\_34

m/z 406.0728

0123E+344.5RT (min)

lcqtof\_t3\_pos\_1\_008\_31

m/z 406.2455

024E+377.5RT (min)

lcqtof\_t3\_pos\_4\_042\_18

m/z 406.6859

01234E+311.5RT (min)

lcqtof\_t3\_pos\_4\_007\_14

m/z 406.7741

02468E+311.5RT (min)

lcqtof\_t3\_pos\_4\_007\_14

m/z 407.0307

024E+388.5RT (min)

lcqtof\_t3\_pos\_4\_088\_59

m/z 407.0750

0123E+311.5RT (min)

lcqtof\_t3\_pos\_1\_014\_14

m/z 407.1847

01234E+355.5RT (min)

lcqtof\_t3\_pos\_2\_055\_18

m/z 407.2079

012E+34.55RT (min)

lcqtof\_t3\_pos\_4\_020\_19

m/z 407.7806

01E+466.5RT (min)

lcqtof\_t3\_pos\_4\_029\_37

m/z 408.0881

01234E+355.5RT (min)

lcqtof\_t3\_pos\_4\_105\_61

m/z 408.0922

012E+317.518RT (min)

lcqtof\_t3\_pos\_2\_005\_17

m/z 408.7992

012E+35.56RT (min)

lcqtof\_t3\_pos\_4\_003\_05

m/z 409.1444

024E+35.56RT (min)

lcqtof\_t3\_pos\_4\_020\_19

m/z 409.1913

0246E+34.55RT (min)

lcqtof\_t3\_pos\_2\_011\_19

m/z 409.2056

01234E+377.5RT (min)

lcqtof\_t3\_pos\_1\_053\_18

m/z 409.8604

01E+317.518RT (min)

lcqtof\_t3\_pos\_2\_011\_19

m/z 410.2759

012E+317.518RT (min)

lcqtof\_t3\_pos\_2\_005\_17

m/z 411.7172

0246E+355.5RT (min)

lcqtof\_t3\_pos\_4\_021\_01

m/z 411.7214

0246E+317.518RT (min)

lcqtof\_t3\_pos\_4\_021\_01

m/z 412.7398

012E+344.5RT (min)

lcqtof\_t3\_pos\_3\_003\_14

m/z 412.7779

0246E+38.59RT (min)

lcqtof\_t3\_pos\_4\_015\_16

m/z 413.6545

01234E+344.5RT (min)

lcqtof\_t3\_pos\_4\_076\_58

m/z 414.1426

024E+377.5RT (min)

lcqtof\_t3\_pos\_4\_021\_01

m/z 414.6124

012E+355.5RT (min)

lcqtof\_t3\_pos\_4\_049\_55

m/z 414.7645

0123E+317.518RT (min)

lcqtof\_t3\_pos\_4\_099\_61

m/z 414.8506

0246E+311.5RT (min)

lcqtof\_t3\_pos\_2\_005\_17

m/z 415.0539

012E+38.59RT (min)

lcqtof\_t3\_pos\_4\_104\_61

m/z 415.1969

01234E+34.55RT (min)

lcqtof\_t3\_pos\_4\_007\_14

m/z 415.2687

02468E+344.5RT (min)

lcqtof\_t3\_pos\_4\_049\_55

m/z 415.3410

012E+36.57RT (min)

lcqtof\_t3\_pos\_2\_020\_27

m/z 415.3436

0123E+355.5RT (min)

lcqtof\_t3\_pos\_4\_029\_37

m/z 415.7769

0123E+35.56RT (min)

lcqtof\_t3\_pos\_2\_008\_31

m/z 415.9826

02468E+366.5RT (min)

lcqtof\_t3\_pos\_4\_029\_37

m/z 416.1531

01234E+34.55RT (min)

lcqtof\_t3\_pos\_2\_020\_27

m/z 416.2876

01234E+35.56RT (min)

lcqtof\_t3\_pos\_2\_027\_38

m/z 416.3142

024E+35.56RT (min)

lcqtof\_t3\_pos\_4\_021\_01

m/z 416.6067

01234E+344.5RT (min)

lcqtof\_t3\_pos\_2\_022\_32

m/z 416.7345

024E+311.5RT (min)

lcqtof\_t3\_pos\_2\_022\_32

m/z 417.2198

01E+344.5RT (min)

lcqtof\_t3\_pos\_2\_050\_14

m/z 417.2539

012E+34.55RT (min)

lcqtof\_t3\_pos\_2\_029\_06

m/z 418.1367

0246E+344.5RT (min)

lcqtof\_t3\_pos\_2\_039\_23

m/z 418.2278

0246E+311.5RT (min)

lcqtof\_t3\_pos\_2\_039\_23

m/z 418.2466

01234E+37.58RT (min)

lcqtof\_t3\_pos\_1\_014\_14

m/z 418.3046

024E+36.57RT (min)

lcqtof\_t3\_pos\_4\_029\_37

m/z 418.6339

0246E+36.57RT (min)

lcqtof\_t3\_pos\_4\_029\_37

m/z 418.9942

0246E+366.5RT (min)

lcqtof\_t3\_pos\_4\_089\_59

m/z 419.0634

0123E+366.5RT (min)

lcqtof\_t3\_pos\_2\_028\_34

m/z 419.1816

012E+366.5RT (min)

lcqtof\_t3\_pos\_2\_052\_36

m/z 420.2707

0246E+344.5RT (min)

lcqtof\_t3\_pos\_4\_014\_24

m/z 420.7972

01234E+311.5RT (min)

lcqtof\_t3\_pos\_4\_007\_14

m/z 420.9662

012E+311.5RT (min)

lcqtof\_t3\_pos\_4\_026\_17

m/z 421.0510

01234E+388.5RT (min)

lcqtof\_t3\_pos\_4\_079\_58

m/z 421.0781

024E+34.55RT (min)

lcqtof\_t3\_pos\_2\_011\_19

m/z 421.1983

0246E+35.56RT (min)

lcqtof\_t3\_pos\_4\_024\_20

m/z 421.5595

0246E+344.5RT (min)

lcqtof\_t3\_pos\_4\_037\_32

m/z 421.8877

0246E+311.5RT (min)

lcqtof\_t3\_pos\_2\_031\_01

m/z 422.1835

0123E+366.5RT (min)

lcqtof\_t3\_pos\_1\_002\_36

m/z 422.7791

0246E+311.5RT (min)

lcqtof\_t3\_pos\_4\_029\_37

m/z 422.8828

012E+317.518RT (min)

lcqtof\_t3\_pos\_3\_017\_51

m/z 423.0641

02468E+311.5RT (min)

lcqtof\_t3\_pos\_4\_047\_13

m/z 423.2742

0246E+34.55RT (min)

lcqtof\_t3\_pos\_4\_029\_37

m/z 423.7767

0123E+317.518RT (min)

lcqtof\_t3\_pos\_4\_090\_60

m/z 424.0704

01234E+355.5RT (min)

lcqtof\_t3\_pos\_3\_105\_61

m/z 424.1478

0123E+344.5RT (min)

lcqtof\_t3\_pos\_4\_005\_30

m/z 424.1659

0246E+344.5RT (min)

lcqtof\_t3\_pos\_2\_037\_46

m/z 426.1216

012E+311.5RT (min)

lcqtof\_t3\_pos\_1\_005\_46

m/z 426.7952

01234E+35.56RT (min)

lcqtof\_t3\_pos\_2\_008\_31

m/z 427.2278

02468E+344.5RT (min)

lcqtof\_t3\_pos\_4\_021\_01

m/z 427.3263

01234E+355.5RT (min)

lcqtof\_t3\_pos\_4\_034\_23

m/z 428.0545

02468E+377.5RT (min)

lcqtof\_t3\_pos\_4\_089\_59

m/z 428.0802

01234E+37.58RT (min)

lcqtof\_t3\_pos\_4\_042\_18

m/z 428.9055

0246E+39.510RT (min)

lcqtof\_t3\_pos\_4\_049\_55

m/z 429.0872

0246E+344.5RT (min)

lcqtof\_t3\_pos\_2\_022\_32

m/z 429.1300

0123E+36.57RT (min)

lcqtof\_t3\_pos\_2\_010\_26

m/z 429.1999

01234E+344.5RT (min)

lcqtof\_t3\_pos\_2\_020\_27

m/z 429.4508

012E+317.518RT (min)

lcqtof\_t3\_pos\_2\_011\_19

m/z 430.2281

0246E+377.5RT (min)

lcqtof\_t3\_pos\_4\_042\_18

m/z 430.2834

0123E+344.5RT (min)

lcqtof\_t3\_pos\_2\_039\_23

m/z 430.6550

024E+35.56RT (min)

lcqtof\_t3\_pos\_4\_029\_37

m/z 430.9037

01E+49.510RT (min)

lcqtof\_t3\_pos\_4\_049\_55

m/z 431.2148

024E+344.5RT (min)

lcqtof\_t3\_pos\_4\_029\_37

m/z 431.2749

02468E+344.5RT (min)

lcqtof\_t3\_pos\_4\_014\_24

m/z 431.9174

0123E+317.518RT (min)

lcqtof\_t3\_pos\_2\_007\_07

m/z 432.1198

0123E+34.55RT (min)

lcqtof\_t3\_pos\_2\_011\_19

m/z 433.0092

0123E+377.5RT (min)

lcqtof\_t3\_pos\_4\_081\_58

m/z 433.2414

01E+317.518RT (min)

lcqtof\_t3\_pos\_2\_001\_49

m/z 434.2377

012E+36.57RT (min)

lcqtof\_t3\_pos\_1\_018\_30

m/z 435.0664

0123E+388.5RT (min)

lcqtof\_t3\_pos\_4\_082\_59

m/z 435.1939

0123E+37.58RT (min)

lcqtof\_t3\_pos\_4\_042\_18

m/z 436.2279

012E+311.5RT (min)

lcqtof\_t3\_pos\_2\_041\_54

m/z 436.2678

0123E+37.58RT (min)

lcqtof\_t3\_pos\_4\_024\_20

m/z 436.6901

0246E+311.5RT (min)

lcqtof\_t3\_pos\_4\_021\_01

m/z 436.7198

01234E+344.5RT (min)

lcqtof\_t3\_pos\_2\_050\_14

m/z 436.7741

0123E+317.518RT (min)

lcqtof\_t3\_pos\_4\_082\_59

m/z 436.8854

012E+411.5RT (min)

lcqtof\_t3\_pos\_4\_049\_55

m/z 437.2920

0246E+311.5RT (min)

lcqtof\_t3\_pos\_2\_034\_24

m/z 438.1504

02468E+311.5RT (min)

lcqtof\_t3\_pos\_4\_014\_24

m/z 439.1008

012E+35.56RT (min)

lcqtof\_t3\_pos\_2\_011\_19

m/z 439.1048

0246E+344.5RT (min)

lcqtof\_t3\_pos\_4\_034\_23

m/z 439.2814

0123E+399.5RT (min)

lcqtof\_t3\_pos\_4\_070\_57

m/z 439.3273

024E+35.56RT (min)

lcqtof\_t3\_pos\_4\_029\_37

m/z 440.2854

012E+35.56RT (min)

lcqtof\_t3\_pos\_4\_038\_21

m/z 440.8274

0123E+36.57RT (min)

lcqtof\_t3\_pos\_4\_029\_37

m/z 440.9882

012E+34.55RT (min)

lcqtof\_t3\_pos\_2\_010\_26

m/z 441.1096

024E+311.5RT (min)

lcqtof\_t3\_pos\_4\_029\_37

m/z 441.3643

0123E+377.5RT (min)

lcqtof\_t3\_pos\_4\_029\_37

m/z 442.1802

02468E+344.5RT (min)

lcqtof\_t3\_pos\_2\_001\_49

m/z 442.8235

012E+317.518RT (min)

lcqtof\_t3\_pos\_1\_012\_42

m/z 443.0702

01234E+388.5RT (min)

lcqtof\_t3\_pos\_4\_083\_59

m/z 443.3505

024E+366.5RT (min)

lcqtof\_t3\_pos\_4\_029\_37

m/z 443.8822

02468E+311.5RT (min)

lcqtof\_t3\_pos\_4\_021\_01

m/z 443.9484

01E+355.5RT (min)

lcqtof\_t3\_pos\_4\_112\_62

m/z 444.0328

01234E+311.5RT (min)

lcqtof\_t3\_pos\_2\_028\_34

m/z 444.6860

02468E+311.5RT (min)

lcqtof\_t3\_pos\_4\_021\_01

m/z 444.8425

01234E+366.5RT (min)

lcqtof\_t3\_pos\_4\_054\_31

m/z 444.8797

02468E+311.5RT (min)

lcqtof\_t3\_pos\_4\_021\_01

m/z 444.8875

0246E+311.5RT (min)

lcqtof\_t3\_pos\_2\_012\_35

m/z 445.1174

024E+35.56RT (min)

lcqtof\_t3\_pos\_2\_011\_19

m/z 445.7900

01234E+317.518RT (min)

lcqtof\_t3\_pos\_4\_106\_62

m/z 446.1602

024E+311.5RT (min)

lcqtof\_t3\_pos\_4\_015\_16

m/z 446.2918

012E+355.5RT (min)

lcqtof\_t3\_pos\_4\_008\_15

m/z 446.3148

01234E+35.56RT (min)

lcqtof\_t3\_pos\_4\_034\_23

m/z 447.0079

012E+344.5RT (min)

lcqtof\_t3\_pos\_2\_030\_33

m/z 447.2637

0123E+34.55RT (min)

lcqtof\_t3\_pos\_4\_020\_19

m/z 447.2697

0246E+344.5RT (min)

lcqtof\_t3\_pos\_2\_022\_32

m/z 448.0239

0123E+37.58RT (min)

lcqtof\_t3\_pos\_3\_101\_61

m/z 448.2147

01234E+37.58RT (min)

lcqtof\_t3\_pos\_4\_015\_16

m/z 448.3249

0246E+377.5RT (min)

lcqtof\_t3\_pos\_4\_021\_01

m/z 448.3496

0246E+344.5RT (min)

lcqtof\_t3\_pos\_2\_022\_32

m/z 448.7562

02468E+311.5RT (min)

lcqtof\_t3\_pos\_4\_021\_01

m/z 448.8393

01234E+36.57RT (min)

lcqtof\_t3\_pos\_4\_029\_37

m/z 449.0775

024E+388.5RT (min)

lcqtof\_t3\_pos\_4\_081\_58

m/z 449.9361

0246E+311.5RT (min)

lcqtof\_t3\_pos\_4\_014\_24

m/z 449.9715

01234E+355.5RT (min)

lcqtof\_t3\_pos\_4\_049\_55

m/z 450.0119

0123E+36.57RT (min)

lcqtof\_t3\_pos\_4\_054\_31

m/z 450.1075

01234E+34.55RT (min)

lcqtof\_t3\_pos\_4\_020\_19

m/z 450.1613

0123E+311.5RT (min)

lcqtof\_t3\_pos\_2\_037\_46

m/z 451.2890

024E+377.5RT (min)

lcqtof\_t3\_pos\_2\_001\_49

m/z 452.0256

012E+344.5RT (min)

lcqtof\_t3\_pos\_2\_007\_07

m/z 452.6626

012E+311.5RT (min)

lcqtof\_t3\_pos\_4\_021\_01

m/z 452.8341

01E+411.5RT (min)

lcqtof\_t3\_pos\_2\_050\_14

m/z 453.3053

012E+37.58RT (min)

lcqtof\_t3\_pos\_4\_070\_57

m/z 453.7067

012E+355.5RT (min)

lcqtof\_t3\_pos\_2\_012\_35

m/z 453.8605

01E+317.518RT (min)

lcqtof\_t3\_pos\_2\_026\_40

m/z 454.0662

024E+311.5RT (min)

lcqtof\_t3\_pos\_3\_092\_60

m/z 454.1953

01234E+36.57RT (min)

lcqtof\_t3\_pos\_4\_020\_19

m/z 454.2782

024E+344.5RT (min)

lcqtof\_t3\_pos\_4\_003\_05

m/z 455.1891

02468E+366.5RT (min)

lcqtof\_t3\_pos\_3\_004\_07

m/z 455.3151

0123E+38.59RT (min)

lcqtof\_t3\_pos\_4\_070\_57

m/z 456.0000

024E+36.57RT (min)

lcqtof\_t3\_pos\_4\_029\_37

m/z 456.1156

01234E+344.5RT (min)

lcqtof\_t3\_pos\_4\_029\_37

m/z 456.1868

0123E+34.55RT (min)

lcqtof\_t3\_pos\_2\_037\_46

m/z 456.2744

0123E+34.55RT (min)

lcqtof\_t3\_pos\_4\_020\_19

m/z 456.6646

0123E+36.57RT (min)

lcqtof\_t3\_pos\_4\_046\_34

m/z 457.0366

02468E+311.5RT (min)

lcqtof\_t3\_pos\_2\_015\_13

m/z 457.3442

02468E+34.55RT (min)

lcqtof\_t3\_pos\_4\_029\_37

m/z 457.3582

0246E+355.5RT (min)

lcqtof\_t3\_pos\_4\_029\_37

m/z 458.0437

01E+344.5RT (min)

lcqtof\_t3\_pos\_2\_011\_19

m/z 458.1463

024E+344.5RT (min)

lcqtof\_t3\_pos\_2\_020\_27

m/z 458.1782

01234E+36.57RT (min)

lcqtof\_t3\_pos\_3\_005\_18

m/z 458.3424

012E+34.55RT (min)

lcqtof\_t3\_pos\_4\_034\_23

m/z 458.7910

0123E+355.5RT (min)

lcqtof\_t3\_pos\_4\_008\_15

m/z 458.7916

012E+317.518RT (min)

lcqtof\_t3\_pos\_4\_083\_59

m/z 459.3763

0123E+377.5RT (min)

lcqtof\_t3\_pos\_4\_029\_37

m/z 460.0770

012E+34.55RT (min)

lcqtof\_t3\_pos\_2\_011\_19

m/z 460.2646

02468E+344.5RT (min)

lcqtof\_t3\_pos\_4\_029\_37

m/z 460.2921

0123E+366.5RT (min)

lcqtof\_t3\_pos\_4\_067\_57

m/z 460.7140

0123E+311.5RT (min)

lcqtof\_t3\_pos\_2\_012\_35

m/z 461.2799

012E+34.55RT (min)

lcqtof\_t3\_pos\_2\_029\_06

m/z 462.2924

02468E+344.5RT (min)

lcqtof\_t3\_pos\_4\_021\_01

m/z 462.6494

0123E+311.5RT (min)

lcqtof\_t3\_pos\_4\_007\_14

m/z 462.7826

0246E+311.5RT (min)

lcqtof\_t3\_pos\_4\_021\_01

m/z 463.8660

012E+311.5RT (min)

lcqtof\_t3\_pos\_2\_050\_14

m/z 463.8780

012E+317.518RT (min)

lcqtof\_t3\_pos\_3\_017\_51

m/z 463.9094

01234E+311.5RT (min)

lcqtof\_t3\_pos\_4\_007\_14

m/z 464.1120

0123E+311.5RT (min)

lcqtof\_t3\_pos\_1\_002\_36

m/z 464.2158

0123E+34.55RT (min)

lcqtof\_t3\_pos\_2\_011\_19

m/z 464.6441

0123E+311.5RT (min)

lcqtof\_t3\_pos\_4\_007\_14

m/z 465.2297

0246E+377.5RT (min)

lcqtof\_t3\_pos\_1\_053\_18

m/z 465.9696

024E+311.5RT (min)

lcqtof\_t3\_pos\_4\_001\_49

m/z 466.1251

024E+344.5RT (min)

lcqtof\_t3\_pos\_2\_025\_52

m/z 466.3442

0123E+355.5RT (min)

lcqtof\_t3\_pos\_4\_029\_37

m/z 466.7385

0246E+311.5RT (min)

lcqtof\_t3\_pos\_4\_029\_37

m/z 467.1602

0246E+311.5RT (min)

lcqtof\_t3\_pos\_4\_072\_57

m/z 467.3027

01E+44.55RT (min)

lcqtof\_t3\_pos\_4\_029\_37

m/z 468.1261

024E+35.56RT (min)

lcqtof\_t3\_pos\_3\_050\_27

m/z 468.1263

024E+35.56RT (min)

lcqtof\_t3\_pos\_3\_053\_19

m/z 468.3048

0123E+355.5RT (min)

lcqtof\_t3\_pos\_4\_008\_15

m/z 469.0321

012E+344.5RT (min)

lcqtof\_t3\_pos\_2\_010\_26

m/z 469.0488

0123E+311.5RT (min)

lcqtof\_t3\_pos\_2\_022\_32

m/z 469.2874

0123E+36.57RT (min)

lcqtof\_t3\_pos\_4\_105\_61

m/z 470.6981

01234E+311.5RT (min)

lcqtof\_t3\_pos\_2\_030\_33

m/z 471.0203

0123E+37.58RT (min)

lcqtof\_t3\_pos\_2\_010\_26

m/z 471.2283

0246E+311.5RT (min)

lcqtof\_t3\_pos\_2\_039\_23

m/z 471.2947

0246E+35.56RT (min)

lcqtof\_t3\_pos\_4\_021\_01

m/z 472.2396

024E+35.56RT (min)

lcqtof\_t3\_pos\_4\_021\_01

m/z 473.1935

0123E+311.5RT (min)

lcqtof\_t3\_pos\_2\_017\_51

m/z 473.2872

012E+355.5RT (min)

lcqtof\_t3\_pos\_4\_105\_61

m/z 473.3995

012E+417.518RT (min)

lcqtof\_t3\_pos\_4\_042\_18

m/z 474.3096

02468E+344.5RT (min)

lcqtof\_t3\_pos\_4\_007\_14

m/z 474.6933

024E+311.5RT (min)

lcqtof\_t3\_pos\_2\_022\_32

m/z 474.9432

012E+317.518RT (min)

lcqtof\_t3\_pos\_2\_051\_11

m/z 475.0717

024E+355.5RT (min)

lcqtof\_t3\_pos\_4\_104\_61

m/z 475.2208

0246E+344.5RT (min)

lcqtof\_t3\_pos\_4\_014\_24

m/z 475.2690

012E+35.56RT (min)

lcqtof\_t3\_pos\_4\_106\_62

m/z 475.9407

0123E+34.55RT (min)

lcqtof\_t3\_pos\_2\_050\_14

m/z 476.7607

0246E+311.5RT (min)

lcqtof\_t3\_pos\_4\_007\_14

m/z 476.9412

02468E+311.5RT (min)

lcqtof\_t3\_pos\_4\_021\_01

m/z 477.1029

024E+366.5RT (min)

lcqtof\_t3\_pos\_4\_013\_29

m/z 477.1097

0123E+388.5RT (min)

lcqtof\_t3\_pos\_4\_088\_59

m/z 477.3400

0246E+344.5RT (min)

lcqtof\_t3\_pos\_2\_022\_32

m/z 478.2386

0123E+37.58RT (min)

lcqtof\_t3\_pos\_1\_047\_26

m/z 478.2716

01E+39.510RT (min)

lcqtof\_t3\_pos\_4\_034\_23

m/z 479.3148

012E+355.5RT (min)

lcqtof\_t3\_pos\_4\_054\_31

m/z 480.1667

01234E+377.5RT (min)

lcqtof\_t3\_pos\_2\_027\_38

m/z 480.7428

0246E+311.5RT (min)

lcqtof\_t3\_pos\_2\_022\_32

m/z 481.1167

0246E+311.5RT (min)

lcqtof\_t3\_pos\_2\_028\_34

m/z 481.6006

01234E+355.5RT (min)

lcqtof\_t3\_pos\_2\_050\_14

m/z 482.3284

012E+377.5RT (min)

lcqtof\_t3\_pos\_4\_044\_38

m/z 482.7370

024E+311.5RT (min)

lcqtof\_t3\_pos\_2\_022\_32

m/z 482.8403

01234E+311.5RT (min)

lcqtof\_t3\_pos\_2\_031\_01

m/z 482.9283

02468E+311.5RT (min)

lcqtof\_t3\_pos\_2\_015\_13

m/z 483.2676

0123E+34.55RT (min)

lcqtof\_t3\_pos\_2\_050\_14

m/z 483.3186

01234E+34.55RT (min)

lcqtof\_t3\_pos\_4\_029\_37

m/z 484.0935

024E+36.57RT (min)

lcqtof\_t3\_pos\_4\_029\_37

m/z 484.2026

01234E+344.5RT (min)

lcqtof\_t3\_pos\_4\_029\_37

m/z 484.3096

0123E+399.5RT (min)

lcqtof\_t3\_pos\_4\_081\_58

m/z 484.4569

012E+313.514RT (min)

lcqtof\_t3\_pos\_4\_111\_62

m/z 484.8037

02468E+311.5RT (min)

lcqtof\_t3\_pos\_2\_052\_36

m/z 485.3556

01E+44.55RT (min)

lcqtof\_t3\_pos\_4\_029\_37

m/z 486.0085

01234E+377.5RT (min)

lcqtof\_t3\_pos\_4\_081\_58

m/z 486.0965

012E+311.5RT (min)

lcqtof\_t3\_pos\_2\_052\_36

m/z 486.1329

01234E+311.5RT (min)

lcqtof\_t3\_pos\_2\_007\_07

m/z 486.3637

0123E+344.5RT (min)

lcqtof\_t3\_pos\_4\_029\_37

m/z 486.7939

02468E+311.5RT (min)

lcqtof\_t3\_pos\_2\_012\_35

m/z 487.3154

01234E+344.5RT (min)

lcqtof\_t3\_pos\_4\_029\_37

m/z 487.3206

01234E+35.56RT (min)

lcqtof\_t3\_pos\_4\_029\_37

m/z 487.3423

01234E+355.5RT (min)

lcqtof\_t3\_pos\_4\_029\_37

m/z 487.4176

02468E+31414.5RT (min)

lcqtof\_t3\_pos\_4\_042\_18

m/z 487.9514

0123E+311.5RT (min)

lcqtof\_t3\_pos\_4\_001\_49

m/z 489.2002

01234E+311.5RT (min)

lcqtof\_t3\_pos\_4\_029\_37

m/z 490.0632

01E+38.59RT (min)

lcqtof\_t3\_pos\_2\_018\_41

m/z 490.3206

01234E+355.5RT (min)

lcqtof\_t3\_pos\_4\_008\_15

m/z 491.0359

02468E+311.5RT (min)

lcqtof\_t3\_pos\_2\_027\_38

m/z 491.2599

024E+344.5RT (min)

lcqtof\_t3\_pos\_1\_053\_18

m/z 491.3001

0123E+355.5RT (min)

lcqtof\_t3\_pos\_2\_039\_23

m/z 491.3016

0123E+355.5RT (min)

lcqtof\_t3\_pos\_4\_029\_37

m/z 492.1033

012E+34.55RT (min)

lcqtof\_t3\_pos\_2\_011\_19

m/z 492.1891

01234E+344.5RT (min)

lcqtof\_t3\_pos\_4\_072\_57

m/z 493.1922

012E+344.5RT (min)

lcqtof\_t3\_pos\_4\_015\_16

m/z 494.1178

0246E+377.5RT (min)

lcqtof\_t3\_pos\_4\_042\_18

m/z 494.6495

02468E+311.5RT (min)

lcqtof\_t3\_pos\_4\_021\_01

m/z 494.8274

01E+317.518RT (min)

lcqtof\_t3\_pos\_2\_035\_44

m/z 494.9765

01234E+311.5RT (min)

lcqtof\_t3\_pos\_4\_035\_45

m/z 495.2842

024E+311.5RT (min)

lcqtof\_t3\_pos\_2\_022\_32

m/z 495.2980

0123E+355.5RT (min)

lcqtof\_t3\_pos\_4\_015\_16

m/z 496.8174

01E+411.5RT (min)

lcqtof\_t3\_pos\_2\_031\_01

m/z 496.9425

02468E+311.5RT (min)

lcqtof\_t3\_pos\_4\_021\_01

m/z 497.0550

02468E+311.5RT (min)

lcqtof\_t3\_pos\_2\_022\_32

m/z 497.2766

012E+313.514RT (min)

lcqtof\_t3\_pos\_4\_034\_23

m/z 497.5402

02468E+314.515RT (min)

lcqtof\_t3\_pos\_4\_014\_24

m/z 498.0792

01234E+311.5RT (min)

lcqtof\_t3\_pos\_4\_001\_49

m/z 498.1101

01234E+377.5RT (min)

lcqtof\_t3\_pos\_4\_040\_28

m/z 498.3679

01E+44.55RT (min)

lcqtof\_t3\_pos\_4\_029\_37

m/z 499.0885

02468E+366.5RT (min)

lcqtof\_t3\_pos\_4\_013\_29

m/z 499.3069

0246E+344.5RT (min)

lcqtof\_t3\_pos\_4\_029\_37

m/z 499.3709

024E+355.5RT (min)

lcqtof\_t3\_pos\_4\_029\_37

m/z 499.9091

012E+317.518RT (min)

lcqtof\_t3\_pos\_2\_011\_19

m/z 500.0698

012E+35.56RT (min)

lcqtof\_t3\_pos\_2\_020\_27

m/z 500.2878

0246E+344.5RT (min)

lcqtof\_t3\_pos\_4\_029\_37

m/z 500.9901

02468E+311.5RT (min)

lcqtof\_t3\_pos\_4\_029\_37

m/z 501.0576

012E+34.55RT (min)

lcqtof\_t3\_pos\_2\_020\_27

m/z 501.3140

0123E+37.58RT (min)

lcqtof\_t3\_pos\_4\_024\_20

m/z 502.1839

0246E+311.5RT (min)

lcqtof\_t3\_pos\_2\_037\_46

m/z 502.3390

0123E+39.510RT (min)

lcqtof\_t3\_pos\_4\_042\_18

m/z 502.3842

01234E+37.58RT (min)

lcqtof\_t3\_pos\_4\_021\_01

m/z 502.6413

024E+311.5RT (min)

lcqtof\_t3\_pos\_4\_007\_14

m/z 503.2972

01234E+344.5RT (min)

lcqtof\_t3\_pos\_2\_008\_31

m/z 504.2750

01234E+35.56RT (min)

lcqtof\_t3\_pos\_4\_107\_62

m/z 504.3214

0246E+366.5RT (min)

lcqtof\_t3\_pos\_4\_021\_01

m/z 505.1852

01234E+355.5RT (min)

lcqtof\_t3\_pos\_4\_001\_49

m/z 505.2630

01234E+311.5RT (min)

lcqtof\_t3\_pos\_4\_088\_59

m/z 505.3031

012E+34.55RT (min)

lcqtof\_t3\_pos\_2\_029\_06

m/z 506.1845

0246E+344.5RT (min)

lcqtof\_t3\_pos\_2\_008\_31

m/z 506.7592

024E+34.55RT (min)

lcqtof\_t3\_pos\_4\_021\_01

m/z 507.3148

02468E+355.5RT (min)

lcqtof\_t3\_pos\_4\_029\_37

m/z 507.3528

0246E+35.56RT (min)

lcqtof\_t3\_pos\_1\_010\_27

m/z 507.3686

024E+344.5RT (min)

lcqtof\_t3\_pos\_4\_037\_32

m/z 507.8890

024E+311.5RT (min)

lcqtof\_t3\_pos\_4\_008\_15

m/z 508.2165

0123E+311.5RT (min)

lcqtof\_t3\_pos\_2\_007\_07

m/z 508.3258

012E+355.5RT (min)

lcqtof\_t3\_pos\_4\_017\_51

m/z 508.8606

012E+317.518RT (min)

lcqtof\_t3\_pos\_2\_011\_19

m/z 508.8608

0246E+311.5RT (min)

lcqtof\_t3\_pos\_4\_007\_14

m/z 509.1942

024E+344.5RT (min)

lcqtof\_t3\_pos\_4\_015\_16

m/z 509.3087

0246E+311.5RT (min)

lcqtof\_t3\_pos\_4\_014\_24

m/z 509.7047

012E+310.511RT (min)

lcqtof\_t3\_pos\_4\_103\_61

m/z 509.9351

02468E+311.5RT (min)

lcqtof\_t3\_pos\_4\_001\_49

m/z 509.9936

0123E+37.58RT (min)

lcqtof\_t3\_pos\_3\_005\_18

m/z 510.3062

012E+39.510RT (min)

lcqtof\_t3\_pos\_4\_076\_58

m/z 510.6251

01234E+311.5RT (min)

lcqtof\_t3\_pos\_4\_007\_14

m/z 510.8141

012E+317.518RT (min)

lcqtof\_t3\_pos\_1\_007\_07

m/z 511.0505

01234E+34.55RT (min)

lcqtof\_t3\_pos\_2\_011\_19

m/z 511.1457

024E+36.57RT (min)

lcqtof\_t3\_pos\_2\_055\_18

m/z 511.3497

0123E+388.5RT (min)

lcqtof\_t3\_pos\_4\_072\_57

m/z 511.5565

0246E+314.515RT (min)

lcqtof\_t3\_pos\_4\_014\_24

m/z 512.1688

0123E+36.57RT (min)

lcqtof\_t3\_pos\_2\_011\_19

m/z 512.3317

0123E+355.5RT (min)

lcqtof\_t3\_pos\_4\_003\_05

m/z 512.7003

024E+311.5RT (min)

lcqtof\_t3\_pos\_4\_021\_01

m/z 513.1061

024E+35.56RT (min)

lcqtof\_t3\_pos\_2\_011\_19

m/z 513.3295

024E+355.5RT (min)

lcqtof\_t3\_pos\_4\_029\_37

m/z 513.3333

024E+344.5RT (min)

lcqtof\_t3\_pos\_4\_029\_37

m/z 514.0931

012E+34.55RT (min)

lcqtof\_t3\_pos\_2\_011\_19

m/z 514.3253

012E+36.57RT (min)

lcqtof\_t3\_pos\_4\_042\_18

m/z 514.6805

02468E+311.5RT (min)

lcqtof\_t3\_pos\_2\_022\_32

m/z 514.9866

012E+37.58RT (min)

lcqtof\_t3\_pos\_2\_055\_18

m/z 515.2410

0123E+36.57RT (min)

lcqtof\_t3\_pos\_1\_010\_27

m/z 515.3646

024E+36.57RT (min)

lcqtof\_t3\_pos\_4\_029\_37

m/z 516.0995

024E+377.5RT (min)

lcqtof\_t3\_pos\_4\_042\_18

m/z 516.2089

0123E+37.58RT (min)

lcqtof\_t3\_pos\_2\_010\_26

m/z 516.6741

0246E+311.5RT (min)

lcqtof\_t3\_pos\_2\_008\_31

m/z 517.2805

01234E+344.5RT (min)

lcqtof\_t3\_pos\_2\_008\_31

m/z 517.3091

0123E+355.5RT (min)

lcqtof\_t3\_pos\_4\_015\_16

m/z 517.8877

0123E+36.57RT (min)

lcqtof\_t3\_pos\_4\_010\_36

m/z 519.7097

012E+31111.5RT (min)

lcqtof\_t3\_pos\_4\_110\_62

m/z 520.0592

01234E+311.5RT (min)

lcqtof\_t3\_pos\_4\_001\_49

m/z 520.2756

0246E+388.5RT (min)

lcqtof\_t3\_pos\_4\_042\_18

m/z 520.6077

024E+311.5RT (min)

lcqtof\_t3\_pos\_4\_007\_14

m/z 520.8228

02468E+311.5RT (min)

lcqtof\_t3\_pos\_2\_050\_14

m/z 521.8489

012E+317.518RT (min)

lcqtof\_t3\_pos\_2\_020\_27

m/z 522.6031

024E+311.5RT (min)

lcqtof\_t3\_pos\_4\_007\_14

m/z 524.6049

01234E+311.5RT (min)

lcqtof\_t3\_pos\_4\_007\_14

m/z 524.7338

0246E+36.57RT (min)

lcqtof\_t3\_pos\_4\_029\_37

m/z 525.3381

0246E+311.512RT (min)

lcqtof\_t3\_pos\_2\_038\_21

m/z 526.3951

012E+34.55RT (min)

lcqtof\_t3\_pos\_3\_005\_18

m/z 528.0607

012E+34.55RT (min)

lcqtof\_t3\_pos\_2\_036\_20

m/z 528.1815

0246E+311.5RT (min)

lcqtof\_t3\_pos\_2\_041\_54

m/z 528.4973

012E+312.513RT (min)

lcqtof\_t3\_pos\_4\_113\_62

m/z 528.8497

024E+311.5RT (min)

lcqtof\_t3\_pos\_2\_022\_32

m/z 529.2372

0246E+344.5RT (min)

lcqtof\_t3\_pos\_4\_015\_16

m/z 529.3029

0123E+311.5RT (min)

lcqtof\_t3\_pos\_2\_052\_36

m/z 530.2756

012E+36.57RT (min)

lcqtof\_t3\_pos\_1\_010\_27

m/z 530.6535

024E+311.5RT (min)

lcqtof\_t3\_pos\_3\_007\_32

m/z 530.8527

012E+317.518RT (min)

lcqtof\_t3\_pos\_2\_046\_30

m/z 531.2914

02468E+317.518RT (min)

lcqtof\_t3\_pos\_4\_021\_01

m/z 531.8368

012E+311.5RT (min)

lcqtof\_t3\_pos\_2\_002\_37

m/z 531.8653

01E+317.518RT (min)

lcqtof\_t3\_pos\_2\_036\_20

m/z 531.9180

01234E+311.5RT (min)

lcqtof\_t3\_pos\_4\_001\_49

m/z 532.3950

024E+35.56RT (min)

lcqtof\_t3\_pos\_4\_029\_37

m/z 532.6493

01234E+311.5RT (min)

lcqtof\_t3\_pos\_4\_001\_49

m/z 534.2989

0123E+317.518RT (min)

lcqtof\_t3\_pos\_2\_053\_15

m/z 534.6458

012E+311.5RT (min)

lcqtof\_t3\_pos\_2\_030\_33

m/z 535.1658

012E+377.5RT (min)

lcqtof\_t3\_pos\_2\_005\_17

m/z 537.0358

024E+311.5RT (min)

lcqtof\_t3\_pos\_4\_037\_32

m/z 537.1946

024E+311.5RT (min)

lcqtof\_t3\_pos\_4\_015\_16

m/z 537.9922

01234E+311.5RT (min)

lcqtof\_t3\_pos\_4\_037\_32

m/z 538.1287

0246E+311.5RT (min)

lcqtof\_t3\_pos\_4\_034\_23

m/z 538.7051

01234E+311.5RT (min)

lcqtof\_t3\_pos\_2\_022\_32

m/z 538.7871

0123E+34.55RT (min)

lcqtof\_t3\_pos\_3\_081\_58

m/z 539.7557

02468E+344.5RT (min)

lcqtof\_t3\_pos\_4\_007\_14

m/z 540.1483

012E+37.58RT (min)

lcqtof\_t3\_pos\_2\_055\_18

m/z 540.6895

01234E+311.5RT (min)

lcqtof\_t3\_pos\_2\_002\_37

m/z 541.4007

01234E+366.5RT (min)

lcqtof\_t3\_pos\_4\_029\_37

m/z 541.7766

012E+317.518RT (min)

lcqtof\_t3\_pos\_2\_052\_36

m/z 542.2799

01E+317.518RT (min)

lcqtof\_t3\_pos\_2\_052\_36

m/z 542.3235

01E+417.518RT (min)

lcqtof\_t3\_pos\_4\_021\_01

m/z 542.3571

012E+34.55RT (min)

lcqtof\_t3\_pos\_2\_039\_23

m/z 542.9321

01E+317.518RT (min)

lcqtof\_t3\_pos\_2\_013\_29

m/z 543.0422

01234E+317.518RT (min)

lcqtof\_t3\_pos\_4\_021\_01

m/z 543.2359

02468E+36.57RT (min)

lcqtof\_t3\_pos\_4\_020\_19

m/z 543.2487

0246E+344.5RT (min)

lcqtof\_t3\_pos\_4\_005\_30

m/z 543.2914

024E+317.518RT (min)

lcqtof\_t3\_pos\_4\_021\_01

m/z 543.4976

02468E+314.515RT (min)

lcqtof\_t3\_pos\_4\_029\_37

m/z 544.6185

02468E+311.5RT (min)

lcqtof\_t3\_pos\_4\_021\_01

m/z 544.7464

024E+311.5RT (min)

lcqtof\_t3\_pos\_4\_007\_14

m/z 545.2576

0123E+36.57RT (min)

lcqtof\_t3\_pos\_3\_014\_38

m/z 545.9732

012E+37.58RT (min)

lcqtof\_t3\_pos\_4\_083\_59

m/z 546.2881

024E+344.5RT (min)

lcqtof\_t3\_pos\_4\_034\_23

m/z 547.2562

012E+366.5RT (min)

lcqtof\_t3\_pos\_3\_003\_14

m/z 547.3550

02468E+317.518RT (min)

lcqtof\_t3\_pos\_4\_021\_01

m/z 548.3455

0246E+366.5RT (min)

lcqtof\_t3\_pos\_4\_021\_01

m/z 548.3598

02468E+377.5RT (min)

lcqtof\_t3\_pos\_1\_001\_49

m/z 549.0639

01E+31111.5RT (min)

lcqtof\_t3\_pos\_4\_081\_58

m/z 549.1163

0123E+34.55RT (min)

lcqtof\_t3\_pos\_4\_020\_19

m/z 549.3395

01E+44.55RT (min)

lcqtof\_t3\_pos\_4\_029\_37

m/z 549.7875

0246E+317.518RT (min)

lcqtof\_t3\_pos\_4\_021\_01

m/z 550.2913

0123E+317.518RT (min)

lcqtof\_t3\_pos\_4\_021\_01

m/z 552.2267

01234E+344.5RT (min)

lcqtof\_t3\_pos\_4\_006\_41

m/z 552.7719

01E+317.518RT (min)

lcqtof\_t3\_pos\_3\_010\_01

m/z 554.2196

0123E+36.57RT (min)

lcqtof\_t3\_pos\_2\_011\_19

m/z 556.4293

02468E+366.5RT (min)

lcqtof\_t3\_pos\_4\_029\_37

m/z 557.3562

024E+317.518RT (min)

lcqtof\_t3\_pos\_2\_005\_17

m/z 558.2412

012E+366.5RT (min)

lcqtof\_t3\_pos\_2\_050\_14

m/z 558.3112

024E+37.58RT (min)

lcqtof\_t3\_pos\_4\_112\_62

m/z 558.3630

0123E+36.57RT (min)

lcqtof\_t3\_pos\_4\_021\_01

m/z 558.8671

0123E+317.518RT (min)

lcqtof\_t3\_pos\_4\_106\_62

m/z 559.0207

024E+311.5RT (min)

lcqtof\_t3\_pos\_2\_012\_35

m/z 560.0892

012E+34.55RT (min)

lcqtof\_t3\_pos\_2\_020\_27

m/z 561.4257

0123E+36.57RT (min)

lcqtof\_t3\_pos\_4\_003\_05

m/z 562.8175

01E+411.5RT (min)

lcqtof\_t3\_pos\_2\_012\_35

m/z 562.8181

01E+317.518RT (min)

lcqtof\_t3\_pos\_2\_006\_04

m/z 563.0453

012E+37.58RT (min)

lcqtof\_t3\_pos\_2\_055\_18

m/z 563.0624

012E+38.59RT (min)

lcqtof\_t3\_pos\_4\_089\_59

m/z 563.0963

0123E+377.5RT (min)

lcqtof\_t3\_pos\_2\_010\_26

m/z 563.4266

012E+39.510RT (min)

lcqtof\_t3\_pos\_4\_069\_57

m/z 563.8309

0123E+355.5RT (min)

lcqtof\_t3\_pos\_2\_050\_14

m/z 563.8516

0123E+311.5RT (min)

lcqtof\_t3\_pos\_4\_013\_29

m/z 565.0416

0246E+311.5RT (min)

lcqtof\_t3\_pos\_2\_022\_32

m/z 565.2598

012E+35.56RT (min)

lcqtof\_t3\_pos\_4\_029\_37

m/z 566.5466

0246E+35.56RT (min)

lcqtof\_t3\_pos\_2\_012\_35

m/z 567.4989

0246E+313.514RT (min)

lcqtof\_t3\_pos\_2\_008\_31

m/z 567.8972

0123E+317.518RT (min)

lcqtof\_t3\_pos\_2\_013\_29

m/z 568.5800

0246E+311.5RT (min)

lcqtof\_t3\_pos\_4\_021\_01

m/z 568.8762

01E+317.518RT (min)

lcqtof\_t3\_pos\_4\_006\_41

m/z 569.3929

012E+388.5RT (min)

lcqtof\_t3\_pos\_4\_068\_57

m/z 569.6152

02468E+31313.5RT (min)

lcqtof\_t3\_pos\_3\_093\_60

m/z 570.3184

0246E+344.5RT (min)

lcqtof\_t3\_pos\_4\_029\_37

m/z 570.3879

0123E+355.5RT (min)

lcqtof\_t3\_pos\_4\_029\_37

m/z 570.4212

01234E+355.5RT (min)

lcqtof\_t3\_pos\_4\_029\_37

m/z 570.4434

012E+35.56RT (min)

lcqtof\_t3\_pos\_4\_003\_05

m/z 570.9113

01E+46.57RT (min)

lcqtof\_t3\_pos\_4\_029\_37

m/z 572.6254

02468E+311.5RT (min)

lcqtof\_t3\_pos\_4\_021\_01

m/z 573.0417

0123E+34.55RT (min)

lcqtof\_t3\_pos\_2\_020\_27

m/z 573.8487

0123E+317.518RT (min)

lcqtof\_t3\_pos\_2\_012\_35

m/z 574.1866

0246E+311.5RT (min)

lcqtof\_t3\_pos\_1\_014\_14

m/z 574.6263

02468E+311.5RT (min)

lcqtof\_t3\_pos\_4\_029\_37

m/z 574.8326

02468E+311.5RT (min)

lcqtof\_t3\_pos\_2\_031\_01

m/z 575.4011

0246E+366.5RT (min)

lcqtof\_t3\_pos\_4\_029\_37

m/z 575.4132

02468E+314.515RT (min)

lcqtof\_t3\_pos\_4\_029\_37

m/z 575.4443

0246E+355.5RT (min)

lcqtof\_t3\_pos\_2\_002\_37

m/z 576.6287

0123E+311.5RT (min)

lcqtof\_t3\_pos\_4\_001\_49

m/z 576.8486

012E+317.518RT (min)

lcqtof\_t3\_pos\_2\_004\_42

m/z 578.5687

024E+311.5RT (min)

lcqtof\_t3\_pos\_4\_007\_14

m/z 578.8002

012E+317.518RT (min)

lcqtof\_t3\_pos\_1\_013\_20

m/z 579.0599

0123E+34.55RT (min)

lcqtof\_t3\_pos\_2\_011\_19

m/z 579.3350

02468E+34.55RT (min)

lcqtof\_t3\_pos\_4\_021\_01

m/z 580.2114

01E+411.5RT (min)

lcqtof\_t3\_pos\_2\_022\_32

m/z 580.5612

0246E+311.5RT (min)

lcqtof\_t3\_pos\_4\_007\_14

m/z 580.8641

02468E+311.5RT (min)

lcqtof\_t3\_pos\_2\_052\_36

m/z 581.0150

012E+311.5RT (min)

lcqtof\_t3\_pos\_4\_052\_06

m/z 581.9297

0123E+35.56RT (min)

lcqtof\_t3\_pos\_4\_003\_05

m/z 582.5582

01234E+311.5RT (min)

lcqtof\_t3\_pos\_4\_007\_14

m/z 583.4166

0246E+366.5RT (min)

lcqtof\_t3\_pos\_4\_029\_37

m/z 583.8525

012E+355.5RT (min)

lcqtof\_t3\_pos\_4\_112\_62

m/z 584.4597

0246E+34.55RT (min)

lcqtof\_t3\_pos\_4\_029\_37

m/z 584.8688

01E+317.518RT (min)

lcqtof\_t3\_pos\_2\_054\_09

m/z 585.0797

01234E+34.55RT (min)

lcqtof\_t3\_pos\_4\_107\_62

m/z 585.1252

0123E+37.58RT (min)

lcqtof\_t3\_pos\_2\_010\_26

m/z 585.2047

0246E+344.5RT (min)

lcqtof\_t3\_pos\_2\_010\_26

m/z 585.3391

0123E+377.5RT (min)

lcqtof\_t3\_pos\_4\_021\_01

m/z 585.8116

0123E+344.5RT (min)

lcqtof\_t3\_pos\_2\_050\_14

m/z 586.6749

0246E+313.514RT (min)

lcqtof\_t3\_pos\_4\_014\_24

m/z 587.2635

012E+34.55RT (min)

lcqtof\_t3\_pos\_3\_081\_58

m/z 588.3950

01234E+366.5RT (min)

lcqtof\_t3\_pos\_4\_029\_37

m/z 588.6126

024E+311.5RT (min)

lcqtof\_t3\_pos\_4\_001\_49

m/z 588.8189

01234E+317.518RT (min)

lcqtof\_t3\_pos\_2\_009\_50

m/z 588.9393

01234E+35.56RT (min)

lcqtof\_t3\_pos\_4\_029\_37

m/z 589.8342

012E+317.518RT (min)

lcqtof\_t3\_pos\_2\_016\_39

m/z 590.0054

02468E+311.5RT (min)

lcqtof\_t3\_pos\_4\_037\_32

m/z 590.6068

01234E+311.5RT (min)

lcqtof\_t3\_pos\_2\_012\_35

m/z 591.4195

01234E+355.5RT (min)

lcqtof\_t3\_pos\_4\_021\_01

m/z 592.3165

01234E+34.55RT (min)

lcqtof\_t3\_pos\_4\_021\_01

m/z 592.6089

0123E+311.5RT (min)

lcqtof\_t3\_pos\_2\_012\_35

m/z 593.6011

0123E+312.513RT (min)

lcqtof\_t3\_pos\_4\_082\_59

m/z 594.2039

024E+344.5RT (min)

lcqtof\_t3\_pos\_2\_013\_29

m/z 595.0297

0123E+37.58RT (min)

lcqtof\_t3\_pos\_4\_088\_59

m/z 595.1695

0246E+311.5RT (min)

lcqtof\_t3\_pos\_2\_050\_14

m/z 596.0251

012E+37.58RT (min)

lcqtof\_t3\_pos\_4\_087\_59

m/z 596.1527

0123E+311.5RT (min)

lcqtof\_t3\_pos\_3\_003\_14

m/z 596.2921

024E+355.5RT (min)

lcqtof\_t3\_pos\_4\_021\_01

m/z 596.6265

0123E+355.5RT (min)

lcqtof\_t3\_pos\_3\_015\_13

m/z 596.8397

0246E+311.5RT (min)

lcqtof\_t3\_pos\_2\_022\_32

m/z 597.2710

0123E+388.5RT (min)

lcqtof\_t3\_pos\_2\_055\_18

m/z 597.4358

01234E+355.5RT (min)

lcqtof\_t3\_pos\_4\_029\_37

m/z 598.6633

024E+311.5RT (min)

lcqtof\_t3\_pos\_2\_022\_32

m/z 598.8378

0123E+317.518RT (min)

lcqtof\_t3\_pos\_2\_011\_19

m/z 599.0070

012E+355.5RT (min)

lcqtof\_t3\_pos\_2\_055\_18

m/z 599.2947

0123E+311.5RT (min)

lcqtof\_t3\_pos\_2\_029\_06

m/z 599.3405

0123E+344.5RT (min)

lcqtof\_t3\_pos\_4\_046\_34

m/z 599.8423

01234E+311.5RT (min)

lcqtof\_t3\_pos\_2\_031\_01

m/z 599.8635

0123E+317.518RT (min)

lcqtof\_t3\_pos\_2\_011\_19

m/z 599.8813

01234E+317.518RT (min)

lcqtof\_t3\_pos\_4\_020\_19

m/z 600.3394

0246E+344.5RT (min)

lcqtof\_t3\_pos\_2\_008\_31

m/z 600.3925

0123E+344.5RT (min)

lcqtof\_t3\_pos\_2\_034\_24

m/z 600.4006

01234E+35.56RT (min)

lcqtof\_t3\_pos\_4\_029\_37

m/z 600.8774

0123E+317.518RT (min)

lcqtof\_t3\_pos\_2\_024\_45

m/z 601.3840

024E+311.5RT (min)

lcqtof\_t3\_pos\_2\_015\_13

m/z 602.1928

0246E+311.5RT (min)

lcqtof\_t3\_pos\_2\_022\_32

m/z 602.3821

012E+36.57RT (min)

lcqtof\_t3\_pos\_4\_005\_30

m/z 602.5792

0246E+311.5RT (min)

lcqtof\_t3\_pos\_4\_021\_01

m/z 603.6855

0246E+313.514RT (min)

lcqtof\_t3\_pos\_3\_092\_60

m/z 605.0316

012E+355.5RT (min)

lcqtof\_t3\_pos\_4\_105\_61

m/z 605.0710

0123E+34.55RT (min)

lcqtof\_t3\_pos\_2\_011\_19

m/z 607.1811

0123E+377.5RT (min)

lcqtof\_t3\_pos\_2\_020\_27

m/z 607.3355

02468E+317.518RT (min)

lcqtof\_t3\_pos\_4\_021\_01

m/z 607.8319

01234E+355.5RT (min)

lcqtof\_t3\_pos\_2\_050\_14

m/z 608.0481

024E+38.59RT (min)

lcqtof\_t3\_pos\_4\_089\_59

m/z 608.0876

02468E+311.5RT (min)

lcqtof\_t3\_pos\_4\_082\_59

m/z 608.8811

012E+317.518RT (min)

lcqtof\_t3\_pos\_2\_045\_10

m/z 609.3695

0123E+311.5RT (min)

lcqtof\_t3\_pos\_2\_010\_26

m/z 609.3850

01234E+37.58RT (min)

lcqtof\_t3\_pos\_4\_029\_37

m/z 610.0742

02468E+313.514RT (min)

lcqtof\_t3\_pos\_3\_105\_61

m/z 610.3890

024E+34.55RT (min)

lcqtof\_t3\_pos\_4\_021\_01

m/z 610.9163

01E+317.518RT (min)

lcqtof\_t3\_pos\_2\_038\_21

m/z 611.0521

012E+317.518RT (min)

lcqtof\_t3\_pos\_1\_001\_49

m/z 611.0882

0123E+34.55RT (min)

lcqtof\_t3\_pos\_2\_011\_19

m/z 611.4464

01234E+355.5RT (min)

lcqtof\_t3\_pos\_4\_029\_37

m/z 612.4168

012E+399.5RT (min)

lcqtof\_t3\_pos\_4\_084\_59

m/z 612.8829

01E+411.5RT (min)

lcqtof\_t3\_pos\_4\_021\_01

m/z 613.2693

02468E+34.55RT (min)

lcqtof\_t3\_pos\_4\_015\_16

m/z 613.3681

012E+399.5RT (min)

lcqtof\_t3\_pos\_4\_086\_59

m/z 615.3436

01234E+35.56RT (min)

lcqtof\_t3\_pos\_4\_113\_62

m/z 616.1442

02468E+311.5RT (min)

lcqtof\_t3\_pos\_2\_010\_26

m/z 616.1634

0246E+311.5RT (min)

lcqtof\_t3\_pos\_4\_034\_23

m/z 616.8582

0246E+311.5RT (min)

lcqtof\_t3\_pos\_2\_012\_35

m/z 617.2630

02468E+344.5RT (min)

lcqtof\_t3\_pos\_2\_050\_14

m/z 617.3215

024E+317.518RT (min)

lcqtof\_t3\_pos\_4\_021\_01

m/z 617.8578

01E+317.518RT (min)

lcqtof\_t3\_pos\_2\_016\_39

m/z 618.4639

02468E+314.515RT (min)

lcqtof\_t3\_pos\_4\_029\_37

m/z 618.9032

02468E+311.5RT (min)

lcqtof\_t3\_pos\_2\_015\_13

m/z 620.2413

024E+344.5RT (min)

lcqtof\_t3\_pos\_4\_015\_16

m/z 620.2512

0246E+366.5RT (min)

lcqtof\_t3\_pos\_4\_007\_14

m/z 621.4421

012E+39.510RT (min)

lcqtof\_t3\_pos\_4\_077\_58

m/z 621.8895

0123E+317.518RT (min)

lcqtof\_t3\_pos\_4\_006\_41

m/z 622.2516

0123E+317.518RT (min)

lcqtof\_t3\_pos\_4\_106\_62

m/z 623.2293

0246E+344.5RT (min)

lcqtof\_t3\_pos\_4\_042\_18

m/z 623.3813

0123E+36.57RT (min)

lcqtof\_t3\_pos\_4\_003\_05

m/z 624.8488

0246E+317.518RT (min)

lcqtof\_t3\_pos\_2\_012\_35

m/z 625.2056

024E+377.5RT (min)

lcqtof\_t3\_pos\_4\_113\_62

m/z 625.2311

0123E+344.5RT (min)

lcqtof\_t3\_pos\_2\_010\_26

m/z 625.8488

01E+411.5RT (min)

lcqtof\_t3\_pos\_2\_015\_13

m/z 626.5354

02468E+311.5RT (min)

lcqtof\_t3\_pos\_4\_021\_01

m/z 626.8504

0123E+317.518RT (min)

lcqtof\_t3\_pos\_2\_005\_17

m/z 626.9489

01234E+36.57RT (min)

lcqtof\_t3\_pos\_4\_029\_37

m/z 627.3841

012E+35.56RT (min)

lcqtof\_t3\_pos\_4\_073\_57

m/z 627.5368

02468E+314.515RT (min)

lcqtof\_t3\_pos\_2\_003\_16

m/z 627.9539

024E+36.57RT (min)

lcqtof\_t3\_pos\_4\_029\_37

m/z 628.3181

02468E+344.5RT (min)

lcqtof\_t3\_pos\_4\_021\_01

m/z 628.8589

0123E+411.5RT (min)

lcqtof\_t3\_pos\_4\_021\_01

m/z 629.8636

01E+411.5RT (min)

lcqtof\_t3\_pos\_2\_015\_13

m/z 630.0694

0246E+311.5RT (min)

lcqtof\_t3\_pos\_4\_085\_59

m/z 630.2199

0246E+311.5RT (min)

lcqtof\_t3\_pos\_4\_014\_24

m/z 630.5953

0246E+311.5RT (min)

lcqtof\_t3\_pos\_4\_021\_01

m/z 630.7926

01E+411.5RT (min)

lcqtof\_t3\_pos\_2\_050\_14

m/z 630.7991

01E+317.518RT (min)

lcqtof\_t3\_pos\_2\_053\_15

m/z 631.0186

01234E+366.5RT (min)

lcqtof\_t3\_pos\_4\_021\_01

m/z 631.0511

01E+38.59RT (min)

lcqtof\_t3\_pos\_4\_088\_59

m/z 631.3396

0123E+366.5RT (min)

lcqtof\_t3\_pos\_4\_021\_01

m/z 632.5922

0246E+311.5RT (min)

lcqtof\_t3\_pos\_4\_029\_37

m/z 633.0268

012E+311.5RT (min)

lcqtof\_t3\_pos\_2\_022\_32

m/z 634.5681

024E+311.5RT (min)

lcqtof\_t3\_pos\_4\_021\_01

m/z 634.8599

0123E+366.5RT (min)

lcqtof\_t3\_pos\_4\_007\_14

m/z 636.3952

0246E+366.5RT (min)

lcqtof\_t3\_pos\_4\_021\_01

m/z 637.1414

02468E+311.5RT (min)

lcqtof\_t3\_pos\_4\_029\_37

m/z 637.2553

0246E+34.55RT (min)

lcqtof\_t3\_pos\_4\_015\_16

m/z 637.8497

01E+411.5RT (min)

lcqtof\_t3\_pos\_2\_031\_01

m/z 637.8505

012E+317.518RT (min)

lcqtof\_t3\_pos\_2\_005\_17

m/z 638.1287

012E+377.5RT (min)

lcqtof\_t3\_pos\_2\_021\_22

m/z 638.5194

024E+311.5RT (min)

lcqtof\_t3\_pos\_4\_007\_14

m/z 639.2246

01234E+34.55RT (min)

lcqtof\_t3\_pos\_4\_015\_16

m/z 639.9591

01234E+366.5RT (min)

lcqtof\_t3\_pos\_4\_029\_37

m/z 640.3763

012E+38.59RT (min)

lcqtof\_t3\_pos\_4\_079\_58

m/z 640.5201

01234E+311.5RT (min)

lcqtof\_t3\_pos\_4\_007\_14

m/z 641.0346

012E+34.55RT (min)

lcqtof\_t3\_pos\_2\_027\_38

m/z 641.2466

024E+34.55RT (min)

lcqtof\_t3\_pos\_4\_015\_16

m/z 641.8369

0123E+317.518RT (min)

lcqtof\_t3\_pos\_2\_006\_04

m/z 642.8511

012E+317.518RT (min)

lcqtof\_t3\_pos\_2\_019\_08

m/z 643.3606

0246E+311.5RT (min)

lcqtof\_t3\_pos\_2\_005\_17

m/z 643.3837

01E+317.518RT (min)

lcqtof\_t3\_pos\_2\_016\_39

m/z 643.4000

01234E+317.518RT (min)

lcqtof\_t3\_pos\_4\_028\_22

m/z 643.8677

0123E+311.5RT (min)

lcqtof\_t3\_pos\_2\_010\_26

m/z 644.3306

012E+37.58RT (min)

lcqtof\_t3\_pos\_2\_011\_19

m/z 644.8386

012E+317.518RT (min)

lcqtof\_t3\_pos\_2\_004\_42

m/z 645.2877

024E+34.55RT (min)

lcqtof\_t3\_pos\_4\_015\_16

m/z 645.8349

01234E+311.5RT (min)

lcqtof\_t3\_pos\_3\_010\_01

m/z 645.8392

012E+317.518RT (min)

lcqtof\_t3\_pos\_2\_005\_17

m/z 646.1432

012E+344.5RT (min)

lcqtof\_t3\_pos\_4\_034\_23

m/z 646.3466

01E+317.518RT (min)

lcqtof\_t3\_pos\_2\_025\_52

m/z 646.5716

0123E+311.5RT (min)

lcqtof\_t3\_pos\_4\_029\_37

m/z 646.7847

012E+317.518RT (min)

lcqtof\_t3\_pos\_1\_013\_20

m/z 647.0443

012E+34.55RT (min)

lcqtof\_t3\_pos\_2\_020\_27

m/z 648.5640

0123E+311.5RT (min)

lcqtof\_t3\_pos\_4\_001\_49

m/z 648.8510

0246E+311.5RT (min)

lcqtof\_t3\_pos\_2\_002\_37

m/z 649.3928

01E+35.56RT (min)

lcqtof\_t3\_pos\_4\_072\_57

m/z 651.1155

024E+311.5RT (min)

lcqtof\_t3\_pos\_2\_039\_23

m/z 651.2262

024E+34.55RT (min)

lcqtof\_t3\_pos\_4\_015\_16

m/z 651.3548

0123E+311.5RT (min)

lcqtof\_t3\_pos\_2\_050\_14

m/z 652.0569

0246E+311.5RT (min)

lcqtof\_t3\_pos\_2\_028\_34

m/z 652.8518

012E+317.518RT (min)

lcqtof\_t3\_pos\_2\_004\_42

m/z 653.2611

024E+34.55RT (min)

lcqtof\_t3\_pos\_4\_015\_16

m/z 654.2037

0246E+311.5RT (min)

lcqtof\_t3\_pos\_4\_037\_32

m/z 654.3087

01234E+34.55RT (min)

lcqtof\_t3\_pos\_4\_029\_37

m/z 654.4215

0123E+34.55RT (min)

lcqtof\_t3\_pos\_4\_021\_01

m/z 654.6351

024E+314.515RT (min)

lcqtof\_t3\_pos\_4\_021\_01

m/z 655.3989

012E+38.59RT (min)

lcqtof\_t3\_pos\_4\_087\_59

m/z 655.4448

0246E+31414.5RT (min)

lcqtof\_t3\_pos\_2\_039\_23

m/z 656.4099

012E+388.5RT (min)

lcqtof\_t3\_pos\_4\_090\_60

m/z 656.6173

0123E+311.5RT (min)

lcqtof\_t3\_pos\_2\_022\_32

m/z 656.8065

02468E+311.5RT (min)

lcqtof\_t3\_pos\_4\_001\_49

m/z 656.8331

01E+411.5RT (min)

lcqtof\_t3\_pos\_2\_022\_32

m/z 657.1266

0246E+311.5RT (min)

lcqtof\_t3\_pos\_4\_029\_37

m/z 657.8192

012E+317.518RT (min)

lcqtof\_t3\_pos\_2\_006\_04

m/z 657.9932

0246E+311.5RT (min)

lcqtof\_t3\_pos\_4\_037\_32

m/z 658.8320

02468E+311.5RT (min)

lcqtof\_t3\_pos\_2\_003\_16

m/z 660.0832

01234E+311.5RT (min)

lcqtof\_t3\_pos\_1\_014\_14

m/z 663.8544

012E+411.5RT (min)

lcqtof\_t3\_pos\_2\_015\_13

m/z 664.8307

0246E+311.5RT (min)

lcqtof\_t3\_pos\_2\_012\_35

m/z 665.4208

012E+317.518RT (min)

lcqtof\_t3\_pos\_1\_004\_05

m/z 666.3906

024E+34.55RT (min)

lcqtof\_t3\_pos\_4\_021\_01

m/z 666.4211

012E+355.5RT (min)

lcqtof\_t3\_pos\_4\_029\_37

m/z 666.9886

0123E+311.5RT (min)

lcqtof\_t3\_pos\_4\_010\_36

m/z 667.1223

01234E+344.5RT (min)

lcqtof\_t3\_pos\_2\_012\_35

m/z 667.1262

024E+311.5RT (min)

lcqtof\_t3\_pos\_4\_029\_37

m/z 667.1523

024E+355.5RT (min)

lcqtof\_t3\_pos\_4\_042\_18

m/z 667.8282

024E+311.5RT (min)

lcqtof\_t3\_pos\_2\_031\_01

m/z 667.8375

012E+317.518RT (min)

lcqtof\_t3\_pos\_2\_013\_29

m/z 668.5184

024E+366.5RT (min)

lcqtof\_t3\_pos\_4\_029\_37

m/z 669.2728

0246E+311.5RT (min)

lcqtof\_t3\_pos\_2\_007\_07

m/z 669.3755

02468E+34.55RT (min)

lcqtof\_t3\_pos\_4\_021\_01

m/z 669.8736

0246E+311.5RT (min)

lcqtof\_t3\_pos\_2\_015\_13

m/z 670.2745

0246E+311.5RT (min)

lcqtof\_t3\_pos\_2\_041\_54

m/z 671.8429

01E+411.5RT (min)

lcqtof\_t3\_pos\_2\_031\_01

m/z 671.8511

012E+317.518RT (min)

lcqtof\_t3\_pos\_2\_004\_42

m/z 672.7784

012E+411.5RT (min)

lcqtof\_t3\_pos\_4\_047\_13

m/z 673.3995

01234E+377.5RT (min)

lcqtof\_t3\_pos\_2\_011\_19

m/z 674.0129

0123E+36.57RT (min)

lcqtof\_t3\_pos\_2\_012\_35

m/z 674.2864

024E+311.5RT (min)

lcqtof\_t3\_pos\_2\_007\_07

m/z 674.7747

0123E+411.5RT (min)

lcqtof\_t3\_pos\_4\_047\_13

m/z 675.2850

024E+311.5RT (min)

lcqtof\_t3\_pos\_2\_007\_07

m/z 675.3610

01234E+344.5RT (min)

lcqtof\_t3\_pos\_4\_029\_37

m/z 676.6741

0123E+35.56RT (min)

lcqtof\_t3\_pos\_4\_021\_01

m/z 676.8575

0123E+317.518RT (min)

lcqtof\_t3\_pos\_2\_021\_22

m/z 677.0104

01234E+35.56RT (min)

lcqtof\_t3\_pos\_4\_021\_01

m/z 677.3674

012E+317.518RT (min)

lcqtof\_t3\_pos\_2\_013\_29

m/z 677.8538

0246E+311.5RT (min)

lcqtof\_t3\_pos\_2\_005\_17

m/z 678.3704

024E+34.55RT (min)

lcqtof\_t3\_pos\_4\_021\_01

m/z 678.5233

02468E+311.5RT (min)

lcqtof\_t3\_pos\_4\_021\_01

m/z 678.6537

02468E+366.5RT (min)

lcqtof\_t3\_pos\_4\_021\_01

m/z 678.8915

01E+317.518RT (min)

lcqtof\_t3\_pos\_2\_004\_42

m/z 679.8307

0246E+311.5RT (min)

lcqtof\_t3\_pos\_2\_050\_14

m/z 679.8347

01E+317.518RT (min)

lcqtof\_t3\_pos\_2\_035\_44

m/z 680.3328

01E+317.518RT (min)

lcqtof\_t3\_pos\_3\_008\_42

m/z 680.4103

012E+355.5RT (min)

lcqtof\_t3\_pos\_4\_017\_51

m/z 680.4276

01234E+366.5RT (min)

lcqtof\_t3\_pos\_4\_021\_01

m/z 681.3836

0123E+34.55RT (min)

lcqtof\_t3\_pos\_2\_050\_14

m/z 681.8851

012E+34.55RT (min)

lcqtof\_t3\_pos\_2\_050\_14

m/z 682.0690

024E+311.5RT (min)

lcqtof\_t3\_pos\_1\_014\_14

m/z 682.2396

01234E+34.55RT (min)

lcqtof\_t3\_pos\_4\_015\_16

m/z 682.3294

0246E+34.55RT (min)

lcqtof\_t3\_pos\_4\_029\_37

m/z 682.4968

024E+31414.5RT (min)

lcqtof\_t3\_pos\_2\_039\_23

m/z 684.0336

01234E+311.5RT (min)

lcqtof\_t3\_pos\_4\_045\_04

m/z 684.4985

0246E+311.5RT (min)

lcqtof\_t3\_pos\_4\_021\_01

m/z 684.8431

012E+317.518RT (min)

lcqtof\_t3\_pos\_2\_007\_07

m/z 685.3455

024E+311.5RT (min)

lcqtof\_t3\_pos\_2\_050\_14

m/z 685.3459

01E+317.518RT (min)

lcqtof\_t3\_pos\_2\_019\_08

m/z 685.8328

024E+311.5RT (min)

lcqtof\_t3\_pos\_2\_012\_35

m/z 686.2234

0123E+344.5RT (min)

lcqtof\_t3\_pos\_2\_054\_09

m/z 686.8934

02468E+311.5RT (min)

lcqtof\_t3\_pos\_2\_015\_13

m/z 687.8168

01234E+311.5RT (min)

lcqtof\_t3\_pos\_2\_050\_14

m/z 688.2953

01234E+34.55RT (min)

lcqtof\_t3\_pos\_4\_015\_16

m/z 689.0859

01234E+311.5RT (min)

lcqtof\_t3\_pos\_2\_050\_14

m/z 689.4734

024E+36.57RT (min)

lcqtof\_t3\_pos\_4\_029\_37

m/z 690.5374

0246E+311.5RT (min)

lcqtof\_t3\_pos\_4\_021\_01

m/z 690.9935

0123E+317.518RT (min)

lcqtof\_t3\_pos\_4\_110\_62

m/z 693.3378

012E+311.5RT (min)

lcqtof\_t3\_pos\_2\_031\_01

m/z 694.2617

024E+34.55RT (min)

lcqtof\_t3\_pos\_4\_015\_16

m/z 694.8394

0123E+317.518RT (min)

lcqtof\_t3\_pos\_2\_013\_29

m/z 695.1863

012E+344.5RT (min)

lcqtof\_t3\_pos\_4\_008\_15

m/z 695.2937

0123E+34.55RT (min)

lcqtof\_t3\_pos\_4\_107\_62

m/z 695.5416

012E+317.518RT (min)

lcqtof\_t3\_pos\_2\_007\_07

m/z 695.8407

024E+315.516RT (min)

lcqtof\_t3\_pos\_2\_050\_14

m/z 696.8142

024E+311.5RT (min)

lcqtof\_t3\_pos\_4\_001\_49

m/z 697.8494

0123E+411.5RT (min)

lcqtof\_t3\_pos\_2\_015\_13

m/z 697.8522

01E+317.518RT (min)

lcqtof\_t3\_pos\_2\_013\_29

m/z 698.0308

012E+311.5RT (min)

lcqtof\_t3\_pos\_1\_014\_14

m/z 698.3491

01234E+35.56RT (min)

lcqtof\_t3\_pos\_4\_021\_01

m/z 698.6604

02468E+312.513RT (min)

lcqtof\_t3\_pos\_2\_005\_17

m/z 698.7809

01E+411.5RT (min)

lcqtof\_t3\_pos\_3\_003\_14

m/z 698.7856

01234E+35.56RT (min)

lcqtof\_t3\_pos\_4\_021\_01

m/z 699.8525

02468E+311.5RT (min)

lcqtof\_t3\_pos\_4\_021\_01

m/z 700.0369

012E+388.5RT (min)

lcqtof\_t3\_pos\_4\_080\_58

m/z 700.3378

024E+344.5RT (min)

lcqtof\_t3\_pos\_4\_007\_14

m/z 700.3380

0123E+311.5RT (min)

lcqtof\_t3\_pos\_4\_007\_14

m/z 700.4456

012E+388.5RT (min)

lcqtof\_t3\_pos\_4\_084\_59

m/z 700.7954

01E+411.5RT (min)

lcqtof\_t3\_pos\_3\_010\_01

m/z 701.2687

01234E+35.56RT (min)

lcqtof\_t3\_pos\_4\_029\_37

m/z 702.6463

01E+411.5RT (min)

lcqtof\_t3\_pos\_4\_021\_01

m/z 702.8072

024E+311.5RT (min)

lcqtof\_t3\_pos\_4\_021\_01

m/z 703.8707

0123E+317.518RT (min)

lcqtof\_t3\_pos\_2\_005\_17

m/z 703.8713

01234E+317.518RT (min)

lcqtof\_t3\_pos\_2\_007\_07

m/z 704.0430

01234E+311.5RT (min)

lcqtof\_t3\_pos\_4\_007\_14

m/z 705.8359

01E+411.5RT (min)

lcqtof\_t3\_pos\_2\_031\_01

m/z 705.8375

012E+317.518RT (min)

lcqtof\_t3\_pos\_2\_017\_51

m/z 706.2275

0123E+344.5RT (min)

lcqtof\_t3\_pos\_4\_007\_14

m/z 706.3465

01E+317.518RT (min)

lcqtof\_t3\_pos\_2\_038\_21

m/z 706.5214

0123E+311.5RT (min)

lcqtof\_t3\_pos\_4\_029\_37

m/z 709.1826

024E+344.5RT (min)

lcqtof\_t3\_pos\_2\_022\_32

m/z 709.8251

0123E+317.518RT (min)

lcqtof\_t3\_pos\_2\_016\_39

m/z 710.7951

024E+31414.5RT (min)

lcqtof\_t3\_pos\_2\_022\_32

m/z 710.8495

01234E+317.518RT (min)

lcqtof\_t3\_pos\_2\_005\_17

m/z 711.3596

012E+317.518RT (min)

lcqtof\_t3\_pos\_2\_020\_27

m/z 711.5080

0123E+355.5RT (min)

lcqtof\_t3\_pos\_4\_003\_05

m/z 711.6844

01E+41313.5RT (min)

lcqtof\_t3\_pos\_3\_093\_60

m/z 711.8496

02468E+311.5RT (min)

lcqtof\_t3\_pos\_2\_010\_26

m/z 712.8191

012E+317.518RT (min)

lcqtof\_t3\_pos\_2\_021\_22

m/z 712.9926

0246E+311.5RT (min)

lcqtof\_t3\_pos\_4\_019\_03

m/z 713.8158

0123E+317.518RT (min)

lcqtof\_t3\_pos\_2\_004\_42

m/z 713.8235

02468E+311.5RT (min)

lcqtof\_t3\_pos\_2\_050\_14

m/z 714.2157

01234E+313.514RT (min)

lcqtof\_t3\_pos\_3\_081\_58

m/z 714.3449

01E+317.518RT (min)

lcqtof\_t3\_pos\_2\_026\_40

m/z 714.7731

012E+317.518RT (min)

lcqtof\_t3\_pos\_2\_033\_53

m/z 716.8401

0246E+311.5RT (min)

lcqtof\_t3\_pos\_2\_027\_38

m/z 717.1658

0246E+355.5RT (min)

lcqtof\_t3\_pos\_2\_055\_18

m/z 718.8391

01E+417.518RT (min)

lcqtof\_t3\_pos\_2\_011\_19

m/z 719.3387

0246E+311.5RT (min)

lcqtof\_t3\_pos\_2\_050\_14

m/z 719.3423

01E+317.518RT (min)

lcqtof\_t3\_pos\_2\_036\_20

m/z 720.4927

024E+311.5RT (min)

lcqtof\_t3\_pos\_4\_021\_01

m/z 720.8521

012E+317.518RT (min)

lcqtof\_t3\_pos\_2\_025\_52

m/z 721.4763

012E+38.59RT (min)

lcqtof\_t3\_pos\_4\_100\_61

m/z 721.8124

024E+311.5RT (min)

lcqtof\_t3\_pos\_3\_010\_01

m/z 722.3993

01234E+355.5RT (min)

lcqtof\_t3\_pos\_2\_050\_14

m/z 723.3686

01E+417.518RT (min)

lcqtof\_t3\_pos\_4\_021\_01

m/z 724.2462

01234E+317.518RT (min)

lcqtof\_t3\_pos\_4\_011\_10

m/z 724.3663

0246E+35.56RT (min)

lcqtof\_t3\_pos\_4\_021\_01

m/z 724.3747

024E+317.518RT (min)

lcqtof\_t3\_pos\_4\_021\_01

m/z 724.7817

01E+411.5RT (min)

lcqtof\_t3\_pos\_2\_050\_14

m/z 724.8185

01E+411.5RT (min)

lcqtof\_t3\_pos\_2\_022\_32

m/z 725.2546

0123E+317.518RT (min)

lcqtof\_t3\_pos\_4\_022\_25

m/z 725.2560

0123E+34.55RT (min)

lcqtof\_t3\_pos\_4\_072\_57

m/z 725.2755

0123E+34.55RT (min)

lcqtof\_t3\_pos\_2\_011\_19

m/z 725.4989

01234E+31313.5RT (min)

lcqtof\_t3\_pos\_2\_047\_02

m/z 725.5359

01234E+317.518RT (min)

lcqtof\_t3\_pos\_4\_021\_01

m/z 725.8160

012E+317.518RT (min)

lcqtof\_t3\_pos\_2\_047\_02

m/z 725.8270

0246E+312.513RT (min)

lcqtof\_t3\_pos\_2\_040\_28

m/z 726.0276

012E+317.518RT (min)

lcqtof\_t3\_pos\_4\_018\_42

m/z 726.0334

024E+311.5RT (min)

lcqtof\_t3\_pos\_4\_007\_14

m/z 726.4818

0246E+311.5RT (min)

lcqtof\_t3\_pos\_4\_021\_01

m/z 726.4899

01234E+313.514RT (min)

lcqtof\_t3\_pos\_2\_039\_23

m/z 727.2729

0123E+34.55RT (min)

lcqtof\_t3\_pos\_2\_020\_27

m/z 727.3264

0123E+311.5RT (min)

lcqtof\_t3\_pos\_2\_050\_14

m/z 727.8259

024E+311.5RT (min)

lcqtof\_t3\_pos\_2\_031\_01

m/z 728.4369

01E+38.59RT (min)

lcqtof\_t3\_pos\_4\_076\_58

m/z 730.3782

024E+344.5RT (min)

lcqtof\_t3\_pos\_4\_021\_01

m/z 731.4544

012E+355.5RT (min)

lcqtof\_t3\_pos\_4\_003\_05

m/z 731.8420

0123E+411.5RT (min)

lcqtof\_t3\_pos\_2\_015\_13

m/z 732.8115

02468E+311.5RT (min)

lcqtof\_t3\_pos\_4\_001\_49

m/z 733.3315

0246E+355.5RT (min)

lcqtof\_t3\_pos\_4\_021\_01

m/z 733.3759

024E+311.512RT (min)

lcqtof\_t3\_pos\_1\_014\_14

m/z 733.8462

02468E+311.5RT (min)

lcqtof\_t3\_pos\_4\_021\_01

m/z 734.9756

0123E+311.5RT (min)

lcqtof\_t3\_pos\_4\_019\_03

m/z 734.9785

012E+311.5RT (min)

lcqtof\_t3\_pos\_4\_010\_36

m/z 735.8125

0246E+311.5RT (min)

lcqtof\_t3\_pos\_2\_031\_01

m/z 735.8219

012E+317.518RT (min)

lcqtof\_t3\_pos\_2\_018\_41

m/z 736.3106

024E+344.5RT (min)

lcqtof\_t3\_pos\_4\_015\_16

m/z 737.2935

012E+344.5RT (min)

lcqtof\_t3\_pos\_4\_074\_58

m/z 738.4727

0246E+311.5RT (min)

lcqtof\_t3\_pos\_4\_021\_01

m/z 738.6259

02468E+313.514RT (min)

lcqtof\_t3\_pos\_4\_042\_18

m/z 739.8295

01E+411.5RT (min)

lcqtof\_t3\_pos\_3\_010\_01

m/z 739.8326

012E+317.518RT (min)

lcqtof\_t3\_pos\_2\_025\_52

m/z 740.4628

01234E+317.518RT (min)

lcqtof\_t3\_pos\_4\_090\_60

m/z 740.4641

0246E+311.5RT (min)

lcqtof\_t3\_pos\_4\_021\_01

m/z 740.7657

0123E+411.5RT (min)

lcqtof\_t3\_pos\_4\_047\_13

m/z 741.4630

024E+34.55RT (min)

lcqtof\_t3\_pos\_4\_021\_01

m/z 742.3527

01234E+34.55RT (min)

lcqtof\_t3\_pos\_4\_007\_14

m/z 742.7804

024E+31414.5RT (min)

lcqtof\_t3\_pos\_4\_014\_24

m/z 743.4422

0123E+388.5RT (min)

lcqtof\_t3\_pos\_4\_015\_16

m/z 745.3503

012E+317.518RT (min)

lcqtof\_t3\_pos\_2\_021\_22

m/z 745.8462

02468E+311.5RT (min)

lcqtof\_t3\_pos\_2\_013\_29

m/z 746.5046

024E+377.5RT (min)

lcqtof\_t3\_pos\_4\_021\_01

m/z 746.8954

01E+317.518RT (min)

lcqtof\_t3\_pos\_2\_013\_29

m/z 747.2599

012E+317.518RT (min)

lcqtof\_t3\_pos\_4\_020\_19

m/z 747.8175

02468E+311.5RT (min)

lcqtof\_t3\_pos\_2\_050\_14

m/z 747.8237

012E+317.518RT (min)

lcqtof\_t3\_pos\_2\_014\_03

m/z 748.3270

01E+317.518RT (min)

lcqtof\_t3\_pos\_2\_003\_16

m/z 748.4985

024E+311.5RT (min)

lcqtof\_t3\_pos\_4\_021\_01

m/z 748.8522

012E+317.518RT (min)

lcqtof\_t3\_pos\_3\_020\_12

m/z 750.5179

0123E+313.514RT (min)

lcqtof\_t3\_pos\_1\_007\_07

m/z 751.2466

01234E+31414.5RT (min)

lcqtof\_t3\_pos\_4\_041\_54

m/z 752.7152

012E+311.5RT (min)

lcqtof\_t3\_pos\_3\_010\_01

m/z 752.8217

0123E+317.518RT (min)

lcqtof\_t3\_pos\_2\_013\_29

m/z 753.3350

01E+317.518RT (min)

lcqtof\_t3\_pos\_2\_011\_19

m/z 753.3356

0246E+311.5RT (min)

lcqtof\_t3\_pos\_2\_050\_14

m/z 753.5343

0123E+315.516RT (min)

lcqtof\_t3\_pos\_2\_050\_14

m/z 753.6591

0246E+31515.5RT (min)

lcqtof\_t3\_pos\_3\_077\_58

m/z 753.6694

012E+311.5RT (min)

lcqtof\_t3\_pos\_2\_031\_01

m/z 753.8314

0246E+311.5RT (min)

lcqtof\_t3\_pos\_3\_003\_14

m/z 753.8441

01E+317.518RT (min)

lcqtof\_t3\_pos\_2\_044\_25

m/z 754.7083

01E+311.5RT (min)

lcqtof\_t3\_pos\_2\_015\_13

m/z 754.8774

0246E+311.5RT (min)

lcqtof\_t3\_pos\_2\_015\_13

m/z 755.5759

01234E+313.514RT (min)

lcqtof\_t3\_pos\_2\_050\_14

m/z 755.6720

024E+311.5RT (min)

lcqtof\_t3\_pos\_2\_031\_01

m/z 756.2158

01234E+34.55RT (min)

lcqtof\_t3\_pos\_2\_027\_38

m/z 756.3270

01E+317.518RT (min)

lcqtof\_t3\_pos\_2\_017\_51

m/z 756.9595

0123E+311.5RT (min)

lcqtof\_t3\_pos\_4\_001\_49

m/z 757.6691

0246E+311.5RT (min)

lcqtof\_t3\_pos\_2\_015\_13

m/z 758.6235

0123E+311.5RT (min)

lcqtof\_t3\_pos\_2\_031\_01

m/z 758.8185

01E+411.5RT (min)

lcqtof\_t3\_pos\_4\_009\_50

m/z 758.8259

012E+317.518RT (min)

lcqtof\_t3\_pos\_2\_042\_12

m/z 759.6624

01234E+311.5RT (min)

lcqtof\_t3\_pos\_2\_031\_01

m/z 760.6257

02468E+311.5RT (min)

lcqtof\_t3\_pos\_2\_015\_13

m/z 760.6503

01E+414.515RT (min)

lcqtof\_t3\_pos\_4\_042\_18

m/z 761.3198

01234E+311.5RT (min)

lcqtof\_t3\_pos\_2\_050\_14

m/z 761.4508

0246E+377.5RT (min)

lcqtof\_t3\_pos\_4\_042\_18

m/z 762.8181

02468E+314.515RT (min)

lcqtof\_t3\_pos\_4\_014\_24

m/z 762.8215

012E+411.5RT (min)

lcqtof\_t3\_pos\_3\_010\_01

m/z 762.8361

012E+317.518RT (min)

lcqtof\_t3\_pos\_2\_007\_07

m/z 763.5807

01234E+311.5RT (min)

lcqtof\_t3\_pos\_2\_015\_13

m/z 764.5070

012E+38.59RT (min)

lcqtof\_t3\_pos\_4\_081\_58

m/z 764.6200

02468E+311.5RT (min)

lcqtof\_t3\_pos\_2\_015\_13

m/z 764.8241

01234E+317.518RT (min)

lcqtof\_t3\_pos\_2\_021\_22

m/z 764.8287

0123E+317.518RT (min)

lcqtof\_t3\_pos\_2\_043\_48

m/z 765.8361

01E+317.518RT (min)

lcqtof\_t3\_pos\_2\_018\_41

m/z 765.8372

0123E+411.5RT (min)

lcqtof\_t3\_pos\_2\_015\_13

m/z 766.2326

01234E+34.55RT (min)

lcqtof\_t3\_pos\_4\_020\_19

m/z 766.6165

0123E+311.5RT (min)

lcqtof\_t3\_pos\_2\_015\_13

m/z 767.4524

024E+37.58RT (min)

lcqtof\_t3\_pos\_4\_050\_48

m/z 767.5323

012E+39.510RT (min)

lcqtof\_t3\_pos\_4\_091\_60

m/z 767.6169

01234E+314.515RT (min)

lcqtof\_t3\_pos\_4\_047\_13

m/z 767.8356

02468E+311.5RT (min)

lcqtof\_t3\_pos\_4\_021\_01

m/z 768.5363

01234E+311.5RT (min)

lcqtof\_t3\_pos\_4\_047\_13

m/z 768.8232

024E+314.515RT (min)

lcqtof\_t3\_pos\_4\_093\_60

m/z 769.5398

01E+41414.5RT (min)

lcqtof\_t3\_pos\_2\_039\_23

m/z 769.5734

02468E+311.5RT (min)

lcqtof\_t3\_pos\_4\_047\_13

m/z 769.8483

0246E+355.5RT (min)

lcqtof\_t3\_pos\_4\_021\_01

m/z 771.4945

0246E+31313.5RT (min)

lcqtof\_t3\_pos\_2\_047\_02

m/z 773.7797

01234E+314.515RT (min)

lcqtof\_t3\_pos\_2\_034\_24

m/z 773.8333

012E+317.518RT (min)

lcqtof\_t3\_pos\_2\_047\_02

m/z 774.5300

0246E+311.5RT (min)

lcqtof\_t3\_pos\_2\_015\_13

m/z 775.4899

02468E+311.5RT (min)

lcqtof\_t3\_pos\_4\_021\_01

m/z 777.8177

0123E+317.518RT (min)

lcqtof\_t3\_pos\_2\_023\_05

m/z 778.5885

01234E+311.5RT (min)

lcqtof\_t3\_pos\_4\_029\_37

m/z 778.8306

01234E+317.518RT (min)

lcqtof\_t3\_pos\_2\_007\_07

m/z 779.3424

012E+317.518RT (min)

lcqtof\_t3\_pos\_2\_013\_29

m/z 779.4866

02468E+311.5RT (min)

lcqtof\_t3\_pos\_4\_021\_01

m/z 779.8387

02468E+311.5RT (min)

lcqtof\_t3\_pos\_2\_031\_01

m/z 780.8065

012E+317.518RT (min)

lcqtof\_t3\_pos\_2\_025\_52

m/z 781.4288

0246E+344.5RT (min)

lcqtof\_t3\_pos\_4\_021\_01

m/z 781.5577

0123E+311.5RT (min)

lcqtof\_t3\_pos\_4\_007\_14

m/z 781.8094

0246E+311.5RT (min)

lcqtof\_t3\_pos\_2\_053\_15

m/z 781.8157

012E+317.518RT (min)

lcqtof\_t3\_pos\_2\_004\_42

m/z 782.3195

0123E+317.518RT (min)

lcqtof\_t3\_pos\_4\_011\_10

m/z 782.3312

0246E+344.5RT (min)

lcqtof\_t3\_pos\_4\_015\_16

m/z 782.7844

0246E+311.5RT (min)

lcqtof\_t3\_pos\_4\_037\_32

m/z 783.4977

02468E+35.56RT (min)

lcqtof\_t3\_pos\_4\_029\_37

m/z 783.5552

01234E+311.5RT (min)

lcqtof\_t3\_pos\_2\_031\_01

m/z 784.5088

0246E+317.518RT (min)

lcqtof\_t3\_pos\_4\_087\_59

m/z 785.0555

0246E+51313.5RT (min)

lcqtof\_t3\_pos\_2\_028\_34

m/z 785.1551

0123E+355.5RT (min)

lcqtof\_t3\_pos\_2\_010\_26

m/z 785.4931

0123E+355.5RT (min)

lcqtof\_t3\_pos\_4\_021\_01

m/z 785.5518

0123E+311.5RT (min)

lcqtof\_t3\_pos\_2\_031\_01

m/z 785.8585

024E+31414.5RT (min)

lcqtof\_t3\_pos\_2\_022\_32

m/z 786.5110

01234E+311.5RT (min)

lcqtof\_t3\_pos\_2\_031\_01

m/z 787.3275

01E+317.518RT (min)

lcqtof\_t3\_pos\_2\_004\_42

m/z 787.4450

024E+377.5RT (min)

lcqtof\_t3\_pos\_4\_024\_20

m/z 788.2676

0123E+366.5RT (min)

lcqtof\_t3\_pos\_2\_027\_38

m/z 788.5074

0246E+311.5RT (min)

lcqtof\_t3\_pos\_2\_031\_01

m/z 788.8289

0246E+311.5RT (min)

lcqtof\_t3\_pos\_3\_003\_14

m/z 789.2516

02468E+388.5RT (min)

lcqtof\_t3\_pos\_4\_042\_18

m/z 789.2693

02468E+344.5RT (min)

lcqtof\_t3\_pos\_2\_010\_26

m/z 789.7976

0246E+311.5RT (min)

lcqtof\_t3\_pos\_3\_003\_14

m/z 790.2462

012E+41414.5RT (min)

lcqtof\_t3\_pos\_4\_041\_54

m/z 790.3036

0123E+366.5RT (min)

lcqtof\_t3\_pos\_2\_011\_19

m/z 790.4988

01234E+311.5RT (min)

lcqtof\_t3\_pos\_2\_046\_30

m/z 791.4590

0246E+311.5RT (min)

lcqtof\_t3\_pos\_4\_021\_01

m/z 792.7811

01234E+317.518RT (min)

lcqtof\_t3\_pos\_2\_018\_41

m/z 792.8049

01E+411.5RT (min)

lcqtof\_t3\_pos\_2\_012\_35

m/z 793.4645

0246E+311.5RT (min)

lcqtof\_t3\_pos\_4\_021\_01

m/z 793.7933

012E+317.518RT (min)

lcqtof\_t3\_pos\_2\_022\_32

m/z 794.4254

02468E+311.5RT (min)

lcqtof\_t3\_pos\_4\_007\_14

m/z 795.3123

01234E+311.5RT (min)

lcqtof\_t3\_pos\_2\_050\_14

m/z 795.4595

0123E+311.5RT (min)

lcqtof\_t3\_pos\_4\_021\_01

m/z 795.8159

024E+311.5RT (min)

lcqtof\_t3\_pos\_2\_010\_26

m/z 797.2879

024E+34.55RT (min)

lcqtof\_t3\_pos\_4\_024\_20

m/z 797.5287

02468E+317.518RT (min)

lcqtof\_t3\_pos\_2\_003\_16

m/z 799.1772

024E+51313.5RT (min)

lcqtof\_t3\_pos\_3\_093\_60

m/z 800.4867

0123E+36.57RT (min)

lcqtof\_t3\_pos\_4\_029\_37

m/z 801.8367

02468E+311.5RT (min)

lcqtof\_t3\_pos\_4\_021\_01

m/z 802.5066

01234E+36.57RT (min)

lcqtof\_t3\_pos\_4\_029\_37

m/z 802.8107

0123E+317.518RT (min)

lcqtof\_t3\_pos\_2\_011\_19

m/z 802.9560

0123E+311.5RT (min)

lcqtof\_t3\_pos\_4\_029\_37

m/z 803.8140

012E+317.518RT (min)

lcqtof\_t3\_pos\_2\_021\_22

m/z 804.4761

01E+411.5RT (min)

lcqtof\_t3\_pos\_4\_021\_01

m/z 804.8393

024E+317.518RT (min)

lcqtof\_t3\_pos\_2\_007\_07

m/z 806.2954

02468E+34.55RT (min)

lcqtof\_t3\_pos\_4\_029\_37

m/z 807.4937

01234E+35.56RT (min)

lcqtof\_t3\_pos\_4\_024\_20

m/z 807.8163

01E+411.5RT (min)

lcqtof\_t3\_pos\_2\_031\_01

m/z 807.8199

012E+317.518RT (min)

lcqtof\_t3\_pos\_2\_017\_51

m/z 808.3017

01234E+344.5RT (min)

lcqtof\_t3\_pos\_2\_054\_09

m/z 809.2860

0123E+317.518RT (min)

lcqtof\_t3\_pos\_4\_016\_40

m/z 810.7498

01E+411.5RT (min)

lcqtof\_t3\_pos\_4\_005\_30

m/z 811.2573

02468E+34.55RT (min)

lcqtof\_t3\_pos\_4\_029\_37

m/z 812.8301

0123E+317.518RT (min)

lcqtof\_t3\_pos\_2\_004\_42

m/z 813.3290

012E+317.518RT (min)

lcqtof\_t3\_pos\_2\_017\_51

m/z 813.8267

02468E+311.5RT (min)

lcqtof\_t3\_pos\_2\_005\_17

m/z 813.9997

0246E+377.5RT (min)

lcqtof\_t3\_pos\_4\_029\_37

m/z 815.8056

01E+411.5RT (min)

lcqtof\_t3\_pos\_2\_050\_14

m/z 815.8081

012E+317.518RT (min)

lcqtof\_t3\_pos\_2\_014\_03

m/z 816.3193

012E+317.518RT (min)

lcqtof\_t3\_pos\_2\_015\_13

m/z 818.0721

0123E+355.5RT (min)

lcqtof\_t3\_pos\_2\_050\_14

m/z 818.3919

0246E+311.5RT (min)

lcqtof\_t3\_pos\_4\_007\_14

m/z 818.5095

02468E+317.518RT (min)

lcqtof\_t3\_pos\_2\_039\_23

m/z 818.5841

01E+399.5RT (min)

lcqtof\_t3\_pos\_4\_106\_62

m/z 820.3025

0123E+317.518RT (min)

lcqtof\_t3\_pos\_4\_023\_39

m/z 820.8150

012E+317.518RT (min)

lcqtof\_t3\_pos\_2\_046\_30

m/z 821.3212

02468E+311.5RT (min)

lcqtof\_t3\_pos\_2\_050\_14

m/z 821.3270

012E+317.518RT (min)

lcqtof\_t3\_pos\_2\_025\_52

m/z 821.5292

01234E+317.518RT (min)

lcqtof\_t3\_pos\_3\_016\_34

m/z 821.8196

02468E+311.5RT (min)

lcqtof\_t3\_pos\_2\_013\_29

m/z 821.8351

012E+317.518RT (min)

lcqtof\_t3\_pos\_2\_026\_40

m/z 822.4658

01234E+36.57RT (min)

lcqtof\_t3\_pos\_4\_029\_37

m/z 822.5433

01E+417.518RT (min)

lcqtof\_t3\_pos\_2\_003\_16

m/z 822.8667

024E+311.5RT (min)

lcqtof\_t3\_pos\_3\_015\_13

m/z 823.5476

0123E+317.518RT (min)

lcqtof\_t3\_pos\_2\_003\_16

m/z 823.5561

01E+412.513RT (min)

lcqtof\_t3\_pos\_3\_093\_60

m/z 823.7946

024E+31414.5RT (min)

lcqtof\_t3\_pos\_2\_022\_32

m/z 824.3026

01E+317.518RT (min)

lcqtof\_t3\_pos\_2\_018\_41

m/z 824.4760

0246E+344.5RT (min)

lcqtof\_t3\_pos\_4\_021\_01

m/z 824.4820

024E+36.57RT (min)

lcqtof\_t3\_pos\_2\_027\_38

m/z 824.4899

024E+36.57RT (min)

lcqtof\_t3\_pos\_4\_044\_38

m/z 824.5561

01E+417.518RT (min)

lcqtof\_t3\_pos\_2\_003\_16

m/z 825.2559

0246E+31414.5RT (min)

lcqtof\_t3\_pos\_4\_041\_54

m/z 825.5584

024E+317.518RT (min)

lcqtof\_t3\_pos\_2\_040\_28

m/z 827.2574

02468E+31414.5RT (min)

lcqtof\_t3\_pos\_4\_041\_54

m/z 829.3095

01234E+311.5RT (min)

lcqtof\_t3\_pos\_2\_050\_14

m/z 829.4808

0246E+312.513RT (min)

lcqtof\_t3\_pos\_3\_018\_37

m/z 829.8126

02468E+311.5RT (min)

lcqtof\_t3\_pos\_3\_010\_01

m/z 830.4326

0123E+366.5RT (min)

lcqtof\_t3\_pos\_2\_050\_14

m/z 831.0975

0123E+366.5RT (min)

lcqtof\_t3\_pos\_2\_050\_14

m/z 831.2318

02468E+34.55RT (min)

lcqtof\_t3\_pos\_4\_024\_20

m/z 831.4321

0123E+366.5RT (min)

lcqtof\_t3\_pos\_4\_021\_01

m/z 831.9703

024E+377.5RT (min)

lcqtof\_t3\_pos\_2\_011\_19

m/z 832.8239

0123E+317.518RT (min)

lcqtof\_t3\_pos\_2\_007\_07

m/z 834.2383

01234E+314.515RT (min)

lcqtof\_t3\_pos\_2\_006\_04

m/z 835.3377

012E+344.5RT (min)

lcqtof\_t3\_pos\_4\_069\_57

m/z 835.8300

02468E+311.5RT (min)

lcqtof\_t3\_pos\_4\_021\_01

m/z 836.3448

02468E+377.5RT (min)

lcqtof\_t3\_pos\_4\_024\_20

m/z 838.4048

024E+311.5RT (min)

lcqtof\_t3\_pos\_4\_047\_13

m/z 839.6519

012E+314.515RT (min)

lcqtof\_t3\_pos\_2\_015\_13

m/z 840.3344

024E+317.518RT (min)

lcqtof\_t3\_pos\_4\_106\_62

m/z 840.3433

0123E+34.55RT (min)

lcqtof\_t3\_pos\_4\_067\_57

m/z 840.4051

0246E+311.5RT (min)

lcqtof\_t3\_pos\_4\_047\_13

m/z 841.3419

012E+34.55RT (min)

lcqtof\_t3\_pos\_4\_072\_57

m/z 841.8134

01E+411.5RT (min)

lcqtof\_t3\_pos\_2\_031\_01

m/z 841.8171

012E+317.518RT (min)

lcqtof\_t3\_pos\_2\_012\_35

m/z 842.3997

0246E+311.5RT (min)

lcqtof\_t3\_pos\_4\_021\_01

m/z 842.4628

0123E+377.5RT (min)

lcqtof\_t3\_pos\_2\_027\_38

m/z 844.3976

0246E+36.57RT (min)

lcqtof\_t3\_pos\_4\_029\_37

m/z 844.4417

0123E+311.5RT (min)

lcqtof\_t3\_pos\_2\_012\_35

m/z 844.5201

012E+317.518RT (min)

lcqtof\_t3\_pos\_2\_020\_27

m/z 844.6674

01234E+417.518RT (min)

lcqtof\_t3\_pos\_4\_014\_24

m/z 845.5211

0246E+377.5RT (min)

lcqtof\_t3\_pos\_2\_002\_37

m/z 845.6788

01E+417.518RT (min)

lcqtof\_t3\_pos\_4\_014\_24

m/z 845.8008

0123E+317.518RT (min)

lcqtof\_t3\_pos\_2\_016\_39

m/z 846.5285

01234E+317.518RT (min)

lcqtof\_t3\_pos\_2\_002\_37

m/z 846.8188

0123E+317.518RT (min)

lcqtof\_t3\_pos\_2\_016\_39

m/z 847.3297

012E+317.518RT (min)

lcqtof\_t3\_pos\_3\_002\_17

m/z 847.8213

02468E+311.5RT (min)

lcqtof\_t3\_pos\_2\_005\_17

m/z 848.1956

024E+31313.5RT (min)

lcqtof\_t3\_pos\_2\_007\_07

m/z 848.3193

012E+311.5RT (min)

lcqtof\_t3\_pos\_2\_005\_17

m/z 848.4790

0123E+35.56RT (min)

lcqtof\_t3\_pos\_2\_020\_27

m/z 849.5847

0123E+31414.5RT (min)

lcqtof\_t3\_pos\_4\_091\_60

m/z 849.7977

0246E+311.5RT (min)

lcqtof\_t3\_pos\_2\_031\_01

m/z 850.3065

01E+317.518RT (min)

lcqtof\_t3\_pos\_2\_010\_26

m/z 851.3215

0123E+317.518RT (min)

lcqtof\_t3\_pos\_4\_016\_40

m/z 851.5669

0123E+317.518RT (min)

lcqtof\_t3\_pos\_2\_020\_27

m/z 852.0627

01234E+35.56RT (min)

lcqtof\_t3\_pos\_4\_007\_14

m/z 854.3786

0123E+311.5RT (min)

lcqtof\_t3\_pos\_4\_008\_15

m/z 855.1896

0123E+344.5RT (min)

lcqtof\_t3\_pos\_2\_022\_32

m/z 855.3153

0246E+311.5RT (min)

lcqtof\_t3\_pos\_2\_050\_14

m/z 855.3276

01E+317.518RT (min)

lcqtof\_t3\_pos\_2\_035\_44

m/z 855.8293

0246E+317.518RT (min)

lcqtof\_t3\_pos\_2\_021\_22

m/z 856.3766

01234E+311.5RT (min)

lcqtof\_t3\_pos\_4\_021\_01

m/z 856.8151

024E+311.5RT (min)

lcqtof\_t3\_pos\_3\_003\_14

m/z 858.3018

01E+317.518RT (min)

lcqtof\_t3\_pos\_2\_012\_35

m/z 858.3742

01234E+311.5RT (min)

lcqtof\_t3\_pos\_4\_021\_01

m/z 859.5624

01234E+31414.5RT (min)

lcqtof\_t3\_pos\_4\_108\_62

m/z 860.3381

01234E+317.518RT (min)

lcqtof\_t3\_pos\_4\_020\_19

m/z 860.7902

02468E+311.5RT (min)

lcqtof\_t3\_pos\_4\_001\_49

m/z 861.7756

012E+317.518RT (min)

lcqtof\_t3\_pos\_2\_021\_22

m/z 862.5698

0246E+313.514RT (min)

lcqtof\_t3\_pos\_4\_006\_41

m/z 863.3034

01234E+311.5RT (min)

lcqtof\_t3\_pos\_2\_050\_14

m/z 864.5161

0123E+317.518RT (min)

lcqtof\_t3\_pos\_2\_020\_27

m/z 865.2730

01E+41414.5RT (min)

lcqtof\_t3\_pos\_4\_041\_54

m/z 866.4911

0123E+377.5RT (min)

lcqtof\_t3\_pos\_2\_020\_27

m/z 866.6638

012E+314.515RT (min)

lcqtof\_t3\_pos\_3\_023\_36

m/z 867.3340

0246E+34.55RT (min)

lcqtof\_t3\_pos\_4\_066\_57

m/z 868.6793

01234E+314.515RT (min)

lcqtof\_t3\_pos\_2\_015\_13

m/z 869.3510

01234E+34.55RT (min)

lcqtof\_t3\_pos\_4\_069\_57

m/z 869.4828

0123E+377.5RT (min)

lcqtof\_t3\_pos\_2\_036\_20

m/z 870.5251

024E+317.518RT (min)

lcqtof\_t3\_pos\_2\_002\_37

m/z 870.7834

0123E+317.518RT (min)

lcqtof\_t3\_pos\_2\_039\_23

m/z 871.3747

01234E+31313.5RT (min)

lcqtof\_t3\_pos\_4\_023\_39

m/z 871.5370

0123E+312.513RT (min)

lcqtof\_t3\_pos\_1\_028\_08

m/z 871.8007

0123E+317.518RT (min)

lcqtof\_t3\_pos\_2\_020\_27

m/z 872.1453

0246E+311.5RT (min)

lcqtof\_t3\_pos\_2\_031\_01

m/z 872.3235

012E+317.518RT (min)

lcqtof\_t3\_pos\_4\_017\_51

m/z 872.5400

0123E+355.5RT (min)

lcqtof\_t3\_pos\_4\_008\_15

m/z 872.7057

0123E+31515.5RT (min)

lcqtof\_t3\_pos\_1\_014\_14

m/z 872.8151

01234E+317.518RT (min)

lcqtof\_t3\_pos\_2\_036\_20

m/z 872.8303

02468E+314.515RT (min)

lcqtof\_t3\_pos\_3\_013\_24

m/z 873.5497

01234E+317.518RT (min)

lcqtof\_t3\_pos\_4\_029\_37

m/z 874.2021

02468E+31414.5RT (min)

lcqtof\_t3\_pos\_4\_012\_12

m/z 874.5585

0246E+317.518RT (min)

lcqtof\_t3\_pos\_4\_021\_01

m/z 874.6566

01234E+31515.5RT (min)

lcqtof\_t3\_pos\_2\_012\_35

m/z 875.5524

012E+317.518RT (min)

lcqtof\_t3\_pos\_2\_003\_16

m/z 875.8008

01E+411.5RT (min)

lcqtof\_t3\_pos\_2\_031\_01

m/z 875.8121

012E+317.518RT (min)

lcqtof\_t3\_pos\_2\_011\_19

m/z 877.2357

024E+34.55RT (min)

lcqtof\_t3\_pos\_4\_024\_20

m/z 881.3464

0123E+317.518RT (min)

lcqtof\_t3\_pos\_4\_098\_61

m/z 881.6635

0246E+314.515RT (min)

lcqtof\_t3\_pos\_2\_039\_23

m/z 881.8214

02468E+311.5RT (min)

lcqtof\_t3\_pos\_3\_010\_01

m/z 882.9008

02468E+314.515RT (min)

lcqtof\_t3\_pos\_4\_014\_24

m/z 883.3733

01234E+34.55RT (min)

lcqtof\_t3\_pos\_4\_015\_16

m/z 883.7919

0246E+311.5RT (min)

lcqtof\_t3\_pos\_2\_020\_27

m/z 884.2961

012E+317.518RT (min)

lcqtof\_t3\_pos\_2\_017\_51

m/z 886.4586

0123E+377.5RT (min)

lcqtof\_t3\_pos\_2\_011\_19

m/z 886.6716

024E+314.515RT (min)

lcqtof\_t3\_pos\_2\_015\_13

m/z 886.7660

012E+411.5RT (min)

lcqtof\_t3\_pos\_3\_010\_01

m/z 888.6564

024E+314.515RT (min)

lcqtof\_t3\_pos\_3\_047\_23

m/z 888.7537

012E+31515.5RT (min)

lcqtof\_t3\_pos\_2\_042\_12

m/z 888.7852

012E+317.518RT (min)

lcqtof\_t3\_pos\_2\_033\_53

m/z 889.3108

0246E+311.5RT (min)

lcqtof\_t3\_pos\_2\_050\_14

m/z 889.3752

0123E+488.5RT (min)

lcqtof\_t3\_pos\_4\_107\_62

m/z 889.6365

0123E+313.514RT (min)

lcqtof\_t3\_pos\_2\_046\_30

m/z 889.8117

02468E+311.5RT (min)

lcqtof\_t3\_pos\_2\_031\_01

m/z 892.2896

012E+317.518RT (min)

lcqtof\_t3\_pos\_1\_010\_27

m/z 893.1990

012E+34.55RT (min)

lcqtof\_t3\_pos\_3\_006\_20

m/z 893.6963

01234E+317.518RT (min)

lcqtof\_t3\_pos\_3\_016\_34

m/z 893.8015

01E+41414.5RT (min)

lcqtof\_t3\_pos\_4\_088\_59

m/z 894.2193

0123E+34.55RT (min)

lcqtof\_t3\_pos\_3\_053\_19

m/z 894.7988

02468E+311.5RT (min)

lcqtof\_t3\_pos\_2\_012\_35

m/z 895.7186

0123E+317.518RT (min)

lcqtof\_t3\_pos\_3\_051\_35

m/z 895.8899

0246E+39.510RT (min)

lcqtof\_t3\_pos\_2\_050\_14

m/z 896.3568

0246E+311.5RT (min)

lcqtof\_t3\_pos\_4\_021\_01

m/z 896.7739

02468E+317.518RT (min)

lcqtof\_t3\_pos\_4\_029\_37

m/z 897.2681

01E+41414.5RT (min)

lcqtof\_t3\_pos\_3\_080\_58

m/z 897.2924

01234E+311.5RT (min)

lcqtof\_t3\_pos\_2\_050\_14

m/z 897.5485

01234E+31313.5RT (min)

lcqtof\_t3\_pos\_3\_032\_02

m/z 897.8014

0246E+311.5RT (min)

lcqtof\_t3\_pos\_3\_010\_01

m/z 898.3587

02468E+311.5RT (min)

lcqtof\_t3\_pos\_4\_021\_01

m/z 898.3979

02468E+39.510RT (min)

lcqtof\_t3\_pos\_2\_050\_14

m/z 899.2684

01E+41414.5RT (min)

lcqtof\_t3\_pos\_3\_104\_61

m/z 900.1353

01234E+311.5RT (min)

lcqtof\_t3\_pos\_2\_031\_01

m/z 900.3571

02468E+311.5RT (min)

lcqtof\_t3\_pos\_4\_021\_01

m/z 900.5182

0123E+377.5RT (min)

lcqtof\_t3\_pos\_2\_028\_34

m/z 900.5729

01234E+317.518RT (min)

lcqtof\_t3\_pos\_4\_021\_01

m/z 902.3575

024E+311.5RT (min)

lcqtof\_t3\_pos\_4\_021\_01

m/z 903.4669

024E+377.5RT (min)

lcqtof\_t3\_pos\_2\_001\_49

m/z 904.7836

0246E+311.5RT (min)

lcqtof\_t3\_pos\_2\_050\_14

m/z 907.8149

012E+317.518RT (min)

lcqtof\_t3\_pos\_4\_019\_03

m/z 908.3690

0123E+34.55RT (min)

lcqtof\_t3\_pos\_4\_066\_57

m/z 909.4682

01234E+377.5RT (min)

lcqtof\_t3\_pos\_3\_014\_38

m/z 909.7425

024E+31414.5RT (min)

lcqtof\_t3\_pos\_4\_023\_39

m/z 909.8002

01E+411.5RT (min)

lcqtof\_t3\_pos\_2\_031\_01

m/z 910.2579

0246E+314.515RT (min)

lcqtof\_t3\_pos\_4\_076\_58

m/z 911.3808

0123E+34.55RT (min)

lcqtof\_t3\_pos\_4\_015\_16

m/z 911.8359

024E+313.514RT (min)

lcqtof\_t3\_pos\_3\_096\_60

m/z 912.1375

02468E+311.5RT (min)

lcqtof\_t3\_pos\_2\_003\_16

m/z 912.3412

01234E+311.5RT (min)

lcqtof\_t3\_pos\_4\_021\_01

m/z 913.3927

0123E+317.518RT (min)

lcqtof\_t3\_pos\_4\_106\_62

m/z 913.7922

0123E+317.518RT (min)

lcqtof\_t3\_pos\_2\_025\_52

m/z 914.3346

0246E+311.5RT (min)

lcqtof\_t3\_pos\_4\_021\_01

m/z 914.7990

012E+317.518RT (min)

lcqtof\_t3\_pos\_2\_006\_04

m/z 915.8052

0246E+311.5RT (min)

lcqtof\_t3\_pos\_2\_003\_16

m/z 916.3334

02468E+311.5RT (min)

lcqtof\_t3\_pos\_4\_021\_01

m/z 916.5515

0123E+317.518RT (min)

lcqtof\_t3\_pos\_4\_021\_01

m/z 917.4662

0246E+311.5RT (min)

lcqtof\_t3\_pos\_2\_005\_17

m/z 917.6966

0246E+317.518RT (min)

lcqtof\_t3\_pos\_2\_039\_23

m/z 917.7880

0246E+311.5RT (min)

lcqtof\_t3\_pos\_2\_031\_01

m/z 917.7888

0246E+31515.5RT (min)

lcqtof\_t3\_pos\_4\_042\_18

m/z 918.3303

01234E+311.5RT (min)

lcqtof\_t3\_pos\_4\_021\_01

m/z 918.7131

01234E+317.518RT (min)

lcqtof\_t3\_pos\_4\_014\_24

m/z 919.7103

0246E+317.518RT (min)

lcqtof\_t3\_pos\_2\_039\_23

m/z 920.7134

01234E+317.518RT (min)

lcqtof\_t3\_pos\_2\_039\_23

m/z 921.6460

01E+412.513RT (min)

lcqtof\_t3\_pos\_3\_013\_24

m/z 923.3058

0246E+311.5RT (min)

lcqtof\_t3\_pos\_4\_017\_51

m/z 923.3278

0123E+317.518RT (min)

lcqtof\_t3\_pos\_2\_003\_16

m/z 923.8135

0246E+317.518RT (min)

lcqtof\_t3\_pos\_2\_024\_45

m/z 924.7951

01E+41515.5RT (min)

lcqtof\_t3\_pos\_4\_042\_18

m/z 928.7772

02468E+311.5RT (min)

lcqtof\_t3\_pos\_4\_017\_51

m/z 929.7645

0123E+317.518RT (min)

lcqtof\_t3\_pos\_2\_042\_12

m/z 931.2853

01234E+311.5RT (min)

lcqtof\_t3\_pos\_4\_017\_51

m/z 931.6090

01234E+31414.5RT (min)

lcqtof\_t3\_pos\_4\_108\_62

m/z 933.8258

0246E+31414.5RT (min)

lcqtof\_t3\_pos\_2\_050\_14

m/z 935.1475

024E+311.5RT (min)

lcqtof\_t3\_pos\_4\_015\_16

m/z 936.7886

0246E+31414.5RT (min)

lcqtof\_t3\_pos\_4\_041\_54

m/z 938.2689

01E+41414.5RT (min)

lcqtof\_t3\_pos\_3\_105\_61

m/z 938.7043

0123E+31515.5RT (min)

lcqtof\_t3\_pos\_2\_008\_31

m/z 938.7652

012E+317.518RT (min)

lcqtof\_t3\_pos\_2\_018\_41

m/z 939.7788

012E+317.518RT (min)

lcqtof\_t3\_pos\_2\_042\_12

m/z 940.1400

0246E+311.5RT (min)

lcqtof\_t3\_pos\_4\_015\_16

m/z 940.4547

0123E+311.5RT (min)

lcqtof\_t3\_pos\_2\_003\_16

m/z 940.8094

0123E+317.518RT (min)

lcqtof\_t3\_pos\_2\_018\_41

m/z 942.8325

02468E+314.515RT (min)

lcqtof\_t3\_pos\_2\_034\_24

m/z 943.7907

01E+411.5RT (min)

lcqtof\_t3\_pos\_2\_031\_01

m/z 943.7936

01E+317.518RT (min)

lcqtof\_t3\_pos\_2\_011\_19

m/z 945.4617

024E+311.5RT (min)

lcqtof\_t3\_pos\_2\_050\_14

m/z 946.0081

012E+366.5RT (min)

lcqtof\_t3\_pos\_2\_050\_14

m/z 946.5169

024E+366.5RT (min)

lcqtof\_t3\_pos\_4\_021\_01

m/z 949.4323

024E+344.5RT (min)

lcqtof\_t3\_pos\_4\_014\_24

m/z 949.8106

0246E+311.5RT (min)

lcqtof\_t3\_pos\_2\_031\_01

m/z 950.3090

012E+311.5RT (min)

lcqtof\_t3\_pos\_2\_031\_01

m/z 950.7464

01E+411.5RT (min)

lcqtof\_t3\_pos\_4\_001\_49

m/z 951.7768

0246E+311.5RT (min)

lcqtof\_t3\_pos\_2\_031\_01

m/z 952.4850

0246E+311.5RT (min)

lcqtof\_t3\_pos\_4\_047\_13

m/z 954.3221

02468E+311.5RT (min)

lcqtof\_t3\_pos\_4\_021\_01

m/z 954.4475

02468E+377.5RT (min)

lcqtof\_t3\_pos\_4\_042\_18

m/z 954.7513

01E+411.5RT (min)

lcqtof\_t3\_pos\_2\_010\_26

m/z 954.7886

0246E+311.5RT (min)

lcqtof\_t3\_pos\_2\_012\_35

m/z 955.6558

01234E+314.515RT (min)

lcqtof\_t3\_pos\_2\_039\_23

m/z 956.7910

012E+317.518RT (min)

lcqtof\_t3\_pos\_2\_005\_17

m/z 957.2969

0246E+311.5RT (min)

lcqtof\_t3\_pos\_4\_017\_51

m/z 957.8028

02468E+311.5RT (min)

lcqtof\_t3\_pos\_2\_013\_29

m/z 958.1483

01234E+311.5RT (min)

lcqtof\_t3\_pos\_4\_047\_13

m/z 958.8187

012E+315.516RT (min)

lcqtof\_t3\_pos\_2\_019\_08

m/z 962.3148

024E+311.5RT (min)

lcqtof\_t3\_pos\_4\_021\_01

m/z 962.8019

02468E+311.5RT (min)

lcqtof\_t3\_pos\_4\_015\_16

m/z 962.8189

0123E+31515.5RT (min)

lcqtof\_t3\_pos\_4\_112\_62

m/z 963.1366

01234E+311.5RT (min)

lcqtof\_t3\_pos\_4\_015\_16

m/z 963.3763

024E+36.57RT (min)

lcqtof\_t3\_pos\_2\_020\_27

m/z 963.4676

0123E+311.5RT (min)

lcqtof\_t3\_pos\_4\_015\_16

m/z 964.7878

0123E+317.518RT (min)

lcqtof\_t3\_pos\_2\_018\_41

m/z 965.2841

0123E+311.5RT (min)

lcqtof\_t3\_pos\_2\_050\_14

m/z 965.7880

0246E+311.5RT (min)

lcqtof\_t3\_pos\_2\_031\_01

m/z 966.7313

01234E+31515.5RT (min)

lcqtof\_t3\_pos\_2\_002\_37

m/z 968.1245

024E+311.5RT (min)

lcqtof\_t3\_pos\_2\_031\_01

m/z 969.7199

0123E+314.515RT (min)

lcqtof\_t3\_pos\_2\_039\_23

m/z 970.3013

0246E+311.5RT (min)

lcqtof\_t3\_pos\_4\_021\_01

m/z 971.0123

01E+41515.5RT (min)

lcqtof\_t3\_pos\_3\_092\_60

m/z 971.4172

024E+344.5RT (min)

lcqtof\_t3\_pos\_2\_034\_24

m/z 971.7274

0123E+31515.5RT (min)

lcqtof\_t3\_pos\_2\_012\_35

m/z 973.4479

0123E+311.5RT (min)

lcqtof\_t3\_pos\_4\_017\_51

m/z 974.8780

02468E+31515.5RT (min)

lcqtof\_t3\_pos\_1\_002\_36

m/z 975.1475

02468E+311.5RT (min)

lcqtof\_t3\_pos\_4\_047\_13

m/z 975.7745

01234E+314.515RT (min)

lcqtof\_t3\_pos\_4\_109\_62

m/z 976.2887

02468E+311.5RT (min)

lcqtof\_t3\_pos\_4\_021\_01

m/z 976.8340

0246E+311.5RT (min)

lcqtof\_t3\_pos\_4\_007\_14

m/z 977.7849

01E+411.5RT (min)

lcqtof\_t3\_pos\_2\_031\_01

m/z 978.2783

01234E+311.5RT (min)

lcqtof\_t3\_pos\_4\_021\_01

m/z 980.4641

0246E+311.5RT (min)

lcqtof\_t3\_pos\_4\_015\_16

m/z 981.7814

012E+317.518RT (min)

lcqtof\_t3\_pos\_2\_020\_27

m/z 982.6762

0123E+314.515RT (min)

lcqtof\_t3\_pos\_2\_039\_23

m/z 982.7914

01E+317.518RT (min)

lcqtof\_t3\_pos\_3\_008\_42

m/z 983.8021

0246E+311.5RT (min)

lcqtof\_t3\_pos\_2\_031\_01

m/z 984.2963

012E+311.5RT (min)

lcqtof\_t3\_pos\_2\_005\_17

m/z 985.4620

02468E+311.5RT (min)

lcqtof\_t3\_pos\_4\_015\_16

m/z 985.7815

02468E+311.5RT (min)

lcqtof\_t3\_pos\_2\_031\_01

m/z 986.1351

01234E+311.5RT (min)

lcqtof\_t3\_pos\_4\_015\_16

m/z 986.7499

012E+317.518RT (min)

lcqtof\_t3\_pos\_2\_006\_04

m/z 987.7021

0123E+314.515RT (min)

lcqtof\_t3\_pos\_2\_002\_37

m/z 988.2637

024E+311.5RT (min)

lcqtof\_t3\_pos\_4\_007\_14

m/z 988.9360

0246E+314.515RT (min)

lcqtof\_t3\_pos\_4\_012\_12

m/z 990.2589

024E+311.5RT (min)

lcqtof\_t3\_pos\_4\_007\_14

m/z 990.3512

0246E+344.5RT (min)

lcqtof\_t3\_pos\_4\_015\_16

m/z 990.7394

01234E+31515.5RT (min)

lcqtof\_t3\_pos\_2\_012\_35

m/z 991.2980

0246E+311.5RT (min)

lcqtof\_t3\_pos\_2\_050\_14

m/z 991.7906

02468E+311.5RT (min)

lcqtof\_t3\_pos\_4\_017\_51

m/z 991.7982

0123E+317.518RT (min)

lcqtof\_t3\_pos\_2\_048\_43

m/z 991.8146

02468E+314.515RT (min)

lcqtof\_t3\_pos\_1\_008\_31

m/z 996.1027

01234E+311.5RT (min)

lcqtof\_t3\_pos\_4\_017\_51

m/z 996.7495

0123E+317.518RT (min)

lcqtof\_t3\_pos\_2\_037\_46

m/z 996.7958

0246E+311.5RT (min)

lcqtof\_t3\_pos\_4\_049\_55

m/z 997.7363

012E+317.518RT (min)

lcqtof\_t3\_pos\_2\_023\_05

m/z 999.2812

0123E+311.5RT (min)

lcqtof\_t3\_pos\_2\_050\_14

m/z 1001.0527

02468E+31515.5RT (min)

lcqtof\_t3\_pos\_3\_090\_60

m/z 1002.9363

0246E+314.515RT (min)

lcqtof\_t3\_pos\_2\_022\_32

m/z 1003.1331

0246E+311.5RT (min)

lcqtof\_t3\_pos\_4\_015\_16

m/z 1003.7941

01E+411.5RT (min)

lcqtof\_t3\_pos\_2\_015\_13

m/z 1004.7934

0246E+311.5RT (min)

lcqtof\_t3\_pos\_3\_015\_13

m/z 1008.1270

02468E+311.5RT (min)

lcqtof\_t3\_pos\_4\_015\_16

m/z 1008.4587

024E+311.5RT (min)

lcqtof\_t3\_pos\_4\_015\_16

m/z 1008.8065

0123E+317.518RT (min)

lcqtof\_t3\_pos\_2\_036\_20

m/z 1008.8271

01234E+314.515RT (min)

lcqtof\_t3\_pos\_2\_002\_37

m/z 1008.9004

012E+315.516RT (min)

lcqtof\_t3\_pos\_4\_028\_22

m/z 1009.7880

012E+313.514RT (min)

lcqtof\_t3\_pos\_2\_012\_35

m/z 1009.8210

0123E+314.515RT (min)

lcqtof\_t3\_pos\_2\_012\_35

m/z 1010.8182

024E+314.515RT (min)

lcqtof\_t3\_pos\_4\_048\_11

m/z 1011.7729

02468E+311.5RT (min)

lcqtof\_t3\_pos\_2\_031\_01

m/z 1011.8071

02468E+314.515RT (min)

lcqtof\_t3\_pos\_4\_041\_54

m/z 1012.2746

0246E+311.5RT (min)

lcqtof\_t3\_pos\_4\_021\_01

m/z 1013.4503

0246E+311.5RT (min)

lcqtof\_t3\_pos\_4\_039\_02

m/z 1013.7829

01234E+311.5RT (min)

lcqtof\_t3\_pos\_4\_007\_14

m/z 1014.1158

0123E+311.5RT (min)

lcqtof\_t3\_pos\_4\_039\_02

m/z 1015.8688

024E+31515.5RT (min)

lcqtof\_t3\_pos\_2\_012\_35

m/z 1017.2957

02468E+311.5RT (min)

lcqtof\_t3\_pos\_2\_031\_01

m/z 1017.7990

024E+311.5RT (min)

lcqtof\_t3\_pos\_3\_002\_17

m/z 1017.8895

01234E+31515.5RT (min)

lcqtof\_t3\_pos\_2\_012\_35

m/z 1018.7379

01E+411.5RT (min)

lcqtof\_t3\_pos\_4\_001\_49

m/z 1019.7716

01234E+311.5RT (min)

lcqtof\_t3\_pos\_2\_053\_15

m/z 1020.2679

0246E+311.5RT (min)

lcqtof\_t3\_pos\_4\_021\_01

m/z 1022.7761

024E+311.5RT (min)

lcqtof\_t3\_pos\_2\_012\_35

m/z 1023.8289

012E+317.518RT (min)

lcqtof\_t3\_pos\_2\_032\_47

m/z 1024.7845

012E+317.518RT (min)

lcqtof\_t3\_pos\_3\_005\_18

m/z 1025.2841

0246E+311.5RT (min)

lcqtof\_t3\_pos\_4\_017\_51

m/z 1025.7896

01E+411.5RT (min)

lcqtof\_t3\_pos\_4\_015\_16

m/z 1026.0331

0246E+31515.5RT (min)

lcqtof\_t3\_pos\_1\_026\_32

m/z 1026.1309

01234E+311.5RT (min)

lcqtof\_t3\_pos\_4\_015\_16

m/z 1028.2743

0246E+311.5RT (min)

lcqtof\_t3\_pos\_4\_021\_01

m/z 1030.2661

02468E+311.5RT (min)

lcqtof\_t3\_pos\_4\_021\_01

m/z 1030.7849

02468E+311.5RT (min)

lcqtof\_t3\_pos\_4\_015\_16

m/z 1031.1217

024E+311.5RT (min)

lcqtof\_t3\_pos\_4\_015\_16

m/z 1031.4579

01234E+311.5RT (min)

lcqtof\_t3\_pos\_4\_015\_16

m/z 1032.7815

012E+317.518RT (min)

lcqtof\_t3\_pos\_4\_013\_29

m/z 1032.7889

02468E+313.514RT (min)

lcqtof\_t3\_pos\_2\_008\_31

m/z 1033.2715

0123E+311.5RT (min)

lcqtof\_t3\_pos\_2\_050\_14

m/z 1033.5040

0123E+37.58RT (min)

lcqtof\_t3\_pos\_3\_029\_26

m/z 1033.7638

024E+311.5RT (min)

lcqtof\_t3\_pos\_2\_005\_17

m/z 1034.2480

02468E+311.5RT (min)

lcqtof\_t3\_pos\_4\_021\_01

m/z 1034.9254

012E+31515.5RT (min)

lcqtof\_t3\_pos\_4\_111\_62

m/z 1036.1144

0246E+311.5RT (min)

lcqtof\_t3\_pos\_4\_039\_02

m/z 1036.2546

01234E+311.5RT (min)

lcqtof\_t3\_pos\_4\_007\_14

m/z 1039.9940

0246E+314.515RT (min)

lcqtof\_t3\_pos\_3\_094\_60

m/z 1041.4390

024E+311.5RT (min)

lcqtof\_t3\_pos\_4\_017\_51

m/z 1042.0148

0246E+314.515RT (min)

lcqtof\_t3\_pos\_3\_093\_60

m/z 1045.7719

0246E+311.5RT (min)

lcqtof\_t3\_pos\_2\_010\_26

m/z 1048.0096

0246E+314.515RT (min)

lcqtof\_t3\_pos\_1\_026\_32

m/z 1048.3031

012E+41414.5RT (min)

lcqtof\_t3\_pos\_3\_105\_61

m/z 1048.4601

0246E+311.5RT (min)

lcqtof\_t3\_pos\_4\_015\_16

m/z 1048.7523

01234E+317.518RT (min)

lcqtof\_t3\_pos\_2\_036\_20

m/z 1048.7579

02468E+311.5RT (min)

lcqtof\_t3\_pos\_4\_029\_37

m/z 1049.3074

01E+41414.5RT (min)

lcqtof\_t3\_pos\_3\_089\_59

m/z 1049.3146

0246E+314.515RT (min)

lcqtof\_t3\_pos\_4\_014\_24

m/z 1051.7876

024E+311.5RT (min)

lcqtof\_t3\_pos\_2\_013\_29

m/z 1053.7631

0246E+311.5RT (min)

lcqtof\_t3\_pos\_2\_031\_01

m/z 1054.1227

01234E+311.5RT (min)

lcqtof\_t3\_pos\_4\_015\_16

m/z 1058.7729

0123E+317.518RT (min)

lcqtof\_t3\_pos\_2\_021\_22

m/z 1059.1085

0123E+311.5RT (min)

lcqtof\_t3\_pos\_4\_039\_02

m/z 1059.2769

024E+311.5RT (min)

lcqtof\_t3\_pos\_2\_050\_14

m/z 1059.4497

0123E+311.5RT (min)

lcqtof\_t3\_pos\_4\_039\_02

m/z 1059.7753

0246E+311.5RT (min)

lcqtof\_t3\_pos\_4\_017\_51

m/z 1059.7909

0123E+317.518RT (min)

lcqtof\_t3\_pos\_2\_016\_39

m/z 1060.9872

012E+31313.5RT (min)

lcqtof\_t3\_pos\_3\_032\_02

m/z 1064.0955

01234E+311.5RT (min)

lcqtof\_t3\_pos\_4\_017\_51

m/z 1064.7352

012E+317.518RT (min)

lcqtof\_t3\_pos\_2\_021\_22

m/z 1064.7573

024E+311.5RT (min)

lcqtof\_t3\_pos\_4\_001\_49

m/z 1065.7457

012E+317.518RT (min)

lcqtof\_t3\_pos\_2\_048\_43

m/z 1065.9689

01E+31616.5RT (min)

lcqtof\_t3\_pos\_4\_111\_62

m/z 1066.7576

02468E+311.5RT (min)

lcqtof\_t3\_pos\_2\_050\_14

m/z 1067.2656

0123E+311.5RT (min)

lcqtof\_t3\_pos\_2\_050\_14

m/z 1067.9781

0123E+315.516RT (min)

lcqtof\_t3\_pos\_4\_092\_60

m/z 1070.7483

01E+411.5RT (min)

lcqtof\_t3\_pos\_4\_001\_49

m/z 1071.1249

0246E+311.5RT (min)

lcqtof\_t3\_pos\_4\_047\_13

m/z 1071.4526

024E+311.5RT (min)

lcqtof\_t3\_pos\_4\_015\_16

m/z 1071.7770

01E+411.5RT (min)

lcqtof\_t3\_pos\_4\_047\_13

m/z 1072.9379

01234E+31515.5RT (min)

lcqtof\_t3\_pos\_4\_087\_59

m/z 1076.4459

024E+311.5RT (min)

lcqtof\_t3\_pos\_4\_015\_16

m/z 1076.7874

012E+317.518RT (min)

lcqtof\_t3\_pos\_2\_006\_04

m/z 1079.7662

0246E+311.5RT (min)

lcqtof\_t3\_pos\_2\_031\_01

m/z 1081.4381

0246E+311.5RT (min)

lcqtof\_t3\_pos\_4\_039\_02

m/z 1082.1016

0123E+311.5RT (min)

lcqtof\_t3\_pos\_4\_039\_02

m/z 1082.9994

012E+315.516RT (min)

lcqtof\_t3\_pos\_4\_098\_61

m/z 1085.2810

02468E+311.5RT (min)

lcqtof\_t3\_pos\_2\_020\_27

m/z 1085.7845

024E+311.5RT (min)

lcqtof\_t3\_pos\_4\_015\_16

m/z 1085.8313

02468E+314.515RT (min)

lcqtof\_t3\_pos\_4\_041\_54

m/z 1086.7216

02468E+311.5RT (min)

lcqtof\_t3\_pos\_4\_001\_49

m/z 1086.7566

0246E+311.5RT (min)

lcqtof\_t3\_pos\_4\_050\_48

m/z 1087.7529

024E+311.5RT (min)

lcqtof\_t3\_pos\_2\_050\_14

m/z 1090.7616

024E+311.5RT (min)

lcqtof\_t3\_pos\_2\_002\_37

m/z 1093.2731

024E+311.5RT (min)

lcqtof\_t3\_pos\_2\_050\_14

m/z 1093.7844

02468E+311.5RT (min)

lcqtof\_t3\_pos\_4\_015\_16

m/z 1094.1160

024E+311.5RT (min)

lcqtof\_t3\_pos\_4\_047\_13

m/z 1095.0022

012E+315.516RT (min)

lcqtof\_t3\_pos\_4\_106\_62

m/z 1095.9618

012E+31313.5RT (min)

lcqtof\_t3\_pos\_3\_096\_60

m/z 1096.0060

012E+315.516RT (min)

lcqtof\_t3\_pos\_4\_086\_59

m/z 1097.2498

0246E+314.515RT (min)

lcqtof\_t3\_pos\_4\_012\_12

m/z 1098.7759

02468E+311.5RT (min)

lcqtof\_t3\_pos\_4\_015\_16

m/z 1099.1068

024E+311.5RT (min)

lcqtof\_t3\_pos\_4\_015\_16

m/z 1099.4424

024E+311.5RT (min)

lcqtof\_t3\_pos\_4\_015\_16

m/z 1100.7819

0123E+317.518RT (min)

lcqtof\_t3\_pos\_4\_023\_39

m/z 1101.0363

012E+31616.5RT (min)

lcqtof\_t3\_pos\_4\_088\_59

m/z 1101.2601

0123E+311.5RT (min)

lcqtof\_t3\_pos\_2\_050\_14

m/z 1102.9680

0123E+31515.5RT (min)

lcqtof\_t3\_pos\_2\_022\_32

m/z 1104.1024

0246E+311.5RT (min)

lcqtof\_t3\_pos\_4\_039\_02

m/z 1106.7473

01234E+311.5RT (min)

lcqtof\_t3\_pos\_2\_005\_17

m/z 1109.4204

01234E+311.5RT (min)

lcqtof\_t3\_pos\_4\_017\_51

m/z 1110.7878

0246E+317.518RT (min)

lcqtof\_t3\_pos\_4\_006\_41

m/z 1111.8906

024E+31515.5RT (min)

lcqtof\_t3\_pos\_4\_084\_59

m/z 1112.8069

024E+311.5RT (min)

lcqtof\_t3\_pos\_4\_008\_15

m/z 1112.9689

01E+31515.5RT (min)

lcqtof\_t3\_pos\_4\_085\_59

m/z 1113.7590

0246E+311.5RT (min)

lcqtof\_t3\_pos\_2\_031\_01

m/z 1114.7572

01234E+311.5RT (min)

lcqtof\_t3\_pos\_2\_020\_27

m/z 1116.4454

0246E+311.5RT (min)

lcqtof\_t3\_pos\_4\_015\_16

m/z 1116.7277

0246E+311.5RT (min)

lcqtof\_t3\_pos\_2\_050\_14

m/z 1116.7561

0123E+317.518RT (min)

lcqtof\_t3\_pos\_2\_028\_34

m/z 1116.8358

01234E+31313.5RT (min)

lcqtof\_t3\_pos\_2\_005\_17

m/z 1119.2688

02468E+311.5RT (min)

lcqtof\_t3\_pos\_2\_005\_17

m/z 1119.7744

024E+311.5RT (min)

lcqtof\_t3\_pos\_2\_031\_01

m/z 1121.4320

02468E+311.5RT (min)

lcqtof\_t3\_pos\_4\_015\_16

m/z 1121.7644

024E+311.5RT (min)

lcqtof\_t3\_pos\_4\_015\_16

m/z 1122.1035

01234E+311.5RT (min)

lcqtof\_t3\_pos\_4\_015\_16

m/z 1122.8291

02468E+314.515RT (min)

lcqtof\_t3\_pos\_3\_105\_61

m/z 1123.3234

01E+41414.5RT (min)

lcqtof\_t3\_pos\_3\_105\_61

m/z 1124.9354

0246E+31414.5RT (min)

lcqtof\_t3\_pos\_2\_002\_37

m/z 1125.9312

01234E+31414.5RT (min)

lcqtof\_t3\_pos\_2\_008\_31

m/z 1126.7549

0123E+317.518RT (min)

lcqtof\_t3\_pos\_3\_010\_01

m/z 1126.9917

012E+31515.5RT (min)

lcqtof\_t3\_pos\_4\_092\_60

m/z 1127.0907

0123E+311.5RT (min)

lcqtof\_t3\_pos\_4\_039\_02

m/z 1127.2611

01234E+311.5RT (min)

lcqtof\_t3\_pos\_2\_010\_26

m/z 1127.4219

0123E+311.5RT (min)

lcqtof\_t3\_pos\_4\_015\_16

m/z 1127.7694

0246E+311.5RT (min)

lcqtof\_t3\_pos\_4\_017\_51

m/z 1127.7832

012E+317.518RT (min)

lcqtof\_t3\_pos\_2\_017\_51

m/z 1130.2025

0246E+311.5RT (min)

lcqtof\_t3\_pos\_4\_021\_01

m/z 1132.0884

01234E+311.5RT (min)

lcqtof\_t3\_pos\_4\_017\_51

m/z 1132.1886

02468E+311.5RT (min)

lcqtof\_t3\_pos\_4\_021\_01

m/z 1132.7039

012E+317.518RT (min)

lcqtof\_t3\_pos\_3\_030\_40

m/z 1132.7373

024E+311.5RT (min)

lcqtof\_t3\_pos\_4\_017\_51

m/z 1133.3131

02468E+31515.5RT (min)

lcqtof\_t3\_pos\_4\_078\_58

m/z 1133.8136

024E+31515.5RT (min)

lcqtof\_t3\_pos\_4\_076\_58

m/z 1134.1855

0246E+311.5RT (min)

lcqtof\_t3\_pos\_4\_021\_01

m/z 1134.7467

0246E+311.5RT (min)

lcqtof\_t3\_pos\_4\_017\_51

m/z 1135.2424

012E+311.5RT (min)

lcqtof\_t3\_pos\_4\_017\_51

m/z 1136.1890

0246E+311.5RT (min)

lcqtof\_t3\_pos\_4\_021\_01

m/z 1136.9839

01E+414.515RT (min)

lcqtof\_t3\_pos\_4\_012\_12

m/z 1139.1100

0246E+311.5RT (min)

lcqtof\_t3\_pos\_4\_047\_13

m/z 1139.4386

024E+311.5RT (min)

lcqtof\_t3\_pos\_4\_047\_13

m/z 1140.7661

024E+311.5RT (min)

lcqtof\_t3\_pos\_4\_047\_13

m/z 1142.7405

02468E+311.5RT (min)

lcqtof\_t3\_pos\_4\_017\_51

m/z 1144.0989

02468E+311.5RT (min)

lcqtof\_t3\_pos\_4\_015\_16

m/z 1144.4353

024E+311.5RT (min)

lcqtof\_t3\_pos\_4\_015\_16

m/z 1144.8370

01234E+31313.5RT (min)

lcqtof\_t3\_pos\_2\_003\_16

m/z 1145.3385

0246E+31313.5RT (min)

lcqtof\_t3\_pos\_2\_013\_29

m/z 1145.7826

02468E+311.5RT (min)

lcqtof\_t3\_pos\_4\_047\_13

m/z 1146.5151

02468E+36.57RT (min)

lcqtof\_t3\_pos\_4\_029\_37

m/z 1147.7520

024E+311.5RT (min)

lcqtof\_t3\_pos\_3\_010\_01

m/z 1148.1798

0246E+311.5RT (min)

lcqtof\_t3\_pos\_4\_021\_01

m/z 1149.4180

024E+311.5RT (min)

lcqtof\_t3\_pos\_4\_039\_02

m/z 1150.0903

0123E+311.5RT (min)

lcqtof\_t3\_pos\_4\_039\_02

m/z 1150.1702

0246E+311.5RT (min)

lcqtof\_t3\_pos\_4\_021\_01

m/z 1152.3591

0246E+31313.5RT (min)

lcqtof\_t3\_pos\_2\_003\_16

m/z 1152.4825

02468E+36.57RT (min)

lcqtof\_t3\_pos\_4\_029\_37

m/z 1152.7498

0246E+313.514RT (min)

lcqtof\_t3\_pos\_2\_039\_23

m/z 1153.2646

0246E+311.5RT (min)

lcqtof\_t3\_pos\_4\_039\_02

m/z 1153.7727

024E+311.5RT (min)

lcqtof\_t3\_pos\_4\_039\_02

m/z 1154.7056

0246E+311.5RT (min)

lcqtof\_t3\_pos\_4\_001\_49

m/z 1155.7433

0123E+311.5RT (min)

lcqtof\_t3\_pos\_2\_031\_01

m/z 1158.7465

0123E+311.5RT (min)

lcqtof\_t3\_pos\_3\_023\_36

m/z 1159.8479

0246E+314.515RT (min)

lcqtof\_t3\_pos\_4\_037\_32

m/z 1160.8269

024E+314.515RT (min)

lcqtof\_t3\_pos\_4\_012\_12

m/z 1161.2529

024E+311.5RT (min)

lcqtof\_t3\_pos\_4\_039\_02

m/z 1161.7231

0246E+377.5RT (min)

lcqtof\_t3\_pos\_4\_029\_37

m/z 1161.7660

02468E+311.5RT (min)

lcqtof\_t3\_pos\_4\_015\_16

m/z 1162.0986

024E+311.5RT (min)

lcqtof\_t3\_pos\_4\_047\_13

m/z 1166.7567

02468E+311.5RT (min)

lcqtof\_t3\_pos\_4\_015\_16

m/z 1166.8591

0246E+31313.5RT (min)

lcqtof\_t3\_pos\_3\_029\_26

m/z 1166.9355

02468E+31414.5RT (min)

lcqtof\_t3\_pos\_4\_014\_24

m/z 1167.0994

024E+311.5RT (min)

lcqtof\_t3\_pos\_4\_015\_16

m/z 1167.3616

01234E+31313.5RT (min)

lcqtof\_t3\_pos\_3\_029\_26

m/z 1167.4299

01234E+311.5RT (min)

lcqtof\_t3\_pos\_4\_015\_16

m/z 1168.7462

012E+317.518RT (min)

lcqtof\_t3\_pos\_3\_012\_16

m/z 1168.7474

01E+411.5RT (min)

lcqtof\_t3\_pos\_4\_047\_13

m/z 1169.2421

0123E+311.5RT (min)

lcqtof\_t3\_pos\_4\_017\_51

m/z 1172.0808

024E+311.5RT (min)

lcqtof\_t3\_pos\_4\_039\_02

m/z 1172.4215

0123E+311.5RT (min)

lcqtof\_t3\_pos\_4\_039\_02

m/z 1177.8826

02468E+31313.5RT (min)

lcqtof\_t3\_pos\_2\_005\_17

m/z 1178.7749

01234E+317.518RT (min)

lcqtof\_t3\_pos\_4\_006\_41

m/z 1179.3660

0123E+31313.5RT (min)

lcqtof\_t3\_pos\_2\_040\_28

m/z 1182.7395

0123E+311.5RT (min)

lcqtof\_t3\_pos\_2\_031\_01

m/z 1184.4287

0246E+311.5RT (min)

lcqtof\_t3\_pos\_4\_047\_13

m/z 1184.7161

01234E+311.5RT (min)

lcqtof\_t3\_pos\_3\_010\_01

m/z 1184.7406

012E+317.518RT (min)

lcqtof\_t3\_pos\_2\_020\_27

m/z 1187.2614

0246E+311.5RT (min)

lcqtof\_t3\_pos\_4\_015\_16

m/z 1187.7692

024E+311.5RT (min)

lcqtof\_t3\_pos\_4\_015\_16

m/z 1189.4155

02468E+311.5RT (min)

lcqtof\_t3\_pos\_4\_015\_16

m/z 1189.7491

024E+311.5RT (min)

lcqtof\_t3\_pos\_4\_015\_16

m/z 1190.0974

01234E+311.5RT (min)

lcqtof\_t3\_pos\_4\_015\_16

m/z 1192.3420

024E+31313.5RT (min)

lcqtof\_t3\_pos\_3\_105\_61

m/z 1193.3682

02468E+313.514RT (min)

lcqtof\_t3\_pos\_3\_087\_59

m/z 1194.7211

012E+317.518RT (min)

lcqtof\_t3\_pos\_2\_045\_10

m/z 1195.0880

012E+311.5RT (min)

lcqtof\_t3\_pos\_4\_019\_03

m/z 1195.2555

01234E+311.5RT (min)

lcqtof\_t3\_pos\_4\_039\_02

m/z 1195.4182

0123E+311.5RT (min)

lcqtof\_t3\_pos\_4\_039\_02

m/z 1195.7527

0246E+311.5RT (min)

lcqtof\_t3\_pos\_4\_017\_51

m/z 1195.7812

01E+317.518RT (min)

lcqtof\_t3\_pos\_3\_006\_20

m/z 1198.3452

0246E+314.515RT (min)

lcqtof\_t3\_pos\_3\_105\_61
