## Supplementary material for "The Everything Bagel Feature Finder: Ultra-fast automated feature finding for untargeted metabolomics": SI Data File: tool_unique_xic_st_EB_ST004504.html

EB-unique features (not detected by the other two tools) - ST004504

### EB-unique features (not detected by the other two tools) - ST004504

91 features · orbitrap · XIC window m/z ± 10 ppm (≥ 0.002 Da) · ± 0.5 min

Every panel is a feature **EB** detected that neither of the other two tools detected — three-way unique across EB (standard mode (min\_detection\_rate=0.1)), Masscube and XCMS, under the standard matching rule (m/z ± 10 ppm; apex within the reference RT window ± 0.2 min). It shows the MS1 extracted-ion chromatogram at the feature's m/z, drawn from the raw mzML named in `alignment_reference_file`, zoomed to ± 0.5 min around the feature's reported apex RT. **Gray solid line** = the feature's reported apex RT; **red dashed line** = EB's assigned MS/MS — `MS2_scan_id` (native scan number) in `MS2_source_file`; the panel widens to keep an off-apex MS/MS on-screen. Features with no assigned MS/MS show only the gray line. y is per-cell autoscaled. Rows keep the table order (m/z ascending).

m/z 85.3949

0123E+513.514RT (min)

lcorbi\_t3\_pos\_1\_032\_26

m/z 119.0807

01E+511.5RT (min)

lcorbi\_t3\_pos\_4\_095\_60

m/z 120.0904

01E+511.5RT (min)

lcorbi\_t3\_pos\_4\_076\_58

m/z 130.0692

012E+511.5RT (min)

lcorbi\_t3\_pos\_4\_080\_58

m/z 134.0188

01E+511.5RT (min)

lcorbi\_t3\_pos\_1\_028\_32

m/z 134.0634

01234E+511.5RT (min)

lcorbi\_t3\_pos\_4\_090\_60

m/z 134.0826

0246E+511.5RT (min)

lcorbi\_t3\_pos\_4\_091\_60

m/z 138.0582

02468E+611.5RT (min)

lcorbi\_t3\_pos\_4\_097\_60

m/z 138.1221

024E+522.5RT (min)

lcorbi\_t3\_pos\_1\_036\_08

m/z 147.0861

0123E+511.5RT (min)

lcorbi\_t3\_pos\_4\_090\_60

m/z 148.0393

01E+522.5RT (min)

lcorbi\_t3\_pos\_1\_034\_34

m/z 148.0733

0123E+511.5RT (min)

lcorbi\_t3\_pos\_4\_085\_59

m/z 148.0757

012E+522.5RT (min)

lcorbi\_t3\_pos\_2\_053\_28

m/z 149.0791

01E+511.5RT (min)

lcorbi\_t3\_pos\_4\_112\_62

m/z 150.0647

012E+511.5RT (min)

lcorbi\_t3\_pos\_4\_089\_59

m/z 154.0434

01E+511.5RT (min)

lcorbi\_t3\_pos\_4\_094\_60

m/z 154.0838

01E+522.5RT (min)

lcorbi\_t3\_pos\_4\_006\_12

m/z 155.0815

01E+511.5RT (min)

lcorbi\_t3\_pos\_4\_096\_60

m/z 157.0486

012E+511.5RT (min)

lcorbi\_t3\_pos\_2\_024\_15

m/z 164.1125

02468E+511.5RT (min)

lcorbi\_t3\_pos\_4\_090\_60

m/z 168.1019

01E+511.5RT (min)

lcorbi\_t3\_pos\_4\_070\_57

m/z 170.0577

01E+511.5RT (min)

lcorbi\_t3\_pos\_4\_083\_59

m/z 175.0390

012E+57.58RT (min)

lcorbi\_t3\_pos\_4\_100\_61

m/z 177.1063

012E+511.5RT (min)

lcorbi\_t3\_pos\_4\_082\_59

m/z 182.0811

01E+511.5RT (min)

lcorbi\_t3\_pos\_2\_020\_22

m/z 187.0788

024E+511.5RT (min)

lcorbi\_t3\_pos\_4\_073\_57

m/z 189.0468

01E+511.5RT (min)

lcorbi\_t3\_pos\_1\_020\_31

m/z 189.0900

0246E+511.5RT (min)

lcorbi\_t3\_pos\_4\_094\_60

m/z 191.0316

01E+611.5RT (min)

lcorbi\_t3\_pos\_1\_034\_34

m/z 197.0271

02468E+411.5RT (min)

lcorbi\_t3\_pos\_1\_008\_36

m/z 203.0339

0123E+588.5RT (min)

lcorbi\_t3\_pos\_3\_034\_26

m/z 218.1387

01E+522.5RT (min)

lcorbi\_t3\_pos\_4\_021\_25

m/z 218.1991

024E+56.57RT (min)

lcorbi\_t3\_pos\_4\_074\_58

m/z 220.0862

012E+511.5RT (min)

lcorbi\_t3\_pos\_1\_034\_34

m/z 221.0185

01234E+511.5RT (min)

lcorbi\_t3\_pos\_4\_075\_58

m/z 230.0983

0123E+511.5RT (min)

lcorbi\_t3\_pos\_4\_090\_60

m/z 240.2322

01E+510.511RT (min)

lcorbi\_t3\_pos\_4\_105\_61

m/z 243.9288

01E+511.5RT (min)

lcorbi\_t3\_pos\_1\_029\_29

m/z 244.8706

01E+511.5RT (min)

lcorbi\_t3\_pos\_1\_036\_08

m/z 248.8705

012E+511.5RT (min)

lcorbi\_t3\_pos\_1\_029\_29

m/z 258.1183

01E+511.5RT (min)

lcorbi\_t3\_pos\_3\_003\_33

m/z 261.9626

01234E+511.5RT (min)

lcorbi\_t3\_pos\_1\_036\_08

m/z 262.9070

012E+511.5RT (min)

lcorbi\_t3\_pos\_1\_036\_08

m/z 268.1410

01E+511.5RT (min)

lcorbi\_t3\_pos\_3\_027\_27

m/z 274.8813

012E+511.5RT (min)

lcorbi\_t3\_pos\_1\_020\_31

m/z 280.8473

01E+511.5RT (min)

lcorbi\_t3\_pos\_1\_034\_34

m/z 284.9102

0123E+511.5RT (min)

lcorbi\_t3\_pos\_1\_020\_31

m/z 290.1347

01E+511.5RT (min)

lcorbi\_t3\_pos\_4\_086\_59

m/z 300.0948

012E+57.58RT (min)

lcorbi\_t3\_pos\_4\_105\_61

m/z 313.0293

01E+511.5RT (min)

lcorbi\_t3\_pos\_1\_034\_34

m/z 319.2268

012E+511.512RT (min)

lcorbi\_t3\_pos\_1\_037\_19

m/z 319.2268

0123E+510.511RT (min)

lcorbi\_t3\_pos\_1\_037\_19

m/z 323.0115

012E+511.5RT (min)

lcorbi\_t3\_pos\_3\_031\_45

m/z 326.1842

01E+54.55RT (min)

lcorbi\_t3\_pos\_4\_112\_62

m/z 326.8869

01E+511.5RT (min)

lcorbi\_t3\_pos\_1\_029\_29

m/z 333.9653

01234E+511.5RT (min)

lcorbi\_t3\_pos\_2\_022\_38

m/z 344.1495

0123E+511.5RT (min)

lcorbi\_t3\_pos\_4\_090\_60

m/z 347.2583

0123E+51111.5RT (min)

lcorbi\_t3\_pos\_4\_108\_62

m/z 354.8949

012E+511.5RT (min)

lcorbi\_t3\_pos\_1\_020\_31

m/z 356.2303

012E+599.5RT (min)

lcorbi\_t3\_pos\_1\_021\_18

m/z 361.0782

01234E+511.5RT (min)

lcorbi\_t3\_pos\_3\_010\_44

m/z 371.1104

012E+57.58RT (min)

lcorbi\_t3\_pos\_4\_105\_61

m/z 380.3373

01E+51010.5RT (min)

lcorbi\_t3\_pos\_3\_021\_39

m/z 382.7671

01E+511.5RT (min)

lcorbi\_t3\_pos\_4\_070\_57

m/z 390.7989

01E+511.5RT (min)

lcorbi\_t3\_pos\_2\_028\_24

m/z 394.8743

0123E+511.5RT (min)

lcorbi\_t3\_pos\_1\_029\_29

m/z 398.7617

01E+511.5RT (min)

lcorbi\_t3\_pos\_3\_024\_07

m/z 400.8275

0123E+511.5RT (min)

lcorbi\_t3\_pos\_2\_028\_24

m/z 409.3104

01234E+513.514RT (min)

lcorbi\_t3\_pos\_4\_111\_62

m/z 412.8534

01234E+511.5RT (min)

lcorbi\_t3\_pos\_2\_028\_24

m/z 419.3222

01E+61414.5RT (min)

lcorbi\_t3\_pos\_1\_008\_36

m/z 422.8821

01234E+511.5RT (min)

lcorbi\_t3\_pos\_3\_012\_31

m/z 432.1476

0123E+522.5RT (min)

lcorbi\_t3\_pos\_4\_004\_14

m/z 488.6553

012E+511.5RT (min)

lcorbi\_t3\_pos\_4\_067\_57

m/z 496.3676

012E+51111.5RT (min)

lcorbi\_t3\_pos\_1\_034\_34

m/z 500.9059

01E+511.5RT (min)

lcorbi\_t3\_pos\_4\_068\_57

m/z 504.8466

0123E+511.5RT (min)

lcorbi\_t3\_pos\_4\_080\_58

m/z 512.4153

01E+51010.5RT (min)

lcorbi\_t3\_pos\_3\_020\_35

m/z 520.3275

01E+51111.5RT (min)

lcorbi\_t3\_pos\_1\_029\_29

m/z 524.4007

01E+511.512RT (min)

lcorbi\_t3\_pos\_1\_029\_29

m/z 598.8370

0123E+511.5RT (min)

lcorbi\_t3\_pos\_4\_075\_58

m/z 613.1595

01E+511.5RT (min)

lcorbi\_t3\_pos\_3\_011\_42

m/z 629.5108

012E+51515.5RT (min)

lcorbi\_t3\_pos\_1\_054\_48

m/z 666.6107

01E+51515.5RT (min)

lcorbi\_t3\_pos\_4\_081\_58

m/z 676.5320

012E+51414.5RT (min)

lcorbi\_t3\_pos\_1\_028\_32

m/z 682.5982

012E+51515.5RT (min)

lcorbi\_t3\_pos\_4\_070\_57

m/z 727.5103

012E+51313.5RT (min)

lcorbi\_t3\_pos\_4\_032\_40

m/z 766.6420

0123E+513.514RT (min)

lcorbi\_t3\_pos\_1\_036\_08

m/z 788.5208

0123E+513.514RT (min)

lcorbi\_t3\_pos\_4\_083\_59

m/z 798.6506

0123E+513.514RT (min)

lcorbi\_t3\_pos\_1\_003\_45

m/z 854.7227

012E+51515.5RT (min)

lcorbi\_t3\_pos\_4\_083\_59
