## Supplementary material for "The Everything Bagel Feature Finder: Ultra-fast automated feature finding for untargeted metabolomics": SI Data File: tool_unique_xic_st_EB_ST004626.html

EB-unique features (not detected by the other two tools) - ST004626

### EB-unique features (not detected by the other two tools) - ST004626

1800 features · qtof · XIC window m/z ± 25 ppm (≥ 0.002 Da) · ± 0.5 min

Every panel is a feature **EB** detected that neither of the other two tools detected — three-way unique across EB (standard mode (min\_detection\_rate=0.1)), Masscube and XCMS, under the standard matching rule (m/z ± 25 ppm; apex within the reference RT window ± 0.2 min). It shows the MS1 extracted-ion chromatogram at the feature's m/z, drawn from the raw mzML named in `alignment_reference_file`, zoomed to ± 0.5 min around the feature's reported apex RT. **Gray solid line** = the feature's reported apex RT; **red dashed line** = EB's assigned MS/MS — `MS2_scan_id` (native scan number) in `MS2_source_file`; the panel widens to keep an off-apex MS/MS on-screen. Features with no assigned MS/MS show only the gray line. y is per-cell autoscaled. Rows keep the table order (m/z ascending).

m/z 74.0227

0246E+311.5RT (min)

lcqtof\_t3\_neg\_1\_055\_32

m/z 74.0233

01234E+311.5RT (min)

lcqtof\_t3\_neg\_4\_040\_24

m/z 79.9559

01234E+355.5RT (min)

lcqtof\_t3\_neg\_4\_046\_33

m/z 80.9630

0123E+311.5RT (min)

lcqtof\_t3\_neg\_4\_019\_05

m/z 86.9939

012E+311.5RT (min)

lcqtof\_t3\_neg\_1\_036\_14

m/z 89.0434

0123E+311.5RT (min)

lcqtof\_t3\_neg\_1\_015\_34

m/z 90.0472

012E+311.5RT (min)

lcqtof\_t3\_neg\_1\_055\_32

m/z 93.0338

0123E+34.55RT (min)

lcqtof\_t3\_neg\_1\_018\_33

m/z 97.9599

0123E+311.5RT (min)

lcqtof\_t3\_neg\_4\_040\_24

m/z 102.9707

012E+311.5RT (min)

lcqtof\_t3\_neg\_1\_036\_14

m/z 103.0583

01234E+311.5RT (min)

lcqtof\_t3\_neg\_3\_006\_23

m/z 104.0346

01234E+311.5RT (min)

lcqtof\_t3\_neg\_1\_015\_34

m/z 107.0446

0123E+311.5RT (min)

lcqtof\_t3\_neg\_1\_055\_32

m/z 108.9014

024E+311.5RT (min)

lcqtof\_t3\_neg\_4\_040\_24

m/z 110.8980

01234E+311.5RT (min)

lcqtof\_t3\_neg\_4\_040\_24

m/z 110.9835

0123E+311.5RT (min)

lcqtof\_t3\_neg\_4\_040\_24

m/z 113.0346

0246E+311.5RT (min)

lcqtof\_t3\_neg\_1\_015\_34

m/z 116.0499

012E+344.5RT (min)

lcqtof\_t3\_neg\_1\_044\_37

m/z 117.0480

02468E+311.5RT (min)

lcqtof\_t3\_neg\_1\_055\_32

m/z 119.0720

0246E+311.5RT (min)

lcqtof\_t3\_neg\_1\_055\_32

m/z 122.0641

0123E+311.5RT (min)

lcqtof\_t3\_neg\_1\_055\_32

m/z 124.0150

01234E+311.5RT (min)

lcqtof\_t3\_neg\_4\_037\_01

m/z 125.0234

0246E+311.5RT (min)

lcqtof\_t3\_neg\_4\_018\_23

m/z 125.0245

01234E+344.5RT (min)

lcqtof\_t3\_neg\_4\_030\_31

m/z 129.0185

01E+344.5RT (min)

lcqtof\_t3\_neg\_1\_009\_50

m/z 129.0839

024E+344.5RT (min)

lcqtof\_t3\_neg\_3\_018\_32

m/z 130.0872

01234E+311.5RT (min)

lcqtof\_t3\_neg\_1\_032\_36

m/z 131.0342

0123E+311.5RT (min)

lcqtof\_t3\_neg\_3\_006\_23

m/z 131.0443

012E+344.5RT (min)

lcqtof\_t3\_neg\_3\_032\_24

m/z 132.0874

0246E+311.5RT (min)

lcqtof\_t3\_neg\_1\_032\_36

m/z 133.0523

024E+344.5RT (min)

lcqtof\_t3\_neg\_3\_032\_24

m/z 134.0169

0246E+311.5RT (min)

lcqtof\_t3\_neg\_4\_018\_23

m/z 134.0337

024E+311.5RT (min)

lcqtof\_t3\_neg\_1\_032\_36

m/z 135.0955

024E+311.5RT (min)

lcqtof\_t3\_neg\_4\_040\_24

m/z 136.0158

0123E+388.5RT (min)

lcqtof\_t3\_neg\_2\_053\_38

m/z 136.0236

0246E+311.5RT (min)

lcqtof\_t3\_neg\_4\_040\_24

m/z 137.0349

012E+344.5RT (min)

lcqtof\_t3\_neg\_1\_029\_44

m/z 137.0437

0246E+311.5RT (min)

lcqtof\_t3\_neg\_1\_055\_32

m/z 138.0263

024E+36.57RT (min)

lcqtof\_t3\_neg\_4\_034\_34

m/z 138.8754

012E+311.5RT (min)

lcqtof\_t3\_neg\_4\_052\_03\_rerun

m/z 139.0063

01E+344.5RT (min)

lcqtof\_t3\_neg\_1\_041\_54

m/z 140.9171

01E+411.5RT (min)

lcqtof\_t3\_neg\_4\_052\_03\_rerun

m/z 142.0498

01234E+311.5RT (min)

lcqtof\_t3\_neg\_2\_030\_23

m/z 142.0658

01E+344.5RT (min)

lcqtof\_t3\_neg\_2\_026\_36

m/z 143.0347

01E+311.5RT (min)

lcqtof\_t3\_neg\_4\_018\_23

m/z 143.0457

0123E+311.5RT (min)

lcqtof\_t3\_neg\_4\_034\_34

m/z 145.0509

01E+34.55RT (min)

lcqtof\_t3\_neg\_3\_087\_59

m/z 146.0623

012E+344.5RT (min)

lcqtof\_t3\_neg\_2\_041\_54

m/z 146.0638

02468E+311.5RT (min)

lcqtof\_t3\_neg\_1\_012\_16

m/z 146.9458

012E+311.5RT (min)

lcqtof\_t3\_neg\_4\_044\_48

m/z 147.0664

012E+344.5RT (min)

lcqtof\_t3\_neg\_2\_005\_34

m/z 147.0664

01234E+344.5RT (min)

lcqtof\_t3\_neg\_4\_034\_34

m/z 149.0238

01E+477.5RT (min)

lcqtof\_t3\_neg\_4\_102\_61

m/z 149.0747

01234E+311.5RT (min)

lcqtof\_t3\_neg\_1\_055\_32

m/z 150.0114

01234E+311.5RT (min)

lcqtof\_t3\_neg\_1\_051\_26

m/z 150.0559

01E+344.5RT (min)

lcqtof\_t3\_neg\_1\_053\_01

m/z 152.9176

012E+311.5RT (min)

lcqtof\_t3\_neg\_1\_018\_33

m/z 153.0670

0123E+311.5RT (min)

lcqtof\_t3\_neg\_2\_028\_24

m/z 155.0032

012E+311.5RT (min)

lcqtof\_t3\_neg\_4\_040\_24

m/z 155.0455

012E+311.5RT (min)

lcqtof\_t3\_neg\_4\_016\_14

m/z 157.0153

024E+311.5RT (min)

lcqtof\_t3\_neg\_4\_105\_61

m/z 158.0822

0246E+311.5RT (min)

lcqtof\_t3\_neg\_2\_028\_24

m/z 158.1002

012E+311.5RT (min)

lcqtof\_t3\_neg\_1\_036\_14

m/z 159.0778

0246E+311.5RT (min)

lcqtof\_t3\_neg\_4\_040\_24

m/z 160.1166

01234E+311.5RT (min)

lcqtof\_t3\_neg\_1\_038\_15

m/z 161.0465

012E+311.5RT (min)

lcqtof\_t3\_neg\_4\_051\_15

m/z 161.0644

012E+316.517RT (min)

lcqtof\_t3\_neg\_4\_046\_33

m/z 161.0650

012E+311.5RT (min)

lcqtof\_t3\_neg\_2\_030\_23

m/z 161.9935

012E+311.5RT (min)

lcqtof\_t3\_neg\_1\_015\_34

m/z 163.0331

0246E+311.5RT (min)

lcqtof\_t3\_neg\_4\_004\_20

m/z 163.0817

012E+311.5RT (min)

lcqtof\_t3\_neg\_1\_055\_32

m/z 164.0374

01234E+311.5RT (min)

lcqtof\_t3\_neg\_1\_032\_36

m/z 164.0634

012E+311.5RT (min)

lcqtof\_t3\_neg\_3\_097\_60

m/z 164.9982

01E+311.5RT (min)

lcqtof\_t3\_neg\_2\_042\_13

m/z 165.0417

01234E+344.5RT (min)

lcqtof\_t3\_neg\_1\_002\_13

m/z 165.0543

0123E+35.56RT (min)

lcqtof\_t3\_neg\_4\_016\_14

m/z 165.8414

02468E+311.5RT (min)

lcqtof\_t3\_neg\_4\_014\_13

m/z 167.0222

012E+344.5RT (min)

lcqtof\_t3\_neg\_1\_046\_17

m/z 169.0883

012E+344.5RT (min)

lcqtof\_t3\_neg\_1\_040\_08

m/z 170.9973

0123E+311.5RT (min)

lcqtof\_t3\_neg\_1\_012\_16

m/z 171.0768

02468E+311.5RT (min)

lcqtof\_t3\_neg\_1\_038\_15

m/z 171.0930

01E+344.5RT (min)

lcqtof\_t3\_neg\_4\_052\_03\_rerun

m/z 171.1013

01E+36.57RT (min)

lcqtof\_t3\_neg\_3\_087\_59

m/z 171.8316

01E+411.5RT (min)

lcqtof\_t3\_neg\_4\_045\_02\_rerun

m/z 173.0464

0123E+344.5RT (min)

lcqtof\_t3\_neg\_4\_018\_23

m/z 173.0466

0123E+344.5RT (min)

lcqtof\_t3\_neg\_1\_044\_37

m/z 173.0559

0123E+311.5RT (min)

lcqtof\_t3\_neg\_1\_022\_28

m/z 173.1009

0246E+344.5RT (min)

lcqtof\_t3\_neg\_2\_019\_32

m/z 173.8302

01E+411.5RT (min)

lcqtof\_t3\_neg\_4\_052\_03\_rerun

m/z 175.0151

0246E+311.5RT (min)

lcqtof\_t3\_neg\_4\_103\_61

m/z 175.0718

01234E+311.5RT (min)

lcqtof\_t3\_neg\_1\_022\_28

m/z 176.0060

012E+316.517RT (min)

lcqtof\_t3\_neg\_2\_019\_32

m/z 176.0092

012E+311.5RT (min)

lcqtof\_t3\_neg\_4\_037\_01

m/z 176.0637

024E+34.55RT (min)

lcqtof\_t3\_neg\_4\_018\_23

m/z 176.9097

01234E+311.5RT (min)

lcqtof\_t3\_neg\_1\_055\_32

m/z 178.0581

02468E+35.56RT (min)

lcqtof\_t3\_neg\_4\_041\_54

m/z 179.0542

012E+44.55RT (min)

lcqtof\_t3\_neg\_4\_037\_01

m/z 180.1225

02468E+311.5RT (min)

lcqtof\_t3\_neg\_4\_040\_24

m/z 181.0700

012E+311.5RT (min)

lcqtof\_t3\_neg\_4\_002\_17

m/z 181.9658

01E+311.5RT (min)

lcqtof\_t3\_neg\_1\_022\_28

m/z 182.0428

0246E+311.5RT (min)

lcqtof\_t3\_neg\_2\_043\_31

m/z 182.0537

02468E+344.5RT (min)

lcqtof\_t3\_neg\_4\_015\_36

m/z 183.8475

02468E+311.5RT (min)

lcqtof\_t3\_neg\_1\_010\_27

m/z 184.9912

012E+344.5RT (min)

lcqtof\_t3\_neg\_3\_031\_19

m/z 185.0557

012E+311.5RT (min)

lcqtof\_t3\_neg\_1\_053\_01

m/z 186.0373

012E+311.5RT (min)

lcqtof\_t3\_neg\_4\_015\_36

m/z 186.0768

0246E+344.5RT (min)

lcqtof\_t3\_neg\_4\_030\_31

m/z 186.9285

01E+411.5RT (min)

lcqtof\_t3\_neg\_3\_018\_32

m/z 188.0428

01234E+311.5RT (min)

lcqtof\_t3\_neg\_4\_030\_31

m/z 188.0505

01234E+344.5RT (min)

lcqtof\_t3\_neg\_4\_018\_23

m/z 188.0567

012E+311.5RT (min)

lcqtof\_t3\_neg\_2\_030\_23

m/z 188.0690

0123E+311.5RT (min)

lcqtof\_t3\_neg\_1\_015\_34

m/z 189.0391

01E+344.5RT (min)

lcqtof\_t3\_neg\_1\_037\_25

m/z 189.0551

01E+316.517RT (min)

lcqtof\_t3\_neg\_4\_010\_46

m/z 189.0641

0123E+344.5RT (min)

lcqtof\_t3\_neg\_4\_027\_41

m/z 190.0063

0123E+311.5RT (min)

lcqtof\_t3\_neg\_1\_051\_26

m/z 190.9828

0123E+344.5RT (min)

lcqtof\_t3\_neg\_1\_044\_37

m/z 191.0662

012E+311.5RT (min)

lcqtof\_t3\_neg\_1\_022\_28

m/z 193.0999

0123E+34.55RT (min)

lcqtof\_t3\_neg\_1\_043\_38

m/z 194.0345

0123E+37.58RT (min)

lcqtof\_t3\_neg\_4\_005\_18

m/z 194.0457

0123E+35.56RT (min)

lcqtof\_t3\_neg\_4\_037\_01

m/z 195.0660

012E+34.55RT (min)

lcqtof\_t3\_neg\_4\_048\_19

m/z 196.8110

0246E+311.5RT (min)

lcqtof\_t3\_neg\_4\_021\_21

m/z 196.9753

01234E+311.5RT (min)

lcqtof\_t3\_neg\_4\_039\_06\_rerun

m/z 197.0207

01E+366.5RT (min)

lcqtof\_t3\_neg\_3\_020\_10

m/z 197.0551

012E+311.5RT (min)

lcqtof\_t3\_neg\_3\_012\_14

m/z 197.0681

024E+344.5RT (min)

lcqtof\_t3\_neg\_1\_002\_13

m/z 198.0785

012E+344.5RT (min)

lcqtof\_t3\_neg\_4\_030\_31

m/z 200.0082

0123E+344.5RT (min)

lcqtof\_t3\_neg\_1\_041\_54

m/z 200.0090

0123E+344.5RT (min)

lcqtof\_t3\_neg\_4\_039\_06\_rerun

m/z 200.0246

012E+316.517RT (min)

lcqtof\_t3\_neg\_4\_004\_20

m/z 200.0593

012E+316.517RT (min)

lcqtof\_t3\_neg\_4\_032\_04

m/z 200.9743

012E+311.5RT (min)

lcqtof\_t3\_neg\_4\_052\_03\_rerun

m/z 200.9994

012E+344.5RT (min)

lcqtof\_t3\_neg\_1\_037\_25

m/z 201.0349

012E+311.5RT (min)

lcqtof\_t3\_neg\_1\_036\_14

m/z 201.0718

012E+355.5RT (min)

lcqtof\_t3\_neg\_1\_051\_26

m/z 201.1245

0123E+344.5RT (min)

lcqtof\_t3\_neg\_3\_012\_14

m/z 203.0017

01E+44.55RT (min)

lcqtof\_t3\_neg\_4\_046\_33

m/z 203.0558

012E+344.5RT (min)

lcqtof\_t3\_neg\_2\_043\_31

m/z 204.0558

01234E+311.5RT (min)

lcqtof\_t3\_neg\_4\_087\_59

m/z 204.0577

0123E+316.517RT (min)

lcqtof\_t3\_neg\_4\_039\_06

m/z 204.1065

024E+311.5RT (min)

lcqtof\_t3\_neg\_1\_036\_14

m/z 204.7909

0123E+311.5RT (min)

lcqtof\_t3\_neg\_4\_105\_61

m/z 205.0525

01E+416.517RT (min)

lcqtof\_t3\_neg\_4\_051\_15

m/z 207.1065

024E+344.5RT (min)

lcqtof\_t3\_neg\_4\_011\_32

m/z 207.1139

01234E+35.56RT (min)

lcqtof\_t3\_neg\_2\_043\_31

m/z 209.0531

01234E+311.5RT (min)

lcqtof\_t3\_neg\_4\_044\_48

m/z 209.0678

012E+311.5RT (min)

lcqtof\_t3\_neg\_4\_037\_01

m/z 209.9746

0123E+311.5RT (min)

lcqtof\_t3\_neg\_4\_040\_24

m/z 210.0347

0123E+311.5RT (min)

lcqtof\_t3\_neg\_1\_055\_32

m/z 211.0177

02468E+311.5RT (min)

lcqtof\_t3\_neg\_1\_055\_32

m/z 214.0185

01E+311.5RT (min)

lcqtof\_t3\_neg\_4\_034\_34

m/z 214.9961

024E+311.5RT (min)

lcqtof\_t3\_neg\_4\_034\_34

m/z 215.0021

01E+344.5RT (min)

lcqtof\_t3\_neg\_1\_046\_17

m/z 215.0021

012E+35.56RT (min)

lcqtof\_t3\_neg\_4\_003\_37

m/z 215.1014

024E+311.5RT (min)

lcqtof\_t3\_neg\_4\_040\_24

m/z 215.8670

024E+311.5RT (min)

lcqtof\_t3\_neg\_1\_002\_13

m/z 216.0703

01234E+311.5RT (min)

lcqtof\_t3\_neg\_1\_036\_14

m/z 216.9082

012E+311.5RT (min)

lcqtof\_t3\_neg\_4\_094\_60

m/z 217.0349

012E+344.5RT (min)

lcqtof\_t3\_neg\_3\_021\_34

m/z 218.0715

012E+316.517RT (min)

lcqtof\_t3\_neg\_4\_004\_20

m/z 219.0354

0123E+311.5RT (min)

lcqtof\_t3\_neg\_4\_037\_01

m/z 219.1078

012E+344.5RT (min)

lcqtof\_t3\_neg\_2\_043\_31

m/z 219.9718

012E+311.5RT (min)

lcqtof\_t3\_neg\_1\_032\_36

m/z 221.0294

02468E+311.5RT (min)

lcqtof\_t3\_neg\_4\_008\_35

m/z 222.9152

024E+311.5RT (min)

lcqtof\_t3\_neg\_4\_011\_32

m/z 223.0711

012E+311.5RT (min)

lcqtof\_t3\_neg\_1\_036\_14

m/z 223.0734

0123E+34.55RT (min)

lcqtof\_t3\_neg\_4\_018\_23

m/z 223.0811

012E+311.5RT (min)

lcqtof\_t3\_neg\_4\_041\_54

m/z 224.0847

0123E+311.5RT (min)

lcqtof\_t3\_neg\_1\_020\_47

m/z 226.0933

012E+311.5RT (min)

lcqtof\_t3\_neg\_4\_016\_14

m/z 226.9858

01234E+311.5RT (min)

lcqtof\_t3\_neg\_4\_055\_38

m/z 227.0278

0123E+344.5RT (min)

lcqtof\_t3\_neg\_1\_035\_31

m/z 227.0919

012E+34.55RT (min)

lcqtof\_t3\_neg\_4\_055\_38

m/z 228.8125

01234E+311.5RT (min)

lcqtof\_t3\_neg\_4\_016\_14

m/z 229.0826

01E+411.5RT (min)

lcqtof\_t3\_neg\_1\_036\_14

m/z 229.1545

01234E+344.5RT (min)

lcqtof\_t3\_neg\_4\_016\_14

m/z 229.9167

01E+316.517RT (min)

lcqtof\_t3\_neg\_1\_015\_34

m/z 229.9989

01234E+311.5RT (min)

lcqtof\_t3\_neg\_2\_030\_23

m/z 230.0125

01E+44.55RT (min)

lcqtof\_t3\_neg\_4\_046\_33

m/z 232.0220

0246E+311.5RT (min)

lcqtof\_t3\_neg\_4\_018\_23

m/z 232.0455

012E+311.5RT (min)

lcqtof\_t3\_neg\_1\_032\_36

m/z 232.0607

0123E+344.5RT (min)

lcqtof\_t3\_neg\_4\_045\_02\_rerun

m/z 232.0963

0123E+34.55RT (min)

lcqtof\_t3\_neg\_4\_044\_48

m/z 233.0646

012E+34.55RT (min)

lcqtof\_t3\_neg\_3\_031\_19

m/z 233.1049

02468E+34.55RT (min)

lcqtof\_t3\_neg\_4\_011\_32

m/z 235.9263

012E+311.5RT (min)

lcqtof\_t3\_neg\_1\_029\_44

m/z 236.0518

012E+344.5RT (min)

lcqtof\_t3\_neg\_2\_028\_24

m/z 237.0391

0123E+37.58RT (min)

lcqtof\_t3\_neg\_4\_026\_30

m/z 237.0877

01E+38.59RT (min)

lcqtof\_t3\_neg\_2\_036\_06

m/z 238.0667

0123E+311.5RT (min)

lcqtof\_t3\_neg\_1\_055\_32

m/z 238.0793

012E+344.5RT (min)

lcqtof\_t3\_neg\_1\_005\_18

m/z 239.9343

0246E+311.5RT (min)

lcqtof\_t3\_neg\_4\_037\_01

m/z 240.0991

012E+344.5RT (min)

lcqtof\_t3\_neg\_1\_041\_54

m/z 241.0587

012E+311.5RT (min)

lcqtof\_t3\_neg\_1\_036\_14

m/z 241.9315

012E+311.5RT (min)

lcqtof\_t3\_neg\_4\_051\_15

m/z 243.0083

012E+34.55RT (min)

lcqtof\_t3\_neg\_1\_015\_34

m/z 243.0269

0123E+311.5RT (min)

lcqtof\_t3\_neg\_4\_018\_23

m/z 243.9558

012E+311.5RT (min)

lcqtof\_t3\_neg\_4\_034\_34

m/z 244.0331

0123E+316.517RT (min)

lcqtof\_t3\_neg\_4\_003\_37

m/z 244.9655

0246E+311.5RT (min)

lcqtof\_t3\_neg\_1\_001\_49

m/z 245.0663

012E+34.55RT (min)

lcqtof\_t3\_neg\_4\_018\_23

m/z 245.9102

0123E+377.5RT (min)

lcqtof\_t3\_neg\_4\_016\_14

m/z 246.0362

0123E+34.55RT (min)

lcqtof\_t3\_neg\_4\_037\_01

m/z 246.0471

0246E+311.5RT (min)

lcqtof\_t3\_neg\_1\_019\_23

m/z 247.1214

01234E+34.55RT (min)

lcqtof\_t3\_neg\_3\_018\_32

m/z 247.9079

0123E+377.5RT (min)

lcqtof\_t3\_neg\_4\_016\_14

m/z 248.0237

0246E+311.5RT (min)

lcqtof\_t3\_neg\_4\_112\_62

m/z 248.0661

012E+316.517RT (min)

lcqtof\_t3\_neg\_4\_033\_53

m/z 248.9001

0123E+311.5RT (min)

lcqtof\_t3\_neg\_2\_026\_36

m/z 249.0406

02468E+311.5RT (min)

lcqtof\_t3\_neg\_4\_045\_02\_rerun

m/z 250.1090

012E+34.55RT (min)

lcqtof\_t3\_neg\_1\_044\_37

m/z 250.1097

0123E+34.55RT (min)

lcqtof\_t3\_neg\_4\_037\_01

m/z 250.8424

01234E+311.5RT (min)

lcqtof\_t3\_neg\_4\_011\_32

m/z 251.0938

012E+311.5RT (min)

lcqtof\_t3\_neg\_2\_026\_36

m/z 252.8274

012E+311.5RT (min)

lcqtof\_t3\_neg\_2\_042\_13

m/z 254.0769

0123E+344.5RT (min)

lcqtof\_t3\_neg\_1\_042\_46

m/z 254.0963

012E+311.5RT (min)

lcqtof\_t3\_neg\_4\_047\_07

m/z 254.8330

012E+311.5RT (min)

lcqtof\_t3\_neg\_4\_046\_33

m/z 254.9825

01E+311.5RT (min)

lcqtof\_t3\_neg\_1\_055\_32

m/z 254.9872

01234E+34.55RT (min)

lcqtof\_t3\_neg\_4\_016\_14

m/z 254.9935

024E+355.5RT (min)

lcqtof\_t3\_neg\_4\_046\_33

m/z 255.0198

024E+311.5RT (min)

lcqtof\_t3\_neg\_4\_024\_26

m/z 255.9292

0123E+311.5RT (min)

lcqtof\_t3\_neg\_1\_055\_32

m/z 256.1108

024E+311.5RT (min)

lcqtof\_t3\_neg\_2\_008\_07

m/z 256.8766

0246E+311.5RT (min)

lcqtof\_t3\_neg\_1\_017\_51

m/z 256.9023

0123E+311.5RT (min)

lcqtof\_t3\_neg\_4\_046\_33

m/z 256.9293

012E+311.5RT (min)

lcqtof\_t3\_neg\_3\_018\_32

m/z 257.0788

01E+344.5RT (min)

lcqtof\_t3\_neg\_1\_044\_37

m/z 257.1503

012E+344.5RT (min)

lcqtof\_t3\_neg\_3\_045\_31

m/z 258.1154

01234E+311.5RT (min)

lcqtof\_t3\_neg\_4\_018\_23

m/z 258.8743

012E+311.5RT (min)

lcqtof\_t3\_neg\_1\_017\_51

m/z 258.9823

01E+311.5RT (min)

lcqtof\_t3\_neg\_4\_045\_02\_rerun

m/z 260.0522

024E+311.5RT (min)

lcqtof\_t3\_neg\_4\_039\_06\_rerun

m/z 260.0788

012E+344.5RT (min)

lcqtof\_t3\_neg\_1\_044\_37

m/z 260.8727

012E+316.517RT (min)

lcqtof\_t3\_neg\_1\_027\_21

m/z 260.8797

0246E+311.5RT (min)

lcqtof\_t3\_neg\_1\_033\_53

m/z 261.1324

012E+366.5RT (min)

lcqtof\_t3\_neg\_4\_005\_18

m/z 261.1371

0123E+34.55RT (min)

lcqtof\_t3\_neg\_1\_044\_37

m/z 262.8672

0123E+311.5RT (min)

lcqtof\_t3\_neg\_4\_030\_31

m/z 262.9787

01E+311.5RT (min)

lcqtof\_t3\_neg\_3\_019\_37

m/z 263.0289

01E+34.55RT (min)

lcqtof\_t3\_neg\_1\_044\_37

m/z 263.1440

0246E+35.56RT (min)

lcqtof\_t3\_neg\_4\_011\_32

m/z 264.0349

01234E+34.55RT (min)

lcqtof\_t3\_neg\_1\_046\_17

m/z 264.0945

0246E+35.56RT (min)

lcqtof\_t3\_neg\_4\_100\_61

m/z 264.1066

012E+34.55RT (min)

lcqtof\_t3\_neg\_4\_037\_01

m/z 265.0919

0246E+344.5RT (min)

lcqtof\_t3\_neg\_1\_002\_13

m/z 265.9403

01E+311.5RT (min)

lcqtof\_t3\_neg\_2\_051\_14

m/z 266.0569

012E+311.5RT (min)

lcqtof\_t3\_neg\_2\_019\_32

m/z 266.9051

0246E+311.5RT (min)

lcqtof\_t3\_neg\_4\_011\_32

m/z 266.9317

012E+311.5RT (min)

lcqtof\_t3\_neg\_1\_017\_51

m/z 267.0710

0123E+311.5RT (min)

lcqtof\_t3\_neg\_4\_055\_38

m/z 268.0371

0123E+388.5RT (min)

lcqtof\_t3\_neg\_4\_055\_38

m/z 268.0383

01234E+36.57RT (min)

lcqtof\_t3\_neg\_4\_004\_20

m/z 269.0876

01234E+311.5RT (min)

lcqtof\_t3\_neg\_4\_010\_46

m/z 269.1271

0246E+344.5RT (min)

lcqtof\_t3\_neg\_1\_031\_24

m/z 270.0817

012E+311.5RT (min)

lcqtof\_t3\_neg\_3\_012\_14

m/z 270.1024

024E+311.5RT (min)

lcqtof\_t3\_neg\_1\_031\_24

m/z 271.0200

02468E+311.5RT (min)

lcqtof\_t3\_neg\_1\_055\_32

m/z 272.0737

024E+311.5RT (min)

lcqtof\_t3\_neg\_2\_019\_32

m/z 272.1704

0123E+34.55RT (min)

lcqtof\_t3\_neg\_1\_032\_36

m/z 272.8981

01E+311.5RT (min)

lcqtof\_t3\_neg\_4\_055\_38

m/z 273.0231

024E+316.517RT (min)

lcqtof\_t3\_neg\_4\_001\_49

m/z 273.0857

02468E+35.56RT (min)

lcqtof\_t3\_neg\_4\_030\_31

m/z 273.9299

012E+388.5RT (min)

lcqtof\_t3\_neg\_1\_030\_42

m/z 273.9837

012E+355.5RT (min)

lcqtof\_t3\_neg\_3\_018\_32

m/z 273.9903

01234E+311.5RT (min)

lcqtof\_t3\_neg\_4\_016\_14

m/z 274.8378

012E+311.5RT (min)

lcqtof\_t3\_neg\_4\_046\_33

m/z 275.0554

012E+311.5RT (min)

lcqtof\_t3\_neg\_4\_016\_14

m/z 275.0778

024E+311.5RT (min)

lcqtof\_t3\_neg\_1\_036\_14

m/z 275.0917

0123E+311.5RT (min)

lcqtof\_t3\_neg\_4\_019\_05

m/z 276.0343

0123E+34.55RT (min)

lcqtof\_t3\_neg\_1\_026\_19

m/z 276.8849

0123E+311.5RT (min)

lcqtof\_t3\_neg\_4\_011\_32

m/z 276.9084

024E+311.5RT (min)

lcqtof\_t3\_neg\_1\_055\_32

m/z 277.0701

012E+36.57RT (min)

lcqtof\_t3\_neg\_4\_005\_18

m/z 277.1567

012E+35.56RT (min)

lcqtof\_t3\_neg\_4\_016\_14

m/z 277.9832

024E+311.5RT (min)

lcqtof\_t3\_neg\_3\_018\_32

m/z 278.0369

0123E+311.5RT (min)

lcqtof\_t3\_neg\_4\_019\_05

m/z 278.1046

012E+34.55RT (min)

lcqtof\_t3\_neg\_4\_037\_01

m/z 278.8783

012E+311.5RT (min)

lcqtof\_t3\_neg\_2\_019\_32

m/z 279.1105

0123E+344.5RT (min)

lcqtof\_t3\_neg\_1\_055\_32

m/z 279.6117

0123E+36.57RT (min)

lcqtof\_t3\_neg\_4\_016\_14

m/z 279.9078

024E+311.5RT (min)

lcqtof\_t3\_neg\_1\_022\_28

m/z 280.1023

024E+311.5RT (min)

lcqtof\_t3\_neg\_3\_006\_23

m/z 280.9433

0246E+311.5RT (min)

lcqtof\_t3\_neg\_1\_001\_49

m/z 281.0089

0246E+36.57RT (min)

lcqtof\_t3\_neg\_4\_002\_17

m/z 281.0742

012E+311.5RT (min)

lcqtof\_t3\_neg\_1\_032\_36

m/z 281.1376

0246E+311.5RT (min)

lcqtof\_t3\_neg\_4\_040\_24

m/z 282.0963

0246E+311.5RT (min)

lcqtof\_t3\_neg\_4\_040\_24

m/z 283.0602

0123E+36.57RT (min)

lcqtof\_t3\_neg\_4\_004\_20

m/z 284.1088

012E+344.5RT (min)

lcqtof\_t3\_neg\_4\_032\_04

m/z 284.1255

012E+344.5RT (min)

lcqtof\_t3\_neg\_1\_041\_54

m/z 284.1371

0123E+311.5RT (min)

lcqtof\_t3\_neg\_1\_055\_32

m/z 284.9139

0246E+311.5RT (min)

lcqtof\_t3\_neg\_4\_011\_32

m/z 285.1600

0123E+377.5RT (min)

lcqtof\_t3\_neg\_4\_055\_38

m/z 285.9086

01234E+311.5RT (min)

lcqtof\_t3\_neg\_1\_015\_34

m/z 286.8187

01E+311.5RT (min)

lcqtof\_t3\_neg\_1\_055\_32

m/z 287.0250

0123E+34.55RT (min)

lcqtof\_t3\_neg\_4\_016\_14

m/z 287.2012

01234E+36.57RT (min)

lcqtof\_t3\_neg\_1\_015\_34

m/z 287.9857

0246E+311.5RT (min)

lcqtof\_t3\_neg\_4\_055\_38

m/z 288.0414

01E+36.57RT (min)

lcqtof\_t3\_neg\_1\_035\_31

m/z 288.0600

01234E+344.5RT (min)

lcqtof\_t3\_neg\_1\_036\_14

m/z 288.8154

01E+311.5RT (min)

lcqtof\_t3\_neg\_3\_018\_32

m/z 288.8763

0123E+31.52RT (min)

lcqtof\_t3\_neg\_1\_044\_37

m/z 288.9275

012E+311.5RT (min)

lcqtof\_t3\_neg\_4\_014\_13

m/z 289.0380

012E+316.517RT (min)

lcqtof\_t3\_neg\_4\_012\_27

m/z 289.0707

0123E+316.517RT (min)

lcqtof\_t3\_neg\_4\_007\_09

m/z 289.1088

012E+316.517RT (min)

lcqtof\_t3\_neg\_4\_030\_31

m/z 289.1145

01234E+311.5RT (min)

lcqtof\_t3\_neg\_2\_030\_23

m/z 289.9390

01E+411.5RT (min)

lcqtof\_t3\_neg\_4\_016\_14

m/z 290.0326

012E+316.517RT (min)

lcqtof\_t3\_neg\_4\_022\_28

m/z 290.0370

024E+311.5RT (min)

lcqtof\_t3\_neg\_3\_088\_59

m/z 290.7652

01E+311.5RT (min)

lcqtof\_t3\_neg\_4\_051\_15

m/z 290.8740

0123E+31.52RT (min)

lcqtof\_t3\_neg\_1\_032\_36

m/z 291.0201

01234E+311.5RT (min)

lcqtof\_t3\_neg\_1\_015\_34

m/z 291.0309

012E+316.517RT (min)

lcqtof\_t3\_neg\_4\_004\_20

m/z 291.2219

01234E+411.5RT (min)

lcqtof\_t3\_neg\_1\_018\_33

m/z 291.9197

0123E+311.5RT (min)

lcqtof\_t3\_neg\_1\_002\_13

m/z 292.2209

0123E+411.5RT (min)

lcqtof\_t3\_neg\_1\_032\_36

m/z 292.2985

0246E+411.5RT (min)

lcqtof\_t3\_neg\_1\_053\_01

m/z 292.9399

01234E+311.5RT (min)

lcqtof\_t3\_neg\_1\_036\_14

m/z 293.1241

01E+34.55RT (min)

lcqtof\_t3\_neg\_2\_054\_29

m/z 294.0296

012E+344.5RT (min)

lcqtof\_t3\_neg\_1\_034\_30

m/z 294.8913

0123E+311.5RT (min)

lcqtof\_t3\_neg\_4\_055\_38

m/z 295.0411

0246E+311.5RT (min)

lcqtof\_t3\_neg\_3\_018\_32

m/z 295.8885

01E+316.517RT (min)

lcqtof\_t3\_neg\_1\_015\_34

m/z 296.0733

0246E+311.5RT (min)

lcqtof\_t3\_neg\_1\_019\_23

m/z 296.0862

01234E+311.5RT (min)

lcqtof\_t3\_neg\_4\_055\_38

m/z 297.0379

0123E+311.5RT (min)

lcqtof\_t3\_neg\_3\_018\_32

m/z 297.1027

0123E+311.5RT (min)

lcqtof\_t3\_neg\_2\_026\_36

m/z 297.8954

024E+311.5RT (min)

lcqtof\_t3\_neg\_4\_037\_01

m/z 298.0777

012E+37.58RT (min)

lcqtof\_t3\_neg\_4\_024\_26

m/z 298.1197

024E+311.5RT (min)

lcqtof\_t3\_neg\_4\_039\_06\_rerun

m/z 298.7508

012E+311.5RT (min)

lcqtof\_t3\_neg\_4\_016\_14

m/z 299.0830

01E+411.5RT (min)

lcqtof\_t3\_neg\_4\_039\_06\_rerun

m/z 299.8904

0123E+311.5RT (min)

lcqtof\_t3\_neg\_4\_051\_15

m/z 300.0433

024E+311.5RT (min)

lcqtof\_t3\_neg\_4\_018\_23

m/z 300.0490

0123E+344.5RT (min)

lcqtof\_t3\_neg\_1\_014\_04

m/z 300.0851

012E+311.5RT (min)

lcqtof\_t3\_neg\_4\_010\_46

m/z 300.7480

0123E+311.5RT (min)

lcqtof\_t3\_neg\_4\_016\_14

m/z 300.9284

0246E+311.5RT (min)

lcqtof\_t3\_neg\_4\_046\_33

m/z 301.1094

012E+344.5RT (min)

lcqtof\_t3\_neg\_1\_031\_24

m/z 302.7473

0123E+311.5RT (min)

lcqtof\_t3\_neg\_4\_016\_14

m/z 306.0232

024E+34.55RT (min)

lcqtof\_t3\_neg\_4\_004\_20

m/z 306.8333

0246E+311.5RT (min)

lcqtof\_t3\_neg\_4\_040\_24

m/z 307.0311

01234E+311.5RT (min)

lcqtof\_t3\_neg\_4\_016\_14

m/z 307.0996

01234E+344.5RT (min)

lcqtof\_t3\_neg\_3\_018\_32

m/z 307.1183

0246E+34.55RT (min)

lcqtof\_t3\_neg\_4\_040\_24

m/z 308.1929

0246E+344.5RT (min)

lcqtof\_t3\_neg\_2\_019\_32

m/z 309.0722

0123E+311.5RT (min)

lcqtof\_t3\_neg\_1\_019\_23

m/z 309.1015

012E+311.5RT (min)

lcqtof\_t3\_neg\_1\_053\_01

m/z 309.9194

0123E+311.5RT (min)

lcqtof\_t3\_neg\_4\_037\_01

m/z 311.0287

024E+311.5RT (min)

lcqtof\_t3\_neg\_4\_030\_31

m/z 312.0179

012E+311.5RT (min)

lcqtof\_t3\_neg\_4\_055\_38

m/z 312.7921

0123E+311.5RT (min)

lcqtof\_t3\_neg\_4\_046\_33

m/z 312.8747

02468E+311.5RT (min)

lcqtof\_t3\_neg\_4\_001\_49

m/z 314.0570

0123E+388.5RT (min)

lcqtof\_t3\_neg\_4\_048\_19

m/z 315.1035

0246E+311.5RT (min)

lcqtof\_t3\_neg\_1\_031\_24

m/z 315.8827

0123E+311.5RT (min)

lcqtof\_t3\_neg\_1\_053\_01

m/z 316.8627

02468E+311.5RT (min)

lcqtof\_t3\_neg\_2\_028\_24

m/z 316.9813

012E+311.5RT (min)

lcqtof\_t3\_neg\_3\_016\_35

m/z 317.0974

012E+344.5RT (min)

lcqtof\_t3\_neg\_1\_044\_37

m/z 318.0496

0123E+311.5RT (min)

lcqtof\_t3\_neg\_4\_046\_33

m/z 318.1035

0123E+344.5RT (min)

lcqtof\_t3\_neg\_1\_031\_24

m/z 318.8366

0246E+311.5RT (min)

lcqtof\_t3\_neg\_1\_015\_34

m/z 318.8764

01234E+311.5RT (min)

lcqtof\_t3\_neg\_4\_039\_06\_rerun

m/z 318.9543

0123E+316.517RT (min)

lcqtof\_t3\_neg\_4\_022\_28

m/z 319.0751

01234E+311.5RT (min)

lcqtof\_t3\_neg\_4\_030\_31

m/z 319.0840

012E+311.5RT (min)

lcqtof\_t3\_neg\_1\_015\_34

m/z 319.1103

0123E+355.5RT (min)

lcqtof\_t3\_neg\_2\_038\_18

m/z 320.0730

0123E+37.58RT (min)

lcqtof\_t3\_neg\_4\_005\_18

m/z 320.0979

0123E+344.5RT (min)

lcqtof\_t3\_neg\_4\_041\_54

m/z 320.8237

01234E+311.5RT (min)

lcqtof\_t3\_neg\_4\_046\_33

m/z 320.9861

012E+34.55RT (min)

lcqtof\_t3\_neg\_1\_015\_34

m/z 321.0590

01E+37.58RT (min)

lcqtof\_t3\_neg\_1\_026\_19

m/z 322.1121

024E+311.5RT (min)

lcqtof\_t3\_neg\_1\_042\_46

m/z 323.1147

024E+311.5RT (min)

lcqtof\_t3\_neg\_4\_030\_31

m/z 323.9172

01234E+311.5RT (min)

lcqtof\_t3\_neg\_4\_055\_38

m/z 324.0818

01234E+311.5RT (min)

lcqtof\_t3\_neg\_1\_053\_01

m/z 324.1178

01E+411.5RT (min)

lcqtof\_t3\_neg\_4\_039\_06\_rerun

m/z 324.8899

02468E+311.5RT (min)

lcqtof\_t3\_neg\_1\_036\_14

m/z 325.0835

01234E+311.5RT (min)

lcqtof\_t3\_neg\_4\_028\_25

m/z 325.8996

01234E+311.5RT (min)

lcqtof\_t3\_neg\_1\_002\_13

m/z 327.0098

0123E+311.5RT (min)

lcqtof\_t3\_neg\_4\_055\_38

m/z 327.0859

01234E+344.5RT (min)

lcqtof\_t3\_neg\_1\_044\_37

m/z 327.1251

0246E+311.5RT (min)

lcqtof\_t3\_neg\_4\_040\_24

m/z 327.8885

0246E+311.5RT (min)

lcqtof\_t3\_neg\_4\_055\_38

m/z 328.0556

0246E+311.5RT (min)

lcqtof\_t3\_neg\_2\_026\_36

m/z 329.1773

0123E+36.57RT (min)

lcqtof\_t3\_neg\_4\_037\_01

m/z 329.8855

0246E+311.5RT (min)

lcqtof\_t3\_neg\_4\_055\_38

m/z 330.0933

0246E+311.5RT (min)

lcqtof\_t3\_neg\_4\_010\_46

m/z 330.1238

0123E+344.5RT (min)

lcqtof\_t3\_neg\_4\_040\_24

m/z 330.8416

0123E+311.5RT (min)

lcqtof\_t3\_neg\_4\_003\_37

m/z 330.8547

024E+311.5RT (min)

lcqtof\_t3\_neg\_4\_046\_33

m/z 331.0893

012E+34.55RT (min)

lcqtof\_t3\_neg\_4\_037\_01

m/z 331.1108

0246E+344.5RT (min)

lcqtof\_t3\_neg\_4\_030\_31

m/z 331.1602

0123E+344.5RT (min)

lcqtof\_t3\_neg\_4\_018\_23

m/z 332.7501

024E+311.5RT (min)

lcqtof\_t3\_neg\_4\_037\_01

m/z 333.0268

0246E+311.5RT (min)

lcqtof\_t3\_neg\_4\_003\_37

m/z 333.0588

02468E+311.5RT (min)

lcqtof\_t3\_neg\_1\_019\_23

m/z 333.1507

0123E+35.56RT (min)

lcqtof\_t3\_neg\_4\_004\_20

m/z 335.0777

02468E+311.5RT (min)

lcqtof\_t3\_neg\_4\_034\_34

m/z 335.1162

0123E+311.5RT (min)

lcqtof\_t3\_neg\_2\_051\_14

m/z 335.1284

02468E+311.5RT (min)

lcqtof\_t3\_neg\_4\_090\_60

m/z 336.8862

02468E+311.5RT (min)

lcqtof\_t3\_neg\_4\_039\_06\_rerun

m/z 337.0028

012E+34.55RT (min)

lcqtof\_t3\_neg\_1\_010\_27

m/z 337.0384

0246E+311.5RT (min)

lcqtof\_t3\_neg\_4\_011\_32

m/z 337.0876

012E+344.5RT (min)

lcqtof\_t3\_neg\_1\_051\_26

m/z 338.0907

01234E+311.5RT (min)

lcqtof\_t3\_neg\_1\_055\_32

m/z 338.1341

012E+344.5RT (min)

lcqtof\_t3\_neg\_2\_043\_31

m/z 338.1361

0123E+35.56RT (min)

lcqtof\_t3\_neg\_4\_048\_19

m/z 339.0629

0123E+344.5RT (min)

lcqtof\_t3\_neg\_4\_037\_01

m/z 340.7354

024E+311.5RT (min)

lcqtof\_t3\_neg\_4\_051\_15

m/z 340.7829

0123E+311.5RT (min)

lcqtof\_t3\_neg\_4\_037\_01

m/z 340.8284

012E+311.5RT (min)

lcqtof\_t3\_neg\_4\_046\_33

m/z 340.8864

012E+311.5RT (min)

lcqtof\_t3\_neg\_3\_018\_32

m/z 341.0974

02468E+344.5RT (min)

lcqtof\_t3\_neg\_4\_045\_02\_rerun

m/z 342.0746

0246E+311.5RT (min)

lcqtof\_t3\_neg\_1\_006\_05

m/z 343.0034

0246E+311.5RT (min)

lcqtof\_t3\_neg\_4\_052\_03\_rerun

m/z 343.0898

012E+311.5RT (min)

lcqtof\_t3\_neg\_1\_042\_46

m/z 343.0961

0123E+34.55RT (min)

lcqtof\_t3\_neg\_2\_019\_32

m/z 343.9000

0246E+311.5RT (min)

lcqtof\_t3\_neg\_4\_030\_31

m/z 344.7773

0123E+311.5RT (min)

lcqtof\_t3\_neg\_4\_041\_54

m/z 345.0397

0123E+355.5RT (min)

lcqtof\_t3\_neg\_1\_027\_21

m/z 345.1783

0123E+344.5RT (min)

lcqtof\_t3\_neg\_1\_044\_37

m/z 345.9453

01234E+311.5RT (min)

lcqtof\_t3\_neg\_4\_055\_38

m/z 346.1525

01234E+344.5RT (min)

lcqtof\_t3\_neg\_4\_023\_45

m/z 346.1870

012E+377.5RT (min)

lcqtof\_t3\_neg\_4\_012\_27

m/z 346.7277

01234E+311.5RT (min)

lcqtof\_t3\_neg\_4\_051\_15

m/z 347.1265

012E+344.5RT (min)

lcqtof\_t3\_neg\_1\_005\_18

m/z 347.1973

0123E+366.5RT (min)

lcqtof\_t3\_neg\_4\_030\_31

m/z 348.7691

012E+311.5RT (min)

lcqtof\_t3\_neg\_4\_041\_54

m/z 348.8952

01E+411.5RT (min)

lcqtof\_t3\_neg\_4\_039\_06\_rerun

m/z 348.9793

0123E+311.5RT (min)

lcqtof\_t3\_neg\_4\_055\_38

m/z 348.9982

0123E+411.5RT (min)

lcqtof\_t3\_neg\_4\_045\_02\_rerun

m/z 349.0618

01234E+311.5RT (min)

lcqtof\_t3\_neg\_4\_024\_26

m/z 350.0312

01E+344.5RT (min)

lcqtof\_t3\_neg\_1\_036\_14

m/z 350.0935

0123E+344.5RT (min)

lcqtof\_t3\_neg\_4\_043\_11

m/z 351.1049

02468E+311.5RT (min)

lcqtof\_t3\_neg\_4\_018\_23

m/z 351.2246

01E+31010.5RT (min)

lcqtof\_t3\_neg\_4\_090\_60

m/z 352.0764

0246E+311.5RT (min)

lcqtof\_t3\_neg\_4\_010\_46

m/z 352.1311

0123E+35.56RT (min)

lcqtof\_t3\_neg\_4\_048\_19

m/z 352.1662

0123E+35.56RT (min)

lcqtof\_t3\_neg\_4\_037\_01

m/z 352.7668

0246E+311.5RT (min)

lcqtof\_t3\_neg\_4\_016\_14

m/z 352.9038

0246E+311.5RT (min)

lcqtof\_t3\_neg\_4\_046\_33

m/z 353.9966

01E+411.5RT (min)

lcqtof\_t3\_neg\_4\_045\_02\_rerun

m/z 354.0714

024E+311.5RT (min)

lcqtof\_t3\_neg\_4\_039\_06\_rerun

m/z 355.0804

01234E+311.5RT (min)

lcqtof\_t3\_neg\_2\_019\_32

m/z 355.1237

012E+411.5RT (min)

lcqtof\_t3\_neg\_4\_045\_02\_rerun

m/z 355.9949

024E+311.5RT (min)

lcqtof\_t3\_neg\_1\_051\_26

m/z 356.0029

02468E+311.5RT (min)

lcqtof\_t3\_neg\_2\_052\_17

m/z 356.0060

01234E+34.55RT (min)

lcqtof\_t3\_neg\_4\_004\_20

m/z 356.1355

01234E+344.5RT (min)

lcqtof\_t3\_neg\_4\_027\_41

m/z 356.2535

0123E+34.55RT (min)

lcqtof\_t3\_neg\_4\_055\_38

m/z 356.7687

0123E+311.5RT (min)

lcqtof\_t3\_neg\_4\_046\_33

m/z 357.0862

01234E+311.5RT (min)

lcqtof\_t3\_neg\_1\_031\_24

m/z 357.2156

01234E+355.5RT (min)

lcqtof\_t3\_neg\_4\_018\_23

m/z 360.1277

012E+344.5RT (min)

lcqtof\_t3\_neg\_3\_018\_32

m/z 360.9739

012E+316.517RT (min)

lcqtof\_t3\_neg\_1\_015\_34

m/z 362.2107

01E+36.57RT (min)

lcqtof\_t3\_neg\_1\_053\_01

m/z 363.1154

0246E+344.5RT (min)

lcqtof\_t3\_neg\_3\_002\_46

m/z 363.1924

012E+377.5RT (min)

lcqtof\_t3\_neg\_4\_004\_20

m/z 363.8774

012E+316.517RT (min)

lcqtof\_t3\_neg\_1\_001\_49

m/z 364.0607

01234E+311.5RT (min)

lcqtof\_t3\_neg\_4\_046\_33

m/z 365.0849

0123E+311.5RT (min)

lcqtof\_t3\_neg\_4\_034\_34

m/z 365.1046

0246E+344.5RT (min)

lcqtof\_t3\_neg\_3\_032\_24

m/z 368.0255

01E+355.5RT (min)

lcqtof\_t3\_neg\_1\_017\_51

m/z 369.0791

012E+344.5RT (min)

lcqtof\_t3\_neg\_1\_005\_18

m/z 369.1016

0246E+311.5RT (min)

lcqtof\_t3\_neg\_4\_033\_53

m/z 369.1310

012E+344.5RT (min)

lcqtof\_t3\_neg\_4\_037\_01

m/z 370.0239

0123E+355.5RT (min)

lcqtof\_t3\_neg\_4\_033\_53

m/z 370.1008

024E+344.5RT (min)

lcqtof\_t3\_neg\_3\_052\_41

m/z 370.1055

0246E+311.5RT (min)

lcqtof\_t3\_neg\_4\_049\_55

m/z 370.1251

0123E+311.5RT (min)

lcqtof\_t3\_neg\_2\_022\_47

m/z 370.2331

01234E+34.55RT (min)

lcqtof\_t3\_neg\_4\_003\_37

m/z 371.1155

012E+344.5RT (min)

lcqtof\_t3\_neg\_1\_005\_18

m/z 371.1471

01E+344.5RT (min)

lcqtof\_t3\_neg\_4\_009\_50

m/z 371.1959

0123E+344.5RT (min)

lcqtof\_t3\_neg\_4\_040\_24

m/z 372.0844

01234E+311.5RT (min)

lcqtof\_t3\_neg\_1\_031\_24

m/z 372.1354

024E+37.58RT (min)

lcqtof\_t3\_neg\_4\_037\_01

m/z 372.1911

012E+344.5RT (min)

lcqtof\_t3\_neg\_1\_044\_37

m/z 372.2500

01234E+35.56RT (min)

lcqtof\_t3\_neg\_4\_018\_23

m/z 372.7537

012E+311.5RT (min)

lcqtof\_t3\_neg\_4\_041\_54

m/z 373.9882

012E+311.5RT (min)

lcqtof\_t3\_neg\_2\_015\_33

m/z 374.1413

0123E+311.5RT (min)

lcqtof\_t3\_neg\_4\_047\_07

m/z 375.9086

0246E+311.5RT (min)

lcqtof\_t3\_neg\_4\_051\_15

m/z 375.9793

02468E+311.5RT (min)

lcqtof\_t3\_neg\_4\_045\_02\_rerun

m/z 376.8000

0246E+311.5RT (min)

lcqtof\_t3\_neg\_1\_055\_32

m/z 376.9750

02468E+311.5RT (min)

lcqtof\_t3\_neg\_2\_055\_16

m/z 377.0244

0123E+37.58RT (min)

lcqtof\_t3\_neg\_4\_024\_26

m/z 378.0735

01234E+311.5RT (min)

lcqtof\_t3\_neg\_4\_051\_15

m/z 378.1296

0123E+344.5RT (min)

lcqtof\_t3\_neg\_2\_005\_34

m/z 378.7839

024E+311.5RT (min)

lcqtof\_t3\_neg\_4\_046\_33

m/z 379.0458

0246E+311.5RT (min)

lcqtof\_t3\_neg\_4\_030\_31

m/z 379.0936

01234E+355.5RT (min)

lcqtof\_t3\_neg\_1\_051\_26

m/z 379.1954

01234E+366.5RT (min)

lcqtof\_t3\_neg\_4\_037\_01

m/z 380.0062

012E+311.5RT (min)

lcqtof\_t3\_neg\_4\_034\_34

m/z 380.0861

02468E+311.5RT (min)

lcqtof\_t3\_neg\_4\_047\_07

m/z 381.0332

0246E+311.5RT (min)

lcqtof\_t3\_neg\_4\_016\_14

m/z 381.1596

0246E+311.5RT (min)

lcqtof\_t3\_neg\_4\_093\_60

m/z 382.8485

02468E+311.5RT (min)

lcqtof\_t3\_neg\_4\_051\_15

m/z 383.0992

0246E+311.5RT (min)

lcqtof\_t3\_neg\_4\_047\_07

m/z 385.1545

024E+366.5RT (min)

lcqtof\_t3\_neg\_4\_016\_14

m/z 385.1556

012E+311.5RT (min)

lcqtof\_t3\_neg\_1\_020\_47

m/z 386.0142

012E+316.517RT (min)

lcqtof\_t3\_neg\_4\_002\_17

m/z 386.0325

012E+311.5RT (min)

lcqtof\_t3\_neg\_1\_055\_32

m/z 386.1483

012E+311.5RT (min)

lcqtof\_t3\_neg\_1\_055\_32

m/z 386.1718

0246E+366.5RT (min)

lcqtof\_t3\_neg\_4\_037\_01

m/z 386.7639

01234E+311.5RT (min)

lcqtof\_t3\_neg\_4\_046\_33

m/z 386.9418

0123E+316.517RT (min)

lcqtof\_t3\_neg\_4\_003\_37

m/z 387.0048

012E+316.517RT (min)

lcqtof\_t3\_neg\_4\_006\_16

m/z 387.2397

0123E+31212.5RT (min)

lcqtof\_t3\_neg\_4\_036\_47

m/z 387.9977

0123E+34.55RT (min)

lcqtof\_t3\_neg\_4\_108\_62

m/z 388.1096

01234E+35.56RT (min)

lcqtof\_t3\_neg\_4\_038\_29

m/z 388.1419

012E+311.5RT (min)

lcqtof\_t3\_neg\_4\_044\_48

m/z 388.8143

024E+311.5RT (min)

lcqtof\_t3\_neg\_4\_046\_33

m/z 388.9772

01234E+311.5RT (min)

lcqtof\_t3\_neg\_4\_016\_14

m/z 389.0586

012E+311.5RT (min)

lcqtof\_t3\_neg\_3\_018\_32

m/z 389.0668

012E+37.58RT (min)

lcqtof\_t3\_neg\_4\_005\_18

m/z 389.2078

02468E+344.5RT (min)

lcqtof\_t3\_neg\_1\_055\_32

m/z 390.0830

02468E+311.5RT (min)

lcqtof\_t3\_neg\_4\_040\_24

m/z 390.7080

01234E+311.5RT (min)

lcqtof\_t3\_neg\_4\_037\_01

m/z 390.7595

01234E+311.5RT (min)

lcqtof\_t3\_neg\_4\_055\_38

m/z 390.8148

024E+311.5RT (min)

lcqtof\_t3\_neg\_2\_028\_24

m/z 391.9009

01234E+311.5RT (min)

lcqtof\_t3\_neg\_4\_055\_38

m/z 392.7513

0246E+311.5RT (min)

lcqtof\_t3\_neg\_4\_037\_01

m/z 392.7951

0123E+311.5RT (min)

lcqtof\_t3\_neg\_3\_018\_32

m/z 392.8748

02468E+311.5RT (min)

lcqtof\_t3\_neg\_4\_046\_33

m/z 393.0385

024E+355.5RT (min)

lcqtof\_t3\_neg\_4\_005\_18

m/z 393.1240

0123E+34.55RT (min)

lcqtof\_t3\_neg\_3\_022\_27

m/z 393.1446

02468E+35.56RT (min)

lcqtof\_t3\_neg\_2\_053\_38

m/z 394.0079

0246E+311.5RT (min)

lcqtof\_t3\_neg\_4\_016\_14

m/z 394.8925

012E+311.5RT (min)

lcqtof\_t3\_neg\_4\_052\_03\_rerun

m/z 396.0964

0123E+34.55RT (min)

lcqtof\_t3\_neg\_4\_055\_38

m/z 396.7443

02468E+311.5RT (min)

lcqtof\_t3\_neg\_4\_037\_01

m/z 398.0042

0123E+311.5RT (min)

lcqtof\_t3\_neg\_2\_019\_32

m/z 398.1416

0246E+311.5RT (min)

lcqtof\_t3\_neg\_4\_039\_06\_rerun

m/z 398.6945

01234E+311.5RT (min)

lcqtof\_t3\_neg\_4\_037\_01

m/z 398.8398

024E+311.5RT (min)

lcqtof\_t3\_neg\_4\_046\_33

m/z 399.1115

024E+344.5RT (min)

lcqtof\_t3\_neg\_4\_018\_23

m/z 399.9890

0246E+37.58RT (min)

lcqtof\_t3\_neg\_4\_005\_18

m/z 401.0302

012E+344.5RT (min)

lcqtof\_t3\_neg\_1\_036\_14

m/z 401.2118

0123E+344.5RT (min)

lcqtof\_t3\_neg\_4\_018\_23

m/z 402.1361

0246E+344.5RT (min)

lcqtof\_t3\_neg\_4\_030\_31

m/z 402.2061

01234E+377.5RT (min)

lcqtof\_t3\_neg\_4\_048\_19

m/z 403.1860

01234E+344.5RT (min)

lcqtof\_t3\_neg\_1\_044\_37

m/z 403.5529

0246E+344.5RT (min)

lcqtof\_t3\_neg\_4\_003\_37

m/z 403.9180

024E+311.5RT (min)

lcqtof\_t3\_neg\_1\_036\_14

m/z 404.1027

01E+316.517RT (min)

lcqtof\_t3\_neg\_4\_053\_12

m/z 404.2078

0123E+377.5RT (min)

lcqtof\_t3\_neg\_4\_004\_20

m/z 404.6874

024E+311.5RT (min)

lcqtof\_t3\_neg\_4\_051\_15

m/z 404.7364

0123E+311.5RT (min)

lcqtof\_t3\_neg\_4\_046\_33

m/z 404.9165

024E+311.5RT (min)

lcqtof\_t3\_neg\_4\_016\_14

m/z 404.9486

012E+311.5RT (min)

lcqtof\_t3\_neg\_1\_015\_34

m/z 406.1323

012E+34.55RT (min)

lcqtof\_t3\_neg\_4\_048\_19

m/z 406.6869

012E+311.5RT (min)

lcqtof\_t3\_neg\_4\_051\_15

m/z 406.7306

012E+311.5RT (min)

lcqtof\_t3\_neg\_4\_041\_54

m/z 406.8656

01E+316.517RT (min)

lcqtof\_t3\_neg\_1\_003\_07

m/z 406.9428

012E+311.5RT (min)

lcqtof\_t3\_neg\_4\_030\_31

m/z 408.2111

0123E+36.57RT (min)

lcqtof\_t3\_neg\_4\_016\_14

m/z 408.8749

012E+311.5RT (min)

lcqtof\_t3\_neg\_2\_019\_32

m/z 409.0820

0246E+355.5RT (min)

lcqtof\_t3\_neg\_4\_005\_18

m/z 409.1267

0123E+311.5RT (min)

lcqtof\_t3\_neg\_4\_081\_58

m/z 409.1536

0123E+344.5RT (min)

lcqtof\_t3\_neg\_2\_034\_19

m/z 410.7220

024E+311.5RT (min)

lcqtof\_t3\_neg\_4\_016\_14

m/z 411.1124

01E+311.5RT (min)

lcqtof\_t3\_neg\_3\_036\_16

m/z 412.0645

01E+344.5RT (min)

lcqtof\_t3\_neg\_4\_053\_12

m/z 412.1200

0123E+344.5RT (min)

lcqtof\_t3\_neg\_1\_051\_26

m/z 413.0153

01E+316.517RT (min)

lcqtof\_t3\_neg\_4\_020\_22

m/z 413.1773

01234E+377.5RT (min)

lcqtof\_t3\_neg\_4\_004\_20

m/z 414.0116

0123E+316.517RT (min)

lcqtof\_t3\_neg\_4\_009\_50

m/z 414.7356

024E+311.5RT (min)

lcqtof\_t3\_neg\_4\_046\_33

m/z 415.0052

012E+316.517RT (min)

lcqtof\_t3\_neg\_4\_011\_32

m/z 415.2026

0246E+377.5RT (min)

lcqtof\_t3\_neg\_4\_016\_14

m/z 416.1291

012E+35.56RT (min)

lcqtof\_t3\_neg\_1\_026\_19

m/z 416.6656

0123E+311.5RT (min)

lcqtof\_t3\_neg\_4\_051\_15

m/z 417.1570

012E+344.5RT (min)

lcqtof\_t3\_neg\_4\_028\_25

m/z 418.6626

0123E+311.5RT (min)

lcqtof\_t3\_neg\_4\_051\_15

m/z 419.9002

0246E+311.5RT (min)

lcqtof\_t3\_neg\_1\_036\_14

m/z 420.1166

0123E+35.56RT (min)

lcqtof\_t3\_neg\_4\_048\_19

m/z 420.6613

012E+311.5RT (min)

lcqtof\_t3\_neg\_4\_016\_14

m/z 420.8949

024E+311.5RT (min)

lcqtof\_t3\_neg\_1\_036\_14

m/z 421.0733

0246E+311.5RT (min)

lcqtof\_t3\_neg\_4\_018\_23

m/z 421.2221

0123E+39.510RT (min)

lcqtof\_t3\_neg\_4\_093\_60

m/z 421.2517

012E+38.59RT (min)

lcqtof\_t3\_neg\_4\_051\_15

m/z 422.0978

01234E+311.5RT (min)

lcqtof\_t3\_neg\_4\_010\_46

m/z 422.1791

012E+36.57RT (min)

lcqtof\_t3\_neg\_1\_010\_27

m/z 422.9622

01E+411.5RT (min)

lcqtof\_t3\_neg\_4\_045\_02\_rerun

m/z 423.0681

024E+311.5RT (min)

lcqtof\_t3\_neg\_2\_019\_32

m/z 424.7211

02468E+311.5RT (min)

lcqtof\_t3\_neg\_4\_055\_38

m/z 424.9686

0123E+311.5RT (min)

lcqtof\_t3\_neg\_4\_001\_49

m/z 425.0837

01234E+311.5RT (min)

lcqtof\_t3\_neg\_4\_018\_23

m/z 426.1250

024E+34.55RT (min)

lcqtof\_t3\_neg\_4\_038\_29

m/z 426.1427

0246E+344.5RT (min)

lcqtof\_t3\_neg\_1\_002\_13

m/z 426.9429

0123E+411.5RT (min)

lcqtof\_t3\_neg\_4\_045\_02\_rerun

m/z 427.0123

02468E+311.5RT (min)

lcqtof\_t3\_neg\_4\_016\_14

m/z 427.1259

02468E+311.5RT (min)

lcqtof\_t3\_neg\_4\_018\_23

m/z 427.2506

024E+377.5RT (min)

lcqtof\_t3\_neg\_4\_005\_18

m/z 428.0380

0123E+311.5RT (min)

lcqtof\_t3\_neg\_4\_044\_48

m/z 428.1539

0246E+344.5RT (min)

lcqtof\_t3\_neg\_1\_009\_50

m/z 429.2009

012E+34.55RT (min)

lcqtof\_t3\_neg\_4\_018\_23

m/z 430.0728

0246E+35.56RT (min)

lcqtof\_t3\_neg\_4\_004\_20

m/z 430.1299

01234E+36.57RT (min)

lcqtof\_t3\_neg\_4\_005\_18

m/z 430.7080

01234E+311.5RT (min)

lcqtof\_t3\_neg\_4\_046\_33

m/z 430.9270

01234E+311.5RT (min)

lcqtof\_t3\_neg\_4\_051\_15

m/z 430.9294

012E+316.517RT (min)

lcqtof\_t3\_neg\_3\_024\_08

m/z 432.2449

0246E+34.55RT (min)

lcqtof\_t3\_neg\_2\_043\_31

m/z 432.7842

0246E+311.5RT (min)

lcqtof\_t3\_neg\_4\_046\_33

m/z 433.1332

01234E+34.55RT (min)

lcqtof\_t3\_neg\_4\_048\_19

m/z 434.0266

0123E+311.5RT (min)

lcqtof\_t3\_neg\_2\_019\_32

m/z 434.7600

0246E+311.5RT (min)

lcqtof\_t3\_neg\_1\_055\_32

m/z 435.0060

01E+311.5RT (min)

lcqtof\_t3\_neg\_2\_005\_34

m/z 435.0513

012E+311.5RT (min)

lcqtof\_t3\_neg\_2\_019\_32

m/z 435.8800

01234E+311.5RT (min)

lcqtof\_t3\_neg\_1\_036\_14

m/z 436.1345

01234E+311.5RT (min)

lcqtof\_t3\_neg\_4\_040\_24

m/z 436.1659

01234E+377.5RT (min)

lcqtof\_t3\_neg\_4\_004\_20

m/z 436.7439

024E+311.5RT (min)

lcqtof\_t3\_neg\_4\_046\_33

m/z 436.8610

02468E+311.5RT (min)

lcqtof\_t3\_neg\_4\_037\_01

m/z 438.1432

01234E+344.5RT (min)

lcqtof\_t3\_neg\_3\_004\_26

m/z 438.1665

0123E+37.58RT (min)

lcqtof\_t3\_neg\_4\_016\_14

m/z 439.0969

0123E+311.5RT (min)

lcqtof\_t3\_neg\_1\_032\_36

m/z 441.2196

0123E+36.57RT (min)

lcqtof\_t3\_neg\_4\_026\_30

m/z 442.0188

012E+311.5RT (min)

lcqtof\_t3\_neg\_4\_030\_31

m/z 442.8127

024E+311.5RT (min)

lcqtof\_t3\_neg\_4\_046\_33

m/z 443.1863

012E+344.5RT (min)

lcqtof\_t3\_neg\_4\_055\_38

m/z 443.8960

02468E+311.5RT (min)

lcqtof\_t3\_neg\_2\_042\_13

m/z 444.1410

02468E+35.56RT (min)

lcqtof\_t3\_neg\_4\_004\_20

m/z 445.0930

0123E+34.55RT (min)

lcqtof\_t3\_neg\_1\_010\_27

m/z 446.1271

012E+344.5RT (min)

lcqtof\_t3\_neg\_1\_022\_28

m/z 447.0881

012E+34.55RT (min)

lcqtof\_t3\_neg\_4\_004\_20

m/z 447.9911

01E+311.5RT (min)

lcqtof\_t3\_neg\_4\_045\_02\_rerun

m/z 448.1654

024E+35.56RT (min)

lcqtof\_t3\_neg\_4\_055\_38

m/z 448.7563

0123E+311.5RT (min)

lcqtof\_t3\_neg\_4\_046\_33

m/z 449.1314

0123E+35.56RT (min)

lcqtof\_t3\_neg\_4\_014\_13

m/z 449.9704

012E+34.55RT (min)

lcqtof\_t3\_neg\_1\_028\_20

m/z 450.1644

024E+35.56RT (min)

lcqtof\_t3\_neg\_4\_055\_38

m/z 450.6657

01234E+311.5RT (min)

lcqtof\_t3\_neg\_4\_037\_01

m/z 451.1411

024E+366.5RT (min)

lcqtof\_t3\_neg\_4\_016\_14

m/z 451.2275

0123E+344.5RT (min)

lcqtof\_t3\_neg\_4\_043\_11

m/z 452.8324

0123E+311.5RT (min)

lcqtof\_t3\_neg\_4\_046\_33

m/z 453.0941

01234E+311.5RT (min)

lcqtof\_t3\_neg\_4\_038\_29

m/z 454.1453

0246E+44.55RT (min)

lcqtof\_t3\_neg\_3\_031\_19

m/z 454.1486

012E+36.57RT (min)

lcqtof\_t3\_neg\_1\_053\_01

m/z 455.0948

0123E+344.5RT (min)

lcqtof\_t3\_neg\_4\_003\_37

m/z 455.1244

012E+36.57RT (min)

lcqtof\_t3\_neg\_4\_034\_34

m/z 455.9307

012E+311.5RT (min)

lcqtof\_t3\_neg\_4\_045\_02\_rerun

m/z 455.9869

01E+34.55RT (min)

lcqtof\_t3\_neg\_1\_028\_20

m/z 456.1382

02468E+311.5RT (min)

lcqtof\_t3\_neg\_1\_031\_24

m/z 456.1606

01234E+344.5RT (min)

lcqtof\_t3\_neg\_4\_004\_20

m/z 456.7004

01E+411.5RT (min)

lcqtof\_t3\_neg\_4\_037\_01

m/z 456.7995

02468E+311.5RT (min)

lcqtof\_t3\_neg\_4\_046\_33

m/z 457.1199

024E+311.5RT (min)

lcqtof\_t3\_neg\_1\_032\_36

m/z 457.1209

012E+36.57RT (min)

lcqtof\_t3\_neg\_4\_005\_18

m/z 457.2590

0246E+344.5RT (min)

lcqtof\_t3\_neg\_4\_016\_14

m/z 458.0384

024E+311.5RT (min)

lcqtof\_t3\_neg\_1\_055\_32

m/z 458.1176

0123E+311.5RT (min)

lcqtof\_t3\_neg\_4\_019\_05

m/z 459.8852

01234E+311.5RT (min)

lcqtof\_t3\_neg\_1\_032\_36

m/z 460.2822

01234E+366.5RT (min)

lcqtof\_t3\_neg\_4\_018\_23

m/z 461.1464

024E+34.55RT (min)

lcqtof\_t3\_neg\_1\_026\_19

m/z 461.1640

012E+377.5RT (min)

lcqtof\_t3\_neg\_4\_004\_20

m/z 461.1659

01E+366.5RT (min)

lcqtof\_t3\_neg\_2\_040\_43

m/z 461.9077

01234E+311.5RT (min)

lcqtof\_t3\_neg\_4\_030\_31

m/z 462.1366

012E+311.5RT (min)

lcqtof\_t3\_neg\_2\_011\_25

m/z 462.1798

01E+366.5RT (min)

lcqtof\_t3\_neg\_4\_051\_15

m/z 463.9549

01E+311.5RT (min)

lcqtof\_t3\_neg\_1\_055\_32

m/z 465.0686

024E+34.55RT (min)

lcqtof\_t3\_neg\_4\_048\_19

m/z 465.2086

024E+36.57RT (min)

lcqtof\_t3\_neg\_4\_099\_61

m/z 466.6397

012E+311.5RT (min)

lcqtof\_t3\_neg\_4\_051\_15

m/z 466.8279

0246E+311.5RT (min)

lcqtof\_t3\_neg\_4\_046\_33

m/z 468.4195

01E+313.514RT (min)

lcqtof\_t3\_neg\_4\_086\_59

m/z 468.6793

01E+411.5RT (min)

lcqtof\_t3\_neg\_4\_037\_01

m/z 469.0903

02468E+34.55RT (min)

lcqtof\_t3\_neg\_1\_010\_27

m/z 469.2177

012E+35.56RT (min)

lcqtof\_t3\_neg\_2\_021\_27

m/z 469.2211

0123E+377.5RT (min)

lcqtof\_t3\_neg\_4\_044\_48

m/z 470.1454

01E+366.5RT (min)

lcqtof\_t3\_neg\_1\_018\_33

m/z 470.6773

01E+411.5RT (min)

lcqtof\_t3\_neg\_4\_037\_01

m/z 470.8992

012E+316.517RT (min)

lcqtof\_t3\_neg\_4\_049\_55

m/z 470.9791

0123E+311.5RT (min)

lcqtof\_t3\_neg\_1\_055\_32

m/z 471.0204

01234E+311.5RT (min)

lcqtof\_t3\_neg\_1\_032\_36

m/z 471.0847

0246E+34.55RT (min)

lcqtof\_t3\_neg\_4\_048\_19

m/z 471.2932

024E+344.5RT (min)

lcqtof\_t3\_neg\_4\_016\_14

m/z 471.9030

012E+311.5RT (min)

lcqtof\_t3\_neg\_4\_045\_02\_rerun

m/z 472.6290

0123E+311.5RT (min)

lcqtof\_t3\_neg\_4\_051\_15

m/z 472.6881

02468E+311.5RT (min)

lcqtof\_t3\_neg\_4\_041\_54

m/z 473.1502

01E+444.5RT (min)

lcqtof\_t3\_neg\_4\_045\_02\_rerun

m/z 474.0903

01234E+34.55RT (min)

lcqtof\_t3\_neg\_4\_004\_20

m/z 474.6831

012E+311.5RT (min)

lcqtof\_t3\_neg\_4\_034\_34

m/z 475.1193

0246E+35.56RT (min)

lcqtof\_t3\_neg\_4\_004\_20

m/z 475.3055

02468E+366.5RT (min)

lcqtof\_t3\_neg\_4\_030\_31

m/z 475.8624

01E+311.5RT (min)

lcqtof\_t3\_neg\_1\_017\_51

m/z 476.8437

0246E+311.5RT (min)

lcqtof\_t3\_neg\_4\_046\_33

m/z 476.9872

01234E+311.5RT (min)

lcqtof\_t3\_neg\_1\_018\_33

m/z 477.0979

012E+35.56RT (min)

lcqtof\_t3\_neg\_1\_028\_20

m/z 477.1901

024E+311.5RT (min)

lcqtof\_t3\_neg\_4\_039\_06\_rerun

m/z 477.4659

0246E+31414.5RT (min)

lcqtof\_t3\_neg\_4\_047\_07

m/z 478.1805

0123E+37.58RT (min)

lcqtof\_t3\_neg\_2\_013\_26

m/z 480.0865

0246E+311.5RT (min)

lcqtof\_t3\_neg\_1\_055\_32

m/z 480.6183

01234E+311.5RT (min)

lcqtof\_t3\_neg\_4\_016\_14

m/z 481.0729

012E+311.5RT (min)

lcqtof\_t3\_neg\_3\_036\_16

m/z 481.1898

0246E+36.57RT (min)

lcqtof\_t3\_neg\_4\_016\_14

m/z 481.2412

012E+36.57RT (min)

lcqtof\_t3\_neg\_4\_037\_01

m/z 482.0894

0123E+35.56RT (min)

lcqtof\_t3\_neg\_1\_026\_19

m/z 482.2067

01E+411.5RT (min)

lcqtof\_t3\_neg\_4\_039\_06\_rerun

m/z 482.2076

01234E+36.57RT (min)

lcqtof\_t3\_neg\_3\_004\_26

m/z 483.3196

01234E+355.5RT (min)

lcqtof\_t3\_neg\_4\_055\_38

m/z 484.0753

0123E+311.5RT (min)

lcqtof\_t3\_neg\_2\_053\_38

m/z 484.8711

012E+316.517RT (min)

lcqtof\_t3\_neg\_1\_001\_49

m/z 485.8759

012E+311.5RT (min)

lcqtof\_t3\_neg\_1\_036\_14

m/z 486.1945

012E+344.5RT (min)

lcqtof\_t3\_neg\_1\_001\_49

m/z 486.2038

012E+38.59RT (min)

lcqtof\_t3\_neg\_4\_088\_59

m/z 486.2594

012E+355.5RT (min)

lcqtof\_t3\_neg\_4\_055\_38

m/z 486.8697

024E+311.5RT (min)

lcqtof\_t3\_neg\_1\_036\_14

m/z 486.8834

0123E+311.5RT (min)

lcqtof\_t3\_neg\_4\_045\_02\_rerun

m/z 487.1046

0123E+377.5RT (min)

lcqtof\_t3\_neg\_4\_037\_01

m/z 487.1314

012E+34.55RT (min)

lcqtof\_t3\_neg\_2\_054\_29

m/z 487.1787

0246E+311.5RT (min)

lcqtof\_t3\_neg\_4\_018\_23

m/z 488.1374

0246E+344.5RT (min)

lcqtof\_t3\_neg\_1\_005\_18

m/z 488.1901

01E+411.5RT (min)

lcqtof\_t3\_neg\_4\_039\_06\_rerun

m/z 488.6698

02468E+311.5RT (min)

lcqtof\_t3\_neg\_4\_041\_54

m/z 489.1870

01E+411.5RT (min)

lcqtof\_t3\_neg\_4\_039\_06\_rerun

m/z 490.0686

0123E+311.5RT (min)

lcqtof\_t3\_neg\_2\_026\_36

m/z 491.0551

01234E+311.5RT (min)

lcqtof\_t3\_neg\_2\_019\_32

m/z 491.0600

01234E+355.5RT (min)

lcqtof\_t3\_neg\_1\_005\_18

m/z 491.1255

0123E+344.5RT (min)

lcqtof\_t3\_neg\_4\_016\_14

m/z 491.3091

024E+355.5RT (min)

lcqtof\_t3\_neg\_4\_018\_23

m/z 492.0861

0123E+311.5RT (min)

lcqtof\_t3\_neg\_1\_055\_32

m/z 492.7183

02468E+311.5RT (min)

lcqtof\_t3\_neg\_4\_046\_33

m/z 492.9548

012E+311.5RT (min)

lcqtof\_t3\_neg\_4\_046\_33

m/z 494.1677

012E+37.58RT (min)

lcqtof\_t3\_neg\_2\_038\_18

m/z 494.7022

02468E+311.5RT (min)

lcqtof\_t3\_neg\_4\_046\_33

m/z 494.9092

01234E+311.5RT (min)

lcqtof\_t3\_neg\_3\_028\_13

m/z 495.2411

0123E+35.56RT (min)

lcqtof\_t3\_neg\_2\_034\_19

m/z 496.6972

024E+311.5RT (min)

lcqtof\_t3\_neg\_4\_041\_54

m/z 496.9482

012E+316.517RT (min)

lcqtof\_t3\_neg\_4\_013\_39

m/z 497.0840

0246E+34.55RT (min)

lcqtof\_t3\_neg\_3\_031\_19

m/z 497.1426

0246E+344.5RT (min)

lcqtof\_t3\_neg\_4\_037\_01

m/z 498.1218

012E+377.5RT (min)

lcqtof\_t3\_neg\_1\_036\_14

m/z 498.9155

0123E+311.5RT (min)

lcqtof\_t3\_neg\_4\_051\_15

m/z 498.9170

012E+316.517RT (min)

lcqtof\_t3\_neg\_3\_111\_62

m/z 499.1359

012E+37.58RT (min)

lcqtof\_t3\_neg\_4\_051\_15

m/z 499.1455

0123E+366.5RT (min)

lcqtof\_t3\_neg\_4\_037\_01

m/z 499.1623

0246E+311.5RT (min)

lcqtof\_t3\_neg\_4\_028\_25

m/z 501.8585

01234E+311.5RT (min)

lcqtof\_t3\_neg\_4\_037\_01

m/z 502.0429

0246E+34.55RT (min)

lcqtof\_t3\_neg\_4\_004\_20

m/z 502.1609

012E+344.5RT (min)

lcqtof\_t3\_neg\_4\_037\_01

m/z 503.3276

012E+366.5RT (min)

lcqtof\_t3\_neg\_1\_044\_37

m/z 504.0804

0246E+311.5RT (min)

lcqtof\_t3\_neg\_4\_055\_38

m/z 504.3302

012E+344.5RT (min)

lcqtof\_t3\_neg\_4\_030\_31

m/z 504.9366

0123E+316.517RT (min)

lcqtof\_t3\_neg\_4\_039\_06\_rerun

m/z 505.1818

0246E+311.5RT (min)

lcqtof\_t3\_neg\_4\_047\_07

m/z 506.1161

0123E+34.55RT (min)

lcqtof\_t3\_neg\_4\_004\_20

m/z 506.2343

0246E+366.5RT (min)

lcqtof\_t3\_neg\_4\_041\_54

m/z 506.7249

024E+311.5RT (min)

lcqtof\_t3\_neg\_4\_046\_33

m/z 507.2040

0123E+311.5RT (min)

lcqtof\_t3\_neg\_2\_022\_47

m/z 508.1696

0123E+377.5RT (min)

lcqtof\_t3\_neg\_1\_026\_19

m/z 508.1768

024E+344.5RT (min)

lcqtof\_t3\_neg\_4\_037\_01

m/z 508.2065

0246E+36.57RT (min)

lcqtof\_t3\_neg\_4\_034\_34

m/z 508.2943

0123E+311.5RT (min)

lcqtof\_t3\_neg\_4\_040\_24

m/z 509.1923

0123E+36.57RT (min)

lcqtof\_t3\_neg\_4\_005\_18

m/z 510.1572

0123E+35.56RT (min)

lcqtof\_t3\_neg\_2\_021\_27

m/z 510.2053

012E+311.5RT (min)

lcqtof\_t3\_neg\_1\_020\_47

m/z 510.2957

024E+34.55RT (min)

lcqtof\_t3\_neg\_4\_016\_14

m/z 510.6632

012E+411.5RT (min)

lcqtof\_t3\_neg\_4\_037\_01

m/z 510.9191

0246E+311.5RT (min)

lcqtof\_t3\_neg\_4\_037\_01

m/z 511.0982

012E+37.58RT (min)

lcqtof\_t3\_neg\_1\_005\_18

m/z 511.1716

012E+377.5RT (min)

lcqtof\_t3\_neg\_4\_044\_48

m/z 511.8844

01E+411.5RT (min)

lcqtof\_t3\_neg\_2\_042\_13

m/z 512.0489

01234E+311.5RT (min)

lcqtof\_t3\_neg\_1\_032\_36

m/z 512.8790

0246E+316.517RT (min)

lcqtof\_t3\_neg\_4\_025\_52

m/z 513.0336

01234E+311.5RT (min)

lcqtof\_t3\_neg\_4\_030\_31

m/z 513.0422

012E+355.5RT (min)

lcqtof\_t3\_neg\_4\_034\_34

m/z 513.0664

01234E+311.5RT (min)

lcqtof\_t3\_neg\_1\_036\_14

m/z 513.0956

012E+37.58RT (min)

lcqtof\_t3\_neg\_1\_005\_18

m/z 513.1683

0123E+377.5RT (min)

lcqtof\_t3\_neg\_1\_016\_48

m/z 514.0684

01234E+311.5RT (min)

lcqtof\_t3\_neg\_1\_036\_14

m/z 514.1081

012E+377.5RT (min)

lcqtof\_t3\_neg\_1\_018\_33

m/z 514.1261

024E+37.58RT (min)

lcqtof\_t3\_neg\_4\_037\_01

m/z 514.3632

012E+344.5RT (min)

lcqtof\_t3\_neg\_4\_018\_23

m/z 514.9543

0123E+311.5RT (min)

lcqtof\_t3\_neg\_4\_046\_33

m/z 517.0645

01234E+311.5RT (min)

lcqtof\_t3\_neg\_1\_036\_14

m/z 517.1474

0246E+35.56RT (min)

lcqtof\_t3\_neg\_4\_055\_38

m/z 517.2000

0123E+36.57RT (min)

lcqtof\_t3\_neg\_4\_055\_38

m/z 517.8712

01234E+311.5RT (min)

lcqtof\_t3\_neg\_4\_026\_30

m/z 518.0470

012E+311.5RT (min)

lcqtof\_t3\_neg\_1\_055\_32

m/z 519.8661

01234E+311.5RT (min)

lcqtof\_t3\_neg\_4\_026\_30

m/z 520.1178

012E+377.5RT (min)

lcqtof\_t3\_neg\_4\_022\_28

m/z 520.8210

0246E+311.5RT (min)

lcqtof\_t3\_neg\_4\_051\_15

m/z 521.1415

01E+411.5RT (min)

lcqtof\_t3\_neg\_4\_045\_02\_rerun

m/z 523.0633

01234E+311.5RT (min)

lcqtof\_t3\_neg\_1\_055\_32

m/z 523.2154

0246E+35.56RT (min)

lcqtof\_t3\_neg\_1\_010\_27

m/z 523.9170

012E+311.5RT (min)

lcqtof\_t3\_neg\_4\_045\_02\_rerun

m/z 523.9743

012E+34.55RT (min)

lcqtof\_t3\_neg\_4\_108\_62

m/z 524.1549

0123E+377.5RT (min)

lcqtof\_t3\_neg\_1\_026\_19

m/z 524.1899

012E+311.5RT (min)

lcqtof\_t3\_neg\_1\_041\_54

m/z 524.8953

024E+311.5RT (min)

lcqtof\_t3\_neg\_4\_016\_14

m/z 525.2138

0123E+35.56RT (min)

lcqtof\_t3\_neg\_1\_010\_27

m/z 526.0649

02468E+311.5RT (min)

lcqtof\_t3\_neg\_4\_055\_38

m/z 526.1622

012E+311.5RT (min)

lcqtof\_t3\_neg\_1\_017\_51

m/z 526.7637

0123E+366.5RT (min)

lcqtof\_t3\_neg\_2\_051\_14

m/z 526.8898

0246E+311.5RT (min)

lcqtof\_t3\_neg\_4\_016\_14

m/z 527.1592

01234E+311.5RT (min)

lcqtof\_t3\_neg\_1\_044\_37

m/z 527.8715

0246E+311.5RT (min)

lcqtof\_t3\_neg\_1\_032\_36

m/z 528.6362

01E+411.5RT (min)

lcqtof\_t3\_neg\_4\_037\_01

m/z 528.9379

012E+311.5RT (min)

lcqtof\_t3\_neg\_4\_041\_54

m/z 530.5854

0123E+311.5RT (min)

lcqtof\_t3\_neg\_4\_051\_15

m/z 530.6304

01E+411.5RT (min)

lcqtof\_t3\_neg\_4\_037\_01

m/z 530.8599

0123E+311.5RT (min)

lcqtof\_t3\_neg\_4\_051\_15

m/z 531.1793

0246E+311.5RT (min)

lcqtof\_t3\_neg\_1\_020\_47

m/z 532.3556

01234E+344.5RT (min)

lcqtof\_t3\_neg\_2\_043\_31

m/z 532.8198

01E+316.517RT (min)

lcqtof\_t3\_neg\_1\_025\_52

m/z 532.8729

024E+311.5RT (min)

lcqtof\_t3\_neg\_1\_036\_14

m/z 534.1889

0123E+36.57RT (min)

lcqtof\_t3\_neg\_1\_010\_27

m/z 534.9387

0246E+311.5RT (min)

lcqtof\_t3\_neg\_4\_046\_33

m/z 535.2606

0246E+355.5RT (min)

lcqtof\_t3\_neg\_4\_018\_23

m/z 536.1756

01234E+34.55RT (min)

lcqtof\_t3\_neg\_4\_003\_37

m/z 537.0955

024E+399.5RT (min)

lcqtof\_t3\_neg\_4\_055\_38

m/z 537.1370

01234E+311.5RT (min)

lcqtof\_t3\_neg\_1\_032\_36

m/z 539.0070

024E+311.5RT (min)

lcqtof\_t3\_neg\_1\_032\_36

m/z 539.1823

02468E+377.5RT (min)

lcqtof\_t3\_neg\_4\_004\_20

m/z 539.2483

012E+35.56RT (min)

lcqtof\_t3\_neg\_2\_053\_38

m/z 539.8860

012E+311.5RT (min)

lcqtof\_t3\_neg\_4\_045\_02\_rerun

m/z 540.2533

012E+36.57RT (min)

lcqtof\_t3\_neg\_4\_037\_01

m/z 540.3294

02468E+316.517RT (min)

lcqtof\_t3\_neg\_4\_016\_14

m/z 540.5747

012E+311.5RT (min)

lcqtof\_t3\_neg\_2\_051\_14

m/z 540.8253

0246E+311.5RT (min)

lcqtof\_t3\_neg\_1\_015\_34

m/z 542.0605

01234E+311.5RT (min)

lcqtof\_t3\_neg\_4\_055\_38

m/z 543.0085

0246E+37.58RT (min)

lcqtof\_t3\_neg\_4\_101\_61

m/z 543.8395

0123E+311.5RT (min)

lcqtof\_t3\_neg\_2\_019\_32

m/z 544.8255

02468E+311.5RT (min)

lcqtof\_t3\_neg\_2\_051\_14

m/z 546.0414

01E+311.5RT (min)

lcqtof\_t3\_neg\_2\_055\_16

m/z 546.1694

012E+311.5RT (min)

lcqtof\_t3\_neg\_1\_053\_01

m/z 546.6285

0246E+311.5RT (min)

lcqtof\_t3\_neg\_4\_055\_38

m/z 547.2335

0246E+35.56RT (min)

lcqtof\_t3\_neg\_4\_048\_19

m/z 548.0457

024E+311.5RT (min)

lcqtof\_t3\_neg\_4\_055\_38

m/z 548.1851

0246E+311.5RT (min)

lcqtof\_t3\_neg\_2\_051\_14

m/z 549.1675

02468E+311.5RT (min)

lcqtof\_t3\_neg\_4\_047\_07

m/z 549.8646

0123E+311.5RT (min)

lcqtof\_t3\_neg\_1\_032\_36

m/z 550.3776

0123E+31313.5RT (min)

lcqtof\_t3\_neg\_2\_052\_17

m/z 550.5544

01234E+311.5RT (min)

lcqtof\_t3\_neg\_4\_016\_14

m/z 551.1942

012E+36.57RT (min)

lcqtof\_t3\_neg\_1\_010\_27

m/z 551.2092

0246E+344.5RT (min)

lcqtof\_t3\_neg\_1\_044\_37

m/z 552.5502

012E+311.5RT (min)

lcqtof\_t3\_neg\_4\_016\_14

m/z 552.9832

0123E+311.5RT (min)

lcqtof\_t3\_neg\_1\_055\_32

m/z 553.8742

0123E+316.517RT (min)

lcqtof\_t3\_neg\_1\_011\_45

m/z 554.1018

0246E+311.5RT (min)

lcqtof\_t3\_neg\_1\_055\_32

m/z 554.1219

012E+311.5RT (min)

lcqtof\_t3\_neg\_4\_018\_23

m/z 554.1572

01E+311.5RT (min)

lcqtof\_t3\_neg\_1\_020\_47

m/z 554.6503

0246E+311.5RT (min)

lcqtof\_t3\_neg\_4\_033\_53

m/z 555.0953

0123E+34.55RT (min)

lcqtof\_t3\_neg\_4\_004\_20

m/z 555.1539

01234E+311.5RT (min)

lcqtof\_t3\_neg\_1\_020\_47

m/z 555.1807

01234E+36.57RT (min)

lcqtof\_t3\_neg\_4\_005\_18

m/z 558.0621

01234E+311.5RT (min)

lcqtof\_t3\_neg\_1\_032\_36

m/z 559.0427

01234E+311.5RT (min)

lcqtof\_t3\_neg\_2\_019\_32

m/z 559.0461

0123E+355.5RT (min)

lcqtof\_t3\_neg\_1\_005\_18

m/z 559.1189

012E+344.5RT (min)

lcqtof\_t3\_neg\_4\_016\_14

m/z 559.2516

0123E+344.5RT (min)

lcqtof\_t3\_neg\_1\_019\_23

m/z 559.9976

01234E+311.5RT (min)

lcqtof\_t3\_neg\_2\_019\_32

m/z 562.6862

01E+411.5RT (min)

lcqtof\_t3\_neg\_4\_046\_33

m/z 562.8953

024E+311.5RT (min)

lcqtof\_t3\_neg\_1\_002\_13

m/z 564.8517

0246E+311.5RT (min)

lcqtof\_t3\_neg\_4\_016\_14

m/z 565.0624

0246E+355.5RT (min)

lcqtof\_t3\_neg\_4\_099\_61

m/z 565.1315

01234E+35.56RT (min)

lcqtof\_t3\_neg\_1\_010\_27

m/z 565.2118

02468E+377.5RT (min)

lcqtof\_t3\_neg\_3\_050\_29

m/z 565.4144

0123E+35.56RT (min)

lcqtof\_t3\_neg\_1\_015\_34

m/z 566.0524

012E+311.5RT (min)

lcqtof\_t3\_neg\_1\_019\_23

m/z 566.8342

0123E+311.5RT (min)

lcqtof\_t3\_neg\_3\_018\_32

m/z 566.9020

01234E+311.5RT (min)

lcqtof\_t3\_neg\_4\_051\_15

m/z 568.3626

024E+316.517RT (min)

lcqtof\_t3\_neg\_4\_016\_14

m/z 568.9949

01E+311.5RT (min)

lcqtof\_t3\_neg\_3\_046\_36

m/z 569.8455

024E+311.5RT (min)

lcqtof\_t3\_neg\_4\_037\_01

m/z 571.8416

0123E+311.5RT (min)

lcqtof\_t3\_neg\_4\_037\_01

m/z 572.1466

0123E+344.5RT (min)

lcqtof\_t3\_neg\_1\_022\_28

m/z 572.2852

012E+344.5RT (min)

lcqtof\_t3\_neg\_1\_036\_14

m/z 572.9235

0123E+316.517RT (min)

lcqtof\_t3\_neg\_4\_045\_02\_rerun

m/z 573.0818

01234E+311.5RT (min)

lcqtof\_t3\_neg\_4\_055\_38

m/z 573.3636

0123E+36.57RT (min)

lcqtof\_t3\_neg\_4\_018\_23

m/z 574.9664

0123E+311.5RT (min)

lcqtof\_t3\_neg\_1\_055\_32

m/z 576.1586

0123E+377.5RT (min)

lcqtof\_t3\_neg\_1\_026\_19

m/z 577.3246

01E+37.58RT (min)

lcqtof\_t3\_neg\_4\_048\_19

m/z 578.7805

01E+411.5RT (min)

lcqtof\_t3\_neg\_4\_051\_15

m/z 579.3140

02468E+37.58RT (min)

lcqtof\_t3\_neg\_4\_110\_62

m/z 579.8723

01E+411.5RT (min)

lcqtof\_t3\_neg\_1\_002\_13

m/z 580.0551

012E+38.59RT (min)

lcqtof\_t3\_neg\_1\_037\_25

m/z 580.8043

0246E+311.5RT (min)

lcqtof\_t3\_neg\_1\_055\_32

m/z 582.1136

01234E+37.58RT (min)

lcqtof\_t3\_neg\_4\_037\_01

m/z 582.8560

02468E+311.5RT (min)

lcqtof\_t3\_neg\_1\_032\_36

m/z 583.1301

024E+311.5RT (min)

lcqtof\_t3\_neg\_4\_088\_59

m/z 586.1558

01234E+311.5RT (min)

lcqtof\_t3\_neg\_2\_053\_38

m/z 588.8093

0246E+311.5RT (min)

lcqtof\_t3\_neg\_4\_051\_15

m/z 591.9061

0123E+311.5RT (min)

lcqtof\_t3\_neg\_4\_045\_02\_rerun

m/z 592.1442

0123E+377.5RT (min)

lcqtof\_t3\_neg\_1\_026\_19

m/z 592.1495

0123E+311.5RT (min)

lcqtof\_t3\_neg\_2\_051\_14

m/z 592.2745

024E+388.5RT (min)

lcqtof\_t3\_neg\_4\_005\_18

m/z 592.7654

01E+411.5RT (min)

lcqtof\_t3\_neg\_4\_046\_33

m/z 592.9011

01234E+311.5RT (min)

lcqtof\_t3\_neg\_4\_055\_38

m/z 593.2708

024E+388.5RT (min)

lcqtof\_t3\_neg\_4\_024\_26

m/z 594.9030

0123E+311.5RT (min)

lcqtof\_t3\_neg\_4\_046\_33

m/z 595.2709

024E+36.57RT (min)

lcqtof\_t3\_neg\_4\_004\_20

m/z 595.8480

0246E+311.5RT (min)

lcqtof\_t3\_neg\_1\_036\_14

m/z 596.0798

01234E+35.56RT (min)

lcqtof\_t3\_neg\_4\_048\_19

m/z 596.3089

0123E+37.58RT (min)

lcqtof\_t3\_neg\_3\_109\_62

m/z 596.4036

01234E+355.5RT (min)

lcqtof\_t3\_neg\_4\_018\_23

m/z 596.7861

01E+411.5RT (min)

lcqtof\_t3\_neg\_4\_046\_33

m/z 598.2766

024E+377.5RT (min)

lcqtof\_t3\_neg\_4\_004\_20

m/z 598.4180

0246E+366.5RT (min)

lcqtof\_t3\_neg\_4\_018\_23

m/z 598.7844

0246E+311.5RT (min)

lcqtof\_t3\_neg\_4\_055\_38

m/z 598.8357

0123E+311.5RT (min)

lcqtof\_t3\_neg\_1\_017\_51

m/z 598.8502

01234E+311.5RT (min)

lcqtof\_t3\_neg\_4\_051\_15

m/z 599.5519

01234E+311.5RT (min)

lcqtof\_t3\_neg\_4\_037\_01

m/z 600.8091

012E+316.517RT (min)

lcqtof\_t3\_neg\_1\_039\_43

m/z 601.0443

01E+38.59RT (min)

lcqtof\_t3\_neg\_4\_104\_61

m/z 601.1875

02468E+311.5RT (min)

lcqtof\_t3\_neg\_4\_097\_60

m/z 601.1896

0246E+36.57RT (min)

lcqtof\_t3\_neg\_4\_004\_20

m/z 601.3312

0123E+344.5RT (min)

lcqtof\_t3\_neg\_4\_016\_14

m/z 602.0959

0246E+35.56RT (min)

lcqtof\_t3\_neg\_4\_048\_19

m/z 602.7852

01E+411.5RT (min)

lcqtof\_t3\_neg\_2\_051\_14

m/z 604.3177

0123E+316.517RT (min)

lcqtof\_t3\_neg\_4\_051\_15

m/z 604.3934

02468E+35.56RT (min)

lcqtof\_t3\_neg\_4\_030\_31

m/z 604.8147

012E+316.517RT (min)

lcqtof\_t3\_neg\_2\_056\_56

m/z 605.0735

01234E+311.5RT (min)

lcqtof\_t3\_neg\_1\_032\_36

m/z 606.5359

0246E+311.5RT (min)

lcqtof\_t3\_neg\_4\_016\_14

m/z 606.9930

024E+311.5RT (min)

lcqtof\_t3\_neg\_2\_026\_36

m/z 607.1357

012E+311.5RT (min)

lcqtof\_t3\_neg\_3\_033\_53

m/z 607.1579

02468E+311.5RT (min)

lcqtof\_t3\_neg\_2\_051\_14

m/z 607.8762

0123E+311.5RT (min)

lcqtof\_t3\_neg\_4\_045\_02\_rerun

m/z 608.5201

0246E+311.5RT (min)

lcqtof\_t3\_neg\_4\_016\_14

m/z 610.2189

02468E+355.5RT (min)

lcqtof\_t3\_neg\_4\_005\_18

m/z 610.5065

01234E+311.5RT (min)

lcqtof\_t3\_neg\_4\_016\_14

m/z 611.0303

0123E+36.57RT (min)

lcqtof\_t3\_neg\_4\_048\_19

m/z 611.8188

0123E+311.5RT (min)

lcqtof\_t3\_neg\_3\_018\_32

m/z 611.8453

012E+316.517RT (min)

lcqtof\_t3\_neg\_1\_003\_07

m/z 612.3140

0123E+355.5RT (min)

lcqtof\_t3\_neg\_4\_003\_37

m/z 612.6047

02468E+311.5RT (min)

lcqtof\_t3\_neg\_4\_046\_33

m/z 612.8139

01E+411.5RT (min)

lcqtof\_t3\_neg\_1\_036\_14

m/z 613.3559

0123E+37.58RT (min)

lcqtof\_t3\_neg\_4\_016\_14

m/z 615.1348

012E+311.5RT (min)

lcqtof\_t3\_neg\_1\_055\_32

m/z 617.7359

024E+366.5RT (min)

lcqtof\_t3\_neg\_4\_016\_14

m/z 617.8512

01234E+311.5RT (min)

lcqtof\_t3\_neg\_1\_043\_38

m/z 624.5504

0246E+311.5RT (min)

lcqtof\_t3\_neg\_4\_014\_13

m/z 624.8157

02468E+311.5RT (min)

lcqtof\_t3\_neg\_1\_055\_32

m/z 625.0985

01E+311.5RT (min)

lcqtof\_t3\_neg\_1\_034\_30

m/z 626.1627

01234E+344.5RT (min)

lcqtof\_t3\_neg\_4\_018\_23

m/z 627.0289

024E+311.5RT (min)

lcqtof\_t3\_neg\_3\_018\_32

m/z 627.0327

012E+355.5RT (min)

lcqtof\_t3\_neg\_4\_103\_61

m/z 627.5327

0246E+311.5RT (min)

lcqtof\_t3\_neg\_4\_037\_01

m/z 627.9832

024E+311.5RT (min)

lcqtof\_t3\_neg\_3\_018\_32

m/z 628.8708

01E+316.517RT (min)

lcqtof\_t3\_neg\_4\_091\_60

m/z 629.0624

0246E+311.5RT (min)

lcqtof\_t3\_neg\_2\_026\_36

m/z 629.1228

0246E+311.5RT (min)

lcqtof\_t3\_neg\_1\_036\_14

m/z 629.1300

02468E+311.5RT (min)

lcqtof\_t3\_neg\_2\_051\_14

m/z 629.2039

0246E+311.5RT (min)

lcqtof\_t3\_neg\_4\_040\_24

m/z 630.4185

024E+35.56RT (min)

lcqtof\_t3\_neg\_1\_048\_35

m/z 630.8817

01234E+311.5RT (min)

lcqtof\_t3\_neg\_2\_042\_13

m/z 634.1243

01234E+311.5RT (min)

lcqtof\_t3\_neg\_4\_018\_23

m/z 634.8213

0123E+311.5RT (min)

lcqtof\_t3\_neg\_1\_055\_32

m/z 634.8890

012E+311.5RT (min)

lcqtof\_t3\_neg\_4\_037\_01

m/z 635.1247

0123E+311.5RT (min)

lcqtof\_t3\_neg\_4\_018\_23

m/z 636.0854

0123E+311.5RT (min)

lcqtof\_t3\_neg\_1\_038\_15

m/z 636.9833

012E+311.5RT (min)

lcqtof\_t3\_neg\_2\_026\_36

m/z 637.8301

024E+311.5RT (min)

lcqtof\_t3\_neg\_2\_050\_01

m/z 640.5167

024E+311.5RT (min)

lcqtof\_t3\_neg\_4\_051\_15

m/z 640.6946

02468E+311.5RT (min)

lcqtof\_t3\_neg\_4\_046\_33

m/z 640.8307

02468E+311.5RT (min)

lcqtof\_t3\_neg\_4\_037\_01

m/z 640.9083

0123E+316.517RT (min)

lcqtof\_t3\_neg\_4\_039\_06\_rerun

m/z 641.0200

012E+38.59RT (min)

lcqtof\_t3\_neg\_1\_040\_08

m/z 642.3553

0123E+344.5RT (min)

lcqtof\_t3\_neg\_4\_055\_38

m/z 642.5721

02468E+311.5RT (min)

lcqtof\_t3\_neg\_4\_055\_38

m/z 644.0269

012E+311.5RT (min)

lcqtof\_t3\_neg\_1\_032\_36

m/z 644.1181

01234E+344.5RT (min)

lcqtof\_t3\_neg\_2\_030\_23

m/z 644.1455

012E+377.5RT (min)

lcqtof\_t3\_neg\_1\_026\_19

m/z 644.2197

02468E+344.5RT (min)

lcqtof\_t3\_neg\_4\_040\_24

m/z 644.4368

0123E+34.55RT (min)

lcqtof\_t3\_neg\_1\_044\_37

m/z 644.5695

0246E+311.5RT (min)

lcqtof\_t3\_neg\_4\_041\_54

m/z 645.1141

0123E+344.5RT (min)

lcqtof\_t3\_neg\_1\_044\_37

m/z 647.5261

02468E+311.5RT (min)

lcqtof\_t3\_neg\_2\_042\_13

m/z 647.8578

01E+411.5RT (min)

lcqtof\_t3\_neg\_2\_042\_13

m/z 647.8627

012E+316.517RT (min)

lcqtof\_t3\_neg\_4\_035\_08

m/z 648.0358

01E+38.59RT (min)

lcqtof\_t3\_neg\_4\_048\_19

m/z 649.0919

0123E+311.5RT (min)

lcqtof\_t3\_neg\_1\_019\_23

m/z 650.3242

0246E+312.513RT (min)

lcqtof\_t3\_neg\_1\_005\_18

m/z 650.7148

01E+411.5RT (min)

lcqtof\_t3\_neg\_4\_051\_15

m/z 650.8335

02468E+311.5RT (min)

lcqtof\_t3\_neg\_1\_032\_36

m/z 651.1060

01234E+311.5RT (min)

lcqtof\_t3\_neg\_2\_051\_14

m/z 651.8704

0246E+311.5RT (min)

lcqtof\_t3\_neg\_2\_042\_13

m/z 652.1919

0123E+311.5RT (min)

lcqtof\_t3\_neg\_4\_040\_24

m/z 652.2984

01234E+34.55RT (min)

lcqtof\_t3\_neg\_4\_055\_38

m/z 654.0552

012E+38.59RT (min)

lcqtof\_t3\_neg\_4\_100\_61

m/z 655.1075

0246E+311.5RT (min)

lcqtof\_t3\_neg\_4\_018\_23

m/z 655.1686

0246E+311.5RT (min)

lcqtof\_t3\_neg\_4\_051\_15

m/z 655.5171

024E+311.5RT (min)

lcqtof\_t3\_neg\_4\_037\_01

m/z 656.7964

0246E+311.5RT (min)

lcqtof\_t3\_neg\_4\_051\_15

m/z 657.3845

024E+311.5RT (min)

lcqtof\_t3\_neg\_2\_042\_13

m/z 657.5145

02468E+311.5RT (min)

lcqtof\_t3\_neg\_4\_037\_01

m/z 658.5140

02468E+311.5RT (min)

lcqtof\_t3\_neg\_4\_037\_01

m/z 658.5438

02468E+311.5RT (min)

lcqtof\_t3\_neg\_4\_041\_54

m/z 659.0750

0123E+311.5RT (min)

lcqtof\_t3\_neg\_2\_051\_14

m/z 662.1802

024E+344.5RT (min)

lcqtof\_t3\_neg\_1\_031\_24

m/z 662.5339

0246E+311.5RT (min)

lcqtof\_t3\_neg\_4\_046\_33

m/z 663.8333

024E+311.5RT (min)

lcqtof\_t3\_neg\_1\_038\_15

m/z 664.0713

01234E+35.56RT (min)

lcqtof\_t3\_neg\_4\_048\_19

m/z 664.4960

02468E+311.5RT (min)

lcqtof\_t3\_neg\_4\_037\_01

m/z 664.7684

01E+411.5RT (min)

lcqtof\_t3\_neg\_4\_046\_33

m/z 665.1141

0123E+311.5RT (min)

lcqtof\_t3\_neg\_4\_055\_38

m/z 665.1154

0246E+344.5RT (min)

lcqtof\_t3\_neg\_1\_044\_37

m/z 665.1876

0246E+344.5RT (min)

lcqtof\_t3\_neg\_2\_028\_24

m/z 665.3342

012E+37.58RT (min)

lcqtof\_t3\_neg\_4\_048\_19

m/z 665.3699

0123E+311.5RT (min)

lcqtof\_t3\_neg\_4\_045\_02\_rerun

m/z 665.4963

024E+311.5RT (min)

lcqtof\_t3\_neg\_4\_051\_15

m/z 666.0928

01234E+311.5RT (min)

lcqtof\_t3\_neg\_3\_012\_14

m/z 666.1440

0123E+377.5RT (min)

lcqtof\_t3\_neg\_1\_026\_19

m/z 666.6063

0246E+311.5RT (min)

lcqtof\_t3\_neg\_4\_037\_01

m/z 666.8087

0246E+311.5RT (min)

lcqtof\_t3\_neg\_1\_055\_32

m/z 666.8380

0123E+311.5RT (min)

lcqtof\_t3\_neg\_4\_051\_15

m/z 667.2094

0246E+344.5RT (min)

lcqtof\_t3\_neg\_4\_040\_24

m/z 667.8518

0246E+316.517RT (min)

lcqtof\_t3\_neg\_4\_015\_36

m/z 668.4756

0123E+311.5RT (min)

lcqtof\_t3\_neg\_4\_016\_14

m/z 668.7751

01E+411.5RT (min)

lcqtof\_t3\_neg\_1\_036\_14

m/z 668.7985

012E+316.517RT (min)

lcqtof\_t3\_neg\_1\_015\_34

m/z 668.8043

01E+411.5RT (min)

lcqtof\_t3\_neg\_3\_018\_32

m/z 669.1573

0246E+311.5RT (min)

lcqtof\_t3\_neg\_1\_031\_24

m/z 670.7741

01E+411.5RT (min)

lcqtof\_t3\_neg\_1\_036\_14

m/z 670.8581

024E+311.5RT (min)

lcqtof\_t3\_neg\_2\_005\_34

m/z 672.4863

024E+311.5RT (min)

lcqtof\_t3\_neg\_4\_051\_15

m/z 673.0999

012E+311.5RT (min)

lcqtof\_t3\_neg\_1\_036\_14

m/z 673.3531

0123E+344.5RT (min)

lcqtof\_t3\_neg\_4\_003\_37

m/z 673.5850

012E+312.513RT (min)

lcqtof\_t3\_neg\_1\_035\_31

m/z 674.9816

024E+311.5RT (min)

lcqtof\_t3\_neg\_2\_043\_31

m/z 675.8661

0123E+311.5RT (min)

lcqtof\_t3\_neg\_4\_045\_02\_rerun

m/z 676.5087

0246E+31313.5RT (min)

lcqtof\_t3\_neg\_4\_019\_05

m/z 676.8629

01234E+316.517RT (min)

lcqtof\_t3\_neg\_4\_005\_18

m/z 679.3709

02468E+316.517RT (min)

lcqtof\_t3\_neg\_4\_055\_38

m/z 679.5007

02468E+311.5RT (min)

lcqtof\_t3\_neg\_2\_042\_13

m/z 679.8282

012E+316.517RT (min)

lcqtof\_t3\_neg\_1\_013\_09

m/z 679.9987

01E+411.5RT (min)

lcqtof\_t3\_neg\_2\_019\_32

m/z 680.0476

0123E+311.5RT (min)

lcqtof\_t3\_neg\_4\_016\_14

m/z 681.1502

01234E+311.5RT (min)

lcqtof\_t3\_neg\_4\_040\_24

m/z 682.1676

0246E+311.5RT (min)

lcqtof\_t3\_neg\_2\_019\_32

m/z 682.3266

012E+37.58RT (min)

lcqtof\_t3\_neg\_2\_047\_28

m/z 683.2817

012E+34.55RT (min)

lcqtof\_t3\_neg\_4\_042\_10

m/z 685.8616

012E+411.5RT (min)

lcqtof\_t3\_neg\_2\_042\_13

m/z 686.0141

02468E+311.5RT (min)

lcqtof\_t3\_neg\_1\_032\_36

m/z 686.8396

01234E+311.5RT (min)

lcqtof\_t3\_neg\_2\_002\_35

m/z 687.4899

0246E+311.5RT (min)

lcqtof\_t3\_neg\_4\_037\_01

m/z 689.0715

012E+311.5RT (min)

lcqtof\_t3\_neg\_3\_012\_14

m/z 689.8353

02468E+311.5RT (min)

lcqtof\_t3\_neg\_1\_036\_14

m/z 691.8796

0246E+311.5RT (min)

lcqtof\_t3\_neg\_2\_042\_13

m/z 693.4756

024E+311.5RT (min)

lcqtof\_t3\_neg\_2\_050\_01

m/z 693.8502

01E+411.5RT (min)

lcqtof\_t3\_neg\_2\_050\_01

m/z 693.8544

012E+316.517RT (min)

lcqtof\_t3\_neg\_4\_054\_44

m/z 694.9562

02468E+316.517RT (min)

lcqtof\_t3\_neg\_4\_011\_32

m/z 695.0170

01234E+311.5RT (min)

lcqtof\_t3\_neg\_3\_018\_32

m/z 695.5060

01234E+355.5RT (min)

lcqtof\_t3\_neg\_4\_018\_23

m/z 695.9716

02468E+311.5RT (min)

lcqtof\_t3\_neg\_2\_019\_32

m/z 697.2684

0123E+35.56RT (min)

lcqtof\_t3\_neg\_4\_016\_14

m/z 698.4736

0246E+311.5RT (min)

lcqtof\_t3\_neg\_4\_037\_01

m/z 698.6488

0246E+311.5RT (min)

lcqtof\_t3\_neg\_4\_051\_15

m/z 699.3630

02468E+311.5RT (min)

lcqtof\_t3\_neg\_4\_045\_02\_rerun

m/z 699.3757

01E+316.517RT (min)

lcqtof\_t3\_neg\_1\_011\_45

m/z 699.8655

0246E+311.5RT (min)

lcqtof\_t3\_neg\_4\_045\_02\_rerun

m/z 700.9101

0123E+316.517RT (min)

lcqtof\_t3\_neg\_4\_039\_06\_rerun

m/z 701.8173

024E+311.5RT (min)

lcqtof\_t3\_neg\_1\_055\_32

m/z 702.5261

0246E+311.5RT (min)

lcqtof\_t3\_neg\_4\_046\_33

m/z 702.8090

024E+311.5RT (min)

lcqtof\_t3\_neg\_1\_032\_36

m/z 702.8743

012E+316.517RT (min)

lcqtof\_t3\_neg\_1\_015\_34

m/z 702.8771

012E+316.517RT (min)

lcqtof\_t3\_neg\_1\_040\_08

m/z 704.5153

02468E+311.5RT (min)

lcqtof\_t3\_neg\_4\_046\_33

m/z 704.9684

0123E+311.5RT (min)

lcqtof\_t3\_neg\_2\_043\_31

m/z 705.8128

024E+311.5RT (min)

lcqtof\_t3\_neg\_3\_048\_01

m/z 705.8217

01E+316.517RT (min)

lcqtof\_t3\_neg\_3\_047\_42

m/z 706.1760

01E+34.55RT (min)

lcqtof\_t3\_neg\_4\_013\_39

m/z 706.4727

02468E+311.5RT (min)

lcqtof\_t3\_neg\_4\_037\_01

m/z 706.8391

012E+316.517RT (min)

lcqtof\_t3\_neg\_4\_047\_07

m/z 707.0947

01234E+377.5RT (min)

lcqtof\_t3\_neg\_4\_016\_14

m/z 707.1706

024E+344.5RT (min)

lcqtof\_t3\_neg\_1\_055\_32

m/z 707.3528

024E+311.5RT (min)

lcqtof\_t3\_neg\_4\_045\_02\_rerun

m/z 707.8446

012E+316.517RT (min)

lcqtof\_t3\_neg\_4\_030\_31

m/z 707.8520

024E+311.5RT (min)

lcqtof\_t3\_neg\_4\_045\_02\_rerun

m/z 708.6541

0123E+31414.5RT (min)

lcqtof\_t3\_neg\_4\_016\_14

m/z 708.6787

02468E+311.5RT (min)

lcqtof\_t3\_neg\_4\_046\_33

m/z 708.8041

024E+311.5RT (min)

lcqtof\_t3\_neg\_4\_038\_29

m/z 708.8990

02468E+311.5RT (min)

lcqtof\_t3\_neg\_2\_042\_13

m/z 709.4701

02468E+316.517RT (min)

lcqtof\_t3\_neg\_4\_011\_32

m/z 709.4874

0123E+355.5RT (min)

lcqtof\_t3\_neg\_4\_030\_31

m/z 710.0578

0123E+311.5RT (min)

lcqtof\_t3\_neg\_2\_051\_14

m/z 711.5033

024E+366.5RT (min)

lcqtof\_t3\_neg\_4\_034\_34

m/z 712.1882

0246E+311.5RT (min)

lcqtof\_t3\_neg\_4\_040\_24

m/z 712.8352

02468E+311.5RT (min)

lcqtof\_t3\_neg\_2\_023\_37

m/z 714.4568

01234E+311.5RT (min)

lcqtof\_t3\_neg\_4\_051\_15

m/z 715.3448

0123E+311.5RT (min)

lcqtof\_t3\_neg\_2\_055\_16

m/z 715.8426

01E+411.5RT (min)

lcqtof\_t3\_neg\_2\_042\_13

m/z 715.8527

012E+316.517RT (min)

lcqtof\_t3\_neg\_4\_042\_10

m/z 716.1448

0246E+31313.5RT (min)

lcqtof\_t3\_neg\_3\_089\_59

m/z 716.4753

02468E+316.517RT (min)

lcqtof\_t3\_neg\_4\_011\_32

m/z 716.5140

024E+311.5RT (min)

lcqtof\_t3\_neg\_4\_046\_33

m/z 717.3502

024E+366.5RT (min)

lcqtof\_t3\_neg\_4\_016\_14

m/z 717.4254

01E+37.58RT (min)

lcqtof\_t3\_neg\_1\_025\_52

m/z 718.5050

02468E+311.5RT (min)

lcqtof\_t3\_neg\_4\_046\_33

m/z 718.7025

02468E+311.5RT (min)

lcqtof\_t3\_neg\_4\_051\_15

m/z 718.8528

0246E+311.5RT (min)

lcqtof\_t3\_neg\_4\_051\_15

m/z 719.8555

0123E+411.5RT (min)

lcqtof\_t3\_neg\_2\_042\_13

m/z 719.8657

012E+316.517RT (min)

lcqtof\_t3\_neg\_4\_046\_33

m/z 720.4512

01234E+31414.5RT (min)

lcqtof\_t3\_neg\_4\_003\_37

m/z 720.5040

0246E+311.5RT (min)

lcqtof\_t3\_neg\_4\_055\_38

m/z 722.0446

01E+38.59RT (min)

lcqtof\_t3\_neg\_4\_088\_59

m/z 722.7928

012E+411.5RT (min)

lcqtof\_t3\_neg\_1\_038\_15

m/z 724.3953

0246E+35.56RT (min)

lcqtof\_t3\_neg\_4\_003\_37

m/z 725.1092

02468E+37.58RT (min)

lcqtof\_t3\_neg\_4\_005\_18

m/z 726.3746

012E+311.5RT (min)

lcqtof\_t3\_neg\_2\_042\_13

m/z 726.7522

01E+411.5RT (min)

lcqtof\_t3\_neg\_4\_055\_38

m/z 727.8469

01234E+316.517RT (min)

lcqtof\_t3\_neg\_4\_043\_11

m/z 728.1149

0123E+377.5RT (min)

lcqtof\_t3\_neg\_1\_026\_19

m/z 728.3443

024E+311.5RT (min)

lcqtof\_t3\_neg\_2\_052\_17

m/z 728.3541

01E+316.517RT (min)

lcqtof\_t3\_neg\_4\_029\_40

m/z 728.4943

01234E+366.5RT (min)

lcqtof\_t3\_neg\_1\_019\_23

m/z 728.8298

0123E+311.5RT (min)

lcqtof\_t3\_neg\_3\_019\_37

m/z 729.8892

0246E+312.513RT (min)

lcqtof\_t3\_neg\_2\_041\_54

m/z 731.8152

02468E+311.5RT (min)

lcqtof\_t3\_neg\_1\_036\_14

m/z 732.0143

012E+38.59RT (min)

lcqtof\_t3\_neg\_1\_024\_10

m/z 732.0581

0123E+35.56RT (min)

lcqtof\_t3\_neg\_4\_048\_19

m/z 732.5938

0246E+311.5RT (min)

lcqtof\_t3\_neg\_4\_037\_01

m/z 732.7625

01E+411.5RT (min)

lcqtof\_t3\_neg\_4\_046\_33

m/z 733.3603

012E+316.517RT (min)

lcqtof\_t3\_neg\_4\_039\_06\_rerun

m/z 734.3577

0123E+311.5RT (min)

lcqtof\_t3\_neg\_4\_045\_02\_rerun

m/z 734.7977

0246E+311.5RT (min)

lcqtof\_t3\_neg\_1\_055\_32

m/z 735.8276

01E+411.5RT (min)

lcqtof\_t3\_neg\_1\_038\_15

m/z 736.3434

01E+316.517RT (min)

lcqtof\_t3\_neg\_1\_054\_41

m/z 736.7812

012E+316.517RT (min)

lcqtof\_t3\_neg\_4\_007\_09

m/z 738.7614

01E+411.5RT (min)

lcqtof\_t3\_neg\_1\_038\_15

m/z 739.1751

0123E+355.5RT (min)

lcqtof\_t3\_neg\_1\_005\_18

m/z 741.3405

01234E+34.55RT (min)

lcqtof\_t3\_neg\_4\_016\_14

m/z 741.3411

01E+316.517RT (min)

lcqtof\_t3\_neg\_4\_039\_06\_rerun

m/z 741.3460

01E+411.5RT (min)

lcqtof\_t3\_neg\_4\_045\_02\_rerun

m/z 742.3452

012E+311.5RT (min)

lcqtof\_t3\_neg\_2\_051\_14

m/z 742.4607

0246E+311.5RT (min)

lcqtof\_t3\_neg\_4\_037\_01

m/z 742.7172

02468E+311.5RT (min)

lcqtof\_t3\_neg\_4\_037\_01

m/z 742.9719

024E+311.5RT (min)

lcqtof\_t3\_neg\_1\_032\_36

m/z 743.8129

02468E+311.5RT (min)

lcqtof\_t3\_neg\_2\_050\_01

m/z 744.8511

024E+316.517RT (min)

lcqtof\_t3\_neg\_4\_052\_03\_rerun

m/z 745.1891

0246E+355.5RT (min)

lcqtof\_t3\_neg\_4\_024\_26

m/z 747.0975

0123E+311.5RT (min)

lcqtof\_t3\_neg\_2\_026\_36

m/z 747.3364

01234E+311.5RT (min)

lcqtof\_t3\_neg\_1\_032\_36

m/z 747.8140

0123E+316.517RT (min)

lcqtof\_t3\_neg\_1\_039\_43

m/z 749.3331

024E+311.5RT (min)

lcqtof\_t3\_neg\_2\_051\_14

m/z 749.8333

02468E+311.5RT (min)

lcqtof\_t3\_neg\_2\_055\_16

m/z 750.6606

02468E+311.5RT (min)

lcqtof\_t3\_neg\_4\_037\_01

m/z 753.7164

012E+314.515RT (min)

lcqtof\_t3\_neg\_4\_032\_04

m/z 753.8481

01234E+316.517RT (min)

lcqtof\_t3\_neg\_4\_045\_02\_rerun

m/z 753.8493

01234E+411.5RT (min)

lcqtof\_t3\_neg\_2\_042\_13

m/z 754.8257

01E+411.5RT (min)

lcqtof\_t3\_neg\_1\_032\_36

m/z 755.7706

01234E+313.514RT (min)

lcqtof\_t3\_neg\_4\_011\_32

m/z 756.4252

024E+377.5RT (min)

lcqtof\_t3\_neg\_4\_004\_20

m/z 756.4286

0246E+311.5RT (min)

lcqtof\_t3\_neg\_4\_037\_01

m/z 758.7516

012E+31414.5RT (min)

lcqtof\_t3\_neg\_4\_033\_53

m/z 759.3749

0123E+316.517RT (min)

lcqtof\_t3\_neg\_4\_023\_45

m/z 760.3680

0123E+311.5RT (min)

lcqtof\_t3\_neg\_2\_042\_13

m/z 760.4888

024E+311.5RT (min)

lcqtof\_t3\_neg\_4\_055\_38

m/z 760.6857

01E+411.5RT (min)

lcqtof\_t3\_neg\_4\_051\_15

m/z 761.4363

0246E+312.513RT (min)

lcqtof\_t3\_neg\_1\_019\_23

m/z 761.6609

0246E+31414.5RT (min)

lcqtof\_t3\_neg\_2\_026\_36

m/z 761.8355

01E+316.517RT (min)

lcqtof\_t3\_neg\_1\_019\_23

m/z 761.8420

01234E+316.517RT (min)

lcqtof\_t3\_neg\_4\_022\_28

m/z 762.3377

0246E+311.5RT (min)

lcqtof\_t3\_neg\_2\_055\_16

m/z 762.3425

012E+316.517RT (min)

lcqtof\_t3\_neg\_1\_009\_50

m/z 762.3942

012E+377.5RT (min)

lcqtof\_t3\_neg\_1\_005\_18

m/z 762.4857

0246E+311.5RT (min)

lcqtof\_t3\_neg\_4\_055\_38

m/z 763.0039

01234E+311.5RT (min)

lcqtof\_t3\_neg\_2\_019\_32

m/z 763.9581

0246E+311.5RT (min)

lcqtof\_t3\_neg\_3\_018\_32

m/z 764.4421

02468E+311.5RT (min)

lcqtof\_t3\_neg\_2\_042\_13

m/z 765.7862

024E+31414.5RT (min)

lcqtof\_t3\_neg\_4\_011\_32

m/z 766.7270

012E+314.515RT (min)

lcqtof\_t3\_neg\_4\_035\_08

m/z 767.3481

0123E+316.517RT (min)

lcqtof\_t3\_neg\_4\_039\_06\_rerun

m/z 767.6792

024E+31414.5RT (min)

lcqtof\_t3\_neg\_2\_019\_32

m/z 767.8339

024E+355.5RT (min)

lcqtof\_t3\_neg\_4\_016\_14

m/z 768.3561

01234E+311.5RT (min)

lcqtof\_t3\_neg\_4\_045\_02\_rerun

m/z 768.8960

01234E+316.517RT (min)

lcqtof\_t3\_neg\_4\_045\_02\_rerun

m/z 769.0643

012E+377.5RT (min)

lcqtof\_t3\_neg\_4\_051\_15

m/z 769.0792

0123E+311.5RT (min)

lcqtof\_t3\_neg\_2\_015\_33

m/z 769.8235

01E+411.5RT (min)

lcqtof\_t3\_neg\_2\_051\_14

m/z 769.8328

012E+316.517RT (min)

lcqtof\_t3\_neg\_4\_029\_40

m/z 770.3337

012E+316.517RT (min)

lcqtof\_t3\_neg\_1\_025\_52

m/z 770.3905

024E+36.57RT (min)

lcqtof\_t3\_neg\_1\_015\_34

m/z 770.8052

0246E+311.5RT (min)

lcqtof\_t3\_neg\_1\_032\_36

m/z 770.8618

012E+316.517RT (min)

lcqtof\_t3\_neg\_1\_017\_51

m/z 772.4325

02468E+311.5RT (min)

lcqtof\_t3\_neg\_4\_037\_01

m/z 772.7835

01234E+31414.5RT (min)

lcqtof\_t3\_neg\_4\_019\_05

m/z 772.9617

012E+311.5RT (min)

lcqtof\_t3\_neg\_2\_002\_35

m/z 773.8110

01E+316.517RT (min)

lcqtof\_t3\_neg\_1\_050\_40

m/z 774.8361

0123E+316.517RT (min)

lcqtof\_t3\_neg\_4\_042\_10

m/z 775.3289

01234E+355.5RT (min)

lcqtof\_t3\_neg\_4\_016\_14

m/z 775.3433

012E+316.517RT (min)

lcqtof\_t3\_neg\_1\_052\_12

m/z 775.4279

01234E+311.5RT (min)

lcqtof\_t3\_neg\_4\_037\_01

m/z 775.8354

012E+316.517RT (min)

lcqtof\_t3\_neg\_4\_042\_10

m/z 776.3410

0123E+311.5RT (min)

lcqtof\_t3\_neg\_4\_045\_02\_rerun

m/z 776.4589

0246E+311.5RT (min)

lcqtof\_t3\_neg\_4\_055\_38

m/z 776.6796

0123E+31414.5RT (min)

lcqtof\_t3\_neg\_2\_026\_36

m/z 776.8867

0246E+311.5RT (min)

lcqtof\_t3\_neg\_2\_042\_13

m/z 778.3201

01E+316.517RT (min)

lcqtof\_t3\_neg\_1\_050\_40

m/z 778.4539

024E+311.5RT (min)

lcqtof\_t3\_neg\_4\_030\_31

m/z 778.7378

01E+314.515RT (min)

lcqtof\_t3\_neg\_4\_001\_49

m/z 778.7451

01E+411.5RT (min)

lcqtof\_t3\_neg\_4\_051\_15

m/z 779.4065

02468E+377.5RT (min)

lcqtof\_t3\_neg\_4\_004\_20

m/z 780.8250

01E+411.5RT (min)

lcqtof\_t3\_neg\_2\_043\_31

m/z 781.7526

01E+314.515RT (min)

lcqtof\_t3\_neg\_4\_029\_40

m/z 782.3708

0123E+31414.5RT (min)

lcqtof\_t3\_neg\_4\_087\_59

m/z 782.4119

0246E+311.5RT (min)

lcqtof\_t3\_neg\_4\_051\_15

m/z 782.5202

0123E+311.5RT (min)

lcqtof\_t3\_neg\_4\_037\_01

m/z 783.3225

01E+316.517RT (min)

lcqtof\_t3\_neg\_1\_001\_49

m/z 783.3265

0246E+311.5RT (min)

lcqtof\_t3\_neg\_2\_051\_14

m/z 783.8342

0123E+316.517RT (min)

lcqtof\_t3\_neg\_4\_039\_06\_rerun

m/z 784.3313

012E+311.5RT (min)

lcqtof\_t3\_neg\_2\_054\_29

m/z 784.5130

0123E+311.5RT (min)

lcqtof\_t3\_neg\_4\_037\_01

m/z 784.8286

02468E+316.517RT (min)

lcqtof\_t3\_neg\_4\_045\_02\_rerun

m/z 784.8295

01E+411.5RT (min)

lcqtof\_t3\_neg\_1\_012\_16

m/z 786.7947

01234E+311.5RT (min)

lcqtof\_t3\_neg\_2\_019\_32

m/z 786.8427

0123E+311.5RT (min)

lcqtof\_t3\_neg\_4\_051\_15

m/z 787.4669

01234E+311.5RT (min)

lcqtof\_t3\_neg\_2\_042\_13

m/z 787.8425

01234E+411.5RT (min)

lcqtof\_t3\_neg\_2\_042\_13

m/z 787.8455

0123E+316.517RT (min)

lcqtof\_t3\_neg\_4\_045\_02\_rerun

m/z 788.7917

012E+316.517RT (min)

lcqtof\_t3\_neg\_4\_015\_36

m/z 789.4648

024E+311.5RT (min)

lcqtof\_t3\_neg\_2\_042\_13

m/z 789.8455

012E+311.5RT (min)

lcqtof\_t3\_neg\_2\_050\_01

m/z 790.7891

012E+411.5RT (min)

lcqtof\_t3\_neg\_1\_038\_15

m/z 792.8018

012E+316.517RT (min)

lcqtof\_t3\_neg\_1\_025\_52

m/z 793.3566

01234E+316.517RT (min)

lcqtof\_t3\_neg\_4\_045\_02\_rerun

m/z 794.3633

01234E+311.5RT (min)

lcqtof\_t3\_neg\_2\_042\_13

m/z 795.8338

01234E+316.517RT (min)

lcqtof\_t3\_neg\_4\_039\_06\_rerun

m/z 796.1000

012E+377.5RT (min)

lcqtof\_t3\_neg\_1\_013\_09

m/z 796.3322

0246E+311.5RT (min)

lcqtof\_t3\_neg\_2\_050\_01

m/z 796.3372

012E+316.517RT (min)

lcqtof\_t3\_neg\_1\_009\_50

m/z 797.5135

02468E+31313.5RT (min)

lcqtof\_t3\_neg\_3\_089\_59

m/z 798.7127

012E+31414.5RT (min)

lcqtof\_t3\_neg\_4\_036\_47

m/z 799.8218

01234E+316.517RT (min)

lcqtof\_t3\_neg\_1\_018\_33

m/z 800.7614

01E+411.5RT (min)

lcqtof\_t3\_neg\_4\_055\_38

m/z 801.3432

01234E+316.517RT (min)

lcqtof\_t3\_neg\_4\_052\_03\_rerun

m/z 802.3466

024E+311.5RT (min)

lcqtof\_t3\_neg\_4\_045\_02\_rerun

m/z 802.4686

0246E+31313.5RT (min)

lcqtof\_t3\_neg\_2\_047\_28

m/z 803.8140

01E+411.5RT (min)

lcqtof\_t3\_neg\_1\_038\_15

m/z 803.8223

012E+316.517RT (min)

lcqtof\_t3\_neg\_4\_045\_02\_rerun

m/z 804.3179

024E+311.5RT (min)

lcqtof\_t3\_neg\_1\_036\_14

m/z 804.3303

012E+316.517RT (min)

lcqtof\_t3\_neg\_1\_050\_40

m/z 804.7780

01E+411.5RT (min)

lcqtof\_t3\_neg\_3\_018\_32

m/z 804.7852

012E+316.517RT (min)

lcqtof\_t3\_neg\_1\_039\_43

m/z 809.1193

024E+344.5RT (min)

lcqtof\_t3\_neg\_4\_003\_37

m/z 809.3374

012E+316.517RT (min)

lcqtof\_t3\_neg\_1\_049\_55

m/z 810.3334

01234E+311.5RT (min)

lcqtof\_t3\_neg\_4\_045\_02\_rerun

m/z 810.7529

012E+314.515RT (min)

lcqtof\_t3\_neg\_4\_037\_01

m/z 810.8337

024E+316.517RT (min)

lcqtof\_t3\_neg\_4\_039\_06\_rerun

m/z 810.9550

0123E+311.5RT (min)

lcqtof\_t3\_neg\_2\_043\_31

m/z 811.2917

01234E+366.5RT (min)

lcqtof\_t3\_neg\_1\_010\_27

m/z 811.8055

01E+411.5RT (min)

lcqtof\_t3\_neg\_2\_051\_14

m/z 812.3091

01E+316.517RT (min)

lcqtof\_t3\_neg\_1\_027\_21

m/z 812.7351

012E+31414.5RT (min)

lcqtof\_t3\_neg\_4\_038\_29

m/z 812.8443

024E+316.517RT (min)

lcqtof\_t3\_neg\_4\_045\_02\_rerun

m/z 815.7935

0123E+316.517RT (min)

lcqtof\_t3\_neg\_1\_017\_51

m/z 815.8085

01234E+31414.5RT (min)

lcqtof\_t3\_neg\_1\_055\_32

m/z 815.9733

0246E+311.5RT (min)

lcqtof\_t3\_neg\_2\_019\_32

m/z 816.8049

0123E+316.517RT (min)

lcqtof\_t3\_neg\_1\_021\_39

m/z 817.3192

02468E+311.5RT (min)

lcqtof\_t3\_neg\_1\_036\_14

m/z 817.3210

01E+316.517RT (min)

lcqtof\_t3\_neg\_1\_050\_40

m/z 817.8186

02468E+311.5RT (min)

lcqtof\_t3\_neg\_1\_036\_14

m/z 818.3184

0123E+311.5RT (min)

lcqtof\_t3\_neg\_1\_036\_14

m/z 818.5901

024E+31414.5RT (min)

lcqtof\_t3\_neg\_4\_003\_37

m/z 820.3204

0246E+34.55RT (min)

lcqtof\_t3\_neg\_4\_002\_17

m/z 820.8326

0123E+31414.5RT (min)

lcqtof\_t3\_neg\_1\_055\_32

m/z 821.2271

012E+34.55RT (min)

lcqtof\_t3\_neg\_4\_004\_20

m/z 821.8365

01234E+411.5RT (min)

lcqtof\_t3\_neg\_2\_042\_13

m/z 821.8380

01234E+316.517RT (min)

lcqtof\_t3\_neg\_4\_052\_03\_rerun

m/z 822.8043

01E+411.5RT (min)

lcqtof\_t3\_neg\_3\_018\_32

m/z 823.4857

0246E+31313.5RT (min)

lcqtof\_t3\_neg\_1\_003\_07

m/z 823.8405

012E+311.5RT (min)

lcqtof\_t3\_neg\_2\_050\_01

m/z 824.9725

024E+311.5RT (min)

lcqtof\_t3\_neg\_1\_032\_36

m/z 825.3071

0123E+311.5RT (min)

lcqtof\_t3\_neg\_1\_036\_14

m/z 826.4548

024E+344.5RT (min)

lcqtof\_t3\_neg\_4\_016\_14

m/z 827.3446

0123E+316.517RT (min)

lcqtof\_t3\_neg\_4\_045\_02\_rerun

m/z 829.8269

024E+316.517RT (min)

lcqtof\_t3\_neg\_4\_052\_03\_rerun

m/z 829.8563

01234E+31414.5RT (min)

lcqtof\_t3\_neg\_4\_011\_32

m/z 830.1920

0246E+344.5RT (min)

lcqtof\_t3\_neg\_4\_011\_32

m/z 830.3256

0246E+311.5RT (min)

lcqtof\_t3\_neg\_1\_053\_01

m/z 830.3302

012E+316.517RT (min)

lcqtof\_t3\_neg\_1\_008\_02

m/z 830.7612

0123E+316.517RT (min)

lcqtof\_t3\_neg\_1\_001\_49

m/z 830.7650

01E+411.5RT (min)

lcqtof\_t3\_neg\_4\_037\_01

m/z 831.5944

024E+316.517RT (min)

lcqtof\_t3\_neg\_4\_003\_37

m/z 831.9496

024E+311.5RT (min)

lcqtof\_t3\_neg\_2\_019\_32

m/z 834.4150

01E+411.5RT (min)

lcqtof\_t3\_neg\_4\_016\_14

m/z 834.4512

0246E+312.513RT (min)

lcqtof\_t3\_neg\_4\_030\_31

m/z 835.3416

01234E+316.517RT (min)

lcqtof\_t3\_neg\_4\_039\_06\_rerun

m/z 836.3430

0246E+311.5RT (min)

lcqtof\_t3\_neg\_4\_045\_02\_rerun

m/z 836.3972

02468E+311.5RT (min)

lcqtof\_t3\_neg\_4\_016\_14

m/z 836.8896

0123E+316.517RT (min)

lcqtof\_t3\_neg\_4\_043\_11

m/z 837.5817

02468E+31313.5RT (min)

lcqtof\_t3\_neg\_1\_006\_05

m/z 837.8088

012E+316.517RT (min)

lcqtof\_t3\_neg\_4\_032\_04

m/z 837.8103

01E+411.5RT (min)

lcqtof\_t3\_neg\_3\_012\_14

m/z 838.3204

012E+316.517RT (min)

lcqtof\_t3\_neg\_1\_042\_46

m/z 838.7938

02468E+311.5RT (min)

lcqtof\_t3\_neg\_1\_032\_36

m/z 838.8331

012E+316.517RT (min)

lcqtof\_t3\_neg\_1\_026\_19

m/z 839.2850

012E+377.5RT (min)

lcqtof\_t3\_neg\_2\_026\_36

m/z 839.3757

0123E+377.5RT (min)

lcqtof\_t3\_neg\_1\_026\_19

m/z 840.7906

01234E+316.517RT (min)

lcqtof\_t3\_neg\_4\_051\_15

m/z 840.9417

012E+311.5RT (min)

lcqtof\_t3\_neg\_2\_026\_36

m/z 841.5790

012E+366.5RT (min)

lcqtof\_t3\_neg\_1\_019\_23

m/z 842.8264

024E+316.517RT (min)

lcqtof\_t3\_neg\_4\_039\_06

m/z 843.3313

012E+316.517RT (min)

lcqtof\_t3\_neg\_1\_008\_02

m/z 843.8304

012E+316.517RT (min)

lcqtof\_t3\_neg\_4\_039\_06\_rerun

m/z 846.3123

01E+316.517RT (min)

lcqtof\_t3\_neg\_1\_012\_16

m/z 846.4915

01234E+36.57RT (min)

lcqtof\_t3\_neg\_4\_048\_19

m/z 846.7327

02468E+311.5RT (min)

lcqtof\_t3\_neg\_4\_051\_15

m/z 848.2430

01234E+34.55RT (min)

lcqtof\_t3\_neg\_1\_026\_19

m/z 848.8152

01E+411.5RT (min)

lcqtof\_t3\_neg\_1\_032\_36

m/z 849.5086

01234E+377.5RT (min)

lcqtof\_t3\_neg\_4\_055\_38

m/z 850.5995

024E+313.514RT (min)

lcqtof\_t3\_neg\_1\_002\_13

m/z 850.9995

012E+38.59RT (min)

lcqtof\_t3\_neg\_4\_106\_62

m/z 851.3141

02468E+311.5RT (min)

lcqtof\_t3\_neg\_1\_036\_14

m/z 851.3240

01E+316.517RT (min)

lcqtof\_t3\_neg\_3\_044\_38

m/z 851.8241

0123E+316.517RT (min)

lcqtof\_t3\_neg\_4\_052\_03\_rerun

m/z 852.3153

0123E+311.5RT (min)

lcqtof\_t3\_neg\_2\_051\_14

m/z 852.8170

0246E+316.517RT (min)

lcqtof\_t3\_neg\_4\_046\_33

m/z 854.1893

0123E+38.59RT (min)

lcqtof\_t3\_neg\_4\_053\_12

m/z 854.2410

02468E+34.55RT (min)

lcqtof\_t3\_neg\_4\_004\_20

m/z 854.5935

01E+416.517RT (min)

lcqtof\_t3\_neg\_4\_016\_14

m/z 854.8294

0123E+316.517RT (min)

lcqtof\_t3\_neg\_1\_010\_27

m/z 855.8304

01234E+316.517RT (min)

lcqtof\_t3\_neg\_4\_039\_06\_rerun

m/z 856.7923

0123E+316.517RT (min)

lcqtof\_t3\_neg\_4\_027\_41

m/z 857.0236

012E+38.59RT (min)

lcqtof\_t3\_neg\_4\_102\_61

m/z 857.4163

0246E+39.510RT (min)

lcqtof\_t3\_neg\_3\_076\_58

m/z 857.8361

012E+311.5RT (min)

lcqtof\_t3\_neg\_2\_050\_01

m/z 858.7785

01E+411.5RT (min)

lcqtof\_t3\_neg\_1\_036\_14

m/z 859.3005

01234E+311.5RT (min)

lcqtof\_t3\_neg\_1\_036\_14

m/z 860.7932

0123E+316.517RT (min)

lcqtof\_t3\_neg\_1\_039\_43

m/z 861.3430

0123E+316.517RT (min)

lcqtof\_t3\_neg\_4\_045\_02\_rerun

m/z 861.7370

0246E+313.514RT (min)

lcqtof\_t3\_neg\_4\_040\_24

m/z 862.6825

01234E+314.515RT (min)

lcqtof\_t3\_neg\_4\_037\_01

m/z 863.8205

01234E+316.517RT (min)

lcqtof\_t3\_neg\_4\_039\_06\_rerun

m/z 864.3201

0246E+311.5RT (min)

lcqtof\_t3\_neg\_3\_048\_01

m/z 864.3267

012E+316.517RT (min)

lcqtof\_t3\_neg\_1\_008\_02

m/z 864.4952

024E+31313.5RT (min)

lcqtof\_t3\_neg\_4\_085\_59

m/z 865.4878

01234E+377.5RT (min)

lcqtof\_t3\_neg\_1\_026\_19

m/z 865.5828

0123E+31313.5RT (min)

lcqtof\_t3\_neg\_2\_050\_01

m/z 866.3542

0246E+311.5RT (min)

lcqtof\_t3\_neg\_4\_051\_15

m/z 867.8145

01234E+316.517RT (min)

lcqtof\_t3\_neg\_4\_001\_49

m/z 868.7500

01E+411.5RT (min)

lcqtof\_t3\_neg\_1\_018\_33

m/z 869.3265

01234E+316.517RT (min)

lcqtof\_t3\_neg\_4\_052\_03\_rerun

m/z 870.3370

0246E+311.5RT (min)

lcqtof\_t3\_neg\_4\_045\_02\_rerun

m/z 870.8103

0123E+316.517RT (min)

lcqtof\_t3\_neg\_4\_014\_13

m/z 871.8030

01E+411.5RT (min)

lcqtof\_t3\_neg\_1\_038\_15

m/z 871.8065

012E+316.517RT (min)

lcqtof\_t3\_neg\_4\_047\_07

m/z 872.3110

012E+316.517RT (min)

lcqtof\_t3\_neg\_1\_032\_36

m/z 872.3450

0246E+311.5RT (min)

lcqtof\_t3\_neg\_4\_037\_01

m/z 872.7523

0123E+316.517RT (min)

lcqtof\_t3\_neg\_3\_032\_24

m/z 877.3264

012E+316.517RT (min)

lcqtof\_t3\_neg\_4\_043\_11

m/z 878.3229

024E+311.5RT (min)

lcqtof\_t3\_neg\_4\_045\_02\_rerun

m/z 878.8250

01234E+316.517RT (min)

lcqtof\_t3\_neg\_4\_031\_42

m/z 880.3024

01E+316.517RT (min)

lcqtof\_t3\_neg\_1\_039\_43

m/z 880.4192

01234E+377.5RT (min)

lcqtof\_t3\_neg\_4\_004\_20

m/z 880.8226

01234E+316.517RT (min)

lcqtof\_t3\_neg\_4\_050\_43

m/z 882.4120

01234E+377.5RT (min)

lcqtof\_t3\_neg\_4\_004\_20

m/z 883.7857

012E+316.517RT (min)

lcqtof\_t3\_neg\_1\_039\_43

m/z 883.9593

01234E+311.5RT (min)

lcqtof\_t3\_neg\_2\_019\_32

m/z 884.1047

01234E+344.5RT (min)

lcqtof\_t3\_neg\_1\_055\_32

m/z 884.3355

024E+311.5RT (min)

lcqtof\_t3\_neg\_4\_037\_01

m/z 884.7927

012E+316.517RT (min)

lcqtof\_t3\_neg\_4\_052\_03\_rerun

m/z 885.3071

01E+411.5RT (min)

lcqtof\_t3\_neg\_4\_045\_02\_rerun

m/z 885.3096

01E+316.517RT (min)

lcqtof\_t3\_neg\_1\_013\_09

m/z 885.8058

01E+411.5RT (min)

lcqtof\_t3\_neg\_1\_036\_14

m/z 886.3084

0123E+311.5RT (min)

lcqtof\_t3\_neg\_3\_012\_14

m/z 886.3349

024E+311.5RT (min)

lcqtof\_t3\_neg\_4\_051\_15

m/z 889.8307

0123E+316.517RT (min)

lcqtof\_t3\_neg\_4\_045\_02\_rerun

m/z 890.7973

01E+411.5RT (min)

lcqtof\_t3\_neg\_1\_032\_36

m/z 892.9547

0123E+311.5RT (min)

lcqtof\_t3\_neg\_2\_002\_35

m/z 893.2992

01234E+311.5RT (min)

lcqtof\_t3\_neg\_2\_051\_14

m/z 893.7941

02468E+311.5RT (min)

lcqtof\_t3\_neg\_1\_036\_14

m/z 893.8045

0123E+316.517RT (min)

lcqtof\_t3\_neg\_1\_039\_43

m/z 895.3364

0123E+316.517RT (min)

lcqtof\_t3\_neg\_4\_052\_03\_rerun

m/z 896.3448

024E+311.5RT (min)

lcqtof\_t3\_neg\_2\_042\_13

m/z 896.4492

024E+35.56RT (min)

lcqtof\_t3\_neg\_4\_004\_20

m/z 897.8101

012E+411.5RT (min)

lcqtof\_t3\_neg\_2\_055\_16

m/z 897.8108

01234E+316.517RT (min)

lcqtof\_t3\_neg\_4\_045\_02\_rerun

m/z 898.3111

0246E+311.5RT (min)

lcqtof\_t3\_neg\_2\_050\_01

m/z 898.3145

012E+316.517RT (min)

lcqtof\_t3\_neg\_4\_025\_52

m/z 898.7462

0123E+316.517RT (min)

lcqtof\_t3\_neg\_1\_015\_34

m/z 898.7968

02468E+311.5RT (min)

lcqtof\_t3\_neg\_1\_032\_36

m/z 899.9343

0123E+311.5RT (min)

lcqtof\_t3\_neg\_2\_019\_32

m/z 900.3252

01234E+311.5RT (min)

lcqtof\_t3\_neg\_4\_051\_15

m/z 902.8253

02468E+316.517RT (min)

lcqtof\_t3\_neg\_4\_045\_02\_rerun

m/z 903.3279

024E+316.517RT (min)

lcqtof\_t3\_neg\_4\_050\_43

m/z 904.3284

0246E+311.5RT (min)

lcqtof\_t3\_neg\_4\_045\_02\_rerun

m/z 904.3946

0123E+355.5RT (min)

lcqtof\_t3\_neg\_4\_016\_14

m/z 904.4512

01234E+377.5RT (min)

lcqtof\_t3\_neg\_4\_004\_20

m/z 905.3707

0246E+34.55RT (min)

lcqtof\_t3\_neg\_4\_037\_01

m/z 905.4499

01234E+377.5RT (min)

lcqtof\_t3\_neg\_4\_004\_20

m/z 905.7978

01E+411.5RT (min)

lcqtof\_t3\_neg\_3\_015\_15

m/z 905.8044

0123E+316.517RT (min)

lcqtof\_t3\_neg\_4\_048\_19

m/z 906.3004

01234E+311.5RT (min)

lcqtof\_t3\_neg\_1\_036\_14

m/z 906.3145

012E+316.517RT (min)

lcqtof\_t3\_neg\_4\_025\_52

m/z 906.3160

024E+38.59RT (min)

lcqtof\_t3\_neg\_4\_012\_27

m/z 906.8279

01E+316.517RT (min)

lcqtof\_t3\_neg\_1\_041\_54

m/z 908.3273

024E+311.5RT (min)

lcqtof\_t3\_neg\_4\_037\_01

m/z 908.7804

0123E+316.517RT (min)

lcqtof\_t3\_neg\_4\_045\_02

m/z 910.8025

024E+316.517RT (min)

lcqtof\_t3\_neg\_4\_045\_02\_rerun

m/z 911.3116

012E+316.517RT (min)

lcqtof\_t3\_neg\_4\_039\_06

m/z 911.4982

0246E+311.5RT (min)

lcqtof\_t3\_neg\_4\_045\_02\_rerun

m/z 911.8081

012E+316.517RT (min)

lcqtof\_t3\_neg\_4\_045\_02

m/z 912.3130

0246E+311.5RT (min)

lcqtof\_t3\_neg\_4\_045\_02\_rerun

m/z 913.7936

01E+316.517RT (min)

lcqtof\_t3\_neg\_1\_005\_18

m/z 914.2959

01E+316.517RT (min)

lcqtof\_t3\_neg\_1\_013\_09

m/z 915.1595

0123E+38.59RT (min)

lcqtof\_t3\_neg\_4\_053\_12

m/z 916.8029

01E+411.5RT (min)

lcqtof\_t3\_neg\_1\_032\_36

m/z 919.3066

01E+316.517RT (min)

lcqtof\_t3\_neg\_1\_005\_18

m/z 919.8124

012E+316.517RT (min)

lcqtof\_t3\_neg\_4\_030\_31

m/z 920.3004

0123E+311.5RT (min)

lcqtof\_t3\_neg\_2\_051\_14

m/z 920.8056

0246E+316.517RT (min)

lcqtof\_t3\_neg\_4\_052\_03\_rerun

m/z 920.8338

01E+411.5RT (min)

lcqtof\_t3\_neg\_2\_042\_13

m/z 922.8005

012E+316.517RT (min)

lcqtof\_t3\_neg\_1\_022\_28

m/z 923.8290

0123E+316.517RT (min)

lcqtof\_t3\_neg\_4\_016\_14

m/z 924.7668

0123E+316.517RT (min)

lcqtof\_t3\_neg\_4\_027\_41

m/z 926.7607

012E+316.517RT (min)

lcqtof\_t3\_neg\_4\_040\_24

m/z 926.7851

01E+411.5RT (min)

lcqtof\_t3\_neg\_1\_036\_14

m/z 927.2865

024E+311.5RT (min)

lcqtof\_t3\_neg\_1\_036\_14

m/z 929.4539

01234E+377.5RT (min)

lcqtof\_t3\_neg\_4\_004\_20

m/z 929.8408

0246E+316.517RT (min)

lcqtof\_t3\_neg\_4\_045\_02\_rerun

m/z 930.3124

0246E+311.5RT (min)

lcqtof\_t3\_neg\_4\_037\_01

m/z 930.3361

024E+311.5RT (min)

lcqtof\_t3\_neg\_3\_028\_13

m/z 930.8370

02468E+311.5RT (min)

lcqtof\_t3\_neg\_2\_042\_13

m/z 931.4697

01234E+377.5RT (min)

lcqtof\_t3\_neg\_1\_026\_19

m/z 931.8125

01234E+316.517RT (min)

lcqtof\_t3\_neg\_4\_045\_02\_rerun

m/z 932.3032

0246E+311.5RT (min)

lcqtof\_t3\_neg\_1\_012\_16

m/z 932.3174

01E+316.517RT (min)

lcqtof\_t3\_neg\_4\_043\_11

m/z 934.1543

02468E+311.5RT (min)

lcqtof\_t3\_neg\_4\_045\_02\_rerun

m/z 934.4847

024E+311.5RT (min)

lcqtof\_t3\_neg\_4\_045\_02\_rerun

m/z 935.8022

0123E+316.517RT (min)

lcqtof\_t3\_neg\_4\_006\_16

m/z 936.7401

01E+411.5RT (min)

lcqtof\_t3\_neg\_1\_032\_36

m/z 936.8134

01E+416.517RT (min)

lcqtof\_t3\_neg\_4\_045\_02\_rerun

m/z 937.3214

01234E+316.517RT (min)

lcqtof\_t3\_neg\_4\_052\_03\_rerun

m/z 938.3224

0246E+311.5RT (min)

lcqtof\_t3\_neg\_4\_045\_02\_rerun

m/z 938.7989

0123E+316.517RT (min)

lcqtof\_t3\_neg\_4\_016\_14

m/z 939.4813

02468E+311.5RT (min)

lcqtof\_t3\_neg\_4\_045\_02\_rerun

m/z 939.7934

012E+316.517RT (min)

lcqtof\_t3\_neg\_1\_025\_52

m/z 940.2939

024E+311.5RT (min)

lcqtof\_t3\_neg\_3\_015\_15

m/z 940.3048

012E+316.517RT (min)

lcqtof\_t3\_neg\_1\_029\_44

m/z 940.4550

0123E+35.56RT (min)

lcqtof\_t3\_neg\_4\_051\_15

m/z 940.7271

012E+316.517RT (min)

lcqtof\_t3\_neg\_4\_011\_32

m/z 942.2828

01234E+311.5RT (min)

lcqtof\_t3\_neg\_4\_051\_15

m/z 943.9950

01234E+366.5RT (min)

lcqtof\_t3\_neg\_4\_016\_14

m/z 944.2828

0246E+311.5RT (min)

lcqtof\_t3\_neg\_4\_051\_15

m/z 944.4951

01234E+366.5RT (min)

lcqtof\_t3\_neg\_4\_016\_14

m/z 945.3056

012E+316.517RT (min)

lcqtof\_t3\_neg\_4\_052\_03

m/z 946.3073

0246E+311.5RT (min)

lcqtof\_t3\_neg\_4\_045\_02\_rerun

m/z 946.8111

01234E+316.517RT (min)

lcqtof\_t3\_neg\_4\_045\_02\_rerun

m/z 946.8114

01E+411.5RT (min)

lcqtof\_t3\_neg\_4\_045\_02\_rerun

m/z 948.2919

012E+316.517RT (min)

lcqtof\_t3\_neg\_1\_036\_14

m/z 948.8096

01234E+316.517RT (min)

lcqtof\_t3\_neg\_4\_045\_02\_rerun

m/z 950.7527

01E+411.5RT (min)

lcqtof\_t3\_neg\_1\_036\_14

m/z 951.2944

024E+311.5RT (min)

lcqtof\_t3\_neg\_1\_044\_37

m/z 951.7703

012E+316.517RT (min)

lcqtof\_t3\_neg\_1\_052\_12

m/z 951.9484

0123E+311.5RT (min)

lcqtof\_t3\_neg\_2\_019\_32

m/z 952.7778

012E+316.517RT (min)

lcqtof\_t3\_neg\_1\_054\_41

m/z 953.2955

01E+316.517RT (min)

lcqtof\_t3\_neg\_1\_039\_43

m/z 953.7941

01E+411.5RT (min)

lcqtof\_t3\_neg\_2\_051\_14

m/z 954.2928

0123E+311.5RT (min)

lcqtof\_t3\_neg\_4\_045\_02\_rerun

m/z 957.1508

0246E+311.5RT (min)

lcqtof\_t3\_neg\_4\_045\_02\_rerun

m/z 957.8121

0123E+411.5RT (min)

lcqtof\_t3\_neg\_1\_002\_13

m/z 958.7830

01E+411.5RT (min)

lcqtof\_t3\_neg\_1\_032\_36

m/z 959.4667

024E+377.5RT (min)

lcqtof\_t3\_neg\_4\_004\_20

m/z 960.9454

012E+311.5RT (min)

lcqtof\_t3\_neg\_2\_050\_01

m/z 961.2845

0246E+311.5RT (min)

lcqtof\_t3\_neg\_1\_036\_14

m/z 961.7996

012E+316.517RT (min)

lcqtof\_t3\_neg\_1\_050\_40

m/z 962.1418

01E+411.5RT (min)

lcqtof\_t3\_neg\_4\_045\_02\_rerun

m/z 962.4520

0246E+366.5RT (min)

lcqtof\_t3\_neg\_4\_016\_14

m/z 962.4755

0246E+311.5RT (min)

lcqtof\_t3\_neg\_4\_045\_02\_rerun

m/z 962.8164

02468E+316.517RT (min)

lcqtof\_t3\_neg\_4\_052\_03\_rerun

m/z 964.3324

0246E+311.5RT (min)

lcqtof\_t3\_neg\_2\_042\_13

m/z 965.8045

01234E+316.517RT (min)

lcqtof\_t3\_neg\_4\_052\_03\_rerun

m/z 966.3140

01E+316.517RT (min)

lcqtof\_t3\_neg\_4\_042\_10

m/z 966.7883

01E+411.5RT (min)

lcqtof\_t3\_neg\_2\_050\_01

m/z 967.4672

02468E+311.5RT (min)

lcqtof\_t3\_neg\_4\_045\_02\_rerun

m/z 967.6929

0246E+313.514RT (min)

lcqtof\_t3\_neg\_4\_034\_34

m/z 967.8035

01234E+311.5RT (min)

lcqtof\_t3\_neg\_4\_045\_02\_rerun

m/z 967.8506

01E+31414.5RT (min)

lcqtof\_t3\_neg\_2\_052\_17

m/z 970.8149

02468E+316.517RT (min)

lcqtof\_t3\_neg\_4\_045\_02\_rerun

m/z 971.3057

0123E+316.517RT (min)

lcqtof\_t3\_neg\_4\_052\_03\_rerun

m/z 972.3189

024E+311.5RT (min)

lcqtof\_t3\_neg\_1\_012\_16

m/z 973.7834

01E+411.5RT (min)

lcqtof\_t3\_neg\_1\_038\_15

m/z 973.8104

012E+316.517RT (min)

lcqtof\_t3\_neg\_4\_039\_06

m/z 974.2869

024E+311.5RT (min)

lcqtof\_t3\_neg\_2\_050\_01

m/z 974.3106

012E+316.517RT (min)

lcqtof\_t3\_neg\_1\_036\_14

m/z 974.7816

02468E+311.5RT (min)

lcqtof\_t3\_neg\_2\_027\_15

m/z 974.8203

02468E+31313.5RT (min)

lcqtof\_t3\_neg\_4\_040\_24

m/z 975.3455

0123E+377.5RT (min)

lcqtof\_t3\_neg\_4\_004\_20

m/z 975.9047

01E+314.515RT (min)

lcqtof\_t3\_neg\_2\_041\_54

m/z 976.2980

024E+311.5RT (min)

lcqtof\_t3\_neg\_4\_037\_01

m/z 976.7953

0123E+316.517RT (min)

lcqtof\_t3\_neg\_4\_031\_42

m/z 978.7937

01234E+316.517RT (min)

lcqtof\_t3\_neg\_4\_052\_03\_rerun

m/z 979.2990

012E+316.517RT (min)

lcqtof\_t3\_neg\_4\_052\_03\_rerun

m/z 979.8047

01E+316.517RT (min)

lcqtof\_t3\_neg\_1\_031\_24

m/z 980.2976

0246E+311.5RT (min)

lcqtof\_t3\_neg\_4\_045\_02\_rerun

m/z 981.7813

01E+316.517RT (min)

lcqtof\_t3\_neg\_1\_042\_46

m/z 982.2806

02468E+311.5RT (min)

lcqtof\_t3\_neg\_4\_037\_01

m/z 982.2993

012E+316.517RT (min)

lcqtof\_t3\_neg\_4\_039\_06\_rerun

m/z 983.4534

02468E+377.5RT (min)

lcqtof\_t3\_neg\_4\_004\_20

m/z 984.7913

01E+411.5RT (min)

lcqtof\_t3\_neg\_1\_032\_36

m/z 985.1377

0246E+311.5RT (min)

lcqtof\_t3\_neg\_4\_045\_02\_rerun

m/z 985.2894

01234E+311.5RT (min)

lcqtof\_t3\_neg\_1\_032\_36

m/z 985.4696

0246E+311.5RT (min)

lcqtof\_t3\_neg\_4\_045\_02\_rerun

m/z 987.2917

012E+316.517RT (min)

lcqtof\_t3\_neg\_1\_032\_36

m/z 987.7877

01E+411.5RT (min)

lcqtof\_t3\_neg\_4\_045\_02\_rerun

m/z 987.8483

01E+314.515RT (min)

lcqtof\_t3\_neg\_2\_032\_39

m/z 988.2632

02468E+311.5RT (min)

lcqtof\_t3\_neg\_4\_051\_15

m/z 988.2849

01234E+311.5RT (min)

lcqtof\_t3\_neg\_3\_012\_14

m/z 988.7899

024E+316.517RT (min)

lcqtof\_t3\_neg\_4\_052\_03\_rerun

m/z 989.1580

0123E+38.59RT (min)

lcqtof\_t3\_neg\_4\_053\_12

m/z 990.1302

01E+411.5RT (min)

lcqtof\_t3\_neg\_4\_045\_02\_rerun

m/z 990.2666

0246E+311.5RT (min)

lcqtof\_t3\_neg\_4\_037\_01

m/z 990.4626

024E+311.5RT (min)

lcqtof\_t3\_neg\_4\_045\_02\_rerun

m/z 990.7467

024E+311.5RT (min)

lcqtof\_t3\_neg\_2\_005\_34

m/z 992.7688

01234E+316.517RT (min)

lcqtof\_t3\_neg\_4\_032\_04

m/z 992.7780

01E+411.5RT (min)

lcqtof\_t3\_neg\_1\_032\_36

m/z 994.7753

01E+411.5RT (min)

lcqtof\_t3\_neg\_1\_036\_14

m/z 995.2802

0246E+311.5RT (min)

lcqtof\_t3\_neg\_1\_036\_14

m/z 995.4550

0246E+311.5RT (min)

lcqtof\_t3\_neg\_2\_051\_14

m/z 995.7811

0246E+311.5RT (min)

lcqtof\_t3\_neg\_2\_051\_14

m/z 998.2527

024E+311.5RT (min)

lcqtof\_t3\_neg\_4\_037\_01

m/z 998.3246

024E+311.5RT (min)

lcqtof\_t3\_neg\_2\_042\_13

m/z 998.8158

02468E+311.5RT (min)

lcqtof\_t3\_neg\_2\_042\_13

m/z 999.7985

0123E+316.517RT (min)

lcqtof\_t3\_neg\_4\_053\_12

m/z 1000.2496

0246E+311.5RT (min)

lcqtof\_t3\_neg\_4\_037\_01

m/z 1000.3101

0123E+316.517RT (min)

lcqtof\_t3\_neg\_4\_052\_03\_rerun

m/z 1002.2446

0246E+311.5RT (min)

lcqtof\_t3\_neg\_4\_037\_01

m/z 1002.4771

02468E+311.5RT (min)

lcqtof\_t3\_neg\_2\_042\_13

m/z 1002.7646

012E+411.5RT (min)

lcqtof\_t3\_neg\_1\_038\_15

m/z 1003.7942

0123E+316.517RT (min)

lcqtof\_t3\_neg\_4\_006\_16

m/z 1003.9714

01234E+366.5RT (min)

lcqtof\_t3\_neg\_4\_037\_01

m/z 1004.2437

024E+311.5RT (min)

lcqtof\_t3\_neg\_4\_037\_01

m/z 1005.3057

01234E+316.517RT (min)

lcqtof\_t3\_neg\_4\_052\_03\_rerun

m/z 1005.7106

024E+313.514RT (min)

lcqtof\_t3\_neg\_4\_018\_23

m/z 1005.8407

01E+313.514RT (min)

lcqtof\_t3\_neg\_2\_027\_15

m/z 1006.2524

0246E+311.5RT (min)

lcqtof\_t3\_neg\_4\_037\_01

m/z 1006.3090

0246E+311.5RT (min)

lcqtof\_t3\_neg\_4\_045\_02\_rerun

m/z 1006.7152

024E+311.5RT (min)

lcqtof\_t3\_neg\_1\_048\_35

m/z 1006.7615

012E+316.517RT (min)

lcqtof\_t3\_neg\_1\_040\_08

m/z 1007.7874

012E+316.517RT (min)

lcqtof\_t3\_neg\_4\_028\_25

m/z 1008.1362

0246E+311.5RT (min)

lcqtof\_t3\_neg\_4\_045\_02\_rerun

m/z 1008.2957

012E+316.517RT (min)

lcqtof\_t3\_neg\_4\_037\_01

m/z 1008.7363

012E+316.517RT (min)

lcqtof\_t3\_neg\_4\_049\_55

m/z 1013.1257

0246E+311.5RT (min)

lcqtof\_t3\_neg\_4\_045\_02\_rerun

m/z 1013.3044

012E+316.517RT (min)

lcqtof\_t3\_neg\_4\_045\_02\_rerun

m/z 1013.4561

0246E+311.5RT (min)

lcqtof\_t3\_neg\_4\_045\_02\_rerun

m/z 1014.2949

02468E+311.5RT (min)

lcqtof\_t3\_neg\_4\_045\_02\_rerun

m/z 1014.7952

01E+411.5RT (min)

lcqtof\_t3\_neg\_4\_045\_02\_rerun

m/z 1014.7952

0123E+316.517RT (min)

lcqtof\_t3\_neg\_4\_028\_25

m/z 1016.2717

01E+316.517RT (min)

lcqtof\_t3\_neg\_1\_021\_39

m/z 1016.9625

01E+31515.5RT (min)

lcqtof\_t3\_neg\_4\_030\_31

m/z 1018.1141

0246E+311.5RT (min)

lcqtof\_t3\_neg\_4\_045\_02\_rerun

m/z 1019.2821

01234E+311.5RT (min)

lcqtof\_t3\_neg\_1\_032\_36

m/z 1019.7714

012E+316.517RT (min)

lcqtof\_t3\_neg\_1\_036\_14

m/z 1020.7714

0123E+316.517RT (min)

lcqtof\_t3\_neg\_4\_031\_42

m/z 1021.2889

01E+316.517RT (min)

lcqtof\_t3\_neg\_1\_010\_27

m/z 1021.7793

01E+411.5RT (min)

lcqtof\_t3\_neg\_4\_045\_02\_rerun

m/z 1022.2803

01234E+311.5RT (min)

lcqtof\_t3\_neg\_4\_045\_02\_rerun

m/z 1024.8053

01E+411.5RT (min)

lcqtof\_t3\_neg\_2\_042\_13

m/z 1025.1385

02468E+311.5RT (min)

lcqtof\_t3\_neg\_4\_045\_02\_rerun

m/z 1025.4725

0246E+311.5RT (min)

lcqtof\_t3\_neg\_2\_042\_13

m/z 1025.8000

0123E+411.5RT (min)

lcqtof\_t3\_neg\_2\_042\_13

m/z 1027.2706

01234E+311.5RT (min)

lcqtof\_t3\_neg\_1\_048\_35

m/z 1028.7644

0246E+316.517RT (min)

lcqtof\_t3\_neg\_4\_039\_06\_rerun

m/z 1028.9347

01E+311.5RT (min)

lcqtof\_t3\_neg\_1\_048\_35

m/z 1029.2716

0246E+311.5RT (min)

lcqtof\_t3\_neg\_1\_036\_14

m/z 1029.6299

0123E+312.513RT (min)

lcqtof\_t3\_neg\_1\_003\_07

m/z 1029.7719

012E+316.517RT (min)

lcqtof\_t3\_neg\_4\_047\_07

m/z 1030.4639

02468E+311.5RT (min)

lcqtof\_t3\_neg\_4\_045\_02\_rerun

m/z 1030.8130

02468E+316.517RT (min)

lcqtof\_t3\_neg\_4\_045\_02\_rerun

m/z 1033.7791

01234E+316.517RT (min)

lcqtof\_t3\_neg\_4\_039\_06\_rerun

m/z 1033.7823

01E+411.5RT (min)

lcqtof\_t3\_neg\_1\_053\_01

m/z 1034.7367

0123E+316.517RT (min)

lcqtof\_t3\_neg\_1\_003\_07

m/z 1034.7616

02468E+311.5RT (min)

lcqtof\_t3\_neg\_2\_043\_31

m/z 1035.4554

01E+411.5RT (min)

lcqtof\_t3\_neg\_4\_045\_02\_rerun

m/z 1035.7891

024E+311.5RT (min)

lcqtof\_t3\_neg\_1\_038\_15

m/z 1036.1213

0246E+311.5RT (min)

lcqtof\_t3\_neg\_4\_045\_02\_rerun

m/z 1036.7357

01E+411.5RT (min)

lcqtof\_t3\_neg\_1\_038\_15

m/z 1038.8008

02468E+316.517RT (min)

lcqtof\_t3\_neg\_4\_039\_06\_rerun

m/z 1039.2961

0123E+316.517RT (min)

lcqtof\_t3\_neg\_4\_039\_06\_rerun

m/z 1040.3044

0246E+311.5RT (min)

lcqtof\_t3\_neg\_4\_045\_02\_rerun

m/z 1041.7700

012E+316.517RT (min)

lcqtof\_t3\_neg\_4\_039\_06\_rerun

m/z 1041.7737

01E+411.5RT (min)

lcqtof\_t3\_neg\_1\_038\_15

m/z 1042.2789

01E+316.517RT (min)

lcqtof\_t3\_neg\_1\_013\_09

m/z 1042.7697

02468E+311.5RT (min)

lcqtof\_t3\_neg\_1\_010\_27

m/z 1043.4756

012E+312.513RT (min)

lcqtof\_t3\_neg\_1\_019\_23

m/z 1044.7716

012E+316.517RT (min)

lcqtof\_t3\_neg\_4\_044\_48

m/z 1044.7725

01E+411.5RT (min)

lcqtof\_t3\_neg\_2\_043\_31

m/z 1046.1066

0123E+311.5RT (min)

lcqtof\_t3\_neg\_2\_051\_14

m/z 1046.7838

01234E+316.517RT (min)

lcqtof\_t3\_neg\_4\_039\_06\_rerun

m/z 1047.2853

012E+316.517RT (min)

lcqtof\_t3\_neg\_4\_039\_06\_rerun

m/z 1048.2887

0246E+311.5RT (min)

lcqtof\_t3\_neg\_4\_045\_02\_rerun

m/z 1050.2189

0246E+311.5RT (min)

lcqtof\_t3\_neg\_4\_037\_01

m/z 1050.2717

01E+316.517RT (min)

lcqtof\_t3\_neg\_1\_009\_50

m/z 1052.7791

01E+411.5RT (min)

lcqtof\_t3\_neg\_1\_032\_36

m/z 1053.1216

02468E+311.5RT (min)

lcqtof\_t3\_neg\_4\_045\_02\_rerun

m/z 1053.2804

024E+311.5RT (min)

lcqtof\_t3\_neg\_1\_032\_36

m/z 1053.4585

02468E+311.5RT (min)

lcqtof\_t3\_neg\_4\_045\_02\_rerun

m/z 1054.7826

01E+416.517RT (min)

lcqtof\_t3\_neg\_4\_045\_02\_rerun

m/z 1055.2848

01E+316.517RT (min)

lcqtof\_t3\_neg\_1\_026\_19

m/z 1056.2250

0246E+311.5RT (min)

lcqtof\_t3\_neg\_4\_037\_01

m/z 1056.2805

024E+311.5RT (min)

lcqtof\_t3\_neg\_4\_045\_02\_rerun

m/z 1056.7848

0123E+316.517RT (min)

lcqtof\_t3\_neg\_4\_027\_41

m/z 1058.1145

01E+411.5RT (min)

lcqtof\_t3\_neg\_4\_045\_02\_rerun

m/z 1058.2146

02468E+311.5RT (min)

lcqtof\_t3\_neg\_4\_037\_01

m/z 1058.2523

01E+316.517RT (min)

lcqtof\_t3\_neg\_3\_014\_39

m/z 1058.4507

0246E+311.5RT (min)

lcqtof\_t3\_neg\_4\_045\_02\_rerun

m/z 1060.4918

024E+34.55RT (min)

lcqtof\_t3\_neg\_4\_016\_14

m/z 1060.7654

01E+411.5RT (min)

lcqtof\_t3\_neg\_1\_032\_36

m/z 1062.2062

01E+411.5RT (min)

lcqtof\_t3\_neg\_4\_037\_01

m/z 1063.2637

0246E+311.5RT (min)

lcqtof\_t3\_neg\_1\_036\_14

m/z 1063.4426

0246E+311.5RT (min)

lcqtof\_t3\_neg\_2\_051\_14

m/z 1064.1088

024E+311.5RT (min)

lcqtof\_t3\_neg\_4\_045\_02\_rerun

m/z 1064.2002

02468E+311.5RT (min)

lcqtof\_t3\_neg\_4\_037\_01

m/z 1065.8075

01234E+316.517RT (min)

lcqtof\_t3\_neg\_4\_045\_02\_rerun

m/z 1066.2117

02468E+311.5RT (min)

lcqtof\_t3\_neg\_4\_037\_01

m/z 1066.3138

024E+311.5RT (min)

lcqtof\_t3\_neg\_1\_002\_13

m/z 1067.7787

0123E+316.517RT (min)

lcqtof\_t3\_neg\_4\_039\_06\_rerun

m/z 1068.7751

01E+411.5RT (min)

lcqtof\_t3\_neg\_1\_053\_01

m/z 1069.2766

0123E+311.5RT (min)

lcqtof\_t3\_neg\_2\_050\_01

m/z 1069.9542

02468E+31414.5RT (min)

lcqtof\_t3\_neg\_4\_040\_24

m/z 1070.4655

01E+411.5RT (min)

lcqtof\_t3\_neg\_2\_042\_13

m/z 1071.7766

012E+316.517RT (min)

lcqtof\_t3\_neg\_4\_042\_10

m/z 1072.7898

0246E+316.517RT (min)

lcqtof\_t3\_neg\_4\_039\_06\_rerun

m/z 1073.2936

0123E+316.517RT (min)

lcqtof\_t3\_neg\_4\_039\_06\_rerun

m/z 1074.0920

0123E+311.5RT (min)

lcqtof\_t3\_neg\_2\_051\_14

m/z 1074.1932

0246E+311.5RT (min)

lcqtof\_t3\_neg\_4\_037\_01

m/z 1074.2970

0246E+311.5RT (min)

lcqtof\_t3\_neg\_4\_045\_02\_rerun

m/z 1074.5076

0123E+31313.5RT (min)

lcqtof\_t3\_neg\_4\_074\_58

m/z 1075.4519

01E+411.5RT (min)

lcqtof\_t3\_neg\_4\_045\_02\_rerun

m/z 1075.7599

0123E+316.517RT (min)

lcqtof\_t3\_neg\_4\_039\_06\_rerun

m/z 1076.1227

02468E+311.5RT (min)

lcqtof\_t3\_neg\_4\_045\_02\_rerun

m/z 1076.2800

012E+316.517RT (min)

lcqtof\_t3\_neg\_4\_045\_02

m/z 1076.7151

012E+316.517RT (min)

lcqtof\_t3\_neg\_4\_053\_12

m/z 1081.1129

0246E+311.5RT (min)

lcqtof\_t3\_neg\_4\_045\_02\_rerun

m/z 1081.2831

012E+316.517RT (min)

lcqtof\_t3\_neg\_4\_024\_26

m/z 1081.4458

0246E+311.5RT (min)

lcqtof\_t3\_neg\_4\_045\_02\_rerun

m/z 1081.7900

024E+316.517RT (min)

lcqtof\_t3\_neg\_4\_052\_03\_rerun

m/z 1082.0203

024E+35.56RT (min)

lcqtof\_t3\_neg\_4\_016\_14

m/z 1082.2866

02468E+311.5RT (min)

lcqtof\_t3\_neg\_4\_045\_02\_rerun

m/z 1082.5220

0123E+35.56RT (min)

lcqtof\_t3\_neg\_2\_051\_14

m/z 1082.7825

02468E+311.5RT (min)

lcqtof\_t3\_neg\_4\_045\_02\_rerun

m/z 1084.2651

01E+316.517RT (min)

lcqtof\_t3\_neg\_1\_024\_10

m/z 1086.1053

02468E+311.5RT (min)

lcqtof\_t3\_neg\_2\_051\_14

m/z 1086.4374

01234E+311.5RT (min)

lcqtof\_t3\_neg\_4\_045\_02\_rerun

m/z 1086.7665

01E+411.5RT (min)

lcqtof\_t3\_neg\_2\_053\_38

m/z 1087.2738

01234E+311.5RT (min)

lcqtof\_t3\_neg\_1\_044\_37

m/z 1088.2092

0246E+311.5RT (min)

lcqtof\_t3\_neg\_4\_037\_01

m/z 1088.7445

0123E+316.517RT (min)

lcqtof\_t3\_neg\_4\_048\_19

m/z 1089.2738

01E+316.517RT (min)

lcqtof\_t3\_neg\_1\_040\_08

m/z 1089.7643

01E+411.5RT (min)

lcqtof\_t3\_neg\_4\_045\_02\_rerun

m/z 1090.2664

01234E+311.5RT (min)

lcqtof\_t3\_neg\_4\_045\_02\_rerun

m/z 1092.7566

01E+411.5RT (min)

lcqtof\_t3\_neg\_1\_032\_36

m/z 1092.7928

01E+411.5RT (min)

lcqtof\_t3\_neg\_2\_042\_13

m/z 1093.1265

01E+411.5RT (min)

lcqtof\_t3\_neg\_2\_042\_13

m/z 1093.4626

0246E+311.5RT (min)

lcqtof\_t3\_neg\_1\_002\_13

m/z 1094.7556

01E+411.5RT (min)

lcqtof\_t3\_neg\_2\_026\_36

m/z 1095.2556

01234E+311.5RT (min)

lcqtof\_t3\_neg\_2\_002\_35

m/z 1096.7560

0246E+316.517RT (min)

lcqtof\_t3\_neg\_4\_045\_02\_rerun

m/z 1097.2584

0246E+311.5RT (min)

lcqtof\_t3\_neg\_3\_012\_14

m/z 1097.7568

01E+411.5RT (min)

lcqtof\_t3\_neg\_2\_051\_14

m/z 1097.7605

012E+316.517RT (min)

lcqtof\_t3\_neg\_3\_034\_04

m/z 1098.2570

012E+311.5RT (min)

lcqtof\_t3\_neg\_3\_012\_14

m/z 1098.4504

01E+411.5RT (min)

lcqtof\_t3\_neg\_4\_045\_02\_rerun

m/z 1098.7906

0246E+316.517RT (min)

lcqtof\_t3\_neg\_4\_045\_02\_rerun

m/z 1100.1936

0246E+311.5RT (min)

lcqtof\_t3\_neg\_4\_037\_01

m/z 1102.1833

02468E+313.514RT (min)

lcqtof\_t3\_neg\_3\_081\_58

m/z 1102.1967

02468E+311.5RT (min)

lcqtof\_t3\_neg\_4\_037\_01

m/z 1102.7606

01E+411.5RT (min)

lcqtof\_t3\_neg\_1\_053\_01

m/z 1103.4409

01E+411.5RT (min)

lcqtof\_t3\_neg\_4\_045\_02\_rerun

m/z 1103.7729

0246E+311.5RT (min)

lcqtof\_t3\_neg\_4\_045\_02\_rerun

m/z 1104.1071

02468E+311.5RT (min)

lcqtof\_t3\_neg\_4\_045\_02\_rerun

m/z 1104.1887

02468E+311.5RT (min)

lcqtof\_t3\_neg\_4\_037\_01

m/z 1104.7439

01E+411.5RT (min)

lcqtof\_t3\_neg\_1\_036\_14

m/z 1105.3036

0246E+312.513RT (min)

lcqtof\_t3\_neg\_1\_055\_32

m/z 1106.7953

0246E+316.517RT (min)

lcqtof\_t3\_neg\_4\_045\_02\_rerun

m/z 1107.2885

0123E+316.517RT (min)

lcqtof\_t3\_neg\_4\_045\_02\_rerun

m/z 1107.7230

01234E+313.514RT (min)

lcqtof\_t3\_neg\_4\_018\_23

m/z 1108.2883

02468E+311.5RT (min)

lcqtof\_t3\_neg\_4\_045\_02\_rerun

m/z 1109.1028

01234E+311.5RT (min)

lcqtof\_t3\_neg\_3\_012\_14

m/z 1109.4352

024E+311.5RT (min)

lcqtof\_t3\_neg\_4\_045\_02\_rerun

m/z 1109.7751

0123E+316.517RT (min)

lcqtof\_t3\_neg\_4\_047\_07

m/z 1110.7622

0246E+311.5RT (min)

lcqtof\_t3\_neg\_1\_034\_30

m/z 1112.7542

012E+316.517RT (min)

lcqtof\_t3\_neg\_4\_049\_55

m/z 1114.0906

024E+311.5RT (min)

lcqtof\_t3\_neg\_3\_012\_14

m/z 1114.7723

01234E+316.517RT (min)

lcqtof\_t3\_neg\_4\_050\_43

m/z 1115.2844

012E+316.517RT (min)

lcqtof\_t3\_neg\_4\_039\_06\_rerun

m/z 1116.1262

02468E+311.5RT (min)

lcqtof\_t3\_neg\_2\_042\_13

m/z 1116.2744

0246E+311.5RT (min)

lcqtof\_t3\_neg\_4\_045\_02\_rerun

m/z 1117.7545

012E+316.517RT (min)

lcqtof\_t3\_neg\_4\_045\_02\_rerun

m/z 1118.2688

01E+316.517RT (min)

lcqtof\_t3\_neg\_4\_045\_02\_rerun

m/z 1121.1121

01E+411.5RT (min)

lcqtof\_t3\_neg\_4\_045\_02\_rerun

m/z 1121.2642

01234E+311.5RT (min)

lcqtof\_t3\_neg\_2\_026\_36

m/z 1121.4456

01E+411.5RT (min)

lcqtof\_t3\_neg\_4\_045\_02\_rerun

m/z 1122.7650

0246E+316.517RT (min)

lcqtof\_t3\_neg\_4\_050\_43

m/z 1123.2552

01E+316.517RT (min)

lcqtof\_t3\_neg\_4\_039\_06

m/z 1123.7588

01E+411.5RT (min)

lcqtof\_t3\_neg\_4\_045\_02\_rerun

m/z 1124.2620

0246E+311.5RT (min)

lcqtof\_t3\_neg\_4\_045\_02\_rerun

m/z 1124.7784

0123E+316.517RT (min)

lcqtof\_t3\_neg\_4\_023\_45

m/z 1126.1034

01E+411.5RT (min)

lcqtof\_t3\_neg\_4\_045\_02\_rerun

m/z 1126.4382

0246E+311.5RT (min)

lcqtof\_t3\_neg\_4\_045\_02\_rerun

m/z 1128.4778

0246E+36.57RT (min)

lcqtof\_t3\_neg\_4\_055\_38

m/z 1128.7554

01E+411.5RT (min)

lcqtof\_t3\_neg\_1\_032\_36

m/z 1129.2823

01234E+311.5RT (min)

lcqtof\_t3\_neg\_1\_002\_13

m/z 1131.2555

02468E+311.5RT (min)

lcqtof\_t3\_neg\_1\_036\_14

m/z 1131.4287

02468E+311.5RT (min)

lcqtof\_t3\_neg\_4\_045\_02\_rerun

m/z 1131.7544

01E+411.5RT (min)

lcqtof\_t3\_neg\_3\_012\_14

m/z 1132.0959

0246E+311.5RT (min)

lcqtof\_t3\_neg\_4\_045\_02\_rerun

m/z 1132.2515

0123E+311.5RT (min)

lcqtof\_t3\_neg\_3\_012\_14

m/z 1135.7710

01E+411.5RT (min)

lcqtof\_t3\_neg\_1\_002\_13

m/z 1136.7628

02468E+311.5RT (min)

lcqtof\_t3\_neg\_1\_012\_16

m/z 1137.2678

0123E+311.5RT (min)

lcqtof\_t3\_neg\_1\_012\_16

m/z 1137.4213

0123E+311.5RT (min)

lcqtof\_t3\_neg\_3\_012\_14

m/z 1138.4548

01E+411.5RT (min)

lcqtof\_t3\_neg\_2\_042\_13

m/z 1138.7479

0246E+316.517RT (min)

lcqtof\_t3\_neg\_4\_038\_29

m/z 1139.7540

012E+316.517RT (min)

lcqtof\_t3\_neg\_1\_013\_09

m/z 1140.7728

0246E+316.517RT (min)

lcqtof\_t3\_neg\_4\_052\_03\_rerun

m/z 1141.2670

012E+316.517RT (min)

lcqtof\_t3\_neg\_4\_039\_06\_rerun

m/z 1142.2804

0246E+311.5RT (min)

lcqtof\_t3\_neg\_4\_045\_02\_rerun

m/z 1143.4358

01E+411.5RT (min)

lcqtof\_t3\_neg\_4\_045\_02\_rerun

m/z 1143.7488

012E+316.517RT (min)

lcqtof\_t3\_neg\_4\_039\_06

m/z 1144.1083

02468E+311.5RT (min)

lcqtof\_t3\_neg\_4\_045\_02\_rerun

m/z 1144.2655

01E+316.517RT (min)

lcqtof\_t3\_neg\_1\_012\_16

m/z 1144.7297

012E+316.517RT (min)

lcqtof\_t3\_neg\_4\_048\_19

m/z 1144.7454

0246E+311.5RT (min)

lcqtof\_t3\_neg\_2\_050\_01

m/z 1148.6057

01234E+312.513RT (min)

lcqtof\_t3\_neg\_4\_094\_60

m/z 1149.1028

02468E+311.5RT (min)

lcqtof\_t3\_neg\_4\_045\_02\_rerun

m/z 1149.2673

012E+316.517RT (min)

lcqtof\_t3\_neg\_4\_052\_03

m/z 1149.4326

02468E+311.5RT (min)

lcqtof\_t3\_neg\_4\_045\_02\_rerun

m/z 1149.7737

01234E+316.517RT (min)

lcqtof\_t3\_neg\_4\_039\_06\_rerun

m/z 1150.2703

0246E+311.5RT (min)

lcqtof\_t3\_neg\_4\_045\_02\_rerun

m/z 1150.7682

0246E+311.5RT (min)

lcqtof\_t3\_neg\_4\_045\_02\_rerun

m/z 1152.2426

01E+316.517RT (min)

lcqtof\_t3\_neg\_4\_048\_19

m/z 1154.0953

02468E+311.5RT (min)

lcqtof\_t3\_neg\_2\_051\_14

m/z 1154.4246

0246E+311.5RT (min)

lcqtof\_t3\_neg\_4\_045\_02\_rerun

m/z 1154.7318

01234E+316.517RT (min)

lcqtof\_t3\_neg\_4\_042\_10

m/z 1154.7614

01E+411.5RT (min)

lcqtof\_t3\_neg\_1\_032\_36

m/z 1155.2623

0123E+311.5RT (min)

lcqtof\_t3\_neg\_1\_032\_36

m/z 1156.7469

012E+316.517RT (min)

lcqtof\_t3\_neg\_4\_024\_26

m/z 1157.2546

01E+411.5RT (min)

lcqtof\_t3\_neg\_4\_045\_02\_rerun

m/z 1157.2581

01E+316.517RT (min)

lcqtof\_t3\_neg\_4\_045\_02

m/z 1157.7549

01E+411.5RT (min)

lcqtof\_t3\_neg\_4\_045\_02\_rerun

m/z 1158.2538

024E+311.5RT (min)

lcqtof\_t3\_neg\_4\_045\_02\_rerun

m/z 1158.8844

024E+314.515RT (min)

lcqtof\_t3\_neg\_4\_003\_37

m/z 1159.4215

0246E+311.5RT (min)

lcqtof\_t3\_neg\_2\_051\_14

m/z 1159.8842

024E+314.515RT (min)

lcqtof\_t3\_neg\_4\_008\_35

m/z 1160.7463

02468E+311.5RT (min)

lcqtof\_t3\_neg\_1\_032\_36

m/z 1161.1138

02468E+311.5RT (min)

lcqtof\_t3\_neg\_2\_055\_16

m/z 1161.4492

0246E+311.5RT (min)

lcqtof\_t3\_neg\_2\_042\_13

m/z 1162.1396

02468E+311.5RT (min)

lcqtof\_t3\_neg\_4\_037\_01

m/z 1162.7437

01E+411.5RT (min)

lcqtof\_t3\_neg\_1\_044\_37

m/z 1164.7493

024E+316.517RT (min)

lcqtof\_t3\_neg\_4\_052\_03\_rerun

m/z 1165.2471

0246E+311.5RT (min)

lcqtof\_t3\_neg\_1\_036\_14

m/z 1165.7434

01E+411.5RT (min)

lcqtof\_t3\_neg\_1\_036\_14

m/z 1165.7526

012E+316.517RT (min)

lcqtof\_t3\_neg\_1\_046\_17

m/z 1166.1011

01E+411.5RT (min)

lcqtof\_t3\_neg\_4\_045\_02\_rerun

m/z 1166.2471

0123E+311.5RT (min)

lcqtof\_t3\_neg\_4\_045\_02\_rerun

m/z 1166.4379

01E+411.5RT (min)

lcqtof\_t3\_neg\_4\_045\_02\_rerun

m/z 1166.7999

0246E+316.517RT (min)

lcqtof\_t3\_neg\_4\_056\_56

m/z 1169.7606

012E+316.517RT (min)

lcqtof\_t3\_neg\_4\_052\_03\_rerun

m/z 1170.7390

0246E+311.5RT (min)

lcqtof\_t3\_neg\_1\_044\_37

m/z 1170.7534

012E+316.517RT (min)

lcqtof\_t3\_neg\_4\_045\_02\_rerun

m/z 1171.2379

0123E+311.5RT (min)

lcqtof\_t3\_neg\_2\_019\_32

m/z 1171.4263

01E+411.5RT (min)

lcqtof\_t3\_neg\_4\_045\_02\_rerun

m/z 1171.7649

024E+311.5RT (min)

lcqtof\_t3\_neg\_2\_048\_30

m/z 1172.0955

02468E+311.5RT (min)

lcqtof\_t3\_neg\_4\_045\_02\_rerun

m/z 1172.7218

02468E+311.5RT (min)

lcqtof\_t3\_neg\_3\_012\_14

m/z 1172.7440

01E+316.517RT (min)

lcqtof\_t3\_neg\_4\_014\_13

m/z 1174.7751

0246E+316.517RT (min)

lcqtof\_t3\_neg\_4\_039\_06\_rerun

m/z 1175.4843

024E+36.57RT (min)

lcqtof\_t3\_neg\_4\_055\_38

m/z 1176.2815

024E+311.5RT (min)

lcqtof\_t3\_neg\_4\_045\_02\_rerun

m/z 1177.0847

024E+311.5RT (min)

lcqtof\_t3\_neg\_4\_045\_02\_rerun

m/z 1177.4170

0246E+311.5RT (min)

lcqtof\_t3\_neg\_4\_045\_02\_rerun

m/z 1177.7482

01E+411.5RT (min)

lcqtof\_t3\_neg\_1\_046\_17

m/z 1177.7534

012E+316.517RT (min)

lcqtof\_t3\_neg\_4\_045\_02

m/z 1178.7484

0246E+311.5RT (min)

lcqtof\_t3\_neg\_1\_046\_17

m/z 1182.0813

024E+311.5RT (min)

lcqtof\_t3\_neg\_3\_012\_14

m/z 1182.7489

01234E+316.517RT (min)

lcqtof\_t3\_neg\_4\_052\_03\_rerun

m/z 1183.2767

0123E+316.517RT (min)

lcqtof\_t3\_neg\_4\_054\_44

m/z 1183.4442

01E+411.5RT (min)

lcqtof\_t3\_neg\_2\_042\_13

m/z 1184.1157

02468E+311.5RT (min)

lcqtof\_t3\_neg\_2\_042\_13

m/z 1184.2651

0246E+311.5RT (min)

lcqtof\_t3\_neg\_4\_045\_02\_rerun

m/z 1186.2341

01E+316.517RT (min)

lcqtof\_t3\_neg\_1\_004\_11

m/z 1188.7643

01E+411.5RT (min)

lcqtof\_t3\_neg\_4\_045\_02\_rerun

m/z 1189.1000

02468E+311.5RT (min)

lcqtof\_t3\_neg\_4\_045\_02\_rerun

m/z 1189.4357

02468E+311.5RT (min)

lcqtof\_t3\_neg\_4\_045\_02\_rerun

m/z 1190.7544

0246E+316.517RT (min)

lcqtof\_t3\_neg\_4\_039\_06\_rerun

m/z 1192.2498

024E+311.5RT (min)

lcqtof\_t3\_neg\_4\_045\_02\_rerun

m/z 1192.7074

02468E+311.5RT (min)

lcqtof\_t3\_neg\_1\_032\_36

m/z 1192.7507

012E+316.517RT (min)

lcqtof\_t3\_neg\_4\_013\_39

m/z 1194.0886

01E+411.5RT (min)

lcqtof\_t3\_neg\_4\_045\_02\_rerun

m/z 1194.4258

02468E+311.5RT (min)

lcqtof\_t3\_neg\_4\_045\_02\_rerun

m/z 1194.7574

0246E+311.5RT (min)

lcqtof\_t3\_neg\_4\_045\_02\_rerun

m/z 1195.0873

0123E+311.5RT (min)

lcqtof\_t3\_neg\_4\_045\_02\_rerun

m/z 1196.7391

02468E+311.5RT (min)

lcqtof\_t3\_neg\_1\_043\_38

m/z 1196.7488

0123E+316.517RT (min)

lcqtof\_t3\_neg\_4\_049\_55

m/z 1197.2357

0123E+311.5RT (min)

lcqtof\_t3\_neg\_2\_043\_31

m/z 1198.7393

01E+411.5RT (min)

lcqtof\_t3\_neg\_2\_051\_14

m/z 1199.2452

0246E+311.5RT (min)

lcqtof\_t3\_neg\_1\_036\_14

m/z 1199.4111

02468E+311.5RT (min)

lcqtof\_t3\_neg\_4\_045\_02\_rerun

m/z 1199.7393

01E+411.5RT (min)

lcqtof\_t3\_neg\_4\_045\_02\_rerun
