## Supplementary material for "The Everything Bagel Feature Finder: Ultra-fast automated feature finding for untargeted metabolomics": SI Data File: tool_unique_xic_st_EB_ST004651.html

 
 
 EB-unique features (not detected by the other two tools) - ST004651 
 
 EB-unique features (not detected by the other two tools) - ST004651 
 4052 features · qtof · XIC window m/z ± 25 ppm (≥ 0.002 Da) · ± 0.5 min 
 Every panel is a feature  EB  detected that neither of the other two tools detected — three-way unique across EB (standard mode (min_detection_rate=0.1)), Masscube and XCMS, under the standard matching rule (m/z ± 25 ppm; apex within the reference RT window ± 0.2 min). It shows the MS1 extracted-ion chromatogram at the feature's m/z, drawn from the raw mzML named in  alignment_reference_file , zoomed to ± 0.5 min around the feature's reported apex RT.  Gray solid line  = the feature's reported apex RT;  red dashed line  = EB's assigned MS/MS —  MS2_scan_id  (native scan number) in  MS2_source_file ; the panel widens to keep an off-apex MS/MS on-screen. Features with no assigned MS/MS show only the gray line. y is per-cell autoscaled. Rows keep the table order (m/z ascending). 
   m/z 71.0136     0   1   2   3E+3   6.5   7  RT (min)      lcqtof_hilic_neg_4_088_59    m/z 71.0148     0   1E+3   5   5.5  RT (min)       lcqtof_hilic_neg_1_015_33    m/z 71.0500     0   1   2   3E+3   1   1.5  RT (min)      lcqtof_hilic_neg_4_055_32    m/z 73.0109     0   1   2   3E+3   4.5   5  RT (min)      lcqtof_hilic_neg_4_039_18    m/z 76.0305     0   2   4E+3   6   6.5  RT (min)      lcqtof_hilic_neg_4_008_24    m/z 76.0310     0   2   4E+3   6   6.5  RT (min)      lcqtof_hilic_neg_4_055_32    m/z 79.0544     0   1   2   3E+3   1   1.5  RT (min)      lcqtof_hilic_neg_4_039_18    m/z 79.9565     0   1   2E+3   4   4.5  RT (min)      lcqtof_hilic_neg_2_016_33    m/z 81.0339     0   1   2   3E+3   1   1.5  RT (min)      lcqtof_hilic_neg_4_039_18    m/z 84.0445     0   2   4E+3   1.5   2  RT (min)      lcqtof_hilic_neg_4_027_19    m/z 84.0451     0   1   2E+3   6   6.5  RT (min)      lcqtof_hilic_neg_4_048_23    m/z 85.0040     0   1   2   3E+3   3   3.5  RT (min)      lcqtof_hilic_neg_4_039_18    m/z 87.0445     0   1   2   3   4E+3   1.5   2  RT (min)       lcqtof_hilic_neg_4_028_36    m/z 89.0426     0   1   2   3   4E+3   1   1.5  RT (min)       lcqtof_hilic_neg_4_050_02    m/z 90.0464     0   1   2E+3   6.5   7  RT (min)      lcqtof_hilic_neg_1_037_24    m/z 92.0471     0   1   2   3E+3   5.5   6  RT (min)      lcqtof_hilic_neg_4_019_03    m/z 93.0449     0   2   4E+3   2   2.5  RT (min)       lcqtof_hilic_neg_4_032_01    m/z 94.0365     0   1   2   3E+3   0.5   1  RT (min)      lcqtof_hilic_neg_4_027_19    m/z 95.9751     0   2   4   6E+3   0.5   1  RT (min)      lcqtof_hilic_neg_3_045_18    m/z 96.0382     0   2   4   6   8E+3   3.5   4  RT (min)       lcqtof_hilic_neg_4_008_24    m/z 99.9489     0   1   2E+3   6   6.5  RT (min)      lcqtof_hilic_neg_2_015_06    m/z 100.0392     0   1E+3   5   5.5  RT (min)      lcqtof_hilic_neg_4_053_14    m/z 101.0280     0   2   4   6E+3   5   5.5  RT (min)       lcqtof_hilic_neg_1_022_36    m/z 104.0433     0   1   2E+3   1   1.5  RT (min)      lcqtof_hilic_neg_4_055_32    m/z 105.0343     0   1   2   3   4E+3   0.5   1  RT (min)      lcqtof_hilic_neg_4_102_61    m/z 105.0849     0   1   2   3   4E+3   6   6.5  RT (min)      lcqtof_hilic_neg_4_008_24    m/z 107.0495     0   2   4E+4   1   1.5  RT (min)      lcqtof_hilic_neg_4_053_14    m/z 110.0244     0   1E+3   4.5   5  RT (min)      lcqtof_hilic_neg_2_029_31    m/z 110.0246     0   1   2E+3   3.5   4  RT (min)      lcqtof_hilic_neg_4_023_15    m/z 110.0419     0   1   2E+3   6   6.5  RT (min)      lcqtof_hilic_neg_4_048_23    m/z 110.9847     0   1   2E+3   5.5   6  RT (min)      lcqtof_hilic_neg_4_096_60    m/z 111.0195     0   1   2   3   4E+3   5   5.5  RT (min)      lcqtof_hilic_neg_4_028_36    m/z 113.0142     0   1   2E+3   6   6.5  RT (min)      lcqtof_hilic_neg_2_015_06    m/z 113.0243     0   2   4E+3   5.5   6  RT (min)       lcqtof_hilic_neg_4_028_36    m/z 113.0551     0   1E+3   3.5   4  RT (min)      lcqtof_hilic_neg_4_003_16    m/z 113.0935     0   2   4E+3   5   5.5  RT (min)      lcqtof_hilic_neg_4_032_01    m/z 113.9890     0   1   2E+3   1   1.5  RT (min)      lcqtof_hilic_neg_1_006_22    m/z 114.0637     0   1   2E+3   3.5   4  RT (min)      lcqtof_hilic_neg_4_013_20    m/z 115.0027     0   1   2   3E+3   1   1.5  RT (min)       lcqtof_hilic_neg_4_004_47    m/z 115.9208     0   1   2   3   4E+3   1   1.5  RT (min)      lcqtof_hilic_neg_1_002_39    m/z 116.0355     0   1E+3   3   3.5  RT (min)      lcqtof_hilic_neg_2_021_17    m/z 116.0716     0   1   2   3   4E+3   6   6.5  RT (min)      lcqtof_hilic_neg_1_023_34    m/z 116.0719     0   1   2   3E+3   3.5   4  RT (min)      lcqtof_hilic_neg_4_026_04    m/z 117.0741     0   1   2   3E+3   5   5.5  RT (min)      lcqtof_hilic_neg_4_006_30    m/z 117.0756     0   1E+3   5   5.5  RT (min)      lcqtof_hilic_neg_2_031_01    m/z 118.0288     0   1   2E+3   0.5   1  RT (min)      lcqtof_hilic_neg_4_034_31    m/z 118.0404     0   2   4   6E+3   5.5   6  RT (min)       lcqtof_hilic_neg_4_008_24    m/z 119.0499     0   1   2E+3   5.5   6  RT (min)      lcqtof_hilic_neg_4_053_14    m/z 120.0120     0   2   4   6   8E+3   4   4.5  RT (min)       lcqtof_hilic_neg_4_052_13    m/z 124.0156     0   2   4   6E+3   0.5   1  RT (min)      lcqtof_hilic_neg_4_039_18    m/z 124.9834     0   1   2   3   4E+3   3.5   4  RT (min)       lcqtof_hilic_neg_4_108_62    m/z 124.9873     0   1   2   3   4E+3   5.5   6  RT (min)       lcqtof_hilic_neg_4_048_23    m/z 125.0101     0   1E+3   5   5.5   6  RT (min)      lcqtof_hilic_neg_1_054_01    m/z 125.0598     0   2   4E+3   1   1.5  RT (min)      lcqtof_hilic_neg_4_048_23    m/z 125.0968     0   1   2   3E+3   1   1.5  RT (min)       lcqtof_hilic_neg_4_039_18    m/z 126.0273     0   2   4E+3   1.5   2  RT (min)      lcqtof_hilic_neg_4_088_59    m/z 126.9659     0   1E+3   6.5   7  RT (min)      lcqtof_hilic_neg_1_035_23    m/z 126.9889     0   1   2E+3   4   4.5  RT (min)      lcqtof_hilic_neg_2_016_33    m/z 128.0542     0   2   4E+3   6   6.5  RT (min)      lcqtof_hilic_neg_4_050_02    m/z 129.0081     0   1   2   3   4E+3   5   5.5  RT (min)      lcqtof_hilic_neg_4_055_32    m/z 129.0185     0   1   2E+4   1   1.5  RT (min)      lcqtof_hilic_neg_2_021_17    m/z 129.0367     0   1   2E+3   1   1.5  RT (min)      lcqtof_hilic_neg_2_047_32    m/z 129.1248     0   1   2   3   4E+3   0.5   1  RT (min)      lcqtof_hilic_neg_4_050_02    m/z 129.9624     0   1   2   3E+3   4.5   5  RT (min)      lcqtof_hilic_neg_4_035_33    m/z 130.0041     0   2   4E+3   5.5   6  RT (min)      lcqtof_hilic_neg_4_008_24    m/z 130.0401     0   2   4   6E+3   1.5   2  RT (min)      lcqtof_hilic_neg_4_093_60    m/z 130.0681     0   1   2   3   4E+3   2   2.5  RT (min)      lcqtof_hilic_neg_4_032_01    m/z 130.0867     0   2   4   6E+3   5.5   6  RT (min)      lcqtof_hilic_neg_1_035_23    m/z 131.0362     0   1E+3   5   5.5  RT (min)      lcqtof_hilic_neg_1_023_34    m/z 131.0636     0   2   4   6E+3   6   6.5  RT (min)      lcqtof_hilic_neg_4_008_24    m/z 133.0334     0   1E+5   1   1.5  RT (min)      lcqtof_hilic_neg_4_050_02    m/z 133.0654     0   2   4   6   8E+3   0.5   1  RT (min)      lcqtof_hilic_neg_4_053_14    m/z 135.0809     0   2   4   6E+3   1   1.5  RT (min)      lcqtof_hilic_neg_4_053_14    m/z 135.0961     0   1   2   3E+3   6.5   7  RT (min)      lcqtof_hilic_neg_4_008_24    m/z 137.0987     0   1   2E+3   7   7.5  RT (min)      lcqtof_hilic_neg_4_055_32    m/z 138.0376     0   2   4   6   8E+3   1   1.5  RT (min)      lcqtof_hilic_neg_4_032_01    m/z 138.0404     0   2   4   6E+3   7.5   8  RT (min)       lcqtof_hilic_neg_4_008_24    m/z 140.0103     0   1   2E+3   4.5   5  RT (min)      lcqtof_hilic_neg_1_003_37    m/z 140.0119     0   1   2E+3   6.5   7  RT (min)      lcqtof_hilic_neg_4_048_23    m/z 142.0124     0   1   2E+3   6   6.5  RT (min)       lcqtof_hilic_neg_4_052_13    m/z 142.0295     0   1E+3   2   2.5  RT (min)      lcqtof_hilic_neg_4_004_47    m/z 142.0652     0   1   2   3   4E+3   4   4.5  RT (min)      lcqtof_hilic_neg_4_052_13    m/z 142.0652     0   1   2   3E+3   5   5.5  RT (min)       lcqtof_hilic_neg_4_053_14    m/z 142.0710     0   2   4   6E+3   7   7.5  RT (min)      lcqtof_hilic_neg_4_008_24    m/z 144.0449     0   1   2E+3   4   4.5  RT (min)      lcqtof_hilic_neg_4_089_59    m/z 144.0456     0   1   2E+3   5   5.5  RT (min)      lcqtof_hilic_neg_4_006_30    m/z 144.0605     0   1   2   3   4E+3   5.5   6  RT (min)      lcqtof_hilic_neg_4_050_02    m/z 144.9761     0   1   2   3E+3   6.5   7  RT (min)      lcqtof_hilic_neg_2_020_23    m/z 145.0791     0   1E+4   1   1.5  RT (min)      lcqtof_hilic_neg_4_008_24    m/z 145.9643     0   1   2   3   4E+3   1   1.5  RT (min)      lcqtof_hilic_neg_4_050_02    m/z 146.0118     0   2   4E+3   2   2.5  RT (min)       lcqtof_hilic_neg_4_079_58    m/z 146.9825     0   1   2   3E+3   6.5   7  RT (min)      lcqtof_hilic_neg_2_052_24    m/z 147.0286     0   1   2E+3   1   1.5  RT (min)      lcqtof_hilic_neg_4_048_23    m/z 147.9612     0   1   2E+3   1   1.5  RT (min)      lcqtof_hilic_neg_4_020_42    m/z 148.0703     0   1   2   3   4E+3   6   6.5  RT (min)      lcqtof_hilic_neg_4_008_24    m/z 148.9554     0   2   4   6E+3   1   1.5  RT (min)       lcqtof_hilic_neg_4_050_02    m/z 149.0450     0   1E+4   5.5   6  RT (min)      lcqtof_hilic_neg_1_027_27    m/z 149.0961     0   1   2   3   4E+3   0.5   1  RT (min)      lcqtof_hilic_neg_4_053_14    m/z 149.9122     0   1   2E+3   1   1.5  RT (min)      lcqtof_hilic_neg_4_038_10    m/z 149.9542     0   1   2E+3   0.5   1  RT (min)      lcqtof_hilic_neg_4_048_23    m/z 151.0617     0   2   4   6E+3   3   3.5  RT (min)      lcqtof_hilic_neg_4_008_24    m/z 151.0702     0   1   2E+4   1   1.5  RT (min)       lcqtof_hilic_neg_4_055_32    m/z 152.0279     0   1   2   3E+3   4   4.5  RT (min)      lcqtof_hilic_neg_4_032_01    m/z 152.0435     0   1   2   3E+3   1   1.5  RT (min)      lcqtof_hilic_neg_4_015_09    m/z 152.0608     0   2   4   6   8E+3   6   6.5  RT (min)      lcqtof_hilic_neg_1_037_24    m/z 152.0821     0   2   4E+3   6   6.5  RT (min)      lcqtof_hilic_neg_4_008_24    m/z 152.0997     0   1   2E+3   5   5.5  RT (min)       lcqtof_hilic_neg_4_013_20    m/z 152.9678     0   1   2   3E+3   0.5   1  RT (min)       lcqtof_hilic_neg_2_034_47    m/z 152.9851     0   2   4   6   8E+3   1   1.5  RT (min)      lcqtof_hilic_neg_4_053_14    m/z 152.9945     0   1E+4   6.5   7  RT (min)      lcqtof_hilic_neg_4_048_23    m/z 153.0201     0   1   2E+4   1   1.5  RT (min)      lcqtof_hilic_neg_4_022_34    m/z 153.0655     0   2   4   6E+3   1   1.5  RT (min)      lcqtof_hilic_neg_4_034_31    m/z 153.0921     0   1E+3   1   1.5  RT (min)      lcqtof_hilic_neg_4_018_27    m/z 153.8688     0   1   2E+3   7.5   8  RT (min)      lcqtof_hilic_neg_1_016_28    m/z 154.0605     0   1   2E+3   5   5.5  RT (min)      lcqtof_hilic_neg_4_031_38    m/z 154.0625     0   2   4   6E+3   0.5   1  RT (min)      lcqtof_hilic_neg_2_045_11    m/z 154.8939     0   1E+3   7.5   8  RT (min)      lcqtof_hilic_neg_2_028_05    m/z 154.9472     0   1   2   3   4E+3   2   2.5  RT (min)      lcqtof_hilic_neg_4_005_29    m/z 155.1431     0   1E+3   0.5   1  RT (min)      lcqtof_hilic_neg_4_034_31    m/z 155.9061     0   1   2   3   4E+3   0.5   1  RT (min)      lcqtof_hilic_neg_2_047_32    m/z 156.9592     0   1   2E+3   4   4.5  RT (min)      lcqtof_hilic_neg_1_022_36    m/z 156.9839     0   1E+3   6   6.5  RT (min)      lcqtof_hilic_neg_2_002_36    m/z 158.0453     0   2   4   6   8E+3   4   4.5  RT (min)      lcqtof_hilic_neg_4_040_05    m/z 158.0601     0   1E+3   1   1.5  RT (min)      lcqtof_hilic_neg_2_051_14    m/z 158.9658     0   1   2E+3   4   4.5  RT (min)      lcqtof_hilic_neg_1_029_06    m/z 159.0378     0   2   4E+3   5.5   6  RT (min)       lcqtof_hilic_neg_4_008_24    m/z 159.0764     0   1   2   3E+3   5.5   6  RT (min)      lcqtof_hilic_neg_1_014_31    m/z 159.0920     0   1   2   3E+3   5   5.5  RT (min)       lcqtof_hilic_neg_4_053_14    m/z 159.9180     0   1   2E+3   1.5   2  RT (min)      lcqtof_hilic_neg_2_045_11    m/z 159.9694     0   1   2   3E+3   4   4.5  RT (min)      lcqtof_hilic_neg_1_029_06    m/z 160.0243     0   1   2   3   4E+3   2   2.5  RT (min)      lcqtof_hilic_neg_4_082_59    m/z 160.0397     0   1E+3   4.5   5  RT (min)      lcqtof_hilic_neg_1_013_38    m/z 161.0187     0   2   4E+3   2   2.5  RT (min)       lcqtof_hilic_neg_4_088_59    m/z 161.0450     0   1   2E+3   6   6.5  RT (min)       lcqtof_hilic_neg_2_004_26    m/z 161.0453     0   2   4   6E+3   5.5   6  RT (min)       lcqtof_hilic_neg_4_028_36    m/z 161.0649     0   2   4E+3   6   6.5  RT (min)      lcqtof_hilic_neg_1_022_36    m/z 161.0806     0   2   4   6E+3   7   7.5  RT (min)      lcqtof_hilic_neg_4_008_24    m/z 162.0245     0   2   4   6E+3   0.5   1  RT (min)       lcqtof_hilic_neg_4_013_20    m/z 162.0419     0   2   4E+3   3.5   4  RT (min)      lcqtof_hilic_neg_4_048_23    m/z 162.9161     0   2   4E+3   1   1.5  RT (min)      lcqtof_hilic_neg_1_032_18    m/z 163.0753     0   2   4   6E+3   1   1.5  RT (min)      lcqtof_hilic_neg_4_053_14    m/z 163.8799     0   1E+3   1   1.5  RT (min)      lcqtof_hilic_neg_4_015_09    m/z 163.9136     0   1   2E+3   1.5   2  RT (min)      lcqtof_hilic_neg_2_045_11    m/z 164.0351     0   1   2   3   4E+3   0.5   1  RT (min)      lcqtof_hilic_neg_4_038_10    m/z 164.0647     0   2   4   6E+3   4   4.5  RT (min)      lcqtof_hilic_neg_4_040_05    m/z 164.0733     0   1E+4   1   1.5  RT (min)       lcqtof_hilic_neg_4_055_32    m/z 164.9764     0   1   2   3E+3   6   6.5  RT (min)      lcqtof_hilic_neg_2_020_23    m/z 164.9856     0   2   4   6E+3   0.5   1  RT (min)       lcqtof_hilic_neg_4_101_61    m/z 165.0980     0   1   2   3   4E+3   5.5   6  RT (min)      lcqtof_hilic_neg_1_005_32    m/z 166.0590     0   1   2   3   4E+3   1   1.5  RT (min)      lcqtof_hilic_neg_4_053_14    m/z 166.9914     0   1   2E+3   6   6.5  RT (min)      lcqtof_hilic_neg_2_020_23    m/z 167.0641     0   2   4E+3   5.5   6  RT (min)       lcqtof_hilic_neg_4_055_32    m/z 167.9749     0   1E+4   6   6.5  RT (min)      lcqtof_hilic_neg_3_030_23    m/z 168.1050     0   1   2   3   4E+3   1   1.5  RT (min)      lcqtof_hilic_neg_1_005_32    m/z 168.9623     0   1   2   3E+3   1   1.5  RT (min)       lcqtof_hilic_neg_4_089_59    m/z 169.0235     0   1E+3   5   5.5  RT (min)      lcqtof_hilic_neg_4_032_01    m/z 169.1245     0   1   2   3E+3   0.5   1  RT (min)      lcqtof_hilic_neg_4_053_14    m/z 169.9553     0   1   2   3E+3   6.5   7  RT (min)      lcqtof_hilic_neg_2_020_23    m/z 170.0439     0   1E+3   1   1.5  RT (min)      lcqtof_hilic_neg_4_104_61    m/z 170.9393     0   1   2E+3   2   2.5  RT (min)      lcqtof_hilic_neg_2_002_36    m/z 170.9757     0   1   2   3   4E+3   0.5   1  RT (min)      lcqtof_hilic_neg_4_004_47    m/z 173.0061     0   1E+3   8   8.5  RT (min)      lcqtof_hilic_neg_2_016_33    m/z 173.1080     0   1   2   3   4E+3   4   4.5  RT (min)      lcqtof_hilic_neg_2_036_34    m/z 173.9615     0   1   2E+3   1   1.5  RT (min)      lcqtof_hilic_neg_3_026_42    m/z 174.0848     0   1   2E+3   1   1.5  RT (min)      lcqtof_hilic_neg_4_039_18    m/z 175.0794     0   2   4   6   8E+3   5.5   6  RT (min)      lcqtof_hilic_neg_1_035_23    m/z 175.0905     0   1   2   3E+3   6.5   7  RT (min)      lcqtof_hilic_neg_1_013_38    m/z 176.0636     0   2   4   6E+3   1   1.5  RT (min)      lcqtof_hilic_neg_1_023_34    m/z 176.8484     0   1   2E+3   3   3.5  RT (min)      lcqtof_hilic_neg_3_043_13    m/z 176.8739     0   2   4   6E+3   1   1.5  RT (min)      lcqtof_hilic_neg_1_004_14    m/z 177.0922     0   2   4   6E+3   0.5   1  RT (min)      lcqtof_hilic_neg_2_015_06    m/z 177.9408     0   1   2E+3   6.5   7  RT (min)      lcqtof_hilic_neg_4_008_24    m/z 178.0052     0   1   2   3E+3   6   6.5  RT (min)      lcqtof_hilic_neg_4_053_14    m/z 178.0513     0   1   2E+3   0.5   1  RT (min)      lcqtof_hilic_neg_4_036_11    m/z 178.9482     0   1   2   3E+3   0.5   1  RT (min)      lcqtof_hilic_neg_4_053_14    m/z 179.0287     0   2   4   6E+3   6.5   7  RT (min)      lcqtof_hilic_neg_3_093_60    m/z 179.0319     0   1E+4   0.5   1  RT (min)      lcqtof_hilic_neg_4_050_02    m/z 179.1063     0   1   2E+3   6.5   7  RT (min)      lcqtof_hilic_neg_1_005_32    m/z 180.0519     0   1   2   3   4E+3   4   4.5  RT (min)      lcqtof_hilic_neg_4_047_06    m/z 180.0671     0   2   4E+3   0.5   1  RT (min)      lcqtof_hilic_neg_4_048_23    m/z 180.1031     0   1   2   3   4E+3   0.5   1  RT (min)      lcqtof_hilic_neg_4_022_34    m/z 180.1243     0   2   4   6   8E+3   1   1.5  RT (min)      lcqtof_hilic_neg_4_008_24    m/z 181.1272     0   1   2E+3   7   7.5  RT (min)      lcqtof_hilic_neg_4_008_24    m/z 181.9542     0   1E+4   1   1.5  RT (min)       lcqtof_hilic_neg_1_032_18    m/z 182.0316     0   2   4   6E+3   3.5   4  RT (min)      lcqtof_hilic_neg_2_031_01    m/z 182.9703     0   2   4E+3   4   4.5  RT (min)      lcqtof_hilic_neg_1_005_32    m/z 183.1397     0   1   2E+3   1   1.5  RT (min)      lcqtof_hilic_neg_4_053_14    m/z 183.9691     0   2   4   6E+3   6   6.5  RT (min)      lcqtof_hilic_neg_3_054_37    m/z 183.9823     0   1   2   3E+3   6   6.5  RT (min)      lcqtof_hilic_neg_1_035_23    m/z 184.0128     0   1   2   3E+4   4   4.5  RT (min)      lcqtof_hilic_neg_4_055_32    m/z 184.0137     0   1   2E+4   1   1.5  RT (min)      lcqtof_hilic_neg_4_047_06    m/z 184.0974     0   1   2E+4   5   5.5  RT (min)      lcqtof_hilic_neg_4_005_29    m/z 184.9556     0   1E+3   1   1.5  RT (min)      lcqtof_hilic_neg_1_014_31    m/z 184.9904     0   1E+4   0.5   1  RT (min)      lcqtof_hilic_neg_4_027_19    m/z 185.0104     0   1E+4   1   1.5  RT (min)      lcqtof_hilic_neg_4_040_05    m/z 185.1004     0   1   2   3E+3   4.5   5  RT (min)      lcqtof_hilic_neg_4_003_16    m/z 186.0726     0   2   4E+3   4.5   5  RT (min)      lcqtof_hilic_neg_1_005_32    m/z 186.1203     0   2   4   6E+3   1   1.5  RT (min)      lcqtof_hilic_neg_4_053_14    m/z 186.9794     0   1   2   3E+3   0.5   1  RT (min)       lcqtof_hilic_neg_4_053_14    m/z 187.0718     0   1   2   3E+3   7   7.5  RT (min)      lcqtof_hilic_neg_2_051_14    m/z 188.0097     0   1   2   3E+3   5.5   6  RT (min)      lcqtof_hilic_neg_4_053_14    m/z 188.0335     0   2   4   6   8E+3   2   2.5  RT (min)      lcqtof_hilic_neg_3_078_58    m/z 188.0414     0   1   2   3E+3   4   4.5  RT (min)      lcqtof_hilic_neg_2_044_37    m/z 188.0630     0   2   4   6E+3   6   6.5  RT (min)       lcqtof_hilic_neg_4_008_24    m/z 188.9864     0   2   4   6   8E+4   1   1.5  RT (min)      lcqtof_hilic_neg_4_022_34    m/z 189.0361     0   1   2   3   4E+3   0.5   1  RT (min)      lcqtof_hilic_neg_4_050_02    m/z 189.0888     0   1   2   3   4E+3   2   2.5  RT (min)      lcqtof_hilic_neg_1_005_32    m/z 189.0963     0   2   4   6   8E+3   5.5   6  RT (min)       lcqtof_hilic_neg_4_008_24    m/z 190.0506     0   1   2   3E+3   5   5.5  RT (min)       lcqtof_hilic_neg_4_050_02    m/z 190.0864     0   1   2   3E+3   3   3.5  RT (min)      lcqtof_hilic_neg_4_054_48    m/z 190.1277     0   2   4E+3   1.5   2  RT (min)      lcqtof_hilic_neg_1_005_32    m/z 191.0662     0   2   4   6E+3   7   7.5  RT (min)       lcqtof_hilic_neg_1_004_14    m/z 191.1056     0   1   2   3   4E+3   1   1.5  RT (min)      lcqtof_hilic_neg_4_032_01    m/z 191.8156     0   1   2   3E+3   4.5   5  RT (min)      lcqtof_hilic_neg_3_067_57    m/z 191.9512     0   1   2   3   4E+3   1   1.5  RT (min)      lcqtof_hilic_neg_4_052_13    m/z 192.0669     0   2   4   6   8E+3   5.5   6  RT (min)      lcqtof_hilic_neg_4_028_36    m/z 192.0805     0   2   4   6E+3   2   2.5  RT (min)      lcqtof_hilic_neg_4_055_32    m/z 192.9102     0   1E+3   0.5   1  RT (min)      lcqtof_hilic_neg_3_068_57    m/z 192.9799     0   2   4   6E+3   1   1.5  RT (min)      lcqtof_hilic_neg_4_053_14    m/z 193.0318     0   1   2E+3   1   1.5  RT (min)      lcqtof_hilic_neg_3_044_09    m/z 193.0502     0   1   2   3   4E+3   1   1.5  RT (min)       lcqtof_hilic_neg_4_053_14    m/z 193.1586     0   1   2E+3   0.5   1  RT (min)      lcqtof_hilic_neg_2_015_06    m/z 194.1006     0   1   2   3E+3   1.5   2  RT (min)      lcqtof_hilic_neg_4_048_23    m/z 194.8549     0   1   2   3   4E+3   0.5   1  RT (min)      lcqtof_hilic_neg_4_084_59    m/z 194.9945     0   2   4E+3   2.5   3  RT (min)       lcqtof_hilic_neg_4_056_56    m/z 194.9964     0   1   2   3E+3   3.5   4  RT (min)      lcqtof_hilic_neg_3_036_17    m/z 195.0382     0   1   2E+3   5.5   6  RT (min)      lcqtof_hilic_neg_4_023_15    m/z 195.0598     0   2   4   6E+3   3   3.5  RT (min)      lcqtof_hilic_neg_2_024_10    m/z 196.0520     0   2   4   6E+3   1.5   2  RT (min)      lcqtof_hilic_neg_4_008_24    m/z 196.1055     0   1   2E+3   1   1.5  RT (min)      lcqtof_hilic_neg_4_018_27    m/z 197.0294     0   1   2   3   4E+3   5   5.5  RT (min)      lcqtof_hilic_neg_4_010_28    m/z 198.0315     0   1   2E+3   1   1.5  RT (min)      lcqtof_hilic_neg_4_042_22    m/z 198.0649     0   1   2E+3   0.5   1  RT (min)      lcqtof_hilic_neg_4_013_20    m/z 198.0687     0   1   2   3E+3   4   4.5  RT (min)      lcqtof_hilic_neg_3_030_23    m/z 198.1124     0   1   2E+3   1   1.5  RT (min)      lcqtof_hilic_neg_4_053_14    m/z 198.9563     0   1   2E+3   6   6.5  RT (min)      lcqtof_hilic_neg_1_013_38    m/z 198.9836     0   1   2E+3   6   6.5  RT (min)      lcqtof_hilic_neg_2_047_32    m/z 199.0746     0   1   2   3E+3   0.5   1  RT (min)      lcqtof_hilic_neg_4_052_13    m/z 200.0380     0   1   2   3E+3   5.5   6  RT (min)      lcqtof_hilic_neg_4_052_13    m/z 200.0422     0   2   4   6   8E+3   2   2.5  RT (min)      lcqtof_hilic_neg_4_018_27    m/z 200.0616     0   1   2E+3   1   1.5  RT (min)      lcqtof_hilic_neg_2_016_33    m/z 200.9755     0   1   2E+3   1   1.5  RT (min)      lcqtof_hilic_neg_2_044_37    m/z 200.9969     0   1   2   3E+3   6   6.5  RT (min)      lcqtof_hilic_neg_1_035_23    m/z 201.0883     0   1   2   3E+3   7   7.5  RT (min)      lcqtof_hilic_neg_2_051_14    m/z 201.9817     0   1   2E+3   6   6.5  RT (min)      lcqtof_hilic_neg_1_045_07    m/z 202.0256     0   1   2   3   4E+3   0.5   1  RT (min)      lcqtof_hilic_neg_4_053_14    m/z 202.0526     0   1   2E+3   6   6.5  RT (min)      lcqtof_hilic_neg_2_020_23    m/z 202.1091     0   1   2   3   4E+3   1   1.5  RT (min)      lcqtof_hilic_neg_4_050_02    m/z 203.0000     0   2   4   6E+4   1   1.5  RT (min)      lcqtof_hilic_neg_4_053_14    m/z 203.0678     0   1   2   3   4E+3   6   6.5  RT (min)      lcqtof_hilic_neg_2_002_36    m/z 203.0686     0   1   2   3E+3   0.5   1  RT (min)      lcqtof_hilic_neg_4_056_56    m/z 203.1311     0   1   2   3   4E+3   1.5   2  RT (min)      lcqtof_hilic_neg_4_055_32    m/z 203.9566     0   1E+3   6   6.5  RT (min)       lcqtof_hilic_neg_4_014_46    m/z 204.0867     0   1   2E+3   4   4.5  RT (min)      lcqtof_hilic_neg_3_034_24    m/z 204.7912     0   1   2   3   4E+3   4.5   5  RT (min)      lcqtof_hilic_neg_4_026_04    m/z 204.9674     0   1E+3   1   1.5  RT (min)      lcqtof_hilic_neg_2_029_31    m/z 205.0870     0   1   2   3E+3   1   1.5  RT (min)      lcqtof_hilic_neg_1_027_27    m/z 205.1227     0   2   4   6   8E+3   0.5   1  RT (min)      lcqtof_hilic_neg_2_021_17    m/z 205.9495     0   1E+3   0.5   1  RT (min)      lcqtof_hilic_neg_1_054_01    m/z 206.0019     0   1   2   3   4E+3   5   5.5  RT (min)      lcqtof_hilic_neg_4_053_14    m/z 206.0146     0   1   2   3   4E+3   2   2.5  RT (min)      lcqtof_hilic_neg_4_108_62    m/z 206.1542     0   1E+4   1   1.5  RT (min)      lcqtof_hilic_neg_1_029_06    m/z 207.1062     0   1   2   3   4E+3   2.5   3  RT (min)      lcqtof_hilic_neg_4_002_25    m/z 207.1386     0   1   2   3E+3   0.5   1  RT (min)      lcqtof_hilic_neg_4_027_19    m/z 207.9938     0   1   2   3   4E+3   2   2.5  RT (min)      lcqtof_hilic_neg_4_075_58    m/z 208.0610     0   2   4E+4   0.5   1  RT (min)      lcqtof_hilic_neg_4_052_13    m/z 208.1146     0   1   2   3   4E+3   1   1.5  RT (min)      lcqtof_hilic_neg_4_048_23    m/z 208.9376     0   1   2E+3   0.5   1  RT (min)      lcqtof_hilic_neg_4_050_02    m/z 208.9840     0   1E+3   5.5   6  RT (min)      lcqtof_hilic_neg_1_025_52    m/z 209.0009     0   2   4E+3   0.5   1  RT (min)      lcqtof_hilic_neg_2_040_20    m/z 209.0460     0   1   2   3E+3   1   1.5  RT (min)      lcqtof_hilic_neg_4_034_31    m/z 209.0743     0   1   2   3   4E+3   2   2.5  RT (min)      lcqtof_hilic_neg_4_026_04    m/z 209.1149     0   1   2   3E+3   3   3.5  RT (min)      lcqtof_hilic_neg_1_005_32    m/z 210.0988     0   1   2   3E+3   3   3.5  RT (min)      lcqtof_hilic_neg_4_068_57    m/z 210.1026     0   1   2   3   4E+3   0.5   1  RT (min)      lcqtof_hilic_neg_4_042_22    m/z 210.9436     0   1   2   3E+3   0.5   1  RT (min)      lcqtof_hilic_neg_4_050_02    m/z 211.0014     0   1   2   3E+3   7   7.5  RT (min)       lcqtof_hilic_neg_1_007_05    m/z 212.9341     0   2   4E+3   0.5   1  RT (min)       lcqtof_hilic_neg_4_004_47    m/z 212.9454     0   1   2E+3   0.5   1  RT (min)      lcqtof_hilic_neg_4_053_14    m/z 213.0067     0   1E+4   1   1.5  RT (min)      lcqtof_hilic_neg_2_051_14    m/z 213.1258     0   1   2   3E+3   5   5.5  RT (min)      lcqtof_hilic_neg_4_028_36    m/z 213.8782     0   1   2   3   4E+3   6   6.5  RT (min)      lcqtof_hilic_neg_1_001_49    m/z 213.9241     0   1   2E+3   4.5   5  RT (min)      lcqtof_hilic_neg_4_053_14    m/z 213.9635     0   1   2   3   4E+3   1   1.5  RT (min)      lcqtof_hilic_neg_4_008_24    m/z 214.0327     0   1   2   3E+3   6   6.5  RT (min)      lcqtof_hilic_neg_4_082_59    m/z 214.0332     0   2   4   6E+3   5   5.5  RT (min)      lcqtof_hilic_neg_4_048_23    m/z 214.0581     0   1   2E+3   1   1.5  RT (min)      lcqtof_hilic_neg_4_051_43    m/z 214.9837     0   1E+3   6   6.5  RT (min)      lcqtof_hilic_neg_4_001_49    m/z 214.9943     0   2   4   6E+3   6   6.5  RT (min)      lcqtof_hilic_neg_3_034_24    m/z 215.0523     0   1   2E+3   4.5   5  RT (min)      lcqtof_hilic_neg_4_053_14    m/z 215.1213     0   1   2   3E+3   4.5   5  RT (min)      lcqtof_hilic_neg_4_005_29    m/z 215.9483     0   1   2E+3   6   6.5  RT (min)      lcqtof_hilic_neg_4_016_37    m/z 215.9548     0   1   2E+3   4   4.5  RT (min)      lcqtof_hilic_neg_1_006_22    m/z 216.0551     0   2   4   6E+3   0.5   1  RT (min)      lcqtof_hilic_neg_4_055_32    m/z 216.1681     0   1   2E+3   1   1.5  RT (min)      lcqtof_hilic_neg_4_053_14    m/z 216.7794     0   1   2E+3   8.5   9  RT (min)       lcqtof_hilic_neg_4_003_16    m/z 216.8543     0   1   2E+3   8   8.5  RT (min)      lcqtof_hilic_neg_3_055_01    m/z 216.9294     0   1   2E+3   0.5   1  RT (min)      lcqtof_hilic_neg_4_099_61    m/z 217.0044     0   1   2   3E+3   1   1.5  RT (min)      lcqtof_hilic_neg_4_056_56    m/z 217.0143     0   2   4E+3   1   1.5  RT (min)      lcqtof_hilic_neg_4_050_02    m/z 217.0166     0   2   4   6   8E+3   1   1.5  RT (min)      lcqtof_hilic_neg_4_022_34    m/z 217.0815     0   1E+3   4.5   5  RT (min)      lcqtof_hilic_neg_4_016_37    m/z 217.1085     0   1   2   3   4E+3   1.5   2  RT (min)       lcqtof_hilic_neg_2_053_18    m/z 217.9408     0   1E+3   2   2.5  RT (min)      lcqtof_hilic_neg_1_045_07    m/z 217.9770     0   2   4   6E+3   1   1.5  RT (min)      lcqtof_hilic_neg_4_052_13    m/z 217.9915     0   1E+3   5.5   6  RT (min)      lcqtof_hilic_neg_2_026_28    m/z 218.0352     0   1   2   3   4E+3   3   3.5  RT (min)      lcqtof_hilic_neg_1_004_14    m/z 218.0673     0   2   4E+3   1.5   2  RT (min)      lcqtof_hilic_neg_4_003_16    m/z 218.0697     0   1   2E+3   3   3.5  RT (min)      lcqtof_hilic_neg_4_032_01    m/z 218.0770     0   1   2   3E+3   7   7.5  RT (min)      lcqtof_hilic_neg_4_023_15    m/z 218.1011     0   2   4   6E+3   4   4.5  RT (min)      lcqtof_hilic_neg_2_036_34    m/z 218.1022     0   2   4E+3   4   4.5  RT (min)      lcqtof_hilic_neg_1_023_34    m/z 219.9475     0   1   2   3   4E+3   1   1.5  RT (min)      lcqtof_hilic_neg_4_023_15    m/z 219.9752     0   1   2   3   4E+3   1   1.5  RT (min)      lcqtof_hilic_neg_4_003_16    m/z 219.9997     0   1   2   3E+3   4.5   5  RT (min)      lcqtof_hilic_neg_2_053_18    m/z 221.0249     0   1   2   3E+3   2   2.5  RT (min)       lcqtof_hilic_neg_4_080_58    m/z 221.0555     0   1   2   3E+3   1   1.5  RT (min)      lcqtof_hilic_neg_2_032_02    m/z 221.0756     0   2   4   6   8E+3   3   3.5  RT (min)      lcqtof_hilic_neg_4_004_47    m/z 221.9450     0   1   2   3   4E+3   0.5   1  RT (min)      lcqtof_hilic_neg_4_053_14    m/z 221.9627     0   1   2   3   4E+3   1   1.5  RT (min)      lcqtof_hilic_neg_4_039_18    m/z 222.0790     0   1   2E+3   1.5   2  RT (min)      lcqtof_hilic_neg_1_007_05    m/z 222.0797     0   2   4   6   8E+3   0.5   1  RT (min)      lcqtof_hilic_neg_4_052_13    m/z 222.0842     0   1   2   3   4E+3   1   1.5  RT (min)      lcqtof_hilic_neg_4_032_01    m/z 222.1148     0   1E+4   0.5   1  RT (min)      lcqtof_hilic_neg_4_045_26    m/z 222.9016     0   1E+3   0.5   1  RT (min)      lcqtof_hilic_neg_1_046_30    m/z 223.0964     0   1   2E+3   1   1.5  RT (min)       lcqtof_hilic_neg_1_004_14    m/z 223.1678     0   2   4   6E+3   0.5   1  RT (min)      lcqtof_hilic_neg_4_032_01    m/z 223.9419     0   1   2E+3   0.5   1  RT (min)      lcqtof_hilic_neg_4_053_14    m/z 224.1370     0   1   2   3E+3   1   1.5  RT (min)      lcqtof_hilic_neg_4_053_14    m/z 224.1653     0   2   4   6   8E+3   1   1.5  RT (min)      lcqtof_hilic_neg_4_026_04    m/z 224.9799     0   1   2E+3   6   6.5  RT (min)      lcqtof_hilic_neg_4_016_37    m/z 225.0623     0   2   4   6E+3   5.5   6  RT (min)      lcqtof_hilic_neg_4_008_24    m/z 225.0858     0   1   2E+3   7   7.5  RT (min)      lcqtof_hilic_neg_3_079_58    m/z 225.1491     0   2   4E+3   0.5   1  RT (min)      lcqtof_hilic_neg_2_015_06    m/z 225.2220     0   2   4   6   8E+3   1   1.5  RT (min)      lcqtof_hilic_neg_4_053_14    m/z 225.2231     0   2   4   6   8E+3   0.5   1  RT (min)      lcqtof_hilic_neg_4_053_14    m/z 225.5624     0   1E+3   3   3.5  RT (min)      lcqtof_hilic_neg_3_055_01    m/z 226.1199     0   1   2E+3   6   6.5  RT (min)      lcqtof_hilic_neg_2_044_37    m/z 226.1454     0   1   2   3   4E+3   1   1.5  RT (min)      lcqtof_hilic_neg_4_020_42    m/z 227.0603     0   1   2   3   4E+3   0.5   1  RT (min)      lcqtof_hilic_neg_4_034_31    m/z 227.0798     0   2   4   6E+3   7.5   8  RT (min)      lcqtof_hilic_neg_4_008_24    m/z 227.1025     0   1   2   3E+3   6   6.5  RT (min)      lcqtof_hilic_neg_4_052_13    m/z 227.1654     0   1   2   3   4E+3   0.5   1  RT (min)      lcqtof_hilic_neg_1_029_06    m/z 228.0307     0   1   2   3E+3   2   2.5  RT (min)       lcqtof_hilic_neg_4_052_13    m/z 228.9405     0   1   2E+3   4   4.5  RT (min)      lcqtof_hilic_neg_1_004_14    m/z 228.9834     0   1   2E+3   6   6.5  RT (min)      lcqtof_hilic_neg_2_002_36    m/z 229.0479     0   2   4   6E+3   3.5   4  RT (min)      lcqtof_hilic_neg_1_019_02    m/z 229.0518     0   1   2   3E+3   5   5.5  RT (min)      lcqtof_hilic_neg_4_048_23    m/z 229.1411     0   2   4   6E+3   0.5   1  RT (min)      lcqtof_hilic_neg_4_050_02    m/z 230.0135     0   1E+5   0.5   1  RT (min)      lcqtof_hilic_neg_4_055_32    m/z 230.0475     0   1   2E+3   4.5   5  RT (min)      lcqtof_hilic_neg_2_046_41    m/z 230.8683     0   1   2   3   4E+3   1   1.5  RT (min)      lcqtof_hilic_neg_1_021_42    m/z 230.9281     0   1   2E+3   0.5   1  RT (min)      lcqtof_hilic_neg_4_053_14    m/z 231.0633     0   1   2E+3   7   7.5  RT (min)       lcqtof_hilic_neg_2_005_13    m/z 231.1372     0   1   2E+3   5   5.5  RT (min)      lcqtof_hilic_neg_1_004_14    m/z 231.9776     0   1   2   3E+3   5.5   6  RT (min)      lcqtof_hilic_neg_1_005_32    m/z 232.0001     0   1   2E+3   0.5   1  RT (min)      lcqtof_hilic_neg_4_035_33    m/z 232.0448     0   1   2E+3   6.5   7  RT (min)      lcqtof_hilic_neg_4_052_13    m/z 232.1015     0   2   4   6E+3   1.5   2  RT (min)      lcqtof_hilic_neg_4_055_32    m/z 232.1190     0   1   2   3E+3   1.5   2  RT (min)      lcqtof_hilic_neg_4_034_31    m/z 232.9562     0   1   2   3E+3   6.5   7  RT (min)      lcqtof_hilic_neg_2_020_23    m/z 233.0104     0   1   2E+3   2   2.5  RT (min)       lcqtof_hilic_neg_3_110_62    m/z 233.0931     0   1   2E+3   4   4.5  RT (min)      lcqtof_hilic_neg_2_036_34    m/z 233.9760     0   1   2   3E+3   4.5   5  RT (min)      lcqtof_hilic_neg_2_004_26    m/z 234.0060     0   1   2E+3   4   4.5  RT (min)      lcqtof_hilic_neg_1_050_12    m/z 234.0625     0   1   2   3   4E+3   1.5   2  RT (min)      lcqtof_hilic_neg_1_053_19    m/z 234.0627     0   1   2   3   4E+3   3.5   4  RT (min)      lcqtof_hilic_neg_4_048_23    m/z 234.1133     0   2   4E+3   1   1.5  RT (min)      lcqtof_hilic_neg_4_048_23    m/z 234.8206     0   1   2E+3   4   4.5  RT (min)      lcqtof_hilic_neg_2_018_16    m/z 235.0929     0   2   4E+3   0.5   1  RT (min)      lcqtof_hilic_neg_4_050_02    m/z 235.1372     0   2   4E+3   1   1.5  RT (min)       lcqtof_hilic_neg_4_032_01    m/z 235.9768     0   1E+3   7   7.5  RT (min)      lcqtof_hilic_neg_2_013_29    m/z 236.1024     0   1   2   3E+3   5   5.5  RT (min)       lcqtof_hilic_neg_4_013_20    m/z 236.8848     0   1E+3   7.5   8  RT (min)      lcqtof_hilic_neg_1_005_32    m/z 236.9685     0   1   2E+3   0.5   1  RT (min)      lcqtof_hilic_neg_4_007_45    m/z 237.0764     0   1   2E+3   0.5   1  RT (min)       lcqtof_hilic_neg_4_022_34    m/z 238.0033     0   1E+3   6   6.5  RT (min)      lcqtof_hilic_neg_1_027_27    m/z 238.1216     0   1   2   3E+3   0.5   1  RT (min)      lcqtof_hilic_neg_4_048_23    m/z 238.1401     0   1   2   3E+3   0.5   1  RT (min)      lcqtof_hilic_neg_4_050_02    m/z 238.9971     0   1   2E+3   4   4.5  RT (min)      lcqtof_hilic_neg_2_015_06    m/z 239.0012     0   2   4   6E+3   0.5   1  RT (min)       lcqtof_hilic_neg_4_053_14    m/z 239.0036     0   2   4E+3   2   2.5  RT (min)      lcqtof_hilic_neg_4_018_27    m/z 239.8951     0   1E+3   8   8.5  RT (min)      lcqtof_hilic_neg_2_019_12    m/z 239.9031     0   1   2E+3   1   1.5  RT (min)      lcqtof_hilic_neg_4_038_10    m/z 239.9113     0   1E+3   2   2.5  RT (min)      lcqtof_hilic_neg_3_007_04    m/z 240.9000     0   2   4E+3   1   1.5  RT (min)      lcqtof_hilic_neg_4_038_10    m/z 240.9765     0   1   2E+3   1   1.5  RT (min)      lcqtof_hilic_neg_4_081_58    m/z 241.1339     0   1E+4   0.5   1  RT (min)      lcqtof_hilic_neg_4_018_27    m/z 241.2175     0   1   2E+3   1   1.5  RT (min)      lcqtof_hilic_neg_4_020_42    m/z 241.7577     0   1   2   3E+3   4.5   5  RT (min)      lcqtof_hilic_neg_4_007_45    m/z 241.9157     0   1   2   3E+3   0.5   1  RT (min)      lcqtof_hilic_neg_4_028_36    m/z 241.9461     0   2   4E+3   4   4.5  RT (min)       lcqtof_hilic_neg_1_003_37    m/z 242.1513     0   1E+3   7   7.5  RT (min)      lcqtof_hilic_neg_4_016_37    m/z 242.1532     0   2   4   6   8E+3   1.5   2  RT (min)       lcqtof_hilic_neg_4_055_32    m/z 243.0173     0   1E+3   5   5.5  RT (min)      lcqtof_hilic_neg_3_043_13    m/z 243.0601     0   1   2E+3   6   6.5  RT (min)       lcqtof_hilic_neg_4_052_13    m/z 243.1645     0   1E+4   1   1.5  RT (min)      lcqtof_hilic_neg_2_016_33    m/z 244.0753     0   2   4   6E+3   2   2.5  RT (min)      lcqtof_hilic_neg_4_056_56    m/z 244.0842     0   2   4E+3   6   6.5  RT (min)      lcqtof_hilic_neg_4_050_02    m/z 244.1299     0   1   2   3E+3   6   6.5  RT (min)      lcqtof_hilic_neg_3_030_23    m/z 244.9612     0   2   4   6E+3   0.5   1  RT (min)       lcqtof_hilic_neg_4_004_47    m/z 244.9978     0   1   2   3E+3   6.5   7  RT (min)      lcqtof_hilic_neg_1_027_27    m/z 245.0198     0   2   4   6   8E+3   4.5   5  RT (min)      lcqtof_hilic_neg_4_104_61    m/z 245.0829     0   2   4   6E+3   0.5   1  RT (min)      lcqtof_hilic_neg_4_055_32    m/z 245.9841     0   2   4   6E+3   4   4.5  RT (min)      lcqtof_hilic_neg_1_022_36    m/z 246.0067     0   2   4   6   8E+3   1   1.5  RT (min)      lcqtof_hilic_neg_4_035_33    m/z 246.0305     0   1   2   3E+3   3   3.5  RT (min)      lcqtof_hilic_neg_3_054_37    m/z 246.0470     0   1E+3   5.5   6  RT (min)      lcqtof_hilic_neg_1_013_38    m/z 246.0999     0   2   4   6E+3   2   2.5  RT (min)       lcqtof_hilic_neg_4_008_24    m/z 246.5744     0   1E+3   3.5   4  RT (min)      lcqtof_hilic_neg_1_004_14    m/z 247.0334     0   1E+4   2   2.5  RT (min)       lcqtof_hilic_neg_4_083_59    m/z 247.0625     0   2   4E+3   0.5   1  RT (min)      lcqtof_hilic_neg_4_031_38    m/z 247.1361     0   2   4   6   8E+3   0.5   1  RT (min)      lcqtof_hilic_neg_4_027_19    m/z 247.1703     0   1   2E+3   0.5   1  RT (min)      lcqtof_hilic_neg_3_025_52    m/z 247.8094     0   2   4E+3   0.5   1  RT (min)      lcqtof_hilic_neg_4_022_34    m/z 247.8939     0   1   2E+3   1   1.5  RT (min)      lcqtof_hilic_neg_4_012_07    m/z 247.9959     0   1   2E+3   1   1.5  RT (min)      lcqtof_hilic_neg_4_053_14    m/z 248.0807     0   2   4E+3   2   2.5  RT (min)      lcqtof_hilic_neg_4_004_47    m/z 248.0863     0   1   2E+4   1   1.5  RT (min)       lcqtof_hilic_neg_4_050_02    m/z 248.9495     0   2   4   6E+3   1   1.5  RT (min)      lcqtof_hilic_neg_2_053_18    m/z 248.9908     0   2   4   6E+3   4   4.5  RT (min)      lcqtof_hilic_neg_1_022_36    m/z 249.0308     0   1   2E+3   4.5   5  RT (min)      lcqtof_hilic_neg_4_032_01    m/z 249.0623     0   1E+3   1.5   2  RT (min)      lcqtof_hilic_neg_2_047_32    m/z 249.0991     0   1   2   3E+3   3.5   4  RT (min)      lcqtof_hilic_neg_4_008_24    m/z 249.1246     0   1   2   3E+3   4   4.5  RT (min)       lcqtof_hilic_neg_4_102_61    m/z 249.9533     0   1E+3   1   1.5  RT (min)      lcqtof_hilic_neg_2_053_18    m/z 249.9550     0   1   2   3E+3   4   4.5  RT (min)      lcqtof_hilic_neg_2_047_32    m/z 250.0313     0   1   2E+3   6   6.5  RT (min)       lcqtof_hilic_neg_2_020_23    m/z 250.0321     0   1   2   3E+3   4.5   5  RT (min)      lcqtof_hilic_neg_4_053_14    m/z 250.0547     0   2   4E+3   5   5.5  RT (min)      lcqtof_hilic_neg_4_008_24    m/z 250.8486     0   1   2E+3   7.5   8  RT (min)      lcqtof_hilic_neg_2_028_05    m/z 250.9999     0   2   4   6   8E+3   4   4.5  RT (min)      lcqtof_hilic_neg_1_005_32    m/z 251.0766     0   2   4   6   8E+3   5   5.5  RT (min)       lcqtof_hilic_neg_1_011_46    m/z 251.1291     0   1   2   3   4E+3   1   1.5  RT (min)       lcqtof_hilic_neg_2_016_33    m/z 251.1957     0   2   4   6   8E+3   0.5   1  RT (min)      lcqtof_hilic_neg_4_008_24    m/z 251.9919     0   1E+3   7   7.5  RT (min)      lcqtof_hilic_neg_2_004_26    m/z 252.0917     0   2   4E+3   5.5   6  RT (min)      lcqtof_hilic_neg_1_005_32    m/z 252.0928     0   2   4   6   8E+3   3.5   4  RT (min)       lcqtof_hilic_neg_4_008_24    m/z 253.0480     0   1   2   3   4E+3   1   1.5  RT (min)      lcqtof_hilic_neg_4_023_15    m/z 253.0562     0   1   2   3   4E+3   3.5   4  RT (min)       lcqtof_hilic_neg_1_037_24    m/z 253.0719     0   1   2E+3   5.5   6  RT (min)      lcqtof_hilic_neg_4_013_20    m/z 253.1012     0   2   4E+3   1   1.5  RT (min)      lcqtof_hilic_neg_4_053_14    m/z 253.1124     0   2   4   6E+3   1.5   2  RT (min)      lcqtof_hilic_neg_4_008_24    m/z 253.2127     0   2   4   6E+3   1   1.5  RT (min)      lcqtof_hilic_neg_4_008_24    m/z 253.2178     0   1   2   3E+3   4   4.5  RT (min)      lcqtof_hilic_neg_4_052_13    m/z 253.9021     0   1E+3   7.5   8  RT (min)      lcqtof_hilic_neg_2_002_36    m/z 254.0948     0   2   4   6E+3   3   3.5  RT (min)      lcqtof_hilic_neg_4_004_47    m/z 254.1486     0   1   2   3   4E+3   0.5   1  RT (min)      lcqtof_hilic_neg_4_048_23    m/z 254.9729     0   1   2   3E+3   4   4.5  RT (min)      lcqtof_hilic_neg_1_045_07    m/z 255.0788     0   1   2E+3   5.5   6  RT (min)      lcqtof_hilic_neg_4_012_07    m/z 255.1062     0   2   4E+3   1   1.5  RT (min)      lcqtof_hilic_neg_4_047_06    m/z 256.1914     0   1E+4   0.5   1  RT (min)      lcqtof_hilic_neg_4_026_04    m/z 256.6299     0   1   2   3E+3   2   2.5  RT (min)      lcqtof_hilic_neg_2_031_01    m/z 256.9601     0   1   2   3   4E+3   0.5   1  RT (min)       lcqtof_hilic_neg_4_077_58    m/z 257.0056     0   1   2   3E+3   6.5   7  RT (min)       lcqtof_hilic_neg_4_048_23    m/z 257.0108     0   1E+3   2   2.5  RT (min)      lcqtof_hilic_neg_1_035_23    m/z 257.0431     0   2   4E+3   4.5   5  RT (min)      lcqtof_hilic_neg_1_035_23    m/z 257.0773     0   1   2E+3   1   1.5  RT (min)      lcqtof_hilic_neg_2_005_13    m/z 257.0775     0   1   2E+3   2   2.5  RT (min)      lcqtof_hilic_neg_1_022_36    m/z 257.0785     0   1   2   3   4E+3   2   2.5  RT (min)      lcqtof_hilic_neg_4_028_36    m/z 257.9923     0   1   2E+3   5   5.5  RT (min)      lcqtof_hilic_neg_4_032_01    m/z 257.9950     0   1   2E+3   6   6.5  RT (min)      lcqtof_hilic_neg_1_003_37    m/z 258.0746     0   1   2   3E+3   6   6.5  RT (min)       lcqtof_hilic_neg_4_053_14    m/z 258.0825     0   2   4E+3   2.5   3  RT (min)      lcqtof_hilic_neg_2_048_35    m/z 258.0829     0   1E+3   7   7.5  RT (min)       lcqtof_hilic_neg_2_005_13    m/z 258.1811     0   1   2E+3   1   1.5  RT (min)      lcqtof_hilic_neg_4_035_33    m/z 259.1195     0   1   2E+3   6.5   7  RT (min)      lcqtof_hilic_neg_1_035_23    m/z 259.9743     0   2   4E+3   0.5   1  RT (min)      lcqtof_hilic_neg_4_004_47    m/z 259.9851     0   1E+3   5   5.5  RT (min)      lcqtof_hilic_neg_1_031_16    m/z 260.0229     0   1E+3   5.5   6  RT (min)      lcqtof_hilic_neg_1_015_33    m/z 260.0556     0   1   2   3E+3   5   5.5  RT (min)       lcqtof_hilic_neg_4_022_34    m/z 260.0871     0   1   2   3E+3   6.5   7  RT (min)      lcqtof_hilic_neg_4_011_35    m/z 261.0303     0   1E+4   2   2.5  RT (min)       lcqtof_hilic_neg_2_045_11    m/z 261.0673     0   1   2   3E+3   1   1.5  RT (min)      lcqtof_hilic_neg_4_036_11    m/z 261.0941     0   2   4E+3   6   6.5  RT (min)      lcqtof_hilic_neg_4_032_01    m/z 261.9883     0   1E+3   6   6.5  RT (min)      lcqtof_hilic_neg_2_047_32    m/z 262.1390     0   2   4E+3   1   1.5  RT (min)      lcqtof_hilic_neg_4_055_32    m/z 262.8152     0   1   2   3E+3   7   7.5  RT (min)      lcqtof_hilic_neg_3_023_08    m/z 262.8398     0   1   2E+3   0.5   1  RT (min)      lcqtof_hilic_neg_4_052_13    m/z 263.0038     0   1   2   3E+3   6.5   7  RT (min)      lcqtof_hilic_neg_1_053_19    m/z 263.0365     0   1   2E+3   5   5.5  RT (min)      lcqtof_hilic_neg_4_052_13    m/z 263.0933     0   1   2   3   4E+4   2   2.5  RT (min)      lcqtof_hilic_neg_1_022_36    m/z 263.1441     0   1   2E+3   2.5   3  RT (min)      lcqtof_hilic_neg_1_005_32    m/z 263.9303     0   2   4   6E+3   8   8.5  RT (min)      lcqtof_hilic_neg_4_053_14    m/z 263.9882     0   1E+3   3.5   4  RT (min)      lcqtof_hilic_neg_1_004_14    m/z 264.0166     0   1E+3   7   7.5  RT (min)      lcqtof_hilic_neg_1_005_32    m/z 264.1286     0   2   4E+3   0.5   1  RT (min)      lcqtof_hilic_neg_4_050_02    m/z 264.8323     0   1   2   3E+3   0.5   1  RT (min)      lcqtof_hilic_neg_4_050_02    m/z 265.0502     0   2   4   6E+3   1   1.5  RT (min)       lcqtof_hilic_neg_3_044_09    m/z 265.0528     0   2   4E+3   0.5   1  RT (min)       lcqtof_hilic_neg_4_053_14    m/z 265.0857     0   1   2   3E+3   6.5   7  RT (min)      lcqtof_hilic_neg_1_045_07    m/z 265.1084     0   1   2   3E+3   2.5   3  RT (min)       lcqtof_hilic_neg_4_029_17    m/z 265.9453     0   1   2   3E+3   1   1.5  RT (min)      lcqtof_hilic_neg_2_051_14    m/z 266.0766     0   1   2   3   4E+3   7   7.5  RT (min)       lcqtof_hilic_neg_4_052_13    m/z 266.0877     0   2   4   6   8E+3   6.5   7  RT (min)      lcqtof_hilic_neg_4_048_23    m/z 266.1191     0   1   2   3   4E+3   4   4.5  RT (min)      lcqtof_hilic_neg_4_055_32    m/z 266.8357     0   1   2E+3   0.5   1  RT (min)      lcqtof_hilic_neg_4_034_31    m/z 266.9630     0   2   4   6E+3   0.5   1  RT (min)      lcqtof_hilic_neg_4_053_14    m/z 267.0736     0   1   2   3E+3   3.5   4  RT (min)      lcqtof_hilic_neg_4_048_23    m/z 267.1446     0   2   4E+3   4   4.5  RT (min)      lcqtof_hilic_neg_4_095_60    m/z 267.1468     0   2   4E+3   6   6.5  RT (min)       lcqtof_hilic_neg_4_053_14    m/z 267.2325     0   1   2   3   4E+3   0.5   1  RT (min)      lcqtof_hilic_neg_2_016_33    m/z 267.9616     0   1   2   3E+3   1   1.5  RT (min)      lcqtof_hilic_neg_2_053_18    m/z 268.0689     0   1   2   3   4E+3   0.5   1  RT (min)      lcqtof_hilic_neg_4_047_06    m/z 268.0864     0   1E+4   5   5.5  RT (min)      lcqtof_hilic_neg_4_028_36    m/z 268.1219     0   1   2E+4   0.5   1  RT (min)      lcqtof_hilic_neg_4_050_02    m/z 268.1505     0   1   2E+3   7.5   8  RT (min)      lcqtof_hilic_neg_2_052_24    m/z 268.8007     0   1   2   3E+3   4   4.5  RT (min)       lcqtof_hilic_neg_4_103_61    m/z 268.8010     0   1   2   3E+3   4   4.5  RT (min)      lcqtof_hilic_neg_4_042_22    m/z 269.0113     0   1   2E+3   0.5   1  RT (min)      lcqtof_hilic_neg_4_016_37    m/z 269.0409     0   1   2   3   4E+3   3.5   4  RT (min)      lcqtof_hilic_neg_3_032_19    m/z 269.0752     0   1E+3   7   7.5  RT (min)      lcqtof_hilic_neg_2_043_15    m/z 269.1190     0   2   4   6   8E+3   6.5   7  RT (min)      lcqtof_hilic_neg_4_008_24    m/z 269.1297     0   2   4   6E+3   1   1.5  RT (min)      lcqtof_hilic_neg_4_053_14    m/z 269.1488     0   1   2   3   4E+3   0.5   1  RT (min)      lcqtof_hilic_neg_4_048_23    m/z 269.1738     0   1   2   3   4E+3   7   7.5  RT (min)       lcqtof_hilic_neg_2_052_24    m/z 269.1788     0   2   4   6E+3   0.5   1  RT (min)      lcqtof_hilic_neg_4_053_14    m/z 269.2477     0   1   2E+3   3.5   4  RT (min)      lcqtof_hilic_neg_4_053_14    m/z 269.2513     0   1   2   3E+3   0.5   1  RT (min)      lcqtof_hilic_neg_2_016_33    m/z 270.1091     0   1E+4   5   5.5  RT (min)      lcqtof_hilic_neg_2_047_32    m/z 270.7900     0   1   2   3   4E+3   4.5   5  RT (min)      lcqtof_hilic_neg_4_050_02    m/z 270.8807     0   1E+3   4.5   5  RT (min)      lcqtof_hilic_neg_4_009_50    m/z 270.9352     0   1   2E+3   4   4.5  RT (min)      lcqtof_hilic_neg_1_020_09    m/z 271.0043     0   1E+3   5   5.5  RT (min)      lcqtof_hilic_neg_2_031_01    m/z 271.8566     0   1E+3   8   8.5  RT (min)       lcqtof_hilic_neg_4_076_58    m/z 271.9756     0   1   2   3   4E+3   0.5   1  RT (min)      lcqtof_hilic_neg_4_004_47    m/z 272.0777     0   2   4E+3   3.5   4  RT (min)       lcqtof_hilic_neg_4_089_59    m/z 272.1288     0   2   4E+3   0.5   1  RT (min)      lcqtof_hilic_neg_4_053_14    m/z 272.1495     0   1   2   3E+3   0.5   1  RT (min)      lcqtof_hilic_neg_1_029_06    m/z 272.9328     0   2   4E+3   1.5   2  RT (min)      lcqtof_hilic_neg_1_032_18    m/z 272.9869     0   2   4   6E+3   1   1.5  RT (min)      lcqtof_hilic_neg_2_004_26    m/z 273.0767     0   2   4   6   8E+3   6   6.5  RT (min)      lcqtof_hilic_neg_4_048_23    m/z 273.1209     0   1   2E+3   6   6.5  RT (min)      lcqtof_hilic_neg_1_003_37    m/z 273.1421     0   1   2   3   4E+3   1   1.5  RT (min)      lcqtof_hilic_neg_4_032_01    m/z 273.9076     0   1   2   3E+3   1   1.5  RT (min)      lcqtof_hilic_neg_4_034_31    m/z 273.9424     0   1   2   3E+3   6   6.5  RT (min)      lcqtof_hilic_neg_2_051_14    m/z 274.0257     0   1E+3   6   6.5  RT (min)       lcqtof_hilic_neg_2_002_36    m/z 274.0759     0   1E+3   5   5.5  RT (min)      lcqtof_hilic_neg_4_027_19    m/z 274.1445     0   2   4   6E+3   0.5   1  RT (min)      lcqtof_hilic_neg_2_015_06    m/z 275.1159     0   2   4   6E+3   5.5   6  RT (min)      lcqtof_hilic_neg_4_003_16    m/z 275.9359     0   1E+3   4.5   5  RT (min)      lcqtof_hilic_neg_3_025_52    m/z 275.9593     0   2   4   6   8E+3   8   8.5  RT (min)       lcqtof_hilic_neg_4_053_14    m/z 275.9997     0   1   2   3   4E+3   1   1.5  RT (min)      lcqtof_hilic_neg_4_035_33    m/z 276.0344     0   2   4E+3   1   1.5  RT (min)      lcqtof_hilic_neg_4_109_62    m/z 276.7141     0   2   4   6E+3   0.5   1  RT (min)      lcqtof_hilic_neg_4_015_09    m/z 276.8447     0   1E+3   8   8.5  RT (min)       lcqtof_hilic_neg_4_111_62    m/z 277.0121     0   1   2   3E+3   5.5   6  RT (min)       lcqtof_hilic_neg_4_048_23    m/z 277.0472     0   2   4   6E+3   2   2.5  RT (min)      lcqtof_hilic_neg_3_077_58    m/z 277.0927     0   1   2E+3   5   5.5  RT (min)      lcqtof_hilic_neg_1_037_24    m/z 277.1013     0   2   4   6   8E+3   7   7.5  RT (min)       lcqtof_hilic_neg_4_008_24    m/z 277.9840     0   1   2   3   4E+3   5.5   6  RT (min)      lcqtof_hilic_neg_1_005_32    m/z 277.9987     0   1   2E+3   2   2.5  RT (min)      lcqtof_hilic_neg_4_078_58    m/z 278.0364     0   1E+4   2   2.5  RT (min)      lcqtof_hilic_neg_4_004_47    m/z 278.0874     0   1   2E+3   6   6.5  RT (min)      lcqtof_hilic_neg_1_003_37    m/z 278.1116     0   1   2E+3   6   6.5  RT (min)       lcqtof_hilic_neg_4_053_14    m/z 278.8735     0   1E+3   8   8.5  RT (min)       lcqtof_hilic_neg_4_080_58    m/z 278.9157     0   1   2   3   4E+3   0.5   1  RT (min)      lcqtof_hilic_neg_4_053_14    m/z 278.9294     0   2   4E+3   0.5   1  RT (min)      lcqtof_hilic_neg_3_066_57    m/z 278.9601     0   2   4E+3   1   1.5  RT (min)      lcqtof_hilic_neg_3_045_18    m/z 279.0117     0   1   2E+3   5   5.5  RT (min)      lcqtof_hilic_neg_4_053_14    m/z 279.0289     0   1   2E+3   2   2.5  RT (min)       lcqtof_hilic_neg_4_074_58    m/z 279.0401     0   1E+3   4.5   5  RT (min)      lcqtof_hilic_neg_2_013_29    m/z 279.1226     0   1E+3   4.5   5  RT (min)      lcqtof_hilic_neg_2_047_32    m/z 279.1261     0   2   4   6   8E+3   2   2.5  RT (min)       lcqtof_hilic_neg_1_014_31    m/z 279.2990     0   1   2   3E+5   0.5   1  RT (min)      lcqtof_hilic_neg_4_050_02    m/z 280.0375     0   1   2   3E+3   1.5   2  RT (min)      lcqtof_hilic_neg_4_021_21    m/z 280.0381     0   1   2   3   4E+3   3   3.5  RT (min)      lcqtof_hilic_neg_1_007_05    m/z 280.2355     0   1   2   3   4E+3   3   3.5  RT (min)      lcqtof_hilic_neg_4_016_37    m/z 281.0431     0   2   4E+3   4.5   5  RT (min)       lcqtof_hilic_neg_1_035_23    m/z 281.0472     0   1   2   3   4E+3   3   3.5  RT (min)      lcqtof_hilic_neg_3_090_60    m/z 281.0723     0   2   4E+3   1   1.5  RT (min)      lcqtof_hilic_neg_1_032_18    m/z 281.1042     0   1   2   3E+3   5   5.5  RT (min)       lcqtof_hilic_neg_4_013_20    m/z 281.1097     0   2   4   6E+3   6.5   7  RT (min)      lcqtof_hilic_neg_1_003_37    m/z 281.2496     0   1E+3   15   15.5  RT (min)       lcqtof_hilic_neg_4_051_43    m/z 282.0052     0   1E+3   5.5   6  RT (min)      lcqtof_hilic_neg_3_024_35    m/z 282.0356     0   1   2   3   4E+3   6.5   7  RT (min)       lcqtof_hilic_neg_4_052_13    m/z 282.8923     0   2   4   6E+3   0.5   1  RT (min)      lcqtof_hilic_neg_4_006_30    m/z 282.9909     0   1E+3   4.5   5  RT (min)      lcqtof_hilic_neg_2_031_01    m/z 283.0909     0   2   4E+3   5.5   6  RT (min)      lcqtof_hilic_neg_4_053_14    m/z 283.8861     0   2   4E+3   1   1.5  RT (min)      lcqtof_hilic_neg_4_018_27    m/z 284.0178     0   1   2E+3   5   5.5  RT (min)       lcqtof_hilic_neg_4_034_31    m/z 284.1266     0   1   2E+3   3.5   4  RT (min)      lcqtof_hilic_neg_1_004_14    m/z 284.2659     0   1   2   3E+3   15   15.5  RT (min)      lcqtof_hilic_neg_4_047_06    m/z 284.2674     0   2   4   6   8E+3   1   1.5  RT (min)      lcqtof_hilic_neg_4_024_40    m/z 286.0266     0   1   2E+3   6   6.5  RT (min)      lcqtof_hilic_neg_1_003_37    m/z 286.0272     0   1E+3   7   7.5  RT (min)      lcqtof_hilic_neg_2_002_36    m/z 286.0572     0   2   4   6E+3   1.5   2  RT (min)      lcqtof_hilic_neg_2_052_24    m/z 286.1293     0   2   4   6E+3   1.5   2  RT (min)      lcqtof_hilic_neg_4_053_14    m/z 286.2484     0   1   2E+3   0.5   1  RT (min)      lcqtof_hilic_neg_4_048_23    m/z 286.9169     0   2   4   6E+3   0.5   1  RT (min)       lcqtof_hilic_neg_4_004_47    m/z 286.9921     0   1E+3   6   6.5  RT (min)      lcqtof_hilic_neg_1_019_02    m/z 287.0463     0   2   4E+3   2   2.5  RT (min)       lcqtof_hilic_neg_4_076_58    m/z 287.0588     0   1   2E+3   2.5   3  RT (min)       lcqtof_hilic_neg_3_055_01    m/z 287.1492     0   2   4E+3   1   1.5  RT (min)      lcqtof_hilic_neg_4_048_23    m/z 287.2014     0   1   2E+3   3.5   4  RT (min)      lcqtof_hilic_neg_3_055_01    m/z 287.2231     0   2   4E+3   0.5   1  RT (min)      lcqtof_hilic_neg_1_029_06    m/z 287.9818     0   1   2E+3   0.5   1  RT (min)      lcqtof_hilic_neg_4_028_36    m/z 287.9828     0   1   2E+3   6   6.5  RT (min)       lcqtof_hilic_neg_4_087_59    m/z 288.0179     0   1E+3   4   4.5  RT (min)      lcqtof_hilic_neg_2_031_01    m/z 288.0491     0   1   2E+3   8   8.5  RT (min)       lcqtof_hilic_neg_4_034_31    m/z 288.0679     0   1   2E+3   4.5   5  RT (min)      lcqtof_hilic_neg_4_008_24    m/z 288.1018     0   2   4   6E+3   5.5   6  RT (min)      lcqtof_hilic_neg_4_008_24    m/z 288.2074     0   1   2E+3   1   1.5  RT (min)      lcqtof_hilic_neg_4_045_26    m/z 288.8953     0   1   2   3E+3   0.5   1  RT (min)       lcqtof_hilic_neg_4_021_21    m/z 288.9312     0   1   2   3E+3   0.5   1  RT (min)       lcqtof_hilic_neg_3_068_57    m/z 288.9850     0   1E+3   3.5   4  RT (min)      lcqtof_hilic_neg_1_027_27    m/z 288.9935     0   1   2E+3   5.5   6  RT (min)      lcqtof_hilic_neg_1_013_38    m/z 289.0398     0   2   4   6E+3   2   2.5  RT (min)       lcqtof_hilic_neg_4_074_58    m/z 289.0531     0   2   4E+3   1   1.5  RT (min)      lcqtof_hilic_neg_4_047_06    m/z 289.0745     0   1E+4   0.5   1  RT (min)       lcqtof_hilic_neg_4_004_47    m/z 289.1060     0   1   2E+3   1   1.5  RT (min)      lcqtof_hilic_neg_4_053_14    m/z 289.1399     0   1   2E+3   7   7.5  RT (min)      lcqtof_hilic_neg_1_003_37    m/z 289.9218     0   1   2E+3   3.5   4  RT (min)      lcqtof_hilic_neg_3_040_14    m/z 290.0027     0   1E+3   6   6.5  RT (min)      lcqtof_hilic_neg_2_047_32    m/z 290.0536     0   1   2E+3   6.5   7  RT (min)      lcqtof_hilic_neg_4_040_05    m/z 290.0816     0   2   4E+3   3.5   4  RT (min)      lcqtof_hilic_neg_1_005_32    m/z 290.0872     0   2   4   6E+4   5.5   6  RT (min)       lcqtof_hilic_neg_1_014_31    m/z 290.1287     0   2   4   6   8E+3   1.5   2  RT (min)       lcqtof_hilic_neg_4_055_32    m/z 290.1765     0   1   2E+3   0.5   1  RT (min)      lcqtof_hilic_neg_2_053_18    m/z 290.8738     0   1   2   3E+3   1   1.5  RT (min)      lcqtof_hilic_neg_2_054_42    m/z 291.0469     0   1   2E+3   1   1.5  RT (min)      lcqtof_hilic_neg_4_053_14    m/z 291.9666     0   1   2E+3   1   1.5  RT (min)      lcqtof_hilic_neg_3_044_09    m/z 292.0790     0   1   2E+3   5.5   6  RT (min)      lcqtof_hilic_neg_1_023_34    m/z 292.0833     0   1   2E+3   3.5   4  RT (min)      lcqtof_hilic_neg_4_039_18    m/z 292.0862     0   1   2   3   4E+3   6   6.5  RT (min)      lcqtof_hilic_neg_2_029_31    m/z 292.1928     0   1   2E+3   0.5   1  RT (min)      lcqtof_hilic_neg_1_032_18    m/z 293.0597     0   1   2E+3   2.5   3  RT (min)       lcqtof_hilic_neg_3_080_58    m/z 293.1166     0   2   4   6E+3   2.5   3  RT (min)      lcqtof_hilic_neg_4_048_23    m/z 293.9713     0   1   2E+3   1   1.5  RT (min)      lcqtof_hilic_neg_2_005_13    m/z 293.9971     0   1E+3   6   6.5  RT (min)      lcqtof_hilic_neg_2_043_15    m/z 294.0208     0   1   2   3   4E+3   4   4.5  RT (min)      lcqtof_hilic_neg_2_047_32    m/z 294.0974     0   1   2   3E+3   0.5   1  RT (min)      lcqtof_hilic_neg_2_015_06    m/z 294.1076     0   1   2   3E+3   3.5   4  RT (min)      lcqtof_hilic_neg_4_013_20    m/z 294.8122     0   1E+3   8.5   9  RT (min)       lcqtof_hilic_neg_4_030_44    m/z 294.8297     0   1   2E+3   7.5   8  RT (min)      lcqtof_hilic_neg_2_031_01    m/z 294.9848     0   2   4E+3   3   3.5  RT (min)      lcqtof_hilic_neg_1_006_22    m/z 294.9929     0   2   4   6E+3   8   8.5  RT (min)      lcqtof_hilic_neg_4_023_15    m/z 295.0047     0   2   4   6   8E+3   3   3.5  RT (min)      lcqtof_hilic_neg_1_005_32    m/z 295.0127     0   1   2   3   4E+3   5.5   6  RT (min)      lcqtof_hilic_neg_4_008_24    m/z 295.0524     0   2   4   6E+3   7   7.5  RT (min)      lcqtof_hilic_neg_4_055_32    m/z 295.0526     0   1   2   3   4E+3   3.5   4  RT (min)       lcqtof_hilic_neg_1_029_06    m/z 295.0721     0   1   2   3E+3   8   8.5  RT (min)       lcqtof_hilic_neg_4_094_60    m/z 295.1884     0   1   2   3   4E+3   7   7.5  RT (min)      lcqtof_hilic_neg_2_052_24    m/z 295.1922     0   2   4E+3   1   1.5  RT (min)      lcqtof_hilic_neg_4_035_33    m/z 295.8387     0   1   2   3E+3   0.5   1  RT (min)      lcqtof_hilic_neg_4_053_14    m/z 296.0072     0   1   2E+3   2   2.5  RT (min)      lcqtof_hilic_neg_3_076_58    m/z 296.0859     0   1   2E+3   6.5   7  RT (min)      lcqtof_hilic_neg_1_035_23    m/z 296.1155     0   1   2   3   4E+3   0.5   1  RT (min)      lcqtof_hilic_neg_4_040_05    m/z 296.1405     0   1   2   3E+3   1   1.5  RT (min)      lcqtof_hilic_neg_4_047_06    m/z 296.9274     0   2   4E+3   2   2.5  RT (min)      lcqtof_hilic_neg_4_003_16    m/z 296.9932     0   2   4   6E+3   4.5   5  RT (min)      lcqtof_hilic_neg_1_004_14    m/z 297.0040     0   1   2E+3   4   4.5  RT (min)      lcqtof_hilic_neg_3_034_24    m/z 297.9100     0   1   2E+3   1   1.5  RT (min)      lcqtof_hilic_neg_1_042_26    m/z 298.0081     0   1   2E+3   4.5   5  RT (min)      lcqtof_hilic_neg_4_053_14    m/z 298.0993     0   1   2   3   4E+3   5.5   6  RT (min)      lcqtof_hilic_neg_1_029_06    m/z 298.1158     0   2   4   6E+3   7   7.5  RT (min)       lcqtof_hilic_neg_4_008_24    m/z 299.0095     0   1E+3   1   1.5  RT (min)      lcqtof_hilic_neg_1_034_08    m/z 299.0145     0   1E+3   6   6.5  RT (min)      lcqtof_hilic_neg_2_002_36    m/z 299.0873     0   2   4E+3   5   5.5  RT (min)       lcqtof_hilic_neg_1_035_23    m/z 299.0966     0   1   2   3E+3   5   5.5  RT (min)       lcqtof_hilic_neg_4_012_07    m/z 299.0976     0   1   2   3   4E+3   5.5   6  RT (min)       lcqtof_hilic_neg_4_013_20    m/z 299.1485     0   1   2   3E+3   7.5   8  RT (min)      lcqtof_hilic_neg_1_037_24    m/z 299.1577     0   1   2E+3   4   4.5  RT (min)      lcqtof_hilic_neg_2_028_05    m/z 299.8864     0   1   2   3   4E+3   4.5   5  RT (min)      lcqtof_hilic_neg_4_053_14    m/z 300.0168     0   2   4   6   8E+3   4.5   5  RT (min)      lcqtof_hilic_neg_1_005_32    m/z 300.1558     0   2   4   6E+3   1.5   2  RT (min)      lcqtof_hilic_neg_4_054_48    m/z 300.7476     0   1E+3   4.5   5  RT (min)      lcqtof_hilic_neg_2_013_29    m/z 300.7489     0   1   2E+3   4   4.5  RT (min)      lcqtof_hilic_neg_4_050_02    m/z 301.0519     0   1   2   3E+3   5   5.5  RT (min)      lcqtof_hilic_neg_4_008_24    m/z 301.0996     0   2   4E+3   3.5   4  RT (min)      lcqtof_hilic_neg_4_008_24    m/z 301.3230     0   2   4   6E+3   0.5   1  RT (min)      lcqtof_hilic_neg_4_040_05    m/z 301.9879     0   1   2   3   4E+3   1   1.5  RT (min)      lcqtof_hilic_neg_4_027_19    m/z 302.0469     0   1   2E+3   0.5   1  RT (min)      lcqtof_hilic_neg_1_041_54    m/z 302.0488     0   1   2   3E+3   1.5   2  RT (min)      lcqtof_hilic_neg_4_005_29    m/z 302.1953     0   2   4E+3   3   3.5  RT (min)      lcqtof_hilic_neg_4_013_20    m/z 302.9948     0   1   2   3   4E+3   5   5.5  RT (min)      lcqtof_hilic_neg_4_032_01    m/z 302.9951     0   1   2   3E+3   5   5.5  RT (min)      lcqtof_hilic_neg_1_054_01    m/z 303.0526     0   2   4   6E+3   3   3.5  RT (min)      lcqtof_hilic_neg_1_022_36    m/z 303.0529     0   1   2   3E+3   0.5   1  RT (min)      lcqtof_hilic_neg_4_014_46    m/z 303.0670     0   1   2E+3   1   1.5  RT (min)      lcqtof_hilic_neg_4_018_27    m/z 303.0864     0   1   2   3E+3   2   2.5  RT (min)      lcqtof_hilic_neg_3_054_37    m/z 304.0381     0   1   2   3E+3   2.5   3  RT (min)      lcqtof_hilic_neg_2_028_05    m/z 304.0906     0   1   2E+3   5   5.5  RT (min)       lcqtof_hilic_neg_4_013_20    m/z 304.1049     0   1   2E+3   6   6.5  RT (min)      lcqtof_hilic_neg_4_003_16    m/z 304.1356     0   2   4   6E+3   0.5   1  RT (min)      lcqtof_hilic_neg_1_032_18    m/z 304.2376     0   1   2E+3   3   3.5  RT (min)      lcqtof_hilic_neg_4_018_27    m/z 305.0215     0   1   2E+3   6   6.5  RT (min)      lcqtof_hilic_neg_1_037_24    m/z 305.0477     0   1   2   3E+3   0.5   1  RT (min)      lcqtof_hilic_neg_2_005_13    m/z 305.0557     0   1E+3   3.5   4  RT (min)      lcqtof_hilic_neg_1_027_27    m/z 305.0652     0   2   4E+3   5.5   6  RT (min)      lcqtof_hilic_neg_4_003_16    m/z 305.2466     0   1   2   3E+3   3   3.5  RT (min)       lcqtof_hilic_neg_4_003_16    m/z 305.8175     0   1   2E+3   8   8.5  RT (min)      lcqtof_hilic_neg_2_027_07    m/z 305.9475     0   1   2   3   4E+3   1   1.5  RT (min)      lcqtof_hilic_neg_2_051_14    m/z 306.0183     0   1   2E+3   5.5   6  RT (min)      lcqtof_hilic_neg_4_032_01    m/z 306.0590     0   2   4   6E+3   5.5   6  RT (min)      lcqtof_hilic_neg_2_020_23    m/z 306.0755     0   1   2   3E+3   8   8.5  RT (min)       lcqtof_hilic_neg_2_020_23    m/z 306.1397     0   1   2   3   4E+3   4   4.5  RT (min)      lcqtof_hilic_neg_1_037_24    m/z 306.9091     0   1   2   3   4E+3   0.5   1  RT (min)      lcqtof_hilic_neg_3_074_58    m/z 306.9650     0   1   2E+3   5.5   6  RT (min)      lcqtof_hilic_neg_2_036_34    m/z 307.0988     0   1   2   3   4E+3   5.5   6  RT (min)       lcqtof_hilic_neg_1_005_32    m/z 307.1254     0   1   2   3E+3   1   1.5  RT (min)      lcqtof_hilic_neg_1_054_01    m/z 307.6788     0   1E+3   3   3.5  RT (min)      lcqtof_hilic_neg_1_001_49    m/z 307.9619     0   1   2E+3   1   1.5  RT (min)      lcqtof_hilic_neg_3_101_61    m/z 309.0760     0   1   2E+3   7   7.5  RT (min)      lcqtof_hilic_neg_4_048_23    m/z 310.0627     0   1   2   3E+3   6.5   7  RT (min)       lcqtof_hilic_neg_4_073_57    m/z 310.1185     0   2   4   6E+3   3   3.5  RT (min)      lcqtof_hilic_neg_4_094_60    m/z 310.1711     0   1   2E+4   1   1.5  RT (min)      lcqtof_hilic_neg_4_039_18    m/z 310.2119     0   1   2   3E+3   1   1.5  RT (min)      lcqtof_hilic_neg_4_053_14    m/z 310.9019     0   1   2E+3   4.5   5  RT (min)      lcqtof_hilic_neg_1_005_32    m/z 311.0242     0   1E+4   3   3.5  RT (min)      lcqtof_hilic_neg_4_098_61    m/z 311.0500     0   1   2   3E+3   6.5   7  RT (min)      lcqtof_hilic_neg_4_008_24    m/z 311.9681     0   1E+3   2   2.5  RT (min)      lcqtof_hilic_neg_4_075_58    m/z 312.0169     0   1   2E+3   1   1.5  RT (min)      lcqtof_hilic_neg_1_054_01    m/z 312.0671     0   2   4   6E+3   5.5   6  RT (min)      lcqtof_hilic_neg_4_034_31    m/z 312.1013     0   1   2E+3   3.5   4  RT (min)      lcqtof_hilic_neg_1_005_32    m/z 312.2237     0   2   4   6E+3   1   1.5  RT (min)      lcqtof_hilic_neg_4_053_14    m/z 312.9644     0   1   2E+3   2   2.5  RT (min)      lcqtof_hilic_neg_4_074_58    m/z 313.0170     0   1   2   3E+3   4   4.5  RT (min)      lcqtof_hilic_neg_1_022_36    m/z 313.1790     0   1   2   3E+3   6   6.5  RT (min)      lcqtof_hilic_neg_4_040_05    m/z 313.2140     0   1   2   3   4E+3   1   1.5  RT (min)      lcqtof_hilic_neg_4_048_23    m/z 314.0314     0   1   2   3E+3   0.5   1  RT (min)      lcqtof_hilic_neg_4_053_14    m/z 314.1675     0   2   4   6E+3   6   6.5  RT (min)      lcqtof_hilic_neg_4_008_24    m/z 314.2410     0   2   4E+3   1   1.5  RT (min)      lcqtof_hilic_neg_4_032_01    m/z 315.0016     0   1   2E+3   4   4.5  RT (min)      lcqtof_hilic_neg_1_014_31    m/z 315.1018     0   1   2   3   4E+3   8   8.5  RT (min)       lcqtof_hilic_neg_2_052_24    m/z 315.1961     0   2   4   6E+3   0.5   1  RT (min)      lcqtof_hilic_neg_4_053_14    m/z 316.0283     0   1   2E+4   1   1.5  RT (min)      lcqtof_hilic_neg_4_005_29    m/z 316.0433     0   1E+4   5.5   6  RT (min)       lcqtof_hilic_neg_1_054_01    m/z 316.0439     0   1E+4   7   7.5  RT (min)      lcqtof_hilic_neg_2_048_35    m/z 316.0439     0   1   2E+4   5.5   6  RT (min)      lcqtof_hilic_neg_1_022_36    m/z 316.0454     0   1   2   3E+3   2   2.5  RT (min)       lcqtof_hilic_neg_4_080_58    m/z 316.0627     0   1   2E+3   0.5   1  RT (min)      lcqtof_hilic_neg_2_043_15    m/z 316.9227     0   2   4   6   8E+3   0.5   1  RT (min)      lcqtof_hilic_neg_4_014_46    m/z 316.9897     0   1   2E+3   1.5   2  RT (min)      lcqtof_hilic_neg_1_031_16    m/z 317.9265     0   1   2   3   4E+3   0.5   1  RT (min)      lcqtof_hilic_neg_4_054_48    m/z 318.0147     0   1E+3   7   7.5  RT (min)      lcqtof_hilic_neg_1_001_49    m/z 318.0528     0   2   4E+3   7.5   8  RT (min)       lcqtof_hilic_neg_4_090_60    m/z 318.1456     0   1   2   3E+3   5   5.5  RT (min)      lcqtof_hilic_neg_4_016_37    m/z 318.3235     0   2   4   6   8E+3   0.5   1  RT (min)      lcqtof_hilic_neg_4_008_24    m/z 318.8282     0   1   2E+3   7.5   8  RT (min)      lcqtof_hilic_neg_1_001_49    m/z 319.1956     0   1   2E+3   0.5   1  RT (min)      lcqtof_hilic_neg_4_009_50    m/z 319.8192     0   1E+3   8   8.5  RT (min)      lcqtof_hilic_neg_2_020_23    m/z 319.8217     0   1E+3   8   8.5  RT (min)      lcqtof_hilic_neg_2_052_24    m/z 320.1080     0   1   2E+3   5.5   6  RT (min)      lcqtof_hilic_neg_1_007_05    m/z 320.1336     0   1   2E+3   6   6.5  RT (min)      lcqtof_hilic_neg_1_037_24    m/z 320.8249     0   1   2E+3   7.5   8  RT (min)      lcqtof_hilic_neg_1_001_49    m/z 320.8257     0   1   2E+3   7.5   8  RT (min)      lcqtof_hilic_neg_1_001_49    m/z 321.0121     0   1E+3   2   2.5  RT (min)      lcqtof_hilic_neg_4_098_61    m/z 321.0303     0   1   2   3E+3   4   4.5  RT (min)      lcqtof_hilic_neg_4_055_32    m/z 321.1012     0   2   4   6   8E+3   6   6.5  RT (min)      lcqtof_hilic_neg_4_047_06    m/z 321.2097     0   1   2   3E+3   15   15.5  RT (min)       lcqtof_hilic_neg_4_019_03    m/z 322.0047     0   1   2E+3   2   2.5  RT (min)      lcqtof_hilic_neg_3_110_62    m/z 322.0452     0   2   4   6E+3   7   7.5  RT (min)       lcqtof_hilic_neg_1_007_05    m/z 322.9858     0   1   2E+3   7.5   8  RT (min)      lcqtof_hilic_neg_1_035_23    m/z 323.0027     0   1E+3   2   2.5  RT (min)      lcqtof_hilic_neg_3_113_62    m/z 323.0323     0   1   2   3   4E+3   5   5.5  RT (min)      lcqtof_hilic_neg_4_055_32    m/z 323.1064     0   1   2   3E+3   2   2.5  RT (min)      lcqtof_hilic_neg_1_030_17    m/z 323.2231     0   1   2   3   4E+3   0.5   1  RT (min)      lcqtof_hilic_neg_4_002_25    m/z 323.8165     0   1E+3   8   8.5  RT (min)      lcqtof_hilic_neg_2_015_06    m/z 324.0312     0   2   4   6   8E+3   6.5   7  RT (min)      lcqtof_hilic_neg_4_048_23    m/z 324.0386     0   1   2E+3   7   7.5  RT (min)      lcqtof_hilic_neg_4_008_24    m/z 324.0877     0   2   4E+3   0.5   1  RT (min)      lcqtof_hilic_neg_4_010_28    m/z 324.0930     0   1   2   3E+3   5   5.5  RT (min)      lcqtof_hilic_neg_4_055_32    m/z 324.1715     0   1   2   3   4E+3   1   1.5  RT (min)      lcqtof_hilic_neg_4_047_06    m/z 325.1355     0   1   2E+3   1   1.5  RT (min)      lcqtof_hilic_neg_4_050_02    m/z 326.1320     0   1E+3   5   5.5  RT (min)      lcqtof_hilic_neg_4_042_22    m/z 326.2701     0   1   2   3   4E+3   1   1.5  RT (min)      lcqtof_hilic_neg_4_003_16    m/z 326.9901     0   2   4   6E+3   6   6.5  RT (min)      lcqtof_hilic_neg_1_023_34    m/z 327.0186     0   1   2   3E+3   1   1.5  RT (min)      lcqtof_hilic_neg_3_003_20    m/z 327.0395     0   2   4   6E+3   1   1.5  RT (min)      lcqtof_hilic_neg_4_004_47    m/z 327.0631     0   1   2   3E+3   5.5   6  RT (min)      lcqtof_hilic_neg_1_022_36    m/z 327.0969     0   2   4E+3   1   1.5  RT (min)      lcqtof_hilic_neg_2_015_06    m/z 327.9523     0   1   2E+3   1.5   2  RT (min)      lcqtof_hilic_neg_1_035_23    m/z 328.0412     0   1   2   3E+3   5.5   6  RT (min)      lcqtof_hilic_neg_4_022_34    m/z 328.1301     0   1   2E+3   0.5   1  RT (min)      lcqtof_hilic_neg_2_013_29    m/z 328.1378     0   2   4   6E+3   3.5   4  RT (min)      lcqtof_hilic_neg_4_008_24    m/z 328.2302     0   1E+5   1   1.5  RT (min)       lcqtof_hilic_neg_4_022_34    m/z 329.0308     0   1   2E+3   6.5   7  RT (min)      lcqtof_hilic_neg_1_014_31    m/z 329.2080     0   2   4   6E+3   4.5   5  RT (min)      lcqtof_hilic_neg_2_003_25    m/z 329.2526     0   1E+4   3.5   4  RT (min)      lcqtof_hilic_neg_4_032_01    m/z 330.0072     0   1   2   3E+3   7.5   8  RT (min)      lcqtof_hilic_neg_2_052_24    m/z 330.0939     0   1E+4   4   4.5  RT (min)       lcqtof_hilic_neg_4_102_61    m/z 330.2512     0   1   2   3E+3   3.5   4  RT (min)      lcqtof_hilic_neg_4_032_01    m/z 330.9792     0   2   4   6E+3   0.5   1  RT (min)      lcqtof_hilic_neg_4_013_20    m/z 331.0095     0   1   2   3E+3   0.5   1  RT (min)      lcqtof_hilic_neg_4_050_02    m/z 331.1081     0   1   2E+3   0.5   1  RT (min)       lcqtof_hilic_neg_4_050_02    m/z 331.2650     0   1E+4   1   1.5  RT (min)      lcqtof_hilic_neg_4_054_48    m/z 331.2985     0   2   4   6   8E+3   0.5   1  RT (min)      lcqtof_hilic_neg_4_008_24    m/z 331.9906     0   1E+3   4.5   5  RT (min)      lcqtof_hilic_neg_4_031_38    m/z 332.0467     0   1E+3   3.5   4  RT (min)      lcqtof_hilic_neg_4_018_27    m/z 332.0855     0   2   4   6   8E+3   0.5   1  RT (min)      lcqtof_hilic_neg_4_039_18    m/z 332.0897     0   2   4   6E+3   6   6.5  RT (min)      lcqtof_hilic_neg_4_048_23    m/z 332.0972     0   1   2   3E+3   5.5   6  RT (min)      lcqtof_hilic_neg_4_031_38    m/z 332.1200     0   1   2   3E+3   5   5.5  RT (min)      lcqtof_hilic_neg_1_005_32    m/z 332.1949     0   1   2E+4   0.5   1  RT (min)      lcqtof_hilic_neg_4_039_18    m/z 333.0384     0   1   2E+3   7   7.5  RT (min)      lcqtof_hilic_neg_1_001_49    m/z 333.0742     0   2   4   6E+3   6   6.5  RT (min)      lcqtof_hilic_neg_4_048_23    m/z 333.0830     0   1E+3   5.5   6  RT (min)      lcqtof_hilic_neg_1_019_02    m/z 333.1212     0   2   4   6   8E+3   4.5   5  RT (min)       lcqtof_hilic_neg_4_053_14    m/z 333.1218     0   1   2   3E+3   6.5   7  RT (min)      lcqtof_hilic_neg_4_047_06    m/z 333.2799     0   1   2   3   4E+3   0.5   1  RT (min)      lcqtof_hilic_neg_2_016_33    m/z 334.0123     0   1   2   3E+3   5.5   6  RT (min)      lcqtof_hilic_neg_4_023_15    m/z 334.1018     0   2   4   6   8E+4   6   6.5  RT (min)      lcqtof_hilic_neg_4_047_06    m/z 334.1354     0   1E+3   0.5   1  RT (min)      lcqtof_hilic_neg_4_047_06    m/z 335.0742     0   2   4   6   8E+3   0.5   1  RT (min)       lcqtof_hilic_neg_4_005_29    m/z 335.1053     0   2   4   6E+3   5.5   6  RT (min)      lcqtof_hilic_neg_4_008_24    m/z 335.1257     0   1   2   3   4E+3   6.5   7  RT (min)      lcqtof_hilic_neg_4_047_06    m/z 335.2574     0   2   4E+3   0.5   1  RT (min)      lcqtof_hilic_neg_4_053_14    m/z 335.9922     0   2   4   6E+3   0.5   1  RT (min)      lcqtof_hilic_neg_4_004_47    m/z 336.0505     0   1E+3   7   7.5  RT (min)       lcqtof_hilic_neg_2_004_26    m/z 337.0324     0   1E+4   0.5   1  RT (min)      lcqtof_hilic_neg_1_053_19    m/z 337.0396     0   1E+3   6.5   7  RT (min)      lcqtof_hilic_neg_2_039_38    m/z 337.0679     0   1   2E+3   5.5   6  RT (min)      lcqtof_hilic_neg_4_008_24    m/z 337.8115     0   1E+3   8   8.5  RT (min)      lcqtof_hilic_neg_4_068_57    m/z 338.1899     0   2   4   6E+3   1   1.5  RT (min)      lcqtof_hilic_neg_4_047_06    m/z 338.2468     0   2   4   6   8E+3   1   1.5  RT (min)      lcqtof_hilic_neg_4_032_01    m/z 338.2482     0   2   4   6   8E+3   0.5   1  RT (min)      lcqtof_hilic_neg_4_032_01    m/z 338.2700     0   2   4   6E+3   1   1.5  RT (min)      lcqtof_hilic_neg_2_016_33    m/z 339.0155     0   1   2E+3   4   4.5  RT (min)      lcqtof_hilic_neg_1_054_01    m/z 339.0280     0   2   4E+3   5   5.5  RT (min)      lcqtof_hilic_neg_4_008_24    m/z 339.0555     0   1E+3   5   5.5  RT (min)       lcqtof_hilic_neg_4_052_13    m/z 340.0690     0   1   2   3E+3   2.5   3  RT (min)      lcqtof_hilic_neg_1_007_05    m/z 340.0826     0   1   2   3   4E+3   6.5   7  RT (min)      lcqtof_hilic_neg_4_097_60    m/z 340.0827     0   2   4E+3   7   7.5  RT (min)       lcqtof_hilic_neg_4_092_60    m/z 340.2864     0   1   2   3E+3   1   1.5  RT (min)      lcqtof_hilic_neg_4_043_12    m/z 341.0805     0   1   2E+3   3.5   4  RT (min)      lcqtof_hilic_neg_4_004_47    m/z 341.1567     0   1   2   3   4E+3   6   6.5  RT (min)      lcqtof_hilic_neg_4_048_23    m/z 341.8688     0   1E+3   8   8.5  RT (min)      lcqtof_hilic_neg_2_046_41    m/z 342.1130     0   1   2   3E+3   7   7.5  RT (min)      lcqtof_hilic_neg_2_028_05    m/z 342.9735     0   2   4E+3   1   1.5  RT (min)       lcqtof_hilic_neg_4_075_58    m/z 342.9879     0   1E+3   3.5   4  RT (min)      lcqtof_hilic_neg_2_055_30    m/z 343.0909     0   2   4   6   8E+3   6.5   7  RT (min)       lcqtof_hilic_neg_4_052_13    m/z 343.1537     0   1   2   3   4E+3   7   7.5  RT (min)      lcqtof_hilic_neg_1_037_24    m/z 344.0685     0   1   2E+3   1   1.5  RT (min)      lcqtof_hilic_neg_1_024_21    m/z 344.2886     0   2   4E+3   0.5   1  RT (min)      lcqtof_hilic_neg_4_018_27    m/z 344.9706     0   1   2E+3   1   1.5  RT (min)      lcqtof_hilic_neg_4_079_58    m/z 345.1284     0   1   2E+3   7   7.5  RT (min)      lcqtof_hilic_neg_2_005_13    m/z 345.1803     0   2   4   6E+3   4   4.5  RT (min)      lcqtof_hilic_neg_4_013_20    m/z 346.9643     0   1   2E+3   6.5   7  RT (min)      lcqtof_hilic_neg_2_042_27    m/z 346.9767     0   1   2   3E+3   4.5   5  RT (min)      lcqtof_hilic_neg_1_032_18    m/z 347.0783     0   2   4E+3   1   1.5  RT (min)      lcqtof_hilic_neg_4_016_37    m/z 347.2678     0   1E+4   0.5   1  RT (min)      lcqtof_hilic_neg_4_055_32    m/z 348.1504     0   1   2   3E+3   0.5   1  RT (min)      lcqtof_hilic_neg_4_034_31    m/z 348.1998     0   1   2   3   4E+3   1   1.5  RT (min)      lcqtof_hilic_neg_1_003_37    m/z 348.2464     0   1   2   3   4E+3   1.5   2  RT (min)      lcqtof_hilic_neg_4_053_14    m/z 348.2665     0   2   4   6   8E+3   0.5   1  RT (min)      lcqtof_hilic_neg_4_008_24    m/z 348.9193     0   1   2   3E+3   1   1.5  RT (min)      lcqtof_hilic_neg_1_032_18    m/z 348.9399     0   1   2E+3   0.5   1  RT (min)      lcqtof_hilic_neg_4_032_01    m/z 349.0595     0   1   2   3E+3   7   7.5  RT (min)       lcqtof_hilic_neg_4_072_57    m/z 349.0937     0   1   2E+3   1   1.5  RT (min)      lcqtof_hilic_neg_1_012_04    m/z 349.1248     0   1   2   3E+3   0.5   1  RT (min)      lcqtof_hilic_neg_4_027_19    m/z 349.1305     0   1   2   3E+3   5   5.5  RT (min)      lcqtof_hilic_neg_4_002_25    m/z 349.2048     0   1   2E+3   4   4.5  RT (min)      lcqtof_hilic_neg_4_041_54    m/z 349.9245     0   1E+3   1   1.5  RT (min)      lcqtof_hilic_neg_3_025_52    m/z 350.0089     0   2   4   6E+3   0.5   1  RT (min)      lcqtof_hilic_neg_4_004_47    m/z 350.0640     0   1E+3   4.5   5  RT (min)      lcqtof_hilic_neg_2_042_27    m/z 350.0921     0   2   4E+3   6   6.5  RT (min)      lcqtof_hilic_neg_1_011_46    m/z 350.8632     0   1E+3   4.5   5  RT (min)      lcqtof_hilic_neg_1_016_28    m/z 351.0067     0   1   2   3   4E+3   0.5   1  RT (min)      lcqtof_hilic_neg_4_004_47    m/z 351.0262     0   1   2   3E+3   1   1.5  RT (min)      lcqtof_hilic_neg_4_039_18    m/z 351.0737     0   1   2   3E+3   3   3.5  RT (min)      lcqtof_hilic_neg_3_054_37    m/z 351.1924     0   2   4   6E+3   1   1.5  RT (min)      lcqtof_hilic_neg_4_011_35    m/z 351.2507     0   1E+3   15   15.5  RT (min)       lcqtof_hilic_neg_4_047_06    m/z 352.0176     0   1   2   3   4E+3   5.5   6  RT (min)      lcqtof_hilic_neg_4_022_34    m/z 352.1212     0   1E+3   0.5   1  RT (min)      lcqtof_hilic_neg_4_019_03    m/z 352.1442     0   2   4   6E+3   7   7.5  RT (min)      lcqtof_hilic_neg_1_037_24    m/z 352.1876     0   1   2E+3   0.5   1  RT (min)      lcqtof_hilic_neg_4_008_24    m/z 352.2843     0   1   2   3E+3   0.5   1  RT (min)      lcqtof_hilic_neg_4_003_16    m/z 353.1391     0   1   2   3E+3   5.5   6  RT (min)      lcqtof_hilic_neg_1_054_01    m/z 353.1405     0   1   2   3E+3   1   1.5  RT (min)      lcqtof_hilic_neg_3_052_22    m/z 354.0153     0   1   2E+3   5.5   6  RT (min)      lcqtof_hilic_neg_4_004_47    m/z 354.0716     0   1   2E+3   4   4.5  RT (min)       lcqtof_hilic_neg_4_101_61    m/z 354.2711     0   1   2   3E+3   0.5   1  RT (min)      lcqtof_hilic_neg_4_050_02    m/z 354.3011     0   1   2   3   4E+3   1   1.5  RT (min)      lcqtof_hilic_neg_4_003_16    m/z 355.0314     0   1   2E+3   5   5.5  RT (min)      lcqtof_hilic_neg_4_053_14    m/z 355.1021     0   1   2   3E+3   6.5   7  RT (min)      lcqtof_hilic_neg_4_047_06    m/z 355.2618     0   1   2   3E+3   3.5   4  RT (min)      lcqtof_hilic_neg_1_035_23    m/z 355.9893     0   2   4E+3   6   6.5  RT (min)      lcqtof_hilic_neg_1_037_24    m/z 356.0600     0   1   2   3   4E+3   1.5   2  RT (min)      lcqtof_hilic_neg_2_020_23    m/z 356.1187     0   1   2   3E+3   0.5   1  RT (min)      lcqtof_hilic_neg_4_053_14    m/z 356.1277     0   1E+3   1   1.5  RT (min)      lcqtof_hilic_neg_1_036_03    m/z 356.9893     0   1   2   3E+3   1   1.5  RT (min)      lcqtof_hilic_neg_4_056_56    m/z 357.0308     0   1   2   3E+3   5   5.5  RT (min)      lcqtof_hilic_neg_1_054_01    m/z 357.1075     0   1   2   3E+3   6   6.5  RT (min)      lcqtof_hilic_neg_2_022_46    m/z 357.1100     0   1   2   3E+3   6.5   7  RT (min)      lcqtof_hilic_neg_4_047_06    m/z 357.1265     0   1   2   3   4E+3   5   5.5  RT (min)       lcqtof_hilic_neg_4_008_24    m/z 357.1497     0   1   2   3   4E+3   3.5   4  RT (min)      lcqtof_hilic_neg_3_032_19    m/z 357.2291     0   1   2E+3   1   1.5  RT (min)      lcqtof_hilic_neg_1_014_31    m/z 357.2816     0   2   4   6   8E+3   1   1.5  RT (min)      lcqtof_hilic_neg_4_004_47    m/z 358.0405     0   1   2   3   4E+3   6.5   7  RT (min)      lcqtof_hilic_neg_3_030_23    m/z 358.0827     0   1   2   3E+3   6   6.5  RT (min)      lcqtof_hilic_neg_1_035_23    m/z 358.1210     0   1   2E+3   1   1.5  RT (min)      lcqtof_hilic_neg_4_026_04    m/z 358.2518     0   2   4E+3   7   7.5  RT (min)      lcqtof_hilic_neg_4_008_24    m/z 359.0013     0   2   4E+3   3.5   4  RT (min)      lcqtof_hilic_neg_1_029_06    m/z 359.0335     0   1   2E+3   4   4.5  RT (min)      lcqtof_hilic_neg_3_011_34    m/z 359.0377     0   1   2   3   4E+3   0.5   1  RT (min)      lcqtof_hilic_neg_4_039_18    m/z 359.0520     0   2   4   6   8E+3   4.5   5  RT (min)       lcqtof_hilic_neg_4_039_18    m/z 359.2948     0   2   4E+3   1   1.5  RT (min)      lcqtof_hilic_neg_4_007_45    m/z 360.0883     0   2   4E+3   3.5   4  RT (min)      lcqtof_hilic_neg_4_005_29    m/z 360.1541     0   1   2E+3   6.5   7  RT (min)      lcqtof_hilic_neg_1_003_37    m/z 360.2931     0   2   4   6   8E+3   0.5   1  RT (min)      lcqtof_hilic_neg_4_052_13    m/z 361.0046     0   2   4   6E+3   0.5   1  RT (min)      lcqtof_hilic_neg_4_053_14    m/z 361.0969     0   1E+3   6   6.5  RT (min)      lcqtof_hilic_neg_1_049_55    m/z 361.0992     0   1E+4   0.5   1  RT (min)      lcqtof_hilic_neg_4_050_02    m/z 361.1617     0   1   2E+3   6   6.5  RT (min)      lcqtof_hilic_neg_1_007_05    m/z 361.9760     0   2   4   6E+3   4   4.5  RT (min)      lcqtof_hilic_neg_1_004_14    m/z 362.0494     0   2   4E+3   7   7.5  RT (min)      lcqtof_hilic_neg_4_048_23    m/z 362.1348     0   1   2   3E+3   0.5   1  RT (min)      lcqtof_hilic_neg_4_048_23    m/z 362.1953     0   1   2   3   4E+3   7   7.5  RT (min)      lcqtof_hilic_neg_4_055_32    m/z 362.7995     0   1E+3   8.5   9  RT (min)       lcqtof_hilic_neg_4_107_62    m/z 362.9750     0   1   2E+3   4   4.5  RT (min)      lcqtof_hilic_neg_1_045_07    m/z 363.0782     0   1   2   3   4E+3   0.5   1  RT (min)      lcqtof_hilic_neg_2_016_33    m/z 363.1229     0   1   2E+3   0.5   1  RT (min)      lcqtof_hilic_neg_1_012_04    m/z 363.1636     0   1E+4   0.5   1  RT (min)      lcqtof_hilic_neg_4_004_47    m/z 363.3517     0   1   2   3   4E+3   1   1.5  RT (min)      lcqtof_hilic_neg_4_008_24    m/z 364.0591     0   1   2   3E+3   1   1.5  RT (min)      lcqtof_hilic_neg_3_020_21    m/z 364.0813     0   1   2   3   4E+3   5.5   6  RT (min)      lcqtof_hilic_neg_4_028_36    m/z 364.1605     0   2   4E+3   4.5   5  RT (min)      lcqtof_hilic_neg_4_032_01    m/z 364.1952     0   2   4E+3   4   4.5  RT (min)      lcqtof_hilic_neg_4_013_20    m/z 364.2617     0   2   4   6   8E+3   1   1.5  RT (min)       lcqtof_hilic_neg_4_048_23    m/z 364.9697     0   1   2   3   4E+3   5.5   6  RT (min)       lcqtof_hilic_neg_1_023_34    m/z 365.2206     0   2   4   6   8E+3   1   1.5  RT (min)      lcqtof_hilic_neg_4_040_05    m/z 366.0105     0   1   2   3   4E+3   6   6.5  RT (min)      lcqtof_hilic_neg_4_010_28    m/z 366.0665     0   1   2E+3   5   5.5  RT (min)       lcqtof_hilic_neg_4_002_25    m/z 366.1372     0   2   4E+3   4   4.5  RT (min)       lcqtof_hilic_neg_4_017_51    m/z 366.2713     0   2   4   6E+3   1   1.5  RT (min)      lcqtof_hilic_neg_4_053_14    m/z 366.2714     0   2   4   6E+3   0.5   1  RT (min)      lcqtof_hilic_neg_4_053_14    m/z 367.0363     0   1E+3   6   6.5  RT (min)      lcqtof_hilic_neg_1_008_13    m/z 367.0403     0   2   4   6E+3   6.5   7  RT (min)      lcqtof_hilic_neg_1_005_32    m/z 368.1201     0   1   2   3   4E+3   1.5   2  RT (min)      lcqtof_hilic_neg_4_042_22    m/z 368.2997     0   2   4   6   8E+3   3.5   4  RT (min)      lcqtof_hilic_neg_4_052_13    m/z 368.3600     0   1   2   3E+3   0.5   1  RT (min)      lcqtof_hilic_neg_4_054_48    m/z 368.9605     0   1E+3   5.5   6  RT (min)      lcqtof_hilic_neg_4_073_57    m/z 368.9624     0   2   4   6   8E+3   0.5   1  RT (min)      lcqtof_hilic_neg_3_066_57    m/z 369.0411     0   2   4E+3   5   5.5  RT (min)       lcqtof_hilic_neg_4_008_24    m/z 369.0973     0   1   2   3E+3   6.5   7  RT (min)      lcqtof_hilic_neg_4_048_23    m/z 369.1294     0   1   2E+3   6.5   7  RT (min)       lcqtof_hilic_neg_4_053_14    m/z 369.1878     0   1E+4   6   6.5  RT (min)      lcqtof_hilic_neg_4_008_24    m/z 369.2708     0   2   4   6E+3   1   1.5  RT (min)      lcqtof_hilic_neg_4_003_16    m/z 370.0435     0   1   2   3E+3   5   5.5  RT (min)      lcqtof_hilic_neg_4_055_32    m/z 370.0443     0   1   2E+3   4   4.5  RT (min)      lcqtof_hilic_neg_1_003_37    m/z 370.0633     0   1   2   3E+3   6.5   7  RT (min)      lcqtof_hilic_neg_4_008_24    m/z 370.0873     0   1   2E+3   6   6.5  RT (min)      lcqtof_hilic_neg_4_003_16    m/z 370.0962     0   2   4   6   8E+3   5.5   6  RT (min)      lcqtof_hilic_neg_4_008_24    m/z 370.1084     0   2   4E+3   6.5   7  RT (min)       lcqtof_hilic_neg_4_052_13    m/z 370.1992     0   2   4   6E+3   7   7.5  RT (min)      lcqtof_hilic_neg_4_048_23    m/z 370.8966     0   1   2   3E+3   0.5   1  RT (min)      lcqtof_hilic_neg_4_035_33    m/z 371.0057     0   1   2   3   4E+3   0.5   1  RT (min)      lcqtof_hilic_neg_1_028_47    m/z 372.0269     0   2   4   6   8E+3   4   4.5  RT (min)      lcqtof_hilic_neg_1_037_24    m/z 372.1004     0   1E+3   6   6.5  RT (min)      lcqtof_hilic_neg_1_005_32    m/z 372.1684     0   1   2E+3   6.5   7  RT (min)      lcqtof_hilic_neg_1_003_37    m/z 372.7913     0   1E+3   7.5   8  RT (min)      lcqtof_hilic_neg_1_009_50    m/z 372.9953     0   1   2   3   4E+3   1   1.5  RT (min)      lcqtof_hilic_neg_4_039_18    m/z 373.0521     0   1   2   3E+3   5.5   6  RT (min)      lcqtof_hilic_neg_4_003_16    m/z 373.1273     0   2   4   6   8E+3   4.5   5  RT (min)      lcqtof_hilic_neg_4_053_14    m/z 373.2003     0   2   4   6   8E+3   1   1.5  RT (min)      lcqtof_hilic_neg_4_053_14    m/z 373.2353     0   1   2   3E+3   0.5   1  RT (min)      lcqtof_hilic_neg_4_053_14    m/z 373.2680     0   1   2   3   4E+3   4.5   5  RT (min)      lcqtof_hilic_neg_4_008_24    m/z 374.0058     0   1   2E+3   7   7.5  RT (min)      lcqtof_hilic_neg_1_016_28    m/z 374.0067     0   1   2E+3   5.5   6  RT (min)       lcqtof_hilic_neg_4_053_14    m/z 374.1566     0   1   2   3E+3   4   4.5  RT (min)      lcqtof_hilic_neg_1_022_36    m/z 374.1614     0   2   4   6E+3   6   6.5  RT (min)      lcqtof_hilic_neg_4_009_50    m/z 374.3085     0   2   4   6   8E+3   1   1.5  RT (min)      lcqtof_hilic_neg_4_005_29    m/z 374.3752     0   2   4   6E+3   0.5   1  RT (min)      lcqtof_hilic_neg_4_008_24    m/z 374.9560     0   1E+3   5.5   6  RT (min)       lcqtof_hilic_neg_1_046_30    m/z 375.1666     0   2   4E+3   5.5   6  RT (min)       lcqtof_hilic_neg_4_008_24    m/z 375.2179     0   2   4   6E+3   0.5   1  RT (min)      lcqtof_hilic_neg_4_053_14    m/z 375.2518     0   1   2E+4   0.5   1  RT (min)      lcqtof_hilic_neg_4_003_16    m/z 375.3238     0   1   2   3E+3   0.5   1  RT (min)      lcqtof_hilic_neg_4_053_14    m/z 376.0361     0   1   2   3   4E+3   4   4.5  RT (min)      lcqtof_hilic_neg_1_032_18    m/z 376.2198     0   1   2E+3   7   7.5  RT (min)      lcqtof_hilic_neg_4_017_51    m/z 376.3222     0   1   2   3   4E+3   1   1.5  RT (min)      lcqtof_hilic_neg_2_013_29    m/z 376.3918     0   2   4   6   8E+3   0.5   1  RT (min)      lcqtof_hilic_neg_4_008_24    m/z 376.8565     0   1E+3   4.5   5  RT (min)      lcqtof_hilic_neg_3_012_27    m/z 377.0559     0   1E+4   1   1.5  RT (min)      lcqtof_hilic_neg_4_021_21    m/z 377.1534     0   1   2   3E+3   3.5   4  RT (min)      lcqtof_hilic_neg_2_031_01    m/z 377.2293     0   1   2E+3   4.5   5  RT (min)      lcqtof_hilic_neg_1_022_36    m/z 377.2414     0   1   2E+3   15   15.5  RT (min)       lcqtof_hilic_neg_4_012_07    m/z 377.3414     0   1   2E+3   0.5   1  RT (min)      lcqtof_hilic_neg_4_053_14    m/z 378.1047     0   1   2E+3   2   2.5  RT (min)      lcqtof_hilic_neg_4_073_57    m/z 378.1998     0   1   2   3E+3   1   1.5  RT (min)      lcqtof_hilic_neg_4_039_18    m/z 378.9061     0   1   2   3   4E+3   1   1.5  RT (min)      lcqtof_hilic_neg_4_006_30    m/z 378.9884     0   2   4   6   8E+3   0.5   1  RT (min)      lcqtof_hilic_neg_1_024_21    m/z 379.0907     0   1   2E+3   4   4.5  RT (min)      lcqtof_hilic_neg_4_013_20    m/z 379.1457     0   2   4   6   8E+3   4.5   5  RT (min)      lcqtof_hilic_neg_4_026_04    m/z 379.3480     0   1   2   3   4E+3   3.5   4  RT (min)      lcqtof_hilic_neg_4_055_32    m/z 379.3566     0   1   2   3E+3   0.5   1  RT (min)      lcqtof_hilic_neg_4_035_33    m/z 380.0535     0   1   2   3E+3   5.5   6  RT (min)      lcqtof_hilic_neg_4_028_36    m/z 380.0876     0   1   2E+3   7   7.5  RT (min)      lcqtof_hilic_neg_1_035_23    m/z 380.1218     0   1   2   3E+3   6   6.5  RT (min)      lcqtof_hilic_neg_1_011_46    m/z 380.2331     0   1   2   3E+3   7   7.5  RT (min)      lcqtof_hilic_neg_4_055_32    m/z 381.1848     0   1   2E+3   1   1.5  RT (min)      lcqtof_hilic_neg_4_077_58    m/z 381.1873     0   1   2E+3   5   5.5  RT (min)      lcqtof_hilic_neg_4_005_29    m/z 382.0347     0   2   4E+3   0.5   1  RT (min)      lcqtof_hilic_neg_4_004_47    m/z 382.0573     0   1   2E+3   4   4.5  RT (min)      lcqtof_hilic_neg_4_054_48    m/z 382.0668     0   1   2   3   4E+3   1   1.5  RT (min)      lcqtof_hilic_neg_1_032_18    m/z 382.2341     0   2   4   6E+3   1.5   2  RT (min)      lcqtof_hilic_neg_1_014_31    m/z 382.3212     0   2   4E+3   0.5   1  RT (min)      lcqtof_hilic_neg_2_015_06    m/z 382.3730     0   1   2E+3   0.5   1  RT (min)      lcqtof_hilic_neg_4_028_36    m/z 382.8887     0   2   4   6E+3   0.5   1  RT (min)      lcqtof_hilic_neg_3_081_58    m/z 382.8891     0   2   4   6   8E+3   0.5   1  RT (min)      lcqtof_hilic_neg_4_004_47    m/z 383.0327     0   1   2   3E+3   0.5   1  RT (min)      lcqtof_hilic_neg_4_004_47    m/z 383.1935     0   1   2   3   4E+3   7   7.5  RT (min)      lcqtof_hilic_neg_1_022_36    m/z 383.8447     0   2   4E+3   4.5   5  RT (min)      lcqtof_hilic_neg_4_053_14    m/z 383.9684     0   1E+3   4.5   5  RT (min)      lcqtof_hilic_neg_4_005_29    m/z 383.9774     0   1   2   3E+3   0.5   1  RT (min)      lcqtof_hilic_neg_3_066_57    m/z 383.9859     0   2   4E+3   0.5   1  RT (min)      lcqtof_hilic_neg_4_052_13    m/z 384.1480     0   2   4   6   8E+3   5   5.5  RT (min)      lcqtof_hilic_neg_4_050_02    m/z 384.2312     0   1   2E+3   4.5   5  RT (min)      lcqtof_hilic_neg_3_040_14    m/z 384.2742     0   1   2   3E+3   1   1.5  RT (min)      lcqtof_hilic_neg_4_034_31    m/z 384.9524     0   1   2E+3   0.5   1  RT (min)      lcqtof_hilic_neg_4_016_37    m/z 385.0703     0   2   4   6E+3   0.5   1  RT (min)      lcqtof_hilic_neg_4_018_27    m/z 385.9836     0   1   2   3E+3   0.5   1  RT (min)      lcqtof_hilic_neg_4_052_13    m/z 386.0452     0   1   2E+3   0.5   1  RT (min)      lcqtof_hilic_neg_2_016_33    m/z 386.1178     0   1E+4   0.5   1  RT (min)      lcqtof_hilic_neg_4_022_34    m/z 386.1349     0   1   2   3E+3   1   1.5  RT (min)      lcqtof_hilic_neg_4_026_04    m/z 386.3376     0   2   4   6E+3   0.5   1  RT (min)      lcqtof_hilic_neg_4_053_14    m/z 387.1639     0   2   4   6E+3   5   5.5  RT (min)      lcqtof_hilic_neg_2_047_32    m/z 387.2536     0   2   4   6E+3   1   1.5  RT (min)      lcqtof_hilic_neg_4_006_30    m/z 387.9422     0   1E+3   5   5.5  RT (min)      lcqtof_hilic_neg_4_004_47    m/z 387.9751     0   2   4   6   8E+3   1   1.5  RT (min)      lcqtof_hilic_neg_2_050_19    m/z 388.0442     0   2   4   6E+3   6   6.5  RT (min)      lcqtof_hilic_neg_1_023_34    m/z 388.4207     0   1   2E+3   0.5   1  RT (min)      lcqtof_hilic_neg_4_013_20    m/z 388.9697     0   2   4E+3   1   1.5  RT (min)      lcqtof_hilic_neg_2_050_19    m/z 389.0262     0   1   2E+3   5.5   6  RT (min)      lcqtof_hilic_neg_4_052_13    m/z 389.0547     0   2   4   6E+3   0.5   1  RT (min)      lcqtof_hilic_neg_4_005_29    m/z 389.1601     0   1E+3   1   1.5  RT (min)       lcqtof_hilic_neg_3_043_13    m/z 389.2068     0   1   2   3E+3   4   4.5  RT (min)      lcqtof_hilic_neg_2_047_32    m/z 389.3505     0   2   4   6E+3   0.5   1  RT (min)      lcqtof_hilic_neg_4_055_32    m/z 390.0578     0   1   2E+3   3.5   4  RT (min)      lcqtof_hilic_neg_4_016_37    m/z 390.1228     0   1   2   3E+3   7   7.5  RT (min)      lcqtof_hilic_neg_2_052_24    m/z 390.2415     0   2   4   6E+3   4   4.5  RT (min)      lcqtof_hilic_neg_4_053_14    m/z 390.9814     0   1   2E+3   8.5   9  RT (min)      lcqtof_hilic_neg_1_003_37    m/z 391.0714     0   1   2   3E+3   6   6.5  RT (min)      lcqtof_hilic_neg_4_031_38    m/z 391.1203     0   1   2   3   4E+3   6.5   7  RT (min)      lcqtof_hilic_neg_1_005_32    m/z 391.1363     0   2   4   6   8E+3   6   6.5  RT (min)       lcqtof_hilic_neg_4_008_24    m/z 392.0069     0   1   2   3E+3   2   2.5  RT (min)       lcqtof_hilic_neg_3_111_62    m/z 392.0409     0   1   2E+3   3.5   4  RT (min)      lcqtof_hilic_neg_2_031_01    m/z 392.2487     0   1   2E+3   5   5.5  RT (min)      lcqtof_hilic_neg_4_034_31    m/z 392.2871     0   2   4   6E+3   0.5   1  RT (min)      lcqtof_hilic_neg_4_053_14    m/z 392.3166     0   2   4E+3   0.5   1  RT (min)      lcqtof_hilic_neg_4_053_14    m/z 392.7786     0   1E+3   7.5   8  RT (min)      lcqtof_hilic_neg_1_027_27    m/z 393.1084     0   1   2E+3   8   8.5  RT (min)       lcqtof_hilic_neg_2_002_36    m/z 393.2448     0   1   2   3E+3   5   5.5  RT (min)      lcqtof_hilic_neg_4_034_31    m/z 393.2847     0   1   2E+3   1   1.5  RT (min)      lcqtof_hilic_neg_4_003_16    m/z 393.6248     0   1   2   3E+3   5   5.5  RT (min)      lcqtof_hilic_neg_3_012_27    m/z 393.8365     0   1E+3   8   8.5  RT (min)      lcqtof_hilic_neg_2_019_12    m/z 393.9682     0   1   2E+3   5   5.5  RT (min)      lcqtof_hilic_neg_4_032_01    m/z 394.1303     0   2   4   6E+3   6   6.5  RT (min)      lcqtof_hilic_neg_4_002_25    m/z 394.2969     0   1   2   3   4E+3   0.5   1  RT (min)      lcqtof_hilic_neg_3_025_52    m/z 395.0604     0   1   2   3   4E+3   6.5   7  RT (min)      lcqtof_hilic_neg_4_040_05    m/z 395.1046     0   2   4E+3   0.5   1  RT (min)      lcqtof_hilic_neg_2_016_33    m/z 395.1338     0   2   4   6E+4   5.5   6  RT (min)      lcqtof_hilic_neg_1_049_55    m/z 395.1373     0   1   2E+3   4.5   5  RT (min)      lcqtof_hilic_neg_4_085_59    m/z 395.3166     0   1   2E+3   0.5   1  RT (min)      lcqtof_hilic_neg_1_004_14    m/z 396.3950     0   1   2   3   4E+3   0.5   1  RT (min)      lcqtof_hilic_neg_2_016_33    m/z 397.0458     0   1   2E+3   0.5   1  RT (min)      lcqtof_hilic_neg_2_007_03    m/z 397.1966     0   2   4   6   8E+3   6   6.5  RT (min)      lcqtof_hilic_neg_4_008_24    m/z 397.2223     0   1   2E+3   7   7.5  RT (min)      lcqtof_hilic_neg_4_055_32    m/z 397.2448     0   1   2   3E+3   4.5   5  RT (min)      lcqtof_hilic_neg_4_048_23    m/z 397.3637     0   1E+3   15   15.5  RT (min)       lcqtof_hilic_neg_4_001_49    m/z 397.9044     0   2   4   6E+3   0.5   1  RT (min)      lcqtof_hilic_neg_4_004_47    m/z 397.9062     0   2   4   6E+3   0.5   1  RT (min)      lcqtof_hilic_neg_4_004_47    m/z 398.0701     0   1   2E+4   6   6.5  RT (min)      lcqtof_hilic_neg_4_014_46    m/z 398.9612     0   1   2   3E+3   4   4.5  RT (min)      lcqtof_hilic_neg_1_005_32    m/z 399.0009     0   1   2E+3   5.5   6  RT (min)      lcqtof_hilic_neg_4_003_16    m/z 399.0832     0   1   2   3E+3   6   6.5  RT (min)      lcqtof_hilic_neg_2_052_24    m/z 399.1151     0   1   2   3   4E+3   1   1.5  RT (min)      lcqtof_hilic_neg_3_042_02    m/z 399.1521     0   2   4E+3   0.5   1  RT (min)       lcqtof_hilic_neg_4_081_58    m/z 399.1861     0   2   4E+3   1   1.5  RT (min)       lcqtof_hilic_neg_4_032_01    m/z 399.2408     0   1E+3   7   7.5  RT (min)      lcqtof_hilic_neg_4_040_05    m/z 400.1452     0   1   2E+3   0.5   1  RT (min)      lcqtof_hilic_neg_1_004_14    m/z 400.9978     0   2   4   6E+3   6.5   7  RT (min)      lcqtof_hilic_neg_1_014_31    m/z 401.0244     0   2   4   6E+3   7   7.5  RT (min)       lcqtof_hilic_neg_4_054_48    m/z 401.1083     0   2   4E+3   2.5   3  RT (min)       lcqtof_hilic_neg_4_013_20    m/z 401.1279     0   1   2   3E+3   0.5   1  RT (min)      lcqtof_hilic_neg_4_053_14    m/z 401.1585     0   1   2   3   4E+3   7   7.5  RT (min)      lcqtof_hilic_neg_2_052_24    m/z 401.1644     0   1   2E+3   15   15.5  RT (min)       lcqtof_hilic_neg_4_012_07    m/z 401.1953     0   1   2   3   4E+3   0.5   1  RT (min)      lcqtof_hilic_neg_4_005_29    m/z 401.2115     0   1   2E+3   5.5   6  RT (min)      lcqtof_hilic_neg_4_011_35    m/z 401.9533     0   1E+3   6   6.5  RT (min)      lcqtof_hilic_neg_2_002_36    m/z 402.1451     0   1E+3   0.5   1  RT (min)      lcqtof_hilic_neg_1_010_15    m/z 402.1993     0   2   4E+3   1   1.5  RT (min)      lcqtof_hilic_neg_4_039_18    m/z 402.1995     0   2   4E+3   5.5   6  RT (min)       lcqtof_hilic_neg_4_048_23    m/z 403.0555     0   2   4E+3   6   6.5  RT (min)      lcqtof_hilic_neg_1_035_23    m/z 403.0765     0   1   2   3E+3   5   5.5  RT (min)      lcqtof_hilic_neg_4_053_14    m/z 403.2673     0   1   2E+3   15   15.5  RT (min)      lcqtof_hilic_neg_1_005_32    m/z 403.2833     0   2   4   6   8E+3   0.5   1  RT (min)      lcqtof_hilic_neg_4_052_13    m/z 403.3062     0   1   2E+3   1   1.5  RT (min)      lcqtof_hilic_neg_4_015_09    m/z 403.3102     0   1   2E+3   0.5   1  RT (min)      lcqtof_hilic_neg_2_055_30    m/z 404.0623     0   1   2E+3   1.5   2  RT (min)      lcqtof_hilic_neg_4_040_05    m/z 404.1034     0   2   4   6E+3   2   2.5  RT (min)       lcqtof_hilic_neg_3_073_57    m/z 404.1698     0   1   2   3E+3   6   6.5  RT (min)      lcqtof_hilic_neg_4_052_13    m/z 404.2824     0   2   4   6   8E+3   1   1.5  RT (min)      lcqtof_hilic_neg_4_038_10    m/z 404.2929     0   2   4   6E+3   0.5   1  RT (min)      lcqtof_hilic_neg_1_012_04    m/z 404.9141     0   1   2E+3   5.5   6  RT (min)      lcqtof_hilic_neg_4_047_06    m/z 404.9368     0   1   2   3   4E+3   1   1.5  RT (min)      lcqtof_hilic_neg_4_004_47    m/z 404.9991     0   1   2E+3   5.5   6  RT (min)      lcqtof_hilic_neg_4_052_13    m/z 405.0803     0   1   2   3   4E+3   4.5   5  RT (min)      lcqtof_hilic_neg_4_048_23    m/z 405.0939     0   1   2   3E+3   0.5   1  RT (min)      lcqtof_hilic_neg_1_001_49    m/z 405.1714     0   1   2   3   4E+3   0.5   1  RT (min)      lcqtof_hilic_neg_4_027_19    m/z 405.1874     0   2   4   6E+3   4.5   5  RT (min)      lcqtof_hilic_neg_1_027_27    m/z 405.2161     0   2   4   6   8E+3   3.5   4  RT (min)      lcqtof_hilic_neg_1_007_05    m/z 405.2615     0   2   4E+3   7   7.5  RT (min)       lcqtof_hilic_neg_4_008_24    m/z 406.0696     0   1   2E+3   0.5   1  RT (min)      lcqtof_hilic_neg_4_050_02    m/z 406.1074     0   1   2   3E+3   7   7.5  RT (min)      lcqtof_hilic_neg_1_035_23    m/z 406.1084     0   2   4   6   8E+3   6   6.5  RT (min)      lcqtof_hilic_neg_3_011_34    m/z 406.1320     0   1   2   3E+3   0.5   1  RT (min)      lcqtof_hilic_neg_2_037_21    m/z 406.1328     0   2   4E+3   0.5   1  RT (min)      lcqtof_hilic_neg_4_053_14    m/z 406.1515     0   1   2   3E+3   8   8.5  RT (min)       lcqtof_hilic_neg_4_008_24    m/z 406.2912     0   1   2E+3   3.5   4  RT (min)      lcqtof_hilic_neg_3_007_04    m/z 406.3671     0   1   2   3   4E+3   1   1.5  RT (min)      lcqtof_hilic_neg_4_008_24    m/z 406.9033     0   1   2E+3   4.5   5  RT (min)      lcqtof_hilic_neg_2_031_01    m/z 407.0168     0   2   4   6E+3   5   5.5  RT (min)      lcqtof_hilic_neg_4_008_24    m/z 407.0820     0   1E+4   1   1.5  RT (min)      lcqtof_hilic_neg_3_012_27    m/z 407.1767     0   2   4   6   8E+3   5.5   6  RT (min)      lcqtof_hilic_neg_4_014_46    m/z 407.2821     0   2   4   6   8E+3   0.5   1  RT (min)      lcqtof_hilic_neg_4_053_14    m/z 407.2974     0   1   2   3   4E+3   0.5   1  RT (min)      lcqtof_hilic_neg_2_020_23    m/z 407.3163     0   1   2   3   4E+3   0.5   1  RT (min)      lcqtof_hilic_neg_4_053_14    m/z 408.0002     0   1   2   3   4E+3   6   6.5  RT (min)      lcqtof_hilic_neg_1_023_34    m/z 408.1506     0   1   2   3   4E+3   1   1.5  RT (min)      lcqtof_hilic_neg_4_003_16    m/z 408.2836     0   2   4E+3   0.5   1  RT (min)      lcqtof_hilic_neg_4_003_16    m/z 409.1642     0   1   2   3E+3   5.5   6  RT (min)       lcqtof_hilic_neg_4_033_53    m/z 409.1897     0   2   4E+3   1   1.5  RT (min)      lcqtof_hilic_neg_4_032_01    m/z 409.2072     0   1   2   3E+3   1.5   2  RT (min)      lcqtof_hilic_neg_4_048_23    m/z 409.9969     0   1   2   3   4E+3   7   7.5  RT (min)      lcqtof_hilic_neg_2_001_49    m/z 410.2377     0   2   4   6E+3   4.5   5  RT (min)      lcqtof_hilic_neg_4_040_05    m/z 411.0768     0   1E+3   4.5   5  RT (min)      lcqtof_hilic_neg_1_013_38    m/z 411.1791     0   1E+4   0.5   1  RT (min)      lcqtof_hilic_neg_4_050_02    m/z 411.2646     0   1   2   3E+3   0.5   1  RT (min)      lcqtof_hilic_neg_4_048_23    m/z 412.0453     0   1   2E+3   1   1.5  RT (min)      lcqtof_hilic_neg_2_007_03    m/z 412.0819     0   1   2E+3   0.5   1  RT (min)      lcqtof_hilic_neg_1_035_23    m/z 412.1447     0   2   4E+3   5   5.5  RT (min)       lcqtof_hilic_neg_4_008_24    m/z 412.2317     0   2   4E+3   0.5   1  RT (min)      lcqtof_hilic_neg_1_036_03    m/z 412.2899     0   1E+4   1   1.5  RT (min)      lcqtof_hilic_neg_4_052_13    m/z 412.3566     0   2   4   6E+3   0.5   1  RT (min)      lcqtof_hilic_neg_2_029_31    m/z 412.3862     0   1E+4   1   1.5  RT (min)      lcqtof_hilic_neg_4_040_05    m/z 413.0414     0   1   2   3   4E+3   6   6.5  RT (min)      lcqtof_hilic_neg_2_022_46    m/z 413.1031     0   1   2E+3   1   1.5  RT (min)      lcqtof_hilic_neg_1_029_06    m/z 413.2224     0   1E+3   15   15.5  RT (min)       lcqtof_hilic_neg_4_012_07    m/z 413.2437     0   2   4   6   8E+3   1   1.5  RT (min)      lcqtof_hilic_neg_4_045_26    m/z 414.1829     0   1   2   3   4E+3   6   6.5  RT (min)      lcqtof_hilic_neg_2_052_24    m/z 414.2054     0   1   2   3E+3   1   1.5  RT (min)      lcqtof_hilic_neg_2_051_14    m/z 414.2887     0   2   4E+3   1   1.5  RT (min)      lcqtof_hilic_neg_4_007_45    m/z 414.3039     0   1   2   3E+3   3.5   4  RT (min)      lcqtof_hilic_neg_2_044_37    m/z 414.3652     0   2   4E+3   0.5   1  RT (min)      lcqtof_hilic_neg_4_053_14    m/z 414.9140     0   2   4   6   8E+3   0.5   1  RT (min)      lcqtof_hilic_neg_3_081_58    m/z 415.0854     0   1   2   3E+3   1   1.5  RT (min)      lcqtof_hilic_neg_2_053_18    m/z 415.2041     0   2   4   6   8E+3   7   7.5  RT (min)       lcqtof_hilic_neg_3_030_23    m/z 415.2463     0   1   2   3   4E+3   0.5   1  RT (min)      lcqtof_hilic_neg_4_053_14    m/z 416.0119     0   1E+4   7   7.5  RT (min)      lcqtof_hilic_neg_4_055_32    m/z 416.1434     0   2   4E+3   6   6.5  RT (min)      lcqtof_hilic_neg_4_008_24    m/z 416.1508     0   1   2E+3   5.5   6  RT (min)      lcqtof_hilic_neg_2_033_53    m/z 416.2075     0   2   4E+3   0.5   1  RT (min)      lcqtof_hilic_neg_2_015_06    m/z 416.2855     0   2   4   6E+3   0.5   1  RT (min)      lcqtof_hilic_neg_4_048_23    m/z 417.1429     0   2   4   6   8E+3   5.5   6  RT (min)      lcqtof_hilic_neg_4_013_20    m/z 417.3284     0   1   2   3E+3   3.5   4  RT (min)      lcqtof_hilic_neg_4_034_31    m/z 417.3705     0   2   4E+3   0.5   1  RT (min)      lcqtof_hilic_neg_4_008_24    m/z 417.9543     0   1E+3   4.5   5  RT (min)      lcqtof_hilic_neg_4_023_15    m/z 418.0662     0   1   2E+3   4   4.5  RT (min)      lcqtof_hilic_neg_1_041_54    m/z 418.1484     0   2   4E+3   7   7.5  RT (min)      lcqtof_hilic_neg_1_037_24    m/z 418.2151     0   1E+3   15   15.5  RT (min)      lcqtof_hilic_neg_4_019_03    m/z 418.3002     0   1   2   3E+3   0.5   1  RT (min)      lcqtof_hilic_neg_4_053_14    m/z 418.3260     0   1   2E+3   1   1.5  RT (min)      lcqtof_hilic_neg_1_032_18    m/z 418.9535     0   1   2   3   4E+3   1   1.5  RT (min)      lcqtof_hilic_neg_4_087_59    m/z 419.1075     0   1   2   3E+3   6   6.5  RT (min)      lcqtof_hilic_neg_1_047_29    m/z 419.1673     0   1E+3   15   15.5  RT (min)       lcqtof_hilic_neg_4_051_43    m/z 419.2133     0   1E+4   1   1.5  RT (min)      lcqtof_hilic_neg_4_047_06    m/z 419.2800     0   2   4   6   8E+3   0.5   1  RT (min)      lcqtof_hilic_neg_4_023_15    m/z 419.4619     0   1   2   3E+3   0.5   1  RT (min)      lcqtof_hilic_neg_4_008_24    m/z 419.9978     0   1   2   3   4E+3   1   1.5  RT (min)      lcqtof_hilic_neg_1_032_18    m/z 420.1380     0   1   2   3   4E+3   6   6.5  RT (min)       lcqtof_hilic_neg_1_037_24    m/z 420.1565     0   1   2   3E+3   0.5   1  RT (min)      lcqtof_hilic_neg_2_051_14    m/z 420.1647     0   2   4E+3   3   3.5  RT (min)       lcqtof_hilic_neg_3_090_60    m/z 420.3208     0   1   2   3   4E+3   1   1.5  RT (min)      lcqtof_hilic_neg_4_040_05    m/z 420.8405     0   1E+3   8   8.5  RT (min)      lcqtof_hilic_neg_2_027_07    m/z 421.0434     0   1   2E+3   1   1.5  RT (min)      lcqtof_hilic_neg_4_015_09    m/z 421.1048     0   2   4   6   8E+3   7   7.5  RT (min)      lcqtof_hilic_neg_1_031_16    m/z 422.0272     0   2   4E+3   6.5   7  RT (min)      lcqtof_hilic_neg_1_037_24    m/z 422.1776     0   1   2   3E+3   6.5   7  RT (min)      lcqtof_hilic_neg_4_028_36    m/z 422.2844     0   2   4E+3   1   1.5  RT (min)      lcqtof_hilic_neg_4_048_23    m/z 423.0072     0   1   2E+3   7   7.5  RT (min)      lcqtof_hilic_neg_4_054_48    m/z 423.0826     0   1E+4   1   1.5  RT (min)      lcqtof_hilic_neg_4_015_09    m/z 423.1595     0   1   2E+3   3.5   4  RT (min)      lcqtof_hilic_neg_1_032_18    m/z 423.1609     0   1   2   3E+3   0.5   1  RT (min)      lcqtof_hilic_neg_4_047_06    m/z 423.3161     0   2   4E+3   0.5   1  RT (min)      lcqtof_hilic_neg_4_003_16    m/z 423.4182     0   1   2E+3   0.5   1  RT (min)      lcqtof_hilic_neg_4_028_36    m/z 423.9912     0   2   4   6E+3   5.5   6  RT (min)      lcqtof_hilic_neg_1_022_36    m/z 424.1332     0   1   2   3   4E+3   3.5   4  RT (min)      lcqtof_hilic_neg_1_053_19    m/z 424.1580     0   2   4   6E+3   6   6.5  RT (min)      lcqtof_hilic_neg_4_040_05    m/z 424.2482     0   1   2E+3   4   4.5  RT (min)      lcqtof_hilic_neg_3_048_05    m/z 425.0682     0   1   2   3   4E+3   5.5   6  RT (min)      lcqtof_hilic_neg_4_003_16    m/z 425.3011     0   2   4   6E+3   2   2.5  RT (min)      lcqtof_hilic_neg_4_003_16    m/z 425.3034     0   2   4   6E+3   1   1.5  RT (min)       lcqtof_hilic_neg_4_020_42    m/z 425.9914     0   1   2E+3   0.5   1  RT (min)      lcqtof_hilic_neg_4_032_01    m/z 427.0021     0   1   2E+3   4   4.5  RT (min)      lcqtof_hilic_neg_1_024_21    m/z 427.2616     0   1   2   3E+3   1   1.5  RT (min)      lcqtof_hilic_neg_2_031_01    m/z 427.3666     0   1E+3   15   15.5  RT (min)      lcqtof_hilic_neg_4_047_06    m/z 427.9789     0   1   2E+3   5.5   6  RT (min)       lcqtof_hilic_neg_4_048_23    m/z 427.9882     0   1   2E+3   0.5   1  RT (min)      lcqtof_hilic_neg_4_028_36    m/z 428.0370     0   1E+3   5.5   6  RT (min)      lcqtof_hilic_neg_2_001_49    m/z 428.0748     0   1E+3   0.5   1  RT (min)      lcqtof_hilic_neg_2_020_23    m/z 429.0818     0   1   2   3   4E+3   3.5   4  RT (min)      lcqtof_hilic_neg_2_050_19    m/z 429.0824     0   1E+3   6   6.5  RT (min)      lcqtof_hilic_neg_2_034_47    m/z 429.0864     0   1   2   3   4E+3   1   1.5  RT (min)      lcqtof_hilic_neg_4_098_61    m/z 429.1248     0   2   4   6   8E+3   3   3.5  RT (min)      lcqtof_hilic_neg_4_027_19    m/z 429.1614     0   2   4E+3   5.5   6  RT (min)      lcqtof_hilic_neg_4_014_46    m/z 429.1624     0   1   2E+3   5   5.5  RT (min)      lcqtof_hilic_neg_2_050_19    m/z 429.2119     0   1   2   3   4E+3   1   1.5  RT (min)      lcqtof_hilic_neg_2_050_19    m/z 429.3116     0   2   4   6   8E+3   0.5   1  RT (min)      lcqtof_hilic_neg_4_003_16    m/z 429.3324     0   2   4   6E+3   0.5   1  RT (min)      lcqtof_hilic_neg_4_053_14    m/z 429.3708     0   1   2E+3   0.5   1  RT (min)      lcqtof_hilic_neg_4_052_13    m/z 429.9292     0   1   2   3   4E+3   0.5   1  RT (min)      lcqtof_hilic_neg_4_004_47    m/z 430.0816     0   2   4   6E+3   5.5   6  RT (min)       lcqtof_hilic_neg_1_005_32    m/z 430.1129     0   2   4E+3   7.5   8  RT (min)      lcqtof_hilic_neg_4_002_25    m/z 430.1137     0   2   4E+3   7.5   8  RT (min)      lcqtof_hilic_neg_3_048_05    m/z 431.0017     0   1   2   3E+3   6.5   7  RT (min)      lcqtof_hilic_neg_1_016_28    m/z 431.1408     0   2   4E+3   7   7.5  RT (min)      lcqtof_hilic_neg_2_020_23    m/z 431.3105     0   1   2E+3   3.5   4  RT (min)      lcqtof_hilic_neg_4_048_23    m/z 431.9716     0   2   4   6E+3   0.5   1  RT (min)      lcqtof_hilic_neg_4_004_47    m/z 432.1873     0   2   4E+3   0.5   1  RT (min)      lcqtof_hilic_neg_4_047_06    m/z 433.0677     0   1   2E+3   6   6.5  RT (min)      lcqtof_hilic_neg_4_066_57    m/z 433.0906     0   1   2   3   4E+3   4   4.5  RT (min)       lcqtof_hilic_neg_1_053_19    m/z 433.1285     0   1   2   3   4E+3   6   6.5  RT (min)      lcqtof_hilic_neg_2_022_46    m/z 433.1738     0   2   4   6E+3   6   6.5  RT (min)      lcqtof_hilic_neg_4_014_46    m/z 433.2256     0   2   4   6E+3   1   1.5  RT (min)      lcqtof_hilic_neg_1_054_01    m/z 433.2951     0   2   4   6   8E+3   0.5   1  RT (min)      lcqtof_hilic_neg_4_003_16    m/z 433.3522     0   1   2E+3   4.5   5  RT (min)      lcqtof_hilic_neg_4_004_47    m/z 433.5818     0   2   4   6E+3   5   5.5  RT (min)      lcqtof_hilic_neg_1_006_22    m/z 434.0535     0   1   2E+3   5   5.5  RT (min)      lcqtof_hilic_neg_4_026_04    m/z 434.0551     0   1   2E+3   5   5.5  RT (min)      lcqtof_hilic_neg_4_012_07    m/z 434.1208     0   1   2   3E+3   1.5   2  RT (min)      lcqtof_hilic_neg_2_026_28    m/z 434.2195     0   2   4   6   8E+3   0.5   1  RT (min)      lcqtof_hilic_neg_2_033_53    m/z 434.2560     0   1   2   3   4E+3   4   4.5  RT (min)       lcqtof_hilic_neg_1_037_24    m/z 434.3335     0   1   2   3   4E+3   0.5   1  RT (min)      lcqtof_hilic_neg_2_051_14    m/z 435.0418     0   1   2   3E+3   4   4.5  RT (min)      lcqtof_hilic_neg_4_002_25    m/z 435.0830     0   1   2   3E+3   0.5   1  RT (min)      lcqtof_hilic_neg_4_044_08    m/z 435.1130     0   2   4E+3   8   8.5  RT (min)      lcqtof_hilic_neg_4_055_32    m/z 435.1611     0   2   4E+3   5   5.5  RT (min)       lcqtof_hilic_neg_2_001_49    m/z 435.2143     0   1   2E+3   0.5   1  RT (min)      lcqtof_hilic_neg_4_006_30    m/z 435.3091     0   1   2   3E+4   0.5   1  RT (min)       lcqtof_hilic_neg_4_003_16    m/z 435.3195     0   1E+3   3.5   4  RT (min)      lcqtof_hilic_neg_4_036_11    m/z 435.9637     0   1   2E+3   1   1.5  RT (min)      lcqtof_hilic_neg_1_040_20    m/z 436.0836     0   2   4E+3   1   1.5  RT (min)      lcqtof_hilic_neg_1_026_10    m/z 436.1325     0   1   2   3   4E+3   1   1.5  RT (min)      lcqtof_hilic_neg_4_053_14    m/z 436.9156     0   2   4   6   8E+3   0.5   1  RT (min)      lcqtof_hilic_neg_4_004_47    m/z 436.9480     0   1   2E+3   5.5   6  RT (min)       lcqtof_hilic_neg_4_026_04    m/z 437.0288     0   2   4E+3   5   5.5  RT (min)      lcqtof_hilic_neg_4_008_24    m/z 437.1278     0   2   4   6   8E+3   1   1.5  RT (min)      lcqtof_hilic_neg_4_053_14    m/z 437.4337     0   1   2E+3   0.5   1  RT (min)      lcqtof_hilic_neg_4_031_38    m/z 438.0314     0   1   2   3E+3   5   5.5  RT (min)      lcqtof_hilic_neg_4_055_32    m/z 438.1354     0   1   2E+3   0.5   1  RT (min)      lcqtof_hilic_neg_1_035_23    m/z 438.2563     0   2   4   6E+3   4   4.5  RT (min)      lcqtof_hilic_neg_1_030_17    m/z 438.2656     0   2   4   6   8E+3   3.5   4  RT (min)      lcqtof_hilic_neg_1_007_05    m/z 439.0709     0   1   2   3   4E+3   1   1.5  RT (min)      lcqtof_hilic_neg_4_035_33    m/z 439.1259     0   2   4E+3   5.5   6  RT (min)       lcqtof_hilic_neg_4_012_07    m/z 439.1531     0   1   2   3E+3   6.5   7  RT (min)      lcqtof_hilic_neg_4_052_13    m/z 439.1633     0   1   2E+3   3.5   4  RT (min)      lcqtof_hilic_neg_1_022_36    m/z 439.3648     0   1   2   3   4E+4   0.5   1  RT (min)      lcqtof_hilic_neg_2_052_24    m/z 440.2360     0   2   4   6E+3   1   1.5  RT (min)      lcqtof_hilic_neg_1_032_18    m/z 440.3437     0   1   2   3E+3   0.5   1  RT (min)      lcqtof_hilic_neg_4_005_29    m/z 441.0338     0   2   4   6   8E+3   6   6.5  RT (min)       lcqtof_hilic_neg_3_011_34    m/z 441.0339     0   1   2   3   4E+3   7   7.5  RT (min)      lcqtof_hilic_neg_2_020_23    m/z 441.0408     0   1   2   3E+3   5.5   6  RT (min)      lcqtof_hilic_neg_4_018_27    m/z 441.1128     0   1   2E+3   6   6.5  RT (min)      lcqtof_hilic_neg_4_048_23    m/z 441.3450     0   1   2E+3   4.5   5  RT (min)       lcqtof_hilic_neg_4_004_47    m/z 441.9754     0   1E+3   7   7.5  RT (min)      lcqtof_hilic_neg_2_026_28    m/z 441.9917     0   1   2E+3   5.5   6  RT (min)       lcqtof_hilic_neg_4_028_36    m/z 442.0883     0   1   2   3E+3   6   6.5  RT (min)      lcqtof_hilic_neg_1_007_05    m/z 442.1366     0   2   4   6E+3   1.5   2  RT (min)      lcqtof_hilic_neg_4_023_15    m/z 442.2075     0   2   4   6   8E+3   3.5   4  RT (min)      lcqtof_hilic_neg_2_020_23    m/z 442.9656     0   1   2   3   4E+3   7   7.5  RT (min)      lcqtof_hilic_neg_2_007_03    m/z 443.0840     0   1   2E+3   5   5.5  RT (min)      lcqtof_hilic_neg_2_039_38    m/z 443.0976     0   1   2E+3   1   1.5  RT (min)      lcqtof_hilic_neg_1_023_34    m/z 443.2037     0   1   2E+3   7   7.5  RT (min)      lcqtof_hilic_neg_4_044_08    m/z 443.3036     0   1   2E+3   15   15.5  RT (min)       lcqtof_hilic_neg_4_047_06    m/z 443.4019     0   2   4   6E+3   0.5   1  RT (min)      lcqtof_hilic_neg_4_053_14    m/z 443.9422     0   2   4   6E+3   0.5   1  RT (min)      lcqtof_hilic_neg_4_004_47    m/z 443.9492     0   2   4   6E+3   0.5   1  RT (min)      lcqtof_hilic_neg_4_004_47    m/z 444.0923     0   1   2   3E+3   6   6.5  RT (min)      lcqtof_hilic_neg_4_040_05    m/z 444.9253     0   1E+3   6   6.5  RT (min)      lcqtof_hilic_neg_1_012_04    m/z 445.0657     0   1   2   3   4E+3   3.5   4  RT (min)       lcqtof_hilic_neg_1_053_19    m/z 445.0782     0   1   2   3   4E+3   4.5   5  RT (min)      lcqtof_hilic_neg_2_050_19    m/z 445.1500     0   2   4   6E+3   6.5   7  RT (min)      lcqtof_hilic_neg_4_040_05    m/z 445.1663     0   1   2E+3   6.5   7  RT (min)      lcqtof_hilic_neg_2_020_23    m/z 445.2418     0   2   4   6E+3   0.5   1  RT (min)      lcqtof_hilic_neg_4_047_06    m/z 445.2572     0   2   4   6E+3   0.5   1  RT (min)      lcqtof_hilic_neg_4_011_35    m/z 445.3580     0   1   2E+3   3.5   4  RT (min)      lcqtof_hilic_neg_4_048_23    m/z 445.9796     0   2   4   6   8E+3   6.5   7  RT (min)      lcqtof_hilic_neg_3_048_05    m/z 446.1238     0   2   4   6   8E+3   7   7.5  RT (min)       lcqtof_hilic_neg_3_066_57    m/z 447.0614     0   1   2E+3   8   8.5  RT (min)      lcqtof_hilic_neg_4_016_37    m/z 447.1180     0   2   4   6E+3   7   7.5  RT (min)       lcqtof_hilic_neg_1_014_31    m/z 447.1725     0   2   4E+3   0.5   1  RT (min)      lcqtof_hilic_neg_1_004_14    m/z 447.2960     0   1   2   3   4E+3   1.5   2  RT (min)      lcqtof_hilic_neg_4_032_01    m/z 448.2781     0   2   4   6E+3   1   1.5  RT (min)      lcqtof_hilic_neg_4_053_14    m/z 448.2956     0   1   2E+4   0.5   1  RT (min)      lcqtof_hilic_neg_4_003_16    m/z 448.9823     0   1   2   3E+3   1   1.5  RT (min)      lcqtof_hilic_neg_4_076_58    m/z 449.0112     0   1E+3   5.5   6  RT (min)      lcqtof_hilic_neg_2_033_53    m/z 449.0861     0   1   2   3   4E+3   6   6.5  RT (min)       lcqtof_hilic_neg_4_072_57    m/z 449.1242     0   2   4E+3   1   1.5  RT (min)      lcqtof_hilic_neg_1_032_18    m/z 449.1526     0   1   2   3E+3   2.5   3  RT (min)      lcqtof_hilic_neg_4_027_19    m/z 449.1761     0   2   4E+3   0.5   1  RT (min)      lcqtof_hilic_neg_4_023_15    m/z 449.2156     0   1   2E+3   0.5   1  RT (min)      lcqtof_hilic_neg_4_025_52    m/z 449.2245     0   1   2   3E+3   7   7.5  RT (min)      lcqtof_hilic_neg_4_044_08    m/z 450.2144     0   1   2   3   4E+3   0.5   1  RT (min)      lcqtof_hilic_neg_4_048_23    m/z 450.2872     0   2   4   6   8E+3   0.5   1  RT (min)      lcqtof_hilic_neg_4_047_06    m/z 451.0719     0   1   2E+3   7   7.5  RT (min)      lcqtof_hilic_neg_1_023_34    m/z 451.2018     0   2   4   6   8E+3   4   4.5  RT (min)      lcqtof_hilic_neg_1_041_54    m/z 451.3229     0   2   4   6E+3   1   1.5  RT (min)      lcqtof_hilic_neg_4_022_34    m/z 451.4514     0   1E+3   15.5   16  RT (min)       lcqtof_hilic_neg_4_074_58    m/z 452.1320     0   2   4   6E+3   6.5   7  RT (min)       lcqtof_hilic_neg_4_002_25    m/z 452.2788     0   1   2E+3   3   3.5  RT (min)      lcqtof_hilic_neg_4_083_59    m/z 452.3316     0   2   4E+3   3   3.5  RT (min)       lcqtof_hilic_neg_4_016_37    m/z 452.8918     0   1   2   3   4E+3   0.5   1  RT (min)      lcqtof_hilic_neg_3_079_58    m/z 453.1151     0   1   2E+3   1   1.5  RT (min)      lcqtof_hilic_neg_1_052_40    m/z 453.9539     0   2   4   6   8E+3   4   4.5  RT (min)      lcqtof_hilic_neg_4_039_18    m/z 454.1274     0   2   4   6E+3   3.5   4  RT (min)       lcqtof_hilic_neg_4_101_61    m/z 454.1372     0   1   2   3E+3   2.5   3  RT (min)      lcqtof_hilic_neg_4_054_48    m/z 454.2349     0   2   4   6E+3   1   1.5  RT (min)      lcqtof_hilic_neg_4_013_20    m/z 454.3242     0   2   4   6E+3   1   1.5  RT (min)      lcqtof_hilic_neg_4_053_14    m/z 454.3465     0   1   2   3   4E+3   4   4.5  RT (min)      lcqtof_hilic_neg_4_001_49    m/z 454.4117     0   2   4   6E+3   1   1.5  RT (min)      lcqtof_hilic_neg_4_008_24    m/z 454.9154     0   1E+3   4   4.5  RT (min)      lcqtof_hilic_neg_2_035_09    m/z 455.0519     0   2   4   6E+3   0.5   1  RT (min)      lcqtof_hilic_neg_1_028_47    m/z 455.8706     0   2   4E+3   0.5   1  RT (min)      lcqtof_hilic_neg_4_054_48    m/z 455.9542     0   2   4   6E+3   4   4.5  RT (min)      lcqtof_hilic_neg_4_039_18    m/z 456.0008     0   1E+3   1   1.5  RT (min)      lcqtof_hilic_neg_2_037_21    m/z 456.0551     0   1   2   3E+3   0.5   1  RT (min)      lcqtof_hilic_neg_1_001_49    m/z 456.0710     0   2   4   6   8E+3   3.5   4  RT (min)      lcqtof_hilic_neg_4_013_20    m/z 456.1046     0   2   4E+3   7   7.5  RT (min)      lcqtof_hilic_neg_2_020_23    m/z 456.2028     0   1   2   3E+3   0.5   1  RT (min)      lcqtof_hilic_neg_2_021_17    m/z 456.3139     0   2   4   6   8E+3   0.5   1  RT (min)      lcqtof_hilic_neg_1_007_05    m/z 456.7388     0   1E+3   5   5.5  RT (min)      lcqtof_hilic_neg_2_036_34    m/z 456.8684     0   2   4   6E+3   0.5   1  RT (min)      lcqtof_hilic_neg_4_004_47    m/z 456.9842     0   1   2   3E+3   1   1.5  RT (min)      lcqtof_hilic_neg_2_024_10    m/z 457.0116     0   1   2E+3   5   5.5  RT (min)      lcqtof_hilic_neg_4_053_14    m/z 457.0160     0   1   2E+3   5.5   6  RT (min)      lcqtof_hilic_neg_4_052_13    m/z 457.1460     0   1   2   3   4E+3   1   1.5  RT (min)      lcqtof_hilic_neg_4_053_14    m/z 457.2589     0   1   2   3   4E+3   6.5   7  RT (min)      lcqtof_hilic_neg_4_053_14    m/z 458.0431     0   1   2E+3   1   1.5  RT (min)      lcqtof_hilic_neg_4_026_04    m/z 458.0467     0   2   4   6E+3   2.5   3  RT (min)      lcqtof_hilic_neg_2_050_19    m/z 458.1368     0   2   4   6E+3   1   1.5  RT (min)      lcqtof_hilic_neg_4_053_14    m/z 458.1416     0   2   4E+3   4.5   5  RT (min)      lcqtof_hilic_neg_4_016_37    m/z 458.2685     0   2   4   6   8E+3   1   1.5  RT (min)      lcqtof_hilic_neg_4_004_47    m/z 458.3383     0   2   4   6   8E+3   4.5   5  RT (min)      lcqtof_hilic_neg_4_008_24    m/z 458.8105     0   1E+3   6.5   7  RT (min)      lcqtof_hilic_neg_3_089_59    m/z 458.9971     0   2   4   6   8E+3   0.5   1  RT (min)      lcqtof_hilic_neg_2_053_18    m/z 459.0449     0   1   2   3   4E+3   6.5   7  RT (min)      lcqtof_hilic_neg_1_035_23    m/z 459.0449     0   1   2   3   4E+3   6   6.5  RT (min)       lcqtof_hilic_neg_1_053_19    m/z 459.0461     0   1   2   3E+3   6   6.5  RT (min)      lcqtof_hilic_neg_1_023_34    m/z 459.0824     0   2   4   6E+3   6   6.5  RT (min)      lcqtof_hilic_neg_3_048_05    m/z 459.1497     0   1   2E+3   4.5   5  RT (min)      lcqtof_hilic_neg_1_054_01    m/z 459.2575     0   1   2E+3   7   7.5  RT (min)      lcqtof_hilic_neg_4_048_23    m/z 459.9983     0   1   2E+3   1   1.5  RT (min)      lcqtof_hilic_neg_2_035_09    m/z 460.3392     0   1   2   3E+3   3.5   4  RT (min)      lcqtof_hilic_neg_4_036_11    m/z 461.9832     0   1   2   3E+3   1   1.5  RT (min)      lcqtof_hilic_neg_4_004_47    m/z 462.2323     0   2   4E+3   0.5   1  RT (min)      lcqtof_hilic_neg_4_013_20    m/z 462.2784     0   1   2   3   4E+3   4.5   5  RT (min)      lcqtof_hilic_neg_1_037_24    m/z 462.9636     0   1   2   3   4E+3   0.5   1  RT (min)      lcqtof_hilic_neg_4_035_33    m/z 463.0381     0   1E+3   5.5   6  RT (min)      lcqtof_hilic_neg_1_027_27    m/z 463.1928     0   2   4   6   8E+4   0.5   1  RT (min)      lcqtof_hilic_neg_1_032_18    m/z 463.1950     0   2   4   6   8E+3   6   6.5  RT (min)       lcqtof_hilic_neg_1_011_46    m/z 464.1136     0   2   4E+3   1   1.5  RT (min)      lcqtof_hilic_neg_1_003_37    m/z 465.1087     0   2   4   6E+3   7.5   8  RT (min)       lcqtof_hilic_neg_4_048_23    m/z 465.1822     0   1   2E+3   3.5   4  RT (min)      lcqtof_hilic_neg_1_042_26    m/z 465.2373     0   2   4   6   8E+3   1   1.5  RT (min)       lcqtof_hilic_neg_4_009_50    m/z 465.9614     0   1   2E+3   5   5.5  RT (min)      lcqtof_hilic_neg_4_015_09    m/z 466.1397     0   1E+3   7   7.5  RT (min)      lcqtof_hilic_neg_2_052_24    m/z 466.2530     0   1   2   3E+3   3   3.5  RT (min)      lcqtof_hilic_neg_4_006_30    m/z 467.1434     0   1   2   3   4E+3   6.5   7  RT (min)       lcqtof_hilic_neg_1_005_32    m/z 467.1755     0   2   4   6E+3   7   7.5  RT (min)       lcqtof_hilic_neg_4_048_23    m/z 467.9298     0   1E+3   4.5   5  RT (min)      lcqtof_hilic_neg_4_023_15    m/z 468.0399     0   2   4   6   8E+3   1.5   2  RT (min)      lcqtof_hilic_neg_4_004_47    m/z 468.0832     0   1   2   3E+3   0.5   1  RT (min)      lcqtof_hilic_neg_4_033_53    m/z 468.0868     0   1   2E+3   1   1.5  RT (min)      lcqtof_hilic_neg_1_054_01    m/z 468.1024     0   1   2E+3   1   1.5  RT (min)      lcqtof_hilic_neg_4_039_18    m/z 468.2722     0   1   2E+3   4   4.5  RT (min)      lcqtof_hilic_neg_2_047_32    m/z 469.0783     0   1   2   3   4E+3   0.5   1  RT (min)      lcqtof_hilic_neg_4_004_47    m/z 469.2631     0   1   2   3E+4   1   1.5  RT (min)      lcqtof_hilic_neg_4_003_16    m/z 469.2652     0   1   2E+3   5   5.5  RT (min)      lcqtof_hilic_neg_1_035_23    m/z 470.0791     0   1   2   3E+3   7   7.5  RT (min)      lcqtof_hilic_neg_1_037_24    m/z 470.1029     0   1   2E+3   1   1.5  RT (min)      lcqtof_hilic_neg_3_055_01    m/z 470.3068     0   1   2   3E+3   3.5   4  RT (min)      lcqtof_hilic_neg_4_048_23    m/z 470.3945     0   1   2   3E+3   0.5   1  RT (min)      lcqtof_hilic_neg_4_053_14    m/z 471.0186     0   2   4E+3   5.5   6  RT (min)      lcqtof_hilic_neg_4_053_14    m/z 471.0743     0   1   2E+3   4.5   5  RT (min)      lcqtof_hilic_neg_2_050_19    m/z 471.1671     0   1E+4   6   6.5  RT (min)      lcqtof_hilic_neg_1_005_32    m/z 471.2924     0   1   2E+3   6.5   7  RT (min)      lcqtof_hilic_neg_4_053_14    m/z 471.8842     0   1   2   3E+3   0.5   1  RT (min)      lcqtof_hilic_neg_4_076_58    m/z 471.9547     0   1E+3   7   7.5  RT (min)      lcqtof_hilic_neg_2_047_32    m/z 472.0104     0   1   2   3   4E+3   0.5   1  RT (min)      lcqtof_hilic_neg_4_036_11    m/z 472.1325     0   2   4   6E+3   7.5   8  RT (min)      lcqtof_hilic_neg_4_040_05    m/z 472.2495     0   1E+3   1   1.5  RT (min)      lcqtof_hilic_neg_1_006_22    m/z 472.7226     0   1   2E+3   7.5   8  RT (min)      lcqtof_hilic_neg_3_030_23    m/z 472.8208     0   1   2E+3   7   7.5  RT (min)       lcqtof_hilic_neg_4_089_59    m/z 472.8836     0   1   2   3   4E+3   0.5   1  RT (min)      lcqtof_hilic_neg_4_014_46    m/z 472.9893     0   1E+3   5.5   6  RT (min)      lcqtof_hilic_neg_4_026_04    m/z 473.0182     0   1E+3   1   1.5  RT (min)      lcqtof_hilic_neg_1_006_22    m/z 473.3999     0   1E+3   15   15.5  RT (min)       lcqtof_hilic_neg_4_019_03    m/z 474.0282     0   2   4   6   8E+3   0.5   1  RT (min)      lcqtof_hilic_neg_4_018_27    m/z 474.2658     0   1   2E+3   3   3.5  RT (min)       lcqtof_hilic_neg_4_083_59    m/z 474.7198     0   1E+3   7.5   8  RT (min)      lcqtof_hilic_neg_3_034_24    m/z 475.0838     0   2   4   6E+3   0.5   1  RT (min)      lcqtof_hilic_neg_4_053_14    m/z 475.3776     0   1   2   3   4E+4   0.5   1  RT (min)      lcqtof_hilic_neg_4_074_58    m/z 476.0627     0   1   2E+3   5.5   6  RT (min)      lcqtof_hilic_neg_1_037_24    m/z 476.0699     0   2   4E+3   0.5   1  RT (min)      lcqtof_hilic_neg_4_005_29    m/z 476.1475     0   2   4E+3   0.5   1  RT (min)      lcqtof_hilic_neg_4_032_01    m/z 476.2046     0   1   2   3   4E+3   1   1.5  RT (min)      lcqtof_hilic_neg_4_045_26    m/z 476.2477     0   1   2   3   4E+3   0.5   1  RT (min)      lcqtof_hilic_neg_4_013_20    m/z 476.3313     0   1E+4   0.5   1  RT (min)      lcqtof_hilic_neg_4_003_16    m/z 477.9533     0   2   4   6E+3   1   1.5  RT (min)      lcqtof_hilic_neg_3_066_57    m/z 478.0138     0   2   4E+3   0.5   1  RT (min)      lcqtof_hilic_neg_1_028_47    m/z 478.0143     0   1   2E+3   4.5   5  RT (min)      lcqtof_hilic_neg_2_040_20    m/z 478.0929     0   1   2E+3   0.5   1  RT (min)      lcqtof_hilic_neg_1_004_14    m/z 478.7909     0   1   2   3   4E+3   7.5   8  RT (min)      lcqtof_hilic_neg_4_003_16    m/z 478.8170     0   1   2   3E+3   5   5.5  RT (min)      lcqtof_hilic_neg_1_005_32    m/z 479.0954     0   1   2   3   4E+3   5.5   6  RT (min)      lcqtof_hilic_neg_1_013_38    m/z 479.1549     0   2   4   6   8E+3   7.5   8  RT (min)       lcqtof_hilic_neg_4_008_24    m/z 479.2094     0   1   2   3E+3   7   7.5  RT (min)      lcqtof_hilic_neg_4_014_46    m/z 479.3372     0   1E+3   15   15.5  RT (min)       lcqtof_hilic_neg_4_012_07    m/z 479.4786     0   1   2E+3   0.5   1  RT (min)      lcqtof_hilic_neg_4_022_34    m/z 479.4817     0   1E+3   15.5   16  RT (min)       lcqtof_hilic_neg_4_032_01    m/z 479.9684     0   1E+3   6   6.5  RT (min)       lcqtof_hilic_neg_1_048_45    m/z 480.0122     0   1E+3   4.5   5  RT (min)      lcqtof_hilic_neg_2_013_29    m/z 480.0920     0   2   4   6E+3   0.5   1  RT (min)      lcqtof_hilic_neg_2_047_32    m/z 480.1018     0   2   4E+3   6   6.5  RT (min)      lcqtof_hilic_neg_2_028_05    m/z 480.1027     0   1   2   3   4E+3   5.5   6  RT (min)      lcqtof_hilic_neg_2_039_38    m/z 480.1985     0   2   4   6E+3   0.5   1  RT (min)      lcqtof_hilic_neg_4_050_02    m/z 480.2205     0   1   2   3E+3   0.5   1  RT (min)       lcqtof_hilic_neg_4_032_01    m/z 480.2300     0   1E+3   4   4.5  RT (min)      lcqtof_hilic_neg_4_032_01    m/z 481.0865     0   1   2   3E+3   0.5   1  RT (min)      lcqtof_hilic_neg_2_047_32    m/z 481.1033     0   2   4   6   8E+3   4   4.5  RT (min)      lcqtof_hilic_neg_1_006_22    m/z 481.2044     0   1E+3   3.5   4  RT (min)      lcqtof_hilic_neg_1_004_14    m/z 481.2701     0   1   2   3E+3   1   1.5  RT (min)      lcqtof_hilic_neg_1_053_19    m/z 481.3122     0   1   2E+5   3.5   4  RT (min)      lcqtof_hilic_neg_4_074_58    m/z 481.5330     0   2   4   6   8E+3   4   4.5  RT (min)      lcqtof_hilic_neg_4_053_14    m/z 482.0338     0   2   4E+3   6.5   7  RT (min)       lcqtof_hilic_neg_4_048_23    m/z 482.3020     0   1   2   3E+3   0.5   1  RT (min)      lcqtof_hilic_neg_4_053_14    m/z 482.3510     0   1   2   3   4E+3   0.5   1  RT (min)      lcqtof_hilic_neg_4_050_02    m/z 483.2578     0   1E+3   4.5   5  RT (min)      lcqtof_hilic_neg_1_014_31    m/z 483.2839     0   2   4   6   8E+3   0.5   1  RT (min)      lcqtof_hilic_neg_2_005_13    m/z 483.3586     0   2   4   6E+3   3.5   4  RT (min)       lcqtof_hilic_neg_4_052_13    m/z 483.4311     0   1E+3   15   15.5  RT (min)      lcqtof_hilic_neg_4_011_35    m/z 484.1024     0   1   2   3   4E+3   5   5.5  RT (min)      lcqtof_hilic_neg_4_054_48    m/z 484.1960     0   1   2   3   4E+3   1   1.5  RT (min)      lcqtof_hilic_neg_1_042_26    m/z 485.1099     0   1   2E+3   6   6.5  RT (min)      lcqtof_hilic_neg_1_035_23    m/z 485.2857     0   1   2   3E+3   4.5   5  RT (min)      lcqtof_hilic_neg_3_070_57    m/z 485.3749     0   2   4   6E+3   3.5   4  RT (min)       lcqtof_hilic_neg_4_052_13    m/z 486.0239     0   1   2   3E+3   1   1.5  RT (min)      lcqtof_hilic_neg_1_023_34    m/z 486.0398     0   1   2   3E+3   4.5   5  RT (min)      lcqtof_hilic_neg_2_050_19    m/z 486.0509     0   2   4E+3   3.5   4  RT (min)      lcqtof_hilic_neg_1_006_22    m/z 486.1082     0   2   4   6E+3   6   6.5  RT (min)      lcqtof_hilic_neg_2_020_23    m/z 486.1930     0   1   2   3E+3   0.5   1  RT (min)      lcqtof_hilic_neg_4_053_14    m/z 486.2574     0   1   2   3E+3   5   5.5  RT (min)      lcqtof_hilic_neg_4_048_23    m/z 486.3509     0   1E+3   15   15.5  RT (min)      lcqtof_hilic_neg_4_025_52    m/z 486.3564     0   1   2   3E+3   3.5   4  RT (min)      lcqtof_hilic_neg_1_035_23    m/z 487.0440     0   1   2   3   4E+3   0.5   1  RT (min)      lcqtof_hilic_neg_4_018_27    m/z 487.1139     0   2   4E+3   4   4.5  RT (min)      lcqtof_hilic_neg_4_021_21    m/z 487.1210     0   2   4   6   8E+3   0.5   1  RT (min)      lcqtof_hilic_neg_1_016_28    m/z 487.2939     0   1E+3   15   15.5  RT (min)       lcqtof_hilic_neg_4_022_34    m/z 487.3088     0   1   2   3   4E+3   6   6.5  RT (min)      lcqtof_hilic_neg_4_053_14    m/z 487.9807     0   1   2   3E+3   5   5.5  RT (min)      lcqtof_hilic_neg_4_052_13    m/z 488.0385     0   1E+3   4.5   5  RT (min)      lcqtof_hilic_neg_1_053_19    m/z 488.0759     0   2   4E+3   1.5   2  RT (min)      lcqtof_hilic_neg_4_042_22    m/z 488.1836     0   1   2E+3   7   7.5  RT (min)      lcqtof_hilic_neg_4_048_23    m/z 488.2957     0   2   4E+3   0.5   1  RT (min)      lcqtof_hilic_neg_1_007_05    m/z 488.8919     0   1   2   3E+3   0.5   1  RT (min)      lcqtof_hilic_neg_4_025_52    m/z 488.9616     0   1   2E+3   5.5   6  RT (min)      lcqtof_hilic_neg_4_026_04    m/z 489.1585     0   1   2   3   4E+3   0.5   1  RT (min)      lcqtof_hilic_neg_4_035_33    m/z 490.8018     0   1   2   3E+3   0.5   1  RT (min)      lcqtof_hilic_neg_3_076_58    m/z 492.1201     0   1   2E+3   6   6.5  RT (min)      lcqtof_hilic_neg_4_003_16    m/z 492.2794     0   1   2   3E+3   0.5   1  RT (min)      lcqtof_hilic_neg_4_032_01    m/z 492.7457     0   1E+3   7.5   8  RT (min)      lcqtof_hilic_neg_1_009_50    m/z 492.8272     0   1   2   3E+3   5   5.5  RT (min)       lcqtof_hilic_neg_2_047_32    m/z 493.0521     0   1   2   3E+3   5.5   6  RT (min)      lcqtof_hilic_neg_4_006_30    m/z 493.1575     0   1E+3   3.5   4  RT (min)      lcqtof_hilic_neg_2_055_30    m/z 493.4636     0   2   4E+3   0.5   1  RT (min)      lcqtof_hilic_neg_2_016_33    m/z 493.9397     0   1E+3   7   7.5  RT (min)      lcqtof_hilic_neg_2_047_32    m/z 494.1160     0   2   4   6E+3   7.5   8  RT (min)      lcqtof_hilic_neg_4_040_05    m/z 494.1571     0   2   4   6E+3   0.5   1  RT (min)      lcqtof_hilic_neg_4_021_21    m/z 494.2888     0   1   2   3E+3   3.5   4  RT (min)      lcqtof_hilic_neg_4_053_14    m/z 494.2934     0   1   2   3   4E+3   0.5   1  RT (min)      lcqtof_hilic_neg_4_032_01    m/z 495.1269     0   1   2   3E+3   1   1.5  RT (min)      lcqtof_hilic_neg_4_006_30    m/z 495.1906     0   2   4   6E+3   0.5   1  RT (min)      lcqtof_hilic_neg_4_013_20    m/z 495.2574     0   1   2   3   4E+3   3.5   4  RT (min)      lcqtof_hilic_neg_4_040_05    m/z 495.4663     0   2   4E+3   0.5   1  RT (min)      lcqtof_hilic_neg_4_032_01    m/z 495.9624     0   1   2E+3   5.5   6  RT (min)       lcqtof_hilic_neg_4_053_14    m/z 496.1412     0   1   2   3   4E+3   6   6.5  RT (min)      lcqtof_hilic_neg_2_015_06    m/z 496.1443     0   2   4   6   8E+3   0.5   1  RT (min)      lcqtof_hilic_neg_1_026_10    m/z 496.3015     0   1   2E+3   3.5   4  RT (min)      lcqtof_hilic_neg_2_031_01    m/z 496.4456     0   1   2E+3   0.5   1  RT (min)      lcqtof_hilic_neg_2_029_31    m/z 496.8693     0   1   2E+3   6.5   7  RT (min)       lcqtof_hilic_neg_4_002_25    m/z 496.8846     0   1E+3   4   4.5  RT (min)      lcqtof_hilic_neg_1_053_19    m/z 497.0455     0   1   2   3E+3   7.5   8  RT (min)      lcqtof_hilic_neg_1_004_14    m/z 497.1365     0   1   2E+3   6   6.5  RT (min)       lcqtof_hilic_neg_4_067_57    m/z 497.1414     0   1   2E+3   5   5.5  RT (min)      lcqtof_hilic_neg_4_052_13    m/z 497.3014     0   1E+3   15   15.5  RT (min)      lcqtof_hilic_neg_1_005_32    m/z 497.3933     0   1   2   3   4E+3   4   4.5  RT (min)      lcqtof_hilic_neg_4_026_04    m/z 497.9476     0   1   2E+3   0.5   1  RT (min)      lcqtof_hilic_neg_4_050_02    m/z 498.2911     0   1   2E+3   4.5   5  RT (min)      lcqtof_hilic_neg_4_023_15    m/z 499.0514     0   1   2E+3   2.5   3  RT (min)      lcqtof_hilic_neg_2_050_19    m/z 499.0746     0   1   2   3   4E+3   4   4.5  RT (min)      lcqtof_hilic_neg_2_036_34    m/z 499.0999     0   1   2E+3   0.5   1  RT (min)      lcqtof_hilic_neg_2_045_11    m/z 499.1873     0   1   2   3   4E+3   4.5   5  RT (min)      lcqtof_hilic_neg_4_027_19    m/z 499.2228     0   1   2E+3   6   6.5  RT (min)      lcqtof_hilic_neg_4_002_25    m/z 500.9346     0   1E+3   4   4.5  RT (min)      lcqtof_hilic_neg_1_032_18    m/z 501.1038     0   1   2   3E+3   8   8.5  RT (min)       lcqtof_hilic_neg_4_094_60    m/z 501.9164     0   1E+3   4.5   5  RT (min)      lcqtof_hilic_neg_4_035_33    m/z 502.0404     0   1   2E+3   5   5.5  RT (min)       lcqtof_hilic_neg_4_012_07    m/z 502.0410     0   2   4   6E+3   6   6.5  RT (min)      lcqtof_hilic_neg_4_035_33    m/z 502.1366     0   1   2   3   4E+3   1   1.5  RT (min)      lcqtof_hilic_neg_4_020_42    m/z 502.2546     0   2   4   6   8E+3   6   6.5  RT (min)      lcqtof_hilic_neg_4_008_24    m/z 502.2818     0   2   4   6E+3   7   7.5  RT (min)       lcqtof_hilic_neg_4_031_38    m/z 502.3409     0   1   2   3E+3   15   15.5  RT (min)       lcqtof_hilic_neg_4_050_02    m/z 502.7022     0   1E+3   7.5   8  RT (min)      lcqtof_hilic_neg_2_052_24    m/z 502.7460     0   1E+3   8   8.5  RT (min)      lcqtof_hilic_neg_3_034_24    m/z 502.9658     0   2   4   6E+3   0.5   1  RT (min)      lcqtof_hilic_neg_4_004_47    m/z 503.0802     0   1   2E+3   1   1.5  RT (min)      lcqtof_hilic_neg_4_018_27    m/z 503.0893     0   1   2E+3   5   5.5  RT (min)       lcqtof_hilic_neg_4_032_01    m/z 503.8973     0   1   2   3E+3   0.5   1  RT (min)      lcqtof_hilic_neg_4_013_20    m/z 504.0224     0   1   2E+3   4.5   5  RT (min)      lcqtof_hilic_neg_4_015_09    m/z 504.0794     0   1   2   3E+3   6   6.5  RT (min)      lcqtof_hilic_neg_1_016_28    m/z 504.2298     0   1   2   3   4E+3   1   1.5  RT (min)      lcqtof_hilic_neg_1_053_19    m/z 504.6984     0   1E+3   7.5   8  RT (min)      lcqtof_hilic_neg_2_015_06    m/z 504.9363     0   1   2E+3   5.5   6  RT (min)       lcqtof_hilic_neg_4_013_20    m/z 505.0164     0   1   2   3   4E+3   5   5.5  RT (min)       lcqtof_hilic_neg_4_008_24    m/z 505.1004     0   1   2   3   4E+3   2.5   3  RT (min)       lcqtof_hilic_neg_4_111_62    m/z 505.1728     0   1E+3   7   7.5  RT (min)      lcqtof_hilic_neg_4_089_59    m/z 506.0213     0   1   2   3E+3   4.5   5  RT (min)       lcqtof_hilic_neg_4_013_20    m/z 506.0703     0   1   2   3E+3   7.5   8  RT (min)      lcqtof_hilic_neg_2_028_05    m/z 506.0772     0   2   4   6   8E+3   1   1.5  RT (min)      lcqtof_hilic_neg_4_006_30    m/z 506.8150     0   1   2   3E+3   0.5   1  RT (min)      lcqtof_hilic_neg_4_054_48    m/z 507.0712     0   1   2   3E+3   7   7.5  RT (min)      lcqtof_hilic_neg_4_048_23    m/z 507.1421     0   1   2   3E+3   6.5   7  RT (min)       lcqtof_hilic_neg_4_052_13    m/z 507.2678     0   2   4   6E+3   3.5   4  RT (min)      lcqtof_hilic_neg_4_032_01    m/z 507.2784     0   2   4E+3   1   1.5  RT (min)      lcqtof_hilic_neg_4_053_14    m/z 507.3700     0   1E+3   15   15.5  RT (min)       lcqtof_hilic_neg_4_044_08    m/z 507.5046     0   1   2   3E+3   0.5   1  RT (min)      lcqtof_hilic_neg_4_048_23    m/z 508.1805     0   1   2E+3   5   5.5  RT (min)      lcqtof_hilic_neg_4_019_03    m/z 508.2585     0   1   2   3   4E+3   0.5   1  RT (min)      lcqtof_hilic_neg_4_005_29    m/z 508.2850     0   2   4   6E+3   0.5   1  RT (min)      lcqtof_hilic_neg_3_055_01    m/z 508.3055     0   1   2   3E+3   7   7.5  RT (min)      lcqtof_hilic_neg_4_054_48    m/z 509.0278     0   1   2E+3   5.5   6  RT (min)      lcqtof_hilic_neg_4_018_27    m/z 509.1854     0   1   2E+3   5   5.5  RT (min)      lcqtof_hilic_neg_4_013_20    m/z 509.9788     0   1   2   3E+3   5.5   6  RT (min)       lcqtof_hilic_neg_4_052_13    m/z 510.1428     0   2   4   6E+3   6   6.5  RT (min)      lcqtof_hilic_neg_4_031_38    m/z 510.2953     0   2   4   6   8E+3   5   5.5  RT (min)      lcqtof_hilic_neg_4_053_14    m/z 510.9288     0   1E+3   5.5   6  RT (min)       lcqtof_hilic_neg_3_098_61    m/z 511.1083     0   2   4   6   8E+3   6.5   7  RT (min)      lcqtof_hilic_neg_4_002_25    m/z 511.1735     0   1   2E+3   6   6.5  RT (min)      lcqtof_hilic_neg_2_034_47    m/z 511.2065     0   1   2E+3   1   1.5  RT (min)      lcqtof_hilic_neg_2_014_08    m/z 511.2869     0   2   4E+3   4.5   5  RT (min)      lcqtof_hilic_neg_4_040_05    m/z 512.0648     0   1   2   3E+3   2.5   3  RT (min)      lcqtof_hilic_neg_2_050_19    m/z 512.1329     0   1   2   3   4E+3   1   1.5  RT (min)      lcqtof_hilic_neg_4_015_09    m/z 512.2317     0   1   2E+3   4   4.5  RT (min)      lcqtof_hilic_neg_4_023_15    m/z 512.2521     0   1   2E+3   0.5   1  RT (min)      lcqtof_hilic_neg_4_002_25    m/z 512.3702     0   1   2   3E+3   15   15.5  RT (min)       lcqtof_hilic_neg_4_050_02    m/z 512.7248     0   1E+3   7.5   8  RT (min)      lcqtof_hilic_neg_2_055_30    m/z 513.0941     0   2   4   6E+3   7.5   8  RT (min)      lcqtof_hilic_neg_1_037_24    m/z 513.3550     0   1   2   3   4E+3   0.5   1  RT (min)      lcqtof_hilic_neg_2_031_01    m/z 513.8239     0   1   2E+3   6.5   7  RT (min)      lcqtof_hilic_neg_4_002_25    m/z 513.9404     0   1E+3   7   7.5  RT (min)      lcqtof_hilic_neg_2_048_35    m/z 514.1183     0   1   2E+3   5   5.5  RT (min)      lcqtof_hilic_neg_4_029_17    m/z 514.1248     0   1   2   3   4E+3   6   6.5  RT (min)      lcqtof_hilic_neg_4_093_60    m/z 514.2355     0   1   2E+4   1   1.5  RT (min)      lcqtof_hilic_neg_4_050_02    m/z 514.3613     0   2   4   6E+3   5   5.5  RT (min)       lcqtof_hilic_neg_4_048_23    m/z 514.6055     0   1   2   3   4E+3   5   5.5  RT (min)      lcqtof_hilic_neg_1_013_38    m/z 514.9619     0   1E+3   4   4.5  RT (min)      lcqtof_hilic_neg_1_048_45    m/z 515.0398     0   2   4   6E+3   0.5   1  RT (min)      lcqtof_hilic_neg_4_013_20    m/z 515.0818     0   1   2E+3   3.5   4  RT (min)      lcqtof_hilic_neg_1_020_09    m/z 515.2043     0   1   2   3E+3   4   4.5  RT (min)      lcqtof_hilic_neg_4_050_02    m/z 516.0488     0   1   2   3E+3   6   6.5  RT (min)      lcqtof_hilic_neg_4_055_32    m/z 516.0989     0   2   4   6E+3   7.5   8  RT (min)      lcqtof_hilic_neg_4_040_05    m/z 516.2341     0   2   4E+3   1   1.5  RT (min)      lcqtof_hilic_neg_4_005_29    m/z 516.3192     0   1E+3   15   15.5  RT (min)      lcqtof_hilic_neg_4_022_34    m/z 517.1540     0   1   2E+3   2.5   3  RT (min)      lcqtof_hilic_neg_3_109_62    m/z 517.2057     0   2   4   6E+3   6.5   7  RT (min)      lcqtof_hilic_neg_4_052_13    m/z 517.3278     0   2   4   6   8E+3   0.5   1  RT (min)      lcqtof_hilic_neg_4_008_24    m/z 518.1451     0   1   2E+3   6   6.5  RT (min)      lcqtof_hilic_neg_2_022_46    m/z 518.2013     0   2   4   6E+3   0.5   1  RT (min)      lcqtof_hilic_neg_4_013_20    m/z 518.3149     0   2   4E+3   4   4.5  RT (min)      lcqtof_hilic_neg_1_010_15    m/z 518.4276     0   2   4E+3   0.5   1  RT (min)      lcqtof_hilic_neg_4_010_28    m/z 518.9163     0   2   4E+3   0.5   1  RT (min)      lcqtof_hilic_neg_4_033_53    m/z 519.0496     0   2   4   6E+3   3.5   4  RT (min)      lcqtof_hilic_neg_4_013_20    m/z 519.0516     0   2   4   6E+3   7.5   8  RT (min)       lcqtof_hilic_neg_4_048_23    m/z 519.0585     0   1   2   3E+3   5   5.5  RT (min)      lcqtof_hilic_neg_4_053_14    m/z 519.0615     0   1   2   3E+3   4   4.5  RT (min)      lcqtof_hilic_neg_1_006_22    m/z 519.1036     0   2   4   6E+3   1   1.5  RT (min)      lcqtof_hilic_neg_4_027_19    m/z 519.1148     0   1   2   3E+3   7   7.5  RT (min)      lcqtof_hilic_neg_4_008_24    m/z 519.1408     0   2   4E+3   0.5   1  RT (min)       lcqtof_hilic_neg_4_018_27    m/z 519.9978     0   1   2   3E+3   4.5   5  RT (min)      lcqtof_hilic_neg_4_013_20    m/z 520.1666     0   1   2   3E+3   3.5   4  RT (min)      lcqtof_hilic_neg_4_054_48    m/z 520.2268     0   1   2E+3   3   3.5  RT (min)      lcqtof_hilic_neg_2_051_14    m/z 520.2635     0   1   2   3E+3   4   4.5  RT (min)      lcqtof_hilic_neg_4_003_16    m/z 520.3043     0   2   4   6E+3   4   4.5  RT (min)      lcqtof_hilic_neg_4_053_14    m/z 520.3729     0   1   2   3   4E+3   3.5   4  RT (min)      lcqtof_hilic_neg_4_003_16    m/z 521.1494     0   1   2   3E+3   0.5   1  RT (min)       lcqtof_hilic_neg_2_002_36    m/z 521.1498     0   2   4   6E+3   6.5   7  RT (min)      lcqtof_hilic_neg_4_002_25    m/z 521.2455     0   1   2   3   4E+3   3.5   4  RT (min)      lcqtof_hilic_neg_4_003_16    m/z 521.2903     0   2   4   6   8E+3   6   6.5  RT (min)      lcqtof_hilic_neg_4_008_24    m/z 521.2932     0   1E+4   1   1.5  RT (min)      lcqtof_hilic_neg_4_005_29    m/z 521.3315     0   1   2E+3   15   15.5  RT (min)      lcqtof_hilic_neg_1_008_13    m/z 521.3810     0   2   4E+3   1   1.5  RT (min)      lcqtof_hilic_neg_4_053_14    m/z 521.3826     0   1E+3   15   15.5  RT (min)       lcqtof_hilic_neg_4_019_03    m/z 521.4932     0   1   2   3   4E+3   0.5   1  RT (min)      lcqtof_hilic_neg_4_053_14    m/z 521.5104     0   1   2   3E+3   0.5   1  RT (min)      lcqtof_hilic_neg_4_034_31    m/z 521.8398     0   1   2E+3   0.5   1  RT (min)      lcqtof_hilic_neg_4_054_48    m/z 522.3193     0   2   4E+3   3.5   4  RT (min)      lcqtof_hilic_neg_4_053_14    m/z 522.4024     0   2   4   6E+3   4   4.5  RT (min)      lcqtof_hilic_neg_4_026_04    m/z 522.8300     0   2   4E+3   0.5   1  RT (min)      lcqtof_hilic_neg_3_079_58    m/z 523.3101     0   2   4   6   8E+3   1   1.5  RT (min)      lcqtof_hilic_neg_4_032_01    m/z 523.3277     0   1   2E+3   15   15.5  RT (min)      lcqtof_hilic_neg_1_010_15    m/z 523.3796     0   1   2   3E+3   0.5   1  RT (min)      lcqtof_hilic_neg_4_014_46    m/z 523.9877     0   1   2   3E+3   6.5   7  RT (min)       lcqtof_hilic_neg_4_086_59    m/z 524.1385     0   1E+3   0.5   1  RT (min)      lcqtof_hilic_neg_2_039_38    m/z 524.4152     0   2   4   6   8E+3   4   4.5  RT (min)      lcqtof_hilic_neg_4_026_04    m/z 524.4810     0   1E+3   15.5   16  RT (min)      lcqtof_hilic_neg_4_002_25    m/z 524.6842     0   1E+3   7.5   8  RT (min)      lcqtof_hilic_neg_2_015_06    m/z 524.9630     0   2   4   6   8E+3   0.5   1  RT (min)      lcqtof_hilic_neg_1_028_47    m/z 525.0391     0   1   2E+3   0.5   1  RT (min)      lcqtof_hilic_neg_4_077_58    m/z 525.1313     0   2   4   6E+3   0.5   1  RT (min)      lcqtof_hilic_neg_1_005_32    m/z 525.2633     0   1   2E+3   0.5   1  RT (min)      lcqtof_hilic_neg_2_007_03    m/z 525.3316     0   1   2E+4   4   4.5  RT (min)       lcqtof_hilic_neg_4_053_14    m/z 525.3351     0   2   4E+3   0.5   1  RT (min)      lcqtof_hilic_neg_4_053_14    m/z 525.3628     0   1   2E+3   1   1.5  RT (min)      lcqtof_hilic_neg_4_026_04    m/z 525.3807     0   2   4E+3   1   1.5  RT (min)      lcqtof_hilic_neg_4_053_14    m/z 526.0416     0   1   2E+3   1   1.5  RT (min)      lcqtof_hilic_neg_4_054_48    m/z 526.0712     0   1E+3   1   1.5  RT (min)      lcqtof_hilic_neg_3_037_11    m/z 526.0858     0   1   2   3   4E+3   6   6.5  RT (min)       lcqtof_hilic_neg_4_045_26    m/z 526.1318     0   1   2   3E+3   0.5   1  RT (min)      lcqtof_hilic_neg_3_044_09    m/z 526.1338     0   2   4   6E+3   0.5   1  RT (min)      lcqtof_hilic_neg_3_040_14    m/z 526.3047     0   1   2E+5   3.5   4  RT (min)      lcqtof_hilic_neg_4_003_16    m/z 526.6771     0   1E+3   7.5   8  RT (min)      lcqtof_hilic_neg_2_015_06    m/z 526.7641     0   1   2E+3   5.5   6  RT (min)      lcqtof_hilic_neg_2_051_14    m/z 527.3445     0   1   2   3   4E+3   0.5   1  RT (min)      lcqtof_hilic_neg_1_046_30    m/z 527.7009     0   1E+3   5   5.5  RT (min)       lcqtof_hilic_neg_4_012_07    m/z 528.0531     0   1   2   3E+3   7.5   8  RT (min)      lcqtof_hilic_neg_2_028_05    m/z 528.0897     0   1   2   3   4E+4   6.5   7  RT (min)      lcqtof_hilic_neg_4_089_59    m/z 528.2175     0   1   2E+3   4   4.5  RT (min)      lcqtof_hilic_neg_2_021_17    m/z 528.3972     0   2   4E+3   0.5   1  RT (min)      lcqtof_hilic_neg_4_040_05    m/z 528.9371     0   2   4   6E+3   7   7.5  RT (min)       lcqtof_hilic_neg_4_084_59    m/z 529.0839     0   2   4   6   8E+3   6   6.5  RT (min)      lcqtof_hilic_neg_4_048_23    m/z 529.0892     0   1   2   3E+3   5   5.5  RT (min)      lcqtof_hilic_neg_4_053_14    m/z 529.2359     0   2   4E+3   0.5   1  RT (min)      lcqtof_hilic_neg_4_045_26    m/z 530.0967     0   1   2E+3   5   5.5  RT (min)      lcqtof_hilic_neg_4_032_01    m/z 530.2563     0   2   4E+3   5.5   6  RT (min)       lcqtof_hilic_neg_4_011_35    m/z 530.3439     0   1   2E+3   0.5   1  RT (min)      lcqtof_hilic_neg_2_049_55    m/z 530.9539     0   1   2E+3   6   6.5  RT (min)      lcqtof_hilic_neg_4_003_16    m/z 531.0397     0   1   2   3   4E+3   6   6.5  RT (min)      lcqtof_hilic_neg_4_018_27    m/z 531.1519     0   2   4   6   8E+3   3   3.5  RT (min)      lcqtof_hilic_neg_2_039_38    m/z 531.1703     0   1   2   3   4E+3   1   1.5  RT (min)       lcqtof_hilic_neg_4_019_03    m/z 531.2050     0   1E+4   0.5   1  RT (min)      lcqtof_hilic_neg_2_016_33    m/z 532.1940     0   2   4   6E+3   2   2.5  RT (min)       lcqtof_hilic_neg_4_038_10    m/z 532.3539     0   2   4   6E+3   5   5.5  RT (min)      lcqtof_hilic_neg_4_048_23    m/z 533.0155     0   1E+3   6   6.5  RT (min)      lcqtof_hilic_neg_4_031_38    m/z 533.1222     0   1   2   3E+3   6.5   7  RT (min)      lcqtof_hilic_neg_4_040_05    m/z 533.1678     0   1E+3   1   1.5  RT (min)      lcqtof_hilic_neg_2_050_19    m/z 533.2413     0   2   4   6   8E+3   2   2.5  RT (min)      lcqtof_hilic_neg_4_003_16    m/z 533.2540     0   1E+3   5.5   6  RT (min)      lcqtof_hilic_neg_4_032_01    m/z 533.7325     0   1E+3   8   8.5  RT (min)      lcqtof_hilic_neg_3_012_27    m/z 534.1051     0   1   2   3E+3   6   6.5  RT (min)      lcqtof_hilic_neg_2_022_46    m/z 534.4897     0   2   4   6E+3   0.5   1  RT (min)      lcqtof_hilic_neg_4_034_31    m/z 535.0762     0   2   4   6E+3   7.5   8  RT (min)      lcqtof_hilic_neg_2_052_24    m/z 535.1653     0   1   2E+3   3   3.5  RT (min)      lcqtof_hilic_neg_2_001_49    m/z 535.1808     0   1   2   3   4E+3   0.5   1  RT (min)       lcqtof_hilic_neg_4_006_30    m/z 535.1880     0   1   2   3   4E+3   5   5.5  RT (min)      lcqtof_hilic_neg_4_026_04    m/z 535.2399     0   1E+3   4   4.5  RT (min)      lcqtof_hilic_neg_1_031_16    m/z 535.3660     0   1   2E+3   3.5   4  RT (min)      lcqtof_hilic_neg_4_053_14    m/z 535.8492     0   1   2   3E+3   0.5   1  RT (min)      lcqtof_hilic_neg_4_076_58    m/z 535.9727     0   1   2E+3   4.5   5  RT (min)      lcqtof_hilic_neg_2_050_19    m/z 536.0896     0   1   2   3   4E+3   1   1.5  RT (min)      lcqtof_hilic_neg_4_050_02    m/z 536.3000     0   1E+3   15   15.5  RT (min)      lcqtof_hilic_neg_1_002_39    m/z 536.3224     0   1   2E+3   2   2.5  RT (min)      lcqtof_hilic_neg_2_021_17    m/z 537.0233     0   1   2E+3   6   6.5  RT (min)      lcqtof_hilic_neg_4_034_31    m/z 537.1292     0   2   4   6E+3   6.5   7  RT (min)       lcqtof_hilic_neg_4_008_24    m/z 537.1387     0   1E+3   6   6.5  RT (min)      lcqtof_hilic_neg_2_016_33    m/z 537.2528     0   1   2E+3   4   4.5  RT (min)      lcqtof_hilic_neg_4_052_13    m/z 537.3506     0   1   2   3   4E+3   7   7.5  RT (min)      lcqtof_hilic_neg_1_037_24    m/z 537.3777     0   2   4   6   8E+3   1   1.5  RT (min)       lcqtof_hilic_neg_4_008_24    m/z 537.8430     0   1   2E+3   0.5   1  RT (min)      lcqtof_hilic_neg_3_078_58    m/z 537.9719     0   1   2E+3   4.5   5  RT (min)      lcqtof_hilic_neg_1_040_20    m/z 538.1395     0   1   2E+3   6   6.5  RT (min)      lcqtof_hilic_neg_4_003_16    m/z 538.2267     0   2   4   6E+3   1.5   2  RT (min)       lcqtof_hilic_neg_4_027_19    m/z 538.2324     0   2   4   6   8E+3   4   4.5  RT (min)       lcqtof_hilic_neg_4_040_05    m/z 538.2691     0   1   2E+3   4   4.5  RT (min)      lcqtof_hilic_neg_1_010_15    m/z 538.3160     0   1   2E+3   15   15.5  RT (min)      lcqtof_hilic_neg_1_009_50    m/z 538.4875     0   1E+3   15.5   16  RT (min)       lcqtof_hilic_neg_4_103_61    m/z 538.7293     0   1E+3   8   8.5  RT (min)       lcqtof_hilic_neg_4_023_15    m/z 539.0292     0   2   4   6E+3   3.5   4  RT (min)      lcqtof_hilic_neg_4_027_19    m/z 539.1216     0   2   4E+3   4.5   5  RT (min)      lcqtof_hilic_neg_4_005_29    m/z 540.0386     0   2   4   6   8E+3   6.5   7  RT (min)       lcqtof_hilic_neg_4_008_24    m/z 540.0801     0   1E+3   5.5   6  RT (min)      lcqtof_hilic_neg_3_012_27    m/z 540.0889     0   1   2   3E+3   7   7.5  RT (min)       lcqtof_hilic_neg_4_032_01    m/z 540.1223     0   1   2E+3   6   6.5  RT (min)      lcqtof_hilic_neg_4_035_33    m/z 540.3859     0   2   4   6E+3   7   7.5  RT (min)      lcqtof_hilic_neg_4_008_24    m/z 540.4715     0   2   4   6   8E+3   0.5   1  RT (min)      lcqtof_hilic_neg_4_053_14    m/z 541.0757     0   1   2E+3   5.5   6  RT (min)      lcqtof_hilic_neg_4_053_14    m/z 541.1119     0   1   2   3E+3   0.5   1  RT (min)      lcqtof_hilic_neg_4_053_14    m/z 541.4359     0   2   4   6E+3   1.5   2  RT (min)      lcqtof_hilic_neg_4_040_05    m/z 541.4534     0   1   2   3E+3   4   4.5  RT (min)      lcqtof_hilic_neg_4_003_16    m/z 541.7875     0   2   4   6E+3   6.5   7  RT (min)      lcqtof_hilic_neg_4_035_33    m/z 542.0380     0   1E+3   6.5   7  RT (min)      lcqtof_hilic_neg_3_035_46    m/z 542.1361     0   2   4   6E+3   6.5   7  RT (min)       lcqtof_hilic_neg_4_094_60    m/z 542.2679     0   1   2   3   4E+3   1   1.5  RT (min)      lcqtof_hilic_neg_4_053_14    m/z 542.2712     0   1E+4   1   1.5  RT (min)      lcqtof_hilic_neg_4_050_02    m/z 543.0541     0   1   2   3E+3   3.5   4  RT (min)      lcqtof_hilic_neg_2_050_19    m/z 543.2951     0   1   2   3E+3   3.5   4  RT (min)      lcqtof_hilic_neg_1_014_31    m/z 544.0524     0   2   4   6   8E+3   6   6.5  RT (min)      lcqtof_hilic_neg_4_082_59    m/z 544.1817     0   1   2   3E+3   1   1.5  RT (min)      lcqtof_hilic_neg_4_027_19    m/z 544.2984     0   1   2   3   4E+3   3   3.5  RT (min)      lcqtof_hilic_neg_4_053_14    m/z 545.0709     0   1   2   3E+3   5.5   6  RT (min)      lcqtof_hilic_neg_4_003_16    m/z 545.2225     0   1   2E+3   6   6.5  RT (min)      lcqtof_hilic_neg_4_011_35    m/z 545.2714     0   1   2   3   4E+3   4   4.5  RT (min)      lcqtof_hilic_neg_1_003_37    m/z 546.0379     0   1E+3   5   5.5  RT (min)      lcqtof_hilic_neg_4_006_30    m/z 546.1614     0   1E+4   1   1.5  RT (min)      lcqtof_hilic_neg_4_053_14    m/z 546.2625     0   1   2   3   4E+3   0.5   1  RT (min)      lcqtof_hilic_neg_1_053_19    m/z 546.2727     0   2   4   6E+3   3.5   4  RT (min)       lcqtof_hilic_neg_4_003_16    m/z 546.2823     0   1   2   3E+3   4.5   5  RT (min)       lcqtof_hilic_neg_4_040_05    m/z 546.3137     0   2   4E+3   3.5   4  RT (min)      lcqtof_hilic_neg_4_034_31    m/z 546.3924     0   1   2   3E+3   3.5   4  RT (min)      lcqtof_hilic_neg_4_053_14    m/z 546.9277     0   2   4E+3   0.5   1  RT (min)      lcqtof_hilic_neg_4_033_53    m/z 547.0993     0   1   2E+3   5.5   6  RT (min)      lcqtof_hilic_neg_4_050_02    m/z 547.2705     0   2   4   6E+3   0.5   1  RT (min)       lcqtof_hilic_neg_1_004_14    m/z 547.3242     0   1E+3   15   15.5  RT (min)       lcqtof_hilic_neg_4_012_07    m/z 547.3293     0   1   2E+3   2   2.5  RT (min)      lcqtof_hilic_neg_3_055_01    m/z 548.0099     0   1   2   3E+3   0.5   1  RT (min)      lcqtof_hilic_neg_4_004_47    m/z 548.2031     0   2   4E+3   0.5   1  RT (min)      lcqtof_hilic_neg_4_016_37    m/z 548.3420     0   1   2   3E+3   4   4.5  RT (min)      lcqtof_hilic_neg_1_010_15    m/z 548.3766     0   1E+3   3.5   4  RT (min)      lcqtof_hilic_neg_4_003_16    m/z 549.1976     0   1   2   3E+3   0.5   1  RT (min)      lcqtof_hilic_neg_4_004_47    m/z 549.2349     0   1   2E+3   2   2.5  RT (min)      lcqtof_hilic_neg_1_054_01    m/z 549.2914     0   1   2   3E+3   0.5   1  RT (min)      lcqtof_hilic_neg_1_004_14    m/z 549.5588     0   1   2   3E+3   0.5   1  RT (min)      lcqtof_hilic_neg_4_047_06    m/z 550.0810     0   1   2E+3   6.5   7  RT (min)      lcqtof_hilic_neg_4_040_05    m/z 550.1537     0   1   2E+3   6   6.5  RT (min)      lcqtof_hilic_neg_4_066_57    m/z 550.4046     0   2   4   6E+3   1   1.5  RT (min)      lcqtof_hilic_neg_4_053_14    m/z 550.4257     0   1   2   3E+3   3.5   4  RT (min)      lcqtof_hilic_neg_4_053_14    m/z 551.2164     0   1E+3   4   4.5  RT (min)      lcqtof_hilic_neg_1_054_01    m/z 551.2521     0   1E+3   3.5   4  RT (min)      lcqtof_hilic_neg_2_055_30    m/z 551.3578     0   1E+3   3   3.5  RT (min)      lcqtof_hilic_neg_2_021_17    m/z 551.3923     0   1   2   3   4E+3   1   1.5  RT (min)      lcqtof_hilic_neg_4_055_32    m/z 551.8942     0   1E+3   4.5   5  RT (min)      lcqtof_hilic_neg_1_054_01    m/z 552.1013     0   1   2   3   4E+3   0.5   1  RT (min)      lcqtof_hilic_neg_4_013_20    m/z 552.5071     0   1E+3   15.5   16  RT (min)      lcqtof_hilic_neg_4_094_60    m/z 552.8585     0   1   2E+3   0.5   1  RT (min)      lcqtof_hilic_neg_3_076_58    m/z 552.9968     0   1   2E+3   4   4.5  RT (min)      lcqtof_hilic_neg_1_020_09    m/z 553.1185     0   1   2   3   4E+3   5.5   6  RT (min)      lcqtof_hilic_neg_4_026_04    m/z 554.0947     0   1   2E+3   1   1.5  RT (min)      lcqtof_hilic_neg_2_021_17    m/z 554.0970     0   1   2E+3   6.5   7  RT (min)      lcqtof_hilic_neg_4_002_25    m/z 554.3679     0   2   4E+3   0.5   1  RT (min)       lcqtof_hilic_neg_4_053_14    m/z 554.6898     0   1E+3   8   8.5  RT (min)      lcqtof_hilic_neg_1_024_21    m/z 555.0071     0   2   4   6E+3   3.5   4  RT (min)      lcqtof_hilic_neg_2_050_19    m/z 555.0643     0   1   2   3   4E+3   5.5   6  RT (min)       lcqtof_hilic_neg_4_052_13    m/z 555.1135     0   2   4   6   8E+3   6.5   7  RT (min)      lcqtof_hilic_neg_4_048_23    m/z 555.9645     0   1   2   3E+3   5   5.5  RT (min)       lcqtof_hilic_neg_4_052_13    m/z 556.1049     0   1   2E+3   1   1.5  RT (min)      lcqtof_hilic_neg_2_031_01    m/z 556.1530     0   1   2E+3   4.5   5  RT (min)      lcqtof_hilic_neg_1_047_29    m/z 556.2795     0   1   2   3E+3   3.5   4  RT (min)      lcqtof_hilic_neg_4_003_16    m/z 556.3284     0   2   4E+3   0.5   1  RT (min)      lcqtof_hilic_neg_1_024_21    m/z 557.0629     0   1   2   3E+3   1   1.5  RT (min)      lcqtof_hilic_neg_2_001_49    m/z 557.1502     0   1   2E+3   4.5   5  RT (min)      lcqtof_hilic_neg_2_013_29    m/z 557.2416     0   2   4   6E+3   7   7.5  RT (min)       lcqtof_hilic_neg_4_052_13    m/z 558.1998     0   1   2   3E+3   6.5   7  RT (min)      lcqtof_hilic_neg_4_047_06    m/z 558.2561     0   1   2   3E+3   0.5   1  RT (min)      lcqtof_hilic_neg_4_040_05    m/z 558.3070     0   2   4E+3   7   7.5  RT (min)      lcqtof_hilic_neg_1_023_34    m/z 559.1027     0   2   4   6   8E+3   6   6.5  RT (min)       lcqtof_hilic_neg_4_090_60    m/z 559.1128     0   2   4   6   8E+3   6.5   7  RT (min)       lcqtof_hilic_neg_4_008_24    m/z 559.2993     0   2   4   6E+3   7   7.5  RT (min)      lcqtof_hilic_neg_1_014_31    m/z 559.3542     0   2   4   6E+3   0.5   1  RT (min)      lcqtof_hilic_neg_4_003_16    m/z 560.1750     0   2   4   6E+3   1   1.5  RT (min)      lcqtof_hilic_neg_4_053_14    m/z 560.2509     0   1   2E+3   6.5   7  RT (min)      lcqtof_hilic_neg_4_048_23    m/z 560.2697     0   1   2E+3   7   7.5  RT (min)       lcqtof_hilic_neg_1_035_23    m/z 560.9888     0   1E+3   6   6.5  RT (min)       lcqtof_hilic_neg_4_029_17    m/z 561.1807     0   1   2   3E+3   6.5   7  RT (min)      lcqtof_hilic_neg_4_001_49    m/z 562.3990     0   2   4   6E+3   0.5   1  RT (min)      lcqtof_hilic_neg_1_046_30    m/z 563.1960     0   2   4   6E+3   0.5   1  RT (min)      lcqtof_hilic_neg_4_034_31    m/z 563.4326     0   1E+4   1   1.5  RT (min)      lcqtof_hilic_neg_4_018_27    m/z 563.9333     0   2   4E+3   0.5   1  RT (min)       lcqtof_hilic_neg_4_004_47    m/z 564.7838     0   1   2E+3   3.5   4  RT (min)      lcqtof_hilic_neg_4_004_47    m/z 565.0762     0   2   4   6E+3   0.5   1  RT (min)       lcqtof_hilic_neg_4_027_19    m/z 565.1723     0   1E+3   15   15.5  RT (min)      lcqtof_hilic_neg_4_051_43    m/z 566.1445     0   2   4   6E+3   6.5   7  RT (min)       lcqtof_hilic_neg_4_048_23    m/z 566.2483     0   2   4   6E+3   0.5   1  RT (min)      lcqtof_hilic_neg_4_052_13    m/z 566.6981     0   1   2   3   4E+3   3.5   4  RT (min)      lcqtof_hilic_neg_1_054_01    m/z 566.8824     0   2   4   6   8E+3   0.5   1  RT (min)      lcqtof_hilic_neg_4_035_33    m/z 566.9081     0   2   4   6   8E+3   3.5   4  RT (min)      lcqtof_hilic_neg_4_052_13    m/z 567.0508     0   2   4E+3   8   8.5  RT (min)      lcqtof_hilic_neg_4_034_31    m/z 567.2043     0   1   2   3   4E+3   6   6.5  RT (min)      lcqtof_hilic_neg_2_029_31    m/z 567.2209     0   1E+3   3.5   4  RT (min)      lcqtof_hilic_neg_1_010_15    m/z 567.9578     0   1   2   3E+3   5.5   6  RT (min)       lcqtof_hilic_neg_4_046_41    m/z 568.0499     0   2   4   6E+3   6.5   7  RT (min)       lcqtof_hilic_neg_4_048_23    m/z 568.2784     0   1   2E+3   4.5   5  RT (min)      lcqtof_hilic_neg_2_021_17    m/z 568.3841     0   1E+4   0.5   1  RT (min)      lcqtof_hilic_neg_4_023_15    m/z 569.0325     0   1   2   3   4E+3   3.5   4  RT (min)      lcqtof_hilic_neg_4_053_14    m/z 569.5515     0   2   4   6E+3   0.5   1  RT (min)      lcqtof_hilic_neg_4_008_24    m/z 569.9076     0   1   2   3   4E+3   5   5.5  RT (min)      lcqtof_hilic_neg_4_032_01    m/z 569.9168     0   1   2   3E+3   1   1.5  RT (min)      lcqtof_hilic_neg_4_054_48    m/z 569.9669     0   1   2   3   4E+3   0.5   1  RT (min)      lcqtof_hilic_neg_4_032_01    m/z 570.0077     0   2   4   6   8E+3   0.5   1  RT (min)      lcqtof_hilic_neg_1_028_47    m/z 570.0650     0   1   2E+3   3.5   4  RT (min)      lcqtof_hilic_neg_1_053_19    m/z 570.2042     0   2   4   6   8E+3   4.5   5  RT (min)      lcqtof_hilic_neg_4_013_20    m/z 570.2112     0   1   2   3   4E+3   1   1.5  RT (min)      lcqtof_hilic_neg_4_026_04    m/z 570.3611     0   2   4   6E+3   4.5   5  RT (min)       lcqtof_hilic_neg_4_008_24    m/z 570.3962     0   1   2E+3   4.5   5  RT (min)      lcqtof_hilic_neg_4_004_47    m/z 570.7954     0   1   2E+3   7   7.5  RT (min)       lcqtof_hilic_neg_4_097_60    m/z 571.0439     0   2   4   6E+3   7   7.5  RT (min)       lcqtof_hilic_neg_4_016_37    m/z 571.0847     0   1   2   3   4E+3   7   7.5  RT (min)      lcqtof_hilic_neg_4_008_24    m/z 572.0078     0   1   2   3   4E+3   0.5   1  RT (min)      lcqtof_hilic_neg_2_034_47    m/z 572.1370     0   2   4   6E+3   1   1.5  RT (min)      lcqtof_hilic_neg_4_010_28    m/z 572.1429     0   1E+3   6   6.5  RT (min)      lcqtof_hilic_neg_2_043_15    m/z 572.3026     0   1   2   3E+3   0.5   1  RT (min)      lcqtof_hilic_neg_4_004_47    m/z 573.0029     0   1   2   3E+3   5   5.5  RT (min)      lcqtof_hilic_neg_4_008_24    m/z 573.0817     0   2   4   6E+3   7   7.5  RT (min)      lcqtof_hilic_neg_4_011_35    m/z 573.0868     0   2   4E+3   3.5   4  RT (min)      lcqtof_hilic_neg_1_053_19    m/z 573.2713     0   2   4   6E+3   1   1.5  RT (min)      lcqtof_hilic_neg_3_003_20    m/z 573.2924     0   1E+3   6   6.5  RT (min)      lcqtof_hilic_neg_2_051_14    m/z 573.2927     0   2   4   6E+3   1   1.5  RT (min)      lcqtof_hilic_neg_4_039_18    m/z 574.1553     0   2   4   6E+3   1   1.5  RT (min)      lcqtof_hilic_neg_4_005_29    m/z 575.2307     0   1   2   3E+3   6   6.5  RT (min)      lcqtof_hilic_neg_4_002_25    m/z 575.2797     0   1E+3   2.5   3  RT (min)      lcqtof_hilic_neg_1_054_01    m/z 575.3026     0   2   4   6E+3   4   4.5  RT (min)      lcqtof_hilic_neg_4_008_24    m/z 576.2089     0   2   4   6E+3   4.5   5  RT (min)       lcqtof_hilic_neg_4_034_31    m/z 576.2864     0   1   2   3E+3   3.5   4  RT (min)      lcqtof_hilic_neg_1_016_28    m/z 576.4918     0   1   2   3E+4   0.5   1  RT (min)      lcqtof_hilic_neg_2_029_31    m/z 576.4929     0   1   2   3E+4   0.5   1  RT (min)      lcqtof_hilic_neg_1_035_23    m/z 577.0479     0   1   2   3E+3   1   1.5  RT (min)      lcqtof_hilic_neg_4_004_47    m/z 577.3570     0   1   2E+3   1   1.5  RT (min)      lcqtof_hilic_neg_3_002_10    m/z 577.4438     0   1E+3   3.5   4  RT (min)      lcqtof_hilic_neg_4_010_28    m/z 578.1178     0   1   2E+3   5.5   6  RT (min)      lcqtof_hilic_neg_4_003_16    m/z 578.2423     0   1   2   3E+3   6   6.5  RT (min)      lcqtof_hilic_neg_4_002_25    m/z 578.2687     0   1   2   3E+4   1   1.5  RT (min)      lcqtof_hilic_neg_1_012_04    m/z 578.3767     0   1E+3   7   7.5  RT (min)      lcqtof_hilic_neg_4_004_47    m/z 578.7386     0   1   2E+3   7   7.5  RT (min)      lcqtof_hilic_neg_1_037_24    m/z 579.1806     0   1   2   3E+3   1.5   2  RT (min)      lcqtof_hilic_neg_1_032_18    m/z 579.1901     0   2   4E+3   1   1.5  RT (min)      lcqtof_hilic_neg_4_013_20    m/z 579.2228     0   1   2   3   4E+3   4   4.5  RT (min)      lcqtof_hilic_neg_1_007_05    m/z 579.3920     0   1   2E+3   2.5   3  RT (min)      lcqtof_hilic_neg_4_032_01    m/z 579.5392     0   1E+3   15.5   16  RT (min)       lcqtof_hilic_neg_4_013_20    m/z 580.2259     0   2   4   6E+3   4.5   5  RT (min)       lcqtof_hilic_neg_4_034_31    m/z 580.2277     0   1   2E+3   4.5   5  RT (min)      lcqtof_hilic_neg_1_014_31    m/z 581.0920     0   2   4   6E+3   6.5   7  RT (min)      lcqtof_hilic_neg_4_008_24    m/z 581.2446     0   1   2E+3   4   4.5  RT (min)      lcqtof_hilic_neg_3_040_14    m/z 581.2836     0   2   4   6   8E+3   0.5   1  RT (min)      lcqtof_hilic_neg_1_042_26    m/z 581.2990     0   2   4   6E+3   1   1.5  RT (min)      lcqtof_hilic_neg_4_045_26    m/z 581.4008     0   2   4   6   8E+3   0.5   1  RT (min)      lcqtof_hilic_neg_4_053_14    m/z 581.4429     0   2   4   6   8E+3   1   1.5  RT (min)      lcqtof_hilic_neg_4_053_14    m/z 581.8294     0   1   2   3E+3   6.5   7  RT (min)       lcqtof_hilic_neg_4_012_07    m/z 581.9560     0   1E+3   5   5.5  RT (min)      lcqtof_hilic_neg_4_042_22    m/z 582.0369     0   1   2   3E+3   6.5   7  RT (min)      lcqtof_hilic_neg_4_034_31    m/z 582.2372     0   1   2E+3   4   4.5  RT (min)      lcqtof_hilic_neg_1_017_51    m/z 582.3471     0   1   2   3   4E+3   5   5.5  RT (min)      lcqtof_hilic_neg_1_035_23    m/z 583.9197     0   2   4E+3   1.5   2  RT (min)      lcqtof_hilic_neg_4_004_47    m/z 584.0246     0   1   2   3   4E+3   0.5   1  RT (min)      lcqtof_hilic_neg_1_028_47    m/z 584.2542     0   1E+3   1   1.5  RT (min)      lcqtof_hilic_neg_2_046_41    m/z 584.9944     0   1E+3   5.5   6  RT (min)      lcqtof_hilic_neg_3_035_46    m/z 585.2715     0   2   4E+3   4   4.5  RT (min)      lcqtof_hilic_neg_1_030_17    m/z 585.3040     0   2   4E+3   4.5   5  RT (min)       lcqtof_hilic_neg_4_040_05    m/z 585.6100     0   2   4   6   8E+3   0.5   1  RT (min)      lcqtof_hilic_neg_4_008_24    m/z 585.8768     0   1E+3   4.5   5  RT (min)      lcqtof_hilic_neg_4_010_28    m/z 586.8533     0   1   2   3E+3   6.5   7  RT (min)       lcqtof_hilic_neg_4_002_25    m/z 588.0310     0   2   4   6E+3   8   8.5  RT (min)      lcqtof_hilic_neg_4_016_37    m/z 589.1284     0   1   2E+3   0.5   1  RT (min)      lcqtof_hilic_neg_1_003_37    m/z 589.2993     0   1E+3   4   4.5  RT (min)      lcqtof_hilic_neg_3_055_01    m/z 589.3782     0   1   2   3   4E+3   3   3.5  RT (min)      lcqtof_hilic_neg_4_011_35    m/z 589.8975     0   1   2   3E+4   4.5   5  RT (min)      lcqtof_hilic_neg_4_026_04    m/z 590.3124     0   1   2E+3   2   2.5  RT (min)      lcqtof_hilic_neg_1_032_18    m/z 590.7094     0   1E+3   7.5   8  RT (min)      lcqtof_hilic_neg_1_037_24    m/z 590.9703     0   1   2E+3   5   5.5  RT (min)      lcqtof_hilic_neg_4_050_02    m/z 591.4229     0   2   4   6   8E+3   0.5   1  RT (min)      lcqtof_hilic_neg_2_043_15    m/z 592.1518     0   1E+3   5   5.5  RT (min)      lcqtof_hilic_neg_4_045_26    m/z 592.2745     0   1   2   3E+3   4.5   5  RT (min)      lcqtof_hilic_neg_1_037_24    m/z 592.9100     0   1   2   3   4E+3   5   5.5  RT (min)      lcqtof_hilic_neg_4_032_01    m/z 592.9607     0   2   4   6   8E+3   0.5   1  RT (min)      lcqtof_hilic_neg_4_054_48    m/z 592.9839     0   1   2E+3   5   5.5  RT (min)      lcqtof_hilic_neg_4_023_15    m/z 593.2739     0   1   2   3E+3   4.5   5  RT (min)      lcqtof_hilic_neg_4_040_05    m/z 593.2747     0   1   2   3   4E+3   4.5   5  RT (min)      lcqtof_hilic_neg_4_048_23    m/z 594.9049     0   2   4   6   8E+3   0.5   1  RT (min)      lcqtof_hilic_neg_4_004_47    m/z 595.0690     0   1   2   3   4E+3   5   5.5  RT (min)      lcqtof_hilic_neg_4_034_31    m/z 595.3008     0   1   2E+3   0.5   1  RT (min)      lcqtof_hilic_neg_4_030_44    m/z 595.3303     0   2   4   6   8E+3   0.5   1  RT (min)      lcqtof_hilic_neg_4_038_10    m/z 595.9339     0   1E+4   0.5   1  RT (min)      lcqtof_hilic_neg_4_053_14    m/z 595.9345     0   1E+3   4.5   5  RT (min)      lcqtof_hilic_neg_4_013_20    m/z 596.2863     0   1   2   3   4E+3   4   4.5  RT (min)      lcqtof_hilic_neg_3_048_05    m/z 596.7727     0   1   2E+3   4   4.5  RT (min)      lcqtof_hilic_neg_4_049_55    m/z 596.9318     0   1E+3   1.5   2  RT (min)      lcqtof_hilic_neg_4_009_50    m/z 597.0774     0   1E+3   5   5.5  RT (min)      lcqtof_hilic_neg_4_006_30    m/z 597.3040     0   2   4E+3   1   1.5  RT (min)      lcqtof_hilic_neg_4_005_29    m/z 597.4020     0   2   4   6E+3   15   15.5  RT (min)       lcqtof_hilic_neg_4_012_07    m/z 598.1693     0   1   2E+3   6.5   7  RT (min)      lcqtof_hilic_neg_4_048_23    m/z 598.3829     0   2   4E+3   0.5   1  RT (min)      lcqtof_hilic_neg_4_053_14    m/z 598.3900     0   1   2E+3   3.5   4  RT (min)      lcqtof_hilic_neg_1_003_37    m/z 598.4190     0   1   2   3E+3   3   3.5  RT (min)       lcqtof_hilic_neg_4_048_23    m/z 598.9250     0   1   2E+3   0.5   1  RT (min)      lcqtof_hilic_neg_4_052_13    m/z 599.0268     0   1E+3   0.5   1  RT (min)      lcqtof_hilic_neg_4_007_45    m/z 600.2906     0   2   4E+3   4   4.5  RT (min)      lcqtof_hilic_neg_4_003_16    m/z 600.6578     0   2   4   6   8E+3   0.5   1  RT (min)      lcqtof_hilic_neg_4_008_24    m/z 600.6607     0   1   2E+3   8   8.5  RT (min)       lcqtof_hilic_neg_4_013_20    m/z 600.9460     0   1   2   3   4E+3   0.5   1  RT (min)      lcqtof_hilic_neg_4_004_47    m/z 600.9811     0   1   2   3   4E+3   0.5   1  RT (min)      lcqtof_hilic_neg_1_001_49    m/z 600.9824     0   2   4E+3   1   1.5  RT (min)      lcqtof_hilic_neg_4_004_47    m/z 601.3331     0   1E+3   7   7.5  RT (min)      lcqtof_hilic_neg_3_040_14    m/z 603.0143     0   1   2   3   4E+3   6.5   7  RT (min)       lcqtof_hilic_neg_4_028_36    m/z 603.1744     0   1   2E+3   5   5.5  RT (min)      lcqtof_hilic_neg_4_012_07    m/z 603.2042     0   1   2   3E+3   0.5   1  RT (min)      lcqtof_hilic_neg_1_025_52    m/z 603.3912     0   1   2   3   4E+3   1.5   2  RT (min)       lcqtof_hilic_neg_4_048_23    m/z 603.8113     0   1   2   3E+3   6.5   7  RT (min)       lcqtof_hilic_neg_4_044_08    m/z 603.8884     0   2   4   6   8E+3   0.5   1  RT (min)      lcqtof_hilic_neg_4_004_47    m/z 604.0777     0   1   2   3   4E+3   0.5   1  RT (min)      lcqtof_hilic_neg_4_044_08    m/z 604.0928     0   1   2   3   4E+3   6   6.5  RT (min)       lcqtof_hilic_neg_4_018_27    m/z 604.1608     0   2   4   6   8E+3   1   1.5  RT (min)      lcqtof_hilic_neg_4_053_14    m/z 604.8898     0   2   4   6   8E+3   0.5   1  RT (min)      lcqtof_hilic_neg_4_004_47    m/z 605.9743     0   1   2   3E+3   5.5   6  RT (min)      lcqtof_hilic_neg_4_032_01    m/z 606.9306     0   1   2E+3   4   4.5  RT (min)      lcqtof_hilic_neg_3_003_20    m/z 607.3317     0   1   2   3E+3   4   4.5  RT (min)      lcqtof_hilic_neg_3_018_33    m/z 607.8309     0   1   2   3   4E+3   0.5   1  RT (min)      lcqtof_hilic_neg_4_077_58    m/z 608.1104     0   1E+3   5   5.5  RT (min)      lcqtof_hilic_neg_4_050_02    m/z 608.2029     0   2   4   6E+3   6.5   7  RT (min)       lcqtof_hilic_neg_1_035_23    m/z 608.2539     0   1   2E+3   0.5   1  RT (min)      lcqtof_hilic_neg_1_027_27    m/z 608.8959     0   2   4E+3   0.5   1  RT (min)      lcqtof_hilic_neg_4_054_48    m/z 609.3025     0   2   4   6E+3   3.5   4  RT (min)      lcqtof_hilic_neg_4_003_16    m/z 609.3558     0   1   2E+3   3.5   4  RT (min)      lcqtof_hilic_neg_2_004_26    m/z 610.0403     0   1   2   3E+3   0.5   1  RT (min)      lcqtof_hilic_neg_1_049_55    m/z 610.2460     0   1   2   3   4E+3   4   4.5  RT (min)      lcqtof_hilic_neg_1_035_23    m/z 610.3096     0   1   2E+3   3.5   4  RT (min)      lcqtof_hilic_neg_4_052_13    m/z 610.3495     0   2   4   6E+3   0.5   1  RT (min)      lcqtof_hilic_neg_4_002_25    m/z 610.4785     0   1E+4   1   1.5  RT (min)      lcqtof_hilic_neg_4_053_14    m/z 610.7892     0   1   2   3E+3   3.5   4  RT (min)      lcqtof_hilic_neg_4_004_47    m/z 611.3785     0   1   2   3E+3   3.5   4  RT (min)      lcqtof_hilic_neg_4_074_58    m/z 611.4391     0   1   2   3E+3   3.5   4  RT (min)      lcqtof_hilic_neg_4_040_05    m/z 612.0366     0   1E+3   5.5   6  RT (min)      lcqtof_hilic_neg_1_037_24    m/z 612.4922     0   1   2   3   4E+3   1   1.5  RT (min)      lcqtof_hilic_neg_4_053_14    m/z 612.9531     0   1   2   3   4E+3   0.5   1  RT (min)      lcqtof_hilic_neg_4_004_47    m/z 613.0551     0   2   4   6E+3   6   6.5  RT (min)       lcqtof_hilic_neg_4_027_19    m/z 613.1028     0   1   2   3   4E+3   1.5   2  RT (min)      lcqtof_hilic_neg_2_035_09    m/z 613.1727     0   1   2E+3   6   6.5  RT (min)      lcqtof_hilic_neg_4_035_33    m/z 613.2481     0   1   2   3E+3   3.5   4  RT (min)      lcqtof_hilic_neg_1_020_09    m/z 613.5070     0   2   4   6   8E+3   1   1.5  RT (min)      lcqtof_hilic_neg_4_053_14    m/z 614.1310     0   2   4E+3   1.5   2  RT (min)      lcqtof_hilic_neg_1_032_18    m/z 614.1437     0   1   2E+3   1.5   2  RT (min)      lcqtof_hilic_neg_1_001_49    m/z 614.2274     0   1   2   3   4E+3   0.5   1  RT (min)      lcqtof_hilic_neg_2_016_33    m/z 614.2549     0   2   4E+3   4.5   5  RT (min)      lcqtof_hilic_neg_2_052_24    m/z 614.4005     0   1   2   3E+3   15   15.5  RT (min)      lcqtof_hilic_neg_4_019_03    m/z 614.4490     0   2   4   6E+3   0.5   1  RT (min)      lcqtof_hilic_neg_1_054_01    m/z 615.0623     0   2   4   6   8E+3   5   5.5  RT (min)       lcqtof_hilic_neg_4_048_23    m/z 615.0850     0   1   2E+3   5.5   6  RT (min)      lcqtof_hilic_neg_4_019_03    m/z 615.1349     0   1   2   3   4E+3   6.5   7  RT (min)      lcqtof_hilic_neg_4_040_05    m/z 615.3026     0   1E+4   0.5   1  RT (min)      lcqtof_hilic_neg_1_053_19    m/z 616.1408     0   1   2   3E+3   8   8.5  RT (min)       lcqtof_hilic_neg_2_052_24    m/z 616.2629     0   2   4   6E+3   0.5   1  RT (min)      lcqtof_hilic_neg_2_006_22    m/z 616.3307     0   1   2   3   4E+3   3.5   4  RT (min)      lcqtof_hilic_neg_4_022_34    m/z 616.4034     0   1   2   3   4E+3   1   1.5  RT (min)      lcqtof_hilic_neg_4_048_23    m/z 617.1353     0   1   2   3   4E+3   6   6.5  RT (min)       lcqtof_hilic_neg_4_026_04    m/z 618.0463     0   1   2   3E+3   6   6.5  RT (min)      lcqtof_hilic_neg_2_047_32    m/z 618.1815     0   1   2   3   4E+3   4.5   5  RT (min)      lcqtof_hilic_neg_2_028_05    m/z 618.2334     0   1   2E+3   5.5   6  RT (min)      lcqtof_hilic_neg_1_004_14    m/z 618.3582     0   2   4E+4   1   1.5  RT (min)      lcqtof_hilic_neg_4_034_31    m/z 618.4756     0   1E+4   0.5   1  RT (min)      lcqtof_hilic_neg_2_043_15    m/z 619.1188     0   2   4E+3   8.5   9  RT (min)      lcqtof_hilic_neg_4_055_32    m/z 619.1442     0   1E+3   6.5   7  RT (min)      lcqtof_hilic_neg_2_015_06    m/z 619.2406     0   1   2E+3   6.5   7  RT (min)      lcqtof_hilic_neg_1_007_05    m/z 620.4315     0   1E+3   15   15.5  RT (min)       lcqtof_hilic_neg_4_001_49    m/z 620.7292     0   1E+3   8   8.5  RT (min)      lcqtof_hilic_neg_4_056_56    m/z 620.8584     0   1   2   3E+3   3   3.5  RT (min)      lcqtof_hilic_neg_2_005_13    m/z 621.1073     0   1   2   3E+3   5.5   6  RT (min)      lcqtof_hilic_neg_4_013_20    m/z 621.2874     0   2   4   6E+3   0.5   1  RT (min)      lcqtof_hilic_neg_1_004_14    m/z 621.3277     0   1E+3   3.5   4  RT (min)      lcqtof_hilic_neg_2_051_14    m/z 621.3651     0   1   2E+3   3   3.5  RT (min)      lcqtof_hilic_neg_2_021_17    m/z 621.8072     0   1   2E+3   6   6.5  RT (min)      lcqtof_hilic_neg_4_031_38    m/z 621.8654     0   1   2E+3   3   3.5  RT (min)      lcqtof_hilic_neg_2_031_01    m/z 621.9512     0   2   4   6   8E+3   3.5   4  RT (min)      lcqtof_hilic_neg_4_008_24    m/z 622.3726     0   1E+4   3.5   4  RT (min)      lcqtof_hilic_neg_4_022_34    m/z 622.4109     0   1   2E+3   3.5   4  RT (min)      lcqtof_hilic_neg_4_035_33    m/z 622.4398     0   2   4E+3   1.5   2  RT (min)      lcqtof_hilic_neg_4_009_50    m/z 622.6529     0   1E+3   7.5   8  RT (min)      lcqtof_hilic_neg_2_015_06    m/z 623.1240     0   1   2E+3   1   1.5  RT (min)      lcqtof_hilic_neg_4_004_47    m/z 623.1323     0   1   2   3E+3   7.5   8  RT (min)      lcqtof_hilic_neg_1_023_34    m/z 623.1982     0   2   4   6E+3   4   4.5  RT (min)      lcqtof_hilic_neg_2_022_46    m/z 623.3688     0   1E+3   4   4.5  RT (min)      lcqtof_hilic_neg_4_001_49    m/z 623.3871     0   1   2E+3   5   5.5  RT (min)      lcqtof_hilic_neg_4_050_02    m/z 623.8464     0   1   2E+3   0.5   1  RT (min)      lcqtof_hilic_neg_3_079_58    m/z 623.9544     0   1   2E+3   5   5.5  RT (min)       lcqtof_hilic_neg_4_052_13    m/z 624.3464     0   1   2   3E+3   1   1.5  RT (min)      lcqtof_hilic_neg_4_012_07    m/z 624.6500     0   1   2E+3   7.5   8  RT (min)      lcqtof_hilic_neg_1_029_06    m/z 624.9753     0   1   2   3   4E+3   3.5   4  RT (min)      lcqtof_hilic_neg_1_037_24    m/z 625.2213     0   1   2   3E+3   1   1.5  RT (min)      lcqtof_hilic_neg_4_005_29    m/z 625.3050     0   1   2   3E+3   6   6.5  RT (min)       lcqtof_hilic_neg_4_053_14    m/z 625.3340     0   2   4   6   8E+3   1   1.5  RT (min)      lcqtof_hilic_neg_4_018_27    m/z 625.3889     0   1   2   3E+3   0.5   1  RT (min)      lcqtof_hilic_neg_4_033_53    m/z 625.9695     0   2   4   6   8E+3   5   5.5  RT (min)       lcqtof_hilic_neg_4_008_24    m/z 626.1943     0   1   2   3   4E+3   1.5   2  RT (min)      lcqtof_hilic_neg_4_006_30    m/z 626.2898     0   2   4   6   8E+3   0.5   1  RT (min)      lcqtof_hilic_neg_4_002_25    m/z 626.9030     0   2   4   6E+3   5   5.5  RT (min)      lcqtof_hilic_neg_4_032_01    m/z 627.0984     0   1   2E+3   6.5   7  RT (min)      lcqtof_hilic_neg_1_045_07    m/z 627.1611     0   1   2E+3   7   7.5  RT (min)      lcqtof_hilic_neg_1_035_23    m/z 627.2015     0   2   4   6E+3   4   4.5  RT (min)      lcqtof_hilic_neg_2_036_34    m/z 627.3441     0   1   2E+3   3   3.5  RT (min)      lcqtof_hilic_neg_1_054_01    m/z 627.4752     0   1   2   3   4E+3   1.5   2  RT (min)       lcqtof_hilic_neg_4_055_32    m/z 627.9611     0   2   4   6E+3   1   1.5  RT (min)      lcqtof_hilic_neg_4_054_48    m/z 628.2342     0   1   2   3   4E+3   3.5   4  RT (min)      lcqtof_hilic_neg_1_012_04    m/z 628.3257     0   2   4   6   8E+3   4   4.5  RT (min)       lcqtof_hilic_neg_4_053_14    m/z 628.8832     0   2   4E+3   1   1.5  RT (min)       lcqtof_hilic_neg_4_071_57    m/z 630.2332     0   1   2   3   4E+3   0.5   1  RT (min)      lcqtof_hilic_neg_2_025_52    m/z 630.3237     0   1   2   3   4E+3   3.5   4  RT (min)      lcqtof_hilic_neg_4_052_13    m/z 630.3582     0   1   2E+3   3   3.5  RT (min)      lcqtof_hilic_neg_2_031_01    m/z 631.2559     0   2   4   6   8E+3   1   1.5  RT (min)      lcqtof_hilic_neg_3_002_10    m/z 631.2921     0   1   2   3E+3   7   7.5  RT (min)      lcqtof_hilic_neg_1_004_14    m/z 631.3932     0   2   4   6   8E+3   4   4.5  RT (min)       lcqtof_hilic_neg_4_048_23    m/z 631.7996     0   1   2E+3   6.5   7  RT (min)      lcqtof_hilic_neg_1_045_07    m/z 632.2309     0   1   2E+3   0.5   1  RT (min)      lcqtof_hilic_neg_1_024_21    m/z 632.2474     0   1   2   3E+3   6   6.5  RT (min)      lcqtof_hilic_neg_1_029_06    m/z 632.3045     0   1E+4   3.5   4  RT (min)      lcqtof_hilic_neg_4_003_16    m/z 632.3466     0   1   2   3   4E+3   7   7.5  RT (min)      lcqtof_hilic_neg_4_040_05    m/z 632.3972     0   2   4   6   8E+3   0.5   1  RT (min)      lcqtof_hilic_neg_4_048_23    m/z 632.4622     0   1   2   3   4E+3   0.5   1  RT (min)      lcqtof_hilic_neg_1_017_51    m/z 632.9727     0   1   2   3E+3   0.5   1  RT (min)      lcqtof_hilic_neg_1_028_47    m/z 633.0995     0   2   4E+3   6.5   7  RT (min)      lcqtof_hilic_neg_2_020_23    m/z 633.2229     0   1   2   3E+3   1   1.5  RT (min)      lcqtof_hilic_neg_4_005_29    m/z 633.3510     0   2   4E+3   0.5   1  RT (min)      lcqtof_hilic_neg_1_006_22    m/z 633.3873     0   2   4E+3   0.5   1  RT (min)      lcqtof_hilic_neg_4_048_23    m/z 633.4481     0   2   4E+3   15   15.5  RT (min)      lcqtof_hilic_neg_4_050_02    m/z 634.0092     0   1   2E+3   6   6.5  RT (min)      lcqtof_hilic_neg_2_029_31    m/z 634.2088     0   1   2   3E+3   8   8.5  RT (min)       lcqtof_hilic_neg_1_037_24    m/z 634.4028     0   2   4   6E+3   4   4.5  RT (min)      lcqtof_hilic_neg_4_008_24    m/z 635.1226     0   2   4   6E+3   6.5   7  RT (min)       lcqtof_hilic_neg_4_096_60    m/z 635.8535     0   1   2E+3   4.5   5  RT (min)      lcqtof_hilic_neg_4_053_14    m/z 636.1268     0   1   2E+3   8   8.5  RT (min)       lcqtof_hilic_neg_2_048_35    m/z 636.2745     0   1   2   3E+3   5.5   6  RT (min)      lcqtof_hilic_neg_4_053_14    m/z 636.3448     0   1   2E+3   6   6.5  RT (min)      lcqtof_hilic_neg_4_016_37    m/z 636.5505     0   2   4   6E+3   0.5   1  RT (min)      lcqtof_hilic_neg_2_002_36    m/z 637.1171     0   1   2E+3   6.5   7  RT (min)      lcqtof_hilic_neg_4_002_25    m/z 637.1387     0   2   4   6E+3   6   6.5  RT (min)      lcqtof_hilic_neg_1_014_31    m/z 637.2664     0   1   2E+3   3.5   4  RT (min)      lcqtof_hilic_neg_2_005_13    m/z 638.3945     0   1   2   3   4E+3   5   5.5  RT (min)      lcqtof_hilic_neg_4_032_01    m/z 638.4305     0   1   2E+3   15   15.5  RT (min)      lcqtof_hilic_neg_4_047_06    m/z 638.5952     0   1E+3   15.5   16  RT (min)      lcqtof_hilic_neg_3_103_61    m/z 638.8578     0   2   4   6E+3   0.5   1  RT (min)      lcqtof_hilic_neg_4_004_47    m/z 638.9130     0   1   2   3   4E+3   1   1.5  RT (min)       lcqtof_hilic_neg_4_071_57    m/z 639.2808     0   1E+3   3.5   4  RT (min)      lcqtof_hilic_neg_1_048_45    m/z 639.3909     0   2   4   6E+3   1   1.5  RT (min)      lcqtof_hilic_neg_4_003_16    m/z 639.5394     0   1   2   3   4E+3   2   2.5  RT (min)      lcqtof_hilic_neg_4_031_38    m/z 640.1384     0   1E+3   6   6.5  RT (min)      lcqtof_hilic_neg_2_015_06    m/z 640.3439     0   1E+3   7   7.5  RT (min)      lcqtof_hilic_neg_1_035_23    m/z 640.3658     0   2   4   6E+3   3.5   4  RT (min)      lcqtof_hilic_neg_4_052_13    m/z 641.2300     0   1   2E+3   0.5   1  RT (min)      lcqtof_hilic_neg_1_010_15    m/z 641.2985     0   2   4   6E+3   0.5   1  RT (min)       lcqtof_hilic_neg_4_003_16    m/z 641.3582     0   1   2   3   4E+3   3   3.5  RT (min)      lcqtof_hilic_neg_2_005_13    m/z 641.8601     0   1   2   3E+3   3   3.5  RT (min)      lcqtof_hilic_neg_2_005_13    m/z 641.8857     0   1E+4   1   1.5  RT (min)      lcqtof_hilic_neg_4_013_20    m/z 642.3641     0   1   2E+3   3   3.5  RT (min)      lcqtof_hilic_neg_2_018_16    m/z 642.3769     0   2   4   6E+3   3.5   4  RT (min)      lcqtof_hilic_neg_4_052_13    m/z 642.4891     0   1   2   3   4E+3   3.5   4  RT (min)       lcqtof_hilic_neg_4_053_14    m/z 642.7024     0   1E+3   8   8.5  RT (min)       lcqtof_hilic_neg_4_011_35    m/z 642.7637     0   1   2   3   4E+3   4   4.5  RT (min)       lcqtof_hilic_neg_4_033_53    m/z 643.1254     0   2   4   6E+3   0.5   1  RT (min)      lcqtof_hilic_neg_4_022_34    m/z 643.2673     0   2   4   6E+3   1   1.5  RT (min)      lcqtof_hilic_neg_4_019_03    m/z 643.4355     0   2   4   6E+3   4   4.5  RT (min)      lcqtof_hilic_neg_4_034_31    m/z 643.8998     0   2   4   6   8E+3   0.5   1  RT (min)      lcqtof_hilic_neg_4_004_47    m/z 644.1704     0   2   4E+3   0.5   1  RT (min)      lcqtof_hilic_neg_4_018_27    m/z 644.2984     0   1   2   3E+3   4   4.5  RT (min)      lcqtof_hilic_neg_4_032_01    m/z 644.4350     0   1   2E+3   4   4.5  RT (min)      lcqtof_hilic_neg_4_022_34    m/z 645.0735     0   1   2   3   4E+3   7.5   8  RT (min)      lcqtof_hilic_neg_4_055_32    m/z 645.1186     0   1   2E+3   7.5   8  RT (min)      lcqtof_hilic_neg_1_022_36    m/z 645.2933     0   1   2   3E+3   1   1.5  RT (min)      lcqtof_hilic_neg_1_031_16    m/z 645.4389     0   2   4E+3   0.5   1  RT (min)       lcqtof_hilic_neg_4_029_17    m/z 645.4407     0   1   2   3   4E+3   5   5.5  RT (min)      lcqtof_hilic_neg_4_048_23    m/z 645.5638     0   2   4   6E+3   0.5   1  RT (min)      lcqtof_hilic_neg_1_010_15    m/z 645.9875     0   2   4E+3   0.5   1  RT (min)      lcqtof_hilic_neg_2_034_47    m/z 646.1788     0   1   2   3E+3   6   6.5  RT (min)       lcqtof_hilic_neg_2_028_05    m/z 646.2033     0   2   4   6   8E+3   6   6.5  RT (min)      lcqtof_hilic_neg_2_020_23    m/z 646.2263     0   2   4   6E+3   7   7.5  RT (min)      lcqtof_hilic_neg_4_008_24    m/z 646.3217     0   1E+3   5   5.5  RT (min)      lcqtof_hilic_neg_2_051_14    m/z 646.3309     0   2   4   6E+3   0.5   1  RT (min)      lcqtof_hilic_neg_4_047_06    m/z 646.4518     0   1   2   3E+3   4   4.5  RT (min)      lcqtof_hilic_neg_4_022_34    m/z 646.5029     0   1E+3   3.5   4  RT (min)      lcqtof_hilic_neg_4_053_14    m/z 646.9845     0   1   2   3E+3   0.5   1  RT (min)      lcqtof_hilic_neg_2_049_55    m/z 647.0644     0   1   2E+3   6.5   7  RT (min)      lcqtof_hilic_neg_2_048_35    m/z 647.3074     0   1   2E+3   7   7.5  RT (min)      lcqtof_hilic_neg_4_002_25    m/z 647.3199     0   2   4   6   8E+3   4.5   5  RT (min)      lcqtof_hilic_neg_4_052_13    m/z 647.3204     0   1   2   3   4E+3   4.5   5  RT (min)      lcqtof_hilic_neg_2_031_01    m/z 647.4040     0   1E+3   4   4.5  RT (min)      lcqtof_hilic_neg_1_035_23    m/z 647.4070     0   1   2   3   4E+3   0.5   1  RT (min)      lcqtof_hilic_neg_4_032_01    m/z 648.0093     0   2   4   6E+3   0.5   1  RT (min)      lcqtof_hilic_neg_4_055_32    m/z 648.0329     0   1   2   3E+3   8.5   9  RT (min)       lcqtof_hilic_neg_4_028_36    m/z 648.1627     0   2   4E+3   6.5   7  RT (min)      lcqtof_hilic_neg_1_035_23    m/z 648.9042     0   1   2   3   4E+3   6   6.5  RT (min)      lcqtof_hilic_neg_4_055_32    m/z 649.2948     0   2   4E+3   3.5   4  RT (min)      lcqtof_hilic_neg_4_050_02    m/z 649.3893     0   1   2   3   4E+3   0.5   1  RT (min)      lcqtof_hilic_neg_4_048_23    m/z 649.4570     0   1   2   3E+3   0.5   1  RT (min)      lcqtof_hilic_neg_1_035_23    m/z 649.6877     0   1   2   3E+3   0.5   1  RT (min)      lcqtof_hilic_neg_4_008_24    m/z 649.8420     0   1   2   3   4E+3   6   6.5  RT (min)      lcqtof_hilic_neg_4_011_35    m/z 649.9395     0   1   2E+3   5   5.5  RT (min)      lcqtof_hilic_neg_4_019_03    m/z 650.3420     0   1   2   3E+3   6   6.5  RT (min)      lcqtof_hilic_neg_4_011_35    m/z 650.3993     0   2   4E+3   4.5   5  RT (min)      lcqtof_hilic_neg_4_008_24    m/z 651.2202     0   1   2E+3   6   6.5  RT (min)      lcqtof_hilic_neg_2_015_06    m/z 651.3088     0   1   2E+3   3.5   4  RT (min)      lcqtof_hilic_neg_1_003_37    m/z 651.3116     0   1   2E+3   7   7.5  RT (min)      lcqtof_hilic_neg_1_045_07    m/z 651.9689     0   1E+3   5.5   6  RT (min)      lcqtof_hilic_neg_4_021_21    m/z 652.1865     0   2   4   6E+3   0.5   1  RT (min)      lcqtof_hilic_neg_1_004_14    m/z 652.2281     0   2   4   6E+3   1   1.5  RT (min)      lcqtof_hilic_neg_4_021_21    m/z 652.2380     0   2   4E+3   1   1.5  RT (min)      lcqtof_hilic_neg_4_026_04    m/z 652.4761     0   1   2E+3   0.5   1  RT (min)      lcqtof_hilic_neg_1_035_23    m/z 652.9800     0   1   2   3   4E+3   6   6.5  RT (min)       lcqtof_hilic_neg_4_026_04    m/z 653.2964     0   1   2   3E+3   5.5   6  RT (min)      lcqtof_hilic_neg_4_048_23    m/z 653.3025     0   1E+4   1   1.5  RT (min)      lcqtof_hilic_neg_4_040_05    m/z 653.6825     0   1E+3   8   8.5  RT (min)      lcqtof_hilic_neg_2_043_15    m/z 653.8642     0   2   4   6   8E+3   0.5   1  RT (min)      lcqtof_hilic_neg_4_004_47    m/z 653.9308     0   2   4   6E+3   0.5   1  RT (min)      lcqtof_hilic_neg_4_004_47    m/z 654.3190     0   1   2   3   4E+3   3.5   4  RT (min)      lcqtof_hilic_neg_1_037_24    m/z 654.4784     0   2   4E+3   15   15.5  RT (min)      lcqtof_hilic_neg_4_023_15    m/z 654.8538     0   1   2   3E+3   0.5   1  RT (min)      lcqtof_hilic_neg_4_077_58    m/z 654.9239     0   2   4E+3   0.5   1  RT (min)      lcqtof_hilic_neg_4_004_47    m/z 654.9328     0   1   2E+3   4.5   5  RT (min)      lcqtof_hilic_neg_1_003_37    m/z 655.1888     0   2   4E+3   5   5.5  RT (min)      lcqtof_hilic_neg_4_038_10    m/z 655.2794     0   2   4   6E+3   0.5   1  RT (min)      lcqtof_hilic_neg_4_047_06    m/z 655.9014     0   2   4   6   8E+3   1   1.5  RT (min)       lcqtof_hilic_neg_4_054_48    m/z 656.2648     0   2   4E+3   0.5   1  RT (min)      lcqtof_hilic_neg_4_007_45    m/z 656.3391     0   1   2   3E+3   4.5   5  RT (min)      lcqtof_hilic_neg_4_013_20    m/z 656.3571     0   2   4   6E+3   6   6.5  RT (min)      lcqtof_hilic_neg_4_028_36    m/z 656.7781     0   2   4E+3   3.5   4  RT (min)       lcqtof_hilic_neg_4_014_46    m/z 657.2482     0   1E+4   0.5   1  RT (min)      lcqtof_hilic_neg_1_006_22    m/z 657.3152     0   2   4E+3   0.5   1  RT (min)      lcqtof_hilic_neg_4_048_23    m/z 657.3224     0   1   2   3   4E+3   7   7.5  RT (min)      lcqtof_hilic_neg_1_037_24    m/z 657.3308     0   1   2E+3   3   3.5  RT (min)      lcqtof_hilic_neg_1_004_14    m/z 657.3851     0   1   2   3E+3   4   4.5  RT (min)      lcqtof_hilic_neg_4_039_18    m/z 658.0417     0   1   2   3   4E+3   6.5   7  RT (min)      lcqtof_hilic_neg_4_007_45    m/z 658.1658     0   1   2   3   4E+3   4.5   5  RT (min)      lcqtof_hilic_neg_4_034_31    m/z 658.2846     0   1   2E+3   4.5   5  RT (min)      lcqtof_hilic_neg_2_004_26    m/z 658.4089     0   2   4E+3   0.5   1  RT (min)      lcqtof_hilic_neg_4_021_21    m/z 658.4444     0   1   2   3E+3   3   3.5  RT (min)      lcqtof_hilic_neg_1_022_36    m/z 658.4545     0   1   2   3E+3   4   4.5  RT (min)      lcqtof_hilic_neg_4_011_35    m/z 658.9604     0   1   2   3   4E+3   5   5.5  RT (min)      lcqtof_hilic_neg_4_053_14    m/z 659.1394     0   2   4   6   8E+3   6   6.5  RT (min)       lcqtof_hilic_neg_4_048_23    m/z 659.2441     0   2   4E+3   6.5   7  RT (min)      lcqtof_hilic_neg_4_054_48    m/z 659.3737     0   2   4   6E+3   0.5   1  RT (min)      lcqtof_hilic_neg_4_029_17    m/z 659.5776     0   2   4   6E+3   0.5   1  RT (min)      lcqtof_hilic_neg_2_043_15    m/z 659.9156     0   1   2   3   4E+3   0.5   1  RT (min)      lcqtof_hilic_neg_4_004_47    m/z 659.9963     0   2   4E+3   7   7.5  RT (min)       lcqtof_hilic_neg_4_008_24    m/z 660.1304     0   1   2   3   4E+3   1   1.5  RT (min)      lcqtof_hilic_neg_4_021_21    m/z 660.2704     0   2   4   6   8E+3   6   6.5  RT (min)      lcqtof_hilic_neg_4_052_13    m/z 660.8961     0   2   4   6E+3   5   5.5  RT (min)      lcqtof_hilic_neg_4_053_14    m/z 660.9249     0   1   2E+3   1   1.5  RT (min)      lcqtof_hilic_neg_1_001_49    m/z 660.9519     0   2   4E+3   0.5   1  RT (min)      lcqtof_hilic_neg_2_012_43    m/z 660.9739     0   1   2E+3   5   5.5  RT (min)      lcqtof_hilic_neg_2_031_01    m/z 661.9957     0   1   2   3E+3   7.5   8  RT (min)      lcqtof_hilic_neg_2_020_23    m/z 662.4795     0   1   2E+3   3.5   4  RT (min)      lcqtof_hilic_neg_4_052_13    m/z 662.5226     0   1   2E+3   3.5   4  RT (min)      lcqtof_hilic_neg_4_032_01    m/z 663.2757     0   1   2E+3   3.5   4  RT (min)      lcqtof_hilic_neg_4_019_03    m/z 663.3040     0   2   4   6   8E+3   4.5   5  RT (min)      lcqtof_hilic_neg_4_013_20    m/z 663.4442     0   2   4   6   8E+3   0.5   1  RT (min)       lcqtof_hilic_neg_4_010_28    m/z 663.4736     0   1   2E+3   3.5   4  RT (min)      lcqtof_hilic_neg_4_032_01    m/z 664.0883     0   1   2E+3   8.5   9  RT (min)      lcqtof_hilic_neg_2_047_32    m/z 664.1819     0   1   2E+3   8   8.5  RT (min)      lcqtof_hilic_neg_1_037_24    m/z 664.2505     0   1   2E+3   5.5   6  RT (min)      lcqtof_hilic_neg_4_113_62    m/z 664.2875     0   1   2   3   4E+3   3.5   4  RT (min)      lcqtof_hilic_neg_4_050_02    m/z 664.3920     0   2   4   6E+3   0.5   1  RT (min)      lcqtof_hilic_neg_4_048_23    m/z 664.4874     0   1   2E+3   3.5   4  RT (min)      lcqtof_hilic_neg_4_032_01    m/z 664.6934     0   1E+3   8   8.5  RT (min)       lcqtof_hilic_neg_4_004_47    m/z 664.9860     0   2   4   6E+3   5   5.5  RT (min)      lcqtof_hilic_neg_4_008_24    m/z 665.0937     0   1   2   3E+3   7   7.5  RT (min)      lcqtof_hilic_neg_2_043_15    m/z 665.2692     0   1   2   3E+3   0.5   1  RT (min)      lcqtof_hilic_neg_4_010_28    m/z 665.4155     0   1   2   3   4E+3   3   3.5  RT (min)      lcqtof_hilic_neg_4_022_34    m/z 666.3941     0   1   2   3   4E+4   1   1.5  RT (min)      lcqtof_hilic_neg_4_047_06    m/z 666.4816     0   1   2   3   4E+3   3   3.5  RT (min)      lcqtof_hilic_neg_4_016_37    m/z 666.5341     0   1   2   3   4E+3   0.5   1  RT (min)      lcqtof_hilic_neg_2_047_32    m/z 666.5519     0   1   2E+3   0.5   1  RT (min)      lcqtof_hilic_neg_4_023_15    m/z 666.8722     0   1E+4   1   1.5  RT (min)      lcqtof_hilic_neg_4_074_58    m/z 667.0622     0   1   2   3E+3   0.5   1  RT (min)      lcqtof_hilic_neg_4_002_25    m/z 667.0654     0   2   4   6   8E+3   8   8.5  RT (min)       lcqtof_hilic_neg_2_020_23    m/z 667.3670     0   1   2   3   4E+3   4   4.5  RT (min)      lcqtof_hilic_neg_4_004_47    m/z 668.2851     0   2   4   6E+3   3.5   4  RT (min)      lcqtof_hilic_neg_4_014_46    m/z 668.3154     0   2   4   6   8E+3   0.5   1  RT (min)      lcqtof_hilic_neg_4_045_26    m/z 668.3672     0   2   4   6E+3   3.5   4  RT (min)      lcqtof_hilic_neg_4_074_58    m/z 668.4377     0   1   2   3   4E+3   0.5   1  RT (min)      lcqtof_hilic_neg_4_048_23    m/z 669.0844     0   1   2E+3   6   6.5  RT (min)       lcqtof_hilic_neg_4_081_58    m/z 669.1215     0   1   2   3E+3   7   7.5  RT (min)      lcqtof_hilic_neg_3_030_23    m/z 669.1725     0   1   2   3E+3   9   9.5  RT (min)       lcqtof_hilic_neg_4_040_05    m/z 669.8383     0   1   2E+3   4.5   5  RT (min)      lcqtof_hilic_neg_4_029_17    m/z 669.9124     0   1   2E+3   5   5.5  RT (min)      lcqtof_hilic_neg_4_008_24    m/z 670.1739     0   1E+3   6   6.5  RT (min)      lcqtof_hilic_neg_2_036_34    m/z 670.8465     0   2   4   6E+3   2   2.5  RT (min)       lcqtof_hilic_neg_4_014_46    m/z 671.1542     0   1   2   3   4E+3   6.5   7  RT (min)      lcqtof_hilic_neg_2_020_23    m/z 671.1637     0   1   2E+3   5   5.5  RT (min)       lcqtof_hilic_neg_4_012_07    m/z 671.3677     0   2   4E+3   0.5   1  RT (min)      lcqtof_hilic_neg_4_050_02    m/z 671.5140     0   1   2   3E+3   3.5   4  RT (min)      lcqtof_hilic_neg_4_032_01    m/z 671.5191     0   2   4   6E+3   0.5   1  RT (min)      lcqtof_hilic_neg_4_013_20    m/z 671.5853     0   1   2   3E+3   0.5   1  RT (min)      lcqtof_hilic_neg_2_042_27    m/z 672.1035     0   1   2E+3   7   7.5  RT (min)      lcqtof_hilic_neg_2_043_15    m/z 672.2132     0   2   4   6E+3   0.5   1  RT (min)      lcqtof_hilic_neg_4_029_17    m/z 672.3897     0   2   4E+3   5   5.5  RT (min)      lcqtof_hilic_neg_4_053_14    m/z 672.9143     0   2   4   6E+3   0.5   1  RT (min)      lcqtof_hilic_neg_4_004_47    m/z 673.0065     0   1   2   3E+3   0.5   1  RT (min)      lcqtof_hilic_neg_4_004_47    m/z 673.3535     0   1   2E+3   6.5   7  RT (min)      lcqtof_hilic_neg_3_030_23    m/z 673.3574     0   2   4E+3   6   6.5  RT (min)      lcqtof_hilic_neg_4_031_38    m/z 673.3759     0   1   2   3E+3   1   1.5  RT (min)      lcqtof_hilic_neg_4_005_29    m/z 673.4236     0   2   4   6   8E+3   0.5   1  RT (min)      lcqtof_hilic_neg_4_016_37    m/z 673.9561     0   1   2E+3   5.5   6  RT (min)       lcqtof_hilic_neg_4_006_30    m/z 674.1438     0   1   2E+3   4.5   5  RT (min)      lcqtof_hilic_neg_1_035_23    m/z 674.4037     0   1   2   3E+3   0.5   1  RT (min)      lcqtof_hilic_neg_4_015_09    m/z 674.9235     0   1   2   3E+3   1   1.5  RT (min)      lcqtof_hilic_neg_4_079_58    m/z 674.9696     0   1   2   3E+3   0.5   1  RT (min)      lcqtof_hilic_neg_2_023_48    m/z 675.0984     0   1   2   3   4E+3   1   1.5  RT (min)      lcqtof_hilic_neg_1_006_22    m/z 675.3519     0   1   2   3E+3   7   7.5  RT (min)      lcqtof_hilic_neg_4_093_60    m/z 675.4854     0   2   4   6   8E+3   4   4.5  RT (min)      lcqtof_hilic_neg_4_011_35    m/z 675.6700     0   1E+3   8   8.5  RT (min)      lcqtof_hilic_neg_2_031_01    m/z 675.8579     0   1   2   3E+3   4.5   5  RT (min)      lcqtof_hilic_neg_4_012_07    m/z 675.9432     0   1E+3   5   5.5  RT (min)      lcqtof_hilic_neg_4_015_09    m/z 676.9968     0   2   4   6E+3   8   8.5  RT (min)      lcqtof_hilic_neg_4_011_35    m/z 677.0537     0   1   2E+3   5.5   6  RT (min)      lcqtof_hilic_neg_4_053_14    m/z 677.0851     0   2   4   6E+3   6   6.5  RT (min)      lcqtof_hilic_neg_4_034_31    m/z 677.1444     0   2   4   6   8E+3   6   6.5  RT (min)       lcqtof_hilic_neg_4_052_13    m/z 677.2008     0   1   2E+3   7   7.5  RT (min)      lcqtof_hilic_neg_3_034_24    m/z 677.2062     0   1   2   3   4E+3   4   4.5  RT (min)       lcqtof_hilic_neg_4_045_26    m/z 677.3689     0   1   2   3E+3   6   6.5  RT (min)      lcqtof_hilic_neg_4_028_36    m/z 678.3969     0   2   4   6E+3   1   1.5  RT (min)      lcqtof_hilic_neg_4_039_18    m/z 678.4160     0   1   2   3E+3   0.5   1  RT (min)      lcqtof_hilic_neg_1_004_14    m/z 678.4799     0   1E+4   3.5   4  RT (min)      lcqtof_hilic_neg_4_008_24    m/z 678.9272     0   1E+3   5   5.5  RT (min)      lcqtof_hilic_neg_1_054_01    m/z 678.9366     0   2   4   6E+3   6   6.5  RT (min)      lcqtof_hilic_neg_4_055_32    m/z 679.0491     0   1   2   3E+3   7.5   8  RT (min)      lcqtof_hilic_neg_3_004_32    m/z 679.0854     0   1   2   3E+3   0.5   1  RT (min)      lcqtof_hilic_neg_1_032_18    m/z 679.3397     0   1   2E+3   7   7.5  RT (min)      lcqtof_hilic_neg_4_008_24    m/z 679.4230     0   1   2   3E+3   3   3.5  RT (min)      lcqtof_hilic_neg_4_002_25    m/z 679.5651     0   1   2E+3   3   3.5  RT (min)      lcqtof_hilic_neg_1_005_32    m/z 679.7367     0   2   4   6   8E+3   4   4.5  RT (min)      lcqtof_hilic_neg_4_041_54    m/z 680.2581     0   1   2   3E+3   1   1.5  RT (min)      lcqtof_hilic_neg_4_021_21    m/z 680.3196     0   2   4   6E+3   3.5   4  RT (min)      lcqtof_hilic_neg_4_052_13    m/z 680.4972     0   2   4   6E+3   3.5   4  RT (min)       lcqtof_hilic_neg_4_097_60    m/z 680.7897     0   1   2E+3   6.5   7  RT (min)       lcqtof_hilic_neg_4_012_07    m/z 681.2365     0   1   2   3E+3   1   1.5  RT (min)      lcqtof_hilic_neg_4_021_21    m/z 681.7360     0   2   4E+3   4   4.5  RT (min)       lcqtof_hilic_neg_4_033_53    m/z 682.0900     0   1   2E+3   1   1.5  RT (min)      lcqtof_hilic_neg_2_033_53    m/z 682.2119     0   2   4   6   8E+3   6   6.5  RT (min)       lcqtof_hilic_neg_4_053_14    m/z 682.2443     0   1   2E+3   4.5   5  RT (min)      lcqtof_hilic_neg_2_052_24    m/z 682.3589     0   1   2   3   4E+3   4.5   5  RT (min)       lcqtof_hilic_neg_4_090_60    m/z 682.4708     0   1   2E+3   3.5   4  RT (min)      lcqtof_hilic_neg_4_048_23    m/z 682.7343     0   1   2   3   4E+3   4   4.5  RT (min)      lcqtof_hilic_neg_4_033_53    m/z 683.0509     0   2   4   6   8E+3   5   5.5  RT (min)       lcqtof_hilic_neg_4_048_23    m/z 683.4645     0   1   2   3   4E+3   4   4.5  RT (min)      lcqtof_hilic_neg_4_084_59    m/z 683.5115     0   1   2E+3   3.5   4  RT (min)      lcqtof_hilic_neg_4_032_01    m/z 684.1545     0   1   2   3E+3   4   4.5  RT (min)      lcqtof_hilic_neg_1_032_18    m/z 684.2634     0   1   2E+3   3.5   4  RT (min)      lcqtof_hilic_neg_1_042_26    m/z 684.2799     0   1   2E+3   5.5   6  RT (min)      lcqtof_hilic_neg_4_092_60    m/z 685.0023     0   2   4   6E+3   8   8.5  RT (min)      lcqtof_hilic_neg_4_016_37    m/z 685.1494     0   2   4E+3   4   4.5  RT (min)      lcqtof_hilic_neg_1_032_18    m/z 685.3179     0   1   2   3   4E+3   2.5   3  RT (min)      lcqtof_hilic_neg_4_027_19    m/z 685.6700     0   2   4   6E+3   0.5   1  RT (min)      lcqtof_hilic_neg_4_008_24    m/z 685.9591     0   2   4   6   8E+3   6   6.5  RT (min)       lcqtof_hilic_neg_4_055_32    m/z 686.3193     0   2   4   6E+3   3.5   4  RT (min)       lcqtof_hilic_neg_4_039_18    m/z 686.3642     0   1   2   3   4E+3   0.5   1  RT (min)      lcqtof_hilic_neg_4_047_06    m/z 686.4410     0   1   2E+4   0.5   1  RT (min)      lcqtof_hilic_neg_4_023_15    m/z 686.4493     0   2   4   6E+3   6   6.5  RT (min)      lcqtof_hilic_neg_4_055_32    m/z 686.6650     0   1E+3   8   8.5  RT (min)      lcqtof_hilic_neg_4_018_27    m/z 686.6683     0   1E+3   8.5   9  RT (min)       lcqtof_hilic_neg_4_052_13    m/z 687.0745     0   1   2E+3   7   7.5  RT (min)      lcqtof_hilic_neg_3_029_29    m/z 687.0797     0   2   4E+3   6   6.5  RT (min)      lcqtof_hilic_neg_4_035_33    m/z 687.1595     0   1   2E+3   8   8.5  RT (min)       lcqtof_hilic_neg_1_037_24    m/z 687.2893     0   1E+4   1   1.5  RT (min)      lcqtof_hilic_neg_4_045_26    m/z 687.4943     0   2   4   6E+3   4   4.5  RT (min)      lcqtof_hilic_neg_3_031_36    m/z 687.6744     0   2   4   6E+3   3.5   4  RT (min)      lcqtof_hilic_neg_4_032_01    m/z 688.2514     0   1E+4   1   1.5  RT (min)      lcqtof_hilic_neg_1_042_26    m/z 688.2546     0   1   2   3E+3   3.5   4  RT (min)      lcqtof_hilic_neg_3_066_57    m/z 688.2862     0   1   2E+3   3.5   4  RT (min)      lcqtof_hilic_neg_1_054_01    m/z 688.3847     0   1   2   3E+3   4   4.5  RT (min)      lcqtof_hilic_neg_4_085_59    m/z 688.5087     0   2   4   6   8E+3   0.5   1  RT (min)      lcqtof_hilic_neg_1_034_08    m/z 689.0363     0   1   2E+3   0.5   1  RT (min)      lcqtof_hilic_neg_1_049_55    m/z 689.0453     0   1   2   3   4E+3   8   8.5  RT (min)       lcqtof_hilic_neg_2_020_23    m/z 689.0911     0   2   4E+3   5.5   6  RT (min)      lcqtof_hilic_neg_4_026_04    m/z 689.1271     0   2   4   6   8E+3   8   8.5  RT (min)      lcqtof_hilic_neg_4_008_24    m/z 689.2276     0   2   4   6   8E+3   0.5   1  RT (min)      lcqtof_hilic_neg_4_018_27    m/z 689.2953     0   1   2   3   4E+3   1   1.5  RT (min)      lcqtof_hilic_neg_4_006_30    m/z 689.3622     0   1E+3   4   4.5  RT (min)      lcqtof_hilic_neg_3_024_35    m/z 689.4834     0   1   2E+3   3.5   4  RT (min)      lcqtof_hilic_neg_4_052_13    m/z 690.6849     0   1E+3   4.5   5  RT (min)      lcqtof_hilic_neg_4_001_49    m/z 691.0051     0   1   2   3E+3   7   7.5  RT (min)       lcqtof_hilic_neg_4_048_23    m/z 691.3172     0   1   2E+3   1.5   2  RT (min)      lcqtof_hilic_neg_1_046_30    m/z 691.3473     0   1   2E+3   4   4.5  RT (min)      lcqtof_hilic_neg_4_013_20    m/z 691.3939     0   2   4   6E+3   0.5   1  RT (min)      lcqtof_hilic_neg_2_051_14    m/z 691.5646     0   1   2E+3   3.5   4  RT (min)      lcqtof_hilic_neg_4_010_28    m/z 691.7791     0   1   2E+3   6.5   7  RT (min)      lcqtof_hilic_neg_4_012_07    m/z 692.0473     0   2   4E+3   8   8.5  RT (min)      lcqtof_hilic_neg_2_047_32    m/z 692.0905     0   2   4   6E+3   0.5   1  RT (min)      lcqtof_hilic_neg_4_013_20    m/z 692.1609     0   2   4   6E+3   1   1.5  RT (min)      lcqtof_hilic_neg_4_026_04    m/z 692.4202     0   2   4   6   8E+3   4.5   5  RT (min)      lcqtof_hilic_neg_4_004_47    m/z 692.9451     0   1   2   3E+3   0.5   1  RT (min)      lcqtof_hilic_neg_4_054_48    m/z 692.9761     0   1   2   3E+3   1   1.5  RT (min)      lcqtof_hilic_neg_1_028_47    m/z 693.0450     0   1   2E+3   6.5   7  RT (min)      lcqtof_hilic_neg_3_023_08    m/z 693.0629     0   1   2   3E+3   8   8.5  RT (min)      lcqtof_hilic_neg_4_016_37    m/z 693.1400     0   2   4   6E+3   6.5   7  RT (min)      lcqtof_hilic_neg_4_040_05    m/z 693.1544     0   2   4   6   8E+3   1   1.5  RT (min)      lcqtof_hilic_neg_4_004_47    m/z 693.1765     0   1   2   3E+3   4   4.5  RT (min)      lcqtof_hilic_neg_2_036_34    m/z 693.2217     0   2   4   6E+3   4.5   5  RT (min)      lcqtof_hilic_neg_4_050_02    m/z 693.2245     0   1   2E+3   4.5   5  RT (min)      lcqtof_hilic_neg_1_008_13    m/z 693.2437     0   2   4E+3   3.5   4  RT (min)       lcqtof_hilic_neg_1_020_09    m/z 693.4379     0   1E+4   0.5   1  RT (min)      lcqtof_hilic_neg_4_004_47    m/z 693.4523     0   1   2   3   4E+3   3.5   4  RT (min)      lcqtof_hilic_neg_4_032_01    m/z 693.5016     0   1E+3   3.5   4  RT (min)      lcqtof_hilic_neg_4_032_01    m/z 693.5331     0   2   4   6E+3   0.5   1  RT (min)      lcqtof_hilic_neg_4_027_19    m/z 693.5684     0   1   2   3E+3   4   4.5  RT (min)      lcqtof_hilic_neg_4_004_47    m/z 694.0350     0   1   2   3   4E+3   8.5   9  RT (min)       lcqtof_hilic_neg_3_031_36    m/z 695.3939     0   1   2E+3   5   5.5  RT (min)      lcqtof_hilic_neg_4_032_01    m/z 696.1546     0   1   2E+3   8   8.5  RT (min)      lcqtof_hilic_neg_4_007_45    m/z 696.2690     0   1   2   3   4E+3   0.5   1  RT (min)      lcqtof_hilic_neg_4_047_06    m/z 696.2936     0   2   4   6E+3   3.5   4  RT (min)      lcqtof_hilic_neg_4_050_02    m/z 696.6421     0   1E+3   15.5   16  RT (min)       lcqtof_hilic_neg_4_103_61    m/z 696.7681     0   1   2E+3   5.5   6  RT (min)      lcqtof_hilic_neg_1_004_14    m/z 697.0566     0   1   2   3E+3   6.5   7  RT (min)      lcqtof_hilic_neg_3_023_08    m/z 697.1506     0   1   2   3   4E+3   6.5   7  RT (min)      lcqtof_hilic_neg_2_028_05    m/z 697.2267     0   1   2E+3   0.5   1  RT (min)      lcqtof_hilic_neg_2_051_14    m/z 697.5094     0   2   4E+3   4.5   5  RT (min)      lcqtof_hilic_neg_4_031_38    m/z 698.3429     0   1   2   3E+3   3.5   4  RT (min)      lcqtof_hilic_neg_4_008_24    m/z 698.5312     0   2   4E+3   0.5   1  RT (min)      lcqtof_hilic_neg_4_048_23    m/z 698.6088     0   1   2   3E+3   0.5   1  RT (min)      lcqtof_hilic_neg_2_042_27    m/z 698.6288     0   2   4E+3   1   1.5  RT (min)      lcqtof_hilic_neg_4_048_23    m/z 698.6451     0   2   4E+3   0.5   1  RT (min)      lcqtof_hilic_neg_4_048_23    m/z 699.0602     0   2   4E+4   1   1.5  RT (min)      lcqtof_hilic_neg_4_021_21    m/z 699.1249     0   2   4E+3   4   4.5  RT (min)      lcqtof_hilic_neg_2_053_18    m/z 699.2419     0   1   2   3E+3   4.5   5  RT (min)      lcqtof_hilic_neg_3_055_01    m/z 699.3570     0   1   2   3   4E+3   1   1.5  RT (min)      lcqtof_hilic_neg_4_039_18    m/z 699.3848     0   1   2E+3   0.5   1  RT (min)      lcqtof_hilic_neg_1_004_14    m/z 699.4001     0   1   2   3   4E+3   4   4.5  RT (min)      lcqtof_hilic_neg_4_032_01    m/z 699.4341     0   2   4   6E+3   3.5   4  RT (min)      lcqtof_hilic_neg_4_048_23    m/z 699.5990     0   2   4   6E+3   0.5   1  RT (min)      lcqtof_hilic_neg_2_028_05    m/z 700.0526     0   1E+4   1   1.5  RT (min)      lcqtof_hilic_neg_1_024_21    m/z 700.2218     0   1   2E+3   4.5   5  RT (min)      lcqtof_hilic_neg_2_047_32    m/z 700.2638     0   2   4   6E+3   4.5   5  RT (min)      lcqtof_hilic_neg_2_050_19    m/z 700.2881     0   2   4   6E+3   0.5   1  RT (min)      lcqtof_hilic_neg_4_047_06    m/z 700.2988     0   1   2E+3   0.5   1  RT (min)      lcqtof_hilic_neg_4_034_31    m/z 700.4362     0   2   4   6E+3   4   4.5  RT (min)      lcqtof_hilic_neg_4_013_20    m/z 700.4860     0   1   2E+3   15   15.5  RT (min)      lcqtof_hilic_neg_1_008_13    m/z 701.1238     0   1   2E+3   4   4.5  RT (min)      lcqtof_hilic_neg_1_032_18    m/z 701.2617     0   1   2   3   4E+3   4.5   5  RT (min)      lcqtof_hilic_neg_4_013_20    m/z 701.3774     0   1   2E+3   4   4.5  RT (min)       lcqtof_hilic_neg_4_039_18    m/z 701.3897     0   2   4E+3   1   1.5  RT (min)      lcqtof_hilic_neg_4_046_41    m/z 702.3187     0   1   2E+3   1   1.5  RT (min)      lcqtof_hilic_neg_4_003_16    m/z 702.3856     0   1   2   3   4E+3   4.5   5  RT (min)      lcqtof_hilic_neg_1_004_14    m/z 702.4563     0   1E+3   7   7.5  RT (min)      lcqtof_hilic_neg_4_054_48    m/z 702.5154     0   1   2   3   4E+3   4   4.5  RT (min)      lcqtof_hilic_neg_3_028_31    m/z 702.5459     0   2   4   6E+3   3   3.5  RT (min)      lcqtof_hilic_neg_4_023_15    m/z 703.0024     0   2   4   6   8E+3   7   7.5  RT (min)      lcqtof_hilic_neg_2_031_01    m/z 703.1360     0   1   2   3E+3   9   9.5   10  RT (min)       lcqtof_hilic_neg_4_055_32    m/z 703.2225     0   2   4E+3   0.5   1  RT (min)      lcqtof_hilic_neg_1_037_24    m/z 703.4685     0   1   2   3   4E+3   1   1.5  RT (min)      lcqtof_hilic_neg_4_006_30    m/z 703.4811     0   1   2   3   4E+3   3.5   4  RT (min)      lcqtof_hilic_neg_4_031_38    m/z 703.4944     0   1   2E+3   15.5   16  RT (min)       lcqtof_hilic_neg_1_003_37    m/z 703.8649     0   2   4   6E+3   0.5   1  RT (min)      lcqtof_hilic_neg_4_004_47    m/z 703.9581     0   2   4   6E+3   5   5.5  RT (min)      lcqtof_hilic_neg_4_050_02    m/z 704.4514     0   1   2E+3   4   4.5  RT (min)      lcqtof_hilic_neg_1_014_31    m/z 704.4924     0   2   4   6   8E+3   0.5   1  RT (min)      lcqtof_hilic_neg_4_047_06    m/z 704.5464     0   1   2   3E+3   3.5   4  RT (min)      lcqtof_hilic_neg_4_053_14    m/z 705.0389     0   1   2E+3   0.5   1  RT (min)      lcqtof_hilic_neg_1_028_47    m/z 705.0750     0   2   4E+3   6   6.5  RT (min)      lcqtof_hilic_neg_4_007_45    m/z 705.2391     0   1   2   3   4E+3   1   1.5  RT (min)      lcqtof_hilic_neg_4_013_20    m/z 705.3936     0   1   2   3E+3   4   4.5  RT (min)      lcqtof_hilic_neg_4_021_21    m/z 705.4103     0   1   2   3E+4   0.5   1  RT (min)       lcqtof_hilic_neg_4_031_38    m/z 705.5724     0   2   4   6   8E+3   0.5   1  RT (min)      lcqtof_hilic_neg_4_008_24    m/z 705.9553     0   1   2   3   4E+3   0.5   1  RT (min)      lcqtof_hilic_neg_4_054_48    m/z 706.1849     0   2   4   6E+3   0.5   1  RT (min)      lcqtof_hilic_neg_4_026_04    m/z 706.3887     0   2   4   6E+3   5   5.5  RT (min)      lcqtof_hilic_neg_4_053_14    m/z 706.4125     0   1   2   3E+3   1   1.5  RT (min)      lcqtof_hilic_neg_2_016_33    m/z 706.4357     0   1   2E+3   0.5   1  RT (min)      lcqtof_hilic_neg_4_040_05    m/z 706.4763     0   1E+3   7   7.5  RT (min)      lcqtof_hilic_neg_4_040_05    m/z 706.9594     0   1   2E+3   0.5   1  RT (min)      lcqtof_hilic_neg_4_054_48    m/z 707.1690     0   2   4   6   8E+3   7   7.5  RT (min)      lcqtof_hilic_neg_3_004_32    m/z 707.2135     0   2   4   6   8E+3   1   1.5  RT (min)      lcqtof_hilic_neg_4_006_30    m/z 707.3663     0   2   4   6E+3   1   1.5  RT (min)      lcqtof_hilic_neg_4_023_15    m/z 707.5248     0   1   2   3E+3   0.5   1  RT (min)      lcqtof_hilic_neg_2_043_15    m/z 707.5429     0   1   2   3   4E+3   0.5   1  RT (min)      lcqtof_hilic_neg_4_027_19    m/z 707.5791     0   1E+4   0.5   1  RT (min)      lcqtof_hilic_neg_4_004_47    m/z 707.9893     0   2   4   6   8E+3   0.5   1  RT (min)      lcqtof_hilic_neg_1_028_47    m/z 708.4671     0   1   2E+3   1   1.5  RT (min)      lcqtof_hilic_neg_1_003_37    m/z 708.6639     0   1E+3   7.5   8  RT (min)      lcqtof_hilic_neg_2_048_35    m/z 709.0584     0   1   2E+3   7   7.5  RT (min)      lcqtof_hilic_neg_2_055_30    m/z 709.0637     0   1   2   3E+3   8   8.5  RT (min)       lcqtof_hilic_neg_3_054_37    m/z 709.3390     0   1   2   3E+3   4.5   5  RT (min)      lcqtof_hilic_neg_2_035_09    m/z 709.3569     0   2   4   6E+3   4.5   5  RT (min)      lcqtof_hilic_neg_4_005_29    m/z 709.4327     0   1   2   3   4E+3   0.5   1  RT (min)      lcqtof_hilic_neg_1_030_17    m/z 709.4784     0   1   2   3   4E+3   1   1.5  RT (min)      lcqtof_hilic_neg_1_013_38    m/z 710.0697     0   1   2E+3   6   6.5  RT (min)      lcqtof_hilic_neg_4_054_48    m/z 710.0867     0   1   2   3E+3   8.5   9  RT (min)       lcqtof_hilic_neg_2_047_32    m/z 710.2315     0   1   2   3E+3   1   1.5  RT (min)      lcqtof_hilic_neg_4_006_30    m/z 710.6598     0   1E+3   7.5   8  RT (min)      lcqtof_hilic_neg_2_002_36    m/z 710.8572     0   1   2   3   4E+3   0.5   1  RT (min)      lcqtof_hilic_neg_4_004_47    m/z 710.9718     0   1   2   3E+3   4   4.5  RT (min)      lcqtof_hilic_neg_4_048_23    m/z 711.1686     0   1   2   3   4E+3   1   1.5  RT (min)      lcqtof_hilic_neg_4_026_04    m/z 711.3643     0   2   4   6   8E+3   0.5   1  RT (min)      lcqtof_hilic_neg_3_040_14    m/z 711.9685     0   1   2   3E+3   4   4.5  RT (min)      lcqtof_hilic_neg_4_035_33    m/z 712.0732     0   1   2   3E+3   8   8.5  RT (min)      lcqtof_hilic_neg_2_020_23    m/z 712.6120     0   1   2   3   4E+3   3   3.5  RT (min)      lcqtof_hilic_neg_1_037_24    m/z 712.6721     0   2   4   6E+3   1   1.5  RT (min)      lcqtof_hilic_neg_4_016_37    m/z 713.3489     0   1   2E+3   3.5   4  RT (min)      lcqtof_hilic_neg_1_051_35    m/z 713.3928     0   1   2E+3   6   6.5  RT (min)      lcqtof_hilic_neg_4_053_14    m/z 713.6440     0   1E+3   15.5   16  RT (min)      lcqtof_hilic_neg_4_007_45    m/z 713.8443     0   2   4   6E+3   0.5   1  RT (min)      lcqtof_hilic_neg_4_054_48    m/z 713.8961     0   2   4   6   8E+3   0.5   1  RT (min)      lcqtof_hilic_neg_4_004_47    m/z 713.9426     0   1   2   3   4E+3   5.5   6  RT (min)       lcqtof_hilic_neg_4_032_01    m/z 713.9854     0   2   4   6   8E+3   0.5   1  RT (min)      lcqtof_hilic_neg_4_004_47    m/z 714.3804     0   1   2E+3   3   3.5  RT (min)      lcqtof_hilic_neg_2_039_38    m/z 714.3990     0   1   2   3E+3   0.5   1  RT (min)      lcqtof_hilic_neg_4_048_23    m/z 714.4604     0   1E+3   15   15.5  RT (min)      lcqtof_hilic_neg_4_043_12    m/z 714.9872     0   2   4E+3   0.5   1  RT (min)      lcqtof_hilic_neg_4_004_47    m/z 715.2970     0   2   4   6   8E+3   0.5   1  RT (min)      lcqtof_hilic_neg_4_038_10    m/z 715.3799     0   1E+3   3.5   4  RT (min)      lcqtof_hilic_neg_2_042_27    m/z 715.9429     0   1E+4   1   1.5  RT (min)      lcqtof_hilic_neg_1_009_50    m/z 716.2633     0   1   2   3E+3   0.5   1  RT (min)      lcqtof_hilic_neg_4_042_22    m/z 716.3375     0   2   4   6E+3   1   1.5  RT (min)      lcqtof_hilic_neg_4_029_17    m/z 716.3477     0   1   2   3   4E+3   5.5   6  RT (min)      lcqtof_hilic_neg_4_050_02    m/z 716.8509     0   1   2   3E+3   6   6.5  RT (min)      lcqtof_hilic_neg_2_051_14    m/z 717.3524     0   1   2E+3   6   6.5  RT (min)      lcqtof_hilic_neg_2_051_14    m/z 717.4280     0   2   4E+3   1   1.5  RT (min)      lcqtof_hilic_neg_4_027_19    m/z 717.4323     0   2   4   6E+3   0.5   1  RT (min)      lcqtof_hilic_neg_4_010_28    m/z 717.4869     0   1   2   3   4E+3   1   1.5  RT (min)      lcqtof_hilic_neg_4_006_30    m/z 717.8795     0   2   4   6E+3   0.5   1  RT (min)      lcqtof_hilic_neg_4_004_47    m/z 718.2252     0   1   2E+3   6   6.5  RT (min)      lcqtof_hilic_neg_4_052_13    m/z 718.3583     0   1E+4   0.5   1  RT (min)      lcqtof_hilic_neg_4_053_14    m/z 718.5738     0   2   4E+3   1   1.5  RT (min)      lcqtof_hilic_neg_1_005_32    m/z 718.6416     0   2   4   6   8E+3   0.5   1  RT (min)      lcqtof_hilic_neg_1_037_24    m/z 718.7447     0   1   2E+3   6.5   7  RT (min)       lcqtof_hilic_neg_4_087_59    m/z 719.1364     0   1   2   3E+3   6.5   7  RT (min)      lcqtof_hilic_neg_2_028_05    m/z 719.4838     0   1E+3   15   15.5  RT (min)       lcqtof_hilic_neg_4_042_22    m/z 719.5289     0   1   2   3E+5   4   4.5  RT (min)      lcqtof_hilic_neg_2_018_16    m/z 719.9504     0   1E+3   5.5   6  RT (min)      lcqtof_hilic_neg_4_012_07    m/z 720.2055     0   1E+3   5.5   6  RT (min)       lcqtof_hilic_neg_4_019_03    m/z 720.6631     0   2   4   6E+3   0.5   1  RT (min)      lcqtof_hilic_neg_4_008_24    m/z 721.0064     0   2   4E+3   0.5   1  RT (min)      lcqtof_hilic_neg_1_028_47    m/z 721.0417     0   1   2E+3   5   5.5  RT (min)       lcqtof_hilic_neg_4_004_47    m/z 721.4678     0   1   2   3   4E+3   0.5   1  RT (min)      lcqtof_hilic_neg_4_003_16    m/z 722.1920     0   2   4   6   8E+3   0.5   1  RT (min)      lcqtof_hilic_neg_1_028_47    m/z 722.2198     0   1   2   3   4E+3   6   6.5  RT (min)      lcqtof_hilic_neg_4_012_07    m/z 722.5728     0   2   4   6E+3   0.5   1  RT (min)      lcqtof_hilic_neg_4_050_02    m/z 722.6867     0   1   2   3   4E+3   4.5   5  RT (min)      lcqtof_hilic_neg_4_053_14    m/z 723.0460     0   1   2   3   4E+3   6.5   7  RT (min)      lcqtof_hilic_neg_2_043_15    m/z 724.0662     0   1   2E+3   6.5   7  RT (min)      lcqtof_hilic_neg_2_029_31    m/z 725.0573     0   1   2   3E+3   7.5   8  RT (min)      lcqtof_hilic_neg_2_047_32    m/z 725.5537     0   2   4   6E+3   3.5   4  RT (min)       lcqtof_hilic_neg_4_094_60    m/z 726.1606     0   2   4   6   8E+3   1   1.5  RT (min)      lcqtof_hilic_neg_4_026_04    m/z 726.3790     0   2   4   6E+3   5.5   6  RT (min)      lcqtof_hilic_neg_4_003_16    m/z 726.6202     0   1   2   3E+3   3   3.5  RT (min)      lcqtof_hilic_neg_1_037_24    m/z 727.0342     0   2   4E+3   7.5   8  RT (min)      lcqtof_hilic_neg_4_016_37    m/z 727.1665     0   2   4   6E+3   1   1.5  RT (min)      lcqtof_hilic_neg_4_026_04    m/z 727.2214     0   1   2   3   4E+3   1   1.5  RT (min)       lcqtof_hilic_neg_4_013_20    m/z 727.3582     0   1   2   3E+3   3.5   4  RT (min)      lcqtof_hilic_neg_3_083_59    m/z 727.6265     0   1   2E+3   3   3.5  RT (min)      lcqtof_hilic_neg_1_005_32    m/z 727.9517     0   1   2   3E+3   6   6.5  RT (min)      lcqtof_hilic_neg_1_005_32    m/z 727.9881     0   1   2   3   4E+3   7.5   8  RT (min)      lcqtof_hilic_neg_4_022_34    m/z 728.1711     0   1E+3   5.5   6  RT (min)      lcqtof_hilic_neg_4_023_15    m/z 728.9626     0   1   2   3   4E+3   5   5.5  RT (min)      lcqtof_hilic_neg_4_053_14    m/z 729.3490     0   2   4   6E+3   1   1.5  RT (min)      lcqtof_hilic_neg_4_003_16    m/z 729.3680     0   2   4   6   8E+3   1   1.5  RT (min)      lcqtof_hilic_neg_4_053_14    m/z 729.3860     0   1   2E+3   5   5.5  RT (min)      lcqtof_hilic_neg_4_053_14    m/z 729.5562     0   1   2   3E+3   0.5   1  RT (min)      lcqtof_hilic_neg_4_029_17    m/z 730.1349     0   1   2   3E+3   8   8.5  RT (min)      lcqtof_hilic_neg_2_052_24    m/z 730.2211     0   2   4   6E+3   3.5   4  RT (min)      lcqtof_hilic_neg_1_053_19    m/z 730.2778     0   2   4   6E+3   0.5   1  RT (min)      lcqtof_hilic_neg_4_026_04    m/z 730.3933     0   1   2   3E+3   0.5   1  RT (min)      lcqtof_hilic_neg_4_048_23    m/z 730.5535     0   1E+4   0.5   1  RT (min)      lcqtof_hilic_neg_4_004_47    m/z 730.5901     0   2   4E+3   3.5   4  RT (min)      lcqtof_hilic_neg_4_006_30    m/z 730.9847     0   1   2E+3   5   5.5   6  RT (min)      lcqtof_hilic_neg_4_026_04    m/z 731.0294     0   1   2   3E+3   0.5   1  RT (min)      lcqtof_hilic_neg_2_051_14    m/z 731.0389     0   1   2   3E+3   5   5.5  RT (min)      lcqtof_hilic_neg_4_017_51    m/z 731.0433     0   2   4   6E+3   5   5.5  RT (min)      lcqtof_hilic_neg_4_011_35    m/z 731.0917     0   2   4E+3   7.5   8  RT (min)      lcqtof_hilic_neg_4_034_31    m/z 731.2081     0   1   2   3   4E+3   4.5   5  RT (min)      lcqtof_hilic_neg_4_045_26    m/z 731.2881     0   1   2   3   4E+3   1   1.5  RT (min)      lcqtof_hilic_neg_4_053_14    m/z 731.3094     0   1   2   3E+3   0.5   1  RT (min)      lcqtof_hilic_neg_4_050_02    m/z 731.4402     0   1   2E+3   1   1.5  RT (min)      lcqtof_hilic_neg_4_013_20    m/z 731.4979     0   2   4   6E+3   1   1.5  RT (min)      lcqtof_hilic_neg_1_007_05    m/z 731.5060     0   1   2   3   4E+3   5   5.5  RT (min)      lcqtof_hilic_neg_4_048_23    m/z 731.5281     0   2   4   6E+3   3.5   4  RT (min)      lcqtof_hilic_neg_4_032_01    m/z 732.1343     0   1   2   3E+3   8   8.5  RT (min)      lcqtof_hilic_neg_2_052_24    m/z 732.2172     0   2   4   6E+3   3.5   4  RT (min)       lcqtof_hilic_neg_1_053_19    m/z 732.3856     0   2   4E+3   1   1.5  RT (min)      lcqtof_hilic_neg_1_054_01    m/z 732.4625     0   2   4   6E+3   0.5   1  RT (min)      lcqtof_hilic_neg_4_032_01    m/z 732.4822     0   2   4E+3   3.5   4  RT (min)      lcqtof_hilic_neg_4_032_01    m/z 732.5843     0   1   2   3E+3   4   4.5  RT (min)      lcqtof_hilic_neg_4_004_47    m/z 733.0276     0   1   2E+3   0.5   1  RT (min)      lcqtof_hilic_neg_2_051_14    m/z 733.1375     0   1   2   3E+3   8   8.5  RT (min)      lcqtof_hilic_neg_1_037_24    m/z 733.2919     0   2   4E+3   1   1.5  RT (min)      lcqtof_hilic_neg_4_053_14    m/z 733.3589     0   1   2E+3   4   4.5  RT (min)       lcqtof_hilic_neg_4_105_61    m/z 733.3970     0   2   4E+3   0.5   1  RT (min)      lcqtof_hilic_neg_4_006_30    m/z 733.4372     0   1E+4   1   1.5  RT (min)      lcqtof_hilic_neg_4_040_05    m/z 733.5782     0   1   2E+3   0.5   1  RT (min)      lcqtof_hilic_neg_4_004_47    m/z 733.5906     0   1   2   3   4E+3   4   4.5  RT (min)      lcqtof_hilic_neg_4_002_25    m/z 734.1719     0   1   2   3E+3   8   8.5  RT (min)       lcqtof_hilic_neg_4_008_24    m/z 734.9903     0   1E+3   5.5   6  RT (min)      lcqtof_hilic_neg_2_026_28    m/z 735.4777     0   1E+3   7   7.5  RT (min)      lcqtof_hilic_neg_4_040_05    m/z 735.5580     0   2   4   6E+3   0.5   1  RT (min)      lcqtof_hilic_neg_4_005_29    m/z 735.5806     0   2   4   6   8E+3   15   15.5  RT (min)       lcqtof_hilic_neg_4_050_02    m/z 735.5966     0   1   2   3E+3   0.5   1  RT (min)      lcqtof_hilic_neg_4_004_47    m/z 737.2614     0   2   4   6E+3   1   1.5  RT (min)      lcqtof_hilic_neg_1_042_26    m/z 737.3530     0   1E+4   4.5   5  RT (min)      lcqtof_hilic_neg_4_011_35    m/z 737.5346     0   2   4   6E+3   1   1.5  RT (min)      lcqtof_hilic_neg_4_053_14    m/z 738.0562     0   1   2E+3   6   6.5  RT (min)      lcqtof_hilic_neg_1_027_27    m/z 738.9800     0   1   2   3E+3   0.5   1  RT (min)      lcqtof_hilic_neg_1_028_47    m/z 739.2720     0   1   2   3   4E+3   5.5   6  RT (min)      lcqtof_hilic_neg_4_035_33    m/z 739.7219     0   2   4   6E+3   0.5   1  RT (min)      lcqtof_hilic_neg_1_005_32    m/z 739.7883     0   2   4   6E+3   0.5   1  RT (min)      lcqtof_hilic_neg_4_001_49    m/z 739.7924     0   2   4   6E+3   1   1.5  RT (min)      lcqtof_hilic_neg_4_001_49    m/z 740.1970     0   1   2E+3   6   6.5  RT (min)      lcqtof_hilic_neg_2_051_14    m/z 740.2144     0   2   4E+3   5.5   6  RT (min)      lcqtof_hilic_neg_4_013_20    m/z 740.3785     0   2   4E+3   5   5.5  RT (min)      lcqtof_hilic_neg_4_053_14    m/z 740.4332     0   2   4E+3   4   4.5  RT (min)      lcqtof_hilic_neg_4_048_23    m/z 740.5288     0   2   4   6   8E+3   5   5.5  RT (min)       lcqtof_hilic_neg_4_048_23    m/z 740.7608     0   2   4   6E+3   0.5   1  RT (min)      lcqtof_hilic_neg_4_008_24    m/z 741.1082     0   1   2   3   4E+3   4   4.5  RT (min)       lcqtof_hilic_neg_1_032_18    m/z 741.3507     0   2   4   6E+3   0.5   1  RT (min)      lcqtof_hilic_neg_4_023_15    m/z 741.3558     0   2   4   6E+3   1   1.5  RT (min)      lcqtof_hilic_neg_4_032_01    m/z 741.3864     0   2   4   6E+3   4.5   5  RT (min)      lcqtof_hilic_neg_3_011_34    m/z 741.7666     0   1   2E+3   4.5   5  RT (min)      lcqtof_hilic_neg_4_032_01    m/z 742.3821     0   1   2   3   4E+3   5.5   6  RT (min)      lcqtof_hilic_neg_4_053_14    m/z 742.4564     0   1   2   3   4E+3   0.5   1  RT (min)      lcqtof_hilic_neg_4_011_35    m/z 742.6067     0   2   4   6   8E+3   0.5   1  RT (min)      lcqtof_hilic_neg_4_002_25    m/z 743.0269     0   2   4   6   8E+3   3   3.5  RT (min)      lcqtof_hilic_neg_4_048_23    m/z 743.1087     0   1   2   3E+3   4   4.5  RT (min)      lcqtof_hilic_neg_1_032_18    m/z 743.3440     0   2   4   6E+3   0.5   1  RT (min)      lcqtof_hilic_neg_4_045_26    m/z 743.3847     0   1   2   3E+3   4   4.5  RT (min)       lcqtof_hilic_neg_4_022_34    m/z 743.6558     0   1   2E+3   15.5   16  RT (min)       lcqtof_hilic_neg_4_101_61    m/z 744.0555     0   1   2   3   4E+3   0.5   1  RT (min)      lcqtof_hilic_neg_4_053_14    m/z 744.1741     0   1   2E+3   6.5   7  RT (min)      lcqtof_hilic_neg_4_054_48    m/z 744.2301     0   2   4   6E+3   0.5   1  RT (min)      lcqtof_hilic_neg_4_010_28    m/z 744.2796     0   2   4   6E+3   6   6.5  RT (min)      lcqtof_hilic_neg_4_052_13    m/z 744.4356     0   2   4E+3   4   4.5  RT (min)      lcqtof_hilic_neg_1_054_01    m/z 744.4644     0   1   2E+3   15   15.5  RT (min)      lcqtof_hilic_neg_1_007_05    m/z 744.5966     0   1E+3   7.5   8  RT (min)      lcqtof_hilic_neg_1_029_06    m/z 744.7544     0   1   2   3   4E+3   0.5   1  RT (min)      lcqtof_hilic_neg_2_047_32    m/z 745.0274     0   2   4E+3   6.5   7  RT (min)      lcqtof_hilic_neg_2_043_15    m/z 745.0398     0   1   2E+3   5.5   6  RT (min)      lcqtof_hilic_neg_4_053_14    m/z 745.3643     0   1   2   3   4E+3   0.5   1  RT (min)      lcqtof_hilic_neg_4_013_20    m/z 745.6868     0   1   2   3E+3   3.5   4  RT (min)      lcqtof_hilic_neg_4_005_29    m/z 746.3678     0   1   2   3E+3   4.5   5  RT (min)       lcqtof_hilic_neg_4_031_38    m/z 746.5601     0   2   4E+3   2   2.5  RT (min)      lcqtof_hilic_neg_4_048_23    m/z 746.6582     0   2   4E+3   0.5   1  RT (min)      lcqtof_hilic_neg_2_052_24    m/z 746.9185     0   1E+3   5   5.5  RT (min)      lcqtof_hilic_neg_2_055_30    m/z 747.0261     0   1   2   3   4E+3   6.5   7  RT (min)      lcqtof_hilic_neg_2_043_15    m/z 747.0449     0   1E+3   6   6.5  RT (min)      lcqtof_hilic_neg_2_013_29    m/z 747.0933     0   1   2E+3   7   7.5  RT (min)      lcqtof_hilic_neg_2_044_37    m/z 747.0953     0   1E+3   6   6.5  RT (min)      lcqtof_hilic_neg_3_030_23    m/z 747.3038     0   2   4   6   8E+3   1   1.5  RT (min)      lcqtof_hilic_neg_4_053_14    m/z 747.3684     0   2   4   6E+3   5.5   6  RT (min)       lcqtof_hilic_neg_4_053_14    m/z 747.4132     0   1   2   3E+3   4   4.5  RT (min)      lcqtof_hilic_neg_4_013_20    m/z 747.4400     0   1   2   3E+3   3.5   4  RT (min)      lcqtof_hilic_neg_4_011_35    m/z 747.6074     0   2   4E+3   15   15.5  RT (min)      lcqtof_hilic_neg_4_050_02    m/z 748.4924     0   1E+4   3   3.5  RT (min)       lcqtof_hilic_neg_4_048_23    m/z 748.6728     0   2   4   6E+3   0.5   1  RT (min)      lcqtof_hilic_neg_2_052_24    m/z 748.7031     0   1   2E+3   3.5   4  RT (min)      lcqtof_hilic_neg_4_005_29    m/z 749.0618     0   2   4E+3   3.5   4  RT (min)      lcqtof_hilic_neg_4_032_01    m/z 749.2538     0   1   2   3   4E+3   1   1.5  RT (min)      lcqtof_hilic_neg_4_026_04    m/z 749.5543     0   2   4   6   8E+3   0.5   1  RT (min)      lcqtof_hilic_neg_4_023_15    m/z 749.6024     0   2   4   6   8E+3   4   4.5  RT (min)      lcqtof_hilic_neg_4_055_32    m/z 750.3793     0   1   2   3E+3   3   3.5  RT (min)      lcqtof_hilic_neg_2_044_37    m/z 750.8413     0   2   4   6   8E+3   1   1.5  RT (min)      lcqtof_hilic_neg_4_004_47    m/z 750.9980     0   1   2   3   4E+3   0.5   1  RT (min)      lcqtof_hilic_neg_2_034_47    m/z 751.0598     0   1   2E+3   5.5   6  RT (min)      lcqtof_hilic_neg_4_050_02    m/z 751.3662     0   2   4   6E+3   4.5   5  RT (min)       lcqtof_hilic_neg_4_022_34    m/z 751.5184     0   1   2   3   4E+3   2   2.5  RT (min)      lcqtof_hilic_neg_4_048_23    m/z 751.6548     0   1E+3   8   8.5  RT (min)      lcqtof_hilic_neg_2_026_28    m/z 751.6560     0   1E+3   8   8.5  RT (min)      lcqtof_hilic_neg_3_029_29    m/z 752.4941     0   1E+3   15.5   16  RT (min)       lcqtof_hilic_neg_4_041_54    m/z 753.0314     0   2   4   6   8E+3   0.5   1  RT (min)      lcqtof_hilic_neg_1_028_47    m/z 753.4872     0   1   2   3E+3   3.5   4  RT (min)      lcqtof_hilic_neg_4_022_34    m/z 753.5286     0   1   2   3   4E+3   3.5   4  RT (min)      lcqtof_hilic_neg_4_004_47    m/z 753.5496     0   1   2E+3   15   15.5  RT (min)       lcqtof_hilic_neg_1_006_22    m/z 753.6037     0   1   2   3   4E+3   0.5   1  RT (min)      lcqtof_hilic_neg_4_010_28    m/z 753.7958     0   1   2E+3   4.5   5  RT (min)      lcqtof_hilic_neg_4_023_15    m/z 754.0361     0   1   2   3E+3   0.5   1  RT (min)      lcqtof_hilic_neg_1_028_47    m/z 754.2402     0   1   2   3E+3   6   6.5  RT (min)      lcqtof_hilic_neg_4_052_13    m/z 754.4442     0   2   4E+3   0.5   1  RT (min)      lcqtof_hilic_neg_4_032_01    m/z 754.5806     0   1   2   3E+3   4   4.5  RT (min)       lcqtof_hilic_neg_4_002_25    m/z 755.5807     0   1   2   3E+3   3   3.5  RT (min)       lcqtof_hilic_neg_4_005_29    m/z 755.7663     0   1   2E+3   0.5   1  RT (min)      lcqtof_hilic_neg_4_008_24    m/z 756.0450     0   1   2   3E+3   7   7.5  RT (min)      lcqtof_hilic_neg_1_014_31    m/z 756.4123     0   1   2E+3   3.5   4  RT (min)      lcqtof_hilic_neg_4_016_37    m/z 757.0827     0   1   2   3E+3   4   4.5  RT (min)      lcqtof_hilic_neg_1_032_18    m/z 757.2390     0   1   2   3   4E+3   3.5   4  RT (min)      lcqtof_hilic_neg_4_013_20    m/z 757.2853     0   2   4   6E+3   7   7.5  RT (min)      lcqtof_hilic_neg_1_023_34    m/z 757.4205     0   2   4E+3   0.5   1  RT (min)      lcqtof_hilic_neg_4_048_23    m/z 758.2204     0   1   2   3   4E+3   4.5   5  RT (min)      lcqtof_hilic_neg_4_013_20    m/z 758.2357     0   2   4   6   8E+3   3.5   4  RT (min)      lcqtof_hilic_neg_4_027_19    m/z 758.3085     0   2   4   6E+3   1   1.5  RT (min)      lcqtof_hilic_neg_4_003_16    m/z 758.3787     0   1   2E+3   4   4.5  RT (min)      lcqtof_hilic_neg_3_054_37    m/z 759.2182     0   2   4   6E+3   4.5   5  RT (min)      lcqtof_hilic_neg_4_013_20    m/z 759.2338     0   1   2E+3   3.5   4  RT (min)       lcqtof_hilic_neg_4_013_20    m/z 759.4142     0   2   4   6   8E+3   0.5   1  RT (min)      lcqtof_hilic_neg_4_022_34    m/z 759.8234     0   1   2   3E+3   4.5   5  RT (min)      lcqtof_hilic_neg_4_012_07    m/z 760.3237     0   2   4   6E+3   1   1.5  RT (min)      lcqtof_hilic_neg_4_023_15    m/z 760.8813     0   1   2   3E+3   5   5.5  RT (min)      lcqtof_hilic_neg_4_029_17    m/z 761.2567     0   2   4   6E+3   0.5   1  RT (min)      lcqtof_hilic_neg_1_031_16    m/z 761.3849     0   2   4   6E+3   4   4.5  RT (min)       lcqtof_hilic_neg_4_102_61    m/z 761.4182     0   2   4   6E+3   0.5   1  RT (min)      lcqtof_hilic_neg_4_045_26    m/z 761.4328     0   2   4   6   8E+3   1   1.5  RT (min)      lcqtof_hilic_neg_4_005_29    m/z 762.3242     0   1   2   3E+3   1   1.5  RT (min)      lcqtof_hilic_neg_4_053_14    m/z 762.3903     0   1   2   3   4E+3   4   4.5  RT (min)       lcqtof_hilic_neg_4_104_61    m/z 763.3442     0   2   4   6E+3   1   1.5  RT (min)      lcqtof_hilic_neg_4_053_14    m/z 763.4272     0   2   4   6E+3   7   7.5  RT (min)       lcqtof_hilic_neg_1_035_23    m/z 763.9214     0   2   4   6E+3   0.5   1  RT (min)      lcqtof_hilic_neg_4_004_47    m/z 765.7728     0   2   4   6E+3   0.5   1  RT (min)      lcqtof_hilic_neg_4_008_24    m/z 766.0001     0   1   2E+3   0.5   1  RT (min)      lcqtof_hilic_neg_4_014_46    m/z 766.8528     0   1   2E+3   5   5.5  RT (min)       lcqtof_hilic_neg_4_104_61    m/z 767.4651     0   1E+3   3.5   4  RT (min)      lcqtof_hilic_neg_1_036_03    m/z 767.5256     0   1   2E+3   15.5   16  RT (min)       lcqtof_hilic_neg_4_032_01    m/z 767.8352     0   1   2   3E+3   7   7.5  RT (min)      lcqtof_hilic_neg_1_004_14    m/z 767.9597     0   1   2   3E+3   0.5   1  RT (min)      lcqtof_hilic_neg_1_028_47    m/z 768.9653     0   1   2E+3   0.5   1  RT (min)      lcqtof_hilic_neg_1_028_47    m/z 769.0797     0   1E+3   7   7.5  RT (min)      lcqtof_hilic_neg_2_039_38    m/z 769.3956     0   2   4E+3   1   1.5  RT (min)      lcqtof_hilic_neg_4_032_01    m/z 769.4868     0   1   2   3   4E+3   0.5   1  RT (min)      lcqtof_hilic_neg_2_032_02    m/z 769.6677     0   1E+4   3   3.5  RT (min)      lcqtof_hilic_neg_1_037_24    m/z 769.9533     0   1   2   3E+3   1.5   2  RT (min)      lcqtof_hilic_neg_4_013_20    m/z 770.3640     0   1   2   3E+3   3.5   4  RT (min)      lcqtof_hilic_neg_4_032_01    m/z 770.9364     0   1E+3   6   6.5  RT (min)      lcqtof_hilic_neg_1_031_16    m/z 771.1507     0   1   2   3E+3   6   6.5  RT (min)      lcqtof_hilic_neg_4_011_35    m/z 771.3918     0   1   2   3E+3   4   4.5  RT (min)      lcqtof_hilic_neg_3_031_36    m/z 771.4823     0   1   2   3E+3   0.5   1  RT (min)      lcqtof_hilic_neg_2_032_02    m/z 771.5425     0   2   4E+3   0.5   1  RT (min)      lcqtof_hilic_neg_2_047_32    m/z 771.6135     0   1   2   3E+3   4   4.5  RT (min)      lcqtof_hilic_neg_4_008_24    m/z 771.6807     0   1E+3   15.5   16  RT (min)       lcqtof_hilic_neg_4_084_59    m/z 772.3939     0   2   4E+3   3   3.5  RT (min)      lcqtof_hilic_neg_1_035_23    m/z 772.6180     0   1   2E+3   4   4.5  RT (min)      lcqtof_hilic_neg_4_055_32    m/z 772.6619     0   1   2E+3   15.5   16  RT (min)       lcqtof_hilic_neg_4_049_55    m/z 772.9515     0   1   2   3E+3   6   6.5  RT (min)      lcqtof_hilic_neg_4_077_58    m/z 773.0815     0   2   4   6   8E+3   1   1.5  RT (min)      lcqtof_hilic_neg_4_004_47    m/z 773.1929     0   2   4E+3   4.5   5  RT (min)      lcqtof_hilic_neg_1_053_19    m/z 773.8615     0   1   2   3   4E+3   3.5   4  RT (min)      lcqtof_hilic_neg_1_054_01    m/z 773.8687     0   2   4   6E+3   0.5   1  RT (min)      lcqtof_hilic_neg_4_004_47    m/z 774.1662     0   2   4   6E+3   6   6.5  RT (min)      lcqtof_hilic_neg_4_035_33    m/z 774.3719     0   1   2   3E+3   3   3.5  RT (min)      lcqtof_hilic_neg_2_048_35    m/z 774.3730     0   2   4   6E+3   5   5.5  RT (min)      lcqtof_hilic_neg_4_032_01    m/z 774.4296     0   1   2   3   4E+3   0.5   1  RT (min)      lcqtof_hilic_neg_4_018_27    m/z 774.4871     0   1   2   3E+4   3.5   4  RT (min)      lcqtof_hilic_neg_1_022_36    m/z 774.5896     0   1   2   3E+3   0.5   1  RT (min)      lcqtof_hilic_neg_4_029_17    m/z 774.8621     0   1   2   3   4E+3   0.5   1  RT (min)      lcqtof_hilic_neg_4_076_58    m/z 775.0634     0   2   4   6E+3   8   8.5  RT (min)      lcqtof_hilic_neg_4_028_36    m/z 775.1647     0   1   2   3E+3   6   6.5  RT (min)       lcqtof_hilic_neg_4_068_57    m/z 775.3329     0   2   4   6   8E+3   1   1.5  RT (min)      lcqtof_hilic_neg_4_053_14    m/z 775.9861     0   1   2E+3   7.5   8  RT (min)      lcqtof_hilic_neg_2_020_23    m/z 776.3612     0   2   4   6   8E+3   3.5   4  RT (min)      lcqtof_hilic_neg_4_003_16    m/z 776.5045     0   1E+4   4   4.5  RT (min)      lcqtof_hilic_neg_4_011_35    m/z 776.5386     0   2   4   6E+3   4   4.5  RT (min)      lcqtof_hilic_neg_1_014_31    m/z 776.6023     0   1   2   3   4E+3   0.5   1  RT (min)      lcqtof_hilic_neg_2_021_17    m/z 776.6816     0   2   4   6E+3   4   4.5  RT (min)      lcqtof_hilic_neg_4_004_47    m/z 776.8458     0   2   4   6E+3   0.5   1  RT (min)      lcqtof_hilic_neg_4_004_47    m/z 777.4661     0   2   4   6   8E+3   0.5   1  RT (min)      lcqtof_hilic_neg_4_034_31    m/z 777.4900     0   1   2E+3   3.5   4  RT (min)      lcqtof_hilic_neg_1_030_17    m/z 778.0182     0   1   2   3   4E+3   0.5   1  RT (min)      lcqtof_hilic_neg_4_025_52    m/z 778.1110     0   1   2   3   4E+3   6.5   7  RT (min)      lcqtof_hilic_neg_4_022_34    m/z 778.3884     0   1   2E+3   4.5   5  RT (min)      lcqtof_hilic_neg_1_024_21    m/z 778.6238     0   1   2E+3   8   8.5  RT (min)      lcqtof_hilic_neg_2_040_20    m/z 779.3312     0   1   2   3E+3   1.5   2  RT (min)       lcqtof_hilic_neg_4_021_21    m/z 779.3850     0   1   2   3   4E+3   4.5   5  RT (min)      lcqtof_hilic_neg_2_053_18    m/z 779.4084     0   2   4   6E+3   3.5   4  RT (min)      lcqtof_hilic_neg_4_016_37    m/z 780.2688     0   1   2   3E+3   7   7.5  RT (min)      lcqtof_hilic_neg_2_044_37    m/z 780.9370     0   2   4   6E+3   0.5   1  RT (min)      lcqtof_hilic_neg_1_028_47    m/z 781.4247     0   2   4E+3   3   3.5  RT (min)      lcqtof_hilic_neg_3_028_31    m/z 781.4670     0   1   2   3   4E+3   0.5   1  RT (min)      lcqtof_hilic_neg_4_011_35    m/z 781.4795     0   2   4   6E+3   3.5   4  RT (min)      lcqtof_hilic_neg_1_011_46    m/z 782.4987     0   2   4   6E+3   3   3.5  RT (min)      lcqtof_hilic_neg_3_055_01    m/z 782.7227     0   1E+3   15.5   16  RT (min)       lcqtof_hilic_neg_4_100_61    m/z 783.0186     0   1   2   3E+3   0.5   1  RT (min)      lcqtof_hilic_neg_1_049_55    m/z 783.1087     0   1   2   3   4E+3   6.5   7  RT (min)       lcqtof_hilic_neg_4_048_23    m/z 784.4419     0   1   2   3E+3   3.5   4  RT (min)      lcqtof_hilic_neg_1_005_32    m/z 784.8277     0   2   4E+3   0.5   1  RT (min)      lcqtof_hilic_neg_4_055_32    m/z 785.5988     0   1E+3   15.5   16  RT (min)      lcqtof_hilic_neg_2_027_07    m/z 786.0132     0   1   2   3   4E+3   7.5   8  RT (min)       lcqtof_hilic_neg_4_012_07    m/z 786.5500     0   1   2   3E+3   0.5   1  RT (min)      lcqtof_hilic_neg_4_023_15    m/z 786.7043     0   1   2   3   4E+3   3   3.5  RT (min)       lcqtof_hilic_neg_4_008_24    m/z 786.9653     0   2   4E+3   0.5   1  RT (min)      lcqtof_hilic_neg_4_004_47    m/z 787.2733     0   1   2   3E+3   3.5   4  RT (min)      lcqtof_hilic_neg_4_013_20    m/z 787.3299     0   2   4   6   8E+3   6   6.5  RT (min)       lcqtof_hilic_neg_4_069_57    m/z 787.4944     0   2   4   6E+3   1   1.5  RT (min)      lcqtof_hilic_neg_4_005_29    m/z 787.5114     0   2   4E+3   3   3.5  RT (min)      lcqtof_hilic_neg_2_047_32    m/z 787.6793     0   1   2   3   4E+3   0.5   1  RT (min)      lcqtof_hilic_neg_1_037_24    m/z 787.8222     0   2   4   6E+3   0.5   1  RT (min)      lcqtof_hilic_neg_4_075_58    m/z 787.9466     0   1E+3   5.5   6  RT (min)      lcqtof_hilic_neg_4_098_61    m/z 788.2573     0   1   2   3E+3   5   5.5  RT (min)      lcqtof_hilic_neg_2_042_27    m/z 788.4643     0   2   4   6E+3   3   3.5   4  RT (min)      lcqtof_hilic_neg_1_035_23    m/z 788.5817     0   1   2   3   4E+4   3.5   4  RT (min)      lcqtof_hilic_neg_1_030_17    m/z 789.1752     0   1   2   3E+3   3.5   4  RT (min)      lcqtof_hilic_neg_1_024_21    m/z 789.3610     0   2   4E+3   1   1.5  RT (min)      lcqtof_hilic_neg_4_003_16    m/z 789.3780     0   1   2   3E+3   4   4.5  RT (min)      lcqtof_hilic_neg_4_004_47    m/z 789.4606     0   2   4   6E+3   1   1.5  RT (min)      lcqtof_hilic_neg_1_013_38    m/z 790.1430     0   1   2E+3   5.5   6  RT (min)      lcqtof_hilic_neg_4_053_14    m/z 790.8422     0   2   4   6   8E+3   0.5   1  RT (min)       lcqtof_hilic_neg_4_004_47    m/z 791.1305     0   1   2   3E+3   6   6.5  RT (min)       lcqtof_hilic_neg_4_029_17    m/z 791.1656     0   1   2   3E+3   4.5   5  RT (min)      lcqtof_hilic_neg_4_045_26    m/z 791.6260     0   2   4   6   8E+3   1   1.5  RT (min)      lcqtof_hilic_neg_4_026_04    m/z 791.8092     0   2   4   6E+3   0.5   1  RT (min)      lcqtof_hilic_neg_4_008_24    m/z 791.8506     0   2   4E+3   0.5   1  RT (min)      lcqtof_hilic_neg_4_028_36    m/z 792.1145     0   2   4E+3   6   6.5  RT (min)      lcqtof_hilic_neg_4_011_35    m/z 792.4673     0   1   2E+3   3.5   4  RT (min)      lcqtof_hilic_neg_2_021_17    m/z 793.1203     0   1   2E+3   6.5   7  RT (min)      lcqtof_hilic_neg_1_055_25    m/z 793.4103     0   2   4   6E+3   6   6.5  RT (min)      lcqtof_hilic_neg_4_016_37    m/z 793.5115     0   2   4   6E+3   0.5   1  RT (min)      lcqtof_hilic_neg_1_004_14    m/z 793.5964     0   1   2E+3   4   4.5  RT (min)      lcqtof_hilic_neg_1_037_24    m/z 794.0482     0   1   2   3   4E+3   0.5   1  RT (min)      lcqtof_hilic_neg_4_014_46    m/z 794.0833     0   2   4   6   8E+3   0.5   1  RT (min)      lcqtof_hilic_neg_4_004_47    m/z 794.6158     0   1   2E+3   4.5   5  RT (min)      lcqtof_hilic_neg_4_014_46    m/z 795.2406     0   1   2   3E+3   6   6.5  RT (min)      lcqtof_hilic_neg_4_003_16    m/z 795.5543     0   1E+3   15.5   16  RT (min)       lcqtof_hilic_neg_4_102_61    m/z 795.6165     0   1E+3   8   8.5  RT (min)      lcqtof_hilic_neg_2_043_15    m/z 795.6784     0   1   2E+3   15.5   16  RT (min)      lcqtof_hilic_neg_4_019_03    m/z 795.8746     0   1   2E+3   5   5.5  RT (min)      lcqtof_hilic_neg_4_001_49    m/z 795.9579     0   1E+3   6   6.5  RT (min)      lcqtof_hilic_neg_2_042_27    m/z 796.0883     0   2   4E+3   6.5   7  RT (min)      lcqtof_hilic_neg_4_028_36    m/z 796.3064     0   1   2   3   4E+3   3.5   4  RT (min)      lcqtof_hilic_neg_4_013_20    m/z 796.4915     0   1   2E+3   1   1.5  RT (min)      lcqtof_hilic_neg_4_026_04    m/z 796.9481     0   1   2E+3   5   5.5  RT (min)      lcqtof_hilic_neg_4_023_15    m/z 797.3191     0   1   2   3   4E+3   4.5   5  RT (min)      lcqtof_hilic_neg_2_002_36    m/z 797.4254     0   1E+4   0.5   1  RT (min)      lcqtof_hilic_neg_1_027_27    m/z 797.7344     0   1E+3   15.5   16  RT (min)       lcqtof_hilic_neg_4_093_60    m/z 797.9680     0   1   2E+3   7.5   8  RT (min)      lcqtof_hilic_neg_2_020_23    m/z 798.0654     0   2   4E+3   7.5   8  RT (min)       lcqtof_hilic_neg_4_031_38    m/z 798.9469     0   2   4E+3   0.5   1  RT (min)      lcqtof_hilic_neg_1_028_47    m/z 799.7011     0   2   4   6E+3   3   3.5  RT (min)       lcqtof_hilic_neg_1_005_32    m/z 799.7620     0   2   4E+3   6.5   7  RT (min)       lcqtof_hilic_neg_4_012_07    m/z 800.4407     0   2   4   6   8E+3   3.5   4  RT (min)      lcqtof_hilic_neg_4_013_20    m/z 800.5915     0   1   2   3   4E+3   3.5   4  RT (min)      lcqtof_hilic_neg_4_005_29    m/z 800.6011     0   1E+3   8   8.5  RT (min)      lcqtof_hilic_neg_2_008_44    m/z 800.8438     0   1E+3   5   5.5  RT (min)       lcqtof_hilic_neg_4_030_44    m/z 801.3904     0   1   2E+3   5   5.5  RT (min)      lcqtof_hilic_neg_4_031_38    m/z 801.4431     0   1   2   3E+3   0.5   1  RT (min)      lcqtof_hilic_neg_4_048_23    m/z 801.4692     0   2   4   6E+3   4.5   5  RT (min)      lcqtof_hilic_neg_4_055_32    m/z 801.6978     0   1E+3   15.5   16  RT (min)       lcqtof_hilic_neg_4_105_61    m/z 801.7995     0   2   4   6E+3   1   1.5  RT (min)      lcqtof_hilic_neg_4_055_32    m/z 802.1807     0   1   2   3   4E+3   5.5   6  RT (min)      lcqtof_hilic_neg_4_026_04    m/z 802.4564     0   1   2E+3   0.5   1  RT (min)      lcqtof_hilic_neg_4_010_28    m/z 802.4648     0   1   2   3   4E+3   3   3.5  RT (min)      lcqtof_hilic_neg_1_023_34    m/z 802.9579     0   1E+3   5   5.5  RT (min)      lcqtof_hilic_neg_4_006_30    m/z 803.1558     0   2   4   6E+3   6.5   7  RT (min)      lcqtof_hilic_neg_1_037_24    m/z 803.4407     0   2   4E+3   4.5   5  RT (min)      lcqtof_hilic_neg_4_039_18    m/z 803.7068     0   1   2E+3   3   3.5  RT (min)      lcqtof_hilic_neg_4_005_29    m/z 803.7761     0   1   2   3E+3   4.5   5  RT (min)      lcqtof_hilic_neg_4_053_14    m/z 803.9886     0   1E+3   3   3.5  RT (min)      lcqtof_hilic_neg_4_003_16    m/z 804.0699     0   2   4E+3   0.5   1  RT (min)      lcqtof_hilic_neg_4_003_16    m/z 804.2352     0   2   4   6   8E+3   0.5   1  RT (min)      lcqtof_hilic_neg_4_003_16    m/z 804.3488     0   1   2   3   4E+3   0.5   1  RT (min)      lcqtof_hilic_neg_4_005_29    m/z 804.3501     0   1   2   3E+3   7   7.5  RT (min)       lcqtof_hilic_neg_4_048_23    m/z 804.4507     0   1   2   3   4E+3   0.5   1  RT (min)      lcqtof_hilic_neg_4_003_16    m/z 804.5804     0   1E+3   15   15.5  RT (min)       lcqtof_hilic_neg_2_047_32    m/z 804.9175     0   2   4E+3   0.5   1  RT (min)      lcqtof_hilic_neg_4_004_47    m/z 805.1473     0   1   2   3   4E+3   3.5   4  RT (min)      lcqtof_hilic_neg_1_006_22    m/z 805.8686     0   1   2E+3   3   3.5  RT (min)      lcqtof_hilic_neg_4_005_29    m/z 806.4899     0   2   4   6E+3   3   3.5  RT (min)      lcqtof_hilic_neg_1_035_23    m/z 806.6451     0   2   4E+3   4.5   5  RT (min)      lcqtof_hilic_neg_4_053_14    m/z 806.7854     0   2   4   6E+3   1   1.5  RT (min)      lcqtof_hilic_neg_2_052_24    m/z 807.3944     0   1E+4   4.5   5  RT (min)      lcqtof_hilic_neg_4_006_30    m/z 807.5250     0   1   2E+3   3.5   4  RT (min)      lcqtof_hilic_neg_1_019_02    m/z 808.2375     0   1   2E+3   5   5.5  RT (min)      lcqtof_hilic_neg_2_042_27    m/z 808.3157     0   1   2   3E+3   3.5   4  RT (min)      lcqtof_hilic_neg_3_044_09    m/z 808.3208     0   2   4   6   8E+3   0.5   1  RT (min)      lcqtof_hilic_neg_4_013_20    m/z 808.3728     0   2   4E+3   5   5.5  RT (min)      lcqtof_hilic_neg_4_053_14    m/z 808.3845     0   1E+3   5   5.5  RT (min)      lcqtof_hilic_neg_1_031_16    m/z 808.4495     0   1   2   3   4E+3   3   3.5  RT (min)      lcqtof_hilic_neg_3_028_31    m/z 808.5116     0   1   2   3   4E+4   3.5   4  RT (min)      lcqtof_hilic_neg_1_014_31    m/z 809.6580     0   1   2E+4   3.5   4  RT (min)      lcqtof_hilic_neg_4_035_33    m/z 809.8730     0   2   4   6E+3   0.5   1  RT (min)      lcqtof_hilic_neg_4_004_47    m/z 810.2016     0   1   2   3E+3   7   7.5  RT (min)      lcqtof_hilic_neg_1_022_36    m/z 810.4546     0   1   2   3   4E+3   3   3.5  RT (min)      lcqtof_hilic_neg_3_028_31    m/z 810.5922     0   1   2   3   4E+3   1   1.5  RT (min)      lcqtof_hilic_neg_4_013_20    m/z 810.8992     0   1   2   3   4E+3   1   1.5  RT (min)      lcqtof_hilic_neg_4_004_47    m/z 810.9298     0   1   2E+3   6   6.5  RT (min)      lcqtof_hilic_neg_1_012_04    m/z 810.9628     0   2   4   6E+3   0.5   1  RT (min)      lcqtof_hilic_neg_4_081_58    m/z 811.0333     0   1   2E+3   0.5   1  RT (min)      lcqtof_hilic_neg_4_014_46    m/z 811.2802     0   1   2E+3   1   1.5  RT (min)      lcqtof_hilic_neg_1_006_22    m/z 811.3347     0   1   2   3E+3   1.5   2  RT (min)       lcqtof_hilic_neg_4_021_21    m/z 812.2693     0   1   2   3   4E+3   1   1.5  RT (min)      lcqtof_hilic_neg_2_050_19    m/z 812.3974     0   2   4   6E+3   0.5   1  RT (min)      lcqtof_hilic_neg_4_015_09    m/z 812.4655     0   2   4   6E+3   4   4.5  RT (min)       lcqtof_hilic_neg_4_048_23    m/z 812.5963     0   1   2   3E+3   1   1.5  RT (min)      lcqtof_hilic_neg_4_013_20    m/z 812.6732     0   1   2E+3   3.5   4  RT (min)      lcqtof_hilic_neg_4_005_29    m/z 813.0148     0   1   2   3   4E+3   6.5   7  RT (min)      lcqtof_hilic_neg_2_043_15    m/z 813.4668     0   2   4   6E+3   3.5   4  RT (min)      lcqtof_hilic_neg_1_012_04    m/z 813.4850     0   2   4   6E+3   0.5   1  RT (min)      lcqtof_hilic_neg_4_010_28    m/z 813.6918     0   1   2E+4   0.5   1  RT (min)      lcqtof_hilic_neg_4_004_47    m/z 813.7233     0   1E+3   15.5   16  RT (min)       lcqtof_hilic_neg_4_107_62    m/z 813.9362     0   1   2   3E+3   5   5.5  RT (min)      lcqtof_hilic_neg_4_045_26    m/z 813.9994     0   1   2   3E+3   0.5   1  RT (min)      lcqtof_hilic_neg_1_028_47    m/z 814.1826     0   1   2E+3   0.5   1  RT (min)      lcqtof_hilic_neg_4_004_47    m/z 814.2675     0   1   2E+3   2.5   3  RT (min)      lcqtof_hilic_neg_2_042_27    m/z 814.4908     0   2   4   6   8E+3   0.5   1  RT (min)      lcqtof_hilic_neg_4_038_10    m/z 814.9054     0   1   2E+3   5   5.5  RT (min)      lcqtof_hilic_neg_4_053_14    m/z 815.0122     0   1   2   3E+3   6.5   7  RT (min)      lcqtof_hilic_neg_2_043_15    m/z 815.1771     0   2   4   6E+3   4.5   5  RT (min)      lcqtof_hilic_neg_4_013_20    m/z 815.4987     0   2   4E+3   3.5   4  RT (min)      lcqtof_hilic_neg_4_013_20    m/z 815.5981     0   1   2   3   4E+3   0.5   1  RT (min)      lcqtof_hilic_neg_1_046_30    m/z 815.6078     0   2   4   6   8E+3   2   2.5  RT (min)       lcqtof_hilic_neg_4_048_23    m/z 815.6718     0   2   4   6   8E+3   3   3.5  RT (min)      lcqtof_hilic_neg_1_037_24    m/z 815.8156     0   2   4   6E+3   1   1.5  RT (min)      lcqtof_hilic_neg_4_008_24    m/z 816.7215     0   1   2   3   4E+3   7   7.5  RT (min)       lcqtof_hilic_neg_4_044_08    m/z 817.1739     0   2   4   6E+3   4.5   5  RT (min)      lcqtof_hilic_neg_4_013_20    m/z 817.5660     0   1   2   3E+4   3.5   4  RT (min)      lcqtof_hilic_neg_4_016_37    m/z 817.7161     0   1   2   3   4E+3   0.5   1  RT (min)      lcqtof_hilic_neg_4_004_47    m/z 818.1851     0   2   4E+3   6   6.5  RT (min)       lcqtof_hilic_neg_4_052_13    m/z 818.2692     0   2   4   6E+3   6   6.5  RT (min)      lcqtof_hilic_neg_4_072_57    m/z 818.6273     0   1   2E+3   1.5   2  RT (min)      lcqtof_hilic_neg_4_005_29    m/z 818.6339     0   1   2   3   4E+3   1   1.5  RT (min)      lcqtof_hilic_neg_4_013_20    m/z 818.6907     0   2   4   6   8E+3   0.5   1  RT (min)      lcqtof_hilic_neg_2_016_33    m/z 818.7468     0   1   2   3E+3   0.5   1  RT (min)      lcqtof_hilic_neg_4_041_54    m/z 819.0513     0   1   2E+3   5.5   6  RT (min)      lcqtof_hilic_neg_4_019_03    m/z 819.1500     0   1   2E+3   7.5   8  RT (min)      lcqtof_hilic_neg_2_047_32    m/z 819.7090     0   1   2   3   4E+3   0.5   1  RT (min)      lcqtof_hilic_neg_4_008_24    m/z 819.7243     0   1   2   3E+3   0.5   1  RT (min)      lcqtof_hilic_neg_4_014_46    m/z 819.9555     0   1   2   3E+3   5.5   6  RT (min)      lcqtof_hilic_neg_4_026_04    m/z 820.7529     0   1   2   3   4E+3   0.5   1  RT (min)      lcqtof_hilic_neg_4_004_47    m/z 821.3359     0   2   4E+3   5.5   6  RT (min)       lcqtof_hilic_neg_4_052_13    m/z 821.7234     0   2   4   6E+3   0.5   1  RT (min)      lcqtof_hilic_neg_4_008_24    m/z 822.0607     0   2   4E+3   0.5   1  RT (min)      lcqtof_hilic_neg_1_011_46    m/z 822.1702     0   2   4E+3   0.5   1  RT (min)      lcqtof_hilic_neg_4_010_28    m/z 822.2791     0   1   2E+3   6   6.5  RT (min)      lcqtof_hilic_neg_1_048_45    m/z 822.2988     0   1   2   3E+3   1   1.5  RT (min)      lcqtof_hilic_neg_4_027_19    m/z 822.4549     0   1   2   3   4E+3   0.5   1  RT (min)      lcqtof_hilic_neg_1_051_35    m/z 822.4689     0   2   4E+3   4   4.5  RT (min)      lcqtof_hilic_neg_1_003_37    m/z 822.5742     0   1E+3   15.5   16  RT (min)       lcqtof_hilic_neg_4_103_61    m/z 823.0609     0   1   2E+3   0.5   1  RT (min)      lcqtof_hilic_neg_1_011_46    m/z 823.1619     0   1   2   3E+3   0.5   1  RT (min)      lcqtof_hilic_neg_4_010_28    m/z 823.2321     0   1E+3   7   7.5  RT (min)      lcqtof_hilic_neg_2_031_01    m/z 823.5520     0   1E+5   3.5   4  RT (min)      lcqtof_hilic_neg_4_016_37    m/z 823.6935     0   1E+3   15.5   16  RT (min)      lcqtof_hilic_neg_4_081_58    m/z 823.7363     0   1   2   3   4E+3   0.5   1  RT (min)      lcqtof_hilic_neg_4_008_24    m/z 824.4045     0   1   2   3E+3   3.5   4  RT (min)      lcqtof_hilic_neg_2_026_28    m/z 824.8894     0   2   4E+3   0.5   1  RT (min)      lcqtof_hilic_neg_4_004_47    m/z 825.0681     0   1   2   3E+3   5.5   6  RT (min)      lcqtof_hilic_neg_4_013_20    m/z 825.2004     0   1   2   3   4E+3   0.5   1  RT (min)      lcqtof_hilic_neg_4_007_45    m/z 825.4849     0   1   2   3   4E+3   4   4.5  RT (min)      lcqtof_hilic_neg_1_003_37    m/z 825.7340     0   1E+3   4.5   5  RT (min)      lcqtof_hilic_neg_4_005_29    m/z 825.9698     0   1   2   3E+3   0.5   1  RT (min)      lcqtof_hilic_neg_4_007_45    m/z 826.3124     0   2   4   6E+3   1   1.5  RT (min)      lcqtof_hilic_neg_4_053_14    m/z 826.3839     0   2   4   6E+3   0.5   1  RT (min)      lcqtof_hilic_neg_4_053_14    m/z 826.4274     0   1   2   3E+3   0.5   1  RT (min)      lcqtof_hilic_neg_2_020_23    m/z 826.4572     0   1E+3   7   7.5  RT (min)      lcqtof_hilic_neg_2_051_14    m/z 826.9754     0   1   2   3E+3   0.5   1  RT (min)      lcqtof_hilic_neg_2_033_53    m/z 827.0740     0   2   4   6   8E+3   0.5   1  RT (min)      lcqtof_hilic_neg_4_003_16    m/z 827.1951     0   2   4   6   8E+3   0.5   1  RT (min)      lcqtof_hilic_neg_4_004_47    m/z 827.2306     0   1   2E+3   0.5   1  RT (min)      lcqtof_hilic_neg_4_005_29    m/z 827.3704     0   1   2E+3   3   3.5  RT (min)      lcqtof_hilic_neg_2_005_13    m/z 827.4932     0   2   4E+3   0.5   1  RT (min)      lcqtof_hilic_neg_2_051_14    m/z 827.8577     0   1   2E+3   0.5   1  RT (min)      lcqtof_hilic_neg_4_011_35    m/z 828.4395     0   1   2   3E+3   0.5   1  RT (min)      lcqtof_hilic_neg_2_020_23    m/z 828.5715     0   1E+3   15   15.5  RT (min)      lcqtof_hilic_neg_2_030_40    m/z 828.6838     0   2   4E+3   0.5   1  RT (min)       lcqtof_hilic_neg_4_004_47    m/z 828.9694     0   1   2E+3   0.5   1  RT (min)      lcqtof_hilic_neg_1_028_47    m/z 829.1040     0   2   4   6   8E+3   0.5   1  RT (min)      lcqtof_hilic_neg_4_004_47    m/z 829.2507     0   1   2   3   4E+3   3   3.5  RT (min)      lcqtof_hilic_neg_1_003_37    m/z 829.2541     0   1   2   3E+3   3   3.5  RT (min)      lcqtof_hilic_neg_1_008_13    m/z 829.4587     0   1   2   3E+3   4   4.5  RT (min)      lcqtof_hilic_neg_3_024_35    m/z 829.4907     0   2   4E+3   0.5   1  RT (min)      lcqtof_hilic_neg_1_004_14    m/z 829.6588     0   1   2E+3   1.5   2  RT (min)      lcqtof_hilic_neg_4_003_16    m/z 829.7084     0   1   2   3E+3   0.5   1  RT (min)      lcqtof_hilic_neg_4_002_25    m/z 830.0938     0   1   2E+3   6.5   7  RT (min)      lcqtof_hilic_neg_1_045_07    m/z 830.1073     0   1E+3   3   3.5  RT (min)      lcqtof_hilic_neg_4_010_28    m/z 830.2621     0   1   2E+3   3   3.5  RT (min)      lcqtof_hilic_neg_1_003_37    m/z 830.3919     0   1   2E+3   3   3.5  RT (min)      lcqtof_hilic_neg_1_013_38    m/z 830.4695     0   1   2E+3   0.5   1  RT (min)      lcqtof_hilic_neg_4_018_27    m/z 830.5107     0   1   2   3E+3   3.5   4  RT (min)      lcqtof_hilic_neg_4_050_02    m/z 831.1033     0   1   2   3E+3   0.5   1  RT (min)      lcqtof_hilic_neg_4_003_16    m/z 831.1538     0   1   2   3   4E+3   4.5   5  RT (min)      lcqtof_hilic_neg_2_050_19    m/z 831.5214     0   2   4   6   8E+3   0.5   1  RT (min)      lcqtof_hilic_neg_4_039_18    m/z 832.1781     0   1   2E+3   7   7.5  RT (min)      lcqtof_hilic_neg_1_014_31    m/z 832.3147     0   2   4   6E+3   6   6.5  RT (min)      lcqtof_hilic_neg_4_012_07    m/z 832.3980     0   1   2   3E+3   3   3.5  RT (min)      lcqtof_hilic_neg_1_010_15    m/z 832.4481     0   1   2   3E+3   3   3.5  RT (min)      lcqtof_hilic_neg_4_011_35    m/z 832.4750     0   1   2   3   4E+3   4   4.5  RT (min)      lcqtof_hilic_neg_4_013_20    m/z 833.1509     0   1   2   3E+3   4.5   5  RT (min)      lcqtof_hilic_neg_2_050_19    m/z 833.6007     0   1   2   3   4E+3   1   1.5  RT (min)      lcqtof_hilic_neg_4_027_19    m/z 834.0808     0   2   4   6   8E+3   0.5   1  RT (min)      lcqtof_hilic_neg_3_015_43    m/z 834.4457     0   1   2   3   4E+3   4.5   5  RT (min)      lcqtof_hilic_neg_2_047_32    m/z 834.4527     0   2   4   6E+3   7   7.5  RT (min)       lcqtof_hilic_neg_4_008_24    m/z 835.0786     0   2   4   6   8E+3   0.5   1  RT (min)      lcqtof_hilic_neg_3_015_43    m/z 835.3444     0   1   2E+3   5   5.5  RT (min)       lcqtof_hilic_neg_4_112_62    m/z 835.4647     0   1E+5   3.5   4  RT (min)      lcqtof_hilic_neg_1_049_55    m/z 835.6264     0   1   2   3E+3   3   3.5  RT (min)      lcqtof_hilic_neg_2_042_27    m/z 836.9847     0   2   4   6   8E+3   0.5   1  RT (min)      lcqtof_hilic_neg_4_004_47    m/z 837.1372     0   1   2   3E+3   6   6.5  RT (min)       lcqtof_hilic_neg_4_023_15    m/z 837.2838     0   2   4   6E+3   3   3.5  RT (min)      lcqtof_hilic_neg_4_013_20    m/z 837.4195     0   1   2   3E+3   3   3.5  RT (min)      lcqtof_hilic_neg_1_013_38    m/z 837.4830     0   1   2   3E+3   3.5   4  RT (min)      lcqtof_hilic_neg_1_036_03    m/z 837.5080     0   1   2E+3   1   1.5  RT (min)      lcqtof_hilic_neg_4_008_24    m/z 837.7071     0   1   2E+3   15.5   16  RT (min)      lcqtof_hilic_neg_4_003_16    m/z 837.7570     0   1   2E+3   4.5   5  RT (min)      lcqtof_hilic_neg_4_023_15    m/z 837.9842     0   2   4   6   8E+3   0.5   1  RT (min)      lcqtof_hilic_neg_4_004_47    m/z 838.5345     0   2   4   6E+3   3.5   4  RT (min)      lcqtof_hilic_neg_4_028_36    m/z 838.6111     0   1E+3   4.5   5  RT (min)      lcqtof_hilic_neg_1_015_33    m/z 838.6163     0   2   4E+3   4.5   5  RT (min)      lcqtof_hilic_neg_4_053_14    m/z 839.1392     0   2   4   6   8E+3   4   4.5  RT (min)      lcqtof_hilic_neg_4_040_05    m/z 839.4202     0   2   4   6   8E+3   0.5   1  RT (min)      lcqtof_hilic_neg_4_010_28    m/z 839.4481     0   2   4   6E+3   4.5   5  RT (min)      lcqtof_hilic_neg_1_032_18    m/z 839.4920     0   1   2   3   4E+3   3.5   4  RT (min)      lcqtof_hilic_neg_1_036_03    m/z 839.6140     0   1E+3   4.5   5  RT (min)      lcqtof_hilic_neg_2_029_31    m/z 839.6638     0   1E+3   15.5   16  RT (min)       lcqtof_hilic_neg_4_105_61    m/z 840.0339     0   2   4E+3   6.5   7  RT (min)       lcqtof_hilic_neg_4_086_59    m/z 840.4379     0   2   4E+3   3.5   4  RT (min)       lcqtof_hilic_neg_1_032_18    m/z 840.4678     0   1   2   3   4E+3   4   4.5  RT (min)      lcqtof_hilic_neg_4_027_19    m/z 840.7449     0   1E+4   0.5   1  RT (min)      lcqtof_hilic_neg_1_051_35    m/z 840.9038     0   1   2   3   4E+3   0.5   1  RT (min)      lcqtof_hilic_neg_2_034_47    m/z 841.3724     0   1   2E+3   1.5   2  RT (min)      lcqtof_hilic_neg_4_039_18    m/z 841.4908     0   1   2   3   4E+3   0.5   1  RT (min)      lcqtof_hilic_neg_3_007_04    m/z 841.6280     0   1   2E+3   1   1.5  RT (min)      lcqtof_hilic_neg_4_035_33    m/z 841.7471     0   2   4   6E+3   0.5   1  RT (min)      lcqtof_hilic_neg_1_051_35    m/z 842.3598     0   2   4E+3   5   5.5  RT (min)      lcqtof_hilic_neg_4_032_01    m/z 842.4613     0   2   4   6E+3   1   1.5  RT (min)      lcqtof_hilic_neg_1_035_23    m/z 842.5777     0   1   2   3   4E+4   3.5   4  RT (min)      lcqtof_hilic_neg_4_031_38    m/z 843.1296     0   1   2   3E+3   4.5   5  RT (min)      lcqtof_hilic_neg_4_053_14    m/z 843.4337     0   2   4   6E+3   0.5   1  RT (min)      lcqtof_hilic_neg_4_054_48    m/z 843.7767     0   1   2   3   4E+3   4.5   5  RT (min)      lcqtof_hilic_neg_4_047_06    m/z 843.9893     0   1   2E+3   0.5   1  RT (min)      lcqtof_hilic_neg_1_028_47    m/z 844.4130     0   1   2   3   4E+3   4   4.5  RT (min)      lcqtof_hilic_neg_2_029_31    m/z 844.4802     0   2   4   6   8E+3   4   4.5  RT (min)      lcqtof_hilic_neg_1_035_23    m/z 844.5724     0   1   2   3   4E+3   4.5   5   5.5  RT (min)      lcqtof_hilic_neg_1_022_36    m/z 844.5855     0   2   4E+3   5   5.5  RT (min)      lcqtof_hilic_neg_4_048_23    m/z 845.1149     0   2   4   6E+3   1   1.5  RT (min)      lcqtof_hilic_neg_4_004_47    m/z 845.4075     0   2   4E+3   3.5   4  RT (min)      lcqtof_hilic_neg_1_020_09    m/z 845.5501     0   1E+3   15.5   16  RT (min)       lcqtof_hilic_neg_4_033_53    m/z 845.5831     0   1   2   3   4E+3   5   5.5  RT (min)      lcqtof_hilic_neg_4_048_23    m/z 846.2892     0   2   4   6   8E+3   1   1.5  RT (min)       lcqtof_hilic_neg_3_003_20    m/z 846.2905     0   2   4   6   8E+3   1   1.5  RT (min)      lcqtof_hilic_neg_3_003_20    m/z 846.3476     0   1   2E+3   5   5.5  RT (min)      lcqtof_hilic_neg_4_048_23    m/z 846.4193     0   1   2E+3   4   4.5  RT (min)      lcqtof_hilic_neg_2_048_35    m/z 846.4354     0   1   2   3   4E+3   2.5   3  RT (min)      lcqtof_hilic_neg_1_035_23    m/z 846.5378     0   2   4   6E+3   0.5   1  RT (min)      lcqtof_hilic_neg_4_015_09    m/z 847.4076     0   1   2   3E+3   3.5   4  RT (min)      lcqtof_hilic_neg_4_005_29    m/z 847.7156     0   1   2   3   4E+3   0.5   1  RT (min)      lcqtof_hilic_neg_2_021_17    m/z 848.5077     0   2   4E+3   4.5   5  RT (min)      lcqtof_hilic_neg_4_105_61    m/z 848.8262     0   2   4E+3   0.5   1  RT (min)      lcqtof_hilic_neg_4_016_37    m/z 848.9338     0   1E+3   6   6.5  RT (min)      lcqtof_hilic_neg_1_012_04    m/z 849.4382     0   2   4   6E+3   4.5   5  RT (min)       lcqtof_hilic_neg_4_042_22    m/z 849.4721     0   2   4   6E+3   0.5   1  RT (min)      lcqtof_hilic_neg_4_039_18    m/z 849.6441     0   1   2   3E+3   4   4.5  RT (min)      lcqtof_hilic_neg_4_055_32    m/z 849.6447     0   1E+4   4   4.5  RT (min)      lcqtof_hilic_neg_1_005_32    m/z 850.4720     0   2   4   6   8E+3   3.5   4  RT (min)      lcqtof_hilic_neg_4_039_18    m/z 850.6531     0   2   4   6   8E+3   3   3.5  RT (min)      lcqtof_hilic_neg_1_037_24    m/z 851.0132     0   1   2E+3   1   1.5  RT (min)      lcqtof_hilic_neg_4_004_47    m/z 852.5953     0   2   4   6E+3   4.5   5  RT (min)      lcqtof_hilic_neg_4_048_23    m/z 852.6626     0   1E+4   3.5   4  RT (min)      lcqtof_hilic_neg_1_005_32    m/z 852.6654     0   1   2   3   4E+3   4.5   5  RT (min)      lcqtof_hilic_neg_4_055_32    m/z 852.6658     0   1   2E+3   4   4.5  RT (min)      lcqtof_hilic_neg_3_034_24    m/z 852.7168     0   1   2E+3   3   3.5  RT (min)      lcqtof_hilic_neg_4_005_29    m/z 852.9282     0   1   2   3   4E+3   6   6.5  RT (min)      lcqtof_hilic_neg_4_026_04    m/z 853.1785     0   2   4E+3   7   7.5  RT (min)      lcqtof_hilic_neg_1_005_32    m/z 853.6127     0   2   4   6   8E+3   5   5.5  RT (min)       lcqtof_hilic_neg_4_048_23    m/z 853.6923     0   1E+3   15.5   16  RT (min)       lcqtof_hilic_neg_4_102_61    m/z 854.2422     0   1   2E+3   5   5.5  RT (min)      lcqtof_hilic_neg_2_042_27    m/z 854.3872     0   1   2   3E+3   1.5   2  RT (min)      lcqtof_hilic_neg_4_021_21    m/z 854.4789     0   1   2   3   4E+3   3.5   4  RT (min)      lcqtof_hilic_neg_4_005_29    m/z 854.9987     0   2   4E+3   0.5   1  RT (min)      lcqtof_hilic_neg_4_014_46    m/z 855.3319     0   1   2   3   4E+3   0.5   1  RT (min)      lcqtof_hilic_neg_4_013_20    m/z 855.9560     0   2   4E+3   3.5   4  RT (min)      lcqtof_hilic_neg_4_053_14    m/z 855.9923     0   1   2   3   4E+3   0.5   1  RT (min)      lcqtof_hilic_neg_4_014_46    m/z 856.1256     0   1   2E+3   3   3.5  RT (min)      lcqtof_hilic_neg_4_032_01    m/z 856.4882     0   1   2E+3   3   3.5  RT (min)      lcqtof_hilic_neg_4_005_29    m/z 856.8949     0   1   2   3E+3   3   3.5  RT (min)      lcqtof_hilic_neg_4_005_29    m/z 857.4418     0   1   2   3   4E+3   4   4.5  RT (min)       lcqtof_hilic_neg_4_013_20    m/z 857.4463     0   2   4E+3   0.5   1  RT (min)      lcqtof_hilic_neg_4_007_45    m/z 857.9160     0   1   2   3   4E+3   0.5   1  RT (min)      lcqtof_hilic_neg_2_034_47    m/z 857.9690     0   1   2   3E+3   3.5   4  RT (min)      lcqtof_hilic_neg_4_053_14    m/z 858.4210     0   2   4E+3   0.5   1  RT (min)      lcqtof_hilic_neg_4_045_26    m/z 858.5618     0   1   2   3   4E+3   3.5   4  RT (min)       lcqtof_hilic_neg_1_003_37    m/z 859.0411     0   1   2E+3   0.5   1  RT (min)      lcqtof_hilic_neg_1_011_46    m/z 859.1301     0   2   4   6   8E+3   1   1.5  RT (min)      lcqtof_hilic_neg_4_004_47    m/z 859.2028     0   2   4   6E+3   6   6.5  RT (min)      lcqtof_hilic_neg_4_023_15    m/z 859.3613     0   2   4E+3   0.5   1  RT (min)      lcqtof_hilic_neg_4_053_14    m/z 859.4477     0   2   4E+3   0.5   1  RT (min)      lcqtof_hilic_neg_2_051_14    m/z 859.5050     0   2   4E+3   4   4.5  RT (min)       lcqtof_hilic_neg_4_013_20    m/z 859.5837     0   2   4   6   8E+3   3   3.5  RT (min)      lcqtof_hilic_neg_4_016_37    m/z 860.5537     0   2   4   6E+3   0.5   1  RT (min)      lcqtof_hilic_neg_3_044_09    m/z 861.3109     0   1   2   3   4E+3   1   1.5  RT (min)      lcqtof_hilic_neg_3_003_20    m/z 861.3693     0   2   4   6   8E+3   3.5   4  RT (min)      lcqtof_hilic_neg_4_019_03    m/z 861.6387     0   1   2   3   4E+3   0.5   1  RT (min)      lcqtof_hilic_neg_1_011_46    m/z 861.6945     0   1   2   3   4E+3   4   4.5  RT (min)      lcqtof_hilic_neg_2_031_01    m/z 862.3055     0   1   2   3   4E+3   1   1.5  RT (min)      lcqtof_hilic_neg_3_003_20    m/z 862.3658     0   2   4   6   8E+3   3.5   4  RT (min)      lcqtof_hilic_neg_4_019_03    m/z 862.4198     0   1   2   3   4E+3   4   4.5  RT (min)      lcqtof_hilic_neg_1_024_21    m/z 863.3600     0   2   4   6E+3   0.5   1  RT (min)      lcqtof_hilic_neg_4_053_14    m/z 863.8580     0   1   2E+3   5   5.5  RT (min)      lcqtof_hilic_neg_4_010_28    m/z 864.1509     0   1   2E+3   5.5   6  RT (min)      lcqtof_hilic_neg_4_050_02    m/z 864.7386     0   1E+3   15.5   16  RT (min)       lcqtof_hilic_neg_4_073_57    m/z 864.9153     0   1   2E+3   5   5.5  RT (min)      lcqtof_hilic_neg_4_035_33    m/z 865.3752     0   1E+3   5   5.5  RT (min)      lcqtof_hilic_neg_1_016_28    m/z 865.8899     0   2   4E+3   0.5   1  RT (min)      lcqtof_hilic_neg_4_014_46    m/z 866.2323     0   1   2   3E+3   3.5   4  RT (min)      lcqtof_hilic_neg_1_053_19    m/z 866.4761     0   2   4E+3   4   4.5  RT (min)      lcqtof_hilic_neg_4_013_20    m/z 866.5780     0   1   2E+3   4   4.5  RT (min)      lcqtof_hilic_neg_2_031_01    m/z 866.9040     0   1   2E+3   6   6.5  RT (min)       lcqtof_hilic_neg_2_018_16    m/z 867.1332     0   1   2   3   4E+3   0.5   1  RT (min)      lcqtof_hilic_neg_4_013_20    m/z 867.4731     0   2   4   6E+3   4   4.5  RT (min)      lcqtof_hilic_neg_4_013_20    m/z 867.6337     0   2   4   6E+3   0.5   1  RT (min)      lcqtof_hilic_neg_4_053_14    m/z 867.6923     0   1E+3   15.5   16  RT (min)       lcqtof_hilic_neg_4_103_61    m/z 867.7143     0   2   4E+3   4.5   5  RT (min)      lcqtof_hilic_neg_4_008_24    m/z 867.8265     0   1   2   3E+3   1   1.5  RT (min)      lcqtof_hilic_neg_4_054_48    m/z 868.3458     0   1   2   3E+3   0.5   1  RT (min)      lcqtof_hilic_neg_4_054_48    m/z 868.8277     0   1   2   3E+3   5   5.5  RT (min)       lcqtof_hilic_neg_4_050_02    m/z 868.9183     0   2   4   6   8E+3   0.5   1  RT (min)      lcqtof_hilic_neg_4_076_58    m/z 869.3348     0   1   2E+3   5   5.5  RT (min)       lcqtof_hilic_neg_4_044_08    m/z 869.3774     0   1   2   3E+3   5   5.5  RT (min)      lcqtof_hilic_neg_4_031_38    m/z 869.6033     0   1   2   3E+3   3.5   4  RT (min)      lcqtof_hilic_neg_4_005_29    m/z 869.9191     0   2   4   6E+3   0.5   1  RT (min)      lcqtof_hilic_neg_4_054_48    m/z 870.4286     0   1   2   3E+3   4   4.5  RT (min)      lcqtof_hilic_neg_2_048_35    m/z 870.4899     0   1   2   3   4E+3   2   2.5  RT (min)      lcqtof_hilic_neg_1_035_23    m/z 870.9548     0   1E+3   5   5.5  RT (min)      lcqtof_hilic_neg_4_005_29    m/z 871.0095     0   1   2E+3   7   7.5   8  RT (min)      lcqtof_hilic_neg_1_030_17    m/z 871.5095     0   2   4E+3   4   4.5  RT (min)      lcqtof_hilic_neg_4_048_23    m/z 871.6037     0   2   4E+3   5   5.5  RT (min)      lcqtof_hilic_neg_4_048_23    m/z 871.6694     0   2   4   6E+3   2   2.5  RT (min)      lcqtof_hilic_neg_4_048_23    m/z 872.6716     0   1   2   3   4E+3   1.5   2  RT (min)       lcqtof_hilic_neg_4_053_14    m/z 873.5428     0   1   2E+3   3.5   4  RT (min)      lcqtof_hilic_neg_1_047_29    m/z 873.6614     0   1E+4   1.5   2  RT (min)      lcqtof_hilic_neg_4_055_32    m/z 873.7007     0   1E+4   0.5   1  RT (min)      lcqtof_hilic_neg_1_014_31    m/z 874.5590     0   1   2E+3   0.5   1  RT (min)      lcqtof_hilic_neg_4_003_16    m/z 875.2241     0   1   2E+3   5   5.5  RT (min)      lcqtof_hilic_neg_2_050_19    m/z 875.5880     0   1E+4   0.5   1  RT (min)      lcqtof_hilic_neg_1_013_38    m/z 876.1898     0   2   4   6E+3   5.5   6  RT (min)      lcqtof_hilic_neg_4_013_20    m/z 876.3773     0   1E+3   5   5.5  RT (min)      lcqtof_hilic_neg_2_002_36    m/z 876.3920     0   2   4   6E+3   4   4.5  RT (min)      lcqtof_hilic_neg_1_032_18    m/z 877.2525     0   1   2E+3   4.5   5  RT (min)       lcqtof_hilic_neg_4_076_58    m/z 877.7506     0   1E+3   16   16.5  RT (min)       lcqtof_hilic_neg_4_105_61    m/z 878.2504     0   1   2E+3   4.5   5  RT (min)       lcqtof_hilic_neg_4_080_58    m/z 878.3906     0   1   2   3   4E+3   4   4.5  RT (min)      lcqtof_hilic_neg_1_032_18    m/z 878.5787     0   2   4   6E+3   4.5   5  RT (min)      lcqtof_hilic_neg_4_050_02    m/z 879.6571     0   1   2E+3   3.5   4  RT (min)      lcqtof_hilic_neg_4_052_13    m/z 880.5823     0   1   2   3E+4   1   1.5  RT (min)      lcqtof_hilic_neg_4_011_35    m/z 880.5919     0   2   4   6E+3   4.5   5  RT (min)       lcqtof_hilic_neg_4_053_14    m/z 881.3403     0   1   2   3   4E+3   1   1.5  RT (min)      lcqtof_hilic_neg_1_042_26    m/z 881.4771     0   1   2   3   4E+3   4   4.5  RT (min)      lcqtof_hilic_neg_4_013_20    m/z 881.5053     0   1   2E+3   7   7.5   8  RT (min)      lcqtof_hilic_neg_1_030_17    m/z 881.5930     0   1E+3   4.5   5  RT (min)      lcqtof_hilic_neg_2_029_31    m/z 881.6757     0   1   2E+3   3.5   4  RT (min)      lcqtof_hilic_neg_4_029_17    m/z 882.2151     0   1   2E+3   0.5   1  RT (min)      lcqtof_hilic_neg_4_040_05    m/z 882.5049     0   1   2E+3   7.5   8  RT (min)      lcqtof_hilic_neg_1_016_28    m/z 882.5800     0   2   4   6   8E+3   1   1.5  RT (min)      lcqtof_hilic_neg_2_044_37    m/z 882.5938     0   1   2E+3   8   8.5  RT (min)      lcqtof_hilic_neg_2_027_07    m/z 882.6008     0   1E+3   8   8.5  RT (min)      lcqtof_hilic_neg_3_038_06    m/z 882.9297     0   2   4   6   8E+3   5   5.5  RT (min)      lcqtof_hilic_neg_4_008_24    m/z 883.0059     0   1   2E+3   7.5   8  RT (min)      lcqtof_hilic_neg_1_016_28    m/z 883.0281     0   2   4E+3   0.5   1  RT (min)      lcqtof_hilic_neg_4_014_46    m/z 883.0339     0   1   2   3   4E+3   6   6.5  RT (min)      lcqtof_hilic_neg_4_055_32    m/z 883.5549     0   2   4   6   8E+3   3   3.5  RT (min)      lcqtof_hilic_neg_1_023_34    m/z 883.5552     0   1   2E+3   1   1.5  RT (min)      lcqtof_hilic_neg_2_005_13    m/z 884.0246     0   1   2   3   4E+3   0.5   1  RT (min)      lcqtof_hilic_neg_4_014_46    m/z 884.1772     0   1   2E+3   5   5.5  RT (min)      lcqtof_hilic_neg_4_011_35    m/z 885.1135     0   1   2   3   4E+3   4.5   5  RT (min)      lcqtof_hilic_neg_4_053_14    m/z 885.1736     0   1   2   3E+3   5   5.5  RT (min)      lcqtof_hilic_neg_2_042_27    m/z 885.5645     0   2   4E+3   2.5   3  RT (min)      lcqtof_hilic_neg_1_037_24    m/z 885.6321     0   1   2E+3   4   4.5  RT (min)      lcqtof_hilic_neg_1_005_32    m/z 885.9338     0   1   2   3   4E+3   0.5   1  RT (min)      lcqtof_hilic_neg_4_004_47    m/z 886.1788     0   1E+3   5   5.5  RT (min)      lcqtof_hilic_neg_2_039_38    m/z 886.3575     0   2   4   6E+3   0.5   1  RT (min)      lcqtof_hilic_neg_4_013_20    m/z 886.3708     0   2   4   6E+3   6   6.5  RT (min)      lcqtof_hilic_neg_4_050_02    m/z 886.3793     0   2   4E+3   0.5   1  RT (min)      lcqtof_hilic_neg_4_053_14    m/z 886.4280     0   1   2   3   4E+3   4   4.5  RT (min)       lcqtof_hilic_neg_2_053_18    m/z 886.4890     0   2   4E+3   1   1.5  RT (min)      lcqtof_hilic_neg_1_035_23    m/z 887.1081     0   2   4E+3   4.5   5  RT (min)      lcqtof_hilic_neg_4_053_14    m/z 887.2590     0   2   4E+3   4   4.5  RT (min)      lcqtof_hilic_neg_2_031_01    m/z 887.4198     0   1   2   3E+3   4   4.5  RT (min)      lcqtof_hilic_neg_4_016_37    m/z 887.5771     0   2   4   6E+3   2.5   3  RT (min)      lcqtof_hilic_neg_1_037_24    m/z 887.7356     0   1   2   3E+3   4.5   5  RT (min)      lcqtof_hilic_neg_4_029_17    m/z 887.8600     0   2   4   6   8E+3   4   4.5  RT (min)      lcqtof_hilic_neg_4_032_01    m/z 888.6062     0   2   4   6E+3   3.5   4  RT (min)      lcqtof_hilic_neg_4_023_15    m/z 888.9403     0   1   2   3E+3   0.5   1  RT (min)      lcqtof_hilic_neg_1_028_47    m/z 890.0099     0   1   2E+3   4.5   5  RT (min)      lcqtof_hilic_neg_4_080_58    m/z 890.5956     0   2   4   6E+3   2   2.5  RT (min)      lcqtof_hilic_neg_4_011_35    m/z 891.2374     0   2   4   6E+3   6   6.5  RT (min)       lcqtof_hilic_neg_4_052_13    m/z 891.2629     0   2   4   6   8E+3   0.5   1  RT (min)      lcqtof_hilic_neg_4_023_15    m/z 892.5414     0   2   4   6E+3   1   1.5  RT (min)      lcqtof_hilic_neg_1_030_17    m/z 892.6329     0   2   4E+3   0.5   1  RT (min)      lcqtof_hilic_neg_4_050_02    m/z 892.9903     0   1   2   3E+3   7   7.5  RT (min)      lcqtof_hilic_neg_1_010_15    m/z 893.5841     0   1   2E+3   8   8.5  RT (min)      lcqtof_hilic_neg_2_031_01    m/z 893.5906     0   1E+3   8   8.5  RT (min)      lcqtof_hilic_neg_3_038_06    m/z 893.8763     0   1   2E+4   0.5   1  RT (min)      lcqtof_hilic_neg_4_004_47    m/z 894.3635     0   2   4   6   8E+3   0.5   1  RT (min)      lcqtof_hilic_neg_1_004_14    m/z 895.3674     0   1   2   3   4E+3   0.5   1  RT (min)      lcqtof_hilic_neg_1_004_14    m/z 895.4449     0   2   4   6   8E+3   3.5   4  RT (min)       lcqtof_hilic_neg_4_013_20    m/z 895.4935     0   1E+3   4   4.5  RT (min)      lcqtof_hilic_neg_1_030_17    m/z 895.5590     0   1E+4   0.5   1  RT (min)      lcqtof_hilic_neg_4_038_10    m/z 895.5602     0   1   2   3E+3   4   4.5  RT (min)      lcqtof_hilic_neg_1_007_05    m/z 896.1896     0   2   4   6   8E+3   0.5   1  RT (min)      lcqtof_hilic_neg_4_010_28    m/z 896.6606     0   1   2   3E+3   0.5   1  RT (min)      lcqtof_hilic_neg_4_050_02    m/z 896.7238     0   2   4E+3   0.5   1  RT (min)      lcqtof_hilic_neg_4_034_31    m/z 897.1900     0   2   4E+3   0.5   1  RT (min)      lcqtof_hilic_neg_4_010_28    m/z 897.3836     0   1   2   3   4E+3   5   5.5  RT (min)      lcqtof_hilic_neg_4_053_14    m/z 897.4255     0   1   2   3   4E+3   4   4.5  RT (min)      lcqtof_hilic_neg_1_020_09    m/z 897.5935     0   1   2E+3   3.5   4  RT (min)      lcqtof_hilic_neg_2_055_30    m/z 898.5981     0   2   4   6   8E+3   3.5   4  RT (min)      lcqtof_hilic_neg_2_013_29    m/z 898.6392     0   1   2   3   4E+3   3.5   4  RT (min)      lcqtof_hilic_neg_4_018_27    m/z 899.0554     0   1   2E+3   15   15.5  RT (min)       lcqtof_hilic_neg_4_050_02    m/z 899.1114     0   2   4   6E+3   6   6.5  RT (min)      lcqtof_hilic_neg_4_031_38    m/z 899.3613     0   1   2   3   4E+3   5   5.5  RT (min)      lcqtof_hilic_neg_4_032_01    m/z 899.3636     0   1E+3   5   5.5  RT (min)      lcqtof_hilic_neg_1_051_35    m/z 899.4088     0   2   4   6   8E+3   4   4.5  RT (min)      lcqtof_hilic_neg_4_013_20    m/z 899.9993     0   1E+3   6   6.5  RT (min)      lcqtof_hilic_neg_4_077_58    m/z 900.0777     0   1   2   3   4E+3   6   6.5  RT (min)      lcqtof_hilic_neg_4_056_56    m/z 900.5098     0   2   4   6E+3   4.5   5  RT (min)      lcqtof_hilic_neg_1_035_23    m/z 900.8701     0   2   4E+3   0.5   1  RT (min)      lcqtof_hilic_neg_4_004_47    m/z 901.1779     0   2   4   6E+3   0.5   1  RT (min)      lcqtof_hilic_neg_4_004_47    m/z 901.7497     0   2   4   6   8E+3   0.5   1  RT (min)      lcqtof_hilic_neg_4_008_24    m/z 902.4107     0   1   2E+3   4   4.5  RT (min)       lcqtof_hilic_neg_1_040_20    m/z 903.3262     0   1   2E+3   5   5.5  RT (min)       lcqtof_hilic_neg_4_105_61    m/z 903.3730     0   1   2   3E+3   5   5.5  RT (min)      lcqtof_hilic_neg_4_028_36    m/z 903.6008     0   2   4   6   8E+3   1.5   2  RT (min)      lcqtof_hilic_neg_4_050_02    m/z 903.6077     0   1   2   3   4E+3   3   3.5  RT (min)      lcqtof_hilic_neg_4_053_14    m/z 903.7693     0   2   4   6E+3   3   3.5  RT (min)      lcqtof_hilic_neg_4_008_24    m/z 903.9576     0   1   2   3E+3   0.5   1  RT (min)      lcqtof_hilic_neg_2_034_47    m/z 904.0245     0   1   2E+3   6   6.5  RT (min)      lcqtof_hilic_neg_4_055_32    m/z 904.3958     0   1E+3   5.5   6  RT (min)      lcqtof_hilic_neg_1_004_14    m/z 904.5840     0   1   2E+3   8   8.5  RT (min)      lcqtof_hilic_neg_2_032_02    m/z 904.5872     0   1   2   3E+3   1   1.5  RT (min)      lcqtof_hilic_neg_4_003_16    m/z 904.6298     0   1   2E+3   4.5   5  RT (min)      lcqtof_hilic_neg_1_035_23    m/z 904.7488     0   2   4   6E+3   1.5   2  RT (min)      lcqtof_hilic_neg_1_005_32    m/z 905.0832     0   2   4   6E+3   1   1.5  RT (min)      lcqtof_hilic_neg_4_004_47    m/z 905.4506     0   2   4   6   8E+3   4   4.5  RT (min)      lcqtof_hilic_neg_4_027_19    m/z 905.5620     0   2   4E+3   4   4.5  RT (min)      lcqtof_hilic_neg_4_053_14    m/z 906.5502     0   1   2E+3   1   1.5  RT (min)      lcqtof_hilic_neg_3_068_57    m/z 906.6231     0   2   4   6E+3   1.5   2  RT (min)      lcqtof_hilic_neg_1_005_32    m/z 907.1720     0   1   2   3E+3   5   5.5  RT (min)      lcqtof_hilic_neg_4_050_02    m/z 907.3177     0   2   4   6E+3   6   6.5  RT (min)      lcqtof_hilic_neg_4_052_13    m/z 907.5936     0   1   2   3   4E+3   2   2.5  RT (min)      lcqtof_hilic_neg_2_052_24    m/z 908.4519     0   1   2   3   4E+3   4   4.5  RT (min)      lcqtof_hilic_neg_1_054_01    m/z 908.8964     0   1   2E+3   6   6.5  RT (min)      lcqtof_hilic_neg_2_038_04    m/z 909.3112     0   2   4   6E+3   1   1.5  RT (min)      lcqtof_hilic_neg_1_032_18    m/z 909.6315     0   1   2   3   4E+3   0.5   1  RT (min)      lcqtof_hilic_neg_4_045_26    m/z 910.3496     0   1E+3   5   5.5  RT (min)      lcqtof_hilic_neg_4_003_16    m/z 910.4626     0   1   2   3E+3   4   4.5  RT (min)      lcqtof_hilic_neg_1_010_15    m/z 910.6003     0   1   2   3E+3   1   1.5  RT (min)      lcqtof_hilic_neg_1_054_01    m/z 911.3353     0   1   2E+3   6   6.5  RT (min)      lcqtof_hilic_neg_2_041_54    m/z 911.5541     0   1   2E+3   1   1.5  RT (min)      lcqtof_hilic_neg_4_026_04    m/z 911.6445     0   2   4   6   8E+3   4.5   5  RT (min)      lcqtof_hilic_neg_4_031_38    m/z 911.6522     0   1   2E+3   3   3.5  RT (min)       lcqtof_hilic_neg_4_032_01    m/z 911.9489     0   2   4   6E+3   6   6.5  RT (min)       lcqtof_hilic_neg_4_001_49    m/z 912.3221     0   2   4   6   8E+3   6   6.5  RT (min)      lcqtof_hilic_neg_4_052_13    m/z 912.4706     0   1E+3   5   5.5  RT (min)      lcqtof_hilic_neg_4_034_31    m/z 913.3895     0   1   2   3E+3   4   4.5  RT (min)      lcqtof_hilic_neg_1_040_20    m/z 914.0518     0   2   4   6E+3   0.5   1  RT (min)      lcqtof_hilic_neg_4_014_46    m/z 914.2977     0   1   2   3   4E+3   0.5   1  RT (min)      lcqtof_hilic_neg_2_034_47    m/z 915.0517     0   1   2   3E+3   0.5   1  RT (min)      lcqtof_hilic_neg_4_014_46    m/z 915.5764     0   1E+3   8   8.5  RT (min)      lcqtof_hilic_neg_2_043_15    m/z 915.8863     0   2   4   6E+3   0.5   1  RT (min)      lcqtof_hilic_neg_4_004_47    m/z 916.1540     0   1   2E+3   6   6.5  RT (min)      lcqtof_hilic_neg_4_029_17    m/z 917.0427     0   1   2E+3   6   6.5  RT (min)      lcqtof_hilic_neg_4_034_31    m/z 917.3747     0   2   4   6E+3   6   6.5  RT (min)      lcqtof_hilic_neg_4_052_13    m/z 917.5054     0   1   2E+3   5   5.5  RT (min)      lcqtof_hilic_neg_1_022_36    m/z 917.5840     0   1   2   3   4E+3   3   3.5  RT (min)      lcqtof_hilic_neg_4_086_59    m/z 917.8256     0   2   4E+3   2   2.5  RT (min)      lcqtof_hilic_neg_4_008_24    m/z 918.5394     0   2   4   6E+3   1   1.5  RT (min)      lcqtof_hilic_neg_4_022_34    m/z 919.3337     0   1E+3   15   15.5  RT (min)      lcqtof_hilic_neg_2_055_30    m/z 919.4756     0   1   2E+3   1   1.5  RT (min)      lcqtof_hilic_neg_2_032_02    m/z 919.7219     0   1   2   3   4E+3   4.5   5  RT (min)      lcqtof_hilic_neg_4_006_30    m/z 919.9205     0   1   2   3E+3   6   6.5  RT (min)      lcqtof_hilic_neg_4_026_04    m/z 920.1904     0   1   2   3   4E+3   0.5   1  RT (min)      lcqtof_hilic_neg_4_004_47    m/z 920.6603     0   1   2   3E+3   4.5   5  RT (min)      lcqtof_hilic_neg_1_035_23    m/z 921.2194     0   2   4   6E+3   7   7.5  RT (min)      lcqtof_hilic_neg_2_051_14    m/z 921.7210     0   1   2E+3   4.5   5  RT (min)      lcqtof_hilic_neg_4_022_34    m/z 922.5186     0   1   2E+3   4   4.5  RT (min)      lcqtof_hilic_neg_1_031_16    m/z 923.5047     0   1E+4   4   4.5  RT (min)      lcqtof_hilic_neg_3_048_05    m/z 923.5577     0   1   2   3E+3   4   4.5  RT (min)      lcqtof_hilic_neg_2_052_24    m/z 923.5712     0   2   4E+3   4.5   5  RT (min)       lcqtof_hilic_neg_4_053_14    m/z 923.7712     0   1   2   3E+3   1   1.5  RT (min)      lcqtof_hilic_neg_1_001_49    m/z 924.6519     0   2   4E+3   3.5   4  RT (min)      lcqtof_hilic_neg_4_005_29    m/z 924.7158     0   2   4   6   8E+3   1   1.5  RT (min)      lcqtof_hilic_neg_2_047_32    m/z 925.5217     0   1E+4   4   4.5  RT (min)      lcqtof_hilic_neg_4_040_05    m/z 925.6230     0   1   2   3E+3   2   2.5  RT (min)       lcqtof_hilic_neg_2_031_01    m/z 926.5207     0   2   4   6E+3   4.5   5  RT (min)       lcqtof_hilic_neg_4_040_05    m/z 926.7724     0   2   4   6   8E+3   4.5   5  RT (min)       lcqtof_hilic_neg_4_040_05    m/z 927.0197     0   1   2   3E+3   5.5   6  RT (min)      lcqtof_hilic_neg_4_018_27    m/z 927.3212     0   2   4E+3   0.5   1  RT (min)      lcqtof_hilic_neg_1_006_22    m/z 927.5906     0   2   4E+3   3.5   4  RT (min)      lcqtof_hilic_neg_2_051_14    m/z 927.6432     0   2   4   6E+3   3.5   4  RT (min)       lcqtof_hilic_neg_4_048_23    m/z 927.7412     0   2   4E+3   4.5   5  RT (min)      lcqtof_hilic_neg_4_047_06    m/z 928.1479     0   1   2   3E+3   4.5   5  RT (min)      lcqtof_hilic_neg_4_035_33    m/z 928.3886     0   2   4E+3   1.5   2  RT (min)      lcqtof_hilic_neg_2_004_26    m/z 928.6318     0   2   4   6E+3   1   1.5  RT (min)      lcqtof_hilic_neg_1_037_24    m/z 928.6661     0   1   2   3   4E+3   4.5   5  RT (min)      lcqtof_hilic_neg_4_048_23    m/z 928.7512     0   2   4   6   8E+3   1   1.5  RT (min)      lcqtof_hilic_neg_2_047_32    m/z 929.0884     0   2   4E+3   4.5   5  RT (min)      lcqtof_hilic_neg_4_053_14    m/z 929.8798     0   1E+4   0.5   1  RT (min)      lcqtof_hilic_neg_4_004_47    m/z 930.3612     0   1   2E+3   15   15.5  RT (min)      lcqtof_hilic_neg_4_019_03    m/z 930.5179     0   1   2   3   4E+3   4   4.5  RT (min)      lcqtof_hilic_neg_4_048_23    m/z 930.5524     0   1   2   3   4E+3   4   4.5  RT (min)      lcqtof_hilic_neg_2_031_01    m/z 930.6229     0   2   4   6   8E+3   1.5   2  RT (min)      lcqtof_hilic_neg_1_005_32    m/z 931.3738     0   1   2   3   4E+3   5   5.5  RT (min)      lcqtof_hilic_neg_4_053_14    m/z 931.5797     0   1   2   3E+3   4   4.5  RT (min)      lcqtof_hilic_neg_4_052_13    m/z 932.1305     0   1   2   3E+3   5.5   6  RT (min)      lcqtof_hilic_neg_4_053_14    m/z 932.4900     0   1   2   3   4E+3   0.5   1  RT (min)      lcqtof_hilic_neg_4_004_47    m/z 932.6104     0   1   2   3E+3   1   1.5  RT (min)      lcqtof_hilic_neg_3_001_49    m/z 932.9233     0   1   2   3E+3   5   5.5  RT (min)      lcqtof_hilic_neg_4_053_14    m/z 933.3516     0   2   4E+3   5   5.5  RT (min)      lcqtof_hilic_neg_4_032_01    m/z 933.3538     0   1E+3   5   5.5  RT (min)      lcqtof_hilic_neg_1_031_16    m/z 933.4841     0   1   2   3E+3   0.5   1  RT (min)      lcqtof_hilic_neg_4_007_45    m/z 933.5649     0   2   4   6E+3   3   3.5  RT (min)      lcqtof_hilic_neg_1_023_34    m/z 933.6389     0   2   4E+3   1   1.5  RT (min)      lcqtof_hilic_neg_1_003_37    m/z 933.6447     0   1   2E+3   4   4.5  RT (min)      lcqtof_hilic_neg_1_005_32    m/z 933.8510     0   2   4   6E+3   2   2.5  RT (min)       lcqtof_hilic_neg_4_008_24    m/z 934.1866     0   1   2   3E+3   6   6.5  RT (min)      lcqtof_hilic_neg_4_011_35    m/z 934.7005     0   2   4   6E+3   4   4.5  RT (min)      lcqtof_hilic_neg_1_007_05    m/z 935.0038     0   1   2   3   4E+3   5   5.5  RT (min)      lcqtof_hilic_neg_4_001_49    m/z 935.3058     0   1   2E+3   0.5   1  RT (min)      lcqtof_hilic_neg_1_009_50    m/z 935.6946     0   2   4   6   8E+3   4   4.5  RT (min)      lcqtof_hilic_neg_1_007_05    m/z 936.2579     0   1   2E+3   1   1.5  RT (min)      lcqtof_hilic_neg_1_040_20    m/z 936.3896     0   2   4   6E+3   4   4.5  RT (min)      lcqtof_hilic_neg_4_013_20    m/z 936.4705     0   1   2E+3   5   5.5  RT (min)      lcqtof_hilic_neg_4_028_36    m/z 936.6162     0   1   2   3E+3   1   1.5  RT (min)      lcqtof_hilic_neg_1_054_01    m/z 936.8172     0   1   2E+3   5   5.5  RT (min)       lcqtof_hilic_neg_4_030_44    m/z 936.8523     0   2   4   6   8E+3   0.5   1  RT (min)      lcqtof_hilic_neg_4_016_37    m/z 937.3196     0   1   2E+3   5   5.5  RT (min)      lcqtof_hilic_neg_4_043_12    m/z 937.3635     0   1   2   3E+3   5   5.5  RT (min)      lcqtof_hilic_neg_4_028_36    m/z 937.3859     0   2   4   6E+3   4   4.5  RT (min)      lcqtof_hilic_neg_4_013_20    m/z 937.4776     0   2   4E+3   5   5.5  RT (min)       lcqtof_hilic_neg_4_028_36    m/z 937.5406     0   1   2   3E+3   7   7.5  RT (min)      lcqtof_hilic_neg_1_035_23    m/z 937.7034     0   2   4E+3   0.5   1  RT (min)      lcqtof_hilic_neg_3_040_14    m/z 937.8044     0   1E+4   0.5   1  RT (min)      lcqtof_hilic_neg_4_004_47    m/z 938.2037     0   2   4E+3   6   6.5  RT (min)      lcqtof_hilic_neg_4_029_17    m/z 938.4040     0   1   2E+3   4   4.5  RT (min)      lcqtof_hilic_neg_2_050_19    m/z 938.4907     0   1   2E+3   5   5.5  RT (min)       lcqtof_hilic_neg_4_011_35    m/z 938.5250     0   1   2E+3   1   1.5  RT (min)      lcqtof_hilic_neg_2_005_13    m/z 938.6263     0   1   2   3   4E+3   1.5   2  RT (min)       lcqtof_hilic_neg_4_016_37    m/z 938.6303     0   1E+4   3.5   4  RT (min)      lcqtof_hilic_neg_4_028_36    m/z 938.6783     0   1   2   3E+3   3   3.5  RT (min)       lcqtof_hilic_neg_4_029_17    m/z 939.5027     0   1   2E+3   5   5.5  RT (min)      lcqtof_hilic_neg_4_034_31    m/z 939.7333     0   2   4   6   8E+3   3   3.5  RT (min)      lcqtof_hilic_neg_4_002_25    m/z 940.6248     0   1   2   3E+3   3.5   4  RT (min)      lcqtof_hilic_neg_1_010_15    m/z 941.2980     0   2   4   6E+3   4   4.5  RT (min)      lcqtof_hilic_neg_1_023_34    m/z 941.3156     0   2   4   6E+3   0.5   1  RT (min)      lcqtof_hilic_neg_4_004_47    m/z 941.4921     0   1   2   3   4E+3   4   4.5  RT (min)      lcqtof_hilic_neg_4_031_38    m/z 942.2113     0   2   4   6   8E+3   7   7.5  RT (min)      lcqtof_hilic_neg_2_043_15    m/z 942.4984     0   1   2E+3   4   4.5  RT (min)      lcqtof_hilic_neg_1_027_27    m/z 942.5695     0   2   4   6E+3   1   1.5  RT (min)      lcqtof_hilic_neg_2_042_27    m/z 942.9946     0   2   4   6   8E+3   0.5   1  RT (min)      lcqtof_hilic_neg_4_014_46    m/z 943.1439     0   1   2   3E+3   4.5   5  RT (min)      lcqtof_hilic_neg_4_022_34    m/z 943.2222     0   1   2E+3   6   6.5  RT (min)      lcqtof_hilic_neg_4_033_53    m/z 943.4753     0   1E+3   1   1.5  RT (min)      lcqtof_hilic_neg_1_035_23    m/z 943.5695     0   2   4E+3   4   4.5  RT (min)      lcqtof_hilic_neg_4_032_01    m/z 943.6448     0   1   2   3E+3   4   4.5  RT (min)      lcqtof_hilic_neg_4_048_23    m/z 943.8368     0   2   4   6   8E+3   5   5.5  RT (min)      lcqtof_hilic_neg_4_053_14    m/z 943.9907     0   2   4   6E+3   0.5   1  RT (min)      lcqtof_hilic_neg_4_014_46    m/z 944.1737     0   2   4   6E+3   5.5   6  RT (min)      lcqtof_hilic_neg_4_013_20    m/z 944.3584     0   1E+3   5   5.5  RT (min)      lcqtof_hilic_neg_1_025_52    m/z 945.7941     0   2   4   6E+3   0.5   1  RT (min)      lcqtof_hilic_neg_1_022_36    m/z 945.8953     0   2   4   6   8E+3   5   5.5  RT (min)       lcqtof_hilic_neg_4_048_23    m/z 946.4728     0   1   2E+3   0.5   1  RT (min)      lcqtof_hilic_neg_1_035_23    m/z 946.5397     0   2   4   6   8E+3   1   1.5  RT (min)      lcqtof_hilic_neg_4_048_23    m/z 946.6726     0   1   2E+3   4   4.5  RT (min)      lcqtof_hilic_neg_2_047_32    m/z 946.7876     0   2   4E+3   0.5   1  RT (min)      lcqtof_hilic_neg_4_016_37    m/z 946.9228     0   1E+3   6   6.5  RT (min)      lcqtof_hilic_neg_1_028_47    m/z 947.4589     0   1   2E+3   4   4.5  RT (min)      lcqtof_hilic_neg_2_044_37    m/z 947.4632     0   2   4   6E+3   0.5   1  RT (min)      lcqtof_hilic_neg_4_047_06    m/z 947.5200     0   1   2   3E+3   4   4.5  RT (min)      lcqtof_hilic_neg_2_031_01    m/z 947.5422     0   1   2   3   4E+3   1   1.5  RT (min)      lcqtof_hilic_neg_4_048_23    m/z 947.5604     0   1   2   3   4E+3   4   4.5  RT (min)      lcqtof_hilic_neg_2_047_32    m/z 947.5864     0   2   4E+3   1   1.5  RT (min)      lcqtof_hilic_neg_2_051_14    m/z 947.6865     0   1   2E+3   3   3.5  RT (min)      lcqtof_hilic_neg_4_032_01    m/z 949.0000     0   1   2   3E+3   0.5   1  RT (min)      lcqtof_hilic_neg_2_022_46    m/z 949.1268     0   2   4   6   8E+3   0.5   1  RT (min)      lcqtof_hilic_neg_4_004_47    m/z 949.5221     0   1   2   3   4E+3   1   1.5  RT (min)      lcqtof_hilic_neg_4_023_15    m/z 949.5803     0   1   2   3E+3   4   4.5  RT (min)      lcqtof_hilic_neg_1_005_32    m/z 949.6007     0   1E+3   15.5   16  RT (min)      lcqtof_hilic_neg_1_005_32    m/z 950.5041     0   1   2   3   4E+3   4.5   5  RT (min)      lcqtof_hilic_neg_4_048_23    m/z 950.5086     0   2   4   6   8E+3   4.5   5  RT (min)      lcqtof_hilic_neg_4_050_02    m/z 950.5184     0   1   2   3E+3   1   1.5  RT (min)      lcqtof_hilic_neg_4_023_15    m/z 950.8820     0   2   4   6E+3   0.5   1  RT (min)      lcqtof_hilic_neg_1_014_31    m/z 950.9148     0   2   4   6E+3   5   5.5  RT (min)      lcqtof_hilic_neg_4_008_24    m/z 951.4979     0   1   2E+3   4.5   5  RT (min)      lcqtof_hilic_neg_3_054_37    m/z 951.5463     0   2   4   6E+3   4   4.5  RT (min)       lcqtof_hilic_neg_4_040_05    m/z 951.6781     0   1   2E+3   4   4.5  RT (min)      lcqtof_hilic_neg_4_016_37    m/z 952.5703     0   1   2   3   4E+3   3.5   4  RT (min)      lcqtof_hilic_neg_2_028_05    m/z 952.5980     0   1   2   3   4E+3   3.5   4  RT (min)      lcqtof_hilic_neg_4_011_35    m/z 952.6229     0   1   2   3   4E+3   1   1.5  RT (min)      lcqtof_hilic_neg_1_008_13    m/z 953.7954     0   1   2   3   4E+3   0.5   1  RT (min)      lcqtof_hilic_neg_3_075_58    m/z 953.8038     0   1   2   3E+3   4.5   5  RT (min)      lcqtof_hilic_neg_1_005_32    m/z 953.8093     0   1   2E+3   1   1.5  RT (min)      lcqtof_hilic_neg_2_033_53    m/z 954.5953     0   1   2   3   4E+3   3.5   4  RT (min)      lcqtof_hilic_neg_2_002_36    m/z 955.3580     0   2   4   6   8E+3   0.5   1  RT (min)      lcqtof_hilic_neg_4_027_19    m/z 955.5493     0   2   4   6E+3   3   3.5  RT (min)      lcqtof_hilic_neg_1_023_34    m/z 955.5752     0   2   4E+3   4   4.5  RT (min)      lcqtof_hilic_neg_4_052_13    m/z 955.5986     0   1E+3   7   7.5  RT (min)      lcqtof_hilic_neg_1_041_54    m/z 956.2318     0   1   2   3   4E+3   0.5   1  RT (min)      lcqtof_hilic_neg_4_040_05    m/z 956.5604     0   2   4   6   8E+3   3   3.5  RT (min)      lcqtof_hilic_neg_1_023_34    m/z 956.5989     0   2   4   6E+3   3.5   4  RT (min)      lcqtof_hilic_neg_2_051_14    m/z 956.6703     0   2   4   6E+3   5   5.5  RT (min)      lcqtof_hilic_neg_4_048_23    m/z 956.7949     0   2   4   6E+3   0.5   1  RT (min)      lcqtof_hilic_neg_4_031_38    m/z 957.6495     0   1   2   3   4E+3   2   2.5  RT (min)      lcqtof_hilic_neg_4_001_49    m/z 957.6714     0   1   2   3E+3   5   5.5  RT (min)      lcqtof_hilic_neg_4_048_23    m/z 957.8014     0   2   4   6E+3   0.5   1  RT (min)      lcqtof_hilic_neg_4_104_61    m/z 957.8266     0   1   2   3E+3   4.5   5  RT (min)      lcqtof_hilic_neg_2_047_32    m/z 958.6024     0   1   2   3E+3   5   5.5  RT (min)       lcqtof_hilic_neg_4_048_23    m/z 958.6168     0   1   2   3E+3   2   2.5  RT (min)      lcqtof_hilic_neg_4_023_15    m/z 958.8121     0   1   2E+3   5   5.5  RT (min)      lcqtof_hilic_neg_4_020_42    m/z 959.4998     0   1   2   3E+3   4.5   5  RT (min)      lcqtof_hilic_neg_4_011_35    m/z 959.6095     0   1   2E+3   1   1.5  RT (min)      lcqtof_hilic_neg_2_036_34    m/z 959.6370     0   1   2   3   4E+3   4   4.5  RT (min)      lcqtof_hilic_neg_4_052_13    m/z 959.6456     0   2   4E+3   0.5   1  RT (min)      lcqtof_hilic_neg_4_052_13    m/z 959.7523     0   1   2   3   4E+3   0.5   1  RT (min)      lcqtof_hilic_neg_4_053_14    m/z 959.7771     0   2   4E+3   2   2.5  RT (min)      lcqtof_hilic_neg_1_005_32    m/z 960.6140     0   1   2   3   4E+3   3   3.5  RT (min)      lcqtof_hilic_neg_2_044_37    m/z 961.5352     0   1E+4   1   1.5  RT (min)      lcqtof_hilic_neg_1_013_38    m/z 961.5770     0   2   4E+3   4   4.5  RT (min)      lcqtof_hilic_neg_2_047_32    m/z 961.6148     0   1   2   3E+3   0.5   1  RT (min)      lcqtof_hilic_neg_4_004_47    m/z 961.6192     0   1E+3   7   7.5  RT (min)      lcqtof_hilic_neg_4_035_33    m/z 961.6503     0   1   2   3   4E+3   4   4.5  RT (min)      lcqtof_hilic_neg_4_053_14    m/z 961.6910     0   1   2   3   4E+3   1   1.5  RT (min)      lcqtof_hilic_neg_4_004_47    m/z 961.7325     0   2   4   6   8E+3   3   3.5  RT (min)      lcqtof_hilic_neg_1_037_24    m/z 962.6003     0   1   2   3E+3   1   1.5  RT (min)      lcqtof_hilic_neg_4_047_06    m/z 963.2877     0   1   2E+3   0.5   1  RT (min)      lcqtof_hilic_neg_1_046_30    m/z 963.5880     0   1   2   3E+3   4   4.5  RT (min)       lcqtof_hilic_neg_2_047_32    m/z 963.9208     0   1   2   3E+3   0.5   1  RT (min)      lcqtof_hilic_neg_2_034_47    m/z 964.2844     0   1   2   3E+3   6   6.5  RT (min)      lcqtof_hilic_neg_4_052_13    m/z 964.5321     0   1   2   3E+3   4.5   5  RT (min)      lcqtof_hilic_neg_4_022_34    m/z 964.5527     0   1E+3   4.5   5  RT (min)       lcqtof_hilic_neg_1_010_15    m/z 964.5555     0   2   4   6   8E+3   1   1.5  RT (min)      lcqtof_hilic_neg_1_013_38    m/z 964.8961     0   2   4   6   8E+3   0.5   1  RT (min)      lcqtof_hilic_neg_2_029_31    m/z 965.1921     0   2   4   6E+3   6   6.5  RT (min)      lcqtof_hilic_neg_4_050_02    m/z 965.3621     0   1   2E+3   5   5.5  RT (min)      lcqtof_hilic_neg_4_005_29    m/z 965.4128     0   1   2E+3   4   4.5  RT (min)      lcqtof_hilic_neg_2_013_29    m/z 965.5114     0   1   2   3E+3   4.5   5  RT (min)      lcqtof_hilic_neg_1_022_36    m/z 965.6918     0   1   2E+3   3   3.5  RT (min)       lcqtof_hilic_neg_2_028_05    m/z 965.9028     0   1   2E+3   5   5.5  RT (min)      lcqtof_hilic_neg_4_001_49    m/z 966.2850     0   2   4   6E+3   0.5   1  RT (min)      lcqtof_hilic_neg_4_023_15    m/z 966.5582     0   2   4   6   8E+3   1   1.5  RT (min)      lcqtof_hilic_neg_1_013_38    m/z 966.6982     0   2   4   6E+3   5   5.5  RT (min)       lcqtof_hilic_neg_4_048_23    m/z 967.3408     0   2   4   6E+3   5   5.5  RT (min)      lcqtof_hilic_neg_4_032_01    m/z 967.4122     0   1   2E+3   4   4.5  RT (min)      lcqtof_hilic_neg_2_043_15    m/z 967.7059     0   2   4   6E+3   4.5   5  RT (min)      lcqtof_hilic_neg_4_048_23    m/z 967.7059     0   1   2   3   4E+3   0.5   1  RT (min)      lcqtof_hilic_neg_4_023_15    m/z 968.3233     0   1   2   3   4E+3   1   1.5  RT (min)      lcqtof_hilic_neg_4_027_19    m/z 968.5314     0   2   4   6   8E+3   7   7.5  RT (min)      lcqtof_hilic_neg_4_048_23    m/z 968.5845     0   1   2   3   4E+3   4   4.5  RT (min)      lcqtof_hilic_neg_2_005_13    m/z 968.5934     0   2   4   6   8E+3   1   1.5  RT (min)      lcqtof_hilic_neg_4_011_35    m/z 968.7915     0   1   2   3   4E+3   4   4.5  RT (min)      lcqtof_hilic_neg_2_052_24    m/z 968.8475     0   1   2   3E+3   5   5.5  RT (min)      lcqtof_hilic_neg_4_052_13    m/z 969.0457     0   2   4   6   8E+3   0.5   1  RT (min)      lcqtof_hilic_neg_4_026_04    m/z 969.0728     0   1   2   3E+3   4.5   5  RT (min)      lcqtof_hilic_neg_4_053_14    m/z 969.3534     0   1   2   3   4E+3   4   4.5  RT (min)      lcqtof_hilic_neg_1_020_09    m/z 969.5624     0   2   4E+3   1   1.5  RT (min)      lcqtof_hilic_neg_2_031_01    m/z 969.5943     0   2   4   6E+3   4   4.5  RT (min)      lcqtof_hilic_neg_2_031_01    m/z 970.0921     0   2   4E+3   6   6.5  RT (min)      lcqtof_hilic_neg_4_045_26    m/z 970.4959     0   1   2   3E+3   1   1.5  RT (min)      lcqtof_hilic_neg_4_048_23    m/z 970.5692     0   1E+4   1   1.5  RT (min)      lcqtof_hilic_neg_2_042_27    m/z 970.8083     0   1   2   3E+3   5   5.5  RT (min)       lcqtof_hilic_neg_4_024_40    m/z 971.0709     0   2   4E+3   4.5   5  RT (min)      lcqtof_hilic_neg_4_053_14    m/z 971.0793     0   1   2E+3   4.5   5  RT (min)      lcqtof_hilic_neg_4_006_30    m/z 971.3134     0   1   2E+3   5   5.5  RT (min)      lcqtof_hilic_neg_4_019_03    m/z 971.5798     0   2   4   6   8E+3   1   1.5  RT (min)      lcqtof_hilic_neg_2_042_27    m/z 971.7058     0   1   2   3   4E+3   3   3.5  RT (min)      lcqtof_hilic_neg_1_037_24    m/z 972.3493     0   2   4   6   8E+3   0.5   1  RT (min)      lcqtof_hilic_neg_4_027_19    m/z 972.5080     0   1E+3   4   4.5  RT (min)      lcqtof_hilic_neg_1_054_01    m/z 972.5176     0   1   2   3   4E+3   1   1.5  RT (min)      lcqtof_hilic_neg_4_032_01    m/z 972.5684     0   2   4   6E+3   4   4.5  RT (min)      lcqtof_hilic_neg_2_047_32    m/z 972.5964     0   1   2   3E+3   1   1.5  RT (min)      lcqtof_hilic_neg_1_010_15    m/z 972.6202     0   2   4   6E+3   1.5   2  RT (min)      lcqtof_hilic_neg_4_005_29    m/z 972.6638     0   1   2   3E+4   0.5   1  RT (min)      lcqtof_hilic_neg_1_020_09    m/z 972.7122     0   2   4   6   8E+3   3   3.5  RT (min)      lcqtof_hilic_neg_1_037_24    m/z 972.7214     0   2   4E+3   1   1.5  RT (min)      lcqtof_hilic_neg_4_008_24    m/z 974.1412     0   1   2   3   4E+3   7   7.5  RT (min)      lcqtof_hilic_neg_1_010_15    m/z 974.3890     0   1   2E+3   0.5   1  RT (min)      lcqtof_hilic_neg_2_031_01    m/z 974.5407     0   2   4   6E+3   1   1.5  RT (min)      lcqtof_hilic_neg_4_032_01    m/z 974.5739     0   1   2   3   4E+3   4.5   5  RT (min)       lcqtof_hilic_neg_4_053_14    m/z 974.6052     0   2   4   6   8E+3   1   1.5  RT (min)      lcqtof_hilic_neg_4_045_26    m/z 974.7366     0   2   4   6   8E+3   1   1.5  RT (min)      lcqtof_hilic_neg_4_008_24    m/z 975.1840     0   1   2   3   4E+3   0.5   1  RT (min)      lcqtof_hilic_neg_4_017_51    m/z 975.3775     0   1   2E+3   0.5   1  RT (min)      lcqtof_hilic_neg_2_055_30    m/z 975.4734     0   1   2   3   4E+3   4   4.5  RT (min)      lcqtof_hilic_neg_2_051_14    m/z 975.4976     0   1   2   3E+3   5   5.5  RT (min)      lcqtof_hilic_neg_4_053_14    m/z 975.5328     0   1   2   3E+3   1   1.5  RT (min)      lcqtof_hilic_neg_4_023_15    m/z 975.5425     0   2   4E+3   4   4.5  RT (min)      lcqtof_hilic_neg_4_052_13    m/z 975.6154     0   1   2   3   4E+3   1   1.5  RT (min)      lcqtof_hilic_neg_4_053_14    m/z 975.6154     0   1   2   3   4E+3   1   1.5  RT (min)      lcqtof_hilic_neg_1_010_15    m/z 976.4761     0   1   2E+3   4   4.5  RT (min)      lcqtof_hilic_neg_2_051_14    m/z 976.5455     0   1   2   3E+3   1   1.5  RT (min)      lcqtof_hilic_neg_4_023_15    m/z 976.7846     0   2   4   6E+3   1   1.5  RT (min)      lcqtof_hilic_neg_1_005_32    m/z 977.6147     0   2   4   6E+3   0.5   1  RT (min)      lcqtof_hilic_neg_4_005_29    m/z 977.6490     0   1   2   3   4E+3   4   4.5  RT (min)      lcqtof_hilic_neg_4_053_14    m/z 977.6859     0   1   2   3   4E+3   3   3.5  RT (min)      lcqtof_hilic_neg_3_024_35    m/z 977.7380     0   1E+3   0.5   1  RT (min)      lcqtof_hilic_neg_4_048_23    m/z 977.8319     0   2   4   6E+3   5   5.5  RT (min)       lcqtof_hilic_neg_4_032_01    m/z 978.5804     0   1   2E+3   3.5   4  RT (min)      lcqtof_hilic_neg_4_023_15    m/z 978.6406     0   2   4   6   8E+3   0.5   1  RT (min)      lcqtof_hilic_neg_4_018_27    m/z 978.6671     0   2   4E+3   3   3.5  RT (min)      lcqtof_hilic_neg_2_002_36    m/z 979.0107     0   1   2   3E+3   6   6.5  RT (min)      lcqtof_hilic_neg_1_023_34    m/z 979.5647     0   1   2   3E+3   7   7.5  RT (min)      lcqtof_hilic_neg_4_004_47    m/z 979.5921     0   1   2   3E+3   2   2.5  RT (min)      lcqtof_hilic_neg_2_031_01    m/z 979.6462     0   1   2E+3   1   1.5  RT (min)      lcqtof_hilic_neg_4_053_14    m/z 979.6802     0   1   2   3   4E+3   3   3.5  RT (min)      lcqtof_hilic_neg_2_002_36    m/z 980.5769     0   1   2   3E+3   4   4.5  RT (min)      lcqtof_hilic_neg_4_052_13    m/z 981.2612     0   1   2E+3   8   8.5  RT (min)      lcqtof_hilic_neg_2_009_50    m/z 981.4466     0   1   2E+3   4   4.5  RT (min)      lcqtof_hilic_neg_1_054_01    m/z 981.5321     0   1   2   3   4E+3   4.5   5  RT (min)      lcqtof_hilic_neg_2_002_36    m/z 981.7109     0   1   2E+3   1   1.5  RT (min)      lcqtof_hilic_neg_4_004_47    m/z 982.8311     0   1   2   3E+3   4.5   5  RT (min)      lcqtof_hilic_neg_2_052_24    m/z 983.5571     0   2   4   6E+3   0.5   1  RT (min)      lcqtof_hilic_neg_1_032_18    m/z 983.6041     0   1   2   3   4E+3   4   4.5  RT (min)      lcqtof_hilic_neg_4_018_27    m/z 983.6276     0   1   2   3E+3   2   2.5  RT (min)      lcqtof_hilic_neg_2_031_01    m/z 983.6457     0   1   2E+3   5   5.5  RT (min)      lcqtof_hilic_neg_4_055_32    m/z 983.6878     0   2   4   6E+3   5   5.5  RT (min)       lcqtof_hilic_neg_4_048_23    m/z 984.3589     0   2   4   6E+3   6   6.5  RT (min)      lcqtof_hilic_neg_4_052_13    m/z 984.5349     0   1   2   3E+3   1   1.5  RT (min)      lcqtof_hilic_neg_4_032_01    m/z 984.5707     0   1   2E+3   4   4.5  RT (min)      lcqtof_hilic_neg_2_031_01    m/z 984.5944     0   2   4   6   8E+3   0.5   1  RT (min)      lcqtof_hilic_neg_1_013_38    m/z 984.6879     0   1   2   3   4E+3   5   5.5  RT (min)      lcqtof_hilic_neg_4_048_23    m/z 984.7596     0   2   4E+3   0.5   1  RT (min)      lcqtof_hilic_neg_4_053_14    m/z 985.1708     0   1   2   3   4E+3   7   7.5  RT (min)      lcqtof_hilic_neg_2_031_01    m/z 985.3151     0   2   4   6   8E+3   6   6.5  RT (min)       lcqtof_hilic_neg_2_005_13    m/z 985.5976     0   2   4E+3   0.5   1  RT (min)      lcqtof_hilic_neg_1_013_38    m/z 985.6642     0   1   2   3   4E+3   5   5.5  RT (min)      lcqtof_hilic_neg_4_055_32    m/z 985.8351     0   1E+4   0.5   1  RT (min)      lcqtof_hilic_neg_4_004_47    m/z 986.5253     0   1   2E+3   4.5   5  RT (min)      lcqtof_hilic_neg_3_030_23    m/z 986.6270     0   1   2   3E+3   4   4.5  RT (min)      lcqtof_hilic_neg_1_029_06    m/z 986.7680     0   1   2   3E+3   0.5   1  RT (min)      lcqtof_hilic_neg_4_004_47    m/z 986.8516     0   1E+4   0.5   1  RT (min)      lcqtof_hilic_neg_4_004_47    m/z 986.9065     0   1   2   3   4E+3   6   6.5  RT (min)       lcqtof_hilic_neg_4_026_04    m/z 987.5682     0   1   2   3   4E+3   1   1.5  RT (min)      lcqtof_hilic_neg_2_042_27    m/z 987.6161     0   2   4   6   8E+3   0.5   1  RT (min)      lcqtof_hilic_neg_1_013_38    m/z 987.6693     0   2   4   6   8E+3   0.5   1  RT (min)      lcqtof_hilic_neg_4_004_47    m/z 987.7629     0   1   2   3E+3   1.5   2  RT (min)      lcqtof_hilic_neg_4_054_48    m/z 987.8253     0   2   4   6   8E+3   0.5   1  RT (min)       lcqtof_hilic_neg_4_052_13    m/z 988.0318     0   2   4   6E+3   0.5   1  RT (min)      lcqtof_hilic_neg_4_014_46    m/z 988.4337     0   1   2   3E+3   1   1.5  RT (min)      lcqtof_hilic_neg_1_036_03    m/z 988.7145     0   2   4   6   8E+3   1   1.5  RT (min)      lcqtof_hilic_neg_2_047_32    m/z 989.0356     0   1   2   3   4E+3   0.5   1  RT (min)      lcqtof_hilic_neg_4_014_46    m/z 989.5454     0   2   4E+3   1   1.5  RT (min)      lcqtof_hilic_neg_4_005_29    m/z 989.5495     0   2   4   6E+3   1   1.5  RT (min)      lcqtof_hilic_neg_2_048_35    m/z 989.6649     0   1E+3   1   1.5  RT (min)      lcqtof_hilic_neg_2_031_01    m/z 990.0620     0   1   2   3   4E+3   6   6.5  RT (min)       lcqtof_hilic_neg_4_045_26    m/z 990.5116     0   2   4E+3   7   7.5  RT (min)      lcqtof_hilic_neg_4_048_23    m/z 991.3542     0   1   2E+3   4   4.5  RT (min)      lcqtof_hilic_neg_1_040_20    m/z 991.5215     0   1   2   3E+3   4   4.5  RT (min)      lcqtof_hilic_neg_2_005_13    m/z 991.5837     0   2   4   6   8E+3   1   1.5  RT (min)      lcqtof_hilic_neg_1_027_27    m/z 991.6746     0   2   4   6E+3   3.5   4  RT (min)      lcqtof_hilic_neg_4_012_07    m/z 991.9177     0   2   4   6E+3   0.5   1  RT (min)      lcqtof_hilic_neg_4_034_31    m/z 992.2405     0   1E+3   8   8.5  RT (min)      lcqtof_hilic_neg_2_016_33    m/z 992.6582     0   1   2   3E+3   1   1.5  RT (min)      lcqtof_hilic_neg_2_005_13    m/z 992.9174     0   1   2   3   4E+3   0.5   1  RT (min)      lcqtof_hilic_neg_4_034_31    m/z 993.1754     0   2   4   6E+3   0.5   1  RT (min)      lcqtof_hilic_neg_4_004_47    m/z 993.4369     0   1   2   3E+3   4   4.5  RT (min)      lcqtof_hilic_neg_1_010_15    m/z 993.5629     0   1   2   3   4E+3   1   1.5  RT (min)      lcqtof_hilic_neg_2_039_38    m/z 994.1721     0   2   4E+3   0.5   1  RT (min)      lcqtof_hilic_neg_4_004_47    m/z 994.6397     0   1   2   3E+3   4.5   5  RT (min)      lcqtof_hilic_neg_4_008_24    m/z 994.6491     0   1   2   3   4E+3   1.5   2  RT (min)      lcqtof_hilic_neg_4_034_31    m/z 994.6704     0   1   2E+3   1   1.5  RT (min)      lcqtof_hilic_neg_2_005_13    m/z 995.0082     0   1   2   3E+3   5.5   6  RT (min)      lcqtof_hilic_neg_4_032_01    m/z 995.1257     0   1   2   3E+3   7   7.5  RT (min)      lcqtof_hilic_neg_1_027_27    m/z 995.1723     0   1   2   3   4E+3   0.5   1  RT (min)      lcqtof_hilic_neg_4_004_47    m/z 995.2871     0   2   4   6   8E+3   1   1.5  RT (min)      lcqtof_hilic_neg_4_026_04    m/z 995.4368     0   1   2E+3   4   4.5  RT (min)      lcqtof_hilic_neg_1_004_14    m/z 995.5795     0   1   2E+3   1   1.5  RT (min)      lcqtof_hilic_neg_1_013_38    m/z 995.8087     0   2   4   6E+3   4   4.5  RT (min)      lcqtof_hilic_neg_4_008_24    m/z 995.9136     0   1   2   3E+3   5   5.5  RT (min)       lcqtof_hilic_neg_4_079_58    m/z 996.6582     0   1   2   3   4E+3   1   1.5  RT (min)      lcqtof_hilic_neg_1_054_01    m/z 997.4745     0   1   2E+3   4.5   5  RT (min)      lcqtof_hilic_neg_4_011_35    m/z 997.6115     0   1   2   3   4E+3   1   1.5  RT (min)      lcqtof_hilic_neg_2_036_34    m/z 997.7004     0   1   2E+3   3   3.5  RT (min)      lcqtof_hilic_neg_1_025_52    m/z 997.8250     0   2   4   6   8E+3   4   4.5  RT (min)      lcqtof_hilic_neg_4_008_24    m/z 998.1603     0   2   4E+3   5   5.5  RT (min)      lcqtof_hilic_neg_4_032_01    m/z 998.2311     0   1E+3   8   8.5  RT (min)      lcqtof_hilic_neg_1_029_06    m/z 998.3350     0   1   2   3   4E+3   1   1.5  RT (min)      lcqtof_hilic_neg_4_007_45    m/z 998.5488     0   1   2E+3   1   1.5  RT (min)      lcqtof_hilic_neg_4_023_15    m/z 998.5652     0   1   2E+3   4   4.5  RT (min)      lcqtof_hilic_neg_2_047_32    m/z 998.6924     0   1E+3   3.5   4  RT (min)      lcqtof_hilic_neg_2_042_27    m/z 998.7870     0   2   4   6E+3   2   2.5  RT (min)      lcqtof_hilic_neg_4_001_49    m/z 999.1987     0   1   2   3   4E+3   4.5   5  RT (min)      lcqtof_hilic_neg_4_035_33    m/z 999.6488     0   2   4   6E+3   1.5   2  RT (min)      lcqtof_hilic_neg_4_005_29    m/z 999.6786     0   1   2E+3   2   2.5  RT (min)      lcqtof_hilic_neg_1_035_23    m/z 999.7047     0   1   2   3   4E+3   4.5   5  RT (min)      lcqtof_hilic_neg_4_035_33    m/z 999.7120     0   1E+3   3.5   4  RT (min)      lcqtof_hilic_neg_4_032_01    m/z 1000.1201     0   1   2   3E+3   5.5   6  RT (min)      lcqtof_hilic_neg_4_023_15    m/z 1000.3635     0   1   2   3   4E+3   6   6.5  RT (min)      lcqtof_hilic_neg_4_012_07    m/z 1000.6450     0   1   2   3E+3   4   4.5  RT (min)      lcqtof_hilic_neg_4_050_02    m/z 1000.7399     0   2   4   6   8E+3   0.5   1  RT (min)      lcqtof_hilic_neg_4_004_47    m/z 1000.7919     0   1E+4   0.5   1  RT (min)      lcqtof_hilic_neg_4_004_47    m/z 1000.8892     0   2   4   6E+3   0.5   1  RT (min)      lcqtof_hilic_neg_4_004_47    m/z 1001.0326     0   2   4   6E+3   0.5   1  RT (min)      lcqtof_hilic_neg_3_036_17    m/z 1001.0345     0   1   2   3E+3   5.5   6  RT (min)      lcqtof_hilic_neg_4_045_26    m/z 1001.3457     0   1E+3   5   5.5  RT (min)      lcqtof_hilic_neg_2_026_28    m/z 1001.4730     0   1   2E+3   4   4.5  RT (min)      lcqtof_hilic_neg_3_040_14    m/z 1001.5913     0   2   4   6   8E+3   0.5   1  RT (min)      lcqtof_hilic_neg_4_013_20    m/z 1001.6311     0   2   4E+3   0.5   1  RT (min)      lcqtof_hilic_neg_2_042_27    m/z 1001.7482     0   2   4   6E+3   0.5   1  RT (min)      lcqtof_hilic_neg_4_026_04    m/z 1001.9352     0   1   2   3   4E+3   5   5.5  RT (min)       lcqtof_hilic_neg_4_019_03    m/z 1001.9380     0   1   2   3   4E+3   5.5   6  RT (min)       lcqtof_hilic_neg_4_019_03    m/z 1002.2928     0   1E+3   0.5   1  RT (min)      lcqtof_hilic_neg_1_047_29    m/z 1002.4816     0   1   2E+3   4   4.5  RT (min)      lcqtof_hilic_neg_3_040_14    m/z 1002.5005     0   1   2E+3   1   1.5  RT (min)      lcqtof_hilic_neg_4_006_30    m/z 1002.7552     0   2   4   6   8E+3   0.5   1  RT (min)      lcqtof_hilic_neg_4_099_61    m/z 1002.9924     0   1   2   3E+3   5   5.5  RT (min)       lcqtof_hilic_neg_4_017_51    m/z 1003.0705     0   2   4   6   8E+3   7   7.5  RT (min)      lcqtof_hilic_neg_1_004_14    m/z 1003.4545     0   1   2E+3   5   5.5  RT (min)      lcqtof_hilic_neg_4_011_35    m/z 1003.6151     0   2   4E+3   5   5.5  RT (min)      lcqtof_hilic_neg_4_048_23    m/z 1003.6473     0   1   2   3E+3   0.5   1  RT (min)      lcqtof_hilic_neg_4_018_27    m/z 1003.6477     0   1   2E+3   0.5   1  RT (min)      lcqtof_hilic_neg_2_032_02    m/z 1003.6667     0   1E+3   1   1.5  RT (min)      lcqtof_hilic_neg_2_033_53    m/z 1003.6842     0   1   2   3E+3   4.5   5  RT (min)      lcqtof_hilic_neg_4_022_34    m/z 1003.6869     0   1   2   3E+3   3   3.5  RT (min)      lcqtof_hilic_neg_2_004_26    m/z 1003.9590     0   2   4   6   8E+3   0.5   1  RT (min)      lcqtof_hilic_neg_4_004_47    m/z 1004.6765     0   1   2E+3   3.5   4  RT (min)      lcqtof_hilic_neg_4_023_15    m/z 1004.9698     0   1   2E+3   6   6.5  RT (min)      lcqtof_hilic_neg_3_055_01    m/z 1005.1856     0   1   2E+3   6   6.5  RT (min)       lcqtof_hilic_neg_4_010_28    m/z 1005.3127     0   1   2E+3   5   5.5  RT (min)       lcqtof_hilic_neg_4_020_42    m/z 1005.3511     0   1   2   3E+3   5   5.5  RT (min)      lcqtof_hilic_neg_4_028_36    m/z 1005.6940     0   1   2   3E+3   1   1.5  RT (min)      lcqtof_hilic_neg_2_041_54    m/z 1005.7368     0   1   2   3   4E+3   0.5   1  RT (min)      lcqtof_hilic_neg_4_053_14    m/z 1006.1454     0   1   2   3E+3   5.5   6  RT (min)      lcqtof_hilic_neg_4_026_04    m/z 1006.6893     0   1   2E+3   3.5   4  RT (min)      lcqtof_hilic_neg_4_005_29    m/z 1006.7160     0   1E+3   1   1.5  RT (min)      lcqtof_hilic_neg_3_048_05    m/z 1007.0507     0   1   2   3   4E+3   5.5   6  RT (min)       lcqtof_hilic_neg_4_012_07    m/z 1007.7054     0   1   2   3   4E+3   1   1.5  RT (min)      lcqtof_hilic_neg_4_056_56    m/z 1007.7325     0   2   4E+3   0.5   1  RT (min)      lcqtof_hilic_neg_3_030_23    m/z 1007.7349     0   2   4   6E+3   0.5   1  RT (min)      lcqtof_hilic_neg_4_048_23    m/z 1007.9639     0   1   2   3   4E+3   0.5   1  RT (min)      lcqtof_hilic_neg_4_075_58    m/z 1008.2707     0   1   2E+3   8   8.5  RT (min)      lcqtof_hilic_neg_1_037_24    m/z 1008.6539     0   2   4   6E+3   5   5.5  RT (min)       lcqtof_hilic_neg_4_095_60    m/z 1008.9671     0   1   2   3   4E+3   0.5   1  RT (min)      lcqtof_hilic_neg_3_080_58    m/z 1009.2262     0   1E+3   8   8.5  RT (min)      lcqtof_hilic_neg_1_055_25    m/z 1009.4736     0   1   2E+3   4   4.5  RT (min)      lcqtof_hilic_neg_2_031_01    m/z 1009.6949     0   1   2   3E+3   1   1.5  RT (min)      lcqtof_hilic_neg_1_005_32    m/z 1009.6982     0   2   4E+3   4.5   5  RT (min)       lcqtof_hilic_neg_4_047_06    m/z 1010.4796     0   1   2E+3   4   4.5  RT (min)      lcqtof_hilic_neg_2_031_01    m/z 1011.0537     0   1   2   3E+3   4.5   5  RT (min)      lcqtof_hilic_neg_4_053_14    m/z 1011.6453     0   2   4   6E+3   4   4.5  RT (min)      lcqtof_hilic_neg_2_051_14    m/z 1011.6931     0   1   2E+3   0.5   1  RT (min)      lcqtof_hilic_neg_2_043_15    m/z 1011.7025     0   1   2   3   4E+3   4.5   5  RT (min)      lcqtof_hilic_neg_4_047_06    m/z 1011.8253     0   2   4E+3   5   5.5  RT (min)      lcqtof_hilic_neg_4_032_01    m/z 1012.1593     0   2   4   6E+3   5.5   6  RT (min)      lcqtof_hilic_neg_4_013_20    m/z 1012.4883     0   1E+3   4   4.5  RT (min)      lcqtof_hilic_neg_1_054_01    m/z 1012.6413     0   1   2E+3   1   1.5  RT (min)      lcqtof_hilic_neg_4_008_24    m/z 1013.0515     0   1   2   3   4E+3   4.5   5  RT (min)      lcqtof_hilic_neg_4_053_14    m/z 1013.0522     0   1   2   3E+3   4.5   5  RT (min)      lcqtof_hilic_neg_4_008_24    m/z 1013.0717     0   1   2   3   4E+3   4.5   5  RT (min)      lcqtof_hilic_neg_4_001_49    m/z 1013.5091     0   1E+3   4   4.5  RT (min)      lcqtof_hilic_neg_2_031_01    m/z 1013.8819     0   2   4   6E+3   5   5.5  RT (min)      lcqtof_hilic_neg_4_048_23    m/z 1014.0517     0   1   2E+3   4.5   5  RT (min)      lcqtof_hilic_neg_4_053_14    m/z 1014.6476     0   1   2   3E+3   1   1.5  RT (min)      lcqtof_hilic_neg_4_008_24    m/z 1015.4661     0   1   2   3   4E+3   4.5   5  RT (min)      lcqtof_hilic_neg_4_028_36    m/z 1015.7122     0   1   2   3   4E+3   3.5   4  RT (min)      lcqtof_hilic_neg_4_053_14    m/z 1015.7831     0   2   4   6   8E+3   0.5   1  RT (min)      lcqtof_hilic_neg_4_004_47    m/z 1016.4381     0   1E+4   0.5   1  RT (min)      lcqtof_hilic_neg_4_018_27    m/z 1016.5885     0   2   4   6E+3   4   4.5  RT (min)       lcqtof_hilic_neg_4_048_23    m/z 1016.6659     0   1   2E+3   1   1.5  RT (min)      lcqtof_hilic_neg_4_008_24    m/z 1017.5022     0   2   4E+3   4.5   5  RT (min)      lcqtof_hilic_neg_3_030_23    m/z 1017.8336     0   2   4E+3   0.5   1  RT (min)      lcqtof_hilic_neg_1_013_38    m/z 1018.2075     0   2   4E+3   6   6.5  RT (min)      lcqtof_hilic_neg_2_028_05    m/z 1018.6314     0   1   2   3E+3   4.5   5  RT (min)      lcqtof_hilic_neg_4_008_24    m/z 1018.9414     0   2   4   6E+3   0.5   1  RT (min)      lcqtof_hilic_neg_4_055_32    m/z 1019.3296     0   2   4   6   8E+3   0.5   1  RT (min)      lcqtof_hilic_neg_4_013_20    m/z 1019.6373     0   1   2E+3   3   3.5  RT (min)      lcqtof_hilic_neg_1_054_01    m/z 1019.9852     0   1   2   3   4E+3   0.5   1  RT (min)      lcqtof_hilic_neg_4_007_45    m/z 1020.8248     0   2   4E+3   5   5.5  RT (min)      lcqtof_hilic_neg_4_053_14    m/z 1020.8553     0   1   2   3   4E+3   4.5   5  RT (min)      lcqtof_hilic_neg_4_008_24    m/z 1021.8696     0   2   4   6   8E+3   0.5   1  RT (min)      lcqtof_hilic_neg_4_016_37    m/z 1021.8967     0   1   2   3   4E+3   0.5   1  RT (min)      lcqtof_hilic_neg_2_034_47    m/z 1022.1732     0   1E+3   5   5.5  RT (min)      lcqtof_hilic_neg_4_006_30    m/z 1022.6080     0   1E+4   4   4.5  RT (min)      lcqtof_hilic_neg_2_048_35    m/z 1022.6693     0   2   4   6E+3   0.5   1  RT (min)      lcqtof_hilic_neg_1_023_34    m/z 1023.5406     0   1   2E+3   4   4.5  RT (min)      lcqtof_hilic_neg_2_010_45    m/z 1023.6508     0   1   2   3E+4   4.5   5  RT (min)      lcqtof_hilic_neg_2_047_32    m/z 1025.0406     0   1   2E+3   0.5   1  RT (min)      lcqtof_hilic_neg_2_047_32    m/z 1026.3027     0   1E+3   5   5.5  RT (min)      lcqtof_hilic_neg_4_106_62    m/z 1026.7979     0   2   4   6   8E+3   2   2.5  RT (min)       lcqtof_hilic_neg_4_008_24    m/z 1027.2509     0   2   4   6E+3   6   6.5  RT (min)      lcqtof_hilic_neg_4_023_15    m/z 1027.3611     0   2   4   6   8E+3   1.5   2  RT (min)       lcqtof_hilic_neg_4_021_21    m/z 1027.6479     0   2   4   6   8E+3   0.5   1  RT (min)      lcqtof_hilic_neg_4_022_34    m/z 1028.3656     0   2   4   6E+3   1.5   2  RT (min)      lcqtof_hilic_neg_4_021_21    m/z 1028.4552     0   2   4   6E+3   4.5   5  RT (min)      lcqtof_hilic_neg_4_055_32    m/z 1028.7440     0   2   4   6   8E+3   0.5   1  RT (min)      lcqtof_hilic_neg_4_022_34    m/z 1028.8281     0   1   2   3   4E+3   0.5   1  RT (min)      lcqtof_hilic_neg_1_003_37    m/z 1029.0203     0   2   4   6   8E+3   0.5   1  RT (min)      lcqtof_hilic_neg_3_036_17    m/z 1029.6256     0   1   2   3   4E+3   0.5   1  RT (min)      lcqtof_hilic_neg_1_032_18    m/z 1029.8313     0   1   2   3   4E+3   0.5   1  RT (min)      lcqtof_hilic_neg_1_030_17    m/z 1030.2487     0   1   2   3E+3   0.5   1  RT (min)      lcqtof_hilic_neg_4_040_05    m/z 1030.5996     0   1E+3   4.5   5  RT (min)      lcqtof_hilic_neg_4_019_03    m/z 1030.6395     0   2   4   6E+3   5   5.5  RT (min)       lcqtof_hilic_neg_4_048_23    m/z 1030.6830     0   1   2   3   4E+3   3.5   4  RT (min)      lcqtof_hilic_neg_2_042_27    m/z 1030.7690     0   1   2   3   4E+3   0.5   1  RT (min)      lcqtof_hilic_neg_1_049_55    m/z 1031.5406     0   1   2E+3   1   1.5  RT (min)      lcqtof_hilic_neg_4_005_29    m/z 1031.6549     0   1   2   3E+3   5   5.5  RT (min)      lcqtof_hilic_neg_4_048_23    m/z 1031.6909     0   1   2E+3   3.5   4  RT (min)      lcqtof_hilic_neg_1_027_27    m/z 1031.7096     0   1   2E+3   3.5   4  RT (min)      lcqtof_hilic_neg_4_053_14    m/z 1032.6410     0   1   2E+3   3   3.5  RT (min)      lcqtof_hilic_neg_2_031_01    m/z 1032.7629     0   1E+4   5   5.5  RT (min)       lcqtof_hilic_neg_4_008_24    m/z 1033.0986     0   2   4   6E+3   7.5   8  RT (min)      lcqtof_hilic_neg_2_002_36    m/z 1033.1489     0   1   2E+3   3   3.5  RT (min)      lcqtof_hilic_neg_2_031_01    m/z 1033.3555     0   1   2   3   4E+3   5   5.5  RT (min)      lcqtof_hilic_neg_4_032_01    m/z 1033.6498     0   1   2   3E+3   3   3.5  RT (min)      lcqtof_hilic_neg_2_005_13    m/z 1033.8990     0   1   2E+3   5   5.5  RT (min)      lcqtof_hilic_neg_3_040_14    m/z 1034.1506     0   1   2   3   4E+3   3   3.5  RT (min)      lcqtof_hilic_neg_2_021_17    m/z 1034.6564     0   1   2E+3   3   3.5  RT (min)      lcqtof_hilic_neg_1_054_01    m/z 1034.6931     0   2   4   6E+3   4.5   5  RT (min)      lcqtof_hilic_neg_2_052_24    m/z 1035.3324     0   2   4   6E+3   5   5.5  RT (min)      lcqtof_hilic_neg_4_053_14    m/z 1035.3427     0   1E+5   1.5   2  RT (min)      lcqtof_hilic_neg_4_013_20    m/z 1035.7612     0   1   2   3   4E+3   5   5.5  RT (min)      lcqtof_hilic_neg_1_003_37    m/z 1035.8207     0   2   4   6E+3   4   4.5  RT (min)      lcqtof_hilic_neg_4_053_14    m/z 1036.2426     0   1   2   3   4E+3   4   4.5  RT (min)      lcqtof_hilic_neg_4_053_14    m/z 1036.8269     0   1   2   3E+3   5   5.5  RT (min)      lcqtof_hilic_neg_4_052_13    m/z 1036.8651     0   1   2   3E+4   0.5   1  RT (min)      lcqtof_hilic_neg_4_022_34    m/z 1036.9948     0   1   2E+3   4   4.5  RT (min)      lcqtof_hilic_neg_4_053_14    m/z 1037.4791     0   2   4E+3   4.5   5  RT (min)      lcqtof_hilic_neg_1_022_36    m/z 1038.4800     0   1   2   3E+3   4.5   5  RT (min)      lcqtof_hilic_neg_1_022_36    m/z 1038.7238     0   2   4   6E+3   4.5   5  RT (min)      lcqtof_hilic_neg_2_052_24    m/z 1038.8353     0   2   4E+3   1   1.5  RT (min)      lcqtof_hilic_neg_4_004_47    m/z 1038.9185     0   1E+4   1   1.5  RT (min)      lcqtof_hilic_neg_2_052_24    m/z 1039.2942     0   1   2E+3   5   5.5  RT (min)      lcqtof_hilic_neg_4_106_62    m/z 1039.3015     0   1   2   3   4E+3   0.5   1  RT (min)      lcqtof_hilic_neg_4_010_28    m/z 1039.9141     0   2   4   6   8E+3   0.5   1  RT (min)      lcqtof_hilic_neg_4_008_24    m/z 1039.9174     0   2   4   6   8E+3   1   1.5  RT (min)      lcqtof_hilic_neg_4_008_24    m/z 1040.1864     0   1   2   3   4E+3   6   6.5  RT (min)      lcqtof_hilic_neg_2_003_25    m/z 1040.8123     0   1   2   3E+3   2   2.5  RT (min)      lcqtof_hilic_neg_1_005_32    m/z 1041.9298     0   2   4   6E+3   1   1.5  RT (min)      lcqtof_hilic_neg_4_008_24    m/z 1041.9373     0   2   4   6E+3   0.5   1  RT (min)      lcqtof_hilic_neg_4_008_24    m/z 1042.6211     0   2   4E+3   4.5   5  RT (min)      lcqtof_hilic_neg_4_040_05    m/z 1042.7462     0   1   2   3   4E+3   4.5   5  RT (min)      lcqtof_hilic_neg_4_055_32    m/z 1043.6357     0   1   2   3E+3   4.5   5  RT (min)      lcqtof_hilic_neg_4_008_24    m/z 1044.7382     0   2   4E+3   1   1.5  RT (min)      lcqtof_hilic_neg_2_036_34    m/z 1044.8129     0   1E+3   5   5.5  RT (min)      lcqtof_hilic_neg_1_047_29    m/z 1045.1559     0   1   2   3E+3   5   5.5  RT (min)      lcqtof_hilic_neg_4_032_01    m/z 1045.6486     0   1   2   3   4E+3   4.5   5  RT (min)      lcqtof_hilic_neg_2_047_32    m/z 1045.8015     0   2   4E+3   4   4.5  RT (min)      lcqtof_hilic_neg_2_047_32    m/z 1045.8195     0   2   4   6E+3   5   5.5  RT (min)      lcqtof_hilic_neg_4_032_01    m/z 1046.2894     0   1   2E+3   1   1.5  RT (min)      lcqtof_hilic_neg_2_013_29    m/z 1046.3811     0   2   4E+3   0.5   1  RT (min)      lcqtof_hilic_neg_4_013_20    m/z 1046.4999     0   1   2   3   4E+3   4.5   5  RT (min)      lcqtof_hilic_neg_4_050_02    m/z 1046.5983     0   2   4E+3   5   5.5  RT (min)       lcqtof_hilic_neg_4_008_24    m/z 1046.6631     0   1   2   3E+3   3   3.5  RT (min)      lcqtof_hilic_neg_2_005_13    m/z 1046.6666     0   2   4   6   8E+3   0.5   1  RT (min)      lcqtof_hilic_neg_1_023_34    m/z 1046.8379     0   2   4   6   8E+3   0.5   1  RT (min)      lcqtof_hilic_neg_4_004_47    m/z 1046.8446     0   2   4   6E+3   0.5   1  RT (min)      lcqtof_hilic_neg_1_003_37    m/z 1047.3860     0   1   2   3E+3   0.5   1  RT (min)      lcqtof_hilic_neg_4_013_20    m/z 1047.6635     0   1   2   3   4E+3   4.5   5  RT (min)      lcqtof_hilic_neg_4_055_32    m/z 1047.7903     0   2   4   6E+3   0.5   1  RT (min)      lcqtof_hilic_neg_1_031_16    m/z 1047.8474     0   1   2   3E+3   0.5   1  RT (min)      lcqtof_hilic_neg_2_029_31    m/z 1048.3877     0   1   2   3E+3   1   1.5  RT (min)      lcqtof_hilic_neg_4_027_19    m/z 1048.8591     0   2   4   6E+3   0.5   1  RT (min)      lcqtof_hilic_neg_4_011_35    m/z 1049.1980     0   1   2   3E+3   0.5   1  RT (min)      lcqtof_hilic_neg_4_040_05    m/z 1049.5142     0   1   2E+3   4.5   5  RT (min)      lcqtof_hilic_neg_4_003_16    m/z 1049.5665     0   1   2   3E+3   4   4.5  RT (min)      lcqtof_hilic_neg_1_031_16    m/z 1049.5692     0   2   4   6   8E+3   0.5   1  RT (min)      lcqtof_hilic_neg_1_031_16    m/z 1050.4351     0   1   2   3   4E+3   4.5   5  RT (min)      lcqtof_hilic_neg_1_003_37    m/z 1051.7073     0   2   4   6   8E+3   0.5   1  RT (min)      lcqtof_hilic_neg_1_051_35    m/z 1052.7656     0   2   4   6E+3   0.5   1  RT (min)      lcqtof_hilic_neg_4_032_01    m/z 1052.8212     0   1   2E+3   1   1.5  RT (min)      lcqtof_hilic_neg_1_022_36    m/z 1052.9489     0   2   4E+3   1   1.5  RT (min)      lcqtof_hilic_neg_4_008_24    m/z 1053.0333     0   1   2E+3   4.5   5  RT (min)      lcqtof_hilic_neg_4_075_58    m/z 1053.3018     0   2   4   6   8E+3   6   6.5  RT (min)      lcqtof_hilic_neg_4_073_57    m/z 1053.4530     0   1   2E+3   4.5   5  RT (min)      lcqtof_hilic_neg_1_003_37    m/z 1053.6652     0   1   2   3E+3   4.5   5  RT (min)      lcqtof_hilic_neg_4_045_26    m/z 1053.7107     0   2   4   6E+3   4.5   5  RT (min)      lcqtof_hilic_neg_2_047_32    m/z 1054.0746     0   1   2   3   4E+3   4.5   5  RT (min)      lcqtof_hilic_neg_4_022_34    m/z 1054.0980     0   2   4   6   8E+3   6   6.5  RT (min)      lcqtof_hilic_neg_4_018_27    m/z 1054.1388     0   1E+4   0.5   1  RT (min)      lcqtof_hilic_neg_4_004_47    m/z 1054.4486     0   1   2   3E+3   4.5   5  RT (min)      lcqtof_hilic_neg_2_048_35    m/z 1054.8979     0   1   2   3   4E+3   0.5   1  RT (min)      lcqtof_hilic_neg_4_011_35    m/z 1055.0742     0   1   2   3   4E+3   4.5   5  RT (min)      lcqtof_hilic_neg_4_022_34    m/z 1055.1343     0   2   4   6   8E+3   0.5   1  RT (min)      lcqtof_hilic_neg_4_004_47    m/z 1055.1625     0   2   4   6E+3   7.5   8  RT (min)       lcqtof_hilic_neg_1_005_32    m/z 1055.6632     0   2   4   6E+3   4.5   5  RT (min)      lcqtof_hilic_neg_4_031_38    m/z 1055.7227     0   2   4E+3   4   4.5  RT (min)       lcqtof_hilic_neg_4_082_59    m/z 1056.0477     0   1   2E+3   4.5   5  RT (min)      lcqtof_hilic_neg_4_001_49    m/z 1056.3390     0   2   4E+3   1   1.5  RT (min)      lcqtof_hilic_neg_4_027_19    m/z 1056.6958     0   2   4E+3   3.5   4  RT (min)      lcqtof_hilic_neg_4_005_29    m/z 1056.7404     0   2   4E+3   4   4.5  RT (min)       lcqtof_hilic_neg_4_048_23    m/z 1057.3048     0   1   2   3E+3   1   1.5  RT (min)      lcqtof_hilic_neg_2_040_20    m/z 1057.5315     0   1   2   3E+3   4.5   5  RT (min)      lcqtof_hilic_neg_4_053_14    m/z 1057.7014     0   1   2E+3   3.5   4  RT (min)      lcqtof_hilic_neg_4_005_29    m/z 1058.8430     0   2   4   6E+3   0.5   1  RT (min)      lcqtof_hilic_neg_1_008_13    m/z 1059.3003     0   1   2   3E+3   1   1.5  RT (min)      lcqtof_hilic_neg_2_013_29    m/z 1059.4579     0   1   2   3E+3   4.5   5  RT (min)      lcqtof_hilic_neg_4_011_35    m/z 1059.6589     0   2   4E+3   2   2.5  RT (min)      lcqtof_hilic_neg_4_011_35    m/z 1060.4025     0   1   2   3E+3   1   1.5  RT (min)      lcqtof_hilic_neg_1_027_27    m/z 1060.4626     0   1   2   3E+3   4.5   5  RT (min)      lcqtof_hilic_neg_4_016_37    m/z 1060.4940     0   1   2   3E+3   6   6.5  RT (min)      lcqtof_hilic_neg_4_053_14    m/z 1060.6780     0   1   2   3E+3   5   5.5  RT (min)      lcqtof_hilic_neg_4_048_23    m/z 1060.8008     0   1E+4   0.5   1  RT (min)      lcqtof_hilic_neg_4_004_47    m/z 1062.6267     0   1   2   3   4E+3   4.5   5  RT (min)      lcqtof_hilic_neg_4_055_32    m/z 1062.6434     0   1E+3   4.5   5  RT (min)      lcqtof_hilic_neg_2_004_26    m/z 1062.8982     0   1E+4   0.5   1  RT (min)      lcqtof_hilic_neg_4_011_35    m/z 1063.3623     0   2   4   6   8E+3   6   6.5  RT (min)      lcqtof_hilic_neg_4_052_13    m/z 1063.8805     0   1   2   3   4E+3   2   2.5  RT (min)      lcqtof_hilic_neg_1_005_32    m/z 1063.9032     0   2   4E+3   1   1.5  RT (min)      lcqtof_hilic_neg_2_052_24    m/z 1064.9130     0   1   2   3   4E+3   1   1.5  RT (min)      lcqtof_hilic_neg_2_052_24    m/z 1065.6373     0   1   2   3E+3   4   4.5  RT (min)      lcqtof_hilic_neg_4_031_38    m/z 1065.9451     0   1   2E+4   0.5   1  RT (min)      lcqtof_hilic_neg_4_008_24    m/z 1066.6542     0   2   4E+3   1   1.5  RT (min)      lcqtof_hilic_neg_2_020_23    m/z 1066.6857     0   1   2   3   4E+3   6.5   7  RT (min)       lcqtof_hilic_neg_4_012_07    m/z 1067.3513     0   2   4E+3   5   5.5  RT (min)      lcqtof_hilic_neg_4_053_14    m/z 1067.4080     0   1   2   3E+3   1   1.5  RT (min)      lcqtof_hilic_neg_1_016_28    m/z 1067.4857     0   1   2E+3   5   5.5  RT (min)      lcqtof_hilic_neg_4_010_28    m/z 1067.9340     0   2   4   6   8E+3   1   1.5  RT (min)      lcqtof_hilic_neg_2_052_24    m/z 1068.4918     0   1E+3   8   8.5  RT (min)      lcqtof_hilic_neg_3_026_42    m/z 1068.6127     0   2   4   6E+3   3.5   4  RT (min)      lcqtof_hilic_neg_4_032_01    m/z 1068.6736     0   1   2E+3   4.5   5  RT (min)      lcqtof_hilic_neg_1_008_13    m/z 1069.0665     0   1   2   3   4E+3   6   6.5  RT (min)      lcqtof_hilic_neg_4_014_46    m/z 1069.7573     0   1   2   3E+3   5   5.5  RT (min)       lcqtof_hilic_neg_1_035_23    m/z 1069.9456     0   2   4E+3   0.5   1  RT (min)      lcqtof_hilic_neg_4_008_24    m/z 1070.9865     0   1E+3   5   5.5  RT (min)      lcqtof_hilic_neg_2_032_02    m/z 1071.6892     0   2   4E+3   4   4.5  RT (min)      lcqtof_hilic_neg_1_007_05    m/z 1071.7743     0   2   4   6E+3   4.5   5  RT (min)      lcqtof_hilic_neg_4_055_32    m/z 1072.5562     0   2   4   6   8E+3   0.5   1  RT (min)      lcqtof_hilic_neg_1_031_16    m/z 1072.6138     0   2   4   6E+3   1   1.5  RT (min)      lcqtof_hilic_neg_1_023_34    m/z 1072.9673     0   1E+3   6   6.5  RT (min)      lcqtof_hilic_neg_1_030_17    m/z 1073.0708     0   2   4   6E+3   0.5   1  RT (min)      lcqtof_hilic_neg_3_036_17    m/z 1073.2950     0   1   2   3E+3   5   5.5  RT (min)       lcqtof_hilic_neg_4_106_62    m/z 1073.3383     0   1   2   3E+3   5   5.5  RT (min)      lcqtof_hilic_neg_4_028_36    m/z 1073.4257     0   1   2E+3   4.5   5  RT (min)      lcqtof_hilic_neg_2_036_34    m/z 1074.4015     0   1   2E+3   1   1.5  RT (min)      lcqtof_hilic_neg_4_004_47    m/z 1074.6833     0   1   2   3E+3   4.5   5  RT (min)      lcqtof_hilic_neg_4_055_32    m/z 1074.6942     0   2   4E+3   3.5   4   4.5  RT (min)      lcqtof_hilic_neg_4_052_13    m/z 1074.7958     0   2   4   6   8E+3   4.5   5  RT (min)      lcqtof_hilic_neg_4_055_32    m/z 1075.0300     0   1   2   3E+3   5.5   6  RT (min)      lcqtof_hilic_neg_4_038_10    m/z 1075.3723     0   1   2   3E+3   6   6.5  RT (min)      lcqtof_hilic_neg_4_052_13    m/z 1075.4359     0   1   2   3   4E+3   4.5   5  RT (min)      lcqtof_hilic_neg_2_002_36    m/z 1075.5238     0   1   2E+3   5.5   6  RT (min)      lcqtof_hilic_neg_2_051_14    m/z 1076.0472     0   2   4   6E+3   6   6.5  RT (min)      lcqtof_hilic_neg_4_056_56    m/z 1076.0780     0   2   4   6   8E+3   6   6.5  RT (min)      lcqtof_hilic_neg_4_018_27    m/z 1076.3444     0   2   4   6   8E+3   3.5   4  RT (min)      lcqtof_hilic_neg_1_020_09    m/z 1076.6895     0   2   4   6E+3   3.5   4  RT (min)      lcqtof_hilic_neg_1_004_14    m/z 1076.7756     0   2   4E+3   0.5   1  RT (min)      lcqtof_hilic_neg_4_053_14    m/z 1077.4370     0   1   2   3E+3   4.5   5  RT (min)      lcqtof_hilic_neg_4_048_23    m/z 1077.6443     0   2   4   6E+3   0.5   1  RT (min)      lcqtof_hilic_neg_4_045_26    m/z 1078.4415     0   1   2   3E+3   4.5   5  RT (min)      lcqtof_hilic_neg_4_048_23    m/z 1078.8441     0   2   4   6E+3   0.5   1  RT (min)      lcqtof_hilic_neg_4_004_47    m/z 1079.3467     0   2   4   6E+3   3.5   4  RT (min)      lcqtof_hilic_neg_1_040_20    m/z 1080.0248     0   2   4E+3   0.5   1  RT (min)      lcqtof_hilic_neg_4_014_46    m/z 1080.1537     0   1   2   3   4E+3   5.5   6  RT (min)      lcqtof_hilic_neg_4_013_20    m/z 1080.3409     0   2   4   6E+3   3.5   4  RT (min)      lcqtof_hilic_neg_1_053_19    m/z 1080.6177     0   2   4E+3   4.5   5  RT (min)      lcqtof_hilic_neg_1_005_32    m/z 1080.8113     0   2   4   6E+3   5   5.5  RT (min)       lcqtof_hilic_neg_4_008_24    m/z 1081.4242     0   2   4   6E+3   1   1.5  RT (min)      lcqtof_hilic_neg_4_013_20    m/z 1082.6284     0   2   4E+3   4.5   5  RT (min)      lcqtof_hilic_neg_1_005_32    m/z 1082.7303     0   1   2   3   4E+3   4   4.5  RT (min)      lcqtof_hilic_neg_3_048_05    m/z 1083.1748     0   1   2E+3   0.5   1  RT (min)      lcqtof_hilic_neg_4_007_45    m/z 1083.7046     0   2   4   6E+3   1   1.5  RT (min)      lcqtof_hilic_neg_4_053_14    m/z 1083.7490     0   2   4   6E+3   0.5   1  RT (min)      lcqtof_hilic_neg_3_040_14    m/z 1083.9614     0   1   2   3E+3   0.5   1  RT (min)      lcqtof_hilic_neg_4_004_47    m/z 1084.1803     0   1   2E+3   0.5   1  RT (min)      lcqtof_hilic_neg_2_033_53    m/z 1084.6537     0   1   2   3   4E+3   4.5   5  RT (min)      lcqtof_hilic_neg_3_055_01    m/z 1085.3472     0   1   2   3   4E+3   1   1.5  RT (min)      lcqtof_hilic_neg_4_005_29    m/z 1085.6602     0   1   2E+3   4.5   5  RT (min)      lcqtof_hilic_neg_3_055_01    m/z 1086.1605     0   2   4   6   8E+3   0.5   1  RT (min)      lcqtof_hilic_neg_4_001_49    m/z 1086.6702     0   1   2   3   4E+3   4.5   5  RT (min)      lcqtof_hilic_neg_3_055_01    m/z 1087.6448     0   1   2E+3   4.5   5  RT (min)      lcqtof_hilic_neg_4_022_34    m/z 1087.6747     0   1   2E+3   4.5   5  RT (min)      lcqtof_hilic_neg_3_055_01    m/z 1088.5641     0   1   2   3   4E+3   4   4.5  RT (min)      lcqtof_hilic_neg_4_014_46    m/z 1088.8839     0   2   4E+3   1   1.5  RT (min)      lcqtof_hilic_neg_2_001_49    m/z 1089.1506     0   1   2   3E+3   5   5.5  RT (min)      lcqtof_hilic_neg_4_053_14    m/z 1089.4957     0   1E+3   4.5   5  RT (min)      lcqtof_hilic_neg_4_053_14    m/z 1089.6511     0   1E+3   4.5   5  RT (min)      lcqtof_hilic_neg_4_006_30    m/z 1089.9523     0   2   4   6   8E+3   0.5   1  RT (min)      lcqtof_hilic_neg_4_011_35    m/z 1090.1425     0   1   2E+3   5   5.5  RT (min)      lcqtof_hilic_neg_4_001_49    m/z 1090.3979     0   2   4E+3   4.5   5  RT (min)      lcqtof_hilic_neg_4_028_36    m/z 1090.4221     0   2   4   6   8E+3   0.5   1  RT (min)      lcqtof_hilic_neg_4_042_22    m/z 1090.4794     0   1   2   3   4E+3   5   5.5  RT (min)      lcqtof_hilic_neg_4_053_14    m/z 1092.4965     0   1   2   3E+3   4.5   5  RT (min)      lcqtof_hilic_neg_4_053_14    m/z 1092.7347     0   2   4   6E+3   0.5   1  RT (min)      lcqtof_hilic_neg_3_040_14    m/z 1092.7800     0   1   2   3E+3   4.5   5  RT (min)      lcqtof_hilic_neg_1_005_32    m/z 1093.3384     0   2   4E+3   7   7.5  RT (min)      lcqtof_hilic_neg_2_044_37    m/z 1093.6029     0   1   2   3   4E+3   4   4.5  RT (min)      lcqtof_hilic_neg_4_013_20    m/z 1093.6616     0   1   2   3   4E+3   4.5   5  RT (min)       lcqtof_hilic_neg_4_047_06    m/z 1094.2384     0   1   2   3E+3   6   6.5  RT (min)      lcqtof_hilic_neg_4_003_16    m/z 1094.3624     0   2   4   6E+3   1   1.5  RT (min)      lcqtof_hilic_neg_4_013_20    m/z 1094.5970     0   1   2   3   4E+3   1   1.5  RT (min)      lcqtof_hilic_neg_2_036_34    m/z 1094.6326     0   1E+3   4   4.5  RT (min)      lcqtof_hilic_neg_4_026_04    m/z 1094.7133     0   2   4   6   8E+3   4.5   5  RT (min)      lcqtof_hilic_neg_4_034_31    m/z 1094.7787     0   1   2   3   4E+3   0.5   1  RT (min)      lcqtof_hilic_neg_1_031_16    m/z 1094.7896     0   1E+3   5   5.5  RT (min)      lcqtof_hilic_neg_4_109_62    m/z 1094.8580     0   2   4   6   8E+3   0.5   1  RT (min)      lcqtof_hilic_neg_4_004_47    m/z 1094.9413     0   1   2E+3   6   6.5  RT (min)      lcqtof_hilic_neg_4_026_04    m/z 1095.0371     0   1   2E+3   4.5   5  RT (min)      lcqtof_hilic_neg_4_009_50    m/z 1095.3726     0   1   2   3   4E+3   1   1.5  RT (min)      lcqtof_hilic_neg_4_013_20    m/z 1095.6151     0   1E+3   4   4.5  RT (min)      lcqtof_hilic_neg_2_007_03    m/z 1095.7880     0   1   2   3E+3   0.5   1  RT (min)      lcqtof_hilic_neg_2_018_16    m/z 1095.8569     0   2   4   6   8E+3   0.5   1  RT (min)      lcqtof_hilic_neg_4_004_47    m/z 1096.5935     0   1   2   3   4E+3   4.5   5  RT (min)      lcqtof_hilic_neg_1_005_32    m/z 1096.7715     0   1   2   3E+3   5   5.5  RT (min)      lcqtof_hilic_neg_4_048_23    m/z 1097.0206     0   1   2   3   4E+3   4.5   5  RT (min)      lcqtof_hilic_neg_4_031_38    m/z 1097.0505     0   1   2   3E+3   4.5   5  RT (min)      lcqtof_hilic_neg_4_004_47    m/z 1097.0557     0   2   4   6E+3   6   6.5  RT (min)      lcqtof_hilic_neg_4_031_38    m/z 1097.0630     0   1   2   3   4E+3   4.5   5  RT (min)      lcqtof_hilic_neg_4_028_36    m/z 1097.3551     0   2   4   6   8E+3   1   1.5  RT (min)      lcqtof_hilic_neg_4_005_29    m/z 1097.6193     0   2   4   6E+3   1   1.5  RT (min)      lcqtof_hilic_neg_2_036_34    m/z 1097.6371     0   2   4   6   8E+3   2.5   3  RT (min)      lcqtof_hilic_neg_1_013_38    m/z 1097.7998     0   1   2E+3   0.5   1  RT (min)      lcqtof_hilic_neg_2_026_28    m/z 1098.0615     0   1   2   3   4E+3   6   6.5  RT (min)      lcqtof_hilic_neg_4_018_27    m/z 1098.9146     0   2   4   6   8E+3   0.5   1  RT (min)      lcqtof_hilic_neg_1_031_16    m/z 1099.5686     0   2   4   6E+3   4   4.5  RT (min)      lcqtof_hilic_neg_1_012_04    m/z 1099.7401     0   1   2   3   4E+3   4.5   5  RT (min)      lcqtof_hilic_neg_4_016_37    m/z 1100.4424     0   1   2E+3   4.5   5  RT (min)      lcqtof_hilic_neg_2_002_36    m/z 1100.9265     0   2   4   6   8E+3   0.5   1  RT (min)      lcqtof_hilic_neg_1_008_13    m/z 1101.3379     0   1   2   3   4E+3   5   5.5  RT (min)      lcqtof_hilic_neg_4_053_14    m/z 1101.5681     0   2   4E+3   4   4.5  RT (min)      lcqtof_hilic_neg_1_012_04    m/z 1101.6428     0   2   4   6E+3   0.5   1  RT (min)      lcqtof_hilic_neg_4_045_26    m/z 1101.6710     0   2   4   6E+3   2.5   3  RT (min)      lcqtof_hilic_neg_1_013_38    m/z 1101.8856     0   1   2   3   4E+3   5   5.5  RT (min)      lcqtof_hilic_neg_4_029_17    m/z 1101.9307     0   2   4E+3   0.5   1  RT (min)      lcqtof_hilic_neg_2_021_17    m/z 1101.9526     0   1E+3   6   6.5  RT (min)      lcqtof_hilic_neg_4_026_04    m/z 1102.6046     0   1   2   3E+3   4.5   5  RT (min)      lcqtof_hilic_neg_3_004_32    m/z 1103.3644     0   1   2   3E+3   3.5   4  RT (min)      lcqtof_hilic_neg_1_053_19    m/z 1103.6616     0   2   4   6   8E+3   0.5   1  RT (min)      lcqtof_hilic_neg_4_045_26    m/z 1103.7606     0   2   4   6   8E+3   4   4.5  RT (min)      lcqtof_hilic_neg_4_048_23    m/z 1104.2599     0   1   2   3E+3   0.5   1  RT (min)      lcqtof_hilic_neg_4_040_05    m/z 1104.4277     0   1E+4   1   1.5  RT (min)      lcqtof_hilic_neg_4_013_20    m/z 1104.6079     0   1   2   3E+3   4.5   5  RT (min)      lcqtof_hilic_neg_2_047_32    m/z 1104.6710     0   1   2   3E+3   1   1.5  RT (min)      lcqtof_hilic_neg_1_046_30    m/z 1104.7037     0   2   4E+3   4.5   5  RT (min)       lcqtof_hilic_neg_4_008_24    m/z 1104.8130     0   1   2   3E+3   5   5.5  RT (min)      lcqtof_hilic_neg_4_052_13    m/z 1104.8687     0   2   4   6   8E+3   0.5   1  RT (min)      lcqtof_hilic_neg_4_055_32    m/z 1105.3663     0   1   2   3   4E+3   3.5   4  RT (min)       lcqtof_hilic_neg_1_053_19    m/z 1106.2393     0   1   2E+3   6   6.5  RT (min)      lcqtof_hilic_neg_4_005_29    m/z 1106.7509     0   2   4   6   8E+3   0.5   1  RT (min)      lcqtof_hilic_neg_3_040_14    m/z 1106.8478     0   1   2   3E+3   0.5   1  RT (min)      lcqtof_hilic_neg_4_033_53    m/z 1107.2909     0   1   2   3E+3   5   5.5  RT (min)      lcqtof_hilic_neg_4_111_62    m/z 1107.7190     0   2   4   6E+3   4.5   5  RT (min)      lcqtof_hilic_neg_4_034_31    m/z 1107.8596     0   2   4   6E+3   4.5   5  RT (min)      lcqtof_hilic_neg_4_055_32    m/z 1108.0348     0   1   2   3E+3   6   6.5  RT (min)       lcqtof_hilic_neg_1_010_15    m/z 1110.2643     0   1   2E+3   6   6.5  RT (min)      lcqtof_hilic_neg_4_041_54    m/z 1110.6042     0   1   2   3E+3   4   4.5  RT (min)      lcqtof_hilic_neg_1_036_03    m/z 1110.7747     0   2   4   6E+3   0.5   1  RT (min)      lcqtof_hilic_neg_3_040_14    m/z 1111.6029     0   1   2   3E+3   4   4.5  RT (min)      lcqtof_hilic_neg_1_012_04    m/z 1111.7468     0   2   4E+3   4.5   5  RT (min)      lcqtof_hilic_neg_4_016_37    m/z 1111.8225     0   1   2   3E+3   5   5.5  RT (min)       lcqtof_hilic_neg_4_053_14    m/z 1112.6625     0   1   2   3   4E+3   4   4.5  RT (min)      lcqtof_hilic_neg_4_031_38    m/z 1112.8158     0   1   2E+3   5   5.5  RT (min)      lcqtof_hilic_neg_4_003_16    m/z 1113.1433     0   1   2   3   4E+3   5   5.5  RT (min)      lcqtof_hilic_neg_4_053_14    m/z 1113.7240     0   1   2   3E+3   5   5.5  RT (min)      lcqtof_hilic_neg_4_002_25    m/z 1113.8406     0   2   4   6E+3   2   2.5  RT (min)      lcqtof_hilic_neg_3_004_32    m/z 1114.5896     0   1   2E+3   5   5.5  RT (min)       lcqtof_hilic_neg_2_052_24    m/z 1114.7620     0   1E+3   1   1.5  RT (min)      lcqtof_hilic_neg_2_041_54    m/z 1115.3282     0   1   2   3E+3   7   7.5  RT (min)      lcqtof_hilic_neg_2_044_37    m/z 1115.7946     0   2   4E+3   3.5   4  RT (min)      lcqtof_hilic_neg_4_055_32    m/z 1116.3060     0   2   4E+3   6   6.5  RT (min)      lcqtof_hilic_neg_4_023_15    m/z 1116.4309     0   1   2   3E+3   4.5   5  RT (min)      lcqtof_hilic_neg_4_050_02    m/z 1116.7323     0   1   2E+3   3.5   4  RT (min)      lcqtof_hilic_neg_4_016_37    m/z 1117.0601     0   2   4   6E+3   0.5   1  RT (min)      lcqtof_hilic_neg_4_054_48    m/z 1117.9772     0   2   4   6E+3   0.5   1  RT (min)      lcqtof_hilic_neg_4_028_36    m/z 1118.4254     0   1   2   3   4E+3   4.5   5  RT (min)      lcqtof_hilic_neg_4_053_14    m/z 1118.5802     0   2   4E+3   4.5   5  RT (min)      lcqtof_hilic_neg_2_047_32    m/z 1119.6826     0   2   4   6   8E+3   0.5   1  RT (min)      lcqtof_hilic_neg_4_026_04    m/z 1119.6881     0   1   2E+3   1   1.5  RT (min)      lcqtof_hilic_neg_2_031_01    m/z 1119.7881     0   1E+4   0.5   1  RT (min)      lcqtof_hilic_neg_4_004_47    m/z 1119.9875     0   1   2   3   4E+3   0.5   1  RT (min)      lcqtof_hilic_neg_4_034_31    m/z 1120.4651     0   2   4   6E+3   0.5   1  RT (min)      lcqtof_hilic_neg_4_015_09    m/z 1120.5862     0   1   2   3   4E+3   4.5   5  RT (min)      lcqtof_hilic_neg_2_047_32    m/z 1120.8456     0   1   2   3   4E+3   0.5   1  RT (min)      lcqtof_hilic_neg_4_047_06    m/z 1122.8708     0   1   2   3   4E+3   0.5   1  RT (min)      lcqtof_hilic_neg_3_023_08    m/z 1123.8944     0   1   2   3E+3   4.5   5  RT (min)      lcqtof_hilic_neg_4_092_60    m/z 1124.6493     0   1   2E+3   1   1.5  RT (min)      lcqtof_hilic_neg_2_055_30    m/z 1124.8011     0   1   2   3   4E+3   0.5   1  RT (min)      lcqtof_hilic_neg_4_004_47    m/z 1124.8910     0   2   4   6E+3   0.5   1  RT (min)      lcqtof_hilic_neg_4_004_47    m/z 1125.7260     0   2   4   6E+3   1   1.5  RT (min)      lcqtof_hilic_neg_2_031_01    m/z 1125.9839     0   1   2   3   4E+3   0.5   1  RT (min)      lcqtof_hilic_neg_4_014_46    m/z 1126.4618     0   2   4E+3   4.5   5  RT (min)       lcqtof_hilic_neg_4_053_14    m/z 1126.4645     0   1   2E+3   4.5   5  RT (min)      lcqtof_hilic_neg_4_003_16    m/z 1126.6614     0   2   4E+3   1   1.5  RT (min)      lcqtof_hilic_neg_1_023_34    m/z 1126.7332     0   1E+4   1   1.5  RT (min)      lcqtof_hilic_neg_2_031_01    m/z 1127.0360     0   2   4   6   8E+3   0.5   1  RT (min)      lcqtof_hilic_neg_4_004_47    m/z 1127.6697     0   1   2   3E+3   1   1.5  RT (min)      lcqtof_hilic_neg_3_006_30    m/z 1127.7443     0   2   4E+3   1   1.5  RT (min)      lcqtof_hilic_neg_3_055_01    m/z 1127.9664     0   1   2   3E+3   0.5   1  RT (min)      lcqtof_hilic_neg_4_014_46    m/z 1128.7153     0   2   4   6E+3   0.5   1   1.5  RT (min)      lcqtof_hilic_neg_4_045_26    m/z 1128.7506     0   1   2   3E+3   1   1.5  RT (min)      lcqtof_hilic_neg_2_031_01    m/z 1130.3824     0   1   2E+3   4.5   5  RT (min)      lcqtof_hilic_neg_2_048_35    m/z 1130.4867     0   1   2E+3   4.5   5  RT (min)      lcqtof_hilic_neg_4_048_23    m/z 1131.5964     0   1   2E+3   4   4.5  RT (min)      lcqtof_hilic_neg_1_035_23    m/z 1132.3914     0   1   2   3   4E+3   4.5   5  RT (min)      lcqtof_hilic_neg_2_002_36    m/z 1132.7227     0   2   4   6E+3   3   3.5  RT (min)      lcqtof_hilic_neg_1_042_26    m/z 1133.4707     0   1E+4   4.5   5  RT (min)      lcqtof_hilic_neg_4_003_16    m/z 1133.6014     0   1   2E+3   3.5   4  RT (min)      lcqtof_hilic_neg_2_007_03    m/z 1134.3953     0   1   2E+3   4.5   5  RT (min)      lcqtof_hilic_neg_1_051_35    m/z 1134.4756     0   1   2   3E+3   5   5.5  RT (min)      lcqtof_hilic_neg_4_053_14    m/z 1134.4896     0   1E+3   4.5   5  RT (min)      lcqtof_hilic_neg_3_005_15    m/z 1135.3348     0   2   4E+3   5   5.5  RT (min)      lcqtof_hilic_neg_4_053_14    m/z 1135.4708     0   2   4E+3   5   5.5  RT (min)      lcqtof_hilic_neg_4_053_14    m/z 1135.8708     0   1E+4   0.5   1  RT (min)      lcqtof_hilic_neg_4_014_46    m/z 1135.9271     0   1   2E+3   6   6.5  RT (min)      lcqtof_hilic_neg_2_031_01    m/z 1136.1514     0   1E+3   5   5.5  RT (min)      lcqtof_hilic_neg_3_043_13    m/z 1136.4741     0   1E+4   1   1.5  RT (min)      lcqtof_hilic_neg_4_013_20    m/z 1136.4957     0   1   2   3   4E+3   1   1.5  RT (min)      lcqtof_hilic_neg_4_001_49    m/z 1136.7583     0   1   2   3   4E+3   4.5   5  RT (min)      lcqtof_hilic_neg_4_031_38    m/z 1136.8593     0   1   2E+3   5   5.5  RT (min)      lcqtof_hilic_neg_4_041_54    m/z 1136.8901     0   2   4   6   8E+3   4.5   5  RT (min)      lcqtof_hilic_neg_4_055_32    m/z 1137.0291     0   1   2E+3   4.5   5  RT (min)      lcqtof_hilic_neg_4_006_30    m/z 1137.4962     0   1   2E+3   1   1.5  RT (min)      lcqtof_hilic_neg_4_001_49    m/z 1137.8468     0   1E+3   5   5.5  RT (min)      lcqtof_hilic_neg_2_034_47    m/z 1137.9862     0   2   4   6E+3   0.5   1  RT (min)      lcqtof_hilic_neg_2_002_36    m/z 1138.7399     0   1   2   3E+3   1   1.5  RT (min)      lcqtof_hilic_neg_4_016_37    m/z 1138.9894     0   1   2   3   4E+3   4.5   5  RT (min)      lcqtof_hilic_neg_4_053_14    m/z 1139.0452     0   1   2   3   4E+3   4.5   5  RT (min)      lcqtof_hilic_neg_4_035_33    m/z 1139.5619     0   2   4   6E+3   0.5   1  RT (min)      lcqtof_hilic_neg_4_006_30    m/z 1139.6954     0   1   2   3E+3   4   4.5  RT (min)      lcqtof_hilic_neg_4_019_03    m/z 1139.7235     0   1   2   3   4E+3   3.5   4  RT (min)      lcqtof_hilic_neg_4_004_47    m/z 1140.0140     0   1   2   3E+3   4.5   5  RT (min)      lcqtof_hilic_neg_4_016_37    m/z 1140.7174     0   2   4   6   8E+3   4   4.5  RT (min)      lcqtof_hilic_neg_4_005_29    m/z 1140.7600     0   1   2   3E+3   0.5   1  RT (min)      lcqtof_hilic_neg_4_054_48    m/z 1141.2816     0   1   2E+3   5   5.5  RT (min)       lcqtof_hilic_neg_4_112_62    m/z 1141.3224     0   1   2   3E+3   5   5.5  RT (min)      lcqtof_hilic_neg_4_028_36    m/z 1141.6350     0   2   4E+3   7   7.5  RT (min)       lcqtof_hilic_neg_4_048_23    m/z 1141.9027     0   1   2   3E+3   0.5   1  RT (min)      lcqtof_hilic_neg_4_068_57    m/z 1142.3734     0   2   4   6E+3   6   6.5  RT (min)      lcqtof_hilic_neg_4_052_13    m/z 1142.7458     0   2   4E+3   0.5   1  RT (min)      lcqtof_hilic_neg_4_053_14    m/z 1142.7810     0   1   2   3E+3   1   1.5  RT (min)      lcqtof_hilic_neg_2_041_54    m/z 1143.1222     0   1   2   3   4E+3   0.5   1  RT (min)      lcqtof_hilic_neg_4_014_46    m/z 1144.0261     0   1   2   3   4E+3   0.5   1  RT (min)      lcqtof_hilic_neg_4_011_35    m/z 1144.1329     0   1   2E+3   0.5   1  RT (min)      lcqtof_hilic_neg_4_014_46    m/z 1144.6335     0   1   2E+3   4   4.5  RT (min)      lcqtof_hilic_neg_1_005_32    m/z 1146.2516     0   1   2E+3   7   7.5  RT (min)      lcqtof_hilic_neg_3_055_01    m/z 1146.6394     0   1   2   3E+3   3.5   4  RT (min)      lcqtof_hilic_neg_2_005_13    m/z 1147.6676     0   1   2   3   4E+3   5   5.5  RT (min)       lcqtof_hilic_neg_4_016_37    m/z 1147.6980     0   1   2   3   4E+3   0.5   1  RT (min)      lcqtof_hilic_neg_4_050_02    m/z 1148.3584     0   1   2   3E+3   4.5   5  RT (min)      lcqtof_hilic_neg_4_016_37    m/z 1148.7284     0   1   2   3   4E+3   1   1.5  RT (min)      lcqtof_hilic_neg_2_029_31    m/z 1148.8815     0   2   4   6E+3   0.5   1  RT (min)      lcqtof_hilic_neg_1_045_07    m/z 1149.7228     0   1   2E+3   1   1.5  RT (min)      lcqtof_hilic_neg_3_055_01    m/z 1149.8528     0   1E+3   1   1.5  RT (min)      lcqtof_hilic_neg_2_048_35    m/z 1150.6661     0   1   2   3E+3   3.5   4  RT (min)      lcqtof_hilic_neg_4_052_13    m/z 1150.6824     0   2   4E+3   3   3.5  RT (min)      lcqtof_hilic_neg_2_004_26    m/z 1150.7399     0   2   4   6E+3   1   1.5  RT (min)      lcqtof_hilic_neg_1_054_01    m/z 1150.8925     0   2   4E+3   0.5   1  RT (min)      lcqtof_hilic_neg_1_045_07    m/z 1151.7433     0   1   2   3E+3   1   1.5  RT (min)      lcqtof_hilic_neg_3_055_01    m/z 1151.8214     0   2   4   6E+3   0.5   1  RT (min)      lcqtof_hilic_neg_1_051_35    m/z 1152.4059     0   2   4   6E+3   6   6.5  RT (min)      lcqtof_hilic_neg_4_052_13    m/z 1152.7510     0   2   4   6E+3   1   1.5  RT (min)      lcqtof_hilic_neg_3_055_01    m/z 1153.6671     0   1   2   3   4E+3   3.5   4  RT (min)      lcqtof_hilic_neg_4_003_16    m/z 1153.6952     0   2   4   6E+3   3   3.5  RT (min)      lcqtof_hilic_neg_4_006_30    m/z 1153.7577     0   2   4E+3   1   1.5  RT (min)      lcqtof_hilic_neg_1_054_01    m/z 1153.8284     0   1   2   3   4E+3   0.5   1  RT (min)      lcqtof_hilic_neg_4_054_48    m/z 1154.7758     0   2   4E+3   1   1.5  RT (min)      lcqtof_hilic_neg_1_054_01    m/z 1155.7694     0   1   2E+3   1   1.5  RT (min)      lcqtof_hilic_neg_2_031_01    m/z 1155.7738     0   2   4E+3   1   1.5  RT (min)      lcqtof_hilic_neg_4_006_30    m/z 1156.7739     0   1   2E+3   1   1.5  RT (min)      lcqtof_hilic_neg_4_003_16    m/z 1156.7859     0   1E+3   1   1.5  RT (min)      lcqtof_hilic_neg_1_054_01    m/z 1157.1417     0   1   2   3E+3   5   5.5  RT (min)      lcqtof_hilic_neg_4_032_01    m/z 1157.4747     0   2   4   6   8E+3   0.5   1  RT (min)      lcqtof_hilic_neg_4_004_47    m/z 1157.4911     0   1   2E+3   5   5.5  RT (min)      lcqtof_hilic_neg_4_012_07    m/z 1157.6068     0   1   2E+3   3.5   4  RT (min)      lcqtof_hilic_neg_2_005_13    m/z 1157.7020     0   2   4   6E+3   5   5.5  RT (min)      lcqtof_hilic_neg_4_048_23    m/z 1158.4736     0   1E+3   5   5.5  RT (min)      lcqtof_hilic_neg_4_003_16    m/z 1158.6813     0   2   4E+3   5   5.5  RT (min)      lcqtof_hilic_neg_4_022_34    m/z 1158.7964     0   1E+3   5   5.5  RT (min)      lcqtof_hilic_neg_4_016_37    m/z 1158.8849     0   2   4   6   8E+3   5   5.5  RT (min)      lcqtof_hilic_neg_4_055_32    m/z 1158.9348     0   1   2E+3   0.5   1  RT (min)      lcqtof_hilic_neg_3_081_58    m/z 1159.1349     0   1   2   3E+3   6   6.5  RT (min)      lcqtof_hilic_neg_4_006_30    m/z 1159.9340     0   2   4   6E+3   0.5   1  RT (min)      lcqtof_hilic_neg_4_023_15    m/z 1160.5894     0   1E+4   0.5   1  RT (min)      lcqtof_hilic_neg_4_018_27    m/z 1160.7263     0   1   2   3E+3   3.5   4  RT (min)      lcqtof_hilic_neg_2_020_23    m/z 1160.9310     0   1   2   3   4E+3   0.5   1  RT (min)      lcqtof_hilic_neg_4_023_15    m/z 1161.0132     0   2   4   6   8E+3   0.5   1  RT (min)      lcqtof_hilic_neg_4_022_34    m/z 1161.6395     0   2   4   6   8E+3   5   5.5  RT (min)       lcqtof_hilic_neg_4_031_38    m/z 1161.8892     0   2   4   6   8E+3   4.5   5  RT (min)      lcqtof_hilic_neg_4_055_32    m/z 1161.9421     0   2   4   6   8E+3   0.5   1  RT (min)      lcqtof_hilic_neg_4_023_15    m/z 1162.6366     0   2   4E+3   5   5.5  RT (min)       lcqtof_hilic_neg_4_016_37    m/z 1162.7438     0   1   2   3E+3   3.5   4  RT (min)      lcqtof_hilic_neg_3_030_23    m/z 1162.8986     0   2   4   6   8E+3   4.5   5  RT (min)      lcqtof_hilic_neg_4_055_32    m/z 1163.6176     0   2   4   6   8E+3   7   7.5  RT (min)      lcqtof_hilic_neg_4_048_23    m/z 1163.6393     0   1   2E+3   3.5   4  RT (min)      lcqtof_hilic_neg_1_004_14    m/z 1164.4209     0   2   4   6E+3   6   6.5  RT (min)      lcqtof_hilic_neg_4_052_13    m/z 1164.7936     0   2   4   6   8E+3   0.5   1  RT (min)      lcqtof_hilic_neg_4_011_35    m/z 1165.0106     0   2   4   6   8E+3   0.5   1  RT (min)      lcqtof_hilic_neg_1_051_35    m/z 1165.6581     0   1   2E+3   3.5   4  RT (min)      lcqtof_hilic_neg_1_010_15    m/z 1165.8113     0   1   2E+3   4.5   5  RT (min)      lcqtof_hilic_neg_4_017_51    m/z 1166.7312     0   2   4   6   8E+3   4   4.5  RT (min)      lcqtof_hilic_neg_4_032_01    m/z 1166.7455     0   1   2   3   4E+3   0.5   1  RT (min)      lcqtof_hilic_neg_4_053_14    m/z 1167.5232     0   2   4   6   8E+3   0.5   1  RT (min)      lcqtof_hilic_neg_2_042_27    m/z 1167.6255     0   2   4   6E+3   3.5   4  RT (min)      lcqtof_hilic_neg_1_008_13    m/z 1167.7478     0   2   4   6E+3   4   4.5  RT (min)      lcqtof_hilic_neg_4_016_37    m/z 1168.5272     0   2   4E+3   0.5   1  RT (min)      lcqtof_hilic_neg_2_042_27    m/z 1169.3235     0   1   2   3   4E+3   5   5.5  RT (min)      lcqtof_hilic_neg_4_053_14    m/z 1169.8607     0   2   4   6E+3   3   3.5  RT (min)      lcqtof_hilic_neg_4_048_23    m/z 1169.8818     0   1   2E+3   5   5.5  RT (min)      lcqtof_hilic_neg_2_004_26    m/z 1170.0375     0   2   4   6E+3   0.5   1  RT (min)      lcqtof_hilic_neg_2_002_36    m/z 1170.4880     0   1E+4   1   1.5  RT (min)      lcqtof_hilic_neg_4_004_47    m/z 1170.8420     0   1   2E+3   3   3.5  RT (min)      lcqtof_hilic_neg_1_041_54    m/z 1171.0369     0   2   4E+3   0.5   1  RT (min)      lcqtof_hilic_neg_4_016_37    m/z 1172.0525     0   2   4   6E+3   0.5   1  RT (min)      lcqtof_hilic_neg_1_013_38    m/z 1172.7264     0   1   2E+3   0.5   1  RT (min)      lcqtof_hilic_neg_4_053_14    m/z 1172.8488     0   1E+3   3   3.5  RT (min)      lcqtof_hilic_neg_2_001_49    m/z 1172.8568     0   1   2E+3   0.5   1  RT (min)      lcqtof_hilic_neg_1_028_47    m/z 1173.2531     0   1   2   3E+3   6   6.5  RT (min)      lcqtof_hilic_neg_4_010_28    m/z 1173.6736     0   1   2   3   4E+3   3.5   4  RT (min)      lcqtof_hilic_neg_4_050_02    m/z 1173.6885     0   2   4   6   8E+3   5   5.5  RT (min)       lcqtof_hilic_neg_4_031_38    m/z 1173.8497     0   1   2   3E+3   0.5   1  RT (min)      lcqtof_hilic_neg_4_048_23    m/z 1173.8506     0   1   2E+3   1   1.5  RT (min)      lcqtof_hilic_neg_3_030_23    m/z 1174.4568     0   1   2   3E+3   4.5   5  RT (min)       lcqtof_hilic_neg_4_032_01    m/z 1174.6741     0   2   4   6E+3   5   5.5  RT (min)      lcqtof_hilic_neg_4_031_38    m/z 1174.7333     0   1   2   3E+3   0.5   1  RT (min)      lcqtof_hilic_neg_4_052_13    m/z 1174.8429     0   1   2E+3   1   1.5  RT (min)      lcqtof_hilic_neg_4_048_23    m/z 1174.8529     0   1   2E+3   0.5   1  RT (min)      lcqtof_hilic_neg_2_034_47    m/z 1175.2761     0   1   2   3E+3   5   5.5  RT (min)      lcqtof_hilic_neg_4_046_41    m/z 1175.4528     0   1   2   3E+3   4.5   5  RT (min)      lcqtof_hilic_neg_4_003_16    m/z 1175.4553     0   1   2   3   4E+3   4.5   5  RT (min)       lcqtof_hilic_neg_4_032_01    m/z 1175.5978     0   1   2E+3   3.5   4  RT (min)      lcqtof_hilic_neg_1_004_14    m/z 1176.4540     0   1E+3   4.5   5  RT (min)      lcqtof_hilic_neg_4_003_16    m/z 1176.6918     0   1   2E+3   3.5   4  RT (min)      lcqtof_hilic_neg_1_004_14    m/z 1177.6705     0   2   4E+3   3.5   4  RT (min)      lcqtof_hilic_neg_4_003_16    m/z 1177.7577     0   1   2   3   4E+3   0.5   1  RT (min)      lcqtof_hilic_neg_4_010_28    m/z 1178.1458     0   1   2   3   4E+3   0.5   1  RT (min)      lcqtof_hilic_neg_4_032_01    m/z 1178.2853     0   1   2   3E+3   0.5   1  RT (min)      lcqtof_hilic_neg_4_040_05    m/z 1178.7693     0   1   2   3E+3   1   1.5  RT (min)      lcqtof_hilic_neg_3_055_01    m/z 1178.7848     0   1E+3   4   4.5  RT (min)      lcqtof_hilic_neg_3_055_01    m/z 1178.8823     0   2   4   6E+3   0.5   1  RT (min)      lcqtof_hilic_neg_4_026_04    m/z 1178.9873     0   1   2E+3   4.5   5  RT (min)      lcqtof_hilic_neg_4_011_35    m/z 1179.1486     0   1   2   3E+3   0.5   1  RT (min)      lcqtof_hilic_neg_4_014_46    m/z 1179.7310     0   1   2E+3   4.5   5  RT (min)      lcqtof_hilic_neg_4_050_02    m/z 1179.8087     0   1   2   3   4E+3   5   5.5  RT (min)      lcqtof_hilic_neg_4_053_14    m/z 1179.8905     0   2   4   6   8E+3   1   1.5  RT (min)      lcqtof_hilic_neg_4_026_04    m/z 1180.1597     0   1   2   3E+3   5   5.5  RT (min)      lcqtof_hilic_neg_4_023_15    m/z 1180.5527     0   2   4   6E+3   0.5   1  RT (min)      lcqtof_hilic_neg_2_016_33    m/z 1180.6669     0   2   4E+3   3.5   4  RT (min)      lcqtof_hilic_neg_4_052_13    m/z 1180.7925     0   2   4E+3   5   5.5  RT (min)      lcqtof_hilic_neg_4_053_14    m/z 1180.7935     0   2   4E+3   0.5   1  RT (min)      lcqtof_hilic_neg_4_045_26    m/z 1180.8938     0   1   2   3E+3   1   1.5  RT (min)      lcqtof_hilic_neg_4_050_02    m/z 1180.9349     0   2   4   6E+3   4.5   5  RT (min)      lcqtof_hilic_neg_4_055_32    m/z 1181.1289     0   1   2   3   4E+3   5   5.5  RT (min)      lcqtof_hilic_neg_4_032_01    m/z 1181.4612     0   1   2E+3   5   5.5  RT (min)      lcqtof_hilic_neg_4_052_13    m/z 1181.5129     0   1E+5   0.5   1  RT (min)      lcqtof_hilic_neg_3_003_20    m/z 1181.6139     0   1E+3   3.5   4  RT (min)      lcqtof_hilic_neg_2_004_26    m/z 1181.6906     0   1E+3   7   7.5  RT (min)      lcqtof_hilic_neg_4_014_46    m/z 1181.9092     0   2   4   6   8E+3   1   1.5  RT (min)      lcqtof_hilic_neg_4_026_04    m/z 1181.9838     0   1   2   3E+3   4.5   5  RT (min)      lcqtof_hilic_neg_4_009_50    m/z 1182.7758     0   2   4   6   8E+3   0.5   1  RT (min)      lcqtof_hilic_neg_4_004_47    m/z 1183.4384     0   1   2   3E+3   0.5   1  RT (min)      lcqtof_hilic_neg_4_027_19    m/z 1183.6182     0   1   2E+3   3.5   4  RT (min)      lcqtof_hilic_neg_1_031_16    m/z 1183.9193     0   1   2E+3   1   1.5  RT (min)      lcqtof_hilic_neg_4_050_02    m/z 1185.5182     0   1   2   3E+3   1   1.5  RT (min)      lcqtof_hilic_neg_2_053_18    m/z 1185.6963     0   2   4   6   8E+3   5   5.5  RT (min)      lcqtof_hilic_neg_4_048_23    m/z 1187.0403     0   1   2E+3   0.5   1  RT (min)      lcqtof_hilic_neg_4_014_46    m/z 1187.7002     0   1   2   3E+3   7   7.5  RT (min)      lcqtof_hilic_neg_4_002_25    m/z 1187.9213     0   2   4   6   8E+3   5   5.5  RT (min)      lcqtof_hilic_neg_4_055_32    m/z 1187.9354     0   1   2   3E+3   0.5   1  RT (min)      lcqtof_hilic_neg_4_014_46    m/z 1188.4897     0   1   2   3   4E+3   0.5   1  RT (min)      lcqtof_hilic_neg_4_005_29    m/z 1188.6666     0   2   4   6   8E+3   5   5.5  RT (min)       lcqtof_hilic_neg_4_016_37    m/z 1188.8472     0   1   2   3E+3   4   4.5  RT (min)      lcqtof_hilic_neg_1_005_32    m/z 1188.9265     0   2   4E+3   5   5.5  RT (min)      lcqtof_hilic_neg_4_055_32    m/z 1189.1049     0   1   2E+3   0.5   1  RT (min)      lcqtof_hilic_neg_1_005_32    m/z 1190.1145     0   1   2E+3   0.5   1  RT (min)      lcqtof_hilic_neg_1_005_32    m/z 1190.8501     0   2   4   6E+3   0.5   1  RT (min)      lcqtof_hilic_neg_4_055_32    m/z 1191.0839     0   1E+4   0.5   1  RT (min)      lcqtof_hilic_neg_4_004_47    m/z 1191.5165     0   1   2   3E+3   0.5   1  RT (min)      lcqtof_hilic_neg_2_042_27    m/z 1192.8668     0   2   4   6E+3   5   5.5  RT (min)       lcqtof_hilic_neg_4_048_23    m/z 1193.0469     0   2   4E+3   0.5   1  RT (min)      lcqtof_hilic_neg_4_011_35    m/z 1193.1342     0   2   4   6   8E+3   0.5   1  RT (min)      lcqtof_hilic_neg_2_047_32    m/z 1193.7206     0   1   2   3E+3   7   7.5  RT (min)      lcqtof_hilic_neg_4_002_25    m/z 1193.8026     0   1   2E+3   4.5   5  RT (min)       lcqtof_hilic_neg_4_056_56    m/z 1193.8693     0   2   4E+3   5   5.5  RT (min)      lcqtof_hilic_neg_4_048_23    m/z 1193.8906     0   2   4   6   8E+3   0.5   1  RT (min)      lcqtof_hilic_neg_3_035_46    m/z 1194.0356     0   2   4   6   8E+3   1   1.5  RT (min)      lcqtof_hilic_neg_4_008_24    m/z 1194.0790     0   2   4   6E+3   0.5   1  RT (min)      lcqtof_hilic_neg_4_004_47    m/z 1194.0964     0   2   4E+3   0.5   1  RT (min)      lcqtof_hilic_neg_4_026_04    m/z 1195.3026     0   1   2   3E+3   6   6.5  RT (min)      lcqtof_hilic_neg_4_010_28    m/z 1195.6593     0   1E+3   3.5   4  RT (min)      lcqtof_hilic_neg_1_009_50    m/z 1195.9099     0   2   4   6E+3   0.5   1  RT (min)      lcqtof_hilic_neg_2_021_17    m/z 1195.9583     0   2   4   6   8E+3   4.5   5  RT (min)       lcqtof_hilic_neg_4_055_32    m/z 1196.0442     0   2   4   6E+3   1   1.5  RT (min)      lcqtof_hilic_neg_4_008_24    m/z 1196.0513     0   2   4   6E+3   0.5   1  RT (min)      lcqtof_hilic_neg_4_048_23    m/z 1196.8480     0   2   4   6E+3   0.5   1  RT (min)      lcqtof_hilic_neg_4_004_47    m/z 1196.8837     0   2   4   6   8E+3   3   3.5  RT (min)      lcqtof_hilic_neg_4_028_36    m/z 1196.9414     0   1E+3   6   6.5  RT (min)      lcqtof_hilic_neg_2_032_02    m/z 1197.0577     0   2   4E+3   0.5   1  RT (min)      lcqtof_hilic_neg_4_016_37    m/z 1197.8451     0   2   4   6E+3   5   5.5  RT (min)      lcqtof_hilic_neg_4_055_32    m/z 1198.0719     0   2   4   6   8E+3   0.5   1  RT (min)      lcqtof_hilic_neg_2_029_31    m/z 1199.0706     0   2   4   6   8E+3   0.5   1  RT (min)      lcqtof_hilic_neg_4_011_35    m/z 1199.6488     0   1   2   3   4E+3   3.5   4  RT (min)      lcqtof_hilic_neg_4_003_16    m/z 1199.8679     0   2   4   6E+3   0.5   1  RT (min)      lcqtof_hilic_neg_4_004_47   
