## Supplementary material for "The Everything Bagel Feature Finder: Ultra-fast automated feature finding for untargeted metabolomics": SI Data File: tool_unique_xic_st_Masscube_ST004500.html

Masscube-unique features (not detected by the other two tools) - ST004500

### Masscube-unique features (not detected by the other two tools) - ST004500

351 features · orbitrap · XIC window m/z ± 10 ppm (≥ 0.002 Da) · ± 0.5 min

Every panel is a feature **Masscube** detected that neither of the other two tools detected — three-way unique across EB (standard mode (min\_detection\_rate=0.1)), Masscube and XCMS, under the standard matching rule (m/z ± 10 ppm; apex within the reference RT window ± 0.2 min). It shows the MS1 extracted-ion chromatogram at the feature's m/z, drawn from the raw mzML named in `alignment_reference_file`, zoomed to ± 0.5 min around the feature's reported apex RT. **Gray solid line** = the feature's reported apex RT; **red dashed line** = masscube's assigned MS/MS — `MS2_scan_id` in `MS2_source_file`; the panel widens to keep an off-apex MS/MS on-screen. Features with no assigned MS/MS show only the gray line. y is per-cell autoscaled. Rows keep the table order (m/z ascending).

m/z 68.9957

01E+511.5RT (min)

lcorbi\_t3\_neg\_4\_110\_62

m/z 89.0244

02468E+51.52RT (min)

lcorbi\_t3\_neg\_4\_063\_15

m/z 89.0244

02468E+51.52RT (min)

lcorbi\_t3\_neg\_4\_063\_15

m/z 96.9601

012E+511.5RT (min)

lcorbi\_t3\_neg\_4\_087\_59

m/z 96.9695

01E+611.5RT (min)

lcorbi\_t3\_neg\_4\_095\_60

m/z 96.9696

01E+611.5RT (min)

lcorbi\_t3\_neg\_4\_085\_59

m/z 100.9335

0246E+51313.5RT (min)

lcorbi\_t3\_neg\_1\_054\_37

m/z 100.9335

0246E+51414.5RT (min)

lcorbi\_t3\_neg\_3\_002\_24

m/z 100.9335

0123E+51515.5RT (min)

lcorbi\_t3\_neg\_1\_060\_39

m/z 100.9335

0246E+514.515RT (min)

lcorbi\_t3\_neg\_3\_002\_24

m/z 101.9336

01E+51414.5RT (min)

lcorbi\_t3\_neg\_1\_048\_12

m/z 112.9855

024E+566.5RT (min)

lcorbi\_t3\_neg\_2\_065\_51

m/z 112.9855

0123E+612.51517.5RT (min)

lcorbi\_t3\_neg\_4\_091\_60

m/z 112.9855

01E+51.52RT (min)

lcorbi\_t3\_neg\_4\_000\_dd

m/z 112.9856

02468E+51010.5RT (min)

lcorbi\_t3\_neg\_2\_058\_39

m/z 112.9856

0123E+61015RT (min)

lcorbi\_t3\_neg\_4\_093\_60

m/z 112.9856

024E+55.56RT (min)

lcorbi\_t3\_neg\_2\_058\_39

m/z 112.9856

024E+54.55RT (min)

lcorbi\_t3\_neg\_1\_057\_49

m/z 112.9856

0123E+533.5RT (min)

lcorbi\_t3\_neg\_2\_065\_51

m/z 112.9856

024E+555.5RT (min)

lcorbi\_t3\_neg\_2\_065\_51

m/z 112.9856

0246E+56.57RT (min)

lcorbi\_t3\_neg\_2\_061\_50

m/z 112.9856

0123E+61313.514RT (min)

lcorbi\_t3\_neg\_4\_111\_62

m/z 112.9856

0123E+611.5RT (min)

lcorbi\_t3\_neg\_4\_052\_26

m/z 112.9856

02468E+599.5RT (min)

lcorbi\_t3\_neg\_1\_065\_51

m/z 112.9856

0123E+611.5RT (min)

lcorbi\_t3\_neg\_4\_026\_46

m/z 112.9856

02468E+577.5RT (min)

lcorbi\_t3\_neg\_1\_061\_50

m/z 112.9856

02468E+599.5RT (min)

lcorbi\_t3\_neg\_2\_061\_50

m/z 112.9856

0246E+422.5RT (min)

lcorbi\_t3\_neg\_2\_048\_27

m/z 112.9857

01E+61212.5RT (min)

lcorbi\_t3\_neg\_2\_061\_50

m/z 112.9876

0123E+67.51012.5RT (min)

lcorbi\_t3\_neg\_4\_108\_62

m/z 112.9877

01E+61515.5RT (min)

lcorbi\_t3\_neg\_1\_057\_49

m/z 121.0295

01E+65.56RT (min)

lcorbi\_t3\_neg\_4\_023\_18

m/z 128.0353

0246E+51.52RT (min)

lcorbi\_t3\_neg\_3\_005\_23

m/z 134.8946

0123E+51414.5RT (min)

lcorbi\_t3\_neg\_1\_056\_56

m/z 134.8946

01E+51313.5RT (min)

lcorbi\_t3\_neg\_1\_054\_37

m/z 134.8946

02468E+41212.5RT (min)

lcorbi\_t3\_neg\_1\_047\_02

m/z 134.8946

012E+51515.5RT (min)

lcorbi\_t3\_neg\_1\_054\_37

m/z 134.8946

01E+513.514RT (min)

lcorbi\_t3\_neg\_1\_054\_37

m/z 134.8946

012E+514.515RT (min)

lcorbi\_t3\_neg\_1\_052\_24

m/z 134.8946

012E+51515.5RT (min)

lcorbi\_t3\_neg\_1\_049\_55

m/z 134.8946

012E+51414.5RT (min)

lcorbi\_t3\_neg\_1\_056\_56

m/z 136.8628

01E+51.52RT (min)

lcorbi\_t3\_neg\_4\_067\_57

m/z 152.0467

0246E+533.5RT (min)

lcorbi\_t3\_neg\_4\_090\_60

m/z 158.0121

01E+513.514RT (min)

lcorbi\_t3\_neg\_4\_042\_14

m/z 158.9754

0123E+513.514RT (min)

lcorbi\_t3\_neg\_3\_010\_15

m/z 158.9754

0246E+514.515RT (min)

lcorbi\_t3\_neg\_1\_052\_24

m/z 158.9754

01E+51212.5RT (min)

lcorbi\_t3\_neg\_2\_036\_35

m/z 158.9755

01E+51111.5RT (min)

lcorbi\_t3\_neg\_2\_045\_30

m/z 158.9755

012E+59.510RT (min)

lcorbi\_t3\_neg\_2\_048\_27

m/z 158.9755

01E+510.511RT (min)

lcorbi\_t3\_neg\_1\_063\_14

m/z 158.9755

01E+510.511RT (min)

lcorbi\_t3\_neg\_2\_043\_33

m/z 158.9755

01E+599.5RT (min)

lcorbi\_t3\_neg\_2\_048\_27

m/z 158.9755

01E+51111.5RT (min)

lcorbi\_t3\_neg\_1\_054\_37

m/z 158.9755

02468E+411.512RT (min)

lcorbi\_t3\_neg\_4\_007\_13

m/z 158.9755

01E+511.512RT (min)

lcorbi\_t3\_neg\_1\_054\_37

m/z 158.9755

0123E+588.5RT (min)

lcorbi\_t3\_neg\_2\_048\_27

m/z 158.9755

0123E+513.514RT (min)

lcorbi\_t3\_neg\_1\_047\_02

m/z 158.9755

012E+515.516RT (min)

lcorbi\_t3\_neg\_3\_028\_35

m/z 158.9755

012E+512.513RT (min)

lcorbi\_t3\_neg\_4\_016\_05

m/z 158.9755

0123E+588.5RT (min)

lcorbi\_t3\_neg\_2\_048\_27

m/z 158.9755

0123E+57.58RT (min)

lcorbi\_t3\_neg\_2\_048\_27

m/z 158.9755

0246E+51414.5RT (min)

lcorbi\_t3\_neg\_1\_052\_24

m/z 158.9755

01234E+51313.5RT (min)

lcorbi\_t3\_neg\_1\_054\_37

m/z 158.9755

01234E+512.513RT (min)

lcorbi\_t3\_neg\_1\_054\_37

m/z 158.9755

0123E+57.58RT (min)

lcorbi\_t3\_neg\_2\_048\_27

m/z 158.9755

01E+51212.5RT (min)

lcorbi\_t3\_neg\_1\_054\_37

m/z 158.9755

01E+51212.5RT (min)

lcorbi\_t3\_neg\_2\_022\_24

m/z 158.9755

01E+51010.5RT (min)

lcorbi\_t3\_neg\_2\_045\_30

m/z 158.9755

01234E+512.513RT (min)

lcorbi\_t3\_neg\_1\_054\_37

m/z 158.9755

01E+51010.5RT (min)

lcorbi\_t3\_neg\_2\_002\_26

m/z 159.0084

01234E+51414.5RT (min)

lcorbi\_t3\_neg\_4\_048\_03

m/z 160.8420

012E+588.5RT (min)

lcorbi\_t3\_neg\_1\_034\_04

m/z 160.8420

012E+51111.5RT (min)

lcorbi\_t3\_neg\_1\_023\_03

m/z 160.8420

012E+511.512RT (min)

lcorbi\_t3\_neg\_1\_001\_49

m/z 160.8420

012E+511.512RT (min)

lcorbi\_t3\_neg\_1\_001\_49

m/z 160.8420

01E+51212.5RT (min)

lcorbi\_t3\_neg\_1\_001\_49

m/z 160.8420

012E+511.512RT (min)

lcorbi\_t3\_neg\_1\_001\_49

m/z 160.8420

012E+510.511RT (min)

lcorbi\_t3\_neg\_1\_023\_03

m/z 160.8420

01E+51111.5RT (min)

lcorbi\_t3\_neg\_1\_023\_03

m/z 160.8420

012E+511.512RT (min)

lcorbi\_t3\_neg\_1\_001\_49

m/z 160.8420

01E+512.513RT (min)

lcorbi\_t3\_neg\_1\_032\_13

m/z 160.8420

012E+510.511RT (min)

lcorbi\_t3\_neg\_1\_023\_03

m/z 160.8420

01E+512.513RT (min)

lcorbi\_t3\_neg\_4\_007\_13

m/z 160.8420

01E+51212.5RT (min)

lcorbi\_t3\_neg\_1\_032\_13

m/z 160.8420

01E+512.513RT (min)

lcorbi\_t3\_neg\_1\_001\_49

m/z 160.8420

012E+510.511RT (min)

lcorbi\_t3\_neg\_1\_023\_03

m/z 160.8420

012E+51212.5RT (min)

lcorbi\_t3\_neg\_1\_001\_49

m/z 160.8420

012E+57.58RT (min)

lcorbi\_t3\_neg\_1\_034\_04

m/z 160.8421

012E+577.5RT (min)

lcorbi\_t3\_neg\_1\_034\_04

m/z 160.8421

012E+53.54RT (min)

lcorbi\_t3\_neg\_1\_023\_03

m/z 160.8421

01E+54.55RT (min)

lcorbi\_t3\_neg\_1\_034\_04

m/z 160.8421

01E+544.5RT (min)

lcorbi\_t3\_neg\_1\_023\_03

m/z 160.8421

01E+533.5RT (min)

lcorbi\_t3\_neg\_1\_034\_04

m/z 160.8421

012E+533.5RT (min)

lcorbi\_t3\_neg\_1\_023\_03

m/z 160.8421

012E+515.516RT (min)

lcorbi\_t3\_neg\_4\_005\_27

m/z 160.8421

01E+555.5RT (min)

lcorbi\_t3\_neg\_1\_034\_04

m/z 160.8421

012E+500.5RT (min)

lcorbi\_t3\_neg\_4\_066\_57

m/z 160.8421

01E+544.5RT (min)

lcorbi\_t3\_neg\_4\_034\_01

m/z 160.8421

02468E+416.517RT (min)

lcorbi\_t3\_neg\_4\_007\_13

m/z 160.8421

012E+500.5RT (min)

lcorbi\_t3\_neg\_4\_068\_57

m/z 160.8421

012E+533.5RT (min)

lcorbi\_t3\_neg\_1\_023\_03

m/z 160.8421

01E+51515.5RT (min)

lcorbi\_t3\_neg\_4\_036\_16

m/z 160.8421

0246E+51.52RT (min)

lcorbi\_t3\_neg\_1\_032\_13

m/z 160.8421

012E+533.5RT (min)

lcorbi\_t3\_neg\_1\_023\_03

m/z 160.8421

01E+51616.5RT (min)

lcorbi\_t3\_neg\_4\_007\_13

m/z 160.8421

01E+516.517RT (min)

lcorbi\_t3\_neg\_4\_007\_13

m/z 160.8421

01E+544.5RT (min)

lcorbi\_t3\_neg\_4\_054\_36

m/z 160.8421

01E+522.5RT (min)

lcorbi\_t3\_neg\_2\_010\_02

m/z 160.8421

012E+533.5RT (min)

lcorbi\_t3\_neg\_1\_023\_03

m/z 160.8421

012E+599.5RT (min)

lcorbi\_t3\_neg\_1\_034\_04

m/z 160.8422

01E+515.516RT (min)

lcorbi\_t3\_neg\_4\_007\_13

m/z 160.8422

012E+515.516RT (min)

lcorbi\_t3\_neg\_4\_038\_15

m/z 160.8422

0123E+61.52RT (min)

lcorbi\_t3\_neg\_1\_010\_17

m/z 160.8422

01234E+600.5RT (min)

lcorbi\_t3\_neg\_2\_047\_48

m/z 160.8422

01E+515.516RT (min)

lcorbi\_t3\_neg\_4\_052\_26

m/z 161.9866

012E+63.54RT (min)

lcorbi\_t3\_neg\_4\_098\_61

m/z 161.9866

012E+633.5RT (min)

lcorbi\_t3\_neg\_4\_100\_61

m/z 161.9867

0123E+522.5RT (min)

lcorbi\_t3\_neg\_2\_051\_34

m/z 162.8391

01E+55.56RT (min)

lcorbi\_t3\_neg\_2\_038\_22

m/z 162.8391

01E+510.511RT (min)

lcorbi\_t3\_neg\_2\_051\_34

m/z 162.8391

02468E+410.511RT (min)

lcorbi\_t3\_neg\_2\_049\_55

m/z 162.8391

02468E+41313.5RT (min)

lcorbi\_t3\_neg\_4\_046\_37

m/z 162.8391

01E+51212.5RT (min)

lcorbi\_t3\_neg\_1\_001\_49

m/z 162.8391

01E+544.5RT (min)

lcorbi\_t3\_neg\_4\_074\_58

m/z 162.8391

01E+512.513RT (min)

lcorbi\_t3\_neg\_1\_032\_13

m/z 162.8391

01E+512.513RT (min)

lcorbi\_t3\_neg\_1\_001\_49

m/z 162.8391

02468E+41111.5RT (min)

lcorbi\_t3\_neg\_4\_039\_32

m/z 162.8391

0123E+600.5RT (min)

lcorbi\_t3\_neg\_3\_024\_26

m/z 162.8391

01E+566.5RT (min)

lcorbi\_t3\_neg\_2\_034\_19

m/z 162.8391

01E+512.513RT (min)

lcorbi\_t3\_neg\_4\_007\_13

m/z 162.8391

012E+511.512RT (min)

lcorbi\_t3\_neg\_1\_001\_49

m/z 162.8391

012E+599.5RT (min)

lcorbi\_t3\_neg\_1\_034\_04

m/z 162.8391

0246E+51.52RT (min)

lcorbi\_t3\_neg\_1\_032\_13

m/z 162.8391

01E+53.54RT (min)

lcorbi\_t3\_neg\_1\_023\_03

m/z 162.8391

01E+52.53RT (min)

lcorbi\_t3\_neg\_4\_036\_16

m/z 162.8391

01E+51212.5RT (min)

lcorbi\_t3\_neg\_1\_001\_49

m/z 162.8391

012E+500.5RT (min)

lcorbi\_t3\_neg\_4\_066\_57

m/z 162.8391

01E+511.512RT (min)

lcorbi\_t3\_neg\_1\_039\_38

m/z 162.8391

01E+55.56RT (min)

lcorbi\_t3\_neg\_1\_034\_04

m/z 162.8391

01E+511.512RT (min)

lcorbi\_t3\_neg\_1\_007\_26

m/z 162.8391

01E+52.53RT (min)

lcorbi\_t3\_neg\_2\_010\_02

m/z 162.8391

012E+51212.5RT (min)

lcorbi\_t3\_neg\_1\_001\_49

m/z 162.8391

01E+544.5RT (min)

lcorbi\_t3\_neg\_1\_023\_03

m/z 162.8391

01E+555.5RT (min)

lcorbi\_t3\_neg\_1\_034\_04

m/z 162.8391

012E+533.5RT (min)

lcorbi\_t3\_neg\_1\_023\_03

m/z 162.8391

01E+54.55RT (min)

lcorbi\_t3\_neg\_1\_034\_04

m/z 162.8391

012E+515.516RT (min)

lcorbi\_t3\_neg\_4\_037\_29

m/z 162.8391

012E+533.5RT (min)

lcorbi\_t3\_neg\_1\_023\_03

m/z 162.8391

012E+533.5RT (min)

lcorbi\_t3\_neg\_1\_023\_03

m/z 162.8391

012E+511.512RT (min)

lcorbi\_t3\_neg\_1\_001\_49

m/z 162.8391

012E+500.5RT (min)

lcorbi\_t3\_neg\_4\_068\_57

m/z 162.8391

02468E+499.5RT (min)

lcorbi\_t3\_neg\_2\_048\_27

m/z 162.8391

01E+544.5RT (min)

lcorbi\_t3\_neg\_1\_023\_03

m/z 162.8391

012E+577.5RT (min)

lcorbi\_t3\_neg\_1\_034\_04

m/z 162.8392

01E+515.516RT (min)

lcorbi\_t3\_neg\_4\_007\_13

m/z 162.8392

01E+515.516RT (min)

lcorbi\_t3\_neg\_4\_007\_13

m/z 162.8392

012E+51515.5RT (min)

lcorbi\_t3\_neg\_4\_037\_29

m/z 162.8392

01E+51616.5RT (min)

lcorbi\_t3\_neg\_4\_007\_13

m/z 162.8392

012E+515.516RT (min)

lcorbi\_t3\_neg\_4\_037\_29

m/z 164.8361

02468E+499.5RT (min)

lcorbi\_t3\_neg\_1\_008\_08

m/z 170.9443

0246E+41717.5RT (min)

lcorbi\_t3\_neg\_2\_013\_08

m/z 174.9561

01E+51.52RT (min)

lcorbi\_t3\_neg\_1\_005\_07

m/z 174.9561

01E+52.53RT (min)

lcorbi\_t3\_neg\_1\_004\_22

m/z 174.9561

01E+51.52RT (min)

lcorbi\_t3\_neg\_1\_005\_07

m/z 174.9561

0246E+466.5RT (min)

lcorbi\_t3\_neg\_2\_048\_27

m/z 174.9562

01E+600.5RT (min)

lcorbi\_t3\_neg\_1\_001\_49

m/z 178.0183

01E+51414.5RT (min)

lcorbi\_t3\_neg\_3\_035\_03

m/z 179.0147

012E+51414.5RT (min)

lcorbi\_t3\_neg\_4\_030\_04

m/z 179.0147

02468E+41313.5RT (min)

lcorbi\_t3\_neg\_4\_090\_60

m/z 180.9730

01E+5910RT (min)

lcorbi\_t3\_neg\_4\_089\_59

m/z 180.9730

01E+5810RT (min)

lcorbi\_t3\_neg\_4\_072\_57

m/z 180.9731

01E+517.518RT (min)

lcorbi\_t3\_neg\_4\_034\_01

m/z 180.9731

02468E+416.517RT (min)

lcorbi\_t3\_neg\_4\_033\_53

m/z 180.9731

02468E+41616.5RT (min)

lcorbi\_t3\_neg\_4\_051\_40

m/z 180.9731

02468E+416.517RT (min)

lcorbi\_t3\_neg\_4\_056\_56

m/z 180.9733

01E+518.519RT (min)

lcorbi\_t3\_neg\_1\_001\_49

m/z 191.0514

012E+411.5RT (min)

lcorbi\_t3\_neg\_4\_034\_01

m/z 195.8109

01E+51313.5RT (min)

lcorbi\_t3\_neg\_4\_043\_11

m/z 195.8109

01E+513.514RT (min)

lcorbi\_t3\_neg\_4\_043\_11

m/z 197.8079

01E+513.514RT (min)

lcorbi\_t3\_neg\_1\_029\_21

m/z 197.8079

01E+513.514RT (min)

lcorbi\_t3\_neg\_1\_042\_18

m/z 197.8079

01E+513.514RT (min)

lcorbi\_t3\_neg\_4\_014\_20

m/z 197.8079

01E+513.514RT (min)

lcorbi\_t3\_neg\_1\_042\_18

m/z 197.8079

01E+51313.5RT (min)

lcorbi\_t3\_neg\_4\_040\_19

m/z 197.8079

01E+51212.5RT (min)

lcorbi\_t3\_neg\_4\_048\_03

m/z 197.8080

012E+55.56RT (min)

lcorbi\_t3\_neg\_2\_048\_27

m/z 197.9632

0246E+41818.5RT (min)

lcorbi\_t3\_neg\_2\_008\_16

m/z 197.9632

0246E+41818.5RT (min)

lcorbi\_t3\_neg\_2\_010\_02

m/z 203.0212

01234E+51414.5RT (min)

lcorbi\_t3\_neg\_4\_048\_03

m/z 203.0212

01234E+51414.5RT (min)

lcorbi\_t3\_neg\_3\_035\_03

m/z 204.0175

01234E+514.515RT (min)

lcorbi\_t3\_neg\_3\_051\_46

m/z 204.0175

0123E+514.515RT (min)

lcorbi\_t3\_neg\_4\_037\_29

m/z 204.0175

0123E+51515.5RT (min)

lcorbi\_t3\_neg\_4\_037\_29

m/z 204.0175

01E+61414.5RT (min)

lcorbi\_t3\_neg\_4\_030\_04

m/z 204.0175

012E+51313.5RT (min)

lcorbi\_t3\_neg\_4\_097\_60

m/z 204.0176

01E+61414.5RT (min)

lcorbi\_t3\_neg\_4\_048\_03

m/z 204.0176

0123E+514.515RT (min)

lcorbi\_t3\_neg\_4\_037\_29

m/z 204.0176

0246E+51414.5RT (min)

lcorbi\_t3\_neg\_4\_020\_44

m/z 205.0138

024E+511.5RT (min)

lcorbi\_t3\_neg\_3\_019\_12

m/z 205.0139

012E+51515.5RT (min)

lcorbi\_t3\_neg\_4\_007\_13

m/z 205.0139

024E+514.515RT (min)

lcorbi\_t3\_neg\_3\_041\_54

m/z 205.0139

012E+514.515RT (min)

lcorbi\_t3\_neg\_4\_023\_18

m/z 205.0139

0123E+51111.5RT (min)

lcorbi\_t3\_neg\_2\_026\_03

m/z 205.0139

012E+51515.5RT (min)

lcorbi\_t3\_neg\_4\_029\_31

m/z 205.0139

012E+61414.5RT (min)

lcorbi\_t3\_neg\_4\_048\_03

m/z 205.0139

012E+512.513RT (min)

lcorbi\_t3\_neg\_1\_005\_07

m/z 205.0139

012E+51313.5RT (min)

lcorbi\_t3\_neg\_2\_019\_07

m/z 205.0139

024E+514.515RT (min)

lcorbi\_t3\_neg\_4\_023\_18

m/z 205.0139

0123E+51313.5RT (min)

lcorbi\_t3\_neg\_4\_097\_60

m/z 205.0139

01234E+514.51515.5RT (min)

lcorbi\_t3\_neg\_4\_001\_49

m/z 205.0139

01234E+514.515RT (min)

lcorbi\_t3\_neg\_4\_009\_50

m/z 205.0139

012E+61414.5RT (min)

lcorbi\_t3\_neg\_4\_030\_04

m/z 205.0139

01E+61414.5RT (min)

lcorbi\_t3\_neg\_4\_020\_44

m/z 205.0524

01E+51818.5RT (min)

lcorbi\_t3\_neg\_3\_023\_07

m/z 205.8397

01E+55.56RT (min)

lcorbi\_t3\_neg\_1\_054\_37

m/z 205.8397

01E+566.5RT (min)

lcorbi\_t3\_neg\_1\_054\_37

m/z 207.8366

01E+54.55RT (min)

lcorbi\_t3\_neg\_3\_022\_27

m/z 207.8366

01E+555.5RT (min)

lcorbi\_t3\_neg\_1\_054\_37

m/z 207.8366

01E+53.54RT (min)

lcorbi\_t3\_neg\_3\_032\_17

m/z 207.8366

01E+55.56RT (min)

lcorbi\_t3\_neg\_1\_054\_37

m/z 212.9107

012E+51515.5RT (min)

lcorbi\_t3\_neg\_1\_047\_02

m/z 212.9108

012E+515.516RT (min)

lcorbi\_t3\_neg\_3\_037\_36

m/z 212.9108

012E+51515.5RT (min)

lcorbi\_t3\_neg\_1\_047\_02

m/z 212.9108

02468E+41818.5RT (min)

lcorbi\_t3\_neg\_3\_037\_36

m/z 215.8684

01E+54.55RT (min)

lcorbi\_t3\_neg\_1\_043\_14

m/z 217.0346

012E+31414.5RT (min)

lcorbi\_t3\_neg\_4\_019\_08

m/z 218.0311

0123E+51414.5RT (min)

lcorbi\_t3\_neg\_2\_055\_18

m/z 218.0312

01E+31414.5RT (min)

lcorbi\_t3\_neg\_4\_019\_08

m/z 219.8453

01E+53.54RT (min)

lcorbi\_t3\_neg\_3\_053\_14

m/z 219.8453

012E+522.5RT (min)

lcorbi\_t3\_neg\_4\_036\_16

m/z 219.8453

01E+533.5RT (min)

lcorbi\_t3\_neg\_4\_010\_28

m/z 219.8453

01E+53.54RT (min)

lcorbi\_t3\_neg\_3\_053\_14

m/z 219.8453

01E+52.53RT (min)

lcorbi\_t3\_neg\_3\_022\_27

m/z 226.9784

0246E+417.518RT (min)

lcorbi\_t3\_neg\_2\_051\_34

m/z 231.9315

01E+51313.5RT (min)

lcorbi\_t3\_neg\_4\_038\_15

m/z 231.9315

01E+513.514RT (min)

lcorbi\_t3\_neg\_4\_054\_36

m/z 231.9315

01E+51313.5RT (min)

lcorbi\_t3\_neg\_1\_055\_05

m/z 231.9315

01E+51313.5RT (min)

lcorbi\_t3\_neg\_3\_030\_33

m/z 235.9259

01E+533.5RT (min)

lcorbi\_t3\_neg\_1\_043\_14

m/z 235.9259

01E+50.51RT (min)

lcorbi\_t3\_neg\_1\_044\_23

m/z 235.9259

012E+544.5RT (min)

lcorbi\_t3\_neg\_1\_043\_14

m/z 235.9259

01E+53.54RT (min)

lcorbi\_t3\_neg\_1\_043\_14

m/z 248.9602

01E+58910RT (min)

lcorbi\_t3\_neg\_4\_032\_23

m/z 248.9602

01E+51313.5RT (min)

lcorbi\_t3\_neg\_3\_039\_29

m/z 248.9602

01E+51313.5RT (min)

lcorbi\_t3\_neg\_2\_008\_16

m/z 248.9602

01E+51313.5RT (min)

lcorbi\_t3\_neg\_3\_024\_26

m/z 248.9602

01E+51111.5RT (min)

lcorbi\_t3\_neg\_3\_029\_02

m/z 248.9602

02468E+41212.5RT (min)

lcorbi\_t3\_neg\_2\_007\_14

m/z 248.9602

01E+511.512RT (min)

lcorbi\_t3\_neg\_3\_051\_46

m/z 248.9602

01E+51111.5RT (min)

lcorbi\_t3\_neg\_4\_034\_01

m/z 248.9602

01E+51212.5RT (min)

lcorbi\_t3\_neg\_3\_044\_01

m/z 248.9603

01E+51616.5RT (min)

lcorbi\_t3\_neg\_4\_029\_31

m/z 248.9603

012E+516.517RT (min)

lcorbi\_t3\_neg\_4\_029\_31

m/z 248.9603

012E+516.517RT (min)

lcorbi\_t3\_neg\_4\_034\_01

m/z 248.9603

012E+516.517RT (min)

lcorbi\_t3\_neg\_4\_036\_16

m/z 262.0378

01E+31414.5RT (min)

lcorbi\_t3\_neg\_4\_019\_08

m/z 291.0831

01E+62.53RT (min)

lcorbi\_t3\_neg\_4\_042\_14

m/z 291.0831

01E+622.5RT (min)

lcorbi\_t3\_neg\_4\_042\_14

m/z 315.0410

012E+52.53RT (min)

lcorbi\_t3\_neg\_2\_007\_14

m/z 315.0410

012E+52.53RT (min)

lcorbi\_t3\_neg\_2\_007\_14

m/z 315.0410

0246E+51.52RT (min)

lcorbi\_t3\_neg\_2\_007\_14

m/z 315.0411

012E+51.52RT (min)

lcorbi\_t3\_neg\_3\_031\_45

m/z 316.9474

01E+512.513RT (min)

lcorbi\_t3\_neg\_3\_035\_03

m/z 316.9474

02468E+41313.5RT (min)

lcorbi\_t3\_neg\_3\_049\_55

m/z 316.9474

02468E+411.512RT (min)

lcorbi\_t3\_neg\_3\_035\_03

m/z 316.9474

0246E+41010.5RT (min)

lcorbi\_t3\_neg\_4\_048\_03

m/z 316.9474

02468E+41313.5RT (min)

lcorbi\_t3\_neg\_3\_018\_13

m/z 316.9474

0246E+410.511RT (min)

lcorbi\_t3\_neg\_4\_030\_04

m/z 316.9475

0246E+49.510RT (min)

lcorbi\_t3\_neg\_4\_030\_04

m/z 316.9475

02468E+49.510RT (min)

lcorbi\_t3\_neg\_4\_040\_19

m/z 316.9475

02468E+41313.5RT (min)

lcorbi\_t3\_neg\_3\_018\_13

m/z 316.9476

02468E+499.5RT (min)

lcorbi\_t3\_neg\_4\_005\_27

m/z 316.9476

01E+599.5RT (min)

lcorbi\_t3\_neg\_4\_040\_19

m/z 317.0663

01E+699.5RT (min)

lcorbi\_t3\_neg\_4\_023\_18

m/z 317.0664

01E+699.5RT (min)

lcorbi\_t3\_neg\_4\_023\_18

m/z 317.0664

01E+699.5RT (min)

lcorbi\_t3\_neg\_4\_023\_18

m/z 325.1297

01E+611.5RT (min)

lcorbi\_t3\_neg\_4\_011\_24

m/z 341.9995

01E+52.53RT (min)

lcorbi\_t3\_neg\_1\_043\_14

m/z 341.9995

01E+522.5RT (min)

lcorbi\_t3\_neg\_1\_043\_14

m/z 343.9943

01E+611.5RT (min)

lcorbi\_t3\_neg\_3\_010\_15

m/z 343.9944

01E+611.5RT (min)

lcorbi\_t3\_neg\_1\_051\_15

m/z 343.9944

01E+62.53RT (min)

lcorbi\_t3\_neg\_4\_042\_14

m/z 343.9944

01E+633.5RT (min)

lcorbi\_t3\_neg\_1\_051\_15

m/z 343.9945

01E+61.52RT (min)

lcorbi\_t3\_neg\_1\_051\_15

m/z 343.9945

01E+63.54RT (min)

lcorbi\_t3\_neg\_3\_053\_14

m/z 343.9945

0246E+51.52RT (min)

lcorbi\_t3\_neg\_1\_047\_02

m/z 343.9945

01E+633.5RT (min)

lcorbi\_t3\_neg\_4\_042\_14

m/z 343.9945

01E+622.5RT (min)

lcorbi\_t3\_neg\_4\_042\_14

m/z 344.9972

012E+533.5RT (min)

lcorbi\_t3\_neg\_4\_042\_14

m/z 344.9973

012E+52.53RT (min)

lcorbi\_t3\_neg\_1\_043\_14

m/z 344.9973

012E+52.53RT (min)

lcorbi\_t3\_neg\_4\_042\_14

m/z 344.9974

012E+51.52RT (min)

lcorbi\_t3\_neg\_1\_043\_14

m/z 344.9975

012E+51.52RT (min)

lcorbi\_t3\_neg\_1\_043\_14

m/z 353.0100

01E+51.52RT (min)

lcorbi\_t3\_neg\_2\_054\_15

m/z 353.0100

01E+533.5RT (min)

lcorbi\_t3\_neg\_4\_038\_15

m/z 353.0100

01E+52.53RT (min)

lcorbi\_t3\_neg\_4\_034\_01

m/z 353.0100

01E+52.53RT (min)

lcorbi\_t3\_neg\_2\_054\_15

m/z 353.0100

01E+53.54RT (min)

lcorbi\_t3\_neg\_4\_038\_15

m/z 353.0100

012E+533.5RT (min)

lcorbi\_t3\_neg\_4\_042\_14

m/z 353.0100

012E+52.53RT (min)

lcorbi\_t3\_neg\_4\_042\_14

m/z 353.0100

01E+51.52RT (min)

lcorbi\_t3\_neg\_3\_044\_01

m/z 367.9060

01E+517.518RT (min)

lcorbi\_t3\_neg\_2\_016\_11

m/z 367.9060

01E+517.518RT (min)

lcorbi\_t3\_neg\_3\_030\_33

m/z 369.0047

01E+511.5RT (min)

lcorbi\_t3\_neg\_2\_048\_27

m/z 369.0050

01E+522.5RT (min)

lcorbi\_t3\_neg\_4\_033\_53

m/z 424.0854

0246E+54.55RT (min)

lcorbi\_t3\_neg\_4\_023\_18

m/z 491.3593

012E+51414.5RT (min)

lcorbi\_t3\_neg\_1\_007\_26

m/z 515.2871

0246E+57.58RT (min)

lcorbi\_t3\_neg\_4\_023\_18

m/z 520.8986

024E+51818.5RT (min)

lcorbi\_t3\_neg\_4\_019\_08

m/z 588.8850

024E+51818.5RT (min)

lcorbi\_t3\_neg\_4\_019\_08

m/z 591.4630

01E+512.513RT (min)

lcorbi\_t3\_neg\_4\_042\_14

m/z 654.2879

02468E+513.514RT (min)

lcorbi\_t3\_neg\_4\_019\_08

m/z 654.2884

02468E+513.514RT (min)

lcorbi\_t3\_neg\_4\_019\_08

m/z 711.6378

024E+51414.5RT (min)

lcorbi\_t3\_neg\_1\_055\_05

m/z 746.4978

01E+61313.5RT (min)

lcorbi\_t3\_neg\_3\_005\_23

m/z 752.6192

02468E+51414.5RT (min)

lcorbi\_t3\_neg\_4\_019\_08

m/z 768.4818

02468E+512.513RT (min)

lcorbi\_t3\_neg\_2\_053\_23

m/z 776.6098

01E+613.514RT (min)

lcorbi\_t3\_neg\_4\_019\_08

m/z 780.6408

024E+51414.5RT (min)

lcorbi\_t3\_neg\_4\_019\_08

m/z 782.4960

01E+61414.5RT (min)

lcorbi\_t3\_neg\_2\_053\_23

m/z 782.6559

02468E+51414.5RT (min)

lcorbi\_t3\_neg\_4\_019\_08

m/z 797.6115

01E+613.514RT (min)

lcorbi\_t3\_neg\_4\_019\_08

m/z 800.5353

01E+61414.5RT (min)

lcorbi\_t3\_neg\_4\_019\_08

m/z 808.6164

024E+512.513RT (min)

lcorbi\_t3\_neg\_1\_052\_24

m/z 811.6352

01E+61313.5RT (min)

lcorbi\_t3\_neg\_3\_002\_24

m/z 822.5110

01E+613.514RT (min)

lcorbi\_t3\_neg\_4\_019\_08

m/z 823.5142

02468E+513.514RT (min)

lcorbi\_t3\_neg\_4\_019\_08

m/z 826.5283

024E+513.514RT (min)

lcorbi\_t3\_neg\_4\_019\_08

m/z 826.5511

012E+61414.5RT (min)

lcorbi\_t3\_neg\_4\_052\_26

m/z 830.5822

01E+61414.5RT (min)

lcorbi\_t3\_neg\_1\_043\_14

m/z 840.6787

0123E+613.514RT (min)

lcorbi\_t3\_neg\_2\_052\_32

m/z 841.6638

024E+51414.5RT (min)

lcorbi\_t3\_neg\_4\_019\_08

m/z 843.6795

024E+51414.5RT (min)

lcorbi\_t3\_neg\_4\_019\_08

m/z 843.7148

0246E+51414.5RT (min)

lcorbi\_t3\_neg\_4\_019\_08

m/z 853.6825

02468E+513.514RT (min)

lcorbi\_t3\_neg\_3\_052\_32

m/z 856.5342

01E+612.513RT (min)

lcorbi\_t3\_neg\_2\_053\_23

m/z 857.5375

024E+512.513RT (min)

lcorbi\_t3\_neg\_2\_053\_23

m/z 858.5495

0246E+512.513RT (min)

lcorbi\_t3\_neg\_4\_032\_23

m/z 858.5498

0246E+51313.5RT (min)

lcorbi\_t3\_neg\_2\_053\_23

m/z 858.6685

01E+514.515RT (min)

lcorbi\_t3\_neg\_2\_054\_15

m/z 860.5654

024E+51313.5RT (min)

lcorbi\_t3\_neg\_4\_032\_23

m/z 865.6542

02468E+513.514RT (min)

lcorbi\_t3\_neg\_4\_019\_08

m/z 866.6573

012E+613.514RT (min)

lcorbi\_t3\_neg\_4\_019\_08

m/z 867.6700

01E+613.514RT (min)

lcorbi\_t3\_neg\_4\_019\_08

m/z 867.6703

01E+613.514RT (min)

lcorbi\_t3\_neg\_4\_019\_08

m/z 869.6852

01E+61414.5RT (min)

lcorbi\_t3\_neg\_4\_019\_08

m/z 871.7012

0246E+51414.5RT (min)

lcorbi\_t3\_neg\_4\_019\_08

m/z 872.6594

024E+512.513RT (min)

lcorbi\_t3\_neg\_1\_052\_24

m/z 876.6907

0246E+51313.5RT (min)

lcorbi\_t3\_neg\_1\_052\_24

m/z 876.6908

0246E+51313.5RT (min)

lcorbi\_t3\_neg\_2\_022\_24

m/z 936.6595

024E+513.514RT (min)

lcorbi\_t3\_neg\_4\_019\_08
