## Supplementary material for "The Everything Bagel Feature Finder: Ultra-fast automated feature finding for untargeted metabolomics": SI Data File: tool_unique_xic_st_Masscube_ST004501.html

Masscube-unique features (not detected by the other two tools) - ST004501

### Masscube-unique features (not detected by the other two tools) - ST004501

178 features · orbitrap · XIC window m/z ± 10 ppm (≥ 0.002 Da) · ± 0.5 min

Every panel is a feature **Masscube** detected that neither of the other two tools detected — three-way unique across EB (standard mode (min\_detection\_rate=0.1)), Masscube and XCMS, under the standard matching rule (m/z ± 10 ppm; apex within the reference RT window ± 0.2 min). It shows the MS1 extracted-ion chromatogram at the feature's m/z, drawn from the raw mzML named in `alignment_reference_file`, zoomed to ± 0.5 min around the feature's reported apex RT. **Gray solid line** = the feature's reported apex RT; **red dashed line** = masscube's assigned MS/MS — `MS2_scan_id` in `MS2_source_file`; the panel widens to keep an off-apex MS/MS on-screen. Features with no assigned MS/MS show only the gray line. y is per-cell autoscaled. Rows keep the table order (m/z ascending).

m/z 60.0444

0123E+522.5RT (min)

lcorbi\_hilic\_pos\_2\_029\_19

m/z 60.0444

0246E+511.5RT (min)

lcorbi\_hilic\_pos\_2\_016\_18

m/z 60.0444

0246E+511.5RT (min)

lcorbi\_hilic\_pos\_2\_014\_20

m/z 60.0444

01E+633.5RT (min)

lcorbi\_hilic\_pos\_1\_006\_11

m/z 74.0964

01E+610RT (min)

lcorbi\_hilic\_pos\_4\_097\_60

m/z 74.0964

01E+67.51012.5RT (min)

lcorbi\_hilic\_pos\_4\_074\_58

m/z 74.0964

0246E+51111.5RT (min)

lcorbi\_hilic\_pos\_4\_057\_49

m/z 74.0964

0246E+510.511RT (min)

lcorbi\_hilic\_pos\_4\_063\_15

m/z 74.0964

0246E+51212.5RT (min)

lcorbi\_hilic\_pos\_4\_061\_50

m/z 74.0964

024E+51010.5RT (min)

lcorbi\_hilic\_pos\_3\_063\_15

m/z 74.0964

024E+59.510RT (min)

lcorbi\_hilic\_pos\_4\_060\_14

m/z 74.0964

0246E+51010.5RT (min)

lcorbi\_hilic\_pos\_4\_061\_50

m/z 74.0964

0246E+51010.5RT (min)

lcorbi\_hilic\_pos\_4\_057\_49

m/z 74.0964

0246E+51111.5RT (min)

lcorbi\_hilic\_pos\_3\_064\_13

m/z 74.0964

0246E+510.511RT (min)

lcorbi\_hilic\_pos\_4\_060\_14

m/z 74.0964

0246E+555.5RT (min)

lcorbi\_hilic\_pos\_4\_059\_13

m/z 74.0964

024E+51111.5RT (min)

lcorbi\_hilic\_pos\_2\_060\_14

m/z 74.0965

024E+599.5RT (min)

lcorbi\_hilic\_pos\_4\_057\_49

m/z 74.0965

024E+58.59RT (min)

lcorbi\_hilic\_pos\_4\_058\_17

m/z 85.0284

01E+622.5RT (min)

lcorbi\_hilic\_pos\_4\_020\_37

m/z 91.0290

01E+533.54RT (min)

lcorbi\_hilic\_pos\_4\_090\_60

m/z 91.0290

01E+544.5RT (min)

lcorbi\_hilic\_pos\_4\_086\_59

m/z 91.0291

0246E+400.5RT (min)

lcorbi\_hilic\_pos\_2\_014\_20

m/z 91.0291

02468E+43.54RT (min)

lcorbi\_hilic\_pos\_4\_077\_58

m/z 91.0291

02468E+41.52RT (min)

lcorbi\_hilic\_pos\_1\_059\_17

m/z 91.0291

02468E+42.53RT (min)

lcorbi\_hilic\_pos\_3\_063\_15

m/z 91.0291

01E+500.5RT (min)

lcorbi\_hilic\_pos\_4\_060\_14

m/z 91.0291

02468E+400.5RT (min)

lcorbi\_hilic\_pos\_4\_083\_59

m/z 91.0291

01E+545RT (min)

lcorbi\_hilic\_pos\_1\_034\_37

m/z 91.0291

01E+500.5RT (min)

lcorbi\_hilic\_pos\_2\_059\_16

m/z 91.0291

02468E+44.55RT (min)

lcorbi\_hilic\_pos\_3\_065\_51

m/z 91.0291

0246E+44.55RT (min)

lcorbi\_hilic\_pos\_4\_080\_58

m/z 92.0368

0246E+502RT (min)

lcorbi\_hilic\_pos\_2\_057\_49

m/z 92.0368

024E+533.5RT (min)

lcorbi\_hilic\_pos\_2\_060\_14

m/z 92.0369

0123E+544.5RT (min)

lcorbi\_hilic\_pos\_3\_062\_39

m/z 92.0369

01234E+545RT (min)

lcorbi\_hilic\_pos\_3\_062\_39

m/z 92.0369

024E+52.53RT (min)

lcorbi\_hilic\_pos\_3\_058\_16

m/z 92.0369

01234E+544.5RT (min)

lcorbi\_hilic\_pos\_2\_058\_39

m/z 103.0390

02468E+522.5RT (min)

lcorbi\_hilic\_pos\_4\_020\_37

m/z 111.0917

01E+55.566.5RT (min)

lcorbi\_hilic\_pos\_4\_072\_57

m/z 111.0917

01E+54.55RT (min)

lcorbi\_hilic\_pos\_4\_072\_57

m/z 111.0917

01E+555.5RT (min)

lcorbi\_hilic\_pos\_4\_072\_57

m/z 111.0917

01E+555.5RT (min)

lcorbi\_hilic\_pos\_4\_066\_57

m/z 111.0917

01E+55.56RT (min)

lcorbi\_hilic\_pos\_1\_001\_49

m/z 113.1073

01E+555.5RT (min)

lcorbi\_hilic\_pos\_4\_018\_31

m/z 118.0863

02468E+65.56RT (min)

lcorbi\_hilic\_pos\_1\_008\_38

m/z 122.0713

02468E+555.5RT (min)

lcorbi\_hilic\_pos\_4\_020\_37

m/z 124.0869

012E+54.55RT (min)

lcorbi\_hilic\_pos\_4\_082\_59

m/z 124.0869

01234E+502RT (min)

lcorbi\_hilic\_pos\_4\_109\_62

m/z 124.0869

012E+544.5RT (min)

lcorbi\_hilic\_pos\_4\_107\_62

m/z 124.0869

012E+54.55RT (min)

lcorbi\_hilic\_pos\_4\_082\_59

m/z 124.0869

012E+50.51RT (min)

lcorbi\_hilic\_pos\_4\_082\_59

m/z 124.0870

01E+500.5RT (min)

lcorbi\_hilic\_pos\_4\_082\_59

m/z 124.0870

012E+53.54RT (min)

lcorbi\_hilic\_pos\_4\_082\_59

m/z 124.0870

012E+50.51RT (min)

lcorbi\_hilic\_pos\_4\_082\_59

m/z 125.9862

0246E+41010.5RT (min)

lcorbi\_hilic\_pos\_3\_030\_39

m/z 125.9862

012E+599.5RT (min)

lcorbi\_hilic\_pos\_4\_020\_37

m/z 125.9862

01234E+599.5RT (min)

lcorbi\_hilic\_pos\_4\_075\_58

m/z 125.9863

01234E+58.59RT (min)

lcorbi\_hilic\_pos\_4\_074\_58

m/z 137.0458

012E+644.5RT (min)

lcorbi\_hilic\_pos\_4\_020\_37

m/z 139.1229

02468E+455.5RT (min)

lcorbi\_hilic\_pos\_4\_077\_58

m/z 139.9879

01234E+57.58RT (min)

lcorbi\_hilic\_pos\_4\_050\_16

m/z 139.9880

0123E+58.59RT (min)

lcorbi\_hilic\_pos\_3\_030\_39

m/z 140.9859

0246E+57.58RT (min)

lcorbi\_hilic\_pos\_1\_007\_16

m/z 143.9968

01E+59.510RT (min)

lcorbi\_hilic\_pos\_4\_078\_58

m/z 143.9968

01234E+5910RT (min)

lcorbi\_hilic\_pos\_4\_078\_58

m/z 143.9968

01E+51010.5RT (min)

lcorbi\_hilic\_pos\_4\_078\_58

m/z 143.9968

01E+59.510RT (min)

lcorbi\_hilic\_pos\_4\_078\_58

m/z 143.9968

01234E+5910RT (min)

lcorbi\_hilic\_pos\_4\_080\_58

m/z 143.9969

01234E+58.59RT (min)

lcorbi\_hilic\_pos\_4\_081\_58

m/z 144.0478

0246E+555.5RT (min)

lcorbi\_hilic\_pos\_4\_020\_37

m/z 144.9569

01E+57.58RT (min)

lcorbi\_hilic\_pos\_1\_007\_16

m/z 144.9821

02468E+533.5RT (min)

lcorbi\_hilic\_pos\_1\_005\_45

m/z 144.9821

01E+633.5RT (min)

lcorbi\_hilic\_pos\_1\_001\_49

m/z 144.9822

01234E+54.55RT (min)

lcorbi\_hilic\_pos\_1\_002\_29

m/z 144.9822

01E+622.5RT (min)

lcorbi\_hilic\_pos\_1\_005\_45

m/z 144.9822

02468E+544.5RT (min)

lcorbi\_hilic\_pos\_1\_002\_29

m/z 144.9822

01E+611.5RT (min)

lcorbi\_hilic\_pos\_1\_002\_29

m/z 144.9822

0123E+54.55RT (min)

lcorbi\_hilic\_pos\_1\_002\_29

m/z 144.9822

01234E+60.51RT (min)

lcorbi\_hilic\_pos\_1\_002\_29

m/z 144.9822

024E+60.51RT (min)

lcorbi\_hilic\_pos\_1\_001\_49

m/z 144.9822

02468E+500.5RT (min)

lcorbi\_hilic\_pos\_1\_001\_49

m/z 144.9822

024E+611.5RT (min)

lcorbi\_hilic\_pos\_1\_001\_49

m/z 144.9822

024E+60.51RT (min)

lcorbi\_hilic\_pos\_1\_001\_49

m/z 144.9822

024E+611.5RT (min)

lcorbi\_hilic\_pos\_1\_001\_49

m/z 144.9822

024E+60.51RT (min)

lcorbi\_hilic\_pos\_1\_001\_49

m/z 144.9822

02468E+533.5RT (min)

lcorbi\_hilic\_pos\_1\_002\_29

m/z 144.9822

01E+61.52RT (min)

lcorbi\_hilic\_pos\_1\_001\_49

m/z 144.9822

01E+62.53RT (min)

lcorbi\_hilic\_pos\_1\_001\_49

m/z 144.9822

01E+622.5RT (min)

lcorbi\_hilic\_pos\_1\_001\_49

m/z 144.9822

01E+62.53RT (min)

lcorbi\_hilic\_pos\_1\_001\_49

m/z 144.9822

01E+62.53RT (min)

lcorbi\_hilic\_pos\_1\_001\_49

m/z 144.9822

01E+62.53RT (min)

lcorbi\_hilic\_pos\_1\_001\_49

m/z 144.9822

01E+622.5RT (min)

lcorbi\_hilic\_pos\_1\_001\_49

m/z 144.9822

0246E+500.5RT (min)

lcorbi\_hilic\_pos\_1\_004\_22

m/z 144.9822

01E+622.5RT (min)

lcorbi\_hilic\_pos\_1\_001\_49

m/z 144.9822

0246E+544.5RT (min)

lcorbi\_hilic\_pos\_1\_008\_38

m/z 144.9822

0246E+544.5RT (min)

lcorbi\_hilic\_pos\_1\_002\_29

m/z 146.9803

01E+51313.5RT (min)

lcorbi\_hilic\_pos\_4\_084\_59

m/z 146.9803

012E+60.51RT (min)

lcorbi\_hilic\_pos\_1\_002\_29

m/z 146.9803

01234E+522.5RT (min)

lcorbi\_hilic\_pos\_1\_004\_22

m/z 146.9803

0123E+500.5RT (min)

lcorbi\_hilic\_pos\_1\_002\_29

m/z 146.9803

024E+544.5RT (min)

lcorbi\_hilic\_pos\_1\_001\_49

m/z 146.9803

012E+60.51RT (min)

lcorbi\_hilic\_pos\_1\_001\_49

m/z 146.9803

012E+611.5RT (min)

lcorbi\_hilic\_pos\_1\_001\_49

m/z 146.9803

01234E+52.53RT (min)

lcorbi\_hilic\_pos\_1\_002\_29

m/z 146.9803

0246E+51.52RT (min)

lcorbi\_hilic\_pos\_1\_001\_49

m/z 146.9803

0123E+600.5RT (min)

lcorbi\_hilic\_pos\_1\_004\_22

m/z 146.9803

012E+60.51RT (min)

lcorbi\_hilic\_pos\_1\_002\_29

m/z 146.9803

012E+60.51RT (min)

lcorbi\_hilic\_pos\_1\_001\_49

m/z 146.9803

024E+52.533.5RT (min)

lcorbi\_hilic\_pos\_1\_001\_49

m/z 146.9803

01234E+500.5RT (min)

lcorbi\_hilic\_pos\_1\_001\_49

m/z 146.9803

02468E+41212.5RT (min)

lcorbi\_hilic\_pos\_4\_071\_57

m/z 146.9803

01E+512.513RT (min)

lcorbi\_hilic\_pos\_4\_069\_57

m/z 146.9804

024E+544.5RT (min)

lcorbi\_hilic\_pos\_1\_001\_49

m/z 146.9804

01234E+511.5RT (min)

lcorbi\_hilic\_pos\_1\_002\_29

m/z 146.9804

024E+52.53RT (min)

lcorbi\_hilic\_pos\_1\_001\_49

m/z 146.9804

012E+611.5RT (min)

lcorbi\_hilic\_pos\_1\_001\_49

m/z 146.9804

01234E+522.5RT (min)

lcorbi\_hilic\_pos\_1\_002\_29

m/z 146.9804

01234E+52.53RT (min)

lcorbi\_hilic\_pos\_1\_002\_29

m/z 146.9804

0246E+522.5RT (min)

lcorbi\_hilic\_pos\_1\_001\_49

m/z 146.9804

012E+544.5RT (min)

lcorbi\_hilic\_pos\_1\_003\_24

m/z 146.9804

0123E+51.52RT (min)

lcorbi\_hilic\_pos\_1\_011\_09

m/z 146.9804

0123E+54.55RT (min)

lcorbi\_hilic\_pos\_1\_001\_49

m/z 146.9804

024E+522.5RT (min)

lcorbi\_hilic\_pos\_1\_005\_45

m/z 146.9804

01234E+52.53RT (min)

lcorbi\_hilic\_pos\_1\_002\_29

m/z 149.0298

02468E+40.51RT (min)

lcorbi\_hilic\_pos\_4\_081\_58

m/z 151.0353

024E+588.5RT (min)

lcorbi\_hilic\_pos\_4\_081\_58

m/z 151.0353

01E+599.5RT (min)

lcorbi\_hilic\_pos\_4\_106\_62

m/z 151.0353

01E+599.5RT (min)

lcorbi\_hilic\_pos\_4\_106\_62

m/z 167.0128

01E+59.510RT (min)

lcorbi\_hilic\_pos\_4\_069\_57

m/z 167.0128

0123E+58.59RT (min)

lcorbi\_hilic\_pos\_4\_080\_58

m/z 167.0128

0123E+599.5RT (min)

lcorbi\_hilic\_pos\_4\_078\_58

m/z 167.0128

01E+59.510RT (min)

lcorbi\_hilic\_pos\_4\_066\_57

m/z 167.0128

01E+59.510RT (min)

lcorbi\_hilic\_pos\_4\_076\_58

m/z 167.0128

0123E+599.5RT (min)

lcorbi\_hilic\_pos\_4\_080\_58

m/z 173.1286

0246E+54.55RT (min)

lcorbi\_hilic\_pos\_4\_020\_37

m/z 173.1287

0246E+54.55RT (min)

lcorbi\_hilic\_pos\_4\_020\_37

m/z 173.1762

01E+666.5RT (min)

lcorbi\_hilic\_pos\_4\_020\_37

m/z 174.1603

02468E+53.54RT (min)

lcorbi\_hilic\_pos\_4\_020\_37

m/z 184.9856

0123E+57.58RT (min)

lcorbi\_hilic\_pos\_2\_019\_16

m/z 188.1029

0246E+566.5RT (min)

lcorbi\_hilic\_pos\_4\_020\_37

m/z 188.1758

0246E+55.56RT (min)

lcorbi\_hilic\_pos\_4\_020\_37

m/z 198.0875

01E+666.5RT (min)

lcorbi\_hilic\_pos\_4\_020\_37

m/z 203.0855

024E+533.5RT (min)

lcorbi\_hilic\_pos\_3\_059\_14

m/z 212.1031

01E+666.5RT (min)

lcorbi\_hilic\_pos\_4\_020\_37

m/z 216.0769

01E+633.5RT (min)

lcorbi\_hilic\_pos\_4\_020\_37

m/z 218.2065

01E+433.5RT (min)

lcorbi\_hilic\_pos\_3\_064\_13

m/z 218.2115

02468E+73.54RT (min)

lcorbi\_hilic\_pos\_4\_039\_15

m/z 218.2115

0123E+63.54RT (min)

lcorbi\_hilic\_pos\_4\_046\_21

m/z 236.0779

02468E+544.5RT (min)

lcorbi\_hilic\_pos\_4\_020\_37

m/z 239.1060

01234E+65.56RT (min)

lcorbi\_hilic\_pos\_4\_020\_37

m/z 247.1666

01E+544.5RT (min)

lcorbi\_hilic\_pos\_4\_076\_58

m/z 247.1666

01E+544.5RT (min)

lcorbi\_hilic\_pos\_4\_105\_61

m/z 249.1234

01E+611.5RT (min)

lcorbi\_hilic\_pos\_4\_113\_62

m/z 250.0712

024E+53.54RT (min)

lcorbi\_hilic\_pos\_4\_020\_37

m/z 256.3000

0246E+522.5RT (min)

lcorbi\_hilic\_pos\_4\_016\_28

m/z 261.0879

0246E+55.56RT (min)

lcorbi\_hilic\_pos\_4\_020\_37

m/z 265.1119

01E+655.5RT (min)

lcorbi\_hilic\_pos\_4\_020\_37

m/z 268.0817

0246E+63.54RT (min)

lcorbi\_hilic\_pos\_4\_020\_37

m/z 269.0850

02468E+53.54RT (min)

lcorbi\_hilic\_pos\_4\_020\_37

m/z 324.1232

01E+811.5RT (min)

lcorbi\_hilic\_pos\_2\_016\_18

m/z 335.1022

012E+60.51RT (min)

lcorbi\_hilic\_pos\_3\_037\_20

m/z 349.0813

0246E+50.51RT (min)

lcorbi\_hilic\_pos\_3\_037\_20

m/z 350.0157

01E+677.5RT (min)

lcorbi\_hilic\_pos\_4\_074\_58

m/z 355.3686

02468E+522.5RT (min)

lcorbi\_hilic\_pos\_4\_017\_51

m/z 358.1286

01E+611.5RT (min)

lcorbi\_hilic\_pos\_4\_020\_37

m/z 379.2230

01234E+622.5RT (min)

lcorbi\_hilic\_pos\_4\_020\_37

m/z 379.2230

01234E+61.52RT (min)

lcorbi\_hilic\_pos\_4\_020\_37

m/z 379.2231

01E+61.52RT (min)

lcorbi\_hilic\_pos\_1\_014\_27

m/z 394.3107

01E+51.52RT (min)

lcorbi\_hilic\_pos\_3\_023\_18

m/z 394.3108

01E+51.52RT (min)

lcorbi\_hilic\_pos\_1\_008\_38

m/z 394.3108

01E+51.52RT (min)

lcorbi\_hilic\_pos\_2\_016\_18

m/z 442.3376

0246E+50.51RT (min)

lcorbi\_hilic\_pos\_4\_018\_31

m/z 487.1260

01234E+60.51RT (min)

lcorbi\_hilic\_pos\_3\_037\_20

m/z 504.1527

01E+60.51RT (min)

lcorbi\_hilic\_pos\_3\_037\_20

m/z 519.4058

024E+50.51RT (min)

lcorbi\_hilic\_pos\_4\_018\_31

m/z 548.4268

024E+50.51RT (min)

lcorbi\_hilic\_pos\_4\_018\_31
