## Supplementary material for "The Everything Bagel Feature Finder: Ultra-fast automated feature finding for untargeted metabolomics": SI Data File: tool_unique_xic_st_Masscube_ST004502.html

Masscube-unique features (not detected by the other two tools) - ST004502

### Masscube-unique features (not detected by the other two tools) - ST004502

1439 features · orbitrap · XIC window m/z ± 10 ppm (≥ 0.002 Da) · ± 0.5 min

Every panel is a feature **Masscube** detected that neither of the other two tools detected — three-way unique across EB (standard mode (min\_detection\_rate=0.1)), Masscube and XCMS, under the standard matching rule (m/z ± 10 ppm; apex within the reference RT window ± 0.2 min). It shows the MS1 extracted-ion chromatogram at the feature's m/z, drawn from the raw mzML named in `alignment_reference_file`, zoomed to ± 0.5 min around the feature's reported apex RT. **Gray solid line** = the feature's reported apex RT; **red dashed line** = masscube's assigned MS/MS — `MS2_scan_id` in `MS2_source_file`; the panel widens to keep an off-apex MS/MS on-screen. Features with no assigned MS/MS show only the gray line. y is per-cell autoscaled. Rows keep the table order (m/z ascending).

m/z 54.5527

01E+62.53RT (min)

lcorbi\_hilic\_neg\_2\_062\_14

m/z 55.1487

01234E+644.5RT (min)

lcorbi\_hilic\_neg\_4\_063\_14

m/z 55.1488

0123E+63.54RT (min)

lcorbi\_hilic\_neg\_3\_064\_16

m/z 57.8154

0246E+544.5RT (min)

lcorbi\_hilic\_neg\_4\_064\_17

m/z 62.9290

012E+600.5RT (min)

lcorbi\_hilic\_neg\_1\_057\_49

m/z 64.9272

0246E+500.5RT (min)

lcorbi\_hilic\_neg\_3\_057\_49

m/z 68.9957

0123E+555.5RT (min)

lcorbi\_hilic\_neg\_1\_011\_01

m/z 72.0807

01E+655.5RT (min)

lcorbi\_hilic\_neg\_2\_059\_15

m/z 72.0807

012E+677.5RT (min)

lcorbi\_hilic\_neg\_4\_061\_50

m/z 72.0807

0246E+58.59RT (min)

lcorbi\_hilic\_neg\_4\_059\_13

m/z 74.0964

01E+633.5RT (min)

lcorbi\_hilic\_neg\_1\_058\_39

m/z 74.0964

024E+51111.5RT (min)

lcorbi\_hilic\_neg\_1\_060\_13

m/z 74.0964

024E+62.53RT (min)

lcorbi\_hilic\_neg\_1\_057\_49

m/z 74.0964

01E+69.510RT (min)

lcorbi\_hilic\_neg\_1\_057\_49

m/z 74.0964

012E+60.51RT (min)

lcorbi\_hilic\_neg\_3\_059\_14

m/z 74.0964

01E+71.52RT (min)

lcorbi\_hilic\_neg\_1\_057\_49

m/z 74.0964

0246E+512.513RT (min)

lcorbi\_hilic\_neg\_1\_058\_39

m/z 74.0964

012E+63.54RT (min)

lcorbi\_hilic\_neg\_2\_057\_49

m/z 74.0964

0123E+62.53RT (min)

lcorbi\_hilic\_neg\_3\_057\_49

m/z 74.0964

0123E+63.54RT (min)

lcorbi\_hilic\_neg\_1\_057\_49

m/z 74.0964

024E+522.5RT (min)

lcorbi\_hilic\_neg\_1\_059\_14

m/z 74.0964

012E+64.55RT (min)

lcorbi\_hilic\_neg\_1\_057\_49

m/z 74.0964

01E+67.58RT (min)

lcorbi\_hilic\_neg\_1\_057\_49

m/z 74.0964

0246E+511.512RT (min)

lcorbi\_hilic\_neg\_2\_058\_16

m/z 75.0997

012E+600.5RT (min)

lcorbi\_hilic\_neg\_1\_057\_49

m/z 75.0997

02468E+511.5RT (min)

lcorbi\_hilic\_neg\_1\_057\_49

m/z 78.9590

012E+566.5RT (min)

lcorbi\_hilic\_neg\_4\_097\_60

m/z 78.9590

012E+57.58RT (min)

lcorbi\_hilic\_neg\_3\_031\_14

m/z 81.0446

01E+644.5RT (min)

lcorbi\_hilic\_neg\_1\_059\_14

m/z 81.0447

01E+622.5RT (min)

lcorbi\_hilic\_neg\_2\_061\_50

m/z 81.0447

02468E+566.5RT (min)

lcorbi\_hilic\_neg\_2\_061\_50

m/z 81.0447

012E+63.54RT (min)

lcorbi\_hilic\_neg\_4\_058\_15

m/z 81.0447

01E+62.53RT (min)

lcorbi\_hilic\_neg\_3\_057\_49

m/z 81.0447

0246E+55.56RT (min)

lcorbi\_hilic\_neg\_1\_058\_39

m/z 81.0447

0246E+56.57RT (min)

lcorbi\_hilic\_neg\_4\_058\_15

m/z 82.5370

0123E+588.5RT (min)

lcorbi\_hilic\_neg\_2\_060\_39

m/z 84.9597

0246E+58.59RT (min)

lcorbi\_hilic\_neg\_1\_059\_14

m/z 84.9597

024E+58.59RT (min)

lcorbi\_hilic\_neg\_4\_062\_39

m/z 85.0296

0246E+53.54RT (min)

lcorbi\_hilic\_neg\_3\_010\_18

m/z 87.0088

01E+51.52RT (min)

lcorbi\_hilic\_neg\_1\_028\_45

m/z 88.1120

0246E+533.5RT (min)

lcorbi\_hilic\_neg\_3\_064\_16

m/z 89.0244

0123E+64.55RT (min)

lcorbi\_hilic\_neg\_1\_051\_33

m/z 89.0244

01234E+655.5RT (min)

lcorbi\_hilic\_neg\_1\_003\_14

m/z 89.0244

01E+70.51RT (min)

lcorbi\_hilic\_neg\_1\_004\_17

m/z 89.0244

01234E+634RT (min)

lcorbi\_hilic\_neg\_4\_036\_36

m/z 89.0244

02468E+53.54RT (min)

lcorbi\_hilic\_neg\_4\_004\_13

m/z 89.0244

02468E+50.51RT (min)

lcorbi\_hilic\_neg\_1\_001\_49

m/z 89.0244

02468E+50.51RT (min)

lcorbi\_hilic\_neg\_1\_002\_26

m/z 89.0244

01E+63.54RT (min)

lcorbi\_hilic\_neg\_4\_023\_34

m/z 89.0244

01E+63.54RT (min)

lcorbi\_hilic\_neg\_4\_031\_14

m/z 89.0244

0123E+6468RT (min)

lcorbi\_hilic\_neg\_4\_013\_46

m/z 89.0244

01E+633.5RT (min)

lcorbi\_hilic\_neg\_4\_031\_14

m/z 89.0244

01234E+633.5RT (min)

lcorbi\_hilic\_neg\_4\_036\_36

m/z 89.0244

02468E+500.5RT (min)

lcorbi\_hilic\_neg\_1\_001\_49

m/z 89.0244

01E+63.54RT (min)

lcorbi\_hilic\_neg\_4\_036\_36

m/z 89.0244

012E+711.5RT (min)

lcorbi\_hilic\_neg\_1\_005\_36

m/z 89.0245

01234E+64RT (min)

lcorbi\_hilic\_neg\_2\_020\_27

m/z 89.0245

01E+63.54RT (min)

lcorbi\_hilic\_neg\_4\_036\_36

m/z 89.0245

01234E+544.5RT (min)

lcorbi\_hilic\_neg\_1\_054\_27

m/z 89.0245

02468E+622.5RT (min)

lcorbi\_hilic\_neg\_4\_048\_37

m/z 89.0245

01E+711.5RT (min)

lcorbi\_hilic\_neg\_1\_004\_17

m/z 89.0245

01E+64.55RT (min)

lcorbi\_hilic\_neg\_1\_048\_35

m/z 89.0245

01234E+655.5RT (min)

lcorbi\_hilic\_neg\_2\_020\_27

m/z 89.0245

012E+633.5RT (min)

lcorbi\_hilic\_neg\_4\_038\_27

m/z 89.0245

0123E+62.53RT (min)

lcorbi\_hilic\_neg\_4\_013\_46

m/z 89.0245

0123E+633.5RT (min)

lcorbi\_hilic\_neg\_4\_037\_31

m/z 89.0245

02468E+61.52RT (min)

lcorbi\_hilic\_neg\_1\_039\_38

m/z 89.0245

01234E+544.5RT (min)

lcorbi\_hilic\_neg\_1\_054\_27

m/z 89.0245

02468E+61.52RT (min)

lcorbi\_hilic\_neg\_2\_040\_35

m/z 89.0245

01E+722.5RT (min)

lcorbi\_hilic\_neg\_2\_048\_38

m/z 89.0245

0123E+62.53RT (min)

lcorbi\_hilic\_neg\_3\_009\_50

m/z 89.0245

02468E+62.53RT (min)

lcorbi\_hilic\_neg\_4\_048\_37

m/z 90.0277

01E+52.53RT (min)

lcorbi\_hilic\_neg\_4\_023\_34

m/z 90.0277

01E+52.53RT (min)

lcorbi\_hilic\_neg\_4\_026\_33

m/z 90.0277

012E+52.53RT (min)

lcorbi\_hilic\_neg\_3\_044\_32

m/z 90.0278

0246E+511.5RT (min)

lcorbi\_hilic\_neg\_1\_005\_36

m/z 90.0549

0123E+666.5RT (min)

lcorbi\_hilic\_neg\_2\_062\_14

m/z 90.5258

024E+58.59RT (min)

lcorbi\_hilic\_neg\_1\_059\_14

m/z 90.9766

0123E+755.5RT (min)

lcorbi\_hilic\_neg\_3\_064\_16

m/z 91.0542

01E+73.54RT (min)

lcorbi\_hilic\_neg\_2\_061\_50

m/z 92.9280

01E+57.58RT (min)

lcorbi\_hilic\_neg\_4\_073\_57

m/z 94.0450

012E+588.5RT (min)

lcorbi\_hilic\_neg\_2\_062\_14

m/z 94.9251

012E+533.5RT (min)

lcorbi\_hilic\_neg\_1\_008\_39

m/z 94.9251

0246E+544.5RT (min)

lcorbi\_hilic\_neg\_1\_007\_10

m/z 96.9601

01E+544.5RT (min)

lcorbi\_hilic\_neg\_4\_082\_59

m/z 96.9601

0123E+50.51RT (min)

lcorbi\_hilic\_neg\_4\_004\_13

m/z 96.9601

01E+53.54RT (min)

lcorbi\_hilic\_neg\_4\_073\_57

m/z 96.9601

012E+53.54RT (min)

lcorbi\_hilic\_neg\_2\_001\_49

m/z 96.9601

012E+50.51RT (min)

lcorbi\_hilic\_neg\_4\_078\_58

m/z 96.9601

024E+50.51RT (min)

lcorbi\_hilic\_neg\_1\_001\_49

m/z 96.9601

024E+50.51RT (min)

lcorbi\_hilic\_neg\_1\_001\_49

m/z 96.9601

012E+511.5RT (min)

lcorbi\_hilic\_neg\_4\_004\_13

m/z 96.9601

0123E+544.5RT (min)

lcorbi\_hilic\_neg\_1\_012\_32

m/z 96.9601

0246E+500.5RT (min)

lcorbi\_hilic\_neg\_2\_005\_24

m/z 96.9601

0246E+500.5RT (min)

lcorbi\_hilic\_neg\_2\_005\_24

m/z 96.9601

01234E+544.5RT (min)

lcorbi\_hilic\_neg\_4\_015\_24

m/z 96.9601

0123E+54.55RT (min)

lcorbi\_hilic\_neg\_2\_056\_56

m/z 96.9601

0246E+500.5RT (min)

lcorbi\_hilic\_neg\_4\_004\_13

m/z 96.9601

01E+534RT (min)

lcorbi\_hilic\_neg\_4\_108\_62

m/z 96.9601

0246E+511.5RT (min)

lcorbi\_hilic\_neg\_1\_003\_14

m/z 96.9601

01234E+54.55RT (min)

lcorbi\_hilic\_neg\_1\_009\_50

m/z 96.9601

0246E+50.51RT (min)

lcorbi\_hilic\_neg\_3\_012\_35

m/z 96.9601

012E+555.5RT (min)

lcorbi\_hilic\_neg\_1\_053\_16

m/z 96.9601

0123E+555.5RT (min)

lcorbi\_hilic\_neg\_1\_015\_30

m/z 96.9601

02468E+500.5RT (min)

lcorbi\_hilic\_neg\_3\_000\_cc

m/z 96.9601

012E+544.5RT (min)

lcorbi\_hilic\_neg\_1\_054\_27

m/z 96.9602

01234E+54.55RT (min)

lcorbi\_hilic\_neg\_1\_011\_01

m/z 96.9603

01234E+54.55RT (min)

lcorbi\_hilic\_neg\_1\_019\_06

m/z 96.9603

01234E+54.55RT (min)

lcorbi\_hilic\_neg\_1\_019\_06

m/z 96.9649

0246E+38.59RT (min)

lcorbi\_hilic\_neg\_4\_027\_47

m/z 96.9687

01E+588.5RT (min)

lcorbi\_hilic\_neg\_1\_003\_14

m/z 97.9913

0123E+68.59RT (min)

lcorbi\_hilic\_neg\_1\_061\_50

m/z 97.9913

0246E+51010.5RT (min)

lcorbi\_hilic\_neg\_1\_065\_51

m/z 97.9913

02468E+59.510RT (min)

lcorbi\_hilic\_neg\_1\_058\_39

m/z 97.9913

024E+510.511RT (min)

lcorbi\_hilic\_neg\_1\_062\_16

m/z 98.9068

02468E+41111.512RT (min)

lcorbi\_hilic\_neg\_4\_104\_61

m/z 98.9068

024E+500.5RT (min)

lcorbi\_hilic\_neg\_2\_044\_46

m/z 98.9068

01E+51112RT (min)

lcorbi\_hilic\_neg\_4\_108\_62

m/z 98.9392

01E+50.51RT (min)

lcorbi\_hilic\_neg\_4\_082\_59

m/z 98.9392

01E+50.51RT (min)

lcorbi\_hilic\_neg\_4\_082\_59

m/z 98.9392

012E+50.51RT (min)

lcorbi\_hilic\_neg\_4\_082\_59

m/z 98.9392

01E+567RT (min)

lcorbi\_hilic\_neg\_4\_071\_57

m/z 98.9392

01E+56RT (min)

lcorbi\_hilic\_neg\_4\_083\_59

m/z 98.9558

01E+58.59RT (min)

lcorbi\_hilic\_neg\_4\_095\_60

m/z 98.9558

01E+577.5RT (min)

lcorbi\_hilic\_neg\_1\_023\_23

m/z 98.9558

0123E+57.58RT (min)

lcorbi\_hilic\_neg\_4\_082\_59

m/z 98.9558

0123E+57.58RT (min)

lcorbi\_hilic\_neg\_4\_087\_59

m/z 98.9559

0123E+57.58RT (min)

lcorbi\_hilic\_neg\_4\_097\_60

m/z 99.9258

012E+588.5RT (min)

lcorbi\_hilic\_neg\_4\_026\_33

m/z 99.9258

0123E+588.5RT (min)

lcorbi\_hilic\_neg\_4\_053\_23

m/z 99.9258

012E+588.5RT (min)

lcorbi\_hilic\_neg\_4\_026\_33

m/z 99.9258

0123E+544.5RT (min)

lcorbi\_hilic\_neg\_3\_012\_35

m/z 99.9258

0123E+57.58RT (min)

lcorbi\_hilic\_neg\_4\_053\_23

m/z 99.9258

012E+53.54RT (min)

lcorbi\_hilic\_neg\_4\_040\_17

m/z 99.9259

012E+53.54RT (min)

lcorbi\_hilic\_neg\_4\_051\_16

m/z 99.9260

01E+600.5RT (min)

lcorbi\_hilic\_neg\_3\_000\_aa

m/z 100.0756

01E+611.5RT (min)

lcorbi\_hilic\_neg\_1\_063\_17

m/z 100.9261

01E+54.55RT (min)

lcorbi\_hilic\_neg\_4\_098\_61

m/z 100.9261

01E+54.55RT (min)

lcorbi\_hilic\_neg\_4\_098\_61

m/z 100.9261

01E+54.55RT (min)

lcorbi\_hilic\_neg\_4\_098\_61

m/z 100.9261

02468E+44.55RT (min)

lcorbi\_hilic\_neg\_4\_104\_61

m/z 100.9335

01E+688.5RT (min)

lcorbi\_hilic\_neg\_4\_082\_59

m/z 100.9336

01E+688.5RT (min)

lcorbi\_hilic\_neg\_4\_090\_60

m/z 100.9336

02468E+58.59RT (min)

lcorbi\_hilic\_neg\_4\_048\_37

m/z 100.9336

012E+61.52RT (min)

lcorbi\_hilic\_neg\_4\_004\_13

m/z 100.9336

012E+60.51RT (min)

lcorbi\_hilic\_neg\_2\_048\_38

m/z 100.9336

02468E+500.5RT (min)

lcorbi\_hilic\_neg\_2\_011\_29

m/z 100.9336

02468E+522.5RT (min)

lcorbi\_hilic\_neg\_2\_011\_29

m/z 100.9336

0246E+57.58RT (min)

lcorbi\_hilic\_neg\_3\_023\_36

m/z 100.9336

02468E+51.52RT (min)

lcorbi\_hilic\_neg\_2\_027\_02

m/z 100.9336

012E+60.51RT (min)

lcorbi\_hilic\_neg\_3\_053\_13

m/z 100.9336

012E+60.51RT (min)

lcorbi\_hilic\_neg\_4\_009\_50

m/z 100.9336

02468E+57.58RT (min)

lcorbi\_hilic\_neg\_4\_048\_37

m/z 100.9336

012E+61.52RT (min)

lcorbi\_hilic\_neg\_3\_017\_51

m/z 100.9336

01E+61.52RT (min)

lcorbi\_hilic\_neg\_4\_016\_29

m/z 100.9336

02468E+57.58RT (min)

lcorbi\_hilic\_neg\_4\_055\_38

m/z 100.9336

01E+633.5RT (min)

lcorbi\_hilic\_neg\_2\_031\_34

m/z 100.9336

01E+62.53RT (min)

lcorbi\_hilic\_neg\_4\_013\_46

m/z 100.9336

02468E+57.58RT (min)

lcorbi\_hilic\_neg\_4\_055\_38

m/z 100.9336

01E+688.5RT (min)

lcorbi\_hilic\_neg\_4\_096\_60

m/z 100.9336

02468E+500.5RT (min)

lcorbi\_hilic\_neg\_2\_011\_29

m/z 100.9336

02468E+58.59RT (min)

lcorbi\_hilic\_neg\_4\_048\_37

m/z 100.9336

012E+600.5RT (min)

lcorbi\_hilic\_neg\_2\_000\_aa

m/z 100.9336

0246E+58.59RT (min)

lcorbi\_hilic\_neg\_3\_039\_31

m/z 100.9336

02468E+58.59RT (min)

lcorbi\_hilic\_neg\_4\_048\_37

m/z 100.9336

02468E+52.53RT (min)

lcorbi\_hilic\_neg\_3\_001\_49

m/z 100.9336

02468E+57.58RT (min)

lcorbi\_hilic\_neg\_4\_055\_38

m/z 100.9337

01E+622.5RT (min)

lcorbi\_hilic\_neg\_4\_016\_29

m/z 100.9337

02468E+533.5RT (min)

lcorbi\_hilic\_neg\_4\_016\_29

m/z 101.0244

012E+65.56RT (min)

lcorbi\_hilic\_neg\_3\_053\_13

m/z 101.9337

01E+555.5RT (min)

lcorbi\_hilic\_neg\_4\_054\_30

m/z 101.9337

01E+54.55RT (min)

lcorbi\_hilic\_neg\_4\_080\_58

m/z 101.9337

012E+555.5RT (min)

lcorbi\_hilic\_neg\_4\_079\_58

m/z 101.9337

01E+54.55RT (min)

lcorbi\_hilic\_neg\_4\_075\_58

m/z 101.9337

01E+57.58RT (min)

lcorbi\_hilic\_neg\_4\_056\_56

m/z 101.9337

01E+500.5RT (min)

lcorbi\_hilic\_neg\_4\_072\_57

m/z 101.9337

01E+544.5RT (min)

lcorbi\_hilic\_neg\_4\_075\_58

m/z 101.9337

01E+51.52RT (min)

lcorbi\_hilic\_neg\_4\_006\_06

m/z 101.9337

01E+52.53RT (min)

lcorbi\_hilic\_neg\_4\_015\_24

m/z 101.9337

012E+555.5RT (min)

lcorbi\_hilic\_neg\_4\_092\_60

m/z 101.9337

012E+555.5RT (min)

lcorbi\_hilic\_neg\_4\_086\_59

m/z 101.9337

01E+500.5RT (min)

lcorbi\_hilic\_neg\_1\_048\_35

m/z 101.9337

0123E+50.51RT (min)

lcorbi\_hilic\_neg\_4\_070\_57

m/z 101.9337

01E+588.5RT (min)

lcorbi\_hilic\_neg\_4\_027\_47

m/z 101.9337

01E+588.5RT (min)

lcorbi\_hilic\_neg\_4\_047\_07

m/z 101.9337

01E+52.53RT (min)

lcorbi\_hilic\_neg\_4\_020\_04

m/z 101.9337

01E+51.52RT (min)

lcorbi\_hilic\_neg\_4\_020\_04

m/z 101.9337

01E+52.53RT (min)

lcorbi\_hilic\_neg\_3\_008\_04

m/z 101.9337

01E+57.58RT (min)

lcorbi\_hilic\_neg\_4\_055\_38

m/z 101.9337

012E+556RT (min)

lcorbi\_hilic\_neg\_4\_085\_59

m/z 101.9337

012E+566.5RT (min)

lcorbi\_hilic\_neg\_4\_075\_58

m/z 101.9338

01E+54.55RT (min)

lcorbi\_hilic\_neg\_4\_074\_58

m/z 101.9338

012E+54.55RT (min)

lcorbi\_hilic\_neg\_4\_095\_60

m/z 101.9338

01E+522.5RT (min)

lcorbi\_hilic\_neg\_4\_050\_26

m/z 101.9338

01E+544.5RT (min)

lcorbi\_hilic\_neg\_4\_095\_60

m/z 101.9338

01E+522.5RT (min)

lcorbi\_hilic\_neg\_4\_000\_cc

m/z 101.9338

01E+522.5RT (min)

lcorbi\_hilic\_neg\_4\_028\_03

m/z 102.0338

01234E+58.59RT (min)

lcorbi\_hilic\_neg\_1\_059\_14

m/z 102.1277

01234E+52.53RT (min)

lcorbi\_hilic\_neg\_1\_061\_50

m/z 102.9568

012E+577.5RT (min)

lcorbi\_hilic\_neg\_4\_069\_57

m/z 102.9568

012E+57.58RT (min)

lcorbi\_hilic\_neg\_4\_069\_57

m/z 102.9568

012E+566.5RT (min)

lcorbi\_hilic\_neg\_2\_012\_14

m/z 102.9568

012E+566.5RT (min)

lcorbi\_hilic\_neg\_3\_031\_14

m/z 102.9568

012E+577.5RT (min)

lcorbi\_hilic\_neg\_4\_068\_57

m/z 102.9568

012E+566.5RT (min)

lcorbi\_hilic\_neg\_1\_011\_01

m/z 102.9702

01E+599.5RT (min)

lcorbi\_hilic\_neg\_4\_064\_17

m/z 103.0400

01234E+5510RT (min)

lcorbi\_hilic\_neg\_4\_095\_60

m/z 103.9552

02468E+600.5RT (min)

lcorbi\_hilic\_neg\_1\_057\_49

m/z 103.9552

01E+61.52RT (min)

lcorbi\_hilic\_neg\_1\_057\_49

m/z 104.1069

0246E+73.54RT (min)

lcorbi\_hilic\_neg\_3\_064\_16

m/z 104.1069

012E+666.5RT (min)

lcorbi\_hilic\_neg\_4\_063\_14

m/z 104.9538

01E+555.5RT (min)

lcorbi\_hilic\_neg\_1\_015\_30

m/z 104.9538

01E+555.5RT (min)

lcorbi\_hilic\_neg\_4\_073\_57

m/z 104.9538

01E+555.5RT (min)

lcorbi\_hilic\_neg\_4\_069\_57

m/z 104.9538

01E+55.56RT (min)

lcorbi\_hilic\_neg\_4\_076\_58

m/z 104.9538

01E+555.5RT (min)

lcorbi\_hilic\_neg\_3\_006\_28

m/z 105.0381

0246E+54.55RT (min)

lcorbi\_hilic\_neg\_1\_012\_32

m/z 105.0412

01234E+444.5RT (min)

lcorbi\_hilic\_neg\_4\_097\_60

m/z 105.9534

024E+51.52RT (min)

lcorbi\_hilic\_neg\_1\_057\_49

m/z 105.9534

01E+611.5RT (min)

lcorbi\_hilic\_neg\_1\_057\_49

m/z 105.9535

01234E+600.5RT (min)

lcorbi\_hilic\_neg\_1\_057\_49

m/z 106.0498

024E+56.57RT (min)

lcorbi\_hilic\_neg\_2\_057\_49

m/z 106.0862

024E+52.53RT (min)

lcorbi\_hilic\_neg\_4\_059\_13

m/z 106.0862

01234E+655.5RT (min)

lcorbi\_hilic\_neg\_2\_060\_39

m/z 108.9019

02468E+41.52RT (min)

lcorbi\_hilic\_neg\_1\_008\_39

m/z 110.0086

012E+588.5RT (min)

lcorbi\_hilic\_neg\_1\_058\_39

m/z 111.0916

01E+600.5RT (min)

lcorbi\_hilic\_neg\_1\_057\_49

m/z 112.9855

012E+755.5RT (min)

lcorbi\_hilic\_neg\_1\_013\_47

m/z 112.9856

01234E+611.5RT (min)

lcorbi\_hilic\_neg\_1\_013\_47

m/z 112.9856

0123E+62.53RT (min)

lcorbi\_hilic\_neg\_1\_009\_50

m/z 112.9856

0123E+61.52RT (min)

lcorbi\_hilic\_neg\_3\_013\_37

m/z 112.9856

01234E+611.5RT (min)

lcorbi\_hilic\_neg\_1\_017\_51

m/z 112.9856

01234E+611.5RT (min)

lcorbi\_hilic\_neg\_1\_009\_50

m/z 112.9856

0123E+60.51RT (min)

lcorbi\_hilic\_neg\_1\_009\_50

m/z 112.9856

012E+66.57RT (min)

lcorbi\_hilic\_neg\_1\_029\_31

m/z 112.9856

0123E+63.54RT (min)

lcorbi\_hilic\_neg\_1\_005\_36

m/z 112.9856

0123E+666.5RT (min)

lcorbi\_hilic\_neg\_1\_009\_50

m/z 112.9856

01E+60.51RT (min)

lcorbi\_hilic\_neg\_2\_025\_52

m/z 112.9856

01234E+611.5RT (min)

lcorbi\_hilic\_neg\_1\_013\_47

m/z 112.9856

0123E+66.57RT (min)

lcorbi\_hilic\_neg\_1\_009\_50

m/z 112.9856

0123E+66.57RT (min)

lcorbi\_hilic\_neg\_1\_005\_36

m/z 112.9856

0123E+60.51RT (min)

lcorbi\_hilic\_neg\_1\_017\_51

m/z 112.9856

0123E+62.53RT (min)

lcorbi\_hilic\_neg\_1\_009\_50

m/z 112.9856

012E+63.54RT (min)

lcorbi\_hilic\_neg\_1\_020\_13

m/z 112.9856

0123E+644.5RT (min)

lcorbi\_hilic\_neg\_1\_005\_36

m/z 112.9856

01234E+64.55RT (min)

lcorbi\_hilic\_neg\_1\_005\_36

m/z 112.9856

01E+600.5RT (min)

lcorbi\_hilic\_neg\_2\_028\_33

m/z 112.9856

012E+63.54RT (min)

lcorbi\_hilic\_neg\_1\_011\_01

m/z 112.9856

01234E+666.5RT (min)

lcorbi\_hilic\_neg\_1\_005\_36

m/z 112.9856

0123E+644.5RT (min)

lcorbi\_hilic\_neg\_1\_009\_50

m/z 112.9856

01234E+60.51RT (min)

lcorbi\_hilic\_neg\_1\_015\_30

m/z 112.9856

0123E+63.54RT (min)

lcorbi\_hilic\_neg\_1\_009\_50

m/z 112.9856

0123E+666.5RT (min)

lcorbi\_hilic\_neg\_1\_005\_36

m/z 112.9856

01234E+60.51RT (min)

lcorbi\_hilic\_neg\_1\_011\_01

m/z 112.9856

0123E+633.5RT (min)

lcorbi\_hilic\_neg\_1\_026\_34

m/z 112.9856

01E+633.5RT (min)

lcorbi\_hilic\_neg\_3\_043\_02

m/z 112.9856

0246E+44.55RT (min)

lcorbi\_hilic\_neg\_1\_028\_45

m/z 112.9856

012E+600.5RT (min)

lcorbi\_hilic\_neg\_1\_022\_48

m/z 112.9856

0123E+666.5RT (min)

lcorbi\_hilic\_neg\_1\_020\_13

m/z 112.9857

0123E+622.5RT (min)

lcorbi\_hilic\_neg\_1\_009\_50

m/z 112.9857

0123E+644.5RT (min)

lcorbi\_hilic\_neg\_1\_005\_36

m/z 112.9857

012E+61.52RT (min)

lcorbi\_hilic\_neg\_1\_015\_30

m/z 112.9857

01234E+64.55RT (min)

lcorbi\_hilic\_neg\_1\_009\_50

m/z 112.9857

012E+63.54RT (min)

lcorbi\_hilic\_neg\_1\_011\_01

m/z 112.9857

012E+677.5RT (min)

lcorbi\_hilic\_neg\_1\_005\_36

m/z 112.9858

0123E+633.5RT (min)

lcorbi\_hilic\_neg\_1\_009\_50

m/z 112.9860

0123E+633.5RT (min)

lcorbi\_hilic\_neg\_1\_009\_50

m/z 113.0243

01234E+52.53RT (min)

lcorbi\_hilic\_neg\_1\_026\_34

m/z 114.0662

01E+83.54RT (min)

lcorbi\_hilic\_neg\_1\_063\_17

m/z 115.0632

0246E+53.54RT (min)

lcorbi\_hilic\_neg\_1\_060\_13

m/z 115.9206

012E+55.56RT (min)

lcorbi\_hilic\_neg\_4\_031\_14

m/z 115.9206

012E+555.5RT (min)

lcorbi\_hilic\_neg\_4\_053\_23

m/z 115.9207

01E+53.54RT (min)

lcorbi\_hilic\_neg\_4\_074\_58

m/z 115.9207

012E+500.5RT (min)

lcorbi\_hilic\_neg\_1\_048\_35

m/z 115.9207

0246E+43.54RT (min)

lcorbi\_hilic\_neg\_4\_006\_06

m/z 115.9207

0123E+51.52RT (min)

lcorbi\_hilic\_neg\_4\_028\_03

m/z 115.9207

01E+522.5RT (min)

lcorbi\_hilic\_neg\_4\_092\_60

m/z 115.9207

0123E+50.51RT (min)

lcorbi\_hilic\_neg\_4\_074\_58

m/z 115.9207

01E+500.5RT (min)

lcorbi\_hilic\_neg\_4\_074\_58

m/z 115.9207

01E+500.5RT (min)

lcorbi\_hilic\_neg\_1\_049\_55

m/z 115.9207

01E+52.53RT (min)

lcorbi\_hilic\_neg\_4\_006\_06

m/z 115.9207

01E+522.5RT (min)

lcorbi\_hilic\_neg\_4\_028\_03

m/z 115.9207

01E+51.52RT (min)

lcorbi\_hilic\_neg\_4\_006\_06

m/z 115.9207

01E+52.53RT (min)

lcorbi\_hilic\_neg\_3\_008\_04

m/z 115.9207

01E+500.5RT (min)

lcorbi\_hilic\_neg\_4\_074\_58

m/z 115.9207

01E+588.5RT (min)

lcorbi\_hilic\_neg\_4\_034\_45

m/z 115.9207

01E+588.5RT (min)

lcorbi\_hilic\_neg\_4\_056\_56

m/z 115.9207

0123E+51.52RT (min)

lcorbi\_hilic\_neg\_4\_004\_13

m/z 115.9207

01E+57.58RT (min)

lcorbi\_hilic\_neg\_4\_041\_54

m/z 115.9207

01E+54.55RT (min)

lcorbi\_hilic\_neg\_4\_074\_58

m/z 115.9207

01E+588.5RT (min)

lcorbi\_hilic\_neg\_4\_033\_53

m/z 115.9207

01E+57.58RT (min)

lcorbi\_hilic\_neg\_4\_049\_55

m/z 115.9207

02468E+43.54RT (min)

lcorbi\_hilic\_neg\_4\_074\_58

m/z 115.9207

01E+533.5RT (min)

lcorbi\_hilic\_neg\_3\_008\_04

m/z 115.9208

01E+522.5RT (min)

lcorbi\_hilic\_neg\_4\_006\_06

m/z 115.9208

01E+58.59RT (min)

lcorbi\_hilic\_neg\_4\_048\_37

m/z 116.0018

01234E+58.59RT (min)

lcorbi\_hilic\_neg\_1\_061\_50

m/z 116.1070

01E+73.54RT (min)

lcorbi\_hilic\_neg\_4\_060\_16

m/z 116.9285

01234E+53.54RT (min)

lcorbi\_hilic\_neg\_2\_027\_02

m/z 116.9285

0246E+555.5RT (min)

lcorbi\_hilic\_neg\_1\_037\_46

m/z 116.9285

02468E+566.5RT (min)

lcorbi\_hilic\_neg\_3\_044\_32

m/z 116.9285

024E+57.58RT (min)

lcorbi\_hilic\_neg\_1\_055\_24

m/z 116.9285

012E+52.53RT (min)

lcorbi\_hilic\_neg\_3\_022\_06

m/z 116.9285

01234E+53.54RT (min)

lcorbi\_hilic\_neg\_2\_027\_02

m/z 116.9285

012E+51.52RT (min)

lcorbi\_hilic\_neg\_3\_055\_26

m/z 116.9285

0123E+51.52RT (min)

lcorbi\_hilic\_neg\_4\_050\_26

m/z 116.9285

02468E+566.5RT (min)

lcorbi\_hilic\_neg\_3\_044\_32

m/z 116.9285

0246E+355.5RT (min)

lcorbi\_hilic\_neg\_2\_009\_50

m/z 116.9285

01E+611.5RT (min)

lcorbi\_hilic\_neg\_4\_006\_06

m/z 116.9285

02468E+555.5RT (min)

lcorbi\_hilic\_neg\_4\_004\_13

m/z 116.9285

0246E+555.5RT (min)

lcorbi\_hilic\_neg\_2\_006\_45

m/z 116.9286

02468E+54.55RT (min)

lcorbi\_hilic\_neg\_2\_054\_13

m/z 116.9286

01E+46.57RT (min)

lcorbi\_hilic\_neg\_4\_005\_32

m/z 116.9286

012E+52.53RT (min)

lcorbi\_hilic\_neg\_3\_022\_06

m/z 116.9286

012E+52.53RT (min)

lcorbi\_hilic\_neg\_4\_045\_25

m/z 116.9286

02468E+54.55RT (min)

lcorbi\_hilic\_neg\_3\_016\_16

m/z 116.9286

0123E+522.5RT (min)

lcorbi\_hilic\_neg\_4\_050\_26

m/z 116.9286

0123E+522.5RT (min)

lcorbi\_hilic\_neg\_4\_050\_26

m/z 116.9286

0246E+544.5RT (min)

lcorbi\_hilic\_neg\_1\_054\_27

m/z 116.9286

012E+52.53RT (min)

lcorbi\_hilic\_neg\_4\_045\_25

m/z 116.9286

0123E+533.5RT (min)

lcorbi\_hilic\_neg\_4\_029\_02

m/z 116.9286

0123E+51.52RT (min)

lcorbi\_hilic\_neg\_4\_051\_16

m/z 116.9287

02468E+51.52RT (min)

lcorbi\_hilic\_neg\_4\_051\_16

m/z 117.0194

024E+622.5RT (min)

lcorbi\_hilic\_neg\_1\_005\_36

m/z 117.0849

01E+83.54RT (min)

lcorbi\_hilic\_neg\_1\_058\_39

m/z 117.9286

01E+57.58RT (min)

lcorbi\_hilic\_neg\_4\_085\_59

m/z 117.9286

01E+57.58RT (min)

lcorbi\_hilic\_neg\_4\_082\_59

m/z 117.9286

01E+577.5RT (min)

lcorbi\_hilic\_neg\_4\_038\_27

m/z 118.0860

0246E+755.5RT (min)

lcorbi\_hilic\_neg\_2\_062\_14

m/z 118.0862

01E+855.5RT (min)

lcorbi\_hilic\_neg\_4\_059\_13

m/z 118.0881

0246E+63.54RT (min)

lcorbi\_hilic\_neg\_1\_058\_39

m/z 118.9925

01E+53.54RT (min)

lcorbi\_hilic\_neg\_1\_021\_28

m/z 119.0171

02468E+36.57RT (min)

lcorbi\_hilic\_neg\_2\_005\_24

m/z 119.0172

02468E+366.5RT (min)

lcorbi\_hilic\_neg\_2\_005\_24

m/z 119.0855

02468E+63.54RT (min)

lcorbi\_hilic\_neg\_2\_061\_50

m/z 119.9075

02468E+499.5RT (min)

lcorbi\_hilic\_neg\_4\_031\_14

m/z 119.9075

01E+58.599.5RT (min)

lcorbi\_hilic\_neg\_4\_074\_58

m/z 119.9076

01E+599.5RT (min)

lcorbi\_hilic\_neg\_4\_091\_60

m/z 119.9076

01E+58.59RT (min)

lcorbi\_hilic\_neg\_4\_082\_59

m/z 121.9661

01E+61212.5RT (min)

lcorbi\_hilic\_neg\_3\_063\_13

m/z 121.9661

01E+611.512RT (min)

lcorbi\_hilic\_neg\_2\_063\_13

m/z 121.9661

01E+611.512RT (min)

lcorbi\_hilic\_neg\_4\_065\_51

m/z 121.9661

01E+612.513RT (min)

lcorbi\_hilic\_neg\_4\_060\_16

m/z 121.9661

01E+61313.5RT (min)

lcorbi\_hilic\_neg\_3\_064\_16

m/z 121.9661

01E+63.54RT (min)

lcorbi\_hilic\_neg\_3\_057\_49

m/z 121.9661

01E+610.511RT (min)

lcorbi\_hilic\_neg\_3\_058\_39

m/z 121.9661

0246E+61.52RT (min)

lcorbi\_hilic\_neg\_1\_057\_49

m/z 121.9661

01E+612.513RT (min)

lcorbi\_hilic\_neg\_4\_065\_51

m/z 121.9661

024E+522.5RT (min)

lcorbi\_hilic\_neg\_4\_059\_13

m/z 121.9661

024E+577.5RT (min)

lcorbi\_hilic\_neg\_1\_057\_49

m/z 121.9661

01E+61313.5RT (min)

lcorbi\_hilic\_neg\_3\_063\_13

m/z 121.9661

01E+61111.5RT (min)

lcorbi\_hilic\_neg\_3\_063\_13

m/z 121.9661

0246E+52.53RT (min)

lcorbi\_hilic\_neg\_3\_065\_51

m/z 121.9661

01E+61212.5RT (min)

lcorbi\_hilic\_neg\_2\_060\_39

m/z 121.9661

0246E+51.52RT (min)

lcorbi\_hilic\_neg\_1\_063\_17

m/z 121.9661

01E+61111.5RT (min)

lcorbi\_hilic\_neg\_3\_061\_50

m/z 121.9661

024E+700.5RT (min)

lcorbi\_hilic\_neg\_1\_057\_49

m/z 122.0964

012E+611.5RT (min)

lcorbi\_hilic\_neg\_1\_060\_13

m/z 122.9695

02468E+500.5RT (min)

lcorbi\_hilic\_neg\_3\_057\_49

m/z 123.0403

024E+588.5RT (min)

lcorbi\_hilic\_neg\_2\_062\_14

m/z 123.0552

01E+611.5RT (min)

lcorbi\_hilic\_neg\_1\_057\_49

m/z 123.9643

0246E+51111.5RT (min)

lcorbi\_hilic\_neg\_2\_061\_50

m/z 123.9643

012E+700.5RT (min)

lcorbi\_hilic\_neg\_1\_057\_49

m/z 123.9643

0246E+51111.5RT (min)

lcorbi\_hilic\_neg\_3\_063\_13

m/z 123.9643

0246E+51212.5RT (min)

lcorbi\_hilic\_neg\_2\_060\_39

m/z 123.9643

0246E+51313.5RT (min)

lcorbi\_hilic\_neg\_3\_064\_16

m/z 123.9643

0246E+51212.5RT (min)

lcorbi\_hilic\_neg\_4\_062\_39

m/z 123.9643

0246E+512.513RT (min)

lcorbi\_hilic\_neg\_4\_065\_51

m/z 123.9643

0123E+61.52RT (min)

lcorbi\_hilic\_neg\_1\_057\_49

m/z 123.9643

0246E+511.512RT (min)

lcorbi\_hilic\_neg\_2\_063\_13

m/z 123.9643

0246E+511.512RT (min)

lcorbi\_hilic\_neg\_3\_063\_13

m/z 124.0867

0246E+54.55RT (min)

lcorbi\_hilic\_neg\_1\_061\_50

m/z 124.0868

01E+600.5RT (min)

lcorbi\_hilic\_neg\_4\_063\_14

m/z 124.0868

01234E+611.5RT (min)

lcorbi\_hilic\_neg\_1\_058\_39

m/z 124.0868

01E+61.52RT (min)

lcorbi\_hilic\_neg\_1\_062\_16

m/z 124.0868

01E+644.5RT (min)

lcorbi\_hilic\_neg\_3\_058\_39

m/z 124.0869

012E+63.54RT (min)

lcorbi\_hilic\_neg\_4\_058\_15

m/z 124.0869

01E+62.53RT (min)

lcorbi\_hilic\_neg\_2\_062\_14

m/z 124.0869

012E+666.5RT (min)

lcorbi\_hilic\_neg\_4\_065\_51

m/z 124.0869

02468E+544.5RT (min)

lcorbi\_hilic\_neg\_3\_063\_13

m/z 125.0709

024E+566.5RT (min)

lcorbi\_hilic\_neg\_4\_061\_50

m/z 125.9862

0246E+51212.5RT (min)

lcorbi\_hilic\_neg\_4\_057\_49

m/z 125.9862

0123E+68.59RT (min)

lcorbi\_hilic\_neg\_4\_062\_39

m/z 125.9862

0246E+511.512RT (min)

lcorbi\_hilic\_neg\_2\_065\_51

m/z 125.9862

0246E+512.513RT (min)

lcorbi\_hilic\_neg\_3\_061\_50

m/z 125.9862

0246E+51313.5RT (min)

lcorbi\_hilic\_neg\_3\_057\_49

m/z 125.9862

0246E+512.513RT (min)

lcorbi\_hilic\_neg\_2\_064\_17

m/z 128.0192

012E+688.5RT (min)

lcorbi\_hilic\_neg\_2\_060\_39

m/z 128.0352

02468E+544.5RT (min)

lcorbi\_hilic\_neg\_1\_047\_05

m/z 128.0353

01E+63.54RT (min)

lcorbi\_hilic\_neg\_2\_022\_37

m/z 128.0353

02468E+577.5RT (min)

lcorbi\_hilic\_neg\_1\_047\_05

m/z 128.0353

01E+73.54RT (min)

lcorbi\_hilic\_neg\_4\_082\_59

m/z 128.0390

0123E+533.5RT (min)

lcorbi\_hilic\_neg\_4\_088\_59

m/z 128.9595

01E+50.51RT (min)

lcorbi\_hilic\_neg\_4\_090\_60

m/z 128.9595

01E+52.53RT (min)

lcorbi\_hilic\_neg\_1\_037\_46

m/z 128.9595

01E+51.52RT (min)

lcorbi\_hilic\_neg\_4\_070\_57

m/z 128.9595

02468E+42.53RT (min)

lcorbi\_hilic\_neg\_3\_021\_20

m/z 128.9595

02468E+40.51RT (min)

lcorbi\_hilic\_neg\_4\_090\_60

m/z 128.9595

02468E+41.52RT (min)

lcorbi\_hilic\_neg\_4\_094\_60

m/z 128.9595

01E+51.52RT (min)

lcorbi\_hilic\_neg\_4\_070\_57

m/z 128.9595

02468E+40.51RT (min)

lcorbi\_hilic\_neg\_4\_090\_60

m/z 128.9595

02468E+433.5RT (min)

lcorbi\_hilic\_neg\_4\_112\_62

m/z 128.9596

01E+52.53RT (min)

lcorbi\_hilic\_neg\_1\_037\_46

m/z 128.9596

02468E+422.5RT (min)

lcorbi\_hilic\_neg\_4\_029\_02

m/z 129.0200

0123E+588.5RT (min)

lcorbi\_hilic\_neg\_4\_061\_50

m/z 129.0385

0246E+53.54RT (min)

lcorbi\_hilic\_neg\_4\_082\_59

m/z 129.0754

02468E+611.5RT (min)

lcorbi\_hilic\_neg\_4\_061\_50

m/z 129.9363

01E+58.59RT (min)

lcorbi\_hilic\_neg\_4\_074\_58

m/z 129.9363

01E+599.5RT (min)

lcorbi\_hilic\_neg\_4\_074\_58

m/z 130.0167

0123E+588.5RT (min)

lcorbi\_hilic\_neg\_4\_061\_50

m/z 130.0862

012E+63.54RT (min)

lcorbi\_hilic\_neg\_2\_060\_39

m/z 130.1590

01E+61.52RT (min)

lcorbi\_hilic\_neg\_4\_065\_51

m/z 130.1590

012E+622.5RT (min)

lcorbi\_hilic\_neg\_4\_065\_51

m/z 130.9836

0246E+41010.5RT (min)

lcorbi\_hilic\_neg\_4\_088\_59

m/z 130.9836

02468E+499.5RT (min)

lcorbi\_hilic\_neg\_4\_111\_62

m/z 130.9836

0246E+49.510RT (min)

lcorbi\_hilic\_neg\_4\_082\_59

m/z 130.9836

0246E+49.510RT (min)

lcorbi\_hilic\_neg\_4\_089\_59

m/z 130.9836

0246E+49.510RT (min)

lcorbi\_hilic\_neg\_4\_108\_62

m/z 132.1018

01E+755.5RT (min)

lcorbi\_hilic\_neg\_1\_063\_17

m/z 132.8678

012E+60.51RT (min)

lcorbi\_hilic\_neg\_3\_054\_29

m/z 132.8678

01234E+55.56RT (min)

lcorbi\_hilic\_neg\_4\_030\_05

m/z 132.8678

012E+600.5RT (min)

lcorbi\_hilic\_neg\_3\_055\_26

m/z 132.8678

01234E+55.56RT (min)

lcorbi\_hilic\_neg\_2\_038\_36

m/z 132.8679

0123E+566.5RT (min)

lcorbi\_hilic\_neg\_4\_038\_27

m/z 132.8679

0123E+56.57RT (min)

lcorbi\_hilic\_neg\_4\_030\_05

m/z 132.8679

012E+601RT (min)

lcorbi\_hilic\_neg\_4\_071\_57

m/z 132.8679

02468E+522.5RT (min)

lcorbi\_hilic\_neg\_3\_042\_05

m/z 132.8679

02468E+51.52RT (min)

lcorbi\_hilic\_neg\_3\_042\_05

m/z 132.8679

0246E+522.5RT (min)

lcorbi\_hilic\_neg\_3\_042\_05

m/z 132.8679

0246E+544.5RT (min)

lcorbi\_hilic\_neg\_4\_055\_38

m/z 132.8679

024E+52.53RT (min)

lcorbi\_hilic\_neg\_2\_044\_46

m/z 132.8679

012E+60.51RT (min)

lcorbi\_hilic\_neg\_3\_054\_29

m/z 132.8679

02468E+53.54RT (min)

lcorbi\_hilic\_neg\_2\_022\_37

m/z 132.8679

0246E+57.58RT (min)

lcorbi\_hilic\_neg\_1\_007\_10

m/z 132.8679

02468E+51.52RT (min)

lcorbi\_hilic\_neg\_3\_042\_05

m/z 132.8679

0246E+51.52RT (min)

lcorbi\_hilic\_neg\_1\_023\_23

m/z 132.8679

0246E+533.5RT (min)

lcorbi\_hilic\_neg\_1\_007\_10

m/z 132.8679

02468E+500.5RT (min)

lcorbi\_hilic\_neg\_3\_033\_53

m/z 132.8679

0246E+611.5RT (min)

lcorbi\_hilic\_neg\_1\_007\_10

m/z 132.8679

0246E+53.54RT (min)

lcorbi\_hilic\_neg\_3\_024\_27

m/z 132.8679

012E+56.57RT (min)

lcorbi\_hilic\_neg\_1\_046\_19

m/z 132.8679

0123E+612.513RT (min)

lcorbi\_hilic\_neg\_4\_026\_33

m/z 132.8679

0246E+53.54RT (min)

lcorbi\_hilic\_neg\_1\_027\_18

m/z 132.8679

02468E+54.55RT (min)

lcorbi\_hilic\_neg\_1\_007\_10

m/z 132.8679

02468E+53.54RT (min)

lcorbi\_hilic\_neg\_1\_046\_19

m/z 132.8680

0123E+612.513RT (min)

lcorbi\_hilic\_neg\_4\_036\_36

m/z 132.8680

02468E+588.5RT (min)

lcorbi\_hilic\_neg\_1\_046\_19

m/z 132.8680

0123E+61212.5RT (min)

lcorbi\_hilic\_neg\_4\_048\_37

m/z 132.8680

012E+613.514RT (min)

lcorbi\_hilic\_neg\_4\_048\_37

m/z 132.8680

0123E+61212.5RT (min)

lcorbi\_hilic\_neg\_4\_031\_14

m/z 132.8680

0123E+610.511RT (min)

lcorbi\_hilic\_neg\_3\_021\_20

m/z 132.8680

0123E+61212.5RT (min)

lcorbi\_hilic\_neg\_4\_034\_45

m/z 132.8680

01234E+61012RT (min)

lcorbi\_hilic\_neg\_4\_086\_59

m/z 132.8680

01234E+6810RT (min)

lcorbi\_hilic\_neg\_4\_085\_59

m/z 132.8680

012E+61010.5RT (min)

lcorbi\_hilic\_neg\_3\_037\_12

m/z 132.8680

01234E+681012RT (min)

lcorbi\_hilic\_neg\_4\_095\_60

m/z 132.8680

01234E+610RT (min)

lcorbi\_hilic\_neg\_4\_071\_57

m/z 132.8680

01234E+61012RT (min)

lcorbi\_hilic\_neg\_4\_077\_58

m/z 132.8680

0246E+544.5RT (min)

lcorbi\_hilic\_neg\_1\_027\_18

m/z 132.8680

02468E+544.5RT (min)

lcorbi\_hilic\_neg\_1\_007\_10

m/z 132.8680

01E+68.59RT (min)

lcorbi\_hilic\_neg\_1\_046\_19

m/z 132.8680

02468E+522.5RT (min)

lcorbi\_hilic\_neg\_2\_056\_56

m/z 132.8680

02468E+544.5RT (min)

lcorbi\_hilic\_neg\_1\_007\_10

m/z 132.8714

02468E+410.511RT (min)

lcorbi\_hilic\_neg\_2\_013\_18

m/z 132.9002

01E+57.58RT (min)

lcorbi\_hilic\_neg\_4\_105\_61

m/z 132.9002

01E+57.58RT (min)

lcorbi\_hilic\_neg\_4\_112\_62

m/z 133.0142

01E+63.54RT (min)

lcorbi\_hilic\_neg\_1\_005\_36

m/z 133.0142

01E+644.5RT (min)

lcorbi\_hilic\_neg\_1\_005\_36

m/z 133.0142

01E+644.5RT (min)

lcorbi\_hilic\_neg\_1\_005\_36

m/z 133.0143

01E+644.5RT (min)

lcorbi\_hilic\_neg\_1\_005\_36

m/z 133.0143

024E+666.5RT (min)

lcorbi\_hilic\_neg\_1\_048\_35

m/z 133.0143

012E+666.5RT (min)

lcorbi\_hilic\_neg\_1\_029\_31

m/z 133.0520

012E+73.54RT (min)

lcorbi\_hilic\_neg\_3\_044\_32

m/z 133.0521

01E+66.57RT (min)

lcorbi\_hilic\_neg\_3\_011\_24

m/z 133.0521

01E+73.54RT (min)

lcorbi\_hilic\_neg\_4\_090\_60

m/z 133.0562

01E+533.5RT (min)

lcorbi\_hilic\_neg\_4\_090\_60

m/z 133.0634

0246E+444.5RT (min)

lcorbi\_hilic\_neg\_3\_057\_49

m/z 134.0447

02468E+577.5RT (min)

lcorbi\_hilic\_neg\_1\_061\_50

m/z 134.0472

02468E+45.56RT (min)

lcorbi\_hilic\_neg\_4\_113\_62

m/z 134.8653

01234E+55.56RT (min)

lcorbi\_hilic\_neg\_1\_029\_31

m/z 134.8653

024E+681012RT (min)

lcorbi\_hilic\_neg\_4\_084\_59

m/z 134.8653

0246E+55.56RT (min)

lcorbi\_hilic\_neg\_3\_056\_56

m/z 134.8653

01E+655.5RT (min)

lcorbi\_hilic\_neg\_1\_055\_24

m/z 134.8653

024E+6810RT (min)

lcorbi\_hilic\_neg\_4\_085\_59

m/z 134.8653

01234E+55.56RT (min)

lcorbi\_hilic\_neg\_4\_030\_05

m/z 134.8654

0123E+6024RT (min)

lcorbi\_hilic\_neg\_4\_092\_60

m/z 134.8654

01E+600.5RT (min)

lcorbi\_hilic\_neg\_3\_031\_14

m/z 134.8654

01E+60.51RT (min)

lcorbi\_hilic\_neg\_1\_003\_14

m/z 134.8654

02468E+522.5RT (min)

lcorbi\_hilic\_neg\_3\_042\_05

m/z 134.8654

01E+61.52RT (min)

lcorbi\_hilic\_neg\_3\_042\_05

m/z 134.8654

024E+55.56RT (min)

lcorbi\_hilic\_neg\_4\_074\_58

m/z 134.8654

024E+566.5RT (min)

lcorbi\_hilic\_neg\_4\_076\_58

m/z 134.8654

012E+600.5RT (min)

lcorbi\_hilic\_neg\_3\_055\_26

m/z 134.8654

024E+681012RT (min)

lcorbi\_hilic\_neg\_4\_083\_59

m/z 134.8654

024E+67.51012.5RT (min)

lcorbi\_hilic\_neg\_4\_077\_58

m/z 134.8654

0123E+566.5RT (min)

lcorbi\_hilic\_neg\_3\_024\_27

m/z 134.8654

01E+61.52RT (min)

lcorbi\_hilic\_neg\_3\_042\_05

m/z 134.8654

0246E+52.53RT (min)

lcorbi\_hilic\_neg\_2\_044\_46

m/z 134.8654

0246E+611.5RT (min)

lcorbi\_hilic\_neg\_1\_027\_18

m/z 134.8654

01E+622.5RT (min)

lcorbi\_hilic\_neg\_1\_005\_36

m/z 134.8655

01E+622.5RT (min)

lcorbi\_hilic\_neg\_3\_042\_05

m/z 134.8655

02468E+588.5RT (min)

lcorbi\_hilic\_neg\_1\_046\_19

m/z 134.8655

01E+622.5RT (min)

lcorbi\_hilic\_neg\_1\_005\_36

m/z 134.8655

0246E+57.58RT (min)

lcorbi\_hilic\_neg\_1\_018\_21

m/z 134.8671

0123E+610.511RT (min)

lcorbi\_hilic\_neg\_1\_046\_19

m/z 134.8688

01E+5101112RT (min)

lcorbi\_hilic\_neg\_4\_082\_59

m/z 134.8689

024E+41313.5RT (min)

lcorbi\_hilic\_neg\_4\_055\_38

m/z 134.8690

0246E+412.513RT (min)

lcorbi\_hilic\_neg\_1\_046\_19

m/z 134.8946

01E+666.5RT (min)

lcorbi\_hilic\_neg\_4\_042\_28

m/z 134.8946

02468E+500.5RT (min)

lcorbi\_hilic\_neg\_4\_074\_58

m/z 134.8946

01E+600.5RT (min)

lcorbi\_hilic\_neg\_3\_000\_aa

m/z 134.8946

012E+65.56RT (min)

lcorbi\_hilic\_neg\_4\_084\_59

m/z 134.8946

01E+65.56RT (min)

lcorbi\_hilic\_neg\_4\_083\_59

m/z 134.8946

01E+60.51RT (min)

lcorbi\_hilic\_neg\_4\_090\_60

m/z 134.8947

01E+68.59RT (min)

lcorbi\_hilic\_neg\_4\_084\_59

m/z 135.0299

02468E+566.5RT (min)

lcorbi\_hilic\_neg\_1\_016\_37

m/z 135.0299

0246E+566.5RT (min)

lcorbi\_hilic\_neg\_1\_029\_31

m/z 135.0299

0246E+566.5RT (min)

lcorbi\_hilic\_neg\_1\_026\_34

m/z 135.0451

012E+61.52RT (min)

lcorbi\_hilic\_neg\_4\_112\_62

m/z 135.0451

0123E+61.52RT (min)

lcorbi\_hilic\_neg\_4\_112\_62

m/z 135.0451

0123E+61.52RT (min)

lcorbi\_hilic\_neg\_4\_106\_62

m/z 135.0452

01E+622.5RT (min)

lcorbi\_hilic\_neg\_4\_038\_27

m/z 135.8947

012E+57.58RT (min)

lcorbi\_hilic\_neg\_4\_090\_60

m/z 135.8948

0246E+544.5RT (min)

lcorbi\_hilic\_neg\_4\_071\_57

m/z 136.0436

012E+67.58RT (min)

lcorbi\_hilic\_neg\_2\_028\_33

m/z 136.0617

01E+61.52RT (min)

lcorbi\_hilic\_neg\_1\_065\_51

m/z 136.0620

0246E+533.5RT (min)

lcorbi\_hilic\_neg\_3\_061\_50

m/z 136.1121

0123E+73.54RT (min)

lcorbi\_hilic\_neg\_4\_057\_49

m/z 136.8626

024E+54.55RT (min)

lcorbi\_hilic\_neg\_1\_019\_06

m/z 136.8626

01E+61313.5RT (min)

lcorbi\_hilic\_neg\_4\_031\_14

m/z 136.8626

01E+612.513RT (min)

lcorbi\_hilic\_neg\_4\_055\_38

m/z 136.8626

012E+6810RT (min)

lcorbi\_hilic\_neg\_4\_085\_59

m/z 136.8626

012E+681012RT (min)

lcorbi\_hilic\_neg\_4\_095\_60

m/z 136.8626

01E+61012RT (min)

lcorbi\_hilic\_neg\_4\_080\_58

m/z 136.8626

012E+681012RT (min)

lcorbi\_hilic\_neg\_4\_093\_60

m/z 136.8627

01E+600.5RT (min)

lcorbi\_hilic\_neg\_4\_111\_62

m/z 136.8627

01E+55.56RT (min)

lcorbi\_hilic\_neg\_1\_029\_31

m/z 136.8627

0123E+500.5RT (min)

lcorbi\_hilic\_neg\_3\_000\_dd

m/z 136.8627

01E+61111.5RT (min)

lcorbi\_hilic\_neg\_1\_046\_19

m/z 136.8627

01E+612.513RT (min)

lcorbi\_hilic\_neg\_4\_055\_38

m/z 136.8628

0123E+51.52RT (min)

lcorbi\_hilic\_neg\_2\_021\_05

m/z 136.8628

01E+61010.5RT (min)

lcorbi\_hilic\_neg\_1\_046\_19

m/z 136.8628

012E+55.56RT (min)

lcorbi\_hilic\_neg\_4\_069\_57

m/z 136.8628

0123E+522.5RT (min)

lcorbi\_hilic\_neg\_2\_021\_05

m/z 136.8628

012E+55.56RT (min)

lcorbi\_hilic\_neg\_3\_056\_56

m/z 136.8628

0123E+522.5RT (min)

lcorbi\_hilic\_neg\_3\_042\_05

m/z 136.8628

024E+500.5RT (min)

lcorbi\_hilic\_neg\_1\_003\_14

m/z 136.8628

012E+52.53RT (min)

lcorbi\_hilic\_neg\_2\_041\_54

m/z 136.8628

012E+611.5RT (min)

lcorbi\_hilic\_neg\_3\_038\_21

m/z 136.8628

0123E+522.5RT (min)

lcorbi\_hilic\_neg\_1\_005\_36

m/z 136.8628

0123E+51.52RT (min)

lcorbi\_hilic\_neg\_3\_042\_05

m/z 136.8628

024E+54.55RT (min)

lcorbi\_hilic\_neg\_1\_001\_49

m/z 136.8628

0123E+51.52RT (min)

lcorbi\_hilic\_neg\_2\_021\_05

m/z 136.8628

01E+56.57RT (min)

lcorbi\_hilic\_neg\_4\_085\_59

m/z 136.8628

0246E+54.55RT (min)

lcorbi\_hilic\_neg\_1\_047\_05

m/z 136.8629

0246E+545RT (min)

lcorbi\_hilic\_neg\_1\_047\_05

m/z 136.8916

012E+500.5RT (min)

lcorbi\_hilic\_neg\_4\_083\_59

m/z 136.8916

012E+500.5RT (min)

lcorbi\_hilic\_neg\_4\_083\_59

m/z 136.8916

01234E+50.51RT (min)

lcorbi\_hilic\_neg\_4\_083\_59

m/z 136.8917

02468E+56.57RT (min)

lcorbi\_hilic\_neg\_4\_093\_60

m/z 136.8917

02468E+56.57RT (min)

lcorbi\_hilic\_neg\_4\_093\_60

m/z 136.8917

01E+66.577.5RT (min)

lcorbi\_hilic\_neg\_4\_083\_59

m/z 136.8917

0246E+57.58RT (min)

lcorbi\_hilic\_neg\_4\_093\_60

m/z 136.8917

0246E+56.57RT (min)

lcorbi\_hilic\_neg\_3\_036\_17

m/z 136.8917

012E+500.5RT (min)

lcorbi\_hilic\_neg\_1\_049\_55

m/z 136.8917

0246E+56.57RT (min)

lcorbi\_hilic\_neg\_3\_036\_17

m/z 136.9400

01E+58.59RT (min)

lcorbi\_hilic\_neg\_4\_074\_58

m/z 136.9400

01E+588.5RT (min)

lcorbi\_hilic\_neg\_4\_103\_61

m/z 136.9400

01E+58.59RT (min)

lcorbi\_hilic\_neg\_4\_068\_57

m/z 136.9400

01E+588.5RT (min)

lcorbi\_hilic\_neg\_4\_066\_57

m/z 137.0274

0246E+53.54RT (min)

lcorbi\_hilic\_neg\_1\_001\_49

m/z 137.0274

0246E+544.5RT (min)

lcorbi\_hilic\_neg\_1\_001\_49

m/z 137.0275

0246E+666.5RT (min)

lcorbi\_hilic\_neg\_1\_055\_24

m/z 137.0275

01E+64.55RT (min)

lcorbi\_hilic\_neg\_1\_012\_32

m/z 137.0276

0123E+666.5RT (min)

lcorbi\_hilic\_neg\_1\_047\_05

m/z 137.0276

0123E+67.58RT (min)

lcorbi\_hilic\_neg\_4\_097\_60

m/z 137.0276

01E+56.57RT (min)

lcorbi\_hilic\_neg\_2\_005\_24

m/z 137.0277

01E+566.5RT (min)

lcorbi\_hilic\_neg\_2\_005\_24

m/z 137.8917

01E+57.58RT (min)

lcorbi\_hilic\_neg\_4\_098\_61

m/z 137.8917

01E+57.58RT (min)

lcorbi\_hilic\_neg\_4\_112\_62

m/z 137.9874

01E+600.5RT (min)

lcorbi\_hilic\_neg\_3\_057\_49

m/z 138.0195

0246E+513.514RT (min)

lcorbi\_hilic\_neg\_1\_005\_36

m/z 138.0195

02468E+51414.5RT (min)

lcorbi\_hilic\_neg\_1\_001\_49

m/z 138.0196

024E+50.51RT (min)

lcorbi\_hilic\_neg\_1\_029\_31

m/z 138.0196

024E+50.51RT (min)

lcorbi\_hilic\_neg\_1\_029\_31

m/z 138.0196

01E+60.51RT (min)

lcorbi\_hilic\_neg\_4\_089\_59

m/z 138.0196

01E+60.51RT (min)

lcorbi\_hilic\_neg\_4\_089\_59

m/z 138.0581

01E+677.5RT (min)

lcorbi\_hilic\_neg\_1\_057\_49

m/z 139.0178

01E+68.59RT (min)

lcorbi\_hilic\_neg\_1\_061\_50

m/z 139.0682

012E+533.5RT (min)

lcorbi\_hilic\_neg\_4\_096\_60

m/z 139.1229

01E+600.5RT (min)

lcorbi\_hilic\_neg\_1\_057\_49

m/z 139.1230

01E+60.51RT (min)

lcorbi\_hilic\_neg\_2\_057\_49

m/z 139.9878

02468E+588.5RT (min)

lcorbi\_hilic\_neg\_4\_064\_17

m/z 139.9878

01E+67.58RT (min)

lcorbi\_hilic\_neg\_4\_057\_49

m/z 139.9879

01234E+688.5RT (min)

lcorbi\_hilic\_neg\_1\_058\_39

m/z 140.9857

01234E+688.5RT (min)

lcorbi\_hilic\_neg\_1\_060\_13

m/z 140.9951

0246E+488.5RT (min)

lcorbi\_hilic\_neg\_4\_064\_17

m/z 141.9586

0246E+500.5RT (min)

lcorbi\_hilic\_neg\_3\_057\_49

m/z 141.9833

01E+688.5RT (min)

lcorbi\_hilic\_neg\_1\_058\_39

m/z 141.9833

024E+588.5RT (min)

lcorbi\_hilic\_neg\_4\_061\_50

m/z 141.9834

02468E+500.5RT (min)

lcorbi\_hilic\_neg\_2\_057\_49

m/z 142.0975

012E+766.5RT (min)

lcorbi\_hilic\_neg\_4\_065\_51

m/z 142.1226

0123E+633.5RT (min)

lcorbi\_hilic\_neg\_1\_059\_14

m/z 142.1338

01E+62.53RT (min)

lcorbi\_hilic\_neg\_4\_065\_51

m/z 142.1338

0246E+53.54RT (min)

lcorbi\_hilic\_neg\_4\_058\_15

m/z 142.1338

0246E+511.5RT (min)

lcorbi\_hilic\_neg\_3\_059\_14

m/z 142.1339

0246E+50.51RT (min)

lcorbi\_hilic\_neg\_4\_062\_39

m/z 142.1339

0246E+51.52RT (min)

lcorbi\_hilic\_neg\_1\_065\_51

m/z 143.9968

01234E+68.59RT (min)

lcorbi\_hilic\_neg\_4\_062\_39

m/z 143.9970

02468E+51313.5RT (min)

lcorbi\_hilic\_neg\_3\_064\_16

m/z 143.9970

01E+610.511RT (min)

lcorbi\_hilic\_neg\_1\_060\_13

m/z 143.9970

01E+61111.5RT (min)

lcorbi\_hilic\_neg\_4\_059\_13

m/z 143.9970

0246E+513.514RT (min)

lcorbi\_hilic\_neg\_3\_062\_17

m/z 143.9970

02468E+511.512RT (min)

lcorbi\_hilic\_neg\_4\_065\_51

m/z 143.9970

02468E+51212.5RT (min)

lcorbi\_hilic\_neg\_4\_062\_39

m/z 144.1019

024E+755.5RT (min)

lcorbi\_hilic\_neg\_3\_064\_16

m/z 144.9234

01E+60.51RT (min)

lcorbi\_hilic\_neg\_4\_068\_57

m/z 144.9234

01E+61.52RT (min)

lcorbi\_hilic\_neg\_4\_068\_57

m/z 144.9234

0246E+52.53RT (min)

lcorbi\_hilic\_neg\_4\_067\_57

m/z 144.9234

01E+666.5RT (min)

lcorbi\_hilic\_neg\_4\_072\_57

m/z 144.9234

0123E+57.58RT (min)

lcorbi\_hilic\_neg\_4\_036\_36

m/z 144.9234

0123E+57.58RT (min)

lcorbi\_hilic\_neg\_4\_048\_37

m/z 144.9234

024E+50.51RT (min)

lcorbi\_hilic\_neg\_4\_068\_57

m/z 144.9234

0246E+500.5RT (min)

lcorbi\_hilic\_neg\_1\_053\_16

m/z 144.9234

01E+60.51RT (min)

lcorbi\_hilic\_neg\_4\_068\_57

m/z 144.9234

01234E+57.58RT (min)

lcorbi\_hilic\_neg\_4\_041\_54

m/z 144.9234

02468E+511.5RT (min)

lcorbi\_hilic\_neg\_4\_040\_17

m/z 144.9235

01E+600.5RT (min)

lcorbi\_hilic\_neg\_2\_000\_aa

m/z 144.9235

0246E+52.53RT (min)

lcorbi\_hilic\_neg\_4\_067\_57

m/z 144.9235

0246E+522.5RT (min)

lcorbi\_hilic\_neg\_4\_068\_57

m/z 144.9235

0123E+533.5RT (min)

lcorbi\_hilic\_neg\_4\_050\_26

m/z 144.9235

0123E+58.59RT (min)

lcorbi\_hilic\_neg\_4\_041\_54

m/z 144.9235

01234E+533.5RT (min)

lcorbi\_hilic\_neg\_4\_000\_bb

m/z 144.9235

0246E+52.53RT (min)

lcorbi\_hilic\_neg\_4\_067\_57

m/z 144.9821

0246E+800.5RT (min)

lcorbi\_hilic\_neg\_1\_057\_49

m/z 144.9821

0246E+622.5RT (min)

lcorbi\_hilic\_neg\_4\_058\_15

m/z 144.9821

0246E+677.5RT (min)

lcorbi\_hilic\_neg\_1\_057\_49

m/z 144.9821

0123E+644.5RT (min)

lcorbi\_hilic\_neg\_3\_063\_13

m/z 144.9821

01E+71111.5RT (min)

lcorbi\_hilic\_neg\_2\_060\_39

m/z 144.9821

01E+73.54RT (min)

lcorbi\_hilic\_neg\_3\_057\_49

m/z 144.9822

01E+710.511RT (min)

lcorbi\_hilic\_neg\_3\_058\_39

m/z 144.9822

01E+712.513RT (min)

lcorbi\_hilic\_neg\_4\_065\_51

m/z 144.9822

01E+711.512RT (min)

lcorbi\_hilic\_neg\_2\_063\_13

m/z 144.9822

01E+71212.5RT (min)

lcorbi\_hilic\_neg\_3\_064\_16

m/z 144.9822

01E+71111.5RT (min)

lcorbi\_hilic\_neg\_3\_058\_39

m/z 144.9822

01E+71313.5RT (min)

lcorbi\_hilic\_neg\_2\_064\_17

m/z 144.9822

01E+712.513RT (min)

lcorbi\_hilic\_neg\_3\_064\_16

m/z 144.9822

01E+711.512RT (min)

lcorbi\_hilic\_neg\_2\_063\_13

m/z 144.9822

02468E+71.52RT (min)

lcorbi\_hilic\_neg\_1\_057\_49

m/z 144.9836

01E+711.512RT (min)

lcorbi\_hilic\_neg\_3\_062\_17

m/z 144.9917

01234E+32.53RT (min)

lcorbi\_hilic\_neg\_1\_059\_14

m/z 145.0143

01E+64.55RT (min)

lcorbi\_hilic\_neg\_1\_005\_36

m/z 145.9235

01E+50.51RT (min)

lcorbi\_hilic\_neg\_4\_074\_58

m/z 145.9235

02468E+40.51RT (min)

lcorbi\_hilic\_neg\_4\_090\_60

m/z 145.9791

0246E+51.52RT (min)

lcorbi\_hilic\_neg\_1\_057\_49

m/z 145.9791

02468E+411.512RT (min)

lcorbi\_hilic\_neg\_2\_057\_49

m/z 145.9791

0246E+411.512RT (min)

lcorbi\_hilic\_neg\_3\_061\_50

m/z 145.9792

024E+600.5RT (min)

lcorbi\_hilic\_neg\_1\_057\_49

m/z 145.9853

024E+51111.5RT (min)

lcorbi\_hilic\_neg\_2\_060\_39

m/z 145.9853

024E+51212.5RT (min)

lcorbi\_hilic\_neg\_2\_060\_39

m/z 145.9853

0123E+700.5RT (min)

lcorbi\_hilic\_neg\_1\_057\_49

m/z 145.9854

01234E+61.52RT (min)

lcorbi\_hilic\_neg\_1\_057\_49

m/z 145.9854

0246E+511.512RT (min)

lcorbi\_hilic\_neg\_2\_063\_13

m/z 145.9854

0246E+511.512RT (min)

lcorbi\_hilic\_neg\_2\_063\_13

m/z 145.9855

024E+53.54RT (min)

lcorbi\_hilic\_neg\_4\_057\_49

m/z 146.0298

024E+588.5RT (min)

lcorbi\_hilic\_neg\_1\_058\_39

m/z 146.0491

01234E+56.57RT (min)

lcorbi\_hilic\_neg\_2\_031\_34

m/z 146.0496

0123E+566.5RT (min)

lcorbi\_hilic\_neg\_4\_087\_59

m/z 146.9387

02468E+50.51RT (min)

lcorbi\_hilic\_neg\_4\_000\_aa

m/z 146.9611

0123E+599.5RT (min)

lcorbi\_hilic\_neg\_4\_088\_59

m/z 146.9612

0123E+5910RT (min)

lcorbi\_hilic\_neg\_4\_088\_59

m/z 146.9612

012E+59.510RT (min)

lcorbi\_hilic\_neg\_4\_088\_59

m/z 146.9612

01E+59.510RT (min)

lcorbi\_hilic\_neg\_4\_088\_59

m/z 146.9659

01E+533.5RT (min)

lcorbi\_hilic\_neg\_3\_004\_41

m/z 146.9659

01E+533.5RT (min)

lcorbi\_hilic\_neg\_3\_004\_41

m/z 146.9801

0123E+61313.5RT (min)

lcorbi\_hilic\_neg\_2\_065\_51

m/z 146.9801

024E+612.513RT (min)

lcorbi\_hilic\_neg\_4\_065\_51

m/z 146.9801

024E+612.513RT (min)

lcorbi\_hilic\_neg\_2\_063\_13

m/z 146.9801

0246E+711.5RT (min)

lcorbi\_hilic\_neg\_3\_057\_49

m/z 146.9802

0246E+61111.5RT (min)

lcorbi\_hilic\_neg\_3\_063\_13

m/z 146.9802

024E+610.511RT (min)

lcorbi\_hilic\_neg\_3\_058\_39

m/z 146.9802

0246E+611.512RT (min)

lcorbi\_hilic\_neg\_2\_063\_13

m/z 146.9802

0123E+677.5RT (min)

lcorbi\_hilic\_neg\_1\_057\_49

m/z 146.9802

0123E+800.5RT (min)

lcorbi\_hilic\_neg\_1\_057\_49

m/z 146.9802

0123E+62.53RT (min)

lcorbi\_hilic\_neg\_1\_059\_14

m/z 146.9802

01234E+71.52RT (min)

lcorbi\_hilic\_neg\_1\_057\_49

m/z 146.9802

0246E+611.512RT (min)

lcorbi\_hilic\_neg\_2\_063\_13

m/z 146.9803

0246E+63.54RT (min)

lcorbi\_hilic\_neg\_3\_057\_49

m/z 146.9840

01234E+60.51RT (min)

lcorbi\_hilic\_neg\_1\_057\_49

m/z 147.9772

012E+600.5RT (min)

lcorbi\_hilic\_neg\_1\_057\_49

m/z 147.9834

01E+700.5RT (min)

lcorbi\_hilic\_neg\_1\_057\_49

m/z 147.9835

01E+61.52RT (min)

lcorbi\_hilic\_neg\_1\_057\_49

m/z 148.0967

01234E+63.54RT (min)

lcorbi\_hilic\_neg\_2\_060\_39

m/z 148.1175

012E+766.5RT (min)

lcorbi\_hilic\_neg\_4\_065\_51

m/z 149.0091

01E+67.58RT (min)

lcorbi\_hilic\_neg\_1\_048\_35

m/z 149.0091

01E+677.5RT (min)

lcorbi\_hilic\_neg\_1\_026\_34

m/z 149.0092

01E+677.5RT (min)

lcorbi\_hilic\_neg\_1\_026\_34

m/z 149.0092

012E+666.5RT (min)

lcorbi\_hilic\_neg\_1\_005\_36

m/z 149.0092

01E+677.5RT (min)

lcorbi\_hilic\_neg\_1\_026\_34

m/z 149.0092

012E+66.57RT (min)

lcorbi\_hilic\_neg\_1\_005\_36

m/z 149.0092

01E+677.5RT (min)

lcorbi\_hilic\_neg\_1\_026\_34

m/z 149.0093

01E+66.57RT (min)

lcorbi\_hilic\_neg\_1\_026\_34

m/z 149.0277

01E+64.55RT (min)

lcorbi\_hilic\_neg\_1\_005\_36

m/z 150.0312

01E+64.55RT (min)

lcorbi\_hilic\_neg\_1\_012\_32

m/z 151.0352

0246E+688.5RT (min)

lcorbi\_hilic\_neg\_1\_058\_39

m/z 151.0579

01E+544.5RT (min)

lcorbi\_hilic\_neg\_4\_082\_59

m/z 151.0764

01E+60.51RT (min)

lcorbi\_hilic\_neg\_3\_000\_cc

m/z 151.0964

0246E+611.5RT (min)

lcorbi\_hilic\_neg\_1\_059\_14

m/z 152.0365

02468E+588.5RT (min)

lcorbi\_hilic\_neg\_1\_058\_39

m/z 152.9176

0246E+410.511RT (min)

lcorbi\_hilic\_neg\_4\_007\_21

m/z 152.9176

0246E+410.511RT (min)

lcorbi\_hilic\_neg\_4\_007\_21

m/z 152.9176

01E+59.510RT (min)

lcorbi\_hilic\_neg\_4\_051\_16

m/z 152.9176

012E+51012RT (min)

lcorbi\_hilic\_neg\_4\_074\_58

m/z 152.9176

02468E+410.511RT (min)

lcorbi\_hilic\_neg\_2\_003\_20

m/z 152.9176

02468E+41010.5RT (min)

lcorbi\_hilic\_neg\_2\_014\_31

m/z 152.9176

02468E+41010.5RT (min)

lcorbi\_hilic\_neg\_2\_003\_20

m/z 152.9176

01E+599.5RT (min)

lcorbi\_hilic\_neg\_4\_048\_37

m/z 152.9176

01E+599.5RT (min)

lcorbi\_hilic\_neg\_4\_013\_46

m/z 152.9470

01E+58.59RT (min)

lcorbi\_hilic\_neg\_1\_059\_14

m/z 153.0326

02468E+588.5RT (min)

lcorbi\_hilic\_neg\_1\_058\_39

m/z 153.0771

01E+766.5RT (min)

lcorbi\_hilic\_neg\_3\_061\_50

m/z 153.8686

01E+522.5RT (min)

lcorbi\_hilic\_neg\_3\_022\_06

m/z 153.8686

01E+500.5RT (min)

lcorbi\_hilic\_neg\_4\_066\_57

m/z 153.8686

01E+500.5RT (min)

lcorbi\_hilic\_neg\_4\_074\_58

m/z 153.8686

012E+50.51RT (min)

lcorbi\_hilic\_neg\_4\_066\_57

m/z 153.8686

012E+50.51RT (min)

lcorbi\_hilic\_neg\_4\_066\_57

m/z 153.8686

01E+500.5RT (min)

lcorbi\_hilic\_neg\_4\_074\_58

m/z 153.8686

01E+522.5RT (min)

lcorbi\_hilic\_neg\_3\_022\_06

m/z 154.9473

01E+566.5RT (min)

lcorbi\_hilic\_neg\_1\_007\_10

m/z 154.9473

0123E+566.5RT (min)

lcorbi\_hilic\_neg\_1\_020\_13

m/z 155.0408

01E+63.54RT (min)

lcorbi\_hilic\_neg\_1\_058\_39

m/z 155.0539

01E+63.54RT (min)

lcorbi\_hilic\_neg\_1\_064\_15

m/z 155.8656

01E+50.51RT (min)

lcorbi\_hilic\_neg\_4\_066\_57

m/z 155.8656

01E+511.5RT (min)

lcorbi\_hilic\_neg\_4\_093\_60

m/z 155.8656

01E+50.51RT (min)

lcorbi\_hilic\_neg\_4\_066\_57

m/z 156.0767

012E+677.5RT (min)

lcorbi\_hilic\_neg\_1\_057\_49

m/z 156.0777

012E+54.55RT (min)

lcorbi\_hilic\_neg\_4\_001\_49

m/z 156.0778

01E+55.56RT (min)

lcorbi\_hilic\_neg\_4\_017\_51

m/z 156.0778

012E+566.5RT (min)

lcorbi\_hilic\_neg\_4\_026\_33

m/z 156.0778

012E+55.56RT (min)

lcorbi\_hilic\_neg\_4\_026\_33

m/z 156.0778

012E+55.56RT (min)

lcorbi\_hilic\_neg\_4\_026\_33

m/z 156.0779

02468E+57.58RT (min)

lcorbi\_hilic\_neg\_2\_058\_16

m/z 156.0779

024E+51.52RT (min)

lcorbi\_hilic\_neg\_4\_092\_60

m/z 156.1382

01E+62.53RT (min)

lcorbi\_hilic\_neg\_2\_064\_17

m/z 156.1746

0246E+533.5RT (min)

lcorbi\_hilic\_neg\_4\_065\_51

m/z 157.0118

01E+52.53RT (min)

lcorbi\_hilic\_neg\_4\_075\_58

m/z 157.0118

01E+52.53RT (min)

lcorbi\_hilic\_neg\_4\_074\_58

m/z 157.0118

01E+52.53RT (min)

lcorbi\_hilic\_neg\_4\_075\_58

m/z 157.0118

01E+52.53RT (min)

lcorbi\_hilic\_neg\_4\_075\_58

m/z 157.0119

01E+51.52RT (min)

lcorbi\_hilic\_neg\_2\_012\_14

m/z 157.0119

01E+522.5RT (min)

lcorbi\_hilic\_neg\_4\_075\_58

m/z 158.8468

01E+59.510RT (min)

lcorbi\_hilic\_neg\_3\_030\_30

m/z 158.8468

01E+51111.5RT (min)

lcorbi\_hilic\_neg\_4\_031\_14

m/z 158.8468

01E+510.511RT (min)

lcorbi\_hilic\_neg\_4\_046\_15

m/z 158.8468

01E+51010.5RT (min)

lcorbi\_hilic\_neg\_3\_055\_26

m/z 158.8468

01E+59.510RT (min)

lcorbi\_hilic\_neg\_3\_030\_30

m/z 158.8468

01E+51010.5RT (min)

lcorbi\_hilic\_neg\_3\_055\_26

m/z 158.8469

01E+510.511RT (min)

lcorbi\_hilic\_neg\_4\_031\_14

m/z 158.9640

0123E+755.5RT (min)

lcorbi\_hilic\_neg\_3\_064\_16

m/z 159.0083

01E+522.5RT (min)

lcorbi\_hilic\_neg\_4\_082\_59

m/z 159.0083

01E+522.5RT (min)

lcorbi\_hilic\_neg\_4\_083\_59

m/z 159.9793

0246E+499.5RT (min)

lcorbi\_hilic\_neg\_4\_090\_60

m/z 160.0806

02468E+577.5RT (min)

lcorbi\_hilic\_neg\_1\_055\_24

m/z 160.8420

012E+57.58RT (min)

lcorbi\_hilic\_neg\_4\_067\_57

m/z 160.8421

012E+51.52RT (min)

lcorbi\_hilic\_neg\_3\_052\_40

m/z 160.8421

01234E+53.54RT (min)

lcorbi\_hilic\_neg\_4\_099\_61

m/z 160.8421

01234E+533.5RT (min)

lcorbi\_hilic\_neg\_4\_099\_61

m/z 160.8421

02468E+53.54RT (min)

lcorbi\_hilic\_neg\_4\_000\_bb

m/z 160.8421

0123E+57.58RT (min)

lcorbi\_hilic\_neg\_1\_036\_09

m/z 160.8421

012E+57.58RT (min)

lcorbi\_hilic\_neg\_2\_035\_22

m/z 160.8421

012E+51.52RT (min)

lcorbi\_hilic\_neg\_3\_052\_40

m/z 160.8421

0246E+511.5RT (min)

lcorbi\_hilic\_neg\_3\_052\_40

m/z 160.8421

012E+511.5RT (min)

lcorbi\_hilic\_neg\_4\_040\_17

m/z 160.8422

012E+61111.5RT (min)

lcorbi\_hilic\_neg\_4\_031\_14

m/z 160.8422

01E+61212.5RT (min)

lcorbi\_hilic\_neg\_2\_034\_01

m/z 160.8422

01E+69.510RT (min)

lcorbi\_hilic\_neg\_3\_030\_30

m/z 160.8422

012E+61111.5RT (min)

lcorbi\_hilic\_neg\_4\_031\_14

m/z 160.8422

01E+611.512RT (min)

lcorbi\_hilic\_neg\_4\_031\_14

m/z 160.8422

012E+610.511RT (min)

lcorbi\_hilic\_neg\_4\_031\_14

m/z 160.8422

012E+61111.5RT (min)

lcorbi\_hilic\_neg\_4\_031\_14

m/z 160.8422

01E+611.512RT (min)

lcorbi\_hilic\_neg\_3\_006\_28

m/z 160.8422

02468E+51212.5RT (min)

lcorbi\_hilic\_neg\_2\_034\_01

m/z 160.8422

0246E+512.513RT (min)

lcorbi\_hilic\_neg\_3\_036\_17

m/z 160.8422

0246E+513.514RT (min)

lcorbi\_hilic\_neg\_4\_031\_14

m/z 160.8422

0246E+513.514RT (min)

lcorbi\_hilic\_neg\_4\_027\_47

m/z 160.8422

012E+610.511RT (min)

lcorbi\_hilic\_neg\_4\_038\_27

m/z 160.8422

01E+69.510RT (min)

lcorbi\_hilic\_neg\_3\_030\_30

m/z 160.8422

0123E+600.5RT (min)

lcorbi\_hilic\_neg\_2\_000\_bb

m/z 160.8422

0246E+51313.5RT (min)

lcorbi\_hilic\_neg\_4\_031\_14

m/z 160.8422

0246E+51414.5RT (min)

lcorbi\_hilic\_neg\_4\_048\_37

m/z 160.8423

024E+666.5RT (min)

lcorbi\_hilic\_neg\_4\_076\_58

m/z 160.9355

012E+52.53RT (min)

lcorbi\_hilic\_neg\_4\_082\_59

m/z 161.9796

01E+50.51RT (min)

lcorbi\_hilic\_neg\_4\_105\_61

m/z 162.0073

01234E+58.59RT (min)

lcorbi\_hilic\_neg\_3\_059\_14

m/z 162.1123

01E+855.5RT (min)

lcorbi\_hilic\_neg\_1\_062\_16

m/z 162.8391

012E+511.5RT (min)

lcorbi\_hilic\_neg\_4\_040\_17

m/z 162.8391

01E+61010.5RT (min)

lcorbi\_hilic\_neg\_4\_042\_28

m/z 162.8391

024E+666.5RT (min)

lcorbi\_hilic\_neg\_4\_076\_58

m/z 162.8391

0123E+61011RT (min)

lcorbi\_hilic\_neg\_4\_081\_58

m/z 162.8391

0123E+50.51RT (min)

lcorbi\_hilic\_neg\_3\_056\_56

m/z 162.8391

01E+60.51RT (min)

lcorbi\_hilic\_neg\_3\_055\_26

m/z 162.8391

012E+61011RT (min)

lcorbi\_hilic\_neg\_4\_074\_58

m/z 162.8391

0123E+511.5RT (min)

lcorbi\_hilic\_neg\_3\_050\_03

m/z 162.8391

012E+610.511RT (min)

lcorbi\_hilic\_neg\_4\_031\_14

m/z 162.8392

0123E+600.5RT (min)

lcorbi\_hilic\_neg\_2\_000\_bb

m/z 162.8392

0246E+51212.5RT (min)

lcorbi\_hilic\_neg\_4\_048\_37

m/z 162.8392

02468E+51212.5RT (min)

lcorbi\_hilic\_neg\_3\_018\_38

m/z 162.8392

01E+69.510RT (min)

lcorbi\_hilic\_neg\_3\_030\_30

m/z 162.8392

01E+69.510RT (min)

lcorbi\_hilic\_neg\_3\_030\_30

m/z 162.8392

01E+611.512RT (min)

lcorbi\_hilic\_neg\_4\_031\_14

m/z 162.8392

024E+512.513RT (min)

lcorbi\_hilic\_neg\_4\_023\_34

m/z 162.8392

01E+611.512RT (min)

lcorbi\_hilic\_neg\_4\_031\_14

m/z 162.8392

0246E+51313.5RT (min)

lcorbi\_hilic\_neg\_4\_031\_14

m/z 162.8392

0246E+51414.5RT (min)

lcorbi\_hilic\_neg\_4\_048\_37

m/z 162.8392

0246E+51212.5RT (min)

lcorbi\_hilic\_neg\_4\_049\_55

m/z 162.8392

01E+611.512RT (min)

lcorbi\_hilic\_neg\_4\_031\_14

m/z 162.8392

01E+69.51010.5RT (min)

lcorbi\_hilic\_neg\_4\_066\_57

m/z 162.8392

0246E+51313.5RT (min)

lcorbi\_hilic\_neg\_4\_044\_01

m/z 162.8392

0246E+513.514RT (min)

lcorbi\_hilic\_neg\_4\_031\_14

m/z 162.8393

0123E+57.58RT (min)

lcorbi\_hilic\_neg\_4\_066\_57

m/z 162.8394

01234E+57.58RT (min)

lcorbi\_hilic\_neg\_1\_027\_18

m/z 162.9674

02468E+588.5RT (min)

lcorbi\_hilic\_neg\_2\_063\_13

m/z 162.9822

01E+544.5RT (min)

lcorbi\_hilic\_neg\_3\_001\_49

m/z 162.9823

01E+544.5RT (min)

lcorbi\_hilic\_neg\_3\_001\_49

m/z 162.9823

01E+555.5RT (min)

lcorbi\_hilic\_neg\_3\_002\_15

m/z 162.9823

0246E+56RT (min)

lcorbi\_hilic\_neg\_1\_011\_01

m/z 162.9823

012E+50.51RT (min)

lcorbi\_hilic\_neg\_2\_031\_34

m/z 162.9824

012E+50.51RT (min)

lcorbi\_hilic\_neg\_3\_055\_26

m/z 162.9824

01E+533.5RT (min)

lcorbi\_hilic\_neg\_3\_002\_15

m/z 162.9824

01E+544.5RT (min)

lcorbi\_hilic\_neg\_3\_002\_15

m/z 162.9824

02468E+512RT (min)

lcorbi\_hilic\_neg\_1\_002\_26

m/z 162.9824

01E+555.5RT (min)

lcorbi\_hilic\_neg\_3\_002\_15

m/z 162.9824

01E+50.51RT (min)

lcorbi\_hilic\_neg\_4\_014\_18

m/z 162.9824

01E+54.55RT (min)

lcorbi\_hilic\_neg\_3\_009\_50

m/z 162.9824

012E+50.51RT (min)

lcorbi\_hilic\_neg\_2\_031\_34

m/z 162.9824

01E+555.5RT (min)

lcorbi\_hilic\_neg\_3\_017\_51

m/z 163.1155

01E+755.5RT (min)

lcorbi\_hilic\_neg\_1\_062\_16

m/z 163.8974

01E+58.59RT (min)

lcorbi\_hilic\_neg\_4\_066\_57

m/z 163.8974

01E+58.59RT (min)

lcorbi\_hilic\_neg\_3\_030\_30

m/z 164.8360

0246E+555.5RT (min)

lcorbi\_hilic\_neg\_4\_000\_aa

m/z 164.8361

024E+510.511RT (min)

lcorbi\_hilic\_neg\_4\_017\_51

m/z 164.8361

0246E+51010.5RT (min)

lcorbi\_hilic\_neg\_4\_044\_01

m/z 164.8361

0246E+511.5RT (min)

lcorbi\_hilic\_neg\_2\_000\_aa

m/z 164.8361

0246E+51010.5RT (min)

lcorbi\_hilic\_neg\_4\_044\_01

m/z 164.8361

01E+63.54RT (min)

lcorbi\_hilic\_neg\_2\_000\_aa

m/z 164.8361

02468E+53.54RT (min)

lcorbi\_hilic\_neg\_2\_000\_aa

m/z 164.8362

01E+610.511RT (min)

lcorbi\_hilic\_neg\_4\_078\_58

m/z 164.8362

0246E+5910RT (min)

lcorbi\_hilic\_neg\_4\_066\_57

m/z 164.8362

01E+600.5RT (min)

lcorbi\_hilic\_neg\_2\_000\_bb

m/z 164.8363

0246E+51.52RT (min)

lcorbi\_hilic\_neg\_2\_000\_aa

m/z 164.9272

02468E+499.510RT (min)

lcorbi\_hilic\_neg\_4\_084\_59

m/z 164.9272

01E+599.5RT (min)

lcorbi\_hilic\_neg\_4\_099\_61

m/z 164.9272

01E+599.5RT (min)

lcorbi\_hilic\_neg\_4\_104\_61

m/z 164.9273

01E+599.5RT (min)

lcorbi\_hilic\_neg\_4\_074\_58

m/z 164.9273

01E+599.5RT (min)

lcorbi\_hilic\_neg\_4\_069\_57

m/z 165.1123

01E+60.51RT (min)

lcorbi\_hilic\_neg\_2\_060\_39

m/z 167.0127

02468E+51212.5RT (min)

lcorbi\_hilic\_neg\_3\_057\_49

m/z 167.0127

01E+611.512RT (min)

lcorbi\_hilic\_neg\_3\_057\_49

m/z 167.0127

01E+610.511RT (min)

lcorbi\_hilic\_neg\_3\_058\_39

m/z 167.0127

024E+68.59RT (min)

lcorbi\_hilic\_neg\_4\_062\_39

m/z 167.0127

02468E+512.513RT (min)

lcorbi\_hilic\_neg\_3\_061\_50

m/z 167.0128

01E+61111.5RT (min)

lcorbi\_hilic\_neg\_3\_058\_39

m/z 167.8352

0246E+58.59RT (min)

lcorbi\_hilic\_neg\_4\_054\_30

m/z 167.8354

01E+68.59RT (min)

lcorbi\_hilic\_neg\_4\_099\_61

m/z 167.8366

0123E+511.5RT (min)

lcorbi\_hilic\_neg\_4\_070\_57

m/z 167.8367

01234E+500.5RT (min)

lcorbi\_hilic\_neg\_1\_010\_22

m/z 167.8367

012E+577.5RT (min)

lcorbi\_hilic\_neg\_4\_074\_58

m/z 167.8367

01234E+50.51RT (min)

lcorbi\_hilic\_neg\_1\_001\_49

m/z 167.8367

01234E+500.5RT (min)

lcorbi\_hilic\_neg\_1\_009\_50

m/z 167.8368

01234E+52.53RT (min)

lcorbi\_hilic\_neg\_4\_083\_59

m/z 167.8368

012E+52.53RT (min)

lcorbi\_hilic\_neg\_1\_004\_17

m/z 168.0160

0123E+58.59RT (min)

lcorbi\_hilic\_neg\_1\_059\_14

m/z 168.1018

024E+522.5RT (min)

lcorbi\_hilic\_neg\_4\_058\_15

m/z 168.1018

0246E+51.52RT (min)

lcorbi\_hilic\_neg\_3\_064\_16

m/z 168.1019

01E+611.5RT (min)

lcorbi\_hilic\_neg\_1\_063\_17

m/z 168.9814

01E+622.5RT (min)

lcorbi\_hilic\_neg\_4\_066\_57

m/z 169.0458

02468E+588.5RT (min)

lcorbi\_hilic\_neg\_1\_058\_39

m/z 169.8340

0246E+511.5RT (min)

lcorbi\_hilic\_neg\_3\_050\_03

m/z 169.8341

024E+50.51RT (min)

lcorbi\_hilic\_neg\_1\_001\_49

m/z 169.8341

024E+500.5RT (min)

lcorbi\_hilic\_neg\_1\_004\_17

m/z 169.8341

02468E+59.510RT (min)

lcorbi\_hilic\_neg\_4\_096\_60

m/z 169.8341

01234E+5789RT (min)

lcorbi\_hilic\_neg\_4\_075\_58

m/z 169.8341

024E+500.5RT (min)

lcorbi\_hilic\_neg\_1\_001\_49

m/z 169.8341

024E+500.5RT (min)

lcorbi\_hilic\_neg\_1\_007\_10

m/z 169.8341

024E+50.51RT (min)

lcorbi\_hilic\_neg\_1\_001\_49

m/z 169.8341

0246E+500.5RT (min)

lcorbi\_hilic\_neg\_1\_010\_22

m/z 169.8342

01234E+52.53RT (min)

lcorbi\_hilic\_neg\_1\_004\_17

m/z 170.0924

02468E+577.5RT (min)

lcorbi\_hilic\_neg\_1\_059\_14

m/z 170.8707

01E+51010.5RT (min)

lcorbi\_hilic\_neg\_4\_000\_aa

m/z 171.8315

012E+52.53RT (min)

lcorbi\_hilic\_neg\_1\_004\_17

m/z 171.8315

0123E+52.53RT (min)

lcorbi\_hilic\_neg\_4\_083\_59

m/z 171.8315

012E+5101214RT (min)

lcorbi\_hilic\_neg\_4\_081\_58

m/z 171.8315

0123E+52.53RT (min)

lcorbi\_hilic\_neg\_4\_083\_59

m/z 171.8315

01234E+589RT (min)

lcorbi\_hilic\_neg\_4\_106\_62

m/z 171.8316

0123E+58.59RT (min)

lcorbi\_hilic\_neg\_4\_082\_59

m/z 171.8317

024E+58.59RT (min)

lcorbi\_hilic\_neg\_2\_024\_11

m/z 173.0042

01234E+544.5RT (min)

lcorbi\_hilic\_neg\_1\_008\_39

m/z 173.4079

01E+600.5RT (min)

lcorbi\_hilic\_neg\_2\_057\_49

m/z 173.8288

02468E+488.5RT (min)

lcorbi\_hilic\_neg\_2\_019\_21

m/z 173.8288

02468E+488.5RT (min)

lcorbi\_hilic\_neg\_1\_040\_44

m/z 173.8288

02468E+47.58RT (min)

lcorbi\_hilic\_neg\_1\_024\_11

m/z 173.9262

01E+599.5RT (min)

lcorbi\_hilic\_neg\_4\_074\_58

m/z 173.9262

01E+58.59RT (min)

lcorbi\_hilic\_neg\_4\_076\_58

m/z 173.9262

01E+599.5RT (min)

lcorbi\_hilic\_neg\_4\_074\_58

m/z 174.1852

012E+622.5RT (min)

lcorbi\_hilic\_neg\_3\_057\_49

m/z 174.1852

01234E+733.5RT (min)

lcorbi\_hilic\_neg\_2\_058\_16

m/z 174.9599

0123E+48.59RT (min)

lcorbi\_hilic\_neg\_4\_008\_35

m/z 175.0154

0246E+50.51RT (min)

lcorbi\_hilic\_neg\_4\_063\_14

m/z 175.1190

01E+777.5RT (min)

lcorbi\_hilic\_neg\_4\_057\_49

m/z 175.1884

01234E+633.5RT (min)

lcorbi\_hilic\_neg\_2\_058\_16

m/z 175.9593

012E+599.5RT (min)

lcorbi\_hilic\_neg\_4\_074\_58

m/z 175.9593

012E+599.5RT (min)

lcorbi\_hilic\_neg\_4\_081\_58

m/z 175.9593

01E+59.510RT (min)

lcorbi\_hilic\_neg\_4\_074\_58

m/z 175.9593

012E+59.510RT (min)

lcorbi\_hilic\_neg\_4\_074\_58

m/z 175.9593

01E+599.5RT (min)

lcorbi\_hilic\_neg\_4\_084\_59

m/z 175.9593

01E+59.510RT (min)

lcorbi\_hilic\_neg\_4\_074\_58

m/z 175.9593

01E+599.5RT (min)

lcorbi\_hilic\_neg\_4\_088\_59

m/z 176.1281

0123E+633.5RT (min)

lcorbi\_hilic\_neg\_3\_063\_13

m/z 176.8258

01E+58.59RT (min)

lcorbi\_hilic\_neg\_4\_090\_60

m/z 176.8258

01E+59.510RT (min)

lcorbi\_hilic\_neg\_4\_090\_60

m/z 176.8258

01E+599.5RT (min)

lcorbi\_hilic\_neg\_4\_090\_60

m/z 176.8258

01E+58.59RT (min)

lcorbi\_hilic\_neg\_4\_090\_60

m/z 176.8258

01E+59.510RT (min)

lcorbi\_hilic\_neg\_4\_090\_60

m/z 177.1023

01E+711.5RT (min)

lcorbi\_hilic\_neg\_4\_059\_13

m/z 177.9029

012E+599.5RT (min)

lcorbi\_hilic\_neg\_3\_031\_14

m/z 177.9029

012E+58.59RT (min)

lcorbi\_hilic\_neg\_4\_034\_45

m/z 177.9029

012E+599.5RT (min)

lcorbi\_hilic\_neg\_3\_031\_14

m/z 177.9030

01E+599.5RT (min)

lcorbi\_hilic\_neg\_3\_031\_14

m/z 177.9031

02468E+400.5RT (min)

lcorbi\_hilic\_neg\_1\_025\_52

m/z 177.9031

024E+589RT (min)

lcorbi\_hilic\_neg\_2\_002\_28

m/z 177.9031

01234E+57.58RT (min)

lcorbi\_hilic\_neg\_1\_044\_03

m/z 177.9031

0123E+577.5RT (min)

lcorbi\_hilic\_neg\_3\_030\_30

m/z 177.9031

01234E+57.58RT (min)

lcorbi\_hilic\_neg\_3\_043\_02

m/z 177.9031

012E+588.59RT (min)

lcorbi\_hilic\_neg\_2\_028\_33

m/z 177.9031

012E+58.59RT (min)

lcorbi\_hilic\_neg\_2\_028\_33

m/z 177.9032

02468E+400.5RT (min)

lcorbi\_hilic\_neg\_1\_011\_01

m/z 177.9032

0123E+589RT (min)

lcorbi\_hilic\_neg\_1\_030\_04

m/z 178.8649

01E+599.5RT (min)

lcorbi\_hilic\_neg\_4\_088\_59

m/z 178.8649

01E+599.5RT (min)

lcorbi\_hilic\_neg\_4\_113\_62

m/z 178.9488

01E+599.5RT (min)

lcorbi\_hilic\_neg\_4\_093\_60

m/z 178.9488

01E+599.5RT (min)

lcorbi\_hilic\_neg\_4\_074\_58

m/z 179.0561

0246E+65.56RT (min)

lcorbi\_hilic\_neg\_2\_020\_27

m/z 179.0562

01E+62.53RT (min)

lcorbi\_hilic\_neg\_4\_014\_18

m/z 180.0413

02468E+444.5RT (min)

lcorbi\_hilic\_neg\_1\_056\_56

m/z 180.0414

01E+556RT (min)

lcorbi\_hilic\_neg\_3\_000\_bb

m/z 180.0414

02468E+46.57RT (min)

lcorbi\_hilic\_neg\_4\_084\_59

m/z 180.0414

0246E+42.53RT (min)

lcorbi\_hilic\_neg\_1\_055\_24

m/z 180.0414

02468E+444.5RT (min)

lcorbi\_hilic\_neg\_1\_056\_56

m/z 180.0414

02468E+43.54RT (min)

lcorbi\_hilic\_neg\_1\_054\_27

m/z 180.0414

01E+555.5RT (min)

lcorbi\_hilic\_neg\_2\_056\_56

m/z 180.0414

02468E+400.5RT (min)

lcorbi\_hilic\_neg\_2\_004\_09

m/z 180.0414

02468E+43.54RT (min)

lcorbi\_hilic\_neg\_1\_047\_05

m/z 180.0414

01E+50.51RT (min)

lcorbi\_hilic\_neg\_1\_011\_01

m/z 180.0414

02468E+45.56RT (min)

lcorbi\_hilic\_neg\_3\_015\_47

m/z 180.0414

01E+555.5RT (min)

lcorbi\_hilic\_neg\_1\_026\_34

m/z 180.0414

02468E+444.5RT (min)

lcorbi\_hilic\_neg\_2\_009\_50

m/z 180.0414

01E+55.56RT (min)

lcorbi\_hilic\_neg\_1\_040\_44

m/z 180.0414

02468E+466.5RT (min)

lcorbi\_hilic\_neg\_3\_041\_54

m/z 180.0414

02468E+489RT (min)

lcorbi\_hilic\_neg\_4\_091\_60

m/z 180.0414

0246E+42.53RT (min)

lcorbi\_hilic\_neg\_1\_055\_24

m/z 180.0415

0246E+42.53RT (min)

lcorbi\_hilic\_neg\_1\_035\_29

m/z 180.0415

02468E+43.54RT (min)

lcorbi\_hilic\_neg\_1\_047\_05

m/z 180.0415

02468E+444.5RT (min)

lcorbi\_hilic\_neg\_1\_038\_15

m/z 180.9730

0123E+55.56RT (min)

lcorbi\_hilic\_neg\_4\_102\_61

m/z 181.0410

0123E+366.5RT (min)

lcorbi\_hilic\_neg\_3\_044\_32

m/z 181.0448

02468E+53.54RT (min)

lcorbi\_hilic\_neg\_1\_012\_32

m/z 181.0449

02468E+544.5RT (min)

lcorbi\_hilic\_neg\_1\_012\_32

m/z 181.1389

0123E+777.5RT (min)

lcorbi\_hilic\_neg\_1\_057\_49

m/z 181.9918

01234E+711.5RT (min)

lcorbi\_hilic\_neg\_2\_020\_27

m/z 181.9918

012E+70.51RT (min)

lcorbi\_hilic\_neg\_4\_113\_62

m/z 182.0458

02468E+477.5RT (min)

lcorbi\_hilic\_neg\_4\_068\_57

m/z 182.0458

02468E+455.5RT (min)

lcorbi\_hilic\_neg\_4\_087\_59

m/z 182.0458

012E+566.5RT (min)

lcorbi\_hilic\_neg\_1\_023\_23

m/z 182.0458

02468E+477.5RT (min)

lcorbi\_hilic\_neg\_4\_073\_57

m/z 182.0458

02468E+466.5RT (min)

lcorbi\_hilic\_neg\_4\_079\_58

m/z 182.0458

02468E+455.5RT (min)

lcorbi\_hilic\_neg\_4\_086\_59

m/z 182.9884

01E+500.5RT (min)

lcorbi\_hilic\_neg\_1\_002\_26

m/z 182.9884

01E+50.51RT (min)

lcorbi\_hilic\_neg\_1\_002\_26

m/z 182.9884

0100.5RT (min)

lcorbi\_hilic\_neg\_3\_057\_49

m/z 182.9884

02468E+42.53RT (min)

lcorbi\_hilic\_neg\_1\_003\_14

m/z 182.9884

01E+50.51RT (min)

lcorbi\_hilic\_neg\_1\_002\_26

m/z 182.9885

02468E+411.5RT (min)

lcorbi\_hilic\_neg\_2\_020\_27

m/z 182.9885

02468E+411.5RT (min)

lcorbi\_hilic\_neg\_2\_020\_27

m/z 182.9885

02468E+411.5RT (min)

lcorbi\_hilic\_neg\_4\_112\_62

m/z 184.0944

02468E+655.5RT (min)

lcorbi\_hilic\_neg\_3\_064\_16

m/z 184.9856

012E+688.5RT (min)

lcorbi\_hilic\_neg\_1\_058\_39

m/z 185.0233

01234E+58.59RT (min)

lcorbi\_hilic\_neg\_1\_059\_14

m/z 185.1284

01E+611.5RT (min)

lcorbi\_hilic\_neg\_1\_060\_13

m/z 185.9834

0123E+688.5RT (min)

lcorbi\_hilic\_neg\_1\_060\_13

m/z 186.1852

01E+633.5RT (min)

lcorbi\_hilic\_neg\_2\_057\_49

m/z 186.9809

02468E+588.5RT (min)

lcorbi\_hilic\_neg\_1\_058\_39

m/z 186.9810

024E+500.5RT (min)

lcorbi\_hilic\_neg\_4\_057\_49

m/z 187.0070

02468E+611.5RT (min)

lcorbi\_hilic\_neg\_3\_024\_27

m/z 187.0336

01E+444.5RT (min)

lcorbi\_hilic\_neg\_4\_089\_59

m/z 187.0380

012E+744.5RT (min)

lcorbi\_hilic\_neg\_1\_005\_36

m/z 187.0381

01E+64.55RT (min)

lcorbi\_hilic\_neg\_1\_029\_31

m/z 189.0042

0246E+577.5RT (min)

lcorbi\_hilic\_neg\_1\_005\_36

m/z 190.1802

02468E+633.5RT (min)

lcorbi\_hilic\_neg\_2\_057\_49

m/z 190.9286

01E+60.51RT (min)

lcorbi\_hilic\_neg\_3\_000\_aa

m/z 191.0357

01E+44.55RT (min)

lcorbi\_hilic\_neg\_2\_005\_24

m/z 192.0525

01E+64.55RT (min)

lcorbi\_hilic\_neg\_2\_005\_24

m/z 192.0545

01E+64.55RT (min)

lcorbi\_hilic\_neg\_2\_005\_24

m/z 192.0547

0123E+64.55RT (min)

lcorbi\_hilic\_neg\_2\_005\_24

m/z 192.1230

0246E+53.54RT (min)

lcorbi\_hilic\_neg\_2\_060\_39

m/z 193.1435

02468E+60.51RT (min)

lcorbi\_hilic\_neg\_2\_060\_39

m/z 194.0514

01E+744.5RT (min)

lcorbi\_hilic\_neg\_1\_012\_32

m/z 194.0517

01E+64.55RT (min)

lcorbi\_hilic\_neg\_2\_005\_24

m/z 194.9248

012E+47.58RT (min)

lcorbi\_hilic\_neg\_1\_039\_38

m/z 194.9389

02468E+577.5RT (min)

lcorbi\_hilic\_neg\_1\_055\_24

m/z 194.9389

01E+677.5RT (min)

lcorbi\_hilic\_neg\_1\_012\_32

m/z 194.9397

02468E+577.5RT (min)

lcorbi\_hilic\_neg\_1\_047\_05

m/z 194.9884

02468E+40.51RT (min)

lcorbi\_hilic\_neg\_4\_088\_59

m/z 194.9884

0246E+42.53RT (min)

lcorbi\_hilic\_neg\_4\_071\_57

m/z 195.1227

012E+711.5RT (min)

lcorbi\_hilic\_neg\_4\_057\_49

m/z 195.1227

012E+611.5RT (min)

lcorbi\_hilic\_neg\_1\_060\_13

m/z 195.8110

024E+511.5RT (min)

lcorbi\_hilic\_neg\_4\_093\_60

m/z 195.8110

024E+511.5RT (min)

lcorbi\_hilic\_neg\_4\_093\_60

m/z 195.8110

024E+51.52RT (min)

lcorbi\_hilic\_neg\_4\_051\_16

m/z 195.8110

01E+60.51RT (min)

lcorbi\_hilic\_neg\_4\_089\_59

m/z 195.8110

0123E+51.52RT (min)

lcorbi\_hilic\_neg\_4\_086\_59

m/z 195.8110

024E+511.5RT (min)

lcorbi\_hilic\_neg\_4\_051\_16

m/z 197.0774

012E+544.5RT (min)

lcorbi\_hilic\_neg\_4\_085\_59

m/z 197.8080

024E+51.52RT (min)

lcorbi\_hilic\_neg\_2\_050\_17

m/z 197.8080

012E+522.5RT (min)

lcorbi\_hilic\_neg\_4\_051\_16

m/z 197.8080

0123E+522.5RT (min)

lcorbi\_hilic\_neg\_3\_049\_55

m/z 197.8080

0246E+51.52RT (min)

lcorbi\_hilic\_neg\_4\_051\_16

m/z 197.8080

0246E+51.52RT (min)

lcorbi\_hilic\_neg\_4\_088\_59

m/z 197.8080

0246E+511.5RT (min)

lcorbi\_hilic\_neg\_4\_051\_16

m/z 197.8080

0246E+511.5RT (min)

lcorbi\_hilic\_neg\_4\_094\_60

m/z 197.8080

01234E+51.52RT (min)

lcorbi\_hilic\_neg\_4\_088\_59

m/z 197.8080

0123E+52.53RT (min)

lcorbi\_hilic\_neg\_2\_050\_17

m/z 197.8080

0123E+51.52RT (min)

lcorbi\_hilic\_neg\_3\_037\_12

m/z 198.1237

01E+66.57RT (min)

lcorbi\_hilic\_neg\_4\_065\_51

m/z 198.1853

0123E+61.52RT (min)

lcorbi\_hilic\_neg\_2\_060\_39

m/z 199.8050

01E+52.53RT (min)

lcorbi\_hilic\_neg\_3\_043\_02

m/z 199.8050

01E+51.52RT (min)

lcorbi\_hilic\_neg\_3\_037\_12

m/z 199.8050

01234E+53.54RT (min)

lcorbi\_hilic\_neg\_4\_112\_62

m/z 199.8050

0123E+51.52RT (min)

lcorbi\_hilic\_neg\_4\_088\_59

m/z 199.8050

0123E+511.5RT (min)

lcorbi\_hilic\_neg\_4\_093\_60

m/z 199.8050

01E+51.52RT (min)

lcorbi\_hilic\_neg\_3\_043\_02

m/z 199.8050

012E+51.52RT (min)

lcorbi\_hilic\_neg\_4\_097\_60

m/z 199.8050

012E+51.52RT (min)

lcorbi\_hilic\_neg\_4\_086\_59

m/z 199.8050

012E+511.5RT (min)

lcorbi\_hilic\_neg\_4\_092\_60

m/z 200.2009

01E+72.53RT (min)

lcorbi\_hilic\_neg\_4\_059\_13

m/z 200.2009

0246E+622.5RT (min)

lcorbi\_hilic\_neg\_2\_057\_49

m/z 201.2042

01E+62.53RT (min)

lcorbi\_hilic\_neg\_4\_059\_13

m/z 202.2165

012E+72.53RT (min)

lcorbi\_hilic\_neg\_4\_057\_49

m/z 202.2165

012E+622.5RT (min)

lcorbi\_hilic\_neg\_4\_059\_13

m/z 202.2165

01E+61.52RT (min)

lcorbi\_hilic\_neg\_2\_062\_14

m/z 202.9971

01E+588.5RT (min)

lcorbi\_hilic\_neg\_3\_064\_16

m/z 203.0526

01E+655.5RT (min)

lcorbi\_hilic\_neg\_3\_060\_15

m/z 203.0810

01E+600.5RT (min)

lcorbi\_hilic\_neg\_1\_058\_39

m/z 203.0811

024E+53.54RT (min)

lcorbi\_hilic\_neg\_1\_064\_15

m/z 203.0814

012E+51313.5RT (min)

lcorbi\_hilic\_neg\_2\_060\_39

m/z 203.0815

01E+51313.5RT (min)

lcorbi\_hilic\_neg\_1\_065\_51

m/z 203.0853

024E+511.512RT (min)

lcorbi\_hilic\_neg\_4\_065\_51

m/z 203.0853

024E+512.513RT (min)

lcorbi\_hilic\_neg\_2\_064\_17

m/z 203.0853

024E+513.514RT (min)

lcorbi\_hilic\_neg\_1\_065\_51

m/z 203.2198

012E+62.53RT (min)

lcorbi\_hilic\_neg\_4\_057\_49

m/z 204.1225

0123E+744.5RT (min)

lcorbi\_hilic\_neg\_1\_060\_13

m/z 204.1957

024E+633.5RT (min)

lcorbi\_hilic\_neg\_2\_057\_49

m/z 205.1991

024E+533.5RT (min)

lcorbi\_hilic\_neg\_4\_059\_13

m/z 208.9345

01234E+55.56RT (min)

lcorbi\_hilic\_neg\_1\_023\_23

m/z 208.9346

0123E+577.5RT (min)

lcorbi\_hilic\_neg\_1\_004\_17

m/z 209.1648

01E+61.52RT (min)

lcorbi\_hilic\_neg\_1\_061\_50

m/z 211.0975

01E+60.51RT (min)

lcorbi\_hilic\_neg\_3\_000\_cc

m/z 213.8798

01E+58.59RT (min)

lcorbi\_hilic\_neg\_4\_066\_57

m/z 213.8798

01E+58.59RT (min)

lcorbi\_hilic\_neg\_4\_075\_58

m/z 213.8798

01E+58.59RT (min)

lcorbi\_hilic\_neg\_4\_067\_57

m/z 213.8798

012E+577.5RT (min)

lcorbi\_hilic\_neg\_4\_076\_58

m/z 213.8798

01E+588.5RT (min)

lcorbi\_hilic\_neg\_4\_066\_57

m/z 213.8798

01E+588.5RT (min)

lcorbi\_hilic\_neg\_4\_066\_57

m/z 214.2529

01E+61.52RT (min)

lcorbi\_hilic\_neg\_4\_065\_51

m/z 214.9175

01E+58.59RT (min)

lcorbi\_hilic\_neg\_2\_062\_14

m/z 215.0318

0123E+644.5RT (min)

lcorbi\_hilic\_neg\_1\_003\_14

m/z 215.0328

01E+74.55RT (min)

lcorbi\_hilic\_neg\_1\_003\_14

m/z 215.0328

0123E+644.5RT (min)

lcorbi\_hilic\_neg\_4\_077\_58

m/z 215.0329

0246E+64.55RT (min)

lcorbi\_hilic\_neg\_1\_003\_14

m/z 216.1958

0246E+533.5RT (min)

lcorbi\_hilic\_neg\_2\_057\_49

m/z 216.1958

01234E+61.52RT (min)

lcorbi\_hilic\_neg\_2\_060\_39

m/z 216.1958

0123E+622.5RT (min)

lcorbi\_hilic\_neg\_3\_061\_50

m/z 217.0504

0123E+522.5RT (min)

lcorbi\_hilic\_neg\_4\_036\_36

m/z 218.2043

024E+52.53RT (min)

lcorbi\_hilic\_neg\_1\_063\_17

m/z 218.2052

01E+622.5RT (min)

lcorbi\_hilic\_neg\_2\_062\_14

m/z 218.2113

0246E+82.53RT (min)

lcorbi\_hilic\_neg\_2\_062\_14

m/z 218.2114

01234E+822.5RT (min)

lcorbi\_hilic\_neg\_2\_057\_49

m/z 218.2114

012E+822.5RT (min)

lcorbi\_hilic\_neg\_2\_062\_14

m/z 218.2177

012E+622.5RT (min)

lcorbi\_hilic\_neg\_2\_057\_49

m/z 218.2180

01234E+62.53RT (min)

lcorbi\_hilic\_neg\_2\_062\_14

m/z 219.0372

0246E+511.5RT (min)

lcorbi\_hilic\_neg\_4\_096\_60

m/z 219.0662

0246E+61.52RT (min)

lcorbi\_hilic\_neg\_4\_109\_62

m/z 219.0662

012E+53.54RT (min)

lcorbi\_hilic\_neg\_4\_038\_27

m/z 219.0662

0246E+61.52RT (min)

lcorbi\_hilic\_neg\_4\_112\_62

m/z 219.0662

0246E+61.52RT (min)

lcorbi\_hilic\_neg\_4\_108\_62

m/z 219.0662

0123E+53.54RT (min)

lcorbi\_hilic\_neg\_4\_038\_27

m/z 219.0663

0123E+53.54RT (min)

lcorbi\_hilic\_neg\_4\_038\_27

m/z 219.0663

01E+622.5RT (min)

lcorbi\_hilic\_neg\_4\_055\_38

m/z 219.0663

012E+622.5RT (min)

lcorbi\_hilic\_neg\_4\_038\_27

m/z 219.0663

01E+622.5RT (min)

lcorbi\_hilic\_neg\_4\_055\_38

m/z 219.0663

0246E+61.52RT (min)

lcorbi\_hilic\_neg\_4\_108\_62

m/z 219.2094

01E+62.53RT (min)

lcorbi\_hilic\_neg\_2\_062\_14

m/z 219.2145

02468E+72.53RT (min)

lcorbi\_hilic\_neg\_2\_062\_14

m/z 219.2146

0246E+722.5RT (min)

lcorbi\_hilic\_neg\_2\_057\_49

m/z 219.2146

0123E+722.5RT (min)

lcorbi\_hilic\_neg\_1\_059\_14

m/z 219.8454

0246E+50.51RT (min)

lcorbi\_hilic\_neg\_3\_000\_aa

m/z 219.8454

01E+511.5RT (min)

lcorbi\_hilic\_neg\_3\_052\_40

m/z 219.8454

01E+51.52RT (min)

lcorbi\_hilic\_neg\_3\_052\_40

m/z 219.8455

02468E+51010.511RT (min)

lcorbi\_hilic\_neg\_4\_074\_58

m/z 219.8456

0246E+58.59RT (min)

lcorbi\_hilic\_neg\_1\_026\_34

m/z 219.8456

024E+588.5RT (min)

lcorbi\_hilic\_neg\_1\_006\_08

m/z 219.8456

02468E+588.59RT (min)

lcorbi\_hilic\_neg\_4\_066\_57

m/z 219.8456

0246E+58.59RT (min)

lcorbi\_hilic\_neg\_1\_016\_37

m/z 220.0696

02468E+51.52RT (min)

lcorbi\_hilic\_neg\_4\_108\_62

m/z 220.0696

01E+61.52RT (min)

lcorbi\_hilic\_neg\_4\_109\_62

m/z 220.2177

01234E+62.53RT (min)

lcorbi\_hilic\_neg\_2\_062\_14

m/z 221.8424

024E+51010.5RT (min)

lcorbi\_hilic\_neg\_4\_076\_58

m/z 221.8424

024E+510.511RT (min)

lcorbi\_hilic\_neg\_4\_074\_58

m/z 221.8424

0246E+413.514RT (min)

lcorbi\_hilic\_neg\_4\_067\_57

m/z 221.8425

024E+510RT (min)

lcorbi\_hilic\_neg\_4\_066\_57

m/z 221.8426

0246E+566.5RT (min)

lcorbi\_hilic\_neg\_4\_076\_58

m/z 221.8426

024E+50.51RT (min)

lcorbi\_hilic\_neg\_3\_000\_aa

m/z 221.8426

024E+58.59RT (min)

lcorbi\_hilic\_neg\_1\_004\_17

m/z 221.8426

024E+588.5RT (min)

lcorbi\_hilic\_neg\_2\_000\_bb

m/z 223.0686

024E+34.55RT (min)

lcorbi\_hilic\_neg\_2\_005\_24

m/z 223.0924

0123E+65.56RT (min)

lcorbi\_hilic\_neg\_3\_063\_13

m/z 225.0616

024E+65.56RT (min)

lcorbi\_hilic\_neg\_2\_020\_27

m/z 226.2166

0246E+511.5RT (min)

lcorbi\_hilic\_neg\_2\_061\_50

m/z 226.9180

01E+599.5RT (min)

lcorbi\_hilic\_neg\_4\_074\_58

m/z 226.9180

01E+599.5RT (min)

lcorbi\_hilic\_neg\_4\_074\_58

m/z 226.9180

01E+599.5RT (min)

lcorbi\_hilic\_neg\_4\_074\_58

m/z 226.9514

02468E+755.5RT (min)

lcorbi\_hilic\_neg\_3\_064\_16

m/z 226.9947

01E+511.5RT (min)

lcorbi\_hilic\_neg\_4\_090\_60

m/z 227.0924

0246E+50.51RT (min)

lcorbi\_hilic\_neg\_3\_000\_cc

m/z 227.1754

012E+71.52RT (min)

lcorbi\_hilic\_neg\_1\_061\_50

m/z 228.1786

0123E+61.52RT (min)

lcorbi\_hilic\_neg\_1\_061\_50

m/z 228.2322

0246E+52.53RT (min)

lcorbi\_hilic\_neg\_4\_059\_13

m/z 229.1546

012E+655.5RT (min)

lcorbi\_hilic\_neg\_4\_058\_15

m/z 230.1439

01234E+73.54RT (min)

lcorbi\_hilic\_neg\_1\_063\_17

m/z 230.2478

0123E+72.53RT (min)

lcorbi\_hilic\_neg\_4\_059\_13

m/z 230.2478

01234E+61.52RT (min)

lcorbi\_hilic\_neg\_4\_059\_13

m/z 231.2511

024E+51.52RT (min)

lcorbi\_hilic\_neg\_4\_059\_13

m/z 231.2511

01234E+62.53RT (min)

lcorbi\_hilic\_neg\_4\_059\_13

m/z 231.9316

01E+511.5RT (min)

lcorbi\_hilic\_neg\_4\_082\_59

m/z 231.9316

02468E+41.52RT (min)

lcorbi\_hilic\_neg\_4\_110\_62

m/z 231.9316

01E+511.5RT (min)

lcorbi\_hilic\_neg\_4\_082\_59

m/z 231.9316

0246E+43.54RT (min)

lcorbi\_hilic\_neg\_4\_106\_62

m/z 231.9316

02468E+42.53RT (min)

lcorbi\_hilic\_neg\_4\_103\_61

m/z 232.1907

01E+63.54RT (min)

lcorbi\_hilic\_neg\_2\_062\_14

m/z 232.2269

012E+62.53RT (min)

lcorbi\_hilic\_neg\_2\_057\_49

m/z 232.9246

012E+555.5RT (min)

lcorbi\_hilic\_neg\_2\_000\_cc

m/z 232.9281

0123E+58.59RT (min)

lcorbi\_hilic\_neg\_3\_059\_14

m/z 233.1546

012E+411.5RT (min)

lcorbi\_hilic\_neg\_4\_061\_50

m/z 233.9148

0246E+500.5RT (min)

lcorbi\_hilic\_neg\_1\_057\_49

m/z 234.2063

0123E+733.5RT (min)

lcorbi\_hilic\_neg\_2\_057\_49

m/z 234.2064

012E+733.5RT (min)

lcorbi\_hilic\_neg\_4\_065\_51

m/z 235.1704

0246E+60.51RT (min)

lcorbi\_hilic\_neg\_3\_000\_cc

m/z 235.2096

01234E+633.5RT (min)

lcorbi\_hilic\_neg\_2\_057\_49

m/z 235.9130

0246E+500.5RT (min)

lcorbi\_hilic\_neg\_1\_057\_49

m/z 236.1736

02468E+50.51RT (min)

lcorbi\_hilic\_neg\_3\_000\_cc

m/z 236.1736

02468E+511.5RT (min)

lcorbi\_hilic\_neg\_2\_000\_bb

m/z 238.9316

01E+555.5RT (min)

lcorbi\_hilic\_neg\_4\_082\_59

m/z 239.1490

024E+611.5RT (min)

lcorbi\_hilic\_neg\_4\_061\_50

m/z 239.9665

0123E+58.59RT (min)

lcorbi\_hilic\_neg\_1\_059\_14

m/z 241.0104

02468E+411.5RT (min)

lcorbi\_hilic\_neg\_4\_090\_60

m/z 243.9680

012E+52.53RT (min)

lcorbi\_hilic\_neg\_4\_074\_58

m/z 244.2271

0246E+511.5RT (min)

lcorbi\_hilic\_neg\_2\_061\_50

m/z 244.2272

01E+61.52RT (min)

lcorbi\_hilic\_neg\_2\_061\_50

m/z 244.2272

0246E+52.53RT (min)

lcorbi\_hilic\_neg\_2\_057\_49

m/z 244.2634

01234E+62.53RT (min)

lcorbi\_hilic\_neg\_4\_065\_51

m/z 244.2634

02468E+51.52RT (min)

lcorbi\_hilic\_neg\_4\_065\_51

m/z 244.9849

0246E+411.5RT (min)

lcorbi\_hilic\_neg\_4\_080\_58

m/z 245.2668

024E+52.53RT (min)

lcorbi\_hilic\_neg\_4\_065\_51

m/z 246.2063

012E+63.54RT (min)

lcorbi\_hilic\_neg\_2\_060\_39

m/z 246.2356

01234E+52.53RT (min)

lcorbi\_hilic\_neg\_4\_059\_13

m/z 246.2426

01E+82.53RT (min)

lcorbi\_hilic\_neg\_2\_057\_49

m/z 246.2426

0123E+722.5RT (min)

lcorbi\_hilic\_neg\_4\_057\_49

m/z 246.2427

0123E+722.5RT (min)

lcorbi\_hilic\_neg\_4\_059\_13

m/z 246.2496

0246E+52.53RT (min)

lcorbi\_hilic\_neg\_4\_059\_13

m/z 247.1665

01E+51.52RT (min)

lcorbi\_hilic\_neg\_1\_057\_49

m/z 247.1665

01234E+544.5RT (min)

lcorbi\_hilic\_neg\_1\_058\_39

m/z 247.2459

01E+72.53RT (min)

lcorbi\_hilic\_neg\_2\_057\_49

m/z 247.2460

024E+622.5RT (min)

lcorbi\_hilic\_neg\_4\_059\_13

m/z 247.2460

01234E+622.5RT (min)

lcorbi\_hilic\_neg\_4\_057\_49

m/z 248.9604

012E+566.5RT (min)

lcorbi\_hilic\_neg\_4\_107\_62

m/z 253.0858

012E+80.51RT (min)

lcorbi\_hilic\_neg\_1\_057\_49

m/z 253.1182

012E+611.5RT (min)

lcorbi\_hilic\_neg\_1\_065\_51

m/z 255.0919

012E+60.51RT (min)

lcorbi\_hilic\_neg\_1\_061\_50

m/z 255.8223

01234E+58.59RT (min)

lcorbi\_hilic\_neg\_4\_066\_57

m/z 255.8223

0123E+58.59RT (min)

lcorbi\_hilic\_neg\_4\_067\_57

m/z 255.9440

01E+68.59RT (min)

lcorbi\_hilic\_neg\_4\_062\_39

m/z 255.9440

01E+68.59RT (min)

lcorbi\_hilic\_neg\_1\_059\_14

m/z 256.2634

012E+522.5RT (min)

lcorbi\_hilic\_neg\_4\_060\_16

m/z 257.8192

01234E+58.59RT (min)

lcorbi\_hilic\_neg\_4\_066\_57

m/z 257.8192

01234E+58.59RT (min)

lcorbi\_hilic\_neg\_4\_000\_cc

m/z 257.8192

01234E+58.59RT (min)

lcorbi\_hilic\_neg\_3\_000\_cc

m/z 258.1099

0246E+766.5RT (min)

lcorbi\_hilic\_neg\_2\_062\_14

m/z 259.0008

02468E+411.5RT (min)

lcorbi\_hilic\_neg\_4\_102\_61

m/z 259.0009

02468E+411.5RT (min)

lcorbi\_hilic\_neg\_4\_090\_60

m/z 260.2583

0246E+52.53RT (min)

lcorbi\_hilic\_neg\_4\_057\_49

m/z 262.2375

01E+72.53RT (min)

lcorbi\_hilic\_neg\_2\_057\_49

m/z 262.2375

0246E+622.5RT (min)

lcorbi\_hilic\_neg\_2\_057\_49

m/z 262.2376

01E+61.52RT (min)

lcorbi\_hilic\_neg\_1\_059\_14

m/z 263.2408

012E+62.53RT (min)

lcorbi\_hilic\_neg\_2\_057\_49

m/z 264.9910

0246E+411.5RT (min)

lcorbi\_hilic\_neg\_4\_091\_60

m/z 272.2332

012E+61.52RT (min)

lcorbi\_hilic\_neg\_1\_057\_49

m/z 272.2583

02468E+511.5RT (min)

lcorbi\_hilic\_neg\_2\_061\_50

m/z 274.2739

0246E+72.53RT (min)

lcorbi\_hilic\_neg\_2\_061\_50

m/z 274.2740

024E+61.52RT (min)

lcorbi\_hilic\_neg\_4\_057\_49

m/z 274.2740

01E+71.52RT (min)

lcorbi\_hilic\_neg\_2\_061\_50

m/z 275.2771

01E+72.53RT (min)

lcorbi\_hilic\_neg\_2\_061\_50

m/z 275.2773

01E+61.52RT (min)

lcorbi\_hilic\_neg\_2\_061\_50

m/z 275.2773

02468E+51.52RT (min)

lcorbi\_hilic\_neg\_4\_057\_49

m/z 277.1264

01E+60.51RT (min)

lcorbi\_hilic\_neg\_4\_081\_58

m/z 281.1393

012E+62.53RT (min)

lcorbi\_hilic\_neg\_4\_105\_61

m/z 281.1394

0123E+53.54RT (min)

lcorbi\_hilic\_neg\_4\_023\_34

m/z 281.1394

01E+633.5RT (min)

lcorbi\_hilic\_neg\_4\_014\_18

m/z 281.1394

012E+62.53RT (min)

lcorbi\_hilic\_neg\_4\_104\_61

m/z 281.1394

012E+633.5RT (min)

lcorbi\_hilic\_neg\_4\_105\_61

m/z 281.1394

01E+61.52RT (min)

lcorbi\_hilic\_neg\_4\_049\_55

m/z 281.1394

0246E+61.52RT (min)

lcorbi\_hilic\_neg\_4\_104\_61

m/z 281.1395

0246E+53.54RT (min)

lcorbi\_hilic\_neg\_3\_055\_26

m/z 281.1395

0123E+622.5RT (min)

lcorbi\_hilic\_neg\_4\_050\_26

m/z 281.1395

024E+622.5RT (min)

lcorbi\_hilic\_neg\_4\_105\_61

m/z 282.1427

01E+61.52RT (min)

lcorbi\_hilic\_neg\_2\_045\_26

m/z 282.1427

01E+61.52RT (min)

lcorbi\_hilic\_neg\_4\_099\_61

m/z 282.1427

0246E+522.5RT (min)

lcorbi\_hilic\_neg\_4\_050\_26

m/z 282.1428

01E+61.52RT (min)

lcorbi\_hilic\_neg\_4\_099\_61

m/z 284.2947

024E+52.53RT (min)

lcorbi\_hilic\_neg\_4\_061\_50

m/z 288.2896

0246E+622.5RT (min)

lcorbi\_hilic\_neg\_2\_062\_14

m/z 288.2896

01E+61.52RT (min)

lcorbi\_hilic\_neg\_2\_062\_14

m/z 288.2896

012E+622.5RT (min)

lcorbi\_hilic\_neg\_4\_065\_51

m/z 288.9364

012E+599.5RT (min)

lcorbi\_hilic\_neg\_4\_080\_58

m/z 288.9364

012E+599.5RT (min)

lcorbi\_hilic\_neg\_4\_081\_58

m/z 288.9364

01E+59.510RT (min)

lcorbi\_hilic\_neg\_4\_074\_58

m/z 288.9364

012E+58.59RT (min)

lcorbi\_hilic\_neg\_4\_080\_58

m/z 288.9364

01E+59.510RT (min)

lcorbi\_hilic\_neg\_4\_074\_58

m/z 288.9364

01E+59.510RT (min)

lcorbi\_hilic\_neg\_4\_074\_58

m/z 288.9364

01E+59.510RT (min)

lcorbi\_hilic\_neg\_4\_074\_58

m/z 288.9364

01E+59.510RT (min)

lcorbi\_hilic\_neg\_4\_081\_58

m/z 288.9364

012E+599.5RT (min)

lcorbi\_hilic\_neg\_4\_080\_58

m/z 288.9364

012E+58.59RT (min)

lcorbi\_hilic\_neg\_4\_066\_57

m/z 288.9364

012E+58.59RT (min)

lcorbi\_hilic\_neg\_4\_080\_58

m/z 288.9364

012E+599.5RT (min)

lcorbi\_hilic\_neg\_4\_074\_58

m/z 288.9364

012E+599.5RT (min)

lcorbi\_hilic\_neg\_4\_074\_58

m/z 288.9364

012E+599.5RT (min)

lcorbi\_hilic\_neg\_4\_078\_58

m/z 289.0893

012E+61.52RT (min)

lcorbi\_hilic\_neg\_4\_059\_13

m/z 289.2930

01E+622.5RT (min)

lcorbi\_hilic\_neg\_2\_062\_14

m/z 290.1386

0246E+62.53RT (min)

lcorbi\_hilic\_neg\_4\_059\_13

m/z 290.2688

01E+72.53RT (min)

lcorbi\_hilic\_neg\_4\_057\_49

m/z 290.2688

012E+61.52RT (min)

lcorbi\_hilic\_neg\_4\_065\_51

m/z 290.2688

01E+722.5RT (min)

lcorbi\_hilic\_neg\_1\_059\_14

m/z 290.2689

01E+61.52RT (min)

lcorbi\_hilic\_neg\_3\_057\_49

m/z 291.1419

01E+62.53RT (min)

lcorbi\_hilic\_neg\_3\_063\_13

m/z 291.2721

012E+622.5RT (min)

lcorbi\_hilic\_neg\_1\_059\_14

m/z 291.2722

012E+62.53RT (min)

lcorbi\_hilic\_neg\_4\_057\_49

m/z 294.1238

0246E+55.56RT (min)

lcorbi\_hilic\_neg\_4\_097\_60

m/z 294.9388

01E+755.5RT (min)

lcorbi\_hilic\_neg\_3\_064\_16

m/z 297.1704

0123E+60.51RT (min)

lcorbi\_hilic\_neg\_3\_000\_cc

m/z 298.1738

0246E+50.51RT (min)

lcorbi\_hilic\_neg\_3\_000\_cc

m/z 298.1739

024E+511.5RT (min)

lcorbi\_hilic\_neg\_2\_000\_bb

m/z 302.3052

012E+61.52RT (min)

lcorbi\_hilic\_neg\_2\_061\_50

m/z 302.3052

0123E+722.5RT (min)

lcorbi\_hilic\_neg\_2\_061\_50

m/z 303.1999

01E+60.51RT (min)

lcorbi\_hilic\_neg\_4\_112\_62

m/z 303.3085

0246E+622.5RT (min)

lcorbi\_hilic\_neg\_2\_061\_50

m/z 304.1444

01E+62.53RT (min)

lcorbi\_hilic\_neg\_4\_057\_49

m/z 304.2845

02468E+52.53RT (min)

lcorbi\_hilic\_neg\_4\_065\_51

m/z 304.9140

02468E+5910RT (min)

lcorbi\_hilic\_neg\_4\_101\_61

m/z 304.9140

01E+699.5RT (min)

lcorbi\_hilic\_neg\_4\_074\_58

m/z 304.9140

01E+699.5RT (min)

lcorbi\_hilic\_neg\_4\_074\_58

m/z 304.9140

0246E+59.510RT (min)

lcorbi\_hilic\_neg\_4\_101\_61

m/z 304.9140

01E+68.59RT (min)

lcorbi\_hilic\_neg\_4\_080\_58

m/z 304.9140

0123E+59.510RT (min)

lcorbi\_hilic\_neg\_4\_101\_61

m/z 304.9140

01E+68.59RT (min)

lcorbi\_hilic\_neg\_4\_080\_58

m/z 304.9140

01E+699.5RT (min)

lcorbi\_hilic\_neg\_4\_081\_58

m/z 304.9140

01E+699.5RT (min)

lcorbi\_hilic\_neg\_4\_074\_58

m/z 309.2570

01E+60.51RT (min)

lcorbi\_hilic\_neg\_4\_095\_60

m/z 310.8703

01E+599.5RT (min)

lcorbi\_hilic\_neg\_4\_074\_58

m/z 310.8703

01E+58.59RT (min)

lcorbi\_hilic\_neg\_4\_079\_58

m/z 310.8703

01E+58.59RT (min)

lcorbi\_hilic\_neg\_4\_079\_58

m/z 310.8704

01E+599.5RT (min)

lcorbi\_hilic\_neg\_4\_074\_58

m/z 310.8704

01E+599.5RT (min)

lcorbi\_hilic\_neg\_4\_074\_58

m/z 312.1551

01E+83.54RT (min)

lcorbi\_hilic\_neg\_4\_057\_49

m/z 313.1581

01E+73.54RT (min)

lcorbi\_hilic\_neg\_1\_065\_51

m/z 315.0412

01234E+55.56RT (min)

lcorbi\_hilic\_neg\_4\_055\_38

m/z 315.0412

0246E+566.5RT (min)

lcorbi\_hilic\_neg\_4\_079\_58

m/z 315.0412

0123E+55.56RT (min)

lcorbi\_hilic\_neg\_1\_044\_03

m/z 315.0412

0123E+55.56RT (min)

lcorbi\_hilic\_neg\_1\_044\_03

m/z 315.0412

0123E+55.56RT (min)

lcorbi\_hilic\_neg\_2\_022\_37

m/z 315.0413

01234E+56.57RT (min)

lcorbi\_hilic\_neg\_1\_016\_37

m/z 316.2481

012E+633.5RT (min)

lcorbi\_hilic\_neg\_4\_059\_13

m/z 318.3001

012E+622.5RT (min)

lcorbi\_hilic\_neg\_2\_061\_50

m/z 318.3002

0246E+63.54RT (min)

lcorbi\_hilic\_neg\_1\_065\_51

m/z 323.1752

01234E+83.54RT (min)

lcorbi\_hilic\_neg\_4\_057\_49

m/z 324.1229

0246E+711.5RT (min)

lcorbi\_hilic\_neg\_4\_065\_51

m/z 324.1782

02468E+73.54RT (min)

lcorbi\_hilic\_neg\_4\_057\_49

m/z 325.1813

02468E+63.54RT (min)

lcorbi\_hilic\_neg\_4\_065\_51

m/z 330.3365

0246E+622.5RT (min)

lcorbi\_hilic\_neg\_2\_061\_50

m/z 330.3366

0246E+51.52RT (min)

lcorbi\_hilic\_neg\_2\_061\_50

m/z 336.3606

012E+63.54RT (min)

lcorbi\_hilic\_neg\_1\_065\_51

m/z 339.3734

01E+655.5RT (min)

lcorbi\_hilic\_neg\_4\_063\_14

m/z 340.2594

01E+71.52RT (min)

lcorbi\_hilic\_neg\_4\_057\_49

m/z 340.2595

02468E+61.52RT (min)

lcorbi\_hilic\_neg\_3\_061\_50

m/z 341.1603

02468E+511.5RT (min)

lcorbi\_hilic\_neg\_2\_000\_bb

m/z 341.2426

024E+40.51RT (min)

lcorbi\_hilic\_neg\_4\_066\_57

m/z 341.2627

012E+61.52RT (min)

lcorbi\_hilic\_neg\_4\_057\_49

m/z 341.2628

01E+61.52RT (min)

lcorbi\_hilic\_neg\_3\_061\_50

m/z 342.1272

0123E+666.5RT (min)

lcorbi\_hilic\_neg\_3\_061\_50

m/z 342.2638

02468E+52.53RT (min)

lcorbi\_hilic\_neg\_2\_063\_13

m/z 343.9946

01234E+66.57RT (min)

lcorbi\_hilic\_neg\_4\_075\_58

m/z 343.9947

01234E+66.57RT (min)

lcorbi\_hilic\_neg\_4\_075\_58

m/z 343.9947

01E+677.5RT (min)

lcorbi\_hilic\_neg\_4\_075\_58

m/z 343.9947

01E+677.5RT (min)

lcorbi\_hilic\_neg\_1\_005\_36

m/z 343.9947

01234E+66.57RT (min)

lcorbi\_hilic\_neg\_4\_074\_58

m/z 343.9948

01E+677.5RT (min)

lcorbi\_hilic\_neg\_4\_074\_58

m/z 343.9948

01E+67.58RT (min)

lcorbi\_hilic\_neg\_4\_074\_58

m/z 343.9948

02468E+57.58RT (min)

lcorbi\_hilic\_neg\_4\_046\_15

m/z 343.9949

01E+67.58RT (min)

lcorbi\_hilic\_neg\_4\_074\_58

m/z 343.9949

02468E+57.58RT (min)

lcorbi\_hilic\_neg\_1\_012\_32

m/z 343.9949

0246E+588.5RT (min)

lcorbi\_hilic\_neg\_1\_012\_32

m/z 343.9949

01E+67.58RT (min)

lcorbi\_hilic\_neg\_4\_075\_58

m/z 346.0094

01E+666.5RT (min)

lcorbi\_hilic\_neg\_4\_061\_50

m/z 346.0094

01E+66.57RT (min)

lcorbi\_hilic\_neg\_2\_062\_14

m/z 346.0094

02468E+56.57RT (min)

lcorbi\_hilic\_neg\_1\_064\_15

m/z 346.0698

01E+62.53RT (min)

lcorbi\_hilic\_neg\_2\_013\_18

m/z 348.0013

0123E+688.5RT (min)

lcorbi\_hilic\_neg\_4\_079\_58

m/z 349.0177

02468E+67.58RT (min)

lcorbi\_hilic\_neg\_1\_059\_14

m/z 350.0155

02468E+67.58RT (min)

lcorbi\_hilic\_neg\_1\_059\_14

m/z 351.0134

01234E+67.58RT (min)

lcorbi\_hilic\_neg\_1\_059\_14

m/z 351.2067

0246E+711.5RT (min)

lcorbi\_hilic\_neg\_1\_061\_50

m/z 351.2067

01234E+71.52RT (min)

lcorbi\_hilic\_neg\_4\_057\_49

m/z 352.2097

01E+711.5RT (min)

lcorbi\_hilic\_neg\_1\_061\_50

m/z 353.0104

02468E+56.57RT (min)

lcorbi\_hilic\_neg\_3\_012\_35

m/z 353.0104

02468E+56.57RT (min)

lcorbi\_hilic\_neg\_3\_039\_31

m/z 353.0104

01E+666.5RT (min)

lcorbi\_hilic\_neg\_4\_074\_58

m/z 353.0104

02468E+56.57RT (min)

lcorbi\_hilic\_neg\_3\_012\_35

m/z 353.0104

01E+68RT (min)

lcorbi\_hilic\_neg\_4\_078\_58

m/z 353.0104

02468E+566.5RT (min)

lcorbi\_hilic\_neg\_4\_023\_34

m/z 353.0104

0246E+366.5RT (min)

lcorbi\_hilic\_neg\_4\_005\_32

m/z 353.2126

0246E+511.5RT (min)

lcorbi\_hilic\_neg\_4\_065\_51

m/z 355.0758

02468E+50.51RT (min)

lcorbi\_hilic\_neg\_4\_065\_51

m/z 355.3682

024E+555.5RT (min)

lcorbi\_hilic\_neg\_4\_063\_14

m/z 356.0985

024E+52.53RT (min)

lcorbi\_hilic\_neg\_4\_014\_18

m/z 369.1333

012E+611.5RT (min)

lcorbi\_hilic\_neg\_1\_057\_49

m/z 370.9996

0246E+57.58RT (min)

lcorbi\_hilic\_neg\_1\_059\_14

m/z 371.9974

02468E+57.58RT (min)

lcorbi\_hilic\_neg\_1\_059\_14

m/z 372.1587

0246E+51.52RT (min)

lcorbi\_hilic\_neg\_4\_059\_13

m/z 377.2081

024E+51.52RT (min)

lcorbi\_hilic\_neg\_2\_020\_27

m/z 379.2228

01E+71.52RT (min)

lcorbi\_hilic\_neg\_4\_065\_51

m/z 379.2229

01234E+61.52RT (min)

lcorbi\_hilic\_neg\_1\_061\_50

m/z 380.1706

02468E+611.5RT (min)

lcorbi\_hilic\_neg\_1\_057\_49

m/z 380.2260

01234E+622.5RT (min)

lcorbi\_hilic\_neg\_4\_065\_51

m/z 380.2262

01E+61.52RT (min)

lcorbi\_hilic\_neg\_1\_061\_50

m/z 380.2262

0123E+61.52RT (min)

lcorbi\_hilic\_neg\_3\_065\_51

m/z 384.1444

01E+711.5RT (min)

lcorbi\_hilic\_neg\_4\_065\_51

m/z 391.1204

0246E+566.5RT (min)

lcorbi\_hilic\_neg\_4\_053\_23

m/z 391.2844

01234E+70.51RT (min)

lcorbi\_hilic\_neg\_2\_062\_14

m/z 394.3104

01E+82.53RT (min)

lcorbi\_hilic\_neg\_3\_065\_51

m/z 394.3104

01E+722.5RT (min)

lcorbi\_hilic\_neg\_4\_065\_51

m/z 394.3104

012E+611.5RT (min)

lcorbi\_hilic\_neg\_4\_061\_50

m/z 394.3105

01E+71.52RT (min)

lcorbi\_hilic\_neg\_3\_061\_50

m/z 395.3134

024E+72.53RT (min)

lcorbi\_hilic\_neg\_3\_065\_51

m/z 395.3136

01234E+622.5RT (min)

lcorbi\_hilic\_neg\_4\_065\_51

m/z 395.3137

0123E+61.52RT (min)

lcorbi\_hilic\_neg\_3\_061\_50

m/z 396.3166

02468E+62.53RT (min)

lcorbi\_hilic\_neg\_3\_065\_51

m/z 398.1531

02468E+52.53RT (min)

lcorbi\_hilic\_neg\_3\_063\_13

m/z 401.2046

0246E+522.5RT (min)

lcorbi\_hilic\_neg\_2\_057\_49

m/z 438.3790

0123E+70.51RT (min)

lcorbi\_hilic\_neg\_4\_057\_49

m/z 452.2905

01E+63.54RT (min)

lcorbi\_hilic\_neg\_1\_047\_05

m/z 452.2912

0246E+544.5RT (min)

lcorbi\_hilic\_neg\_4\_030\_05

m/z 453.3432

01E+622.5RT (min)

lcorbi\_hilic\_neg\_3\_065\_51

m/z 453.3433

01E+61.52RT (min)

lcorbi\_hilic\_neg\_3\_061\_50

m/z 453.3433

0246E+62.53RT (min)

lcorbi\_hilic\_neg\_3\_061\_50

m/z 454.3467

01E+633.5RT (min)

lcorbi\_hilic\_neg\_4\_057\_49

m/z 468.3084

01234E+644.5RT (min)

lcorbi\_hilic\_neg\_3\_064\_16

m/z 480.3448

01E+644.5RT (min)

lcorbi\_hilic\_neg\_2\_062\_14

m/z 482.3252

012E+644.5RT (min)

lcorbi\_hilic\_neg\_2\_062\_14

m/z 482.3600

024E+644.5RT (min)

lcorbi\_hilic\_neg\_2\_062\_14

m/z 482.4050

0246E+70.51RT (min)

lcorbi\_hilic\_neg\_4\_065\_51

m/z 496.3396

012E+83.54RT (min)

lcorbi\_hilic\_neg\_3\_062\_17

m/z 497.3428

024E+73.54RT (min)

lcorbi\_hilic\_neg\_3\_062\_17

m/z 498.3454

01E+73.54RT (min)

lcorbi\_hilic\_neg\_3\_062\_17

m/z 498.4011

02468E+52.53RT (min)

lcorbi\_hilic\_neg\_1\_057\_49

m/z 508.3757

0123E+644.5RT (min)

lcorbi\_hilic\_neg\_4\_063\_14

m/z 510.3552

01234E+644.5RT (min)

lcorbi\_hilic\_neg\_4\_063\_14

m/z 513.1768

012E+611.5RT (min)

lcorbi\_hilic\_neg\_4\_101\_61

m/z 513.1770

012E+60.51RT (min)

lcorbi\_hilic\_neg\_2\_019\_21

m/z 518.3222

0246E+53.54RT (min)

lcorbi\_hilic\_neg\_4\_059\_13

m/z 519.3264

024E+544.5RT (min)

lcorbi\_hilic\_neg\_2\_061\_50

m/z 520.3396

012E+73.54RT (min)

lcorbi\_hilic\_neg\_2\_058\_16

m/z 520.3396

0246E+744.5RT (min)

lcorbi\_hilic\_neg\_3\_064\_16

m/z 521.3428

012E+744.5RT (min)

lcorbi\_hilic\_neg\_3\_064\_16

m/z 521.3430

0246E+63.54RT (min)

lcorbi\_hilic\_neg\_2\_058\_16

m/z 522.3543

012E+744.5RT (min)

lcorbi\_hilic\_neg\_4\_063\_14

m/z 522.3548

012E+73.54RT (min)

lcorbi\_hilic\_neg\_1\_063\_17

m/z 523.3582

0246E+644.5RT (min)

lcorbi\_hilic\_neg\_4\_063\_14

m/z 524.3696

0123E+744.5RT (min)

lcorbi\_hilic\_neg\_4\_057\_49

m/z 524.3704

01E+73.54RT (min)

lcorbi\_hilic\_neg\_1\_063\_17

m/z 524.3708

0123E+633.5RT (min)

lcorbi\_hilic\_neg\_2\_062\_14

m/z 525.3734

02468E+644.5RT (min)

lcorbi\_hilic\_neg\_4\_057\_49

m/z 525.3740

0123E+63.54RT (min)

lcorbi\_hilic\_neg\_1\_063\_17

m/z 526.3760

01E+644.5RT (min)

lcorbi\_hilic\_neg\_3\_057\_49

m/z 526.4311

01234E+70.51RT (min)

lcorbi\_hilic\_neg\_2\_057\_49

m/z 534.2951

01E+644.5RT (min)

lcorbi\_hilic\_neg\_4\_063\_14

m/z 540.3308

0246E+63.54RT (min)

lcorbi\_hilic\_neg\_2\_020\_27

m/z 542.3230

01E+644.5RT (min)

lcorbi\_hilic\_neg\_3\_064\_16

m/z 544.3370

024E+555.5RT (min)

lcorbi\_hilic\_neg\_2\_005\_24

m/z 544.3392

01E+73.54RT (min)

lcorbi\_hilic\_neg\_1\_063\_17

m/z 544.4106

01E+644.5RT (min)

lcorbi\_hilic\_neg\_2\_061\_50

m/z 545.3424

0246E+644.5RT (min)

lcorbi\_hilic\_neg\_1\_061\_50

m/z 545.3428

0123E+63.54RT (min)

lcorbi\_hilic\_neg\_1\_063\_17

m/z 546.3527

0123E+644.5RT (min)

lcorbi\_hilic\_neg\_1\_061\_50

m/z 546.3542

01E+63.54RT (min)

lcorbi\_hilic\_neg\_1\_063\_17

m/z 546.4262

024E+544.5RT (min)

lcorbi\_hilic\_neg\_1\_061\_50

m/z 547.3576

024E+544.5RT (min)

lcorbi\_hilic\_neg\_4\_059\_13

m/z 568.3394

024E+53.54RT (min)

lcorbi\_hilic\_neg\_1\_063\_17

m/z 568.3504

024E+544.5RT (min)

lcorbi\_hilic\_neg\_2\_041\_54

m/z 651.5659

024E+53.54RT (min)

lcorbi\_hilic\_neg\_4\_062\_39

m/z 651.5661

0246E+544.5RT (min)

lcorbi\_hilic\_neg\_4\_061\_50

m/z 690.5057

02468E+52.53RT (min)

lcorbi\_hilic\_neg\_3\_057\_49

m/z 708.5108

024E+50.51RT (min)

lcorbi\_hilic\_neg\_2\_057\_49

m/z 715.0628

024E+54.55RT (min)

lcorbi\_hilic\_neg\_2\_062\_14

m/z 744.5528

01E+61.52RT (min)

lcorbi\_hilic\_neg\_1\_057\_49

m/z 747.5655

01E+63.54RT (min)

lcorbi\_hilic\_neg\_4\_067\_57

m/z 758.4814

024E+544.5RT (min)

lcorbi\_hilic\_neg\_1\_051\_33

m/z 758.5680

02468E+62.53RT (min)

lcorbi\_hilic\_neg\_4\_059\_13

m/z 758.5682

01E+61.52RT (min)

lcorbi\_hilic\_neg\_1\_057\_49

m/z 758.5684

012E+60.51RT (min)

lcorbi\_hilic\_neg\_2\_065\_51

m/z 758.5688

024E+633.5RT (min)

lcorbi\_hilic\_neg\_3\_062\_17

m/z 759.5716

0123E+62.53RT (min)

lcorbi\_hilic\_neg\_4\_059\_13

m/z 759.5721

012E+633.5RT (min)

lcorbi\_hilic\_neg\_2\_062\_14

m/z 760.5833

012E+62.53RT (min)

lcorbi\_hilic\_neg\_4\_059\_13

m/z 761.5869

01E+62.53RT (min)

lcorbi\_hilic\_neg\_4\_059\_13

m/z 782.5680

012E+611.5RT (min)

lcorbi\_hilic\_neg\_1\_057\_49

m/z 782.5680

01E+62.53RT (min)

lcorbi\_hilic\_neg\_4\_059\_13

m/z 782.5690

012E+633.5RT (min)

lcorbi\_hilic\_neg\_1\_059\_14

m/z 783.6761

024E+51.52RT (min)

lcorbi\_hilic\_neg\_1\_057\_49

m/z 785.6908

01E+61.52RT (min)

lcorbi\_hilic\_neg\_1\_057\_49

m/z 786.5986

01E+633.5RT (min)

lcorbi\_hilic\_neg\_3\_063\_13

m/z 786.5992

012E+62.53RT (min)

lcorbi\_hilic\_neg\_4\_059\_13
