## Supplementary material for "The Everything Bagel Feature Finder: Ultra-fast automated feature finding for untargeted metabolomics": SI Data File: tool_unique_xic_st_Masscube_ST004504.html

 
 
 Masscube-unique features (not detected by the other two tools) - ST004504 
 
 Masscube-unique features (not detected by the other two tools) - ST004504 
 13133 features · orbitrap · XIC window m/z ± 10 ppm (≥ 0.002 Da) · ± 0.5 min 
 Every panel is a feature  Masscube  detected that neither of the other two tools detected — three-way unique across EB (standard mode (min_detection_rate=0.1)), Masscube and XCMS, under the standard matching rule (m/z ± 10 ppm; apex within the reference RT window ± 0.2 min). It shows the MS1 extracted-ion chromatogram at the feature's m/z, drawn from the raw mzML named in  alignment_reference_file , zoomed to ± 0.5 min around the feature's reported apex RT.  Gray solid line  = the feature's reported apex RT;  red dashed line  = masscube's assigned MS/MS —  MS2_scan_id  in  MS2_source_file ; the panel widens to keep an off-apex MS/MS on-screen. Features with no assigned MS/MS show only the gray line. y is per-cell autoscaled. Rows keep the table order (m/z ascending). 
   m/z 52.5998     0   2   4   6E+5   14   14.5  RT (min)      lcorbi_t3_pos_3_034_26    m/z 52.5999     0   1   15.5   16  RT (min)      lcorbi_t3_pos_1_038_30    m/z 52.5999     0   1   2   3   4E+5   15   15.5  RT (min)      lcorbi_t3_pos_4_019_47    m/z 52.5999     0   1   13.5   14  RT (min)      lcorbi_t3_pos_2_039_30    m/z 52.5999     0   1   16.5   17  RT (min)    lcorbi_t3_pos_1_051_09    m/z 52.5999     0   1   15.5   16  RT (min)      lcorbi_t3_pos_1_028_32    m/z 52.5999     0   1E+4   13.5   14  RT (min)      lcorbi_t3_pos_2_047_31    m/z 52.5999     0   1   17.5   18  RT (min)    lcorbi_t3_pos_1_023_02    m/z 52.5999     0   1   2E+5   14   14.5  RT (min)      lcorbi_t3_pos_2_016_23    m/z 52.5999     0   1   13.5   14  RT (min)      lcorbi_t3_pos_2_032_39    m/z 54.4917     0   1E+4   13.5   14  RT (min)      lcorbi_t3_pos_2_039_30    m/z 54.4917     0   1   16.5   17  RT (min)    lcorbi_t3_pos_2_052_48    m/z 54.4917     0   1   16.5   17  RT (min)    lcorbi_t3_pos_1_051_09    m/z 54.4917     0   1   2E+4   15.5   16  RT (min)      lcorbi_t3_pos_1_028_32    m/z 54.4917     0   1   13.5   14  RT (min)      lcorbi_t3_pos_2_032_39    m/z 54.4917     0   1E+6   14   14.5  RT (min)      lcorbi_t3_pos_3_034_26    m/z 54.4917     0   2   4   6   8E+5   14   14.5  RT (min)      lcorbi_t3_pos_3_053_48    m/z 54.4917     0   1   17.5   18  RT (min)    lcorbi_t3_pos_1_023_02    m/z 54.4917     0   2   4   6   8E+5   15   15.5  RT (min)      lcorbi_t3_pos_4_019_47    m/z 54.4917     0   1   16   16.5  RT (min)      lcorbi_t3_pos_1_034_34    m/z 54.4917     0   2   4   6   8E+5   14   14.5  RT (min)      lcorbi_t3_pos_3_053_48    m/z 54.4917     0   1E+4   14   14.5  RT (min)      lcorbi_t3_pos_2_016_23    m/z 54.4917     0   1   16.5   17  RT (min)    lcorbi_t3_pos_1_051_09    m/z 54.4917     0   1   15.5   16  RT (min)      lcorbi_t3_pos_1_055_33    m/z 54.4917     0   1   15.5   16  RT (min)      lcorbi_t3_pos_3_044_46    m/z 54.6031     0   1   2   3E+5   14   14.5  RT (min)      lcorbi_t3_pos_3_034_26    m/z 54.6031     0   1   16.5   17  RT (min)    lcorbi_t3_pos_1_051_09    m/z 54.6031     0   1   15.5   16  RT (min)      lcorbi_t3_pos_1_028_32    m/z 54.6032     0   1   14   14.5  RT (min)      lcorbi_t3_pos_2_016_23    m/z 55.1488     0   1   2E+6   11.5   12  RT (min)      lcorbi_t3_pos_1_008_36    m/z 55.1488     0   1   12   12.5  RT (min)      lcorbi_t3_pos_2_009_50    m/z 55.1488     0   1   2E+6   10.5   11  RT (min)      lcorbi_t3_pos_1_002_01    m/z 55.1488     0   1   12.5   13  RT (min)      lcorbi_t3_pos_2_012_01    m/z 55.1488     0   1E+4   12.5   13  RT (min)      lcorbi_t3_pos_1_023_02    m/z 55.1488     0   2   4   6   8E+3   12   12.5  RT (min)      lcorbi_t3_pos_1_034_34    m/z 55.1488     0   1E+5   11.5   12  RT (min)      lcorbi_t3_pos_1_028_32    m/z 55.1488     0   1   12   12.5  RT (min)      lcorbi_t3_pos_1_031_40    m/z 55.1488     0   1E+6   10.5   11  RT (min)      lcorbi_t3_pos_3_031_45    m/z 55.1489     0   1   2E+6   10.5   11  RT (min)      lcorbi_t3_pos_1_002_01    m/z 55.1489     0   2   4   6E+5   11.5   12  RT (min)      lcorbi_t3_pos_2_022_38    m/z 55.2603     0   2   4   6E+5   10.5   11  RT (min)      lcorbi_t3_pos_1_002_01    m/z 55.2603     0   1E+4   12   12.5  RT (min)      lcorbi_t3_pos_1_031_40    m/z 55.2603     0   1   12   12.5  RT (min)      lcorbi_t3_pos_1_034_34    m/z 55.2603     0   1E+4   12.5   13  RT (min)      lcorbi_t3_pos_1_023_02    m/z 55.2603     0   2   4   6E+5   11.5   12  RT (min)      lcorbi_t3_pos_1_008_36    m/z 57.0698     0   2   4   6   8E+4   15.5   16  RT (min)      lcorbi_t3_pos_1_023_02    m/z 57.0699     0   2   4   6E+4   14.5   15  RT (min)      lcorbi_t3_pos_1_034_34    m/z 57.0699     0   2   4E+4   15   15.5  RT (min)      lcorbi_t3_pos_2_012_01    m/z 57.0699     0   2   4   6   8E+5   13.5   14  RT (min)      lcorbi_t3_pos_3_047_40    m/z 57.0699     0   1   2E+4   12   12.5  RT (min)      lcorbi_t3_pos_2_032_39    m/z 57.0699     0   1E+5   13.5   14  RT (min)      lcorbi_t3_pos_3_056_56    m/z 57.0699     0   1   2E+5   13.5   14  RT (min)      lcorbi_t3_pos_1_008_36    m/z 57.0699     0   2   4E+4   14   14.5  RT (min)      lcorbi_t3_pos_2_022_38    m/z 57.0699     0   1   2   3   4E+5   12.5   13  RT (min)      lcorbi_t3_pos_2_054_29    m/z 57.0699     0   2   4E+4   14.5   15  RT (min)      lcorbi_t3_pos_2_049_55    m/z 57.0699     0   1   2E+4   12.5   13  RT (min)      lcorbi_t3_pos_3_031_45    m/z 57.0699     0   1   2   3   4E+4   14   14.5  RT (min)      lcorbi_t3_pos_3_003_33    m/z 57.8155     0   1   12   12.5  RT (min)      lcorbi_t3_pos_1_023_02    m/z 57.8155     0   1   12   12.5  RT (min)      lcorbi_t3_pos_1_051_09    m/z 57.8155     0   2   4   6E+5   11   11.5  RT (min)      lcorbi_t3_pos_1_008_36    m/z 57.8155     0   1   12   12.5  RT (min)      lcorbi_t3_pos_1_023_02    m/z 57.8155     0   1   11.5   12  RT (min)      lcorbi_t3_pos_1_034_34    m/z 57.8155     0   2   4   6E+5   10.5   11  RT (min)      lcorbi_t3_pos_3_031_45    m/z 57.8155     0   2   4   6E+4   11.5   12  RT (min)      lcorbi_t3_pos_1_028_32    m/z 57.8155     0   1   11.5   12  RT (min)      lcorbi_t3_pos_1_034_34    m/z 58.0395     0   1   2   3E+5   10.5   11  RT (min)      lcorbi_t3_pos_3_024_07    m/z 58.0395     0   1   13   13.5  RT (min)      lcorbi_t3_pos_1_023_02    m/z 58.0395     0   1   2   3   4E+5   11.5   12  RT (min)      lcorbi_t3_pos_1_008_36    m/z 58.0395     0   1   12.5   13  RT (min)      lcorbi_t3_pos_2_012_01    m/z 58.0395     0   1E+4   12.5   13  RT (min)      lcorbi_t3_pos_1_031_40    m/z 58.0395     0   1E+4   12   12.5  RT (min)      lcorbi_t3_pos_3_003_33    m/z 58.0395     0   2   4   6E+3   12   12.5  RT (min)      lcorbi_t3_pos_3_013_19    m/z 58.0395     0   1   12   12.5  RT (min)      lcorbi_t3_pos_1_034_34    m/z 58.0395     0   2   4E+5   11   11.5  RT (min)      lcorbi_t3_pos_1_002_01    m/z 58.2634     0   1E+4   13   13.5  RT (min)      lcorbi_t3_pos_1_031_40    m/z 58.2634     0   1   2E+4   13.5   14  RT (min)      lcorbi_t3_pos_1_023_02    m/z 58.2634     0   1E+6   11.5   12  RT (min)      lcorbi_t3_pos_2_054_29    m/z 58.2634     0   2   4   6   8E+3   12.5   13  RT (min)      lcorbi_t3_pos_2_022_38    m/z 58.2634     0   1E+6   12   12.5  RT (min)      lcorbi_t3_pos_1_008_36    m/z 58.2634     0   1E+4   12.5   13  RT (min)      lcorbi_t3_pos_1_034_34    m/z 58.2634     0   1   2E+4   13.5   14  RT (min)      lcorbi_t3_pos_1_023_02    m/z 58.2634     0   2   4   6   8E+5   11   11.5  RT (min)      lcorbi_t3_pos_2_013_33    m/z 58.2634     0   1   13   13.5  RT (min)      lcorbi_t3_pos_2_009_50    m/z 58.2634     0   1E+4   13   13.5  RT (min)      lcorbi_t3_pos_1_051_09    m/z 58.2635     0   1   2E+4   13.5   14  RT (min)      lcorbi_t3_pos_1_023_02    m/z 59.0491     0   1E+6   15   16   17  RT (min)       lcorbi_t3_pos_1_019_16    m/z 59.0491     0   1   2E+6   15   16   17  RT (min)       lcorbi_t3_pos_1_055_33    m/z 59.0491     0   1E+6   16  RT (min)       lcorbi_t3_pos_1_058_14    m/z 59.0491     0   1E+6   14   16  RT (min)       lcorbi_t3_pos_1_060_39    m/z 59.0491     0   1   17.5   18  RT (min)     lcorbi_t3_pos_1_049_55    m/z 59.0491     0   1   2E+6   16  RT (min)       lcorbi_t3_pos_1_050_13    m/z 59.0491     0   1E+6   16.5   17  RT (min)       lcorbi_t3_pos_4_083_59    m/z 59.0492     0   1E+5   6.5   7  RT (min)      lcorbi_t3_pos_3_001_49    m/z 59.5739     0   1   2   3   4E+5   14   14.5  RT (min)      lcorbi_t3_pos_3_000_cc    m/z 60.4899     0   1   2   3E+5   10.5   11  RT (min)      lcorbi_t3_pos_1_050_13    m/z 60.4901     0   1   2   3   4E+5   10.5   11  RT (min)      lcorbi_t3_pos_3_027_27    m/z 61.0647     0   1   2E+6   17   18  RT (min)       lcorbi_t3_pos_4_082_59    m/z 61.0648     0   2   4   6   8E+5   10   10.5  RT (min)      lcorbi_t3_pos_1_057_49    m/z 61.0648     0   1   2   3E+6   14   16  RT (min)       lcorbi_t3_pos_1_030_21    m/z 61.0648     0   1   2   3E+6   15.5   16   16.5  RT (min)       lcorbi_t3_pos_1_008_36    m/z 61.0648     0   2   4   6E+5   1.5   2  RT (min)      lcorbi_t3_pos_4_057_49    m/z 61.0648     0   1   17.5   18  RT (min)     lcorbi_t3_pos_1_027_43    m/z 61.0648     0   1   2E+6   13   13.5  RT (min)       lcorbi_t3_pos_1_059_17    m/z 61.0648     0   1E+6   10.5   11  RT (min)      lcorbi_t3_pos_1_058_14    m/z 61.0648     0   1   17   17.5  RT (min)     lcorbi_t3_pos_1_025_52    m/z 61.0648     0   1   16.5   17  RT (min)     lcorbi_t3_pos_1_005_42    m/z 61.0648     0   1   2   3   4E+6   15   16  RT (min)       lcorbi_t3_pos_1_012_14    m/z 61.0648     0   1   2E+6   13   13.5   14  RT (min)       lcorbi_t3_pos_1_058_14    m/z 61.0648     0   1   2   3   4E+6   16   17  RT (min)       lcorbi_t3_pos_1_015_11    m/z 61.0648     0   2   4   6E+5   9.5   10  RT (min)      lcorbi_t3_pos_1_058_14    m/z 61.0648     0   2   4   6E+5   9   9.5  RT (min)      lcorbi_t3_pos_1_057_49    m/z 61.0648     0   2   4   6E+5   9   9.5  RT (min)      lcorbi_t3_pos_1_058_14    m/z 61.0648     0   2   4   6   8E+5   11   11.5  RT (min)      lcorbi_t3_pos_1_061_50    m/z 61.0648     0   1   2E+6   16   17  RT (min)       lcorbi_t3_pos_4_098_61    m/z 61.0648     0   1E+6   11.5   12  RT (min)      lcorbi_t3_pos_1_057_49    m/z 61.0648     0   1E+6   11   11.5  RT (min)      lcorbi_t3_pos_1_057_49    m/z 61.0648     0   1E+6   0.5   1  RT (min)      lcorbi_t3_pos_4_057_49    m/z 61.0648     0   1   2   3   4E+6   16   18  RT (min)       lcorbi_t3_pos_1_001_49    m/z 67.7982     0   2   4   6E+5   14.5   15  RT (min)      lcorbi_t3_pos_3_000_cc    m/z 67.7982     0   1   2   3   4E+5   14   14.5  RT (min)      lcorbi_t3_pos_4_105_61    m/z 67.7982     0   2   4E+5   14.5   15  RT (min)      lcorbi_t3_pos_4_113_62    m/z 67.7982     0   1   2   3   4E+5   14   14.5  RT (min)      lcorbi_t3_pos_4_105_61    m/z 69.0447     0   1E+5   6.5   7  RT (min)      lcorbi_t3_pos_4_010_43    m/z 69.0698     0   1   2   3E+5   13   13.5  RT (min)      lcorbi_t3_pos_1_058_14    m/z 69.0698     0   1   2E+5   12.5   13  RT (min)      lcorbi_t3_pos_1_061_50    m/z 71.0127     0   2   4   6E+5   15  RT (min)       lcorbi_t3_pos_1_022_20    m/z 71.0127     0   2   4   6E+5   8   9  RT (min)       lcorbi_t3_pos_1_039_23    m/z 71.0127     0   2   4   6E+5   8   9  RT (min)       lcorbi_t3_pos_1_016_05    m/z 71.0127     0   1   2E+6   12  RT (min)       lcorbi_t3_pos_1_047_41    m/z 71.0127     0   2   4E+5   18   18.5  RT (min)       lcorbi_t3_pos_4_068_57    m/z 71.0127     0   2   4   6E+5   8   10  RT (min)       lcorbi_t3_pos_4_108_62    m/z 71.0127     0   2   4   6   8E+5   8   10  RT (min)       lcorbi_t3_pos_1_018_35    m/z 71.0127     0   2   4   6E+5   10   15  RT (min)       lcorbi_t3_pos_4_075_58    m/z 71.0127     0   1   2E+6   6.5   7  RT (min)       lcorbi_t3_pos_1_046_22    m/z 71.0127     0   2   4E+5   18.5   19  RT (min)       lcorbi_t3_pos_4_081_58    m/z 71.0127     0   2   4   6E+5   10   15  RT (min)       lcorbi_t3_pos_4_099_61    m/z 71.0854     0   1E+6   12.5   13  RT (min)      lcorbi_t3_pos_4_046_44    m/z 71.0855     0   1E+5   14.5   15  RT (min)      lcorbi_t3_pos_4_033_53    m/z 71.0855     0   1   2   3E+4   12.5   13  RT (min)      lcorbi_t3_pos_4_012_37    m/z 71.0855     0   2   4E+6   12.5   13  RT (min)      lcorbi_t3_pos_4_016_15    m/z 71.0855     0   1E+5   15.5   16  RT (min)      lcorbi_t3_pos_1_023_02    m/z 71.0855     0   1E+5   14.5   15  RT (min)      lcorbi_t3_pos_2_049_55    m/z 71.0855     0   1E+5   14   14.5  RT (min)      lcorbi_t3_pos_3_013_19    m/z 71.0855     0   1E+5   14   14.5  RT (min)      lcorbi_t3_pos_1_047_41    m/z 71.0855     0   1   2E+4   12.5   13  RT (min)      lcorbi_t3_pos_3_024_07    m/z 71.0855     0   1   2E+5   14   14.5  RT (min)      lcorbi_t3_pos_4_028_04    m/z 71.0855     0   1   2   3E+5   13.5   14  RT (min)      lcorbi_t3_pos_3_015_06    m/z 71.0855     0   1E+5   14.5   15  RT (min)      lcorbi_t3_pos_1_034_34    m/z 71.0855     0   1E+6   13.5   14  RT (min)      lcorbi_t3_pos_3_025_52    m/z 71.0855     0   1   2   3   4E+6   13.5   14  RT (min)      lcorbi_t3_pos_3_047_40    m/z 72.0807     0   1E+5   14   14.5  RT (min)      lcorbi_t3_pos_1_028_32    m/z 72.0807     0   1E+6   18.5   19  RT (min)      lcorbi_t3_pos_4_076_58    m/z 72.0807     0   1E+6   1   1.5  RT (min)       lcorbi_t3_pos_4_079_58    m/z 72.0807     0   1   2   3   4E+5   10  RT (min)       lcorbi_t3_pos_4_104_61    m/z 72.0807     0   1   2   3E+5   8   8.5  RT (min)      lcorbi_t3_pos_1_057_49    m/z 72.0807     0   1E+5   17   17.5  RT (min)       lcorbi_t3_pos_4_074_58    m/z 72.0807     0   1E+5   0.5   1  RT (min)      lcorbi_t3_pos_2_037_27    m/z 72.0807     0   1E+5   15.5   16  RT (min)      lcorbi_t3_pos_1_023_02    m/z 72.0807     0   1E+6   18   18.5  RT (min)      lcorbi_t3_pos_4_076_58    m/z 72.0807     0   1E+6   17.5   18  RT (min)      lcorbi_t3_pos_4_075_58    m/z 72.0807     0   1E+6   18   18.5  RT (min)      lcorbi_t3_pos_4_076_58    m/z 72.0807     0   1E+5   17   17.5  RT (min)      lcorbi_t3_pos_4_074_58    m/z 72.0807     0   1E+5   0.5   1  RT (min)      lcorbi_t3_pos_2_037_27    m/z 72.0807     0   1E+5   0   0.5  RT (min)      lcorbi_t3_pos_2_037_27    m/z 72.9371     0   2   4   6E+5   0   0.5  RT (min)      lcorbi_t3_pos_1_005_42    m/z 73.0648     0   1   17.5   18   18.5  RT (min)     lcorbi_t3_pos_1_056_56    m/z 73.0648     0   2   4E+5   15.5   16   16.5  RT (min)       lcorbi_t3_pos_1_052_38    m/z 73.0648     0   2   4E+5   15   16   17  RT (min)       lcorbi_t3_pos_1_053_39    m/z 73.0648     0   2   4E+5   16   17  RT (min)       lcorbi_t3_pos_1_054_48    m/z 73.0648     0   1   16.5   17  RT (min)     lcorbi_t3_pos_1_055_33    m/z 74.0600     0   2   4   6   8E+5   15.5   16  RT (min)       lcorbi_t3_pos_1_044_25    m/z 74.0964     0   1   17   17.5   18  RT (min)     lcorbi_t3_pos_1_001_49    m/z 74.0964     0   1   2E+6   17.5   18   18.5  RT (min)       lcorbi_t3_pos_4_092_60    m/z 74.0964     0   2   4   6   8E+5   12   12.5  RT (min)      lcorbi_t3_pos_1_059_17    m/z 74.0964     0   1E+6   12.5   13  RT (min)      lcorbi_t3_pos_1_057_49    m/z 74.0964     0   2   4   6E+5   10   10.5  RT (min)      lcorbi_t3_pos_1_057_49    m/z 74.0964     0   2   4   6E+5   9   9.5  RT (min)      lcorbi_t3_pos_1_057_49    m/z 74.0964     0   1   2   3   4E+5   0.5   1  RT (min)      lcorbi_t3_pos_4_057_49    m/z 74.0964     0   1   2E+6   13   14  RT (min)       lcorbi_t3_pos_1_008_36    m/z 74.0964     0   1   2E+6   17.5   18  RT (min)       lcorbi_t3_pos_4_105_61    m/z 74.0964     0   1E+6   7.5   10   12.5  RT (min)       lcorbi_t3_pos_1_032_26    m/z 74.0964     0   2   4   6   8E+5   13   13.5   14  RT (min)       lcorbi_t3_pos_4_066_57    m/z 75.0440     0   1   2E+5   15   16  RT (min)       lcorbi_t3_pos_4_081_58    m/z 75.0764     0   1   14.5   15  RT (min)       lcorbi_t3_pos_1_060_39    m/z 75.0804     0   1   2E+6   15   15.5  RT (min)       lcorbi_t3_pos_2_062_13    m/z 75.0804     0   1   2E+6   12.5   13  RT (min)       lcorbi_t3_pos_1_057_49    m/z 75.0804     0   1   17   18  RT (min)     lcorbi_t3_pos_1_034_34    m/z 75.0804     0   1   2   3E+6   13.5   14  RT (min)       lcorbi_t3_pos_1_028_32    m/z 75.0804     0   1   2E+6   13.5   14  RT (min)       lcorbi_t3_pos_4_080_58    m/z 75.0804     0   1   2E+5   6.5   7  RT (min)      lcorbi_t3_pos_2_025_52    m/z 75.0804     0   1E+6   13   13.5  RT (min)       lcorbi_t3_pos_2_027_05    m/z 75.0804     0   1E+6   12.5   13  RT (min)       lcorbi_t3_pos_2_061_50    m/z 75.0804     0   2   4   6   8E+5   1.5   2  RT (min)       lcorbi_t3_pos_4_057_49    m/z 75.0804     0   1   2E+6   12   12.5  RT (min)       lcorbi_t3_pos_1_057_49    m/z 75.0804     0   2   4   6E+5   9   9.5  RT (min)       lcorbi_t3_pos_1_057_49    m/z 75.0804     0   1   2E+6   14   14.5  RT (min)       lcorbi_t3_pos_2_016_23    m/z 75.0804     0   1   2   3   4E+6   15.5   16  RT (min)       lcorbi_t3_pos_1_005_42    m/z 75.0804     0   1   2   3   4E+6   15.5   16  RT (min)       lcorbi_t3_pos_1_035_28    m/z 75.0804     0   1   2   3   4E+6   17   18  RT (min)       lcorbi_t3_pos_4_092_60    m/z 75.0804     0   1E+6   11.5   12  RT (min)       lcorbi_t3_pos_1_058_14    m/z 75.0804     0   1   2E+6   14   14.5  RT (min)       lcorbi_t3_pos_1_058_14    m/z 75.0804     0   1   2   3   4E+6   16   17   18  RT (min)       lcorbi_t3_pos_1_033_53    m/z 75.0804     0   2   4   6   8E+5   10   10.5  RT (min)       lcorbi_t3_pos_1_064_16    m/z 75.0804     0   1E+6   13.5   14  RT (min)       lcorbi_t3_pos_2_064_17    m/z 75.0804     0   1   2   3   4E+6   14   16  RT (min)       lcorbi_t3_pos_1_030_21    m/z 75.0804     0   1E+6   11.5   12  RT (min)       lcorbi_t3_pos_1_063_15    m/z 75.0804     0   1   16.5   17  RT (min)     lcorbi_t3_pos_1_038_30    m/z 75.0804     0   2   4   6E+5   9.5   10  RT (min)       lcorbi_t3_pos_1_059_17    m/z 75.0804     0   2   4   6   8E+5   10   10.5  RT (min)       lcorbi_t3_pos_1_058_14    m/z 75.0804     0   1   2   3   4E+6   14.5   15  RT (min)       lcorbi_t3_pos_1_002_01    m/z 75.0804     0   1E+6   10.5   11  RT (min)       lcorbi_t3_pos_1_058_14    m/z 75.0804     0   1   17  RT (min)     lcorbi_t3_pos_1_031_40    m/z 75.0804     0   1E+6   0.5   1  RT (min)       lcorbi_t3_pos_4_057_49    m/z 75.0804     0   2   4   6   8E+5   9.5   10  RT (min)       lcorbi_t3_pos_1_058_14    m/z 75.0804     0   1   2   3E+6   13   13.5  RT (min)       lcorbi_t3_pos_1_036_08    m/z 75.0804     0   1   2E+6   13   13.5  RT (min)       lcorbi_t3_pos_1_057_49    m/z 75.0804     0   1   2E+6   12   12.5  RT (min)       lcorbi_t3_pos_1_057_49    m/z 75.0804     0   1E+6   12   12.5  RT (min)       lcorbi_t3_pos_2_013_33    m/z 75.0804     0   1E+6   1   1.5  RT (min)       lcorbi_t3_pos_4_057_49    m/z 75.0804     0   1E+6   11   11.5  RT (min)       lcorbi_t3_pos_1_058_14    m/z 75.0804     0   1   2   3   4E+6   17.5   18  RT (min)       lcorbi_t3_pos_4_098_61    m/z 76.0226     0   2   4   6E+5   14.5   15  RT (min)      lcorbi_t3_pos_3_000_cc    m/z 78.1750     0   1   15   15.5  RT (min)      lcorbi_t3_pos_1_034_34    m/z 78.1750     0   1   2   3   4E+5   14   14.5  RT (min)      lcorbi_t3_pos_4_019_47    m/z 78.1750     0   1   16   16.5  RT (min)      lcorbi_t3_pos_1_023_02    m/z 78.1751     0   1   14.5   15  RT (min)      lcorbi_t3_pos_3_013_19    m/z 78.1751     0   1   12.5   13  RT (min)      lcorbi_t3_pos_2_032_39    m/z 78.1751     0   2   4   6E+4   13   13.5  RT (min)      lcorbi_t3_pos_2_054_29    m/z 78.1751     0   1   2   3   4E+5   13   13.5  RT (min)      lcorbi_t3_pos_3_034_26    m/z 78.2802     0   2   4   6   8E+5   14   14.5  RT (min)      lcorbi_t3_pos_3_000_aa    m/z 79.0212     0   2   4   6E+4   0   0.5  RT (min)       lcorbi_t3_pos_1_002_01    m/z 79.0212     0   1E+5   0.5   1  RT (min)      lcorbi_t3_pos_1_009_50    m/z 79.0753     0   1   17   17.5  RT (min)     lcorbi_t3_pos_1_001_49    m/z 79.0753     0   1   17   18  RT (min)     lcorbi_t3_pos_1_025_52    m/z 79.0753     0   1E+6   14   16  RT (min)       lcorbi_t3_pos_1_021_18    m/z 79.0753     0   1E+6   15.5   16  RT (min)       lcorbi_t3_pos_1_014_03    m/z 79.0753     0   2   4   6   8E+5   16   17  RT (min)       lcorbi_t3_pos_1_033_53    m/z 79.0753     0   1E+6   16   17  RT (min)       lcorbi_t3_pos_1_005_42    m/z 79.0753     0   1E+6   14.5   15  RT (min)       lcorbi_t3_pos_1_018_35    m/z 79.0753     0   1   17   17.5  RT (min)     lcorbi_t3_pos_1_047_41    m/z 79.0753     0   2   4   6E+5   17   18  RT (min)       lcorbi_t3_pos_4_098_61    m/z 79.0753     0   1   2   3   4E+5   13.5   14  RT (min)       lcorbi_t3_pos_2_021_09    m/z 79.0753     0   2   4   6E+5   8   10   12  RT (min)       lcorbi_t3_pos_1_032_26    m/z 80.0495     0   1   2   3E+4   14   16  RT (min)       lcorbi_t3_pos_3_000_aa    m/z 80.9478     0   2   4   6E+5   17.5   18  RT (min)       lcorbi_t3_pos_4_095_60    m/z 80.9478     0   2   4   6E+5   16   17   18  RT (min)       lcorbi_t3_pos_4_106_62    m/z 81.0310     0   1E+5   18   18.5  RT (min)      lcorbi_t3_pos_4_083_59    m/z 81.0310     0   1E+5   18   18.5  RT (min)      lcorbi_t3_pos_4_099_61    m/z 81.0698     0   1   2   3E+6   12   14  RT (min)       lcorbi_t3_pos_1_013_10    m/z 81.0698     0   1   2E+6   12.5   13  RT (min)       lcorbi_t3_pos_1_032_26    m/z 81.0698     0   1   2E+6   12   13  RT (min)       lcorbi_t3_pos_1_016_05    m/z 81.0698     0   1   2E+6   10   12   14  RT (min)       lcorbi_t3_pos_1_031_40    m/z 81.0699     0   1   2E+6   14   16   18  RT (min)       lcorbi_t3_pos_1_034_34    m/z 81.0699     0   1   2E+6   10   12.5   15  RT (min)       lcorbi_t3_pos_1_038_30    m/z 81.0699     0   1   2E+6   12   12.5   13  RT (min)       lcorbi_t3_pos_1_036_08    m/z 81.0699     0   1   2E+6   12.5   15   17.5  RT (min)       lcorbi_t3_pos_1_031_40    m/z 81.0699     0   1   2E+6   15  RT (min)       lcorbi_t3_pos_1_001_49    m/z 81.0699     0   1   2E+6   13   14   15  RT (min)       lcorbi_t3_pos_1_026_17    m/z 81.0699     0   1   2E+6   16   18  RT (min)       lcorbi_t3_pos_1_019_16    m/z 81.0699     0   1   2E+6   12   14  RT (min)       lcorbi_t3_pos_1_012_14    m/z 81.0699     0   1   2E+6   14   16  RT (min)       lcorbi_t3_pos_1_005_42    m/z 82.7283     0   2   4   6   8E+4   14   14.5  RT (min)      lcorbi_t3_pos_2_013_33    m/z 82.7325     0   1   16   16.5  RT (min)      lcorbi_t3_pos_2_022_38    m/z 82.7325     0   1   17.5   18  RT (min)    lcorbi_t3_pos_2_012_01    m/z 82.7326     0   1   16.5   17  RT (min)    lcorbi_t3_pos_1_028_32    m/z 82.7326     0   1   17.5   18  RT (min)    lcorbi_t3_pos_1_031_40    m/z 82.7326     0   1   16.5   17  RT (min)    lcorbi_t3_pos_1_034_34    m/z 82.7326     0   2   4E+5   15   15.5  RT (min)      lcorbi_t3_pos_2_023_26    m/z 82.7326     0   1   16.5   17  RT (min)    lcorbi_t3_pos_1_034_34    m/z 83.0467     0   1E+6   1   1.5  RT (min)      lcorbi_t3_pos_4_057_49    m/z 83.0491     0   1E+5   11.5   12  RT (min)      lcorbi_t3_pos_3_024_07    m/z 83.0855     0   1E+7   16   17  RT (min)       lcorbi_t3_pos_1_017_51    m/z 83.0855     0   2   4   6E+6   12   12.5  RT (min)       lcorbi_t3_pos_1_058_14    m/z 83.0855     0   2   4E+6   9   9.5  RT (min)      lcorbi_t3_pos_1_058_14    m/z 83.0855     0   1E+7   16   17  RT (min)       lcorbi_t3_pos_1_027_43    m/z 83.0855     0   1   17   18  RT (min)     lcorbi_t3_pos_1_038_30    m/z 83.0855     0   2   4E+6   11.5   12  RT (min)      lcorbi_t3_pos_1_059_17    m/z 83.0855     0   1   2   3E+6   7.5   8  RT (min)      lcorbi_t3_pos_1_058_14    m/z 83.0855     0   1   2   3   4E+6   8.5   9  RT (min)      lcorbi_t3_pos_1_059_17    m/z 83.0855     0   2   4   6E+6   14   16  RT (min)       lcorbi_t3_pos_4_076_58    m/z 83.0855     0   1E+7   15   16  RT (min)       lcorbi_t3_pos_1_045_12    m/z 83.0855     0   1   2E+6   7   7.5  RT (min)      lcorbi_t3_pos_1_059_17    m/z 83.0855     0   1   2   3E+6   8   8.5  RT (min)      lcorbi_t3_pos_1_058_14    m/z 83.0855     0   1E+6   8   8.5  RT (min)      lcorbi_t3_pos_3_061_50    m/z 83.0855     0   1E+7   16   17   18  RT (min)       lcorbi_t3_pos_1_025_52    m/z 83.0855     0   1E+7   14   16  RT (min)       lcorbi_t3_pos_1_005_42    m/z 83.0855     0   1E+7   15   16  RT (min)       lcorbi_t3_pos_1_021_18    m/z 83.0855     0   1   2E+6   0.5   1  RT (min)      lcorbi_t3_pos_4_057_49    m/z 83.0855     0   2   4   6   8E+6   17   17.5  RT (min)       lcorbi_t3_pos_4_099_61    m/z 83.0855     0   1E+6   12   12.5  RT (min)      lcorbi_t3_pos_4_064_39    m/z 83.0855     0   1   2E+6   1.5   2  RT (min)      lcorbi_t3_pos_4_057_49    m/z 83.0855     0   1   16.5   17  RT (min)     lcorbi_t3_pos_1_033_53    m/z 83.0855     0   2   4   6E+6   16   16.5  RT (min)       lcorbi_t3_pos_4_077_58    m/z 83.0855     0   1   2E+6   11.5   12  RT (min)      lcorbi_t3_pos_3_062_15    m/z 83.8377     0   1   2   3   4E+5   13.5   14  RT (min)      lcorbi_t3_pos_1_005_42    m/z 83.8377     0   1   14.5   15  RT (min)      lcorbi_t3_pos_1_028_32    m/z 83.8377     0   2   4   6   8E+4   14   14.5  RT (min)      lcorbi_t3_pos_1_008_36    m/z 83.8377     0   1   14.5   15  RT (min)      lcorbi_t3_pos_1_034_34    m/z 83.8378     0   1   12.5   13  RT (min)      lcorbi_t3_pos_2_013_33    m/z 83.8378     0   1E+4   14   14.5  RT (min)      lcorbi_t3_pos_1_008_36    m/z 83.8378     0   2   4   6E+5   13   13.5  RT (min)      lcorbi_t3_pos_3_011_42    m/z 83.8378     0   1   2E+4   15   15.5  RT (min)      lcorbi_t3_pos_1_031_40    m/z 83.9492     0   1E+4   12.5   13  RT (min)      lcorbi_t3_pos_2_013_33    m/z 83.9492     0   1   14.5   15  RT (min)      lcorbi_t3_pos_1_028_32    m/z 83.9492     0   1   2   3E+4   14   14.5  RT (min)      lcorbi_t3_pos_1_008_36    m/z 83.9492     0   1   14   14.5  RT (min)      lcorbi_t3_pos_1_008_36    m/z 83.9492     0   1   2E+4   15   15.5  RT (min)      lcorbi_t3_pos_1_031_40    m/z 83.9492     0   1   15.5   16  RT (min)      lcorbi_t3_pos_2_012_01    m/z 83.9493     0   1   2E+4   14.5   15  RT (min)      lcorbi_t3_pos_1_034_34    m/z 84.0444     0   2   4E+5   5   5.5  RT (min)      lcorbi_t3_pos_1_057_49    m/z 84.0444     0   2   4   6   8E+5   1.5   2  RT (min)      lcorbi_t3_pos_1_057_49    m/z 84.0616     0   1E+4   13   13.5  RT (min)      lcorbi_t3_pos_2_013_33    m/z 84.0616     0   1   2   3E+4   13   13.5  RT (min)      lcorbi_t3_pos_2_013_33    m/z 84.0617     0   1   2   3   4E+5   14   14.5  RT (min)      lcorbi_t3_pos_3_040_41    m/z 84.0617     0   1   15   15.5  RT (min)      lcorbi_t3_pos_1_034_34    m/z 84.0617     0   1   2E+5   14   14.5  RT (min)      lcorbi_t3_pos_1_008_36    m/z 84.0617     0   1   14.5   15  RT (min)      lcorbi_t3_pos_1_028_32    m/z 84.0617     0   1   15   15.5  RT (min)      lcorbi_t3_pos_1_034_34    m/z 84.0617     0   1   15.5   16  RT (min)      lcorbi_t3_pos_2_012_01    m/z 84.0617     0   1   15   15.5  RT (min)      lcorbi_t3_pos_1_034_34    m/z 84.0617     0   1   15.5   16  RT (min)      lcorbi_t3_pos_1_031_40    m/z 84.0617     0   1   14.5   15  RT (min)      lcorbi_t3_pos_1_047_41    m/z 84.0617     0   1   15.5   16  RT (min)      lcorbi_t3_pos_1_031_40    m/z 84.0617     0   1   14.5   15  RT (min)      lcorbi_t3_pos_1_028_32    m/z 84.0617     0   1   15.5   16  RT (min)      lcorbi_t3_pos_2_012_01    m/z 84.0617     0   2   4   6E+5   14   14.5  RT (min)      lcorbi_t3_pos_1_005_42    m/z 84.0618     0   1   15.5   16  RT (min)      lcorbi_t3_pos_1_031_40    m/z 84.0807     0   1E+5   16.5   17  RT (min)       lcorbi_t3_pos_4_077_58    m/z 84.0808     0   1   2   3   4E+5   16   17   18  RT (min)       lcorbi_t3_pos_1_001_49    m/z 84.0848     0   1   16   17  RT (min)       lcorbi_t3_pos_1_021_18    m/z 84.0888     0   2   4   6   8E+5   14   16  RT (min)       lcorbi_t3_pos_1_036_08    m/z 84.0888     0   1E+6   15.5   16  RT (min)       lcorbi_t3_pos_1_012_14    m/z 84.0888     0   1E+6   15   16  RT (min)       lcorbi_t3_pos_1_013_10    m/z 84.0888     0   1   18  RT (min)     lcorbi_t3_pos_1_031_40    m/z 84.0888     0   1   17   17.5  RT (min)     lcorbi_t3_pos_1_038_30    m/z 84.0888     0   1E+6   15.5   16  RT (min)       lcorbi_t3_pos_1_017_51    m/z 84.0888     0   2   4   6E+5   16   17  RT (min)       lcorbi_t3_pos_1_028_32    m/z 84.0888     0   2   4   6   8E+5   16   16.5  RT (min)       lcorbi_t3_pos_1_005_42    m/z 84.0888     0   2   4   6   8E+5   14   16  RT (min)       lcorbi_t3_pos_1_008_36    m/z 84.1730     0   2   4E+4   14   14.5  RT (min)      lcorbi_t3_pos_1_008_36    m/z 84.1731     0   1   13   13.5  RT (min)      lcorbi_t3_pos_2_013_33    m/z 84.1731     0   1   14.5   15  RT (min)      lcorbi_t3_pos_1_028_32    m/z 84.1731     0   1   2E+4   15.5   16  RT (min)      lcorbi_t3_pos_1_031_40    m/z 84.1731     0   1   14.5   15  RT (min)      lcorbi_t3_pos_1_028_32    m/z 84.1731     0   1   13   13.5  RT (min)      lcorbi_t3_pos_2_013_33    m/z 84.1731     0   1   15   15.5  RT (min)      lcorbi_t3_pos_1_034_34    m/z 84.1732     0   1   15.5   16  RT (min)      lcorbi_t3_pos_2_012_01    m/z 84.1732     0   1   15   15.5  RT (min)      lcorbi_t3_pos_1_034_34    m/z 84.1732     0   1   15.5   16  RT (min)      lcorbi_t3_pos_2_012_01    m/z 84.2466     0   2   4E+5   14.5   15  RT (min)      lcorbi_t3_pos_3_000_cc    m/z 84.2855     0   1   13   13.5  RT (min)      lcorbi_t3_pos_1_002_01    m/z 84.2856     0   2   4   6   8E+3   12.5   13  RT (min)      lcorbi_t3_pos_2_032_39    m/z 84.2856     0   1   16   16.5  RT (min)      lcorbi_t3_pos_1_023_02    m/z 84.2856     0   1   15   15.5  RT (min)      lcorbi_t3_pos_1_028_32    m/z 84.2856     0   1   14.5   15  RT (min)      lcorbi_t3_pos_1_047_41    m/z 84.2856     0   1   16   16.5  RT (min)      lcorbi_t3_pos_1_023_02    m/z 84.2856     0   1   15.5   16  RT (min)      lcorbi_t3_pos_1_031_40    m/z 84.2856     0   1   13   13.5  RT (min)      lcorbi_t3_pos_3_024_07    m/z 84.2856     0   1   2   3E+5   14.5   15  RT (min)      lcorbi_t3_pos_1_008_36    m/z 84.2856     0   1   2   3E+6   13   13.5  RT (min)      lcorbi_t3_pos_1_002_01    m/z 84.2856     0   1   2   3E+6   14   14.5  RT (min)      lcorbi_t3_pos_1_005_42    m/z 84.2856     0   1   15   15.5  RT (min)      lcorbi_t3_pos_1_034_34    m/z 84.2856     0   1E+5   14.5   15  RT (min)      lcorbi_t3_pos_1_008_36    m/z 84.2856     0   1   2   3E+6   14   14.5  RT (min)      lcorbi_t3_pos_1_005_42    m/z 84.2857     0   1   15   15.5  RT (min)      lcorbi_t3_pos_1_028_32    m/z 84.2857     0   1   15   15.5  RT (min)      lcorbi_t3_pos_1_034_34    m/z 84.2857     0   1   15   15.5  RT (min)      lcorbi_t3_pos_1_028_32    m/z 84.2857     0   1   16.5   17  RT (min)    lcorbi_t3_pos_1_023_02    m/z 84.2857     0   1   2E+6   13   13.5  RT (min)      lcorbi_t3_pos_1_002_01    m/z 84.2857     0   1   16   16.5  RT (min)      lcorbi_t3_pos_1_001_49    m/z 84.2857     0   1E+4   15.5   16  RT (min)      lcorbi_t3_pos_1_051_09    m/z 84.3971     0   1E+6   14   14.5  RT (min)      lcorbi_t3_pos_1_005_42    m/z 84.3971     0   1   14.5   15  RT (min)      lcorbi_t3_pos_2_006_47    m/z 84.3971     0   2   4   6   8E+3   13   13.5  RT (min)      lcorbi_t3_pos_2_032_39    m/z 84.3971     0   1   13   13.5  RT (min)      lcorbi_t3_pos_1_002_01    m/z 84.3971     0   1E+6   13   13.5  RT (min)      lcorbi_t3_pos_1_002_01    m/z 84.3971     0   1E+5   14.5   15  RT (min)      lcorbi_t3_pos_1_008_36    m/z 84.3971     0   1   15   15.5  RT (min)      lcorbi_t3_pos_1_028_32    m/z 84.3971     0   1   16.5   17  RT (min)    lcorbi_t3_pos_1_023_02    m/z 84.3971     0   1   14.5   15  RT (min)      lcorbi_t3_pos_1_030_21    m/z 84.3971     0   1   2E+4   13   13.5  RT (min)      lcorbi_t3_pos_1_002_01    m/z 84.3971     0   1   15.5   16  RT (min)      lcorbi_t3_pos_1_031_40    m/z 84.3971     0   1   2   3   4E+4   14.5   15  RT (min)      lcorbi_t3_pos_1_008_36    m/z 84.3971     0   1E+6   13   13.5  RT (min)      lcorbi_t3_pos_1_002_01    m/z 84.3972     0   1   15   15.5  RT (min)      lcorbi_t3_pos_1_028_32    m/z 84.3972     0   1   16   16.5  RT (min)      lcorbi_t3_pos_1_023_02    m/z 84.3972     0   1   2E+4   15   15.5  RT (min)      lcorbi_t3_pos_1_034_34    m/z 84.3972     0   1   16   16.5  RT (min)      lcorbi_t3_pos_1_023_02    m/z 84.3972     0   1   2E+4   15   15.5  RT (min)      lcorbi_t3_pos_1_034_34    m/z 84.3972     0   1   15.5   16  RT (min)      lcorbi_t3_pos_1_025_52    m/z 84.5015     0   1   2E+4   12.5   13  RT (min)      lcorbi_t3_pos_3_024_07    m/z 84.5089     0   2   4E+4   14.5   15  RT (min)      lcorbi_t3_pos_1_008_36    m/z 84.5090     0   1   13   13.5  RT (min)      lcorbi_t3_pos_2_032_39    m/z 84.5094     0   1   15.5   16  RT (min)      lcorbi_t3_pos_2_009_50    m/z 84.5095     0   1   13   13.5  RT (min)      lcorbi_t3_pos_3_024_07    m/z 84.5095     0   1   16   16.5  RT (min)      lcorbi_t3_pos_2_056_56    m/z 84.5095     0   1E+6   14.5   15  RT (min)      lcorbi_t3_pos_1_005_42    m/z 84.5096     0   1E+4   15.5   16  RT (min)      lcorbi_t3_pos_1_034_34    m/z 84.5096     0   1   16   16.5  RT (min)      lcorbi_t3_pos_1_051_09    m/z 84.5096     0   1   16.5   17  RT (min)    lcorbi_t3_pos_1_023_02    m/z 84.5096     0   1   15   15.5  RT (min)      lcorbi_t3_pos_2_025_52    m/z 84.5096     0   1   16.5   17  RT (min)    lcorbi_t3_pos_1_023_02    m/z 84.5096     0   1   15   15.5  RT (min)      lcorbi_t3_pos_1_033_53    m/z 84.5096     0   1   16.5   17  RT (min)    lcorbi_t3_pos_1_023_02    m/z 84.5096     0   1   16   16.5  RT (min)      lcorbi_t3_pos_1_051_09    m/z 84.5096     0   1   16.5   17  RT (min)    lcorbi_t3_pos_1_023_02    m/z 84.5096     0   1   15   15.5  RT (min)      lcorbi_t3_pos_2_006_47    m/z 84.5096     0   1E+6   14   14.5  RT (min)      lcorbi_t3_pos_1_005_42    m/z 84.6129     0   1E+4   13.5   14  RT (min)      lcorbi_t3_pos_1_002_01    m/z 84.6211     0   1   14.5   15  RT (min)      lcorbi_t3_pos_1_008_36    m/z 84.6211     0   1   2E+4   15   15.5  RT (min)      lcorbi_t3_pos_1_033_53    m/z 84.6211     0   1   16.5   17  RT (min)    lcorbi_t3_pos_1_023_02    m/z 84.6211     0   1   16   16.5  RT (min)      lcorbi_t3_pos_1_051_09    m/z 84.6211     0   1   14.5   15  RT (min)      lcorbi_t3_pos_2_046_07    m/z 84.6211     0   1   16   16.5  RT (min)      lcorbi_t3_pos_1_051_09    m/z 84.6211     0   2   4   6E+5   14.5   15  RT (min)      lcorbi_t3_pos_1_005_42    m/z 84.6211     0   1   16.5   17  RT (min)    lcorbi_t3_pos_1_023_02    m/z 84.6211     0   1   2E+4   15.5   16  RT (min)      lcorbi_t3_pos_1_034_34    m/z 84.6211     0   1   16   16.5  RT (min)      lcorbi_t3_pos_1_051_09    m/z 84.6211     0   1   15   15.5  RT (min)      lcorbi_t3_pos_1_030_21    m/z 84.9596     0   2   4E+5   17.5   18  RT (min)      lcorbi_t3_pos_4_093_60    m/z 84.9596     0   2   4E+5   0   2  RT (min)       lcorbi_t3_pos_4_083_59    m/z 84.9596     0   1   2   3   4E+5   18   18.5  RT (min)      lcorbi_t3_pos_4_066_57    m/z 84.9596     0   1   2   3   4E+5   18   18.5  RT (min)      lcorbi_t3_pos_4_072_57    m/z 84.9596     0   1   2   3   4E+5   18   18.5  RT (min)      lcorbi_t3_pos_4_084_59    m/z 84.9597     0   2   4   6E+5   1   2   3  RT (min)       lcorbi_t3_pos_4_082_59    m/z 84.9597     0   1   2   3   4E+6   0   0.5  RT (min)      lcorbi_t3_pos_1_007_07    m/z 84.9597     0   1   2   3   4E+5   2   2.5  RT (min)      lcorbi_t3_pos_1_016_05    m/z 84.9597     0   2   4E+5   3.5   4  RT (min)       lcorbi_t3_pos_4_106_62    m/z 85.1011     0   1E+6   8   8.5  RT (min)      lcorbi_t3_pos_1_063_15    m/z 85.1011     0   1   2   3   4E+6   16   17   18  RT (min)       lcorbi_t3_pos_4_090_60    m/z 85.1011     0   1E+6   0.5   1  RT (min)      lcorbi_t3_pos_4_057_49    m/z 85.1011     0   1E+6   7   7.5  RT (min)      lcorbi_t3_pos_1_061_50    m/z 85.1012     0   1   2E+6   10   12  RT (min)       lcorbi_t3_pos_1_062_13    m/z 85.1012     0   1   16.5   17   17.5  RT (min)     lcorbi_t3_pos_1_008_36    m/z 85.1012     0   1   2   3E+6   13.5   14  RT (min)       lcorbi_t3_pos_1_058_14    m/z 85.1012     0   2   4   6   8E+6   15.5   16  RT (min)       lcorbi_t3_pos_1_013_10    m/z 85.1012     0   1   2   3E+6   14   14.5  RT (min)      lcorbi_t3_pos_1_058_14    m/z 85.1012     0   2   4E+6   13   13.5  RT (min)       lcorbi_t3_pos_1_036_08    m/z 85.1012     0   2   4E+5   7   7.5  RT (min)      lcorbi_t3_pos_2_057_49    m/z 85.1012     0   1   2E+6   11.5   12  RT (min)      lcorbi_t3_pos_1_059_17    m/z 85.1012     0   1E+6   8   8.5  RT (min)      lcorbi_t3_pos_1_057_49    m/z 85.1012     0   1   2E+6   9.5   10  RT (min)      lcorbi_t3_pos_1_058_14    m/z 85.1012     0   1   17.5   18  RT (min)     lcorbi_t3_pos_1_001_49    m/z 85.1012     0   2   4E+6   16  RT (min)       lcorbi_t3_pos_1_050_13    m/z 85.1012     0   2   4   6E+6   10   12.5   15  RT (min)       lcorbi_t3_pos_1_007_07    m/z 85.1012     0   1E+6   8.5   9  RT (min)      lcorbi_t3_pos_2_057_49    m/z 85.1012     0   1   2E+6   9   9.5  RT (min)      lcorbi_t3_pos_1_058_14    m/z 85.1012     0   1   2E+6   8.5   9  RT (min)      lcorbi_t3_pos_1_057_49    m/z 85.1012     0   1E+6   10.5   11  RT (min)      lcorbi_t3_pos_1_063_15    m/z 85.1012     0   1   2E+6   10   10.5  RT (min)      lcorbi_t3_pos_1_057_49    m/z 85.1012     0   1E+6   1.5   2  RT (min)      lcorbi_t3_pos_4_057_49    m/z 85.1012     0   2   4   6   8E+6   15   15.5  RT (min)       lcorbi_t3_pos_1_012_14    m/z 85.1012     0   2   4   6E+6   16   16.5   17  RT (min)       lcorbi_t3_pos_1_005_42    m/z 85.1012     0   1E+6   7   7.5  RT (min)      lcorbi_t3_pos_1_057_49    m/z 86.0600     0   2   4   6   8E+4   4.5   5  RT (min)      lcorbi_t3_pos_4_081_58    m/z 86.0600     0   1E+5   4.5   5  RT (min)      lcorbi_t3_pos_4_106_62    m/z 86.0600     0   1   2E+5   4.5   5  RT (min)       lcorbi_t3_pos_1_006_15    m/z 86.0600     0   1E+5   2.5   3  RT (min)      lcorbi_t3_pos_4_091_60    m/z 86.0600     0   1E+5   4.5   5  RT (min)      lcorbi_t3_pos_1_058_14    m/z 86.0600     0   1   2   3   4E+5   1   1.5  RT (min)       lcorbi_t3_pos_4_087_59    m/z 86.0600     0   1E+5   4.5   5  RT (min)      lcorbi_t3_pos_4_106_62    m/z 86.0964     0   1   17   17.5   18  RT (min)     lcorbi_t3_pos_1_027_43    m/z 86.0964     0   2   4   6E+5   2.5   3  RT (min)       lcorbi_t3_pos_4_075_58    m/z 86.0964     0   2   4   6E+5   3   3.5  RT (min)       lcorbi_t3_pos_4_076_58    m/z 86.0964     0   1   2E+5   15.5   16   16.5  RT (min)       lcorbi_t3_pos_1_006_15    m/z 86.1045     0   1   17   18  RT (min)     lcorbi_t3_pos_1_001_49    m/z 86.1045     0   2   4   6E+5   16   18  RT (min)       lcorbi_t3_pos_1_012_14    m/z 86.1045     0   1   17  RT (min)     lcorbi_t3_pos_1_030_21    m/z 86.5044     0   1   12.5   13  RT (min)      lcorbi_t3_pos_2_013_33    m/z 86.5044     0   1   14.5   15  RT (min)      lcorbi_t3_pos_1_034_34    m/z 86.5044     0   1   14.5   15  RT (min)      lcorbi_t3_pos_1_028_32    m/z 86.5044     0   1   14.5   15  RT (min)      lcorbi_t3_pos_1_028_32    m/z 86.5044     0   1   15   15.5  RT (min)      lcorbi_t3_pos_1_031_40    m/z 86.5044     0   1   14.5   15  RT (min)      lcorbi_t3_pos_1_034_34    m/z 86.5044     0   1   12.5   13  RT (min)      lcorbi_t3_pos_3_024_07    m/z 86.5044     0   1   2   3   4E+5   14   14.5  RT (min)      lcorbi_t3_pos_1_008_36    m/z 86.5044     0   1   14.5   15  RT (min)      lcorbi_t3_pos_3_003_33    m/z 86.5044     0   2   4   6   8E+3   12.5   13  RT (min)      lcorbi_t3_pos_3_031_45    m/z 86.5044     0   1   15   15.5  RT (min)      lcorbi_t3_pos_2_012_01    m/z 86.5044     0   1   12.5   13  RT (min)      lcorbi_t3_pos_2_013_33    m/z 86.5044     0   2   4   6   8E+5   13.5   14  RT (min)      lcorbi_t3_pos_1_005_42    m/z 86.5044     0   1   15   15.5  RT (min)      lcorbi_t3_pos_1_031_40    m/z 86.5044     0   1E+6   13   13.5  RT (min)      lcorbi_t3_pos_3_011_42    m/z 86.5044     0   1   14.5   15  RT (min)      lcorbi_t3_pos_1_034_34    m/z 86.5044     0   1   14   14.5  RT (min)      lcorbi_t3_pos_2_022_38    m/z 86.5045     0   1   15.5   16  RT (min)      lcorbi_t3_pos_2_012_01    m/z 86.6157     0   1   15   15.5  RT (min)      lcorbi_t3_pos_1_031_40    m/z 86.6157     0   1   14.5   15  RT (min)      lcorbi_t3_pos_1_034_34    m/z 86.6158     0   1   2   3   4E+5   14   14.5  RT (min)      lcorbi_t3_pos_1_008_36    m/z 86.6158     0   1   14   14.5  RT (min)      lcorbi_t3_pos_2_022_38    m/z 86.6158     0   1   15   15.5  RT (min)      lcorbi_t3_pos_1_031_40    m/z 86.6158     0   1   14.5   15  RT (min)      lcorbi_t3_pos_1_034_34    m/z 86.6159     0   1   12.5   13  RT (min)      lcorbi_t3_pos_3_031_45    m/z 86.6159     0   1   12.5   13  RT (min)      lcorbi_t3_pos_2_013_33    m/z 86.6159     0   1E+4   15   15.5  RT (min)      lcorbi_t3_pos_2_012_01    m/z 86.6159     0   1   12.5   13  RT (min)      lcorbi_t3_pos_2_013_33    m/z 86.6159     0   1   14.5   15  RT (min)      lcorbi_t3_pos_1_028_32    m/z 86.6159     0   1   14.5   15  RT (min)      lcorbi_t3_pos_1_034_34    m/z 86.7283     0   1E+5   13   13.5  RT (min)      lcorbi_t3_pos_2_013_33    m/z 86.7284     0   1   14.5   15  RT (min)      lcorbi_t3_pos_2_022_38    m/z 86.7284     0   1   15   15.5  RT (min)      lcorbi_t3_pos_1_031_40    m/z 86.7284     0   1   2   3   4E+4   12.5   13  RT (min)      lcorbi_t3_pos_2_013_33    m/z 86.7284     0   1E+5   13   13.5  RT (min)      lcorbi_t3_pos_2_013_33    m/z 86.7284     0   1   15.5   16  RT (min)      lcorbi_t3_pos_1_031_40    m/z 86.7284     0   1   15.5   16  RT (min)      lcorbi_t3_pos_2_012_01    m/z 86.7284     0   1   15   15.5  RT (min)      lcorbi_t3_pos_1_034_34    m/z 86.7284     0   1   15.5   16  RT (min)      lcorbi_t3_pos_2_012_01    m/z 86.7284     0   1   14.5   15  RT (min)      lcorbi_t3_pos_1_028_32    m/z 86.7284     0   2   4   6E+5   14   14.5  RT (min)      lcorbi_t3_pos_1_008_36    m/z 86.7284     0   1   14.5   15  RT (min)      lcorbi_t3_pos_2_022_38    m/z 86.7284     0   1   15.5   16  RT (min)      lcorbi_t3_pos_1_031_40    m/z 86.7284     0   1E+6   14   14.5  RT (min)      lcorbi_t3_pos_1_005_42    m/z 86.7284     0   1   15   15.5  RT (min)      lcorbi_t3_pos_1_041_54    m/z 86.7284     0   1   15   15.5  RT (min)      lcorbi_t3_pos_1_049_55    m/z 86.7284     0   1   15   15.5  RT (min)      lcorbi_t3_pos_1_034_34    m/z 86.7284     0   1   15   15.5  RT (min)      lcorbi_t3_pos_1_034_34    m/z 86.8398     0   2   4   6E+4   14   14.5  RT (min)      lcorbi_t3_pos_1_008_36    m/z 86.8398     0   1   14.5   15  RT (min)      lcorbi_t3_pos_2_022_38    m/z 86.8398     0   1   15.5   16  RT (min)      lcorbi_t3_pos_1_031_40    m/z 86.8398     0   1   15   15.5  RT (min)      lcorbi_t3_pos_1_034_34    m/z 86.8398     0   1   15   15.5  RT (min)      lcorbi_t3_pos_1_034_34    m/z 86.8398     0   1   15.5   16  RT (min)      lcorbi_t3_pos_1_031_40    m/z 86.8398     0   1   15   15.5  RT (min)      lcorbi_t3_pos_1_056_56    m/z 86.8398     0   1   2E+4   14.5   15  RT (min)      lcorbi_t3_pos_1_028_32    m/z 86.8398     0   1   2E+4   14.5   15  RT (min)      lcorbi_t3_pos_1_028_32    m/z 86.8398     0   1   15   15.5  RT (min)      lcorbi_t3_pos_1_031_40    m/z 86.8399     0   1   14.5   15  RT (min)      lcorbi_t3_pos_4_049_55    m/z 86.8399     0   2   4   6   8E+3   12.5   13  RT (min)      lcorbi_t3_pos_2_013_33    m/z 86.8399     0   1E+4   15.5   16  RT (min)      lcorbi_t3_pos_2_012_01    m/z 86.8399     0   2   4   6   8E+3   13   13.5  RT (min)      lcorbi_t3_pos_2_013_33    m/z 86.9512     0   1E+6   13   13.5  RT (min)      lcorbi_t3_pos_1_002_01    m/z 86.9522     0   1   13   13.5  RT (min)      lcorbi_t3_pos_2_013_33    m/z 86.9522     0   1   2   3E+5   13   13.5  RT (min)      lcorbi_t3_pos_1_002_01    m/z 86.9522     0   1   2   3   4E+4   13   13.5  RT (min)      lcorbi_t3_pos_2_013_33    m/z 86.9523     0   2   4   6   8E+5   13   13.5  RT (min)      lcorbi_t3_pos_1_002_01    m/z 86.9523     0   1E+5   14.5   15  RT (min)      lcorbi_t3_pos_1_008_36    m/z 86.9523     0   1E+6   14   14.5  RT (min)      lcorbi_t3_pos_1_008_36    m/z 86.9523     0   1   2E+4   15   15.5  RT (min)      lcorbi_t3_pos_1_034_34    m/z 86.9523     0   1   2E+4   14.5   15  RT (min)      lcorbi_t3_pos_1_028_32    m/z 86.9523     0   1   15.5   16  RT (min)      lcorbi_t3_pos_1_041_54    m/z 86.9523     0   1   16   16.5  RT (min)      lcorbi_t3_pos_1_023_02    m/z 86.9523     0   1   16   16.5  RT (min)      lcorbi_t3_pos_1_023_02    m/z 86.9523     0   1   16.5   17  RT (min)    lcorbi_t3_pos_1_023_02    m/z 86.9523     0   1   2E+4   15   15.5  RT (min)      lcorbi_t3_pos_1_034_34    m/z 86.9523     0   1   16   16.5  RT (min)      lcorbi_t3_pos_1_051_09    m/z 86.9523     0   1   13   13.5  RT (min)      lcorbi_t3_pos_3_031_45    m/z 86.9523     0   1E+4   15.5   16  RT (min)      lcorbi_t3_pos_2_012_01    m/z 86.9523     0   2   4   6E+4   14.5   15  RT (min)      lcorbi_t3_pos_1_008_36    m/z 86.9523     0   1   15.5   16  RT (min)      lcorbi_t3_pos_1_031_40    m/z 86.9523     0   1   15.5   16  RT (min)      lcorbi_t3_pos_1_031_40    m/z 86.9523     0   1E+6   14   14.5  RT (min)      lcorbi_t3_pos_1_005_42    m/z 86.9523     0   1E+4   15.5   16  RT (min)      lcorbi_t3_pos_2_012_01    m/z 86.9523     0   1E+6   14   14.5  RT (min)      lcorbi_t3_pos_1_008_36    m/z 86.9524     0   1   16.5   17  RT (min)    lcorbi_t3_pos_1_023_02    m/z 86.9524     0   1   2E+4   15   15.5  RT (min)      lcorbi_t3_pos_1_034_34    m/z 86.9524     0   1   15   15.5  RT (min)      lcorbi_t3_pos_2_025_52    m/z 86.9524     0   1   14.5   15  RT (min)      lcorbi_t3_pos_1_030_21    m/z 86.9524     0   1   15.5   16  RT (min)      lcorbi_t3_pos_1_049_55    m/z 87.0635     0   2   4   6E+4   13   13.5  RT (min)      lcorbi_t3_pos_1_002_01    m/z 87.0636     0   2   4   6E+5   14   14.5  RT (min)      lcorbi_t3_pos_1_008_36    m/z 87.0636     0   2   4E+5   14   14.5  RT (min)      lcorbi_t3_pos_1_005_42    m/z 87.0636     0   1   2   3E+5   13   13.5  RT (min)      lcorbi_t3_pos_1_002_01    m/z 87.0636     0   1   14.5   15  RT (min)      lcorbi_t3_pos_1_028_32    m/z 87.0638     0   1   2E+4   14.5   15  RT (min)      lcorbi_t3_pos_1_008_36    m/z 87.0638     0   1   16.5   17  RT (min)    lcorbi_t3_pos_1_023_02    m/z 87.0638     0   1   15.5   16  RT (min)      lcorbi_t3_pos_2_012_01    m/z 87.0638     0   1   15.5   16  RT (min)      lcorbi_t3_pos_2_012_01    m/z 87.0638     0   1E+5   14   14.5  RT (min)      lcorbi_t3_pos_1_008_36    m/z 87.0638     0   1   16.5   17  RT (min)    lcorbi_t3_pos_1_023_02    m/z 87.0638     0   1   2E+4   15   15.5  RT (min)      lcorbi_t3_pos_1_034_34    m/z 87.0638     0   1   2E+4   15   15.5  RT (min)      lcorbi_t3_pos_1_034_34    m/z 87.0638     0   1   15.5   16  RT (min)      lcorbi_t3_pos_1_031_40    m/z 87.0638     0   1   15.5   16  RT (min)      lcorbi_t3_pos_1_031_40    m/z 87.0638     0   1   2   3E+4   14.5   15  RT (min)      lcorbi_t3_pos_1_008_36    m/z 87.0638     0   1   2E+4   15   15.5  RT (min)      lcorbi_t3_pos_1_034_34    m/z 87.0638     0   1   16   16.5  RT (min)      lcorbi_t3_pos_1_023_02    m/z 87.0777     0   1   14   14.5  RT (min)      lcorbi_t3_pos_1_058_14    m/z 87.0804     0   1   2   3   4E+5   13   13.5  RT (min)      lcorbi_t3_pos_1_064_16    m/z 87.0804     0   2   4   6E+5   15   15.5  RT (min)      lcorbi_t3_pos_1_058_14    m/z 87.0917     0   2   4E+6   0   0.5  RT (min)      lcorbi_t3_pos_1_002_01    m/z 87.1761     0   1   2E+4   13   13.5  RT (min)      lcorbi_t3_pos_1_002_01    m/z 87.1762     0   1   16   16.5  RT (min)      lcorbi_t3_pos_1_051_09    m/z 87.1762     0   1   15   15.5  RT (min)      lcorbi_t3_pos_1_034_34    m/z 87.1762     0   1   14.5   15  RT (min)      lcorbi_t3_pos_2_017_51    m/z 87.1762     0   1   15.5   16  RT (min)      lcorbi_t3_pos_2_049_55    m/z 87.1762     0   1   15.5   16  RT (min)      lcorbi_t3_pos_2_049_55    m/z 87.1762     0   1   16.5   17  RT (min)    lcorbi_t3_pos_1_023_02    m/z 87.1762     0   1   15   15.5  RT (min)      lcorbi_t3_pos_1_030_21    m/z 87.1762     0   1E+6   14   14.5  RT (min)      lcorbi_t3_pos_1_005_42    m/z 87.1762     0   1   15   15.5  RT (min)      lcorbi_t3_pos_1_041_54    m/z 87.1762     0   1   16.5   17  RT (min)    lcorbi_t3_pos_1_023_02    m/z 87.1762     0   1   15   15.5  RT (min)      lcorbi_t3_pos_1_028_32    m/z 87.1762     0   1E+6   14   14.5  RT (min)      lcorbi_t3_pos_1_005_42    m/z 87.1762     0   1   15.5   16  RT (min)      lcorbi_t3_pos_1_031_40    m/z 87.1762     0   1   2E+4   15.5   16  RT (min)      lcorbi_t3_pos_1_025_52    m/z 87.1763     0   1   15   15.5  RT (min)      lcorbi_t3_pos_1_033_53    m/z 87.1763     0   1   16   16.5  RT (min)      lcorbi_t3_pos_2_056_56    m/z 87.1763     0   1   15   15.5  RT (min)      lcorbi_t3_pos_1_033_53    m/z 87.2875     0   2   4E+5   14   14.5  RT (min)      lcorbi_t3_pos_1_005_42    m/z 87.2877     0   1   16.5   17  RT (min)    lcorbi_t3_pos_1_023_02    m/z 87.2877     0   1   15   15.5  RT (min)      lcorbi_t3_pos_1_028_32    m/z 87.2877     0   1E+4   15.5   16  RT (min)      lcorbi_t3_pos_2_049_55    m/z 87.2877     0   1   15.5   16  RT (min)      lcorbi_t3_pos_1_031_40    m/z 87.2877     0   1E+4   13   13.5  RT (min)      lcorbi_t3_pos_1_002_01    m/z 87.2877     0   1   15   15.5  RT (min)      lcorbi_t3_pos_1_030_21    m/z 87.2877     0   1   14.5   15  RT (min)      lcorbi_t3_pos_2_046_07    m/z 87.2878     0   1   15.5   16  RT (min)      lcorbi_t3_pos_1_049_55    m/z 87.2878     0   1   16.5   17  RT (min)    lcorbi_t3_pos_1_023_02    m/z 87.3988     0   1   2E+6   14   14.5  RT (min)      lcorbi_t3_pos_1_046_22    m/z 87.4001     0   1   13   13.5  RT (min)      lcorbi_t3_pos_2_032_39    m/z 87.4002     0   1   2E+4   13.5   14  RT (min)      lcorbi_t3_pos_2_054_29    m/z 87.4002     0   2   4E+5   13.5   14  RT (min)      lcorbi_t3_pos_4_016_15    m/z 87.4002     0   1   13   13.5  RT (min)      lcorbi_t3_pos_2_032_39    m/z 87.4002     0   1   16.5   17  RT (min)    lcorbi_t3_pos_1_023_02    m/z 87.4002     0   1   2   3   4E+5   13.5   14  RT (min)      lcorbi_t3_pos_1_002_01    m/z 87.4002     0   2   4   6E+4   13.5   14  RT (min)      lcorbi_t3_pos_1_002_01    m/z 87.4003     0   1   16.5   17  RT (min)    lcorbi_t3_pos_1_023_02    m/z 87.4003     0   1   16   16.5  RT (min)      lcorbi_t3_pos_1_051_09    m/z 87.4003     0   1   16   16.5  RT (min)      lcorbi_t3_pos_1_031_40    m/z 87.4003     0   1   15.5   16  RT (min)      lcorbi_t3_pos_3_013_19    m/z 87.4003     0   2   4E+4   14.5   15  RT (min)      lcorbi_t3_pos_1_008_36    m/z 87.4003     0   1   16   16.5  RT (min)      lcorbi_t3_pos_1_051_09    m/z 87.4003     0   1   15   15.5  RT (min)      lcorbi_t3_pos_1_028_32    m/z 87.4003     0   1   2E+6   14.5   15  RT (min)      lcorbi_t3_pos_1_005_42    m/z 87.4003     0   1   2E+6   14.5   15  RT (min)      lcorbi_t3_pos_1_005_42    m/z 87.4003     0   1   16.5   17  RT (min)    lcorbi_t3_pos_1_023_02    m/z 87.4003     0   1   15   15.5  RT (min)      lcorbi_t3_pos_1_028_32    m/z 87.4003     0   1   15.5   16  RT (min)      lcorbi_t3_pos_1_034_34    m/z 87.4003     0   1   2E+5   14.5   15  RT (min)      lcorbi_t3_pos_1_008_36    m/z 87.4003     0   1   15.5   16  RT (min)      lcorbi_t3_pos_1_034_34    m/z 87.4003     0   1   15.5   16  RT (min)      lcorbi_t3_pos_1_034_34    m/z 87.4003     0   1   14.5   15  RT (min)      lcorbi_t3_pos_1_008_36    m/z 87.4004     0   1   15   15.5  RT (min)      lcorbi_t3_pos_1_030_21    m/z 87.5116     0   1   15   15.5  RT (min)      lcorbi_t3_pos_1_030_21    m/z 87.5117     0   2   4   6   8E+4   13.5   14  RT (min)      lcorbi_t3_pos_1_002_01    m/z 87.5117     0   1   13.5   14  RT (min)      lcorbi_t3_pos_1_002_01    m/z 87.5117     0   1   2E+4   15   15.5  RT (min)      lcorbi_t3_pos_1_028_32    m/z 87.5117     0   1   15.5   16  RT (min)      lcorbi_t3_pos_1_034_34    m/z 87.5117     0   1   16   16.5  RT (min)      lcorbi_t3_pos_1_051_09    m/z 87.5117     0   1   16.5   17  RT (min)    lcorbi_t3_pos_1_023_02    m/z 87.5117     0   1   2   3E+4   14.5   15  RT (min)      lcorbi_t3_pos_1_008_36    m/z 87.5117     0   1E+6   14.5   15  RT (min)      lcorbi_t3_pos_1_005_42    m/z 87.5118     0   1E+6   14.5   15  RT (min)      lcorbi_t3_pos_1_005_42    m/z 87.5118     0   1   2   3   4E+4   14.5   15  RT (min)      lcorbi_t3_pos_1_008_36    m/z 87.5118     0   1E+4   14.5   15   15.5  RT (min)      lcorbi_t3_pos_2_017_51    m/z 87.5118     0   1   2E+4   15   15.5  RT (min)      lcorbi_t3_pos_1_028_32    m/z 87.5118     0   1   16   16.5  RT (min)      lcorbi_t3_pos_1_051_09    m/z 87.5118     0   1   15.5   16  RT (min)      lcorbi_t3_pos_1_034_34    m/z 87.5118     0   1   16.5   17  RT (min)    lcorbi_t3_pos_1_023_02    m/z 87.5118     0   1   15.5   16  RT (min)      lcorbi_t3_pos_2_049_55    m/z 87.5118     0   1   16.5   17  RT (min)    lcorbi_t3_pos_2_012_01    m/z 87.5118     0   1   2E+4   15   15.5  RT (min)      lcorbi_t3_pos_1_028_32    m/z 87.5118     0   1   16.5   17  RT (min)    lcorbi_t3_pos_1_023_02    m/z 87.6241     0   1   15.5   16  RT (min)      lcorbi_t3_pos_1_034_34    m/z 87.6242     0   1   17   17.5  RT (min)    lcorbi_t3_pos_1_023_02    m/z 87.6242     0   1   14.5   15  RT (min)      lcorbi_t3_pos_1_008_36    m/z 87.6242     0   2   4   6E+4   13.5   14  RT (min)      lcorbi_t3_pos_2_054_29    m/z 87.6242     0   1   16   16.5  RT (min)      lcorbi_t3_pos_1_031_40    m/z 88.0757     0   1E+5   0.5   1  RT (min)      lcorbi_t3_pos_2_011_10    m/z 88.0757     0   1   2   3   4E+5   18.5   19  RT (min)      lcorbi_t3_pos_4_092_60    m/z 88.0757     0   1E+5   0.5   1  RT (min)      lcorbi_t3_pos_2_014_37    m/z 88.0757     0   1   2   3   4E+5   18.5   19  RT (min)      lcorbi_t3_pos_4_092_60    m/z 88.0757     0   1E+5   1   1.5  RT (min)      lcorbi_t3_pos_2_021_09    m/z 88.1086     0   2   4E+5   16   17   18  RT (min)       lcorbi_t3_pos_1_027_43    m/z 88.1089     0   2   4   6E+5   17.5   18  RT (min)       lcorbi_t3_pos_4_105_61    m/z 88.1091     0   2   4E+4   10   10.5  RT (min)      lcorbi_t3_pos_1_064_16    m/z 88.1091     0   2   4E+4   9.5   10  RT (min)      lcorbi_t3_pos_1_061_50    m/z 88.1094     0   1   17.5   18  RT (min)     lcorbi_t3_pos_1_025_52    m/z 88.1094     0   1   2   3E+5   14   14.5  RT (min)      lcorbi_t3_pos_1_057_49    m/z 88.1097     0   1E+5   12.5   13  RT (min)      lcorbi_t3_pos_1_059_17    m/z 88.1105     0   2   4   6E+6   9   9.5  RT (min)      lcorbi_t3_pos_3_059_17    m/z 88.1105     0   2   4   6E+6   9   9.5  RT (min)      lcorbi_t3_pos_3_059_17    m/z 88.1107     0   1E+7   14   14.5  RT (min)      lcorbi_t3_pos_4_062_13    m/z 88.1120     0   2   4   6   8E+5   0   0.5  RT (min)      lcorbi_t3_pos_2_061_50    m/z 88.1120     0   1   2   3E+6   7   7.5  RT (min)      lcorbi_t3_pos_2_063_16    m/z 88.1120     0   2   4E+6   7.5   8  RT (min)      lcorbi_t3_pos_2_057_49    m/z 88.1120     0   1   2E+6   6   6.5  RT (min)      lcorbi_t3_pos_3_057_49    m/z 88.1120     0   2   4   6   8E+5   5.5   6  RT (min)      lcorbi_t3_pos_4_064_39    m/z 88.1120     0   2   4   6   8E+6   8.5   9  RT (min)      lcorbi_t3_pos_1_062_13    m/z 88.1120     0   2   4E+6   8   8.5  RT (min)      lcorbi_t3_pos_3_057_49    m/z 88.1121     0   1E+7   8.5   9  RT (min)      lcorbi_t3_pos_1_059_17    m/z 88.1121     0   1   2E+7   11.5   12  RT (min)      lcorbi_t3_pos_1_057_49    m/z 88.1121     0   1   2E+7   12   12.5  RT (min)      lcorbi_t3_pos_1_059_17    m/z 88.1121     0   1E+7   10.5   11  RT (min)      lcorbi_t3_pos_1_059_17    m/z 88.1121     0   2   4   6   8E+7   5   10   15  RT (min)       lcorbi_t3_pos_4_090_60    m/z 88.1121     0   1   2   3E+7   13   14  RT (min)       lcorbi_t3_pos_1_059_17    m/z 88.1121     0   1E+7   1.5   2  RT (min)      lcorbi_t3_pos_4_057_49    m/z 88.1121     0   1E+7   11   11.5  RT (min)      lcorbi_t3_pos_1_060_39    m/z 88.1121     0   1   2   3E+7   13   13.5  RT (min)       lcorbi_t3_pos_1_057_49    m/z 88.1121     0   1E+7   10   10.5  RT (min)      lcorbi_t3_pos_1_062_13    m/z 88.1121     0   1E+7   1   1.5  RT (min)      lcorbi_t3_pos_4_057_49    m/z 88.5200     0   1E+4   14   14.5  RT (min)      lcorbi_t3_pos_2_017_51    m/z 88.5200     0   1   14.5   15  RT (min)      lcorbi_t3_pos_1_030_21    m/z 88.5200     0   1   15   15.5  RT (min)      lcorbi_t3_pos_1_056_56    m/z 88.5201     0   1   14.5   15  RT (min)      lcorbi_t3_pos_1_041_54    m/z 88.5201     0   1E+4   14   14.5  RT (min)      lcorbi_t3_pos_2_017_51    m/z 88.5201     0   1   14.5   15  RT (min)      lcorbi_t3_pos_1_021_18    m/z 88.7440     0   1   15.5   16  RT (min)      lcorbi_t3_pos_1_025_52    m/z 88.7440     0   1   15.5   16  RT (min)      lcorbi_t3_pos_1_025_52    m/z 88.7440     0   1   15   15.5  RT (min)      lcorbi_t3_pos_2_009_50    m/z 88.7440     0   1   15.5   16  RT (min)      lcorbi_t3_pos_1_049_55    m/z 88.7440     0   1   14.5   15  RT (min)      lcorbi_t3_pos_1_030_21    m/z 88.7440     0   1E+4   14.5   15  RT (min)      lcorbi_t3_pos_2_046_07    m/z 88.7441     0   1   14.5   15   15.5  RT (min)      lcorbi_t3_pos_2_025_52    m/z 88.7441     0   2   4   6   8E+5   14   14.5  RT (min)      lcorbi_t3_pos_1_005_42    m/z 88.7441     0   1   15   15.5  RT (min)      lcorbi_t3_pos_1_041_54    m/z 89.1073     0   2   4   6   8E+4   1   1.5  RT (min)      lcorbi_t3_pos_4_098_61    m/z 89.1080     0   1   17   18  RT (min)     lcorbi_t3_pos_1_025_52    m/z 89.1091     0   1E+5   13   13.5  RT (min)      lcorbi_t3_pos_2_057_49    m/z 89.1153     0   2   4   6   8E+4   0   1  RT (min)       lcorbi_t3_pos_4_084_59    m/z 89.1153     0   1   2   3   4E+6   17   18  RT (min)       lcorbi_t3_pos_4_083_59    m/z 89.1153     0   1   2   3E+6   16   18  RT (min)       lcorbi_t3_pos_4_105_61    m/z 89.1153     0   1E+5   0   0.5  RT (min)      lcorbi_t3_pos_1_031_40    m/z 89.1153     0   1   2   3   4E+6   14   16   18  RT (min)       lcorbi_t3_pos_4_093_60    m/z 89.1154     0   1   2E+6   14   14.5  RT (min)      lcorbi_t3_pos_1_057_49    m/z 89.1154     0   2   4   6E+5   9   9.5  RT (min)      lcorbi_t3_pos_1_062_13    m/z 89.1154     0   1E+6   12   12.5  RT (min)      lcorbi_t3_pos_1_058_14    m/z 89.1154     0   1E+6   11   11.5  RT (min)      lcorbi_t3_pos_1_061_50    m/z 89.1154     0   2   4   6E+5   9.5   10  RT (min)      lcorbi_t3_pos_1_061_50    m/z 89.1154     0   1E+6   10.5   11  RT (min)      lcorbi_t3_pos_1_057_49    m/z 89.1154     0   1E+5   0   0.5  RT (min)      lcorbi_t3_pos_1_031_40    m/z 89.1154     0   1   2E+6   13.5   14  RT (min)      lcorbi_t3_pos_1_058_14    m/z 89.1154     0   2   4   6E+5   11   11.5  RT (min)      lcorbi_t3_pos_3_058_16    m/z 89.1154     0   2   4   6   8E+5   1.5   2  RT (min)      lcorbi_t3_pos_4_057_49    m/z 89.1154     0   1E+6   10   10.5  RT (min)      lcorbi_t3_pos_1_059_17    m/z 89.1154     0   2   4   6E+5   8.5   9  RT (min)      lcorbi_t3_pos_1_059_17    m/z 89.1154     0   1E+6   11.5   12  RT (min)      lcorbi_t3_pos_1_057_49    m/z 89.1154     0   1E+6   12   12.5  RT (min)      lcorbi_t3_pos_2_057_49    m/z 89.1154     0   1E+6   14   14.5  RT (min)      lcorbi_t3_pos_2_058_15    m/z 89.1154     0   1   2E+6   13   14  RT (min)       lcorbi_t3_pos_4_066_57    m/z 89.1154     0   1E+5   0.5   1  RT (min)      lcorbi_t3_pos_1_024_04    m/z 89.1154     0   2   4   6E+5   9   9.5  RT (min)      lcorbi_t3_pos_1_064_16    m/z 89.1154     0   1   2E+5   0   0.5  RT (min)      lcorbi_t3_pos_1_002_01    m/z 89.1154     0   1E+6   11.5   12  RT (min)      lcorbi_t3_pos_1_058_14    m/z 89.1155     0   1   2   3   4E+6   15  RT (min)       lcorbi_t3_pos_1_027_43    m/z 89.8430     0   1   16.5   17  RT (min)    lcorbi_t3_pos_1_023_02    m/z 90.0669     0   1   16.5   17  RT (min)    lcorbi_t3_pos_1_023_02    m/z 90.0669     0   1   16   16.5  RT (min)      lcorbi_t3_pos_2_056_56    m/z 90.0669     0   1   15   15.5  RT (min)      lcorbi_t3_pos_1_047_41    m/z 90.0669     0   1   15   15.5  RT (min)      lcorbi_t3_pos_1_047_41    m/z 90.0669     0   1   16   16.5  RT (min)      lcorbi_t3_pos_1_051_09    m/z 90.0669     0   2   4   6   8E+5   13.5   14  RT (min)      lcorbi_t3_pos_1_002_01    m/z 90.0669     0   2   4   6   8E+4   13.5   14  RT (min)      lcorbi_t3_pos_1_002_01    m/z 90.0669     0   1   16.5   17  RT (min)    lcorbi_t3_pos_1_023_02    m/z 90.0669     0   1   2E+4   13.5   14  RT (min)      lcorbi_t3_pos_1_002_01    m/z 90.0669     0   1   15.5   16  RT (min)      lcorbi_t3_pos_1_034_34    m/z 90.0669     0   1   16   16.5  RT (min)      lcorbi_t3_pos_1_051_09    m/z 90.0669     0   1E+6   14.5   15  RT (min)      lcorbi_t3_pos_1_005_42    m/z 90.0670     0   1   2E+4   15   15.5  RT (min)      lcorbi_t3_pos_1_028_32    m/z 90.0670     0   1   16.5   17  RT (min)    lcorbi_t3_pos_1_023_02    m/z 90.0670     0   1   2   3E+4   14.5   15  RT (min)      lcorbi_t3_pos_1_008_36    m/z 90.0670     0   1   15.5   16  RT (min)      lcorbi_t3_pos_1_034_34    m/z 90.0670     0   2   4   6   8E+4   14.5   15  RT (min)      lcorbi_t3_pos_1_008_36    m/z 90.0670     0   1   2E+4   15   15.5  RT (min)      lcorbi_t3_pos_1_028_32    m/z 90.0670     0   1   16   16.5  RT (min)      lcorbi_t3_pos_1_049_55    m/z 90.0670     0   1   16   16.5  RT (min)      lcorbi_t3_pos_1_031_40    m/z 90.1783     0   1   16   16.5  RT (min)      lcorbi_t3_pos_1_051_09    m/z 90.1783     0   1   16   16.5  RT (min)      lcorbi_t3_pos_1_031_40    m/z 90.1784     0   1   16.5   17  RT (min)    lcorbi_t3_pos_1_023_02    m/z 90.1784     0   1   15   15.5  RT (min)      lcorbi_t3_pos_1_028_32    m/z 90.1784     0   1   14.5   15  RT (min)      lcorbi_t3_pos_1_008_36    m/z 90.1784     0   1   15.5   16  RT (min)      lcorbi_t3_pos_1_034_34    m/z 90.1784     0   1   16.5   17  RT (min)    lcorbi_t3_pos_1_023_02    m/z 90.1852     0   1   2E+4   14.5   15  RT (min)      lcorbi_t3_pos_1_005_42    m/z 90.2908     0   1   14.5   15  RT (min)      lcorbi_t3_pos_1_008_36    m/z 90.2908     0   1   15.5   16  RT (min)      lcorbi_t3_pos_1_028_32    m/z 90.2908     0   1   16.5   17  RT (min)    lcorbi_t3_pos_1_023_02    m/z 90.9477     0   1   2E+6   0   0.5  RT (min)       lcorbi_t3_pos_1_005_42    m/z 90.9766     0   1E+5   13  RT (min)       lcorbi_t3_pos_1_022_20    m/z 90.9766     0   2   4   6E+5   17.5   18  RT (min)       lcorbi_t3_pos_4_105_61    m/z 91.4097     0   1   2E+4   15   15.5  RT (min)      lcorbi_t3_pos_1_056_56    m/z 91.4097     0   1   2E+4   14.5   15  RT (min)      lcorbi_t3_pos_1_033_53    m/z 91.4097     0   1   15   15.5  RT (min)      lcorbi_t3_pos_1_025_52    m/z 91.4097     0   2   4   6   8E+5   13.5   14  RT (min)      lcorbi_t3_pos_3_040_41    m/z 91.4098     0   1   14.5   15  RT (min)      lcorbi_t3_pos_1_041_54    m/z 91.4098     0   1   14.5   15  RT (min)      lcorbi_t3_pos_2_049_55    m/z 91.4098     0   1E+4   14   14.5  RT (min)      lcorbi_t3_pos_1_047_41    m/z 91.4098     0   1E+6   13.5   14  RT (min)      lcorbi_t3_pos_1_005_42    m/z 91.4098     0   1   14   14.5  RT (min)      lcorbi_t3_pos_1_030_21    m/z 91.5222     0   1   14.5   15  RT (min)      lcorbi_t3_pos_1_033_53    m/z 91.5222     0   1   15.5   16  RT (min)      lcorbi_t3_pos_1_001_49    m/z 91.5222     0   1   2E+4   15   15.5  RT (min)      lcorbi_t3_pos_1_041_54    m/z 91.5222     0   1   2   3   4E+5   14   14.5  RT (min)      lcorbi_t3_pos_1_005_42    m/z 91.5222     0   1   14.5   15  RT (min)      lcorbi_t3_pos_1_030_21    m/z 91.6337     0   1   14.5   15  RT (min)      lcorbi_t3_pos_1_030_21    m/z 91.6337     0   1   15   15.5  RT (min)      lcorbi_t3_pos_1_041_54    m/z 91.6337     0   1   15.5   16  RT (min)      lcorbi_t3_pos_1_056_56    m/z 91.6337     0   1E+6   14   14.5  RT (min)      lcorbi_t3_pos_3_040_41    m/z 91.6337     0   1   15   15.5  RT (min)      lcorbi_t3_pos_1_041_54    m/z 91.6337     0   1   15   15.5  RT (min)      lcorbi_t3_pos_1_041_54    m/z 91.6337     0   1   14.5   15  RT (min)      lcorbi_t3_pos_1_021_18    m/z 91.6337     0   1   15   15.5  RT (min)      lcorbi_t3_pos_1_049_55    m/z 91.6337     0   1   14   14.5  RT (min)      lcorbi_t3_pos_2_046_07    m/z 91.6337     0   1   14.5   15  RT (min)      lcorbi_t3_pos_2_025_52    m/z 91.6337     0   1   14.5   15  RT (min)      lcorbi_t3_pos_1_021_18    m/z 91.6337     0   1   15   15.5  RT (min)      lcorbi_t3_pos_1_056_56    m/z 91.6337     0   1   15.5   16  RT (min)      lcorbi_t3_pos_1_001_49    m/z 91.6337     0   1   15   15.5  RT (min)      lcorbi_t3_pos_1_041_54    m/z 91.6337     0   1   2E+6   14   14.5  RT (min)      lcorbi_t3_pos_1_005_42    m/z 91.6338     0   1   14   14.5  RT (min)      lcorbi_t3_pos_2_046_07    m/z 91.8577     0   1   15.5   16  RT (min)      lcorbi_t3_pos_1_025_52    m/z 91.8577     0   1E+4   14.5   15  RT (min)      lcorbi_t3_pos_2_017_51    m/z 91.8577     0   1E+6   14   14.5  RT (min)      lcorbi_t3_pos_1_005_42    m/z 91.8577     0   1   2   3E+4   15   15.5  RT (min)      lcorbi_t3_pos_1_041_54    m/z 91.8577     0   1   14.5   15  RT (min)      lcorbi_t3_pos_1_021_18    m/z 91.8577     0   1   15.5   16  RT (min)      lcorbi_t3_pos_1_025_52    m/z 91.8577     0   1   15   15.5  RT (min)      lcorbi_t3_pos_2_038_03    m/z 91.8577     0   1   2E+3   14   14.5  RT (min)      lcorbi_t3_pos_4_010_43    m/z 91.8577     0   1E+4   14.5   15  RT (min)      lcorbi_t3_pos_2_017_51    m/z 91.8577     0   1   14.5   15  RT (min)      lcorbi_t3_pos_1_030_21    m/z 92.0815     0   1   15.5   16  RT (min)      lcorbi_t3_pos_1_056_56    m/z 92.0815     0   1   15.5   16  RT (min)      lcorbi_t3_pos_1_056_56    m/z 92.0816     0   1   15.5   16  RT (min)      lcorbi_t3_pos_1_025_52    m/z 93.0650     0   1E+5   9.5   10  RT (min)      lcorbi_t3_pos_1_034_34    m/z 95.0491     0   1E+5   15.5   16  RT (min)       lcorbi_t3_pos_4_110_62    m/z 95.0491     0   1E+5   15   15.5  RT (min)       lcorbi_t3_pos_4_111_62    m/z 97.0284     0   1E+5   0   0.5  RT (min)      lcorbi_t3_pos_1_051_09    m/z 97.0648     0   1   18  RT (min)     lcorbi_t3_pos_1_056_56    m/z 97.0648     0   2   4   6   8E+4   0   0.5  RT (min)      lcorbi_t3_pos_1_051_09    m/z 97.0648     0   2   4   6E+5   16   18  RT (min)       lcorbi_t3_pos_1_053_39    m/z 97.1011     0   2   4   6E+5   17   18  RT (min)       lcorbi_t3_pos_4_106_62    m/z 97.1012     0   2   4   6   8E+5   15   16  RT (min)       lcorbi_t3_pos_4_075_58    m/z 97.1012     0   1E+6   15.5   16  RT (min)       lcorbi_t3_pos_1_007_07    m/z 97.1012     0   2   4   6   8E+5   16   17  RT (min)       lcorbi_t3_pos_1_051_09    m/z 97.1012     0   2   4   6E+5   12.5   13   13.5  RT (min)       lcorbi_t3_pos_1_052_38    m/z 97.1012     0   1E+6   16   17  RT (min)       lcorbi_t3_pos_1_022_20    m/z 97.1012     0   1E+6   16   17  RT (min)       lcorbi_t3_pos_1_025_52    m/z 97.1012     0   1E+6   15.5   16   16.5  RT (min)       lcorbi_t3_pos_1_044_25    m/z 97.1012     0   1E+6   16   16.5  RT (min)       lcorbi_t3_pos_1_021_18    m/z 97.1012     0   1E+6   14   15  RT (min)       lcorbi_t3_pos_1_034_34    m/z 97.1012     0   1E+6   16   17   18  RT (min)       lcorbi_t3_pos_1_027_43    m/z 97.1012     0   2   4   6E+5   17   18  RT (min)       lcorbi_t3_pos_4_094_60    m/z 97.1012     0   2   4E+5   12   12.5  RT (min)      lcorbi_t3_pos_1_059_17    m/z 97.1012     0   1E+6   15   16   17  RT (min)       lcorbi_t3_pos_1_012_14    m/z 97.1012     0   2   4   6   8E+5   13   13.5  RT (min)       lcorbi_t3_pos_1_032_26    m/z 97.8210     0   2   4E+5   13.5   14  RT (min)      lcorbi_t3_pos_3_047_40    m/z 97.8210     0   1   2E+4   15   15.5  RT (min)      lcorbi_t3_pos_2_012_01    m/z 97.8210     0   1E+5   13.5   14  RT (min)      lcorbi_t3_pos_1_008_36    m/z 97.8210     0   1   2   3E+4   14.5   15  RT (min)      lcorbi_t3_pos_1_034_34    m/z 97.8210     0   1   2   3E+4   15   15.5  RT (min)      lcorbi_t3_pos_1_051_09    m/z 97.8210     0   2   4E+5   12.5   13  RT (min)      lcorbi_t3_pos_4_016_15    m/z 97.8210     0   1E+4   14   14.5  RT (min)      lcorbi_t3_pos_3_003_33    m/z 97.8210     0   1   2E+5   12.5   13  RT (min)      lcorbi_t3_pos_1_002_01    m/z 97.8210     0   1E+4   12   12.5  RT (min)      lcorbi_t3_pos_2_032_39    m/z 97.8210     0   1   2   3E+4   15.5   16  RT (min)      lcorbi_t3_pos_1_023_02    m/z 98.9841     0   1E+6   2   2.5  RT (min)      lcorbi_t3_pos_4_057_49    m/z 98.9842     0   1E+5   10.5   11  RT (min)      lcorbi_t3_pos_3_000_aa    m/z 98.9842     0   1E+6   1.5   2  RT (min)      lcorbi_t3_pos_4_057_49    m/z 99.0033     0   2   4E+5   0   0.5  RT (min)      lcorbi_t3_pos_1_001_49    m/z 99.0441     0   1   2E+5   4.5   5  RT (min)       lcorbi_t3_pos_1_004_24    m/z 100.0195     0   2   4   6   8E+5   17.5   18  RT (min)      lcorbi_t3_pos_4_105_61    m/z 100.0756     0   2   4   6E+5   5   10   15  RT (min)       lcorbi_t3_pos_4_075_58    m/z 100.0756     0   2   4   6E+5   5   10   15  RT (min)       lcorbi_t3_pos_4_081_58    m/z 100.9989     0   2   4   6E+5   0   0.5  RT (min)      lcorbi_t3_pos_1_001_49    m/z 101.0961     0   1   2E+6   10   12  RT (min)       lcorbi_t3_pos_1_057_49    m/z 101.0961     0   1   2   3E+6   15   16  RT (min)       lcorbi_t3_pos_1_053_39    m/z 101.0961     0   1E+6   10   10.5  RT (min)      lcorbi_t3_pos_1_057_49    m/z 101.0961     0   1   2   3E+6   16  RT (min)       lcorbi_t3_pos_1_054_48    m/z 101.0961     0   2   4   6   8E+5   1.5   2  RT (min)      lcorbi_t3_pos_4_057_49    m/z 101.0961     0   1E+6   9.5   10  RT (min)      lcorbi_t3_pos_1_062_13    m/z 101.0961     0   1   2   3E+6   16   16.5  RT (min)       lcorbi_t3_pos_1_055_33    m/z 101.0961     0   1E+6   8.5   9  RT (min)      lcorbi_t3_pos_1_058_14    m/z 101.0961     0   2   4   6   8E+5   8   8.5  RT (min)      lcorbi_t3_pos_1_057_49    m/z 101.0961     0   1   2   3E+6   15   16  RT (min)       lcorbi_t3_pos_1_058_14    m/z 101.0961     0   1   2   3E+5   0   0.5  RT (min)      lcorbi_t3_pos_1_007_07    m/z 101.0961     0   1E+6   9.5   10  RT (min)      lcorbi_t3_pos_1_059_17    m/z 101.0961     0   1E+6   10.5   11  RT (min)      lcorbi_t3_pos_1_058_14    m/z 101.0961     0   1   2E+6   13.5   14  RT (min)       lcorbi_t3_pos_1_058_14    m/z 101.0961     0   1   17  RT (min)     lcorbi_t3_pos_1_027_43    m/z 101.0961     0   2   4   6   8E+5   8.5   9  RT (min)      lcorbi_t3_pos_1_060_39    m/z 101.0961     0   1E+6   11   11.5  RT (min)      lcorbi_t3_pos_1_058_14    m/z 101.0961     0   1   17.5   18  RT (min)     lcorbi_t3_pos_1_025_52    m/z 101.0961     0   1E+6   12.5   13  RT (min)      lcorbi_t3_pos_1_060_39    m/z 101.0961     0   1   17.5   18  RT (min)     lcorbi_t3_pos_1_049_55    m/z 101.0961     0   1   2E+6   16.5   17  RT (min)       lcorbi_t3_pos_4_084_59    m/z 101.0961     0   1   16.5   17   17.5  RT (min)     lcorbi_t3_pos_1_021_18    m/z 101.0961     0   2   4   6   8E+5   8   8.5  RT (min)      lcorbi_t3_pos_1_058_14    m/z 101.0961     0   1E+6   11.5   12  RT (min)      lcorbi_t3_pos_1_057_49    m/z 101.0961     0   2   4   6E+5   12   12.5  RT (min)      lcorbi_t3_pos_4_060_16    m/z 101.0961     0   1   2   3E+6   15.5   16  RT (min)       lcorbi_t3_pos_1_052_38    m/z 101.0961     0   1E+6   11   11.5  RT (min)      lcorbi_t3_pos_1_060_39    m/z 102.0550     0   1E+6   1   1.5  RT (min)      lcorbi_t3_pos_4_087_59    m/z 102.0913     0   2   4   6   8E+4   0   0.5  RT (min)      lcorbi_t3_pos_3_000_aa    m/z 102.0914     0   1E+5   9   9.5  RT (min)      lcorbi_t3_pos_1_002_01    m/z 102.0914     0   1E+5   8.5   9  RT (min)      lcorbi_t3_pos_3_000_aa    m/z 102.0914     0   2   4   6   8E+4   9.5   10  RT (min)      lcorbi_t3_pos_1_009_50    m/z 102.0995     0   2   4   6E+4   9   9.5  RT (min)      lcorbi_t3_pos_1_002_01    m/z 102.0995     0   1E+5   10   10.5  RT (min)      lcorbi_t3_pos_1_006_15    m/z 102.1231     0   2   4E+5   17.5   18  RT (min)      lcorbi_t3_pos_4_099_61    m/z 102.1232     0   2   4E+5   16.5   17  RT (min)      lcorbi_t3_pos_4_099_61    m/z 102.1234     0   2   4E+5   16.5   17  RT (min)      lcorbi_t3_pos_4_099_61    m/z 102.1239     0   2   4   6E+5   17.5   18  RT (min)      lcorbi_t3_pos_4_106_62    m/z 102.1240     0   2   4   6E+5   18   18.5  RT (min)       lcorbi_t3_pos_4_106_62    m/z 102.1240     0   2   4   6E+5   18   18.5  RT (min)      lcorbi_t3_pos_4_084_59    m/z 102.1241     0   2   4E+4   8.5   9  RT (min)      lcorbi_t3_pos_2_063_16    m/z 102.1241     0   2   4   6E+5   18   18.5  RT (min)       lcorbi_t3_pos_4_091_60    m/z 102.1242     0   1   18   18.5  RT (min)     lcorbi_t3_pos_1_025_52    m/z 102.1252     0   1   17.5   18  RT (min)     lcorbi_t3_pos_1_001_49    m/z 102.1253     0   1   2E+5   13   13.5  RT (min)      lcorbi_t3_pos_1_057_49    m/z 102.1254     0   1E+5   12   12.5  RT (min)      lcorbi_t3_pos_1_057_49    m/z 102.1255     0   1   2E+5   12   12.5  RT (min)      lcorbi_t3_pos_4_089_59    m/z 102.1255     0   2   4   6   8E+4   9.5   10  RT (min)      lcorbi_t3_pos_1_062_13    m/z 102.1255     0   1E+5   12.5   13  RT (min)      lcorbi_t3_pos_1_065_51    m/z 102.1255     0   1E+5   12   12.5  RT (min)      lcorbi_t3_pos_1_058_14    m/z 102.1256     0   1E+5   11   11.5  RT (min)      lcorbi_t3_pos_1_061_50    m/z 102.1257     0   1E+5   11.5   12  RT (min)      lcorbi_t3_pos_1_060_39    m/z 102.1257     0   2   4   6E+4   10   10.5  RT (min)      lcorbi_t3_pos_1_064_16    m/z 102.1268     0   1   2   3   4E+7   12.5   13  RT (min)      lcorbi_t3_pos_4_096_60    m/z 102.1277     0   1E+7   10   10.5  RT (min)      lcorbi_t3_pos_1_062_13    m/z 102.1277     0   2   4   6E+6   4.5   5  RT (min)      lcorbi_t3_pos_1_057_49    m/z 102.1277     0   1E+6   0   0.5  RT (min)      lcorbi_t3_pos_1_061_50    m/z 102.1277     0   1E+7   9   9.5  RT (min)      lcorbi_t3_pos_3_059_17    m/z 102.1277     0   1   2E+6   2   2.5  RT (min)      lcorbi_t3_pos_1_061_50    m/z 102.1277     0   1E+6   6   6.5  RT (min)      lcorbi_t3_pos_4_065_51    m/z 102.1277     0   2   4   6   8E+6   7   7.5  RT (min)      lcorbi_t3_pos_1_061_50    m/z 102.1277     0   1E+6   5.5   6  RT (min)      lcorbi_t3_pos_4_064_39    m/z 102.1278     0   1   2E+7   1   1.5  RT (min)      lcorbi_t3_pos_4_057_49    m/z 102.1278     0   1E+7   11.5   12  RT (min)      lcorbi_t3_pos_4_057_49    m/z 102.1278     0   1   2   3   4E+7   13   13.5  RT (min)      lcorbi_t3_pos_1_058_14    m/z 102.1278     0   1   2   3   4E+7   13.5   14  RT (min)      lcorbi_t3_pos_1_059_17    m/z 102.1278     0   1E+8   18   18.5  RT (min)       lcorbi_t3_pos_4_090_60    m/z 102.1278     0   1   2E+7   10.5   11  RT (min)      lcorbi_t3_pos_1_059_17    m/z 102.1278     0   1   2E+7   1.5   2  RT (min)      lcorbi_t3_pos_4_057_49    m/z 102.1278     0   1   17.5   18  RT (min)     lcorbi_t3_pos_1_034_34    m/z 102.1282     0   1   2   3   4E+7   13   13.5  RT (min)      lcorbi_t3_pos_4_089_59    m/z 102.1292     0   2   4   6   8E+7   17.5   18  RT (min)       lcorbi_t3_pos_4_093_60    m/z 102.1294     0   2   4   6E+6   8   8.5  RT (min)      lcorbi_t3_pos_2_060_14    m/z 102.1306     0   1E+6   17.5   18  RT (min)       lcorbi_t3_pos_4_105_61    m/z 102.1307     0   1   2   3E+5   12.5   13  RT (min)      lcorbi_t3_pos_3_057_49    m/z 102.1308     0   1E+6   13   13.5  RT (min)      lcorbi_t3_pos_4_099_61    m/z 102.1308     0   1E+6   14   14.5  RT (min)      lcorbi_t3_pos_4_106_62    m/z 102.1308     0   1   2E+6   15.5   16  RT (min)      lcorbi_t3_pos_4_089_59    m/z 102.1308     0   1   2E+6   16.5   17  RT (min)      lcorbi_t3_pos_4_076_58    m/z 102.1308     0   1   2E+6   15   15.5  RT (min)      lcorbi_t3_pos_4_082_59    m/z 102.1308     0   1E+6   14   14.5  RT (min)      lcorbi_t3_pos_4_111_62    m/z 102.1308     0   1   2E+6   16   16.5  RT (min)      lcorbi_t3_pos_4_077_58    m/z 102.1308     0   1   2E+6   16.5   17  RT (min)      lcorbi_t3_pos_4_078_58    m/z 102.1308     0   1   2E+6   14.5   15  RT (min)      lcorbi_t3_pos_4_090_60    m/z 102.1308     0   1E+5   8.5   9  RT (min)      lcorbi_t3_pos_1_062_13    m/z 102.1308     0   1E+6   13.5   14  RT (min)      lcorbi_t3_pos_4_099_61    m/z 102.1308     0   1E+6   14   14.5  RT (min)      lcorbi_t3_pos_4_083_59    m/z 102.1308     0   1E+6   14   14.5  RT (min)      lcorbi_t3_pos_4_083_59    m/z 102.1308     0   1   2E+6   17   17.5  RT (min)      lcorbi_t3_pos_4_075_58    m/z 102.1308     0   1   2E+6   17   17.5  RT (min)      lcorbi_t3_pos_4_099_61    m/z 102.1309     0   1   2E+6   15   15.5  RT (min)      lcorbi_t3_pos_4_098_61    m/z 102.1309     0   1E+6   14   14.5  RT (min)      lcorbi_t3_pos_4_108_62    m/z 102.1309     0   1   2E+6   15   15.5  RT (min)      lcorbi_t3_pos_4_090_60    m/z 102.1309     0   1   2E+6   15.5   16  RT (min)      lcorbi_t3_pos_4_083_59    m/z 102.1309     0   1E+6   14.5   15  RT (min)      lcorbi_t3_pos_4_091_60    m/z 102.1309     0   1E+6   14   14.5  RT (min)      lcorbi_t3_pos_4_099_61    m/z 102.1309     0   1E+6   14   14.5  RT (min)      lcorbi_t3_pos_4_108_62    m/z 102.1309     0   1E+6   13.5   14  RT (min)      lcorbi_t3_pos_4_102_61    m/z 102.1309     0   1E+6   14.5   15  RT (min)      lcorbi_t3_pos_4_082_59    m/z 102.1309     0   1E+6   14.5   15  RT (min)      lcorbi_t3_pos_4_105_61    m/z 102.1309     0   1   2E+6   16   16.5  RT (min)      lcorbi_t3_pos_4_104_61    m/z 102.1310     0   1E+6   15   15.5  RT (min)      lcorbi_t3_pos_4_101_61    m/z 102.1312     0   1E+5   7.5   8  RT (min)      lcorbi_t3_pos_2_057_49    m/z 102.1313     0   1E+5   8.5   9  RT (min)      lcorbi_t3_pos_2_057_49    m/z 102.1314     0   2   4   6   8E+4   9   9.5  RT (min)      lcorbi_t3_pos_3_059_17    m/z 102.9701     0   1E+5   18   18.5  RT (min)      lcorbi_t3_pos_4_099_61    m/z 102.9702     0   2   4   6   8E+5   1   1.5  RT (min)       lcorbi_t3_pos_4_074_58    m/z 103.1117     0   1   2   3E+7   16   17  RT (min)       lcorbi_t3_pos_1_011_44    m/z 103.1117     0   1E+7   11   12   13  RT (min)       lcorbi_t3_pos_1_058_14    m/z 103.1117     0   1   2   3E+7   15.5   16  RT (min)       lcorbi_t3_pos_1_003_45    m/z 103.1117     0   1E+7   12   12.5  RT (min)      lcorbi_t3_pos_1_059_17    m/z 103.1117     0   2   4   6E+6   1.5   2  RT (min)      lcorbi_t3_pos_4_057_49    m/z 103.1117     0   1E+7   12.5   13  RT (min)      lcorbi_t3_pos_1_061_50    m/z 103.1117     0   1   2E+6   7   7.5  RT (min)      lcorbi_t3_pos_1_061_50    m/z 103.1117     0   1   2E+7   13.5   14  RT (min)      lcorbi_t3_pos_1_058_14    m/z 103.1117     0   1   2   3   4E+6   8.5   9  RT (min)      lcorbi_t3_pos_1_060_39    m/z 103.1117     0   1   2E+6   9   9.5  RT (min)      lcorbi_t3_pos_4_060_16    m/z 103.1117     0   2   4   6   8E+6   10.5   11  RT (min)      lcorbi_t3_pos_1_060_39    m/z 103.1117     0   2   4   6E+6   9   9.5  RT (min)      lcorbi_t3_pos_1_059_17    m/z 103.1117     0   1   2E+6   7.5   8  RT (min)      lcorbi_t3_pos_1_063_15    m/z 103.1117     0   2   4   6E+6   10   10.5  RT (min)      lcorbi_t3_pos_1_065_51    m/z 103.1117     0   1E+7   10.5   11  RT (min)      lcorbi_t3_pos_1_058_14    m/z 103.1117     0   1E+7   13   13.5  RT (min)       lcorbi_t3_pos_1_060_39    m/z 103.1117     0   2   4   6E+6   10   10.5  RT (min)      lcorbi_t3_pos_1_057_49    m/z 103.1117     0   1E+7   12   14  RT (min)       lcorbi_t3_pos_1_058_14    m/z 103.1117     0   2   4   6E+6   9.5   10  RT (min)      lcorbi_t3_pos_1_058_14    m/z 103.1117     0   1   2   3E+7   16   17  RT (min)       lcorbi_t3_pos_1_022_20    m/z 103.1117     0   1   2   3E+7   16   17   18  RT (min)       lcorbi_t3_pos_1_004_24    m/z 103.1117     0   1E+6   7.5   8  RT (min)      lcorbi_t3_pos_2_057_49    m/z 103.1117     0   2   4E+6   11   11.5  RT (min)      lcorbi_t3_pos_2_061_50    m/z 103.1118     0   1   2   3E+7   16   17  RT (min)       lcorbi_t3_pos_1_001_49    m/z 103.1118     0   1   17.5   18  RT (min)     lcorbi_t3_pos_1_027_43    m/z 103.1118     0   1   2   3E+7   16   17  RT (min)       lcorbi_t3_pos_1_033_53    m/z 103.1118     0   2   4   6   8E+6   0.5   1  RT (min)      lcorbi_t3_pos_4_057_49    m/z 103.1118     0   1   17   18  RT (min)     lcorbi_t3_pos_1_008_36    m/z 103.1118     0   1   2E+7   16.5   17   17.5  RT (min)       lcorbi_t3_pos_4_080_58    m/z 103.1118     0   1   17   18  RT (min)     lcorbi_t3_pos_1_030_21    m/z 103.1131     0   1   2E+7   18   18.5  RT (min)       lcorbi_t3_pos_4_078_58    m/z 103.1132     0   1   2E+7   17.5   18  RT (min)       lcorbi_t3_pos_4_082_59    m/z 103.1146     0   1   2   3   4E+5   15   15.5  RT (min)      lcorbi_t3_pos_4_101_61    m/z 103.1146     0   1   2   3E+5   13.5   14  RT (min)      lcorbi_t3_pos_4_066_57    m/z 103.1247     0   1E+5   13   13.5  RT (min)      lcorbi_t3_pos_1_062_13    m/z 103.1247     0   1E+5   12   12.5  RT (min)      lcorbi_t3_pos_1_060_39    m/z 103.1247     0   1E+5   14   14.5  RT (min)      lcorbi_t3_pos_4_057_49    m/z 103.1278     0   1E+4   12   12.5  RT (min)      lcorbi_t3_pos_1_058_14    m/z 103.1279     0   1E+4   15   15.5  RT (min)      lcorbi_t3_pos_4_058_14    m/z 103.1310     0   2   4   6   8E+5   8   8.5  RT (min)      lcorbi_t3_pos_1_062_13    m/z 103.1310     0   1E+5   0.5   1  RT (min)      lcorbi_t3_pos_2_026_44    m/z 103.1310     0   1E+6   1.5   2  RT (min)      lcorbi_t3_pos_4_057_49    m/z 103.1310     0   1E+6   11   11.5  RT (min)      lcorbi_t3_pos_1_061_50    m/z 103.1310     0   1E+5   0.5   1  RT (min)      lcorbi_t3_pos_2_016_23    m/z 103.1310     0   1E+6   10   10.5  RT (min)      lcorbi_t3_pos_1_062_13    m/z 103.1310     0   1E+6   10.5   11  RT (min)      lcorbi_t3_pos_1_063_15    m/z 103.1310     0   1   2E+6   11.5   12  RT (min)      lcorbi_t3_pos_1_060_39    m/z 103.1310     0   1E+6   9   9.5  RT (min)      lcorbi_t3_pos_1_064_16    m/z 103.1310     0   1E+6   12.5   13  RT (min)      lcorbi_t3_pos_4_059_15    m/z 103.1310     0   1E+5   0.5   1  RT (min)      lcorbi_t3_pos_1_060_39    m/z 103.1310     0   1E+6   1   1.5  RT (min)      lcorbi_t3_pos_4_057_49    m/z 103.1310     0   1E+6   9.5   10  RT (min)      lcorbi_t3_pos_1_061_50    m/z 103.1310     0   1E+6   10.5   11  RT (min)      lcorbi_t3_pos_1_058_14    m/z 103.1310     0   1E+5   0   0.5  RT (min)      lcorbi_t3_pos_1_057_49    m/z 103.1310     0   1E+6   10   10.5  RT (min)      lcorbi_t3_pos_1_057_49    m/z 103.1310     0   2   4   6   8E+5   8.5   9  RT (min)      lcorbi_t3_pos_1_061_50    m/z 103.1310     0   1E+5   1   1.5  RT (min)      lcorbi_t3_pos_2_028_24    m/z 103.1310     0   1E+5   0   0.5  RT (min)      lcorbi_t3_pos_2_016_23    m/z 103.1310     0   1   2E+5   2.5   3  RT (min)      lcorbi_t3_pos_4_108_62    m/z 103.1325     0   1   2   3   4E+6   14   14.5  RT (min)      lcorbi_t3_pos_1_058_14    m/z 104.1069     0   1   2   3E+5   12   12.5  RT (min)      lcorbi_t3_pos_1_031_40    m/z 104.1069     0   2   4   6   8E+4   12.5   13  RT (min)      lcorbi_t3_pos_1_023_02    m/z 104.1069     0   1   2   3E+5   12   12.5  RT (min)      lcorbi_t3_pos_1_031_40    m/z 104.1150     0   1   2E+6   16   16.5  RT (min)       lcorbi_t3_pos_1_008_36    m/z 104.1150     0   2   4   6E+5   0.5   1  RT (min)      lcorbi_t3_pos_4_057_49    m/z 104.1150     0   1   17.5   18  RT (min)     lcorbi_t3_pos_1_051_09    m/z 104.1150     0   1E+6   16   17   18  RT (min)       lcorbi_t3_pos_4_094_60    m/z 104.1150     0   1   2E+6   16   17  RT (min)       lcorbi_t3_pos_1_011_44    m/z 104.1150     0   1E+6   12.5   13  RT (min)      lcorbi_t3_pos_1_057_49    m/z 104.1150     0   1   17  RT (min)     lcorbi_t3_pos_1_038_30    m/z 104.1150     0   1   2E+6   15   16  RT (min)       lcorbi_t3_pos_1_004_24    m/z 104.1150     0   1E+6   11   12   13  RT (min)       lcorbi_t3_pos_1_036_08    m/z 104.1150     0   1   17   18  RT (min)     lcorbi_t3_pos_1_021_18    m/z 104.1150     0   1   17  RT (min)     lcorbi_t3_pos_1_028_32    m/z 104.1150     0   2   4E+5   10   10.5  RT (min)      lcorbi_t3_pos_1_057_49    m/z 104.1150     0   2   4   6E+5   0.5   1  RT (min)      lcorbi_t3_pos_4_057_49    m/z 104.1150     0   1   17   17.5   18  RT (min)     lcorbi_t3_pos_1_001_49    m/z 105.0032     0   1E+5   2   2.5  RT (min)      lcorbi_t3_pos_2_050_18    m/z 105.0032     0   1   2E+5   3.5   4  RT (min)       lcorbi_t3_pos_4_106_62    m/z 105.0032     0   1E+5   2   2.5  RT (min)      lcorbi_t3_pos_3_024_07    m/z 105.0032     0   1E+5   2   2.5  RT (min)      lcorbi_t3_pos_1_064_16    m/z 105.0033     0   1E+5   2   2.5  RT (min)      lcorbi_t3_pos_2_050_18    m/z 105.1160     0   1E+5   15.5   16  RT (min)      lcorbi_t3_pos_1_024_04    m/z 107.0491     0   1E+5   9.5   10  RT (min)      lcorbi_t3_pos_3_030_22    m/z 107.0491     0   2   4E+5   13  RT (min)       lcorbi_t3_pos_1_029_29    m/z 107.0491     0   1E+5   10   10.5  RT (min)      lcorbi_t3_pos_3_033_53    m/z 107.0491     0   1   2   3   4E+5   12   14  RT (min)       lcorbi_t3_pos_4_087_59    m/z 107.0491     0   2   4E+5   16   17  RT (min)       lcorbi_t3_pos_1_013_10    m/z 107.0702     0   1E+5   0   0.5  RT (min)      lcorbi_t3_pos_1_031_40    m/z 107.0702     0   1   2E+5   6.5   7  RT (min)      lcorbi_t3_pos_3_000_aa    m/z 107.0702     0   1   2E+5   12.5   13  RT (min)      lcorbi_t3_pos_3_020_35    m/z 107.0855     0   1E+5   15.5   16  RT (min)      lcorbi_t3_pos_1_023_02    m/z 107.0855     0   1E+5   14.5   15  RT (min)      lcorbi_t3_pos_1_034_34    m/z 111.0440     0   2   4   6E+4   14   14.5  RT (min)      lcorbi_t3_pos_2_022_38    m/z 111.0440     0   1   2   3   4E+5   12.5   13  RT (min)      lcorbi_t3_pos_1_002_01    m/z 111.0440     0   2   4E+4   12   12.5  RT (min)      lcorbi_t3_pos_2_032_39    m/z 111.0440     0   1E+5   0   0.5  RT (min)      lcorbi_t3_pos_1_045_12    m/z 111.0440     0   1E+5   15.5   16  RT (min)      lcorbi_t3_pos_1_023_02    m/z 111.0440     0   1   2   3   4E+5   13.5   14  RT (min)      lcorbi_t3_pos_1_008_36    m/z 111.0440     0   1E+5   0.5   1  RT (min)      lcorbi_t3_pos_1_044_25    m/z 111.0440     0   1E+5   0.5   1  RT (min)      lcorbi_t3_pos_1_044_25    m/z 111.0440     0   1E+5   0   0.5  RT (min)      lcorbi_t3_pos_1_044_25    m/z 111.0440     0   2   4E+4   12.5   13  RT (min)      lcorbi_t3_pos_2_013_33    m/z 111.0440     0   1E+5   14   14.5  RT (min)      lcorbi_t3_pos_1_028_32    m/z 111.0440     0   1   2   3   4E+5   13.5   14  RT (min)      lcorbi_t3_pos_4_019_47    m/z 111.0440     0   2   4   6   8E+4   14   14.5  RT (min)      lcorbi_t3_pos_1_030_21    m/z 111.0440     0   2   4   6E+5   13.5   14  RT (min)       lcorbi_t3_pos_1_005_42    m/z 111.0440     0   1   2   3E+5   10   10.5  RT (min)       lcorbi_t3_pos_4_074_58    m/z 111.0440     0   1E+5   0.5   1  RT (min)      lcorbi_t3_pos_2_009_50    m/z 111.0440     0   1E+5   15   15.5  RT (min)      lcorbi_t3_pos_1_031_40    m/z 111.0440     0   1E+5   0.5   1  RT (min)      lcorbi_t3_pos_1_047_41    m/z 111.0440     0   2   4   6   8E+4   14.5   15  RT (min)      lcorbi_t3_pos_1_034_34    m/z 111.0440     0   2   4   6E+4   15   15.5  RT (min)      lcorbi_t3_pos_2_012_01    m/z 111.0440     0   2   4   6   8E+4   14.5   15  RT (min)      lcorbi_t3_pos_1_034_34    m/z 111.0440     0   1E+5   0.5   1  RT (min)      lcorbi_t3_pos_1_047_41    m/z 111.0917     0   1   2E+6   0   0.5  RT (min)      lcorbi_t3_pos_1_001_49    m/z 111.1168     0   1   2   3E+5   10   10.5  RT (min)      lcorbi_t3_pos_2_022_38    m/z 111.1168     0   1E+5   10   10.5  RT (min)      lcorbi_t3_pos_2_014_37    m/z 112.0505     0   1   2   3   4E+5   13.5   14  RT (min)       lcorbi_t3_pos_1_001_49    m/z 112.0505     0   2   4E+5   10   12  RT (min)       lcorbi_t3_pos_1_035_28    m/z 112.0505     0   2   4E+5   5   10  RT (min)       lcorbi_t3_pos_1_008_36    m/z 112.0505     0   1   2   3   4E+5   5   10  RT (min)       lcorbi_t3_pos_1_002_01    m/z 112.0505     0   2   4E+5   10   12  RT (min)       lcorbi_t3_pos_1_032_26    m/z 112.0505     0   1   2E+5   15   16  RT (min)       lcorbi_t3_pos_1_025_52    m/z 113.0597     0   2   4   6E+5   8   9  RT (min)       lcorbi_t3_pos_1_006_15    m/z 113.0597     0   1E+6   11   12  RT (min)       lcorbi_t3_pos_1_044_25    m/z 113.0597     0   2   4   6E+5   6   8   10  RT (min)       lcorbi_t3_pos_1_010_46    m/z 113.0597     0   2   4   6E+5   10   12  RT (min)       lcorbi_t3_pos_1_051_09    m/z 113.0597     0   2   4E+5   18   18.5  RT (min)       lcorbi_t3_pos_4_080_58    m/z 113.0597     0   2   4   6   8E+5   6   8  RT (min)       lcorbi_t3_pos_1_002_01    m/z 113.0597     0   2   4   6E+5   7.5   8   8.5  RT (min)       lcorbi_t3_pos_1_012_14    m/z 113.0597     0   2   4   6E+5   9.5   10  RT (min)       lcorbi_t3_pos_1_028_32    m/z 113.1324     0   2   4E+6   12.5   13  RT (min)      lcorbi_t3_pos_4_016_15    m/z 113.1324     0   1E+5   14   14.5  RT (min)      lcorbi_t3_pos_2_038_03    m/z 113.1324     0   1E+5   14.5   15  RT (min)      lcorbi_t3_pos_1_034_34    m/z 113.1324     0   2   4E+6   13.5   14  RT (min)      lcorbi_t3_pos_3_047_40    m/z 113.1324     0   1   2   3   4E+4   12.5   13  RT (min)      lcorbi_t3_pos_4_012_37    m/z 113.1324     0   1   2   3E+6   13.5   14  RT (min)      lcorbi_t3_pos_4_001_49    m/z 113.1324     0   1E+5   15.5   16  RT (min)      lcorbi_t3_pos_1_023_02    m/z 113.1324     0   1E+5   14.5   15  RT (min)      lcorbi_t3_pos_1_031_40    m/z 113.1324     0   1E+5   15   15.5  RT (min)      lcorbi_t3_pos_2_012_01    m/z 113.1324     0   1E+6   13.5   14  RT (min)      lcorbi_t3_pos_3_025_52    m/z 113.1324     0   1E+5   14   14.5  RT (min)      lcorbi_t3_pos_3_003_33    m/z 113.1325     0   1E+5   15   15.5  RT (min)      lcorbi_t3_pos_2_056_56    m/z 113.1325     0   2   4   6E+5   12.5   13  RT (min)      lcorbi_t3_pos_2_054_29    m/z 113.1325     0   1E+6   12.5   13  RT (min)      lcorbi_t3_pos_1_002_01    m/z 113.1325     0   1E+5   14   14.5  RT (min)      lcorbi_t3_pos_2_006_47    m/z 114.0913     0   1E+6   5   10  RT (min)       lcorbi_t3_pos_3_000_aa    m/z 114.1358     0   1   2E+5   12.5   13  RT (min)      lcorbi_t3_pos_2_054_29    m/z 114.1358     0   2   4   6   8E+3   12.5   13  RT (min)      lcorbi_t3_pos_4_012_37    m/z 115.0066     0   2   4   6   8E+4   7   7.5  RT (min)       lcorbi_t3_pos_4_090_60    m/z 115.0754     0   1   2   3E+5   15.5   16  RT (min)       lcorbi_t3_pos_1_057_49    m/z 115.0754     0   1E+5   0   0.5  RT (min)       lcorbi_t3_pos_1_055_33    m/z 116.0706     0   2   4   6   8E+4   4.5   5  RT (min)       lcorbi_t3_pos_4_066_57    m/z 116.1070     0   1E+5   2   2.5  RT (min)      lcorbi_t3_pos_4_073_57    m/z 116.1429     0   1E+7   16.5   17  RT (min)      lcorbi_t3_pos_4_069_57    m/z 116.1433     0   2   4   6E+6   1   1.5  RT (min)      lcorbi_t3_pos_4_057_49    m/z 116.1434     0   1   2   3E+7   18   18.5  RT (min)       lcorbi_t3_pos_4_074_58    m/z 116.1434     0   1   2E+6   7   7.5  RT (min)      lcorbi_t3_pos_1_061_50    m/z 116.1434     0   1   2   3   4E+6   9.5   10  RT (min)      lcorbi_t3_pos_1_061_50    m/z 116.1434     0   1   2E+6   9   9.5  RT (min)      lcorbi_t3_pos_3_059_17    m/z 116.1434     0   1   2E+6   4.5   5  RT (min)      lcorbi_t3_pos_1_057_49    m/z 116.1434     0   1   2E+6   8.5   9  RT (min)      lcorbi_t3_pos_1_062_13    m/z 116.1434     0   1   2E+6   8   8.5  RT (min)      lcorbi_t3_pos_3_057_49    m/z 116.1434     0   2   4   6E+6   1.5   2  RT (min)      lcorbi_t3_pos_4_057_49    m/z 116.1434     0   1   2   3E+6   8.5   9  RT (min)      lcorbi_t3_pos_2_057_49    m/z 116.1434     0   2   4   6E+5   6   6.5  RT (min)      lcorbi_t3_pos_4_061_50    m/z 116.1434     0   1E+7   12   14  RT (min)       lcorbi_t3_pos_1_059_17    m/z 116.1434     0   1E+6   7.5   8  RT (min)      lcorbi_t3_pos_2_057_49    m/z 116.1449     0   1   2   3E+7   17   18  RT (min)       lcorbi_t3_pos_4_075_58    m/z 116.1454     0   2   4E+4   10   10.5  RT (min)      lcorbi_t3_pos_1_062_13    m/z 116.1454     0   2   4   6E+4   11   11.5  RT (min)      lcorbi_t3_pos_1_061_50    m/z 116.1455     0   2   4   6   8E+4   10.5   11  RT (min)      lcorbi_t3_pos_1_057_49    m/z 116.1458     0   1   2E+5   14   14.5  RT (min)      lcorbi_t3_pos_1_058_14    m/z 116.1465     0   1   2E+5   13   13.5   14  RT (min)       lcorbi_t3_pos_4_066_57    m/z 116.1466     0   1   2   3   4E+4   9.5   10  RT (min)      lcorbi_t3_pos_2_059_39    m/z 116.1466     0   2   4   6E+5   18   18.5  RT (min)       lcorbi_t3_pos_4_091_60    m/z 116.1466     0   1   2   3   4E+5   14.5   15  RT (min)      lcorbi_t3_pos_4_108_62    m/z 116.1466     0   1   2E+5   12   13  RT (min)       lcorbi_t3_pos_4_067_57    m/z 116.1466     0   2   4   6   8E+4   11.5   12  RT (min)      lcorbi_t3_pos_1_060_39    m/z 116.1466     0   1E+5   12   12.5  RT (min)      lcorbi_t3_pos_1_058_14    m/z 116.1466     0   1   2E+5   12.5   13  RT (min)      lcorbi_t3_pos_4_105_61    m/z 116.1467     0   1   2E+5   12.5   13  RT (min)      lcorbi_t3_pos_4_104_61    m/z 116.1467     0   1   2   3   4E+5   10   12   14  RT (min)       lcorbi_t3_pos_4_108_62    m/z 116.1467     0   1   2E+5   13   13.5  RT (min)       lcorbi_t3_pos_4_069_57    m/z 116.1467     0   1   2   3   4E+5   17.5   18  RT (min)      lcorbi_t3_pos_4_069_57    m/z 116.1467     0   2   4E+5   18   18.5  RT (min)       lcorbi_t3_pos_4_076_58    m/z 116.1468     0   1   2   3E+5   13.5   14  RT (min)      lcorbi_t3_pos_4_101_61    m/z 116.1468     0   2   4E+5   15.5   16  RT (min)      lcorbi_t3_pos_4_107_62    m/z 116.1468     0   2   4E+5   17.5   18  RT (min)       lcorbi_t3_pos_4_105_61    m/z 116.1469     0   2   4E+5   16.5   17  RT (min)      lcorbi_t3_pos_4_075_58    m/z 116.1469     0   2   4E+5   16   16.5  RT (min)      lcorbi_t3_pos_4_075_58    m/z 116.1469     0   2   4E+5   17   17.5  RT (min)      lcorbi_t3_pos_4_078_58    m/z 116.1470     0   1   2   3   4E+5   16.5   17  RT (min)      lcorbi_t3_pos_4_069_57    m/z 116.9661     0   1   2   3E+5   0   0.5  RT (min)      lcorbi_t3_pos_1_031_40    m/z 116.9661     0   1   2   3E+5   1   1.5  RT (min)      lcorbi_t3_pos_1_019_16    m/z 116.9661     0   1   2   3E+5   0   0.5  RT (min)      lcorbi_t3_pos_1_029_29    m/z 116.9662     0   1   2E+5   3.5   4  RT (min)      lcorbi_t3_pos_1_057_49    m/z 116.9662     0   1   2   3E+5   0.5   1  RT (min)      lcorbi_t3_pos_1_016_05    m/z 117.0222     0   1   2E+5   5.5   6  RT (min)      lcorbi_t3_pos_1_058_14    m/z 117.0222     0   1E+5   0   0.5  RT (min)      lcorbi_t3_pos_1_008_36    m/z 117.0222     0   2   4E+5   18   18.5  RT (min)      lcorbi_t3_pos_4_090_60    m/z 117.0222     0   2   4E+5   18   18.5  RT (min)      lcorbi_t3_pos_4_075_58    m/z 117.0222     0   1   2E+5   0.5   1  RT (min)      lcorbi_t3_pos_1_008_36    m/z 117.0222     0   2   4E+5   17.5   18  RT (min)      lcorbi_t3_pos_4_093_60    m/z 117.0222     0   1   2E+5   0   1  RT (min)       lcorbi_t3_pos_1_006_15    m/z 117.0222     0   1E+5   0   0.5  RT (min)      lcorbi_t3_pos_1_008_36    m/z 117.0223     0   1E+5   1   1.5  RT (min)      lcorbi_t3_pos_1_007_07    m/z 117.0223     0   1E+5   1.5   2  RT (min)       lcorbi_t3_pos_1_007_07    m/z 117.0223     0   1   2   3E+5   1.5   2  RT (min)       lcorbi_t3_pos_4_075_58    m/z 117.0223     0   2   4   6   8E+4   2.5   3  RT (min)      lcorbi_t3_pos_2_003_19    m/z 117.0223     0   1E+5   1.5   2  RT (min)       lcorbi_t3_pos_1_007_07    m/z 117.0223     0   2   4   6   8E+4   2.5   3  RT (min)      lcorbi_t3_pos_2_000_cc    m/z 117.0223     0   1E+5   1   1.5  RT (min)       lcorbi_t3_pos_4_071_57    m/z 117.0223     0   1   2   3E+5   1   1.5  RT (min)       lcorbi_t3_pos_4_077_58    m/z 117.1404     0   1E+5   17.5   18   18.5  RT (min)       lcorbi_t3_pos_4_082_59    m/z 117.1404     0   1E+5   17.5   18  RT (min)       lcorbi_t3_pos_4_093_60    m/z 117.1466     0   1   2E+6   18   18.5  RT (min)       lcorbi_t3_pos_4_090_60    m/z 117.1467     0   1   2E+6   16   18  RT (min)       lcorbi_t3_pos_4_098_61    m/z 117.1467     0   2   4   6E+5   12.5   13  RT (min)      lcorbi_t3_pos_1_062_13    m/z 117.1467     0   1E+6   13.5   14  RT (min)       lcorbi_t3_pos_1_008_36    m/z 117.1467     0   2   4   6E+5   13.5   14  RT (min)      lcorbi_t3_pos_4_057_49    m/z 117.1467     0   2   4   6E+5   12   12.5  RT (min)      lcorbi_t3_pos_1_063_15    m/z 117.1467     0   1   2E+6   18   18.5  RT (min)       lcorbi_t3_pos_4_092_60    m/z 117.1467     0   2   4E+5   11.5   12  RT (min)      lcorbi_t3_pos_1_057_49    m/z 117.1467     0   2   4E+5   1   1.5  RT (min)      lcorbi_t3_pos_4_057_49    m/z 117.9586     0   1E+6   0   0.5  RT (min)      lcorbi_t3_pos_1_011_44    m/z 118.0862     0   1E+7   1.5   2  RT (min)       lcorbi_t3_pos_4_074_58    m/z 118.9426     0   2   4E+6   0   0.5  RT (min)      lcorbi_t3_pos_1_005_42    m/z 119.1066     0   1   16.5   17  RT (min)     lcorbi_t3_pos_1_055_33    m/z 119.1066     0   1E+6   14   14.5  RT (min)      lcorbi_t3_pos_1_057_49    m/z 119.1066     0   1   17  RT (min)     lcorbi_t3_pos_1_038_30    m/z 119.1066     0   1   17.5   18  RT (min)     lcorbi_t3_pos_2_014_37    m/z 119.1066     0   1E+6   14.5   15  RT (min)       lcorbi_t3_pos_1_006_15    m/z 119.1066     0   1   16.5   17   17.5  RT (min)     lcorbi_t3_pos_1_034_34    m/z 119.1066     0   1E+6   13.5   14  RT (min)       lcorbi_t3_pos_1_058_14    m/z 119.1066     0   1   2E+6   16   17   18  RT (min)       lcorbi_t3_pos_1_050_13    m/z 119.1066     0   1E+6   15   16  RT (min)       lcorbi_t3_pos_1_059_17    m/z 119.1066     0   1   2E+6   16  RT (min)       lcorbi_t3_pos_1_053_39    m/z 119.1066     0   1   17   18  RT (min)     lcorbi_t3_pos_1_051_09    m/z 119.1067     0   2   4   6E+5   9.5   10  RT (min)      lcorbi_t3_pos_1_059_17    m/z 119.1067     0   1E+6   12   12.5  RT (min)      lcorbi_t3_pos_1_059_17    m/z 119.1067     0   1E+6   11   11.5  RT (min)      lcorbi_t3_pos_1_058_14    m/z 119.1067     0   1E+6   13   13.5  RT (min)       lcorbi_t3_pos_1_058_14    m/z 119.1067     0   1E+6   11.5   12  RT (min)      lcorbi_t3_pos_1_057_49    m/z 119.1067     0   1E+6   11.5   12  RT (min)      lcorbi_t3_pos_1_060_39    m/z 119.1067     0   1E+6   10.5   11  RT (min)      lcorbi_t3_pos_1_058_14    m/z 119.1067     0   1E+6   12.5   13  RT (min)      lcorbi_t3_pos_1_057_49    m/z 119.1067     0   1E+6   10   12  RT (min)       lcorbi_t3_pos_1_060_39    m/z 119.1067     0   2   4   6E+5   9   9.5  RT (min)      lcorbi_t3_pos_1_057_49    m/z 119.1067     0   2   4   6   8E+5   10   10.5  RT (min)      lcorbi_t3_pos_1_057_49    m/z 120.0808     0   2   4   6   8E+5   0   0.5  RT (min)      lcorbi_t3_pos_1_001_49    m/z 120.9808     0   1   2   3   4E+5   0.5   1  RT (min)      lcorbi_t3_pos_4_076_58    m/z 120.9808     0   1   2E+5   0   0.5  RT (min)      lcorbi_t3_pos_4_079_58    m/z 121.0284     0   1   2   3   4E+6   12.5   13  RT (min)      lcorbi_t3_pos_1_002_01    m/z 121.0284     0   1   2E+6   13.5   14  RT (min)      lcorbi_t3_pos_4_001_49    m/z 121.0284     0   1E+6   13.5   14  RT (min)      lcorbi_t3_pos_1_008_36    m/z 121.0284     0   1   2   3E+5   16   16.5  RT (min)      lcorbi_t3_pos_1_023_02    m/z 121.0284     0   1   2   3E+5   15.5   16  RT (min)      lcorbi_t3_pos_1_023_02    m/z 121.0284     0   1   2   3E+5   15   15.5  RT (min)      lcorbi_t3_pos_1_051_09    m/z 121.0284     0   2   4E+4   12.5   13  RT (min)      lcorbi_t3_pos_2_013_33    m/z 121.0284     0   1   2   3E+6   13.5   14  RT (min)      lcorbi_t3_pos_1_008_36    m/z 121.0284     0   1   2   3E+5   14.5   15  RT (min)      lcorbi_t3_pos_1_034_34    m/z 121.0284     0   1E+6   10   10.5  RT (min)      lcorbi_t3_pos_1_002_01    m/z 121.0284     0   1E+6   10.5   11  RT (min)      lcorbi_t3_pos_1_008_36    m/z 121.0284     0   1   2E+5   11   11.5  RT (min)      lcorbi_t3_pos_1_028_32    m/z 121.0284     0   1   2   3E+5   15.5   16  RT (min)      lcorbi_t3_pos_1_023_02    m/z 121.0284     0   1   2   3E+5   11   11.5  RT (min)      lcorbi_t3_pos_1_031_40    m/z 121.0284     0   1   2   3E+5   14.5   15  RT (min)      lcorbi_t3_pos_1_041_54    m/z 121.0284     0   1   2   3   4E+5   14   14.5  RT (min)      lcorbi_t3_pos_1_030_21    m/z 121.0284     0   1   2   3E+5   14   14.5  RT (min)      lcorbi_t3_pos_1_028_32    m/z 121.0284     0   1   2   3E+5   14.5   15  RT (min)      lcorbi_t3_pos_1_031_40    m/z 121.1223     0   1   2E+6   7.5   8  RT (min)      lcorbi_t3_pos_2_057_49    m/z 121.1223     0   1   2   3E+6   7   7.5  RT (min)      lcorbi_t3_pos_1_059_17    m/z 121.1223     0   1E+6   6.5   7  RT (min)      lcorbi_t3_pos_2_061_50    m/z 121.1223     0   2   4E+6   8   8.5  RT (min)      lcorbi_t3_pos_1_059_17    m/z 121.1223     0   2   4   6E+6   8.5   9  RT (min)      lcorbi_t3_pos_1_060_39    m/z 121.1223     0   1E+7   1.5   2  RT (min)      lcorbi_t3_pos_4_057_49    m/z 121.1223     0   1   2   3E+6   7.5   8  RT (min)      lcorbi_t3_pos_1_063_15    m/z 121.1224     0   1E+7   0.5   1  RT (min)      lcorbi_t3_pos_4_057_49    m/z 121.1224     0   2   4   6E+7   15.5   16  RT (min)       lcorbi_t3_pos_1_003_45    m/z 121.1243     0   2   4   6   8E+4   9   9.5  RT (min)      lcorbi_t3_pos_1_059_17    m/z 121.1244     0   1E+5   9   9.5  RT (min)      lcorbi_t3_pos_1_057_49    m/z 121.1245     0   2   4   6E+4   9.5   10  RT (min)      lcorbi_t3_pos_2_057_49    m/z 121.1245     0   1E+6   15.5   16  RT (min)       lcorbi_t3_pos_1_004_24    m/z 121.1246     0   1   19   19.5  RT (min)    lcorbi_t3_pos_1_023_02    m/z 121.1246     0   1E+6   14   15  RT (min)       lcorbi_t3_pos_1_002_01    m/z 121.1246     0   2   4E+5   13.5   14  RT (min)      lcorbi_t3_pos_1_058_14    m/z 121.1247     0   1   2   3   4E+5   12.5   13  RT (min)      lcorbi_t3_pos_1_058_14    m/z 121.1247     0   1E+6   16   16.5  RT (min)       lcorbi_t3_pos_1_006_15    m/z 121.1247     0   1   2E+5   12.5   13  RT (min)      lcorbi_t3_pos_1_060_39    m/z 121.1248     0   1   2E+5   12   12.5  RT (min)      lcorbi_t3_pos_1_059_17    m/z 121.1250     0   1   19   19.5  RT (min)    lcorbi_t3_pos_1_023_02    m/z 121.1255     0   1E+5   10.5   11  RT (min)      lcorbi_t3_pos_1_058_14    m/z 121.1257     0   1E+5   10   10.5  RT (min)      lcorbi_t3_pos_1_057_49    m/z 121.1257     0   2   4E+5   13   13.5  RT (min)      lcorbi_t3_pos_4_096_60    m/z 121.1257     0   1   2   3   4E+5   11   11.5  RT (min)      lcorbi_t3_pos_4_090_60    m/z 121.1257     0   1E+5   9.5   10  RT (min)      lcorbi_t3_pos_1_060_39    m/z 121.1258     0   1E+5   10.5   11  RT (min)      lcorbi_t3_pos_1_060_39    m/z 121.1258     0   2   4   6   8E+4   11   11.5  RT (min)      lcorbi_t3_pos_4_057_49    m/z 121.1258     0   1   2   3   4E+5   11   11.5  RT (min)      lcorbi_t3_pos_4_090_60    m/z 121.1258     0   1   2   3   4E+5   11   11.5  RT (min)      lcorbi_t3_pos_4_090_60    m/z 121.1258     0   1   2   3E+5   11.5   12  RT (min)      lcorbi_t3_pos_4_095_60    m/z 121.1258     0   1   2E+5   11.5   12  RT (min)      lcorbi_t3_pos_1_060_39    m/z 121.1258     0   1   2E+5   12   12.5  RT (min)      lcorbi_t3_pos_1_062_13    m/z 121.1259     0   1   2   3E+5   11   11.5  RT (min)      lcorbi_t3_pos_4_079_58    m/z 121.1259     0   1E+5   11   11.5  RT (min)      lcorbi_t3_pos_2_061_50    m/z 121.1259     0   2   4   6   8E+4   10   10.5  RT (min)      lcorbi_t3_pos_1_065_51    m/z 121.1259     0   2   4   6E+5   17.5   18  RT (min)       lcorbi_t3_pos_4_107_62    m/z 121.1259     0   1E+6   15   15.5  RT (min)       lcorbi_t3_pos_1_012_14    m/z 121.1259     0   1   2   3   4E+5   11.5   12  RT (min)      lcorbi_t3_pos_4_094_60    m/z 121.1259     0   1   2   3   4E+5   11.5   12  RT (min)      lcorbi_t3_pos_4_094_60    m/z 121.1260     0   2   4   6   8E+5   17   17.5  RT (min)      lcorbi_t3_pos_4_075_58    m/z 121.1260     0   2   4E+5   12   12.5  RT (min)      lcorbi_t3_pos_4_091_60    m/z 121.1260     0   2   4E+5   12.5   13  RT (min)      lcorbi_t3_pos_4_091_60    m/z 121.1260     0   2   4E+5   16   17   18  RT (min)       lcorbi_t3_pos_2_009_50    m/z 121.1260     0   2   4   6E+5   14   16  RT (min)       lcorbi_t3_pos_4_098_61    m/z 121.1260     0   1   16.5   17  RT (min)     lcorbi_t3_pos_1_055_33    m/z 121.1260     0   2   4   6E+5   12   14   16  RT (min)       lcorbi_t3_pos_1_048_47    m/z 121.1260     0   1   2   3E+5   12.5   13  RT (min)      lcorbi_t3_pos_4_103_61    m/z 121.1260     0   2   4   6   8E+5   16   16.5  RT (min)      lcorbi_t3_pos_4_076_58    m/z 121.1260     0   1   2   3E+5   10.5   11  RT (min)      lcorbi_t3_pos_4_091_60    m/z 121.1260     0   2   4   6E+5   16.5   17  RT (min)      lcorbi_t3_pos_4_110_62    m/z 121.1260     0   2   4   6E+5   16.5   17  RT (min)       lcorbi_t3_pos_4_094_60    m/z 121.1260     0   2   4   6   8E+5   17   17.5   18  RT (min)       lcorbi_t3_pos_4_076_58    m/z 121.1260     0   2   4   6   8E+5   17   17.5  RT (min)      lcorbi_t3_pos_4_076_58    m/z 121.1260     0   2   4   6   8E+5   17   17.5  RT (min)      lcorbi_t3_pos_4_078_58    m/z 121.1260     0   1   2   3   4E+5   14   14.5  RT (min)      lcorbi_t3_pos_1_063_15    m/z 121.1260     0   2   4   6E+5   15   16   17  RT (min)       lcorbi_t3_pos_4_066_57    m/z 121.1260     0   2   4   6   8E+5   15   16  RT (min)       lcorbi_t3_pos_4_091_60    m/z 121.1260     0   2   4   6E+5   14   15   16  RT (min)       lcorbi_t3_pos_4_099_61    m/z 121.1260     0   2   4   6E+5   17.5   18  RT (min)      lcorbi_t3_pos_4_111_62    m/z 121.1260     0   2   4   6E+5   15.5   16  RT (min)      lcorbi_t3_pos_4_081_58    m/z 121.1260     0   1   17   17.5  RT (min)     lcorbi_t3_pos_1_001_49    m/z 121.1260     0   1   2   3   4E+5   12   12.5  RT (min)      lcorbi_t3_pos_4_097_60    m/z 121.1260     0   1   17   17.5  RT (min)     lcorbi_t3_pos_1_028_32    m/z 121.1260     0   1   17.5   18  RT (min)     lcorbi_t3_pos_1_027_43    m/z 121.1260     0   2   4   6E+5   18   18.5  RT (min)      lcorbi_t3_pos_4_094_60    m/z 121.1260     0   2   4   6E+5   16.5   17  RT (min)      lcorbi_t3_pos_4_081_58    m/z 121.1260     0   2   4   6E+5   14   14.5  RT (min)      lcorbi_t3_pos_4_088_59    m/z 121.1260     0   1   2   3E+5   11.5   12  RT (min)      lcorbi_t3_pos_4_095_60    m/z 121.1261     0   2   4E+5   13.5   14  RT (min)      lcorbi_t3_pos_4_074_58    m/z 121.1261     0   2   4   6E+5   14.5   15  RT (min)      lcorbi_t3_pos_4_084_59    m/z 121.1261     0   2   4   6E+5   14.5   15  RT (min)      lcorbi_t3_pos_4_082_59    m/z 121.1261     0   1   2   3   4E+5   13   13.5  RT (min)      lcorbi_t3_pos_4_102_61    m/z 121.1261     0   2   4   6   8E+5   17   17.5  RT (min)       lcorbi_t3_pos_4_097_60    m/z 121.1261     0   2   4E+5   13   13.5  RT (min)      lcorbi_t3_pos_4_070_57    m/z 121.1261     0   2   4   6E+5   16   16.5  RT (min)      lcorbi_t3_pos_4_092_60    m/z 121.1261     0   1E+6   16  RT (min)       lcorbi_t3_pos_1_035_28    m/z 121.1261     0   2   4   6E+5   12   14  RT (min)       lcorbi_t3_pos_4_090_60    m/z 121.1261     0   2   4   6   8E+5   16  RT (min)       lcorbi_t3_pos_4_093_60    m/z 121.1262     0   2   4E+5   15   15.5  RT (min)      lcorbi_t3_pos_4_066_57    m/z 121.1262     0   2   4   6E+5   17   17.5  RT (min)      lcorbi_t3_pos_4_090_60    m/z 121.1262     0   2   4E+5   14   14.5  RT (min)      lcorbi_t3_pos_4_111_62    m/z 121.1262     0   2   4   6E+5   14.5   15  RT (min)      lcorbi_t3_pos_4_090_60    m/z 121.1262     0   1   2   3   4E+5   13.5   14  RT (min)      lcorbi_t3_pos_4_078_58    m/z 121.1263     0   2   4E+5   14   14.5  RT (min)      lcorbi_t3_pos_4_112_62    m/z 121.1272     0   1   2   3E+5   18   18.5  RT (min)      lcorbi_t3_pos_4_094_60    m/z 122.0317     0   1   2   3   4E+4   15   15.5  RT (min)      lcorbi_t3_pos_1_051_09    m/z 122.0317     0   2   4E+4   15.5   16  RT (min)      lcorbi_t3_pos_1_023_02    m/z 122.0317     0   1   2   3   4E+5   13.5   14  RT (min)      lcorbi_t3_pos_1_005_42    m/z 122.0317     0   1   2   3   4E+4   14   14.5  RT (min)      lcorbi_t3_pos_1_028_32    m/z 122.0317     0   1   2   3   4E+4   14   14.5  RT (min)      lcorbi_t3_pos_1_030_21    m/z 122.0317     0   1   2   3   4E+4   14.5   15  RT (min)      lcorbi_t3_pos_1_034_34    m/z 122.0317     0   1   2E+5   13.5   14  RT (min)      lcorbi_t3_pos_1_008_36    m/z 122.0317     0   1   2   3   4E+5   12.5   13  RT (min)      lcorbi_t3_pos_1_002_01    m/z 122.0318     0   1   12   12.5  RT (min)      lcorbi_t3_pos_2_032_39    m/z 122.0362     0   2   4E+5   10   10.5  RT (min)      lcorbi_t3_pos_1_028_32    m/z 122.0964     0   1E+5   2.5   3  RT (min)       lcorbi_t3_pos_1_001_49    m/z 122.0964     0   1   2   3E+6   0   0.5  RT (min)      lcorbi_t3_pos_1_001_49    m/z 122.0964     0   2   4   6E+5   5   5.5  RT (min)      lcorbi_t3_pos_3_000_aa    m/z 122.1256     0   1   2E+6   13   13.5  RT (min)       lcorbi_t3_pos_1_052_38    m/z 122.1256     0   1   2   3   4E+6   16   17  RT (min)       lcorbi_t3_pos_1_007_07    m/z 122.1256     0   2   4   6   8E+5   10.5   11  RT (min)      lcorbi_t3_pos_1_059_17    m/z 122.1256     0   1   17.5   18  RT (min)     lcorbi_t3_pos_1_056_56    m/z 122.1256     0   2   4   6E+5   9   9.5  RT (min)      lcorbi_t3_pos_1_057_49    m/z 122.1256     0   2   4   6   8E+5   0.5   1  RT (min)      lcorbi_t3_pos_4_057_49    m/z 122.1256     0   1   2E+6   14   14.5  RT (min)      lcorbi_t3_pos_1_057_49    m/z 122.1256     0   1E+6   12   12.5  RT (min)      lcorbi_t3_pos_1_059_17    m/z 122.1256     0   1   2   3   4E+6   16   17  RT (min)       lcorbi_t3_pos_1_003_45    m/z 122.1256     0   2   4   6E+5   9   9.5  RT (min)      lcorbi_t3_pos_1_059_17    m/z 122.1256     0   1E+6   13.5   14  RT (min)       lcorbi_t3_pos_2_021_09    m/z 122.1256     0   1   17   17.5  RT (min)     lcorbi_t3_pos_1_038_30    m/z 122.1256     0   1   17.5   18  RT (min)     lcorbi_t3_pos_1_051_09    m/z 122.1256     0   1   2   3E+6   16   17  RT (min)       lcorbi_t3_pos_1_033_53    m/z 122.1256     0   1   2   3   4E+6   16   17   18  RT (min)       lcorbi_t3_pos_1_005_42    m/z 122.1256     0   1   17  RT (min)     lcorbi_t3_pos_1_008_36    m/z 122.1256     0   2   4   6E+5   1.5   2  RT (min)      lcorbi_t3_pos_4_057_49    m/z 122.1256     0   1   17   17.5  RT (min)     lcorbi_t3_pos_1_001_49    m/z 122.1269     0   1   2   3E+6   15   16   17  RT (min)       lcorbi_t3_pos_1_039_23    m/z 122.1269     0   2   4   6   8E+5   13   13.5  RT (min)      lcorbi_t3_pos_2_061_50    m/z 123.0138     0   1   2E+5   3.5   4  RT (min)       lcorbi_t3_pos_4_082_59    m/z 123.0552     0   1E+6   2.5   3  RT (min)      lcorbi_t3_pos_4_088_59    m/z 123.0552     0   1   2E+6   2.5   3  RT (min)       lcorbi_t3_pos_4_087_59    m/z 123.0552     0   1E+5   0.5   1  RT (min)      lcorbi_t3_pos_1_034_34    m/z 123.0552     0   1E+6   3   3.5  RT (min)       lcorbi_t3_pos_4_082_59    m/z 123.0552     0   1   2E+5   0.5   1  RT (min)      lcorbi_t3_pos_1_031_40    m/z 123.0553     0   1E+6   3   3.5  RT (min)       lcorbi_t3_pos_4_082_59    m/z 123.0553     0   1E+5   4.5   5  RT (min)       lcorbi_t3_pos_1_012_14    m/z 123.0553     0   1   2E+5   1   1.5  RT (min)       lcorbi_t3_pos_4_088_59    m/z 123.0553     0   1   2E+5   1   1.5  RT (min)      lcorbi_t3_pos_1_054_48    m/z 123.0916     0   1   18   18.5  RT (min)     lcorbi_t3_pos_1_027_43    m/z 123.0917     0   2   4E+5   5   5.5  RT (min)      lcorbi_t3_pos_3_000_aa    m/z 123.1265     0   1E+5   15   16  RT (min)       lcorbi_t3_pos_4_094_60    m/z 123.1265     0   1E+5   14.5   15  RT (min)       lcorbi_t3_pos_1_053_39    m/z 123.1265     0   1   2   3E+5   16  RT (min)       lcorbi_t3_pos_1_005_42    m/z 123.1265     0   1   2E+5   15   16   17  RT (min)       lcorbi_t3_pos_1_011_44    m/z 125.0709     0   1E+5   1   1.5  RT (min)       lcorbi_t3_pos_1_032_26    m/z 126.0549     0   2   4   6E+4   18.5   19  RT (min)       lcorbi_t3_pos_4_075_58    m/z 126.0549     0   2   4   6E+4   19   19.5  RT (min)       lcorbi_t3_pos_4_081_58    m/z 126.0549     0   2   4   6E+4   18.5   19  RT (min)       lcorbi_t3_pos_4_082_59    m/z 126.0549     0   2   4   6E+4   18.5   19  RT (min)       lcorbi_t3_pos_4_083_59    m/z 126.0550     0   2   4   6   8E+4   0.5   1  RT (min)       lcorbi_t3_pos_3_000_aa    m/z 126.0550     0   1E+5   1   2   3  RT (min)       lcorbi_t3_pos_4_077_58    m/z 126.0550     0   1E+5   2  RT (min)       lcorbi_t3_pos_4_071_57    m/z 126.0550     0   1E+5   4.5   5  RT (min)       lcorbi_t3_pos_2_014_37    m/z 127.0390     0   1   2   3E+5   1   1.5  RT (min)      lcorbi_t3_pos_4_079_58    m/z 127.0390     0   1   2E+5   1   1.5  RT (min)       lcorbi_t3_pos_4_074_58    m/z 128.0706     0   1E+5   1   1.5  RT (min)      lcorbi_t3_pos_1_028_32    m/z 129.0754     0   1E+4   1   1.5  RT (min)      lcorbi_t3_pos_4_079_58    m/z 129.0754     0   1   2E+6   3   3.5  RT (min)       lcorbi_t3_pos_4_090_60    m/z 129.1274     0   1   2E+5   10   10.5  RT (min)      lcorbi_t3_pos_1_034_34    m/z 129.1274     0   1E+5   10   10.5  RT (min)      lcorbi_t3_pos_2_052_48    m/z 129.1274     0   1   2   3   4E+4   10.5   11  RT (min)      lcorbi_t3_pos_1_023_02    m/z 130.0498     0   2   4   6E+4   18.5   19  RT (min)      lcorbi_t3_pos_4_090_60    m/z 130.0498     0   2   4   6E+4   18.5   19  RT (min)      lcorbi_t3_pos_4_082_59    m/z 130.0499     0   1E+5   2  RT (min)       lcorbi_t3_pos_1_006_15    m/z 130.0499     0   1   2   3   4E+5   2   4  RT (min)       lcorbi_t3_pos_1_014_03    m/z 130.0499     0   1E+6   1.5   2  RT (min)       lcorbi_t3_pos_1_032_26    m/z 130.0499     0   1   2E+5   3   4  RT (min)       lcorbi_t3_pos_1_004_24    m/z 130.0499     0   1E+6   1.5   2  RT (min)       lcorbi_t3_pos_4_084_59    m/z 130.1590     0   1   2E+6   15   15.5  RT (min)      lcorbi_t3_pos_1_057_49    m/z 130.1590     0   2   4   6E+5   8.5   9  RT (min)      lcorbi_t3_pos_1_058_14    m/z 130.1590     0   2   4   6   8E+4   3.5   4  RT (min)      lcorbi_t3_pos_4_098_61    m/z 130.1590     0   1E+6   12   12.5  RT (min)      lcorbi_t3_pos_2_058_15    m/z 130.1590     0   1E+6   12.5   13  RT (min)      lcorbi_t3_pos_1_059_17    m/z 130.1590     0   1   2E+6   14   14.5  RT (min)      lcorbi_t3_pos_1_057_49    m/z 130.1590     0   1E+6   12   12.5  RT (min)      lcorbi_t3_pos_1_058_14    m/z 130.1590     0   2   4   6   8E+5   11   11.5  RT (min)      lcorbi_t3_pos_2_059_39    m/z 130.1590     0   2   4   6E+5   8.5   9  RT (min)      lcorbi_t3_pos_1_060_39    m/z 130.1590     0   1E+5   0.5   1  RT (min)      lcorbi_t3_pos_1_059_17    m/z 130.1590     0   1   2   3   4E+6   18   18.5  RT (min)       lcorbi_t3_pos_4_106_62    m/z 130.1590     0   2   4   6E+5   11.5   12  RT (min)      lcorbi_t3_pos_4_062_13    m/z 130.1590     0   1   2   3E+6   16   18  RT (min)       lcorbi_t3_pos_4_105_61    m/z 130.1590     0   1E+6   10.5   11  RT (min)      lcorbi_t3_pos_1_058_14    m/z 130.1590     0   1E+5   2.5   3  RT (min)       lcorbi_t3_pos_4_079_58    m/z 130.1590     0   2   4   6   8E+4   0.5   1  RT (min)      lcorbi_t3_pos_3_024_07    m/z 130.1590     0   2   4   6   8E+5   4.5   5  RT (min)      lcorbi_t3_pos_3_057_49    m/z 130.1590     0   2   4   6E+5   9.5   10  RT (min)      lcorbi_t3_pos_1_058_14    m/z 130.1590     0   2   4   6E+5   9   9.5  RT (min)      lcorbi_t3_pos_1_057_49    m/z 130.1590     0   1E+5   2   2.5  RT (min)      lcorbi_t3_pos_3_016_21    m/z 130.1590     0   2   4   6   8E+6   6   8  RT (min)       lcorbi_t3_pos_3_000_aa    m/z 130.1590     0   1   2   3E+6   17.5   18  RT (min)       lcorbi_t3_pos_4_075_58    m/z 130.1590     0   1E+6   10  RT (min)       lcorbi_t3_pos_1_059_17    m/z 130.9818     0   1E+5   1.5   2  RT (min)       lcorbi_t3_pos_1_022_20    m/z 131.1624     0   1   2   3   4E+5   18   18.5  RT (min)       lcorbi_t3_pos_4_098_61    m/z 131.1624     0   1E+5   13   13.5  RT (min)       lcorbi_t3_pos_4_080_58    m/z 131.1624     0   2   4   6E+5   5   5.5  RT (min)      lcorbi_t3_pos_3_000_aa    m/z 132.1019     0   1E+5   1   1.5  RT (min)      lcorbi_t3_pos_1_041_54    m/z 132.1019     0   1E+5   1   1.5  RT (min)      lcorbi_t3_pos_1_019_16    m/z 132.1019     0   2   4   6E+5   3.5   4  RT (min)       lcorbi_t3_pos_4_074_58    m/z 132.1019     0   2   4   6E+5   3.5   4  RT (min)       lcorbi_t3_pos_4_066_57    m/z 132.1019     0   1E+6   3   3.5  RT (min)      lcorbi_t3_pos_4_081_58    m/z 132.9049     0   2   4   6E+6   0   0.5  RT (min)      lcorbi_t3_pos_1_002_01    m/z 135.0328     0   1   2E+5   0   0.5  RT (min)       lcorbi_t3_pos_1_035_28    m/z 135.0328     0   1   2E+5   0.5   1  RT (min)       lcorbi_t3_pos_1_058_14    m/z 135.0328     0   1   2E+5   1   1.5  RT (min)       lcorbi_t3_pos_2_058_15    m/z 135.0328     0   2   4   6E+5   2   4   6  RT (min)       lcorbi_t3_pos_1_038_30    m/z 135.0328     0   2   4   6E+5   4.5   5  RT (min)       lcorbi_t3_pos_2_029_43    m/z 135.0328     0   2   4   6   8E+5   18   18.5  RT (min)       lcorbi_t3_pos_4_095_60    m/z 135.0328     0   2   4E+5   2   2.5  RT (min)       lcorbi_t3_pos_4_057_49    m/z 135.0328     0   2   4   6E+5   2   4   6  RT (min)       lcorbi_t3_pos_1_014_03    m/z 135.0328     0   2   4   6   8E+5   1   1.5  RT (min)       lcorbi_t3_pos_4_077_58    m/z 135.0666     0   1E+6   2   2.5  RT (min)       lcorbi_t3_pos_4_090_60    m/z 135.0666     0   1E+6   1.5   2  RT (min)       lcorbi_t3_pos_4_091_60    m/z 135.0804     0   2   4   6E+5   15   15.5  RT (min)       lcorbi_t3_pos_1_065_51    m/z 135.0804     0   1   2   3   4E+5   13.5   14  RT (min)       lcorbi_t3_pos_1_058_14    m/z 135.0804     0   2   4   6   8E+5   15   15.5  RT (min)       lcorbi_t3_pos_1_037_19    m/z 135.0804     0   1E+5   8   8.5  RT (min)      lcorbi_t3_pos_2_024_15    m/z 135.0804     0   1E+5   8.5   9  RT (min)      lcorbi_t3_pos_2_033_53    m/z 135.0804     0   2   4   6   8E+5   15.5   16  RT (min)       lcorbi_t3_pos_1_029_29    m/z 135.0804     0   1E+5   8.5   9  RT (min)      lcorbi_t3_pos_3_000_cc    m/z 135.0804     0   2   4   6   8E+5   16.5   17  RT (min)       lcorbi_t3_pos_4_088_59    m/z 135.0804     0   2   4   6   8E+5   15.5   16  RT (min)       lcorbi_t3_pos_1_022_20    m/z 135.0804     0   2   4E+5   14.5   15  RT (min)       lcorbi_t3_pos_2_031_20    m/z 135.0804     0   2   4   6E+5   16   17   18  RT (min)       lcorbi_t3_pos_1_051_09    m/z 135.0804     0   2   4   6   8E+5   16   17  RT (min)       lcorbi_t3_pos_1_038_30    m/z 135.0804     0   1   2   3   4E+5   10   10.5  RT (min)       lcorbi_t3_pos_4_081_58    m/z 135.0804     0   1   17   17.5  RT (min)     lcorbi_t3_pos_1_041_54    m/z 135.0804     0   2   4   6   8E+5   16  RT (min)       lcorbi_t3_pos_4_082_59    m/z 135.0804     0   2   4   6   8E+5   15   15.5  RT (min)       lcorbi_t3_pos_1_012_14    m/z 135.0804     0   1   2   3   4E+5   13   13.5  RT (min)       lcorbi_t3_pos_4_072_57    m/z 135.9692     0   1E+6   0   0.5  RT (min)      lcorbi_t3_pos_1_011_44    m/z 136.0215     0   1E+5   0   0.5  RT (min)      lcorbi_t3_pos_2_023_26    m/z 136.0215     0   1E+5   0   0.5  RT (min)      lcorbi_t3_pos_2_022_38    m/z 136.0215     0   1E+5   7.5   8  RT (min)      lcorbi_t3_pos_3_025_52    m/z 136.0618     0   1E+5   0.5   1  RT (min)       lcorbi_t3_pos_4_106_62    m/z 136.0618     0   1   2E+5   2.5   3  RT (min)       lcorbi_t3_pos_4_086_59    m/z 136.0618     0   1E+5   0   0.5  RT (min)       lcorbi_t3_pos_4_075_58    m/z 136.0618     0   1E+5   2.5   3  RT (min)      lcorbi_t3_pos_3_056_56    m/z 136.0618     0   1E+5   0   0.5  RT (min)       lcorbi_t3_pos_4_110_62    m/z 136.0618     0   1   2E+5   18   18.5  RT (min)       lcorbi_t3_pos_4_090_60    m/z 136.0618     0   1   2   3   4E+5   0.5   1  RT (min)       lcorbi_t3_pos_4_091_60    m/z 136.0618     0   1E+5   0.5   1  RT (min)       lcorbi_t3_pos_4_106_62    m/z 136.0618     0   1E+6   1.5   2  RT (min)       lcorbi_t3_pos_4_082_59    m/z 136.0618     0   1E+5   0   0.5  RT (min)       lcorbi_t3_pos_4_112_62    m/z 136.0618     0   1E+5   0   0.5  RT (min)       lcorbi_t3_pos_4_080_58    m/z 136.0618     0   1E+5   0   0.5  RT (min)       lcorbi_t3_pos_4_112_62    m/z 136.0618     0   1E+5   0.5   1  RT (min)       lcorbi_t3_pos_4_098_61    m/z 136.0618     0   1E+5   0.5   1  RT (min)       lcorbi_t3_pos_4_090_60    m/z 136.0618     0   1E+5   0.5   1  RT (min)       lcorbi_t3_pos_4_109_62    m/z 137.0548     0   2   4   6   8E+4   9.5   10  RT (min)      lcorbi_t3_pos_1_002_01    m/z 137.0587     0   1   2E+4   1   1.5  RT (min)      lcorbi_t3_pos_4_070_57    m/z 137.0597     0   1E+5   10   10.5  RT (min)      lcorbi_t3_pos_1_051_09    m/z 137.0597     0   1   2   3E+4   10.5   11  RT (min)      lcorbi_t3_pos_1_023_02    m/z 137.0597     0   2   4   6E+6   10   10.5  RT (min)      lcorbi_t3_pos_1_028_32    m/z 137.0903     0   2   4   6   8E+5   3.5   4  RT (min)       lcorbi_t3_pos_4_090_60    m/z 137.0903     0   2   4   6   8E+5   3   3.5  RT (min)       lcorbi_t3_pos_4_092_60    m/z 137.1074     0   2   4   6E+5   0   0.5  RT (min)      lcorbi_t3_pos_1_001_49    m/z 138.0526     0   1   2E+5   1   1.5  RT (min)      lcorbi_t3_pos_1_030_21    m/z 138.0630     0   1E+4   10   10.5  RT (min)      lcorbi_t3_pos_1_051_09    m/z 138.0630     0   2   4   6E+5   10   10.5  RT (min)      lcorbi_t3_pos_1_028_32    m/z 138.0914     0   2   4   6E+5   0   0.5  RT (min)      lcorbi_t3_pos_1_001_49    m/z 138.1278     0   2   4   6   8E+5   5   5.5  RT (min)      lcorbi_t3_pos_3_000_aa    m/z 139.0389     0   1   2E+5   14   14.5  RT (min)      lcorbi_t3_pos_1_047_41    m/z 139.0390     0   1   2   3E+4   12   12.5  RT (min)      lcorbi_t3_pos_2_032_39    m/z 139.0390     0   1   2E+5   15.5   16  RT (min)       lcorbi_t3_pos_1_023_02    m/z 139.0390     0   1   2   3E+6   12.5   13  RT (min)      lcorbi_t3_pos_4_016_15    m/z 139.0390     0   1   2E+5   14.5   15  RT (min)       lcorbi_t3_pos_1_031_40    m/z 139.0390     0   1   2E+5   15   15.5  RT (min)      lcorbi_t3_pos_1_051_09    m/z 139.0390     0   1   2E+5   15   16  RT (min)       lcorbi_t3_pos_1_034_34    m/z 139.0390     0   1E+5   11   11.5  RT (min)      lcorbi_t3_pos_1_031_40    m/z 139.0390     0   1   2E+5   14   14.5  RT (min)      lcorbi_t3_pos_1_028_32    m/z 139.0390     0   1   2E+6   13.5   14  RT (min)      lcorbi_t3_pos_1_008_36    m/z 139.0390     0   1   2   3   4E+4   12.5   13  RT (min)      lcorbi_t3_pos_2_013_33    m/z 139.0502     0   2   4E+5   1   1.5  RT (min)       lcorbi_t3_pos_3_021_39    m/z 139.1230     0   1E+7   0   0.5  RT (min)      lcorbi_t3_pos_1_001_49    m/z 140.0423     0   1   2   3E+5   13.5   14  RT (min)      lcorbi_t3_pos_1_005_42    m/z 140.1263     0   1E+6   0   0.5  RT (min)      lcorbi_t3_pos_1_001_49    m/z 141.0725     0   1E+5   1.5   2  RT (min)      lcorbi_t3_pos_4_093_60    m/z 141.0785     0   2   4   6E+6   1.5   2  RT (min)       lcorbi_t3_pos_4_090_60    m/z 141.0886     0   1   2E+5   1   1.5  RT (min)      lcorbi_t3_pos_1_016_05    m/z 141.9586     0   1   2E+6   0   0.5  RT (min)      lcorbi_t3_pos_1_002_01    m/z 142.9386     0   1   2   3E+5   13   13.5  RT (min)       lcorbi_t3_pos_1_011_44    m/z 142.9386     0   1E+5   1   1.5  RT (min)       lcorbi_t3_pos_1_031_40    m/z 143.0015     0   1   2E+5   1   1.5  RT (min)      lcorbi_t3_pos_4_080_58    m/z 143.0015     0   1   2   3   4E+5   18   18.5  RT (min)      lcorbi_t3_pos_4_091_60    m/z 143.0015     0   2   4E+5   18   18.5  RT (min)      lcorbi_t3_pos_4_107_62    m/z 143.0015     0   2   4E+5   17.5   18  RT (min)      lcorbi_t3_pos_4_096_60    m/z 143.0015     0   2   4E+5   17.5   18  RT (min)      lcorbi_t3_pos_4_095_60    m/z 143.0015     0   1   2   3   4E+5   18   18.5  RT (min)      lcorbi_t3_pos_4_076_58    m/z 143.0015     0   2   4E+5   18   18.5  RT (min)      lcorbi_t3_pos_4_097_60    m/z 143.0016     0   1   2E+5   0   0.5  RT (min)      lcorbi_t3_pos_1_038_30    m/z 143.0016     0   1E+5   0   0.5  RT (min)      lcorbi_t3_pos_1_047_41    m/z 144.0655     0   1E+6   3   3.5  RT (min)       lcorbi_t3_pos_4_085_59    m/z 144.0655     0   1E+5   1   1.5  RT (min)      lcorbi_t3_pos_2_012_01    m/z 144.0655     0   1E+5   1   1.5  RT (min)      lcorbi_t3_pos_1_031_40    m/z 144.0655     0   1E+6   3.5   4  RT (min)       lcorbi_t3_pos_4_082_59    m/z 144.0656     0   1   2   3E+5   1   1.5  RT (min)       lcorbi_t3_pos_4_057_49    m/z 145.0171     0   2   4   6   8E+5   17.5   18  RT (min)       lcorbi_t3_pos_4_105_61    m/z 145.0172     0   2   4   6   8E+5   17.5   18  RT (min)      lcorbi_t3_pos_4_105_61    m/z 145.0172     0   2   4   6   8E+5   2   4  RT (min)       lcorbi_t3_pos_4_078_58    m/z 145.0346     0   2   4   6E+4   2.5   3  RT (min)       lcorbi_t3_pos_4_104_61    m/z 145.0346     0   1E+5   18   18.5  RT (min)      lcorbi_t3_pos_4_111_62    m/z 145.0346     0   2   4   6E+4   3   3.5  RT (min)       lcorbi_t3_pos_4_100_61    m/z 145.0346     0   1E+5   0   0.5  RT (min)      lcorbi_t3_pos_1_056_56    m/z 147.0502     0   1E+5   18.5   19  RT (min)      lcorbi_t3_pos_4_101_61    m/z 147.0502     0   1   2   3   4E+5   18   18.5  RT (min)       lcorbi_t3_pos_4_084_59    m/z 147.0502     0   1   2   3E+5   18   18.5  RT (min)      lcorbi_t3_pos_4_098_61    m/z 147.0502     0   1   2E+5   2   3  RT (min)       lcorbi_t3_pos_4_110_62    m/z 147.0502     0   1   2   3   4E+5   18.5   19  RT (min)      lcorbi_t3_pos_4_095_60    m/z 148.1121     0   1E+6   0   0.5  RT (min)      lcorbi_t3_pos_1_001_49    m/z 148.9757     0   1E+5   2   2.5  RT (min)      lcorbi_t3_pos_1_020_31    m/z 148.9757     0   1E+5   2   2.5  RT (min)      lcorbi_t3_pos_1_011_44    m/z 148.9757     0   1E+5   2   2.5  RT (min)      lcorbi_t3_pos_1_015_11    m/z 148.9757     0   1E+5   1.5   2  RT (min)      lcorbi_t3_pos_1_006_15    m/z 148.9758     0   1E+5   2   2.5  RT (min)      lcorbi_t3_pos_1_018_35    m/z 149.0168     0   2   4   6E+5   13   13.5  RT (min)      lcorbi_t3_pos_4_075_58    m/z 149.0197     0   2   4   6   8E+3   12.5   13  RT (min)      lcorbi_t3_pos_4_012_37    m/z 149.0215     0   1   2E+6   15   15.5  RT (min)      lcorbi_t3_pos_4_086_59    m/z 149.0233     0   2   4   6E+6   10   11  RT (min)       lcorbi_t3_pos_1_008_36    m/z 149.0233     0   1   2E+7   12.5   13  RT (min)      lcorbi_t3_pos_1_002_01    m/z 149.0233     0   1E+6   11   11.5  RT (min)      lcorbi_t3_pos_1_031_40    m/z 149.0233     0   1E+6   11.5   12  RT (min)      lcorbi_t3_pos_1_023_02    m/z 149.0233     0   1E+5   5   5.5  RT (min)      lcorbi_t3_pos_2_040_25    m/z 149.0233     0   1   2E+6   10.5   11  RT (min)      lcorbi_t3_pos_3_000_aa    m/z 149.0233     0   1E+6   14.5   15  RT (min)      lcorbi_t3_pos_2_009_50    m/z 149.0233     0   1E+5   2   4  RT (min)       lcorbi_t3_pos_4_079_58    m/z 149.0233     0   1   2E+7   13.5   14  RT (min)      lcorbi_t3_pos_4_001_49    m/z 149.0233     0   1E+5   7.5   8  RT (min)      lcorbi_t3_pos_2_040_25    m/z 149.0233     0   1E+5   1   1.5  RT (min)       lcorbi_t3_pos_4_077_58    m/z 149.0233     0   1E+6   14.5   15  RT (min)      lcorbi_t3_pos_2_009_50    m/z 149.0233     0   1E+6   14.5   15  RT (min)      lcorbi_t3_pos_2_049_55    m/z 149.0233     0   1   2E+6   15   16  RT (min)       lcorbi_t3_pos_1_034_34    m/z 149.0233     0   1   2E+6   14.5   15  RT (min)      lcorbi_t3_pos_1_031_40    m/z 149.0233     0   1   2E+6   14   16  RT (min)       lcorbi_t3_pos_1_028_32    m/z 149.0233     0   1E+6   14.5   15  RT (min)      lcorbi_t3_pos_1_056_56    m/z 149.0233     0   1   16.5   17  RT (min)     lcorbi_t3_pos_1_001_49    m/z 149.0233     0   2   4   6   8E+4   0.5   1  RT (min)      lcorbi_t3_pos_1_050_13    m/z 149.0233     0   1E+6   15   16   17  RT (min)       lcorbi_t3_pos_1_051_09    m/z 149.0233     0   1   2E+7   13.5   14  RT (min)      lcorbi_t3_pos_1_008_36    m/z 149.0233     0   1E+6   0.5   1  RT (min)      lcorbi_t3_pos_4_057_49    m/z 149.0233     0   1E+6   14   14.5  RT (min)      lcorbi_t3_pos_2_022_38    m/z 149.0233     0   1E+6   16   17   18  RT (min)       lcorbi_t3_pos_4_082_59    m/z 149.0233     0   1   2E+6   11   11.5  RT (min)      lcorbi_t3_pos_1_028_32    m/z 149.0233     0   1   2E+6   14.5   15  RT (min)      lcorbi_t3_pos_1_041_54    m/z 149.0233     0   1   2E+6   14   14.5  RT (min)      lcorbi_t3_pos_2_006_47    m/z 149.0233     0   1   2   3   4E+6   14   14.5  RT (min)      lcorbi_t3_pos_2_017_51    m/z 149.0233     0   1   2E+6   1   1.5  RT (min)       lcorbi_t3_pos_4_057_49    m/z 149.0233     0   2   4   6E+6   10   10.5  RT (min)      lcorbi_t3_pos_1_002_01    m/z 149.0233     0   1   2   3   4E+7   13.5   14  RT (min)       lcorbi_t3_pos_3_047_40    m/z 149.0233     0   1E+6   14   14.5  RT (min)      lcorbi_t3_pos_3_003_33    m/z 149.0233     0   1   2   3   4E+5   12.5   13  RT (min)      lcorbi_t3_pos_2_013_33    m/z 149.0233     0   1E+6   15.5   16  RT (min)      lcorbi_t3_pos_1_023_02    m/z 149.0233     0   2   4E+5   0   0.5  RT (min)      lcorbi_t3_pos_1_050_13    m/z 149.0233     0   2   4   6E+4   2   3  RT (min)       lcorbi_t3_pos_1_050_13    m/z 149.0233     0   2   4E+5   0.5   1  RT (min)      lcorbi_t3_pos_1_050_13    m/z 149.0233     0   2   4E+5   0   0.5  RT (min)      lcorbi_t3_pos_1_050_13    m/z 149.0234     0   1E+6   14   14.5  RT (min)      lcorbi_t3_pos_2_025_52    m/z 149.0234     0   1   2   3   4E+7   12.5   13  RT (min)      lcorbi_t3_pos_4_016_15    m/z 149.0260     0   1   2   3E+5   13.5   14  RT (min)       lcorbi_t3_pos_4_019_47    m/z 149.0264     0   2   4   6E+3   12.5   13  RT (min)      lcorbi_t3_pos_3_031_45    m/z 149.0265     0   1E+4   12   12.5  RT (min)      lcorbi_t3_pos_2_032_39    m/z 149.0478     0   1E+5   1   1.5  RT (min)      lcorbi_t3_pos_4_078_58    m/z 149.0484     0   1E+5   18   18.5  RT (min)      lcorbi_t3_pos_4_076_58    m/z 150.0266     0   1E+6   13.5   14  RT (min)      lcorbi_t3_pos_4_010_43    m/z 150.0267     0   1   2E+6   12.5   13  RT (min)      lcorbi_t3_pos_1_002_01    m/z 150.0267     0   1E+5   11   11.5  RT (min)      lcorbi_t3_pos_1_031_40    m/z 150.0267     0   1E+5   14.5   15  RT (min)      lcorbi_t3_pos_1_034_34    m/z 150.0267     0   1E+5   15.5   16  RT (min)      lcorbi_t3_pos_1_023_02    m/z 150.0267     0   1   2   3   4E+6   13.5   14  RT (min)      lcorbi_t3_pos_3_047_40    m/z 150.0267     0   1   2E+5   14   14.5  RT (min)      lcorbi_t3_pos_1_030_21    m/z 150.0267     0   1   2E+5   14   14.5  RT (min)      lcorbi_t3_pos_1_028_32    m/z 150.0267     0   1E+5   15   15.5  RT (min)      lcorbi_t3_pos_1_051_09    m/z 150.0267     0   2   4   6E+5   10   10.5  RT (min)      lcorbi_t3_pos_1_002_01    m/z 150.0267     0   1   2E+5   11   11.5  RT (min)      lcorbi_t3_pos_1_028_32    m/z 150.0267     0   1   2E+5   14   14.5  RT (min)      lcorbi_t3_pos_1_030_21    m/z 150.0267     0   1E+5   15   16  RT (min)       lcorbi_t3_pos_1_056_56    m/z 150.0267     0   2   4E+4   11   11.5  RT (min)      lcorbi_t3_pos_1_034_34    m/z 150.0267     0   1   2   3E+4   12   12.5  RT (min)      lcorbi_t3_pos_2_032_39    m/z 150.1277     0   1   2   3E+6   0   0.5  RT (min)      lcorbi_t3_pos_1_001_49    m/z 151.0275     0   1   2E+4   15.5   16  RT (min)      lcorbi_t3_pos_1_023_02    m/z 151.0965     0   2   4   6   8E+5   3   3.5  RT (min)       lcorbi_t3_pos_4_113_62    m/z 151.0965     0   1   2   3   4E+5   11.5   12  RT (min)       lcorbi_t3_pos_4_082_59    m/z 151.0965     0   1E+5   4.5   5  RT (min)      lcorbi_t3_pos_1_060_39    m/z 151.0965     0   2   4   6   8E+5   3.5   4  RT (min)       lcorbi_t3_pos_4_066_57    m/z 152.9471     0   2   4   6E+5   17.5   18  RT (min)      lcorbi_t3_pos_4_105_61    m/z 152.9471     0   2   4   6E+5   18   18.5  RT (min)      lcorbi_t3_pos_4_091_60    m/z 153.0434     0   1E+5   5.5   6  RT (min)       lcorbi_t3_pos_4_099_61    m/z 153.0434     0   1E+5   5   5.5  RT (min)       lcorbi_t3_pos_4_111_62    m/z 153.0658     0   2   4   6E+5   3   3.5  RT (min)       lcorbi_t3_pos_4_069_57    m/z 153.0658     0   2   4E+5   3.5   4  RT (min)       lcorbi_t3_pos_4_066_57    m/z 155.1179     0   2   4   6E+5   0   0.5  RT (min)      lcorbi_t3_pos_1_001_49    m/z 158.0812     0   2   4   6   8E+5   3.5   4  RT (min)       lcorbi_t3_pos_4_085_59    m/z 158.0812     0   1E+6   4.5   5  RT (min)       lcorbi_t3_pos_4_082_59    m/z 158.1539     0   2   4   6   8E+5   9   9.5  RT (min)      lcorbi_t3_pos_3_044_46    m/z 158.1539     0   1E+5   11   11.5  RT (min)      lcorbi_t3_pos_4_103_61    m/z 158.1539     0   1E+5   13   13.5  RT (min)       lcorbi_t3_pos_1_057_49    m/z 158.1539     0   1E+5   12   12.5  RT (min)       lcorbi_t3_pos_4_088_59    m/z 158.1539     0   2   4   6   8E+4   13.5   14  RT (min)      lcorbi_t3_pos_4_106_62    m/z 158.1539     0   1E+5   12.5   13  RT (min)       lcorbi_t3_pos_4_106_62    m/z 158.1539     0   1E+5   15   15.5  RT (min)       lcorbi_t3_pos_2_003_19    m/z 158.1539     0   1E+5   14.5   15  RT (min)      lcorbi_t3_pos_1_057_49    m/z 158.1540     0   1   2E+5   9.5   10  RT (min)      lcorbi_t3_pos_3_003_33    m/z 158.1540     0   1E+5   16   17  RT (min)       lcorbi_t3_pos_4_083_59    m/z 158.1540     0   1E+5   12   12.5  RT (min)      lcorbi_t3_pos_4_090_60    m/z 158.1540     0   1E+5   9.5   10  RT (min)      lcorbi_t3_pos_2_046_07    m/z 158.1540     0   2   4   6   8E+4   0   0.5  RT (min)      lcorbi_t3_pos_1_055_33    m/z 158.1540     0   2   4   6   8E+4   10.5   11  RT (min)      lcorbi_t3_pos_4_082_59    m/z 158.1540     0   1E+5   11.5   12  RT (min)      lcorbi_t3_pos_4_097_60    m/z 158.1540     0   2   4   6   8E+4   10   10.5  RT (min)      lcorbi_t3_pos_4_077_58    m/z 158.9739     0   2   4E+5   0   0.5  RT (min)      lcorbi_t3_pos_1_012_14    m/z 159.0692     0   1E+5   18   18.5  RT (min)      lcorbi_t3_pos_4_082_59    m/z 159.0692     0   1E+5   18   18.5  RT (min)      lcorbi_t3_pos_4_107_62    m/z 159.1573     0   1E+5   9   9.5  RT (min)      lcorbi_t3_pos_1_041_54    m/z 159.9692     0   1   2   3E+6   0   0.5  RT (min)      lcorbi_t3_pos_1_002_01    m/z 160.1808     0   1   2   3   4E+5   18.5   19  RT (min)      lcorbi_t3_pos_4_091_60    m/z 160.1808     0   1   2   3E+5   18.5   19  RT (min)      lcorbi_t3_pos_4_069_57    m/z 161.0121     0   1   2   3E+5   0   0.5  RT (min)      lcorbi_t3_pos_4_014_48    m/z 161.0121     0   1E+6   18   18.5  RT (min)       lcorbi_t3_pos_4_069_57    m/z 161.0121     0   1E+6   18   18.5  RT (min)       lcorbi_t3_pos_4_077_58    m/z 161.0121     0   1   2   3   4E+5   18.5   19  RT (min)       lcorbi_t3_pos_4_070_57    m/z 161.0121     0   1E+6   17.5   18  RT (min)      lcorbi_t3_pos_4_105_61    m/z 161.0121     0   1E+6   17.5   18  RT (min)      lcorbi_t3_pos_4_095_60    m/z 161.0121     0   1E+6   18   18.5  RT (min)      lcorbi_t3_pos_4_093_60    m/z 161.0121     0   1E+6   18   18.5  RT (min)       lcorbi_t3_pos_4_081_58    m/z 161.0121     0   1E+6   17.5   18  RT (min)      lcorbi_t3_pos_4_105_61    m/z 161.0121     0   1E+6   18   18.5  RT (min)      lcorbi_t3_pos_4_094_60    m/z 161.0121     0   1E+6   17.5   18  RT (min)      lcorbi_t3_pos_4_093_60    m/z 161.0121     0   1E+6   18   18.5  RT (min)      lcorbi_t3_pos_4_107_62    m/z 161.0121     0   1E+6   18   18.5  RT (min)      lcorbi_t3_pos_4_106_62    m/z 161.0122     0   2   4   6   8E+5   1   1.5  RT (min)       lcorbi_t3_pos_4_077_58    m/z 161.0122     0   1E+6   17.5   18  RT (min)      lcorbi_t3_pos_4_105_61    m/z 162.0583     0   2   4E+5   3.5   4  RT (min)       lcorbi_t3_pos_4_074_58    m/z 162.0584     0   2   4   6E+5   3   3.5  RT (min)       lcorbi_t3_pos_4_081_58    m/z 162.1278     0   1E+6   5   5.5  RT (min)      lcorbi_t3_pos_3_000_aa    m/z 163.0277     0   1E+6   4.5   5  RT (min)      lcorbi_t3_pos_2_062_13    m/z 163.0277     0   1   2   3   4E+6   18.5   19  RT (min)      lcorbi_t3_pos_4_069_57    m/z 163.0277     0   1   2   3   4E+6   17.5   18  RT (min)      lcorbi_t3_pos_4_105_61    m/z 163.0277     0   1   2   3   4E+6   18.5   19  RT (min)       lcorbi_t3_pos_4_069_57    m/z 163.0277     0   1E+6   1   1.5  RT (min)       lcorbi_t3_pos_4_070_57    m/z 163.0277     0   1   2   3   4E+6   18   18.5  RT (min)       lcorbi_t3_pos_4_069_57    m/z 163.0277     0   1E+6   2.5   3  RT (min)      lcorbi_t3_pos_1_060_39    m/z 163.0277     0   1   2   3E+6   18   18.5  RT (min)       lcorbi_t3_pos_4_081_58    m/z 163.0277     0   1   2E+6   18.5   19  RT (min)      lcorbi_t3_pos_4_076_58    m/z 163.0277     0   1E+6   19   19.5  RT (min)       lcorbi_t3_pos_4_092_60    m/z 163.0278     0   1E+6   4.5   5  RT (min)       lcorbi_t3_pos_4_112_62    m/z 163.0278     0   1E+6   2.5   3  RT (min)      lcorbi_t3_pos_1_063_15    m/z 163.0278     0   1   2   3   4E+6   17.5   18  RT (min)      lcorbi_t3_pos_4_105_61    m/z 163.0278     0   2   4   6E+5   10   10.5  RT (min)      lcorbi_t3_pos_2_062_13    m/z 163.0278     0   2   4   6   8E+5   3.5   4  RT (min)      lcorbi_t3_pos_4_059_15    m/z 163.0278     0   2   4   6E+5   9.5   10  RT (min)      lcorbi_t3_pos_1_062_13    m/z 163.0278     0   2   4   6E+5   8   8.5  RT (min)      lcorbi_t3_pos_1_061_50    m/z 163.0278     0   2   4   6E+5   9   9.5  RT (min)      lcorbi_t3_pos_1_059_17    m/z 163.0278     0   1E+6   2   4  RT (min)       lcorbi_t3_pos_1_065_51    m/z 163.0278     0   2   4   6   8E+5   6   6.5  RT (min)      lcorbi_t3_pos_1_059_17    m/z 163.0278     0   2   4   6   8E+5   7   7.5  RT (min)      lcorbi_t3_pos_1_060_39    m/z 163.0278     0   2   4   6E+5   9   9.5  RT (min)      lcorbi_t3_pos_1_064_16    m/z 163.0278     0   2   4   6   8E+5   3.5   4  RT (min)      lcorbi_t3_pos_1_060_39    m/z 163.0278     0   1E+6   2.5   5  RT (min)       lcorbi_t3_pos_4_096_60    m/z 163.0278     0   1   2   3   4E+6   17.5   18  RT (min)      lcorbi_t3_pos_4_095_60    m/z 163.0278     0   2   4E+5   10.5   11  RT (min)      lcorbi_t3_pos_2_062_13    m/z 163.0278     0   1E+6   2   2.5  RT (min)      lcorbi_t3_pos_1_057_49    m/z 163.0278     0   2   4E+5   10   10.5  RT (min)      lcorbi_t3_pos_3_059_17    m/z 163.0278     0   2   4   6E+5   9.5   10  RT (min)      lcorbi_t3_pos_3_058_16    m/z 163.0278     0   1E+6   7   7.5  RT (min)      lcorbi_t3_pos_1_057_49    m/z 163.0389     0   1   2E+5   6.5   7  RT (min)      lcorbi_t3_pos_2_053_28    m/z 163.0389     0   1E+6   12.5   13  RT (min)      lcorbi_t3_pos_1_002_01    m/z 163.0389     0   2   4   6E+4   15   15.5  RT (min)      lcorbi_t3_pos_1_031_40    m/z 163.0390     0   2   4   6   8E+4   14.5   15  RT (min)      lcorbi_t3_pos_1_034_34    m/z 163.0390     0   1   2   3   4E+5   13.5   14  RT (min)      lcorbi_t3_pos_1_008_36    m/z 163.0390     0   2   4   6   8E+4   15.5   16  RT (min)      lcorbi_t3_pos_1_023_02    m/z 163.0390     0   2   4   6E+4   15   15.5  RT (min)      lcorbi_t3_pos_1_051_09    m/z 163.0390     0   2   4   6E+4   14   14.5  RT (min)      lcorbi_t3_pos_1_047_41    m/z 163.0390     0   2   4   6E+5   12.5   13  RT (min)      lcorbi_t3_pos_1_002_01    m/z 163.0390     0   2   4   6E+4   14.5   15  RT (min)      lcorbi_t3_pos_1_041_54    m/z 163.0390     0   1E+4   12.5   13  RT (min)      lcorbi_t3_pos_3_024_07    m/z 163.0390     0   2   4   6E+4   14   14.5  RT (min)      lcorbi_t3_pos_1_028_32    m/z 163.0390     0   1E+4   12.5   13  RT (min)      lcorbi_t3_pos_2_013_33    m/z 163.0390     0   2   4   6E+4   14.5   15  RT (min)      lcorbi_t3_pos_1_041_54    m/z 163.0390     0   2   4   6E+4   14.5   15  RT (min)      lcorbi_t3_pos_1_031_40    m/z 163.0753     0   2   4E+5   7.5   8  RT (min)      lcorbi_t3_pos_3_000_aa    m/z 163.0753     0   2   4E+5   7.5   8  RT (min)      lcorbi_t3_pos_3_000_aa    m/z 164.0311     0   1   2E+5   18   18.5  RT (min)      lcorbi_t3_pos_4_091_60    m/z 164.0311     0   1   2   3E+5   0   0.5  RT (min)      lcorbi_t3_pos_1_007_07    m/z 164.0311     0   1E+5   18   18.5  RT (min)      lcorbi_t3_pos_4_077_58    m/z 164.0311     0   1   2E+5   18   18.5  RT (min)      lcorbi_t3_pos_4_107_62    m/z 164.1165     0   1E+4   1   1.5  RT (min)       lcorbi_t3_pos_4_090_60    m/z 164.9205     0   1   2   3   4E+5   13   13.5  RT (min)       lcorbi_t3_pos_1_011_44    m/z 164.9205     0   2   4E+5   14   16   18  RT (min)       lcorbi_t3_pos_4_105_61    m/z 165.0607     0   1E+5   0.5   1   1.5  RT (min)       lcorbi_t3_pos_4_098_61    m/z 165.0607     0   1   2   3   4E+5   0   0.5  RT (min)      lcorbi_t3_pos_3_054_15    m/z 165.0608     0   1   2   3   4E+5   1   1.5  RT (min)       lcorbi_t3_pos_4_074_58    m/z 165.0910     0   1   2E+4   15.5   16  RT (min)      lcorbi_t3_pos_1_028_32    m/z 165.0910     0   1   16.5   17  RT (min)    lcorbi_t3_pos_1_051_09    m/z 165.0910     0   1E+4   13.5   14  RT (min)      lcorbi_t3_pos_2_031_20    m/z 165.0910     0   2   4   6E+5   14   14.5  RT (min)      lcorbi_t3_pos_4_105_61    m/z 165.0910     0   1   17.5   18  RT (min)    lcorbi_t3_pos_1_023_02    m/z 165.0910     0   1E+4   13.5   14  RT (min)      lcorbi_t3_pos_2_032_39    m/z 165.0910     0   2   4   6E+4   14   14.5  RT (min)      lcorbi_t3_pos_2_016_23    m/z 165.1386     0   1   2   3   4E+6   0   0.5  RT (min)      lcorbi_t3_pos_1_002_01    m/z 166.0498     0   2   4   6E+5   3.5   4  RT (min)       lcorbi_t3_pos_2_000_aa    m/z 166.0498     0   2   4   6   8E+5   3.5   4  RT (min)       lcorbi_t3_pos_4_098_61    m/z 166.0498     0   2   4   6E+5   3   3.5  RT (min)       lcorbi_t3_pos_4_111_62    m/z 166.0499     0   2   4   6E+5   4.5   5  RT (min)      lcorbi_t3_pos_4_000_aa    m/z 166.0862     0   2   4   6E+5   4   4.5  RT (min)       lcorbi_t3_pos_4_074_58    m/z 166.0862     0   2   4   6   8E+5   3.5   4  RT (min)       lcorbi_t3_pos_4_081_58    m/z 166.1226     0   1E+6   0   0.5  RT (min)      lcorbi_t3_pos_1_001_49    m/z 166.9863     0   1E+5   2   3  RT (min)       lcorbi_t3_pos_4_071_57    m/z 166.9863     0   2   4   6   8E+4   2.5   3  RT (min)       lcorbi_t3_pos_4_107_62    m/z 167.0206     0   1E+5   18   18.5  RT (min)       lcorbi_t3_pos_4_076_58    m/z 167.0339     0   1   2   3E+5   14   14.5  RT (min)      lcorbi_t3_pos_1_028_32    m/z 167.0339     0   1E+6   12.5   13  RT (min)      lcorbi_t3_pos_4_030_05    m/z 167.0339     0   1   2E+5   11   11.5  RT (min)      lcorbi_t3_pos_1_031_40    m/z 167.0339     0   2   4   6E+4   12   12.5  RT (min)      lcorbi_t3_pos_2_032_39    m/z 167.0339     0   1   2   3E+5   16   18  RT (min)       lcorbi_t3_pos_1_031_40    m/z 167.0339     0   1   2   3E+5   14   14.5  RT (min)      lcorbi_t3_pos_1_047_41    m/z 167.0339     0   2   4   6E+6   12.5   13  RT (min)      lcorbi_t3_pos_2_054_29    m/z 167.0339     0   1   2   3E+5   14.5   15  RT (min)      lcorbi_t3_pos_2_009_50    m/z 167.0339     0   1   2   3E+5   14.5   15  RT (min)      lcorbi_t3_pos_4_028_04    m/z 167.0339     0   1   2E+5   11.5   12  RT (min)       lcorbi_t3_pos_1_023_02    m/z 167.0339     0   1E+6   11   11.5  RT (min)      lcorbi_t3_pos_1_028_32    m/z 167.0339     0   1   2   3E+5   14   14.5  RT (min)      lcorbi_t3_pos_4_028_04    m/z 167.0339     0   1   2   3E+5   15.5   16  RT (min)      lcorbi_t3_pos_1_023_02    m/z 167.0339     0   1   2   3E+5   15   15.5  RT (min)      lcorbi_t3_pos_2_052_48    m/z 167.0339     0   1   2   3E+5   16  RT (min)       lcorbi_t3_pos_1_034_34    m/z 167.0339     0   1   2E+5   15.5   16  RT (min)      lcorbi_t3_pos_2_052_48    m/z 167.0339     0   2   4   6E+5   11.5   12  RT (min)      lcorbi_t3_pos_4_075_58    m/z 167.0339     0   1   2E+6   1   1.5  RT (min)      lcorbi_t3_pos_4_057_49    m/z 168.0372     0   1   2E+5   13.5   14  RT (min)      lcorbi_t3_pos_1_008_36    m/z 168.0372     0   1E+4   12   12.5  RT (min)      lcorbi_t3_pos_2_032_39    m/z 168.0372     0   2   4   6E+3   12.5   13  RT (min)      lcorbi_t3_pos_3_024_07    m/z 168.0372     0   1   2   3E+4   15   15.5  RT (min)      lcorbi_t3_pos_2_012_01    m/z 168.0372     0   2   4E+4   15.5   16  RT (min)      lcorbi_t3_pos_1_023_02    m/z 168.0372     0   2   4   6   8E+5   12.5   13  RT (min)      lcorbi_t3_pos_4_016_15    m/z 168.0372     0   1   2   3   4E+4   15   15.5  RT (min)      lcorbi_t3_pos_1_031_40    m/z 168.0372     0   2   4   6E+5   13.5   14  RT (min)      lcorbi_t3_pos_3_047_40    m/z 168.0372     0   1   2   3   4E+4   14   14.5  RT (min)      lcorbi_t3_pos_1_028_32    m/z 168.0372     0   1   2   3   4E+4   14   14.5  RT (min)      lcorbi_t3_pos_1_028_32    m/z 168.0372     0   1   2   3E+4   14   14.5  RT (min)      lcorbi_t3_pos_2_022_38    m/z 170.9577     0   2   4   6E+5   18   18.5  RT (min)      lcorbi_t3_pos_4_069_57    m/z 170.9577     0   1E+5   0   0.5  RT (min)      lcorbi_t3_pos_1_029_29    m/z 170.9577     0   1E+5   1.5   2  RT (min)      lcorbi_t3_pos_4_057_49    m/z 170.9577     0   2   4   6   8E+5   17.5   18  RT (min)      lcorbi_t3_pos_4_105_61    m/z 170.9577     0   2   4   6   8E+4   5   5.5  RT (min)      lcorbi_t3_pos_4_015_45    m/z 170.9577     0   1E+5   4.5   5  RT (min)      lcorbi_t3_pos_3_004_24    m/z 170.9577     0   2   4   6E+4   2   2.5  RT (min)      lcorbi_t3_pos_3_058_16    m/z 170.9577     0   2   4   6E+4   0.5   1  RT (min)      lcorbi_t3_pos_4_082_59    m/z 170.9577     0   1   2E+5   0.5   1  RT (min)      lcorbi_t3_pos_4_066_57    m/z 170.9577     0   2   4   6   8E+4   0.5   1  RT (min)      lcorbi_t3_pos_4_082_59    m/z 170.9577     0   1   2E+5   0.5   1  RT (min)      lcorbi_t3_pos_4_066_57    m/z 172.1696     0   1   2   3   4E+5   9.5   10  RT (min)      lcorbi_t3_pos_3_044_46    m/z 172.1696     0   1E+5   10   10.5  RT (min)      lcorbi_t3_pos_2_052_48    m/z 173.0961     0   1E+5   16   16.5  RT (min)       lcorbi_t3_pos_4_099_61    m/z 173.4111     0   1E+6   17.5   18  RT (min)       lcorbi_t3_pos_4_106_62    m/z 173.4116     0   2   4E+5   8.5   9  RT (min)       lcorbi_t3_pos_4_093_60    m/z 173.4120     0   1E+6   12   12.5  RT (min)       lcorbi_t3_pos_4_107_62    m/z 173.4121     0   2   4   6   8E+5   8.5   9  RT (min)      lcorbi_t3_pos_4_101_61    m/z 173.4121     0   1E+6   17.5   18  RT (min)       lcorbi_t3_pos_4_106_62    m/z 173.4121     0   2   4   6E+5   7   7.5  RT (min)       lcorbi_t3_pos_4_101_61    m/z 173.4122     0   1E+6   17.5   18  RT (min)       lcorbi_t3_pos_4_093_60    m/z 173.4122     0   1E+6   17   17.5  RT (min)       lcorbi_t3_pos_4_096_60    m/z 173.4122     0   1E+6   18   18.5  RT (min)       lcorbi_t3_pos_4_076_58    m/z 173.4123     0   2   4   6   8E+5   11   11.5  RT (min)       lcorbi_t3_pos_4_101_61    m/z 173.4123     0   1E+6   12   12.5  RT (min)       lcorbi_t3_pos_4_098_61    m/z 173.4123     0   2   4   6   8E+5   10   10.5  RT (min)       lcorbi_t3_pos_4_104_61    m/z 173.4123     0   1E+6   15.5   16  RT (min)       lcorbi_t3_pos_4_089_59    m/z 173.4123     0   1   2E+6   16.5   17  RT (min)       lcorbi_t3_pos_4_092_60    m/z 173.4123     0   2   4   6E+5   10   10.5  RT (min)       lcorbi_t3_pos_4_091_60    m/z 173.4123     0   1   18.5   19  RT (min)    lcorbi_t3_pos_1_023_02    m/z 173.4123     0   2   4   6E+5   10   10.5  RT (min)      lcorbi_t3_pos_4_094_60    m/z 173.4123     0   1E+6   11   11.5  RT (min)       lcorbi_t3_pos_4_086_59    m/z 173.4123     0   1   18.5   19  RT (min)    lcorbi_t3_pos_1_023_02    m/z 173.4123     0   2   4   6E+5   9   9.5  RT (min)       lcorbi_t3_pos_4_084_59    m/z 173.4123     0   1E+6   17   17.5  RT (min)       lcorbi_t3_pos_4_076_58    m/z 173.4124     0   2   4   6E+5   10   10.5  RT (min)      lcorbi_t3_pos_4_085_59    m/z 173.4124     0   1   2E+6   15   15.5  RT (min)       lcorbi_t3_pos_4_085_59    m/z 173.4124     0   1E+6   17   17.5  RT (min)       lcorbi_t3_pos_4_101_61    m/z 173.4124     0   1E+6   13.5   14  RT (min)       lcorbi_t3_pos_4_104_61    m/z 173.4124     0   1E+6   11.5   12  RT (min)       lcorbi_t3_pos_4_102_61    m/z 173.4124     0   1E+6   16.5   17  RT (min)       lcorbi_t3_pos_4_074_58    m/z 173.4124     0   2   4   6E+5   10   10.5  RT (min)       lcorbi_t3_pos_4_083_59    m/z 173.4124     0   2   4   6E+5   10.5   11  RT (min)       lcorbi_t3_pos_4_085_59    m/z 173.4124     0   1E+6   16   16.5  RT (min)       lcorbi_t3_pos_4_074_58    m/z 173.4124     0   2   4   6   8E+5   10.5   11  RT (min)       lcorbi_t3_pos_4_081_58    m/z 173.4124     0   2   4   6E+5   9.5   10  RT (min)       lcorbi_t3_pos_4_100_61    m/z 173.4124     0   1E+6   17.5   18  RT (min)       lcorbi_t3_pos_4_100_61    m/z 173.4124     0   1   2   3   4E+6   13   13.5  RT (min)       lcorbi_t3_pos_4_080_58    m/z 173.4124     0   1E+6   17   17.5  RT (min)       lcorbi_t3_pos_4_080_58    m/z 173.4124     0   1E+6   16.5   17  RT (min)       lcorbi_t3_pos_4_097_60    m/z 173.4124     0   2   4   6E+5   9.5   10  RT (min)       lcorbi_t3_pos_4_085_59    m/z 173.4124     0   2   4E+5   9   9.5  RT (min)       lcorbi_t3_pos_4_093_60    m/z 173.4124     0   1E+6   15   15.5  RT (min)       lcorbi_t3_pos_4_103_61    m/z 173.4124     0   2   4   6E+5   9.5   10  RT (min)      lcorbi_t3_pos_4_083_59    m/z 173.4124     0   2   4   6E+5   9.5   10  RT (min)       lcorbi_t3_pos_4_076_58    m/z 173.4124     0   2   4   6   8E+5   10.5   11  RT (min)       lcorbi_t3_pos_4_103_61    m/z 173.4124     0   1E+6   15.5   16  RT (min)       lcorbi_t3_pos_4_089_59    m/z 173.4124     0   1E+6   9   9.5  RT (min)       lcorbi_t3_pos_4_104_61    m/z 173.4124     0   2   4   6E+5   10.5   11  RT (min)       lcorbi_t3_pos_4_106_62    m/z 173.4124     0   1E+6   12.5   13  RT (min)       lcorbi_t3_pos_4_098_61    m/z 173.4124     0   1E+6   12   12.5  RT (min)       lcorbi_t3_pos_4_111_62    m/z 173.4124     0   1E+6   18   18.5  RT (min)      lcorbi_t3_pos_4_079_58    m/z 173.4124     0   1   2E+6   13   13.5  RT (min)       lcorbi_t3_pos_4_075_58    m/z 173.4124     0   1E+6   12.5   13  RT (min)       lcorbi_t3_pos_4_105_61    m/z 173.4124     0   1   18   18.5  RT (min)     lcorbi_t3_pos_1_023_02    m/z 173.4124     0   1E+6   18   18.5  RT (min)       lcorbi_t3_pos_4_072_57    m/z 173.4124     0   2   4   6   8E+5   10   10.5  RT (min)       lcorbi_t3_pos_4_102_61    m/z 173.4124     0   1   2E+6   18   18.5  RT (min)       lcorbi_t3_pos_4_090_60    m/z 173.4124     0   2   4   6   8E+5   11   11.5  RT (min)       lcorbi_t3_pos_4_070_57    m/z 173.4124     0   1   2E+6   13   13.5  RT (min)       lcorbi_t3_pos_4_063_17    m/z 173.4124     0   1E+6   11.5   12  RT (min)       lcorbi_t3_pos_4_074_58    m/z 173.4124     0   2   4   6   8E+5   11   11.5  RT (min)       lcorbi_t3_pos_4_098_61    m/z 173.4124     0   1   2E+6   12.5   13  RT (min)       lcorbi_t3_pos_4_074_58    m/z 173.4124     0   1E+6   17.5   18  RT (min)       lcorbi_t3_pos_4_101_61    m/z 173.4124     0   1E+6   13   13.5  RT (min)       lcorbi_t3_pos_4_087_59    m/z 173.4124     0   1E+6   14   14.5  RT (min)       lcorbi_t3_pos_4_105_61    m/z 173.4124     0   2   4   6E+5   8   8.5  RT (min)       lcorbi_t3_pos_4_108_62    m/z 173.4125     0   1E+6   17   17.5  RT (min)       lcorbi_t3_pos_4_094_60    m/z 173.4125     0   1E+6   14   14.5  RT (min)      lcorbi_t3_pos_4_088_59    m/z 173.4125     0   2   4   6   8E+5   10.5   11  RT (min)       lcorbi_t3_pos_4_073_57    m/z 173.4125     0   2   4   6   8E+5   11   11.5  RT (min)       lcorbi_t3_pos_4_107_62    m/z 173.4125     0   2   4   6E+5   9.5   10  RT (min)       lcorbi_t3_pos_4_074_58    m/z 173.4125     0   1E+6   18   18.5  RT (min)       lcorbi_t3_pos_4_081_58    m/z 173.4125     0   1E+6   12   12.5  RT (min)       lcorbi_t3_pos_4_105_61    m/z 173.4125     0   1E+6   14   14.5  RT (min)      lcorbi_t3_pos_4_101_61    m/z 173.4125     0   2   4   6E+5   10.5   11  RT (min)       lcorbi_t3_pos_4_089_59    m/z 173.4125     0   1E+6   12.5   13  RT (min)       lcorbi_t3_pos_4_082_59    m/z 173.4125     0   1E+6   14.5   15  RT (min)       lcorbi_t3_pos_4_101_61    m/z 173.4125     0   1E+6   16   16.5  RT (min)       lcorbi_t3_pos_4_071_57    m/z 173.4125     0   1E+6   15.5   16  RT (min)       lcorbi_t3_pos_4_096_60    m/z 173.4125     0   1E+6   15   15.5  RT (min)       lcorbi_t3_pos_4_110_62    m/z 173.4125     0   1E+6   15.5   16  RT (min)       lcorbi_t3_pos_4_101_61    m/z 173.4125     0   1E+6   12   12.5  RT (min)       lcorbi_t3_pos_4_107_62    m/z 173.4125     0   2   4   6   8E+5   11.5   12  RT (min)       lcorbi_t3_pos_4_109_62    m/z 173.4125     0   1E+6   14   14.5  RT (min)       lcorbi_t3_pos_4_101_61    m/z 173.4125     0   1E+6   14   14.5   15  RT (min)       lcorbi_t3_pos_4_087_59    m/z 173.4125     0   1E+6   12.5   13  RT (min)       lcorbi_t3_pos_4_089_59    m/z 173.4125     0   1   2E+6   15   15.5  RT (min)       lcorbi_t3_pos_4_091_60    m/z 173.4125     0   1E+6   16   16.5  RT (min)       lcorbi_t3_pos_4_089_59    m/z 173.4125     0   1E+6   17.5   18  RT (min)       lcorbi_t3_pos_4_078_58    m/z 173.4125     0   1E+6   13.5   14  RT (min)       lcorbi_t3_pos_4_109_62    m/z 173.4125     0   1E+6   13   13.5  RT (min)       lcorbi_t3_pos_4_100_61    m/z 173.4125     0   1E+6   14   14.5  RT (min)       lcorbi_t3_pos_4_104_61    m/z 173.4125     0   1E+6   13.5   14  RT (min)      lcorbi_t3_pos_4_080_58    m/z 173.4125     0   1E+6   16   16.5  RT (min)       lcorbi_t3_pos_4_090_60    m/z 173.4125     0   1   2   3   4E+6   13   13.5  RT (min)       lcorbi_t3_pos_4_080_58    m/z 173.4125     0   1E+6   15.5   16  RT (min)       lcorbi_t3_pos_4_082_59    m/z 173.4125     0   1E+6   16   16.5  RT (min)       lcorbi_t3_pos_4_083_59    m/z 173.4125     0   1E+6   15.5   16  RT (min)       lcorbi_t3_pos_4_111_62    m/z 173.4125     0   1E+6   11.5   12  RT (min)       lcorbi_t3_pos_4_102_61    m/z 173.4125     0   2   4   6   8E+5   11   11.5  RT (min)       lcorbi_t3_pos_4_070_57    m/z 173.4125     0   1E+6   12.5   13  RT (min)       lcorbi_t3_pos_4_088_59    m/z 173.4125     0   1E+6   11.5   12  RT (min)       lcorbi_t3_pos_4_071_57    m/z 173.4125     0   1E+6   12.5   13  RT (min)       lcorbi_t3_pos_4_099_61    m/z 173.4125     0   2   4   6E+5   10   10.5  RT (min)       lcorbi_t3_pos_4_106_62    m/z 173.4125     0   2   4   6   8E+5   11   11.5  RT (min)       lcorbi_t3_pos_4_083_59    m/z 173.4125     0   1E+6   15   15.5  RT (min)      lcorbi_t3_pos_4_083_59    m/z 173.4126     0   1E+6   15.5   16  RT (min)       lcorbi_t3_pos_4_111_62    m/z 173.4126     0   2   4E+5   9   9.5  RT (min)       lcorbi_t3_pos_4_091_60    m/z 173.4126     0   1E+6   17.5   18  RT (min)       lcorbi_t3_pos_4_100_61    m/z 173.4126     0   1E+6   17.5   18  RT (min)       lcorbi_t3_pos_4_096_60    m/z 173.4126     0   1E+6   12   12.5  RT (min)       lcorbi_t3_pos_4_078_58    m/z 173.4126     0   1E+6   12   12.5  RT (min)       lcorbi_t3_pos_4_100_61    m/z 173.4126     0   1E+6   12   12.5  RT (min)       lcorbi_t3_pos_4_100_61    m/z 173.4126     0   1E+6   16   17  RT (min)       lcorbi_t3_pos_4_109_62    m/z 173.4126     0   1E+6   11.5   12  RT (min)       lcorbi_t3_pos_4_104_61    m/z 173.4126     0   1   2E+6   16.5   17  RT (min)       lcorbi_t3_pos_4_093_60    m/z 173.4126     0   1E+6   15   15.5  RT (min)       lcorbi_t3_pos_4_105_61    m/z 173.4126     0   1E+6   18   18.5  RT (min)       lcorbi_t3_pos_4_110_62    m/z 173.4127     0   1E+6   11.5   12  RT (min)      lcorbi_t3_pos_4_099_61    m/z 173.4127     0   1   2E+6   17   17.5  RT (min)       lcorbi_t3_pos_4_103_61    m/z 173.4127     0   1E+6   14.5   15  RT (min)      lcorbi_t3_pos_4_094_60    m/z 173.4127     0   1   19   19.5  RT (min)    lcorbi_t3_pos_1_023_02    m/z 173.4128     0   1   18.5   19  RT (min)    lcorbi_t3_pos_1_001_49    m/z 173.4128     0   1E+6   18   18.5  RT (min)       lcorbi_t3_pos_4_076_58    m/z 173.4132     0   1E+6   17   17.5  RT (min)       lcorbi_t3_pos_4_106_62    m/z 174.1853     0   1   2   3   4E+6   1   1.5  RT (min)      lcorbi_t3_pos_4_057_49    m/z 175.0609     0   1   2   3E+4   1.5   2  RT (min)       lcorbi_t3_pos_4_057_49    m/z 175.0642     0   1   2   3E+5   10   12  RT (min)       lcorbi_t3_pos_4_076_58    m/z 175.0642     0   1   2   3E+5   10   12  RT (min)       lcorbi_t3_pos_4_093_60    m/z 175.0642     0   1   2   3E+5   10   12  RT (min)       lcorbi_t3_pos_4_083_59    m/z 175.0642     0   1   2   3E+5   10   12  RT (min)       lcorbi_t3_pos_4_075_58    m/z 175.0642     0   1   2   3E+5   8.5   9  RT (min)       lcorbi_t3_pos_4_096_60    m/z 175.0642     0   1   2   3E+6   18.5   19  RT (min)      lcorbi_t3_pos_4_076_58    m/z 176.0836     0   1E+5   1   1.5  RT (min)       lcorbi_t3_pos_4_085_59    m/z 176.9845     0   1E+6   0   0.5  RT (min)      lcorbi_t3_pos_1_007_07    m/z 177.0048     0   1E+6   0   0.5  RT (min)      lcorbi_t3_pos_1_005_42    m/z 177.0797     0   2   4E+6   17.5   18  RT (min)      lcorbi_t3_pos_4_105_61    m/z 177.0798     0   1E+6   8  RT (min)       lcorbi_t3_pos_1_028_32    m/z 177.0798     0   1E+6   7   8  RT (min)       lcorbi_t3_pos_4_109_62    m/z 177.0798     0   2   4E+6   18.5   19  RT (min)       lcorbi_t3_pos_4_076_58    m/z 177.0798     0   2   4   6   8E+5   10   10.5  RT (min)      lcorbi_t3_pos_1_064_16    m/z 177.0798     0   2   4E+6   17.5   18  RT (min)      lcorbi_t3_pos_4_105_61    m/z 177.0798     0   1   2   3E+6   18.5   19  RT (min)       lcorbi_t3_pos_4_072_57    m/z 177.0798     0   2   4E+5   5.5   6  RT (min)      lcorbi_t3_pos_3_060_39    m/z 177.0798     0   2   4E+6   18   18.5  RT (min)       lcorbi_t3_pos_4_084_59    m/z 177.0798     0   1E+6   7   8  RT (min)       lcorbi_t3_pos_1_009_50    m/z 177.0798     0   2   4E+5   11.5   12  RT (min)      lcorbi_t3_pos_3_057_49    m/z 177.0798     0   2   4   6E+5   11   11.5  RT (min)      lcorbi_t3_pos_1_061_50    m/z 177.0798     0   1   2   3E+6   1.5   2  RT (min)      lcorbi_t3_pos_4_057_49    m/z 177.0798     0   2   4   6E+5   6   6.5  RT (min)      lcorbi_t3_pos_1_062_13    m/z 177.0798     0   2   4   6E+5   5.5   6  RT (min)      lcorbi_t3_pos_4_057_49    m/z 177.0798     0   2   4E+6   17.5   18  RT (min)       lcorbi_t3_pos_4_095_60    m/z 177.0798     0   2   4E+6   18.5   19  RT (min)       lcorbi_t3_pos_4_077_58    m/z 177.0798     0   1E+6   7   8   9  RT (min)       lcorbi_t3_pos_4_098_61    m/z 177.0798     0   1E+6   10   15  RT (min)       lcorbi_t3_pos_1_036_08    m/z 177.0798     0   2   4   6E+5   10.5   11  RT (min)      lcorbi_t3_pos_2_063_16    m/z 177.0798     0   2   4E+6   18.5   19  RT (min)       lcorbi_t3_pos_4_077_58    m/z 177.0798     0   2   4E+6   18   18.5  RT (min)       lcorbi_t3_pos_4_080_58    m/z 177.0798     0   2   4   6E+5   9.5   10  RT (min)      lcorbi_t3_pos_3_057_49    m/z 177.0798     0   1E+6   7   8   9  RT (min)       lcorbi_t3_pos_4_070_57    m/z 177.0798     0   1E+6   8   9   10  RT (min)       lcorbi_t3_pos_1_008_36    m/z 177.0798     0   1   2   3   4E+6   18.5   19  RT (min)       lcorbi_t3_pos_4_077_58    m/z 177.1022     0   2   4   6   8E+5   3   3.5  RT (min)       lcorbi_t3_pos_4_069_57    m/z 177.1022     0   2   4   6   8E+5   3.5   4  RT (min)       lcorbi_t3_pos_4_066_57    m/z 178.0161     0   1E+6   0   0.5  RT (min)      lcorbi_t3_pos_1_005_42    m/z 178.0831     0   1   2   3E+5   18   18.5  RT (min)       lcorbi_t3_pos_4_066_57    m/z 178.0831     0   2   4   6E+4   9.5   10  RT (min)      lcorbi_t3_pos_1_031_40    m/z 178.0831     0   1   2   3E+5   18.5   19  RT (min)       lcorbi_t3_pos_4_069_57    m/z 178.0832     0   2   4   6E+4   8   9  RT (min)       lcorbi_t3_pos_1_002_01    m/z 179.0001     0   1   2E+5   0.5   1  RT (min)      lcorbi_t3_pos_1_032_26    m/z 179.0001     0   1   2E+5   1   1.5  RT (min)       lcorbi_t3_pos_1_020_31    m/z 179.0227     0   1E+5   18   18.5  RT (min)      lcorbi_t3_pos_4_094_60    m/z 179.0928     0   2   4   6E+5   3.5   4  RT (min)       lcorbi_t3_pos_4_090_60    m/z 179.0928     0   2   4   6E+5   3   3.5  RT (min)       lcorbi_t3_pos_4_094_60    m/z 180.0034     0   2   4   6   8E+5   0   0.5  RT (min)      lcorbi_t3_pos_1_005_42    m/z 180.0888     0   1   2E+5   9.5   10  RT (min)      lcorbi_t3_pos_1_028_32    m/z 181.0383     0   1E+6   18   19  RT (min)       lcorbi_t3_pos_4_093_60    m/z 181.0383     0   2   4   6E+5   4.5   5  RT (min)       lcorbi_t3_pos_2_027_05    m/z 181.0384     0   1E+6   18   18.5  RT (min)      lcorbi_t3_pos_4_107_62    m/z 181.0495     0   2   4E+4   14   14.5  RT (min)      lcorbi_t3_pos_1_028_32    m/z 181.0495     0   1   2   3   4E+5   13.5   14  RT (min)       lcorbi_t3_pos_3_047_40    m/z 181.0495     0   2   4E+4   15.5   16  RT (min)      lcorbi_t3_pos_1_023_02    m/z 181.0495     0   1E+5   12.5   13  RT (min)      lcorbi_t3_pos_1_002_01    m/z 181.0495     0   1   2   3   4E+4   14.5   15  RT (min)      lcorbi_t3_pos_1_034_34    m/z 181.0495     0   1E+4   12   12.5  RT (min)      lcorbi_t3_pos_2_032_39    m/z 181.0496     0   2   4E+4   14.5   15  RT (min)      lcorbi_t3_pos_1_031_40    m/z 181.0496     0   1   2E+5   13.5   14  RT (min)      lcorbi_t3_pos_1_008_36    m/z 181.0859     0   2   4E+4   15   15.5  RT (min)      lcorbi_t3_pos_1_051_09    m/z 181.0859     0   2   4E+4   15   15.5  RT (min)      lcorbi_t3_pos_1_051_09    m/z 181.0859     0   1   2   3E+5   13.5   14  RT (min)       lcorbi_t3_pos_3_047_40    m/z 181.0859     0   1E+5   13.5   14  RT (min)      lcorbi_t3_pos_1_008_36    m/z 181.0859     0   2   4   6E+4   14.5   15  RT (min)      lcorbi_t3_pos_1_034_34    m/z 181.0859     0   2   4E+4   15.5   16  RT (min)      lcorbi_t3_pos_1_023_02    m/z 181.0859     0   1   2   3   4E+4   14   14.5  RT (min)      lcorbi_t3_pos_1_028_32    m/z 181.1335     0   2   4   6E+5   0   0.5  RT (min)      lcorbi_t3_pos_1_001_49    m/z 182.1652     0   1   2E+6   0   0.5  RT (min)      lcorbi_t3_pos_1_001_49    m/z 183.1492     0   1   2   3   4E+6   0   0.5  RT (min)      lcorbi_t3_pos_1_001_49    m/z 187.0815     0   1   2E+5   0   0.5  RT (min)      lcorbi_t3_pos_2_013_33    m/z 187.0815     0   1E+5   1.5   2  RT (min)      lcorbi_t3_pos_4_070_57    m/z 187.0815     0   2   4   6   8E+5   18   18.5  RT (min)       lcorbi_t3_pos_4_076_58    m/z 187.0815     0   2   4   6   8E+5   18   18.5  RT (min)       lcorbi_t3_pos_4_066_57    m/z 187.0815     0   1   2   3   4E+5   1   1.5  RT (min)      lcorbi_t3_pos_4_077_58    m/z 187.0815     0   2   4   6   8E+5   18   18.5  RT (min)       lcorbi_t3_pos_4_066_57    m/z 187.0815     0   1   2   3   4E+5   18.5   19  RT (min)       lcorbi_t3_pos_4_100_61    m/z 187.0815     0   1   2E+5   0   0.5  RT (min)      lcorbi_t3_pos_2_019_13    m/z 187.0815     0   1   2   3E+5   0.5   1  RT (min)      lcorbi_t3_pos_4_080_58    m/z 187.0815     0   1   2   3E+5   18.5   19  RT (min)       lcorbi_t3_pos_4_070_57    m/z 187.0815     0   2   4   6   8E+5   17.5   18  RT (min)       lcorbi_t3_pos_4_105_61    m/z 187.0816     0   1E+6   17.5   18  RT (min)      lcorbi_t3_pos_4_095_60    m/z 189.0792     0   1E+5   18   18.5  RT (min)      lcorbi_t3_pos_4_097_60    m/z 189.0870     0   1   2   3E+4   1   1.5  RT (min)      lcorbi_t3_pos_4_090_60    m/z 189.0971     0   1E+6   19   19.5  RT (min)      lcorbi_t3_pos_4_069_57    m/z 189.0971     0   1   2   3E+6   18   18.5  RT (min)      lcorbi_t3_pos_4_072_57    m/z 189.0971     0   1   2E+6   0   1  RT (min)       lcorbi_t3_pos_1_006_15    m/z 189.0971     0   2   4   6E+5   5   5.5  RT (min)      lcorbi_t3_pos_4_060_16    m/z 189.0971     0   1   2   3E+6   18   18.5  RT (min)      lcorbi_t3_pos_4_072_57    m/z 189.0971     0   1E+6   18.5   19  RT (min)       lcorbi_t3_pos_4_072_57    m/z 189.0971     0   1   2   3E+6   18.5   19  RT (min)       lcorbi_t3_pos_4_072_57    m/z 189.0971     0   1E+6   5   10  RT (min)       lcorbi_t3_pos_4_080_58    m/z 189.0971     0   1E+6   18.5   19  RT (min)       lcorbi_t3_pos_4_072_57    m/z 189.0972     0   2   4   6E+5   9.5   10  RT (min)      lcorbi_t3_pos_2_060_14    m/z 189.0972     0   2   4   6E+5   7.5   8  RT (min)      lcorbi_t3_pos_3_059_17    m/z 189.0972     0   2   4   6   8E+5   6   8  RT (min)       lcorbi_t3_pos_4_067_57    m/z 189.0972     0   1   2   3E+6   10   15  RT (min)       lcorbi_t3_pos_4_105_61    m/z 189.0972     0   2   4   6E+5   8.5   9  RT (min)      lcorbi_t3_pos_2_060_14    m/z 189.0972     0   2   4E+5   10   10.5  RT (min)      lcorbi_t3_pos_4_057_49    m/z 189.0972     0   2   4E+5   10.5   11  RT (min)      lcorbi_t3_pos_2_060_14    m/z 189.0972     0   1E+6   6.5   7   7.5  RT (min)       lcorbi_t3_pos_1_046_22    m/z 189.0972     0   2   4   6E+5   5   5.5  RT (min)      lcorbi_t3_pos_2_060_14    m/z 189.0972     0   2   4   6E+5   7.5   8  RT (min)      lcorbi_t3_pos_4_057_49    m/z 190.0980     0   1E+5   6.5   7  RT (min)       lcorbi_t3_pos_1_019_16    m/z 190.0980     0   1   2   3E+5   18   18.5  RT (min)      lcorbi_t3_pos_4_081_58    m/z 190.0980     0   1E+5   18.5   19  RT (min)      lcorbi_t3_pos_4_069_57    m/z 190.0980     0   1   2   3E+5   18.5   19  RT (min)      lcorbi_t3_pos_4_076_58    m/z 190.0980     0   1   2   3E+5   18   18.5  RT (min)      lcorbi_t3_pos_4_076_58    m/z 190.0980     0   1E+5   0   0.5  RT (min)      lcorbi_t3_pos_1_008_36    m/z 190.0980     0   1E+5   0   0.5  RT (min)      lcorbi_t3_pos_1_006_15    m/z 190.0980     0   1   2E+5   0   0.5  RT (min)      lcorbi_t3_pos_1_029_29    m/z 190.0980     0   1   2   3E+5   18.5   19  RT (min)      lcorbi_t3_pos_4_076_58    m/z 190.1074     0   1E+5   1   1.5  RT (min)       lcorbi_t3_pos_3_029_13    m/z 190.1590     0   1E+6   5   5.5  RT (min)      lcorbi_t3_pos_3_000_aa    m/z 190.1590     0   1E+6   5   5.5  RT (min)      lcorbi_t3_pos_3_000_aa    m/z 190.1590     0   1E+6   5   5.5  RT (min)      lcorbi_t3_pos_3_000_aa    m/z 191.0226     0   1E+5   0   2  RT (min)       lcorbi_t3_pos_1_006_15    m/z 191.0226     0   1   2E+5   0   2  RT (min)       lcorbi_t3_pos_1_002_01    m/z 191.0226     0   1E+5   0   0.5  RT (min)      lcorbi_t3_pos_1_035_28    m/z 191.0226     0   1E+5   2   3  RT (min)       lcorbi_t3_pos_1_027_43    m/z 191.0227     0   1E+5   0   0.5  RT (min)      lcorbi_t3_pos_1_008_36    m/z 191.0227     0   1   2E+5   1   1.5  RT (min)       lcorbi_t3_pos_1_031_40    m/z 191.0227     0   1E+5   1   1.5  RT (min)      lcorbi_t3_pos_2_027_05    m/z 191.0227     0   1   2E+5   1   2   3  RT (min)       lcorbi_t3_pos_1_034_34    m/z 191.0702     0   1E+6   13.5   14  RT (min)      lcorbi_t3_pos_1_008_36    m/z 191.0702     0   1   2E+6   13.5   14  RT (min)      lcorbi_t3_pos_3_047_40    m/z 191.0702     0   2   4   6E+5   12.5   13  RT (min)      lcorbi_t3_pos_1_002_01    m/z 191.0702     0   1E+5   15   15.5  RT (min)      lcorbi_t3_pos_2_012_01    m/z 191.0702     0   1E+5   15.5   16  RT (min)      lcorbi_t3_pos_1_023_02    m/z 191.0702     0   1E+5   14.5   15  RT (min)      lcorbi_t3_pos_1_034_34    m/z 191.0703     0   1E+5   14   14.5  RT (min)      lcorbi_t3_pos_1_028_32    m/z 191.0703     0   1E+5   14   14.5  RT (min)      lcorbi_t3_pos_1_047_41    m/z 191.0703     0   1   2E+4   12   12.5  RT (min)      lcorbi_t3_pos_2_032_39    m/z 191.0703     0   1E+5   14.5   15  RT (min)      lcorbi_t3_pos_1_034_34    m/z 191.0703     0   1E+5   14.5   15  RT (min)      lcorbi_t3_pos_1_031_40    m/z 191.0703     0   1E+5   11.5   12  RT (min)      lcorbi_t3_pos_1_023_02    m/z 191.0703     0   2   4   6   8E+4   14   14.5  RT (min)      lcorbi_t3_pos_3_003_33    m/z 191.0901     0   1   11.5   12  RT (min)      lcorbi_t3_pos_1_038_30    m/z 191.0947     0   1E+5   18.5   19  RT (min)      lcorbi_t3_pos_4_077_58    m/z 191.0947     0   2   4E+5   18   18.5  RT (min)      lcorbi_t3_pos_4_082_59    m/z 191.0947     0   1   2E+5   0   0.5  RT (min)      lcorbi_t3_pos_1_029_29    m/z 191.0947     0   1   2E+5   0   0.5  RT (min)      lcorbi_t3_pos_1_022_20    m/z 191.0947     0   1   2E+5   1   1.5  RT (min)      lcorbi_t3_pos_4_077_58    m/z 191.0947     0   1   2E+5   0   0.5  RT (min)      lcorbi_t3_pos_1_022_20    m/z 191.0947     0   1   2   3E+5   18.5   19  RT (min)      lcorbi_t3_pos_4_084_59    m/z 191.0947     0   2   4E+5   18.5   19  RT (min)      lcorbi_t3_pos_4_069_57    m/z 191.0947     0   1E+5   0.5   1  RT (min)      lcorbi_t3_pos_1_005_42    m/z 191.0947     0   1E+5   0.5   1  RT (min)      lcorbi_t3_pos_1_023_02    m/z 191.0947     0   1E+5   6   7  RT (min)       lcorbi_t3_pos_4_090_60    m/z 191.0947     0   2   4   6E+4   1   1.5  RT (min)      lcorbi_t3_pos_1_006_15    m/z 191.0947     0   1   2E+5   18.5   19  RT (min)      lcorbi_t3_pos_4_084_59    m/z 191.0947     0   1E+5   5.5   6  RT (min)       lcorbi_t3_pos_4_099_61    m/z 191.0948     0   2   4   6E+5   17.5   18  RT (min)      lcorbi_t3_pos_4_105_61    m/z 191.1066     0   2   4E+5   11   12   13  RT (min)       lcorbi_t3_pos_1_032_26    m/z 191.1066     0   2   4   6   8E+5   14   15   16  RT (min)       lcorbi_t3_pos_1_040_27    m/z 191.1066     0   2   4   6   8E+5   14   15  RT (min)       lcorbi_t3_pos_4_092_60    m/z 191.1066     0   2   4   6   8E+4   8.5   9  RT (min)      lcorbi_t3_pos_1_021_18    m/z 191.1066     0   1E+5   8.5   9  RT (min)      lcorbi_t3_pos_4_078_58    m/z 191.1066     0   1E+6   12.5   15  RT (min)       lcorbi_t3_pos_4_098_61    m/z 191.1066     0   2   4   6   8E+5   16  RT (min)       lcorbi_t3_pos_4_095_60    m/z 191.1067     0   1   17   17.5  RT (min)     lcorbi_t3_pos_1_031_40    m/z 191.1067     0   2   4   6   8E+5   17  RT (min)       lcorbi_t3_pos_4_094_60    m/z 191.1067     0   2   4   6   8E+5   12.5   15  RT (min)       lcorbi_t3_pos_4_113_62    m/z 191.1178     0   2   4E+6   3.5   4  RT (min)       lcorbi_t3_pos_4_106_62    m/z 191.1178     0   1   2   3E+6   2.5   3  RT (min)       lcorbi_t3_pos_4_113_62    m/z 191.1543     0   1E+6   0   0.5  RT (min)      lcorbi_t3_pos_1_001_49    m/z 191.1794     0   1   2   3E+4   12   12.5  RT (min)      lcorbi_t3_pos_1_023_02    m/z 191.1794     0   1   2   3E+5   11   11.5  RT (min)      lcorbi_t3_pos_1_055_33    m/z 191.1794     0   1E+6   10.5   11  RT (min)      lcorbi_t3_pos_3_037_32    m/z 191.1794     0   1   2E+4   12   12.5  RT (min)      lcorbi_t3_pos_2_012_01    m/z 191.1794     0   1   2   3E+4   11.5   12  RT (min)      lcorbi_t3_pos_2_014_37    m/z 191.1794     0   1E+6   11   11.5  RT (min)      lcorbi_t3_pos_1_008_36    m/z 191.1794     0   1   2   3E+4   12   12.5  RT (min)      lcorbi_t3_pos_1_023_02    m/z 191.1794     0   1E+6   10.5   11  RT (min)      lcorbi_t3_pos_2_054_29    m/z 192.1100     0   1E+5   15   16   17  RT (min)       lcorbi_t3_pos_4_103_61    m/z 192.1211     0   2   4   6E+5   3.5   4  RT (min)      lcorbi_t3_pos_4_106_62    m/z 193.1699     0   1E+7   0   0.5  RT (min)      lcorbi_t3_pos_1_002_01    m/z 194.1732     0   1   2E+6   0   0.5  RT (min)      lcorbi_t3_pos_1_002_01    m/z 195.0876     0   2   4E+5   3.5   4  RT (min)       lcorbi_t3_pos_1_006_15    m/z 195.0877     0   1E+4   1.5   2  RT (min)       lcorbi_t3_pos_4_057_49    m/z 195.0877     0   1   2E+5   14.5   15   15.5  RT (min)       lcorbi_t3_pos_1_004_24    m/z 195.0903     0   1E+6   18.5   19  RT (min)       lcorbi_t3_pos_4_076_58    m/z 195.0904     0   1E+6   17.5   18  RT (min)      lcorbi_t3_pos_4_075_58    m/z 195.0904     0   1E+5   5   5.5  RT (min)       lcorbi_t3_pos_4_012_37    m/z 195.0904     0   1   2   3E+5   10   12  RT (min)       lcorbi_t3_pos_4_030_05    m/z 195.0904     0   1   2E+5   6  RT (min)       lcorbi_t3_pos_2_016_23    m/z 195.0904     0   2   4   6   8E+4   5   5.5  RT (min)      lcorbi_t3_pos_2_032_39    m/z 195.0904     0   1E+5   12.5   13  RT (min)       lcorbi_t3_pos_1_026_17    m/z 195.0904     0   1E+5   13   13.5  RT (min)       lcorbi_t3_pos_1_018_35    m/z 195.0904     0   1   2   3E+5   7   7.5  RT (min)       lcorbi_t3_pos_3_034_26    m/z 195.1227     0   2   4   6E+5   4.5   5  RT (min)      lcorbi_t3_pos_4_000_aa    m/z 195.1492     0   2   4E+5   5.5   6  RT (min)      lcorbi_t3_pos_4_003_39    m/z 196.1194     0   2   4   6   8E+5   1   1.5  RT (min)      lcorbi_t3_pos_4_092_60    m/z 197.0271     0   2   4E+5   17.5   18  RT (min)      lcorbi_t3_pos_4_095_60    m/z 197.0272     0   2   4E+5   17.5   18  RT (min)      lcorbi_t3_pos_4_105_61    m/z 197.0808     0   1   2E+4   14   14.5  RT (min)      lcorbi_t3_pos_2_013_33    m/z 197.0808     0   1   16.5   17  RT (min)    lcorbi_t3_pos_1_034_34    m/z 197.0809     0   1   15.5   16  RT (min)      lcorbi_t3_pos_1_008_36    m/z 197.0809     0   1   16.5   17  RT (min)    lcorbi_t3_pos_1_028_32    m/z 197.0809     0   1   17.5   18  RT (min)    lcorbi_t3_pos_1_031_40    m/z 198.1237     0   1E+5   1   1.5  RT (min)       lcorbi_t3_pos_4_088_59    m/z 198.1237     0   1E+5   1   1.5  RT (min)       lcorbi_t3_pos_4_082_59    m/z 199.1692     0   1   2E+5   10   10.5  RT (min)      lcorbi_t3_pos_3_047_40    m/z 199.1692     0   1   2E+5   10   10.5  RT (min)      lcorbi_t3_pos_1_038_30    m/z 200.2009     0   1   2E+5   10   10.5  RT (min)      lcorbi_t3_pos_1_002_01    m/z 201.0434     0   1   2E+5   18   18.5  RT (min)      lcorbi_t3_pos_4_069_57    m/z 202.0161     0   2   4   6E+6   0   0.5  RT (min)      lcorbi_t3_pos_1_002_01    m/z 202.0396     0   1E+5   4   5  RT (min)       lcorbi_t3_pos_4_098_61    m/z 202.0777     0   2   4   6   8E+5   0   0.5  RT (min)      lcorbi_t3_pos_2_057_49    m/z 202.1802     0   2   4   6   8E+5   9   9.5  RT (min)      lcorbi_t3_pos_2_052_48    m/z 202.2165     0   1   2   3E+6   1   1.5  RT (min)      lcorbi_t3_pos_4_057_49    m/z 203.0590     0   1E+6   17.5   18  RT (min)      lcorbi_t3_pos_4_105_61    m/z 203.0591     0   1E+5   0   0.5  RT (min)      lcorbi_t3_pos_2_025_52    m/z 203.0591     0   1E+5   0.5   1  RT (min)      lcorbi_t3_pos_1_049_55    m/z 203.0591     0   2   4   6   8E+5   1.5   2  RT (min)      lcorbi_t3_pos_4_057_49    m/z 203.0591     0   1   2   3E+5   0   0.5  RT (min)       lcorbi_t3_pos_1_038_30    m/z 203.0591     0   1   2E+5   0   0.5   1  RT (min)       lcorbi_t3_pos_1_047_41    m/z 203.0591     0   1E+6   17.5   18  RT (min)      lcorbi_t3_pos_4_105_61    m/z 203.0829     0   1E+6   0   1  RT (min)       lcorbi_t3_pos_4_099_61    m/z 203.0855     0   2   4   6E+5   2.5   5   7.5  RT (min)       lcorbi_t3_pos_1_061_50    m/z 203.0855     0   2   4   6E+5   2.5   5   7.5  RT (min)       lcorbi_t3_pos_4_059_15    m/z 203.0855     0   2   4   6E+5   1   1.5  RT (min)       lcorbi_t3_pos_4_059_15    m/z 204.0624     0   1E+5   18   18.5  RT (min)      lcorbi_t3_pos_4_091_60    m/z 204.0624     0   1E+5   18   18.5  RT (min)      lcorbi_t3_pos_4_104_61    m/z 204.0625     0   1E+5   18   18.5  RT (min)      lcorbi_t3_pos_4_073_57    m/z 204.1230     0   2   4E+6   1.5   2  RT (min)       lcorbi_t3_pos_4_066_57    m/z 205.0746     0   1   2   3E+6   18.5   19  RT (min)      lcorbi_t3_pos_4_076_58    m/z 205.0747     0   1   2   3E+6   18.5   19  RT (min)      lcorbi_t3_pos_4_076_58    m/z 205.0747     0   2   4   6E+5   11   11.5  RT (min)      lcorbi_t3_pos_2_058_15    m/z 205.0747     0   2   4   6   8E+6   18.5   19  RT (min)       lcorbi_t3_pos_4_091_60    m/z 205.0747     0   2   4   6   8E+6   18   18.5  RT (min)       lcorbi_t3_pos_4_091_60    m/z 205.0747     0   2   4   6   8E+5   7.5   8  RT (min)      lcorbi_t3_pos_4_064_39    m/z 205.0747     0   2   4   6E+6   18.5   19  RT (min)      lcorbi_t3_pos_4_072_57    m/z 205.0747     0   1E+6   10   15  RT (min)       lcorbi_t3_pos_1_030_21    m/z 205.0747     0   2   4E+5   11.5   12  RT (min)      lcorbi_t3_pos_3_064_14    m/z 205.0747     0   2   4E+6   18.5   19  RT (min)      lcorbi_t3_pos_4_079_58    m/z 205.0747     0   1   2   3   4E+6   1.5   2  RT (min)      lcorbi_t3_pos_4_057_49    m/z 205.0747     0   2   4   6   8E+6   17.5   18  RT (min)      lcorbi_t3_pos_4_105_61    m/z 205.0747     0   2   4   6   8E+5   10.5   11  RT (min)      lcorbi_t3_pos_2_058_15    m/z 205.0747     0   2   4   6   8E+6   18   18.5  RT (min)       lcorbi_t3_pos_4_104_61    m/z 205.0747     0   1E+6   7   7.5   8  RT (min)       lcorbi_t3_pos_1_037_19    m/z 205.0747     0   2   4   6   8E+5   5.5   6  RT (min)      lcorbi_t3_pos_2_058_15    m/z 205.0747     0   2   4   6   8E+5   4.5   5  RT (min)      lcorbi_t3_pos_4_057_49    m/z 205.0747     0   2   4   6E+5   10.5   11  RT (min)      lcorbi_t3_pos_3_064_14    m/z 205.0747     0   2   4   6   8E+6   17.5   18  RT (min)      lcorbi_t3_pos_4_105_61    m/z 205.0747     0   2   4E+6   18.5   19  RT (min)      lcorbi_t3_pos_4_079_58    m/z 205.0747     0   2   4   6   8E+6   10   15  RT (min)       lcorbi_t3_pos_4_075_58    m/z 205.0859     0   1E+5   10   10.5  RT (min)      lcorbi_t3_pos_3_024_07    m/z 206.0781     0   1E+5   7   7.5  RT (min)       lcorbi_t3_pos_4_105_61    m/z 206.0781     0   1E+5   7   7.5  RT (min)       lcorbi_t3_pos_4_016_15    m/z 206.0781     0   2   4E+5   17.5   18  RT (min)      lcorbi_t3_pos_4_105_61    m/z 207.1077     0   2   4   6E+5   0   0.5  RT (min)      lcorbi_t3_pos_1_027_43    m/z 207.1077     0   1   2   3   4E+5   1   1.5  RT (min)       lcorbi_t3_pos_4_077_58    m/z 207.1077     0   1E+5   14.5   15  RT (min)       lcorbi_t3_pos_4_100_61    m/z 207.1078     0   1E+6   17.5   18  RT (min)      lcorbi_t3_pos_4_075_58    m/z 207.1078     0   1   2   3   4E+5   6  RT (min)       lcorbi_t3_pos_3_039_03    m/z 208.1808     0   1E+6   0   0.5  RT (min)      lcorbi_t3_pos_1_001_49    m/z 209.0808     0   1   2E+6   12.5   13  RT (min)      lcorbi_t3_pos_4_046_44    m/z 209.0808     0   2   4   6   8E+5   10   10.5  RT (min)      lcorbi_t3_pos_1_002_01    m/z 209.0808     0   1   2E+5   11   11.5  RT (min)      lcorbi_t3_pos_1_028_32    m/z 209.0808     0   1   2   3E+5   11   11.5  RT (min)      lcorbi_t3_pos_1_031_40    m/z 209.0808     0   1   2   3E+5   11   11.5  RT (min)      lcorbi_t3_pos_1_031_40    m/z 209.0808     0   2   4   6E+6   12.5   13  RT (min)      lcorbi_t3_pos_2_054_29    m/z 209.0809     0   1   2   3E+5   14   14.5  RT (min)      lcorbi_t3_pos_1_028_32    m/z 209.0809     0   1   2   3   4E+5   14   14.5  RT (min)      lcorbi_t3_pos_3_044_46    m/z 209.0809     0   1   2   3E+5   15.5   16  RT (min)      lcorbi_t3_pos_1_023_02    m/z 209.0809     0   1   2   3   4E+6   13.5   14  RT (min)      lcorbi_t3_pos_4_001_49    m/z 209.0809     0   1   2   3E+6   13.5   14  RT (min)      lcorbi_t3_pos_1_039_23    m/z 209.0809     0   1   2E+5   15   15.5  RT (min)      lcorbi_t3_pos_2_052_48    m/z 209.0809     0   1   2   3   4E+5   14   14.5  RT (min)      lcorbi_t3_pos_1_033_53    m/z 209.0809     0   1   2   3E+5   15   15.5  RT (min)      lcorbi_t3_pos_1_051_09    m/z 209.0809     0   1   2   3E+5   14.5   15  RT (min)      lcorbi_t3_pos_1_034_34    m/z 209.0809     0   1   2   3E+6   13.5   14  RT (min)      lcorbi_t3_pos_1_008_36    m/z 209.0809     0   1   2   3   4E+5   15   15.5  RT (min)      lcorbi_t3_pos_1_001_49    m/z 209.0809     0   1   2   3   4E+4   12   12.5  RT (min)      lcorbi_t3_pos_2_032_39    m/z 209.1172     0   1E+6   10   12.5   15  RT (min)       lcorbi_t3_pos_4_083_59    m/z 209.1172     0   1E+6   14   16  RT (min)       lcorbi_t3_pos_4_111_62    m/z 209.1172     0   1E+5   7.5   8  RT (min)       lcorbi_t3_pos_4_066_57    m/z 209.1172     0   1E+5   7.5   8  RT (min)      lcorbi_t3_pos_3_000_aa    m/z 209.1172     0   2   4   6E+5   10   11  RT (min)       lcorbi_t3_pos_4_073_57    m/z 209.1172     0   2   4   6E+5   13.5   14  RT (min)       lcorbi_t3_pos_1_057_49    m/z 209.1172     0   1E+6   15   16   17  RT (min)       lcorbi_t3_pos_4_074_58    m/z 209.1172     0   1E+6   10   15  RT (min)       lcorbi_t3_pos_4_092_60    m/z 209.1172     0   1E+6   15.5   16  RT (min)       lcorbi_t3_pos_1_010_46    m/z 209.1172     0   2   4   6   8E+5   14   14.5  RT (min)      lcorbi_t3_pos_1_057_49    m/z 209.1172     0   1E+6   12.5   15  RT (min)       lcorbi_t3_pos_4_112_62    m/z 209.1172     0   1E+6   16  RT (min)       lcorbi_t3_pos_4_070_57    m/z 209.1649     0   1   2E+6   0   0.5  RT (min)      lcorbi_t3_pos_1_002_01    m/z 210.0842     0   1E+4   12   12.5  RT (min)      lcorbi_t3_pos_2_032_39    m/z 210.0842     0   2   4   6E+4   14.5   15  RT (min)      lcorbi_t3_pos_1_041_54    m/z 210.0842     0   1E+4   12.5   13  RT (min)      lcorbi_t3_pos_2_013_33    m/z 210.0842     0   2   4   6E+4   14   14.5  RT (min)      lcorbi_t3_pos_1_033_53    m/z 210.0842     0   2   4   6   8E+5   13.5   14  RT (min)      lcorbi_t3_pos_1_005_42    m/z 210.0842     0   2   4   6   8E+5   12.5   13  RT (min)      lcorbi_t3_pos_2_054_29    m/z 210.0842     0   1E+5   15.5   16  RT (min)      lcorbi_t3_pos_1_023_02    m/z 210.0842     0   2   4   6E+4   14.5   15  RT (min)      lcorbi_t3_pos_1_034_34    m/z 210.0842     0   1   2   3E+5   13.5   14  RT (min)      lcorbi_t3_pos_1_008_36    m/z 210.0842     0   1   2   3   4E+4   14   14.5  RT (min)      lcorbi_t3_pos_3_003_33    m/z 210.0842     0   2   4   6E+4   14.5   15  RT (min)      lcorbi_t3_pos_1_031_40    m/z 210.0842     0   1   2   3   4E+4   14   14.5  RT (min)      lcorbi_t3_pos_2_022_38    m/z 210.0842     0   2   4   6   8E+4   14   14.5  RT (min)      lcorbi_t3_pos_2_017_51    m/z 210.0842     0   2   4E+4   15   15.5  RT (min)      lcorbi_t3_pos_2_012_01    m/z 210.1965     0   2   4   6E+6   0   0.5  RT (min)      lcorbi_t3_pos_1_002_01    m/z 211.1936     0   1   2E+5   16   18  RT (min)       lcorbi_t3_pos_4_099_61    m/z 213.0046     0   2   4   6E+5   17.5   18  RT (min)      lcorbi_t3_pos_4_105_61    m/z 216.9228     0   2   4E+5   18   18.5  RT (min)      lcorbi_t3_pos_4_080_58    m/z 216.9228     0   2   4   6E+5   18   18.5  RT (min)      lcorbi_t3_pos_4_111_62    m/z 217.1090     0   2   4   6   8E+5   17.5   18  RT (min)      lcorbi_t3_pos_4_105_61    m/z 218.1954     0   1E+5   7   7.5  RT (min)      lcorbi_t3_pos_4_080_58    m/z 218.2114     0   2   4   6E+7   1   1.5  RT (min)      lcorbi_t3_pos_4_057_49    m/z 219.0315     0   1E+6   0   0.5  RT (min)      lcorbi_t3_pos_1_007_07    m/z 219.0518     0   1E+6   0   0.5  RT (min)      lcorbi_t3_pos_1_005_42    m/z 219.1267     0   1   2E+6   17.5   18  RT (min)      lcorbi_t3_pos_4_105_61    m/z 219.1267     0   1E+6   1.5   2  RT (min)      lcorbi_t3_pos_4_057_49    m/z 219.1268     0   1   2E+5   8  RT (min)       lcorbi_t3_pos_4_100_61    m/z 219.1268     0   1E+6   18.5   19  RT (min)       lcorbi_t3_pos_4_076_58    m/z 219.1744     0   1   2   3   4E+6   10.5   11  RT (min)      lcorbi_t3_pos_1_055_33    m/z 219.1744     0   2   4   6   8E+4   11   11.5  RT (min)      lcorbi_t3_pos_1_027_43    m/z 219.1744     0   2   4   6E+6   10   10.5  RT (min)      lcorbi_t3_pos_3_035_08    m/z 219.1744     0   2   4   6E+6   10.5   11  RT (min)      lcorbi_t3_pos_1_055_33    m/z 219.1744     0   2   4   6E+6   9.5   10  RT (min)      lcorbi_t3_pos_2_047_31    m/z 219.1744     0   1E+5   11   11.5  RT (min)      lcorbi_t3_pos_2_014_37    m/z 219.2147     0   2   4   6   8E+6   1   1.5  RT (min)      lcorbi_t3_pos_4_057_49    m/z 220.0392     0   2   4   6E+5   0   0.5  RT (min)      lcorbi_t3_pos_1_008_36    m/z 220.0392     0   2   4E+5   0.5   1  RT (min)      lcorbi_t3_pos_1_009_50    m/z 220.0392     0   2   4   6E+5   0   0.5  RT (min)       lcorbi_t3_pos_1_008_36    m/z 220.0392     0   2   4   6E+5   0   0.5  RT (min)       lcorbi_t3_pos_1_008_36    m/z 220.0392     0   2   4   6   8E+4   18.5   19  RT (min)      lcorbi_t3_pos_4_075_58    m/z 220.0392     0   2   4   6E+5   0   0.5  RT (min)      lcorbi_t3_pos_1_008_36    m/z 220.0392     0   2   4   6E+5   0   0.5  RT (min)      lcorbi_t3_pos_1_008_36    m/z 220.0392     0   2   4   6E+5   0   1  RT (min)       lcorbi_t3_pos_1_012_14    m/z 220.0393     0   1   2   3E+5   0.5   1  RT (min)      lcorbi_t3_pos_1_008_36    m/z 220.0502     0   2   4E+5   0   0.5  RT (min)      lcorbi_t3_pos_1_007_07    m/z 220.1777     0   1   2   3E+4   11   11.5  RT (min)      lcorbi_t3_pos_2_014_37    m/z 220.1777     0   1E+6   10.5   11  RT (min)      lcorbi_t3_pos_1_055_33    m/z 220.1777     0   2   4   6E+5   10.5   11  RT (min)      lcorbi_t3_pos_1_055_33    m/z 220.1777     0   1E+6   9.5   10  RT (min)      lcorbi_t3_pos_2_047_31    m/z 220.1777     0   1E+6   10   10.5  RT (min)      lcorbi_t3_pos_3_035_08    m/z 220.9346     0   2   4   6E+5   17.5   18  RT (min)      lcorbi_t3_pos_4_105_61    m/z 221.0471     0   2   4   6   8E+5   15   15.5  RT (min)       lcorbi_t3_pos_1_006_15    m/z 222.0504     0   1E+6   0   0.5  RT (min)      lcorbi_t3_pos_1_005_42    m/z 223.0853     0   2   4E+5   1.5   2  RT (min)      lcorbi_t3_pos_4_057_49    m/z 223.0853     0   1E+6   17.5   18  RT (min)       lcorbi_t3_pos_4_105_61    m/z 223.0853     0   1   2E+5   5   5.5  RT (min)      lcorbi_t3_pos_4_034_42    m/z 223.0853     0   1E+6   18   18.5  RT (min)       lcorbi_t3_pos_4_076_58    m/z 223.0853     0   1E+6   18   18.5  RT (min)      lcorbi_t3_pos_4_076_58    m/z 223.0853     0   1E+6   18   18.5  RT (min)       lcorbi_t3_pos_4_078_58    m/z 223.0853     0   1   2   3E+5   5.5   6  RT (min)      lcorbi_t3_pos_4_003_39    m/z 223.0853     0   1E+5   6   6.5  RT (min)      lcorbi_t3_pos_3_004_24    m/z 225.1653     0   1E+6   1   1.5  RT (min)       lcorbi_t3_pos_4_097_60    m/z 225.1961     0   1E+6   9   9.5  RT (min)      lcorbi_t3_pos_1_002_01    m/z 225.1962     0   2   4   6   8E+4   10   10.5  RT (min)      lcorbi_t3_pos_1_023_02    m/z 225.1962     0   1E+6   9.5   10  RT (min)      lcorbi_t3_pos_1_034_34    m/z 225.1962     0   1E+6   9.5   10  RT (min)      lcorbi_t3_pos_1_008_36    m/z 225.1962     0   1E+6   10   10.5  RT (min)      lcorbi_t3_pos_1_034_34    m/z 225.2074     0   1   2   3   4E+6   0   0.5  RT (min)      lcorbi_t3_pos_1_002_01    m/z 226.1914     0   2   4   6E+6   0   0.5  RT (min)      lcorbi_t3_pos_1_002_01    m/z 226.8935     0   1   2   3E+6   0   0.5  RT (min)      lcorbi_t3_pos_1_007_07    m/z 226.9515     0   2   4   6   8E+5   17.5   18  RT (min)      lcorbi_t3_pos_4_105_61    m/z 227.1754     0   1E+6   0   0.5  RT (min)      lcorbi_t3_pos_1_001_49    m/z 230.2115     0   1   2   3E+4   10.5   11  RT (min)      lcorbi_t3_pos_2_052_48    m/z 230.2115     0   1E+6   9.5   10  RT (min)      lcorbi_t3_pos_3_052_47    m/z 230.2479     0   1E+6   9.5   10  RT (min)      lcorbi_t3_pos_2_056_56    m/z 231.0288     0   1   2   3E+6   8.5   9  RT (min)       lcorbi_t3_pos_1_051_09    m/z 231.0288     0   1E+6   8.5   9  RT (min)       lcorbi_t3_pos_1_023_02    m/z 231.2512     0   1   2E+5   9.5   10  RT (min)      lcorbi_t3_pos_2_056_56    m/z 232.9282     0   2   4   6   8E+5   0   0.5  RT (min)      lcorbi_t3_pos_1_007_07    m/z 233.0445     0   1E+6   8.5   9  RT (min)      lcorbi_t3_pos_1_023_02    m/z 233.0445     0   1   2E+6   8.5   9  RT (min)       lcorbi_t3_pos_1_051_09    m/z 233.1900     0   1   2E+4   12   12.5  RT (min)      lcorbi_t3_pos_2_012_01    m/z 233.1900     0   2   4   6E+5   11   11.5  RT (min)      lcorbi_t3_pos_1_008_36    m/z 233.1900     0   1   2E+4   11.5   12  RT (min)      lcorbi_t3_pos_2_014_37    m/z 233.1900     0   2   4   6   8E+5   10.5   11  RT (min)       lcorbi_t3_pos_3_000_aa    m/z 234.2000     0   2   4   6E+4   7   7.5  RT (min)      lcorbi_t3_pos_2_044_21    m/z 234.2064     0   2   4E+6   1   1.5  RT (min)      lcorbi_t3_pos_4_057_49    m/z 235.1137     0   1E+5   15   16  RT (min)       lcorbi_t3_pos_4_093_60    m/z 235.1137     0   1E+5   14   14.5  RT (min)       lcorbi_t3_pos_1_041_54    m/z 235.1137     0   1E+5   15   15.5  RT (min)       lcorbi_t3_pos_4_113_62    m/z 235.1137     0   1E+5   15   15.5  RT (min)       lcorbi_t3_pos_4_087_59    m/z 235.1178     0   1   17.5   18  RT (min)      lcorbi_t3_pos_4_105_61    m/z 235.1217     0   1E+5   11   12  RT (min)       lcorbi_t3_pos_4_100_61    m/z 235.1217     0   1E+5   10   11  RT (min)       lcorbi_t3_pos_1_052_38    m/z 237.1373     0   1   2   3E+6   18   18.5  RT (min)       lcorbi_t3_pos_4_106_62    m/z 237.1373     0   1   2   3E+6   18   18.5  RT (min)      lcorbi_t3_pos_4_095_60    m/z 237.1373     0   1   2   3E+6   18.5   19  RT (min)       lcorbi_t3_pos_4_076_58    m/z 237.1373     0   1E+6   1.5   2  RT (min)      lcorbi_t3_pos_4_057_49    m/z 237.1373     0   1   2   3E+6   18.5   19  RT (min)       lcorbi_t3_pos_4_076_58    m/z 237.1373     0   1   2   3E+6   17.5   18  RT (min)      lcorbi_t3_pos_4_095_60    m/z 237.1373     0   1   2   3E+6   18   18.5  RT (min)       lcorbi_t3_pos_4_091_60    m/z 237.1373     0   1   2   3E+6   18   18.5  RT (min)       lcorbi_t3_pos_4_076_58    m/z 237.1374     0   1   2   3E+6   17.5   18  RT (min)      lcorbi_t3_pos_4_105_61    m/z 237.1374     0   1E+5   5.5   6  RT (min)      lcorbi_t3_pos_4_003_39    m/z 237.1374     0   1   2   3   4E+5   11   11.5  RT (min)       lcorbi_t3_pos_1_001_49    m/z 237.1374     0   2   4E+5   8   10  RT (min)       lcorbi_t3_pos_1_021_18    m/z 237.1374     0   2   4E+5   10   12   14  RT (min)       lcorbi_t3_pos_2_044_21    m/z 237.1374     0   1   2   3   4E+6   18   18.5  RT (min)       lcorbi_t3_pos_4_098_61    m/z 237.1374     0   1   2   3E+6   17.5   18  RT (min)       lcorbi_t3_pos_4_075_58    m/z 237.1374     0   1   2   3E+6   18   18.5  RT (min)       lcorbi_t3_pos_4_090_60    m/z 237.1374     0   2   4E+5   9   10  RT (min)       lcorbi_t3_pos_1_040_27    m/z 238.0498     0   2   4   6E+5   0   0.5  RT (min)      lcorbi_t3_pos_1_005_42    m/z 239.1490     0   2   4E+5   5   5.5  RT (min)      lcorbi_t3_pos_3_000_aa    m/z 239.1618     0   2   4   6   8E+5   9   9.5  RT (min)      lcorbi_t3_pos_2_047_31    m/z 239.1618     0   1   2E+4   10.5   11  RT (min)      lcorbi_t3_pos_1_023_02    m/z 239.1618     0   1   2   3E+5   10   10.5  RT (min)      lcorbi_t3_pos_1_034_34    m/z 239.2249     0   1E+5   17   18  RT (min)       lcorbi_t3_pos_4_083_59    m/z 240.2322     0   1E+4   10   10.5  RT (min)      lcorbi_t3_pos_2_012_01    m/z 240.2322     0   2   4   6E+5   9.5   10  RT (min)      lcorbi_t3_pos_2_022_38    m/z 240.2323     0   1   2E+4   10   10.5  RT (min)      lcorbi_t3_pos_1_031_40    m/z 240.2686     0   1E+5   13.5   14  RT (min)      lcorbi_t3_pos_3_013_19    m/z 240.2686     0   2   4   6   8E+4   14   14.5  RT (min)       lcorbi_t3_pos_1_023_02    m/z 240.2686     0   2   4   6   8E+4   14.5   15  RT (min)      lcorbi_t3_pos_1_023_02    m/z 240.2686     0   1E+5   13.5   14  RT (min)      lcorbi_t3_pos_3_013_19    m/z 240.2686     0   2   4   6E+5   16.5   17  RT (min)      lcorbi_t3_pos_4_075_58    m/z 240.2686     0   1E+5   13.5   14  RT (min)      lcorbi_t3_pos_3_013_19    m/z 240.2686     0   2   4   6   8E+4   14.5   15  RT (min)      lcorbi_t3_pos_1_023_02    m/z 240.2686     0   2   4   6   8E+5   11.5   12  RT (min)      lcorbi_t3_pos_4_012_37    m/z 240.2686     0   1   2   3   4E+5   11.5   12  RT (min)      lcorbi_t3_pos_2_032_39    m/z 240.2686     0   1   2E+5   13.5   14  RT (min)       lcorbi_t3_pos_1_051_09    m/z 240.2686     0   1   2E+5   13.5   14  RT (min)      lcorbi_t3_pos_1_051_09    m/z 240.2686     0   2   4   6   8E+4   14   14.5  RT (min)      lcorbi_t3_pos_1_051_09    m/z 240.2686     0   1E+5   14   14.5  RT (min)      lcorbi_t3_pos_1_023_02    m/z 240.2686     0   2   4   6   8E+5   17   17.5  RT (min)       lcorbi_t3_pos_4_080_58    m/z 240.2686     0   2   4   6E+5   16.5   17  RT (min)      lcorbi_t3_pos_4_075_58    m/z 240.2686     0   2   4   6   8E+4   14.5   15  RT (min)      lcorbi_t3_pos_1_023_02    m/z 240.2686     0   1E+5   13   13.5  RT (min)       lcorbi_t3_pos_3_013_19    m/z 240.2686     0   1E+5   11.5   12  RT (min)      lcorbi_t3_pos_2_032_39    m/z 240.2686     0   1   2E+5   14   14.5  RT (min)      lcorbi_t3_pos_1_051_09    m/z 240.2686     0   1E+5   13   13.5  RT (min)       lcorbi_t3_pos_3_013_19    m/z 240.2686     0   1   2E+5   14   14.5  RT (min)      lcorbi_t3_pos_1_051_09    m/z 240.2686     0   1E+5   13   13.5  RT (min)       lcorbi_t3_pos_3_013_19    m/z 240.2686     0   2   4   6E+5   16.5   17  RT (min)       lcorbi_t3_pos_4_075_58    m/z 240.2687     0   2   4   6   8E+5   17   17.5  RT (min)       lcorbi_t3_pos_4_077_58    m/z 241.0591     0   1   2E+5   18   18.5  RT (min)       lcorbi_t3_pos_4_101_61    m/z 241.1733     0   1   2E+5   10   10.5  RT (min)       lcorbi_t3_pos_1_008_36    m/z 241.1733     0   1E+5   10   10.5  RT (min)       lcorbi_t3_pos_1_041_54    m/z 241.1733     0   1E+5   10   10.5  RT (min)      lcorbi_t3_pos_4_028_04    m/z 241.1733     0   2   4   6   8E+4   10.5   11  RT (min)      lcorbi_t3_pos_1_025_52    m/z 241.1734     0   1E+5   15   16   17  RT (min)       lcorbi_t3_pos_4_086_59    m/z 241.1734     0   1E+5   16   16.5  RT (min)       lcorbi_t3_pos_4_083_59    m/z 241.2720     0   1   2E+5   12.5   13  RT (min)       lcorbi_t3_pos_3_047_40    m/z 243.0628     0   1   2   3E+5   6.5   7  RT (min)      lcorbi_t3_pos_2_053_28    m/z 244.1908     0   2   4   6   8E+5   12.5   13  RT (min)      lcorbi_t3_pos_2_002_06    m/z 244.2635     0   2   4   6E+4   10   10.5  RT (min)      lcorbi_t3_pos_1_023_02    m/z 244.2635     0   1   2E+5   9.5   10  RT (min)      lcorbi_t3_pos_2_056_56    m/z 246.2427     0   1E+7   8   8.5  RT (min)      lcorbi_t3_pos_1_023_02    m/z 246.2428     0   2   4   6E+6   1   1.5  RT (min)      lcorbi_t3_pos_4_057_49    m/z 247.0601     0   1E+7   1   1.5  RT (min)      lcorbi_t3_pos_4_057_49    m/z 248.9006     0   1   2   3   4E+6   0   0.5  RT (min)      lcorbi_t3_pos_1_007_07    m/z 249.1234     0   2   4E+5   5   5.5  RT (min)       lcorbi_t3_pos_4_112_62    m/z 249.1850     0   1E+5   11   11.5  RT (min)      lcorbi_t3_pos_1_033_53    m/z 249.1850     0   1   2   3E+5   10   10.5  RT (min)      lcorbi_t3_pos_4_105_61    m/z 249.1850     0   1   2   3E+5   10.5   11  RT (min)      lcorbi_t3_pos_3_025_52    m/z 249.1850     0   1   2   3E+6   10.5   11  RT (min)       lcorbi_t3_pos_3_000_aa    m/z 250.1187     0   1   2   3E+5   9.5   10  RT (min)      lcorbi_t3_pos_1_034_34    m/z 251.0468     0   1E+4   13   13.5  RT (min)      lcorbi_t3_pos_2_012_01    m/z 251.0468     0   1   2   3   4E+4   13   13.5  RT (min)      lcorbi_t3_pos_1_031_40    m/z 251.0468     0   2   4   6E+6   11.5   12  RT (min)      lcorbi_t3_pos_3_011_42    m/z 251.0468     0   2   4   6   8E+4   12.5   13  RT (min)      lcorbi_t3_pos_1_028_32    m/z 251.0468     0   1E+5   12.5   13  RT (min)      lcorbi_t3_pos_1_034_34    m/z 251.0468     0   2   4   6   8E+3   11   11.5  RT (min)      lcorbi_t3_pos_2_032_39    m/z 251.0468     0   1   2E+4   13.5   14  RT (min)      lcorbi_t3_pos_1_023_02    m/z 251.0468     0   2   4   6E+6   12   12.5  RT (min)       lcorbi_t3_pos_1_008_36    m/z 251.0469     0   2   4   6   8E+4   12.5   13  RT (min)       lcorbi_t3_pos_1_033_53    m/z 251.1642     0   1   2   3E+5   14   15  RT (min)       lcorbi_t3_pos_4_108_62    m/z 251.2006     0   2   4   6   8E+5   11   11.5  RT (min)      lcorbi_t3_pos_1_055_33    m/z 251.2006     0   1   2   3   4E+4   12   12.5  RT (min)      lcorbi_t3_pos_1_023_02    m/z 251.2006     0   2   4   6   8E+5   11   11.5  RT (min)      lcorbi_t3_pos_2_017_51    m/z 251.2006     0   1   2   3   4E+4   12   12.5  RT (min)      lcorbi_t3_pos_2_012_01    m/z 251.2006     0   2   4E+6   11   11.5  RT (min)      lcorbi_t3_pos_1_008_36    m/z 251.2006     0   2   4E+4   11.5   12  RT (min)      lcorbi_t3_pos_2_014_37    m/z 251.2006     0   1   2   3E+4   11.5   12  RT (min)      lcorbi_t3_pos_1_041_54    m/z 251.2006     0   1   2   3E+4   11.5   12  RT (min)      lcorbi_t3_pos_1_041_54    m/z 252.0501     0   1   13   13.5  RT (min)      lcorbi_t3_pos_2_012_01    m/z 252.0502     0   1   12.5   13  RT (min)      lcorbi_t3_pos_1_034_34    m/z 252.0502     0   1   11   11.5  RT (min)      lcorbi_t3_pos_2_032_39    m/z 252.0502     0   1   13.5   14  RT (min)      lcorbi_t3_pos_1_023_02    m/z 252.0502     0   1   2E+4   12.5   13  RT (min)      lcorbi_t3_pos_1_028_32    m/z 252.0502     0   1   13   13.5  RT (min)      lcorbi_t3_pos_1_031_40    m/z 252.0502     0   2   4   6   8E+5   12   12.5  RT (min)      lcorbi_t3_pos_3_047_40    m/z 252.0502     0   1   2E+4   12.5   13  RT (min)      lcorbi_t3_pos_1_030_21    m/z 252.0502     0   1   13.5   14  RT (min)      lcorbi_t3_pos_1_023_02    m/z 252.0502     0   2   4   6   8E+5   11   11.5  RT (min)      lcorbi_t3_pos_2_054_29    m/z 252.2039     0   1E+4   11.5   12  RT (min)      lcorbi_t3_pos_2_014_37    m/z 252.2039     0   1E+5   11   11.5  RT (min)      lcorbi_t3_pos_1_055_33    m/z 252.2040     0   1E+4   12   12.5  RT (min)      lcorbi_t3_pos_2_012_01    m/z 252.2040     0   2   4   6   8E+5   11   11.5  RT (min)      lcorbi_t3_pos_1_008_36    m/z 252.2040     0   2   4   6E+5   10.5   11  RT (min)      lcorbi_t3_pos_2_054_29    m/z 252.2040     0   2   4   6E+5   10.5   11  RT (min)      lcorbi_t3_pos_3_037_32    m/z 253.0703     0   2   4   6   8E+4   9   9.5  RT (min)      lcorbi_t3_pos_4_003_39    m/z 253.0787     0   1   2E+5   9   9.5  RT (min)      lcorbi_t3_pos_1_051_09    m/z 253.0788     0   1   2   3   4E+5   9   9.5  RT (min)      lcorbi_t3_pos_3_013_19    m/z 253.0858     0   1   2   3   4E+6   9.5   10  RT (min)      lcorbi_t3_pos_1_023_02    m/z 253.0860     0   2   4E+7   1   1.5  RT (min)      lcorbi_t3_pos_4_057_49    m/z 253.0893     0   2   4   6   8E+3   9.5   10  RT (min)      lcorbi_t3_pos_1_023_02    m/z 253.0930     0   2   4E+5   9   9.5  RT (min)      lcorbi_t3_pos_1_051_09    m/z 253.0931     0   1E+5   9   9.5  RT (min)      lcorbi_t3_pos_1_034_34    m/z 253.0981     0   1E+5   8   8.5  RT (min)      lcorbi_t3_pos_3_037_32    m/z 253.1060     0   1   2E+5   8.5   9  RT (min)      lcorbi_t3_pos_4_101_61    m/z 253.2386     0   1E+6   0   0.5  RT (min)      lcorbi_t3_pos_1_001_49    m/z 254.0892     0   2   4   6E+5   9.5   10  RT (min)      lcorbi_t3_pos_1_023_02    m/z 254.0892     0   2   4   6   8E+6   1   1.5  RT (min)      lcorbi_t3_pos_4_057_49    m/z 254.0892     0   1E+7   9   9.5  RT (min)      lcorbi_t3_pos_1_051_09    m/z 254.2227     0   1E+6   0   0.5  RT (min)      lcorbi_t3_pos_1_001_49    m/z 254.2478     0   1   2E+5   9.5   10  RT (min)      lcorbi_t3_pos_2_052_48    m/z 255.0924     0   1E+6   9   9.5  RT (min)      lcorbi_t3_pos_1_051_09    m/z 255.1510     0   1E+4   10.5   11  RT (min)      lcorbi_t3_pos_2_014_37    m/z 255.1510     0   2   4E+5   10.5   11  RT (min)      lcorbi_t3_pos_1_055_33    m/z 255.1510     0   1E+4   10.5   11  RT (min)      lcorbi_t3_pos_2_014_37    m/z 255.1510     0   2   4E+5   9.5   10  RT (min)      lcorbi_t3_pos_2_039_30    m/z 255.1591     0   2   4E+5   8.5   9  RT (min)       lcorbi_t3_pos_3_000_aa    m/z 256.2635     0   2   4   6E+4   11.5   12  RT (min)      lcorbi_t3_pos_3_024_07    m/z 256.2635     0   1   2E+5   12.5   13  RT (min)      lcorbi_t3_pos_3_025_52    m/z 256.2636     0   1E+5   12.5   13  RT (min)      lcorbi_t3_pos_3_049_55    m/z 256.2636     0   2   4   6E+5   11.5   12  RT (min)      lcorbi_t3_pos_2_013_33    m/z 256.2636     0   2   4E+5   11.5   12  RT (min)      lcorbi_t3_pos_4_037_26    m/z 256.2636     0   2   4   6E+5   12.5   13  RT (min)      lcorbi_t3_pos_3_047_40    m/z 256.2636     0   1E+5   13   13.5  RT (min)       lcorbi_t3_pos_1_034_34    m/z 256.2636     0   1E+5   15   16  RT (min)       lcorbi_t3_pos_1_035_28    m/z 256.2636     0   1E+5   13   13.5  RT (min)      lcorbi_t3_pos_2_049_55    m/z 256.2636     0   1E+5   13   13.5  RT (min)      lcorbi_t3_pos_2_041_54    m/z 256.2636     0   1E+5   12.5   13  RT (min)      lcorbi_t3_pos_4_049_55    m/z 256.2636     0   2   4E+5   12   12.5  RT (min)      lcorbi_t3_pos_3_033_53    m/z 256.2636     0   1   2   3E+5   12.5   13  RT (min)      lcorbi_t3_pos_2_017_51    m/z 256.2636     0   2   4   6E+5   12   12.5  RT (min)       lcorbi_t3_pos_3_041_54    m/z 256.2636     0   2   4   6   8E+5   12.5   13  RT (min)       lcorbi_t3_pos_1_008_36    m/z 256.2998     0   2   4E+5   10.5   11  RT (min)      lcorbi_t3_pos_1_001_49    m/z 256.2999     0   1E+6   10   10.5  RT (min)      lcorbi_t3_pos_1_001_49    m/z 256.2999     0   2   4   6   8E+5   10.5   11  RT (min)      lcorbi_t3_pos_1_001_49    m/z 256.2999     0   2   4E+5   10   10.5  RT (min)      lcorbi_t3_pos_1_034_34    m/z 256.2999     0   2   4E+5   9   9.5  RT (min)      lcorbi_t3_pos_2_047_31    m/z 257.0366     0   2   4   6E+5   17.5   18  RT (min)      lcorbi_t3_pos_4_105_61    m/z 257.0509     0   1E+6   1   1.5  RT (min)       lcorbi_t3_pos_4_093_60    m/z 257.3032     0   1E+5   10.5   11  RT (min)      lcorbi_t3_pos_1_001_49    m/z 257.3032     0   1E+6   10   10.5  RT (min)      lcorbi_t3_pos_1_008_36    m/z 257.3032     0   2   4   6   8E+4   10.5   11  RT (min)      lcorbi_t3_pos_1_001_49    m/z 257.3032     0   1E+5   10.5   11  RT (min)      lcorbi_t3_pos_1_001_49    m/z 257.3033     0   1   2E+5   10   10.5  RT (min)      lcorbi_t3_pos_1_001_49    m/z 258.1102     0   1E+5   18   18.5  RT (min)      lcorbi_t3_pos_4_101_61    m/z 258.2428     0   1   2E+4   12   12.5  RT (min)      lcorbi_t3_pos_1_023_02    m/z 258.2428     0   1   2E+6   10   10.5  RT (min)      lcorbi_t3_pos_3_052_47    m/z 258.2428     0   1E+6   11   11.5  RT (min)      lcorbi_t3_pos_3_044_46    m/z 258.2428     0   1   2E+4   11.5   12  RT (min)      lcorbi_t3_pos_1_034_34    m/z 258.2428     0   1   2E+4   11.5   12  RT (min)      lcorbi_t3_pos_2_052_48    m/z 258.9976     0   2   4E+5   17.5   18  RT (min)      lcorbi_t3_pos_4_105_61    m/z 260.1622     0   1   2   3   4E+6   1   1.5  RT (min)       lcorbi_t3_pos_2_037_27    m/z 260.2221     0   1   2E+5   8   8.5  RT (min)       lcorbi_t3_pos_4_000_dd    m/z 261.1486     0   1   2   3E+6   12.5   13  RT (min)      lcorbi_t3_pos_4_016_15    m/z 261.1486     0   1E+6   12.5   13  RT (min)      lcorbi_t3_pos_1_002_01    m/z 261.1486     0   1E+4   12   12.5  RT (min)      lcorbi_t3_pos_2_032_39    m/z 261.1486     0   1E+6   12.5   13  RT (min)      lcorbi_t3_pos_2_054_29    m/z 261.1486     0   1E+5   14   14.5  RT (min)      lcorbi_t3_pos_2_038_03    m/z 261.1486     0   1E+5   15   16  RT (min)       lcorbi_t3_pos_2_012_01    m/z 261.1486     0   1E+5   15   15.5  RT (min)      lcorbi_t3_pos_1_051_09    m/z 261.1486     0   1E+5   15   16  RT (min)       lcorbi_t3_pos_2_049_55    m/z 261.1486     0   2   4   6   8E+4   14   14.5  RT (min)      lcorbi_t3_pos_4_049_55    m/z 261.1486     0   2   4   6   8E+4   15.5   16  RT (min)      lcorbi_t3_pos_1_023_02    m/z 261.1486     0   2   4   6   8E+4   14   14.5  RT (min)      lcorbi_t3_pos_3_049_55    m/z 261.1486     0   1E+6   13.5   14  RT (min)       lcorbi_t3_pos_3_047_40    m/z 261.1486     0   1E+5   15   15.5  RT (min)      lcorbi_t3_pos_1_031_40    m/z 261.1486     0   1   2E+6   13.5   14  RT (min)       lcorbi_t3_pos_4_019_47    m/z 261.1486     0   1   2E+4   12.5   13  RT (min)      lcorbi_t3_pos_4_012_37    m/z 261.1486     0   1E+5   14   14.5  RT (min)      lcorbi_t3_pos_2_046_07    m/z 261.1486     0   1   2E+4   12.5   13  RT (min)      lcorbi_t3_pos_2_032_39    m/z 261.1486     0   2   4   6   8E+4   14   14.5  RT (min)      lcorbi_t3_pos_3_013_19    m/z 261.1486     0   1E+4   12.5   13  RT (min)      lcorbi_t3_pos_3_031_45    m/z 261.1486     0   1E+5   16  RT (min)       lcorbi_t3_pos_1_034_34    m/z 261.2213     0   2   4   6   8E+5   10   10.5  RT (min)       lcorbi_t3_pos_3_000_aa    m/z 262.1518     0   2   4   6   8E+3   12.5   13  RT (min)      lcorbi_t3_pos_3_024_07    m/z 262.1519     0   1   2E+4   14   14.5  RT (min)      lcorbi_t3_pos_3_003_33    m/z 262.1520     0   2   4E+5   13.5   14  RT (min)      lcorbi_t3_pos_4_019_47    m/z 262.1520     0   2   4E+5   12.5   13  RT (min)      lcorbi_t3_pos_4_016_15    m/z 262.2278     0   2   4   6   8E+5   0   0.5  RT (min)      lcorbi_t3_pos_1_001_49    m/z 262.2377     0   1   2E+6   8.5   9  RT (min)      lcorbi_t3_pos_2_056_56    m/z 264.8729     0   2   4   6E+6   0   0.5  RT (min)      lcorbi_t3_pos_1_007_07    m/z 266.2843     0   2   4   6E+5   17   17.5  RT (min)       lcorbi_t3_pos_4_077_58    m/z 266.2843     0   2   4E+5   17   17.5  RT (min)      lcorbi_t3_pos_4_081_58    m/z 266.2843     0   2   4E+5   16.5   17  RT (min)       lcorbi_t3_pos_4_074_58    m/z 268.2272     0   2   4   6   8E+5   10.5   11  RT (min)      lcorbi_t3_pos_4_046_44    m/z 268.2272     0   1   2E+6   10.5   11  RT (min)      lcorbi_t3_pos_1_002_01    m/z 268.2272     0   1E+4   12   12.5  RT (min)      lcorbi_t3_pos_2_012_01    m/z 268.2272     0   1E+5   11   11.5  RT (min)      lcorbi_t3_pos_3_003_33    m/z 268.2272     0   1   2E+4   12   12.5  RT (min)      lcorbi_t3_pos_1_023_02    m/z 268.2272     0   1   2E+4   11.5   12  RT (min)      lcorbi_t3_pos_2_014_37    m/z 268.2272     0   1E+4   11.5   12  RT (min)      lcorbi_t3_pos_1_041_54    m/z 268.2272     0   1   2   3E+6   11   11.5  RT (min)      lcorbi_t3_pos_1_008_36    m/z 268.2999     0   2   4E+4   12   12.5  RT (min)      lcorbi_t3_pos_2_032_39    m/z 268.2999     0   2   4   6E+5   12.5   13  RT (min)      lcorbi_t3_pos_4_012_37    m/z 268.2999     0   1E+5   14   14.5  RT (min)      lcorbi_t3_pos_3_013_19    m/z 268.2999     0   1   2E+4   15   15.5  RT (min)      lcorbi_t3_pos_1_023_02    m/z 268.2999     0   1   2   3E+4   14.5   15  RT (min)      lcorbi_t3_pos_1_051_09    m/z 268.2999     0   1   2   3E+4   14.5   15  RT (min)      lcorbi_t3_pos_1_051_09    m/z 268.2999     0   2   4E+5   13.5   14  RT (min)      lcorbi_t3_pos_3_047_40    m/z 268.3000     0   2   4   6E+5   12   12.5  RT (min)      lcorbi_t3_pos_2_054_29    m/z 268.3000     0   1   2   3E+4   14.5   15  RT (min)      lcorbi_t3_pos_1_051_09    m/z 268.3000     0   1   2   3E+4   14.5   15  RT (min)      lcorbi_t3_pos_1_051_09    m/z 268.3000     0   2   4E+4   15   15.5  RT (min)      lcorbi_t3_pos_1_023_02    m/z 268.3000     0   2   4E+4   15   15.5  RT (min)      lcorbi_t3_pos_1_023_02    m/z 268.3000     0   1   2E+5   12   12.5  RT (min)      lcorbi_t3_pos_4_012_37    m/z 268.3000     0   1   2   3E+4   15   15.5  RT (min)      lcorbi_t3_pos_1_023_02    m/z 268.3000     0   2   4E+4   12   12.5  RT (min)      lcorbi_t3_pos_2_032_39    m/z 268.3000     0   1E+5   13.5   14  RT (min)       lcorbi_t3_pos_3_013_19    m/z 269.0809     0   1E+6   9   9.5  RT (min)      lcorbi_t3_pos_1_002_01    m/z 269.0809     0   1   2   3E+5   10.5   11  RT (min)       lcorbi_t3_pos_1_031_40    m/z 269.0809     0   2   4   6E+5   10   10.5  RT (min)      lcorbi_t3_pos_1_034_34    m/z 269.0809     0   1E+5   10.5   11  RT (min)       lcorbi_t3_pos_1_023_02    m/z 269.0809     0   1E+5   10.5   11  RT (min)       lcorbi_t3_pos_1_023_02    m/z 269.0809     0   1   2E+6   1   1.5  RT (min)      lcorbi_t3_pos_4_057_49    m/z 269.0809     0   1   2E+6   10   10.5  RT (min)       lcorbi_t3_pos_1_028_32    m/z 269.0809     0   1E+5   10.5   11  RT (min)      lcorbi_t3_pos_2_012_01    m/z 269.0809     0   1   2E+5   10.5   11  RT (min)      lcorbi_t3_pos_1_031_40    m/z 269.0809     0   1E+6   9   9.5  RT (min)      lcorbi_t3_pos_3_037_32    m/z 269.2305     0   2   4E+5   11   11.5  RT (min)      lcorbi_t3_pos_1_008_36    m/z 270.0842     0   1   2   3E+5   10   10.5  RT (min)      lcorbi_t3_pos_1_038_30    m/z 272.2584     0   1   2   3E+5   9.5   10  RT (min)      lcorbi_t3_pos_1_027_43    m/z 273.1825     0   2   4E+5   10   10.5  RT (min)      lcorbi_t3_pos_2_047_31    m/z 273.1826     0   2   4E+5   10.5   11  RT (min)      lcorbi_t3_pos_3_037_32    m/z 273.1826     0   1E+4   11.5   12  RT (min)      lcorbi_t3_pos_2_014_37    m/z 273.1826     0   2   4   6E+5   11   11.5  RT (min)      lcorbi_t3_pos_1_008_36    m/z 274.1414     0   2   4   6   8E+5   1   1.5  RT (min)       lcorbi_t3_pos_4_093_60    m/z 274.1890     0   2   4E+5   1   1.5  RT (min)      lcorbi_t3_pos_4_097_60    m/z 274.2741     0   2   4E+5   9.5   10  RT (min)      lcorbi_t3_pos_1_001_49    m/z 274.2741     0   1E+6   9.5   10  RT (min)      lcorbi_t3_pos_2_056_56    m/z 274.2741     0   1   2   3   4E+6   9.5   10  RT (min)      lcorbi_t3_pos_1_051_09    m/z 274.9752     0   1E+6   17.5   18  RT (min)      lcorbi_t3_pos_4_105_61    m/z 275.2581     0   1E+6   12   12.5  RT (min)      lcorbi_t3_pos_3_025_52    m/z 275.2582     0   1   2   3E+4   12   12.5  RT (min)      lcorbi_t3_pos_3_003_33    m/z 275.2582     0   1   2E+4   12   12.5  RT (min)      lcorbi_t3_pos_4_048_24    m/z 275.2582     0   1E+6   11   11.5  RT (min)      lcorbi_t3_pos_2_013_33    m/z 275.2582     0   1E+6   11   11.5  RT (min)      lcorbi_t3_pos_3_011_42    m/z 275.2582     0   1E+6   12   12.5  RT (min)      lcorbi_t3_pos_1_008_36    m/z 275.2582     0   1E+4   12.5   13  RT (min)      lcorbi_t3_pos_2_049_55    m/z 275.2582     0   1   2E+4   12.5   13  RT (min)      lcorbi_t3_pos_1_031_40    m/z 275.2582     0   1E+4   12.5   13  RT (min)      lcorbi_t3_pos_1_034_34    m/z 275.2774     0   2   4   6E+5   9.5   10  RT (min)      lcorbi_t3_pos_1_051_09    m/z 277.0082     0   1E+6   17.5   18  RT (min)      lcorbi_t3_pos_4_105_61    m/z 279.1591     0   2   4   6   8E+4   12   12.5  RT (min)      lcorbi_t3_pos_1_023_02    m/z 279.1591     0   1   2E+6   1   1.5  RT (min)      lcorbi_t3_pos_4_057_49    m/z 279.1592     0   2   4   6   8E+4   11.5   12  RT (min)      lcorbi_t3_pos_1_023_02    m/z 279.1592     0   1   2E+6   11   11.5  RT (min)      lcorbi_t3_pos_1_028_32    m/z 279.1592     0   2   4   6E+6   13.5   14  RT (min)      lcorbi_t3_pos_3_047_40    m/z 279.1592     0   1E+5   11   11.5  RT (min)      lcorbi_t3_pos_1_031_40    m/z 279.1592     0   1   2   3   4E+5   15.5   16  RT (min)      lcorbi_t3_pos_1_023_02    m/z 279.1592     0   1   2   3E+5   15   15.5   16  RT (min)       lcorbi_t3_pos_2_012_01    m/z 279.1592     0   1   2   3E+5   15   16  RT (min)       lcorbi_t3_pos_1_051_09    m/z 279.1592     0   2   4   6E+6   10   10.5  RT (min)      lcorbi_t3_pos_1_002_01    m/z 279.1592     0   2   4   6   8E+4   11   11.5  RT (min)      lcorbi_t3_pos_1_051_09    m/z 279.1592     0   1   2E+5   14   14.5  RT (min)      lcorbi_t3_pos_3_003_33    m/z 279.1592     0   2   4   6E+5   13.5   14  RT (min)      lcorbi_t3_pos_1_005_42    m/z 279.1592     0   1E+5   11   11.5  RT (min)      lcorbi_t3_pos_1_051_09    m/z 279.1592     0   1   2   3E+5   14   14.5  RT (min)      lcorbi_t3_pos_1_047_41    m/z 279.1592     0   2   4   6   8E+4   11.5   12  RT (min)      lcorbi_t3_pos_1_031_40    m/z 279.1592     0   2   4   6E+4   12   12.5  RT (min)      lcorbi_t3_pos_2_032_39    m/z 279.1592     0   2   4   6E+5   11   11.5  RT (min)      lcorbi_t3_pos_1_030_21    m/z 279.1592     0   2   4   6E+6   12.5   13  RT (min)      lcorbi_t3_pos_2_054_29    m/z 279.1592     0   2   4   6   8E+4   12   12.5  RT (min)      lcorbi_t3_pos_1_023_02    m/z 279.1592     0   1E+7   13   14  RT (min)       lcorbi_t3_pos_4_016_15    m/z 279.1592     0   2   4   6   8E+4   11.5   12  RT (min)      lcorbi_t3_pos_1_031_40    m/z 279.2319     0   2   4   6E+4   10.5   11  RT (min)      lcorbi_t3_pos_2_014_37    m/z 279.2319     0   1E+5   10.5   11  RT (min)      lcorbi_t3_pos_2_038_03    m/z 279.2319     0   1   2   3   4E+5   10   10.5  RT (min)      lcorbi_t3_pos_3_047_40    m/z 279.2319     0   2   4   6E+5   10.5   11  RT (min)      lcorbi_t3_pos_2_017_51    m/z 279.2319     0   2   4   6E+5   10.5   11  RT (min)      lcorbi_t3_pos_2_006_47    m/z 279.2319     0   2   4   6E+4   11   11.5  RT (min)      lcorbi_t3_pos_1_049_55    m/z 279.2320     0   1   2   3E+5   12   12.5  RT (min)      lcorbi_t3_pos_4_078_58    m/z 279.2320     0   1   2   3   4E+5   11   11.5  RT (min)      lcorbi_t3_pos_3_000_aa    m/z 279.2543     0   1E+6   0   0.5  RT (min)      lcorbi_t3_pos_1_001_49    m/z 280.1625     0   1   2E+4   11.5   12  RT (min)      lcorbi_t3_pos_1_023_02    m/z 280.1625     0   1   2   3E+4   11   11.5  RT (min)      lcorbi_t3_pos_1_031_40    m/z 280.1625     0   1E+5   12.5   13  RT (min)      lcorbi_t3_pos_2_054_29    m/z 280.1625     0   2   4   6   8E+4   14.5   15  RT (min)      lcorbi_t3_pos_2_049_55    m/z 280.1625     0   2   4   6E+4   14   14.5  RT (min)      lcorbi_t3_pos_1_047_41    m/z 280.1625     0   1E+6   12.5   13  RT (min)      lcorbi_t3_pos_2_054_29    m/z 280.1625     0   2   4   6E+4   14   14.5  RT (min)      lcorbi_t3_pos_3_013_19    m/z 280.1625     0   2   4   6   8E+4   14.5   15  RT (min)      lcorbi_t3_pos_1_031_40    m/z 280.1625     0   1E+6   10   10.5  RT (min)      lcorbi_t3_pos_1_002_01    m/z 280.1625     0   2   4   6E+4   14   14.5  RT (min)      lcorbi_t3_pos_4_048_24    m/z 280.1625     0   2   4   6   8E+4   15   15.5  RT (min)      lcorbi_t3_pos_2_012_01    m/z 280.1625     0   1E+4   12   12.5  RT (min)      lcorbi_t3_pos_2_032_39    m/z 280.1625     0   1   2   3E+5   13.5   14  RT (min)      lcorbi_t3_pos_1_008_36    m/z 280.1626     0   1E+6   13.5   14  RT (min)      lcorbi_t3_pos_3_047_40    m/z 280.1626     0   2   4   6   8E+4   15.5   16  RT (min)      lcorbi_t3_pos_1_023_02    m/z 280.1626     0   2   4   6E+4   14.5   15  RT (min)      lcorbi_t3_pos_1_034_34    m/z 280.1626     0   2   4   6   8E+5   13.5   14  RT (min)      lcorbi_t3_pos_3_017_51    m/z 280.2353     0   2   4   6E+5   9.5   10  RT (min)       lcorbi_t3_pos_3_000_aa    m/z 280.2635     0   2   4E+5   10   10.5  RT (min)      lcorbi_t3_pos_3_000_aa    m/z 281.1496     0   2   4   6E+5   5   5.5  RT (min)       lcorbi_t3_pos_3_000_aa    m/z 282.2218     0   1   2E+5   13   13.5  RT (min)       lcorbi_t3_pos_3_033_53    m/z 282.2792     0   2   4   6E+5   11.5   12  RT (min)      lcorbi_t3_pos_2_054_29    m/z 282.2792     0   2   4E+4   13   13.5  RT (min)      lcorbi_t3_pos_2_022_38    m/z 282.2792     0   2   4E+5   12.5   13  RT (min)      lcorbi_t3_pos_3_047_40    m/z 282.2792     0   2   4E+4   13.5   14  RT (min)      lcorbi_t3_pos_1_051_09    m/z 282.2792     0   1   2   3E+4   11.5   12  RT (min)      lcorbi_t3_pos_2_032_39    m/z 282.2792     0   2   4   6E+4   14   14.5  RT (min)      lcorbi_t3_pos_1_023_02    m/z 282.2793     0   2   4   6E+4   13.5   14  RT (min)      lcorbi_t3_pos_1_034_34    m/z 282.2793     0   2   4   6E+4   13   13.5  RT (min)      lcorbi_t3_pos_1_034_34    m/z 282.8834     0   2   4E+6   0   0.5  RT (min)      lcorbi_t3_pos_1_007_07    m/z 282.9051     0   2   4   6E+5   17.5   18  RT (min)      lcorbi_t3_pos_4_105_61    m/z 283.2633     0   1E+5   13   13.5  RT (min)       lcorbi_t3_pos_3_047_40    m/z 283.2745     0   1   14   14.5  RT (min)      lcorbi_t3_pos_1_023_02    m/z 283.2745     0   2   4   6E+5   11.5   12  RT (min)      lcorbi_t3_pos_4_012_37    m/z 283.2745     0   2   4   6   8E+4   14   14.5  RT (min)      lcorbi_t3_pos_1_051_09    m/z 283.2745     0   2   4   6E+5   11.5   12  RT (min)      lcorbi_t3_pos_4_012_37    m/z 283.2745     0   2   4   6E+4   13   13.5  RT (min)      lcorbi_t3_pos_3_013_19    m/z 283.2745     0   1E+5   13.5   14  RT (min)      lcorbi_t3_pos_1_051_09    m/z 283.2745     0   1   2   3E+4   14   14.5  RT (min)      lcorbi_t3_pos_1_023_02    m/z 283.2745     0   1E+5   11.5   12  RT (min)      lcorbi_t3_pos_2_032_39    m/z 283.2745     0   2   4   6E+4   13   13.5  RT (min)      lcorbi_t3_pos_3_013_19    m/z 283.2745     0   1   14.5   15  RT (min)      lcorbi_t3_pos_1_023_02    m/z 283.2745     0   1   14.5   15  RT (min)      lcorbi_t3_pos_1_023_02    m/z 283.2745     0   2   4   6E+4   13.5   14  RT (min)      lcorbi_t3_pos_3_013_19    m/z 284.1341     0   2   4E+5   5   5.5  RT (min)      lcorbi_t3_pos_3_000_aa    m/z 284.2948     0   2   4   6   8E+4   14.5   15  RT (min)      lcorbi_t3_pos_2_056_56    m/z 284.2948     0   2   4   6E+4   14   14.5  RT (min)      lcorbi_t3_pos_2_009_50    m/z 284.2948     0   2   4   6   8E+4   14   14.5  RT (min)      lcorbi_t3_pos_1_031_40    m/z 284.2948     0   1   2   3   4E+4   12   12.5  RT (min)      lcorbi_t3_pos_2_047_31    m/z 284.2949     0   2   4   6E+5   13   13.5  RT (min)      lcorbi_t3_pos_3_033_53    m/z 284.2949     0   1   2   3   4E+4   12   12.5  RT (min)      lcorbi_t3_pos_3_031_45    m/z 284.2949     0   2   4E+5   12   12.5  RT (min)      lcorbi_t3_pos_4_037_26    m/z 284.2949     0   1E+5   13.5   14  RT (min)      lcorbi_t3_pos_1_030_21    m/z 284.2949     0   2   4   6E+4   13.5   14  RT (min)      lcorbi_t3_pos_2_041_54    m/z 284.2949     0   2   4   6E+5   13   13.5  RT (min)      lcorbi_t3_pos_1_008_36    m/z 284.2949     0   2   4   6E+5   13   13.5  RT (min)      lcorbi_t3_pos_3_026_04    m/z 284.2949     0   2   4   6   8E+4   13.5   14  RT (min)      lcorbi_t3_pos_1_034_34    m/z 284.2949     0   2   4   6   8E+4   13.5   14  RT (min)      lcorbi_t3_pos_1_028_32    m/z 284.2949     0   2   4   6   8E+5   12   12.5  RT (min)      lcorbi_t3_pos_3_011_42    m/z 284.2949     0   1E+5   12   12.5  RT (min)      lcorbi_t3_pos_2_013_33    m/z 284.2949     0   2   4   6   8E+4   14   14.5  RT (min)      lcorbi_t3_pos_1_056_56    m/z 284.2949     0   1   2   3E+5   13   13.5  RT (min)      lcorbi_t3_pos_3_056_56    m/z 284.2949     0   1E+5   14.5   15  RT (min)      lcorbi_t3_pos_1_023_02    m/z 285.1540     0   1E+5   3.5   4  RT (min)      lcorbi_t3_pos_1_058_14    m/z 285.1540     0   1   2E+5   0.5   1  RT (min)       lcorbi_t3_pos_1_058_14    m/z 285.1540     0   1   2E+5   19   19.5  RT (min)       lcorbi_t3_pos_4_072_57    m/z 285.1540     0   1   2   3E+5   0   0.5  RT (min)       lcorbi_t3_pos_1_028_32    m/z 285.1540     0   1E+5   2   3  RT (min)       lcorbi_t3_pos_1_058_14    m/z 287.1352     0   2   4E+5   1   1.5  RT (min)      lcorbi_t3_pos_2_030_14    m/z 287.2006     0   2   4E+4   12.5   13  RT (min)      lcorbi_t3_pos_1_038_30    m/z 287.2006     0   1E+5   13   13.5  RT (min)      lcorbi_t3_pos_1_051_09    m/z 287.2006     0   2   4   6   8E+4   13   13.5  RT (min)      lcorbi_t3_pos_1_056_56    m/z 287.2006     0   1   2   3E+4   12.5   13  RT (min)      lcorbi_t3_pos_1_041_54    m/z 287.2006     0   1E+4   12.5   13  RT (min)      lcorbi_t3_pos_2_049_55    m/z 287.2006     0   1   2E+5   12   12.5  RT (min)      lcorbi_t3_pos_1_051_09    m/z 287.2006     0   1   2   3E+6   11   11.5  RT (min)       lcorbi_t3_pos_3_047_40    m/z 287.2006     0   1   2   3E+4   12   12.5  RT (min)       lcorbi_t3_pos_1_031_40    m/z 287.2006     0   2   4E+4   12.5   13  RT (min)      lcorbi_t3_pos_1_023_02    m/z 288.2040     0   1   12   12.5  RT (min)      lcorbi_t3_pos_1_031_40    m/z 288.2040     0   1   12   12.5  RT (min)      lcorbi_t3_pos_1_031_40    m/z 288.2897     0   1E+5   9.5   10  RT (min)      lcorbi_t3_pos_1_023_02    m/z 288.2898     0   1E+5   9.5   10  RT (min)      lcorbi_t3_pos_1_023_02    m/z 289.0894     0   2   4   6   8E+5   5   5.5  RT (min)      lcorbi_t3_pos_3_000_aa    m/z 289.2738     0   1E+6   12   12.5  RT (min)       lcorbi_t3_pos_3_020_35    m/z 289.2738     0   1E+6   11.5   12  RT (min)       lcorbi_t3_pos_1_032_26    m/z 289.2738     0   1E+6   12   12.5  RT (min)      lcorbi_t3_pos_4_084_59    m/z 290.2690     0   1   2   3   4E+4   10   10.5  RT (min)      lcorbi_t3_pos_1_023_02    m/z 290.2690     0   1   2   3   4E+6   9.5   10  RT (min)      lcorbi_t3_pos_2_052_48    m/z 290.2690     0   1   2   3E+5   9.5   10  RT (min)      lcorbi_t3_pos_1_049_55    m/z 290.2690     0   1   2E+6   9.5   10  RT (min)      lcorbi_t3_pos_1_034_34    m/z 290.9475     0   1E+6   0   0.5  RT (min)      lcorbi_t3_pos_1_007_07    m/z 291.2724     0   2   4   6   8E+5   9.5   10  RT (min)      lcorbi_t3_pos_2_052_48    m/z 292.1520     0   1E+6   1   1.5  RT (min)       lcorbi_t3_pos_4_090_60    m/z 292.2846     0   1   12.5   13  RT (min)      lcorbi_t3_pos_1_031_40    m/z 292.2847     0   1   12.5   13  RT (min)      lcorbi_t3_pos_1_034_34    m/z 292.2847     0   1   13   13.5  RT (min)      lcorbi_t3_pos_1_031_40    m/z 292.2847     0   1E+6   12   12.5  RT (min)      lcorbi_t3_pos_1_008_36    m/z 292.2847     0   1E+4   12   12.5  RT (min)      lcorbi_t3_pos_1_028_32    m/z 292.2847     0   1   2   3E+4   12   12.5  RT (min)      lcorbi_t3_pos_1_028_32    m/z 292.2847     0   2   4E+4   12   12.5  RT (min)      lcorbi_t3_pos_4_041_54    m/z 292.2847     0   2   4   6   8E+5   11   11.5  RT (min)      lcorbi_t3_pos_2_013_33    m/z 292.2847     0   1   13   13.5  RT (min)      lcorbi_t3_pos_2_012_01    m/z 292.9858     0   1E+6   17.5   18  RT (min)      lcorbi_t3_pos_4_105_61    m/z 293.0979     0   1E+6   1   1.5  RT (min)       lcorbi_t3_pos_4_074_58    m/z 293.0980     0   1E+6   1   1.5  RT (min)       lcorbi_t3_pos_4_074_58    m/z 293.0980     0   1E+6   1   1.5  RT (min)       lcorbi_t3_pos_4_074_58    m/z 293.0980     0   2   4   6   8E+5   1   1.5  RT (min)      lcorbi_t3_pos_4_078_58    m/z 299.0914     0   2   4   6E+5   8.5   9  RT (min)       lcorbi_t3_pos_1_051_09    m/z 300.2898     0   1   2E+5   10   10.5  RT (min)      lcorbi_t3_pos_1_032_26    m/z 301.1410     0   1   2E+6   10   10.5  RT (min)      lcorbi_t3_pos_1_002_01    m/z 301.1411     0   1E+4   11.5   12  RT (min)      lcorbi_t3_pos_1_023_02    m/z 301.1411     0   1   2E+4   11   11.5  RT (min)      lcorbi_t3_pos_1_051_09    m/z 301.1411     0   1   2   3   4E+4   11   11.5  RT (min)      lcorbi_t3_pos_1_031_40    m/z 301.1411     0   1E+5   11   11.5  RT (min)      lcorbi_t3_pos_3_013_19    m/z 302.1445     0   1E+5   11   11.5  RT (min)      lcorbi_t3_pos_1_028_32    m/z 302.1445     0   1   11   11.5  RT (min)      lcorbi_t3_pos_1_031_40    m/z 302.3054     0   2   4E+5   10   10.5  RT (min)      lcorbi_t3_pos_2_013_33    m/z 302.3054     0   1E+4   11   11.5  RT (min)      lcorbi_t3_pos_1_034_34    m/z 302.3054     0   2   4E+5   11   11.5  RT (min)      lcorbi_t3_pos_1_008_36    m/z 302.3054     0   1E+4   11.5   12  RT (min)      lcorbi_t3_pos_1_031_40    m/z 303.2319     0   1   2   3E+5   11   11.5  RT (min)       lcorbi_t3_pos_1_029_29    m/z 303.2319     0   2   4   6E+5   11   11.5  RT (min)       lcorbi_t3_pos_2_013_33    m/z 303.2319     0   1   2   3E+4   12.5   13  RT (min)      lcorbi_t3_pos_1_028_32    m/z 303.2320     0   2   4   6   8E+5   11.5   12  RT (min)      lcorbi_t3_pos_3_011_42    m/z 303.2320     0   2   4   6E+5   12   12.5  RT (min)       lcorbi_t3_pos_1_008_36    m/z 303.2320     0   1   2   3E+5   11   11.5  RT (min)       lcorbi_t3_pos_1_029_29    m/z 303.2320     0   1   2E+4   13   13.5  RT (min)      lcorbi_t3_pos_1_031_40    m/z 303.2894     0   1   2E+5   11.5   12  RT (min)      lcorbi_t3_pos_2_013_33    m/z 303.2894     0   2   4   6   8E+3   11.5   12  RT (min)      lcorbi_t3_pos_3_031_45    m/z 303.2894     0   2   4   6   8E+5   12.5   13  RT (min)      lcorbi_t3_pos_1_008_36    m/z 303.2894     0   1   13.5   14  RT (min)      lcorbi_t3_pos_1_031_40    m/z 303.2894     0   1E+4   13   13.5  RT (min)      lcorbi_t3_pos_3_003_33    m/z 303.2895     0   1   2   3E+3   14   14.5  RT (min)      lcorbi_t3_pos_2_012_01    m/z 303.2895     0   1   13.5   14  RT (min)      lcorbi_t3_pos_1_034_34    m/z 303.2895     0   1   13.5   14  RT (min)      lcorbi_t3_pos_4_033_53    m/z 304.2999     0   1   2E+5   10   10.5  RT (min)      lcorbi_t3_pos_1_027_43    m/z 304.2999     0   1E+6   9.5   10  RT (min)      lcorbi_t3_pos_1_021_18    m/z 304.3000     0   2   4E+4   10   10.5  RT (min)      lcorbi_t3_pos_1_023_02    m/z 305.1478     0   1E+5   9.5   10  RT (min)      lcorbi_t3_pos_1_051_09    m/z 305.1478     0   2   4   6   8E+5   9.5   10  RT (min)      lcorbi_t3_pos_1_028_32    m/z 306.3004     0   1E+6   12   12.5  RT (min)       lcorbi_t3_pos_1_032_26    m/z 306.3004     0   1   2   3E+5   12   12.5  RT (min)       lcorbi_t3_pos_1_008_36    m/z 306.9198     0   1   2E+6   0   0.5  RT (min)      lcorbi_t3_pos_1_009_50    m/z 308.2220     0   1   2   3E+4   10.5   11  RT (min)      lcorbi_t3_pos_1_023_02    m/z 308.2220     0   1   2E+5   10   10.5  RT (min)      lcorbi_t3_pos_1_051_09    m/z 308.2220     0   2   4   6E+6   10   10.5  RT (min)      lcorbi_t3_pos_1_055_33    m/z 308.2220     0   1E+7   10   10.5  RT (min)      lcorbi_t3_pos_1_028_32    m/z 308.2221     0   1   2E+6   1   1.5  RT (min)      lcorbi_t3_pos_4_057_49    m/z 309.2254     0   1E+6   10   10.5  RT (min)      lcorbi_t3_pos_1_028_32    m/z 309.2254     0   1E+6   9   9.5  RT (min)      lcorbi_t3_pos_3_024_07    m/z 309.2254     0   2   4   6E+4   10   10.5  RT (min)      lcorbi_t3_pos_2_052_48    m/z 309.2254     0   2   4   6   8E+5   10   10.5  RT (min)      lcorbi_t3_pos_1_030_21    m/z 309.2254     0   1   2   3   4E+4   10   10.5  RT (min)      lcorbi_t3_pos_1_051_09    m/z 309.2255     0   1E+4   10.5   11  RT (min)      lcorbi_t3_pos_1_023_02    m/z 309.2901     0   2   4E+5   12   12.5  RT (min)      lcorbi_t3_pos_2_054_29    m/z 309.2901     0   2   4   6E+5   12   12.5  RT (min)       lcorbi_t3_pos_4_075_58    m/z 309.2902     0   1   14.5   15  RT (min)      lcorbi_t3_pos_1_023_02    m/z 309.2902     0   1   14.5   15  RT (min)      lcorbi_t3_pos_1_023_02    m/z 310.3106     0   2   4   6E+5   12   12.5  RT (min)      lcorbi_t3_pos_2_054_29    m/z 310.3106     0   1   2E+4   12   12.5  RT (min)      lcorbi_t3_pos_3_052_47    m/z 311.3058     0   1   2   3E+4   15   15.5  RT (min)      lcorbi_t3_pos_1_023_02    m/z 311.3058     0   2   4   6E+5   12   12.5  RT (min)      lcorbi_t3_pos_2_054_29    m/z 311.3058     0   1   14.5   15  RT (min)      lcorbi_t3_pos_1_051_09    m/z 311.3058     0   1   2   3E+4   15   15.5  RT (min)      lcorbi_t3_pos_1_023_02    m/z 311.3058     0   2   4   6E+5   12.5   13  RT (min)      lcorbi_t3_pos_1_002_01    m/z 311.3058     0   1   2E+5   12   12.5  RT (min)      lcorbi_t3_pos_4_012_37    m/z 311.3059     0   2   4   6   8E+4   12   12.5  RT (min)      lcorbi_t3_pos_2_032_39    m/z 312.1554     0   1   2E+7   1   1.5  RT (min)      lcorbi_t3_pos_4_057_49    m/z 312.3262     0   1E+5   13.5   14  RT (min)      lcorbi_t3_pos_3_044_46    m/z 314.0918     0   2   4E+5   1   1.5  RT (min)       lcorbi_t3_pos_3_029_13    m/z 315.2319     0   1E+6   12   12.5  RT (min)      lcorbi_t3_pos_1_008_36    m/z 315.2319     0   1E+5   13.5   14  RT (min)      lcorbi_t3_pos_1_023_02    m/z 315.2320     0   1   2   3E+4   14   14.5  RT (min)      lcorbi_t3_pos_1_025_52    m/z 315.2320     0   1   2   3   4E+4   13   13.5  RT (min)      lcorbi_t3_pos_2_012_01    m/z 315.2320     0   1   2   3E+6   11   11.5  RT (min)       lcorbi_t3_pos_2_013_33    m/z 315.2320     0   2   4   6E+4   13   13.5  RT (min)      lcorbi_t3_pos_1_031_40    m/z 315.2320     0   1   2   3   4E+5   12   12.5  RT (min)      lcorbi_t3_pos_2_006_47    m/z 315.2320     0   2   4E+6   12   12.5  RT (min)       lcorbi_t3_pos_1_008_36    m/z 315.2320     0   1   2   3   4E+4   12.5   13  RT (min)      lcorbi_t3_pos_1_034_34    m/z 315.2320     0   1   2   3E+6   11.5   12  RT (min)      lcorbi_t3_pos_4_015_45    m/z 315.2320     0   2   4   6E+4   14   14.5  RT (min)      lcorbi_t3_pos_1_023_02    m/z 315.2320     0   1   2   3E+6   11   11.5  RT (min)       lcorbi_t3_pos_4_015_45    m/z 315.2320     0   1   2   3   4E+4   12.5   13  RT (min)       lcorbi_t3_pos_1_034_34    m/z 315.2320     0   2   4E+6   12   12.5  RT (min)       lcorbi_t3_pos_1_008_36    m/z 315.2320     0   2   4   6E+5   12.5   13  RT (min)       lcorbi_t3_pos_2_014_37    m/z 315.2320     0   1E+5   12.5   13  RT (min)       lcorbi_t3_pos_3_003_33    m/z 315.2320     0   1E+5   12.5   13  RT (min)       lcorbi_t3_pos_3_003_33    m/z 315.2320     0   1   2   3E+6   11   11.5  RT (min)       lcorbi_t3_pos_3_037_32    m/z 316.2353     0   1E+5   12.5   13  RT (min)      lcorbi_t3_pos_2_014_37    m/z 316.2353     0   1E+4   13   13.5  RT (min)      lcorbi_t3_pos_2_012_01    m/z 316.2353     0   1   2   3E+4   12.5   13  RT (min)      lcorbi_t3_pos_1_034_34    m/z 316.2353     0   1E+6   12   12.5  RT (min)      lcorbi_t3_pos_1_008_36    m/z 316.2353     0   2   4   6E+5   11   11.5  RT (min)      lcorbi_t3_pos_2_054_29    m/z 316.2354     0   1E+6   12   12.5  RT (min)      lcorbi_t3_pos_1_008_36    m/z 316.2354     0   2   4   6E+5   11   11.5  RT (min)      lcorbi_t3_pos_3_037_32    m/z 316.2354     0   1   2   3E+4   12.5   13  RT (min)      lcorbi_t3_pos_3_003_33    m/z 316.2354     0   2   4   6E+5   11   11.5  RT (min)      lcorbi_t3_pos_4_015_45    m/z 316.2354     0   2   4E+4   13.5   14  RT (min)      lcorbi_t3_pos_1_023_02    m/z 317.1020     0   1   2   3E+6   8.5   9  RT (min)       lcorbi_t3_pos_1_051_09    m/z 317.1021     0   1E+6   8.5   9  RT (min)       lcorbi_t3_pos_1_023_02    m/z 317.2476     0   1E+5   12.5   13  RT (min)       lcorbi_t3_pos_3_013_19    m/z 317.2476     0   1E+4   12.5   13  RT (min)      lcorbi_t3_pos_1_028_32    m/z 318.1054     0   2   4E+5   8.5   9  RT (min)      lcorbi_t3_pos_1_051_09    m/z 318.3003     0   2   4   6   8E+6   10.5   11  RT (min)      lcorbi_t3_pos_1_008_36    m/z 318.3003     0   2   4   6E+6   10.5   11  RT (min)      lcorbi_t3_pos_2_022_38    m/z 318.3003     0   2   4   6E+5   11   11.5  RT (min)      lcorbi_t3_pos_1_034_34    m/z 318.3004     0   1   2E+5   11   11.5  RT (min)      lcorbi_t3_pos_1_031_40    m/z 318.3004     0   2   4   6   8E+6   10.5   11  RT (min)       lcorbi_t3_pos_1_008_36    m/z 318.3004     0   1   2E+5   11   11.5  RT (min)      lcorbi_t3_pos_2_012_01    m/z 319.2843     0   1E+6   11   11.5  RT (min)      lcorbi_t3_pos_3_011_42    m/z 319.2844     0   1   2   3   4E+4   12   12.5  RT (min)      lcorbi_t3_pos_3_003_33    m/z 319.2844     0   1E+4   12.5   13  RT (min)      lcorbi_t3_pos_1_034_34    m/z 319.2844     0   1E+4   12.5   13  RT (min)      lcorbi_t3_pos_1_031_40    m/z 319.2844     0   1   12.5   13  RT (min)      lcorbi_t3_pos_1_025_52    m/z 319.2844     0   2   4   6   8E+3   12.5   13  RT (min)      lcorbi_t3_pos_2_009_50    m/z 319.2844     0   1E+6   11   11.5  RT (min)      lcorbi_t3_pos_2_013_33    m/z 319.2844     0   1E+6   12   12.5  RT (min)      lcorbi_t3_pos_1_008_36    m/z 319.3037     0   1E+6   10.5   11  RT (min)      lcorbi_t3_pos_2_022_38    m/z 319.3037     0   1E+4   11   11.5  RT (min)      lcorbi_t3_pos_2_012_01    m/z 319.3038     0   1   2   3E+4   11   11.5  RT (min)      lcorbi_t3_pos_2_012_01    m/z 319.3038     0   1E+6   10   10.5  RT (min)      lcorbi_t3_pos_3_024_07    m/z 319.3038     0   2   4E+4   11   11.5  RT (min)      lcorbi_t3_pos_1_031_40    m/z 319.3038     0   1E+6   10.5   11  RT (min)      lcorbi_t3_pos_1_008_36    m/z 319.3038     0   1E+5   11   11.5  RT (min)      lcorbi_t3_pos_1_034_34    m/z 320.3161     0   1E+4   13   13.5  RT (min)      lcorbi_t3_pos_2_022_38    m/z 320.3161     0   1   14   14.5  RT (min)      lcorbi_t3_pos_2_012_01    m/z 320.3161     0   2   4   6E+4   11.5   12  RT (min)      lcorbi_t3_pos_2_013_33    m/z 320.3161     0   2   4   6E+5   12.5   13  RT (min)      lcorbi_t3_pos_1_008_36    m/z 320.3161     0   1   13.5   14  RT (min)      lcorbi_t3_pos_1_034_34    m/z 320.3161     0   1E+4   13   13.5  RT (min)      lcorbi_t3_pos_1_033_53    m/z 320.3161     0   1   13.5   14  RT (min)      lcorbi_t3_pos_1_056_56    m/z 320.3161     0   1   13.5   14  RT (min)      lcorbi_t3_pos_1_031_40    m/z 320.3162     0   2   4E+5   11.5   12  RT (min)      lcorbi_t3_pos_2_013_33    m/z 321.3154     0   2   4   6E+4   15   15.5  RT (min)      lcorbi_t3_pos_1_025_52    m/z 321.3154     0   1E+5   14   14.5  RT (min)      lcorbi_t3_pos_3_044_46    m/z 321.3154     0   1E+4   12.5   13  RT (min)      lcorbi_t3_pos_3_052_47    m/z 321.3154     0   2   4E+4   14.5   15  RT (min)      lcorbi_t3_pos_2_038_03    m/z 321.3154     0   2   4   6E+4   14.5   15  RT (min)      lcorbi_t3_pos_1_041_54    m/z 321.3154     0   2   4   6E+4   14   14.5  RT (min)      lcorbi_t3_pos_2_017_51    m/z 321.3154     0   2   4E+4   14.5   15  RT (min)      lcorbi_t3_pos_2_052_48    m/z 321.3154     0   2   4E+4   14   14.5  RT (min)      lcorbi_t3_pos_1_033_53    m/z 321.3154     0   2   4   6E+4   15.5   16  RT (min)      lcorbi_t3_pos_1_023_02    m/z 321.3154     0   2   4E+5   13.5   14  RT (min)       lcorbi_t3_pos_1_005_42    m/z 323.1754     0   1E+7   5   5.5  RT (min)      lcorbi_t3_pos_4_048_24    m/z 323.1754     0   1E+7   1   1.5  RT (min)      lcorbi_t3_pos_4_057_49    m/z 324.1241     0   1   2E+7   1   1.5  RT (min)       lcorbi_t3_pos_4_057_49    m/z 324.1521     0   1E+5   5.5   6  RT (min)      lcorbi_t3_pos_4_103_61    m/z 324.1788     0   1   2   3   4E+6   5   5.5  RT (min)      lcorbi_t3_pos_4_048_24    m/z 324.1789     0   1   2   3E+6   1   1.5  RT (min)      lcorbi_t3_pos_4_057_49    m/z 325.1287     0   2   4   6E+5   3   3.5  RT (min)       lcorbi_t3_pos_4_092_60    m/z 325.1288     0   2   4   6E+5   3.5   4  RT (min)       lcorbi_t3_pos_4_090_60    m/z 325.2528     0   2   4E+4   15   15.5  RT (min)      lcorbi_t3_pos_1_008_36    m/z 325.2528     0   1   16   16.5  RT (min)      lcorbi_t3_pos_1_034_34    m/z 325.2528     0   1   15.5   16  RT (min)      lcorbi_t3_pos_1_055_33    m/z 325.2528     0   1E+4   13.5   14  RT (min)      lcorbi_t3_pos_3_037_32    m/z 325.2528     0   1   16.5   17  RT (min)    lcorbi_t3_pos_2_012_01    m/z 325.2528     0   1   16   16.5  RT (min)      lcorbi_t3_pos_1_034_34    m/z 325.2528     0   1   2E+4   15   15.5  RT (min)      lcorbi_t3_pos_1_008_36    m/z 325.2561     0   1E+6   12   12.5  RT (min)      lcorbi_t3_pos_3_000_aa    m/z 326.1265     0   1   2E+5   9.5   10  RT (min)      lcorbi_t3_pos_1_051_09    m/z 326.1265     0   1E+6   9.5   10  RT (min)      lcorbi_t3_pos_1_028_32    m/z 327.0782     0   2   4E+4   11   11.5  RT (min)      lcorbi_t3_pos_1_025_52    m/z 327.0782     0   2   4E+5   10.5   11  RT (min)      lcorbi_t3_pos_1_008_36    m/z 327.0782     0   2   4   6E+5   9.5   10  RT (min)      lcorbi_t3_pos_2_013_33    m/z 327.0782     0   2   4E+5   10.5   11  RT (min)      lcorbi_t3_pos_1_033_53    m/z 327.0782     0   2   4E+4   11   11.5  RT (min)      lcorbi_t3_pos_1_031_40    m/z 327.0782     0   1   2   3   4E+5   10.5   11  RT (min)      lcorbi_t3_pos_1_028_32    m/z 327.0783     0   2   4   6E+4   11   11.5  RT (min)      lcorbi_t3_pos_1_023_02    m/z 327.1264     0   1   2   3   4E+5   1   1.5  RT (min)       lcorbi_t3_pos_1_031_40    m/z 328.1603     0   2   4   6   8E+5   5   5.5  RT (min)      lcorbi_t3_pos_3_000_aa    m/z 329.1239     0   2   4   6E+5   1   1.5  RT (min)       lcorbi_t3_pos_3_055_10    m/z 329.1596     0   1E+5   10.5   11  RT (min)      lcorbi_t3_pos_2_047_31    m/z 329.1596     0   2   4   6E+4   12   12.5  RT (min)      lcorbi_t3_pos_3_013_19    m/z 329.1596     0   1   2   3E+4   12   12.5  RT (min)      lcorbi_t3_pos_2_052_48    m/z 329.1597     0   2   4   6   8E+5   10.5   11  RT (min)      lcorbi_t3_pos_3_011_42    m/z 329.1597     0   1E+6   11.5   12  RT (min)      lcorbi_t3_pos_3_044_46    m/z 329.1597     0   1E+5   11.5   12  RT (min)      lcorbi_t3_pos_4_028_04    m/z 329.1597     0   1   2E+4   12   12.5  RT (min)      lcorbi_t3_pos_2_056_56    m/z 329.1597     0   1E+6   10.5   11  RT (min)      lcorbi_t3_pos_3_052_47    m/z 329.1598     0   1   2E+4   12.5   13  RT (min)      lcorbi_t3_pos_2_056_56    m/z 330.2040     0   1E+4   10.5   11  RT (min)      lcorbi_t3_pos_1_023_02    m/z 330.2040     0   1   2E+6   10   10.5  RT (min)      lcorbi_t3_pos_2_022_38    m/z 330.2041     0   1   2E+6   10   10.5  RT (min)      lcorbi_t3_pos_1_028_32    m/z 330.2041     0   1   2E+6   9   9.5  RT (min)      lcorbi_t3_pos_2_047_31    m/z 330.2041     0   2   4   6E+4   10   10.5  RT (min)      lcorbi_t3_pos_1_051_09    m/z 330.3004     0   2   4   6   8E+4   11.5   12  RT (min)      lcorbi_t3_pos_1_028_32    m/z 330.3004     0   2   4E+4   11.5   12  RT (min)      lcorbi_t3_pos_1_028_32    m/z 330.3005     0   1E+4   12   12.5  RT (min)      lcorbi_t3_pos_2_012_01    m/z 330.3005     0   2   4   6E+5   11   11.5  RT (min)      lcorbi_t3_pos_1_008_36    m/z 330.3005     0   2   4E+5   10.5   11  RT (min)      lcorbi_t3_pos_3_024_07    m/z 330.3005     0   2   4E+5   10.5   11  RT (min)      lcorbi_t3_pos_2_013_33    m/z 331.1905     0   1   2E+6   11   11.5  RT (min)      lcorbi_t3_pos_2_054_29    m/z 331.1906     0   1   13   13.5  RT (min)      lcorbi_t3_pos_2_012_01    m/z 331.1906     0   1E+4   13   13.5  RT (min)      lcorbi_t3_pos_1_031_40    m/z 331.1906     0   1   2E+6   12   12.5  RT (min)      lcorbi_t3_pos_3_047_40    m/z 331.1906     0   1   2   3E+4   12.5   13  RT (min)      lcorbi_t3_pos_1_028_32    m/z 331.1906     0   1E+4   12.5   13  RT (min)      lcorbi_t3_pos_1_034_34    m/z 331.1906     0   1   2E+4   13   13.5  RT (min)      lcorbi_t3_pos_1_023_02    m/z 331.1906     0   1   2   3E+4   12   12.5  RT (min)      lcorbi_t3_pos_1_028_32    m/z 331.2074     0   2   4E+5   10   10.5  RT (min)      lcorbi_t3_pos_1_028_32    m/z 332.1939     0   1   13   13.5  RT (min)      lcorbi_t3_pos_1_023_02    m/z 332.2797     0   2   4E+5   10.5   11  RT (min)      lcorbi_t3_pos_2_006_47    m/z 332.3313     0   1E+6   10.5   11  RT (min)      lcorbi_t3_pos_2_046_07    m/z 332.3313     0   2   4   6   8E+5   9.5   10  RT (min)      lcorbi_t3_pos_3_024_07    m/z 332.3313     0   1   2   3E+5   10.5   11  RT (min)      lcorbi_t3_pos_2_038_03    m/z 332.3313     0   2   4   6   8E+4   10.5   11  RT (min)      lcorbi_t3_pos_2_052_48    m/z 332.3314     0   2   4   6E+5   9.5   10  RT (min)      lcorbi_t3_pos_3_035_08    m/z 333.1158     0   2   4   6E+5   5   5.5  RT (min)      lcorbi_t3_pos_3_000_aa    m/z 333.2426     0   2   4   6E+5   11.5   12  RT (min)       lcorbi_t3_pos_1_054_48    m/z 333.2637     0   1E+5   9.5   10  RT (min)      lcorbi_t3_pos_4_028_04    m/z 333.3000     0   1E+5   12   12.5  RT (min)      lcorbi_t3_pos_1_028_32    m/z 333.3001     0   1   2   3E+5   12   12.5  RT (min)      lcorbi_t3_pos_2_022_38    m/z 333.3001     0   1E+5   12   12.5  RT (min)      lcorbi_t3_pos_1_028_32    m/z 333.3001     0   2   4   6   8E+4   12   12.5  RT (min)      lcorbi_t3_pos_1_028_32    m/z 333.3001     0   1E+5   12   12.5  RT (min)      lcorbi_t3_pos_2_022_38    m/z 333.3002     0   1E+4   12.5   13  RT (min)      lcorbi_t3_pos_1_049_55    m/z 333.3002     0   1   2   3E+5   11   11.5  RT (min)      lcorbi_t3_pos_3_031_45    m/z 333.3002     0   2   4E+5   12   12.5  RT (min)      lcorbi_t3_pos_1_008_36    m/z 333.3002     0   1E+4   13   13.5  RT (min)      lcorbi_t3_pos_2_012_01    m/z 333.3002     0   1E+4   12.5   13  RT (min)      lcorbi_t3_pos_2_009_50    m/z 333.3002     0   1   2   3   4E+5   12   12.5  RT (min)      lcorbi_t3_pos_2_017_51    m/z 335.0682     0   2   4   6E+5   8.5   9  RT (min)      lcorbi_t3_pos_1_051_09    m/z 336.3025     0   1   12.5   13  RT (min)       lcorbi_t3_pos_1_034_34    m/z 336.3025     0   1   2   3E+4   12.5   13  RT (min)      lcorbi_t3_pos_2_056_56    m/z 336.3025     0   1E+5   12.5   13  RT (min)      lcorbi_t3_pos_2_009_50    m/z 336.3026     0   1E+4   12.5   13  RT (min)       lcorbi_t3_pos_1_031_40    m/z 336.3068     0   1E+4   11.5   12  RT (min)       lcorbi_t3_pos_4_088_59    m/z 336.3111     0   1E+6   11   11.5  RT (min)      lcorbi_t3_pos_2_013_33    m/z 336.3111     0   2   4   6   8E+5   11   11.5  RT (min)      lcorbi_t3_pos_3_011_42    m/z 336.3608     0   2   4   6   8E+4   11   11.5  RT (min)      lcorbi_t3_pos_2_038_03    m/z 336.3608     0   1   2   3   4E+4   11   11.5  RT (min)      lcorbi_t3_pos_1_041_54    m/z 336.3608     0   2   4   6E+5   10.5   11  RT (min)      lcorbi_t3_pos_1_047_41    m/z 336.3608     0   1E+4   11   11.5  RT (min)      lcorbi_t3_pos_1_001_49    m/z 336.3608     0   1E+4   11   11.5  RT (min)      lcorbi_t3_pos_1_056_56    m/z 337.2351     0   2   4E+5   10.5   11  RT (min)      lcorbi_t3_pos_2_017_51    m/z 337.2739     0   2   4   6E+4   15   15.5  RT (min)      lcorbi_t3_pos_1_008_36    m/z 338.3419     0   1   2E+6   14   14.5  RT (min)      lcorbi_t3_pos_1_055_33    m/z 338.3419     0   2   4   6E+4   11.5   12  RT (min)      lcorbi_t3_pos_2_039_30    m/z 338.3419     0   1E+6   15.5   16  RT (min)      lcorbi_t3_pos_1_023_02    m/z 338.3419     0   1E+7   13   13.5  RT (min)      lcorbi_t3_pos_4_105_61    m/z 338.3419     0   1   2E+6   14   14.5  RT (min)      lcorbi_t3_pos_2_017_51    m/z 338.3420     0   1   2E+7   13.5   14  RT (min)      lcorbi_t3_pos_1_020_31    m/z 338.3420     0   1E+7   13.5   14  RT (min)       lcorbi_t3_pos_2_033_53    m/z 338.3420     0   1E+6   15   15.5  RT (min)      lcorbi_t3_pos_1_051_09    m/z 338.3420     0   1   2   3   4E+7   13   13.5  RT (min)       lcorbi_t3_pos_2_002_06    m/z 338.3420     0   1E+6   15   15.5  RT (min)      lcorbi_t3_pos_2_056_56    m/z 338.3420     0   1E+6   12.5   13  RT (min)       lcorbi_t3_pos_3_052_47    m/z 338.3420     0   2   4   6E+5   12   12.5  RT (min)      lcorbi_t3_pos_2_032_39    m/z 338.3420     0   1E+6   16   16.5  RT (min)      lcorbi_t3_pos_1_023_02    m/z 338.3420     0   1E+6   15.5   16  RT (min)      lcorbi_t3_pos_1_051_09    m/z 338.3420     0   1E+8   13   13.5  RT (min)       lcorbi_t3_pos_3_000_aa    m/z 338.3420     0   2   4E+5   12.5   13  RT (min)      lcorbi_t3_pos_2_047_31    m/z 338.3420     0   1E+6   16   16.5  RT (min)      lcorbi_t3_pos_1_023_02    m/z 338.3420     0   1E+6   14.5   15  RT (min)      lcorbi_t3_pos_2_052_48    m/z 338.3420     0   2   4   6E+4   11.5   12  RT (min)      lcorbi_t3_pos_2_032_39    m/z 338.3420     0   1E+6   16   16.5  RT (min)      lcorbi_t3_pos_1_023_02    m/z 338.3420     0   1E+6   15   15.5  RT (min)      lcorbi_t3_pos_2_056_56    m/z 338.3420     0   1E+7   13   13.5  RT (min)       lcorbi_t3_pos_3_053_48    m/z 338.3420     0   2   4   6E+6   13.5   14  RT (min)      lcorbi_t3_pos_4_010_43    m/z 338.3420     0   2   4E+5   12   12.5  RT (min)      lcorbi_t3_pos_2_047_31    m/z 338.3420     0   1E+6   12.5   13  RT (min)       lcorbi_t3_pos_3_052_47    m/z 338.3420     0   1E+7   13   13.5  RT (min)       lcorbi_t3_pos_4_105_61    m/z 338.3420     0   1   2   3E+6   14   14.5  RT (min)       lcorbi_t3_pos_3_044_46    m/z 338.3420     0   1E+6   15.5   16  RT (min)      lcorbi_t3_pos_1_023_02    m/z 338.3420     0   1E+8   13   13.5  RT (min)       lcorbi_t3_pos_3_000_aa    m/z 338.3420     0   1E+6   15   15.5  RT (min)       lcorbi_t3_pos_1_027_43    m/z 338.3420     0   1   2E+7   13   13.5  RT (min)       lcorbi_t3_pos_1_020_31    m/z 338.3420     0   1E+6   14.5   15  RT (min)       lcorbi_t3_pos_2_052_48    m/z 338.3420     0   1   2E+6   14   14.5  RT (min)       lcorbi_t3_pos_1_038_30    m/z 338.3420     0   1E+7   13.5   14  RT (min)      lcorbi_t3_pos_1_005_42    m/z 338.3420     0   1E+7   13   13.5  RT (min)       lcorbi_t3_pos_4_105_61    m/z 338.3420     0   1E+6   15   15.5  RT (min)      lcorbi_t3_pos_1_051_09    m/z 338.3420     0   1E+6   14   14.5   15  RT (min)       lcorbi_t3_pos_1_055_33    m/z 338.3420     0   1E+6   14   14.5   15  RT (min)       lcorbi_t3_pos_1_028_32    m/z 338.3420     0   1E+6   12.5   13  RT (min)       lcorbi_t3_pos_3_052_47    m/z 338.3421     0   1E+6   0.5   1  RT (min)      lcorbi_t3_pos_4_057_49    m/z 339.2880     0   1   2   3E+4   13.5   14  RT (min)      lcorbi_t3_pos_2_013_33    m/z 339.2895     0   1   17   17.5  RT (min)    lcorbi_t3_pos_2_012_01    m/z 339.2895     0   1   2   3E+4   16   16.5  RT (min)      lcorbi_t3_pos_1_028_32    m/z 339.2895     0   1E+5   15   15.5  RT (min)      lcorbi_t3_pos_1_008_36    m/z 339.2896     0   1   2E+5   14   14.5  RT (min)      lcorbi_t3_pos_3_011_42    m/z 339.2896     0   1   2   3E+4   16   16.5  RT (min)      lcorbi_t3_pos_1_028_32    m/z 339.2896     0   1E+4   15.5   16  RT (min)      lcorbi_t3_pos_3_003_33    m/z 339.2896     0   1   2E+4   16   16.5  RT (min)      lcorbi_t3_pos_1_034_34    m/z 339.2896     0   1E+4   15.5   16  RT (min)      lcorbi_t3_pos_2_022_38    m/z 339.2896     0   2   4   6E+4   15   15.5  RT (min)      lcorbi_t3_pos_1_008_36    m/z 339.2896     0   1E+4   15.5   16  RT (min)      lcorbi_t3_pos_2_022_38    m/z 339.2896     0   1E+4   13.5   14  RT (min)      lcorbi_t3_pos_3_031_45    m/z 339.2896     0   1E+4   13.5   14  RT (min)      lcorbi_t3_pos_3_031_45    m/z 339.2896     0   1   2E+5   14   14.5  RT (min)      lcorbi_t3_pos_3_011_42    m/z 339.2896     0   1   16.5   17  RT (min)    lcorbi_t3_pos_1_031_40    m/z 339.3452     0   1E+6   14   14.5  RT (min)      lcorbi_t3_pos_3_047_40    m/z 339.3452     0   1   2   3E+5   12.5   13  RT (min)      lcorbi_t3_pos_1_002_01    m/z 339.3453     0   1   2E+5   14.5   15  RT (min)       lcorbi_t3_pos_2_052_48    m/z 339.3453     0   1   2   3E+5   14   14.5  RT (min)       lcorbi_t3_pos_1_033_53    m/z 339.3453     0   1   2E+6   13.5   14  RT (min)      lcorbi_t3_pos_3_047_40    m/z 339.3453     0   1   2   3E+6   12.5   13  RT (min)      lcorbi_t3_pos_1_002_01    m/z 339.3453     0   1   2   3E+5   14.5   15  RT (min)      lcorbi_t3_pos_1_055_33    m/z 339.3453     0   1   2E+5   15   15.5  RT (min)      lcorbi_t3_pos_2_012_01    m/z 339.3453     0   1   2E+5   15   15.5  RT (min)      lcorbi_t3_pos_2_014_37    m/z 339.3453     0   1   2   3   4E+5   12.5   13  RT (min)       lcorbi_t3_pos_3_052_47    m/z 339.3453     0   1   2   3E+5   15.5   16  RT (min)      lcorbi_t3_pos_1_023_02    m/z 339.3453     0   1   2E+5   15   15.5  RT (min)      lcorbi_t3_pos_2_012_01    m/z 339.3453     0   1   2   3E+5   14   14.5  RT (min)      lcorbi_t3_pos_1_030_21    m/z 339.3453     0   1   2   3E+5   14   14.5  RT (min)      lcorbi_t3_pos_1_033_53    m/z 339.3453     0   1   2   3E+6   13.5   14  RT (min)      lcorbi_t3_pos_3_047_40    m/z 339.3453     0   1   2   3E+5   14.5   15  RT (min)      lcorbi_t3_pos_2_038_03    m/z 339.3453     0   1E+5   12.5   13  RT (min)       lcorbi_t3_pos_2_047_31    m/z 339.3453     0   1   2   3E+5   14.5   15  RT (min)      lcorbi_t3_pos_1_033_53    m/z 339.3453     0   1   2   3   4E+5   12.5   13  RT (min)      lcorbi_t3_pos_3_052_47    m/z 339.3453     0   1E+6   13.5   14  RT (min)       lcorbi_t3_pos_1_008_36    m/z 339.3453     0   1   2   3E+5   15   15.5  RT (min)      lcorbi_t3_pos_1_049_55    m/z 339.3453     0   1   2   3E+5   14   14.5  RT (min)      lcorbi_t3_pos_2_046_07    m/z 339.3453     0   1   2   3   4E+5   14   14.5  RT (min)       lcorbi_t3_pos_2_017_51    m/z 339.3453     0   1   2   3E+5   14.5   15  RT (min)      lcorbi_t3_pos_2_014_37    m/z 339.3453     0   1   2   3E+5   15   15.5  RT (min)      lcorbi_t3_pos_1_025_52    m/z 339.3453     0   1   2E+5   15.5   16  RT (min)      lcorbi_t3_pos_1_051_09    m/z 339.3453     0   1   2   3E+6   12.5   13  RT (min)       lcorbi_t3_pos_2_016_23    m/z 339.3453     0   1   2   3E+5   15   15.5  RT (min)       lcorbi_t3_pos_1_027_43    m/z 339.3453     0   1   2   3E+5   15.5   16  RT (min)      lcorbi_t3_pos_1_051_09    m/z 339.3454     0   1   2   3E+5   14.5   15  RT (min)      lcorbi_t3_pos_1_041_54    m/z 340.3486     0   1   2   3   4E+4   12.5   13  RT (min)      lcorbi_t3_pos_3_052_47    m/z 340.3486     0   2   4   6E+5   13   13.5  RT (min)      lcorbi_t3_pos_3_052_47    m/z 340.3487     0   2   4   6E+4   15   15.5  RT (min)      lcorbi_t3_pos_1_025_52    m/z 340.3487     0   2   4   6E+4   15   15.5  RT (min)      lcorbi_t3_pos_1_027_43    m/z 340.3576     0   1   2   3E+5   14   14.5  RT (min)      lcorbi_t3_pos_2_017_51    m/z 341.1750     0   1E+5   16  RT (min)       lcorbi_t3_pos_4_085_59    m/z 343.2956     0   2   4   6E+4   10   10.5  RT (min)      lcorbi_t3_pos_1_031_40    m/z 343.2957     0   1E+6   9.5   10  RT (min)      lcorbi_t3_pos_1_028_32    m/z 343.2957     0   1   2   3E+4   10   10.5  RT (min)      lcorbi_t3_pos_1_031_40    m/z 343.2957     0   1E+6   9   9.5  RT (min)      lcorbi_t3_pos_3_024_07    m/z 344.3161     0   2   4E+5   10.5   11  RT (min)      lcorbi_t3_pos_2_013_33    m/z 344.3161     0   2   4   6   8E+4   11.5   12  RT (min)      lcorbi_t3_pos_1_028_32    m/z 344.3161     0   1E+4   12   12.5  RT (min)      lcorbi_t3_pos_1_031_40    m/z 344.3161     0   1   2   3E+4   11.5   12  RT (min)      lcorbi_t3_pos_1_034_34    m/z 344.3161     0   2   4   6E+5   11   11.5  RT (min)      lcorbi_t3_pos_1_008_36    m/z 344.3161     0   2   4   6E+5   11   11.5  RT (min)      lcorbi_t3_pos_1_008_36    m/z 344.3161     0   2   4E+5   10   10.5  RT (min)      lcorbi_t3_pos_3_024_07    m/z 344.3162     0   1E+4   11.5   12  RT (min)      lcorbi_t3_pos_1_031_40    m/z 346.0095     0   2   4   6E+5   1   1.5  RT (min)       lcorbi_t3_pos_4_080_58    m/z 346.1780     0   1E+4   10   10.5  RT (min)      lcorbi_t3_pos_1_051_09    m/z 346.1780     0   2   4E+5   10   10.5  RT (min)      lcorbi_t3_pos_1_028_32    m/z 347.3158     0   1   13.5   14  RT (min)      lcorbi_t3_pos_2_056_56    m/z 347.3158     0   1   13.5   14  RT (min)      lcorbi_t3_pos_1_031_40    m/z 347.3158     0   2   4   6E+5   11.5   12  RT (min)      lcorbi_t3_pos_2_013_33    m/z 347.3158     0   2   4   6   8E+5   12.5   13  RT (min)      lcorbi_t3_pos_1_008_36    m/z 347.3158     0   1   13   13.5  RT (min)      lcorbi_t3_pos_1_034_34    m/z 347.3158     0   1E+4   13   13.5  RT (min)      lcorbi_t3_pos_1_028_32    m/z 347.3158     0   1   13   13.5  RT (min)      lcorbi_t3_pos_2_049_55    m/z 350.3266     0   1E+4   12.5   13  RT (min)      lcorbi_t3_pos_1_049_55    m/z 350.3266     0   1E+5   12   12.5  RT (min)      lcorbi_t3_pos_1_028_32    m/z 350.3266     0   1   12.5   13  RT (min)      lcorbi_t3_pos_1_031_40    m/z 350.3266     0   2   4   6   8E+5   12   12.5  RT (min)      lcorbi_t3_pos_1_008_36    m/z 350.3266     0   1E+4   12.5   13  RT (min)      lcorbi_t3_pos_1_049_55    m/z 350.3266     0   1   2   3E+4   12.5   13  RT (min)      lcorbi_t3_pos_1_041_54    m/z 350.3266     0   1   2E+4   12.5   13  RT (min)      lcorbi_t3_pos_1_034_34    m/z 350.3266     0   1E+4   12.5   13  RT (min)      lcorbi_t3_pos_2_009_50    m/z 350.3266     0   2   4   6   8E+5   12   12.5  RT (min)      lcorbi_t3_pos_1_008_36    m/z 350.3266     0   2   4   6   8E+5   12   12.5  RT (min)      lcorbi_t3_pos_1_008_36    m/z 350.3267     0   1   2   3   4E+5   11   11.5  RT (min)      lcorbi_t3_pos_3_031_45    m/z 350.3267     0   1E+4   13   13.5  RT (min)      lcorbi_t3_pos_2_012_01    m/z 350.3267     0   2   4   6   8E+5   12   12.5  RT (min)      lcorbi_t3_pos_1_008_36    m/z 350.3267     0   2   4   6E+5   11   11.5  RT (min)      lcorbi_t3_pos_2_013_33    m/z 350.3267     0   1   2E+5   12   12.5  RT (min)      lcorbi_t3_pos_2_022_38    m/z 350.3267     0   1   2E+6   11.5   12  RT (min)       lcorbi_t3_pos_4_084_59    m/z 350.3267     0   2   4E+5   12   12.5  RT (min)      lcorbi_t3_pos_2_022_38    m/z 350.3267     0   1   13   13.5  RT (min)      lcorbi_t3_pos_1_031_40    m/z 350.3268     0   1E+4   12.5   13  RT (min)      lcorbi_t3_pos_2_009_50    m/z 350.3268     0   2   4   6E+5   11   11.5  RT (min)       lcorbi_t3_pos_2_013_33    m/z 350.3268     0   2   4   6E+5   11   11.5  RT (min)       lcorbi_t3_pos_2_013_33    m/z 350.8926     0   2   4E+5   17.5   18  RT (min)      lcorbi_t3_pos_4_105_61    m/z 351.2069     0   1E+7   1   1.5  RT (min)      lcorbi_t3_pos_4_057_49    m/z 353.0787     0   2   4   6E+6   8.5   9  RT (min)      lcorbi_t3_pos_1_051_09    m/z 353.1725     0   1   12.5   13  RT (min)      lcorbi_t3_pos_1_034_34    m/z 353.1725     0   2   4E+5   12   12.5  RT (min)      lcorbi_t3_pos_1_008_36    m/z 353.1725     0   2   4   6E+5   12   12.5  RT (min)      lcorbi_t3_pos_3_047_40    m/z 353.1725     0   2   4   6E+5   11   11.5  RT (min)      lcorbi_t3_pos_2_054_29    m/z 353.1725     0   1   13   13.5  RT (min)      lcorbi_t3_pos_1_051_09    m/z 353.1725     0   2   4E+5   11   11.5  RT (min)      lcorbi_t3_pos_4_012_37    m/z 353.1726     0   1   2E+4   12   12.5  RT (min)      lcorbi_t3_pos_2_022_38    m/z 353.1726     0   1   13   13.5  RT (min)      lcorbi_t3_pos_1_023_02    m/z 353.1726     0   1   12.5   13  RT (min)      lcorbi_t3_pos_1_028_32    m/z 353.2664     0   1   13.5   14  RT (min)      lcorbi_t3_pos_2_012_01    m/z 353.2664     0   1   13.5   14  RT (min)      lcorbi_t3_pos_2_012_01    m/z 353.2665     0   1   2E+4   13   13.5  RT (min)      lcorbi_t3_pos_3_003_33    m/z 353.2665     0   1   2   3E+4   13   13.5  RT (min)      lcorbi_t3_pos_1_034_34    m/z 354.0820     0   1E+6   8.5   9  RT (min)      lcorbi_t3_pos_1_051_09    m/z 355.0253     0   1   2E+5   1   1.5  RT (min)       lcorbi_t3_pos_4_081_58    m/z 355.0758     0   1E+6   8.5   9  RT (min)      lcorbi_t3_pos_1_023_02    m/z 355.0758     0   1   2E+6   8.5   9  RT (min)      lcorbi_t3_pos_1_051_09    m/z 355.2269     0   1   2   3   4E+5   9.5   10  RT (min)      lcorbi_t3_pos_1_056_56    m/z 358.2014     0   2   4   6E+6   9   9.5  RT (min)       lcorbi_t3_pos_1_043_06    m/z 358.2015     0   1E+6   12   12.5  RT (min)      lcorbi_t3_pos_2_057_49    m/z 358.2015     0   1   2E+6   13   13.5  RT (min)      lcorbi_t3_pos_1_064_16    m/z 358.2015     0   1E+6   12   12.5  RT (min)      lcorbi_t3_pos_3_057_49    m/z 358.2016     0   1E+6   1.5   2  RT (min)      lcorbi_t3_pos_4_057_49    m/z 358.2016     0   1   2E+6   13.5   14  RT (min)      lcorbi_t3_pos_1_059_17    m/z 358.2016     0   1   2   3E+6   14   14.5  RT (min)      lcorbi_t3_pos_1_058_14    m/z 358.2016     0   1   2   3   4E+6   14.5   15  RT (min)      lcorbi_t3_pos_1_062_13    m/z 358.9966     0   1   2E+5   1   1.5  RT (min)      lcorbi_t3_pos_1_029_29    m/z 359.2048     0   1E+6   10   12.5   15  RT (min)       lcorbi_t3_pos_4_090_60    m/z 359.2049     0   1E+6   16   17  RT (min)       lcorbi_t3_pos_4_071_57    m/z 359.2049     0   1E+6   16   17  RT (min)       lcorbi_t3_pos_4_069_57    m/z 359.2049     0   2   4   6E+5   13.5   14  RT (min)      lcorbi_t3_pos_1_058_14    m/z 359.2049     0   2   4   6   8E+5   16   17  RT (min)       lcorbi_t3_pos_4_070_57    m/z 359.2049     0   1E+6   17.5   18  RT (min)       lcorbi_t3_pos_4_068_57    m/z 359.2050     0   2   4   6   8E+5   14.5   15  RT (min)      lcorbi_t3_pos_1_062_13    m/z 360.3239     0   2   4   6E+4   12.5   13  RT (min)      lcorbi_t3_pos_3_052_47    m/z 360.3239     0   2   4   6E+4   15   15.5  RT (min)      lcorbi_t3_pos_1_051_09    m/z 360.3240     0   1E+4   12.5   13  RT (min)      lcorbi_t3_pos_2_047_31    m/z 360.3240     0   1E+6   13   13.5  RT (min)      lcorbi_t3_pos_3_053_48    m/z 360.3240     0   1E+5   14   14.5  RT (min)       lcorbi_t3_pos_1_038_30    m/z 360.3240     0   1E+5   14   14.5  RT (min)      lcorbi_t3_pos_1_038_30    m/z 360.3240     0   2   4   6   8E+4   14.5   15  RT (min)      lcorbi_t3_pos_2_052_48    m/z 360.3240     0   1   2E+5   14   14.5  RT (min)      lcorbi_t3_pos_3_044_46    m/z 360.3240     0   2   4   6   8E+4   14.5   15  RT (min)      lcorbi_t3_pos_1_041_54    m/z 360.3240     0   2   4   6   8E+4   14.5   15  RT (min)      lcorbi_t3_pos_2_052_48    m/z 360.3240     0   2   4   6E+4   12.5   13  RT (min)      lcorbi_t3_pos_3_052_47    m/z 360.3240     0   2   4   6E+4   15   15.5  RT (min)      lcorbi_t3_pos_2_052_48    m/z 360.3240     0   2   4   6E+4   12.5   13  RT (min)      lcorbi_t3_pos_3_052_47    m/z 360.3240     0   1   2E+5   16   17   18  RT (min)       lcorbi_t3_pos_4_086_59    m/z 360.3240     0   2   4   6E+4   15   15.5  RT (min)      lcorbi_t3_pos_1_051_09    m/z 360.3240     0   2   4   6E+4   12.5   13  RT (min)      lcorbi_t3_pos_3_052_47    m/z 360.3240     0   2   4   6E+4   15   15.5  RT (min)      lcorbi_t3_pos_1_025_52    m/z 360.3240     0   2   4   6   8E+4   15   15.5  RT (min)      lcorbi_t3_pos_1_027_43    m/z 360.3240     0   2   4   6E+4   15.5   16  RT (min)      lcorbi_t3_pos_1_023_02    m/z 360.3240     0   1   2   3E+5   13.5   14  RT (min)      lcorbi_t3_pos_3_025_52    m/z 360.3240     0   1E+5   14   14.5  RT (min)      lcorbi_t3_pos_1_055_33    m/z 360.3240     0   2   4   6   8E+4   14   14.5  RT (min)      lcorbi_t3_pos_2_017_51    m/z 360.3240     0   2   4   6   8E+4   15.5   16  RT (min)      lcorbi_t3_pos_1_023_02    m/z 361.2221     0   2   4E+5   10.5   11  RT (min)      lcorbi_t3_pos_3_011_42    m/z 361.2222     0   2   4   6E+5   10   10.5  RT (min)      lcorbi_t3_pos_2_039_30    m/z 361.2222     0   2   4E+5   10   10.5  RT (min)      lcorbi_t3_pos_2_047_31    m/z 361.2223     0   1   2E+5   11.5   12  RT (min)      lcorbi_t3_pos_2_014_37    m/z 361.2223     0   1   2E+4   12   12.5  RT (min)      lcorbi_t3_pos_2_052_48    m/z 361.2223     0   1   2   3   4E+4   11.5   12  RT (min)      lcorbi_t3_pos_2_012_01    m/z 361.2711     0   2   4   6E+5   1   1.5  RT (min)       lcorbi_t3_pos_4_096_60    m/z 363.1721     0   1E+4   12.5   13  RT (min)      lcorbi_t3_pos_1_028_32    m/z 363.1721     0   1   13   13.5  RT (min)      lcorbi_t3_pos_1_031_40    m/z 363.1721     0   2   4   6E+5   12   12.5  RT (min)      lcorbi_t3_pos_1_008_36    m/z 363.1722     0   2   4E+5   11   11.5  RT (min)      lcorbi_t3_pos_1_002_01    m/z 363.1722     0   1   2E+4   12.5   13  RT (min)      lcorbi_t3_pos_1_034_34    m/z 363.1722     0   1   13.5   14  RT (min)      lcorbi_t3_pos_1_023_02    m/z 363.1722     0   2   4E+5   11   11.5  RT (min)      lcorbi_t3_pos_1_002_01    m/z 363.3106     0   1E+4   12   12.5  RT (min)      lcorbi_t3_pos_1_041_54    m/z 363.3107     0   2   4   6   8E+5   11.5   12  RT (min)      lcorbi_t3_pos_2_017_51    m/z 363.3107     0   1E+4   12   12.5  RT (min)      lcorbi_t3_pos_1_034_34    m/z 363.3107     0   1E+6   11.5   12  RT (min)      lcorbi_t3_pos_1_008_36    m/z 363.3107     0   1E+4   12.5   13  RT (min)      lcorbi_t3_pos_2_056_56    m/z 363.3107     0   1E+4   12.5   13  RT (min)      lcorbi_t3_pos_1_031_40    m/z 363.3107     0   1   2   3   4E+4   12   12.5  RT (min)      lcorbi_t3_pos_3_003_33    m/z 363.3108     0   1E+6   10.5   11  RT (min)      lcorbi_t3_pos_2_013_33    m/z 363.3108     0   2   4   6E+5   10.5   11  RT (min)      lcorbi_t3_pos_3_031_45    m/z 363.3109     0   1E+6   10.5   11  RT (min)      lcorbi_t3_pos_3_031_45    m/z 364.1602     0   1   2E+5   9.5   10  RT (min)      lcorbi_t3_pos_1_030_21    m/z 364.1602     0   1   2E+5   10   10.5  RT (min)      lcorbi_t3_pos_1_041_54    m/z 364.1602     0   2   4   6   8E+4   10   10.5  RT (min)      lcorbi_t3_pos_1_025_52    m/z 364.1604     0   1   2E+5   17   18  RT (min)       lcorbi_t3_pos_4_098_61    m/z 364.3424     0   1   13   13.5  RT (min)      lcorbi_t3_pos_1_034_34    m/z 364.3424     0   1   13   13.5  RT (min)      lcorbi_t3_pos_1_034_34    m/z 364.3424     0   1E+4   13   13.5  RT (min)      lcorbi_t3_pos_2_022_38    m/z 364.3424     0   1   2   3   4E+5   11.5   12  RT (min)      lcorbi_t3_pos_2_013_33    m/z 364.3424     0   2   4   6E+5   12.5   13  RT (min)      lcorbi_t3_pos_1_008_36    m/z 364.3424     0   1   13.5   14  RT (min)      lcorbi_t3_pos_1_031_40    m/z 364.3424     0   1   13.5   14  RT (min)      lcorbi_t3_pos_2_012_01    m/z 365.3140     0   1   2E+5   13   13.5  RT (min)       lcorbi_t3_pos_4_072_57    m/z 366.3732     0   1E+5   14.5   15  RT (min)      lcorbi_t3_pos_1_028_32    m/z 366.3732     0   1E+5   15   15.5  RT (min)      lcorbi_t3_pos_4_033_53    m/z 366.3732     0   1   2E+4   13   13.5  RT (min)      lcorbi_t3_pos_2_054_29    m/z 366.3732     0   2   4   6   8E+5   13   13.5  RT (min)      lcorbi_t3_pos_4_012_37    m/z 366.3732     0   2   4   6   8E+5   14   14.5  RT (min)      lcorbi_t3_pos_3_047_40    m/z 366.3732     0   2   4E+5   14   14.5  RT (min)      lcorbi_t3_pos_3_025_52    m/z 366.3733     0   1E+5   16  RT (min)       lcorbi_t3_pos_1_038_30    m/z 366.3733     0   2   4   6E+5   14   14.5  RT (min)      lcorbi_t3_pos_3_047_40    m/z 366.3733     0   1   2   3E+4   12.5   13  RT (min)      lcorbi_t3_pos_4_012_37    m/z 366.3733     0   1E+5   14.5   15  RT (min)      lcorbi_t3_pos_4_041_54    m/z 366.3733     0   1   2   3E+4   12.5   13  RT (min)      lcorbi_t3_pos_4_012_37    m/z 366.3733     0   1E+5   15.5   16  RT (min)      lcorbi_t3_pos_2_012_01    m/z 366.3734     0   1E+5   14.5   15  RT (min)      lcorbi_t3_pos_3_013_19    m/z 367.3361     0   1   18   18.5  RT (min)    lcorbi_t3_pos_1_051_09    m/z 367.3361     0   1   17.5   18  RT (min)    lcorbi_t3_pos_2_012_01    m/z 367.3362     0   1   18.5   19  RT (min)    lcorbi_t3_pos_1_051_09    m/z 367.3362     0   1   19   19.5  RT (min)    lcorbi_t3_pos_1_023_02    m/z 367.3362     0   1   2E+4   14.5   15  RT (min)      lcorbi_t3_pos_2_054_29    m/z 367.3362     0   1   16.5   17  RT (min)     lcorbi_t3_pos_1_028_32    m/z 367.3362     0   1   19   19.5  RT (min)    lcorbi_t3_pos_1_023_02    m/z 367.3362     0   1   2E+4   14.5   15  RT (min)      lcorbi_t3_pos_3_011_42    m/z 367.3362     0   1   16.5   17  RT (min)    lcorbi_t3_pos_1_028_32    m/z 367.3362     0   1   2E+4   14.5   15  RT (min)      lcorbi_t3_pos_3_011_42    m/z 367.3362     0   1   17   17.5  RT (min)     lcorbi_t3_pos_1_034_34    m/z 367.3362     0   1   17.5   18  RT (min)    lcorbi_t3_pos_1_031_40    m/z 367.3362     0   1E+4   14   14.5  RT (min)      lcorbi_t3_pos_3_024_07    m/z 367.3362     0   1   16.5   17  RT (min)     lcorbi_t3_pos_2_022_38    m/z 367.3767     0   1   2E+5   14   14.5  RT (min)      lcorbi_t3_pos_3_047_40    m/z 368.3394     0   1E+4   14.5   15  RT (min)      lcorbi_t3_pos_3_011_42    m/z 368.3394     0   1   14   14.5  RT (min)      lcorbi_t3_pos_3_024_07    m/z 368.3395     0   1   16.5   17  RT (min)    lcorbi_t3_pos_1_028_32    m/z 368.3395     0   1   17   17.5  RT (min)    lcorbi_t3_pos_1_034_34    m/z 368.3395     0   1   17.5   18  RT (min)    lcorbi_t3_pos_1_031_40    m/z 368.3395     0   1   16.5   17  RT (min)    lcorbi_t3_pos_2_022_38    m/z 368.3396     0   1   16.5   17  RT (min)    lcorbi_t3_pos_2_022_38    m/z 369.1334     0   2   4   6E+5   9   9.5  RT (min)      lcorbi_t3_pos_1_051_09    m/z 369.3518     0   1E+6   14.5   15  RT (min)       lcorbi_t3_pos_4_016_15    m/z 369.3518     0   1   2   3E+6   16   16.5  RT (min)       lcorbi_t3_pos_4_068_57    m/z 369.3518     0   1   18   18.5  RT (min)    lcorbi_t3_pos_1_023_02    m/z 369.3518     0   1E+5   14   14.5  RT (min)       lcorbi_t3_pos_3_024_07    m/z 369.3518     0   2   4   6   8E+4   14   14.5  RT (min)      lcorbi_t3_pos_4_012_37    m/z 369.3518     0   1   17.5   18  RT (min)    lcorbi_t3_pos_1_031_40    m/z 369.3518     0   1E+5   14   14.5  RT (min)      lcorbi_t3_pos_2_032_39    m/z 369.3518     0   1   17.5   18  RT (min)    lcorbi_t3_pos_2_012_01    m/z 369.3518     0   1   2   3E+6   16.5   17  RT (min)      lcorbi_t3_pos_4_072_57    m/z 369.3518     0   1   19   19.5  RT (min)    lcorbi_t3_pos_1_023_02    m/z 369.3518     0   1   17   17.5  RT (min)     lcorbi_t3_pos_1_034_34    m/z 369.3518     0   1   18   18.5  RT (min)    lcorbi_t3_pos_1_023_02    m/z 369.3518     0   1   19   19.5  RT (min)    lcorbi_t3_pos_1_023_02    m/z 369.3519     0   1   2   3E+6   16.5   17  RT (min)       lcorbi_t3_pos_4_079_58    m/z 369.3519     0   1E+5   14.5   15  RT (min)      lcorbi_t3_pos_3_011_42    m/z 369.3519     0   1   2   3E+6   16   16.5  RT (min)       lcorbi_t3_pos_4_066_57    m/z 369.3519     0   1   16.5   17  RT (min)     lcorbi_t3_pos_1_028_32    m/z 369.3519     0   1   17.5   18  RT (min)    lcorbi_t3_pos_2_025_52    m/z 369.3519     0   1   17   17.5   18  RT (min)     lcorbi_t3_pos_1_031_40    m/z 369.3519     0   1   16.5   17  RT (min)     lcorbi_t3_pos_2_022_38    m/z 369.3519     0   1   2   3E+6   16.5   17  RT (min)       lcorbi_t3_pos_4_068_57    m/z 369.3519     0   1   17.5   18  RT (min)     lcorbi_t3_pos_2_012_01    m/z 369.3522     0   1   2   3E+6   16.5   17  RT (min)      lcorbi_t3_pos_4_072_57    m/z 370.1053     0   1   2   3E+6   8.5   9  RT (min)      lcorbi_t3_pos_1_051_09    m/z 370.1053     0   1E+6   8.5   9  RT (min)      lcorbi_t3_pos_1_023_02    m/z 370.3551     0   1   17.5   18  RT (min)    lcorbi_t3_pos_2_012_01    m/z 370.3551     0   1   17   17.5  RT (min)    lcorbi_t3_pos_1_034_34    m/z 370.3552     0   2   4   6E+4   14.5   15  RT (min)      lcorbi_t3_pos_3_011_42    m/z 370.3552     0   1   17.5   18  RT (min)    lcorbi_t3_pos_1_031_40    m/z 370.3552     0   1E+6   16   16.5  RT (min)      lcorbi_t3_pos_4_068_57    m/z 370.3552     0   1   17.5   18  RT (min)    lcorbi_t3_pos_1_031_40    m/z 370.3552     0   1   16.5   17  RT (min)    lcorbi_t3_pos_1_028_32    m/z 370.3552     0   2   4   6E+4   14   14.5  RT (min)      lcorbi_t3_pos_2_013_33    m/z 370.3552     0   2   4E+4   14   14.5  RT (min)      lcorbi_t3_pos_3_024_07    m/z 370.3553     0   2   4   6   8E+5   16   16.5  RT (min)      lcorbi_t3_pos_4_067_57    m/z 370.3553     0   1E+6   16.5   17  RT (min)       lcorbi_t3_pos_4_069_57    m/z 370.3554     0   1E+6   16   16.5  RT (min)      lcorbi_t3_pos_4_070_57    m/z 371.0202     0   1E+5   1   1.5  RT (min)       lcorbi_t3_pos_2_040_25    m/z 371.3157     0   1E+5   1   1.5  RT (min)      lcorbi_t3_pos_3_003_33    m/z 371.3159     0   2   4   6E+5   12  RT (min)       lcorbi_t3_pos_4_105_61    m/z 371.3269     0   1E+4   11   11.5  RT (min)      lcorbi_t3_pos_1_034_34    m/z 371.3270     0   2   4   6   8E+3   11   11.5  RT (min)      lcorbi_t3_pos_1_031_40    m/z 371.3270     0   1   11   11.5  RT (min)      lcorbi_t3_pos_2_012_01    m/z 371.3270     0   2   4E+5   10   10.5  RT (min)      lcorbi_t3_pos_2_013_33    m/z 372.1023     0   2   4   6   8E+5   8.5   9  RT (min)      lcorbi_t3_pos_1_051_09    m/z 372.1866     0   1E+6   5   5.5  RT (min)      lcorbi_t3_pos_3_000_aa    m/z 372.3193     0   1E+5   13   13.5  RT (min)       lcorbi_t3_pos_2_008_16    m/z 372.3193     0   1   2   3E+5   13   13.5  RT (min)       lcorbi_t3_pos_1_013_10    m/z 373.2375     0   2   4   6   8E+5   8.5   9  RT (min)      lcorbi_t3_pos_2_056_56    m/z 376.2120     0   1   2E+6   16   18  RT (min)       lcorbi_t3_pos_1_027_43    m/z 376.2120     0   1E+6   10   12  RT (min)       lcorbi_t3_pos_4_072_57    m/z 376.2120     0   1   2E+6   15   16   17  RT (min)       lcorbi_t3_pos_1_021_18    m/z 376.2120     0   2   4   6   8E+5   13   13.5  RT (min)      lcorbi_t3_pos_1_064_16    m/z 376.2120     0   2   4   6E+5   12   12.5  RT (min)      lcorbi_t3_pos_1_059_17    m/z 376.2120     0   2   4E+5   11.5   12  RT (min)      lcorbi_t3_pos_2_059_39    m/z 376.2121     0   1   2E+6   14   15   16  RT (min)       lcorbi_t3_pos_1_018_35    m/z 376.2121     0   1   2E+6   15   16  RT (min)       lcorbi_t3_pos_1_011_44    m/z 376.2121     0   1   2E+6   15   15.5  RT (min)       lcorbi_t3_pos_1_040_27    m/z 376.2121     0   1   2E+6   16   17  RT (min)       lcorbi_t3_pos_1_038_30    m/z 376.2121     0   1   2E+6   15.5   16   16.5  RT (min)       lcorbi_t3_pos_4_067_57    m/z 376.2121     0   2   4   6E+5   13.5   14  RT (min)      lcorbi_t3_pos_3_065_51    m/z 376.2121     0   2   4   6   8E+5   13.5   14  RT (min)      lcorbi_t3_pos_1_059_17    m/z 376.2121     0   1E+6   14   14.5  RT (min)      lcorbi_t3_pos_1_058_14    m/z 376.2122     0   1   2E+6   12   14   16  RT (min)       lcorbi_t3_pos_4_086_59    m/z 376.2122     0   1   2E+6   16   18  RT (min)       lcorbi_t3_pos_4_096_60    m/z 377.1420     0   2   4E+5   5   5.5  RT (min)      lcorbi_t3_pos_3_000_aa    m/z 377.2153     0   1   2   3E+5   8.5   9  RT (min)      lcorbi_t3_pos_1_033_53    m/z 377.2154     0   2   4E+5   14   16  RT (min)       lcorbi_t3_pos_4_083_59    m/z 377.2154     0   1   2   3E+5   8.5   9  RT (min)      lcorbi_t3_pos_1_033_53    m/z 377.2154     0   2   4   6E+5   16  RT (min)       lcorbi_t3_pos_4_106_62    m/z 377.2154     0   2   4   6E+5   14   16  RT (min)       lcorbi_t3_pos_1_013_10    m/z 377.2154     0   2   4   6E+5   16  RT (min)       lcorbi_t3_pos_1_011_44    m/z 377.2154     0   1   2   3E+5   8.5   9  RT (min)      lcorbi_t3_pos_2_009_50    m/z 377.2154     0   2   4   6E+5   14   16  RT (min)       lcorbi_t3_pos_4_067_57    m/z 377.2155     0   2   4   6E+5   14   16  RT (min)       lcorbi_t3_pos_4_092_60    m/z 377.2155     0   2   4   6E+5   14   16  RT (min)       lcorbi_t3_pos_1_038_30    m/z 377.2156     0   2   4E+5   16   17  RT (min)       lcorbi_t3_pos_4_085_59    m/z 377.2156     0   1   17.5   18  RT (min)     lcorbi_t3_pos_1_025_52    m/z 377.2156     0   2   4E+5   14   16  RT (min)       lcorbi_t3_pos_4_078_58    m/z 377.3206     0   2   4   6E+5   13.5   14  RT (min)       lcorbi_t3_pos_4_082_59    m/z 377.3206     0   2   4   6E+5   13.5   14  RT (min)      lcorbi_t3_pos_4_089_59    m/z 377.3262     0   1   2   3   4E+4   12   12.5  RT (min)       lcorbi_t3_pos_1_034_34    m/z 377.3263     0   1   2E+5   12   12.5  RT (min)      lcorbi_t3_pos_1_028_32    m/z 377.3263     0   1   2   3   4E+5   12   12.5  RT (min)       lcorbi_t3_pos_3_003_33    m/z 377.3263     0   1   12.5   13  RT (min)      lcorbi_t3_pos_1_031_40    m/z 377.3263     0   1E+5   12   12.5  RT (min)      lcorbi_t3_pos_2_025_52    m/z 377.3263     0   1   12.5   13  RT (min)      lcorbi_t3_pos_1_031_40    m/z 377.3264     0   1E+4   12.5   13  RT (min)      lcorbi_t3_pos_2_012_01    m/z 377.3264     0   1E+6   11   11.5  RT (min)      lcorbi_t3_pos_3_024_07    m/z 377.3264     0   1E+6   11   11.5  RT (min)       lcorbi_t3_pos_2_013_33    m/z 377.3264     0   2   4   6E+5   12   12.5  RT (min)      lcorbi_t3_pos_2_017_51    m/z 377.3264     0   1   2E+4   12.5   13  RT (min)      lcorbi_t3_pos_1_034_34    m/z 377.3264     0   1   2   3   4E+5   10.5   11  RT (min)      lcorbi_t3_pos_3_031_45    m/z 377.3264     0   2   4   6   8E+5   12   12.5  RT (min)       lcorbi_t3_pos_2_017_51    m/z 378.3298     0   2   4   6E+5   11.5   12  RT (min)       lcorbi_t3_pos_3_020_35    m/z 379.1310     0   1E+6   17   18  RT (min)       lcorbi_t3_pos_4_075_58    m/z 379.2820     0   1   13   13.5  RT (min)      lcorbi_t3_pos_1_028_32    m/z 379.2820     0   1   14   14.5  RT (min)      lcorbi_t3_pos_1_023_02    m/z 379.2821     0   1   14   14.5  RT (min)      lcorbi_t3_pos_1_023_02    m/z 379.2821     0   1   2   3E+4   11.5   12  RT (min)      lcorbi_t3_pos_2_013_33    m/z 379.2821     0   1   13.5   14  RT (min)      lcorbi_t3_pos_1_034_34    m/z 379.2821     0   1   2   3   4E+5   12.5   13  RT (min)      lcorbi_t3_pos_1_008_36    m/z 379.2821     0   1   14   14.5  RT (min)      lcorbi_t3_pos_2_012_01    m/z 380.1341     0   1   2   3E+5   9.5   10  RT (min)      lcorbi_t3_pos_1_034_34    m/z 380.1341     0   1   2   3E+5   9.5   10  RT (min)      lcorbi_t3_pos_1_030_21    m/z 380.1341     0   1   2E+5   9.5   10  RT (min)      lcorbi_t3_pos_2_017_51    m/z 380.1343     0   1   2   3E+5   18   18.5  RT (min)      lcorbi_t3_pos_4_082_59    m/z 380.3372     0   2   4   6   8E+5   11.5   12  RT (min)      lcorbi_t3_pos_2_017_51    m/z 380.3372     0   1E+4   12   12.5  RT (min)      lcorbi_t3_pos_1_041_54    m/z 380.3372     0   2   4   6E+4   12   12.5  RT (min)      lcorbi_t3_pos_1_028_32    m/z 380.3372     0   1E+4   12.5   13  RT (min)      lcorbi_t3_pos_2_012_01    m/z 380.3372     0   1   2E+4   12.5   13  RT (min)      lcorbi_t3_pos_1_031_40    m/z 380.3372     0   1E+6   11.5   12  RT (min)      lcorbi_t3_pos_1_008_36    m/z 380.3373     0   1E+4   12   12.5  RT (min)      lcorbi_t3_pos_1_041_54    m/z 380.3373     0   2   4   6   8E+5   10.5   11  RT (min)       lcorbi_t3_pos_3_024_07    m/z 380.3373     0   1E+6   10.5   11  RT (min)      lcorbi_t3_pos_2_013_33    m/z 381.1673     0   2   4E+5   9   10  RT (min)       lcorbi_t3_pos_2_050_18    m/z 381.1674     0   1E+6   17   18  RT (min)       lcorbi_t3_pos_4_095_60    m/z 381.1674     0   1E+6   17.5   18  RT (min)       lcorbi_t3_pos_4_067_57    m/z 381.1675     0   1E+6   17.5   18  RT (min)       lcorbi_t3_pos_4_111_62    m/z 381.1675     0   1E+6   18  RT (min)       lcorbi_t3_pos_4_076_58    m/z 381.1675     0   1E+6   16   17   18  RT (min)       lcorbi_t3_pos_4_066_57    m/z 381.1675     0   1E+6   16   17  RT (min)       lcorbi_t3_pos_4_079_58    m/z 381.2977     0   1   2E+4   14.5   15  RT (min)      lcorbi_t3_pos_2_012_01    m/z 381.2978     0   1E+4   13.5   14  RT (min)      lcorbi_t3_pos_3_003_33    m/z 381.2978     0   1   2E+4   14   14.5  RT (min)      lcorbi_t3_pos_1_034_34    m/z 381.2978     0   1   2E+4   14   14.5  RT (min)      lcorbi_t3_pos_1_034_34    m/z 382.1707     0   1E+5   8.5   9  RT (min)      lcorbi_t3_pos_1_033_53    m/z 383.2041     0   2   4   6   8E+5   10   10.5  RT (min)      lcorbi_t3_pos_2_047_31    m/z 383.2041     0   1E+6   10   10.5  RT (min)      lcorbi_t3_pos_2_039_30    m/z 383.2042     0   1   2E+4   11.5   12  RT (min)      lcorbi_t3_pos_2_012_01    m/z 383.2042     0   1   2   3   4E+4   11.5   12  RT (min)      lcorbi_t3_pos_2_009_50    m/z 383.2042     0   2   4   6E+4   11   11.5  RT (min)      lcorbi_t3_pos_2_025_52    m/z 383.2042     0   1   2E+4   11.5   12  RT (min)      lcorbi_t3_pos_2_009_50    m/z 383.2042     0   2   4E+4   11.5   12  RT (min)      lcorbi_t3_pos_2_025_52    m/z 383.2042     0   1   2   3   4E+5   11   11.5  RT (min)      lcorbi_t3_pos_2_022_38    m/z 383.2042     0   2   4   6   8E+5   11   11.5  RT (min)      lcorbi_t3_pos_2_017_51    m/z 384.1444     0   1   2E+6   1   1.5  RT (min)      lcorbi_t3_pos_4_057_49    m/z 387.1804     0   2   4   6E+5   9   9.5  RT (min)      lcorbi_t3_pos_2_012_01    m/z 387.1804     0   1   2   3   4E+5   9   9.5  RT (min)      lcorbi_t3_pos_2_012_01    m/z 387.1804     0   2   4   6   8E+5   9   9.5  RT (min)      lcorbi_t3_pos_1_031_40    m/z 388.2542     0   2   4E+5   5   5.5  RT (min)      lcorbi_t3_pos_3_000_aa    m/z 388.3425     0   1E+5   13.5   14  RT (min)       lcorbi_t3_pos_1_019_16    m/z 389.1232     0   1E+6   8   8.5  RT (min)      lcorbi_t3_pos_1_023_02    m/z 391.2455     0   2   4   6   8E+5   13   13.5  RT (min)      lcorbi_t3_pos_4_076_58    m/z 391.2647     0   1   2   3E+4   15   15.5  RT (min)      lcorbi_t3_pos_1_051_09    m/z 391.2649     0   1E+4   14   14.5  RT (min)      lcorbi_t3_pos_2_022_38    m/z 391.2652     0   1   2   3E+4   14   14.5  RT (min)      lcorbi_t3_pos_4_049_55    m/z 391.2669     0   1   2   3   4E+4   14.5   15  RT (min)      lcorbi_t3_pos_2_009_50    m/z 391.2671     0   2   4   6E+5   13.5   14  RT (min)      lcorbi_t3_pos_3_040_41    m/z 391.2691     0   1E+6   12.5   13  RT (min)      lcorbi_t3_pos_4_016_15    m/z 391.2691     0   1   2   3   4E+4   14   14.5  RT (min)      lcorbi_t3_pos_1_028_32    m/z 391.2691     0   1   12.5   13  RT (min)      lcorbi_t3_pos_3_024_07    m/z 391.2691     0   2   4E+4   14   14.5  RT (min)      lcorbi_t3_pos_2_025_52    m/z 391.2692     0   2   4E+4   14   14.5  RT (min)      lcorbi_t3_pos_2_022_38    m/z 391.2692     0   1   2   3   4E+4   14.5   15  RT (min)      lcorbi_t3_pos_2_049_55    m/z 391.2692     0   1   2   3   4E+4   15   15.5  RT (min)      lcorbi_t3_pos_2_012_01    m/z 391.2692     0   2   4   6   8E+5   13.5   14  RT (min)      lcorbi_t3_pos_3_047_40    m/z 391.2692     0   1E+6   12.5   13  RT (min)      lcorbi_t3_pos_3_011_42    m/z 391.2692     0   2   4E+4   14.5   15  RT (min)      lcorbi_t3_pos_1_034_34    m/z 391.2692     0   2   4   6E+4   14.5   15  RT (min)      lcorbi_t3_pos_1_041_54    m/z 391.2692     0   1   2   3E+5   13.5   14  RT (min)      lcorbi_t3_pos_1_008_36    m/z 391.2692     0   1   2   3   4E+4   14   14.5  RT (min)      lcorbi_t3_pos_1_028_32    m/z 391.2692     0   1E+5   14   14.5  RT (min)      lcorbi_t3_pos_2_017_51    m/z 391.2692     0   2   4   6   8E+3   12.5   13  RT (min)      lcorbi_t3_pos_3_031_45    m/z 391.2692     0   2   4E+4   15   15.5  RT (min)      lcorbi_t3_pos_1_031_40    m/z 391.2692     0   1   12   12.5  RT (min)      lcorbi_t3_pos_2_032_39    m/z 391.2692     0   2   4E+4   14.5   15  RT (min)      lcorbi_t3_pos_1_056_56    m/z 391.2693     0   2   4E+4   15.5   16  RT (min)      lcorbi_t3_pos_1_023_02    m/z 391.2694     0   1   2   3   4E+4   15   15.5  RT (min)      lcorbi_t3_pos_2_012_01    m/z 391.2842     0   1E+8   12.5   13  RT (min)      lcorbi_t3_pos_1_002_01    m/z 391.2843     0   2   4   6   8E+5   12   12.5  RT (min)      lcorbi_t3_pos_2_032_39    m/z 391.2843     0   1E+5   5.5   6  RT (min)      lcorbi_t3_pos_4_003_39    m/z 391.2843     0   1E+8   12.5   13  RT (min)      lcorbi_t3_pos_2_054_29    m/z 391.2844     0   1   18   18.5  RT (min)    lcorbi_t3_pos_2_052_48    m/z 391.2844     0   1E+5   4.5   5  RT (min)      lcorbi_t3_pos_4_008_30    m/z 391.2844     0   1E+5   6.5   7  RT (min)      lcorbi_t3_pos_3_017_51    m/z 391.2844     0   1   17.5   18  RT (min)    lcorbi_t3_pos_2_052_48    m/z 391.2844     0   1E+8   13.5   14  RT (min)      lcorbi_t3_pos_3_040_41    m/z 391.2844     0   1E+5   2   4  RT (min)       lcorbi_t3_pos_4_078_58    m/z 391.2844     0   1E+6   1.5   2  RT (min)      lcorbi_t3_pos_4_057_49    m/z 391.2844     0   1   17   17.5  RT (min)    lcorbi_t3_pos_1_027_43    m/z 391.2844     0   2   4   6E+6   10   10.5  RT (min)      lcorbi_t3_pos_4_057_49    m/z 391.2844     0   1   18   18.5  RT (min)    lcorbi_t3_pos_1_027_43    m/z 391.2844     0   2   4   6E+6   0.5   1  RT (min)      lcorbi_t3_pos_4_057_49    m/z 391.2844     0   1   16.5   17  RT (min)    lcorbi_t3_pos_1_055_33    m/z 391.2844     0   1   2   3   4E+7   12.5   13  RT (min)      lcorbi_t3_pos_3_053_48    m/z 391.2844     0   2   4   6E+6   16   16.5  RT (min)      lcorbi_t3_pos_1_027_43    m/z 391.2844     0   1E+7   14   14.5  RT (min)      lcorbi_t3_pos_1_013_10    m/z 391.2844     0   1E+7   14   14.5  RT (min)      lcorbi_t3_pos_1_053_39    m/z 391.2844     0   2   4   6E+6   14.5   15  RT (min)      lcorbi_t3_pos_2_038_03    m/z 391.2844     0   2   4   6E+6   15   15.5  RT (min)      lcorbi_t3_pos_2_012_01    m/z 391.2844     0   2   4   6E+6   14   14.5  RT (min)      lcorbi_t3_pos_2_038_03    m/z 391.2844     0   1   2E+8   13.5   14  RT (min)      lcorbi_t3_pos_3_047_40    m/z 391.2844     0   2   4   6E+6   15.5   16  RT (min)      lcorbi_t3_pos_2_000_aa    m/z 391.2844     0   2   4E+6   14   14.5  RT (min)      lcorbi_t3_pos_3_003_33    m/z 391.2844     0   2   4   6E+7   13.5   14  RT (min)      lcorbi_t3_pos_1_008_36    m/z 391.2845     0   2   4   6   8E+6   15   15.5  RT (min)      lcorbi_t3_pos_1_027_43    m/z 391.2845     0   2   4   6E+6   15   15.5  RT (min)      lcorbi_t3_pos_1_055_33    m/z 391.2845     0   1   2E+7   13.5   14  RT (min)      lcorbi_t3_pos_1_039_23    m/z 391.2845     0   2   4   6   8E+6   15   15.5  RT (min)      lcorbi_t3_pos_1_027_43    m/z 391.2845     0   1   17   17.5  RT (min)    lcorbi_t3_pos_1_027_43    m/z 391.2845     0   2   4   6E+6   14.5   15  RT (min)      lcorbi_t3_pos_2_052_48    m/z 391.2845     0   2   4   6   8E+6   14.5   15  RT (min)      lcorbi_t3_pos_1_034_34    m/z 391.2845     0   1   16.5   17  RT (min)    lcorbi_t3_pos_1_055_33    m/z 391.2845     0   1   17.5   18  RT (min)    lcorbi_t3_pos_2_052_48    m/z 391.2845     0   2   4   6   8E+6   15   15.5  RT (min)      lcorbi_t3_pos_1_051_09    m/z 391.2845     0   2   4   6   8E+6   15.5   16  RT (min)      lcorbi_t3_pos_1_023_02    m/z 391.2845     0   2   4   6   8E+6   14.5   15  RT (min)      lcorbi_t3_pos_1_034_34    m/z 391.2846     0   1   16.5   17  RT (min)    lcorbi_t3_pos_1_055_33    m/z 391.2846     0   2   4   6E+6   16   16.5  RT (min)      lcorbi_t3_pos_2_052_48    m/z 391.2846     0   1   17   17.5  RT (min)    lcorbi_t3_pos_1_027_43    m/z 391.2993     0   1   12   12.5  RT (min)      lcorbi_t3_pos_2_032_39    m/z 391.2995     0   1E+6   12.5   13  RT (min)      lcorbi_t3_pos_2_054_29    m/z 391.2995     0   2   4E+4   15   15.5  RT (min)      lcorbi_t3_pos_1_051_09    m/z 391.2995     0   2   4   6E+4   15.5   16  RT (min)      lcorbi_t3_pos_1_023_02    m/z 391.2995     0   2   4   6E+4   15.5   16  RT (min)      lcorbi_t3_pos_1_023_02    m/z 391.2995     0   2   4   6E+4   14.5   15  RT (min)      lcorbi_t3_pos_1_041_54    m/z 391.2995     0   1   2   3E+4   14   14.5  RT (min)      lcorbi_t3_pos_3_013_19    m/z 391.2995     0   1E+6   12.5   13  RT (min)      lcorbi_t3_pos_4_016_15    m/z 391.2996     0   1E+4   12.5   13  RT (min)      lcorbi_t3_pos_2_013_33    m/z 391.2996     0   2   4E+4   14.5   15  RT (min)      lcorbi_t3_pos_1_034_34    m/z 391.2996     0   1   2   3E+5   13.5   14  RT (min)      lcorbi_t3_pos_1_008_36    m/z 391.2996     0   2   4E+4   15   15.5  RT (min)      lcorbi_t3_pos_1_051_09    m/z 391.2996     0   1E+4   12.5   13  RT (min)      lcorbi_t3_pos_3_031_45    m/z 391.2996     0   1E+6   13.5   14  RT (min)      lcorbi_t3_pos_3_047_40    m/z 391.2996     0   1E+6   13.5   14  RT (min)      lcorbi_t3_pos_4_019_47    m/z 391.2996     0   1   2   3   4E+4   14.5   15  RT (min)      lcorbi_t3_pos_2_009_50    m/z 391.2996     0   2   4E+4   14   14.5  RT (min)      lcorbi_t3_pos_1_021_18    m/z 391.2996     0   2   4E+4   14.5   15  RT (min)      lcorbi_t3_pos_1_031_40    m/z 391.2997     0   1   2   3   4E+4   14   14.5  RT (min)      lcorbi_t3_pos_2_022_38    m/z 391.2997     0   1   2   3E+4   14   14.5  RT (min)      lcorbi_t3_pos_3_003_33    m/z 391.3038     0   1   2   3E+4   15   15.5  RT (min)      lcorbi_t3_pos_2_012_01    m/z 391.3120     0   1   12   12.5  RT (min)      lcorbi_t3_pos_2_032_39    m/z 391.3419     0   2   4E+4   12.5   13  RT (min)      lcorbi_t3_pos_2_022_38    m/z 391.3420     0   2   4   6E+4   13   13.5  RT (min)      lcorbi_t3_pos_1_028_32    m/z 391.3420     0   2   4   6E+4   13   13.5  RT (min)      lcorbi_t3_pos_1_034_34    m/z 391.3420     0   2   4E+5   12.5   13  RT (min)      lcorbi_t3_pos_1_008_36    m/z 391.3420     0   1   2E+5   11.5   12  RT (min)      lcorbi_t3_pos_2_013_33    m/z 391.3420     0   2   4   6E+4   13.5   14  RT (min)      lcorbi_t3_pos_2_056_56    m/z 391.3420     0   2   4   6E+4   13   13.5  RT (min)      lcorbi_t3_pos_1_034_34    m/z 391.3420     0   2   4   6E+4   12.5   13  RT (min)      lcorbi_t3_pos_3_003_33    m/z 392.2574     0   2   4   6E+5   13   13.5  RT (min)      lcorbi_t3_pos_4_075_58    m/z 392.2575     0   2   4   6E+5   13   13.5  RT (min)      lcorbi_t3_pos_4_074_58    m/z 392.2876     0   1   2E+5   12   12.5  RT (min)      lcorbi_t3_pos_2_032_39    m/z 392.2876     0   1   2   3   4E+7   12.5   13  RT (min)      lcorbi_t3_pos_2_054_29    m/z 392.2877     0   1E+7   12.5   13  RT (min)      lcorbi_t3_pos_3_053_48    m/z 392.2877     0   1   2   3   4E+7   12.5   13  RT (min)      lcorbi_t3_pos_1_002_01    m/z 392.2878     0   1E+7   13.5   14  RT (min)      lcorbi_t3_pos_1_008_36    m/z 392.2878     0   2   4E+7   13.5   14  RT (min)      lcorbi_t3_pos_3_047_40    m/z 392.2878     0   1   17   17.5  RT (min)    lcorbi_t3_pos_1_055_33    m/z 392.2878     0   1   17.5   18  RT (min)    lcorbi_t3_pos_2_052_48    m/z 392.2878     0   1   2   3   4E+7   12.5   13  RT (min)      lcorbi_t3_pos_1_002_01    m/z 392.2878     0   1   2E+6   10   10.5  RT (min)      lcorbi_t3_pos_4_057_49    m/z 392.2879     0   1E+6   15   15.5  RT (min)      lcorbi_t3_pos_2_012_01    m/z 392.2879     0   1E+6   14   14.5  RT (min)      lcorbi_t3_pos_3_003_33    m/z 392.2879     0   1E+6   15   15.5  RT (min)      lcorbi_t3_pos_2_056_56    m/z 392.2879     0   1   17.5   18  RT (min)    lcorbi_t3_pos_2_052_48    m/z 392.2879     0   1   17.5   18  RT (min)    lcorbi_t3_pos_2_052_48    m/z 392.2879     0   1   2E+6   15.5   16  RT (min)      lcorbi_t3_pos_1_023_02    m/z 392.2879     0   1   2E+6   14.5   15  RT (min)      lcorbi_t3_pos_1_034_34    m/z 392.2879     0   1   17.5   18  RT (min)    lcorbi_t3_pos_2_052_48    m/z 392.2879     0   1E+6   0.5   1  RT (min)      lcorbi_t3_pos_4_057_49    m/z 392.2879     0   1E+6   15.5   16  RT (min)      lcorbi_t3_pos_1_050_13    m/z 392.2879     0   1   2E+6   14   14.5  RT (min)      lcorbi_t3_pos_1_047_41    m/z 392.2879     0   1   17.5   18  RT (min)    lcorbi_t3_pos_2_052_48    m/z 392.2879     0   1E+6   14.5   15  RT (min)      lcorbi_t3_pos_2_038_03    m/z 392.2879     0   1E+6   14   14.5  RT (min)      lcorbi_t3_pos_2_038_03    m/z 392.2880     0   1E+6   16   16.5  RT (min)      lcorbi_t3_pos_1_055_33    m/z 392.2880     0   1E+6   16   16.5  RT (min)      lcorbi_t3_pos_1_055_33    m/z 393.2905     0   1   2   3   4E+4   12   12.5  RT (min)      lcorbi_t3_pos_2_032_39    m/z 393.2907     0   1   2E+5   14   14.5  RT (min)      lcorbi_t3_pos_1_047_41    m/z 393.2907     0   2   4   6   8E+5   12.5   13  RT (min)      lcorbi_t3_pos_2_054_29    m/z 393.2908     0   1E+5   14   14.5  RT (min)      lcorbi_t3_pos_3_003_33    m/z 393.2908     0   2   4   6E+6   12.5   13  RT (min)      lcorbi_t3_pos_1_002_01    m/z 393.2908     0   1   2   3E+5   14   14.5  RT (min)      lcorbi_t3_pos_1_021_18    m/z 393.2909     0   1   2   3E+5   16  RT (min)       lcorbi_t3_pos_1_023_02    m/z 393.2909     0   1   2E+5   15   15.5  RT (min)      lcorbi_t3_pos_2_056_56    m/z 393.2909     0   1   2   3E+5   15   15.5  RT (min)      lcorbi_t3_pos_1_051_09    m/z 393.2909     0   1   2E+6   13.5   14  RT (min)      lcorbi_t3_pos_1_008_36    m/z 393.2909     0   1   2E+5   15   15.5  RT (min)      lcorbi_t3_pos_2_012_01    m/z 393.2910     0   2   4   6E+6   13.5   14  RT (min)      lcorbi_t3_pos_3_047_40    m/z 393.2910     0   1   2   3E+5   14.5   15  RT (min)      lcorbi_t3_pos_1_034_34    m/z 393.2910     0   1   2E+5   14.5   15  RT (min)       lcorbi_t3_pos_1_034_34    m/z 393.2910     0   1   2E+5   14.5   15  RT (min)      lcorbi_t3_pos_1_034_34    m/z 393.2910     0   1   2E+5   14.5   15  RT (min)      lcorbi_t3_pos_2_038_03    m/z 393.3155     0   2   4   6E+5   12.5   13  RT (min)      lcorbi_t3_pos_1_054_48    m/z 393.3156     0   2   4E+5   12   12.5  RT (min)      lcorbi_t3_pos_2_005_11    m/z 394.1780     0   1E+5   10   10.5  RT (min)      lcorbi_t3_pos_1_030_21    m/z 394.2224     0   2   4   6E+5   13   13.5  RT (min)      lcorbi_t3_pos_3_059_17    m/z 394.2225     0   1E+6   14   16  RT (min)       lcorbi_t3_pos_1_033_53    m/z 394.2225     0   1E+6   14   16  RT (min)       lcorbi_t3_pos_4_098_61    m/z 394.2225     0   1E+6   14   16  RT (min)       lcorbi_t3_pos_4_094_60    m/z 394.2225     0   1E+6   16   17  RT (min)       lcorbi_t3_pos_1_021_18    m/z 394.2225     0   1   2E+6   10   15  RT (min)       lcorbi_t3_pos_1_024_04    m/z 394.2225     0   1E+6   14   16   18  RT (min)       lcorbi_t3_pos_4_083_59    m/z 394.2225     0   2   4   6   8E+5   7.5   10   12.5  RT (min)       lcorbi_t3_pos_4_066_57    m/z 394.2225     0   1   2E+6   14   16  RT (min)       lcorbi_t3_pos_1_038_30    m/z 394.2225     0   1   2E+6   16   18  RT (min)       lcorbi_t3_pos_1_043_06    m/z 394.2930     0   2   4E+4   14.5   15  RT (min)      lcorbi_t3_pos_1_034_34    m/z 394.2930     0   1E+4   12   12.5  RT (min)      lcorbi_t3_pos_2_032_39    m/z 394.2932     0   1   2   3   4E+4   15.5   16  RT (min)      lcorbi_t3_pos_1_023_02    m/z 394.2932     0   2   4   6E+5   12.5   13  RT (min)      lcorbi_t3_pos_4_016_15    m/z 394.2932     0   1   2   3E+4   14   14.5  RT (min)      lcorbi_t3_pos_2_022_38    m/z 394.2932     0   1   2   3   4E+4   14   14.5  RT (min)      lcorbi_t3_pos_1_028_32    m/z 394.2933     0   2   4   6E+5   13.5   14  RT (min)      lcorbi_t3_pos_3_047_40    m/z 394.2933     0   1   12.5   13  RT (min)      lcorbi_t3_pos_2_013_33    m/z 394.2933     0   2   4   6   8E+3   12.5   13  RT (min)      lcorbi_t3_pos_3_024_07    m/z 394.2933     0   1   2   3E+4   15   15.5  RT (min)      lcorbi_t3_pos_2_012_01    m/z 394.2934     0   1   2   3E+4   15   15.5  RT (min)      lcorbi_t3_pos_1_051_09    m/z 394.2935     0   2   4   6E+5   13.5   14  RT (min)      lcorbi_t3_pos_4_019_47    m/z 394.3103     0   1E+7   1   1.5  RT (min)      lcorbi_t3_pos_4_057_49    m/z 394.3469     0   1E+4   13   13.5  RT (min)      lcorbi_t3_pos_2_047_31    m/z 394.3469     0   2   4   6E+5   14.5   15  RT (min)       lcorbi_t3_pos_3_015_06    m/z 394.3469     0   2   4   6E+4   16   16.5  RT (min)      lcorbi_t3_pos_1_027_43    m/z 394.3469     0   2   4   6E+5   13.5   14  RT (min)      lcorbi_t3_pos_3_053_48    m/z 394.3469     0   1   2E+5   14.5   15  RT (min)      lcorbi_t3_pos_4_025_52    m/z 394.3469     0   1   2   3   4E+5   14.5   15  RT (min)      lcorbi_t3_pos_3_041_54    m/z 394.3469     0   2   4E+4   15   15.5  RT (min)      lcorbi_t3_pos_4_028_04    m/z 394.3469     0   1   2E+4   13.5   14  RT (min)      lcorbi_t3_pos_2_039_30    m/z 394.3469     0   1E+4   13.5   14  RT (min)      lcorbi_t3_pos_2_031_20    m/z 394.3469     0   2   4   6   8E+4   15.5   16  RT (min)       lcorbi_t3_pos_1_038_30    m/z 394.3469     0   1E+4   13.5   14  RT (min)      lcorbi_t3_pos_3_052_47    m/z 394.3470     0   2   4   6E+5   13.5   14  RT (min)      lcorbi_t3_pos_2_016_23    m/z 394.3470     0   2   4   6E+4   16   16.5  RT (min)      lcorbi_t3_pos_1_027_43    m/z 394.3470     0   2   4E+4   16   16.5  RT (min)       lcorbi_t3_pos_2_052_48    m/z 394.3470     0   2   4   6E+5   13.5   14  RT (min)      lcorbi_t3_pos_3_053_48    m/z 394.3470     0   2   4   6   8E+4   15   15.5  RT (min)      lcorbi_t3_pos_3_044_46    m/z 394.3470     0   2   4   6   8E+4   15.5   16  RT (min)      lcorbi_t3_pos_1_038_30    m/z 394.3470     0   2   4E+4   16   16.5  RT (min)      lcorbi_t3_pos_2_014_37    m/z 394.3527     0   1   2E+4   13   13.5  RT (min)      lcorbi_t3_pos_2_047_31    m/z 394.3527     0   1E+6   12   12.5  RT (min)      lcorbi_t3_pos_2_022_38    m/z 394.3527     0   2   4   6E+5   12   12.5  RT (min)      lcorbi_t3_pos_1_033_53    m/z 394.3527     0   1E+5   12   12.5  RT (min)      lcorbi_t3_pos_2_025_52    m/z 394.3527     0   1   2   3   4E+4   12.5   13  RT (min)      lcorbi_t3_pos_1_031_40    m/z 394.3528     0   2   4   6E+5   10.5   11  RT (min)      lcorbi_t3_pos_2_013_33    m/z 394.3528     0   2   4   6E+4   12   12.5  RT (min)      lcorbi_t3_pos_1_034_34    m/z 394.3528     0   1E+6   11   11.5  RT (min)       lcorbi_t3_pos_2_013_33    m/z 394.3528     0   1   2   3E+4   13   13.5  RT (min)      lcorbi_t3_pos_2_012_01    m/z 394.3528     0   1   2   3E+4   12.5   13  RT (min)      lcorbi_t3_pos_2_012_01    m/z 394.3528     0   1E+6   11   11.5  RT (min)      lcorbi_t3_pos_3_024_07    m/z 394.3528     0   1   2   3   4E+4   12.5   13  RT (min)      lcorbi_t3_pos_1_031_40    m/z 394.3528     0   2   4   6E+4   12   12.5  RT (min)      lcorbi_t3_pos_1_041_54    m/z 394.3528     0   2   4E+5   12   12.5  RT (min)      lcorbi_t3_pos_1_028_32    m/z 394.3529     0   2   4   6   8E+5   12   12.5  RT (min)      lcorbi_t3_pos_4_025_52    m/z 394.3529     0   1E+6   11   11.5  RT (min)       lcorbi_t3_pos_2_013_33    m/z 395.2258     0   2   4E+5   14   16  RT (min)       lcorbi_t3_pos_4_092_60    m/z 395.2258     0   1E+5   7.5   8  RT (min)      lcorbi_t3_pos_1_049_55    m/z 395.2258     0   1   2E+5   7.5   8  RT (min)      lcorbi_t3_pos_4_066_57    m/z 395.2258     0   1E+5   7.5   8  RT (min)      lcorbi_t3_pos_4_071_57    m/z 395.2259     0   2   4E+5   15.5   16  RT (min)       lcorbi_t3_pos_1_018_35    m/z 395.2259     0   2   4E+5   16   16.5  RT (min)       lcorbi_t3_pos_4_094_60    m/z 395.2259     0   2   4E+5   15.5   16  RT (min)       lcorbi_t3_pos_1_010_46    m/z 395.2260     0   2   4E+5   14   16   18  RT (min)       lcorbi_t3_pos_1_025_52    m/z 395.2261     0   2   4E+5   16   17  RT (min)       lcorbi_t3_pos_4_085_59    m/z 395.3503     0   1   2E+5   14.5   15  RT (min)       lcorbi_t3_pos_3_015_06    m/z 395.3562     0   1E+6   11   11.5  RT (min)       lcorbi_t3_pos_4_086_59    m/z 395.3562     0   1E+6   11   11.5  RT (min)      lcorbi_t3_pos_4_086_59    m/z 395.3562     0   1   2   3E+5   11.5   12  RT (min)       lcorbi_t3_pos_1_008_36    m/z 397.1412     0   1   2E+5   8.5   9  RT (min)       lcorbi_t3_pos_1_025_52    m/z 397.1414     0   1   2E+5   17   18  RT (min)       lcorbi_t3_pos_4_107_62    m/z 399.1777     0   1   2   3   4E+5   8   10  RT (min)       lcorbi_t3_pos_4_066_57    m/z 399.1778     0   1   2   3E+5   7.5   8  RT (min)      lcorbi_t3_pos_4_040_21    m/z 399.1778     0   1   2E+5   7.5   8  RT (min)      lcorbi_t3_pos_1_023_02    m/z 399.1779     0   1E+6   12.5   15   17.5  RT (min)       lcorbi_t3_pos_4_107_62    m/z 399.1779     0   1E+6   16   17  RT (min)       lcorbi_t3_pos_4_068_57    m/z 399.1779     0   1E+6   15.5   16  RT (min)      lcorbi_t3_pos_1_064_16    m/z 399.1780     0   1   2   3E+5   13.5   14  RT (min)       lcorbi_t3_pos_2_040_25    m/z 399.1780     0   1   2E+6   16   18  RT (min)       lcorbi_t3_pos_4_088_59    m/z 399.1780     0   1E+6   16   17   18  RT (min)       lcorbi_t3_pos_4_076_58    m/z 399.1780     0   1E+6   16   17   18  RT (min)       lcorbi_t3_pos_4_111_62    m/z 400.1813     0   2   4E+5   16   18  RT (min)       lcorbi_t3_pos_4_107_62    m/z 400.1815     0   2   4E+5   16   17  RT (min)       lcorbi_t3_pos_4_076_58    m/z 401.2120     0   1E+5   15   15.5  RT (min)       lcorbi_t3_pos_3_026_04    m/z 401.2120     0   1E+5   15   15.5  RT (min)       lcorbi_t3_pos_4_091_60    m/z 402.2084     0   1E+5   12.5   13  RT (min)      lcorbi_t3_pos_2_013_33    m/z 402.2084     0   2   4E+5   14.5   15  RT (min)       lcorbi_t3_pos_4_095_60    m/z 402.2084     0   2   4E+5   15   15.5  RT (min)       lcorbi_t3_pos_1_036_08    m/z 403.2118     0   1E+5   15   16  RT (min)       lcorbi_t3_pos_4_094_60    m/z 403.2326     0   1   2E+6   10.5   11  RT (min)      lcorbi_t3_pos_2_032_39    m/z 403.2327     0   1   2   3E+5   11.5   12  RT (min)      lcorbi_t3_pos_3_013_19    m/z 403.2327     0   1   2   3E+5   11.5   12  RT (min)      lcorbi_t3_pos_1_047_41    m/z 403.2327     0   2   4   6E+6   10.5   11  RT (min)      lcorbi_t3_pos_3_052_47    m/z 403.2327     0   1E+5   12   12.5  RT (min)      lcorbi_t3_pos_2_052_48    m/z 403.2328     0   2   4   6E+6   11.5   12  RT (min)      lcorbi_t3_pos_3_044_46    m/z 403.2328     0   1   2   3E+5   12   12.5  RT (min)      lcorbi_t3_pos_3_013_19    m/z 403.2328     0   1   2   3   4E+6   10.5   11  RT (min)      lcorbi_t3_pos_3_011_42    m/z 403.2328     0   1E+5   12   12.5  RT (min)      lcorbi_t3_pos_2_056_56    m/z 403.2328     0   2   4   6E+5   11.5   12  RT (min)      lcorbi_t3_pos_2_046_07    m/z 403.2328     0   2   4E+4   12   12.5  RT (min)      lcorbi_t3_pos_1_041_54    m/z 403.2328     0   1   2   3E+4   12.5   13  RT (min)      lcorbi_t3_pos_1_023_02    m/z 403.2329     0   1   2   3   4E+4   12   12.5  RT (min)      lcorbi_t3_pos_1_031_40    m/z 404.2069     0   1E+5   15   16   17  RT (min)       lcorbi_t3_pos_4_102_61    m/z 404.2070     0   1E+5   16   16.5  RT (min)       lcorbi_t3_pos_4_112_62    m/z 404.2362     0   1E+6   10.5   11  RT (min)      lcorbi_t3_pos_3_052_47    m/z 404.2362     0   1E+5   10.5   11  RT (min)      lcorbi_t3_pos_2_047_31    m/z 404.2362     0   1   2   3   4E+4   12   12.5  RT (min)      lcorbi_t3_pos_2_052_48    m/z 404.2362     0   1E+6   11.5   12  RT (min)      lcorbi_t3_pos_3_044_46    m/z 404.2362     0   1   2   3E+4   12   12.5  RT (min)      lcorbi_t3_pos_2_056_56    m/z 404.2363     0   2   4   6   8E+4   11.5   12  RT (min)      lcorbi_t3_pos_3_013_19    m/z 405.4140     0   2   4   6E+5   12.5   13  RT (min)       lcorbi_t3_pos_4_094_60    m/z 408.3107     0   2   4   6E+5   12.5   13  RT (min)      lcorbi_t3_pos_2_002_06    m/z 408.3108     0   2   4   6E+5   13.5   14  RT (min)       lcorbi_t3_pos_4_019_47    m/z 408.3108     0   1   2E+4   12.5   13  RT (min)      lcorbi_t3_pos_3_031_45    m/z 408.3108     0   1   2   3E+4   12   12.5  RT (min)      lcorbi_t3_pos_2_032_39    m/z 408.3684     0   2   4   6E+5   12.5   13  RT (min)      lcorbi_t3_pos_3_025_52    m/z 408.3684     0   2   4   6   8E+5   11.5   12  RT (min)      lcorbi_t3_pos_2_013_33    m/z 408.3684     0   1   2   3E+4   13   13.5  RT (min)      lcorbi_t3_pos_1_028_32    m/z 408.3684     0   1   2   3E+4   12.5   13  RT (min)      lcorbi_t3_pos_2_022_38    m/z 408.3685     0   1   2E+4   13.5   14  RT (min)      lcorbi_t3_pos_1_031_40    m/z 408.3685     0   1E+4   13.5   14  RT (min)      lcorbi_t3_pos_2_012_01    m/z 408.3685     0   2   4   6   8E+5   12.5   13  RT (min)      lcorbi_t3_pos_1_008_36    m/z 408.3685     0   1   2   3E+4   13   13.5  RT (min)      lcorbi_t3_pos_1_034_34    m/z 410.3139     0   1E+4   12.5   13  RT (min)      lcorbi_t3_pos_2_047_31    m/z 410.3180     0   1   2   3   4E+4   14.5   15  RT (min)      lcorbi_t3_pos_1_038_30    m/z 410.3180     0   1   16   16.5  RT (min)      lcorbi_t3_pos_1_023_02    m/z 410.3181     0   1   2E+4   14.5   15  RT (min)      lcorbi_t3_pos_1_038_30    m/z 410.3181     0   1   2   3E+4   15   15.5  RT (min)      lcorbi_t3_pos_1_027_43    m/z 410.3181     0   1E+4   15   15.5  RT (min)      lcorbi_t3_pos_2_052_48    m/z 410.3181     0   1E+4   13   13.5  RT (min)      lcorbi_t3_pos_3_037_32    m/z 410.3181     0   1   2   3   4E+4   14   14.5  RT (min)      lcorbi_t3_pos_3_044_46    m/z 410.3182     0   1   16   16.5  RT (min)      lcorbi_t3_pos_1_023_02    m/z 410.3182     0   1   2E+4   14.5   15  RT (min)      lcorbi_t3_pos_1_055_33    m/z 410.3182     0   1   15   15.5  RT (min)      lcorbi_t3_pos_2_014_37    m/z 410.3182     0   2   4   6   8E+5   14   14.5  RT (min)       lcorbi_t3_pos_3_047_40    m/z 410.3182     0   1   2   3   4E+4   14   14.5  RT (min)      lcorbi_t3_pos_3_044_46    m/z 410.3182     0   2   4   6E+5   11.5   12  RT (min)      lcorbi_t3_pos_4_057_49    m/z 410.3182     0   1   15.5   16  RT (min)      lcorbi_t3_pos_1_051_09    m/z 411.3215     0   1   16   16.5  RT (min)      lcorbi_t3_pos_1_023_02    m/z 411.3215     0   1   14.5   15  RT (min)      lcorbi_t3_pos_1_055_33    m/z 411.3216     0   1   14.5   15  RT (min)      lcorbi_t3_pos_1_055_33    m/z 411.3216     0   1   16   16.5  RT (min)      lcorbi_t3_pos_1_023_02    m/z 411.3216     0   1   12.5   13  RT (min)      lcorbi_t3_pos_2_039_30    m/z 411.3260     0   1E+4   12.5   13  RT (min)       lcorbi_t3_pos_2_047_31    m/z 413.2337     0   1E+5   13   13.5  RT (min)      lcorbi_t3_pos_4_080_58    m/z 413.2662     0   1   2E+5   12   12.5  RT (min)      lcorbi_t3_pos_2_032_39    m/z 413.2662     0   2   4   6E+6   12.5   13  RT (min)      lcorbi_t3_pos_2_054_29    m/z 413.2662     0   1   2   3E+5   12.5   13  RT (min)      lcorbi_t3_pos_4_012_37    m/z 413.2662     0   1   2E+7   12.5   13  RT (min)      lcorbi_t3_pos_4_016_15    m/z 413.2662     0   1   2E+5   12.5   13  RT (min)      lcorbi_t3_pos_3_024_07    m/z 413.2663     0   1   2E+7   12.5   13  RT (min)      lcorbi_t3_pos_4_012_37    m/z 413.2663     0   1E+6   16   16.5  RT (min)      lcorbi_t3_pos_1_027_43    m/z 413.2663     0   2   4   6E+5   14   14.5  RT (min)      lcorbi_t3_pos_1_028_32    m/z 413.2663     0   1E+6   15   15.5  RT (min)      lcorbi_t3_pos_1_031_40    m/z 413.2663     0   1   2   3   4E+6   10   10.5  RT (min)      lcorbi_t3_pos_4_057_49    m/z 413.2663     0   1   2E+6   14   14.5  RT (min)      lcorbi_t3_pos_1_038_30    m/z 413.2663     0   2   4   6   8E+5   15   15.5  RT (min)      lcorbi_t3_pos_2_056_56    m/z 413.2663     0   2   4   6   8E+5   14.5   15  RT (min)      lcorbi_t3_pos_1_031_40    m/z 413.2664     0   1E+6   15.5   16  RT (min)      lcorbi_t3_pos_1_023_02    m/z 413.2664     0   2   4   6E+5   15   15.5  RT (min)      lcorbi_t3_pos_1_001_49    m/z 413.2664     0   1E+6   15.5   16  RT (min)      lcorbi_t3_pos_1_023_02    m/z 413.2664     0   2   4   6E+6   13.5   14  RT (min)      lcorbi_t3_pos_3_015_06    m/z 413.2664     0   1E+6   14   14.5  RT (min)      lcorbi_t3_pos_2_046_07    m/z 413.2664     0   2   4   6E+5   14   14.5  RT (min)      lcorbi_t3_pos_2_022_38    m/z 413.2664     0   1E+6   15.5   16  RT (min)      lcorbi_t3_pos_1_023_02    m/z 413.2664     0   1   2E+7   13.5   14  RT (min)      lcorbi_t3_pos_3_047_40    m/z 413.2664     0   1   2   3E+6   18   18.5  RT (min)       lcorbi_t3_pos_4_070_57    m/z 413.2664     0   1   2   3E+6   18   18.5  RT (min)      lcorbi_t3_pos_4_077_58    m/z 413.2664     0   2   4   6   8E+5   14   14.5  RT (min)      lcorbi_t3_pos_3_013_19    m/z 413.2664     0   1E+6   15.5   16  RT (min)      lcorbi_t3_pos_1_023_02    m/z 413.2664     0   1E+7   13.5   14  RT (min)      lcorbi_t3_pos_4_001_49    m/z 413.2664     0   1E+6   15   15.5  RT (min)      lcorbi_t3_pos_1_051_09    m/z 413.2664     0   1E+6   15.5   16  RT (min)      lcorbi_t3_pos_1_023_02    m/z 413.2664     0   2   4   6   8E+5   15.5   16  RT (min)      lcorbi_t3_pos_2_030_14    m/z 413.2664     0   1E+6   14   14.5  RT (min)      lcorbi_t3_pos_2_046_07    m/z 413.2664     0   2   4   6E+5   15   15.5  RT (min)      lcorbi_t3_pos_1_001_49    m/z 413.2664     0   1   2E+7   12.5   13  RT (min)      lcorbi_t3_pos_4_012_37    m/z 413.2665     0   2   4   6   8E+5   0.5   1  RT (min)      lcorbi_t3_pos_4_057_49    m/z 414.2695     0   2   4   6   8E+4   12   12.5  RT (min)      lcorbi_t3_pos_2_032_39    m/z 414.2696     0   2   4   6E+4   12.5   13  RT (min)      lcorbi_t3_pos_3_024_07    m/z 414.2696     0   2   4E+6   12.5   13  RT (min)      lcorbi_t3_pos_4_016_15    m/z 414.2697     0   1   2E+5   15   15.5  RT (min)      lcorbi_t3_pos_1_031_40    m/z 414.2697     0   1E+6   10   10.5  RT (min)      lcorbi_t3_pos_4_057_49    m/z 414.2697     0   1   2   3E+5   16   16.5  RT (min)      lcorbi_t3_pos_1_023_02    m/z 414.2697     0   1E+5   15   15.5  RT (min)      lcorbi_t3_pos_1_001_49    m/z 414.2697     0   1   2E+5   14   14.5  RT (min)      lcorbi_t3_pos_1_021_18    m/z 414.2697     0   1E+6   12.5   13  RT (min)      lcorbi_t3_pos_2_054_29    m/z 414.2697     0   2   4   6E+4   12.5   13  RT (min)      lcorbi_t3_pos_3_024_07    m/z 414.2697     0   2   4   6E+6   12.5   13  RT (min)      lcorbi_t3_pos_2_054_29    m/z 414.2698     0   1E+5   14   14.5  RT (min)      lcorbi_t3_pos_2_022_38    m/z 414.2698     0   1E+5   14.5   15  RT (min)      lcorbi_t3_pos_1_034_34    m/z 414.2698     0   1E+5   14   14.5  RT (min)      lcorbi_t3_pos_4_048_24    m/z 414.2698     0   1   2   3E+5   16   16.5  RT (min)      lcorbi_t3_pos_1_023_02    m/z 414.2698     0   1   2E+5   15   15.5  RT (min)      lcorbi_t3_pos_1_051_09    m/z 414.2698     0   1   2E+5   14   14.5  RT (min)      lcorbi_t3_pos_3_013_19    m/z 414.2698     0   1   2   3E+5   15.5   16  RT (min)      lcorbi_t3_pos_1_023_02    m/z 414.2698     0   1   2E+5   15   15.5  RT (min)      lcorbi_t3_pos_2_056_56    m/z 414.2698     0   1   2E+5   14.5   15  RT (min)      lcorbi_t3_pos_2_049_55    m/z 414.2698     0   1   2E+5   15   15.5  RT (min)      lcorbi_t3_pos_2_056_56    m/z 414.2698     0   1   2   3E+5   15.5   16  RT (min)      lcorbi_t3_pos_1_023_02    m/z 414.2698     0   1   2   3E+5   15.5   16  RT (min)      lcorbi_t3_pos_1_023_02    m/z 414.2698     0   1   2E+5   14.5   15  RT (min)      lcorbi_t3_pos_1_031_40    m/z 414.2698     0   2   4   6E+6   13.5   14  RT (min)      lcorbi_t3_pos_3_047_40    m/z 414.2698     0   2   4   6   8E+5   17.5   18  RT (min)       lcorbi_t3_pos_4_079_58    m/z 414.2699     0   1   2E+5   14.5   15  RT (min)      lcorbi_t3_pos_1_041_54    m/z 414.2699     0   1E+5   14.5   15  RT (min)      lcorbi_t3_pos_1_034_34    m/z 414.2699     0   1   2E+5   15   15.5  RT (min)      lcorbi_t3_pos_1_051_09    m/z 414.2699     0   1E+5   15   15.5  RT (min)      lcorbi_t3_pos_1_001_49    m/z 415.2726     0   1E+4   12   12.5  RT (min)      lcorbi_t3_pos_2_032_39    m/z 415.2727     0   1   2E+4   12.5   13  RT (min)      lcorbi_t3_pos_4_012_37    m/z 415.2727     0   2   4   6E+5   12.5   13  RT (min)      lcorbi_t3_pos_2_054_29    m/z 415.2727     0   1E+4   12.5   13  RT (min)      lcorbi_t3_pos_3_024_07    m/z 415.2727     0   2   4E+4   14.5   15  RT (min)      lcorbi_t3_pos_2_009_50    m/z 415.2728     0   1   2   3   4E+4   14.5   15  RT (min)      lcorbi_t3_pos_2_049_55    m/z 415.2728     0   1   2   3E+4   14   14.5  RT (min)      lcorbi_t3_pos_4_048_24    m/z 415.2728     0   2   4   6E+4   14   14.5  RT (min)      lcorbi_t3_pos_2_017_51    m/z 415.2728     0   2   4   6E+5   12.5   13  RT (min)      lcorbi_t3_pos_4_016_15    m/z 415.2728     0   1   2   3E+4   14   14.5  RT (min)      lcorbi_t3_pos_3_013_19    m/z 415.2728     0   1E+5   18   18.5  RT (min)       lcorbi_t3_pos_4_085_59    m/z 415.2728     0   2   4   6E+5   13.5   14  RT (min)       lcorbi_t3_pos_3_047_40    m/z 415.2728     0   1   2   3   4E+4   14   14.5  RT (min)      lcorbi_t3_pos_2_038_03    m/z 415.2728     0   1   2E+5   13.5   14  RT (min)      lcorbi_t3_pos_3_025_52    m/z 415.2728     0   2   4   6E+5   13.5   14  RT (min)      lcorbi_t3_pos_4_019_47    m/z 415.2728     0   1   2   3   4E+4   15   15.5  RT (min)      lcorbi_t3_pos_2_056_56    m/z 415.2728     0   1   2   3E+4   14   14.5  RT (min)      lcorbi_t3_pos_3_013_19    m/z 415.2728     0   2   4E+4   15   15.5  RT (min)      lcorbi_t3_pos_1_051_09    m/z 415.2728     0   2   4E+4   15   15.5  RT (min)      lcorbi_t3_pos_1_051_09    m/z 415.2729     0   2   4E+5   13.5   14  RT (min)      lcorbi_t3_pos_4_001_49    m/z 415.2729     0   2   4E+4   15.5   16  RT (min)      lcorbi_t3_pos_1_023_02    m/z 415.2730     0   2   4E+4   15.5   16  RT (min)      lcorbi_t3_pos_1_023_02    m/z 415.3208     0   2   4   6   8E+5   12   12.5  RT (min)      lcorbi_t3_pos_1_002_01    m/z 415.3208     0   1   13.5   14  RT (min)      lcorbi_t3_pos_1_034_34    m/z 415.3209     0   1   14   14.5  RT (min)      lcorbi_t3_pos_1_031_40    m/z 415.3209     0   1   14   14.5  RT (min)      lcorbi_t3_pos_2_012_01    m/z 416.2127     0   1E+6   5   5.5  RT (min)      lcorbi_t3_pos_3_000_aa    m/z 416.3074     0   1E+5   15   15.5  RT (min)      lcorbi_t3_pos_1_008_36    m/z 416.3075     0   1   2E+4   13   13.5  RT (min)      lcorbi_t3_pos_2_047_31    m/z 416.3076     0   1   2E+4   15.5   16  RT (min)      lcorbi_t3_pos_1_055_33    m/z 416.3076     0   1   2E+4   15.5   16  RT (min)      lcorbi_t3_pos_1_055_33    m/z 417.3152     0   1   16.5   17  RT (min)    lcorbi_t3_pos_1_031_40    m/z 417.3152     0   1   13.5   14  RT (min)      lcorbi_t3_pos_3_052_47    m/z 417.3153     0   1   15.5   16  RT (min)      lcorbi_t3_pos_2_038_03    m/z 417.3153     0   1   16.5   17  RT (min)    lcorbi_t3_pos_1_031_40    m/z 417.3153     0   1   13   13.5  RT (min)      lcorbi_t3_pos_2_047_31    m/z 417.3153     0   1   13.5   14  RT (min)      lcorbi_t3_pos_2_013_33    m/z 417.3153     0   2   4   6   8E+3   13.5   14  RT (min)      lcorbi_t3_pos_3_037_32    m/z 417.3153     0   1   2E+4   16   16.5  RT (min)      lcorbi_t3_pos_2_014_37    m/z 417.3153     0   2   4   6E+4   15   15.5  RT (min)      lcorbi_t3_pos_3_044_46    m/z 417.3153     0   1   16.5   17  RT (min)    lcorbi_t3_pos_2_012_01    m/z 417.3154     0   2   4   6   8E+6   14   14.5  RT (min)      lcorbi_t3_pos_4_037_26    m/z 417.3154     0   1   2E+4   16   16.5  RT (min)      lcorbi_t3_pos_1_034_34    m/z 417.3154     0   2   4   6E+4   15.5   16  RT (min)      lcorbi_t3_pos_1_055_33    m/z 417.3154     0   1   17   17.5  RT (min)    lcorbi_t3_pos_1_023_02    m/z 417.3154     0   2   4   6E+4   15.5   16  RT (min)      lcorbi_t3_pos_1_055_33    m/z 417.3154     0   1E+6   15   15.5  RT (min)      lcorbi_t3_pos_1_008_36    m/z 417.3154     0   2   4   6E+5   15   15.5  RT (min)      lcorbi_t3_pos_4_001_49    m/z 417.3154     0   1   15.5   16  RT (min)      lcorbi_t3_pos_2_038_03    m/z 418.2588     0   2   4   6   8E+5   10.5   11  RT (min)      lcorbi_t3_pos_4_056_56    m/z 418.2588     0   1   2E+6   10   12   14  RT (min)       lcorbi_t3_pos_1_039_23    m/z 418.2589     0   2   4   6E+5   12.5   13  RT (min)      lcorbi_t3_pos_1_060_39    m/z 418.2589     0   2   4   6   8E+5   13   13.5  RT (min)      lcorbi_t3_pos_1_064_16    m/z 418.2590     0   2   4   6   8E+5   14   14.5  RT (min)      lcorbi_t3_pos_3_057_49    m/z 418.2590     0   1E+6   15   16   17  RT (min)       lcorbi_t3_pos_4_070_57    m/z 418.2590     0   1E+6   16   16.5  RT (min)       lcorbi_t3_pos_4_071_57    m/z 418.2590     0   1E+6   16   16.5  RT (min)       lcorbi_t3_pos_4_068_57    m/z 418.2590     0   2   4   6E+5   13.5   14  RT (min)      lcorbi_t3_pos_1_057_49    m/z 418.2592     0   1E+6   16   17  RT (min)       lcorbi_t3_pos_4_073_57    m/z 418.3186     0   1   15.5   16  RT (min)      lcorbi_t3_pos_1_021_18    m/z 418.3187     0   1E+4   13.5   14  RT (min)      lcorbi_t3_pos_2_013_33    m/z 418.3187     0   1   13.5   14  RT (min)      lcorbi_t3_pos_3_035_08    m/z 418.3187     0   1   13   13.5  RT (min)      lcorbi_t3_pos_2_047_31    m/z 418.3187     0   1   13.5   14  RT (min)      lcorbi_t3_pos_3_035_08    m/z 418.3187     0   1   2   3E+6   13.5   14  RT (min)      lcorbi_t3_pos_4_015_45    m/z 418.3187     0   1   2E+5   15   15.5  RT (min)      lcorbi_t3_pos_4_001_49    m/z 418.3187     0   2   4   6E+5   15   15.5  RT (min)      lcorbi_t3_pos_3_047_40    m/z 418.3187     0   1   16.5   17  RT (min)    lcorbi_t3_pos_2_012_01    m/z 418.3187     0   1E+4   15.5   16  RT (min)      lcorbi_t3_pos_3_003_33    m/z 418.3187     0   1   17   17.5  RT (min)    lcorbi_t3_pos_1_023_02    m/z 418.3188     0   1E+4   15.5   16  RT (min)      lcorbi_t3_pos_4_028_04    m/z 418.3188     0   1E+6   14   14.5  RT (min)      lcorbi_t3_pos_4_016_15    m/z 418.3188     0   1   13.5   14  RT (min)      lcorbi_t3_pos_3_037_32    m/z 418.3188     0   1   2   3   4E+5   15   15.5  RT (min)      lcorbi_t3_pos_1_008_36    m/z 418.3188     0   1   17   17.5  RT (min)    lcorbi_t3_pos_1_023_02    m/z 418.3188     0   1E+4   16   16.5  RT (min)      lcorbi_t3_pos_2_014_37    m/z 418.3188     0   1   16.5   17  RT (min)    lcorbi_t3_pos_1_031_40    m/z 418.3188     0   2   4   6   8E+5   14.5   15  RT (min)      lcorbi_t3_pos_1_005_42    m/z 418.3188     0   1   16   16.5  RT (min)      lcorbi_t3_pos_1_034_34    m/z 418.3188     0   1   16   16.5  RT (min)      lcorbi_t3_pos_2_049_55    m/z 418.3188     0   1   16.5   17  RT (min)    lcorbi_t3_pos_1_031_40    m/z 418.3188     0   1   2E+4   15   15.5  RT (min)      lcorbi_t3_pos_3_044_46    m/z 418.3188     0   2   4E+4   15   15.5  RT (min)      lcorbi_t3_pos_3_025_52    m/z 419.2428     0   1   2E+5   9   9.5  RT (min)      lcorbi_t3_pos_3_013_19    m/z 419.3157     0   1E+5   15   15.5  RT (min)      lcorbi_t3_pos_2_049_55    m/z 419.3157     0   1E+5   12.5   13  RT (min)       lcorbi_t3_pos_2_013_33    m/z 419.3157     0   1E+5   12.5   13  RT (min)      lcorbi_t3_pos_3_024_07    m/z 419.3157     0   1E+6   13   13.5  RT (min)      lcorbi_t3_pos_3_011_42    m/z 419.3157     0   1   16.5   17  RT (min)    lcorbi_t3_pos_2_012_01    m/z 419.3157     0   2   4   6E+5   13   13.5  RT (min)      lcorbi_t3_pos_3_011_42    m/z 419.3158     0   1E+5   14.5   15  RT (min)      lcorbi_t3_pos_2_041_54    m/z 419.3158     0   1   2E+5   16   16.5  RT (min)      lcorbi_t3_pos_1_034_34    m/z 419.3158     0   1E+6   14   14.5  RT (min)      lcorbi_t3_pos_4_025_52    m/z 419.3158     0   2   4   6   8E+4   15   15.5  RT (min)      lcorbi_t3_pos_4_033_53    m/z 419.3158     0   1E+5   14.5   15  RT (min)      lcorbi_t3_pos_2_041_54    m/z 419.3158     0   1E+5   15.5   16  RT (min)      lcorbi_t3_pos_2_012_01    m/z 419.3159     0   1   2   3E+5   14.5   15  RT (min)       lcorbi_t3_pos_1_028_32    m/z 419.3159     0   1   16.5   17  RT (min)    lcorbi_t3_pos_1_031_40    m/z 419.3189     0   1   16.5   17  RT (min)    lcorbi_t3_pos_1_031_40    m/z 420.2190     0   1   2   3E+5   16   17  RT (min)       lcorbi_t3_pos_1_015_11    m/z 420.2190     0   1   2E+5   15   16  RT (min)       lcorbi_t3_pos_1_016_05    m/z 420.2190     0   1E+5   13.5   14  RT (min)      lcorbi_t3_pos_2_003_19    m/z 420.2190     0   1   2   3E+5   14.5   15   15.5  RT (min)       lcorbi_t3_pos_1_006_15    m/z 420.3192     0   1E+5   14.5   15  RT (min)      lcorbi_t3_pos_1_028_32    m/z 420.3192     0   1E+5   14.5   15  RT (min)      lcorbi_t3_pos_1_034_34    m/z 420.3192     0   1E+5   15   15.5  RT (min)      lcorbi_t3_pos_1_031_40    m/z 421.3525     0   1E+6   11.5   12  RT (min)       lcorbi_t3_pos_1_032_26    m/z 422.3266     0   1E+4   12   12.5  RT (min)      lcorbi_t3_pos_2_032_39    m/z 422.3266     0   1E+4   12.5   13  RT (min)      lcorbi_t3_pos_3_024_07    m/z 422.3266     0   1   2   3   4E+4   14   14.5  RT (min)      lcorbi_t3_pos_3_003_33    m/z 422.3266     0   2   4   6   8E+4   15   15.5  RT (min)      lcorbi_t3_pos_1_051_09    m/z 422.3266     0   1E+4   12.5   13  RT (min)      lcorbi_t3_pos_2_013_33    m/z 422.3267     0   2   4E+5   13.5   14  RT (min)      lcorbi_t3_pos_3_047_40    m/z 422.3267     0   2   4E+5   13.5   14  RT (min)      lcorbi_t3_pos_4_019_47    m/z 422.3267     0   1   2E+5   13.5   14  RT (min)      lcorbi_t3_pos_1_008_36    m/z 422.3267     0   1   2   3E+5   13.5   14  RT (min)      lcorbi_t3_pos_3_047_40    m/z 422.3267     0   2   4   6   8E+4   14   14.5  RT (min)      lcorbi_t3_pos_1_047_41    m/z 422.3268     0   1E+4   12.5   13  RT (min)      lcorbi_t3_pos_3_024_07    m/z 422.3268     0   2   4   6   8E+4   14   14.5  RT (min)      lcorbi_t3_pos_1_047_41    m/z 423.2143     0   1E+6   17   17.5  RT (min)       lcorbi_t3_pos_4_105_61    m/z 423.2144     0   1E+6   17   18  RT (min)       lcorbi_t3_pos_4_107_62    m/z 423.2144     0   1E+6   17.5   18  RT (min)       lcorbi_t3_pos_4_109_62    m/z 423.2145     0   1E+6   17.5   18  RT (min)       lcorbi_t3_pos_4_113_62    m/z 423.4250     0   2   4   6E+5   12   12.5  RT (min)      lcorbi_t3_pos_3_055_10    m/z 423.4250     0   2   4   6   8E+5   12   12.5  RT (min)      lcorbi_t3_pos_4_095_60    m/z 423.4250     0   2   4E+5   13.5   14  RT (min)       lcorbi_t3_pos_4_091_60    m/z 423.4250     0   2   4   6   8E+5   12   12.5  RT (min)      lcorbi_t3_pos_2_037_27    m/z 424.2176     0   2   4   6   8E+4   10.5   11  RT (min)      lcorbi_t3_pos_1_031_40    m/z 424.2176     0   1   2   3E+5   10   10.5  RT (min)      lcorbi_t3_pos_1_008_36    m/z 424.2176     0   1   2   3E+5   10   10.5  RT (min)      lcorbi_t3_pos_1_033_53    m/z 424.2176     0   2   4   6E+4   10.5   11  RT (min)      lcorbi_t3_pos_1_027_43    m/z 424.2178     0   1E+5   13   13.5  RT (min)       lcorbi_t3_pos_4_068_57    m/z 424.3634     0   2   4E+5   10.5   11  RT (min)      lcorbi_t3_pos_1_036_08    m/z 424.3634     0   1E+6   10.5   11  RT (min)       lcorbi_t3_pos_3_031_45    m/z 424.3634     0   1E+6   10.5   11  RT (min)       lcorbi_t3_pos_3_031_45    m/z 424.3634     0   1   2E+4   12.5   13  RT (min)      lcorbi_t3_pos_2_012_01    m/z 424.3634     0   1E+6   11.5   12  RT (min)      lcorbi_t3_pos_1_008_36    m/z 424.3634     0   1   2E+4   12   12.5  RT (min)      lcorbi_t3_pos_1_031_40    m/z 424.3634     0   2   4   6   8E+4   11.5   12  RT (min)      lcorbi_t3_pos_3_003_33    m/z 424.3634     0   1   2E+4   12.5   13  RT (min)      lcorbi_t3_pos_2_012_01    m/z 424.3634     0   1   2   3E+4   12.5   13  RT (min)      lcorbi_t3_pos_1_025_52    m/z 424.3634     0   1   2E+4   12   12.5  RT (min)      lcorbi_t3_pos_1_034_34    m/z 424.3635     0   1   2   3E+4   12   12.5  RT (min)      lcorbi_t3_pos_1_034_34    m/z 424.3635     0   2   4   6   8E+5   11.5   12  RT (min)      lcorbi_t3_pos_2_017_51    m/z 425.1744     0   1   2E+5   16   17  RT (min)       lcorbi_t3_pos_4_109_62    m/z 425.1744     0   1E+5   16   17  RT (min)       lcorbi_t3_pos_4_069_57    m/z 425.2146     0   2   4   6E+5   10.5   11  RT (min)      lcorbi_t3_pos_2_047_31    m/z 425.2147     0   2   4   6E+4   12   12.5  RT (min)      lcorbi_t3_pos_2_014_37    m/z 425.2147     0   1   2   3   4E+6   10.5   11  RT (min)      lcorbi_t3_pos_3_052_47    m/z 425.2147     0   2   4E+5   11.5   12  RT (min)      lcorbi_t3_pos_4_048_24    m/z 425.2147     0   2   4   6E+4   12   12.5  RT (min)      lcorbi_t3_pos_2_056_56    m/z 425.2148     0   1   2   3E+4   12   12.5  RT (min)      lcorbi_t3_pos_1_056_56    m/z 425.2148     0   1   2   3   4E+6   11.5   12  RT (min)      lcorbi_t3_pos_3_056_56    m/z 425.2148     0   2   4E+4   11.5   12  RT (min)      lcorbi_t3_pos_1_038_30    m/z 425.2148     0   2   4E+4   12.5   13  RT (min)      lcorbi_t3_pos_2_012_01    m/z 425.2148     0   1E+5   12   12.5  RT (min)      lcorbi_t3_pos_2_052_48    m/z 425.2148     0   2   4   6E+4   12.5   13  RT (min)      lcorbi_t3_pos_2_056_56    m/z 425.2148     0   1E+5   12   12.5  RT (min)      lcorbi_t3_pos_2_038_03    m/z 425.2148     0   1   2   3E+6   10.5   11  RT (min)      lcorbi_t3_pos_2_031_20    m/z 426.2181     0   1   2E+4   12   12.5  RT (min)      lcorbi_t3_pos_2_052_48    m/z 426.2181     0   2   4   6E+5   10.5   11  RT (min)      lcorbi_t3_pos_2_039_30    m/z 426.2182     0   2   4   6   8E+5   11.5   12  RT (min)      lcorbi_t3_pos_3_056_56    m/z 426.2182     0   2   4   6   8E+5   11.5   12  RT (min)      lcorbi_t3_pos_3_044_46    m/z 426.2182     0   1E+5   11.5   12  RT (min)      lcorbi_t3_pos_3_049_55    m/z 426.2182     0   1   2E+4   12   12.5  RT (min)      lcorbi_t3_pos_2_056_56    m/z 426.2182     0   2   4   6E+4   12   12.5  RT (min)      lcorbi_t3_pos_3_013_19    m/z 426.2182     0   1E+5   11.5   12  RT (min)      lcorbi_t3_pos_4_028_04    m/z 426.2182     0   1   2   3E+4   12   12.5  RT (min)      lcorbi_t3_pos_2_038_03    m/z 426.2182     0   1E+6   10.5   11  RT (min)      lcorbi_t3_pos_3_037_32    m/z 426.2183     0   1   2E+4   12.5   13  RT (min)      lcorbi_t3_pos_2_056_56    m/z 426.3662     0   1E+6   14   14.5  RT (min)      lcorbi_t3_pos_1_024_04    m/z 426.3662     0   1E+6   14   14.5  RT (min)      lcorbi_t3_pos_1_024_04    m/z 427.3208     0   2   4E+5   12.5   13  RT (min)       lcorbi_t3_pos_3_029_13    m/z 427.3208     0   2   4   6E+5   12   12.5   13  RT (min)       lcorbi_t3_pos_1_054_48    m/z 429.2401     0   2   4   6E+4   12   12.5  RT (min)      lcorbi_t3_pos_2_032_39    m/z 429.2402     0   1E+5   15.5   16  RT (min)      lcorbi_t3_pos_1_023_02    m/z 429.2402     0   1   2   3   4E+6   12.5   13  RT (min)      lcorbi_t3_pos_1_002_01    m/z 429.2402     0   1E+5   14.5   15  RT (min)      lcorbi_t3_pos_1_031_40    m/z 429.2402     0   2   4   6E+4   12.5   13  RT (min)      lcorbi_t3_pos_4_012_37    m/z 429.2402     0   1E+5   14.5   15  RT (min)      lcorbi_t3_pos_2_049_55    m/z 429.2402     0   1E+5   15.5   16  RT (min)      lcorbi_t3_pos_1_023_02    m/z 429.2402     0   1E+5   15   15.5  RT (min)      lcorbi_t3_pos_1_001_49    m/z 429.2402     0   1E+5   14   14.5  RT (min)      lcorbi_t3_pos_2_022_38    m/z 429.2402     0   1E+5   15   15.5  RT (min)      lcorbi_t3_pos_1_001_49    m/z 429.2402     0   1E+6   13.5   14  RT (min)      lcorbi_t3_pos_4_010_43    m/z 429.2402     0   2   4   6   8E+5   17.5   18  RT (min)       lcorbi_t3_pos_4_075_58    m/z 429.2402     0   1   2   3E+5   14   14.5  RT (min)      lcorbi_t3_pos_1_055_33    m/z 429.2402     0   1   2   3   4E+6   13.5   14  RT (min)      lcorbi_t3_pos_4_001_49    m/z 429.2402     0   1E+5   15.5   16  RT (min)      lcorbi_t3_pos_1_023_02    m/z 429.2402     0   1E+5   15   15.5  RT (min)      lcorbi_t3_pos_1_001_49    m/z 429.2402     0   2   4   6E+6   12.5   13  RT (min)      lcorbi_t3_pos_2_054_29    m/z 429.2402     0   1E+5   15   15.5  RT (min)      lcorbi_t3_pos_1_051_09    m/z 429.2402     0   2   4E+4   12   12.5  RT (min)      lcorbi_t3_pos_2_047_31    m/z 429.2402     0   1E+5   15.5   16  RT (min)      lcorbi_t3_pos_1_023_02    m/z 429.2402     0   1E+5   14.5   15  RT (min)      lcorbi_t3_pos_1_034_34    m/z 429.2402     0   1E+6   13.5   14  RT (min)      lcorbi_t3_pos_3_056_56    m/z 429.2402     0   2   4E+4   12.5   13  RT (min)      lcorbi_t3_pos_3_024_07    m/z 429.2402     0   1   2E+5   14   14.5  RT (min)      lcorbi_t3_pos_3_013_19    m/z 429.2402     0   1E+6   10   10.5  RT (min)      lcorbi_t3_pos_4_057_49    m/z 429.2402     0   1E+6   12.5   13  RT (min)      lcorbi_t3_pos_2_054_29    m/z 429.2403     0   2   4E+5   14   14.5  RT (min)      lcorbi_t3_pos_2_046_07    m/z 429.2403     0   1E+5   15   15.5  RT (min)      lcorbi_t3_pos_1_001_49    m/z 429.2403     0   1   2E+6   13.5   14  RT (min)      lcorbi_t3_pos_2_002_06    m/z 429.2403     0   1E+5   14.5   15  RT (min)      lcorbi_t3_pos_1_034_34    m/z 429.2403     0   1E+5   15   15.5  RT (min)      lcorbi_t3_pos_2_012_01    m/z 429.2403     0   1   2E+6   12.5   13  RT (min)      lcorbi_t3_pos_4_015_45    m/z 429.2404     0   1   2E+6   12.5   13  RT (min)      lcorbi_t3_pos_4_015_45    m/z 430.2435     0   1   2   3E+4   12   12.5  RT (min)      lcorbi_t3_pos_2_032_39    m/z 430.2436     0   1   2E+5   18   18.5  RT (min)      lcorbi_t3_pos_4_106_62    m/z 430.2436     0   2   4E+4   14.5   15  RT (min)      lcorbi_t3_pos_1_034_34    m/z 430.2436     0   1E+6   13.5   14  RT (min)      lcorbi_t3_pos_3_047_40    m/z 430.2436     0   1E+6   13.5   14  RT (min)      lcorbi_t3_pos_3_047_40    m/z 430.2436     0   2   4E+4   15.5   16  RT (min)      lcorbi_t3_pos_1_023_02    m/z 430.2436     0   2   4E+4   14.5   15  RT (min)      lcorbi_t3_pos_2_049_55    m/z 430.2436     0   1   2   3   4E+4   15   15.5  RT (min)      lcorbi_t3_pos_2_012_01    m/z 430.2436     0   2   4   6E+4   14   14.5  RT (min)      lcorbi_t3_pos_3_013_19    m/z 430.2436     0   1E+6   12.5   13  RT (min)      lcorbi_t3_pos_1_002_01    m/z 430.2436     0   1E+6   12.5   13  RT (min)      lcorbi_t3_pos_4_016_15    m/z 430.2436     0   2   4E+4   15.5   16  RT (min)      lcorbi_t3_pos_1_023_02    m/z 430.2436     0   1   2   3E+4   14   14.5  RT (min)      lcorbi_t3_pos_2_022_38    m/z 430.2436     0   2   4E+4   15   15.5  RT (min)      lcorbi_t3_pos_1_051_09    m/z 430.2436     0   2   4E+4   14.5   15  RT (min)      lcorbi_t3_pos_1_031_40    m/z 430.2436     0   1   2   3   4E+4   15   15.5  RT (min)      lcorbi_t3_pos_1_001_49    m/z 430.2436     0   1   2E+5   18   18.5  RT (min)      lcorbi_t3_pos_4_073_57    m/z 430.2436     0   1   2   3E+4   14   14.5  RT (min)      lcorbi_t3_pos_2_022_38    m/z 430.2436     0   1E+5   14   14.5  RT (min)      lcorbi_t3_pos_2_046_07    m/z 430.2436     0   2   4   6   8E+5   12.5   13  RT (min)      lcorbi_t3_pos_3_011_42    m/z 430.2436     0   1E+4   12.5   13  RT (min)      lcorbi_t3_pos_3_024_07    m/z 430.2436     0   2   4   6   8E+4   14   14.5  RT (min)      lcorbi_t3_pos_2_046_07    m/z 430.2436     0   1   2E+4   12.5   13  RT (min)      lcorbi_t3_pos_4_012_37    m/z 430.2437     0   2   4E+4   15.5   16  RT (min)      lcorbi_t3_pos_1_023_02    m/z 430.2437     0   1E+6   12.5   13  RT (min)      lcorbi_t3_pos_1_002_01    m/z 430.2437     0   1   2E+4   12.5   13  RT (min)      lcorbi_t3_pos_2_039_30    m/z 430.2438     0   2   4E+4   15.5   16  RT (min)      lcorbi_t3_pos_1_023_02    m/z 430.2438     0   2   4E+4   14.5   15  RT (min)      lcorbi_t3_pos_1_041_54    m/z 430.2588     0   1   2E+5   11   11.5  RT (min)      lcorbi_t3_pos_3_033_53    m/z 430.2589     0   1E+5   11.5   12  RT (min)      lcorbi_t3_pos_1_055_33    m/z 430.2589     0   1E+5   11.5   12  RT (min)      lcorbi_t3_pos_1_047_41    m/z 430.2590     0   1E+5   16   17  RT (min)       lcorbi_t3_pos_2_011_10    m/z 430.3804     0   1   17   17.5  RT (min)    lcorbi_t3_pos_1_023_02    m/z 430.3805     0   1E+6   14.5   15  RT (min)      lcorbi_t3_pos_1_005_42    m/z 430.3805     0   1   15.5   16  RT (min)      lcorbi_t3_pos_1_021_18    m/z 430.3805     0   1   16.5   17  RT (min)    lcorbi_t3_pos_1_001_49    m/z 430.3805     0   1   15   15.5  RT (min)      lcorbi_t3_pos_1_030_21    m/z 430.3805     0   1   16   16.5  RT (min)      lcorbi_t3_pos_1_001_49    m/z 430.3805     0   1   15.5   16  RT (min)      lcorbi_t3_pos_1_028_32    m/z 430.3806     0   2   4   6E+4   15   15.5  RT (min)      lcorbi_t3_pos_1_008_36    m/z 430.3806     0   1   15.5   16  RT (min)      lcorbi_t3_pos_1_034_34    m/z 430.3806     0   1   16.5   17  RT (min)    lcorbi_t3_pos_1_001_49    m/z 430.3806     0   1E+5   15   15.5  RT (min)      lcorbi_t3_pos_1_008_36    m/z 430.3806     0   1   15   15.5  RT (min)      lcorbi_t3_pos_1_047_41    m/z 430.3806     0   1   16   16.5  RT (min)      lcorbi_t3_pos_1_056_56    m/z 430.3806     0   1   15.5   16  RT (min)      lcorbi_t3_pos_1_041_54    m/z 430.3806     0   1   2E+4   15.5   16  RT (min)      lcorbi_t3_pos_1_021_18    m/z 430.3807     0   1   16   16.5  RT (min)      lcorbi_t3_pos_1_041_54    m/z 430.3808     0   1   17   17.5  RT (min)    lcorbi_t3_pos_1_023_02    m/z 431.2384     0   1   2   3   4E+5   13.5   14  RT (min)       lcorbi_t3_pos_4_019_47    m/z 431.2428     0   1E+5   15.5   16  RT (min)      lcorbi_t3_pos_1_023_02    m/z 431.2428     0   2   4   6   8E+4   15   15.5  RT (min)      lcorbi_t3_pos_2_012_01    m/z 431.3840     0   1   15.5   16  RT (min)      lcorbi_t3_pos_1_021_18    m/z 431.3840     0   1   13.5   14  RT (min)      lcorbi_t3_pos_1_002_01    m/z 431.3840     0   1   17   17.5  RT (min)    lcorbi_t3_pos_1_023_02    m/z 431.3841     0   1   17   17.5  RT (min)    lcorbi_t3_pos_1_023_02    m/z 433.3313     0   1E+5   14   14.5  RT (min)       lcorbi_t3_pos_1_008_36    m/z 433.3314     0   2   4   6E+4   14   14.5  RT (min)      lcorbi_t3_pos_1_008_36    m/z 433.3314     0   1   2   3   4E+4   14.5   15  RT (min)      lcorbi_t3_pos_1_028_32    m/z 433.3314     0   1   2E+4   14.5   15  RT (min)      lcorbi_t3_pos_2_017_51    m/z 433.3314     0   1   13   13.5  RT (min)      lcorbi_t3_pos_2_013_33    m/z 433.3314     0   1   2   3E+4   15.5   16  RT (min)      lcorbi_t3_pos_1_031_40    m/z 433.3314     0   1   2   3E+4   15   15.5  RT (min)      lcorbi_t3_pos_1_041_54    m/z 433.3314     0   1   2   3   4E+4   14.5   15  RT (min)      lcorbi_t3_pos_1_028_32    m/z 433.3314     0   1   2   3E+4   15   15.5  RT (min)      lcorbi_t3_pos_2_009_50    m/z 433.3314     0   1   2   3E+4   15   15.5  RT (min)      lcorbi_t3_pos_1_034_34    m/z 436.2693     0   1E+6   10   15  RT (min)       lcorbi_t3_pos_1_020_31    m/z 436.2694     0   1   17   17.5  RT (min)     lcorbi_t3_pos_1_031_40    m/z 436.2694     0   1   17.5   18  RT (min)     lcorbi_t3_pos_1_025_52    m/z 436.2694     0   1E+6   15   16  RT (min)       lcorbi_t3_pos_1_029_29    m/z 436.2694     0   1E+6   16.5   17  RT (min)       lcorbi_t3_pos_4_067_57    m/z 436.2694     0   2   4   6E+5   13   13.5  RT (min)      lcorbi_t3_pos_1_064_16    m/z 436.2694     0   1E+6   16  RT (min)       lcorbi_t3_pos_1_037_19    m/z 436.2695     0   1E+6   15.5   16  RT (min)       lcorbi_t3_pos_4_109_62    m/z 436.2695     0   1E+6   17  RT (min)       lcorbi_t3_pos_4_083_59    m/z 436.2695     0   1E+6   16   16.5  RT (min)       lcorbi_t3_pos_4_092_60    m/z 436.3420     0   1   2E+4   12   12.5  RT (min)      lcorbi_t3_pos_2_032_39    m/z 436.3421     0   1   2E+4   12.5   13  RT (min)      lcorbi_t3_pos_3_024_07    m/z 436.3422     0   1   18.5   19  RT (min)    lcorbi_t3_pos_1_001_49    m/z 436.3422     0   1   2E+5   14.5   15  RT (min)      lcorbi_t3_pos_1_049_55    m/z 436.3422     0   1   2   3E+4   12.5   13  RT (min)      lcorbi_t3_pos_3_024_07    m/z 436.3422     0   2   4E+4   12.5   13  RT (min)      lcorbi_t3_pos_2_032_39    m/z 436.3422     0   1E+6   15   15.5  RT (min)      lcorbi_t3_pos_1_001_49    m/z 436.3422     0   1   2E+5   14.5   15  RT (min)      lcorbi_t3_pos_1_034_34    m/z 436.3422     0   1   17   17.5  RT (min)    lcorbi_t3_pos_1_001_49    m/z 436.3422     0   1   17   17.5  RT (min)    lcorbi_t3_pos_1_001_49    m/z 436.3422     0   1E+6   12.5   13  RT (min)      lcorbi_t3_pos_2_054_29    m/z 436.3422     0   1E+5   14   14.5  RT (min)      lcorbi_t3_pos_2_006_47    m/z 436.3422     0   1   2E+5   15.5   16  RT (min)      lcorbi_t3_pos_1_023_02    m/z 436.3422     0   1   2E+5   14.5   15  RT (min)      lcorbi_t3_pos_1_034_34    m/z 436.3422     0   1   17   17.5  RT (min)    lcorbi_t3_pos_1_001_49    m/z 436.3422     0   1E+6   15   15.5  RT (min)      lcorbi_t3_pos_1_001_49    m/z 436.3422     0   1   2E+5   14   14.5  RT (min)      lcorbi_t3_pos_1_021_18    m/z 436.3422     0   1   2E+5   14.5   15  RT (min)      lcorbi_t3_pos_2_049_55    m/z 436.3422     0   2   4   6E+5   13.5   14  RT (min)      lcorbi_t3_pos_1_008_36    m/z 436.3422     0   1   2E+5   15.5   16  RT (min)      lcorbi_t3_pos_1_023_02    m/z 436.3422     0   1   2E+5   14.5   15  RT (min)      lcorbi_t3_pos_1_049_55    m/z 436.3423     0   1   2E+5   15.5   16  RT (min)      lcorbi_t3_pos_1_023_02    m/z 436.3423     0   1   18   18.5  RT (min)    lcorbi_t3_pos_1_001_49    m/z 436.3423     0   1E+6   12.5   13  RT (min)      lcorbi_t3_pos_4_016_15    m/z 436.3423     0   1   17   17.5  RT (min)    lcorbi_t3_pos_1_001_49    m/z 436.3423     0   1   2E+5   15.5   16  RT (min)      lcorbi_t3_pos_1_023_02    m/z 436.3423     0   1E+6   13.5   14  RT (min)      lcorbi_t3_pos_1_005_42    m/z 436.3424     0   1   2E+6   12.5   13  RT (min)      lcorbi_t3_pos_2_054_29    m/z 436.3424     0   1E+6   15   15.5  RT (min)      lcorbi_t3_pos_1_001_49    m/z 437.2728     0   1   2   3E+5   9   9.5  RT (min)      lcorbi_t3_pos_2_022_38    m/z 437.2728     0   1   2E+5   9   9.5  RT (min)      lcorbi_t3_pos_1_055_33    m/z 437.2728     0   1   2   3   4E+5   16   16.5   17  RT (min)       lcorbi_t3_pos_4_075_58    m/z 437.2728     0   1   2   3   4E+5   15   15.5   16  RT (min)       lcorbi_t3_pos_4_113_62    m/z 437.2728     0   1   2   3E+5   9   9.5  RT (min)      lcorbi_t3_pos_1_051_09    m/z 437.2728     0   1   2E+5   9   9.5  RT (min)      lcorbi_t3_pos_2_049_55    m/z 437.2729     0   1   2   3   4E+5   15.5   16   16.5  RT (min)       lcorbi_t3_pos_4_094_60    m/z 437.3455     0   1   2   3E+5   15   15.5  RT (min)      lcorbi_t3_pos_1_001_49    m/z 437.3455     0   1   2   3E+5   15   15.5  RT (min)      lcorbi_t3_pos_1_001_49    m/z 437.3455     0   1E+4   12.5   13  RT (min)      lcorbi_t3_pos_2_013_33    m/z 437.3455     0   1   2   3E+5   15   15.5  RT (min)      lcorbi_t3_pos_1_001_49    m/z 437.3456     0   1E+5   14.5   15  RT (min)      lcorbi_t3_pos_1_034_34    m/z 437.3456     0   1E+5   14.5   15  RT (min)      lcorbi_t3_pos_1_034_34    m/z 437.3456     0   2   4   6E+5   13.5   14  RT (min)      lcorbi_t3_pos_1_008_36    m/z 437.3456     0   2   4   6   8E+5   13.5   14  RT (min)      lcorbi_t3_pos_1_008_36    m/z 437.3456     0   2   4E+5   12.5   13  RT (min)      lcorbi_t3_pos_4_016_15    m/z 437.3456     0   2   4   6   8E+4   15.5   16  RT (min)      lcorbi_t3_pos_1_023_02    m/z 437.3456     0   1   2   3E+5   15   15.5  RT (min)      lcorbi_t3_pos_1_001_49    m/z 437.3456     0   1   12   12.5  RT (min)      lcorbi_t3_pos_2_032_39    m/z 437.3456     0   1E+5   14   14.5  RT (min)      lcorbi_t3_pos_1_008_36    m/z 437.3456     0   1E+6   13.5   14  RT (min)      lcorbi_t3_pos_1_005_42    m/z 437.3458     0   2   4   6   8E+3   12.5   13  RT (min)      lcorbi_t3_pos_3_024_07    m/z 438.3789     0   1E+5   12   12.5  RT (min)      lcorbi_t3_pos_1_034_34    m/z 438.3789     0   2   4E+4   12.5   13  RT (min)      lcorbi_t3_pos_1_031_40    m/z 438.3790     0   2   4E+4   12.5   13  RT (min)      lcorbi_t3_pos_1_031_40    m/z 438.3790     0   1   2E+6   10.5   11  RT (min)       lcorbi_t3_pos_2_013_33    m/z 438.3790     0   1   2   3   4E+4   12.5   13  RT (min)      lcorbi_t3_pos_2_012_01    m/z 438.3790     0   1E+6   12   12.5  RT (min)      lcorbi_t3_pos_1_028_32    m/z 438.3790     0   2   4E+4   12.5   13  RT (min)      lcorbi_t3_pos_1_031_40    m/z 438.3790     0   1   2E+6   11.5   12  RT (min)      lcorbi_t3_pos_1_008_36    m/z 438.3790     0   1   2   3   4E+4   12.5   13  RT (min)      lcorbi_t3_pos_2_012_01    m/z 438.3790     0   2   4   6E+5   10.5   11  RT (min)      lcorbi_t3_pos_3_031_45    m/z 438.3790     0   1E+6   10.5   11  RT (min)      lcorbi_t3_pos_3_031_45    m/z 438.3790     0   1   2E+6   10.5   11  RT (min)      lcorbi_t3_pos_3_024_07    m/z 438.3790     0   1   2   3   4E+4   12.5   13  RT (min)      lcorbi_t3_pos_2_012_01    m/z 438.3790     0   1   2E+5   12   12.5  RT (min)      lcorbi_t3_pos_1_034_34    m/z 438.3790     0   2   4   6   8E+5   12   12.5  RT (min)      lcorbi_t3_pos_1_028_32    m/z 438.3790     0   2   4   6E+5   12   12.5  RT (min)      lcorbi_t3_pos_2_041_54    m/z 438.3790     0   1E+6   12   12.5  RT (min)      lcorbi_t3_pos_2_017_51    m/z 438.3790     0   1   2   3   4E+4   13   13.5  RT (min)      lcorbi_t3_pos_2_012_01    m/z 438.3791     0   1   2   3   4E+4   12.5   13  RT (min)      lcorbi_t3_pos_2_012_01    m/z 439.1881     0   2   4   6   8E+4   10.5   11  RT (min)      lcorbi_t3_pos_1_025_52    m/z 439.1881     0   1   2E+5   10   10.5  RT (min)      lcorbi_t3_pos_3_025_52    m/z 439.1882     0   1   2E+5   10   10.5  RT (min)      lcorbi_t3_pos_1_030_21    m/z 439.1882     0   1   2E+5   10   10.5  RT (min)      lcorbi_t3_pos_2_025_52    m/z 439.1884     0   1   2E+5   17   18  RT (min)       lcorbi_t3_pos_4_086_59    m/z 439.3823     0   1E+4   12.5   13  RT (min)      lcorbi_t3_pos_2_012_01    m/z 439.3823     0   1E+4   12.5   13  RT (min)      lcorbi_t3_pos_2_012_01    m/z 439.3824     0   1   2E+6   11   11.5  RT (min)       lcorbi_t3_pos_4_088_59    m/z 439.3824     0   1   2E+4   12.5   13  RT (min)      lcorbi_t3_pos_1_031_40    m/z 439.3824     0   1   2E+5   12   12.5  RT (min)      lcorbi_t3_pos_1_028_32    m/z 439.3824     0   2   4   6E+5   10.5   11  RT (min)      lcorbi_t3_pos_2_013_33    m/z 439.3824     0   1   2   3E+4   12   12.5  RT (min)      lcorbi_t3_pos_1_041_54    m/z 439.3824     0   1   2E+6   11   11.5  RT (min)      lcorbi_t3_pos_4_087_59    m/z 439.3824     0   1   2E+4   12.5   13  RT (min)      lcorbi_t3_pos_1_031_40    m/z 439.3824     0   2   4E+5   11.5   12  RT (min)      lcorbi_t3_pos_2_017_51    m/z 439.3824     0   2   4   6E+5   10.5   11  RT (min)      lcorbi_t3_pos_2_013_33    m/z 439.3824     0   2   4   6E+5   11.5   12  RT (min)      lcorbi_t3_pos_1_008_36    m/z 439.3824     0   1   2E+5   12   12.5  RT (min)      lcorbi_t3_pos_1_028_32    m/z 439.4198     0   2   4   6E+5   12.5   13  RT (min)       lcorbi_t3_pos_4_097_60    m/z 441.1885     0   2   4   6E+5   11.5   12  RT (min)      lcorbi_t3_pos_4_001_49    m/z 441.1886     0   1   2   3E+4   12   12.5  RT (min)      lcorbi_t3_pos_2_052_48    m/z 441.1886     0   2   4   6E+4   12   12.5  RT (min)      lcorbi_t3_pos_3_013_19    m/z 441.1886     0   1   2   3E+4   12   12.5  RT (min)      lcorbi_t3_pos_2_038_03    m/z 441.1886     0   2   4   6   8E+5   11.5   12  RT (min)      lcorbi_t3_pos_3_056_56    m/z 441.1886     0   1   2E+4   12   12.5  RT (min)      lcorbi_t3_pos_2_056_56    m/z 441.1886     0   2   4   6E+5   10.5   11  RT (min)      lcorbi_t3_pos_3_031_45    m/z 441.1886     0   2   4   6   8E+5   10.5   11  RT (min)      lcorbi_t3_pos_3_052_47    m/z 441.2248     0   1E+6   12.5   15   17.5  RT (min)       lcorbi_t3_pos_4_084_59    m/z 441.2249     0   1E+6   16   18  RT (min)       lcorbi_t3_pos_4_086_59    m/z 441.2249     0   2   4E+5   15   15.5  RT (min)      lcorbi_t3_pos_2_059_39    m/z 441.2249     0   1E+6   16   18  RT (min)       lcorbi_t3_pos_4_091_60    m/z 441.2249     0   1E+6   16   17   18  RT (min)       lcorbi_t3_pos_4_112_62    m/z 442.2282     0   1E+5   9   9.5  RT (min)      lcorbi_t3_pos_1_051_09    m/z 442.2282     0   1   2E+5   9   9.5  RT (min)      lcorbi_t3_pos_2_022_38    m/z 442.2282     0   1   2E+5   9   9.5  RT (min)      lcorbi_t3_pos_1_041_54    m/z 443.2438     0   2   4   6E+5   1   1.5  RT (min)       lcorbi_t3_pos_3_055_10    m/z 444.2553     0   1E+5   14.5   15   15.5  RT (min)       lcorbi_t3_pos_4_092_60    m/z 444.2554     0   1E+5   15   15.5  RT (min)       lcorbi_t3_pos_4_099_61    m/z 445.1202     0   1E+6   13.5   14  RT (min)      lcorbi_t3_pos_4_105_61    m/z 446.1207     0   2   4E+5   14   14.5  RT (min)      lcorbi_t3_pos_4_112_62    m/z 446.1208     0   2   4   6   8E+5   14   14.5  RT (min)       lcorbi_t3_pos_4_113_62    m/z 446.1209     0   2   4   6   8E+5   14   14.5  RT (min)      lcorbi_t3_pos_4_113_62    m/z 446.2154     0   1E+5   15   15.5  RT (min)       lcorbi_t3_pos_2_003_19    m/z 447.1177     0   2   4E+5   14   14.5  RT (min)      lcorbi_t3_pos_4_113_62    m/z 447.3470     0   1E+5   14   14.5  RT (min)      lcorbi_t3_pos_3_025_52    m/z 447.3470     0   2   4   6E+5   14   14.5  RT (min)       lcorbi_t3_pos_3_002_16    m/z 447.3470     0   1E+5   14.5   15  RT (min)      lcorbi_t3_pos_3_056_56    m/z 447.3470     0   1E+5   14.5   15  RT (min)      lcorbi_t3_pos_3_044_46    m/z 447.3470     0   2   4E+5   13.5   14  RT (min)      lcorbi_t3_pos_4_105_61    m/z 447.3471     0   2   4   6E+5   14   14.5  RT (min)       lcorbi_t3_pos_3_002_16    m/z 447.3471     0   2   4   6E+5   14   14.5  RT (min)      lcorbi_t3_pos_3_002_16    m/z 448.2694     0   1   2E+5   16   17  RT (min)       lcorbi_t3_pos_2_002_06    m/z 448.2694     0   1   2E+5   10   10.5  RT (min)      lcorbi_t3_pos_1_038_30    m/z 448.2694     0   1   2E+5   15   15.5  RT (min)       lcorbi_t3_pos_4_101_61    m/z 448.2694     0   1   2E+5   10   10.5  RT (min)      lcorbi_t3_pos_1_008_36    m/z 448.2694     0   1E+5   10.5   11  RT (min)      lcorbi_t3_pos_1_041_54    m/z 448.3504     0   1   2E+5   14   14.5  RT (min)      lcorbi_t3_pos_3_002_16    m/z 450.3576     0   1E+4   12   12.5  RT (min)      lcorbi_t3_pos_2_032_39    m/z 450.3578     0   1   2E+5   12.5   13  RT (min)      lcorbi_t3_pos_2_054_29    m/z 450.3578     0   1E+5   14   14.5  RT (min)      lcorbi_t3_pos_1_033_53    m/z 450.3579     0   2   4   6E+5   12.5   13  RT (min)      lcorbi_t3_pos_3_011_42    m/z 450.3579     0   1E+5   15.5   16  RT (min)      lcorbi_t3_pos_1_023_02    m/z 450.3579     0   2   4   6   8E+4   14   14.5  RT (min)      lcorbi_t3_pos_2_025_52    m/z 450.3579     0   1E+5   14.5   15  RT (min)      lcorbi_t3_pos_1_034_34    m/z 450.3579     0   2   4   6E+4   14   14.5  RT (min)      lcorbi_t3_pos_2_022_38    m/z 450.3579     0   1E+5   15   16  RT (min)       lcorbi_t3_pos_1_023_02    m/z 450.3579     0   1E+5   16  RT (min)       lcorbi_t3_pos_1_031_40    m/z 450.3579     0   2   4   6E+4   14   14.5  RT (min)      lcorbi_t3_pos_2_022_38    m/z 450.3579     0   1E+5   15   16  RT (min)       lcorbi_t3_pos_2_012_01    m/z 450.3580     0   1   2   3   4E+5   13.5   14  RT (min)      lcorbi_t3_pos_1_008_36    m/z 450.3580     0   1   2E+5   13.5   14  RT (min)      lcorbi_t3_pos_3_056_56    m/z 450.3580     0   2   4   6E+5   13.5   14  RT (min)       lcorbi_t3_pos_3_047_40    m/z 450.3580     0   2   4   6E+5   13.5   14  RT (min)      lcorbi_t3_pos_4_019_47    m/z 450.3580     0   1E+5   14   14.5  RT (min)      lcorbi_t3_pos_1_028_32    m/z 450.3580     0   1   2   3   4E+5   13.5   14  RT (min)      lcorbi_t3_pos_4_001_49    m/z 451.1017     0   2   4E+5   1   1.5  RT (min)       lcorbi_t3_pos_4_095_60    m/z 451.3614     0   1E+5   13.5   14  RT (min)      lcorbi_t3_pos_4_019_47    m/z 452.3946     0   1   2   3   4E+4   13   13.5  RT (min)      lcorbi_t3_pos_1_041_54    m/z 452.3946     0   1   2   3   4E+4   12.5   13  RT (min)      lcorbi_t3_pos_1_028_32    m/z 452.3946     0   1   2E+4   12.5   13  RT (min)      lcorbi_t3_pos_2_025_52    m/z 452.3947     0   2   4   6E+5   11.5   12  RT (min)      lcorbi_t3_pos_2_013_33    m/z 452.3947     0   1   2   3E+4   13   13.5  RT (min)      lcorbi_t3_pos_1_031_40    m/z 452.3947     0   2   4E+4   12.5   13  RT (min)      lcorbi_t3_pos_2_022_38    m/z 452.3947     0   2   4   6E+4   11   11.5  RT (min)      lcorbi_t3_pos_3_024_07    m/z 452.3948     0   2   4   6E+5   12.5   13  RT (min)      lcorbi_t3_pos_1_008_36    m/z 452.3948     0   2   4   6E+5   12.5   13  RT (min)      lcorbi_t3_pos_1_008_36    m/z 453.2248     0   1   2E+5   10.5   11  RT (min)      lcorbi_t3_pos_1_033_53    m/z 455.4148     0   2   4   6E+5   12.5   13  RT (min)      lcorbi_t3_pos_1_043_06    m/z 455.4148     0   2   4   6   8E+5   12.5   13  RT (min)       lcorbi_t3_pos_4_096_60    m/z 457.1366     0   1   2E+5   15.5   16   16.5  RT (min)       lcorbi_t3_pos_4_105_61    m/z 457.1988     0   1E+5   9   9.5  RT (min)      lcorbi_t3_pos_2_022_38    m/z 461.2696     0   1E+5   15   15.5  RT (min)       lcorbi_t3_pos_4_091_60    m/z 462.1470     0   1E+6   13.5   14  RT (min)      lcorbi_t3_pos_4_105_61    m/z 462.2658     0   2   4   6   8E+4   13   13.5  RT (min)      lcorbi_t3_pos_3_011_42    m/z 462.2660     0   1E+5   13.5   14  RT (min)      lcorbi_t3_pos_1_052_38    m/z 462.2660     0   1E+5   13   13.5  RT (min)      lcorbi_t3_pos_2_054_29    m/z 462.2660     0   1   2   3   4E+5   14   15  RT (min)       lcorbi_t3_pos_3_000_aa    m/z 462.2660     0   1   2   3E+5   15   15.5  RT (min)       lcorbi_t3_pos_4_098_61    m/z 462.2660     0   1   2   3   4E+5   14.5   15   15.5  RT (min)       lcorbi_t3_pos_4_095_60    m/z 462.2660     0   2   4   6   8E+4   12.5   13  RT (min)      lcorbi_t3_pos_3_031_45    m/z 463.1474     0   2   4   6   8E+5   14   14.5  RT (min)       lcorbi_t3_pos_4_113_62    m/z 463.1474     0   2   4   6E+5   14   14.5  RT (min)       lcorbi_t3_pos_4_112_62    m/z 463.1475     0   2   4   6E+5   14   14.5  RT (min)      lcorbi_t3_pos_4_112_62    m/z 463.1477     0   2   4E+5   13.5   14  RT (min)      lcorbi_t3_pos_4_105_61    m/z 463.2694     0   1E+5   15   15.5  RT (min)       lcorbi_t3_pos_4_091_60    m/z 464.1443     0   2   4E+5   14   14.5  RT (min)      lcorbi_t3_pos_4_113_62    m/z 464.3733     0   1   2   3   4E+4   12.5   13  RT (min)      lcorbi_t3_pos_3_024_07    m/z 464.3734     0   1   2E+5   12.5   13  RT (min)      lcorbi_t3_pos_1_002_01    m/z 464.3734     0   1   2   3   4E+4   12   12.5  RT (min)      lcorbi_t3_pos_2_032_39    m/z 464.3734     0   1E+5   14.5   15  RT (min)      lcorbi_t3_pos_1_031_40    m/z 464.3734     0   2   4   6E+4   15   15.5  RT (min)      lcorbi_t3_pos_1_051_09    m/z 464.3735     0   2   4   6   8E+4   14.5   15  RT (min)      lcorbi_t3_pos_1_034_34    m/z 464.3735     0   2   4   6   8E+4   15.5   16  RT (min)      lcorbi_t3_pos_1_023_02    m/z 464.3736     0   2   4E+4   14   14.5  RT (min)      lcorbi_t3_pos_3_003_33    m/z 464.3736     0   2   4E+5   12.5   13  RT (min)      lcorbi_t3_pos_4_016_15    m/z 464.3736     0   2   4E+5   13.5   14  RT (min)      lcorbi_t3_pos_4_019_47    m/z 464.3736     0   1   2   3E+5   13.5   14  RT (min)      lcorbi_t3_pos_3_047_40    m/z 464.3736     0   1   2E+5   13.5   14  RT (min)      lcorbi_t3_pos_1_008_36    m/z 464.3736     0   1   2   3E+4   12.5   13  RT (min)      lcorbi_t3_pos_4_012_37    m/z 466.2587     0   1E+6   10   12.5   15  RT (min)       lcorbi_t3_pos_4_074_58    m/z 466.2589     0   2   4   6   8E+5   16   16.5  RT (min)       lcorbi_t3_pos_4_067_57    m/z 466.2590     0   2   4   6   8E+5   16   16.5  RT (min)       lcorbi_t3_pos_4_070_57    m/z 467.2621     0   1   2   3E+5   10.5   11  RT (min)      lcorbi_t3_pos_3_044_46    m/z 467.2621     0   1   2   3E+5   10.5   11  RT (min)      lcorbi_t3_pos_4_019_47    m/z 468.3083     0   1   11.5   12  RT (min)      lcorbi_t3_pos_1_023_02    m/z 468.3084     0   1E+4   11   11.5  RT (min)      lcorbi_t3_pos_1_031_40    m/z 468.3084     0   2   4   6   8E+5   10.5   11  RT (min)      lcorbi_t3_pos_1_008_36    m/z 468.3084     0   1   2   3   4E+4   11   11.5  RT (min)      lcorbi_t3_pos_1_034_34    m/z 468.3084     0   1   2   3E+5   11   11.5  RT (min)      lcorbi_t3_pos_1_028_32    m/z 468.3084     0   1   11.5   12  RT (min)      lcorbi_t3_pos_1_023_02    m/z 468.3084     0   1E+4   11   11.5  RT (min)      lcorbi_t3_pos_1_031_40    m/z 468.3085     0   2   4   6   8E+5   10   10.5  RT (min)      lcorbi_t3_pos_2_032_39    m/z 468.3895     0   2   4E+5   10.5   11  RT (min)       lcorbi_t3_pos_3_031_45    m/z 468.3896     0   2   4E+5   10.5   11  RT (min)       lcorbi_t3_pos_2_013_33    m/z 468.3896     0   2   4   6   8E+4   11.5   12  RT (min)      lcorbi_t3_pos_1_033_53    m/z 468.3896     0   2   4   6E+5   11.5   12  RT (min)      lcorbi_t3_pos_1_008_36    m/z 468.3897     0   1   2   3   4E+4   12   12.5  RT (min)      lcorbi_t3_pos_1_056_56    m/z 468.3897     0   2   4E+4   12   12.5  RT (min)      lcorbi_t3_pos_1_041_54    m/z 471.2143     0   2   4E+5   17.5   18   18.5  RT (min)       lcorbi_t3_pos_4_074_58    m/z 473.3988     0   1   16   16.5  RT (min)      lcorbi_t3_pos_2_014_37    m/z 473.3989     0   1   2E+5   14   14.5  RT (min)      lcorbi_t3_pos_2_016_23    m/z 473.3989     0   2   4   6   8E+5   15   15.5  RT (min)      lcorbi_t3_pos_4_001_49    m/z 473.3990     0   1   2   3   4E+4   15.5   16  RT (min)      lcorbi_t3_pos_1_028_32    m/z 473.3990     0   1   13.5   14  RT (min)      lcorbi_t3_pos_2_039_30    m/z 473.3990     0   1   16.5   17  RT (min)    lcorbi_t3_pos_1_027_43    m/z 473.3990     0   1   16.5   17  RT (min)    lcorbi_t3_pos_2_052_48    m/z 473.3990     0   1   16.5   17  RT (min)    lcorbi_t3_pos_1_051_09    m/z 473.3990     0   1E+7   15   15.5  RT (min)      lcorbi_t3_pos_4_019_47    m/z 473.3990     0   1E+7   14   14.5  RT (min)      lcorbi_t3_pos_2_016_23    m/z 473.3990     0   2   4E+4   15.5   16  RT (min)      lcorbi_t3_pos_3_044_46    m/z 473.3990     0   1   16   16.5  RT (min)      lcorbi_t3_pos_1_034_34    m/z 473.3990     0   1   17.5   18  RT (min)    lcorbi_t3_pos_1_023_02    m/z 473.3990     0   1   13.5   14  RT (min)      lcorbi_t3_pos_2_047_31    m/z 473.3990     0   1   13.5   14  RT (min)      lcorbi_t3_pos_2_032_39    m/z 473.3990     0   1   2E+4   16   16.5  RT (min)      lcorbi_t3_pos_3_013_19    m/z 473.3991     0   1   2E+7   14   14.5  RT (min)      lcorbi_t3_pos_3_034_26    m/z 474.4023     0   1   17.5   18  RT (min)    lcorbi_t3_pos_1_023_02    m/z 474.4024     0   1   17   17.5  RT (min)    lcorbi_t3_pos_2_012_01    m/z 474.4024     0   1   15.5   16  RT (min)      lcorbi_t3_pos_1_030_21    m/z 474.4024     0   1   16.5   17  RT (min)    lcorbi_t3_pos_1_049_55    m/z 474.4024     0   1   15.5   16  RT (min)      lcorbi_t3_pos_1_028_32    m/z 474.4024     0   2   4   6E+6   15   15.5  RT (min)      lcorbi_t3_pos_4_019_47    m/z 474.4024     0   1   2   3E+6   14   14.5  RT (min)      lcorbi_t3_pos_2_016_23    m/z 474.4024     0   2   4   6   8E+4   14   14.5  RT (min)      lcorbi_t3_pos_2_016_23    m/z 474.4024     0   1   16   16.5  RT (min)      lcorbi_t3_pos_2_014_37    m/z 474.4024     0   1   16   16.5  RT (min)      lcorbi_t3_pos_1_041_54    m/z 474.4024     0   1   2E+5   15   15.5  RT (min)      lcorbi_t3_pos_4_001_49    m/z 474.4024     0   1E+4   15.5   16  RT (min)      lcorbi_t3_pos_3_044_46    m/z 474.4024     0   1   16.5   17  RT (min)    lcorbi_t3_pos_1_051_09    m/z 474.4024     0   1   13.5   14  RT (min)      lcorbi_t3_pos_2_032_39    m/z 474.4025     0   2   4   6E+6   15   15.5  RT (min)      lcorbi_t3_pos_4_019_47    m/z 474.4026     0   1   2E+4   15.5   16  RT (min)      lcorbi_t3_pos_3_044_46    m/z 474.4026     0   1   13.5   14  RT (min)      lcorbi_t3_pos_2_047_31    m/z 474.4026     0   1   16   16.5  RT (min)      lcorbi_t3_pos_1_034_34    m/z 474.4026     0   1   16.5   17  RT (min)    lcorbi_t3_pos_2_052_48    m/z 475.4056     0   1   15.5   16  RT (min)      lcorbi_t3_pos_1_055_33    m/z 475.4056     0   1   15.5   16  RT (min)      lcorbi_t3_pos_1_038_30    m/z 475.4056     0   1   17.5   18  RT (min)    lcorbi_t3_pos_1_023_02    m/z 475.4056     0   1   16.5   17  RT (min)    lcorbi_t3_pos_1_027_43    m/z 475.4056     0   1   16.5   17  RT (min)    lcorbi_t3_pos_1_051_09    m/z 475.4056     0   1   16   16.5  RT (min)      lcorbi_t3_pos_2_014_37    m/z 475.4057     0   1   13.5   14  RT (min)      lcorbi_t3_pos_2_013_33    m/z 475.4057     0   1   15.5   16  RT (min)      lcorbi_t3_pos_1_028_32    m/z 475.4057     0   1   16.5   17  RT (min)    lcorbi_t3_pos_1_051_09    m/z 475.4057     0   1   13.5   14  RT (min)      lcorbi_t3_pos_2_032_39    m/z 475.4057     0   1   2E+4   14   14.5  RT (min)      lcorbi_t3_pos_2_016_23    m/z 475.4058     0   1   16   16.5  RT (min)      lcorbi_t3_pos_1_034_34    m/z 475.4058     0   1   13.5   14  RT (min)      lcorbi_t3_pos_2_047_31    m/z 475.4058     0   1E+6   15   15.5  RT (min)      lcorbi_t3_pos_4_019_47    m/z 475.4058     0   1   15.5   16  RT (min)      lcorbi_t3_pos_3_044_46    m/z 477.3967     0   2   4E+5   12.5   13  RT (min)      lcorbi_t3_pos_4_096_60    m/z 478.2929     0   2   4E+5   11.5   12  RT (min)      lcorbi_t3_pos_1_008_36    m/z 478.2930     0   2   4E+4   11.5   12  RT (min)      lcorbi_t3_pos_1_021_18    m/z 478.2930     0   1E+4   11.5   12  RT (min)      lcorbi_t3_pos_1_034_34    m/z 478.3161     0   1E+6   10.5   11  RT (min)      lcorbi_t3_pos_4_056_56    m/z 478.3162     0   2   4   6E+5   10.5   11  RT (min)      lcorbi_t3_pos_4_056_56    m/z 478.3162     0   2   4   6   8E+5   10.5   11  RT (min)      lcorbi_t3_pos_4_056_56    m/z 478.3164     0   2   4E+5   13.5   14  RT (min)      lcorbi_t3_pos_1_057_49    m/z 478.3164     0   2   4   6   8E+5   15   15.5  RT (min)       lcorbi_t3_pos_4_073_57    m/z 478.3164     0   2   4   6   8E+5   16   16.5  RT (min)       lcorbi_t3_pos_4_069_57    m/z 478.3165     0   2   4   6   8E+5   16   16.5  RT (min)       lcorbi_t3_pos_4_072_57    m/z 478.3890     0   1E+7   13   13.5  RT (min)      lcorbi_t3_pos_4_076_58    m/z 478.3890     0   2   4   6   8E+4   12.5   13  RT (min)      lcorbi_t3_pos_3_024_07    m/z 478.3890     0   1E+7   12.5   13  RT (min)      lcorbi_t3_pos_4_016_15    m/z 478.3890     0   2   4   6   8E+4   12   12.5  RT (min)      lcorbi_t3_pos_2_032_39    m/z 478.3891     0   1   17.5   18  RT (min)    lcorbi_t3_pos_1_031_40    m/z 478.3891     0   1   18   18.5  RT (min)    lcorbi_t3_pos_1_031_40    m/z 478.3891     0   2   4   6E+5   16   16.5  RT (min)      lcorbi_t3_pos_4_049_55    m/z 478.3891     0   2   4E+6   12.5   13  RT (min)      lcorbi_t3_pos_1_002_01    m/z 478.3891     0   1E+5   12.5   13  RT (min)      lcorbi_t3_pos_4_012_37    m/z 478.3891     0   2   4   6E+5   15.5   16  RT (min)      lcorbi_t3_pos_3_005_25    m/z 478.3891     0   1   2E+7   13   13.5  RT (min)       lcorbi_t3_pos_4_063_17    m/z 478.3892     0   2   4   6   8E+6   13.5   14  RT (min)      lcorbi_t3_pos_4_019_47    m/z 478.3892     0   1   17   17.5  RT (min)    lcorbi_t3_pos_1_031_40    m/z 478.3892     0   2   4   6   8E+5   16   16.5  RT (min)      lcorbi_t3_pos_1_008_36    m/z 478.3892     0   2   4   6E+5   14   14.5  RT (min)      lcorbi_t3_pos_2_022_38    m/z 478.3892     0   2   4   6   8E+5   14   14.5  RT (min)      lcorbi_t3_pos_1_028_32    m/z 478.3892     0   1E+6   17.5   18  RT (min)      lcorbi_t3_pos_4_075_58    m/z 478.3892     0   1   16.5   17  RT (min)    lcorbi_t3_pos_1_008_36    m/z 478.3892     0   1   2   3E+6   13.5   14  RT (min)      lcorbi_t3_pos_1_008_36    m/z 478.3892     0   1E+6   16   17  RT (min)       lcorbi_t3_pos_4_075_58    m/z 478.3892     0   2   4   6E+5   15   15.5  RT (min)      lcorbi_t3_pos_1_051_09    m/z 478.3892     0   2   4   6E+5   16   16.5  RT (min)      lcorbi_t3_pos_4_051_34    m/z 478.3892     0   2   4   6   8E+5   14.5   15  RT (min)      lcorbi_t3_pos_1_034_34    m/z 478.3892     0   2   4   6E+5   15   15.5  RT (min)      lcorbi_t3_pos_1_051_09    m/z 478.3892     0   1   16.5   17  RT (min)    lcorbi_t3_pos_1_031_40    m/z 478.3892     0   1E+7   13   13.5  RT (min)       lcorbi_t3_pos_4_080_58    m/z 478.3892     0   2   4   6   8E+5   14.5   15  RT (min)      lcorbi_t3_pos_1_034_34    m/z 478.3892     0   1E+7   13   13.5  RT (min)       lcorbi_t3_pos_4_075_58    m/z 478.3892     0   1E+6   15.5   16  RT (min)      lcorbi_t3_pos_1_008_36    m/z 478.3892     0   1   17   17.5  RT (min)    lcorbi_t3_pos_1_031_40    m/z 478.3892     0   1   17   17.5  RT (min)    lcorbi_t3_pos_1_031_40    m/z 478.3892     0   1E+6   14   14.5  RT (min)      lcorbi_t3_pos_1_008_36    m/z 478.3892     0   1E+6   15.5   16  RT (min)      lcorbi_t3_pos_1_008_36    m/z 478.3892     0   1E+7   13   13.5  RT (min)      lcorbi_t3_pos_4_077_58    m/z 478.3892     0   2   4   6E+6   13.5   14  RT (min)      lcorbi_t3_pos_3_047_40    m/z 478.3893     0   2   4   6E+5   14   14.5  RT (min)      lcorbi_t3_pos_3_049_55    m/z 478.3893     0   1E+6   16   16.5   17  RT (min)       lcorbi_t3_pos_4_078_58    m/z 478.3893     0   1E+6   16   16.5  RT (min)       lcorbi_t3_pos_4_081_58    m/z 478.3893     0   2   4   6E+5   15   15.5  RT (min)      lcorbi_t3_pos_1_051_09    m/z 478.3893     0   1E+6   17.5   18  RT (min)      lcorbi_t3_pos_4_075_58    m/z 478.3893     0   1   2   3E+6   13.5   14  RT (min)      lcorbi_t3_pos_4_071_57    m/z 478.3893     0   2   4   6E+6   12.5   13  RT (min)      lcorbi_t3_pos_1_002_01    m/z 478.3893     0   1   17   17.5  RT (min)    lcorbi_t3_pos_1_031_40    m/z 478.3894     0   2   4   6E+5   0.5   1  RT (min)      lcorbi_t3_pos_4_057_49    m/z 478.3894     0   2   4   6   8E+5   16   16.5  RT (min)      lcorbi_t3_pos_1_008_36    m/z 478.3894     0   1   18   18.5  RT (min)    lcorbi_t3_pos_1_031_40    m/z 478.3894     0   1   18.5   19  RT (min)    lcorbi_t3_pos_1_051_09    m/z 479.3196     0   1   2   3E+5   10.5   11  RT (min)      lcorbi_t3_pos_3_025_52    m/z 479.3924     0   1   2E+4   12   12.5  RT (min)      lcorbi_t3_pos_2_032_39    m/z 479.3925     0   2   4E+4   12.5   13  RT (min)      lcorbi_t3_pos_4_012_37    m/z 479.3925     0   1   2E+6   12.5   13  RT (min)      lcorbi_t3_pos_4_016_15    m/z 479.3925     0   1   2E+6   12.5   13  RT (min)      lcorbi_t3_pos_2_054_29    m/z 479.3926     0   1   2E+6   13.5   14  RT (min)      lcorbi_t3_pos_3_047_40    m/z 479.3926     0   2   4   6E+5   13.5   14  RT (min)      lcorbi_t3_pos_1_008_36    m/z 479.3926     0   2   4   6   8E+5   13.5   14  RT (min)      lcorbi_t3_pos_1_008_36    m/z 479.3926     0   1   2E+6   13.5   14  RT (min)      lcorbi_t3_pos_4_019_47    m/z 479.3926     0   1   2   3E+5   15   15.5  RT (min)      lcorbi_t3_pos_1_031_40    m/z 479.3926     0   1   2E+5   15   15.5  RT (min)      lcorbi_t3_pos_2_012_01    m/z 479.3926     0   1   2E+5   15   15.5  RT (min)      lcorbi_t3_pos_1_051_09    m/z 479.3926     0   1   2E+5   14.5   15  RT (min)      lcorbi_t3_pos_1_034_34    m/z 479.3926     0   1   2   3   4E+5   16   16.5  RT (min)       lcorbi_t3_pos_4_075_58    m/z 479.3926     0   1   2E+5   14.5   15  RT (min)      lcorbi_t3_pos_1_034_34    m/z 479.3927     0   1   2E+5   15   15.5  RT (min)      lcorbi_t3_pos_1_051_09    m/z 480.3954     0   1E+4   12.5   13  RT (min)       lcorbi_t3_pos_4_012_37    m/z 480.3957     0   1   2   3   4E+5   13.5   14  RT (min)      lcorbi_t3_pos_3_047_40    m/z 480.3958     0   2   4   6   8E+4   12.5   13  RT (min)      lcorbi_t3_pos_2_054_29    m/z 482.3242     0   2   4   6E+4   11   11.5  RT (min)      lcorbi_t3_pos_1_028_32    m/z 482.3242     0   2   4E+5   10   10.5  RT (min)       lcorbi_t3_pos_1_002_01    m/z 482.3242     0   1E+5   11.5   12  RT (min)      lcorbi_t3_pos_1_034_34    m/z 482.3242     0   2   4   6E+5   11   11.5  RT (min)      lcorbi_t3_pos_1_008_36    m/z 482.3605     0   1E+5   12   12.5  RT (min)      lcorbi_t3_pos_2_022_38    m/z 482.3606     0   1   2E+4   12.5   13  RT (min)      lcorbi_t3_pos_1_034_34    m/z 482.3606     0   1   12.5   13  RT (min)      lcorbi_t3_pos_2_012_01    m/z 482.3606     0   2   4   6E+4   12   12.5  RT (min)      lcorbi_t3_pos_3_003_33    m/z 482.3606     0   1   12.5   13  RT (min)      lcorbi_t3_pos_1_031_40    m/z 482.3606     0   1   2E+4   12   12.5  RT (min)      lcorbi_t3_pos_1_034_34    m/z 482.3606     0   1   2E+4   13   13.5  RT (min)      lcorbi_t3_pos_1_023_02    m/z 482.3688     0   1E+4   11.5   12  RT (min)      lcorbi_t3_pos_1_044_25    m/z 482.4051     0   1   2   3   4E+4   12.5   13  RT (min)      lcorbi_t3_pos_2_012_01    m/z 482.4052     0   1   2   3   4E+4   12.5   13  RT (min)      lcorbi_t3_pos_2_012_01    m/z 482.4052     0   1   2   3E+5   12   12.5  RT (min)      lcorbi_t3_pos_2_025_52    m/z 482.4052     0   2   4E+4   12   12.5   13  RT (min)      lcorbi_t3_pos_1_025_52    m/z 482.4052     0   1E+5   12   12.5  RT (min)      lcorbi_t3_pos_1_034_34    m/z 482.4052     0   2   4   6E+4   12   12.5  RT (min)      lcorbi_t3_pos_2_009_50    m/z 482.4052     0   2   4E+4   12   12.5  RT (min)      lcorbi_t3_pos_1_025_52    m/z 482.4052     0   1E+5   12   12.5  RT (min)      lcorbi_t3_pos_1_034_34    m/z 482.4052     0   1E+6   11.5   12  RT (min)      lcorbi_t3_pos_2_017_51    m/z 482.4052     0   1   2E+6   10.5   11  RT (min)      lcorbi_t3_pos_2_013_33    m/z 482.4052     0   1E+6   11.5   12  RT (min)      lcorbi_t3_pos_2_022_38    m/z 482.4052     0   2   4E+5   12   12.5  RT (min)      lcorbi_t3_pos_2_025_52    m/z 482.4053     0   1   2   3   4E+4   12.5   13  RT (min)      lcorbi_t3_pos_2_012_01    m/z 482.4053     0   1   2E+6   10.5   11  RT (min)       lcorbi_t3_pos_3_031_45    m/z 482.4053     0   1   2   3   4E+4   12.5   13  RT (min)      lcorbi_t3_pos_2_012_01    m/z 482.4053     0   2   4   6   8E+5   10.5   11  RT (min)      lcorbi_t3_pos_3_031_45    m/z 482.4053     0   2   4   6E+5   12   12.5  RT (min)      lcorbi_t3_pos_1_028_32    m/z 482.4053     0   1E+6   10.5   11  RT (min)      lcorbi_t3_pos_2_013_33    m/z 482.4053     0   1E+6   1   1.5  RT (min)      lcorbi_t3_pos_4_057_49    m/z 482.4054     0   1   2E+6   10.5   11  RT (min)       lcorbi_t3_pos_2_013_33    m/z 483.2719     0   2   4E+5   17   18  RT (min)       lcorbi_t3_pos_4_088_59    m/z 483.2719     0   2   4E+5   17   17.5  RT (min)       lcorbi_t3_pos_4_073_57    m/z 483.2722     0   2   4E+5   17   18  RT (min)       lcorbi_t3_pos_4_076_58    m/z 483.4086     0   1   2   3E+5   11.5   12  RT (min)      lcorbi_t3_pos_1_033_53    m/z 483.4086     0   1   2E+4   12   12.5   13  RT (min)      lcorbi_t3_pos_1_025_52    m/z 483.4086     0   1   2E+4   12   12.5  RT (min)      lcorbi_t3_pos_1_025_52    m/z 483.4086     0   1   2E+4   12   12.5  RT (min)      lcorbi_t3_pos_1_034_34    m/z 483.4086     0   1E+4   12.5   13  RT (min)      lcorbi_t3_pos_2_012_01    m/z 483.4086     0   1   2E+6   11   11.5  RT (min)      lcorbi_t3_pos_4_082_59    m/z 483.4086     0   1   2E+4   12   12.5  RT (min)      lcorbi_t3_pos_2_009_50    m/z 483.4086     0   1E+4   12.5   13  RT (min)      lcorbi_t3_pos_2_012_01    m/z 483.4087     0   1   2E+6   11   11.5  RT (min)       lcorbi_t3_pos_4_082_59    m/z 483.4087     0   2   4E+4   12   12.5  RT (min)      lcorbi_t3_pos_1_034_34    m/z 483.4088     0   2   4E+5   11.5   12  RT (min)      lcorbi_t3_pos_2_017_51    m/z 483.4088     0   2   4E+5   10.5   11  RT (min)      lcorbi_t3_pos_2_013_33    m/z 483.4088     0   2   4E+5   10.5   11  RT (min)      lcorbi_t3_pos_2_013_33    m/z 483.4088     0   1   2   3E+5   11.5   12  RT (min)       lcorbi_t3_pos_4_041_54    m/z 483.4088     0   2   4E+5   10.5   11  RT (min)      lcorbi_t3_pos_3_024_07    m/z 483.4088     0   1   2   3   4E+4   12   12.5  RT (min)      lcorbi_t3_pos_1_034_34    m/z 484.2693     0   1E+6   9.5   10  RT (min)      lcorbi_t3_pos_1_033_53    m/z 484.2693     0   1E+6   9   9.5  RT (min)      lcorbi_t3_pos_2_013_33    m/z 484.2693     0   1E+6   9.5   10  RT (min)      lcorbi_t3_pos_1_055_33    m/z 484.2693     0   1E+6   9.5   10  RT (min)      lcorbi_t3_pos_1_028_32    m/z 484.2693     0   1E+6   9   9.5  RT (min)      lcorbi_t3_pos_1_026_17    m/z 484.2693     0   1E+6   10   15  RT (min)       lcorbi_t3_pos_4_074_58    m/z 484.2693     0   2   4   6   8E+5   9.5   10  RT (min)      lcorbi_t3_pos_2_046_07    m/z 484.2694     0   1   2E+5   10.5   11  RT (min)      lcorbi_t3_pos_1_033_53    m/z 484.2694     0   1E+6   9   9.5  RT (min)      lcorbi_t3_pos_2_013_33    m/z 484.2694     0   1E+5   10   10.5  RT (min)      lcorbi_t3_pos_1_001_49    m/z 484.2696     0   1E+6   17   18  RT (min)       lcorbi_t3_pos_4_086_59    m/z 484.2696     0   1E+6   15.5   16   16.5  RT (min)       lcorbi_t3_pos_4_105_61    m/z 484.2696     0   1E+6   15   16  RT (min)       lcorbi_t3_pos_4_113_62    m/z 484.2696     0   1E+6   16   16.5  RT (min)       lcorbi_t3_pos_4_067_57    m/z 484.2696     0   1E+6   16   16.5  RT (min)       lcorbi_t3_pos_4_091_60    m/z 484.2696     0   1E+6   16   17  RT (min)       lcorbi_t3_pos_4_098_61    m/z 484.2697     0   1E+6   16.5   17  RT (min)       lcorbi_t3_pos_4_083_59    m/z 484.2700     0   1E+6   16   17  RT (min)       lcorbi_t3_pos_4_085_59    m/z 484.2700     0   1E+6   16   17   18  RT (min)       lcorbi_t3_pos_4_112_62    m/z 485.2727     0   1   2   3E+5   9.5   10  RT (min)      lcorbi_t3_pos_1_008_36    m/z 485.2727     0   1   2   3E+5   9.5   10  RT (min)      lcorbi_t3_pos_1_033_53    m/z 485.2728     0   2   4   6   8E+4   10   10.5  RT (min)      lcorbi_t3_pos_1_025_52    m/z 485.2728     0   1   2E+5   9.5   10  RT (min)      lcorbi_t3_pos_3_025_52    m/z 485.2728     0   1E+5   10   10.5  RT (min)      lcorbi_t3_pos_1_049_55    m/z 485.2729     0   1   2   3E+5   15   16   17  RT (min)       lcorbi_t3_pos_4_066_57    m/z 485.2730     0   1   2   3E+5   15.5   16  RT (min)       lcorbi_t3_pos_4_075_58    m/z 487.1881     0   1   2E+5   10.5   11  RT (min)      lcorbi_t3_pos_3_047_40    m/z 487.1884     0   1E+5   18   18.5  RT (min)       lcorbi_t3_pos_4_113_62    m/z 489.2250     0   2   4   6   8E+5   14   16   18  RT (min)       lcorbi_t3_pos_4_098_61    m/z 490.2281     0   1E+5   9.5   10  RT (min)      lcorbi_t3_pos_2_041_54    m/z 490.3166     0   1E+5   13   13.5  RT (min)       lcorbi_t3_pos_4_075_58    m/z 490.4255     0   1   2   3E+7   15   15.5  RT (min)      lcorbi_t3_pos_4_019_47    m/z 490.4255     0   1E+4   13.5   14  RT (min)      lcorbi_t3_pos_2_039_30    m/z 490.4255     0   2   4E+5   14   14.5  RT (min)      lcorbi_t3_pos_2_016_23    m/z 490.4255     0   1   16.5   17  RT (min)    lcorbi_t3_pos_1_051_09    m/z 490.4255     0   1   13.5   14  RT (min)      lcorbi_t3_pos_2_047_31    m/z 490.4255     0   1   16.5   17  RT (min)     lcorbi_t3_pos_2_052_48    m/z 490.4255     0   1   16.5   17  RT (min)    lcorbi_t3_pos_1_051_09    m/z 490.4256     0   1   2   3E+7   14   14.5  RT (min)      lcorbi_t3_pos_3_053_48    m/z 490.4256     0   1   2   3   4E+4   15.5   16  RT (min)      lcorbi_t3_pos_1_028_32    m/z 490.4256     0   1   2E+4   16   16.5  RT (min)       lcorbi_t3_pos_1_034_34    m/z 490.4256     0   1   2E+4   15.5   16  RT (min)      lcorbi_t3_pos_1_055_33    m/z 490.4256     0   2   4   6   8E+4   15.5   16  RT (min)      lcorbi_t3_pos_3_044_46    m/z 490.4256     0   2   4E+5   15   15.5  RT (min)      lcorbi_t3_pos_3_025_52    m/z 490.4256     0   1   13.5   14  RT (min)      lcorbi_t3_pos_2_032_39    m/z 490.4256     0   1   2E+7   14   14.5  RT (min)      lcorbi_t3_pos_2_016_23    m/z 490.4260     0   1   17.5   18  RT (min)     lcorbi_t3_pos_1_023_02    m/z 491.4288     0   1E+5   14   14.5  RT (min)      lcorbi_t3_pos_2_016_23    m/z 491.4289     0   1   16.5   17  RT (min)    lcorbi_t3_pos_1_051_09    m/z 491.4289     0   1E+7   15   15.5  RT (min)      lcorbi_t3_pos_4_019_47    m/z 491.4289     0   1   13.5   14  RT (min)      lcorbi_t3_pos_2_039_30    m/z 491.4290     0   1   16.5   17  RT (min)    lcorbi_t3_pos_2_052_48    m/z 491.4290     0   1   13.5   14  RT (min)      lcorbi_t3_pos_2_047_31    m/z 491.4290     0   2   4   6E+6   14   14.5  RT (min)      lcorbi_t3_pos_2_016_23    m/z 491.4290     0   1   2   3   4E+5   15   15.5  RT (min)      lcorbi_t3_pos_4_001_49    m/z 491.4290     0   1   2E+3   13.5   14  RT (min)      lcorbi_t3_pos_2_032_39    m/z 491.4290     0   1   16   16.5  RT (min)      lcorbi_t3_pos_1_034_34    m/z 491.4290     0   1E+7   14   14.5  RT (min)      lcorbi_t3_pos_3_034_26    m/z 491.4290     0   1   15.5   16  RT (min)      lcorbi_t3_pos_1_055_33    m/z 491.4290     0   1   2   3   4E+4   15.5   16  RT (min)      lcorbi_t3_pos_3_044_46    m/z 491.4290     0   1   2E+4   15.5   16  RT (min)      lcorbi_t3_pos_1_028_32    m/z 491.4291     0   1   17.5   18  RT (min)    lcorbi_t3_pos_1_023_02    m/z 491.4291     0   1   16   16.5  RT (min)      lcorbi_t3_pos_2_014_37    m/z 491.4291     0   1   2E+4   15   15.5  RT (min)      lcorbi_t3_pos_1_008_36    m/z 492.4046     0   1   2   3   4E+4   12   12.5  RT (min)      lcorbi_t3_pos_2_032_39    m/z 492.4047     0   1   2   3E+5   15   15.5  RT (min)      lcorbi_t3_pos_1_051_09    m/z 492.4048     0   2   4   6E+4   12.5   13  RT (min)      lcorbi_t3_pos_4_012_37    m/z 492.4048     0   1   2   3E+6   12.5   13  RT (min)      lcorbi_t3_pos_4_016_15    m/z 492.4048     0   2   4E+4   12.5   13  RT (min)      lcorbi_t3_pos_2_032_39    m/z 492.4049     0   1   2E+5   14   14.5  RT (min)      lcorbi_t3_pos_3_049_55    m/z 492.4049     0   1   2   3E+5   14.5   15  RT (min)      lcorbi_t3_pos_4_033_53    m/z 492.4049     0   1   2   3E+5   14.5   15  RT (min)      lcorbi_t3_pos_1_034_34    m/z 492.4049     0   1   2   3E+5   15   15.5  RT (min)      lcorbi_t3_pos_1_051_09    m/z 492.4049     0   1   2   3E+5   15   15.5  RT (min)      lcorbi_t3_pos_1_051_09    m/z 492.4049     0   2   4   6E+5   16.5   17  RT (min)      lcorbi_t3_pos_4_075_58    m/z 492.4049     0   1   2E+5   14   14.5  RT (min)      lcorbi_t3_pos_2_022_38    m/z 492.4049     0   1   2   3   4E+5   15   16  RT (min)       lcorbi_t3_pos_1_023_02    m/z 492.4049     0   2   4E+5   14   14.5  RT (min)      lcorbi_t3_pos_3_047_40    m/z 492.4049     0   2   4E+5   17   17.5  RT (min)       lcorbi_t3_pos_4_071_57    m/z 492.4049     0   2   4   6E+5   12.5   13  RT (min)      lcorbi_t3_pos_2_054_29    m/z 492.4050     0   1   2   3E+5   14.5   15  RT (min)      lcorbi_t3_pos_1_034_34    m/z 492.4050     0   2   4E+5   17  RT (min)       lcorbi_t3_pos_4_075_58    m/z 492.4050     0   1   2   3E+6   13.5   14  RT (min)      lcorbi_t3_pos_3_047_40    m/z 492.4050     0   2   4E+5   13.5   14  RT (min)      lcorbi_t3_pos_1_008_36    m/z 492.4050     0   2   4E+5   16   17  RT (min)       lcorbi_t3_pos_4_070_57    m/z 492.4050     0   1   2   3E+6   12.5   13  RT (min)      lcorbi_t3_pos_2_013_33    m/z 492.4050     0   2   4   6E+5   15.5   16  RT (min)       lcorbi_t3_pos_4_075_58    m/z 492.4050     0   2   4   6E+5   13.5   14  RT (min)      lcorbi_t3_pos_3_047_40    m/z 492.4050     0   1   2E+6   13.5   14  RT (min)      lcorbi_t3_pos_3_033_53    m/z 492.4050     0   2   4   6E+5   16   16.5  RT (min)       lcorbi_t3_pos_4_069_57    m/z 492.4050     0   1   2   3E+5   15   15.5  RT (min)      lcorbi_t3_pos_1_051_09    m/z 492.4050     0   1E+6   13.5   14  RT (min)      lcorbi_t3_pos_1_008_36    m/z 492.4051     0   2   4E+5   16.5   17  RT (min)      lcorbi_t3_pos_4_076_58    m/z 492.4052     0   2   4E+5   17   17.5  RT (min)      lcorbi_t3_pos_4_079_58    m/z 492.4053     0   2   4E+5   17   17.5  RT (min)       lcorbi_t3_pos_4_074_58    m/z 492.4321     0   1   16.5   17  RT (min)    lcorbi_t3_pos_1_031_40    m/z 492.4322     0   1   13.5   14  RT (min)      lcorbi_t3_pos_2_032_39    m/z 492.4322     0   1   15.5   16  RT (min)      lcorbi_t3_pos_1_030_21    m/z 492.4322     0   2   4   6E+4   14   14.5  RT (min)      lcorbi_t3_pos_2_016_23    m/z 492.4322     0   1   16.5   17  RT (min)    lcorbi_t3_pos_1_051_09    m/z 492.4322     0   1   13.5   14  RT (min)      lcorbi_t3_pos_2_031_20    m/z 492.4322     0   1   16.5   17  RT (min)    lcorbi_t3_pos_1_051_09    m/z 492.4322     0   1   16   16.5  RT (min)      lcorbi_t3_pos_1_041_54    m/z 492.4322     0   2   4   6   8E+4   15   15.5  RT (min)      lcorbi_t3_pos_4_001_49    m/z 492.4322     0   1   13.5   14  RT (min)      lcorbi_t3_pos_2_047_31    m/z 492.4322     0   1E+6   15   15.5  RT (min)      lcorbi_t3_pos_4_019_47    m/z 492.4322     0   1E+6   15   15.5  RT (min)      lcorbi_t3_pos_4_019_47    m/z 492.4322     0   1   15.5   16  RT (min)      lcorbi_t3_pos_1_055_33    m/z 492.4322     0   1   2   3E+6   14   14.5  RT (min)      lcorbi_t3_pos_4_105_61    m/z 492.4322     0   1   17   17.5  RT (min)    lcorbi_t3_pos_1_023_02    m/z 492.4322     0   1   15.5   16  RT (min)      lcorbi_t3_pos_3_044_46    m/z 492.4322     0   1   16   16.5  RT (min)      lcorbi_t3_pos_1_034_34    m/z 492.4323     0   1   15.5   16  RT (min)      lcorbi_t3_pos_1_028_32    m/z 492.4324     0   1   16   16.5  RT (min)      lcorbi_t3_pos_2_041_54    m/z 493.4082     0   1   2E+4   12   12.5  RT (min)      lcorbi_t3_pos_2_032_39    m/z 493.4082     0   1   2E+4   12.5   13  RT (min)      lcorbi_t3_pos_4_012_37    m/z 493.4082     0   1E+6   12.5   13  RT (min)      lcorbi_t3_pos_4_016_15    m/z 493.4082     0   2   4   6   8E+5   12.5   13  RT (min)      lcorbi_t3_pos_1_002_01    m/z 493.4082     0   2   4   6   8E+4   14   14.5  RT (min)      lcorbi_t3_pos_3_013_19    m/z 493.4082     0   1E+5   14.5   15  RT (min)      lcorbi_t3_pos_1_031_40    m/z 493.4083     0   1E+6   12.5   13  RT (min)      lcorbi_t3_pos_4_016_15    m/z 493.4083     0   1E+5   15.5   16  RT (min)      lcorbi_t3_pos_1_023_02    m/z 493.4083     0   2   4   6E+5   12.5   13  RT (min)      lcorbi_t3_pos_2_054_29    m/z 493.4083     0   1   2   3E+5   13.5   14  RT (min)      lcorbi_t3_pos_1_008_36    m/z 493.4083     0   1E+5   14   14.5  RT (min)      lcorbi_t3_pos_1_047_41    m/z 493.4084     0   1E+5   14   14.5  RT (min)      lcorbi_t3_pos_2_025_52    m/z 493.4084     0   1E+5   15.5   16  RT (min)      lcorbi_t3_pos_1_023_02    m/z 493.4084     0   1   2E+5   13.5   14  RT (min)      lcorbi_t3_pos_1_008_36    m/z 493.4084     0   1E+5   14.5   15  RT (min)      lcorbi_t3_pos_1_034_34    m/z 493.4084     0   2   4E+5   13.5   14  RT (min)      lcorbi_t3_pos_1_008_36    m/z 493.4084     0   1E+5   15   15.5  RT (min)      lcorbi_t3_pos_1_051_09    m/z 493.4084     0   1E+5   15   15.5  RT (min)      lcorbi_t3_pos_2_012_01    m/z 493.4084     0   1E+5   14.5   15  RT (min)      lcorbi_t3_pos_1_034_34    m/z 494.3195     0   1   13   13.5  RT (min)      lcorbi_t3_pos_2_031_20    m/z 494.3195     0   2   4E+5   13   13.5  RT (min)      lcorbi_t3_pos_2_016_23    m/z 494.3196     0   1   2   3   4E+5   14   14.5  RT (min)       lcorbi_t3_pos_3_015_06    m/z 494.3196     0   2   4   6   8E+4   14.5   15  RT (min)      lcorbi_t3_pos_1_038_30    m/z 494.3196     0   1E+4   12.5   13  RT (min)      lcorbi_t3_pos_2_047_31    m/z 494.3196     0   1   13   13.5  RT (min)      lcorbi_t3_pos_2_039_30    m/z 494.3196     0   2   4   6   8E+4   15   15.5  RT (min)      lcorbi_t3_pos_1_038_30    m/z 494.3196     0   1   2   3   4E+4   15   15.5  RT (min)      lcorbi_t3_pos_2_014_37    m/z 494.3196     0   2   4   6E+4   15.5   16  RT (min)      lcorbi_t3_pos_1_027_43    m/z 494.3196     0   2   4E+5   13.5   14  RT (min)      lcorbi_t3_pos_3_053_48    m/z 494.3196     0   2   4   6E+4   15.5   16  RT (min)      lcorbi_t3_pos_2_052_48    m/z 494.3196     0   1   2   3   4E+4   15.5   16  RT (min)      lcorbi_t3_pos_2_052_48    m/z 494.3197     0   1   2   3   4E+5   14   14.5  RT (min)       lcorbi_t3_pos_3_015_06    m/z 494.3197     0   2   4   6E+4   15.5   16  RT (min)      lcorbi_t3_pos_1_027_43    m/z 494.3240     0   1E+6   10   10.5  RT (min)      lcorbi_t3_pos_3_024_07    m/z 494.3240     0   1E+6   10   10.5  RT (min)      lcorbi_t3_pos_2_032_39    m/z 494.3242     0   1E+5   11   11.5  RT (min)      lcorbi_t3_pos_1_034_34    m/z 494.3242     0   1E+4   11.5   12  RT (min)      lcorbi_t3_pos_2_012_01    m/z 494.3242     0   1   12   12.5  RT (min)      lcorbi_t3_pos_1_023_02    m/z 494.3242     0   1E+6   11   11.5  RT (min)      lcorbi_t3_pos_1_008_36    m/z 494.3242     0   1E+4   12   12.5  RT (min)      lcorbi_t3_pos_1_023_02    m/z 494.3242     0   2   4   6E+5   11   11.5  RT (min)      lcorbi_t3_pos_1_028_32    m/z 494.3242     0   1E+4   11.5   12  RT (min)      lcorbi_t3_pos_1_031_40    m/z 494.3242     0   1E+4   11.5   12  RT (min)      lcorbi_t3_pos_1_031_40    m/z 494.3242     0   1E+6   11   11.5  RT (min)      lcorbi_t3_pos_2_022_38    m/z 494.3242     0   1E+4   12   12.5  RT (min)      lcorbi_t3_pos_1_023_02    m/z 496.3052     0   2   4E+5   11   11.5  RT (min)      lcorbi_t3_pos_4_075_58    m/z 496.3176     0   1   2   3E+5   10.5   11  RT (min)      lcorbi_t3_pos_2_032_39    m/z 496.3177     0   2   4E+5   10.5   11  RT (min)      lcorbi_t3_pos_1_002_01    m/z 496.3180     0   2   4E+5   10.5   11  RT (min)      lcorbi_t3_pos_1_002_01    m/z 496.3180     0   1   12   12.5  RT (min)      lcorbi_t3_pos_1_031_40    m/z 496.3180     0   2   4E+5   11.5   12  RT (min)      lcorbi_t3_pos_1_008_36    m/z 496.3180     0   1   12   12.5  RT (min)      lcorbi_t3_pos_1_034_34    m/z 496.3180     0   1   11.5   12  RT (min)      lcorbi_t3_pos_1_028_32    m/z 496.3180     0   1   12.5   13  RT (min)      lcorbi_t3_pos_1_023_02    m/z 496.3182     0   1   12   12.5  RT (min)      lcorbi_t3_pos_1_051_09    m/z 496.3395     0   1   2   3E+7   1   1.5  RT (min)      lcorbi_t3_pos_4_057_49    m/z 496.3396     0   1E+6   12   12.5  RT (min)      lcorbi_t3_pos_3_013_19    m/z 496.3396     0   2   4   6   8E+7   10.5   11  RT (min)       lcorbi_t3_pos_1_002_01    m/z 496.3397     0   1E+5   12.5   13  RT (min)      lcorbi_t3_pos_1_023_02    m/z 496.3397     0   2   4   6E+6   11.5   12  RT (min)      lcorbi_t3_pos_1_028_32    m/z 496.3398     0   2   4   6E+4   12.5   13  RT (min)      lcorbi_t3_pos_2_012_01    m/z 496.3398     0   2   4   6   8E+7   11.5   12  RT (min)      lcorbi_t3_pos_1_008_36    m/z 496.3398     0   2   4   6   8E+6   10.5   11  RT (min)      lcorbi_t3_pos_3_024_07    m/z 496.3398     0   1E+5   12   12.5  RT (min)      lcorbi_t3_pos_1_001_49    m/z 496.3398     0   1E+5   12   12.5  RT (min)      lcorbi_t3_pos_1_031_40    m/z 496.3398     0   2   4E+5   12   12.5  RT (min)      lcorbi_t3_pos_1_034_34    m/z 496.3398     0   1   2E+7   11.5   12  RT (min)      lcorbi_t3_pos_2_022_38    m/z 496.3398     0   2   4   6E+7   10.5   11  RT (min)      lcorbi_t3_pos_3_031_45    m/z 496.3613     0   2   4E+5   10.5   11  RT (min)      lcorbi_t3_pos_1_002_01    m/z 497.2665     0   1E+6   11.5   12  RT (min)      lcorbi_t3_pos_1_054_48    m/z 497.2666     0   2   4   6E+5   11.5   12  RT (min)       lcorbi_t3_pos_4_083_59    m/z 497.2667     0   1E+6   12   12.5  RT (min)       lcorbi_t3_pos_1_054_48    m/z 497.3430     0   2   4   6   8E+6   1   1.5  RT (min)      lcorbi_t3_pos_4_057_49    m/z 497.3431     0   1E+6   11.5   12  RT (min)      lcorbi_t3_pos_1_028_32    m/z 497.3432     0   1E+7   10.5   11  RT (min)      lcorbi_t3_pos_3_031_45    m/z 497.3432     0   1   2   3E+4   12   12.5  RT (min)      lcorbi_t3_pos_1_031_40    m/z 497.3432     0   2   4   6E+6   11.5   12  RT (min)      lcorbi_t3_pos_2_022_38    m/z 497.3432     0   1   2E+7   11.5   12  RT (min)      lcorbi_t3_pos_1_008_36    m/z 497.3432     0   2   4   6   8E+6   11.5   12  RT (min)      lcorbi_t3_pos_3_049_55    m/z 497.3432     0   1E+5   12   12.5  RT (min)      lcorbi_t3_pos_1_034_34    m/z 497.3432     0   1   2E+7   10.5   11  RT (min)      lcorbi_t3_pos_1_002_01    m/z 497.3432     0   1   2   3E+4   12.5   13  RT (min)      lcorbi_t3_pos_1_023_02    m/z 497.3433     0   1   2   3   4E+4   12   12.5  RT (min)      lcorbi_t3_pos_1_049_55    m/z 497.3433     0   1E+4   12   12.5  RT (min)      lcorbi_t3_pos_1_025_52    m/z 497.3433     0   1   2E+5   12   12.5  RT (min)      lcorbi_t3_pos_3_013_19    m/z 497.3434     0   1   2E+6   10.5   11  RT (min)      lcorbi_t3_pos_2_013_33    m/z 498.3457     0   1   12.5   13  RT (min)      lcorbi_t3_pos_1_023_02    m/z 498.3458     0   1   12.5   13  RT (min)      lcorbi_t3_pos_1_023_02    m/z 498.3458     0   2   4E+4   12   12.5  RT (min)      lcorbi_t3_pos_3_013_19    m/z 498.3458     0   1E+4   12   12.5  RT (min)      lcorbi_t3_pos_1_031_40    m/z 498.3459     0   1E+5   11.5   12  RT (min)      lcorbi_t3_pos_1_028_32    m/z 498.3459     0   1   12   12.5  RT (min)      lcorbi_t3_pos_1_051_09    m/z 498.3460     0   2   4   6E+5   11.5   12  RT (min)      lcorbi_t3_pos_2_022_38    m/z 498.3460     0   1   2E+6   10.5   11  RT (min)      lcorbi_t3_pos_3_031_45    m/z 498.3460     0   1   2   3   4E+6   11.5   12  RT (min)      lcorbi_t3_pos_1_008_36    m/z 498.3460     0   1   2   3E+6   10.5   11  RT (min)      lcorbi_t3_pos_1_002_01    m/z 498.3460     0   1E+4   12   12.5  RT (min)      lcorbi_t3_pos_1_034_34    m/z 502.2929     0   1E+6   10.5   11  RT (min)      lcorbi_t3_pos_2_054_29    m/z 502.2929     0   2   4E+5   10.5   11  RT (min)      lcorbi_t3_pos_3_024_07    m/z 502.2930     0   1E+5   11.5   12  RT (min)      lcorbi_t3_pos_3_013_19    m/z 502.2930     0   1   12.5   13  RT (min)      lcorbi_t3_pos_1_023_02    m/z 502.2930     0   1   12   12.5  RT (min)      lcorbi_t3_pos_1_023_02    m/z 502.2930     0   1   12   12.5  RT (min)      lcorbi_t3_pos_1_051_09    m/z 502.2930     0   1E+4   12   12.5  RT (min)      lcorbi_t3_pos_1_051_09    m/z 502.2930     0   1   2E+4   11.5   12  RT (min)      lcorbi_t3_pos_1_034_34    m/z 505.1988     0   1E+5   9.5   10  RT (min)       lcorbi_t3_pos_2_017_51    m/z 508.3760     0   2   4   6   8E+5   12   12.5  RT (min)       lcorbi_t3_pos_1_008_36    m/z 508.3760     0   2   4   6   8E+5   11   11.5  RT (min)      lcorbi_t3_pos_2_013_33    m/z 508.3760     0   1E+6   11   11.5  RT (min)      lcorbi_t3_pos_1_002_01    m/z 508.3760     0   2   4   6   8E+4   12.5   13  RT (min)      lcorbi_t3_pos_1_028_32    m/z 508.3760     0   1   13   13.5  RT (min)      lcorbi_t3_pos_2_012_01    m/z 508.3760     0   1   13   13.5  RT (min)      lcorbi_t3_pos_1_051_09    m/z 508.3760     0   1E+4   13   13.5  RT (min)      lcorbi_t3_pos_1_023_02    m/z 508.3761     0   2   4   6   8E+4   12   12.5  RT (min)      lcorbi_t3_pos_1_028_32    m/z 508.3761     0   1E+4   11   11.5  RT (min)      lcorbi_t3_pos_2_032_39    m/z 508.3761     0   2   4   6   8E+4   12.5   13  RT (min)      lcorbi_t3_pos_1_028_32    m/z 508.3762     0   1   13   13.5  RT (min)      lcorbi_t3_pos_1_051_09    m/z 509.3864     0   1E+6   12   12.5  RT (min)      lcorbi_t3_pos_2_008_16    m/z 510.2658     0   1E+5   15   16  RT (min)       lcorbi_t3_pos_4_093_60    m/z 510.2658     0   1E+5   15   15.5  RT (min)       lcorbi_t3_pos_4_102_61    m/z 510.3553     0   1E+4   12   12.5  RT (min)      lcorbi_t3_pos_1_028_32    m/z 510.3553     0   1   12.5   13  RT (min)      lcorbi_t3_pos_2_012_01    m/z 510.3554     0   1   12.5   13  RT (min)      lcorbi_t3_pos_1_034_34    m/z 510.3554     0   1E+6   11   11.5  RT (min)      lcorbi_t3_pos_2_054_29    m/z 510.3554     0   1E+6   12   12.5  RT (min)      lcorbi_t3_pos_1_008_36    m/z 510.3554     0   1   2   3   4E+5   11   11.5  RT (min)      lcorbi_t3_pos_3_031_45    m/z 510.3554     0   1   12.5   13  RT (min)      lcorbi_t3_pos_1_031_40    m/z 510.3554     0   1   13   13.5  RT (min)      lcorbi_t3_pos_1_023_02    m/z 510.3554     0   1   2   3E+4   12   12.5  RT (min)      lcorbi_t3_pos_3_003_33    m/z 510.3554     0   1E+4   12.5   13  RT (min)      lcorbi_t3_pos_1_034_34    m/z 510.3555     0   1   13   13.5  RT (min)      lcorbi_t3_pos_1_023_02    m/z 512.4157     0   1   2   3E+5   10.5   11  RT (min)      lcorbi_t3_pos_2_013_33    m/z 514.3057     0   1   12.5   13  RT (min)      lcorbi_t3_pos_1_041_54    m/z 514.3057     0   1   2   3E+5   12   12.5  RT (min)      lcorbi_t3_pos_2_017_51    m/z 514.3058     0   1   13   13.5  RT (min)      lcorbi_t3_pos_1_025_52    m/z 514.3058     0   1   2E+4   12   12.5  RT (min)      lcorbi_t3_pos_1_030_21    m/z 514.3058     0   1   13   13.5  RT (min)      lcorbi_t3_pos_1_049_55    m/z 514.3059     0   1E+4   12.5   13  RT (min)      lcorbi_t3_pos_2_025_52    m/z 514.3059     0   2   4E+5   12   12.5  RT (min)      lcorbi_t3_pos_3_025_52    m/z 515.1910     0   2   4   6E+4   9.5   10  RT (min)      lcorbi_t3_pos_1_023_02    m/z 518.3217     0   1   12   12.5  RT (min)      lcorbi_t3_pos_1_031_40    m/z 518.3217     0   1   12.5   13  RT (min)      lcorbi_t3_pos_1_023_02    m/z 518.3218     0   1   2E+6   1   1.5  RT (min)      lcorbi_t3_pos_4_057_49    m/z 518.3218     0   2   4   6E+4   11.5   12  RT (min)      lcorbi_t3_pos_3_013_19    m/z 518.3218     0   1   12.5   13  RT (min)      lcorbi_t3_pos_1_023_02    m/z 518.3218     0   1E+5   11.5   12  RT (min)      lcorbi_t3_pos_1_028_32    m/z 518.3219     0   1   2E+6   11.5   12  RT (min)      lcorbi_t3_pos_3_047_40    m/z 518.3220     0   1   2E+4   12   12.5  RT (min)      lcorbi_t3_pos_1_034_34    m/z 518.3221     0   1   2   3E+6   11.5   12  RT (min)      lcorbi_t3_pos_1_008_36    m/z 518.3229     0   1E+4   12   12.5  RT (min)      lcorbi_t3_pos_2_025_52    m/z 518.3240     0   1E+6   10   10.5  RT (min)      lcorbi_t3_pos_3_024_07    m/z 518.3240     0   1E+6   10   10.5  RT (min)      lcorbi_t3_pos_2_013_33    m/z 518.4566     0   1   17.5   18  RT (min)    lcorbi_t3_pos_1_023_02    m/z 518.4568     0   2   4   6E+5   15   15.5  RT (min)      lcorbi_t3_pos_1_005_42    m/z 518.4569     0   1   14   14.5  RT (min)      lcorbi_t3_pos_1_002_01    m/z 518.4569     0   1   16.5   17  RT (min)    lcorbi_t3_pos_1_001_49    m/z 518.4569     0   1   13.5   14  RT (min)      lcorbi_t3_pos_2_032_39    m/z 518.4569     0   1   15.5   16  RT (min)      lcorbi_t3_pos_1_038_30    m/z 518.4570     0   1   15.5   16  RT (min)      lcorbi_t3_pos_1_028_32    m/z 518.4570     0   1E+6   14   14.5  RT (min)      lcorbi_t3_pos_4_105_61    m/z 519.3250     0   1E+4   12.5   13  RT (min)      lcorbi_t3_pos_1_023_02    m/z 519.3250     0   2   4   6E+5   11.5   12  RT (min)      lcorbi_t3_pos_3_047_40    m/z 519.3250     0   1   12   12.5  RT (min)      lcorbi_t3_pos_1_051_09    m/z 519.3252     0   2   4   6   8E+5   11.5   12  RT (min)      lcorbi_t3_pos_1_008_36    m/z 519.3252     0   1   12   12.5  RT (min)      lcorbi_t3_pos_1_034_34    m/z 519.3252     0   1   12   12.5  RT (min)      lcorbi_t3_pos_1_031_40    m/z 519.3263     0   1E+5   11.5   12  RT (min)      lcorbi_t3_pos_2_022_38    m/z 520.3350     0   2   4   6E+6   10   10.5  RT (min)      lcorbi_t3_pos_3_031_45    m/z 520.3396     0   1   2E+7   11   11.5  RT (min)      lcorbi_t3_pos_1_008_36    m/z 520.3396     0   1   2E+5   11.5   12  RT (min)      lcorbi_t3_pos_1_034_34    m/z 520.3396     0   2   4   6E+6   1   1.5  RT (min)      lcorbi_t3_pos_4_057_49    m/z 520.3396     0   1   2   3   4E+4   12   12.5  RT (min)      lcorbi_t3_pos_1_051_09    m/z 520.3396     0   1   2E+7   10.5   11  RT (min)      lcorbi_t3_pos_3_031_45    m/z 520.3396     0   1   2   3E+6   11.5   12  RT (min)      lcorbi_t3_pos_1_028_32    m/z 520.3397     0   1   2E+5   11.5   12  RT (min)      lcorbi_t3_pos_1_034_34    m/z 520.3397     0   1   2   3E+4   12   12.5  RT (min)      lcorbi_t3_pos_2_012_01    m/z 520.3397     0   1   2   3   4E+6   11.5   12  RT (min)      lcorbi_t3_pos_3_003_33    m/z 520.3397     0   1   2E+4   12   12.5  RT (min)      lcorbi_t3_pos_1_023_02    m/z 520.3398     0   1E+7   11   11.5  RT (min)      lcorbi_t3_pos_1_033_53    m/z 521.3430     0   1   2E+6   10   10.5  RT (min)      lcorbi_t3_pos_2_032_39    m/z 521.3431     0   2   4   6E+6   10.5   11  RT (min)      lcorbi_t3_pos_3_031_45    m/z 521.3431     0   2   4   6E+4   11.5   12  RT (min)      lcorbi_t3_pos_1_034_34    m/z 521.3431     0   1E+4   12   12.5  RT (min)      lcorbi_t3_pos_1_031_40    m/z 521.3431     0   2   4   6E+5   11.5   12  RT (min)      lcorbi_t3_pos_1_028_32    m/z 521.3432     0   2   4   6E+4   11.5   12  RT (min)      lcorbi_t3_pos_1_034_34    m/z 521.3432     0   1E+4   12   12.5  RT (min)      lcorbi_t3_pos_1_023_02    m/z 521.3432     0   1E+4   12   12.5  RT (min)      lcorbi_t3_pos_1_023_02    m/z 521.3432     0   2   4   6E+6   11   11.5  RT (min)       lcorbi_t3_pos_1_008_36    m/z 521.3432     0   1E+6   11.5   12  RT (min)      lcorbi_t3_pos_3_003_33    m/z 521.3432     0   1E+4   12   12.5  RT (min)      lcorbi_t3_pos_1_051_09    m/z 522.3459     0   2   4   6   8E+5   10.5   11  RT (min)      lcorbi_t3_pos_3_031_45    m/z 522.3460     0   1   11.5   12  RT (min)      lcorbi_t3_pos_3_056_56    m/z 522.3460     0   2   4   6   8E+5   10.5   11  RT (min)      lcorbi_t3_pos_2_013_33    m/z 522.3460     0   2   4   6   8E+5   10.5   11  RT (min)      lcorbi_t3_pos_2_013_33    m/z 522.3551     0   2   4   6E+6   1   1.5  RT (min)      lcorbi_t3_pos_4_057_49    m/z 522.3552     0   2   4   6   8E+5   12   12.5  RT (min)      lcorbi_t3_pos_1_028_32    m/z 522.3553     0   1   2   3   4E+6   12   12.5  RT (min)      lcorbi_t3_pos_2_022_38    m/z 522.3553     0   1E+7   11.5   12  RT (min)      lcorbi_t3_pos_1_008_36    m/z 522.3553     0   1   2E+4   12   12.5  RT (min)      lcorbi_t3_pos_2_009_50    m/z 522.3553     0   2   4   6   8E+4   12   12.5  RT (min)      lcorbi_t3_pos_1_034_34    m/z 522.3554     0   1   2E+4   12.5   13  RT (min)      lcorbi_t3_pos_2_012_01    m/z 522.3554     0   1   2   3E+5   12   12.5  RT (min)      lcorbi_t3_pos_3_013_19    m/z 522.3554     0   1   2E+4   12.5   13  RT (min)      lcorbi_t3_pos_1_051_09    m/z 522.3554     0   1   2E+4   12.5   13  RT (min)      lcorbi_t3_pos_1_031_40    m/z 522.3554     0   1E+7   11.5   12  RT (min)      lcorbi_t3_pos_1_008_36    m/z 522.3595     0   2   4   6E+4   13   13.5  RT (min)      lcorbi_t3_pos_1_023_02    m/z 522.3634     0   1   13   13.5  RT (min)      lcorbi_t3_pos_1_023_02    m/z 523.3587     0   1   2E+4   13   13.5  RT (min)      lcorbi_t3_pos_1_023_02    m/z 523.3587     0   2   4   6   8E+4   12   12.5  RT (min)      lcorbi_t3_pos_3_013_19    m/z 523.3588     0   1   2   3   4E+6   10.5   11  RT (min)      lcorbi_t3_pos_2_032_39    m/z 523.3588     0   1E+4   12   12.5  RT (min)      lcorbi_t3_pos_1_041_54    m/z 523.3588     0   1   2   3   4E+6   11   11.5  RT (min)       lcorbi_t3_pos_2_054_29    m/z 523.3588     0   1   2E+4   13   13.5  RT (min)      lcorbi_t3_pos_1_023_02    m/z 523.3588     0   1   2   3   4E+6   11.5   12  RT (min)      lcorbi_t3_pos_1_008_36    m/z 523.3588     0   1   12.5   13  RT (min)      lcorbi_t3_pos_2_012_01    m/z 523.3588     0   1   2E+5   12   12.5  RT (min)      lcorbi_t3_pos_1_028_32    m/z 523.3588     0   1E+6   12   12.5  RT (min)      lcorbi_t3_pos_2_022_38    m/z 523.3588     0   1   2E+4   12   12.5  RT (min)      lcorbi_t3_pos_1_034_34    m/z 523.3588     0   1   12.5   13  RT (min)      lcorbi_t3_pos_1_031_40    m/z 523.3588     0   1   12.5   13  RT (min)      lcorbi_t3_pos_2_012_01    m/z 523.3589     0   1   2   3   4E+6   11.5   12  RT (min)      lcorbi_t3_pos_1_008_36    m/z 523.3589     0   1E+6   11.5   12  RT (min)      lcorbi_t3_pos_2_017_51    m/z 523.3590     0   2   4   6   8E+5   10.5   11  RT (min)      lcorbi_t3_pos_2_032_39    m/z 524.3578     0   1   2E+4   12   12.5  RT (min)       lcorbi_t3_pos_3_041_54    m/z 524.3616     0   2   4   6E+5   11   11.5  RT (min)      lcorbi_t3_pos_2_054_29    m/z 524.3616     0   2   4   6E+5   11.5   12  RT (min)       lcorbi_t3_pos_2_044_21    m/z 524.3708     0   1E+7   1.5   2  RT (min)      lcorbi_t3_pos_4_057_49    m/z 524.3708     0   2   4E+4   13.5   14  RT (min)      lcorbi_t3_pos_1_023_02    m/z 524.3708     0   1E+6   12.5   13  RT (min)      lcorbi_t3_pos_2_022_38    m/z 524.3708     0   1   2E+5   12.5   13  RT (min)      lcorbi_t3_pos_1_028_32    m/z 524.3709     0   2   4   6E+4   12.5   13  RT (min)      lcorbi_t3_pos_1_034_34    m/z 524.3709     0   1   2E+4   13   13.5  RT (min)      lcorbi_t3_pos_2_012_01    m/z 524.3709     0   1   2   3   4E+7   12   12.5  RT (min)      lcorbi_t3_pos_1_008_36    m/z 524.3709     0   1   2   3E+4   13   13.5  RT (min)      lcorbi_t3_pos_1_031_40    m/z 524.3710     0   1   2   3E+4   13   13.5  RT (min)      lcorbi_t3_pos_1_051_09    m/z 524.3710     0   2   4E+4   13.5   14  RT (min)      lcorbi_t3_pos_1_023_02    m/z 524.3710     0   1   2   3E+4   13   13.5  RT (min)      lcorbi_t3_pos_2_012_01    m/z 524.3711     0   1   2   3   4E+4   12.5   13  RT (min)      lcorbi_t3_pos_2_009_50    m/z 525.3604     0   2   4   6   8E+5   11.5   12  RT (min)      lcorbi_t3_pos_4_096_60    m/z 525.3604     0   1E+6   11.5   12  RT (min)       lcorbi_t3_pos_2_037_27    m/z 525.3604     0   1E+6   11.5   12  RT (min)       lcorbi_t3_pos_4_091_60    m/z 525.3604     0   2   4   6   8E+5   11.5   12  RT (min)      lcorbi_t3_pos_4_092_60    m/z 525.3604     0   1E+6   11.5   12  RT (min)      lcorbi_t3_pos_1_003_45    m/z 525.3609     0   1E+4   12   12.5  RT (min)       lcorbi_t3_pos_1_008_36    m/z 525.3742     0   2   4   6   8E+6   12   12.5  RT (min)      lcorbi_t3_pos_3_047_40    m/z 525.3742     0   2   4E+5   12.5   13  RT (min)      lcorbi_t3_pos_2_022_38    m/z 525.3743     0   1   2E+4   13.5   14  RT (min)      lcorbi_t3_pos_1_023_02    m/z 525.3744     0   1E+5   12.5   13  RT (min)      lcorbi_t3_pos_1_028_32    m/z 525.3744     0   1   2E+4   13.5   14  RT (min)      lcorbi_t3_pos_1_023_02    m/z 525.3744     0   2   4   6   8E+6   11.5   12  RT (min)      lcorbi_t3_pos_3_034_26    m/z 525.3744     0   1   2E+4   13   13.5  RT (min)      lcorbi_t3_pos_1_051_09    m/z 525.3744     0   1E+4   13   13.5  RT (min)      lcorbi_t3_pos_1_031_40    m/z 525.3744     0   1   2E+4   13   13.5  RT (min)      lcorbi_t3_pos_1_051_09    m/z 525.3744     0   2   4   6E+6   11   11.5  RT (min)      lcorbi_t3_pos_4_012_37    m/z 525.3744     0   1   2E+4   12.5   13  RT (min)      lcorbi_t3_pos_1_034_34    m/z 525.3744     0   1E+4   13   13.5  RT (min)      lcorbi_t3_pos_2_009_50    m/z 525.3745     0   1E+4   13   13.5  RT (min)      lcorbi_t3_pos_1_041_54    m/z 525.3745     0   1   2E+4   13   13.5  RT (min)      lcorbi_t3_pos_1_051_09    m/z 525.3745     0   1E+7   12   12.5  RT (min)      lcorbi_t3_pos_1_008_36    m/z 525.3745     0   1   11   11.5  RT (min)      lcorbi_t3_pos_2_032_39    m/z 526.2928     0   1   12.5   13  RT (min)      lcorbi_t3_pos_1_023_02    m/z 526.3164     0   1   2   3E+5   15   16  RT (min)       lcorbi_t3_pos_2_003_19    m/z 526.3772     0   1E+4   12.5   13  RT (min)      lcorbi_t3_pos_1_028_32    m/z 526.3772     0   1E+6   12   12.5  RT (min)      lcorbi_t3_pos_1_008_36    m/z 526.3772     0   1   13   13.5  RT (min)      lcorbi_t3_pos_1_031_40    m/z 526.3772     0   1E+6   12   12.5  RT (min)      lcorbi_t3_pos_3_047_40    m/z 526.3772     0   1   13   13.5  RT (min)      lcorbi_t3_pos_1_031_40    m/z 526.3772     0   1E+6   11.5   12  RT (min)      lcorbi_t3_pos_2_054_29    m/z 526.3772     0   2   4   6E+5   12   12.5  RT (min)      lcorbi_t3_pos_3_025_52    m/z 526.3772     0   1   13.5   14  RT (min)      lcorbi_t3_pos_1_023_02    m/z 526.3772     0   2   4   6E+4   12.5   13  RT (min)      lcorbi_t3_pos_2_022_38    m/z 526.3772     0   1E+5   11   11.5  RT (min)      lcorbi_t3_pos_2_032_39    m/z 526.3772     0   1   12.5   13  RT (min)      lcorbi_t3_pos_1_034_34    m/z 526.3773     0   1   13.5   14  RT (min)      lcorbi_t3_pos_1_023_02    m/z 526.4312     0   1   2   3   4E+4   12.5   13  RT (min)      lcorbi_t3_pos_2_012_01    m/z 526.4312     0   1   2E+5   12   12.5  RT (min)      lcorbi_t3_pos_1_034_34    m/z 526.4312     0   1   2   3   4E+4   12.5   13  RT (min)      lcorbi_t3_pos_1_025_52    m/z 526.4312     0   2   4   6E+5   10.5   11  RT (min)      lcorbi_t3_pos_3_024_07    m/z 526.4312     0   2   4   6   8E+4   12   12.5  RT (min)      lcorbi_t3_pos_1_028_32    m/z 526.4312     0   1   2   3   4E+4   12   12.5  RT (min)      lcorbi_t3_pos_2_056_56    m/z 526.4312     0   1E+6   10.5   11  RT (min)      lcorbi_t3_pos_3_031_45    m/z 526.4313     0   1E+6   10.5   11  RT (min)      lcorbi_t3_pos_2_013_33    m/z 526.4314     0   1   2   3   4E+4   12   12.5  RT (min)      lcorbi_t3_pos_1_056_56    m/z 526.4314     0   1E+6   11.5   12  RT (min)      lcorbi_t3_pos_2_022_38    m/z 526.4314     0   1   2   3   4E+5   11.5   12  RT (min)      lcorbi_t3_pos_1_033_53    m/z 526.4314     0   1   2   3   4E+4   12   12.5   13  RT (min)      lcorbi_t3_pos_1_025_52    m/z 526.4314     0   1   2   3   4E+4   12   12.5  RT (min)      lcorbi_t3_pos_1_025_52    m/z 526.4314     0   2   4E+5   11.5   12  RT (min)      lcorbi_t3_pos_2_041_54    m/z 526.4314     0   1E+6   11.5   12  RT (min)      lcorbi_t3_pos_1_033_53    m/z 526.4314     0   1E+6   11.5   12  RT (min)      lcorbi_t3_pos_1_008_36    m/z 526.4314     0   1E+6   10.5   11  RT (min)       lcorbi_t3_pos_2_013_33    m/z 526.4314     0   1   2   3   4E+4   12.5   13  RT (min)      lcorbi_t3_pos_2_056_56    m/z 527.3196     0   1E+5   11   11.5  RT (min)      lcorbi_t3_pos_3_047_40    m/z 527.4347     0   1   2E+4   12   12.5  RT (min)      lcorbi_t3_pos_1_056_56    m/z 527.4347     0   2   4E+5   10.5   11  RT (min)      lcorbi_t3_pos_3_031_45    m/z 527.4348     0   1   2E+4   12   12.5  RT (min)      lcorbi_t3_pos_1_025_52    m/z 527.4348     0   1E+5   11.5   12  RT (min)      lcorbi_t3_pos_2_041_54    m/z 527.4348     0   2   4   6E+4   12   12.5  RT (min)      lcorbi_t3_pos_1_034_34    m/z 527.4348     0   1   2E+6   11   11.5  RT (min)       lcorbi_t3_pos_4_082_59    m/z 531.2717     0   1E+5   11   11.5  RT (min)       lcorbi_t3_pos_4_001_49    m/z 531.2718     0   1   2E+5   11   11.5  RT (min)      lcorbi_t3_pos_1_008_36    m/z 531.4076     0   1   2   3E+4   13   13.5  RT (min)      lcorbi_t3_pos_2_039_30    m/z 531.4077     0   1   2E+4   12.5   13  RT (min)      lcorbi_t3_pos_2_032_39    m/z 531.4078     0   2   4E+6   13   13.5   14  RT (min)       lcorbi_t3_pos_3_053_48    m/z 531.4078     0   1   2E+5   15   15.5  RT (min)      lcorbi_t3_pos_2_049_55    m/z 531.4078     0   1   2E+4   13   13.5  RT (min)      lcorbi_t3_pos_2_039_30    m/z 531.4078     0   2   4E+6   13   13.5  RT (min)      lcorbi_t3_pos_2_016_23    m/z 531.4078     0   2   4E+4   13   13.5  RT (min)      lcorbi_t3_pos_2_054_29    m/z 531.4078     0   1   2E+5   16   16.5  RT (min)      lcorbi_t3_pos_1_023_02    m/z 531.4078     0   2   4E+6   13   13.5  RT (min)      lcorbi_t3_pos_4_015_45    m/z 531.4079     0   1   2E+5   15   15.5  RT (min)      lcorbi_t3_pos_2_052_48    m/z 531.4079     0   1   2E+5   15.5   16  RT (min)      lcorbi_t3_pos_1_027_43    m/z 531.4079     0   1   2E+5   14.5   15  RT (min)      lcorbi_t3_pos_1_038_30    m/z 531.4079     0   1   2E+5   15.5   16  RT (min)      lcorbi_t3_pos_1_025_52    m/z 531.4079     0   1E+5   15.5   16  RT (min)      lcorbi_t3_pos_2_012_01    m/z 531.4079     0   2   4E+6   13   13.5  RT (min)      lcorbi_t3_pos_3_053_48    m/z 531.4080     0   1   2E+5   15   15.5  RT (min)       lcorbi_t3_pos_2_041_54    m/z 531.4080     0   1   2E+5   14.5   15  RT (min)      lcorbi_t3_pos_1_055_33    m/z 531.4080     0   1   2E+5   16   16.5  RT (min)      lcorbi_t3_pos_1_023_02    m/z 531.4080     0   1   2E+5   15   15.5  RT (min)      lcorbi_t3_pos_2_025_52    m/z 531.4080     0   1   2E+5   15.5   16  RT (min)      lcorbi_t3_pos_1_001_49    m/z 531.4080     0   1   2E+5   14.5   15  RT (min)       lcorbi_t3_pos_1_038_30    m/z 532.4112     0   1E+5   14.5   15  RT (min)       lcorbi_t3_pos_1_055_33    m/z 532.4112     0   1E+4   12.5   13  RT (min)      lcorbi_t3_pos_2_032_39    m/z 532.4112     0   1E+5   14   14.5  RT (min)      lcorbi_t3_pos_3_056_56    m/z 532.4112     0   2   4   6   8E+4   15.5   16  RT (min)      lcorbi_t3_pos_1_027_43    m/z 532.4112     0   1E+5   14   14.5  RT (min)      lcorbi_t3_pos_3_025_52    m/z 532.4112     0   1E+6   13   13.5  RT (min)      lcorbi_t3_pos_2_016_23    m/z 532.4112     0   2   4   6   8E+4   15.5   16  RT (min)      lcorbi_t3_pos_1_027_43    m/z 532.4112     0   1E+4   13   13.5  RT (min)      lcorbi_t3_pos_2_031_20    m/z 532.4112     0   1E+5   14.5   15  RT (min)      lcorbi_t3_pos_3_044_46    m/z 532.4113     0   1E+6   14   14.5  RT (min)       lcorbi_t3_pos_3_047_40    m/z 532.4113     0   1E+6   14   14.5  RT (min)       lcorbi_t3_pos_1_005_42    m/z 532.4113     0   1E+6   13   13.5  RT (min)      lcorbi_t3_pos_3_053_48    m/z 532.4113     0   1E+5   14.5   15  RT (min)      lcorbi_t3_pos_1_038_30    m/z 532.4113     0   2   4   6   8E+4   15   15.5  RT (min)       lcorbi_t3_pos_2_014_37    m/z 532.4113     0   1E+4   12.5   13  RT (min)      lcorbi_t3_pos_2_047_31    m/z 532.4113     0   1E+5   15   15.5  RT (min)      lcorbi_t3_pos_2_041_54    m/z 532.4113     0   2   4   6E+4   16   16.5  RT (min)       lcorbi_t3_pos_1_023_02    m/z 532.4113     0   2   4   6   8E+4   15   15.5  RT (min)      lcorbi_t3_pos_2_052_48    m/z 532.4113     0   2   4   6   8E+4   15.5   16  RT (min)      lcorbi_t3_pos_2_056_56    m/z 532.4113     0   2   4   6   8E+4   15.5   16  RT (min)      lcorbi_t3_pos_2_056_56    m/z 532.4114     0   2   4   6   8E+4   15   15.5  RT (min)      lcorbi_t3_pos_2_025_52    m/z 532.4114     0   2   4   6   8E+4   15   15.5  RT (min)      lcorbi_t3_pos_2_038_03    m/z 532.4114     0   1E+5   14.5   15  RT (min)      lcorbi_t3_pos_1_047_41    m/z 532.4624     0   2   4E+5   11.5   12  RT (min)       lcorbi_t3_pos_1_003_45    m/z 536.1652     0   1   17   17.5  RT (min)    lcorbi_t3_pos_1_023_02    m/z 536.1654     0   1   16   16.5  RT (min)      lcorbi_t3_pos_1_049_55    m/z 536.1654     0   1   17   17.5  RT (min)    lcorbi_t3_pos_1_023_02    m/z 536.1656     0   1   15   15.5  RT (min)       lcorbi_t3_pos_4_028_04    m/z 536.1656     0   2   4E+5   14.5   15  RT (min)      lcorbi_t3_pos_4_051_34    m/z 536.1656     0   1   15.5   16  RT (min)      lcorbi_t3_pos_1_034_34    m/z 536.1657     0   1E+7   14   14.5  RT (min)      lcorbi_t3_pos_4_105_61    m/z 537.1665     0   2   4E+6   14   14.5  RT (min)      lcorbi_t3_pos_4_105_61    m/z 537.5353     0   1E+5   15.5   16  RT (min)      lcorbi_t3_pos_2_025_52    m/z 537.5354     0   1   2E+4   13   13.5  RT (min)      lcorbi_t3_pos_2_047_31    m/z 537.5354     0   1   16.5   17  RT (min)    lcorbi_t3_pos_2_012_01    m/z 537.5354     0   1   2   3E+4   13.5   14  RT (min)      lcorbi_t3_pos_2_039_30    m/z 537.5354     0   1   2E+5   15.5   16  RT (min)      lcorbi_t3_pos_1_041_54    m/z 537.5354     0   1   2   3E+5   15   15.5  RT (min)      lcorbi_t3_pos_3_044_46    m/z 537.5354     0   1   2   3   4E+6   0.5   1  RT (min)      lcorbi_t3_pos_4_057_49    m/z 537.5354     0   1   16.5   17  RT (min)    lcorbi_t3_pos_2_056_56    m/z 537.5354     0   1   2   3E+6   13.5   14  RT (min)      lcorbi_t3_pos_2_016_23    m/z 537.5354     0   1E+5   16   16.5  RT (min)      lcorbi_t3_pos_1_025_52    m/z 537.5354     0   1   2   3   4E+6   13.5   14  RT (min)      lcorbi_t3_pos_2_054_29    m/z 537.5354     0   1   2E+5   15   16   17  RT (min)       lcorbi_t3_pos_1_055_33    m/z 537.5354     0   1E+6   14.5   15  RT (min)      lcorbi_t3_pos_4_025_52    m/z 537.5354     0   1   17   17.5  RT (min)     lcorbi_t3_pos_1_023_02    m/z 537.5354     0   1   16.5   17  RT (min)    lcorbi_t3_pos_2_012_01    m/z 537.5354     0   1E+5   16   16.5  RT (min)      lcorbi_t3_pos_1_027_43    m/z 537.5354     0   1   17   17.5  RT (min)    lcorbi_t3_pos_1_023_02    m/z 537.5354     0   1   17   17.5  RT (min)    lcorbi_t3_pos_1_023_02    m/z 537.5354     0   1   16.5   17  RT (min)    lcorbi_t3_pos_2_012_01    m/z 537.5354     0   1   2E+5   15.5   16  RT (min)      lcorbi_t3_pos_2_038_03    m/z 537.5354     0   2   4   6E+6   13.5   14  RT (min)      lcorbi_t3_pos_2_016_23    m/z 537.5354     0   1   2E+6   14.5   15  RT (min)      lcorbi_t3_pos_4_025_52    m/z 537.5355     0   2   4   6E+5   15   16  RT (min)       lcorbi_t3_pos_1_008_36    m/z 537.5355     0   1   2E+4   13.5   14  RT (min)      lcorbi_t3_pos_4_012_37    m/z 537.5355     0   1E+5   16   16.5  RT (min)      lcorbi_t3_pos_2_052_48    m/z 537.5355     0   2   4   6E+6   15   16  RT (min)       lcorbi_t3_pos_1_005_42    m/z 537.5355     0   1   2   3E+5   15   15.5  RT (min)      lcorbi_t3_pos_2_017_51    m/z 537.5356     0   1   17   17.5  RT (min)    lcorbi_t3_pos_1_023_02    m/z 537.5356     0   1E+5   16   16.5  RT (min)      lcorbi_t3_pos_1_031_40    m/z 537.5356     0   1   2E+5   15.5   16  RT (min)      lcorbi_t3_pos_1_021_18    m/z 538.1642     0   1   2   3E+6   14   14.5  RT (min)       lcorbi_t3_pos_4_105_61    m/z 538.5195     0   1   16   16.5  RT (min)      lcorbi_t3_pos_1_031_40    m/z 538.5196     0   1   2   3E+4   14.5   15  RT (min)      lcorbi_t3_pos_1_008_36    m/z 538.5196     0   1   15   15.5  RT (min)      lcorbi_t3_pos_1_028_32    m/z 538.5198     0   1   15.5   16  RT (min)      lcorbi_t3_pos_1_034_34    m/z 538.5387     0   1   13   13.5  RT (min)      lcorbi_t3_pos_2_047_31    m/z 538.5387     0   1E+4   13.5   14  RT (min)      lcorbi_t3_pos_3_052_47    m/z 538.5388     0   2   4   6E+4   16   16.5  RT (min)      lcorbi_t3_pos_1_027_43    m/z 538.5388     0   1   2   3   4E+4   16   16.5  RT (min)      lcorbi_t3_pos_1_025_52    m/z 538.5388     0   1E+6   0.5   1  RT (min)      lcorbi_t3_pos_4_057_49    m/z 538.5388     0   2   4   6E+4   15.5   16  RT (min)      lcorbi_t3_pos_1_041_54    m/z 538.5388     0   1   2E+6   13.5   14  RT (min)      lcorbi_t3_pos_3_053_48    m/z 538.5388     0   1   17   17.5  RT (min)    lcorbi_t3_pos_1_023_02    m/z 538.5388     0   1   16.5   17  RT (min)    lcorbi_t3_pos_1_025_52    m/z 538.5388     0   2   4   6   8E+4   15.5   16  RT (min)      lcorbi_t3_pos_2_025_52    m/z 538.5388     0   1   2E+6   13.5   14  RT (min)      lcorbi_t3_pos_2_016_23    m/z 538.5388     0   1E+5   15.5   16  RT (min)      lcorbi_t3_pos_1_038_30    m/z 538.5389     0   1E+6   13.5   14  RT (min)      lcorbi_t3_pos_2_016_23    m/z 538.5389     0   2   4   6E+4   16   16.5  RT (min)      lcorbi_t3_pos_2_052_48    m/z 538.5389     0   1   2E+6   14.5   15  RT (min)       lcorbi_t3_pos_1_005_42    m/z 538.5389     0   1   16.5   17  RT (min)    lcorbi_t3_pos_2_012_01    m/z 538.5389     0   1E+5   15   15.5  RT (min)      lcorbi_t3_pos_3_044_46    m/z 538.5389     0   2   4   6   8E+5   14.5   15  RT (min)       lcorbi_t3_pos_4_025_52    m/z 538.5389     0   2   4   6   8E+4   15.5   16  RT (min)      lcorbi_t3_pos_1_028_32    m/z 538.5389     0   1E+4   13   13.5  RT (min)      lcorbi_t3_pos_2_047_31    m/z 538.5389     0   2   4   6   8E+4   15.5   16  RT (min)      lcorbi_t3_pos_2_041_54    m/z 538.5389     0   1   17   17.5  RT (min)    lcorbi_t3_pos_1_023_02    m/z 538.5389     0   1   2   3E+5   14.5   15  RT (min)      lcorbi_t3_pos_3_025_52    m/z 538.5389     0   1   2   3E+3   13.5   14  RT (min)      lcorbi_t3_pos_2_013_33    m/z 538.5389     0   1E+6   14.5   15  RT (min)      lcorbi_t3_pos_4_051_34    m/z 538.5389     0   2   4   6   8E+4   15.5   16  RT (min)      lcorbi_t3_pos_1_041_54    m/z 538.5389     0   1   16.5   17  RT (min)    lcorbi_t3_pos_2_012_01    m/z 538.5390     0   1E+5   15   15.5  RT (min)      lcorbi_t3_pos_1_055_33    m/z 538.5390     0   2   4   6   8E+4   15   15.5  RT (min)      lcorbi_t3_pos_2_022_38    m/z 538.5390     0   2   4   6E+4   16   16.5  RT (min)      lcorbi_t3_pos_2_014_37    m/z 538.5390     0   1E+5   15   15.5  RT (min)      lcorbi_t3_pos_2_017_51    m/z 538.5390     0   1E+6   14.5   15  RT (min)      lcorbi_t3_pos_3_033_53    m/z 538.5390     0   1   2   3E+5   15   15.5  RT (min)      lcorbi_t3_pos_1_008_36    m/z 538.5390     0   1   17   17.5  RT (min)     lcorbi_t3_pos_1_023_02    m/z 538.5390     0   1   17   17.5  RT (min)    lcorbi_t3_pos_1_023_02    m/z 539.1642     0   1E+6   14   14.5  RT (min)      lcorbi_t3_pos_4_105_61    m/z 539.5230     0   2   4E+5   13.5   14  RT (min)      lcorbi_t3_pos_4_069_57    m/z 539.5230     0   2   4E+5   13.5   14  RT (min)      lcorbi_t3_pos_4_071_57    m/z 539.5420     0   1   2   3   4E+5   14.5   15  RT (min)      lcorbi_t3_pos_1_005_42    m/z 540.1620     0   2   4E+5   14   14.5  RT (min)       lcorbi_t3_pos_4_113_62    m/z 542.3216     0   1E+4   11.5   12  RT (min)       lcorbi_t3_pos_1_034_34    m/z 542.3219     0   1E+4   12   12.5  RT (min)      lcorbi_t3_pos_1_051_09    m/z 542.3220     0   1   2E+4   12   12.5  RT (min)      lcorbi_t3_pos_1_023_02    m/z 542.3220     0   2   4   6E+4   11.5   12  RT (min)      lcorbi_t3_pos_1_028_32    m/z 542.3221     0   1   12   12.5  RT (min)      lcorbi_t3_pos_2_012_01    m/z 542.3221     0   2   4   6   8E+4   11.5   12  RT (min)      lcorbi_t3_pos_3_003_33    m/z 542.3222     0   2   4E+5   11   11.5  RT (min)      lcorbi_t3_pos_1_008_36    m/z 543.3274     0   1E+5   10.5   11  RT (min)      lcorbi_t3_pos_2_054_29    m/z 543.3275     0   1   2E+5   10.5   11  RT (min)      lcorbi_t3_pos_4_067_57    m/z 544.3376     0   1   2   3E+4   13   13.5  RT (min)      lcorbi_t3_pos_1_023_02    m/z 544.3378     0   1   2   3E+6   11.5   12  RT (min)       lcorbi_t3_pos_1_047_41    m/z 544.3379     0   1   2   3   4E+5   11.5   12  RT (min)       lcorbi_t3_pos_3_013_19    m/z 544.3380     0   2   4E+5   11.5   12  RT (min)      lcorbi_t3_pos_1_028_32    m/z 544.3387     0   2   4E+4   12   12.5  RT (min)      lcorbi_t3_pos_1_031_40    m/z 544.3395     0   2   4   6   8E+6   10   10.5  RT (min)      lcorbi_t3_pos_2_032_39    m/z 544.3396     0   1   2E+5   12   12.5  RT (min)      lcorbi_t3_pos_1_051_09    m/z 544.3396     0   1E+7   10.5   11  RT (min)      lcorbi_t3_pos_2_054_29    m/z 544.3396     0   1   2   3E+5   11.5   12  RT (min)      lcorbi_t3_pos_1_051_09    m/z 544.3397     0   1   2E+5   11.5   12  RT (min)      lcorbi_t3_pos_1_041_54    m/z 544.3398     0   1   2   3   4E+6   1   1.5  RT (min)      lcorbi_t3_pos_4_057_49    m/z 544.4112     0   1   2E+4   11   11.5  RT (min)      lcorbi_t3_pos_1_041_54    m/z 544.4112     0   1   2   3E+5   11   11.5  RT (min)      lcorbi_t3_pos_1_021_18    m/z 544.4112     0   2   4   6   8E+5   11   11.5  RT (min)      lcorbi_t3_pos_4_048_24    m/z 544.4112     0   1   11.5   12  RT (min)      lcorbi_t3_pos_1_049_55    m/z 544.4112     0   2   4E+5   11   11.5  RT (min)      lcorbi_t3_pos_1_033_53    m/z 544.4113     0   1E+4   11   11.5  RT (min)      lcorbi_t3_pos_2_009_50    m/z 544.4114     0   1   11.5   12  RT (min)      lcorbi_t3_pos_2_056_56    m/z 545.3410     0   1E+4   12.5   13  RT (min)      lcorbi_t3_pos_1_023_02    m/z 545.3414     0   1E+6   11.5   12  RT (min)       lcorbi_t3_pos_3_013_19    m/z 545.3416     0   1E+5   11.5   12  RT (min)      lcorbi_t3_pos_3_013_19    m/z 545.3430     0   1E+5   11.5   12  RT (min)      lcorbi_t3_pos_1_028_32    m/z 545.3431     0   1   2   3E+6   10.5   11  RT (min)      lcorbi_t3_pos_2_054_29    m/z 545.3431     0   1E+4   12   12.5  RT (min)      lcorbi_t3_pos_1_023_02    m/z 545.3431     0   2   4   6E+4   12   12.5  RT (min)      lcorbi_t3_pos_1_051_09    m/z 545.3432     0   1   2   3   4E+5   11.5   12  RT (min)      lcorbi_t3_pos_3_003_33    m/z 545.3432     0   1E+4   12   12.5  RT (min)      lcorbi_t3_pos_1_056_56    m/z 545.3432     0   1   2   3E+6   10.5   11  RT (min)      lcorbi_t3_pos_2_054_29    m/z 545.3432     0   1   2E+4   12   12.5  RT (min)      lcorbi_t3_pos_1_031_40    m/z 545.3432     0   1   2E+6   10   10.5  RT (min)      lcorbi_t3_pos_2_032_39    m/z 545.3432     0   2   4   6   8E+4   11.5   12  RT (min)      lcorbi_t3_pos_1_051_09    m/z 546.3530     0   1E+4   13   13.5  RT (min)      lcorbi_t3_pos_1_031_40    m/z 546.3530     0   1   2E+4   13.5   14  RT (min)      lcorbi_t3_pos_1_023_02    m/z 546.3530     0   1   13   13.5  RT (min)      lcorbi_t3_pos_1_051_09    m/z 546.3530     0   1   12.5   13  RT (min)      lcorbi_t3_pos_1_034_34    m/z 546.3531     0   1   12.5   13  RT (min)      lcorbi_t3_pos_1_023_02    m/z 546.3531     0   1   2E+4   13.5   14  RT (min)      lcorbi_t3_pos_1_023_02    m/z 546.3531     0   2   4   6E+4   12.5   13  RT (min)      lcorbi_t3_pos_2_012_01    m/z 546.3531     0   1   2E+4   12.5   13  RT (min)      lcorbi_t3_pos_1_023_02    m/z 546.3532     0   1   2E+5   12.5   13  RT (min)      lcorbi_t3_pos_1_051_09    m/z 546.3532     0   2   4E+5   12   12.5  RT (min)      lcorbi_t3_pos_1_031_40    m/z 546.3537     0   2   4   6   8E+5   1.5   2  RT (min)      lcorbi_t3_pos_4_057_49    m/z 546.4270     0   2   4   6E+3   11.5   12  RT (min)      lcorbi_t3_pos_2_009_50    m/z 546.4270     0   1E+6   11   11.5  RT (min)      lcorbi_t3_pos_4_010_43    m/z 546.4270     0   1   2   3E+5   11.5   12  RT (min)      lcorbi_t3_pos_1_021_18    m/z 546.4270     0   1   12   12.5  RT (min)      lcorbi_t3_pos_2_056_56    m/z 546.4271     0   1E+4   12   12.5  RT (min)      lcorbi_t3_pos_1_049_55    m/z 546.4271     0   1E+4   11.5   12  RT (min)      lcorbi_t3_pos_1_041_54    m/z 546.4881     0   1   12   12.5  RT (min)      lcorbi_t3_pos_2_039_30    m/z 546.4881     0   1   12   12.5  RT (min)      lcorbi_t3_pos_2_047_31    m/z 546.4882     0   1   2   3   4E+4   14   14.5  RT (min)      lcorbi_t3_pos_1_033_53    m/z 546.4882     0   1   2   3   4E+4   13.5   14  RT (min)      lcorbi_t3_pos_4_041_54    m/z 546.4882     0   2   4   6E+5   12.5   13  RT (min)      lcorbi_t3_pos_3_053_48    m/z 546.4882     0   1   2   3E+4   15   15.5  RT (min)      lcorbi_t3_pos_1_023_02    m/z 546.4882     0   2   4   6   8E+4   14   14.5  RT (min)      lcorbi_t3_pos_2_038_03    m/z 546.4882     0   2   4   6   8E+5   12.5   13  RT (min)      lcorbi_t3_pos_4_015_45    m/z 546.4882     0   1   2E+5   13.5   14  RT (min)      lcorbi_t3_pos_3_056_56    m/z 546.4882     0   1E+5   12.5   13  RT (min)      lcorbi_t3_pos_3_037_32    m/z 547.3566     0   2   4   6   8E+4   12.5   13  RT (min)      lcorbi_t3_pos_1_051_09    m/z 547.3566     0   1E+4   12.5   13  RT (min)      lcorbi_t3_pos_1_023_02    m/z 547.3588     0   2   4E+5   10.5   11  RT (min)       lcorbi_t3_pos_1_002_01    m/z 547.3590     0   1E+5   12   12.5  RT (min)      lcorbi_t3_pos_1_031_40    m/z 550.3405     0   2   4   6E+5   8.5   9  RT (min)      lcorbi_t3_pos_3_056_56    m/z 550.3406     0   1E+6   8   8.5  RT (min)      lcorbi_t3_pos_3_052_47    m/z 550.3407     0   2   4   6   8E+5   8   8.5  RT (min)      lcorbi_t3_pos_3_053_48    m/z 553.3898     0   2   4E+4   15.5   16  RT (min)      lcorbi_t3_pos_1_027_43    m/z 553.3898     0   1   2   3E+5   14   14.5  RT (min)      lcorbi_t3_pos_3_047_40    m/z 553.3898     0   1E+6   13   13.5  RT (min)      lcorbi_t3_pos_4_015_45    m/z 553.3898     0   2   4   6E+4   14.5   15  RT (min)      lcorbi_t3_pos_1_038_30    m/z 553.3898     0   1   2   3   4E+4   16   16.5  RT (min)      lcorbi_t3_pos_1_023_02    m/z 553.3898     0   1E+4   13   13.5  RT (min)      lcorbi_t3_pos_2_039_30    m/z 553.3898     0   2   4E+4   15.5   16  RT (min)      lcorbi_t3_pos_1_027_43    m/z 553.3898     0   1E+6   13   13.5  RT (min)      lcorbi_t3_pos_3_053_48    m/z 553.3898     0   1   2   3   4E+4   16   16.5  RT (min)      lcorbi_t3_pos_1_023_02    m/z 553.3899     0   1E+4   13   13.5  RT (min)      lcorbi_t3_pos_2_031_20    m/z 553.3899     0   2   4E+4   14.5   15  RT (min)      lcorbi_t3_pos_1_055_33    m/z 553.3899     0   1E+6   13   13.5  RT (min)       lcorbi_t3_pos_3_053_48    m/z 553.3899     0   2   4   6   8E+4   14.5   15  RT (min)       lcorbi_t3_pos_3_044_46    m/z 553.3899     0   2   4E+5   14   14.5  RT (min)      lcorbi_t3_pos_4_019_47    m/z 553.3899     0   2   4   6E+4   14.5   15  RT (min)      lcorbi_t3_pos_1_038_30    m/z 553.3899     0   1   2   3   4E+4   15   15.5  RT (min)       lcorbi_t3_pos_2_052_48    m/z 553.3900     0   1E+4   12.5   13  RT (min)      lcorbi_t3_pos_2_047_31    m/z 560.5038     0   1E+6   15   15.5  RT (min)      lcorbi_t3_pos_3_026_04    m/z 560.5039     0   1   2   3   4E+6   14   14.5  RT (min)       lcorbi_t3_pos_3_034_26    m/z 560.5039     0   1   2   3E+6   14   14.5  RT (min)       lcorbi_t3_pos_2_016_23    m/z 560.5039     0   1E+5   13.5   14  RT (min)       lcorbi_t3_pos_3_024_07    m/z 560.5039     0   1   2   3E+6   15   15.5  RT (min)      lcorbi_t3_pos_4_019_47    m/z 560.5039     0   1   2   3   4E+4   14   14.5  RT (min)      lcorbi_t3_pos_2_016_23    m/z 560.5039     0   1   16.5   17  RT (min)    lcorbi_t3_pos_1_051_09    m/z 560.5039     0   1   16.5   17  RT (min)    lcorbi_t3_pos_1_051_09    m/z 560.5040     0   1   16.5   17  RT (min)    lcorbi_t3_pos_2_052_48    m/z 560.5040     0   2   4   6E+4   13.5   14  RT (min)      lcorbi_t3_pos_3_052_47    m/z 560.5040     0   1   17   17.5  RT (min)    lcorbi_t3_pos_1_051_09    m/z 560.5040     0   1   2   3E+5   13.5   14  RT (min)       lcorbi_t3_pos_2_032_39    m/z 560.5040     0   1   16   16.5  RT (min)      lcorbi_t3_pos_1_028_32    m/z 560.5040     0   1E+4   15.5   16  RT (min)      lcorbi_t3_pos_2_006_47    m/z 560.5040     0   1   15.5   16  RT (min)      lcorbi_t3_pos_1_028_32    m/z 560.5040     0   1   16   16.5  RT (min)      lcorbi_t3_pos_1_034_34    m/z 560.5040     0   1   16.5   17  RT (min)    lcorbi_t3_pos_2_052_48    m/z 560.5041     0   1   2E+4   15.5   16  RT (min)      lcorbi_t3_pos_3_044_46    m/z 560.5041     0   1   16.5   17  RT (min)    lcorbi_t3_pos_1_051_09    m/z 560.5041     0   1   2   3   4E+4   13.5   14  RT (min)      lcorbi_t3_pos_3_035_08    m/z 560.5041     0   1   15.5   16  RT (min)      lcorbi_t3_pos_1_030_21    m/z 560.5041     0   1   2   3E+4   15   15.5  RT (min)      lcorbi_t3_pos_4_010_43    m/z 560.5042     0   1   2   3E+5   15   15.5  RT (min)      lcorbi_t3_pos_3_047_40    m/z 560.5042     0   1   17.5   18  RT (min)    lcorbi_t3_pos_1_023_02    m/z 560.5042     0   1   15.5   16  RT (min)      lcorbi_t3_pos_1_028_32    m/z 561.5072     0   2   4   6   8E+5   14   14.5  RT (min)      lcorbi_t3_pos_4_037_26    m/z 561.5072     0   1   16.5   17  RT (min)    lcorbi_t3_pos_1_051_09    m/z 561.5073     0   1E+5   15   15.5  RT (min)      lcorbi_t3_pos_3_047_40    m/z 561.5073     0   1   2E+4   14   14.5  RT (min)      lcorbi_t3_pos_2_016_23    m/z 561.5073     0   1   16.5   17  RT (min)    lcorbi_t3_pos_1_051_09    m/z 561.5073     0   1E+6   15   15.5  RT (min)      lcorbi_t3_pos_4_019_47    m/z 561.5074     0   1   2   3E+4   13.5   14  RT (min)      lcorbi_t3_pos_3_024_07    m/z 561.5074     0   1E+4   15.5   16  RT (min)      lcorbi_t3_pos_3_044_46    m/z 561.5074     0   1   16.5   17  RT (min)    lcorbi_t3_pos_2_009_50    m/z 561.5074     0   1   16   16.5  RT (min)      lcorbi_t3_pos_1_034_34    m/z 561.5074     0   2   4   6E+3   14   14.5  RT (min)      lcorbi_t3_pos_3_011_42    m/z 561.5074     0   1   15.5   16  RT (min)      lcorbi_t3_pos_1_028_32    m/z 561.5075     0   1E+6   14   14.5  RT (min)      lcorbi_t3_pos_3_034_26    m/z 561.5075     0   1   15.5   16  RT (min)      lcorbi_t3_pos_1_028_32    m/z 561.5075     0   1   17.5   18  RT (min)    lcorbi_t3_pos_1_023_02    m/z 561.5075     0   1   15.5   16  RT (min)      lcorbi_t3_pos_1_030_21    m/z 561.5075     0   1   16   16.5  RT (min)      lcorbi_t3_pos_1_041_54    m/z 561.5075     0   1   16.5   17  RT (min)    lcorbi_t3_pos_1_051_09    m/z 561.5076     0   1   15.5   16  RT (min)      lcorbi_t3_pos_2_046_07    m/z 562.4830     0   1E+4   12   12.5  RT (min)      lcorbi_t3_pos_2_047_31    m/z 562.4831     0   1E+6   12   12.5  RT (min)      lcorbi_t3_pos_2_031_20    m/z 562.4831     0   1   2   3   4E+4   14   14.5  RT (min)      lcorbi_t3_pos_1_051_09    m/z 562.4831     0   2   4   6E+4   14   14.5  RT (min)      lcorbi_t3_pos_2_052_48    m/z 562.4831     0   1E+6   12   12.5  RT (min)      lcorbi_t3_pos_2_031_20    m/z 562.4831     0   1   2   3E+4   14.5   15  RT (min)       lcorbi_t3_pos_2_056_56    m/z 562.4831     0   1   2   3   4E+4   14   14.5  RT (min)      lcorbi_t3_pos_1_001_49    m/z 562.4831     0   1   2E+5   13.5   14  RT (min)      lcorbi_t3_pos_1_047_41    m/z 562.4832     0   1   2   3   4E+4   14.5   15  RT (min)      lcorbi_t3_pos_1_023_02    m/z 562.4832     0   1   12   12.5  RT (min)      lcorbi_t3_pos_2_047_31    m/z 562.4832     0   1   2   3   4E+4   14.5   15  RT (min)      lcorbi_t3_pos_1_023_02    m/z 562.4832     0   1   2E+4   12   12.5  RT (min)      lcorbi_t3_pos_2_039_30    m/z 562.4832     0   1   2E+5   13.5   14  RT (min)       lcorbi_t3_pos_1_038_30    m/z 562.4832     0   1E+6   13   13.5  RT (min)      lcorbi_t3_pos_3_025_52    m/z 562.4832     0   2   4   6   8E+4   13.5   14  RT (min)       lcorbi_t3_pos_2_025_52    m/z 562.4832     0   1E+5   13.5   14  RT (min)      lcorbi_t3_pos_1_038_30    m/z 562.4832     0   1   2E+6   12   12.5  RT (min)      lcorbi_t3_pos_4_015_45    m/z 562.4832     0   1E+6   13   13.5  RT (min)       lcorbi_t3_pos_3_056_56    m/z 562.4832     0   2   4   6E+4   14   14.5  RT (min)       lcorbi_t3_pos_2_014_37    m/z 562.4832     0   1   2   3E+4   14   14.5  RT (min)      lcorbi_t3_pos_1_049_55    m/z 562.4832     0   1   2   3   4E+4   14.5   15  RT (min)      lcorbi_t3_pos_1_001_49    m/z 562.4832     0   2   4   6   8E+5   13   13.5  RT (min)      lcorbi_t3_pos_3_025_52    m/z 562.4833     0   1E+5   13.5   14  RT (min)       lcorbi_t3_pos_3_003_33    m/z 562.4833     0   2   4   6E+5   13   13.5  RT (min)       lcorbi_t3_pos_2_017_51    m/z 562.4833     0   1   2E+6   12   12.5  RT (min)      lcorbi_t3_pos_2_013_33    m/z 563.4862     0   1   2   3   4E+4   14.5   15  RT (min)      lcorbi_t3_pos_1_023_02    m/z 563.4864     0   1   2   3   4E+4   14   14.5  RT (min)      lcorbi_t3_pos_1_027_43    m/z 563.4864     0   1   12   12.5  RT (min)      lcorbi_t3_pos_2_047_31    m/z 563.4864     0   1   2E+4   14   14.5  RT (min)      lcorbi_t3_pos_2_049_55    m/z 563.4864     0   2   4   6   8E+5   13   13.5  RT (min)       lcorbi_t3_pos_3_047_40    m/z 563.4864     0   1   12   12.5  RT (min)      lcorbi_t3_pos_2_047_31    m/z 563.4864     0   2   4   6E+4   13.5   14  RT (min)      lcorbi_t3_pos_3_003_33    m/z 563.4864     0   1   2   3E+4   13.5   14  RT (min)      lcorbi_t3_pos_2_025_52    m/z 563.4864     0   1   2   3E+5   12   12.5  RT (min)      lcorbi_t3_pos_2_031_20    m/z 563.4864     0   1   2   3E+4   14   14.5  RT (min)      lcorbi_t3_pos_2_014_37    m/z 563.4864     0   2   4   6E+5   12   12.5  RT (min)      lcorbi_t3_pos_2_031_20    m/z 563.4865     0   2   4   6E+4   13.5   14  RT (min)      lcorbi_t3_pos_1_038_30    m/z 563.4865     0   1   12   12.5  RT (min)      lcorbi_t3_pos_2_039_30    m/z 563.4865     0   1   2   3   4E+4   14.5   15  RT (min)      lcorbi_t3_pos_1_023_02    m/z 563.4865     0   2   4   6E+5   13   13.5  RT (min)       lcorbi_t3_pos_3_056_56    m/z 563.4865     0   1E+4   14.5   15  RT (min)      lcorbi_t3_pos_2_056_56    m/z 563.4865     0   1   2E+4   14   14.5  RT (min)      lcorbi_t3_pos_1_051_09    m/z 563.4865     0   1   2   3E+4   13.5   14  RT (min)      lcorbi_t3_pos_2_025_52    m/z 563.4865     0   1   2E+5   13   13.5  RT (min)      lcorbi_t3_pos_2_017_51    m/z 563.4866     0   2   4   6E+5   12   12.5  RT (min)      lcorbi_t3_pos_4_015_45    m/z 563.4867     0   1   2E+4   14   14.5  RT (min)      lcorbi_t3_pos_1_056_56    m/z 563.5032     0   1   16   16.5  RT (min)      lcorbi_t3_pos_1_023_02    m/z 563.5036     0   2   4   6E+5   13   13.5  RT (min)      lcorbi_t3_pos_4_015_45    m/z 563.5036     0   1E+5   15   15.5  RT (min)      lcorbi_t3_pos_2_025_52    m/z 563.5036     0   1E+4   12.5   13  RT (min)      lcorbi_t3_pos_2_047_31    m/z 563.5036     0   1   2   3   4E+4   15.5   16  RT (min)      lcorbi_t3_pos_2_052_48    m/z 563.5036     0   1   2   3E+4   15.5   16  RT (min)      lcorbi_t3_pos_1_027_43    m/z 563.5036     0   1   2   3E+4   15   15.5  RT (min)       lcorbi_t3_pos_2_014_37    m/z 563.5036     0   2   4   6   8E+5   13.5   14  RT (min)      lcorbi_t3_pos_3_053_48    m/z 563.5036     0   1   2E+4   13   13.5  RT (min)      lcorbi_t3_pos_3_052_47    m/z 563.5036     0   1   15.5   16  RT (min)      lcorbi_t3_pos_1_027_43    m/z 563.5036     0   2   4   6   8E+5   14.5   15  RT (min)       lcorbi_t3_pos_3_000_aa    m/z 563.5036     0   2   4   6   8E+4   14.5   15  RT (min)      lcorbi_t3_pos_1_038_30    m/z 563.5036     0   1   2   3E+4   15   15.5  RT (min)      lcorbi_t3_pos_2_014_37    m/z 563.5036     0   1E+5   15   15.5  RT (min)      lcorbi_t3_pos_2_041_54    m/z 563.5036     0   2   4   6   8E+4   14.5   15  RT (min)       lcorbi_t3_pos_4_028_04    m/z 563.5037     0   2   4   6   8E+4   14.5   15  RT (min)       lcorbi_t3_pos_4_028_04    m/z 563.5037     0   1   2   3   4E+5   14   14.5  RT (min)       lcorbi_t3_pos_4_025_52    m/z 564.5071     0   1   2   3E+5   14   14.5  RT (min)       lcorbi_t3_pos_3_017_51    m/z 564.5349     0   1   13.5   14  RT (min)      lcorbi_t3_pos_2_013_33    m/z 564.5351     0   1   15.5   16  RT (min)      lcorbi_t3_pos_1_034_34    m/z 564.5352     0   1   15.5   16  RT (min)      lcorbi_t3_pos_1_028_32    m/z 564.5352     0   2   4   6E+3   13   13.5  RT (min)      lcorbi_t3_pos_3_031_45    m/z 564.5353     0   1   16.5   17  RT (min)    lcorbi_t3_pos_2_012_01    m/z 564.5353     0   1   16   16.5  RT (min)      lcorbi_t3_pos_1_031_40    m/z 564.5353     0   1   16.5   17  RT (min)    lcorbi_t3_pos_2_012_01    m/z 564.5353     0   1   15   15.5   16  RT (min)      lcorbi_t3_pos_1_028_32    m/z 564.5354     0   1   2   3E+3   13.5   14  RT (min)       lcorbi_t3_pos_3_011_42    m/z 564.5354     0   2   4   6E+4   14.5   15  RT (min)      lcorbi_t3_pos_1_008_36    m/z 564.5354     0   1   15   15.5  RT (min)      lcorbi_t3_pos_3_003_33    m/z 565.5385     0   1   16.5   17  RT (min)    lcorbi_t3_pos_2_012_01    m/z 565.5387     0   1   2   3   4E+4   14.5   15  RT (min)      lcorbi_t3_pos_1_008_36    m/z 565.5390     0   1   16   16.5  RT (min)      lcorbi_t3_pos_1_031_40    m/z 565.5667     0   1E+5   15.5   16  RT (min)      lcorbi_t3_pos_1_055_33    m/z 565.5668     0   1   2E+5   15   15.5  RT (min)      lcorbi_t3_pos_3_044_46    m/z 565.5668     0   1E+5   15.5   16  RT (min)      lcorbi_t3_pos_2_038_03    m/z 565.5668     0   1E+5   16   16.5  RT (min)       lcorbi_t3_pos_2_052_48    m/z 565.5668     0   1   16.5   17  RT (min)     lcorbi_t3_pos_1_027_43    m/z 565.5668     0   1   17   17.5  RT (min)     lcorbi_t3_pos_1_023_02    m/z 565.5668     0   1   2   3   4E+6   14   14.5  RT (min)      lcorbi_t3_pos_3_053_48    m/z 565.5668     0   1   2   3E+4   13.5   14  RT (min)      lcorbi_t3_pos_2_054_29    m/z 565.5668     0   1   2   3   4E+6   14   15  RT (min)       lcorbi_t3_pos_3_053_48    m/z 565.5668     0   1E+5   16   16.5  RT (min)      lcorbi_t3_pos_1_034_34    m/z 565.5668     0   1E+5   15.5   16  RT (min)      lcorbi_t3_pos_1_038_30    m/z 565.5668     0   1   16.5   17  RT (min)    lcorbi_t3_pos_1_056_56    m/z 565.5668     0   1   2   3   4E+5   14   14.5  RT (min)      lcorbi_t3_pos_2_016_23    m/z 565.5668     0   1   2E+4   13.5   14  RT (min)      lcorbi_t3_pos_2_047_31    m/z 565.5668     0   1E+4   13.5   14  RT (min)      lcorbi_t3_pos_2_032_39    m/z 565.5668     0   1   2   3   4E+5   15   15.5  RT (min)      lcorbi_t3_pos_1_008_36    m/z 565.5669     0   2   4E+5   15   15.5  RT (min)       lcorbi_t3_pos_3_025_52    m/z 565.5669     0   1   16.5   17  RT (min)    lcorbi_t3_pos_2_012_01    m/z 565.5669     0   1   17   17.5  RT (min)     lcorbi_t3_pos_1_023_02    m/z 565.5669     0   1   2   3   4E+6   14.5   15   15.5  RT (min)       lcorbi_t3_pos_3_041_54    m/z 565.5669     0   1E+5   15.5   16  RT (min)      lcorbi_t3_pos_1_030_21    m/z 565.5669     0   1   2   3E+6   0.5   1  RT (min)      lcorbi_t3_pos_4_057_49    m/z 565.5670     0   2   4   6   8E+4   16   16.5  RT (min)      lcorbi_t3_pos_2_009_50    m/z 565.5670     0   1   2   3E+6   14   14.5  RT (min)      lcorbi_t3_pos_4_016_15    m/z 565.5670     0   1E+5   15.5   16  RT (min)      lcorbi_t3_pos_4_048_24    m/z 565.5672     0   1E+5   15.5   16  RT (min)      lcorbi_t3_pos_4_048_24    m/z 566.5507     0   1   15.5   16  RT (min)      lcorbi_t3_pos_1_028_32    m/z 566.5508     0   1   13.5   14  RT (min)      lcorbi_t3_pos_2_013_33    m/z 566.5508     0   1   15.5   16  RT (min)      lcorbi_t3_pos_1_028_32    m/z 566.5509     0   1   16   16.5  RT (min)      lcorbi_t3_pos_1_034_34    m/z 566.5509     0   2   4E+4   15   15.5  RT (min)      lcorbi_t3_pos_1_008_36    m/z 566.5509     0   1   16.5   17  RT (min)    lcorbi_t3_pos_2_012_01    m/z 566.5510     0   1   16   16.5  RT (min)      lcorbi_t3_pos_1_031_40    m/z 566.5510     0   1   13.5   14  RT (min)      lcorbi_t3_pos_3_011_42    m/z 566.5510     0   1   15.5   16  RT (min)      lcorbi_t3_pos_1_034_34    m/z 566.5512     0   1   16.5   17  RT (min)    lcorbi_t3_pos_2_012_01    m/z 566.5700     0   1E+4   13.5   14  RT (min)      lcorbi_t3_pos_2_047_31    m/z 566.5701     0   1E+5   15   15.5  RT (min)       lcorbi_t3_pos_1_008_36    m/z 566.5701     0   1   17   17.5  RT (min)    lcorbi_t3_pos_1_023_02    m/z 566.5702     0   1E+5   15   15.5  RT (min)      lcorbi_t3_pos_3_025_52    m/z 566.5702     0   2   4   6   8E+4   15.5   16  RT (min)      lcorbi_t3_pos_1_055_33    m/z 566.5702     0   1   2   3   4E+4   16   16.5  RT (min)      lcorbi_t3_pos_2_049_55    m/z 566.5702     0   2   4E+4   16   16.5  RT (min)      lcorbi_t3_pos_2_052_48    m/z 566.5702     0   2   4E+4   16   16.5  RT (min)      lcorbi_t3_pos_2_014_37    m/z 566.5702     0   1   16.5   17  RT (min)    lcorbi_t3_pos_2_056_56    m/z 566.5702     0   1E+6   14   14.5  RT (min)      lcorbi_t3_pos_3_053_48    m/z 566.5702     0   1E+6   14.5   15  RT (min)      lcorbi_t3_pos_3_041_54    m/z 566.5702     0   1E+5   15.5   16  RT (min)      lcorbi_t3_pos_1_038_30    m/z 566.5702     0   2   4   6   8E+4   15.5   16  RT (min)      lcorbi_t3_pos_1_030_21    m/z 566.5702     0   1E+4   13.5   14  RT (min)      lcorbi_t3_pos_3_035_08    m/z 566.5702     0   1   16.5   17  RT (min)    lcorbi_t3_pos_1_025_52    m/z 566.5702     0   1E+5   15   15.5  RT (min)      lcorbi_t3_pos_2_017_51    m/z 566.5702     0   2   4   6E+4   15.5   16  RT (min)      lcorbi_t3_pos_1_028_32    m/z 566.5702     0   2   4   6E+4   15.5   16  RT (min)      lcorbi_t3_pos_2_038_03    m/z 566.5703     0   1   16.5   17  RT (min)    lcorbi_t3_pos_1_031_40    m/z 566.5703     0   1   17   17.5  RT (min)    lcorbi_t3_pos_1_023_02    m/z 566.5703     0   2   4E+3   13.5   14  RT (min)      lcorbi_t3_pos_2_054_29    m/z 566.5703     0   1   16.5   17  RT (min)    lcorbi_t3_pos_1_025_52    m/z 566.5703     0   2   4E+5   15   15.5  RT (min)       lcorbi_t3_pos_4_025_52    m/z 566.5704     0   1E+6   14   14.5  RT (min)      lcorbi_t3_pos_2_016_23    m/z 568.3398     0   1E+6   10.5   11  RT (min)      lcorbi_t3_pos_2_032_39    m/z 568.3398     0   1E+4   12   12.5  RT (min)      lcorbi_t3_pos_1_051_09    m/z 568.3398     0   1E+4   12   12.5  RT (min)      lcorbi_t3_pos_1_051_09    m/z 568.3398     0   1E+6   10.5   11  RT (min)      lcorbi_t3_pos_2_054_29    m/z 568.3398     0   1E+4   12   12.5  RT (min)      lcorbi_t3_pos_1_023_02    m/z 568.3398     0   2   4   6E+3   12   12.5  RT (min)      lcorbi_t3_pos_2_012_01    m/z 568.3399     0   2   4   6   8E+5   10   10.5  RT (min)      lcorbi_t3_pos_2_032_39    m/z 568.3399     0   1   2E+4   12.5   13  RT (min)      lcorbi_t3_pos_1_023_02    m/z 568.3399     0   2   4E+4   11.5   12  RT (min)      lcorbi_t3_pos_3_013_19    m/z 568.3399     0   1   2E+5   11.5   12  RT (min)      lcorbi_t3_pos_2_022_38    m/z 568.3399     0   1   2   3E+4   11.5   12  RT (min)      lcorbi_t3_pos_1_034_34    m/z 568.3400     0   1   12   12.5  RT (min)      lcorbi_t3_pos_2_012_01    m/z 568.5665     0   1   16.5   17  RT (min)    lcorbi_t3_pos_1_031_40    m/z 568.5666     0   1   15.5   16  RT (min)      lcorbi_t3_pos_1_028_32    m/z 570.3556     0   1   12.5   13  RT (min)      lcorbi_t3_pos_1_023_02    m/z 570.3556     0   1   12.5   13  RT (min)      lcorbi_t3_pos_1_023_02    m/z 570.3556     0   1   12.5   13  RT (min)      lcorbi_t3_pos_1_023_02    m/z 570.3556     0   2   4E+5   10.5   11  RT (min)      lcorbi_t3_pos_1_002_01    m/z 570.3556     0   1   2E+5   11.5   12  RT (min)      lcorbi_t3_pos_3_013_19    m/z 570.4575     0   1E+6   10.5   11  RT (min)       lcorbi_t3_pos_2_013_33    m/z 570.4576     0   1   2   3E+4   12   12.5  RT (min)      lcorbi_t3_pos_2_009_50    m/z 570.4576     0   2   4E+4   12   12.5  RT (min)      lcorbi_t3_pos_2_009_50    m/z 570.4576     0   1   2   3E+5   11.5   12  RT (min)      lcorbi_t3_pos_2_025_52    m/z 570.4576     0   1   2   3   4E+4   12   12.5   13  RT (min)      lcorbi_t3_pos_1_025_52    m/z 570.4577     0   1   2   3E+4   12   12.5  RT (min)      lcorbi_t3_pos_2_056_56    m/z 570.4577     0   1E+5   11.5   12  RT (min)      lcorbi_t3_pos_1_041_54    m/z 570.4577     0   2   4   6   8E+4   12   12.5  RT (min)      lcorbi_t3_pos_1_041_54    m/z 570.4577     0   1   2   3E+4   12   12.5  RT (min)      lcorbi_t3_pos_1_056_56    m/z 570.4577     0   1E+6   11.5   12  RT (min)       lcorbi_t3_pos_2_017_51    m/z 570.4577     0   2   4   6E+5   11.5   12  RT (min)      lcorbi_t3_pos_1_028_32    m/z 570.4578     0   1   2   3   4E+4   12   12.5  RT (min)      lcorbi_t3_pos_1_025_52    m/z 570.4578     0   1E+6   10.5   11  RT (min)      lcorbi_t3_pos_3_031_45    m/z 571.4610     0   1E+4   12   12.5  RT (min)      lcorbi_t3_pos_1_056_56    m/z 571.4611     0   1   2   3   4E+5   11.5   12  RT (min)      lcorbi_t3_pos_1_008_36    m/z 571.4612     0   1   2E+6   10.5   11  RT (min)      lcorbi_t3_pos_4_083_59    m/z 572.3711     0   1   12.5   13  RT (min)      lcorbi_t3_pos_1_051_09    m/z 572.3712     0   2   4   6E+5   11   11.5  RT (min)      lcorbi_t3_pos_2_054_29    m/z 572.3712     0   1   13   13.5  RT (min)      lcorbi_t3_pos_1_023_02    m/z 572.3715     0   1   2   3   4E+4   10.5   11  RT (min)      lcorbi_t3_pos_2_032_39    m/z 574.3166     0   1E+5   15   16  RT (min)       lcorbi_t3_pos_4_083_59    m/z 574.3374     0   1   2   3E+5   9   9.5  RT (min)      lcorbi_t3_pos_1_023_02    m/z 574.5195     0   1   16   16.5  RT (min)      lcorbi_t3_pos_1_041_54    m/z 574.5195     0   1   16.5   17  RT (min)    lcorbi_t3_pos_1_051_09    m/z 574.5196     0   2   4   6   8E+5   15   15.5  RT (min)      lcorbi_t3_pos_4_019_47    m/z 574.5196     0   1   16.5   17  RT (min)    lcorbi_t3_pos_1_031_40    m/z 574.5196     0   1   16   16.5  RT (min)      lcorbi_t3_pos_1_028_32    m/z 574.5196     0   2   4   6   8E+5   14   14.5  RT (min)      lcorbi_t3_pos_2_016_23    m/z 574.5196     0   1   15.5   16  RT (min)      lcorbi_t3_pos_3_044_46    m/z 574.5196     0   1   13.5   14  RT (min)      lcorbi_t3_pos_3_024_07    m/z 574.5196     0   1   16.5   17  RT (min)    lcorbi_t3_pos_2_052_48    m/z 574.5196     0   1   15.5   16  RT (min)      lcorbi_t3_pos_1_028_32    m/z 574.5196     0   1   16.5   17  RT (min)    lcorbi_t3_pos_1_051_09    m/z 574.5196     0   1   13.5   14  RT (min)      lcorbi_t3_pos_2_032_39    m/z 574.5196     0   2   4   6   8E+4   14   14.5  RT (min)      lcorbi_t3_pos_2_016_23    m/z 574.5196     0   1   17.5   18  RT (min)    lcorbi_t3_pos_1_023_02    m/z 574.5197     0   1   13.5   14  RT (min)      lcorbi_t3_pos_3_052_47    m/z 574.5197     0   1E+6   14   14.5  RT (min)      lcorbi_t3_pos_4_105_61    m/z 574.5198     0   1   15.5   16  RT (min)      lcorbi_t3_pos_1_038_30    m/z 575.5034     0   2   4   6E+4   16   16.5  RT (min)      lcorbi_t3_pos_1_034_34    m/z 575.5034     0   2   4   6E+4   16   16.5  RT (min)      lcorbi_t3_pos_1_034_34    m/z 575.5035     0   1   17   17.5  RT (min)    lcorbi_t3_pos_2_012_01    m/z 575.5035     0   1   16.5   17  RT (min)    lcorbi_t3_pos_2_012_01    m/z 575.5036     0   1E+6   14.5   15  RT (min)      lcorbi_t3_pos_1_034_34    m/z 575.5036     0   2   4   6   8E+5   13.5   14  RT (min)       lcorbi_t3_pos_1_052_38    m/z 575.5036     0   1   12.5   13  RT (min)      lcorbi_t3_pos_3_024_07    m/z 575.5036     0   1   17   17.5  RT (min)    lcorbi_t3_pos_1_023_02    m/z 575.5036     0   1   17   17.5  RT (min)    lcorbi_t3_pos_1_023_02    m/z 575.5036     0   2   4   6E+4   15.5   16  RT (min)      lcorbi_t3_pos_1_028_32    m/z 575.5036     0   2   4   6E+4   15   15.5  RT (min)      lcorbi_t3_pos_1_008_36    m/z 575.5036     0   2   4   6E+4   15.5   16  RT (min)      lcorbi_t3_pos_1_028_32    m/z 575.5036     0   2   4E+4   15.5   16  RT (min)      lcorbi_t3_pos_2_012_01    m/z 575.5036     0   2   4   6   8E+5   13   13.5  RT (min)       lcorbi_t3_pos_2_023_26    m/z 575.5036     0   2   4E+4   15   15.5  RT (min)      lcorbi_t3_pos_1_031_40    m/z 575.5036     0   1   16.5   17  RT (min)    lcorbi_t3_pos_1_031_40    m/z 575.5231     0   2   4   6E+5   14   14.5  RT (min)      lcorbi_t3_pos_4_105_61    m/z 576.4657     0   1   2   3E+4   13   13.5  RT (min)      lcorbi_t3_pos_1_002_01    m/z 576.4986     0   1   12   12.5  RT (min)      lcorbi_t3_pos_2_032_39    m/z 576.4987     0   2   4   6E+5   12.5   13  RT (min)      lcorbi_t3_pos_4_015_45    m/z 576.4987     0   1   12   12.5  RT (min)      lcorbi_t3_pos_2_039_30    m/z 576.4987     0   1E+4   12   12.5  RT (min)      lcorbi_t3_pos_2_013_33    m/z 576.4987     0   1   12   12.5  RT (min)      lcorbi_t3_pos_2_047_31    m/z 576.4987     0   1E+5   12   12.5  RT (min)      lcorbi_t3_pos_2_031_20    m/z 576.4990     0   1   2   3   4E+4   15   15.5  RT (min)      lcorbi_t3_pos_1_023_02    m/z 576.4990     0   2   4E+4   13.5   14  RT (min)       lcorbi_t3_pos_1_030_21    m/z 576.4990     0   1   2   3   4E+4   13.5   14  RT (min)       lcorbi_t3_pos_2_022_38    m/z 576.4990     0   1   2   3   4E+4   15   15.5  RT (min)      lcorbi_t3_pos_1_023_02    m/z 576.4990     0   2   4   6E+5   13   13.5  RT (min)      lcorbi_t3_pos_3_047_40    m/z 576.4991     0   1   2   3   4E+4   13.5   14  RT (min)       lcorbi_t3_pos_1_021_18    m/z 576.4992     0   1   2E+5   14   14.5  RT (min)       lcorbi_t3_pos_2_038_03    m/z 576.4992     0   1   2E+5   13.5   14  RT (min)       lcorbi_t3_pos_2_041_54    m/z 576.4994     0   1E+5   13.5   14  RT (min)       lcorbi_t3_pos_3_044_46    m/z 576.4996     0   1   2   3   4E+4   13.5   14  RT (min)       lcorbi_t3_pos_2_022_38    m/z 576.5028     0   1   2   3   4E+4   15   15.5  RT (min)      lcorbi_t3_pos_1_023_02    m/z 576.5031     0   1   2   3E+4   16   16.5  RT (min)      lcorbi_t3_pos_1_034_34    m/z 576.5067     0   1   2E+4   15.5   16  RT (min)      lcorbi_t3_pos_1_028_32    m/z 576.5069     0   1   16.5   17  RT (min)    lcorbi_t3_pos_1_031_40    m/z 577.3372     0   1   2   3E+6   8.5   9  RT (min)      lcorbi_t3_pos_2_012_01    m/z 577.5191     0   1   17   17.5  RT (min)    lcorbi_t3_pos_2_012_01    m/z 577.5192     0   1E+5   14   14.5  RT (min)      lcorbi_t3_pos_3_011_42    m/z 577.5192     0   1   16.5   17  RT (min)    lcorbi_t3_pos_1_031_40    m/z 577.5192     0   1   17   17.5  RT (min)    lcorbi_t3_pos_2_012_01    m/z 577.5192     0   2   4E+5   15   15.5  RT (min)      lcorbi_t3_pos_4_019_47    m/z 577.5193     0   1E+6   15   15.5  RT (min)      lcorbi_t3_pos_1_034_34    m/z 577.5193     0   1   2   3   4E+4   15   15.5  RT (min)      lcorbi_t3_pos_1_008_36    m/z 577.5193     0   1   2   3E+4   15.5   16  RT (min)      lcorbi_t3_pos_2_022_38    m/z 577.5193     0   1   2   3   4E+4   16   16.5  RT (min)      lcorbi_t3_pos_1_028_32    m/z 577.5194     0   1   17.5   18  RT (min)    lcorbi_t3_pos_1_023_02    m/z 577.5194     0   1   2   3E+4   16   16.5  RT (min)      lcorbi_t3_pos_1_034_34    m/z 577.5194     0   1   2E+4   15.5   16  RT (min)      lcorbi_t3_pos_3_003_33    m/z 577.5194     0   1   2   3   4E+4   15   15.5  RT (min)      lcorbi_t3_pos_1_008_36    m/z 578.3405     0   1E+6   8.5   9  RT (min)      lcorbi_t3_pos_2_012_01    m/z 578.4166     0   2   4   6E+5   9.5   10  RT (min)       lcorbi_t3_pos_4_092_60    m/z 578.4168     0   2   4   6E+5   9   9.5  RT (min)      lcorbi_t3_pos_4_095_60    m/z 578.4168     0   2   4   6E+5   9   9.5  RT (min)       lcorbi_t3_pos_4_092_60    m/z 578.5226     0   1   2E+5   13.5   14  RT (min)       lcorbi_t3_pos_2_013_33    m/z 578.5228     0   1   16   16.5  RT (min)      lcorbi_t3_pos_1_034_34    m/z 578.5229     0   1   15   15.5  RT (min)      lcorbi_t3_pos_1_008_36    m/z 580.2962     0   2   4   6   8E+5   10.5   11  RT (min)      lcorbi_t3_pos_4_079_58    m/z 581.4539     0   1E+6   13.5   14  RT (min)      lcorbi_t3_pos_2_016_23    m/z 581.4540     0   1   17   17.5  RT (min)    lcorbi_t3_pos_1_023_02    m/z 581.4541     0   1   16   16.5  RT (min)      lcorbi_t3_pos_1_041_54    m/z 581.4541     0   1   13.5   14  RT (min)      lcorbi_t3_pos_2_039_30    m/z 581.4542     0   1E+6   14   14.5  RT (min)      lcorbi_t3_pos_2_016_23    m/z 581.4542     0   1E+6   14   14.5  RT (min)      lcorbi_t3_pos_3_053_48    m/z 581.4542     0   1   13   13.5  RT (min)      lcorbi_t3_pos_2_047_31    m/z 581.4542     0   1   16   16.5  RT (min)      lcorbi_t3_pos_2_014_37    m/z 581.4542     0   1E+4   15.5   16  RT (min)      lcorbi_t3_pos_4_028_04    m/z 581.4542     0   1   2E+4   15   15.5  RT (min)      lcorbi_t3_pos_2_046_07    m/z 581.4542     0   1   2E+5   13.5   14  RT (min)      lcorbi_t3_pos_4_015_45    m/z 581.4542     0   1   16.5   17  RT (min)    lcorbi_t3_pos_1_001_49    m/z 581.4543     0   1   16.5   17  RT (min)    lcorbi_t3_pos_1_027_43    m/z 581.4543     0   1   2   3E+4   15   15.5  RT (min)      lcorbi_t3_pos_3_044_46    m/z 581.4543     0   1   13.5   14  RT (min)       lcorbi_t3_pos_2_031_20    m/z 581.4543     0   1   2   3   4E+5   15   15.5  RT (min)      lcorbi_t3_pos_4_025_52    m/z 581.4543     0   1   15.5   16  RT (min)      lcorbi_t3_pos_1_038_30    m/z 581.4543     0   1E+4   16   16.5  RT (min)      lcorbi_t3_pos_2_052_48    m/z 581.4543     0   1   15.5   16  RT (min)      lcorbi_t3_pos_1_038_30    m/z 581.4544     0   1   15.5   16  RT (min)      lcorbi_t3_pos_2_025_52    m/z 581.4544     0   1   17   17.5  RT (min)    lcorbi_t3_pos_1_023_02    m/z 581.4546     0   1   17   17.5  RT (min)    lcorbi_t3_pos_1_023_02    m/z 582.4576     0   1   16.5   17  RT (min)    lcorbi_t3_pos_1_027_43    m/z 582.4576     0   1   13.5   14  RT (min)      lcorbi_t3_pos_2_031_20    m/z 582.4576     0   1   15.5   16  RT (min)      lcorbi_t3_pos_1_038_30    m/z 582.4576     0   1   2E+4   15   15.5  RT (min)      lcorbi_t3_pos_3_044_46    m/z 582.4576     0   1   13.5   14  RT (min)      lcorbi_t3_pos_2_039_30    m/z 582.4577     0   1   2   3E+5   13.5   14  RT (min)      lcorbi_t3_pos_2_016_23    m/z 582.4577     0   2   4   6E+5   14   14.5  RT (min)      lcorbi_t3_pos_4_105_61    m/z 582.5091     0   1   16   16.5  RT (min)      lcorbi_t3_pos_1_031_40    m/z 582.5092     0   1   13   13.5  RT (min)      lcorbi_t3_pos_3_024_07    m/z 582.5092     0   1   2   3E+4   15   15.5  RT (min)      lcorbi_t3_pos_1_008_36    m/z 582.5092     0   2   4   6E+4   15   15.5  RT (min)      lcorbi_t3_pos_1_008_36    m/z 582.5093     0   1   15   15.5  RT (min)      lcorbi_t3_pos_2_022_38    m/z 582.5094     0   1   15.5   16  RT (min)      lcorbi_t3_pos_1_028_32    m/z 582.5094     0   1   16   16.5  RT (min)      lcorbi_t3_pos_1_031_40    m/z 582.5094     0   1   13.5   14  RT (min)      lcorbi_t3_pos_3_011_42    m/z 582.5094     0   1   15.5   16  RT (min)      lcorbi_t3_pos_1_034_34    m/z 582.5094     0   1   15.5   16  RT (min)      lcorbi_t3_pos_1_034_34    m/z 582.5094     0   1   13.5   14  RT (min)      lcorbi_t3_pos_3_011_42    m/z 582.5094     0   1   15.5   16  RT (min)      lcorbi_t3_pos_1_028_32    m/z 582.5095     0   1   16   16.5  RT (min)      lcorbi_t3_pos_1_031_40    m/z 582.5455     0   1   2   3E+3   13.5   14  RT (min)      lcorbi_t3_pos_2_013_33    m/z 582.5456     0   1   13   13.5  RT (min)      lcorbi_t3_pos_3_024_07    m/z 582.5457     0   1   15.5   16  RT (min)      lcorbi_t3_pos_1_028_32    m/z 582.5458     0   1   15.5   16  RT (min)      lcorbi_t3_pos_1_028_32    m/z 582.5458     0   1   15.5   16  RT (min)      lcorbi_t3_pos_1_034_34    m/z 582.5458     0   1   15   15.5  RT (min)      lcorbi_t3_pos_2_022_38    m/z 582.5458     0   1   16.5   17  RT (min)    lcorbi_t3_pos_2_012_01    m/z 582.5459     0   1   16   16.5  RT (min)      lcorbi_t3_pos_1_031_40    m/z 582.5459     0   1   2   3E+4   15   15.5  RT (min)      lcorbi_t3_pos_1_008_36    m/z 582.5459     0   2   4   6E+4   14.5   15  RT (min)      lcorbi_t3_pos_1_008_36    m/z 583.5492     0   1   15.5   16  RT (min)      lcorbi_t3_pos_1_028_32    m/z 583.5492     0   1   16   16.5  RT (min)      lcorbi_t3_pos_1_031_40    m/z 583.5492     0   1   16.5   17  RT (min)    lcorbi_t3_pos_2_012_01    m/z 583.5493     0   1   15   15.5  RT (min)      lcorbi_t3_pos_2_022_38    m/z 583.5494     0   1   2   3   4E+4   14.5   15  RT (min)      lcorbi_t3_pos_1_008_36    m/z 584.5250     0   1   16.5   17  RT (min)    lcorbi_t3_pos_1_031_40    m/z 584.5250     0   1   16.5   17  RT (min)    lcorbi_t3_pos_2_012_01    m/z 584.5250     0   1   16.5   17  RT (min)    lcorbi_t3_pos_1_031_40    m/z 584.5250     0   2   4   6   8E+4   15   15.5  RT (min)      lcorbi_t3_pos_1_008_36    m/z 584.5250     0   1   15.5   16  RT (min)      lcorbi_t3_pos_2_022_38    m/z 584.5250     0   1   17   17.5  RT (min)    lcorbi_t3_pos_1_023_02    m/z 584.5250     0   1   17   17.5  RT (min)    lcorbi_t3_pos_1_023_02    m/z 584.5251     0   1   15.5   16  RT (min)      lcorbi_t3_pos_2_022_38    m/z 584.5251     0   1   16   16.5  RT (min)      lcorbi_t3_pos_1_034_34    m/z 584.5251     0   1   15.5   16  RT (min)      lcorbi_t3_pos_1_028_32    m/z 584.5252     0   1   13.5   14  RT (min)      lcorbi_t3_pos_2_013_33    m/z 584.5252     0   1   13.5   14  RT (min)      lcorbi_t3_pos_2_013_33    m/z 593.5981     0   1   2   3E+4   15.5   16  RT (min)      lcorbi_t3_pos_1_030_21    m/z 593.5981     0   1   17.5   18  RT (min)    lcorbi_t3_pos_1_023_02    m/z 593.5982     0   1   2   3E+4   16   16.5  RT (min)      lcorbi_t3_pos_2_041_54    m/z 593.5982     0   2   4   6E+5   15   15.5  RT (min)      lcorbi_t3_pos_1_005_42    m/z 593.5982     0   1   17.5   18  RT (min)    lcorbi_t3_pos_1_023_02    m/z 593.5982     0   2   4E+4   15.5   16  RT (min)      lcorbi_t3_pos_2_017_51    m/z 593.5982     0   1   16.5   17  RT (min)    lcorbi_t3_pos_1_025_52    m/z 593.5982     0   1E+4   14   14.5  RT (min)      lcorbi_t3_pos_4_015_45    m/z 593.5982     0   1   2E+4   16   16.5  RT (min)      lcorbi_t3_pos_2_025_52    m/z 593.5982     0   1   16.5   17  RT (min)    lcorbi_t3_pos_2_052_48    m/z 593.5982     0   1   14   14.5  RT (min)      lcorbi_t3_pos_2_054_29    m/z 593.5982     0   1   16.5   17  RT (min)    lcorbi_t3_pos_2_014_37    m/z 593.5982     0   1   2E+4   15.5   16  RT (min)      lcorbi_t3_pos_1_033_53    m/z 593.5982     0   1E+5   15   15.5  RT (min)      lcorbi_t3_pos_4_001_49    m/z 593.5982     0   1   13.5   14  RT (min)      lcorbi_t3_pos_2_047_31    m/z 593.5982     0   2   4E+4   15.5   16  RT (min)      lcorbi_t3_pos_3_044_46    m/z 593.5983     0   1   16.5   17  RT (min)    lcorbi_t3_pos_1_056_56    m/z 593.5983     0   1   14   14.5  RT (min)      lcorbi_t3_pos_3_037_32    m/z 593.5983     0   1   16.5   17  RT (min)    lcorbi_t3_pos_1_031_40    m/z 593.5984     0   1   2E+5   15   15.5  RT (min)      lcorbi_t3_pos_3_047_40    m/z 593.5984     0   1   13.5   14  RT (min)      lcorbi_t3_pos_3_031_45    m/z 593.5984     0   1E+5   15   15.5  RT (min)       lcorbi_t3_pos_1_008_36    m/z 598.5406     0   1   16.5   17  RT (min)    lcorbi_t3_pos_1_031_40    m/z 598.5406     0   1   15.5   16  RT (min)      lcorbi_t3_pos_2_022_38    m/z 598.5406     0   1   16   16.5  RT (min)      lcorbi_t3_pos_1_034_34    m/z 598.5407     0   1   16   16.5  RT (min)      lcorbi_t3_pos_1_034_34    m/z 598.5407     0   1   16.5   17  RT (min)    lcorbi_t3_pos_2_012_01    m/z 598.5407     0   1   16   16.5  RT (min)      lcorbi_t3_pos_2_052_48    m/z 598.5408     0   1   13.5   14  RT (min)      lcorbi_t3_pos_3_011_42    m/z 598.5408     0   2   4   6   8E+3   13.5   14  RT (min)      lcorbi_t3_pos_3_024_07    m/z 598.5408     0   1   15.5   16  RT (min)      lcorbi_t3_pos_1_028_32    m/z 598.5408     0   1   2   3E+4   15   15.5  RT (min)      lcorbi_t3_pos_1_008_36    m/z 599.3191     0   1E+6   8.5   9  RT (min)      lcorbi_t3_pos_1_023_02    m/z 601.5191     0   1   16.5   17  RT (min)    lcorbi_t3_pos_2_012_01    m/z 601.5192     0   1   2   3E+4   15.5   16  RT (min)      lcorbi_t3_pos_2_022_38    m/z 601.5192     0   2   4   6   8E+4   15.5   16  RT (min)      lcorbi_t3_pos_1_028_32    m/z 601.5192     0   2   4   6   8E+4   15.5   16  RT (min)      lcorbi_t3_pos_1_028_32    m/z 601.5192     0   1   16.5   17  RT (min)    lcorbi_t3_pos_1_031_40    m/z 601.5193     0   2   4E+4   16   16.5  RT (min)      lcorbi_t3_pos_1_034_34    m/z 601.5194     0   1   16.5   17  RT (min)    lcorbi_t3_pos_1_031_40    m/z 601.5195     0   1   2   3   4E+4   13.5   14  RT (min)      lcorbi_t3_pos_3_024_07    m/z 601.5196     0   1   2   3E+4   13.5   14  RT (min)       lcorbi_t3_pos_3_011_42    m/z 603.5348     0   1   16.5   17  RT (min)    lcorbi_t3_pos_2_009_50    m/z 603.5348     0   1   16.5   17  RT (min)    lcorbi_t3_pos_1_031_40    m/z 603.5348     0   2   4   6E+4   15.5   16  RT (min)      lcorbi_t3_pos_1_008_36    m/z 603.5348     0   2   4   6E+4   15.5   16  RT (min)      lcorbi_t3_pos_1_008_36    m/z 603.5349     0   1   2   3E+4   15.5   16  RT (min)      lcorbi_t3_pos_3_003_33    m/z 603.5349     0   2   4E+4   16   16.5  RT (min)      lcorbi_t3_pos_1_034_34    m/z 603.5349     0   2   4E+4   16   16.5  RT (min)      lcorbi_t3_pos_1_028_32    m/z 603.5349     0   2   4   6E+4   15   15.5  RT (min)      lcorbi_t3_pos_1_008_36    m/z 603.5349     0   1   17   17.5  RT (min)    lcorbi_t3_pos_2_012_01    m/z 603.5349     0   2   4E+4   16   16.5  RT (min)      lcorbi_t3_pos_1_034_34    m/z 603.5349     0   2   4   6E+5   15   15.5  RT (min)      lcorbi_t3_pos_1_005_42    m/z 603.5349     0   2   4E+4   16   16.5  RT (min)      lcorbi_t3_pos_1_028_32    m/z 603.5349     0   2   4E+4   16   16.5  RT (min)      lcorbi_t3_pos_1_028_32    m/z 603.5350     0   1   17.5   18  RT (min)    lcorbi_t3_pos_1_023_02    m/z 603.5350     0   1   2E+4   15.5   16  RT (min)      lcorbi_t3_pos_1_047_41    m/z 603.5350     0   2   4   6E+5   15   15.5  RT (min)      lcorbi_t3_pos_3_040_41    m/z 603.5350     0   1   17.5   18  RT (min)    lcorbi_t3_pos_1_023_02    m/z 604.5278     0   1   15   15.5  RT (min)      lcorbi_t3_pos_1_008_36    m/z 604.5381     0   1   17   17.5  RT (min)    lcorbi_t3_pos_2_012_01    m/z 604.5382     0   1   16.5   17  RT (min)    lcorbi_t3_pos_1_031_40    m/z 604.5382     0   1   16   16.5  RT (min)      lcorbi_t3_pos_1_034_34    m/z 604.5384     0   1   16   16.5  RT (min)      lcorbi_t3_pos_1_028_32    m/z 604.5384     0   1   2E+4   15.5   16  RT (min)      lcorbi_t3_pos_3_003_33    m/z 606.5093     0   1E+4   13.5   14  RT (min)      lcorbi_t3_pos_2_013_33    m/z 606.5094     0   1   15   15.5  RT (min)      lcorbi_t3_pos_2_022_38    m/z 606.5094     0   1   16   16.5  RT (min)      lcorbi_t3_pos_1_031_40    m/z 606.5095     0   1   2E+4   15.5   16  RT (min)      lcorbi_t3_pos_1_028_32    m/z 606.5095     0   1   15.5   16  RT (min)      lcorbi_t3_pos_1_034_34    m/z 606.5096     0   1   15.5   16  RT (min)      lcorbi_t3_pos_1_034_34    m/z 606.5096     0   1   2   3   4E+4   14.5   15  RT (min)      lcorbi_t3_pos_1_008_36    m/z 606.5096     0   1   16   16.5  RT (min)      lcorbi_t3_pos_1_031_40    m/z 606.5097     0   1   2E+4   15.5   16  RT (min)      lcorbi_t3_pos_1_028_32    m/z 608.5249     0   2   4   6   8E+5   14.5   15  RT (min)      lcorbi_t3_pos_3_040_41    m/z 608.5250     0   1   15.5   16  RT (min)      lcorbi_t3_pos_2_022_38    m/z 608.5250     0   1   16.5   17  RT (min)    lcorbi_t3_pos_2_056_56    m/z 608.5250     0   1   13.5   14  RT (min)      lcorbi_t3_pos_3_011_42    m/z 608.5250     0   2   4   6   8E+5   14.5   15  RT (min)      lcorbi_t3_pos_1_005_42    m/z 608.5250     0   1   15.5   16  RT (min)      lcorbi_t3_pos_2_022_38    m/z 608.5250     0   1   2   3E+3   13.5   14  RT (min)      lcorbi_t3_pos_3_031_45    m/z 608.5250     0   1   2   3E+3   13   13.5  RT (min)      lcorbi_t3_pos_3_031_45    m/z 608.5250     0   1   16   16.5  RT (min)      lcorbi_t3_pos_1_056_56    m/z 608.5251     0   1   15.5   16  RT (min)      lcorbi_t3_pos_1_028_32    m/z 608.5251     0   1   15.5   16  RT (min)      lcorbi_t3_pos_2_022_38    m/z 608.5251     0   1   2E+5   15   15.5  RT (min)      lcorbi_t3_pos_1_008_36    m/z 608.5252     0   1   17   17.5  RT (min)    lcorbi_t3_pos_1_023_02    m/z 608.5252     0   1   16.5   17  RT (min)    lcorbi_t3_pos_1_031_40    m/z 608.5252     0   1   16.5   17  RT (min)    lcorbi_t3_pos_1_031_40    m/z 608.5252     0   1   16   16.5  RT (min)      lcorbi_t3_pos_1_034_34    m/z 608.5252     0   1E+4   13.5   14  RT (min)      lcorbi_t3_pos_2_013_33    m/z 609.5284     0   1   13.5   14  RT (min)      lcorbi_t3_pos_3_031_45    m/z 609.5284     0   1   16.5   17  RT (min)    lcorbi_t3_pos_2_012_01    m/z 609.5284     0   1   16.5   17  RT (min)    lcorbi_t3_pos_1_031_40    m/z 609.5285     0   1   16   16.5  RT (min)      lcorbi_t3_pos_1_034_34    m/z 609.5285     0   1   15.5   16  RT (min)      lcorbi_t3_pos_1_034_34    m/z 609.5286     0   1   15.5   16  RT (min)      lcorbi_t3_pos_1_028_32    m/z 609.5286     0   1E+4   13.5   14  RT (min)      lcorbi_t3_pos_2_013_33    m/z 609.5286     0   2   4   6   8E+4   15   15.5  RT (min)      lcorbi_t3_pos_1_008_36    m/z 609.5287     0   1   16.5   17  RT (min)    lcorbi_t3_pos_1_031_40    m/z 609.5288     0   1   13.5   14  RT (min)      lcorbi_t3_pos_3_011_42    m/z 610.1842     0   1   17   17.5  RT (min)    lcorbi_t3_pos_1_023_02    m/z 610.1844     0   1   13.5   14  RT (min)      lcorbi_t3_pos_4_046_44    m/z 610.1844     0   1   16.5   17  RT (min)    lcorbi_t3_pos_1_027_43    m/z 610.1844     0   1   16   16.5  RT (min)      lcorbi_t3_pos_1_034_34    m/z 610.1844     0   1   16.5   17  RT (min)    lcorbi_t3_pos_1_049_55    m/z 610.1844     0   1E+7   14   14.5  RT (min)      lcorbi_t3_pos_4_105_61    m/z 610.1845     0   1   2E+4   15   15.5  RT (min)      lcorbi_t3_pos_4_010_43    m/z 610.1845     0   1E+4   15.5   16  RT (min)      lcorbi_t3_pos_4_028_04    m/z 610.1846     0   1E+6   14.5   15  RT (min)      lcorbi_t3_pos_4_051_34    m/z 610.5360     0   1   16   16.5  RT (min)      lcorbi_t3_pos_2_009_50    m/z 610.5363     0   1   16.5   17  RT (min)    lcorbi_t3_pos_2_009_50    m/z 610.5363     0   1   13.5   14  RT (min)      lcorbi_t3_pos_2_032_39    m/z 610.5404     0   1   16.5   17  RT (min)    lcorbi_t3_pos_1_031_40    m/z 610.5407     0   1   2   3E+4   16   16.5  RT (min)      lcorbi_t3_pos_1_034_34    m/z 610.5407     0   1   15.5   16  RT (min)      lcorbi_t3_pos_2_006_47    m/z 610.5408     0   1   13.5   14  RT (min)      lcorbi_t3_pos_3_024_07    m/z 610.5408     0   1   16.5   17  RT (min)    lcorbi_t3_pos_2_012_01    m/z 610.5408     0   1   2E+4   15   15.5  RT (min)      lcorbi_t3_pos_2_017_51    m/z 610.5408     0   1   16   16.5  RT (min)      lcorbi_t3_pos_2_009_50    m/z 610.5408     0   1   13.5   14  RT (min)      lcorbi_t3_pos_3_011_42    m/z 610.5408     0   1   2   3E+5   15   15.5  RT (min)      lcorbi_t3_pos_1_008_36    m/z 610.5408     0   1   2E+6   15   15.5  RT (min)      lcorbi_t3_pos_1_005_42    m/z 610.5408     0   1   2   3E+4   15.5   16  RT (min)      lcorbi_t3_pos_1_028_32    m/z 610.5408     0   1   2E+6   14.5   15  RT (min)      lcorbi_t3_pos_3_040_41    m/z 610.5408     0   1   17   17.5  RT (min)    lcorbi_t3_pos_1_023_02    m/z 610.5408     0   1   2E+6   14   14.5  RT (min)      lcorbi_t3_pos_3_034_26    m/z 610.5409     0   1   13.5   14  RT (min)      lcorbi_t3_pos_3_011_42    m/z 611.1849     0   1   15.5   16  RT (min)      lcorbi_t3_pos_4_041_54    m/z 611.1850     0   1   17   17.5  RT (min)    lcorbi_t3_pos_1_023_02    m/z 611.1850     0   2   4   6   8E+6   14   14.5  RT (min)      lcorbi_t3_pos_4_105_61    m/z 611.1851     0   1   16   16.5  RT (min)      lcorbi_t3_pos_1_034_34    m/z 611.1851     0   1   16.5   17  RT (min)    lcorbi_t3_pos_1_049_55    m/z 611.1852     0   2   4   6E+5   14.5   15  RT (min)      lcorbi_t3_pos_4_051_34    m/z 611.5439     0   1   16.5   17  RT (min)    lcorbi_t3_pos_1_031_40    m/z 611.5440     0   1E+6   15   15.5  RT (min)      lcorbi_t3_pos_1_005_42    m/z 611.5442     0   1   16   16.5  RT (min)      lcorbi_t3_pos_1_034_34    m/z 611.5442     0   1E+5   15   15.5  RT (min)      lcorbi_t3_pos_1_008_36    m/z 611.5442     0   1   17   17.5  RT (min)    lcorbi_t3_pos_1_023_02    m/z 611.5442     0   1   15.5   16  RT (min)      lcorbi_t3_pos_1_047_41    m/z 611.5442     0   1   15.5   16  RT (min)      lcorbi_t3_pos_1_028_32    m/z 611.5442     0   1   16   16.5  RT (min)      lcorbi_t3_pos_2_009_50    m/z 611.5442     0   1E+4   13.5   14  RT (min)      lcorbi_t3_pos_2_032_39    m/z 611.5442     0   1   15.5   16  RT (min)      lcorbi_t3_pos_2_041_54    m/z 611.5442     0   1   13.5   14  RT (min)      lcorbi_t3_pos_3_024_07    m/z 611.5442     0   1E+6   14.5   15  RT (min)      lcorbi_t3_pos_3_040_41    m/z 611.5443     0   1E+5   15   15.5  RT (min)      lcorbi_t3_pos_4_001_49    m/z 611.5443     0   1   16.5   17  RT (min)    lcorbi_t3_pos_2_012_01    m/z 611.5443     0   1   13.5   14  RT (min)      lcorbi_t3_pos_3_011_42    m/z 611.5444     0   1   13.5   14  RT (min)      lcorbi_t3_pos_3_011_42    m/z 612.1822     0   1   16   16.5  RT (min)      lcorbi_t3_pos_1_034_34    m/z 612.1826     0   2   4E+6   14   14.5  RT (min)      lcorbi_t3_pos_4_105_61    m/z 612.5472     0   1   16.5   17  RT (min)    lcorbi_t3_pos_2_009_50    m/z 612.5472     0   1E+5   14   14.5  RT (min)      lcorbi_t3_pos_4_016_15    m/z 612.5473     0   1   13.5   14  RT (min)      lcorbi_t3_pos_3_024_07    m/z 612.5474     0   1   15.5   16  RT (min)      lcorbi_t3_pos_2_006_47    m/z 612.5476     0   1   15.5   16  RT (min)      lcorbi_t3_pos_1_030_21    m/z 612.5520     0   2   4   6   8E+5   14   14.5  RT (min)       lcorbi_t3_pos_3_011_42    m/z 612.5521     0   1   15.5   16  RT (min)      lcorbi_t3_pos_2_006_47    m/z 612.5562     0   1   2E+5   15   15.5  RT (min)      lcorbi_t3_pos_1_008_36    m/z 612.5564     0   1   17   17.5  RT (min)    lcorbi_t3_pos_2_012_01    m/z 612.5564     0   1   16   16.5  RT (min)      lcorbi_t3_pos_2_025_52    m/z 612.5564     0   1   17.5   18  RT (min)    lcorbi_t3_pos_1_023_02    m/z 612.5564     0   1   17.5   18  RT (min)    lcorbi_t3_pos_1_023_02    m/z 612.5564     0   1   13.5   14  RT (min)      lcorbi_t3_pos_3_031_45    m/z 612.5564     0   1   16   16.5  RT (min)      lcorbi_t3_pos_1_034_34    m/z 612.5564     0   1E+6   15   15.5  RT (min)      lcorbi_t3_pos_1_005_42    m/z 612.5564     0   1   16.5   17  RT (min)    lcorbi_t3_pos_1_031_40    m/z 612.5565     0   1   16   16.5  RT (min)      lcorbi_t3_pos_1_028_32    m/z 612.5565     0   1E+5   15   15.5  RT (min)      lcorbi_t3_pos_4_001_49    m/z 612.5565     0   1   13.5   14  RT (min)      lcorbi_t3_pos_2_013_33    m/z 612.5566     0   1   17.5   18  RT (min)    lcorbi_t3_pos_1_023_02    m/z 612.5566     0   1   17   17.5  RT (min)    lcorbi_t3_pos_2_012_01    m/z 612.6277     0   1   16   16.5  RT (min)      lcorbi_t3_pos_1_041_54    m/z 612.6278     0   1   15.5   16  RT (min)      lcorbi_t3_pos_1_033_53    m/z 612.6278     0   1E+6   13.5   14  RT (min)      lcorbi_t3_pos_1_036_08    m/z 612.6278     0   1E+4   15.5   16  RT (min)      lcorbi_t3_pos_1_047_41    m/z 612.6280     0   2   4E+5   15   15.5  RT (min)      lcorbi_t3_pos_1_005_42    m/z 612.6280     0   1E+6   13.5   14  RT (min)      lcorbi_t3_pos_3_010_44    m/z 613.1826     0   1   2E+6   14   14.5  RT (min)      lcorbi_t3_pos_4_105_61    m/z 613.2718     0   2   4E+5   12   12.5  RT (min)      lcorbi_t3_pos_4_057_49    m/z 613.2719     0   1   15.5   16  RT (min)      lcorbi_t3_pos_1_025_52    m/z 613.2719     0   1   2   3E+5   14   14.5  RT (min)      lcorbi_t3_pos_3_047_40    m/z 613.2720     0   1   15   15.5  RT (min)      lcorbi_t3_pos_1_041_54    m/z 613.2720     0   1   12.5   13  RT (min)      lcorbi_t3_pos_2_047_31    m/z 613.2720     0   1   14.5   15  RT (min)      lcorbi_t3_pos_1_038_30    m/z 613.2720     0   1   14.5   15  RT (min)      lcorbi_t3_pos_1_038_30    m/z 613.2720     0   2   4   6   8E+5   14   14.5  RT (min)      lcorbi_t3_pos_3_041_54    m/z 613.2720     0   1   13   13.5  RT (min)      lcorbi_t3_pos_2_031_20    m/z 613.2720     0   1   15.5   16  RT (min)      lcorbi_t3_pos_1_025_52    m/z 613.2720     0   1   16   16.5  RT (min)      lcorbi_t3_pos_1_023_02    m/z 613.2721     0   1   15   15.5  RT (min)      lcorbi_t3_pos_1_041_54    m/z 613.2721     0   1   14.5   15  RT (min)      lcorbi_t3_pos_2_006_47    m/z 613.2722     0   1   15.5   16  RT (min)      lcorbi_t3_pos_1_027_43    m/z 613.2722     0   1   15   15.5  RT (min)      lcorbi_t3_pos_1_041_54    m/z 613.4804     0   1   2E+4   15.5   16  RT (min)      lcorbi_t3_pos_1_034_34    m/z 613.4804     0   1   2E+4   15   15.5  RT (min)      lcorbi_t3_pos_1_008_36    m/z 613.4804     0   1   16.5   17  RT (min)    lcorbi_t3_pos_1_031_40    m/z 613.4805     0   1   15.5   16  RT (min)      lcorbi_t3_pos_2_022_38    m/z 613.4805     0   1   16.5   17  RT (min)    lcorbi_t3_pos_2_012_01    m/z 613.4806     0   2   4E+4   15   15.5  RT (min)      lcorbi_t3_pos_1_008_36    m/z 613.4806     0   1   16.5   17  RT (min)    lcorbi_t3_pos_1_031_40    m/z 613.4806     0   1E+4   13.5   14  RT (min)      lcorbi_t3_pos_2_013_33    m/z 613.4806     0   1   2E+4   16   16.5  RT (min)      lcorbi_t3_pos_1_034_34    m/z 613.4806     0   1   15.5   16  RT (min)      lcorbi_t3_pos_1_028_32    m/z 613.4806     0   2   4   6   8E+3   15.5   16  RT (min)      lcorbi_t3_pos_3_003_33    m/z 613.5595     0   1   17.5   18  RT (min)    lcorbi_t3_pos_1_023_02    m/z 613.5598     0   1   17   17.5  RT (min)    lcorbi_t3_pos_2_012_01    m/z 613.5598     0   1   15.5   16  RT (min)      lcorbi_t3_pos_2_006_47    m/z 613.5599     0   1   2   3   4E+3   13.5   14  RT (min)      lcorbi_t3_pos_3_024_07    m/z 613.5599     0   1E+5   15   15.5  RT (min)      lcorbi_t3_pos_1_008_36    m/z 613.5599     0   1   16   16.5  RT (min)      lcorbi_t3_pos_1_028_32    m/z 613.5599     0   2   4   6E+5   15   15.5  RT (min)      lcorbi_t3_pos_1_005_42    m/z 613.5599     0   1   16.5   17  RT (min)    lcorbi_t3_pos_1_031_40    m/z 613.5599     0   1   16.5   17  RT (min)    lcorbi_t3_pos_1_031_40    m/z 613.5600     0   2   4   6E+5   14   14.5  RT (min)      lcorbi_t3_pos_4_016_15    m/z 613.5600     0   1   16   16.5  RT (min)      lcorbi_t3_pos_1_034_34    m/z 613.5600     0   1   13.5   14  RT (min)      lcorbi_t3_pos_2_013_33    m/z 613.5600     0   1   17.5   18  RT (min)    lcorbi_t3_pos_1_023_02    m/z 613.5600     0   2   4E+5   15   15.5  RT (min)      lcorbi_t3_pos_3_040_41    m/z 613.5600     0   1   15.5   16  RT (min)      lcorbi_t3_pos_1_033_53    m/z 613.5600     0   1   16.5   17  RT (min)    lcorbi_t3_pos_2_009_50    m/z 613.5601     0   2   4   6   8E+3   14   14.5  RT (min)      lcorbi_t3_pos_3_011_42    m/z 614.1816     0   2   4   6   8E+5   14   14.5  RT (min)      lcorbi_t3_pos_4_105_61    m/z 614.4838     0   1   2E+4   12   12.5  RT (min)      lcorbi_t3_pos_1_025_52    m/z 614.4840     0   1   2   3   4E+6   10.5   11  RT (min)       lcorbi_t3_pos_4_082_59    m/z 614.4840     0   1   2E+5   11.5   12  RT (min)      lcorbi_t3_pos_2_025_52    m/z 614.4840     0   2   4E+5   11.5   12  RT (min)      lcorbi_t3_pos_1_033_53    m/z 614.4840     0   2   4E+5   11.5   12  RT (min)      lcorbi_t3_pos_4_049_55    m/z 614.4840     0   1   2   3E+4   12   12.5  RT (min)      lcorbi_t3_pos_2_012_01    m/z 614.4840     0   2   4   6   8E+5   11.5   12  RT (min)       lcorbi_t3_pos_2_017_51    m/z 614.4840     0   1   2   3   4E+4   11.5   12  RT (min)      lcorbi_t3_pos_2_009_50    m/z 614.4840     0   1   2   3   4E+6   10.5   11  RT (min)      lcorbi_t3_pos_4_082_59    m/z 614.4840     0   1   2   3   4E+6   10.5   11  RT (min)       lcorbi_t3_pos_4_082_59    m/z 614.4840     0   1   2E+4   12   12.5  RT (min)      lcorbi_t3_pos_1_056_56    m/z 614.4840     0   2   4E+5   10.5   11  RT (min)      lcorbi_t3_pos_3_024_07    m/z 614.4840     0   1   2E+4   12   12.5  RT (min)      lcorbi_t3_pos_1_025_52    m/z 614.4840     0   1   2   3   4E+6   10.5   11  RT (min)       lcorbi_t3_pos_4_082_59    m/z 614.4841     0   1   2E+5   11.5   12  RT (min)      lcorbi_t3_pos_2_025_52    m/z 614.4842     0   1   2E+4   12   12.5  RT (min)      lcorbi_t3_pos_1_049_55    m/z 614.6436     0   1E+6   13.5   14  RT (min)       lcorbi_t3_pos_4_095_60    m/z 614.6436     0   1E+6   13.5   14  RT (min)       lcorbi_t3_pos_4_090_60    m/z 614.6436     0   1E+6   13.5   14  RT (min)       lcorbi_t3_pos_4_094_60    m/z 614.6439     0   1E+6   13.5   14  RT (min)      lcorbi_t3_pos_4_095_60    m/z 615.4874     0   1E+6   10.5   11  RT (min)      lcorbi_t3_pos_4_082_59    m/z 615.4960     0   1   16.5   17  RT (min)    lcorbi_t3_pos_1_031_40    m/z 615.4961     0   1   15.5   16  RT (min)      lcorbi_t3_pos_1_028_32    m/z 615.4962     0   1   17   17.5  RT (min)    lcorbi_t3_pos_1_023_02    m/z 615.4962     0   1   16   16.5  RT (min)      lcorbi_t3_pos_2_049_55    m/z 615.4963     0   2   4E+4   15   15.5  RT (min)      lcorbi_t3_pos_1_008_36    m/z 615.4963     0   1   16   16.5  RT (min)      lcorbi_t3_pos_1_034_34    m/z 615.4963     0   1   2E+4   13.5   14  RT (min)      lcorbi_t3_pos_3_024_07    m/z 615.4963     0   2   4   6E+5   14.5   15  RT (min)      lcorbi_t3_pos_3_026_04    m/z 615.4963     0   1   17   17.5  RT (min)    lcorbi_t3_pos_1_023_02    m/z 615.4963     0   1   2E+5   13.5   14  RT (min)      lcorbi_t3_pos_3_011_42    m/z 615.4964     0   1   16.5   17  RT (min)    lcorbi_t3_pos_1_049_55    m/z 615.4964     0   1   16.5   17  RT (min)    lcorbi_t3_pos_2_012_01    m/z 615.4964     0   1   15.5   16  RT (min)      lcorbi_t3_pos_2_006_47    m/z 615.4964     0   1   16   16.5  RT (min)      lcorbi_t3_pos_1_034_34    m/z 616.4995     0   1   16   16.5  RT (min)      lcorbi_t3_pos_1_034_34    m/z 616.4996     0   1   15.5   16  RT (min)      lcorbi_t3_pos_2_022_38    m/z 616.4996     0   1   15.5   16  RT (min)      lcorbi_t3_pos_1_028_32    m/z 616.4996     0   1E+4   13.5   14  RT (min)      lcorbi_t3_pos_2_013_33    m/z 616.4996     0   1   16.5   17  RT (min)    lcorbi_t3_pos_2_012_01    m/z 616.4996     0   1E+4   13.5   14  RT (min)      lcorbi_t3_pos_3_024_07    m/z 616.4996     0   1E+5   13.5   14  RT (min)       lcorbi_t3_pos_3_011_42    m/z 616.4996     0   1   16.5   17  RT (min)    lcorbi_t3_pos_1_031_40    m/z 616.4998     0   1   2   3   4E+4   15   15.5  RT (min)      lcorbi_t3_pos_1_008_36    m/z 617.5118     0   1   16.5   17  RT (min)    lcorbi_t3_pos_1_049_55    m/z 617.5118     0   1   15.5   16  RT (min)      lcorbi_t3_pos_1_021_18    m/z 617.5118     0   1   17.5   18  RT (min)    lcorbi_t3_pos_1_023_02    m/z 617.5119     0   1   17.5   18  RT (min)    lcorbi_t3_pos_1_023_02    m/z 617.5120     0   2   4   6E+4   15   15.5  RT (min)      lcorbi_t3_pos_1_008_36    m/z 617.5120     0   1   16.5   17  RT (min)    lcorbi_t3_pos_1_031_40    m/z 617.5120     0   1   15.5   16  RT (min)      lcorbi_t3_pos_2_022_38    m/z 617.5121     0   1E+4   13.5   14  RT (min)      lcorbi_t3_pos_2_013_33    m/z 617.5122     0   1   2   3   4E+5   14   14.5  RT (min)      lcorbi_t3_pos_3_011_42    m/z 617.5122     0   1   16   16.5  RT (min)      lcorbi_t3_pos_1_034_34    m/z 617.5131     0   1   17   17.5  RT (min)    lcorbi_t3_pos_2_012_01    m/z 617.5138     0   1   2   3E+4   15   15.5  RT (min)      lcorbi_t3_pos_1_008_36    m/z 617.5139     0   1   16.5   17  RT (min)    lcorbi_t3_pos_1_031_40    m/z 617.5141     0   2   4   6   8E+3   13.5   14  RT (min)      lcorbi_t3_pos_3_031_45    m/z 617.5142     0   1   15   15.5  RT (min)      lcorbi_t3_pos_1_008_36    m/z 617.5143     0   1   15.5   16  RT (min)      lcorbi_t3_pos_1_028_32    m/z 618.2273     0   1   14.5   15  RT (min)      lcorbi_t3_pos_1_038_30    m/z 618.2274     0   1   13   13.5  RT (min)      lcorbi_t3_pos_2_031_20    m/z 618.5129     0   2   4   6E+4   13   13.5  RT (min)      lcorbi_t3_pos_4_046_44    m/z 618.5129     0   1   2   3   4E+5   14   14.5  RT (min)       lcorbi_t3_pos_4_019_47    m/z 618.5130     0   1   12.5   13  RT (min)      lcorbi_t3_pos_2_047_31    m/z 618.5130     0   1   13   13.5  RT (min)      lcorbi_t3_pos_3_035_08    m/z 618.5142     0   1   2   3   4E+4   16   16.5  RT (min)       lcorbi_t3_pos_1_034_34    m/z 618.5148     0   1   2   3E+4   15   15.5  RT (min)      lcorbi_t3_pos_2_014_37    m/z 618.5150     0   1   2   3E+5   14.5   15  RT (min)      lcorbi_t3_pos_4_079_58    m/z 618.5151     0   2   4E+5   14.5   15  RT (min)       lcorbi_t3_pos_1_032_26    m/z 618.5152     0   1   2   3   4E+5   14.5   15  RT (min)      lcorbi_t3_pos_2_020_22    m/z 618.5155     0   2   4E+4   14.5   15  RT (min)       lcorbi_t3_pos_3_044_46    m/z 620.5254     0   1   16   16.5  RT (min)      lcorbi_t3_pos_1_031_40    m/z 622.5406     0   1   15.5   16  RT (min)      lcorbi_t3_pos_2_022_38    m/z 622.5407     0   1   16   16.5  RT (min)      lcorbi_t3_pos_1_034_34    m/z 622.5407     0   1   15.5   16  RT (min)      lcorbi_t3_pos_1_028_32    m/z 622.5408     0   1   15.5   16  RT (min)      lcorbi_t3_pos_1_028_32    m/z 622.5408     0   1   16.5   17  RT (min)    lcorbi_t3_pos_2_012_01    m/z 622.5408     0   2   4E+3   13.5   14  RT (min)      lcorbi_t3_pos_2_013_33    m/z 622.5409     0   1   16.5   17  RT (min)    lcorbi_t3_pos_1_031_40    m/z 622.5409     0   2   4E+3   13.5   14  RT (min)      lcorbi_t3_pos_2_013_33    m/z 622.5409     0   1   16.5   17  RT (min)    lcorbi_t3_pos_1_031_40    m/z 622.5410     0   1   15   15.5  RT (min)      lcorbi_t3_pos_1_008_36    m/z 622.5410     0   1   16   16.5  RT (min)      lcorbi_t3_pos_1_034_34    m/z 622.5902     0   2   4E+5   14.5   15  RT (min)      lcorbi_t3_pos_2_030_14    m/z 624.5564     0   1   16.5   17  RT (min)    lcorbi_t3_pos_1_031_40    m/z 624.5564     0   1   16   16.5  RT (min)      lcorbi_t3_pos_1_034_34    m/z 624.5565     0   1   15.5   16  RT (min)      lcorbi_t3_pos_1_028_32    m/z 624.5565     0   1   16   16.5  RT (min)      lcorbi_t3_pos_1_034_34    m/z 624.5565     0   1   16   16.5  RT (min)      lcorbi_t3_pos_1_028_32    m/z 624.5565     0   1   2   3E+4   15   15.5  RT (min)      lcorbi_t3_pos_1_008_36    m/z 624.5566     0   1   17   17.5  RT (min)    lcorbi_t3_pos_2_012_01    m/z 624.5566     0   1   13.5   14  RT (min)      lcorbi_t3_pos_2_013_33    m/z 624.5566     0   1   16.5   17  RT (min)    lcorbi_t3_pos_1_031_40    m/z 624.5567     0   1   13.5   14  RT (min)      lcorbi_t3_pos_3_024_07    m/z 624.5568     0   1   17   17.5  RT (min)    lcorbi_t3_pos_2_012_01    m/z 624.6402     0   2   4   6   8E+5   0.5   1  RT (min)      lcorbi_t3_pos_4_057_49    m/z 624.6402     0   1   17   17.5  RT (min)     lcorbi_t3_pos_1_023_02    m/z 624.6403     0   1   2   3   4E+4   16   16.5  RT (min)      lcorbi_t3_pos_4_033_53    m/z 624.6403     0   2   4E+4   15.5   16  RT (min)      lcorbi_t3_pos_2_041_54    m/z 624.6403     0   1   2   3   4E+4   15.5   16  RT (min)       lcorbi_t3_pos_2_049_55    m/z 624.6404     0   2   4   6E+4   15.5   16  RT (min)      lcorbi_t3_pos_1_021_18    m/z 624.6404     0   1E+4   13   13.5  RT (min)      lcorbi_t3_pos_2_047_31    m/z 624.6404     0   2   4E+4   15.5   16  RT (min)      lcorbi_t3_pos_2_038_03    m/z 624.6404     0   1   13   13.5  RT (min)      lcorbi_t3_pos_2_032_39    m/z 624.6404     0   1E+4   13.5   14  RT (min)      lcorbi_t3_pos_3_037_32    m/z 624.6404     0   1   16.5   17  RT (min)     lcorbi_t3_pos_2_056_56    m/z 624.6404     0   2   4E+4   15   15.5  RT (min)      lcorbi_t3_pos_2_017_51    m/z 624.6404     0   2   4   6E+4   15   15.5  RT (min)       lcorbi_t3_pos_1_030_21    m/z 624.6404     0   2   4   6E+4   15   15.5  RT (min)      lcorbi_t3_pos_1_047_41    m/z 624.6404     0   1   2   3E+4   16   16.5  RT (min)       lcorbi_t3_pos_1_051_09    m/z 624.6404     0   2   4   6E+5   14.5   15  RT (min)       lcorbi_t3_pos_3_026_04    m/z 624.6404     0   1   2   3   4E+4   16   16.5  RT (min)      lcorbi_t3_pos_1_027_43    m/z 624.6405     0   1   2   3   4E+4   15.5   16  RT (min)       lcorbi_t3_pos_2_025_52    m/z 624.6405     0   1   16.5   17  RT (min)    lcorbi_t3_pos_1_025_52    m/z 624.6405     0   1   2   3E+4   16   16.5  RT (min)      lcorbi_t3_pos_2_049_55    m/z 624.6405     0   1   2   3E+5   13.5   14  RT (min)      lcorbi_t3_pos_2_016_23    m/z 624.6405     0   1E+5   15   15.5  RT (min)      lcorbi_t3_pos_3_056_56    m/z 624.6406     0   1   13.5   14  RT (min)      lcorbi_t3_pos_3_052_47    m/z 624.6406     0   1   2   3   4E+5   14.5   15  RT (min)       lcorbi_t3_pos_4_025_52    m/z 624.6406     0   2   4   6E+4   15   15.5  RT (min)      lcorbi_t3_pos_3_044_46    m/z 624.6407     0   2   4   6   8E+5   13.5   14  RT (min)      lcorbi_t3_pos_3_053_48    m/z 624.6407     0   1   2   3E+4   16   16.5  RT (min)       lcorbi_t3_pos_1_049_55    m/z 630.5058     0   1   14.5   15  RT (min)      lcorbi_t3_pos_1_008_36    m/z 630.5094     0   1   2E+4   15.5   16  RT (min)      lcorbi_t3_pos_1_028_32    m/z 630.5094     0   1   16.5   17  RT (min)    lcorbi_t3_pos_2_012_01    m/z 630.5094     0   1   2E+4   15   15.5  RT (min)      lcorbi_t3_pos_1_028_32    m/z 630.5095     0   1   15.5   16  RT (min)      lcorbi_t3_pos_1_034_34    m/z 630.5095     0   1   16   16.5  RT (min)      lcorbi_t3_pos_1_031_40    m/z 630.5095     0   1   13.5   14  RT (min)      lcorbi_t3_pos_2_013_33    m/z 630.5096     0   1   16   16.5  RT (min)      lcorbi_t3_pos_1_031_40    m/z 630.5144     0   1E+5   14.5   15  RT (min)      lcorbi_t3_pos_1_008_36    m/z 632.5251     0   1   2   3E+4   15.5   16  RT (min)      lcorbi_t3_pos_2_025_52    m/z 632.5251     0   1   2   3   4E+4   15.5   16  RT (min)      lcorbi_t3_pos_1_034_34    m/z 632.5252     0   1   2E+4   15.5   16  RT (min)      lcorbi_t3_pos_3_003_33    m/z 632.5252     0   1   2   3E+4   15   15.5  RT (min)       lcorbi_t3_pos_2_022_38    m/z 632.5252     0   1   2   3   4E+4   15.5   16  RT (min)      lcorbi_t3_pos_1_034_34    m/z 632.5252     0   1   2   3   4E+4   15.5   16  RT (min)      lcorbi_t3_pos_1_028_32    m/z 632.5252     0   1E+4   16   16.5  RT (min)      lcorbi_t3_pos_4_033_53    m/z 632.5252     0   1   2E+5   15   15.5  RT (min)      lcorbi_t3_pos_1_008_36    m/z 632.5253     0   1   16   16.5  RT (min)      lcorbi_t3_pos_1_031_40    m/z 632.5253     0   1   2E+5   14.5   15  RT (min)      lcorbi_t3_pos_4_001_49    m/z 632.5254     0   1E+6   14.5   15  RT (min)      lcorbi_t3_pos_1_005_42    m/z 632.5254     0   1E+5   15   15.5  RT (min)      lcorbi_t3_pos_1_008_36    m/z 632.5255     0   1   16.5   17  RT (min)     lcorbi_t3_pos_2_012_01    m/z 632.5255     0   2   4   6   8E+5   14.5   15  RT (min)      lcorbi_t3_pos_3_040_41    m/z 632.5283     0   1   2E+4   13.5   14  RT (min)      lcorbi_t3_pos_3_031_45    m/z 632.5283     0   1   2E+4   13   13.5  RT (min)      lcorbi_t3_pos_3_024_07    m/z 633.2264     0   2   4   6   8E+5   2   2.5  RT (min)       lcorbi_t3_pos_4_096_60    m/z 633.2266     0   2   4   6E+5   2.5   3  RT (min)       lcorbi_t3_pos_4_090_60    m/z 633.5283     0   1   15   15.5  RT (min)      lcorbi_t3_pos_2_022_38    m/z 633.5285     0   1   15.5   16  RT (min)      lcorbi_t3_pos_2_022_38    m/z 633.5285     0   1   2   3   4E+4   15   15.5  RT (min)      lcorbi_t3_pos_1_008_36    m/z 633.5285     0   1   15.5   16  RT (min)      lcorbi_t3_pos_1_034_34    m/z 633.5286     0   1   15.5   16  RT (min)      lcorbi_t3_pos_1_028_32    m/z 633.5286     0   1   16   16.5  RT (min)      lcorbi_t3_pos_1_031_40    m/z 633.5286     0   2   4E+5   14.5   15  RT (min)      lcorbi_t3_pos_1_005_42    m/z 633.5286     0   1   16.5   17  RT (min)    lcorbi_t3_pos_2_012_01    m/z 633.5287     0   1   15.5   16  RT (min)      lcorbi_t3_pos_1_034_34    m/z 633.5287     0   1   2   3   4E+4   15   15.5  RT (min)      lcorbi_t3_pos_1_008_36    m/z 633.5288     0   1E+5   15   15.5  RT (min)      lcorbi_t3_pos_1_008_36    m/z 633.5288     0   1   16   16.5  RT (min)      lcorbi_t3_pos_1_031_40    m/z 633.5315     0   1E+4   13   13.5  RT (min)      lcorbi_t3_pos_3_024_07    m/z 634.5319     0   1   2   3E+3   13.5   14  RT (min)      lcorbi_t3_pos_3_031_45    m/z 634.5406     0   1   17   17.5  RT (min)    lcorbi_t3_pos_1_023_02    m/z 634.5407     0   1E+6   14   14.5  RT (min)      lcorbi_t3_pos_4_016_15    m/z 634.5407     0   1   16.5   17  RT (min)    lcorbi_t3_pos_1_031_40    m/z 634.5408     0   1   16.5   17  RT (min)    lcorbi_t3_pos_2_012_01    m/z 634.5408     0   1   16.5   17  RT (min)    lcorbi_t3_pos_1_031_40    m/z 634.5408     0   1   16   16.5  RT (min)      lcorbi_t3_pos_1_034_34    m/z 634.5408     0   2   4   6E+3   13.5   14  RT (min)      lcorbi_t3_pos_3_011_42    m/z 634.5408     0   1   2E+6   14.5   15  RT (min)      lcorbi_t3_pos_1_005_42    m/z 634.5408     0   2   4   6   8E+3   13.5   14  RT (min)      lcorbi_t3_pos_2_013_33    m/z 634.5408     0   2   4   6   8E+4   15   15.5  RT (min)      lcorbi_t3_pos_1_008_36    m/z 634.5408     0   1   15.5   16  RT (min)      lcorbi_t3_pos_1_028_32    m/z 634.5408     0   1E+4   15.5   16  RT (min)      lcorbi_t3_pos_2_022_38    m/z 634.5408     0   2   4E+5   15   15.5  RT (min)      lcorbi_t3_pos_1_008_36    m/z 634.5408     0   1   16.5   17  RT (min)    lcorbi_t3_pos_2_012_01    m/z 634.5408     0   1   15.5   16  RT (min)      lcorbi_t3_pos_1_028_32    m/z 634.5409     0   2   4   6   8E+3   13.5   14  RT (min)      lcorbi_t3_pos_3_031_45    m/z 634.5410     0   1E+6   14.5   15  RT (min)      lcorbi_t3_pos_3_033_53    m/z 634.5410     0   1   16   16.5  RT (min)      lcorbi_t3_pos_1_034_34    m/z 634.5411     0   1E+4   13.5   14  RT (min)      lcorbi_t3_pos_2_054_29    m/z 635.5440     0   1   17   17.5  RT (min)    lcorbi_t3_pos_1_023_02    m/z 635.5442     0   1   2   3   4E+3   13.5   14  RT (min)      lcorbi_t3_pos_3_031_45    m/z 635.5442     0   2   4E+5   14   14.5  RT (min)      lcorbi_t3_pos_4_016_15    m/z 635.5442     0   1   16   16.5  RT (min)      lcorbi_t3_pos_1_034_34    m/z 635.5442     0   1   2   3   4E+3   13.5   14  RT (min)      lcorbi_t3_pos_2_013_33    m/z 635.5442     0   1   15.5   16  RT (min)      lcorbi_t3_pos_1_028_32    m/z 635.5442     0   1E+6   14.5   15  RT (min)      lcorbi_t3_pos_1_005_42    m/z 635.5442     0   1   16.5   17  RT (min)    lcorbi_t3_pos_1_031_40    m/z 635.5442     0   1   2E+5   15   15.5  RT (min)      lcorbi_t3_pos_1_008_36    m/z 635.5442     0   1   15.5   16  RT (min)      lcorbi_t3_pos_2_022_38    m/z 635.5442     0   1   15.5   16  RT (min)      lcorbi_t3_pos_1_028_32    m/z 635.5442     0   1   16.5   17  RT (min)    lcorbi_t3_pos_2_012_01    m/z 635.5443     0   1   16.5   17  RT (min)    lcorbi_t3_pos_2_012_01    m/z 635.5443     0   1   17   17.5  RT (min)    lcorbi_t3_pos_1_023_02    m/z 635.5444     0   2   4   6E+5   14   14.5  RT (min)      lcorbi_t3_pos_3_034_26    m/z 635.5444     0   1   16.5   17  RT (min)    lcorbi_t3_pos_1_031_40    m/z 635.5444     0   1   15.5   16  RT (min)      lcorbi_t3_pos_2_022_38    m/z 635.5444     0   1   15   15.5  RT (min)      lcorbi_t3_pos_2_046_07    m/z 635.5445     0   1   13.5   14  RT (min)      lcorbi_t3_pos_1_002_01    m/z 636.5471     0   1   16   16.5  RT (min)      lcorbi_t3_pos_2_009_50    m/z 636.5475     0   1   13.5   14  RT (min)      lcorbi_t3_pos_3_031_45    m/z 636.5476     0   1   13.5   14  RT (min)      lcorbi_t3_pos_3_031_45    m/z 636.5476     0   1   15.5   16  RT (min)      lcorbi_t3_pos_4_041_54    m/z 636.5478     0   1   15.5   16  RT (min)      lcorbi_t3_pos_2_017_51    m/z 636.5481     0   1   15.5   16  RT (min)      lcorbi_t3_pos_1_030_21    m/z 636.5522     0   1   16   16.5  RT (min)      lcorbi_t3_pos_1_034_34    m/z 636.5562     0   1E+4   13.5   14  RT (min)      lcorbi_t3_pos_3_031_45    m/z 636.5564     0   1   17   17.5  RT (min)    lcorbi_t3_pos_1_023_02    m/z 636.5564     0   2   4E+4   13.5   14  RT (min)      lcorbi_t3_pos_3_011_42    m/z 636.5564     0   1   16.5   17  RT (min)    lcorbi_t3_pos_1_031_40    m/z 636.5564     0   1   2   3   4E+5   15   15.5  RT (min)      lcorbi_t3_pos_4_001_49    m/z 636.5564     0   1E+4   13.5   14  RT (min)      lcorbi_t3_pos_3_031_45    m/z 636.5565     0   1   2   3E+6   15   15.5  RT (min)      lcorbi_t3_pos_1_005_42    m/z 636.5565     0   1   15.5   16  RT (min)      lcorbi_t3_pos_1_028_32    m/z 636.5565     0   1   2   3   4E+4   13.5   14  RT (min)      lcorbi_t3_pos_2_013_33    m/z 636.5565     0   1   2E+6   15   15.5  RT (min)      lcorbi_t3_pos_3_040_41    m/z 636.5565     0   1   2   3E+4   15.5   16  RT (min)      lcorbi_t3_pos_1_030_21    m/z 636.5566     0   1   2   3E+5   15   15.5  RT (min)      lcorbi_t3_pos_1_008_36    m/z 636.5566     0   1   2E+6   14   14.5  RT (min)      lcorbi_t3_pos_4_016_15    m/z 636.5566     0   1   16.5   17  RT (min)    lcorbi_t3_pos_2_012_01    m/z 636.5568     0   1   16.5   17  RT (min)    lcorbi_t3_pos_1_051_09    m/z 637.3056     0   1   13.5   14  RT (min)      lcorbi_t3_pos_2_012_01    m/z 637.3056     0   1   2E+4   13   13.5  RT (min)      lcorbi_t3_pos_1_034_34    m/z 637.3056     0   2   4   6   8E+5   12.5   13  RT (min)      lcorbi_t3_pos_1_008_36    m/z 637.3056     0   1   2E+4   13   13.5  RT (min)      lcorbi_t3_pos_1_028_32    m/z 637.3056     0   1   11.5   12  RT (min)      lcorbi_t3_pos_3_024_07    m/z 637.3057     0   1   2E+4   13   13.5  RT (min)      lcorbi_t3_pos_3_003_33    m/z 637.3058     0   2   4   6E+5   12.5   13  RT (min)      lcorbi_t3_pos_1_008_36    m/z 637.3058     0   1E+6   11.5   12  RT (min)      lcorbi_t3_pos_3_011_42    m/z 637.3058     0   1   13.5   14  RT (min)      lcorbi_t3_pos_1_031_40    m/z 637.4804     0   1   15.5   16  RT (min)      lcorbi_t3_pos_1_034_34    m/z 637.4804     0   1   15.5   16  RT (min)      lcorbi_t3_pos_1_028_32    m/z 637.5596     0   1   17   17.5  RT (min)    lcorbi_t3_pos_1_023_02    m/z 637.5598     0   1E+6   15   15.5  RT (min)      lcorbi_t3_pos_1_005_42    m/z 637.5598     0   1E+4   13.5   14  RT (min)      lcorbi_t3_pos_2_013_33    m/z 637.5599     0   1   15.5   16  RT (min)      lcorbi_t3_pos_1_030_21    m/z 637.5599     0   1E+6   15   15.5  RT (min)      lcorbi_t3_pos_3_026_04    m/z 637.5599     0   1   16.5   17  RT (min)    lcorbi_t3_pos_1_031_40    m/z 637.5600     0   1   13.5   14  RT (min)      lcorbi_t3_pos_2_032_39    m/z 637.5600     0   1   16.5   17  RT (min)    lcorbi_t3_pos_2_012_01    m/z 637.5600     0   1   2E+4   13.5   14  RT (min)      lcorbi_t3_pos_3_011_42    m/z 637.5600     0   1   16.5   17  RT (min)    lcorbi_t3_pos_1_025_52    m/z 637.5600     0   1E+4   15.5   16  RT (min)      lcorbi_t3_pos_2_046_07    m/z 637.5600     0   1   15.5   16  RT (min)      lcorbi_t3_pos_1_028_32    m/z 637.5600     0   1E+4   13.5   14  RT (min)      lcorbi_t3_pos_2_013_33    m/z 637.5600     0   1E+5   15   15.5  RT (min)      lcorbi_t3_pos_1_008_36    m/z 637.5600     0   1E+6   14.5   15  RT (min)      lcorbi_t3_pos_3_026_04    m/z 637.5600     0   1   15.5   16  RT (min)      lcorbi_t3_pos_4_048_24    m/z 637.5600     0   1E+6   14   14.5  RT (min)      lcorbi_t3_pos_4_016_15    m/z 637.5600     0   1   16   16.5  RT (min)      lcorbi_t3_pos_1_034_34    m/z 637.5602     0   1E+6   14   14.5  RT (min)      lcorbi_t3_pos_3_034_26    m/z 637.5602     0   1   16   16.5  RT (min)      lcorbi_t3_pos_1_034_34    m/z 638.5630     0   1   13.5   14  RT (min)      lcorbi_t3_pos_3_031_45    m/z 638.5632     0   1   16.5   17  RT (min)    lcorbi_t3_pos_1_041_54    m/z 638.5633     0   1   13.5   14  RT (min)      lcorbi_t3_pos_3_031_45    m/z 638.5633     0   1   15.5   16  RT (min)      lcorbi_t3_pos_2_017_51    m/z 638.5677     0   2   4   6   8E+5   15   15.5  RT (min)      lcorbi_t3_pos_4_025_52    m/z 638.5677     0   1   16.5   17  RT (min)    lcorbi_t3_pos_1_056_56    m/z 638.5718     0   1   16   16.5  RT (min)      lcorbi_t3_pos_2_038_03    m/z 638.5718     0   1   13.5   14  RT (min)      lcorbi_t3_pos_2_013_33    m/z 638.5719     0   1   2E+6   14   14.5  RT (min)      lcorbi_t3_pos_4_016_15    m/z 638.5720     0   1   13.5   14  RT (min)      lcorbi_t3_pos_3_031_45    m/z 638.5720     0   1   17   17.5  RT (min)    lcorbi_t3_pos_2_012_01    m/z 638.5720     0   1   16.5   17  RT (min)    lcorbi_t3_pos_1_025_52    m/z 638.5720     0   1E+4   15.5   16  RT (min)      lcorbi_t3_pos_3_003_33    m/z 638.5721     0   1   2   3E+4   15.5   16  RT (min)      lcorbi_t3_pos_2_022_38    m/z 638.5721     0   1E+5   15.5   16  RT (min)      lcorbi_t3_pos_1_008_36    m/z 638.5721     0   1   2   3E+6   15   15.5  RT (min)      lcorbi_t3_pos_1_005_42    m/z 638.5721     0   1   16   16.5  RT (min)      lcorbi_t3_pos_1_034_34    m/z 638.5721     0   1   2   3E+6   15   15.5  RT (min)      lcorbi_t3_pos_3_040_41    m/z 638.5721     0   1   16   16.5  RT (min)      lcorbi_t3_pos_1_034_34    m/z 638.5722     0   1   2   3   4E+5   15   15.5  RT (min)      lcorbi_t3_pos_1_008_36    m/z 638.5722     0   2   4   6E+4   14   14.5  RT (min)      lcorbi_t3_pos_3_011_42    m/z 638.5722     0   1   16.5   17  RT (min)    lcorbi_t3_pos_1_031_40    m/z 638.5722     0   1   16   16.5  RT (min)      lcorbi_t3_pos_1_028_32    m/z 638.5722     0   1   13.5   14  RT (min)      lcorbi_t3_pos_3_024_07    m/z 638.5722     0   1   17.5   18  RT (min)    lcorbi_t3_pos_1_023_02    m/z 638.6089     0   2   4   6E+5   13.5   14  RT (min)      lcorbi_t3_pos_4_050_31    m/z 638.6560     0   1   2   3   4E+4   16   16.5  RT (min)      lcorbi_t3_pos_1_049_55    m/z 638.6560     0   2   4E+5   13.5   14  RT (min)      lcorbi_t3_pos_3_053_48    m/z 638.6560     0   1   13.5   14  RT (min)      lcorbi_t3_pos_2_039_30    m/z 638.6560     0   1   2   3E+4   16   16.5  RT (min)      lcorbi_t3_pos_1_056_56    m/z 638.6562     0   1   2   3E+4   15.5   16  RT (min)      lcorbi_t3_pos_2_041_54    m/z 638.6562     0   1   2   3E+4   15.5   16  RT (min)      lcorbi_t3_pos_2_025_52    m/z 638.6562     0   1   2   3   4E+4   15   15.5  RT (min)       lcorbi_t3_pos_1_030_21    m/z 638.6562     0   1   2   3   4E+4   15.5   16  RT (min)      lcorbi_t3_pos_1_021_18    m/z 638.6562     0   1   2   3E+5   13.5   14  RT (min)      lcorbi_t3_pos_2_016_23    m/z 638.6562     0   2   4   6   8E+4   15   15.5  RT (min)      lcorbi_t3_pos_3_056_56    m/z 638.6562     0   1   16   16.5  RT (min)       lcorbi_t3_pos_1_025_52    m/z 638.6562     0   1   2   3E+4   16   16.5  RT (min)      lcorbi_t3_pos_2_052_48    m/z 638.6562     0   1   13.5   14  RT (min)      lcorbi_t3_pos_3_035_08    m/z 638.6562     0   1   13   13.5  RT (min)      lcorbi_t3_pos_2_047_31    m/z 638.6562     0   2   4E+4   15   15.5  RT (min)      lcorbi_t3_pos_2_046_07    m/z 638.6562     0   2   4E+5   13.5   14  RT (min)      lcorbi_t3_pos_3_053_48    m/z 639.4961     0   1   16   16.5  RT (min)      lcorbi_t3_pos_1_034_34    m/z 639.4962     0   1   15.5   16  RT (min)      lcorbi_t3_pos_2_022_38    m/z 639.4962     0   1   16.5   17  RT (min)    lcorbi_t3_pos_2_012_01    m/z 639.4962     0   1   15.5   16  RT (min)      lcorbi_t3_pos_2_022_38    m/z 639.4962     0   2   4   6E+5   14.5   15  RT (min)      lcorbi_t3_pos_1_005_42    m/z 639.4962     0   1   13.5   14  RT (min)      lcorbi_t3_pos_2_013_33    m/z 639.4962     0   1   16.5   17  RT (min)    lcorbi_t3_pos_2_012_01    m/z 639.4963     0   1E+5   15   15.5  RT (min)      lcorbi_t3_pos_1_008_36    m/z 639.4963     0   1   16   16.5  RT (min)      lcorbi_t3_pos_1_034_34    m/z 639.4963     0   1   16.5   17  RT (min)    lcorbi_t3_pos_1_031_40    m/z 639.4964     0   2   4E+4   15   15.5  RT (min)      lcorbi_t3_pos_1_008_36    m/z 639.4964     0   1   15.5   16  RT (min)      lcorbi_t3_pos_1_028_32    m/z 639.4964     0   1   2   3E+4   15   15.5  RT (min)      lcorbi_t3_pos_1_008_36    m/z 639.4966     0   1   2   3E+3   13.5   14  RT (min)      lcorbi_t3_pos_3_011_42    m/z 639.5754     0   1E+5   14   14.5  RT (min)      lcorbi_t3_pos_1_002_01    m/z 639.5755     0   1E+4   15.5   16  RT (min)      lcorbi_t3_pos_2_022_38    m/z 639.5755     0   1   17.5   18  RT (min)    lcorbi_t3_pos_1_023_02    m/z 639.5755     0   1E+6   15   15.5  RT (min)      lcorbi_t3_pos_1_005_42    m/z 639.5755     0   1   16   16.5  RT (min)      lcorbi_t3_pos_2_025_52    m/z 639.5755     0   1   16.5   17  RT (min)    lcorbi_t3_pos_1_031_40    m/z 639.5755     0   1   13.5   14  RT (min)      lcorbi_t3_pos_2_013_33    m/z 639.5756     0   2   4   6   8E+5   14   14.5  RT (min)      lcorbi_t3_pos_3_034_26    m/z 639.5756     0   1   16   16.5  RT (min)      lcorbi_t3_pos_1_034_34    m/z 639.5756     0   1   16.5   17  RT (min)    lcorbi_t3_pos_2_009_50    m/z 639.5756     0   1   16.5   17  RT (min)    lcorbi_t3_pos_2_009_50    m/z 639.5756     0   1   15.5   16  RT (min)      lcorbi_t3_pos_1_047_41    m/z 639.5756     0   1E+5   15   15.5  RT (min)      lcorbi_t3_pos_1_008_36    m/z 639.5756     0   2   4   6E+4   14   14.5  RT (min)      lcorbi_t3_pos_1_002_01    m/z 639.5756     0   1   17   17.5  RT (min)    lcorbi_t3_pos_2_012_01    m/z 639.5756     0   1   17   17.5  RT (min)    lcorbi_t3_pos_2_012_01    m/z 639.5756     0   1   16   16.5  RT (min)      lcorbi_t3_pos_1_028_32    m/z 639.5756     0   1   17.5   18  RT (min)    lcorbi_t3_pos_1_023_02    m/z 639.5757     0   1E+6   15   15.5  RT (min)      lcorbi_t3_pos_3_040_41    m/z 639.5757     0   1   16.5   17  RT (min)    lcorbi_t3_pos_1_056_56    m/z 639.5757     0   1   15.5   16  RT (min)      lcorbi_t3_pos_2_017_51    m/z 639.5757     0   1   13.5   14  RT (min)      lcorbi_t3_pos_3_024_07    m/z 639.5758     0   1   15.5   16  RT (min)      lcorbi_t3_pos_2_017_51    m/z 639.5758     0   1E+6   14   14.5  RT (min)      lcorbi_t3_pos_4_016_15    m/z 639.5759     0   1   13.5   14  RT (min)      lcorbi_t3_pos_2_032_39    m/z 640.4995     0   1   16.5   17  RT (min)    lcorbi_t3_pos_1_031_40    m/z 640.4995     0   1   2   3   4E+4   15   15.5  RT (min)      lcorbi_t3_pos_1_008_36    m/z 640.4996     0   1   13.5   14  RT (min)      lcorbi_t3_pos_3_024_07    m/z 640.4996     0   1   15.5   16  RT (min)      lcorbi_t3_pos_2_022_38    m/z 640.4997     0   1   15.5   16  RT (min)      lcorbi_t3_pos_1_028_32    m/z 640.5788     0   1   13.5   14  RT (min)      lcorbi_t3_pos_3_031_45    m/z 640.5788     0   1   14   14.5  RT (min)      lcorbi_t3_pos_2_013_33    m/z 640.5789     0   1   16   16.5  RT (min)      lcorbi_t3_pos_1_028_32    m/z 640.5789     0   1   13.5   14  RT (min)      lcorbi_t3_pos_3_031_45    m/z 640.5789     0   1   15.5   16  RT (min)      lcorbi_t3_pos_2_022_38    m/z 640.5790     0   1   13.5   14  RT (min)      lcorbi_t3_pos_2_013_33    m/z 640.5791     0   1   16.5   17  RT (min)    lcorbi_t3_pos_2_009_50    m/z 640.5791     0   1   16   16.5  RT (min)      lcorbi_t3_pos_1_034_34    m/z 640.5791     0   1   17   17.5  RT (min)    lcorbi_t3_pos_1_031_40    m/z 640.5793     0   1   15   15.5  RT (min)      lcorbi_t3_pos_1_008_36    m/z 640.5838     0   1   2   3   4E+5   14   14.5  RT (min)      lcorbi_t3_pos_3_011_42    m/z 640.5875     0   2   4   6   8E+5   15   15.5  RT (min)      lcorbi_t3_pos_1_008_36    m/z 640.5875     0   2   4   6E+4   15.5   16  RT (min)      lcorbi_t3_pos_1_008_36    m/z 640.5876     0   1   16.5   17  RT (min)    lcorbi_t3_pos_1_031_40    m/z 640.5877     0   1   16.5   17  RT (min)    lcorbi_t3_pos_1_034_34    m/z 640.5878     0   1   17   17.5  RT (min)    lcorbi_t3_pos_2_012_01    m/z 640.5878     0   1   16   16.5  RT (min)      lcorbi_t3_pos_1_028_32    m/z 640.5878     0   2   4   6E+5   15   15.5  RT (min)      lcorbi_t3_pos_3_026_04    m/z 640.5879     0   1   17   17.5  RT (min)    lcorbi_t3_pos_2_012_01    m/z 640.5879     0   1   17   17.5  RT (min)    lcorbi_t3_pos_2_012_01    m/z 641.3030     0   1E+4   13   13.5  RT (min)      lcorbi_t3_pos_3_052_47    m/z 641.3031     0   1E+4   15   15.5  RT (min)      lcorbi_t3_pos_2_025_52    m/z 641.3032     0   1   2E+4   13.5   14  RT (min)      lcorbi_t3_pos_4_046_44    m/z 641.3032     0   2   4   6E+4   14.5   15  RT (min)      lcorbi_t3_pos_3_056_56    m/z 641.3032     0   1E+6   14.5   15  RT (min)       lcorbi_t3_pos_1_005_42    m/z 641.3032     0   1   16   16.5  RT (min)      lcorbi_t3_pos_1_025_52    m/z 641.3032     0   1   15.5   16  RT (min)      lcorbi_t3_pos_1_041_54    m/z 641.3033     0   1   2   3   4E+6   13.5   14  RT (min)       lcorbi_t3_pos_4_105_61    m/z 641.3033     0   1   16.5   17  RT (min)    lcorbi_t3_pos_1_023_02    m/z 641.3033     0   1   2E+4   15   15.5  RT (min)      lcorbi_t3_pos_1_038_30    m/z 641.3033     0   1   2   3E+4   15   15.5  RT (min)      lcorbi_t3_pos_1_030_21    m/z 641.3034     0   1E+4   15.5   16  RT (min)      lcorbi_t3_pos_2_009_50    m/z 641.3034     0   1   15   15.5  RT (min)      lcorbi_t3_pos_1_038_30    m/z 641.3034     0   1   2E+4   13   13.5  RT (min)      lcorbi_t3_pos_2_031_20    m/z 641.3034     0   1   13   13.5  RT (min)      lcorbi_t3_pos_2_047_31    m/z 641.3034     0   1   16   16.5  RT (min)      lcorbi_t3_pos_1_027_43    m/z 641.3034     0   1   16   16.5  RT (min)      lcorbi_t3_pos_1_056_56    m/z 641.3034     0   1   16   16.5  RT (min)      lcorbi_t3_pos_1_025_52    m/z 641.3034     0   1   2   3   4E+4   14.5   15  RT (min)      lcorbi_t3_pos_3_044_46    m/z 641.3034     0   1   2   3   4E+5   13.5   14  RT (min)      lcorbi_t3_pos_4_015_45    m/z 641.3034     0   2   4   6E+5   14.5   15  RT (min)      lcorbi_t3_pos_4_025_52    m/z 641.3036     0   1   16.5   17  RT (min)    lcorbi_t3_pos_1_023_02    m/z 641.3039     0   1   16   16.5  RT (min)      lcorbi_t3_pos_2_056_56    m/z 641.5075     0   1   13.5   14  RT (min)      lcorbi_t3_pos_3_031_45    m/z 641.5117     0   1E+4   15.5   16  RT (min)      lcorbi_t3_pos_2_017_51    m/z 641.5118     0   1   16   16.5  RT (min)      lcorbi_t3_pos_1_034_34    m/z 641.5118     0   1   17   17.5  RT (min)    lcorbi_t3_pos_1_023_02    m/z 641.5119     0   1   16.5   17  RT (min)    lcorbi_t3_pos_1_031_40    m/z 641.5119     0   1   15.5   16  RT (min)      lcorbi_t3_pos_1_028_32    m/z 641.5119     0   1   17   17.5  RT (min)    lcorbi_t3_pos_2_012_01    m/z 641.5120     0   1   15.5   16  RT (min)      lcorbi_t3_pos_2_022_38    m/z 641.5120     0   2   4   6E+5   15   15.5  RT (min)      lcorbi_t3_pos_1_005_42    m/z 641.5120     0   2   4   6E+5   14   14.5  RT (min)      lcorbi_t3_pos_3_034_26    m/z 641.5120     0   2   4   6E+5   15   15.5  RT (min)      lcorbi_t3_pos_3_040_41    m/z 641.5120     0   2   4   6E+4   15   15.5  RT (min)      lcorbi_t3_pos_1_008_36    m/z 641.5120     0   1   16   16.5  RT (min)      lcorbi_t3_pos_1_034_34    m/z 641.5120     0   1   13.5   14  RT (min)      lcorbi_t3_pos_2_013_33    m/z 641.5120     0   1   16.5   17  RT (min)    lcorbi_t3_pos_2_012_01    m/z 641.5121     0   1   16   16.5  RT (min)      lcorbi_t3_pos_2_025_52    m/z 641.5121     0   2   4   6E+5   14   14.5  RT (min)      lcorbi_t3_pos_4_016_15    m/z 641.5821     0   1   16   16.5  RT (min)      lcorbi_t3_pos_3_003_33    m/z 641.5911     0   1   17   17.5  RT (min)    lcorbi_t3_pos_1_031_40    m/z 641.5911     0   1   17   17.5  RT (min)    lcorbi_t3_pos_1_031_40    m/z 641.5912     0   1   14   14.5  RT (min)      lcorbi_t3_pos_2_013_33    m/z 641.5912     0   1   16.5   17  RT (min)    lcorbi_t3_pos_1_034_34    m/z 641.5912     0   1   16   16.5  RT (min)      lcorbi_t3_pos_1_028_32    m/z 641.5912     0   1   16   16.5  RT (min)      lcorbi_t3_pos_2_022_38    m/z 641.5912     0   1   13.5   14  RT (min)      lcorbi_t3_pos_3_024_07    m/z 641.5913     0   1   14   14.5  RT (min)      lcorbi_t3_pos_2_013_33    m/z 642.3065     0   1E+4   13   13.5  RT (min)      lcorbi_t3_pos_2_031_20    m/z 642.3066     0   1   13.5   14  RT (min)      lcorbi_t3_pos_4_046_44    m/z 642.3066     0   1   15   15.5  RT (min)      lcorbi_t3_pos_1_038_30    m/z 642.3066     0   1   15   15.5  RT (min)      lcorbi_t3_pos_1_038_30    m/z 642.3066     0   2   4E+5   14.5   15  RT (min)      lcorbi_t3_pos_1_005_42    m/z 642.3066     0   1   15.5   16  RT (min)      lcorbi_t3_pos_1_041_54    m/z 642.3066     0   1   13   13.5  RT (min)      lcorbi_t3_pos_2_047_31    m/z 642.3066     0   1   16   16.5  RT (min)      lcorbi_t3_pos_1_056_56    m/z 642.3067     0   1   16   16.5  RT (min)      lcorbi_t3_pos_1_025_52    m/z 642.3068     0   1E+6   13.5   14  RT (min)      lcorbi_t3_pos_4_105_61    m/z 642.3428     0   1E+5   15   16  RT (min)       lcorbi_t3_pos_4_080_58    m/z 642.3428     0   1   2E+5   11   11.5  RT (min)      lcorbi_t3_pos_1_008_36    m/z 642.5152     0   1   15.5   16  RT (min)      lcorbi_t3_pos_1_028_32    m/z 642.5152     0   1   15.5   16  RT (min)      lcorbi_t3_pos_2_022_38    m/z 642.5153     0   1   16   16.5  RT (min)      lcorbi_t3_pos_1_034_34    m/z 642.5154     0   1   16.5   17  RT (min)    lcorbi_t3_pos_1_031_40    m/z 642.5154     0   1   13.5   14  RT (min)      lcorbi_t3_pos_2_013_33    m/z 642.5154     0   2   4E+4   15   15.5  RT (min)       lcorbi_t3_pos_1_008_36    m/z 642.5155     0   1   16.5   17  RT (min)    lcorbi_t3_pos_2_012_01    m/z 642.5155     0   1   13.5   14  RT (min)      lcorbi_t3_pos_3_011_42    m/z 642.6656     0   1   15.5   16  RT (min)      lcorbi_t3_pos_1_030_21    m/z 642.6658     0   1   2E+4   16   16.5  RT (min)      lcorbi_t3_pos_1_021_18    m/z 642.6658     0   1   2   3E+4   15   15.5  RT (min)      lcorbi_t3_pos_3_025_52    m/z 642.6658     0   1   16.5   17  RT (min)    lcorbi_t3_pos_1_056_56    m/z 642.6658     0   1   17   17.5  RT (min)    lcorbi_t3_pos_2_056_56    m/z 642.6659     0   1E+4   15.5   16  RT (min)      lcorbi_t3_pos_2_017_51    m/z 642.6659     0   1   16.5   17  RT (min)    lcorbi_t3_pos_1_056_56    m/z 642.6659     0   1E+4   16   16.5  RT (min)      lcorbi_t3_pos_2_041_54    m/z 642.6659     0   1   2E+4   16   16.5  RT (min)      lcorbi_t3_pos_1_021_18    m/z 642.6659     0   2   4   6E+5   15   15.5  RT (min)      lcorbi_t3_pos_1_005_42    m/z 642.6660     0   1E+5   15   15.5  RT (min)      lcorbi_t3_pos_4_025_52    m/z 642.6660     0   1   16   16.5  RT (min)      lcorbi_t3_pos_1_041_54    m/z 642.6662     0   2   4E+5   15   15.5  RT (min)      lcorbi_t3_pos_3_040_41    m/z 643.5274     0   1   13.5   14  RT (min)      lcorbi_t3_pos_2_013_33    m/z 643.5274     0   1   17   17.5  RT (min)    lcorbi_t3_pos_2_012_01    m/z 643.5274     0   1   17.5   18  RT (min)    lcorbi_t3_pos_1_023_02    m/z 643.5274     0   1   16.5   17  RT (min)    lcorbi_t3_pos_1_056_56    m/z 643.5275     0   1   2E+4   15.5   16  RT (min)      lcorbi_t3_pos_1_008_36    m/z 643.5276     0   1   16   16.5  RT (min)      lcorbi_t3_pos_1_034_34    m/z 643.5276     0   1E+5   15   15.5  RT (min)      lcorbi_t3_pos_1_008_36    m/z 643.5276     0   1   16   16.5  RT (min)      lcorbi_t3_pos_1_034_34    m/z 643.5276     0   1   17.5   18  RT (min)    lcorbi_t3_pos_1_023_02    m/z 643.5276     0   1   2   3E+3   14   14.5  RT (min)      lcorbi_t3_pos_4_012_37    m/z 643.5276     0   1   16   16.5  RT (min)      lcorbi_t3_pos_1_028_32    m/z 643.5276     0   1   15.5   16  RT (min)      lcorbi_t3_pos_3_003_33    m/z 643.5276     0   1   13.5   14  RT (min)      lcorbi_t3_pos_3_031_45    m/z 643.5276     0   1   15.5   16  RT (min)      lcorbi_t3_pos_2_022_38    m/z 643.5276     0   1   16.5   17  RT (min)    lcorbi_t3_pos_1_031_40    m/z 643.5276     0   2   4   6E+5   15   15.5  RT (min)      lcorbi_t3_pos_1_005_42    m/z 643.5277     0   2   4   6E+5   14   14.5  RT (min)      lcorbi_t3_pos_4_016_15    m/z 643.6336     0   1   16   16.5  RT (min)      lcorbi_t3_pos_2_056_56    m/z 643.6336     0   1   16   16.5  RT (min)      lcorbi_t3_pos_1_049_55    m/z 643.6337     0   1   15   15.5  RT (min)      lcorbi_t3_pos_1_021_18    m/z 643.6338     0   1   15   15.5  RT (min)      lcorbi_t3_pos_1_030_21    m/z 644.5308     0   1   17   17.5  RT (min)    lcorbi_t3_pos_2_012_01    m/z 644.5309     0   1   16.5   17  RT (min)    lcorbi_t3_pos_1_031_40    m/z 644.5310     0   1   15.5   16  RT (min)      lcorbi_t3_pos_2_022_38    m/z 644.5310     0   1   13.5   14  RT (min)      lcorbi_t3_pos_2_013_33    m/z 644.5310     0   1   15.5   16  RT (min)      lcorbi_t3_pos_1_008_36    m/z 644.5310     0   1   16   16.5  RT (min)      lcorbi_t3_pos_1_028_32    m/z 644.5310     0   1   16.5   17  RT (min)    lcorbi_t3_pos_1_031_40    m/z 644.5310     0   1   16   16.5  RT (min)      lcorbi_t3_pos_1_028_32    m/z 644.5310     0   1   13.5   14  RT (min)      lcorbi_t3_pos_3_031_45    m/z 644.5310     0   1   13.5   14  RT (min)      lcorbi_t3_pos_3_024_07    m/z 644.5310     0   1   2   3E+4   15   15.5  RT (min)      lcorbi_t3_pos_1_008_36    m/z 644.5310     0   1   16   16.5  RT (min)      lcorbi_t3_pos_1_034_34    m/z 644.5312     0   1   16   16.5  RT (min)      lcorbi_t3_pos_1_034_34    m/z 645.5432     0   1   14   14.5  RT (min)      lcorbi_t3_pos_2_013_33    m/z 645.6492     0   1   16   16.5  RT (min)      lcorbi_t3_pos_1_056_56    m/z 645.6493     0   1E+6   14.5   15  RT (min)      lcorbi_t3_pos_3_026_04    m/z 645.6493     0   1   16   16.5  RT (min)      lcorbi_t3_pos_1_041_54    m/z 645.6494     0   1   15   15.5  RT (min)      lcorbi_t3_pos_1_047_41    m/z 645.6494     0   1   15.5   16  RT (min)      lcorbi_t3_pos_1_021_18    m/z 645.6494     0   1   15   15.5  RT (min)      lcorbi_t3_pos_2_046_07    m/z 645.6494     0   1   16   16.5  RT (min)      lcorbi_t3_pos_2_049_55    m/z 645.6495     0   1E+4   15   15.5  RT (min)      lcorbi_t3_pos_3_025_52    m/z 645.6495     0   1   15.5   16  RT (min)      lcorbi_t3_pos_1_030_21    m/z 645.6495     0   1   15.5   16  RT (min)      lcorbi_t3_pos_2_025_52    m/z 645.6495     0   1   16.5   17  RT (min)    lcorbi_t3_pos_2_056_56    m/z 645.6496     0   1   16.5   17  RT (min)    lcorbi_t3_pos_1_025_52    m/z 645.6496     0   1   16   16.5  RT (min)      lcorbi_t3_pos_1_056_56    m/z 645.6497     0   1E+6   14.5   15  RT (min)      lcorbi_t3_pos_3_026_04    m/z 645.6572     0   2   4   6E+4   14.5   15  RT (min)      lcorbi_t3_pos_1_005_42    m/z 646.2586     0   1   15.5   16  RT (min)      lcorbi_t3_pos_2_052_48    m/z 646.2586     0   1   2E+4   13   13.5  RT (min)      lcorbi_t3_pos_2_031_20    m/z 646.2586     0   1   2E+6   13.5   14  RT (min)      lcorbi_t3_pos_4_105_61    m/z 646.2586     0   1   2E+4   13   13.5  RT (min)      lcorbi_t3_pos_2_031_20    m/z 646.2586     0   1   15   15.5  RT (min)      lcorbi_t3_pos_1_038_30    m/z 646.2586     0   1   15.5   16  RT (min)      lcorbi_t3_pos_2_052_48    m/z 646.2586     0   1   16   16.5  RT (min)      lcorbi_t3_pos_1_027_43    m/z 646.2588     0   1   15.5   16  RT (min)      lcorbi_t3_pos_2_014_37    m/z 646.2588     0   1E+4   14.5   15  RT (min)      lcorbi_t3_pos_3_044_46    m/z 646.2588     0   1   15   15.5  RT (min)      lcorbi_t3_pos_1_038_30    m/z 646.2588     0   1   2   3E+4   13   13.5  RT (min)      lcorbi_t3_pos_2_047_31    m/z 646.2590     0   1E+4   14.5   15  RT (min)      lcorbi_t3_pos_3_044_46    m/z 646.6615     0   1   16.5   17  RT (min)    lcorbi_t3_pos_1_049_55    m/z 646.6617     0   1   16   16.5  RT (min)      lcorbi_t3_pos_2_009_50    m/z 646.6617     0   1   15.5   16  RT (min)      lcorbi_t3_pos_2_006_47    m/z 646.6617     0   2   4   6E+5   14.5   15  RT (min)      lcorbi_t3_pos_3_040_41    m/z 646.6617     0   1   15.5   16  RT (min)      lcorbi_t3_pos_2_041_54    m/z 646.6617     0   1   15.5   16  RT (min)      lcorbi_t3_pos_1_030_21    m/z 646.6618     0   1   16.5   17  RT (min)    lcorbi_t3_pos_1_001_49    m/z 646.6618     0   2   4E+4   15   15.5  RT (min)      lcorbi_t3_pos_4_001_49    m/z 646.6618     0   1   16.5   17  RT (min)    lcorbi_t3_pos_2_056_56    m/z 646.6618     0   1   2E+4   15   15.5  RT (min)      lcorbi_t3_pos_3_056_56    m/z 646.6618     0   2   4   6E+5   15   15.5  RT (min)      lcorbi_t3_pos_1_005_42    m/z 647.4588     0   1E+4   16   16.5  RT (min)      lcorbi_t3_pos_2_046_07    m/z 647.5766     0   1   16.5   17  RT (min)    lcorbi_t3_pos_2_022_38    m/z 647.5767     0   1   17.5   18  RT (min)    lcorbi_t3_pos_1_031_40    m/z 647.6648     0   1   15.5   16  RT (min)      lcorbi_t3_pos_2_041_54    m/z 647.6649     0   1   16   16.5  RT (min)      lcorbi_t3_pos_2_049_55    m/z 647.6649     0   1   16.5   17  RT (min)    lcorbi_t3_pos_4_033_53    m/z 647.6649     0   1   15.5   16  RT (min)      lcorbi_t3_pos_1_021_18    m/z 647.6649     0   1   16.5   17  RT (min)    lcorbi_t3_pos_1_049_55    m/z 647.6650     0   1   2E+5   15   15.5  RT (min)      lcorbi_t3_pos_4_001_49    m/z 647.6650     0   1   16.5   17  RT (min)    lcorbi_t3_pos_1_025_52    m/z 647.6650     0   1   2E+6   14.5   15  RT (min)      lcorbi_t3_pos_3_026_04    m/z 647.6651     0   1   16   16.5  RT (min)      lcorbi_t3_pos_1_041_54    m/z 647.6652     0   1   15.5   16  RT (min)      lcorbi_t3_pos_1_047_41    m/z 647.6652     0   1   16.5   17  RT (min)    lcorbi_t3_pos_2_056_56    m/z 647.6652     0   2   4   6E+4   15   15.5  RT (min)      lcorbi_t3_pos_4_010_43    m/z 649.6804     0   1   17   17.5  RT (min)    lcorbi_t3_pos_2_056_56    m/z 649.6806     0   1   16.5   17  RT (min)    lcorbi_t3_pos_1_049_55    m/z 649.6807     0   1E+4   15.5   16  RT (min)      lcorbi_t3_pos_2_046_07    m/z 649.6808     0   1E+6   15   15.5  RT (min)      lcorbi_t3_pos_1_005_42    m/z 649.6808     0   1   15.5   16  RT (min)      lcorbi_t3_pos_1_030_21    m/z 649.6808     0   2   4   6   8E+5   15   15.5  RT (min)      lcorbi_t3_pos_1_005_42    m/z 649.6808     0   1   16.5   17  RT (min)    lcorbi_t3_pos_2_009_50    m/z 649.6808     0   1   15.5   16  RT (min)      lcorbi_t3_pos_1_033_53    m/z 649.6808     0   1   16   16.5  RT (min)      lcorbi_t3_pos_1_041_54    m/z 649.6808     0   1   15.5   16  RT (min)      lcorbi_t3_pos_1_033_53    m/z 652.6203     0   1   16   16.5  RT (min)      lcorbi_t3_pos_2_009_50    m/z 652.6204     0   2   4   6E+5   14.5   15  RT (min)      lcorbi_t3_pos_3_026_04    m/z 652.6204     0   1   15.5   16  RT (min)      lcorbi_t3_pos_1_021_18    m/z 652.6204     0   1   15.5   16  RT (min)      lcorbi_t3_pos_1_030_21    m/z 652.6204     0   1   16   16.5  RT (min)      lcorbi_t3_pos_2_038_03    m/z 652.6204     0   1   16.5   17  RT (min)    lcorbi_t3_pos_1_025_52    m/z 652.6204     0   1   16.5   17  RT (min)    lcorbi_t3_pos_1_056_56    m/z 652.6204     0   1   15   15.5  RT (min)      lcorbi_t3_pos_2_017_51    m/z 652.6205     0   1E+4   15   15.5  RT (min)      lcorbi_t3_pos_3_056_56    m/z 652.6205     0   1   16   16.5  RT (min)      lcorbi_t3_pos_1_041_54    m/z 652.6205     0   1   16.5   17  RT (min)    lcorbi_t3_pos_2_056_56    m/z 652.6206     0   2   4E+4   15   15.5  RT (min)      lcorbi_t3_pos_4_001_49    m/z 652.6716     0   2   4E+4   15.5   16  RT (min)      lcorbi_t3_pos_4_049_55    m/z 652.6717     0   1   2   3   4E+4   16   16.5  RT (min)      lcorbi_t3_pos_2_014_37    m/z 652.6717     0   1   2   3E+4   15.5   16  RT (min)      lcorbi_t3_pos_2_041_54    m/z 652.6717     0   1   17   17.5  RT (min)    lcorbi_t3_pos_1_023_02    m/z 652.6718     0   1   2   3   4E+4   15.5   16  RT (min)      lcorbi_t3_pos_2_025_52    m/z 652.6718     0   2   4E+4   15   15.5  RT (min)      lcorbi_t3_pos_3_044_46    m/z 652.6718     0   1   16.5   17  RT (min)    lcorbi_t3_pos_1_031_40    m/z 652.6718     0   1   13.5   14  RT (min)      lcorbi_t3_pos_2_039_30    m/z 652.6718     0   1   16.5   17  RT (min)    lcorbi_t3_pos_2_056_56    m/z 652.6718     0   1   2   3E+4   15.5   16  RT (min)       lcorbi_t3_pos_2_038_03    m/z 652.6719     0   2   4E+4   14   14.5  RT (min)      lcorbi_t3_pos_2_016_23    m/z 652.6719     0   1   13.5   14  RT (min)      lcorbi_t3_pos_3_031_45    m/z 652.6719     0   1   13.5   14  RT (min)      lcorbi_t3_pos_2_032_39    m/z 652.6719     0   1   2E+5   15   15.5  RT (min)       lcorbi_t3_pos_3_047_40    m/z 652.6720     0   2   4   6   8E+5   14.5   15  RT (min)       lcorbi_t3_pos_4_051_34    m/z 652.6720     0   1   2   3   4E+4   15   15.5  RT (min)      lcorbi_t3_pos_2_046_07    m/z 652.6720     0   1E+4   13.5   14  RT (min)      lcorbi_t3_pos_2_054_29    m/z 652.6720     0   2   4   6E+4   15   15.5  RT (min)      lcorbi_t3_pos_3_056_56    m/z 652.6720     0   2   4   6E+5   14   14.5  RT (min)      lcorbi_t3_pos_3_053_48    m/z 652.6721     0   1   2   3E+4   16   16.5  RT (min)       lcorbi_t3_pos_2_052_48    m/z 653.6752     0   1   2   3E+5   14.5   15  RT (min)      lcorbi_t3_pos_3_033_53    m/z 654.3320     0   1   2E+4   13.5   14  RT (min)      lcorbi_t3_pos_1_049_55    m/z 654.3321     0   1   13.5   14  RT (min)      lcorbi_t3_pos_1_041_54    m/z 654.3322     0   2   4   6   8E+5   12.5   13  RT (min)      lcorbi_t3_pos_1_008_36    m/z 654.3322     0   1   13.5   14  RT (min)      lcorbi_t3_pos_1_031_40    m/z 654.3322     0   1   11.5   12  RT (min)      lcorbi_t3_pos_3_024_07    m/z 654.3322     0   1E+6   11.5   12  RT (min)      lcorbi_t3_pos_3_011_42    m/z 654.3322     0   1   13   13.5  RT (min)      lcorbi_t3_pos_1_034_34    m/z 654.3323     0   1E+6   12.5   13  RT (min)      lcorbi_t3_pos_1_008_36    m/z 654.3324     0   1   2E+4   13   13.5  RT (min)      lcorbi_t3_pos_1_028_32    m/z 654.3324     0   1   14   14.5  RT (min)      lcorbi_t3_pos_2_012_01    m/z 654.3324     0   1E+6   11.5   12  RT (min)      lcorbi_t3_pos_2_013_33    m/z 654.5478     0   1   16.5   17  RT (min)    lcorbi_t3_pos_1_031_40    m/z 655.3354     0   1   11.5   12  RT (min)      lcorbi_t3_pos_3_024_07    m/z 655.3355     0   1   13.5   14  RT (min)      lcorbi_t3_pos_1_031_40    m/z 655.3356     0   1   13   13.5  RT (min)      lcorbi_t3_pos_1_034_34    m/z 655.3356     0   1   2E+4   13   13.5  RT (min)      lcorbi_t3_pos_1_028_32    m/z 655.3356     0   2   4E+5   12.5   13  RT (min)      lcorbi_t3_pos_1_008_36    m/z 655.3356     0   2   4   6E+5   11.5   12  RT (min)      lcorbi_t3_pos_3_011_42    m/z 655.3357     0   1   13.5   14  RT (min)      lcorbi_t3_pos_2_012_01    m/z 658.5102     0   1E+4   12   12.5  RT (min)      lcorbi_t3_pos_1_025_52    m/z 658.5102     0   2   4   6   8E+4   11.5   12  RT (min)      lcorbi_t3_pos_1_041_54    m/z 658.5102     0   1   2E+6   10.5   11  RT (min)       lcorbi_t3_pos_4_089_59    m/z 658.5406     0   1   17   17.5  RT (min)    lcorbi_t3_pos_1_023_02    m/z 658.5408     0   1   15.5   16  RT (min)      lcorbi_t3_pos_1_028_32    m/z 658.5410     0   1   13.5   14  RT (min)      lcorbi_t3_pos_2_013_33    m/z 658.5410     0   1   16.5   17  RT (min)    lcorbi_t3_pos_1_031_40    m/z 658.5411     0   1   16   16.5  RT (min)      lcorbi_t3_pos_1_034_34    m/z 660.5564     0   1   17   17.5  RT (min)    lcorbi_t3_pos_1_023_02    m/z 660.5564     0   1   16   16.5  RT (min)      lcorbi_t3_pos_1_034_34    m/z 660.5564     0   2   4   6E+4   13.5   14  RT (min)      lcorbi_t3_pos_1_002_01    m/z 660.5564     0   1   15.5   16  RT (min)      lcorbi_t3_pos_1_028_32    m/z 660.5565     0   1   16.5   17  RT (min)    lcorbi_t3_pos_1_031_40    m/z 660.5566     0   1   15   15.5  RT (min)      lcorbi_t3_pos_1_008_36    m/z 660.5566     0   1   15.5   16  RT (min)      lcorbi_t3_pos_1_028_32    m/z 660.5566     0   1   16   16.5  RT (min)      lcorbi_t3_pos_1_034_34    m/z 662.2326     0   1   15   15.5  RT (min)      lcorbi_t3_pos_1_038_30    m/z 662.2327     0   2   4   6E+5   13.5   14  RT (min)      lcorbi_t3_pos_4_105_61    m/z 662.5006     0   2   4   6E+4   14.5   15  RT (min)      lcorbi_t3_pos_3_044_46    m/z 662.5006     0   1   2   3   4E+5   14   14.5  RT (min)       lcorbi_t3_pos_1_005_42    m/z 662.5008     0   2   4   6E+5   13.5   14  RT (min)      lcorbi_t3_pos_4_015_45    m/z 662.5008     0   1   13   13.5  RT (min)      lcorbi_t3_pos_2_047_31    m/z 662.5008     0   1   2E+4   15.5   16  RT (min)      lcorbi_t3_pos_2_052_48    m/z 662.5008     0   1E+5   14.5   15  RT (min)      lcorbi_t3_pos_3_044_46    m/z 662.5008     0   1   2E+4   15.5   16  RT (min)      lcorbi_t3_pos_1_027_43    m/z 662.5008     0   1   13   13.5  RT (min)      lcorbi_t3_pos_3_052_47    m/z 662.5008     0   1   2   3   4E+4   15   15.5  RT (min)      lcorbi_t3_pos_1_055_33    m/z 662.5008     0   1   2E+4   16   16.5  RT (min)      lcorbi_t3_pos_1_027_43    m/z 662.5008     0   1   2   3   4E+4   15   15.5  RT (min)      lcorbi_t3_pos_1_055_33    m/z 662.5008     0   1E+6   13.5   14  RT (min)      lcorbi_t3_pos_4_015_45    m/z 662.5011     0   1   2E+4   16   16.5  RT (min)      lcorbi_t3_pos_1_027_43    m/z 663.4534     0   1   17   17.5  RT (min)     lcorbi_t3_pos_2_012_01    m/z 663.4538     0   1E+5   15   15.5  RT (min)      lcorbi_t3_pos_2_017_51    m/z 663.4538     0   1E+5   15.5   16  RT (min)      lcorbi_t3_pos_1_055_33    m/z 663.4539     0   2   4   6   8E+4   16   16.5  RT (min)       lcorbi_t3_pos_2_052_48    m/z 663.4540     0   1E+5   16   16.5  RT (min)       lcorbi_t3_pos_1_034_34    m/z 663.4540     0   1   17.5   18  RT (min)    lcorbi_t3_pos_1_023_02    m/z 663.4540     0   1   16.5   17  RT (min)    lcorbi_t3_pos_1_049_55    m/z 663.4540     0   1E+5   15.5   16  RT (min)       lcorbi_t3_pos_1_030_21    m/z 663.4540     0   2   4   6   8E+4   16   16.5   17  RT (min)       lcorbi_t3_pos_2_049_55    m/z 663.4540     0   1E+5   16   17   18  RT (min)       lcorbi_t3_pos_1_028_32    m/z 663.4540     0   1   16.5   17  RT (min)     lcorbi_t3_pos_1_031_40    m/z 663.4540     0   1   16.5   17  RT (min)    lcorbi_t3_pos_2_056_56    m/z 663.4540     0   1   17   17.5  RT (min)    lcorbi_t3_pos_1_023_02    m/z 663.4540     0   2   4E+5   15   15.5  RT (min)       lcorbi_t3_pos_1_008_36    m/z 663.4541     0   2   4   6E+4   13.5   14  RT (min)      lcorbi_t3_pos_2_039_30    m/z 663.4541     0   1   17   17.5  RT (min)     lcorbi_t3_pos_2_012_01    m/z 663.4541     0   1   2   3E+6   14   14.5  RT (min)      lcorbi_t3_pos_4_030_05    m/z 663.4541     0   1E+6   15   15.5  RT (min)      lcorbi_t3_pos_4_025_52    m/z 663.4541     0   1   2E+4   13.5   14  RT (min)      lcorbi_t3_pos_3_011_42    m/z 663.4541     0   1E+5   16   16.5  RT (min)       lcorbi_t3_pos_1_034_34    m/z 663.4542     0   1   17   17.5  RT (min)    lcorbi_t3_pos_2_012_01    m/z 663.4542     0   1   16.5   17  RT (min)    lcorbi_t3_pos_4_033_53    m/z 663.4542     0   2   4E+4   13.5   14  RT (min)      lcorbi_t3_pos_2_013_33    m/z 663.4542     0   1   17   17.5  RT (min)     lcorbi_t3_pos_1_023_02    m/z 663.4543     0   2   4   6E+4   13.5   14  RT (min)       lcorbi_t3_pos_3_024_07    m/z 663.4544     0   1   2E+6   0.5   1  RT (min)      lcorbi_t3_pos_4_057_49    m/z 663.4544     0   1E+5   16   16.5  RT (min)      lcorbi_t3_pos_2_025_52    m/z 663.4966     0   1E+4   14.5   15  RT (min)      lcorbi_t3_pos_3_044_46    m/z 663.5039     0   2   4E+5   13.5   14  RT (min)      lcorbi_t3_pos_2_016_23    m/z 663.5039     0   1   15.5   16  RT (min)      lcorbi_t3_pos_1_027_43    m/z 663.5040     0   1   13   13.5  RT (min)      lcorbi_t3_pos_3_035_08    m/z 663.5040     0   1   2E+4   15   15.5  RT (min)      lcorbi_t3_pos_1_055_33    m/z 663.5040     0   1   16   16.5  RT (min)      lcorbi_t3_pos_1_027_43    m/z 663.5041     0   1   2E+4   15   15.5  RT (min)      lcorbi_t3_pos_1_055_33    m/z 663.5041     0   1   13   13.5  RT (min)      lcorbi_t3_pos_3_052_47    m/z 663.5041     0   1E+4   15.5   16  RT (min)      lcorbi_t3_pos_2_014_37    m/z 663.5042     0   1   2E+4   15   15.5  RT (min)      lcorbi_t3_pos_1_038_30    m/z 663.5754     0   1E+5   14.5   15  RT (min)       lcorbi_t3_pos_1_002_01    m/z 664.1168     0   2   4   6E+5   1   1.5  RT (min)      lcorbi_t3_pos_2_027_05    m/z 664.4573     0   1   17   17.5  RT (min)    lcorbi_t3_pos_2_012_01    m/z 664.4573     0   1   17   17.5  RT (min)    lcorbi_t3_pos_1_023_02    m/z 664.4573     0   1E+6   15   15.5  RT (min)      lcorbi_t3_pos_3_026_04    m/z 664.4574     0   2   4   6E+4   15.5   16  RT (min)      lcorbi_t3_pos_2_041_54    m/z 664.4574     0   2   4E+4   15.5   16  RT (min)      lcorbi_t3_pos_2_022_38    m/z 664.4574     0   1E+6   14   14.5  RT (min)      lcorbi_t3_pos_4_016_15    m/z 664.4574     0   2   4   6E+4   15.5   16  RT (min)      lcorbi_t3_pos_1_028_32    m/z 664.4574     0   1   16.5   17  RT (min)    lcorbi_t3_pos_1_049_55    m/z 664.4574     0   1   2   3   4E+4   16   16.5  RT (min)      lcorbi_t3_pos_2_049_55    m/z 664.4575     0   2   4   6   8E+4   15.5   16  RT (min)      lcorbi_t3_pos_1_030_21    m/z 664.4575     0   2   4   6E+4   15.5   16  RT (min)      lcorbi_t3_pos_1_028_32    m/z 664.4575     0   1   17   17.5  RT (min)    lcorbi_t3_pos_2_012_01    m/z 664.4575     0   1E+4   13.5   14  RT (min)      lcorbi_t3_pos_3_011_42    m/z 664.4575     0   1   2   3   4E+4   16   16.5  RT (min)       lcorbi_t3_pos_1_034_34    m/z 664.4575     0   1   16.5   17  RT (min)    lcorbi_t3_pos_1_031_40    m/z 664.4575     0   1   2E+4   13.5   14  RT (min)      lcorbi_t3_pos_2_013_33    m/z 664.4576     0   1   2   3E+4   13.5   14  RT (min)      lcorbi_t3_pos_2_039_30    m/z 664.4576     0   1   16.5   17  RT (min)    lcorbi_t3_pos_2_012_01    m/z 664.4576     0   1   2E+5   15   15.5  RT (min)      lcorbi_t3_pos_1_008_36    m/z 664.4576     0   1   16.5   17  RT (min)    lcorbi_t3_pos_4_033_53    m/z 664.4576     0   1   2   3   4E+4   14   14.5  RT (min)      lcorbi_t3_pos_2_016_23    m/z 664.4576     0   1   2   3   4E+5   15   15.5  RT (min)      lcorbi_t3_pos_4_025_52    m/z 664.4576     0   1   17   17.5  RT (min)    lcorbi_t3_pos_1_023_02    m/z 664.4576     0   1   2E+4   13.5   14  RT (min)      lcorbi_t3_pos_3_024_07    m/z 664.4576     0   1   2E+4   13.5   14  RT (min)      lcorbi_t3_pos_2_013_33    m/z 664.4577     0   1E+6   15   15.5  RT (min)       lcorbi_t3_pos_3_026_04    m/z 664.4579     0   1   17.5   18  RT (min)     lcorbi_t3_pos_1_023_02    m/z 664.6028     0   1   17   17.5  RT (min)    lcorbi_t3_pos_1_034_34    m/z 664.6028     0   1   17   17.5  RT (min)    lcorbi_t3_pos_1_034_34    m/z 664.6029     0   2   4   6E+5   16   16.5  RT (min)      lcorbi_t3_pos_4_074_58    m/z 664.6030     0   2   4E+4   16   16.5  RT (min)       lcorbi_t3_pos_1_008_36    m/z 664.6030     0   1   2E+3   14   14.5  RT (min)      lcorbi_t3_pos_3_024_07    m/z 664.6030     0   1   17.5   18  RT (min)    lcorbi_t3_pos_1_031_40    m/z 664.6031     0   2   4E+4   16   16.5  RT (min)       lcorbi_t3_pos_1_008_36    m/z 664.6031     0   1   17.5   18  RT (min)    lcorbi_t3_pos_2_012_01    m/z 664.6032     0   2   4   6   8E+5   16   16.5  RT (min)       lcorbi_t3_pos_4_075_58    m/z 664.6032     0   2   4   6   8E+5   16.5   17  RT (min)       lcorbi_t3_pos_4_078_58    m/z 664.6032     0   1   16.5   17  RT (min)    lcorbi_t3_pos_1_028_32    m/z 664.6032     0   2   4   6   8E+5   16.5   17  RT (min)       lcorbi_t3_pos_4_078_58    m/z 664.6033     0   1   14   14.5  RT (min)      lcorbi_t3_pos_2_013_33    m/z 664.6034     0   1   17.5   18  RT (min)    lcorbi_t3_pos_1_031_40    m/z 665.4606     0   2   4   6   8E+4   15   15.5  RT (min)      lcorbi_t3_pos_1_008_36    m/z 665.4606     0   1   16.5   17  RT (min)    lcorbi_t3_pos_1_031_40    m/z 665.4607     0   1   2E+4   16   16.5  RT (min)      lcorbi_t3_pos_1_034_34    m/z 665.4607     0   1   2E+4   16   16.5  RT (min)      lcorbi_t3_pos_1_034_34    m/z 665.4607     0   1   2E+4   15.5   16  RT (min)      lcorbi_t3_pos_2_022_38    m/z 665.4608     0   1   2   3   4E+4   15.5   16  RT (min)      lcorbi_t3_pos_1_028_32    m/z 666.6189     0   1   2   3E+6   16.5   17  RT (min)       lcorbi_t3_pos_4_069_57    m/z 666.6190     0   1   2   3   4E+6   16.5   17  RT (min)       lcorbi_t3_pos_4_072_57    m/z 666.6190     0   1   2   3   4E+6   16.5   17  RT (min)       lcorbi_t3_pos_4_071_57    m/z 666.6192     0   1   2   3E+6   16   16.5  RT (min)       lcorbi_t3_pos_4_075_58    m/z 666.6874     0   1   13.5   14  RT (min)      lcorbi_t3_pos_2_013_33    m/z 666.6874     0   1E+5   15   15.5  RT (min)      lcorbi_t3_pos_3_047_40    m/z 666.6874     0   1   2   3E+4   15.5   16  RT (min)      lcorbi_t3_pos_1_030_21    m/z 666.6875     0   1   16.5   17  RT (min)    lcorbi_t3_pos_1_031_40    m/z 666.6875     0   1   2E+4   16   16.5  RT (min)      lcorbi_t3_pos_2_052_48    m/z 666.6875     0   1   13.5   14  RT (min)      lcorbi_t3_pos_3_024_07    m/z 666.6875     0   1   13.5   14  RT (min)      lcorbi_t3_pos_2_039_30    m/z 666.6876     0   1   16.5   17  RT (min)    lcorbi_t3_pos_2_056_56    m/z 666.6877     0   1   16.5   17  RT (min)    lcorbi_t3_pos_1_001_49    m/z 667.6224     0   1E+6   16   16.5  RT (min)      lcorbi_t3_pos_4_075_58    m/z 667.6224     0   1   2E+6   16   16.5  RT (min)       lcorbi_t3_pos_4_066_57    m/z 667.6910     0   1   2E+5   14.5   15  RT (min)      lcorbi_t3_pos_4_051_34    m/z 668.6258     0   2   4E+5   16.5   17  RT (min)       lcorbi_t3_pos_4_071_57    m/z 668.6259     0   2   4E+5   16.5   17  RT (min)       lcorbi_t3_pos_4_072_57    m/z 669.3344     0   1   15.5   16  RT (min)      lcorbi_t3_pos_1_041_54    m/z 669.3344     0   2   4   6E+4   13.5   14  RT (min)      lcorbi_t3_pos_2_031_20    m/z 669.3344     0   1E+6   14.5   15  RT (min)      lcorbi_t3_pos_4_025_52    m/z 669.3344     0   1   16   16.5  RT (min)      lcorbi_t3_pos_1_025_52    m/z 669.3344     0   1   16   16.5  RT (min)      lcorbi_t3_pos_1_056_56    m/z 669.3344     0   1E+5   15   15.5  RT (min)       lcorbi_t3_pos_3_056_56    m/z 669.3344     0   1   2E+4   15   15.5  RT (min)      lcorbi_t3_pos_4_041_54    m/z 669.3344     0   1E+4   15.5   16  RT (min)      lcorbi_t3_pos_2_041_54    m/z 669.3346     0   2   4E+6   13.5   14  RT (min)      lcorbi_t3_pos_3_053_48    m/z 669.3346     0   1   13   13.5  RT (min)      lcorbi_t3_pos_2_047_31    m/z 669.3346     0   1   2   3   4E+4   15   15.5  RT (min)      lcorbi_t3_pos_2_017_51    m/z 669.3346     0   2   4   6E+4   13.5   14  RT (min)      lcorbi_t3_pos_4_015_45    m/z 669.3346     0   1   2E+4   15.5   16  RT (min)      lcorbi_t3_pos_2_025_52    m/z 669.3346     0   1   13.5   14  RT (min)      lcorbi_t3_pos_3_035_08    m/z 669.3346     0   1   13.5   14  RT (min)      lcorbi_t3_pos_2_054_29    m/z 669.3347     0   1   2   3E+6   13.5   14  RT (min)      lcorbi_t3_pos_4_015_45    m/z 669.3348     0   1   16.5   17  RT (min)    lcorbi_t3_pos_2_012_01    m/z 669.3348     0   2   4   6E+6   14   14.5  RT (min)      lcorbi_t3_pos_4_105_61    m/z 669.3349     0   1   17   17.5  RT (min)    lcorbi_t3_pos_1_023_02    m/z 669.3350     0   2   4   6E+5   14.5   15  RT (min)      lcorbi_t3_pos_3_047_40    m/z 669.6404     0   1   16   16.5  RT (min)      lcorbi_t3_pos_2_009_50    m/z 669.6404     0   1   16   16.5  RT (min)      lcorbi_t3_pos_1_025_52    m/z 669.6405     0   1   2   3   4E+4   14.5   15  RT (min)      lcorbi_t3_pos_3_025_52    m/z 669.6406     0   1   16   16.5  RT (min)      lcorbi_t3_pos_1_025_52    m/z 669.6406     0   1   2E+4   15   15.5  RT (min)      lcorbi_t3_pos_3_049_55    m/z 669.6406     0   1   2E+4   15.5   16  RT (min)      lcorbi_t3_pos_2_041_54    m/z 669.6406     0   2   4   6   8E+5   14.5   15  RT (min)       lcorbi_t3_pos_3_026_04    m/z 669.6406     0   1   15.5   16  RT (min)      lcorbi_t3_pos_1_041_54    m/z 669.6406     0   1   2E+4   15   15.5  RT (min)      lcorbi_t3_pos_1_047_41    m/z 669.6408     0   2   4   6   8E+5   14.5   15  RT (min)      lcorbi_t3_pos_1_005_42    m/z 669.6408     0   1   2   3E+4   15   15.5  RT (min)       lcorbi_t3_pos_1_021_18    m/z 669.6409     0   1E+4   15   15.5  RT (min)      lcorbi_t3_pos_2_046_07    m/z 669.6411     0   1   15.5   16  RT (min)      lcorbi_t3_pos_1_041_54    m/z 670.3376     0   1   15.5   16  RT (min)      lcorbi_t3_pos_4_048_24    m/z 670.3376     0   1   2E+4   15   15.5  RT (min)      lcorbi_t3_pos_2_017_51    m/z 670.3377     0   1E+4   15   15.5  RT (min)      lcorbi_t3_pos_4_041_54    m/z 670.3378     0   1   13   13.5  RT (min)      lcorbi_t3_pos_2_047_31    m/z 670.3378     0   1   13.5   14  RT (min)      lcorbi_t3_pos_3_035_08    m/z 670.3378     0   1   13.5   14  RT (min)      lcorbi_t3_pos_2_013_33    m/z 670.3378     0   1E+4   13.5   14  RT (min)      lcorbi_t3_pos_2_031_20    m/z 670.3379     0   1E+5   14.5   15  RT (min)      lcorbi_t3_pos_4_001_49    m/z 670.3379     0   1   13.5   14  RT (min)      lcorbi_t3_pos_1_002_01    m/z 670.3379     0   1   15.5   16  RT (min)      lcorbi_t3_pos_1_041_54    m/z 670.3379     0   1   2E+6   14.5   15  RT (min)      lcorbi_t3_pos_3_041_54    m/z 670.3379     0   1   15.5   16  RT (min)      lcorbi_t3_pos_1_041_54    m/z 670.3379     0   1   15.5   16  RT (min)      lcorbi_t3_pos_2_041_54    m/z 670.3380     0   1E+4   13.5   14  RT (min)      lcorbi_t3_pos_4_015_45    m/z 670.3380     0   1   16   16.5  RT (min)      lcorbi_t3_pos_1_056_56    m/z 670.3380     0   1   17   17.5  RT (min)    lcorbi_t3_pos_1_023_02    m/z 670.3380     0   2   4   6   8E+4   15   15.5  RT (min)      lcorbi_t3_pos_3_025_52    m/z 670.3381     0   1   16.5   17  RT (min)    lcorbi_t3_pos_1_025_52    m/z 670.3383     0   1   17   17.5  RT (min)    lcorbi_t3_pos_1_023_02    m/z 670.6526     0   2   4   6E+5   14.5   15  RT (min)      lcorbi_t3_pos_1_005_42    m/z 670.6526     0   1   15   15.5  RT (min)      lcorbi_t3_pos_2_017_51    m/z 670.6526     0   1   15.5   16  RT (min)      lcorbi_t3_pos_1_041_54    m/z 670.6527     0   1   16   16.5  RT (min)    lcorbi_t3_pos_1_025_52    m/z 670.6527     0   1   15.5   16  RT (min)      lcorbi_t3_pos_1_033_53    m/z 670.6528     0   1   15   15.5  RT (min)      lcorbi_t3_pos_1_030_21    m/z 670.6530     0   1   16   16.5  RT (min)      lcorbi_t3_pos_1_049_55    m/z 671.5760     0   1E+6   16   16.5  RT (min)       lcorbi_t3_pos_4_075_58    m/z 671.5760     0   1   16.5   17  RT (min)     lcorbi_t3_pos_1_028_32    m/z 671.5761     0   1E+6   16.5   17  RT (min)       lcorbi_t3_pos_4_073_57    m/z 671.5765     0   1   17.5   18  RT (min)    lcorbi_t3_pos_1_031_40    m/z 671.5765     0   1   16.5   17  RT (min)     lcorbi_t3_pos_1_028_32    m/z 671.5765     0   1   17.5   18  RT (min)    lcorbi_t3_pos_1_031_40    m/z 671.5766     0   1   14   14.5  RT (min)      lcorbi_t3_pos_2_032_39    m/z 671.5766     0   1   2   3E+4   14   14.5  RT (min)      lcorbi_t3_pos_2_013_33    m/z 671.5766     0   1   17.5   18  RT (min)    lcorbi_t3_pos_2_012_01    m/z 671.5766     0   1E+4   14.5   15  RT (min)      lcorbi_t3_pos_3_011_42    m/z 671.5766     0   1   14.5   15  RT (min)      lcorbi_t3_pos_1_002_01    m/z 671.5766     0   1   2E+4   14   14.5  RT (min)      lcorbi_t3_pos_3_024_07    m/z 671.5766     0   1   17   17.5  RT (min)    lcorbi_t3_pos_1_034_34    m/z 671.5767     0   1   18   18.5  RT (min)    lcorbi_t3_pos_1_023_02    m/z 671.6557     0   1   16   16.5  RT (min)      lcorbi_t3_pos_1_056_56    m/z 671.6559     0   1   15.5   16  RT (min)      lcorbi_t3_pos_1_041_54    m/z 671.6560     0   1   15   15.5  RT (min)      lcorbi_t3_pos_2_017_51    m/z 671.6560     0   1   2E+6   14.5   15  RT (min)      lcorbi_t3_pos_1_005_42    m/z 671.6560     0   1   15.5   16  RT (min)      lcorbi_t3_pos_2_025_52    m/z 671.6560     0   1   2   3E+4   15   15.5  RT (min)      lcorbi_t3_pos_3_056_56    m/z 671.6560     0   1   2   3E+4   15.5   16  RT (min)      lcorbi_t3_pos_1_021_18    m/z 671.6562     0   1   16.5   17  RT (min)    lcorbi_t3_pos_1_025_52    m/z 671.6562     0   1   15   15.5  RT (min)      lcorbi_t3_pos_1_030_21    m/z 672.5778     0   1   16.5   17  RT (min)     lcorbi_t3_pos_2_022_38    m/z 672.5779     0   2   4   6E+5   16.5   17  RT (min)      lcorbi_t3_pos_4_077_58    m/z 672.5784     0   1   16.5   17  RT (min)    lcorbi_t3_pos_1_028_32    m/z 672.5797     0   1   17.5   18  RT (min)    lcorbi_t3_pos_1_031_40    m/z 672.5798     0   1   17   17.5  RT (min)    lcorbi_t3_pos_1_034_34    m/z 672.5799     0   1   14   14.5  RT (min)      lcorbi_t3_pos_3_031_45    m/z 672.5800     0   1   14.5   15  RT (min)      lcorbi_t3_pos_3_011_42    m/z 672.5800     0   1   17.5   18  RT (min)    lcorbi_t3_pos_2_012_01    m/z 672.5800     0   1   14.5   15  RT (min)      lcorbi_t3_pos_3_011_42    m/z 672.5800     0   2   4   6E+5   16.5   17  RT (min)      lcorbi_t3_pos_4_068_57    m/z 672.5801     0   1   14   14.5  RT (min)      lcorbi_t3_pos_3_024_07    m/z 672.5801     0   1   17   17.5  RT (min)    lcorbi_t3_pos_1_034_34    m/z 672.6682     0   1   16.5   17  RT (min)    lcorbi_t3_pos_1_001_49    m/z 672.6684     0   1   16   16.5  RT (min)      lcorbi_t3_pos_2_049_55    m/z 672.6684     0   1   16   16.5  RT (min)      lcorbi_t3_pos_1_041_54    m/z 672.6684     0   1   15.5   16  RT (min)      lcorbi_t3_pos_2_025_52    m/z 672.6684     0   1   15.5   16  RT (min)      lcorbi_t3_pos_4_041_54    m/z 672.6684     0   1   16.5   17  RT (min)    lcorbi_t3_pos_1_049_55    m/z 672.6684     0   1   15   15.5  RT (min)      lcorbi_t3_pos_2_017_51    m/z 672.6685     0   1   16.5   17  RT (min)    lcorbi_t3_pos_1_025_52    m/z 672.6685     0   1   15.5   16  RT (min)      lcorbi_t3_pos_1_021_18    m/z 672.6686     0   1   15.5   16  RT (min)      lcorbi_t3_pos_1_030_21    m/z 672.6686     0   2   4   6   8E+5   14.5   15  RT (min)      lcorbi_t3_pos_1_005_42    m/z 672.6686     0   1   2E+4   15   15.5  RT (min)      lcorbi_t3_pos_4_010_43    m/z 673.6715     0   1   16.5   17  RT (min)    lcorbi_t3_pos_1_049_55    m/z 673.6715     0   1   16   16.5  RT (min)      lcorbi_t3_pos_1_041_54    m/z 673.6716     0   1   16.5   17  RT (min)    lcorbi_t3_pos_1_025_52    m/z 673.6716     0   1   2   3E+4   15   15.5  RT (min)      lcorbi_t3_pos_3_056_56    m/z 673.6716     0   1   15.5   16  RT (min)      lcorbi_t3_pos_2_006_47    m/z 673.6717     0   1   2   3E+4   15.5   16  RT (min)      lcorbi_t3_pos_1_021_18    m/z 673.6717     0   1   16   16.5  RT (min)      lcorbi_t3_pos_2_049_55    m/z 673.6717     0   1   2   3E+6   14.5   15  RT (min)      lcorbi_t3_pos_3_026_04    m/z 673.6717     0   1   15.5   16  RT (min)      lcorbi_t3_pos_2_041_54    m/z 673.6717     0   1   2   3   4E+5   15   15.5  RT (min)      lcorbi_t3_pos_4_001_49    m/z 673.6718     0   1   2   3E+6   14.5   15  RT (min)      lcorbi_t3_pos_1_005_42    m/z 674.2874     0   1   2   3   4E+5   12.5   13  RT (min)       lcorbi_t3_pos_4_105_61    m/z 674.2876     0   1   2   3   4E+5   13   13.5  RT (min)       lcorbi_t3_pos_2_032_39    m/z 674.2876     0   1   2   3   4E+5   13   13.5  RT (min)       lcorbi_t3_pos_2_047_31    m/z 674.2876     0   1   2   3E+5   13.5   14  RT (min)       lcorbi_t3_pos_2_039_30    m/z 674.2878     0   1   2E+5   13.5   14  RT (min)       lcorbi_t3_pos_2_031_20    m/z 674.2879     0   2   4   6   8E+4   15   15.5  RT (min)      lcorbi_t3_pos_3_025_52    m/z 674.2880     0   2   4E+5   14.5   15  RT (min)      lcorbi_t3_pos_4_025_52    m/z 674.2896     0   1   15.5   16  RT (min)      lcorbi_t3_pos_2_038_03    m/z 674.2896     0   1E+4   15   15.5  RT (min)      lcorbi_t3_pos_4_041_54    m/z 674.2896     0   1   15.5   16  RT (min)      lcorbi_t3_pos_1_041_54    m/z 674.2898     0   1   15.5   16  RT (min)      lcorbi_t3_pos_1_038_30    m/z 674.2898     0   1   16   16.5  RT (min)      lcorbi_t3_pos_1_025_52    m/z 674.2899     0   1   16   16.5  RT (min)      lcorbi_t3_pos_1_056_56    m/z 674.2899     0   1   15.5   16  RT (min)      lcorbi_t3_pos_1_033_53    m/z 674.2901     0   1   16.5   17  RT (min)    lcorbi_t3_pos_2_056_56    m/z 674.2902     0   1   17   17.5  RT (min)    lcorbi_t3_pos_1_023_02    m/z 674.2904     0   1   17   17.5  RT (min)    lcorbi_t3_pos_1_023_02    m/z 674.6838     0   1   16   16.5  RT (min)      lcorbi_t3_pos_2_038_03    m/z 674.6838     0   1   15.5   16  RT (min)      lcorbi_t3_pos_2_006_47    m/z 674.6839     0   1   16   16.5  RT (min)      lcorbi_t3_pos_1_041_54    m/z 674.6839     0   1   16.5   17  RT (min)    lcorbi_t3_pos_1_025_52    m/z 674.6839     0   1   16   16.5  RT (min)      lcorbi_t3_pos_2_009_50    m/z 674.6839     0   2   4   6   8E+5   15   15.5  RT (min)       lcorbi_t3_pos_3_040_41    m/z 674.6839     0   1   16.5   17  RT (min)    lcorbi_t3_pos_1_001_49    m/z 674.6840     0   1   15.5   16  RT (min)      lcorbi_t3_pos_1_021_18    m/z 674.6840     0   1   15.5   16  RT (min)      lcorbi_t3_pos_2_017_51    m/z 674.6840     0   1   15.5   16  RT (min)      lcorbi_t3_pos_1_030_21    m/z 674.6840     0   1   2   3E+4   15   15.5  RT (min)      lcorbi_t3_pos_3_025_52    m/z 674.6842     0   2   4   6   8E+5   15   15.5  RT (min)      lcorbi_t3_pos_1_005_42    m/z 674.6842     0   1   16.5   17  RT (min)    lcorbi_t3_pos_1_049_55    m/z 675.2910     0   1E+5   13   13.5  RT (min)       lcorbi_t3_pos_2_047_31    m/z 675.2912     0   2   4   6E+4   13.5   14  RT (min)      lcorbi_t3_pos_2_031_20    m/z 675.2930     0   1   15.5   16  RT (min)      lcorbi_t3_pos_1_041_54    m/z 675.5437     0   1   2E+4   14.5   15  RT (min)      lcorbi_t3_pos_1_034_34    m/z 675.5438     0   1   15   15.5  RT (min)      lcorbi_t3_pos_1_051_09    m/z 675.5438     0   1   15.5   16  RT (min)      lcorbi_t3_pos_1_023_02    m/z 675.5438     0   1   14   14.5  RT (min)      lcorbi_t3_pos_1_028_32    m/z 675.5439     0   1E+6   13.5   14  RT (min)      lcorbi_t3_pos_3_047_40    m/z 675.5439     0   1   2E+4   14   14.5  RT (min)      lcorbi_t3_pos_2_022_38    m/z 675.5439     0   1   14.5   15  RT (min)      lcorbi_t3_pos_1_041_54    m/z 675.5439     0   1   15   15.5  RT (min)      lcorbi_t3_pos_1_031_40    m/z 675.5439     0   1   15   15.5  RT (min)      lcorbi_t3_pos_1_051_09    m/z 675.5439     0   2   4E+5   12.5   13  RT (min)      lcorbi_t3_pos_3_011_42    m/z 675.5439     0   2   4E+5   13.5   14  RT (min)      lcorbi_t3_pos_1_008_36    m/z 675.5439     0   1   12.5   13  RT (min)      lcorbi_t3_pos_3_024_07    m/z 675.5439     0   1   15   15.5  RT (min)      lcorbi_t3_pos_1_031_40    m/z 675.5440     0   1E+6   13.5   14  RT (min)      lcorbi_t3_pos_3_047_40    m/z 675.5440     0   1   2E+5   14   14.5  RT (min)      lcorbi_t3_pos_1_008_36    m/z 675.5440     0   1   15.5   16  RT (min)      lcorbi_t3_pos_1_023_02    m/z 675.5441     0   1   12.5   13  RT (min)      lcorbi_t3_pos_2_032_39    m/z 675.6766     0   1   12.5   13  RT (min)      lcorbi_t3_pos_3_052_47    m/z 675.6872     0   1   16.5   17  RT (min)    lcorbi_t3_pos_1_001_49    m/z 675.6872     0   1   15.5   16  RT (min)      lcorbi_t3_pos_1_033_53    m/z 675.6872     0   1   2E+4   15.5   16  RT (min)      lcorbi_t3_pos_2_017_51    m/z 675.6872     0   1   15.5   16  RT (min)      lcorbi_t3_pos_1_030_21    m/z 675.6872     0   2   4   6E+5   15   15.5  RT (min)      lcorbi_t3_pos_4_025_52    m/z 675.6872     0   2   4   6   8E+4   15   15.5  RT (min)      lcorbi_t3_pos_3_025_52    m/z 675.6873     0   1   16   16.5  RT (min)      lcorbi_t3_pos_2_049_55    m/z 675.6873     0   1E+4   15.5   16  RT (min)      lcorbi_t3_pos_2_046_07    m/z 675.6873     0   1   16.5   17  RT (min)    lcorbi_t3_pos_1_049_55    m/z 675.6873     0   1   2   3E+6   15   15.5  RT (min)      lcorbi_t3_pos_1_005_42    m/z 675.6874     0   1   2   3E+6   15   15.5  RT (min)      lcorbi_t3_pos_3_026_04    m/z 675.6874     0   1   2E+4   16   16.5  RT (min)      lcorbi_t3_pos_1_041_54    m/z 675.6874     0   1   16   16.5  RT (min)      lcorbi_t3_pos_2_025_52    m/z 675.6875     0   1   16.5   17  RT (min)    lcorbi_t3_pos_4_033_53    m/z 676.3032     0   1   2   3E+5   13.5   14  RT (min)       lcorbi_t3_pos_2_044_21    m/z 676.5471     0   1   15.5   16  RT (min)      lcorbi_t3_pos_1_023_02    m/z 676.5472     0   1   15   15.5  RT (min)      lcorbi_t3_pos_1_031_40    m/z 676.5472     0   1   14   14.5  RT (min)      lcorbi_t3_pos_3_003_33    m/z 676.5473     0   1   2E+5   14   14.5  RT (min)      lcorbi_t3_pos_1_008_36    m/z 676.5473     0   2   4   6E+4   12.5   13  RT (min)      lcorbi_t3_pos_1_002_01    m/z 676.5473     0   1   14   14.5  RT (min)      lcorbi_t3_pos_1_028_32    m/z 676.5473     0   1E+5   14   14.5  RT (min)      lcorbi_t3_pos_1_008_36    m/z 676.5473     0   1   2   3E+5   12.5   13  RT (min)      lcorbi_t3_pos_1_002_01    m/z 676.5473     0   1   12.5   13  RT (min)      lcorbi_t3_pos_3_024_07    m/z 676.5473     0   1   14.5   15  RT (min)      lcorbi_t3_pos_1_034_34    m/z 676.5474     0   1   15   15.5  RT (min)      lcorbi_t3_pos_1_031_40    m/z 676.5474     0   1   14.5   15  RT (min)      lcorbi_t3_pos_1_028_32    m/z 676.5476     0   1   12.5   13  RT (min)      lcorbi_t3_pos_2_032_39    m/z 676.6112     0   1   15.5   16  RT (min)      lcorbi_t3_pos_1_041_54    m/z 676.6995     0   1   17   17.5  RT (min)    lcorbi_t3_pos_2_056_56    m/z 676.6995     0   1   16   16.5  RT (min)      lcorbi_t3_pos_1_041_54    m/z 676.6996     0   1E+6   15   15.5  RT (min)      lcorbi_t3_pos_1_005_42    m/z 676.6996     0   1   15.5   16  RT (min)      lcorbi_t3_pos_1_021_18    m/z 676.6996     0   1E+5   15   15.5  RT (min)      lcorbi_t3_pos_4_001_49    m/z 676.6997     0   1   16.5   17  RT (min)    lcorbi_t3_pos_4_033_53    m/z 676.6997     0   1E+6   15   15.5  RT (min)      lcorbi_t3_pos_3_040_41    m/z 676.6997     0   1   15.5   16  RT (min)      lcorbi_t3_pos_1_030_21    m/z 676.6997     0   1   16   16.5  RT (min)      lcorbi_t3_pos_2_025_52    m/z 676.6997     0   1   16.5   17  RT (min)    lcorbi_t3_pos_1_056_56    m/z 676.6997     0   1   16.5   17  RT (min)    lcorbi_t3_pos_1_025_52    m/z 676.6998     0   1   15.5   16  RT (min)      lcorbi_t3_pos_1_030_21    m/z 676.6998     0   1E+4   15.5   16  RT (min)      lcorbi_t3_pos_3_056_56    m/z 676.6998     0   1   15.5   16  RT (min)      lcorbi_t3_pos_2_046_07    m/z 677.4754     0   2   4E+5   13   13.5  RT (min)      lcorbi_t3_pos_4_037_26    m/z 677.4758     0   1   12.5   13  RT (min)      lcorbi_t3_pos_3_037_32    m/z 677.7028     0   1   16.5   17  RT (min)    lcorbi_t3_pos_1_056_56    m/z 677.7029     0   1   2   3   4E+6   15   15.5  RT (min)      lcorbi_t3_pos_1_005_42    m/z 677.7029     0   1   16.5   17  RT (min)    lcorbi_t3_pos_2_009_50    m/z 677.7029     0   1   15.5   16  RT (min)      lcorbi_t3_pos_1_030_21    m/z 677.7030     0   1   2   3   4E+5   15   15.5  RT (min)      lcorbi_t3_pos_4_001_49    m/z 677.7030     0   1   2   3   4E+6   15   15.5  RT (min)      lcorbi_t3_pos_3_040_41    m/z 677.7030     0   1   15.5   16  RT (min)      lcorbi_t3_pos_1_047_41    m/z 677.7030     0   2   4E+4   15.5   16  RT (min)      lcorbi_t3_pos_3_056_56    m/z 677.7030     0   1   16   16.5  RT (min)      lcorbi_t3_pos_2_041_54    m/z 677.7030     0   1   17   17.5  RT (min)    lcorbi_t3_pos_2_056_56    m/z 677.7030     0   1   16   16.5  RT (min)      lcorbi_t3_pos_1_041_54    m/z 677.7030     0   1   2E+4   15.5   16  RT (min)      lcorbi_t3_pos_2_046_07    m/z 677.7030     0   1   16   16.5  RT (min)      lcorbi_t3_pos_2_041_54    m/z 677.7031     0   1   16.5   17  RT (min)    lcorbi_t3_pos_1_056_56    m/z 678.5070     0   1   2   3E+4   15.5   16  RT (min)      lcorbi_t3_pos_2_014_37    m/z 678.5071     0   1E+4   13   13.5  RT (min)      lcorbi_t3_pos_3_052_47    m/z 678.5072     0   1E+5   14.5   15  RT (min)       lcorbi_t3_pos_4_028_04    m/z 678.5072     0   2   4   6E+4   15   15.5  RT (min)      lcorbi_t3_pos_2_041_54    m/z 678.5072     0   1   2   3   4E+4   15.5   16  RT (min)      lcorbi_t3_pos_1_027_43    m/z 678.5072     0   2   4E+4   15   15.5  RT (min)      lcorbi_t3_pos_1_034_34    m/z 678.5072     0   1   2   3E+4   15.5   16  RT (min)      lcorbi_t3_pos_1_001_49    m/z 678.5072     0   1   2   3E+4   15   15.5  RT (min)      lcorbi_t3_pos_2_014_37    m/z 678.5072     0   1E+5   15   15.5  RT (min)       lcorbi_t3_pos_1_038_30    m/z 678.5072     0   1   2E+5   14   14.5  RT (min)       lcorbi_t3_pos_4_025_52    m/z 678.5072     0   2   4   6   8E+3   13   13.5  RT (min)      lcorbi_t3_pos_3_035_08    m/z 678.5073     0   1   16   16.5  RT (min)      lcorbi_t3_pos_1_023_02    m/z 678.5073     0   1   16   16.5  RT (min)      lcorbi_t3_pos_1_023_02    m/z 678.5073     0   2   4E+4   15.5   16  RT (min)       lcorbi_t3_pos_2_052_48    m/z 678.5073     0   1   2   3   4E+4   15.5   16  RT (min)      lcorbi_t3_pos_1_027_43    m/z 678.5075     0   1E+5   14.5   15  RT (min)       lcorbi_t3_pos_3_056_56    m/z 678.5076     0   1   2E+5   14.5   15  RT (min)       lcorbi_t3_pos_3_044_46    m/z 678.5077     0   1   2E+5   14.5   15  RT (min)       lcorbi_t3_pos_3_044_46    m/z 678.6267     0   1   16.5   17  RT (min)     lcorbi_t3_pos_1_025_52    m/z 678.6269     0   1   16.5   17  RT (min)    lcorbi_t3_pos_1_056_56    m/z 678.6269     0   1   15.5   16  RT (min)      lcorbi_t3_pos_2_041_54    m/z 678.6270     0   2   4   6E+5   14.5   15  RT (min)      lcorbi_t3_pos_3_026_04    m/z 678.6270     0   1   16   16.5  RT (min)      lcorbi_t3_pos_1_041_54    m/z 678.6270     0   2   4   6E+5   14.5   15  RT (min)      lcorbi_t3_pos_3_026_04    m/z 678.6270     0   1   16.5   17  RT (min)    lcorbi_t3_pos_1_025_52    m/z 678.6270     0   1   15   15.5  RT (min)      lcorbi_t3_pos_2_046_07    m/z 678.6271     0   2   4   6   8E+4   15   15.5  RT (min)      lcorbi_t3_pos_4_001_49    m/z 678.6271     0   1   15.5   16  RT (min)      lcorbi_t3_pos_1_047_41    m/z 678.6271     0   1   15.5   16  RT (min)      lcorbi_t3_pos_1_021_18    m/z 678.6272     0   1   2   3E+4   15   15.5  RT (min)      lcorbi_t3_pos_4_010_43    m/z 678.6272     0   2   4   6E+5   14.5   15  RT (min)      lcorbi_t3_pos_1_005_42    m/z 679.5103     0   2   4   6   8E+5   13   13.5  RT (min)      lcorbi_t3_pos_4_015_45    m/z 679.5104     0   1E+6   13   13.5  RT (min)      lcorbi_t3_pos_4_037_26    m/z 679.5104     0   1   16   16.5  RT (min)      lcorbi_t3_pos_1_023_02    m/z 679.5105     0   1   13   13.5  RT (min)      lcorbi_t3_pos_3_052_47    m/z 679.5105     0   2   4   6   8E+3   13   13.5  RT (min)      lcorbi_t3_pos_3_035_08    m/z 679.5106     0   2   4   6   8E+4   14.5   15  RT (min)      lcorbi_t3_pos_3_044_46    m/z 679.5106     0   2   4E+4   14.5   15  RT (min)      lcorbi_t3_pos_1_038_30    m/z 679.5106     0   2   4   6E+4   14.5   15  RT (min)      lcorbi_t3_pos_3_056_56    m/z 679.5107     0   2   4E+4   15   15.5  RT (min)      lcorbi_t3_pos_1_038_30    m/z 679.5107     0   2   4   6   8E+5   14   14.5  RT (min)       lcorbi_t3_pos_3_015_06    m/z 679.5107     0   1   2   3E+4   15.5   16  RT (min)      lcorbi_t3_pos_1_027_43    m/z 679.5107     0   1   2E+4   15   15.5  RT (min)      lcorbi_t3_pos_2_014_37    m/z 679.5107     0   1   2E+4   15.5   16  RT (min)      lcorbi_t3_pos_2_052_48    m/z 679.5108     0   1   2   3   4E+4   14.5   15  RT (min)       lcorbi_t3_pos_4_028_04    m/z 679.5108     0   1E+5   14.5   15  RT (min)      lcorbi_t3_pos_3_044_46    m/z 679.5108     0   1   2   3E+4   15   15.5  RT (min)      lcorbi_t3_pos_2_025_52    m/z 679.5108     0   1   2E+4   15.5   16  RT (min)      lcorbi_t3_pos_2_052_48    m/z 679.6550     0   2   4E+5   13.5   14  RT (min)       lcorbi_t3_pos_4_097_60    m/z 679.6550     0   2   4   6E+5   13.5   14  RT (min)       lcorbi_t3_pos_4_095_60    m/z 679.6551     0   2   4   6E+5   13.5   14  RT (min)       lcorbi_t3_pos_4_092_60    m/z 679.7181     0   1   16.5   17  RT (min)    lcorbi_t3_pos_2_049_55    m/z 679.7184     0   1   16   16.5  RT (min)      lcorbi_t3_pos_4_048_24    m/z 679.7185     0   1   15.5   16  RT (min)      lcorbi_t3_pos_2_006_47    m/z 679.7185     0   1   16   16.5  RT (min)      lcorbi_t3_pos_2_025_52    m/z 679.7186     0   1   17   17.5  RT (min)    lcorbi_t3_pos_1_056_56    m/z 679.7186     0   1   2   3E+4   15.5   16  RT (min)      lcorbi_t3_pos_3_025_52    m/z 679.7186     0   1   17   17.5  RT (min)    lcorbi_t3_pos_1_056_56    m/z 679.7186     0   1   17   17.5  RT (min)    lcorbi_t3_pos_1_025_52    m/z 679.7187     0   1   16   16.5  RT (min)      lcorbi_t3_pos_2_006_47    m/z 679.7187     0   1   16.5   17  RT (min)    lcorbi_t3_pos_1_041_54    m/z 679.7188     0   1   15.5   16  RT (min)      lcorbi_t3_pos_2_046_07    m/z 680.4804     0   2   4   6E+4   15.5   16  RT (min)      lcorbi_t3_pos_2_006_47    m/z 680.4804     0   1E+6   15   15.5  RT (min)       lcorbi_t3_pos_3_040_41    m/z 680.4804     0   2   4E+4   15.5   16  RT (min)      lcorbi_t3_pos_3_003_33    m/z 680.4804     0   1   16.5   17  RT (min)     lcorbi_t3_pos_2_012_01    m/z 680.4804     0   1E+5   15   15.5  RT (min)       lcorbi_t3_pos_3_056_56    m/z 680.4804     0   1E+5   15.5   16  RT (min)       lcorbi_t3_pos_1_028_32    m/z 680.4804     0   2   4   6E+4   16   16.5  RT (min)       lcorbi_t3_pos_2_049_55    m/z 680.4804     0   1   17   17.5  RT (min)     lcorbi_t3_pos_2_012_01    m/z 680.4804     0   1   17   17.5  RT (min)     lcorbi_t3_pos_1_023_02    m/z 680.4804     0   2   4   6E+4   16   16.5  RT (min)       lcorbi_t3_pos_1_034_34    m/z 680.4805     0   2   4   6E+4   16   16.5  RT (min)      lcorbi_t3_pos_2_049_55    m/z 680.4806     0   1E+6   15   15.5  RT (min)       lcorbi_t3_pos_1_005_42    m/z 680.4806     0   1E+5   15.5   16  RT (min)       lcorbi_t3_pos_2_046_07    m/z 680.4806     0   1   2   3E+5   15   15.5  RT (min)       lcorbi_t3_pos_1_008_36    m/z 680.4806     0   1E+5   13.5   14  RT (min)      lcorbi_t3_pos_2_039_30    m/z 680.4806     0   1   2   3   4E+4   14   14.5  RT (min)      lcorbi_t3_pos_1_002_01    m/z 680.4806     0   2   4E+5   15   15.5  RT (min)       lcorbi_t3_pos_4_001_49    m/z 680.4806     0   1E+6   14   14.5  RT (min)      lcorbi_t3_pos_4_016_15    m/z 680.4806     0   1   2E+4   13.5   14  RT (min)      lcorbi_t3_pos_3_011_42    m/z 680.4806     0   2   4   6E+4   15.5   16  RT (min)      lcorbi_t3_pos_2_041_54    m/z 680.4806     0   1   16.5   17  RT (min)    lcorbi_t3_pos_1_001_49    m/z 680.4806     0   1E+5   15.5   16  RT (min)       lcorbi_t3_pos_1_030_21    m/z 680.4806     0   1   16.5   17  RT (min)    lcorbi_t3_pos_4_033_53    m/z 680.4806     0   2   4   6   8E+4   13.5   14  RT (min)      lcorbi_t3_pos_2_031_20    m/z 680.4806     0   1   16.5   17  RT (min)     lcorbi_t3_pos_1_031_40    m/z 680.4860     0   2   4E+4   13.5   14  RT (min)       lcorbi_t3_pos_3_031_45    m/z 680.6344     0   1   16   16.5  RT (min)       lcorbi_t3_pos_2_049_55    m/z 680.6357     0   1   16.5   17  RT (min)     lcorbi_t3_pos_1_056_56    m/z 680.6424     0   1   15.5   16  RT (min)      lcorbi_t3_pos_1_047_41    m/z 680.6425     0   1   16   16.5  RT (min)      lcorbi_t3_pos_2_025_52    m/z 680.6425     0   1   16.5   17  RT (min)     lcorbi_t3_pos_1_025_52    m/z 680.6426     0   1   16.5   17  RT (min)    lcorbi_t3_pos_1_025_52    m/z 680.6427     0   1   2E+4   15   15.5  RT (min)      lcorbi_t3_pos_3_025_52    m/z 680.6427     0   1   15.5   16  RT (min)      lcorbi_t3_pos_1_030_21    m/z 680.6427     0   1E+5   15   15.5  RT (min)      lcorbi_t3_pos_4_001_49    m/z 680.6428     0   1E+4   15   15.5  RT (min)       lcorbi_t3_pos_3_056_56    m/z 680.6428     0   1   16   16.5  RT (min)      lcorbi_t3_pos_1_041_54    m/z 680.6428     0   2   4   6E+5   15   15.5  RT (min)       lcorbi_t3_pos_3_026_04    m/z 680.6428     0   1   15.5   16  RT (min)      lcorbi_t3_pos_1_047_41    m/z 680.6428     0   1   15.5   16  RT (min)      lcorbi_t3_pos_1_033_53    m/z 680.6429     0   2   4   6E+5   15   15.5  RT (min)      lcorbi_t3_pos_1_005_42    m/z 681.4837     0   1   16   16.5  RT (min)      lcorbi_t3_pos_2_009_50    m/z 681.4838     0   1   16.5   17  RT (min)    lcorbi_t3_pos_1_031_40    m/z 681.4839     0   1E+5   15   15.5  RT (min)      lcorbi_t3_pos_1_008_36    m/z 681.4839     0   1   17   17.5  RT (min)    lcorbi_t3_pos_1_023_02    m/z 681.4839     0   1   2E+4   16   16.5  RT (min)      lcorbi_t3_pos_2_049_55    m/z 681.4840     0   2   4E+4   15.5   16  RT (min)      lcorbi_t3_pos_1_033_53    m/z 681.4840     0   1E+5   15   15.5  RT (min)      lcorbi_t3_pos_1_008_36    m/z 681.4840     0   1   17   17.5  RT (min)    lcorbi_t3_pos_2_012_01    m/z 681.4840     0   1   2   3   4E+4   15.5   16  RT (min)      lcorbi_t3_pos_4_049_55    m/z 681.4840     0   1   2   3E+5   15   15.5  RT (min)      lcorbi_t3_pos_4_025_52    m/z 681.4840     0   1   2   3   4E+4   16   16.5  RT (min)      lcorbi_t3_pos_1_034_34    m/z 681.4840     0   1   2E+4   13.5   14  RT (min)      lcorbi_t3_pos_3_011_42    m/z 681.4840     0   1   2   3   4E+4   15.5   16  RT (min)      lcorbi_t3_pos_1_028_32    m/z 681.4840     0   2   4E+4   15.5   16  RT (min)      lcorbi_t3_pos_1_030_21    m/z 681.4840     0   1E+4   13.5   14  RT (min)      lcorbi_t3_pos_3_024_07    m/z 681.4840     0   1   16.5   17  RT (min)    lcorbi_t3_pos_1_025_52    m/z 681.4840     0   1   2E+4   16   16.5  RT (min)      lcorbi_t3_pos_2_025_52    m/z 681.4840     0   1   16.5   17  RT (min)    lcorbi_t3_pos_1_025_52    m/z 681.4840     0   1   2E+4   14   14.5  RT (min)      lcorbi_t3_pos_2_016_23    m/z 681.4840     0   2   4   6   8E+5   14   14.5  RT (min)      lcorbi_t3_pos_3_034_26    m/z 681.4841     0   2   4   6E+4   15   15.5  RT (min)      lcorbi_t3_pos_3_044_46    m/z 682.2902     0   1   15   15.5  RT (min)      lcorbi_t3_pos_1_038_30    m/z 682.2903     0   1E+4   13   13.5  RT (min)      lcorbi_t3_pos_2_047_31    m/z 682.5468     0   2   4E+5   11.5   12  RT (min)      lcorbi_t3_pos_4_094_60    m/z 682.5469     0   2   4E+5   11.5   12  RT (min)      lcorbi_t3_pos_4_095_60    m/z 682.5470     0   2   4   6E+5   11.5   12  RT (min)       lcorbi_t3_pos_1_003_45    m/z 682.5470     0   2   4   6E+5   11.5   12  RT (min)       lcorbi_t3_pos_1_003_45    m/z 682.5472     0   2   4E+5   11.5   12  RT (min)       lcorbi_t3_pos_4_096_60    m/z 682.6582     0   1   16.5   17  RT (min)    lcorbi_t3_pos_1_056_56    m/z 682.6583     0   1E+4   15.5   16  RT (min)      lcorbi_t3_pos_2_046_07    m/z 682.6583     0   1   16.5   17  RT (min)    lcorbi_t3_pos_2_049_55    m/z 682.6584     0   1   17   17.5  RT (min)    lcorbi_t3_pos_2_056_56    m/z 682.6584     0   1   16   16.5  RT (min)      lcorbi_t3_pos_1_021_18    m/z 682.6584     0   2   4   6   8E+4   15   15.5  RT (min)      lcorbi_t3_pos_4_001_49    m/z 682.6584     0   1   16.5   17  RT (min)    lcorbi_t3_pos_1_056_56    m/z 682.6584     0   1   15.5   16  RT (min)      lcorbi_t3_pos_4_049_55    m/z 682.6584     0   1   16   16.5  RT (min)      lcorbi_t3_pos_1_041_54    m/z 682.6584     0   1   17   17.5  RT (min)    lcorbi_t3_pos_2_056_56    m/z 682.6584     0   1   15.5   16  RT (min)      lcorbi_t3_pos_1_047_41    m/z 682.6585     0   2   4   6   8E+5   15   15.5  RT (min)      lcorbi_t3_pos_3_040_41    m/z 682.6585     0   1   16   16.5  RT (min)      lcorbi_t3_pos_2_038_03    m/z 682.6585     0   2   4   6   8E+5   15   15.5  RT (min)      lcorbi_t3_pos_3_040_41    m/z 683.4624     0   2   4   6E+5   13   13.5  RT (min)      lcorbi_t3_pos_4_015_45    m/z 683.4624     0   1   2   3E+4   15   15.5  RT (min)      lcorbi_t3_pos_2_014_37    m/z 683.4624     0   1   13   13.5  RT (min)      lcorbi_t3_pos_3_035_08    m/z 683.4624     0   1   13   13.5  RT (min)      lcorbi_t3_pos_3_035_08    m/z 683.4624     0   2   4   6   8E+5   13.5   14  RT (min)      lcorbi_t3_pos_3_053_48    m/z 683.4625     0   2   4   6E+5   13   13.5  RT (min)      lcorbi_t3_pos_4_015_45    m/z 683.4625     0   1   2   3   4E+4   15   15.5  RT (min)      lcorbi_t3_pos_1_038_30    m/z 683.4625     0   1   2   3   4E+4   14.5   15  RT (min)      lcorbi_t3_pos_1_038_30    m/z 683.4626     0   1   15.5   16  RT (min)      lcorbi_t3_pos_1_027_43    m/z 683.4626     0   1   2E+4   15.5   16  RT (min)      lcorbi_t3_pos_2_052_48    m/z 683.4626     0   2   4   6E+5   14   14.5  RT (min)       lcorbi_t3_pos_3_015_06    m/z 683.4626     0   1E+5   14.5   15  RT (min)      lcorbi_t3_pos_3_044_46    m/z 683.4627     0   2   4   6E+5   14   14.5  RT (min)       lcorbi_t3_pos_3_015_06    m/z 683.4628     0   1   2   3E+4   15.5   16  RT (min)      lcorbi_t3_pos_2_014_37    m/z 683.4628     0   1   2   3   4E+4   14.5   15  RT (min)      lcorbi_t3_pos_4_028_04    m/z 683.4862     0   2   4E+5   11   11.5  RT (min)       lcorbi_t3_pos_4_091_60    m/z 683.4866     0   2   4   6E+5   11   11.5  RT (min)       lcorbi_t3_pos_1_003_45    m/z 684.2030     0   1   16.5   17  RT (min)    lcorbi_t3_pos_1_049_55    m/z 684.2030     0   1   16   16.5  RT (min)      lcorbi_t3_pos_1_034_34    m/z 684.2030     0   1   16.5   17  RT (min)    lcorbi_t3_pos_2_014_37    m/z 684.2032     0   1   17.5   18  RT (min)    lcorbi_t3_pos_1_023_02    m/z 684.2032     0   2   4   6E+5   15   15.5  RT (min)      lcorbi_t3_pos_1_005_42    m/z 684.2032     0   1   16   16.5  RT (min)    lcorbi_t3_pos_2_014_37    m/z 684.2033     0   1E+4   15.5   16  RT (min)      lcorbi_t3_pos_4_049_55    m/z 684.2033     0   2   4   6E+4   14   14.5  RT (min)      lcorbi_t3_pos_1_002_01    m/z 684.2034     0   1E+7   14   14.5  RT (min)      lcorbi_t3_pos_4_105_61    m/z 684.2034     0   1   2E+4   15   15.5  RT (min)      lcorbi_t3_pos_4_010_43    m/z 684.2034     0   1   17.5   18  RT (min)    lcorbi_t3_pos_1_023_02    m/z 684.2034     0   1E+6   15   15.5  RT (min)       lcorbi_t3_pos_4_051_34    m/z 684.2034     0   1E+4   15.5   16  RT (min)      lcorbi_t3_pos_4_028_04    m/z 684.2034     0   1   14   14.5  RT (min)      lcorbi_t3_pos_4_046_44    m/z 684.2036     0   1   16.5   17  RT (min)    lcorbi_t3_pos_1_025_52    m/z 684.6502     0   2   4E+5   15   15.5  RT (min)      lcorbi_t3_pos_1_005_42    m/z 684.6502     0   1   16.5   17  RT (min)    lcorbi_t3_pos_2_009_50    m/z 684.6503     0   1   16   16.5  RT (min)      lcorbi_t3_pos_1_034_34    m/z 684.6504     0   2   4E+4   15   15.5  RT (min)      lcorbi_t3_pos_1_008_36    m/z 684.6504     0   1   17   17.5  RT (min)    lcorbi_t3_pos_2_012_01    m/z 684.6504     0   1   13.5   14  RT (min)      lcorbi_t3_pos_3_031_45    m/z 684.6504     0   1   15.5   16  RT (min)      lcorbi_t3_pos_2_017_51    m/z 684.6504     0   1   16   16.5  RT (min)      lcorbi_t3_pos_2_025_52    m/z 684.6504     0   1E+4   15.5   16  RT (min)      lcorbi_t3_pos_2_022_38    m/z 684.6504     0   1   16.5   17  RT (min)    lcorbi_t3_pos_1_031_40    m/z 684.6504     0   1   16   16.5  RT (min)      lcorbi_t3_pos_1_028_32    m/z 684.6505     0   1   15.5   16  RT (min)      lcorbi_t3_pos_2_017_51    m/z 684.6505     0   1   17   17.5  RT (min)    lcorbi_t3_pos_2_012_01    m/z 684.6505     0   1   14   14.5  RT (min)      lcorbi_t3_pos_3_011_42    m/z 684.6507     0   1   14   14.5  RT (min)      lcorbi_t3_pos_2_013_33    m/z 685.2035     0   1   16   16.5  RT (min)      lcorbi_t3_pos_1_034_34    m/z 685.2038     0   1   15.5   16  RT (min)      lcorbi_t3_pos_4_028_04    m/z 685.2038     0   1   16.5   17  RT (min)    lcorbi_t3_pos_1_049_55    m/z 685.2038     0   1   16   16.5  RT (min)      lcorbi_t3_pos_1_034_34    m/z 685.2038     0   1   16.5   17  RT (min)    lcorbi_t3_pos_1_049_55    m/z 685.2039     0   2   4   6E+5   15   15.5  RT (min)      lcorbi_t3_pos_4_051_34    m/z 685.2040     0   1   16   16.5  RT (min)      lcorbi_t3_pos_1_034_34    m/z 685.2040     0   1   15.5   16  RT (min)      lcorbi_t3_pos_4_049_55    m/z 685.2040     0   1   17.5   18  RT (min)    lcorbi_t3_pos_1_023_02    m/z 685.2042     0   2   4   6   8E+6   14   14.5  RT (min)      lcorbi_t3_pos_4_105_61    m/z 685.6534     0   1   14   14.5  RT (min)      lcorbi_t3_pos_3_011_42    m/z 685.6536     0   1   2   3   4E+4   15.5   16  RT (min)      lcorbi_t3_pos_1_008_36    m/z 685.6536     0   1   2   3   4E+4   15.5   16  RT (min)      lcorbi_t3_pos_1_008_36    m/z 685.6537     0   1   16   16.5  RT (min)      lcorbi_t3_pos_1_034_34    m/z 685.6537     0   2   4E+4   15   15.5  RT (min)      lcorbi_t3_pos_1_008_36    m/z 685.6537     0   1   17   17.5  RT (min)    lcorbi_t3_pos_2_012_01    m/z 685.6537     0   1   15.5   16  RT (min)      lcorbi_t3_pos_2_022_38    m/z 685.6537     0   1   16   16.5  RT (min)      lcorbi_t3_pos_1_028_32    m/z 685.6538     0   1   16   16.5  RT (min)      lcorbi_t3_pos_1_028_32    m/z 685.6538     0   1   16.5   17  RT (min)    lcorbi_t3_pos_1_031_40    m/z 685.6538     0   1   17   17.5  RT (min)    lcorbi_t3_pos_2_012_01    m/z 685.6538     0   1   2   3E+3   13.5   14  RT (min)      lcorbi_t3_pos_3_031_45    m/z 685.6539     0   1   14   14.5  RT (min)      lcorbi_t3_pos_3_011_42    m/z 686.2011     0   1   16   16.5  RT (min)      lcorbi_t3_pos_2_014_37    m/z 686.2011     0   1   17   17.5  RT (min)    lcorbi_t3_pos_1_023_02    m/z 686.2012     0   1   16   16.5  RT (min)      lcorbi_t3_pos_1_034_34    m/z 686.2014     0   1   16.5   17  RT (min)    lcorbi_t3_pos_1_049_55    m/z 686.2020     0   2   4   6E+6   14   14.5  RT (min)      lcorbi_t3_pos_4_105_61    m/z 686.5417     0   2   4   6E+5   11   11.5  RT (min)       lcorbi_t3_pos_1_003_45    m/z 687.2022     0   1   2E+6   14   14.5  RT (min)      lcorbi_t3_pos_4_105_61    m/z 687.5688     0   1   16.5   17  RT (min)    lcorbi_t3_pos_1_028_32    m/z 687.5688     0   1   17.5   18  RT (min)    lcorbi_t3_pos_1_031_40    m/z 687.5689     0   1   17   17.5  RT (min)    lcorbi_t3_pos_1_034_34    m/z 687.5689     0   1   16.5   17  RT (min)    lcorbi_t3_pos_2_022_38    m/z 687.5691     0   1   16.5   17  RT (min)    lcorbi_t3_pos_1_028_32    m/z 687.5692     0   2   4   6E+4   14   14.5  RT (min)      lcorbi_t3_pos_3_024_07    m/z 687.5692     0   2   4   6E+4   14   14.5  RT (min)      lcorbi_t3_pos_2_013_33    m/z 688.2008     0   1E+6   14   14.5  RT (min)      lcorbi_t3_pos_4_105_61    m/z 689.5594     0   1   15   15.5  RT (min)      lcorbi_t3_pos_1_031_40    m/z 689.5594     0   1   15.5   16  RT (min)      lcorbi_t3_pos_2_012_01    m/z 689.5594     0   1   15.5   16  RT (min)      lcorbi_t3_pos_1_023_02    m/z 689.5595     0   1   16   16.5  RT (min)      lcorbi_t3_pos_1_023_02    m/z 689.5596     0   1   14.5   15  RT (min)      lcorbi_t3_pos_1_028_32    m/z 689.5596     0   1   14.5   15  RT (min)      lcorbi_t3_pos_1_030_21    m/z 689.5596     0   1E+5   14   14.5  RT (min)      lcorbi_t3_pos_1_008_36    m/z 689.5596     0   1   2E+4   14.5   15  RT (min)      lcorbi_t3_pos_3_003_33    m/z 689.5596     0   2   4   6E+5   14   14.5  RT (min)       lcorbi_t3_pos_3_047_40    m/z 689.5597     0   1   12.5   13  RT (min)      lcorbi_t3_pos_2_032_39    m/z 689.5598     0   1   14.5   15  RT (min)      lcorbi_t3_pos_1_034_34    m/z 689.5598     0   1   12.5   13  RT (min)      lcorbi_t3_pos_3_024_07    m/z 689.5600     0   1   15   15.5  RT (min)      lcorbi_t3_pos_1_031_40    m/z 689.5600     0   2   4   6E+5   13   13.5  RT (min)      lcorbi_t3_pos_4_016_15    m/z 690.2636     0   1   13.5   14  RT (min)      lcorbi_t3_pos_2_031_20    m/z 690.2636     0   1   15   15.5  RT (min)      lcorbi_t3_pos_2_017_51    m/z 690.2636     0   1   15   15.5  RT (min)      lcorbi_t3_pos_3_056_56    m/z 690.2637     0   1   15.5   16  RT (min)      lcorbi_t3_pos_1_041_54    m/z 690.2638     0   1E+4   15.5   16  RT (min)      lcorbi_t3_pos_2_025_52    m/z 690.2638     0   1   15   15.5  RT (min)      lcorbi_t3_pos_2_006_47    m/z 690.2638     0   1   15.5   16  RT (min)      lcorbi_t3_pos_2_041_54    m/z 690.2638     0   1E+4   13   13.5  RT (min)      lcorbi_t3_pos_2_047_31    m/z 690.2638     0   1   16   16.5  RT (min)      lcorbi_t3_pos_1_025_52    m/z 690.2640     0   2   4   6   8E+5   14   14.5  RT (min)      lcorbi_t3_pos_4_105_61    m/z 690.5071     0   2   4E+4   14.5   15  RT (min)      lcorbi_t3_pos_1_028_32    m/z 690.5071     0   1E+4   13   13.5  RT (min)      lcorbi_t3_pos_2_013_33    m/z 690.5071     0   1   2   3E+4   15   15.5  RT (min)      lcorbi_t3_pos_1_034_34    m/z 690.5072     0   2   4   6E+4   14.5   15  RT (min)      lcorbi_t3_pos_1_008_36    m/z 690.5072     0   1   2E+4   15   15.5  RT (min)      lcorbi_t3_pos_2_038_03    m/z 690.5072     0   1   2   3E+4   15   15.5  RT (min)      lcorbi_t3_pos_1_028_32    m/z 690.5072     0   1   2   3   4E+4   14.5   15  RT (min)      lcorbi_t3_pos_2_022_38    m/z 690.5072     0   1   2   3E+4   15.5   16  RT (min)      lcorbi_t3_pos_1_031_40    m/z 690.5072     0   1E+4   15.5   16  RT (min)      lcorbi_t3_pos_2_012_01    m/z 690.5072     0   1   2   3   4E+5   14   14.5  RT (min)       lcorbi_t3_pos_1_008_36    m/z 690.5072     0   1   2   3E+4   15.5   16  RT (min)      lcorbi_t3_pos_1_031_40    m/z 690.5630     0   1   2   3   4E+4   14   14.5  RT (min)      lcorbi_t3_pos_1_008_36    m/z 690.5630     0   1   14.5   15  RT (min)      lcorbi_t3_pos_1_034_34    m/z 690.5630     0   1   14.5   15  RT (min)      lcorbi_t3_pos_1_028_32    m/z 690.5630     0   1   2   3E+4   13   13.5  RT (min)      lcorbi_t3_pos_3_011_42    m/z 690.5630     0   1   15   15.5  RT (min)      lcorbi_t3_pos_1_031_40    m/z 690.6187     0   1   2E+6   16.5   17  RT (min)       lcorbi_t3_pos_4_079_58    m/z 690.6189     0   1   2E+6   16   16.5  RT (min)       lcorbi_t3_pos_4_071_57    m/z 690.6190     0   1   2E+6   16.5   17  RT (min)      lcorbi_t3_pos_4_079_58    m/z 691.3141     0   1   15.5   16  RT (min)      lcorbi_t3_pos_1_027_43    m/z 691.3142     0   2   4   6E+4   13   13.5  RT (min)       lcorbi_t3_pos_3_037_32    m/z 691.3146     0   1   15   15.5  RT (min)      lcorbi_t3_pos_1_038_30    m/z 691.3147     0   1   15.5   16  RT (min)      lcorbi_t3_pos_2_014_37    m/z 691.6222     0   1E+6   16   16.5  RT (min)       lcorbi_t3_pos_4_075_58    m/z 691.6223     0   1E+6   16   16.5  RT (min)       lcorbi_t3_pos_4_073_57    m/z 691.6224     0   1E+6   16.5   17  RT (min)       lcorbi_t3_pos_4_069_57    m/z 691.6225     0   1E+6   16   16.5  RT (min)       lcorbi_t3_pos_4_073_57    m/z 694.5019     0   1   12.5   13  RT (min)      lcorbi_t3_pos_2_047_31    m/z 694.5022     0   1   12.5   13  RT (min)      lcorbi_t3_pos_3_037_32    m/z 695.5744     0   2   4   6E+5   16   16.5  RT (min)       lcorbi_t3_pos_4_075_58    m/z 695.5744     0   2   4   6E+5   16   16.5  RT (min)       lcorbi_t3_pos_4_075_58    m/z 695.5744     0   2   4   6   8E+5   16   16.5  RT (min)       lcorbi_t3_pos_4_071_57    m/z 695.5744     0   2   4   6   8E+5   16.5   17  RT (min)       lcorbi_t3_pos_4_069_57    m/z 696.5538     0   2   4   6E+5   11.5   12  RT (min)       lcorbi_t3_pos_1_003_45    m/z 697.3656     0   1   2   3   4E+4   15   15.5  RT (min)      lcorbi_t3_pos_2_017_51    m/z 697.3656     0   1E+5   15   15.5  RT (min)      lcorbi_t3_pos_3_056_56    m/z 697.3656     0   2   4   6   8E+6   15   15.5  RT (min)      lcorbi_t3_pos_1_005_42    m/z 697.3656     0   1   16.5   17  RT (min)    lcorbi_t3_pos_1_025_52    m/z 697.3656     0   1   16.5   17  RT (min)    lcorbi_t3_pos_1_031_40    m/z 697.3656     0   1   13.5   14  RT (min)      lcorbi_t3_pos_2_032_39    m/z 697.3657     0   1   2E+5   13.5   14  RT (min)      lcorbi_t3_pos_1_002_01    m/z 697.3657     0   1   16.5   17  RT (min)    lcorbi_t3_pos_2_056_56    m/z 697.3657     0   1   2   3E+4   16   16.5  RT (min)      lcorbi_t3_pos_1_041_54    m/z 697.3657     0   2   4E+6   14   14.5  RT (min)       lcorbi_t3_pos_3_053_48    m/z 697.3657     0   1E+4   15.5   16  RT (min)      lcorbi_t3_pos_2_041_54    m/z 697.3657     0   1E+4   15.5   16  RT (min)      lcorbi_t3_pos_4_041_54    m/z 697.3657     0   1   2   3   4E+6   14.5   15  RT (min)      lcorbi_t3_pos_4_019_47    m/z 697.3658     0   1E+4   13.5   14  RT (min)      lcorbi_t3_pos_2_039_30    m/z 697.3658     0   1   2E+5   15   15.5  RT (min)      lcorbi_t3_pos_4_010_43    m/z 697.3658     0   1   17   17.5  RT (min)    lcorbi_t3_pos_1_023_02    m/z 697.3658     0   1   2   3E+4   16   16.5  RT (min)      lcorbi_t3_pos_1_041_54    m/z 697.3658     0   2   4   6   8E+6   14   14.5  RT (min)       lcorbi_t3_pos_4_105_61    m/z 697.3658     0   1E+4   15.5   16  RT (min)      lcorbi_t3_pos_4_041_54    m/z 697.3659     0   1   2   3E+4   15.5   16  RT (min)      lcorbi_t3_pos_2_017_51    m/z 698.3690     0   2   4   6E+4   15   15.5  RT (min)      lcorbi_t3_pos_4_010_43    m/z 698.3690     0   1E+4   13.5   14  RT (min)      lcorbi_t3_pos_2_054_29    m/z 698.3690     0   1   16.5   17  RT (min)    lcorbi_t3_pos_1_049_55    m/z 698.3690     0   1   16   16.5  RT (min)      lcorbi_t3_pos_1_041_54    m/z 698.3690     0   1   16.5   17  RT (min)    lcorbi_t3_pos_4_033_53    m/z 698.3690     0   1E+6   14.5   15  RT (min)      lcorbi_t3_pos_4_054_18    m/z 698.3690     0   1   15.5   16  RT (min)      lcorbi_t3_pos_2_041_54    m/z 698.3690     0   1E+4   13.5   14  RT (min)      lcorbi_t3_pos_2_032_39    m/z 698.3690     0   1   15.5   16  RT (min)      lcorbi_t3_pos_4_041_54    m/z 698.3691     0   1E+6   14   14.5  RT (min)      lcorbi_t3_pos_4_037_26    m/z 698.3691     0   1   2   3E+6   14   14.5  RT (min)      lcorbi_t3_pos_4_105_61    m/z 698.3691     0   1   13.5   14  RT (min)      lcorbi_t3_pos_2_047_31    m/z 698.3691     0   1   16.5   17  RT (min)    lcorbi_t3_pos_2_056_56    m/z 698.3691     0   1   2E+4   13.5   14  RT (min)      lcorbi_t3_pos_2_031_20    m/z 698.3692     0   1E+6   14   14.5  RT (min)      lcorbi_t3_pos_3_053_48    m/z 698.3692     0   1   2E+6   15   15.5  RT (min)      lcorbi_t3_pos_1_005_42    m/z 698.3692     0   1   17   17.5  RT (min)    lcorbi_t3_pos_1_023_02    m/z 698.3692     0   1   16   16.5  RT (min)      lcorbi_t3_pos_1_041_54    m/z 698.3692     0   1   2   3   4E+4   15   15.5  RT (min)      lcorbi_t3_pos_3_056_56    m/z 698.3692     0   1   15.5   16  RT (min)      lcorbi_t3_pos_2_006_47    m/z 698.3693     0   1E+6   14.5   15   15.5  RT (min)      lcorbi_t3_pos_3_017_51    m/z 698.3694     0   1   13.5   14  RT (min)      lcorbi_t3_pos_2_013_33    m/z 698.6656     0   1   17   17.5  RT (min)    lcorbi_t3_pos_1_031_40    m/z 698.6658     0   1   17   17.5  RT (min)    lcorbi_t3_pos_1_031_40    m/z 698.6658     0   1   17   17.5  RT (min)    lcorbi_t3_pos_2_012_01    m/z 698.6660     0   1   16   16.5  RT (min)      lcorbi_t3_pos_1_028_32    m/z 698.6660     0   1   16.5   17  RT (min)    lcorbi_t3_pos_1_034_34    m/z 698.6660     0   1   17   17.5  RT (min)    lcorbi_t3_pos_2_012_01    m/z 698.6661     0   2   4   6   8E+3   13.5   14  RT (min)      lcorbi_t3_pos_3_024_07    m/z 698.6662     0   1   14   14.5  RT (min)      lcorbi_t3_pos_3_011_42    m/z 698.6663     0   1   16   16.5  RT (min)      lcorbi_t3_pos_1_034_34    m/z 699.3723     0   2   4   6   8E+5   14   14.5  RT (min)      lcorbi_t3_pos_4_105_61    m/z 699.6690     0   1   17   17.5  RT (min)    lcorbi_t3_pos_1_031_40    m/z 699.6695     0   1   16   16.5  RT (min)      lcorbi_t3_pos_1_028_32    m/z 699.6695     0   1   15.5   16  RT (min)      lcorbi_t3_pos_1_008_36    m/z 699.6695     0   1   16   16.5  RT (min)      lcorbi_t3_pos_1_034_34    m/z 701.5594     0   1   15.5   16  RT (min)      lcorbi_t3_pos_1_023_02    m/z 701.5595     0   1   15   15.5  RT (min)      lcorbi_t3_pos_1_051_09    m/z 701.5595     0   1   2   3   4E+6   13.5   14  RT (min)      lcorbi_t3_pos_4_019_47    m/z 701.5596     0   2   4   6   8E+3   14   14.5  RT (min)      lcorbi_t3_pos_4_049_55    m/z 701.5596     0   1   14.5   15  RT (min)      lcorbi_t3_pos_1_028_32    m/z 701.5596     0   1   14.5   15  RT (min)      lcorbi_t3_pos_1_041_54    m/z 701.5596     0   1   2   3   4E+6   13   13.5  RT (min)      lcorbi_t3_pos_3_011_42    m/z 701.5596     0   1E+6   14   14.5  RT (min)      lcorbi_t3_pos_1_008_36    m/z 701.5596     0   2   4   6E+4   12.5   13  RT (min)      lcorbi_t3_pos_2_054_29    m/z 701.5596     0   1   2   3   4E+6   13.5   14  RT (min)      lcorbi_t3_pos_3_047_40    m/z 701.5596     0   1   14.5   15  RT (min)      lcorbi_t3_pos_1_028_32    m/z 701.5596     0   1   14.5   15  RT (min)      lcorbi_t3_pos_1_034_34    m/z 701.5596     0   1   14.5   15  RT (min)      lcorbi_t3_pos_1_034_34    m/z 701.5597     0   1   15.5   16  RT (min)      lcorbi_t3_pos_2_012_01    m/z 701.5597     0   1   15   15.5  RT (min)      lcorbi_t3_pos_1_031_40    m/z 701.5597     0   1E+4   14.5   15  RT (min)      lcorbi_t3_pos_3_003_33    m/z 701.5597     0   1   15   15.5  RT (min)      lcorbi_t3_pos_2_012_01    m/z 701.5597     0   1   2   3E+5   14   14.5  RT (min)      lcorbi_t3_pos_1_008_36    m/z 702.3211     0   1   13.5   14  RT (min)      lcorbi_t3_pos_2_047_31    m/z 702.3211     0   1E+6   14   14.5  RT (min)      lcorbi_t3_pos_3_053_48    m/z 702.3211     0   1   15.5   16  RT (min)      lcorbi_t3_pos_2_041_54    m/z 702.3211     0   1   16.5   17  RT (min)    lcorbi_t3_pos_1_025_52    m/z 702.3211     0   1   16.5   17  RT (min)    lcorbi_t3_pos_1_056_56    m/z 702.3212     0   1   2E+6   14   14.5  RT (min)       lcorbi_t3_pos_4_105_61    m/z 702.3212     0   1E+4   15.5   16  RT (min)      lcorbi_t3_pos_4_041_54    m/z 702.3212     0   1   17   17.5  RT (min)    lcorbi_t3_pos_1_023_02    m/z 702.3212     0   1   15.5   16  RT (min)      lcorbi_t3_pos_4_049_55    m/z 702.3212     0   1   16   16.5  RT (min)      lcorbi_t3_pos_2_049_55    m/z 702.3212     0   1E+6   14.5   15  RT (min)      lcorbi_t3_pos_3_041_54    m/z 702.3212     0   1   13.5   14  RT (min)      lcorbi_t3_pos_1_002_01    m/z 702.3212     0   1   13.5   14  RT (min)      lcorbi_t3_pos_2_031_20    m/z 702.3212     0   1   16.5   17  RT (min)    lcorbi_t3_pos_1_025_52    m/z 702.3212     0   1   2E+4   15   15.5  RT (min)      lcorbi_t3_pos_3_056_56    m/z 702.3213     0   1   13.5   14  RT (min)      lcorbi_t3_pos_2_032_39    m/z 702.3213     0   1   16   16.5  RT (min)      lcorbi_t3_pos_1_041_54    m/z 702.3213     0   1E+5   15   15.5  RT (min)      lcorbi_t3_pos_4_001_49    m/z 702.5362     0   2   4   6   8E+5   10.5   11  RT (min)       lcorbi_t3_pos_1_032_26    m/z 702.5362     0   1E+6   10.5   11  RT (min)       lcorbi_t3_pos_4_083_59    m/z 702.5628     0   1   15.5   16  RT (min)      lcorbi_t3_pos_1_023_02    m/z 702.5628     0   1   15.5   16  RT (min)      lcorbi_t3_pos_1_023_02    m/z 702.5629     0   1   14.5   15  RT (min)      lcorbi_t3_pos_1_041_54    m/z 702.5629     0   1   2   3E+4   12.5   13  RT (min)      lcorbi_t3_pos_2_054_29    m/z 702.5630     0   1   2   3E+4   14.5   15  RT (min)      lcorbi_t3_pos_1_034_34    m/z 702.5630     0   1   14.5   15  RT (min)      lcorbi_t3_pos_1_028_32    m/z 702.5630     0   1E+6   13.5   14  RT (min)      lcorbi_t3_pos_4_019_47    m/z 702.5630     0   1   15   15.5  RT (min)      lcorbi_t3_pos_1_056_56    m/z 702.5630     0   1E+6   13.5   14  RT (min)      lcorbi_t3_pos_3_047_40    m/z 702.5630     0   1   2E+4   15   15.5  RT (min)      lcorbi_t3_pos_1_051_09    m/z 702.5630     0   1   15   15.5  RT (min)      lcorbi_t3_pos_1_031_40    m/z 702.5630     0   2   4   6E+5   14   14.5  RT (min)      lcorbi_t3_pos_1_008_36    m/z 702.5630     0   1E+6   13   13.5  RT (min)      lcorbi_t3_pos_1_002_01    m/z 702.5630     0   2   4E+5   14   14.5  RT (min)      lcorbi_t3_pos_4_001_49    m/z 702.5630     0   1E+6   14   14.5  RT (min)      lcorbi_t3_pos_3_047_40    m/z 702.5630     0   1   14   14.5  RT (min)      lcorbi_t3_pos_2_022_38    m/z 702.5631     0   1   2   3E+4   14.5   15  RT (min)      lcorbi_t3_pos_1_034_34    m/z 702.5631     0   1   12.5   13  RT (min)      lcorbi_t3_pos_2_032_39    m/z 702.5631     0   1   14.5   15  RT (min)      lcorbi_t3_pos_1_041_54    m/z 702.5631     0   1   14   14.5  RT (min)      lcorbi_t3_pos_2_022_38    m/z 703.3243     0   1   15.5   16  RT (min)      lcorbi_t3_pos_2_041_54    m/z 703.3244     0   1   15   15.5  RT (min)      lcorbi_t3_pos_3_056_56    m/z 703.3244     0   1   16.5   17  RT (min)    lcorbi_t3_pos_1_056_56    m/z 703.3244     0   1   15.5   16  RT (min)      lcorbi_t3_pos_4_041_54    m/z 703.3244     0   1   13.5   14  RT (min)      lcorbi_t3_pos_2_047_31    m/z 703.3246     0   1   15.5   16  RT (min)      lcorbi_t3_pos_1_038_30    m/z 703.3246     0   2   4   6   8E+5   14   14.5  RT (min)      lcorbi_t3_pos_4_105_61    m/z 703.3246     0   1   16.5   17  RT (min)    lcorbi_t3_pos_1_025_52    m/z 703.5661     0   1   2E+4   14.5   15  RT (min)      lcorbi_t3_pos_1_047_41    m/z 703.5664     0   1   15   15.5   16  RT (min)      lcorbi_t3_pos_1_056_56    m/z 703.5664     0   1E+4   12.5   13  RT (min)      lcorbi_t3_pos_2_032_39    m/z 703.5670     0   1   2   3   4E+4   14   14.5  RT (min)      lcorbi_t3_pos_1_008_36    m/z 703.5744     0   1   2E+4   13   13.5  RT (min)      lcorbi_t3_pos_2_054_29    m/z 703.5746     0   1E+4   12.5   13  RT (min)      lcorbi_t3_pos_2_032_39    m/z 703.5748     0   1   2   3E+6   14   14.5  RT (min)      lcorbi_t3_pos_1_008_36    m/z 703.5748     0   1E+6   11.5   12  RT (min)      lcorbi_t3_pos_4_057_49    m/z 703.5750     0   1E+7   14   14.5  RT (min)      lcorbi_t3_pos_4_019_47    m/z 703.5750     0   1   2E+4   14.5   15  RT (min)      lcorbi_t3_pos_2_022_38    m/z 703.5750     0   1   2   3E+4   15.5   16  RT (min)      lcorbi_t3_pos_1_031_40    m/z 703.5750     0   1   2   3E+4   15.5   16  RT (min)      lcorbi_t3_pos_1_051_09    m/z 703.5750     0   1   2E+4   14.5   15  RT (min)       lcorbi_t3_pos_3_013_19    m/z 703.5750     0   1   2   3E+4   15   15.5  RT (min)      lcorbi_t3_pos_1_034_34    m/z 703.5751     0   1   2   3E+4   15   15.5  RT (min)      lcorbi_t3_pos_1_034_34    m/z 703.5751     0   1   2   3E+5   13   13.5  RT (min)       lcorbi_t3_pos_1_002_01    m/z 703.5752     0   1E+4   14.5   15  RT (min)      lcorbi_t3_pos_2_038_03    m/z 703.5752     0   1E+4   15.5   16  RT (min)      lcorbi_t3_pos_2_012_01    m/z 703.5752     0   1   15   15.5  RT (min)      lcorbi_t3_pos_1_041_54    m/z 703.5753     0   1   2   3E+4   15.5   16  RT (min)      lcorbi_t3_pos_1_051_09    m/z 703.5852     0   1   16   16.5  RT (min)      lcorbi_t3_pos_1_023_02    m/z 703.5854     0   1   16   16.5  RT (min)      lcorbi_t3_pos_1_023_02    m/z 703.5860     0   1   15.5   16  RT (min)      lcorbi_t3_pos_2_012_01    m/z 704.5227     0   2   4   6E+4   14.5   15  RT (min)      lcorbi_t3_pos_3_044_46    m/z 704.5227     0   1   16.5   17  RT (min)    lcorbi_t3_pos_1_023_02    m/z 704.5228     0   1   2E+6   14   14.5  RT (min)       lcorbi_t3_pos_1_005_42    m/z 704.5228     0   1E+4   16   16.5  RT (min)      lcorbi_t3_pos_2_056_56    m/z 704.5228     0   1   2   3E+4   15.5   16  RT (min)      lcorbi_t3_pos_1_041_54    m/z 704.5228     0   1   2   3   4E+5   14.5   15  RT (min)       lcorbi_t3_pos_4_025_52    m/z 704.5228     0   1   2   3E+4   15.5   16  RT (min)      lcorbi_t3_pos_1_041_54    m/z 704.5228     0   1E+4   13   13.5  RT (min)      lcorbi_t3_pos_3_035_08    m/z 704.5229     0   1E+5   14.5   15  RT (min)       lcorbi_t3_pos_3_044_46    m/z 704.5229     0   1E+4   15.5   16  RT (min)      lcorbi_t3_pos_2_014_37    m/z 704.5229     0   1   2   3   4E+4   15.5   16  RT (min)      lcorbi_t3_pos_1_027_43    m/z 704.5230     0   2   4   6E+4   15   15.5  RT (min)      lcorbi_t3_pos_1_055_33    m/z 704.5230     0   1   2   3   4E+4   16   16.5  RT (min)      lcorbi_t3_pos_1_027_43    m/z 704.5230     0   1E+4   16   16.5  RT (min)      lcorbi_t3_pos_2_056_56    m/z 704.5230     0   1   2   3   4E+4   15   15.5  RT (min)      lcorbi_t3_pos_1_033_53    m/z 704.5230     0   1   2   3E+4   15   15.5  RT (min)      lcorbi_t3_pos_2_038_03    m/z 704.5230     0   1E+4   15.5   16  RT (min)      lcorbi_t3_pos_2_052_48    m/z 704.5231     0   1   16.5   17  RT (min)    lcorbi_t3_pos_1_023_02    m/z 704.5231     0   1E+6   14.5   15  RT (min)      lcorbi_t3_pos_1_005_42    m/z 704.5232     0   2   4   6E+5   14   14.5  RT (min)       lcorbi_t3_pos_3_040_41    m/z 704.5232     0   1   16.5   17  RT (min)    lcorbi_t3_pos_1_023_02    m/z 704.5778     0   1   12.5   13  RT (min)      lcorbi_t3_pos_2_032_39    m/z 704.5783     0   1   2E+4   13   13.5  RT (min)      lcorbi_t3_pos_2_054_29    m/z 704.5783     0   2   4   6E+5   11.5   12  RT (min)      lcorbi_t3_pos_4_057_49    m/z 704.5783     0   1   16   16.5  RT (min)      lcorbi_t3_pos_1_023_02    m/z 704.5784     0   1E+4   14.5   15  RT (min)      lcorbi_t3_pos_3_013_19    m/z 704.5784     0   1E+6   13   13.5  RT (min)      lcorbi_t3_pos_2_054_29    m/z 704.5785     0   1   15.5   16  RT (min)      lcorbi_t3_pos_1_031_40    m/z 704.5785     0   1E+4   14.5   15  RT (min)      lcorbi_t3_pos_2_022_38    m/z 704.5786     0   1   2E+4   15.5   16  RT (min)      lcorbi_t3_pos_1_051_09    m/z 704.5786     0   1   2E+4   15   15.5  RT (min)      lcorbi_t3_pos_1_034_34    m/z 704.5786     0   1   2E+4   15.5   16  RT (min)      lcorbi_t3_pos_1_051_09    m/z 704.5786     0   1   16   16.5  RT (min)      lcorbi_t3_pos_1_023_02    m/z 704.5786     0   1E+6   14   14.5  RT (min)      lcorbi_t3_pos_1_008_36    m/z 704.5786     0   2   4   6E+6   14   14.5  RT (min)      lcorbi_t3_pos_4_019_47    m/z 704.5786     0   2   4   6E+6   14   14.5  RT (min)      lcorbi_t3_pos_4_019_47    m/z 704.5786     0   1   2E+4   15   15.5  RT (min)      lcorbi_t3_pos_1_034_34    m/z 704.5786     0   1   14.5   15  RT (min)      lcorbi_t3_pos_1_047_41    m/z 704.5787     0   1   15   15.5  RT (min)      lcorbi_t3_pos_2_009_50    m/z 704.5787     0   2   4E+5   14   14.5  RT (min)      lcorbi_t3_pos_1_008_36    m/z 705.5262     0   1   2   3   4E+4   15   15.5  RT (min)      lcorbi_t3_pos_1_033_53    m/z 705.5262     0   1E+4   15.5   16  RT (min)      lcorbi_t3_pos_2_049_55    m/z 705.5262     0   1E+6   14   14.5  RT (min)      lcorbi_t3_pos_1_005_42    m/z 705.5262     0   1   13   13.5  RT (min)      lcorbi_t3_pos_3_035_08    m/z 705.5263     0   1   16   16.5  RT (min)      lcorbi_t3_pos_1_025_52    m/z 705.5264     0   1E+4   15.5   16  RT (min)      lcorbi_t3_pos_2_014_37    m/z 705.5264     0   1   2   3E+4   15   15.5  RT (min)      lcorbi_t3_pos_1_055_33    m/z 705.5264     0   1   16.5   17  RT (min)    lcorbi_t3_pos_1_023_02    m/z 705.5264     0   1   2   3E+4   15   15.5  RT (min)      lcorbi_t3_pos_1_038_30    m/z 705.5264     0   1   2   3   4E+4   14.5   15  RT (min)      lcorbi_t3_pos_3_044_46    m/z 705.5264     0   1   15.5   16  RT (min)      lcorbi_t3_pos_2_049_55    m/z 705.5264     0   1   16   16.5  RT (min)      lcorbi_t3_pos_1_027_43    m/z 705.5265     0   1   16.5   17  RT (min)    lcorbi_t3_pos_1_023_02    m/z 705.5265     0   1   2   3   4E+4   14.5   15  RT (min)      lcorbi_t3_pos_3_044_46    m/z 705.5265     0   1   16   16.5  RT (min)      lcorbi_t3_pos_1_001_49    m/z 705.5265     0   1   15.5   16  RT (min)      lcorbi_t3_pos_1_049_55    m/z 705.5266     0   1   15.5   16  RT (min)      lcorbi_t3_pos_1_027_43    m/z 705.5815     0   1   13   13.5  RT (min)      lcorbi_t3_pos_4_012_37    m/z 705.5816     0   1   15   15.5  RT (min)      lcorbi_t3_pos_1_034_34    m/z 705.5816     0   1   2   3E+5   13   13.5  RT (min)      lcorbi_t3_pos_1_002_01    m/z 705.5817     0   1   13   13.5  RT (min)      lcorbi_t3_pos_2_054_29    m/z 705.5818     0   1   13   13.5  RT (min)      lcorbi_t3_pos_1_002_01    m/z 705.5818     0   1   14.5   15  RT (min)      lcorbi_t3_pos_2_022_38    m/z 705.5818     0   1   15.5   16  RT (min)      lcorbi_t3_pos_1_051_09    m/z 705.5818     0   1   15   15.5  RT (min)      lcorbi_t3_pos_1_034_34    m/z 705.5818     0   1   16   16.5  RT (min)      lcorbi_t3_pos_1_023_02    m/z 705.5818     0   1   12.5   13  RT (min)      lcorbi_t3_pos_2_032_39    m/z 705.5818     0   1   14.5   15  RT (min)      lcorbi_t3_pos_1_028_32    m/z 705.5818     0   1   14.5   15  RT (min)      lcorbi_t3_pos_1_028_32    m/z 705.5818     0   1   15.5   16  RT (min)      lcorbi_t3_pos_1_051_09    m/z 705.5818     0   1   2   3E+5   14   14.5  RT (min)      lcorbi_t3_pos_1_008_36    m/z 705.5820     0   1E+4   14   14.5  RT (min)      lcorbi_t3_pos_2_017_51    m/z 705.5824     0   1   15.5   16  RT (min)      lcorbi_t3_pos_1_031_40    m/z 705.5902     0   1E+5   14   14.5  RT (min)      lcorbi_t3_pos_1_008_36    m/z 705.6618     0   2   4   6E+5   13.5   14  RT (min)       lcorbi_t3_pos_1_003_45    m/z 705.6618     0   2   4   6E+5   13.5   14  RT (min)      lcorbi_t3_pos_4_093_60    m/z 706.5295     0   1   16   16.5  RT (min)      lcorbi_t3_pos_1_023_02    m/z 706.5300     0   1   2E+4   13.5   14  RT (min)      lcorbi_t3_pos_3_034_26    m/z 706.5341     0   1   15   15.5  RT (min)      lcorbi_t3_pos_1_034_34    m/z 706.5379     0   1   13   13.5  RT (min)      lcorbi_t3_pos_2_054_29    m/z 706.5383     0   1   15.5   16  RT (min)      lcorbi_t3_pos_1_051_09    m/z 706.5383     0   1   15.5   16  RT (min)      lcorbi_t3_pos_1_051_09    m/z 706.5384     0   1   16   16.5  RT (min)      lcorbi_t3_pos_1_023_02    m/z 706.5384     0   1E+4   13   13.5  RT (min)      lcorbi_t3_pos_4_012_37    m/z 706.5385     0   1   12.5   13  RT (min)      lcorbi_t3_pos_2_032_39    m/z 706.5385     0   1E+4   14.5   15  RT (min)      lcorbi_t3_pos_3_013_19    m/z 706.6134     0   1   16.5   17  RT (min)    lcorbi_t3_pos_3_003_33    m/z 706.6136     0   1   17   17.5  RT (min)    lcorbi_t3_pos_1_034_34    m/z 706.6136     0   1   16.5   17  RT (min)    lcorbi_t3_pos_1_028_32    m/z 706.6136     0   1   16.5   17  RT (min)    lcorbi_t3_pos_3_003_33    m/z 706.6136     0   1E+4   14   14.5  RT (min)      lcorbi_t3_pos_3_024_07    m/z 706.6136     0   1   17.5   18  RT (min)    lcorbi_t3_pos_2_012_01    m/z 706.6137     0   1   2   3E+4   16   16.5  RT (min)      lcorbi_t3_pos_1_008_36    m/z 706.6138     0   1   17.5   18  RT (min)    lcorbi_t3_pos_1_031_40    m/z 706.6323     0   1   16.5   17  RT (min)    lcorbi_t3_pos_1_031_40    m/z 707.6775     0   2   4   6   8E+5   13.5   14  RT (min)       lcorbi_t3_pos_4_092_60    m/z 707.6776     0   2   4   6E+5   13.5   14  RT (min)       lcorbi_t3_pos_4_090_60    m/z 707.6776     0   2   4   6E+5   13.5   14  RT (min)      lcorbi_t3_pos_4_093_60    m/z 707.6776     0   2   4   6   8E+5   13.5   14  RT (min)      lcorbi_t3_pos_4_095_60    m/z 708.5119     0   1   16.5   17  RT (min)    lcorbi_t3_pos_1_001_49    m/z 708.5536     0   2   4   6E+5   11   11.5  RT (min)      lcorbi_t3_pos_1_003_45    m/z 708.5537     0   2   4   6E+5   11   11.5  RT (min)       lcorbi_t3_pos_1_003_45    m/z 709.5570     0   1   2E+6   11.5   12  RT (min)      lcorbi_t3_pos_4_091_60    m/z 709.6928     0   2   4   6E+5   13.5   14  RT (min)       lcorbi_t3_pos_4_091_60    m/z 709.6929     0   2   4E+5   13.5   14  RT (min)       lcorbi_t3_pos_4_093_60    m/z 709.6930     0   2   4E+5   14   14.5  RT (min)      lcorbi_t3_pos_4_092_60    m/z 710.5692     0   2   4   6E+5   11.5   12  RT (min)       lcorbi_t3_pos_4_093_60    m/z 710.5694     0   2   4   6   8E+5   11.5   12  RT (min)       lcorbi_t3_pos_1_003_45    m/z 710.5694     0   2   4   6   8E+5   11.5   12  RT (min)       lcorbi_t3_pos_4_091_60    m/z 710.5695     0   2   4   6E+5   11.5   12  RT (min)      lcorbi_t3_pos_4_095_60    m/z 711.5687     0   1E+4   14.5   15  RT (min)      lcorbi_t3_pos_3_011_42    m/z 711.5688     0   1   17.5   18  RT (min)    lcorbi_t3_pos_1_031_40    m/z 711.5689     0   1E+4   14   14.5  RT (min)      lcorbi_t3_pos_3_024_07    m/z 711.5690     0   1   16   16.5  RT (min)      lcorbi_t3_pos_2_022_38    m/z 711.5691     0   2   4E+5   15   15.5  RT (min)      lcorbi_t3_pos_2_023_26    m/z 711.5691     0   1   16.5   17  RT (min)    lcorbi_t3_pos_1_028_32    m/z 711.5692     0   1   17.5   18  RT (min)    lcorbi_t3_pos_2_012_01    m/z 712.4916     0   1   14.5   15  RT (min)      lcorbi_t3_pos_1_028_32    m/z 712.4918     0   2   4   6   8E+4   14   14.5  RT (min)      lcorbi_t3_pos_1_008_36    m/z 712.4920     0   1   15   15.5  RT (min)      lcorbi_t3_pos_1_031_40    m/z 712.5487     0   2   4   6E+5   11   11.5  RT (min)       lcorbi_t3_pos_1_003_45    m/z 712.5489     0   2   4   6E+5   11   11.5  RT (min)      lcorbi_t3_pos_1_003_45    m/z 712.6814     0   1   14   14.5  RT (min)      lcorbi_t3_pos_3_031_45    m/z 712.6815     0   1   2E+4   15.5   16  RT (min)      lcorbi_t3_pos_1_008_36    m/z 712.6815     0   1   16.5   17  RT (min)    lcorbi_t3_pos_1_034_34    m/z 712.6816     0   1   16   16.5  RT (min)      lcorbi_t3_pos_2_022_38    m/z 712.6816     0   1   14   14.5  RT (min)      lcorbi_t3_pos_2_013_33    m/z 712.6816     0   1   17   17.5  RT (min)    lcorbi_t3_pos_1_031_40    m/z 712.6816     0   1   13.5   14  RT (min)      lcorbi_t3_pos_3_024_07    m/z 712.6816     0   1   17   17.5  RT (min)    lcorbi_t3_pos_1_031_40    m/z 712.6816     0   1   16   16.5  RT (min)      lcorbi_t3_pos_2_022_38    m/z 712.6816     0   1   17   17.5  RT (min)    lcorbi_t3_pos_2_012_01    m/z 712.6816     0   2   4E+5   15   15.5  RT (min)      lcorbi_t3_pos_1_005_42    m/z 712.6816     0   2   4E+4   15.5   16  RT (min)      lcorbi_t3_pos_1_008_36    m/z 712.6817     0   1   16   16.5  RT (min)      lcorbi_t3_pos_1_028_32    m/z 712.6817     0   1   17   17.5  RT (min)    lcorbi_t3_pos_2_012_01    m/z 712.6818     0   1   16   16.5  RT (min)      lcorbi_t3_pos_1_028_32    m/z 712.6818     0   1   16.5   17  RT (min)    lcorbi_t3_pos_1_034_34    m/z 712.6820     0   1   17   17.5  RT (min)    lcorbi_t3_pos_1_031_40    m/z 713.5880     0   2   4E+5   12   12.5  RT (min)       lcorbi_t3_pos_4_096_60    m/z 713.5882     0   2   4E+5   12   12.5  RT (min)       lcorbi_t3_pos_1_003_45    m/z 713.6849     0   1   16   16.5  RT (min)      lcorbi_t3_pos_2_022_38    m/z 713.6849     0   1   16.5   17  RT (min)    lcorbi_t3_pos_1_034_34    m/z 713.6850     0   1   2   3E+4   15.5   16  RT (min)      lcorbi_t3_pos_1_008_36    m/z 713.6850     0   1   16   16.5  RT (min)      lcorbi_t3_pos_1_028_32    m/z 713.6850     0   1   17   17.5  RT (min)    lcorbi_t3_pos_2_012_01    m/z 713.6851     0   1   13.5   14  RT (min)      lcorbi_t3_pos_3_024_07    m/z 713.6851     0   1   17   17.5  RT (min)    lcorbi_t3_pos_1_031_40    m/z 713.6852     0   1   14   14.5  RT (min)      lcorbi_t3_pos_2_013_33    m/z 713.6853     0   1   15.5   16  RT (min)      lcorbi_t3_pos_1_008_36    m/z 714.5066     0   1   2   3   4E+4   13   13.5  RT (min)      lcorbi_t3_pos_2_013_33    m/z 714.5071     0   1   15.5   16  RT (min)      lcorbi_t3_pos_2_012_01    m/z 714.5072     0   1   2   3E+4   14   14.5  RT (min)      lcorbi_t3_pos_1_008_36    m/z 714.5072     0   1   2   3E+4   14.5   15  RT (min)      lcorbi_t3_pos_1_008_36    m/z 714.5072     0   1   15.5   16  RT (min)      lcorbi_t3_pos_1_031_40    m/z 714.5072     0   1   2   3E+4   15   15.5  RT (min)      lcorbi_t3_pos_1_034_34    m/z 714.5072     0   1   2   3E+4   15   15.5  RT (min)      lcorbi_t3_pos_1_034_34    m/z 714.5073     0   2   4   6E+5   14   14.5  RT (min)      lcorbi_t3_pos_1_005_42    m/z 714.5073     0   1   2   3   4E+4   13   13.5  RT (min)       lcorbi_t3_pos_2_013_33    m/z 714.5074     0   1   15   15.5  RT (min)      lcorbi_t3_pos_1_028_32    m/z 714.5074     0   1   2E+5   14   14.5  RT (min)      lcorbi_t3_pos_1_008_36    m/z 714.5074     0   1   15.5   16  RT (min)      lcorbi_t3_pos_1_031_40    m/z 714.5074     0   1   14.5   15  RT (min)      lcorbi_t3_pos_2_022_38    m/z 714.5075     0   1   15.5   16  RT (min)      lcorbi_t3_pos_1_031_40    m/z 714.5642     0   2   4   6   8E+5   11.5   12  RT (min)       lcorbi_t3_pos_1_003_45    m/z 714.5644     0   2   4   6   8E+5   11.5   12  RT (min)       lcorbi_t3_pos_1_003_45    m/z 715.5106     0   1   15   15.5  RT (min)      lcorbi_t3_pos_1_034_34    m/z 715.5107     0   1   15.5   16  RT (min)      lcorbi_t3_pos_1_031_40    m/z 715.5107     0   1E+5   14   14.5  RT (min)      lcorbi_t3_pos_1_008_36    m/z 715.5108     0   1   2E+4   14.5   15  RT (min)      lcorbi_t3_pos_1_028_32    m/z 715.5109     0   1   15.5   16  RT (min)      lcorbi_t3_pos_1_031_40    m/z 715.5110     0   1E+4   14.5   15  RT (min)      lcorbi_t3_pos_1_008_36    m/z 715.5110     0   1   15   15.5  RT (min)      lcorbi_t3_pos_1_034_34    m/z 715.5112     0   1   2E+4   15   15.5  RT (min)      lcorbi_t3_pos_1_028_32    m/z 716.5226     0   2   4   6E+4   13   13.5  RT (min)      lcorbi_t3_pos_2_013_33    m/z 716.5226     0   1   2   3E+4   13   13.5  RT (min)      lcorbi_t3_pos_3_024_07    m/z 716.5227     0   1E+6   12.5   13  RT (min)      lcorbi_t3_pos_4_057_49    m/z 716.5228     0   2   4   6E+4   13   13.5  RT (min)       lcorbi_t3_pos_2_013_33    m/z 716.5229     0   1   2   3   4E+4   14.5   15  RT (min)      lcorbi_t3_pos_2_017_51    m/z 716.5229     0   2   4E+4   15.5   16  RT (min)      lcorbi_t3_pos_1_034_34    m/z 716.5230     0   2   4   6E+5   14.5   15  RT (min)      lcorbi_t3_pos_1_008_36    m/z 716.5230     0   1   2E+4   14.5   15  RT (min)      lcorbi_t3_pos_1_047_41    m/z 716.5230     0   2   4E+6   14   14.5  RT (min)      lcorbi_t3_pos_1_005_42    m/z 716.5230     0   1   2   3E+4   16   16.5  RT (min)      lcorbi_t3_pos_2_012_01    m/z 716.5230     0   1E+6   14.5   15  RT (min)      lcorbi_t3_pos_1_008_36    m/z 716.5230     0   1   2E+4   15   15.5  RT (min)      lcorbi_t3_pos_1_041_54    m/z 716.5230     0   2   4   6   8E+5   14   14.5  RT (min)      lcorbi_t3_pos_3_040_41    m/z 716.5230     0   1   16   16.5  RT (min)      lcorbi_t3_pos_1_001_49    m/z 716.5230     0   1   2   3E+4   15   15.5  RT (min)      lcorbi_t3_pos_1_030_21    m/z 716.5230     0   1   15.5   16  RT (min)      lcorbi_t3_pos_1_049_55    m/z 716.5230     0   2   4E+4   15.5   16  RT (min)      lcorbi_t3_pos_1_031_40    m/z 716.5230     0   2   4E+4   15   15.5  RT (min)      lcorbi_t3_pos_1_028_32    m/z 716.5231     0   1E+6   14.5   15  RT (min)      lcorbi_t3_pos_1_008_36    m/z 716.5232     0   2   4E+4   15   15.5  RT (min)      lcorbi_t3_pos_1_034_34    m/z 717.5256     0   1E+4   13   13.5  RT (min)      lcorbi_t3_pos_3_024_07    m/z 717.5259     0   1   2E+6   14   14.5  RT (min)      lcorbi_t3_pos_1_005_42    m/z 717.5262     0   1   2   3   4E+4   13   13.5  RT (min)      lcorbi_t3_pos_2_013_33    m/z 717.5262     0   1   2E+4   15.5   16  RT (min)      lcorbi_t3_pos_1_049_55    m/z 717.5262     0   1   16   16.5  RT (min)      lcorbi_t3_pos_1_001_49    m/z 717.5262     0   1   2E+4   15.5   16  RT (min)      lcorbi_t3_pos_1_049_55    m/z 717.5263     0   1   2E+4   14.5   15  RT (min)      lcorbi_t3_pos_1_047_41    m/z 717.5263     0   2   4   6E+5   14.5   15  RT (min)      lcorbi_t3_pos_1_008_36    m/z 717.5263     0   1   15   15.5  RT (min)      lcorbi_t3_pos_1_041_54    m/z 717.5263     0   2   4   6   8E+5   14.5   15  RT (min)      lcorbi_t3_pos_1_008_36    m/z 717.5263     0   1   2   3E+4   14.5   15  RT (min)      lcorbi_t3_pos_2_017_51    m/z 717.5264     0   1   2   3E+4   15   15.5  RT (min)      lcorbi_t3_pos_1_028_32    m/z 717.5264     0   1   2E+4   15.5   16  RT (min)      lcorbi_t3_pos_1_031_40    m/z 717.5264     0   1   15   15.5  RT (min)      lcorbi_t3_pos_1_030_21    m/z 717.5265     0   1   2   3E+4   15   15.5  RT (min)      lcorbi_t3_pos_1_034_34    m/z 717.5266     0   1   2   3E+5   14.5   15  RT (min)      lcorbi_t3_pos_1_008_36    m/z 717.5268     0   1   15.5   16  RT (min)      lcorbi_t3_pos_1_041_54    m/z 718.2949     0   1   16.5   17  RT (min)    lcorbi_t3_pos_1_049_55    m/z 718.2950     0   1   16   16.5  RT (min)      lcorbi_t3_pos_1_041_54    m/z 718.2952     0   1   13.5   14  RT (min)      lcorbi_t3_pos_1_002_01    m/z 718.2952     0   1   16.5   17  RT (min)    lcorbi_t3_pos_1_025_52    m/z 718.2952     0   2   4E+5   14.5   15  RT (min)      lcorbi_t3_pos_3_041_54    m/z 718.2953     0   1   15.5   16  RT (min)      lcorbi_t3_pos_2_041_54    m/z 718.2953     0   1   15.5   16  RT (min)      lcorbi_t3_pos_2_041_54    m/z 718.2953     0   2   4   6   8E+5   14   14.5  RT (min)      lcorbi_t3_pos_4_105_61    m/z 718.2953     0   1   16.5   17  RT (min)    lcorbi_t3_pos_2_056_56    m/z 718.2953     0   1   15   15.5  RT (min)      lcorbi_t3_pos_3_056_56    m/z 718.2954     0   1   13.5   14  RT (min)      lcorbi_t3_pos_2_032_39    m/z 718.5384     0   1E+6   13.5   14  RT (min)       lcorbi_t3_pos_3_053_48    m/z 718.5384     0   1E+4   13   13.5  RT (min)      lcorbi_t3_pos_3_037_32    m/z 718.5384     0   1   16   16.5  RT (min)      lcorbi_t3_pos_2_012_01    m/z 718.5384     0   2   4   6   8E+5   0   0.5  RT (min)      lcorbi_t3_pos_4_057_49    m/z 718.5384     0   1   2   3E+4   15   15.5  RT (min)      lcorbi_t3_pos_2_038_03    m/z 718.5384     0   1   16.5   17  RT (min)    lcorbi_t3_pos_1_023_02    m/z 718.5384     0   1   2   3E+4   16   16.5  RT (min)      lcorbi_t3_pos_1_031_40    m/z 718.5384     0   2   4   6   8E+4   15   15.5  RT (min)      lcorbi_t3_pos_1_033_53    m/z 718.5384     0   1E+6   14.5   15  RT (min)       lcorbi_t3_pos_4_025_52    m/z 718.5384     0   1E+4   13   13.5  RT (min)      lcorbi_t3_pos_3_037_32    m/z 718.5385     0   2   4E+4   15   15.5  RT (min)      lcorbi_t3_pos_1_028_32    m/z 718.5385     0   1E+6   14.5   15  RT (min)       lcorbi_t3_pos_1_008_36    m/z 718.5385     0   1   16.5   17  RT (min)    lcorbi_t3_pos_1_023_02    m/z 718.5385     0   1   2   3E+4   16   16.5  RT (min)      lcorbi_t3_pos_1_031_40    m/z 718.5385     0   1   2E+6   14.5   15  RT (min)      lcorbi_t3_pos_1_005_42    m/z 718.5385     0   2   4   6E+4   15   15.5  RT (min)      lcorbi_t3_pos_1_047_41    m/z 718.5385     0   1   2   3   4E+4   15   15.5  RT (min)      lcorbi_t3_pos_3_049_55    m/z 718.5385     0   2   4E+4   15   15.5  RT (min)      lcorbi_t3_pos_1_028_32    m/z 718.5385     0   1   2   3   4E+5   14.5   15  RT (min)       lcorbi_t3_pos_1_008_36    m/z 718.5385     0   1   2   3E+4   15.5   16  RT (min)      lcorbi_t3_pos_1_034_34    m/z 718.5385     0   1   2E+5   14.5   15  RT (min)       lcorbi_t3_pos_1_008_36    m/z 718.5386     0   1   2   3E+4   15.5   16  RT (min)      lcorbi_t3_pos_1_034_34    m/z 718.5386     0   1E+6   13.5   14  RT (min)       lcorbi_t3_pos_4_015_45    m/z 718.5386     0   2   4   6E+4   15   15.5  RT (min)      lcorbi_t3_pos_2_017_51    m/z 718.5387     0   1   16.5   17  RT (min)    lcorbi_t3_pos_1_023_02    m/z 718.5387     0   1   2   3E+4   16   16.5  RT (min)      lcorbi_t3_pos_1_027_43    m/z 718.5389     0   1   2   3E+4   15.5   16  RT (min)      lcorbi_t3_pos_2_049_55    m/z 719.5418     0   1   13   13.5  RT (min)      lcorbi_t3_pos_2_047_31    m/z 719.5418     0   1   2   3E+4   15.5   16  RT (min)      lcorbi_t3_pos_1_034_34    m/z 719.5418     0   1   2   3E+4   15   15.5  RT (min)      lcorbi_t3_pos_1_038_30    m/z 719.5418     0   1   13   13.5  RT (min)      lcorbi_t3_pos_2_039_30    m/z 719.5418     0   2   4   6E+5   13.5   14  RT (min)      lcorbi_t3_pos_4_037_26    m/z 719.5418     0   1   13   13.5  RT (min)      lcorbi_t3_pos_3_037_32    m/z 719.5419     0   1   2E+4   15   15.5  RT (min)      lcorbi_t3_pos_3_049_55    m/z 719.5419     0   1E+4   15.5   16  RT (min)      lcorbi_t3_pos_2_049_55    m/z 719.5420     0   1   2E+4   16   16.5  RT (min)      lcorbi_t3_pos_1_031_40    m/z 719.5420     0   1E+5   14.5   15  RT (min)       lcorbi_t3_pos_1_008_36    m/z 719.5420     0   1   2   3   4E+4   14.5   15  RT (min)      lcorbi_t3_pos_2_017_51    m/z 719.5420     0   1   2   3E+4   15.5   16  RT (min)      lcorbi_t3_pos_1_034_34    m/z 719.5420     0   1   2E+4   15   15.5  RT (min)      lcorbi_t3_pos_2_041_54    m/z 719.5420     0   1   2   3   4E+4   15   15.5  RT (min)      lcorbi_t3_pos_1_028_32    m/z 719.5420     0   1   2   3   4E+4   15   15.5  RT (min)      lcorbi_t3_pos_1_028_32    m/z 719.5420     0   1   16   16.5  RT (min)      lcorbi_t3_pos_1_025_52    m/z 719.5420     0   1   2E+4   16   16.5  RT (min)      lcorbi_t3_pos_1_031_40    m/z 719.5421     0   1E+5   14.5   15  RT (min)      lcorbi_t3_pos_1_008_36    m/z 719.5421     0   2   4E+5   14.5   15  RT (min)      lcorbi_t3_pos_1_008_36    m/z 719.5421     0   1E+6   14.5   15  RT (min)      lcorbi_t3_pos_1_005_42    m/z 719.5422     0   1E+6   14.5   15  RT (min)      lcorbi_t3_pos_1_005_42    m/z 719.5422     0   1   13   13.5  RT (min)      lcorbi_t3_pos_2_047_31    m/z 720.5174     0   1E+4   13   13.5  RT (min)      lcorbi_t3_pos_3_035_08    m/z 720.5176     0   1   2E+4   15   15.5  RT (min)      lcorbi_t3_pos_1_034_34    m/z 720.5176     0   1   12.5   13  RT (min)      lcorbi_t3_pos_2_047_31    m/z 720.5176     0   1   2   3E+4   14.5   15  RT (min)      lcorbi_t3_pos_1_055_33    m/z 720.5177     0   1   13   13.5  RT (min)      lcorbi_t3_pos_2_039_30    m/z 720.5177     0   1   2E+6   13   13.5  RT (min)      lcorbi_t3_pos_4_015_45    m/z 720.5178     0   1   16   16.5  RT (min)      lcorbi_t3_pos_1_023_02    m/z 720.5178     0   1   2   3   4E+4   14.5   15  RT (min)      lcorbi_t3_pos_1_038_30    m/z 720.5178     0   2   4E+4   13   13.5  RT (min)      lcorbi_t3_pos_1_002_01    m/z 720.5178     0   1   15.5   16  RT (min)      lcorbi_t3_pos_1_056_56    m/z 720.5178     0   1E+6   14   14.5  RT (min)      lcorbi_t3_pos_4_019_47    m/z 720.5178     0   1   2   3   4E+4   14.5   15  RT (min)      lcorbi_t3_pos_1_038_30    m/z 720.5178     0   1   15.5   16  RT (min)      lcorbi_t3_pos_1_027_43    m/z 720.5179     0   1   2E+4   15   15.5  RT (min)      lcorbi_t3_pos_2_052_48    m/z 720.5179     0   1   2E+4   15   15.5  RT (min)      lcorbi_t3_pos_2_049_55    m/z 720.5179     0   1   2   3   4E+4   14.5   15  RT (min)      lcorbi_t3_pos_1_038_30    m/z 720.5179     0   1E+5   14   14.5  RT (min)      lcorbi_t3_pos_3_044_46    m/z 720.5180     0   1   15.5   16  RT (min)      lcorbi_t3_pos_1_051_09    m/z 720.5180     0   2   4   6   8E+5   14   14.5  RT (min)      lcorbi_t3_pos_3_047_40    m/z 720.5180     0   2   4   6   8E+5   14   14.5  RT (min)      lcorbi_t3_pos_3_040_41    m/z 720.5180     0   1   16   16.5  RT (min)      lcorbi_t3_pos_1_023_02    m/z 720.5180     0   1   2E+4   15.5   16  RT (min)      lcorbi_t3_pos_1_025_52    m/z 720.5180     0   1   2E+4   15   15.5  RT (min)      lcorbi_t3_pos_2_052_48    m/z 720.5453     0   1   2   3E+4   15   15.5  RT (min)      lcorbi_t3_pos_1_028_32    m/z 720.5453     0   2   4   6E+4   14.5   15  RT (min)      lcorbi_t3_pos_1_008_36    m/z 721.5208     0   1E+4   13   13.5  RT (min)      lcorbi_t3_pos_2_031_20    m/z 721.5210     0   1   15   15.5  RT (min)      lcorbi_t3_pos_2_038_03    m/z 721.5210     0   1   12.5   13  RT (min)      lcorbi_t3_pos_2_047_31    m/z 721.5211     0   2   4   6   8E+5   13   13.5  RT (min)      lcorbi_t3_pos_3_053_48    m/z 721.5212     0   1E+6   13   13.5  RT (min)      lcorbi_t3_pos_4_015_45    m/z 721.5212     0   2   4   6E+4   14   14.5  RT (min)      lcorbi_t3_pos_3_044_46    m/z 721.5212     0   1   13   13.5  RT (min)      lcorbi_t3_pos_2_039_30    m/z 721.5212     0   1   16   16.5  RT (min)      lcorbi_t3_pos_1_023_02    m/z 721.5212     0   1   15.5   16  RT (min)      lcorbi_t3_pos_1_051_09    m/z 721.5213     0   1   2E+4   14.5   15  RT (min)      lcorbi_t3_pos_1_038_30    m/z 721.5213     0   1E+5   14   14.5  RT (min)      lcorbi_t3_pos_1_008_36    m/z 721.5213     0   1   2   3E+4   15   15.5  RT (min)      lcorbi_t3_pos_1_034_34    m/z 721.5213     0   1   2E+4   14.5   15  RT (min)      lcorbi_t3_pos_1_055_33    m/z 721.5213     0   1   15.5   16  RT (min)      lcorbi_t3_pos_1_001_49    m/z 721.5213     0   1   16   16.5  RT (min)      lcorbi_t3_pos_1_023_02    m/z 721.5214     0   2   4   6E+5   14   14.5  RT (min)      lcorbi_t3_pos_4_019_47    m/z 721.5214     0   2   4E+5   14   14.5  RT (min)      lcorbi_t3_pos_1_005_42    m/z 721.5214     0   1E+4   15   15.5  RT (min)      lcorbi_t3_pos_2_052_48    m/z 721.5214     0   2   4   6   8E+3   13   13.5  RT (min)      lcorbi_t3_pos_3_035_08    m/z 721.5216     0   1   15.5   16  RT (min)      lcorbi_t3_pos_1_027_43    m/z 723.4937     0   1   2E+4   15   15.5  RT (min)      lcorbi_t3_pos_4_028_04    m/z 723.4939     0   1   13   13.5  RT (min)      lcorbi_t3_pos_2_047_31    m/z 723.4939     0   2   4E+4   14.5   15  RT (min)      lcorbi_t3_pos_3_044_46    m/z 723.4939     0   1   13   13.5  RT (min)      lcorbi_t3_pos_3_035_08    m/z 723.4940     0   1   13   13.5  RT (min)      lcorbi_t3_pos_2_031_20    m/z 723.4940     0   2   4E+5   13.5   14  RT (min)      lcorbi_t3_pos_4_037_26    m/z 723.4940     0   2   4   6E+5   13.5   14  RT (min)      lcorbi_t3_pos_4_030_05    m/z 723.5369     0   2   4E+4   14   14.5  RT (min)       lcorbi_t3_pos_1_008_36    m/z 725.3969     0   1E+4   13.5   14  RT (min)      lcorbi_t3_pos_2_047_31    m/z 725.3970     0   2   4   6   8E+3   14   14.5  RT (min)      lcorbi_t3_pos_4_012_37    m/z 725.3970     0   2   4   6E+6   14   14.5  RT (min)       lcorbi_t3_pos_4_105_61    m/z 725.3970     0   1   2   3   4E+5   14   14.5  RT (min)      lcorbi_t3_pos_1_002_01    m/z 725.3970     0   1   16.5   17  RT (min)    lcorbi_t3_pos_1_056_56    m/z 725.3970     0   1   16.5   17  RT (min)    lcorbi_t3_pos_2_009_50    m/z 725.3970     0   1   2E+5   15   15.5  RT (min)      lcorbi_t3_pos_3_025_52    m/z 725.3971     0   1   16   16.5  RT (min)      lcorbi_t3_pos_1_033_53    m/z 725.3971     0   1   17   17.5  RT (min)    lcorbi_t3_pos_1_025_52    m/z 725.3971     0   1   15.5   16  RT (min)      lcorbi_t3_pos_1_030_21    m/z 725.3971     0   1   16   16.5  RT (min)      lcorbi_t3_pos_2_025_52    m/z 725.3971     0   2   4   6   8E+6   15   15.5  RT (min)       lcorbi_t3_pos_3_041_54    m/z 725.3972     0   1E+4   15.5   16  RT (min)      lcorbi_t3_pos_2_017_51    m/z 725.3972     0   2   4   6E+4   14   14.5  RT (min)      lcorbi_t3_pos_2_054_29    m/z 725.3972     0   1   17.5   18  RT (min)    lcorbi_t3_pos_1_023_02    m/z 725.3972     0   1   17   17.5  RT (min)    lcorbi_t3_pos_2_056_56    m/z 725.3972     0   1   17   17.5  RT (min)    lcorbi_t3_pos_2_012_01    m/z 725.3972     0   1   16   16.5  RT (min)      lcorbi_t3_pos_2_038_03    m/z 725.3972     0   1   17   17.5  RT (min)    lcorbi_t3_pos_2_012_01    m/z 725.3972     0   1   16   16.5  RT (min)      lcorbi_t3_pos_1_041_54    m/z 725.3973     0   1   13.5   14  RT (min)      lcorbi_t3_pos_2_032_39    m/z 725.3973     0   1   17.5   18  RT (min)    lcorbi_t3_pos_1_023_02    m/z 725.3973     0   1   16.5   17  RT (min)    lcorbi_t3_pos_1_049_55    m/z 725.5509     0   2   4   6   8E+4   14.5   15  RT (min)       lcorbi_t3_pos_1_030_21    m/z 725.5509     0   2   4   6E+5   14   14.5  RT (min)       lcorbi_t3_pos_3_047_40    m/z 725.5511     0   2   4   6E+4   14   14.5  RT (min)       lcorbi_t3_pos_1_008_36    m/z 725.5512     0   2   4   6   8E+4   14.5   15  RT (min)       lcorbi_t3_pos_3_013_19    m/z 725.5513     0   1   16   16.5  RT (min)      lcorbi_t3_pos_1_023_02    m/z 725.5568     0   1   2E+4   15.5   16  RT (min)      lcorbi_t3_pos_1_051_09    m/z 725.5572     0   1   2E+4   15.5   16  RT (min)      lcorbi_t3_pos_1_051_09    m/z 725.5572     0   1E+6   14   14.5  RT (min)      lcorbi_t3_pos_4_019_47    m/z 725.5572     0   1   2   3   4E+4   13   13.5  RT (min)      lcorbi_t3_pos_1_002_01    m/z 725.5572     0   1E+4   13   13.5  RT (min)      lcorbi_t3_pos_2_054_29    m/z 725.5573     0   1   12.5   13  RT (min)      lcorbi_t3_pos_2_032_39    m/z 726.4003     0   1   17   17.5  RT (min)    lcorbi_t3_pos_2_056_56    m/z 726.4004     0   1   2   3E+6   15   15.5  RT (min)      lcorbi_t3_pos_3_040_41    m/z 726.4004     0   1   16.5   17  RT (min)    lcorbi_t3_pos_2_009_50    m/z 726.4004     0   2   4   6E+4   15   15.5  RT (min)      lcorbi_t3_pos_3_025_52    m/z 726.4004     0   1   17   17.5  RT (min)    lcorbi_t3_pos_1_025_52    m/z 726.4004     0   1   16   16.5  RT (min)      lcorbi_t3_pos_1_041_54    m/z 726.4004     0   1E+6   14   14.5  RT (min)      lcorbi_t3_pos_3_053_48    m/z 726.4004     0   1   16.5   17  RT (min)    lcorbi_t3_pos_1_056_56    m/z 726.4005     0   1   16   16.5  RT (min)      lcorbi_t3_pos_1_033_53    m/z 726.4005     0   1   2   3   4E+4   14   14.5  RT (min)      lcorbi_t3_pos_2_054_29    m/z 726.4005     0   1   17.5   18  RT (min)    lcorbi_t3_pos_1_023_02    m/z 726.4005     0   1   16   16.5  RT (min)      lcorbi_t3_pos_2_038_03    m/z 726.4005     0   1   15.5   16  RT (min)      lcorbi_t3_pos_2_017_51    m/z 726.4005     0   1   15.5   16  RT (min)      lcorbi_t3_pos_1_030_21    m/z 726.4005     0   1   16.5   17  RT (min)    lcorbi_t3_pos_1_056_56    m/z 726.4006     0   1   15.5   16  RT (min)      lcorbi_t3_pos_2_017_51    m/z 726.4006     0   1E+5   14   14.5  RT (min)      lcorbi_t3_pos_1_002_01    m/z 726.4006     0   1   13.5   14  RT (min)      lcorbi_t3_pos_2_032_39    m/z 726.4006     0   1   2E+6   14   14.5  RT (min)      lcorbi_t3_pos_4_105_61    m/z 726.4006     0   1   2E+4   15.5   16  RT (min)      lcorbi_t3_pos_2_006_47    m/z 726.4006     0   1   16   16.5  RT (min)      lcorbi_t3_pos_2_025_52    m/z 726.4006     0   1   14   14.5  RT (min)      lcorbi_t3_pos_4_012_37    m/z 726.4008     0   1   13.5   14  RT (min)      lcorbi_t3_pos_2_047_31    m/z 726.5047     0   1   15   15.5  RT (min)      lcorbi_t3_pos_1_038_30    m/z 726.5047     0   1   15   15.5  RT (min)      lcorbi_t3_pos_1_055_33    m/z 726.5048     0   1   16   16.5  RT (min)      lcorbi_t3_pos_1_027_43    m/z 726.5050     0   1   15.5   16  RT (min)      lcorbi_t3_pos_1_027_43    m/z 726.5057     0   1E+4   15.5   16  RT (min)      lcorbi_t3_pos_2_052_48    m/z 726.5073     0   1   2E+4   14.5   15  RT (min)      lcorbi_t3_pos_3_044_46    m/z 726.5602     0   1   16   16.5  RT (min)      lcorbi_t3_pos_1_023_02    m/z 726.5604     0   1   12.5   13  RT (min)      lcorbi_t3_pos_2_032_39    m/z 726.5605     0   2   4   6E+4   15.5   16  RT (min)      lcorbi_t3_pos_1_051_09    m/z 726.5608     0   1   2E+4   13   13.5  RT (min)      lcorbi_t3_pos_2_054_29    m/z 726.7914     0   1   16   16.5  RT (min)      lcorbi_t3_pos_1_033_53    m/z 727.4038     0   2   4   6E+5   14   14.5  RT (min)      lcorbi_t3_pos_4_105_61    m/z 727.5643     0   2   4   6E+4   14.5   15  RT (min)      lcorbi_t3_pos_2_016_23    m/z 727.5662     0   1   18   18.5  RT (min)    lcorbi_t3_pos_1_023_02    m/z 727.5662     0   1   16.5   17  RT (min)    lcorbi_t3_pos_1_034_34    m/z 727.5662     0   1   17.5   18  RT (min)    lcorbi_t3_pos_2_012_01    m/z 727.5663     0   1   16   16.5  RT (min)      lcorbi_t3_pos_1_047_41    m/z 727.5663     0   1   17   17.5  RT (min)    lcorbi_t3_pos_2_009_50    m/z 727.5664     0   1   16.5   17  RT (min)    lcorbi_t3_pos_1_028_32    m/z 727.5664     0   1   17.5   18  RT (min)    lcorbi_t3_pos_1_031_40    m/z 727.5664     0   1   18   18.5  RT (min)    lcorbi_t3_pos_1_023_02    m/z 727.5664     0   1   16.5   17  RT (min)    lcorbi_t3_pos_1_034_34    m/z 727.5664     0   1   2E+6   15.5   16  RT (min)      lcorbi_t3_pos_1_005_42    m/z 727.5664     0   1   16.5   17  RT (min)    lcorbi_t3_pos_2_025_52    m/z 727.5664     0   1   17.5   18  RT (min)    lcorbi_t3_pos_1_031_40    m/z 727.5664     0   2   4   6   8E+4   15.5   16  RT (min)      lcorbi_t3_pos_1_008_36    m/z 727.5664     0   1   16   16.5  RT (min)      lcorbi_t3_pos_2_022_38    m/z 727.5664     0   2   4   6E+4   14   14.5  RT (min)      lcorbi_t3_pos_2_013_33    m/z 727.5666     0   1   16   16.5  RT (min)      lcorbi_t3_pos_2_046_07    m/z 727.5666     0   2   4   6E+4   14   14.5  RT (min)      lcorbi_t3_pos_2_013_33    m/z 727.5675     0   2   4   6E+5   11.5   12  RT (min)       lcorbi_t3_pos_1_003_45    m/z 727.5676     0   2   4E+5   11.5   12  RT (min)       lcorbi_t3_pos_4_090_60    m/z 727.5677     0   2   4   6E+5   11.5   12  RT (min)       lcorbi_t3_pos_4_092_60    m/z 727.5678     0   2   4   6E+5   11.5   12  RT (min)      lcorbi_t3_pos_4_093_60    m/z 727.5678     0   2   4   6E+5   11   11.5  RT (min)       lcorbi_t3_pos_4_095_60    m/z 727.5708     0   2   4E+4   15.5   16  RT (min)      lcorbi_t3_pos_4_001_49    m/z 727.5750     0   1   2E+4   14.5   15  RT (min)       lcorbi_t3_pos_3_053_48    m/z 727.5751     0   1   14.5   15  RT (min)      lcorbi_t3_pos_1_002_01    m/z 727.5752     0   1   14   14.5  RT (min)      lcorbi_t3_pos_2_047_31    m/z 728.5226     0   1   2E+4   14.5   15  RT (min)      lcorbi_t3_pos_1_034_34    m/z 728.5228     0   1   2   3   4E+4   14   14.5  RT (min)      lcorbi_t3_pos_3_003_33    m/z 728.5228     0   1   2E+4   14.5   15  RT (min)      lcorbi_t3_pos_1_034_34    m/z 728.5229     0   1   15   15.5  RT (min)      lcorbi_t3_pos_1_031_40    m/z 728.5230     0   1   15   15.5  RT (min)      lcorbi_t3_pos_2_012_01    m/z 728.5230     0   1E+6   12.5   13  RT (min)      lcorbi_t3_pos_3_011_42    m/z 728.5230     0   2   4   6E+4   14   14.5  RT (min)      lcorbi_t3_pos_2_022_38    m/z 728.5230     0   2   4   6E+5   14   14.5  RT (min)       lcorbi_t3_pos_1_008_36    m/z 728.5230     0   2   4   6E+4   14.5   15  RT (min)      lcorbi_t3_pos_1_028_32    m/z 728.5230     0   1   2E+4   14.5   15  RT (min)      lcorbi_t3_pos_1_034_34    m/z 728.5231     0   1   15   15.5  RT (min)      lcorbi_t3_pos_1_031_40    m/z 728.5232     0   1   12.5   13  RT (min)      lcorbi_t3_pos_3_024_07    m/z 728.5232     0   1   2E+6   13.5   14  RT (min)       lcorbi_t3_pos_4_085_59    m/z 728.5232     0   1   2E+4   12.5   13  RT (min)      lcorbi_t3_pos_2_013_33    m/z 728.5232     0   1E+6   13   13.5  RT (min)      lcorbi_t3_pos_3_011_42    m/z 728.5232     0   1E+6   12.5   13  RT (min)      lcorbi_t3_pos_3_011_42    m/z 728.5233     0   1   12.5   13  RT (min)      lcorbi_t3_pos_3_024_07    m/z 728.5696     0   1   16.5   17  RT (min)    lcorbi_t3_pos_2_038_03    m/z 728.5699     0   1   16.5   17  RT (min)    lcorbi_t3_pos_1_033_53    m/z 728.5699     0   1   16   16.5  RT (min)      lcorbi_t3_pos_2_022_38    m/z 728.5699     0   2   4   6   8E+5   15.5   16  RT (min)      lcorbi_t3_pos_3_040_41    m/z 728.5700     0   1   16   16.5  RT (min)      lcorbi_t3_pos_2_046_07    m/z 728.5700     0   1   2   3   4E+4   14   14.5  RT (min)      lcorbi_t3_pos_2_013_33    m/z 728.5700     0   1   17.5   18  RT (min)    lcorbi_t3_pos_1_031_40    m/z 728.5700     0   1   16.5   17  RT (min)    lcorbi_t3_pos_1_034_34    m/z 728.5700     0   1   16.5   17  RT (min)    lcorbi_t3_pos_1_028_32    m/z 728.5700     0   1   17.5   18  RT (min)    lcorbi_t3_pos_1_031_40    m/z 728.5700     0   2   4   6   8E+4   15.5   16  RT (min)      lcorbi_t3_pos_1_008_36    m/z 728.5701     0   1   17   17.5  RT (min)    lcorbi_t3_pos_2_009_50    m/z 728.5702     0   1   17.5   18  RT (min)    lcorbi_t3_pos_2_012_01    m/z 728.5702     0   1   16.5   17  RT (min)    lcorbi_t3_pos_1_034_34    m/z 728.5702     0   1E+6   15.5   16  RT (min)      lcorbi_t3_pos_1_005_42    m/z 728.5719     0   2   4   6E+4   14   14.5  RT (min)      lcorbi_t3_pos_3_031_45    m/z 729.5260     0   1   2   3   4E+4   14.5   15  RT (min)      lcorbi_t3_pos_1_034_34    m/z 729.5262     0   1   15   15.5  RT (min)      lcorbi_t3_pos_2_012_01    m/z 729.5262     0   1   2   3E+4   14   14.5  RT (min)      lcorbi_t3_pos_2_022_38    m/z 729.5263     0   1   15   15.5  RT (min)      lcorbi_t3_pos_1_031_40    m/z 729.5264     0   1   15   15.5  RT (min)      lcorbi_t3_pos_1_031_40    m/z 729.5264     0   1   12.5   13  RT (min)      lcorbi_t3_pos_3_024_07    m/z 729.5264     0   1   12.5   13  RT (min)      lcorbi_t3_pos_2_013_33    m/z 729.5264     0   1   15   15.5  RT (min)      lcorbi_t3_pos_1_031_40    m/z 729.5265     0   1   2   3   4E+4   14.5   15  RT (min)      lcorbi_t3_pos_1_034_34    m/z 729.5265     0   1   12.5   13  RT (min)      lcorbi_t3_pos_2_013_33    m/z 729.5265     0   1   2   3E+5   14   14.5  RT (min)       lcorbi_t3_pos_1_008_36    m/z 729.5265     0   1   2   3   4E+4   14.5   15  RT (min)      lcorbi_t3_pos_1_034_34    m/z 729.5266     0   1   15   15.5  RT (min)      lcorbi_t3_pos_1_031_40    m/z 729.5732     0   1E+4   14   14.5  RT (min)      lcorbi_t3_pos_2_013_33    m/z 729.5732     0   1   16.5   17  RT (min)    lcorbi_t3_pos_1_034_34    m/z 729.5733     0   1   17.5   18  RT (min)    lcorbi_t3_pos_1_031_40    m/z 729.5735     0   1   16   16.5  RT (min)      lcorbi_t3_pos_2_022_38    m/z 729.5737     0   1   2   3   4E+4   15.5   16  RT (min)      lcorbi_t3_pos_1_008_36    m/z 729.5737     0   1   16.5   17  RT (min)    lcorbi_t3_pos_1_028_32    m/z 729.5738     0   1   16   16.5  RT (min)      lcorbi_t3_pos_1_008_36    m/z 729.5907     0   1   12.5   13  RT (min)      lcorbi_t3_pos_2_032_39    m/z 729.5908     0   1   13   13.5  RT (min)      lcorbi_t3_pos_1_002_01    m/z 729.5908     0   1   12.5   13  RT (min)      lcorbi_t3_pos_2_032_39    m/z 729.5910     0   1   15.5   16  RT (min)      lcorbi_t3_pos_1_051_09    m/z 729.5910     0   1   16   16.5  RT (min)      lcorbi_t3_pos_1_023_02    m/z 729.5910     0   1   2   3   4E+4   14.5   15  RT (min)      lcorbi_t3_pos_1_008_36    m/z 729.5910     0   1   16   16.5  RT (min)      lcorbi_t3_pos_1_023_02    m/z 729.5911     0   1E+4   15   15.5   16  RT (min)      lcorbi_t3_pos_2_009_50    m/z 729.5912     0   1   15.5   16  RT (min)      lcorbi_t3_pos_1_031_40    m/z 729.5912     0   1   15   15.5  RT (min)      lcorbi_t3_pos_1_034_34    m/z 729.5912     0   1   2E+5   14   14.5  RT (min)      lcorbi_t3_pos_1_008_36    m/z 729.5912     0   1   15   15.5  RT (min)      lcorbi_t3_pos_1_034_34    m/z 729.5912     0   2   4   6E+5   13   13.5  RT (min)      lcorbi_t3_pos_1_002_01    m/z 729.5912     0   1   2E+6   14   14.5  RT (min)      lcorbi_t3_pos_3_047_40    m/z 729.5912     0   1   15.5   16  RT (min)      lcorbi_t3_pos_1_051_09    m/z 729.5912     0   1E+5   15   15.5  RT (min)       lcorbi_t3_pos_3_013_19    m/z 729.5913     0   1   14.5   15  RT (min)      lcorbi_t3_pos_1_047_41    m/z 729.5915     0   1E+5   14.5   15  RT (min)      lcorbi_t3_pos_3_013_19    m/z 730.3524     0   1   16   16.5  RT (min)      lcorbi_t3_pos_1_041_54    m/z 730.3524     0   1   16.5   17  RT (min)    lcorbi_t3_pos_1_041_54    m/z 730.3525     0   1   16.5   17  RT (min)    lcorbi_t3_pos_1_049_55    m/z 730.3526     0   1   14   14.5  RT (min)      lcorbi_t3_pos_2_054_29    m/z 730.3526     0   1E+4   15.5   16  RT (min)      lcorbi_t3_pos_3_056_56    m/z 730.3526     0   1   15.5   16  RT (min)      lcorbi_t3_pos_1_030_21    m/z 730.3526     0   1   15.5   16  RT (min)      lcorbi_t3_pos_2_006_47    m/z 730.3526     0   2   4E+4   15   15.5  RT (min)      lcorbi_t3_pos_1_008_36    m/z 730.3526     0   1   2E+4   16   16.5  RT (min)      lcorbi_t3_pos_1_033_53    m/z 730.3526     0   1   15.5   16  RT (min)      lcorbi_t3_pos_2_017_51    m/z 730.3526     0   1   2E+5   15   15.5  RT (min)      lcorbi_t3_pos_3_047_40    m/z 730.3526     0   1E+6   14.5   15  RT (min)      lcorbi_t3_pos_4_105_61    m/z 730.3526     0   1   16   16.5  RT (min)      lcorbi_t3_pos_2_038_03    m/z 730.3526     0   1   17   17.5  RT (min)    lcorbi_t3_pos_2_012_01    m/z 730.3527     0   1   17.5   18  RT (min)    lcorbi_t3_pos_1_023_02    m/z 730.3527     0   1   15.5   16  RT (min)      lcorbi_t3_pos_2_017_51    m/z 730.3527     0   1   17   17.5  RT (min)    lcorbi_t3_pos_2_056_56    m/z 730.3528     0   1   17.5   18  RT (min)    lcorbi_t3_pos_1_023_02    m/z 730.3530     0   1   13.5   14  RT (min)      lcorbi_t3_pos_2_047_31    m/z 730.5343     0   1   12.5   13  RT (min)      lcorbi_t3_pos_3_031_45    m/z 730.5385     0   1E+4   13   13.5  RT (min)      lcorbi_t3_pos_3_011_42    m/z 730.5385     0   1   16   16.5  RT (min)      lcorbi_t3_pos_1_023_02    m/z 730.5386     0   1   16   16.5  RT (min)      lcorbi_t3_pos_1_023_02    m/z 730.5386     0   1   2E+4   12.5   13  RT (min)      lcorbi_t3_pos_2_013_33    m/z 730.5386     0   1   2E+5   14.5   15  RT (min)      lcorbi_t3_pos_1_028_32    m/z 730.5387     0   1   16   16.5  RT (min)      lcorbi_t3_pos_1_023_02    m/z 730.5387     0   1E+6   14   14.5  RT (min)      lcorbi_t3_pos_1_008_36    m/z 730.5387     0   1   2   3E+6   13   13.5  RT (min)      lcorbi_t3_pos_3_011_42    m/z 730.5387     0   1E+4   15.5   16  RT (min)      lcorbi_t3_pos_2_012_01    m/z 730.5387     0   1   15   15.5  RT (min)      lcorbi_t3_pos_1_056_56    m/z 730.5388     0   1   12.5   13  RT (min)      lcorbi_t3_pos_3_031_45    m/z 730.5388     0   1   15.5   16  RT (min)      lcorbi_t3_pos_1_031_40    m/z 730.5388     0   1   15   15.5  RT (min)      lcorbi_t3_pos_1_031_40    m/z 730.5388     0   1E+6   14   14.5  RT (min)       lcorbi_t3_pos_4_019_47    m/z 730.5388     0   1   2   3E+4   15   15.5  RT (min)      lcorbi_t3_pos_1_034_34    m/z 730.5388     0   1E+6   14   14.5  RT (min)       lcorbi_t3_pos_1_008_36    m/z 730.5388     0   1   2   3E+4   14.5   15  RT (min)      lcorbi_t3_pos_1_034_34    m/z 730.5388     0   1   2   3E+4   15   15.5  RT (min)      lcorbi_t3_pos_1_034_34    m/z 730.5389     0   1   2   3   4E+6   14   14.5  RT (min)      lcorbi_t3_pos_4_086_59    m/z 730.5389     0   1   2   3E+5   14.5   15  RT (min)      lcorbi_t3_pos_2_022_38    m/z 730.5390     0   1   16   16.5  RT (min)      lcorbi_t3_pos_1_023_02    m/z 730.5772     0   1   2   3E+5   15   15.5  RT (min)       lcorbi_t3_pos_3_040_41    m/z 730.5941     0   1   12.5   13  RT (min)      lcorbi_t3_pos_2_032_39    m/z 730.5942     0   1   16   16.5  RT (min)      lcorbi_t3_pos_1_023_02    m/z 730.5943     0   1   12.5   13  RT (min)      lcorbi_t3_pos_2_032_39    m/z 730.5944     0   1   13   13.5  RT (min)      lcorbi_t3_pos_2_054_29    m/z 730.5944     0   1   15.5   16  RT (min)      lcorbi_t3_pos_2_012_01    m/z 730.5944     0   1   16   16.5  RT (min)      lcorbi_t3_pos_1_023_02    m/z 730.5945     0   1E+4   14   14.5  RT (min)      lcorbi_t3_pos_1_008_36    m/z 730.5945     0   1   15.5   16  RT (min)      lcorbi_t3_pos_1_051_09    m/z 730.5945     0   1   2   3E+5   13   13.5  RT (min)      lcorbi_t3_pos_1_002_01    m/z 730.5946     0   1   15.5   16  RT (min)      lcorbi_t3_pos_1_031_40    m/z 730.5946     0   1   2   3   4E+4   14.5   15  RT (min)       lcorbi_t3_pos_3_013_19    m/z 730.5946     0   1E+6   14   14.5  RT (min)      lcorbi_t3_pos_3_047_40    m/z 730.5946     0   1E+5   14   14.5  RT (min)      lcorbi_t3_pos_1_008_36    m/z 730.5946     0   1   15   15.5  RT (min)      lcorbi_t3_pos_1_034_34    m/z 730.5946     0   1   15.5   16  RT (min)      lcorbi_t3_pos_1_051_09    m/z 730.5946     0   1E+4   14.5   15  RT (min)      lcorbi_t3_pos_2_006_47    m/z 730.5946     0   1   15   15.5  RT (min)      lcorbi_t3_pos_1_034_34    m/z 730.5948     0   1E+4   14.5   15  RT (min)      lcorbi_t3_pos_1_008_36    m/z 731.3560     0   1   16   16.5  RT (min)      lcorbi_t3_pos_1_033_53    m/z 731.3562     0   2   4   6E+5   14.5   15  RT (min)      lcorbi_t3_pos_4_105_61    m/z 731.5420     0   1E+4   15.5   16  RT (min)      lcorbi_t3_pos_2_012_01    m/z 731.5420     0   2   4E+4   15   15.5  RT (min)      lcorbi_t3_pos_1_034_34    m/z 731.5420     0   1E+4   13   13.5  RT (min)      lcorbi_t3_pos_3_011_42    m/z 731.5420     0   1   15.5   16  RT (min)      lcorbi_t3_pos_1_031_40    m/z 731.5420     0   2   4E+4   15   15.5  RT (min)      lcorbi_t3_pos_1_034_34    m/z 731.5421     0   1   2   3   4E+5   14   14.5  RT (min)      lcorbi_t3_pos_1_008_36    m/z 731.5421     0   1E+5   14.5   15  RT (min)      lcorbi_t3_pos_1_028_32    m/z 731.5421     0   1   16   16.5  RT (min)      lcorbi_t3_pos_1_023_02    m/z 731.5421     0   1   15   15.5  RT (min)      lcorbi_t3_pos_1_031_40    m/z 731.5421     0   1E+4   15.5   16  RT (min)      lcorbi_t3_pos_2_012_01    m/z 731.5421     0   2   4   6   8E+5   13   13.5  RT (min)      lcorbi_t3_pos_4_016_15    m/z 731.5421     0   2   4E+4   14.5   15  RT (min)      lcorbi_t3_pos_1_034_34    m/z 731.5421     0   1   16   16.5  RT (min)      lcorbi_t3_pos_1_023_02    m/z 731.5421     0   1   2E+5   14.5   15  RT (min)      lcorbi_t3_pos_1_030_21    m/z 731.5423     0   1   15   15.5  RT (min)      lcorbi_t3_pos_1_056_56    m/z 731.5423     0   1   12.5   13  RT (min)      lcorbi_t3_pos_3_031_45    m/z 731.5423     0   1E+5   14.5   15  RT (min)      lcorbi_t3_pos_1_047_41    m/z 731.5426     0   1   16   16.5  RT (min)      lcorbi_t3_pos_1_023_02    m/z 731.6056     0   1E+5   13   13.5  RT (min)      lcorbi_t3_pos_1_002_01    m/z 731.6062     0   1   2   3E+4   13   13.5  RT (min)      lcorbi_t3_pos_1_002_01    m/z 731.6063     0   1   15.5   16  RT (min)      lcorbi_t3_pos_1_049_55    m/z 731.6064     0   1   16.5   17  RT (min)    lcorbi_t3_pos_1_023_02    m/z 731.6065     0   1   15.5   16  RT (min)      lcorbi_t3_pos_1_051_09    m/z 731.6065     0   2   4   6E+5   14   14.5  RT (min)      lcorbi_t3_pos_1_005_42    m/z 731.6065     0   1   2E+4   14.5   15  RT (min)      lcorbi_t3_pos_1_047_41    m/z 731.6066     0   1   15   15.5  RT (min)      lcorbi_t3_pos_1_030_21    m/z 731.6067     0   1   15.5   16  RT (min)      lcorbi_t3_pos_1_031_40    m/z 731.6068     0   1   15.5   16  RT (min)       lcorbi_t3_pos_1_041_54    m/z 731.6068     0   1   16.5   17  RT (min)    lcorbi_t3_pos_1_023_02    m/z 731.6074     0   1   15   15.5  RT (min)       lcorbi_t3_pos_1_033_53    m/z 732.5454     0   1   12.5   13  RT (min)      lcorbi_t3_pos_3_031_45    m/z 732.5536     0   1E+5   13   13.5  RT (min)      lcorbi_t3_pos_1_002_01    m/z 732.5536     0   1   13   13.5  RT (min)      lcorbi_t3_pos_3_031_45    m/z 732.5537     0   1   12.5   13  RT (min)      lcorbi_t3_pos_2_032_39    m/z 732.5539     0   1   15   15.5  RT (min)      lcorbi_t3_pos_2_009_50    m/z 732.5540     0   1   13   13.5  RT (min)      lcorbi_t3_pos_2_054_29    m/z 732.5540     0   1   16   16.5  RT (min)      lcorbi_t3_pos_1_023_02    m/z 732.5541     0   1   2   3   4E+4   15   15.5  RT (min)      lcorbi_t3_pos_1_034_34    m/z 732.5542     0   1E+4   15.5   16  RT (min)      lcorbi_t3_pos_2_012_01    m/z 732.5542     0   2   4E+4   15   15.5  RT (min)      lcorbi_t3_pos_1_028_32    m/z 732.5543     0   1   15.5   16  RT (min)      lcorbi_t3_pos_1_051_09    m/z 732.5543     0   1   2   3   4E+4   15   15.5  RT (min)      lcorbi_t3_pos_1_034_34    m/z 732.5543     0   1   15.5   16  RT (min)      lcorbi_t3_pos_1_051_09    m/z 732.5543     0   2   4E+4   14.5   15  RT (min)      lcorbi_t3_pos_1_028_32    m/z 732.5544     0   2   4E+4   14.5   15  RT (min)      lcorbi_t3_pos_1_033_53    m/z 732.5544     0   1   15.5   16  RT (min)      lcorbi_t3_pos_2_056_56    m/z 732.5544     0   1   2E+5   14.5   15  RT (min)      lcorbi_t3_pos_2_046_07    m/z 732.5546     0   1   2   3E+4   15.5   16  RT (min)      lcorbi_t3_pos_1_031_40    m/z 732.6100     0   1   16.5   17  RT (min)    lcorbi_t3_pos_1_023_02    m/z 733.5574     0   1   13   13.5  RT (min)      lcorbi_t3_pos_3_024_07    m/z 733.5574     0   1   2E+6   14   14.5  RT (min)      lcorbi_t3_pos_1_005_42    m/z 733.5575     0   1   15   15.5  RT (min)      lcorbi_t3_pos_1_034_34    m/z 733.5576     0   1   2   3E+6   14   14.5  RT (min)      lcorbi_t3_pos_3_047_40    m/z 733.5576     0   1   12.5   13  RT (min)      lcorbi_t3_pos_2_032_39    m/z 733.5576     0   1   13   13.5  RT (min)      lcorbi_t3_pos_4_012_37    m/z 733.5576     0   1   2E+5   14   14.5  RT (min)      lcorbi_t3_pos_1_008_36    m/z 733.5577     0   1E+4   15   15.5  RT (min)      lcorbi_t3_pos_3_013_19    m/z 733.5577     0   2   4E+4   13   13.5  RT (min)      lcorbi_t3_pos_1_002_01    m/z 733.5577     0   1   15.5   16  RT (min)      lcorbi_t3_pos_1_051_09    m/z 733.5577     0   1   15.5   16  RT (min)      lcorbi_t3_pos_1_031_40    m/z 733.5577     0   1   2   3E+6   13   13.5  RT (min)      lcorbi_t3_pos_4_016_15    m/z 733.5578     0   2   4E+5   14   14.5  RT (min)      lcorbi_t3_pos_1_008_36    m/z 733.5578     0   1   2   3   4E+4   14.5   15  RT (min)      lcorbi_t3_pos_1_028_32    m/z 733.5578     0   1   16   16.5  RT (min)      lcorbi_t3_pos_1_023_02    m/z 733.5578     0   1   2E+5   14.5   15  RT (min)      lcorbi_t3_pos_1_008_36    m/z 733.5578     0   1   2E+6   14   14.5  RT (min)      lcorbi_t3_pos_1_005_42    m/z 733.5578     0   1   15.5   16  RT (min)      lcorbi_t3_pos_1_051_09    m/z 733.5579     0   1   16   16.5  RT (min)      lcorbi_t3_pos_1_023_02    m/z 733.5579     0   1E+5   14.5   15  RT (min)      lcorbi_t3_pos_1_047_41    m/z 734.5607     0   1   12.5   13  RT (min)      lcorbi_t3_pos_2_032_39    m/z 734.5609     0   1   13   13.5  RT (min)      lcorbi_t3_pos_2_054_29    m/z 734.5651     0   1   2E+4   15.5   16  RT (min)      lcorbi_t3_pos_1_041_54    m/z 734.5651     0   1   13   13.5  RT (min)      lcorbi_t3_pos_4_012_37    m/z 734.5655     0   1E+6   13.5   14  RT (min)      lcorbi_t3_pos_1_002_01    m/z 734.5657     0   1   15.5   16  RT (min)      lcorbi_t3_pos_1_051_09    m/z 734.5696     0   1   15   15.5  RT (min)      lcorbi_t3_pos_1_028_32    m/z 734.5696     0   1   2E+5   13   13.5  RT (min)      lcorbi_t3_pos_1_002_01    m/z 734.5696     0   1E+6   14   14.5  RT (min)       lcorbi_t3_pos_1_005_42    m/z 734.5696     0   1E+5   14.5   15  RT (min)      lcorbi_t3_pos_1_008_36    m/z 734.5696     0   1   15   15.5  RT (min)      lcorbi_t3_pos_1_028_32    m/z 734.5697     0   1   15.5   16  RT (min)      lcorbi_t3_pos_2_049_55    m/z 734.5700     0   1   15.5   16  RT (min)      lcorbi_t3_pos_1_049_55    m/z 734.5700     0   1   16   16.5  RT (min)      lcorbi_t3_pos_1_051_09    m/z 734.5700     0   1   16   16.5  RT (min)      lcorbi_t3_pos_1_023_02    m/z 734.5700     0   1   16.5   17  RT (min)    lcorbi_t3_pos_1_023_02    m/z 734.5700     0   1   2E+4   15   15.5  RT (min)      lcorbi_t3_pos_1_033_53    m/z 734.5700     0   1   15   15.5  RT (min)      lcorbi_t3_pos_1_033_53    m/z 734.5702     0   1E+6   14   14.5  RT (min)      lcorbi_t3_pos_1_005_42    m/z 734.5703     0   1   16.5   17  RT (min)    lcorbi_t3_pos_1_023_02    m/z 734.5703     0   1   16.5   17  RT (min)    lcorbi_t3_pos_1_023_02    m/z 735.5684     0   2   4E+5   14   14.5  RT (min)      lcorbi_t3_pos_1_005_42    m/z 735.5729     0   2   4   6   8E+4   13   13.5  RT (min)      lcorbi_t3_pos_1_002_01    m/z 735.5732     0   1   15.5   16  RT (min)      lcorbi_t3_pos_1_056_56    m/z 735.5733     0   1   16.5   17  RT (min)    lcorbi_t3_pos_1_023_02    m/z 735.5733     0   1   2E+4   15   15.5  RT (min)      lcorbi_t3_pos_1_033_53    m/z 735.5736     0   1   16   16.5  RT (min)      lcorbi_t3_pos_1_051_09    m/z 736.4914     0   1   14.5   15  RT (min)      lcorbi_t3_pos_1_034_34    m/z 736.4916     0   1   15   15.5  RT (min)      lcorbi_t3_pos_1_031_40    m/z 736.4918     0   1   14.5   15  RT (min)      lcorbi_t3_pos_1_034_34    m/z 736.4918     0   1E+5   14   14.5  RT (min)      lcorbi_t3_pos_1_008_36    m/z 736.4918     0   1   14.5   15  RT (min)      lcorbi_t3_pos_3_003_33    m/z 736.4920     0   1E+4   13   13.5  RT (min)      lcorbi_t3_pos_3_011_42    m/z 736.4920     0   1E+4   12.5   13  RT (min)      lcorbi_t3_pos_3_024_07    m/z 736.4920     0   1   15.5   16  RT (min)      lcorbi_t3_pos_2_012_01    m/z 736.4922     0   1   2   3   4E+5   13   13.5  RT (min)      lcorbi_t3_pos_3_011_42    m/z 737.4950     0   2   4   6E+4   14   14.5  RT (min)      lcorbi_t3_pos_1_008_36    m/z 737.4952     0   1   15   15.5  RT (min)      lcorbi_t3_pos_1_031_40    m/z 737.4953     0   1   14.5   15  RT (min)      lcorbi_t3_pos_1_028_32    m/z 737.4953     0   1   14.5   15  RT (min)      lcorbi_t3_pos_1_034_34    m/z 737.5794     0   2   4E+5   11.5   12  RT (min)       lcorbi_t3_pos_1_003_45    m/z 738.5054     0   1   2   3   4E+4   15.5   16  RT (min)      lcorbi_t3_pos_2_012_01    m/z 738.5059     0   1   2   3   4E+4   15   15.5  RT (min)      lcorbi_t3_pos_1_034_34    m/z 738.5066     0   1   15   15.5   16  RT (min)      lcorbi_t3_pos_1_056_56    m/z 738.5067     0   1   2   3   4E+4   14.5   15  RT (min)      lcorbi_t3_pos_1_047_41    m/z 738.5068     0   1   2   3   4E+4   15   15.5  RT (min)      lcorbi_t3_pos_1_034_34    m/z 738.5070     0   1   2   3   4E+4   15.5   16  RT (min)      lcorbi_t3_pos_1_031_40    m/z 738.5070     0   2   4   6   8E+5   11.5   12  RT (min)      lcorbi_t3_pos_4_057_49    m/z 738.5070     0   1E+5   13   13.5  RT (min)       lcorbi_t3_pos_2_013_33    m/z 738.5071     0   1E+5   13   13.5  RT (min)      lcorbi_t3_pos_2_013_33    m/z 738.5071     0   1   2   3E+6   13   13.5  RT (min)      lcorbi_t3_pos_3_011_42    m/z 738.5072     0   1   2   3   4E+4   15.5   16  RT (min)      lcorbi_t3_pos_2_012_01    m/z 738.5072     0   1   2E+4   14.5   15  RT (min)      lcorbi_t3_pos_1_030_21    m/z 738.5072     0   1E+6   14   14.5  RT (min)      lcorbi_t3_pos_1_008_36    m/z 738.5072     0   2   4E+4   14.5   15  RT (min)      lcorbi_t3_pos_1_028_32    m/z 738.5072     0   1E+5   12.5   13  RT (min)      lcorbi_t3_pos_3_031_45    m/z 738.5072     0   2   4E+4   14.5   15  RT (min)      lcorbi_t3_pos_1_028_32    m/z 738.5072     0   1   2   3   4E+4   15.5   16  RT (min)      lcorbi_t3_pos_1_031_40    m/z 738.5073     0   2   4E+4   15   15.5  RT (min)      lcorbi_t3_pos_1_028_32    m/z 739.5090     0   1E+6   14   14.5  RT (min)       lcorbi_t3_pos_3_040_41    m/z 739.5092     0   1   2E+6   14   14.5  RT (min)       lcorbi_t3_pos_1_005_42    m/z 739.5096     0   1   2E+4   15.5   16  RT (min)      lcorbi_t3_pos_2_012_01    m/z 739.5098     0   1   2E+4   15.5   16  RT (min)      lcorbi_t3_pos_2_012_01    m/z 739.5100     0   1   2   3E+4   15   15.5  RT (min)      lcorbi_t3_pos_1_034_34    m/z 739.5100     0   1   2E+4   15.5   16  RT (min)      lcorbi_t3_pos_2_012_01    m/z 739.5104     0   1   2E+4   14.5   15  RT (min)      lcorbi_t3_pos_1_028_32    m/z 739.5104     0   1   2   3E+4   15   15.5  RT (min)      lcorbi_t3_pos_1_034_34    m/z 739.5105     0   2   4   6   8E+4   13   13.5  RT (min)      lcorbi_t3_pos_2_013_33    m/z 739.5105     0   2   4   6   8E+4   13   13.5  RT (min)      lcorbi_t3_pos_2_013_33    m/z 739.5106     0   1E+5   14   14.5  RT (min)      lcorbi_t3_pos_3_025_52    m/z 739.5106     0   1   2   3E+4   15   15.5  RT (min)      lcorbi_t3_pos_1_034_34    m/z 739.5106     0   1   2   3E+4   15.5   16  RT (min)      lcorbi_t3_pos_1_031_40    m/z 739.5106     0   2   4   6E+5   14   14.5  RT (min)      lcorbi_t3_pos_1_008_36    m/z 739.5107     0   1   2   3E+4   15   15.5  RT (min)      lcorbi_t3_pos_1_034_34    m/z 739.5107     0   1   2E+4   15   15.5   16  RT (min)      lcorbi_t3_pos_1_056_56    m/z 739.5108     0   1   2   3E+4   15.5   16  RT (min)      lcorbi_t3_pos_1_031_40    m/z 739.5108     0   1   2E+4   14.5   15  RT (min)      lcorbi_t3_pos_1_028_32    m/z 739.5108     0   1   2E+4   14.5   15  RT (min)      lcorbi_t3_pos_1_030_21    m/z 739.5109     0   1   14.5   15  RT (min)      lcorbi_t3_pos_1_047_41    m/z 739.5950     0   2   4E+5   12   12.5  RT (min)       lcorbi_t3_pos_1_003_45    m/z 739.5954     0   2   4E+5   11.5   12  RT (min)       lcorbi_t3_pos_1_003_45    m/z 740.5142     0   1E+5   14.5   15  RT (min)      lcorbi_t3_pos_2_022_38    m/z 740.5183     0   2   4   6E+5   13   13.5  RT (min)      lcorbi_t3_pos_2_013_33    m/z 740.5185     0   2   4   6E+5   13   13.5  RT (min)      lcorbi_t3_pos_3_031_45    m/z 740.5211     0   1E+6   14   14.5  RT (min)       lcorbi_t3_pos_4_054_18    m/z 740.5213     0   1   2   3E+4   14.5   15  RT (min)      lcorbi_t3_pos_2_006_47    m/z 740.5213     0   1   2E+6   14   14.5  RT (min)       lcorbi_t3_pos_1_005_42    m/z 740.5214     0   1   2E+4   15   15.5  RT (min)      lcorbi_t3_pos_2_049_55    m/z 740.5216     0   1   2E+4   15   15.5  RT (min)      lcorbi_t3_pos_1_041_54    m/z 740.5223     0   1   2   3E+4   15.5   16  RT (min)      lcorbi_t3_pos_1_031_40    m/z 740.5224     0   1   2   3E+4   15   15.5  RT (min)      lcorbi_t3_pos_1_034_34    m/z 740.5225     0   1   2   3E+5   14.5   15  RT (min)      lcorbi_t3_pos_1_008_36    m/z 740.5227     0   1   2   3E+5   12.5   13  RT (min)      lcorbi_t3_pos_3_011_42    m/z 740.5229     0   1   2   3E+4   15.5   16  RT (min)      lcorbi_t3_pos_1_031_40    m/z 740.5230     0   1   2   3   4E+5   14.5   15  RT (min)       lcorbi_t3_pos_1_008_36    m/z 740.5230     0   1E+4   15.5   16  RT (min)      lcorbi_t3_pos_2_012_01    m/z 740.5230     0   1   16   16.5  RT (min)      lcorbi_t3_pos_2_056_56    m/z 740.5230     0   1   2   3E+4   15   15.5  RT (min)      lcorbi_t3_pos_1_034_34    m/z 740.5230     0   1   2E+6   14   14.5  RT (min)      lcorbi_t3_pos_1_005_42    m/z 740.5230     0   1   2   3E+4   15   15.5  RT (min)      lcorbi_t3_pos_1_028_32    m/z 740.5231     0   1   2   3E+4   15   15.5  RT (min)      lcorbi_t3_pos_1_028_32    m/z 740.5231     0   2   4   6   8E+5   12.5   13  RT (min)      lcorbi_t3_pos_4_057_49    m/z 740.5231     0   1E+5   14.5   15  RT (min)      lcorbi_t3_pos_3_056_56    m/z 740.5232     0   1   2   3E+4   15   15.5  RT (min)      lcorbi_t3_pos_1_028_32    m/z 740.5233     0   1   15   15.5  RT (min)      lcorbi_t3_pos_2_038_03    m/z 740.5233     0   1   14.5   15  RT (min)      lcorbi_t3_pos_1_030_21    m/z 741.5177     0   1   14.5   15  RT (min)      lcorbi_t3_pos_1_047_41    m/z 741.5254     0   2   4   6   8E+5   14   14.5  RT (min)       lcorbi_t3_pos_1_005_42    m/z 741.5262     0   1   15.5   16  RT (min)      lcorbi_t3_pos_2_009_50    m/z 741.5263     0   1   15.5   16  RT (min)      lcorbi_t3_pos_1_056_56    m/z 741.5264     0   1   2E+4   15.5   16  RT (min)      lcorbi_t3_pos_1_031_40    m/z 741.5264     0   1   2   3E+4   15   15.5  RT (min)      lcorbi_t3_pos_1_034_34    m/z 741.5265     0   1E+5   14.5   15  RT (min)      lcorbi_t3_pos_1_008_36    m/z 741.5266     0   1   2   3E+4   15   15.5  RT (min)      lcorbi_t3_pos_1_034_34    m/z 741.5266     0   1   2E+4   14.5   15  RT (min)      lcorbi_t3_pos_1_047_41    m/z 741.5266     0   1   2   3E+4   15   15.5  RT (min)      lcorbi_t3_pos_1_028_32    m/z 741.5266     0   1E+5   14.5   15  RT (min)      lcorbi_t3_pos_1_008_36    m/z 741.5266     0   1   15   15.5  RT (min)      lcorbi_t3_pos_1_028_32    m/z 741.5266     0   1   2E+5   14.5   15  RT (min)       lcorbi_t3_pos_1_008_36    m/z 741.5268     0   1   15   15.5  RT (min)      lcorbi_t3_pos_1_028_32    m/z 741.5268     0   1   15.5   16   16.5  RT (min)      lcorbi_t3_pos_2_012_01    m/z 741.5269     0   1   15.5   16  RT (min)      lcorbi_t3_pos_1_056_56    m/z 741.5274     0   1   2E+5   13   13.5  RT (min)      lcorbi_t3_pos_3_031_45    m/z 741.5465     0   2   4   6E+5   9.5   10  RT (min)       lcorbi_t3_pos_1_003_45    m/z 742.4996     0   1   15.5   16  RT (min)      lcorbi_t3_pos_1_027_43    m/z 742.4996     0   1   15   15.5  RT (min)      lcorbi_t3_pos_2_052_48    m/z 742.4998     0   1   14.5   15  RT (min)      lcorbi_t3_pos_1_038_30    m/z 742.4998     0   1   14.5   15  RT (min)      lcorbi_t3_pos_1_055_33    m/z 742.4999     0   1   13   13.5  RT (min)      lcorbi_t3_pos_3_035_08    m/z 742.5347     0   1   2E+4   14.5   15  RT (min)      lcorbi_t3_pos_2_046_07    m/z 742.5379     0   1E+6   13   13.5  RT (min)      lcorbi_t3_pos_2_013_33    m/z 742.5380     0   1E+6   13   13.5  RT (min)      lcorbi_t3_pos_3_024_07    m/z 742.5384     0   1   2   3E+5   13   13.5  RT (min)      lcorbi_t3_pos_3_011_42    m/z 742.5384     0   1   16   16.5  RT (min)      lcorbi_t3_pos_2_056_56    m/z 742.5386     0   1   12.5   13  RT (min)      lcorbi_t3_pos_2_013_33    m/z 742.5386     0   1E+6   14.5   15  RT (min)      lcorbi_t3_pos_2_022_38    m/z 742.5386     0   1   15.5   16  RT (min)      lcorbi_t3_pos_1_056_56    m/z 742.5386     0   1   2E+4   15.5   16  RT (min)      lcorbi_t3_pos_2_012_01    m/z 742.5387     0   1   15.5   16  RT (min)      lcorbi_t3_pos_1_041_54    m/z 742.5387     0   1   16   16.5  RT (min)      lcorbi_t3_pos_1_025_52    m/z 742.5387     0   1   15   15.5  RT (min)      lcorbi_t3_pos_1_030_21    m/z 742.5387     0   1   15   15.5  RT (min)      lcorbi_t3_pos_1_021_18    m/z 742.5387     0   1   2   3   4E+4   14.5   15  RT (min)      lcorbi_t3_pos_1_034_34    m/z 742.5388     0   1   15.5   16  RT (min)      lcorbi_t3_pos_1_049_55    m/z 742.5388     0   1   12.5   13  RT (min)      lcorbi_t3_pos_3_031_45    m/z 742.5388     0   1E+4   15.5   16  RT (min)      lcorbi_t3_pos_2_009_50    m/z 742.5388     0   1   15   15.5  RT (min)      lcorbi_t3_pos_2_025_52    m/z 742.5388     0   1   16   16.5  RT (min)      lcorbi_t3_pos_1_031_40    m/z 742.5388     0   1   15   15.5  RT (min)      lcorbi_t3_pos_1_031_40    m/z 742.5389     0   1E+5   14.5   15  RT (min)      lcorbi_t3_pos_2_022_38    m/z 742.5389     0   1   2E+4   15   15.5  RT (min)      lcorbi_t3_pos_1_034_34    m/z 742.5390     0   1   12.5   13  RT (min)      lcorbi_t3_pos_2_013_33    m/z 742.5390     0   2   4E+5   14   14.5  RT (min)      lcorbi_t3_pos_3_040_41    m/z 742.5750     0   1   16   16.5  RT (min)      lcorbi_t3_pos_1_031_40    m/z 742.5830     0   2   4   6   8E+4   14.5   15  RT (min)      lcorbi_t3_pos_1_008_36    m/z 743.5420     0   2   4   6   8E+5   13   13.5  RT (min)      lcorbi_t3_pos_2_023_26    m/z 743.5420     0   2   4   6   8E+4   14.5   15  RT (min)      lcorbi_t3_pos_1_008_36    m/z 743.5420     0   1   2   3   4E+5   13   13.5  RT (min)      lcorbi_t3_pos_2_013_33    m/z 743.5420     0   1   2   3E+4   16   16.5  RT (min)      lcorbi_t3_pos_1_031_40    m/z 743.5421     0   1   2E+4   12.5   13  RT (min)      lcorbi_t3_pos_2_013_33    m/z 743.5421     0   1   2E+4   14.5   15  RT (min)      lcorbi_t3_pos_2_017_51    m/z 743.5421     0   1   2E+4   14.5   15  RT (min)      lcorbi_t3_pos_2_017_51    m/z 743.5421     0   1   15   15.5  RT (min)      lcorbi_t3_pos_1_033_53    m/z 743.5421     0   1   15   15.5  RT (min)      lcorbi_t3_pos_1_030_21    m/z 743.5422     0   1   2   3E+6   13.5   14  RT (min)       lcorbi_t3_pos_4_084_59    m/z 743.5422     0   2   4   6   8E+5   13   13.5  RT (min)      lcorbi_t3_pos_2_023_26    m/z 743.5422     0   1   2   3   4E+6   13.5   14  RT (min)      lcorbi_t3_pos_4_086_59    m/z 743.5422     0   1   2   3   4E+5   14.5   15  RT (min)      lcorbi_t3_pos_1_008_36    m/z 743.5422     0   1   2   3E+6   13.5   14  RT (min)      lcorbi_t3_pos_4_087_59    m/z 743.5423     0   1E+4   12.5   13  RT (min)      lcorbi_t3_pos_2_013_33    m/z 743.5423     0   1   2   3E+4   15   15.5  RT (min)      lcorbi_t3_pos_1_031_40    m/z 743.5423     0   1   15.5   16  RT (min)      lcorbi_t3_pos_1_041_54    m/z 743.5424     0   1   12.5   13  RT (min)      lcorbi_t3_pos_3_024_07    m/z 743.5424     0   2   4   6   8E+5   14   14.5  RT (min)      lcorbi_t3_pos_1_005_42    m/z 743.5425     0   1   16   16.5  RT (min)      lcorbi_t3_pos_1_025_52    m/z 743.5425     0   1   2   3E+4   15.5   16  RT (min)      lcorbi_t3_pos_1_031_40    m/z 743.5425     0   1   15.5   16  RT (min)      lcorbi_t3_pos_1_056_56    m/z 744.5538     0   1   13   13.5  RT (min)      lcorbi_t3_pos_4_012_37    m/z 744.5539     0   1   16   16.5  RT (min)      lcorbi_t3_pos_1_025_52    m/z 744.5540     0   1   2   3E+6   13   13.5  RT (min)      lcorbi_t3_pos_2_013_33    m/z 744.5541     0   1   2   3   4E+4   13   13.5  RT (min)      lcorbi_t3_pos_2_013_33    m/z 744.5542     0   1   2   3E+4   15   15.5  RT (min)      lcorbi_t3_pos_1_047_41    m/z 744.5542     0   1E+4   13   13.5  RT (min)      lcorbi_t3_pos_3_035_08    m/z 744.5542     0   2   4   6E+4   15   15.5  RT (min)      lcorbi_t3_pos_2_046_07    m/z 744.5542     0   1   2   3   4E+4   16   16.5  RT (min)      lcorbi_t3_pos_1_031_40    m/z 744.5542     0   1   16.5   17  RT (min)    lcorbi_t3_pos_2_012_01    m/z 744.5542     0   1   12.5   13  RT (min)      lcorbi_t3_pos_2_032_39    m/z 744.5542     0   1E+6   13   13.5  RT (min)      lcorbi_t3_pos_3_024_07    m/z 744.5542     0   2   4E+4   15   15.5  RT (min)      lcorbi_t3_pos_2_046_07    m/z 744.5543     0   2   4   6E+5   14.5   15  RT (min)       lcorbi_t3_pos_1_008_36    m/z 744.5543     0   1   2E+4   15.5   16  RT (min)      lcorbi_t3_pos_2_009_50    m/z 744.5543     0   1E+5   15   15.5  RT (min)      lcorbi_t3_pos_1_028_32    m/z 744.5543     0   1   2   3   4E+4   16   16.5  RT (min)      lcorbi_t3_pos_1_031_40    m/z 744.5543     0   2   4E+6   14.5   15  RT (min)       lcorbi_t3_pos_1_005_42    m/z 744.5543     0   1E+5   15.5   16  RT (min)      lcorbi_t3_pos_1_028_32    m/z 744.5544     0   1E+5   15.5   16  RT (min)      lcorbi_t3_pos_1_034_34    m/z 744.5544     0   1E+5   15   15.5  RT (min)      lcorbi_t3_pos_1_028_32    m/z 744.5545     0   1   2E+4   15.5   16  RT (min)      lcorbi_t3_pos_2_009_50    m/z 744.5545     0   1E+6   14.5   15  RT (min)       lcorbi_t3_pos_1_008_36    m/z 744.5546     0   1   16.5   17  RT (min)    lcorbi_t3_pos_1_023_02    m/z 744.5548     0   1E+6   13   13.5  RT (min)      lcorbi_t3_pos_2_032_39    m/z 744.5904     0   1E+5   13.5   14  RT (min)      lcorbi_t3_pos_1_002_01    m/z 744.5906     0   1E+4   13   13.5  RT (min)      lcorbi_t3_pos_2_013_33    m/z 744.5906     0   1   13   13.5  RT (min)      lcorbi_t3_pos_1_002_01    m/z 744.5906     0   1   13   13.5  RT (min)      lcorbi_t3_pos_2_032_39    m/z 744.5909     0   1   13.5   14  RT (min)      lcorbi_t3_pos_1_002_01    m/z 744.5910     0   1   16.5   17  RT (min)    lcorbi_t3_pos_1_023_02    m/z 744.5910     0   1   14   14.5  RT (min)      lcorbi_t3_pos_2_031_20    m/z 744.5912     0   1   2E+4   14   14.5  RT (min)      lcorbi_t3_pos_3_037_32    m/z 744.5912     0   1   2E+6   14.5   15  RT (min)      lcorbi_t3_pos_1_002_01    m/z 744.5917     0   1E+5   15.5   16  RT (min)      lcorbi_t3_pos_4_001_49    m/z 744.5919     0   1   16   16.5  RT (min)      lcorbi_t3_pos_3_049_55    m/z 744.5919     0   2   4   6   8E+5   14.5   15  RT (min)      lcorbi_t3_pos_3_053_48    m/z 744.5919     0   2   4   6E+5   14.5   15  RT (min)      lcorbi_t3_pos_2_016_23    m/z 744.5923     0   1   18   18.5  RT (min)    lcorbi_t3_pos_1_023_02    m/z 744.5926     0   1   17   17.5  RT (min)    lcorbi_t3_pos_1_027_43    m/z 744.5926     0   1   18   18.5  RT (min)    lcorbi_t3_pos_1_023_02    m/z 744.5926     0   1   16.5   17  RT (min)    lcorbi_t3_pos_2_041_54    m/z 744.5927     0   1   16.5   17  RT (min)    lcorbi_t3_pos_1_028_32    m/z 744.5927     0   1   17.5   18  RT (min)    lcorbi_t3_pos_2_056_56    m/z 744.5927     0   1   17.5   18  RT (min)    lcorbi_t3_pos_1_031_40    m/z 744.5927     0   1E+4   16   16.5  RT (min)      lcorbi_t3_pos_2_022_38    m/z 744.5928     0   1   16.5   17  RT (min)    lcorbi_t3_pos_2_025_52    m/z 744.5928     0   1E+7   15.5   16  RT (min)       lcorbi_t3_pos_1_005_42    m/z 744.5928     0   1   2E+6   14.5   15  RT (min)      lcorbi_t3_pos_4_016_15    m/z 744.5928     0   1   16.5   17  RT (min)    lcorbi_t3_pos_1_034_34    m/z 744.5928     0   1   17   17.5  RT (min)    lcorbi_t3_pos_2_014_37    m/z 744.5928     0   1   2   3E+4   16   16.5  RT (min)      lcorbi_t3_pos_1_047_41    m/z 744.5928     0   2   4E+5   15.5   16  RT (min)      lcorbi_t3_pos_1_008_36    m/z 744.5928     0   1   17.5   18  RT (min)    lcorbi_t3_pos_2_012_01    m/z 744.5928     0   1   17   17.5  RT (min)    lcorbi_t3_pos_1_041_54    m/z 744.5929     0   1   17.5   18  RT (min)    lcorbi_t3_pos_2_056_56    m/z 744.5929     0   1   18   18.5  RT (min)    lcorbi_t3_pos_1_023_02    m/z 744.5930     0   1   16.5   17  RT (min)    lcorbi_t3_pos_1_034_34    m/z 744.5930     0   1   17.5   18  RT (min)    lcorbi_t3_pos_2_056_56    m/z 744.5932     0   1   17.5   18  RT (min)    lcorbi_t3_pos_2_056_56    m/z 745.5573     0   1   13   13.5  RT (min)      lcorbi_t3_pos_3_035_08    m/z 745.5574     0   1   16   16.5  RT (min)      lcorbi_t3_pos_2_056_56    m/z 745.5574     0   1   12.5   13  RT (min)      lcorbi_t3_pos_2_032_39    m/z 745.5575     0   1   16.5   17  RT (min)    lcorbi_t3_pos_1_023_02    m/z 745.5575     0   1E+6   13   13.5  RT (min)      lcorbi_t3_pos_2_013_33    m/z 745.5576     0   1   13   13.5  RT (min)      lcorbi_t3_pos_2_013_33    m/z 745.5576     0   1   16   16.5  RT (min)      lcorbi_t3_pos_1_001_49    m/z 745.5576     0   1   2   3   4E+4   15.5   16  RT (min)      lcorbi_t3_pos_1_028_32    m/z 745.5576     0   1   17   17.5  RT (min)    lcorbi_t3_pos_1_023_02    m/z 745.5576     0   1   2E+4   13   13.5  RT (min)      lcorbi_t3_pos_1_002_01    m/z 745.5577     0   1E+6   13.5   14  RT (min)      lcorbi_t3_pos_2_013_33    m/z 745.5578     0   1   2   3   4E+5   14.5   15  RT (min)      lcorbi_t3_pos_4_025_52    m/z 745.5578     0   1E+6   14.5   15  RT (min)      lcorbi_t3_pos_4_025_52    m/z 745.5578     0   1   2   3   4E+4   15   15.5  RT (min)      lcorbi_t3_pos_1_028_32    m/z 745.5578     0   1   2   3   4E+4   15   15.5  RT (min)      lcorbi_t3_pos_1_028_32    m/z 745.5578     0   1   2   3   4E+4   15   15.5  RT (min)      lcorbi_t3_pos_2_046_07    m/z 745.5578     0   1   2   3E+5   14.5   15  RT (min)       lcorbi_t3_pos_1_008_36    m/z 745.5578     0   1   16   16.5  RT (min)      lcorbi_t3_pos_1_031_40    m/z 745.5578     0   1   15.5   16  RT (min)      lcorbi_t3_pos_2_009_50    m/z 745.5578     0   1   2   3   4E+4   15   15.5  RT (min)      lcorbi_t3_pos_1_034_34    m/z 745.5578     0   1   2   3   4E+4   15.5   16  RT (min)      lcorbi_t3_pos_1_034_34    m/z 745.5579     0   1   2E+6   14.5   15  RT (min)      lcorbi_t3_pos_1_005_42    m/z 745.5579     0   1   15.5   16  RT (min)      lcorbi_t3_pos_1_031_40    m/z 745.5580     0   1   16   16.5  RT (min)      lcorbi_t3_pos_1_049_55    m/z 745.5580     0   2   4   6E+5   14.5   15  RT (min)      lcorbi_t3_pos_1_008_36    m/z 745.5580     0   1   2   3   4E+4   15   15.5  RT (min)      lcorbi_t3_pos_2_046_07    m/z 745.5580     0   1   16.5   17  RT (min)    lcorbi_t3_pos_1_023_02    m/z 745.5941     0   1   2   3   4E+4   13.5   14  RT (min)      lcorbi_t3_pos_1_002_01    m/z 745.5942     0   1   13.5   14  RT (min)      lcorbi_t3_pos_1_002_01    m/z 745.5945     0   2   4E+4   16   16.5  RT (min)      lcorbi_t3_pos_1_031_40    m/z 745.5949     0   1   2E+5   14.5   15  RT (min)      lcorbi_t3_pos_3_011_42    m/z 745.5954     0   1   2   3   4E+5   15.5   16  RT (min)      lcorbi_t3_pos_4_025_52    m/z 745.5956     0   1   18   18.5  RT (min)    lcorbi_t3_pos_1_023_02    m/z 745.5960     0   1   16.5   17  RT (min)    lcorbi_t3_pos_2_038_03    m/z 745.5960     0   1E+6   14.5   15  RT (min)      lcorbi_t3_pos_3_053_48    m/z 745.5960     0   1   17.5   18  RT (min)    lcorbi_t3_pos_1_031_40    m/z 745.5960     0   1E+6   14.5   15  RT (min)      lcorbi_t3_pos_4_105_61    m/z 745.5962     0   1   2   3E+5   15.5   16  RT (min)      lcorbi_t3_pos_1_008_36    m/z 745.5962     0   1E+6   14.5   15  RT (min)      lcorbi_t3_pos_1_002_01    m/z 745.5962     0   1   16.5   17  RT (min)    lcorbi_t3_pos_1_028_32    m/z 745.5962     0   2   4E+6   15.5   16  RT (min)      lcorbi_t3_pos_1_005_42    m/z 745.5962     0   1   16   16.5  RT (min)      lcorbi_t3_pos_1_047_41    m/z 745.5962     0   1   18   18.5  RT (min)    lcorbi_t3_pos_1_023_02    m/z 745.5962     0   1   16   16.5  RT (min)      lcorbi_t3_pos_2_022_38    m/z 745.5962     0   1E+4   16   16.5  RT (min)      lcorbi_t3_pos_4_049_55    m/z 745.5962     0   1   17.5   18  RT (min)    lcorbi_t3_pos_2_012_01    m/z 745.5962     0   1   16.5   17  RT (min)    lcorbi_t3_pos_2_041_54    m/z 745.5962     0   1   17   17.5  RT (min)    lcorbi_t3_pos_1_056_56    m/z 745.5963     0   1   17.5   18  RT (min)    lcorbi_t3_pos_2_056_56    m/z 745.5963     0   1   17   17.5  RT (min)    lcorbi_t3_pos_2_009_50    m/z 745.5963     0   1   17   17.5  RT (min)    lcorbi_t3_pos_2_049_55    m/z 745.5963     0   1   16.5   17  RT (min)    lcorbi_t3_pos_1_034_34    m/z 745.5964     0   1   2   3E+5   14.5   15  RT (min)      lcorbi_t3_pos_4_015_45    m/z 745.5964     0   1   16.5   17  RT (min)    lcorbi_t3_pos_1_034_34    m/z 745.5965     0   1   17.5   18  RT (min)    lcorbi_t3_pos_2_056_56    m/z 746.3268     0   2   4E+5   14.5   15  RT (min)      lcorbi_t3_pos_4_105_61    m/z 746.5612     0   1   2E+5   13   13.5  RT (min)      lcorbi_t3_pos_1_002_01    m/z 746.5614     0   1   2   3E+4   16   16.5  RT (min)      lcorbi_t3_pos_1_031_40    m/z 746.5615     0   1E+5   13   13.5  RT (min)      lcorbi_t3_pos_3_024_07    m/z 746.5617     0   1   2   3E+4   16   16.5  RT (min)      lcorbi_t3_pos_1_031_40    m/z 746.5655     0   1   2E+4   15   15.5  RT (min)      lcorbi_t3_pos_2_046_07    m/z 746.5689     0   1E+5   13   13.5  RT (min)      lcorbi_t3_pos_1_002_01    m/z 746.5692     0   2   4E+4   15   15.5  RT (min)      lcorbi_t3_pos_1_028_32    m/z 746.5693     0   2   4E+4   15   15.5  RT (min)      lcorbi_t3_pos_1_028_32    m/z 746.5693     0   1   2   3E+4   15.5   16  RT (min)      lcorbi_t3_pos_1_031_40    m/z 746.5694     0   2   4E+4   15.5   16  RT (min)      lcorbi_t3_pos_1_028_32    m/z 746.5694     0   1   16   16.5  RT (min)      lcorbi_t3_pos_1_031_40    m/z 746.5695     0   2   4E+4   15   15.5  RT (min)      lcorbi_t3_pos_1_028_32    m/z 746.5695     0   1   16   16.5  RT (min)      lcorbi_t3_pos_1_031_40    m/z 746.5698     0   1   2   3E+4   16   16.5  RT (min)      lcorbi_t3_pos_1_031_40    m/z 746.5702     0   1   16.5   17  RT (min)    lcorbi_t3_pos_2_012_01    m/z 746.5992     0   1   18   18.5  RT (min)    lcorbi_t3_pos_1_023_02    m/z 746.5992     0   1   18   18.5  RT (min)    lcorbi_t3_pos_1_023_02    m/z 746.5996     0   1   17.5   18  RT (min)    lcorbi_t3_pos_2_012_01    m/z 746.5997     0   1   17   17.5  RT (min)    lcorbi_t3_pos_1_049_55    m/z 746.5998     0   1   16.5   17  RT (min)    lcorbi_t3_pos_1_041_54    m/z 746.5998     0   1   2E+6   15.5   16  RT (min)      lcorbi_t3_pos_1_008_36    m/z 746.5998     0   1   16.5   17  RT (min)    lcorbi_t3_pos_1_034_34    m/z 746.5999     0   1   16.5   17  RT (min)    lcorbi_t3_pos_1_034_34    m/z 746.6028     0   1   2   3E+4   16   16.5  RT (min)      lcorbi_t3_pos_1_047_41    m/z 746.6032     0   1   17   17.5  RT (min)    lcorbi_t3_pos_1_031_40    m/z 746.6058     0   1   16   16.5  RT (min)      lcorbi_t3_pos_2_006_47    m/z 746.6060     0   1   2   3E+6   14.5   15  RT (min)      lcorbi_t3_pos_3_026_04    m/z 746.6060     0   1   2   3   4E+3   13.5   14  RT (min)      lcorbi_t3_pos_2_013_33    m/z 746.6061     0   1   2E+6   15.5   16  RT (min)      lcorbi_t3_pos_1_008_36    m/z 746.6061     0   1E+4   13   13.5  RT (min)      lcorbi_t3_pos_2_032_39    m/z 746.6062     0   1E+5   13.5   14  RT (min)      lcorbi_t3_pos_1_002_01    m/z 746.6062     0   1   16.5   17  RT (min)    lcorbi_t3_pos_1_034_34    m/z 746.6062     0   1   16   16.5  RT (min)      lcorbi_t3_pos_1_028_32    m/z 746.6064     0   1E+4   16   16.5  RT (min)      lcorbi_t3_pos_3_003_33    m/z 746.6065     0   1   13.5   14  RT (min)      lcorbi_t3_pos_1_002_01    m/z 746.6074     0   1   17   17.5  RT (min)    lcorbi_t3_pos_2_012_01    m/z 746.6081     0   1   16   16.5  RT (min)      lcorbi_t3_pos_2_022_38    m/z 746.6084     0   1   17   17.5  RT (min)    lcorbi_t3_pos_1_031_40    m/z 747.5734     0   1   2E+4   15   15.5  RT (min)      lcorbi_t3_pos_1_028_32    m/z 747.5734     0   1   15.5   16  RT (min)      lcorbi_t3_pos_1_031_40    m/z 747.5734     0   1   2E+4   15   15.5  RT (min)      lcorbi_t3_pos_1_028_32    m/z 747.6028     0   1   16.5   17  RT (min)    lcorbi_t3_pos_1_034_34    m/z 747.6032     0   1   16.5   17  RT (min)    lcorbi_t3_pos_1_034_34    m/z 747.6069     0   1   16.5   17  RT (min)    lcorbi_t3_pos_4_033_53    m/z 747.6090     0   1   16.5   17  RT (min)    lcorbi_t3_pos_1_034_34    m/z 747.6096     0   2   4E+5   15.5   16  RT (min)      lcorbi_t3_pos_1_008_36    m/z 747.6098     0   2   4E+5   15.5   16  RT (min)      lcorbi_t3_pos_1_008_36    m/z 747.6099     0   1   13.5   14  RT (min)      lcorbi_t3_pos_1_002_01    m/z 747.6099     0   1   16   16.5  RT (min)      lcorbi_t3_pos_1_028_32    m/z 747.6100     0   1   13.5   14  RT (min)      lcorbi_t3_pos_2_013_33    m/z 747.6101     0   1   16   16.5  RT (min)      lcorbi_t3_pos_2_022_38    m/z 747.6102     0   1   2   3E+5   13.5   14  RT (min)      lcorbi_t3_pos_3_024_07    m/z 747.6104     0   1   13   13.5  RT (min)      lcorbi_t3_pos_2_032_39    m/z 747.6110     0   1   16   16.5  RT (min)      lcorbi_t3_pos_2_022_38    m/z 747.6110     0   1   15.5   16  RT (min)      lcorbi_t3_pos_2_046_07    m/z 747.6112     0   1   17   17.5  RT (min)    lcorbi_t3_pos_1_031_40    m/z 747.6119     0   1   17   17.5  RT (min)    lcorbi_t3_pos_2_012_01    m/z 747.6120     0   1   17   17.5  RT (min)    lcorbi_t3_pos_1_031_40    m/z 748.5124     0   2   4E+5   12   12.5  RT (min)      lcorbi_t3_pos_2_005_11    m/z 748.5125     0   2   4E+5   12   12.5  RT (min)       lcorbi_t3_pos_1_054_48    m/z 748.5130     0   1   15   15.5  RT (min)      lcorbi_t3_pos_1_031_40    m/z 748.5488     0   2   4   6E+3   13   13.5  RT (min)       lcorbi_t3_pos_3_052_47    m/z 748.5488     0   1   13   13.5  RT (min)      lcorbi_t3_pos_2_039_30    m/z 748.5490     0   1E+4   13   13.5  RT (min)      lcorbi_t3_pos_2_047_31    m/z 748.5490     0   2   4   6   8E+5   13.5   14  RT (min)      lcorbi_t3_pos_4_030_05    m/z 748.5492     0   1   15.5   16  RT (min)      lcorbi_t3_pos_1_027_43    m/z 748.5492     0   1E+4   15.5   16  RT (min)      lcorbi_t3_pos_2_052_48    m/z 748.5493     0   1   16   16.5  RT (min)      lcorbi_t3_pos_1_027_43    m/z 748.5493     0   1   2E+4   15   15.5  RT (min)      lcorbi_t3_pos_1_055_33    m/z 748.5493     0   1   2   3E+4   14.5   15  RT (min)      lcorbi_t3_pos_3_044_46    m/z 748.6134     0   2   4E+5   15   15.5  RT (min)      lcorbi_t3_pos_3_040_41    m/z 748.6140     0   1   16.5   17  RT (min)    lcorbi_t3_pos_1_034_34    m/z 748.6146     0   1   16.5   17  RT (min)    lcorbi_t3_pos_1_034_34    m/z 748.6149     0   1   17   17.5  RT (min)    lcorbi_t3_pos_2_012_01    m/z 748.6149     0   1   16   16.5  RT (min)      lcorbi_t3_pos_1_028_32    m/z 748.6149     0   2   4E+4   15.5   16  RT (min)      lcorbi_t3_pos_1_008_36    m/z 748.6150     0   1E+5   13.5   14  RT (min)      lcorbi_t3_pos_3_024_07    m/z 748.6150     0   1   16   16.5  RT (min)      lcorbi_t3_pos_1_028_32    m/z 748.6151     0   1   17   17.5  RT (min)    lcorbi_t3_pos_1_031_40    m/z 748.6153     0   2   4E+4   15.5   16  RT (min)      lcorbi_t3_pos_1_008_36    m/z 749.5478     0   1   16.5   17  RT (min)    lcorbi_t3_pos_1_033_53    m/z 749.5480     0   2   4   6   8E+4   14.5   15  RT (min)      lcorbi_t3_pos_2_016_23    m/z 749.5480     0   1   17.5   18  RT (min)    lcorbi_t3_pos_2_012_01    m/z 749.5480     0   1   16   16.5  RT (min)      lcorbi_t3_pos_2_022_38    m/z 749.5480     0   1   16.5   17  RT (min)    lcorbi_t3_pos_1_034_34    m/z 749.5480     0   1   17.5   18  RT (min)    lcorbi_t3_pos_2_012_01    m/z 749.5480     0   1   16.5   17  RT (min)    lcorbi_t3_pos_2_025_52    m/z 749.5481     0   1   2   3E+4   16   16.5  RT (min)      lcorbi_t3_pos_1_008_36    m/z 749.5482     0   1   17   17.5  RT (min)    lcorbi_t3_pos_2_009_50    m/z 749.5482     0   2   4   6E+5   15.5   16  RT (min)      lcorbi_t3_pos_1_005_42    m/z 749.5482     0   1   16.5   17  RT (min)    lcorbi_t3_pos_1_028_32    m/z 749.5482     0   1   17   17.5  RT (min)    lcorbi_t3_pos_1_034_34    m/z 749.5482     0   1   17.5   18  RT (min)    lcorbi_t3_pos_1_031_40    m/z 749.5483     0   1   2   3E+4   14   14.5  RT (min)      lcorbi_t3_pos_2_013_33    m/z 749.5483     0   2   4   6   8E+5   15.5   16  RT (min)      lcorbi_t3_pos_1_005_42    m/z 749.5483     0   1   16   16.5  RT (min)      lcorbi_t3_pos_2_022_38    m/z 749.5483     0   1   16   16.5  RT (min)      lcorbi_t3_pos_2_046_07    m/z 749.5484     0   1   18   18.5  RT (min)    lcorbi_t3_pos_1_023_02    m/z 749.5488     0   1   2   3E+4   15.5   16  RT (min)      lcorbi_t3_pos_1_008_36    m/z 750.5067     0   1E+5   13.5   14  RT (min)      lcorbi_t3_pos_1_008_36    m/z 750.5068     0   2   4   6   8E+5   12.5   13  RT (min)      lcorbi_t3_pos_3_011_42    m/z 750.5069     0   1   2   3   4E+3   14   14.5  RT (min)       lcorbi_t3_pos_3_003_33    m/z 750.5069     0   1   15   15.5  RT (min)      lcorbi_t3_pos_2_012_01    m/z 750.5069     0   1   14.5   15  RT (min)      lcorbi_t3_pos_1_031_40    m/z 750.5070     0   1   2   3E+4   14   14.5  RT (min)      lcorbi_t3_pos_1_028_32    m/z 750.5070     0   1   2   3E+5   13.5   14  RT (min)       lcorbi_t3_pos_1_008_36    m/z 750.5070     0   2   4   6E+3   14   14.5  RT (min)      lcorbi_t3_pos_1_034_34    m/z 750.5070     0   1   2   3E+4   14   14.5  RT (min)      lcorbi_t3_pos_1_028_32    m/z 750.5072     0   1   12.5   13  RT (min)      lcorbi_t3_pos_2_013_33    m/z 750.5072     0   1   12   12.5  RT (min)      lcorbi_t3_pos_3_024_07    m/z 750.5525     0   1   2   3E+5   15.5   16  RT (min)      lcorbi_t3_pos_2_017_51    m/z 750.5526     0   1   17.5   18  RT (min)    lcorbi_t3_pos_1_023_02    m/z 750.5529     0   2   4E+4   16   16.5  RT (min)       lcorbi_t3_pos_2_049_55    m/z 750.5547     0   1E+5   15   15.5  RT (min)       lcorbi_t3_pos_3_044_46    m/z 750.5569     0   1   17.5   18  RT (min)     lcorbi_t3_pos_2_012_01    m/z 750.5569     0   1   16.5   17  RT (min)     lcorbi_t3_pos_1_031_40    m/z 750.5574     0   1   16.5   17  RT (min)     lcorbi_t3_pos_1_031_40    m/z 750.5580     0   2   4   6E+5   15.5   16  RT (min)       lcorbi_t3_pos_2_022_38    m/z 750.5583     0   1   17   17.5  RT (min)     lcorbi_t3_pos_1_023_02    m/z 750.5583     0   1E+5   15.5   16  RT (min)      lcorbi_t3_pos_1_033_53    m/z 750.5584     0   1   16.5   17  RT (min)     lcorbi_t3_pos_2_012_01    m/z 750.5584     0   1   2   3E+5   15.5   16  RT (min)       lcorbi_t3_pos_1_030_21    m/z 750.5586     0   2   4E+4   16   16.5  RT (min)       lcorbi_t3_pos_1_034_34    m/z 750.5586     0   1E+6   14   14.5  RT (min)      lcorbi_t3_pos_4_105_61    m/z 750.5586     0   1   16.5   17  RT (min)    lcorbi_t3_pos_1_001_49    m/z 750.5586     0   1   16.5   17  RT (min)    lcorbi_t3_pos_1_001_49    m/z 750.5587     0   1E+6   14.5   15  RT (min)       lcorbi_t3_pos_4_054_18    m/z 750.5587     0   1   16.5   17  RT (min)    lcorbi_t3_pos_4_033_53    m/z 750.5587     0   2   4E+4   15.5   16  RT (min)      lcorbi_t3_pos_2_041_54    m/z 750.5588     0   1E+5   13.5   14  RT (min)      lcorbi_t3_pos_2_047_31    m/z 750.5588     0   2   4   6E+4   15.5   16  RT (min)       lcorbi_t3_pos_1_028_32    m/z 750.5589     0   1   2   3E+4   13.5   14  RT (min)      lcorbi_t3_pos_2_032_39    m/z 750.5589     0   1   2   3E+4   13.5   14  RT (min)      lcorbi_t3_pos_2_032_39    m/z 750.5594     0   2   4   6E+5   0.5   1  RT (min)      lcorbi_t3_pos_4_057_49    m/z 751.5103     0   1E+5   13.5   14  RT (min)      lcorbi_t3_pos_1_008_36    m/z 751.5104     0   2   4   6E+3   14   14.5  RT (min)      lcorbi_t3_pos_1_034_34    m/z 751.5622     0   1   16.5   17  RT (min)    lcorbi_t3_pos_2_056_56    m/z 751.5622     0   1   2E+5   14   14.5  RT (min)      lcorbi_t3_pos_2_013_33    m/z 751.5623     0   2   4E+4   15.5   16  RT (min)      lcorbi_t3_pos_1_028_32    m/z 751.5623     0   1   2E+4   13.5   14  RT (min)      lcorbi_t3_pos_3_037_32    m/z 751.5623     0   1E+4   13.5   14  RT (min)      lcorbi_t3_pos_3_052_47    m/z 751.5624     0   2   4   6E+5   14   14.5  RT (min)      lcorbi_t3_pos_3_011_42    m/z 751.5624     0   2   4   6   8E+4   15.5   16  RT (min)      lcorbi_t3_pos_1_030_21    m/z 751.5624     0   1E+5   15.5   16  RT (min)      lcorbi_t3_pos_2_022_38    m/z 751.5624     0   1   16.5   17  RT (min)     lcorbi_t3_pos_1_056_56    m/z 751.5625     0   1   13.5   14  RT (min)      lcorbi_t3_pos_3_024_07    m/z 751.5625     0   1   13.5   14  RT (min)       lcorbi_t3_pos_3_024_07    m/z 751.5625     0   1   2   3   4E+4   16   16.5  RT (min)      lcorbi_t3_pos_2_022_38    m/z 751.5625     0   1   2   3E+4   15.5   16  RT (min)      lcorbi_t3_pos_2_041_54    m/z 751.5625     0   1   16.5   17  RT (min)    lcorbi_t3_pos_4_033_53    m/z 751.5626     0   1   2   3E+4   16   16.5  RT (min)      lcorbi_t3_pos_2_049_55    m/z 751.5627     0   1E+5   15.5   16  RT (min)      lcorbi_t3_pos_1_047_41    m/z 751.5630     0   1   2   3E+4   16   16.5  RT (min)      lcorbi_t3_pos_1_034_34    m/z 751.5634     0   1   17   17.5  RT (min)    lcorbi_t3_pos_2_012_01    m/z 751.5636     0   1   2E+4   16   16.5  RT (min)      lcorbi_t3_pos_1_028_32    m/z 752.5223     0   1   12.5   13  RT (min)      lcorbi_t3_pos_3_024_07    m/z 752.5225     0   1E+5   14   14.5  RT (min)       lcorbi_t3_pos_2_022_38    m/z 752.5226     0   1   12.5   13  RT (min)      lcorbi_t3_pos_2_013_33    m/z 752.5226     0   1   2   3E+4   14.5   15  RT (min)      lcorbi_t3_pos_1_034_34    m/z 752.5226     0   1E+6   13.5   14  RT (min)       lcorbi_t3_pos_1_008_36    m/z 752.5227     0   2   4E+4   14   14.5  RT (min)      lcorbi_t3_pos_1_028_32    m/z 752.5227     0   1   2E+4   15   15.5  RT (min)      lcorbi_t3_pos_1_031_40    m/z 752.5227     0   1   2E+4   14   14.5  RT (min)       lcorbi_t3_pos_2_022_38    m/z 752.5227     0   1   2E+4   15   15.5  RT (min)      lcorbi_t3_pos_2_012_01    m/z 752.5228     0   1   2E+4   15   15.5  RT (min)      lcorbi_t3_pos_1_031_40    m/z 752.5228     0   1   2   3   4E+6   12.5   13  RT (min)      lcorbi_t3_pos_3_011_42    m/z 752.5228     0   1   2   3E+4   14.5   15  RT (min)      lcorbi_t3_pos_1_034_34    m/z 752.5228     0   2   4E+4   14   14.5  RT (min)      lcorbi_t3_pos_1_028_32    m/z 752.5228     0   2   4   6   8E+5   14   14.5  RT (min)       lcorbi_t3_pos_1_008_36    m/z 752.5229     0   1   2E+4   15   15.5  RT (min)      lcorbi_t3_pos_1_031_40    m/z 752.5232     0   1   14.5   15  RT (min)      lcorbi_t3_pos_1_041_54    m/z 752.5655     0   2   4E+4   16   16.5  RT (min)      lcorbi_t3_pos_2_022_38    m/z 752.5658     0   1   16   16.5  RT (min)      lcorbi_t3_pos_1_031_40    m/z 753.4281     0   1E+4   14   14.5  RT (min)      lcorbi_t3_pos_3_031_45    m/z 753.4282     0   1   16   16.5  RT (min)      lcorbi_t3_pos_2_025_52    m/z 753.4282     0   1   16.5   17  RT (min)    lcorbi_t3_pos_1_041_54    m/z 753.4282     0   2   4   6   8E+6   14.5   15  RT (min)       lcorbi_t3_pos_4_105_61    m/z 753.4282     0   1   16   16.5  RT (min)      lcorbi_t3_pos_2_041_54    m/z 753.4282     0   1E+5   15.5   16  RT (min)      lcorbi_t3_pos_3_025_52    m/z 753.4282     0   1E+4   13.5   14  RT (min)      lcorbi_t3_pos_2_032_39    m/z 753.4282     0   1   17   17.5  RT (min)    lcorbi_t3_pos_1_025_52    m/z 753.4282     0   1   17.5   18  RT (min)    lcorbi_t3_pos_1_023_02    m/z 753.4283     0   2   4   6E+6   15   15.5  RT (min)      lcorbi_t3_pos_3_041_54    m/z 753.4283     0   1E+4   16   16.5  RT (min)      lcorbi_t3_pos_4_041_54    m/z 753.4283     0   2   4   6E+6   15   15.5  RT (min)      lcorbi_t3_pos_1_005_42    m/z 753.4283     0   1   16   16.5  RT (min)      lcorbi_t3_pos_1_033_53    m/z 753.4284     0   2   4E+4   15.5   16  RT (min)      lcorbi_t3_pos_3_056_56    m/z 753.4284     0   1   2   3E+4   15.5   16  RT (min)      lcorbi_t3_pos_3_056_56    m/z 753.4284     0   1   17   17.5  RT (min)    lcorbi_t3_pos_1_001_49    m/z 753.4284     0   1   16.5   17  RT (min)    lcorbi_t3_pos_2_009_50    m/z 753.4284     0   1   17   17.5  RT (min)    lcorbi_t3_pos_2_056_56    m/z 753.4284     0   1   17   17.5  RT (min)    lcorbi_t3_pos_2_012_01    m/z 753.4284     0   1   13.5   14  RT (min)      lcorbi_t3_pos_2_047_31    m/z 753.4284     0   1   17   17.5  RT (min)    lcorbi_t3_pos_1_056_56    m/z 753.4284     0   1   2   3   4E+6   14.5   15  RT (min)      lcorbi_t3_pos_3_053_48    m/z 753.4284     0   1   2   3E+4   14   14.5  RT (min)       lcorbi_t3_pos_4_046_44    m/z 753.4285     0   1E+4   15.5   16  RT (min)      lcorbi_t3_pos_3_044_46    m/z 753.4286     0   1   17.5   18  RT (min)    lcorbi_t3_pos_1_023_02    m/z 753.5260     0   1   2E+4   12.5   13  RT (min)      lcorbi_t3_pos_2_013_33    m/z 753.5261     0   2   4   6   8E+3   12.5   13  RT (min)      lcorbi_t3_pos_3_024_07    m/z 753.5261     0   1   2E+4   14.5   15  RT (min)      lcorbi_t3_pos_1_034_34    m/z 753.5261     0   1   2   3   4E+4   14   14.5  RT (min)       lcorbi_t3_pos_1_028_32    m/z 753.5261     0   2   4   6   8E+4   14   14.5  RT (min)      lcorbi_t3_pos_2_022_38    m/z 753.5262     0   1E+4   15   15.5  RT (min)      lcorbi_t3_pos_2_012_01    m/z 753.5262     0   2   4   6E+5   13.5   14  RT (min)       lcorbi_t3_pos_1_008_36    m/z 753.5262     0   1   15   15.5  RT (min)      lcorbi_t3_pos_1_031_40    m/z 753.5262     0   1   2E+4   14.5   15  RT (min)      lcorbi_t3_pos_1_034_34    m/z 753.5263     0   1E+6   12.5   13  RT (min)      lcorbi_t3_pos_3_011_42    m/z 753.5264     0   2   4   6E+5   13.5   14  RT (min)      lcorbi_t3_pos_1_005_42    m/z 753.5265     0   1   15   15.5  RT (min)      lcorbi_t3_pos_1_031_40    m/z 753.5265     0   1   2   3   4E+5   14   14.5  RT (min)      lcorbi_t3_pos_1_008_36    m/z 753.6382     0   1   14.5   15  RT (min)      lcorbi_t3_pos_1_021_18    m/z 753.6384     0   1   15.5   16  RT (min)      lcorbi_t3_pos_1_001_49    m/z 753.6385     0   1   15.5   16  RT (min)      lcorbi_t3_pos_1_001_49    m/z 753.6427     0   1   15   15.5  RT (min)      lcorbi_t3_pos_1_041_54    m/z 753.6434     0   1   15   15.5  RT (min)      lcorbi_t3_pos_2_038_03    m/z 753.6460     0   1   15   15.5  RT (min)      lcorbi_t3_pos_1_041_54    m/z 754.4315     0   1   16   16.5  RT (min)      lcorbi_t3_pos_2_041_54    m/z 754.4315     0   1   16   16.5  RT (min)      lcorbi_t3_pos_1_021_18    m/z 754.4315     0   1   2E+6   15   15.5  RT (min)      lcorbi_t3_pos_3_041_54    m/z 754.4315     0   1   2   3   4E+4   15.5   16  RT (min)      lcorbi_t3_pos_3_025_52    m/z 754.4316     0   1   17   17.5  RT (min)    lcorbi_t3_pos_2_056_56    m/z 754.4316     0   1   16   16.5  RT (min)      lcorbi_t3_pos_2_041_54    m/z 754.4316     0   1   17   17.5  RT (min)    lcorbi_t3_pos_2_012_01    m/z 754.4316     0   1   17   17.5  RT (min)    lcorbi_t3_pos_1_049_55    m/z 754.4316     0   1E+4   16   16.5  RT (min)      lcorbi_t3_pos_4_049_55    m/z 754.4316     0   1   17.5   18  RT (min)    lcorbi_t3_pos_1_023_02    m/z 754.4316     0   1   16   16.5  RT (min)      lcorbi_t3_pos_4_041_54    m/z 754.4317     0   1   16.5   17  RT (min)    lcorbi_t3_pos_1_041_54    m/z 754.4317     0   1E+4   14   14.5  RT (min)      lcorbi_t3_pos_3_035_08    m/z 754.4317     0   1   2E+6   15   15.5  RT (min)      lcorbi_t3_pos_3_041_54    m/z 754.4317     0   1   2   3   4E+3   13.5   14  RT (min)      lcorbi_t3_pos_2_032_39    m/z 754.4317     0   1   14   14.5  RT (min)      lcorbi_t3_pos_2_054_29    m/z 754.4317     0   1   2E+4   15.5   16  RT (min)      lcorbi_t3_pos_3_056_56    m/z 754.4317     0   1   2   3E+6   14.5   15  RT (min)      lcorbi_t3_pos_4_105_61    m/z 754.4320     0   1E+6   14.5   15  RT (min)      lcorbi_t3_pos_4_016_15    m/z 754.5370     0   1E+7   13.5   14  RT (min)       lcorbi_t3_pos_1_005_42    m/z 754.5370     0   1   15.5   16  RT (min)      lcorbi_t3_pos_1_051_09    m/z 754.5375     0   1E+4   15   15.5  RT (min)      lcorbi_t3_pos_2_049_55    m/z 754.5376     0   1   2E+7   13   13.5  RT (min)      lcorbi_t3_pos_3_011_42    m/z 754.5379     0   2   4   6   8E+5   13   13.5  RT (min)      lcorbi_t3_pos_1_002_01    m/z 754.5382     0   2   4   6E+4   14.5   15  RT (min)      lcorbi_t3_pos_1_034_34    m/z 754.5382     0   1E+4   12.5   13  RT (min)      lcorbi_t3_pos_2_013_33    m/z 754.5382     0   2   4   6E+6   14   14.5  RT (min)      lcorbi_t3_pos_1_008_36    m/z 754.5382     0   2   4E+5   10   10.5  RT (min)      lcorbi_t3_pos_4_057_49    m/z 754.5382     0   1   2   3E+4   15   15.5  RT (min)      lcorbi_t3_pos_1_049_55    m/z 754.5384     0   2   4E+4   14   14.5  RT (min)      lcorbi_t3_pos_1_047_41    m/z 754.5384     0   1   2   3E+6   14   14.5  RT (min)      lcorbi_t3_pos_1_008_36    m/z 754.5384     0   1   2   3E+7   13   13.5  RT (min)      lcorbi_t3_pos_3_011_42    m/z 754.5384     0   2   4   6E+4   14.5   15  RT (min)      lcorbi_t3_pos_1_028_32    m/z 754.5385     0   2   4E+5   10.5   11  RT (min)      lcorbi_t3_pos_4_057_49    m/z 754.5386     0   1   15.5   16  RT (min)      lcorbi_t3_pos_1_051_09    m/z 754.5386     0   1E+4   14.5   15  RT (min)      lcorbi_t3_pos_3_013_19    m/z 754.5386     0   1   16   16.5  RT (min)      lcorbi_t3_pos_1_023_02    m/z 754.5388     0   1   12.5   13  RT (min)      lcorbi_t3_pos_2_032_39    m/z 754.5389     0   1   2   3   4E+4   15   15.5  RT (min)      lcorbi_t3_pos_1_031_40    m/z 754.6505     0   1E+6   14   14.5  RT (min)      lcorbi_t3_pos_1_005_42    m/z 754.6506     0   1   16   16.5  RT (min)      lcorbi_t3_pos_1_001_49    m/z 754.6506     0   1   15.5   16  RT (min)      lcorbi_t3_pos_1_049_55    m/z 754.6506     0   1E+4   14.5   15  RT (min)      lcorbi_t3_pos_2_017_51    m/z 754.6507     0   1   15.5   16  RT (min)      lcorbi_t3_pos_1_025_52    m/z 754.6508     0   1   15.5   16  RT (min)      lcorbi_t3_pos_1_041_54    m/z 754.6509     0   1   15   15.5  RT (min)      lcorbi_t3_pos_1_030_21    m/z 754.6509     0   1   15   15.5  RT (min)      lcorbi_t3_pos_1_041_54    m/z 755.4346     0   2   4   6   8E+5   14.5   15  RT (min)      lcorbi_t3_pos_4_105_61    m/z 755.5403     0   2   4   6E+6   13.5   14  RT (min)       lcorbi_t3_pos_1_005_42    m/z 755.5417     0   1   2E+4   14.5   15  RT (min)      lcorbi_t3_pos_1_034_34    m/z 755.5417     0   1   12.5   13  RT (min)      lcorbi_t3_pos_2_013_33    m/z 755.5418     0   1   2   3E+6   14   14.5  RT (min)      lcorbi_t3_pos_1_008_36    m/z 755.5418     0   1   2   3E+4   15   15.5  RT (min)      lcorbi_t3_pos_1_031_40    m/z 755.5418     0   1   15   15.5  RT (min)      lcorbi_t3_pos_2_049_55    m/z 755.5419     0   1E+7   13   13.5  RT (min)      lcorbi_t3_pos_3_011_42    m/z 755.5420     0   2   4   6   8E+5   14   14.5  RT (min)      lcorbi_t3_pos_1_008_36    m/z 755.5420     0   1E+4   14.5   15  RT (min)      lcorbi_t3_pos_3_049_55    m/z 755.5420     0   1   2   3   4E+4   14.5   15  RT (min)      lcorbi_t3_pos_1_028_32    m/z 755.5420     0   1E+7   13   13.5  RT (min)      lcorbi_t3_pos_3_011_42    m/z 755.5421     0   1   2   3E+4   14   14.5  RT (min)      lcorbi_t3_pos_1_047_41    m/z 755.5424     0   1   15.5   16  RT (min)      lcorbi_t3_pos_1_001_49    m/z 755.5900     0   2   4E+5   11.5   12  RT (min)      lcorbi_t3_pos_1_003_45    m/z 755.6537     0   1E+6   12.5   13  RT (min)      lcorbi_t3_pos_4_057_49    m/z 755.6538     0   1   2   3   4E+5   14.5   15  RT (min)      lcorbi_t3_pos_4_025_52    m/z 755.6538     0   1   2   3   4E+6   14   14.5  RT (min)      lcorbi_t3_pos_1_005_42    m/z 755.6539     0   1   2E+4   15.5   16  RT (min)      lcorbi_t3_pos_1_049_55    m/z 755.6540     0   1E+6   14   14.5  RT (min)      lcorbi_t3_pos_3_040_41    m/z 755.6540     0   1   2   3E+4   15.5   16  RT (min)      lcorbi_t3_pos_1_041_54    m/z 755.6540     0   1   15   15.5  RT (min)      lcorbi_t3_pos_1_033_53    m/z 755.6540     0   1   2   3E+4   15   15.5  RT (min)      lcorbi_t3_pos_1_041_54    m/z 755.6540     0   1   15   15.5  RT (min)      lcorbi_t3_pos_1_033_53    m/z 755.6540     0   1E+5   14.5   15  RT (min)      lcorbi_t3_pos_3_056_56    m/z 755.6540     0   1   2E+4   15   15.5  RT (min)      lcorbi_t3_pos_1_030_21    m/z 755.6540     0   1   15.5   16  RT (min)      lcorbi_t3_pos_1_025_52    m/z 755.6541     0   1   2   3E+4   15   15.5  RT (min)      lcorbi_t3_pos_1_041_54    m/z 755.6541     0   1   2   3   4E+4   14.5   15  RT (min)      lcorbi_t3_pos_2_046_07    m/z 755.6542     0   2   4E+4   14.5   15  RT (min)      lcorbi_t3_pos_2_017_51    m/z 755.6542     0   1   2   3E+4   15.5   16  RT (min)      lcorbi_t3_pos_1_041_54    m/z 756.5450     0   1   2   3E+4   12.5   13  RT (min)      lcorbi_t3_pos_3_031_45    m/z 756.5452     0   1   2   3   4E+6   13   13.5  RT (min)      lcorbi_t3_pos_3_011_42    m/z 756.5487     0   1   2   3   4E+4   12.5   13  RT (min)      lcorbi_t3_pos_3_031_45    m/z 756.5494     0   1E+7   14   14.5  RT (min)      lcorbi_t3_pos_1_008_36    m/z 756.5534     0   2   4E+6   12   12.5  RT (min)      lcorbi_t3_pos_4_057_49    m/z 756.5535     0   1   2   3   4E+4   12.5   13  RT (min)      lcorbi_t3_pos_3_031_45    m/z 756.5536     0   2   4   6E+4   15.5   16  RT (min)      lcorbi_t3_pos_2_012_01    m/z 756.5536     0   1   2E+5   14.5   15  RT (min)      lcorbi_t3_pos_2_046_07    m/z 756.5536     0   1E+5   14.5   15  RT (min)      lcorbi_t3_pos_1_028_32    m/z 756.5536     0   2   4   6E+4   15.5   16  RT (min)      lcorbi_t3_pos_2_012_01    m/z 756.5537     0   1   2E+5   14.5   15  RT (min)      lcorbi_t3_pos_2_046_07    m/z 756.5537     0   1   2   3   4E+4   15   15.5  RT (min)      lcorbi_t3_pos_2_009_50    m/z 756.5537     0   2   4   6   8E+4   15.5   16  RT (min)      lcorbi_t3_pos_1_031_40    m/z 756.5537     0   1E+5   15   15.5  RT (min)      lcorbi_t3_pos_1_034_34    m/z 756.5537     0   2   4   6   8E+4   15.5   16  RT (min)      lcorbi_t3_pos_1_031_40    m/z 756.5538     0   2   4E+4   13   13.5  RT (min)      lcorbi_t3_pos_2_013_33    m/z 756.5538     0   1   16   16.5  RT (min)      lcorbi_t3_pos_1_023_02    m/z 756.5538     0   1E+5   14.5   15  RT (min)      lcorbi_t3_pos_1_028_32    m/z 756.5538     0   2   4E+4   13   13.5  RT (min)       lcorbi_t3_pos_2_013_33    m/z 756.5539     0   1E+5   15   15.5  RT (min)      lcorbi_t3_pos_1_034_34    m/z 756.5539     0   1   16   16.5  RT (min)      lcorbi_t3_pos_1_023_02    m/z 756.5540     0   2   4   6E+4   15.5   16  RT (min)      lcorbi_t3_pos_2_012_01    m/z 756.5541     0   1   2E+7   14   14.5  RT (min)      lcorbi_t3_pos_1_005_42    m/z 756.5542     0   1   16   16.5  RT (min)      lcorbi_t3_pos_1_023_02    m/z 756.5543     0   2   4   6E+6   13   13.5  RT (min)      lcorbi_t3_pos_2_054_29    m/z 756.5990     0   1   2E+5   14.5   15  RT (min)      lcorbi_t3_pos_1_008_36    m/z 756.5991     0   1   15   15.5  RT (min)      lcorbi_t3_pos_1_028_32    m/z 756.5992     0   1   15.5   16  RT (min)      lcorbi_t3_pos_1_031_40    m/z 756.6660     0   1   16   16.5  RT (min)      lcorbi_t3_pos_1_025_52    m/z 756.6660     0   1   16   16.5  RT (min)      lcorbi_t3_pos_1_025_52    m/z 756.6661     0   1   16   16.5  RT (min)      lcorbi_t3_pos_1_025_52    m/z 756.6662     0   2   4   6E+5   14.5   15  RT (min)      lcorbi_t3_pos_1_005_42    m/z 756.6662     0   2   4   6E+5   14.5   15  RT (min)      lcorbi_t3_pos_1_005_42    m/z 756.6662     0   1E+4   14.5   15  RT (min)      lcorbi_t3_pos_2_017_51    m/z 756.6662     0   1   15   15.5  RT (min)      lcorbi_t3_pos_1_033_53    m/z 756.6664     0   1   15.5   16  RT (min)      lcorbi_t3_pos_1_041_54    m/z 756.6664     0   1   15   15.5  RT (min)      lcorbi_t3_pos_1_030_21    m/z 757.5483     0   1E+4   12.5   13  RT (min)      lcorbi_t3_pos_3_031_45    m/z 757.5525     0   1   2E+4   12.5   13  RT (min)      lcorbi_t3_pos_3_031_45    m/z 757.5570     0   1   2E+4   13   13.5  RT (min)      lcorbi_t3_pos_3_011_42    m/z 757.5572     0   1   2E+6   12   12.5  RT (min)      lcorbi_t3_pos_4_057_49    m/z 757.5572     0   2   4   6E+4   15.5   16  RT (min)      lcorbi_t3_pos_1_031_40    m/z 757.5572     0   2   4E+4   13   13.5  RT (min)      lcorbi_t3_pos_2_013_33    m/z 757.5572     0   2   4E+4   14.5   15  RT (min)      lcorbi_t3_pos_1_028_32    m/z 757.5572     0   2   4   6E+4   14.5   15  RT (min)      lcorbi_t3_pos_1_028_32    m/z 757.5572     0   2   4E+4   13   13.5  RT (min)      lcorbi_t3_pos_2_013_33    m/z 757.5573     0   2   4E+4   14.5   15  RT (min)      lcorbi_t3_pos_1_047_41    m/z 757.5573     0   1   2   3   4E+4   14.5   15  RT (min)      lcorbi_t3_pos_1_047_41    m/z 757.5574     0   1   2E+6   14   14.5  RT (min)      lcorbi_t3_pos_3_025_52    m/z 757.5574     0   2   4   6E+4   15   15.5  RT (min)       lcorbi_t3_pos_1_034_34    m/z 757.5574     0   1E+4   15   15.5  RT (min)      lcorbi_t3_pos_2_009_50    m/z 757.5574     0   2   4   6E+4   15.5   16  RT (min)      lcorbi_t3_pos_1_031_40    m/z 757.5574     0   2   4E+6   14   14.5  RT (min)      lcorbi_t3_pos_1_008_36    m/z 757.5575     0   2   4   6E+6   13   13.5  RT (min)      lcorbi_t3_pos_3_011_42    m/z 757.5575     0   1   2   3   4E+4   15.5   16  RT (min)      lcorbi_t3_pos_2_012_01    m/z 757.5575     0   1   2   3   4E+4   15.5   16  RT (min)      lcorbi_t3_pos_2_012_01    m/z 757.5575     0   2   4   6E+4   15   15.5  RT (min)      lcorbi_t3_pos_1_034_34    m/z 757.5575     0   1   16   16.5  RT (min)      lcorbi_t3_pos_1_023_02    m/z 757.5576     0   1   16   16.5  RT (min)      lcorbi_t3_pos_1_023_02    m/z 757.5577     0   1E+7   14   14.5  RT (min)       lcorbi_t3_pos_1_005_42    m/z 757.6026     0   1   15.5   16  RT (min)      lcorbi_t3_pos_1_031_40    m/z 757.6026     0   1   15   15.5  RT (min)      lcorbi_t3_pos_1_028_32    m/z 757.6220     0   1   15.5   16  RT (min)      lcorbi_t3_pos_1_034_34    m/z 757.6222     0   1   2E+4   13.5   14  RT (min)      lcorbi_t3_pos_1_002_01    m/z 757.6224     0   1   15   15.5  RT (min)      lcorbi_t3_pos_1_030_21    m/z 757.6224     0   2   4   6E+4   14.5   15  RT (min)      lcorbi_t3_pos_1_008_36    m/z 757.6224     0   1   15   15.5  RT (min)      lcorbi_t3_pos_1_021_18    m/z 757.6224     0   1   2E+4   16   16.5  RT (min)      lcorbi_t3_pos_1_051_09    m/z 757.6225     0   1   2E+4   16   16.5  RT (min)      lcorbi_t3_pos_1_051_09    m/z 757.6226     0   1   16.5   17  RT (min)    lcorbi_t3_pos_1_023_02    m/z 757.6226     0   1   15   15.5  RT (min)      lcorbi_t3_pos_1_028_32    m/z 757.6228     0   1   16.5   17  RT (min)    lcorbi_t3_pos_1_023_02    m/z 757.6693     0   1   15.5   16  RT (min)      lcorbi_t3_pos_2_049_55    m/z 757.6693     0   1E+6   13   13.5  RT (min)      lcorbi_t3_pos_4_057_49    m/z 757.6694     0   1   16   16.5  RT (min)      lcorbi_t3_pos_1_001_49    m/z 757.6695     0   1   15.5   16  RT (min)      lcorbi_t3_pos_1_041_54    m/z 757.6695     0   1   15.5   16  RT (min)      lcorbi_t3_pos_1_041_54    m/z 757.6695     0   1   16   16.5  RT (min)      lcorbi_t3_pos_2_056_56    m/z 757.6695     0   1   2   3E+4   15   15.5  RT (min)      lcorbi_t3_pos_1_021_18    m/z 757.6696     0   1   2   3E+4   14.5   15  RT (min)      lcorbi_t3_pos_2_017_51    m/z 757.6696     0   1   2   3E+4   15   15.5  RT (min)      lcorbi_t3_pos_1_021_18    m/z 757.6696     0   1   2E+6   14.5   15  RT (min)      lcorbi_t3_pos_1_005_42    m/z 757.6697     0   1E+5   14.5   15  RT (min)      lcorbi_t3_pos_3_025_52    m/z 757.6697     0   1   16   16.5  RT (min)      lcorbi_t3_pos_1_056_56    m/z 757.6698     0   1   2E+6   14.5   15  RT (min)       lcorbi_t3_pos_1_005_42    m/z 757.6698     0   1   15.5   16  RT (min)      lcorbi_t3_pos_2_049_55    m/z 757.6698     0   1E+4   15   15.5  RT (min)      lcorbi_t3_pos_3_049_55    m/z 757.6699     0   1E+4   15   15.5  RT (min)      lcorbi_t3_pos_4_049_55    m/z 758.2218     0   1   17.5   18  RT (min)    lcorbi_t3_pos_1_023_02    m/z 758.2218     0   1   17.5   18  RT (min)    lcorbi_t3_pos_1_023_02    m/z 758.2219     0   1   16   16.5  RT (min)      lcorbi_t3_pos_1_034_34    m/z 758.2219     0   1   17   17.5  RT (min)    lcorbi_t3_pos_2_056_56    m/z 758.2219     0   1   13.5   14  RT (min)      lcorbi_t3_pos_3_052_47    m/z 758.2219     0   1E+6   15   15.5  RT (min)      lcorbi_t3_pos_4_051_34    m/z 758.2220     0   1   16.5   17  RT (min)    lcorbi_t3_pos_1_049_55    m/z 758.2220     0   1   16   16.5  RT (min)      lcorbi_t3_pos_2_041_54    m/z 758.2220     0   2   4   6   8E+5   14   14.5  RT (min)      lcorbi_t3_pos_4_016_15    m/z 758.2220     0   1   2E+4   15.5   16  RT (min)      lcorbi_t3_pos_4_010_43    m/z 758.2220     0   1E+7   14.5   15  RT (min)       lcorbi_t3_pos_4_105_61    m/z 758.2220     0   1   15.5   16  RT (min)      lcorbi_t3_pos_4_049_55    m/z 758.2221     0   1   17   17.5  RT (min)    lcorbi_t3_pos_1_001_49    m/z 758.2222     0   1   14   14.5  RT (min)      lcorbi_t3_pos_4_046_44    m/z 758.3836     0   1   16   16.5  RT (min)      lcorbi_t3_pos_4_041_54    m/z 758.3836     0   1   17   17.5  RT (min)    lcorbi_t3_pos_1_025_52    m/z 758.3836     0   2   4   6   8E+5   14.5   15  RT (min)      lcorbi_t3_pos_3_053_48    m/z 758.3837     0   1E+5   15.5   16  RT (min)      lcorbi_t3_pos_3_047_40    m/z 758.3838     0   1E+6   14.5   15  RT (min)       lcorbi_t3_pos_4_105_61    m/z 758.3838     0   1   2E+4   15.5   16  RT (min)      lcorbi_t3_pos_3_025_52    m/z 758.3838     0   1   16.5   17  RT (min)    lcorbi_t3_pos_2_049_55    m/z 758.3838     0   1   16.5   17  RT (min)    lcorbi_t3_pos_1_041_54    m/z 758.3838     0   1   16   16.5  RT (min)      lcorbi_t3_pos_1_038_30    m/z 758.3838     0   1   2   3E+4   14   14.5  RT (min)      lcorbi_t3_pos_1_002_01    m/z 758.3838     0   1   16   16.5  RT (min)      lcorbi_t3_pos_2_025_52    m/z 758.3838     0   2   4E+5   14.5   15  RT (min)      lcorbi_t3_pos_4_016_15    m/z 758.3838     0   1   17   17.5  RT (min)    lcorbi_t3_pos_1_027_43    m/z 758.3838     0   1   14   14.5  RT (min)      lcorbi_t3_pos_4_046_44    m/z 758.3840     0   1   2E+4   13.5   14  RT (min)      lcorbi_t3_pos_2_032_39    m/z 758.3840     0   1   13.5   14  RT (min)      lcorbi_t3_pos_2_047_31    m/z 758.3844     0   1   17.5   18  RT (min)    lcorbi_t3_pos_1_023_02    m/z 758.4969     0   1E+4   15   15.5  RT (min)      lcorbi_t3_pos_2_012_01    m/z 758.4970     0   1   14.5   15  RT (min)      lcorbi_t3_pos_1_034_34    m/z 758.4970     0   1   2E+4   14   14.5  RT (min)      lcorbi_t3_pos_2_022_38    m/z 758.4970     0   1E+4   12.5   13  RT (min)      lcorbi_t3_pos_2_013_33    m/z 758.4970     0   1   15   15.5  RT (min)      lcorbi_t3_pos_1_031_40    m/z 758.4970     0   1   2E+4   14   14.5  RT (min)      lcorbi_t3_pos_2_022_38    m/z 758.4970     0   1E+4   12.5   13  RT (min)       lcorbi_t3_pos_3_024_07    m/z 758.4971     0   1   15   15.5  RT (min)      lcorbi_t3_pos_1_031_40    m/z 758.4972     0   2   4   6E+4   14   14.5  RT (min)      lcorbi_t3_pos_1_028_32    m/z 758.4972     0   1   14.5   15  RT (min)      lcorbi_t3_pos_1_034_34    m/z 758.4973     0   1   15   15.5  RT (min)      lcorbi_t3_pos_1_031_40    m/z 758.4977     0   1E+4   15   15.5  RT (min)      lcorbi_t3_pos_2_012_01    m/z 758.5280     0   1   16   16.5  RT (min)      lcorbi_t3_pos_1_023_02    m/z 758.5283     0   1   15.5   16  RT (min)      lcorbi_t3_pos_1_031_40    m/z 758.5596     0   1E+6   13   13.5  RT (min)      lcorbi_t3_pos_3_011_42    m/z 758.5597     0   2   4   6   8E+3   12.5   13  RT (min)      lcorbi_t3_pos_3_024_07    m/z 758.5645     0   1   2   3   4E+5   14.5   15  RT (min)      lcorbi_t3_pos_4_048_24    m/z 758.5650     0   1E+8   13   13.5  RT (min)      lcorbi_t3_pos_1_002_01    m/z 758.5652     0   1   2   3E+5   15.5   16  RT (min)      lcorbi_t3_pos_1_034_34    m/z 758.5684     0   1   13   13.5  RT (min)       lcorbi_t3_pos_2_032_39    m/z 758.5685     0   1E+4   13   13.5  RT (min)      lcorbi_t3_pos_3_031_45    m/z 758.5686     0   2   4E+5   18   18.5  RT (min)       lcorbi_t3_pos_4_072_57    m/z 758.5687     0   2   4   6   8E+5   13   13.5  RT (min)      lcorbi_t3_pos_1_002_01    m/z 758.5688     0   1   2   3E+7   12.5   13  RT (min)      lcorbi_t3_pos_4_057_49    m/z 758.5690     0   2   4E+7   14   14.5  RT (min)      lcorbi_t3_pos_3_033_53    m/z 758.5691     0   1   2   3   4E+7   13   13.5  RT (min)       lcorbi_t3_pos_1_002_01    m/z 758.5692     0   1   16.5   17  RT (min)    lcorbi_t3_pos_1_023_02    m/z 758.5692     0   1   2   3   4E+5   15   15.5  RT (min)      lcorbi_t3_pos_1_028_32    m/z 758.5693     0   1   2E+5   15.5   16  RT (min)      lcorbi_t3_pos_1_031_40    m/z 758.5693     0   1   2   3E+5   15.5   16  RT (min)      lcorbi_t3_pos_1_051_09    m/z 758.5694     0   1E+8   14   14.5  RT (min)      lcorbi_t3_pos_1_005_42    m/z 758.5694     0   1   2   3E+5   15   15.5  RT (min)      lcorbi_t3_pos_1_034_34    m/z 758.5694     0   1   2   3   4E+5   15   15.5  RT (min)      lcorbi_t3_pos_1_028_32    m/z 758.5694     0   1   2   3E+5   15   15.5  RT (min)      lcorbi_t3_pos_1_034_34    m/z 758.5694     0   2   4E+5   15   15.5  RT (min)      lcorbi_t3_pos_1_028_32    m/z 758.5694     0   1E+8   13   13.5  RT (min)      lcorbi_t3_pos_1_002_01    m/z 758.5696     0   1E+7   14   14.5  RT (min)      lcorbi_t3_pos_4_025_52    m/z 758.5697     0   2   4   6   8E+4   16   16.5  RT (min)      lcorbi_t3_pos_2_056_56    m/z 758.5698     0   1E+8   14   14.5  RT (min)       lcorbi_t3_pos_1_005_42    m/z 758.5699     0   1   2E+5   16   16.5  RT (min)      lcorbi_t3_pos_1_031_40    m/z 758.5700     0   1E+5   16   16.5  RT (min)      lcorbi_t3_pos_1_023_02    m/z 758.5706     0   1   16.5   17  RT (min)    lcorbi_t3_pos_1_023_02    m/z 758.6097     0   2   4   6   8E+4   14.5   15  RT (min)      lcorbi_t3_pos_1_008_36    m/z 758.6099     0   1   13   13.5  RT (min)      lcorbi_t3_pos_1_002_01    m/z 758.6100     0   2   4   6E+5   14   14.5  RT (min)      lcorbi_t3_pos_1_005_42    m/z 758.6100     0   1   13   13.5  RT (min)      lcorbi_t3_pos_1_002_01    m/z 758.6104     0   1   15.5   16  RT (min)      lcorbi_t3_pos_1_001_49    m/z 758.6217     0   1   15.5   16  RT (min)      lcorbi_t3_pos_1_001_49    m/z 759.2218     0   1E+4   14   14.5  RT (min)      lcorbi_t3_pos_4_046_44    m/z 759.2224     0   1   16   16.5  RT (min)      lcorbi_t3_pos_1_034_34    m/z 759.2224     0   1   17.5   18  RT (min)    lcorbi_t3_pos_1_023_02    m/z 759.2226     0   1   16.5   17  RT (min)    lcorbi_t3_pos_1_049_55    m/z 759.2226     0   1   16.5   17  RT (min)    lcorbi_t3_pos_1_034_34    m/z 759.2227     0   2   4E+5   15   15.5  RT (min)      lcorbi_t3_pos_1_005_42    m/z 759.2227     0   1   15.5   16  RT (min)      lcorbi_t3_pos_4_028_04    m/z 759.2227     0   2   4   6   8E+6   14.5   15  RT (min)      lcorbi_t3_pos_4_105_61    m/z 759.2228     0   1   16.5   17  RT (min)    lcorbi_t3_pos_1_049_55    m/z 759.2228     0   1   15.5   16  RT (min)      lcorbi_t3_pos_4_041_54    m/z 759.2228     0   1   2   3E+4   15.5   16  RT (min)      lcorbi_t3_pos_1_008_36    m/z 759.3872     0   2   4E+5   14.5   15  RT (min)      lcorbi_t3_pos_4_105_61    m/z 759.4997     0   1   15   15.5  RT (min)      lcorbi_t3_pos_1_031_40    m/z 759.5003     0   2   4   6   8E+3   14   14.5  RT (min)      lcorbi_t3_pos_2_022_38    m/z 759.5004     0   1   2   3E+5   12.5   13  RT (min)      lcorbi_t3_pos_3_011_42    m/z 759.5004     0   1   15   15.5  RT (min)      lcorbi_t3_pos_2_012_01    m/z 759.5005     0   1   12.5   13  RT (min)      lcorbi_t3_pos_3_024_07    m/z 759.5006     0   1   15   15.5  RT (min)      lcorbi_t3_pos_1_031_40    m/z 759.5006     0   1E+4   14   14.5  RT (min)      lcorbi_t3_pos_2_022_38    m/z 759.5006     0   1   14.5   15  RT (min)      lcorbi_t3_pos_1_034_34    m/z 759.5006     0   1   15   15.5  RT (min)      lcorbi_t3_pos_1_031_40    m/z 759.5007     0   1   2   3   4E+4   14   14.5  RT (min)      lcorbi_t3_pos_1_028_32    m/z 759.5680     0   1E+7   12.5   13  RT (min)      lcorbi_t3_pos_4_057_49    m/z 759.5681     0   2   4E+7   13   13.5  RT (min)      lcorbi_t3_pos_1_002_01    m/z 759.5716     0   2   4E+7   13   13.5  RT (min)      lcorbi_t3_pos_1_002_01    m/z 759.5716     0   1   13   13.5  RT (min)      lcorbi_t3_pos_2_032_39    m/z 759.5718     0   1   13   13.5  RT (min)      lcorbi_t3_pos_3_031_45    m/z 759.5720     0   1   2   3E+4   13   13.5  RT (min)      lcorbi_t3_pos_1_002_01    m/z 759.5724     0   2   4   6E+5   14.5   15  RT (min)      lcorbi_t3_pos_2_017_51    m/z 759.5725     0   1   2E+5   14.5   15  RT (min)      lcorbi_t3_pos_2_006_47    m/z 759.5725     0   1E+5   16   16.5  RT (min)      lcorbi_t3_pos_1_023_02    m/z 759.5726     0   1   16.5   17  RT (min)    lcorbi_t3_pos_1_023_02    m/z 759.5726     0   1E+5   15.5   16  RT (min)      lcorbi_t3_pos_1_051_09    m/z 759.5726     0   1E+5   15   15.5  RT (min)      lcorbi_t3_pos_1_034_34    m/z 759.5728     0   1E+5   15.5   16  RT (min)      lcorbi_t3_pos_1_031_40    m/z 759.5728     0   1   2E+5   15   15.5  RT (min)      lcorbi_t3_pos_1_028_32    m/z 759.5728     0   1E+5   15   15.5  RT (min)      lcorbi_t3_pos_1_028_32    m/z 759.5729     0   1E+5   15   15.5  RT (min)      lcorbi_t3_pos_1_028_32    m/z 759.5729     0   2   4   6E+4   15.5   16  RT (min)      lcorbi_t3_pos_1_049_55    m/z 759.5730     0   1   2   3E+5   14.5   15  RT (min)      lcorbi_t3_pos_2_006_47    m/z 759.5730     0   1E+5   15   15.5  RT (min)      lcorbi_t3_pos_1_034_34    m/z 759.5731     0   1   2   3   4E+6   14.5   15  RT (min)      lcorbi_t3_pos_1_008_36    m/z 759.5732     0   1E+5   16   16.5  RT (min)       lcorbi_t3_pos_1_023_02    m/z 759.6376     0   2   4   6E+4   13.5   14  RT (min)      lcorbi_t3_pos_1_002_01    m/z 759.6378     0   1   16.5   17  RT (min)    lcorbi_t3_pos_1_023_02    m/z 759.6378     0   1   15.5   16  RT (min)      lcorbi_t3_pos_1_028_32    m/z 759.6378     0   1   16   16.5  RT (min)      lcorbi_t3_pos_1_051_09    m/z 759.6380     0   1   15.5   16  RT (min)      lcorbi_t3_pos_1_034_34    m/z 759.6380     0   1E+4   13.5   14  RT (min)      lcorbi_t3_pos_2_013_33    m/z 759.6380     0   2   4E+4   14.5   15  RT (min)      lcorbi_t3_pos_1_008_36    m/z 759.6380     0   1   16   16.5  RT (min)      lcorbi_t3_pos_1_031_40    m/z 759.6380     0   1   15   15.5  RT (min)      lcorbi_t3_pos_1_028_32    m/z 759.6380     0   1   15.5   16  RT (min)      lcorbi_t3_pos_1_034_34    m/z 759.6381     0   1   17   17.5  RT (min)    lcorbi_t3_pos_1_023_02    m/z 759.6384     0   1   13   13.5  RT (min)      lcorbi_t3_pos_2_032_39    m/z 760.2204     0   2   4   6E+5   15   15.5  RT (min)      lcorbi_t3_pos_4_051_34    m/z 760.2204     0   1   16.5   17  RT (min)    lcorbi_t3_pos_1_049_55    m/z 760.2205     0   1   15.5   16  RT (min)      lcorbi_t3_pos_4_049_55    m/z 760.2206     0   1   16   16.5  RT (min)      lcorbi_t3_pos_1_034_34    m/z 760.2207     0   2   4   6E+6   14.5   15  RT (min)      lcorbi_t3_pos_4_105_61    m/z 760.2209     0   1   16   16.5  RT (min)      lcorbi_t3_pos_1_034_34    m/z 760.2209     0   1   17.5   18  RT (min)    lcorbi_t3_pos_1_023_02    m/z 760.2210     0   1   16.5   17  RT (min)    lcorbi_t3_pos_1_049_55    m/z 760.4891     0   2   4   6   8E+4   14   14.5  RT (min)      lcorbi_t3_pos_1_008_36    m/z 760.4892     0   1   15.5   16  RT (min)      lcorbi_t3_pos_1_031_40    m/z 760.4893     0   1   15.5   16  RT (min)      lcorbi_t3_pos_1_031_40    m/z 760.4894     0   1   2E+4   15.5   16  RT (min)      lcorbi_t3_pos_2_012_01    m/z 760.4894     0   1   14.5   15  RT (min)      lcorbi_t3_pos_1_028_32    m/z 760.4895     0   1   15   15.5  RT (min)      lcorbi_t3_pos_1_034_34    m/z 760.5079     0   1E+7   13   13.5  RT (min)      lcorbi_t3_pos_3_011_42    m/z 760.5083     0   1   2   3E+4   15   15.5  RT (min)      lcorbi_t3_pos_1_041_54    m/z 760.5124     0   1   12.5   13  RT (min)      lcorbi_t3_pos_3_024_07    m/z 760.5124     0   1   2E+4   12.5   13  RT (min)      lcorbi_t3_pos_2_013_33    m/z 760.5126     0   1   2   3E+4   15.5   16  RT (min)      lcorbi_t3_pos_1_001_49    m/z 760.5126     0   2   4   6   8E+4   14.5   15  RT (min)      lcorbi_t3_pos_1_028_32    m/z 760.5126     0   2   4E+4   15   15.5  RT (min)       lcorbi_t3_pos_1_031_40    m/z 760.5126     0   2   4   6   8E+4   14.5   15  RT (min)      lcorbi_t3_pos_1_034_34    m/z 760.5126     0   2   4E+4   15   15.5  RT (min)      lcorbi_t3_pos_1_031_40    m/z 760.5126     0   2   4   6E+4   14.5   15  RT (min)      lcorbi_t3_pos_1_034_34    m/z 760.5126     0   1E+4   14.5   15  RT (min)      lcorbi_t3_pos_4_048_24    m/z 760.5126     0   2   4   6E+4   14.5   15  RT (min)      lcorbi_t3_pos_2_022_38    m/z 760.5127     0   1   2E+6   13   13.5  RT (min)      lcorbi_t3_pos_3_011_42    m/z 760.5127     0   2   4E+4   15.5   16  RT (min)       lcorbi_t3_pos_2_012_01    m/z 760.5128     0   1E+4   14.5   15  RT (min)      lcorbi_t3_pos_4_048_24    m/z 760.5129     0   1   2E+4   15   15.5  RT (min)      lcorbi_t3_pos_1_049_55    m/z 760.5129     0   2   4E+4   15.5   16  RT (min)      lcorbi_t3_pos_1_031_40    m/z 760.5466     0   1   15.5   16  RT (min)      lcorbi_t3_pos_2_014_37    m/z 760.5484     0   1   13   13.5  RT (min)      lcorbi_t3_pos_2_047_31    m/z 760.5488     0   1   13   13.5  RT (min)      lcorbi_t3_pos_2_039_30    m/z 760.5489     0   2   4E+5   13   13.5  RT (min)      lcorbi_t3_pos_2_016_23    m/z 760.5490     0   2   4E+5   13.5   14  RT (min)      lcorbi_t3_pos_3_053_48    m/z 760.5495     0   1   15.5   16  RT (min)      lcorbi_t3_pos_1_027_43    m/z 760.5753     0   1E+4   13   13.5  RT (min)      lcorbi_t3_pos_2_032_39    m/z 760.5753     0   1   2   3E+4   14.5   15  RT (min)      lcorbi_t3_pos_1_047_41    m/z 760.5754     0   1   2   3   4E+4   14.5   15  RT (min)      lcorbi_t3_pos_3_003_33    m/z 760.5758     0   1E+4   13   13.5  RT (min)      lcorbi_t3_pos_2_032_39    m/z 760.5797     0   1E+4   13   13.5  RT (min)      lcorbi_t3_pos_2_032_39    m/z 760.5835     0   1   2   3E+7   13   13.5  RT (min)      lcorbi_t3_pos_4_057_49    m/z 760.5846     0   1   2   3E+4   16   16.5  RT (min)       lcorbi_t3_pos_2_056_56    m/z 760.5846     0   2   4E+4   16   16.5  RT (min)      lcorbi_t3_pos_1_049_55    m/z 760.5846     0   1E+5   16   16.5  RT (min)      lcorbi_t3_pos_1_051_09    m/z 760.5846     0   2   4E+4   15.5   16  RT (min)      lcorbi_t3_pos_2_009_50    m/z 760.5848     0   2   4   6   8E+4   15   15.5  RT (min)      lcorbi_t3_pos_2_041_54    m/z 760.5848     0   1E+5   16   16.5  RT (min)       lcorbi_t3_pos_1_051_09    m/z 760.5848     0   1E+5   15   15.5  RT (min)      lcorbi_t3_pos_2_022_38    m/z 760.5849     0   1E+5   15.5   16  RT (min)       lcorbi_t3_pos_1_034_34    m/z 760.5850     0   1E+5   15   15.5  RT (min)       lcorbi_t3_pos_1_028_32    m/z 760.5851     0   1   2   3   4E+5   14.5   15  RT (min)       lcorbi_t3_pos_2_017_51    m/z 760.5852     0   1   2E+5   15   15.5  RT (min)       lcorbi_t3_pos_1_033_53    m/z 760.5852     0   1   2E+5   15   15.5  RT (min)      lcorbi_t3_pos_1_047_41    m/z 760.5853     0   1   16.5   17  RT (min)    lcorbi_t3_pos_1_023_02    m/z 760.5854     0   1   16.5   17  RT (min)    lcorbi_t3_pos_1_023_02    m/z 760.5854     0   1   16.5   17  RT (min)    lcorbi_t3_pos_1_023_02    m/z 760.5854     0   2   4   6E+4   16   16.5  RT (min)      lcorbi_t3_pos_1_023_02    m/z 760.5855     0   1   16.5   17  RT (min)    lcorbi_t3_pos_1_023_02    m/z 760.5856     0   2   4   6E+4   16   16.5  RT (min)      lcorbi_t3_pos_1_023_02    m/z 760.5856     0   1   2E+4   13   13.5  RT (min)      lcorbi_t3_pos_1_002_01    m/z 760.5857     0   1E+7   13.5   14  RT (min)      lcorbi_t3_pos_1_002_01    m/z 760.5858     0   1   16.5   17  RT (min)    lcorbi_t3_pos_1_023_02    m/z 760.5863     0   1   2   3E+4   16   16.5  RT (min)      lcorbi_t3_pos_2_056_56    m/z 760.5869     0   1   2E+5   15   15.5  RT (min)       lcorbi_t3_pos_1_047_41    m/z 760.5874     0   2   4   6E+6   14.5   15  RT (min)       lcorbi_t3_pos_1_008_36    m/z 760.5874     0   1E+7   14.5   15  RT (min)       lcorbi_t3_pos_1_008_36    m/z 760.5874     0   1   2E+4   13.5   14  RT (min)      lcorbi_t3_pos_1_002_01    m/z 760.5876     0   2   4   6E+7   13.5   14  RT (min)      lcorbi_t3_pos_1_002_01    m/z 760.5878     0   1   2   3   4E+5   15   15.5  RT (min)      lcorbi_t3_pos_4_001_49    m/z 760.6253     0   1   2   3E+4   13.5   14  RT (min)      lcorbi_t3_pos_1_002_01    m/z 760.6410     0   1   16.5   17  RT (min)    lcorbi_t3_pos_1_023_02    m/z 761.2207     0   1   2   3E+6   14.5   15  RT (min)      lcorbi_t3_pos_4_105_61    m/z 761.5158     0   1   12.5   13  RT (min)      lcorbi_t3_pos_3_024_07    m/z 761.5158     0   1   2E+4   14   14.5  RT (min)      lcorbi_t3_pos_1_047_41    m/z 761.5160     0   1   2E+4   14.5   15  RT (min)      lcorbi_t3_pos_2_022_38    m/z 761.5161     0   1   2   3E+4   15   15.5  RT (min)      lcorbi_t3_pos_1_031_40    m/z 761.5161     0   1   2   3   4E+4   14.5   15  RT (min)      lcorbi_t3_pos_1_034_34    m/z 761.5161     0   1   2   3   4E+4   14.5   15  RT (min)      lcorbi_t3_pos_1_034_34    m/z 761.5161     0   1   2E+4   12.5   13  RT (min)      lcorbi_t3_pos_2_013_33    m/z 761.5162     0   1   2   3E+4   15.5   16  RT (min)      lcorbi_t3_pos_2_012_01    m/z 761.5162     0   2   4   6E+4   14.5   15  RT (min)      lcorbi_t3_pos_1_028_32    m/z 761.5162     0   1E+4   14.5   15  RT (min)      lcorbi_t3_pos_2_041_54    m/z 761.5163     0   1E+6   13   13.5  RT (min)      lcorbi_t3_pos_3_011_42    m/z 761.5163     0   1   2   3E+4   15.5   16  RT (min)      lcorbi_t3_pos_1_031_40    m/z 761.5163     0   1   2   3E+4   15   15.5  RT (min)      lcorbi_t3_pos_1_031_40    m/z 761.5164     0   2   4E+6   13   13.5  RT (min)      lcorbi_t3_pos_3_011_42    m/z 761.5782     0   1   13   13.5  RT (min)      lcorbi_t3_pos_2_032_39    m/z 761.5788     0   1   14.5   15  RT (min)      lcorbi_t3_pos_1_047_41    m/z 761.5788     0   1   2   3E+5   14   14.5  RT (min)      lcorbi_t3_pos_3_047_40    m/z 761.5789     0   1   16   16.5  RT (min)      lcorbi_t3_pos_1_023_02    m/z 761.5790     0   1   2E+4   15.5   16  RT (min)      lcorbi_t3_pos_1_051_09    m/z 761.5792     0   1   13   13.5  RT (min)      lcorbi_t3_pos_2_032_39    m/z 761.5873     0   1   13   13.5  RT (min)      lcorbi_t3_pos_2_032_39    m/z 761.5877     0   1E+7   13   13.5  RT (min)      lcorbi_t3_pos_4_057_49    m/z 761.5878     0   1E+4   16   16.5  RT (min)      lcorbi_t3_pos_2_056_56    m/z 761.5882     0   1   2   3E+6   0   0.5  RT (min)      lcorbi_t3_pos_4_057_49    m/z 761.5882     0   2   4E+4   16   16.5  RT (min)      lcorbi_t3_pos_1_051_09    m/z 761.5883     0   1   2   3E+4   15.5   16  RT (min)       lcorbi_t3_pos_1_049_55    m/z 761.5885     0   2   4E+4   16   16.5  RT (min)       lcorbi_t3_pos_1_051_09    m/z 761.5885     0   2   4   6   8E+4   15   15.5  RT (min)      lcorbi_t3_pos_1_028_32    m/z 761.5885     0   1   2   3   4E+4   15.5   16  RT (min)      lcorbi_t3_pos_2_009_50    m/z 761.5885     0   1E+5   15   15.5  RT (min)      lcorbi_t3_pos_1_030_21    m/z 761.5887     0   1   16.5   17  RT (min)    lcorbi_t3_pos_1_023_02    m/z 761.5888     0   1   2   3E+7   13.5   14  RT (min)      lcorbi_t3_pos_1_046_22    m/z 761.5888     0   2   4E+4   15.5   16  RT (min)      lcorbi_t3_pos_1_034_34    m/z 761.5888     0   1   2   3E+4   15   15.5  RT (min)      lcorbi_t3_pos_2_025_52    m/z 761.5890     0   1E+4   16   16.5  RT (min)       lcorbi_t3_pos_2_056_56    m/z 761.5890     0   2   4E+6   13.5   14  RT (min)      lcorbi_t3_pos_1_002_01    m/z 761.5890     0   2   4   6   8E+4   15   15.5  RT (min)       lcorbi_t3_pos_1_033_53    m/z 761.5891     0   1   16.5   17  RT (min)    lcorbi_t3_pos_1_023_02    m/z 761.5891     0   1   16.5   17  RT (min)    lcorbi_t3_pos_1_023_02    m/z 761.5892     0   1   13.5   14  RT (min)      lcorbi_t3_pos_1_002_01    m/z 761.5894     0   1   2E+7   14   14.5  RT (min)       lcorbi_t3_pos_1_005_42    m/z 761.5894     0   2   4   6   8E+6   13.5   14  RT (min)      lcorbi_t3_pos_3_034_26    m/z 761.5897     0   1   16.5   17  RT (min)    lcorbi_t3_pos_1_023_02    m/z 761.5903     0   1   16.5   17  RT (min)    lcorbi_t3_pos_1_023_02    m/z 761.5905     0   1   2E+6   14.5   15  RT (min)       lcorbi_t3_pos_1_008_36    m/z 761.5906     0   1E+5   15   15.5  RT (min)       lcorbi_t3_pos_1_047_41    m/z 761.5906     0   1E+5   15   15.5  RT (min)       lcorbi_t3_pos_1_047_41    m/z 761.5906     0   1   2E+7   14.5   15  RT (min)       lcorbi_t3_pos_1_005_42    m/z 761.5907     0   2   4E+6   14.5   15  RT (min)       lcorbi_t3_pos_1_008_36    m/z 761.5908     0   1   16.5   17  RT (min)    lcorbi_t3_pos_2_012_01    m/z 762.2192     0   1E+6   14.5   15  RT (min)      lcorbi_t3_pos_4_105_61    m/z 762.5187     0   1   15.5   16  RT (min)      lcorbi_t3_pos_1_001_49    m/z 762.5191     0   1   12.5   13  RT (min)      lcorbi_t3_pos_3_024_07    m/z 762.5195     0   1   2   3   4E+4   14   14.5  RT (min)      lcorbi_t3_pos_1_008_36    m/z 762.5237     0   2   4   6E+5   14   14.5  RT (min)      lcorbi_t3_pos_1_008_36    m/z 762.5274     0   1   2   3E+4   15   15.5  RT (min)      lcorbi_t3_pos_1_034_34    m/z 762.5276     0   2   4   6E+4   13   13.5  RT (min)      lcorbi_t3_pos_2_013_33    m/z 762.5280     0   2   4   6E+4   14.5   15  RT (min)      lcorbi_t3_pos_1_028_32    m/z 762.5281     0   1   15.5   16  RT (min)      lcorbi_t3_pos_1_031_40    m/z 762.5281     0   2   4   6E+4   14.5   15  RT (min)      lcorbi_t3_pos_1_028_32    m/z 762.5281     0   1   2E+4   14.5   15  RT (min)      lcorbi_t3_pos_1_033_53    m/z 762.5281     0   1   2   3E+4   15   15.5  RT (min)      lcorbi_t3_pos_1_031_40    m/z 762.5281     0   1   2E+4   14.5   15  RT (min)      lcorbi_t3_pos_2_046_07    m/z 762.5282     0   1   2   3E+4   15   15.5  RT (min)      lcorbi_t3_pos_1_034_34    m/z 762.5282     0   2   4E+4   15   15.5  RT (min)      lcorbi_t3_pos_1_028_32    m/z 762.5282     0   1   15.5   16  RT (min)      lcorbi_t3_pos_1_031_40    m/z 762.5282     0   1   2   3   4E+6   14   14.5  RT (min)      lcorbi_t3_pos_1_008_36    m/z 762.5283     0   1E+4   15.5   16  RT (min)      lcorbi_t3_pos_2_012_01    m/z 762.5283     0   1E+4   15.5   16  RT (min)      lcorbi_t3_pos_2_012_01    m/z 762.5283     0   1   15   15.5  RT (min)      lcorbi_t3_pos_2_009_50    m/z 762.5284     0   1   15.5   16  RT (min)      lcorbi_t3_pos_1_049_55    m/z 762.5909     0   1   2   3E+6   13   13.5  RT (min)      lcorbi_t3_pos_4_057_49    m/z 762.5910     0   1   13   13.5  RT (min)      lcorbi_t3_pos_2_032_39    m/z 762.5915     0   1   13.5   14  RT (min)      lcorbi_t3_pos_1_002_01    m/z 762.5916     0   1   16   16.5  RT (min)      lcorbi_t3_pos_1_049_55    m/z 762.5918     0   1   16.5   17  RT (min)    lcorbi_t3_pos_1_023_02    m/z 762.5919     0   1   2E+4   16   16.5  RT (min)      lcorbi_t3_pos_1_051_09    m/z 762.5919     0   1   2E+4   16   16.5  RT (min)      lcorbi_t3_pos_1_051_09    m/z 762.5919     0   2   4   6E+4   14.5   15  RT (min)      lcorbi_t3_pos_2_046_07    m/z 762.5920     0   1   16.5   17  RT (min)    lcorbi_t3_pos_1_023_02    m/z 762.5920     0   1   2   3E+4   15   15.5  RT (min)      lcorbi_t3_pos_1_033_53    m/z 762.5920     0   1   2E+4   15.5   16  RT (min)      lcorbi_t3_pos_2_009_50    m/z 762.5920     0   1E+4   15   15.5  RT (min)      lcorbi_t3_pos_2_025_52    m/z 762.5921     0   1   15   15.5  RT (min)      lcorbi_t3_pos_2_038_03    m/z 762.5921     0   2   4   6E+4   15   15.5  RT (min)      lcorbi_t3_pos_1_030_21    m/z 762.5923     0   1   16.5   17  RT (min)    lcorbi_t3_pos_1_023_02    m/z 762.5923     0   1   16.5   17  RT (min)    lcorbi_t3_pos_1_023_02    m/z 762.5924     0   2   4E+4   15   15.5  RT (min)      lcorbi_t3_pos_1_047_41    m/z 762.5924     0   1   2E+4   16   16.5  RT (min)      lcorbi_t3_pos_1_051_09    m/z 762.5926     0   1   2E+4   15.5   16  RT (min)      lcorbi_t3_pos_1_034_34    m/z 762.5926     0   1E+6   13.5   14  RT (min)      lcorbi_t3_pos_1_002_01    m/z 762.5928     0   2   4   6E+6   13.5   14  RT (min)      lcorbi_t3_pos_1_002_01    m/z 762.5930     0   2   4E+6   14.5   15  RT (min)       lcorbi_t3_pos_1_005_42    m/z 762.5930     0   2   4E+5   14.5   15  RT (min)      lcorbi_t3_pos_4_001_49    m/z 762.5930     0   2   4E+5   14.5   15  RT (min)       lcorbi_t3_pos_1_008_36    m/z 762.5932     0   2   4E+6   14.5   15  RT (min)      lcorbi_t3_pos_1_005_42    m/z 762.5932     0   1E+4   16   16.5  RT (min)      lcorbi_t3_pos_2_056_56    m/z 762.5937     0   1   16.5   17  RT (min)    lcorbi_t3_pos_1_023_02    m/z 763.5308     0   1   2   3E+4   13   13.5  RT (min)      lcorbi_t3_pos_2_013_33    m/z 763.5314     0   1   15.5   16  RT (min)      lcorbi_t3_pos_1_031_40    m/z 763.5315     0   1   2   3E+4   14.5   15  RT (min)      lcorbi_t3_pos_1_028_32    m/z 763.5316     0   1   14.5   15  RT (min)      lcorbi_t3_pos_1_030_21    m/z 763.5316     0   1   15.5   16  RT (min)      lcorbi_t3_pos_1_031_40    m/z 763.5316     0   1   2   3   4E+4   15   15.5  RT (min)      lcorbi_t3_pos_1_034_34    m/z 763.5317     0   2   4   6E+5   14   14.5  RT (min)      lcorbi_t3_pos_1_005_42    m/z 763.5317     0   1   2   3   4E+4   15   15.5  RT (min)      lcorbi_t3_pos_1_034_34    m/z 763.5318     0   1   15.5   16  RT (min)      lcorbi_t3_pos_2_012_01    m/z 763.5318     0   1   2E+5   14   14.5  RT (min)      lcorbi_t3_pos_1_008_36    m/z 763.5319     0   1   2   3   4E+4   15   15.5  RT (min)      lcorbi_t3_pos_1_034_34    m/z 763.5943     0   2   4   6E+4   13   13.5  RT (min)      lcorbi_t3_pos_1_002_01    m/z 763.5946     0   1   2   3E+4   13.5   14  RT (min)      lcorbi_t3_pos_1_002_01    m/z 763.5947     0   1E+4   15   15.5  RT (min)      lcorbi_t3_pos_1_047_41    m/z 763.5947     0   1E+5   14.5   15  RT (min)       lcorbi_t3_pos_4_025_52    m/z 763.5947     0   1   16   16.5  RT (min)      lcorbi_t3_pos_1_051_09    m/z 763.5948     0   1E+6   14.5   15  RT (min)      lcorbi_t3_pos_1_005_42    m/z 763.5948     0   1   16   16.5  RT (min)      lcorbi_t3_pos_1_051_09    m/z 763.5948     0   1   15   15.5  RT (min)      lcorbi_t3_pos_1_028_32    m/z 763.5948     0   1   15   15.5  RT (min)      lcorbi_t3_pos_1_021_18    m/z 763.5948     0   1   16   16.5  RT (min)      lcorbi_t3_pos_1_049_55    m/z 763.5949     0   1   15.5   16  RT (min)      lcorbi_t3_pos_1_034_34    m/z 763.5949     0   1   15   15.5  RT (min)      lcorbi_t3_pos_2_041_54    m/z 763.5949     0   1   16.5   17  RT (min)    lcorbi_t3_pos_1_023_02    m/z 763.5951     0   1   15.5   16  RT (min)      lcorbi_t3_pos_2_049_55    m/z 763.5954     0   1   15   15.5  RT (min)      lcorbi_t3_pos_2_025_52    m/z 763.5993     0   1   16.5   17  RT (min)    lcorbi_t3_pos_1_023_02    m/z 764.5224     0   1   15   15.5  RT (min)      lcorbi_t3_pos_2_012_01    m/z 764.5224     0   1   15   15.5  RT (min)      lcorbi_t3_pos_2_012_01    m/z 764.5226     0   1   2   3   4E+4   14.5   15  RT (min)      lcorbi_t3_pos_1_034_34    m/z 764.5226     0   1   14.5   15  RT (min)      lcorbi_t3_pos_4_033_53    m/z 764.5227     0   1   15   15.5  RT (min)      lcorbi_t3_pos_2_012_01    m/z 764.5228     0   1   12.5   13  RT (min)      lcorbi_t3_pos_2_013_33    m/z 764.5228     0   1E+5   14.5   15  RT (min)      lcorbi_t3_pos_1_034_34    m/z 764.5228     0   1   12.5   13  RT (min)      lcorbi_t3_pos_3_024_07    m/z 764.5228     0   1   2   3   4E+4   14.5   15  RT (min)      lcorbi_t3_pos_1_034_34    m/z 764.5228     0   1   12.5   13  RT (min)      lcorbi_t3_pos_2_013_33    m/z 764.5228     0   1   15   15.5  RT (min)      lcorbi_t3_pos_1_031_40    m/z 764.5229     0   1   15   15.5  RT (min)      lcorbi_t3_pos_1_031_40    m/z 764.5230     0   1   15.5   16  RT (min)      lcorbi_t3_pos_2_012_01    m/z 764.5230     0   2   4   6   8E+4   12.5   13  RT (min)      lcorbi_t3_pos_2_013_33    m/z 764.5232     0   1   15   15.5  RT (min)      lcorbi_t3_pos_2_012_01    m/z 764.5232     0   1   2E+6   12.5   13  RT (min)      lcorbi_t3_pos_2_013_33    m/z 764.5232     0   1   2E+6   12.5   13  RT (min)      lcorbi_t3_pos_2_013_33    m/z 764.5354     0   1   2   3E+5   14   14.5  RT (min)      lcorbi_t3_pos_1_008_36    m/z 764.5740     0   1   17   17.5  RT (min)     lcorbi_t3_pos_1_023_02    m/z 764.5743     0   1   16.5   17  RT (min)    lcorbi_t3_pos_1_001_49    m/z 764.5743     0   2   4E+4   15.5   16  RT (min)       lcorbi_t3_pos_2_046_07    m/z 764.5744     0   2   4E+4   15.5   16  RT (min)      lcorbi_t3_pos_2_022_38    m/z 764.5744     0   1E+4   13.5   14  RT (min)      lcorbi_t3_pos_2_047_31    m/z 764.5744     0   1   2   3   4E+4   15.5   16  RT (min)      lcorbi_t3_pos_2_041_54    m/z 764.5744     0   1   2E+5   15   15.5  RT (min)      lcorbi_t3_pos_3_047_40    m/z 764.5744     0   1E+5   15   15.5  RT (min)       lcorbi_t3_pos_4_025_52    m/z 764.5744     0   1   16.5   17  RT (min)     lcorbi_t3_pos_2_012_01    m/z 764.5744     0   1   2   3E+4   16   16.5  RT (min)       lcorbi_t3_pos_2_038_03    m/z 764.5744     0   1   2   3E+4   16   16.5  RT (min)       lcorbi_t3_pos_1_034_34    m/z 764.5744     0   1E+4   13.5   14  RT (min)      lcorbi_t3_pos_3_052_47    m/z 764.5744     0   1E+6   14   14.5  RT (min)      lcorbi_t3_pos_4_105_61    m/z 764.5744     0   2   4   6E+5   14   14.5  RT (min)      lcorbi_t3_pos_3_053_48    m/z 764.5744     0   1   16.5   17  RT (min)     lcorbi_t3_pos_1_031_40    m/z 764.5745     0   2   4E+4   15.5   16  RT (min)      lcorbi_t3_pos_1_038_30    m/z 764.5745     0   1   13.5   14  RT (min)      lcorbi_t3_pos_3_031_45    m/z 764.5745     0   1E+5   15   15.5  RT (min)       lcorbi_t3_pos_3_025_52    m/z 764.5745     0   1E+6   14.5   15  RT (min)       lcorbi_t3_pos_3_033_53    m/z 764.5746     0   1   2   3   4E+4   15.5   16  RT (min)      lcorbi_t3_pos_2_041_54    m/z 764.5746     0   2   4   6   8E+5   14.5   15  RT (min)       lcorbi_t3_pos_4_054_18    m/z 764.5746     0   1   2E+4   14   14.5  RT (min)      lcorbi_t3_pos_2_016_23    m/z 764.5746     0   1   16.5   17  RT (min)     lcorbi_t3_pos_4_033_53    m/z 764.5746     0   1   2   3   4E+3   13.5   14  RT (min)      lcorbi_t3_pos_2_032_39    m/z 764.5747     0   1   16.5   17  RT (min)    lcorbi_t3_pos_4_033_53    m/z 764.5752     0   1   17   17.5  RT (min)    lcorbi_t3_pos_1_023_02    m/z 765.5244     0   1   2   3E+5   14   14.5  RT (min)      lcorbi_t3_pos_1_008_36    m/z 765.5248     0   1   2E+4   14   14.5  RT (min)       lcorbi_t3_pos_3_003_33    m/z 765.5260     0   2   4E+5   13.5   14  RT (min)       lcorbi_t3_pos_1_008_36    m/z 765.5261     0   2   4   6E+4   14   14.5  RT (min)      lcorbi_t3_pos_1_028_32    m/z 765.5261     0   1   12.5   13  RT (min)      lcorbi_t3_pos_2_013_33    m/z 765.5262     0   1   2   3E+5   14.5   15  RT (min)      lcorbi_t3_pos_1_034_34    m/z 765.5262     0   2   4   6E+4   14.5   15  RT (min)      lcorbi_t3_pos_1_034_34    m/z 765.5262     0   1   12.5   13  RT (min)      lcorbi_t3_pos_2_013_33    m/z 765.5264     0   1   12.5   13  RT (min)      lcorbi_t3_pos_3_024_07    m/z 765.5776     0   1   16.5   17  RT (min)    lcorbi_t3_pos_1_031_40    m/z 765.5777     0   1   2   3E+4   15.5   16  RT (min)      lcorbi_t3_pos_2_022_38    m/z 765.5778     0   1   16.5   17  RT (min)    lcorbi_t3_pos_2_012_01    m/z 765.5779     0   2   4E+5   14.5   15  RT (min)       lcorbi_t3_pos_3_033_53    m/z 765.5779     0   1   2   3   4E+4   15   15.5  RT (min)      lcorbi_t3_pos_1_008_36    m/z 765.5779     0   1   16   16.5  RT (min)      lcorbi_t3_pos_1_034_34    m/z 765.5779     0   1   13.5   14  RT (min)       lcorbi_t3_pos_3_052_47    m/z 765.5779     0   1   2   3E+4   15.5   16  RT (min)      lcorbi_t3_pos_1_038_30    m/z 765.5780     0   1   2E+4   16   16.5  RT (min)      lcorbi_t3_pos_2_038_03    m/z 765.5781     0   2   4E+5   14   14.5  RT (min)      lcorbi_t3_pos_4_105_61    m/z 765.5782     0   1   2   3E+3   13.5   14  RT (min)      lcorbi_t3_pos_3_031_45    m/z 766.5362     0   1   2E+4   15.5   16  RT (min)      lcorbi_t3_pos_2_012_01    m/z 766.5362     0   1   2E+4   15   15.5  RT (min)      lcorbi_t3_pos_2_012_01    m/z 766.5363     0   1   16   16.5  RT (min)      lcorbi_t3_pos_1_027_43    m/z 766.5366     0   1   2E+4   15   15.5  RT (min)      lcorbi_t3_pos_1_031_40    m/z 766.5366     0   1   15.5   16  RT (min)      lcorbi_t3_pos_1_025_52    m/z 766.5366     0   1   2   3E+4   15   15.5  RT (min)      lcorbi_t3_pos_1_034_34    m/z 766.5367     0   1E+4   15   15.5  RT (min)      lcorbi_t3_pos_4_033_53    m/z 766.5367     0   1   2   3   4E+5   14.5   15  RT (min)      lcorbi_t3_pos_1_028_32    m/z 766.5367     0   1   15   15.5  RT (min)      lcorbi_t3_pos_1_049_55    m/z 766.5368     0   1   16   16.5  RT (min)      lcorbi_t3_pos_2_052_48    m/z 766.5368     0   1   2E+4   15.5   16  RT (min)      lcorbi_t3_pos_2_012_01    m/z 766.5370     0   1   2E+4   15.5   16  RT (min)      lcorbi_t3_pos_2_012_01    m/z 766.5377     0   2   4   6   8E+4   14.5   15  RT (min)      lcorbi_t3_pos_1_034_34    m/z 766.5381     0   1   2E+4   15   15.5  RT (min)      lcorbi_t3_pos_1_031_40    m/z 766.5381     0   1   2E+5   14.5   15  RT (min)      lcorbi_t3_pos_1_034_34    m/z 766.5382     0   1E+5   12.5   13  RT (min)      lcorbi_t3_pos_3_011_42    m/z 766.5382     0   1   15   15.5  RT (min)      lcorbi_t3_pos_1_025_52    m/z 766.5384     0   1E+5   12.5   13  RT (min)      lcorbi_t3_pos_2_013_33    m/z 766.5384     0   1   15   15.5  RT (min)      lcorbi_t3_pos_1_041_54    m/z 766.5385     0   1   2   3   4E+6   13   13.5  RT (min)      lcorbi_t3_pos_3_011_42    m/z 766.5385     0   1   2   3   4E+6   13   13.5  RT (min)      lcorbi_t3_pos_3_011_42    m/z 766.5747     0   1E+6   14   14.5  RT (min)       lcorbi_t3_pos_1_016_05    m/z 766.5750     0   2   4   6E+5   14   14.5  RT (min)       lcorbi_t3_pos_1_029_29    m/z 766.5807     0   1   16   16.5  RT (min)      lcorbi_t3_pos_1_031_40    m/z 766.5807     0   1   2   3   4E+4   15.5   16  RT (min)      lcorbi_t3_pos_1_034_34    m/z 767.5398     0   1   15.5   16  RT (min)      lcorbi_t3_pos_2_012_01    m/z 767.5400     0   1E+5   15   15.5  RT (min)      lcorbi_t3_pos_2_012_01    m/z 767.5401     0   1   2E+6   13.5   14  RT (min)       lcorbi_t3_pos_2_027_05    m/z 767.5402     0   1   2   3E+4   15.5   16  RT (min)      lcorbi_t3_pos_2_012_01    m/z 767.5406     0   1   15.5   16  RT (min)      lcorbi_t3_pos_2_012_01    m/z 767.5406     0   1   2E+5   15   15.5  RT (min)      lcorbi_t3_pos_1_031_40    m/z 767.5416     0   1   2E+5   15   15.5  RT (min)      lcorbi_t3_pos_1_031_40    m/z 767.5416     0   1   2E+6   13   13.5  RT (min)      lcorbi_t3_pos_3_011_42    m/z 767.5418     0   1E+6   13.5   14  RT (min)       lcorbi_t3_pos_1_005_42    m/z 767.5418     0   1   2E+4   12.5   13  RT (min)      lcorbi_t3_pos_2_013_33    m/z 767.5421     0   1E+6   13   13.5  RT (min)      lcorbi_t3_pos_2_013_33    m/z 767.5422     0   1   2E+4   12.5   13  RT (min)      lcorbi_t3_pos_2_013_33    m/z 768.5453     0   1   12.5   13  RT (min)      lcorbi_t3_pos_3_031_45    m/z 768.5454     0   2   4   6E+5   13.5   14  RT (min)       lcorbi_t3_pos_2_023_26    m/z 768.5454     0   1E+4   12.5   13  RT (min)      lcorbi_t3_pos_3_024_07    m/z 768.5538     0   1   2   3   4E+4   13   13.5  RT (min)      lcorbi_t3_pos_3_011_42    m/z 768.5538     0   1E+7   13   13.5  RT (min)      lcorbi_t3_pos_3_011_42    m/z 768.5538     0   1   2   3E+4   15.5   16  RT (min)      lcorbi_t3_pos_1_031_40    m/z 768.5539     0   1E+4   13   13.5  RT (min)      lcorbi_t3_pos_2_013_33    m/z 768.5539     0   1   2   3E+4   15   15.5  RT (min)      lcorbi_t3_pos_1_034_34    m/z 768.5539     0   2   4   6E+5   11.5   12  RT (min)      lcorbi_t3_pos_4_057_49    m/z 768.5539     0   1E+4   13   13.5  RT (min)      lcorbi_t3_pos_2_013_33    m/z 768.5540     0   1   2   3   4E+5   14.5   15  RT (min)      lcorbi_t3_pos_1_028_32    m/z 768.5540     0   1   2E+4   15.5   16  RT (min)      lcorbi_t3_pos_2_012_01    m/z 768.5540     0   1   15   15.5  RT (min)      lcorbi_t3_pos_1_056_56    m/z 768.5540     0   1   2   3   4E+4   15   15.5  RT (min)      lcorbi_t3_pos_1_034_34    m/z 768.5541     0   1   16   16.5  RT (min)      lcorbi_t3_pos_1_031_40    m/z 768.5542     0   2   4E+5   14.5   15  RT (min)      lcorbi_t3_pos_1_047_41    m/z 768.5542     0   2   4E+5   14.5   15  RT (min)      lcorbi_t3_pos_1_047_41    m/z 768.5542     0   1   16.5   17  RT (min)    lcorbi_t3_pos_1_023_02    m/z 768.5542     0   1   2   3E+4   15   15.5  RT (min)      lcorbi_t3_pos_1_034_34    m/z 768.5542     0   1   2E+4   15.5   16  RT (min)      lcorbi_t3_pos_2_012_01    m/z 768.5542     0   1   2E+4   15.5   16  RT (min)      lcorbi_t3_pos_2_012_01    m/z 768.5543     0   2   4E+4   14.5   15  RT (min)      lcorbi_t3_pos_2_025_52    m/z 768.5544     0   1   16.5   17  RT (min)    lcorbi_t3_pos_1_023_02    m/z 768.5897     0   1   13   13.5  RT (min)      lcorbi_t3_pos_2_032_39    m/z 768.5898     0   1   2E+4   13   13.5  RT (min)      lcorbi_t3_pos_3_024_07    m/z 768.5899     0   2   4   6E+3   13   13.5  RT (min)      lcorbi_t3_pos_4_012_37    m/z 768.5902     0   1   2E+6   14   14.5  RT (min)       lcorbi_t3_pos_3_040_41    m/z 768.5903     0   1   16.5   17  RT (min)    lcorbi_t3_pos_1_023_02    m/z 768.5904     0   1   2   3E+4   16   16.5  RT (min)      lcorbi_t3_pos_1_051_09    m/z 768.5904     0   2   4   6E+6   13.5   14  RT (min)      lcorbi_t3_pos_1_002_01    m/z 768.5904     0   1   2   3   4E+4   15   15.5  RT (min)      lcorbi_t3_pos_1_028_32    m/z 768.5904     0   1   16   16.5  RT (min)      lcorbi_t3_pos_1_049_55    m/z 768.5904     0   1   15.5   16  RT (min)      lcorbi_t3_pos_1_034_34    m/z 768.5904     0   1   2   3   4E+4   14.5   15  RT (min)      lcorbi_t3_pos_2_017_51    m/z 768.5905     0   1   2   3   4E+6   14.5   15  RT (min)       lcorbi_t3_pos_1_005_42    m/z 768.5905     0   1   2E+4   15.5   16  RT (min)      lcorbi_t3_pos_1_034_34    m/z 768.5905     0   1   2   3   4E+4   13.5   14  RT (min)      lcorbi_t3_pos_1_002_01    m/z 768.5906     0   1   15   15.5  RT (min)      lcorbi_t3_pos_1_030_21    m/z 768.5906     0   1   15   15.5  RT (min)      lcorbi_t3_pos_1_030_21    m/z 768.5906     0   1   16   16.5  RT (min)      lcorbi_t3_pos_1_051_09    m/z 768.5906     0   2   4   6   8E+5   14.5   15  RT (min)       lcorbi_t3_pos_1_008_36    m/z 768.5906     0   1   2E+4   15   15.5  RT (min)      lcorbi_t3_pos_1_047_41    m/z 768.5906     0   1   2   3E+4   15   15.5  RT (min)      lcorbi_t3_pos_1_021_18    m/z 768.5906     0   1   15   15.5  RT (min)      lcorbi_t3_pos_2_038_03    m/z 768.5906     0   2   4   6   8E+4   13.5   14  RT (min)      lcorbi_t3_pos_1_002_01    m/z 768.5906     0   1   17   17.5  RT (min)    lcorbi_t3_pos_1_023_02    m/z 768.5907     0   1   16.5   17  RT (min)    lcorbi_t3_pos_1_023_02    m/z 768.5908     0   1   16.5   17  RT (min)    lcorbi_t3_pos_1_023_02    m/z 768.5909     0   1   2E+5   14.5   15  RT (min)      lcorbi_t3_pos_1_008_36    m/z 768.5910     0   1   16.5   17  RT (min)    lcorbi_t3_pos_2_012_01    m/z 768.5910     0   1   16.5   17  RT (min)    lcorbi_t3_pos_1_023_02    m/z 768.5910     0   1   16.5   17  RT (min)    lcorbi_t3_pos_1_023_02    m/z 768.6571     0   1   15   15.5  RT (min)      lcorbi_t3_pos_2_056_56    m/z 768.6572     0   1   2E+4   15   15.5  RT (min)      lcorbi_t3_pos_1_056_56    m/z 768.6572     0   2   4   6E+4   14   14.5  RT (min)       lcorbi_t3_pos_1_047_41    m/z 768.6573     0   2   4   6E+4   14   14.5  RT (min)      lcorbi_t3_pos_1_021_18    m/z 768.6573     0   2   4   6   8E+4   14   14.5  RT (min)      lcorbi_t3_pos_1_030_21    m/z 768.6573     0   2   4E+4   14   14.5  RT (min)       lcorbi_t3_pos_2_046_07    m/z 768.6573     0   1   14.5   15  RT (min)      lcorbi_t3_pos_1_041_54    m/z 768.6573     0   1   15   15.5  RT (min)      lcorbi_t3_pos_1_049_55    m/z 768.6574     0   1E+4   14.5   15  RT (min)      lcorbi_t3_pos_2_041_54    m/z 768.6575     0   1   14.5   15  RT (min)      lcorbi_t3_pos_1_041_54    m/z 768.6577     0   1   2   3E+5   13.5   14  RT (min)       lcorbi_t3_pos_4_001_49    m/z 769.5574     0   1   15   15.5  RT (min)      lcorbi_t3_pos_2_038_03    m/z 769.5575     0   1   2E+5   14.5   15  RT (min)      lcorbi_t3_pos_2_022_38    m/z 769.5575     0   1   2E+4   13   13.5  RT (min)      lcorbi_t3_pos_3_011_42    m/z 769.5575     0   1   2   3   4E+4   13   13.5  RT (min)      lcorbi_t3_pos_2_013_33    m/z 769.5576     0   1   15.5   16  RT (min)      lcorbi_t3_pos_1_031_40    m/z 769.5576     0   1   13   13.5  RT (min)      lcorbi_t3_pos_2_013_33    m/z 769.5576     0   1   2E+5   14.5   15  RT (min)      lcorbi_t3_pos_2_022_38    m/z 769.5577     0   1   2E+4   15.5   16  RT (min)      lcorbi_t3_pos_2_012_01    m/z 769.5577     0   1E+5   14.5   15  RT (min)      lcorbi_t3_pos_1_028_32    m/z 769.5577     0   1   2E+5   14.5   15  RT (min)      lcorbi_t3_pos_2_022_38    m/z 769.5577     0   1   2E+4   14.5   15  RT (min)      lcorbi_t3_pos_2_038_03    m/z 769.5577     0   1   2E+4   15.5   16  RT (min)      lcorbi_t3_pos_2_012_01    m/z 769.5577     0   1   15   15.5  RT (min)      lcorbi_t3_pos_1_034_34    m/z 769.5578     0   1   15   15.5  RT (min)      lcorbi_t3_pos_1_034_34    m/z 769.5579     0   1   15.5   16  RT (min)      lcorbi_t3_pos_1_001_49    m/z 769.5580     0   1   15.5   16  RT (min)      lcorbi_t3_pos_2_012_01    m/z 769.5580     0   1   2   3E+4   15   15.5  RT (min)      lcorbi_t3_pos_1_034_34    m/z 769.5580     0   1   2   3E+4   15.5   16  RT (min)      lcorbi_t3_pos_1_025_52    m/z 769.5932     0   1   13   13.5  RT (min)      lcorbi_t3_pos_4_012_37    m/z 769.5932     0   1   13   13.5  RT (min)      lcorbi_t3_pos_2_032_39    m/z 769.5938     0   1   16   16.5  RT (min)      lcorbi_t3_pos_1_049_55    m/z 769.5938     0   2   4   6E+4   13.5   14  RT (min)      lcorbi_t3_pos_1_002_01    m/z 769.5938     0   1   2   3E+4   13.5   14  RT (min)      lcorbi_t3_pos_1_002_01    m/z 769.5939     0   1   2E+4   15   15.5  RT (min)      lcorbi_t3_pos_1_028_32    m/z 769.5939     0   1   2E+4   15   15.5  RT (min)      lcorbi_t3_pos_1_047_41    m/z 769.5939     0   1   16   16.5  RT (min)      lcorbi_t3_pos_1_049_55    m/z 769.5940     0   1   16.5   17  RT (min)    lcorbi_t3_pos_1_023_02    m/z 769.5940     0   1   15.5   16  RT (min)      lcorbi_t3_pos_1_034_34    m/z 769.5940     0   1   2E+6   14.5   15  RT (min)      lcorbi_t3_pos_1_005_42    m/z 769.5940     0   1   2   3E+5   14.5   15  RT (min)      lcorbi_t3_pos_1_008_36    m/z 769.5940     0   1   16.5   17  RT (min)    lcorbi_t3_pos_1_023_02    m/z 769.5940     0   1   16.5   17  RT (min)    lcorbi_t3_pos_1_023_02    m/z 769.5941     0   1E+4   15   15.5  RT (min)      lcorbi_t3_pos_2_038_03    m/z 769.5941     0   1   16.5   17  RT (min)    lcorbi_t3_pos_1_023_02    m/z 769.5941     0   1   16   16.5  RT (min)      lcorbi_t3_pos_1_051_09    m/z 769.5942     0   1E+4   14.5   15  RT (min)      lcorbi_t3_pos_2_017_51    m/z 769.5942     0   1   15.5   16  RT (min)      lcorbi_t3_pos_2_049_55    m/z 769.5945     0   1   2   3E+6   13.5   14  RT (min)      lcorbi_t3_pos_1_002_01    m/z 769.5946     0   1   16.5   17  RT (min)    lcorbi_t3_pos_1_023_02    m/z 769.6695     0   2   4   6   8E+4   14   14.5  RT (min)       lcorbi_t3_pos_3_025_52    m/z 769.6696     0   1   2E+4   15.5   16  RT (min)      lcorbi_t3_pos_1_001_49    m/z 769.6698     0   2   4   6E+4   14.5   15  RT (min)      lcorbi_t3_pos_1_030_21    m/z 770.4970     0   2   4   6E+5   12   12.5  RT (min)       lcorbi_t3_pos_1_054_48    m/z 770.4970     0   2   4E+5   11.5   12  RT (min)       lcorbi_t3_pos_2_005_11    m/z 770.4970     0   2   4E+5   11.5   12  RT (min)       lcorbi_t3_pos_2_005_11    m/z 770.5608     0   1E+4   15.5   16  RT (min)      lcorbi_t3_pos_2_049_55    m/z 770.5608     0   1   12.5   13  RT (min)      lcorbi_t3_pos_3_031_45    m/z 770.5614     0   1E+6   14   14.5  RT (min)       lcorbi_t3_pos_4_084_59    m/z 770.5655     0   1E+5   14.5   15  RT (min)      lcorbi_t3_pos_2_046_07    m/z 770.5677     0   1   2E+6   13.5   14  RT (min)      lcorbi_t3_pos_2_023_26    m/z 770.5686     0   2   4   6   8E+5   13   13.5  RT (min)      lcorbi_t3_pos_3_011_42    m/z 770.5689     0   1E+6   14   14.5  RT (min)      lcorbi_t3_pos_1_008_36    m/z 770.5689     0   1E+5   14.5   15  RT (min)      lcorbi_t3_pos_2_046_07    m/z 770.5690     0   2   4   6   8E+3   13   13.5  RT (min)      lcorbi_t3_pos_3_031_45    m/z 770.5690     0   1   13   13.5  RT (min)      lcorbi_t3_pos_2_013_33    m/z 770.5692     0   1   15.5   16  RT (min)      lcorbi_t3_pos_2_009_50    m/z 770.5692     0   1   2   3   4E+4   14.5   15  RT (min)      lcorbi_t3_pos_1_030_21    m/z 770.5694     0   2   4   6E+4   14.5   15  RT (min)      lcorbi_t3_pos_2_006_47    m/z 770.5694     0   2   4   6   8E+5   14   14.5  RT (min)      lcorbi_t3_pos_4_025_52    m/z 770.5694     0   1E+7   14   14.5  RT (min)       lcorbi_t3_pos_4_084_59    m/z 770.5694     0   1E+6   12.5   13  RT (min)      lcorbi_t3_pos_4_057_49    m/z 770.5694     0   1   15.5   16  RT (min)      lcorbi_t3_pos_1_049_55    m/z 770.5696     0   1   2   3E+4   15   15.5  RT (min)      lcorbi_t3_pos_1_034_34    m/z 770.5696     0   1   2   3E+4   15   15.5  RT (min)      lcorbi_t3_pos_1_034_34    m/z 770.5696     0   1   15   15.5  RT (min)      lcorbi_t3_pos_2_038_03    m/z 770.5697     0   1   2   3E+4   15.5   16  RT (min)      lcorbi_t3_pos_1_031_40    m/z 770.5698     0   1   16   16.5  RT (min)      lcorbi_t3_pos_1_001_49    m/z 770.5698     0   1   2   3E+4   15   15.5  RT (min)      lcorbi_t3_pos_1_028_32    m/z 770.5698     0   1   15   15.5  RT (min)      lcorbi_t3_pos_2_038_03    m/z 770.5699     0   1   2E+4   15.5   16  RT (min)      lcorbi_t3_pos_2_012_01    m/z 770.5699     0   1   16   16.5  RT (min)      lcorbi_t3_pos_2_056_56    m/z 770.5971     0   1   16.5   17  RT (min)    lcorbi_t3_pos_1_023_02    m/z 770.6054     0   1E+4   15   15.5  RT (min)      lcorbi_t3_pos_2_006_47    m/z 770.6056     0   1   2E+4   15.5   16  RT (min)      lcorbi_t3_pos_1_034_34    m/z 770.6058     0   1   2E+4   16   16.5  RT (min)      lcorbi_t3_pos_1_051_09    m/z 770.6059     0   1   2E+5   14.5   15  RT (min)      lcorbi_t3_pos_1_008_36    m/z 770.6059     0   1   15.5   16  RT (min)      lcorbi_t3_pos_1_034_34    m/z 770.6059     0   2   4   6E+5   13.5   14  RT (min)      lcorbi_t3_pos_1_002_01    m/z 770.6060     0   1E+6   14.5   15  RT (min)      lcorbi_t3_pos_1_005_42    m/z 770.6060     0   1   15.5   16  RT (min)      lcorbi_t3_pos_2_049_55    m/z 770.6060     0   1   2   3   4E+5   14.5   15  RT (min)      lcorbi_t3_pos_1_008_36    m/z 770.6060     0   1   16   16.5  RT (min)      lcorbi_t3_pos_1_056_56    m/z 770.6061     0   1E+4   15   15.5  RT (min)      lcorbi_t3_pos_2_046_07    m/z 770.6061     0   1   16.5   17  RT (min)    lcorbi_t3_pos_1_023_02    m/z 770.6062     0   1   16   16.5  RT (min)      lcorbi_t3_pos_1_001_49    m/z 770.6066     0   1   2E+4   13.5   14  RT (min)       lcorbi_t3_pos_1_002_01    m/z 770.6068     0   1   16.5   17  RT (min)    lcorbi_t3_pos_1_023_02    m/z 770.6083     0   1   2E+4   13.5   14  RT (min)      lcorbi_t3_pos_1_002_01    m/z 770.6726     0   1   2   3E+4   15   15.5  RT (min)      lcorbi_t3_pos_1_041_54    m/z 770.6726     0   1   2   3E+4   15.5   16  RT (min)      lcorbi_t3_pos_1_001_49    m/z 770.6728     0   1E+5   14.5   15  RT (min)      lcorbi_t3_pos_1_021_18    m/z 770.6728     0   1   2   3E+5   14   14.5  RT (min)       lcorbi_t3_pos_3_025_52    m/z 770.6728     0   1   2   3E+4   15   15.5  RT (min)      lcorbi_t3_pos_1_041_54    m/z 770.6728     0   1E+4   14.5   15  RT (min)      lcorbi_t3_pos_2_041_54    m/z 770.6728     0   1E+6   14   14.5  RT (min)      lcorbi_t3_pos_1_005_42    m/z 770.6729     0   1E+6   14   14.5  RT (min)       lcorbi_t3_pos_1_005_42    m/z 770.6729     0   1   15   15.5  RT (min)      lcorbi_t3_pos_4_033_53    m/z 770.6729     0   1   2E+5   14.5   15  RT (min)      lcorbi_t3_pos_1_047_41    m/z 770.6729     0   1   2   3E+4   15.5   16  RT (min)      lcorbi_t3_pos_1_001_49    m/z 770.6730     0   1   2E+5   14.5   15  RT (min)      lcorbi_t3_pos_1_030_21    m/z 770.6732     0   1   2E+4   14.5   15  RT (min)      lcorbi_t3_pos_2_038_03    m/z 770.6734     0   1   2E+4   14.5   15  RT (min)      lcorbi_t3_pos_2_038_03    m/z 771.5722     0   2   4   6E+4   13   13.5  RT (min)      lcorbi_t3_pos_3_011_42    m/z 771.5728     0   2   4E+5   12.5   13  RT (min)      lcorbi_t3_pos_4_057_49    m/z 771.5728     0   1   16   16.5  RT (min)      lcorbi_t3_pos_1_031_40    m/z 771.5728     0   1E+4   13   13.5  RT (min)      lcorbi_t3_pos_2_013_33    m/z 771.5729     0   1   13   13.5  RT (min)      lcorbi_t3_pos_3_031_45    m/z 771.5729     0   1E+4   15   15.5  RT (min)      lcorbi_t3_pos_2_041_54    m/z 771.5729     0   1   2   3   4E+5   14   14.5  RT (min)      lcorbi_t3_pos_4_025_52    m/z 771.5730     0   1   15.5   16  RT (min)      lcorbi_t3_pos_2_009_50    m/z 771.5730     0   1   2   3E+4   14.5   15  RT (min)      lcorbi_t3_pos_1_047_41    m/z 771.5730     0   1   15.5   16  RT (min)      lcorbi_t3_pos_1_049_55    m/z 771.5731     0   2   4   6E+5   14   14.5  RT (min)      lcorbi_t3_pos_1_005_42    m/z 771.5732     0   1   2   3E+4   14.5   15  RT (min)      lcorbi_t3_pos_1_030_21    m/z 771.5732     0   1   2   3   4E+4   14.5   15   15.5  RT (min)      lcorbi_t3_pos_1_028_32    m/z 771.5732     0   1   16   16.5  RT (min)      lcorbi_t3_pos_2_056_56    m/z 771.5732     0   2   4   6E+5   14   14.5  RT (min)      lcorbi_t3_pos_1_008_36    m/z 771.5732     0   1   16   16.5  RT (min)      lcorbi_t3_pos_1_001_49    m/z 771.5733     0   1   15.5   16  RT (min)      lcorbi_t3_pos_2_012_01    m/z 771.5733     0   1   2   3E+4   15   15.5  RT (min)      lcorbi_t3_pos_1_034_34    m/z 771.5733     0   1   15.5   16  RT (min)      lcorbi_t3_pos_1_031_40    m/z 771.5734     0   1   2   3E+4   15   15.5  RT (min)      lcorbi_t3_pos_1_034_34    m/z 771.6092     0   1   16.5   17  RT (min)    lcorbi_t3_pos_1_023_02    m/z 771.6094     0   1   16   16.5  RT (min)      lcorbi_t3_pos_1_051_09    m/z 771.6095     0   2   4   6   8E+4   15   15.5  RT (min)       lcorbi_t3_pos_1_028_32    m/z 771.6096     0   1   2   3E+5   14.5   15  RT (min)      lcorbi_t3_pos_1_008_36    m/z 771.6097     0   1   16   16.5  RT (min)      lcorbi_t3_pos_1_051_09    m/z 771.6097     0   1   16.5   17  RT (min)    lcorbi_t3_pos_1_023_02    m/z 771.6098     0   1E+5   14.5   15  RT (min)      lcorbi_t3_pos_1_008_36    m/z 771.6099     0   2   4   6   8E+4   15.5   16  RT (min)      lcorbi_t3_pos_1_034_34    m/z 772.5126     0   1E+6   12   12.5  RT (min)       lcorbi_t3_pos_1_054_48    m/z 772.5126     0   2   4   6   8E+5   12   12.5  RT (min)       lcorbi_t3_pos_3_029_13    m/z 772.5489     0   2   4   6E+5   13   13.5  RT (min)       lcorbi_t3_pos_1_054_48    m/z 772.5490     0   2   4   6E+5   13   13.5  RT (min)       lcorbi_t3_pos_1_054_48    m/z 772.5492     0   2   4   6E+5   13   13.5  RT (min)       lcorbi_t3_pos_2_005_11    m/z 772.5840     0   1   13   13.5  RT (min)      lcorbi_t3_pos_2_013_33    m/z 772.5845     0   1   13   13.5  RT (min)      lcorbi_t3_pos_3_024_07    m/z 772.5848     0   1E+6   13   13.5  RT (min)      lcorbi_t3_pos_4_057_49    m/z 772.5848     0   2   4   6E+4   15   15.5  RT (min)      lcorbi_t3_pos_2_006_47    m/z 772.5851     0   1E+5   14.5   15  RT (min)      lcorbi_t3_pos_2_017_51    m/z 772.5851     0   1E+6   13.5   14  RT (min)      lcorbi_t3_pos_1_002_01    m/z 772.5852     0   2   4   6   8E+4   13   13.5  RT (min)      lcorbi_t3_pos_1_002_01    m/z 772.5852     0   1   2E+4   15   15.5  RT (min)      lcorbi_t3_pos_1_030_21    m/z 772.5852     0   1   16   16.5  RT (min)      lcorbi_t3_pos_1_001_49    m/z 772.5852     0   1   2   3E+5   14.5   15  RT (min)      lcorbi_t3_pos_3_025_52    m/z 772.5852     0   1   15.5   16  RT (min)      lcorbi_t3_pos_1_041_54    m/z 772.5853     0   1   2   3E+4   15   15.5  RT (min)      lcorbi_t3_pos_1_033_53    m/z 772.5854     0   1   16   16.5  RT (min)      lcorbi_t3_pos_1_001_49    m/z 772.5854     0   1   16   16.5  RT (min)      lcorbi_t3_pos_1_001_49    m/z 772.5854     0   1   2   3E+6   14   14.5  RT (min)      lcorbi_t3_pos_1_005_42    m/z 772.5854     0   1   2   3E+6   14.5   15  RT (min)      lcorbi_t3_pos_1_005_42    m/z 772.5854     0   1   15.5   16  RT (min)      lcorbi_t3_pos_2_009_50    m/z 772.5854     0   1   16   16.5  RT (min)      lcorbi_t3_pos_1_051_09    m/z 772.5854     0   1   15.5   16  RT (min)      lcorbi_t3_pos_2_009_50    m/z 772.5854     0   1   2E+6   14.5   15  RT (min)       lcorbi_t3_pos_1_008_36    m/z 772.5854     0   1   2E+4   15.5   16  RT (min)      lcorbi_t3_pos_2_049_55    m/z 772.5854     0   1   2E+4   15   15.5  RT (min)      lcorbi_t3_pos_1_030_21    m/z 772.5855     0   1   16   16.5  RT (min)      lcorbi_t3_pos_1_001_49    m/z 772.5855     0   1   16.5   17  RT (min)    lcorbi_t3_pos_1_023_02    m/z 772.5855     0   1E+4   15   15.5  RT (min)      lcorbi_t3_pos_2_025_52    m/z 772.5856     0   1   16.5   17  RT (min)    lcorbi_t3_pos_1_023_02    m/z 772.5856     0   1   16.5   17  RT (min)    lcorbi_t3_pos_2_012_01    m/z 772.5856     0   2   4   6   8E+4   14.5   15  RT (min)      lcorbi_t3_pos_2_017_51    m/z 772.5858     0   1   16.5   17  RT (min)    lcorbi_t3_pos_1_023_02    m/z 772.6220     0   2   4   6   8E+5   14.5   15  RT (min)      lcorbi_t3_pos_4_019_47    m/z 772.6220     0   2   4   6   8E+5   14.5   15  RT (min)      lcorbi_t3_pos_4_019_47    m/z 772.6222     0   1   16   16.5  RT (min)      lcorbi_t3_pos_1_031_40    m/z 772.6222     0   1E+5   15.5   16  RT (min)      lcorbi_t3_pos_1_034_34    m/z 772.6222     0   1   2   3E+5   13.5   14  RT (min)      lcorbi_t3_pos_1_002_01    m/z 772.6223     0   1   2   3E+4   15.5   16  RT (min)      lcorbi_t3_pos_1_034_34    m/z 772.6238     0   1   2   3   4E+5   15.5   16  RT (min)       lcorbi_t3_pos_1_028_32    m/z 772.6238     0   1   17   17.5  RT (min)    lcorbi_t3_pos_1_023_02    m/z 772.6241     0   1   16   16.5  RT (min)      lcorbi_t3_pos_1_051_09    m/z 772.6242     0   1   17.5   18  RT (min)    lcorbi_t3_pos_1_001_49    m/z 772.6885     0   1   2   3E+6   14   14.5  RT (min)      lcorbi_t3_pos_4_092_60    m/z 772.6886     0   2   4   6   8E+5   14   14.5  RT (min)      lcorbi_t3_pos_1_005_42    m/z 772.6886     0   1   15.5   16  RT (min)      lcorbi_t3_pos_1_001_49    m/z 772.6886     0   1E+5   14.5   15  RT (min)      lcorbi_t3_pos_2_046_07    m/z 772.6887     0   1   2   3E+4   14.5   15  RT (min)      lcorbi_t3_pos_1_021_18    m/z 772.6887     0   1   2E+4   15   15.5  RT (min)      lcorbi_t3_pos_1_041_54    m/z 772.6888     0   1   2   3   4E+4   14.5   15  RT (min)      lcorbi_t3_pos_1_030_21    m/z 772.6888     0   1   15.5   16  RT (min)      lcorbi_t3_pos_1_049_55    m/z 772.6889     0   1   2E+4   15   15.5  RT (min)      lcorbi_t3_pos_1_041_54    m/z 773.5883     0   2   4   6E+5   13   13.5  RT (min)      lcorbi_t3_pos_4_057_49    m/z 773.5886     0   1   2   3   4E+4   13   13.5  RT (min)      lcorbi_t3_pos_1_002_01    m/z 773.5887     0   2   4   6   8E+5   13.5   14  RT (min)      lcorbi_t3_pos_1_002_01    m/z 773.5888     0   1E+6   14.5   15  RT (min)      lcorbi_t3_pos_1_008_36    m/z 773.5888     0   1   2E+4   15   15.5  RT (min)      lcorbi_t3_pos_2_006_47    m/z 773.5889     0   1   15   15.5  RT (min)      lcorbi_t3_pos_1_028_32    m/z 773.5889     0   1   2   3E+4   15   15.5  RT (min)      lcorbi_t3_pos_1_033_53    m/z 773.5889     0   1   15.5   16  RT (min)      lcorbi_t3_pos_1_056_56    m/z 773.5890     0   1   15.5   16  RT (min)      lcorbi_t3_pos_1_041_54    m/z 773.5890     0   1   16   16.5  RT (min)      lcorbi_t3_pos_1_051_09    m/z 773.5890     0   1   2   3   4E+4   14.5   15  RT (min)      lcorbi_t3_pos_2_017_51    m/z 773.5891     0   1   15.5   16  RT (min)      lcorbi_t3_pos_1_031_40    m/z 773.5892     0   1   16.5   17  RT (min)    lcorbi_t3_pos_1_023_02    m/z 773.5892     0   1   16.5   17  RT (min)    lcorbi_t3_pos_1_023_02    m/z 773.5892     0   2   4   6E+4   14.5   15  RT (min)      lcorbi_t3_pos_2_017_51    m/z 773.5896     0   1   15.5   16  RT (min)      lcorbi_t3_pos_1_034_34    m/z 773.6250     0   1   16   16.5  RT (min)      lcorbi_t3_pos_1_031_40    m/z 773.6251     0   1   16   16.5  RT (min)      lcorbi_t3_pos_1_051_09    m/z 773.6251     0   2   4   6E+3   13.5   14  RT (min)      lcorbi_t3_pos_2_054_29    m/z 773.6253     0   1   13.5   14  RT (min)      lcorbi_t3_pos_1_002_01    m/z 773.6254     0   2   4   6E+4   15.5   16  RT (min)      lcorbi_t3_pos_1_034_34    m/z 773.6255     0   1   2E+5   15.5   16  RT (min)      lcorbi_t3_pos_1_028_32    m/z 773.6255     0   1   2E+5   14.5   15  RT (min)      lcorbi_t3_pos_1_008_36    m/z 773.6258     0   1   16   16.5  RT (min)      lcorbi_t3_pos_1_051_09    m/z 773.6268     0   1   17   17.5  RT (min)    lcorbi_t3_pos_1_023_02    m/z 774.3578     0   1   2   3   4E+5   14.5   15  RT (min)       lcorbi_t3_pos_4_105_61    m/z 774.4920     0   2   4E+5   12.5   13  RT (min)       lcorbi_t3_pos_1_054_48    m/z 774.5282     0   2   4E+5   12.5   13  RT (min)       lcorbi_t3_pos_1_054_48    m/z 774.5282     0   1   2E+4   13   13.5  RT (min)      lcorbi_t3_pos_2_013_33    m/z 774.5284     0   1E+4   13   13.5  RT (min)      lcorbi_t3_pos_2_013_33    m/z 774.5286     0   1E+4   15.5   16  RT (min)      lcorbi_t3_pos_2_012_01    m/z 774.5286     0   2   4E+5   12.5   13  RT (min)       lcorbi_t3_pos_1_054_48    m/z 774.5286     0   1   15   15.5  RT (min)      lcorbi_t3_pos_1_034_34    m/z 774.5287     0   1   15.5   16  RT (min)      lcorbi_t3_pos_1_031_40    m/z 774.5288     0   2   4   6E+4   14   14.5  RT (min)      lcorbi_t3_pos_1_008_36    m/z 774.5288     0   1   2E+5   14   14.5  RT (min)      lcorbi_t3_pos_1_008_36    m/z 774.5288     0   1   14.5   15  RT (min)      lcorbi_t3_pos_1_028_32    m/z 774.5644     0   2   4   6   8E+4   12   12.5  RT (min)      lcorbi_t3_pos_2_013_33    m/z 774.5646     0   1   2E+4   14   14.5  RT (min)       lcorbi_t3_pos_1_034_34    m/z 774.5646     0   1E+4   14.5   15  RT (min)      lcorbi_t3_pos_2_012_01    m/z 774.5646     0   1E+4   14.5   15  RT (min)      lcorbi_t3_pos_2_012_01    m/z 774.5646     0   2   4   6   8E+3   12   12.5  RT (min)      lcorbi_t3_pos_3_031_45    m/z 774.5647     0   1E+5   13.5   14  RT (min)       lcorbi_t3_pos_3_003_33    m/z 774.5648     0   1   2E+4   14.5   15  RT (min)      lcorbi_t3_pos_1_031_40    m/z 774.5654     0   1   2   3   4E+4   14   14.5  RT (min)      lcorbi_t3_pos_1_034_34    m/z 774.5919     0   1   13   13.5  RT (min)      lcorbi_t3_pos_2_047_31    m/z 774.5923     0   1   15   15.5  RT (min)      lcorbi_t3_pos_1_047_41    m/z 774.5932     0   2   4   6E+5   13.5   14  RT (min)       lcorbi_t3_pos_3_053_48    m/z 774.6008     0   1E+6   14.5   15  RT (min)      lcorbi_t3_pos_1_005_42    m/z 774.6008     0   1   15.5   16  RT (min)      lcorbi_t3_pos_1_034_34    m/z 774.6008     0   1E+4   13   13.5  RT (min)      lcorbi_t3_pos_3_052_47    m/z 774.6008     0   1   13   13.5  RT (min)      lcorbi_t3_pos_2_047_31    m/z 774.6008     0   1   2E+5   14.5   15  RT (min)      lcorbi_t3_pos_1_008_36    m/z 774.6009     0   1   16   16.5  RT (min)      lcorbi_t3_pos_1_051_09    m/z 774.6009     0   1E+5   14.5   15  RT (min)      lcorbi_t3_pos_1_008_36    m/z 774.6010     0   1   16.5   17  RT (min)    lcorbi_t3_pos_2_012_01    m/z 774.6010     0   1   16   16.5  RT (min)      lcorbi_t3_pos_1_031_40    m/z 774.6010     0   1   2   3E+4   15   15.5  RT (min)      lcorbi_t3_pos_1_030_21    m/z 774.6010     0   1   16.5   17  RT (min)    lcorbi_t3_pos_1_023_02    m/z 774.6010     0   1   16   16.5  RT (min)      lcorbi_t3_pos_1_001_49    m/z 774.6010     0   1   2   3E+4   15   15.5  RT (min)      lcorbi_t3_pos_2_046_07    m/z 774.6010     0   1   2   3E+5   13.5   14  RT (min)      lcorbi_t3_pos_1_002_01    m/z 774.6010     0   1   15.5   16  RT (min)      lcorbi_t3_pos_2_049_55    m/z 774.6011     0   1   15.5   16  RT (min)      lcorbi_t3_pos_2_009_50    m/z 774.6011     0   1   16.5   17  RT (min)    lcorbi_t3_pos_1_023_02    m/z 774.6011     0   1   15   15.5  RT (min)      lcorbi_t3_pos_1_028_32    m/z 774.6011     0   1   16   16.5  RT (min)      lcorbi_t3_pos_2_056_56    m/z 774.6012     0   1   15   15.5  RT (min)      lcorbi_t3_pos_1_028_32    m/z 774.6012     0   1   17   17.5  RT (min)    lcorbi_t3_pos_1_023_02    m/z 774.6012     0   1   15.5   16  RT (min)      lcorbi_t3_pos_2_025_52    m/z 774.6012     0   1E+6   14.5   15  RT (min)      lcorbi_t3_pos_1_005_42    m/z 774.6012     0   2   4E+5   14.5   15  RT (min)      lcorbi_t3_pos_1_008_36    m/z 775.6043     0   1   15.5   16  RT (min)      lcorbi_t3_pos_1_034_34    m/z 775.6044     0   1   16.5   17  RT (min)    lcorbi_t3_pos_1_023_02    m/z 775.6046     0   2   4   6E+5   14.5   15  RT (min)       lcorbi_t3_pos_1_005_42    m/z 775.6046     0   1   16   16.5  RT (min)      lcorbi_t3_pos_1_031_40    m/z 775.6046     0   1E+5   13.5   14  RT (min)      lcorbi_t3_pos_1_002_01    m/z 775.6047     0   1   2E+5   14.5   15  RT (min)      lcorbi_t3_pos_1_008_36    m/z 775.6047     0   1   2E+4   15   15.5  RT (min)      lcorbi_t3_pos_1_028_32    m/z 775.6048     0   1   15.5   16  RT (min)      lcorbi_t3_pos_1_034_34    m/z 775.6048     0   1   16.5   17  RT (min)    lcorbi_t3_pos_1_023_02    m/z 775.6048     0   2   4   6E+5   14.5   15  RT (min)       lcorbi_t3_pos_1_005_42    m/z 775.6048     0   1   2E+4   15   15.5  RT (min)      lcorbi_t3_pos_1_028_32    m/z 775.6048     0   1E+5   14.5   15  RT (min)      lcorbi_t3_pos_1_008_36    m/z 775.6048     0   1   16   16.5  RT (min)      lcorbi_t3_pos_1_001_49    m/z 775.6048     0   1   16   16.5  RT (min)      lcorbi_t3_pos_1_031_40    m/z 775.6049     0   2   4   6   8E+4   14.5   15  RT (min)      lcorbi_t3_pos_1_008_36    m/z 775.7274     0   1   17   17.5  RT (min)    lcorbi_t3_pos_1_041_54    m/z 775.7277     0   1   17   17.5  RT (min)    lcorbi_t3_pos_1_049_55    m/z 776.5205     0   1   14.5   15  RT (min)      lcorbi_t3_pos_1_034_34    m/z 776.5206     0   1   14.5   15  RT (min)      lcorbi_t3_pos_1_028_32    m/z 776.5206     0   1   2   3E+5   13   13.5  RT (min)      lcorbi_t3_pos_3_011_42    m/z 776.5206     0   1   14   14.5  RT (min)      lcorbi_t3_pos_2_022_38    m/z 776.5206     0   1   15   15.5  RT (min)      lcorbi_t3_pos_1_031_40    m/z 776.5207     0   1   15   15.5  RT (min)      lcorbi_t3_pos_1_031_40    m/z 776.5208     0   1   15.5   16  RT (min)      lcorbi_t3_pos_2_012_01    m/z 776.5208     0   1E+4   12.5   13  RT (min)      lcorbi_t3_pos_3_024_07    m/z 776.5208     0   1   14.5   15  RT (min)      lcorbi_t3_pos_1_034_34    m/z 776.5208     0   1   14.5   15  RT (min)      lcorbi_t3_pos_1_034_34    m/z 776.5210     0   1E+5   14   14.5  RT (min)      lcorbi_t3_pos_1_008_36    m/z 776.7306     0   1   16   16.5  RT (min)      lcorbi_t3_pos_1_030_21    m/z 776.7306     0   1   16.5   17  RT (min)    lcorbi_t3_pos_2_041_54    m/z 776.7306     0   1   17   17.5  RT (min)    lcorbi_t3_pos_1_041_54    m/z 776.7307     0   1   17   17.5  RT (min)    lcorbi_t3_pos_1_049_55    m/z 776.7307     0   2   4   6   8E+3   15.5   16  RT (min)      lcorbi_t3_pos_4_010_43    m/z 776.7308     0   1   17.5   18  RT (min)    lcorbi_t3_pos_1_025_52    m/z 776.7308     0   1E+6   15.5   16  RT (min)      lcorbi_t3_pos_1_005_42    m/z 776.7308     0   1   17.5   18  RT (min)    lcorbi_t3_pos_2_056_56    m/z 776.7308     0   1   17   17.5  RT (min)    lcorbi_t3_pos_1_041_54    m/z 776.7308     0   1   16   16.5  RT (min)      lcorbi_t3_pos_2_046_07    m/z 776.7309     0   1   16.5   17  RT (min)    lcorbi_t3_pos_1_033_53    m/z 776.7309     0   1   16   16.5  RT (min)      lcorbi_t3_pos_1_047_41    m/z 777.5238     0   1   15   15.5  RT (min)      lcorbi_t3_pos_1_031_40    m/z 777.5241     0   1   14.5   15  RT (min)      lcorbi_t3_pos_1_034_34    m/z 777.5243     0   1   14.5   15  RT (min)      lcorbi_t3_pos_1_028_32    m/z 777.6295     0   1E+5   14.5   15  RT (min)      lcorbi_t3_pos_1_030_21    m/z 777.6352     0   1   2E+4   15   15.5  RT (min)      lcorbi_t3_pos_1_049_55    m/z 777.6357     0   1E+6   14   14.5  RT (min)       lcorbi_t3_pos_4_091_60    m/z 777.6358     0   2   4E+5   14   14.5  RT (min)      lcorbi_t3_pos_1_043_06    m/z 777.6360     0   2   4   6   8E+5   14   14.5  RT (min)       lcorbi_t3_pos_4_092_60    m/z 777.6361     0   1   14.5   15  RT (min)      lcorbi_t3_pos_1_041_54    m/z 777.6370     0   1E+5   14   14.5  RT (min)      lcorbi_t3_pos_1_047_41    m/z 778.4538     0   2   4   6E+5   13   13.5  RT (min)      lcorbi_t3_pos_4_087_59    m/z 778.5372     0   1   15.5   16  RT (min)      lcorbi_t3_pos_1_001_49    m/z 778.5376     0   1E+7   14   14.5  RT (min)       lcorbi_t3_pos_1_008_36    m/z 778.5378     0   1E+4   12.5   13  RT (min)      lcorbi_t3_pos_2_013_33    m/z 778.5378     0   1   12.5   13  RT (min)      lcorbi_t3_pos_3_031_45    m/z 778.5380     0   1E+5   14.5   15  RT (min)      lcorbi_t3_pos_1_034_34    m/z 778.5380     0   1   12.5   13  RT (min)      lcorbi_t3_pos_3_024_07    m/z 778.5380     0   1E+5   14.5   15  RT (min)      lcorbi_t3_pos_1_034_34    m/z 778.5380     0   2   4   6   8E+4   15   15.5  RT (min)      lcorbi_t3_pos_2_012_01    m/z 778.5380     0   1E+4   12.5   13  RT (min)      lcorbi_t3_pos_2_013_33    m/z 778.5380     0   2   4   6   8E+4   15.5   16  RT (min)      lcorbi_t3_pos_2_012_01    m/z 778.5380     0   2   4   6E+4   14   14.5  RT (min)      lcorbi_t3_pos_2_022_38    m/z 778.5380     0   1   2   3   4E+7   13   13.5  RT (min)      lcorbi_t3_pos_3_011_42    m/z 778.5381     0   1E+5   15   15.5  RT (min)      lcorbi_t3_pos_1_031_40    m/z 778.5381     0   1E+5   14.5   15  RT (min)      lcorbi_t3_pos_1_028_32    m/z 778.5382     0   1E+5   15   15.5  RT (min)      lcorbi_t3_pos_1_031_40    m/z 778.5382     0   1E+5   14.5   15  RT (min)      lcorbi_t3_pos_1_034_34    m/z 778.5382     0   1E+5   14.5   15  RT (min)      lcorbi_t3_pos_1_028_32    m/z 778.5383     0   1   2   3E+4   15   15.5  RT (min)      lcorbi_t3_pos_2_009_50    m/z 778.5757     0   2   4   6   8E+4   14.5   15  RT (min)      lcorbi_t3_pos_1_028_32    m/z 778.5804     0   1   15   15.5  RT (min)      lcorbi_t3_pos_1_031_40    m/z 778.6231     0   2   4E+5   13   13.5  RT (min)      lcorbi_t3_pos_4_082_59    m/z 778.6415     0   1E+4   15   15.5  RT (min)      lcorbi_t3_pos_1_041_54    m/z 778.6416     0   1   2E+4   14.5   15  RT (min)      lcorbi_t3_pos_1_033_53    m/z 778.6416     0   1   15.5   16  RT (min)      lcorbi_t3_pos_1_001_49    m/z 778.6416     0   1   2E+4   14.5   15  RT (min)      lcorbi_t3_pos_1_047_41    m/z 778.6416     0   1   15   15.5  RT (min)      lcorbi_t3_pos_1_041_54    m/z 778.6417     0   1   2   3E+4   15.5   16  RT (min)      lcorbi_t3_pos_1_025_52    m/z 778.6417     0   1   2E+4   14.5   15  RT (min)      lcorbi_t3_pos_1_021_18    m/z 778.6418     0   1   2E+4   15   15.5   16  RT (min)      lcorbi_t3_pos_1_056_56    m/z 778.6418     0   1   2E+4   14.5   15  RT (min)      lcorbi_t3_pos_1_021_18    m/z 778.6418     0   2   4E+4   14.5   15  RT (min)      lcorbi_t3_pos_2_017_51    m/z 779.5402     0   1   2E+7   13   13.5  RT (min)      lcorbi_t3_pos_3_011_42    m/z 779.5404     0   1   2   3   4E+6   14   14.5  RT (min)       lcorbi_t3_pos_4_001_49    m/z 779.5406     0   1E+7   13.5   14  RT (min)       lcorbi_t3_pos_1_005_42    m/z 779.5409     0   1   2E+4   12.5   13  RT (min)      lcorbi_t3_pos_2_013_33    m/z 779.5410     0   2   4   6E+6   14   14.5  RT (min)       lcorbi_t3_pos_1_008_36    m/z 779.5412     0   1E+4   12.5   13  RT (min)      lcorbi_t3_pos_3_031_45    m/z 779.5412     0   1   2E+4   12.5   13  RT (min)      lcorbi_t3_pos_2_013_33    m/z 779.5413     0   2   4   6   8E+3   12.5   13  RT (min)      lcorbi_t3_pos_3_024_07    m/z 779.5413     0   1   2   3   4E+4   14.5   15  RT (min)      lcorbi_t3_pos_3_003_33    m/z 779.5416     0   1   2   3   4E+4   15   15.5  RT (min)      lcorbi_t3_pos_2_012_01    m/z 779.5416     0   1E+4   14.5   15  RT (min)      lcorbi_t3_pos_2_049_55    m/z 779.5416     0   1   2   3   4E+4   14   14.5  RT (min)      lcorbi_t3_pos_2_022_38    m/z 779.5416     0   2   4E+4   14.5   15  RT (min)      lcorbi_t3_pos_1_034_34    m/z 779.5417     0   2   4   6E+4   15   15.5  RT (min)      lcorbi_t3_pos_1_031_40    m/z 779.5417     0   2   4E+4   14.5   15  RT (min)      lcorbi_t3_pos_1_034_34    m/z 779.5417     0   2   4E+4   14.5   15  RT (min)      lcorbi_t3_pos_1_034_34    m/z 779.5417     0   2   4   6E+4   15   15.5  RT (min)      lcorbi_t3_pos_1_031_40    m/z 779.5418     0   1   2   3   4E+4   15.5   16  RT (min)      lcorbi_t3_pos_2_012_01    m/z 779.5418     0   2   4   6E+4   14.5   15  RT (min)      lcorbi_t3_pos_1_028_32    m/z 779.5420     0   1E+4   15   15.5  RT (min)      lcorbi_t3_pos_2_049_55    m/z 779.6446     0   1E+5   14.5   15  RT (min)      lcorbi_t3_pos_1_047_41    m/z 779.6447     0   1   2E+4   15   15.5   16  RT (min)      lcorbi_t3_pos_1_056_56    m/z 779.6448     0   1E+6   11.5   12  RT (min)      lcorbi_t3_pos_4_057_49    m/z 779.6448     0   1   14.5   15  RT (min)      lcorbi_t3_pos_1_021_18    m/z 779.6449     0   1   2   3   4E+4   14.5   15  RT (min)      lcorbi_t3_pos_1_033_53    m/z 779.6450     0   1   14.5   15  RT (min)      lcorbi_t3_pos_1_021_18    m/z 779.6450     0   1   2   3   4E+4   15   15.5  RT (min)      lcorbi_t3_pos_1_041_54    m/z 779.6452     0   1   15.5   16  RT (min)      lcorbi_t3_pos_1_001_49    m/z 779.6456     0   1   2E+4   15.5   16  RT (min)      lcorbi_t3_pos_1_056_56    m/z 779.6501     0   1E+5   14.5   15  RT (min)       lcorbi_t3_pos_2_017_51    m/z 779.6510     0   1   2E+5   14   14.5  RT (min)       lcorbi_t3_pos_3_056_56    m/z 779.6510     0   1   2   3E+4   15   15.5  RT (min)      lcorbi_t3_pos_1_041_54    m/z 780.4690     0   2   4E+5   13.5   14  RT (min)      lcorbi_t3_pos_4_082_59    m/z 780.4790     0   1   14   14.5  RT (min)      lcorbi_t3_pos_1_028_32    m/z 780.5449     0   1E+4   12.5   13  RT (min)      lcorbi_t3_pos_3_031_45    m/z 780.5450     0   1E+4   12.5   13  RT (min)      lcorbi_t3_pos_3_031_45    m/z 780.5483     0   1   2   3E+4   12.5   13  RT (min)      lcorbi_t3_pos_3_031_45    m/z 780.5489     0   1E+6   14   14.5  RT (min)      lcorbi_t3_pos_2_046_07    m/z 780.5522     0   1   2E+7   14   14.5  RT (min)      lcorbi_t3_pos_4_025_52    m/z 780.5523     0   1   2   3   4E+4   16   16.5  RT (min)      lcorbi_t3_pos_2_056_56    m/z 780.5525     0   1   2E+5   15   15.5  RT (min)      lcorbi_t3_pos_1_034_34    m/z 780.5526     0   1   2E+7   14   14.5  RT (min)      lcorbi_t3_pos_1_008_36    m/z 780.5526     0   2   4E+7   14   14.5  RT (min)      lcorbi_t3_pos_1_005_42    m/z 780.5526     0   2   4   6E+4   15   15.5  RT (min)      lcorbi_t3_pos_1_049_55    m/z 780.5526     0   1   16.5   17  RT (min)    lcorbi_t3_pos_1_023_02    m/z 780.5528     0   1   2E+5   14.5   15  RT (min)      lcorbi_t3_pos_2_022_38    m/z 780.5529     0   1E+5   15.5   16  RT (min)      lcorbi_t3_pos_2_012_01    m/z 780.5532     0   1   2E+5   15   15.5  RT (min)       lcorbi_t3_pos_1_034_34    m/z 780.5533     0   1   2E+4   16   16.5  RT (min)      lcorbi_t3_pos_1_023_02    m/z 780.5533     0   1E+5   15.5   16  RT (min)      lcorbi_t3_pos_2_012_01    m/z 780.5534     0   1   2E+5   15   15.5  RT (min)      lcorbi_t3_pos_1_034_34    m/z 780.5534     0   1E+5   15.5   16  RT (min)      lcorbi_t3_pos_1_031_40    m/z 780.5534     0   1   2E+5   14.5   15  RT (min)      lcorbi_t3_pos_1_028_32    m/z 780.5535     0   2   4   6E+4   14.5   15  RT (min)      lcorbi_t3_pos_2_038_03    m/z 780.5536     0   2   4   6E+6   11.5   12  RT (min)      lcorbi_t3_pos_4_057_49    m/z 780.5536     0   1   2   3E+7   13   13.5  RT (min)      lcorbi_t3_pos_3_011_42    m/z 780.5537     0   1E+5   15   15.5  RT (min)      lcorbi_t3_pos_1_031_40    m/z 780.5537     0   1   2E+5   14.5   15  RT (min)      lcorbi_t3_pos_2_022_38    m/z 780.5537     0   1E+5   15.5   16  RT (min)       lcorbi_t3_pos_1_031_40    m/z 780.5538     0   2   4E+4   12.5   13  RT (min)       lcorbi_t3_pos_2_013_33    m/z 780.5539     0   2   4   6E+4   13   13.5  RT (min)       lcorbi_t3_pos_3_011_42    m/z 780.5962     0   2   4E+4   14.5   15  RT (min)      lcorbi_t3_pos_1_028_32    m/z 780.5962     0   1   2E+5   14   14.5  RT (min)      lcorbi_t3_pos_1_008_36    m/z 780.5964     0   1   15.5   16  RT (min)      lcorbi_t3_pos_1_031_40    m/z 780.5964     0   1   15   15.5  RT (min)      lcorbi_t3_pos_1_031_40    m/z 780.5965     0   2   4E+4   14.5   15  RT (min)      lcorbi_t3_pos_1_028_32    m/z 780.5965     0   1   2   3E+4   15   15.5  RT (min)      lcorbi_t3_pos_1_034_34    m/z 780.5966     0   2   4   6   8E+4   15   15.5  RT (min)      lcorbi_t3_pos_1_034_34    m/z 780.6391     0   2   4   6E+5   13.5   14  RT (min)      lcorbi_t3_pos_4_082_59    m/z 780.6510     0   1E+4   14.5   15  RT (min)      lcorbi_t3_pos_2_017_51    m/z 780.6534     0   1E+4   15.5   16  RT (min)      lcorbi_t3_pos_2_009_50    m/z 780.6570     0   1   14.5   15  RT (min)      lcorbi_t3_pos_1_033_53    m/z 780.6572     0   1   15   15.5  RT (min)      lcorbi_t3_pos_1_021_18    m/z 780.6572     0   1   15   15.5  RT (min)      lcorbi_t3_pos_1_041_54    m/z 780.6572     0   1   15.5   16  RT (min)      lcorbi_t3_pos_1_049_55    m/z 780.6572     0   1   15.5   16  RT (min)      lcorbi_t3_pos_1_001_49    m/z 780.6573     0   2   4   6E+5   14   14.5  RT (min)      lcorbi_t3_pos_1_005_42    m/z 780.6574     0   2   4   6E+5   14   14.5  RT (min)       lcorbi_t3_pos_1_005_42    m/z 781.4594     0   1   17.5   18  RT (min)    lcorbi_t3_pos_2_056_56    m/z 781.4595     0   1E+4   13.5   14  RT (min)      lcorbi_t3_pos_2_047_31    m/z 781.4596     0   1   17.5   18  RT (min)    lcorbi_t3_pos_2_056_56    m/z 781.4596     0   1   18   18.5  RT (min)    lcorbi_t3_pos_1_023_02    m/z 781.4596     0   1   16.5   17  RT (min)    lcorbi_t3_pos_2_041_54    m/z 781.4596     0   1   16.5   17  RT (min)    lcorbi_t3_pos_1_041_54    m/z 781.4596     0   1   17   17.5  RT (min)    lcorbi_t3_pos_1_049_55    m/z 781.4596     0   1   16.5   17  RT (min)    lcorbi_t3_pos_2_025_52    m/z 781.4597     0   1E+4   16   16.5  RT (min)      lcorbi_t3_pos_4_041_54    m/z 781.4597     0   1   2E+6   14.5   15  RT (min)       lcorbi_t3_pos_3_053_48    m/z 781.4597     0   1   2   3E+4   14   14.5  RT (min)      lcorbi_t3_pos_3_052_47    m/z 781.4597     0   1   2E+4   14   14.5  RT (min)       lcorbi_t3_pos_2_054_29    m/z 781.4597     0   1   2E+6   14.5   15  RT (min)      lcorbi_t3_pos_3_053_48    m/z 781.4598     0   1   17   17.5  RT (min)    lcorbi_t3_pos_1_025_52    m/z 781.4598     0   1   2   3   4E+6   15.5   16  RT (min)      lcorbi_t3_pos_1_005_42    m/z 781.4598     0   1   17   17.5  RT (min)    lcorbi_t3_pos_2_009_50    m/z 781.4598     0   2   4   6E+4   15.5   16  RT (min)      lcorbi_t3_pos_3_025_52    m/z 781.4598     0   1   2   3   4E+6   15   15.5  RT (min)       lcorbi_t3_pos_3_041_54    m/z 781.4598     0   1   2E+4   14   14.5  RT (min)      lcorbi_t3_pos_2_032_39    m/z 781.4598     0   1   18   18.5  RT (min)    lcorbi_t3_pos_1_023_02    m/z 781.4600     0   1   18   18.5  RT (min)    lcorbi_t3_pos_1_023_02    m/z 781.5515     0   1E+4   12.5   13  RT (min)      lcorbi_t3_pos_3_024_07    m/z 781.5564     0   1   16   16.5  RT (min)      lcorbi_t3_pos_1_023_02    m/z 781.5564     0   1   16.5   17  RT (min)    lcorbi_t3_pos_1_023_02    m/z 781.5564     0   1   2   3E+4   15.5   16  RT (min)      lcorbi_t3_pos_1_051_09    m/z 781.5564     0   2   4   6E+4   15.5   16  RT (min)      lcorbi_t3_pos_2_012_01    m/z 781.5566     0   1   2   3E+4   16   16.5  RT (min)      lcorbi_t3_pos_1_023_02    m/z 781.5567     0   2   4   6   8E+4   12.5   13  RT (min)      lcorbi_t3_pos_2_013_33    m/z 781.5567     0   2   4   6E+4   15.5   16  RT (min)      lcorbi_t3_pos_2_012_01    m/z 781.5568     0   1E+5   15   15.5  RT (min)      lcorbi_t3_pos_1_034_34    m/z 781.5570     0   1E+7   14   14.5  RT (min)       lcorbi_t3_pos_1_008_36    m/z 781.5572     0   1E+5   14.5   15  RT (min)      lcorbi_t3_pos_2_022_38    m/z 781.5572     0   1   2   3E+6   11.5   12  RT (min)      lcorbi_t3_pos_4_057_49    m/z 781.5572     0   1E+5   15.5   16  RT (min)      lcorbi_t3_pos_1_031_40    m/z 781.5572     0   2   4   6E+4   15.5   16  RT (min)      lcorbi_t3_pos_2_012_01    m/z 781.5572     0   2   4   6E+4   14.5   15  RT (min)      lcorbi_t3_pos_1_047_41    m/z 781.5572     0   1E+5   15   15.5  RT (min)      lcorbi_t3_pos_1_034_34    m/z 781.5572     0   1E+5   14.5   15  RT (min)      lcorbi_t3_pos_1_028_32    m/z 781.5573     0   1E+5   14.5   15  RT (min)      lcorbi_t3_pos_2_022_38    m/z 781.5573     0   1   2   3   4E+4   14.5   15  RT (min)      lcorbi_t3_pos_2_038_03    m/z 781.5573     0   1E+5   15   15.5  RT (min)      lcorbi_t3_pos_1_034_34    m/z 781.5573     0   1E+5   15   15.5  RT (min)      lcorbi_t3_pos_1_031_40    m/z 781.5574     0   2   4E+4   14.5   15  RT (min)      lcorbi_t3_pos_4_048_24    m/z 781.5574     0   1   2   3   4E+4   15   15.5  RT (min)      lcorbi_t3_pos_1_049_55    m/z 781.5575     0   1   2E+4   13   13.5  RT (min)      lcorbi_t3_pos_2_013_33    m/z 781.5575     0   1   2E+4   13   13.5  RT (min)      lcorbi_t3_pos_2_013_33    m/z 781.5576     0   1   2   3E+4   15   15.5  RT (min)      lcorbi_t3_pos_1_041_54    m/z 781.6602     0   1E+6   12.5   13  RT (min)      lcorbi_t3_pos_4_057_49    m/z 781.6605     0   1E+4   14.5   15  RT (min)      lcorbi_t3_pos_2_006_47    m/z 781.6605     0   1   2E+6   14   14.5  RT (min)      lcorbi_t3_pos_1_005_42    m/z 781.6606     0   1   14.5   15  RT (min)      lcorbi_t3_pos_1_033_53    m/z 781.6606     0   1   15.5   16  RT (min)      lcorbi_t3_pos_1_025_52    m/z 781.6606     0   2   4   6   8E+4   14.5   15  RT (min)      lcorbi_t3_pos_3_056_56    m/z 781.6606     0   1   15.5   16  RT (min)      lcorbi_t3_pos_1_001_49    m/z 781.6607     0   1   2   3E+4   14.5   15  RT (min)      lcorbi_t3_pos_2_017_51    m/z 781.6607     0   1   16   16.5  RT (min)      lcorbi_t3_pos_2_056_56    m/z 781.6607     0   1   15.5   16  RT (min)      lcorbi_t3_pos_1_056_56    m/z 781.6608     0   2   4   6   8E+6   13   13.5  RT (min)       lcorbi_t3_pos_1_036_08    m/z 781.6608     0   1   15   15.5  RT (min)      lcorbi_t3_pos_1_041_54    m/z 781.6608     0   1   15   15.5  RT (min)      lcorbi_t3_pos_1_033_53    m/z 781.6608     0   1   15   15.5  RT (min)      lcorbi_t3_pos_1_033_53    m/z 781.6610     0   1E+4   15   15.5  RT (min)      lcorbi_t3_pos_2_049_55    m/z 782.4606     0   1   13.5   14  RT (min)      lcorbi_t3_pos_2_047_31    m/z 782.4628     0   1   16.5   17  RT (min)    lcorbi_t3_pos_2_025_52    m/z 782.4628     0   1   17   17.5  RT (min)    lcorbi_t3_pos_1_049_55    m/z 782.4629     0   1   16   16.5  RT (min)      lcorbi_t3_pos_1_028_32    m/z 782.4629     0   1E+4   14   14.5  RT (min)      lcorbi_t3_pos_2_032_39    m/z 782.4629     0   1E+5   15.5   16  RT (min)      lcorbi_t3_pos_4_025_52    m/z 782.4630     0   1   17   17.5  RT (min)    lcorbi_t3_pos_2_009_50    m/z 782.4630     0   1E+6   14.5   15  RT (min)      lcorbi_t3_pos_3_053_48    m/z 782.4630     0   1   16   16.5  RT (min)      lcorbi_t3_pos_1_030_21    m/z 782.4630     0   1E+6   15.5   16  RT (min)      lcorbi_t3_pos_1_005_42    m/z 782.4630     0   1E+4   14   14.5  RT (min)      lcorbi_t3_pos_2_054_29    m/z 782.4630     0   1   16   16.5  RT (min)      lcorbi_t3_pos_4_041_54    m/z 782.4630     0   1   17.5   18  RT (min)    lcorbi_t3_pos_2_056_56    m/z 782.4630     0   1   17   17.5  RT (min)    lcorbi_t3_pos_1_025_52    m/z 782.4631     0   1E+6   14.5   15  RT (min)      lcorbi_t3_pos_3_053_48    m/z 782.4632     0   1   2E+4   15.5   16  RT (min)      lcorbi_t3_pos_3_025_52    m/z 782.4632     0   1   16.5   17  RT (min)    lcorbi_t3_pos_1_041_54    m/z 782.4632     0   1E+6   15.5   16  RT (min)      lcorbi_t3_pos_3_041_54    m/z 782.4632     0   1   18   18.5  RT (min)    lcorbi_t3_pos_1_023_02    m/z 782.4633     0   1   16   16.5  RT (min)      lcorbi_t3_pos_2_017_51    m/z 782.4635     0   1   2E+6   14.5   15  RT (min)      lcorbi_t3_pos_4_105_61    m/z 782.4946     0   2   4   6E+5   13.5   14  RT (min)      lcorbi_t3_pos_1_005_42    m/z 782.4947     0   1   2   3   4E+4   15   15.5  RT (min)      lcorbi_t3_pos_1_031_40    m/z 782.4947     0   1   2E+4   15.5   16  RT (min)      lcorbi_t3_pos_2_012_01    m/z 782.4948     0   1   2   3E+4   14.5   15  RT (min)      lcorbi_t3_pos_1_028_32    m/z 782.4948     0   1   2E+4   15.5   16  RT (min)      lcorbi_t3_pos_2_012_01    m/z 782.4948     0   2   4E+4   14.5   15  RT (min)      lcorbi_t3_pos_1_034_34    m/z 782.4948     0   1   2E+4   14.5   15  RT (min)      lcorbi_t3_pos_2_041_54    m/z 782.4948     0   1   2   3E+4   14.5   15  RT (min)      lcorbi_t3_pos_1_028_32    m/z 782.4948     0   1   2   3E+5   12.5   13  RT (min)      lcorbi_t3_pos_3_024_07    m/z 782.4949     0   1   2   3   4E+4   15   15.5  RT (min)      lcorbi_t3_pos_1_031_40    m/z 782.4951     0   2   4E+4   14.5   15  RT (min)      lcorbi_t3_pos_1_034_34    m/z 782.4952     0   1   2   3   4E+4   14   14.5  RT (min)      lcorbi_t3_pos_1_047_41    m/z 782.4954     0   1E+5   14   14.5  RT (min)      lcorbi_t3_pos_1_008_36    m/z 782.4956     0   1   2   3   4E+5   13   13.5  RT (min)      lcorbi_t3_pos_3_011_42    m/z 782.4962     0   1   2   3   4E+5   13   13.5  RT (min)      lcorbi_t3_pos_3_011_42    m/z 782.5551     0   1   12.5   13  RT (min)      lcorbi_t3_pos_3_031_45    m/z 782.5557     0   1   12.5   13  RT (min)      lcorbi_t3_pos_3_031_45    m/z 782.5599     0   1   2   3E+4   12.5   13  RT (min)      lcorbi_t3_pos_2_013_33    m/z 782.5606     0   2   4   6   8E+5   11.5   12  RT (min)      lcorbi_t3_pos_4_057_49    m/z 782.5608     0   1E+6   13   13.5  RT (min)      lcorbi_t3_pos_1_002_01    m/z 782.5608     0   1E+5   15   15.5  RT (min)      lcorbi_t3_pos_1_041_54    m/z 782.5678     0   2   4   6   8E+6   13   13.5  RT (min)      lcorbi_t3_pos_3_011_42    m/z 782.5684     0   1   2E+6   13   13.5  RT (min)      lcorbi_t3_pos_2_013_33    m/z 782.5684     0   2   4   6E+4   13   13.5  RT (min)       lcorbi_t3_pos_2_013_33    m/z 782.5684     0   1   2E+4   13   13.5  RT (min)      lcorbi_t3_pos_3_031_45    m/z 782.5685     0   1E+7   13   13.5  RT (min)      lcorbi_t3_pos_1_002_01    m/z 782.5685     0   2   4   6E+6   14.5   15  RT (min)      lcorbi_t3_pos_1_008_36    m/z 782.5686     0   1   2   3E+7   13   13.5  RT (min)      lcorbi_t3_pos_1_002_01    m/z 782.5690     0   1   2   3   4E+5   14.5   15  RT (min)      lcorbi_t3_pos_1_028_32    m/z 782.5691     0   1   2   3   4E+7   14   14.5  RT (min)      lcorbi_t3_pos_1_005_42    m/z 782.5691     0   1   2E+5   15   15.5  RT (min)      lcorbi_t3_pos_1_034_34    m/z 782.5692     0   1   2E+5   15.5   16  RT (min)      lcorbi_t3_pos_1_031_40    m/z 782.5692     0   1   16.5   17  RT (min)    lcorbi_t3_pos_1_023_02    m/z 782.5692     0   1   2E+5   15.5   16  RT (min)      lcorbi_t3_pos_1_031_40    m/z 782.5692     0   1E+5   15.5   16  RT (min)      lcorbi_t3_pos_1_049_55    m/z 782.5692     0   1   2E+5   15   15.5  RT (min)      lcorbi_t3_pos_1_034_34    m/z 782.5692     0   1E+7   12   12.5  RT (min)      lcorbi_t3_pos_4_057_49    m/z 782.5692     0   2   4E+5   14.5   15  RT (min)      lcorbi_t3_pos_1_047_41    m/z 782.5694     0   2   4   6E+4   16   16.5  RT (min)      lcorbi_t3_pos_2_056_56    m/z 782.5694     0   1   2E+5   16   16.5  RT (min)      lcorbi_t3_pos_1_051_09    m/z 782.5694     0   1E+5   15.5   16  RT (min)       lcorbi_t3_pos_2_012_01    m/z 782.5694     0   1   2E+5   15   15.5  RT (min)      lcorbi_t3_pos_1_034_34    m/z 782.5695     0   1E+5   15.5   16  RT (min)      lcorbi_t3_pos_2_012_01    m/z 782.5696     0   1E+5   16   16.5  RT (min)      lcorbi_t3_pos_1_023_02    m/z 782.5698     0   2   4E+7   14   14.5  RT (min)       lcorbi_t3_pos_1_008_36    m/z 782.5698     0   2   4   6E+4   16   16.5  RT (min)      lcorbi_t3_pos_1_023_02    m/z 782.6120     0   1   14.5   15  RT (min)      lcorbi_t3_pos_1_028_32    m/z 782.6544     0   2   4E+5   13.5   14  RT (min)      lcorbi_t3_pos_4_087_59    m/z 782.6727     0   1   15.5   16  RT (min)      lcorbi_t3_pos_2_009_50    m/z 782.6728     0   1   14.5   15  RT (min)      lcorbi_t3_pos_2_017_51    m/z 782.6729     0   1   15   15.5  RT (min)      lcorbi_t3_pos_1_021_18    m/z 782.6729     0   2   4   6E+5   13.5   14  RT (min)       lcorbi_t3_pos_1_036_08    m/z 782.6729     0   2   4   6E+5   14   14.5  RT (min)      lcorbi_t3_pos_1_005_42    m/z 782.6730     0   1   16   16.5  RT (min)      lcorbi_t3_pos_1_001_49    m/z 782.6730     0   1   15.5   16  RT (min)      lcorbi_t3_pos_1_041_54    m/z 782.6730     0   1   15   15.5  RT (min)      lcorbi_t3_pos_1_030_21    m/z 782.6730     0   2   4   6E+5   14.5   15  RT (min)      lcorbi_t3_pos_1_005_42    m/z 782.6731     0   1   2E+4   16   16.5  RT (min)      lcorbi_t3_pos_1_025_52    m/z 782.6732     0   1   15.5   16  RT (min)      lcorbi_t3_pos_1_056_56    m/z 783.4650     0   2   4   6E+5   14.5   15  RT (min)      lcorbi_t3_pos_4_105_61    m/z 783.4979     0   1E+5   12.5   13  RT (min)      lcorbi_t3_pos_3_024_07    m/z 783.4980     0   1   15.5   16  RT (min)      lcorbi_t3_pos_2_012_01    m/z 783.4981     0   1   2   3   4E+4   15   15.5  RT (min)      lcorbi_t3_pos_1_031_40    m/z 783.4981     0   1E+5   12.5   13  RT (min)      lcorbi_t3_pos_2_013_33    m/z 783.4982     0   1   14.5   15  RT (min)      lcorbi_t3_pos_1_028_32    m/z 783.4982     0   1   14.5   15  RT (min)      lcorbi_t3_pos_1_034_34    m/z 783.4982     0   1   2   3   4E+4   15   15.5  RT (min)      lcorbi_t3_pos_1_031_40    m/z 783.4982     0   1   14.5   15  RT (min)      lcorbi_t3_pos_1_028_32    m/z 783.4986     0   1   14.5   15  RT (min)      lcorbi_t3_pos_1_034_34    m/z 783.4987     0   1E+5   14   14.5  RT (min)      lcorbi_t3_pos_1_008_36    m/z 783.5628     0   1E+4   12.5   13  RT (min)      lcorbi_t3_pos_3_031_45    m/z 783.5632     0   1E+4   12.5   13  RT (min)      lcorbi_t3_pos_3_031_45    m/z 783.5633     0   2   4E+4   15.5   16  RT (min)      lcorbi_t3_pos_1_031_40    m/z 783.5633     0   1E+4   12.5   13  RT (min)      lcorbi_t3_pos_3_031_45    m/z 783.5717     0   2   4   6E+4   13   13.5  RT (min)      lcorbi_t3_pos_3_031_45    m/z 783.5718     0   2   4   6E+6   13   13.5  RT (min)      lcorbi_t3_pos_1_002_01    m/z 783.5718     0   1E+5   13   13.5  RT (min)      lcorbi_t3_pos_2_013_33    m/z 783.5722     0   2   4   6E+4   13   13.5  RT (min)       lcorbi_t3_pos_2_013_33    m/z 783.5724     0   1E+7   13   13.5  RT (min)      lcorbi_t3_pos_1_002_01    m/z 783.5726     0   1E+7   14   14.5  RT (min)      lcorbi_t3_pos_3_041_54    m/z 783.5726     0   1E+5   15   15.5  RT (min)      lcorbi_t3_pos_1_034_34    m/z 783.5727     0   1   2   3E+4   16   16.5  RT (min)      lcorbi_t3_pos_2_056_56    m/z 783.5727     0   1   16.5   17  RT (min)    lcorbi_t3_pos_1_023_02    m/z 783.5727     0   2   4   6E+6   12   12.5  RT (min)      lcorbi_t3_pos_4_057_49    m/z 783.5728     0   2   4   6E+4   16   16.5  RT (min)      lcorbi_t3_pos_1_023_02    m/z 783.5728     0   1E+5   15.5   16  RT (min)      lcorbi_t3_pos_1_031_40    m/z 783.5728     0   1   2E+5   14.5   15  RT (min)      lcorbi_t3_pos_1_028_32    m/z 783.5728     0   1   2E+7   14   14.5  RT (min)       lcorbi_t3_pos_1_005_42    m/z 783.5729     0   1E+5   15.5   16  RT (min)      lcorbi_t3_pos_1_031_40    m/z 783.5729     0   1   2   3E+6   14.5   15  RT (min)      lcorbi_t3_pos_1_008_36    m/z 783.5729     0   1E+5   15.5   16  RT (min)      lcorbi_t3_pos_2_012_01    m/z 783.5730     0   2   4E+4   15   15.5  RT (min)      lcorbi_t3_pos_2_025_52    m/z 783.5730     0   1   2   3E+6   14.5   15  RT (min)      lcorbi_t3_pos_1_008_36    m/z 783.5730     0   1E+5   15.5   16  RT (min)      lcorbi_t3_pos_2_012_01    m/z 783.5730     0   2   4   6   8E+4   16   16.5  RT (min)      lcorbi_t3_pos_1_051_09    m/z 783.5730     0   1E+5   15   15.5  RT (min)      lcorbi_t3_pos_1_034_34    m/z 783.5730     0   1   2E+7   14   14.5  RT (min)      lcorbi_t3_pos_1_008_36    m/z 783.5730     0   2   4   6E+4   16   16.5  RT (min)      lcorbi_t3_pos_1_023_02    m/z 783.5730     0   1E+5   15   15.5  RT (min)      lcorbi_t3_pos_1_034_34    m/z 783.5730     0   2   4E+4   15   15.5  RT (min)      lcorbi_t3_pos_2_038_03    m/z 783.5732     0   2   4E+6   14   14.5  RT (min)       lcorbi_t3_pos_1_008_36    m/z 783.5741     0   2   4   6E+6   13   13.5  RT (min)       lcorbi_t3_pos_1_002_01    m/z 783.6758     0   1E+6   13   13.5  RT (min)      lcorbi_t3_pos_4_057_49    m/z 783.6760     0   1E+6   13   13.5  RT (min)      lcorbi_t3_pos_4_095_60    m/z 783.6760     0   1   2   3E+4   15.5   16  RT (min)      lcorbi_t3_pos_1_025_52    m/z 783.6760     0   1   2E+6   14   14.5  RT (min)      lcorbi_t3_pos_1_005_42    m/z 783.6762     0   2   4   6E+6   13.5   14  RT (min)       lcorbi_t3_pos_4_091_60    m/z 783.6762     0   1   2E+6   14.5   15  RT (min)      lcorbi_t3_pos_1_005_42    m/z 783.6762     0   1   16   16.5  RT (min)      lcorbi_t3_pos_1_001_49    m/z 783.6762     0   1   15.5   16  RT (min)      lcorbi_t3_pos_1_041_54    m/z 783.6762     0   1E+4   15.5   16  RT (min)      lcorbi_t3_pos_2_009_50    m/z 783.6762     0   1   16   16.5  RT (min)      lcorbi_t3_pos_1_001_49    m/z 783.6762     0   1   2E+6   14.5   15  RT (min)      lcorbi_t3_pos_1_005_42    m/z 783.6762     0   1   2   3E+4   14.5   15  RT (min)      lcorbi_t3_pos_2_017_51    m/z 783.6762     0   2   4E+5   14   14.5  RT (min)      lcorbi_t3_pos_3_040_41    m/z 783.6762     0   1   2E+4   14.5   15  RT (min)      lcorbi_t3_pos_2_017_51    m/z 783.6763     0   1E+6   13.5   14  RT (min)      lcorbi_t3_pos_4_093_60    m/z 783.6763     0   1   15   15.5  RT (min)      lcorbi_t3_pos_2_041_54    m/z 783.6763     0   1E+4   15   15.5  RT (min)      lcorbi_t3_pos_1_047_41    m/z 783.6763     0   1   2   3E+4   16   16.5  RT (min)      lcorbi_t3_pos_1_025_52    m/z 783.6764     0   1E+4   15   15.5  RT (min)      lcorbi_t3_pos_2_038_03    m/z 783.6764     0   1   15   15.5  RT (min)      lcorbi_t3_pos_1_030_21    m/z 783.6764     0   1   15   15.5  RT (min)      lcorbi_t3_pos_1_033_53    m/z 783.6764     0   2   4   6E+4   14.5   15  RT (min)      lcorbi_t3_pos_1_030_21    m/z 783.6764     0   1E+4   15   15.5  RT (min)      lcorbi_t3_pos_2_006_47    m/z 783.6764     0   1   2E+6   13.5   14  RT (min)       lcorbi_t3_pos_1_036_08    m/z 783.6764     0   1   15.5   16  RT (min)      lcorbi_t3_pos_1_056_56    m/z 784.5108     0   1   2   3E+5   14   14.5  RT (min)      lcorbi_t3_pos_1_008_36    m/z 784.5111     0   1   2E+4   14.5   15  RT (min)      lcorbi_t3_pos_1_034_34    m/z 784.5117     0   1   2E+4   14.5   15  RT (min)      lcorbi_t3_pos_1_034_34    m/z 784.5121     0   1E+4   15   15.5  RT (min)      lcorbi_t3_pos_2_012_01    m/z 784.5122     0   1   15   15.5  RT (min)      lcorbi_t3_pos_1_031_40    m/z 784.5124     0   1E+4   14   14.5  RT (min)      lcorbi_t3_pos_2_022_38    m/z 784.5125     0   2   4E+4   12.5   13  RT (min)      lcorbi_t3_pos_2_013_33    m/z 784.5126     0   1   15   15.5  RT (min)      lcorbi_t3_pos_1_031_40    m/z 784.5129     0   1   2   3E+5   14   14.5  RT (min)      lcorbi_t3_pos_1_008_36    m/z 784.5664     0   1   13   13.5  RT (min)      lcorbi_t3_pos_3_031_45    m/z 784.5665     0   1   14.5   15  RT (min)      lcorbi_t3_pos_3_049_55    m/z 784.5753     0   1   2   3E+4   13   13.5  RT (min)      lcorbi_t3_pos_2_032_39    m/z 784.5764     0   1E+7   13   13.5  RT (min)      lcorbi_t3_pos_4_057_49    m/z 784.5798     0   1   2E+6   13   13.5  RT (min)      lcorbi_t3_pos_2_013_33    m/z 784.5834     0   1   2   3   4E+4   15.5   16  RT (min)      lcorbi_t3_pos_4_033_53    m/z 784.5843     0   1   2E+6   13   13.5  RT (min)       lcorbi_t3_pos_1_002_01    m/z 784.5843     0   2   4E+4   15.5   16  RT (min)      lcorbi_t3_pos_2_049_55    m/z 784.5846     0   1   2   3   4E+4   16   16.5  RT (min)      lcorbi_t3_pos_2_056_56    m/z 784.5846     0   2   4   6   8E+4   15   15.5  RT (min)      lcorbi_t3_pos_1_033_53    m/z 784.5846     0   1E+5   15.5   16  RT (min)      lcorbi_t3_pos_1_031_40    m/z 784.5846     0   2   4   6   8E+6   13.5   14  RT (min)      lcorbi_t3_pos_4_016_15    m/z 784.5847     0   2   4   6E+4   15.5   16  RT (min)      lcorbi_t3_pos_1_041_54    m/z 784.5847     0   1   2E+5   15   15.5  RT (min)      lcorbi_t3_pos_2_006_47    m/z 784.5847     0   1   2   3   4E+4   16   16.5  RT (min)      lcorbi_t3_pos_2_056_56    m/z 784.5847     0   1E+5   16   16.5  RT (min)      lcorbi_t3_pos_1_051_09    m/z 784.5847     0   1   16.5   17  RT (min)    lcorbi_t3_pos_1_023_02    m/z 784.5848     0   1E+6   14.5   15  RT (min)      lcorbi_t3_pos_4_010_43    m/z 784.5848     0   1   2   3   4E+5   14.5   15  RT (min)      lcorbi_t3_pos_2_017_51    m/z 784.5849     0   1E+5   15   15.5  RT (min)      lcorbi_t3_pos_1_030_21    m/z 784.5850     0   1   16.5   17  RT (min)     lcorbi_t3_pos_1_023_02    m/z 784.5850     0   1   16.5   17  RT (min)    lcorbi_t3_pos_1_023_02    m/z 784.5850     0   1E+5   15   15.5  RT (min)      lcorbi_t3_pos_1_028_32    m/z 784.5850     0   2   4   6   8E+5   14.5   15  RT (min)      lcorbi_t3_pos_2_046_07    m/z 784.5850     0   2   4   6E+4   15   15.5  RT (min)      lcorbi_t3_pos_1_041_54    m/z 784.5850     0   1E+5   13   13.5  RT (min)      lcorbi_t3_pos_1_002_01    m/z 784.5851     0   1   2   3   4E+7   14   14.5  RT (min)      lcorbi_t3_pos_1_005_42    m/z 784.5851     0   1   2   3E+5   14.5   15  RT (min)      lcorbi_t3_pos_2_046_07    m/z 784.5852     0   2   4   6E+6   14.5   15  RT (min)      lcorbi_t3_pos_3_047_40    m/z 784.5852     0   1   2   3   4E+7   14   15  RT (min)       lcorbi_t3_pos_1_005_42    m/z 784.5852     0   1   16.5   17  RT (min)    lcorbi_t3_pos_2_012_01    m/z 784.5856     0   1   16.5   17  RT (min)    lcorbi_t3_pos_1_023_02    m/z 784.6880     0   1   15.5   16  RT (min)      lcorbi_t3_pos_2_041_54    m/z 784.6881     0   1   15.5   16  RT (min)      lcorbi_t3_pos_2_009_50    m/z 784.6884     0   1   15   15.5  RT (min)      lcorbi_t3_pos_1_047_41    m/z 784.6884     0   1   15.5   16  RT (min)      lcorbi_t3_pos_1_041_54    m/z 784.6884     0   1   2   3E+5   14.5   15  RT (min)      lcorbi_t3_pos_4_025_52    m/z 784.6884     0   1   2   3   4E+4   14.5   15  RT (min)      lcorbi_t3_pos_3_056_56    m/z 784.6885     0   1   15.5   16  RT (min)      lcorbi_t3_pos_1_041_54    m/z 784.6885     0   1   2   3E+6   13.5   14  RT (min)      lcorbi_t3_pos_1_003_45    m/z 784.6885     0   1E+6   14.5   15  RT (min)      lcorbi_t3_pos_1_005_42    m/z 784.6885     0   1   2   3   4E+6   13.5   14  RT (min)       lcorbi_t3_pos_4_093_60    m/z 784.6886     0   1   15   15.5  RT (min)      lcorbi_t3_pos_1_047_41    m/z 784.6886     0   1   15   15.5  RT (min)      lcorbi_t3_pos_1_047_41    m/z 784.6886     0   1   16   16.5  RT (min)      lcorbi_t3_pos_1_001_49    m/z 784.6886     0   1   15   15.5  RT (min)      lcorbi_t3_pos_1_021_18    m/z 784.6887     0   1   15.5   16  RT (min)      lcorbi_t3_pos_2_009_50    m/z 784.6887     0   1E+6   14.5   15  RT (min)      lcorbi_t3_pos_1_005_42    m/z 785.5790     0   1   2E+4   13   13.5  RT (min)      lcorbi_t3_pos_2_032_39    m/z 785.5794     0   1   2E+5   13   13.5  RT (min)      lcorbi_t3_pos_4_057_49    m/z 785.5874     0   1   2   3E+4   15.5   16  RT (min)      lcorbi_t3_pos_4_033_53    m/z 785.5876     0   1E+6   13   13.5  RT (min)      lcorbi_t3_pos_1_002_01    m/z 785.5879     0   1   2   3E+4   13   13.5  RT (min)      lcorbi_t3_pos_1_002_01    m/z 785.5882     0   1E+5   15   15.5  RT (min)      lcorbi_t3_pos_2_006_47    m/z 785.5883     0   2   4   6   8E+4   15.5   16  RT (min)      lcorbi_t3_pos_1_031_40    m/z 785.5884     0   2   4   6   8E+4   15   15.5  RT (min)      lcorbi_t3_pos_1_030_21    m/z 785.5884     0   1E+5   15   15.5  RT (min)       lcorbi_t3_pos_1_028_32    m/z 785.5885     0   1   16.5   17  RT (min)    lcorbi_t3_pos_1_023_02    m/z 785.5886     0   1E+6   14.5   15  RT (min)      lcorbi_t3_pos_1_008_36    m/z 785.5886     0   1   2E+5   14.5   15  RT (min)      lcorbi_t3_pos_2_017_51    m/z 785.5886     0   2   4E+4   16   16.5  RT (min)      lcorbi_t3_pos_1_051_09    m/z 785.5886     0   2   4E+4   15   15.5  RT (min)      lcorbi_t3_pos_1_033_53    m/z 785.5886     0   1   2   3E+4   15.5   16  RT (min)      lcorbi_t3_pos_2_049_55    m/z 785.5887     0   1   16.5   17  RT (min)    lcorbi_t3_pos_1_023_02    m/z 785.5887     0   1   2   3   4E+6   13.5   14  RT (min)      lcorbi_t3_pos_4_016_15    m/z 785.5888     0   1   2E+7   14   14.5  RT (min)      lcorbi_t3_pos_1_005_42    m/z 785.5888     0   1E+4   16   16.5  RT (min)      lcorbi_t3_pos_2_056_56    m/z 785.5888     0   1   2   3E+6   14.5   15  RT (min)      lcorbi_t3_pos_1_008_36    m/z 785.5888     0   1   2E+7   14   14.5  RT (min)       lcorbi_t3_pos_1_005_42    m/z 785.5888     0   1   2E+4   16   16.5  RT (min)      lcorbi_t3_pos_2_056_56    m/z 785.5890     0   1   16.5   17  RT (min)    lcorbi_t3_pos_1_023_02    m/z 785.5891     0   2   4   6E+4   15   15.5  RT (min)      lcorbi_t3_pos_1_034_34    m/z 785.5891     0   1   16.5   17  RT (min)    lcorbi_t3_pos_1_023_02    m/z 785.6277     0   2   4E+5   12   12.5  RT (min)       lcorbi_t3_pos_2_037_27    m/z 785.6278     0   2   4E+5   12   12.5  RT (min)       lcorbi_t3_pos_4_092_60    m/z 785.6280     0   2   4E+5   12   12.5  RT (min)       lcorbi_t3_pos_4_090_60    m/z 785.6281     0   2   4E+5   12   12.5  RT (min)      lcorbi_t3_pos_4_093_60    m/z 785.6532     0   1   16   16.5  RT (min)      lcorbi_t3_pos_1_049_55    m/z 785.6533     0   1   16   16.5  RT (min)      lcorbi_t3_pos_4_033_53    m/z 785.6533     0   1   15.5   16  RT (min)      lcorbi_t3_pos_1_041_54    m/z 785.6534     0   1   16   16.5  RT (min)      lcorbi_t3_pos_1_031_40    m/z 785.6534     0   1   15.5   16  RT (min)      lcorbi_t3_pos_1_034_34    m/z 785.6534     0   1   2E+6   13.5   14  RT (min)      lcorbi_t3_pos_4_016_15    m/z 785.6534     0   2   4   6   8E+3   13.5   14  RT (min)      lcorbi_t3_pos_2_013_33    m/z 785.6535     0   1   15.5   16  RT (min)      lcorbi_t3_pos_1_028_32    m/z 785.6536     0   1   13   13.5  RT (min)      lcorbi_t3_pos_3_024_07    m/z 785.6536     0   1   2E+4   15   15.5  RT (min)      lcorbi_t3_pos_3_056_56    m/z 785.6536     0   1   2E+4   15   15.5  RT (min)      lcorbi_t3_pos_2_022_38    m/z 785.6536     0   1   15.5   16  RT (min)      lcorbi_t3_pos_1_028_32    m/z 785.6536     0   2   4   6   8E+3   13.5   14  RT (min)      lcorbi_t3_pos_2_054_29    m/z 785.6536     0   1   2E+4   15   15.5  RT (min)      lcorbi_t3_pos_2_017_51    m/z 785.6537     0   1   15.5   16  RT (min)      lcorbi_t3_pos_1_028_32    m/z 785.6538     0   1   17   17.5  RT (min)    lcorbi_t3_pos_1_023_02    m/z 785.6538     0   1E+6   14.5   15  RT (min)      lcorbi_t3_pos_3_047_40    m/z 785.6538     0   1   2E+6   14.5   15  RT (min)      lcorbi_t3_pos_4_019_47    m/z 785.6538     0   1   15.5   16  RT (min)      lcorbi_t3_pos_1_034_34    m/z 785.6538     0   1   15.5   16  RT (min)      lcorbi_t3_pos_1_028_32    m/z 785.6538     0   1   16.5   17  RT (min)    lcorbi_t3_pos_2_012_01    m/z 785.6538     0   1   16   16.5  RT (min)      lcorbi_t3_pos_1_051_09    m/z 785.6540     0   1   2E+5   14.5   15  RT (min)      lcorbi_t3_pos_1_008_36    m/z 785.6543     0   1   13   13.5  RT (min)      lcorbi_t3_pos_2_032_39    m/z 785.6914     0   1   2   3   4E+4   15   15.5  RT (min)      lcorbi_t3_pos_2_046_07    m/z 785.6914     0   2   4   6E+5   12.5   13  RT (min)      lcorbi_t3_pos_4_057_49    m/z 785.6914     0   1   2   3E+4   16   16.5  RT (min)      lcorbi_t3_pos_1_025_52    m/z 785.6916     0   1   15.5   16  RT (min)      lcorbi_t3_pos_2_025_52    m/z 785.6916     0   1E+4   16   16.5  RT (min)      lcorbi_t3_pos_4_033_53    m/z 785.6916     0   1   2E+4   15.5   16  RT (min)      lcorbi_t3_pos_1_041_54    m/z 785.6917     0   1   2E+4   15   15.5  RT (min)      lcorbi_t3_pos_2_046_07    m/z 785.6917     0   1   2   3   4E+5   14.5   15  RT (min)      lcorbi_t3_pos_4_025_52    m/z 785.6918     0   1E+4   15   15.5  RT (min)      lcorbi_t3_pos_2_041_54    m/z 785.6918     0   1E+5   14.5   15  RT (min)      lcorbi_t3_pos_3_056_56    m/z 785.6918     0   1   15   15.5  RT (min)      lcorbi_t3_pos_1_030_21    m/z 785.6918     0   1   2   3   4E+6   14.5   15  RT (min)      lcorbi_t3_pos_1_005_42    m/z 785.6918     0   1   2E+4   16   16.5  RT (min)      lcorbi_t3_pos_1_049_55    m/z 785.6919     0   1   2E+4   15   15.5  RT (min)      lcorbi_t3_pos_1_021_18    m/z 785.6919     0   1E+7   13.5   14  RT (min)       lcorbi_t3_pos_4_093_60    m/z 785.6919     0   1E+7   13.5   14  RT (min)       lcorbi_t3_pos_1_003_45    m/z 785.6920     0   1   16   16.5  RT (min)      lcorbi_t3_pos_1_001_49    m/z 785.6921     0   1E+7   13.5   14  RT (min)      lcorbi_t3_pos_4_096_60    m/z 786.4148     0   1   17   17.5  RT (min)    lcorbi_t3_pos_1_049_55    m/z 786.4149     0   1   16.5   17  RT (min)    lcorbi_t3_pos_2_041_54    m/z 786.4150     0   1   17.5   18  RT (min)    lcorbi_t3_pos_2_056_56    m/z 786.4150     0   1E+4   15.5   16  RT (min)      lcorbi_t3_pos_3_025_52    m/z 786.4151     0   1   17.5   18  RT (min)    lcorbi_t3_pos_2_056_56    m/z 786.4152     0   1   16   16.5  RT (min)      lcorbi_t3_pos_4_041_54    m/z 786.4152     0   1   16.5   17  RT (min)    lcorbi_t3_pos_2_025_52    m/z 786.4152     0   2   4   6E+5   15.5   16  RT (min)      lcorbi_t3_pos_1_005_42    m/z 786.4153     0   2   4   6E+5   14.5   15  RT (min)      lcorbi_t3_pos_4_105_61    m/z 786.4153     0   1   16.5   17  RT (min)    lcorbi_t3_pos_1_041_54    m/z 786.5278     0   1   15.5   16  RT (min)      lcorbi_t3_pos_1_031_40    m/z 786.5282     0   1   15   15.5  RT (min)      lcorbi_t3_pos_1_031_40    m/z 786.5283     0   1E+6   12.5   13  RT (min)       lcorbi_t3_pos_1_054_48    m/z 786.5284     0   2   4   6   8E+5   12   12.5  RT (min)       lcorbi_t3_pos_2_005_11    m/z 786.5284     0   1E+5   14   14.5  RT (min)      lcorbi_t3_pos_1_008_36    m/z 786.5284     0   2   4   6   8E+5   12   12.5  RT (min)       lcorbi_t3_pos_2_005_11    m/z 786.5285     0   2   4   6   8E+4   14.5   15  RT (min)      lcorbi_t3_pos_1_028_32    m/z 786.5286     0   2   4   6   8E+5   12   12.5  RT (min)       lcorbi_t3_pos_2_005_11    m/z 786.5911     0   1   2E+6   13   13.5  RT (min)      lcorbi_t3_pos_4_057_49    m/z 786.5919     0   2   4   6   8E+4   15   15.5  RT (min)      lcorbi_t3_pos_1_047_41    m/z 786.5920     0   1   16.5   17  RT (min)    lcorbi_t3_pos_1_023_02    m/z 786.5957     0   1E+4   13   13.5  RT (min)      lcorbi_t3_pos_2_032_39    m/z 786.5961     0   1E+5   15.5   16  RT (min)      lcorbi_t3_pos_3_013_19    m/z 786.5998     0   1E+4   13   13.5  RT (min)      lcorbi_t3_pos_2_032_39    m/z 786.6002     0   1   2E+5   15   15.5  RT (min)      lcorbi_t3_pos_1_028_32    m/z 786.6004     0   1   2E+5   15   15.5  RT (min)      lcorbi_t3_pos_1_030_21    m/z 786.6004     0   1E+4   13   13.5  RT (min)      lcorbi_t3_pos_4_012_37    m/z 786.6004     0   1E+5   16   16.5  RT (min)      lcorbi_t3_pos_1_051_09    m/z 786.6004     0   1E+7   13.5   14  RT (min)       lcorbi_t3_pos_4_016_15    m/z 786.6004     0   1E+5   15.5   16  RT (min)      lcorbi_t3_pos_1_034_34    m/z 786.6004     0   1E+5   16   16.5  RT (min)      lcorbi_t3_pos_1_031_40    m/z 786.6005     0   2   4   6   8E+4   16   16.5  RT (min)      lcorbi_t3_pos_1_049_55    m/z 786.6005     0   1   2   3E+6   14.5   15  RT (min)      lcorbi_t3_pos_1_008_36    m/z 786.6005     0   1   2E+5   15   15.5  RT (min)      lcorbi_t3_pos_1_028_32    m/z 786.6005     0   1E+7   13.5   14  RT (min)      lcorbi_t3_pos_1_002_01    m/z 786.6005     0   2   4   6E+6   14.5   15  RT (min)      lcorbi_t3_pos_1_008_36    m/z 786.6006     0   1   16.5   17  RT (min)    lcorbi_t3_pos_1_023_02    m/z 786.6006     0   1E+5   15.5   16  RT (min)      lcorbi_t3_pos_1_034_34    m/z 786.6006     0   1E+5   15.5   16  RT (min)      lcorbi_t3_pos_2_009_50    m/z 786.6006     0   1   2E+5   15   15.5  RT (min)      lcorbi_t3_pos_1_028_32    m/z 786.6007     0   1E+5   16   16.5  RT (min)      lcorbi_t3_pos_1_051_09    m/z 786.6007     0   1   16.5   17  RT (min)    lcorbi_t3_pos_1_023_02    m/z 786.6008     0   1E+7   14.5   15  RT (min)       lcorbi_t3_pos_1_008_36    m/z 786.6008     0   1   16.5   17  RT (min)     lcorbi_t3_pos_1_023_02    m/z 786.6009     0   1   2   3E+4   13.5   14  RT (min)      lcorbi_t3_pos_1_002_01    m/z 786.6010     0   2   4   6   8E+7   14.5   15  RT (min)       lcorbi_t3_pos_1_005_42    m/z 786.6010     0   1   2   3E+4   13.5   14  RT (min)      lcorbi_t3_pos_2_054_29    m/z 786.6402     0   1   16.5   17  RT (min)    lcorbi_t3_pos_1_023_02    m/z 786.6435     0   1E+5   14.5   15  RT (min)      lcorbi_t3_pos_1_008_36    m/z 786.6438     0   1E+5   15.5   16  RT (min)      lcorbi_t3_pos_1_034_34    m/z 786.6562     0   1   16   16.5  RT (min)      lcorbi_t3_pos_4_033_53    m/z 786.6564     0   1   13.5   14  RT (min)      lcorbi_t3_pos_2_013_33    m/z 786.6564     0   1E+6   13.5   14  RT (min)      lcorbi_t3_pos_4_016_15    m/z 786.6565     0   1E+4   15   15.5  RT (min)      lcorbi_t3_pos_1_047_41    m/z 786.6567     0   1   16   16.5  RT (min)      lcorbi_t3_pos_1_051_09    m/z 786.6567     0   1   15.5   16  RT (min)      lcorbi_t3_pos_1_034_34    m/z 786.6567     0   2   4E+5   14.5   15  RT (min)      lcorbi_t3_pos_4_054_18    m/z 786.6567     0   1   13   13.5  RT (min)      lcorbi_t3_pos_3_024_07    m/z 786.6568     0   1E+5   14.5   15  RT (min)      lcorbi_t3_pos_1_008_36    m/z 786.6568     0   1   2   3E+4   16   16.5  RT (min)      lcorbi_t3_pos_1_031_40    m/z 786.6568     0   1   15.5   16  RT (min)      lcorbi_t3_pos_1_028_32    m/z 786.6568     0   2   4   6E+5   14.5   15  RT (min)      lcorbi_t3_pos_3_047_40    m/z 786.6568     0   2   4   6   8E+5   14.5   15  RT (min)      lcorbi_t3_pos_1_005_42    m/z 786.6568     0   1E+5   13.5   14  RT (min)      lcorbi_t3_pos_1_002_01    m/z 786.6569     0   1   2   3E+4   15   15.5  RT (min)      lcorbi_t3_pos_1_008_36    m/z 786.6570     0   1   17   17.5  RT (min)    lcorbi_t3_pos_1_023_02    m/z 786.6570     0   1   2E+4   15   15.5  RT (min)      lcorbi_t3_pos_2_022_38    m/z 786.6570     0   1   15.5   16  RT (min)      lcorbi_t3_pos_1_034_34    m/z 786.6572     0   1   13   13.5  RT (min)      lcorbi_t3_pos_2_032_39    m/z 786.7031     0   2   4   6E+5   13.5   14   14.5  RT (min)      lcorbi_t3_pos_4_095_60    m/z 787.5996     0   1   2   3E+4   16   16.5  RT (min)      lcorbi_t3_pos_4_033_53    m/z 787.6032     0   1   13   13.5  RT (min)      lcorbi_t3_pos_2_032_39    m/z 787.6032     0   1   13   13.5  RT (min)      lcorbi_t3_pos_2_032_39    m/z 787.6036     0   2   4   6   8E+4   13.5   14  RT (min)      lcorbi_t3_pos_2_054_29    m/z 787.6039     0   2   4   6   8E+6   13.5   14  RT (min)      lcorbi_t3_pos_4_016_15    m/z 787.6039     0   1E+5   15.5   16  RT (min)      lcorbi_t3_pos_1_034_34    m/z 787.6039     0   1E+5   15.5   16  RT (min)      lcorbi_t3_pos_3_013_19    m/z 787.6039     0   1E+4   13   13.5  RT (min)      lcorbi_t3_pos_4_012_37    m/z 787.6040     0   1E+5   15   15.5  RT (min)      lcorbi_t3_pos_1_028_32    m/z 787.6040     0   1   16.5   17  RT (min)    lcorbi_t3_pos_1_023_02    m/z 787.6040     0   2   4   6E+6   0   0.5  RT (min)      lcorbi_t3_pos_4_057_49    m/z 787.6040     0   2   4   6E+4   16   16.5  RT (min)       lcorbi_t3_pos_1_051_09    m/z 787.6040     0   1E+5   15.5   16  RT (min)       lcorbi_t3_pos_1_034_34    m/z 787.6040     0   1E+5   15   15.5  RT (min)      lcorbi_t3_pos_1_028_32    m/z 787.6041     0   2   4   6   8E+6   13.5   14  RT (min)      lcorbi_t3_pos_1_002_01    m/z 787.6042     0   1E+5   15   15.5  RT (min)      lcorbi_t3_pos_1_028_32    m/z 787.6042     0   1   13.5   14  RT (min)      lcorbi_t3_pos_1_002_01    m/z 787.6042     0   2   4   6E+4   15   15.5  RT (min)      lcorbi_t3_pos_1_030_21    m/z 787.6042     0   1   2   3   4E+4   16   16.5  RT (min)      lcorbi_t3_pos_1_001_49    m/z 787.6042     0   1   16.5   17  RT (min)    lcorbi_t3_pos_1_023_02    m/z 787.6042     0   1E+5   15.5   16  RT (min)      lcorbi_t3_pos_1_034_34    m/z 787.6042     0   2   4   6E+4   16   16.5  RT (min)      lcorbi_t3_pos_1_031_40    m/z 787.6043     0   1E+5   15   15.5  RT (min)      lcorbi_t3_pos_1_047_41    m/z 787.6043     0   1   16.5   17  RT (min)    lcorbi_t3_pos_1_023_02    m/z 787.6043     0   2   4   6E+4   16   16.5  RT (min)      lcorbi_t3_pos_1_051_09    m/z 787.6044     0   2   4E+6   14.5   15  RT (min)       lcorbi_t3_pos_1_008_36    m/z 787.6044     0   1   2E+6   14.5   15  RT (min)      lcorbi_t3_pos_1_008_36    m/z 787.6044     0   1E+6   14.5   15  RT (min)      lcorbi_t3_pos_1_008_36    m/z 787.6046     0   1   2   3E+7   14.5   15  RT (min)      lcorbi_t3_pos_1_005_42    m/z 787.6436     0   2   4   6E+5   13   13.5  RT (min)       lcorbi_t3_pos_4_092_60    m/z 787.6436     0   2   4   6   8E+5   13   13.5  RT (min)      lcorbi_t3_pos_4_097_60    m/z 787.6437     0   2   4   6E+5   13   13.5  RT (min)      lcorbi_t3_pos_4_094_60    m/z 787.6437     0   2   4   6E+5   12.5   13  RT (min)       lcorbi_t3_pos_4_095_60    m/z 787.6441     0   2   4   6E+5   12   12.5  RT (min)       lcorbi_t3_pos_4_096_60    m/z 787.6604     0   2   4   6E+5   14   14.5  RT (min)      lcorbi_t3_pos_4_016_15    m/z 787.6604     0   1   16   16.5  RT (min)      lcorbi_t3_pos_1_031_40    m/z 787.6605     0   1   2   3   4E+4   15   15.5  RT (min)      lcorbi_t3_pos_1_008_36    m/z 787.6606     0   1   13.5   14  RT (min)      lcorbi_t3_pos_3_031_45    m/z 787.6607     0   1E+4   15.5   16  RT (min)      lcorbi_t3_pos_2_022_38    m/z 787.6687     0   1   16   16.5  RT (min)      lcorbi_t3_pos_1_034_34    m/z 787.6688     0   1   16   16.5  RT (min)      lcorbi_t3_pos_1_034_34    m/z 787.6688     0   1   16.5   17  RT (min)    lcorbi_t3_pos_2_012_01    m/z 787.6688     0   2   4   6E+3   13.5   14  RT (min)      lcorbi_t3_pos_2_013_33    m/z 787.6688     0   1   16.5   17  RT (min)    lcorbi_t3_pos_1_031_40    m/z 787.6689     0   1   13.5   14  RT (min)      lcorbi_t3_pos_2_032_39    m/z 787.6690     0   1   17   17.5  RT (min)    lcorbi_t3_pos_1_023_02    m/z 787.6690     0   1E+6   14.5   15  RT (min)      lcorbi_t3_pos_4_019_47    m/z 787.6690     0   1   17   17.5  RT (min)    lcorbi_t3_pos_1_023_02    m/z 787.6690     0   1   13.5   14  RT (min)      lcorbi_t3_pos_2_013_33    m/z 787.6690     0   1   16.5   17  RT (min)    lcorbi_t3_pos_1_051_09    m/z 787.6690     0   1   15.5   16  RT (min)      lcorbi_t3_pos_1_028_32    m/z 787.6690     0   2   4   6E+3   13.5   14  RT (min)      lcorbi_t3_pos_2_054_29    m/z 787.6691     0   1   16.5   17  RT (min)    lcorbi_t3_pos_2_012_01    m/z 787.6692     0   1E+5   15   15.5  RT (min)      lcorbi_t3_pos_1_008_36    m/z 787.6692     0   1   15.5   16  RT (min)      lcorbi_t3_pos_1_033_53    m/z 787.6693     0   1   16.5   17  RT (min)    lcorbi_t3_pos_2_012_01    m/z 787.7069     0   1E+4   16   16.5  RT (min)      lcorbi_t3_pos_1_056_56    m/z 787.7071     0   1   15.5   16  RT (min)      lcorbi_t3_pos_1_041_54    m/z 787.7072     0   1   15   15.5  RT (min)      lcorbi_t3_pos_2_006_47    m/z 787.7072     0   1E+4   15   15.5  RT (min)      lcorbi_t3_pos_3_056_56    m/z 787.7072     0   1   16   16.5  RT (min)      lcorbi_t3_pos_2_009_50    m/z 787.7074     0   1   16.5   17  RT (min)    lcorbi_t3_pos_2_056_56    m/z 787.7074     0   2   4   6E+5   14.5   15  RT (min)      lcorbi_t3_pos_1_005_42    m/z 787.7074     0   2   4   6E+5   14.5   15  RT (min)      lcorbi_t3_pos_1_005_42    m/z 787.7074     0   1   15   15.5  RT (min)      lcorbi_t3_pos_1_030_21    m/z 787.7074     0   1   16   16.5  RT (min)      lcorbi_t3_pos_1_025_52    m/z 787.7074     0   1   15.5   16  RT (min)      lcorbi_t3_pos_2_038_03    m/z 787.7074     0   1   15.5   16  RT (min)      lcorbi_t3_pos_1_021_18    m/z 787.7074     0   1   2E+6   13.5   14  RT (min)      lcorbi_t3_pos_4_093_60    m/z 787.7076     0   1   2E+6   13.5   14  RT (min)       lcorbi_t3_pos_4_095_60    m/z 788.5437     0   1   2E+4   14.5   15   15.5  RT (min)      lcorbi_t3_pos_1_028_32    m/z 788.5438     0   1E+5   14   14.5  RT (min)      lcorbi_t3_pos_1_008_36    m/z 788.5438     0   1   15.5   16  RT (min)      lcorbi_t3_pos_1_031_40    m/z 788.5439     0   1   2E+6   12.5   13  RT (min)      lcorbi_t3_pos_2_005_11    m/z 788.5440     0   1   2E+6   12.5   13  RT (min)       lcorbi_t3_pos_1_054_48    m/z 788.5440     0   1   2E+6   12.5   13  RT (min)       lcorbi_t3_pos_2_005_11    m/z 788.5440     0   1   15.5   16  RT (min)      lcorbi_t3_pos_1_031_40    m/z 788.5440     0   1   2E+6   12.5   13  RT (min)       lcorbi_t3_pos_2_005_11    m/z 788.5440     0   1   15   15.5  RT (min)      lcorbi_t3_pos_1_034_34    m/z 788.5442     0   1   2E+6   12.5   13  RT (min)      lcorbi_t3_pos_1_054_48    m/z 788.6023     0   1   2E+4   15.5   16  RT (min)      lcorbi_t3_pos_1_041_54    m/z 788.6062     0   2   4   6E+4   15   15.5  RT (min)      lcorbi_t3_pos_2_022_38    m/z 788.6066     0   1E+4   13   13.5  RT (min)      lcorbi_t3_pos_2_032_39    m/z 788.6068     0   1E+4   13   13.5  RT (min)      lcorbi_t3_pos_2_032_39    m/z 788.6073     0   1   2   3E+6   13.5   14  RT (min)      lcorbi_t3_pos_4_016_15    m/z 788.6074     0   1   2E+6   13.5   14  RT (min)      lcorbi_t3_pos_1_002_01    m/z 788.6075     0   1   2   3E+4   15   15.5  RT (min)      lcorbi_t3_pos_2_006_47    m/z 788.6076     0   1   2E+4   16   16.5  RT (min)      lcorbi_t3_pos_1_051_09    m/z 788.6076     0   1   2   3E+4   16   16.5  RT (min)      lcorbi_t3_pos_1_051_09    m/z 788.6076     0   1   2   3E+4   15   15.5  RT (min)      lcorbi_t3_pos_2_022_38    m/z 788.6076     0   1   2   3   4E+4   15   15.5  RT (min)      lcorbi_t3_pos_1_030_21    m/z 788.6110     0   1E+4   13   13.5  RT (min)      lcorbi_t3_pos_2_032_39    m/z 788.6117     0   1   13.5   14  RT (min)      lcorbi_t3_pos_2_013_33    m/z 788.6119     0   1   16.5   17  RT (min)    lcorbi_t3_pos_1_023_02    m/z 788.6119     0   2   4   6   8E+6   14.5   15  RT (min)      lcorbi_t3_pos_1_005_42    m/z 788.6121     0   1   15.5   16  RT (min)      lcorbi_t3_pos_1_041_54    m/z 788.6155     0   1   2E+4   15.5   16  RT (min)      lcorbi_t3_pos_2_038_03    m/z 788.6156     0   1   16.5   17  RT (min)    lcorbi_t3_pos_2_012_01    m/z 788.6156     0   1   16   16.5  RT (min)      lcorbi_t3_pos_1_031_40    m/z 788.6156     0   1   2   3E+4   15.5   16  RT (min)      lcorbi_t3_pos_1_028_32    m/z 788.6157     0   1E+6   14.5   15  RT (min)      lcorbi_t3_pos_1_008_36    m/z 788.6157     0   1   2   3E+4   15.5   16  RT (min)      lcorbi_t3_pos_1_028_32    m/z 788.6158     0   2   4   6   8E+6   14.5   15  RT (min)      lcorbi_t3_pos_1_008_36    m/z 788.6158     0   1   17   17.5  RT (min)    lcorbi_t3_pos_1_023_02    m/z 788.6158     0   1   2   3E+4   15.5   16  RT (min)      lcorbi_t3_pos_1_034_34    m/z 788.6160     0   1E+7   14.5   15  RT (min)       lcorbi_t3_pos_4_019_47    m/z 788.6161     0   2   4   6   8E+3   13.5   14  RT (min)      lcorbi_t3_pos_2_054_29    m/z 788.6161     0   1   2   3E+4   15.5   16  RT (min)      lcorbi_t3_pos_1_028_32    m/z 788.6162     0   1   2E+4   16   16.5  RT (min)      lcorbi_t3_pos_1_051_09    m/z 788.6164     0   1   2   3E+4   15.5   16  RT (min)      lcorbi_t3_pos_1_034_34    m/z 788.6164     0   1   2   3E+6   0.5   1  RT (min)      lcorbi_t3_pos_4_057_49    m/z 788.6165     0   1   16.5   17  RT (min)    lcorbi_t3_pos_2_012_01    m/z 788.6165     0   2   4E+6   13.5   14  RT (min)      lcorbi_t3_pos_1_002_01    m/z 788.6720     0   1   16.5   17  RT (min)    lcorbi_t3_pos_1_031_40    m/z 788.6722     0   1   2   3E+3   13.5   14  RT (min)      lcorbi_t3_pos_4_012_37    m/z 788.6722     0   1   17   17.5  RT (min)    lcorbi_t3_pos_1_023_02    m/z 788.6723     0   1   16   16.5  RT (min)      lcorbi_t3_pos_1_034_34    m/z 788.6723     0   1   17   17.5  RT (min)    lcorbi_t3_pos_1_023_02    m/z 788.6724     0   2   4E+4   15.5   16  RT (min)      lcorbi_t3_pos_2_022_38    m/z 788.6724     0   1   2   3E+5   15   15.5  RT (min)      lcorbi_t3_pos_3_047_40    m/z 788.6725     0   1   13.5   14  RT (min)      lcorbi_t3_pos_3_024_07    m/z 788.6726     0   1   16.5   17  RT (min)    lcorbi_t3_pos_1_051_09    m/z 788.6726     0   1   13.5   14  RT (min)      lcorbi_t3_pos_2_032_39    m/z 788.6726     0   2   4E+4   15.5   16  RT (min)      lcorbi_t3_pos_1_028_32    m/z 788.6726     0   2   4E+5   14.5   15  RT (min)      lcorbi_t3_pos_4_019_47    m/z 788.6728     0   1   13.5   14  RT (min)      lcorbi_t3_pos_1_002_01    m/z 788.6728     0   2   4   6E+5   14   14.5  RT (min)      lcorbi_t3_pos_3_034_26    m/z 789.5472     0   1E+6   12.5   13  RT (min)      lcorbi_t3_pos_2_005_11    m/z 789.5473     0   1   15.5   16  RT (min)      lcorbi_t3_pos_1_031_40    m/z 789.5474     0   1   14.5   15  RT (min)      lcorbi_t3_pos_1_028_32    m/z 789.5474     0   1E+6   12.5   13  RT (min)      lcorbi_t3_pos_1_054_48    m/z 789.5474     0   1E+6   12.5   13  RT (min)      lcorbi_t3_pos_2_005_11    m/z 789.5475     0   1E+6   12.5   13  RT (min)      lcorbi_t3_pos_2_005_11    m/z 789.6098     0   1   13   13.5  RT (min)      lcorbi_t3_pos_2_032_39    m/z 789.6103     0   1   16.5   17  RT (min)    lcorbi_t3_pos_1_023_02    m/z 789.6106     0   1   15   15.5  RT (min)      lcorbi_t3_pos_3_049_55    m/z 789.6107     0   1   16   16.5  RT (min)      lcorbi_t3_pos_1_051_09    m/z 789.6108     0   1   2E+4   15   15.5  RT (min)      lcorbi_t3_pos_2_046_07    m/z 789.6109     0   1E+4   15   15.5  RT (min)      lcorbi_t3_pos_2_006_47    m/z 789.6109     0   1   2E+4   15.5   16  RT (min)      lcorbi_t3_pos_1_041_54    m/z 789.6151     0   1   15.5   16  RT (min)      lcorbi_t3_pos_2_038_03    m/z 789.6154     0   1   2   3E+6   14.5   15  RT (min)      lcorbi_t3_pos_3_040_41    m/z 789.6154     0   1   16.5   17  RT (min)    lcorbi_t3_pos_1_023_02    m/z 789.6155     0   1   15.5   16  RT (min)      lcorbi_t3_pos_1_034_34    m/z 789.6190     0   1   13   13.5  RT (min)      lcorbi_t3_pos_2_032_39    m/z 789.6195     0   1   16   16.5  RT (min)      lcorbi_t3_pos_1_049_55    m/z 789.6196     0   2   4E+6   14.5   15  RT (min)      lcorbi_t3_pos_4_019_47    m/z 789.6196     0   1E+4   13.5   14  RT (min)      lcorbi_t3_pos_2_054_29    m/z 789.6197     0   1E+5   13.5   14  RT (min)      lcorbi_t3_pos_1_002_01    m/z 789.6198     0   1   2E+4   15.5   16  RT (min)      lcorbi_t3_pos_1_028_32    m/z 789.6198     0   1   2   3E+6   14.5   15  RT (min)      lcorbi_t3_pos_3_047_40    m/z 789.6198     0   1   2E+4   15.5   16  RT (min)      lcorbi_t3_pos_1_028_32    m/z 789.6198     0   1   13   13.5  RT (min)      lcorbi_t3_pos_3_024_07    m/z 789.6198     0   1   15   15.5  RT (min)      lcorbi_t3_pos_1_021_18    m/z 789.6198     0   1   13.5   14  RT (min)      lcorbi_t3_pos_2_013_33    m/z 789.6199     0   1   15.5   16  RT (min)      lcorbi_t3_pos_1_034_34    m/z 789.6199     0   2   4   6E+5   14.5   15  RT (min)      lcorbi_t3_pos_1_008_36    m/z 789.6200     0   1   16.5   17  RT (min)    lcorbi_t3_pos_2_056_56    m/z 789.6200     0   1E+6   0.5   1  RT (min)      lcorbi_t3_pos_4_057_49    m/z 789.6200     0   1   2E+4   15.5   16  RT (min)      lcorbi_t3_pos_1_028_32    m/z 789.6201     0   1   17   17.5  RT (min)    lcorbi_t3_pos_1_023_02    m/z 789.6201     0   1   16   16.5   17  RT (min)    lcorbi_t3_pos_2_056_56    m/z 789.6203     0   1   16   16.5  RT (min)      lcorbi_t3_pos_1_051_09    m/z 790.5595     0   1   2E+6   13   13.5  RT (min)       lcorbi_t3_pos_1_054_48    m/z 790.5595     0   1   2E+6   12.5   13  RT (min)       lcorbi_t3_pos_1_054_48    m/z 790.5596     0   1   2E+6   13   13.5  RT (min)       lcorbi_t3_pos_1_054_48    m/z 790.5597     0   1   2E+6   12.5   13  RT (min)       lcorbi_t3_pos_2_005_11    m/z 790.5598     0   1   2E+6   12.5   13  RT (min)       lcorbi_t3_pos_3_029_13    m/z 790.6140     0   1   15   15.5  RT (min)      lcorbi_t3_pos_2_046_07    m/z 790.6181     0   1   15.5   16  RT (min)      lcorbi_t3_pos_1_028_32    m/z 790.6182     0   1   16.5   17  RT (min)    lcorbi_t3_pos_2_012_01    m/z 790.6184     0   1   15.5   16  RT (min)      lcorbi_t3_pos_1_034_34    m/z 790.6186     0   1   2E+4   15   15.5  RT (min)      lcorbi_t3_pos_2_022_38    m/z 790.6229     0   1   15.5   16  RT (min)      lcorbi_t3_pos_1_028_32    m/z 790.6230     0   1   16   16.5  RT (min)      lcorbi_t3_pos_1_051_09    m/z 790.6230     0   1E+4   15   15.5  RT (min)      lcorbi_t3_pos_2_046_07    m/z 790.6231     0   1   2E+5   14.5   15  RT (min)      lcorbi_t3_pos_1_008_36    m/z 790.6232     0   1   13.5   14  RT (min)      lcorbi_t3_pos_1_002_01    m/z 790.6232     0   1   13.5   14  RT (min)      lcorbi_t3_pos_2_013_33    m/z 790.6233     0   1E+6   14.5   15  RT (min)      lcorbi_t3_pos_1_005_42    m/z 790.6233     0   1   15.5   16  RT (min)      lcorbi_t3_pos_1_028_32    m/z 790.6233     0   1   13   13.5  RT (min)      lcorbi_t3_pos_2_032_39    m/z 790.6234     0   1   15.5   16  RT (min)      lcorbi_t3_pos_1_034_34    m/z 790.6234     0   1   17   17.5  RT (min)    lcorbi_t3_pos_1_023_02    m/z 790.6234     0   1   16   16.5  RT (min)      lcorbi_t3_pos_1_051_09    m/z 790.6235     0   1   13   13.5  RT (min)      lcorbi_t3_pos_3_024_07    m/z 790.6235     0   1   2   3E+5   14.5   15  RT (min)      lcorbi_t3_pos_4_001_49    m/z 790.6236     0   1   16.5   17  RT (min)    lcorbi_t3_pos_2_012_01    m/z 791.5630     0   1E+6   12.5   13  RT (min)      lcorbi_t3_pos_2_005_11    m/z 791.5630     0   1E+6   12.5   13  RT (min)      lcorbi_t3_pos_3_029_13    m/z 791.5630     0   1E+6   13   13.5  RT (min)      lcorbi_t3_pos_1_054_48    m/z 791.7500     0   1   17   17.5  RT (min)    lcorbi_t3_pos_1_049_55    m/z 791.7505     0   1   16.5   17  RT (min)    lcorbi_t3_pos_1_021_18    m/z 791.7506     0   1   17.5   18  RT (min)    lcorbi_t3_pos_1_025_52    m/z 791.7507     0   2   4E+5   15.5   16  RT (min)      lcorbi_t3_pos_1_005_42    m/z 791.7508     0   1   17   17.5  RT (min)    lcorbi_t3_pos_2_009_50    m/z 792.5525     0   2   4   6E+5   14   14.5  RT (min)       lcorbi_t3_pos_4_084_59    m/z 792.5526     0   2   4   6E+5   14   14.5  RT (min)       lcorbi_t3_pos_4_084_59    m/z 792.5898     0   1E+5   13.5   14  RT (min)      lcorbi_t3_pos_1_002_01    m/z 792.5898     0   1   13   13.5  RT (min)      lcorbi_t3_pos_1_002_01    m/z 792.5902     0   1   15   15.5  RT (min)      lcorbi_t3_pos_1_030_21    m/z 792.5903     0   1   16.5   17  RT (min)    lcorbi_t3_pos_1_023_02    m/z 792.5904     0   1   15.5   16  RT (min)      lcorbi_t3_pos_1_034_34    m/z 792.5904     0   1   16   16.5  RT (min)      lcorbi_t3_pos_1_031_40    m/z 792.5905     0   1   16   16.5  RT (min)      lcorbi_t3_pos_1_051_09    m/z 792.5905     0   1   15   15.5  RT (min)      lcorbi_t3_pos_1_021_18    m/z 792.5906     0   1   15   15.5  RT (min)      lcorbi_t3_pos_1_028_32    m/z 792.5906     0   1E+5   14.5   15  RT (min)      lcorbi_t3_pos_1_008_36    m/z 792.5906     0   1   16.5   17  RT (min)    lcorbi_t3_pos_1_023_02    m/z 792.6482     0   1   2   3E+4   13.5   14  RT (min)       lcorbi_t3_pos_2_017_51    m/z 792.6482     0   1   14.5   15  RT (min)      lcorbi_t3_pos_1_025_52    m/z 792.6482     0   2   4   6E+4   14   14.5  RT (min)      lcorbi_t3_pos_2_006_47    m/z 792.7538     0   1   16.5   17  RT (min)    lcorbi_t3_pos_2_038_03    m/z 792.7539     0   1   17   17.5  RT (min)    lcorbi_t3_pos_1_041_54    m/z 792.7539     0   1   17.5   18  RT (min)    lcorbi_t3_pos_2_056_56    m/z 792.7540     0   1   16   16.5  RT (min)      lcorbi_t3_pos_2_017_51    m/z 792.7540     0   1   16.5   17  RT (min)    lcorbi_t3_pos_1_021_18    m/z 792.7540     0   1   16   16.5  RT (min)      lcorbi_t3_pos_1_030_21    m/z 792.7540     0   1   17   17.5  RT (min)    lcorbi_t3_pos_1_049_55    m/z 792.7540     0   1   17.5   18  RT (min)    lcorbi_t3_pos_1_025_52    m/z 792.7540     0   1   17.5   18  RT (min)    lcorbi_t3_pos_2_056_56    m/z 792.7540     0   1   2E+6   15.5   16  RT (min)      lcorbi_t3_pos_1_005_42    m/z 792.7540     0   1   2E+4   15.5   16  RT (min)      lcorbi_t3_pos_4_010_43    m/z 792.7540     0   1   17   17.5  RT (min)    lcorbi_t3_pos_2_009_50    m/z 792.7541     0   1   2E+4   16   16.5  RT (min)      lcorbi_t3_pos_4_010_43    m/z 793.5581     0   1   2E+5   14   14.5  RT (min)       lcorbi_t3_pos_4_084_59    m/z 793.6605     0   2   4   6   8E+4   14   14.5  RT (min)      lcorbi_t3_pos_1_047_41    m/z 793.6606     0   1   15   15.5  RT (min)      lcorbi_t3_pos_1_049_55    m/z 793.6607     0   1   15   15.5  RT (min)      lcorbi_t3_pos_1_001_49    m/z 793.6607     0   1   14.5   15  RT (min)      lcorbi_t3_pos_1_041_54    m/z 793.6607     0   1   14   14.5  RT (min)      lcorbi_t3_pos_1_033_53    m/z 793.6608     0   1   14   14.5  RT (min)      lcorbi_t3_pos_1_033_53    m/z 793.6611     0   1   15   15.5  RT (min)      lcorbi_t3_pos_1_001_49    m/z 793.6676     0   1   14.5   15  RT (min)      lcorbi_t3_pos_1_041_54    m/z 793.7572     0   1   16   16.5  RT (min)      lcorbi_t3_pos_1_030_21    m/z 793.7572     0   1   17   17.5  RT (min)    lcorbi_t3_pos_1_049_55    m/z 793.7573     0   2   4   6   8E+6   15.5   16  RT (min)      lcorbi_t3_pos_1_005_42    m/z 793.7573     0   1   16.5   17  RT (min)    lcorbi_t3_pos_1_021_18    m/z 793.7573     0   1   17.5   18  RT (min)    lcorbi_t3_pos_2_056_56    m/z 793.7574     0   1   17.5   18  RT (min)    lcorbi_t3_pos_2_056_56    m/z 793.7611     0   1   16.5   17  RT (min)    lcorbi_t3_pos_2_038_03    m/z 793.7614     0   1   16.5   17  RT (min)    lcorbi_t3_pos_2_041_54    m/z 793.7648     0   1   17   17.5  RT (min)    lcorbi_t3_pos_2_009_50    m/z 793.7652     0   1   16   16.5  RT (min)      lcorbi_t3_pos_2_046_07    m/z 793.7654     0   2   4E+4   16   16.5  RT (min)      lcorbi_t3_pos_3_056_56    m/z 793.7656     0   1   2E+5   15.5   16  RT (min)      lcorbi_t3_pos_4_001_49    m/z 793.7657     0   1   17   17.5  RT (min)    lcorbi_t3_pos_1_041_54    m/z 794.6054     0   1   13   13.5  RT (min)      lcorbi_t3_pos_2_032_39    m/z 794.6058     0   1   15   15.5  RT (min)      lcorbi_t3_pos_1_028_32    m/z 794.6059     0   1   2E+4   15   15.5  RT (min)      lcorbi_t3_pos_2_006_47    m/z 794.6060     0   1   2   3E+4   13.5   14  RT (min)      lcorbi_t3_pos_1_002_01    m/z 794.6060     0   1   16   16.5  RT (min)      lcorbi_t3_pos_1_051_09    m/z 794.6060     0   1   16   16.5  RT (min)      lcorbi_t3_pos_1_051_09    m/z 794.6060     0   1E+4   13   13.5  RT (min)      lcorbi_t3_pos_2_032_39    m/z 794.6060     0   1   15   15.5  RT (min)      lcorbi_t3_pos_1_030_21    m/z 794.6060     0   1   15.5   16  RT (min)      lcorbi_t3_pos_1_028_32    m/z 794.6060     0   1   16   16.5   17  RT (min)    lcorbi_t3_pos_2_056_56    m/z 794.6061     0   2   4   6   8E+5   13.5   14  RT (min)      lcorbi_t3_pos_3_011_42    m/z 794.6061     0   1E+6   14.5   15  RT (min)      lcorbi_t3_pos_1_008_36    m/z 794.6061     0   1   15   15.5  RT (min)      lcorbi_t3_pos_1_028_32    m/z 794.6061     0   2   4   6E+5   14.5   15  RT (min)      lcorbi_t3_pos_1_008_36    m/z 794.6061     0   1   2E+4   16   16.5  RT (min)      lcorbi_t3_pos_1_051_09    m/z 794.6061     0   1   2   3   4E+4   15.5   16  RT (min)      lcorbi_t3_pos_1_034_34    m/z 794.6061     0   1   2   3E+6   13.5   14  RT (min)      lcorbi_t3_pos_1_002_01    m/z 794.6062     0   1   16   16.5  RT (min)      lcorbi_t3_pos_1_051_09    m/z 794.6062     0   2   4   6E+4   14.5   15  RT (min)      lcorbi_t3_pos_2_017_51    m/z 794.6062     0   1   2   3E+6   14.5   15  RT (min)       lcorbi_t3_pos_1_005_42    m/z 794.6062     0   1   16.5   17  RT (min)    lcorbi_t3_pos_1_023_02    m/z 794.6062     0   1   16.5   17  RT (min)    lcorbi_t3_pos_1_023_02    m/z 794.6062     0   1   2E+4   15.5   16  RT (min)      lcorbi_t3_pos_1_034_34    m/z 794.6062     0   1   15   15.5  RT (min)      lcorbi_t3_pos_1_028_32    m/z 794.6064     0   1   16   16.5  RT (min)      lcorbi_t3_pos_1_049_55    m/z 794.6065     0   1   16   16.5  RT (min)      lcorbi_t3_pos_1_051_09    m/z 794.6066     0   1   16.5   17  RT (min)    lcorbi_t3_pos_1_023_02    m/z 794.6640     0   2   4E+4   14   14.5  RT (min)      lcorbi_t3_pos_1_033_53    m/z 794.6641     0   1   15   15.5  RT (min)      lcorbi_t3_pos_1_056_56    m/z 794.6641     0   1   14.5   15  RT (min)      lcorbi_t3_pos_2_049_55    m/z 794.6644     0   1   2   3E+5   13.5   14  RT (min)       lcorbi_t3_pos_3_025_52    m/z 794.6644     0   1   2E+4   14.5   15  RT (min)      lcorbi_t3_pos_1_041_54    m/z 794.6646     0   1E+6   13.5   14  RT (min)       lcorbi_t3_pos_1_005_42    m/z 794.6684     0   1   2E+5   14   14.5  RT (min)      lcorbi_t3_pos_2_017_51    m/z 794.6686     0   1   15   15.5  RT (min)      lcorbi_t3_pos_1_001_49    m/z 794.6720     0   2   4   6E+5   13.5   14  RT (min)      lcorbi_t3_pos_3_026_04    m/z 794.6720     0   1   14.5   15  RT (min)      lcorbi_t3_pos_1_030_21    m/z 794.6721     0   1   15   15.5  RT (min)      lcorbi_t3_pos_1_025_52    m/z 794.6724     0   1   14.5   15  RT (min)      lcorbi_t3_pos_1_041_54    m/z 794.6726     0   1   15   15.5  RT (min)      lcorbi_t3_pos_1_025_52    m/z 794.6731     0   1   14.5   15  RT (min)      lcorbi_t3_pos_2_041_54    m/z 794.7632     0   1   2E+6   15   15.5  RT (min)      lcorbi_t3_pos_3_040_41    m/z 794.7633     0   1E+6   15   15.5  RT (min)      lcorbi_t3_pos_3_041_54    m/z 794.7634     0   1   16.5   17  RT (min)    lcorbi_t3_pos_1_041_54    m/z 794.7664     0   1   16.5   17  RT (min)    lcorbi_t3_pos_4_033_53    m/z 794.7665     0   1   16   16.5  RT (min)      lcorbi_t3_pos_2_041_54    m/z 794.7667     0   1   16   16.5  RT (min)      lcorbi_t3_pos_2_041_54    m/z 794.7693     0   1   16.5   17  RT (min)    lcorbi_t3_pos_1_041_54    m/z 794.7694     0   1E+5   15.5   16  RT (min)      lcorbi_t3_pos_4_001_49    m/z 794.7694     0   1   17   17.5  RT (min)    lcorbi_t3_pos_2_056_56    m/z 794.7695     0   1   16   16.5  RT (min)      lcorbi_t3_pos_4_041_54    m/z 794.7695     0   1   15.5   16  RT (min)      lcorbi_t3_pos_4_049_55    m/z 794.7695     0   2   4   6   8E+4   15.5   16  RT (min)      lcorbi_t3_pos_3_025_52    m/z 794.7696     0   1   17   17.5  RT (min)    lcorbi_t3_pos_1_049_55    m/z 794.7696     0   1   16   16.5  RT (min)      lcorbi_t3_pos_2_041_54    m/z 794.7697     0   1   17   17.5  RT (min)    lcorbi_t3_pos_1_001_49    m/z 794.7697     0   1   2   3   4E+4   15.5   16  RT (min)      lcorbi_t3_pos_2_046_07    m/z 794.7698     0   1   16   16.5  RT (min)      lcorbi_t3_pos_4_041_54    m/z 795.6093     0   2   4   6E+4   15   15.5  RT (min)      lcorbi_t3_pos_1_028_32    m/z 795.6094     0   1   16   16.5  RT (min)      lcorbi_t3_pos_1_051_09    m/z 795.6094     0   1E+6   14.5   15  RT (min)       lcorbi_t3_pos_1_005_42    m/z 795.6095     0   1   15.5   16  RT (min)      lcorbi_t3_pos_1_041_54    m/z 795.6095     0   1   15   15.5  RT (min)      lcorbi_t3_pos_2_025_52    m/z 795.6095     0   1   2   3E+4   15   15.5  RT (min)      lcorbi_t3_pos_1_030_21    m/z 795.6095     0   1   16.5   17  RT (min)    lcorbi_t3_pos_1_023_02    m/z 795.6095     0   2   4E+4   15.5   16  RT (min)      lcorbi_t3_pos_1_034_34    m/z 795.6095     0   1   16.5   17  RT (min)    lcorbi_t3_pos_1_023_02    m/z 795.6095     0   1   16   16.5  RT (min)      lcorbi_t3_pos_1_051_09    m/z 795.6095     0   2   4   6E+4   15   15.5  RT (min)      lcorbi_t3_pos_1_028_32    m/z 795.6095     0   1   13.5   14  RT (min)      lcorbi_t3_pos_1_002_01    m/z 795.6096     0   1   16   16.5  RT (min)      lcorbi_t3_pos_1_001_49    m/z 795.6096     0   1E+6   14.5   15  RT (min)      lcorbi_t3_pos_1_005_42    m/z 795.6096     0   1E+6   13.5   14  RT (min)      lcorbi_t3_pos_1_002_01    m/z 795.6098     0   1   2E+4   14.5   15  RT (min)      lcorbi_t3_pos_2_017_51    m/z 795.6098     0   1   2E+5   14.5   15  RT (min)      lcorbi_t3_pos_1_008_36    m/z 795.6098     0   1   16.5   17  RT (min)    lcorbi_t3_pos_1_023_02    m/z 795.6760     0   1   2   3E+4   15.5   16  RT (min)      lcorbi_t3_pos_1_001_49    m/z 795.6762     0   1E+4   14.5   15  RT (min)      lcorbi_t3_pos_2_038_03    m/z 795.6762     0   1   14.5   15  RT (min)      lcorbi_t3_pos_1_041_54    m/z 795.6762     0   2   4   6E+4   14   14.5  RT (min)      lcorbi_t3_pos_2_017_51    m/z 795.6762     0   1   2E+4   14.5   15  RT (min)      lcorbi_t3_pos_1_021_18    m/z 795.6762     0   1   15   15.5  RT (min)      lcorbi_t3_pos_2_049_55    m/z 795.6763     0   1   2   3E+4   15   15.5  RT (min)      lcorbi_t3_pos_1_025_52    m/z 795.6764     0   2   4   6E+4   14   14.5  RT (min)      lcorbi_t3_pos_2_017_51    m/z 795.6764     0   1   14.5   15  RT (min)      lcorbi_t3_pos_1_030_21    m/z 795.6764     0   1E+4   14.5   15  RT (min)      lcorbi_t3_pos_3_049_55    m/z 795.7727     0   2   4E+6   15   15.5  RT (min)       lcorbi_t3_pos_3_033_53    m/z 795.7727     0   2   4   6E+4   15.5   16  RT (min)      lcorbi_t3_pos_4_010_43    m/z 795.7728     0   2   4   6E+6   15   15.5  RT (min)      lcorbi_t3_pos_3_040_41    m/z 795.7728     0   1   15.5   16  RT (min)      lcorbi_t3_pos_4_049_55    m/z 795.7729     0   1   16   16.5  RT (min)      lcorbi_t3_pos_4_048_24    m/z 795.7729     0   2   4   6E+6   15   15.5  RT (min)      lcorbi_t3_pos_3_017_51    m/z 795.7729     0   1   16.5   17  RT (min)    lcorbi_t3_pos_4_033_53    m/z 795.7730     0   1   16   16.5  RT (min)      lcorbi_t3_pos_4_041_54    m/z 795.7730     0   1E+5   15.5   16  RT (min)      lcorbi_t3_pos_3_025_52    m/z 795.7730     0   1   16   16.5  RT (min)      lcorbi_t3_pos_2_025_52    m/z 795.7730     0   2   4E+5   15.5   16  RT (min)      lcorbi_t3_pos_4_001_49    m/z 795.7730     0   1   17   17.5  RT (min)    lcorbi_t3_pos_1_049_55    m/z 795.7730     0   1   17   17.5  RT (min)    lcorbi_t3_pos_2_056_56    m/z 795.7731     0   1   16.5   17  RT (min)    lcorbi_t3_pos_2_049_55    m/z 795.7732     0   1   17   17.5  RT (min)    lcorbi_t3_pos_2_056_56    m/z 796.5843     0   1E+6   14   14.5  RT (min)      lcorbi_t3_pos_1_005_42    m/z 796.5846     0   1E+6   13.5   14  RT (min)      lcorbi_t3_pos_1_002_01    m/z 796.5846     0   1   2   3E+4   15   15.5  RT (min)      lcorbi_t3_pos_1_047_41    m/z 796.5846     0   1   15.5   16  RT (min)      lcorbi_t3_pos_1_041_54    m/z 796.5847     0   2   4E+5   13   13.5  RT (min)      lcorbi_t3_pos_4_057_49    m/z 796.5848     0   1   15.5   16  RT (min)      lcorbi_t3_pos_1_056_56    m/z 796.5849     0   1E+6   14.5   15  RT (min)      lcorbi_t3_pos_1_005_42    m/z 796.5849     0   1E+6   14.5   15  RT (min)      lcorbi_t3_pos_1_005_42    m/z 796.5850     0   1   15   15.5  RT (min)      lcorbi_t3_pos_2_025_52    m/z 796.5851     0   1   15   15.5  RT (min)      lcorbi_t3_pos_1_030_21    m/z 796.5852     0   1   16.5   17  RT (min)    lcorbi_t3_pos_1_023_02    m/z 796.5852     0   1   16   16.5  RT (min)      lcorbi_t3_pos_1_051_09    m/z 796.5852     0   1   15   15.5  RT (min)      lcorbi_t3_pos_1_028_32    m/z 796.5852     0   1   2   3E+4   14.5   15  RT (min)      lcorbi_t3_pos_2_017_51    m/z 796.5852     0   1   16   16.5  RT (min)      lcorbi_t3_pos_1_001_49    m/z 796.5854     0   1   15.5   16  RT (min)      lcorbi_t3_pos_1_041_54    m/z 796.5858     0   1   16.5   17  RT (min)    lcorbi_t3_pos_1_023_02    m/z 796.5885     0   1   2   3E+4   13   13.5  RT (min)      lcorbi_t3_pos_1_002_01    m/z 796.6131     0   1   16   16.5  RT (min)      lcorbi_t3_pos_4_033_53    m/z 796.6206     0   1E+6   14.5   15  RT (min)      lcorbi_t3_pos_3_017_51    m/z 796.6209     0   1   13   13.5  RT (min)      lcorbi_t3_pos_2_032_39    m/z 796.6210     0   1   16   16.5  RT (min)      lcorbi_t3_pos_1_049_55    m/z 796.6213     0   1   16.5   17  RT (min)    lcorbi_t3_pos_1_023_02    m/z 796.6214     0   1   15.5   16  RT (min)      lcorbi_t3_pos_1_034_34    m/z 796.6214     0   1   16   16.5  RT (min)      lcorbi_t3_pos_1_051_09    m/z 796.6215     0   1E+4   15   15.5  RT (min)      lcorbi_t3_pos_2_022_38    m/z 796.6215     0   2   4   6   8E+5   0.5   1  RT (min)      lcorbi_t3_pos_4_057_49    m/z 796.6216     0   1   2E+4   15   15.5  RT (min)      lcorbi_t3_pos_2_046_07    m/z 796.6216     0   1   15.5   16  RT (min)      lcorbi_t3_pos_1_028_32    m/z 796.6216     0   1   2   3E+3   13.5   14  RT (min)      lcorbi_t3_pos_2_054_29    m/z 796.6216     0   1   2E+6   14.5   15  RT (min)      lcorbi_t3_pos_3_047_40    m/z 796.6216     0   1   15.5   16  RT (min)      lcorbi_t3_pos_1_028_32    m/z 796.6216     0   2   4E+5   14.5   15  RT (min)      lcorbi_t3_pos_1_008_36    m/z 796.6216     0   1   15.5   16  RT (min)      lcorbi_t3_pos_1_034_34    m/z 796.6216     0   1   16.5   17  RT (min)    lcorbi_t3_pos_1_051_09    m/z 796.6216     0   1   16   16.5  RT (min)      lcorbi_t3_pos_1_041_54    m/z 796.6216     0   1E+4   15   15.5  RT (min)      lcorbi_t3_pos_2_022_38    m/z 796.6217     0   1   16   16.5  RT (min)      lcorbi_t3_pos_1_051_09    m/z 796.6218     0   1   13   13.5  RT (min)      lcorbi_t3_pos_2_032_39    m/z 796.6218     0   1   15.5   16  RT (min)      lcorbi_t3_pos_2_038_03    m/z 796.6220     0   1   16.5   17  RT (min)    lcorbi_t3_pos_1_051_09    m/z 796.6220     0   1   17   17.5  RT (min)    lcorbi_t3_pos_1_023_02    m/z 796.6220     0   1   2   3   4E+5   13.5   14  RT (min)      lcorbi_t3_pos_1_002_01    m/z 796.6224     0   1   17   17.5  RT (min)    lcorbi_t3_pos_1_023_02    m/z 796.6790     0   1   2E+5   14   14.5  RT (min)      lcorbi_t3_pos_2_017_51    m/z 796.6790     0   1   2E+4   14.5   15  RT (min)      lcorbi_t3_pos_4_048_24    m/z 796.6790     0   1   2E+4   15   15.5  RT (min)      lcorbi_t3_pos_1_025_52    m/z 796.6791     0   1   2   3   4E+4   14.5   15  RT (min)      lcorbi_t3_pos_1_021_18    m/z 796.6791     0   1E+4   15   15.5  RT (min)      lcorbi_t3_pos_4_033_53    m/z 796.6791     0   1   2E+4   15   15.5  RT (min)      lcorbi_t3_pos_2_049_55    m/z 796.6791     0   1   2E+5   14   14.5  RT (min)      lcorbi_t3_pos_2_017_51    m/z 796.6792     0   2   4   6   8E+5   10   10.5  RT (min)      lcorbi_t3_pos_4_057_49    m/z 796.6792     0   2   4   6E+4   14.5   15  RT (min)      lcorbi_t3_pos_1_030_21    m/z 796.6792     0   1   2   3   4E+4   14.5   15  RT (min)      lcorbi_t3_pos_1_041_54    m/z 796.6793     0   2   4   6   8E+5   10.5   11  RT (min)      lcorbi_t3_pos_4_057_49    m/z 796.6796     0   1E+4   15   15.5  RT (min)      lcorbi_t3_pos_4_033_53    m/z 796.6797     0   1   2   3E+4   15.5   16  RT (min)      lcorbi_t3_pos_1_001_49    m/z 797.5884     0   1   2   3E+4   13   13.5  RT (min)      lcorbi_t3_pos_1_002_01    m/z 797.5888     0   2   4   6   8E+4   13.5   14  RT (min)      lcorbi_t3_pos_1_002_01    m/z 797.5888     0   1   15   15.5  RT (min)      lcorbi_t3_pos_1_021_18    m/z 797.5888     0   1   16.5   17  RT (min)    lcorbi_t3_pos_1_023_02    m/z 797.5890     0   2   4E+5   14   14.5  RT (min)      lcorbi_t3_pos_1_005_42    m/z 797.5891     0   1   16   16.5  RT (min)      lcorbi_t3_pos_1_051_09    m/z 797.6250     0   1   15.5   16  RT (min)      lcorbi_t3_pos_1_028_32    m/z 797.6250     0   1   15.5   16  RT (min)      lcorbi_t3_pos_1_034_34    m/z 797.6250     0   1   15   15.5  RT (min)      lcorbi_t3_pos_2_022_38    m/z 797.6251     0   1   17   17.5  RT (min)    lcorbi_t3_pos_1_023_02    m/z 797.6251     0   1   15.5   16  RT (min)      lcorbi_t3_pos_1_034_34    m/z 797.6251     0   1   16   16.5  RT (min)      lcorbi_t3_pos_1_051_09    m/z 797.6251     0   1   16.5   17  RT (min)    lcorbi_t3_pos_2_012_01    m/z 797.6252     0   1E+6   14.5   15  RT (min)      lcorbi_t3_pos_4_019_47    m/z 797.6252     0   1E+6   13.5   14  RT (min)      lcorbi_t3_pos_4_016_15    m/z 797.6252     0   1   15   15.5  RT (min)      lcorbi_t3_pos_2_022_38    m/z 797.6253     0   1   2E+4   15.5   16  RT (min)      lcorbi_t3_pos_1_028_32    m/z 797.6253     0   1   2E+5   14.5   15  RT (min)      lcorbi_t3_pos_1_008_36    m/z 797.6253     0   1   16   16.5  RT (min)      lcorbi_t3_pos_1_031_40    m/z 797.6253     0   1   16   16.5  RT (min)      lcorbi_t3_pos_1_051_09    m/z 797.6255     0   1   17   17.5  RT (min)    lcorbi_t3_pos_1_023_02    m/z 797.6256     0   1   17   17.5  RT (min)    lcorbi_t3_pos_1_023_02    m/z 797.6256     0   1E+6   14.5   15  RT (min)      lcorbi_t3_pos_4_019_47    m/z 797.6257     0   1   15   15.5  RT (min)      lcorbi_t3_pos_2_006_47    m/z 797.6278     0   1   2E+5   13.5   14  RT (min)       lcorbi_t3_pos_1_002_01    m/z 797.6848     0   1E+5   14.5   15  RT (min)      lcorbi_t3_pos_2_046_07    m/z 797.6913     0   1E+6   12   12.5  RT (min)      lcorbi_t3_pos_4_057_49    m/z 797.6914     0   1   2   3   4E+5   14   14.5  RT (min)      lcorbi_t3_pos_3_025_52    m/z 797.6916     0   1   2E+4   14.5   15   15.5  RT (min)      lcorbi_t3_pos_2_025_52    m/z 797.6916     0   1   15   15.5  RT (min)      lcorbi_t3_pos_2_009_50    m/z 797.6917     0   1   15.5   16  RT (min)      lcorbi_t3_pos_1_049_55    m/z 797.6917     0   1   2E+4   14.5   15  RT (min)      lcorbi_t3_pos_1_030_21    m/z 797.6917     0   1   15.5   16  RT (min)      lcorbi_t3_pos_1_025_52    m/z 797.6917     0   1E+4   14.5   15  RT (min)      lcorbi_t3_pos_1_030_21    m/z 797.6917     0   1   15   15.5  RT (min)      lcorbi_t3_pos_1_041_54    m/z 797.6918     0   1   15.5   16  RT (min)      lcorbi_t3_pos_1_025_52    m/z 797.6920     0   2   4   6E+6   14   14.5  RT (min)      lcorbi_t3_pos_1_005_42    m/z 797.6925     0   2   4E+6   14   14.5  RT (min)      lcorbi_t3_pos_3_026_04    m/z 798.5643     0   1   14.5   15  RT (min)      lcorbi_t3_pos_1_031_40    m/z 798.5644     0   1E+5   12   12.5  RT (min)      lcorbi_t3_pos_3_024_07    m/z 798.5644     0   2   4   6E+5   12.5   13  RT (min)      lcorbi_t3_pos_3_011_42    m/z 798.5644     0   1E+6   12.5   13  RT (min)       lcorbi_t3_pos_4_075_58    m/z 798.5645     0   1   2   3E+4   14   14.5  RT (min)      lcorbi_t3_pos_1_028_32    m/z 798.5645     0   1   2E+4   14   14.5  RT (min)      lcorbi_t3_pos_1_028_32    m/z 798.5646     0   2   4   6E+4   14   14.5  RT (min)      lcorbi_t3_pos_1_028_32    m/z 798.5646     0   2   4E+5   13.5   14  RT (min)       lcorbi_t3_pos_1_008_36    m/z 798.5646     0   1E+5   12   12.5  RT (min)      lcorbi_t3_pos_2_013_33    m/z 798.5646     0   2   4   6E+5   12.5   13  RT (min)       lcorbi_t3_pos_3_011_42    m/z 798.5647     0   1   14.5   15  RT (min)      lcorbi_t3_pos_1_031_40    m/z 798.5647     0   1E+4   15   15.5  RT (min)      lcorbi_t3_pos_2_012_01    m/z 798.5648     0   1E+5   13.5   14  RT (min)      lcorbi_t3_pos_1_008_36    m/z 798.5875     0   2   4E+4   15.5   16  RT (min)      lcorbi_t3_pos_1_023_02    m/z 798.5880     0   1   12.5   13  RT (min)      lcorbi_t3_pos_2_013_33    m/z 798.5880     0   1   2   3E+4   14.5   15  RT (min)      lcorbi_t3_pos_2_049_55    m/z 798.5881     0   1   2   3E+4   14.5   15  RT (min)      lcorbi_t3_pos_4_033_53    m/z 798.5882     0   1   12.5   13  RT (min)      lcorbi_t3_pos_3_024_07    m/z 798.5882     0   2   4   6E+6   12.5   13  RT (min)      lcorbi_t3_pos_2_054_29    m/z 798.5882     0   2   4E+4   15   15.5  RT (min)      lcorbi_t3_pos_1_031_40    m/z 798.5882     0   1   2   3   4E+4   14.5   15  RT (min)      lcorbi_t3_pos_1_034_34    m/z 798.5882     0   1   2   3E+4   15   15.5  RT (min)       lcorbi_t3_pos_2_012_01    m/z 798.5883     0   2   4   6E+4   14   14.5  RT (min)       lcorbi_t3_pos_1_028_32    m/z 798.5883     0   2   4E+4   15   15.5  RT (min)      lcorbi_t3_pos_1_001_49    m/z 798.5883     0   1E+7   13.5   14  RT (min)       lcorbi_t3_pos_4_019_47    m/z 798.5884     0   2   4   6   8E+5   12.5   13  RT (min)      lcorbi_t3_pos_4_015_45    m/z 798.5884     0   2   4   6   8E+4   14   14.5  RT (min)      lcorbi_t3_pos_2_006_47    m/z 798.5884     0   1E+7   13.5   14  RT (min)      lcorbi_t3_pos_3_047_40    m/z 798.5884     0   1   2E+6   13.5   14  RT (min)       lcorbi_t3_pos_1_008_36    m/z 798.5886     0   1   2   3E+4   14   14.5  RT (min)       lcorbi_t3_pos_3_003_33    m/z 798.5887     0   2   4   6   8E+5   12.5   13  RT (min)      lcorbi_t3_pos_4_015_45    m/z 798.5888     0   2   4   6   8E+4   14   14.5  RT (min)      lcorbi_t3_pos_1_047_41    m/z 798.6004     0   1E+5   14.5   15  RT (min)      lcorbi_t3_pos_1_008_36    m/z 798.6005     0   1   16   16.5  RT (min)      lcorbi_t3_pos_1_031_40    m/z 798.6006     0   1   15   15.5  RT (min)      lcorbi_t3_pos_1_028_32    m/z 798.6006     0   1   16.5   17  RT (min)    lcorbi_t3_pos_1_023_02    m/z 798.6006     0   1   15.5   16  RT (min)      lcorbi_t3_pos_1_034_34    m/z 798.6007     0   2   4E+5   14.5   15  RT (min)      lcorbi_t3_pos_1_005_42    m/z 798.6007     0   1   16   16.5  RT (min)      lcorbi_t3_pos_1_031_40    m/z 798.6008     0   1   2   3E+4   13.5   14  RT (min)      lcorbi_t3_pos_1_002_01    m/z 798.6010     0   1   16.5   17  RT (min)    lcorbi_t3_pos_1_023_02    m/z 798.6295     0   1   16   16.5  RT (min)      lcorbi_t3_pos_1_031_40    m/z 798.6297     0   1   17   17.5  RT (min)    lcorbi_t3_pos_1_023_02    m/z 798.6320     0   2   4E+5   12.5   13  RT (min)       lcorbi_t3_pos_4_090_60    m/z 798.6331     0   1   15   15.5  RT (min)      lcorbi_t3_pos_1_001_49    m/z 798.6337     0   1   2E+4   15   15.5  RT (min)      lcorbi_t3_pos_1_049_55    m/z 798.6344     0   1   15.5   16  RT (min)      lcorbi_t3_pos_1_028_32    m/z 798.6352     0   1   16   16.5  RT (min)      lcorbi_t3_pos_1_031_40    m/z 798.6360     0   1   2E+4   15   15.5  RT (min)      lcorbi_t3_pos_1_049_55    m/z 798.6362     0   1   15.5   16  RT (min)      lcorbi_t3_pos_1_028_32    m/z 798.6363     0   1   15.5   16  RT (min)      lcorbi_t3_pos_1_034_34    m/z 798.6366     0   1   2E+5   14.5   15  RT (min)      lcorbi_t3_pos_1_041_54    m/z 798.6942     0   2   4   6E+6   12   12.5  RT (min)      lcorbi_t3_pos_4_057_49    m/z 798.6946     0   1   2   3E+5   14.5   15  RT (min)      lcorbi_t3_pos_2_046_07    m/z 798.6946     0   1   2   3E+4   15.5   16  RT (min)       lcorbi_t3_pos_1_025_52    m/z 798.6947     0   2   4   6E+5   14   14.5  RT (min)      lcorbi_t3_pos_3_056_56    m/z 798.6947     0   2   4E+4   14.5   15  RT (min)      lcorbi_t3_pos_1_021_18    m/z 798.6947     0   2   4E+4   15   15.5  RT (min)      lcorbi_t3_pos_2_009_50    m/z 798.6947     0   2   4   6E+4   14.5   15  RT (min)      lcorbi_t3_pos_4_049_55    m/z 798.6948     0   1   2   3E+4   15   15.5  RT (min)      lcorbi_t3_pos_2_038_03    m/z 798.6948     0   1   2   3E+4   15.5   16  RT (min)      lcorbi_t3_pos_1_025_52    m/z 798.6949     0   1   2   3   4E+4   15   15.5  RT (min)      lcorbi_t3_pos_1_041_54    m/z 798.6950     0   2   4   6E+4   14.5   15  RT (min)      lcorbi_t3_pos_1_030_21    m/z 798.6950     0   1   2   3E+7   14   14.5  RT (min)      lcorbi_t3_pos_1_005_42    m/z 798.6952     0   1   2   3E+4   15.5   16  RT (min)      lcorbi_t3_pos_1_025_52    m/z 798.7125     0   1   17   17.5  RT (min)    lcorbi_t3_pos_1_041_54    m/z 798.7126     0   1   17   17.5  RT (min)    lcorbi_t3_pos_1_049_55    m/z 798.7128     0   1   17   17.5  RT (min)    lcorbi_t3_pos_2_009_50    m/z 798.7129     0   1   17.5   18  RT (min)    lcorbi_t3_pos_1_025_52    m/z 798.7129     0   1   16.5   17  RT (min)    lcorbi_t3_pos_2_038_03    m/z 798.7129     0   1   16   16.5  RT (min)      lcorbi_t3_pos_1_047_41    m/z 798.7129     0   1   16.5   17  RT (min)    lcorbi_t3_pos_1_021_18    m/z 798.7130     0   1   17.5   18  RT (min)    lcorbi_t3_pos_1_025_52    m/z 798.7131     0   2   4E+5   15.5   16  RT (min)      lcorbi_t3_pos_3_040_41    m/z 799.5910     0   1   12   12.5  RT (min)      lcorbi_t3_pos_2_047_31    m/z 799.5913     0   1   2   3E+4   15   15.5  RT (min)      lcorbi_t3_pos_2_012_01    m/z 799.5914     0   1   2   3E+4   15.5   16  RT (min)      lcorbi_t3_pos_1_023_02    m/z 799.5914     0   1   2   3E+4   14.5   15  RT (min)      lcorbi_t3_pos_1_041_54    m/z 799.5914     0   1   2E+4   14.5   15  RT (min)      lcorbi_t3_pos_2_049_55    m/z 799.5914     0   1   2   3E+4   15   15.5  RT (min)      lcorbi_t3_pos_2_056_56    m/z 799.5914     0   1   12.5   13  RT (min)      lcorbi_t3_pos_2_013_33    m/z 799.5915     0   1   2   3E+6   12.5   13  RT (min)      lcorbi_t3_pos_2_054_29    m/z 799.5915     0   1   12.5   13  RT (min)      lcorbi_t3_pos_3_024_07    m/z 799.5915     0   2   4   6E+6   13.5   14  RT (min)      lcorbi_t3_pos_3_047_40    m/z 799.5915     0   1   2   3E+4   15   15.5  RT (min)      lcorbi_t3_pos_1_031_40    m/z 799.5915     0   1   2E+4   14.5   15  RT (min)      lcorbi_t3_pos_2_009_50    m/z 799.5915     0   1   14.5   15  RT (min)      lcorbi_t3_pos_1_034_34    m/z 799.5915     0   1E+4   14   14.5  RT (min)      lcorbi_t3_pos_3_003_33    m/z 799.5916     0   1   14.5   15  RT (min)      lcorbi_t3_pos_1_034_34    m/z 799.5916     0   1   2   3   4E+6   12.5   13  RT (min)      lcorbi_t3_pos_1_002_01    m/z 799.5916     0   1E+6   13.5   14  RT (min)       lcorbi_t3_pos_1_008_36    m/z 799.5916     0   1   2   3E+4   14   14.5  RT (min)      lcorbi_t3_pos_1_028_32    m/z 799.5918     0   2   4   6E+6   13.5   14  RT (min)       lcorbi_t3_pos_4_019_47    m/z 799.5918     0   1   2E+4   14   14.5  RT (min)      lcorbi_t3_pos_2_006_47    m/z 799.5920     0   1   2   3E+4   15   15.5  RT (min)      lcorbi_t3_pos_2_012_01    m/z 799.6408     0   1   15.5   16  RT (min)      lcorbi_t3_pos_1_025_52    m/z 799.6410     0   1   15   15.5  RT (min)      lcorbi_t3_pos_1_025_52    m/z 799.6419     0   1   2E+4   15   15.5  RT (min)      lcorbi_t3_pos_1_041_54    m/z 799.6431     0   2   4E+4   14.5   15  RT (min)      lcorbi_t3_pos_1_041_54    m/z 799.6431     0   1   2   3E+4   15   15.5  RT (min)      lcorbi_t3_pos_1_041_54    m/z 799.6435     0   1   2E+4   15   15.5  RT (min)      lcorbi_t3_pos_1_049_55    m/z 799.6436     0   1E+6   13.5   14  RT (min)      lcorbi_t3_pos_1_005_42    m/z 799.6437     0   1   15   15.5  RT (min)      lcorbi_t3_pos_1_056_56    m/z 799.6438     0   1E+6   13.5   14  RT (min)      lcorbi_t3_pos_1_005_42    m/z 799.6689     0   1   13.5   14  RT (min)      lcorbi_t3_pos_1_002_01    m/z 799.6690     0   2   4E+5   14.5   15  RT (min)      lcorbi_t3_pos_3_047_40    m/z 799.6691     0   1E+6   14   14.5  RT (min)      lcorbi_t3_pos_4_016_15    m/z 799.6692     0   1   15   15.5  RT (min)      lcorbi_t3_pos_2_022_38    m/z 799.6692     0   1   17   17.5  RT (min)    lcorbi_t3_pos_1_023_02    m/z 799.6692     0   1E+6   14.5   15  RT (min)      lcorbi_t3_pos_4_019_47    m/z 799.6692     0   1E+4   15.5   16  RT (min)      lcorbi_t3_pos_3_013_19    m/z 799.6692     0   1   16   16.5  RT (min)      lcorbi_t3_pos_2_009_50    m/z 799.6693     0   1   16   16.5  RT (min)      lcorbi_t3_pos_1_031_40    m/z 799.6694     0   1   15.5   16  RT (min)      lcorbi_t3_pos_1_034_34    m/z 799.6694     0   1   13.5   14  RT (min)      lcorbi_t3_pos_2_013_33    m/z 799.6694     0   1   16.5   17  RT (min)    lcorbi_t3_pos_1_051_09    m/z 799.6696     0   1   15   15.5  RT (min)      lcorbi_t3_pos_1_008_36    m/z 799.6697     0   1   17   17.5  RT (min)    lcorbi_t3_pos_1_023_02    m/z 799.6713     0   1   13   13.5  RT (min)      lcorbi_t3_pos_2_032_39    m/z 799.7007     0   1E+5   14.5   15  RT (min)      lcorbi_t3_pos_4_010_43    m/z 799.7009     0   2   4   6   8E+4   14.5   15  RT (min)      lcorbi_t3_pos_2_017_51    m/z 799.7039     0   1E+4   15.5   16  RT (min)      lcorbi_t3_pos_2_009_50    m/z 799.7039     0   1   2   3E+6   14   14.5  RT (min)      lcorbi_t3_pos_1_005_42    m/z 799.7042     0   1   2E+4   14.5   15  RT (min)      lcorbi_t3_pos_2_006_47    m/z 799.7064     0   1   2E+5   14.5   15  RT (min)      lcorbi_t3_pos_3_056_56    m/z 799.7064     0   1   2E+5   14.5   15  RT (min)      lcorbi_t3_pos_3_056_56    m/z 799.7070     0   1E+6   12.5   13  RT (min)      lcorbi_t3_pos_4_057_49    m/z 799.7070     0   1   2E+6   14   14.5  RT (min)      lcorbi_t3_pos_1_005_42    m/z 799.7071     0   1   2   3E+6   14   14.5  RT (min)      lcorbi_t3_pos_1_005_42    m/z 799.7071     0   1   15   15.5  RT (min)      lcorbi_t3_pos_2_049_55    m/z 799.7071     0   1   2   3E+4   14.5   15  RT (min)      lcorbi_t3_pos_1_030_21    m/z 799.7072     0   1   15.5   16  RT (min)      lcorbi_t3_pos_1_025_52    m/z 799.7072     0   1   15.5   16  RT (min)      lcorbi_t3_pos_1_025_52    m/z 799.7072     0   1   15   15.5  RT (min)      lcorbi_t3_pos_2_038_03    m/z 799.7072     0   1   15   15.5  RT (min)      lcorbi_t3_pos_1_021_18    m/z 799.7074     0   1   16   16.5  RT (min)      lcorbi_t3_pos_1_025_52    m/z 799.7074     0   1   15.5   16  RT (min)      lcorbi_t3_pos_1_056_56    m/z 799.7074     0   1   15   15.5  RT (min)      lcorbi_t3_pos_1_041_54    m/z 799.7430     0   1   15.5   16  RT (min)      lcorbi_t3_pos_1_056_56    m/z 799.7430     0   1   14.5   15  RT (min)      lcorbi_t3_pos_1_047_41    m/z 799.7430     0   1   15   15.5  RT (min)      lcorbi_t3_pos_1_030_21    m/z 799.7431     0   1   15.5   16  RT (min)      lcorbi_t3_pos_1_025_52    m/z 799.7433     0   1   15   15.5  RT (min)      lcorbi_t3_pos_1_041_54    m/z 799.7433     0   1   15.5   16  RT (min)      lcorbi_t3_pos_1_041_54    m/z 799.7434     0   1   16   16.5  RT (min)      lcorbi_t3_pos_1_001_49    m/z 800.5204     0   1   14.5   15  RT (min)      lcorbi_t3_pos_1_034_34    m/z 800.5204     0   1   14   14.5  RT (min)      lcorbi_t3_pos_1_047_41    m/z 800.5205     0   1   14.5   15  RT (min)      lcorbi_t3_pos_1_028_32    m/z 800.5206     0   1E+6   13.5   14  RT (min)      lcorbi_t3_pos_1_005_42    m/z 800.5206     0   2   4   6E+5   14   14.5  RT (min)      lcorbi_t3_pos_1_008_36    m/z 800.5206     0   1   15   15.5  RT (min)      lcorbi_t3_pos_1_031_40    m/z 800.5206     0   1   2E+4   14   14.5  RT (min)      lcorbi_t3_pos_3_056_56    m/z 800.5206     0   1   15   15.5  RT (min)      lcorbi_t3_pos_1_056_56    m/z 800.5206     0   1   12.5   13  RT (min)      lcorbi_t3_pos_2_013_33    m/z 800.5206     0   1   12.5   13  RT (min)      lcorbi_t3_pos_3_031_45    m/z 800.5206     0   1   14.5   15  RT (min)      lcorbi_t3_pos_1_028_32    m/z 800.5207     0   1   15.5   16  RT (min)      lcorbi_t3_pos_2_012_01    m/z 800.5207     0   1   15   15.5  RT (min)      lcorbi_t3_pos_2_012_01    m/z 800.5208     0   1   2E+6   13   13.5  RT (min)      lcorbi_t3_pos_3_011_42    m/z 800.5208     0   1   14.5   15  RT (min)      lcorbi_t3_pos_1_028_32    m/z 800.5208     0   1E+6   12.5   13  RT (min)      lcorbi_t3_pos_3_011_42    m/z 800.5208     0   2   4   6   8E+5   13.5   14  RT (min)      lcorbi_t3_pos_3_040_41    m/z 800.5944     0   1   2   3E+4   15   15.5  RT (min)      lcorbi_t3_pos_1_031_40    m/z 800.5945     0   2   4   6   8E+5   12.5   13  RT (min)      lcorbi_t3_pos_2_054_29    m/z 800.5945     0   1E+4   14.5   15  RT (min)      lcorbi_t3_pos_2_049_55    m/z 800.5945     0   1   12.5   13  RT (min)      lcorbi_t3_pos_2_013_33    m/z 800.5945     0   1   15.5   16  RT (min)      lcorbi_t3_pos_1_023_02    m/z 800.5946     0   1   14   14.5  RT (min)      lcorbi_t3_pos_3_013_19    m/z 800.5946     0   1   12   12.5  RT (min)      lcorbi_t3_pos_2_032_39    m/z 800.5946     0   1E+4   14   14.5  RT (min)      lcorbi_t3_pos_3_003_33    m/z 800.5947     0   1   2E+4   14   14.5  RT (min)      lcorbi_t3_pos_2_046_07    m/z 800.5947     0   1   2   3   4E+5   13.5   14  RT (min)      lcorbi_t3_pos_1_008_36    m/z 800.5948     0   1E+6   13.5   14  RT (min)       lcorbi_t3_pos_4_019_47    m/z 800.5948     0   1   15   15.5  RT (min)      lcorbi_t3_pos_2_056_56    m/z 800.5948     0   1E+4   14.5   15  RT (min)      lcorbi_t3_pos_1_034_34    m/z 800.5948     0   2   4   6E+5   13.5   14  RT (min)      lcorbi_t3_pos_3_047_40    m/z 800.6166     0   1   16   16.5  RT (min)      lcorbi_t3_pos_1_031_40    m/z 800.6166     0   1   16   16.5  RT (min)      lcorbi_t3_pos_1_031_40    m/z 800.6166     0   1   15   15.5  RT (min)      lcorbi_t3_pos_1_047_41    m/z 800.6168     0   1   15.5   16  RT (min)      lcorbi_t3_pos_1_034_34    m/z 800.6168     0   1   2E+4   15   15.5  RT (min)      lcorbi_t3_pos_1_028_32    m/z 800.6168     0   1   2E+4   15.5   16  RT (min)      lcorbi_t3_pos_1_028_32    m/z 800.6169     0   1   15.5   16  RT (min)      lcorbi_t3_pos_1_034_34    m/z 800.6169     0   1   13.5   14  RT (min)      lcorbi_t3_pos_2_013_33    m/z 800.6170     0   1   15   15.5  RT (min)      lcorbi_t3_pos_2_006_47    m/z 800.6170     0   2   4   6E+5   14.5   15  RT (min)      lcorbi_t3_pos_3_033_53    m/z 800.6170     0   2   4   6   8E+4   14.5   15  RT (min)      lcorbi_t3_pos_1_008_36    m/z 800.6170     0   2   4E+4   15   15.5  RT (min)      lcorbi_t3_pos_1_008_36    m/z 800.6170     0   1   16   16.5  RT (min)      lcorbi_t3_pos_2_056_56    m/z 800.6170     0   1   16.5   17  RT (min)    lcorbi_t3_pos_1_023_02    m/z 800.6172     0   1   17   17.5  RT (min)    lcorbi_t3_pos_1_023_02    m/z 800.6173     0   1   15.5   16  RT (min)      lcorbi_t3_pos_2_025_52    m/z 800.6468     0   2   4   6   8E+4   15   15.5  RT (min)      lcorbi_t3_pos_1_041_54    m/z 800.6468     0   2   4E+4   15.5   16  RT (min)      lcorbi_t3_pos_1_001_49    m/z 800.6469     0   2   4   6   8E+4   14.5   15  RT (min)      lcorbi_t3_pos_1_041_54    m/z 800.6470     0   2   4E+4   15.5   16  RT (min)      lcorbi_t3_pos_1_001_49    m/z 800.6470     0   1E+6   14   14.5  RT (min)      lcorbi_t3_pos_3_025_52    m/z 800.6470     0   2   4   6E+6   13.5   14  RT (min)      lcorbi_t3_pos_1_005_42    m/z 800.6471     0   1E+5   14   14.5  RT (min)      lcorbi_t3_pos_2_017_51    m/z 800.6472     0   2   4   6E+6   13.5   14   14.5  RT (min)      lcorbi_t3_pos_1_005_42    m/z 800.6474     0   2   4E+4   15.5   16  RT (min)      lcorbi_t3_pos_1_001_49    m/z 800.6494     0   1   2   3E+4   15   15.5  RT (min)       lcorbi_t3_pos_1_056_56    m/z 800.6518     0   1   2   3   4E+4   15   15.5  RT (min)       lcorbi_t3_pos_1_025_52    m/z 800.6722     0   1   13.5   14  RT (min)      lcorbi_t3_pos_1_002_01    m/z 800.6722     0   1   15.5   16  RT (min)      lcorbi_t3_pos_1_028_32    m/z 800.6722     0   1   16.5   17  RT (min)    lcorbi_t3_pos_1_031_40    m/z 800.6724     0   1   15   15.5  RT (min)      lcorbi_t3_pos_2_022_38    m/z 800.6724     0   1   13.5   14  RT (min)      lcorbi_t3_pos_3_031_45    m/z 800.6725     0   2   4   6E+4   15   15.5  RT (min)      lcorbi_t3_pos_1_008_36    m/z 800.6725     0   2   4E+5   14   14.5  RT (min)      lcorbi_t3_pos_4_016_15    m/z 800.6726     0   1   16.5   17  RT (min)    lcorbi_t3_pos_1_051_09    m/z 800.6726     0   1   13   13.5  RT (min)      lcorbi_t3_pos_2_032_39    m/z 800.6728     0   1   15   15.5  RT (min)      lcorbi_t3_pos_1_008_36    m/z 800.6728     0   1   15.5   16  RT (min)      lcorbi_t3_pos_1_034_34    m/z 800.6728     0   2   4E+5   14.5   15  RT (min)      lcorbi_t3_pos_4_019_47    m/z 800.6729     0   1   17   17.5  RT (min)    lcorbi_t3_pos_1_023_02    m/z 800.7102     0   1   2   3   4E+6   12.5   13  RT (min)      lcorbi_t3_pos_4_057_49    m/z 800.7104     0   2   4   6E+4   14.5   15  RT (min)      lcorbi_t3_pos_2_006_47    m/z 800.7106     0   1   2   3E+4   15.5   16  RT (min)      lcorbi_t3_pos_1_025_52    m/z 800.7106     0   1   2E+4   15.5   16  RT (min)      lcorbi_t3_pos_1_025_52    m/z 800.7106     0   2   4   6E+4   15   15.5  RT (min)      lcorbi_t3_pos_1_021_18    m/z 800.7107     0   1E+5   14.5   15  RT (min)      lcorbi_t3_pos_1_047_41    m/z 800.7107     0   1E+7   14   14.5  RT (min)      lcorbi_t3_pos_1_005_42    m/z 800.7107     0   1   2   3E+4   15.5   16  RT (min)      lcorbi_t3_pos_1_001_49    m/z 800.7108     0   1   2   3E+6   14   14.5  RT (min)      lcorbi_t3_pos_4_051_34    m/z 800.7108     0   2   4   6E+4   14.5   15  RT (min)      lcorbi_t3_pos_1_030_21    m/z 800.7108     0   1   2E+4   15.5   16  RT (min)      lcorbi_t3_pos_1_056_56    m/z 800.7108     0   1   2E+4   16   16.5  RT (min)      lcorbi_t3_pos_2_056_56    m/z 800.7109     0   1   2E+4   15   15.5  RT (min)      lcorbi_t3_pos_2_049_55    m/z 800.7109     0   1   2   3E+4   15.5   16  RT (min)      lcorbi_t3_pos_2_009_50    m/z 800.7109     0   2   4   6   8E+6   14   14.5  RT (min)      lcorbi_t3_pos_1_005_42    m/z 800.7109     0   2   4E+4   15   15.5  RT (min)      lcorbi_t3_pos_1_041_54    m/z 800.7110     0   1   2E+4   15   15.5  RT (min)      lcorbi_t3_pos_2_038_03    m/z 800.7110     0   2   4   6E+5   14.5   15  RT (min)      lcorbi_t3_pos_3_056_56    m/z 800.7110     0   1   2E+4   16   16.5  RT (min)      lcorbi_t3_pos_2_056_56    m/z 800.7111     0   1   2E+4   16   16.5  RT (min)      lcorbi_t3_pos_2_056_56    m/z 800.7117     0   1   2E+4   15.5   16  RT (min)      lcorbi_t3_pos_2_049_55    m/z 800.7283     0   1   16.5   17  RT (min)    lcorbi_t3_pos_1_041_54    m/z 800.7284     0   1E+4   15.5   16  RT (min)      lcorbi_t3_pos_3_025_52    m/z 800.7285     0   2   4E+5   15   15.5  RT (min)      lcorbi_t3_pos_3_040_41    m/z 801.5238     0   1   12.5   13  RT (min)      lcorbi_t3_pos_2_013_33    m/z 801.5238     0   1   14.5   15  RT (min)      lcorbi_t3_pos_1_034_34    m/z 801.5240     0   1   15   15.5  RT (min)      lcorbi_t3_pos_1_031_40    m/z 801.5240     0   1   2E+4   14.5   15  RT (min)      lcorbi_t3_pos_1_028_32    m/z 801.5240     0   1   2   3E+5   14   14.5  RT (min)      lcorbi_t3_pos_1_008_36    m/z 801.5240     0   1   14.5   15  RT (min)      lcorbi_t3_pos_1_034_34    m/z 801.5240     0   1   14   14.5  RT (min)      lcorbi_t3_pos_2_022_38    m/z 801.5240     0   1   15   15.5  RT (min)      lcorbi_t3_pos_1_031_40    m/z 801.5241     0   2   4E+5   13.5   14  RT (min)      lcorbi_t3_pos_1_005_42    m/z 801.5241     0   2   4E+5   12.5   13  RT (min)      lcorbi_t3_pos_3_011_42    m/z 801.5242     0   1   12.5   13  RT (min)      lcorbi_t3_pos_3_024_07    m/z 801.5242     0   1   15   15.5  RT (min)      lcorbi_t3_pos_2_012_01    m/z 801.5242     0   1   12.5   13  RT (min)      lcorbi_t3_pos_3_031_45    m/z 801.5974     0   1   12.5   13  RT (min)      lcorbi_t3_pos_2_013_33    m/z 801.5980     0   1   2E+4   15.5   16  RT (min)      lcorbi_t3_pos_1_023_02    m/z 801.6202     0   1   15.5   16  RT (min)      lcorbi_t3_pos_1_025_52    m/z 801.6202     0   1   16   16.5  RT (min)      lcorbi_t3_pos_1_031_40    m/z 801.6202     0   1   2   3   4E+5   14.5   15  RT (min)      lcorbi_t3_pos_1_005_42    m/z 801.6204     0   1   15.5   16  RT (min)      lcorbi_t3_pos_1_025_52    m/z 801.6209     0   1   2   3E+5   14.5   15  RT (min)      lcorbi_t3_pos_1_033_53    m/z 801.6230     0   1   2E+5   14.5   15  RT (min)      lcorbi_t3_pos_1_021_18    m/z 801.6257     0   2   4E+4   14.5   15  RT (min)      lcorbi_t3_pos_1_008_36    m/z 801.6555     0   1   15   15.5  RT (min)      lcorbi_t3_pos_1_041_54    m/z 801.6555     0   1   15   15.5  RT (min)      lcorbi_t3_pos_1_041_54    m/z 801.6590     0   2   4E+4   14.5   15  RT (min)      lcorbi_t3_pos_1_030_21    m/z 801.6591     0   1   2   3   4E+4   14.5   15  RT (min)      lcorbi_t3_pos_1_033_53    m/z 801.6594     0   1   2E+4   15.5   16  RT (min)      lcorbi_t3_pos_1_049_55    m/z 801.6596     0   1   2   3   4E+4   15.5   16  RT (min)      lcorbi_t3_pos_1_001_49    m/z 801.7197     0   1   16   16.5  RT (min)      lcorbi_t3_pos_1_049_55    m/z 801.7199     0   1   15   15.5  RT (min)      lcorbi_t3_pos_1_030_21    m/z 801.7226     0   1   15   15.5  RT (min)      lcorbi_t3_pos_1_033_53    m/z 801.7226     0   1   15.5   16  RT (min)      lcorbi_t3_pos_1_041_54    m/z 801.7229     0   1   16   16.5  RT (min)      lcorbi_t3_pos_1_025_52    m/z 802.5272     0   1E+4   14.5   15  RT (min)      lcorbi_t3_pos_1_047_41    m/z 802.5276     0   1   12.5   13  RT (min)      lcorbi_t3_pos_3_031_45    m/z 802.5315     0   1   12.5   13  RT (min)      lcorbi_t3_pos_3_031_45    m/z 802.5358     0   2   4E+5   13   13.5  RT (min)      lcorbi_t3_pos_3_011_42    m/z 802.5360     0   2   4   6E+4   12.5   13  RT (min)      lcorbi_t3_pos_2_013_33    m/z 802.5360     0   1   14.5   15  RT (min)      lcorbi_t3_pos_2_041_54    m/z 802.5362     0   1E+5   13   13.5  RT (min)      lcorbi_t3_pos_3_011_42    m/z 802.5362     0   1   15   15.5  RT (min)      lcorbi_t3_pos_1_034_34    m/z 802.5363     0   1   2   3   4E+4   15.5   16  RT (min)      lcorbi_t3_pos_1_031_40    m/z 802.5363     0   1   14.5   15  RT (min)      lcorbi_t3_pos_1_028_32    m/z 802.5363     0   1E+4   14.5   15  RT (min)      lcorbi_t3_pos_2_022_38    m/z 802.5363     0   2   4   6   8E+5   14   14.5  RT (min)      lcorbi_t3_pos_1_008_36    m/z 802.5364     0   1   2E+4   15.5   16  RT (min)      lcorbi_t3_pos_2_012_01    m/z 802.5364     0   1   2E+4   15.5   16  RT (min)      lcorbi_t3_pos_2_012_01    m/z 802.5364     0   1   2   3   4E+5   14   14.5  RT (min)      lcorbi_t3_pos_1_008_36    m/z 802.5364     0   1   15   15.5  RT (min)      lcorbi_t3_pos_1_034_34    m/z 802.5365     0   1   2   3   4E+4   15   15.5  RT (min)      lcorbi_t3_pos_1_031_40    m/z 802.5594     0   2   4   6E+5   12.5   13  RT (min)       lcorbi_t3_pos_2_005_11    m/z 802.5598     0   2   4   6E+5   12.5   13  RT (min)       lcorbi_t3_pos_1_054_48    m/z 802.5598     0   2   4   6E+5   12.5   13  RT (min)       lcorbi_t3_pos_2_005_11    m/z 802.6621     0   1E+5   14   14.5  RT (min)      lcorbi_t3_pos_3_056_56    m/z 802.6623     0   1   2E+4   15   15.5  RT (min)      lcorbi_t3_pos_2_038_03    m/z 802.6625     0   1   15.5   16  RT (min)      lcorbi_t3_pos_1_049_55    m/z 802.6626     0   1   2   3E+4   15   15.5  RT (min)      lcorbi_t3_pos_1_041_54    m/z 802.6626     0   1   2   3E+4   15   15.5  RT (min)      lcorbi_t3_pos_1_041_54    m/z 802.6626     0   1   15.5   16  RT (min)      lcorbi_t3_pos_1_001_49    m/z 802.6626     0   1   2   3E+4   14.5   15  RT (min)      lcorbi_t3_pos_2_046_07    m/z 802.6626     0   1   2E+6   14   14.5  RT (min)      lcorbi_t3_pos_1_005_42    m/z 802.6627     0   1   2   3E+4   15   15.5  RT (min)      lcorbi_t3_pos_1_041_54    m/z 802.6627     0   1   2   3   4E+4   14.5   15  RT (min)      lcorbi_t3_pos_1_047_41    m/z 802.6627     0   1   2E+4   14.5   15  RT (min)      lcorbi_t3_pos_2_041_54    m/z 802.6627     0   1   2   3E+4   14.5   15  RT (min)      lcorbi_t3_pos_1_021_18    m/z 802.6628     0   1   2E+6   14   14.5  RT (min)       lcorbi_t3_pos_1_005_42    m/z 802.7256     0   1   16   16.5  RT (min)      lcorbi_t3_pos_2_056_56    m/z 802.7262     0   1   2E+4   15   15.5  RT (min)      lcorbi_t3_pos_1_047_41    m/z 802.7263     0   1E+6   14.5   15  RT (min)      lcorbi_t3_pos_1_005_42    m/z 802.7264     0   1   15.5   16  RT (min)      lcorbi_t3_pos_1_041_54    m/z 802.7265     0   1   16   16.5  RT (min)      lcorbi_t3_pos_1_001_49    m/z 802.7265     0   1   15   15.5  RT (min)      lcorbi_t3_pos_1_030_21    m/z 802.7265     0   2   4   6   8E+4   14.5   15  RT (min)      lcorbi_t3_pos_3_056_56    m/z 802.7266     0   1   15.5   16  RT (min)      lcorbi_t3_pos_2_009_50    m/z 802.7266     0   1E+6   14.5   15  RT (min)      lcorbi_t3_pos_1_005_42    m/z 802.7266     0   1   15.5   16  RT (min)      lcorbi_t3_pos_2_009_50    m/z 802.7266     0   1   15   15.5  RT (min)      lcorbi_t3_pos_1_021_18    m/z 802.7266     0   1   15   15.5  RT (min)      lcorbi_t3_pos_1_033_53    m/z 802.7267     0   1   16   16.5  RT (min)      lcorbi_t3_pos_1_056_56    m/z 803.5398     0   1   2E+4   15.5   16  RT (min)      lcorbi_t3_pos_2_012_01    m/z 803.5399     0   2   4E+4   15.5   16  RT (min)      lcorbi_t3_pos_1_023_02    m/z 803.5400     0   1   2   3E+4   15   15.5  RT (min)      lcorbi_t3_pos_1_031_40    m/z 803.5401     0   2   4E+4   15.5   16  RT (min)      lcorbi_t3_pos_1_023_02    m/z 803.5401     0   1   2   3E+4   14.5   15  RT (min)      lcorbi_t3_pos_2_049_55    m/z 803.5401     0   1   2   3   4E+4   14.5   15  RT (min)      lcorbi_t3_pos_1_034_34    m/z 803.5402     0   1   2   3E+4   15   15.5  RT (min)      lcorbi_t3_pos_2_056_56    m/z 803.5407     0   1E+5   14   14.5  RT (min)      lcorbi_t3_pos_3_013_19    m/z 803.5418     0   1   2   3   4E+4   14.5   15  RT (min)      lcorbi_t3_pos_1_031_40    m/z 803.5421     0   1E+5   14   14.5  RT (min)      lcorbi_t3_pos_3_044_46    m/z 803.5429     0   2   4E+4   15   15.5  RT (min)      lcorbi_t3_pos_1_051_09    m/z 803.5429     0   2   4   6E+5   14   14.5  RT (min)       lcorbi_t3_pos_2_022_38    m/z 803.5430     0   1   2   3   4E+4   14.5   15  RT (min)      lcorbi_t3_pos_1_034_34    m/z 803.5432     0   1   12.5   13  RT (min)      lcorbi_t3_pos_4_012_37    m/z 803.5432     0   1   2   3E+4   14   14.5  RT (min)      lcorbi_t3_pos_2_038_03    m/z 803.5434     0   1   2E+4   15   15.5  RT (min)      lcorbi_t3_pos_2_012_01    m/z 803.5435     0   1   2E+6   13   13.5  RT (min)      lcorbi_t3_pos_3_053_48    m/z 803.5435     0   1E+7   12.5   13  RT (min)      lcorbi_t3_pos_4_016_15    m/z 803.5435     0   1   12   12.5  RT (min)      lcorbi_t3_pos_2_032_39    m/z 804.5392     0   1   12   12.5  RT (min)       lcorbi_t3_pos_2_047_31    m/z 804.5467     0   1E+4   12.5   13  RT (min)      lcorbi_t3_pos_4_012_37    m/z 804.5468     0   1   12   12.5  RT (min)       lcorbi_t3_pos_2_032_39    m/z 804.5469     0   2   4   6   8E+6   12.5   13  RT (min)      lcorbi_t3_pos_4_016_15    m/z 804.5470     0   1E+6   12.5   13  RT (min)      lcorbi_t3_pos_3_053_48    m/z 804.5475     0   2   4   6E+4   14.5   15  RT (min)      lcorbi_t3_pos_1_034_34    m/z 804.5476     0   2   4   6E+6   12.5   13  RT (min)      lcorbi_t3_pos_2_032_39    m/z 804.5505     0   1   2   3   4E+4   14.5   15  RT (min)      lcorbi_t3_pos_2_049_55    m/z 804.5510     0   1   2   3E+4   15   15.5  RT (min)      lcorbi_t3_pos_1_051_09    m/z 804.5513     0   1   2   3E+4   14.5   15  RT (min)      lcorbi_t3_pos_1_031_40    m/z 804.5513     0   1   2   3E+4   15   15.5  RT (min)      lcorbi_t3_pos_2_012_01    m/z 804.5515     0   1   16.5   17  RT (min)    lcorbi_t3_pos_1_023_02    m/z 804.5518     0   1   2   3E+4   15.5   16  RT (min)      lcorbi_t3_pos_1_025_52    m/z 804.5518     0   2   4E+5   14   14.5  RT (min)       lcorbi_t3_pos_2_022_38    m/z 804.5520     0   1   2   3E+4   15   15.5  RT (min)      lcorbi_t3_pos_1_041_54    m/z 804.5520     0   2   4E+4   14   14.5  RT (min)      lcorbi_t3_pos_1_038_30    m/z 804.5520     0   2   4   6E+5   14   14.5  RT (min)      lcorbi_t3_pos_3_013_19    m/z 804.5520     0   1   2E+4   16   16.5  RT (min)      lcorbi_t3_pos_1_051_09    m/z 804.5520     0   1   2   3   4E+4   15.5   16  RT (min)      lcorbi_t3_pos_1_023_02    m/z 804.5520     0   1   2   3E+4   15   15.5  RT (min)      lcorbi_t3_pos_1_034_34    m/z 804.5521     0   1   2   3   4E+4   15.5   16  RT (min)      lcorbi_t3_pos_1_023_02    m/z 804.5521     0   1   2E+4   15   15.5  RT (min)      lcorbi_t3_pos_2_056_56    m/z 804.5521     0   2   4   6   8E+4   14   14.5  RT (min)      lcorbi_t3_pos_1_008_36    m/z 804.5521     0   2   4   6E+4   14.5   15  RT (min)      lcorbi_t3_pos_1_034_34    m/z 804.5522     0   2   4   6E+4   14.5   15  RT (min)      lcorbi_t3_pos_1_034_34    m/z 804.5522     0   1   16.5   17  RT (min)    lcorbi_t3_pos_1_023_02    m/z 804.5523     0   1   2   3E+4   15.5   16  RT (min)      lcorbi_t3_pos_2_012_01    m/z 804.5524     0   1   2E+4   16   16.5  RT (min)      lcorbi_t3_pos_1_023_02    m/z 804.5526     0   1   2E+4   15.5   16  RT (min)      lcorbi_t3_pos_1_056_56    m/z 805.5500     0   1E+4   12   12.5  RT (min)      lcorbi_t3_pos_2_032_39    m/z 805.5500     0   1E+6   12.5   13  RT (min)      lcorbi_t3_pos_2_054_29    m/z 805.5501     0   2   4   6E+3   12.5   13  RT (min)      lcorbi_t3_pos_4_012_37    m/z 805.5545     0   1   2   3E+5   14   14.5  RT (min)      lcorbi_t3_pos_2_038_03    m/z 805.5546     0   2   4   6E+4   14.5   15  RT (min)      lcorbi_t3_pos_1_034_34    m/z 805.5550     0   1   15   15.5  RT (min)      lcorbi_t3_pos_2_056_56    m/z 805.5550     0   1   2   3E+4   14.5   15  RT (min)      lcorbi_t3_pos_1_031_40    m/z 805.5553     0   1   2E+4   14.5   15  RT (min)      lcorbi_t3_pos_2_049_55    m/z 805.5553     0   1   15   15.5  RT (min)      lcorbi_t3_pos_1_034_34    m/z 805.5553     0   2   4   6E+4   14.5   15  RT (min)      lcorbi_t3_pos_1_034_34    m/z 805.5554     0   1E+5   14   14.5  RT (min)       lcorbi_t3_pos_3_003_33    m/z 805.5554     0   1   16.5   17  RT (min)    lcorbi_t3_pos_1_023_02    m/z 805.5555     0   1   15   15.5  RT (min)      lcorbi_t3_pos_1_034_34    m/z 805.5555     0   1   2E+4   14.5   15  RT (min)      lcorbi_t3_pos_1_049_55    m/z 805.5556     0   1   16.5   17  RT (min)    lcorbi_t3_pos_1_023_02    m/z 805.5556     0   1   15.5   16  RT (min)      lcorbi_t3_pos_1_031_40    m/z 805.5556     0   1   2E+5   14   14.5  RT (min)       lcorbi_t3_pos_4_048_24    m/z 805.5557     0   1   15.5   16  RT (min)      lcorbi_t3_pos_1_023_02    m/z 805.5558     0   1   15.5   16  RT (min)      lcorbi_t3_pos_1_023_02    m/z 805.5559     0   1E+4   15.5   16  RT (min)      lcorbi_t3_pos_2_012_01    m/z 806.5544     0   2   4   6E+5   12.5   13  RT (min)       lcorbi_t3_pos_1_054_48    m/z 806.5544     0   2   4   6E+5   12.5   13  RT (min)      lcorbi_t3_pos_1_054_48    m/z 806.5688     0   1   2E+4   16   16.5  RT (min)      lcorbi_t3_pos_2_056_56    m/z 806.5690     0   1E+7   13   13.5  RT (min)      lcorbi_t3_pos_1_002_01    m/z 806.5690     0   1   16.5   17  RT (min)    lcorbi_t3_pos_1_023_02    m/z 806.5692     0   1E+6   14.5   15  RT (min)      lcorbi_t3_pos_1_008_36    m/z 806.5692     0   1   15.5   16  RT (min)      lcorbi_t3_pos_1_025_52    m/z 806.5693     0   1E+7   14   14.5  RT (min)      lcorbi_t3_pos_4_081_58    m/z 806.5693     0   2   4E+4   15.5   16  RT (min)      lcorbi_t3_pos_1_031_40    m/z 806.5693     0   1   2   3   4E+4   16   16.5  RT (min)       lcorbi_t3_pos_1_023_02    m/z 806.5694     0   1   2   3E+4   15.5   16  RT (min)      lcorbi_t3_pos_1_001_49    m/z 806.5694     0   1   2   3   4E+4   15   15.5  RT (min)      lcorbi_t3_pos_1_033_53    m/z 806.5694     0   1   16   16.5  RT (min)       lcorbi_t3_pos_1_023_02    m/z 806.5694     0   2   4   6E+4   14.5   15  RT (min)      lcorbi_t3_pos_1_030_21    m/z 806.5694     0   1   2   3E+4   15   15.5  RT (min)      lcorbi_t3_pos_1_041_54    m/z 806.5694     0   1   2   3E+6   12.5   13  RT (min)      lcorbi_t3_pos_4_057_49    m/z 806.5694     0   1   2E+4   16   16.5  RT (min)      lcorbi_t3_pos_2_056_56    m/z 806.5694     0   1   2E+6   13   13.5  RT (min)      lcorbi_t3_pos_1_002_01    m/z 806.5694     0   1   2   3   4E+4   15.5   16  RT (min)      lcorbi_t3_pos_1_051_09    m/z 806.5694     0   1E+4   12.5   13  RT (min)      lcorbi_t3_pos_2_032_39    m/z 806.5694     0   1   2   3   4E+4   15   15.5  RT (min)       lcorbi_t3_pos_1_034_34    m/z 806.5695     0   1E+6   14.5   15  RT (min)      lcorbi_t3_pos_1_008_36    m/z 806.5695     0   1   16.5   17  RT (min)    lcorbi_t3_pos_1_023_02    m/z 806.5695     0   1E+7   14   14.5  RT (min)       lcorbi_t3_pos_1_005_42    m/z 806.5695     0   1E+4   12.5   13  RT (min)      lcorbi_t3_pos_2_032_39    m/z 806.5695     0   1E+7   13   13.5  RT (min)      lcorbi_t3_pos_1_002_01    m/z 806.5696     0   2   4   6E+4   14.5   15  RT (min)      lcorbi_t3_pos_2_006_47    m/z 806.5696     0   1   2   3   4E+4   15   15.5  RT (min)       lcorbi_t3_pos_1_021_18    m/z 806.5696     0   1   2   3E+4   15.5   16  RT (min)      lcorbi_t3_pos_2_009_50    m/z 806.5696     0   1   16.5   17  RT (min)    lcorbi_t3_pos_1_023_02    m/z 806.5697     0   1   2   3   4E+4   15   15.5  RT (min)      lcorbi_t3_pos_1_034_34    m/z 806.6638     0   2   4   6   8E+5   13   13.5  RT (min)       lcorbi_t3_pos_3_010_44    m/z 806.6702     0   1   2   3   4E+5   14   14.5  RT (min)      lcorbi_t3_pos_4_092_60    m/z 807.5721     0   2   4   6E+6   13   13.5  RT (min)      lcorbi_t3_pos_1_002_01    m/z 807.5724     0   1   16.5   17  RT (min)    lcorbi_t3_pos_1_023_02    m/z 807.5728     0   2   4E+5   14.5   15  RT (min)       lcorbi_t3_pos_1_008_36    m/z 807.5729     0   1   15   15.5  RT (min)      lcorbi_t3_pos_1_041_54    m/z 807.5729     0   2   4E+5   14.5   15  RT (min)      lcorbi_t3_pos_1_008_36    m/z 807.5729     0   1   16   16.5  RT (min)      lcorbi_t3_pos_1_023_02    m/z 807.5729     0   1E+6   12.5   13  RT (min)      lcorbi_t3_pos_4_057_49    m/z 807.5729     0   1E+6   13   13.5  RT (min)      lcorbi_t3_pos_1_002_01    m/z 807.5730     0   1   2   3   4E+4   15.5   16  RT (min)      lcorbi_t3_pos_1_031_40    m/z 807.5730     0   1   2   3E+4   15   15.5  RT (min)      lcorbi_t3_pos_1_028_32    m/z 807.5730     0   1   2   3E+4   15   15.5  RT (min)      lcorbi_t3_pos_1_034_34    m/z 807.5730     0   1   2   3E+4   15   15.5  RT (min)       lcorbi_t3_pos_1_021_18    m/z 807.5730     0   1   15.5   16  RT (min)      lcorbi_t3_pos_1_001_49    m/z 807.5730     0   2   4   6E+6   13   13.5  RT (min)      lcorbi_t3_pos_1_002_01    m/z 807.5730     0   1   2E+4   14.5   15  RT (min)      lcorbi_t3_pos_1_030_21    m/z 807.5730     0   1E+4   15.5   16  RT (min)      lcorbi_t3_pos_2_009_50    m/z 807.5731     0   1   12.5   13  RT (min)      lcorbi_t3_pos_2_032_39    m/z 807.5731     0   1   2   3   4E+4   14.5   15  RT (min)      lcorbi_t3_pos_2_006_47    m/z 807.5731     0   1   2   3E+4   15.5   16  RT (min)      lcorbi_t3_pos_1_051_09    m/z 807.5731     0   1   12.5   13  RT (min)      lcorbi_t3_pos_2_032_39    m/z 807.5732     0   1   16   16.5  RT (min)      lcorbi_t3_pos_1_023_02    m/z 807.5732     0   1   2   3E+4   15   15.5  RT (min)      lcorbi_t3_pos_1_034_34    m/z 807.5732     0   1   16.5   17  RT (min)    lcorbi_t3_pos_1_023_02    m/z 807.5732     0   1   13   13.5  RT (min)      lcorbi_t3_pos_1_002_01    m/z 807.5733     0   1   16.5   17  RT (min)    lcorbi_t3_pos_1_023_02    m/z 807.6676     0   1   15   15.5  RT (min)      lcorbi_t3_pos_1_049_55    m/z 807.6679     0   1   2E+5   14.5   15  RT (min)      lcorbi_t3_pos_2_049_55    m/z 807.6679     0   1   2E+6   13.5   14  RT (min)      lcorbi_t3_pos_1_036_08    m/z 807.6731     0   1   2   3E+5   13.5   14  RT (min)      lcorbi_t3_pos_3_025_52    m/z 807.6731     0   1   15   15.5  RT (min)      lcorbi_t3_pos_1_025_52    m/z 807.6736     0   1   2E+5   14.5   15  RT (min)      lcorbi_t3_pos_1_041_54    m/z 808.5737     0   1   12.5   13  RT (min)      lcorbi_t3_pos_2_032_39    m/z 808.5772     0   2   4E+6   13   13.5  RT (min)      lcorbi_t3_pos_4_057_49    m/z 808.5834     0   1   2   3E+5   13   13.5  RT (min)      lcorbi_t3_pos_1_002_01    m/z 808.5838     0   1E+6   13   13.5  RT (min)       lcorbi_t3_pos_1_002_01    m/z 808.5838     0   1   16.5   17  RT (min)    lcorbi_t3_pos_1_023_02    m/z 808.5839     0   1E+6   13   13.5  RT (min)      lcorbi_t3_pos_1_002_01    m/z 808.5839     0   1   2   3E+4   15.5   16  RT (min)       lcorbi_t3_pos_2_009_50    m/z 808.5840     0   1   16   16.5  RT (min)      lcorbi_t3_pos_1_001_49    m/z 808.5840     0   1   2   3   4E+4   15.5   16  RT (min)       lcorbi_t3_pos_1_041_54    m/z 808.5842     0   1   2E+4   16   16.5  RT (min)      lcorbi_t3_pos_1_025_52    m/z 808.5842     0   1   2   3   4E+4   15   15.5  RT (min)      lcorbi_t3_pos_1_041_54    m/z 808.5842     0   1E+7   13.5   14  RT (min)      lcorbi_t3_pos_1_002_01    m/z 808.5843     0   2   4   6E+4   15   15.5  RT (min)       lcorbi_t3_pos_1_033_53    m/z 808.5843     0   1   2E+4   13   13.5  RT (min)      lcorbi_t3_pos_2_032_39    m/z 808.5844     0   1   16.5   17  RT (min)     lcorbi_t3_pos_1_023_02    m/z 808.5844     0   1E+7   14.5   15  RT (min)       lcorbi_t3_pos_1_005_42    m/z 808.5844     0   1   2E+5   14.5   15  RT (min)      lcorbi_t3_pos_2_017_51    m/z 808.5844     0   1   2   3E+5   14.5   15  RT (min)       lcorbi_t3_pos_2_017_51    m/z 808.5844     0   1   16.5   17  RT (min)    lcorbi_t3_pos_1_023_02    m/z 808.5844     0   2   4E+6   14.5   15  RT (min)       lcorbi_t3_pos_1_008_36    m/z 808.5844     0   1   2   3E+6   14   14.5  RT (min)      lcorbi_t3_pos_4_072_57    m/z 808.5844     0   1   16.5   17  RT (min)    lcorbi_t3_pos_1_023_02    m/z 808.5845     0   1E+7   14   14.5  RT (min)      lcorbi_t3_pos_1_005_42    m/z 808.5846     0   2   4   6   8E+5   14.5   15  RT (min)      lcorbi_t3_pos_2_017_51    m/z 808.5846     0   2   4E+4   15   15.5  RT (min)      lcorbi_t3_pos_1_033_53    m/z 808.5846     0   1   2E+4   13   13.5  RT (min)      lcorbi_t3_pos_2_032_39    m/z 808.5846     0   2   4   6E+4   15   15.5  RT (min)      lcorbi_t3_pos_1_030_21    m/z 808.5847     0   1   2   3   4E+4   15.5   16  RT (min)      lcorbi_t3_pos_1_031_40    m/z 808.5848     0   1   16.5   17  RT (min)    lcorbi_t3_pos_1_023_02    m/z 808.5848     0   1   2   3E+4   16   16.5  RT (min)      lcorbi_t3_pos_1_025_52    m/z 808.5848     0   2   4   6E+4   15.5   16  RT (min)      lcorbi_t3_pos_1_051_09    m/z 808.5849     0   1   16.5   17  RT (min)    lcorbi_t3_pos_1_023_02    m/z 808.5850     0   1   16   16.5  RT (min)      lcorbi_t3_pos_1_001_49    m/z 808.6795     0   2   4E+4   14.5   15  RT (min)      lcorbi_t3_pos_1_041_54    m/z 808.6795     0   1   15   15.5  RT (min)      lcorbi_t3_pos_1_001_49    m/z 808.6796     0   1   15   15.5  RT (min)      lcorbi_t3_pos_1_041_54    m/z 808.6797     0   1   15.5   16  RT (min)      lcorbi_t3_pos_1_001_49    m/z 808.6797     0   1E+6   13.5   14  RT (min)       lcorbi_t3_pos_1_036_08    m/z 808.6797     0   1   15   15.5  RT (min)      lcorbi_t3_pos_2_056_56    m/z 808.6797     0   1   2   3E+4   14.5   15  RT (min)      lcorbi_t3_pos_1_041_54    m/z 808.6798     0   1E+6   13.5   14  RT (min)      lcorbi_t3_pos_1_036_08    m/z 809.4906     0   1   18   18.5  RT (min)    lcorbi_t3_pos_1_023_02    m/z 809.4908     0   1   18   18.5  RT (min)    lcorbi_t3_pos_1_023_02    m/z 809.4909     0   1   16.5   17  RT (min)    lcorbi_t3_pos_2_038_03    m/z 809.4910     0   1E+4   16   16.5  RT (min)      lcorbi_t3_pos_3_056_56    m/z 809.4910     0   1   17.5   18  RT (min)    lcorbi_t3_pos_1_056_56    m/z 809.4910     0   1   14   14.5  RT (min)      lcorbi_t3_pos_2_047_31    m/z 809.4910     0   1   2E+4   16   16.5  RT (min)      lcorbi_t3_pos_2_017_51    m/z 809.4910     0   1   17   17.5  RT (min)    lcorbi_t3_pos_1_041_54    m/z 809.4910     0   1E+6   15.5   16  RT (min)       lcorbi_t3_pos_3_041_54    m/z 809.4910     0   1   2   3E+4   14.5   15  RT (min)       lcorbi_t3_pos_1_002_01    m/z 809.4911     0   1   16   16.5  RT (min)      lcorbi_t3_pos_1_030_21    m/z 809.4912     0   1   16.5   17  RT (min)    lcorbi_t3_pos_1_033_53    m/z 809.4912     0   1   17.5   18  RT (min)    lcorbi_t3_pos_1_025_52    m/z 809.4912     0   2   4   6   8E+4   15.5   16  RT (min)      lcorbi_t3_pos_4_001_49    m/z 809.4912     0   1   14   14.5  RT (min)      lcorbi_t3_pos_3_031_45    m/z 809.4912     0   1   17.5   18  RT (min)    lcorbi_t3_pos_2_012_01    m/z 809.4912     0   1   2E+4   14   14.5  RT (min)      lcorbi_t3_pos_3_035_08    m/z 809.4912     0   1   2E+6   15.5   16  RT (min)      lcorbi_t3_pos_1_005_42    m/z 809.4912     0   1   16.5   17  RT (min)    lcorbi_t3_pos_2_038_03    m/z 809.4913     0   1   2E+6   14.5   15  RT (min)       lcorbi_t3_pos_4_105_61    m/z 809.4914     0   1   16   16.5  RT (min)      lcorbi_t3_pos_2_017_51    m/z 809.5873     0   2   4   6   8E+6   13.5   14  RT (min)      lcorbi_t3_pos_1_002_01    m/z 809.5876     0   1   16.5   17  RT (min)    lcorbi_t3_pos_1_023_02    m/z 809.5876     0   1   2E+5   13   13.5  RT (min)      lcorbi_t3_pos_1_002_01    m/z 809.5877     0   1   16.5   17  RT (min)    lcorbi_t3_pos_1_023_02    m/z 809.5878     0   1   16   16.5  RT (min)      lcorbi_t3_pos_1_001_49    m/z 809.5878     0   2   4E+5   13   13.5  RT (min)      lcorbi_t3_pos_1_002_01    m/z 809.5878     0   1E+4   13   13.5  RT (min)      lcorbi_t3_pos_2_013_33    m/z 809.5879     0   1   2   3E+4   15   15.5  RT (min)      lcorbi_t3_pos_1_033_53    m/z 809.5879     0   2   4E+6   14.5   15  RT (min)      lcorbi_t3_pos_1_005_42    m/z 809.5879     0   1   15.5   16  RT (min)      lcorbi_t3_pos_1_041_54    m/z 809.5879     0   2   4E+6   14   14.5  RT (min)       lcorbi_t3_pos_1_005_42    m/z 809.5879     0   1   16   16.5  RT (min)      lcorbi_t3_pos_2_056_56    m/z 809.5879     0   1   15   15.5  RT (min)      lcorbi_t3_pos_1_041_54    m/z 809.5880     0   2   4E+6   13.5   14  RT (min)      lcorbi_t3_pos_1_002_01    m/z 809.5880     0   1   15.5   16  RT (min)      lcorbi_t3_pos_1_041_54    m/z 809.5880     0   2   4   6   8E+6   14   14.5  RT (min)       lcorbi_t3_pos_1_014_03    m/z 809.5880     0   1   2   3E+6   14.5   15  RT (min)       lcorbi_t3_pos_1_008_36    m/z 809.5880     0   1   2E+5   14.5   15  RT (min)       lcorbi_t3_pos_2_017_51    m/z 809.5881     0   1   2   3   4E+5   14.5   15  RT (min)      lcorbi_t3_pos_2_017_51    m/z 809.5882     0   1   16.5   17  RT (min)    lcorbi_t3_pos_1_023_02    m/z 809.5882     0   1   17   17.5  RT (min)    lcorbi_t3_pos_1_023_02    m/z 809.5882     0   1   2   3E+6   14.5   15  RT (min)       lcorbi_t3_pos_1_005_42    m/z 809.5882     0   1   2E+6   13   13.5  RT (min)      lcorbi_t3_pos_4_057_49    m/z 809.5882     0   1   16.5   17  RT (min)    lcorbi_t3_pos_1_023_02    m/z 809.5882     0   1   2   3E+4   15.5   16  RT (min)      lcorbi_t3_pos_1_031_40    m/z 809.5883     0   1   16.5   17  RT (min)    lcorbi_t3_pos_1_023_02    m/z 809.5884     0   2   4   6   8E+4   14.5   15  RT (min)      lcorbi_t3_pos_2_046_07    m/z 809.5884     0   1   16.5   17  RT (min)    lcorbi_t3_pos_1_023_02    m/z 809.5884     0   1E+4   13   13.5  RT (min)      lcorbi_t3_pos_2_032_39    m/z 809.5885     0   1   2   3E+4   15   15.5  RT (min)      lcorbi_t3_pos_1_033_53    m/z 809.5885     0   1   2   3E+4   15   15.5  RT (min)       lcorbi_t3_pos_1_033_53    m/z 809.5886     0   1E+4   13   13.5  RT (min)      lcorbi_t3_pos_2_032_39    m/z 809.5887     0   1E+4   13   13.5  RT (min)      lcorbi_t3_pos_2_032_39    m/z 809.5887     0   1   2   3   4E+4   15.5   16  RT (min)      lcorbi_t3_pos_1_051_09    m/z 809.6524     0   1   16.5   17  RT (min)    lcorbi_t3_pos_1_023_02    m/z 809.6532     0   1   13.5   14  RT (min)      lcorbi_t3_pos_1_002_01    m/z 809.6535     0   1   16   16.5  RT (min)      lcorbi_t3_pos_1_051_09    m/z 809.6535     0   1   15   15.5  RT (min)      lcorbi_t3_pos_1_028_32    m/z 809.6536     0   1E+5   14.5   15  RT (min)      lcorbi_t3_pos_1_008_36    m/z 809.6536     0   1   2E+4   15.5   16  RT (min)      lcorbi_t3_pos_1_034_34    m/z 809.6538     0   1E+5   14.5   15  RT (min)      lcorbi_t3_pos_1_008_36    m/z 809.6828     0   1   15   15.5  RT (min)      lcorbi_t3_pos_4_033_53    m/z 809.6830     0   1   15   15.5  RT (min)      lcorbi_t3_pos_1_001_49    m/z 809.6830     0   2   4E+4   14.5   15  RT (min)      lcorbi_t3_pos_1_041_54    m/z 809.6831     0   1   15   15.5  RT (min)      lcorbi_t3_pos_1_041_54    m/z 809.6832     0   2   4E+6   13.5   14  RT (min)       lcorbi_t3_pos_1_036_08    m/z 809.6832     0   1E+5   14.5   15  RT (min)      lcorbi_t3_pos_1_021_18    m/z 809.6833     0   2   4E+6   13.5   14  RT (min)      lcorbi_t3_pos_1_036_08    m/z 809.6833     0   2   4E+6   13.5   14  RT (min)       lcorbi_t3_pos_1_036_08    m/z 809.6834     0   1   2   3E+4   15.5   16  RT (min)      lcorbi_t3_pos_1_025_52    m/z 809.6834     0   1   2   3E+4   15   15.5  RT (min)      lcorbi_t3_pos_1_025_52    m/z 809.6834     0   1E+5   14.5   15  RT (min)      lcorbi_t3_pos_1_041_54    m/z 809.6834     0   2   4   6   8E+5   14   14.5  RT (min)      lcorbi_t3_pos_3_041_54    m/z 809.6834     0   2   4   6   8E+5   14   14.5  RT (min)       lcorbi_t3_pos_3_025_52    m/z 809.6834     0   1   15   15.5  RT (min)      lcorbi_t3_pos_1_041_54    m/z 809.6834     0   1   15.5   16  RT (min)      lcorbi_t3_pos_1_001_49    m/z 809.7522     0   2   4   6E+5   14.5   15  RT (min)       lcorbi_t3_pos_3_055_10    m/z 809.7523     0   2   4   6E+5   14.5   15  RT (min)       lcorbi_t3_pos_4_093_60    m/z 809.7523     0   2   4   6E+5   14.5   15  RT (min)       lcorbi_t3_pos_2_037_27    m/z 809.7525     0   2   4   6E+5   14.5   15  RT (min)      lcorbi_t3_pos_4_094_60    m/z 810.4940     0   1   14   14.5  RT (min)      lcorbi_t3_pos_2_032_39    m/z 810.4942     0   1E+4   14   14.5  RT (min)      lcorbi_t3_pos_3_035_08    m/z 810.4943     0   1   16   16.5  RT (min)      lcorbi_t3_pos_2_006_47    m/z 810.4943     0   1   2   3   4E+4   14.5   15  RT (min)      lcorbi_t3_pos_1_002_01    m/z 810.4944     0   1   17.5   18  RT (min)    lcorbi_t3_pos_1_056_56    m/z 810.4944     0   1   16.5   17  RT (min)    lcorbi_t3_pos_1_033_53    m/z 810.4944     0   1   16   16.5  RT (min)      lcorbi_t3_pos_3_025_52    m/z 810.4944     0   2   4   6   8E+5   15   15.5  RT (min)      lcorbi_t3_pos_4_105_61    m/z 810.4944     0   1   17   17.5  RT (min)    lcorbi_t3_pos_2_009_50    m/z 810.4944     0   1   17.5   18  RT (min)    lcorbi_t3_pos_1_025_52    m/z 810.4945     0   2   4   6   8E+5   15.5   16  RT (min)      lcorbi_t3_pos_1_005_42    m/z 810.4945     0   1   16.5   17  RT (min)    lcorbi_t3_pos_2_025_52    m/z 810.4945     0   2   4   6E+5   15.5   16  RT (min)      lcorbi_t3_pos_3_033_53    m/z 810.4946     0   1   2E+4   14.5   15  RT (min)      lcorbi_t3_pos_2_054_29    m/z 810.4948     0   1   14   14.5  RT (min)      lcorbi_t3_pos_2_047_31    m/z 810.5996     0   1   2   3   4E+4   13   13.5  RT (min)      lcorbi_t3_pos_2_032_39    m/z 810.6004     0   2   4   6E+4   16   16.5  RT (min)       lcorbi_t3_pos_1_051_09    m/z 810.6005     0   1   16.5   17  RT (min)     lcorbi_t3_pos_1_023_02    m/z 810.6005     0   2   4   6E+6   0   0.5  RT (min)      lcorbi_t3_pos_4_057_49    m/z 810.6005     0   2   4E+5   13.5   14  RT (min)       lcorbi_t3_pos_2_054_29    m/z 810.6005     0   1E+5   15   15.5  RT (min)      lcorbi_t3_pos_1_028_32    m/z 810.6006     0   2   4   6E+4   15.5   16  RT (min)      lcorbi_t3_pos_1_034_34    m/z 810.6006     0   2   4   6   8E+4   15   15.5  RT (min)      lcorbi_t3_pos_1_030_21    m/z 810.6006     0   2   4   6E+4   15   15.5  RT (min)      lcorbi_t3_pos_1_033_53    m/z 810.6006     0   1   16.5   17  RT (min)    lcorbi_t3_pos_2_012_01    m/z 810.6007     0   2   4   6E+4   15.5   16  RT (min)      lcorbi_t3_pos_1_034_34    m/z 810.6007     0   2   4   6E+4   15.5   16  RT (min)      lcorbi_t3_pos_1_034_34    m/z 810.6008     0   2   4   6   8E+6   13.5   14  RT (min)       lcorbi_t3_pos_1_002_01    m/z 810.6008     0   1   2   3E+6   14.5   15  RT (min)      lcorbi_t3_pos_1_008_36    m/z 810.6008     0   1   2   3E+4   16   16.5  RT (min)      lcorbi_t3_pos_1_001_49    m/z 810.6008     0   1E+5   15   15.5  RT (min)      lcorbi_t3_pos_1_028_32    m/z 810.6008     0   1   2   3   4E+5   15   15.5  RT (min)      lcorbi_t3_pos_2_046_07    m/z 810.6009     0   2   4   6E+6   14.5   15  RT (min)       lcorbi_t3_pos_1_008_36    m/z 810.6009     0   2   4   6E+4   16   16.5  RT (min)      lcorbi_t3_pos_1_051_09    m/z 810.6009     0   1   16.5   17  RT (min)    lcorbi_t3_pos_1_023_02    m/z 810.6009     0   1   16.5   17  RT (min)     lcorbi_t3_pos_1_023_02    m/z 810.6010     0   1   2   3E+4   16   16.5  RT (min)      lcorbi_t3_pos_1_049_55    m/z 810.6010     0   1   2   3   4E+4   13   13.5  RT (min)       lcorbi_t3_pos_2_032_39    m/z 810.6010     0   1E+5   15   15.5  RT (min)      lcorbi_t3_pos_1_047_41    m/z 810.6010     0   1   2   3   4E+7   14.5   15  RT (min)       lcorbi_t3_pos_1_005_42    m/z 810.6010     0   1   17   17.5  RT (min)    lcorbi_t3_pos_1_023_02    m/z 810.6010     0   1E+6   13.5   14  RT (min)       lcorbi_t3_pos_1_002_01    m/z 810.6949     0   2   4   6E+4   14   14.5  RT (min)      lcorbi_t3_pos_2_017_51    m/z 810.6950     0   1   15   15.5   16  RT (min)      lcorbi_t3_pos_1_056_56    m/z 810.6950     0   1   14.5   15  RT (min)      lcorbi_t3_pos_1_030_21    m/z 810.6951     0   1   15.5   16  RT (min)      lcorbi_t3_pos_1_025_52    m/z 810.6951     0   1   15   15.5  RT (min)      lcorbi_t3_pos_1_041_54    m/z 810.6952     0   1   15.5   16  RT (min)      lcorbi_t3_pos_1_001_49    m/z 810.6952     0   1E+4   14.5   15  RT (min)      lcorbi_t3_pos_1_047_41    m/z 810.6952     0   1   15   15.5  RT (min)      lcorbi_t3_pos_1_041_54    m/z 810.6952     0   1   14.5   15  RT (min)      lcorbi_t3_pos_1_021_18    m/z 810.6952     0   1   15   15.5  RT (min)      lcorbi_t3_pos_1_041_54    m/z 810.6952     0   2   4   6   8E+5   14   14.5  RT (min)       lcorbi_t3_pos_1_005_42    m/z 810.6954     0   1   15.5   16  RT (min)      lcorbi_t3_pos_1_025_52    m/z 810.6955     0   1   14.5   15  RT (min)      lcorbi_t3_pos_1_030_21    m/z 811.6032     0   1   2E+4   13   13.5  RT (min)      lcorbi_t3_pos_2_032_39    m/z 811.6034     0   1   2E+5   13.5   14  RT (min)      lcorbi_t3_pos_2_054_29    m/z 811.6040     0   1   16.5   17  RT (min)    lcorbi_t3_pos_1_023_02    m/z 811.6040     0   2   4   6   8E+5   13.5   14  RT (min)      lcorbi_t3_pos_1_002_01    m/z 811.6040     0   1   2   3   4E+6   13.5   14  RT (min)      lcorbi_t3_pos_4_016_15    m/z 811.6042     0   1   2   3E+6   0   0.5  RT (min)      lcorbi_t3_pos_4_057_49    m/z 811.6042     0   1   2   3E+6   13.5   14  RT (min)      lcorbi_t3_pos_4_012_37    m/z 811.6042     0   1   16.5   17  RT (min)    lcorbi_t3_pos_1_023_02    m/z 811.6042     0   2   4E+4   15   15.5  RT (min)      lcorbi_t3_pos_1_028_32    m/z 811.6042     0   2   4   6E+4   15.5   16  RT (min)       lcorbi_t3_pos_1_034_34    m/z 811.6042     0   1   2   3E+4   16   16.5  RT (min)      lcorbi_t3_pos_1_051_09    m/z 811.6042     0   2   4E+4   15.5   16  RT (min)      lcorbi_t3_pos_1_034_34    m/z 811.6042     0   1   2   3   4E+4   15   15.5  RT (min)      lcorbi_t3_pos_1_033_53    m/z 811.6043     0   1   16.5   17  RT (min)    lcorbi_t3_pos_2_012_01    m/z 811.6043     0   2   4E+4   15   15.5  RT (min)      lcorbi_t3_pos_1_030_21    m/z 811.6043     0   1   2E+7   14.5   15  RT (min)       lcorbi_t3_pos_1_005_42    m/z 811.6044     0   1   2   3   4E+4   16   16.5  RT (min)      lcorbi_t3_pos_1_031_40    m/z 811.6044     0   1   2   3E+6   14.5   15  RT (min)       lcorbi_t3_pos_1_008_36    m/z 811.6044     0   1   2   3   4E+6   13.5   14  RT (min)      lcorbi_t3_pos_1_002_01    m/z 811.6044     0   2   4   6E+4   15   15.5  RT (min)      lcorbi_t3_pos_1_028_32    m/z 811.6044     0   1   2   3E+5   15   15.5  RT (min)      lcorbi_t3_pos_2_046_07    m/z 811.6044     0   1E+6   14.5   15  RT (min)      lcorbi_t3_pos_1_008_36    m/z 811.6046     0   1   17   17.5  RT (min)    lcorbi_t3_pos_1_023_02    m/z 811.6046     0   1   16.5   17  RT (min)    lcorbi_t3_pos_1_023_02    m/z 811.6046     0   1   2E+4   16   16.5  RT (min)      lcorbi_t3_pos_1_049_55    m/z 811.6047     0   2   4   6E+4   15   15.5  RT (min)      lcorbi_t3_pos_1_047_41    m/z 811.6047     0   1   2   3   4E+5   14.5   15  RT (min)      lcorbi_t3_pos_4_010_43    m/z 811.6344     0   2   4   6   8E+5   12   12.5  RT (min)      lcorbi_t3_pos_2_037_27    m/z 811.6349     0   2   4   6   8E+5   11.5   12  RT (min)       lcorbi_t3_pos_4_093_60    m/z 811.6349     0   2   4   6E+5   11.5   12  RT (min)       lcorbi_t3_pos_4_090_60    m/z 811.6682     0   1   13   13.5  RT (min)      lcorbi_t3_pos_2_032_39    m/z 811.6686     0   1   16   16.5  RT (min)      lcorbi_t3_pos_4_033_53    m/z 811.6686     0   1   2E+4   13.5   14  RT (min)      lcorbi_t3_pos_2_054_29    m/z 811.6688     0   1   16   16.5  RT (min)      lcorbi_t3_pos_1_031_40    m/z 811.6688     0   1   16.5   17  RT (min)    lcorbi_t3_pos_1_051_09    m/z 811.6688     0   1   15.5   16  RT (min)      lcorbi_t3_pos_1_041_54    m/z 811.6690     0   1   16   16.5  RT (min)      lcorbi_t3_pos_4_033_53    m/z 811.6690     0   1   2E+4   15   15.5  RT (min)      lcorbi_t3_pos_3_003_33    m/z 811.6690     0   1E+5   13.5   14  RT (min)      lcorbi_t3_pos_1_002_01    m/z 811.6690     0   1   2   3   4E+6   14.5   15  RT (min)      lcorbi_t3_pos_4_019_47    m/z 811.6690     0   1   15.5   16  RT (min)      lcorbi_t3_pos_1_028_32    m/z 811.6690     0   1   2   3   4E+6   14.5   15  RT (min)      lcorbi_t3_pos_4_019_47    m/z 811.6690     0   1   2   3   4E+5   15   15.5  RT (min)      lcorbi_t3_pos_1_008_36    m/z 811.6690     0   1   15.5   16  RT (min)      lcorbi_t3_pos_1_028_32    m/z 811.6692     0   1   15.5   16  RT (min)      lcorbi_t3_pos_1_034_34    m/z 811.6692     0   1   16.5   17  RT (min)    lcorbi_t3_pos_1_051_09    m/z 811.6692     0   1E+5   15   15.5  RT (min)      lcorbi_t3_pos_1_008_36    m/z 811.6692     0   1   13   13.5  RT (min)      lcorbi_t3_pos_3_024_07    m/z 811.6692     0   1   13   13.5  RT (min)      lcorbi_t3_pos_2_032_39    m/z 811.6693     0   1   16   16.5  RT (min)      lcorbi_t3_pos_1_051_09    m/z 811.6694     0   2   4   6E+5   14.5   15  RT (min)      lcorbi_t3_pos_4_001_49    m/z 811.6698     0   1   17   17.5  RT (min)    lcorbi_t3_pos_1_023_02    m/z 811.6698     0   1   17   17.5  RT (min)    lcorbi_t3_pos_1_023_02    m/z 811.6977     0   1   2E+4   15   15.5  RT (min)      lcorbi_t3_pos_2_038_03    m/z 811.6984     0   2   4E+5   11.5   12  RT (min)      lcorbi_t3_pos_4_057_49    m/z 811.6986     0   1   15   15.5  RT (min)      lcorbi_t3_pos_2_009_50    m/z 811.6986     0   1   15   15.5  RT (min)      lcorbi_t3_pos_1_041_54    m/z 811.6986     0   1   2   3   4E+4   14.5   15  RT (min)      lcorbi_t3_pos_1_021_18    m/z 811.6986     0   1   15   15.5  RT (min)      lcorbi_t3_pos_2_009_50    m/z 811.6986     0   2   4E+4   14.5   15  RT (min)      lcorbi_t3_pos_1_030_21    m/z 811.6986     0   1   2E+5   14   14.5  RT (min)      lcorbi_t3_pos_2_017_51    m/z 811.6987     0   2   4   6E+4   14.5   15  RT (min)      lcorbi_t3_pos_1_047_41    m/z 811.6987     0   1   15   15.5   16  RT (min)      lcorbi_t3_pos_1_056_56    m/z 811.6988     0   1   15.5   16  RT (min)      lcorbi_t3_pos_1_001_49    m/z 811.6988     0   1   2E+4   14.5   15  RT (min)      lcorbi_t3_pos_2_038_03    m/z 811.6988     0   1   2   3E+4   15.5   16  RT (min)      lcorbi_t3_pos_1_025_52    m/z 811.6989     0   1   15   15.5  RT (min)      lcorbi_t3_pos_1_041_54    m/z 811.6990     0   2   4   6   8E+4   14.5   15  RT (min)      lcorbi_t3_pos_2_017_51    m/z 811.6992     0   1   2   3E+6   14   14.5  RT (min)       lcorbi_t3_pos_1_005_42    m/z 812.6081     0   1   2   3E+4   16   16.5  RT (min)      lcorbi_t3_pos_1_051_09    m/z 812.6082     0   1E+4   16   16.5  RT (min)      lcorbi_t3_pos_4_033_53    m/z 812.6083     0   1E+4   15.5   16  RT (min)      lcorbi_t3_pos_2_049_55    m/z 812.6119     0   1   2   3   4E+4   15   15.5  RT (min)      lcorbi_t3_pos_2_006_47    m/z 812.6124     0   1   16.5   17  RT (min)    lcorbi_t3_pos_1_023_02    m/z 812.6150     0   1   2   3   4E+4   15.5   16  RT (min)       lcorbi_t3_pos_1_034_34    m/z 812.6150     0   1   2E+4   13   13.5  RT (min)      lcorbi_t3_pos_2_032_39    m/z 812.6150     0   1   2E+4   13   13.5  RT (min)      lcorbi_t3_pos_2_032_39    m/z 812.6154     0   1   2   3E+4   13.5   14  RT (min)      lcorbi_t3_pos_2_013_33    m/z 812.6154     0   1   16.5   17  RT (min)    lcorbi_t3_pos_1_023_02    m/z 812.6156     0   1E+5   15   15.5  RT (min)      lcorbi_t3_pos_2_046_07    m/z 812.6156     0   1   2   3E+4   15.5   16  RT (min)       lcorbi_t3_pos_1_028_32    m/z 812.6157     0   1   2   3   4E+4   16   16.5  RT (min)      lcorbi_t3_pos_1_031_40    m/z 812.6157     0   1   2   3E+4   16   16.5  RT (min)      lcorbi_t3_pos_1_051_09    m/z 812.6158     0   2   4   6E+6   0   0.5  RT (min)      lcorbi_t3_pos_4_057_49    m/z 812.6158     0   1   2   3E+6   13.5   14  RT (min)      lcorbi_t3_pos_1_002_01    m/z 812.6159     0   1   2   3   4E+4   15   15.5  RT (min)      lcorbi_t3_pos_1_028_32    m/z 812.6160     0   1   17   17.5  RT (min)    lcorbi_t3_pos_1_023_02    m/z 812.6160     0   1E+6   14.5   15  RT (min)      lcorbi_t3_pos_1_008_36    m/z 812.6160     0   1E+7   14.5   15  RT (min)      lcorbi_t3_pos_1_005_42    m/z 812.6162     0   2   4   6   8E+6   13.5   14  RT (min)      lcorbi_t3_pos_4_016_15    m/z 812.6162     0   1E+7   14.5   15  RT (min)       lcorbi_t3_pos_3_026_04    m/z 812.6162     0   2   4   6   8E+4   15   15.5  RT (min)      lcorbi_t3_pos_2_046_07    m/z 812.6166     0   1E+7   14.5   15  RT (min)      lcorbi_t3_pos_1_005_42    m/z 812.6721     0   1   2E+6   14   14.5  RT (min)      lcorbi_t3_pos_3_034_26    m/z 812.6722     0   1   13   13.5  RT (min)      lcorbi_t3_pos_3_024_07    m/z 812.6722     0   1   13   13.5  RT (min)      lcorbi_t3_pos_2_032_39    m/z 812.6722     0   1   13   13.5  RT (min)      lcorbi_t3_pos_2_032_39    m/z 812.6723     0   1   13.5   14  RT (min)      lcorbi_t3_pos_1_002_01    m/z 812.6723     0   1   15.5   16  RT (min)      lcorbi_t3_pos_1_034_34    m/z 812.6723     0   1   2E+6   14.5   15  RT (min)      lcorbi_t3_pos_4_019_47    m/z 812.6723     0   1   2E+5   15   15.5  RT (min)      lcorbi_t3_pos_1_008_36    m/z 812.6724     0   1   16   16.5  RT (min)      lcorbi_t3_pos_4_033_53    m/z 812.6724     0   1   16   16.5  RT (min)      lcorbi_t3_pos_1_051_09    m/z 812.6724     0   1   2   3   4E+4   15.5   16  RT (min)      lcorbi_t3_pos_1_028_32    m/z 812.6724     0   1   16.5   17  RT (min)    lcorbi_t3_pos_1_051_09    m/z 812.6725     0   1   2   3E+4   15   15.5  RT (min)      lcorbi_t3_pos_1_047_41    m/z 812.6726     0   1   2   3E+4   15   15.5  RT (min)      lcorbi_t3_pos_2_022_38    m/z 812.6726     0   1   2E+6   14.5   15  RT (min)      lcorbi_t3_pos_4_019_47    m/z 812.6726     0   1   15.5   16  RT (min)      lcorbi_t3_pos_1_034_34    m/z 812.6727     0   1E+4   15.5   16  RT (min)      lcorbi_t3_pos_3_013_19    m/z 812.6727     0   1   16   16.5  RT (min)      lcorbi_t3_pos_1_031_40    m/z 812.6728     0   1   17   17.5  RT (min)    lcorbi_t3_pos_1_023_02    m/z 812.6728     0   1   16.5   17  RT (min)    lcorbi_t3_pos_1_051_09    m/z 812.6728     0   2   4   6   8E+4   13.5   14  RT (min)      lcorbi_t3_pos_1_002_01    m/z 812.6731     0   1   17   17.5  RT (min)    lcorbi_t3_pos_1_023_02    m/z 812.6746     0   2   4   6E+5   12.5   13  RT (min)       lcorbi_t3_pos_4_091_60    m/z 812.6750     0   2   4   6E+5   12.5   13  RT (min)       lcorbi_t3_pos_4_093_60    m/z 812.6753     0   2   4   6E+5   12.5   13  RT (min)       lcorbi_t3_pos_4_093_60    m/z 812.7093     0   1   15   15.5  RT (min)       lcorbi_t3_pos_1_041_54    m/z 812.7102     0   1   14.5   15  RT (min)      lcorbi_t3_pos_1_021_18    m/z 812.7103     0   1   2E+4   15   15.5  RT (min)       lcorbi_t3_pos_1_030_21    m/z 812.7107     0   1   15.5   16  RT (min)      lcorbi_t3_pos_1_041_54    m/z 812.7108     0   1   2E+4   14.5   15  RT (min)       lcorbi_t3_pos_1_030_21    m/z 812.7109     0   1   15.5   16  RT (min)      lcorbi_t3_pos_1_025_52    m/z 812.7109     0   2   4   6E+5   14   14.5  RT (min)      lcorbi_t3_pos_1_005_42    m/z 812.7109     0   1   15.5   16  RT (min)      lcorbi_t3_pos_1_056_56    m/z 812.7110     0   1   16   16.5  RT (min)      lcorbi_t3_pos_1_001_49    m/z 812.7110     0   1   2E+4   15   15.5  RT (min)      lcorbi_t3_pos_2_049_55    m/z 812.7110     0   1   15.5   16  RT (min)      lcorbi_t3_pos_1_041_54    m/z 813.6158     0   1   15.5   16  RT (min)      lcorbi_t3_pos_1_041_54    m/z 813.6194     0   1   16   16.5  RT (min)      lcorbi_t3_pos_4_033_53    m/z 813.6194     0   1   16   16.5  RT (min)      lcorbi_t3_pos_2_009_50    m/z 813.6196     0   1   16.5   17  RT (min)    lcorbi_t3_pos_1_023_02    m/z 813.6197     0   1   2E+4   15   15.5  RT (min)      lcorbi_t3_pos_2_006_47    m/z 813.6198     0   1   2E+4   15   15.5  RT (min)      lcorbi_t3_pos_1_028_32    m/z 813.6198     0   1   16   16.5  RT (min)      lcorbi_t3_pos_1_051_09    m/z 813.6198     0   2   4   6E+4   15   15.5  RT (min)      lcorbi_t3_pos_2_017_51    m/z 813.6198     0   2   4E+6   14.5   15  RT (min)      lcorbi_t3_pos_1_005_42    m/z 813.6198     0   1   16   16.5  RT (min)      lcorbi_t3_pos_1_031_40    m/z 813.6198     0   2   4E+6   14.5   15  RT (min)      lcorbi_t3_pos_1_005_42    m/z 813.6199     0   1   2E+4   15.5   16  RT (min)      lcorbi_t3_pos_1_028_32    m/z 813.6200     0   1E+4   13   13.5  RT (min)      lcorbi_t3_pos_2_032_39    m/z 813.6200     0   1   2   3E+4   15.5   16  RT (min)      lcorbi_t3_pos_1_034_34    m/z 813.6200     0   2   4   6   8E+5   14.5   15  RT (min)      lcorbi_t3_pos_1_008_36    m/z 813.6200     0   1   2   3E+6   13.5   14  RT (min)      lcorbi_t3_pos_4_016_15    m/z 813.6200     0   1   17   17.5  RT (min)    lcorbi_t3_pos_1_023_02    m/z 813.6201     0   2   4E+6   14.5   15  RT (min)      lcorbi_t3_pos_1_005_42    m/z 813.6205     0   1   16.5   17  RT (min)    lcorbi_t3_pos_2_012_01    m/z 813.6501     0   2   4   6   8E+5   12.5   13  RT (min)      lcorbi_t3_pos_4_095_60    m/z 813.6502     0   1E+6   12   12.5  RT (min)       lcorbi_t3_pos_4_093_60    m/z 813.6504     0   2   4E+5   12.5   13  RT (min)       lcorbi_t3_pos_4_092_60    m/z 813.6504     0   1E+6   12   12.5  RT (min)       lcorbi_t3_pos_4_094_60    m/z 813.6505     0   1E+6   12   12.5  RT (min)      lcorbi_t3_pos_4_095_60    m/z 813.6508     0   1E+6   12   12.5  RT (min)      lcorbi_t3_pos_4_093_60    m/z 813.6756     0   1   13   13.5  RT (min)      lcorbi_t3_pos_2_032_39    m/z 813.6759     0   2   4   6   8E+5   14   14.5  RT (min)      lcorbi_t3_pos_4_016_15    m/z 813.6760     0   1   13.5   14  RT (min)      lcorbi_t3_pos_2_032_39    m/z 813.6761     0   1   2   3E+4   15.5   16  RT (min)      lcorbi_t3_pos_2_022_38    m/z 813.6765     0   1   2   3E+4   15   15.5  RT (min)      lcorbi_t3_pos_2_017_51    m/z 813.6839     0   1   2E+5   15   15.5  RT (min)       lcorbi_t3_pos_1_008_36    m/z 813.6841     0   1E+5   15   15.5  RT (min)      lcorbi_t3_pos_1_008_36    m/z 813.6844     0   1   16   16.5  RT (min)      lcorbi_t3_pos_1_034_34    m/z 813.6844     0   1   16   16.5  RT (min)      lcorbi_t3_pos_1_034_34    m/z 813.6845     0   1   17   17.5  RT (min)    lcorbi_t3_pos_1_023_02    m/z 813.6846     0   1   13.5   14  RT (min)      lcorbi_t3_pos_2_013_33    m/z 813.6846     0   1   2E+4   15.5   16  RT (min)      lcorbi_t3_pos_1_028_32    m/z 813.6846     0   1   2   3   4E+4   15.5   16  RT (min)      lcorbi_t3_pos_1_033_53    m/z 813.6846     0   1   17   17.5  RT (min)    lcorbi_t3_pos_1_023_02    m/z 813.6846     0   1   17   17.5  RT (min)    lcorbi_t3_pos_1_023_02    m/z 813.6847     0   1   16.5   17  RT (min)    lcorbi_t3_pos_1_049_55    m/z 813.6847     0   1   16.5   17  RT (min)    lcorbi_t3_pos_1_051_09    m/z 813.6847     0   1   2E+4   15.5   16  RT (min)      lcorbi_t3_pos_1_028_32    m/z 813.6848     0   1   16.5   17  RT (min)    lcorbi_t3_pos_2_012_01    m/z 813.6848     0   1   2E+4   13.5   14  RT (min)      lcorbi_t3_pos_2_054_29    m/z 813.6848     0   1E+4   15.5   16  RT (min)      lcorbi_t3_pos_2_041_54    m/z 813.6848     0   1   16   16.5  RT (min)      lcorbi_t3_pos_1_034_34    m/z 813.6848     0   1   16.5   17  RT (min)    lcorbi_t3_pos_1_031_40    m/z 813.6848     0   1   16   16.5  RT (min)      lcorbi_t3_pos_1_034_34    m/z 813.6849     0   1   16.5   17  RT (min)    lcorbi_t3_pos_1_056_56    m/z 813.6849     0   1   2   3E+6   15   15.5  RT (min)      lcorbi_t3_pos_3_047_40    m/z 813.7136     0   1   16   16.5  RT (min)      lcorbi_t3_pos_2_056_56    m/z 813.7137     0   1E+6   12.5   13  RT (min)      lcorbi_t3_pos_4_057_49    m/z 813.7139     0   1   2E+4   15   15.5  RT (min)      lcorbi_t3_pos_1_033_53    m/z 813.7140     0   1E+4   15   15.5  RT (min)      lcorbi_t3_pos_2_038_03    m/z 813.7141     0   1   15   15.5  RT (min)      lcorbi_t3_pos_2_025_52    m/z 813.7142     0   1E+6   13   13.5  RT (min)      lcorbi_t3_pos_4_057_49    m/z 813.7142     0   1E+5   14.5   15  RT (min)      lcorbi_t3_pos_2_017_51    m/z 813.7142     0   2   4E+4   14.5   15  RT (min)      lcorbi_t3_pos_1_047_41    m/z 813.7142     0   1   15   15.5  RT (min)      lcorbi_t3_pos_1_030_21    m/z 813.7142     0   1   16   16.5  RT (min)      lcorbi_t3_pos_1_001_49    m/z 813.7143     0   1   2   3E+4   14.5   15  RT (min)      lcorbi_t3_pos_1_030_21    m/z 813.7143     0   1   2   3   4E+4   14.5   15  RT (min)      lcorbi_t3_pos_1_021_18    m/z 813.7144     0   1   15.5   16  RT (min)      lcorbi_t3_pos_1_056_56    m/z 813.7144     0   1   2E+6   14   14.5  RT (min)      lcorbi_t3_pos_1_005_42    m/z 813.7144     0   1   2E+4   15   15.5  RT (min)      lcorbi_t3_pos_1_041_54    m/z 813.7144     0   1   15.5   16  RT (min)      lcorbi_t3_pos_1_025_52    m/z 813.7145     0   1   15.5   16  RT (min)      lcorbi_t3_pos_1_056_56    m/z 813.7146     0   1   15.5   16  RT (min)      lcorbi_t3_pos_1_056_56    m/z 813.7147     0   1   2   3   4E+4   14.5   15  RT (min)      lcorbi_t3_pos_2_006_47    m/z 814.5510     0   1E+6   12.5   13  RT (min)      lcorbi_t3_pos_3_029_13    m/z 814.6231     0   1   16.5   17  RT (min)    lcorbi_t3_pos_1_023_02    m/z 814.6242     0   1E+6   14.5   15  RT (min)      lcorbi_t3_pos_1_005_42    m/z 814.6262     0   1E+6   12.5   13  RT (min)       lcorbi_t3_pos_4_097_60    m/z 814.6263     0   1E+6   12.5   13  RT (min)       lcorbi_t3_pos_4_093_60    m/z 814.6263     0   1E+6   12.5   13  RT (min)      lcorbi_t3_pos_4_094_60    m/z 814.6263     0   1E+6   12.5   13  RT (min)       lcorbi_t3_pos_4_092_60    m/z 814.6264     0   2   4   6   8E+5   12.5   13  RT (min)       lcorbi_t3_pos_4_096_60    m/z 814.6264     0   2   4   6   8E+5   12   12.5  RT (min)       lcorbi_t3_pos_4_095_60    m/z 814.6273     0   1   13   13.5  RT (min)      lcorbi_t3_pos_2_032_39    m/z 814.6275     0   1   16.5   17  RT (min)    lcorbi_t3_pos_1_023_02    m/z 814.6275     0   1   2   3E+4   15   15.5  RT (min)      lcorbi_t3_pos_1_030_21    m/z 814.6311     0   1   16   16.5  RT (min)      lcorbi_t3_pos_1_051_09    m/z 814.6312     0   1   2   3   4E+4   15.5   16  RT (min)      lcorbi_t3_pos_1_028_32    m/z 814.6313     0   1   2   3   4E+4   15   15.5  RT (min)      lcorbi_t3_pos_1_028_32    m/z 814.6314     0   1   15.5   16  RT (min)      lcorbi_t3_pos_1_041_54    m/z 814.6315     0   1   2E+4   15.5   16  RT (min)      lcorbi_t3_pos_1_034_34    m/z 814.6316     0   1   16.5   17  RT (min)    lcorbi_t3_pos_1_051_09    m/z 814.6317     0   1E+6   14.5   15  RT (min)      lcorbi_t3_pos_1_008_36    m/z 814.6318     0   1   2   3   4E+4   15.5   16  RT (min)      lcorbi_t3_pos_1_028_32    m/z 814.6318     0   1E+5   15   15.5  RT (min)      lcorbi_t3_pos_1_008_36    m/z 814.6318     0   1   16   16.5  RT (min)      lcorbi_t3_pos_1_031_40    m/z 814.6318     0   1   13.5   14  RT (min)      lcorbi_t3_pos_3_011_42    m/z 814.6319     0   1   17   17.5  RT (min)    lcorbi_t3_pos_1_023_02    m/z 814.6319     0   1E+4   13.5   14  RT (min)      lcorbi_t3_pos_2_054_29    m/z 814.6320     0   1   13.5   14  RT (min)      lcorbi_t3_pos_1_002_01    m/z 814.6320     0   1   13   13.5  RT (min)      lcorbi_t3_pos_2_032_39    m/z 814.6320     0   1   17   17.5  RT (min)    lcorbi_t3_pos_1_023_02    m/z 814.6321     0   1   2E+4   15.5   16  RT (min)      lcorbi_t3_pos_1_034_34    m/z 814.6322     0   1   16   16.5  RT (min)      lcorbi_t3_pos_1_031_40    m/z 814.6323     0   1E+6   14   14.5  RT (min)      lcorbi_t3_pos_3_034_26    m/z 814.6324     0   1   13.5   14  RT (min)      lcorbi_t3_pos_1_002_01    m/z 814.6710     0   1   16.5   17  RT (min)    lcorbi_t3_pos_1_034_34    m/z 814.6710     0   1   2   3E+4   16   16.5  RT (min)      lcorbi_t3_pos_1_008_36    m/z 814.6710     0   1   17   17.5   18  RT (min)    lcorbi_t3_pos_1_031_40    m/z 814.6710     0   1   16.5   17  RT (min)    lcorbi_t3_pos_1_028_32    m/z 814.6711     0   1   17   17.5  RT (min)    lcorbi_t3_pos_1_034_34    m/z 814.6712     0   2   4   6E+5   14.5   15  RT (min)      lcorbi_t3_pos_2_023_26    m/z 814.6712     0   1   2   3   4E+4   15.5   16  RT (min)      lcorbi_t3_pos_1_008_36    m/z 814.6712     0   1   16.5   17  RT (min)    lcorbi_t3_pos_1_034_34    m/z 814.6712     0   1   16.5   17  RT (min)    lcorbi_t3_pos_1_028_32    m/z 814.6712     0   2   4E+5   15.5   16  RT (min)      lcorbi_t3_pos_1_005_42    m/z 814.6712     0   1   16   16.5  RT (min)      lcorbi_t3_pos_2_022_38    m/z 814.6714     0   1   17.5   18  RT (min)    lcorbi_t3_pos_2_012_01    m/z 814.6790     0   1   2   3E+5   14.5   15  RT (min)       lcorbi_t3_pos_4_072_57    m/z 814.6832     0   1E+5   13.5   14  RT (min)      lcorbi_t3_pos_2_054_29    m/z 814.6878     0   1   16   16.5  RT (min)      lcorbi_t3_pos_1_034_34    m/z 814.6878     0   1   17   17.5  RT (min)    lcorbi_t3_pos_1_023_02    m/z 814.6878     0   1   16.5   17  RT (min)    lcorbi_t3_pos_2_012_01    m/z 814.6880     0   1E+5   15.5   16  RT (min)      lcorbi_t3_pos_3_003_33    m/z 814.6880     0   1   16   16.5  RT (min)      lcorbi_t3_pos_1_034_34    m/z 814.6881     0   1   16   16.5  RT (min)      lcorbi_t3_pos_1_034_34    m/z 814.6881     0   1   13.5   14  RT (min)      lcorbi_t3_pos_2_054_29    m/z 814.6881     0   1   13.5   14  RT (min)      lcorbi_t3_pos_2_013_33    m/z 814.6882     0   1   13.5   14  RT (min)      lcorbi_t3_pos_2_032_39    m/z 814.6882     0   1   16.5   17  RT (min)    lcorbi_t3_pos_1_031_40    m/z 814.6883     0   1   2E+6   14.5   15  RT (min)      lcorbi_t3_pos_4_019_47    m/z 814.6884     0   1   17   17.5  RT (min)    lcorbi_t3_pos_1_023_02    m/z 814.6884     0   1E+5   15.5   16  RT (min)      lcorbi_t3_pos_1_028_32    m/z 814.6887     0   1   16.5   17  RT (min)    lcorbi_t3_pos_1_031_40    m/z 814.6901     0   1E+6   13   13.5  RT (min)       lcorbi_t3_pos_4_094_60    m/z 814.6901     0   2   4   6   8E+5   13   13.5  RT (min)      lcorbi_t3_pos_4_091_60    m/z 814.6902     0   1E+6   13   13.5  RT (min)       lcorbi_t3_pos_1_003_45    m/z 814.6906     0   1E+6   13   13.5  RT (min)      lcorbi_t3_pos_4_097_60    m/z 814.6907     0   2   4   6   8E+5   12.5   13  RT (min)       lcorbi_t3_pos_4_093_60    m/z 814.6907     0   1   17   17.5  RT (min)    lcorbi_t3_pos_1_023_02    m/z 815.6309     0   1   16.5   17  RT (min)    lcorbi_t3_pos_2_012_01    m/z 815.6349     0   1   15.5   16  RT (min)      lcorbi_t3_pos_1_034_34    m/z 815.6350     0   2   4E+5   14.5   15  RT (min)      lcorbi_t3_pos_3_047_40    m/z 815.6352     0   1   15.5   16  RT (min)      lcorbi_t3_pos_1_034_34    m/z 815.6352     0   1   15   15.5  RT (min)      lcorbi_t3_pos_2_022_38    m/z 815.6352     0   1   13   13.5  RT (min)      lcorbi_t3_pos_3_031_45    m/z 815.6353     0   1   16.5   17  RT (min)    lcorbi_t3_pos_1_051_09    m/z 815.6353     0   1E+5   13.5   14  RT (min)      lcorbi_t3_pos_3_011_42    m/z 815.6353     0   1   16   16.5  RT (min)      lcorbi_t3_pos_2_009_50    m/z 815.6354     0   1   17   17.5  RT (min)    lcorbi_t3_pos_1_023_02    m/z 815.6354     0   1   16.5   17  RT (min)    lcorbi_t3_pos_1_051_09    m/z 815.6355     0   1   13.5   14  RT (min)      lcorbi_t3_pos_2_013_33    m/z 815.6356     0   2   4   6   8E+5   14   14.5  RT (min)      lcorbi_t3_pos_4_016_15    m/z 815.6356     0   2   4   6E+5   14.5   15  RT (min)      lcorbi_t3_pos_1_005_42    m/z 815.6356     0   1E+4   15   15.5  RT (min)      lcorbi_t3_pos_2_017_51    m/z 815.6357     0   1   15.5   16  RT (min)      lcorbi_t3_pos_1_028_32    m/z 815.6357     0   1   15.5   16  RT (min)      lcorbi_t3_pos_1_028_32    m/z 815.6358     0   1E+5   15   15.5  RT (min)      lcorbi_t3_pos_1_008_36    m/z 815.6358     0   1   16   16.5  RT (min)      lcorbi_t3_pos_1_031_40    m/z 815.6358     0   1E+4   13.5   14  RT (min)      lcorbi_t3_pos_2_054_29    m/z 815.6360     0   1E+4   13.5   14  RT (min)      lcorbi_t3_pos_2_054_29    m/z 815.6360     0   1   17   17.5  RT (min)    lcorbi_t3_pos_1_023_02    m/z 815.6657     0   2   4   6   8E+5   12   12.5  RT (min)      lcorbi_t3_pos_4_095_60    m/z 815.6658     0   2   4   6E+5   12   12.5  RT (min)      lcorbi_t3_pos_4_096_60    m/z 815.6661     0   2   4   6   8E+5   12.5   13  RT (min)       lcorbi_t3_pos_4_095_60    m/z 815.6662     0   2   4   6   8E+5   12.5   13  RT (min)       lcorbi_t3_pos_4_092_60    m/z 815.6662     0   2   4   6   8E+5   12   12.5  RT (min)       lcorbi_t3_pos_4_095_60    m/z 815.6663     0   2   4   6   8E+5   12.5   13  RT (min)       lcorbi_t3_pos_4_092_60    m/z 815.6745     0   1   16.5   17  RT (min)    lcorbi_t3_pos_1_034_34    m/z 815.6745     0   1   16.5   17  RT (min)    lcorbi_t3_pos_1_034_34    m/z 815.6746     0   1   16.5   17  RT (min)    lcorbi_t3_pos_1_028_32    m/z 815.6746     0   1   2E+4   15.5   16  RT (min)      lcorbi_t3_pos_1_008_36    m/z 815.6748     0   1   2E+4   14   14.5  RT (min)      lcorbi_t3_pos_2_013_33    m/z 815.6748     0   1   17.5   18  RT (min)    lcorbi_t3_pos_1_031_40    m/z 815.6914     0   1   13.5   14  RT (min)      lcorbi_t3_pos_2_032_39    m/z 815.6916     0   1   16.5   17  RT (min)    lcorbi_t3_pos_1_031_40    m/z 815.6917     0   1   17   17.5  RT (min)    lcorbi_t3_pos_1_023_02    m/z 815.6919     0   2   4   6E+4   15.5   16  RT (min)      lcorbi_t3_pos_1_028_32    m/z 815.6920     0   1   13.5   14  RT (min)      lcorbi_t3_pos_2_013_33    m/z 815.6922     0   1   13.5   14  RT (min)      lcorbi_t3_pos_3_024_07    m/z 815.6922     0   2   4E+4   15.5   16  RT (min)      lcorbi_t3_pos_2_022_38    m/z 815.6926     0   1   16.5   17  RT (min)    lcorbi_t3_pos_1_031_40    m/z 815.6958     0   1   17   17.5  RT (min)    lcorbi_t3_pos_1_023_02    m/z 815.6958     0   1   16.5   17  RT (min)    lcorbi_t3_pos_2_012_01    m/z 815.6972     0   1   17.5   18  RT (min)    lcorbi_t3_pos_1_023_02    m/z 815.6991     0   2   4   6E+4   15.5   16  RT (min)      lcorbi_t3_pos_1_028_32    m/z 815.6994     0   1   2   3   4E+5   15   15.5  RT (min)      lcorbi_t3_pos_1_008_36    m/z 815.6998     0   1   16   16.5  RT (min)      lcorbi_t3_pos_1_034_34    m/z 815.6998     0   1   16   16.5  RT (min)      lcorbi_t3_pos_1_034_34    m/z 815.6999     0   1   2   3   4E+5   15   15.5  RT (min)      lcorbi_t3_pos_3_047_40    m/z 815.6999     0   1   13.5   14  RT (min)      lcorbi_t3_pos_2_054_29    m/z 815.6999     0   1   16   16.5  RT (min)      lcorbi_t3_pos_1_034_34    m/z 815.7006     0   1   16.5   17  RT (min)    lcorbi_t3_pos_1_031_40    m/z 815.7296     0   1   16   16.5  RT (min)      lcorbi_t3_pos_2_056_56    m/z 815.7296     0   1   15.5   16  RT (min)      lcorbi_t3_pos_1_041_54    m/z 815.7296     0   1   15.5   16  RT (min)      lcorbi_t3_pos_2_041_54    m/z 815.7296     0   1   16   16.5  RT (min)      lcorbi_t3_pos_1_001_49    m/z 815.7297     0   1   16   16.5  RT (min)      lcorbi_t3_pos_1_049_55    m/z 815.7298     0   1   15   15.5  RT (min)      lcorbi_t3_pos_1_030_21    m/z 815.7298     0   1   15   15.5  RT (min)      lcorbi_t3_pos_2_046_07    m/z 815.7298     0   2   4E+4   14.5   15  RT (min)      lcorbi_t3_pos_3_025_52    m/z 815.7298     0   1   15   15.5  RT (min)      lcorbi_t3_pos_1_021_18    m/z 815.7298     0   1E+6   14.5   15  RT (min)      lcorbi_t3_pos_1_005_42    m/z 815.7299     0   1   15.5   16  RT (min)      lcorbi_t3_pos_1_041_54    m/z 815.7299     0   1   2E+4   14.5   15  RT (min)      lcorbi_t3_pos_3_056_56    m/z 815.7299     0   1   15   15.5  RT (min)      lcorbi_t3_pos_1_033_53    m/z 815.7300     0   1   16   16.5  RT (min)      lcorbi_t3_pos_1_025_52    m/z 816.4952     0   1   15   15.5  RT (min)      lcorbi_t3_pos_1_031_40    m/z 816.5665     0   2   4   6E+5   12.5   13  RT (min)      lcorbi_t3_pos_1_054_48    m/z 818.5180     0   2   4E+5   11   11.5  RT (min)       lcorbi_t3_pos_1_054_48    m/z 818.6046     0   1   13.5   14  RT (min)      lcorbi_t3_pos_1_002_01    m/z 818.6046     0   1   2   3E+5   14.5   15  RT (min)       lcorbi_t3_pos_1_005_42    m/z 818.6614     0   1   14.5   15  RT (min)      lcorbi_t3_pos_1_041_54    m/z 818.6616     0   1   15   15.5  RT (min)      lcorbi_t3_pos_1_056_56    m/z 818.6616     0   1   2E+4   14.5   15  RT (min)      lcorbi_t3_pos_1_030_21    m/z 818.6616     0   1   2E+4   14.5   15  RT (min)      lcorbi_t3_pos_1_021_18    m/z 818.6617     0   2   4   6E+5   13.5   14  RT (min)       lcorbi_t3_pos_1_005_42    m/z 818.6620     0   1   2E+4   14   14.5  RT (min)      lcorbi_t3_pos_2_017_51    m/z 818.7233     0   1   17   17.5  RT (min)    lcorbi_t3_pos_1_028_32    m/z 818.7234     0   1   16.5   17  RT (min)    lcorbi_t3_pos_1_028_32    m/z 818.7236     0   2   4E+5   15   15.5  RT (min)      lcorbi_t3_pos_4_034_42    m/z 818.7236     0   1   16   16.5  RT (min)      lcorbi_t3_pos_1_008_36    m/z 818.7236     0   1   16   16.5  RT (min)      lcorbi_t3_pos_1_008_36    m/z 818.7236     0   1   14.5   15  RT (min)      lcorbi_t3_pos_3_011_42    m/z 818.7237     0   1   14.5   15  RT (min)      lcorbi_t3_pos_3_024_07    m/z 818.7238     0   1   17.5   18  RT (min)    lcorbi_t3_pos_1_031_40    m/z 818.7238     0   1   17   17.5  RT (min)    lcorbi_t3_pos_1_034_34    m/z 819.5146     0   1   15.5   16  RT (min)      lcorbi_t3_pos_1_023_02    m/z 819.5172     0   1   12.5   13  RT (min)      lcorbi_t3_pos_2_013_33    m/z 819.5172     0   1   15   15.5  RT (min)      lcorbi_t3_pos_1_051_09    m/z 819.5172     0   1   2   3E+5   13.5   14  RT (min)      lcorbi_t3_pos_1_008_36    m/z 819.5172     0   2   4E+3   14   14.5  RT (min)      lcorbi_t3_pos_3_003_33    m/z 819.5172     0   1   12.5   13  RT (min)      lcorbi_t3_pos_4_012_37    m/z 819.5172     0   1   14.5   15  RT (min)      lcorbi_t3_pos_1_034_34    m/z 819.5172     0   1   15   15.5  RT (min)      lcorbi_t3_pos_1_031_40    m/z 819.5173     0   1   2   3E+4   14   14.5  RT (min)      lcorbi_t3_pos_1_028_32    m/z 819.5174     0   1E+6   13.5   14  RT (min)       lcorbi_t3_pos_3_047_40    m/z 819.5175     0   1   14.5   15  RT (min)      lcorbi_t3_pos_2_049_55    m/z 819.5176     0   1   15   15.5  RT (min)      lcorbi_t3_pos_2_056_56    m/z 819.5177     0   1E+6   12.5   13  RT (min)      lcorbi_t3_pos_2_054_29    m/z 819.5179     0   1   14.5   15  RT (min)      lcorbi_t3_pos_1_034_34    m/z 819.6081     0   1E+5   14   14.5  RT (min)       lcorbi_t3_pos_1_032_26    m/z 819.6744     0   1   2   3E+5   13.5   14  RT (min)       lcorbi_t3_pos_4_092_60    m/z 820.5207     0   2   4   6E+3   12.5   13  RT (min)      lcorbi_t3_pos_4_012_37    m/z 820.5207     0   1   12.5   13  RT (min)      lcorbi_t3_pos_2_013_33    m/z 820.5207     0   2   4   6   8E+5   13.5   14  RT (min)      lcorbi_t3_pos_4_019_47    m/z 820.5208     0   1E+4   12   12.5  RT (min)      lcorbi_t3_pos_2_032_39    m/z 820.5208     0   1   15   15.5  RT (min)      lcorbi_t3_pos_1_031_40    m/z 820.5208     0   2   4   6   8E+5   12.5   13  RT (min)      lcorbi_t3_pos_4_016_15    m/z 820.5210     0   1   15   15.5  RT (min)      lcorbi_t3_pos_1_051_09    m/z 820.5211     0   1   2E+5   14   14.5  RT (min)      lcorbi_t3_pos_1_028_32    m/z 820.5212     0   1   14.5   15  RT (min)      lcorbi_t3_pos_1_034_34    m/z 820.5214     0   2   4E+4   14   14.5  RT (min)      lcorbi_t3_pos_2_022_38    m/z 820.5214     0   1   15   15.5  RT (min)      lcorbi_t3_pos_2_012_01    m/z 820.5223     0   1   14.5   15  RT (min)      lcorbi_t3_pos_1_034_34    m/z 820.5252     0   1   2E+5   13.5   14  RT (min)       lcorbi_t3_pos_1_008_36    m/z 820.5257     0   1   15.5   16  RT (min)      lcorbi_t3_pos_1_023_02    m/z 820.5338     0   2   4   6E+5   11.5   12  RT (min)       lcorbi_t3_pos_1_054_48    m/z 820.6214     0   1   2E+5   13.5   14  RT (min)      lcorbi_t3_pos_1_002_01    m/z 820.6214     0   1   15.5   16  RT (min)      lcorbi_t3_pos_1_034_34    m/z 820.6215     0   1   16   16.5  RT (min)      lcorbi_t3_pos_1_031_40    m/z 820.6215     0   1   15.5   16  RT (min)      lcorbi_t3_pos_1_028_32    m/z 820.6215     0   1   16   16.5  RT (min)      lcorbi_t3_pos_1_051_09    m/z 820.6215     0   1   16   16.5  RT (min)      lcorbi_t3_pos_1_051_09    m/z 820.6215     0   2   4   6E+4   15   15.5  RT (min)      lcorbi_t3_pos_1_008_36    m/z 820.6216     0   1   15   15.5  RT (min)      lcorbi_t3_pos_1_028_32    m/z 820.6217     0   1   17   17.5  RT (min)    lcorbi_t3_pos_1_023_02    m/z 820.6217     0   1   15.5   16  RT (min)      lcorbi_t3_pos_1_034_34    m/z 820.6219     0   2   4E+5   13.5   14  RT (min)      lcorbi_t3_pos_1_002_01    m/z 820.6219     0   1   16.5   17  RT (min)    lcorbi_t3_pos_1_023_02    m/z 820.6767     0   1   15.5   16  RT (min)      lcorbi_t3_pos_2_056_56    m/z 820.6773     0   1   15.5   16  RT (min)      lcorbi_t3_pos_1_025_52    m/z 820.6774     0   1   14.5   15  RT (min)      lcorbi_t3_pos_1_041_54    m/z 820.6774     0   1   14.5   15  RT (min)      lcorbi_t3_pos_1_041_54    m/z 820.6774     0   1   14.5   15  RT (min)      lcorbi_t3_pos_1_041_54    m/z 820.6775     0   1   15.5   16  RT (min)      lcorbi_t3_pos_1_056_56    m/z 820.6776     0   2   4   6E+5   14   14.5  RT (min)       lcorbi_t3_pos_4_001_49    m/z 820.6778     0   1E+4   14.5   15  RT (min)      lcorbi_t3_pos_2_025_52    m/z 820.6779     0   1   2   3   4E+5   14   14.5  RT (min)      lcorbi_t3_pos_2_046_07    m/z 820.6780     0   1E+5   14   14.5  RT (min)      lcorbi_t3_pos_1_047_41    m/z 820.6787     0   2   4   6   8E+3   14   14.5  RT (min)      lcorbi_t3_pos_4_048_24    m/z 820.6791     0   1   15   15.5  RT (min)      lcorbi_t3_pos_1_056_56    m/z 820.6794     0   1   2   3E+4   14   14.5  RT (min)      lcorbi_t3_pos_1_030_21    m/z 820.6794     0   1   15   15.5  RT (min)      lcorbi_t3_pos_1_025_52    m/z 820.6794     0   1   2   3E+4   14.5   15  RT (min)      lcorbi_t3_pos_1_021_18    m/z 820.6796     0   1   2   3E+4   14.5   15  RT (min)      lcorbi_t3_pos_1_033_53    m/z 820.7388     0   1   18   18.5  RT (min)    lcorbi_t3_pos_1_031_40    m/z 820.7388     0   1   17   17.5  RT (min)    lcorbi_t3_pos_1_034_34    m/z 820.7389     0   1   18.5   19  RT (min)    lcorbi_t3_pos_1_023_02    m/z 820.7389     0   1   17.5   18  RT (min)    lcorbi_t3_pos_1_034_34    m/z 820.7390     0   1   17   17.5  RT (min)    lcorbi_t3_pos_1_028_32    m/z 820.7390     0   1   14.5   15  RT (min)      lcorbi_t3_pos_3_024_07    m/z 820.7391     0   1   16.5   17  RT (min)    lcorbi_t3_pos_2_022_38    m/z 820.7392     0   1   17   17.5  RT (min)    lcorbi_t3_pos_1_028_32    m/z 820.7392     0   1   18   18.5  RT (min)    lcorbi_t3_pos_1_031_40    m/z 820.7392     0   1   14.5   15  RT (min)      lcorbi_t3_pos_2_013_33    m/z 820.7392     0   1   16.5   17  RT (min)    lcorbi_t3_pos_2_022_38    m/z 820.7392     0   1   16   16.5  RT (min)      lcorbi_t3_pos_1_008_36    m/z 820.7393     0   1   2E+4   16   16.5  RT (min)      lcorbi_t3_pos_3_047_40    m/z 820.7394     0   1   18   18.5  RT (min)    lcorbi_t3_pos_2_012_01    m/z 820.7396     0   1   14.5   15  RT (min)      lcorbi_t3_pos_3_011_42    m/z 821.6249     0   1   16   16.5  RT (min)      lcorbi_t3_pos_1_031_40    m/z 821.6827     0   1   15   15.5  RT (min)      lcorbi_t3_pos_1_001_49    m/z 821.6827     0   1   2E+4   14.5   15  RT (min)      lcorbi_t3_pos_2_049_55    m/z 821.6828     0   2   4E+4   14   14.5  RT (min)      lcorbi_t3_pos_1_047_41    m/z 821.6828     0   1   15   15.5  RT (min)      lcorbi_t3_pos_1_001_49    m/z 821.6829     0   1   15   15.5  RT (min)      lcorbi_t3_pos_1_025_52    m/z 821.6829     0   1E+4   14.5   15  RT (min)      lcorbi_t3_pos_2_038_03    m/z 821.6829     0   1   15   15.5  RT (min)      lcorbi_t3_pos_1_056_56    m/z 821.6829     0   2   4   6E+4   14   14.5  RT (min)      lcorbi_t3_pos_1_030_21    m/z 821.6830     0   1   2E+4   15   15.5  RT (min)      lcorbi_t3_pos_1_049_55    m/z 821.6830     0   1   14.5   15  RT (min)      lcorbi_t3_pos_1_041_54    m/z 821.6832     0   1E+4   14.5   15  RT (min)      lcorbi_t3_pos_4_048_24    m/z 821.6832     0   1   2   3E+5   14   14.5  RT (min)      lcorbi_t3_pos_3_056_56    m/z 821.6832     0   2   4   6   8E+6   13.5   14  RT (min)       lcorbi_t3_pos_3_040_41    m/z 821.6853     0   1E+5   14   14.5  RT (min)      lcorbi_t3_pos_2_046_07    m/z 821.6862     0   1E+7   13.5   14  RT (min)       lcorbi_t3_pos_1_005_42    m/z 821.7888     0   1   17.5   18  RT (min)    lcorbi_t3_pos_1_001_49    m/z 822.5645     0   2   4   6E+5   13   13.5  RT (min)       lcorbi_t3_pos_4_081_58    m/z 822.6290     0   1E+6   14   14.5  RT (min)      lcorbi_t3_pos_4_016_15    m/z 822.6335     0   1E+6   14   14.5  RT (min)      lcorbi_t3_pos_4_016_15    m/z 822.6369     0   1   15.5   16  RT (min)      lcorbi_t3_pos_1_034_34    m/z 822.6370     0   1   15.5   16  RT (min)      lcorbi_t3_pos_1_034_34    m/z 822.6370     0   1   15   15.5  RT (min)      lcorbi_t3_pos_2_022_38    m/z 822.6371     0   1   15.5   16  RT (min)      lcorbi_t3_pos_1_034_34    m/z 822.6371     0   1   15.5   16  RT (min)      lcorbi_t3_pos_1_028_32    m/z 822.6372     0   1   15   15.5  RT (min)      lcorbi_t3_pos_2_022_38    m/z 822.6372     0   1E+5   15   15.5  RT (min)      lcorbi_t3_pos_1_008_36    m/z 822.6372     0   2   4E+5   14.5   15  RT (min)      lcorbi_t3_pos_3_047_40    m/z 822.6372     0   1   16   16.5  RT (min)      lcorbi_t3_pos_1_031_40    m/z 822.6372     0   1   15.5   16  RT (min)      lcorbi_t3_pos_1_028_32    m/z 822.6373     0   1   16   16.5  RT (min)      lcorbi_t3_pos_1_031_40    m/z 822.6374     0   1   16.5   17  RT (min)    lcorbi_t3_pos_1_051_09    m/z 822.6374     0   1   16   16.5  RT (min)      lcorbi_t3_pos_1_049_55    m/z 822.6376     0   1   17   17.5  RT (min)    lcorbi_t3_pos_1_023_02    m/z 822.6378     0   1   17   17.5  RT (min)    lcorbi_t3_pos_1_023_02    m/z 822.6854     0   1   2   3E+4   14.5   15  RT (min)      lcorbi_t3_pos_2_049_55    m/z 822.6856     0   1   2   3E+4   14.5   15  RT (min)      lcorbi_t3_pos_4_048_24    m/z 822.6857     0   1   2   3   4E+4   15   15.5  RT (min)      lcorbi_t3_pos_1_025_52    m/z 822.6858     0   1   2   3   4E+4   15   15.5  RT (min)       lcorbi_t3_pos_1_056_56    m/z 822.6858     0   1   2   3E+4   15   15.5  RT (min)      lcorbi_t3_pos_1_001_49    m/z 822.6858     0   1E+5   14.5   15  RT (min)      lcorbi_t3_pos_1_021_18    m/z 822.6858     0   1   2E+5   14   14.5  RT (min)      lcorbi_t3_pos_1_030_21    m/z 822.6860     0   1   2   3E+4   15   15.5  RT (min)      lcorbi_t3_pos_2_049_55    m/z 822.6860     0   1   2   3   4E+4   14.5   15  RT (min)      lcorbi_t3_pos_2_041_54    m/z 822.6861     0   2   4E+4   14.5   15  RT (min)      lcorbi_t3_pos_1_041_54    m/z 822.6862     0   1   2   3E+4   15   15.5  RT (min)      lcorbi_t3_pos_2_049_55    m/z 822.6885     0   1E+5   14   14.5  RT (min)      lcorbi_t3_pos_1_047_41    m/z 822.6919     0   1E+7   14   14.5  RT (min)       lcorbi_t3_pos_4_091_60    m/z 822.6924     0   1   2   3   4E+7   13.5   14  RT (min)       lcorbi_t3_pos_1_005_42    m/z 822.6928     0   1   2   3   4E+7   13.5   14  RT (min)      lcorbi_t3_pos_1_005_42    m/z 822.6930     0   1   2   3E+4   15   15.5  RT (min)      lcorbi_t3_pos_1_001_49    m/z 822.6934     0   1   2   3E+4   15.5   16  RT (min)      lcorbi_t3_pos_1_001_49    m/z 822.6948     0   2   4   6E+5   11.5   12  RT (min)      lcorbi_t3_pos_4_057_49    m/z 823.6404     0   1   17   17.5  RT (min)    lcorbi_t3_pos_1_023_02    m/z 823.6404     0   1E+5   15   15.5  RT (min)      lcorbi_t3_pos_1_008_36    m/z 823.6405     0   1   13   13.5  RT (min)      lcorbi_t3_pos_2_032_39    m/z 823.6406     0   1   16   16.5  RT (min)      lcorbi_t3_pos_1_031_40    m/z 823.6406     0   1   16   16.5  RT (min)      lcorbi_t3_pos_1_031_40    m/z 823.6407     0   1   13.5   14  RT (min)      lcorbi_t3_pos_2_013_33    m/z 823.6407     0   1   15.5   16  RT (min)      lcorbi_t3_pos_1_028_32    m/z 823.6408     0   1E+4   15   15.5  RT (min)      lcorbi_t3_pos_2_022_38    m/z 823.6409     0   1   2E+5   14.5   15  RT (min)      lcorbi_t3_pos_3_047_40    m/z 823.6409     0   1   15.5   16  RT (min)      lcorbi_t3_pos_1_028_32    m/z 823.6409     0   2   4E+5   14.5   15  RT (min)      lcorbi_t3_pos_4_019_47    m/z 823.6410     0   1   15.5   16  RT (min)      lcorbi_t3_pos_1_034_34    m/z 823.6417     0   1   13.5   14  RT (min)      lcorbi_t3_pos_2_054_29    m/z 823.6946     0   1   2   3E+4   15   15.5  RT (min)      lcorbi_t3_pos_1_041_54    m/z 823.6946     0   1   2   3E+4   15   15.5  RT (min)      lcorbi_t3_pos_1_041_54    m/z 823.6982     0   1   2   3E+4   15   15.5  RT (min)      lcorbi_t3_pos_1_041_54    m/z 823.6982     0   2   4E+4   14.5   15  RT (min)      lcorbi_t3_pos_1_030_21    m/z 823.6982     0   1   2   3E+6   11.5   12  RT (min)      lcorbi_t3_pos_4_057_49    m/z 823.6983     0   2   4   6   8E+4   14   14.5  RT (min)      lcorbi_t3_pos_2_017_51    m/z 823.6983     0   1   2   3   4E+4   15   15.5  RT (min)      lcorbi_t3_pos_1_049_55    m/z 823.6984     0   2   4   6E+4   14.5   15  RT (min)      lcorbi_t3_pos_1_047_41    m/z 823.6984     0   1   2   3   4E+4   14.5   15  RT (min)      lcorbi_t3_pos_1_021_18    m/z 823.6984     0   1   2E+6   14   14.5  RT (min)      lcorbi_t3_pos_4_010_43    m/z 823.6984     0   1   2   3   4E+4   15   15.5  RT (min)      lcorbi_t3_pos_1_049_55    m/z 823.6984     0   1   2E+4   15.5   16  RT (min)      lcorbi_t3_pos_2_056_56    m/z 823.6984     0   1   2   3   4E+4   14.5   15  RT (min)      lcorbi_t3_pos_1_021_18    m/z 823.6985     0   1   2   3E+4   15.5   16  RT (min)      lcorbi_t3_pos_1_001_49    m/z 823.6985     0   1   2E+4   15.5   16  RT (min)      lcorbi_t3_pos_2_056_56    m/z 823.6985     0   1   2   3E+4   14.5   15  RT (min)      lcorbi_t3_pos_1_033_53    m/z 823.6986     0   1   2E+4   14.5   15  RT (min)      lcorbi_t3_pos_2_025_52    m/z 823.6986     0   1   2E+7   14   14.5  RT (min)       lcorbi_t3_pos_1_005_42    m/z 823.6986     0   1   2E+7   14   14.5  RT (min)      lcorbi_t3_pos_1_005_42    m/z 824.6085     0   2   4   6E+5   13.5   14  RT (min)      lcorbi_t3_pos_4_090_60    m/z 824.6446     0   1   2   3E+5   14   14.5  RT (min)       lcorbi_t3_pos_3_034_26    m/z 824.6476     0   1E+5   15   15.5  RT (min)      lcorbi_t3_pos_3_047_40    m/z 824.6524     0   1   17   17.5  RT (min)    lcorbi_t3_pos_1_023_02    m/z 824.6524     0   1   15.5   16  RT (min)      lcorbi_t3_pos_3_003_33    m/z 824.6526     0   1   16   16.5  RT (min)      lcorbi_t3_pos_1_034_34    m/z 824.6526     0   1   15.5   16  RT (min)      lcorbi_t3_pos_1_028_32    m/z 824.6526     0   1   2   3   4E+4   15   15.5  RT (min)      lcorbi_t3_pos_1_008_36    m/z 824.6527     0   1   15.5   16  RT (min)      lcorbi_t3_pos_2_022_38    m/z 824.6528     0   1   16   16.5  RT (min)      lcorbi_t3_pos_1_034_34    m/z 824.6529     0   1   15.5   16  RT (min)      lcorbi_t3_pos_1_028_32    m/z 824.6530     0   1   16.5   17  RT (min)    lcorbi_t3_pos_1_051_09    m/z 824.6531     0   1   16.5   17  RT (min)    lcorbi_t3_pos_2_012_01    m/z 824.6532     0   1   16.5   17  RT (min)    lcorbi_t3_pos_1_031_40    m/z 824.6905     0   1   14.5   15  RT (min)      lcorbi_t3_pos_1_041_54    m/z 824.6906     0   2   4   6E+5   13.5   14  RT (min)      lcorbi_t3_pos_1_005_42    m/z 824.6906     0   1   14   14.5  RT (min)      lcorbi_t3_pos_1_047_41    m/z 824.6906     0   1   2E+4   15   15.5  RT (min)      lcorbi_t3_pos_1_056_56    m/z 824.6906     0   1   2   3E+4   14.5   15  RT (min)      lcorbi_t3_pos_1_033_53    m/z 824.6909     0   1   2   3E+5   13.5   14   14.5  RT (min)      lcorbi_t3_pos_3_033_53    m/z 824.7010     0   2   4   6   8E+4   14.5   15  RT (min)      lcorbi_t3_pos_1_021_18    m/z 824.7011     0   1E+5   14.5   15  RT (min)      lcorbi_t3_pos_1_030_21    m/z 824.7011     0   1E+7   11.5   12  RT (min)      lcorbi_t3_pos_4_057_49    m/z 824.7012     0   2   4   6   8E+4   15   15.5  RT (min)      lcorbi_t3_pos_1_049_55    m/z 824.7012     0   2   4   6   8E+4   15   15.5  RT (min)      lcorbi_t3_pos_1_041_54    m/z 824.7012     0   2   4   6   8E+4   15   15.5  RT (min)      lcorbi_t3_pos_1_041_54    m/z 824.7012     0   2   4E+4   15.5   16  RT (min)      lcorbi_t3_pos_2_056_56    m/z 824.7013     0   1E+5   14.5   15  RT (min)      lcorbi_t3_pos_1_021_18    m/z 824.7013     0   2   4   6E+7   14   14.5  RT (min)       lcorbi_t3_pos_1_005_42    m/z 824.7014     0   2   4   6E+4   15.5   16  RT (min)      lcorbi_t3_pos_1_001_49    m/z 824.7014     0   2   4   6   8E+4   15   15.5  RT (min)      lcorbi_t3_pos_1_049_55    m/z 824.7014     0   1E+5   14.5   15  RT (min)      lcorbi_t3_pos_1_033_53    m/z 824.7015     0   2   4E+4   15.5   16  RT (min)      lcorbi_t3_pos_2_056_56    m/z 824.7016     0   1   2E+5   14   14.5  RT (min)      lcorbi_t3_pos_2_046_07    m/z 824.7017     0   2   4   6   8E+4   15   15.5  RT (min)      lcorbi_t3_pos_1_041_54    m/z 825.7070     0   1   2   3E+4   15.5   16  RT (min)      lcorbi_t3_pos_2_009_50    m/z 825.7072     0   1   2   3E+4   15   15.5  RT (min)      lcorbi_t3_pos_1_041_54    m/z 825.7072     0   1   2   3E+4   15   15.5  RT (min)      lcorbi_t3_pos_2_009_50    m/z 825.7072     0   2   4   6   8E+4   14.5   15  RT (min)      lcorbi_t3_pos_3_056_56    m/z 825.7075     0   1   2   3E+4   15   15.5  RT (min)      lcorbi_t3_pos_1_041_54    m/z 825.7105     0   2   4   6E+4   14.5   15  RT (min)      lcorbi_t3_pos_1_047_41    m/z 825.7106     0   1   2   3   4E+4   14.5   15  RT (min)      lcorbi_t3_pos_1_030_21    m/z 825.7136     0   1E+5   14.5   15  RT (min)      lcorbi_t3_pos_2_017_51    m/z 825.7138     0   1   2   3   4E+6   12   12.5  RT (min)      lcorbi_t3_pos_4_057_49    m/z 825.7138     0   1E+5   14.5   15  RT (min)      lcorbi_t3_pos_3_056_56    m/z 825.7140     0   1E+5   14   14.5  RT (min)      lcorbi_t3_pos_3_025_52    m/z 825.7140     0   1   2E+4   15   15.5  RT (min)      lcorbi_t3_pos_2_041_54    m/z 825.7141     0   1   15.5   16  RT (min)      lcorbi_t3_pos_1_025_52    m/z 825.7141     0   1E+7   14   14.5  RT (min)      lcorbi_t3_pos_1_005_42    m/z 825.7142     0   1   15.5   16  RT (min)      lcorbi_t3_pos_1_025_52    m/z 825.7142     0   1   2   3   4E+4   14.5   15  RT (min)      lcorbi_t3_pos_1_021_18    m/z 825.7143     0   1E+4   15.5   16  RT (min)      lcorbi_t3_pos_2_056_56    m/z 825.7146     0   1E+4   15   15.5  RT (min)      lcorbi_t3_pos_2_038_03    m/z 826.5945     0   2   4   6   8E+5   13   13.5  RT (min)       lcorbi_t3_pos_4_075_58    m/z 826.5962     0   1   14.5   15  RT (min)      lcorbi_t3_pos_1_028_32    m/z 826.6248     0   2   4E+5   13.5   14  RT (min)      lcorbi_t3_pos_4_092_60    m/z 826.7062     0   1   2E+4   15   15.5  RT (min)      lcorbi_t3_pos_1_056_56    m/z 826.7063     0   2   4   6   8E+5   14   14.5  RT (min)      lcorbi_t3_pos_1_005_42    m/z 826.7063     0   1   14.5   15  RT (min)      lcorbi_t3_pos_1_030_21    m/z 826.7064     0   1   2E+4   14.5   15  RT (min)      lcorbi_t3_pos_1_047_41    m/z 826.7064     0   1   14.5   15  RT (min)      lcorbi_t3_pos_1_030_21    m/z 826.7166     0   1E+7   12   12.5  RT (min)      lcorbi_t3_pos_4_057_49    m/z 826.7169     0   1   2   3   4E+4   15.5   16  RT (min)      lcorbi_t3_pos_1_025_52    m/z 826.7169     0   2   4   6   8E+4   14.5   15  RT (min)      lcorbi_t3_pos_1_021_18    m/z 826.7169     0   2   4E+4   15   15.5  RT (min)      lcorbi_t3_pos_2_038_03    m/z 826.7170     0   2   4   6E+4   15   15.5  RT (min)      lcorbi_t3_pos_1_041_54    m/z 826.7170     0   1   2   3   4E+5   14.5   15  RT (min)      lcorbi_t3_pos_3_056_56    m/z 826.7170     0   2   4E+4   15.5   16  RT (min)      lcorbi_t3_pos_1_001_49    m/z 826.7170     0   1   2   3   4E+5   14.5   15  RT (min)      lcorbi_t3_pos_3_056_56    m/z 826.7170     0   1   2   3   4E+4   15.5   16  RT (min)      lcorbi_t3_pos_1_025_52    m/z 826.7172     0   1   2   3   4E+7   14   14.5  RT (min)      lcorbi_t3_pos_1_005_42    m/z 826.7172     0   2   4   6E+4   15   15.5  RT (min)      lcorbi_t3_pos_1_041_54    m/z 826.7172     0   1   2   3   4E+5   14.5   15  RT (min)      lcorbi_t3_pos_2_017_51    m/z 826.7177     0   1E+5   14.5   15  RT (min)      lcorbi_t3_pos_1_030_21    m/z 826.7180     0   2   4   6E+4   15.5   16  RT (min)      lcorbi_t3_pos_2_009_50    m/z 826.7186     0   1   2   3   4E+4   16   16.5  RT (min)      lcorbi_t3_pos_2_056_56    m/z 826.7196     0   1   2   3   4E+4   16   16.5  RT (min)      lcorbi_t3_pos_2_056_56    m/z 826.7196     0   1   2   3   4E+4   16   16.5  RT (min)      lcorbi_t3_pos_2_056_56    m/z 826.7198     0   1   2   3   4E+4   16   16.5  RT (min)      lcorbi_t3_pos_2_056_56    m/z 826.8095     0   2   4E+5   13.5   14  RT (min)      lcorbi_t3_pos_4_090_60    m/z 826.8095     0   2   4E+5   13.5   14  RT (min)      lcorbi_t3_pos_4_093_60    m/z 827.5994     0   2   4E+5   13   13.5  RT (min)      lcorbi_t3_pos_4_075_58    m/z 827.6662     0   2   4E+5   12   12.5  RT (min)       lcorbi_t3_pos_4_095_60    m/z 827.7232     0   1E+5   14.5   15  RT (min)      lcorbi_t3_pos_2_006_47    m/z 827.7292     0   1   2E+4   14.5   15  RT (min)      lcorbi_t3_pos_2_006_47    m/z 827.7292     0   1   2E+4   15.5   16  RT (min)      lcorbi_t3_pos_1_041_54    m/z 827.7292     0   1E+4   16   16.5  RT (min)      lcorbi_t3_pos_2_056_56    m/z 827.7294     0   1   2E+4   15   15.5  RT (min)      lcorbi_t3_pos_1_041_54    m/z 827.7295     0   1   15.5   16  RT (min)      lcorbi_t3_pos_1_056_56    m/z 827.7295     0   2   4E+6   14.5   15  RT (min)      lcorbi_t3_pos_1_005_42    m/z 827.7296     0   1   15.5   16  RT (min)      lcorbi_t3_pos_2_009_50    m/z 827.7296     0   1   2E+6   13   13.5  RT (min)      lcorbi_t3_pos_4_057_49    m/z 827.7296     0   1   2E+4   15   15.5  RT (min)       lcorbi_t3_pos_1_030_21    m/z 827.7297     0   2   4   6   8E+4   14.5   15  RT (min)      lcorbi_t3_pos_2_017_51    m/z 827.7297     0   1   16   16.5  RT (min)      lcorbi_t3_pos_1_001_49    m/z 827.7297     0   2   4   6E+6   14   14.5  RT (min)      lcorbi_t3_pos_1_005_42    m/z 827.7298     0   1   2E+4   15   15.5  RT (min)      lcorbi_t3_pos_2_025_52    m/z 827.7298     0   2   4   6E+6   14   14.5  RT (min)      lcorbi_t3_pos_1_005_42    m/z 827.7298     0   1   2E+4   15.5   16  RT (min)      lcorbi_t3_pos_1_041_54    m/z 827.7298     0   1   2E+4   15   15.5  RT (min)      lcorbi_t3_pos_1_041_54    m/z 827.7298     0   1   15.5   16  RT (min)      lcorbi_t3_pos_1_025_52    m/z 827.7299     0   1E+4   16   16.5  RT (min)      lcorbi_t3_pos_2_056_56    m/z 827.7304     0   1   16   16.5  RT (min)      lcorbi_t3_pos_1_001_49    m/z 828.5386     0   2   4   6E+5   12   12.5  RT (min)       lcorbi_t3_pos_1_054_48    m/z 828.6692     0   1   14.5   15  RT (min)      lcorbi_t3_pos_1_033_53    m/z 828.6693     0   1   15.5   16  RT (min)      lcorbi_t3_pos_1_001_49    m/z 828.6693     0   1E+6   12   12.5  RT (min)       lcorbi_t3_pos_4_093_60    m/z 828.6693     0   1E+6   12.5   13  RT (min)       lcorbi_t3_pos_4_091_60    m/z 828.6694     0   1E+6   12.5   13  RT (min)       lcorbi_t3_pos_4_091_60    m/z 828.6695     0   1   15   15.5  RT (min)      lcorbi_t3_pos_1_056_56    m/z 828.6695     0   1   14.5   15  RT (min)      lcorbi_t3_pos_1_030_21    m/z 828.6695     0   1   15.5   16  RT (min)      lcorbi_t3_pos_1_001_49    m/z 828.6698     0   1   15   15.5  RT (min)      lcorbi_t3_pos_1_041_54    m/z 828.6698     0   1E+6   12   12.5  RT (min)      lcorbi_t3_pos_4_095_60    m/z 828.6744     0   1E+6   12.5   13  RT (min)      lcorbi_t3_pos_4_091_60    m/z 828.7218     0   2   4   6E+5   14   14.5  RT (min)      lcorbi_t3_pos_1_005_42    m/z 828.7220     0   1   2E+4   15.5   16  RT (min)      lcorbi_t3_pos_1_001_49    m/z 828.7222     0   2   4E+4   15.5   16  RT (min)      lcorbi_t3_pos_1_049_55    m/z 828.7222     0   1   2   3E+4   14.5   15  RT (min)      lcorbi_t3_pos_1_021_18    m/z 828.7222     0   1   2   3E+4   15.5   16  RT (min)      lcorbi_t3_pos_1_025_52    m/z 828.7222     0   1   2   3   4E+4   15   15.5  RT (min)      lcorbi_t3_pos_2_038_03    m/z 828.7223     0   2   4E+4   15   15.5  RT (min)      lcorbi_t3_pos_1_041_54    m/z 828.7224     0   2   4E+4   14.5   15  RT (min)      lcorbi_t3_pos_1_030_21    m/z 828.7320     0   2   4E+4   15   15.5  RT (min)      lcorbi_t3_pos_1_041_54    m/z 828.7325     0   1E+7   13   13.5  RT (min)      lcorbi_t3_pos_4_057_49    m/z 828.7326     0   2   4E+4   15   15.5  RT (min)      lcorbi_t3_pos_1_041_54    m/z 828.7326     0   1   2   3E+4   16   16.5  RT (min)      lcorbi_t3_pos_2_056_56    m/z 828.7329     0   1   2   3   4E+4   15.5   16  RT (min)      lcorbi_t3_pos_1_056_56    m/z 828.7329     0   1   2   3   4E+4   16   16.5  RT (min)      lcorbi_t3_pos_1_001_49    m/z 828.7329     0   1   2   3E+4   16   16.5  RT (min)      lcorbi_t3_pos_2_056_56    m/z 828.7329     0   2   4   6E+4   15   15.5  RT (min)      lcorbi_t3_pos_1_030_21    m/z 828.7330     0   2   4E+4   15.5   16  RT (min)      lcorbi_t3_pos_1_041_54    m/z 828.7330     0   1   2E+4   15.5   16  RT (min)      lcorbi_t3_pos_1_056_56    m/z 828.7332     0   1   2E+6   14.5   15  RT (min)      lcorbi_t3_pos_4_025_52    m/z 828.7332     0   1   2E+7   14   14.5  RT (min)      lcorbi_t3_pos_1_005_42    m/z 828.7332     0   2   4E+4   15.5   16  RT (min)      lcorbi_t3_pos_1_041_54    m/z 828.7332     0   1   2E+7   14   14.5  RT (min)      lcorbi_t3_pos_1_005_42    m/z 828.7332     0   2   4E+4   15.5   16  RT (min)      lcorbi_t3_pos_1_041_54    m/z 828.7332     0   1   2   3E+4   15.5   16  RT (min)      lcorbi_t3_pos_1_025_52    m/z 828.7334     0   1   2   3E+4   16   16.5  RT (min)      lcorbi_t3_pos_2_056_56    m/z 828.7334     0   1   2E+5   14.5   15  RT (min)      lcorbi_t3_pos_2_017_51    m/z 828.7334     0   2   4E+5   14.5   15  RT (min)      lcorbi_t3_pos_3_056_56    m/z 829.6818     0   1E+6   12.5   13  RT (min)       lcorbi_t3_pos_4_091_60    m/z 829.6818     0   1E+6   12.5   13  RT (min)       lcorbi_t3_pos_4_092_60    m/z 829.6819     0   2   4   6   8E+5   12.5   13  RT (min)       lcorbi_t3_pos_4_095_60    m/z 829.6820     0   1E+6   12.5   13  RT (min)       lcorbi_t3_pos_4_096_60    m/z 829.6824     0   1E+6   12.5   13  RT (min)      lcorbi_t3_pos_4_096_60    m/z 829.7351     0   1E+5   15.5   16  RT (min)      lcorbi_t3_pos_1_041_54    m/z 829.7380     0   1   2E+4   15.5   16  RT (min)      lcorbi_t3_pos_1_025_52    m/z 829.7383     0   1   16   16.5  RT (min)      lcorbi_t3_pos_1_056_56    m/z 829.7448     0   2   4E+4   15   15.5  RT (min)      lcorbi_t3_pos_1_021_18    m/z 829.7449     0   1   15   15.5  RT (min)      lcorbi_t3_pos_1_047_41    m/z 829.7451     0   1   2   3   4E+4   15.5   16  RT (min)      lcorbi_t3_pos_1_041_54    m/z 829.7452     0   1   16   16.5  RT (min)      lcorbi_t3_pos_1_025_52    m/z 829.7452     0   1   16   16.5  RT (min)      lcorbi_t3_pos_2_056_56    m/z 829.7452     0   1   16   16.5  RT (min)      lcorbi_t3_pos_1_049_55    m/z 829.7455     0   1E+6   14.5   15  RT (min)      lcorbi_t3_pos_1_005_42    m/z 829.7455     0   1   15   15.5  RT (min)      lcorbi_t3_pos_2_025_52    m/z 829.7456     0   1E+5   14.5   15  RT (min)      lcorbi_t3_pos_3_056_56    m/z 829.7458     0   1   16   16.5  RT (min)      lcorbi_t3_pos_2_056_56    m/z 830.5680     0   2   4   6E+5   14   14.5  RT (min)       lcorbi_t3_pos_4_078_58    m/z 830.5682     0   1   2   3E+5   14   14.5  RT (min)       lcorbi_t3_pos_3_047_40    m/z 830.5688     0   1   15.5   16  RT (min)      lcorbi_t3_pos_1_051_09    m/z 830.5695     0   1   16   16.5  RT (min)      lcorbi_t3_pos_1_023_02    m/z 830.5700     0   1   2E+4   12.5   13  RT (min)      lcorbi_t3_pos_2_032_39    m/z 830.5700     0   1   15.5   16  RT (min)      lcorbi_t3_pos_1_051_09    m/z 830.6850     0   1   2E+6   12.5   13  RT (min)       lcorbi_t3_pos_4_024_27    m/z 830.6850     0   1   2   3E+6   12.5   13  RT (min)       lcorbi_t3_pos_4_090_60    m/z 830.6850     0   1   2   3E+6   12.5   13  RT (min)       lcorbi_t3_pos_4_092_60    m/z 830.6850     0   1   2E+4   14.5   15  RT (min)      lcorbi_t3_pos_2_046_07    m/z 830.6850     0   1   2   3E+4   15.5   16  RT (min)      lcorbi_t3_pos_1_049_55    m/z 830.6850     0   1   2   3E+6   12.5   13  RT (min)       lcorbi_t3_pos_4_097_60    m/z 830.6850     0   1   2   3E+6   12.5   13  RT (min)       lcorbi_t3_pos_4_091_60    m/z 830.6851     0   1   15.5   16  RT (min)      lcorbi_t3_pos_1_001_49    m/z 830.6851     0   1   2   3E+4   14.5   15  RT (min)      lcorbi_t3_pos_2_006_47    m/z 830.6852     0   1   2   3   4E+4   14.5   15  RT (min)      lcorbi_t3_pos_1_033_53    m/z 830.6852     0   1   2   3E+6   12.5   13  RT (min)       lcorbi_t3_pos_4_093_60    m/z 830.6852     0   1   2   3   4E+6   12.5   13  RT (min)       lcorbi_t3_pos_4_096_60    m/z 830.6853     0   1   2   3E+6   12   12.5  RT (min)       lcorbi_t3_pos_4_095_60    m/z 830.6853     0   1   2   3   4E+4   14.5   15  RT (min)      lcorbi_t3_pos_1_033_53    m/z 830.6854     0   1   15   15.5  RT (min)      lcorbi_t3_pos_2_049_55    m/z 830.6855     0   2   4   6   8E+5   14   14.5  RT (min)      lcorbi_t3_pos_1_005_42    m/z 830.6873     0   1   2E+6   12.5   13  RT (min)       lcorbi_t3_pos_4_024_27    m/z 830.6876     0   1   2   3E+6   13   13.5  RT (min)       lcorbi_t3_pos_4_094_60    m/z 830.7486     0   1   15.5   16  RT (min)      lcorbi_t3_pos_1_041_54    m/z 830.7486     0   1   16   16.5  RT (min)      lcorbi_t3_pos_1_001_49    m/z 830.7487     0   1   16   16.5  RT (min)      lcorbi_t3_pos_1_001_49    m/z 830.7487     0   1   16   16.5  RT (min)      lcorbi_t3_pos_1_049_55    m/z 830.7487     0   1   15.5   16  RT (min)      lcorbi_t3_pos_1_041_54    m/z 830.7487     0   1   2   3   4E+4   15   15.5  RT (min)      lcorbi_t3_pos_1_033_53    m/z 830.7488     0   2   4E+4   15   15.5  RT (min)      lcorbi_t3_pos_2_006_47    m/z 830.7488     0   1   2   3   4E+6   14.5   15  RT (min)      lcorbi_t3_pos_1_005_42    m/z 830.7489     0   2   4E+5   14.5   15  RT (min)      lcorbi_t3_pos_4_025_52    m/z 830.7489     0   1   2   3E+4   15   15.5  RT (min)      lcorbi_t3_pos_1_030_21    m/z 830.7489     0   1   15.5   16  RT (min)      lcorbi_t3_pos_1_041_54    m/z 830.7489     0   1   2   3   4E+6   14.5   15  RT (min)      lcorbi_t3_pos_1_005_42    m/z 830.7489     0   1E+6   13   13.5  RT (min)      lcorbi_t3_pos_4_057_49    m/z 830.7490     0   2   4   6E+4   14.5   15  RT (min)      lcorbi_t3_pos_2_017_51    m/z 830.7494     0   1   16   16.5  RT (min)      lcorbi_t3_pos_1_001_49    m/z 831.6968     0   1E+6   12.5   13  RT (min)      lcorbi_t3_pos_4_097_60    m/z 831.6970     0   1E+6   12.5   13  RT (min)       lcorbi_t3_pos_4_095_60    m/z 831.6972     0   1E+6   13   13.5  RT (min)       lcorbi_t3_pos_4_092_60    m/z 831.6973     0   1E+6   12.5   13  RT (min)       lcorbi_t3_pos_4_090_60    m/z 831.6973     0   1E+6   13   13.5  RT (min)       lcorbi_t3_pos_4_092_60    m/z 831.6978     0   1E+6   13   13.5  RT (min)       lcorbi_t3_pos_4_092_60    m/z 831.7539     0   1E+6   14   14.5  RT (min)       lcorbi_t3_pos_4_093_60    m/z 831.7540     0   2   4   6   8E+5   14   14.5  RT (min)       lcorbi_t3_pos_4_095_60    m/z 831.7544     0   1E+6   14   14.5  RT (min)      lcorbi_t3_pos_4_093_60    m/z 832.2405     0   1   17.5   18  RT (min)    lcorbi_t3_pos_1_023_02    m/z 832.2406     0   1   16.5   17  RT (min)    lcorbi_t3_pos_1_034_34    m/z 832.2406     0   1   14   14.5  RT (min)      lcorbi_t3_pos_3_011_42    m/z 832.2406     0   2   4   6E+5   15   15.5  RT (min)      lcorbi_t3_pos_1_005_42    m/z 832.2407     0   1   2   3   4E+4   15.5   16  RT (min)      lcorbi_t3_pos_1_008_36    m/z 832.2407     0   1E+6   15   15.5  RT (min)      lcorbi_t3_pos_4_051_34    m/z 832.2408     0   1   17   17.5  RT (min)    lcorbi_t3_pos_1_031_40    m/z 832.2409     0   1   17   17.5  RT (min)    lcorbi_t3_pos_2_012_01    m/z 832.2409     0   1   16   16.5  RT (min)      lcorbi_t3_pos_2_022_38    m/z 832.2409     0   2   4   6E+5   14   14.5  RT (min)      lcorbi_t3_pos_4_016_15    m/z 832.2409     0   1   16   16.5  RT (min)      lcorbi_t3_pos_2_025_52    m/z 832.2409     0   1   16   16.5  RT (min)      lcorbi_t3_pos_2_022_38    m/z 832.2410     0   1   16.5   17  RT (min)    lcorbi_t3_pos_2_009_50    m/z 832.2410     0   1   16.5   17  RT (min)    lcorbi_t3_pos_1_034_34    m/z 832.2410     0   2   4   6   8E+4   15.5   16  RT (min)      lcorbi_t3_pos_1_008_36    m/z 832.2410     0   1   17   17.5  RT (min)    lcorbi_t3_pos_1_031_40    m/z 832.2411     0   1   13.5   14  RT (min)      lcorbi_t3_pos_3_024_07    m/z 832.2411     0   1   17.5   18  RT (min)    lcorbi_t3_pos_1_023_02    m/z 832.2411     0   1   16   16.5  RT (min)      lcorbi_t3_pos_1_028_32    m/z 832.2412     0   2   4   6   8E+5   15   15.5  RT (min)      lcorbi_t3_pos_1_005_42    m/z 832.2412     0   2   4   6E+6   14.5   15  RT (min)       lcorbi_t3_pos_4_105_61    m/z 832.5704     0   2   4E+5   12.5   13  RT (min)       lcorbi_t3_pos_2_005_11    m/z 832.5706     0   2   4   6E+5   12.5   13  RT (min)       lcorbi_t3_pos_1_054_48    m/z 832.5809     0   2   4   6E+4   13   13.5  RT (min)      lcorbi_t3_pos_2_032_39    m/z 832.5825     0   1E+5   13.5   14  RT (min)      lcorbi_t3_pos_2_054_29    m/z 832.5830     0   1   15   15.5  RT (min)      lcorbi_t3_pos_1_028_32    m/z 832.5832     0   1   16   16.5  RT (min)      lcorbi_t3_pos_1_025_52    m/z 832.5834     0   1   16.5   17  RT (min)    lcorbi_t3_pos_1_023_02    m/z 832.5834     0   1   16.5   17  RT (min)    lcorbi_t3_pos_1_023_02    m/z 832.5834     0   1   15   15.5  RT (min)      lcorbi_t3_pos_1_047_41    m/z 832.5835     0   1   16.5   17  RT (min)    lcorbi_t3_pos_1_023_02    m/z 832.5835     0   1   2E+5   14.5   15  RT (min)      lcorbi_t3_pos_1_008_36    m/z 832.5836     0   1   17   17.5  RT (min)    lcorbi_t3_pos_1_023_02    m/z 832.5836     0   1   15   15.5  RT (min)      lcorbi_t3_pos_1_047_41    m/z 832.5836     0   1   15   15.5  RT (min)      lcorbi_t3_pos_1_030_21    m/z 832.5836     0   2   4   6   8E+5   14.5   15  RT (min)      lcorbi_t3_pos_1_005_42    m/z 832.5837     0   1   15.5   16  RT (min)      lcorbi_t3_pos_2_049_55    m/z 832.5839     0   1   15.5   16  RT (min)      lcorbi_t3_pos_1_025_52    m/z 832.5839     0   1E+5   14.5   15  RT (min)      lcorbi_t3_pos_1_008_36    m/z 832.5842     0   1   16.5   17  RT (min)    lcorbi_t3_pos_1_023_02    m/z 832.5842     0   1   2   3E+4   15   15.5  RT (min)      lcorbi_t3_pos_1_033_53    m/z 832.5846     0   1   16.5   17  RT (min)    lcorbi_t3_pos_1_023_02    m/z 832.6372     0   2   4   6E+5   12.5   13  RT (min)       lcorbi_t3_pos_4_090_60    m/z 832.6372     0   2   4   6E+5   12.5   13  RT (min)       lcorbi_t3_pos_4_096_60    m/z 832.7003     0   1   2   3   4E+6   12.5   13  RT (min)       lcorbi_t3_pos_4_093_60    m/z 832.7006     0   1   2   3   4E+6   12.5   13  RT (min)       lcorbi_t3_pos_4_090_60    m/z 832.7006     0   1   2   3   4E+6   13   13.5  RT (min)       lcorbi_t3_pos_4_092_60    m/z 832.7007     0   1   2   3   4E+6   13   13.5  RT (min)       lcorbi_t3_pos_4_091_60    m/z 832.7007     0   1   2   3   4E+6   12.5   13  RT (min)       lcorbi_t3_pos_4_095_60    m/z 832.7008     0   1   2   3   4E+6   12.5   13  RT (min)      lcorbi_t3_pos_4_097_60    m/z 832.7008     0   1   2   3   4E+6   13   13.5  RT (min)       lcorbi_t3_pos_4_092_60    m/z 832.7643     0   1   16   16.5  RT (min)      lcorbi_t3_pos_1_049_55    m/z 832.7644     0   1   2E+4   15   15.5  RT (min)      lcorbi_t3_pos_1_033_53    m/z 832.7644     0   1   15   15.5  RT (min)      lcorbi_t3_pos_2_046_07    m/z 832.7644     0   1   2E+4   15   15.5  RT (min)      lcorbi_t3_pos_1_033_53    m/z 832.7644     0   1   15.5   16  RT (min)      lcorbi_t3_pos_1_041_54    m/z 832.7645     0   2   4   6E+5   14.5   15  RT (min)      lcorbi_t3_pos_1_005_42    m/z 832.7645     0   1   2   3   4E+4   14.5   15  RT (min)      lcorbi_t3_pos_3_025_52    m/z 832.7645     0   1   16   16.5  RT (min)      lcorbi_t3_pos_1_001_49    m/z 832.7646     0   1   16   16.5   17  RT (min)    lcorbi_t3_pos_2_056_56    m/z 833.2414     0   1   2   3E+4   15.5   16  RT (min)      lcorbi_t3_pos_1_008_36    m/z 833.2414     0   2   4   6E+4   15.5   16  RT (min)      lcorbi_t3_pos_1_008_36    m/z 833.2416     0   1   16   16.5  RT (min)      lcorbi_t3_pos_3_003_33    m/z 833.2416     0   2   4E+6   14.5   15  RT (min)      lcorbi_t3_pos_4_105_61    m/z 833.2416     0   2   4   6   8E+5   15   15.5  RT (min)      lcorbi_t3_pos_4_051_34    m/z 833.2416     0   1   17   17.5  RT (min)    lcorbi_t3_pos_1_031_40    m/z 833.2416     0   1   16   16.5  RT (min)      lcorbi_t3_pos_2_022_38    m/z 833.2416     0   1   16.5   17  RT (min)    lcorbi_t3_pos_1_034_34    m/z 833.2416     0   1   16   16.5  RT (min)      lcorbi_t3_pos_1_028_32    m/z 833.2416     0   1   16.5   17  RT (min)    lcorbi_t3_pos_1_034_34    m/z 833.2417     0   1   14   14.5  RT (min)      lcorbi_t3_pos_3_011_42    m/z 833.2417     0   1   13.5   14  RT (min)      lcorbi_t3_pos_3_024_07    m/z 833.2417     0   1   17   17.5  RT (min)    lcorbi_t3_pos_2_012_01    m/z 833.2419     0   1   17.5   18  RT (min)    lcorbi_t3_pos_1_023_02    m/z 833.5870     0   1   16.5   17  RT (min)    lcorbi_t3_pos_1_023_02    m/z 833.5894     0   2   4   6E+5   13.5   14  RT (min)      lcorbi_t3_pos_4_075_58    m/z 834.2394     0   1   2   3E+4   15.5   16  RT (min)      lcorbi_t3_pos_1_008_36    m/z 834.2395     0   2   4E+4   15.5   16  RT (min)      lcorbi_t3_pos_1_008_36    m/z 834.2396     0   1   16.5   17  RT (min)    lcorbi_t3_pos_1_034_34    m/z 834.2396     0   1   13.5   14  RT (min)      lcorbi_t3_pos_3_024_07    m/z 834.2396     0   1   16.5   17  RT (min)    lcorbi_t3_pos_1_034_34    m/z 834.2396     0   1   16   16.5  RT (min)      lcorbi_t3_pos_3_003_33    m/z 834.2397     0   1   13.5   14  RT (min)      lcorbi_t3_pos_3_031_45    m/z 834.2398     0   1   14   14.5  RT (min)      lcorbi_t3_pos_3_011_42    m/z 834.2398     0   1   16   16.5  RT (min)      lcorbi_t3_pos_2_022_38    m/z 834.2398     0   1   16   16.5  RT (min)      lcorbi_t3_pos_1_028_32    m/z 834.2398     0   2   4   6E+5   15   15.5  RT (min)      lcorbi_t3_pos_4_051_34    m/z 834.2400     0   1   17   17.5  RT (min)    lcorbi_t3_pos_1_031_40    m/z 834.2401     0   1   16   16.5  RT (min)      lcorbi_t3_pos_1_028_32    m/z 834.2401     0   1   2   3   4E+6   14.5   15  RT (min)       lcorbi_t3_pos_4_105_61    m/z 834.2401     0   1   17   17.5  RT (min)    lcorbi_t3_pos_1_031_40    m/z 834.2402     0   1   17   17.5  RT (min)    lcorbi_t3_pos_2_012_01    m/z 834.5999     0   2   4   6   8E+5   14.5   15  RT (min)      lcorbi_t3_pos_3_047_40    m/z 834.6001     0   1E+4   13   13.5  RT (min)      lcorbi_t3_pos_2_032_39    m/z 834.6002     0   2   4   6E+6   13.5   14  RT (min)      lcorbi_t3_pos_1_002_01    m/z 834.6002     0   2   4E+6   13.5   14  RT (min)      lcorbi_t3_pos_1_002_01    m/z 834.6004     0   1E+6   14.5   15  RT (min)      lcorbi_t3_pos_1_008_36    m/z 834.6004     0   1   2   3   4E+6   14.5   15  RT (min)       lcorbi_t3_pos_1_005_42    m/z 834.6004     0   1   2E+4   15   15.5  RT (min)      lcorbi_t3_pos_1_030_21    m/z 834.6004     0   1   2   3   4E+4   15   15.5  RT (min)      lcorbi_t3_pos_1_047_41    m/z 834.6005     0   2   4   6E+5   13.5   14  RT (min)      lcorbi_t3_pos_1_002_01    m/z 834.6005     0   1   15.5   16  RT (min)      lcorbi_t3_pos_2_049_55    m/z 834.6006     0   1   16   16.5  RT (min)      lcorbi_t3_pos_2_056_56    m/z 834.6006     0   1   2   3E+4   15.5   16  RT (min)      lcorbi_t3_pos_1_034_34    m/z 834.6006     0   1   16   16.5  RT (min)      lcorbi_t3_pos_1_001_49    m/z 834.6006     0   1E+4   15   15.5  RT (min)      lcorbi_t3_pos_2_025_52    m/z 834.6007     0   1E+4   13   13.5  RT (min)      lcorbi_t3_pos_2_032_39    m/z 834.6007     0   1E+5   14.5   15  RT (min)      lcorbi_t3_pos_2_017_51    m/z 834.6007     0   1   2   3E+4   15.5   16  RT (min)      lcorbi_t3_pos_1_034_34    m/z 834.6007     0   1   2   3E+4   15.5   16  RT (min)      lcorbi_t3_pos_1_034_34    m/z 834.6008     0   1   16.5   17  RT (min)    lcorbi_t3_pos_1_023_02    m/z 834.6008     0   1   2   3   4E+5   14.5   15  RT (min)      lcorbi_t3_pos_1_008_36    m/z 834.6008     0   1   2   3E+6   13.5   14  RT (min)      lcorbi_t3_pos_1_046_22    m/z 834.6008     0   1   2E+6   13.5   14  RT (min)      lcorbi_t3_pos_1_002_01    m/z 834.6009     0   1   2E+4   15   15.5  RT (min)      lcorbi_t3_pos_1_028_32    m/z 834.6010     0   1   2E+4   15   15.5  RT (min)      lcorbi_t3_pos_1_028_32    m/z 834.6010     0   1   16   16.5  RT (min)      lcorbi_t3_pos_1_051_09    m/z 834.6012     0   1   16.5   17  RT (min)    lcorbi_t3_pos_1_023_02    m/z 834.6012     0   1   16.5   17  RT (min)    lcorbi_t3_pos_1_023_02    m/z 835.2392     0   1   14   14.5  RT (min)      lcorbi_t3_pos_3_011_42    m/z 835.2393     0   1   16   16.5  RT (min)      lcorbi_t3_pos_2_022_38    m/z 835.2394     0   1   16   16.5  RT (min)      lcorbi_t3_pos_1_028_32    m/z 835.2394     0   1   13.5   14  RT (min)      lcorbi_t3_pos_3_024_07    m/z 835.2394     0   1   17   17.5  RT (min)    lcorbi_t3_pos_1_031_40    m/z 835.2394     0   1   16.5   17  RT (min)    lcorbi_t3_pos_1_034_34    m/z 835.2395     0   1   15.5   16  RT (min)      lcorbi_t3_pos_1_008_36    m/z 835.2396     0   2   4   6E+4   15.5   16  RT (min)      lcorbi_t3_pos_1_008_36    m/z 835.2396     0   1   17   17.5  RT (min)    lcorbi_t3_pos_2_012_01    m/z 835.2397     0   1   16.5   17  RT (min)    lcorbi_t3_pos_1_034_34    m/z 835.2401     0   1   2E+6   14.5   15  RT (min)      lcorbi_t3_pos_4_105_61    m/z 835.6037     0   1   15.5   16  RT (min)      lcorbi_t3_pos_2_049_55    m/z 835.6038     0   1   16   16.5  RT (min)      lcorbi_t3_pos_2_056_56    m/z 835.6038     0   1E+6   13.5   14  RT (min)      lcorbi_t3_pos_1_002_01    m/z 835.6040     0   1   2   3E+5   13.5   14  RT (min)      lcorbi_t3_pos_1_002_01    m/z 835.6040     0   1   16   16.5  RT (min)      lcorbi_t3_pos_2_056_56    m/z 835.6041     0   1   15   15.5  RT (min)      lcorbi_t3_pos_1_030_21    m/z 835.6041     0   1   16.5   17  RT (min)    lcorbi_t3_pos_1_023_02    m/z 835.6042     0   1   15.5   16  RT (min)      lcorbi_t3_pos_1_034_34    m/z 835.6042     0   1   2   3E+4   15   15.5  RT (min)      lcorbi_t3_pos_1_028_32    m/z 835.6042     0   1   2   3E+5   14.5   15  RT (min)      lcorbi_t3_pos_1_008_36    m/z 835.6043     0   1   16   16.5  RT (min)      lcorbi_t3_pos_1_051_09    m/z 835.6043     0   1   2E+4   15   15.5  RT (min)      lcorbi_t3_pos_2_006_47    m/z 835.6044     0   1   2   3E+4   15   15.5  RT (min)      lcorbi_t3_pos_1_028_32    m/z 835.6044     0   1   16.5   17  RT (min)    lcorbi_t3_pos_1_023_02    m/z 835.6046     0   1   16.5   17  RT (min)    lcorbi_t3_pos_1_023_02    m/z 835.6046     0   1E+5   14.5   15  RT (min)      lcorbi_t3_pos_1_008_36    m/z 835.6046     0   1   16   16.5  RT (min)      lcorbi_t3_pos_1_051_09    m/z 835.6046     0   1   15.5   16  RT (min)      lcorbi_t3_pos_1_034_34    m/z 835.6964     0   2   4E+5   14   14.5  RT (min)       lcorbi_t3_pos_4_094_60    m/z 836.2382     0   1E+6   14.5   15  RT (min)      lcorbi_t3_pos_4_105_61    m/z 836.6152     0   1   16.5   17  RT (min)    lcorbi_t3_pos_1_023_02    m/z 836.6156     0   1E+4   13.5   14  RT (min)      lcorbi_t3_pos_2_013_33    m/z 836.6156     0   1   15.5   16  RT (min)      lcorbi_t3_pos_1_034_34    m/z 836.6157     0   1   16   16.5  RT (min)      lcorbi_t3_pos_2_009_50    m/z 836.6157     0   1   16.5   17  RT (min)    lcorbi_t3_pos_1_023_02    m/z 836.6159     0   2   4   6   8E+5   13.5   14  RT (min)      lcorbi_t3_pos_1_002_01    m/z 836.6159     0   1   16   16.5  RT (min)      lcorbi_t3_pos_1_031_40    m/z 836.6160     0   1   16.5   17  RT (min)    lcorbi_t3_pos_2_012_01    m/z 836.6160     0   1   2   3E+5   14.5   15  RT (min)      lcorbi_t3_pos_1_008_36    m/z 836.6160     0   1   15.5   16  RT (min)      lcorbi_t3_pos_1_028_32    m/z 836.6160     0   1   15.5   16  RT (min)      lcorbi_t3_pos_1_034_34    m/z 836.6160     0   1E+6   13.5   14  RT (min)      lcorbi_t3_pos_4_016_15    m/z 836.6160     0   1   15   15.5  RT (min)      lcorbi_t3_pos_1_030_21    m/z 836.6161     0   1   2E+6   13.5   14  RT (min)      lcorbi_t3_pos_1_002_01    m/z 836.6161     0   1   2E+6   14.5   15  RT (min)      lcorbi_t3_pos_1_005_42    m/z 836.6162     0   1   15   15.5  RT (min)      lcorbi_t3_pos_1_028_32    m/z 836.6162     0   1   16   16.5  RT (min)      lcorbi_t3_pos_1_051_09    m/z 836.6162     0   1   15.5   16  RT (min)      lcorbi_t3_pos_2_025_52    m/z 836.6162     0   1   15.5   16  RT (min)      lcorbi_t3_pos_2_049_55    m/z 836.6163     0   1   16   16.5  RT (min)      lcorbi_t3_pos_1_031_40    m/z 836.6163     0   1E+5   13.5   14  RT (min)      lcorbi_t3_pos_1_002_01    m/z 836.6164     0   1   2E+6   14.5   15  RT (min)      lcorbi_t3_pos_1_005_42    m/z 836.6164     0   1   17   17.5  RT (min)    lcorbi_t3_pos_1_023_02    m/z 836.6164     0   1E+6   14.5   15  RT (min)      lcorbi_t3_pos_1_008_36    m/z 836.6165     0   1   13   13.5  RT (min)      lcorbi_t3_pos_2_032_39    m/z 836.6165     0   1   15.5   16  RT (min)      lcorbi_t3_pos_2_025_52    m/z 837.5219     0   1   2   3E+4   16   16.5  RT (min)      lcorbi_t3_pos_4_001_49    m/z 837.5219     0   1   17.5   18  RT (min)    lcorbi_t3_pos_1_025_52    m/z 837.5219     0   2   4   6   8E+5   15.5   16  RT (min)      lcorbi_t3_pos_1_005_42    m/z 837.5220     0   1   2   3E+5   14.5   15  RT (min)       lcorbi_t3_pos_4_037_26    m/z 837.5220     0   1   17.5   18  RT (min)    lcorbi_t3_pos_1_056_56    m/z 837.5220     0   1   16.5   17  RT (min)    lcorbi_t3_pos_1_033_53    m/z 837.5221     0   1   16   16.5  RT (min)      lcorbi_t3_pos_2_017_51    m/z 837.5221     0   1   14   14.5  RT (min)      lcorbi_t3_pos_2_032_39    m/z 837.5221     0   1   17   17.5  RT (min)    lcorbi_t3_pos_1_041_54    m/z 837.5221     0   2   4   6   8E+5   15   15.5  RT (min)       lcorbi_t3_pos_4_105_61    m/z 837.5221     0   1   2E+4   14.5   15  RT (min)      lcorbi_t3_pos_2_013_33    m/z 837.5222     0   1   16   16.5  RT (min)      lcorbi_t3_pos_1_008_36    m/z 837.5222     0   1   17.5   18  RT (min)    lcorbi_t3_pos_1_025_52    m/z 837.5223     0   1   16.5   17  RT (min)    lcorbi_t3_pos_1_030_21    m/z 837.5223     0   1   17   17.5  RT (min)    lcorbi_t3_pos_2_038_03    m/z 837.5224     0   1   17   17.5  RT (min)    lcorbi_t3_pos_1_041_54    m/z 837.6194     0   2   4   6   8E+5   13.5   14  RT (min)      lcorbi_t3_pos_4_016_15    m/z 837.6195     0   1E+4   13.5   14  RT (min)      lcorbi_t3_pos_2_013_33    m/z 837.6196     0   1   16.5   17  RT (min)    lcorbi_t3_pos_2_012_01    m/z 837.6196     0   1   13.5   14  RT (min)      lcorbi_t3_pos_1_002_01    m/z 837.6196     0   1   2E+5   14.5   15  RT (min)      lcorbi_t3_pos_1_008_36    m/z 837.6197     0   1E+5   13.5   14  RT (min)      lcorbi_t3_pos_1_002_01    m/z 837.6198     0   2   4   6E+5   14.5   15  RT (min)      lcorbi_t3_pos_1_008_36    m/z 837.6198     0   1   15   15.5  RT (min)      lcorbi_t3_pos_2_006_47    m/z 837.6198     0   1   16   16.5  RT (min)      lcorbi_t3_pos_1_031_40    m/z 837.6198     0   1E+6   14.5   15  RT (min)      lcorbi_t3_pos_1_005_42    m/z 837.6198     0   1   15.5   16  RT (min)      lcorbi_t3_pos_1_034_34    m/z 837.6198     0   1   15.5   16  RT (min)      lcorbi_t3_pos_1_034_34    m/z 837.6199     0   1   15.5   16  RT (min)      lcorbi_t3_pos_1_041_54    m/z 837.6199     0   1   2   3E+4   15.5   16  RT (min)      lcorbi_t3_pos_1_028_32    m/z 837.6199     0   1E+4   13   13.5  RT (min)      lcorbi_t3_pos_2_032_39    m/z 837.6200     0   1   16   16.5  RT (min)      lcorbi_t3_pos_1_031_40    m/z 837.6200     0   1   2   3E+4   15   15.5  RT (min)      lcorbi_t3_pos_1_028_32    m/z 837.6200     0   1   17   17.5  RT (min)    lcorbi_t3_pos_1_023_02    m/z 837.6200     0   1   15   15.5  RT (min)      lcorbi_t3_pos_1_047_41    m/z 837.6200     0   1   16.5   17  RT (min)    lcorbi_t3_pos_1_023_02    m/z 837.6201     0   1   2   3E+4   15.5   16  RT (min)      lcorbi_t3_pos_1_028_32    m/z 838.5253     0   1   17.5   18  RT (min)    lcorbi_t3_pos_1_056_56    m/z 838.6237     0   1   16   16.5  RT (min)      lcorbi_t3_pos_2_009_50    m/z 838.6240     0   1   16.5   17  RT (min)    lcorbi_t3_pos_1_023_02    m/z 838.6271     0   1   15   15.5  RT (min)      lcorbi_t3_pos_1_021_18    m/z 838.6308     0   1   16   16.5  RT (min)      lcorbi_t3_pos_1_051_09    m/z 838.6315     0   1   2E+4   15   15.5  RT (min)      lcorbi_t3_pos_2_046_07    m/z 838.6316     0   1   16   16.5  RT (min)      lcorbi_t3_pos_1_031_40    m/z 838.6316     0   1   2   3E+4   15.5   16  RT (min)      lcorbi_t3_pos_1_034_34    m/z 838.6316     0   1   2   3   4E+5   14.5   15  RT (min)      lcorbi_t3_pos_1_008_36    m/z 838.6316     0   1   15   15.5  RT (min)      lcorbi_t3_pos_2_022_38    m/z 838.6316     0   1   2E+4   15   15.5  RT (min)      lcorbi_t3_pos_2_046_07    m/z 838.6316     0   1   13.5   14  RT (min)      lcorbi_t3_pos_2_013_33    m/z 838.6316     0   1E+4   15   15.5  RT (min)      lcorbi_t3_pos_3_056_56    m/z 838.6317     0   1   15.5   16  RT (min)      lcorbi_t3_pos_1_028_32    m/z 838.6317     0   1   16   16.5  RT (min)      lcorbi_t3_pos_1_049_55    m/z 838.6317     0   1E+6   14.5   15  RT (min)      lcorbi_t3_pos_3_047_40    m/z 838.6317     0   1E+6   14.5   15  RT (min)      lcorbi_t3_pos_4_019_47    m/z 838.6318     0   1   2   3E+4   15.5   16  RT (min)      lcorbi_t3_pos_1_034_34    m/z 838.6318     0   1   15.5   16  RT (min)      lcorbi_t3_pos_1_028_32    m/z 838.6318     0   1   2   3E+5   13.5   14  RT (min)      lcorbi_t3_pos_1_002_01    m/z 838.6318     0   1   16.5   17  RT (min)    lcorbi_t3_pos_1_051_09    m/z 838.6318     0   1   15.5   16  RT (min)      lcorbi_t3_pos_1_028_32    m/z 838.6319     0   1E+5   15   15.5  RT (min)      lcorbi_t3_pos_1_008_36    m/z 838.6320     0   1   13   13.5  RT (min)      lcorbi_t3_pos_2_032_39    m/z 838.6320     0   1   16.5   17  RT (min)    lcorbi_t3_pos_1_051_09    m/z 838.6320     0   1   15.5   16  RT (min)      lcorbi_t3_pos_1_034_34    m/z 838.6320     0   1   13   13.5  RT (min)      lcorbi_t3_pos_2_032_39    m/z 838.6322     0   1   17   17.5  RT (min)    lcorbi_t3_pos_1_023_02    m/z 838.6322     0   1   17   17.5  RT (min)    lcorbi_t3_pos_1_023_02    m/z 838.6812     0   2   4   6E+5   12.5   13  RT (min)       lcorbi_t3_pos_4_094_60    m/z 838.6812     0   2   4   6E+5   12.5   13  RT (min)       lcorbi_t3_pos_4_090_60    m/z 838.6813     0   2   4   6E+5   12.5   13  RT (min)       lcorbi_t3_pos_4_091_60    m/z 838.6815     0   2   4   6E+5   12.5   13  RT (min)      lcorbi_t3_pos_4_093_60    m/z 839.6352     0   1   15.5   16  RT (min)      lcorbi_t3_pos_1_034_34    m/z 839.6353     0   1   16   16.5  RT (min)      lcorbi_t3_pos_1_051_09    m/z 839.6353     0   1   2   3E+4   13.5   14  RT (min)      lcorbi_t3_pos_1_002_01    m/z 839.6353     0   1   16.5   17  RT (min)    lcorbi_t3_pos_2_012_01    m/z 839.6354     0   1E+5   15   15.5  RT (min)      lcorbi_t3_pos_1_008_36    m/z 839.6354     0   1   15.5   16  RT (min)      lcorbi_t3_pos_1_028_32    m/z 839.6354     0   1E+6   14   14.5  RT (min)      lcorbi_t3_pos_4_016_15    m/z 839.6354     0   1   2   3E+5   14.5   15  RT (min)      lcorbi_t3_pos_3_047_40    m/z 839.6354     0   1   16   16.5  RT (min)      lcorbi_t3_pos_1_051_09    m/z 839.6354     0   1   15.5   16  RT (min)      lcorbi_t3_pos_1_028_32    m/z 839.6356     0   1   16   16.5  RT (min)      lcorbi_t3_pos_1_049_55    m/z 839.6356     0   2   4   6   8E+5   14.5   15  RT (min)      lcorbi_t3_pos_4_019_47    m/z 839.6356     0   1E+4   13.5   14  RT (min)      lcorbi_t3_pos_2_013_33    m/z 839.6356     0   1   15.5   16  RT (min)      lcorbi_t3_pos_1_034_34    m/z 839.6356     0   1   15   15.5  RT (min)      lcorbi_t3_pos_2_022_38    m/z 839.6357     0   1   15.5   16  RT (min)      lcorbi_t3_pos_1_034_34    m/z 839.6357     0   1   15.5   16  RT (min)      lcorbi_t3_pos_1_028_32    m/z 839.6358     0   2   4E+4   15   15.5  RT (min)      lcorbi_t3_pos_1_008_36    m/z 839.6358     0   2   4   6   8E+5   14   14.5  RT (min)      lcorbi_t3_pos_3_034_26    m/z 839.6362     0   1   13   13.5  RT (min)      lcorbi_t3_pos_2_032_39    m/z 839.6362     0   1   17   17.5  RT (min)    lcorbi_t3_pos_1_023_02    m/z 840.6966     0   1E+6   12.5   13  RT (min)      lcorbi_t3_pos_4_095_60    m/z 841.7090     0   2   4E+5   13   13.5  RT (min)       lcorbi_t3_pos_4_090_60    m/z 842.5176     0   2   4   6E+5   11   11.5  RT (min)      lcorbi_t3_pos_1_043_06    m/z 842.5177     0   2   4   6E+5   11   11.5  RT (min)      lcorbi_t3_pos_1_043_06    m/z 842.5178     0   1E+6   11   11.5  RT (min)       lcorbi_t3_pos_1_054_48    m/z 842.5179     0   2   4   6E+5   11   11.5  RT (min)      lcorbi_t3_pos_1_043_06    m/z 842.7117     0   1E+6   13   13.5  RT (min)      lcorbi_t3_pos_4_097_60    m/z 842.7120     0   1E+6   13   13.5  RT (min)       lcorbi_t3_pos_4_093_60    m/z 842.7122     0   1E+6   13   13.5  RT (min)       lcorbi_t3_pos_4_093_60    m/z 842.7123     0   1E+6   13   13.5  RT (min)       lcorbi_t3_pos_1_003_45    m/z 842.7123     0   1E+6   13   13.5  RT (min)       lcorbi_t3_pos_4_090_60    m/z 842.7128     0   1E+6   13   13.5  RT (min)      lcorbi_t3_pos_4_091_60    m/z 842.7230     0   1E+4   14   14.5  RT (min)      lcorbi_t3_pos_3_024_07    m/z 842.7233     0   1   17   17.5  RT (min)    lcorbi_t3_pos_1_034_34    m/z 842.7233     0   1   16   16.5  RT (min)      lcorbi_t3_pos_1_008_36    m/z 842.7233     0   1   14.5   15  RT (min)      lcorbi_t3_pos_3_011_42    m/z 842.7233     0   1   17   17.5  RT (min)    lcorbi_t3_pos_1_034_34    m/z 842.7233     0   1   17.5   18  RT (min)    lcorbi_t3_pos_1_031_40    m/z 842.7234     0   1   16.5   17  RT (min)    lcorbi_t3_pos_2_022_38    m/z 842.7234     0   1   18   18.5  RT (min)    lcorbi_t3_pos_2_012_01    m/z 842.7234     0   1   17   17.5  RT (min)    lcorbi_t3_pos_1_028_32    m/z 842.7235     0   1   17.5   18  RT (min)    lcorbi_t3_pos_1_031_40    m/z 842.7238     0   1   14.5   15  RT (min)      lcorbi_t3_pos_3_011_42    m/z 843.5212     0   2   4E+5   11   11.5  RT (min)      lcorbi_t3_pos_2_005_11    m/z 843.5214     0   2   4   6E+5   11   11.5  RT (min)       lcorbi_t3_pos_1_054_48    m/z 843.6648     0   1   2   3E+4   14   14.5  RT (min)      lcorbi_t3_pos_1_030_21    m/z 843.6650     0   2   4E+5   13.5   14  RT (min)      lcorbi_t3_pos_1_005_42    m/z 843.7263     0   1   17   17.5  RT (min)    lcorbi_t3_pos_1_034_34    m/z 843.7263     0   1   17.5   18  RT (min)    lcorbi_t3_pos_1_031_40    m/z 843.7265     0   1   18   18.5  RT (min)    lcorbi_t3_pos_2_012_01    m/z 843.7265     0   1   16.5   17  RT (min)    lcorbi_t3_pos_1_028_32    m/z 843.7266     0   1   14   14.5  RT (min)      lcorbi_t3_pos_3_024_07    m/z 843.7266     0   1   17.5   18  RT (min)    lcorbi_t3_pos_1_031_40    m/z 843.7268     0   1   14.5   15  RT (min)      lcorbi_t3_pos_2_013_33    m/z 843.7269     0   1   16.5   17  RT (min)    lcorbi_t3_pos_2_022_38    m/z 843.7269     0   1   2   3   4E+4   16   16.5  RT (min)      lcorbi_t3_pos_1_008_36    m/z 844.5332     0   2   4   6   8E+5   11   11.5  RT (min)      lcorbi_t3_pos_1_043_06    m/z 844.5337     0   1E+6   11   11.5  RT (min)       lcorbi_t3_pos_1_054_48    m/z 844.5338     0   2   4   6   8E+5   11.5   12  RT (min)      lcorbi_t3_pos_1_043_06    m/z 844.6678     0   1   14.5   15  RT (min)      lcorbi_t3_pos_2_049_55    m/z 844.6680     0   1   14.5   15  RT (min)      lcorbi_t3_pos_1_041_54    m/z 844.6680     0   1E+5   14   14.5  RT (min)      lcorbi_t3_pos_3_025_52    m/z 844.6681     0   1   14   14.5  RT (min)      lcorbi_t3_pos_1_047_41    m/z 844.6682     0   1E+4   14.5   15  RT (min)      lcorbi_t3_pos_2_025_52    m/z 844.6682     0   1   14   14.5  RT (min)      lcorbi_t3_pos_1_030_21    m/z 844.6682     0   1   14   14.5  RT (min)      lcorbi_t3_pos_1_030_21    m/z 844.6683     0   1   15   15.5  RT (min)      lcorbi_t3_pos_1_025_52    m/z 844.6685     0   1   15   15.5  RT (min)      lcorbi_t3_pos_1_049_55    m/z 844.7276     0   2   4E+5   13   13.5  RT (min)      lcorbi_t3_pos_4_095_60    m/z 844.7280     0   2   4   6E+5   13   13.5  RT (min)       lcorbi_t3_pos_1_003_45    m/z 844.7287     0   2   4E+5   13   13.5  RT (min)       lcorbi_t3_pos_4_095_60    m/z 844.7342     0   1   14   14.5  RT (min)      lcorbi_t3_pos_2_032_39    m/z 844.7387     0   1   2   3E+4   16   16.5  RT (min)      lcorbi_t3_pos_4_025_52    m/z 844.7388     0   2   4   6E+5   16   16.5  RT (min)      lcorbi_t3_pos_1_005_42    m/z 844.7388     0   1   17.5   18  RT (min)    lcorbi_t3_pos_2_049_55    m/z 844.7388     0   1   17   17.5  RT (min)    lcorbi_t3_pos_1_028_32    m/z 844.7389     0   1   17.5   18  RT (min)    lcorbi_t3_pos_2_009_50    m/z 844.7389     0   1   18   18.5  RT (min)    lcorbi_t3_pos_1_031_40    m/z 844.7389     0   1   14.5   15  RT (min)      lcorbi_t3_pos_3_024_07    m/z 844.7389     0   1   14.5   15  RT (min)      lcorbi_t3_pos_2_013_33    m/z 844.7389     0   1   18   18.5  RT (min)    lcorbi_t3_pos_2_012_01    m/z 844.7390     0   1   16.5   17  RT (min)    lcorbi_t3_pos_2_006_47    m/z 844.7390     0   2   4   6E+5   15.5   16  RT (min)      lcorbi_t3_pos_4_051_34    m/z 844.7390     0   2   4   6E+5   16   16.5  RT (min)      lcorbi_t3_pos_3_040_41    m/z 844.7390     0   1   16.5   17  RT (min)    lcorbi_t3_pos_2_022_38    m/z 844.7390     0   1   16.5   17  RT (min)    lcorbi_t3_pos_2_017_51    m/z 844.7391     0   1   2   3   4E+4   16   16.5  RT (min)      lcorbi_t3_pos_1_008_36    m/z 844.7392     0   1   2   3E+5   15   15.5  RT (min)      lcorbi_t3_pos_4_016_15    m/z 844.7392     0   1   17   17.5  RT (min)    lcorbi_t3_pos_1_034_34    m/z 844.7393     0   1   18   18.5  RT (min)    lcorbi_t3_pos_2_012_01    m/z 844.7394     0   1   17   17.5  RT (min)    lcorbi_t3_pos_1_034_34    m/z 844.7394     0   1   17.5   18  RT (min)    lcorbi_t3_pos_1_056_56    m/z 844.7396     0   1   18.5   19  RT (min)    lcorbi_t3_pos_1_023_02    m/z 845.6806     0   1   14.5   15  RT (min)      lcorbi_t3_pos_1_030_21    m/z 845.6807     0   1   15.5   16  RT (min)      lcorbi_t3_pos_1_025_52    m/z 845.7039     0   2   4   6E+5   12.5   13  RT (min)      lcorbi_t3_pos_4_094_60    m/z 845.7045     0   2   4E+5   13   13.5  RT (min)       lcorbi_t3_pos_4_091_60    m/z 845.7046     0   2   4   6E+5   12.5   13  RT (min)      lcorbi_t3_pos_4_095_60    m/z 845.7422     0   1   14.5   15  RT (min)      lcorbi_t3_pos_2_013_33    m/z 845.7422     0   1   18   18.5  RT (min)    lcorbi_t3_pos_2_012_01    m/z 845.7423     0   1   18   18.5  RT (min)    lcorbi_t3_pos_2_012_01    m/z 845.7424     0   1   17   17.5  RT (min)    lcorbi_t3_pos_1_034_34    m/z 845.7424     0   1   17   17.5  RT (min)    lcorbi_t3_pos_1_034_34    m/z 845.7424     0   1   17   17.5  RT (min)    lcorbi_t3_pos_1_028_32    m/z 845.7425     0   1   16   16.5  RT (min)      lcorbi_t3_pos_1_008_36    m/z 845.7425     0   1   14.5   15  RT (min)      lcorbi_t3_pos_3_024_07    m/z 845.7426     0   1   17   17.5  RT (min)    lcorbi_t3_pos_1_028_32    m/z 845.7427     0   1   18   18.5  RT (min)    lcorbi_t3_pos_1_031_40    m/z 846.5489     0   1E+6   11.5   12  RT (min)       lcorbi_t3_pos_1_054_48    m/z 846.6795     0   2   4E+5   12   12.5  RT (min)      lcorbi_t3_pos_3_055_10    m/z 846.6795     0   2   4   6E+5   12   12.5  RT (min)      lcorbi_t3_pos_1_003_45    m/z 846.6799     0   2   4   6E+5   12   12.5  RT (min)       lcorbi_t3_pos_1_003_45    m/z 846.6835     0   1   15.5   16  RT (min)      lcorbi_t3_pos_1_025_52    m/z 846.6836     0   1   14.5   15  RT (min)      lcorbi_t3_pos_1_030_21    m/z 846.6838     0   1   14.5   15  RT (min)      lcorbi_t3_pos_1_021_18    m/z 846.6838     0   1   15.5   16  RT (min)      lcorbi_t3_pos_1_025_52    m/z 846.6838     0   1   15   15.5  RT (min)      lcorbi_t3_pos_1_041_54    m/z 846.6838     0   1   14   14.5  RT (min)      lcorbi_t3_pos_2_046_07    m/z 846.6839     0   1   15   15.5  RT (min)      lcorbi_t3_pos_1_041_54    m/z 846.6839     0   1   15   15.5  RT (min)      lcorbi_t3_pos_1_049_55    m/z 846.6839     0   1   2   3E+4   15.5   16  RT (min)      lcorbi_t3_pos_1_001_49    m/z 846.6839     0   1E+6   14   14.5  RT (min)      lcorbi_t3_pos_1_005_42    m/z 846.6839     0   1   14.5   15  RT (min)      lcorbi_t3_pos_1_021_18    m/z 846.6840     0   1   15   15.5  RT (min)      lcorbi_t3_pos_1_056_56    m/z 846.6840     0   1   14   14.5  RT (min)      lcorbi_t3_pos_2_046_07    m/z 846.6841     0   1E+6   14   14.5  RT (min)      lcorbi_t3_pos_1_005_42    m/z 846.6841     0   1   15   15.5  RT (min)      lcorbi_t3_pos_1_041_54    m/z 846.6844     0   1E+4   14.5   15  RT (min)      lcorbi_t3_pos_2_038_03    m/z 846.7457     0   1   2E+5   15   15.5  RT (min)      lcorbi_t3_pos_4_016_15    m/z 846.7471     0   1   16.5   17  RT (min)    lcorbi_t3_pos_2_022_38    m/z 846.7500     0   1   17.5   18  RT (min)    lcorbi_t3_pos_2_009_50    m/z 846.7501     0   1   17.5   18  RT (min)    lcorbi_t3_pos_4_033_53    m/z 846.7503     0   1   16.5   17  RT (min)    lcorbi_t3_pos_2_046_07    m/z 846.7537     0   1   18.5   19  RT (min)    lcorbi_t3_pos_1_023_02    m/z 846.7542     0   1   17   17.5  RT (min)    lcorbi_t3_pos_1_033_53    m/z 846.7543     0   1   14.5   15  RT (min)      lcorbi_t3_pos_3_011_42    m/z 846.7544     0   1   18   18.5  RT (min)    lcorbi_t3_pos_1_031_40    m/z 846.7545     0   2   4   6   8E+5   15   15.5  RT (min)      lcorbi_t3_pos_4_016_15    m/z 846.7545     0   1   2E+4   16   16.5  RT (min)      lcorbi_t3_pos_4_001_49    m/z 846.7545     0   1   14.5   15  RT (min)      lcorbi_t3_pos_3_024_07    m/z 846.7545     0   1   2   3E+4   16   16.5  RT (min)      lcorbi_t3_pos_1_008_36    m/z 846.7546     0   1   18.5   19  RT (min)    lcorbi_t3_pos_1_023_02    m/z 846.7546     0   1E+6   16   16.5  RT (min)      lcorbi_t3_pos_4_019_47    m/z 846.7546     0   1E+6   16   16.5  RT (min)      lcorbi_t3_pos_1_005_42    m/z 846.7546     0   1E+6   16   16.5  RT (min)      lcorbi_t3_pos_3_026_04    m/z 846.7546     0   1   18   18.5  RT (min)    lcorbi_t3_pos_2_012_01    m/z 846.7547     0   1   17.5   18  RT (min)    lcorbi_t3_pos_1_034_34    m/z 846.7548     0   1   14.5   15  RT (min)      lcorbi_t3_pos_2_013_33    m/z 846.7548     0   1   16.5   17  RT (min)    lcorbi_t3_pos_1_008_36    m/z 846.7548     0   1   18   18.5  RT (min)    lcorbi_t3_pos_2_012_01    m/z 846.7548     0   1   17   17.5  RT (min)    lcorbi_t3_pos_2_038_03    m/z 846.7548     0   1   16.5   17  RT (min)    lcorbi_t3_pos_2_022_38    m/z 846.7549     0   1   17   17.5  RT (min)    lcorbi_t3_pos_1_028_32    m/z 846.7550     0   1   17.5   18  RT (min)    lcorbi_t3_pos_1_034_34    m/z 847.7575     0   1   18   18.5  RT (min)    lcorbi_t3_pos_1_031_40    m/z 847.7579     0   1   17.5   18  RT (min)    lcorbi_t3_pos_2_009_50    m/z 847.7579     0   1   2E+5   15   15.5  RT (min)      lcorbi_t3_pos_4_016_15    m/z 847.7579     0   1   16.5   17  RT (min)    lcorbi_t3_pos_2_022_38    m/z 847.7581     0   1   18.5   19  RT (min)    lcorbi_t3_pos_1_023_02    m/z 847.7581     0   1   18   18.5  RT (min)    lcorbi_t3_pos_2_012_01    m/z 847.7581     0   1   2E+4   16   16.5  RT (min)      lcorbi_t3_pos_1_008_36    m/z 847.7581     0   1   14.5   15  RT (min)      lcorbi_t3_pos_3_011_42    m/z 847.7581     0   1   16.5   17  RT (min)    lcorbi_t3_pos_2_017_51    m/z 847.7581     0   1   16.5   17  RT (min)    lcorbi_t3_pos_2_022_38    m/z 847.7581     0   2   4   6E+5   16   16.5  RT (min)       lcorbi_t3_pos_1_005_42    m/z 847.7581     0   2   4   6E+5   16   16.5  RT (min)      lcorbi_t3_pos_3_040_41    m/z 847.7581     0   1E+6   15   15.5  RT (min)      lcorbi_t3_pos_2_023_26    m/z 847.7582     0   1   14.5   15  RT (min)      lcorbi_t3_pos_2_032_39    m/z 847.7582     0   1   17   17.5  RT (min)    lcorbi_t3_pos_1_028_32    m/z 847.7582     0   1   2   3   4E+5   16   16.5  RT (min)      lcorbi_t3_pos_1_005_42    m/z 847.7582     0   1   16.5   17  RT (min)    lcorbi_t3_pos_1_008_36    m/z 847.7582     0   2   4   6E+5   15   15.5  RT (min)      lcorbi_t3_pos_3_034_26    m/z 847.7583     0   1   18   18.5  RT (min)    lcorbi_t3_pos_1_031_40    m/z 847.7584     0   1   14.5   15  RT (min)      lcorbi_t3_pos_2_013_33    m/z 847.7584     0   1   18.5   19  RT (min)    lcorbi_t3_pos_1_023_02    m/z 847.7585     0   1   17.5   18  RT (min)    lcorbi_t3_pos_1_034_34    m/z 848.6530     0   1   2E+5   14.5   15  RT (min)      lcorbi_t3_pos_3_047_40    m/z 848.6954     0   2   4   6   8E+5   12   12.5  RT (min)      lcorbi_t3_pos_4_096_60    m/z 848.6956     0   2   4   6   8E+5   12.5   13  RT (min)       lcorbi_t3_pos_4_092_60    m/z 848.6957     0   1E+6   12.5   13  RT (min)       lcorbi_t3_pos_1_003_45    m/z 848.6960     0   2   4   6   8E+5   12   12.5  RT (min)       lcorbi_t3_pos_4_095_60    m/z 848.6993     0   1   15   15.5  RT (min)      lcorbi_t3_pos_1_041_54    m/z 848.6994     0   1   14.5   15  RT (min)      lcorbi_t3_pos_1_021_18    m/z 848.6995     0   1   15.5   16  RT (min)      lcorbi_t3_pos_1_025_52    m/z 848.6996     0   1E+4   14.5   15  RT (min)      lcorbi_t3_pos_2_017_51    m/z 848.6996     0   1   15   15.5  RT (min)      lcorbi_t3_pos_2_038_03    m/z 848.6996     0   1E+6   14   14.5  RT (min)      lcorbi_t3_pos_1_005_42    m/z 848.6996     0   1   14.5   15  RT (min)      lcorbi_t3_pos_1_033_53    m/z 848.6997     0   1   15.5   16  RT (min)      lcorbi_t3_pos_1_025_52    m/z 848.6998     0   1   15   15.5   16  RT (min)      lcorbi_t3_pos_2_009_50    m/z 848.6999     0   1   14.5   15  RT (min)      lcorbi_t3_pos_1_033_53    m/z 848.7618     0   1E+6   15.5   16  RT (min)      lcorbi_t3_pos_3_034_26    m/z 848.7621     0   1   17   17.5  RT (min)    lcorbi_t3_pos_2_006_47    m/z 848.7626     0   1   18   18.5  RT (min)    lcorbi_t3_pos_1_049_55    m/z 848.7659     0   1   14.5   15  RT (min)      lcorbi_t3_pos_2_013_33    m/z 848.7660     0   1   17.5   18  RT (min)    lcorbi_t3_pos_1_034_34    m/z 848.7660     0   1   14.5   15  RT (min)      lcorbi_t3_pos_3_031_45    m/z 848.7660     0   1   17.5   18  RT (min)    lcorbi_t3_pos_2_038_03    m/z 848.7660     0   1   16.5   17  RT (min)    lcorbi_t3_pos_2_046_07    m/z 848.7698     0   1   19   19.5  RT (min)    lcorbi_t3_pos_1_023_02    m/z 848.7699     0   1   18   18.5  RT (min)    lcorbi_t3_pos_1_031_40    m/z 848.7699     0   1   19   19.5  RT (min)    lcorbi_t3_pos_1_023_02    m/z 848.7700     0   1   18.5   19  RT (min)    lcorbi_t3_pos_2_012_01    m/z 848.7701     0   1   15   15.5  RT (min)      lcorbi_t3_pos_3_011_42    m/z 848.7701     0   1   18.5   19  RT (min)    lcorbi_t3_pos_2_012_01    m/z 848.7701     0   1   17   17.5  RT (min)    lcorbi_t3_pos_1_028_32    m/z 848.7702     0   1   17   17.5  RT (min)    lcorbi_t3_pos_1_028_32    m/z 848.7702     0   1   17   17.5  RT (min)    lcorbi_t3_pos_2_022_38    m/z 848.7702     0   1   18   18.5  RT (min)    lcorbi_t3_pos_4_033_53    m/z 848.7703     0   1   17.5   18  RT (min)    lcorbi_t3_pos_2_009_50    m/z 848.7703     0   1   17.5   18  RT (min)    lcorbi_t3_pos_1_034_34    m/z 848.7703     0   1   16.5   17  RT (min)    lcorbi_t3_pos_1_008_36    m/z 848.7704     0   1   16.5   17  RT (min)    lcorbi_t3_pos_1_008_36    m/z 848.7704     0   1   17   17.5  RT (min)    lcorbi_t3_pos_2_022_38    m/z 848.7704     0   2   4   6   8E+5   16   16.5  RT (min)      lcorbi_t3_pos_4_019_47    m/z 848.7704     0   2   4   6   8E+5   16   16.5  RT (min)      lcorbi_t3_pos_1_005_42    m/z 848.7704     0   1   19   19.5  RT (min)    lcorbi_t3_pos_1_023_02    m/z 848.7705     0   1   18   18.5  RT (min)    lcorbi_t3_pos_1_051_09    m/z 848.7706     0   1   15   15.5  RT (min)      lcorbi_t3_pos_1_002_01    m/z 849.7734     0   1   18.5   19  RT (min)    lcorbi_t3_pos_2_012_01    m/z 849.7735     0   1   19   19.5  RT (min)    lcorbi_t3_pos_1_023_02    m/z 849.7738     0   1   17.5   18  RT (min)    lcorbi_t3_pos_1_034_34    m/z 849.7738     0   1   14.5   15  RT (min)      lcorbi_t3_pos_2_013_33    m/z 849.7738     0   1   14.5   15  RT (min)      lcorbi_t3_pos_2_032_39    m/z 849.7738     0   1   17   17.5  RT (min)    lcorbi_t3_pos_1_028_32    m/z 849.7739     0   1   16.5   17  RT (min)    lcorbi_t3_pos_1_008_36    m/z 849.7739     0   1   18   18.5  RT (min)    lcorbi_t3_pos_1_031_40    m/z 849.7739     0   1   17   17.5  RT (min)    lcorbi_t3_pos_2_022_38    m/z 849.7739     0   1   18.5   19  RT (min)    lcorbi_t3_pos_2_012_01    m/z 849.7740     0   2   4E+5   16   16.5  RT (min)      lcorbi_t3_pos_3_040_41    m/z 849.7740     0   1   2   3E+5   15   15.5  RT (min)      lcorbi_t3_pos_4_016_15    m/z 849.7740     0   1   14.5   15  RT (min)      lcorbi_t3_pos_3_024_07    m/z 849.7742     0   1   17.5   18  RT (min)    lcorbi_t3_pos_1_034_34    m/z 849.7742     0   1   15   15.5  RT (min)      lcorbi_t3_pos_1_002_01    m/z 850.6667     0   1   13.5   14  RT (min)       lcorbi_t3_pos_2_032_39    m/z 850.6682     0   1   2   3E+4   15   15.5  RT (min)      lcorbi_t3_pos_1_008_36    m/z 850.6683     0   1   16.5   17  RT (min)    lcorbi_t3_pos_1_051_09    m/z 850.6685     0   1   16   16.5  RT (min)      lcorbi_t3_pos_1_034_34    m/z 850.6686     0   1   15.5   16  RT (min)      lcorbi_t3_pos_3_003_33    m/z 850.6687     0   1   13.5   14  RT (min)      lcorbi_t3_pos_2_032_39    m/z 850.6687     0   1   17   17.5  RT (min)    lcorbi_t3_pos_1_023_02    m/z 850.6687     0   1   16.5   17  RT (min)    lcorbi_t3_pos_2_012_01    m/z 850.6687     0   1   15.5   16  RT (min)      lcorbi_t3_pos_2_022_38    m/z 850.6687     0   1   2E+5   15   15.5  RT (min)      lcorbi_t3_pos_3_047_40    m/z 850.6687     0   1   16.5   17  RT (min)    lcorbi_t3_pos_1_031_40    m/z 850.6687     0   1   13.5   14  RT (min)      lcorbi_t3_pos_2_013_33    m/z 850.6688     0   2   4   6E+5   14   14.5  RT (min)      lcorbi_t3_pos_4_016_15    m/z 850.6688     0   2   4   6E+5   14.5   15  RT (min)      lcorbi_t3_pos_4_019_47    m/z 850.6688     0   1   15.5   16  RT (min)      lcorbi_t3_pos_1_028_32    m/z 850.7815     0   1   17   17.5  RT (min)    lcorbi_t3_pos_2_022_38    m/z 850.7853     0   1   18   18.5  RT (min)    lcorbi_t3_pos_1_031_40    m/z 850.7854     0   1   17.5   18  RT (min)    lcorbi_t3_pos_1_034_34    m/z 850.7856     0   1   15   15.5  RT (min)      lcorbi_t3_pos_2_054_29    m/z 850.7858     0   1   18.5   19  RT (min)    lcorbi_t3_pos_2_012_01    m/z 850.7864     0   1   19   19.5  RT (min)    lcorbi_t3_pos_1_023_02    m/z 852.5601     0   2   4   6   8E+4   14   14.5  RT (min)      lcorbi_t3_pos_1_008_36    m/z 852.5609     0   1   15   15.5  RT (min)      lcorbi_t3_pos_1_031_40    m/z 852.7303     0   2   4E+5   14   14.5  RT (min)      lcorbi_t3_pos_4_093_60    m/z 852.7305     0   2   4E+5   14   14.5  RT (min)       lcorbi_t3_pos_4_091_60    m/z 852.7307     0   2   4   6E+5   14   14.5  RT (min)       lcorbi_t3_pos_4_090_60    m/z 854.3384     0   2   4   6   8E+5   9   9.5  RT (min)      lcorbi_t3_pos_1_034_34    m/z 854.3385     0   2   4   6   8E+5   9   9.5  RT (min)      lcorbi_t3_pos_1_034_34    m/z 854.6821     0   1   2E+6   12.5   13  RT (min)       lcorbi_t3_pos_4_092_60    m/z 855.3416     0   2   4E+4   9   9.5  RT (min)      lcorbi_t3_pos_1_023_02    m/z 855.7440     0   1   15   15.5  RT (min)       lcorbi_t3_pos_1_033_53    m/z 856.7383     0   1   17   17.5  RT (min)    lcorbi_t3_pos_1_034_34    m/z 856.7387     0   1   14.5   15  RT (min)      lcorbi_t3_pos_3_024_07    m/z 856.7391     0   1   17   17.5  RT (min)    lcorbi_t3_pos_1_028_32    m/z 856.7392     0   2   4   6E+4   14.5   15  RT (min)      lcorbi_t3_pos_2_013_33    m/z 856.7396     0   1   17.5   18  RT (min)    lcorbi_t3_pos_1_031_40    m/z 856.7551     0   1   15.5   16  RT (min)      lcorbi_t3_pos_1_041_54    m/z 856.7553     0   1   2E+4   16   16.5  RT (min)      lcorbi_t3_pos_1_049_55    m/z 856.7554     0   1   15   15.5  RT (min)      lcorbi_t3_pos_2_025_52    m/z 856.7555     0   1   16   16.5  RT (min)      lcorbi_t3_pos_1_056_56    m/z 856.7556     0   1   15   15.5  RT (min)      lcorbi_t3_pos_1_033_53    m/z 856.7556     0   1   16   16.5  RT (min)      lcorbi_t3_pos_1_025_52    m/z 856.7557     0   1   15.5   16  RT (min)      lcorbi_t3_pos_1_041_54    m/z 856.7558     0   1   15   15.5  RT (min)      lcorbi_t3_pos_1_047_41    m/z 856.7558     0   2   4   6   8E+5   14.5   15  RT (min)       lcorbi_t3_pos_1_005_42    m/z 856.7558     0   1   15   15.5  RT (min)      lcorbi_t3_pos_1_021_18    m/z 857.7058     0   1   2E+6   13   13.5  RT (min)       lcorbi_t3_pos_4_094_60    m/z 858.7542     0   1   18   18.5  RT (min)    lcorbi_t3_pos_1_031_40    m/z 858.7543     0   1   17   17.5  RT (min)    lcorbi_t3_pos_1_034_34    m/z 858.7545     0   1   16.5   17  RT (min)    lcorbi_t3_pos_2_022_38    m/z 858.7547     0   1   16   16.5  RT (min)      lcorbi_t3_pos_1_008_36    m/z 858.7547     0   1   14.5   15  RT (min)      lcorbi_t3_pos_3_011_42    m/z 858.7548     0   1   17   17.5  RT (min)    lcorbi_t3_pos_3_003_33    m/z 858.7548     0   1   14.5   15  RT (min)      lcorbi_t3_pos_3_024_07    m/z 858.7549     0   1   17   17.5  RT (min)    lcorbi_t3_pos_1_028_32    m/z 858.7549     0   1   17   17.5  RT (min)    lcorbi_t3_pos_1_028_32    m/z 858.7550     0   1   17.5   18  RT (min)    lcorbi_t3_pos_1_034_34    m/z 858.7551     0   1   18   18.5  RT (min)    lcorbi_t3_pos_1_031_40    m/z 858.7708     0   2   4E+5   14.5   15  RT (min)      lcorbi_t3_pos_1_005_42    m/z 858.7713     0   1   15.5   16  RT (min)      lcorbi_t3_pos_1_041_54    m/z 858.7713     0   1   16   16.5  RT (min)      lcorbi_t3_pos_1_001_49    m/z 860.6593     0   2   4   6   8E+5   11   11.5  RT (min)       lcorbi_t3_pos_4_093_60    m/z 860.7698     0   1   18.5   19  RT (min)    lcorbi_t3_pos_2_012_01    m/z 860.7701     0   1   17   17.5  RT (min)    lcorbi_t3_pos_1_028_32    m/z 860.7702     0   1   18   18.5  RT (min)    lcorbi_t3_pos_1_031_40    m/z 860.7703     0   1   17   17.5  RT (min)    lcorbi_t3_pos_3_003_33    m/z 860.7703     0   1   18   18.5  RT (min)    lcorbi_t3_pos_1_031_40    m/z 860.7703     0   1   17.5   18  RT (min)    lcorbi_t3_pos_1_034_34    m/z 860.7703     0   1   14.5   15  RT (min)      lcorbi_t3_pos_3_031_45    m/z 860.7704     0   1   17.5   18  RT (min)    lcorbi_t3_pos_1_034_34    m/z 860.7705     0   1   15   15.5  RT (min)      lcorbi_t3_pos_3_011_42    m/z 860.7706     0   1   16.5   17  RT (min)    lcorbi_t3_pos_1_008_36    m/z 860.7707     0   1   17   17.5  RT (min)    lcorbi_t3_pos_2_022_38    m/z 862.6747     0   1E+6   11.5   12  RT (min)       lcorbi_t3_pos_4_090_60    m/z 862.7347     0   2   4   6E+5   13   13.5  RT (min)       lcorbi_t3_pos_4_095_60    m/z 862.7379     0   2   4   6   8E+5   13   13.5  RT (min)       lcorbi_t3_pos_4_093_60    m/z 862.7382     0   2   4   6E+5   13   13.5  RT (min)      lcorbi_t3_pos_4_090_60    m/z 862.7385     0   2   4   6E+5   13   13.5  RT (min)       lcorbi_t3_pos_4_092_60    m/z 862.7386     0   2   4   6   8E+5   13   13.5  RT (min)       lcorbi_t3_pos_4_091_60    m/z 866.7230     0   2   4E+4   16   16.5  RT (min)      lcorbi_t3_pos_1_008_36    m/z 866.7230     0   1   18   18.5  RT (min)    lcorbi_t3_pos_2_012_01    m/z 866.7231     0   1   17   17.5  RT (min)    lcorbi_t3_pos_1_034_34    m/z 866.7231     0   1   17.5   18  RT (min)    lcorbi_t3_pos_1_031_40    m/z 866.7232     0   1   16.5   17  RT (min)    lcorbi_t3_pos_1_028_32    m/z 866.7233     0   1   17   17.5  RT (min)    lcorbi_t3_pos_1_034_34    m/z 866.7233     0   1   14.5   15  RT (min)      lcorbi_t3_pos_3_011_42    m/z 866.7233     0   1   16.5   17  RT (min)    lcorbi_t3_pos_1_028_32    m/z 866.7233     0   1   16.5   17  RT (min)    lcorbi_t3_pos_2_022_38    m/z 866.7234     0   1   14   14.5  RT (min)      lcorbi_t3_pos_3_024_07    m/z 866.7236     0   1   17.5   18  RT (min)    lcorbi_t3_pos_1_031_40    m/z 866.7238     0   2   4E+4   16   16.5  RT (min)      lcorbi_t3_pos_1_008_36    m/z 866.7239     0   1   18   18.5  RT (min)    lcorbi_t3_pos_2_012_01    m/z 867.7256     0   1   16   16.5  RT (min)      lcorbi_t3_pos_1_008_36    m/z 867.7263     0   1   17   17.5  RT (min)    lcorbi_t3_pos_1_034_34    m/z 867.7264     0   1   17.5   18  RT (min)    lcorbi_t3_pos_1_031_40    m/z 867.7265     0   1   17.5   18  RT (min)    lcorbi_t3_pos_1_031_40    m/z 867.7267     0   1   18   18.5  RT (min)    lcorbi_t3_pos_2_012_01    m/z 867.7267     0   1   16   16.5  RT (min)      lcorbi_t3_pos_1_008_36    m/z 867.7267     0   1   16.5   17  RT (min)    lcorbi_t3_pos_3_003_33    m/z 867.7267     0   1   14.5   15  RT (min)      lcorbi_t3_pos_3_011_42    m/z 867.7269     0   1   18   18.5  RT (min)    lcorbi_t3_pos_2_012_01    m/z 867.7269     0   1   16.5   17  RT (min)    lcorbi_t3_pos_1_028_32    m/z 867.7269     0   1   14   14.5  RT (min)      lcorbi_t3_pos_3_024_07    m/z 867.7271     0   2   4   6   8E+5   15   15.5  RT (min)      lcorbi_t3_pos_2_023_26    m/z 868.7385     0   1   16.5   17  RT (min)    lcorbi_t3_pos_4_049_55    m/z 868.7386     0   1   18   18.5  RT (min)    lcorbi_t3_pos_2_012_01    m/z 868.7386     0   1   17.5   18  RT (min)    lcorbi_t3_pos_2_009_50    m/z 868.7386     0   1   17   17.5  RT (min)    lcorbi_t3_pos_1_034_34    m/z 868.7388     0   1   17.5   18  RT (min)    lcorbi_t3_pos_1_031_40    m/z 868.7388     0   1   17   17.5  RT (min)    lcorbi_t3_pos_1_034_34    m/z 868.7389     0   1   17.5   18  RT (min)    lcorbi_t3_pos_1_049_55    m/z 868.7389     0   1   16.5   17  RT (min)    lcorbi_t3_pos_3_003_33    m/z 868.7390     0   1   14   14.5  RT (min)      lcorbi_t3_pos_3_024_07    m/z 868.7390     0   1   16.5   17  RT (min)    lcorbi_t3_pos_2_022_38    m/z 868.7390     0   1E+6   15.5   16  RT (min)      lcorbi_t3_pos_3_040_41    m/z 868.7391     0   1   14.5   15  RT (min)      lcorbi_t3_pos_3_011_42    m/z 868.7391     0   1   2   3E+4   16   16.5  RT (min)      lcorbi_t3_pos_1_008_36    m/z 868.7391     0   2   4   6   8E+4   16   16.5  RT (min)      lcorbi_t3_pos_4_025_52    m/z 868.7391     0   2   4   6   8E+5   16   16.5  RT (min)       lcorbi_t3_pos_1_005_42    m/z 868.7391     0   1   16.5   17  RT (min)    lcorbi_t3_pos_1_028_32    m/z 868.7391     0   1   2   3E+4   16   16.5  RT (min)      lcorbi_t3_pos_1_008_36    m/z 868.7393     0   1   14.5   15  RT (min)      lcorbi_t3_pos_3_011_42    m/z 868.7394     0   1   18   18.5  RT (min)    lcorbi_t3_pos_2_012_01    m/z 869.7420     0   1   17.5   18  RT (min)    lcorbi_t3_pos_1_031_40    m/z 869.7421     0   1   14   14.5  RT (min)      lcorbi_t3_pos_3_024_07    m/z 869.7422     0   1   16.5   17  RT (min)    lcorbi_t3_pos_2_022_38    m/z 869.7423     0   1   16   16.5  RT (min)      lcorbi_t3_pos_1_008_36    m/z 869.7423     0   1   17   17.5  RT (min)    lcorbi_t3_pos_1_034_34    m/z 869.7424     0   1   17.5   18  RT (min)    lcorbi_t3_pos_2_009_50    m/z 869.7424     0   1   18   18.5  RT (min)    lcorbi_t3_pos_2_012_01    m/z 869.7424     0   2   4   6E+5   16   16.5  RT (min)      lcorbi_t3_pos_1_005_42    m/z 869.7424     0   1   17   17.5  RT (min)    lcorbi_t3_pos_1_028_32    m/z 869.7424     0   1   17.5   18  RT (min)    lcorbi_t3_pos_1_031_40    m/z 869.7424     0   1   17   17.5  RT (min)    lcorbi_t3_pos_1_028_32    m/z 869.7424     0   1   17   17.5  RT (min)    lcorbi_t3_pos_1_034_34    m/z 869.7425     0   2   4E+5   15.5   16  RT (min)      lcorbi_t3_pos_4_051_34    m/z 869.7425     0   1   16.5   17  RT (min)    lcorbi_t3_pos_2_017_51    m/z 869.7425     0   1   16.5   17  RT (min)    lcorbi_t3_pos_1_028_32    m/z 869.7426     0   1   14.5   15  RT (min)      lcorbi_t3_pos_3_011_42    m/z 869.7426     0   1   14.5   15  RT (min)      lcorbi_t3_pos_2_013_33    m/z 870.6709     0   2   4E+5   11.5   12  RT (min)       lcorbi_t3_pos_1_003_45    m/z 870.7457     0   1   14   14.5  RT (min)      lcorbi_t3_pos_3_024_07    m/z 870.7542     0   1E+6   16   16.5  RT (min)      lcorbi_t3_pos_1_005_42    m/z 870.7543     0   1   17.5   18  RT (min)    lcorbi_t3_pos_1_034_34    m/z 870.7543     0   1   17.5   18  RT (min)    lcorbi_t3_pos_1_049_55    m/z 870.7544     0   1   18.5   19  RT (min)    lcorbi_t3_pos_1_023_02    m/z 870.7544     0   1   18   18.5  RT (min)    lcorbi_t3_pos_1_031_40    m/z 870.7545     0   2   4E+4   16   16.5  RT (min)      lcorbi_t3_pos_4_001_49    m/z 870.7545     0   1   16.5   17  RT (min)    lcorbi_t3_pos_2_022_38    m/z 870.7545     0   1   17   17.5  RT (min)    lcorbi_t3_pos_2_038_03    m/z 870.7545     0   1   14.5   15  RT (min)      lcorbi_t3_pos_3_024_07    m/z 870.7545     0   1   17   17.5  RT (min)    lcorbi_t3_pos_1_028_32    m/z 870.7545     0   1   17   17.5  RT (min)    lcorbi_t3_pos_1_034_34    m/z 870.7545     0   1   18   18.5  RT (min)    lcorbi_t3_pos_2_012_01    m/z 870.7546     0   1   16.5   17  RT (min)    lcorbi_t3_pos_2_046_07    m/z 870.7546     0   2   4   6   8E+5   15   15.5  RT (min)      lcorbi_t3_pos_3_034_26    m/z 870.7546     0   1   17   17.5  RT (min)    lcorbi_t3_pos_1_028_32    m/z 870.7547     0   2   4E+4   16   16.5  RT (min)      lcorbi_t3_pos_1_008_36    m/z 870.7547     0   1E+4   14.5   15  RT (min)      lcorbi_t3_pos_3_011_42    m/z 870.7547     0   1   17.5   18  RT (min)    lcorbi_t3_pos_2_009_50    m/z 870.7548     0   1   14.5   15  RT (min)      lcorbi_t3_pos_3_031_45    m/z 870.7548     0   1E+4   14.5   15  RT (min)      lcorbi_t3_pos_3_011_42    m/z 870.7548     0   1   18.5   19  RT (min)    lcorbi_t3_pos_1_023_02    m/z 870.7548     0   1   16.5   17  RT (min)    lcorbi_t3_pos_3_003_33    m/z 870.7548     0   1   2E+6   16   16.5  RT (min)      lcorbi_t3_pos_3_026_04    m/z 870.7548     0   1   17   17.5  RT (min)    lcorbi_t3_pos_1_034_34    m/z 870.7550     0   1   2E+6   16   16.5  RT (min)      lcorbi_t3_pos_3_040_41    m/z 871.7576     0   1   17.5   18  RT (min)    lcorbi_t3_pos_1_034_34    m/z 871.7577     0   1   18.5   19  RT (min)    lcorbi_t3_pos_1_023_02    m/z 871.7577     0   1   17   17.5  RT (min)    lcorbi_t3_pos_2_038_03    m/z 871.7579     0   1   16.5   17  RT (min)    lcorbi_t3_pos_2_046_07    m/z 871.7579     0   1   2   3   4E+4   16   16.5  RT (min)      lcorbi_t3_pos_4_001_49    m/z 871.7579     0   1   17.5   18  RT (min)    lcorbi_t3_pos_2_009_50    m/z 871.7579     0   1   14.5   15  RT (min)      lcorbi_t3_pos_2_032_39    m/z 871.7579     0   1   18   18.5  RT (min)    lcorbi_t3_pos_1_031_40    m/z 871.7580     0   1   16.5   17  RT (min)    lcorbi_t3_pos_2_046_07    m/z 871.7580     0   1E+6   16   16.5  RT (min)      lcorbi_t3_pos_1_005_42    m/z 871.7580     0   1   18   18.5  RT (min)    lcorbi_t3_pos_2_012_01    m/z 871.7581     0   1   18   18.5  RT (min)    lcorbi_t3_pos_2_056_56    m/z 871.7581     0   1   14.5   15  RT (min)      lcorbi_t3_pos_3_024_07    m/z 871.7581     0   1   16.5   17  RT (min)    lcorbi_t3_pos_1_047_41    m/z 871.7582     0   1   14.5   15  RT (min)      lcorbi_t3_pos_3_011_42    m/z 871.7582     0   1   17   17.5  RT (min)    lcorbi_t3_pos_1_028_32    m/z 871.7582     0   2   4   6E+4   16   16.5  RT (min)      lcorbi_t3_pos_1_008_36    m/z 871.7582     0   1   14.5   15  RT (min)      lcorbi_t3_pos_3_011_42    m/z 871.7582     0   1E+6   16   16.5  RT (min)      lcorbi_t3_pos_3_040_41    m/z 871.7582     0   1   16.5   17  RT (min)    lcorbi_t3_pos_3_003_33    m/z 871.7582     0   1   17   17.5  RT (min)    lcorbi_t3_pos_1_028_32    m/z 871.7582     0   1   16.5   17  RT (min)    lcorbi_t3_pos_2_022_38    m/z 871.7582     0   1   17.5   18  RT (min)    lcorbi_t3_pos_2_009_50    m/z 871.7583     0   1   17   17.5  RT (min)    lcorbi_t3_pos_1_034_34    m/z 871.7584     0   1   17   17.5  RT (min)    lcorbi_t3_pos_1_034_34    m/z 871.7584     0   1   14.5   15  RT (min)      lcorbi_t3_pos_3_011_42    m/z 871.7585     0   2   4E+5   15   15.5  RT (min)      lcorbi_t3_pos_3_034_26    m/z 872.6865     0   2   4   6   8E+5   12   12.5  RT (min)       lcorbi_t3_pos_4_092_60    m/z 872.7610     0   1   17   17.5  RT (min)    lcorbi_t3_pos_1_034_34    m/z 872.7612     0   2   4E+5   16   16.5  RT (min)      lcorbi_t3_pos_1_008_36    m/z 872.7612     0   1   18   18.5  RT (min)    lcorbi_t3_pos_4_033_53    m/z 872.7614     0   1   16.5   17  RT (min)    lcorbi_t3_pos_2_006_47    m/z 872.7618     0   1   14.5   15  RT (min)      lcorbi_t3_pos_3_024_07    m/z 872.7659     0   1   2E+6   15   15.5  RT (min)       lcorbi_t3_pos_3_034_26    m/z 872.7698     0   1   18.5   19  RT (min)    lcorbi_t3_pos_1_023_02    m/z 872.7699     0   1   16.5   17  RT (min)    lcorbi_t3_pos_2_022_38    m/z 872.7699     0   1E+4   14.5   15  RT (min)      lcorbi_t3_pos_3_031_45    m/z 872.7699     0   1   18   18.5  RT (min)    lcorbi_t3_pos_1_031_40    m/z 872.7700     0   1   17   17.5  RT (min)    lcorbi_t3_pos_3_049_55    m/z 872.7701     0   1   16.5   17  RT (min)    lcorbi_t3_pos_1_008_36    m/z 872.7701     0   1   17   17.5  RT (min)    lcorbi_t3_pos_2_038_03    m/z 872.7701     0   1   16.5   17  RT (min)    lcorbi_t3_pos_1_008_36    m/z 872.7702     0   1   17   17.5  RT (min)    lcorbi_t3_pos_1_028_32    m/z 872.7702     0   1   14.5   15  RT (min)      lcorbi_t3_pos_3_024_07    m/z 872.7702     0   1   17.5   18  RT (min)    lcorbi_t3_pos_1_034_34    m/z 872.7702     0   1   14.5   15  RT (min)      lcorbi_t3_pos_2_013_33    m/z 872.7702     0   1   18   18.5  RT (min)    lcorbi_t3_pos_2_012_01    m/z 872.7703     0   1   17.5   18  RT (min)    lcorbi_t3_pos_2_009_50    m/z 872.7703     0   1   17   17.5  RT (min)    lcorbi_t3_pos_3_003_33    m/z 872.7703     0   1   2E+6   16   16.5  RT (min)      lcorbi_t3_pos_1_005_42    m/z 872.7703     0   1   18   18.5  RT (min)    lcorbi_t3_pos_2_012_01    m/z 872.7703     0   1E+4   15   15.5  RT (min)      lcorbi_t3_pos_3_011_42    m/z 872.7703     0   1   17.5   18  RT (min)    lcorbi_t3_pos_1_034_34    m/z 872.7704     0   1   16.5   17  RT (min)    lcorbi_t3_pos_1_008_36    m/z 872.7704     0   1   2E+6   16   16.5  RT (min)      lcorbi_t3_pos_3_026_04    m/z 872.7704     0   1E+4   14.5   15  RT (min)      lcorbi_t3_pos_3_011_42    m/z 872.7705     0   1   14.5   15  RT (min)      lcorbi_t3_pos_4_012_37    m/z 873.7733     0   1   18.5   19  RT (min)    lcorbi_t3_pos_1_023_02    m/z 873.7736     0   1   17.5   18  RT (min)    lcorbi_t3_pos_1_034_34    m/z 873.7736     0   1   16.5   17  RT (min)    lcorbi_t3_pos_1_008_36    m/z 873.7736     0   1   17   17.5  RT (min)    lcorbi_t3_pos_1_030_21    m/z 873.7736     0   1   16.5   17  RT (min)    lcorbi_t3_pos_1_008_36    m/z 873.7736     0   1   18   18.5  RT (min)    lcorbi_t3_pos_1_031_40    m/z 873.7736     0   1   15   15.5  RT (min)      lcorbi_t3_pos_3_011_42    m/z 873.7736     0   1   16.5   17  RT (min)    lcorbi_t3_pos_2_046_07    m/z 873.7736     0   1   18   18.5  RT (min)    lcorbi_t3_pos_1_049_55    m/z 873.7737     0   1   16.5   17  RT (min)    lcorbi_t3_pos_2_022_38    m/z 873.7737     0   1   17.5   18  RT (min)    lcorbi_t3_pos_1_034_34    m/z 873.7737     0   1   17.5   18  RT (min)    lcorbi_t3_pos_1_034_34    m/z 873.7738     0   1   16.5   17  RT (min)    lcorbi_t3_pos_1_008_36    m/z 873.7738     0   1E+6   15   15.5  RT (min)      lcorbi_t3_pos_3_034_26    m/z 873.7738     0   1   18   18.5  RT (min)    lcorbi_t3_pos_1_031_40    m/z 873.7738     0   1   17.5   18  RT (min)    lcorbi_t3_pos_2_009_50    m/z 873.7738     0   1   17   17.5  RT (min)    lcorbi_t3_pos_3_003_33    m/z 873.7738     0   1   15   15.5  RT (min)      lcorbi_t3_pos_1_002_01    m/z 873.7739     0   1   17   17.5  RT (min)    lcorbi_t3_pos_2_038_03    m/z 873.7739     0   1   14.5   15  RT (min)      lcorbi_t3_pos_3_011_42    m/z 873.7739     0   1   18   18.5  RT (min)    lcorbi_t3_pos_2_012_01    m/z 873.7739     0   1E+6   16   16.5  RT (min)      lcorbi_t3_pos_1_005_42    m/z 873.7740     0   1E+6   16   16.5  RT (min)      lcorbi_t3_pos_3_026_04    m/z 873.7741     0   1   14.5   15  RT (min)      lcorbi_t3_pos_3_024_07    m/z 873.7742     0   1   19   19.5  RT (min)    lcorbi_t3_pos_1_023_02    m/z 873.7744     0   1   18   18.5  RT (min)    lcorbi_t3_pos_1_051_09    m/z 874.7019     0   2   4   6   8E+5   12   12.5  RT (min)      lcorbi_t3_pos_3_055_10    m/z 874.7021     0   2   4   6E+5   12   12.5  RT (min)      lcorbi_t3_pos_4_024_27    m/z 874.7021     0   1E+6   12   12.5  RT (min)       lcorbi_t3_pos_4_091_60    m/z 874.7764     0   1   15   15.5  RT (min)      lcorbi_t3_pos_1_002_01    m/z 874.7772     0   1   18   18.5  RT (min)    lcorbi_t3_pos_1_031_40    m/z 874.7775     0   1   16.5   17  RT (min)    lcorbi_t3_pos_2_046_07    m/z 874.7775     0   1   2   3E+6   15.5   16  RT (min)      lcorbi_t3_pos_3_034_26    m/z 874.7815     0   1   14.5   15  RT (min)      lcorbi_t3_pos_3_031_45    m/z 874.7817     0   1   14.5   15  RT (min)      lcorbi_t3_pos_3_031_45    m/z 874.7817     0   1   18   18.5  RT (min)    lcorbi_t3_pos_2_009_50    m/z 874.7856     0   1   18   18.5  RT (min)    lcorbi_t3_pos_1_031_40    m/z 874.7856     0   1   17   17.5  RT (min)    lcorbi_t3_pos_2_022_38    m/z 874.7856     0   1   2   3   4E+4   15   15.5  RT (min)      lcorbi_t3_pos_1_002_01    m/z 874.7857     0   1   19   19.5  RT (min)    lcorbi_t3_pos_1_023_02    m/z 874.7858     0   1   18.5   19  RT (min)    lcorbi_t3_pos_2_012_01    m/z 874.7858     0   1   15   15.5  RT (min)      lcorbi_t3_pos_1_002_01    m/z 874.7858     0   1   2E+6   16   16.5  RT (min)      lcorbi_t3_pos_4_051_34    m/z 874.7858     0   1   2   3E+6   16   16.5  RT (min)      lcorbi_t3_pos_4_019_47    m/z 874.7858     0   1   17.5   18  RT (min)    lcorbi_t3_pos_1_034_34    m/z 874.7859     0   1   17   17.5  RT (min)    lcorbi_t3_pos_3_003_33    m/z 874.7859     0   1   18   18.5  RT (min)    lcorbi_t3_pos_4_033_53    m/z 874.7859     0   1   16.5   17  RT (min)    lcorbi_t3_pos_1_008_36    m/z 874.7860     0   1   14.5   15  RT (min)      lcorbi_t3_pos_2_013_33    m/z 874.7860     0   1   17   17.5  RT (min)    lcorbi_t3_pos_1_047_41    m/z 874.7861     0   1   17   17.5  RT (min)    lcorbi_t3_pos_1_028_32    m/z 874.7863     0   1   15   15.5  RT (min)      lcorbi_t3_pos_3_011_42    m/z 874.7863     0   1   19   19.5  RT (min)    lcorbi_t3_pos_1_023_02    m/z 875.7098     0   1   16   16.5  RT (min)      lcorbi_t3_pos_1_008_36    m/z 875.7099     0   1   17   17.5  RT (min)    lcorbi_t3_pos_1_028_32    m/z 875.7100     0   1   16.5   17  RT (min)    lcorbi_t3_pos_2_022_38    m/z 875.7100     0   1E+4   14.5   15  RT (min)      lcorbi_t3_pos_3_024_07    m/z 875.7101     0   1   17   17.5  RT (min)    lcorbi_t3_pos_1_028_32    m/z 875.7102     0   1   2E+4   14.5   15  RT (min)      lcorbi_t3_pos_3_031_45    m/z 875.7102     0   1   18   18.5  RT (min)    lcorbi_t3_pos_2_012_01    m/z 875.7103     0   1   18   18.5  RT (min)    lcorbi_t3_pos_1_031_40    m/z 875.7104     0   1E+4   14.5   15  RT (min)      lcorbi_t3_pos_3_011_42    m/z 875.7105     0   1   17   17.5  RT (min)    lcorbi_t3_pos_1_034_34    m/z 875.7849     0   1   14.5   15  RT (min)      lcorbi_t3_pos_3_031_45    m/z 875.7890     0   1   19   19.5  RT (min)    lcorbi_t3_pos_1_023_02    m/z 875.7891     0   1   18.5   19  RT (min)    lcorbi_t3_pos_2_012_01    m/z 875.7892     0   1   15   15.5  RT (min)      lcorbi_t3_pos_1_002_01    m/z 875.7892     0   1   17.5   18  RT (min)    lcorbi_t3_pos_1_034_34    m/z 875.7892     0   1   18   18.5  RT (min)    lcorbi_t3_pos_1_031_40    m/z 875.7892     0   1E+6   16   16.5  RT (min)      lcorbi_t3_pos_1_005_42    m/z 875.7892     0   1   17.5   18  RT (min)    lcorbi_t3_pos_1_034_34    m/z 875.7893     0   1   16.5   17  RT (min)    lcorbi_t3_pos_1_008_36    m/z 875.7893     0   1   14.5   15  RT (min)      lcorbi_t3_pos_2_013_33    m/z 875.7894     0   1E+6   16   16.5  RT (min)      lcorbi_t3_pos_4_019_47    m/z 875.7894     0   1   16.5   17  RT (min)    lcorbi_t3_pos_2_046_07    m/z 875.7894     0   1   17   17.5  RT (min)    lcorbi_t3_pos_2_022_38    m/z 875.7894     0   1   16.5   17  RT (min)    lcorbi_t3_pos_1_008_36    m/z 875.7895     0   1   17   17.5  RT (min)    lcorbi_t3_pos_1_028_32    m/z 875.7896     0   1   15   15.5  RT (min)      lcorbi_t3_pos_3_011_42    m/z 875.7896     0   1   14.5   15  RT (min)      lcorbi_t3_pos_3_031_45    m/z 875.7897     0   1   18   18.5  RT (min)    lcorbi_t3_pos_2_009_50    m/z 875.7897     0   1   2   3   4E+5   15   15.5  RT (min)      lcorbi_t3_pos_4_016_15    m/z 875.7897     0   1   19   19.5  RT (min)    lcorbi_t3_pos_1_023_02    m/z 875.7898     0   1   18   18.5  RT (min)    lcorbi_t3_pos_1_056_56    m/z 876.6837     0   1   17   17.5  RT (min)    lcorbi_t3_pos_1_023_02    m/z 876.6840     0   1   16   16.5  RT (min)      lcorbi_t3_pos_1_034_34    m/z 876.6843     0   1   15.5   16  RT (min)      lcorbi_t3_pos_2_022_38    m/z 876.6843     0   1   13.5   14  RT (min)      lcorbi_t3_pos_2_032_39    m/z 876.6845     0   1   16.5   17  RT (min)    lcorbi_t3_pos_1_031_40    m/z 876.6846     0   1   16   16.5  RT (min)      lcorbi_t3_pos_1_034_34    m/z 876.6848     0   1E+4   14   14.5  RT (min)      lcorbi_t3_pos_1_002_01    m/z 876.6849     0   1   13.5   14  RT (min)      lcorbi_t3_pos_2_054_29    m/z 876.7176     0   1E+6   12   12.5  RT (min)      lcorbi_t3_pos_4_095_60    m/z 876.7178     0   1E+6   12.5   13  RT (min)       lcorbi_t3_pos_4_091_60    m/z 876.7179     0   1E+6   12.5   13  RT (min)       lcorbi_t3_pos_1_003_45    m/z 876.7179     0   1E+6   12.5   13  RT (min)       lcorbi_t3_pos_4_093_60    m/z 876.7181     0   1E+6   12.5   13  RT (min)       lcorbi_t3_pos_4_094_60    m/z 876.7922     0   1   19   19.5  RT (min)    lcorbi_t3_pos_1_023_02    m/z 876.7927     0   1   14.5   15  RT (min)      lcorbi_t3_pos_3_031_45    m/z 876.7927     0   1   17   17.5  RT (min)    lcorbi_t3_pos_2_022_38    m/z 876.7927     0   1   18   18.5  RT (min)    lcorbi_t3_pos_1_031_40    m/z 876.7928     0   1E+6   15.5   16  RT (min)      lcorbi_t3_pos_3_034_26    m/z 876.7929     0   1   17   17.5  RT (min)    lcorbi_t3_pos_3_013_19    m/z 876.7929     0   1   14.5   15  RT (min)      lcorbi_t3_pos_3_031_45    m/z 876.7929     0   1   15   15.5  RT (min)      lcorbi_t3_pos_2_013_33    m/z 876.7930     0   1   17.5   18  RT (min)    lcorbi_t3_pos_1_034_34    m/z 876.7930     0   1   18   18.5  RT (min)    lcorbi_t3_pos_1_051_09    m/z 876.7931     0   1   16.5   17  RT (min)    lcorbi_t3_pos_1_008_36    m/z 876.7932     0   1   14.5   15  RT (min)      lcorbi_t3_pos_2_013_33    m/z 876.7932     0   1   18   18.5  RT (min)    lcorbi_t3_pos_1_031_40    m/z 876.7935     0   1   14.5   15  RT (min)      lcorbi_t3_pos_2_032_39    m/z 876.7937     0   2   4   6E+5   15   15.5  RT (min)      lcorbi_t3_pos_4_016_15    m/z 876.7971     0   1   17   17.5  RT (min)    lcorbi_t3_pos_2_022_38    m/z 876.8002     0   1E+6   16   16.5  RT (min)      lcorbi_t3_pos_4_019_47    m/z 876.8004     0   1   18.5   19  RT (min)    lcorbi_t3_pos_2_012_01    m/z 876.8005     0   1   17.5   18  RT (min)    lcorbi_t3_pos_1_034_34    m/z 876.8007     0   1E+6   16   16.5  RT (min)      lcorbi_t3_pos_3_033_53    m/z 876.8007     0   1   18.5   19  RT (min)    lcorbi_t3_pos_2_012_01    m/z 876.8009     0   1   18   18.5  RT (min)    lcorbi_t3_pos_1_031_40    m/z 876.8011     0   1   16.5   17  RT (min)    lcorbi_t3_pos_1_008_36    m/z 876.8011     0   1   15   15.5  RT (min)      lcorbi_t3_pos_3_011_42    m/z 876.8013     0   1   17   17.5  RT (min)    lcorbi_t3_pos_1_028_32    m/z 876.8015     0   1   19   19.5  RT (min)    lcorbi_t3_pos_1_023_02    m/z 877.7252     0   1   16.5   17  RT (min)    lcorbi_t3_pos_1_008_36    m/z 877.7254     0   1   17   17.5  RT (min)    lcorbi_t3_pos_2_022_38    m/z 877.7254     0   1   17   17.5  RT (min)    lcorbi_t3_pos_1_028_32    m/z 877.7255     0   1   18   18.5  RT (min)    lcorbi_t3_pos_1_031_40    m/z 877.7257     0   1   16.5   17  RT (min)    lcorbi_t3_pos_1_008_36    m/z 877.7257     0   1   18   18.5  RT (min)    lcorbi_t3_pos_2_012_01    m/z 877.7257     0   1   2   3   4E+4   15   15.5  RT (min)      lcorbi_t3_pos_3_011_42    m/z 877.7258     0   1   2   3   4E+4   14.5   15  RT (min)       lcorbi_t3_pos_2_013_33    m/z 877.7258     0   1   18   18.5  RT (min)    lcorbi_t3_pos_1_031_40    m/z 877.7259     0   1   17.5   18  RT (min)    lcorbi_t3_pos_1_034_34    m/z 877.7259     0   1   18   18.5  RT (min)    lcorbi_t3_pos_2_012_01    m/z 877.7260     0   1   17.5   18  RT (min)    lcorbi_t3_pos_1_034_34    m/z 877.7260     0   1   2   3   4E+4   14.5   15  RT (min)      lcorbi_t3_pos_3_024_07    m/z 877.7264     0   1   2   3   4E+4   14.5   15  RT (min)      lcorbi_t3_pos_3_011_42    m/z 877.7963     0   1E+5   15   15.5  RT (min)      lcorbi_t3_pos_4_016_15    m/z 877.7965     0   1   16.5   17  RT (min)    lcorbi_t3_pos_1_005_42    m/z 877.8005     0   1   15   15.5  RT (min)      lcorbi_t3_pos_4_012_37    m/z 877.8006     0   1   14.5   15  RT (min)      lcorbi_t3_pos_3_031_45    m/z 877.8006     0   1   17   17.5  RT (min)    lcorbi_t3_pos_2_046_07    m/z 877.8008     0   1   17   17.5  RT (min)    lcorbi_t3_pos_3_003_33    m/z 877.8043     0   1   18.5   19  RT (min)    lcorbi_t3_pos_2_012_01    m/z 877.8044     0   1   17.5   18  RT (min)    lcorbi_t3_pos_1_034_34    m/z 877.8044     0   1   18   18.5  RT (min)    lcorbi_t3_pos_1_031_40    m/z 877.8045     0   1   17   17.5  RT (min)    lcorbi_t3_pos_2_022_38    m/z 877.8046     0   1   15   15.5  RT (min)      lcorbi_t3_pos_2_013_33    m/z 877.8047     0   1   18   18.5  RT (min)    lcorbi_t3_pos_4_033_53    m/z 877.8049     0   1   16.5   17  RT (min)    lcorbi_t3_pos_1_008_36    m/z 877.8049     0   2   4   6   8E+5   16   16.5  RT (min)      lcorbi_t3_pos_4_019_47    m/z 877.8051     0   1   15   15.5  RT (min)      lcorbi_t3_pos_3_011_42    m/z 877.8055     0   1   19   19.5  RT (min)    lcorbi_t3_pos_1_023_02    m/z 878.6953     0   1   13.5   14  RT (min)      lcorbi_t3_pos_2_032_39    m/z 878.6995     0   1   16   16.5  RT (min)      lcorbi_t3_pos_1_034_34    m/z 878.6996     0   1   17.5   18  RT (min)    lcorbi_t3_pos_1_023_02    m/z 878.6997     0   1   15.5   16  RT (min)      lcorbi_t3_pos_1_008_36    m/z 878.6998     0   1   16.5   17  RT (min)    lcorbi_t3_pos_1_031_40    m/z 878.6998     0   1   16.5   17  RT (min)    lcorbi_t3_pos_1_031_40    m/z 878.6999     0   1   2E+5   15   15.5  RT (min)      lcorbi_t3_pos_3_047_40    m/z 878.6999     0   1   15.5   16  RT (min)      lcorbi_t3_pos_2_022_38    m/z 878.6999     0   1E+4   15.5   16  RT (min)      lcorbi_t3_pos_3_003_33    m/z 878.7000     0   1   14   14.5  RT (min)      lcorbi_t3_pos_2_054_29    m/z 878.7001     0   1   13.5   14  RT (min)      lcorbi_t3_pos_2_013_33    m/z 878.7001     0   1   17.5   18  RT (min)    lcorbi_t3_pos_1_023_02    m/z 878.7002     0   1   16   16.5  RT (min)      lcorbi_t3_pos_1_034_34    m/z 878.7003     0   1   17   17.5  RT (min)    lcorbi_t3_pos_2_012_01    m/z 878.7003     0   1   16   16.5  RT (min)      lcorbi_t3_pos_1_028_32    m/z 878.7285     0   1   2   3E+4   14.5   15  RT (min)      lcorbi_t3_pos_3_024_07    m/z 878.7286     0   1   18   18.5  RT (min)    lcorbi_t3_pos_1_031_40    m/z 878.7292     0   1   2   3E+5   14.5   15  RT (min)      lcorbi_t3_pos_3_011_42    m/z 878.7294     0   1   18   18.5  RT (min)    lcorbi_t3_pos_2_012_01    m/z 878.8084     0   1   17   17.5  RT (min)    lcorbi_t3_pos_2_022_38    m/z 878.8085     0   1   16.5   17  RT (min)    lcorbi_t3_pos_1_008_36    m/z 879.7407     0   1   17.5   18  RT (min)    lcorbi_t3_pos_1_034_34    m/z 879.7407     0   1   18.5   19  RT (min)    lcorbi_t3_pos_2_012_01    m/z 879.7407     0   2   4   6E+4   15   15.5  RT (min)      lcorbi_t3_pos_3_011_42    m/z 879.7411     0   1   16.5   17  RT (min)    lcorbi_t3_pos_1_008_36    m/z 879.7412     0   2   4   6E+4   14.5   15  RT (min)      lcorbi_t3_pos_3_031_45    m/z 879.7413     0   1   17   17.5  RT (min)    lcorbi_t3_pos_3_003_33    m/z 879.7414     0   2   4   6E+4   14.5   15  RT (min)      lcorbi_t3_pos_3_024_07    m/z 879.7415     0   1   17.5   18  RT (min)    lcorbi_t3_pos_1_034_34    m/z 879.7416     0   1   17   17.5  RT (min)    lcorbi_t3_pos_1_028_32    m/z 879.7416     0   1   18.5   19  RT (min)    lcorbi_t3_pos_2_012_01    m/z 879.7419     0   1E+5   15   15.5  RT (min)       lcorbi_t3_pos_4_016_15    m/z 880.7386     0   1   14   14.5  RT (min)      lcorbi_t3_pos_3_024_07    m/z 880.7390     0   1   2   3E+4   14.5   15  RT (min)      lcorbi_t3_pos_2_013_33    m/z 880.7391     0   1   16.5   17  RT (min)    lcorbi_t3_pos_1_028_32    m/z 880.7396     0   1   16.5   17  RT (min)    lcorbi_t3_pos_3_003_33    m/z 880.7398     0   1   17.5   18  RT (min)    lcorbi_t3_pos_1_031_40    m/z 880.7445     0   1   17   17.5  RT (min)    lcorbi_t3_pos_1_034_34    m/z 880.7446     0   1   17   17.5  RT (min)    lcorbi_t3_pos_1_034_34    m/z 880.7447     0   1   18   18.5  RT (min)    lcorbi_t3_pos_2_012_01    m/z 880.7448     0   1   2   3   4E+4   14.5   15  RT (min)      lcorbi_t3_pos_3_011_42    m/z 880.7451     0   1   2   3   4E+5   16   16.5  RT (min)      lcorbi_t3_pos_1_008_36    m/z 881.7423     0   1   16.5   17  RT (min)    lcorbi_t3_pos_2_022_38    m/z 882.7542     0   1   18   18.5  RT (min)    lcorbi_t3_pos_2_012_01    m/z 882.7543     0   1   14.5   15  RT (min)      lcorbi_t3_pos_3_024_07    m/z 882.7545     0   1   16.5   17  RT (min)    lcorbi_t3_pos_3_003_33    m/z 882.7545     0   1   2   3   4E+4   14.5   15  RT (min)      lcorbi_t3_pos_3_011_42    m/z 882.7545     0   1   2E+5   16   16.5  RT (min)      lcorbi_t3_pos_1_008_36    m/z 882.7546     0   1   18   18.5  RT (min)    lcorbi_t3_pos_1_031_40    m/z 882.7546     0   1   2   3   4E+4   14.5   15  RT (min)      lcorbi_t3_pos_2_013_33    m/z 882.7547     0   1   16.5   17  RT (min)    lcorbi_t3_pos_2_022_38    m/z 882.7549     0   1   17   17.5  RT (min)    lcorbi_t3_pos_1_028_32    m/z 882.7599     0   1   17   17.5  RT (min)    lcorbi_t3_pos_1_034_34    m/z 883.7578     0   1   17   17.5  RT (min)    lcorbi_t3_pos_1_028_32    m/z 883.7578     0   1   17   17.5  RT (min)    lcorbi_t3_pos_1_034_34    m/z 883.7579     0   1   14.5   15  RT (min)      lcorbi_t3_pos_3_024_07    m/z 883.7579     0   2   4   6E+4   16   16.5  RT (min)      lcorbi_t3_pos_1_008_36    m/z 883.7579     0   2   4   6E+5   15   15.5  RT (min)      lcorbi_t3_pos_2_023_26    m/z 883.7581     0   1   17   17.5  RT (min)    lcorbi_t3_pos_1_034_34    m/z 883.7581     0   1   18   18.5  RT (min)    lcorbi_t3_pos_2_012_01    m/z 883.7582     0   1   16.5   17  RT (min)    lcorbi_t3_pos_2_022_38    m/z 883.7582     0   1   2E+4   14.5   15  RT (min)      lcorbi_t3_pos_2_013_33    m/z 883.7585     0   1   18   18.5  RT (min)    lcorbi_t3_pos_1_031_40    m/z 884.7700     0   1   18   18.5  RT (min)    lcorbi_t3_pos_1_031_40    m/z 884.7700     0   1   14.5   15  RT (min)      lcorbi_t3_pos_3_011_42    m/z 884.7701     0   1   16.5   17  RT (min)    lcorbi_t3_pos_2_022_38    m/z 884.7702     0   1   14.5   15  RT (min)      lcorbi_t3_pos_3_024_07    m/z 884.7703     0   1   16   16.5  RT (min)      lcorbi_t3_pos_1_008_36    m/z 884.7703     0   1   16.5   17  RT (min)    lcorbi_t3_pos_2_022_38    m/z 884.7703     0   1   14.5   15  RT (min)      lcorbi_t3_pos_2_013_33    m/z 884.7704     0   1   17   17.5  RT (min)    lcorbi_t3_pos_1_028_32    m/z 884.7704     0   1   16.5   17  RT (min)    lcorbi_t3_pos_1_008_36    m/z 884.7706     0   1   18   18.5  RT (min)    lcorbi_t3_pos_2_012_01    m/z 884.7706     0   1   17.5   18  RT (min)    lcorbi_t3_pos_1_034_34    m/z 884.7706     0   1   2E+4   15   15.5  RT (min)      lcorbi_t3_pos_3_011_42    m/z 885.3675     0   1   15   15.5  RT (min)      lcorbi_t3_pos_1_023_02    m/z 885.3676     0   1   14   14.5  RT (min)      lcorbi_t3_pos_1_034_34    m/z 885.3677     0   1   14.5   15  RT (min)      lcorbi_t3_pos_1_031_40    m/z 885.3677     0   1   14.5   15  RT (min)      lcorbi_t3_pos_1_051_09    m/z 885.3678     0   1   14   14.5  RT (min)      lcorbi_t3_pos_1_034_34    m/z 885.3678     0   1   14.5   15  RT (min)      lcorbi_t3_pos_1_031_40    m/z 885.3678     0   2   4   6E+5   13   13.5  RT (min)      lcorbi_t3_pos_4_019_47    m/z 885.3680     0   1   12   12.5  RT (min)      lcorbi_t3_pos_2_013_33    m/z 885.3680     0   1   2   3   4E+5   12.5   13  RT (min)      lcorbi_t3_pos_1_002_01    m/z 885.3680     0   1   14   14.5  RT (min)      lcorbi_t3_pos_3_003_33    m/z 885.3680     0   1   13.5   14  RT (min)      lcorbi_t3_pos_2_022_38    m/z 885.3680     0   1   14   14.5  RT (min)      lcorbi_t3_pos_1_028_32    m/z 885.3682     0   1   2   3E+5   13.5   14  RT (min)      lcorbi_t3_pos_1_008_36    m/z 885.3682     0   1   15   15.5  RT (min)      lcorbi_t3_pos_1_023_02    m/z 885.3683     0   1   12   12.5  RT (min)      lcorbi_t3_pos_2_013_33    m/z 885.3805     0   1   2E+5   9.5   10  RT (min)      lcorbi_t3_pos_1_031_40    m/z 885.3805     0   2   4   6E+5   9   9.5  RT (min)      lcorbi_t3_pos_1_034_34    m/z 885.6625     0   2   4E+5   11   11.5  RT (min)       lcorbi_t3_pos_4_091_60    m/z 885.7734     0   1   16.5   17  RT (min)    lcorbi_t3_pos_2_022_38    m/z 885.7735     0   1   18   18.5  RT (min)    lcorbi_t3_pos_1_031_40    m/z 885.7736     0   1   16.5   17  RT (min)    lcorbi_t3_pos_2_022_38    m/z 885.7736     0   1   14.5   15  RT (min)      lcorbi_t3_pos_3_011_42    m/z 885.7736     0   1   14.5   15  RT (min)      lcorbi_t3_pos_3_024_07    m/z 885.7736     0   1   18   18.5  RT (min)    lcorbi_t3_pos_2_012_01    m/z 885.7737     0   1   17.5   18  RT (min)    lcorbi_t3_pos_1_034_34    m/z 885.7737     0   1   17   17.5  RT (min)    lcorbi_t3_pos_1_028_32    m/z 885.7738     0   1   14.5   15  RT (min)      lcorbi_t3_pos_2_013_33    m/z 885.7740     0   1   16   16.5  RT (min)      lcorbi_t3_pos_1_008_36    m/z 885.7740     0   1   18   18.5  RT (min)    lcorbi_t3_pos_1_031_40    m/z 886.3708     0   1   14   14.5  RT (min)      lcorbi_t3_pos_1_034_34    m/z 886.3711     0   1   12   12.5  RT (min)      lcorbi_t3_pos_2_013_33    m/z 886.3712     0   1   13.5   14  RT (min)      lcorbi_t3_pos_2_022_38    m/z 886.3715     0   1   12   12.5  RT (min)      lcorbi_t3_pos_2_013_33    m/z 886.6655     0   1E+6   11   11.5  RT (min)       lcorbi_t3_pos_4_091_60    m/z 886.6655     0   1E+6   11   11.5  RT (min)      lcorbi_t3_pos_2_037_27    m/z 886.6658     0   1E+6   11   11.5  RT (min)      lcorbi_t3_pos_2_037_27    m/z 886.6664     0   1E+6   11   11.5  RT (min)      lcorbi_t3_pos_2_037_27    m/z 886.6664     0   1E+6   11   11.5  RT (min)      lcorbi_t3_pos_2_037_27    m/z 886.7820     0   1   14.5   15  RT (min)      lcorbi_t3_pos_2_013_33    m/z 886.7856     0   1   17   17.5  RT (min)    lcorbi_t3_pos_2_022_38    m/z 886.7856     0   1   18.5   19  RT (min)    lcorbi_t3_pos_2_012_01    m/z 886.7857     0   1   16.5   17  RT (min)    lcorbi_t3_pos_1_008_36    m/z 886.7857     0   1   17   17.5  RT (min)    lcorbi_t3_pos_1_028_32    m/z 886.7858     0   1   14.5   15  RT (min)      lcorbi_t3_pos_2_013_33    m/z 886.7858     0   1   14.5   15  RT (min)      lcorbi_t3_pos_3_024_07    m/z 886.7858     0   1   18   18.5  RT (min)    lcorbi_t3_pos_1_031_40    m/z 886.7858     0   1   18   18.5  RT (min)    lcorbi_t3_pos_1_031_40    m/z 886.7858     0   1   17   17.5  RT (min)    lcorbi_t3_pos_1_028_32    m/z 886.7859     0   1   15   15.5  RT (min)      lcorbi_t3_pos_3_011_42    m/z 886.7860     0   1   17.5   18  RT (min)    lcorbi_t3_pos_1_034_34    m/z 887.6783     0   2   4   6   8E+5   11   11.5  RT (min)       lcorbi_t3_pos_4_090_60    m/z 887.7889     0   1   16.5   17  RT (min)    lcorbi_t3_pos_1_008_36    m/z 887.7894     0   1   14.5   15  RT (min)      lcorbi_t3_pos_2_013_33    m/z 887.7894     0   1   17   17.5  RT (min)    lcorbi_t3_pos_1_028_32    m/z 887.7895     0   1   17.5   18  RT (min)    lcorbi_t3_pos_1_034_34    m/z 887.7895     0   1E+4   15   15.5  RT (min)      lcorbi_t3_pos_3_011_42    m/z 887.7895     0   1   18.5   19  RT (min)    lcorbi_t3_pos_2_012_01    m/z 887.7896     0   1   17   17.5  RT (min)    lcorbi_t3_pos_1_028_32    m/z 887.7897     0   1   17.5   18  RT (min)    lcorbi_t3_pos_1_034_34    m/z 887.7899     0   1   18   18.5  RT (min)    lcorbi_t3_pos_1_031_40    m/z 887.7900     0   1   17.5   18  RT (min)    lcorbi_t3_pos_1_034_34    m/z 888.6814     0   1   2E+6   11   11.5  RT (min)       lcorbi_t3_pos_4_090_60    m/z 888.7646     0   1   17   17.5  RT (min)    lcorbi_t3_pos_1_031_40    m/z 888.7646     0   1   17.5   18  RT (min)    lcorbi_t3_pos_2_012_01    m/z 888.7648     0   1   16   16.5  RT (min)      lcorbi_t3_pos_2_022_38    m/z 888.7648     0   1   16   16.5  RT (min)      lcorbi_t3_pos_1_028_32    m/z 888.7650     0   1   14   14.5  RT (min)      lcorbi_t3_pos_2_013_33    m/z 888.7650     0   1   17.5   18  RT (min)    lcorbi_t3_pos_2_012_01    m/z 888.7650     0   1   16.5   17  RT (min)    lcorbi_t3_pos_1_034_34    m/z 888.7651     0   2   4   6E+4   15.5   16  RT (min)      lcorbi_t3_pos_1_008_36    m/z 888.7652     0   1   14   14.5  RT (min)      lcorbi_t3_pos_3_024_07    m/z 888.7653     0   1   16.5   17  RT (min)    lcorbi_t3_pos_1_034_34    m/z 888.7655     0   1E+5   15.5   16  RT (min)      lcorbi_t3_pos_1_008_36    m/z 888.8009     0   1   18   18.5  RT (min)    lcorbi_t3_pos_1_031_40    m/z 888.8010     0   1   17   17.5  RT (min)    lcorbi_t3_pos_2_022_38    m/z 888.8011     0   1   14.5   15  RT (min)      lcorbi_t3_pos_3_024_07    m/z 888.8011     0   1   18.5   19  RT (min)    lcorbi_t3_pos_2_012_01    m/z 888.8012     0   1   17   17.5  RT (min)    lcorbi_t3_pos_1_028_32    m/z 888.8012     0   1   17.5   18  RT (min)    lcorbi_t3_pos_1_034_34    m/z 888.8013     0   1   17.5   18  RT (min)    lcorbi_t3_pos_1_034_34    m/z 888.8015     0   1   14.5   15  RT (min)      lcorbi_t3_pos_2_013_33    m/z 888.8016     0   1   16.5   17  RT (min)    lcorbi_t3_pos_1_008_36    m/z 888.8019     0   1E+4   15   15.5  RT (min)      lcorbi_t3_pos_3_011_42    m/z 889.6935     0   2   4   6E+5   11.5   12  RT (min)       lcorbi_t3_pos_4_090_60    m/z 889.7685     0   1   17   17.5  RT (min)    lcorbi_t3_pos_1_031_40    m/z 889.7687     0   1   2   3   4E+4   15.5   16  RT (min)      lcorbi_t3_pos_1_008_36    m/z 889.8048     0   1   17   17.5  RT (min)    lcorbi_t3_pos_1_028_32    m/z 889.8050     0   1   17.5   18  RT (min)    lcorbi_t3_pos_1_034_34    m/z 889.8055     0   1   15   15.5  RT (min)      lcorbi_t3_pos_3_011_42    m/z 890.6972     0   1E+6   11.5   12  RT (min)       lcorbi_t3_pos_4_090_60    m/z 890.7230     0   1   17.5   18   18.5  RT (min)    lcorbi_t3_pos_2_012_01    m/z 890.7230     0   1   18   18.5  RT (min)    lcorbi_t3_pos_2_012_01    m/z 890.7231     0   1   16.5   17  RT (min)    lcorbi_t3_pos_2_022_38    m/z 890.7231     0   1E+5   16   16.5  RT (min)      lcorbi_t3_pos_1_008_36    m/z 890.7233     0   1   16.5   17  RT (min)    lcorbi_t3_pos_1_028_32    m/z 890.7233     0   1   17.5   18  RT (min)    lcorbi_t3_pos_1_031_40    m/z 890.7233     0   1   17.5   18  RT (min)    lcorbi_t3_pos_1_031_40    m/z 890.7234     0   1   16.5   17  RT (min)    lcorbi_t3_pos_3_003_33    m/z 890.7234     0   1   14.5   15  RT (min)      lcorbi_t3_pos_3_011_42    m/z 890.7234     0   1   14   14.5  RT (min)      lcorbi_t3_pos_3_024_07    m/z 890.7235     0   1   14.5   15  RT (min)      lcorbi_t3_pos_2_013_33    m/z 890.7236     0   1   17   17.5  RT (min)    lcorbi_t3_pos_1_034_34    m/z 890.7799     0   1   17.5   18  RT (min)    lcorbi_t3_pos_2_012_01    m/z 890.7803     0   2   4   6   8E+4   15.5   16  RT (min)      lcorbi_t3_pos_1_008_36    m/z 890.7803     0   1   17   17.5  RT (min)    lcorbi_t3_pos_1_031_40    m/z 890.7803     0   1   14   14.5  RT (min)      lcorbi_t3_pos_3_024_07    m/z 890.7804     0   1   16   16.5  RT (min)      lcorbi_t3_pos_1_028_32    m/z 890.7804     0   1   16.5   17  RT (min)    lcorbi_t3_pos_1_034_34    m/z 890.7806     0   1   16   16.5  RT (min)      lcorbi_t3_pos_3_003_33    m/z 890.7806     0   1   14   14.5  RT (min)      lcorbi_t3_pos_3_011_42    m/z 890.7808     0   1   14   14.5  RT (min)      lcorbi_t3_pos_3_011_42    m/z 890.7811     0   1   16   16.5  RT (min)      lcorbi_t3_pos_3_003_33    m/z 891.7261     0   1   17.5   18  RT (min)    lcorbi_t3_pos_1_031_40    m/z 891.7264     0   1   18   18.5  RT (min)    lcorbi_t3_pos_2_012_01    m/z 891.7266     0   1   16.5   17  RT (min)    lcorbi_t3_pos_2_022_38    m/z 891.7266     0   1   16.5   17  RT (min)    lcorbi_t3_pos_1_028_32    m/z 891.7267     0   1   17.5   18  RT (min)    lcorbi_t3_pos_2_012_01    m/z 891.7267     0   1   17.5   18  RT (min)    lcorbi_t3_pos_1_031_40    m/z 891.7267     0   1   2   3   4E+4   16   16.5  RT (min)      lcorbi_t3_pos_1_008_36    m/z 891.7268     0   1   17   17.5  RT (min)    lcorbi_t3_pos_1_034_34    m/z 891.7269     0   1   14.5   15  RT (min)      lcorbi_t3_pos_2_013_33    m/z 891.7269     0   1   17   17.5  RT (min)    lcorbi_t3_pos_1_034_34    m/z 891.7271     0   1   14.5   15  RT (min)      lcorbi_t3_pos_3_011_42    m/z 892.7384     0   1   17   17.5  RT (min)    lcorbi_t3_pos_1_034_34    m/z 892.7384     0   1   16.5   17  RT (min)    lcorbi_t3_pos_1_047_41    m/z 892.7385     0   2   4   6E+5   16   16.5  RT (min)      lcorbi_t3_pos_1_005_42    m/z 892.7386     0   1   14.5   15  RT (min)      lcorbi_t3_pos_3_011_42    m/z 892.7387     0   1   16.5   17  RT (min)    lcorbi_t3_pos_2_022_38    m/z 892.7387     0   1   14   14.5  RT (min)      lcorbi_t3_pos_3_024_07    m/z 892.7388     0   1   16.5   17  RT (min)    lcorbi_t3_pos_1_028_32    m/z 892.7388     0   1   16   16.5  RT (min)      lcorbi_t3_pos_2_046_07    m/z 892.7388     0   1   18   18.5  RT (min)    lcorbi_t3_pos_2_012_01    m/z 892.7389     0   1   16.5   17  RT (min)    lcorbi_t3_pos_1_047_41    m/z 892.7390     0   1   17.5   18  RT (min)    lcorbi_t3_pos_1_031_40    m/z 892.7390     0   2   4   6   8E+4   16   16.5  RT (min)      lcorbi_t3_pos_1_008_36    m/z 892.7390     0   1   17   17.5  RT (min)    lcorbi_t3_pos_1_034_34    m/z 892.7391     0   2   4   6   8E+4   16   16.5  RT (min)      lcorbi_t3_pos_1_008_36    m/z 892.7391     0   1   17   17.5  RT (min)    lcorbi_t3_pos_2_038_03    m/z 892.7391     0   1   17.5   18  RT (min)    lcorbi_t3_pos_2_009_50    m/z 892.7392     0   1   17   17.5  RT (min)    lcorbi_t3_pos_1_034_34    m/z 892.7394     0   1   18   18.5  RT (min)    lcorbi_t3_pos_2_012_01    m/z 893.7420     0   1   17   17.5  RT (min)    lcorbi_t3_pos_1_034_34    m/z 893.7421     0   2   4   6E+4   16   16.5  RT (min)      lcorbi_t3_pos_1_008_36    m/z 893.7421     0   1   18   18.5  RT (min)    lcorbi_t3_pos_2_012_01    m/z 893.7421     0   1   16.5   17  RT (min)    lcorbi_t3_pos_2_022_38    m/z 893.7422     0   1   16.5   17  RT (min)    lcorbi_t3_pos_1_028_32    m/z 893.7422     0   1   17.5   18  RT (min)    lcorbi_t3_pos_1_025_52    m/z 893.7422     0   2   4   6E+5   15.5   16   16.5  RT (min)      lcorbi_t3_pos_1_005_42    m/z 893.7422     0   1   17   17.5  RT (min)    lcorbi_t3_pos_1_034_34    m/z 893.7422     0   1   16.5   17  RT (min)    lcorbi_t3_pos_2_006_47    m/z 893.7423     0   1E+6   15.5   16  RT (min)       lcorbi_t3_pos_3_026_04    m/z 893.7423     0   1   14   14.5  RT (min)      lcorbi_t3_pos_3_024_07    m/z 893.7424     0   1   17.5   18  RT (min)    lcorbi_t3_pos_1_031_40    m/z 893.7424     0   1   14.5   15  RT (min)      lcorbi_t3_pos_3_011_42    m/z 893.7424     0   1   17.5   18  RT (min)    lcorbi_t3_pos_2_009_50    m/z 893.7426     0   1   17   17.5  RT (min)    lcorbi_t3_pos_1_034_34    m/z 893.7427     0   2   4   6E+4   16   16.5  RT (min)      lcorbi_t3_pos_1_008_36    m/z 893.7428     0   1   18   18.5  RT (min)    lcorbi_t3_pos_2_012_01    m/z 893.7428     0   1   17   17.5  RT (min)    lcorbi_t3_pos_1_034_34    m/z 893.7428     0   2   4   6   8E+4   16   16.5  RT (min)      lcorbi_t3_pos_4_025_52    m/z 894.7455     0   1   16.5   17  RT (min)    lcorbi_t3_pos_3_003_33    m/z 894.7458     0   1   17   17.5  RT (min)    lcorbi_t3_pos_1_034_34    m/z 894.7460     0   1   14   14.5  RT (min)      lcorbi_t3_pos_3_024_07    m/z 894.7498     0   1   16.5   17  RT (min)    lcorbi_t3_pos_2_046_07    m/z 894.7499     0   1   17   17.5  RT (min)    lcorbi_t3_pos_2_038_03    m/z 894.7540     0   1   17   17.5  RT (min)    lcorbi_t3_pos_1_034_34    m/z 894.7541     0   1   14.5   15  RT (min)      lcorbi_t3_pos_3_024_07    m/z 894.7541     0   1   2E+6   16   16.5  RT (min)      lcorbi_t3_pos_3_040_41    m/z 894.7542     0   1   17.5   18  RT (min)    lcorbi_t3_pos_1_031_40    m/z 894.7542     0   1   16.5   17  RT (min)    lcorbi_t3_pos_2_017_51    m/z 894.7543     0   1   17.5   18  RT (min)    lcorbi_t3_pos_2_049_55    m/z 894.7543     0   1E+6   16   16.5  RT (min)      lcorbi_t3_pos_1_005_42    m/z 894.7543     0   1   16.5   17  RT (min)    lcorbi_t3_pos_2_022_38    m/z 894.7544     0   1   17   17.5  RT (min)    lcorbi_t3_pos_1_028_32    m/z 894.7544     0   1   18   18.5  RT (min)    lcorbi_t3_pos_2_012_01    m/z 894.7545     0   1   14.5   15  RT (min)      lcorbi_t3_pos_3_024_07    m/z 894.7545     0   1   2E+4   14.5   15  RT (min)      lcorbi_t3_pos_2_013_33    m/z 894.7545     0   1   17.5   18  RT (min)    lcorbi_t3_pos_2_009_50    m/z 894.7546     0   1   14.5   15  RT (min)      lcorbi_t3_pos_3_011_42    m/z 894.7546     0   1E+5   16   16.5  RT (min)      lcorbi_t3_pos_1_008_36    m/z 894.7547     0   1   2   3E+4   16   16.5  RT (min)      lcorbi_t3_pos_4_010_43    m/z 894.7547     0   1   2E+6   16   16.5  RT (min)      lcorbi_t3_pos_3_026_04    m/z 894.7548     0   1   18   18.5  RT (min)    lcorbi_t3_pos_2_056_56    m/z 895.7043     0   1   14   14.5  RT (min)      lcorbi_t3_pos_1_030_21    m/z 895.7043     0   1   15   15.5  RT (min)      lcorbi_t3_pos_1_025_52    m/z 895.7045     0   1   2   3E+4   14.5   15  RT (min)      lcorbi_t3_pos_1_041_54    m/z 895.7045     0   1   2E+4   15   15.5  RT (min)      lcorbi_t3_pos_1_025_52    m/z 895.7047     0   1   14.5   15  RT (min)      lcorbi_t3_pos_1_033_53    m/z 895.7049     0   1   14   14.5  RT (min)      lcorbi_t3_pos_1_047_41    m/z 895.7050     0   2   4   6E+5   13.5   14  RT (min)      lcorbi_t3_pos_1_005_42    m/z 895.7576     0   1   14.5   15  RT (min)      lcorbi_t3_pos_3_024_07    m/z 895.7580     0   1   17.5   18  RT (min)    lcorbi_t3_pos_1_031_40    m/z 895.7580     0   1   17   17.5  RT (min)    lcorbi_t3_pos_1_034_34    m/z 895.7580     0   1   18   18.5  RT (min)    lcorbi_t3_pos_2_012_01    m/z 895.7580     0   1   17   17.5  RT (min)    lcorbi_t3_pos_1_034_34    m/z 895.7581     0   1   14.5   15  RT (min)      lcorbi_t3_pos_2_013_33    m/z 895.7581     0   1   16.5   17  RT (min)    lcorbi_t3_pos_3_003_33    m/z 895.7581     0   1E+6   16   16.5  RT (min)       lcorbi_t3_pos_3_026_04    m/z 895.7581     0   1   17.5   18  RT (min)    lcorbi_t3_pos_2_009_50    m/z 895.7581     0   1   17   17.5  RT (min)    lcorbi_t3_pos_1_028_32    m/z 895.7582     0   1   16.5   17  RT (min)    lcorbi_t3_pos_2_017_51    m/z 895.7582     0   1E+6   16   16.5  RT (min)      lcorbi_t3_pos_3_040_41    m/z 895.7582     0   2   4E+4   16   16.5  RT (min)      lcorbi_t3_pos_1_008_36    m/z 895.7584     0   1   17   17.5  RT (min)    lcorbi_t3_pos_1_028_32    m/z 896.7614     0   1   16.5   17  RT (min)    lcorbi_t3_pos_2_046_07    m/z 896.7615     0   1   14   14.5   15  RT (min)      lcorbi_t3_pos_3_031_45    m/z 896.7621     0   1   16.5   17  RT (min)    lcorbi_t3_pos_2_022_38    m/z 896.7658     0   1   18   18.5  RT (min)    lcorbi_t3_pos_1_056_56    m/z 896.7660     0   1E+5   16   16.5  RT (min)      lcorbi_t3_pos_4_001_49    m/z 896.7697     0   1   18   18.5  RT (min)    lcorbi_t3_pos_1_031_40    m/z 896.7697     0   1   18.5   19  RT (min)    lcorbi_t3_pos_1_023_02    m/z 896.7697     0   1   17   17.5  RT (min)    lcorbi_t3_pos_1_034_34    m/z 896.7698     0   1   18.5   19  RT (min)    lcorbi_t3_pos_1_023_02    m/z 896.7699     0   1   14.5   15  RT (min)      lcorbi_t3_pos_3_011_42    m/z 896.7699     0   1   14.5   15  RT (min)      lcorbi_t3_pos_3_011_42    m/z 896.7700     0   1   18   18.5  RT (min)    lcorbi_t3_pos_1_031_40    m/z 896.7700     0   1   17   17.5  RT (min)    lcorbi_t3_pos_1_028_32    m/z 896.7700     0   1   17   17.5  RT (min)    lcorbi_t3_pos_3_003_33    m/z 896.7701     0   1   17   17.5  RT (min)    lcorbi_t3_pos_2_025_52    m/z 896.7701     0   1E+7   15   15.5  RT (min)      lcorbi_t3_pos_2_023_26    m/z 896.7702     0   1E+4   14.5   15  RT (min)      lcorbi_t3_pos_2_013_33    m/z 896.7702     0   1   2   3E+6   16   16.5  RT (min)      lcorbi_t3_pos_3_040_41    m/z 896.7702     0   1   17.5   18  RT (min)    lcorbi_t3_pos_2_049_55    m/z 896.7702     0   1   17.5   18  RT (min)    lcorbi_t3_pos_1_034_34    m/z 896.7703     0   1   17   17.5  RT (min)    lcorbi_t3_pos_2_038_03    m/z 896.7703     0   1   16.5   17  RT (min)    lcorbi_t3_pos_1_008_36    m/z 896.7703     0   1   16.5   17  RT (min)    lcorbi_t3_pos_2_006_47    m/z 896.7703     0   1   14.5   15  RT (min)      lcorbi_t3_pos_3_024_07    m/z 896.7703     0   1   18   18.5  RT (min)    lcorbi_t3_pos_2_056_56    m/z 896.7703     0   1   18   18.5  RT (min)    lcorbi_t3_pos_2_012_01    m/z 896.7705     0   1   18.5   19  RT (min)    lcorbi_t3_pos_1_023_02    m/z 897.6937     0   1   17.5   18  RT (min)    lcorbi_t3_pos_1_031_40    m/z 897.6941     0   1   14.5   15  RT (min)      lcorbi_t3_pos_3_011_42    m/z 897.6942     0   1   14   14.5  RT (min)      lcorbi_t3_pos_3_024_07    m/z 897.6942     0   1   16   16.5  RT (min)      lcorbi_t3_pos_1_008_36    m/z 897.6943     0   1   16.5   17  RT (min)    lcorbi_t3_pos_1_028_32    m/z 897.6943     0   1   16.5   17  RT (min)    lcorbi_t3_pos_2_022_38    m/z 897.6945     0   1   17   17.5  RT (min)    lcorbi_t3_pos_1_034_34    m/z 897.7732     0   1   18   18.5  RT (min)    lcorbi_t3_pos_2_056_56    m/z 897.7734     0   1   14.5   15  RT (min)      lcorbi_t3_pos_3_011_42    m/z 897.7735     0   1   2E+4   15   15.5  RT (min)      lcorbi_t3_pos_3_011_42    m/z 897.7736     0   1   17   17.5  RT (min)    lcorbi_t3_pos_1_034_34    m/z 897.7736     0   1   18   18.5  RT (min)    lcorbi_t3_pos_2_012_01    m/z 897.7736     0   1   17.5   18  RT (min)    lcorbi_t3_pos_1_034_34    m/z 897.7736     0   1   14.5   15  RT (min)      lcorbi_t3_pos_3_024_07    m/z 897.7736     0   1   17   17.5  RT (min)    lcorbi_t3_pos_1_028_32    m/z 897.7736     0   1   18   18.5  RT (min)    lcorbi_t3_pos_1_031_40    m/z 897.7736     0   2   4E+5   15   15.5  RT (min)      lcorbi_t3_pos_3_034_26    m/z 897.7737     0   1   17   17.5  RT (min)    lcorbi_t3_pos_2_025_52    m/z 897.7737     0   1   16.5   17  RT (min)    lcorbi_t3_pos_2_046_07    m/z 897.7737     0   1   16.5   17  RT (min)    lcorbi_t3_pos_2_006_47    m/z 897.7737     0   1E+5   16   16.5  RT (min)      lcorbi_t3_pos_1_008_36    m/z 897.7737     0   1   16.5   17  RT (min)    lcorbi_t3_pos_2_022_38    m/z 897.7738     0   1   17   17.5  RT (min)    lcorbi_t3_pos_1_028_32    m/z 897.7738     0   1   14.5   15  RT (min)      lcorbi_t3_pos_3_011_42    m/z 897.7738     0   1   2E+6   16   16.5  RT (min)      lcorbi_t3_pos_3_040_41    m/z 897.7739     0   1   17   17.5  RT (min)    lcorbi_t3_pos_1_030_21    m/z 897.7739     0   1E+4   16   16.5  RT (min)      lcorbi_t3_pos_3_025_52    m/z 897.7739     0   1   17.5   18  RT (min)    lcorbi_t3_pos_2_009_50    m/z 897.7742     0   1   18.5   19  RT (min)    lcorbi_t3_pos_1_023_02    m/z 898.6978     0   1   16   16.5  RT (min)      lcorbi_t3_pos_1_008_36    m/z 898.7767     0   1   14.5   15  RT (min)      lcorbi_t3_pos_3_031_45    m/z 898.7769     0   1   18   18.5  RT (min)    lcorbi_t3_pos_1_056_56    m/z 898.7772     0   1   17   17.5  RT (min)    lcorbi_t3_pos_1_047_41    m/z 898.7775     0   1   17   17.5  RT (min)    lcorbi_t3_pos_1_034_34    m/z 898.7775     0   1   16.5   17  RT (min)    lcorbi_t3_pos_2_017_51    m/z 898.7811     0   2   4   6   8E+6   15   15.5  RT (min)      lcorbi_t3_pos_4_034_42    m/z 898.7813     0   1   14.5   15  RT (min)      lcorbi_t3_pos_3_031_45    m/z 898.7817     0   1   18   18.5  RT (min)    lcorbi_t3_pos_1_056_56    m/z 898.7846     0   1   19   19.5  RT (min)    lcorbi_t3_pos_1_023_02    m/z 898.7854     0   1   14.5   15  RT (min)      lcorbi_t3_pos_3_031_45    m/z 898.7854     0   1   18.5   19  RT (min)    lcorbi_t3_pos_2_012_01    m/z 898.7854     0   1   2   3E+6   16   16.5  RT (min)      lcorbi_t3_pos_3_040_41    m/z 898.7855     0   1   15   15.5  RT (min)      lcorbi_t3_pos_1_002_01    m/z 898.7855     0   1   18   18.5  RT (min)    lcorbi_t3_pos_1_031_40    m/z 898.7855     0   1   18   18.5  RT (min)    lcorbi_t3_pos_1_031_40    m/z 898.7855     0   1   19   19.5  RT (min)    lcorbi_t3_pos_1_023_02    m/z 898.7856     0   1   17.5   18  RT (min)    lcorbi_t3_pos_2_009_50    m/z 898.7856     0   1   17   17.5  RT (min)    lcorbi_t3_pos_2_022_38    m/z 898.7856     0   1E+4   15   15.5  RT (min)      lcorbi_t3_pos_3_011_42    m/z 898.7856     0   1   19   19.5  RT (min)    lcorbi_t3_pos_1_023_02    m/z 898.7856     0   1   17.5   18  RT (min)    lcorbi_t3_pos_1_034_34    m/z 898.7857     0   1   2   3E+6   16   16.5  RT (min)      lcorbi_t3_pos_1_005_42    m/z 898.7857     0   1   14.5   15  RT (min)      lcorbi_t3_pos_2_013_33    m/z 898.7858     0   1   17   17.5  RT (min)    lcorbi_t3_pos_2_022_38    m/z 898.7858     0   1   17   17.5  RT (min)    lcorbi_t3_pos_1_028_32    m/z 898.7858     0   1   16.5   17  RT (min)    lcorbi_t3_pos_1_008_36    m/z 898.7858     0   1   17.5   18  RT (min)    lcorbi_t3_pos_1_034_34    m/z 898.7858     0   1   16.5   17  RT (min)    lcorbi_t3_pos_1_008_36    m/z 898.7859     0   1   17   17.5  RT (min)    lcorbi_t3_pos_3_013_19    m/z 898.7859     0   1   19   19.5  RT (min)    lcorbi_t3_pos_1_023_02    m/z 898.7860     0   1   2   3E+6   16   16.5  RT (min)      lcorbi_t3_pos_3_026_04    m/z 898.7860     0   1   14.5   15  RT (min)      lcorbi_t3_pos_3_031_45    m/z 899.7097     0   1   17   17.5  RT (min)    lcorbi_t3_pos_1_034_34    m/z 899.7097     0   1   14.5   15  RT (min)      lcorbi_t3_pos_3_024_07    m/z 899.7097     0   1   17   17.5  RT (min)    lcorbi_t3_pos_1_034_34    m/z 899.7098     0   1   16.5   17  RT (min)    lcorbi_t3_pos_2_022_38    m/z 899.7098     0   1   17   17.5  RT (min)    lcorbi_t3_pos_1_028_32    m/z 899.7098     0   1   16   16.5  RT (min)      lcorbi_t3_pos_1_008_36    m/z 899.7099     0   1   14.5   15  RT (min)      lcorbi_t3_pos_3_011_42    m/z 899.7100     0   1   14.5   15  RT (min)      lcorbi_t3_pos_2_013_33    m/z 899.7101     0   1   16   16.5  RT (min)      lcorbi_t3_pos_1_008_36    m/z 899.7101     0   1   16.5   17  RT (min)    lcorbi_t3_pos_3_003_33    m/z 899.7102     0   1   18   18.5  RT (min)    lcorbi_t3_pos_2_012_01    m/z 899.7106     0   1   17.5   18  RT (min)    lcorbi_t3_pos_1_031_40    m/z 899.7885     0   1   19   19.5  RT (min)    lcorbi_t3_pos_1_023_02    m/z 899.7889     0   1   19   19.5  RT (min)    lcorbi_t3_pos_1_023_02    m/z 899.7890     0   1E+6   16   16.5  RT (min)      lcorbi_t3_pos_1_005_42    m/z 899.7891     0   1   16.5   17  RT (min)    lcorbi_t3_pos_1_008_36    m/z 899.7891     0   1   17   17.5  RT (min)    lcorbi_t3_pos_2_025_52    m/z 899.7891     0   1   17.5   18  RT (min)    lcorbi_t3_pos_1_034_34    m/z 899.7891     0   1   14.5   15  RT (min)      lcorbi_t3_pos_2_013_33    m/z 899.7892     0   1   16.5   17  RT (min)    lcorbi_t3_pos_2_046_07    m/z 899.7892     0   1   14.5   15  RT (min)      lcorbi_t3_pos_3_024_07    m/z 899.7892     0   1   18   18.5  RT (min)    lcorbi_t3_pos_1_031_40    m/z 899.7892     0   1   14.5   15  RT (min)      lcorbi_t3_pos_2_032_39    m/z 899.7892     0   1   18   18.5  RT (min)    lcorbi_t3_pos_1_031_40    m/z 899.7892     0   1   17   17.5  RT (min)    lcorbi_t3_pos_1_028_32    m/z 899.7893     0   1   17.5   18  RT (min)    lcorbi_t3_pos_2_009_50    m/z 899.7893     0   1   16.5   17  RT (min)    lcorbi_t3_pos_1_008_36    m/z 899.7893     0   1   17   17.5  RT (min)    lcorbi_t3_pos_2_022_38    m/z 899.7894     0   1   18   18.5  RT (min)    lcorbi_t3_pos_1_025_52    m/z 899.7894     0   1   18.5   19  RT (min)    lcorbi_t3_pos_2_012_01    m/z 899.7894     0   1   17   17.5  RT (min)    lcorbi_t3_pos_3_003_33    m/z 899.7894     0   1   2E+6   16   16.5  RT (min)      lcorbi_t3_pos_3_040_41    m/z 899.7894     0   1   17.5   18  RT (min)    lcorbi_t3_pos_1_034_34    m/z 899.7894     0   1   17   17.5  RT (min)    lcorbi_t3_pos_2_025_52    m/z 899.7895     0   1   15   15.5  RT (min)      lcorbi_t3_pos_3_011_42    m/z 899.7895     0   2   4   6E+5   15   15.5  RT (min)      lcorbi_t3_pos_3_034_26    m/z 899.7895     0   1   2E+6   16   16.5  RT (min)      lcorbi_t3_pos_3_026_04    m/z 899.7897     0   1   17.5   18  RT (min)    lcorbi_t3_pos_1_034_34    m/z 899.7898     0   1   19   19.5  RT (min)    lcorbi_t3_pos_1_023_02    m/z 900.7133     0   1   14.5   15  RT (min)      lcorbi_t3_pos_3_024_07    m/z 900.7133     0   1   16   16.5  RT (min)      lcorbi_t3_pos_1_008_36    m/z 900.7133     0   1   17   17.5  RT (min)    lcorbi_t3_pos_1_028_32    m/z 900.7134     0   1   17   17.5  RT (min)    lcorbi_t3_pos_1_034_34    m/z 900.7134     0   1   16.5   17  RT (min)    lcorbi_t3_pos_2_022_38    m/z 900.7134     0   2   4E+5   15   15.5  RT (min)      lcorbi_t3_pos_2_023_26    m/z 900.7135     0   1   17.5   18  RT (min)    lcorbi_t3_pos_1_031_40    m/z 900.7925     0   1   16.5   17  RT (min)    lcorbi_t3_pos_2_017_51    m/z 900.7925     0   1   14.5   15  RT (min)      lcorbi_t3_pos_3_031_45    m/z 900.7927     0   1   17.5   18  RT (min)    lcorbi_t3_pos_1_034_34    m/z 900.7929     0   1   17.5   18  RT (min)    lcorbi_t3_pos_1_034_34    m/z 900.7929     0   1   17   17.5  RT (min)    lcorbi_t3_pos_1_047_41    m/z 900.7935     0   1   18   18.5  RT (min)    lcorbi_t3_pos_1_049_55    m/z 900.7937     0   1   18   18.5  RT (min)    lcorbi_t3_pos_2_009_50    m/z 900.7970     0   1   17   17.5  RT (min)    lcorbi_t3_pos_2_022_38    m/z 900.7983     0   2   4   6E+6   15.5   16  RT (min)       lcorbi_t3_pos_2_023_26    m/z 900.8006     0   1   19   19.5  RT (min)    lcorbi_t3_pos_1_023_02    m/z 900.8008     0   1   17.5   18  RT (min)    lcorbi_t3_pos_1_034_34    m/z 900.8010     0   1   19   19.5  RT (min)    lcorbi_t3_pos_1_023_02    m/z 900.8010     0   1   16.5   17  RT (min)    lcorbi_t3_pos_2_017_51    m/z 900.8010     0   1   18   18.5  RT (min)    lcorbi_t3_pos_1_031_40    m/z 900.8011     0   1   16.5   17  RT (min)    lcorbi_t3_pos_1_008_36    m/z 900.8011     0   1   18   18.5  RT (min)    lcorbi_t3_pos_4_033_53    m/z 900.8012     0   1   14.5   15  RT (min)      lcorbi_t3_pos_3_024_07    m/z 900.8012     0   1   17   17.5  RT (min)    lcorbi_t3_pos_1_028_32    m/z 900.8012     0   1E+6   16   16.5  RT (min)      lcorbi_t3_pos_1_005_42    m/z 900.8013     0   1   14.5   15  RT (min)      lcorbi_t3_pos_2_013_33    m/z 900.8013     0   1E+4   15   15.5  RT (min)      lcorbi_t3_pos_3_011_42    m/z 900.8013     0   1   2E+6   16   16.5  RT (min)      lcorbi_t3_pos_3_026_04    m/z 900.8013     0   1   18.5   19  RT (min)    lcorbi_t3_pos_2_012_01    m/z 900.8014     0   1   17   17.5  RT (min)    lcorbi_t3_pos_1_028_32    m/z 900.8018     0   1   15   15.5  RT (min)      lcorbi_t3_pos_1_002_01    m/z 900.8018     0   1   18   18.5  RT (min)    lcorbi_t3_pos_1_051_09    m/z 900.8019     0   1E+6   15.5   16  RT (min)      lcorbi_t3_pos_3_034_26    m/z 901.7164     0   1   14.5   15  RT (min)      lcorbi_t3_pos_3_031_45    m/z 901.7251     0   1E+4   14.5   15  RT (min)      lcorbi_t3_pos_3_024_07    m/z 901.7252     0   1   17   17.5  RT (min)    lcorbi_t3_pos_1_034_34    m/z 901.7253     0   1   18   18.5  RT (min)    lcorbi_t3_pos_2_012_01    m/z 901.7254     0   1   16.5   17  RT (min)    lcorbi_t3_pos_2_022_38    m/z 901.7255     0   1   18   18.5  RT (min)    lcorbi_t3_pos_1_031_40    m/z 901.7255     0   2   4E+5   16   16.5  RT (min)      lcorbi_t3_pos_3_040_41    m/z 901.7255     0   1   17.5   18  RT (min)    lcorbi_t3_pos_1_034_34    m/z 901.7256     0   1   17   17.5  RT (min)    lcorbi_t3_pos_1_028_32    m/z 901.7256     0   1   14.5   15  RT (min)      lcorbi_t3_pos_3_011_42    m/z 901.7256     0   1   16   16.5  RT (min)      lcorbi_t3_pos_1_008_36    m/z 901.7258     0   1   14.5   15  RT (min)      lcorbi_t3_pos_2_013_33    m/z 901.7260     0   1   18   18.5  RT (min)    lcorbi_t3_pos_1_031_40    m/z 901.8043     0   1   19   19.5  RT (min)    lcorbi_t3_pos_1_023_02    m/z 901.8045     0   1   17   17.5  RT (min)    lcorbi_t3_pos_3_013_19    m/z 901.8046     0   1   14.5   15  RT (min)      lcorbi_t3_pos_2_032_39    m/z 901.8046     0   1   17.5   18  RT (min)    lcorbi_t3_pos_1_034_34    m/z 901.8046     0   1   14.5   15  RT (min)      lcorbi_t3_pos_3_031_45    m/z 901.8047     0   1   17   17.5  RT (min)    lcorbi_t3_pos_2_022_38    m/z 901.8047     0   1   16.5   17  RT (min)    lcorbi_t3_pos_2_017_51    m/z 901.8047     0   1   16.5   17  RT (min)    lcorbi_t3_pos_1_008_36    m/z 901.8047     0   1   18   18.5  RT (min)    lcorbi_t3_pos_1_031_40    m/z 901.8048     0   1   18   18.5  RT (min)    lcorbi_t3_pos_2_009_50    m/z 901.8048     0   1E+6   16   16.5  RT (min)      lcorbi_t3_pos_1_005_42    m/z 901.8048     0   1   17   17.5  RT (min)    lcorbi_t3_pos_1_028_32    m/z 901.8048     0   2   4   6   8E+4   15   15.5  RT (min)      lcorbi_t3_pos_4_016_15    m/z 901.8048     0   1   18.5   19  RT (min)    lcorbi_t3_pos_2_012_01    m/z 901.8049     0   1   17   17.5  RT (min)    lcorbi_t3_pos_2_022_38    m/z 901.8049     0   2   4   6E+5   15.5   16  RT (min)      lcorbi_t3_pos_3_034_26    m/z 901.8049     0   1   14.5   15  RT (min)      lcorbi_t3_pos_2_013_33    m/z 901.8049     0   1E+6   16   16.5  RT (min)      lcorbi_t3_pos_3_026_04    m/z 901.8055     0   1   18   18.5  RT (min)    lcorbi_t3_pos_4_033_53    m/z 902.3943     0   1   14.5   15  RT (min)      lcorbi_t3_pos_1_031_40    m/z 902.3944     0   1   14   14.5  RT (min)      lcorbi_t3_pos_1_034_34    m/z 902.3944     0   1   13.5   14  RT (min)      lcorbi_t3_pos_2_017_51    m/z 902.3944     0   1   12   12.5  RT (min)      lcorbi_t3_pos_2_013_33    m/z 902.3945     0   1   2   3   4E+5   13.5   14  RT (min)      lcorbi_t3_pos_1_008_36    m/z 902.3945     0   1   15   15.5  RT (min)      lcorbi_t3_pos_1_023_02    m/z 902.3945     0   1   14   14.5  RT (min)      lcorbi_t3_pos_1_030_21    m/z 902.3945     0   1   14   14.5  RT (min)      lcorbi_t3_pos_1_034_34    m/z 902.3945     0   1   12   12.5  RT (min)      lcorbi_t3_pos_3_024_07    m/z 902.3945     0   1   15   15.5  RT (min)      lcorbi_t3_pos_1_023_02    m/z 902.3945     0   1   14.5   15  RT (min)      lcorbi_t3_pos_1_031_40    m/z 902.3947     0   1   14   14.5  RT (min)      lcorbi_t3_pos_1_028_32    m/z 902.3947     0   1E+6   12.5   13  RT (min)      lcorbi_t3_pos_1_002_01    m/z 902.7284     0   1E+4   14.5   15  RT (min)      lcorbi_t3_pos_3_011_42    m/z 902.7285     0   1   17   17.5  RT (min)    lcorbi_t3_pos_1_034_34    m/z 902.7289     0   1E+4   14.5   15  RT (min)      lcorbi_t3_pos_3_024_07    m/z 902.7289     0   1   17   17.5  RT (min)    lcorbi_t3_pos_3_003_33    m/z 902.7289     0   1   17   17.5  RT (min)    lcorbi_t3_pos_1_028_32    m/z 902.7291     0   1   18   18.5  RT (min)    lcorbi_t3_pos_2_012_01    m/z 902.7291     0   1   18   18.5  RT (min)    lcorbi_t3_pos_1_031_40    m/z 902.7292     0   1   18   18.5  RT (min)    lcorbi_t3_pos_1_031_40    m/z 902.7295     0   1   17.5   18  RT (min)    lcorbi_t3_pos_1_034_34    m/z 902.8080     0   1   14.5   15  RT (min)      lcorbi_t3_pos_3_031_45    m/z 902.8081     0   1   18   18.5  RT (min)    lcorbi_t3_pos_1_031_40    m/z 902.8082     0   1   16.5   17  RT (min)    lcorbi_t3_pos_1_008_36    m/z 902.8085     0   2   4E+5   16   16.5  RT (min)      lcorbi_t3_pos_3_017_51    m/z 902.8085     0   1E+4   15   15.5  RT (min)      lcorbi_t3_pos_3_011_42    m/z 902.8085     0   1   17   17.5  RT (min)    lcorbi_t3_pos_2_022_38    m/z 902.8086     0   1   17   17.5  RT (min)    lcorbi_t3_pos_2_022_38    m/z 902.8086     0   1   18   18.5  RT (min)    lcorbi_t3_pos_1_031_40    m/z 902.8087     0   1   17.5   18  RT (min)    lcorbi_t3_pos_1_034_34    m/z 902.8087     0   1   16.5   17  RT (min)    lcorbi_t3_pos_1_008_36    m/z 902.8087     0   1   14.5   15  RT (min)      lcorbi_t3_pos_2_013_33    m/z 902.8088     0   1   17   17.5  RT (min)    lcorbi_t3_pos_2_022_38    m/z 902.8089     0   1   15   15.5  RT (min)      lcorbi_t3_pos_2_013_33    m/z 902.8101     0   1   14.5   15  RT (min)      lcorbi_t3_pos_3_031_45    m/z 902.8130     0   1   2E+5   15   15.5  RT (min)      lcorbi_t3_pos_4_016_15    m/z 902.8158     0   1   17   17.5  RT (min)    lcorbi_t3_pos_1_028_32    m/z 902.8162     0   1E+6   16   16.5  RT (min)       lcorbi_t3_pos_4_051_34    m/z 902.8170     0   1E+4   15   15.5  RT (min)      lcorbi_t3_pos_3_011_42    m/z 903.3973     0   1   14   14.5  RT (min)      lcorbi_t3_pos_1_034_34    m/z 903.3973     0   2   4   6E+5   13.5   14  RT (min)      lcorbi_t3_pos_3_047_40    m/z 903.3976     0   2   4   6E+5   13   13.5  RT (min)      lcorbi_t3_pos_4_019_47    m/z 903.3976     0   2   4E+5   13.5   14  RT (min)      lcorbi_t3_pos_1_008_36    m/z 903.3977     0   1   14.5   15  RT (min)      lcorbi_t3_pos_1_031_40    m/z 903.3977     0   1   14.5   15  RT (min)      lcorbi_t3_pos_1_031_40    m/z 903.3978     0   1   14   14.5  RT (min)      lcorbi_t3_pos_1_034_34    m/z 903.3979     0   1   15   15.5  RT (min)      lcorbi_t3_pos_1_023_02    m/z 903.3979     0   2   4   6   8E+3   14   14.5  RT (min)      lcorbi_t3_pos_3_013_19    m/z 903.3979     0   2   4   6E+5   12.5   13  RT (min)      lcorbi_t3_pos_4_016_15    m/z 903.3979     0   1   12   12.5  RT (min)      lcorbi_t3_pos_2_013_33    m/z 903.3979     0   1   14.5   15  RT (min)      lcorbi_t3_pos_2_012_01    m/z 903.3980     0   1   14   14.5  RT (min)      lcorbi_t3_pos_1_028_32    m/z 903.3981     0   1   12   12.5  RT (min)      lcorbi_t3_pos_2_013_33    m/z 903.3983     0   2   4   6E+5   12.5   13  RT (min)      lcorbi_t3_pos_1_002_01    m/z 903.3983     0   1   15   15.5  RT (min)      lcorbi_t3_pos_1_023_02    m/z 903.7410     0   1   17   17.5  RT (min)    lcorbi_t3_pos_3_003_33    m/z 903.7410     0   1E+4   14.5   15  RT (min)      lcorbi_t3_pos_3_031_45    m/z 903.7411     0   1   17   17.5  RT (min)    lcorbi_t3_pos_1_028_32    m/z 903.7411     0   1   17   17.5  RT (min)    lcorbi_t3_pos_1_028_32    m/z 903.7411     0   1   18.5   19  RT (min)    lcorbi_t3_pos_2_012_01    m/z 903.7411     0   1   18.5   19  RT (min)    lcorbi_t3_pos_2_012_01    m/z 903.7412     0   1   16.5   17  RT (min)    lcorbi_t3_pos_1_008_36    m/z 903.7412     0   1   16.5   17  RT (min)    lcorbi_t3_pos_1_008_36    m/z 903.7412     0   1   17.5   18  RT (min)    lcorbi_t3_pos_1_034_34    m/z 903.7412     0   1   18   18.5  RT (min)    lcorbi_t3_pos_1_031_40    m/z 903.7413     0   1   17   17.5  RT (min)    lcorbi_t3_pos_2_022_38    m/z 903.7414     0   1   17.5   18  RT (min)    lcorbi_t3_pos_1_034_34    m/z 903.7414     0   1   2E+4   14.5   15  RT (min)       lcorbi_t3_pos_2_013_33    m/z 903.7414     0   1   17.5   18  RT (min)    lcorbi_t3_pos_1_034_34    m/z 903.7415     0   1E+4   15   15.5  RT (min)      lcorbi_t3_pos_3_011_42    m/z 903.7416     0   1   18   18.5  RT (min)    lcorbi_t3_pos_1_031_40    m/z 903.8123     0   1   17   17.5  RT (min)    lcorbi_t3_pos_4_048_24    m/z 903.8123     0   1   17   17.5  RT (min)    lcorbi_t3_pos_2_022_38    m/z 903.8158     0   1   18   18.5  RT (min)    lcorbi_t3_pos_1_031_40    m/z 903.8169     0   1   17.5   18  RT (min)    lcorbi_t3_pos_2_041_54    m/z 903.8195     0   1   17.5   18  RT (min)    lcorbi_t3_pos_1_034_34    m/z 903.8199     0   1   16.5   17  RT (min)    lcorbi_t3_pos_1_008_36    m/z 903.8199     0   1E+4   15   15.5  RT (min)      lcorbi_t3_pos_2_013_33    m/z 903.8202     0   1   14.5   15  RT (min)      lcorbi_t3_pos_3_031_45    m/z 903.8206     0   2   4   6E+5   16   16.5  RT (min)      lcorbi_t3_pos_4_051_34    m/z 903.8207     0   1   15   15.5  RT (min)      lcorbi_t3_pos_3_011_42    m/z 904.5905     0   1   16   16.5  RT (min)      lcorbi_t3_pos_1_023_02    m/z 904.5917     0   1   2E+4   13   13.5  RT (min)      lcorbi_t3_pos_1_002_01    m/z 904.7441     0   1   18.5   19  RT (min)    lcorbi_t3_pos_2_012_01    m/z 904.7443     0   1   18   18.5  RT (min)    lcorbi_t3_pos_1_031_40    m/z 904.7444     0   1E+4   14.5   15  RT (min)      lcorbi_t3_pos_2_013_33    m/z 904.7446     0   1   17   17.5  RT (min)    lcorbi_t3_pos_2_022_38    m/z 904.7447     0   1   17   17.5  RT (min)    lcorbi_t3_pos_1_028_32    m/z 904.7448     0   1   17   17.5  RT (min)    lcorbi_t3_pos_1_028_32    m/z 904.7448     0   1   16.5   17  RT (min)    lcorbi_t3_pos_1_008_36    m/z 904.7448     0   1   17.5   18  RT (min)    lcorbi_t3_pos_1_034_34    m/z 904.7449     0   1   17.5   18  RT (min)    lcorbi_t3_pos_1_034_34    m/z 904.7449     0   1   18   18.5  RT (min)    lcorbi_t3_pos_1_031_40    m/z 904.8236     0   1   18   18.5  RT (min)    lcorbi_t3_pos_1_031_40    m/z 904.8240     0   1   14.5   15  RT (min)      lcorbi_t3_pos_3_031_45    m/z 904.8242     0   1E+4   15   15.5  RT (min)      lcorbi_t3_pos_3_011_42    m/z 904.8243     0   1   16.5   17  RT (min)    lcorbi_t3_pos_1_008_36    m/z 904.8322     0   1E+6   16   16.5  RT (min)       lcorbi_t3_pos_4_084_59    m/z 905.7565     0   1   18.5   19  RT (min)    lcorbi_t3_pos_2_012_01    m/z 905.7566     0   1   16.5   17  RT (min)    lcorbi_t3_pos_1_008_36    m/z 905.7566     0   1   17   17.5  RT (min)    lcorbi_t3_pos_1_028_32    m/z 905.7568     0   1   17.5   18  RT (min)    lcorbi_t3_pos_1_034_34    m/z 905.7568     0   1   17   17.5  RT (min)    lcorbi_t3_pos_2_022_38    m/z 905.7568     0   1   2E+4   14.5   15  RT (min)      lcorbi_t3_pos_3_031_45    m/z 905.7568     0   1   17.5   18  RT (min)    lcorbi_t3_pos_1_034_34    m/z 905.7570     0   1   18   18.5  RT (min)    lcorbi_t3_pos_1_031_40    m/z 905.7571     0   1   16.5   17  RT (min)    lcorbi_t3_pos_1_008_36    m/z 905.7578     0   1E+4   15   15.5  RT (min)       lcorbi_t3_pos_3_011_42    m/z 905.7623     0   1E+4   15   15.5  RT (min)      lcorbi_t3_pos_3_011_42    m/z 905.7634     0   1E+4   14.5   15  RT (min)      lcorbi_t3_pos_3_024_07    m/z 905.8363     0   2   4E+5   16   16.5  RT (min)      lcorbi_t3_pos_4_084_59    m/z 905.8367     0   2   4   6E+5   16   16.5  RT (min)       lcorbi_t3_pos_4_082_59    m/z 906.2592     0   1   17   17.5  RT (min)    lcorbi_t3_pos_4_033_53    m/z 906.2593     0   1   16.5   17  RT (min)    lcorbi_t3_pos_2_009_50    m/z 906.2593     0   1E+5   15.5   16  RT (min)      lcorbi_t3_pos_1_008_36    m/z 906.2593     0   1   17.5   18  RT (min)    lcorbi_t3_pos_2_012_01    m/z 906.2594     0   1   17.5   18  RT (min)    lcorbi_t3_pos_2_012_01    m/z 906.2594     0   1   16.5   17  RT (min)    lcorbi_t3_pos_1_034_34    m/z 906.2594     0   1   17.5   18  RT (min)    lcorbi_t3_pos_2_012_01    m/z 906.2594     0   1   16   16.5  RT (min)      lcorbi_t3_pos_2_022_38    m/z 906.2595     0   1   2   3   4E+4   15.5   16  RT (min)      lcorbi_t3_pos_1_008_36    m/z 906.2595     0   1   14   14.5  RT (min)      lcorbi_t3_pos_2_013_33    m/z 906.2595     0   1   17   17.5  RT (min)    lcorbi_t3_pos_1_031_40    m/z 906.2595     0   1   16.5   17  RT (min)    lcorbi_t3_pos_2_025_52    m/z 906.2595     0   2   4   6   8E+5   15   15.5  RT (min)      lcorbi_t3_pos_4_051_34    m/z 906.2597     0   1   17.5   18  RT (min)    lcorbi_t3_pos_2_012_01    m/z 906.2597     0   2   4   6E+5   15   15.5  RT (min)      lcorbi_t3_pos_3_040_41    m/z 906.2598     0   1   14   14.5  RT (min)      lcorbi_t3_pos_3_031_45    m/z 906.2600     0   1   2   3   4E+6   14.5   15  RT (min)       lcorbi_t3_pos_4_105_61    m/z 906.6922     0   2   4   6E+5   11   11.5  RT (min)       lcorbi_t3_pos_4_094_60    m/z 906.7604     0   1E+4   14.5   15  RT (min)      lcorbi_t3_pos_3_031_45    m/z 906.7605     0   1   17   17.5  RT (min)    lcorbi_t3_pos_1_028_32    m/z 907.2592     0   1   16.5   17  RT (min)    lcorbi_t3_pos_2_025_52    m/z 907.2598     0   1   14   14.5  RT (min)      lcorbi_t3_pos_3_024_07    m/z 907.2599     0   1   2E+4   15.5   16  RT (min)      lcorbi_t3_pos_1_008_36    m/z 907.2599     0   1   17.5   18  RT (min)    lcorbi_t3_pos_2_012_01    m/z 907.2600     0   1   16   16.5  RT (min)      lcorbi_t3_pos_4_041_54    m/z 907.2600     0   1   17.5   18  RT (min)    lcorbi_t3_pos_2_012_01    m/z 907.2600     0   2   4   6E+5   15   15.5  RT (min)      lcorbi_t3_pos_3_017_51    m/z 907.2601     0   2   4   6E+5   15.5   16  RT (min)      lcorbi_t3_pos_1_005_42    m/z 907.2601     0   2   4E+4   15.5   16  RT (min)      lcorbi_t3_pos_1_008_36    m/z 907.2601     0   1   16.5   17  RT (min)    lcorbi_t3_pos_2_009_50    m/z 907.2602     0   1   17   17.5  RT (min)    lcorbi_t3_pos_1_031_40    m/z 907.2602     0   1   16.5   17  RT (min)    lcorbi_t3_pos_1_034_34    m/z 907.2603     0   1   14   14.5  RT (min)      lcorbi_t3_pos_3_011_42    m/z 907.2604     0   1   16   16.5  RT (min)      lcorbi_t3_pos_2_022_38    m/z 907.2607     0   1   15.5   16  RT (min)      lcorbi_t3_pos_2_017_51    m/z 907.2609     0   1   2   3   4E+6   14.5   15  RT (min)      lcorbi_t3_pos_4_105_61    m/z 908.2580     0   1   14   14.5  RT (min)      lcorbi_t3_pos_3_011_42    m/z 908.2581     0   1   16   16.5  RT (min)      lcorbi_t3_pos_4_049_55    m/z 908.2581     0   2   4E+5   15.5   16  RT (min)      lcorbi_t3_pos_1_005_42    m/z 908.2582     0   2   4E+4   15.5   16  RT (min)      lcorbi_t3_pos_1_008_36    m/z 908.2584     0   1   16   16.5  RT (min)      lcorbi_t3_pos_2_022_38    m/z 908.2584     0   1   17   17.5  RT (min)    lcorbi_t3_pos_1_031_40    m/z 908.2584     0   1   14   14.5  RT (min)      lcorbi_t3_pos_3_024_07    m/z 908.2584     0   1   15.5   16  RT (min)      lcorbi_t3_pos_1_008_36    m/z 908.2585     0   1   2   3E+6   14.5   15  RT (min)       lcorbi_t3_pos_4_105_61    m/z 908.2585     0   2   4   6E+5   15   15.5  RT (min)      lcorbi_t3_pos_3_033_53    m/z 908.2586     0   1   16.5   17  RT (min)    lcorbi_t3_pos_2_009_50    m/z 908.2586     0   1   17.5   18  RT (min)    lcorbi_t3_pos_2_012_01    m/z 908.2586     0   1   16.5   17  RT (min)    lcorbi_t3_pos_1_034_34    m/z 908.2587     0   1   15.5   16  RT (min)      lcorbi_t3_pos_2_017_51    m/z 908.2589     0   1   17.5   18  RT (min)    lcorbi_t3_pos_2_012_01    m/z 909.2581     0   1   16.5   17  RT (min)    lcorbi_t3_pos_1_034_34    m/z 909.2582     0   1   17   17.5  RT (min)    lcorbi_t3_pos_1_031_40    m/z 909.2582     0   1   15.5   16  RT (min)      lcorbi_t3_pos_1_008_36    m/z 909.2582     0   1   16   16.5  RT (min)      lcorbi_t3_pos_2_022_38    m/z 909.2583     0   1   17.5   18  RT (min)    lcorbi_t3_pos_2_012_01    m/z 909.2583     0   1   16   16.5  RT (min)      lcorbi_t3_pos_4_049_55    m/z 909.2584     0   1   14   14.5  RT (min)      lcorbi_t3_pos_3_024_07    m/z 909.2584     0   1   16.5   17  RT (min)    lcorbi_t3_pos_2_025_52    m/z 909.2585     0   1   2E+6   14.5   15  RT (min)      lcorbi_t3_pos_4_105_61    m/z 909.2587     0   1   14   14.5  RT (min)      lcorbi_t3_pos_3_011_42    m/z 910.2566     0   1   16.5   17  RT (min)    lcorbi_t3_pos_1_034_34    m/z 910.2566     0   1   16.5   17  RT (min)    lcorbi_t3_pos_1_034_34    m/z 910.2571     0   1   17.5   18  RT (min)    lcorbi_t3_pos_2_012_01    m/z 910.2572     0   1   16   16.5  RT (min)      lcorbi_t3_pos_2_022_38    m/z 910.2573     0   1E+6   14.5   15  RT (min)      lcorbi_t3_pos_4_105_61    m/z 910.2573     0   1   14   14.5  RT (min)      lcorbi_t3_pos_3_024_07    m/z 910.2573     0   1   14   14.5  RT (min)      lcorbi_t3_pos_3_011_42    m/z 914.7800     0   1   17   17.5  RT (min)    lcorbi_t3_pos_1_031_40    m/z 914.7801     0   1   16.5   17  RT (min)    lcorbi_t3_pos_1_034_34    m/z 914.7803     0   1   14   14.5  RT (min)      lcorbi_t3_pos_2_013_33    m/z 914.7805     0   1   14   14.5  RT (min)      lcorbi_t3_pos_3_024_07    m/z 914.7806     0   2   4   6E+4   15.5   16  RT (min)      lcorbi_t3_pos_1_008_36    m/z 914.7808     0   1   14   14.5  RT (min)      lcorbi_t3_pos_3_011_42    m/z 914.7809     0   1E+4   16   16.5  RT (min)      lcorbi_t3_pos_2_022_38    m/z 914.7812     0   1   17.5   18  RT (min)    lcorbi_t3_pos_2_012_01    m/z 916.7952     0   1   17.5   18  RT (min)    lcorbi_t3_pos_2_012_01    m/z 916.7955     0   1   16   16.5  RT (min)      lcorbi_t3_pos_2_022_38    m/z 916.7955     0   1   14   14.5  RT (min)      lcorbi_t3_pos_3_024_07    m/z 916.7959     0   1   16.5   17  RT (min)    lcorbi_t3_pos_1_034_34    m/z 920.6710     0   1E+6   10   10.5  RT (min)       lcorbi_t3_pos_4_094_60    m/z 923.9232     0   1   17.5   18  RT (min)    lcorbi_t3_pos_1_049_55    m/z 923.9232     0   1   17.5   18  RT (min)    lcorbi_t3_pos_2_009_50    m/z 923.9233     0   1   17   17.5  RT (min)    lcorbi_t3_pos_1_041_54    m/z 923.9235     0   1   16.5   17  RT (min)    lcorbi_t3_pos_1_033_53    m/z 923.9236     0   1   18   18.5  RT (min)    lcorbi_t3_pos_2_056_56    m/z 923.9236     0   2   4   6E+5   15.5   16  RT (min)      lcorbi_t3_pos_3_026_04    m/z 923.9236     0   2   4   6E+4   16   16.5  RT (min)      lcorbi_t3_pos_4_025_52    m/z 923.9236     0   1E+4   16   16.5  RT (min)      lcorbi_t3_pos_3_025_52    m/z 923.9237     0   1   16.5   17  RT (min)    lcorbi_t3_pos_4_048_24    m/z 923.9238     0   2   4E+5   15.5   16  RT (min)      lcorbi_t3_pos_3_041_54    m/z 923.9238     0   1   17   17.5  RT (min)    lcorbi_t3_pos_2_025_52    m/z 923.9238     0   1   17.5   18  RT (min)    lcorbi_t3_pos_1_025_52    m/z 923.9239     0   1   16.5   17  RT (min)    lcorbi_t3_pos_2_017_51    m/z 925.9388     0   1   16.5   17  RT (min)    lcorbi_t3_pos_2_006_47    m/z 925.9389     0   1   17.5   18  RT (min)    lcorbi_t3_pos_1_041_54    m/z 925.9391     0   1   16.5   17  RT (min)    lcorbi_t3_pos_4_049_55    m/z 925.9392     0   1   18   18.5  RT (min)    lcorbi_t3_pos_1_025_52    m/z 925.9393     0   1E+4   16   16.5  RT (min)      lcorbi_t3_pos_3_025_52    m/z 925.9393     0   1   16.5   17  RT (min)    lcorbi_t3_pos_3_049_55    m/z 925.9393     0   1   17.5   18  RT (min)    lcorbi_t3_pos_2_009_50    m/z 925.9393     0   1   18   18.5  RT (min)    lcorbi_t3_pos_1_056_56    m/z 925.9394     0   1   2E+4   16   16.5  RT (min)      lcorbi_t3_pos_4_001_49    m/z 925.9394     0   2   4   6E+5   16   16.5  RT (min)      lcorbi_t3_pos_4_054_18    m/z 925.9395     0   2   4   6   8E+5   16   16.5  RT (min)      lcorbi_t3_pos_3_040_41    m/z 925.9395     0   1   16.5   17  RT (min)    lcorbi_t3_pos_2_017_51    m/z 925.9395     0   1   17.5   18  RT (min)    lcorbi_t3_pos_2_009_50    m/z 925.9395     0   1   17   17.5  RT (min)    lcorbi_t3_pos_2_038_03    m/z 925.9396     0   1   18   18.5  RT (min)    lcorbi_t3_pos_2_056_56    m/z 927.9541     0   1   17.5   18  RT (min)    lcorbi_t3_pos_1_041_54    m/z 927.9544     0   1   17   17.5  RT (min)    lcorbi_t3_pos_2_025_52    m/z 927.9548     0   1   16.5   17  RT (min)    lcorbi_t3_pos_4_049_55    m/z 927.9548     0   1   17   17.5  RT (min)    lcorbi_t3_pos_1_033_53    m/z 927.9549     0   2   4E+5   16   16.5  RT (min)      lcorbi_t3_pos_3_040_41    m/z 927.9549     0   1   16.5   17  RT (min)    lcorbi_t3_pos_3_025_52    m/z 927.9551     0   1   17.5   18  RT (min)    lcorbi_t3_pos_1_041_54    m/z 927.9551     0   1   17.5   18  RT (min)    lcorbi_t3_pos_2_009_50    m/z 927.9551     0   2   4   6E+5   16   16.5  RT (min)      lcorbi_t3_pos_1_005_42    m/z 927.9552     0   1   16.5   17  RT (min)    lcorbi_t3_pos_3_056_56    m/z 927.9553     0   1   18   18.5  RT (min)    lcorbi_t3_pos_1_049_55    m/z 927.9556     0   1   18   18.5  RT (min)    lcorbi_t3_pos_1_001_49    m/z 929.9705     0   1   17   17.5  RT (min)    lcorbi_t3_pos_1_021_18    m/z 929.9707     0   1   17   17.5  RT (min)    lcorbi_t3_pos_4_049_55    m/z 949.9299     0   1   17   17.5  RT (min)    lcorbi_t3_pos_1_041_54    m/z 949.9301     0   1   17.5   18  RT (min)    lcorbi_t3_pos_1_025_52    m/z 949.9304     0   1   18   18.5  RT (min)    lcorbi_t3_pos_2_056_56    m/z 949.9305     0   1   17   17.5  RT (min)    lcorbi_t3_pos_1_041_54    m/z 949.9306     0   1   16.5   17  RT (min)    lcorbi_t3_pos_2_041_54    m/z 949.9306     0   2   4   6   8E+5   15.5   16  RT (min)      lcorbi_t3_pos_3_041_54    m/z 949.9306     0   1   17.5   18  RT (min)    lcorbi_t3_pos_2_009_50    m/z 949.9306     0   1   16   16.5  RT (min)      lcorbi_t3_pos_3_025_52    m/z 949.9306     0   2   4   6E+5   15.5   16  RT (min)      lcorbi_t3_pos_3_040_41    m/z 949.9307     0   1   16.5   17  RT (min)    lcorbi_t3_pos_2_006_47    m/z 949.9307     0   1   16.5   17  RT (min)    lcorbi_t3_pos_1_021_18    m/z 949.9308     0   1   17.5   18  RT (min)    lcorbi_t3_pos_1_056_56    m/z 949.9309     0   1   17   17.5  RT (min)    lcorbi_t3_pos_2_025_52    m/z 950.9426     0   1   18   18.5  RT (min)    lcorbi_t3_pos_1_025_52    m/z 950.9426     0   1   17.5   18  RT (min)    lcorbi_t3_pos_1_049_55    m/z 950.9428     0   2   4E+5   15.5   16  RT (min)      lcorbi_t3_pos_3_041_54    m/z 950.9429     0   1   16.5   17  RT (min)    lcorbi_t3_pos_2_017_51    m/z 950.9431     0   1   16.5   17  RT (min)    lcorbi_t3_pos_1_030_21    m/z 951.9459     0   1   17.5   18  RT (min)    lcorbi_t3_pos_1_049_55    m/z 951.9460     0   1   16   16.5  RT (min)      lcorbi_t3_pos_3_056_56    m/z 951.9460     0   1   16.5   17  RT (min)    lcorbi_t3_pos_4_048_24    m/z 951.9461     0   1   16.5   17  RT (min)    lcorbi_t3_pos_1_033_53    m/z 951.9462     0   1   16.5   17  RT (min)    lcorbi_t3_pos_2_017_51    m/z 951.9462     0   1   17.5   18  RT (min)    lcorbi_t3_pos_2_049_55    m/z 951.9463     0   1   2   3E+4   16   16.5  RT (min)      lcorbi_t3_pos_4_001_49    m/z 951.9464     0   1   18   18.5  RT (min)    lcorbi_t3_pos_1_025_52    m/z 951.9464     0   1   16.5   17  RT (min)    lcorbi_t3_pos_2_006_47    m/z 951.9465     0   1E+6   15.5   16  RT (min)      lcorbi_t3_pos_3_041_54    m/z 951.9466     0   1   17.5   18  RT (min)    lcorbi_t3_pos_2_049_55    m/z 951.9470     0   1   17   17.5  RT (min)    lcorbi_t3_pos_2_025_52    m/z 951.9474     0   1   17   17.5  RT (min)    lcorbi_t3_pos_2_038_03    m/z 952.9578     0   1   17.5   18  RT (min)    lcorbi_t3_pos_1_041_54    m/z 952.9583     0   2   4E+5   16   16.5  RT (min)      lcorbi_t3_pos_3_040_41    m/z 952.9587     0   1   17.5   18  RT (min)    lcorbi_t3_pos_2_049_55    m/z 953.9614     0   1   17.5   18  RT (min)    lcorbi_t3_pos_1_041_54    m/z 953.9615     0   1   18   18.5  RT (min)    lcorbi_t3_pos_1_056_56    m/z 953.9616     0   1   18   18.5  RT (min)    lcorbi_t3_pos_1_001_49    m/z 953.9617     0   1   17   17.5  RT (min)    lcorbi_t3_pos_2_041_54    m/z 953.9620     0   1   17.5   18  RT (min)    lcorbi_t3_pos_1_041_54    m/z 953.9620     0   1   17   17.5  RT (min)    lcorbi_t3_pos_3_049_55    m/z 953.9620     0   1   17   17.5  RT (min)    lcorbi_t3_pos_1_033_53    m/z 953.9620     0   1E+6   16   16.5  RT (min)       lcorbi_t3_pos_3_040_41    m/z 953.9620     0   1   17   17.5  RT (min)    lcorbi_t3_pos_1_033_53    m/z 953.9620     0   1   16.5   17  RT (min)    lcorbi_t3_pos_2_006_47    m/z 953.9622     0   1   17.5   18  RT (min)    lcorbi_t3_pos_2_049_55    m/z 953.9622     0   1   2E+4   16   16.5  RT (min)      lcorbi_t3_pos_3_025_52    m/z 953.9623     0   1   17   17.5  RT (min)    lcorbi_t3_pos_2_038_03    m/z 953.9624     0   1   18   18.5  RT (min)    lcorbi_t3_pos_1_025_52    m/z 953.9625     0   1   16.5   17  RT (min)    lcorbi_t3_pos_2_046_07    m/z 953.9626     0   2   4   6   8E+5   16   16.5  RT (min)      lcorbi_t3_pos_3_033_53    m/z 953.9631     0   1   16.5   17  RT (min)    lcorbi_t3_pos_2_046_07    m/z 953.9631     0   2   4   6E+4   16   16.5  RT (min)      lcorbi_t3_pos_4_025_52    m/z 954.9737     0   1   17.5   18  RT (min)    lcorbi_t3_pos_1_041_54    m/z 954.9739     0   1   17   17.5  RT (min)    lcorbi_t3_pos_1_033_53    m/z 955.9769     0   1   18   18.5  RT (min)    lcorbi_t3_pos_1_001_49    m/z 955.9773     0   1   17   17.5  RT (min)    lcorbi_t3_pos_2_041_54    m/z 955.9774     0   1   18   18.5  RT (min)    lcorbi_t3_pos_2_056_56    m/z 955.9774     0   1   17.5   18  RT (min)    lcorbi_t3_pos_1_041_54    m/z 955.9775     0   1   17   17.5  RT (min)    lcorbi_t3_pos_2_038_03    m/z 955.9775     0   1   17   17.5  RT (min)    lcorbi_t3_pos_1_030_21    m/z 955.9776     0   1   16.5   17  RT (min)    lcorbi_t3_pos_3_025_52    m/z 955.9776     0   1E+6   16   16.5  RT (min)      lcorbi_t3_pos_3_041_54    m/z 955.9776     0   1   17   17.5  RT (min)    lcorbi_t3_pos_3_049_55    m/z 955.9777     0   1   17.5   18  RT (min)    lcorbi_t3_pos_2_009_50    m/z 955.9777     0   1E+6   16   16.5  RT (min)      lcorbi_t3_pos_3_040_41    m/z 955.9777     0   1   17.5   18  RT (min)    lcorbi_t3_pos_1_041_54    m/z 955.9778     0   1   16.5   17  RT (min)    lcorbi_t3_pos_1_047_41    m/z 955.9780     0   1   18   18.5  RT (min)    lcorbi_t3_pos_1_049_55    m/z 955.9785     0   1E+6   16   16.5  RT (min)      lcorbi_t3_pos_3_040_41    m/z 957.9922     0   1   17.5   18  RT (min)    lcorbi_t3_pos_1_041_54    m/z 957.9928     0   1   16.5   17  RT (min)    lcorbi_t3_pos_4_025_52    m/z 957.9928     0   1   18   18.5  RT (min)    lcorbi_t3_pos_1_056_56    m/z 957.9930     0   1   17   17.5  RT (min)    lcorbi_t3_pos_1_030_21    m/z 957.9931     0   2   4   6E+5   16   16.5  RT (min)      lcorbi_t3_pos_3_040_41    m/z 957.9931     0   1   17   17.5  RT (min)    lcorbi_t3_pos_2_041_54    m/z 957.9932     0   1   17.5   18  RT (min)    lcorbi_t3_pos_1_041_54    m/z 957.9932     0   2   4   6   8E+5   16   16.5  RT (min)      lcorbi_t3_pos_3_017_51    m/z 957.9932     0   1   18.5   19  RT (min)    lcorbi_t3_pos_2_056_56    m/z 957.9933     0   1   16.5   17  RT (min)    lcorbi_t3_pos_2_017_51    m/z 957.9935     0   1   16.5   17  RT (min)    lcorbi_t3_pos_3_025_52    m/z 957.9935     0   1   17   17.5  RT (min)    lcorbi_t3_pos_1_047_41    m/z 957.9937     0   1   18   18.5  RT (min)    lcorbi_t3_pos_1_049_55    m/z 962.8174     0   2   4   6E+5   14   14.5  RT (min)      lcorbi_t3_pos_4_105_61    m/z 962.8178     0   1   16.5   17  RT (min)    lcorbi_t3_pos_1_051_09    m/z 980.2780     0   1   17.5   18  RT (min)    lcorbi_t3_pos_2_012_01    m/z 980.2781     0   1   17   17.5  RT (min)    lcorbi_t3_pos_1_031_40    m/z 980.2781     0   1   2   3E+4   15.5   16  RT (min)      lcorbi_t3_pos_1_008_36    m/z 980.2781     0   1   14   14.5  RT (min)      lcorbi_t3_pos_3_024_07    m/z 980.2781     0   1   17.5   18  RT (min)    lcorbi_t3_pos_2_012_01    m/z 980.2783     0   1   16   16.5  RT (min)      lcorbi_t3_pos_1_028_32    m/z 980.2783     0   1   2   3E+4   15.5   16  RT (min)      lcorbi_t3_pos_1_008_36    m/z 980.2784     0   1   16.5   17  RT (min)    lcorbi_t3_pos_1_034_34    m/z 980.2784     0   1   16   16.5  RT (min)      lcorbi_t3_pos_3_049_55    m/z 980.2784     0   1   16   16.5  RT (min)      lcorbi_t3_pos_2_022_38    m/z 980.2784     0   2   4   6E+5   15   15.5  RT (min)      lcorbi_t3_pos_4_051_34    m/z 980.2785     0   2   4   6E+5   15   15.5  RT (min)      lcorbi_t3_pos_3_033_53    m/z 980.2787     0   2   4E+4   15.5   16  RT (min)      lcorbi_t3_pos_4_025_52    m/z 980.2787     0   1   14   14.5  RT (min)      lcorbi_t3_pos_3_011_42    m/z 980.2789     0   1   2   3E+6   14.5   15  RT (min)      lcorbi_t3_pos_4_105_61    m/z 981.2786     0   1   17.5   18  RT (min)    lcorbi_t3_pos_2_012_01    m/z 981.2788     0   1   16.5   17  RT (min)    lcorbi_t3_pos_1_034_34    m/z 981.2788     0   1   16   16.5  RT (min)      lcorbi_t3_pos_2_017_51    m/z 981.2789     0   1   17   17.5  RT (min)    lcorbi_t3_pos_2_009_50    m/z 981.2789     0   1   17.5   18  RT (min)    lcorbi_t3_pos_2_012_01    m/z 981.2789     0   1   2   3   4E+4   15.5   16  RT (min)      lcorbi_t3_pos_1_008_36    m/z 981.2791     0   1   2   3E+4   15.5   16  RT (min)      lcorbi_t3_pos_1_008_36    m/z 981.2791     0   1   14   14.5  RT (min)      lcorbi_t3_pos_3_024_07    m/z 981.2791     0   1   17   17.5  RT (min)    lcorbi_t3_pos_1_031_40    m/z 981.2792     0   1   16   16.5  RT (min)      lcorbi_t3_pos_1_028_32    m/z 981.2792     0   1   16   16.5  RT (min)      lcorbi_t3_pos_4_041_54    m/z 981.2793     0   1E+4   16   16.5  RT (min)      lcorbi_t3_pos_2_022_38    m/z 981.2793     0   2   4   6E+5   15   15.5  RT (min)       lcorbi_t3_pos_4_051_34    m/z 981.2794     0   1   14   14.5  RT (min)      lcorbi_t3_pos_3_011_42    m/z 981.2794     0   1   2   3E+6   14.5   15  RT (min)      lcorbi_t3_pos_4_105_61    m/z 982.2771     0   1   17   17.5  RT (min)    lcorbi_t3_pos_2_009_50    m/z 982.2773     0   1   17.5   18  RT (min)    lcorbi_t3_pos_2_012_01    m/z 982.2774     0   1   16   16.5  RT (min)      lcorbi_t3_pos_1_028_32    m/z 982.2774     0   1   16.5   17  RT (min)    lcorbi_t3_pos_1_034_34    m/z 982.2775     0   1   2   3   4E+4   15.5   16  RT (min)      lcorbi_t3_pos_1_008_36    m/z 982.2775     0   1   15.5   16  RT (min)      lcorbi_t3_pos_1_008_36    m/z 982.2776     0   1   14   14.5  RT (min)      lcorbi_t3_pos_3_024_07    m/z 982.2776     0   1   17.5   18  RT (min)    lcorbi_t3_pos_2_012_01    m/z 982.2776     0   1   14   14.5  RT (min)      lcorbi_t3_pos_3_011_42    m/z 982.2777     0   1   16   16.5  RT (min)      lcorbi_t3_pos_4_041_54    m/z 982.2778     0   1   16   16.5  RT (min)      lcorbi_t3_pos_2_017_51    m/z 982.2779     0   1   2E+6   14.5   15  RT (min)      lcorbi_t3_pos_4_105_61    m/z 982.2779     0   1   17   17.5  RT (min)    lcorbi_t3_pos_1_031_40    m/z 982.2780     0   2   4E+5   15   15.5  RT (min)      lcorbi_t3_pos_3_017_51    m/z 983.2765     0   1   17.5   18  RT (min)    lcorbi_t3_pos_2_012_01    m/z 983.2766     0   1   17.5   18  RT (min)    lcorbi_t3_pos_2_012_01    m/z 983.2767     0   1   17   17.5  RT (min)    lcorbi_t3_pos_1_031_40    m/z 983.2769     0   1   16.5   17  RT (min)    lcorbi_t3_pos_1_034_34    m/z 983.2770     0   1   14   14.5  RT (min)      lcorbi_t3_pos_3_024_07    m/z 983.2770     0   1   16   16.5  RT (min)      lcorbi_t3_pos_1_028_32    m/z 983.2772     0   1   2   3   4E+4   15.5   16  RT (min)      lcorbi_t3_pos_1_008_36    m/z 983.2773     0   1   16   16.5  RT (min)      lcorbi_t3_pos_3_003_33    m/z 983.2774     0   1E+6   14.5   15  RT (min)      lcorbi_t3_pos_4_105_61    m/z 983.2775     0   1   14   14.5  RT (min)      lcorbi_t3_pos_3_011_42    m/z 983.2776     0   1   2   3   4E+4   15.5   16  RT (min)      lcorbi_t3_pos_1_008_36    m/z 983.2777     0   1   16.5   17  RT (min)    lcorbi_t3_pos_1_034_34    m/z 984.2759     0   1   16   16.5  RT (min)      lcorbi_t3_pos_3_003_33    m/z 984.2761     0   1   17.5   18  RT (min)    lcorbi_t3_pos_2_012_01    m/z 984.2763     0   1   16.5   17  RT (min)    lcorbi_t3_pos_1_034_34    m/z 984.2766     0   2   4   6   8E+5   14.5   15  RT (min)      lcorbi_t3_pos_4_105_61    m/z 1054.2965     0   1E+6   14.5   15  RT (min)      lcorbi_t3_pos_4_105_61    m/z 1054.2966     0   1   14   14.5  RT (min)      lcorbi_t3_pos_3_031_45    m/z 1054.2967     0   1   15.5   16  RT (min)      lcorbi_t3_pos_1_008_36    m/z 1054.2968     0   1   17   17.5  RT (min)    lcorbi_t3_pos_1_031_40    m/z 1054.2969     0   1   15.5   16  RT (min)      lcorbi_t3_pos_1_008_36    m/z 1054.2969     0   1   16.5   17  RT (min)    lcorbi_t3_pos_1_028_32    m/z 1054.2969     0   1   16   16.5   17  RT (min)    lcorbi_t3_pos_3_003_33    m/z 1054.2971     0   1   14   14.5  RT (min)      lcorbi_t3_pos_3_011_42    m/z 1054.2971     0   1   17.5   18  RT (min)    lcorbi_t3_pos_2_012_01    m/z 1054.2971     0   1   16.5   17  RT (min)    lcorbi_t3_pos_1_034_34    m/z 1054.2972     0   1   16   16.5  RT (min)      lcorbi_t3_pos_2_022_38    m/z 1054.2973     0   1E+6   14.5   15  RT (min)      lcorbi_t3_pos_4_105_61    m/z 1054.2975     0   1   16.5   17  RT (min)    lcorbi_t3_pos_1_028_32    m/z 1054.2975     0   1   17.5   18  RT (min)    lcorbi_t3_pos_1_031_40    m/z 1054.2976     0   1   17.5   18  RT (min)    lcorbi_t3_pos_2_012_01    m/z 1055.2972     0   1   16.5   17  RT (min)    lcorbi_t3_pos_1_034_34    m/z 1055.2974     0   1   17.5   18  RT (min)    lcorbi_t3_pos_1_031_40    m/z 1055.2975     0   1   17.5   18  RT (min)    lcorbi_t3_pos_2_012_01    m/z 1055.2976     0   1   15.5   16  RT (min)      lcorbi_t3_pos_1_008_36    m/z 1055.2976     0   1   16.5   17  RT (min)    lcorbi_t3_pos_1_028_32    m/z 1055.2976     0   1   14   14.5  RT (min)      lcorbi_t3_pos_3_031_45    m/z 1055.2977     0   1   14   14.5  RT (min)      lcorbi_t3_pos_3_024_07    m/z 1055.2980     0   1   16   16.5   17  RT (min)    lcorbi_t3_pos_3_003_33    m/z 1055.2980     0   1   17   17.5  RT (min)    lcorbi_t3_pos_1_031_40    m/z 1055.2980     0   1   14   14.5  RT (min)      lcorbi_t3_pos_3_011_42    m/z 1055.2980     0   1   2E+6   14.5   15  RT (min)      lcorbi_t3_pos_4_105_61    m/z 1055.2981     0   1   16   16.5  RT (min)      lcorbi_t3_pos_3_003_33    m/z 1055.2983     0   1   17.5   18  RT (min)    lcorbi_t3_pos_2_012_01    m/z 1055.2983     0   1   17   17.5  RT (min)    lcorbi_t3_pos_1_034_34    m/z 1056.2955     0   1   17.5   18  RT (min)    lcorbi_t3_pos_1_031_40    m/z 1056.2959     0   1   16.5   17  RT (min)    lcorbi_t3_pos_1_034_34    m/z 1056.2961     0   1   15.5   16  RT (min)      lcorbi_t3_pos_1_008_36    m/z 1056.2961     0   1   16.5   17  RT (min)    lcorbi_t3_pos_1_028_32    m/z 1056.2962     0   1   16   16.5  RT (min)      lcorbi_t3_pos_2_022_38    m/z 1056.2962     0   1   17.5   18  RT (min)    lcorbi_t3_pos_2_012_01    m/z 1056.2963     0   1   14   14.5  RT (min)      lcorbi_t3_pos_3_011_42    m/z 1056.2965     0   1   16   16.5  RT (min)      lcorbi_t3_pos_3_003_33    m/z 1056.2969     0   1   17   17.5  RT (min)    lcorbi_t3_pos_1_031_40    m/z 1056.2969     0   1E+6   14.5   15  RT (min)      lcorbi_t3_pos_4_105_61    m/z 1056.2971     0   1E+6   14.5   15  RT (min)      lcorbi_t3_pos_4_105_61    m/z 1056.2972     0   1   17.5   18  RT (min)    lcorbi_t3_pos_2_012_01    m/z 1057.2955     0   1   17   17.5   18  RT (min)    lcorbi_t3_pos_1_031_40    m/z 1057.2959     0   1   14   14.5  RT (min)      lcorbi_t3_pos_3_024_07    m/z 1057.2960     0   1   15.5   16  RT (min)      lcorbi_t3_pos_1_008_36    m/z 1057.2962     0   1   14   14.5  RT (min)      lcorbi_t3_pos_3_011_42    m/z 1057.2964     0   1   17   17.5  RT (min)    lcorbi_t3_pos_1_031_40    m/z 1057.2965     0   1   16   16.5   17  RT (min)    lcorbi_t3_pos_3_003_33    m/z 1057.2968     0   1E+6   14.5   15  RT (min)      lcorbi_t3_pos_4_105_61    m/z 1057.2970     0   1E+6   14.5   15  RT (min)      lcorbi_t3_pos_4_105_61    m/z 1058.2952     0   2   4   6E+5   14.5   15  RT (min)      lcorbi_t3_pos_4_105_61    m/z 1128.3164     0   1E+6   15   15.5  RT (min)       lcorbi_t3_pos_4_105_61    m/z 1128.3166     0   1E+6   14.5   15  RT (min)      lcorbi_t3_pos_4_105_61    m/z 1128.3167     0   1   17   17.5  RT (min)    lcorbi_t3_pos_1_034_34    m/z 1129.3164     0   1   17.5   18  RT (min)    lcorbi_t3_pos_1_031_40    m/z 1129.3165     0   1E+6   14.5   15  RT (min)      lcorbi_t3_pos_4_105_61    m/z 1129.3170     0   1   16   16.5  RT (min)      lcorbi_t3_pos_2_022_38    m/z 1129.3170     0   1   14   14.5  RT (min)      lcorbi_t3_pos_2_013_33    m/z 1129.3173     0   1E+6   15   15.5  RT (min)      lcorbi_t3_pos_4_105_61    m/z 1130.3152     0   1E+6   14.5   15  RT (min)      lcorbi_t3_pos_4_105_61    m/z 1130.3152     0   1   17.5   18  RT (min)    lcorbi_t3_pos_1_031_40    m/z 1130.3162     0   1   17   17.5  RT (min)    lcorbi_t3_pos_1_034_34    m/z 1131.3154     0   2   4   6   8E+5   15   15.5  RT (min)      lcorbi_t3_pos_4_105_61    m/z 1131.3157     0   2   4   6   8E+5   14.5   15  RT (min)      lcorbi_t3_pos_4_105_61   
