## Supplementary material for "The Everything Bagel Feature Finder: Ultra-fast automated feature finding for untargeted metabolomics": SI Data File: tool_unique_xic_st_Masscube_ST004626.html

 
 
 Masscube-unique features (not detected by the other two tools) - ST004626 
 
 Masscube-unique features (not detected by the other two tools) - ST004626 
 9990 features · qtof · XIC window m/z ± 25 ppm (≥ 0.002 Da) · ± 0.5 min 
 Every panel is a feature  Masscube  detected that neither of the other two tools detected — three-way unique across EB (standard mode (min_detection_rate=0.1)), Masscube and XCMS, under the standard matching rule (m/z ± 25 ppm; apex within the reference RT window ± 0.2 min). It shows the MS1 extracted-ion chromatogram at the feature's m/z, drawn from the raw mzML named in  alignment_reference_file , zoomed to ± 0.5 min around the feature's reported apex RT.  Gray solid line  = the feature's reported apex RT;  red dashed line  = masscube's assigned MS/MS —  MS2_scan_id  in  MS2_source_file ; the panel widens to keep an off-apex MS/MS on-screen. Features with no assigned MS/MS show only the gray line. y is per-cell autoscaled. Rows keep the table order (m/z ascending). 
   m/z 68.9954     0   1   2   3   4E+3   1   1.5  RT (min)      lcqtof_t3_neg_4_021_21    m/z 79.0487     0   1E+4   1   1.5  RT (min)      lcqtof_t3_neg_4_019_05    m/z 84.9381     0   2   4E+3   16.5   17  RT (min)      lcqtof_t3_neg_4_004_20    m/z 87.0083     0   1E+4   4   4.5  RT (min)      lcqtof_t3_neg_4_051_15    m/z 96.9595     0   2   4   6   8E+3   9   9.5  RT (min)       lcqtof_t3_neg_4_000_ee    m/z 96.9595     0   1   2E+3   17   17.5  RT (min)       lcqtof_t3_neg_3_103_61    m/z 96.9596     0   2   4   6   8E+3   9   9.5  RT (min)      lcqtof_t3_neg_4_000_ee    m/z 96.9597     0   2   4   6E+3   7   7.5  RT (min)       lcqtof_t3_neg_4_016_14    m/z 96.9597     0   1   2   3E+3   17   17.5  RT (min)       lcqtof_t3_neg_4_035_08    m/z 96.9598     0   1   2E+3   17   17.5  RT (min)       lcqtof_t3_neg_3_086_59    m/z 96.9598     0   1   2E+3   17.5   18  RT (min)       lcqtof_t3_neg_2_065_51    m/z 96.9604     0   2   4   6E+4   1   1.5  RT (min)      lcqtof_t3_neg_4_041_54    m/z 101.9377     0   1   2   3E+3   16.5   17  RT (min)      lcqtof_t3_neg_4_007_09    m/z 102.9352     0   1E+4   0.5   1  RT (min)      lcqtof_t3_neg_4_000_ee    m/z 102.9495     0   1   2E+4   0.5   1  RT (min)      lcqtof_t3_neg_4_000_ee    m/z 112.0117     0   1E+4   4   4.5  RT (min)      lcqtof_t3_neg_4_051_15    m/z 112.9720     0   2   4E+3   1   1.5  RT (min)      lcqtof_t3_neg_4_037_01    m/z 112.9844     0   1E+4   9   9.5  RT (min)       lcqtof_t3_neg_4_074_58    m/z 112.9846     0   2   4   6E+4   4   4.5  RT (min)      lcqtof_t3_neg_4_068_57    m/z 112.9846     0   1E+4   5.5   6  RT (min)       lcqtof_t3_neg_4_069_57    m/z 112.9846     0   1E+4   7   7.5  RT (min)       lcqtof_t3_neg_4_069_57    m/z 112.9846     0   1E+4   13.5   14  RT (min)       lcqtof_t3_neg_4_075_58    m/z 112.9846     0   1   2E+4   13   13.5  RT (min)       lcqtof_t3_neg_4_072_57    m/z 112.9847     0   1E+4   6.5   7  RT (min)       lcqtof_t3_neg_4_078_58    m/z 112.9847     0   1E+4   6   6.5  RT (min)       lcqtof_t3_neg_4_076_58    m/z 112.9847     0   1E+4   9   9.5  RT (min)       lcqtof_t3_neg_4_068_57    m/z 112.9847     0   1E+4   12.5   13  RT (min)       lcqtof_t3_neg_4_078_58    m/z 112.9847     0   1E+4   5   5.5  RT (min)      lcqtof_t3_neg_4_072_57    m/z 112.9847     0   1   2   3E+4   14.5   15  RT (min)       lcqtof_t3_neg_4_074_58    m/z 112.9847     0   1E+4   12   12.5  RT (min)       lcqtof_t3_neg_4_068_57    m/z 112.9847     0   1   2E+4   13   13.5  RT (min)       lcqtof_t3_neg_4_072_57    m/z 112.9847     0   1E+4   6.5   7  RT (min)       lcqtof_t3_neg_4_068_57    m/z 112.9848     0   1E+4   8   8.5  RT (min)       lcqtof_t3_neg_4_067_57    m/z 112.9848     0   1E+4   8.5   9  RT (min)       lcqtof_t3_neg_4_066_57    m/z 112.9848     0   1   2E+4   10   10.5  RT (min)       lcqtof_t3_neg_4_068_57    m/z 112.9848     0   1E+4   12.5   13  RT (min)       lcqtof_t3_neg_4_069_57    m/z 112.9848     0   1   2E+4   10.5   11  RT (min)       lcqtof_t3_neg_4_073_57    m/z 112.9848     0   1   2E+5   1   1.5  RT (min)      lcqtof_t3_neg_4_067_57    m/z 112.9848     0   1E+4   6   6.5  RT (min)       lcqtof_t3_neg_4_076_58    m/z 112.9848     0   1E+4   8.5   9  RT (min)       lcqtof_t3_neg_4_068_57    m/z 112.9848     0   1   2E+4   10   10.5  RT (min)       lcqtof_t3_neg_4_079_58    m/z 112.9848     0   1E+4   12   12.5  RT (min)       lcqtof_t3_neg_4_073_57    m/z 112.9848     0   1   2E+4   7   7.5  RT (min)       lcqtof_t3_neg_4_079_58    m/z 112.9848     0   1E+4   11   11.5  RT (min)       lcqtof_t3_neg_4_081_58    m/z 112.9848     0   1   2E+4   13.5   14  RT (min)       lcqtof_t3_neg_4_073_57    m/z 112.9848     0   1E+4   6   6.5  RT (min)       lcqtof_t3_neg_4_078_58    m/z 112.9848     0   1   2E+4   10.5   11  RT (min)       lcqtof_t3_neg_4_072_57    m/z 112.9848     0   1E+4   9.5   10  RT (min)       lcqtof_t3_neg_4_068_57    m/z 112.9848     0   1E+4   5.5   6  RT (min)       lcqtof_t3_neg_4_067_57    m/z 112.9848     0   1E+4   6   6.5  RT (min)      lcqtof_t3_neg_4_067_57    m/z 112.9848     0   1E+4   5   5.5  RT (min)       lcqtof_t3_neg_4_067_57    m/z 112.9848     0   1E+5   1.5   2  RT (min)      lcqtof_t3_neg_4_072_57    m/z 112.9848     0   1E+4   11   11.5  RT (min)       lcqtof_t3_neg_4_071_57    m/z 112.9848     0   1   2   3   4E+4   15   15.5  RT (min)       lcqtof_t3_neg_4_000_ee    m/z 112.9848     0   1E+4   12   12.5  RT (min)       lcqtof_t3_neg_4_078_58    m/z 112.9848     0   1E+4   14   14.5  RT (min)       lcqtof_t3_neg_4_076_58    m/z 112.9848     0   1   2E+4   10   10.5  RT (min)       lcqtof_t3_neg_4_073_57    m/z 112.9849     0   1   2E+4   9.5   10  RT (min)       lcqtof_t3_neg_4_074_58    m/z 112.9849     0   1   2E+4   14   14.5  RT (min)       lcqtof_t3_neg_4_074_58    m/z 112.9849     0   1E+4   4.5   5  RT (min)       lcqtof_t3_neg_4_073_57    m/z 112.9849     0   1E+4   9.5   10  RT (min)       lcqtof_t3_neg_4_072_57    m/z 112.9849     0   1E+5   5   10   15  RT (min)       lcqtof_t3_neg_4_005_18    m/z 112.9850     0   1E+4   11.5   12  RT (min)       lcqtof_t3_neg_4_074_58    m/z 112.9850     0   2   4E+4   4.5   5  RT (min)      lcqtof_t3_neg_4_072_57    m/z 112.9850     0   1   2   3E+4   14.5   15  RT (min)       lcqtof_t3_neg_4_074_58    m/z 112.9850     0   1   2   3E+4   15.5   16  RT (min)      lcqtof_t3_neg_4_076_58    m/z 112.9850     0   1   2   3E+4   15.5   16  RT (min)       lcqtof_t3_neg_4_068_57    m/z 112.9851     0   2   4   6   8E+4   16   16.5  RT (min)       lcqtof_t3_neg_4_018_23    m/z 112.9851     0   1E+5   16.5   17  RT (min)       lcqtof_t3_neg_4_019_05    m/z 112.9851     0   1   2   3   4E+4   15   16   17  RT (min)       lcqtof_t3_neg_4_091_60    m/z 112.9852     0   2   4   6   8E+4   14   16  RT (min)       lcqtof_t3_neg_4_106_62    m/z 112.9852     0   1   2   3   4E+3   4   4.5  RT (min)      lcqtof_t3_neg_3_004_26    m/z 113.9883     0   1   2   3E+3   4   4.5  RT (min)      lcqtof_t3_neg_4_000_aa    m/z 114.9884     0   1   2E+3   17.5   18  RT (min)      lcqtof_t3_neg_3_096_60    m/z 115.0326     0   1E+4   1   1.5  RT (min)      lcqtof_t3_neg_4_040_24    m/z 116.9280     0   1   2   3   4E+3   16.5   17  RT (min)       lcqtof_t3_neg_4_006_16    m/z 124.9635     0   1   2   3   4E+4   1   1.5  RT (min)       lcqtof_t3_neg_4_019_05    m/z 128.0346     0   1   2   3   4E+4   2   2.5  RT (min)      lcqtof_t3_neg_4_085_59    m/z 128.0346     0   2   4   6E+4   1.5   2  RT (min)      lcqtof_t3_neg_4_055_38    m/z 128.0346     0   2   4   6   8E+4   1.5   2  RT (min)      lcqtof_t3_neg_4_030_31    m/z 128.0346     0   2   4   6   8E+4   1.5   2  RT (min)      lcqtof_t3_neg_4_030_31    m/z 128.0347     0   2   4E+4   2   2.5  RT (min)      lcqtof_t3_neg_4_030_31    m/z 128.0348     0   1   2   3   4E+4   2   2.5  RT (min)      lcqtof_t3_neg_4_089_59    m/z 128.0524     0   1E+4   5.5   6  RT (min)       lcqtof_t3_neg_4_040_24    m/z 129.9999     0   2   4E+3   12   12.5  RT (min)      lcqtof_t3_neg_3_077_58    m/z 130.9657     0   2   4   6   8E+3   0.5   1  RT (min)      lcqtof_t3_neg_1_008_02    m/z 130.9658     0   1   2E+3   0   0.5  RT (min)      lcqtof_t3_neg_1_038_15    m/z 130.9658     0   1   2E+3   0   0.5  RT (min)      lcqtof_t3_neg_1_048_35    m/z 130.9658     0   1   2E+4   0.5   1  RT (min)      lcqtof_t3_neg_1_051_26    m/z 130.9659     0   1E+4   0.5   1  RT (min)      lcqtof_t3_neg_1_022_28    m/z 130.9661     0   2   4   6E+3   1   1.5  RT (min)      lcqtof_t3_neg_3_030_03    m/z 132.8671     0   2   4   6   8E+3   1.5   2  RT (min)      lcqtof_t3_neg_4_014_13    m/z 132.8695     0   2   4   6E+4   0.5   1  RT (min)      lcqtof_t3_neg_4_000_ee    m/z 133.0106     0   1   2E+3   9.5   10  RT (min)      lcqtof_t3_neg_1_036_14    m/z 133.0287     0   1E+4   6   6.5  RT (min)      lcqtof_t3_neg_4_109_62    m/z 133.0320     0   1E+4   4.5   5  RT (min)      lcqtof_t3_neg_4_062_13    m/z 133.0514     0   2   4   6E+4   2   2.5  RT (min)      lcqtof_t3_neg_4_097_60    m/z 133.0514     0   2   4   6E+4   2   2.5  RT (min)      lcqtof_t3_neg_4_011_32    m/z 133.0515     0   1   2E+4   2   2.5  RT (min)      lcqtof_t3_neg_1_055_32    m/z 133.0515     0   2   4   6   8E+4   1.5   2  RT (min)      lcqtof_t3_neg_4_011_32    m/z 133.0516     0   2   4   6   8E+4   1.5   2  RT (min)      lcqtof_t3_neg_4_011_32    m/z 133.0516     0   2   4   6E+4   2   2.5  RT (min)      lcqtof_t3_neg_4_011_32    m/z 134.0465     0   1E+4   2   2.5  RT (min)       lcqtof_t3_neg_4_019_05    m/z 134.0467     0   1E+4   1.5   2  RT (min)      lcqtof_t3_neg_4_018_23    m/z 134.0467     0   1E+4   2   2.5  RT (min)      lcqtof_t3_neg_4_018_23    m/z 134.8647     0   2   4   6   8E+3   1.5   2  RT (min)      lcqtof_t3_neg_4_014_13    m/z 134.8649     0   2   4   6E+4   0.5   1  RT (min)      lcqtof_t3_neg_4_000_ee    m/z 134.9874     0   2   4   6   8E+3   1   1.5  RT (min)      lcqtof_t3_neg_1_028_20    m/z 134.9877     0   1E+4   0.5   1  RT (min)      lcqtof_t3_neg_4_000_ee    m/z 135.0435     0   1   2   3   4E+3   9.5   10  RT (min)      lcqtof_t3_neg_4_059_14    m/z 136.0476     0   1E+4   6.5   7  RT (min)      lcqtof_t3_neg_4_004_20    m/z 136.0480     0   1E+4   6.5   7  RT (min)      lcqtof_t3_neg_4_004_20    m/z 136.1614     0   1   2   3   4E+4   1   1.5  RT (min)      lcqtof_t3_neg_1_055_32    m/z 136.8623     0   1E+4   1   1.5  RT (min)      lcqtof_t3_neg_4_040_24    m/z 136.8644     0   1   2E+4   0.5   1  RT (min)      lcqtof_t3_neg_4_000_ee    m/z 139.0066     0   1   2   3E+3   14   14.5  RT (min)       lcqtof_t3_neg_4_011_32    m/z 139.0633     0   1E+4   2   2.5  RT (min)       lcqtof_t3_neg_3_032_24    m/z 139.0634     0   1   2   3E+4   2   2.5  RT (min)       lcqtof_t3_neg_1_031_24    m/z 142.0501     0   2   4   6   8E+1   3.5   4  RT (min)      lcqtof_t3_neg_3_038_06    m/z 142.9430     0   1   2E+3   16.5   17  RT (min)      lcqtof_t3_neg_4_000_bb    m/z 143.8961     0   1   2E+3   16.5   17  RT (min)      lcqtof_t3_neg_1_000_aa    m/z 144.8969     0   1   2E+3   16.5   17  RT (min)      lcqtof_t3_neg_4_000_bb    m/z 145.8940     0   1   2E+3   16.5   17  RT (min)      lcqtof_t3_neg_4_000_aa    m/z 145.9421     0   1E+4   0.5   1  RT (min)      lcqtof_t3_neg_4_000_ee    m/z 146.1683     0   1E+4   1   1.5  RT (min)      lcqtof_t3_neg_1_032_36    m/z 146.8986     0   1   2   3E+3   16.5   17  RT (min)      lcqtof_t3_neg_4_000_aa    m/z 146.9379     0   1   2E+4   18   18.5  RT (min)      lcqtof_t3_neg_4_062_13    m/z 146.9380     0   1E+4   18   18.5  RT (min)      lcqtof_t3_neg_4_112_62    m/z 146.9380     0   1   2E+4   18.5   19  RT (min)      lcqtof_t3_neg_4_061_50    m/z 146.9382     0   1   2   3E+4   17.5   18  RT (min)      lcqtof_t3_neg_4_061_50    m/z 146.9382     0   1   2E+4   12   12.5  RT (min)      lcqtof_t3_neg_4_062_13    m/z 146.9382     0   1   2   3E+4   17   17.5  RT (min)      lcqtof_t3_neg_4_060_15    m/z 146.9382     0   1   2   3   4E+4   1   1.5  RT (min)      lcqtof_t3_neg_4_000_ee    m/z 146.9382     0   1E+4   17   17.5  RT (min)      lcqtof_t3_neg_4_072_57    m/z 146.9382     0   1E+5   0.5   1  RT (min)      lcqtof_t3_neg_4_000_ee    m/z 146.9384     0   1E+4   18.5   19  RT (min)      lcqtof_t3_neg_4_112_62    m/z 146.9386     0   1E+4   12   12.5  RT (min)      lcqtof_t3_neg_4_027_41    m/z 146.9388     0   1E+4   18.5   19  RT (min)      lcqtof_t3_neg_4_103_61    m/z 146.9390     0   1E+4   13.5   14  RT (min)      lcqtof_t3_neg_4_027_41    m/z 146.9392     0   1E+5   1   1.5  RT (min)      lcqtof_t3_neg_4_000_ee    m/z 146.9613     0   1   2   3E+4   7   8  RT (min)       lcqtof_t3_neg_1_008_02    m/z 146.9644     0   1E+4   1   1.5  RT (min)      lcqtof_t3_neg_1_051_26    m/z 146.9644     0   1   2E+4   8   9  RT (min)       lcqtof_t3_neg_1_032_36    m/z 146.9645     0   1E+4   0   0.5  RT (min)      lcqtof_t3_neg_1_036_14    m/z 146.9645     0   1   2E+4   16.5   17  RT (min)       lcqtof_t3_neg_1_043_38    m/z 146.9645     0   1E+4   0.5   1  RT (min)       lcqtof_t3_neg_1_056_56    m/z 146.9646     0   1E+4   0   0.5  RT (min)      lcqtof_t3_neg_1_038_15    m/z 146.9646     0   1E+4   0   0.5  RT (min)       lcqtof_t3_neg_1_037_25    m/z 146.9646     0   1E+4   0   0.5  RT (min)      lcqtof_t3_neg_1_036_14    m/z 146.9646     0   1E+4   0.5   1  RT (min)      lcqtof_t3_neg_1_051_26    m/z 146.9647     0   1E+4   0   0.5  RT (min)      lcqtof_t3_neg_1_036_14    m/z 146.9649     0   1   2   3E+4   5.5   6   6.5  RT (min)       lcqtof_t3_neg_1_059_17    m/z 146.9649     0   1   2E+4   7.5   8  RT (min)       lcqtof_t3_neg_1_000_dd    m/z 146.9650     0   1   2   3E+4   7   7.5  RT (min)       lcqtof_t3_neg_1_064_14    m/z 146.9651     0   1   2   3E+4   6.5   7  RT (min)       lcqtof_t3_neg_1_007_03    m/z 146.9651     0   1   2   3E+4   6.5   7  RT (min)       lcqtof_t3_neg_1_030_42    m/z 146.9652     0   1   2   3E+4   6   7  RT (min)       lcqtof_t3_neg_1_036_14    m/z 148.0703     0   1E+4   4   4.5  RT (min)      lcqtof_t3_neg_3_032_24    m/z 152.9780     0   1   2E+3   17   17.5  RT (min)      lcqtof_t3_neg_3_109_62    m/z 153.0752     0   1   2E+4   6   6.5  RT (min)      lcqtof_t3_neg_3_032_24    m/z 154.9620     0   1   2   3E+3   17.5   18  RT (min)       lcqtof_t3_neg_1_035_31    m/z 154.9632     0   1   2E+3   17.5   18  RT (min)       lcqtof_t3_neg_3_016_35    m/z 154.9644     0   1   2   3   4E+3   17   17.5  RT (min)       lcqtof_t3_neg_1_037_25    m/z 154.9660     0   1   2   3E+3   17   17.5  RT (min)       lcqtof_t3_neg_1_014_04    m/z 154.9663     0   1   2   3E+3   17   18  RT (min)       lcqtof_t3_neg_1_019_23    m/z 156.0657     0   1   2E+2   3.5   4  RT (min)      lcqtof_t3_neg_3_038_06    m/z 156.2045     0   1E+4   1   1.5  RT (min)      lcqtof_t3_neg_1_055_32    m/z 156.9452     0   1   2E+4   1.5   2  RT (min)      lcqtof_t3_neg_3_000_aa    m/z 157.0104     0   1E+3   16   16.5  RT (min)      lcqtof_t3_neg_2_030_23    m/z 157.0108     0   2   4   6E+3   16   16.5  RT (min)      lcqtof_t3_neg_4_041_54    m/z 157.0109     0   2   4   6   8E+3   16   16.5  RT (min)      lcqtof_t3_neg_4_018_23    m/z 157.0153     0   1E+4   17   17.5  RT (min)      lcqtof_t3_neg_4_035_08    m/z 158.0109     0   1   2E+4   13   13.5  RT (min)      lcqtof_t3_neg_4_106_62    m/z 158.0111     0   1   2E+4   10.5   11  RT (min)      lcqtof_t3_neg_4_106_62    m/z 158.0112     0   1E+4   9.5   10  RT (min)      lcqtof_t3_neg_4_106_62    m/z 158.0113     0   1E+2   16   16.5  RT (min)      lcqtof_t3_neg_2_030_23    m/z 158.0114     0   1E+4   8.5   9  RT (min)      lcqtof_t3_neg_4_106_62    m/z 158.0114     0   1E+4   17   17.5  RT (min)      lcqtof_t3_neg_4_035_08    m/z 158.0114     0   2   4   6   8E+4   16   16.5  RT (min)      lcqtof_t3_neg_4_018_23    m/z 158.0115     0   1   2   3   4E+3   13.5   14  RT (min)      lcqtof_t3_neg_3_038_06    m/z 158.0115     0   1   2E+4   10.5   11  RT (min)      lcqtof_t3_neg_4_106_62    m/z 158.0116     0   2   4   6   8E+4   16   16.5  RT (min)      lcqtof_t3_neg_4_018_23    m/z 158.0116     0   2   4   6E+4   16.5   17  RT (min)      lcqtof_t3_neg_4_045_02    m/z 158.0117     0   1   2E+2   15   15.5  RT (min)      lcqtof_t3_neg_3_032_24    m/z 158.0117     0   2   4   6E+4   17   17.5  RT (min)      lcqtof_t3_neg_4_035_08    m/z 158.0118     0   1   2E+4   17   17.5  RT (min)      lcqtof_t3_neg_4_035_08    m/z 158.8918     0   1   2   3E+3   16.5   17  RT (min)      lcqtof_t3_neg_3_018_32    m/z 158.8920     0   1   2   3   4E+3   16.5   17  RT (min)      lcqtof_t3_neg_4_000_aa    m/z 158.8922     0   1   2   3E+3   16.5   17  RT (min)      lcqtof_t3_neg_1_031_24    m/z 158.9781     0   1   2   3   4E+3   17.5   18  RT (min)      lcqtof_t3_neg_3_113_62    m/z 159.0074     0   1E+4   7.5   8  RT (min)      lcqtof_t3_neg_4_085_59    m/z 159.0075     0   1   2E+4   9   9.5  RT (min)      lcqtof_t3_neg_4_111_62    m/z 159.0077     0   1   2E+4   14   14.5  RT (min)      lcqtof_t3_neg_4_095_60    m/z 159.0077     0   1   2   3E+4   11.5   12  RT (min)      lcqtof_t3_neg_4_091_60    m/z 159.0077     0   1   2   3E+4   9.5   10  RT (min)      lcqtof_t3_neg_4_095_60    m/z 159.0077     0   1   2   3E+4   13.5   14  RT (min)      lcqtof_t3_neg_4_107_62    m/z 159.0077     0   1   2E+4   9   9.5  RT (min)       lcqtof_t3_neg_4_068_57    m/z 159.0078     0   1E+2   16   16.5  RT (min)      lcqtof_t3_neg_2_030_23    m/z 159.0078     0   1   2   3E+4   10   10.5  RT (min)      lcqtof_t3_neg_4_091_60    m/z 159.0078     0   1E+4   8.5   9  RT (min)      lcqtof_t3_neg_4_107_62    m/z 159.0079     0   1E+4   8.5   9  RT (min)       lcqtof_t3_neg_4_095_60    m/z 159.0079     0   2   4   6E+3   13.5   14  RT (min)      lcqtof_t3_neg_3_038_06    m/z 159.0079     0   1   2   3E+4   10   10.5  RT (min)      lcqtof_t3_neg_4_068_57    m/z 159.0079     0   1   2   3   4E+4   11   12  RT (min)       lcqtof_t3_neg_4_000_cc    m/z 159.0080     0   1E+2   15   15.5  RT (min)      lcqtof_t3_neg_3_032_24    m/z 159.0080     0   1   2   3   4E+4   11   12  RT (min)       lcqtof_t3_neg_4_005_18    m/z 159.0080     0   1E+5   16   16.5  RT (min)      lcqtof_t3_neg_4_018_23    m/z 159.0080     0   1   2E+4   13   13.5  RT (min)      lcqtof_t3_neg_4_085_59    m/z 159.0080     0   2   4E+4   12   14  RT (min)       lcqtof_t3_neg_4_006_16    m/z 159.0081     0   1E+5   17   17.5  RT (min)      lcqtof_t3_neg_4_035_08    m/z 159.0081     0   1E+4   17   17.5  RT (min)      lcqtof_t3_neg_4_035_08    m/z 159.0082     0   1   2   3E+4   17   17.5  RT (min)      lcqtof_t3_neg_4_035_08    m/z 159.0526     0   1   2   3E+3   4   4.5  RT (min)      lcqtof_t3_neg_4_062_13    m/z 159.0767     0   1E+4   15   15.5  RT (min)       lcqtof_t3_neg_4_108_62    m/z 159.0767     0   1E+4   14.5   15   15.5  RT (min)       lcqtof_t3_neg_4_101_61    m/z 159.0767     0   1   2E+4   13   14  RT (min)       lcqtof_t3_neg_4_082_59    m/z 159.0768     0   1   2E+4   16.5   17  RT (min)       lcqtof_t3_neg_4_000_ff    m/z 159.8926     0   1   2E+3   15   15.5  RT (min)      lcqtof_t3_neg_3_099_61    m/z 160.0254     0   1E+4   16   16.5  RT (min)      lcqtof_t3_neg_4_018_23    m/z 160.0254     0   1E+4   17   17.5  RT (min)      lcqtof_t3_neg_4_035_08    m/z 160.8412     0   1E+4   4.5   5  RT (min)      lcqtof_t3_neg_4_001_49    m/z 160.8423     0   2   4E+4   1.5   2  RT (min)      lcqtof_t3_neg_1_059_17    m/z 160.8918     0   1   2E+3   15   15.5  RT (min)      lcqtof_t3_neg_3_080_58    m/z 161.0214     0   2   4   6   8E+3   7.5   10   12.5  RT (min)       lcqtof_t3_neg_4_016_14    m/z 161.0217     0   2   4   6   8E+3   8   10   12  RT (min)       lcqtof_t3_neg_4_039_06    m/z 161.0221     0   1E+4   16   16.5  RT (min)      lcqtof_t3_neg_4_018_23    m/z 161.0224     0   1E+4   17   17.5  RT (min)      lcqtof_t3_neg_4_035_08    m/z 161.0253     0   1E+4   6.5   7  RT (min)       lcqtof_t3_neg_4_004_20    m/z 161.8925     0   1   2   3   4E+3   16.5   17  RT (min)      lcqtof_t3_neg_2_002_35    m/z 161.8926     0   1   2   3   4E+3   16.5   17  RT (min)      lcqtof_t3_neg_3_018_32    m/z 161.9857     0   2   4   6E+4   3.5   4  RT (min)      lcqtof_t3_neg_4_008_35    m/z 161.9857     0   1E+5   3.5   4  RT (min)      lcqtof_t3_neg_4_005_18    m/z 161.9858     0   2   4   6E+4   3.5   4  RT (min)      lcqtof_t3_neg_4_008_35    m/z 161.9858     0   1E+5   3   3.5  RT (min)       lcqtof_t3_neg_4_005_18    m/z 161.9858     0   1E+5   3   3.5  RT (min)       lcqtof_t3_neg_4_005_18    m/z 161.9858     0   1E+5   3.5   4  RT (min)       lcqtof_t3_neg_4_005_18    m/z 161.9858     0   2   4   6E+4   3   3.5  RT (min)      lcqtof_t3_neg_4_005_18    m/z 161.9860     0   1   2   3E+4   3.5   4  RT (min)      lcqtof_t3_neg_4_049_55    m/z 162.8385     0   2   4E+4   1.5   2  RT (min)      lcqtof_t3_neg_1_059_17    m/z 162.8387     0   2   4E+3   14   14.5  RT (min)       lcqtof_t3_neg_4_045_02    m/z 162.8928     0   2   4   6   8E+3   16.5   17  RT (min)       lcqtof_t3_neg_4_000_aa    m/z 162.8928     0   1   2   3   4E+3   16   16.5  RT (min)       lcqtof_t3_neg_4_018_23    m/z 162.8928     0   1   2   3E+3   15.5   16  RT (min)       lcqtof_t3_neg_3_072_57    m/z 162.8930     0   1   2   3E+3   15   15.5  RT (min)       lcqtof_t3_neg_3_112_62    m/z 162.8931     0   1   2   3E+3   15.5   16  RT (min)       lcqtof_t3_neg_3_112_62    m/z 162.8932     0   1   2E+3   12.5   13  RT (min)       lcqtof_t3_neg_4_032_04    m/z 163.0424     0   1   2   3E+2   4   4.5  RT (min)      lcqtof_t3_neg_4_018_23    m/z 163.0897     0   1   2E+4   4   4.5  RT (min)      lcqtof_t3_neg_3_032_24    m/z 164.8949     0   1   2   3E+3   16.5   17  RT (min)      lcqtof_t3_neg_3_037_05    m/z 164.8951     0   1   2   3   4E+3   16.5   17  RT (min)      lcqtof_t3_neg_4_000_aa    m/z 165.0403     0   1   2   3   4E+4   13.5   14  RT (min)       lcqtof_t3_neg_4_064_17    m/z 165.0404     0   2   4   6E+4   13  RT (min)       lcqtof_t3_neg_4_064_17    m/z 165.0405     0   2   4   6E+4   12.5   13  RT (min)      lcqtof_t3_neg_4_064_17    m/z 165.0405     0   2   4   6E+4   12   13  RT (min)       lcqtof_t3_neg_4_064_17    m/z 165.0405     0   1   2E+4   12   12.5  RT (min)      lcqtof_t3_neg_4_064_17    m/z 165.0406     0   1   2E+4   12   12.5  RT (min)       lcqtof_t3_neg_4_064_17    m/z 165.0406     0   2   4   6E+4   12.5   13  RT (min)      lcqtof_t3_neg_4_064_17    m/z 165.0407     0   2   4   6E+4   13   13.5  RT (min)       lcqtof_t3_neg_4_064_17    m/z 165.0407     0   1   2E+4   12   12.5  RT (min)      lcqtof_t3_neg_4_064_17    m/z 165.0407     0   2   4   6E+4   13.5   14  RT (min)      lcqtof_t3_neg_4_064_17    m/z 165.0407     0   2   4   6E+4   13   13.5  RT (min)       lcqtof_t3_neg_4_064_17    m/z 165.0408     0   1   2E+4   13.5   14  RT (min)      lcqtof_t3_neg_4_064_17    m/z 165.0409     0   2   4   6E+4   12.5   13  RT (min)      lcqtof_t3_neg_4_064_17    m/z 165.0409     0   2   4   6E+4   13   13.5  RT (min)      lcqtof_t3_neg_4_064_17    m/z 165.0413     0   2   4   6E+4   12.5   13  RT (min)      lcqtof_t3_neg_4_064_17    m/z 165.0413     0   1   2E+4   14   14.5  RT (min)      lcqtof_t3_neg_4_064_17    m/z 167.0453     0   1E+4   4   4.5  RT (min)      lcqtof_t3_neg_4_062_13    m/z 167.0650     0   1   2   3E+4   4   4.5  RT (min)      lcqtof_t3_neg_3_032_24    m/z 167.1005     0   1   2E+4   7.5   8  RT (min)      lcqtof_t3_neg_4_005_18    m/z 167.2659     0   1E+4   7.5   8  RT (min)      lcqtof_t3_neg_4_005_18    m/z 168.0226     0   1E+4   1.5   2  RT (min)      lcqtof_t3_neg_4_037_01    m/z 168.0231     0   1   2E+3   2   2.5  RT (min)      lcqtof_t3_neg_1_053_01    m/z 170.0913     0   1E+2   3.5   4  RT (min)      lcqtof_t3_neg_3_038_06    m/z 171.0517     0   2   4E+3   4   4.5  RT (min)      lcqtof_t3_neg_4_029_40    m/z 172.8302     0   2   4   6E+3   1.5   2  RT (min)      lcqtof_t3_neg_4_016_14    m/z 173.0086     0   1   2   3   4E+4   1.5   2  RT (min)      lcqtof_t3_neg_4_059_14    m/z 174.0120     0   1   2   3   4E+3   13.5   14  RT (min)      lcqtof_t3_neg_4_000_cc    m/z 174.0145     0   1   2   3   4E+3   13.5   14  RT (min)      lcqtof_t3_neg_4_000_cc    m/z 174.9552     0   1   2   3   4E+4   17.5   18  RT (min)       lcqtof_t3_neg_1_036_14    m/z 174.9552     0   1E+4   4.5   5  RT (min)      lcqtof_t3_neg_1_064_14    m/z 174.9552     0   1   2E+4   17.5   18  RT (min)       lcqtof_t3_neg_1_036_14    m/z 174.9552     0   1E+4   5   5.5  RT (min)       lcqtof_t3_neg_1_036_14    m/z 174.9552     0   1   2E+4   8   8.5  RT (min)       lcqtof_t3_neg_4_099_61    m/z 174.9552     0   1E+4   4.5   5  RT (min)      lcqtof_t3_neg_1_064_14    m/z 174.9552     0   1E+4   3   3.5  RT (min)       lcqtof_t3_neg_1_064_14    m/z 174.9552     0   1   2E+4   17.5   18  RT (min)       lcqtof_t3_neg_1_036_14    m/z 174.9553     0   1E+4   10.5   11  RT (min)       lcqtof_t3_neg_1_036_14    m/z 174.9553     0   1E+4   12   12.5  RT (min)       lcqtof_t3_neg_3_064_15    m/z 174.9553     0   1E+4   0   0.5  RT (min)       lcqtof_t3_neg_2_062_17    m/z 174.9553     0   1E+4   18.5   19  RT (min)       lcqtof_t3_neg_2_058_15    m/z 174.9553     0   1E+4   5.5   6  RT (min)       lcqtof_t3_neg_3_079_58    m/z 174.9553     0   1   2E+5   1   1.5  RT (min)       lcqtof_t3_neg_1_036_14    m/z 174.9553     0   1E+4   6   6.5  RT (min)       lcqtof_t3_neg_4_074_58    m/z 174.9553     0   1E+4   0   0.5  RT (min)       lcqtof_t3_neg_3_075_58    m/z 174.9554     0   1   2E+4   7   7.5  RT (min)       lcqtof_t3_neg_1_036_14    m/z 174.9554     0   1E+4   4.5   5  RT (min)       lcqtof_t3_neg_1_064_14    m/z 174.9554     0   1E+4   4   4.5  RT (min)      lcqtof_t3_neg_1_064_14    m/z 174.9554     0   1E+5   0.5   1  RT (min)      lcqtof_t3_neg_2_058_15    m/z 174.9554     0   1   2E+4   8.5   9  RT (min)       lcqtof_t3_neg_1_064_14    m/z 174.9554     0   1E+4   10.5   11  RT (min)       lcqtof_t3_neg_4_081_58    m/z 174.9554     0   1E+4   5.5   6  RT (min)       lcqtof_t3_neg_1_064_14    m/z 174.9554     0   1E+4   11   11.5  RT (min)       lcqtof_t3_neg_1_064_14    m/z 174.9554     0   1E+4   18.5   19  RT (min)       lcqtof_t3_neg_1_064_14    m/z 174.9554     0   1E+4   6.5   7  RT (min)       lcqtof_t3_neg_4_080_58    m/z 174.9554     0   1   2E+4   9   9.5  RT (min)       lcqtof_t3_neg_1_064_14    m/z 174.9554     0   1E+4   18.5   19  RT (min)       lcqtof_t3_neg_2_058_15    m/z 174.9554     0   1E+4   2.5   3  RT (min)       lcqtof_t3_neg_1_064_14    m/z 174.9554     0   1   2   3E+4   17.5   18  RT (min)       lcqtof_t3_neg_1_036_14    m/z 174.9554     0   1   2E+4   18   18.5  RT (min)       lcqtof_t3_neg_3_079_58    m/z 174.9554     0   1   2E+4   8   8.5  RT (min)       lcqtof_t3_neg_1_064_14    m/z 174.9554     0   1   2E+4   18   18.5  RT (min)       lcqtof_t3_neg_3_081_58    m/z 174.9554     0   1E+4   12   12.5  RT (min)       lcqtof_t3_neg_3_077_58    m/z 174.9554     0   1   2E+4   7   8  RT (min)       lcqtof_t3_neg_1_036_14    m/z 174.9554     0   1E+4   1.5   2  RT (min)       lcqtof_t3_neg_1_036_14    m/z 174.9554     0   1E+4   10   10.5  RT (min)       lcqtof_t3_neg_1_064_14    m/z 174.9554     0   1E+4   6.5   7  RT (min)       lcqtof_t3_neg_4_076_58    m/z 174.9554     0   1   2E+4   8.5   9  RT (min)       lcqtof_t3_neg_1_064_14    m/z 174.9554     0   1   2E+4   7   7.5  RT (min)       lcqtof_t3_neg_1_064_14    m/z 174.9554     0   1E+4   18.5   19  RT (min)       lcqtof_t3_neg_3_078_58    m/z 174.9554     0   1E+4   0   0.5  RT (min)       lcqtof_t3_neg_2_059_14    m/z 174.9554     0   1E+4   5.5   6  RT (min)       lcqtof_t3_neg_2_062_17    m/z 174.9555     0   1   2E+4   9.5   10  RT (min)       lcqtof_t3_neg_1_036_14    m/z 174.9555     0   1E+4   8   8.5  RT (min)      lcqtof_t3_neg_4_058_16    m/z 174.9555     0   1   2E+4   10   10.5  RT (min)       lcqtof_t3_neg_1_036_14    m/z 174.9555     0   1   2E+4   8.5   9  RT (min)       lcqtof_t3_neg_1_064_14    m/z 174.9555     0   2   4E+4   1.5   2  RT (min)       lcqtof_t3_neg_3_000_aa    m/z 174.9555     0   1E+4   10.5   11  RT (min)       lcqtof_t3_neg_1_064_14    m/z 174.9555     0   1E+4   0   0.5  RT (min)       lcqtof_t3_neg_2_059_14    m/z 174.9555     0   1E+4   8   8.5  RT (min)      lcqtof_t3_neg_4_058_16    m/z 174.9555     0   1E+4   5   5.5  RT (min)      lcqtof_t3_neg_2_059_14    m/z 174.9555     0   1E+4   1.5   2  RT (min)       lcqtof_t3_neg_1_036_14    m/z 174.9555     0   1E+4   11   11.5  RT (min)       lcqtof_t3_neg_1_064_14    m/z 174.9555     0   1E+4   2.5   3  RT (min)       lcqtof_t3_neg_3_076_58    m/z 174.9555     0   1E+4   12.5   13  RT (min)       lcqtof_t3_neg_1_036_14    m/z 174.9555     0   1E+4   2   2.5  RT (min)       lcqtof_t3_neg_2_062_17    m/z 174.9555     0   1   2E+4   9   9.5  RT (min)       lcqtof_t3_neg_1_036_14    m/z 174.9555     0   1E+4   5   5.5  RT (min)       lcqtof_t3_neg_2_059_14    m/z 174.9555     0   1E+4   5.5   6  RT (min)       lcqtof_t3_neg_1_064_14    m/z 174.9555     0   1E+5   1.5   2  RT (min)       lcqtof_t3_neg_1_036_14    m/z 174.9555     0   1E+4   11.5   12  RT (min)       lcqtof_t3_neg_1_064_14    m/z 174.9555     0   1E+4   2   2.5  RT (min)       lcqtof_t3_neg_1_036_14    m/z 174.9555     0   1E+4   5.5   6  RT (min)       lcqtof_t3_neg_1_064_14    m/z 174.9555     0   1E+4   0   0.5  RT (min)       lcqtof_t3_neg_2_058_15    m/z 174.9556     0   1   2E+4   9.5   10  RT (min)       lcqtof_t3_neg_1_036_14    m/z 174.9556     0   1E+4   12   12.5  RT (min)       lcqtof_t3_neg_3_012_14    m/z 174.9556     0   1   2E+5   1.5   2  RT (min)      lcqtof_t3_neg_1_036_14    m/z 174.9556     0   1E+4   0   0.5  RT (min)       lcqtof_t3_neg_2_062_17    m/z 174.9556     0   1E+4   11.5   12  RT (min)       lcqtof_t3_neg_1_064_14    m/z 174.9556     0   1   2E+4   17.5   18  RT (min)       lcqtof_t3_neg_1_036_14    m/z 174.9556     0   1E+4   18.5   19  RT (min)       lcqtof_t3_neg_2_059_14    m/z 174.9556     0   1E+4   6.5   7  RT (min)       lcqtof_t3_neg_4_080_58    m/z 174.9556     0   1E+4   2   2.5  RT (min)       lcqtof_t3_neg_4_064_17    m/z 174.9556     0   1   2E+4   7   7.5  RT (min)       lcqtof_t3_neg_1_064_14    m/z 174.9556     0   1E+4   8.5   9  RT (min)      lcqtof_t3_neg_4_058_16    m/z 174.9556     0   1E+4   5   5.5  RT (min)       lcqtof_t3_neg_4_081_58    m/z 174.9556     0   1   2E+4   6   6.5  RT (min)       lcqtof_t3_neg_1_036_14    m/z 174.9556     0   1   2E+4   8   8.5  RT (min)      lcqtof_t3_neg_1_064_14    m/z 174.9556     0   1   2E+4   10   10.5  RT (min)       lcqtof_t3_neg_1_036_14    m/z 174.9556     0   1E+4   18.5   19  RT (min)       lcqtof_t3_neg_2_058_15    m/z 174.9556     0   1E+4   5.5   6  RT (min)      lcqtof_t3_neg_4_058_16    m/z 174.9556     0   1E+4   6   6.5  RT (min)       lcqtof_t3_neg_4_058_16    m/z 174.9556     0   1   2   3E+4   0.5   1  RT (min)       lcqtof_t3_neg_2_062_17    m/z 174.9557     0   1E+4   11   11.5  RT (min)       lcqtof_t3_neg_1_064_14    m/z 174.9557     0   1E+4   2.5   3  RT (min)       lcqtof_t3_neg_2_062_17    m/z 174.9557     0   1E+4   11   11.5  RT (min)      lcqtof_t3_neg_1_064_14    m/z 174.9557     0   1   2E+4   7   7.5  RT (min)       lcqtof_t3_neg_1_064_14    m/z 174.9557     0   1E+4   6   6.5  RT (min)       lcqtof_t3_neg_4_058_16    m/z 174.9557     0   1E+4   2   2.5  RT (min)       lcqtof_t3_neg_3_074_58    m/z 174.9558     0   1E+4   8.5   9  RT (min)      lcqtof_t3_neg_4_058_16    m/z 174.9559     0   1E+4   9.5   10  RT (min)      lcqtof_t3_neg_3_058_14    m/z 175.0088     0   1   2   3   4E+3   13.5   14  RT (min)      lcqtof_t3_neg_4_108_62    m/z 175.0113     0   1E+4   16.5   17  RT (min)      lcqtof_t3_neg_4_000_bb    m/z 175.0118     0   2   4   6E+3   12.5   13  RT (min)      lcqtof_t3_neg_4_045_02    m/z 175.0118     0   2   4   6E+3   12   12.5  RT (min)      lcqtof_t3_neg_4_000_ff    m/z 175.0121     0   2   4   6E+3   12   12.5  RT (min)      lcqtof_t3_neg_4_000_ff    m/z 175.0134     0   2   4   6E+3   10.5   11  RT (min)      lcqtof_t3_neg_4_000_ff    m/z 177.0045     0   2   4E+3   13.5   14  RT (min)       lcqtof_t3_neg_4_000_aa    m/z 177.0066     0   2   4   6E+3   12   13   14  RT (min)       lcqtof_t3_neg_4_046_33    m/z 177.0086     0   2   4E+3   12.5   13  RT (min)       lcqtof_t3_neg_4_032_04    m/z 177.0094     0   2   4E+3   11   12   13  RT (min)       lcqtof_t3_neg_4_056_56    m/z 177.0096     0   2   4E+3   12   13   14  RT (min)       lcqtof_t3_neg_4_040_24    m/z 177.0096     0   2   4   6E+3   12   13  RT (min)       lcqtof_t3_neg_4_000_ff    m/z 177.0096     0   1   2   3   4E+3   10.5   11  RT (min)      lcqtof_t3_neg_4_038_29    m/z 177.0097     0   2   4E+3   13   14  RT (min)       lcqtof_t3_neg_4_037_01    m/z 177.0098     0   2   4E+3   11.5   12  RT (min)      lcqtof_t3_neg_4_078_58    m/z 177.0106     0   1   2   3E+3   10   10.5  RT (min)      lcqtof_t3_neg_4_040_24    m/z 177.0927     0   1E+4   8.5   9  RT (min)      lcqtof_t3_neg_4_000_ee    m/z 178.1020     0   1E+4   8.5   9  RT (min)      lcqtof_t3_neg_4_000_ee    m/z 178.1020     0   1E+4   8.5   9  RT (min)      lcqtof_t3_neg_4_000_ee    m/z 178.8145     0   2   4   6   8E+3   1.5   2  RT (min)       lcqtof_t3_neg_4_007_09    m/z 178.9769     0   1   2   3E+4   16.5   17  RT (min)       lcqtof_t3_neg_1_008_02    m/z 178.9769     0   1   2E+4   17   17.5  RT (min)       lcqtof_t3_neg_1_002_13    m/z 178.9769     0   1   2   3E+4   17   17.5  RT (min)       lcqtof_t3_neg_1_014_04    m/z 178.9769     0   1   2   3E+4   16.5   17  RT (min)       lcqtof_t3_neg_1_059_17    m/z 178.9770     0   1   2E+4   17   17.5  RT (min)       lcqtof_t3_neg_3_059_13    m/z 178.9770     0   1E+4   18.5   19  RT (min)       lcqtof_t3_neg_1_060_15    m/z 178.9770     0   1   2E+4   17.5   18  RT (min)       lcqtof_t3_neg_1_062_13    m/z 178.9770     0   1   2E+4   17.5   18  RT (min)       lcqtof_t3_neg_3_104_61    m/z 178.9770     0   2   4   6   8E+3   18.5   19  RT (min)       lcqtof_t3_neg_1_059_17    m/z 178.9770     0   2   4   6   8E+3   0   0.5  RT (min)      lcqtof_t3_neg_1_025_52    m/z 178.9770     0   1   2   3E+4   17   17.5  RT (min)       lcqtof_t3_neg_1_059_17    m/z 178.9770     0   1   2   3E+4   17   17.5  RT (min)      lcqtof_t3_neg_1_059_17    m/z 178.9770     0   1E+4   0   0.5  RT (min)      lcqtof_t3_neg_1_015_34    m/z 178.9770     0   1E+4   18   18.5  RT (min)       lcqtof_t3_neg_1_060_15    m/z 178.9770     0   1   2E+4   17.5   18  RT (min)       lcqtof_t3_neg_1_062_13    m/z 178.9770     0   1E+4   0   0.5  RT (min)      lcqtof_t3_neg_1_000_dd    m/z 178.9770     0   1   2   3E+4   16.5   17  RT (min)       lcqtof_t3_neg_1_011_45    m/z 178.9770     0   1E+4   0   0.5  RT (min)      lcqtof_t3_neg_1_004_11    m/z 178.9771     0   2   4   6   8E+3   0.5   1  RT (min)      lcqtof_t3_neg_1_029_44    m/z 178.9771     0   1E+4   0   0.5  RT (min)      lcqtof_t3_neg_1_018_33    m/z 178.9771     0   1E+4   0   0.5  RT (min)      lcqtof_t3_neg_1_018_33    m/z 178.9771     0   1   2   3E+4   0.5   1  RT (min)      lcqtof_t3_neg_1_008_02    m/z 178.9771     0   2   4   6   8E+3   18.5   19  RT (min)       lcqtof_t3_neg_1_059_17    m/z 178.9771     0   1   2   3   4E+4   1   1.5  RT (min)      lcqtof_t3_neg_1_000_dd    m/z 178.9771     0   1E+4   0.5   1  RT (min)      lcqtof_t3_neg_1_009_50    m/z 178.9771     0   1E+4   0   0.5  RT (min)       lcqtof_t3_neg_1_018_33    m/z 178.9771     0   1   2E+4   17   17.5  RT (min)       lcqtof_t3_neg_3_094_60    m/z 178.9771     0   1E+4   18   18.5  RT (min)       lcqtof_t3_neg_1_060_15    m/z 178.9771     0   2   4   6E+4   2.5   5   7.5  RT (min)       lcqtof_t3_neg_1_028_20    m/z 178.9771     0   1E+4   18.5   19  RT (min)       lcqtof_t3_neg_1_064_14    m/z 178.9772     0   2   4   6E+3   18.5   19  RT (min)       lcqtof_t3_neg_3_107_62    m/z 178.9772     0   1   2   3E+4   0.5   1  RT (min)      lcqtof_t3_neg_1_011_45    m/z 178.9772     0   1   2   3E+4   0.5   1  RT (min)      lcqtof_t3_neg_1_011_45    m/z 178.9772     0   2   4   6E+4   0.5   1  RT (min)      lcqtof_t3_neg_4_000_ee    m/z 178.9772     0   2   4   6E+3   18.5   19  RT (min)       lcqtof_t3_neg_3_078_58    m/z 179.1393     0   1E+4   8   8.5  RT (min)      lcqtof_t3_neg_2_053_38    m/z 179.2071     0   1   2   3E+4   8   8.5  RT (min)      lcqtof_t3_neg_2_053_38    m/z 180.0661     0   2   4E+3   2.5   3  RT (min)      lcqtof_t3_neg_1_012_16    m/z 180.9219     0   1   2   3   4E+3   17.5   18  RT (min)      lcqtof_t3_neg_4_091_60    m/z 180.9301     0   1E+4   1   1.5  RT (min)      lcqtof_t3_neg_3_032_24    m/z 180.9720     0   2   4   6E+4   16.5   17  RT (min)      lcqtof_t3_neg_4_016_14    m/z 180.9724     0   2   4   6E+3   15   15.5  RT (min)      lcqtof_t3_neg_4_106_62    m/z 180.9726     0   2   4   6E+4   17   17.5  RT (min)      lcqtof_t3_neg_4_016_14    m/z 180.9726     0   2   4   6E+4   17   17.5  RT (min)      lcqtof_t3_neg_4_016_14    m/z 180.9726     0   2   4   6E+4   16.5   17  RT (min)      lcqtof_t3_neg_4_016_14    m/z 180.9728     0   1   2E+4   17   17.5  RT (min)       lcqtof_t3_neg_1_065_51    m/z 180.9729     0   1   2E+4   16.5   17  RT (min)       lcqtof_t3_neg_1_016_48    m/z 180.9729     0   2   4   6E+4   16.5   17  RT (min)       lcqtof_t3_neg_4_046_33    m/z 180.9729     0   1   2   3E+4   0.5   1  RT (min)      lcqtof_t3_neg_4_000_ee    m/z 180.9730     0   1   2   3E+4   0.5   1  RT (min)      lcqtof_t3_neg_4_000_ee    m/z 180.9730     0   2   4E+4   10   15  RT (min)       lcqtof_t3_neg_4_106_62    m/z 180.9731     0   1E+4   17   17.5  RT (min)      lcqtof_t3_neg_4_016_14    m/z 180.9732     0   1E+5   1.5   2  RT (min)      lcqtof_t3_neg_1_035_31    m/z 180.9732     0   1   2E+5   1.5   2  RT (min)      lcqtof_t3_neg_1_036_14    m/z 180.9745     0   2   4   6   8E+4   2   2.5  RT (min)      lcqtof_t3_neg_4_064_17    m/z 182.0459     0   2   4   6E+3   4   4.5  RT (min)       lcqtof_t3_neg_4_037_01    m/z 182.0600     0   1   2   3E+4   4   4.5  RT (min)      lcqtof_t3_neg_4_004_20    m/z 182.0839     0   1E+4   4.5   5  RT (min)      lcqtof_t3_neg_3_032_24    m/z 182.1286     0   2   4   6E+4   4   4.5  RT (min)      lcqtof_t3_neg_4_004_20    m/z 182.2327     0   1   2E+4   4   4.5  RT (min)      lcqtof_t3_neg_4_004_20    m/z 182.9204     0   1   2   3E+3   17   17.5  RT (min)       lcqtof_t3_neg_4_050_43    m/z 182.9540     0   2   4E+3   1   1.5  RT (min)      lcqtof_t3_neg_4_062_13    m/z 182.9541     0   2   4   6E+3   1   1.5  RT (min)      lcqtof_t3_neg_1_027_21    m/z 183.0113     0   1E+4   10   10.5  RT (min)       lcqtof_t3_neg_4_000_ee    m/z 183.0113     0   2   4E+3   9.5   10  RT (min)       lcqtof_t3_neg_4_007_09    m/z 183.0116     0   2   4   6E+3   14.5   15  RT (min)       lcqtof_t3_neg_4_000_dd    m/z 186.0863     0   1   2   3E+3   2.5   3  RT (min)      lcqtof_t3_neg_4_059_14    m/z 186.0865     0   1   2E+3   3   3.5  RT (min)      lcqtof_t3_neg_1_028_20    m/z 187.0462     0   1   2E+4   2   2.5  RT (min)      lcqtof_t3_neg_3_045_31    m/z 187.0973     0   1   2   3E+2   5.5   6  RT (min)      lcqtof_t3_neg_3_038_06    m/z 187.1459     0   1E+4   5   5.5  RT (min)      lcqtof_t3_neg_4_016_14    m/z 187.8886     0   1   2   3E+3   16.5   17  RT (min)      lcqtof_t3_neg_4_000_aa    m/z 187.9406     0   1   2   3   4E+3   15.5   16  RT (min)      lcqtof_t3_neg_4_105_61    m/z 187.9407     0   1   2E+4   16.5   17  RT (min)       lcqtof_t3_neg_4_000_cc    m/z 187.9410     0   1   2   3   4E+3   13   13.5  RT (min)      lcqtof_t3_neg_4_089_59    m/z 187.9442     0   1E+4   0.5   1  RT (min)      lcqtof_t3_neg_4_000_ee    m/z 187.9446     0   1E+4   0.5   1  RT (min)      lcqtof_t3_neg_4_000_ee    m/z 188.8877     0   1   2   3E+3   16.5   17  RT (min)      lcqtof_t3_neg_4_000_aa    m/z 188.9420     0   1   2   3E+3   16.5   17  RT (min)      lcqtof_t3_neg_4_002_17    m/z 189.0132     0   1E+3   16.5   17  RT (min)      lcqtof_t3_neg_4_010_46    m/z 189.0242     0   2   4   6   8E+3   16.5   17  RT (min)      lcqtof_t3_neg_4_028_25    m/z 190.0184     0   1E+4   13   13.5   14  RT (min)       lcqtof_t3_neg_4_000_ff    m/z 190.0185     0   2   4   6   8E+3   11   12   13  RT (min)       lcqtof_t3_neg_4_113_62    m/z 190.1945     0   1E+4   4.5   5  RT (min)      lcqtof_t3_neg_4_004_20    m/z 190.8880     0   1   2   3   4E+3   16.5   17  RT (min)      lcqtof_t3_neg_4_000_ee    m/z 190.8881     0   1   2   3E+3   16.5   17  RT (min)      lcqtof_t3_neg_3_041_54    m/z 190.9280     0   1E+4   18.5   19  RT (min)       lcqtof_t3_neg_4_063_39    m/z 190.9281     0   2   4   6E+3   17   17.5  RT (min)       lcqtof_t3_neg_4_107_62    m/z 190.9282     0   1E+4   17.5   18  RT (min)       lcqtof_t3_neg_4_000_ee    m/z 190.9282     0   2   4   6E+4   1   1.5  RT (min)      lcqtof_t3_neg_4_000_ee    m/z 190.9286     0   2   4   6E+4   1   1.5  RT (min)       lcqtof_t3_neg_4_000_ee    m/z 190.9286     0   1E+4   17.5   18  RT (min)       lcqtof_t3_neg_4_061_50    m/z 190.9288     0   2   4   6E+3   17.5   18  RT (min)       lcqtof_t3_neg_4_072_57    m/z 190.9292     0   1E+4   1   1.5  RT (min)       lcqtof_t3_neg_4_000_ee    m/z 190.9300     0   1E+4   17   17.5  RT (min)       lcqtof_t3_neg_4_060_15    m/z 190.9300     0   1E+4   6   7  RT (min)       lcqtof_t3_neg_4_060_15    m/z 190.9302     0   1E+4   4   6   8  RT (min)       lcqtof_t3_neg_4_064_17    m/z 190.9303     0   1E+4   5   5.5  RT (min)       lcqtof_t3_neg_4_000_ff    m/z 190.9304     0   1E+4   4   4.5  RT (min)       lcqtof_t3_neg_4_063_39    m/z 190.9310     0   2   4   6E+4   0.5   1  RT (min)       lcqtof_t3_neg_4_000_ee    m/z 191.0071     0   2   4   6E+3   14   14.5  RT (min)      lcqtof_t3_neg_4_012_27    m/z 191.0148     0   2   4   6   8E+3   12   13  RT (min)       lcqtof_t3_neg_4_104_61    m/z 191.0149     0   1E+4   12   13  RT (min)       lcqtof_t3_neg_4_000_ff    m/z 191.0151     0   1E+4   12   14  RT (min)       lcqtof_t3_neg_4_000_dd    m/z 191.0156     0   1E+4   12  RT (min)       lcqtof_t3_neg_4_041_54    m/z 191.0190     0   2   4   6   8E+5   1.5   2  RT (min)       lcqtof_t3_neg_4_059_14    m/z 191.0191     0   1   2   3E+5   1.5   2  RT (min)      lcqtof_t3_neg_2_043_31    m/z 191.0193     0   1   2E+5   1.5   2  RT (min)      lcqtof_t3_neg_1_033_53    m/z 191.0900     0   1E+4   1   1.5  RT (min)      lcqtof_t3_neg_4_016_14    m/z 191.9456     0   1   2   3   4E+3   1   1.5  RT (min)      lcqtof_t3_neg_1_036_14    m/z 191.9459     0   1E+4   8   8.5  RT (min)       lcqtof_t3_neg_1_014_04    m/z 192.1167     0   1   2E+4   9   9.5  RT (min)      lcqtof_t3_neg_4_000_ee    m/z 192.1183     0   1   2E+4   9   9.5  RT (min)      lcqtof_t3_neg_4_000_ee    m/z 194.0455     0   1   2E+4   1   1.5  RT (min)      lcqtof_t3_neg_1_031_24    m/z 194.0812     0   1E+2   7.5   8  RT (min)      lcqtof_t3_neg_3_038_06    m/z 194.0817     0   1E+2   7.5   8  RT (min)      lcqtof_t3_neg_3_038_06    m/z 194.0820     0   2   4   6E+2   8.5   9  RT (min)      lcqtof_t3_neg_3_032_24    m/z 194.0824     0   1   2   3E+4   9   9.5  RT (min)      lcqtof_t3_neg_4_000_ee    m/z 194.0833     0   2   4   6E+2   8.5   9  RT (min)      lcqtof_t3_neg_3_032_24    m/z 195.0539     0   1   2   3E+4   0.5   1  RT (min)      lcqtof_t3_neg_4_000_ee    m/z 195.8101     0   1E+4   14.5   15  RT (min)       lcqtof_t3_neg_4_074_58    m/z 195.8102     0   1E+4   14   14.5  RT (min)       lcqtof_t3_neg_4_107_62    m/z 195.8103     0   2   4   6E+3   12.5   13  RT (min)       lcqtof_t3_neg_3_113_62    m/z 195.8103     0   1E+4   14   14.5  RT (min)       lcqtof_t3_neg_4_018_23    m/z 195.8104     0   2   4   6E+3   13   13.5  RT (min)       lcqtof_t3_neg_3_101_61    m/z 195.8105     0   1   2   3E+3   12   12.5  RT (min)       lcqtof_t3_neg_4_076_58    m/z 195.8106     0   1   2   3E+3   11.5   12  RT (min)       lcqtof_t3_neg_4_034_34    m/z 195.8106     0   2   4   6E+3   12.5   13  RT (min)       lcqtof_t3_neg_3_100_61    m/z 195.8106     0   1E+4   14.5   15  RT (min)       lcqtof_t3_neg_4_029_40    m/z 195.8106     0   2   4   6E+3   13   13.5  RT (min)       lcqtof_t3_neg_3_106_62    m/z 197.0392     0   1E+5   1.5   2  RT (min)       lcqtof_t3_neg_4_046_33    m/z 197.8072     0   2   4   6   8E+3   13   13.5  RT (min)       lcqtof_t3_neg_3_106_62    m/z 197.8072     0   1E+4   14.5   15  RT (min)       lcqtof_t3_neg_4_029_40    m/z 197.8072     0   2   4   6   8E+3   13   13.5  RT (min)       lcqtof_t3_neg_3_107_62    m/z 197.8073     0   1E+4   14   14.5  RT (min)       lcqtof_t3_neg_4_059_14    m/z 197.8073     0   1E+4   14   14.5  RT (min)       lcqtof_t3_neg_4_034_34    m/z 197.8074     0   1   2   3   4E+3   11.5   12  RT (min)       lcqtof_t3_neg_3_076_58    m/z 197.8074     0   2   4E+3   12   12.5  RT (min)       lcqtof_t3_neg_3_080_58    m/z 197.8074     0   2   4   6   8E+3   12.5   13  RT (min)       lcqtof_t3_neg_3_100_61    m/z 197.8075     0   2   4   6   8E+3   12.5   13  RT (min)       lcqtof_t3_neg_3_101_61    m/z 197.8075     0   1E+4   14   14.5  RT (min)       lcqtof_t3_neg_4_059_14    m/z 197.8075     0   1E+4   14.5   15  RT (min)       lcqtof_t3_neg_4_059_14    m/z 197.8076     0   1E+4   14   14.5  RT (min)       lcqtof_t3_neg_4_059_14    m/z 197.9627     0   2   4E+3   16   16.5  RT (min)       lcqtof_t3_neg_4_011_32    m/z 197.9627     0   1   2   3   4E+3   15   15.5  RT (min)       lcqtof_t3_neg_3_088_59    m/z 197.9627     0   1   2   3   4E+3   15   16  RT (min)       lcqtof_t3_neg_4_096_60    m/z 197.9628     0   2   4   6E+3   12   14   16  RT (min)       lcqtof_t3_neg_1_054_41    m/z 198.9357     0   1   2E+3   2   4  RT (min)       lcqtof_t3_neg_4_000_ff    m/z 198.9360     0   1   2E+3   18.5   19  RT (min)       lcqtof_t3_neg_4_088_59    m/z 198.9361     0   1   2E+3   18   18.5  RT (min)       lcqtof_t3_neg_4_104_61    m/z 198.9362     0   1   2E+3   0   0.5  RT (min)       lcqtof_t3_neg_4_073_57    m/z 199.0058     0   1   2E+4   5   5.5  RT (min)      lcqtof_t3_neg_1_000_aa    m/z 199.0063     0   1E+2   3.5   4  RT (min)      lcqtof_t3_neg_3_038_06    m/z 199.0065     0   1   2E+4   4   4.5  RT (min)      lcqtof_t3_neg_3_032_24    m/z 199.8046     0   2   4   6   8E+3   14   14.5  RT (min)       lcqtof_t3_neg_4_031_42    m/z 199.8046     0   2   4   6   8E+3   14   14.5  RT (min)       lcqtof_t3_neg_4_034_34    m/z 199.8048     0   1   2   3   4E+3   12.5   13  RT (min)       lcqtof_t3_neg_3_100_61    m/z 199.8049     0   1   2   3   4E+3   12.5   13  RT (min)      lcqtof_t3_neg_3_107_62    m/z 199.8050     0   2   4   6E+3   14.5   15  RT (min)       lcqtof_t3_neg_4_105_61    m/z 201.0212     0   1E+4   17   17.5  RT (min)      lcqtof_t3_neg_4_035_08    m/z 201.0227     0   1E+4   16   16.5  RT (min)      lcqtof_t3_neg_4_018_23    m/z 202.0193     0   1   2   3E+4   17   17.5  RT (min)      lcqtof_t3_neg_4_035_08    m/z 202.0199     0   1   2   3E+4   16   16.5  RT (min)      lcqtof_t3_neg_4_018_23    m/z 202.0269     0   1   2   3   4E+3   12   12.5  RT (min)      lcqtof_t3_neg_1_061_50    m/z 202.0269     0   1   2   3   4E+3   10.5   11  RT (min)      lcqtof_t3_neg_3_022_27    m/z 202.0270     0   1   2   3   4E+3   12.5   13  RT (min)      lcqtof_t3_neg_1_061_50    m/z 202.0277     0   2   4   6   8E+3   14   14.5  RT (min)      lcqtof_t3_neg_4_035_08    m/z 202.0277     0   1   2   3E+3   13   13.5  RT (min)      lcqtof_t3_neg_1_014_04    m/z 202.0292     0   2   4   6   8E+3   14   14.5  RT (min)      lcqtof_t3_neg_4_000_aa    m/z 202.9395     0   1   2   3E+3   16.5   17  RT (min)      lcqtof_t3_neg_4_000_bb    m/z 203.0183     0   1   2E+4   11.5   12  RT (min)      lcqtof_t3_neg_4_090_60    m/z 203.0184     0   1E+4   9.5   10  RT (min)      lcqtof_t3_neg_4_090_60    m/z 203.0184     0   2   4   6E+4   16   16.5  RT (min)      lcqtof_t3_neg_4_018_23    m/z 203.0188     0   1   2E+4   10.5   11  RT (min)      lcqtof_t3_neg_4_090_60    m/z 203.0190     0   1E+4   10   10.5  RT (min)      lcqtof_t3_neg_4_090_60    m/z 203.0192     0   1E+4   9   9.5  RT (min)      lcqtof_t3_neg_4_090_60    m/z 203.0194     0   1E+2   15   15.5  RT (min)      lcqtof_t3_neg_3_032_24    m/z 203.0195     0   1E+2   16   16.5  RT (min)      lcqtof_t3_neg_2_030_23    m/z 203.0200     0   1E+4   17   17.5  RT (min)      lcqtof_t3_neg_4_035_08    m/z 203.0200     0   2   4   6E+4   17   17.5  RT (min)      lcqtof_t3_neg_4_035_08    m/z 203.0204     0   2   4   6E+4   16.5   17  RT (min)      lcqtof_t3_neg_4_031_42    m/z 203.0233     0   1E+4   13   13.5  RT (min)      lcqtof_t3_neg_4_090_60    m/z 203.0252     0   1   2   3   4E+3   13.5   14  RT (min)      lcqtof_t3_neg_3_038_06    m/z 203.0263     0   1E+4   14   14.5  RT (min)      lcqtof_t3_neg_4_090_60    m/z 203.0272     0   1   2   3E+3   14   14.5  RT (min)      lcqtof_t3_neg_1_007_03    m/z 204.0165     0   1   2E+2   15   15.5  RT (min)      lcqtof_t3_neg_3_032_24    m/z 204.0168     0   1   2   3   4E+4   14  RT (min)       lcqtof_t3_neg_4_084_59    m/z 204.0168     0   2   4   6   8E+1   16   16.5  RT (min)      lcqtof_t3_neg_2_030_23    m/z 204.0168     0   1   2   3   4E+4   13.5   14  RT (min)      lcqtof_t3_neg_4_109_62    m/z 204.0169     0   1E+4   7.5   8  RT (min)       lcqtof_t3_neg_4_083_59    m/z 204.0169     0   1   2E+4   8   8.5   9  RT (min)       lcqtof_t3_neg_4_112_62    m/z 204.0169     0   1   2   3   4E+4   9   10  RT (min)       lcqtof_t3_neg_4_096_60    m/z 204.0170     0   1   2   3E+4   9.5   10  RT (min)      lcqtof_t3_neg_4_084_59    m/z 204.0170     0   1E+5   16.5   17  RT (min)      lcqtof_t3_neg_4_043_11    m/z 204.0170     0   2   4E+4   7.5   10   12.5  RT (min)       lcqtof_t3_neg_4_080_58    m/z 204.0170     0   2   4E+4   10.5   11  RT (min)      lcqtof_t3_neg_4_068_57    m/z 204.0170     0   1E+4   7   7.5  RT (min)       lcqtof_t3_neg_4_111_62    m/z 204.0170     0   1   2   3   4E+4   13.5   14  RT (min)      lcqtof_t3_neg_4_099_61    m/z 204.0170     0   1   2   3   4E+4   9.5   10  RT (min)      lcqtof_t3_neg_4_093_60    m/z 204.0170     0   1   2   3E+4   9   9.5  RT (min)      lcqtof_t3_neg_4_068_57    m/z 204.0170     0   1   2E+4   8   8.5  RT (min)      lcqtof_t3_neg_4_068_57    m/z 204.0170     0   1E+4   10   10.5  RT (min)       lcqtof_t3_neg_3_096_60    m/z 204.0170     0   1   2   3E+4   14   14.5  RT (min)       lcqtof_t3_neg_4_093_60    m/z 204.0171     0   2   4   6E+4   12   14  RT (min)       lcqtof_t3_neg_4_016_14    m/z 204.0171     0   2   4   6E+4   7.5   10   12.5  RT (min)       lcqtof_t3_neg_4_059_14    m/z 204.0171     0   1E+4   7.5   8  RT (min)       lcqtof_t3_neg_4_099_61    m/z 204.0171     0   2   4   6E+4   8   10   12  RT (min)       lcqtof_t3_neg_4_004_20    m/z 204.0171     0   2   4   6E+4   12   14  RT (min)       lcqtof_t3_neg_4_005_18    m/z 204.0172     0   2   4E+4   12.5   13  RT (min)      lcqtof_t3_neg_4_084_59    m/z 204.0172     0   2   4E+4   10   11  RT (min)       lcqtof_t3_neg_4_078_58    m/z 204.0172     0   1   2   3   4E+4   13   13.5  RT (min)      lcqtof_t3_neg_4_106_62    m/z 204.0172     0   1   2   3   4E+4   12.5   13  RT (min)      lcqtof_t3_neg_4_107_62    m/z 204.0172     0   1E+5   17   17.5  RT (min)      lcqtof_t3_neg_4_035_08    m/z 204.0174     0   1   2   3   4E+4   17   17.5  RT (min)      lcqtof_t3_neg_4_035_08    m/z 204.0176     0   1E+5   16   16.5  RT (min)      lcqtof_t3_neg_4_018_23    m/z 204.9389     0   1   2   3E+3   16.5   17  RT (min)       lcqtof_t3_neg_4_000_aa    m/z 204.9393     0   1   2E+3   16   16.5  RT (min)       lcqtof_t3_neg_3_002_46    m/z 204.9694     0   1   2   3   4E+3   15   15.5  RT (min)       lcqtof_t3_neg_4_077_58    m/z 204.9731     0   1   2E+3   16.5   17  RT (min)      lcqtof_t3_neg_4_076_58    m/z 204.9736     0   1E+4   16.5   17  RT (min)       lcqtof_t3_neg_4_000_dd    m/z 205.0090     0   2   4   6E+4   17   17.5  RT (min)      lcqtof_t3_neg_4_035_08    m/z 205.0130     0   1E+4   10.5   11  RT (min)       lcqtof_t3_neg_3_076_58    m/z 205.0131     0   1E+4   7   7.5  RT (min)       lcqtof_t3_neg_4_082_59    m/z 205.0131     0   1   2E+4   7   7.5  RT (min)       lcqtof_t3_neg_4_082_59    m/z 205.0131     0   1   2   3E+4   8.5   9  RT (min)       lcqtof_t3_neg_4_095_60    m/z 205.0131     0   1   2   3E+4   8   8.5  RT (min)       lcqtof_t3_neg_4_105_61    m/z 205.0132     0   1E+4   7   7.5  RT (min)       lcqtof_t3_neg_4_096_60    m/z 205.0132     0   2   4   6E+4   9.5   10  RT (min)       lcqtof_t3_neg_4_092_60    m/z 205.0132     0   2   4E+4   9   9.5  RT (min)       lcqtof_t3_neg_4_085_59    m/z 205.0132     0   1   2E+4   7.5   8  RT (min)       lcqtof_t3_neg_4_085_59    m/z 205.0132     0   1   2   3E+4   8.5   9  RT (min)       lcqtof_t3_neg_4_109_62    m/z 205.0132     0   1E+2   16   16.5  RT (min)      lcqtof_t3_neg_1_019_23    m/z 205.0132     0   2   4   6   8E+4   12   12.5  RT (min)       lcqtof_t3_neg_4_111_62    m/z 205.0133     0   2   4E+4   9   9.5  RT (min)       lcqtof_t3_neg_4_085_59    m/z 205.0133     0   1   2E+5   16   16.5  RT (min)      lcqtof_t3_neg_4_018_23    m/z 205.0133     0   1E+5   17   17.5  RT (min)       lcqtof_t3_neg_4_035_08    m/z 205.0133     0   2   4   6E+4   10   10.5  RT (min)       lcqtof_t3_neg_4_096_60    m/z 205.0133     0   1   2   3   4E+4   9   9.5  RT (min)       lcqtof_t3_neg_4_077_58    m/z 205.0133     0   2   4   6   8E+4   11.5   12  RT (min)       lcqtof_t3_neg_4_092_60    m/z 205.0134     0   2   4   6E+4   10   10.5  RT (min)       lcqtof_t3_neg_4_090_60    m/z 205.0134     0   2   4   6E+4   13   13.5  RT (min)      lcqtof_t3_neg_4_110_62    m/z 205.0134     0   2   4   6E+4   10.5   11  RT (min)       lcqtof_t3_neg_4_073_57    m/z 205.0134     0   2   4   6E+4   13.5   14  RT (min)       lcqtof_t3_neg_4_089_59    m/z 205.0134     0   2   4E+4   13.5   14  RT (min)       lcqtof_t3_neg_4_067_57    m/z 205.0134     0   2   4   6   8E+4   12   12.5  RT (min)       lcqtof_t3_neg_4_059_14    m/z 205.0134     0   1   2   3E+4   8   8.5  RT (min)       lcqtof_t3_neg_4_071_57    m/z 205.0134     0   1   2   3E+4   8.5   9  RT (min)       lcqtof_t3_neg_4_098_61    m/z 205.0134     0   2   4   6E+4   12.5   13  RT (min)       lcqtof_t3_neg_4_071_57    m/z 205.0134     0   2   4   6E+4   13   13.5  RT (min)       lcqtof_t3_neg_4_104_61    m/z 205.0134     0   1   2E+4   7.5   8  RT (min)       lcqtof_t3_neg_4_099_61    m/z 205.0134     0   2   4   6   8E+4   12   13  RT (min)       lcqtof_t3_neg_4_070_57    m/z 205.0134     0   2   4   6E+4   10.5   11  RT (min)       lcqtof_t3_neg_4_073_57    m/z 205.0134     0   2   4   6   8E+4   12   12.5  RT (min)       lcqtof_t3_neg_4_082_59    m/z 205.0134     0   2   4   6E+4   10   10.5  RT (min)       lcqtof_t3_neg_4_073_57    m/z 205.0135     0   1   2E+2   15   15.5  RT (min)      lcqtof_t3_neg_3_032_24    m/z 205.0135     0   2   4   6E+4   13   13.5  RT (min)       lcqtof_t3_neg_4_106_62    m/z 205.0135     0   1   2   3E+4   8.5   9  RT (min)      lcqtof_t3_neg_4_113_62    m/z 205.0135     0   1   2   3E+4   7.5   8  RT (min)       lcqtof_t3_neg_4_067_57    m/z 205.0135     0   1E+4   6.5   7  RT (min)       lcqtof_t3_neg_4_092_60    m/z 205.0135     0   2   4   6   8E+4   11   11.5  RT (min)       lcqtof_t3_neg_4_006_16    m/z 205.0135     0   2   4   6   8E+4   11.5   12  RT (min)       lcqtof_t3_neg_4_000_bb    m/z 205.0135     0   2   4   6   8E+4   12   12.5  RT (min)       lcqtof_t3_neg_4_000_dd    m/z 205.0136     0   2   4E+4   9.5   10  RT (min)      lcqtof_t3_neg_4_103_61    m/z 205.0136     0   2   4   6E+4   13.5   14  RT (min)       lcqtof_t3_neg_4_099_61    m/z 205.0137     0   1E+4   10.5   11  RT (min)       lcqtof_t3_neg_3_076_58    m/z 205.0137     0   2   4   6E+4   11.5   12  RT (min)       lcqtof_t3_neg_4_090_60    m/z 205.0138     0   2   4E+4   14   14.5  RT (min)       lcqtof_t3_neg_4_110_62    m/z 205.0138     0   2   4E+4   9.5   10  RT (min)       lcqtof_t3_neg_4_087_59    m/z 205.0349     0   1E+4   1   1.5  RT (min)      lcqtof_t3_neg_4_003_37    m/z 205.0518     0   1   2   3   4E+3   13   14  RT (min)       lcqtof_t3_neg_4_016_14    m/z 205.0520     0   1E+4   16   16.5  RT (min)      lcqtof_t3_neg_4_018_23    m/z 205.0526     0   1E+4   17   17.5  RT (min)      lcqtof_t3_neg_4_035_08    m/z 206.0164     0   1E+4   17   17.5  RT (min)      lcqtof_t3_neg_4_035_08    m/z 206.0173     0   1E+4   16   16.5  RT (min)      lcqtof_t3_neg_4_018_23    m/z 206.9718     0   1   2   3E+4   17.5   18  RT (min)       lcqtof_t3_neg_4_035_08    m/z 206.9719     0   1E+4   17.5   18  RT (min)       lcqtof_t3_neg_2_064_13    m/z 206.9719     0   1E+4   18   18.5  RT (min)       lcqtof_t3_neg_4_067_57    m/z 206.9719     0   1   2E+4   17   17.5  RT (min)       lcqtof_t3_neg_2_060_39    m/z 206.9719     0   2   4   6   8E+3   18.5   19  RT (min)       lcqtof_t3_neg_4_068_57    m/z 206.9720     0   1E+4   17.5   18  RT (min)       lcqtof_t3_neg_4_091_60    m/z 206.9720     0   1   2E+4   16.5   17  RT (min)      lcqtof_t3_neg_2_060_39    m/z 206.9720     0   1   2E+4   16.5   17  RT (min)       lcqtof_t3_neg_3_090_60    m/z 206.9721     0   2   4E+4   16.5   17  RT (min)       lcqtof_t3_neg_4_018_23    m/z 206.9721     0   2   4E+4   0.5   1  RT (min)       lcqtof_t3_neg_4_011_32    m/z 206.9721     0   1   2   3E+4   17   17.5  RT (min)       lcqtof_t3_neg_4_044_48    m/z 206.9721     0   1E+4   0   0.5  RT (min)       lcqtof_t3_neg_4_011_32    m/z 206.9721     0   2   4   6E+4   1   1.5  RT (min)       lcqtof_t3_neg_1_028_20    m/z 206.9721     0   1E+4   0   0.5  RT (min)       lcqtof_t3_neg_4_011_32    m/z 206.9721     0   2   4   6   8E+4   10  RT (min)       lcqtof_t3_neg_4_000_bb    m/z 206.9721     0   2   4E+4   1   1.5  RT (min)      lcqtof_t3_neg_1_026_19    m/z 206.9721     0   1E+4   0   0.5  RT (min)       lcqtof_t3_neg_4_011_32    m/z 206.9722     0   1E+4   0   0.5  RT (min)      lcqtof_t3_neg_4_011_32    m/z 206.9722     0   2   4E+4   0.5   1  RT (min)       lcqtof_t3_neg_4_011_32    m/z 206.9722     0   1E+4   0   0.5   1  RT (min)       lcqtof_t3_neg_4_011_32    m/z 206.9722     0   2   4   6   8E+3   0.5   1  RT (min)      lcqtof_t3_neg_4_033_53    m/z 206.9722     0   2   4E+4   0.5   1  RT (min)       lcqtof_t3_neg_4_011_32    m/z 206.9722     0   1E+4   0   0.5  RT (min)       lcqtof_t3_neg_4_011_32    m/z 206.9723     0   2   4E+4   0.5   1  RT (min)       lcqtof_t3_neg_4_000_ee    m/z 206.9726     0   1E+4   4   4.5  RT (min)       lcqtof_t3_neg_4_000_aa    m/z 207.0388     0   1E+4   4   4.5  RT (min)      lcqtof_t3_neg_4_051_15    m/z 207.1024     0   1E+4   8   8.5  RT (min)      lcqtof_t3_neg_4_000_ee    m/z 207.1527     0   1E+4   6.5   7  RT (min)      lcqtof_t3_neg_4_000_ee    m/z 207.1527     0   1E+4   7   7.5  RT (min)      lcqtof_t3_neg_4_000_ee    m/z 207.1582     0   1E+4   8   8.5  RT (min)      lcqtof_t3_neg_4_000_ee    m/z 207.9754     0   2   4E+3   1   1.5  RT (min)      lcqtof_t3_neg_4_014_13    m/z 208.0447     0   1E+4   4   4.5  RT (min)      lcqtof_t3_neg_4_006_16    m/z 208.1370     0   1E+4   7   7.5  RT (min)      lcqtof_t3_neg_4_000_ee    m/z 208.1371     0   1E+4   7   7.5  RT (min)      lcqtof_t3_neg_4_000_ee    m/z 208.1372     0   1E+4   7   7.5  RT (min)      lcqtof_t3_neg_4_000_ee    m/z 208.9361     0   1   2   3E+3   10   10.5  RT (min)      lcqtof_t3_neg_1_036_14    m/z 209.0827     0   1E+4   6   6.5  RT (min)      lcqtof_t3_neg_4_000_ee    m/z 210.9490     0   1   2E+4   1   1.5  RT (min)      lcqtof_t3_neg_4_000_ee    m/z 210.9491     0   1   2E+4   1   1.5  RT (min)       lcqtof_t3_neg_4_062_13    m/z 210.9492     0   1E+4   4.5   5  RT (min)       lcqtof_t3_neg_4_006_16    m/z 210.9492     0   2   4E+3   1.5   2  RT (min)      lcqtof_t3_neg_1_010_27    m/z 210.9494     0   2   4E+3   1.5   2  RT (min)      lcqtof_t3_neg_1_009_50    m/z 210.9494     0   1   2E+4   0.5   1  RT (min)      lcqtof_t3_neg_4_000_ee    m/z 210.9495     0   1E+4   1   1.5  RT (min)      lcqtof_t3_neg_1_005_18    m/z 212.0585     0   1E+4   1   1.5  RT (min)      lcqtof_t3_neg_4_011_32    m/z 213.0144     0   1   2   3E+4   6   6.5  RT (min)      lcqtof_t3_neg_3_032_24    m/z 213.0145     0   1   2E+2   5   5.5  RT (min)      lcqtof_t3_neg_3_038_06    m/z 213.0150     0   1   2   3   4E+4   6.5   7  RT (min)       lcqtof_t3_neg_1_048_35    m/z 214.1189     0   1E+4   4   4.5  RT (min)      lcqtof_t3_neg_3_032_24    m/z 214.1192     0   2   4   6   8E+1   3.5   4  RT (min)      lcqtof_t3_neg_3_038_06    m/z 215.0746     0   1   2   3   4E+3   16.5   17  RT (min)      lcqtof_t3_neg_4_000_aa    m/z 215.1477     0   1E+4   6.5   7  RT (min)      lcqtof_t3_neg_4_000_ee    m/z 215.1496     0   1E+4   6.5   7  RT (min)      lcqtof_t3_neg_4_000_ee    m/z 217.0358     0   1E+4   16   16.5  RT (min)      lcqtof_t3_neg_4_018_23    m/z 217.0654     0   1E+4   16   16.5  RT (min)      lcqtof_t3_neg_4_018_23    m/z 218.0315     0   1E+4   12   12.5   13  RT (min)       lcqtof_t3_neg_4_046_33    m/z 218.0318     0   1E+4   13   13.5  RT (min)       lcqtof_t3_neg_4_005_18    m/z 218.0321     0   1E+4   17   17.5  RT (min)      lcqtof_t3_neg_4_035_08    m/z 218.0324     0   2   4   6   8E+3   12.5   13   13.5  RT (min)       lcqtof_t3_neg_4_078_58    m/z 219.0283     0   2   4   6   8E+3   1   1.5  RT (min)      lcqtof_t3_neg_1_039_43    m/z 219.0291     0   1E+4   13  RT (min)       lcqtof_t3_neg_4_000_ff    m/z 219.0292     0   1E+4   12   13  RT (min)       lcqtof_t3_neg_4_046_33    m/z 219.0295     0   1   2E+4   16.5   17  RT (min)      lcqtof_t3_neg_4_091_60    m/z 219.0314     0   1E+4   13.5   14  RT (min)       lcqtof_t3_neg_4_000_dd    m/z 219.0315     0   1E+4   13   13.5  RT (min)       lcqtof_t3_neg_4_107_62    m/z 219.0323     0   1E+4   13   13.5  RT (min)       lcqtof_t3_neg_4_000_cc    m/z 219.0348     0   1   2E+3   13.5   14  RT (min)      lcqtof_t3_neg_2_031_02    m/z 219.1534     0   1E+4   6.5   7  RT (min)      lcqtof_t3_neg_4_000_ee    m/z 220.0598     0   1E+4   1   1.5  RT (min)      lcqtof_t3_neg_2_028_24    m/z 220.8136     0   1E+4   1   1.5  RT (min)      lcqtof_t3_neg_1_010_27    m/z 221.0242     0   1   2   3E+3   17   17.5  RT (min)       lcqtof_t3_neg_4_035_08    m/z 221.0245     0   1   2   3E+3   16.5   17  RT (min)       lcqtof_t3_neg_1_004_11    m/z 221.0811     0   1E+4   7.5   8  RT (min)      lcqtof_t3_neg_4_000_ee    m/z 221.1299     0   1E+4   8   8.5  RT (min)      lcqtof_t3_neg_4_000_ee    m/z 221.1564     0   1   2E+5   8.5   9  RT (min)      lcqtof_t3_neg_4_000_ee    m/z 222.1589     0   1   2   3E+4   9   9.5  RT (min)      lcqtof_t3_neg_4_000_ee    m/z 222.7645     0   1   2   3E+3   17   17.5  RT (min)       lcqtof_t3_neg_4_013_39    m/z 223.0276     0   2   4E+4   12   12.5  RT (min)       lcqtof_t3_neg_4_064_17    m/z 223.0276     0   1   2   3   4E+4   12   12.5  RT (min)      lcqtof_t3_neg_4_064_17    m/z 223.0277     0   2   4E+4   12   12.5  RT (min)      lcqtof_t3_neg_4_064_17    m/z 223.0277     0   2   4   6   8E+4   13.5   14  RT (min)      lcqtof_t3_neg_4_064_17    m/z 223.0278     0   2   4E+4   12   12.5  RT (min)      lcqtof_t3_neg_4_064_17    m/z 223.0278     0   2   4   6E+4   12.5   13  RT (min)      lcqtof_t3_neg_4_064_17    m/z 223.0279     0   1E+5   13   13.5  RT (min)       lcqtof_t3_neg_4_064_17    m/z 223.0279     0   1E+5   12.5   13  RT (min)      lcqtof_t3_neg_4_064_17    m/z 223.0279     0   2   4   6E+4   14   14.5  RT (min)      lcqtof_t3_neg_4_064_17    m/z 223.0279     0   1E+5   13   13.5   14  RT (min)       lcqtof_t3_neg_4_064_17    m/z 223.0279     0   2   4E+4   12.5   13  RT (min)      lcqtof_t3_neg_2_064_13    m/z 223.0279     0   1E+5   13   14  RT (min)       lcqtof_t3_neg_4_064_17    m/z 223.0279     0   1E+5   13.5   14  RT (min)      lcqtof_t3_neg_4_064_17    m/z 223.0279     0   1E+5   13   13.5  RT (min)       lcqtof_t3_neg_4_064_17    m/z 223.0279     0   2   4   6E+4   14   14.5  RT (min)      lcqtof_t3_neg_4_064_17    m/z 223.0280     0   1E+5   13.5   14  RT (min)      lcqtof_t3_neg_4_064_17    m/z 223.0280     0   2   4E+4   13   13.5  RT (min)      lcqtof_t3_neg_2_064_13    m/z 223.0280     0   1E+5   13.5   14  RT (min)      lcqtof_t3_neg_4_064_17    m/z 223.0280     0   1   2   3E+4   13   13.5  RT (min)      lcqtof_t3_neg_1_062_13    m/z 223.0281     0   2   4   6   8E+4   13.5   14  RT (min)      lcqtof_t3_neg_4_064_17    m/z 223.0282     0   2   4   6E+4   14   14.5  RT (min)      lcqtof_t3_neg_4_064_17    m/z 223.0283     0   1   2E+4   12.5   13  RT (min)      lcqtof_t3_neg_2_064_13    m/z 223.1094     0   1E+4   8   8.5  RT (min)      lcqtof_t3_neg_4_000_ee    m/z 225.0070     0   1   2E+3   15   15.5  RT (min)       lcqtof_t3_neg_4_103_61    m/z 225.0073     0   1   2   3E+3   16.5   17  RT (min)       lcqtof_t3_neg_4_091_60    m/z 225.0080     0   1   2   3E+3   15.5   16  RT (min)       lcqtof_t3_neg_4_000_ff    m/z 225.0706     0   1E+4   4   4.5  RT (min)      lcqtof_t3_neg_4_040_24    m/z 225.0954     0   1   2E+4   9   9.5  RT (min)      lcqtof_t3_neg_4_000_ee    m/z 225.1641     0   1   2E+4   5.5   6  RT (min)      lcqtof_t3_neg_4_000_ee    m/z 226.0126     0   1E+4   1   1.5  RT (min)      lcqtof_t3_neg_4_011_32    m/z 226.1460     0   1   2   3E+4   8.5   9  RT (min)      lcqtof_t3_neg_4_000_ee    m/z 226.9636     0   1   2E+3   17   17.5  RT (min)       lcqtof_t3_neg_4_035_08    m/z 226.9640     0   1   2   3   4E+3   16.5   17  RT (min)      lcqtof_t3_neg_1_058_16    m/z 227.1303     0   1   2E+4   8.5   9  RT (min)      lcqtof_t3_neg_4_000_ee    m/z 227.1314     0   1   2E+4   8.5   9  RT (min)      lcqtof_t3_neg_4_000_ee    m/z 227.1330     0   1   2E+4   4   4.5  RT (min)      lcqtof_t3_neg_3_032_24    m/z 227.1331     0   2   4   6E+1   3.5   4  RT (min)      lcqtof_t3_neg_3_038_06    m/z 228.0723     0   1E+4   1   1.5  RT (min)      lcqtof_t3_neg_4_011_32    m/z 228.9593     0   2   4   6E+3   17   17.5  RT (min)       lcqtof_t3_neg_1_021_39    m/z 228.9594     0   1   2   3E+3   5.5   6  RT (min)       lcqtof_t3_neg_1_028_20    m/z 228.9594     0   2   4E+3   5   6   7  RT (min)       lcqtof_t3_neg_1_013_09    m/z 228.9594     0   1   2   3E+3   5   5.5  RT (min)       lcqtof_t3_neg_1_051_26    m/z 228.9595     0   2   4   6E+3   17.5   18  RT (min)       lcqtof_t3_neg_1_039_43    m/z 228.9596     0   1   2   3E+3   2   2.5  RT (min)       lcqtof_t3_neg_1_058_16    m/z 228.9596     0   2   4   6E+3   1.5   2  RT (min)       lcqtof_t3_neg_1_005_18    m/z 228.9596     0   2   4   6   8E+3   1   1.5  RT (min)       lcqtof_t3_neg_1_047_22    m/z 228.9598     0   1   2   3E+3   1.5   2  RT (min)       lcqtof_t3_neg_1_010_27    m/z 228.9598     0   1   2   3   4E+3   4.5   5  RT (min)       lcqtof_t3_neg_1_028_20    m/z 228.9600     0   1   2   3E+3   2.5   3   3.5  RT (min)       lcqtof_t3_neg_1_015_34    m/z 228.9600     0   1   2E+3   6   6.5  RT (min)       lcqtof_t3_neg_4_002_17    m/z 228.9600     0   1E+4   0.5   1  RT (min)       lcqtof_t3_neg_4_000_ee    m/z 230.0312     0   1E+4   8.5   9  RT (min)      lcqtof_t3_neg_4_012_27    m/z 230.8823     0   1E+4   1   1.5  RT (min)       lcqtof_t3_neg_4_106_62    m/z 230.9555     0   1   2E+3   4.5   5  RT (min)       lcqtof_t3_neg_1_013_09    m/z 230.9556     0   1   2   3   4E+3   17   17.5  RT (min)       lcqtof_t3_neg_1_039_43    m/z 230.9556     0   1   2E+3   17   17.5  RT (min)       lcqtof_t3_neg_3_093_60    m/z 231.0310     0   1   2E+4   4   4.5  RT (min)      lcqtof_t3_neg_4_060_15    m/z 231.0410     0   1   2   3E+3   16.5   17  RT (min)      lcqtof_t3_neg_4_002_17    m/z 231.9308     0   2   4   6E+3   14   14.5  RT (min)       lcqtof_t3_neg_4_095_60    m/z 231.9308     0   1E+4   16.5   17  RT (min)       lcqtof_t3_neg_4_035_08    m/z 231.9309     0   2   4   6E+3   14   14.5  RT (min)       lcqtof_t3_neg_4_094_60    m/z 231.9311     0   1   2   3   4E+3   15.5   16  RT (min)       lcqtof_t3_neg_4_105_61    m/z 231.9311     0   2   4E+3   15   15.5  RT (min)       lcqtof_t3_neg_4_106_62    m/z 231.9311     0   2   4E+3   13.5   14  RT (min)       lcqtof_t3_neg_4_077_58    m/z 231.9312     0   1   2   3   4E+3   13   13.5  RT (min)       lcqtof_t3_neg_4_096_60    m/z 231.9312     0   1   2   3   4E+3   15.5   16  RT (min)       lcqtof_t3_neg_4_084_59    m/z 231.9313     0   1   2   3E+3   12.5   13  RT (min)       lcqtof_t3_neg_4_087_59    m/z 231.9313     0   1   2   3   4E+3   13   13.5  RT (min)       lcqtof_t3_neg_4_090_60    m/z 231.9313     0   2   4   6E+3   14   14.5  RT (min)       lcqtof_t3_neg_4_097_60    m/z 231.9314     0   1E+4   16   16.5  RT (min)       lcqtof_t3_neg_4_018_23    m/z 231.9316     0   1   2   3E+3   12.5   13  RT (min)       lcqtof_t3_neg_4_097_60    m/z 231.9322     0   1   2E+4   16.5   17  RT (min)       lcqtof_t3_neg_4_000_cc    m/z 231.9345     0   1E+4   0.5   1  RT (min)      lcqtof_t3_neg_4_000_ee    m/z 231.9357     0   1E+4   0.5   1  RT (min)      lcqtof_t3_neg_4_000_ee    m/z 232.0122     0   1   2E+4   17   17.5  RT (min)       lcqtof_t3_neg_4_035_08    m/z 232.0133     0   1   2   3E+3   11.5   12  RT (min)       lcqtof_t3_neg_4_050_43    m/z 232.0483     0   1   2   3   4E+3   16.5   17  RT (min)      lcqtof_t3_neg_4_000_bb    m/z 232.7674     0   1E+4   0.5   1  RT (min)      lcqtof_t3_neg_4_000_ee    m/z 232.9323     0   1   2   3   4E+3   16.5   17  RT (min)      lcqtof_t3_neg_4_000_bb    m/z 232.9998     0   1   2   3   4E+3   17   17.5  RT (min)      lcqtof_t3_neg_4_035_08    m/z 233.0000     0   2   4   6   8E+3   16.5   17  RT (min)      lcqtof_t3_neg_4_000_dd    m/z 233.0095     0   1   2   3E+3   12.5   13  RT (min)       lcqtof_t3_neg_4_000_dd    m/z 233.0106     0   1   2   3   4E+3   12.5   13  RT (min)      lcqtof_t3_neg_4_000_dd    m/z 233.0160     0   2   4E+3   11.5   12   12.5  RT (min)       lcqtof_t3_neg_4_110_62    m/z 233.0160     0   1   2   3   4E+3   12   12.5  RT (min)       lcqtof_t3_neg_4_029_40    m/z 233.0185     0   2   4   6E+3   12   12.5  RT (min)       lcqtof_t3_neg_4_100_61    m/z 233.0195     0   2   4   6E+3   12   13  RT (min)       lcqtof_t3_neg_4_000_ff    m/z 233.0259     0   2   4E+3   13.5   14  RT (min)      lcqtof_t3_neg_4_000_dd    m/z 233.0308     0   1   2E+3   12   12.5  RT (min)       lcqtof_t3_neg_4_024_26    m/z 233.0340     0   1   2E+3   12   12.5  RT (min)      lcqtof_t3_neg_4_040_24    m/z 233.1552     0   1E+5   16   16.5  RT (min)      lcqtof_t3_neg_4_018_23    m/z 233.1556     0   1   2   3   4E+4   16.5   17  RT (min)      lcqtof_t3_neg_4_043_11    m/z 233.1579     0   1   2E+5   17   17.5  RT (min)      lcqtof_t3_neg_4_106_62    m/z 234.0202     0   2   4E+3   16.5   17  RT (min)      lcqtof_t3_neg_4_000_cc    m/z 234.1892     0   1   2E+4   9.5   10  RT (min)      lcqtof_t3_neg_4_000_ee    m/z 234.1896     0   1   2E+4   10   10.5  RT (min)      lcqtof_t3_neg_4_000_ee    m/z 234.7651     0   1E+4   0.5   1  RT (min)      lcqtof_t3_neg_4_000_ee    m/z 235.1244     0   1E+4   8.5   9  RT (min)      lcqtof_t3_neg_4_000_ee    m/z 235.1252     0   1E+4   8.5   9  RT (min)      lcqtof_t3_neg_4_000_ee    m/z 235.1734     0   1E+4   10.5   11  RT (min)      lcqtof_t3_neg_4_000_ee    m/z 235.1734     0   1E+4   10.5   11  RT (min)      lcqtof_t3_neg_4_000_ee    m/z 235.1740     0   1E+4   10.5   11  RT (min)      lcqtof_t3_neg_4_000_ee    m/z 236.0439     0   2   4   6E+4   1   1.5  RT (min)      lcqtof_t3_neg_4_019_05    m/z 236.1449     0   1   2E+5   8   8.5  RT (min)      lcqtof_t3_neg_4_000_ee    m/z 236.1909     0   1E+4   1   1.5  RT (min)      lcqtof_t3_neg_1_031_24    m/z 237.0381     0   1E+4   1   1.5  RT (min)      lcqtof_t3_neg_4_040_24    m/z 237.1139     0   1E+4   7.5   8  RT (min)      lcqtof_t3_neg_4_000_ee    m/z 237.1508     0   1   2   3   4E+4   8   8.5  RT (min)      lcqtof_t3_neg_4_000_ee    m/z 238.9311     0   1E+4   10   15  RT (min)       lcqtof_t3_neg_4_005_18    m/z 238.9312     0   2   4   6E+3   15   16  RT (min)       lcqtof_t3_neg_4_063_39    m/z 238.9312     0   2   4   6E+3   14.5   15  RT (min)       lcqtof_t3_neg_4_093_60    m/z 238.9312     0   2   4   6   8E+3   15   16  RT (min)       lcqtof_t3_neg_4_018_23    m/z 238.9315     0   1   2E+3   12   12.5  RT (min)       lcqtof_t3_neg_4_012_27    m/z 239.0587     0   1E+4   11   11.5  RT (min)      lcqtof_t3_neg_4_064_17    m/z 239.0588     0   1E+4   11   11.5  RT (min)       lcqtof_t3_neg_4_064_17    m/z 239.0588     0   2   4   6E+4   14   14.5  RT (min)      lcqtof_t3_neg_4_064_17    m/z 239.0588     0   1E+4   10.5   11  RT (min)      lcqtof_t3_neg_4_064_17    m/z 239.0588     0   1   2E+5   13.5   14  RT (min)      lcqtof_t3_neg_4_064_17    m/z 239.0592     0   1E+4   10.5   11  RT (min)      lcqtof_t3_neg_4_064_17    m/z 239.0593     0   1E+5   12   12.5  RT (min)       lcqtof_t3_neg_4_064_17    m/z 239.0593     0   1   2E+5   13.5   14  RT (min)       lcqtof_t3_neg_4_064_17    m/z 239.0594     0   1E+5   12.5   13   13.5  RT (min)       lcqtof_t3_neg_4_036_47    m/z 239.0594     0   1E+5   13.5   14  RT (min)      lcqtof_t3_neg_4_064_17    m/z 239.0594     0   1   2E+5   13   13.5  RT (min)      lcqtof_t3_neg_4_064_17    m/z 239.0595     0   2   4E+4   12.5   13  RT (min)      lcqtof_t3_neg_3_060_16    m/z 239.0595     0   1E+5   12   12.5  RT (min)      lcqtof_t3_neg_4_064_17    m/z 239.0596     0   1   2E+5   13.5   14  RT (min)       lcqtof_t3_neg_4_064_17    m/z 239.0596     0   1E+5   12   12.5  RT (min)      lcqtof_t3_neg_4_064_17    m/z 239.0596     0   1E+5   14   14.5  RT (min)      lcqtof_t3_neg_4_064_17    m/z 239.0597     0   1E+5   12   12.5  RT (min)       lcqtof_t3_neg_4_064_17    m/z 239.0597     0   1E+5   14   14.5  RT (min)       lcqtof_t3_neg_4_064_17    m/z 239.0598     0   1E+5   13.5   14  RT (min)      lcqtof_t3_neg_4_064_17    m/z 239.0598     0   1   2E+5   12.5   13  RT (min)       lcqtof_t3_neg_4_064_17    m/z 239.0598     0   1   2E+5   13.5   14  RT (min)       lcqtof_t3_neg_4_064_17    m/z 239.0599     0   1E+5   12   12.5  RT (min)      lcqtof_t3_neg_4_064_17    m/z 239.0601     0   1E+5   14   14.5  RT (min)      lcqtof_t3_neg_4_064_17    m/z 239.1796     0   1E+4   6   6.5  RT (min)      lcqtof_t3_neg_4_000_ee    m/z 239.7600     0   1   2   3E+3   14   14.5  RT (min)       lcqtof_t3_neg_4_031_42    m/z 239.7605     0   1   2   3E+3   13.5   14  RT (min)       lcqtof_t3_neg_4_020_22    m/z 240.0598     0   1   2   3E+4   12   12.5  RT (min)      lcqtof_t3_neg_4_064_17    m/z 240.0598     0   1   2   3E+4   12   12.5  RT (min)      lcqtof_t3_neg_4_064_17    m/z 240.0600     0   2   4E+4   13   13.5  RT (min)      lcqtof_t3_neg_4_064_17    m/z 240.0600     0   1   2E+4   14   14.5  RT (min)      lcqtof_t3_neg_4_064_17    m/z 240.0602     0   2   4E+4   13.5   14  RT (min)      lcqtof_t3_neg_4_064_17    m/z 240.0602     0   2   4E+4   13.5   14  RT (min)      lcqtof_t3_neg_4_064_17    m/z 240.0602     0   2   4E+4   13.5   14  RT (min)      lcqtof_t3_neg_4_064_17    m/z 240.0602     0   2   4E+4   12.5   13  RT (min)      lcqtof_t3_neg_4_064_17    m/z 240.0602     0   1   2   3E+4   14   14.5  RT (min)      lcqtof_t3_neg_4_064_17    m/z 240.0603     0   2   4E+4   13   13.5  RT (min)      lcqtof_t3_neg_4_064_17    m/z 240.0603     0   2   4E+4   13.5   14  RT (min)      lcqtof_t3_neg_4_064_17    m/z 240.0604     0   2   4E+4   13   13.5  RT (min)      lcqtof_t3_neg_4_064_17    m/z 240.0604     0   2   4E+4   12.5   13  RT (min)      lcqtof_t3_neg_4_064_17    m/z 240.0606     0   2   4E+4   13   13.5  RT (min)      lcqtof_t3_neg_4_064_17    m/z 240.1336     0   1E+4   8.5   9  RT (min)      lcqtof_t3_neg_4_000_ee    m/z 240.1356     0   1E+4   8.5   9  RT (min)      lcqtof_t3_neg_4_000_ee    m/z 241.7574     0   2   4E+3   14   14.5  RT (min)       lcqtof_t3_neg_4_028_25    m/z 241.7576     0   2   4   6E+3   14   14.5  RT (min)       lcqtof_t3_neg_4_029_40    m/z 241.7576     0   1   2   3   4E+3   12.5   13  RT (min)       lcqtof_t3_neg_4_032_04    m/z 241.7577     0   1   2E+3   12   12.5  RT (min)      lcqtof_t3_neg_4_112_62    m/z 242.1754     0   1E+2   6.5   7  RT (min)      lcqtof_t3_neg_3_038_06    m/z 242.1758     0   1E+4   7   7.5  RT (min)      lcqtof_t3_neg_3_032_24    m/z 242.1760     0   1   2E+3   13   13.5  RT (min)      lcqtof_t3_neg_1_002_13    m/z 242.3367     0   1   2E+4   7.5   8  RT (min)      lcqtof_t3_neg_4_000_ee    m/z 242.9430     0   2   4   6   8E+3   17   17.5  RT (min)       lcqtof_t3_neg_4_091_60    m/z 242.9432     0   1E+4   8   9   10  RT (min)       lcqtof_t3_neg_1_000_dd    m/z 242.9432     0   2   4   6   8E+3   16.5   17  RT (min)       lcqtof_t3_neg_4_059_14    m/z 242.9433     0   2   4   6   8E+3   16.5   17  RT (min)       lcqtof_t3_neg_4_063_39    m/z 242.9433     0   1E+4   10   11  RT (min)       lcqtof_t3_neg_1_000_bb    m/z 242.9434     0   2   4   6   8E+3   8   9  RT (min)       lcqtof_t3_neg_1_029_44    m/z 242.9436     0   2   4   6   8E+3   17   17.5  RT (min)      lcqtof_t3_neg_4_059_14    m/z 243.0468     0   1E+4   6   6.5  RT (min)      lcqtof_t3_neg_4_107_62    m/z 243.7548     0   1   2   3   4E+3   13.5   14  RT (min)       lcqtof_t3_neg_4_052_03    m/z 243.7549     0   1   2   3E+3   14   14.5  RT (min)       lcqtof_t3_neg_4_039_06    m/z 243.7551     0   1   2   3E+3   14   14.5  RT (min)       lcqtof_t3_neg_4_035_08    m/z 244.0668     0   1   2   3   4E+3   16.5   17  RT (min)      lcqtof_t3_neg_4_000_aa    m/z 244.1545     0   1E+4   6.5   7  RT (min)      lcqtof_t3_neg_4_000_ee    m/z 245.0301     0   1   2E+3   11.5   12  RT (min)      lcqtof_t3_neg_4_008_35    m/z 245.0626     0   1E+4   7   7.5  RT (min)      lcqtof_t3_neg_4_055_38    m/z 245.9837     0   1   2   3   4E+3   16.5   17  RT (min)      lcqtof_t3_neg_4_016_14    m/z 246.0255     0   1E+4   17   17.5  RT (min)      lcqtof_t3_neg_4_035_08    m/z 246.0267     0   1E+4   16   16.5  RT (min)      lcqtof_t3_neg_4_018_23    m/z 246.0643     0   1E+4   16   16.5  RT (min)      lcqtof_t3_neg_4_018_23    m/z 246.0646     0   1E+4   17   17.5  RT (min)      lcqtof_t3_neg_4_035_08    m/z 246.1900     0   1   2E+4   10   10.5  RT (min)      lcqtof_t3_neg_4_000_ee    m/z 246.1900     0   1   2E+4   10   10.5  RT (min)      lcqtof_t3_neg_4_000_ee    m/z 246.1900     0   1   2E+4   10   10.5  RT (min)      lcqtof_t3_neg_4_000_ee    m/z 247.0249     0   1E+4   16   16.5  RT (min)      lcqtof_t3_neg_4_018_23    m/z 247.0251     0   1E+4   17   17.5  RT (min)      lcqtof_t3_neg_4_035_08    m/z 247.0263     0   1   2   3E+3   13.5   14  RT (min)      lcqtof_t3_neg_4_049_55    m/z 247.0602     0   2   4   6E+3   13.5   14  RT (min)       lcqtof_t3_neg_4_028_25    m/z 247.0603     0   2   4   6E+3   13   14  RT (min)       lcqtof_t3_neg_4_006_16    m/z 247.0607     0   1E+4   17   17.5  RT (min)      lcqtof_t3_neg_4_035_08    m/z 248.0200     0   1E+4   17   17.5  RT (min)      lcqtof_t3_neg_4_035_08    m/z 248.0242     0   2   4   6E+3   11  RT (min)       lcqtof_t3_neg_4_018_23    m/z 248.0245     0   1E+4   16   16.5  RT (min)      lcqtof_t3_neg_4_018_23    m/z 248.0249     0   2   4   6E+3   11   12  RT (min)       lcqtof_t3_neg_4_081_58    m/z 248.9012     0   1   2E+3   16.5   17  RT (min)      lcqtof_t3_neg_1_018_33    m/z 248.9597     0   1E+4   6.5   7  RT (min)       lcqtof_t3_neg_4_108_62    m/z 248.9598     0   1   2   3E+5   16.5   17  RT (min)       lcqtof_t3_neg_4_016_14    m/z 248.9598     0   1E+4   6.5   7  RT (min)       lcqtof_t3_neg_4_110_62    m/z 248.9598     0   1E+4   9   9.5  RT (min)       lcqtof_t3_neg_4_104_61    m/z 248.9599     0   2   4   6E+4   16   16.5  RT (min)      lcqtof_t3_neg_1_042_46    m/z 248.9599     0   1E+4   7   7.5  RT (min)       lcqtof_t3_neg_4_111_62    m/z 248.9599     0   1E+4   9.5   10  RT (min)       lcqtof_t3_neg_4_106_62    m/z 248.9599     0   1E+4   11   11.5  RT (min)       lcqtof_t3_neg_4_109_62    m/z 248.9599     0   1E+4   6   6.5  RT (min)       lcqtof_t3_neg_4_107_62    m/z 248.9599     0   1E+4   7.5   8  RT (min)       lcqtof_t3_neg_4_102_61    m/z 248.9599     0   1E+4   12.5   13  RT (min)       lcqtof_t3_neg_4_099_61    m/z 248.9599     0   1E+4   6.5   7  RT (min)       lcqtof_t3_neg_4_108_62    m/z 248.9600     0   1E+4   5   6  RT (min)       lcqtof_t3_neg_4_111_62    m/z 248.9600     0   1E+4   5.5   6  RT (min)       lcqtof_t3_neg_4_098_61    m/z 248.9600     0   1E+4   10  RT (min)       lcqtof_t3_neg_4_112_62    m/z 248.9600     0   1E+4   7.5   8  RT (min)       lcqtof_t3_neg_4_063_39    m/z 248.9600     0   1E+4   7   7.5  RT (min)       lcqtof_t3_neg_4_106_62    m/z 248.9600     0   1   2E+5   16   16.5  RT (min)       lcqtof_t3_neg_4_018_23    m/z 248.9600     0   2   4   6   8E+3   15   15.5  RT (min)       lcqtof_t3_neg_3_032_24    m/z 248.9600     0   1   2E+4   14.5   15  RT (min)       lcqtof_t3_neg_4_041_54    m/z 248.9600     0   1   2E+4   15.5   16  RT (min)      lcqtof_t3_neg_4_038_29    m/z 248.9600     0   1E+4   12   12.5  RT (min)       lcqtof_t3_neg_4_111_62    m/z 248.9600     0   1   2E+4   15   15.5  RT (min)       lcqtof_t3_neg_4_101_61    m/z 248.9600     0   1E+4   14   14.5  RT (min)       lcqtof_t3_neg_4_065_51    m/z 248.9600     0   1E+4   11.5   12  RT (min)       lcqtof_t3_neg_4_099_61    m/z 248.9600     0   1E+4   11.5   12  RT (min)       lcqtof_t3_neg_4_108_62    m/z 248.9600     0   2   4   6E+4   16   16.5  RT (min)      lcqtof_t3_neg_4_041_54    m/z 248.9600     0   1   2E+4   15   15.5  RT (min)       lcqtof_t3_neg_4_038_29    m/z 248.9600     0   1   2E+4   15   15.5  RT (min)       lcqtof_t3_neg_4_057_49    m/z 248.9600     0   1E+4   10   10.5  RT (min)       lcqtof_t3_neg_4_108_62    m/z 248.9600     0   1   2E+4   14.5   15  RT (min)       lcqtof_t3_neg_4_109_62    m/z 248.9600     0   1   2E+5   16   16.5  RT (min)       lcqtof_t3_neg_4_018_23    m/z 248.9601     0   1E+4   11   11.5  RT (min)       lcqtof_t3_neg_4_104_61    m/z 248.9601     0   1E+4   10.5   11  RT (min)       lcqtof_t3_neg_4_109_62    m/z 248.9601     0   1E+4   14   14.5  RT (min)       lcqtof_t3_neg_4_063_39    m/z 248.9601     0   1E+4   12   12.5  RT (min)       lcqtof_t3_neg_4_036_47    m/z 248.9601     0   1E+4   12.5   13  RT (min)       lcqtof_t3_neg_4_107_62    m/z 248.9601     0   1   2E+4   15   15.5  RT (min)       lcqtof_t3_neg_4_099_61    m/z 248.9601     0   1E+4   7.5   8  RT (min)       lcqtof_t3_neg_4_108_62    m/z 248.9602     0   1E+4   8   8.5  RT (min)      lcqtof_t3_neg_4_110_62    m/z 248.9602     0   1E+5   16.5   17  RT (min)      lcqtof_t3_neg_4_047_07    m/z 248.9602     0   1E+4   14   14.5  RT (min)       lcqtof_t3_neg_4_099_61    m/z 248.9602     0   1E+4   8.5   9  RT (min)       lcqtof_t3_neg_4_063_39    m/z 248.9602     0   1E+4   10.5   11  RT (min)       lcqtof_t3_neg_4_102_61    m/z 248.9602     0   1E+4   10   10.5  RT (min)       lcqtof_t3_neg_4_102_61    m/z 248.9602     0   1E+4   12.5   13  RT (min)       lcqtof_t3_neg_4_113_62    m/z 248.9603     0   1E+4   13   13.5  RT (min)       lcqtof_t3_neg_4_112_62    m/z 248.9603     0   1   2E+4   14.5   15  RT (min)       lcqtof_t3_neg_4_041_54    m/z 248.9604     0   1   2   3   4E+3   13.5   14  RT (min)       lcqtof_t3_neg_3_038_06    m/z 248.9604     0   1E+4   5.5   6  RT (min)       lcqtof_t3_neg_4_099_61    m/z 248.9604     0   1E+4   12   12.5  RT (min)       lcqtof_t3_neg_4_073_57    m/z 248.9604     0   1E+5   16.5   17  RT (min)      lcqtof_t3_neg_4_047_07    m/z 248.9604     0   1E+4   10   10.5  RT (min)       lcqtof_t3_neg_4_103_61    m/z 248.9604     0   1E+4   9.5   10  RT (min)       lcqtof_t3_neg_4_110_62    m/z 248.9604     0   1E+4   10   12  RT (min)       lcqtof_t3_neg_4_113_62    m/z 248.9605     0   1   2E+5   17   17.5  RT (min)       lcqtof_t3_neg_4_106_62    m/z 248.9605     0   1E+4   7   7.5  RT (min)       lcqtof_t3_neg_4_099_61    m/z 248.9607     0   1E+4   5   5.5  RT (min)       lcqtof_t3_neg_4_112_62    m/z 248.9607     0   1E+4   9.5   10  RT (min)      lcqtof_t3_neg_4_107_62    m/z 248.9611     0   2   4   6   8E+4   17   17.5  RT (min)       lcqtof_t3_neg_4_035_08    m/z 248.9615     0   1E+4   5   5.5  RT (min)       lcqtof_t3_neg_4_110_62    m/z 248.9628     0   1   2   3E+5   2   2.5  RT (min)      lcqtof_t3_neg_4_064_17    m/z 248.9629     0   1E+5   3   3.5  RT (min)      lcqtof_t3_neg_4_063_39    m/z 248.9697     0   1E+4   4.5   5  RT (min)       lcqtof_t3_neg_4_028_25    m/z 248.9721     0   2   4   6E+3   16.5   17  RT (min)      lcqtof_t3_neg_4_047_07    m/z 248.9740     0   2   4   6E+3   16.5   17  RT (min)      lcqtof_t3_neg_4_044_48    m/z 249.0082     0   2   4E+3   17   17.5  RT (min)      lcqtof_t3_neg_4_005_18    m/z 249.0108     0   1   2   3E+3   17   17.5  RT (min)      lcqtof_t3_neg_4_035_08    m/z 249.0128     0   2   4E+3   16.5   17  RT (min)      lcqtof_t3_neg_4_003_37    m/z 249.0207     0   1E+4   16   16.5  RT (min)      lcqtof_t3_neg_4_018_23    m/z 249.0280     0   2   4E+3   6   6.5  RT (min)      lcqtof_t3_neg_4_016_14    m/z 249.0762     0   1   2E+4   6   6.5  RT (min)      lcqtof_t3_neg_4_107_62    m/z 249.1549     0   2   4E+4   8.5   9  RT (min)      lcqtof_t3_neg_4_000_ee    m/z 249.9673     0   1   2E+3   15   15.5  RT (min)      lcqtof_t3_neg_4_011_32    m/z 251.1356     0   1E+4   8.5   9  RT (min)      lcqtof_t3_neg_4_000_ee    m/z 253.0566     0   1E+4   1   1.5  RT (min)       lcqtof_t3_neg_4_040_24    m/z 253.1437     0   1E+4   8   8.5  RT (min)      lcqtof_t3_neg_4_000_ee    m/z 254.1991     0   1   2   3E+4   6.5   7  RT (min)      lcqtof_t3_neg_4_000_ee    m/z 255.0664     0   1E+4   8.5   9  RT (min)      lcqtof_t3_neg_2_013_26    m/z 255.2329     0   2   4   6   8E+3   12.5   13  RT (min)      lcqtof_t3_neg_1_035_31    m/z 256.1595     0   1E+4   7.5   8  RT (min)      lcqtof_t3_neg_4_000_ee    m/z 256.9537     0   1E+4   17.5   18  RT (min)       lcqtof_t3_neg_3_099_61    m/z 256.9542     0   1E+4   17.5   18  RT (min)       lcqtof_t3_neg_1_001_49    m/z 256.9544     0   1   2   3E+4   0.5   1  RT (min)       lcqtof_t3_neg_4_000_ee    m/z 256.9544     0   1E+4   17.5   18  RT (min)       lcqtof_t3_neg_1_059_17    m/z 256.9545     0   1   2   3   4E+4   1   1.5  RT (min)       lcqtof_t3_neg_4_062_13    m/z 256.9546     0   2   4   6   8E+3   0   0.5  RT (min)       lcqtof_t3_neg_1_015_34    m/z 256.9546     0   1E+4   17.5   18  RT (min)       lcqtof_t3_neg_1_005_18    m/z 256.9546     0   1   2E+4   1   1.5  RT (min)       lcqtof_t3_neg_1_027_21    m/z 256.9546     0   1   2E+4   1   1.5  RT (min)      lcqtof_t3_neg_1_027_21    m/z 256.9546     0   1E+4   17   17.5  RT (min)       lcqtof_t3_neg_4_046_33    m/z 256.9546     0   1   2E+4   17   17.5  RT (min)       lcqtof_t3_neg_4_000_bb    m/z 256.9546     0   1E+4   4.5   5  RT (min)       lcqtof_t3_neg_4_000_dd    m/z 256.9547     0   2   4   6E+3   1.5   2  RT (min)       lcqtof_t3_neg_1_010_27    m/z 256.9547     0   1   2   3E+4   0.5   1  RT (min)       lcqtof_t3_neg_4_000_ee    m/z 256.9548     0   1   2   3E+4   0.5   1  RT (min)       lcqtof_t3_neg_4_000_ee    m/z 256.9548     0   2   4   6   8E+3   1.5   2  RT (min)       lcqtof_t3_neg_1_008_02    m/z 256.9548     0   2   4   6E+3   0   0.5  RT (min)       lcqtof_t3_neg_1_047_22    m/z 256.9548     0   2   4   6E+3   0   0.5  RT (min)       lcqtof_t3_neg_1_004_11    m/z 256.9548     0   2   4   6E+3   0   0.5  RT (min)      lcqtof_t3_neg_1_047_22    m/z 256.9550     0   1E+4   16.5   17  RT (min)       lcqtof_t3_neg_4_018_23    m/z 256.9550     0   2   4   6E+3   0   0.5  RT (min)      lcqtof_t3_neg_4_008_35    m/z 257.0230     0   1E+4   1   1.5  RT (min)       lcqtof_t3_neg_4_083_59    m/z 257.1762     0   1E+4   9   9.5  RT (min)      lcqtof_t3_neg_4_000_ee    m/z 258.0405     0   1   2   3   4E+3   16.5   17  RT (min)      lcqtof_t3_neg_4_003_37    m/z 258.1738     0   2   4   6   8E+4   6.5   7  RT (min)      lcqtof_t3_neg_4_000_ee    m/z 259.1758     0   1E+4   7   7.5  RT (min)      lcqtof_t3_neg_4_000_ee    m/z 261.0392     0   1   2E+4   16   16.5  RT (min)      lcqtof_t3_neg_4_018_23    m/z 261.0419     0   1   2   3E+3   12   12.5  RT (min)       lcqtof_t3_neg_4_016_14    m/z 261.0744     0   1   2   3E+4   6   6.5  RT (min)       lcqtof_t3_neg_4_099_61    m/z 262.1601     0   1E+4   6.5   7  RT (min)      lcqtof_t3_neg_4_000_ee    m/z 263.0852     0   1E+4   1   1.5  RT (min)       lcqtof_t3_neg_4_097_60    m/z 263.1518     0   1E+4   7.5   8  RT (min)      lcqtof_t3_neg_4_000_ee    m/z 264.1596     0   1E+2   9   9.5  RT (min)      lcqtof_t3_neg_3_038_06    m/z 264.1599     0   1E+2   9   9.5  RT (min)      lcqtof_t3_neg_3_038_06    m/z 264.1604     0   1E+2   9   9.5  RT (min)      lcqtof_t3_neg_3_038_06    m/z 265.1474     0   1E+4   15.5   16  RT (min)       lcqtof_t3_neg_4_058_16    m/z 265.1475     0   1E+4   10   10.5  RT (min)      lcqtof_t3_neg_4_034_34    m/z 265.1476     0   1   2E+4   12   12.5  RT (min)      lcqtof_t3_neg_4_061_50    m/z 265.1476     0   1   2   3E+4   9.5   10  RT (min)      lcqtof_t3_neg_4_018_23    m/z 265.1476     0   1   2E+4   16   16.5  RT (min)       lcqtof_t3_neg_4_060_15    m/z 265.1477     0   2   4E+3   13.5   14  RT (min)       lcqtof_t3_neg_1_019_23    m/z 265.1477     0   1E+4   11   11.5  RT (min)      lcqtof_t3_neg_4_046_33    m/z 265.1477     0   2   4   6E+3   15   15.5  RT (min)       lcqtof_t3_neg_3_108_62    m/z 265.1477     0   1E+4   14.5   15  RT (min)       lcqtof_t3_neg_4_086_59    m/z 265.1478     0   1   2E+4   16   16.5  RT (min)      lcqtof_t3_neg_4_061_50    m/z 265.1481     0   1E+4   14   14.5  RT (min)       lcqtof_t3_neg_4_062_13    m/z 265.1481     0   1E+4   14.5   15  RT (min)       lcqtof_t3_neg_4_005_18    m/z 265.1482     0   1E+4   12   12.5  RT (min)      lcqtof_t3_neg_4_062_13    m/z 265.1484     0   1E+4   10   10.5  RT (min)      lcqtof_t3_neg_4_004_20    m/z 265.1486     0   1E+4   13   13.5  RT (min)      lcqtof_t3_neg_4_034_34    m/z 265.1487     0   1E+4   11   11.5  RT (min)      lcqtof_t3_neg_4_047_07    m/z 265.1489     0   1E+4   11.5   12  RT (min)      lcqtof_t3_neg_4_066_57    m/z 265.1496     0   1E+4   10   10.5  RT (min)      lcqtof_t3_neg_4_060_15    m/z 265.1498     0   1   2E+2   7.5   8  RT (min)      lcqtof_t3_neg_3_038_06    m/z 265.1499     0   1E+4   14   14.5  RT (min)       lcqtof_t3_neg_4_066_57    m/z 265.1500     0   1E+4   13   13.5  RT (min)      lcqtof_t3_neg_4_061_50    m/z 265.1508     0   1E+4   12.5   13  RT (min)      lcqtof_t3_neg_4_084_59    m/z 265.1519     0   1   2E+2   7.5   8  RT (min)      lcqtof_t3_neg_3_038_06    m/z 265.3172     0   1E+4   9   9.5  RT (min)      lcqtof_t3_neg_4_000_ee    m/z 265.9498     0   1E+3   15   15.5  RT (min)      lcqtof_t3_neg_3_032_24    m/z 265.9503     0   2   4   6   8E+3   16.5   17  RT (min)      lcqtof_t3_neg_1_065_51    m/z 266.0726     0   1   2   3   4E+3   1   1.5  RT (min)      lcqtof_t3_neg_1_042_46    m/z 266.1080     0   1E+4   9.5   10  RT (min)      lcqtof_t3_neg_4_000_ee    m/z 266.1086     0   1E+4   9.5   10  RT (min)      lcqtof_t3_neg_4_000_ee    m/z 266.1182     0   1   2E+4   8.5   9  RT (min)      lcqtof_t3_neg_4_000_ee    m/z 266.1184     0   1   2E+4   8.5   9  RT (min)      lcqtof_t3_neg_4_000_ee    m/z 267.9440     0   1   2E+4   1   1.5  RT (min)      lcqtof_t3_neg_4_014_13    m/z 269.0770     0   1   2   3   4E+2   3.5   4  RT (min)      lcqtof_t3_neg_3_038_06    m/z 270.1859     0   1E+4   5.5   6  RT (min)      lcqtof_t3_neg_4_000_ee    m/z 271.2864     0   1E+4   12   12.5  RT (min)      lcqtof_t3_neg_1_006_05    m/z 272.8983     0   2   4   6   8E+3   17   17.5  RT (min)       lcqtof_t3_neg_4_013_39    m/z 274.0876     0   1   2   3E+3   16.5   17  RT (min)      lcqtof_t3_neg_4_024_26    m/z 275.1695     0   1E+4   9   9.5  RT (min)      lcqtof_t3_neg_4_000_ee    m/z 277.1011     0   1E+4   1   1.5  RT (min)       lcqtof_t3_neg_3_032_24    m/z 277.1442     0   2   4E+4   16.5   17  RT (min)      lcqtof_t3_neg_4_016_14    m/z 277.2165     0   1   2E+4   11   11.5  RT (min)      lcqtof_t3_neg_2_016_05    m/z 277.2170     0   1   2E+4   11   11.5  RT (min)      lcqtof_t3_neg_2_016_05    m/z 277.2767     0   1   2E+4   9.5   10  RT (min)      lcqtof_t3_neg_4_059_14    m/z 277.3476     0   2   4   6E+4   9.5   10  RT (min)      lcqtof_t3_neg_4_059_14    m/z 277.4983     0   1E+4   9.5   10  RT (min)      lcqtof_t3_neg_4_059_14    m/z 277.6130     0   1E+4   9.5   10  RT (min)      lcqtof_t3_neg_4_059_14    m/z 278.1550     0   1E+4   7   7.5  RT (min)      lcqtof_t3_neg_4_000_ee    m/z 278.2318     0   2   4E+4   9.5   10  RT (min)      lcqtof_t3_neg_4_059_14    m/z 278.2822     0   1E+4   9.5   10  RT (min)      lcqtof_t3_neg_2_051_14    m/z 278.2824     0   1   2E+4   9.5   10  RT (min)      lcqtof_t3_neg_4_059_14    m/z 278.3554     0   1   2   3E+4   9.5   10  RT (min)      lcqtof_t3_neg_4_059_14    m/z 278.3962     0   1   2E+4   9.5   10  RT (min)      lcqtof_t3_neg_4_059_14    m/z 278.4447     0   1   2   3E+4   9.5   10  RT (min)      lcqtof_t3_neg_4_059_14    m/z 279.0077     0   1E+4   17   17.5  RT (min)       lcqtof_t3_neg_4_000_bb    m/z 279.1377     0   1   2E+4   7   7.5  RT (min)      lcqtof_t3_neg_4_000_ee    m/z 279.2325     0   2   4E+4   11.5   12  RT (min)      lcqtof_t3_neg_2_016_05    m/z 279.2328     0   2   4E+3   13   13.5  RT (min)      lcqtof_t3_neg_1_010_27    m/z 279.2328     0   2   4E+4   11.5   12  RT (min)      lcqtof_t3_neg_2_016_05    m/z 279.3102     0   1E+4   9.5   10  RT (min)      lcqtof_t3_neg_4_059_14    m/z 280.1143     0   1E+2   3.5   4  RT (min)      lcqtof_t3_neg_3_038_06    m/z 280.1144     0   1   2   3E+4   4   4.5  RT (min)      lcqtof_t3_neg_3_032_24    m/z 281.2480     0   2   4   6E+3   10.5   11  RT (min)      lcqtof_t3_neg_1_035_31    m/z 281.2480     0   2   4   6E+3   13   13.5  RT (min)      lcqtof_t3_neg_1_043_38    m/z 281.2482     0   2   4E+3   10.5   11  RT (min)      lcqtof_t3_neg_1_012_16    m/z 281.2482     0   1   2   3   4E+3   10.5   11  RT (min)      lcqtof_t3_neg_1_018_33    m/z 281.2482     0   1   2   3   4E+3   10.5   11  RT (min)      lcqtof_t3_neg_4_058_16    m/z 281.2482     0   2   4E+3   13   13.5  RT (min)      lcqtof_t3_neg_4_034_34    m/z 281.2484     0   2   4   6   8E+3   13   13.5  RT (min)      lcqtof_t3_neg_1_032_36    m/z 281.3438     0   1E+4   9   9.5  RT (min)      lcqtof_t3_neg_2_038_18    m/z 281.3440     0   1E+4   9   9.5  RT (min)      lcqtof_t3_neg_4_005_18    m/z 281.3872     0   1   2E+4   9   9.5  RT (min)      lcqtof_t3_neg_2_038_18    m/z 281.3877     0   1E+4   9   9.5  RT (min)      lcqtof_t3_neg_4_005_18    m/z 282.1306     0   1E+4   6.5   7  RT (min)      lcqtof_t3_neg_4_000_ee    m/z 282.3127     0   1E+4   9   9.5  RT (min)      lcqtof_t3_neg_2_038_18    m/z 282.3130     0   1E+4   9   9.5  RT (min)      lcqtof_t3_neg_4_005_18    m/z 283.2560     0   2   4E+3   12   12.5  RT (min)      lcqtof_t3_neg_1_019_23    m/z 283.2563     0   2   4   6   8E+3   12   12.5  RT (min)      lcqtof_t3_neg_1_019_23    m/z 284.0924     0   2   4   6E+3   8   8.5  RT (min)      lcqtof_t3_neg_4_065_51    m/z 284.0926     0   2   4E+3   8   8.5  RT (min)      lcqtof_t3_neg_4_025_52    m/z 284.1664     0   1E+4   7   7.5  RT (min)      lcqtof_t3_neg_4_000_ee    m/z 284.1720     0   1E+4   7   7.5  RT (min)      lcqtof_t3_neg_4_000_ee    m/z 285.0398     0   1   2E+4   6.5   7  RT (min)      lcqtof_t3_neg_4_005_18    m/z 285.0411     0   1E+4   7   7.5  RT (min)      lcqtof_t3_neg_2_038_18    m/z 285.1208     0   1   2   3E+4   8.5   9  RT (min)      lcqtof_t3_neg_4_000_ee    m/z 285.1259     0   1   2   3E+4   8.5   9  RT (min)      lcqtof_t3_neg_4_000_ee    m/z 286.2047     0   1E+4   8   8.5  RT (min)      lcqtof_t3_neg_4_000_ee    m/z 286.8730     0   1   2E+4   1   1.5  RT (min)      lcqtof_t3_neg_1_006_05    m/z 288.0048     0   1E+4   6   6.5  RT (min)      lcqtof_t3_neg_4_064_17    m/z 288.1196     0   2   4E+3   2   2.5  RT (min)       lcqtof_t3_neg_4_089_59    m/z 288.1198     0   1   2   3   4E+3   2.5   3  RT (min)      lcqtof_t3_neg_1_035_31    m/z 288.1394     0   1   2E+3   16   16.5  RT (min)      lcqtof_t3_neg_1_020_47    m/z 288.1396     0   1   2   3E+3   16.5   17  RT (min)      lcqtof_t3_neg_4_007_09    m/z 288.9356     0   2   4   6E+3   16   16.5  RT (min)      lcqtof_t3_neg_1_019_23    m/z 288.9356     0   1E+3   16   16.5  RT (min)      lcqtof_t3_neg_1_019_23    m/z 288.9360     0   1   2E+2   16   16.5  RT (min)      lcqtof_t3_neg_1_019_23    m/z 289.0372     0   1   2   3   4E+4   1.5   2  RT (min)      lcqtof_t3_neg_4_059_14    m/z 291.0680     0   1E+4   6.5   7  RT (min)       lcqtof_t3_neg_4_004_20    m/z 292.2030     0   1E+4   9.5   10  RT (min)      lcqtof_t3_neg_4_064_17    m/z 292.4903     0   1   2E+4   1   1.5  RT (min)      lcqtof_t3_neg_1_036_14    m/z 292.8071     0   1E+4   1   1.5  RT (min)      lcqtof_t3_neg_4_059_14    m/z 293.1061     0   1   2   3   4E+4   5   5.5  RT (min)      lcqtof_t3_neg_1_045_29    m/z 293.1749     0   1   2E+2   7.5   8  RT (min)      lcqtof_t3_neg_3_038_06    m/z 293.1756     0   2   4   6E+2   7.5   8  RT (min)      lcqtof_t3_neg_3_038_06    m/z 293.1757     0   2   4   6E+2   8   8.5  RT (min)      lcqtof_t3_neg_3_038_06    m/z 293.1776     0   1E+4   16   16.5  RT (min)      lcqtof_t3_neg_4_064_17    m/z 293.1780     0   1E+4   15   15.5  RT (min)       lcqtof_t3_neg_4_065_51    m/z 293.1780     0   1   2   3E+3   8.5   9  RT (min)      lcqtof_t3_neg_3_038_06    m/z 293.1782     0   1E+4   14.5   15  RT (min)       lcqtof_t3_neg_2_000_aa    m/z 293.2574     0   1   2   3   4E+4   1   1.5  RT (min)      lcqtof_t3_neg_1_036_14    m/z 293.2970     0   1   2   3E+4   1   1.5  RT (min)      lcqtof_t3_neg_1_036_14    m/z 293.3446     0   2   4E+4   1   1.5  RT (min)      lcqtof_t3_neg_1_036_14    m/z 293.3551     0   1   2E+4   9   9.5  RT (min)      lcqtof_t3_neg_4_000_ee    m/z 293.4004     0   2   4E+4   1   1.5  RT (min)      lcqtof_t3_neg_1_036_14    m/z 294.4103     0   2   4E+4   1   1.5  RT (min)      lcqtof_t3_neg_1_036_14    m/z 294.4559     0   1   2   3E+4   1   1.5  RT (min)      lcqtof_t3_neg_1_038_15    m/z 294.8730     0   1E+4   0.5   1  RT (min)      lcqtof_t3_neg_4_000_ee    m/z 294.8911     0   1   2E+3   16.5   17  RT (min)      lcqtof_t3_neg_1_040_08    m/z 294.9932     0   1E+4   1.5   2  RT (min)      lcqtof_t3_neg_4_016_14    m/z 295.1044     0   1E+4   1   1.5  RT (min)      lcqtof_t3_neg_4_011_32    m/z 295.1807     0   1   2E+4   9   9.5  RT (min)      lcqtof_t3_neg_4_000_ee    m/z 295.1820     0   1   2E+4   9   9.5  RT (min)      lcqtof_t3_neg_4_000_ee    m/z 295.1861     0   1   2E+4   9   9.5  RT (min)      lcqtof_t3_neg_4_000_ee    m/z 295.2764     0   2   4   6E+2   10.5   11  RT (min)      lcqtof_t3_neg_3_032_24    m/z 295.2768     0   2   4E+4   11   11.5  RT (min)      lcqtof_t3_neg_2_016_05    m/z 295.2770     0   1   2   3   4E+4   11   11.5  RT (min)      lcqtof_t3_neg_2_016_05    m/z 296.2893     0   1E+4   11.5   12  RT (min)      lcqtof_t3_neg_1_006_05    m/z 296.8011     0   1E+4   1   1.5  RT (min)      lcqtof_t3_neg_4_059_14    m/z 296.8895     0   1   2E+3   15   15.5  RT (min)      lcqtof_t3_neg_4_107_62    m/z 297.0467     0   1   2E+4   13   13.5  RT (min)       lcqtof_t3_neg_4_064_17    m/z 297.1516     0   1   2E+4   16.5   17  RT (min)      lcqtof_t3_neg_4_091_60    m/z 297.1520     0   1E+4   12   12.5  RT (min)      lcqtof_t3_neg_4_074_58    m/z 297.1521     0   1E+4   12   12.5  RT (min)      lcqtof_t3_neg_4_095_60    m/z 297.1523     0   1   2E+4   14.5   15  RT (min)       lcqtof_t3_neg_4_103_61    m/z 297.1524     0   1   2E+4   13.5   14  RT (min)       lcqtof_t3_neg_4_094_60    m/z 297.1524     0   1   2E+4   14.5   15  RT (min)       lcqtof_t3_neg_4_074_58    m/z 297.1524     0   1   2E+4   14   14.5  RT (min)       lcqtof_t3_neg_4_090_60    m/z 297.1524     0   1   2E+4   14.5   15  RT (min)       lcqtof_t3_neg_4_106_62    m/z 297.1524     0   1   2E+4   14   14.5  RT (min)       lcqtof_t3_neg_4_091_60    m/z 297.1524     0   1   2E+4   16   16.5  RT (min)       lcqtof_t3_neg_4_084_59    m/z 297.1524     0   1   2E+4   16   16.5  RT (min)       lcqtof_t3_neg_4_098_61    m/z 297.1524     0   1   2E+4   15   15.5  RT (min)       lcqtof_t3_neg_4_080_58    m/z 297.1525     0   1   2E+4   15.5   16  RT (min)       lcqtof_t3_neg_4_000_ee    m/z 297.1525     0   1E+4   12   12.5  RT (min)      lcqtof_t3_neg_4_075_58    m/z 297.1525     0   1E+4   12.5   13  RT (min)      lcqtof_t3_neg_4_085_59    m/z 297.1526     0   1E+4   12   12.5  RT (min)      lcqtof_t3_neg_4_097_60    m/z 297.1526     0   2   4   6   8E+3   10.5   11  RT (min)      lcqtof_t3_neg_4_107_62    m/z 297.1526     0   1   2E+4   15   15.5  RT (min)       lcqtof_t3_neg_4_068_57    m/z 297.1526     0   1   2E+4   14   14.5  RT (min)       lcqtof_t3_neg_4_090_60    m/z 297.1526     0   1E+4   13   13.5  RT (min)       lcqtof_t3_neg_4_107_62    m/z 297.1526     0   1   2E+4   15.5   16  RT (min)      lcqtof_t3_neg_4_098_61    m/z 297.2428     0   2   4   6E+3   10   10.5  RT (min)       lcqtof_t3_neg_2_025_52    m/z 297.2930     0   2   4   6E+4   11.5   12  RT (min)      lcqtof_t3_neg_2_016_05    m/z 297.2932     0   2   4   6E+4   11.5   12  RT (min)      lcqtof_t3_neg_2_016_05    m/z 297.2933     0   2   4   6   8E+3   13   13.5  RT (min)      lcqtof_t3_neg_1_031_24    m/z 298.1564     0   2   4E+3   14.5   15  RT (min)      lcqtof_t3_neg_4_070_57    m/z 298.1830     0   1E+4   7.5   8  RT (min)      lcqtof_t3_neg_4_000_ee    m/z 298.8870     0   1   2E+3   15.5   16  RT (min)      lcqtof_t3_neg_4_105_61    m/z 299.1563     0   2   4   6E+3   2   2.5  RT (min)       lcqtof_t3_neg_4_096_60    m/z 299.1567     0   1E+4   1   1.5  RT (min)      lcqtof_t3_neg_4_040_24    m/z 299.2053     0   1   2   3   4E+4   6.5   7  RT (min)      lcqtof_t3_neg_4_000_ee    m/z 299.2587     0   2   4   6   8E+3   10.5   11  RT (min)      lcqtof_t3_neg_4_055_38    m/z 299.2588     0   2   4   6E+3   10.5   11  RT (min)      lcqtof_t3_neg_2_030_23    m/z 299.2588     0   2   4   6E+3   10.5   11  RT (min)      lcqtof_t3_neg_1_043_38    m/z 299.2590     0   1E+4   11.5   12  RT (min)      lcqtof_t3_neg_4_003_37    m/z 299.2592     0   1E+4   11   11.5  RT (min)      lcqtof_t3_neg_4_018_23    m/z 299.2593     0   1E+4   11   11.5  RT (min)      lcqtof_t3_neg_2_030_23    m/z 299.3080     0   1   2   3E+4   11.5   12  RT (min)      lcqtof_t3_neg_2_016_05    m/z 299.3084     0   1   2   3E+4   11.5   12  RT (min)      lcqtof_t3_neg_2_016_05    m/z 299.3085     0   1   2   3E+4   11.5   12  RT (min)      lcqtof_t3_neg_2_016_05    m/z 299.3087     0   2   4   6E+3   13   13.5  RT (min)      lcqtof_t3_neg_4_040_24    m/z 299.3088     0   2   4   6E+3   13   13.5  RT (min)      lcqtof_t3_neg_4_040_24    m/z 301.0393     0   2   4   6E+3   15   15.5  RT (min)       lcqtof_t3_neg_4_108_62    m/z 301.0394     0   1   2   3   4E+4   4   4.5  RT (min)       lcqtof_t3_neg_4_005_18    m/z 301.0396     0   2   4E+3   15.5   16  RT (min)       lcqtof_t3_neg_4_073_57    m/z 301.0396     0   2   4   6E+3   15   15.5  RT (min)       lcqtof_t3_neg_4_087_59    m/z 301.0398     0   2   4E+3   1   1.5  RT (min)       lcqtof_t3_neg_4_007_09    m/z 301.1654     0   1E+4   7.5   8  RT (min)      lcqtof_t3_neg_3_000_aa    m/z 301.1656     0   1E+4   7.5   8  RT (min)      lcqtof_t3_neg_4_000_ee    m/z 301.1673     0   1E+4   8   8.5  RT (min)      lcqtof_t3_neg_4_000_ee    m/z 302.1094     0   1E+4   4   4.5  RT (min)      lcqtof_t3_neg_3_032_24    m/z 303.2327     0   1   2E+3   13   13.5  RT (min)      lcqtof_t3_neg_1_060_15    m/z 304.2358     0   1   2   3   4E+3   10   10.5  RT (min)      lcqtof_t3_neg_4_037_01    m/z 304.9129     0   1E+4   16   16.5  RT (min)      lcqtof_t3_neg_1_019_23    m/z 304.9132     0   1E+2   15.5   16  RT (min)      lcqtof_t3_neg_3_032_24    m/z 304.9132     0   1E+4   17.5   18  RT (min)       lcqtof_t3_neg_1_049_55    m/z 304.9132     0   1E+4   8   8.5  RT (min)       lcqtof_t3_neg_1_017_51    m/z 304.9132     0   2   4   6   8E+3   17.5   18  RT (min)       lcqtof_t3_neg_1_006_05    m/z 304.9132     0   1E+2   15.5   16  RT (min)      lcqtof_t3_neg_3_032_24    m/z 304.9134     0   1E+4   7   8   9  RT (min)       lcqtof_t3_neg_1_003_07    m/z 304.9136     0   1E+4   7   8  RT (min)       lcqtof_t3_neg_1_004_11    m/z 304.9136     0   1   2E+2   15.5   16  RT (min)      lcqtof_t3_neg_3_032_24    m/z 304.9138     0   1E+4   7   8   9  RT (min)       lcqtof_t3_neg_1_012_16    m/z 304.9235     0   1E+4   1   1.5  RT (min)      lcqtof_t3_neg_4_014_13    m/z 305.0512     0   1E+4   1   1.5  RT (min)      lcqtof_t3_neg_4_018_23    m/z 305.0762     0   2   4   6E+2   1   1.5  RT (min)      lcqtof_t3_neg_4_030_31    m/z 305.0983     0   1   2E+4   2   2.5  RT (min)      lcqtof_t3_neg_4_051_15    m/z 305.0983     0   1E+4   2   2.5  RT (min)      lcqtof_t3_neg_4_002_17    m/z 305.0986     0   1   2E+4   2   2.5  RT (min)       lcqtof_t3_neg_4_060_15    m/z 305.0988     0   1   2E+4   2   2.5  RT (min)      lcqtof_t3_neg_4_051_15    m/z 306.0760     0   1E+4   2   2.5  RT (min)       lcqtof_t3_neg_1_035_31    m/z 306.0761     0   2   4E+4   1.5   2  RT (min)      lcqtof_t3_neg_1_019_23    m/z 306.0836     0   1   2   3E+3   6.5   7  RT (min)       lcqtof_t3_neg_4_048_19    m/z 306.0874     0   1E+4   1   1.5  RT (min)      lcqtof_t3_neg_4_057_49    m/z 306.1932     0   1E+4   10   10.5  RT (min)      lcqtof_t3_neg_4_000_ee    m/z 306.2048     0   1E+4   10   10.5  RT (min)      lcqtof_t3_neg_4_000_ee    m/z 306.2181     0   1E+4   10   10.5  RT (min)      lcqtof_t3_neg_4_064_17    m/z 306.9184     0   2   4E+3   15   15.5  RT (min)       lcqtof_t3_neg_4_079_58    m/z 306.9185     0   1E+4   16   16.5  RT (min)       lcqtof_t3_neg_4_011_32    m/z 306.9186     0   1E+4   10   15  RT (min)       lcqtof_t3_neg_4_000_ee    m/z 306.9187     0   1   2E+3   11.5   12  RT (min)       lcqtof_t3_neg_4_040_24    m/z 306.9222     0   1   2   3E+3   8  RT (min)       lcqtof_t3_neg_4_035_08    m/z 306.9264     0   1E+4   16.5   17  RT (min)       lcqtof_t3_neg_4_044_48    m/z 306.9265     0   1   2   3E+3   7.5   8  RT (min)       lcqtof_t3_neg_4_001_49    m/z 306.9287     0   1   2   3E+3   7   8  RT (min)       lcqtof_t3_neg_4_037_01    m/z 306.9312     0   1   2   3E+3   4.5   5  RT (min)      lcqtof_t3_neg_4_000_aa    m/z 306.9351     0   1   2   3E+3   17   17.5  RT (min)       lcqtof_t3_neg_1_014_04    m/z 307.0712     0   2   4E+3   2   2.5  RT (min)      lcqtof_t3_neg_1_035_31    m/z 307.0802     0   2   4   6E+3   6.5   7  RT (min)       lcqtof_t3_neg_4_048_19    m/z 307.1577     0   2   4   6E+3   14   14.5  RT (min)       lcqtof_t3_neg_4_008_35    m/z 307.1578     0   2   4   6E+3   14.5   15  RT (min)       lcqtof_t3_neg_4_005_18    m/z 307.1582     0   2   4   6E+3   14.5   15  RT (min)       lcqtof_t3_neg_4_000_bb    m/z 308.8690     0   1E+4   0.5   1  RT (min)      lcqtof_t3_neg_4_000_ee    m/z 308.9160     0   1   2E+3   15.5   16  RT (min)      lcqtof_t3_neg_4_113_62    m/z 309.1622     0   1   2E+3   14.5   15  RT (min)      lcqtof_t3_neg_4_000_dd    m/z 309.1702     0   2   4   6   8E+3   10   10.5  RT (min)      lcqtof_t3_neg_2_038_18    m/z 309.1703     0   2   4   6   8E+3   10   10.5  RT (min)      lcqtof_t3_neg_1_034_30    m/z 309.1734     0   1   2E+2   8   8.5  RT (min)      lcqtof_t3_neg_3_038_06    m/z 310.0958     0   1E+4   1   1.5  RT (min)      lcqtof_t3_neg_1_055_32    m/z 310.1403     0   1E+5   5   5.5  RT (min)       lcqtof_t3_neg_2_053_38    m/z 310.1403     0   1E+5   5   5.5  RT (min)       lcqtof_t3_neg_2_053_38    m/z 310.1404     0   1E+5   5   5.5  RT (min)       lcqtof_t3_neg_1_043_38    m/z 310.1405     0   1   2E+4   5.5   6  RT (min)       lcqtof_t3_neg_1_013_09    m/z 310.9302     0   2   4E+3   17   17.5  RT (min)       lcqtof_t3_neg_4_035_08    m/z 310.9302     0   2   4   6E+3   16.5   17  RT (min)      lcqtof_t3_neg_4_113_62    m/z 310.9304     0   2   4E+3   10  RT (min)       lcqtof_t3_neg_1_005_18    m/z 310.9305     0   2   4E+3   9.5   10  RT (min)       lcqtof_t3_neg_1_029_44    m/z 310.9306     0   2   4   6E+3   16   16.5  RT (min)      lcqtof_t3_neg_1_009_50    m/z 311.0962     0   1   2E+3   16.5   17  RT (min)      lcqtof_t3_neg_4_003_37    m/z 311.0972     0   1   2   3E+3   16   16.5  RT (min)      lcqtof_t3_neg_4_065_51    m/z 311.0974     0   1E+4   4   4.5  RT (min)      lcqtof_t3_neg_4_062_13    m/z 311.0981     0   1   2   3E+3   15   15.5  RT (min)      lcqtof_t3_neg_4_030_31    m/z 311.1669     0   1E+4   7.5   8  RT (min)      lcqtof_t3_neg_4_000_ee    m/z 311.1677     0   1   2E+4   13.5   14  RT (min)      lcqtof_t3_neg_3_065_51    m/z 311.1680     0   2   4   6E+4   15.5   16  RT (min)       lcqtof_t3_neg_4_106_62    m/z 311.1680     0   1   2   3   4E+4   11.5   12  RT (min)       lcqtof_t3_neg_4_059_14    m/z 311.1680     0   1   2   3   4E+4   12   13  RT (min)       lcqtof_t3_neg_4_099_61    m/z 311.1680     0   2   4   6E+4   13.5   14   14.5  RT (min)       lcqtof_t3_neg_4_065_51    m/z 311.1680     0   1   2   3   4E+4   12   12.5  RT (min)       lcqtof_t3_neg_4_059_14    m/z 311.1680     0   1   2   3E+4   11.5   12  RT (min)       lcqtof_t3_neg_4_065_51    m/z 311.1680     0   2   4   6E+4   15.5   16  RT (min)       lcqtof_t3_neg_4_065_51    m/z 311.1680     0   1   2E+4   11   11.5  RT (min)       lcqtof_t3_neg_4_100_61    m/z 311.1680     0   2   4   6E+4   14   14.5  RT (min)       lcqtof_t3_neg_4_097_60    m/z 311.1680     0   2   4   6E+4   14.5   15  RT (min)       lcqtof_t3_neg_4_062_13    m/z 311.1680     0   2   4   6   8E+4   14.5   15  RT (min)       lcqtof_t3_neg_4_099_61    m/z 311.1680     0   1   2   3E+4   12.5   13  RT (min)       lcqtof_t3_neg_4_106_62    m/z 311.1680     0   1   2   3   4E+4   13   13.5  RT (min)       lcqtof_t3_neg_4_104_61    m/z 311.1680     0   2   4   6E+4   14.5   15  RT (min)       lcqtof_t3_neg_4_113_62    m/z 311.1680     0   1   2   3   4E+4   11   11.5  RT (min)       lcqtof_t3_neg_4_000_ee    m/z 311.1680     0   1E+4   13.5   14  RT (min)       lcqtof_t3_neg_3_100_61    m/z 311.1681     0   1   2   3   4E+4   12   12.5  RT (min)       lcqtof_t3_neg_4_059_14    m/z 311.1681     0   2   4   6E+4   16   16.5  RT (min)       lcqtof_t3_neg_4_087_59    m/z 311.1681     0   2   4   6E+4   16   16.5  RT (min)       lcqtof_t3_neg_4_059_14    m/z 311.1681     0   2   4E+4   13.5   14  RT (min)       lcqtof_t3_neg_4_089_59    m/z 311.1681     0   2   4   6E+4   16   16.5  RT (min)       lcqtof_t3_neg_4_065_51    m/z 311.1681     0   2   4   6E+4   15.5   16  RT (min)       lcqtof_t3_neg_4_098_61    m/z 311.1681     0   2   4   6E+4   16.5   17  RT (min)      lcqtof_t3_neg_4_091_60    m/z 311.1682     0   1   2   3   4E+4   11.5   12  RT (min)       lcqtof_t3_neg_4_059_14    m/z 311.1682     0   2   4   6E+4   13.5   14  RT (min)       lcqtof_t3_neg_4_100_61    m/z 311.1682     0   2   4   6   8E+4   14.5   15  RT (min)       lcqtof_t3_neg_4_000_ee    m/z 311.1682     0   1   2   3E+4   10.5   11  RT (min)       lcqtof_t3_neg_4_096_60    m/z 311.1682     0   2   4   6E+4   14.5   15  RT (min)       lcqtof_t3_neg_4_107_62    m/z 311.1682     0   2   4   6E+4   10   10.5  RT (min)       lcqtof_t3_neg_4_100_61    m/z 311.1682     0   1   2E+4   11   11.5  RT (min)       lcqtof_t3_neg_4_065_51    m/z 311.1682     0   1   2E+4   10.5   11  RT (min)       lcqtof_t3_neg_4_073_57    m/z 311.1682     0   2   4   6E+4   14   14.5  RT (min)       lcqtof_t3_neg_4_062_13    m/z 311.1683     0   2   4   6E+4   14   14.5  RT (min)       lcqtof_t3_neg_4_057_49    m/z 311.1683     0   1   2E+4   10.5   11  RT (min)       lcqtof_t3_neg_4_082_59    m/z 311.1684     0   1   2   3   4E+4   12   12.5  RT (min)       lcqtof_t3_neg_4_059_14    m/z 311.1685     0   2   4   6E+4   16   16.5  RT (min)       lcqtof_t3_neg_4_065_51    m/z 311.2041     0   1E+4   1   1.5  RT (min)      lcqtof_t3_neg_1_019_23    m/z 311.2940     0   1   2   3   4E+4   1   1.5  RT (min)      lcqtof_t3_neg_1_019_23    m/z 311.3525     0   1E+4   9.5   10  RT (min)      lcqtof_t3_neg_4_000_ee    m/z 312.1712     0   1E+4   13.5   14  RT (min)      lcqtof_t3_neg_4_104_61    m/z 312.1713     0   1E+4   14.5   15  RT (min)      lcqtof_t3_neg_4_100_61    m/z 312.1714     0   1E+4   16   16.5  RT (min)      lcqtof_t3_neg_4_087_59    m/z 313.0778     0   1E+5   12   12.5  RT (min)       lcqtof_t3_neg_4_064_17    m/z 313.0778     0   1   2E+5   13.5   14  RT (min)       lcqtof_t3_neg_4_064_17    m/z 313.0779     0   1   2   3   4E+4   12   12.5  RT (min)       lcqtof_t3_neg_4_064_17    m/z 313.0780     0   1   2E+5   13.5   14  RT (min)       lcqtof_t3_neg_4_064_17    m/z 313.0780     0   1   2E+5   13   13.5  RT (min)       lcqtof_t3_neg_4_064_17    m/z 313.0780     0   1E+5   12.5   13  RT (min)       lcqtof_t3_neg_4_036_47    m/z 313.0780     0   1E+5   13.5   14  RT (min)       lcqtof_t3_neg_4_064_17    m/z 313.0780     0   1   2   3   4E+4   11.5   12  RT (min)       lcqtof_t3_neg_4_064_17    m/z 313.0781     0   1   2E+5   13   13.5  RT (min)      lcqtof_t3_neg_4_064_17    m/z 313.0781     0   1E+5   14   14.5  RT (min)      lcqtof_t3_neg_4_064_17    m/z 313.0781     0   1E+5   13.5   14  RT (min)       lcqtof_t3_neg_4_064_17    m/z 313.0782     0   1   2E+4   14   14.5  RT (min)      lcqtof_t3_neg_4_036_47    m/z 313.0782     0   1E+5   13   13.5  RT (min)      lcqtof_t3_neg_4_036_47    m/z 313.0783     0   1E+5   13.5   14  RT (min)       lcqtof_t3_neg_4_064_17    m/z 313.0783     0   1E+5   13.5   14  RT (min)      lcqtof_t3_neg_4_064_17    m/z 313.0787     0   1E+5   13.5   14  RT (min)      lcqtof_t3_neg_4_064_17    m/z 313.0789     0   1E+5   12   12.5  RT (min)      lcqtof_t3_neg_4_064_17    m/z 313.0793     0   2   4   6   8E+4   12   12.5  RT (min)      lcqtof_t3_neg_4_064_17    m/z 313.0904     0   1   2   3   4E+4   1   1.5  RT (min)       lcqtof_t3_neg_4_040_24    m/z 313.2450     0   1   2E+4   1   1.5  RT (min)      lcqtof_t3_neg_4_059_14    m/z 314.0787     0   1   2   3E+4   13   13.5  RT (min)      lcqtof_t3_neg_3_089_59    m/z 314.0789     0   1   2   3E+4   12.5   13  RT (min)      lcqtof_t3_neg_3_089_59    m/z 314.0790     0   1   2   3E+4   13   13.5  RT (min)      lcqtof_t3_neg_3_089_59    m/z 314.2415     0   1   2   3   4E+4   9.5   10  RT (min)      lcqtof_t3_neg_4_000_ee    m/z 314.9043     0   1E+4   1   1.5  RT (min)      lcqtof_t3_neg_2_016_05    m/z 315.0567     0   1   2E+4   1   1.5  RT (min)      lcqtof_t3_neg_2_028_24    m/z 315.0759     0   1   2E+4   13   13.5  RT (min)      lcqtof_t3_neg_3_089_59    m/z 315.0763     0   1   2E+4   12.5   13  RT (min)      lcqtof_t3_neg_3_089_59    m/z 315.1427     0   1   2E+3   14.5   15  RT (min)      lcqtof_t3_neg_4_000_ff    m/z 316.1095     0   2   4   6   8E+3   2   2.5  RT (min)      lcqtof_t3_neg_1_055_32    m/z 316.1772     0   1E+4   6.5   7  RT (min)      lcqtof_t3_neg_4_000_ee    m/z 316.1924     0   1E+4   5.5   6  RT (min)      lcqtof_t3_neg_4_000_ee    m/z 316.8874     0   1   2E+3   16.5   17  RT (min)      lcqtof_t3_neg_1_000_bb    m/z 316.9468     0   1E+4   10.5   11  RT (min)       lcqtof_t3_neg_4_108_62    m/z 316.9469     0   1E+4   7.5   8  RT (min)       lcqtof_t3_neg_4_101_61    m/z 316.9469     0   1E+4   9   9.5  RT (min)       lcqtof_t3_neg_4_105_61    m/z 316.9469     0   1E+4   10   10.5  RT (min)       lcqtof_t3_neg_4_106_62    m/z 316.9470     0   2   4   6E+3   14   14.5  RT (min)       lcqtof_t3_neg_4_107_62    m/z 316.9470     0   1E+4   7.5   8  RT (min)       lcqtof_t3_neg_4_112_62    m/z 316.9470     0   1   2E+3   13.5   14  RT (min)       lcqtof_t3_neg_3_038_06    m/z 316.9470     0   1E+4   10   10.5  RT (min)       lcqtof_t3_neg_4_106_62    m/z 316.9470     0   1E+4   8.5   9  RT (min)       lcqtof_t3_neg_4_109_62    m/z 316.9470     0   2   4E+3   15   15.5  RT (min)       lcqtof_t3_neg_3_032_24    m/z 316.9470     0   2   4   6   8E+3   12   12.5  RT (min)       lcqtof_t3_neg_4_107_62    m/z 316.9470     0   1E+4   15   15.5  RT (min)       lcqtof_t3_neg_4_098_61    m/z 316.9470     0   1E+4   8.5   9  RT (min)       lcqtof_t3_neg_4_101_61    m/z 316.9471     0   1E+5   16   16.5  RT (min)       lcqtof_t3_neg_4_018_23    m/z 316.9471     0   1E+4   10   12  RT (min)       lcqtof_t3_neg_4_101_61    m/z 316.9471     0   1E+4   9.5   10  RT (min)       lcqtof_t3_neg_4_113_62    m/z 316.9471     0   2   4   6   8E+3   12   12.5  RT (min)       lcqtof_t3_neg_4_113_62    m/z 316.9471     0   1E+5   16   16.5  RT (min)       lcqtof_t3_neg_4_018_23    m/z 316.9471     0   1E+4   8   8.5  RT (min)       lcqtof_t3_neg_4_106_62    m/z 316.9472     0   2   4   6   8E+3   11   11.5  RT (min)       lcqtof_t3_neg_4_104_61    m/z 316.9472     0   1E+4   9   9.5  RT (min)       lcqtof_t3_neg_4_106_62    m/z 316.9472     0   1E+4   15   15.5  RT (min)       lcqtof_t3_neg_4_041_54    m/z 316.9472     0   1E+4   15.5   16  RT (min)       lcqtof_t3_neg_4_059_14    m/z 316.9472     0   1E+4   9.5   10  RT (min)       lcqtof_t3_neg_4_107_62    m/z 316.9473     0   2   4   6E+4   17   17.5  RT (min)       lcqtof_t3_neg_4_035_08    m/z 316.9473     0   2   4   6   8E+3   14.5   15  RT (min)       lcqtof_t3_neg_4_101_61    m/z 316.9473     0   2   4   6   8E+3   12   12.5  RT (min)       lcqtof_t3_neg_4_111_62    m/z 316.9473     0   2   4   6   8E+3   12   12.5  RT (min)       lcqtof_t3_neg_4_106_62    m/z 316.9474     0   1E+4   10.5   11  RT (min)       lcqtof_t3_neg_4_107_62    m/z 316.9474     0   1E+4   11   11.5  RT (min)       lcqtof_t3_neg_4_106_62    m/z 316.9474     0   2   4   6E+3   14   14.5  RT (min)       lcqtof_t3_neg_4_065_51    m/z 316.9474     0   1E+4   10   10.5  RT (min)       lcqtof_t3_neg_4_113_62    m/z 316.9475     0   2   4E+3   9.5   10  RT (min)       lcqtof_t3_neg_4_000_bb    m/z 316.9478     0   2   4   6E+3   10   10.5  RT (min)       lcqtof_t3_neg_4_051_15    m/z 316.9522     0   1   2E+5   17   17.5  RT (min)       lcqtof_t3_neg_4_106_62    m/z 317.1536     0   1E+4   7.5   8  RT (min)      lcqtof_t3_neg_4_000_ee    m/z 317.1890     0   1   2E+4   10   10.5  RT (min)      lcqtof_t3_neg_4_000_ee    m/z 317.1922     0   1   2E+4   10   10.5  RT (min)      lcqtof_t3_neg_4_000_ee    m/z 317.3186     0   1   2   3   4E+3   11   11.5  RT (min)       lcqtof_t3_neg_3_032_24    m/z 317.3190     0   2   4   6   8E+3   10.5   11  RT (min)       lcqtof_t3_neg_1_031_24    m/z 317.9357     0   1   2   3   4E+4   1   1.5  RT (min)      lcqtof_t3_neg_1_008_02    m/z 319.1008     0   1   2   3   4E+4   1   1.5  RT (min)       lcqtof_t3_neg_4_082_59    m/z 319.9332     0   1   2E+4   1   1.5  RT (min)      lcqtof_t3_neg_1_008_02    m/z 320.1176     0   1E+4   4   4.5  RT (min)      lcqtof_t3_neg_4_058_16    m/z 320.2756     0   1   2E+4   6.5   7  RT (min)      lcqtof_t3_neg_4_000_ee    m/z 320.8863     0   1   2   3E+3   16.5   17  RT (min)       lcqtof_t3_neg_4_000_ee    m/z 320.8864     0   1   2E+3   17   17.5  RT (min)       lcqtof_t3_neg_4_107_62    m/z 320.9759     0   1E+4   6.5   7  RT (min)      lcqtof_t3_neg_2_013_26    m/z 320.9763     0   1E+4   6.5   7  RT (min)      lcqtof_t3_neg_4_024_26    m/z 321.1789     0   1E+4   1   1.5  RT (min)      lcqtof_t3_neg_2_028_24    m/z 321.2100     0   2   4   6   8E+3   9.5   10  RT (min)       lcqtof_t3_neg_4_031_42    m/z 321.2102     0   2   4   6   8E+3   9.5   10  RT (min)      lcqtof_t3_neg_4_031_42    m/z 321.2399     0   1   2   3E+4   1   1.5  RT (min)      lcqtof_t3_neg_1_031_24    m/z 321.4696     0   1   2E+4   1   1.5  RT (min)      lcqtof_t3_neg_1_031_24    m/z 321.9036     0   2   4E+3   16.5   17  RT (min)       lcqtof_t3_neg_1_037_25    m/z 322.1611     0   1E+4   7   7.5  RT (min)      lcqtof_t3_neg_4_000_ee    m/z 322.9062     0   2   4   6E+3   17   17.5  RT (min)      lcqtof_t3_neg_4_035_08    m/z 322.9741     0   1E+4   6.5   7  RT (min)      lcqtof_t3_neg_4_024_26    m/z 322.9810     0   1E+4   5   5.5  RT (min)       lcqtof_t3_neg_4_059_14    m/z 323.0116     0   1   2E+4   17   17.5  RT (min)       lcqtof_t3_neg_4_000_bb    m/z 323.0118     0   1   2E+4   16.5   17  RT (min)       lcqtof_t3_neg_4_000_bb    m/z 323.0119     0   1   2E+4   17   17.5  RT (min)       lcqtof_t3_neg_4_000_bb    m/z 323.0121     0   1   2E+4   16.5   17  RT (min)      lcqtof_t3_neg_4_000_bb    m/z 323.0122     0   1   2E+4   17   17.5  RT (min)       lcqtof_t3_neg_4_000_bb    m/z 323.0130     0   1   2E+4   17   17.5  RT (min)      lcqtof_t3_neg_4_000_bb    m/z 323.1064     0   1   2   3   4E+3   9   9.5  RT (min)       lcqtof_t3_neg_4_000_ee    m/z 323.1784     0   1E+3   9.5   10  RT (min)      lcqtof_t3_neg_1_025_52    m/z 325.1834     0   2   4   6   8E+4   15.5   16  RT (min)       lcqtof_t3_neg_4_106_62    m/z 325.1834     0   1   2   3   4E+4   13   13.5  RT (min)       lcqtof_t3_neg_4_099_61    m/z 325.1835     0   1   2   3E+4   11.5   12  RT (min)       lcqtof_t3_neg_4_065_51    m/z 325.1835     0   2   4E+4   13.5   14  RT (min)       lcqtof_t3_neg_4_082_59    m/z 325.1835     0   2   4   6   8E+4   16   16.5  RT (min)       lcqtof_t3_neg_4_057_49    m/z 325.1836     0   2   4   6   8E+4   16   16.5  RT (min)       lcqtof_t3_neg_4_057_49    m/z 325.1836     0   2   4   6   8E+4   15   15.5  RT (min)       lcqtof_t3_neg_4_057_49    m/z 325.1836     0   2   4   6E+4   13.5   14  RT (min)       lcqtof_t3_neg_4_091_60    m/z 325.1836     0   2   4   6E+4   14   14.5  RT (min)       lcqtof_t3_neg_4_097_60    m/z 325.1836     0   1   2   3E+4   12   12.5  RT (min)       lcqtof_t3_neg_4_065_51    m/z 325.1836     0   1   2   3E+4   12.5   13  RT (min)       lcqtof_t3_neg_4_100_61    m/z 325.1836     0   1E+5   14.5   15  RT (min)       lcqtof_t3_neg_4_000_dd    m/z 325.1836     0   2   4   6   8E+4   15   15.5  RT (min)       lcqtof_t3_neg_4_057_49    m/z 325.1837     0   1E+5   10.5   11  RT (min)       lcqtof_t3_neg_4_097_60    m/z 325.1837     0   2   4   6   8E+4   16   16.5  RT (min)       lcqtof_t3_neg_4_057_49    m/z 325.1837     0   2   4   6E+4   13.5   14   14.5  RT (min)       lcqtof_t3_neg_4_057_49    m/z 325.1838     0   2   4   6   8E+4   16   16.5  RT (min)       lcqtof_t3_neg_4_057_49    m/z 325.1838     0   2   4   6   8E+4   15.5   16  RT (min)       lcqtof_t3_neg_4_057_49    m/z 325.1838     0   2   4   6   8E+4   14.5   15  RT (min)       lcqtof_t3_neg_4_068_57    m/z 325.1838     0   2   4   6   8E+4   15   15.5  RT (min)      lcqtof_t3_neg_4_065_51    m/z 325.1838     0   1   2   3E+4   11.5   12  RT (min)       lcqtof_t3_neg_4_057_49    m/z 325.1838     0   2   4   6E+4   14   14.5  RT (min)       lcqtof_t3_neg_4_090_60    m/z 325.1839     0   1   2   3E+4   13   13.5  RT (min)       lcqtof_t3_neg_4_104_61    m/z 325.1839     0   1   2   3E+4   11.5   12  RT (min)       lcqtof_t3_neg_4_057_49    m/z 325.3724     0   1   2E+4   10   10.5  RT (min)      lcqtof_t3_neg_4_000_ee    m/z 326.1615     0   1E+4   7   7.5  RT (min)      lcqtof_t3_neg_4_000_ee    m/z 326.1860     0   1E+4   16.5   17  RT (min)      lcqtof_t3_neg_4_091_60    m/z 326.1867     0   1E+4   14.5   15  RT (min)      lcqtof_t3_neg_4_107_62    m/z 326.1867     0   1E+4   13.5   14  RT (min)      lcqtof_t3_neg_4_093_60    m/z 326.1868     0   1E+4   16   16.5  RT (min)      lcqtof_t3_neg_4_086_59    m/z 326.1870     0   1E+4   14   14.5  RT (min)      lcqtof_t3_neg_4_093_60    m/z 326.1870     0   1E+4   13.5   14  RT (min)      lcqtof_t3_neg_4_094_60    m/z 326.1870     0   1E+4   15.5   16  RT (min)      lcqtof_t3_neg_4_074_58    m/z 326.1872     0   1E+4   15   15.5  RT (min)      lcqtof_t3_neg_4_077_58    m/z 326.9040     0   1E+4   17   17.5  RT (min)       lcqtof_t3_neg_4_112_62    m/z 327.1346     0   1E+4   7.5   8  RT (min)      lcqtof_t3_neg_4_062_13    m/z 327.1347     0   1E+4   8   8.5  RT (min)      lcqtof_t3_neg_4_062_13    m/z 327.2323     0   1   2   3E+3   10   10.5  RT (min)      lcqtof_t3_neg_1_002_13    m/z 327.2325     0   2   4E+3   10   10.5  RT (min)      lcqtof_t3_neg_4_037_01    m/z 328.1091     0   1E+4   1   1.5  RT (min)      lcqtof_t3_neg_4_011_32    m/z 328.1845     0   2   4   6   8E+3   10   10.5  RT (min)      lcqtof_t3_neg_4_000_ee    m/z 328.9024     0   1   2   3E+3   16.5   17  RT (min)      lcqtof_t3_neg_4_000_cc    m/z 329.0594     0   1E+4   1   1.5  RT (min)      lcqtof_t3_neg_4_032_04    m/z 330.1079     0   1   2   3E+4   4   4.5  RT (min)       lcqtof_t3_neg_4_018_23    m/z 330.1318     0   1E+4   7.5   8  RT (min)      lcqtof_t3_neg_4_000_ee    m/z 330.2364     0   1   2   3   4E+3   8.5   9  RT (min)      lcqtof_t3_neg_4_000_ee    m/z 330.8779     0   1   2   3E+4   1   1.5  RT (min)       lcqtof_t3_neg_2_016_05    m/z 331.0504     0   1E+4   0.5   1  RT (min)      lcqtof_t3_neg_4_000_aa    m/z 332.1118     0   1E+4   4   4.5  RT (min)      lcqtof_t3_neg_4_018_23    m/z 333.0751     0   2   4   6E+3   6.5   7  RT (min)       lcqtof_t3_neg_4_004_20    m/z 333.0755     0   1   2   3   4E+3   6.5   7  RT (min)       lcqtof_t3_neg_4_107_62    m/z 333.1570     0   1E+4   6.5   7  RT (min)      lcqtof_t3_neg_4_000_ee    m/z 333.1581     0   1E+4   7   7.5  RT (min)      lcqtof_t3_neg_4_000_ee    m/z 333.1690     0   1   2   3E+3   14.5   15  RT (min)       lcqtof_t3_neg_4_056_56    m/z 333.1698     0   1   2   3E+3   16   16.5  RT (min)       lcqtof_t3_neg_4_000_ee    m/z 333.1700     0   1   2   3E+3   16   16.5  RT (min)      lcqtof_t3_neg_4_012_27    m/z 333.1702     0   1   2   3E+3   14.5   15  RT (min)       lcqtof_t3_neg_4_036_47    m/z 333.1705     0   1   2E+3   16   16.5  RT (min)      lcqtof_t3_neg_4_000_bb    m/z 333.1705     0   1   2   3E+3   15.5   16  RT (min)       lcqtof_t3_neg_4_000_ee    m/z 333.1705     0   1   2   3E+3   15.5   16  RT (min)       lcqtof_t3_neg_4_000_aa    m/z 333.1707     0   1   2   3E+3   15   15.5  RT (min)       lcqtof_t3_neg_4_000_ee    m/z 333.1712     0   1   2   3E+3   15   15.5  RT (min)       lcqtof_t3_neg_4_039_06    m/z 333.1713     0   1   2   3E+3   15.5   16  RT (min)       lcqtof_t3_neg_4_024_26    m/z 333.1721     0   1   2   3E+3   15   15.5  RT (min)       lcqtof_t3_neg_4_065_51    m/z 333.9374     0   1   2E+3   15   15.5  RT (min)       lcqtof_t3_neg_4_092_60    m/z 333.9374     0   1   2E+3   15.5   16  RT (min)       lcqtof_t3_neg_1_042_46    m/z 333.9374     0   1E+3   16   16.5  RT (min)       lcqtof_t3_neg_2_016_05    m/z 333.9374     0   1   2   3E+3   15.5   16  RT (min)       lcqtof_t3_neg_4_099_61    m/z 333.9375     0   1   2E+3   15.5   16  RT (min)       lcqtof_t3_neg_1_019_23    m/z 333.9375     0   1E+4   16   16.5  RT (min)      lcqtof_t3_neg_1_041_54    m/z 333.9376     0   1E+3   15   15.5  RT (min)       lcqtof_t3_neg_3_032_24    m/z 333.9376     0   1E+4   16   16.5  RT (min)      lcqtof_t3_neg_1_042_46    m/z 333.9376     0   1E+4   16   16.5  RT (min)       lcqtof_t3_neg_1_011_45    m/z 335.0488     0   2   4   6   8E+3   2   2.5  RT (min)       lcqtof_t3_neg_4_037_01    m/z 335.1620     0   1E+4   7.5   8  RT (min)      lcqtof_t3_neg_4_000_ee    m/z 335.1696     0   1E+4   10.5   11  RT (min)      lcqtof_t3_neg_4_000_ee    m/z 335.1720     0   1E+4   10.5   11  RT (min)      lcqtof_t3_neg_4_000_ee    m/z 335.1972     0   1E+4   6.5   7  RT (min)      lcqtof_t3_neg_4_000_ee    m/z 335.8673     0   1E+4   0.5   1  RT (min)      lcqtof_t3_neg_4_000_ee    m/z 336.8644     0   2   4E+4   0.5   1  RT (min)      lcqtof_t3_neg_4_000_ee    m/z 336.9293     0   1   2   3E+3   16.5   17  RT (min)      lcqtof_t3_neg_4_000_ee    m/z 337.1871     0   1   2   3   4E+3   10   10.5  RT (min)      lcqtof_t3_neg_4_014_13    m/z 337.1887     0   1E+4   9   9.5  RT (min)      lcqtof_t3_neg_4_000_ee    m/z 337.2490     0   1   2   3   4E+3   9.5   10  RT (min)       lcqtof_t3_neg_3_077_58    m/z 337.8675     0   1E+4   0.5   1  RT (min)      lcqtof_t3_neg_4_000_ee    m/z 338.1972     0   1   2   3E+4   8.5   9  RT (min)      lcqtof_t3_neg_4_000_ee    m/z 338.1987     0   1   2   3E+4   8.5   9  RT (min)      lcqtof_t3_neg_4_000_ee    m/z 338.8640     0   1   2   3   4E+4   0.5   1  RT (min)      lcqtof_t3_neg_4_000_ee    m/z 339.0746     0   1   2E+4   1   1.5  RT (min)       lcqtof_t3_neg_4_082_59    m/z 339.1992     0   2   4E+4   14   14.5  RT (min)       lcqtof_t3_neg_4_083_59    m/z 339.1992     0   1E+4   14   14.5  RT (min)      lcqtof_t3_neg_3_084_59    m/z 339.1992     0   1   2   3E+4   13   13.5  RT (min)       lcqtof_t3_neg_4_100_61    m/z 339.1993     0   1E+4   14.5   15  RT (min)      lcqtof_t3_neg_3_098_61    m/z 339.1993     0   1E+5   14.5   15  RT (min)       lcqtof_t3_neg_4_000_aa    m/z 339.1993     0   2   4   6E+4   14.5   15  RT (min)       lcqtof_t3_neg_4_034_34    m/z 339.1993     0   2   4   6E+4   14.5   15  RT (min)       lcqtof_t3_neg_4_036_47    m/z 339.1994     0   2   4E+4   15.5   16  RT (min)       lcqtof_t3_neg_4_102_61    m/z 339.1994     0   2   4E+4   14   14.5  RT (min)      lcqtof_t3_neg_4_070_57    m/z 339.1994     0   1   2E+4   12   12.5  RT (min)      lcqtof_t3_neg_4_080_58    m/z 339.1994     0   2   4E+4   13.5   14  RT (min)       lcqtof_t3_neg_4_094_60    m/z 339.1995     0   2   4E+4   16   16.5  RT (min)       lcqtof_t3_neg_4_090_60    m/z 339.1995     0   1   2E+4   12   12.5  RT (min)       lcqtof_t3_neg_4_070_57    m/z 339.1995     0   1   2   3   4E+4   13.5   14  RT (min)       lcqtof_t3_neg_4_091_60    m/z 339.1995     0   1   2E+4   13   13.5  RT (min)      lcqtof_t3_neg_4_099_61    m/z 339.1997     0   1   2E+4   11   11.5  RT (min)      lcqtof_t3_neg_4_003_37    m/z 339.1997     0   2   4   6E+4   11   11.5  RT (min)       lcqtof_t3_neg_4_000_ee    m/z 339.1997     0   1   2E+4   11.5   12  RT (min)      lcqtof_t3_neg_4_037_01    m/z 339.1997     0   1   2E+4   11   11.5  RT (min)      lcqtof_t3_neg_4_008_35    m/z 339.1998     0   1   2   3E+4   11.5   12  RT (min)      lcqtof_t3_neg_4_003_37    m/z 339.8638     0   1E+4   0.5   1  RT (min)      lcqtof_t3_neg_4_000_ee    m/z 340.1740     0   1E+4   7.5   8  RT (min)      lcqtof_t3_neg_4_000_ee    m/z 340.2020     0   1E+4   15   15.5  RT (min)      lcqtof_t3_neg_4_105_61    m/z 340.2027     0   1E+4   14   14.5  RT (min)      lcqtof_t3_neg_4_113_62    m/z 340.2029     0   1E+4   16.5   17  RT (min)      lcqtof_t3_neg_4_091_60    m/z 340.2044     0   2   4E+3   12.5   13  RT (min)      lcqtof_t3_neg_4_109_62    m/z 340.2055     0   2   4E+3   12   12.5  RT (min)      lcqtof_t3_neg_4_077_58    m/z 340.8619     0   1   2   3E+4   0.5   1  RT (min)      lcqtof_t3_neg_4_000_ee    m/z 341.1087     0   1   2   3E+3   16.5   17  RT (min)      lcqtof_t3_neg_4_027_41    m/z 341.8943     0   1   2   3E+4   1   1.5  RT (min)       lcqtof_t3_neg_4_014_13    m/z 342.0883     0   1   2E+4   1   1.5  RT (min)      lcqtof_t3_neg_1_006_05    m/z 342.1120     0   1   2   3   4E+4   1   1.5  RT (min)      lcqtof_t3_neg_4_047_07    m/z 342.8599     0   1E+4   0.5   1  RT (min)      lcqtof_t3_neg_4_000_ee    m/z 343.1146     0   1E+4   1   1.5  RT (min)      lcqtof_t3_neg_4_047_07    m/z 343.9919     0   1E+4   7.5   8  RT (min)      lcqtof_t3_neg_4_024_26    m/z 344.0901     0   1E+4   1   1.5  RT (min)      lcqtof_t3_neg_4_028_25    m/z 344.1208     0   1   2E+4   4   4.5  RT (min)      lcqtof_t3_neg_4_018_23    m/z 344.1208     0   1   2E+4   4   4.5  RT (min)       lcqtof_t3_neg_4_018_23    m/z 344.1833     0   1E+4   1   1.5  RT (min)      lcqtof_t3_neg_4_030_31    m/z 345.1027     0   1   2   3   4E+3   15   15.5  RT (min)       lcqtof_t3_neg_4_064_17    m/z 345.1029     0   1   2   3E+3   15.5   16  RT (min)       lcqtof_t3_neg_4_071_57    m/z 345.2243     0   1   2E+4   6.5   7  RT (min)      lcqtof_t3_neg_4_000_ee    m/z 345.9900     0   1E+4   7.5   8  RT (min)      lcqtof_t3_neg_4_024_26    m/z 346.1920     0   1E+4   6.5   7  RT (min)      lcqtof_t3_neg_4_000_ee    m/z 347.0807     0   1   2   3E+3   4   4.5  RT (min)       lcqtof_t3_neg_4_000_aa    m/z 347.0910     0   1   2E+4   1   1.5  RT (min)       lcqtof_t3_neg_1_031_24    m/z 348.0974     0   1E+4   4   4.5  RT (min)      lcqtof_t3_neg_4_062_13    m/z 349.0995     0   1E+4   4   4.5  RT (min)      lcqtof_t3_neg_4_062_13    m/z 349.1062     0   1   2   3   4E+4   1   1.5  RT (min)       lcqtof_t3_neg_4_090_60    m/z 350.1251     0   1E+4   8.5   9  RT (min)      lcqtof_t3_neg_1_019_23    m/z 350.1554     0   1E+4   7   7.5  RT (min)      lcqtof_t3_neg_4_000_ee    m/z 350.8237     0   1E+4   1   1.5  RT (min)      lcqtof_t3_neg_2_028_24    m/z 351.1206     0   1   2   3E+3   15.5   16  RT (min)       lcqtof_t3_neg_4_000_cc    m/z 351.1344     0   1E+4   9   9.5  RT (min)      lcqtof_t3_neg_4_000_ee    m/z 351.1558     0   1E+4   7.5   8  RT (min)      lcqtof_t3_neg_4_000_ee    m/z 351.1562     0   1E+4   7.5   8  RT (min)      lcqtof_t3_neg_4_000_ee    m/z 352.1746     0   1E+4   6.5   7  RT (min)      lcqtof_t3_neg_4_000_ee    m/z 352.8600     0   1   2E+4   1   1.5  RT (min)       lcqtof_t3_neg_4_094_60    m/z 353.1238     0   1E+4   4   4.5  RT (min)      lcqtof_t3_neg_3_032_24    m/z 353.1504     0   1E+4   4   4.5  RT (min)      lcqtof_t3_neg_4_000_ee    m/z 353.1799     0   1E+4   10   10.5  RT (min)      lcqtof_t3_neg_4_000_ee    m/z 353.1809     0   1E+4   9.5   10  RT (min)      lcqtof_t3_neg_4_000_ee    m/z 353.1815     0   1E+4   10   10.5  RT (min)      lcqtof_t3_neg_4_000_ee    m/z 353.1817     0   1E+4   10   10.5  RT (min)      lcqtof_t3_neg_4_000_ee    m/z 353.1857     0   1E+4   4   4.5  RT (min)      lcqtof_t3_neg_3_032_24    m/z 353.1998     0   1E+4   9   9.5  RT (min)      lcqtof_t3_neg_3_006_23    m/z 354.1173     0   1E+4   4   4.5  RT (min)      lcqtof_t3_neg_4_027_41    m/z 354.1396     0   1   2E+4   1   1.5  RT (min)      lcqtof_t3_neg_1_019_23    m/z 354.1747     0   1   2   3   4E+4   1   1.5  RT (min)      lcqtof_t3_neg_4_040_24    m/z 354.2815     0   1   2E+4   11   11.5  RT (min)      lcqtof_t3_neg_4_000_ee    m/z 355.2508     0   1E+4   9.5   10  RT (min)      lcqtof_t3_neg_4_000_ee    m/z 355.2510     0   1E+4   9.5   10  RT (min)      lcqtof_t3_neg_4_000_ee    m/z 355.2540     0   1E+4   9.5   10  RT (min)      lcqtof_t3_neg_4_000_ee    m/z 355.2551     0   1E+4   9.5   10  RT (min)      lcqtof_t3_neg_4_000_ee    m/z 355.2747     0   1E+4   11   11.5  RT (min)      lcqtof_t3_neg_4_000_ee    m/z 355.2786     0   1E+4   11   11.5  RT (min)      lcqtof_t3_neg_4_000_ee    m/z 355.2849     0   1   2   3   4E+3   12.5   13  RT (min)      lcqtof_t3_neg_4_037_01    m/z 356.0886     0   1E+5   1.5   2  RT (min)      lcqtof_t3_neg_4_097_60    m/z 356.0887     0   2   4   6   8E+3   2   2.5  RT (min)      lcqtof_t3_neg_3_092_60    m/z 356.0887     0   2   4   6   8E+3   2   2.5  RT (min)      lcqtof_t3_neg_3_032_24    m/z 356.0889     0   1E+5   1.5   2  RT (min)      lcqtof_t3_neg_4_097_60    m/z 356.1731     0   1E+4   1   1.5  RT (min)      lcqtof_t3_neg_4_040_24    m/z 356.2670     0   1   2   3   4E+4   11   11.5  RT (min)      lcqtof_t3_neg_4_000_ee    m/z 356.2676     0   1   2   3   4E+4   11   11.5  RT (min)      lcqtof_t3_neg_4_000_ee    m/z 356.9237     0   1   2E+3   16.5   17  RT (min)      lcqtof_t3_neg_1_042_46    m/z 357.2063     0   1E+4   16   17  RT (min)       lcqtof_t3_neg_4_090_60    m/z 357.2066     0   1E+4   15   15.5  RT (min)       lcqtof_t3_neg_4_028_25    m/z 357.2073     0   1E+4   15   16  RT (min)       lcqtof_t3_neg_4_045_02    m/z 357.2479     0   1   2E+4   9.5   10  RT (min)      lcqtof_t3_neg_4_000_ee    m/z 358.0099     0   1E+4   2   2.5  RT (min)      lcqtof_t3_neg_4_060_15    m/z 358.0107     0   1E+4   2   2.5  RT (min)      lcqtof_t3_neg_4_051_15    m/z 358.0607     0   1E+4   0.5   1  RT (min)      lcqtof_t3_neg_4_000_aa    m/z 358.2099     0   1E+4   10.5   11  RT (min)      lcqtof_t3_neg_4_000_ee    m/z 358.2100     0   1E+4   10.5   11  RT (min)      lcqtof_t3_neg_4_000_ee    m/z 359.0472     0   1E+4   4   4.5  RT (min)      lcqtof_t3_neg_4_000_aa    m/z 359.1440     0   2   4E+3   5   5.5  RT (min)      lcqtof_t3_neg_4_041_54    m/z 361.1518     0   1E+4   7   7.5  RT (min)      lcqtof_t3_neg_4_000_ee    m/z 361.1655     0   2   4   6E+3   10   10.5  RT (min)       lcqtof_t3_neg_4_055_38    m/z 361.1660     0   1   2   3   4E+3   10   10.5  RT (min)      lcqtof_t3_neg_4_015_36    m/z 362.2638     0   1E+4   11   11.5  RT (min)      lcqtof_t3_neg_4_000_ee    m/z 362.2642     0   1E+4   11   11.5  RT (min)      lcqtof_t3_neg_4_000_ee    m/z 363.0690     0   1   2   3E+4   1   1.5  RT (min)       lcqtof_t3_neg_1_019_23    m/z 363.1457     0   1E+4   1   1.5  RT (min)       lcqtof_t3_neg_4_092_60    m/z 363.2212     0   1E+4   10.5   11  RT (min)      lcqtof_t3_neg_4_019_05    m/z 363.2309     0   1E+4   10   10.5  RT (min)      lcqtof_t3_neg_4_016_14    m/z 364.2511     0   1E+4   11   11.5  RT (min)      lcqtof_t3_neg_4_000_ee    m/z 364.2530     0   1E+4   11   11.5  RT (min)      lcqtof_t3_neg_4_000_ee    m/z 365.1390     0   1   2   3   4E+3   14.5   15  RT (min)       lcqtof_t3_neg_4_090_60    m/z 365.1391     0   1   2   3   4E+3   14.5   15  RT (min)       lcqtof_t3_neg_4_007_09    m/z 365.1392     0   1   2   3   4E+3   15.5   16   16.5  RT (min)       lcqtof_t3_neg_4_001_49    m/z 365.1803     0   1E+4   9   9.5  RT (min)      lcqtof_t3_neg_4_000_ee    m/z 365.1862     0   1E+4   9   9.5  RT (min)      lcqtof_t3_neg_4_000_ee    m/z 365.1938     0   1E+4   9   9.5  RT (min)      lcqtof_t3_neg_4_000_ee    m/z 366.2795     0   1   2   3E+4   6.5   7  RT (min)      lcqtof_t3_neg_4_000_ee    m/z 367.1570     0   1E+4   1   1.5  RT (min)      lcqtof_t3_neg_4_040_24    m/z 367.3533     0   1E+4   7   7.5  RT (min)      lcqtof_t3_neg_4_059_14    m/z 368.2358     0   1   2   3   4E+3   12   12.5  RT (min)      lcqtof_t3_neg_4_037_01    m/z 368.8338     0   1   2E+4   1   1.5  RT (min)       lcqtof_t3_neg_2_028_24    m/z 369.1250     0   1   2   3   4E+3   13.5   14  RT (min)       lcqtof_t3_neg_4_025_52    m/z 369.1265     0   2   4   6E+3   14   14.5  RT (min)       lcqtof_t3_neg_4_102_61    m/z 369.1265     0   2   4   6E+3   14.5   15  RT (min)       lcqtof_t3_neg_4_074_58    m/z 369.1265     0   2   4E+3   16   16.5  RT (min)       lcqtof_t3_neg_4_094_60    m/z 369.1265     0   2   4   6   8E+3   15   15.5  RT (min)       lcqtof_t3_neg_4_011_32    m/z 369.1267     0   2   4   6E+3   14.5   15  RT (min)       lcqtof_t3_neg_4_043_11    m/z 369.1268     0   1E+4   14.5   15  RT (min)       lcqtof_t3_neg_4_000_dd    m/z 369.1268     0   2   4   6E+3   9.5   10  RT (min)       lcqtof_t3_neg_4_000_ee    m/z 369.1330     0   1   2E+4   7   7.5  RT (min)       lcqtof_t3_neg_4_000_ee    m/z 369.3004     0   1   2E+3   12   12.5  RT (min)      lcqtof_t3_neg_1_033_53    m/z 370.0510     0   1   2   3E+4   4   4.5  RT (min)       lcqtof_t3_neg_4_004_20    m/z 370.0599     0   1E+4   0.5   1  RT (min)      lcqtof_t3_neg_4_000_aa    m/z 370.0607     0   1E+4   2   2.5  RT (min)      lcqtof_t3_neg_4_000_aa    m/z 370.8234     0   1E+4   1   1.5  RT (min)      lcqtof_t3_neg_4_059_14    m/z 371.0988     0   1E+4   4   4.5  RT (min)      lcqtof_t3_neg_3_052_41    m/z 371.1322     0   1E+4   7   7.5  RT (min)      lcqtof_t3_neg_4_000_ee    m/z 371.1888     0   2   4   6   8E+3   7.5   8  RT (min)      lcqtof_t3_neg_4_016_14    m/z 371.2439     0   1   2   3E+4   9.5   10  RT (min)      lcqtof_t3_neg_4_000_ee    m/z 372.1007     0   1E+4   1   1.5  RT (min)      lcqtof_t3_neg_4_040_24    m/z 372.2922     0   2   4   6   8E+4   11   11.5  RT (min)      lcqtof_t3_neg_4_000_ee    m/z 373.0666     0   1E+4   0.5   1  RT (min)      lcqtof_t3_neg_4_000_aa    m/z 373.0875     0   1   2   3E+4   4   4.5  RT (min)      lcqtof_t3_neg_1_005_18    m/z 373.2058     0   1   2   3E+4   9   9.5  RT (min)      lcqtof_t3_neg_4_000_ee    m/z 373.2074     0   1   2   3E+4   9   9.5  RT (min)      lcqtof_t3_neg_4_000_ee    m/z 373.2938     0   1   2E+4   11   11.5  RT (min)      lcqtof_t3_neg_4_000_ee    m/z 373.2951     0   1   2E+4   11   11.5  RT (min)      lcqtof_t3_neg_4_000_ee    m/z 374.1812     0   1E+4   6.5   7  RT (min)      lcqtof_t3_neg_4_000_ee    m/z 374.2206     0   1   2E+2   4   4.5  RT (min)      lcqtof_t3_neg_3_038_06    m/z 374.3257     0   1E+4   9.5   10  RT (min)      lcqtof_t3_neg_4_000_ee    m/z 374.3307     0   1E+4   9.5   10  RT (min)      lcqtof_t3_neg_4_000_ee    m/z 374.3325     0   1E+4   9.5   10  RT (min)      lcqtof_t3_neg_4_000_ee    m/z 374.3330     0   1E+4   9.5   10  RT (min)      lcqtof_t3_neg_4_000_ee    m/z 374.9053     0   1   2   3   4E+3   15   15.5  RT (min)       lcqtof_t3_neg_4_075_58    m/z 374.9055     0   2   4   6E+3   16   16.5  RT (min)       lcqtof_t3_neg_4_018_23    m/z 374.9058     0   2   4   6   8E+3   16.5   17  RT (min)       lcqtof_t3_neg_4_000_ff    m/z 375.0617     0   1   2E+4   1   1.5  RT (min)      lcqtof_t3_neg_4_032_04    m/z 375.0784     0   1E+4   1   1.5  RT (min)      lcqtof_t3_neg_2_028_24    m/z 375.1144     0   1E+4   1   1.5  RT (min)      lcqtof_t3_neg_4_040_24    m/z 375.9410     0   1   2   3   4E+4   1   1.5  RT (min)      lcqtof_t3_neg_4_014_13    m/z 377.2717     0   1   2   3E+3   11   11.5  RT (min)       lcqtof_t3_neg_4_000_ff    m/z 377.2718     0   1   2   3E+3   10.5   11  RT (min)       lcqtof_t3_neg_4_003_37    m/z 377.4056     0   1E+4   7   7.5  RT (min)      lcqtof_t3_neg_4_004_20    m/z 377.9392     0   1   2E+4   1   1.5  RT (min)      lcqtof_t3_neg_4_014_13    m/z 378.1084     0   1   2   3E+4   4.5   5  RT (min)       lcqtof_t3_neg_4_004_20    m/z 378.1388     0   1E+4   6.5   7  RT (min)      lcqtof_t3_neg_4_000_ee    m/z 378.1564     0   2   4   6E+3   10   10.5  RT (min)       lcqtof_t3_neg_1_032_36    m/z 378.1564     0   2   4   6E+3   10   10.5  RT (min)      lcqtof_t3_neg_1_032_36    m/z 378.1564     0   2   4   6E+3   10   10.5  RT (min)      lcqtof_t3_neg_1_044_37    m/z 378.9171     0   1   2   3   4E+3   17.5   18  RT (min)       lcqtof_t3_neg_4_106_62    m/z 378.9172     0   1E+4   8   9  RT (min)       lcqtof_t3_neg_4_000_ff    m/z 378.9173     0   2   4   6E+3   17   17.5  RT (min)      lcqtof_t3_neg_4_065_51    m/z 378.9174     0   2   4   6E+3   17   17.5  RT (min)      lcqtof_t3_neg_4_057_49    m/z 378.9174     0   2   4   6   8E+3   17   17.5  RT (min)       lcqtof_t3_neg_4_107_62    m/z 378.9175     0   2   4   6E+3   9.5   10  RT (min)       lcqtof_t3_neg_1_003_07    m/z 378.9176     0   2   4   6E+3   17   17.5  RT (min)       lcqtof_t3_neg_4_057_49    m/z 378.9176     0   2   4   6   8E+3   16.5   17  RT (min)       lcqtof_t3_neg_4_100_61    m/z 378.9176     0   2   4   6E+3   10   10.5  RT (min)       lcqtof_t3_neg_1_005_18    m/z 378.9180     0   2   4   6   8E+3   17   17.5  RT (min)       lcqtof_t3_neg_4_035_08    m/z 379.0580     0   1   2   3   4E+3   9.5   10  RT (min)      lcqtof_t3_neg_4_016_14    m/z 379.1551     0   2   4   6E+3   13.5   14  RT (min)       lcqtof_t3_neg_4_080_58    m/z 379.1554     0   1E+4   14.5   15  RT (min)       lcqtof_t3_neg_4_109_62    m/z 379.1554     0   1E+4   16   16.5  RT (min)       lcqtof_t3_neg_4_046_33    m/z 379.1554     0   1E+4   14.5   15  RT (min)       lcqtof_t3_neg_4_008_35    m/z 379.1554     0   1E+4   15   15.5  RT (min)       lcqtof_t3_neg_4_011_32    m/z 379.1556     0   1E+4   15.5   16  RT (min)       lcqtof_t3_neg_4_036_47    m/z 379.1558     0   1E+4   14.5   15  RT (min)       lcqtof_t3_neg_4_006_16    m/z 379.1558     0   1E+4   15.5   16  RT (min)       lcqtof_t3_neg_4_025_52    m/z 379.1563     0   1   2   3   4E+3   12   12.5  RT (min)       lcqtof_t3_neg_4_044_48    m/z 379.1564     0   2   4   6E+3   11.5   12  RT (min)       lcqtof_t3_neg_4_016_14    m/z 379.1569     0   1   2   3   4E+3   12   12.5  RT (min)       lcqtof_t3_neg_4_082_59    m/z 379.1574     0   1   2   3E+3   11.5   12  RT (min)      lcqtof_t3_neg_1_015_34    m/z 380.1374     0   1E+4   6.5   7  RT (min)      lcqtof_t3_neg_4_000_ee    m/z 380.1702     0   1E+4   7   7.5  RT (min)      lcqtof_t3_neg_4_000_ee    m/z 380.1986     0   1E+4   9   9.5  RT (min)      lcqtof_t3_neg_4_000_ee    m/z 381.1046     0   1E+2   7   7.5  RT (min)      lcqtof_t3_neg_3_038_06    m/z 381.1051     0   2   4   6   8E+2   8   8.5  RT (min)      lcqtof_t3_neg_3_032_24    m/z 381.1056     0   1   2E+4   8   8.5  RT (min)      lcqtof_t3_neg_1_019_23    m/z 381.1942     0   1   2E+4   9   9.5  RT (min)      lcqtof_t3_neg_4_000_ee    m/z 381.2285     0   1   2E+4   9.5   10  RT (min)      lcqtof_t3_neg_4_000_ee    m/z 381.2317     0   1   2E+4   9.5   10  RT (min)      lcqtof_t3_neg_4_000_ee    m/z 381.3732     0   1   2   3   4E+3   13.5   14  RT (min)      lcqtof_t3_neg_4_047_07    m/z 383.1341     0   1E+4   7   7.5  RT (min)      lcqtof_t3_neg_4_000_ee    m/z 383.1414     0   2   4   6E+3   15.5   16  RT (min)       lcqtof_t3_neg_4_066_57    m/z 383.1416     0   2   4   6E+3   15   15.5  RT (min)       lcqtof_t3_neg_4_079_58    m/z 383.1417     0   1E+4   14.5   15  RT (min)       lcqtof_t3_neg_4_018_23    m/z 383.1420     0   1E+4   14.5   15  RT (min)       lcqtof_t3_neg_4_000_dd    m/z 383.1422     0   1E+4   14.5   15  RT (min)       lcqtof_t3_neg_4_025_52    m/z 383.1429     0   1E+4   10   10.5  RT (min)       lcqtof_t3_neg_4_000_ee    m/z 383.1431     0   1E+4   10   10.5  RT (min)      lcqtof_t3_neg_4_000_ee    m/z 383.3165     0   1   2   3E+3   12.5   13  RT (min)      lcqtof_t3_neg_1_035_31    m/z 384.2496     0   1E+2   4   4.5  RT (min)      lcqtof_t3_neg_3_038_06    m/z 384.8748     0   1   2E+3   16.5   17  RT (min)      lcqtof_t3_neg_1_008_02    m/z 384.8752     0   1   2E+3   16.5   17  RT (min)      lcqtof_t3_neg_1_000_dd    m/z 384.9343     0   2   4   6   8E+3   15.5   16  RT (min)       lcqtof_t3_neg_4_107_62    m/z 384.9344     0   1   2   3E+3   14   14.5  RT (min)       lcqtof_t3_neg_4_065_51    m/z 384.9344     0   1   2   3E+3   15   15.5  RT (min)       lcqtof_t3_neg_3_032_24    m/z 384.9344     0   1   2   3   4E+3   15   15.5  RT (min)      lcqtof_t3_neg_1_019_23    m/z 384.9344     0   1E+5   16   16.5  RT (min)       lcqtof_t3_neg_4_018_23    m/z 384.9345     0   2   4   6E+3   15.5   16  RT (min)      lcqtof_t3_neg_1_006_05    m/z 384.9346     0   1E+3   13.5   14  RT (min)       lcqtof_t3_neg_3_038_06    m/z 384.9346     0   2   4   6   8E+3   15   15.5  RT (min)       lcqtof_t3_neg_4_099_61    m/z 384.9346     0   2   4   6   8E+3   15.5   16  RT (min)      lcqtof_t3_neg_4_058_16    m/z 384.9346     0   2   4   6E+3   15   15.5  RT (min)       lcqtof_t3_neg_4_113_62    m/z 384.9347     0   2   4   6E+3   14.5   15  RT (min)       lcqtof_t3_neg_4_106_62    m/z 384.9348     0   2   4   6E+3   10.5   11  RT (min)       lcqtof_t3_neg_4_103_61    m/z 384.9349     0   2   4E+3   11.5   12  RT (min)       lcqtof_t3_neg_4_099_61    m/z 384.9349     0   1   2   3   4E+3   12.5   13  RT (min)       lcqtof_t3_neg_4_101_61    m/z 384.9350     0   2   4E+3   11   12  RT (min)       lcqtof_t3_neg_4_085_59    m/z 384.9350     0   2   4   6E+3   14.5   15  RT (min)       lcqtof_t3_neg_4_098_61    m/z 384.9350     0   2   4   6E+3   9   9.5  RT (min)       lcqtof_t3_neg_4_108_62    m/z 384.9350     0   2   4   6E+3   9.5   10  RT (min)       lcqtof_t3_neg_4_102_61    m/z 384.9362     0   2   4E+4   17   17.5  RT (min)      lcqtof_t3_neg_4_035_08    m/z 384.9374     0   1E+5   17   17.5  RT (min)       lcqtof_t3_neg_4_106_62    m/z 386.0949     0   1E+4   4   4.5  RT (min)      lcqtof_t3_neg_1_002_13    m/z 386.2496     0   1E+4   7.5   8  RT (min)      lcqtof_t3_neg_4_000_ee    m/z 387.0076     0   1E+4   1   1.5  RT (min)      lcqtof_t3_neg_4_018_23    m/z 387.0961     0   2   4E+4   12.5   13  RT (min)      lcqtof_t3_neg_4_064_17    m/z 387.0962     0   1E+4   12   12.5  RT (min)      lcqtof_t3_neg_4_064_17    m/z 387.0966     0   2   4E+4   12.5   13  RT (min)      lcqtof_t3_neg_4_064_17    m/z 387.0967     0   2   4E+4   13   13.5  RT (min)      lcqtof_t3_neg_4_064_17    m/z 387.0968     0   2   4E+4   13.5   14  RT (min)      lcqtof_t3_neg_4_064_17    m/z 387.0968     0   1E+4   12   12.5  RT (min)       lcqtof_t3_neg_4_064_17    m/z 387.0969     0   1E+4   12   12.5  RT (min)      lcqtof_t3_neg_4_064_17    m/z 387.0969     0   1E+4   12   12.5  RT (min)       lcqtof_t3_neg_4_064_17    m/z 387.0970     0   2   4E+4   13   13.5  RT (min)      lcqtof_t3_neg_4_064_17    m/z 387.0970     0   2   4E+4   12.5   13   13.5  RT (min)       lcqtof_t3_neg_4_064_17    m/z 387.0971     0   2   4E+4   12.5   13  RT (min)      lcqtof_t3_neg_4_064_17    m/z 387.0971     0   2   4E+4   13   13.5  RT (min)      lcqtof_t3_neg_4_064_17    m/z 387.0976     0   2   4E+4   13   13.5  RT (min)      lcqtof_t3_neg_4_064_17    m/z 387.0977     0   2   4E+4   13   13.5  RT (min)      lcqtof_t3_neg_4_064_17    m/z 387.1139     0   1   2   3E+3   16.5   17  RT (min)      lcqtof_t3_neg_4_048_19    m/z 387.2202     0   2   4   6   8E+3   10   10.5  RT (min)       lcqtof_t3_neg_4_059_14    m/z 387.2858     0   1E+4   8.5   9  RT (min)      lcqtof_t3_neg_2_030_23    m/z 388.0968     0   1   2E+4   13   13.5  RT (min)      lcqtof_t3_neg_4_064_17    m/z 388.0976     0   1   2E+4   13   13.5  RT (min)      lcqtof_t3_neg_4_064_17    m/z 388.0977     0   1   2E+4   13   13.5  RT (min)      lcqtof_t3_neg_4_064_17    m/z 389.1991     0   1E+5   8.5   9  RT (min)      lcqtof_t3_neg_4_000_ee    m/z 389.2047     0   1E+5   8.5   9  RT (min)      lcqtof_t3_neg_4_000_ee    m/z 390.2056     0   1   2   3   4E+4   8.5   9  RT (min)      lcqtof_t3_neg_4_000_ee    m/z 390.2058     0   1   2   3   4E+4   8.5   9  RT (min)      lcqtof_t3_neg_4_000_ee    m/z 390.8034     0   1   2   3E+4   0.5   1  RT (min)      lcqtof_t3_neg_4_000_ee    m/z 391.8959     0   1   2E+3   16   16.5  RT (min)      lcqtof_t3_neg_1_006_05    m/z 393.0789     0   1   2E+4   1   1.5  RT (min)       lcqtof_t3_neg_1_019_23    m/z 393.1068     0   1E+4   9   9.5  RT (min)      lcqtof_t3_neg_4_000_ee    m/z 393.1706     0   1E+4   15   15.5  RT (min)       lcqtof_t3_neg_4_074_58    m/z 393.1707     0   1   2   3   4E+3   12   12.5  RT (min)       lcqtof_t3_neg_4_110_62    m/z 393.1708     0   1E+4   15   15.5  RT (min)       lcqtof_t3_neg_4_073_57    m/z 393.1708     0   1   2   3   4E+3   11.5   12  RT (min)       lcqtof_t3_neg_4_082_59    m/z 393.1710     0   1E+4   16   16.5  RT (min)       lcqtof_t3_neg_4_111_62    m/z 393.1710     0   1E+4   14.5   15  RT (min)       lcqtof_t3_neg_4_006_16    m/z 393.1711     0   1E+4   14.5   15  RT (min)       lcqtof_t3_neg_4_078_58    m/z 393.1711     0   1E+4   15.5   16  RT (min)       lcqtof_t3_neg_4_027_41    m/z 393.1712     0   1E+4   16   16.5  RT (min)       lcqtof_t3_neg_4_091_60    m/z 393.1712     0   1E+4   14   14.5  RT (min)       lcqtof_t3_neg_4_075_58    m/z 393.1712     0   2   4   6   8E+3   14   14.5  RT (min)       lcqtof_t3_neg_4_099_61    m/z 393.1712     0   1E+4   14.5   15  RT (min)       lcqtof_t3_neg_4_013_39    m/z 393.1716     0   1   2   3   4E+3   12.5   13  RT (min)       lcqtof_t3_neg_4_099_61    m/z 393.9866     0   1E+4   2   2.5  RT (min)      lcqtof_t3_neg_4_060_15    m/z 394.8898     0   1   2   3E+3   17.5   18  RT (min)       lcqtof_t3_neg_4_035_08    m/z 394.8908     0   1   2E+4   17   17.5  RT (min)       lcqtof_t3_neg_4_112_62    m/z 394.8920     0   1E+4   16.5   17  RT (min)       lcqtof_t3_neg_4_017_51    m/z 395.1706     0   2   4   6E+3   10   10.5  RT (min)      lcqtof_t3_neg_4_000_ee    m/z 395.1717     0   1   2   3   4E+3   15   15.5  RT (min)      lcqtof_t3_neg_4_023_45    m/z 395.1722     0   1   2   3   4E+3   14.5   15  RT (min)      lcqtof_t3_neg_4_000_ee    m/z 395.1728     0   1   2   3E+3   15.5   16  RT (min)      lcqtof_t3_neg_4_035_08    m/z 395.2289     0   1   2   3E+3   12   12.5  RT (min)      lcqtof_t3_neg_1_002_13    m/z 395.9072     0   1   2E+3   9   10  RT (min)       lcqtof_t3_neg_1_000_bb    m/z 395.9080     0   1   2   3E+3   8   9  RT (min)       lcqtof_t3_neg_1_021_39    m/z 396.1457     0   1E+3   16   16.5  RT (min)      lcqtof_t3_neg_3_041_54    m/z 396.1460     0   2   4   6E+3   14.5   15  RT (min)       lcqtof_t3_neg_4_005_18    m/z 396.8897     0   1   2   3E+3   16.5   17  RT (min)       lcqtof_t3_neg_4_013_39    m/z 396.8905     0   1   2   3   4E+3   16.5   17  RT (min)       lcqtof_t3_neg_4_000_cc    m/z 397.1584     0   2   4   6   8E+3   10   10.5  RT (min)       lcqtof_t3_neg_4_000_ee    m/z 397.2268     0   1E+4   9   9.5  RT (min)      lcqtof_t3_neg_3_006_23    m/z 397.3684     0   1   2   3   4E+3   12   12.5  RT (min)      lcqtof_t3_neg_2_051_14    m/z 397.3686     0   1   2   3   4E+3   12   12.5  RT (min)      lcqtof_t3_neg_4_059_14    m/z 398.0947     0   1   2E+2   7   7.5  RT (min)      lcqtof_t3_neg_3_038_06    m/z 398.0948     0   1   2E+2   7   7.5  RT (min)      lcqtof_t3_neg_3_038_06    m/z 398.0958     0   1   2   3E+4   8   8.5  RT (min)      lcqtof_t3_neg_1_019_23    m/z 398.0959     0   1E+2   8   8.5  RT (min)      lcqtof_t3_neg_3_032_24    m/z 399.1307     0   1E+4   1   1.5  RT (min)       lcqtof_t3_neg_4_018_23    m/z 399.1444     0   1E+4   7   7.5  RT (min)      lcqtof_t3_neg_4_000_ee    m/z 400.2804     0   1E+4   11   11.5  RT (min)      lcqtof_t3_neg_4_000_ee    m/z 400.2808     0   1E+4   11   11.5  RT (min)      lcqtof_t3_neg_4_000_ee    m/z 400.2835     0   1E+4   11   11.5  RT (min)      lcqtof_t3_neg_4_000_ee    m/z 401.0671     0   1   2   3   4E+3   4   4.5  RT (min)      lcqtof_t3_neg_4_007_09    m/z 401.1512     0   1   2E+4   6.5   7  RT (min)      lcqtof_t3_neg_4_000_ee    m/z 401.2396     0   1E+2   4   4.5  RT (min)      lcqtof_t3_neg_3_038_06    m/z 401.9245     0   1E+3   15   15.5  RT (min)       lcqtof_t3_neg_3_032_24    m/z 401.9246     0   1E+3   15.5   16  RT (min)       lcqtof_t3_neg_1_019_23    m/z 401.9246     0   1   2E+3   15.5   16  RT (min)       lcqtof_t3_neg_4_110_62    m/z 402.1485     0   2   4E+3   4   4.5  RT (min)      lcqtof_t3_neg_4_018_23    m/z 402.2426     0   2   4   6E+1   4   4.5  RT (min)      lcqtof_t3_neg_3_038_06    m/z 402.8637     0   1   2   3   4E+3   16.5   17  RT (min)       lcqtof_t3_neg_4_019_05    m/z 404.0152     0   2   4   6E+3   2.5   3  RT (min)       lcqtof_t3_neg_1_046_17    m/z 404.0154     0   1E+4   4   4.5  RT (min)      lcqtof_t3_neg_4_058_16    m/z 404.0155     0   1   2E+4   1   1.5  RT (min)       lcqtof_t3_neg_4_058_16    m/z 404.0155     0   2   4   6E+3   2.5   3  RT (min)       lcqtof_t3_neg_1_060_15    m/z 404.9168     0   1   2   3E+3   16.5   17  RT (min)      lcqtof_t3_neg_4_000_ee    m/z 405.0191     0   1E+4   2   2.5  RT (min)      lcqtof_t3_neg_4_060_15    m/z 405.2165     0   1   2   3E+3   15   15.5  RT (min)      lcqtof_t3_neg_4_000_ff    m/z 406.1956     0   1E+4   9   9.5  RT (min)      lcqtof_t3_neg_4_000_ee    m/z 406.1960     0   1E+4   9   9.5  RT (min)      lcqtof_t3_neg_4_000_ee    m/z 407.1055     0   1E+4   4   4.5  RT (min)      lcqtof_t3_neg_4_006_16    m/z 407.1840     0   2   4   6E+3   11.5   12  RT (min)       lcqtof_t3_neg_1_006_05    m/z 407.1864     0   1E+4   14.5   15  RT (min)       lcqtof_t3_neg_4_034_34    m/z 407.1866     0   1E+4   14   14.5  RT (min)       lcqtof_t3_neg_4_018_23    m/z 407.1868     0   2   4   6E+3   16   16.5  RT (min)       lcqtof_t3_neg_4_113_62    m/z 407.1870     0   2   4   6   8E+3   15   15.5  RT (min)       lcqtof_t3_neg_4_057_49    m/z 407.1870     0   2   4   6E+3   15.5   16  RT (min)       lcqtof_t3_neg_4_082_59    m/z 407.1870     0   1E+4   14.5   15  RT (min)       lcqtof_t3_neg_4_000_aa    m/z 407.1893     0   1   2E+4   9   9.5  RT (min)      lcqtof_t3_neg_4_000_ee    m/z 407.1896     0   1   2E+4   9   9.5  RT (min)      lcqtof_t3_neg_4_000_ee    m/z 407.4328     0   1E+4   13   13.5  RT (min)      lcqtof_t3_neg_4_019_05    m/z 407.4335     0   1E+4   13   13.5  RT (min)      lcqtof_t3_neg_4_019_05    m/z 408.1903     0   1E+4   10   10.5  RT (min)      lcqtof_t3_neg_4_000_ee    m/z 408.1909     0   1E+4   10.5   11  RT (min)      lcqtof_t3_neg_4_000_ee    m/z 409.1729     0   1   2E+4   7   7.5  RT (min)      lcqtof_t3_neg_4_000_ee    m/z 409.2356     0   1E+4   10   10.5  RT (min)      lcqtof_t3_neg_2_016_05    m/z 409.2356     0   1E+4   10   10.5  RT (min)       lcqtof_t3_neg_2_016_05    m/z 409.2358     0   1E+4   10   10.5  RT (min)      lcqtof_t3_neg_2_016_05    m/z 409.2360     0   1   2   3E+4   10   10.5  RT (min)      lcqtof_t3_neg_4_019_05    m/z 410.1596     0   1   2E+4   1   1.5  RT (min)       lcqtof_t3_neg_4_091_60    m/z 410.1612     0   2   4   6   8E+3   14.5   15  RT (min)       lcqtof_t3_neg_4_018_23    m/z 410.1614     0   2   4   6E+3   15   15.5  RT (min)       lcqtof_t3_neg_4_035_08    m/z 410.1615     0   2   4   6   8E+3   14.5   15  RT (min)       lcqtof_t3_neg_4_000_dd    m/z 410.1634     0   2   4   6E+3   9.5   10  RT (min)      lcqtof_t3_neg_1_036_14    m/z 410.1655     0   2   4E+3   9.5   10  RT (min)      lcqtof_t3_neg_4_059_14    m/z 410.2429     0   1E+4   7   7.5  RT (min)      lcqtof_t3_neg_4_000_ee    m/z 410.3151     0   1E+4   13   13.5  RT (min)      lcqtof_t3_neg_4_057_49    m/z 411.1656     0   1   2   3E+3   14.5   15  RT (min)       lcqtof_t3_neg_4_000_dd    m/z 411.1658     0   1   2   3E+3   14.5   15  RT (min)       lcqtof_t3_neg_4_000_dd    m/z 411.3479     0   1   2   3E+3   12.5   13  RT (min)      lcqtof_t3_neg_2_000_aa    m/z 411.3485     0   1   2E+3   13   13.5  RT (min)      lcqtof_t3_neg_1_051_26    m/z 411.3837     0   2   4   6E+3   12.5   13  RT (min)      lcqtof_t3_neg_4_059_14    m/z 411.9196     0   1   2E+4   1   1.5  RT (min)      lcqtof_t3_neg_4_014_13    m/z 412.2112     0   1   2   3E+3   12   12.5  RT (min)      lcqtof_t3_neg_1_010_27    m/z 412.2796     0   1E+4   1   1.5  RT (min)      lcqtof_t3_neg_3_032_24    m/z 413.0331     0   1E+4   1   1.5  RT (min)       lcqtof_t3_neg_2_028_24    m/z 413.0615     0   1   2E+3   15.5   16  RT (min)      lcqtof_t3_neg_4_006_16    m/z 413.0624     0   1   2E+3   15.5   16  RT (min)      lcqtof_t3_neg_4_018_23    m/z 413.0647     0   1   2E+3   15   15.5  RT (min)       lcqtof_t3_neg_4_002_17    m/z 413.2154     0   1E+4   6.5   7  RT (min)      lcqtof_t3_neg_4_000_ee    m/z 413.9160     0   1E+4   1   1.5  RT (min)      lcqtof_t3_neg_4_014_13    m/z 416.3384     0   1E+4   10   10.5  RT (min)      lcqtof_t3_neg_4_000_ee    m/z 416.3402     0   1E+4   10   10.5  RT (min)      lcqtof_t3_neg_4_000_ee    m/z 416.3444     0   1E+4   10   10.5  RT (min)      lcqtof_t3_neg_4_000_ee    m/z 417.1354     0   2   4E+4   1   1.5  RT (min)       lcqtof_t3_neg_1_019_23    m/z 417.2104     0   1E+4   10   12   14  RT (min)       lcqtof_t3_neg_4_035_08    m/z 417.2108     0   1E+4   10   10.5  RT (min)      lcqtof_t3_neg_4_047_07    m/z 418.1431     0   1E+4   1   1.5  RT (min)       lcqtof_t3_neg_4_092_60    m/z 418.1714     0   1E+2   4   4.5  RT (min)      lcqtof_t3_neg_3_038_06    m/z 419.1800     0   1E+4   10   10.5  RT (min)      lcqtof_t3_neg_4_000_ee    m/z 420.1684     0   2   4   6E+1   4   4.5  RT (min)      lcqtof_t3_neg_3_038_06    m/z 421.2812     0   2   4E+3   10.5   11  RT (min)       lcqtof_t3_neg_1_032_36    m/z 422.1519     0   1E+4   1   1.5  RT (min)      lcqtof_t3_neg_1_023_06    m/z 423.1742     0   1   2   3E+3   14.5   15  RT (min)       lcqtof_t3_neg_4_091_60    m/z 423.1745     0   1   2   3E+3   14.5   15  RT (min)       lcqtof_t3_neg_4_056_56    m/z 423.1753     0   1   2E+3   15.5   16  RT (min)       lcqtof_t3_neg_4_042_10    m/z 423.1757     0   1   2E+3   16   16.5  RT (min)       lcqtof_t3_neg_4_056_56    m/z 423.1761     0   1   2E+3   15   15.5  RT (min)       lcqtof_t3_neg_4_024_26    m/z 423.3303     0   1E+4   8   8.5  RT (min)      lcqtof_t3_neg_4_000_ee    m/z 423.4206     0   1   2   3E+3   14   14.5  RT (min)       lcqtof_t3_neg_4_046_33    m/z 424.0932     0   1E+4   1   1.5  RT (min)      lcqtof_t3_neg_1_019_23    m/z 424.1764     0   2   4E+3   14.5   15  RT (min)       lcqtof_t3_neg_4_031_42    m/z 424.1770     0   2   4   6E+3   14.5   15  RT (min)       lcqtof_t3_neg_4_005_18    m/z 424.1770     0   1   2   3E+3   16   16.5  RT (min)       lcqtof_t3_neg_4_024_26    m/z 425.0158     0   1E+4   1   1.5  RT (min)      lcqtof_t3_neg_4_016_14    m/z 425.1806     0   1   2E+3   15.5   16  RT (min)       lcqtof_t3_neg_4_002_17    m/z 425.1816     0   1   2E+3   15.5   16  RT (min)       lcqtof_t3_neg_4_001_49    m/z 425.1827     0   1   2E+3   15.5   16  RT (min)       lcqtof_t3_neg_4_025_52    m/z 425.2341     0   1E+4   11.5   12  RT (min)      lcqtof_t3_neg_1_006_05    m/z 425.2344     0   1E+4   11.5   12  RT (min)      lcqtof_t3_neg_1_006_05    m/z 425.2740     0   1E+4   11   11.5  RT (min)      lcqtof_t3_neg_4_000_ee    m/z 426.2720     0   1E+4   11   11.5  RT (min)      lcqtof_t3_neg_4_000_ee    m/z 427.1090     0   2   4E+3   9   9.5  RT (min)       lcqtof_t3_neg_4_000_ee    m/z 427.2507     0   1E+4   11.5   12  RT (min)      lcqtof_t3_neg_1_006_05    m/z 427.2512     0   1E+4   11.5   12  RT (min)      lcqtof_t3_neg_1_006_05    m/z 427.3284     0   1E+4   10   10.5  RT (min)       lcqtof_t3_neg_3_071_57    m/z 428.2990     0   1   2   3E+4   10   10.5  RT (min)      lcqtof_t3_neg_4_019_05    m/z 428.2992     0   1   2E+4   10   10.5  RT (min)       lcqtof_t3_neg_2_016_05    m/z 428.2993     0   1E+4   10   10.5  RT (min)      lcqtof_t3_neg_3_037_05    m/z 428.2993     0   1   2E+4   10   10.5  RT (min)      lcqtof_t3_neg_2_016_05    m/z 428.2996     0   1   2   3E+4   10   10.5  RT (min)       lcqtof_t3_neg_4_019_05    m/z 428.2998     0   1E+4   10   10.5  RT (min)      lcqtof_t3_neg_4_028_25    m/z 428.3005     0   1E+4   10   10.5  RT (min)      lcqtof_t3_neg_4_028_25    m/z 428.7950     0   1E+4   1   1.5  RT (min)      lcqtof_t3_neg_4_046_33    m/z 428.8730     0   1E+4   9.5   10  RT (min)      lcqtof_t3_neg_1_045_29    m/z 429.0464     0   1E+4   9.5   10  RT (min)      lcqtof_t3_neg_4_002_17    m/z 429.0470     0   1E+4   9.5   10  RT (min)      lcqtof_t3_neg_4_002_17    m/z 429.0899     0   1   2E+3   14.5   15  RT (min)       lcqtof_t3_neg_4_000_dd    m/z 429.1360     0   1   2   3E+3   15   15.5  RT (min)       lcqtof_t3_neg_4_064_17    m/z 429.1362     0   1   2   3E+3   15   15.5  RT (min)      lcqtof_t3_neg_4_056_56    m/z 429.1367     0   1   2E+3   15.5   16  RT (min)       lcqtof_t3_neg_4_051_15    m/z 429.1369     0   1   2E+3   16.5   17  RT (min)       lcqtof_t3_neg_4_007_09    m/z 429.1372     0   1   2E+3   15   15.5  RT (min)       lcqtof_t3_neg_4_069_57    m/z 429.1381     0   1   2E+3   15.5   16  RT (min)       lcqtof_t3_neg_4_039_06    m/z 429.1389     0   1   2E+3   15.5   16  RT (min)       lcqtof_t3_neg_4_088_59    m/z 430.1786     0   1   2   3E+4   1   1.5  RT (min)      lcqtof_t3_neg_1_031_24    m/z 430.3233     0   1E+4   9   9.5  RT (min)      lcqtof_t3_neg_4_000_ee    m/z 430.3236     0   1E+4   9.5   10  RT (min)      lcqtof_t3_neg_4_000_ee    m/z 430.7782     0   1E+4   1   1.5  RT (min)      lcqtof_t3_neg_4_059_14    m/z 430.7904     0   1E+4   1   1.5  RT (min)      lcqtof_t3_neg_4_046_33    m/z 431.1628     0   1E+4   4   4.5  RT (min)      lcqtof_t3_neg_3_032_24    m/z 431.2192     0   1E+4   9.5   10  RT (min)      lcqtof_t3_neg_4_028_25    m/z 431.2192     0   1   2   3   4E+4   9.5   10  RT (min)       lcqtof_t3_neg_4_019_05    m/z 431.2198     0   1   2   3E+4   9.5   10  RT (min)      lcqtof_t3_neg_2_016_05    m/z 431.2202     0   1   2E+4   9.5   10  RT (min)      lcqtof_t3_neg_3_037_05    m/z 431.3032     0   1E+4   12   12.5  RT (min)       lcqtof_t3_neg_4_019_05    m/z 431.3042     0   1E+4   12   12.5  RT (min)       lcqtof_t3_neg_4_019_05    m/z 432.2241     0   1E+4   9.5   10  RT (min)      lcqtof_t3_neg_4_019_05    m/z 432.3606     0   1E+4   12.5   13  RT (min)      lcqtof_t3_neg_4_062_13    m/z 432.3611     0   1E+4   12.5   13  RT (min)      lcqtof_t3_neg_4_062_13    m/z 432.8644     0   1   2E+3   16.5   17  RT (min)       lcqtof_t3_neg_4_057_49    m/z 432.8646     0   1   2E+3   15.5   16  RT (min)       lcqtof_t3_neg_4_041_54    m/z 432.8654     0   1   2E+3   15.5   16  RT (min)       lcqtof_t3_neg_4_054_44    m/z 433.1274     0   1   2E+3   16   16.5  RT (min)       lcqtof_t3_neg_4_029_40    m/z 433.1281     0   1   2E+3   15.5   16  RT (min)       lcqtof_t3_neg_4_000_ee    m/z 433.2356     0   2   4   6E+4   9.5   10  RT (min)      lcqtof_t3_neg_2_016_05    m/z 433.2358     0   2   4   6E+4   9.5   10  RT (min)       lcqtof_t3_neg_2_016_05    m/z 434.2376     0   1   2E+4   10   10.5  RT (min)      lcqtof_t3_neg_4_019_05    m/z 434.2383     0   1E+4   9.5   10  RT (min)      lcqtof_t3_neg_2_016_05    m/z 434.2387     0   1   2E+4   10   10.5  RT (min)      lcqtof_t3_neg_4_019_05    m/z 434.2388     0   1E+4   9.5   10  RT (min)      lcqtof_t3_neg_2_016_05    m/z 434.2391     0   1   2E+4   10   10.5  RT (min)      lcqtof_t3_neg_4_019_05    m/z 434.2392     0   1   2E+4   10   10.5  RT (min)      lcqtof_t3_neg_4_019_05    m/z 434.2392     0   1   2E+4   10   10.5  RT (min)      lcqtof_t3_neg_4_019_05    m/z 434.2398     0   1   2E+4   10   10.5  RT (min)      lcqtof_t3_neg_4_019_05    m/z 434.2404     0   1   2E+4   10   10.5  RT (min)      lcqtof_t3_neg_4_019_05    m/z 434.8714     0   2   4   6E+3   16.5   17  RT (min)      lcqtof_t3_neg_1_011_45    m/z 435.1636     0   1E+4   7   7.5  RT (min)      lcqtof_t3_neg_4_000_ee    m/z 435.8934     0   1   2   3   4E+3   16.5   17  RT (min)       lcqtof_t3_neg_4_000_ff    m/z 435.8941     0   1   2E+3   15.5   16  RT (min)       lcqtof_t3_neg_4_040_24    m/z 436.1553     0   1E+4   1   1.5  RT (min)      lcqtof_t3_neg_4_019_05    m/z 436.2367     0   1E+4   11.5   12  RT (min)       lcqtof_t3_neg_4_019_05    m/z 437.2334     0   1E+4   11.5   12  RT (min)      lcqtof_t3_neg_4_019_05    m/z 437.2335     0   1E+4   11   11.5  RT (min)      lcqtof_t3_neg_4_019_05    m/z 437.2338     0   1E+4   11.5   12  RT (min)       lcqtof_t3_neg_4_019_05    m/z 437.2906     0   1   2E+3   9.5   10  RT (min)       lcqtof_t3_neg_2_050_01    m/z 437.2908     0   1   2   3   4E+3   9.5   10  RT (min)       lcqtof_t3_neg_4_059_14    m/z 437.3434     0   1E+4   13   13.5  RT (min)      lcqtof_t3_neg_4_064_17    m/z 438.2162     0   1E+4   6.5   7  RT (min)      lcqtof_t3_neg_4_034_34    m/z 438.8111     0   1E+4   1   1.5  RT (min)      lcqtof_t3_neg_4_059_14    m/z 439.0855     0   1E+4   1   1.5  RT (min)       lcqtof_t3_neg_2_030_23    m/z 440.1748     0   1E+4   1   1.5  RT (min)      lcqtof_t3_neg_4_040_24    m/z 440.3126     0   1E+4   11   11.5  RT (min)      lcqtof_t3_neg_4_019_05    m/z 440.8070     0   1E+4   1   1.5  RT (min)      lcqtof_t3_neg_4_059_14    m/z 440.8877     0   1   2   3E+3   7   8   9  RT (min)       lcqtof_t3_neg_1_055_32    m/z 440.8880     0   1   2   3E+3   7   7.5   8  RT (min)       lcqtof_t3_neg_1_029_44    m/z 440.8882     0   1   2   3   4E+3   8   8.5   9  RT (min)       lcqtof_t3_neg_1_039_43    m/z 440.8884     0   1   2   3   4E+3   9   10  RT (min)       lcqtof_t3_neg_1_005_18    m/z 440.8884     0   1   2E+3   7.5   8  RT (min)       lcqtof_t3_neg_4_065_51    m/z 441.1044     0   1   2   3E+3   14.5   15  RT (min)       lcqtof_t3_neg_4_007_09    m/z 441.1048     0   1   2E+3   15.5   16  RT (min)      lcqtof_t3_neg_4_005_18    m/z 441.1059     0   1   2E+3   15.5   16  RT (min)      lcqtof_t3_neg_4_001_49    m/z 441.1727     0   2   4E+4   1   1.5  RT (min)       lcqtof_t3_neg_4_040_24    m/z 442.3204     0   1E+2   8   8.5  RT (min)      lcqtof_t3_neg_3_038_06    m/z 442.3212     0   1E+2   8   8.5  RT (min)      lcqtof_t3_neg_3_038_06    m/z 442.3216     0   1   2E+4   9   9.5  RT (min)      lcqtof_t3_neg_1_019_23    m/z 442.3216     0   1   2   3E+2   8.5   9  RT (min)      lcqtof_t3_neg_3_032_24    m/z 442.3216     0   1E+2   8   8.5  RT (min)      lcqtof_t3_neg_3_038_06    m/z 442.3230     0   1   2   3E+2   8.5   9  RT (min)      lcqtof_t3_neg_3_032_24    m/z 442.8927     0   1   2E+3   8   9  RT (min)       lcqtof_t3_neg_4_088_59    m/z 442.8929     0   1   2E+3   9   9.5  RT (min)       lcqtof_t3_neg_4_075_58    m/z 442.8929     0   1   2   3   4E+3   15   15.5  RT (min)       lcqtof_t3_neg_4_107_62    m/z 442.8932     0   1E+4   16   16.5  RT (min)       lcqtof_t3_neg_4_018_23    m/z 442.8934     0   1   2   3   4E+3   15   15.5  RT (min)       lcqtof_t3_neg_4_100_61    m/z 443.0760     0   1E+4   8   8.5  RT (min)      lcqtof_t3_neg_1_019_23    m/z 443.3101     0   2   4   6   8E+3   11.5   12  RT (min)      lcqtof_t3_neg_4_059_14    m/z 443.3145     0   2   4   6   8E+3   11.5   12  RT (min)      lcqtof_t3_neg_4_059_14    m/z 446.9052     0   2   4E+3   17   17.5  RT (min)       lcqtof_t3_neg_4_035_08    m/z 446.9054     0   1   2E+3   10   10.5  RT (min)      lcqtof_t3_neg_1_036_14    m/z 446.9054     0   1   2   3   4E+3   9.5   10  RT (min)       lcqtof_t3_neg_4_027_41    m/z 446.9054     0   2   4   6E+3   8.5   9   9.5  RT (min)       lcqtof_t3_neg_4_110_62    m/z 446.9056     0   1   2   3E+3   10   11   12  RT (min)       lcqtof_t3_neg_1_012_16    m/z 446.9056     0   1   2   3E+3   10   10.5  RT (min)       lcqtof_t3_neg_1_000_bb    m/z 446.9057     0   1   2   3   4E+3   17   17.5  RT (min)       lcqtof_t3_neg_4_035_08    m/z 446.9058     0   1   2   3E+3   10   11  RT (min)       lcqtof_t3_neg_1_028_20    m/z 447.0928     0   1   2E+4   5.5   6  RT (min)      lcqtof_t3_neg_1_005_18    m/z 447.0940     0   1   2E+4   5.5   6  RT (min)      lcqtof_t3_neg_4_005_18    m/z 447.1341     0   1E+4   1   1.5  RT (min)      lcqtof_t3_neg_4_047_07    m/z 447.1424     0   2   4E+3   15   16  RT (min)       lcqtof_t3_neg_4_006_16    m/z 447.1432     0   2   4E+3   16   16.5  RT (min)       lcqtof_t3_neg_4_023_45    m/z 447.1434     0   2   4E+3   15.5   16  RT (min)       lcqtof_t3_neg_4_016_14    m/z 447.3454     0   1   2   3E+4   11.5   12  RT (min)      lcqtof_t3_neg_4_016_14    m/z 447.3456     0   1   2   3E+4   11.5   12  RT (min)      lcqtof_t3_neg_4_016_14    m/z 448.3500     0   2   4   6   8E+3   11.5   12  RT (min)      lcqtof_t3_neg_4_059_14    m/z 449.2199     0   1E+4   9.5   10  RT (min)      lcqtof_t3_neg_4_000_ee    m/z 449.3109     0   1   2E+4   13   13.5  RT (min)      lcqtof_t3_neg_1_000_aa    m/z 449.3110     0   1   2E+4   12.5   13  RT (min)      lcqtof_t3_neg_1_000_aa    m/z 449.3111     0   1   2E+4   12.5   13  RT (min)      lcqtof_t3_neg_1_000_aa    m/z 449.3115     0   1   2E+4   12.5   13  RT (min)      lcqtof_t3_neg_1_000_aa    m/z 449.3116     0   1   2E+4   12.5   13  RT (min)      lcqtof_t3_neg_1_000_aa    m/z 449.3125     0   1   2E+4   13   13.5  RT (min)      lcqtof_t3_neg_1_000_aa    m/z 450.1458     0   1E+4   1   1.5  RT (min)      lcqtof_t3_neg_4_047_07    m/z 450.1685     0   1E+4   1   1.5  RT (min)      lcqtof_t3_neg_4_040_24    m/z 450.1905     0   1E+4   1   1.5  RT (min)      lcqtof_t3_neg_1_006_05    m/z 450.3076     0   1E+4   9.5   10  RT (min)      lcqtof_t3_neg_4_000_ee    m/z 450.9331     0   1E+4   7.5   8  RT (min)      lcqtof_t3_neg_4_005_18    m/z 451.1249     0   1E+4   1   1.5  RT (min)      lcqtof_t3_neg_2_028_24    m/z 451.1291     0   1   2E+3   14.5   15  RT (min)       lcqtof_t3_neg_4_052_03    m/z 451.4514     0   1   2   3   4E+3   14   14.5  RT (min)       lcqtof_t3_neg_4_000_ee    m/z 452.1034     0   1E+4   1   1.5  RT (min)      lcqtof_t3_neg_4_018_23    m/z 452.2895     0   2   4E+4   9.5   10  RT (min)      lcqtof_t3_neg_2_016_05    m/z 452.2904     0   2   4   6E+4   9.5   10  RT (min)       lcqtof_t3_neg_4_019_05    m/z 452.2918     0   1E+4   9.5   10  RT (min)      lcqtof_t3_neg_3_053_25    m/z 452.8613     0   1   2E+3   16.5   17  RT (min)      lcqtof_t3_neg_1_000_bb    m/z 452.9217     0   1E+4   15.5   16  RT (min)       lcqtof_t3_neg_4_107_62    m/z 452.9220     0   2   4   6E+3   15.5   16  RT (min)       lcqtof_t3_neg_1_019_23    m/z 452.9220     0   2   4   6E+3   15.5   16  RT (min)       lcqtof_t3_neg_1_011_45    m/z 452.9220     0   2   4   6   8E+3   15.5   16  RT (min)       lcqtof_t3_neg_4_099_61    m/z 452.9221     0   2   4   6E+4   17   17.5  RT (min)       lcqtof_t3_neg_4_035_08    m/z 452.9222     0   1E+3   13.5   14  RT (min)       lcqtof_t3_neg_3_038_06    m/z 452.9222     0   2   4   6   8E+3   15.5   16  RT (min)       lcqtof_t3_neg_4_057_49    m/z 452.9222     0   2   4   6E+3   15   15.5  RT (min)       lcqtof_t3_neg_4_092_60    m/z 452.9222     0   1   2   3E+3   15   15.5  RT (min)       lcqtof_t3_neg_3_032_24    m/z 452.9222     0   2   4   6   8E+3   9   9.5  RT (min)       lcqtof_t3_neg_4_112_62    m/z 452.9224     0   2   4   6E+3   10.5   11  RT (min)      lcqtof_t3_neg_4_103_61    m/z 452.9225     0   2   4   6E+3   11   11.5  RT (min)       lcqtof_t3_neg_4_000_ee    m/z 452.9226     0   2   4E+3   11.5   12  RT (min)       lcqtof_t3_neg_4_098_61    m/z 452.9226     0   2   4E+3   12.5   13  RT (min)       lcqtof_t3_neg_4_109_62    m/z 452.9238     0   1E+4   7.5   8  RT (min)       lcqtof_t3_neg_4_005_18    m/z 452.9246     0   1   2E+5   17   17.5  RT (min)       lcqtof_t3_neg_4_106_62    m/z 453.2976     0   1   2E+4   10   10.5  RT (min)      lcqtof_t3_neg_4_019_05    m/z 453.3020     0   1E+4   9.5   10  RT (min)      lcqtof_t3_neg_2_016_05    m/z 453.3023     0   1E+4   9.5   10  RT (min)      lcqtof_t3_neg_2_016_05    m/z 453.3023     0   1   2E+4   10   10.5  RT (min)      lcqtof_t3_neg_4_019_05    m/z 453.3030     0   1   2E+4   10   10.5  RT (min)      lcqtof_t3_neg_4_019_05    m/z 453.3038     0   1   2E+4   10   10.5  RT (min)      lcqtof_t3_neg_4_019_05    m/z 453.3040     0   1   2E+4   10   10.5  RT (min)      lcqtof_t3_neg_4_019_05    m/z 453.3051     0   1   2E+4   10   10.5  RT (min)      lcqtof_t3_neg_4_019_05    m/z 454.2932     0   1E+4   11   11.5  RT (min)      lcqtof_t3_neg_4_019_05    m/z 454.2957     0   1E+4   11.5   12  RT (min)       lcqtof_t3_neg_4_019_05    m/z 454.2968     0   1E+4   11.5   12  RT (min)      lcqtof_t3_neg_4_019_05    m/z 454.3049     0   1E+4   9.5   10  RT (min)      lcqtof_t3_neg_4_040_24    m/z 454.3056     0   1   2E+4   10   10.5  RT (min)      lcqtof_t3_neg_3_053_25    m/z 454.3057     0   1E+5   9.5   10  RT (min)       lcqtof_t3_neg_2_016_05    m/z 454.3057     0   1   2   3   4E+4   10   10.5  RT (min)      lcqtof_t3_neg_2_011_25    m/z 454.3058     0   1E+5   10   10.5  RT (min)       lcqtof_t3_neg_4_019_05    m/z 454.3059     0   1E+5   10   10.5  RT (min)       lcqtof_t3_neg_4_019_05    m/z 454.3062     0   1   2   3E+4   10   10.5  RT (min)      lcqtof_t3_neg_4_028_25    m/z 454.9287     0   1E+4   7.5   8  RT (min)      lcqtof_t3_neg_4_005_18    m/z 455.1857     0   1E+4   4   4.5  RT (min)       lcqtof_t3_neg_4_000_ee    m/z 455.1866     0   1E+4   4.5   5  RT (min)      lcqtof_t3_neg_4_000_ee    m/z 455.2074     0   1   2   3E+4   1   1.5  RT (min)       lcqtof_t3_neg_1_031_24    m/z 455.2925     0   1   2E+4   11   11.5  RT (min)      lcqtof_t3_neg_4_019_05    m/z 455.2926     0   1   2E+4   11.5   12  RT (min)      lcqtof_t3_neg_4_019_05    m/z 455.2928     0   1   2E+4   11.5   12  RT (min)      lcqtof_t3_neg_4_019_05    m/z 455.2931     0   1   2E+4   11.5   12  RT (min)       lcqtof_t3_neg_4_019_05    m/z 455.2935     0   1   2E+4   11   11.5  RT (min)      lcqtof_t3_neg_4_019_05    m/z 455.2970     0   1E+4   11.5   12  RT (min)      lcqtof_t3_neg_4_040_24    m/z 455.3050     0   1   2E+4   1   1.5  RT (min)      lcqtof_t3_neg_1_031_24    m/z 455.3133     0   1E+4   11.5   12  RT (min)      lcqtof_t3_neg_4_019_05    m/z 456.1040     0   1   2E+4   1   1.5  RT (min)      lcqtof_t3_neg_4_047_07    m/z 456.3110     0   1   2E+4   11.5   12  RT (min)      lcqtof_t3_neg_4_019_05    m/z 456.3112     0   1E+4   11.5   12  RT (min)      lcqtof_t3_neg_4_028_25    m/z 456.3114     0   1   2E+4   11.5   12  RT (min)       lcqtof_t3_neg_4_019_05    m/z 456.3117     0   1   2E+4   12   12.5  RT (min)      lcqtof_t3_neg_4_019_05    m/z 456.3217     0   1   2   3E+4   10.5   11  RT (min)      lcqtof_t3_neg_3_037_05    m/z 456.3218     0   1   2   3   4E+4   10   10.5  RT (min)       lcqtof_t3_neg_2_016_05    m/z 456.3220     0   1   2   3E+4   10   10.5  RT (min)      lcqtof_t3_neg_3_037_05    m/z 456.3220     0   2   4   6E+4   10.5   11  RT (min)       lcqtof_t3_neg_4_019_05    m/z 456.3228     0   1   2   3E+4   10   10.5  RT (min)      lcqtof_t3_neg_3_037_05    m/z 456.3248     0   1E+4   10.5   11  RT (min)      lcqtof_t3_neg_4_028_25    m/z 456.3508     0   1E+4   8   8.5  RT (min)      lcqtof_t3_neg_4_000_ee    m/z 456.3520     0   1E+4   8   8.5  RT (min)      lcqtof_t3_neg_4_000_ee    m/z 457.3050     0   1E+4   11.5   12  RT (min)      lcqtof_t3_neg_4_028_25    m/z 458.3194     0   1   2E+4   9.5   10  RT (min)      lcqtof_t3_neg_4_000_ee    m/z 458.3202     0   1   2E+4   9.5   10  RT (min)      lcqtof_t3_neg_4_000_ee    m/z 459.3262     0   2   4E+3   12.5   13  RT (min)       lcqtof_t3_neg_4_097_60    m/z 459.3262     0   1   2   3E+3   15   15.5  RT (min)       lcqtof_t3_neg_4_013_39    m/z 459.3266     0   1E+4   13  RT (min)       lcqtof_t3_neg_4_032_04    m/z 459.3435     0   2   4   6E+3   12   12.5  RT (min)      lcqtof_t3_neg_4_064_17    m/z 459.8835     0   1   2E+3   16   16.5  RT (min)      lcqtof_t3_neg_1_016_48    m/z 459.8836     0   1   2   3E+3   16.5   17  RT (min)      lcqtof_t3_neg_1_015_34    m/z 460.3306     0   1   2   3   4E+3   13   13.5  RT (min)      lcqtof_t3_neg_4_032_04    m/z 460.3306     0   1   2   3E+3   12.5   13  RT (min)      lcqtof_t3_neg_4_057_49    m/z 460.3350     0   1   2   3E+4   9.5   10  RT (min)      lcqtof_t3_neg_4_000_ee    m/z 461.1157     0   1   2   3E+4   13   13.5  RT (min)      lcqtof_t3_neg_4_064_17    m/z 461.1162     0   1   2   3   4E+4   11.5   12  RT (min)       lcqtof_t3_neg_4_064_17    m/z 461.1173     0   1   2   3   4E+4   12   12.5  RT (min)      lcqtof_t3_neg_4_064_17    m/z 461.1382     0   1E+4   1   1.5  RT (min)       lcqtof_t3_neg_4_084_59    m/z 461.1580     0   1   2   3E+3   12   12.5  RT (min)       lcqtof_t3_neg_4_032_04    m/z 461.1586     0   1   2E+3   11.5   12  RT (min)       lcqtof_t3_neg_4_039_06    m/z 461.1588     0   2   4   6E+3   15   15.5  RT (min)       lcqtof_t3_neg_4_058_16    m/z 461.1590     0   2   4   6E+3   14.5   15  RT (min)       lcqtof_t3_neg_4_005_18    m/z 461.1590     0   2   4   6E+3   16   16.5  RT (min)       lcqtof_t3_neg_4_050_43    m/z 461.2336     0   1E+4   7   7.5  RT (min)      lcqtof_t3_neg_4_000_ee    m/z 462.1163     0   1   2E+4   11.5   12  RT (min)      lcqtof_t3_neg_4_064_17    m/z 462.1164     0   1   2E+4   11.5   12  RT (min)      lcqtof_t3_neg_4_064_17    m/z 462.1170     0   1E+4   13   13.5  RT (min)      lcqtof_t3_neg_4_064_17    m/z 462.2857     0   1E+4   7   7.5  RT (min)      lcqtof_t3_neg_4_005_18    m/z 463.1138     0   1E+4   11.5   12  RT (min)      lcqtof_t3_neg_4_064_17    m/z 463.1152     0   1E+4   11.5   12  RT (min)      lcqtof_t3_neg_4_064_17    m/z 464.3153     0   1E+4   10.5   11  RT (min)      lcqtof_t3_neg_4_019_05    m/z 464.3154     0   1E+4   11   11.5  RT (min)      lcqtof_t3_neg_4_059_14    m/z 464.3159     0   1E+4   10.5   11  RT (min)      lcqtof_t3_neg_1_006_05    m/z 465.3560     0   2   4E+3   11   11.5  RT (min)       lcqtof_t3_neg_4_016_14    m/z 466.3089     0   1E+4   8   8.5  RT (min)      lcqtof_t3_neg_4_005_18    m/z 466.7276     0   1E+4   1   1.5  RT (min)      lcqtof_t3_neg_4_037_01    m/z 467.3738     0   1E+4   11.5   12  RT (min)      lcqtof_t3_neg_4_003_37    m/z 468.8964     0   1E+4   16   16.5  RT (min)      lcqtof_t3_neg_1_011_45    m/z 468.8967     0   1E+4   17   17.5  RT (min)      lcqtof_t3_neg_4_035_08    m/z 469.1574     0   2   4   6   8E+3   10   10.5  RT (min)       lcqtof_t3_neg_4_000_ee    m/z 469.9117     0   1   2   3E+3   16   16.5  RT (min)      lcqtof_t3_neg_2_016_05    m/z 469.9119     0   1E+3   15.5   16  RT (min)       lcqtof_t3_neg_1_019_23    m/z 469.9120     0   1   2   3E+3   16   16.5  RT (min)      lcqtof_t3_neg_4_015_36    m/z 469.9122     0   1E+3   15   15.5  RT (min)       lcqtof_t3_neg_3_032_24    m/z 469.9125     0   1   2   3E+3   15.5   16  RT (min)       lcqtof_t3_neg_4_105_61    m/z 471.1856     0   1   2E+4   1   1.5  RT (min)      lcqtof_t3_neg_4_040_24    m/z 471.2757     0   1E+4   11   11.5  RT (min)      lcqtof_t3_neg_4_000_ee    m/z 471.2759     0   1E+4   11   11.5  RT (min)      lcqtof_t3_neg_4_000_ee    m/z 471.2814     0   1E+4   11   11.5  RT (min)      lcqtof_t3_neg_4_000_ee    m/z 471.3326     0   1E+4   10   10.5  RT (min)      lcqtof_t3_neg_4_019_05    m/z 471.3328     0   1E+4   10   10.5  RT (min)       lcqtof_t3_neg_4_019_05    m/z 472.1730     0   1   2E+4   1   1.5  RT (min)      lcqtof_t3_neg_1_019_23    m/z 472.2787     0   1   2E+4   11   11.5  RT (min)      lcqtof_t3_neg_4_000_ee    m/z 472.2798     0   1   2E+4   11   11.5  RT (min)      lcqtof_t3_neg_4_000_ee    m/z 472.3812     0   1E+4   9.5   10  RT (min)      lcqtof_t3_neg_4_000_ee    m/z 472.9042     0   1   2   3E+3   16.5   17  RT (min)      lcqtof_t3_neg_4_000_ee    m/z 473.0542     0   1E+4   1   1.5  RT (min)      lcqtof_t3_neg_2_028_24    m/z 473.0865     0   1E+4   4   4.5  RT (min)      lcqtof_t3_neg_1_005_18    m/z 473.1652     0   1   2E+3   15   15.5  RT (min)       lcqtof_t3_neg_4_051_15    m/z 473.1669     0   1   2E+3   15.5   16  RT (min)       lcqtof_t3_neg_4_000_ee    m/z 473.2821     0   2   4   6E+4   11   11.5  RT (min)      lcqtof_t3_neg_4_000_ee    m/z 473.2822     0   2   4   6   8E+3   11   11.5  RT (min)      lcqtof_t3_neg_4_037_01    m/z 473.2846     0   1   2   3E+3   13.5   14  RT (min)       lcqtof_t3_neg_4_044_48    m/z 473.2856     0   1   2E+3   14   14.5  RT (min)       lcqtof_t3_neg_4_018_23    m/z 473.2874     0   1   2   3   4E+3   12   12.5   13  RT (min)       lcqtof_t3_neg_4_030_31    m/z 473.2884     0   1   2   3   4E+3   12   12.5  RT (min)      lcqtof_t3_neg_4_041_54    m/z 473.2890     0   1   2E+3   15   15.5  RT (min)       lcqtof_t3_neg_4_038_29    m/z 473.2890     0   1   2E+3   16   16.5  RT (min)      lcqtof_t3_neg_4_019_05    m/z 473.2899     0   2   4   6E+3   16.5   17  RT (min)      lcqtof_t3_neg_4_000_bb    m/z 473.2984     0   2   4   6E+3   12   12.5  RT (min)       lcqtof_t3_neg_4_019_05    m/z 473.3098     0   2   4   6E+3   11.5   12  RT (min)       lcqtof_t3_neg_4_058_16    m/z 473.3482     0   1   2E+2   9.5   10  RT (min)      lcqtof_t3_neg_3_032_24    m/z 473.3484     0   1E+3   9.5   10  RT (min)      lcqtof_t3_neg_3_032_24    m/z 474.1092     0   1E+4   1   1.5  RT (min)      lcqtof_t3_neg_4_018_23    m/z 474.2622     0   1   2E+4   9.5   10  RT (min)      lcqtof_t3_neg_2_016_05    m/z 474.2629     0   1E+4   9.5   10  RT (min)      lcqtof_t3_neg_2_030_23    m/z 474.2737     0   1E+4   7.5   8  RT (min)      lcqtof_t3_neg_4_000_ee    m/z 474.2839     0   1   2E+4   11   11.5  RT (min)      lcqtof_t3_neg_4_000_ee    m/z 474.2856     0   1   2E+4   11   11.5  RT (min)      lcqtof_t3_neg_4_000_ee    m/z 474.2887     0   1   2E+4   11   11.5  RT (min)      lcqtof_t3_neg_4_000_ee    m/z 474.2912     0   1   2E+4   11   11.5  RT (min)      lcqtof_t3_neg_4_000_ee    m/z 474.3600     0   1   2   3   4E+3   13   13.5  RT (min)      lcqtof_t3_neg_1_002_13    m/z 474.3658     0   2   4   6   8E+3   11.5   12  RT (min)      lcqtof_t3_neg_4_059_14    m/z 475.1746     0   2   4   6E+3   14.5   15  RT (min)       lcqtof_t3_neg_4_000_ee    m/z 475.1748     0   1   2   3   4E+3   15   15.5   16  RT (min)       lcqtof_t3_neg_4_060_15    m/z 475.1750     0   1   2E+4   10   10.5  RT (min)       lcqtof_t3_neg_4_000_ee    m/z 475.2888     0   2   4   6E+3   11   11.5  RT (min)      lcqtof_t3_neg_4_036_47    m/z 476.1262     0   1E+4   4   4.5  RT (min)      lcqtof_t3_neg_1_002_13    m/z 476.2590     0   1   2E+4   6   6.5  RT (min)      lcqtof_t3_neg_4_000_ee    m/z 476.2747     0   1E+2   8.5   9  RT (min)      lcqtof_t3_neg_3_038_06    m/z 476.2768     0   1E+2   8.5   9  RT (min)      lcqtof_t3_neg_3_038_06    m/z 476.2769     0   1E+2   8.5   9  RT (min)      lcqtof_t3_neg_3_038_06    m/z 476.7117     0   1E+4   1   1.5  RT (min)      lcqtof_t3_neg_4_059_14    m/z 476.7576     0   1E+4   1   1.5  RT (min)      lcqtof_t3_neg_4_037_01    m/z 478.1602     0   1E+4   1   1.5  RT (min)       lcqtof_t3_neg_4_040_24    m/z 478.2940     0   1E+4   10   10.5  RT (min)       lcqtof_t3_neg_4_062_13    m/z 478.2941     0   1E+4   10   10.5  RT (min)       lcqtof_t3_neg_4_037_01    m/z 478.2944     0   1E+4   9.5   10  RT (min)      lcqtof_t3_neg_4_000_ee    m/z 478.2944     0   1E+4   9.5   10  RT (min)      lcqtof_t3_neg_4_000_ee    m/z 478.7071     0   1E+4   1   1.5  RT (min)      lcqtof_t3_neg_4_060_15    m/z 479.1560     0   1   2   3   4E+4   1   1.5  RT (min)       lcqtof_t3_neg_3_082_59    m/z 479.1782     0   2   4   6   8E+3   9.5   10  RT (min)      lcqtof_t3_neg_4_059_14    m/z 479.2988     0   1   2   3   4E+3   9.5   10  RT (min)      lcqtof_t3_neg_4_058_16    m/z 479.3116     0   1   2   3E+3   10   10.5  RT (min)      lcqtof_t3_neg_4_016_14    m/z 479.3212     0   1   2   3   4E+3   11.5   12  RT (min)      lcqtof_t3_neg_4_034_34    m/z 479.3530     0   1E+4   10.5   11  RT (min)      lcqtof_t3_neg_4_000_ee    m/z 480.1825     0   1E+4   1   1.5  RT (min)      lcqtof_t3_neg_1_042_46    m/z 480.1840     0   1E+4   1   1.5  RT (min)      lcqtof_t3_neg_4_010_46    m/z 480.3564     0   1E+4   10.5   11  RT (min)      lcqtof_t3_neg_4_019_05    m/z 480.3568     0   1E+4   10   10.5   11  RT (min)      lcqtof_t3_neg_4_019_05    m/z 480.3570     0   1E+4   10.5   11  RT (min)       lcqtof_t3_neg_4_019_05    m/z 480.3572     0   1E+4   10   10.5  RT (min)      lcqtof_t3_neg_2_016_05    m/z 480.3574     0   1E+4   10.5   11  RT (min)      lcqtof_t3_neg_4_019_05    m/z 480.3576     0   1E+4   10.5   11  RT (min)      lcqtof_t3_neg_4_019_05    m/z 481.2291     0   1E+4   9   9.5  RT (min)      lcqtof_t3_neg_4_000_ee    m/z 481.3124     0   1E+4   11   11.5  RT (min)      lcqtof_t3_neg_4_037_01    m/z 481.3124     0   1   2   3   4E+3   11   11.5  RT (min)      lcqtof_t3_neg_1_036_14    m/z 481.3125     0   2   4   6   8E+4   11   11.5  RT (min)      lcqtof_t3_neg_4_059_14    m/z 481.3125     0   2   4E+4   11   11.5  RT (min)      lcqtof_t3_neg_4_059_14    m/z 481.3329     0   1   2E+3   12   12.5  RT (min)      lcqtof_t3_neg_2_043_31    m/z 481.3898     0   2   4E+3   12   12.5  RT (min)      lcqtof_t3_neg_4_037_01    m/z 482.3714     0   1E+4   10.5   11  RT (min)      lcqtof_t3_neg_2_016_05    m/z 482.3728     0   1   2E+4   11   11.5  RT (min)      lcqtof_t3_neg_4_019_05    m/z 482.3728     0   1   2E+4   10.5   11  RT (min)      lcqtof_t3_neg_1_006_05    m/z 482.3728     0   1   2E+4   10.5   11  RT (min)       lcqtof_t3_neg_4_019_05    m/z 482.3732     0   1E+4   10.5   11  RT (min)      lcqtof_t3_neg_2_016_05    m/z 482.3732     0   1E+4   10.5   11  RT (min)      lcqtof_t3_neg_2_016_05    m/z 482.9754     0   1E+4   1   1.5  RT (min)      lcqtof_t3_neg_4_016_14    m/z 483.3170     0   1   2   3   4E+3   10.5   11  RT (min)      lcqtof_t3_neg_4_037_01    m/z 483.3508     0   1E+4   8.5   9  RT (min)      lcqtof_t3_neg_4_000_ee    m/z 484.3832     0   2   4   6   8E+4   8.5   9  RT (min)      lcqtof_t3_neg_4_000_ee    m/z 484.3885     0   1E+4   11   11.5  RT (min)       lcqtof_t3_neg_4_019_05    m/z 484.3888     0   1E+4   11   11.5  RT (min)      lcqtof_t3_neg_1_006_05    m/z 484.7974     0   1E+4   0.5   1  RT (min)      lcqtof_t3_neg_4_000_ee    m/z 485.2112     0   1   2   3   4E+3   12   12.5  RT (min)      lcqtof_t3_neg_1_002_13    m/z 485.3858     0   1   2E+4   8.5   9  RT (min)      lcqtof_t3_neg_4_000_ee    m/z 485.3866     0   1   2E+4   8.5   9  RT (min)      lcqtof_t3_neg_4_000_ee    m/z 485.3868     0   1   2E+4   8.5   9  RT (min)      lcqtof_t3_neg_4_000_ee    m/z 486.7948     0   1E+4   0.5   1  RT (min)      lcqtof_t3_neg_4_000_ee    m/z 488.1282     0   1E+4   1   1.5  RT (min)       lcqtof_t3_neg_4_018_23    m/z 488.3304     0   1   2   3E+4   9.5   10  RT (min)      lcqtof_t3_neg_4_000_ee    m/z 488.7927     0   1E+4   1   1.5  RT (min)       lcqtof_t3_neg_4_095_60    m/z 488.7928     0   1E+4   0.5   1  RT (min)      lcqtof_t3_neg_4_000_ee    m/z 489.1598     0   1   2E+4   1   1.5  RT (min)      lcqtof_t3_neg_4_040_24    m/z 489.3154     0   2   4E+3   11   11.5  RT (min)      lcqtof_t3_neg_3_073_57    m/z 489.3163     0   2   4   6E+3   11   11.5  RT (min)       lcqtof_t3_neg_4_058_16    m/z 490.1411     0   1E+4   4   4.5  RT (min)      lcqtof_t3_neg_1_008_02    m/z 491.3590     0   1   2   3   4E+3   15   15.5  RT (min)       lcqtof_t3_neg_4_000_aa    m/z 491.3590     0   1E+5   13   13.5  RT (min)      lcqtof_t3_neg_1_000_aa    m/z 491.3590     0   1   2   3E+3   14.5   15  RT (min)       lcqtof_t3_neg_4_000_cc    m/z 491.3592     0   1   2   3E+3   15.5   16  RT (min)       lcqtof_t3_neg_4_000_cc    m/z 491.3592     0   2   4E+4   12.5   13  RT (min)      lcqtof_t3_neg_1_000_aa    m/z 491.3662     0   1   2   3   4E+3   12   12.5  RT (min)       lcqtof_t3_neg_3_080_58    m/z 491.3691     0   2   4   6   8E+3   12   12.5  RT (min)       lcqtof_t3_neg_4_059_14    m/z 492.3606     0   1   2   3E+4   13   13.5  RT (min)      lcqtof_t3_neg_1_000_aa    m/z 492.3611     0   1   2   3E+4   13   13.5  RT (min)      lcqtof_t3_neg_1_000_aa    m/z 492.3612     0   1   2   3E+4   13   13.5  RT (min)      lcqtof_t3_neg_1_000_aa    m/z 492.3619     0   1   2   3E+4   13   13.5  RT (min)      lcqtof_t3_neg_1_000_aa    m/z 492.3620     0   1   2   3E+4   13   13.5  RT (min)      lcqtof_t3_neg_1_000_aa    m/z 493.3594     0   1E+4   13   13.5  RT (min)      lcqtof_t3_neg_1_000_aa    m/z 493.3608     0   1E+4   13.5   14  RT (min)      lcqtof_t3_neg_1_000_aa    m/z 493.3835     0   1E+4   7   7.5  RT (min)      lcqtof_t3_neg_4_000_ee    m/z 493.3891     0   1E+4   12   12.5  RT (min)      lcqtof_t3_neg_4_059_14    m/z 493.3894     0   1E+4   12   12.5  RT (min)      lcqtof_t3_neg_4_059_14    m/z 493.3896     0   1E+4   12   12.5  RT (min)       lcqtof_t3_neg_4_059_14    m/z 493.3897     0   1E+4   11.5   12  RT (min)       lcqtof_t3_neg_4_059_14    m/z 496.3356     0   1E+4   10   10.5  RT (min)      lcqtof_t3_neg_4_019_05    m/z 496.3357     0   1E+4   10   10.5  RT (min)      lcqtof_t3_neg_4_019_05    m/z 496.3365     0   1E+4   10   10.5  RT (min)      lcqtof_t3_neg_4_019_05    m/z 496.8802     0   1E+4   7.5   8  RT (min)      lcqtof_t3_neg_4_005_18    m/z 497.1616     0   1   2   3   4E+3   8.5   9  RT (min)      lcqtof_t3_neg_4_053_12    m/z 497.1666     0   2   4   6   8E+3   14.5   15  RT (min)       lcqtof_t3_neg_4_027_41    m/z 497.1667     0   1E+4   14.5   15  RT (min)       lcqtof_t3_neg_4_000_aa    m/z 497.2711     0   2   4   6   8E+3   9.5   10  RT (min)      lcqtof_t3_neg_2_016_05    m/z 497.3346     0   1E+2   4.5   5  RT (min)      lcqtof_t3_neg_3_038_06    m/z 497.3390     0   1   2   3   4E+4   9.5   10  RT (min)      lcqtof_t3_neg_2_016_05    m/z 497.3391     0   1   2E+3   9   9.5  RT (min)      lcqtof_t3_neg_3_032_24    m/z 497.3393     0   1   2   3E+2   9   9.5  RT (min)      lcqtof_t3_neg_3_032_24    m/z 497.3394     0   1   2E+4   9.5   10  RT (min)      lcqtof_t3_neg_3_032_24    m/z 497.3400     0   1   2   3   4E+4   9.5   10  RT (min)      lcqtof_t3_neg_2_016_05    m/z 497.3400     0   2   4   6   8E+4   10   10.5  RT (min)      lcqtof_t3_neg_4_019_05    m/z 498.3514     0   1   2E+3   9.5   10  RT (min)      lcqtof_t3_neg_3_032_24    m/z 498.3516     0   1E+4   9.5   10  RT (min)      lcqtof_t3_neg_3_032_24    m/z 498.3518     0   2   4   6   8E+2   9.5   10  RT (min)      lcqtof_t3_neg_3_032_24    m/z 499.2197     0   1E+4   9.5   10  RT (min)       lcqtof_t3_neg_4_019_05    m/z 499.2228     0   1E+4   9.5   10  RT (min)      lcqtof_t3_neg_4_019_05    m/z 499.2236     0   2   4   6E+3   14   14.5  RT (min)       lcqtof_t3_neg_4_011_32    m/z 499.2927     0   1E+4   8   8.5  RT (min)      lcqtof_t3_neg_4_005_18    m/z 499.3538     0   1E+2   8.5   9  RT (min)      lcqtof_t3_neg_3_038_06    m/z 499.3539     0   1E+2   8.5   9  RT (min)      lcqtof_t3_neg_3_038_06    m/z 499.3544     0   2   4   6   8E+4   9.5   10  RT (min)      lcqtof_t3_neg_3_032_24    m/z 499.3550     0   2   4   6   8E+2   9.5   10  RT (min)      lcqtof_t3_neg_3_032_24    m/z 499.3552     0   1E+2   8.5   9  RT (min)      lcqtof_t3_neg_3_038_06    m/z 499.3571     0   1E+4   9.5   10  RT (min)      lcqtof_t3_neg_3_032_24    m/z 499.3778     0   2   4   6E+3   12   12.5  RT (min)      lcqtof_t3_neg_4_059_14    m/z 500.1300     0   1E+4   1   1.5  RT (min)      lcqtof_t3_neg_4_018_23    m/z 500.8510     0   1   2E+3   15   15.5  RT (min)       lcqtof_t3_neg_4_043_11    m/z 500.8520     0   1   2   3E+3   16.5   17  RT (min)       lcqtof_t3_neg_4_016_14    m/z 500.8521     0   1   2E+3   15.5   16  RT (min)       lcqtof_t3_neg_4_041_54    m/z 500.8524     0   1   2E+3   15.5   16  RT (min)       lcqtof_t3_neg_4_109_62    m/z 501.1262     0   1   2   3   4E+4   1   1.5  RT (min)       lcqtof_t3_neg_4_018_23    m/z 501.1920     0   1E+4   4.5   5  RT (min)      lcqtof_t3_neg_4_000_ee    m/z 501.1925     0   1E+4   1   1.5  RT (min)      lcqtof_t3_neg_4_018_23    m/z 501.1933     0   1   2   3E+3   15   15.5  RT (min)       lcqtof_t3_neg_4_098_61    m/z 501.1937     0   1   2E+3   14.5   15  RT (min)       lcqtof_t3_neg_4_093_60    m/z 501.1938     0   1   2E+3   16.5   17  RT (min)       lcqtof_t3_neg_4_048_19    m/z 501.1942     0   1   2   3   4E+3   15   15.5  RT (min)       lcqtof_t3_neg_4_000_ee    m/z 501.1943     0   1   2   3E+3   16   16.5  RT (min)       lcqtof_t3_neg_4_000_ee    m/z 501.1944     0   1   2   3E+3   15.5   16  RT (min)       lcqtof_t3_neg_4_040_24    m/z 501.2215     0   1E+4   10   10.5  RT (min)      lcqtof_t3_neg_4_019_05    m/z 501.2236     0   1E+4   10   10.5  RT (min)      lcqtof_t3_neg_4_019_05    m/z 501.3706     0   1   2   3E+4   10   10.5  RT (min)       lcqtof_t3_neg_3_032_24    m/z 501.3708     0   2   4E+3   10   10.5  RT (min)      lcqtof_t3_neg_3_032_24    m/z 501.3714     0   1   2   3   4E+3   10   10.5  RT (min)       lcqtof_t3_neg_3_032_24    m/z 501.3949     0   1E+4   12   12.5  RT (min)      lcqtof_t3_neg_4_016_14    m/z 501.3950     0   1E+4   12   12.5  RT (min)      lcqtof_t3_neg_4_016_14    m/z 501.3951     0   1E+4   12   12.5  RT (min)      lcqtof_t3_neg_4_016_14    m/z 502.3981     0   2   4   6E+3   12   12.5  RT (min)      lcqtof_t3_neg_4_059_14    m/z 502.8498     0   1   2   3E+3   16   16.5  RT (min)       lcqtof_t3_neg_1_019_23    m/z 502.8580     0   2   4   6E+3   16.5   17  RT (min)       lcqtof_t3_neg_1_011_45    m/z 503.0670     0   1E+4   10   10.5  RT (min)      lcqtof_t3_neg_4_002_17    m/z 503.0671     0   1E+4   10   10.5  RT (min)      lcqtof_t3_neg_4_002_17    m/z 503.0676     0   1E+4   10   10.5  RT (min)      lcqtof_t3_neg_4_002_17    m/z 503.2958     0   2   4   6   8E+3   10.5   11  RT (min)       lcqtof_t3_neg_4_037_01    m/z 503.2967     0   2   4   6   8E+3   10.5   11  RT (min)      lcqtof_t3_neg_4_037_01    m/z 503.2971     0   1E+4   9.5   10  RT (min)      lcqtof_t3_neg_4_019_05    m/z 503.3861     0   1E+4   10.5   11  RT (min)      lcqtof_t3_neg_4_019_05    m/z 503.3863     0   1E+4   10.5   11  RT (min)       lcqtof_t3_neg_4_019_05    m/z 503.3864     0   1E+4   10.5   11  RT (min)       lcqtof_t3_neg_4_019_05    m/z 503.8808     0   1   2   3E+3   16.5   17  RT (min)       lcqtof_t3_neg_4_008_35    m/z 504.2333     0   1E+4   1   1.5  RT (min)      lcqtof_t3_neg_4_040_24    m/z 504.4136     0   2   4E+3   12.5   13  RT (min)      lcqtof_t3_neg_4_059_14    m/z 504.7664     0   1E+4   1   1.5  RT (min)       lcqtof_t3_neg_2_028_24    m/z 505.0573     0   2   4   6   8E+3   13   13.5  RT (min)      lcqtof_t3_neg_4_002_17    m/z 505.0583     0   1E+4   13   13.5  RT (min)      lcqtof_t3_neg_4_036_47    m/z 505.1177     0   1   2   3E+3   15.5   16  RT (min)       lcqtof_t3_neg_4_024_26    m/z 505.1179     0   1   2   3   4E+3   15   15.5  RT (min)       lcqtof_t3_neg_4_004_20    m/z 505.1180     0   1   2   3E+3   16   16.5  RT (min)       lcqtof_t3_neg_4_007_09    m/z 505.1183     0   1   2   3E+3   16   16.5  RT (min)       lcqtof_t3_neg_4_004_20    m/z 505.1184     0   1   2   3   4E+3   15.5   16  RT (min)       lcqtof_t3_neg_4_012_27    m/z 505.1187     0   1   2   3   4E+3   15   15.5  RT (min)       lcqtof_t3_neg_4_004_20    m/z 505.1195     0   1   2   3   4E+3   14.5   15  RT (min)       lcqtof_t3_neg_4_010_46    m/z 505.1212     0   1   2   3E+3   14.5   15  RT (min)       lcqtof_t3_neg_4_022_28    m/z 505.1489     0   2   4   6E+3   7.5   8  RT (min)       lcqtof_t3_neg_4_059_14    m/z 505.3318     0   2   4E+3   12.5   13  RT (min)       lcqtof_t3_neg_4_018_23    m/z 505.3319     0   1   2   3E+3   15   15.5  RT (min)       lcqtof_t3_neg_4_032_04    m/z 505.3320     0   2   4   6E+3   13   13.5  RT (min)       lcqtof_t3_neg_4_032_04    m/z 505.3320     0   1   2E+3   16   16.5  RT (min)       lcqtof_t3_neg_4_008_35    m/z 505.3321     0   2   4   6E+3   13   13.5  RT (min)       lcqtof_t3_neg_4_040_24    m/z 505.3328     0   1   2   3E+3   15.5   16  RT (min)       lcqtof_t3_neg_4_000_ee    m/z 505.9881     0   2   4   6   8E+3   1   1.5  RT (min)      lcqtof_t3_neg_1_015_34    m/z 506.1422     0   1E+4   1   1.5  RT (min)      lcqtof_t3_neg_4_028_25    m/z 506.1800     0   1   2E+4   1   1.5  RT (min)      lcqtof_t3_neg_4_047_07    m/z 506.1830     0   1   2E+4   1   1.5  RT (min)      lcqtof_t3_neg_4_047_07    m/z 507.0548     0   2   4   6   8E+3   13   13.5  RT (min)      lcqtof_t3_neg_4_036_47    m/z 507.1204     0   2   4E+4   11.5   12  RT (min)      lcqtof_t3_neg_4_064_17    m/z 507.1206     0   1   2E+4   13   13.5  RT (min)      lcqtof_t3_neg_4_064_17    m/z 507.1209     0   2   4E+4   11.5   12  RT (min)       lcqtof_t3_neg_4_064_17    m/z 507.3459     0   1E+4   7.5   8  RT (min)      lcqtof_t3_neg_4_000_ee    m/z 508.1211     0   1E+4   11.5   12  RT (min)      lcqtof_t3_neg_4_002_17    m/z 508.1217     0   1   2E+4   11.5   12  RT (min)      lcqtof_t3_neg_4_064_17    m/z 508.1224     0   1   2E+4   11.5   12  RT (min)      lcqtof_t3_neg_4_064_17    m/z 508.1230     0   1E+4   13   13.5  RT (min)      lcqtof_t3_neg_4_036_47    m/z 508.8752     0   1   2   3   4E+3   8   9   10  RT (min)       lcqtof_t3_neg_1_012_16    m/z 508.8752     0   1   2   3   4E+3   8   8.5  RT (min)       lcqtof_t3_neg_1_014_04    m/z 508.8752     0   1   2   3E+3   8   10  RT (min)       lcqtof_t3_neg_1_020_47    m/z 508.8753     0   1   2   3   4E+3   8   9  RT (min)       lcqtof_t3_neg_1_004_11    m/z 508.8759     0   1   2   3E+3   8.5   9  RT (min)       lcqtof_t3_neg_1_010_27    m/z 509.1200     0   1E+4   11.5   12  RT (min)      lcqtof_t3_neg_4_064_17    m/z 509.1217     0   1E+4   12   12.5  RT (min)      lcqtof_t3_neg_4_064_17    m/z 509.2944     0   1   2E+4   9   9.5  RT (min)      lcqtof_t3_neg_4_000_ee    m/z 509.2946     0   1   2E+4   9   9.5  RT (min)      lcqtof_t3_neg_4_000_ee    m/z 509.3399     0   1   2E+3   13   13.5  RT (min)      lcqtof_t3_neg_4_033_53    m/z 509.3485     0   1E+4   10.5   11  RT (min)       lcqtof_t3_neg_2_043_31    m/z 510.8803     0   2   4E+3   15.5   16  RT (min)       lcqtof_t3_neg_4_099_61    m/z 510.8805     0   2   4E+3   15.5   16  RT (min)       lcqtof_t3_neg_4_100_61    m/z 510.8807     0   1E+4   16   16.5  RT (min)       lcqtof_t3_neg_4_018_23    m/z 510.8807     0   1   2   3   4E+3   15   15.5  RT (min)       lcqtof_t3_neg_4_108_62    m/z 510.8808     0   1   2E+3   8.5   9   9.5  RT (min)       lcqtof_t3_neg_4_000_ff    m/z 510.8808     0   1   2E+3   15   15.5  RT (min)       lcqtof_t3_neg_3_032_24    m/z 511.0102     0   1E+4   1   1.5  RT (min)      lcqtof_t3_neg_2_028_24    m/z 511.2752     0   1E+4   9   9.5  RT (min)      lcqtof_t3_neg_4_000_ee    m/z 511.2756     0   1E+4   9   9.5  RT (min)      lcqtof_t3_neg_4_000_ee    m/z 512.1761     0   1   2E+4   1   1.5  RT (min)      lcqtof_t3_neg_4_040_24    m/z 512.1772     0   1   2E+4   1   1.5  RT (min)       lcqtof_t3_neg_4_040_24    m/z 512.2368     0   1E+4   6   6.5  RT (min)      lcqtof_t3_neg_4_000_ee    m/z 512.2686     0   1   2   3E+4   6.5   7  RT (min)      lcqtof_t3_neg_4_005_18    m/z 512.7624     0   1E+4   1   1.5  RT (min)      lcqtof_t3_neg_4_046_33    m/z 513.1974     0   1   2E+4   1   1.5  RT (min)      lcqtof_t3_neg_4_018_23    m/z 513.2378     0   2   4   6E+3   14   14.5  RT (min)       lcqtof_t3_neg_4_011_32    m/z 513.2388     0   2   4   6   8E+3   10   10.5  RT (min)      lcqtof_t3_neg_4_019_05    m/z 513.2394     0   2   4   6   8E+3   10   10.5  RT (min)       lcqtof_t3_neg_4_019_05    m/z 513.3103     0   2   4   6E+3   10   10.5  RT (min)      lcqtof_t3_neg_4_055_38    m/z 513.4070     0   1E+4   8   8.5  RT (min)      lcqtof_t3_neg_4_005_18    m/z 514.1941     0   1E+5   1   1.5  RT (min)      lcqtof_t3_neg_4_018_23    m/z 514.2840     0   1   2   3E+5   7   7.5  RT (min)      lcqtof_t3_neg_4_005_18    m/z 514.3240     0   1E+2   4.5   5  RT (min)      lcqtof_t3_neg_3_038_06    m/z 514.4264     0   1   2   3   4E+3   13   13.5  RT (min)       lcqtof_t3_neg_4_000_ee    m/z 514.5544     0   1   2E+4   5   5.5  RT (min)      lcqtof_t3_neg_1_019_23    m/z 514.7540     0   1E+4   1   1.5  RT (min)      lcqtof_t3_neg_4_046_33    m/z 514.8916     0   1   2   3   4E+3   17   17.5  RT (min)       lcqtof_t3_neg_4_106_62    m/z 514.8922     0   1   2   3E+3   17.5   18  RT (min)       lcqtof_t3_neg_4_112_62    m/z 514.8924     0   1   2E+3   10   10.5  RT (min)      lcqtof_t3_neg_1_008_02    m/z 514.8924     0   2   4   6E+3   17   17.5  RT (min)       lcqtof_t3_neg_4_035_08    m/z 514.8925     0   2   4   6E+3   8   9  RT (min)       lcqtof_t3_neg_4_103_61    m/z 514.8925     0   1   2   3E+3   10.5   11  RT (min)       lcqtof_t3_neg_1_004_11    m/z 514.8926     0   2   4E+3   9  RT (min)       lcqtof_t3_neg_4_090_60    m/z 514.8928     0   1   2   3E+3   9   9.5   10  RT (min)       lcqtof_t3_neg_1_013_09    m/z 514.8932     0   1   2   3E+3   10   10.5  RT (min)       lcqtof_t3_neg_1_003_07    m/z 514.8932     0   2   4E+3   9   10  RT (min)       lcqtof_t3_neg_4_053_12    m/z 514.8932     0   1   2   3   4E+3   9   9.5  RT (min)       lcqtof_t3_neg_4_040_24    m/z 514.8935     0   1   2E+3   11.5   12  RT (min)       lcqtof_t3_neg_1_008_02    m/z 514.8937     0   1   2   3E+3   11   11.5  RT (min)       lcqtof_t3_neg_4_031_42    m/z 514.8937     0   1   2E+3   11.5   12  RT (min)       lcqtof_t3_neg_3_034_04    m/z 515.0968     0   1E+4   1   1.5  RT (min)       lcqtof_t3_neg_4_040_24    m/z 515.1302     0   1   2E+3   14.5   15  RT (min)       lcqtof_t3_neg_4_042_10    m/z 515.1307     0   1   2   3E+3   14.5   15  RT (min)       lcqtof_t3_neg_4_021_21    m/z 515.1312     0   1   2   3   4E+3   16   16.5  RT (min)       lcqtof_t3_neg_4_000_ee    m/z 515.1922     0   2   4   6   8E+3   7   7.5  RT (min)       lcqtof_t3_neg_4_109_62    m/z 515.1982     0   1   2E+4   1   1.5  RT (min)      lcqtof_t3_neg_4_018_23    m/z 515.2876     0   1E+5   7   7.5  RT (min)      lcqtof_t3_neg_4_005_18    m/z 515.3269     0   1E+2   4.5   5  RT (min)      lcqtof_t3_neg_3_038_06    m/z 515.3706     0   2   4   6E+3   11.5   12  RT (min)       lcqtof_t3_neg_4_059_14    m/z 516.1812     0   1E+4   1   1.5  RT (min)      lcqtof_t3_neg_4_040_24    m/z 516.2859     0   1   2   3E+4   7   7.5  RT (min)      lcqtof_t3_neg_4_005_18    m/z 516.3294     0   2   4   6   8E+1   4.5   5  RT (min)      lcqtof_t3_neg_3_038_06    m/z 516.8257     0   1E+4   1   1.5  RT (min)      lcqtof_t3_neg_4_026_30    m/z 517.1802     0   1   2   3E+4   1   1.5  RT (min)       lcqtof_t3_neg_4_090_60    m/z 517.3882     0   1E+4   12   12.5  RT (min)       lcqtof_t3_neg_4_059_14    m/z 517.3895     0   1   2   3   4E+3   11.5   12  RT (min)       lcqtof_t3_neg_4_011_32    m/z 518.2511     0   1E+4   9.5   10  RT (min)       lcqtof_t3_neg_4_019_05    m/z 518.3407     0   2   4E+4   9.5   10  RT (min)      lcqtof_t3_neg_2_016_05    m/z 518.3408     0   1   2   3E+4   9.5   10  RT (min)      lcqtof_t3_neg_3_032_24    m/z 518.3412     0   2   4E+4   9.5   10  RT (min)      lcqtof_t3_neg_2_016_05    m/z 518.3418     0   1   2   3E+2   9   9.5  RT (min)      lcqtof_t3_neg_3_032_24    m/z 518.3419     0   1   2E+3   9   9.5  RT (min)      lcqtof_t3_neg_3_032_24    m/z 519.1375     0   1   2   3   4E+3   10   10.5  RT (min)      lcqtof_t3_neg_4_000_ee    m/z 519.1467     0   2   4   6   8E+3   14   14.5  RT (min)       lcqtof_t3_neg_4_074_58    m/z 519.1468     0   1E+4   15   16  RT (min)       lcqtof_t3_neg_4_005_18    m/z 519.1468     0   1E+4   14.5   15  RT (min)       lcqtof_t3_neg_4_000_cc    m/z 519.1468     0   1E+4   15   16  RT (min)       lcqtof_t3_neg_4_035_08    m/z 519.1470     0   1E+4   16   16.5  RT (min)       lcqtof_t3_neg_4_001_49    m/z 519.1470     0   1E+4   15.5   16  RT (min)       lcqtof_t3_neg_4_034_34    m/z 519.1473     0   1   2   3E+3   11.5   12  RT (min)       lcqtof_t3_neg_4_032_04    m/z 519.1475     0   1   2   3E+3   10.5   11  RT (min)       lcqtof_t3_neg_4_000_ff    m/z 519.1476     0   1   2   3E+3   11   11.5  RT (min)      lcqtof_t3_neg_4_000_aa    m/z 519.1479     0   2   4   6   8E+3   14   14.5  RT (min)      lcqtof_t3_neg_4_067_57    m/z 519.1480     0   1E+4   16   16.5  RT (min)       lcqtof_t3_neg_4_000_ee    m/z 519.1486     0   1   2   3   4E+3   12   12.5  RT (min)       lcqtof_t3_neg_4_032_04    m/z 519.1488     0   1   2   3   4E+3   12.5   13  RT (min)       lcqtof_t3_neg_4_000_ff    m/z 519.2120     0   1E+4   1   1.5  RT (min)      lcqtof_t3_neg_4_028_25    m/z 520.1518     0   1   2   3E+3   8   8.5  RT (min)      lcqtof_t3_neg_4_000_ee    m/z 520.1530     0   1   2   3E+3   8   8.5  RT (min)      lcqtof_t3_neg_4_000_ee    m/z 520.2662     0   2   4E+4   10.5   11  RT (min)       lcqtof_t3_neg_4_019_05    m/z 520.2663     0   1   2E+4   10.5   11  RT (min)      lcqtof_t3_neg_4_019_05    m/z 520.2668     0   2   4   6   8E+4   10.5   11  RT (min)       lcqtof_t3_neg_4_019_05    m/z 520.2675     0   1   2E+4   9   9.5  RT (min)      lcqtof_t3_neg_1_019_23    m/z 520.4090     0   2   4   6E+3   12   12.5  RT (min)      lcqtof_t3_neg_4_059_14    m/z 520.9090     0   2   4   6E+3   15.5   16  RT (min)       lcqtof_t3_neg_4_111_62    m/z 520.9094     0   1   2   3E+3   15   15.5  RT (min)       lcqtof_t3_neg_3_032_24    m/z 520.9095     0   2   4   6   8E+4   16   16.5  RT (min)      lcqtof_t3_neg_1_011_45    m/z 520.9095     0   1   2E+5   16   16.5  RT (min)       lcqtof_t3_neg_4_018_23    m/z 520.9095     0   1E+3   13.5   14  RT (min)      lcqtof_t3_neg_3_038_06    m/z 520.9096     0   1   2   3E+3   11.5   12  RT (min)       lcqtof_t3_neg_4_098_61    m/z 520.9096     0   2   4   6E+3   9   9.5  RT (min)       lcqtof_t3_neg_4_113_62    m/z 520.9097     0   2   4   6E+3   8.5   9   9.5  RT (min)       lcqtof_t3_neg_4_108_62    m/z 520.9097     0   2   4E+3   15.5   16  RT (min)       lcqtof_t3_neg_1_019_23    m/z 520.9098     0   1   2   3   4E+3   10.5   11  RT (min)       lcqtof_t3_neg_4_105_61    m/z 520.9098     0   1   2E+5   16   16.5  RT (min)       lcqtof_t3_neg_4_018_23    m/z 520.9099     0   1E+4   6   8  RT (min)       lcqtof_t3_neg_4_000_ff    m/z 520.9100     0   1   2   3   4E+3   12   12.5  RT (min)       lcqtof_t3_neg_4_000_ee    m/z 520.9100     0   2   4   6   8E+3   15.5   16  RT (min)       lcqtof_t3_neg_4_107_62    m/z 520.9102     0   1   2   3E+3   11   11.5  RT (min)       lcqtof_t3_neg_4_090_60    m/z 520.9104     0   2   4E+3   9.5   10  RT (min)      lcqtof_t3_neg_4_098_61    m/z 520.9111     0   2   4   6   8E+4   17   17.5  RT (min)       lcqtof_t3_neg_4_035_08    m/z 520.9129     0   1   2   3E+5   17   17.5  RT (min)       lcqtof_t3_neg_4_106_62    m/z 520.9190     0   1   2E+5   17   17.5  RT (min)       lcqtof_t3_neg_4_108_62    m/z 521.0819     0   1E+4   13   13.5  RT (min)      lcqtof_t3_neg_4_036_47    m/z 521.0823     0   1E+4   12   12.5  RT (min)      lcqtof_t3_neg_4_064_17    m/z 521.0824     0   1E+4   13   13.5  RT (min)      lcqtof_t3_neg_4_064_17    m/z 521.0825     0   1E+4   12   12.5  RT (min)      lcqtof_t3_neg_4_064_17    m/z 521.1423     0   1   2   3E+3   15   15.5  RT (min)      lcqtof_t3_neg_4_025_52    m/z 521.1424     0   1   2   3E+3   14.5   15  RT (min)      lcqtof_t3_neg_4_007_09    m/z 521.1823     0   1   2E+3   10   10.5  RT (min)      lcqtof_t3_neg_1_001_49    m/z 521.1832     0   1   2E+3   10.5   11  RT (min)      lcqtof_t3_neg_1_036_14    m/z 521.2710     0   1   2E+4   9.5   10  RT (min)      lcqtof_t3_neg_2_016_05    m/z 522.0827     0   1E+4   12   12.5  RT (min)      lcqtof_t3_neg_4_064_17    m/z 522.2658     0   1   2E+4   6   6.5  RT (min)      lcqtof_t3_neg_4_000_ee    m/z 522.2816     0   1   2E+4   9.5   10  RT (min)      lcqtof_t3_neg_2_016_05    m/z 522.2844     0   1   2E+4   10   10.5  RT (min)      lcqtof_t3_neg_2_016_05    m/z 522.5338     0   1   2E+3   14.5   15  RT (min)      lcqtof_t3_neg_4_011_32    m/z 522.5340     0   1   2E+3   14.5   15  RT (min)      lcqtof_t3_neg_4_000_bb    m/z 523.2888     0   1E+4   10   10.5  RT (min)      lcqtof_t3_neg_4_019_05    m/z 523.3930     0   1E+4   8.5   9  RT (min)      lcqtof_t3_neg_4_000_ee    m/z 524.2272     0   1   2E+4   1   1.5  RT (min)      lcqtof_t3_neg_4_028_25    m/z 524.2690     0   1E+4   7.5   8  RT (min)      lcqtof_t3_neg_4_058_16    m/z 524.6858     0   1E+4   1   1.5  RT (min)      lcqtof_t3_neg_4_037_01    m/z 525.3174     0   1E+4   9   9.5  RT (min)      lcqtof_t3_neg_4_000_ee    m/z 525.3182     0   1E+4   9   9.5  RT (min)      lcqtof_t3_neg_4_000_ee    m/z 525.3214     0   1E+4   9   9.5  RT (min)      lcqtof_t3_neg_4_000_ee    m/z 525.3920     0   2   4   6E+3   12   12.5  RT (min)      lcqtof_t3_neg_4_059_14    m/z 526.2853     0   1E+4   8   8.5  RT (min)      lcqtof_t3_neg_4_062_13    m/z 526.3011     0   1E+4   9.5   10  RT (min)      lcqtof_t3_neg_4_062_13    m/z 526.3729     0   1E+4   10   10.5  RT (min)      lcqtof_t3_neg_4_019_05    m/z 526.3745     0   1E+4   9.5   10  RT (min)      lcqtof_t3_neg_4_019_05    m/z 526.4018     0   1E+4   8.5   9  RT (min)      lcqtof_t3_neg_4_000_ee    m/z 526.4046     0   1E+4   8.5   9  RT (min)      lcqtof_t3_neg_4_000_ee    m/z 526.4065     0   1E+4   8.5   9  RT (min)      lcqtof_t3_neg_4_000_ee    m/z 526.6390     0   1E+4   1   1.5  RT (min)      lcqtof_t3_neg_4_059_14    m/z 526.8522     0   2   4   6E+3   16.5   17  RT (min)      lcqtof_t3_neg_4_000_ee    m/z 527.2191     0   1   2E+4   4   4.5  RT (min)      lcqtof_t3_neg_4_000_ee    m/z 527.2193     0   1   2E+4   4   4.5  RT (min)      lcqtof_t3_neg_4_000_ee    m/z 527.2301     0   1   2E+4   4.5   5  RT (min)      lcqtof_t3_neg_4_000_ee    m/z 527.2384     0   1E+4   1   1.5  RT (min)      lcqtof_t3_neg_4_028_25    m/z 527.3766     0   2   4   6E+2   9   9.5  RT (min)      lcqtof_t3_neg_3_032_24    m/z 527.3769     0   1   2E+4   9   9.5  RT (min)      lcqtof_t3_neg_3_032_24    m/z 527.3786     0   1   2E+3   9   9.5  RT (min)      lcqtof_t3_neg_3_032_24    m/z 527.8707     0   1   2   3   4E+3   16   16.5  RT (min)       lcqtof_t3_neg_1_009_50    m/z 527.8712     0   1   2E+3   15.5   16  RT (min)       lcqtof_t3_neg_4_000_ee    m/z 528.3090     0   1   2E+4   10.5   11  RT (min)      lcqtof_t3_neg_4_002_17    m/z 528.3110     0   1E+4   9   9.5  RT (min)      lcqtof_t3_neg_4_000_ee    m/z 528.3888     0   1E+4   10   10.5  RT (min)      lcqtof_t3_neg_4_019_05    m/z 528.3891     0   1E+4   10   10.5  RT (min)      lcqtof_t3_neg_4_019_05    m/z 528.3892     0   1E+4   10   10.5  RT (min)      lcqtof_t3_neg_4_019_05    m/z 529.1463     0   1   2   3   4E+3   15.5   16  RT (min)       lcqtof_t3_neg_4_024_26    m/z 529.1464     0   2   4E+3   15   16  RT (min)       lcqtof_t3_neg_4_017_51    m/z 529.3124     0   1E+4   10.5   11  RT (min)       lcqtof_t3_neg_4_037_01    m/z 529.3128     0   1E+4   10.5   11  RT (min)      lcqtof_t3_neg_4_037_01    m/z 529.3922     0   1E+3   9.5   10  RT (min)      lcqtof_t3_neg_3_032_24    m/z 529.3927     0   1   2E+4   9.5   10  RT (min)      lcqtof_t3_neg_3_032_24    m/z 529.3928     0   2   4   6E+2   9.5   10  RT (min)      lcqtof_t3_neg_3_032_24    m/z 529.3930     0   1   2   3   4E+4   10   10.5  RT (min)      lcqtof_t3_neg_2_016_05    m/z 529.3932     0   1   2   3   4E+4   10   10.5  RT (min)      lcqtof_t3_neg_2_016_05    m/z 529.3935     0   1   2   3   4E+4   10   10.5  RT (min)      lcqtof_t3_neg_2_016_05    m/z 530.4295     0   2   4   6E+3   12.5   13  RT (min)      lcqtof_t3_neg_4_059_14    m/z 531.2554     0   1   2   3E+2   4.5   5  RT (min)      lcqtof_t3_neg_3_038_06    m/z 531.2571     0   1E+4   1   1.5  RT (min)      lcqtof_t3_neg_4_040_24    m/z 531.4084     0   1E+4   10   10.5  RT (min)      lcqtof_t3_neg_2_016_05    m/z 531.4091     0   1E+4   10.5   11  RT (min)      lcqtof_t3_neg_2_016_05    m/z 531.4092     0   1E+4   10   10.5  RT (min)      lcqtof_t3_neg_2_016_05    m/z 532.2549     0   2   4E+4   1   1.5  RT (min)      lcqtof_t3_neg_4_040_24    m/z 532.2896     0   1   2   3   4E+4   8   8.5  RT (min)      lcqtof_t3_neg_4_005_18    m/z 533.1444     0   1   2   3E+3   16   16.5  RT (min)      lcqtof_t3_neg_4_028_25    m/z 533.1449     0   1   2   3E+3   16   16.5  RT (min)      lcqtof_t3_neg_4_005_18    m/z 533.1451     0   1   2   3E+3   14.5   15  RT (min)       lcqtof_t3_neg_4_006_16    m/z 533.1453     0   1   2   3E+3   15.5   16  RT (min)       lcqtof_t3_neg_4_005_18    m/z 533.1456     0   1   2   3E+3   15   15.5  RT (min)       lcqtof_t3_neg_4_007_09    m/z 533.1723     0   1   2E+4   1   1.5  RT (min)      lcqtof_t3_neg_4_047_07    m/z 533.2520     0   1E+2   4.5   5  RT (min)      lcqtof_t3_neg_3_038_06    m/z 533.2927     0   1E+4   8   8.5  RT (min)      lcqtof_t3_neg_4_005_18    m/z 533.3077     0   1E+4   11.5   12  RT (min)      lcqtof_t3_neg_4_041_54    m/z 533.3099     0   1E+4   11.5   12  RT (min)      lcqtof_t3_neg_4_065_51    m/z 534.6958     0   1E+4   1   1.5  RT (min)      lcqtof_t3_neg_4_046_33    m/z 534.7143     0   1E+4   1   1.5  RT (min)      lcqtof_t3_neg_4_062_13    m/z 535.0984     0   1   2E+4   12.5   13  RT (min)      lcqtof_t3_neg_4_064_17    m/z 536.0984     0   1E+4   12.5   13  RT (min)      lcqtof_t3_neg_4_064_17    m/z 536.3618     0   1E+4   5   5.5  RT (min)       lcqtof_t3_neg_4_018_23    m/z 536.5494     0   1   2E+3   14.5   15  RT (min)      lcqtof_t3_neg_4_031_42    m/z 536.6644     0   1E+4   1   1.5  RT (min)      lcqtof_t3_neg_4_060_15    m/z 536.8824     0   1   2E+4   17   17.5  RT (min)      lcqtof_t3_neg_4_035_08    m/z 536.8825     0   2   4E+2   13.5   14  RT (min)      lcqtof_t3_neg_3_038_06    m/z 536.8833     0   1E+3   15   15.5  RT (min)      lcqtof_t3_neg_3_032_24    m/z 536.8841     0   1E+3   15.5   16  RT (min)      lcqtof_t3_neg_1_019_23    m/z 536.8842     0   1   2E+4   16   16.5  RT (min)      lcqtof_t3_neg_1_011_45    m/z 537.0928     0   1E+4   12.5   13  RT (min)      lcqtof_t3_neg_4_064_17    m/z 537.8988     0   1E+3   15   15.5  RT (min)       lcqtof_t3_neg_3_032_24    m/z 537.8993     0   1E+3   15.5   16  RT (min)       lcqtof_t3_neg_1_019_23    m/z 537.8994     0   1   2E+3   15.5   16  RT (min)       lcqtof_t3_neg_4_042_10    m/z 537.8995     0   1E+4   16   16.5  RT (min)       lcqtof_t3_neg_1_011_45    m/z 538.4056     0   1E+4   8.5   9  RT (min)      lcqtof_t3_neg_4_000_ee    m/z 539.2188     0   1   2   3E+4   4.5   5  RT (min)      lcqtof_t3_neg_4_000_ee    m/z 539.4307     0   1E+4   11.5   12  RT (min)      lcqtof_t3_neg_4_016_14    m/z 540.0531     0   1E+4   3.5   4  RT (min)      lcqtof_t3_neg_1_032_36    m/z 540.0533     0   1E+4   3.5   4  RT (min)      lcqtof_t3_neg_1_048_35    m/z 540.0539     0   1E+4   2   2.5  RT (min)      lcqtof_t3_neg_4_055_38    m/z 540.0554     0   1E+4   2   2.5  RT (min)      lcqtof_t3_neg_4_055_38    m/z 540.8915     0   1   2   3   4E+3   16.5   17  RT (min)      lcqtof_t3_neg_4_000_ee    m/z 541.3126     0   1E+4   7.5   8  RT (min)      lcqtof_t3_neg_4_000_ee    m/z 541.3128     0   1E+4   7.5   8  RT (min)      lcqtof_t3_neg_4_000_ee    m/z 541.3333     0   1E+4   11   11.5  RT (min)      lcqtof_t3_neg_1_034_30    m/z 541.3861     0   2   4   6E+3   11.5   12  RT (min)       lcqtof_t3_neg_4_016_14    m/z 542.2501     0   1E+4   9.5   10  RT (min)      lcqtof_t3_neg_1_019_23    m/z 542.2501     0   1   2E+4   9.5   10  RT (min)      lcqtof_t3_neg_4_018_23    m/z 542.2502     0   1   2E+4   9.5   10  RT (min)      lcqtof_t3_neg_4_018_23    m/z 543.1614     0   1   2E+3   14.5   15  RT (min)       lcqtof_t3_neg_4_019_05    m/z 543.1619     0   1E+4   10.5   11  RT (min)       lcqtof_t3_neg_4_000_ee    m/z 543.1624     0   1E+4   10   10.5  RT (min)       lcqtof_t3_neg_4_000_ee    m/z 543.1626     0   1   2   3E+3   15.5   16   16.5  RT (min)       lcqtof_t3_neg_4_048_19    m/z 543.1626     0   1   2   3E+3   15   16  RT (min)       lcqtof_t3_neg_4_041_54    m/z 543.1631     0   1   2   3E+3   15   15.5  RT (min)       lcqtof_t3_neg_4_052_03    m/z 543.3370     0   1   2E+4   8.5   9  RT (min)      lcqtof_t3_neg_4_000_ee    m/z 543.3370     0   1   2E+4   8.5   9  RT (min)      lcqtof_t3_neg_4_000_ee    m/z 543.3374     0   1   2E+4   8.5   9  RT (min)      lcqtof_t3_neg_4_000_ee    m/z 543.3440     0   1   2E+4   10   10.5  RT (min)      lcqtof_t3_neg_4_040_24    m/z 543.3440     0   1   2E+4   10   10.5  RT (min)      lcqtof_t3_neg_4_019_05    m/z 543.3445     0   1   2E+4   9.5   10  RT (min)      lcqtof_t3_neg_4_040_24    m/z 543.3447     0   1   2E+4   9.5   10  RT (min)      lcqtof_t3_neg_4_019_05    m/z 543.3448     0   1   2E+4   10   10.5  RT (min)      lcqtof_t3_neg_4_019_05    m/z 543.3450     0   1E+4   9.5   10  RT (min)      lcqtof_t3_neg_2_016_05    m/z 543.3451     0   1   2E+4   9.5   10  RT (min)      lcqtof_t3_neg_4_040_24    m/z 544.2663     0   1E+4   9.5   10  RT (min)      lcqtof_t3_neg_1_019_23    m/z 544.3478     0   1E+4   9   9.5  RT (min)      lcqtof_t3_neg_3_032_24    m/z 544.3480     0   1E+3   9   9.5  RT (min)      lcqtof_t3_neg_3_032_24    m/z 544.3482     0   2   4   6E+4   9   9.5  RT (min)      lcqtof_t3_neg_3_032_24    m/z 544.7010     0   1E+4   1   1.5  RT (min)      lcqtof_t3_neg_4_037_01    m/z 545.2720     0   1   2E+4   9   9.5  RT (min)      lcqtof_t3_neg_4_000_ee    m/z 545.2725     0   1   2E+4   9   9.5  RT (min)      lcqtof_t3_neg_4_000_ee    m/z 546.3634     0   1E+3   9.5   10  RT (min)      lcqtof_t3_neg_3_032_24    m/z 546.3638     0   1   2   3   4E+4   10   10.5  RT (min)      lcqtof_t3_neg_2_016_05    m/z 546.3639     0   1   2E+4   9.5   10  RT (min)      lcqtof_t3_neg_3_032_24    m/z 546.3644     0   1   2   3   4E+4   9.5   10  RT (min)      lcqtof_t3_neg_2_016_05    m/z 546.3644     0   1   2   3   4E+3   9.5   10  RT (min)      lcqtof_t3_neg_3_032_24    m/z 548.2957     0   1   2   3   4E+3   10.5   11  RT (min)       lcqtof_t3_neg_4_030_31    m/z 548.2958     0   1   2E+3   10.5   11  RT (min)      lcqtof_t3_neg_4_038_29    m/z 549.4520     0   2   4   6E+3   12.5   13  RT (min)       lcqtof_t3_neg_4_059_14    m/z 550.2464     0   2   4   6   8E+3   7.5   8  RT (min)      lcqtof_t3_neg_4_059_14    m/z 550.5650     0   1   2E+3   14.5   15  RT (min)      lcqtof_t3_neg_4_000_dd    m/z 551.0548     0   1   2E+4   13   13.5  RT (min)      lcqtof_t3_neg_4_036_47    m/z 551.0564     0   1E+4   12   12.5  RT (min)      lcqtof_t3_neg_4_064_17    m/z 551.0579     0   1E+4   13   13.5  RT (min)      lcqtof_t3_neg_4_002_17    m/z 551.0597     0   1E+4   13   13.5  RT (min)      lcqtof_t3_neg_4_002_17    m/z 552.0554     0   1E+4   13   13.5  RT (min)      lcqtof_t3_neg_4_036_47    m/z 553.3109     0   1   2   3E+4   9   9.5  RT (min)      lcqtof_t3_neg_4_000_ee    m/z 553.3217     0   1   2   3E+4   9   9.5  RT (min)      lcqtof_t3_neg_4_000_ee    m/z 553.3746     0   2   4   6E+3   10   10.5  RT (min)       lcqtof_t3_neg_4_083_59    m/z 553.4062     0   1E+4   8   8.5  RT (min)      lcqtof_t3_neg_4_000_ee    m/z 553.4065     0   1E+4   8   8.5  RT (min)      lcqtof_t3_neg_4_000_ee    m/z 554.3023     0   1E+4   10   10.5  RT (min)      lcqtof_t3_neg_4_062_13    m/z 554.4508     0   2   4E+3   12   12.5  RT (min)      lcqtof_t3_neg_4_037_01    m/z 555.2995     0   2   4E+4   9   9.5  RT (min)       lcqtof_t3_neg_4_000_ee    m/z 556.3034     0   1E+4   9   9.5  RT (min)      lcqtof_t3_neg_4_000_ee    m/z 556.3044     0   1E+4   9   9.5  RT (min)      lcqtof_t3_neg_4_000_ee    m/z 560.2962     0   2   4E+3   10.5   11  RT (min)      lcqtof_t3_neg_4_033_53    m/z 560.7050     0   1E+4   1   1.5  RT (min)      lcqtof_t3_neg_4_046_33    m/z 560.8256     0   1   2E+3   16.5   17  RT (min)      lcqtof_t3_neg_1_003_07    m/z 561.1318     0   1E+4   11   11.5  RT (min)      lcqtof_t3_neg_4_064_17    m/z 561.3053     0   1   2E+4   9   9.5  RT (min)      lcqtof_t3_neg_4_000_ee    m/z 561.3056     0   1   2E+4   9   9.5  RT (min)      lcqtof_t3_neg_4_000_ee    m/z 562.3130     0   1   2E+2   8   8.5  RT (min)      lcqtof_t3_neg_3_038_06    m/z 562.3130     0   1   2E+2   8   8.5  RT (min)      lcqtof_t3_neg_3_038_06    m/z 562.3951     0   2   4E+2   9   9.5  RT (min)      lcqtof_t3_neg_3_032_24    m/z 562.3952     0   1E+4   9.5   10  RT (min)      lcqtof_t3_neg_3_032_24    m/z 562.3963     0   1   2E+4   9.5   10  RT (min)      lcqtof_t3_neg_2_016_05    m/z 563.1502     0   1   2E+4   5   5.5  RT (min)      lcqtof_t3_neg_4_000_ee    m/z 563.5041     0   1   2   3   4E+3   13   13.5  RT (min)      lcqtof_t3_neg_4_016_14    m/z 564.3302     0   1   2E+2   8.5   9  RT (min)      lcqtof_t3_neg_3_038_06    m/z 564.3308     0   1   2E+2   8.5   9  RT (min)      lcqtof_t3_neg_3_038_06    m/z 564.3312     0   1   2E+2   8.5   9  RT (min)      lcqtof_t3_neg_3_038_06    m/z 564.4107     0   2   4   6E+2   9.5   10  RT (min)      lcqtof_t3_neg_3_032_24    m/z 564.4108     0   1E+3   9.5   10  RT (min)      lcqtof_t3_neg_3_032_24    m/z 564.4109     0   2   4E+2   9.5   10  RT (min)      lcqtof_t3_neg_3_032_24    m/z 564.4112     0   1E+4   9.5   10  RT (min)      lcqtof_t3_neg_3_032_24    m/z 564.5719     0   1   2E+4   10   10.5  RT (min)      lcqtof_t3_neg_4_062_13    m/z 564.5723     0   1   2E+4   10   10.5  RT (min)      lcqtof_t3_neg_4_062_13    m/z 565.4451     0   2   4   6E+3   11.5   12  RT (min)       lcqtof_t3_neg_4_016_14    m/z 566.3302     0   1   2   3   4E+4   9.5   10  RT (min)      lcqtof_t3_neg_4_040_24    m/z 566.3316     0   2   4   6E+2   9   9.5  RT (min)      lcqtof_t3_neg_3_032_24    m/z 566.3318     0   1E+4   9.5   10  RT (min)      lcqtof_t3_neg_3_053_25    m/z 566.3327     0   1   2E+4   9.5   10  RT (min)      lcqtof_t3_neg_1_031_24    m/z 567.0868     0   1   2E+4   13   13.5  RT (min)      lcqtof_t3_neg_4_064_17    m/z 567.0875     0   1   2E+4   13   13.5  RT (min)      lcqtof_t3_neg_4_064_17    m/z 567.0877     0   2   4   6E+4   12   12.5  RT (min)       lcqtof_t3_neg_4_064_17    m/z 567.0879     0   1   2E+4   12   12.5  RT (min)      lcqtof_t3_neg_4_002_17    m/z 567.0882     0   2   4   6E+4   12   12.5  RT (min)      lcqtof_t3_neg_4_064_17    m/z 567.0897     0   1   2E+4   13   13.5  RT (min)      lcqtof_t3_neg_4_036_47    m/z 567.3410     0   2   4E+2   9.5   10  RT (min)      lcqtof_t3_neg_3_032_24    m/z 567.3415     0   1E+4   9.5   10  RT (min)      lcqtof_t3_neg_3_032_24    m/z 567.3421     0   1E+4   10   10.5  RT (min)      lcqtof_t3_neg_2_016_05    m/z 567.3428     0   1   2E+3   9.5   10  RT (min)      lcqtof_t3_neg_3_032_24    m/z 567.4626     0   2   4   6   8E+3   11.5   12  RT (min)       lcqtof_t3_neg_4_059_14    m/z 568.0882     0   1   2   3E+4   12   12.5  RT (min)      lcqtof_t3_neg_4_064_17    m/z 568.0891     0   1E+4   12   12.5  RT (min)      lcqtof_t3_neg_4_002_17    m/z 568.0894     0   1E+4   12   12.5  RT (min)      lcqtof_t3_neg_4_002_17    m/z 568.0895     0   1E+4   13   13.5  RT (min)      lcqtof_t3_neg_4_064_17    m/z 568.0896     0   1E+4   13   13.5  RT (min)      lcqtof_t3_neg_4_036_47    m/z 568.0898     0   1E+4   12   12.5  RT (min)      lcqtof_t3_neg_4_002_17    m/z 568.0899     0   1   2   3E+4   12   12.5  RT (min)      lcqtof_t3_neg_4_064_17    m/z 568.3454     0   1E+4   10   10.5  RT (min)      lcqtof_t3_neg_4_040_24    m/z 568.3456     0   1E+4   10   10.5  RT (min)       lcqtof_t3_neg_4_040_24    m/z 568.3456     0   1E+4   10   10.5  RT (min)      lcqtof_t3_neg_4_040_24    m/z 568.3456     0   1E+4   10   10.5  RT (min)      lcqtof_t3_neg_4_040_24    m/z 568.3459     0   1E+4   10   10.5  RT (min)       lcqtof_t3_neg_4_040_24    m/z 568.3462     0   1E+4   10   10.5  RT (min)      lcqtof_t3_neg_4_040_24    m/z 568.6036     0   1   2E+4   11   11.5  RT (min)      lcqtof_t3_neg_4_037_01    m/z 568.6038     0   1E+4   11   11.5  RT (min)      lcqtof_t3_neg_4_059_14    m/z 568.6040     0   1   2E+4   11   11.5  RT (min)      lcqtof_t3_neg_4_037_01    m/z 568.8397     0   1   2E+3   15.5   16  RT (min)       lcqtof_t3_neg_4_048_19    m/z 568.8398     0   1   2   3E+3   16   16.5  RT (min)       lcqtof_t3_neg_4_000_ee    m/z 568.8400     0   1   2E+3   15.5   16  RT (min)       lcqtof_t3_neg_4_026_30    m/z 568.8400     0   1   2   3E+3   16.5   17  RT (min)       lcqtof_t3_neg_4_051_15    m/z 568.8400     0   1   2E+3   15   15.5  RT (min)       lcqtof_t3_neg_4_061_50    m/z 568.8405     0   1   2E+3   16   16.5  RT (min)      lcqtof_t3_neg_4_041_54    m/z 569.0864     0   1   2E+4   12   12.5  RT (min)      lcqtof_t3_neg_4_064_17    m/z 569.0865     0   1   2E+4   12   12.5  RT (min)      lcqtof_t3_neg_4_064_17    m/z 569.0873     0   1E+4   13   13.5  RT (min)      lcqtof_t3_neg_4_036_47    m/z 569.0875     0   1E+4   12   12.5  RT (min)      lcqtof_t3_neg_4_002_17    m/z 569.0878     0   1   2E+4   12   12.5  RT (min)      lcqtof_t3_neg_4_064_17    m/z 569.3528     0   1E+4   8.5   9  RT (min)      lcqtof_t3_neg_4_000_ee    m/z 569.3548     0   1E+4   9   9.5  RT (min)      lcqtof_t3_neg_4_000_ee    m/z 570.3258     0   1E+4   7.5   8  RT (min)      lcqtof_t3_neg_4_000_ee    m/z 570.8449     0   2   4   6E+3   16.5   17  RT (min)       lcqtof_t3_neg_1_011_45    m/z 571.1245     0   1E+4   4   4.5  RT (min)      lcqtof_t3_neg_4_000_ee    m/z 572.2973     0   1E+4   9   9.5  RT (min)      lcqtof_t3_neg_4_000_ee    m/z 572.3020     0   1E+4   9   9.5  RT (min)      lcqtof_t3_neg_4_000_ee    m/z 572.3358     0   1   2   3E+4   9   9.5  RT (min)      lcqtof_t3_neg_4_000_ee    m/z 572.3365     0   1   2   3E+4   9   9.5  RT (min)      lcqtof_t3_neg_4_000_ee    m/z 572.6139     0   1   2E+4   1   1.5  RT (min)      lcqtof_t3_neg_4_062_13    m/z 573.1820     0   2   4   6   8E+3   9   9.5  RT (min)       lcqtof_t3_neg_4_062_13    m/z 573.2246     0   1E+5   4.5   5  RT (min)       lcqtof_t3_neg_4_000_ee    m/z 573.2251     0   1   2   3   4E+4   4.5   5  RT (min)      lcqtof_t3_neg_4_000_ee    m/z 573.4510     0   1   2E+4   12.5   13  RT (min)      lcqtof_t3_neg_4_059_14    m/z 573.4514     0   1   2E+4   12.5   13  RT (min)      lcqtof_t3_neg_4_059_14    m/z 573.4524     0   1   2E+4   11.5   12  RT (min)      lcqtof_t3_neg_4_059_14    m/z 575.4665     0   1   2E+4   12   12.5  RT (min)      lcqtof_t3_neg_4_059_14    m/z 575.4672     0   1   2E+4   11.5   12  RT (min)      lcqtof_t3_neg_4_059_14    m/z 576.4708     0   1E+4   12.5   13  RT (min)      lcqtof_t3_neg_4_059_14    m/z 576.4711     0   1E+4   12.5   13  RT (min)      lcqtof_t3_neg_4_059_14    m/z 576.4719     0   1E+4   12.5   13  RT (min)      lcqtof_t3_neg_4_059_14    m/z 576.6522     0   1E+4   1   1.5  RT (min)      lcqtof_t3_neg_4_037_01    m/z 576.8615     0   1   2E+3   6.5   7  RT (min)       lcqtof_t3_neg_1_039_43    m/z 576.8617     0   1   2   3E+3   6.5   7  RT (min)       lcqtof_t3_neg_4_000_ff    m/z 576.8624     0   1   2E+3   7.5   8  RT (min)       lcqtof_t3_neg_4_112_62    m/z 576.8624     0   1   2   3E+3   7.5   8  RT (min)       lcqtof_t3_neg_4_000_ff    m/z 576.8632     0   1   2   3E+3   8.5   9  RT (min)       lcqtof_t3_neg_1_029_44    m/z 576.8638     0   1   2E+3   8   8.5  RT (min)       lcqtof_t3_neg_1_019_23    m/z 577.0862     0   1E+4   10.5   11  RT (min)      lcqtof_t3_neg_4_002_17    m/z 577.0875     0   1E+4   10.5   11  RT (min)       lcqtof_t3_neg_4_002_17    m/z 577.0885     0   1E+4   10.5   11  RT (min)      lcqtof_t3_neg_4_002_17    m/z 577.0900     0   1E+4   10.5   11  RT (min)      lcqtof_t3_neg_4_002_17    m/z 577.1007     0   1   2   3   4E+3   9.5   10  RT (min)       lcqtof_t3_neg_4_000_ee    m/z 577.3338     0   1E+4   8.5   9  RT (min)      lcqtof_t3_neg_4_000_ee    m/z 577.4833     0   1E+4   12   12.5  RT (min)      lcqtof_t3_neg_4_059_14    m/z 577.4841     0   1E+4   12   12.5  RT (min)      lcqtof_t3_neg_4_059_14    m/z 577.5224     0   1E+4   9.5   10  RT (min)      lcqtof_t3_neg_4_059_14    m/z 577.5852     0   1   2   3E+3   14   14.5  RT (min)      lcqtof_t3_neg_4_030_31    m/z 578.5966     0   1   2E+3   14.5   15  RT (min)       lcqtof_t3_neg_4_000_ee    m/z 578.8678     0   1   2   3   4E+3   15.5   16  RT (min)       lcqtof_t3_neg_4_108_62    m/z 578.8680     0   1   2E+3   15   15.5  RT (min)       lcqtof_t3_neg_3_032_24    m/z 578.8682     0   1E+4   16   16.5  RT (min)       lcqtof_t3_neg_4_018_23    m/z 578.8685     0   1E+4   16   16.5  RT (min)      lcqtof_t3_neg_1_011_45    m/z 578.8686     0   1   2   3   4E+3   15   15.5  RT (min)       lcqtof_t3_neg_4_106_62    m/z 579.0706     0   1E+4   13   13.5  RT (min)      lcqtof_t3_neg_4_064_17    m/z 579.4984     0   2   4   6E+3   12   12.5  RT (min)       lcqtof_t3_neg_4_059_14    m/z 580.5304     0   1   2   3E+4   12.5   13  RT (min)      lcqtof_t3_neg_1_019_23    m/z 580.5307     0   1   2   3E+4   13   13.5  RT (min)      lcqtof_t3_neg_1_019_23    m/z 580.5319     0   1   2E+4   13   13.5  RT (min)      lcqtof_t3_neg_4_018_23    m/z 580.7389     0   1E+4   1   1.5  RT (min)      lcqtof_t3_neg_4_041_54    m/z 580.8662     0   1   2   3   4E+3   16   16.5  RT (min)      lcqtof_t3_neg_1_011_45    m/z 581.0993     0   1   2E+4   12.5   13  RT (min)      lcqtof_t3_neg_4_064_17    m/z 581.1028     0   1   2E+4   12.5   13  RT (min)      lcqtof_t3_neg_4_064_17    m/z 581.5247     0   1   2   3   4E+2   12.5   13  RT (min)      lcqtof_t3_neg_3_032_24    m/z 581.5256     0   1E+2   11   11.5  RT (min)      lcqtof_t3_neg_3_038_06    m/z 581.5256     0   1E+2   11   11.5  RT (min)      lcqtof_t3_neg_3_038_06    m/z 581.5267     0   1   2E+3   14.5   15  RT (min)      lcqtof_t3_neg_1_000_cc    m/z 581.5268     0   1   2   3E+3   14   14.5  RT (min)      lcqtof_t3_neg_2_011_25    m/z 581.5269     0   1   2   3   4E+4   13   13.5  RT (min)      lcqtof_t3_neg_2_030_23    m/z 581.5271     0   1   2   3   4E+2   12.5   13  RT (min)      lcqtof_t3_neg_3_032_24    m/z 582.0997     0   1E+4   12.5   13  RT (min)      lcqtof_t3_neg_4_064_17    m/z 582.2727     0   1   2   3E+4   7   7.5  RT (min)      lcqtof_t3_neg_4_005_18    m/z 582.5094     0   2   4   6E+4   12.5   13  RT (min)      lcqtof_t3_neg_2_030_23    m/z 582.5102     0   1E+4   13   13.5  RT (min)      lcqtof_t3_neg_2_016_05    m/z 582.5105     0   1E+2   11   11.5  RT (min)      lcqtof_t3_neg_3_038_06    m/z 582.5302     0   1E+4   13   13.5  RT (min)      lcqtof_t3_neg_2_030_23    m/z 582.5323     0   1E+4   13   13.5  RT (min)      lcqtof_t3_neg_2_016_05    m/z 582.6438     0   1   2E+4   1   1.5  RT (min)      lcqtof_t3_neg_4_062_13    m/z 582.8790     0   1   2   3E+3   5.5   6  RT (min)      lcqtof_t3_neg_4_099_61    m/z 582.8793     0   2   4E+3   8   9  RT (min)       lcqtof_t3_neg_4_111_62    m/z 582.8794     0   2   4   6E+3   17   17.5  RT (min)       lcqtof_t3_neg_4_035_08    m/z 582.8794     0   2   4   6E+3   17   17.5  RT (min)       lcqtof_t3_neg_4_035_08    m/z 582.8798     0   2   4E+3   7.5   8  RT (min)       lcqtof_t3_neg_4_110_62    m/z 582.8801     0   2   4E+3   9   9.5   10  RT (min)       lcqtof_t3_neg_4_113_62    m/z 582.8804     0   1   2   3   4E+3   8   9  RT (min)       lcqtof_t3_neg_4_094_60    m/z 582.8808     0   2   4E+3   8   10  RT (min)       lcqtof_t3_neg_4_107_62    m/z 582.8812     0   1   2   3E+3   9   9.5  RT (min)       lcqtof_t3_neg_4_063_39    m/z 583.1012     0   1E+4   12.5   13  RT (min)      lcqtof_t3_neg_4_064_17    m/z 583.1176     0   1   2   3E+3   15   15.5  RT (min)       lcqtof_t3_neg_4_013_39    m/z 583.1182     0   1   2   3E+3   15.5   16  RT (min)       lcqtof_t3_neg_4_016_14    m/z 583.1186     0   1   2   3E+3   16   16.5  RT (min)       lcqtof_t3_neg_4_028_25    m/z 583.2743     0   1E+4   7   7.5  RT (min)      lcqtof_t3_neg_4_005_18    m/z 583.5131     0   1E+4   12.5   13  RT (min)      lcqtof_t3_neg_1_019_23    m/z 583.5142     0   1   2E+4   12.5   13  RT (min)      lcqtof_t3_neg_2_030_23    m/z 583.8723     0   1E+4   10   10.5  RT (min)       lcqtof_t3_neg_1_045_29    m/z 583.8728     0   1E+4   10   10.5  RT (min)      lcqtof_t3_neg_1_045_29    m/z 584.3426     0   1E+4   10   10.5  RT (min)       lcqtof_t3_neg_1_031_24    m/z 584.3433     0   1E+4   10   10.5  RT (min)       lcqtof_t3_neg_1_031_24    m/z 584.5243     0   1E+4   12.5   13  RT (min)      lcqtof_t3_neg_2_030_23    m/z 585.2241     0   1   2E+4   4.5   5  RT (min)      lcqtof_t3_neg_4_000_ee    m/z 585.2246     0   1   2E+4   4.5   5  RT (min)       lcqtof_t3_neg_4_000_ee    m/z 585.2352     0   1   2E+4   4.5   5  RT (min)      lcqtof_t3_neg_4_000_ee    m/z 585.3266     0   1E+4   11   11.5  RT (min)       lcqtof_t3_neg_4_034_34    m/z 585.3304     0   1   2E+4   11   11.5  RT (min)      lcqtof_t3_neg_1_051_26    m/z 585.3608     0   1   2   3E+3   13   13.5  RT (min)      lcqtof_t3_neg_2_000_aa    m/z 585.8680     0   1E+4   10   10.5  RT (min)      lcqtof_t3_neg_1_045_29    m/z 585.8696     0   1E+4   10   10.5  RT (min)      lcqtof_t3_neg_1_045_29    m/z 586.3158     0   2   4   6E+3   11   11.5  RT (min)      lcqtof_t3_neg_1_005_18    m/z 586.3282     0   1   2   3   4E+3   11.5   12  RT (min)       lcqtof_t3_neg_4_058_16    m/z 587.1200     0   1   2   3E+3   16   16.5  RT (min)      lcqtof_t3_neg_4_021_21    m/z 587.1207     0   1   2   3E+3   15   15.5  RT (min)       lcqtof_t3_neg_4_032_04    m/z 587.1218     0   1   2   3E+3   15   15.5  RT (min)      lcqtof_t3_neg_4_009_50    m/z 587.1218     0   1   2   3E+3   14.5   15  RT (min)       lcqtof_t3_neg_4_026_30    m/z 587.1226     0   1   2   3E+3   14.5   15  RT (min)       lcqtof_t3_neg_4_027_41    m/z 587.1261     0   1   2   3E+3   14   14.5  RT (min)       lcqtof_t3_neg_4_018_23    m/z 587.3991     0   1   2E+3   9   9.5  RT (min)      lcqtof_t3_neg_3_032_24    m/z 587.3992     0   1E+4   9   9.5  RT (min)      lcqtof_t3_neg_3_032_24    m/z 587.3993     0   1E+3   9   9.5  RT (min)      lcqtof_t3_neg_3_032_24    m/z 588.3248     0   1E+4   9   9.5  RT (min)      lcqtof_t3_neg_4_000_ee    m/z 588.3676     0   1   2E+4   8.5   9  RT (min)      lcqtof_t3_neg_4_000_ee    m/z 588.4013     0   1E+2   8.5   9  RT (min)      lcqtof_t3_neg_3_038_06    m/z 588.4014     0   1E+2   8   8.5  RT (min)      lcqtof_t3_neg_3_038_06    m/z 588.4020     0   1E+5   9   9.5  RT (min)      lcqtof_t3_neg_3_032_24    m/z 588.4021     0   2   4   6   8E+3   9   9.5  RT (min)      lcqtof_t3_neg_3_032_24    m/z 588.4032     0   1E+2   8   8.5  RT (min)      lcqtof_t3_neg_3_038_06    m/z 588.4733     0   1E+4   12.5   13  RT (min)      lcqtof_t3_neg_2_030_23    m/z 588.8962     0   2   4E+2   13.5   14  RT (min)      lcqtof_t3_neg_3_038_06    m/z 588.8966     0   1   2   3   4E+3   8   9  RT (min)       lcqtof_t3_neg_4_093_60    m/z 588.8968     0   2   4   6   8E+4   16   16.5  RT (min)      lcqtof_t3_neg_1_011_45    m/z 588.8968     0   1   2E+3   15   15.5  RT (min)       lcqtof_t3_neg_3_032_24    m/z 588.8969     0   1   2   3E+3   15   15.5  RT (min)       lcqtof_t3_neg_4_083_59    m/z 588.8970     0   1   2E+5   16   16.5  RT (min)       lcqtof_t3_neg_4_018_23    m/z 588.8971     0   1   2   3   4E+3   9   9.5   10  RT (min)       lcqtof_t3_neg_4_087_59    m/z 588.8972     0   2   4E+3   15.5   16  RT (min)       lcqtof_t3_neg_4_099_61    m/z 588.8972     0   1   2   3   4E+3   15.5   16  RT (min)      lcqtof_t3_neg_1_019_23    m/z 588.8975     0   1E+5   17   17.5  RT (min)       lcqtof_t3_neg_4_035_08    m/z 588.8975     0   1   2E+5   16   16.5  RT (min)      lcqtof_t3_neg_4_018_23    m/z 588.8978     0   2   4E+3   7   8  RT (min)       lcqtof_t3_neg_4_103_61    m/z 588.9067     0   1   2   3E+5   17   17.5  RT (min)       lcqtof_t3_neg_4_106_62    m/z 588.9074     0   2   4   6E+4   17   17.5  RT (min)      lcqtof_t3_neg_4_113_62    m/z 589.2988     0   1   2E+4   9   9.5  RT (min)      lcqtof_t3_neg_4_000_ee    m/z 589.4043     0   2   4   6E+3   9.5   10  RT (min)      lcqtof_t3_neg_3_032_24    m/z 589.4068     0   1   2E+4   9.5   10  RT (min)      lcqtof_t3_neg_3_032_24    m/z 589.4134     0   1   2   3E+4   10   10.5  RT (min)      lcqtof_t3_neg_2_016_05    m/z 589.4138     0   1   2   3E+3   9.5   10  RT (min)      lcqtof_t3_neg_3_032_24    m/z 589.4454     0   2   4   6   8E+3   11.5   12  RT (min)      lcqtof_t3_neg_4_059_14    m/z 589.9006     0   1E+4   17   17.5  RT (min)      lcqtof_t3_neg_4_035_08    m/z 590.4160     0   2   4   6   8E+1   8.5   9  RT (min)      lcqtof_t3_neg_3_038_06    m/z 590.4167     0   1E+4   9.5   10  RT (min)      lcqtof_t3_neg_3_032_24    m/z 590.4168     0   1E+2   8.5   9  RT (min)      lcqtof_t3_neg_3_038_06    m/z 590.4176     0   1E+2   8.5   9  RT (min)      lcqtof_t3_neg_3_038_06    m/z 590.4176     0   1   2E+3   9.5   10  RT (min)      lcqtof_t3_neg_3_032_24    m/z 590.4178     0   1E+5   9.5   10  RT (min)      lcqtof_t3_neg_3_032_24    m/z 591.2395     0   1E+4   4   4.5  RT (min)      lcqtof_t3_neg_4_000_ee    m/z 591.3003     0   1E+4   9   9.5  RT (min)      lcqtof_t3_neg_4_000_ee    m/z 591.4293     0   1E+4   10.5   11  RT (min)      lcqtof_t3_neg_4_019_05    m/z 591.4296     0   1E+4   10.5   11  RT (min)      lcqtof_t3_neg_4_019_05    m/z 591.4297     0   1E+4   10.5   11  RT (min)      lcqtof_t3_neg_2_028_24    m/z 591.4632     0   1   2E+4   12.5   13  RT (min)      lcqtof_t3_neg_4_059_14    m/z 592.2051     0   1E+4   5.5   6  RT (min)      lcqtof_t3_neg_4_047_07    m/z 592.4219     0   1   2   3E+4   10   10.5  RT (min)      lcqtof_t3_neg_3_032_24    m/z 592.4324     0   1   2E+3   9.5   10  RT (min)      lcqtof_t3_neg_3_032_24    m/z 592.4329     0   1   2   3   4E+3   10   10.5  RT (min)      lcqtof_t3_neg_3_032_24    m/z 592.4656     0   1E+4   11.5   12  RT (min)      lcqtof_t3_neg_4_059_14    m/z 592.4666     0   1E+4   11.5   12  RT (min)      lcqtof_t3_neg_4_059_14    m/z 592.5622     0   1E+4   1   1.5  RT (min)      lcqtof_t3_neg_4_062_13    m/z 592.6112     0   1   2E+3   14.5   15  RT (min)      lcqtof_t3_neg_4_000_ee    m/z 592.6711     0   1   2E+4   1   1.5  RT (min)      lcqtof_t3_neg_4_037_01    m/z 592.6817     0   1   2E+4   1   1.5  RT (min)      lcqtof_t3_neg_4_062_13    m/z 593.1220     0   1E+5   13.5   14  RT (min)      lcqtof_t3_neg_4_002_17    m/z 593.1222     0   1   2E+4   13.5   14  RT (min)      lcqtof_t3_neg_4_036_47    m/z 593.1243     0   1E+4   13.5   14  RT (min)      lcqtof_t3_neg_4_002_17    m/z 593.2606     0   1E+4   4   4.5  RT (min)      lcqtof_t3_neg_4_000_ee    m/z 593.4752     0   2   4   6E+3   12.5   13  RT (min)      lcqtof_t3_neg_2_048_30    m/z 593.4787     0   1   2E+4   12.5   13  RT (min)      lcqtof_t3_neg_4_059_14    m/z 594.1226     0   1E+4   13.5   14  RT (min)      lcqtof_t3_neg_4_002_17    m/z 594.1234     0   1E+4   13.5   14  RT (min)      lcqtof_t3_neg_4_036_47    m/z 594.2822     0   1   2   3   4E+3   10.5   11  RT (min)       lcqtof_t3_neg_4_037_01    m/z 594.8426     0   1E+4   17   17.5  RT (min)      lcqtof_t3_neg_4_035_08    m/z 594.8510     0   1E+4   16   16.5  RT (min)      lcqtof_t3_neg_1_011_45    m/z 595.1002     0   1   2E+4   12   12.5  RT (min)      lcqtof_t3_neg_4_002_17    m/z 595.1010     0   2   4E+4   12.5   13  RT (min)      lcqtof_t3_neg_4_002_17    m/z 595.1017     0   1   2   3   4E+4   12   12.5  RT (min)      lcqtof_t3_neg_4_064_17    m/z 595.1065     0   2   4   6   8E+4   12.5   13  RT (min)      lcqtof_t3_neg_4_002_17    m/z 595.1102     0   1E+4   13.5   14  RT (min)      lcqtof_t3_neg_4_064_17    m/z 595.1110     0   1E+4   13.5   14  RT (min)      lcqtof_t3_neg_4_064_17    m/z 595.2630     0   2   4   6E+3   11.5   12  RT (min)       lcqtof_t3_neg_1_002_13    m/z 595.3683     0   1E+4   10   10.5  RT (min)      lcqtof_t3_neg_4_040_24    m/z 595.3684     0   1E+4   10   10.5  RT (min)      lcqtof_t3_neg_4_019_05    m/z 595.4296     0   2   4   6   8E+3   12   12.5  RT (min)      lcqtof_t3_neg_4_059_14    m/z 595.4868     0   2   4   6E+3   13   13.5  RT (min)      lcqtof_t3_neg_2_054_29    m/z 595.8577     0   2   4E+3   16.5   17  RT (min)       lcqtof_t3_neg_1_000_bb    m/z 595.8582     0   1   2E+3   16   16.5  RT (min)       lcqtof_t3_neg_4_040_24    m/z 595.8588     0   1   2E+3   16   16.5  RT (min)       lcqtof_t3_neg_4_000_ff    m/z 596.1012     0   1   2E+4   12   12.5  RT (min)      lcqtof_t3_neg_4_064_17    m/z 596.1020     0   1E+4   12   12.5  RT (min)      lcqtof_t3_neg_4_002_17    m/z 596.1032     0   1E+4   12.5   13  RT (min)      lcqtof_t3_neg_4_002_17    m/z 596.2908     0   2   4   6E+3   9.5   10  RT (min)      lcqtof_t3_neg_4_014_13    m/z 596.2909     0   2   4   6   8E+3   9.5   10  RT (min)      lcqtof_t3_neg_4_062_13    m/z 596.3730     0   1   2E+4   9.5   10  RT (min)      lcqtof_t3_neg_3_032_24    m/z 596.3734     0   2   4   6   8E+2   9   9.5  RT (min)      lcqtof_t3_neg_3_032_24    m/z 596.3936     0   1   2   3   4E+3   11.5   12  RT (min)      lcqtof_t3_neg_4_037_01    m/z 596.5256     0   2   4   6   8E+3   13   13.5  RT (min)      lcqtof_t3_neg_2_030_23    m/z 597.0977     0   1   2E+4   12   12.5  RT (min)      lcqtof_t3_neg_4_064_17    m/z 597.1336     0   1   2   3E+3   15   15.5  RT (min)       lcqtof_t3_neg_4_030_31    m/z 597.1338     0   1   2   3E+3   15.5   16   16.5  RT (min)       lcqtof_t3_neg_4_031_42    m/z 597.1338     0   1   2   3E+3   15.5   16   16.5  RT (min)       lcqtof_t3_neg_4_013_39    m/z 597.1343     0   1   2   3E+3   14.5   15  RT (min)       lcqtof_t3_neg_4_031_42    m/z 597.1344     0   1   2   3E+3   15.5   16  RT (min)       lcqtof_t3_neg_4_034_34    m/z 597.3495     0   1E+4   9   9.5  RT (min)      lcqtof_t3_neg_4_000_ee    m/z 597.3496     0   1E+4   9   9.5  RT (min)      lcqtof_t3_neg_4_000_ee    m/z 597.4011     0   2   4   6E+3   10.5   11  RT (min)       lcqtof_t3_neg_4_030_31    m/z 597.4016     0   2   4   6E+3   10   10.5  RT (min)      lcqtof_t3_neg_4_086_59    m/z 597.4486     0   1E+4   12.5   13  RT (min)      lcqtof_t3_neg_4_059_14    m/z 597.4495     0   1E+4   12   12.5  RT (min)       lcqtof_t3_neg_4_059_14    m/z 598.1010     0   1   2E+4   13   13.5  RT (min)      lcqtof_t3_neg_4_036_47    m/z 598.1023     0   1E+4   13   13.5  RT (min)      lcqtof_t3_neg_4_002_17    m/z 598.5035     0   1   2E+4   12.5   13  RT (min)      lcqtof_t3_neg_2_030_23    m/z 598.5212     0   1E+4   13   13.5  RT (min)      lcqtof_t3_neg_2_030_23    m/z 599.1092     0   1   2   3E+4   4   4.5  RT (min)      lcqtof_t3_neg_4_000_ee    m/z 599.2975     0   2   4   6E+3   10.5   11  RT (min)      lcqtof_t3_neg_4_015_36    m/z 599.3256     0   2   4   6E+4   9   9.5  RT (min)       lcqtof_t3_neg_4_000_ee    m/z 599.5270     0   1   2   3E+2   12.5   13  RT (min)      lcqtof_t3_neg_3_032_24    m/z 599.5270     0   1   2   3   4E+3   14   14.5  RT (min)      lcqtof_t3_neg_1_000_dd    m/z 599.5271     0   1E+4   14   14.5  RT (min)      lcqtof_t3_neg_1_000_dd    m/z 599.5276     0   1   2E+3   14.5   15  RT (min)      lcqtof_t3_neg_1_004_11    m/z 599.5277     0   1   2E+3   14.5   15  RT (min)      lcqtof_t3_neg_2_034_19    m/z 599.5278     0   1   2E+3   14   14.5  RT (min)      lcqtof_t3_neg_1_021_39    m/z 599.5288     0   1   2E+2   11.5   12  RT (min)      lcqtof_t3_neg_3_038_06    m/z 599.5291     0   1   2E+2   11.5   12  RT (min)      lcqtof_t3_neg_3_038_06    m/z 599.5294     0   1   2   3E+2   12.5   13  RT (min)      lcqtof_t3_neg_3_032_24    m/z 600.2756     0   1   2E+4   8   8.5  RT (min)      lcqtof_t3_neg_4_005_18    m/z 600.2878     0   2   4E+3   10   10.5  RT (min)      lcqtof_t3_neg_4_034_34    m/z 600.3306     0   1   2E+4   9   9.5  RT (min)      lcqtof_t3_neg_4_000_ee    m/z 600.3910     0   2   4   6E+1   4.5   5  RT (min)      lcqtof_t3_neg_3_038_06    m/z 600.5151     0   1E+4   13   13.5  RT (min)      lcqtof_t3_neg_2_030_23    m/z 600.7873     0   1E+4   1   1.5  RT (min)      lcqtof_t3_neg_4_051_15    m/z 601.5211     0   2   4   6E+3   13   13.5  RT (min)      lcqtof_t3_neg_1_019_23    m/z 601.5274     0   1E+4   13   13.5  RT (min)      lcqtof_t3_neg_1_019_23    m/z 602.2838     0   2   4   6E+3   11.5   12  RT (min)      lcqtof_t3_neg_4_058_16    m/z 602.7022     0   1E+4   1   1.5  RT (min)      lcqtof_t3_neg_4_037_01    m/z 602.7443     0   1E+4   1   1.5  RT (min)      lcqtof_t3_neg_2_028_24    m/z 603.2890     0   2   4E+3   11.5   12  RT (min)       lcqtof_t3_neg_4_059_14    m/z 603.3624     0   1E+4   8   8.5  RT (min)      lcqtof_t3_neg_4_000_ee    m/z 603.5391     0   1E+4   13.5   14  RT (min)      lcqtof_t3_neg_4_030_31    m/z 604.2476     0   1E+4   4.5   5  RT (min)      lcqtof_t3_neg_4_000_ee    m/z 604.2492     0   1E+4   4.5   5  RT (min)      lcqtof_t3_neg_4_000_ee    m/z 604.8684     0   1   2   3   4E+2   13.5   14  RT (min)      lcqtof_t3_neg_3_038_06    m/z 604.8694     0   2   4   6   8E+2   15   15.5  RT (min)      lcqtof_t3_neg_3_032_24    m/z 604.8707     0   1E+3   15.5   16  RT (min)      lcqtof_t3_neg_1_019_23    m/z 604.8711     0   1   2E+4   17   17.5  RT (min)      lcqtof_t3_neg_4_035_08    m/z 604.8715     0   1   2E+4   16   16.5  RT (min)      lcqtof_t3_neg_4_018_23    m/z 605.0469     0   1   2E+3   9.5   10  RT (min)      lcqtof_t3_neg_1_010_27    m/z 605.3294     0   2   4   6E+4   8.5   9  RT (min)      lcqtof_t3_neg_4_000_ee    m/z 605.3922     0   1E+4   9.5   10  RT (min)       lcqtof_t3_neg_1_031_24    m/z 605.4771     0   1E+4   12   12.5  RT (min)      lcqtof_t3_neg_4_037_01    m/z 605.4778     0   1E+4   12   12.5  RT (min)      lcqtof_t3_neg_4_037_01    m/z 605.4782     0   1E+4   12   12.5  RT (min)      lcqtof_t3_neg_4_037_01    m/z 605.4790     0   1E+4   12   12.5  RT (min)      lcqtof_t3_neg_4_037_01    m/z 605.4795     0   1E+4   12   12.5  RT (min)      lcqtof_t3_neg_4_051_15    m/z 605.6225     0   1   2   3E+3   15   15.5  RT (min)      lcqtof_t3_neg_4_063_39    m/z 605.6239     0   1   2   3   4E+3   14.5   15  RT (min)       lcqtof_t3_neg_4_022_28    m/z 605.8864     0   2   4   6   8E+2   15   15.5  RT (min)       lcqtof_t3_neg_3_032_24    m/z 605.8872     0   1E+4   16   16.5  RT (min)       lcqtof_t3_neg_1_011_45    m/z 605.8877     0   1   2E+3   15.5   16  RT (min)       lcqtof_t3_neg_4_098_61    m/z 605.8878     0   1E+3   15.5   16  RT (min)       lcqtof_t3_neg_1_019_23    m/z 606.3382     0   1E+4   8.5   9  RT (min)      lcqtof_t3_neg_4_000_ee    m/z 606.5280     0   2   4   6   8E+3   13   13.5  RT (min)      lcqtof_t3_neg_2_030_23    m/z 607.3296     0   1   2E+4   8.5   9  RT (min)      lcqtof_t3_neg_4_000_ee    m/z 607.3296     0   1E+4   13   13.5  RT (min)      lcqtof_t3_neg_1_000_aa    m/z 607.4081     0   1E+4   10   10.5  RT (min)       lcqtof_t3_neg_1_031_24    m/z 607.4085     0   1E+4   10   10.5  RT (min)      lcqtof_t3_neg_1_031_24    m/z 607.4938     0   1   2   3E+4   12   12.5  RT (min)      lcqtof_t3_neg_4_002_17    m/z 608.1586     0   1E+4   8.5   9  RT (min)      lcqtof_t3_neg_4_102_61    m/z 608.3168     0   2   4E+3   11   11.5  RT (min)       lcqtof_t3_neg_1_005_18    m/z 608.9876     0   1E+4   1   1.5  RT (min)      lcqtof_t3_neg_2_028_24    m/z 609.1162     0   1E+4   12   12.5  RT (min)      lcqtof_t3_neg_4_057_49    m/z 609.2542     0   1   2   3   4E+3   10   10.5  RT (min)      lcqtof_t3_neg_1_002_13    m/z 609.3461     0   2   4E+4   9   9.5  RT (min)      lcqtof_t3_neg_4_000_ee    m/z 609.5564     0   2   4E+2   12.5   13  RT (min)      lcqtof_t3_neg_3_032_24    m/z 609.5570     0   2   4   6E+4   13   13.5  RT (min)      lcqtof_t3_neg_2_030_23    m/z 609.5572     0   1   2E+2   11.5   12  RT (min)      lcqtof_t3_neg_3_038_06    m/z 610.2369     0   1E+4   9.5   10  RT (min)      lcqtof_t3_neg_4_018_23    m/z 610.2387     0   1E+4   9.5   10  RT (min)      lcqtof_t3_neg_4_018_23    m/z 610.3553     0   1   2E+4   9   9.5  RT (min)      lcqtof_t3_neg_4_000_ee    m/z 610.4184     0   2   4   6   8E+1   4.5   5  RT (min)      lcqtof_t3_neg_3_038_06    m/z 610.5404     0   1E+2   11   11.5  RT (min)      lcqtof_t3_neg_3_038_06    m/z 610.5405     0   2   4E+4   13   13.5  RT (min)      lcqtof_t3_neg_2_030_23    m/z 610.5592     0   1E+2   11.5   12  RT (min)      lcqtof_t3_neg_3_038_06    m/z 610.5596     0   1E+4   13   13.5  RT (min)      lcqtof_t3_neg_1_019_23    m/z 610.5597     0   1   2E+4   13   13.5  RT (min)      lcqtof_t3_neg_2_030_23    m/z 610.5606     0   1   2E+2   12.5   13  RT (min)      lcqtof_t3_neg_3_032_24    m/z 610.5610     0   1   2E+2   12.5   13  RT (min)      lcqtof_t3_neg_3_032_24    m/z 610.5612     0   1   2E+4   14   14.5  RT (min)      lcqtof_t3_neg_1_017_51    m/z 610.5617     0   1E+2   11.5   12  RT (min)      lcqtof_t3_neg_3_038_06    m/z 610.6131     0   1E+4   1   1.5  RT (min)      lcqtof_t3_neg_4_046_33    m/z 611.1440     0   1   2   3E+4   2.5   3  RT (min)      lcqtof_t3_neg_4_018_23    m/z 611.1442     0   1   2   3   4E+4   3   3.5  RT (min)       lcqtof_t3_neg_4_088_59    m/z 611.1491     0   1E+4   10   10.5  RT (min)       lcqtof_t3_neg_4_000_ee    m/z 611.1495     0   1E+4   10   10.5  RT (min)       lcqtof_t3_neg_4_000_ee    m/z 611.1496     0   1   2E+3   16   16.5  RT (min)       lcqtof_t3_neg_4_042_10    m/z 611.1499     0   1   2E+3   16   16.5  RT (min)       lcqtof_t3_neg_4_031_42    m/z 611.1504     0   1   2E+3   16   16.5  RT (min)       lcqtof_t3_neg_4_034_34    m/z 611.1505     0   1   2E+3   15   15.5  RT (min)       lcqtof_t3_neg_4_042_10    m/z 611.1508     0   1   2E+3   15.5   16  RT (min)       lcqtof_t3_neg_4_000_ff    m/z 611.2515     0   2   4   6   8E+4   4.5   5  RT (min)      lcqtof_t3_neg_4_000_ee    m/z 611.3927     0   1E+4   7.5   8  RT (min)      lcqtof_t3_neg_4_000_ee    m/z 611.5444     0   1   2E+4   13   13.5  RT (min)      lcqtof_t3_neg_2_030_23    m/z 612.2539     0   1   2E+4   4.5   5  RT (min)      lcqtof_t3_neg_4_000_ee    m/z 612.5198     0   1   2E+4   12.5   13  RT (min)      lcqtof_t3_neg_2_030_23    m/z 612.5202     0   1   2E+4   12.5   13  RT (min)      lcqtof_t3_neg_1_019_23    m/z 612.5549     0   1   2E+4   13   13.5  RT (min)      lcqtof_t3_neg_2_030_23    m/z 612.5557     0   1   2   3E+4   13   13.5  RT (min)      lcqtof_t3_neg_4_018_23    m/z 612.7602     0   1E+4   1   1.5  RT (min)      lcqtof_t3_neg_4_046_33    m/z 613.3733     0   1E+4   9   9.5  RT (min)      lcqtof_t3_neg_4_000_ee    m/z 613.3772     0   1E+4   8.5   9  RT (min)      lcqtof_t3_neg_4_000_ee    m/z 613.3786     0   1E+4   8.5   9  RT (min)      lcqtof_t3_neg_4_000_ee    m/z 613.4430     0   1E+4   11.5   12  RT (min)      lcqtof_t3_neg_4_059_14    m/z 613.4432     0   1E+4   11.5   12  RT (min)      lcqtof_t3_neg_4_059_14    m/z 613.4436     0   1E+4   11.5   12  RT (min)      lcqtof_t3_neg_4_059_14    m/z 613.5234     0   1E+4   12.5   13  RT (min)      lcqtof_t3_neg_2_030_23    m/z 613.5240     0   1E+4   13   13.5  RT (min)      lcqtof_t3_neg_4_018_23    m/z 613.5598     0   2   4   6   8E+3   13   13.5  RT (min)      lcqtof_t3_neg_2_030_23    m/z 614.3436     0   1   2   3E+4   10   10.5  RT (min)      lcqtof_t3_neg_4_014_13    m/z 614.4956     0   2   4   6   8E+3   12.5   13  RT (min)      lcqtof_t3_neg_1_019_23    m/z 614.4959     0   1E+4   12.5   13  RT (min)      lcqtof_t3_neg_2_030_23    m/z 615.2392     0   1E+4   4   4.5  RT (min)      lcqtof_t3_neg_4_000_ee    m/z 615.4300     0   1E+4   11   11.5  RT (min)      lcqtof_t3_neg_4_024_26    m/z 615.4314     0   1E+4   11   11.5  RT (min)       lcqtof_t3_neg_4_034_34    m/z 615.4322     0   1E+4   11   11.5  RT (min)      lcqtof_t3_neg_4_024_26    m/z 615.4940     0   1E+4   12.5   13  RT (min)      lcqtof_t3_neg_2_030_23    m/z 616.2948     0   1   2   3E+3   10   10.5  RT (min)      lcqtof_t3_neg_4_024_26    m/z 616.3220     0   1E+4   9   9.5  RT (min)      lcqtof_t3_neg_4_000_ee    m/z 616.4991     0   2   4   6E+4   13   13.5  RT (min)      lcqtof_t3_neg_1_019_23    m/z 616.5061     0   1E+2   11   11.5  RT (min)      lcqtof_t3_neg_3_038_06    m/z 616.5063     0   2   4   6E+4   13   13.5  RT (min)      lcqtof_t3_neg_1_019_23    m/z 616.5077     0   1E+2   11   11.5  RT (min)      lcqtof_t3_neg_3_038_06    m/z 616.6228     0   1   2E+2   12   12.5  RT (min)      lcqtof_t3_neg_3_032_24    m/z 616.6308     0   1   2E+2   12.5   13  RT (min)      lcqtof_t3_neg_3_032_24    m/z 616.8396     0   1   2E+3   16.5   17  RT (min)      lcqtof_t3_neg_1_013_09    m/z 617.3664     0   1E+4   8.5   9  RT (min)      lcqtof_t3_neg_4_000_ee    m/z 617.4460     0   2   4   6   8E+3   13.5   14  RT (min)      lcqtof_t3_neg_2_000_aa    m/z 617.5098     0   1E+4   13   13.5  RT (min)      lcqtof_t3_neg_4_018_23    m/z 617.5100     0   1   2E+4   13   13.5  RT (min)      lcqtof_t3_neg_1_019_23    m/z 617.5101     0   1   2E+4   13   13.5  RT (min)      lcqtof_t3_neg_1_019_23    m/z 618.5063     0   1   2E+4   13   13.5  RT (min)      lcqtof_t3_neg_1_019_23    m/z 618.5086     0   1   2E+4   13   13.5  RT (min)      lcqtof_t3_neg_2_030_23    m/z 618.6748     0   1E+4   1   1.5  RT (min)      lcqtof_t3_neg_4_051_15    m/z 619.4184     0   2   4   6E+3   11.5   12  RT (min)       lcqtof_t3_neg_4_037_01    m/z 619.4810     0   2   4   6   8E+3   11.5   12  RT (min)      lcqtof_t3_neg_4_059_14    m/z 619.6386     0   1   2   3   4E+3   15   15.5  RT (min)       lcqtof_t3_neg_4_106_62    m/z 620.3447     0   2   4   6   8E+3   11   11.5  RT (min)      lcqtof_t3_neg_4_037_01    m/z 621.2680     0   1E+4   4.5   5  RT (min)      lcqtof_t3_neg_4_000_ee    m/z 621.3610     0   1E+4   8.5   9  RT (min)      lcqtof_t3_neg_4_000_ee    m/z 621.3612     0   1E+4   8.5   9  RT (min)      lcqtof_t3_neg_4_000_ee    m/z 621.3626     0   1E+4   8.5   9  RT (min)      lcqtof_t3_neg_4_000_ee    m/z 621.4926     0   2   4   6E+3   12   12.5  RT (min)      lcqtof_t3_neg_4_059_14    m/z 622.2892     0   1E+4   10   10.5  RT (min)      lcqtof_t3_neg_4_062_13    m/z 622.3791     0   1   2   3   4E+4   9   9.5  RT (min)      lcqtof_t3_neg_3_032_24    m/z 622.3792     0   1   2E+3   9   9.5  RT (min)      lcqtof_t3_neg_3_032_24    m/z 622.3796     0   2   4   6E+3   9   9.5  RT (min)      lcqtof_t3_neg_3_032_24    m/z 622.8559     0   1   2   3E+3   16.5   17  RT (min)      lcqtof_t3_neg_1_042_46    m/z 622.8564     0   1   2   3E+3   16.5   17  RT (min)      lcqtof_t3_neg_1_046_17    m/z 623.1676     0   1   2E+4   12.5   13  RT (min)      lcqtof_t3_neg_4_064_17    m/z 623.1689     0   1E+4   12.5   13  RT (min)      lcqtof_t3_neg_4_036_47    m/z 623.2521     0   1E+4   4   4.5  RT (min)      lcqtof_t3_neg_4_000_ee    m/z 623.3914     0   1E+4   10   10.5  RT (min)      lcqtof_t3_neg_4_019_05    m/z 624.1689     0   1E+4   12.5   13  RT (min)      lcqtof_t3_neg_4_064_17    m/z 624.3934     0   1E+4   9.5   10  RT (min)      lcqtof_t3_neg_3_032_24    m/z 624.3937     0   1E+3   9   9.5  RT (min)      lcqtof_t3_neg_3_032_24    m/z 624.3942     0   1   2E+4   9.5   10  RT (min)      lcqtof_t3_neg_3_032_24    m/z 624.3946     0   1   2   3   4E+3   9.5   10  RT (min)      lcqtof_t3_neg_3_032_24    m/z 624.5200     0   1   2   3E+4   12.5   13  RT (min)      lcqtof_t3_neg_2_030_23    m/z 624.5205     0   1   2E+4   12.5   13  RT (min)      lcqtof_t3_neg_1_019_23    m/z 625.1648     0   1E+4   12.5   13  RT (min)      lcqtof_t3_neg_4_064_17    m/z 625.5222     0   1E+4   12.5   13  RT (min)      lcqtof_t3_neg_2_030_23    m/z 626.5354     0   1   2E+2   11   11.5  RT (min)      lcqtof_t3_neg_3_038_06    m/z 626.5356     0   1   2E+5   13   13.5  RT (min)      lcqtof_t3_neg_2_030_23    m/z 626.5359     0   1   2E+2   11   11.5  RT (min)      lcqtof_t3_neg_3_038_06    m/z 626.5461     0   1   2   3   4E+2   12.5   13  RT (min)      lcqtof_t3_neg_3_032_24    m/z 626.5467     0   1   2   3   4E+2   12.5   13  RT (min)      lcqtof_t3_neg_3_032_24    m/z 626.5471     0   1   2   3   4E+3   16.5   17  RT (min)      lcqtof_t3_neg_2_023_37    m/z 626.5474     0   1   2   3   4E+3   16.5   17  RT (min)      lcqtof_t3_neg_1_018_33    m/z 626.5474     0   1   2E+3   16   16.5  RT (min)      lcqtof_t3_neg_2_056_56    m/z 626.5475     0   1E+2   11.5   12  RT (min)      lcqtof_t3_neg_3_038_06    m/z 626.5476     0   1   2E+3   14.5   15  RT (min)      lcqtof_t3_neg_2_034_19    m/z 626.5481     0   1   2   3E+4   14   14.5  RT (min)      lcqtof_t3_neg_1_000_aa    m/z 626.5484     0   1E+2   11.5   12  RT (min)      lcqtof_t3_neg_3_038_06    m/z 627.2520     0   1   2E+4   4   4.5  RT (min)      lcqtof_t3_neg_4_000_ee    m/z 627.4084     0   1E+2   4.5   5  RT (min)      lcqtof_t3_neg_3_038_06    m/z 627.5293     0   2   4   6   8E+4   13   13.5  RT (min)      lcqtof_t3_neg_2_030_23    m/z 627.5387     0   1E+2   11   11.5  RT (min)      lcqtof_t3_neg_3_038_06    m/z 627.5394     0   1E+2   11   11.5  RT (min)      lcqtof_t3_neg_3_038_06    m/z 627.5395     0   2   4   6   8E+4   13   13.5  RT (min)      lcqtof_t3_neg_2_030_23    m/z 627.5566     0   1E+4   14   14.5  RT (min)      lcqtof_t3_neg_1_022_28    m/z 627.5567     0   1E+4   14   14.5  RT (min)      lcqtof_t3_neg_1_000_dd    m/z 627.5569     0   1   2   3E+3   14.5   15  RT (min)      lcqtof_t3_neg_2_042_13    m/z 627.5570     0   1E+4   13.5   14  RT (min)      lcqtof_t3_neg_2_030_23    m/z 627.5574     0   1   2   3E+2   13   13.5  RT (min)      lcqtof_t3_neg_3_032_24    m/z 627.5596     0   1   2E+2   11.5   12  RT (min)      lcqtof_t3_neg_3_038_06    m/z 628.4110     0   1E+2   4.5   5  RT (min)      lcqtof_t3_neg_3_038_06    m/z 628.5298     0   1E+4   13   13.5  RT (min)      lcqtof_t3_neg_2_016_05    m/z 628.5414     0   1   2E+4   13   13.5  RT (min)      lcqtof_t3_neg_2_030_23    m/z 629.4019     0   1E+4   7.5   8  RT (min)      lcqtof_t3_neg_4_000_ee    m/z 629.5099     0   2   4   6   8E+3   12.5   13  RT (min)      lcqtof_t3_neg_1_019_23    m/z 629.5544     0   1E+4   13   13.5  RT (min)      lcqtof_t3_neg_4_018_23    m/z 630.2454     0   1   2E+4   4   4.5  RT (min)      lcqtof_t3_neg_4_000_ee    m/z 630.7944     0   1E+4   1   1.5  RT (min)      lcqtof_t3_neg_4_037_01    m/z 631.2106     0   1E+4   2.5   3  RT (min)      lcqtof_t3_neg_4_095_60    m/z 631.2108     0   1   2   3   4E+4   3   3.5  RT (min)       lcqtof_t3_neg_1_031_24    m/z 631.2110     0   1   2   3E+4   3   3.5  RT (min)       lcqtof_t3_neg_4_041_54    m/z 631.2110     0   1   2E+4   3   3.5  RT (min)       lcqtof_t3_neg_3_032_24    m/z 631.2110     0   1   2   3E+4   3   3.5  RT (min)       lcqtof_t3_neg_4_028_25    m/z 631.2112     0   1E+4   2.5   3  RT (min)       lcqtof_t3_neg_4_001_49    m/z 631.2113     0   1E+5   3   3.5  RT (min)       lcqtof_t3_neg_4_096_60    m/z 631.2115     0   1E+4   3   3.5  RT (min)       lcqtof_t3_neg_3_038_06    m/z 631.3866     0   1E+4   10   10.5  RT (min)      lcqtof_t3_neg_4_000_ee    m/z 631.6139     0   1   2   3   4E+3   13   13.5  RT (min)      lcqtof_t3_neg_2_028_24    m/z 632.3518     0   1   2   3E+4   8.5   9  RT (min)      lcqtof_t3_neg_4_000_ee    m/z 632.3694     0   1   2   3   4E+4   8.5   9  RT (min)      lcqtof_t3_neg_4_000_ee    m/z 632.3946     0   1   2   3   4E+4   8.5   9  RT (min)      lcqtof_t3_neg_4_000_ee    m/z 633.3276     0   1   2E+4   9   9.5  RT (min)      lcqtof_t3_neg_4_000_ee    m/z 633.3539     0   1E+4   8.5   9  RT (min)      lcqtof_t3_neg_4_000_ee    m/z 633.3868     0   1E+4   8.5   9  RT (min)      lcqtof_t3_neg_4_000_ee    m/z 633.3987     0   1E+4   8.5   9  RT (min)      lcqtof_t3_neg_4_000_ee    m/z 633.6544     0   1   2   3   4E+3   15   15.5  RT (min)      lcqtof_t3_neg_4_000_bb    m/z 633.6548     0   1   2   3   4E+3   15   15.5  RT (min)      lcqtof_t3_neg_4_058_16    m/z 633.6550     0   2   4E+3   15   15.5  RT (min)      lcqtof_t3_neg_4_108_62    m/z 634.2582     0   1E+4   4.5   5  RT (min)      lcqtof_t3_neg_4_000_ee    m/z 634.3178     0   1   2E+4   9.5   10  RT (min)       lcqtof_t3_neg_4_040_24    m/z 634.3181     0   1   2E+4   9.5   10  RT (min)      lcqtof_t3_neg_4_040_24    m/z 635.1546     0   1E+4   11.5   12  RT (min)      lcqtof_t3_neg_4_064_17    m/z 635.2272     0   1   2   3   4E+3   8   8.5  RT (min)      lcqtof_t3_neg_4_004_20    m/z 635.3302     0   1E+4   10   10.5  RT (min)      lcqtof_t3_neg_4_028_25    m/z 636.3336     0   1E+4   10   10.5  RT (min)       lcqtof_t3_neg_4_091_60    m/z 636.4607     0   1   2   3   4E+3   12.5   13  RT (min)      lcqtof_t3_neg_4_014_13    m/z 636.8267     0   1   2   3   4E+3   16.5   17  RT (min)       lcqtof_t3_neg_4_000_ff    m/z 636.8272     0   1   2E+3   15.5   16  RT (min)       lcqtof_t3_neg_4_042_10    m/z 636.8278     0   1   2   3E+3   16   16.5  RT (min)       lcqtof_t3_neg_4_056_56    m/z 637.5876     0   2   4   6E+4   13.5   14  RT (min)      lcqtof_t3_neg_2_016_05    m/z 637.5878     0   1   2   3   4E+4   13.5   14  RT (min)      lcqtof_t3_neg_2_030_23    m/z 637.5880     0   1E+2   11.5   12  RT (min)      lcqtof_t3_neg_3_038_06    m/z 637.5886     0   1E+3   13   13.5  RT (min)      lcqtof_t3_neg_3_032_24    m/z 637.5888     0   1E+3   13   13.5  RT (min)      lcqtof_t3_neg_3_032_24    m/z 638.3548     0   2   4   6   8E+3   11   11.5  RT (min)      lcqtof_t3_neg_4_016_14    m/z 638.5360     0   2   4   6   8E+3   13   13.5  RT (min)      lcqtof_t3_neg_2_030_23    m/z 638.5910     0   1E+4   13.5   14  RT (min)      lcqtof_t3_neg_2_030_23    m/z 638.5910     0   1   2   3E+2   13   13.5  RT (min)      lcqtof_t3_neg_3_032_24    m/z 638.5915     0   1   2   3E+4   13.5   14  RT (min)      lcqtof_t3_neg_2_016_05    m/z 638.5919     0   1   2E+2   11.5   12  RT (min)      lcqtof_t3_neg_3_038_06    m/z 638.8324     0   2   4E+3   16.5   17  RT (min)       lcqtof_t3_neg_1_013_09    m/z 638.8324     0   2   4E+3   16.5   17  RT (min)      lcqtof_t3_neg_1_005_18    m/z 639.1270     0   1   2E+5   12.5   13  RT (min)      lcqtof_t3_neg_4_064_17    m/z 639.1271     0   1E+5   13.5   14  RT (min)      lcqtof_t3_neg_4_002_17    m/z 639.1275     0   1   2E+4   13.5   14  RT (min)      lcqtof_t3_neg_4_002_17    m/z 639.1276     0   1   2   3E+4   13.5   14  RT (min)      lcqtof_t3_neg_4_036_47    m/z 639.1291     0   1   2E+5   12.5   13  RT (min)      lcqtof_t3_neg_4_064_17    m/z 639.5994     0   1   2E+4   5   5.5  RT (min)      lcqtof_t3_neg_4_003_37    m/z 640.1287     0   1E+4   13.5   14  RT (min)      lcqtof_t3_neg_4_036_47    m/z 640.1302     0   1E+4   13.5   14  RT (min)      lcqtof_t3_neg_4_002_17    m/z 641.1252     0   1E+4   13.5   14  RT (min)      lcqtof_t3_neg_4_002_17    m/z 641.1262     0   1E+4   13.5   14  RT (min)      lcqtof_t3_neg_4_036_47    m/z 641.2595     0   1   2E+3   6.5   7  RT (min)      lcqtof_t3_neg_4_016_14    m/z 641.2600     0   2   4   6E+4   5   5.5  RT (min)      lcqtof_t3_neg_4_043_11    m/z 641.2606     0   1   2E+3   8.5   9  RT (min)      lcqtof_t3_neg_4_027_41    m/z 641.2606     0   2   4E+3   9.5   10  RT (min)      lcqtof_t3_neg_1_036_14    m/z 641.2606     0   1   2E+3   8   8.5  RT (min)      lcqtof_t3_neg_4_000_cc    m/z 641.2610     0   2   4   6E+3   9.5   10  RT (min)      lcqtof_t3_neg_3_058_14    m/z 641.2610     0   2   4E+3   16.5   17  RT (min)      lcqtof_t3_neg_4_000_cc    m/z 641.2611     0   1   2E+3   6   6.5  RT (min)      lcqtof_t3_neg_4_049_55    m/z 641.2611     0   2   4E+3   7   7.5  RT (min)       lcqtof_t3_neg_4_062_13    m/z 641.2612     0   2   4   6E+3   9.5   10  RT (min)      lcqtof_t3_neg_3_058_14    m/z 641.2614     0   1   2   3   4E+3   15   15.5  RT (min)       lcqtof_t3_neg_4_012_27    m/z 641.2615     0   1   2E+3   8   8.5  RT (min)      lcqtof_t3_neg_4_032_04    m/z 641.2616     0   1   2   3E+3   14.5   15  RT (min)      lcqtof_t3_neg_4_000_dd    m/z 641.2621     0   1   2E+3   7   7.5  RT (min)      lcqtof_t3_neg_2_064_13    m/z 641.2625     0   1   2   3E+3   9   9.5  RT (min)      lcqtof_t3_neg_4_000_ee    m/z 641.4276     0   2   4   6E+3   10   10.5  RT (min)       lcqtof_t3_neg_4_030_31    m/z 641.5104     0   1E+4   12.5   13  RT (min)      lcqtof_t3_neg_2_030_23    m/z 641.5291     0   1   2E+4   4.5   5  RT (min)      lcqtof_t3_neg_4_000_ee    m/z 641.5838     0   2   4E+3   14   14.5  RT (min)      lcqtof_t3_neg_2_052_17    m/z 642.1250     0   2   4E+4   13   13.5  RT (min)      lcqtof_t3_neg_4_036_47    m/z 642.1280     0   1   2   3E+4   13   13.5  RT (min)      lcqtof_t3_neg_4_002_17    m/z 642.1282     0   1   2   3E+4   13   13.5  RT (min)      lcqtof_t3_neg_4_002_17    m/z 643.1226     0   1   2E+4   13   13.5  RT (min)      lcqtof_t3_neg_4_036_47    m/z 643.1231     0   1E+4   13   13.5  RT (min)      lcqtof_t3_neg_4_002_17    m/z 643.3089     0   1   2E+3   11   11.5  RT (min)      lcqtof_t3_neg_1_022_28    m/z 643.3517     0   1   2   3   4E+4   9   9.5  RT (min)      lcqtof_t3_neg_4_000_ee    m/z 643.3518     0   1   2   3   4E+4   9   9.5  RT (min)      lcqtof_t3_neg_4_000_ee    m/z 643.4054     0   1E+4   11   11.5  RT (min)       lcqtof_t3_neg_4_034_34    m/z 643.4376     0   1   2E+4   8   8.5  RT (min)      lcqtof_t3_neg_4_000_ee    m/z 643.4380     0   1   2E+4   8   8.5  RT (min)      lcqtof_t3_neg_4_000_ee    m/z 643.5240     0   1E+4   13   13.5  RT (min)      lcqtof_t3_neg_2_016_05    m/z 643.5254     0   1   2E+2   11   11.5  RT (min)      lcqtof_t3_neg_3_038_06    m/z 643.5257     0   2   4   6E+4   13   13.5  RT (min)      lcqtof_t3_neg_2_030_23    m/z 643.5260     0   1   2E+2   11   11.5  RT (min)      lcqtof_t3_neg_3_038_06    m/z 644.2774     0   1E+4   4.5   5  RT (min)      lcqtof_t3_neg_4_000_ee    m/z 644.3590     0   1E+4   9   9.5  RT (min)      lcqtof_t3_neg_4_000_ee    m/z 644.5284     0   1   2E+4   13   13.5  RT (min)      lcqtof_t3_neg_2_030_23    m/z 644.8480     0   1   2   3E+3   7.5   8  RT (min)       lcqtof_t3_neg_4_000_ff    m/z 644.8492     0   1   2E+3   9   9.5  RT (min)       lcqtof_t3_neg_4_000_ff    m/z 644.8498     0   1   2   3E+3   6   7  RT (min)       lcqtof_t3_neg_4_000_ff    m/z 645.2569     0   1   2E+3   5.5   6  RT (min)       lcqtof_t3_neg_4_005_18    m/z 645.2576     0   1   2   3E+3   7   7.5  RT (min)      lcqtof_t3_neg_4_005_18    m/z 645.2579     0   1   2E+3   6   6.5  RT (min)       lcqtof_t3_neg_4_043_11    m/z 645.2580     0   1   2E+3   8   8.5  RT (min)      lcqtof_t3_neg_4_004_20    m/z 645.4491     0   1E+4   12.5   13  RT (min)      lcqtof_t3_neg_4_034_34    m/z 645.4494     0   2   4E+3   12.5   13  RT (min)      lcqtof_t3_neg_1_015_34    m/z 646.2890     0   2   4E+3   10   10.5  RT (min)       lcqtof_t3_neg_4_037_01    m/z 646.6610     0   1E+2   12.5   13  RT (min)      lcqtof_t3_neg_3_032_24    m/z 646.6636     0   1E+2   12.5   13  RT (min)      lcqtof_t3_neg_3_032_24    m/z 646.7652     0   1E+4   1   1.5  RT (min)      lcqtof_t3_neg_4_037_01    m/z 646.8553     0   1E+4   17   17.5  RT (min)      lcqtof_t3_neg_4_035_08    m/z 646.8558     0   1E+4   16   16.5  RT (min)      lcqtof_t3_neg_4_048_19    m/z 646.8558     0   1   2   3E+3   15.5   16  RT (min)       lcqtof_t3_neg_4_015_36    m/z 646.8560     0   1E+4   16   16.5  RT (min)       lcqtof_t3_neg_4_018_23    m/z 646.8560     0   1E+3   16   16.5  RT (min)      lcqtof_t3_neg_2_016_05    m/z 646.8562     0   1E+3   15   15.5  RT (min)       lcqtof_t3_neg_3_032_24    m/z 646.8562     0   1   2E+3   15.5   16  RT (min)       lcqtof_t3_neg_1_019_23    m/z 646.8562     0   1   2   3E+3   15.5   16  RT (min)       lcqtof_t3_neg_4_042_10    m/z 647.3689     0   1E+4   8   8.5  RT (min)      lcqtof_t3_neg_4_000_ee    m/z 648.4450     0   1E+4   5.5   6  RT (min)      lcqtof_t3_neg_4_019_05    m/z 648.4453     0   1E+4   5   5.5  RT (min)      lcqtof_t3_neg_4_018_23    m/z 648.6740     0   1   2E+3   15   15.5  RT (min)      lcqtof_t3_neg_4_001_49    m/z 649.2820     0   1E+4   8.5   9  RT (min)      lcqtof_t3_neg_4_000_ee    m/z 649.3592     0   2   4   6E+4   8.5   9  RT (min)      lcqtof_t3_neg_4_000_ee    m/z 650.2586     0   1E+4   7   7.5  RT (min)      lcqtof_t3_neg_4_005_18    m/z 650.3064     0   1   2   3E+3   10.5   11  RT (min)      lcqtof_t3_neg_1_002_13    m/z 650.3102     0   1   2E+3   10   10.5  RT (min)      lcqtof_t3_neg_1_062_13    m/z 650.3598     0   1   2E+4   8.5   9  RT (min)      lcqtof_t3_neg_4_000_ee    m/z 650.6126     0   1E+2   12.5   13  RT (min)      lcqtof_t3_neg_3_032_24    m/z 650.6304     0   1E+4   1   1.5  RT (min)      lcqtof_t3_neg_4_037_01    m/z 650.8652     0   1   2   3   4E+3   7.5   8  RT (min)       lcqtof_t3_neg_4_109_62    m/z 650.8662     0   2   4E+3   17   17.5  RT (min)       lcqtof_t3_neg_4_035_08    m/z 650.8662     0   1   2   3   4E+3   6.5   7  RT (min)       lcqtof_t3_neg_4_113_62    m/z 650.8664     0   1   2   3E+3   8  RT (min)       lcqtof_t3_neg_4_105_61    m/z 650.8669     0   1   2   3   4E+3   7   8  RT (min)       lcqtof_t3_neg_4_112_62    m/z 650.8674     0   2   4   6E+3   8   9  RT (min)       lcqtof_t3_neg_4_000_ff    m/z 650.8678     0   1   2   3E+3   6.5   7  RT (min)       lcqtof_t3_neg_4_096_60    m/z 651.1047     0   1   2E+3   15.5   16  RT (min)       lcqtof_t3_neg_4_030_31    m/z 651.1054     0   1E+4   11   11.5  RT (min)      lcqtof_t3_neg_4_002_17    m/z 651.1054     0   1E+4   11   11.5  RT (min)      lcqtof_t3_neg_4_002_17    m/z 651.1056     0   1   2E+3   15.5   16  RT (min)       lcqtof_t3_neg_4_017_51    m/z 651.1060     0   1E+4   11   11.5  RT (min)      lcqtof_t3_neg_4_002_17    m/z 651.1068     0   1E+4   11   11.5  RT (min)      lcqtof_t3_neg_4_002_17    m/z 651.1076     0   1   2E+3   16   16.5  RT (min)       lcqtof_t3_neg_4_000_ff    m/z 651.1094     0   1E+4   11   11.5  RT (min)      lcqtof_t3_neg_4_064_17    m/z 651.2780     0   1E+4   8.5   9  RT (min)      lcqtof_t3_neg_4_000_ee    m/z 651.2787     0   1E+4   8.5   9  RT (min)      lcqtof_t3_neg_4_000_ee    m/z 651.3407     0   1   2E+3   11.5   12  RT (min)      lcqtof_t3_neg_4_036_47    m/z 651.3564     0   1   2E+4   8.5   9  RT (min)      lcqtof_t3_neg_4_000_ee    m/z 652.3264     0   1   2   3   4E+3   10.5   11  RT (min)      lcqtof_t3_neg_1_002_13    m/z 652.3286     0   1   2   3   4E+3   10   10.5  RT (min)       lcqtof_t3_neg_1_031_24    m/z 652.3303     0   1   2   3E+3   10.5   11  RT (min)      lcqtof_t3_neg_4_041_54    m/z 652.5327     0   1   2E+4   13   13.5  RT (min)      lcqtof_t3_neg_4_018_23    m/z 652.6248     0   1E+2   12.5   13  RT (min)      lcqtof_t3_neg_3_032_24    m/z 652.6272     0   1E+2   12.5   13  RT (min)      lcqtof_t3_neg_3_032_24    m/z 652.6280     0   1E+4   13   13.5  RT (min)      lcqtof_t3_neg_2_016_05    m/z 652.6281     0   1E+4   13   13.5  RT (min)      lcqtof_t3_neg_2_016_05    m/z 653.0865     0   1   2   3E+4   12.5   13  RT (min)      lcqtof_t3_neg_4_064_17    m/z 653.0888     0   1   2   3E+4   13.5   14  RT (min)      lcqtof_t3_neg_4_064_17    m/z 653.1927     0   1E+4   3   3.5  RT (min)      lcqtof_t3_neg_4_019_05    m/z 653.3592     0   1E+4   11.5   12  RT (min)       lcqtof_t3_neg_4_037_01    m/z 653.5176     0   1E+4   13   13.5  RT (min)      lcqtof_t3_neg_2_030_23    m/z 653.5194     0   1E+4   13   13.5  RT (min)      lcqtof_t3_neg_4_018_23    m/z 654.0885     0   1E+4   13   13.5  RT (min)      lcqtof_t3_neg_4_036_47    m/z 654.0893     0   1E+4   13.5   14  RT (min)      lcqtof_t3_neg_4_002_17    m/z 654.0910     0   1   2E+4   13   13.5  RT (min)      lcqtof_t3_neg_4_064_17    m/z 654.4986     0   1   2E+3   13.5   14  RT (min)      lcqtof_t3_neg_4_023_45    m/z 654.5660     0   1E+4   13.5   14  RT (min)       lcqtof_t3_neg_2_030_23    m/z 654.5781     0   1   2   3E+2   13   13.5  RT (min)      lcqtof_t3_neg_3_032_24    m/z 654.5786     0   1   2E+2   11.5   12  RT (min)      lcqtof_t3_neg_3_038_06    m/z 655.0684     0   1   2   3E+4   13   13.5  RT (min)      lcqtof_t3_neg_4_064_17    m/z 655.0692     0   1E+4   12.5   13  RT (min)      lcqtof_t3_neg_4_064_17    m/z 655.0698     0   1   2E+4   12.5   13  RT (min)      lcqtof_t3_neg_4_064_17    m/z 655.0707     0   1   2   3E+4   13   13.5  RT (min)      lcqtof_t3_neg_4_064_17    m/z 655.0710     0   1   2   3E+4   12.5   13  RT (min)      lcqtof_t3_neg_4_064_17    m/z 655.0736     0   1   2E+4   13.5   14  RT (min)      lcqtof_t3_neg_4_064_17    m/z 655.0741     0   1   2E+4   13.5   14  RT (min)      lcqtof_t3_neg_4_064_17    m/z 655.0804     0   1   2   3E+4   13   13.5  RT (min)      lcqtof_t3_neg_4_064_17    m/z 655.0841     0   1   2   3E+4   13   13.5  RT (min)      lcqtof_t3_neg_4_064_17    m/z 655.1034     0   1   2E+3   15.5   16  RT (min)       lcqtof_t3_neg_4_039_06    m/z 655.1036     0   1   2E+3   15   15.5  RT (min)       lcqtof_t3_neg_4_011_32    m/z 655.1038     0   1   2E+3   16   16.5  RT (min)      lcqtof_t3_neg_4_050_43    m/z 655.1062     0   1   2E+3   16   16.5  RT (min)      lcqtof_t3_neg_4_024_26    m/z 656.3886     0   1E+3   9   9.5  RT (min)      lcqtof_t3_neg_3_032_24    m/z 656.3888     0   1E+4   9.5   10  RT (min)      lcqtof_t3_neg_2_016_05    m/z 656.3892     0   1E+4   9   9.5  RT (min)      lcqtof_t3_neg_3_032_24    m/z 656.3892     0   1   2   3E+4   9.5   10  RT (min)      lcqtof_t3_neg_4_019_05    m/z 656.3893     0   1E+4   9.5   10  RT (min)      lcqtof_t3_neg_2_016_05    m/z 656.3900     0   1   2E+3   9   9.5  RT (min)      lcqtof_t3_neg_3_032_24    m/z 656.3963     0   1   2E+4   8   8.5  RT (min)      lcqtof_t3_neg_4_000_ee    m/z 656.8814     0   2   4   6   8E+4   17   17.5  RT (min)      lcqtof_t3_neg_4_035_08    m/z 656.8826     0   1   2   3   4E+2   13.5   14  RT (min)      lcqtof_t3_neg_3_038_06    m/z 656.8840     0   1E+5   16   16.5  RT (min)       lcqtof_t3_neg_4_018_23    m/z 656.8841     0   1   2E+3   15.5   16  RT (min)       lcqtof_t3_neg_1_019_23    m/z 656.8844     0   1E+5   16   16.5  RT (min)      lcqtof_t3_neg_4_018_23    m/z 656.8846     0   1   2   3E+3   8   8.5  RT (min)       lcqtof_t3_neg_4_111_62    m/z 656.8849     0   1E+3   15   15.5  RT (min)       lcqtof_t3_neg_3_032_24    m/z 656.8858     0   1   2E+3   15   15.5  RT (min)       lcqtof_t3_neg_4_055_38    m/z 656.8950     0   1E+5   17   17.5  RT (min)       lcqtof_t3_neg_4_112_62    m/z 657.0605     0   1E+4   13   13.5  RT (min)      lcqtof_t3_neg_4_064_17    m/z 657.0642     0   1E+4   12.5   13  RT (min)      lcqtof_t3_neg_4_064_17    m/z 657.0647     0   1E+4   12.5   13  RT (min)      lcqtof_t3_neg_4_064_17    m/z 657.0652     0   1E+4   13   13.5  RT (min)      lcqtof_t3_neg_4_064_17    m/z 657.2567     0   2   4   6E+4   4.5   5  RT (min)       lcqtof_t3_neg_4_000_ee    m/z 657.4497     0   1E+4   12.5   13  RT (min)      lcqtof_t3_neg_4_003_37    m/z 657.4502     0   1E+4   12   12.5  RT (min)      lcqtof_t3_neg_4_034_34    m/z 658.2598     0   1E+4   4.5   5  RT (min)      lcqtof_t3_neg_4_000_ee    m/z 658.3322     0   2   4   6E+3   10   10.5  RT (min)      lcqtof_t3_neg_4_060_15    m/z 658.3332     0   1E+4   10   10.5  RT (min)      lcqtof_t3_neg_4_058_16    m/z 658.4010     0   1   2E+4   8   8.5  RT (min)      lcqtof_t3_neg_4_000_ee    m/z 658.4022     0   1   2E+4   8   8.5  RT (min)      lcqtof_t3_neg_4_000_ee    m/z 658.4032     0   1E+4   10   10.5  RT (min)      lcqtof_t3_neg_2_016_05    m/z 658.4040     0   1E+4   9.5   10  RT (min)      lcqtof_t3_neg_3_032_24    m/z 659.3407     0   1   2   3E+3   14.5   15  RT (min)      lcqtof_t3_neg_4_012_27    m/z 659.3412     0   1   2   3   4E+3   15   15.5  RT (min)      lcqtof_t3_neg_4_005_18    m/z 660.3534     0   1E+4   9   9.5  RT (min)      lcqtof_t3_neg_4_000_ee    m/z 660.3544     0   1E+4   9   9.5  RT (min)      lcqtof_t3_neg_4_000_ee    m/z 660.3858     0   2   4   6E+4   8.5   9  RT (min)      lcqtof_t3_neg_4_000_ee    m/z 660.4655     0   1E+4   13   13.5  RT (min)      lcqtof_t3_neg_2_030_23    m/z 660.4683     0   2   4   6E+3   12.5   13  RT (min)      lcqtof_t3_neg_4_034_34    m/z 660.6398     0   1E+2   12   12.5  RT (min)      lcqtof_t3_neg_3_032_24    m/z 660.6404     0   1E+2   12.5   13  RT (min)      lcqtof_t3_neg_3_032_24    m/z 660.6406     0   1   2   3E+4   13   13.5  RT (min)      lcqtof_t3_neg_2_016_05    m/z 660.6422     0   1E+2   11   11.5  RT (min)      lcqtof_t3_neg_3_038_06    m/z 660.6438     0   1E+2   11   11.5  RT (min)      lcqtof_t3_neg_3_038_06    m/z 661.3917     0   1   2E+4   9   9.5  RT (min)      lcqtof_t3_neg_4_000_ee    m/z 661.3928     0   1   2E+4   8.5   9  RT (min)      lcqtof_t3_neg_4_000_ee    m/z 661.3936     0   1   2E+4   8.5   9  RT (min)      lcqtof_t3_neg_4_000_ee    m/z 661.4026     0   1   2E+4   8.5   9  RT (min)      lcqtof_t3_neg_4_000_ee    m/z 661.4052     0   1   2E+4   8.5   9  RT (min)      lcqtof_t3_neg_4_000_ee    m/z 661.6518     0   1E+2   12.5   13  RT (min)      lcqtof_t3_neg_3_032_24    m/z 661.6530     0   1E+4   13   13.5  RT (min)      lcqtof_t3_neg_2_016_05    m/z 661.6537     0   1E+2   12.5   13  RT (min)      lcqtof_t3_neg_3_032_24    m/z 662.2786     0   1   2E+4   4   4.5  RT (min)      lcqtof_t3_neg_4_000_ee    m/z 662.4552     0   1E+4   13   13.5  RT (min)      lcqtof_t3_neg_2_030_23    m/z 662.6159     0   1E+4   1   1.5  RT (min)      lcqtof_t3_neg_4_037_01    m/z 662.6558     0   1E+2   12.5   13  RT (min)      lcqtof_t3_neg_3_032_24    m/z 662.6560     0   1E+2   12.5   13  RT (min)      lcqtof_t3_neg_3_032_24    m/z 662.6566     0   1E+2   11   11.5  RT (min)      lcqtof_t3_neg_3_038_06    m/z 662.6567     0   1E+2   11   11.5  RT (min)      lcqtof_t3_neg_3_038_06    m/z 662.6569     0   1   2   3   4E+4   13   13.5  RT (min)      lcqtof_t3_neg_2_016_05    m/z 662.6914     0   1   2E+3   15   15.5  RT (min)      lcqtof_t3_neg_4_032_04    m/z 662.8351     0   1E+4   16.5   17  RT (min)      lcqtof_t3_neg_4_091_60    m/z 662.8378     0   1E+4   16   16.5  RT (min)      lcqtof_t3_neg_4_018_23    m/z 663.1817     0   1E+2   3.5   4  RT (min)      lcqtof_t3_neg_3_038_06    m/z 663.1840     0   1   2   3E+4   4   4.5  RT (min)      lcqtof_t3_neg_3_032_24    m/z 663.8449     0   2   4   6   8E+4   16   16.5  RT (min)       lcqtof_t3_neg_4_018_23    m/z 663.8564     0   2   4   6   8E+2   16   16.5  RT (min)      lcqtof_t3_neg_2_030_23    m/z 663.8568     0   2   4E+2   10.5   11  RT (min)      lcqtof_t3_neg_3_038_06    m/z 663.8570     0   1   2   3   4E+4   12.5   13  RT (min)      lcqtof_t3_neg_1_019_23    m/z 663.8571     0   2   4E+4   12.5   13  RT (min)      lcqtof_t3_neg_2_030_23    m/z 663.8572     0   2   4   6   8E+4   16   16.5  RT (min)      lcqtof_t3_neg_4_018_23    m/z 663.8636     0   2   4   6E+2   13.5   14  RT (min)      lcqtof_t3_neg_3_038_06    m/z 664.3581     0   1   2   3E+4   12.5   13  RT (min)      lcqtof_t3_neg_2_030_23    m/z 664.3584     0   1   2   3E+4   12.5   13  RT (min)      lcqtof_t3_neg_2_030_23    m/z 664.3585     0   1   2E+2   16   16.5  RT (min)      lcqtof_t3_neg_2_030_23    m/z 664.3591     0   1   2E+4   12.5   13  RT (min)      lcqtof_t3_neg_1_019_23    m/z 664.3611     0   2   4E+4   16   16.5  RT (min)      lcqtof_t3_neg_4_018_23    m/z 664.4193     0   1E+4   10.5   11  RT (min)      lcqtof_t3_neg_4_037_01    m/z 664.6972     0   1E+4   1   1.5  RT (min)      lcqtof_t3_neg_4_037_01    m/z 664.8600     0   2   4   6E+2   16   16.5  RT (min)      lcqtof_t3_neg_2_030_23    m/z 664.8604     0   1E+4   12.5   13  RT (min)      lcqtof_t3_neg_2_030_23    m/z 665.1211     0   2   4   6   8E+3   10   10.5  RT (min)       lcqtof_t3_neg_4_000_ee    m/z 665.1214     0   1   2E+3   15   15.5  RT (min)      lcqtof_t3_neg_4_040_24    m/z 665.1218     0   1   2E+3   16   16.5  RT (min)       lcqtof_t3_neg_4_040_24    m/z 665.1220     0   1   2E+3   15.5   16  RT (min)       lcqtof_t3_neg_4_038_29    m/z 665.1222     0   1   2E+3   15.5   16  RT (min)       lcqtof_t3_neg_4_000_ee    m/z 665.1226     0   1   2E+3   16   16.5  RT (min)       lcqtof_t3_neg_4_033_53    m/z 665.3540     0   2   4   6E+3   12.5   13  RT (min)      lcqtof_t3_neg_2_030_23    m/z 665.3546     0   1   2   3E+3   14.5   15  RT (min)       lcqtof_t3_neg_4_000_aa    m/z 665.3548     0   2   4   6E+3   12   12.5  RT (min)       lcqtof_t3_neg_4_000_ff    m/z 665.3551     0   2   4   6E+3   11.5   12  RT (min)       lcqtof_t3_neg_4_033_53    m/z 665.3553     0   1   2   3   4E+3   15.5   16  RT (min)       lcqtof_t3_neg_4_000_bb    m/z 665.3554     0   1   2   3   4E+3   14.5   15  RT (min)       lcqtof_t3_neg_4_003_37    m/z 665.3555     0   2   4   6E+3   16   16.5  RT (min)      lcqtof_t3_neg_4_018_23    m/z 665.3556     0   1   2   3   4E+3   15   15.5   16  RT (min)       lcqtof_t3_neg_4_000_bb    m/z 665.3558     0   2   4   6E+3   12   12.5  RT (min)      lcqtof_t3_neg_4_000_ff    m/z 665.3559     0   1   2   3   4E+3   14.5   15  RT (min)       lcqtof_t3_neg_4_000_bb    m/z 665.3560     0   2   4   6E+2   16   16.5  RT (min)      lcqtof_t3_neg_2_030_23    m/z 665.3560     0   1E+4   16   16.5  RT (min)      lcqtof_t3_neg_4_034_34    m/z 665.3593     0   1E+4   16.5   17  RT (min)      lcqtof_t3_neg_4_015_36    m/z 665.3880     0   1   2E+4   8.5   9  RT (min)      lcqtof_t3_neg_4_000_ee    m/z 665.3892     0   1   2E+4   8.5   9  RT (min)      lcqtof_t3_neg_4_000_ee    m/z 666.3591     0   1   2   3E+3   11.5   12  RT (min)      lcqtof_t3_neg_4_035_08    m/z 666.7901     0   1E+4   1   1.5  RT (min)      lcqtof_t3_neg_4_059_14    m/z 667.1407     0   1E+4   13   13.5  RT (min)      lcqtof_t3_neg_4_036_47    m/z 667.4317     0   1E+4   12   12.5  RT (min)      lcqtof_t3_neg_1_019_23    m/z 667.4358     0   1E+4   12   12.5  RT (min)      lcqtof_t3_neg_2_030_23    m/z 667.5122     0   2   4   6E+3   14   14.5  RT (min)      lcqtof_t3_neg_2_053_38    m/z 667.5146     0   2   4   6   8E+3   13   13.5  RT (min)      lcqtof_t3_neg_2_030_23    m/z 668.1418     0   1E+4   13   13.5  RT (min)      lcqtof_t3_neg_3_089_59    m/z 668.1426     0   1E+4   13   13.5  RT (min)      lcqtof_t3_neg_4_036_47    m/z 668.2623     0   1E+4   8   8.5  RT (min)      lcqtof_t3_neg_4_005_18    m/z 669.1208     0   1E+4   13   13.5  RT (min)      lcqtof_t3_neg_4_036_47    m/z 669.1234     0   1E+4   13   13.5  RT (min)      lcqtof_t3_neg_4_064_17    m/z 669.1371     0   1E+4   13   13.5  RT (min)      lcqtof_t3_neg_4_036_47    m/z 669.2567     0   1   2   3E+3   15   15.5  RT (min)       lcqtof_t3_neg_4_018_23    m/z 669.2567     0   1   2E+3   14.5   15  RT (min)       lcqtof_t3_neg_4_057_49    m/z 669.2567     0   1   2   3E+3   14.5   15  RT (min)       lcqtof_t3_neg_4_006_16    m/z 669.2571     0   1   2   3   4E+3   15.5   16  RT (min)       lcqtof_t3_neg_4_000_ee    m/z 669.2572     0   1   2   3E+3   15.5   16  RT (min)       lcqtof_t3_neg_4_038_29    m/z 669.2578     0   1   2   3E+3   16.5   17  RT (min)       lcqtof_t3_neg_4_000_cc    m/z 669.2578     0   1   2   3E+3   16   16.5  RT (min)       lcqtof_t3_neg_4_000_aa    m/z 669.4356     0   1E+4   8   8.5  RT (min)      lcqtof_t3_neg_4_000_ee    m/z 669.4380     0   1E+4   8   8.5  RT (min)      lcqtof_t3_neg_4_000_ee    m/z 669.4490     0   2   4E+4   12   12.5  RT (min)      lcqtof_t3_neg_2_030_23    m/z 669.4575     0   2   4E+4   12   12.5  RT (min)      lcqtof_t3_neg_4_018_23    m/z 669.4580     0   2   4   6E+3   11.5   12  RT (min)      lcqtof_t3_neg_4_079_58    m/z 669.4586     0   1   2E+4   12   12.5  RT (min)      lcqtof_t3_neg_2_030_23    m/z 669.4618     0   1   2   3E+3   13.5   14  RT (min)      lcqtof_t3_neg_4_088_59    m/z 669.4629     0   1   2   3E+3   13.5   14  RT (min)      lcqtof_t3_neg_4_008_35    m/z 669.5232     0   1E+4   13   13.5  RT (min)      lcqtof_t3_neg_2_030_23    m/z 669.6152     0   2   4E+3   14   14.5  RT (min)      lcqtof_t3_neg_2_062_17    m/z 670.2785     0   1   2   3E+4   4   4.5  RT (min)      lcqtof_t3_neg_4_000_ee    m/z 670.4528     0   1   2E+4   12   12.5  RT (min)      lcqtof_t3_neg_4_018_23    m/z 670.4534     0   1   2E+4   12   12.5  RT (min)      lcqtof_t3_neg_4_034_34    m/z 670.8635     0   1E+4   12.5   13  RT (min)      lcqtof_t3_neg_1_019_23    m/z 670.8643     0   1   2E+2   16   16.5  RT (min)      lcqtof_t3_neg_2_030_23    m/z 670.8643     0   1E+4   12.5   13  RT (min)      lcqtof_t3_neg_4_018_23    m/z 670.8646     0   1E+4   12.5   13  RT (min)      lcqtof_t3_neg_2_030_23    m/z 670.8647     0   1E+4   12.5   13  RT (min)      lcqtof_t3_neg_2_030_23    m/z 671.0969     0   1E+4   11.5   12  RT (min)      lcqtof_t3_neg_4_064_17    m/z 671.0984     0   1E+4   11.5   12  RT (min)      lcqtof_t3_neg_4_064_17    m/z 671.0984     0   1   2   3E+4   12   12.5  RT (min)      lcqtof_t3_neg_4_064_17    m/z 671.0984     0   1   2   3E+4   12   12.5  RT (min)      lcqtof_t3_neg_4_064_17    m/z 671.0990     0   1   2   3E+4   12.5   13  RT (min)      lcqtof_t3_neg_4_064_17    m/z 671.0993     0   1E+4   13   13.5  RT (min)      lcqtof_t3_neg_4_036_47    m/z 671.1000     0   1   2E+4   13.5   14  RT (min)      lcqtof_t3_neg_4_064_17    m/z 671.1001     0   1   2   3E+4   12.5   13  RT (min)      lcqtof_t3_neg_4_064_17    m/z 671.1008     0   1   2   3E+4   12.5   13  RT (min)      lcqtof_t3_neg_4_064_17    m/z 671.1017     0   1   2   3E+4   12   12.5  RT (min)      lcqtof_t3_neg_4_064_17    m/z 671.3674     0   1E+4   16   16.5  RT (min)      lcqtof_t3_neg_4_018_23    m/z 671.4323     0   1E+4   8.5   9  RT (min)      lcqtof_t3_neg_4_000_ee    m/z 671.4332     0   1E+4   8.5   9  RT (min)      lcqtof_t3_neg_4_000_ee    m/z 671.4349     0   1E+4   8.5   9  RT (min)      lcqtof_t3_neg_4_000_ee    m/z 671.4644     0   1   2E+2   10.5   11  RT (min)      lcqtof_t3_neg_3_038_06    m/z 671.4646     0   1   2E+5   12   12.5  RT (min)      lcqtof_t3_neg_1_019_23    m/z 671.4648     0   1   2E+2   10.5   11  RT (min)      lcqtof_t3_neg_3_038_06    m/z 671.4650     0   1   2E+2   10.5   11  RT (min)      lcqtof_t3_neg_3_038_06    m/z 671.4650     0   1   2E+2   10.5   11  RT (min)      lcqtof_t3_neg_3_038_06    m/z 672.0979     0   1E+4   12.5   13  RT (min)      lcqtof_t3_neg_4_036_47    m/z 672.1000     0   1E+4   13   13.5  RT (min)      lcqtof_t3_neg_4_002_17    m/z 672.4678     0   1E+2   10.5   11  RT (min)      lcqtof_t3_neg_3_038_06    m/z 672.4679     0   2   4   6E+4   12   12.5  RT (min)      lcqtof_t3_neg_1_019_23    m/z 672.4686     0   1E+2   10.5   11  RT (min)      lcqtof_t3_neg_3_038_06    m/z 672.4690     0   1E+2   10.5   11  RT (min)      lcqtof_t3_neg_3_038_06    m/z 672.4691     0   1   2E+2   10.5   11  RT (min)      lcqtof_t3_neg_3_038_06    m/z 672.4696     0   1E+2   10.5   11  RT (min)      lcqtof_t3_neg_3_038_06    m/z 672.4818     0   1   2   3   4E+3   13.5   14  RT (min)      lcqtof_t3_neg_4_008_35    m/z 672.8556     0   2   4   6E+2   15   15.5  RT (min)      lcqtof_t3_neg_3_032_24    m/z 672.8564     0   1   2   3E+2   13.5   14  RT (min)      lcqtof_t3_neg_3_038_06    m/z 672.8583     0   1   2   3E+4   17   17.5  RT (min)      lcqtof_t3_neg_4_035_08    m/z 672.8587     0   1E+3   15.5   16  RT (min)      lcqtof_t3_neg_1_019_23    m/z 673.0973     0   1E+4   12.5   13  RT (min)      lcqtof_t3_neg_4_064_17    m/z 673.0978     0   1E+4   12.5   13  RT (min)      lcqtof_t3_neg_4_064_17    m/z 673.0978     0   1E+4   12.5   13  RT (min)      lcqtof_t3_neg_4_064_17    m/z 673.0979     0   1E+4   13   13.5  RT (min)      lcqtof_t3_neg_4_064_17    m/z 673.0984     0   1E+4   12.5   13  RT (min)      lcqtof_t3_neg_4_064_17    m/z 673.0989     0   1E+4   12.5   13  RT (min)      lcqtof_t3_neg_4_064_17    m/z 673.2255     0   1   2   3   4E+3   11.5   12  RT (min)      lcqtof_t3_neg_4_012_27    m/z 673.3225     0   1   2   3E+3   11.5   12  RT (min)       lcqtof_t3_neg_4_046_33    m/z 673.3415     0   1   2   3E+3   16   16.5  RT (min)       lcqtof_t3_neg_4_000_dd    m/z 673.3416     0   1   2   3E+3   16   16.5  RT (min)      lcqtof_t3_neg_4_007_09    m/z 673.3458     0   2   4   6E+3   15   15.5  RT (min)      lcqtof_t3_neg_4_000_dd    m/z 673.3466     0   2   4   6E+3   14   14.5  RT (min)      lcqtof_t3_neg_4_000_dd    m/z 673.3478     0   2   4E+3   14.5   15  RT (min)      lcqtof_t3_neg_4_000_cc    m/z 673.3500     0   2   4E+3   14.5   15  RT (min)      lcqtof_t3_neg_4_000_cc    m/z 673.3587     0   1   2   3E+4   10   10.5  RT (min)      lcqtof_t3_neg_4_000_ee    m/z 673.4660     0   1E+4   12   12.5  RT (min)      lcqtof_t3_neg_1_019_23    m/z 673.4669     0   1E+4   12   12.5  RT (min)      lcqtof_t3_neg_3_006_23    m/z 673.4721     0   2   4   6E+4   12   12.5  RT (min)      lcqtof_t3_neg_2_030_23    m/z 673.4780     0   1   2E+2   10.5   11  RT (min)      lcqtof_t3_neg_3_038_06    m/z 673.4810     0   1   2E+2   11   11.5  RT (min)      lcqtof_t3_neg_3_038_06    m/z 673.4812     0   1   2E+2   10.5   11  RT (min)      lcqtof_t3_neg_3_038_06    m/z 673.4819     0   2   4E+4   13.5   14  RT (min)      lcqtof_t3_neg_4_003_37    m/z 674.3710     0   1E+4   10   10.5  RT (min)      lcqtof_t3_neg_4_000_ee    m/z 674.4820     0   1   2E+4   12   12.5  RT (min)      lcqtof_t3_neg_2_030_23    m/z 674.4833     0   1   2   3   4E+4   12.5   13  RT (min)      lcqtof_t3_neg_4_018_23    m/z 674.4833     0   1   2   3   4E+4   12.5   13  RT (min)       lcqtof_t3_neg_4_055_38    m/z 674.4834     0   1   2E+4   12.5   13  RT (min)      lcqtof_t3_neg_2_030_23    m/z 674.4836     0   2   4   6E+4   12.5   13  RT (min)       lcqtof_t3_neg_4_034_34    m/z 674.5118     0   1E+4   13   13.5  RT (min)      lcqtof_t3_neg_1_006_05    m/z 674.5122     0   1E+4   13   13.5  RT (min)      lcqtof_t3_neg_2_016_05    m/z 674.7985     0   1   2E+4   1   1.5  RT (min)      lcqtof_t3_neg_4_055_38    m/z 675.1747     0   1E+4   3   3.5  RT (min)      lcqtof_t3_neg_4_096_60    m/z 675.1750     0   1E+4   3   3.5  RT (min)      lcqtof_t3_neg_4_040_24    m/z 675.2692     0   1   2   3E+4   4.5   5  RT (min)      lcqtof_t3_neg_4_000_ee    m/z 675.4057     0   1E+5   8.5   9  RT (min)      lcqtof_t3_neg_4_000_ee    m/z 675.4812     0   1E+4   12.5   13  RT (min)      lcqtof_t3_neg_4_060_15    m/z 675.4827     0   1E+4   12   12.5  RT (min)      lcqtof_t3_neg_4_051_15    m/z 675.4860     0   1E+4   12.5   13  RT (min)      lcqtof_t3_neg_4_034_34    m/z 675.4864     0   1E+4   12.5   13  RT (min)      lcqtof_t3_neg_4_030_31    m/z 675.4928     0   2   4   6E+3   13   13.5  RT (min)      lcqtof_t3_neg_4_038_29    m/z 675.6578     0   1E+4   13   13.5  RT (min)       lcqtof_t3_neg_2_016_05    m/z 675.6588     0   1   2E+2   12.5   13  RT (min)      lcqtof_t3_neg_3_032_24    m/z 675.6594     0   1   2E+2   12.5   13  RT (min)      lcqtof_t3_neg_3_032_24    m/z 676.3844     0   2   4E+4   8.5   9  RT (min)      lcqtof_t3_neg_4_000_ee    m/z 676.4066     0   2   4E+4   8.5   9  RT (min)      lcqtof_t3_neg_4_000_ee    m/z 676.4226     0   2   4E+4   8.5   9  RT (min)      lcqtof_t3_neg_4_000_ee    m/z 676.7060     0   1   2   3E+3   15   15.5  RT (min)      lcqtof_t3_neg_4_049_55    m/z 676.7064     0   1   2E+3   15   15.5  RT (min)      lcqtof_t3_neg_4_003_37    m/z 676.7065     0   1   2E+3   15   15.5  RT (min)      lcqtof_t3_neg_4_011_32    m/z 677.1423     0   1E+4   11   11.5  RT (min)      lcqtof_t3_neg_4_064_17    m/z 677.2379     0   1E+4   5.5   6  RT (min)       lcqtof_t3_neg_4_012_27    m/z 677.2379     0   2   4   6   8E+3   6.5   7  RT (min)       lcqtof_t3_neg_4_016_14    m/z 677.2379     0   2   4   6   8E+3   8.5   9  RT (min)       lcqtof_t3_neg_4_013_39    m/z 677.2380     0   2   4   6   8E+3   12   12.5  RT (min)       lcqtof_t3_neg_4_007_09    m/z 677.2380     0   2   4   6E+3   6   6.5  RT (min)       lcqtof_t3_neg_4_059_14    m/z 677.2381     0   1E+4   14.5   15  RT (min)       lcqtof_t3_neg_4_007_09    m/z 677.2381     0   2   4   6   8E+3   8   8.5  RT (min)       lcqtof_t3_neg_4_006_16    m/z 677.2382     0   1E+4   7.5   8  RT (min)       lcqtof_t3_neg_4_024_26    m/z 677.2383     0   2   4   6   8E+3   6.5   7  RT (min)       lcqtof_t3_neg_4_013_39    m/z 677.2383     0   2   4E+3   10.5   11  RT (min)       lcqtof_t3_neg_4_070_57    m/z 677.2384     0   2   4   6   8E+3   16.5   17  RT (min)       lcqtof_t3_neg_4_000_ff    m/z 677.2384     0   2   4   6   8E+3   9.5   10  RT (min)       lcqtof_t3_neg_4_027_41    m/z 677.2385     0   1E+4   14.5   15  RT (min)       lcqtof_t3_neg_4_005_18    m/z 677.2385     0   2   4   6   8E+3   7.5   8  RT (min)       lcqtof_t3_neg_4_027_41    m/z 677.2386     0   2   4   6E+3   12   12.5  RT (min)       lcqtof_t3_neg_4_030_31    m/z 677.2388     0   2   4   6E+3   15   16   17  RT (min)       lcqtof_t3_neg_4_091_60    m/z 677.3462     0   1   2E+4   9   9.5  RT (min)      lcqtof_t3_neg_4_000_ee    m/z 677.3462     0   1   2E+4   9   9.5  RT (min)      lcqtof_t3_neg_4_000_ee    m/z 677.3822     0   1E+4   8.5   9  RT (min)      lcqtof_t3_neg_4_000_ee    m/z 677.3968     0   1E+4   8.5   9  RT (min)      lcqtof_t3_neg_4_000_ee    m/z 677.4244     0   1E+4   8.5   9  RT (min)      lcqtof_t3_neg_4_000_ee    m/z 677.4961     0   2   4   6   8E+3   12.5   13  RT (min)      lcqtof_t3_neg_4_051_15    m/z 677.6290     0   1E+2   12   12.5  RT (min)      lcqtof_t3_neg_3_032_24    m/z 677.6313     0   1E+4   13   13.5  RT (min)      lcqtof_t3_neg_2_016_05    m/z 677.6327     0   1E+2   12.5   13  RT (min)      lcqtof_t3_neg_3_032_24    m/z 677.8714     0   1   2E+4   12.5   13  RT (min)      lcqtof_t3_neg_2_030_23    m/z 677.8720     0   1   2   3E+2   16   16.5  RT (min)      lcqtof_t3_neg_2_030_23    m/z 677.8725     0   1   2   3E+4   16   16.5  RT (min)      lcqtof_t3_neg_4_018_23    m/z 678.3732     0   1E+4   13   13.5  RT (min)      lcqtof_t3_neg_4_018_23    m/z 678.3734     0   1E+2   16   16.5  RT (min)      lcqtof_t3_neg_2_030_23    m/z 678.3763     0   1E+4   12.5   13  RT (min)      lcqtof_t3_neg_3_006_23    m/z 678.8731     0   1   2   3E+2   16   16.5  RT (min)      lcqtof_t3_neg_2_030_23    m/z 678.8780     0   1E+4   16   16.5  RT (min)      lcqtof_t3_neg_4_018_23    m/z 679.1371     0   2   4   6E+3   10   10.5  RT (min)       lcqtof_t3_neg_4_000_ee    m/z 679.1572     0   1E+4   11.5   12  RT (min)      lcqtof_t3_neg_4_064_17    m/z 679.4974     0   1E+4   13   13.5  RT (min)      lcqtof_t3_neg_2_030_23    m/z 679.6446     0   1   2E+2   12.5   13  RT (min)      lcqtof_t3_neg_3_032_24    m/z 679.6466     0   1E+4   13   13.5  RT (min)      lcqtof_t3_neg_2_016_05    m/z 679.6470     0   1   2E+2   12.5   13  RT (min)      lcqtof_t3_neg_3_032_24    m/z 679.6472     0   1E+4   13   13.5  RT (min)      lcqtof_t3_neg_2_016_05    m/z 680.2406     0   2   4   6E+3   10   10.5  RT (min)      lcqtof_t3_neg_4_018_23    m/z 680.2416     0   2   4   6E+3   10   10.5  RT (min)       lcqtof_t3_neg_4_019_05    m/z 680.3583     0   1   2   3   4E+3   11   11.5  RT (min)       lcqtof_t3_neg_4_051_15    m/z 680.4511     0   1E+4   11   11.5  RT (min)      lcqtof_t3_neg_4_037_01    m/z 680.5019     0   1   2E+4   13   13.5  RT (min)      lcqtof_t3_neg_4_034_34    m/z 680.5153     0   1E+4   13   13.5  RT (min)      lcqtof_t3_neg_4_018_23    m/z 680.5669     0   2   4   6   8E+3   12.5   13  RT (min)      lcqtof_t3_neg_4_040_24    m/z 681.1573     0   1E+4   11.5   12  RT (min)      lcqtof_t3_neg_4_064_17    m/z 681.5133     0   1   2E+4   13   13.5  RT (min)      lcqtof_t3_neg_1_019_23    m/z 681.5202     0   1   2E+2   11.5   12  RT (min)      lcqtof_t3_neg_3_038_06    m/z 681.5283     0   1E+4   13   13.5  RT (min)      lcqtof_t3_neg_2_030_23    m/z 681.6137     0   1E+4   12   12.5  RT (min)      lcqtof_t3_neg_2_016_05    m/z 681.6138     0   1E+4   12   12.5  RT (min)      lcqtof_t3_neg_2_030_23    m/z 681.6194     0   1   2E+3   14   14.5  RT (min)      lcqtof_t3_neg_4_037_01    m/z 682.1406     0   1E+4   12   12.5  RT (min)      lcqtof_t3_neg_4_057_49    m/z 682.3429     0   1   2   3   4E+3   11   11.5  RT (min)       lcqtof_t3_neg_4_037_01    m/z 682.5165     0   1E+4   13   13.5  RT (min)      lcqtof_t3_neg_2_030_23    m/z 682.5330     0   1   2E+4   13.5   14  RT (min)      lcqtof_t3_neg_4_018_23    m/z 682.6341     0   1   2   3E+4   13.5   14  RT (min)      lcqtof_t3_neg_1_019_23    m/z 683.1352     0   1   2   3   4E+4   12   12.5  RT (min)      lcqtof_t3_neg_4_057_49    m/z 683.1714     0   1E+4   13   13.5  RT (min)      lcqtof_t3_neg_4_036_47    m/z 683.2242     0   1   2E+4   1   1.5  RT (min)      lcqtof_t3_neg_4_047_07    m/z 683.2979     0   1   2   3E+4   4.5   5  RT (min)      lcqtof_t3_neg_4_000_ee    m/z 683.3074     0   1E+4   9.5   10  RT (min)       lcqtof_t3_neg_4_059_14    m/z 683.3477     0   1E+4   8.5   9  RT (min)      lcqtof_t3_neg_4_000_ee    m/z 683.4230     0   1E+4   13.5   14  RT (min)      lcqtof_t3_neg_1_000_aa    m/z 683.4238     0   1E+4   13   13.5  RT (min)      lcqtof_t3_neg_1_000_aa    m/z 683.4241     0   1E+4   13.5   14  RT (min)      lcqtof_t3_neg_1_000_aa    m/z 683.4248     0   1E+4   13.5   14  RT (min)      lcqtof_t3_neg_1_000_aa    m/z 683.4249     0   1E+4   13.5   14  RT (min)      lcqtof_t3_neg_1_000_aa    m/z 683.4464     0   1E+4   12   12.5  RT (min)      lcqtof_t3_neg_4_059_14    m/z 683.4478     0   1E+4   11.5   12  RT (min)       lcqtof_t3_neg_4_059_14    m/z 683.4479     0   1E+4   11.5   12  RT (min)      lcqtof_t3_neg_4_059_14    m/z 683.4624     0   1E+4   12.5   13  RT (min)       lcqtof_t3_neg_4_034_34    m/z 683.4630     0   2   4   6E+3   12   12.5  RT (min)      lcqtof_t3_neg_2_030_23    m/z 683.5166     0   1E+4   13.5   14  RT (min)      lcqtof_t3_neg_2_016_05    m/z 683.5222     0   1E+4   13   13.5  RT (min)      lcqtof_t3_neg_2_030_23    m/z 683.5336     0   1   2E+4   13.5   14  RT (min)      lcqtof_t3_neg_4_018_23    m/z 683.5959     0   1   2   3E+3   14   14.5  RT (min)      lcqtof_t3_neg_4_059_14    m/z 683.6095     0   1   2   3   4E+3   12.5   13  RT (min)      lcqtof_t3_neg_1_002_13    m/z 683.6374     0   1E+4   13.5   14  RT (min)      lcqtof_t3_neg_1_019_23    m/z 684.1355     0   1E+4   12.5   13  RT (min)      lcqtof_t3_neg_4_001_49    m/z 684.1358     0   1   2E+4   12.5   13  RT (min)      lcqtof_t3_neg_4_057_49    m/z 684.3484     0   2   4E+3   10.5   11  RT (min)      lcqtof_t3_neg_4_014_13    m/z 684.3490     0   2   4E+3   10.5   11  RT (min)      lcqtof_t3_neg_4_062_13    m/z 684.4070     0   1E+4   10.5   11  RT (min)      lcqtof_t3_neg_4_037_01    m/z 684.4078     0   1E+4   10.5   11  RT (min)      lcqtof_t3_neg_4_037_01    m/z 684.5322     0   1E+4   13.5   14  RT (min)      lcqtof_t3_neg_4_018_23    m/z 684.8246     0   1   2E+3   16.5   17  RT (min)      lcqtof_t3_neg_1_016_48    m/z 685.1136     0   1E+4   13.5   14  RT (min)      lcqtof_t3_neg_4_002_17    m/z 685.1142     0   1   2   3E+4   13.5   14  RT (min)      lcqtof_t3_neg_4_064_17    m/z 685.1144     0   1E+4   13   13.5  RT (min)      lcqtof_t3_neg_4_002_17    m/z 685.1157     0   1   2   3E+4   13   13.5  RT (min)      lcqtof_t3_neg_4_064_17    m/z 685.1330     0   1   2E+4   12   12.5  RT (min)      lcqtof_t3_neg_4_057_49    m/z 685.1587     0   1E+4   4   4.5  RT (min)      lcqtof_t3_neg_4_000_ee    m/z 685.3016     0   1E+4   9.5   10  RT (min)      lcqtof_t3_neg_4_059_14    m/z 685.4798     0   1E+4   12   12.5  RT (min)      lcqtof_t3_neg_1_019_23    m/z 685.4798     0   1   2E+4   12.5   13  RT (min)      lcqtof_t3_neg_4_018_23    m/z 685.4804     0   1   2   3E+4   12.5   13  RT (min)       lcqtof_t3_neg_4_034_34    m/z 685.4806     0   1E+4   12   12.5  RT (min)      lcqtof_t3_neg_2_030_23    m/z 685.4816     0   1   2E+4   12.5   13  RT (min)      lcqtof_t3_neg_4_055_38    m/z 685.4914     0   1E+4   12.5   13  RT (min)      lcqtof_t3_neg_4_049_55    m/z 685.5098     0   1   2   3E+4   12.5   13  RT (min)      lcqtof_t3_neg_4_003_37    m/z 685.5878     0   1E+4   13   13.5  RT (min)      lcqtof_t3_neg_2_030_23    m/z 685.6025     0   1   2E+3   14   14.5  RT (min)      lcqtof_t3_neg_4_038_29    m/z 686.1136     0   1E+4   13.5   14  RT (min)      lcqtof_t3_neg_4_002_17    m/z 686.1155     0   1E+4   13   13.5  RT (min)      lcqtof_t3_neg_4_064_17    m/z 686.1155     0   1E+4   13   13.5  RT (min)      lcqtof_t3_neg_4_064_17    m/z 686.1198     0   1E+4   13   13.5  RT (min)      lcqtof_t3_neg_4_064_17    m/z 686.4777     0   1   2E+4   12   12.5  RT (min)      lcqtof_t3_neg_2_030_23    m/z 687.0357     0   1E+4   13   13.5  RT (min)      lcqtof_t3_neg_4_064_17    m/z 687.1145     0   1E+4   13   13.5  RT (min)      lcqtof_t3_neg_4_064_17    m/z 687.2656     0   1E+4   5.5   6  RT (min)      lcqtof_t3_neg_4_007_09    m/z 687.2658     0   1E+4   14.5   15  RT (min)       lcqtof_t3_neg_4_106_62    m/z 687.2666     0   1E+4   15.5   16  RT (min)       lcqtof_t3_neg_4_058_16    m/z 687.2666     0   1E+4   15.5   16  RT (min)       lcqtof_t3_neg_4_058_16    m/z 687.2668     0   1E+4   15.5   16  RT (min)       lcqtof_t3_neg_4_098_61    m/z 687.2672     0   1E+4   16   16.5  RT (min)       lcqtof_t3_neg_4_018_23    m/z 687.2672     0   1E+4   5.5   6  RT (min)      lcqtof_t3_neg_4_032_04    m/z 687.2672     0   1E+4   15.5   16  RT (min)       lcqtof_t3_neg_4_058_16    m/z 687.2678     0   1E+4   16.5   17  RT (min)       lcqtof_t3_neg_4_091_60    m/z 687.2681     0   1E+4   16   16.5  RT (min)       lcqtof_t3_neg_4_058_16    m/z 687.2694     0   1E+4   16   16.5  RT (min)      lcqtof_t3_neg_4_058_16    m/z 687.3380     0   2   4   6   8E+3   12   12.5  RT (min)      lcqtof_t3_neg_4_000_ff    m/z 687.3460     0   1   2   3   4E+3   16   16.5  RT (min)      lcqtof_t3_neg_4_005_18    m/z 687.3468     0   2   4E+3   14   14.5  RT (min)      lcqtof_t3_neg_4_000_dd    m/z 687.3489     0   1   2   3   4E+3   15.5   16  RT (min)      lcqtof_t3_neg_4_004_20    m/z 687.3498     0   2   4E+3   15   15.5  RT (min)      lcqtof_t3_neg_4_000_dd    m/z 687.3502     0   1   2   3   4E+3   15.5   16  RT (min)      lcqtof_t3_neg_4_001_49    m/z 687.3822     0   1   2E+4   9   9.5  RT (min)      lcqtof_t3_neg_4_000_ee    m/z 687.3824     0   1   2E+4   9   9.5  RT (min)      lcqtof_t3_neg_4_000_ee    m/z 687.4104     0   2   4   6E+4   4.5   5  RT (min)      lcqtof_t3_neg_4_000_ee    m/z 687.4435     0   1   2E+4   8   8.5  RT (min)      lcqtof_t3_neg_4_000_ee    m/z 687.4806     0   2   4   6E+3   12.5   13  RT (min)      lcqtof_t3_neg_3_086_59    m/z 687.4937     0   1E+4   12.5   13  RT (min)      lcqtof_t3_neg_4_084_59    m/z 687.5236     0   1   2E+4   12.5   13  RT (min)      lcqtof_t3_neg_4_034_34    m/z 687.6715     0   2   4   6E+4   4.5   5  RT (min)      lcqtof_t3_neg_4_000_ee    m/z 687.7523     0   2   4   6   8E+4   4.5   5  RT (min)      lcqtof_t3_neg_4_000_ee    m/z 687.8980     0   2   4   6E+4   4.5   5  RT (min)      lcqtof_t3_neg_4_000_ee    m/z 688.2696     0   1   2   3E+3   12   12.5  RT (min)      lcqtof_t3_neg_4_011_32    m/z 688.2710     0   1   2   3   4E+3   9.5   10  RT (min)      lcqtof_t3_neg_4_058_16    m/z 688.2718     0   1   2   3   4E+3   11   11.5  RT (min)      lcqtof_t3_neg_4_008_35    m/z 688.2745     0   2   4E+3   11   11.5  RT (min)      lcqtof_t3_neg_4_011_32    m/z 688.3051     0   2   4   6E+3   11   11.5  RT (min)      lcqtof_t3_neg_4_002_17    m/z 688.3315     0   1   2E+4   10   10.5  RT (min)      lcqtof_t3_neg_4_037_01    m/z 688.3396     0   1   2   3   4E+3   11.5   12  RT (min)      lcqtof_t3_neg_4_000_ff    m/z 688.3561     0   1   2E+3   14.5   15  RT (min)      lcqtof_t3_neg_4_000_dd    m/z 688.4910     0   2   4   6E+4   12.5   13  RT (min)      lcqtof_t3_neg_2_030_23    m/z 688.4918     0   2   4E+4   12.5   13  RT (min)      lcqtof_t3_neg_1_019_23    m/z 688.4928     0   1   2E+2   11   11.5  RT (min)      lcqtof_t3_neg_3_038_06    m/z 688.5476     0   1   2E+4   4.5   5  RT (min)      lcqtof_t3_neg_4_000_ee    m/z 688.7523     0   1E+4   1   1.5  RT (min)      lcqtof_t3_neg_4_037_01    m/z 688.8324     0   2   4   6   8E+3   16   16.5  RT (min)      lcqtof_t3_neg_1_011_45    m/z 688.9010     0   1   2   3   4E+4   4.5   5  RT (min)      lcqtof_t3_neg_4_000_ee    m/z 689.0282     0   1   2   3   4E+4   4.5   5  RT (min)      lcqtof_t3_neg_4_000_ee    m/z 689.1658     0   1E+4   4.5   5  RT (min)      lcqtof_t3_neg_4_000_ee    m/z 689.2958     0   1   2   3E+5   4   4.5  RT (min)      lcqtof_t3_neg_4_000_ee    m/z 689.4947     0   1   2E+4   12.5   13  RT (min)      lcqtof_t3_neg_2_030_23    m/z 689.4952     0   1   2E+4   12.5   13  RT (min)      lcqtof_t3_neg_4_018_23    m/z 689.4957     0   1E+4   12.5   13  RT (min)      lcqtof_t3_neg_1_019_23    m/z 690.2780     0   2   4E+3   10   10.5  RT (min)       lcqtof_t3_neg_4_062_13    m/z 690.2942     0   2   4E+4   4   4.5  RT (min)      lcqtof_t3_neg_4_000_ee    m/z 690.3640     0   1E+4   9.5   10  RT (min)      lcqtof_t3_neg_4_040_24    m/z 690.3661     0   1E+4   9.5   10  RT (min)      lcqtof_t3_neg_4_040_24    m/z 690.3661     0   1E+4   9.5   10  RT (min)       lcqtof_t3_neg_4_040_24    m/z 691.4345     0   1   2E+4   12   12.5  RT (min)      lcqtof_t3_neg_1_019_23    m/z 691.5022     0   1E+4   13   13.5  RT (min)      lcqtof_t3_neg_1_006_05    m/z 691.5518     0   1   2E+4   13   13.5  RT (min)      lcqtof_t3_neg_1_002_13    m/z 691.5525     0   1   2   3   4E+3   13   13.5  RT (min)      lcqtof_t3_neg_2_051_14    m/z 691.5528     0   1E+4   12.5   13  RT (min)      lcqtof_t3_neg_1_010_27    m/z 691.5533     0   1E+4   13   13.5  RT (min)      lcqtof_t3_neg_1_010_27    m/z 691.5536     0   1E+4   12   12.5  RT (min)      lcqtof_t3_neg_1_019_23    m/z 691.5548     0   1E+4   12   12.5  RT (min)      lcqtof_t3_neg_2_030_23    m/z 692.4086     0   1E+4   8.5   9  RT (min)      lcqtof_t3_neg_4_000_ee    m/z 692.4360     0   1E+4   11.5   12  RT (min)      lcqtof_t3_neg_1_019_23    m/z 693.3079     0   1E+4   8.5   9  RT (min)      lcqtof_t3_neg_4_000_ee    m/z 693.3843     0   1   2   3   4E+4   8.5   9  RT (min)      lcqtof_t3_neg_4_000_ee    m/z 693.4452     0   1E+4   12   12.5  RT (min)      lcqtof_t3_neg_2_016_05    m/z 693.4463     0   1   2   3E+4   12   12.5  RT (min)      lcqtof_t3_neg_3_006_23    m/z 693.4469     0   1   2   3   4E+4   12   12.5  RT (min)      lcqtof_t3_neg_2_030_23    m/z 693.4478     0   1   2   3   4E+4   12   12.5  RT (min)      lcqtof_t3_neg_1_019_23    m/z 693.4565     0   1   2   3E+2   15   15.5  RT (min)      lcqtof_t3_neg_3_032_24    m/z 693.4568     0   2   4   6E+3   16   16.5  RT (min)      lcqtof_t3_neg_2_016_05    m/z 693.4570     0   2   4   6   8E+2   13.5   14  RT (min)      lcqtof_t3_neg_3_038_06    m/z 693.9568     0   1   2E+2   12   12.5  RT (min)      lcqtof_t3_neg_3_032_24    m/z 693.9569     0   2   4E+2   10.5   11  RT (min)      lcqtof_t3_neg_3_038_06    m/z 693.9572     0   1   2E+2   15   15.5  RT (min)      lcqtof_t3_neg_3_032_24    m/z 693.9572     0   2   4E+2   10.5   11  RT (min)      lcqtof_t3_neg_3_038_06    m/z 693.9572     0   1E+3   13.5   14  RT (min)      lcqtof_t3_neg_3_038_06    m/z 693.9578     0   1   2E+4   16   16.5  RT (min)      lcqtof_t3_neg_2_016_05    m/z 693.9580     0   1   2E+2   12   12.5  RT (min)      lcqtof_t3_neg_3_032_24    m/z 693.9583     0   1   2E+2   12   12.5  RT (min)      lcqtof_t3_neg_3_032_24    m/z 693.9586     0   1   2E+4   12.5   13  RT (min)      lcqtof_t3_neg_2_016_05    m/z 694.2956     0   1   2   3   4E+3   10.5   11  RT (min)      lcqtof_t3_neg_1_033_53    m/z 694.3874     0   1E+4   8.5   9  RT (min)      lcqtof_t3_neg_4_000_ee    m/z 694.4518     0   1   2E+4   12   12.5  RT (min)      lcqtof_t3_neg_4_018_23    m/z 694.4520     0   1E+4   12   12.5  RT (min)      lcqtof_t3_neg_2_030_23    m/z 694.4522     0   1E+4   12   12.5  RT (min)      lcqtof_t3_neg_1_019_23    m/z 694.5236     0   1   2E+4   13   13.5  RT (min)      lcqtof_t3_neg_2_030_23    m/z 694.5258     0   1E+4   13   13.5  RT (min)      lcqtof_t3_neg_4_018_23    m/z 695.2941     0   1E+4   4   4.5  RT (min)      lcqtof_t3_neg_4_000_ee    m/z 695.3079     0   1E+4   8.5   9  RT (min)      lcqtof_t3_neg_4_000_ee    m/z 695.3822     0   1E+4   8.5   9  RT (min)      lcqtof_t3_neg_4_000_ee    m/z 695.3823     0   1E+4   8.5   9  RT (min)      lcqtof_t3_neg_4_000_ee    m/z 695.4610     0   1E+4   12   12.5  RT (min)      lcqtof_t3_neg_2_030_23    m/z 695.4611     0   1E+4   12   12.5  RT (min)      lcqtof_t3_neg_2_030_23    m/z 695.4625     0   1   2   3E+4   12.5   13  RT (min)      lcqtof_t3_neg_4_018_23    m/z 695.4632     0   1   2   3   4E+4   12   12.5  RT (min)       lcqtof_t3_neg_2_030_23    m/z 695.4644     0   1   2   3E+4   12   12.5  RT (min)      lcqtof_t3_neg_1_019_23    m/z 695.5291     0   1E+4   13   13.5  RT (min)      lcqtof_t3_neg_1_019_23    m/z 695.5305     0   1E+4   13   13.5  RT (min)      lcqtof_t3_neg_2_016_05    m/z 695.5343     0   1E+4   13   13.5  RT (min)      lcqtof_t3_neg_2_030_23    m/z 695.5412     0   1E+4   13   13.5  RT (min)      lcqtof_t3_neg_2_030_23    m/z 695.5453     0   1   2E+2   11.5   12  RT (min)      lcqtof_t3_neg_3_038_06    m/z 695.5462     0   1   2   3   4E+2   12.5   13  RT (min)      lcqtof_t3_neg_3_032_24    m/z 696.4652     0   1E+4   12   12.5  RT (min)      lcqtof_t3_neg_4_018_23    m/z 696.4674     0   1E+4   12   12.5  RT (min)      lcqtof_t3_neg_4_018_23    m/z 696.4680     0   1E+4   12   12.5  RT (min)      lcqtof_t3_neg_2_030_23    m/z 696.4680     0   1E+4   12   12.5  RT (min)      lcqtof_t3_neg_2_030_23    m/z 696.5480     0   2   4   6E+3   13   13.5  RT (min)      lcqtof_t3_neg_2_016_05    m/z 696.6503     0   2   4   6   8E+3   13.5   14  RT (min)      lcqtof_t3_neg_1_019_23    m/z 696.6505     0   1E+4   13.5   14  RT (min)      lcqtof_t3_neg_4_018_23    m/z 697.1568     0   1E+4   3   3.5  RT (min)      lcqtof_t3_neg_4_096_60    m/z 697.1851     0   1   2E+4   12.5   13  RT (min)      lcqtof_t3_neg_4_036_47    m/z 697.1856     0   1   2E+4   12.5   13  RT (min)      lcqtof_t3_neg_4_064_17    m/z 697.1859     0   1E+4   13   13.5  RT (min)      lcqtof_t3_neg_4_002_17    m/z 697.4780     0   1E+4   12   12.5  RT (min)      lcqtof_t3_neg_2_030_23    m/z 697.4787     0   2   4E+4   12.5   13  RT (min)      lcqtof_t3_neg_4_018_23    m/z 697.4788     0   1   2   3E+4   12.5   13  RT (min)      lcqtof_t3_neg_3_006_23    m/z 697.4794     0   2   4E+4   12.5   13  RT (min)      lcqtof_t3_neg_4_018_23    m/z 697.4795     0   2   4   6E+4   12.5   13  RT (min)      lcqtof_t3_neg_2_030_23    m/z 697.4798     0   2   4   6E+4   12.5   13  RT (min)      lcqtof_t3_neg_2_030_23    m/z 697.4811     0   1   2   3   4E+2   10.5   11  RT (min)      lcqtof_t3_neg_3_038_06    m/z 698.1872     0   1E+4   12.5   13  RT (min)      lcqtof_t3_neg_4_064_17    m/z 698.4789     0   1   2E+4   12.5   13  RT (min)      lcqtof_t3_neg_4_018_23    m/z 698.4819     0   1   2E+4   12.5   13  RT (min)      lcqtof_t3_neg_2_030_23    m/z 698.4826     0   1   2E+4   12.5   13  RT (min)      lcqtof_t3_neg_2_030_23    m/z 698.6060     0   1E+4   12   12.5  RT (min)      lcqtof_t3_neg_2_030_23    m/z 699.0934     0   1   2   3   4E+4   12.5   13  RT (min)      lcqtof_t3_neg_4_064_17    m/z 699.0938     0   1   2E+4   12.5   13  RT (min)      lcqtof_t3_neg_4_064_17    m/z 699.0940     0   1   2E+4   12.5   13  RT (min)      lcqtof_t3_neg_4_064_17    m/z 699.0946     0   2   4E+4   13.5   14  RT (min)      lcqtof_t3_neg_4_002_17    m/z 699.0948     0   2   4   6E+4   13.5   14  RT (min)      lcqtof_t3_neg_4_064_17    m/z 699.0948     0   2   4E+4   13   13.5  RT (min)      lcqtof_t3_neg_4_002_17    m/z 699.0956     0   1E+4   12.5   13  RT (min)      lcqtof_t3_neg_4_002_17    m/z 699.0957     0   2   4   6E+4   13   13.5  RT (min)      lcqtof_t3_neg_4_064_17    m/z 699.1865     0   1E+4   12.5   13  RT (min)      lcqtof_t3_neg_4_064_17    m/z 699.2845     0   1E+4   4   4.5  RT (min)      lcqtof_t3_neg_4_000_ee    m/z 699.4927     0   1   2E+4   12.5   13  RT (min)      lcqtof_t3_neg_1_019_23    m/z 699.4940     0   2   4   6E+4   12.5   13  RT (min)      lcqtof_t3_neg_2_030_23    m/z 699.4946     0   1E+4   12.5   13  RT (min)      lcqtof_t3_neg_1_019_23    m/z 699.4970     0   1   2   3E+2   11   11.5  RT (min)      lcqtof_t3_neg_3_038_06    m/z 699.4977     0   2   4   6E+4   13.5   14  RT (min)       lcqtof_t3_neg_4_018_23    m/z 699.4987     0   1   2   3   4E+4   13.5   14  RT (min)      lcqtof_t3_neg_4_055_38    m/z 699.4989     0   1   2   3E+2   11   11.5  RT (min)      lcqtof_t3_neg_3_038_06    m/z 699.5086     0   1   2   3E+4   13.5   14  RT (min)      lcqtof_t3_neg_4_034_34    m/z 699.5100     0   2   4   6E+4   13.5   14  RT (min)      lcqtof_t3_neg_4_018_23    m/z 699.6037     0   1   2   3   4E+3   11.5   12  RT (min)      lcqtof_t3_neg_1_002_13    m/z 699.6038     0   1   2   3   4E+3   11.5   12  RT (min)      lcqtof_t3_neg_1_038_15    m/z 699.6042     0   1   2   3E+3   11.5   12  RT (min)      lcqtof_t3_neg_1_009_50    m/z 699.6052     0   1E+3   12   12.5  RT (min)      lcqtof_t3_neg_3_032_24    m/z 699.6060     0   2   4   6   8E+2   11.5   12  RT (min)      lcqtof_t3_neg_3_032_24    m/z 699.6086     0   1E+4   12   12.5  RT (min)      lcqtof_t3_neg_3_006_23    m/z 699.6153     0   2   4E+3   13.5   14  RT (min)      lcqtof_t3_neg_1_015_34    m/z 699.6154     0   1   2E+4   13   13.5  RT (min)      lcqtof_t3_neg_2_059_14    m/z 699.6164     0   1E+2   10.5   11  RT (min)      lcqtof_t3_neg_3_038_06    m/z 700.0936     0   1   2   3E+4   12.5   13  RT (min)      lcqtof_t3_neg_4_064_17    m/z 700.0940     0   1   2   3E+4   13.5   14  RT (min)      lcqtof_t3_neg_4_036_47    m/z 700.0958     0   1   2   3   4E+4   13   13.5  RT (min)      lcqtof_t3_neg_4_064_17    m/z 700.6049     0   2   4E+3   14   14.5  RT (min)      lcqtof_t3_neg_4_019_05    m/z 700.6190     0   1E+4   12   12.5  RT (min)      lcqtof_t3_neg_2_030_23    m/z 700.6191     0   2   4   6E+3   13   13.5  RT (min)      lcqtof_t3_neg_1_006_05    m/z 700.6194     0   1E+4   12   12.5  RT (min)      lcqtof_t3_neg_2_030_23    m/z 701.0797     0   1   2   3E+4   13   13.5  RT (min)      lcqtof_t3_neg_4_064_17    m/z 701.0874     0   1E+4   12.5   13  RT (min)      lcqtof_t3_neg_4_064_17    m/z 701.0883     0   1   2E+4   12.5   13  RT (min)      lcqtof_t3_neg_4_064_17    m/z 701.0900     0   1E+4   12.5   13  RT (min)      lcqtof_t3_neg_4_064_17    m/z 701.0901     0   1   2   3E+4   12.5   13  RT (min)      lcqtof_t3_neg_4_064_17    m/z 701.0913     0   1   2   3E+4   13.5   14  RT (min)      lcqtof_t3_neg_4_002_17    m/z 701.0918     0   1   2   3E+4   13   13.5  RT (min)      lcqtof_t3_neg_4_064_17    m/z 701.2462     0   1E+4   4.5   5  RT (min)      lcqtof_t3_neg_4_000_ee    m/z 701.3919     0   1E+4   10   10.5  RT (min)      lcqtof_t3_neg_4_000_ee    m/z 701.4020     0   1E+4   10   10.5  RT (min)      lcqtof_t3_neg_4_000_ee    m/z 701.4057     0   1E+4   8   8.5  RT (min)      lcqtof_t3_neg_4_000_ee    m/z 701.4640     0   1E+2   15   15.5  RT (min)      lcqtof_t3_neg_3_032_24    m/z 701.4803     0   1   2E+4   8.5   9  RT (min)      lcqtof_t3_neg_4_000_ee    m/z 701.4808     0   1   2E+4   8.5   9  RT (min)      lcqtof_t3_neg_4_000_ee    m/z 701.4828     0   1E+4   12.5   13  RT (min)      lcqtof_t3_neg_2_016_05    m/z 701.4963     0   1   2E+4   12.5   13  RT (min)      lcqtof_t3_neg_4_018_23    m/z 701.4972     0   1E+4   12.5   13  RT (min)      lcqtof_t3_neg_2_030_23    m/z 701.4983     0   1   2E+4   12.5   13  RT (min)      lcqtof_t3_neg_4_018_23    m/z 701.5816     0   1   2   3E+2   11.5   12  RT (min)      lcqtof_t3_neg_3_032_24    m/z 701.5824     0   1E+2   10.5   11  RT (min)      lcqtof_t3_neg_3_038_06    m/z 701.5824     0   2   4   6E+4   12   12.5  RT (min)      lcqtof_t3_neg_2_030_23    m/z 701.5836     0   1E+2   10.5   11  RT (min)      lcqtof_t3_neg_3_038_06    m/z 701.5878     0   1E+4   13.5   14  RT (min)      lcqtof_t3_neg_2_021_27    m/z 701.5878     0   1E+4   13   13.5  RT (min)      lcqtof_t3_neg_1_002_13    m/z 701.5880     0   1E+4   13   13.5  RT (min)      lcqtof_t3_neg_3_037_05    m/z 701.5888     0   2   4   6E+3   16   16.5  RT (min)      lcqtof_t3_neg_1_064_14    m/z 701.5892     0   2   4   6E+3   13.5   14  RT (min)      lcqtof_t3_neg_2_053_38    m/z 701.5903     0   1   2   3E+2   11.5   12  RT (min)      lcqtof_t3_neg_3_032_24    m/z 701.5915     0   1   2   3E+2   12   12.5  RT (min)      lcqtof_t3_neg_3_032_24    m/z 701.5958     0   1E+2   10.5   11  RT (min)      lcqtof_t3_neg_3_038_06    m/z 701.5972     0   1E+2   10.5   11  RT (min)      lcqtof_t3_neg_3_038_06    m/z 701.5988     0   1E+4   13   13.5  RT (min)      lcqtof_t3_neg_3_037_05    m/z 701.6034     0   1E+2   10.5   11  RT (min)      lcqtof_t3_neg_3_038_06    m/z 702.0902     0   1E+4   13   13.5  RT (min)      lcqtof_t3_neg_4_036_47    m/z 702.0902     0   1E+4   13   13.5  RT (min)      lcqtof_t3_neg_4_064_17    m/z 702.3043     0   1E+4   9.5   10  RT (min)      lcqtof_t3_neg_4_040_24    m/z 702.3043     0   1E+4   9.5   10  RT (min)       lcqtof_t3_neg_4_090_60    m/z 702.5092     0   1   2E+2   11   11.5  RT (min)      lcqtof_t3_neg_3_038_06    m/z 702.5812     0   1E+4   13.5   14  RT (min)      lcqtof_t3_neg_2_000_aa    m/z 702.5851     0   1   2E+2   11.5   12  RT (min)      lcqtof_t3_neg_3_032_24    m/z 702.5854     0   1E+4   12   12.5  RT (min)      lcqtof_t3_neg_1_019_23    m/z 702.5860     0   1E+2   10.5   11  RT (min)      lcqtof_t3_neg_3_038_06    m/z 702.5861     0   1   2   3E+4   12   12.5  RT (min)      lcqtof_t3_neg_2_030_23    m/z 702.5867     0   1   2   3E+4   12   12.5  RT (min)      lcqtof_t3_neg_2_030_23    m/z 702.5885     0   2   4   6E+3   13.5   14  RT (min)      lcqtof_t3_neg_2_010_08    m/z 702.5887     0   1   2   3E+3   16   16.5  RT (min)      lcqtof_t3_neg_2_000_aa    m/z 702.5892     0   2   4   6E+3   14   14.5  RT (min)      lcqtof_t3_neg_1_035_31    m/z 702.5898     0   1   2   3E+4   13   13.5  RT (min)      lcqtof_t3_neg_2_030_23    m/z 702.5905     0   2   4   6E+3   13   13.5  RT (min)      lcqtof_t3_neg_2_050_01    m/z 702.5908     0   1   2   3E+3   14.5   15  RT (min)      lcqtof_t3_neg_1_008_02    m/z 702.5909     0   2   4   6E+3   13   13.5  RT (min)      lcqtof_t3_neg_2_033_53    m/z 702.5915     0   1   2E+2   11.5   12  RT (min)      lcqtof_t3_neg_3_032_24    m/z 703.4046     0   1E+5   8.5   9  RT (min)      lcqtof_t3_neg_4_000_ee    m/z 703.5104     0   1   2E+4   12.5   13  RT (min)      lcqtof_t3_neg_2_030_23    m/z 703.5836     0   1E+4   13.5   14  RT (min)      lcqtof_t3_neg_1_006_05    m/z 703.5846     0   1E+4   13   13.5  RT (min)      lcqtof_t3_neg_2_016_05    m/z 703.5920     0   1   2E+3   15.5   16  RT (min)      lcqtof_t3_neg_2_051_14    m/z 703.5922     0   1E+4   13   13.5  RT (min)      lcqtof_t3_neg_2_030_23    m/z 703.6145     0   1E+4   13   13.5  RT (min)      lcqtof_t3_neg_1_006_05    m/z 703.7328     0   1   2   3   4E+3   15   15.5  RT (min)      lcqtof_t3_neg_4_048_19    m/z 704.2416     0   2   4   6   8E+3   10   10.5  RT (min)      lcqtof_t3_neg_4_051_15    m/z 704.2572     0   1   2E+3   16   16.5  RT (min)       lcqtof_t3_neg_4_000_ee    m/z 704.2576     0   1   2E+3   15.5   16  RT (min)      lcqtof_t3_neg_4_024_26    m/z 704.2582     0   1   2E+3   15.5   16  RT (min)      lcqtof_t3_neg_4_000_ff    m/z 704.3796     0   1E+4   9   9.5  RT (min)      lcqtof_t3_neg_4_000_ee    m/z 704.4174     0   2   4E+4   8.5   9  RT (min)      lcqtof_t3_neg_4_000_ee    m/z 704.7252     0   1E+4   1   1.5  RT (min)      lcqtof_t3_neg_4_037_01    m/z 704.7368     0   1   2E+3   15.5   16  RT (min)      lcqtof_t3_neg_4_023_45    m/z 704.8344     0   1E+4   1   1.5  RT (min)      lcqtof_t3_neg_2_026_36    m/z 705.4112     0   1E+4   8.5   9  RT (min)      lcqtof_t3_neg_4_000_ee    m/z 705.4144     0   1E+4   8.5   9  RT (min)      lcqtof_t3_neg_4_000_ee    m/z 705.4170     0   1E+4   8.5   9  RT (min)      lcqtof_t3_neg_4_000_ee    m/z 705.4197     0   1E+4   8.5   9  RT (min)      lcqtof_t3_neg_4_000_ee    m/z 705.5111     0   1   2E+4   13   13.5  RT (min)      lcqtof_t3_neg_2_030_23    m/z 706.5144     0   1E+4   13   13.5  RT (min)      lcqtof_t3_neg_2_030_23    m/z 706.5740     0   2   4E+4   12.5   13  RT (min)      lcqtof_t3_neg_4_040_24    m/z 706.5748     0   1   2E+3   11.5   12  RT (min)      lcqtof_t3_neg_3_032_24    m/z 706.5749     0   1   2E+4   12   12.5  RT (min)      lcqtof_t3_neg_4_019_05    m/z 706.5761     0   2   4   6   8E+3   12   12.5  RT (min)      lcqtof_t3_neg_3_032_24    m/z 706.8130     0   1   2E+4   1   1.5  RT (min)      lcqtof_t3_neg_4_037_01    m/z 706.8184     0   1   2   3E+3   16.5   17  RT (min)      lcqtof_t3_neg_1_005_18    m/z 706.8186     0   1   2   3E+3   16.5   17  RT (min)       lcqtof_t3_neg_1_031_24    m/z 707.5270     0   1   2   3E+4   13   13.5  RT (min)      lcqtof_t3_neg_2_030_23    m/z 707.5846     0   1E+2   11.5   12  RT (min)      lcqtof_t3_neg_3_032_24    m/z 707.5849     0   1E+4   12   12.5  RT (min)      lcqtof_t3_neg_2_016_05    m/z 707.5854     0   1   2   3E+2   12   12.5  RT (min)      lcqtof_t3_neg_3_032_24    m/z 707.5855     0   1E+2   11.5   12  RT (min)      lcqtof_t3_neg_3_032_24    m/z 707.5861     0   1   2   3E+2   12   12.5  RT (min)      lcqtof_t3_neg_3_032_24    m/z 707.5877     0   1E+2   10.5   11  RT (min)      lcqtof_t3_neg_3_038_06    m/z 707.6346     0   1E+4   13.5   14  RT (min)      lcqtof_t3_neg_4_018_23    m/z 707.6852     0   1E+4   13   13.5  RT (min)      lcqtof_t3_neg_4_019_05    m/z 707.6854     0   1E+4   13   13.5  RT (min)      lcqtof_t3_neg_4_019_05    m/z 708.4615     0   1E+4   12   12.5  RT (min)      lcqtof_t3_neg_4_018_23    m/z 708.5302     0   1E+4   13   13.5  RT (min)      lcqtof_t3_neg_2_030_23    m/z 708.5313     0   1E+4   13.5   14  RT (min)      lcqtof_t3_neg_4_018_23    m/z 708.5875     0   1E+2   11.5   12  RT (min)      lcqtof_t3_neg_3_032_24    m/z 708.5878     0   1   2   3   4E+4   12   12.5  RT (min)      lcqtof_t3_neg_3_032_24    m/z 708.5880     0   1   2E+2   11.5   12  RT (min)      lcqtof_t3_neg_3_032_24    m/z 708.5882     0   1   2E+2   10.5   11  RT (min)      lcqtof_t3_neg_3_038_06    m/z 708.5885     0   1   2   3E+2   12   12.5  RT (min)      lcqtof_t3_neg_3_032_24    m/z 708.5893     0   1   2E+2   10.5   11  RT (min)      lcqtof_t3_neg_3_038_06    m/z 708.5899     0   1   2E+2   10.5   11  RT (min)      lcqtof_t3_neg_3_038_06    m/z 708.6016     0   1E+4   13.5   14  RT (min)      lcqtof_t3_neg_4_019_05    m/z 708.6284     0   1   2   3   4E+4   13.5   14  RT (min)      lcqtof_t3_neg_4_018_23    m/z 708.9748     0   2   4E+2   15   15.5  RT (min)      lcqtof_t3_neg_3_032_24    m/z 708.9762     0   1E+4   16   16.5  RT (min)      lcqtof_t3_neg_2_016_05    m/z 708.9767     0   1   2   3E+2   12   12.5  RT (min)      lcqtof_t3_neg_3_032_24    m/z 708.9770     0   2   4   6E+3   13.5   14  RT (min)      lcqtof_t3_neg_3_038_06    m/z 708.9772     0   1   2   3E+2   11   11.5  RT (min)      lcqtof_t3_neg_3_038_06    m/z 709.1673     0   1   2E+4   12   12.5  RT (min)      lcqtof_t3_neg_4_064_17    m/z 709.1682     0   1   2E+4   12   12.5  RT (min)      lcqtof_t3_neg_4_064_17    m/z 709.4153     0   1E+4   8.5   9  RT (min)      lcqtof_t3_neg_4_000_ee    m/z 709.5568     0   1E+4   13   13.5  RT (min)      lcqtof_t3_neg_2_030_23    m/z 709.5969     0   2   4E+4   12.5   13  RT (min)      lcqtof_t3_neg_4_040_24    m/z 709.6016     0   1   2   3E+4   13   13.5  RT (min)      lcqtof_t3_neg_4_011_32    m/z 709.6028     0   1   2   3E+4   13   13.5  RT (min)      lcqtof_t3_neg_4_011_32    m/z 709.6120     0   1   2E+3   12   12.5  RT (min)      lcqtof_t3_neg_3_032_24    m/z 709.6206     0   1E+3   12   12.5  RT (min)      lcqtof_t3_neg_3_032_24    m/z 709.6228     0   1   2   3E+4   13   13.5  RT (min)      lcqtof_t3_neg_2_019_32    m/z 709.6240     0   1   2E+4   12.5   13  RT (min)      lcqtof_t3_neg_2_016_05    m/z 709.6267     0   1   2E+2   11   11.5  RT (min)      lcqtof_t3_neg_3_038_06    m/z 709.6442     0   1   2E+2   10.5   11  RT (min)      lcqtof_t3_neg_3_038_06    m/z 709.6444     0   1   2E+2   11.5   12  RT (min)      lcqtof_t3_neg_3_032_24    m/z 709.6449     0   1E+5   12   12.5  RT (min)      lcqtof_t3_neg_2_030_23    m/z 709.6450     0   1   2E+2   10.5   11  RT (min)      lcqtof_t3_neg_3_038_06    m/z 709.6454     0   1   2E+2   12   12.5  RT (min)      lcqtof_t3_neg_3_032_24    m/z 709.6457     0   2   4E+4   13.5   14  RT (min)      lcqtof_t3_neg_1_019_23    m/z 709.6460     0   2   4   6   8E+4   13   13.5  RT (min)      lcqtof_t3_neg_2_050_01    m/z 709.6464     0   2   4   6E+4   13   13.5  RT (min)      lcqtof_t3_neg_2_055_16    m/z 710.1682     0   1E+4   12   12.5  RT (min)      lcqtof_t3_neg_4_064_17    m/z 710.1684     0   1E+4   12   12.5  RT (min)      lcqtof_t3_neg_4_064_17    m/z 710.2773     0   1   2E+3   10.5   11  RT (min)      lcqtof_t3_neg_1_038_15    m/z 710.4754     0   1   2   3E+4   12   12.5  RT (min)      lcqtof_t3_neg_1_019_23    m/z 710.4770     0   1   2   3   4E+4   12   12.5  RT (min)      lcqtof_t3_neg_2_030_23    m/z 710.5466     0   1E+4   13.5   14  RT (min)      lcqtof_t3_neg_1_019_23    m/z 710.6031     0   2   4   6E+4   12.5   13  RT (min)      lcqtof_t3_neg_2_016_05    m/z 710.6045     0   2   4   6   8E+2   12   12.5  RT (min)      lcqtof_t3_neg_3_032_24    m/z 710.6054     0   2   4   6   8E+2   12   12.5  RT (min)      lcqtof_t3_neg_3_032_24    m/z 710.6061     0   1E+2   11   11.5  RT (min)      lcqtof_t3_neg_3_038_06    m/z 710.6064     0   1E+5   12.5   13  RT (min)       lcqtof_t3_neg_4_040_24    m/z 710.6088     0   1E+2   10.5   11  RT (min)      lcqtof_t3_neg_3_038_06    m/z 710.6236     0   1   2E+2   12   12.5  RT (min)      lcqtof_t3_neg_3_032_24    m/z 710.6249     0   1   2E+2   11.5   12  RT (min)      lcqtof_t3_neg_3_032_24    m/z 710.6297     0   1   2E+4   13.5   14  RT (min)      lcqtof_t3_neg_4_034_34    m/z 710.6304     0   1E+4   13.5   14  RT (min)      lcqtof_t3_neg_2_016_05    m/z 710.6386     0   1   2E+2   10.5   11  RT (min)      lcqtof_t3_neg_3_038_06    m/z 710.6408     0   1   2E+2   10.5   11  RT (min)      lcqtof_t3_neg_3_038_06    m/z 710.6467     0   2   4   6E+4   12.5   13  RT (min)      lcqtof_t3_neg_2_016_05    m/z 710.6469     0   1   2E+4   13.5   14  RT (min)      lcqtof_t3_neg_1_019_23    m/z 710.6473     0   2   4   6E+4   12.5   13  RT (min)      lcqtof_t3_neg_2_016_05    m/z 710.6488     0   2   4E+4   12   12.5  RT (min)      lcqtof_t3_neg_2_030_23    m/z 710.6491     0   1E+4   13   13.5  RT (min)      lcqtof_t3_neg_1_006_05    m/z 710.6510     0   1E+4   14.5   15  RT (min)      lcqtof_t3_neg_1_016_48    m/z 710.6648     0   1   2   3E+4   14   14.5  RT (min)      lcqtof_t3_neg_4_034_34    m/z 711.1297     0   1E+4   12.5   13  RT (min)      lcqtof_t3_neg_4_001_49    m/z 711.1299     0   1   2E+4   12   12.5  RT (min)      lcqtof_t3_neg_4_057_49    m/z 711.1654     0   1E+4   12   12.5  RT (min)      lcqtof_t3_neg_4_064_17    m/z 711.1656     0   1E+4   12   12.5  RT (min)      lcqtof_t3_neg_4_064_17    m/z 711.1692     0   1E+4   12   12.5  RT (min)      lcqtof_t3_neg_4_064_17    m/z 711.2922     0   1E+4   4   4.5  RT (min)      lcqtof_t3_neg_4_000_ee    m/z 711.2929     0   1E+4   4   4.5  RT (min)      lcqtof_t3_neg_4_000_ee    m/z 711.4806     0   1E+4   12   12.5  RT (min)      lcqtof_t3_neg_2_030_23    m/z 711.5466     0   1E+4   13.5   14  RT (min)      lcqtof_t3_neg_4_018_23    m/z 711.5483     0   1E+4   13.5   14  RT (min)      lcqtof_t3_neg_4_018_23    m/z 711.5562     0   2   4E+3   13.5   14  RT (min)      lcqtof_t3_neg_2_030_23    m/z 711.6404     0   1E+4   13.5   14  RT (min)      lcqtof_t3_neg_4_018_23    m/z 711.6434     0   1   2   3E+4   13   13.5  RT (min)      lcqtof_t3_neg_1_006_05    m/z 711.6452     0   1   2   3E+4   13.5   14  RT (min)      lcqtof_t3_neg_2_016_05    m/z 711.6512     0   1E+4   12   12.5  RT (min)      lcqtof_t3_neg_2_030_23    m/z 711.6524     0   1E+4   12.5   13  RT (min)      lcqtof_t3_neg_2_016_05    m/z 711.6532     0   1E+4   12   12.5  RT (min)      lcqtof_t3_neg_2_030_23    m/z 711.6663     0   2   4   6E+3   13.5   14  RT (min)      lcqtof_t3_neg_1_019_23    m/z 712.1309     0   1E+4   12   12.5  RT (min)      lcqtof_t3_neg_4_057_49    m/z 712.4420     0   1E+4   11   11.5  RT (min)      lcqtof_t3_neg_4_037_01    m/z 712.4892     0   1   2   3E+5   12.5   13  RT (min)      lcqtof_t3_neg_2_030_23    m/z 712.4910     0   1E+2   10.5   11  RT (min)      lcqtof_t3_neg_3_038_06    m/z 712.5404     0   2   4   6   8E+3   14   14.5  RT (min)      lcqtof_t3_neg_4_030_31    m/z 713.0547     0   2   4   6   8E+4   13.5   14  RT (min)      lcqtof_t3_neg_4_002_17    m/z 713.0553     0   2   4   6E+4   14   14.5  RT (min)      lcqtof_t3_neg_4_002_17    m/z 713.0553     0   1E+5   13   13.5  RT (min)      lcqtof_t3_neg_4_064_17    m/z 713.0554     0   2   4   6   8E+4   13   13.5  RT (min)      lcqtof_t3_neg_4_036_47    m/z 713.0554     0   1E+5   13   13.5  RT (min)      lcqtof_t3_neg_4_064_17    m/z 713.0561     0   2   4   6E+4   14   14.5  RT (min)      lcqtof_t3_neg_4_002_17    m/z 713.0563     0   1E+5   13.5   14  RT (min)      lcqtof_t3_neg_4_064_17    m/z 713.0578     0   2   4   6   8E+4   13   13.5  RT (min)      lcqtof_t3_neg_4_002_17    m/z 713.1282     0   1E+4   12   12.5  RT (min)      lcqtof_t3_neg_4_057_49    m/z 713.1534     0   1E+4   4   4.5  RT (min)      lcqtof_t3_neg_4_000_ee    m/z 713.1697     0   1E+4   12   12.5  RT (min)      lcqtof_t3_neg_4_064_17    m/z 713.1783     0   1E+4   12   12.5  RT (min)      lcqtof_t3_neg_4_064_17    m/z 713.4936     0   1   2E+2   10.5   11  RT (min)      lcqtof_t3_neg_3_038_06    m/z 713.4937     0   1E+5   12.5   13  RT (min)      lcqtof_t3_neg_2_030_23    m/z 713.4942     0   1   2E+2   10.5   11  RT (min)      lcqtof_t3_neg_3_038_06    m/z 713.4942     0   1E+5   12.5   13  RT (min)      lcqtof_t3_neg_2_030_23    m/z 713.6164     0   1   2E+3   16   16.5  RT (min)      lcqtof_t3_neg_1_008_02    m/z 713.6166     0   1   2E+3   15   15.5  RT (min)      lcqtof_t3_neg_1_017_51    m/z 713.6168     0   1   2E+3   15.5   16  RT (min)      lcqtof_t3_neg_2_052_17    m/z 713.6175     0   1   2   3E+3   14.5   15  RT (min)      lcqtof_t3_neg_1_016_48    m/z 713.6175     0   1   2E+2   12.5   13  RT (min)      lcqtof_t3_neg_3_032_24    m/z 713.6190     0   1   2   3   4E+4   13   13.5  RT (min)      lcqtof_t3_neg_2_030_23    m/z 713.6194     0   2   4E+2   11.5   12  RT (min)      lcqtof_t3_neg_3_038_06    m/z 713.6309     0   2   4E+4   13   13.5  RT (min)      lcqtof_t3_neg_1_006_05    m/z 714.0554     0   2   4E+4   13   13.5  RT (min)      lcqtof_t3_neg_4_036_47    m/z 714.0558     0   2   4   6   8E+4   13.5   14  RT (min)      lcqtof_t3_neg_4_064_17    m/z 714.0568     0   2   4   6   8E+4   13   13.5  RT (min)      lcqtof_t3_neg_4_064_17    m/z 714.1480     0   1E+4   13   13.5  RT (min)      lcqtof_t3_neg_4_064_17    m/z 714.1482     0   1E+4   13   13.5  RT (min)      lcqtof_t3_neg_4_036_47    m/z 714.1489     0   1E+4   13   13.5  RT (min)      lcqtof_t3_neg_4_036_47    m/z 714.2764     0   1   2E+4   4.5   5  RT (min)      lcqtof_t3_neg_4_000_ee    m/z 714.2782     0   1   2E+4   4.5   5  RT (min)      lcqtof_t3_neg_4_000_ee    m/z 714.2860     0   1   2E+4   4   4.5  RT (min)      lcqtof_t3_neg_4_000_ee    m/z 714.4449     0   1E+4   8.5   9  RT (min)      lcqtof_t3_neg_4_000_ee    m/z 714.4450     0   1E+4   8.5   9  RT (min)      lcqtof_t3_neg_4_000_ee    m/z 714.5072     0   1   2   3E+2   11   11.5  RT (min)      lcqtof_t3_neg_3_038_06    m/z 714.5082     0   1   2   3E+2   11   11.5  RT (min)      lcqtof_t3_neg_3_038_06    m/z 714.6220     0   1E+2   12.5   13  RT (min)      lcqtof_t3_neg_3_032_24    m/z 714.6223     0   1   2E+4   14   14.5  RT (min)      lcqtof_t3_neg_1_035_31    m/z 714.6224     0   1E+2   12.5   13  RT (min)      lcqtof_t3_neg_3_032_24    m/z 714.6244     0   1   2   3E+2   11.5   12  RT (min)      lcqtof_t3_neg_3_038_06    m/z 714.6386     0   1   2E+4   12.5   13  RT (min)      lcqtof_t3_neg_2_030_23    m/z 714.6397     0   1   2E+4   12.5   13  RT (min)      lcqtof_t3_neg_2_030_23    m/z 714.7772     0   2   4   6E+4   12.5   13  RT (min)      lcqtof_t3_neg_2_030_23    m/z 714.8408     0   1E+4   17   17.5  RT (min)      lcqtof_t3_neg_4_035_08    m/z 714.8433     0   1E+4   16   16.5  RT (min)      lcqtof_t3_neg_4_018_23    m/z 714.8433     0   1E+4   16   16.5  RT (min)       lcqtof_t3_neg_4_018_23    m/z 714.8433     0   1E+3   15   15.5  RT (min)       lcqtof_t3_neg_3_032_24    m/z 714.8434     0   1E+3   15.5   16  RT (min)       lcqtof_t3_neg_1_019_23    m/z 714.8434     0   1E+4   16   16.5  RT (min)      lcqtof_t3_neg_4_053_12    m/z 714.8436     0   1   2E+3   15.5   16  RT (min)       lcqtof_t3_neg_4_109_62    m/z 714.8436     0   1E+4   16   16.5  RT (min)      lcqtof_t3_neg_4_106_62    m/z 714.9053     0   1E+4   13   13.5  RT (min)      lcqtof_t3_neg_4_034_34    m/z 714.9870     0   1   2E+4   12.5   13  RT (min)      lcqtof_t3_neg_2_030_23    m/z 714.9870     0   1   2E+4   12.5   13  RT (min)      lcqtof_t3_neg_2_030_23    m/z 715.0528     0   2   4   6E+4   12.5   13  RT (min)      lcqtof_t3_neg_4_064_17    m/z 715.0534     0   2   4   6E+4   13   13.5  RT (min)      lcqtof_t3_neg_4_064_17    m/z 715.0536     0   2   4   6E+4   13   13.5  RT (min)      lcqtof_t3_neg_4_064_17    m/z 715.0541     0   1   2   3   4E+4   13.5   14  RT (min)      lcqtof_t3_neg_4_036_47    m/z 715.0542     0   2   4   6E+4   13.5   14  RT (min)      lcqtof_t3_neg_4_064_17    m/z 715.1364     0   1E+4   12.5   13  RT (min)      lcqtof_t3_neg_2_030_23    m/z 715.4411     0   2   4   6E+4   8.5   9  RT (min)      lcqtof_t3_neg_4_000_ee    m/z 715.4809     0   1E+4   12.5   13  RT (min)      lcqtof_t3_neg_1_019_23    m/z 715.4817     0   1E+4   12   12.5  RT (min)      lcqtof_t3_neg_1_019_23    m/z 715.5102     0   1   2   3E+2   11   11.5  RT (min)      lcqtof_t3_neg_3_038_06    m/z 715.5116     0   1   2   3E+2   11   11.5  RT (min)      lcqtof_t3_neg_3_038_06    m/z 715.5651     0   1E+4   13   13.5  RT (min)      lcqtof_t3_neg_4_006_16    m/z 715.5675     0   1E+4   13   13.5  RT (min)      lcqtof_t3_neg_4_024_26    m/z 715.5751     0   1   2   3E+4   13   13.5  RT (min)       lcqtof_t3_neg_4_037_01    m/z 715.5765     0   1E+4   13   13.5  RT (min)      lcqtof_t3_neg_4_006_16    m/z 715.6087     0   1E+4   12.5   13  RT (min)      lcqtof_t3_neg_1_043_38    m/z 715.6114     0   1E+4   16   16.5  RT (min)      lcqtof_t3_neg_3_080_58    m/z 716.0528     0   1   2E+4   13   13.5  RT (min)      lcqtof_t3_neg_4_002_17    m/z 716.0528     0   1   2E+4   13.5   14  RT (min)      lcqtof_t3_neg_4_002_17    m/z 716.0529     0   1   2   3E+4   13   13.5  RT (min)      lcqtof_t3_neg_4_064_17    m/z 716.0530     0   1   2E+4   13.5   14  RT (min)      lcqtof_t3_neg_4_002_17    m/z 716.0531     0   1   2E+4   13.5   14  RT (min)      lcqtof_t3_neg_4_002_17    m/z 716.0532     0   1   2   3E+4   13.5   14  RT (min)      lcqtof_t3_neg_4_064_17    m/z 716.0536     0   1   2E+4   13   13.5  RT (min)      lcqtof_t3_neg_4_036_47    m/z 716.4462     0   1   2E+4   8.5   9  RT (min)      lcqtof_t3_neg_4_000_ee    m/z 716.5134     0   1   2E+2   11   11.5  RT (min)      lcqtof_t3_neg_3_038_06    m/z 716.5228     0   1   2E+2   11   11.5  RT (min)      lcqtof_t3_neg_3_038_06    m/z 716.5229     0   1   2E+2   11   11.5  RT (min)      lcqtof_t3_neg_3_038_06    m/z 716.5760     0   1E+4   13   13.5  RT (min)      lcqtof_t3_neg_4_037_01    m/z 716.6086     0   2   4   6   8E+3   12.5   13  RT (min)      lcqtof_t3_neg_1_034_30    m/z 716.6135     0   1   2   3   4E+3   12.5   13  RT (min)      lcqtof_t3_neg_2_036_06    m/z 716.6138     0   1   2   3   4E+3   12.5   13  RT (min)      lcqtof_t3_neg_2_036_06    m/z 716.6150     0   1E+4   12   12.5  RT (min)      lcqtof_t3_neg_1_036_14    m/z 716.6153     0   1E+4   12   12.5  RT (min)      lcqtof_t3_neg_2_059_14    m/z 716.6182     0   1   2E+4   13.5   14  RT (min)       lcqtof_t3_neg_4_040_24    m/z 716.6221     0   1   2E+4   13.5   14  RT (min)      lcqtof_t3_neg_4_040_24    m/z 716.6287     0   1   2E+4   13.5   14  RT (min)      lcqtof_t3_neg_4_040_24    m/z 716.7941     0   1   2E+4   12.5   13  RT (min)      lcqtof_t3_neg_2_030_23    m/z 717.0512     0   1   2E+4   13.5   14  RT (min)      lcqtof_t3_neg_4_064_17    m/z 717.0521     0   1   2E+4   13   13.5  RT (min)      lcqtof_t3_neg_4_064_17    m/z 717.0528     0   1   2E+4   13   13.5  RT (min)      lcqtof_t3_neg_4_064_17    m/z 717.2730     0   1E+4   1   1.5  RT (min)      lcqtof_t3_neg_4_018_23    m/z 717.2898     0   1   2E+4   4   4.5  RT (min)       lcqtof_t3_neg_4_000_ee    m/z 717.3002     0   1   2E+4   4.5   5  RT (min)      lcqtof_t3_neg_4_000_ee    m/z 717.3020     0   1   2E+4   4   4.5  RT (min)      lcqtof_t3_neg_4_000_ee    m/z 717.3024     0   1   2E+4   4.5   5  RT (min)      lcqtof_t3_neg_4_000_ee    m/z 717.5141     0   1   2   3   4E+4   12.5   13  RT (min)      lcqtof_t3_neg_2_030_23    m/z 717.5258     0   1   2   3E+2   11   11.5  RT (min)      lcqtof_t3_neg_3_038_06    m/z 717.5264     0   1   2   3E+2   11   11.5  RT (min)      lcqtof_t3_neg_3_038_06    m/z 717.5816     0   2   4E+3   13   13.5  RT (min)      lcqtof_t3_neg_4_037_01    m/z 717.6042     0   2   4   6E+3   13   13.5  RT (min)      lcqtof_t3_neg_1_065_51    m/z 717.6057     0   2   4   6   8E+3   13   13.5  RT (min)      lcqtof_t3_neg_1_008_02    m/z 717.6074     0   2   4E+3   12.5   13  RT (min)      lcqtof_t3_neg_1_043_38    m/z 717.6074     0   1E+4   13   13.5  RT (min)      lcqtof_t3_neg_2_027_15    m/z 717.6076     0   2   4   6E+3   13   13.5  RT (min)      lcqtof_t3_neg_1_065_51    m/z 717.6082     0   2   4   6E+3   14   14.5  RT (min)      lcqtof_t3_neg_1_000_aa    m/z 717.6110     0   1   2   3   4E+3   12.5   13  RT (min)      lcqtof_t3_neg_2_033_53    m/z 717.7463     0   1   2   3E+3   15   15.5  RT (min)       lcqtof_t3_neg_4_018_23    m/z 717.7472     0   1   2   3E+3   15.5   16  RT (min)      lcqtof_t3_neg_4_000_ff    m/z 717.7482     0   1   2   3E+3   15.5   16  RT (min)       lcqtof_t3_neg_4_000_ee    m/z 718.2957     0   2   4   6E+3   10.5   11  RT (min)      lcqtof_t3_neg_1_002_13    m/z 718.5322     0   1   2E+2   11   11.5  RT (min)      lcqtof_t3_neg_3_038_06    m/z 718.5346     0   1   2E+2   11   11.5  RT (min)      lcqtof_t3_neg_3_038_06    m/z 718.5537     0   1   2E+4   13   13.5  RT (min)      lcqtof_t3_neg_1_001_49    m/z 718.5718     0   1   2   3E+2   11.5   12  RT (min)      lcqtof_t3_neg_3_032_24    m/z 718.5720     0   1E+4   13   13.5  RT (min)      lcqtof_t3_neg_2_001_49    m/z 718.5722     0   1   2   3E+2   11.5   12  RT (min)      lcqtof_t3_neg_3_032_24    m/z 718.5724     0   1   2E+2   10.5   11  RT (min)      lcqtof_t3_neg_3_038_06    m/z 718.5724     0   1   2   3E+2   11.5   12  RT (min)      lcqtof_t3_neg_3_032_24    m/z 718.5726     0   2   4   6E+4   12   12.5  RT (min)      lcqtof_t3_neg_2_030_23    m/z 718.5728     0   2   4E+4   12   12.5  RT (min)      lcqtof_t3_neg_2_016_05    m/z 718.5738     0   1   2E+2   10.5   11  RT (min)      lcqtof_t3_neg_3_038_06    m/z 718.5738     0   1   2   3   4E+4   13   13.5  RT (min)      lcqtof_t3_neg_2_028_24    m/z 718.5739     0   1E+4   13   13.5  RT (min)      lcqtof_t3_neg_2_049_55    m/z 718.5744     0   2   4   6   8E+3   16.5   17  RT (min)      lcqtof_t3_neg_2_029_12    m/z 718.5780     0   2   4   6E+3   13.5   14  RT (min)      lcqtof_t3_neg_1_019_23    m/z 718.5781     0   1   2   3   4E+3   13.5   14  RT (min)      lcqtof_t3_neg_1_055_32    m/z 718.5800     0   1E+4   13   13.5  RT (min)      lcqtof_t3_neg_2_051_14    m/z 718.5804     0   2   4   6   8E+3   13.5   14  RT (min)      lcqtof_t3_neg_2_019_32    m/z 718.5811     0   2   4   6E+3   14   14.5  RT (min)      lcqtof_t3_neg_2_030_23    m/z 718.5815     0   2   4   6E+3   13.5   14  RT (min)      lcqtof_t3_neg_2_053_38    m/z 718.5818     0   2   4   6E+3   14   14.5  RT (min)      lcqtof_t3_neg_2_051_14    m/z 718.5830     0   2   4   6   8E+3   16   16.5  RT (min)      lcqtof_t3_neg_2_057_49    m/z 718.5838     0   2   4   6   8E+3   16   16.5  RT (min)      lcqtof_t3_neg_2_020_46    m/z 718.5849     0   2   4   6E+3   14   14.5  RT (min)      lcqtof_t3_neg_3_017_51    m/z 718.5850     0   2   4   6E+3   15.5   16  RT (min)      lcqtof_t3_neg_2_030_23    m/z 718.5855     0   2   4   6E+3   16   16.5  RT (min)      lcqtof_t3_neg_2_016_05    m/z 718.5858     0   2   4   6E+3   14.5   15  RT (min)      lcqtof_t3_neg_2_030_23    m/z 718.5863     0   2   4   6E+3   15   15.5  RT (min)      lcqtof_t3_neg_1_056_56    m/z 718.6100     0   1E+5   13.5   14  RT (min)      lcqtof_t3_neg_1_019_23    m/z 718.6326     0   1   2E+3   11.5   12  RT (min)      lcqtof_t3_neg_3_038_06    m/z 718.6331     0   1   2   3E+4   13   13.5  RT (min)      lcqtof_t3_neg_2_016_05    m/z 718.6339     0   2   4   6E+2   12.5   13  RT (min)      lcqtof_t3_neg_3_032_24    m/z 718.7522     0   1   2E+3   15.5   16  RT (min)       lcqtof_t3_neg_4_000_ee    m/z 718.8531     0   2   4E+3   7.5   8  RT (min)       lcqtof_t3_neg_4_000_ff    m/z 718.8534     0   2   4E+3   6.5   7  RT (min)       lcqtof_t3_neg_4_000_ff    m/z 719.0935     0   1   2   3   4E+3   9.5   10  RT (min)       lcqtof_t3_neg_4_053_12    m/z 719.2018     0   1E+4   1   1.5  RT (min)      lcqtof_t3_neg_4_047_07    m/z 719.4442     0   1E+5   8.5   9  RT (min)      lcqtof_t3_neg_4_000_ee    m/z 719.5339     0   1   2E+4   13   13.5  RT (min)      lcqtof_t3_neg_4_018_23    m/z 719.5362     0   1E+4   13   13.5  RT (min)      lcqtof_t3_neg_4_030_31    m/z 719.5366     0   1   2E+4   13   13.5  RT (min)       lcqtof_t3_neg_4_018_23    m/z 719.5630     0   1   2   3E+4   12.5   13  RT (min)      lcqtof_t3_neg_1_007_03    m/z 719.5748     0   1   2E+4   12   12.5  RT (min)      lcqtof_t3_neg_2_030_23    m/z 719.5757     0   1E+5   12.5   13  RT (min)      lcqtof_t3_neg_2_030_23    m/z 719.5760     0   1E+5   12   12.5  RT (min)      lcqtof_t3_neg_2_030_23    m/z 719.5835     0   1   2E+4   15   15.5  RT (min)      lcqtof_t3_neg_1_014_04    m/z 719.5840     0   1   2E+4   14   14.5  RT (min)      lcqtof_t3_neg_1_010_27    m/z 719.5842     0   1E+5   13   13.5  RT (min)       lcqtof_t3_neg_2_059_14    m/z 719.5843     0   1   2E+4   14.5   15  RT (min)      lcqtof_t3_neg_1_014_04    m/z 719.5843     0   1   2E+4   15   15.5  RT (min)      lcqtof_t3_neg_1_007_03    m/z 719.5844     0   1   2E+4   16   16.5  RT (min)      lcqtof_t3_neg_1_000_cc    m/z 719.5845     0   1   2   3E+4   14.5   15  RT (min)      lcqtof_t3_neg_2_059_14    m/z 719.5845     0   1   2   3E+4   13.5   14  RT (min)       lcqtof_t3_neg_1_010_27    m/z 719.5846     0   1   2   3E+4   15.5   16  RT (min)      lcqtof_t3_neg_2_059_14    m/z 719.5846     0   1   2E+4   15   15.5  RT (min)      lcqtof_t3_neg_1_030_42    m/z 719.5846     0   1   2E+4   14.5   15  RT (min)      lcqtof_t3_neg_1_000_cc    m/z 719.5847     0   1   2   3   4E+4   13.5   14  RT (min)      lcqtof_t3_neg_1_001_49    m/z 720.1490     0   1E+4   12   12.5  RT (min)      lcqtof_t3_neg_4_064_17    m/z 720.3123     0   1   2   3E+3   10.5   11  RT (min)      lcqtof_t3_neg_4_046_33    m/z 720.3130     0   1   2   3E+3   10.5   11  RT (min)       lcqtof_t3_neg_4_076_58    m/z 720.3140     0   1   2   3E+3   10   10.5  RT (min)       lcqtof_t3_neg_1_031_24    m/z 720.3156     0   1   2   3E+3   10.5   11  RT (min)       lcqtof_t3_neg_4_056_56    m/z 720.4055     0   1   2   3E+4   8.5   9  RT (min)      lcqtof_t3_neg_4_000_ee    m/z 720.4059     0   1   2   3E+4   8.5   9  RT (min)      lcqtof_t3_neg_4_000_ee    m/z 720.4348     0   1   2   3E+4   8.5   9  RT (min)      lcqtof_t3_neg_4_000_ee    m/z 720.4474     0   1   2   3E+4   8.5   9  RT (min)      lcqtof_t3_neg_4_000_ee    m/z 720.4486     0   1   2   3E+4   8.5   9  RT (min)      lcqtof_t3_neg_4_000_ee    m/z 720.5356     0   1E+4   14   14.5  RT (min)      lcqtof_t3_neg_4_018_23    m/z 720.5394     0   1   2E+3   13.5   14  RT (min)      lcqtof_t3_neg_1_019_23    m/z 720.5866     0   1   2   3E+4   12   12.5  RT (min)      lcqtof_t3_neg_3_006_23    m/z 720.5873     0   1E+2   10.5   11  RT (min)      lcqtof_t3_neg_3_038_06    m/z 720.5874     0   1E+2   10.5   11  RT (min)      lcqtof_t3_neg_3_038_06    m/z 720.5874     0   1E+4   14.5   15  RT (min)      lcqtof_t3_neg_2_062_17    m/z 720.5877     0   1E+4   14.5   15  RT (min)      lcqtof_t3_neg_2_065_51    m/z 720.5877     0   2   4   6E+4   13   13.5  RT (min)      lcqtof_t3_neg_2_059_14    m/z 720.5879     0   1E+4   16   16.5  RT (min)      lcqtof_t3_neg_2_061_50    m/z 720.5880     0   1E+4   16   16.5  RT (min)      lcqtof_t3_neg_1_003_07    m/z 720.5880     0   1E+4   16   16.5  RT (min)      lcqtof_t3_neg_1_065_51    m/z 720.5880     0   2   4   6E+4   13   13.5  RT (min)      lcqtof_t3_neg_2_065_51    m/z 720.5881     0   1E+4   15   15.5  RT (min)      lcqtof_t3_neg_2_058_15    m/z 720.5881     0   1E+4   15   15.5  RT (min)      lcqtof_t3_neg_2_061_50    m/z 720.5883     0   1E+4   14.5   15  RT (min)      lcqtof_t3_neg_2_062_17    m/z 720.5886     0   1E+4   15   15.5  RT (min)      lcqtof_t3_neg_2_065_51    m/z 720.5886     0   1E+4   15   15.5  RT (min)      lcqtof_t3_neg_2_060_39    m/z 720.5888     0   1E+4   15   15.5  RT (min)      lcqtof_t3_neg_1_064_14    m/z 720.5888     0   2   4E+4   14   14.5  RT (min)      lcqtof_t3_neg_1_018_33    m/z 720.5893     0   1E+4   13.5   14  RT (min)      lcqtof_t3_neg_2_016_05    m/z 720.6056     0   1E+2   11.5   12  RT (min)      lcqtof_t3_neg_3_038_06    m/z 720.6078     0   2   4E+4   13.5   14  RT (min)      lcqtof_t3_neg_1_019_23    m/z 720.6498     0   1   2   3   4E+1   13   13.5  RT (min)      lcqtof_t3_neg_3_032_24    m/z 720.6522     0   1E+4   13.5   14  RT (min)      lcqtof_t3_neg_2_016_05    m/z 721.1456     0   1E+4   12   12.5  RT (min)      lcqtof_t3_neg_4_064_17    m/z 721.1479     0   1E+4   12   12.5  RT (min)      lcqtof_t3_neg_4_064_17    m/z 721.3900     0   1E+4   8.5   9  RT (min)      lcqtof_t3_neg_4_000_ee    m/z 721.4001     0   1E+4   8.5   9  RT (min)      lcqtof_t3_neg_4_000_ee    m/z 721.4074     0   1E+4   8.5   9  RT (min)      lcqtof_t3_neg_4_000_ee    m/z 721.5409     0   1E+4   14   14.5  RT (min)      lcqtof_t3_neg_4_018_23    m/z 721.5683     0   1E+4   14   14.5  RT (min)      lcqtof_t3_neg_2_005_34    m/z 721.5837     0   1   2E+4   12   12.5  RT (min)      lcqtof_t3_neg_3_006_23    m/z 721.5840     0   1E+4   13.5   14  RT (min)      lcqtof_t3_neg_1_019_23    m/z 721.5845     0   1E+4   14.5   15  RT (min)      lcqtof_t3_neg_2_061_50    m/z 721.5846     0   1E+4   14.5   15  RT (min)      lcqtof_t3_neg_2_063_16    m/z 721.5846     0   2   4E+4   13   13.5  RT (min)      lcqtof_t3_neg_2_059_14    m/z 721.5847     0   1E+4   15   15.5  RT (min)      lcqtof_t3_neg_1_040_08    m/z 721.5848     0   1E+4   14   14.5  RT (min)      lcqtof_t3_neg_1_035_31    m/z 721.5848     0   2   4   6E+4   13   13.5  RT (min)      lcqtof_t3_neg_1_001_49    m/z 721.5854     0   1   2   3   4E+4   12.5   13  RT (min)      lcqtof_t3_neg_1_019_23    m/z 721.5856     0   1E+4   14.5   15  RT (min)      lcqtof_t3_neg_2_063_16    m/z 721.6052     0   1E+4   13.5   14  RT (min)      lcqtof_t3_neg_1_019_23    m/z 721.6122     0   1E+4   13.5   14  RT (min)      lcqtof_t3_neg_4_018_23    m/z 722.3000     0   1   2E+3   10.5   11  RT (min)       lcqtof_t3_neg_4_036_47    m/z 722.4754     0   1   2E+4   12   12.5  RT (min)      lcqtof_t3_neg_3_006_23    m/z 722.5844     0   1   2   3E+3   14   14.5  RT (min)      lcqtof_t3_neg_1_035_31    m/z 722.5864     0   1   2E+4   13   13.5  RT (min)      lcqtof_t3_neg_2_051_14    m/z 722.5866     0   1E+4   12.5   13  RT (min)      lcqtof_t3_neg_2_030_23    m/z 722.5868     0   1   2E+4   13   13.5  RT (min)      lcqtof_t3_neg_1_001_49    m/z 722.5874     0   1E+4   12   12.5  RT (min)      lcqtof_t3_neg_2_030_23    m/z 722.5874     0   1   2   3E+2   12   12.5  RT (min)      lcqtof_t3_neg_3_032_24    m/z 722.5884     0   1   2   3E+2   10.5   11  RT (min)      lcqtof_t3_neg_3_038_06    m/z 722.5892     0   1   2   3E+2   10.5   11  RT (min)      lcqtof_t3_neg_3_038_06    m/z 722.5906     0   1   2   3E+2   12   12.5  RT (min)      lcqtof_t3_neg_3_032_24    m/z 722.6396     0   1E+4   13.5   14  RT (min)      lcqtof_t3_neg_4_018_23    m/z 723.1580     0   1E+4   11.5   12  RT (min)      lcqtof_t3_neg_4_064_17    m/z 723.4782     0   1E+4   12   12.5  RT (min)      lcqtof_t3_neg_2_030_23    m/z 723.4789     0   1E+4   12   12.5  RT (min)      lcqtof_t3_neg_2_030_23    m/z 723.5976     0   1E+4   12.5   13  RT (min)      lcqtof_t3_neg_2_016_05    m/z 723.5988     0   2   4   6E+2   12   12.5  RT (min)      lcqtof_t3_neg_3_032_24    m/z 723.5997     0   2   4   6E+2   12   12.5  RT (min)      lcqtof_t3_neg_3_032_24    m/z 723.6018     0   1E+3   11.5   12  RT (min)      lcqtof_t3_neg_3_032_24    m/z 724.1590     0   1E+4   11.5   12  RT (min)      lcqtof_t3_neg_4_064_17    m/z 724.4369     0   1E+4   11   11.5  RT (min)      lcqtof_t3_neg_4_037_01    m/z 724.4408     0   1E+4   11   11.5  RT (min)      lcqtof_t3_neg_4_037_01    m/z 724.4894     0   1   2   3E+4   12   12.5  RT (min)      lcqtof_t3_neg_3_006_23    m/z 724.4897     0   1   2   3   4E+4   12.5   13  RT (min)       lcqtof_t3_neg_2_030_23    m/z 724.6244     0   1   2E+3   11.5   12  RT (min)      lcqtof_t3_neg_1_001_49    m/z 724.6260     0   1   2E+3   11.5   12  RT (min)      lcqtof_t3_neg_2_059_14    m/z 724.8700     0   1   2   3E+2   13.5   14  RT (min)      lcqtof_t3_neg_3_038_06    m/z 724.8714     0   2   4E+4   16   16.5  RT (min)      lcqtof_t3_neg_1_011_45    m/z 724.8717     0   2   4   6E+4   17   17.5  RT (min)      lcqtof_t3_neg_4_035_08    m/z 724.8723     0   1E+3   15.5   16  RT (min)       lcqtof_t3_neg_1_019_23    m/z 724.8727     0   1   2E+3   15.5   16  RT (min)      lcqtof_t3_neg_4_058_16    m/z 724.8748     0   1E+3   15   15.5  RT (min)      lcqtof_t3_neg_3_032_24    m/z 724.8844     0   1E+5   17   17.5  RT (min)      lcqtof_t3_neg_4_109_62    m/z 725.1293     0   1E+4   11.5   12  RT (min)       lcqtof_t3_neg_4_064_17    m/z 725.1340     0   1E+4   11.5   12  RT (min)      lcqtof_t3_neg_4_064_17    m/z 725.3858     0   1E+4   8   8.5  RT (min)      lcqtof_t3_neg_4_000_ee    m/z 725.3884     0   1E+4   8   8.5  RT (min)      lcqtof_t3_neg_4_000_ee    m/z 725.4954     0   1E+4   12.5   13  RT (min)      lcqtof_t3_neg_2_030_23    m/z 725.6389     0   1E+4   12.5   13  RT (min)       lcqtof_t3_neg_1_035_31    m/z 725.6389     0   1E+4   13.5   14  RT (min)      lcqtof_t3_neg_1_035_31    m/z 725.6394     0   1E+4   12.5   13  RT (min)       lcqtof_t3_neg_1_035_31    m/z 725.6394     0   1E+4   12.5   13  RT (min)      lcqtof_t3_neg_3_065_51    m/z 725.6396     0   1E+4   16   16.5  RT (min)      lcqtof_t3_neg_1_003_07    m/z 725.6396     0   2   4   6   8E+4   11.5   12  RT (min)       lcqtof_t3_neg_1_035_31    m/z 725.6397     0   1E+5   11.5   12  RT (min)       lcqtof_t3_neg_1_001_49    m/z 725.6398     0   1E+4   15   15.5  RT (min)       lcqtof_t3_neg_1_030_42    m/z 725.6398     0   2   4   6   8E+4   11.5   12  RT (min)       lcqtof_t3_neg_2_016_05    m/z 725.6399     0   1E+4   14.5   15  RT (min)       lcqtof_t3_neg_1_061_50    m/z 725.6400     0   1   2E+4   13   13.5  RT (min)       lcqtof_t3_neg_1_062_13    m/z 725.6402     0   1   2E+4   12.5   13  RT (min)       lcqtof_t3_neg_3_065_51    m/z 725.6402     0   1E+4   15   15.5  RT (min)      lcqtof_t3_neg_1_003_07    m/z 725.6402     0   1E+4   13.5   14  RT (min)       lcqtof_t3_neg_1_035_31    m/z 725.6403     0   1E+4   16   16.5  RT (min)       lcqtof_t3_neg_1_003_07    m/z 725.6403     0   1   2E+4   12.5   13  RT (min)      lcqtof_t3_neg_1_003_07    m/z 725.6403     0   1E+4   15   15.5  RT (min)       lcqtof_t3_neg_1_061_50    m/z 725.6404     0   1E+4   15   15.5  RT (min)       lcqtof_t3_neg_1_059_17    m/z 725.6404     0   1E+4   16   16.5  RT (min)       lcqtof_t3_neg_1_003_07    m/z 725.6404     0   1E+4   13.5   14  RT (min)       lcqtof_t3_neg_1_035_31    m/z 725.6404     0   1E+4   13.5   14  RT (min)       lcqtof_t3_neg_1_035_31    m/z 725.6404     0   1E+4   14.5   15  RT (min)       lcqtof_t3_neg_1_058_16    m/z 725.6405     0   1E+4   16   16.5  RT (min)      lcqtof_t3_neg_1_003_07    m/z 725.6406     0   1E+4   14.5   15  RT (min)       lcqtof_t3_neg_1_063_39    m/z 725.6406     0   1E+4   14   14.5  RT (min)       lcqtof_t3_neg_1_001_49    m/z 725.6406     0   1E+4   16   16.5  RT (min)      lcqtof_t3_neg_1_030_42    m/z 725.6406     0   1E+4   14   14.5  RT (min)      lcqtof_t3_neg_1_062_13    m/z 725.6407     0   1E+4   14.5   15  RT (min)       lcqtof_t3_neg_1_059_17    m/z 725.6408     0   1E+4   14   14.5  RT (min)      lcqtof_t3_neg_1_061_50    m/z 725.6408     0   1E+4   14   14.5  RT (min)       lcqtof_t3_neg_1_061_50    m/z 725.6410     0   1E+4   14.5   15  RT (min)       lcqtof_t3_neg_1_059_17    m/z 725.6414     0   1   2E+4   12.5   13  RT (min)      lcqtof_t3_neg_1_003_07    m/z 726.1343     0   1E+4   11.5   12  RT (min)      lcqtof_t3_neg_4_064_17    m/z 726.3216     0   2   4E+3   10.5   11  RT (min)      lcqtof_t3_neg_4_062_13    m/z 726.3910     0   1E+4   10   10.5  RT (min)      lcqtof_t3_neg_4_028_25    m/z 726.3912     0   1   2E+4   10   10.5  RT (min)      lcqtof_t3_neg_4_040_24    m/z 726.3914     0   1   2E+4   10   10.5  RT (min)      lcqtof_t3_neg_4_040_24    m/z 726.3916     0   1   2E+4   10   10.5  RT (min)      lcqtof_t3_neg_4_040_24    m/z 726.3917     0   1   2E+4   10   10.5  RT (min)       lcqtof_t3_neg_4_040_24    m/z 726.5057     0   1   2   3E+4   12.5   13  RT (min)      lcqtof_t3_neg_2_016_05    m/z 726.5071     0   2   4   6   8E+4   12.5   13  RT (min)      lcqtof_t3_neg_1_019_23    m/z 726.5082     0   1   2E+2   11   11.5  RT (min)      lcqtof_t3_neg_3_038_06    m/z 726.5088     0   1   2E+2   10.5   11  RT (min)      lcqtof_t3_neg_3_038_06    m/z 726.5106     0   2   4   6   8E+4   12.5   13  RT (min)      lcqtof_t3_neg_1_019_23    m/z 726.6398     0   1E+5   13   13.5  RT (min)      lcqtof_t3_neg_2_049_55    m/z 726.6398     0   2   4   6E+4   13   13.5  RT (min)      lcqtof_t3_neg_2_063_16    m/z 726.6401     0   1E+4   16.5   17  RT (min)      lcqtof_t3_neg_2_063_16    m/z 726.6402     0   2   4   6E+4   13   13.5  RT (min)      lcqtof_t3_neg_2_059_14    m/z 726.6408     0   2   4E+4   13   13.5  RT (min)      lcqtof_t3_neg_1_064_14    m/z 726.6410     0   1   2   3E+4   13.5   14  RT (min)      lcqtof_t3_neg_2_041_54    m/z 726.6410     0   1E+4   16   16.5  RT (min)      lcqtof_t3_neg_1_003_07    m/z 726.6412     0   2   4   6E+4   12.5   13  RT (min)      lcqtof_t3_neg_1_019_23    m/z 726.6413     0   1E+4   16   16.5  RT (min)      lcqtof_t3_neg_2_046_45    m/z 726.6416     0   1E+4   15   15.5  RT (min)      lcqtof_t3_neg_1_064_14    m/z 726.6417     0   1E+4   14.5   15  RT (min)      lcqtof_t3_neg_1_061_50    m/z 726.6427     0   2   4   6E+4   12.5   13  RT (min)      lcqtof_t3_neg_1_019_23    m/z 726.6428     0   1   2   3E+4   11.5   12  RT (min)      lcqtof_t3_neg_2_046_45    m/z 726.6430     0   1   2   3E+4   12   12.5  RT (min)      lcqtof_t3_neg_2_026_36    m/z 726.6434     0   1   2   3E+4   12   12.5  RT (min)      lcqtof_t3_neg_2_059_14    m/z 726.6436     0   1   2E+4   11   11.5  RT (min)      lcqtof_t3_neg_1_019_23    m/z 726.6452     0   1   2E+4   11   11.5  RT (min)      lcqtof_t3_neg_1_019_23    m/z 726.6523     0   1E+4   1   1.5  RT (min)      lcqtof_t3_neg_4_062_13    m/z 727.1084     0   1E+4   13   13.5  RT (min)      lcqtof_t3_neg_4_064_17    m/z 727.1090     0   1E+4   13   13.5  RT (min)      lcqtof_t3_neg_4_064_17    m/z 727.3758     0   1E+4   8.5   9  RT (min)      lcqtof_t3_neg_4_000_ee    m/z 727.5104     0   1   2   3E+4   12.5   13  RT (min)      lcqtof_t3_neg_1_019_23    m/z 727.5114     0   1   2   3E+4   12.5   13  RT (min)      lcqtof_t3_neg_1_019_23    m/z 727.6024     0   1E+4   12   12.5  RT (min)      lcqtof_t3_neg_3_006_23    m/z 727.6042     0   1E+4   12   12.5  RT (min)      lcqtof_t3_neg_2_016_05    m/z 727.6362     0   1E+4   13.5   14  RT (min)      lcqtof_t3_neg_2_030_23    m/z 727.6364     0   1   2   3   4E+4   13   13.5  RT (min)      lcqtof_t3_neg_2_036_06    m/z 727.6370     0   1   2E+2   10.5   11  RT (min)      lcqtof_t3_neg_3_038_06    m/z 727.6377     0   1   2E+4   13   13.5  RT (min)      lcqtof_t3_neg_2_047_28    m/z 727.6380     0   1E+4   13.5   14  RT (min)      lcqtof_t3_neg_2_016_05    m/z 727.6382     0   1   2   3   4E+4   13   13.5  RT (min)      lcqtof_t3_neg_2_022_47    m/z 727.6386     0   1   2   3E+4   13   13.5  RT (min)      lcqtof_t3_neg_2_028_24    m/z 727.6387     0   1   2   3E+3   14   14.5  RT (min)      lcqtof_t3_neg_2_036_06    m/z 727.6389     0   2   4E+3   13.5   14  RT (min)      lcqtof_t3_neg_1_015_34    m/z 727.6392     0   2   4E+3   13.5   14  RT (min)      lcqtof_t3_neg_1_035_31    m/z 727.6407     0   1   2   3   4E+3   14.5   15  RT (min)      lcqtof_t3_neg_1_015_34    m/z 727.6412     0   1   2   3E+3   15   15.5  RT (min)      lcqtof_t3_neg_1_056_56    m/z 727.6413     0   1   2   3E+3   15.5   16  RT (min)      lcqtof_t3_neg_1_040_08    m/z 727.6414     0   1   2   3E+4   12.5   13  RT (min)      lcqtof_t3_neg_2_016_05    m/z 727.6414     0   1   2   3   4E+3   16   16.5  RT (min)      lcqtof_t3_neg_1_000_dd    m/z 727.6415     0   2   4E+3   16   16.5  RT (min)      lcqtof_t3_neg_2_054_29    m/z 727.6418     0   1   2   3E+3   14.5   15  RT (min)      lcqtof_t3_neg_1_017_51    m/z 727.6422     0   1   2   3   4E+3   15   15.5  RT (min)      lcqtof_t3_neg_1_000_dd    m/z 727.6429     0   2   4E+3   14   14.5  RT (min)      lcqtof_t3_neg_2_026_36    m/z 727.6452     0   1   2   3   4E+4   12   12.5  RT (min)      lcqtof_t3_neg_2_030_23    m/z 727.6468     0   1   2   3   4E+4   12   12.5  RT (min)      lcqtof_t3_neg_2_030_23    m/z 727.6468     0   2   4E+3   12   12.5  RT (min)      lcqtof_t3_neg_3_032_24    m/z 727.6472     0   2   4E+3   12   12.5  RT (min)      lcqtof_t3_neg_3_032_24    m/z 727.6477     0   2   4   6   8E+3   11.5   12  RT (min)      lcqtof_t3_neg_2_016_05    m/z 728.2442     0   1   2E+3   10   10.5  RT (min)       lcqtof_t3_neg_4_051_15    m/z 728.5168     0   1E+4   12.5   13  RT (min)      lcqtof_t3_neg_1_019_23    m/z 728.5183     0   2   4E+4   12.5   13  RT (min)      lcqtof_t3_neg_1_019_23    m/z 728.5221     0   1   2   3E+2   11   11.5  RT (min)      lcqtof_t3_neg_3_038_06    m/z 728.5240     0   1   2   3E+2   11   11.5  RT (min)      lcqtof_t3_neg_3_038_06    m/z 728.6222     0   1E+4   13   13.5  RT (min)      lcqtof_t3_neg_2_016_05    m/z 728.6261     0   2   4   6   8E+3   12.5   13  RT (min)      lcqtof_t3_neg_2_026_36    m/z 728.6377     0   1E+4   12.5   13  RT (min)      lcqtof_t3_neg_2_065_51    m/z 728.6378     0   2   4   6   8E+3   13   13.5  RT (min)      lcqtof_t3_neg_2_026_36    m/z 728.6381     0   1E+4   12.5   13  RT (min)      lcqtof_t3_neg_2_046_45    m/z 728.6510     0   2   4   6   8E+2   11.5   12  RT (min)      lcqtof_t3_neg_3_038_06    m/z 728.9146     0   1E+4   13.5   14  RT (min)      lcqtof_t3_neg_4_034_34    m/z 729.0828     0   1E+4   13   13.5  RT (min)      lcqtof_t3_neg_4_064_17    m/z 729.0879     0   1E+4   12.5   13  RT (min)      lcqtof_t3_neg_4_064_17    m/z 729.0918     0   1E+4   13   13.5  RT (min)      lcqtof_t3_neg_4_064_17    m/z 729.0932     0   1E+4   13   13.5  RT (min)      lcqtof_t3_neg_4_064_17    m/z 729.1769     0   1E+4   12.5   13  RT (min)      lcqtof_t3_neg_4_036_47    m/z 729.1781     0   1   2E+4   13   13.5  RT (min)      lcqtof_t3_neg_4_064_17    m/z 729.1783     0   1   2E+4   13   13.5  RT (min)      lcqtof_t3_neg_4_002_17    m/z 729.2295     0   1   2E+4   1   1.5  RT (min)      lcqtof_t3_neg_4_047_07    m/z 729.4704     0   1E+4   8.5   9  RT (min)      lcqtof_t3_neg_4_000_ee    m/z 729.5241     0   1   2E+4   12.5   13  RT (min)      lcqtof_t3_neg_1_019_23    m/z 729.5257     0   1   2   3   4E+2   11   11.5  RT (min)      lcqtof_t3_neg_3_038_06    m/z 729.5274     0   1   2   3   4E+2   11   11.5  RT (min)      lcqtof_t3_neg_3_038_06    m/z 729.6126     0   1   2E+5   12   12.5  RT (min)      lcqtof_t3_neg_3_032_24    m/z 729.6128     0   2   4   6E+2   12   12.5  RT (min)      lcqtof_t3_neg_3_032_24    m/z 729.6129     0   1E+3   12   12.5  RT (min)      lcqtof_t3_neg_3_032_24    m/z 729.6134     0   2   4   6   8E+2   12   12.5  RT (min)      lcqtof_t3_neg_3_032_24    m/z 729.6134     0   2   4E+2   12   12.5  RT (min)      lcqtof_t3_neg_3_032_24    m/z 729.6137     0   2   4E+5   12   12.5  RT (min)      lcqtof_t3_neg_2_030_23    m/z 729.6138     0   1   2   3   4E+2   10.5   11  RT (min)      lcqtof_t3_neg_3_038_06    m/z 729.6139     0   1   2   3E+4   15.5   16  RT (min)       lcqtof_t3_neg_1_009_50    m/z 729.6140     0   1   2   3E+4   16.5   17  RT (min)      lcqtof_t3_neg_1_015_34    m/z 729.6140     0   1   2   3E+4   15   15.5  RT (min)       lcqtof_t3_neg_1_009_50    m/z 729.6140     0   1   2   3E+2   10.5   11  RT (min)      lcqtof_t3_neg_3_038_06    m/z 729.6140     0   2   4E+4   13.5   14  RT (min)       lcqtof_t3_neg_1_006_05    m/z 729.6140     0   1   2   3E+4   16.5   17  RT (min)      lcqtof_t3_neg_1_048_35    m/z 729.6141     0   1   2   3E+4   15.5   16  RT (min)       lcqtof_t3_neg_1_009_50    m/z 729.6142     0   1   2   3E+4   15.5   16  RT (min)       lcqtof_t3_neg_2_000_aa    m/z 729.6142     0   1   2   3E+4   15   15.5  RT (min)      lcqtof_t3_neg_1_048_35    m/z 729.6142     0   1   2   3E+4   16   16.5  RT (min)      lcqtof_t3_neg_1_007_03    m/z 729.6142     0   1   2   3E+4   15.5   16  RT (min)       lcqtof_t3_neg_2_062_17    m/z 729.6142     0   2   4   6E+4   14   14.5   15  RT (min)       lcqtof_t3_neg_1_018_33    m/z 729.6142     0   1   2   3E+4   15.5   16  RT (min)       lcqtof_t3_neg_1_009_50    m/z 729.6142     0   1   2   3E+4   15.5   16  RT (min)       lcqtof_t3_neg_1_009_50    m/z 729.6142     0   1   2   3E+4   15   15.5  RT (min)       lcqtof_t3_neg_1_009_50    m/z 729.6142     0   1   2   3E+4   15   15.5  RT (min)      lcqtof_t3_neg_1_054_41    m/z 729.6143     0   1   2   3E+4   15   15.5  RT (min)       lcqtof_t3_neg_1_016_48    m/z 729.6144     0   1   2   3E+4   14.5   15  RT (min)      lcqtof_t3_neg_1_053_01    m/z 729.6144     0   2   4E+4   16.5   17  RT (min)       lcqtof_t3_neg_1_018_33    m/z 729.6144     0   1   2   3E+4   14.5   15  RT (min)       lcqtof_t3_neg_1_008_02    m/z 729.6145     0   2   4   6E+4   14   14.5  RT (min)      lcqtof_t3_neg_1_018_33    m/z 729.6146     0   1   2   3E+4   14.5   15  RT (min)       lcqtof_t3_neg_1_006_05    m/z 729.6146     0   1   2   3E+4   14.5   15  RT (min)       lcqtof_t3_neg_1_004_11    m/z 729.6146     0   2   4   6E+4   13.5   14  RT (min)       lcqtof_t3_neg_1_035_31    m/z 729.6153     0   2   4   6   8E+4   14   14.5  RT (min)      lcqtof_t3_neg_1_018_33    m/z 729.6352     0   2   4E+2   11.5   12  RT (min)      lcqtof_t3_neg_3_038_06    m/z 729.6370     0   1E+5   13.5   14  RT (min)      lcqtof_t3_neg_1_019_23    m/z 730.1776     0   1E+4   13   13.5  RT (min)      lcqtof_t3_neg_4_002_17    m/z 730.1776     0   1E+4   13   13.5  RT (min)      lcqtof_t3_neg_4_002_17    m/z 730.1778     0   1E+4   13   13.5  RT (min)      lcqtof_t3_neg_4_002_17    m/z 730.4650     0   1   2   3E+4   11.5   12  RT (min)      lcqtof_t3_neg_1_019_23    m/z 730.4662     0   1   2   3E+4   12   12.5  RT (min)      lcqtof_t3_neg_2_030_23    m/z 730.5326     0   1   2   3E+4   12.5   13  RT (min)      lcqtof_t3_neg_1_019_23    m/z 730.5329     0   1   2   3E+4   12.5   13  RT (min)      lcqtof_t3_neg_1_019_23    m/z 730.5379     0   1   2E+2   11   11.5  RT (min)      lcqtof_t3_neg_3_038_06    m/z 730.5399     0   1   2E+2   11   11.5  RT (min)      lcqtof_t3_neg_3_038_06    m/z 730.5743     0   1E+4   13.5   14  RT (min)      lcqtof_t3_neg_4_037_01    m/z 730.6057     0   1   2   3   4E+4   13.5   14  RT (min)      lcqtof_t3_neg_1_038_15    m/z 730.6126     0   1   2   3E+2   11.5   12  RT (min)      lcqtof_t3_neg_3_038_06    m/z 730.6164     0   2   4E+2   12   12.5  RT (min)      lcqtof_t3_neg_3_032_24    m/z 730.6164     0   1E+4   15.5   16  RT (min)      lcqtof_t3_neg_1_012_16    m/z 730.6166     0   1E+4   16.5   17  RT (min)      lcqtof_t3_neg_1_012_16    m/z 730.6166     0   1E+4   16   16.5  RT (min)      lcqtof_t3_neg_1_054_41    m/z 730.6168     0   1   2E+5   12   12.5  RT (min)      lcqtof_t3_neg_2_030_23    m/z 730.6168     0   1   2E+4   16.5   17  RT (min)      lcqtof_t3_neg_2_023_37    m/z 730.6168     0   1E+4   15   15.5  RT (min)      lcqtof_t3_neg_1_061_50    m/z 730.6169     0   1E+4   14.5   15  RT (min)      lcqtof_t3_neg_1_055_32    m/z 730.6170     0   1E+4   14.5   15  RT (min)      lcqtof_t3_neg_1_004_11    m/z 730.6170     0   2   4E+2   12   12.5  RT (min)      lcqtof_t3_neg_3_032_24    m/z 730.6170     0   1E+4   16   16.5  RT (min)      lcqtof_t3_neg_2_047_28    m/z 730.6171     0   1E+4   16   16.5  RT (min)      lcqtof_t3_neg_1_000_cc    m/z 730.6172     0   2   4   6   8E+4   13.5   14  RT (min)      lcqtof_t3_neg_1_017_51    m/z 730.6172     0   1E+4   16   16.5  RT (min)      lcqtof_t3_neg_2_047_28    m/z 730.6172     0   1   2E+5   13   13.5  RT (min)      lcqtof_t3_neg_1_002_13    m/z 730.6173     0   1   2E+5   13   13.5  RT (min)      lcqtof_t3_neg_2_050_01    m/z 730.6173     0   1   2   3E+4   14   14.5  RT (min)      lcqtof_t3_neg_1_061_50    m/z 730.6174     0   1   2   3E+2   10.5   11  RT (min)      lcqtof_t3_neg_3_038_06    m/z 730.6175     0   1E+4   14.5   15  RT (min)      lcqtof_t3_neg_1_055_32    m/z 730.6176     0   1E+4   15   15.5  RT (min)      lcqtof_t3_neg_1_014_04    m/z 730.6176     0   1E+4   15.5   16  RT (min)      lcqtof_t3_neg_1_012_16    m/z 730.6178     0   1E+5   13   13.5  RT (min)      lcqtof_t3_neg_2_013_26    m/z 730.6182     0   1   2   3E+2   10.5   11  RT (min)      lcqtof_t3_neg_3_038_06    m/z 730.6270     0   1   2   3E+2   11.5   12  RT (min)      lcqtof_t3_neg_3_038_06    m/z 730.6288     0   1E+5   13.5   14  RT (min)      lcqtof_t3_neg_3_090_60    m/z 730.6460     0   1   2   3E+4   13.5   14  RT (min)       lcqtof_t3_neg_2_030_23    m/z 730.6476     0   1E+4   13   13.5  RT (min)      lcqtof_t3_neg_4_040_24    m/z 730.8190     0   1E+4   17   17.5  RT (min)       lcqtof_t3_neg_4_035_08    m/z 730.8251     0   1E+3   15.5   16  RT (min)      lcqtof_t3_neg_1_019_23    m/z 731.0672     0   2   4   6   8E+4   13.5   14  RT (min)      lcqtof_t3_neg_4_064_17    m/z 731.0677     0   2   4   6   8E+4   13   13.5  RT (min)      lcqtof_t3_neg_4_064_17    m/z 731.0683     0   2   4   6E+4   13   13.5  RT (min)      lcqtof_t3_neg_4_002_17    m/z 731.0688     0   1   2E+4   12.5   13  RT (min)      lcqtof_t3_neg_4_064_17    m/z 731.0698     0   1   2   3E+4   12.5   13  RT (min)      lcqtof_t3_neg_4_036_47    m/z 731.0705     0   2   4   6   8E+4   12.5   13  RT (min)      lcqtof_t3_neg_4_064_17    m/z 731.0716     0   1   2E+4   12.5   13  RT (min)      lcqtof_t3_neg_4_064_17    m/z 731.0841     0   1E+4   12.5   13  RT (min)      lcqtof_t3_neg_4_036_47    m/z 731.1646     0   1   2   3E+4   4   4.5  RT (min)      lcqtof_t3_neg_4_000_ee    m/z 731.1762     0   1E+4   13   13.5  RT (min)      lcqtof_t3_neg_4_064_17    m/z 731.3094     0   1E+4   4   4.5  RT (min)      lcqtof_t3_neg_4_000_ee    m/z 731.4072     0   1E+4   8.5   9  RT (min)      lcqtof_t3_neg_4_000_ee    m/z 731.4114     0   1E+4   9   9.5  RT (min)      lcqtof_t3_neg_4_000_ee    m/z 731.4692     0   1E+4   11.5   12  RT (min)      lcqtof_t3_neg_1_019_23    m/z 731.4706     0   1E+4   12   12.5  RT (min)      lcqtof_t3_neg_2_030_23    m/z 731.5376     0   1   2   3   4E+4   13   13.5  RT (min)      lcqtof_t3_neg_4_018_23    m/z 731.5402     0   1   2E+4   12.5   13  RT (min)      lcqtof_t3_neg_1_019_23    m/z 731.5406     0   1   2E+4   13   13.5  RT (min)      lcqtof_t3_neg_2_030_23    m/z 731.5414     0   1   2   3   4E+4   13   13.5  RT (min)      lcqtof_t3_neg_4_018_23    m/z 731.6187     0   2   4   6E+4   13   13.5  RT (min)      lcqtof_t3_neg_2_050_01    m/z 731.6206     0   2   4   6E+4   13   13.5  RT (min)      lcqtof_t3_neg_2_030_23    m/z 731.6208     0   2   4   6E+4   13   13.5  RT (min)      lcqtof_t3_neg_1_002_13    m/z 731.6210     0   2   4   6E+4   12   12.5  RT (min)      lcqtof_t3_neg_2_030_23    m/z 731.6367     0   1E+4   14   14.5  RT (min)      lcqtof_t3_neg_2_023_37    m/z 731.6522     0   1E+4   13.5   14  RT (min)      lcqtof_t3_neg_4_034_34    m/z 731.6525     0   1E+4   13.5   14   14.5  RT (min)      lcqtof_t3_neg_4_055_38    m/z 731.7644     0   1   2   3E+3   15.5   16  RT (min)      lcqtof_t3_neg_4_037_01    m/z 731.7652     0   1   2   3E+3   15.5   16  RT (min)      lcqtof_t3_neg_4_062_13    m/z 732.0655     0   2   4E+4   13   13.5  RT (min)      lcqtof_t3_neg_4_064_17    m/z 732.0657     0   1   2   3   4E+4   13   13.5  RT (min)      lcqtof_t3_neg_4_002_17    m/z 732.0667     0   1   2   3   4E+4   13.5   14  RT (min)      lcqtof_t3_neg_4_002_17    m/z 732.0677     0   2   4E+4   13.5   14  RT (min)      lcqtof_t3_neg_4_064_17    m/z 732.0688     0   1E+4   12.5   13  RT (min)      lcqtof_t3_neg_4_064_17    m/z 732.0699     0   1E+4   12.5   13  RT (min)      lcqtof_t3_neg_4_064_17    m/z 732.0726     0   1   2   3   4E+4   12.5   13  RT (min)      lcqtof_t3_neg_4_064_17    m/z 732.4802     0   2   4   6   8E+4   12   12.5  RT (min)      lcqtof_t3_neg_1_019_23    m/z 732.4808     0   1   2E+2   10.5   11  RT (min)      lcqtof_t3_neg_3_038_06    m/z 732.4811     0   1   2   3E+2   10.5   11  RT (min)      lcqtof_t3_neg_3_038_06    m/z 732.5470     0   1E+4   13   13.5  RT (min)      lcqtof_t3_neg_4_018_23    m/z 732.5797     0   1E+4   12   12.5  RT (min)      lcqtof_t3_neg_4_040_24    m/z 732.6201     0   1   2E+4   13.5   14  RT (min)      lcqtof_t3_neg_2_030_23    m/z 732.6272     0   1   2E+2   11.5   12  RT (min)      lcqtof_t3_neg_3_038_06    m/z 732.7703     0   1   2E+3   15.5   16  RT (min)      lcqtof_t3_neg_4_000_ee    m/z 733.0639     0   1   2   3   4E+4   13.5   14  RT (min)      lcqtof_t3_neg_4_064_17    m/z 733.0645     0   1   2   3   4E+4   13   13.5  RT (min)      lcqtof_t3_neg_4_064_17    m/z 733.0645     0   1   2E+4   13   13.5  RT (min)      lcqtof_t3_neg_4_036_47    m/z 733.0655     0   1   2   3   4E+4   13   13.5  RT (min)      lcqtof_t3_neg_4_064_17    m/z 733.0663     0   1E+4   12.5   13  RT (min)      lcqtof_t3_neg_4_064_17    m/z 733.0663     0   1   2   3   4E+4   12.5   13  RT (min)      lcqtof_t3_neg_4_064_17    m/z 733.0676     0   1E+4   12.5   13  RT (min)      lcqtof_t3_neg_4_064_17    m/z 733.0689     0   1   2   3E+4   12.5   13  RT (min)      lcqtof_t3_neg_4_064_17    m/z 733.4845     0   1   2E+4   12   12.5  RT (min)      lcqtof_t3_neg_3_006_23    m/z 733.5206     0   1   2   3E+4   12.5   13  RT (min)      lcqtof_t3_neg_4_055_38    m/z 733.5744     0   1E+4   12.5   13  RT (min)      lcqtof_t3_neg_4_011_32    m/z 733.5824     0   1E+4   12.5   13  RT (min)      lcqtof_t3_neg_4_011_32    m/z 733.5904     0   1E+4   12.5   13  RT (min)      lcqtof_t3_neg_4_040_24    m/z 733.6132     0   1   2E+4   13.5   14  RT (min)      lcqtof_t3_neg_4_018_23    m/z 733.6282     0   2   4   6   8E+3   13.5   14  RT (min)      lcqtof_t3_neg_2_030_23    m/z 734.0642     0   1E+4   13.5   14  RT (min)      lcqtof_t3_neg_4_064_17    m/z 734.0646     0   1E+4   13   13.5  RT (min)      lcqtof_t3_neg_4_064_17    m/z 734.0679     0   1E+4   13   13.5  RT (min)      lcqtof_t3_neg_4_002_17    m/z 734.4756     0   2   4   6   8E+4   12   12.5  RT (min)      lcqtof_t3_neg_3_006_23    m/z 734.4759     0   1   2   3   4E+2   10.5   11  RT (min)      lcqtof_t3_neg_3_038_06    m/z 734.4759     0   1   2   3   4E+2   10.5   11  RT (min)      lcqtof_t3_neg_3_038_06    m/z 734.5942     0   1   2   3   4E+4   12.5   13  RT (min)       lcqtof_t3_neg_4_040_24    m/z 734.5950     0   1E+4   12.5   13  RT (min)      lcqtof_t3_neg_4_028_25    m/z 734.6137     0   1   2E+3   13.5   14  RT (min)      lcqtof_t3_neg_2_030_23    m/z 735.3076     0   1E+4   4   4.5  RT (min)      lcqtof_t3_neg_4_000_ee    m/z 735.4673     0   1E+4   1   1.5  RT (min)      lcqtof_t3_neg_4_062_13    m/z 735.4796     0   2   4E+4   12   12.5  RT (min)      lcqtof_t3_neg_1_019_23    m/z 735.4810     0   2   4E+4   12   12.5  RT (min)      lcqtof_t3_neg_1_019_23    m/z 735.4818     0   1E+2   10.5   11  RT (min)      lcqtof_t3_neg_3_038_06    m/z 735.4836     0   1E+2   10.5   11  RT (min)      lcqtof_t3_neg_3_038_06    m/z 735.4868     0   1E+2   10.5   11  RT (min)      lcqtof_t3_neg_3_038_06    m/z 735.5012     0   1   2   3E+3   13.5   14  RT (min)      lcqtof_t3_neg_1_027_21    m/z 735.5782     0   1   2E+4   12.5   13  RT (min)      lcqtof_t3_neg_2_052_17    m/z 735.5782     0   1   2E+4   12.5   13  RT (min)       lcqtof_t3_neg_2_052_17    m/z 735.5792     0   2   4E+4   13   13.5  RT (min)       lcqtof_t3_neg_3_074_58    m/z 735.5793     0   1   2   3E+4   14   14.5  RT (min)       lcqtof_t3_neg_3_084_59    m/z 735.5793     0   1   2   3   4E+4   14   14.5   15  RT (min)       lcqtof_t3_neg_3_097_60    m/z 735.5794     0   1   2E+4   13.5   14  RT (min)      lcqtof_t3_neg_3_090_60    m/z 735.5796     0   1   2   3   4E+4   16   16.5  RT (min)      lcqtof_t3_neg_3_100_61    m/z 735.5796     0   1   2   3   4E+4   15   15.5  RT (min)      lcqtof_t3_neg_3_111_62    m/z 735.5797     0   1   2   3   4E+4   14   14.5  RT (min)       lcqtof_t3_neg_3_091_60    m/z 735.5798     0   1E+5   12   12.5  RT (min)      lcqtof_t3_neg_2_052_17    m/z 735.5798     0   1   2   3   4E+4   14.5   15  RT (min)      lcqtof_t3_neg_3_078_58    m/z 735.5800     0   1   2   3E+4   14   14.5  RT (min)       lcqtof_t3_neg_3_082_59    m/z 735.5802     0   1E+5   12   12.5  RT (min)      lcqtof_t3_neg_2_052_17    m/z 735.5805     0   1   2E+4   13   13.5  RT (min)      lcqtof_t3_neg_3_092_60    m/z 735.5806     0   1E+5   12   12.5  RT (min)      lcqtof_t3_neg_2_052_17    m/z 735.5810     0   1   2   3E+4   13.5   14  RT (min)      lcqtof_t3_neg_3_092_60    m/z 735.6084     0   1E+4   13   13.5  RT (min)      lcqtof_t3_neg_4_040_24    m/z 735.6086     0   1E+4   12.5   13  RT (min)      lcqtof_t3_neg_4_040_24    m/z 735.6086     0   1E+4   12.5   13  RT (min)      lcqtof_t3_neg_4_040_24    m/z 735.6096     0   1E+4   13   13.5  RT (min)      lcqtof_t3_neg_4_011_32    m/z 735.6098     0   1E+4   13   13.5  RT (min)      lcqtof_t3_neg_4_040_24    m/z 735.6244     0   1E+4   13.5   14  RT (min)       lcqtof_t3_neg_4_034_34    m/z 735.6277     0   1E+4   13.5   14  RT (min)      lcqtof_t3_neg_4_040_24    m/z 736.4908     0   1   2   3E+2   10.5   11  RT (min)      lcqtof_t3_neg_3_038_06    m/z 736.4912     0   1   2   3E+2   10.5   11  RT (min)      lcqtof_t3_neg_3_038_06    m/z 736.4925     0   1E+6   12   12.5  RT (min)       lcqtof_t3_neg_2_030_23    m/z 736.5821     0   1   2E+4   12.5   13  RT (min)      lcqtof_t3_neg_2_065_51    m/z 736.5824     0   1   2   3E+4   14   14.5  RT (min)       lcqtof_t3_neg_2_030_23    m/z 736.5825     0   1E+4   12.5   13  RT (min)      lcqtof_t3_neg_2_053_38    m/z 736.5825     0   1E+4   12.5   13  RT (min)      lcqtof_t3_neg_1_048_35    m/z 736.5827     0   1E+4   12   12.5  RT (min)      lcqtof_t3_neg_2_019_32    m/z 736.5828     0   2   4   6E+4   12   12.5  RT (min)      lcqtof_t3_neg_1_043_38    m/z 736.5828     0   1E+4   16   16.5  RT (min)      lcqtof_t3_neg_3_079_58    m/z 736.5828     0   1   2   3   4E+4   12   12.5  RT (min)      lcqtof_t3_neg_1_019_23    m/z 736.5830     0   2   4   6E+4   12   12.5  RT (min)      lcqtof_t3_neg_1_015_34    m/z 736.5831     0   1   2   3   4E+4   14.5   15  RT (min)       lcqtof_t3_neg_2_061_50    m/z 736.5832     0   1   2E+4   16   16.5  RT (min)      lcqtof_t3_neg_3_076_58    m/z 736.5832     0   1   2E+4   15   15.5  RT (min)      lcqtof_t3_neg_3_108_62    m/z 736.5833     0   1   2E+4   13   13.5  RT (min)      lcqtof_t3_neg_3_074_58    m/z 736.5834     0   1   2E+4   14.5   15  RT (min)      lcqtof_t3_neg_3_097_60    m/z 736.5834     0   2   4   6E+4   12   12.5  RT (min)      lcqtof_t3_neg_1_044_37    m/z 736.5838     0   1E+4   16   16.5  RT (min)      lcqtof_t3_neg_3_066_57    m/z 736.5838     0   1E+4   13.5   14  RT (min)      lcqtof_t3_neg_3_090_60    m/z 736.5839     0   1   2   3   4E+4   11   11.5  RT (min)      lcqtof_t3_neg_1_019_23    m/z 736.5839     0   1   2   3   4E+4   11   11.5  RT (min)      lcqtof_t3_neg_1_019_23    m/z 736.5840     0   1E+4   13.5   14  RT (min)      lcqtof_t3_neg_3_093_60    m/z 736.5840     0   1E+4   14   14.5  RT (min)      lcqtof_t3_neg_3_074_58    m/z 736.5846     0   1   2E+4   14.5   15  RT (min)      lcqtof_t3_neg_3_081_58    m/z 736.6255     0   1   2   3E+4   13.5   14  RT (min)       lcqtof_t3_neg_4_034_34    m/z 736.6256     0   1   2E+4   13.5   14  RT (min)      lcqtof_t3_neg_4_018_23    m/z 736.6263     0   1   2E+4   13.5   14  RT (min)      lcqtof_t3_neg_4_003_37    m/z 736.6270     0   1   2E+4   12   12.5  RT (min)      lcqtof_t3_neg_2_030_23    m/z 736.6466     0   2   4   6   8E+3   13.5   14  RT (min)      lcqtof_t3_neg_1_019_23    m/z 736.6598     0   1   2   3E+3   13.5   14  RT (min)      lcqtof_t3_neg_1_019_23    m/z 736.6617     0   1   2E+4   13.5   14  RT (min)      lcqtof_t3_neg_4_018_23    m/z 736.7656     0   2   4E+4   12.5   13  RT (min)      lcqtof_t3_neg_1_019_23    m/z 736.7658     0   2   4   6E+4   12   12.5  RT (min)      lcqtof_t3_neg_2_030_23    m/z 736.8954     0   1E+4   12.5   13  RT (min)      lcqtof_t3_neg_4_034_34    m/z 736.9794     0   1E+4   12   12.5  RT (min)      lcqtof_t3_neg_2_030_23    m/z 737.1328     0   1E+4   12.5   13  RT (min)      lcqtof_t3_neg_4_034_34    m/z 737.2547     0   2   4   6E+3   11.5   12  RT (min)       lcqtof_t3_neg_4_058_16    m/z 737.4088     0   1   2E+4   8.5   9  RT (min)      lcqtof_t3_neg_4_000_ee    m/z 737.4140     0   1   2E+4   8.5   9  RT (min)      lcqtof_t3_neg_4_000_ee    m/z 737.4476     0   2   4   6E+3   12.5   13  RT (min)      lcqtof_t3_neg_1_015_34    m/z 737.4647     0   1E+4   1   1.5  RT (min)      lcqtof_t3_neg_4_062_13    m/z 737.4942     0   1   2   3   4E+2   10.5   11  RT (min)      lcqtof_t3_neg_3_038_06    m/z 737.4942     0   1   2   3   4E+2   10.5   11  RT (min)      lcqtof_t3_neg_3_038_06    m/z 737.5042     0   1   2   3   4E+3   13.5   14  RT (min)      lcqtof_t3_neg_4_033_53    m/z 737.5767     0   1E+4   12.5   13  RT (min)      lcqtof_t3_neg_2_057_49    m/z 737.5767     0   2   4   6   8E+3   12.5   13  RT (min)      lcqtof_t3_neg_3_046_36    m/z 737.5786     0   1   2   3E+4   11.5   12  RT (min)      lcqtof_t3_neg_2_016_05    m/z 737.5789     0   2   4   6E+4   12   12.5  RT (min)      lcqtof_t3_neg_1_036_14    m/z 737.5791     0   1   2   3E+2   11   11.5  RT (min)      lcqtof_t3_neg_3_032_24    m/z 737.5791     0   1   2E+2   10.5   11  RT (min)      lcqtof_t3_neg_3_032_24    m/z 737.5792     0   1E+4   16   16.5  RT (min)      lcqtof_t3_neg_3_095_60    m/z 737.5792     0   1   2E+2   10.5   11  RT (min)      lcqtof_t3_neg_3_032_24    m/z 737.5794     0   1   2   3E+4   12   12.5  RT (min)      lcqtof_t3_neg_2_030_23    m/z 737.5794     0   1   2   3E+4   11.5   12  RT (min)      lcqtof_t3_neg_2_016_05    m/z 737.5795     0   2   4E+4   12   12.5  RT (min)      lcqtof_t3_neg_1_018_33    m/z 737.5795     0   1   2E+4   13   13.5  RT (min)      lcqtof_t3_neg_2_049_55    m/z 737.5796     0   1E+4   14.5   15  RT (min)      lcqtof_t3_neg_3_098_61    m/z 737.5797     0   1E+4   14   14.5  RT (min)      lcqtof_t3_neg_3_090_60    m/z 737.5798     0   1   2   3E+4   14   14.5  RT (min)       lcqtof_t3_neg_2_030_23    m/z 737.5801     0   1   2   3E+2   11   11.5  RT (min)      lcqtof_t3_neg_3_032_24    m/z 737.5802     0   1   2   3E+2   11   11.5  RT (min)      lcqtof_t3_neg_3_032_24    m/z 737.5929     0   1E+4   13   13.5  RT (min)      lcqtof_t3_neg_1_023_06    m/z 737.5935     0   1E+4   13   13.5  RT (min)      lcqtof_t3_neg_1_023_06    m/z 737.6206     0   1E+4   12.5   13  RT (min)      lcqtof_t3_neg_4_028_25    m/z 737.6227     0   1   2   3E+4   13   13.5  RT (min)      lcqtof_t3_neg_4_040_24    m/z 737.6229     0   1E+4   13.5   14  RT (min)      lcqtof_t3_neg_4_008_35    m/z 737.6254     0   1E+4   13.5   14  RT (min)      lcqtof_t3_neg_4_034_34    m/z 737.6331     0   1E+4   13.5   14  RT (min)      lcqtof_t3_neg_4_040_24    m/z 737.6659     0   1E+4   13.5   14  RT (min)      lcqtof_t3_neg_4_018_23    m/z 738.3058     0   1   2   3E+3   11   11.5  RT (min)      lcqtof_t3_neg_1_010_27    m/z 738.4986     0   1   2   3E+2   10.5   11  RT (min)      lcqtof_t3_neg_3_038_06    m/z 738.5064     0   1E+5   12   12.5  RT (min)      lcqtof_t3_neg_3_006_23    m/z 738.5066     0   1   2   3E+2   10.5   11  RT (min)      lcqtof_t3_neg_3_038_06    m/z 738.5073     0   1   2E+5   12   12.5  RT (min)      lcqtof_t3_neg_2_030_23    m/z 738.5076     0   1   2   3   4E+4   12   12.5  RT (min)      lcqtof_t3_neg_1_019_23    m/z 738.5080     0   1E+6   12.5   13  RT (min)       lcqtof_t3_neg_2_030_23    m/z 738.5084     0   1   2   3E+2   11   11.5  RT (min)      lcqtof_t3_neg_3_038_06    m/z 738.5793     0   2   4   6E+3   13.5   14  RT (min)      lcqtof_t3_neg_2_016_05    m/z 738.5801     0   1   2   3E+3   12.5   13  RT (min)      lcqtof_t3_neg_1_011_45    m/z 738.5801     0   1   2   3   4E+3   12.5   13  RT (min)      lcqtof_t3_neg_2_043_31    m/z 738.5806     0   1E+4   15   15.5  RT (min)       lcqtof_t3_neg_2_031_02    m/z 738.5806     0   1E+4   11   11.5  RT (min)      lcqtof_t3_neg_2_030_23    m/z 738.5807     0   2   4   6E+3   13   13.5  RT (min)      lcqtof_t3_neg_2_056_56    m/z 738.5808     0   1   2   3   4E+3   12.5   13  RT (min)      lcqtof_t3_neg_1_033_53    m/z 738.5812     0   1   2E+4   12   12.5  RT (min)      lcqtof_t3_neg_1_043_38    m/z 738.6222     0   1E+2   12   12.5  RT (min)      lcqtof_t3_neg_3_032_24    m/z 738.6228     0   1E+2   12   12.5  RT (min)      lcqtof_t3_neg_3_032_24    m/z 738.6247     0   1E+2   12   12.5  RT (min)      lcqtof_t3_neg_3_032_24    m/z 738.6254     0   1   2E+4   12.5   13  RT (min)      lcqtof_t3_neg_2_016_05    m/z 738.6258     0   1E+2   12   12.5  RT (min)      lcqtof_t3_neg_3_032_24    m/z 738.6267     0   1   2E+2   11   11.5  RT (min)      lcqtof_t3_neg_3_038_06    m/z 738.6275     0   1   2E+2   11   11.5  RT (min)      lcqtof_t3_neg_3_038_06    m/z 738.6344     0   1E+4   13.5   14  RT (min)      lcqtof_t3_neg_4_011_32    m/z 738.6374     0   1   2E+4   13.5   14  RT (min)       lcqtof_t3_neg_4_040_24    m/z 738.6598     0   1E+4   14   14.5  RT (min)      lcqtof_t3_neg_4_018_23    m/z 738.6830     0   1   2   3E+3   13   13.5  RT (min)      lcqtof_t3_neg_2_050_01    m/z 738.7822     0   1   2   3E+4   12.5   13  RT (min)      lcqtof_t3_neg_2_030_23    m/z 739.3338     0   1E+4   8.5   9  RT (min)      lcqtof_t3_neg_4_000_ee    m/z 739.4540     0   1E+4   12.5   13  RT (min)      lcqtof_t3_neg_1_001_49    m/z 739.4762     0   1E+4   12.5   13  RT (min)      lcqtof_t3_neg_1_003_07    m/z 739.4858     0   1E+4   12.5   13  RT (min)      lcqtof_t3_neg_2_056_56    m/z 739.5098     0   1   2   3E+2   10.5   11  RT (min)      lcqtof_t3_neg_3_038_06    m/z 739.5110     0   1   2E+2   11   11.5  RT (min)      lcqtof_t3_neg_3_038_06    m/z 739.5111     0   2   4E+5   12.5   13  RT (min)      lcqtof_t3_neg_2_030_23    m/z 740.3058     0   1E+4   4   4.5  RT (min)      lcqtof_t3_neg_4_000_ee    m/z 740.4702     0   1   2   3   4E+4   11   11.5  RT (min)       lcqtof_t3_neg_4_037_01    m/z 740.4707     0   1   2   3   4E+4   11   11.5  RT (min)      lcqtof_t3_neg_4_037_01    m/z 740.4719     0   1   2   3   4E+4   11   11.5  RT (min)      lcqtof_t3_neg_4_037_01    m/z 740.4915     0   1E+2   4.5   5  RT (min)      lcqtof_t3_neg_3_038_06    m/z 740.5168     0   1   2E+5   12.5   13  RT (min)      lcqtof_t3_neg_2_030_23    m/z 740.5174     0   1   2E+5   12.5   13  RT (min)      lcqtof_t3_neg_2_030_23    m/z 740.5185     0   1   2   3E+2   10.5   11  RT (min)      lcqtof_t3_neg_3_038_06    m/z 740.5203     0   1E+5   12.5   13  RT (min)      lcqtof_t3_neg_2_030_23    m/z 740.5223     0   1   2   3E+2   11   11.5  RT (min)      lcqtof_t3_neg_3_038_06    m/z 740.5224     0   1E+6   12.5   13  RT (min)       lcqtof_t3_neg_2_030_23    m/z 740.5226     0   1   2   3E+2   11   11.5  RT (min)      lcqtof_t3_neg_3_038_06    m/z 740.6366     0   1E+4   12.5   13  RT (min)      lcqtof_t3_neg_2_016_05    m/z 740.6380     0   1   2E+4   12.5   13  RT (min)      lcqtof_t3_neg_2_016_05    m/z 740.6388     0   2   4   6E+2   12   12.5  RT (min)      lcqtof_t3_neg_3_032_24    m/z 740.6389     0   1E+4   12.5   13  RT (min)      lcqtof_t3_neg_2_016_05    m/z 740.6394     0   2   4   6E+2   12.5   13  RT (min)      lcqtof_t3_neg_3_032_24    m/z 740.6408     0   2   4E+2   11   11.5  RT (min)      lcqtof_t3_neg_3_038_06    m/z 740.6430     0   1   2E+2   11   11.5  RT (min)      lcqtof_t3_neg_3_038_06    m/z 740.6771     0   1E+4   14   14.5  RT (min)      lcqtof_t3_neg_4_018_23    m/z 740.7973     0   1   2E+4   12.5   13  RT (min)      lcqtof_t3_neg_2_030_23    m/z 740.8430     0   1   2E+2   13.5   14  RT (min)      lcqtof_t3_neg_3_038_06    m/z 740.8438     0   2   4   6E+2   15   15.5  RT (min)      lcqtof_t3_neg_3_032_24    m/z 740.8451     0   2   4   6   8E+2   15.5   16  RT (min)      lcqtof_t3_neg_1_019_23    m/z 740.8466     0   1   2   3E+4   17   17.5  RT (min)      lcqtof_t3_neg_4_035_08    m/z 741.2068     0   1E+4   4.5   5  RT (min)      lcqtof_t3_neg_4_000_ee    m/z 741.4751     0   1E+4   11   11.5  RT (min)      lcqtof_t3_neg_4_037_01    m/z 741.4954     0   1E+2   4.5   5  RT (min)      lcqtof_t3_neg_3_038_06    m/z 741.5210     0   2   4   6E+4   12.5   13  RT (min)      lcqtof_t3_neg_1_019_23    m/z 741.5230     0   1   2   3E+2   10.5   11  RT (min)      lcqtof_t3_neg_3_038_06    m/z 741.5240     0   1   2   3E+2   10.5   11  RT (min)      lcqtof_t3_neg_3_038_06    m/z 741.5246     0   1   2   3   4E+5   12.5   13  RT (min)       lcqtof_t3_neg_2_030_23    m/z 741.5248     0   2   4   6E+4   12.5   13  RT (min)      lcqtof_t3_neg_2_030_23    m/z 741.5257     0   1   2   3E+2   11   11.5  RT (min)      lcqtof_t3_neg_3_038_06    m/z 741.5258     0   1   2   3E+2   11   11.5  RT (min)      lcqtof_t3_neg_3_038_06    m/z 742.2812     0   1   2   3   4E+3   10   10.5  RT (min)       lcqtof_t3_neg_4_062_13    m/z 742.4517     0   1E+4   10.5   11  RT (min)      lcqtof_t3_neg_4_037_01    m/z 742.5354     0   1   2   3E+5   12.5   13  RT (min)      lcqtof_t3_neg_2_030_23    m/z 742.5368     0   1   2E+2   11   11.5  RT (min)      lcqtof_t3_neg_3_038_06    m/z 742.5376     0   1   2E+2   11   11.5  RT (min)      lcqtof_t3_neg_3_038_06    m/z 742.5390     0   1   2E+2   11   11.5  RT (min)      lcqtof_t3_neg_3_038_06    m/z 742.6292     0   1E+4   14.5   15  RT (min)      lcqtof_t3_neg_2_061_50    m/z 742.6298     0   1   2   3   4E+4   11   11.5  RT (min)      lcqtof_t3_neg_1_019_23    m/z 742.6300     0   1   2   3E+4   12.5   13  RT (min)      lcqtof_t3_neg_2_043_31    m/z 742.6301     0   1   2E+4   12.5   13  RT (min)      lcqtof_t3_neg_1_015_34    m/z 742.6302     0   1   2   3E+4   12.5   13  RT (min)      lcqtof_t3_neg_2_030_23    m/z 742.6302     0   1   2   3E+4   12   12.5  RT (min)      lcqtof_t3_neg_1_019_23    m/z 742.6304     0   2   4E+4   12   12.5  RT (min)      lcqtof_t3_neg_2_030_23    m/z 742.6305     0   1   2   3   4E+4   12   12.5  RT (min)      lcqtof_t3_neg_2_030_23    m/z 742.6306     0   1E+4   15.5   16  RT (min)      lcqtof_t3_neg_2_030_23    m/z 742.6306     0   1   2   3E+4   11   11.5  RT (min)      lcqtof_t3_neg_3_006_23    m/z 742.6307     0   2   4E+4   11   11.5  RT (min)      lcqtof_t3_neg_2_030_23    m/z 742.6307     0   1   2E+4   12   12.5  RT (min)      lcqtof_t3_neg_1_019_23    m/z 742.6308     0   1   2   3E+4   13   13.5  RT (min)      lcqtof_t3_neg_2_005_34    m/z 742.6308     0   1   2   3E+4   12.5   13  RT (min)      lcqtof_t3_neg_2_043_31    m/z 742.6308     0   1E+4   16.5   17  RT (min)      lcqtof_t3_neg_2_043_31    m/z 742.6310     0   1E+4   16   16.5  RT (min)      lcqtof_t3_neg_1_003_07    m/z 742.6311     0   1E+4   15.5   16  RT (min)      lcqtof_t3_neg_2_030_23    m/z 742.6311     0   1E+4   15   15.5  RT (min)      lcqtof_t3_neg_2_061_50    m/z 742.6312     0   2   4E+4   12   12.5  RT (min)      lcqtof_t3_neg_3_039_33    m/z 742.6312     0   1E+4   16   16.5  RT (min)      lcqtof_t3_neg_1_003_07    m/z 742.6312     0   1   2   3E+4   13.5   14  RT (min)      lcqtof_t3_neg_2_015_33    m/z 742.6313     0   1   2   3   4E+4   14.5   15  RT (min)      lcqtof_t3_neg_2_061_50    m/z 742.6313     0   1E+4   16   16.5  RT (min)      lcqtof_t3_neg_2_015_33    m/z 742.6314     0   1E+4   16   16.5  RT (min)      lcqtof_t3_neg_2_023_37    m/z 742.6315     0   1   2   3   4E+4   11   11.5  RT (min)      lcqtof_t3_neg_1_019_23    m/z 742.6316     0   1   2   3E+4   13.5   14  RT (min)      lcqtof_t3_neg_1_015_34    m/z 742.6316     0   1   2   3   4E+4   14   14.5  RT (min)      lcqtof_t3_neg_2_061_50    m/z 742.6333     0   2   4E+4   14.5   15  RT (min)      lcqtof_t3_neg_2_061_50    m/z 742.6345     0   1E+4   16.5   17  RT (min)      lcqtof_t3_neg_2_023_37    m/z 742.6361     0   1E+5   13.5   14  RT (min)      lcqtof_t3_neg_1_043_38    m/z 742.6484     0   1   2   3E+4   12.5   13  RT (min)      lcqtof_t3_neg_2_030_23    m/z 742.6540     0   2   4E+4   13.5   14  RT (min)      lcqtof_t3_neg_2_030_23    m/z 742.6703     0   1E+4   13   13.5  RT (min)      lcqtof_t3_neg_4_018_23    m/z 742.6742     0   1E+4   13   13.5  RT (min)      lcqtof_t3_neg_4_034_34    m/z 743.0296     0   1E+4   13   13.5  RT (min)      lcqtof_t3_neg_4_018_23    m/z 743.1380     0   2   4   6E+4   12.5   13  RT (min)      lcqtof_t3_neg_4_002_17    m/z 743.1382     0   1E+5   12.5   13  RT (min)      lcqtof_t3_neg_4_064_17    m/z 743.1385     0   2   4   6E+4   14   14.5  RT (min)      lcqtof_t3_neg_4_002_17    m/z 743.1386     0   1   2   3   4E+5   13   13.5  RT (min)      lcqtof_t3_neg_4_002_17    m/z 743.1388     0   2   4   6E+4   12.5   13  RT (min)      lcqtof_t3_neg_4_002_17    m/z 743.1389     0   1   2E+5   12.5   13  RT (min)      lcqtof_t3_neg_4_064_17    m/z 743.1390     0   1E+5   14   14.5  RT (min)      lcqtof_t3_neg_4_002_17    m/z 743.1390     0   1E+5   14   14.5  RT (min)      lcqtof_t3_neg_4_002_17    m/z 743.1396     0   1E+5   14   14.5  RT (min)      lcqtof_t3_neg_4_002_17    m/z 743.1405     0   1   2   3   4E+5   13   13.5  RT (min)      lcqtof_t3_neg_4_036_47    m/z 743.2566     0   1E+4   4.5   5  RT (min)      lcqtof_t3_neg_4_000_ee    m/z 743.3618     0   1E+4   1   1.5  RT (min)      lcqtof_t3_neg_4_040_24    m/z 743.4836     0   1   2E+4   12   12.5  RT (min)      lcqtof_t3_neg_1_019_23    m/z 743.5401     0   1E+5   12.5   13  RT (min)      lcqtof_t3_neg_2_030_23    m/z 743.5408     0   1E+2   11   11.5  RT (min)      lcqtof_t3_neg_3_038_06    m/z 743.5416     0   1E+2   11   11.5  RT (min)      lcqtof_t3_neg_3_038_06    m/z 743.5429     0   1E+2   11   11.5  RT (min)      lcqtof_t3_neg_3_038_06    m/z 743.6012     0   1   2E+2   10   10.5  RT (min)      lcqtof_t3_neg_3_038_06    m/z 743.6056     0   1E+3   11   11.5  RT (min)      lcqtof_t3_neg_3_032_24    m/z 743.6070     0   1   2E+4   11.5   12  RT (min)      lcqtof_t3_neg_3_006_23    m/z 743.6161     0   1E+4   16   16.5  RT (min)      lcqtof_t3_neg_1_011_45    m/z 743.6163     0   2   4   6   8E+3   13.5   14  RT (min)      lcqtof_t3_neg_1_019_23    m/z 743.6164     0   1E+4   16   16.5  RT (min)      lcqtof_t3_neg_2_056_56    m/z 743.6166     0   1E+4   16   16.5  RT (min)      lcqtof_t3_neg_2_046_45    m/z 743.6166     0   1E+4   12.5   13  RT (min)      lcqtof_t3_neg_2_023_37    m/z 743.6167     0   1E+4   16   16.5  RT (min)      lcqtof_t3_neg_2_016_05    m/z 743.6175     0   1E+4   14   14.5  RT (min)      lcqtof_t3_neg_2_053_38    m/z 743.6176     0   1E+4   15   15.5  RT (min)      lcqtof_t3_neg_2_054_29    m/z 743.6180     0   1   2E+4   12   12.5  RT (min)      lcqtof_t3_neg_2_030_23    m/z 743.6182     0   2   4   6   8E+3   14.5   15  RT (min)      lcqtof_t3_neg_1_035_31    m/z 743.6186     0   1E+4   14   14.5  RT (min)      lcqtof_t3_neg_1_009_50    m/z 743.6186     0   1E+4   14   14.5  RT (min)      lcqtof_t3_neg_2_030_23    m/z 743.6186     0   1E+4   16   16.5  RT (min)      lcqtof_t3_neg_1_016_48    m/z 743.6191     0   1E+4   15   15.5  RT (min)      lcqtof_t3_neg_2_030_23    m/z 743.6194     0   1E+4   15   15.5  RT (min)      lcqtof_t3_neg_2_045_04    m/z 743.6198     0   1E+4   16   16.5  RT (min)      lcqtof_t3_neg_2_056_56    m/z 743.6198     0   1E+4   15   15.5  RT (min)      lcqtof_t3_neg_2_044_44    m/z 743.6198     0   1   2E+4   14   14.5  RT (min)      lcqtof_t3_neg_1_018_33    m/z 743.6200     0   1   2   3   4E+2   11   11.5  RT (min)      lcqtof_t3_neg_3_032_24    m/z 743.6212     0   1E+4   14   14.5  RT (min)      lcqtof_t3_neg_1_044_37    m/z 743.6212     0   1E+4   14.5   15  RT (min)      lcqtof_t3_neg_1_008_02    m/z 743.6216     0   1E+4   15   15.5  RT (min)      lcqtof_t3_neg_1_008_02    m/z 743.6222     0   1E+4   14.5   15  RT (min)      lcqtof_t3_neg_1_009_50    m/z 743.6238     0   1E+4   14.5   15  RT (min)      lcqtof_t3_neg_2_039_22    m/z 743.6262     0   1   2E+2   9.5   10  RT (min)      lcqtof_t3_neg_3_038_06    m/z 743.6271     0   2   4   6E+2   11   11.5  RT (min)      lcqtof_t3_neg_3_032_24    m/z 743.6290     0   1E+4   16.5   17  RT (min)      lcqtof_t3_neg_2_054_29    m/z 743.6328     0   1   2E+2   9.5   10  RT (min)      lcqtof_t3_neg_3_038_06    m/z 743.6330     0   1   2   3   4E+4   11   11.5  RT (min)      lcqtof_t3_neg_2_016_05    m/z 743.6338     0   1   2   3E+2   10.5   11  RT (min)      lcqtof_t3_neg_3_032_24    m/z 743.6361     0   1   2   3   4E+4   13.5   14  RT (min)       lcqtof_t3_neg_4_040_24    m/z 743.6426     0   1   2E+4   13.5   14  RT (min)      lcqtof_t3_neg_4_028_25    m/z 743.6566     0   1   2   3E+4   13.5   14  RT (min)      lcqtof_t3_neg_2_030_23    m/z 743.6574     0   1   2   3E+2   11.5   12  RT (min)      lcqtof_t3_neg_3_038_06    m/z 743.6595     0   2   4   6   8E+4   13.5   14  RT (min)      lcqtof_t3_neg_4_018_23    m/z 744.1386     0   2   4   6   8E+4   14   14.5  RT (min)      lcqtof_t3_neg_4_036_47    m/z 744.1388     0   1E+5   12.5   13  RT (min)      lcqtof_t3_neg_4_064_17    m/z 744.1389     0   1   2   3E+4   14   14.5  RT (min)      lcqtof_t3_neg_4_064_17    m/z 744.1390     0   1   2   3   4E+4   12.5   13  RT (min)      lcqtof_t3_neg_4_002_17    m/z 744.1390     0   1E+5   12.5   13  RT (min)      lcqtof_t3_neg_4_064_17    m/z 744.1397     0   2   4   6   8E+4   14   14.5  RT (min)      lcqtof_t3_neg_4_002_17    m/z 744.1398     0   1   2   3   4E+4   12.5   13  RT (min)      lcqtof_t3_neg_4_002_17    m/z 744.1398     0   1   2   3   4E+4   12.5   13  RT (min)      lcqtof_t3_neg_4_002_17    m/z 744.1399     0   1   2   3E+5   12.5   13  RT (min)      lcqtof_t3_neg_4_064_17    m/z 744.1404     0   1   2   3   4E+4   12.5   13  RT (min)      lcqtof_t3_neg_4_002_17    m/z 744.1411     0   1E+5   14   14.5  RT (min)      lcqtof_t3_neg_4_036_47    m/z 744.4600     0   1E+4   1   1.5  RT (min)      lcqtof_t3_neg_4_037_01    m/z 744.4813     0   1   2   3E+2   10.5   11  RT (min)      lcqtof_t3_neg_3_038_06    m/z 744.4824     0   1   2   3E+2   10.5   11  RT (min)      lcqtof_t3_neg_3_038_06    m/z 744.4824     0   1E+5   12   12.5  RT (min)      lcqtof_t3_neg_1_019_23    m/z 744.4828     0   1   2   3E+2   10   10.5  RT (min)      lcqtof_t3_neg_3_038_06    m/z 744.5454     0   1   2   3E+4   12.5   13  RT (min)      lcqtof_t3_neg_2_030_23    m/z 744.5468     0   1   2E+2   11   11.5  RT (min)      lcqtof_t3_neg_3_038_06    m/z 744.5468     0   1E+4   12.5   13  RT (min)      lcqtof_t3_neg_2_016_05    m/z 744.5480     0   1   2E+2   11   11.5  RT (min)      lcqtof_t3_neg_3_038_06    m/z 744.5548     0   1E+2   11   11.5  RT (min)      lcqtof_t3_neg_3_038_06    m/z 744.5554     0   1E+2   11   11.5  RT (min)      lcqtof_t3_neg_3_038_06    m/z 744.5809     0   1E+4   12   12.5  RT (min)       lcqtof_t3_neg_2_030_23    m/z 744.5882     0   1E+4   13   13.5  RT (min)      lcqtof_t3_neg_1_007_03    m/z 744.5883     0   1   2E+2   10.5   11  RT (min)      lcqtof_t3_neg_3_038_06    m/z 744.5982     0   1E+4   13.5   14  RT (min)      lcqtof_t3_neg_1_010_27    m/z 744.6000     0   1E+4   13.5   14  RT (min)      lcqtof_t3_neg_1_036_14    m/z 744.6018     0   2   4E+3   15.5   16  RT (min)      lcqtof_t3_neg_2_063_16    m/z 744.6020     0   2   4E+3   16   16.5  RT (min)      lcqtof_t3_neg_1_006_05    m/z 744.6022     0   2   4E+3   16   16.5  RT (min)      lcqtof_t3_neg_2_048_30    m/z 744.6024     0   1E+4   13.5   14  RT (min)      lcqtof_t3_neg_4_018_23    m/z 744.6024     0   2   4   6E+3   16.5   17  RT (min)      lcqtof_t3_neg_2_063_16    m/z 744.6030     0   1E+4   11.5   12  RT (min)      lcqtof_t3_neg_2_016_05    m/z 744.6042     0   1   2   3   4E+3   14.5   15  RT (min)      lcqtof_t3_neg_1_015_34    m/z 744.6042     0   1   2   3   4E+3   14.5   15  RT (min)      lcqtof_t3_neg_1_013_09    m/z 744.6045     0   2   4   6E+3   14   14.5  RT (min)      lcqtof_t3_neg_1_023_06    m/z 744.6046     0   2   4E+3   15.5   16  RT (min)      lcqtof_t3_neg_2_065_51    m/z 744.6065     0   2   4E+3   14.5   15  RT (min)      lcqtof_t3_neg_1_017_51    m/z 744.6112     0   1E+4   12   12.5  RT (min)      lcqtof_t3_neg_1_019_23    m/z 744.6120     0   1E+4   12   12.5  RT (min)      lcqtof_t3_neg_1_019_23    m/z 744.6163     0   1E+4   11.5   12  RT (min)      lcqtof_t3_neg_2_016_05    m/z 744.6178     0   2   4   6   8E+3   12.5   13  RT (min)      lcqtof_t3_neg_2_041_54    m/z 744.6355     0   2   4   6E+3   11   11.5  RT (min)      lcqtof_t3_neg_3_038_06    m/z 744.6377     0   1   2   3E+4   13   13.5  RT (min)      lcqtof_t3_neg_2_016_05    m/z 744.6400     0   1   2E+3   12.5   13  RT (min)      lcqtof_t3_neg_3_032_24    m/z 744.6614     0   1   2E+4   13.5   14  RT (min)      lcqtof_t3_neg_4_018_23    m/z 745.1173     0   2   4   6   8E+4   12   12.5  RT (min)      lcqtof_t3_neg_4_064_17    m/z 745.1323     0   1   2   3E+4   12.5   13  RT (min)      lcqtof_t3_neg_4_036_47    m/z 745.1326     0   1   2   3E+5   12.5   13  RT (min)      lcqtof_t3_neg_4_064_17    m/z 745.1338     0   1   2   3E+4   12.5   13  RT (min)      lcqtof_t3_neg_4_036_47    m/z 745.1344     0   2   4   6   8E+4   12.5   13  RT (min)      lcqtof_t3_neg_4_064_17    m/z 745.1344     0   1   2E+5   13   13.5  RT (min)      lcqtof_t3_neg_4_002_17    m/z 745.1362     0   2   4   6E+4   14   14.5  RT (min)      lcqtof_t3_neg_4_002_17    m/z 745.1362     0   2   4   6E+4   14   14.5  RT (min)      lcqtof_t3_neg_4_002_17    m/z 745.1366     0   1   2E+4   14   14.5  RT (min)      lcqtof_t3_neg_4_002_17    m/z 745.1368     0   2   4   6E+4   14   14.5  RT (min)      lcqtof_t3_neg_4_002_17    m/z 745.1371     0   1E+5   14   14.5  RT (min)      lcqtof_t3_neg_4_064_17    m/z 745.1383     0   2   4   6   8E+4   14   14.5  RT (min)      lcqtof_t3_neg_4_036_47    m/z 745.4840     0   1   2   3   4E+4   12   12.5  RT (min)      lcqtof_t3_neg_1_019_23    m/z 745.4864     0   1E+4   8.5   9  RT (min)      lcqtof_t3_neg_4_000_ee    m/z 745.4882     0   1E+4   8.5   9  RT (min)      lcqtof_t3_neg_4_000_ee    m/z 745.4990     0   1   2E+2   10.5   11  RT (min)      lcqtof_t3_neg_3_038_06    m/z 745.5007     0   1E+5   12   12.5  RT (min)      lcqtof_t3_neg_1_019_23    m/z 745.5024     0   1   2E+2   10.5   11  RT (min)      lcqtof_t3_neg_3_038_06    m/z 745.5578     0   1E+5   13.5   14  RT (min)      lcqtof_t3_neg_4_003_37    m/z 745.5580     0   2   4   6E+4   13.5   14  RT (min)      lcqtof_t3_neg_4_001_49    m/z 745.5584     0   1E+5   13.5   14  RT (min)       lcqtof_t3_neg_1_035_31    m/z 745.5944     0   2   4   6E+4   13.5   14  RT (min)       lcqtof_t3_neg_1_006_05    m/z 745.6059     0   2   4   6E+4   12.5   13  RT (min)       lcqtof_t3_neg_2_053_38    m/z 745.6059     0   2   4   6   8E+4   12.5   13  RT (min)       lcqtof_t3_neg_2_065_51    m/z 745.6068     0   2   4   6E+4   12.5   13  RT (min)      lcqtof_t3_neg_2_023_37    m/z 745.6069     0   2   4   6E+4   12.5   13  RT (min)      lcqtof_t3_neg_2_002_35    m/z 745.6070     0   1E+5   13   13.5  RT (min)      lcqtof_t3_neg_1_008_02    m/z 745.6078     0   2   4   6E+4   14.5   15  RT (min)      lcqtof_t3_neg_1_002_13    m/z 745.6082     0   2   4   6   8E+4   16   16.5  RT (min)       lcqtof_t3_neg_1_000_cc    m/z 745.6083     0   2   4   6E+4   15.5   16  RT (min)       lcqtof_t3_neg_1_006_05    m/z 745.6083     0   2   4   6   8E+4   16   16.5  RT (min)       lcqtof_t3_neg_1_000_cc    m/z 745.6083     0   2   4   6   8E+4   16   16.5  RT (min)       lcqtof_t3_neg_1_000_cc    m/z 745.6084     0   2   4   6E+4   14.5   15  RT (min)       lcqtof_t3_neg_1_006_05    m/z 745.6084     0   2   4   6E+4   15.5   16  RT (min)       lcqtof_t3_neg_1_004_11    m/z 745.6084     0   2   4   6E+4   14.5   15  RT (min)       lcqtof_t3_neg_1_006_05    m/z 745.6084     0   2   4   6E+4   15   15.5  RT (min)       lcqtof_t3_neg_1_005_18    m/z 745.6084     0   2   4   6E+4   14.5   15  RT (min)       lcqtof_t3_neg_1_007_03    m/z 745.6084     0   2   4   6E+4   15   15.5  RT (min)      lcqtof_t3_neg_1_005_18    m/z 745.6084     0   2   4   6   8E+4   16   16.5  RT (min)       lcqtof_t3_neg_1_000_cc    m/z 745.6084     0   2   4   6E+4   15   15.5  RT (min)       lcqtof_t3_neg_1_005_18    m/z 745.6084     0   2   4   6E+4   14   14.5  RT (min)       lcqtof_t3_neg_1_016_48    m/z 745.6084     0   2   4   6E+4   15   15.5  RT (min)       lcqtof_t3_neg_1_001_49    m/z 745.6085     0   2   4   6E+4   14.5   15  RT (min)       lcqtof_t3_neg_1_006_05    m/z 745.6086     0   2   4   6E+4   16   16.5  RT (min)      lcqtof_t3_neg_1_004_11    m/z 745.6086     0   2   4   6E+4   15   15.5  RT (min)      lcqtof_t3_neg_1_028_20    m/z 745.6086     0   2   4   6E+4   16   16.5  RT (min)      lcqtof_t3_neg_1_013_09    m/z 745.6086     0   2   4   6E+4   14.5   15  RT (min)       lcqtof_t3_neg_1_006_05    m/z 745.6088     0   2   4E+4   14   14.5  RT (min)       lcqtof_t3_neg_1_041_54    m/z 745.6088     0   2   4   6E+4   14   14.5  RT (min)       lcqtof_t3_neg_1_016_48    m/z 745.6089     0   2   4   6   8E+4   12.5   13  RT (min)       lcqtof_t3_neg_1_055_32    m/z 745.6092     0   2   4E+4   12.5   13  RT (min)      lcqtof_t3_neg_3_001_49    m/z 745.6092     0   2   4   6E+4   14.5   15  RT (min)      lcqtof_t3_neg_1_002_13    m/z 745.6096     0   1   2   3E+5   12   12.5  RT (min)      lcqtof_t3_neg_1_029_44    m/z 745.6456     0   1   2   3E+4   14   14.5  RT (min)      lcqtof_t3_neg_4_046_33    m/z 745.6568     0   1   2   3   4E+4   13.5   14  RT (min)      lcqtof_t3_neg_4_011_32    m/z 745.6572     0   2   4E+4   13.5   14  RT (min)      lcqtof_t3_neg_4_040_24    m/z 745.7790     0   1   2E+3   15.5   16  RT (min)      lcqtof_t3_neg_4_031_42    m/z 745.7797     0   1   2   3E+3   15.5   16  RT (min)       lcqtof_t3_neg_4_000_ee    m/z 746.1286     0   1E+5   13   13.5  RT (min)      lcqtof_t3_neg_4_036_47    m/z 746.1317     0   1E+5   13   13.5  RT (min)      lcqtof_t3_neg_4_036_47    m/z 746.1322     0   1E+4   12.5   13  RT (min)      lcqtof_t3_neg_4_036_47    m/z 746.1326     0   2   4   6   8E+4   12.5   13  RT (min)      lcqtof_t3_neg_4_064_17    m/z 746.1330     0   1E+4   12.5   13  RT (min)      lcqtof_t3_neg_4_036_47    m/z 746.1332     0   1E+4   12.5   13  RT (min)      lcqtof_t3_neg_4_036_47    m/z 746.1356     0   1   2   3E+4   14   14.5  RT (min)      lcqtof_t3_neg_4_002_17    m/z 746.1360     0   1   2   3   4E+4   14   14.5  RT (min)      lcqtof_t3_neg_4_064_17    m/z 746.1362     0   1E+4   12.5   13  RT (min)      lcqtof_t3_neg_4_036_47    m/z 746.1366     0   1   2   3E+4   14   14.5  RT (min)      lcqtof_t3_neg_4_002_17    m/z 746.4894     0   1E+4   12   12.5  RT (min)      lcqtof_t3_neg_1_019_23    m/z 746.4996     0   1   2   3E+2   10.5   11  RT (min)      lcqtof_t3_neg_3_038_06    m/z 746.5012     0   1   2E+2   10.5   11  RT (min)      lcqtof_t3_neg_3_038_06    m/z 746.5261     0   2   4   6   8E+4   12.5   13  RT (min)      lcqtof_t3_neg_4_033_53    m/z 746.5832     0   2   4   6E+4   12.5   13  RT (min)      lcqtof_t3_neg_2_019_32    m/z 746.5935     0   2   4   6   8E+4   13.5   14  RT (min)       lcqtof_t3_neg_3_083_59    m/z 746.6045     0   2   4   6E+2   10.5   11  RT (min)      lcqtof_t3_neg_3_038_06    m/z 746.6050     0   2   4E+2   10.5   11  RT (min)      lcqtof_t3_neg_3_038_06    m/z 746.6059     0   1   2   3   4E+5   13   13.5  RT (min)       lcqtof_t3_neg_2_050_01    m/z 746.6065     0   2   4   6   8E+4   16.5   17  RT (min)       lcqtof_t3_neg_2_058_15    m/z 746.6068     0   2   4E+5   13   13.5  RT (min)      lcqtof_t3_neg_2_046_45    m/z 746.6070     0   2   4   6E+4   16.5   17  RT (min)      lcqtof_t3_neg_2_023_37    m/z 746.6075     0   1   2   3E+5   13.5   14  RT (min)       lcqtof_t3_neg_1_019_23    m/z 746.6078     0   1E+5   13.5   14  RT (min)      lcqtof_t3_neg_2_053_38    m/z 746.6079     0   2   4   6E+4   14   14.5  RT (min)      lcqtof_t3_neg_2_016_05    m/z 746.6080     0   2   4   6E+4   16   16.5  RT (min)       lcqtof_t3_neg_2_046_45    m/z 746.6080     0   2   4   6E+4   14.5   15  RT (min)       lcqtof_t3_neg_1_014_04    m/z 746.6081     0   2   4   6E+4   16   16.5  RT (min)      lcqtof_t3_neg_2_033_53    m/z 746.6081     0   2   4   6E+4   14.5   15  RT (min)       lcqtof_t3_neg_1_012_16    m/z 746.6081     0   2   4   6E+4   16   16.5  RT (min)      lcqtof_t3_neg_2_046_45    m/z 746.6081     0   2   4   6E+4   14.5   15  RT (min)       lcqtof_t3_neg_1_011_45    m/z 746.6082     0   2   4E+4   13.5   14  RT (min)      lcqtof_t3_neg_1_039_43    m/z 746.6082     0   2   4   6E+4   16   16.5  RT (min)       lcqtof_t3_neg_1_011_45    m/z 746.6082     0   2   4   6E+4   14.5   15  RT (min)      lcqtof_t3_neg_1_012_16    m/z 746.6083     0   2   4   6E+4   15   15.5  RT (min)      lcqtof_t3_neg_2_059_14    m/z 746.6083     0   2   4   6E+4   15   15.5  RT (min)       lcqtof_t3_neg_1_014_04    m/z 746.6084     0   2   4   6E+4   16   16.5  RT (min)       lcqtof_t3_neg_1_011_45    m/z 746.6084     0   2   4   6E+4   15   15.5  RT (min)       lcqtof_t3_neg_2_051_14    m/z 746.6084     0   2   4   6E+4   14   15  RT (min)       lcqtof_t3_neg_2_016_05    m/z 746.6084     0   2   4   6E+4   14.5   15  RT (min)       lcqtof_t3_neg_2_061_50    m/z 746.6084     0   2   4   6E+4   15   15.5  RT (min)       lcqtof_t3_neg_1_008_02    m/z 746.6084     0   2   4   6E+4   14   14.5  RT (min)      lcqtof_t3_neg_1_011_45    m/z 746.6088     0   1E+5   13.5   14  RT (min)       lcqtof_t3_neg_2_056_56    m/z 746.6090     0   2   4   6E+4   16   16.5  RT (min)      lcqtof_t3_neg_1_009_50    m/z 746.6092     0   2   4E+4   15   15.5  RT (min)      lcqtof_t3_neg_1_019_23    m/z 746.6092     0   2   4E+4   14.5   15  RT (min)      lcqtof_t3_neg_1_045_29    m/z 746.6096     0   2   4E+5   12.5   13  RT (min)      lcqtof_t3_neg_2_030_23    m/z 746.6110     0   1E+5   12   12.5  RT (min)      lcqtof_t3_neg_3_015_15    m/z 746.6112     0   1   2   3   4E+5   12   12.5  RT (min)      lcqtof_t3_neg_2_030_23    m/z 746.6118     0   1   2   3E+2   10   10.5  RT (min)      lcqtof_t3_neg_3_038_06    m/z 746.6118     0   1E+5   12   12.5  RT (min)      lcqtof_t3_neg_2_042_13    m/z 746.6128     0   1   2   3E+2   9.5   10  RT (min)      lcqtof_t3_neg_3_038_06    m/z 746.6130     0   1   2E+2   9.5   10  RT (min)      lcqtof_t3_neg_3_038_06    m/z 746.6213     0   1E+5   13.5   14  RT (min)      lcqtof_t3_neg_2_061_50    m/z 746.6289     0   2   4   6E+4   13   13.5  RT (min)      lcqtof_t3_neg_2_016_05    m/z 746.6308     0   2   4   6E+4   13.5   14  RT (min)      lcqtof_t3_neg_2_030_23    m/z 746.6439     0   1E+5   13.5   14  RT (min)       lcqtof_t3_neg_2_019_32    m/z 746.6575     0   1E+5   13.5   14  RT (min)       lcqtof_t3_neg_4_040_24    m/z 747.0106     0   1E+4   1   1.5  RT (min)      lcqtof_t3_neg_4_059_14    m/z 747.1158     0   2   4   6E+4   13   13.5  RT (min)      lcqtof_t3_neg_4_064_17    m/z 747.1277     0   2   4   6E+4   12.5   13  RT (min)      lcqtof_t3_neg_4_064_17    m/z 747.4409     0   2   4   6E+4   8.5   9  RT (min)      lcqtof_t3_neg_4_000_ee    m/z 747.4410     0   2   4   6E+4   8.5   9  RT (min)      lcqtof_t3_neg_4_000_ee    m/z 747.5023     0   1   2   3   4E+4   12   12.5  RT (min)      lcqtof_t3_neg_1_019_23    m/z 747.5108     0   2   4   6   8E+3   13   13.5  RT (min)      lcqtof_t3_neg_2_019_32    m/z 747.5130     0   1   2E+2   10.5   11  RT (min)      lcqtof_t3_neg_3_038_06    m/z 747.5154     0   1   2E+5   12   12.5  RT (min)      lcqtof_t3_neg_1_019_23    m/z 747.5160     0   1   2E+2   10.5   11  RT (min)      lcqtof_t3_neg_3_038_06    m/z 747.5900     0   1   2E+4   12.5   13  RT (min)      lcqtof_t3_neg_4_040_24    m/z 747.6056     0   1   2E+5   12   12.5  RT (min)      lcqtof_t3_neg_2_030_23    m/z 747.6084     0   1   2E+5   12.5   13  RT (min)      lcqtof_t3_neg_2_030_23    m/z 747.6086     0   1   2   3E+4   16.5   17  RT (min)      lcqtof_t3_neg_2_028_24    m/z 747.6090     0   1   2   3E+4   16.5   17  RT (min)      lcqtof_t3_neg_2_036_06    m/z 747.6092     0   1   2   3   4E+5   13   13.5  RT (min)      lcqtof_t3_neg_2_049_55    m/z 747.6093     0   2   4   6E+4   13   13.5  RT (min)       lcqtof_t3_neg_2_050_01    m/z 747.6096     0   1   2   3E+5   13   13.5  RT (min)       lcqtof_t3_neg_2_017_51    m/z 747.6097     0   1   2E+5   13   13.5  RT (min)      lcqtof_t3_neg_2_036_06    m/z 747.6098     0   1   2E+4   14.5   15  RT (min)      lcqtof_t3_neg_1_033_53    m/z 747.6101     0   1   2E+4   13.5   14  RT (min)      lcqtof_t3_neg_1_055_32    m/z 747.6103     0   1   2E+4   13.5   14  RT (min)      lcqtof_t3_neg_1_031_24    m/z 747.6104     0   1   2E+4   15.5   16  RT (min)      lcqtof_t3_neg_2_030_23    m/z 747.6104     0   1   2E+4   16   16.5  RT (min)      lcqtof_t3_neg_1_011_45    m/z 747.6104     0   1   2E+4   15.5   16  RT (min)      lcqtof_t3_neg_2_030_23    m/z 747.6106     0   1   2E+4   15.5   16  RT (min)      lcqtof_t3_neg_2_030_23    m/z 747.6106     0   1   2E+4   15   15.5  RT (min)      lcqtof_t3_neg_1_014_04    m/z 747.6106     0   1   2E+4   16   16.5  RT (min)      lcqtof_t3_neg_1_011_45    m/z 747.6110     0   1   2E+4   15   15.5  RT (min)      lcqtof_t3_neg_2_061_50    m/z 747.6110     0   1   2   3E+4   14   14.5  RT (min)      lcqtof_t3_neg_2_049_55    m/z 747.6112     0   1   2E+4   15   15.5  RT (min)      lcqtof_t3_neg_1_014_04    m/z 747.6119     0   1   2E+5   12.5   13  RT (min)      lcqtof_t3_neg_2_030_23    m/z 747.6162     0   1   2   3E+4   11   11.5  RT (min)      lcqtof_t3_neg_1_019_23    m/z 747.6261     0   1   2   3   4E+4   13.5   14  RT (min)      lcqtof_t3_neg_3_039_33    m/z 747.6383     0   1   2E+4   13.5   14  RT (min)      lcqtof_t3_neg_2_030_23    m/z 747.6665     0   1   2E+4   13.5   14  RT (min)      lcqtof_t3_neg_2_016_05    m/z 747.6712     0   1   2E+2   13   13.5  RT (min)      lcqtof_t3_neg_3_032_24    m/z 748.1136     0   1   2E+4   13   13.5  RT (min)      lcqtof_t3_neg_4_064_17    m/z 748.1250     0   1   2E+4   12.5   13  RT (min)      lcqtof_t3_neg_4_064_17    m/z 748.1254     0   1E+4   12.5   13  RT (min)      lcqtof_t3_neg_4_064_17    m/z 748.1338     0   1   2E+4   13.5   14  RT (min)      lcqtof_t3_neg_4_002_17    m/z 748.1354     0   1   2   3E+4   13   13.5  RT (min)      lcqtof_t3_neg_4_064_17    m/z 748.1355     0   1   2E+4   13   13.5  RT (min)      lcqtof_t3_neg_4_036_47    m/z 748.3003     0   2   4E+3   10   10.5  RT (min)      lcqtof_t3_neg_4_062_13    m/z 748.3024     0   2   4E+3   10   10.5  RT (min)       lcqtof_t3_neg_4_014_13    m/z 748.4448     0   1   2E+4   8.5   9  RT (min)      lcqtof_t3_neg_4_000_ee    m/z 748.5163     0   1E+2   10.5   11  RT (min)      lcqtof_t3_neg_3_038_06    m/z 748.5192     0   2   4   6   8E+4   12   12.5  RT (min)      lcqtof_t3_neg_1_019_23    m/z 748.5407     0   1   2E+4   12.5   13  RT (min)      lcqtof_t3_neg_4_014_13    m/z 748.5432     0   1   2E+5   12.5   13  RT (min)      lcqtof_t3_neg_4_034_34    m/z 748.6095     0   1   2E+2   10.5   11  RT (min)      lcqtof_t3_neg_3_038_06    m/z 748.6095     0   1   2E+2   10.5   11  RT (min)      lcqtof_t3_neg_3_038_06    m/z 748.6107     0   1   2   3E+4   12   12.5  RT (min)      lcqtof_t3_neg_3_006_23    m/z 748.6118     0   2   4   6E+3   16.5   17  RT (min)      lcqtof_t3_neg_2_051_14    m/z 748.6136     0   1E+4   13   13.5  RT (min)      lcqtof_t3_neg_2_063_16    m/z 748.6140     0   2   4   6E+3   13.5   14  RT (min)      lcqtof_t3_neg_1_006_05    m/z 748.6142     0   2   4E+3   15   15.5  RT (min)      lcqtof_t3_neg_2_060_39    m/z 748.6144     0   2   4   6   8E+3   14   14.5  RT (min)      lcqtof_t3_neg_1_016_48    m/z 748.6155     0   2   4E+3   13.5   14  RT (min)      lcqtof_t3_neg_1_031_24    m/z 748.6206     0   2   4E+3   12   12.5  RT (min)      lcqtof_t3_neg_3_032_24    m/z 748.6714     0   1E+2   13   13.5  RT (min)      lcqtof_t3_neg_3_032_24    m/z 748.6728     0   1   2   3   4E+4   13.5   14  RT (min)      lcqtof_t3_neg_2_016_05    m/z 748.6736     0   1E+2   11.5   12  RT (min)      lcqtof_t3_neg_3_038_06    m/z 748.8266     0   1   2E+4   1   1.5  RT (min)      lcqtof_t3_neg_1_036_14    m/z 749.1336     0   1E+4   13   13.5  RT (min)      lcqtof_t3_neg_4_064_17    m/z 749.4538     0   1E+4   8.5   9  RT (min)      lcqtof_t3_neg_4_000_ee    m/z 749.5240     0   1E+4   12   12.5  RT (min)      lcqtof_t3_neg_1_019_23    m/z 749.5476     0   1   2   3   4E+4   12.5   13  RT (min)      lcqtof_t3_neg_4_003_37    m/z 749.5520     0   1   2   3E+4   12.5   13  RT (min)      lcqtof_t3_neg_4_030_31    m/z 749.6060     0   2   4   6E+2   11.5   12  RT (min)      lcqtof_t3_neg_3_032_24    m/z 749.6067     0   2   4   6E+2   12   12.5  RT (min)      lcqtof_t3_neg_3_032_24    m/z 749.6085     0   1E+2   10.5   11  RT (min)      lcqtof_t3_neg_3_038_06    m/z 749.6118     0   1   2E+4   12   12.5  RT (min)      lcqtof_t3_neg_2_016_05    m/z 749.6123     0   1   2E+4   12   12.5  RT (min)      lcqtof_t3_neg_2_016_05    m/z 749.6145     0   1   2E+4   12.5   13  RT (min)      lcqtof_t3_neg_2_016_05    m/z 749.6416     0   2   4E+3   13.5   14  RT (min)      lcqtof_t3_neg_2_030_23    m/z 749.6431     0   1E+4   13.5   14  RT (min)      lcqtof_t3_neg_4_018_23    m/z 750.3318     0   1   2   3E+3   11   11.5  RT (min)      lcqtof_t3_neg_4_051_15    m/z 750.5102     0   1   2E+4   12.5   13  RT (min)      lcqtof_t3_neg_4_018_23    m/z 750.5105     0   1   2E+4   12   12.5  RT (min)      lcqtof_t3_neg_2_030_23    m/z 750.5112     0   1   2E+4   12   12.5  RT (min)      lcqtof_t3_neg_2_030_23    m/z 750.5114     0   1E+4   12   12.5  RT (min)      lcqtof_t3_neg_3_006_23    m/z 750.5146     0   2   4   6   8E+3   13.5   14  RT (min)       lcqtof_t3_neg_2_053_38    m/z 750.6162     0   1   2   3E+2   12   12.5  RT (min)      lcqtof_t3_neg_3_032_24    m/z 750.6182     0   2   4E+2   12   12.5  RT (min)      lcqtof_t3_neg_3_032_24    m/z 750.6182     0   2   4E+2   12   12.5  RT (min)      lcqtof_t3_neg_3_032_24    m/z 750.6192     0   1   2E+2   10.5   11  RT (min)      lcqtof_t3_neg_3_038_06    m/z 750.6210     0   1   2E+4   12.5   13  RT (min)      lcqtof_t3_neg_2_016_05    m/z 750.6415     0   1E+4   13.5   14  RT (min)       lcqtof_t3_neg_4_034_34    m/z 750.6460     0   1E+4   14   14.5  RT (min)      lcqtof_t3_neg_4_003_37    m/z 750.6826     0   1E+4   13.5   14  RT (min)      lcqtof_t3_neg_2_016_05    m/z 750.6892     0   1E+2   13   13.5  RT (min)      lcqtof_t3_neg_3_032_24    m/z 751.1598     0   1E+4   11.5   12  RT (min)      lcqtof_t3_neg_4_064_17    m/z 751.1604     0   1E+4   11.5   12  RT (min)      lcqtof_t3_neg_4_064_17    m/z 751.5166     0   1E+4   12.5   13  RT (min)      lcqtof_t3_neg_4_034_34    m/z 751.5312     0   1   2   3E+4   14   14.5  RT (min)      lcqtof_t3_neg_2_043_31    m/z 751.6212     0   1   2E+3   12   12.5  RT (min)      lcqtof_t3_neg_3_032_24    m/z 751.6217     0   1E+2   10.5   11  RT (min)      lcqtof_t3_neg_3_038_06    m/z 751.6220     0   1   2E+3   12   12.5  RT (min)      lcqtof_t3_neg_3_032_24    m/z 751.6258     0   1   2E+3   12   12.5  RT (min)      lcqtof_t3_neg_3_032_24    m/z 751.6397     0   1   2E+3   11.5   12  RT (min)      lcqtof_t3_neg_1_001_49    m/z 751.7454     0   1   2E+4   4   4.5  RT (min)      lcqtof_t3_neg_4_000_ee    m/z 751.7702     0   1E+4   14   14.5  RT (min)      lcqtof_t3_neg_4_019_05    m/z 752.1620     0   1E+4   11.5   12  RT (min)      lcqtof_t3_neg_4_064_17    m/z 752.2482     0   1E+4   4   4.5  RT (min)      lcqtof_t3_neg_4_000_ee    m/z 752.3140     0   1   2E+4   4   4.5  RT (min)      lcqtof_t3_neg_4_000_ee    m/z 752.4732     0   1E+4   11.5   12  RT (min)      lcqtof_t3_neg_4_037_01    m/z 752.5190     0   2   4   6E+3   13.5   14  RT (min)      lcqtof_t3_neg_2_053_38    m/z 752.5214     0   1E+4   12   12.5  RT (min)      lcqtof_t3_neg_2_030_23    m/z 752.5232     0   1   2   3E+4   12.5   13  RT (min)      lcqtof_t3_neg_4_018_23    m/z 752.5327     0   2   4   6E+3   14   14.5  RT (min)      lcqtof_t3_neg_2_053_38    m/z 752.6047     0   1   2E+4   13   13.5  RT (min)      lcqtof_t3_neg_4_097_60    m/z 752.6100     0   1E+4   12.5   13  RT (min)      lcqtof_t3_neg_4_061_50    m/z 752.6324     0   1E+3   12   12.5  RT (min)      lcqtof_t3_neg_3_032_24    m/z 752.6336     0   2   4   6E+2   12   12.5  RT (min)      lcqtof_t3_neg_3_032_24    m/z 752.6358     0   1E+2   11   11.5  RT (min)      lcqtof_t3_neg_3_038_06    m/z 752.6361     0   1E+2   11   11.5  RT (min)      lcqtof_t3_neg_3_038_06    m/z 752.6398     0   1   2E+5   12.5   13  RT (min)      lcqtof_t3_neg_2_016_05    m/z 752.7435     0   1E+4   1   1.5  RT (min)      lcqtof_t3_neg_4_037_01    m/z 752.7708     0   2   4   6E+2   13   13.5  RT (min)      lcqtof_t3_neg_3_032_24    m/z 752.7712     0   1   2E+4   13.5   14  RT (min)      lcqtof_t3_neg_2_016_05    m/z 753.1318     0   1E+4   12   12.5  RT (min)      lcqtof_t3_neg_4_057_49    m/z 753.1751     0   1E+4   12   12.5  RT (min)      lcqtof_t3_neg_4_064_17    m/z 753.1754     0   1E+4   12   12.5  RT (min)      lcqtof_t3_neg_4_064_17    m/z 753.5260     0   1E+4   12.5   13  RT (min)      lcqtof_t3_neg_4_034_34    m/z 753.5261     0   2   4   6   8E+3   13   13.5  RT (min)      lcqtof_t3_neg_2_043_31    m/z 753.5277     0   1E+4   12.5   13  RT (min)      lcqtof_t3_neg_4_030_31    m/z 753.5285     0   1E+4   12.5   13  RT (min)      lcqtof_t3_neg_4_034_34    m/z 753.5808     0   1E+4   14.5   15  RT (min)      lcqtof_t3_neg_4_016_14    m/z 753.6362     0   1   2   3E+3   12   12.5  RT (min)      lcqtof_t3_neg_3_032_24    m/z 753.6366     0   2   4   6   8E+2   12   12.5  RT (min)      lcqtof_t3_neg_3_032_24    m/z 753.6382     0   1   2E+2   11   11.5  RT (min)      lcqtof_t3_neg_3_038_06    m/z 753.6389     0   1E+2   11   11.5  RT (min)      lcqtof_t3_neg_3_038_06    m/z 753.9134     0   1E+2   12   12.5  RT (min)      lcqtof_t3_neg_3_032_24    m/z 753.9148     0   1E+2   12   12.5  RT (min)      lcqtof_t3_neg_3_032_24    m/z 753.9150     0   1E+4   12.5   13  RT (min)      lcqtof_t3_neg_2_016_05    m/z 753.9159     0   1E+4   12.5   13  RT (min)      lcqtof_t3_neg_2_016_05    m/z 754.0248     0   1E+4   1   1.5  RT (min)      lcqtof_t3_neg_4_059_14    m/z 754.0465     0   1E+4   13   13.5  RT (min)      lcqtof_t3_neg_4_040_24    m/z 754.1327     0   1   2   3E+4   13   13.5  RT (min)      lcqtof_t3_neg_4_040_24    m/z 754.1748     0   1E+4   12   12.5  RT (min)      lcqtof_t3_neg_4_064_17    m/z 754.1776     0   1E+4   12   12.5  RT (min)      lcqtof_t3_neg_4_064_17    m/z 754.2868     0   1   2E+4   13   13.5  RT (min)      lcqtof_t3_neg_4_040_24    m/z 754.3112     0   1E+4   4   4.5  RT (min)      lcqtof_t3_neg_4_000_ee    m/z 754.4647     0   1   2E+4   12   12.5  RT (min)      lcqtof_t3_neg_4_018_23    m/z 754.4654     0   1E+4   11.5   12  RT (min)      lcqtof_t3_neg_2_030_23    m/z 754.5375     0   1   2   3E+4   12.5   13  RT (min)      lcqtof_t3_neg_2_030_23    m/z 754.5399     0   1   2   3   4E+4   12.5   13  RT (min)      lcqtof_t3_neg_2_030_23    m/z 754.5448     0   1   2   3E+2   11   11.5  RT (min)      lcqtof_t3_neg_3_038_06    m/z 754.5564     0   1   2   3   4E+4   12.5   13  RT (min)      lcqtof_t3_neg_1_019_23    m/z 754.5641     0   1   2   3   4E+4   13   13.5  RT (min)      lcqtof_t3_neg_1_032_36    m/z 754.6412     0   1E+5   13   13.5  RT (min)      lcqtof_t3_neg_2_016_05    m/z 754.6424     0   1E+5   12.5   13  RT (min)      lcqtof_t3_neg_2_016_05    m/z 754.6430     0   1   2E+4   12.5   13  RT (min)      lcqtof_t3_neg_2_016_05    m/z 754.6470     0   1   2E+2   11   11.5  RT (min)      lcqtof_t3_neg_3_038_06    m/z 754.6480     0   1   2   3   4E+2   12   12.5  RT (min)      lcqtof_t3_neg_3_032_24    m/z 754.6494     0   1   2   3   4E+2   12.5   13  RT (min)      lcqtof_t3_neg_3_032_24    m/z 754.6509     0   1   2E+2   11   11.5  RT (min)      lcqtof_t3_neg_3_038_06    m/z 754.6510     0   1   2E+2   11   11.5  RT (min)      lcqtof_t3_neg_3_038_06    m/z 755.1336     0   1E+4   12   12.5  RT (min)      lcqtof_t3_neg_4_057_49    m/z 755.1725     0   1E+4   12   12.5  RT (min)      lcqtof_t3_neg_4_002_17    m/z 755.1748     0   1E+4   13   13.5  RT (min)      lcqtof_t3_neg_4_036_47    m/z 755.1750     0   1   2E+4   12   12.5  RT (min)       lcqtof_t3_neg_4_064_17    m/z 755.1758     0   1E+4   12   12.5  RT (min)      lcqtof_t3_neg_4_002_17    m/z 755.2544     0   1   2E+3   11   11.5  RT (min)       lcqtof_t3_neg_4_058_16    m/z 755.2544     0   1   2   3   4E+3   6.5   7  RT (min)       lcqtof_t3_neg_4_004_20    m/z 755.2545     0   1   2   3E+3   9   9.5  RT (min)       lcqtof_t3_neg_4_019_05    m/z 755.2547     0   1   2   3   4E+3   9   9.5  RT (min)       lcqtof_t3_neg_4_010_46    m/z 755.2547     0   1   2   3   4E+3   8.5   9  RT (min)       lcqtof_t3_neg_4_045_02    m/z 755.2549     0   1   2   3E+3   10.5   11  RT (min)       lcqtof_t3_neg_4_005_18    m/z 755.2550     0   2   4E+3   7   7.5  RT (min)       lcqtof_t3_neg_4_040_24    m/z 755.2550     0   1   2   3E+3   12   12.5  RT (min)       lcqtof_t3_neg_4_005_18    m/z 755.2550     0   1   2   3E+3   11.5   12  RT (min)       lcqtof_t3_neg_4_048_19    m/z 755.2550     0   2   4   6   8E+3   9.5   10  RT (min)       lcqtof_t3_neg_4_062_13    m/z 755.2552     0   2   4   6E+3   16   16.5  RT (min)       lcqtof_t3_neg_4_041_54    m/z 755.2552     0   2   4E+3   17   17.5  RT (min)       lcqtof_t3_neg_4_035_08    m/z 755.2552     0   1   2   3E+3   14.5   15  RT (min)       lcqtof_t3_neg_4_021_21    m/z 755.2554     0   1   2   3   4E+3   6.5   7   7.5  RT (min)       lcqtof_t3_neg_4_030_31    m/z 755.2554     0   2   4   6   8E+3   16.5   17  RT (min)       lcqtof_t3_neg_4_000_cc    m/z 755.2554     0   1   2   3E+3   14   15  RT (min)       lcqtof_t3_neg_4_032_04    m/z 755.2555     0   1   2   3E+3   15.5   16  RT (min)       lcqtof_t3_neg_4_006_16    m/z 755.2556     0   1   2   3   4E+3   15.5   16  RT (min)       lcqtof_t3_neg_4_023_45    m/z 755.2556     0   2   4   6E+3   16.5   17  RT (min)       lcqtof_t3_neg_4_040_24    m/z 755.2556     0   1   2E+3   13.5   14  RT (min)       lcqtof_t3_neg_4_042_10    m/z 755.2556     0   1   2   3   4E+3   8   8.5  RT (min)       lcqtof_t3_neg_4_032_04    m/z 755.2557     0   1   2   3E+3   14.5   15  RT (min)       lcqtof_t3_neg_4_024_26    m/z 755.2558     0   2   4E+3   16.5   17  RT (min)      lcqtof_t3_neg_4_044_48    m/z 755.2559     0   1   2   3   4E+3   8   9  RT (min)       lcqtof_t3_neg_4_016_14    m/z 755.2559     0   1   2   3   4E+3   8   8.5  RT (min)       lcqtof_t3_neg_4_000_aa    m/z 755.2560     0   2   4   6E+3   16.5   17  RT (min)       lcqtof_t3_neg_4_010_46    m/z 755.2561     0   1   2   3E+3   14   14.5  RT (min)       lcqtof_t3_neg_4_000_ff    m/z 755.2562     0   1   2E+3   13.5   14  RT (min)       lcqtof_t3_neg_4_029_40    m/z 755.2564     0   1   2   3E+3   11.5   12  RT (min)       lcqtof_t3_neg_4_043_11    m/z 755.2564     0   1   2   3E+3   14.5   15  RT (min)       lcqtof_t3_neg_4_045_02    m/z 755.2567     0   2   4E+3   9.5   10  RT (min)       lcqtof_t3_neg_4_058_16    m/z 755.2574     0   1   2   3E+3   14   14.5  RT (min)      lcqtof_t3_neg_4_000_ee    m/z 755.2576     0   2   4   6   8E+3   9.5   10  RT (min)       lcqtof_t3_neg_4_062_13    m/z 755.2597     0   1   2   3E+3   10.5   11  RT (min)       lcqtof_t3_neg_4_000_dd    m/z 755.3242     0   1E+4   4   4.5  RT (min)      lcqtof_t3_neg_4_000_ee    m/z 755.3277     0   1E+4   4   4.5  RT (min)      lcqtof_t3_neg_4_000_ee    m/z 755.3301     0   1E+4   4.5   5  RT (min)      lcqtof_t3_neg_4_000_ee    m/z 755.4027     0   1E+4   8   8.5  RT (min)      lcqtof_t3_neg_4_000_ee    m/z 755.4095     0   1E+4   8   8.5  RT (min)      lcqtof_t3_neg_4_000_ee    m/z 755.5416     0   1E+4   12.5   13  RT (min)      lcqtof_t3_neg_3_006_23    m/z 755.5427     0   1E+4   12.5   13  RT (min)      lcqtof_t3_neg_1_019_23    m/z 755.5557     0   1E+4   12.5   13  RT (min)      lcqtof_t3_neg_1_019_23    m/z 755.5666     0   1E+4   13   13.5  RT (min)      lcqtof_t3_neg_1_035_31    m/z 755.5727     0   1E+4   13   13.5  RT (min)      lcqtof_t3_neg_1_015_34    m/z 755.6507     0   1E+2   12   12.5  RT (min)      lcqtof_t3_neg_3_032_24    m/z 755.6523     0   1E+2   12.5   13  RT (min)      lcqtof_t3_neg_3_032_24    m/z 755.6541     0   2   4E+5   13   13.5  RT (min)      lcqtof_t3_neg_2_016_05    m/z 755.6542     0   1E+2   11   11.5  RT (min)      lcqtof_t3_neg_3_038_06    m/z 755.6543     0   1E+2   11   11.5  RT (min)      lcqtof_t3_neg_3_038_06    m/z 755.7904     0   1E+4   13   13.5  RT (min)      lcqtof_t3_neg_4_040_24    m/z 756.1480     0   1E+4   13   13.5  RT (min)      lcqtof_t3_neg_4_040_24    m/z 756.1576     0   1E+4   12.5   13  RT (min)      lcqtof_t3_neg_4_057_49    m/z 756.1764     0   1E+4   12   12.5  RT (min)      lcqtof_t3_neg_4_064_17    m/z 756.1766     0   1E+4   12   12.5  RT (min)      lcqtof_t3_neg_4_064_17    m/z 756.4808     0   1   2E+2   10.5   11  RT (min)      lcqtof_t3_neg_3_038_06    m/z 756.4810     0   2   4   6E+5   12   12.5  RT (min)      lcqtof_t3_neg_1_019_23    m/z 756.4816     0   1   2E+2   10   10.5  RT (min)      lcqtof_t3_neg_3_038_06    m/z 756.5504     0   1E+4   12.5   13  RT (min)      lcqtof_t3_neg_2_030_23    m/z 756.5513     0   1   2E+4   12.5   13  RT (min)      lcqtof_t3_neg_2_030_23    m/z 756.5530     0   1   2E+4   12.5   13  RT (min)      lcqtof_t3_neg_2_030_23    m/z 756.6558     0   2   4   6   8E+1   12   12.5  RT (min)      lcqtof_t3_neg_3_032_24    m/z 756.6566     0   2   4   6   8E+1   12.5   13  RT (min)      lcqtof_t3_neg_3_032_24    m/z 756.6584     0   1E+4   12.5   13  RT (min)      lcqtof_t3_neg_2_016_05    m/z 756.6596     0   1E+4   13   13.5  RT (min)      lcqtof_t3_neg_2_016_05    m/z 756.6708     0   1   2E+2   11.5   12  RT (min)      lcqtof_t3_neg_3_038_06    m/z 756.6718     0   1   2   3E+5   13.5   14  RT (min)      lcqtof_t3_neg_4_018_23    m/z 756.7600     0   1E+4   12.5   13  RT (min)      lcqtof_t3_neg_4_055_38    m/z 756.7607     0   1   2E+4   12   12.5  RT (min)      lcqtof_t3_neg_4_034_34    m/z 757.1529     0   1   2   3E+4   12.5   13  RT (min)      lcqtof_t3_neg_4_001_49    m/z 757.1538     0   2   4   6E+4   12.5   13  RT (min)      lcqtof_t3_neg_4_057_49    m/z 757.1727     0   1E+4   12   12.5  RT (min)      lcqtof_t3_neg_4_064_17    m/z 757.4842     0   1   2E+2   10.5   11  RT (min)      lcqtof_t3_neg_3_038_06    m/z 757.4846     0   1   2E+2   10   10.5  RT (min)      lcqtof_t3_neg_3_038_06    m/z 757.4846     0   1   2E+5   12   12.5  RT (min)      lcqtof_t3_neg_1_019_23    m/z 757.5562     0   1E+4   12.5   13  RT (min)      lcqtof_t3_neg_1_019_23    m/z 757.5566     0   1E+4   12.5   13  RT (min)      lcqtof_t3_neg_2_030_23    m/z 757.6409     0   1E+4   13.5   14  RT (min)      lcqtof_t3_neg_1_001_49    m/z 757.6446     0   1E+4   12.5   13  RT (min)      lcqtof_t3_neg_2_030_23    m/z 757.6536     0   1E+2   12   12.5  RT (min)      lcqtof_t3_neg_3_032_24    m/z 757.6546     0   1E+2   12.5   13  RT (min)      lcqtof_t3_neg_3_032_24    m/z 757.6707     0   1E+5   13.5   14  RT (min)      lcqtof_t3_neg_4_018_23    m/z 757.6734     0   1   2E+4   13.5   14  RT (min)      lcqtof_t3_neg_2_030_23    m/z 757.6745     0   1E+2   11.5   12  RT (min)      lcqtof_t3_neg_3_038_06    m/z 758.1553     0   1   2   3   4E+4   12.5   13  RT (min)      lcqtof_t3_neg_4_057_49    m/z 758.1553     0   1   2E+4   12.5   13  RT (min)      lcqtof_t3_neg_4_001_49    m/z 758.3094     0   1E+4   7   7.5  RT (min)      lcqtof_t3_neg_4_059_14    m/z 758.4954     0   1   2   3E+6   12   12.5  RT (min)      lcqtof_t3_neg_3_006_23    m/z 758.4959     0   2   4E+2   16   16.5  RT (min)      lcqtof_t3_neg_2_030_23    m/z 758.4964     0   1   2   3   4E+2   10.5   11  RT (min)      lcqtof_t3_neg_3_038_06    m/z 758.4970     0   1   2   3   4E+2   10.5   11  RT (min)      lcqtof_t3_neg_3_038_06    m/z 758.4979     0   1   2   3E+4   16   16.5  RT (min)      lcqtof_t3_neg_4_018_23    m/z 758.5341     0   1E+4   12   12.5  RT (min)      lcqtof_t3_neg_4_059_14    m/z 758.5345     0   1E+4   12   12.5  RT (min)      lcqtof_t3_neg_4_059_14    m/z 758.5350     0   1E+4   12   12.5  RT (min)      lcqtof_t3_neg_4_059_14    m/z 758.6339     0   2   4   6E+4   12   12.5  RT (min)      lcqtof_t3_neg_1_019_23    m/z 758.6428     0   1   2   3E+3   14   14.5  RT (min)      lcqtof_t3_neg_1_029_44    m/z 758.6429     0   2   4E+3   13.5   14  RT (min)      lcqtof_t3_neg_4_034_34    m/z 758.6775     0   2   4   6   8E+3   13.5   14  RT (min)      lcqtof_t3_neg_2_030_23    m/z 758.6786     0   1   2   3   4E+4   13.5   14  RT (min)      lcqtof_t3_neg_4_018_23    m/z 758.7752     0   1   2   3   4E+2   10.5   11  RT (min)      lcqtof_t3_neg_3_038_06    m/z 758.7754     0   1   2   3   4E+2   10.5   11  RT (min)      lcqtof_t3_neg_3_038_06    m/z 758.9060     0   2   4   6E+4   12   12.5  RT (min)      lcqtof_t3_neg_2_030_23    m/z 758.9062     0   2   4   6E+4   12   12.5  RT (min)      lcqtof_t3_neg_1_019_23    m/z 759.0502     0   1   2   3   4E+3   8.5   9  RT (min)      lcqtof_t3_neg_3_058_14    m/z 759.0592     0   1   2E+4   13.5   14  RT (min)      lcqtof_t3_neg_4_004_20    m/z 759.0602     0   2   4   6   8E+4   14   14.5  RT (min)      lcqtof_t3_neg_4_036_47    m/z 759.0603     0   1   2E+5   13.5   14  RT (min)      lcqtof_t3_neg_4_064_17    m/z 759.0605     0   1E+5   14   14.5  RT (min)      lcqtof_t3_neg_4_036_47    m/z 759.0605     0   1E+5   14   14.5  RT (min)      lcqtof_t3_neg_4_036_47    m/z 759.0616     0   1E+5   14   14.5  RT (min)      lcqtof_t3_neg_4_036_47    m/z 759.0620     0   1E+5   12.5   13  RT (min)      lcqtof_t3_neg_4_036_47    m/z 759.1515     0   1   2E+4   4   4.5  RT (min)      lcqtof_t3_neg_4_000_ee    m/z 759.4989     0   1   2E+6   12   12.5  RT (min)      lcqtof_t3_neg_1_019_23    m/z 759.4999     0   1   2   3   4E+2   10.5   11  RT (min)      lcqtof_t3_neg_3_038_06    m/z 759.5000     0   1   2E+2   16   16.5  RT (min)      lcqtof_t3_neg_2_030_23    m/z 759.5001     0   1   2   3   4E+2   10.5   11  RT (min)      lcqtof_t3_neg_3_038_06    m/z 759.5028     0   1E+4   16   16.5  RT (min)      lcqtof_t3_neg_4_018_23    m/z 759.5620     0   2   4E+3   14   14.5  RT (min)      lcqtof_t3_neg_1_035_31    m/z 759.6102     0   1E+4   13.5   14  RT (min)      lcqtof_t3_neg_2_028_24    m/z 759.6164     0   1   2   3E+3   11.5   12  RT (min)      lcqtof_t3_neg_1_036_14    m/z 759.6164     0   1E+4   14.5   15  RT (min)      lcqtof_t3_neg_1_000_aa    m/z 759.6170     0   1E+4   14.5   15  RT (min)      lcqtof_t3_neg_1_000_aa    m/z 759.6228     0   1E+4   13.5   14  RT (min)      lcqtof_t3_neg_2_028_24    m/z 759.6399     0   1E+4   13.5   14  RT (min)       lcqtof_t3_neg_2_030_23    m/z 759.6500     0   1   2   3   4E+3   13   13.5  RT (min)      lcqtof_t3_neg_2_000_aa    m/z 759.6504     0   1   2   3E+3   13   13.5  RT (min)      lcqtof_t3_neg_1_002_13    m/z 759.7786     0   1   2   3E+4   12   12.5  RT (min)      lcqtof_t3_neg_1_019_23    m/z 759.7790     0   1   2   3E+4   12   12.5  RT (min)      lcqtof_t3_neg_2_030_23    m/z 759.9978     0   1   2E+4   12   12.5  RT (min)      lcqtof_t3_neg_1_019_23    m/z 760.0608     0   2   4   6E+4   14   14.5  RT (min)      lcqtof_t3_neg_4_002_17    m/z 760.0608     0   1E+5   14   14.5  RT (min)      lcqtof_t3_neg_4_002_17    m/z 760.0611     0   1E+5   13.5   14  RT (min)      lcqtof_t3_neg_4_064_17    m/z 760.0616     0   1E+5   14   14.5  RT (min)      lcqtof_t3_neg_4_002_17    m/z 760.0620     0   2   4   6   8E+4   14   14.5  RT (min)      lcqtof_t3_neg_4_002_17    m/z 760.0620     0   1E+4   13.5   14  RT (min)      lcqtof_t3_neg_4_004_20    m/z 760.1499     0   1   2   3   4E+4   12   12.5  RT (min)      lcqtof_t3_neg_2_030_23    m/z 760.1507     0   1   2   3   4E+4   12   12.5  RT (min)      lcqtof_t3_neg_1_019_23    m/z 760.1508     0   2   4E+4   12   12.5  RT (min)      lcqtof_t3_neg_4_018_23    m/z 760.2783     0   1E+4   12   12.5  RT (min)      lcqtof_t3_neg_1_019_23    m/z 760.2794     0   1   2E+4   12   12.5  RT (min)      lcqtof_t3_neg_4_018_23    m/z 760.2807     0   1   2E+4   12   12.5  RT (min)      lcqtof_t3_neg_2_030_23    m/z 760.3315     0   1E+4   12.5   13  RT (min)      lcqtof_t3_neg_4_018_23    m/z 760.3320     0   1E+4   12   12.5  RT (min)      lcqtof_t3_neg_4_018_23    m/z 760.3328     0   1E+4   12.5   13  RT (min)      lcqtof_t3_neg_4_018_23    m/z 760.3331     0   1E+4   12   12.5  RT (min)      lcqtof_t3_neg_1_019_23    m/z 760.4977     0   2   4E+5   12   12.5  RT (min)      lcqtof_t3_neg_1_019_23    m/z 760.5032     0   1   2   3E+2   10.5   11  RT (min)      lcqtof_t3_neg_3_038_06    m/z 760.5103     0   1   16   16.5  RT (min)      lcqtof_t3_neg_2_030_23    m/z 760.5103     0   1   2   3E+2   10.5   11  RT (min)      lcqtof_t3_neg_3_038_06    m/z 760.5104     0   1E+6   12   12.5  RT (min)      lcqtof_t3_neg_1_019_23    m/z 760.5105     0   1E+4   16   16.5  RT (min)      lcqtof_t3_neg_4_018_23    m/z 760.5124     0   1   2   3E+2   10.5   11  RT (min)      lcqtof_t3_neg_3_038_06    m/z 760.5787     0   1E+4   11.5   12  RT (min)      lcqtof_t3_neg_2_016_05    m/z 760.5795     0   2   4   6E+3   14   14.5  RT (min)      lcqtof_t3_neg_2_056_56    m/z 760.5804     0   1E+4   11.5   12  RT (min)      lcqtof_t3_neg_2_016_05    m/z 760.5810     0   2   4E+3   14   14.5  RT (min)      lcqtof_t3_neg_1_018_33    m/z 760.5816     0   2   4   6   8E+3   12.5   13  RT (min)      lcqtof_t3_neg_2_016_05    m/z 760.5820     0   2   4   6   8E+3   12.5   13  RT (min)      lcqtof_t3_neg_2_041_54    m/z 760.5820     0   1E+4   11   11.5  RT (min)      lcqtof_t3_neg_1_019_23    m/z 760.5821     0   1   2E+4   12   12.5  RT (min)      lcqtof_t3_neg_1_043_38    m/z 760.5824     0   1   2E+4   12   12.5  RT (min)      lcqtof_t3_neg_2_051_14    m/z 760.5829     0   2   4   6E+3   13.5   14  RT (min)      lcqtof_t3_neg_1_055_32    m/z 760.5829     0   1E+4   11.5   12  RT (min)      lcqtof_t3_neg_2_016_05    m/z 760.5830     0   2   4   6   8E+3   12.5   13  RT (min)      lcqtof_t3_neg_2_053_38    m/z 760.5832     0   1E+4   12.5   13  RT (min)      lcqtof_t3_neg_1_031_24    m/z 760.5832     0   1E+4   12.5   13  RT (min)      lcqtof_t3_neg_1_031_24    m/z 760.5855     0   2   4E+3   13.5   14  RT (min)      lcqtof_t3_neg_2_025_52    m/z 760.5856     0   2   4   6E+3   13   13.5  RT (min)      lcqtof_t3_neg_2_065_51    m/z 760.5860     0   2   4E+3   13   13.5  RT (min)      lcqtof_t3_neg_2_061_50    m/z 760.5966     0   2   4   6   8E+3   13.5   14  RT (min)      lcqtof_t3_neg_2_019_32    m/z 760.5973     0   2   4   6   8E+3   13   13.5  RT (min)      lcqtof_t3_neg_2_046_45    m/z 760.5973     0   2   4E+3   13   13.5  RT (min)      lcqtof_t3_neg_2_000_aa    m/z 760.6146     0   1E+4   13.5   14  RT (min)      lcqtof_t3_neg_2_028_24    m/z 760.6248     0   1E+4   13.5   14  RT (min)      lcqtof_t3_neg_2_019_32    m/z 760.6346     0   2   4   6   8E+3   13.5   14  RT (min)       lcqtof_t3_neg_2_030_23    m/z 760.6399     0   1E+4   12.5   13  RT (min)      lcqtof_t3_neg_4_018_23    m/z 760.6478     0   1E+4   12.5   13  RT (min)      lcqtof_t3_neg_4_034_34    m/z 760.6490     0   1E+4   12.5   13  RT (min)      lcqtof_t3_neg_4_003_37    m/z 760.7504     0   1E+2   13   13.5  RT (min)      lcqtof_t3_neg_3_032_24    m/z 760.7511     0   1E+4   13.5   14  RT (min)      lcqtof_t3_neg_2_016_05    m/z 760.7532     0   1   2   3   4E+2   11.5   12  RT (min)      lcqtof_t3_neg_3_038_06    m/z 760.7905     0   1   2   3   4E+4   12.5   13  RT (min)      lcqtof_t3_neg_4_018_23    m/z 760.7911     0   1   2   3   4E+4   12.5   13  RT (min)      lcqtof_t3_neg_4_018_23    m/z 760.7918     0   1   2   3E+4   12   12.5  RT (min)      lcqtof_t3_neg_1_019_23    m/z 761.0085     0   1E+4   12.5   13  RT (min)      lcqtof_t3_neg_4_034_34    m/z 761.0576     0   1E+5   12.5   13  RT (min)      lcqtof_t3_neg_4_064_17    m/z 761.0588     0   1E+5   13   13.5  RT (min)      lcqtof_t3_neg_4_064_17    m/z 761.0590     0   1E+4   13   13.5  RT (min)      lcqtof_t3_neg_4_022_28    m/z 761.0594     0   2   4   6   8E+4   13   13.5  RT (min)      lcqtof_t3_neg_4_036_47    m/z 761.0595     0   1E+5   13.5   14  RT (min)      lcqtof_t3_neg_4_064_17    m/z 761.0597     0   1E+5   12.5   13  RT (min)      lcqtof_t3_neg_4_064_17    m/z 761.0611     0   2   4   6   8E+4   14   14.5  RT (min)      lcqtof_t3_neg_4_002_17    m/z 761.5062     0   2   4   6   8E+4   12   12.5  RT (min)      lcqtof_t3_neg_1_019_23    m/z 761.5142     0   1   2E+2   10.5   11  RT (min)      lcqtof_t3_neg_3_038_06    m/z 761.5150     0   2   4E+5   12   12.5  RT (min)      lcqtof_t3_neg_1_019_23    m/z 761.5160     0   1   2   3E+2   10.5   11  RT (min)      lcqtof_t3_neg_3_038_06    m/z 761.5660     0   1E+4   16   16.5  RT (min)      lcqtof_t3_neg_1_035_31    m/z 761.5691     0   1E+4   16   16.5  RT (min)      lcqtof_t3_neg_1_035_31    m/z 761.5717     0   1E+4   14   14.5  RT (min)      lcqtof_t3_neg_2_062_17    m/z 761.5737     0   2   4   6   8E+3   12.5   13  RT (min)      lcqtof_t3_neg_1_043_38    m/z 761.5747     0   1E+4   12.5   13  RT (min)      lcqtof_t3_neg_1_061_50    m/z 761.5751     0   2   4   6   8E+3   14   14.5  RT (min)      lcqtof_t3_neg_2_017_51    m/z 761.5759     0   1E+4   12.5   13  RT (min)      lcqtof_t3_neg_2_019_32    m/z 761.5771     0   1E+4   11   11.5  RT (min)      lcqtof_t3_neg_2_030_23    m/z 761.5782     0   1   2E+4   11.5   12  RT (min)      lcqtof_t3_neg_2_000_aa    m/z 761.5799     0   2   4   6   8E+3   13.5   14  RT (min)      lcqtof_t3_neg_2_053_38    m/z 761.5820     0   1E+4   11   11.5  RT (min)      lcqtof_t3_neg_2_016_05    m/z 761.5821     0   2   4   6E+3   13.5   14  RT (min)      lcqtof_t3_neg_1_037_25    m/z 761.5831     0   1E+4   13   13.5  RT (min)      lcqtof_t3_neg_2_062_17    m/z 761.5844     0   2   4   6E+3   13.5   14  RT (min)      lcqtof_t3_neg_1_009_50    m/z 761.5846     0   1E+4   12   12.5  RT (min)      lcqtof_t3_neg_2_054_29    m/z 761.5846     0   1E+4   12   12.5  RT (min)      lcqtof_t3_neg_2_016_05    m/z 761.5855     0   1E+4   13   13.5  RT (min)      lcqtof_t3_neg_2_055_16    m/z 761.5864     0   2   4E+3   13   13.5  RT (min)      lcqtof_t3_neg_1_019_23    m/z 761.5919     0   1E+4   14   14.5  RT (min)      lcqtof_t3_neg_4_011_32    m/z 761.6278     0   1E+4   13   13.5  RT (min)      lcqtof_t3_neg_2_016_05    m/z 762.0588     0   2   4   6E+4   13.5   14  RT (min)      lcqtof_t3_neg_4_064_17    m/z 762.0592     0   2   4   6E+4   12.5   13  RT (min)      lcqtof_t3_neg_4_064_17    m/z 762.0593     0   1   2   3   4E+4   13   13.5  RT (min)      lcqtof_t3_neg_4_002_17    m/z 762.0599     0   2   4   6E+4   13   13.5  RT (min)      lcqtof_t3_neg_4_064_17    m/z 762.0608     0   2   4   6E+4   13   13.5  RT (min)      lcqtof_t3_neg_4_064_17    m/z 762.5066     0   2   4   6   8E+4   12   12.5  RT (min)      lcqtof_t3_neg_3_006_23    m/z 762.5072     0   2   4   6E+2   10.5   11  RT (min)      lcqtof_t3_neg_3_038_06    m/z 762.5075     0   1E+5   12   12.5  RT (min)      lcqtof_t3_neg_2_030_23    m/z 762.5076     0   2   4   6E+2   10.5   11  RT (min)      lcqtof_t3_neg_3_038_06    m/z 762.5086     0   2   4   6E+2   10.5   11  RT (min)      lcqtof_t3_neg_3_038_06    m/z 762.5166     0   2   4   6E+2   10.5   11  RT (min)      lcqtof_t3_neg_3_038_06    m/z 762.5188     0   2   4   6E+2   10.5   11  RT (min)      lcqtof_t3_neg_3_038_06    m/z 762.5375     0   1   2E+4   12.5   13  RT (min)       lcqtof_t3_neg_1_002_13    m/z 762.5634     0   1E+4   13.5   14  RT (min)      lcqtof_t3_neg_4_034_34    m/z 762.5636     0   1E+4   13.5   14  RT (min)      lcqtof_t3_neg_4_018_23    m/z 762.5646     0   1E+4   14   14.5  RT (min)      lcqtof_t3_neg_4_055_38    m/z 762.5869     0   1   2   3E+4   14   14.5  RT (min)      lcqtof_t3_neg_3_085_59    m/z 762.5978     0   2   4   6E+4   15   15.5  RT (min)       lcqtof_t3_neg_2_057_49    m/z 762.5978     0   2   4   6   8E+4   14   15  RT (min)       lcqtof_t3_neg_2_063_16    m/z 762.5978     0   2   4   6   8E+4   16   16.5  RT (min)      lcqtof_t3_neg_1_000_cc    m/z 762.5978     0   1E+5   16   16.5  RT (min)      lcqtof_t3_neg_2_058_15    m/z 762.5978     0   2   4   6E+4   15   15.5  RT (min)       lcqtof_t3_neg_2_059_14    m/z 762.5978     0   2   4   6   8E+4   16.5   17  RT (min)      lcqtof_t3_neg_2_000_aa    m/z 762.5979     0   2   4   6   8E+4   16   16.5  RT (min)       lcqtof_t3_neg_1_003_07    m/z 762.5979     0   2   4   6E+4   15   15.5  RT (min)       lcqtof_t3_neg_2_062_17    m/z 762.5979     0   2   4   6E+4   14   15  RT (min)       lcqtof_t3_neg_2_060_39    m/z 762.5980     0   2   4   6E+4   14.5   15  RT (min)      lcqtof_t3_neg_1_007_03    m/z 762.5980     0   2   4   6   8E+4   16   16.5  RT (min)       lcqtof_t3_neg_2_059_14    m/z 762.5981     0   2   4   6E+4   14.5   15  RT (min)      lcqtof_t3_neg_2_059_14    m/z 762.5982     0   2   4   6E+4   14.5   15  RT (min)       lcqtof_t3_neg_2_062_17    m/z 762.5982     0   2   4   6E+4   14.5   15  RT (min)       lcqtof_t3_neg_2_062_17    m/z 762.5982     0   2   4   6   8E+4   13   13.5  RT (min)       lcqtof_t3_neg_1_006_05    m/z 762.5982     0   1E+5   16.5   17  RT (min)       lcqtof_t3_neg_2_058_15    m/z 762.5982     0   2   4E+4   14   14.5  RT (min)      lcqtof_t3_neg_1_041_54    m/z 762.5985     0   2   4   6   8E+4   13.5   14  RT (min)      lcqtof_t3_neg_1_044_37    m/z 762.5985     0   1E+5   13   13.5  RT (min)       lcqtof_t3_neg_2_057_49    m/z 762.5987     0   2   4   6   8E+4   13   13.5  RT (min)       lcqtof_t3_neg_1_044_37    m/z 762.5987     0   2   4   6   8E+4   14   14.5  RT (min)       lcqtof_t3_neg_1_032_36    m/z 762.5991     0   2   4   6   8E+4   13.5   14  RT (min)       lcqtof_t3_neg_1_031_24    m/z 762.5995     0   2   4   6   8E+4   14   14.5  RT (min)       lcqtof_t3_neg_1_015_34    m/z 762.5998     0   2   4E+4   13.5   14  RT (min)      lcqtof_t3_neg_1_016_48    m/z 762.7963     0   1E+4   1   1.5  RT (min)      lcqtof_t3_neg_1_018_33    m/z 763.0568     0   1   2E+4   13.5   14  RT (min)      lcqtof_t3_neg_4_002_17    m/z 763.0574     0   1   2E+4   13   13.5  RT (min)      lcqtof_t3_neg_4_002_17    m/z 763.0578     0   1   2   3E+4   13   13.5  RT (min)      lcqtof_t3_neg_4_064_17    m/z 763.0584     0   1   2E+4   13   13.5  RT (min)      lcqtof_t3_neg_4_002_17    m/z 763.0594     0   1   2   3E+4   13.5   14  RT (min)      lcqtof_t3_neg_4_064_17    m/z 763.4720     0   2   4E+4   8.5   9  RT (min)      lcqtof_t3_neg_4_000_ee    m/z 763.5095     0   2   4E+4   12   12.5  RT (min)      lcqtof_t3_neg_2_030_23    m/z 763.5102     0   2   4E+4   12   12.5  RT (min)      lcqtof_t3_neg_2_030_23    m/z 763.5302     0   1   2   3E+4   12   12.5  RT (min)      lcqtof_t3_neg_3_006_23    m/z 763.5318     0   2   4E+4   12   12.5  RT (min)       lcqtof_t3_neg_2_030_23    m/z 763.5330     0   2   4   6   8E+3   13.5   14  RT (min)      lcqtof_t3_neg_1_037_25    m/z 763.5337     0   1E+2   12   12.5  RT (min)      lcqtof_t3_neg_3_032_24    m/z 763.5343     0   1   12   12.5  RT (min)      lcqtof_t3_neg_3_032_24    m/z 763.5968     0   2   4   6E+4   12.5   13  RT (min)      lcqtof_t3_neg_1_001_49    m/z 763.5982     0   1   2   3   4E+4   12.5   13  RT (min)      lcqtof_t3_neg_1_043_38    m/z 763.5994     0   1   2   3   4E+4   12.5   13  RT (min)      lcqtof_t3_neg_1_065_51    m/z 763.5996     0   2   4   6E+4   12.5   13  RT (min)      lcqtof_t3_neg_1_001_49    m/z 763.5998     0   1   2   3E+4   13   13.5  RT (min)      lcqtof_t3_neg_1_003_07    m/z 763.6002     0   1   2   3E+4   14   14.5  RT (min)      lcqtof_t3_neg_1_035_31    m/z 763.6003     0   1   2   3E+4   16   16.5  RT (min)      lcqtof_t3_neg_1_033_53    m/z 763.6004     0   1   2   3E+4   12.5   13  RT (min)      lcqtof_t3_neg_1_001_49    m/z 763.6008     0   2   4E+4   12.5   13  RT (min)      lcqtof_t3_neg_1_061_50    m/z 763.6010     0   1   2   3   4E+4   12.5   13  RT (min)      lcqtof_t3_neg_1_043_38    m/z 763.6011     0   2   4   6E+4   13   13.5  RT (min)      lcqtof_t3_neg_1_001_49    m/z 763.6012     0   1   2   3E+4   15   15.5  RT (min)      lcqtof_t3_neg_2_059_14    m/z 763.6012     0   1   2   3   4E+4   16   16.5  RT (min)      lcqtof_t3_neg_1_003_07    m/z 763.6012     0   1   2   3   4E+4   12.5   13  RT (min)      lcqtof_t3_neg_1_035_31    m/z 763.6013     0   1   2   3E+4   14.5   15  RT (min)      lcqtof_t3_neg_2_060_39    m/z 763.6014     0   1   2   3E+4   14.5   15  RT (min)      lcqtof_t3_neg_2_058_15    m/z 763.6014     0   1   2   3E+4   14.5   15  RT (min)      lcqtof_t3_neg_2_059_14    m/z 763.6014     0   1   2   3   4E+4   16   16.5  RT (min)      lcqtof_t3_neg_2_060_39    m/z 763.6014     0   1   2   3E+4   15   15.5  RT (min)      lcqtof_t3_neg_2_059_14    m/z 763.6015     0   1   2   3E+4   13.5   14  RT (min)      lcqtof_t3_neg_1_043_38    m/z 763.6015     0   2   4   6E+4   13   13.5  RT (min)      lcqtof_t3_neg_1_001_49    m/z 763.6016     0   1   2E+4   14.5   15  RT (min)      lcqtof_t3_neg_1_053_01    m/z 763.6016     0   2   4E+4   16   16.5  RT (min)      lcqtof_t3_neg_2_058_15    m/z 763.6017     0   1   2   3   4E+4   16   16.5  RT (min)      lcqtof_t3_neg_1_003_07    m/z 763.6018     0   2   4   6E+4   13   13.5  RT (min)      lcqtof_t3_neg_1_001_49    m/z 763.6018     0   1   2   3E+4   15.5   16  RT (min)      lcqtof_t3_neg_1_000_dd    m/z 763.6018     0   1   2   3E+4   13.5   14  RT (min)      lcqtof_t3_neg_1_035_31    m/z 763.6022     0   1   2   3   4E+4   16   16.5  RT (min)      lcqtof_t3_neg_1_035_31    m/z 763.6022     0   1   2   3E+4   14.5   15  RT (min)      lcqtof_t3_neg_2_059_14    m/z 763.6027     0   1   2   3E+4   12   12.5  RT (min)      lcqtof_t3_neg_1_033_53    m/z 763.6028     0   1   2   3E+4   13   13.5  RT (min)      lcqtof_t3_neg_1_003_07    m/z 763.6032     0   1   2   3E+4   16.5   17  RT (min)      lcqtof_t3_neg_1_032_36    m/z 763.6118     0   1   2E+4   14   14.5  RT (min)      lcqtof_t3_neg_1_001_49    m/z 763.6245     0   1E+4   12.5   13  RT (min)      lcqtof_t3_neg_4_040_24    m/z 764.0576     0   1E+4   13   13.5  RT (min)      lcqtof_t3_neg_4_064_17    m/z 764.4320     0   1E+4   8.5   9  RT (min)      lcqtof_t3_neg_4_000_ee    m/z 764.4336     0   1E+4   8.5   9  RT (min)      lcqtof_t3_neg_4_000_ee    m/z 764.4490     0   1   2E+4   8.5   9  RT (min)      lcqtof_t3_neg_4_000_ee    m/z 764.4666     0   1   2E+4   8.5   9  RT (min)      lcqtof_t3_neg_4_000_ee    m/z 764.4743     0   1   2E+4   8.5   9  RT (min)      lcqtof_t3_neg_4_000_ee    m/z 764.5240     0   1   2   3   4E+2   10.5   11  RT (min)      lcqtof_t3_neg_3_038_06    m/z 764.5242     0   1   2   3   4E+2   10.5   11  RT (min)      lcqtof_t3_neg_3_038_06    m/z 764.5250     0   1   2E+5   12   12.5  RT (min)      lcqtof_t3_neg_2_030_23    m/z 764.5370     0   2   4   6E+3   13.5   14  RT (min)      lcqtof_t3_neg_2_016_05    m/z 764.5380     0   2   4E+3   14.5   15  RT (min)      lcqtof_t3_neg_2_045_04    m/z 764.5382     0   2   4   6   8E+3   13.5   14  RT (min)      lcqtof_t3_neg_4_033_53    m/z 764.5592     0   1   2E+4   13.5   14  RT (min)      lcqtof_t3_neg_2_030_23    m/z 764.5850     0   1E+4   14   14.5  RT (min)      lcqtof_t3_neg_1_000_aa    m/z 764.5876     0   1E+4   14   14.5  RT (min)      lcqtof_t3_neg_1_000_aa    m/z 764.6030     0   1E+4   14   14.5  RT (min)      lcqtof_t3_neg_1_000_aa    m/z 764.6039     0   2   4   6E+3   13.5   14  RT (min)      lcqtof_t3_neg_1_055_32    m/z 764.6042     0   2   4   6E+3   14   14.5  RT (min)      lcqtof_t3_neg_2_046_45    m/z 764.6046     0   1   2E+4   13   13.5  RT (min)      lcqtof_t3_neg_2_057_49    m/z 764.6054     0   1E+4   12.5   13  RT (min)      lcqtof_t3_neg_1_043_38    m/z 764.6056     0   1E+4   12.5   13  RT (min)      lcqtof_t3_neg_1_061_50    m/z 764.6059     0   1   2E+4   11   11.5  RT (min)      lcqtof_t3_neg_1_019_23    m/z 764.6068     0   1   2   3   4E+4   12   12.5  RT (min)      lcqtof_t3_neg_1_043_38    m/z 765.4118     0   1E+4   8.5   9  RT (min)      lcqtof_t3_neg_4_000_ee    m/z 765.5272     0   1E+5   12   12.5  RT (min)      lcqtof_t3_neg_2_030_23    m/z 765.5273     0   2   4   6   8E+4   12   12.5  RT (min)      lcqtof_t3_neg_3_006_23    m/z 765.5336     0   1E+4   14.5   15  RT (min)       lcqtof_t3_neg_2_065_51    m/z 765.5339     0   1E+4   16.5   17  RT (min)      lcqtof_t3_neg_2_065_51    m/z 765.5350     0   1E+4   13.5   14  RT (min)      lcqtof_t3_neg_2_051_14    m/z 765.5414     0   1E+4   14   14.5  RT (min)      lcqtof_t3_neg_2_061_50    m/z 765.5497     0   1   2   3E+4   13.5   14  RT (min)      lcqtof_t3_neg_1_019_23    m/z 765.5544     0   1E+4   13.5   14  RT (min)      lcqtof_t3_neg_4_003_37    m/z 765.5616     0   1E+4   14   14.5  RT (min)      lcqtof_t3_neg_4_003_37    m/z 765.6202     0   2   4   6E+3   14   14.5  RT (min)      lcqtof_t3_neg_4_011_32    m/z 765.6226     0   2   4   6E+3   14   14.5  RT (min)      lcqtof_t3_neg_4_041_54    m/z 765.6612     0   2   4   6   8E+2   13   13.5  RT (min)      lcqtof_t3_neg_3_032_24    m/z 766.4681     0   1E+4   12   12.5  RT (min)      lcqtof_t3_neg_1_019_23    m/z 766.5338     0   1   2E+5   13.5   14  RT (min)      lcqtof_t3_neg_4_006_16    m/z 766.5344     0   1E+5   13.5   14  RT (min)      lcqtof_t3_neg_4_038_29    m/z 766.5375     0   1E+2   10.5   11  RT (min)      lcqtof_t3_neg_3_038_06    m/z 766.5377     0   1   2E+5   12.5   13  RT (min)      lcqtof_t3_neg_2_030_23    m/z 766.5388     0   1E+2   11   11.5  RT (min)      lcqtof_t3_neg_3_038_06    m/z 766.6399     0   1   2   3E+2   12   12.5  RT (min)      lcqtof_t3_neg_3_032_24    m/z 766.6413     0   1   2E+4   12.5   13  RT (min)      lcqtof_t3_neg_2_016_05    m/z 766.6421     0   1   2   3E+2   12   12.5  RT (min)      lcqtof_t3_neg_3_032_24    m/z 766.6432     0   1E+2   11   11.5  RT (min)      lcqtof_t3_neg_3_038_06    m/z 766.6441     0   1E+2   10.5   11  RT (min)      lcqtof_t3_neg_3_038_06    m/z 767.1599     0   1E+4   11.5   12  RT (min)      lcqtof_t3_neg_4_064_17    m/z 767.5400     0   1   2E+2   10.5   11  RT (min)      lcqtof_t3_neg_3_038_06    m/z 767.5415     0   1   2E+2   10.5   11  RT (min)      lcqtof_t3_neg_3_038_06    m/z 767.6521     0   1E+2   12   12.5  RT (min)      lcqtof_t3_neg_3_032_24    m/z 767.6524     0   1E+2   12   12.5  RT (min)      lcqtof_t3_neg_3_032_24    m/z 767.6542     0   1E+4   12.5   13  RT (min)      lcqtof_t3_neg_2_016_05    m/z 768.3185     0   1   2   3E+4   4   4.5  RT (min)      lcqtof_t3_neg_4_000_ee    m/z 768.4797     0   1E+4   12   12.5  RT (min)      lcqtof_t3_neg_1_019_23    m/z 768.4812     0   1E+4   12   12.5  RT (min)      lcqtof_t3_neg_2_030_23    m/z 768.5026     0   1   2E+4   11.5   12  RT (min)      lcqtof_t3_neg_4_037_01    m/z 768.5430     0   1   2E+4   12.5   13  RT (min)      lcqtof_t3_neg_2_030_23    m/z 768.5432     0   1   2E+4   12.5   13  RT (min)      lcqtof_t3_neg_1_019_23    m/z 768.5437     0   1   2E+4   12.5   13  RT (min)      lcqtof_t3_neg_2_030_23    m/z 768.5443     0   1E+5   12.5   13  RT (min)      lcqtof_t3_neg_3_006_23    m/z 768.5535     0   1   2E+5   12.5   13  RT (min)      lcqtof_t3_neg_2_030_23    m/z 768.5543     0   1   2E+2   11   11.5  RT (min)      lcqtof_t3_neg_3_038_06    m/z 768.5930     0   1E+3   11.5   12  RT (min)      lcqtof_t3_neg_3_032_24    m/z 768.5930     0   1E+3   11.5   12  RT (min)      lcqtof_t3_neg_3_032_24    m/z 768.5937     0   1E+3   11.5   12  RT (min)      lcqtof_t3_neg_3_032_24    m/z 768.5944     0   1E+4   12   12.5  RT (min)      lcqtof_t3_neg_2_016_05    m/z 768.5948     0   1E+4   12   12.5  RT (min)      lcqtof_t3_neg_2_016_05    m/z 768.6542     0   1   2   3E+2   12   12.5  RT (min)      lcqtof_t3_neg_3_032_24    m/z 768.6554     0   1   2E+2   12   12.5  RT (min)      lcqtof_t3_neg_3_032_24    m/z 768.6565     0   1E+2   11   11.5  RT (min)      lcqtof_t3_neg_3_038_06    m/z 768.6571     0   2   4E+2   12   12.5  RT (min)      lcqtof_t3_neg_3_032_24    m/z 768.6573     0   2   4   6E+2   11   11.5  RT (min)      lcqtof_t3_neg_3_038_06    m/z 768.7882     0   1   2E+3   16.5   17  RT (min)      lcqtof_t3_neg_1_005_18    m/z 768.9960     0   1E+4   1   1.5  RT (min)      lcqtof_t3_neg_4_059_14    m/z 769.1587     0   1E+4   11.5   12  RT (min)      lcqtof_t3_neg_4_064_17    m/z 769.5083     0   1E+4   11.5   12  RT (min)      lcqtof_t3_neg_4_037_01    m/z 769.5516     0   2   4   6E+4   12.5   13  RT (min)      lcqtof_t3_neg_3_006_23    m/z 769.5583     0   1   2   3E+2   11   11.5  RT (min)      lcqtof_t3_neg_3_038_06    m/z 769.5596     0   1   2   3E+2   11   11.5  RT (min)      lcqtof_t3_neg_3_038_06    m/z 769.6185     0   1   2   3   4E+3   13   13.5  RT (min)      lcqtof_t3_neg_1_036_14    m/z 769.6222     0   1E+4   11   11.5  RT (min)      lcqtof_t3_neg_2_016_05    m/z 769.6244     0   1E+4   11   11.5  RT (min)      lcqtof_t3_neg_1_019_23    m/z 769.6394     0   1E+4   11.5   12  RT (min)      lcqtof_t3_neg_1_033_53    m/z 769.6659     0   1   2E+4   13   13.5  RT (min)      lcqtof_t3_neg_4_040_24    m/z 769.6663     0   1   2E+4   13   13.5  RT (min)      lcqtof_t3_neg_4_040_24    m/z 769.7778     0   1E+4   13.5   14  RT (min)      lcqtof_t3_neg_2_016_05    m/z 769.7780     0   2   4   6E+2   13   13.5  RT (min)      lcqtof_t3_neg_3_032_24    m/z 769.7805     0   2   4   6E+2   13   13.5  RT (min)      lcqtof_t3_neg_3_032_24    m/z 770.4958     0   2   4   6E+4   12   12.5  RT (min)      lcqtof_t3_neg_1_019_23    m/z 770.4978     0   1   2   3E+2   10.5   11  RT (min)      lcqtof_t3_neg_3_038_06    m/z 770.5594     0   1   2E+4   12.5   13  RT (min)      lcqtof_t3_neg_2_030_23    m/z 770.5616     0   2   4   6   8E+4   12.5   13  RT (min)      lcqtof_t3_neg_2_030_23    m/z 770.5700     0   1   2E+2   11   11.5  RT (min)      lcqtof_t3_neg_3_038_06    m/z 770.5930     0   1E+5   13.5   14  RT (min)      lcqtof_t3_neg_4_016_14    m/z 770.6078     0   1   2E+4   12   12.5  RT (min)      lcqtof_t3_neg_2_016_05    m/z 770.6083     0   1   2   3E+2   10.5   11  RT (min)      lcqtof_t3_neg_3_038_06    m/z 770.6106     0   1   2   3E+2   10.5   11  RT (min)      lcqtof_t3_neg_3_038_06    m/z 770.6292     0   2   4   6E+3   11.5   12  RT (min)      lcqtof_t3_neg_2_016_05    m/z 770.6684     0   1E+3   12   12.5  RT (min)      lcqtof_t3_neg_3_032_24    m/z 770.6712     0   1E+3   12.5   13  RT (min)      lcqtof_t3_neg_3_032_24    m/z 770.6714     0   1E+3   12.5   13  RT (min)      lcqtof_t3_neg_3_032_24    m/z 770.6718     0   1E+4   13   13.5  RT (min)      lcqtof_t3_neg_2_016_05    m/z 770.6880     0   1   2E+4   13.5   14  RT (min)      lcqtof_t3_neg_4_018_23    m/z 770.7800     0   1   2E+2   13   13.5  RT (min)      lcqtof_t3_neg_3_032_24    m/z 770.7804     0   1   2   3E+2   11.5   12  RT (min)      lcqtof_t3_neg_3_038_06    m/z 770.7808     0   1   2   3   4E+4   13.5   14  RT (min)      lcqtof_t3_neg_2_016_05    m/z 770.8510     0   1E+4   13   13.5  RT (min)      lcqtof_t3_neg_4_037_01    m/z 771.2046     0   1E+4   13   13.5  RT (min)      lcqtof_t3_neg_4_002_17    m/z 771.2065     0   1E+4   13   13.5  RT (min)      lcqtof_t3_neg_4_064_17    m/z 771.2068     0   1E+4   13   13.5  RT (min)      lcqtof_t3_neg_4_064_17    m/z 771.2077     0   1E+4   13   13.5  RT (min)      lcqtof_t3_neg_4_002_17    m/z 771.3014     0   1   2   3E+4   4.5   5  RT (min)      lcqtof_t3_neg_4_000_ee    m/z 771.3034     0   1   2   3E+4   4.5   5  RT (min)      lcqtof_t3_neg_4_000_ee    m/z 771.3139     0   1   2   3E+4   4.5   5  RT (min)      lcqtof_t3_neg_4_000_ee    m/z 771.3246     0   1E+4   4.5   5  RT (min)      lcqtof_t3_neg_4_011_32    m/z 771.4991     0   1   2E+4   12   12.5  RT (min)      lcqtof_t3_neg_1_019_23    m/z 771.6162     0   1E+4   13.5   14  RT (min)      lcqtof_t3_neg_1_010_27    m/z 771.6918     0   1E+4   13.5   14  RT (min)      lcqtof_t3_neg_4_018_23    m/z 772.2079     0   1E+4   13   13.5  RT (min)      lcqtof_t3_neg_4_064_17    m/z 772.4660     0   1E+4   13   13.5  RT (min)      lcqtof_t3_neg_4_035_08    m/z 772.4696     0   1E+4   13   13.5  RT (min)      lcqtof_t3_neg_4_035_08    m/z 772.5099     0   1   2   3E+4   12   12.5  RT (min)      lcqtof_t3_neg_2_030_23    m/z 772.5110     0   1   2E+5   12   12.5  RT (min)      lcqtof_t3_neg_2_030_23    m/z 772.5124     0   1   2   3E+2   10.5   11  RT (min)      lcqtof_t3_neg_3_038_06    m/z 772.5140     0   1   2E+2   10.5   11  RT (min)      lcqtof_t3_neg_3_038_06    m/z 772.5282     0   1   2E+5   13   13.5  RT (min)      lcqtof_t3_neg_2_053_38    m/z 772.5701     0   1   2E+4   13   13.5  RT (min)      lcqtof_t3_neg_4_018_23    m/z 772.5752     0   1   2E+4   13   13.5  RT (min)      lcqtof_t3_neg_4_018_23    m/z 772.5869     0   1E+4   13   13.5  RT (min)      lcqtof_t3_neg_1_019_23    m/z 772.6092     0   2   4   6E+4   13.5   14  RT (min)      lcqtof_t3_neg_4_016_14    m/z 772.6654     0   1E+4   13.5   14  RT (min)      lcqtof_t3_neg_4_018_23    m/z 773.0212     0   1   2   3   4E+4   13.5   14  RT (min)      lcqtof_t3_neg_4_002_17    m/z 773.0214     0   2   4E+4   13   13.5  RT (min)      lcqtof_t3_neg_4_064_17    m/z 773.0215     0   2   4E+4   12.5   13  RT (min)      lcqtof_t3_neg_4_064_17    m/z 773.0222     0   1   2   3E+4   14   14.5  RT (min)      lcqtof_t3_neg_4_002_17    m/z 773.0222     0   2   4E+4   12.5   13  RT (min)      lcqtof_t3_neg_4_064_17    m/z 773.0224     0   2   4E+4   13   13.5  RT (min)      lcqtof_t3_neg_4_064_17    m/z 773.0227     0   2   4E+4   13.5   14  RT (min)      lcqtof_t3_neg_4_064_17    m/z 773.0228     0   1   2   3E+4   14   14.5  RT (min)      lcqtof_t3_neg_4_002_17    m/z 773.0230     0   2   4E+4   13.5   14  RT (min)      lcqtof_t3_neg_4_064_17    m/z 773.0236     0   1   2E+4   14   14.5  RT (min)      lcqtof_t3_neg_4_002_17    m/z 773.0239     0   1   2   3   4E+4   13   13.5  RT (min)      lcqtof_t3_neg_4_002_17    m/z 773.0242     0   1   2   3   4E+4   13   13.5  RT (min)      lcqtof_t3_neg_4_002_17    m/z 773.1126     0   1   2   3   4E+4   13.5   14  RT (min)      lcqtof_t3_neg_4_064_17    m/z 773.1132     0   1   2   3   4E+4   13   13.5  RT (min)      lcqtof_t3_neg_4_064_17    m/z 773.1139     0   1   2   3   4E+4   13   13.5  RT (min)      lcqtof_t3_neg_4_064_17    m/z 773.1153     0   1   2   3   4E+4   13   13.5  RT (min)      lcqtof_t3_neg_4_064_17    m/z 773.2044     0   1E+4   13   13.5  RT (min)      lcqtof_t3_neg_4_064_17    m/z 773.3508     0   1E+4   4.5   5  RT (min)      lcqtof_t3_neg_4_000_ee    m/z 773.4848     0   1E+4   8.5   9  RT (min)      lcqtof_t3_neg_4_000_ee    m/z 773.5163     0   1   2E+2   10.5   11  RT (min)      lcqtof_t3_neg_3_038_06    m/z 773.5163     0   1   2E+2   10.5   11  RT (min)      lcqtof_t3_neg_3_038_06    m/z 773.5164     0   1E+5   12   12.5  RT (min)      lcqtof_t3_neg_2_030_23    m/z 773.5185     0   1   2E+2   10.5   11  RT (min)      lcqtof_t3_neg_3_038_06    m/z 773.6274     0   1   2   3E+4   13   13.5  RT (min)      lcqtof_t3_neg_1_000_dd    m/z 773.6278     0   1   2E+4   13   13.5  RT (min)      lcqtof_t3_neg_1_004_11    m/z 773.6305     0   1   2   3   4E+3   13.5   14  RT (min)      lcqtof_t3_neg_1_019_23    m/z 773.6322     0   2   4E+3   14   14.5  RT (min)      lcqtof_t3_neg_1_008_02    m/z 773.6326     0   2   4E+3   14   14.5  RT (min)      lcqtof_t3_neg_1_008_02    m/z 773.6336     0   2   4E+3   14   14.5  RT (min)      lcqtof_t3_neg_1_032_36    m/z 773.6338     0   2   4E+3   14   14.5  RT (min)      lcqtof_t3_neg_1_016_48    m/z 773.6566     0   1   2   3   4E+3   11.5   12  RT (min)      lcqtof_t3_neg_3_038_06    m/z 773.6591     0   1   2E+4   13.5   14  RT (min)      lcqtof_t3_neg_4_018_23    m/z 773.6597     0   1E+4   13.5   14  RT (min)      lcqtof_t3_neg_2_030_23    m/z 773.8110     0   1   2E+3   15.5   16  RT (min)      lcqtof_t3_neg_4_015_36    m/z 773.8111     0   1   2E+3   15.5   16  RT (min)      lcqtof_t3_neg_4_000_cc    m/z 774.0218     0   1   2   3E+4   13   13.5  RT (min)      lcqtof_t3_neg_4_064_17    m/z 774.0220     0   1   2   3E+4   13.5   14  RT (min)      lcqtof_t3_neg_4_064_17    m/z 774.0222     0   1   2   3E+4   13.5   14  RT (min)      lcqtof_t3_neg_4_064_17    m/z 774.0230     0   1   2   3E+4   13   13.5  RT (min)      lcqtof_t3_neg_4_064_17    m/z 774.0250     0   1   2   3E+4   13.5   14  RT (min)      lcqtof_t3_neg_4_064_17    m/z 774.0264     0   1   2   3E+4   12.5   13  RT (min)      lcqtof_t3_neg_4_064_17    m/z 774.0266     0   1E+4   14   14.5  RT (min)      lcqtof_t3_neg_4_064_17    m/z 774.1112     0   1   2   3E+4   13.5   14  RT (min)      lcqtof_t3_neg_4_064_17    m/z 774.1132     0   1E+4   13   13.5  RT (min)      lcqtof_t3_neg_4_036_47    m/z 774.1141     0   1   2   3E+4   13   13.5  RT (min)      lcqtof_t3_neg_4_064_17    m/z 774.2456     0   1E+4   4   4.5  RT (min)      lcqtof_t3_neg_4_000_ee    m/z 774.3280     0   1E+4   4   4.5  RT (min)      lcqtof_t3_neg_4_000_ee    m/z 774.4822     0   2   4   6   8E+3   12.5   13  RT (min)      lcqtof_t3_neg_1_019_23    m/z 774.4961     0   1E+4   13   13.5  RT (min)      lcqtof_t3_neg_4_018_23    m/z 774.5241     0   2   4   6   8E+4   12   12.5  RT (min)      lcqtof_t3_neg_2_030_23    m/z 774.5274     0   1   2E+2   10.5   11  RT (min)      lcqtof_t3_neg_3_038_06    m/z 774.5276     0   1   2   3   4E+2   10.5   11  RT (min)      lcqtof_t3_neg_3_038_06    m/z 774.5292     0   1   2   3   4E+2   10.5   11  RT (min)      lcqtof_t3_neg_3_038_06    m/z 774.6072     0   1E+2   11.5   12  RT (min)      lcqtof_t3_neg_3_032_24    m/z 774.6086     0   1E+2   11.5   12  RT (min)      lcqtof_t3_neg_3_032_24    m/z 774.6097     0   1E+4   12   12.5  RT (min)      lcqtof_t3_neg_2_016_05    m/z 774.6129     0   1   2   3E+2   11.5   12  RT (min)      lcqtof_t3_neg_3_032_24    m/z 774.6280     0   2   4   6   8E+3   13.5   14  RT (min)      lcqtof_t3_neg_1_011_45    m/z 774.6283     0   1E+4   12   12.5  RT (min)      lcqtof_t3_neg_3_032_24    m/z 774.6383     0   1   2E+3   16.5   17  RT (min)      lcqtof_t3_neg_2_002_35    m/z 774.6388     0   2   4   6E+3   13.5   14  RT (min)      lcqtof_t3_neg_4_040_24    m/z 774.6389     0   1   2   3E+3   14   14.5  RT (min)      lcqtof_t3_neg_1_047_22    m/z 774.6394     0   1E+4   13.5   14  RT (min)      lcqtof_t3_neg_1_031_24    m/z 774.6397     0   1   2   3   4E+3   13.5   14  RT (min)      lcqtof_t3_neg_1_031_24    m/z 774.6397     0   2   4   6   8E+3   14   14.5  RT (min)      lcqtof_t3_neg_1_023_06    m/z 774.6510     0   1   2E+4   13.5   14  RT (min)       lcqtof_t3_neg_2_030_23    m/z 774.6576     0   2   4   6E+3   13.5   14  RT (min)      lcqtof_t3_neg_1_019_23    m/z 774.8195     0   1   2   3E+3   5   5.5  RT (min)      lcqtof_t3_neg_1_064_14    m/z 775.0172     0   1   2   3E+4   12.5   13  RT (min)      lcqtof_t3_neg_4_064_17    m/z 775.0212     0   1   2   3E+4   13.5   14  RT (min)      lcqtof_t3_neg_4_064_17    m/z 775.0214     0   1   2   3E+4   13   13.5  RT (min)      lcqtof_t3_neg_4_064_17    m/z 775.0216     0   1   2   3E+4   13   13.5  RT (min)      lcqtof_t3_neg_4_064_17    m/z 775.0218     0   1   2   3E+4   13.5   14  RT (min)      lcqtof_t3_neg_4_064_17    m/z 775.0223     0   1   2   3E+4   13.5   14  RT (min)      lcqtof_t3_neg_4_064_17    m/z 775.0226     0   1   2   3E+4   14   14.5  RT (min)      lcqtof_t3_neg_4_064_17    m/z 775.1070     0   1   2E+4   13.5   14  RT (min)      lcqtof_t3_neg_4_064_17    m/z 775.1104     0   1   2E+4   13   13.5  RT (min)      lcqtof_t3_neg_4_064_17    m/z 775.1112     0   1   2E+4   13.5   14  RT (min)      lcqtof_t3_neg_4_064_17    m/z 775.1133     0   1   2E+4   13   13.5  RT (min)      lcqtof_t3_neg_4_064_17    m/z 775.1142     0   1   2E+4   13.5   14  RT (min)      lcqtof_t3_neg_4_064_17    m/z 775.4370     0   1E+4   8.5   9  RT (min)      lcqtof_t3_neg_4_000_ee    m/z 775.4564     0   1E+4   9   9.5  RT (min)      lcqtof_t3_neg_4_000_ee    m/z 775.4566     0   1E+4   9   9.5  RT (min)      lcqtof_t3_neg_4_000_ee    m/z 775.5303     0   1E+4   12   12.5  RT (min)      lcqtof_t3_neg_1_019_23    m/z 775.5311     0   1   2E+4   12   12.5  RT (min)      lcqtof_t3_neg_2_030_23    m/z 775.5321     0   1   2   3E+4   12.5   13  RT (min)      lcqtof_t3_neg_2_030_23    m/z 775.5321     0   1E+4   12   12.5  RT (min)      lcqtof_t3_neg_3_006_23    m/z 775.5767     0   1   2   3E+4   12.5   13  RT (min)      lcqtof_t3_neg_4_037_01    m/z 775.5857     0   1   2   3   4E+4   12.5   13  RT (min)      lcqtof_t3_neg_4_016_14    m/z 775.5889     0   1   2   3   4E+4   12.5   13  RT (min)      lcqtof_t3_neg_4_059_14    m/z 775.6116     0   2   4   6   8E+2   11.5   12  RT (min)      lcqtof_t3_neg_3_032_24    m/z 775.6122     0   1   2   3E+2   10.5   11  RT (min)      lcqtof_t3_neg_3_038_06    m/z 775.6126     0   1   2E+2   10.5   11  RT (min)      lcqtof_t3_neg_3_038_06    m/z 775.6126     0   2   4   6   8E+2   11.5   12  RT (min)      lcqtof_t3_neg_3_032_24    m/z 775.6150     0   2   4   6   8E+2   11.5   12  RT (min)      lcqtof_t3_neg_3_032_24    m/z 775.6152     0   2   4   6   8E+2   12   12.5  RT (min)      lcqtof_t3_neg_3_032_24    m/z 775.6162     0   1   2   3E+2   10.5   11  RT (min)      lcqtof_t3_neg_3_038_06    m/z 775.6225     0   1   2   3E+2   10.5   11  RT (min)      lcqtof_t3_neg_3_038_06    m/z 775.6369     0   1E+4   12.5   13  RT (min)      lcqtof_t3_neg_2_030_23    m/z 776.0203     0   1E+4   13   13.5  RT (min)      lcqtof_t3_neg_4_064_17    m/z 776.0221     0   1E+4   13   13.5  RT (min)      lcqtof_t3_neg_4_064_17    m/z 776.0232     0   1E+4   13.5   14  RT (min)      lcqtof_t3_neg_4_064_17    m/z 776.0233     0   1E+4   13   13.5  RT (min)      lcqtof_t3_neg_4_064_17    m/z 776.1104     0   1E+4   13   13.5  RT (min)      lcqtof_t3_neg_4_064_17    m/z 776.1120     0   1E+4   13.5   14  RT (min)      lcqtof_t3_neg_4_064_17    m/z 776.2806     0   1   2   3   4E+3   4   4.5  RT (min)      lcqtof_t3_neg_4_037_01    m/z 776.2812     0   1   2   3   4E+3   4   4.5  RT (min)      lcqtof_t3_neg_4_002_17    m/z 776.5416     0   2   4   6E+4   12.5   13  RT (min)      lcqtof_t3_neg_2_030_23    m/z 776.5799     0   2   4E+3   13.5   14  RT (min)      lcqtof_t3_neg_2_030_23    m/z 776.5800     0   1E+4   12.5   13  RT (min)       lcqtof_t3_neg_4_098_61    m/z 776.6194     0   1E+4   13.5   14  RT (min)      lcqtof_t3_neg_4_018_23    m/z 776.6240     0   1E+3   12   12.5  RT (min)      lcqtof_t3_neg_3_032_24    m/z 776.6246     0   1E+2   10.5   11  RT (min)      lcqtof_t3_neg_3_038_06    m/z 776.6252     0   1E+2   10.5   11  RT (min)      lcqtof_t3_neg_3_038_06    m/z 776.6254     0   1E+3   11.5   12  RT (min)      lcqtof_t3_neg_3_032_24    m/z 776.6256     0   2   4   6   8E+5   12   12.5  RT (min)      lcqtof_t3_neg_4_040_24    m/z 776.9087     0   1E+4   12.5   13  RT (min)      lcqtof_t3_neg_4_040_24    m/z 777.3360     0   1E+4   5   5.5  RT (min)      lcqtof_t3_neg_4_000_ee    m/z 777.5174     0   1   2E+3   16   16.5  RT (min)      lcqtof_t3_neg_2_054_29    m/z 777.5176     0   1   2E+3   16   16.5  RT (min)      lcqtof_t3_neg_1_016_48    m/z 777.5178     0   1   2E+3   15.5   16  RT (min)      lcqtof_t3_neg_2_012_41    m/z 777.5182     0   1   2E+3   16   16.5  RT (min)      lcqtof_t3_neg_2_050_01    m/z 777.5189     0   1   2E+3   15   15.5  RT (min)      lcqtof_t3_neg_2_005_34    m/z 777.5464     0   1   2E+4   12.5   13  RT (min)      lcqtof_t3_neg_2_030_23    m/z 777.5607     0   1E+4   13.5   14  RT (min)      lcqtof_t3_neg_4_011_32    m/z 777.6292     0   1E+6   12   12.5  RT (min)      lcqtof_t3_neg_4_049_55    m/z 777.6295     0   2   4   6E+3   12   12.5  RT (min)      lcqtof_t3_neg_3_032_24    m/z 777.6296     0   1   2E+2   10.5   11  RT (min)      lcqtof_t3_neg_3_038_06    m/z 777.6297     0   1   2E+2   10.5   11  RT (min)      lcqtof_t3_neg_3_038_06    m/z 777.6307     0   2   4   6E+3   12   12.5  RT (min)      lcqtof_t3_neg_3_032_24    m/z 777.7705     0   2   4E+3   14   14.5  RT (min)      lcqtof_t3_neg_4_011_32    m/z 777.9104     0   1   2E+4   12.5   13  RT (min)      lcqtof_t3_neg_2_016_05    m/z 777.9109     0   1   2E+4   12.5   13  RT (min)      lcqtof_t3_neg_2_016_05    m/z 777.9110     0   1   2E+2   12   12.5  RT (min)      lcqtof_t3_neg_3_032_24    m/z 777.9114     0   1   2E+2   11.5   12  RT (min)      lcqtof_t3_neg_3_032_24    m/z 778.0440     0   1E+4   12.5   13  RT (min)      lcqtof_t3_neg_4_040_24    m/z 778.2679     0   1   2E+3   10   10.5   11  RT (min)      lcqtof_t3_neg_1_033_53    m/z 778.2695     0   1   2E+3   10.5   11  RT (min)      lcqtof_t3_neg_1_010_27    m/z 778.2868     0   1E+4   12.5   13  RT (min)      lcqtof_t3_neg_2_028_24    m/z 778.2875     0   1E+4   12.5   13  RT (min)      lcqtof_t3_neg_2_028_24    m/z 778.2875     0   1   2   3E+4   12.5   13  RT (min)      lcqtof_t3_neg_4_040_24    m/z 778.4632     0   1   2E+2   10   10.5  RT (min)      lcqtof_t3_neg_3_038_06    m/z 778.4638     0   1   2E+2   10.5   11  RT (min)      lcqtof_t3_neg_3_038_06    m/z 778.4676     0   1E+5   12   12.5  RT (min)      lcqtof_t3_neg_1_019_23    m/z 778.5162     0   1   2E+4   12   12.5  RT (min)      lcqtof_t3_neg_4_046_33    m/z 778.5170     0   1   2E+4   12   12.5  RT (min)      lcqtof_t3_neg_4_046_33    m/z 778.5406     0   1E+4   14   14.5  RT (min)      lcqtof_t3_neg_2_053_38    m/z 778.5411     0   1E+4   13.5   14  RT (min)      lcqtof_t3_neg_2_053_38    m/z 778.5588     0   1E+4   12.5   13  RT (min)      lcqtof_t3_neg_1_019_23    m/z 778.5625     0   1E+4   12.5   13  RT (min)      lcqtof_t3_neg_1_019_23    m/z 778.5748     0   1E+4   14   14.5  RT (min)      lcqtof_t3_neg_3_039_33    m/z 778.5772     0   1   2E+4   14   14.5  RT (min)       lcqtof_t3_neg_2_026_36    m/z 778.5861     0   1   2E+3   11.5   12  RT (min)      lcqtof_t3_neg_3_038_06    m/z 778.5871     0   2   4E+3   11.5   12  RT (min)      lcqtof_t3_neg_1_003_07    m/z 778.5894     0   1E+4   14.5   15  RT (min)      lcqtof_t3_neg_4_059_14    m/z 778.5946     0   1E+4   13   13.5  RT (min)       lcqtof_t3_neg_4_098_61    m/z 778.5996     0   1   2E+3   15.5   16  RT (min)      lcqtof_t3_neg_2_030_23    m/z 778.6336     0   2   4   6E+1   11.5   12  RT (min)      lcqtof_t3_neg_3_032_24    m/z 778.6348     0   2   4E+2   10.5   11  RT (min)      lcqtof_t3_neg_3_038_06    m/z 778.6391     0   2   4   6E+2   12   12.5  RT (min)      lcqtof_t3_neg_3_032_24    m/z 778.6401     0   1   2E+2   10.5   11  RT (min)      lcqtof_t3_neg_3_038_06    m/z 778.6402     0   2   4   6E+3   12   12.5  RT (min)      lcqtof_t3_neg_3_032_24    m/z 778.6410     0   1   2E+2   11   11.5  RT (min)      lcqtof_t3_neg_3_038_06    m/z 779.4678     0   2   4E+4   12   12.5  RT (min)      lcqtof_t3_neg_1_019_23    m/z 779.5282     0   1E+4   14.5   15  RT (min)      lcqtof_t3_neg_3_076_58    m/z 779.5282     0   1   2   3   4E+4   16   16.5  RT (min)       lcqtof_t3_neg_2_051_14    m/z 779.5284     0   1   2   3   4E+4   16   16.5  RT (min)       lcqtof_t3_neg_2_056_56    m/z 779.5286     0   1   2   3E+4   15.5   16  RT (min)       lcqtof_t3_neg_2_016_05    m/z 779.5286     0   1   2   3E+4   15   15.5  RT (min)       lcqtof_t3_neg_2_000_aa    m/z 779.5286     0   1   2E+4   15   15.5  RT (min)       lcqtof_t3_neg_3_094_60    m/z 779.5289     0   1   2   3E+4   15   15.5  RT (min)       lcqtof_t3_neg_2_030_23    m/z 779.5290     0   1   2E+4   14.5   15  RT (min)      lcqtof_t3_neg_3_108_62    m/z 779.5294     0   1   2E+4   14.5   15  RT (min)      lcqtof_t3_neg_3_090_60    m/z 779.5296     0   1   2E+2   10.5   11  RT (min)      lcqtof_t3_neg_3_032_24    m/z 779.5313     0   1E+4   12.5   13  RT (min)      lcqtof_t3_neg_2_036_06    m/z 779.5314     0   1E+4   12.5   13  RT (min)      lcqtof_t3_neg_2_036_06    m/z 779.5314     0   1E+4   12.5   13  RT (min)      lcqtof_t3_neg_2_025_52    m/z 779.5320     0   1E+4   12   12.5  RT (min)      lcqtof_t3_neg_1_035_31    m/z 779.5326     0   1E+4   13.5   14  RT (min)      lcqtof_t3_neg_3_006_23    m/z 779.5326     0   1E+4   12.5   13  RT (min)      lcqtof_t3_neg_2_030_23    m/z 779.5326     0   1   2E+2   10.5   11  RT (min)      lcqtof_t3_neg_3_032_24    m/z 779.5328     0   1E+4   13   13.5  RT (min)      lcqtof_t3_neg_1_014_04    m/z 779.5334     0   1   2   3E+4   12   12.5  RT (min)      lcqtof_t3_neg_1_018_33    m/z 779.5336     0   1   2   3E+4   12   12.5  RT (min)      lcqtof_t3_neg_1_018_33    m/z 779.5336     0   1   2E+4   11   11.5  RT (min)      lcqtof_t3_neg_2_030_23    m/z 779.5340     0   1E+4   14   14.5  RT (min)       lcqtof_t3_neg_3_090_60    m/z 779.5344     0   1   2   3E+4   14   14.5  RT (min)       lcqtof_t3_neg_2_053_38    m/z 779.5347     0   1   2E+4   11   11.5  RT (min)      lcqtof_t3_neg_1_019_23    m/z 779.5350     0   2   4   6E+3   13   13.5  RT (min)      lcqtof_t3_neg_3_053_25    m/z 779.5357     0   1E+4   12.5   13  RT (min)      lcqtof_t3_neg_1_019_23    m/z 779.5357     0   1   2   3E+4   13.5   14  RT (min)      lcqtof_t3_neg_2_053_38    m/z 779.5358     0   2   4   6E+3   13   13.5  RT (min)      lcqtof_t3_neg_1_022_28    m/z 779.5359     0   1   2E+4   11   11.5  RT (min)      lcqtof_t3_neg_2_030_23    m/z 779.5359     0   1   2E+4   13.5   14  RT (min)      lcqtof_t3_neg_1_018_33    m/z 779.5359     0   1   2   3E+4   13.5   14  RT (min)      lcqtof_t3_neg_2_053_38    m/z 779.5362     0   1E+2   10   10.5  RT (min)      lcqtof_t3_neg_3_038_06    m/z 779.5365     0   1E+4   13   13.5  RT (min)      lcqtof_t3_neg_2_036_06    m/z 779.5370     0   1   2   3E+2   11   11.5  RT (min)      lcqtof_t3_neg_3_032_24    m/z 779.5374     0   1E+4   12.5   13  RT (min)      lcqtof_t3_neg_1_019_23    m/z 779.5380     0   1   2E+4   13.5   14  RT (min)      lcqtof_t3_neg_1_018_33    m/z 779.5399     0   1E+2   9.5   10  RT (min)      lcqtof_t3_neg_3_038_06    m/z 779.5874     0   2   4   6E+3   12.5   13  RT (min)      lcqtof_t3_neg_2_041_54    m/z 779.5964     0   2   4   6E+3   11.5   12  RT (min)      lcqtof_t3_neg_2_033_53    m/z 779.5973     0   1   2   3   4E+3   11   11.5  RT (min)      lcqtof_t3_neg_1_001_49    m/z 779.5973     0   1   2E+3   11   11.5  RT (min)      lcqtof_t3_neg_1_031_24    m/z 779.6392     0   1   2   3E+1   11.5   12  RT (min)      lcqtof_t3_neg_3_032_24    m/z 779.6410     0   1   2E+2   10.5   11  RT (min)      lcqtof_t3_neg_3_038_06    m/z 779.6430     0   1   2   3   4E+5   12.5   13  RT (min)      lcqtof_t3_neg_4_009_50    m/z 779.6440     0   2   4   6E+5   12.5   13  RT (min)      lcqtof_t3_neg_4_028_25    m/z 779.6442     0   2   4   6E+5   12.5   13  RT (min)      lcqtof_t3_neg_2_016_05    m/z 779.6443     0   1E+4   12   12.5  RT (min)      lcqtof_t3_neg_3_032_24    m/z 779.6444     0   1   2E+2   10.5   11  RT (min)      lcqtof_t3_neg_3_038_06    m/z 779.6446     0   1   2   3E+4   12   12.5  RT (min)      lcqtof_t3_neg_3_032_24    m/z 779.6448     0   1   2E+2   10.5   11  RT (min)      lcqtof_t3_neg_3_038_06    m/z 779.6452     0   1   2   3E+2   11.5   12  RT (min)      lcqtof_t3_neg_3_032_24    m/z 779.6461     0   1   2E+2   11   11.5  RT (min)      lcqtof_t3_neg_3_038_06    m/z 779.7844     0   1E+4   13   13.5  RT (min)      lcqtof_t3_neg_4_090_60    m/z 780.1468     0   1E+4   13   13.5  RT (min)      lcqtof_t3_neg_4_040_24    m/z 780.3310     0   1   2   3E+4   4   4.5  RT (min)      lcqtof_t3_neg_4_000_ee    m/z 780.4782     0   1   2   3E+2   10.5   11  RT (min)      lcqtof_t3_neg_3_038_06    m/z 780.4790     0   1   2   3E+2   10.5   11  RT (min)      lcqtof_t3_neg_3_038_06    m/z 780.4801     0   2   4   6   8E+4   12   12.5  RT (min)      lcqtof_t3_neg_2_030_23    m/z 780.4808     0   1   2   3E+2   10   10.5  RT (min)      lcqtof_t3_neg_3_038_06    m/z 780.4811     0   2   4   6   8E+4   11.5   12  RT (min)      lcqtof_t3_neg_1_019_23    m/z 780.4812     0   1E+5   12   12.5  RT (min)       lcqtof_t3_neg_1_019_23    m/z 780.4820     0   1   2E+2   10   10.5  RT (min)      lcqtof_t3_neg_3_038_06    m/z 780.5009     0   1E+4   11.5   12  RT (min)      lcqtof_t3_neg_4_037_01    m/z 780.5320     0   2   4   6E+3   12.5   13  RT (min)      lcqtof_t3_neg_1_031_24    m/z 780.5322     0   1E+4   13.5   14  RT (min)      lcqtof_t3_neg_1_018_33    m/z 780.5334     0   1E+4   15   15.5  RT (min)      lcqtof_t3_neg_3_111_62    m/z 780.5352     0   1E+4   14   14.5  RT (min)      lcqtof_t3_neg_1_010_27    m/z 780.5365     0   1E+4   12   12.5  RT (min)      lcqtof_t3_neg_2_019_32    m/z 780.5367     0   1E+4   14   14.5  RT (min)      lcqtof_t3_neg_2_026_36    m/z 780.5367     0   2   4   6   8E+3   13.5   14  RT (min)      lcqtof_t3_neg_1_031_24    m/z 780.5368     0   1E+4   14   14.5  RT (min)      lcqtof_t3_neg_2_005_34    m/z 780.5376     0   1E+4   13.5   14  RT (min)      lcqtof_t3_neg_1_018_33    m/z 780.5394     0   1E+4   11   11.5  RT (min)      lcqtof_t3_neg_1_019_23    m/z 780.5416     0   1E+4   11   11.5  RT (min)      lcqtof_t3_neg_2_030_23    m/z 780.5425     0   1E+4   11   11.5  RT (min)      lcqtof_t3_neg_1_019_23    m/z 780.5628     0   1E+4   12.5   13  RT (min)      lcqtof_t3_neg_1_019_23    m/z 780.6502     0   1E+2   12   12.5  RT (min)      lcqtof_t3_neg_3_032_24    m/z 780.6505     0   1E+2   12   12.5  RT (min)      lcqtof_t3_neg_3_032_24    m/z 780.6506     0   2   4E+4   12.5   13  RT (min)      lcqtof_t3_neg_2_016_05    m/z 780.6512     0   1   2   3E+2   10.5   11  RT (min)      lcqtof_t3_neg_3_038_06    m/z 780.6522     0   1   2E+4   12.5   13  RT (min)      lcqtof_t3_neg_2_016_05    m/z 780.6524     0   1E+5   12.5   13  RT (min)      lcqtof_t3_neg_2_016_05    m/z 780.6541     0   1   2   3E+2   10.5   11  RT (min)      lcqtof_t3_neg_3_038_06    m/z 780.6552     0   2   4   6E+3   12   12.5  RT (min)      lcqtof_t3_neg_3_032_24    m/z 780.6559     0   1E+2   12   12.5  RT (min)      lcqtof_t3_neg_3_032_24    m/z 780.6562     0   1   2E+2   11   11.5  RT (min)      lcqtof_t3_neg_3_038_06    m/z 780.6565     0   1E+2   12   12.5  RT (min)      lcqtof_t3_neg_3_032_24    m/z 780.6567     0   1   2E+2   11   11.5  RT (min)      lcqtof_t3_neg_3_038_06    m/z 780.8227     0   1   2   3E+3   16.5   17  RT (min)       lcqtof_t3_neg_4_000_ee    m/z 781.4386     0   1E+4   8.5   9  RT (min)      lcqtof_t3_neg_4_000_ee    m/z 781.4845     0   2   4   6E+4   12   12.5  RT (min)      lcqtof_t3_neg_1_019_23    m/z 781.4847     0   1   2   3   4E+4   11.5   12  RT (min)      lcqtof_t3_neg_1_019_23    m/z 781.4856     0   2   4   6E+4   12   12.5  RT (min)       lcqtof_t3_neg_1_019_23    m/z 781.4890     0   1   2   3   4E+4   12   12.5  RT (min)      lcqtof_t3_neg_2_030_23    m/z 781.4948     0   1   2E+2   10.5   11  RT (min)      lcqtof_t3_neg_3_038_06    m/z 781.4948     0   1   2E+4   12   12.5  RT (min)      lcqtof_t3_neg_3_006_23    m/z 781.4982     0   1   2   3E+2   10.5   11  RT (min)      lcqtof_t3_neg_3_038_06    m/z 781.5004     0   1   2E+4   12   12.5  RT (min)      lcqtof_t3_neg_2_016_05    m/z 781.5072     0   1   2E+4   12   12.5  RT (min)      lcqtof_t3_neg_2_016_05    m/z 781.5083     0   1   2   3E+2   10.5   11  RT (min)      lcqtof_t3_neg_3_038_06    m/z 781.5265     0   1   2E+4   12.5   13  RT (min)      lcqtof_t3_neg_2_004_48    m/z 781.5268     0   1   2E+4   15   15.5  RT (min)       lcqtof_t3_neg_3_112_62    m/z 781.5270     0   1   2   3E+2   9.5   10  RT (min)      lcqtof_t3_neg_3_038_06    m/z 781.5277     0   1E+4   12.5   13  RT (min)      lcqtof_t3_neg_1_031_24    m/z 781.5277     0   1E+4   14.5   15  RT (min)       lcqtof_t3_neg_3_066_57    m/z 781.5282     0   1   2   3   4E+4   16   16.5  RT (min)       lcqtof_t3_neg_2_065_51    m/z 781.5282     0   1   2   3   4E+4   15   16  RT (min)       lcqtof_t3_neg_2_043_31    m/z 781.5282     0   2   4   6E+3   13   13.5  RT (min)      lcqtof_t3_neg_2_010_08    m/z 781.5282     0   1   2   3   4E+4   15   15.5  RT (min)       lcqtof_t3_neg_2_029_12    m/z 781.5282     0   1   2   3   4E+2   10.5   11  RT (min)      lcqtof_t3_neg_3_032_24    m/z 781.5283     0   1   2   3E+4   15   15.5  RT (min)       lcqtof_t3_neg_2_030_23    m/z 781.5283     0   1E+4   12.5   13  RT (min)      lcqtof_t3_neg_1_031_24    m/z 781.5283     0   1   2   3E+4   16   16.5  RT (min)       lcqtof_t3_neg_2_009_50    m/z 781.5283     0   1   2E+4   14.5   15  RT (min)       lcqtof_t3_neg_1_023_06    m/z 781.5287     0   1   2   3E+2   9.5   10  RT (min)      lcqtof_t3_neg_3_038_06    m/z 781.5288     0   1E+4   13.5   14  RT (min)      lcqtof_t3_neg_2_046_45    m/z 781.5288     0   1E+4   14.5   15  RT (min)       lcqtof_t3_neg_3_075_58    m/z 781.5290     0   2   4   6E+3   13   13.5  RT (min)      lcqtof_t3_neg_2_010_08    m/z 781.5290     0   1E+4   12.5   13  RT (min)      lcqtof_t3_neg_2_020_46    m/z 781.5291     0   1E+4   14   14.5  RT (min)       lcqtof_t3_neg_3_092_60    m/z 781.5294     0   1E+4   12.5   13  RT (min)      lcqtof_t3_neg_1_032_36    m/z 781.5297     0   1E+4   14   14.5  RT (min)       lcqtof_t3_neg_3_090_60    m/z 781.5298     0   1   2   3   4E+2   11   11.5  RT (min)      lcqtof_t3_neg_3_032_24    m/z 781.5300     0   1   2   3   4E+2   11   11.5  RT (min)      lcqtof_t3_neg_3_032_24    m/z 781.5301     0   1E+4   11   11.5  RT (min)      lcqtof_t3_neg_3_006_23    m/z 781.5302     0   1   2   3E+2   9.5   10  RT (min)      lcqtof_t3_neg_3_038_06    m/z 781.5302     0   1   2E+4   11   11.5  RT (min)      lcqtof_t3_neg_2_030_23    m/z 781.5302     0   1E+4   11   11.5  RT (min)      lcqtof_t3_neg_3_006_23    m/z 781.5303     0   1E+4   12   12.5  RT (min)      lcqtof_t3_neg_1_055_32    m/z 781.5312     0   1E+4   13.5   14  RT (min)      lcqtof_t3_neg_2_046_45    m/z 781.5314     0   1E+4   13   13.5  RT (min)      lcqtof_t3_neg_2_056_56    m/z 781.5322     0   2   4   6   8E+3   13   13.5  RT (min)      lcqtof_t3_neg_2_020_46    m/z 781.5401     0   1   2E+4   13   13.5  RT (min)      lcqtof_t3_neg_2_051_14    m/z 781.5520     0   1   2E+4   13   13.5  RT (min)      lcqtof_t3_neg_4_059_14    m/z 781.5581     0   1   2E+4   13   13.5  RT (min)       lcqtof_t3_neg_4_059_14    m/z 781.5959     0   1E+4   12   12.5  RT (min)      lcqtof_t3_neg_4_040_24    m/z 781.5965     0   1E+4   12   12.5  RT (min)       lcqtof_t3_neg_4_040_24    m/z 781.6579     0   2   4   6   8E+1   12   12.5  RT (min)      lcqtof_t3_neg_3_032_24    m/z 781.6580     0   2   4   6   8E+1   12   12.5  RT (min)      lcqtof_t3_neg_3_032_24    m/z 781.6590     0   2   4   6   8E+1   12   12.5  RT (min)      lcqtof_t3_neg_3_032_24    m/z 781.6594     0   2   4   6E+4   12.5   13  RT (min)      lcqtof_t3_neg_2_016_05    m/z 781.6597     0   1   2   3E+4   12   12.5  RT (min)      lcqtof_t3_neg_3_032_24    m/z 781.6598     0   1   2E+2   11   11.5  RT (min)      lcqtof_t3_neg_3_038_06    m/z 781.6600     0   1   2E+2   11   11.5  RT (min)      lcqtof_t3_neg_3_038_06    m/z 781.6602     0   2   4   6   8E+1   12   12.5  RT (min)      lcqtof_t3_neg_3_032_24    m/z 781.6602     0   1   2E+2   10.5   11  RT (min)      lcqtof_t3_neg_3_038_06    m/z 781.6613     0   1   2E+2   10.5   11  RT (min)      lcqtof_t3_neg_3_038_06    m/z 781.8000     0   1E+4   13   13.5  RT (min)      lcqtof_t3_neg_1_031_24    m/z 781.9414     0   1   2E+2   12   12.5  RT (min)      lcqtof_t3_neg_3_032_24    m/z 782.1624     0   1E+4   13   13.5  RT (min)      lcqtof_t3_neg_4_040_24    m/z 782.1631     0   1E+4   13   13.5  RT (min)      lcqtof_t3_neg_1_031_24    m/z 782.4921     0   1   2   3E+2   11   11.5  RT (min)      lcqtof_t3_neg_3_038_06    m/z 782.4931     0   1   2E+4   13   13.5  RT (min)      lcqtof_t3_neg_2_043_31    m/z 782.4947     0   1E+5   13   13.5  RT (min)       lcqtof_t3_neg_4_084_59    m/z 782.4950     0   1E+5   13   13.5  RT (min)       lcqtof_t3_neg_4_086_59    m/z 782.4960     0   1   2E+2   11   11.5  RT (min)      lcqtof_t3_neg_3_038_06    m/z 782.4972     0   1   2E+5   12   12.5  RT (min)      lcqtof_t3_neg_1_019_23    m/z 782.4975     0   1   2E+2   10.5   11  RT (min)      lcqtof_t3_neg_3_038_06    m/z 782.4998     0   1   2E+2   10.5   11  RT (min)      lcqtof_t3_neg_3_038_06    m/z 782.5302     0   1   2E+4   12.5   13  RT (min)      lcqtof_t3_neg_4_016_14    m/z 782.5304     0   1E+4   16   16.5  RT (min)      lcqtof_t3_neg_3_110_62    m/z 782.5312     0   1   2E+4   12   12.5  RT (min)      lcqtof_t3_neg_4_016_14    m/z 782.5324     0   1   2E+4   12   12.5  RT (min)      lcqtof_t3_neg_4_059_14    m/z 782.5327     0   1   2E+4   12   12.5  RT (min)      lcqtof_t3_neg_4_059_14    m/z 782.5336     0   2   4   6   8E+3   13.5   14  RT (min)      lcqtof_t3_neg_1_055_32    m/z 782.5351     0   2   4   6E+3   13   13.5  RT (min)      lcqtof_t3_neg_2_028_24    m/z 782.5373     0   2   4E+3   13.5   14  RT (min)      lcqtof_t3_neg_1_033_53    m/z 782.6670     0   1E+2   12   12.5  RT (min)      lcqtof_t3_neg_3_032_24    m/z 782.6675     0   1   2E+2   12   12.5  RT (min)      lcqtof_t3_neg_3_032_24    m/z 782.6690     0   1   2   3   4E+4   12.5   13  RT (min)      lcqtof_t3_neg_2_016_05    m/z 782.6700     0   1   2E+2   11   11.5  RT (min)      lcqtof_t3_neg_3_038_06    m/z 782.6712     0   1   2E+5   13   13.5  RT (min)      lcqtof_t3_neg_2_016_05    m/z 782.6718     0   1E+3   12.5   13  RT (min)      lcqtof_t3_neg_3_032_24    m/z 782.6718     0   1   2   3   4E+4   12.5   13  RT (min)      lcqtof_t3_neg_2_016_05    m/z 782.6725     0   1   2   3E+2   11   11.5  RT (min)      lcqtof_t3_neg_3_038_06    m/z 782.6726     0   1   2   3E+2   11   11.5  RT (min)      lcqtof_t3_neg_3_038_06    m/z 782.7070     0   1E+4   1   1.5  RT (min)      lcqtof_t3_neg_4_059_14    m/z 782.8299     0   1E+4   17   17.5  RT (min)      lcqtof_t3_neg_4_035_08    m/z 782.8311     0   1E+4   16   16.5  RT (min)       lcqtof_t3_neg_4_018_23    m/z 782.9565     0   1E+4   13   13.5  RT (min)      lcqtof_t3_neg_4_040_24    m/z 783.1607     0   1E+4   4.5   5  RT (min)      lcqtof_t3_neg_4_055_38    m/z 783.1823     0   1E+4   12   12.5  RT (min)      lcqtof_t3_neg_4_002_17    m/z 783.1855     0   1E+4   12.5   13  RT (min)      lcqtof_t3_neg_4_002_17    m/z 783.1875     0   1E+4   12   12.5  RT (min)      lcqtof_t3_neg_4_002_17    m/z 783.1894     0   1   2E+4   12   12.5  RT (min)      lcqtof_t3_neg_4_064_17    m/z 783.4308     0   1E+4   8.5   9  RT (min)      lcqtof_t3_neg_4_000_ee    m/z 783.4320     0   1E+4   8.5   9  RT (min)      lcqtof_t3_neg_4_000_ee    m/z 783.4968     0   1   2   3E+2   11   11.5  RT (min)      lcqtof_t3_neg_3_038_06    m/z 783.4995     0   1E+5   12   12.5  RT (min)      lcqtof_t3_neg_1_019_23    m/z 783.4999     0   1   2   3E+2   10.5   11  RT (min)      lcqtof_t3_neg_3_038_06    m/z 783.5009     0   1   2   3E+2   11   11.5  RT (min)      lcqtof_t3_neg_3_038_06    m/z 783.5010     0   1   2   3E+2   10.5   11  RT (min)      lcqtof_t3_neg_3_038_06    m/z 783.5016     0   1   2   3E+2   10.5   11  RT (min)      lcqtof_t3_neg_3_038_06    m/z 783.5125     0   1E+5   12   12.5  RT (min)      lcqtof_t3_neg_1_019_23    m/z 783.5340     0   1   2   3E+3   14.5   15  RT (min)      lcqtof_t3_neg_1_000_dd    m/z 783.5364     0   1E+4   12   12.5  RT (min)      lcqtof_t3_neg_4_059_14    m/z 783.6116     0   1   2   3   4E+2   11.5   12  RT (min)      lcqtof_t3_neg_3_032_24    m/z 783.6122     0   1   2   3   4E+2   11.5   12  RT (min)      lcqtof_t3_neg_3_032_24    m/z 783.6134     0   1   2E+4   12   12.5  RT (min)      lcqtof_t3_neg_2_016_05    m/z 783.6138     0   1   2E+2   10   10.5  RT (min)      lcqtof_t3_neg_3_038_06    m/z 783.6147     0   1   2   3   4E+2   11.5   12  RT (min)      lcqtof_t3_neg_3_032_24    m/z 783.6148     0   1   2   3   4E+2   11.5   12  RT (min)      lcqtof_t3_neg_3_032_24    m/z 783.6158     0   1   2   3E+2   10.5   11  RT (min)      lcqtof_t3_neg_3_038_06    m/z 783.6740     0   1   2E+2   12   12.5  RT (min)      lcqtof_t3_neg_3_032_24    m/z 783.6742     0   1   2E+2   12.5   13  RT (min)      lcqtof_t3_neg_3_032_24    m/z 783.6750     0   1E+5   12.5   13  RT (min)      lcqtof_t3_neg_2_016_05    m/z 783.6750     0   1   2E+2   12   12.5  RT (min)      lcqtof_t3_neg_3_032_24    m/z 783.6752     0   1   2E+2   12   12.5  RT (min)      lcqtof_t3_neg_3_032_24    m/z 783.6754     0   1   2   3E+2   11   11.5  RT (min)      lcqtof_t3_neg_3_038_06    m/z 783.6755     0   1   2   3E+2   11   11.5  RT (min)      lcqtof_t3_neg_3_038_06    m/z 783.6758     0   1E+6   13   13.5  RT (min)       lcqtof_t3_neg_2_016_05    m/z 783.6765     0   1   2   3E+2   11   11.5  RT (min)      lcqtof_t3_neg_3_038_06    m/z 783.9576     0   1   2E+2   12.5   13  RT (min)      lcqtof_t3_neg_3_032_24    m/z 783.9582     0   1   2E+4   13   13.5  RT (min)      lcqtof_t3_neg_2_016_05    m/z 784.0932     0   1   2E+4   13   13.5  RT (min)      lcqtof_t3_neg_4_040_24    m/z 784.1794     0   1   2   3   4E+4   13   13.5  RT (min)      lcqtof_t3_neg_4_040_24    m/z 784.1797     0   1E+4   13   13.5  RT (min)      lcqtof_t3_neg_4_011_32    m/z 784.1872     0   1   2E+4   12   12.5  RT (min)      lcqtof_t3_neg_4_064_17    m/z 784.1876     0   1   2E+4   12   12.5  RT (min)      lcqtof_t3_neg_4_064_17    m/z 784.1915     0   1E+4   12.5   13  RT (min)      lcqtof_t3_neg_4_002_17    m/z 784.3382     0   1   2   3E+4   13   13.5  RT (min)      lcqtof_t3_neg_4_040_24    m/z 784.5039     0   1   2   3   4E+4   12   12.5  RT (min)      lcqtof_t3_neg_2_030_23    m/z 784.5056     0   1   2   3   4E+4   12   12.5  RT (min)      lcqtof_t3_neg_1_019_23    m/z 784.5060     0   1   2E+5   12   12.5  RT (min)      lcqtof_t3_neg_2_030_23    m/z 784.5100     0   1   2   3E+2   10.5   11  RT (min)      lcqtof_t3_neg_3_038_06    m/z 784.5108     0   1   2E+5   12   12.5  RT (min)      lcqtof_t3_neg_1_019_23    m/z 784.5113     0   1   2   3E+2   10.5   11  RT (min)      lcqtof_t3_neg_3_038_06    m/z 784.5114     0   1   2E+2   11   11.5  RT (min)      lcqtof_t3_neg_3_038_06    m/z 784.5117     0   1   2E+2   11   11.5  RT (min)      lcqtof_t3_neg_3_038_06    m/z 784.6292     0   1   2   3   4E+2   11.5   12  RT (min)      lcqtof_t3_neg_3_032_24    m/z 784.6295     0   1   2   3   4E+2   11.5   12  RT (min)      lcqtof_t3_neg_3_032_24    m/z 784.6299     0   1E+4   12   12.5  RT (min)      lcqtof_t3_neg_2_016_05    m/z 784.6308     0   1E+4   12   12.5  RT (min)      lcqtof_t3_neg_2_016_05    m/z 784.6794     0   1E+2   12   12.5  RT (min)      lcqtof_t3_neg_3_032_24    m/z 784.6798     0   1E+2   12.5   13  RT (min)      lcqtof_t3_neg_3_032_24    m/z 784.6822     0   1   2E+2   11   11.5  RT (min)      lcqtof_t3_neg_3_038_06    m/z 784.6826     0   1   2E+2   11   11.5  RT (min)      lcqtof_t3_neg_3_038_06    m/z 784.6858     0   1   2E+3   12.5   13  RT (min)      lcqtof_t3_neg_3_032_24    m/z 784.6864     0   1   2E+3   12.5   13  RT (min)      lcqtof_t3_neg_3_032_24    m/z 784.6872     0   1   2E+4   13   13.5  RT (min)      lcqtof_t3_neg_2_016_05    m/z 784.6890     0   1   2E+2   11   11.5  RT (min)      lcqtof_t3_neg_3_038_06    m/z 784.6892     0   1   2E+2   11   11.5  RT (min)      lcqtof_t3_neg_3_038_06    m/z 784.7036     0   1E+4   14   14.5  RT (min)      lcqtof_t3_neg_4_018_23    m/z 785.1508     0   1   2E+4   12.5   13  RT (min)      lcqtof_t3_neg_4_057_49    m/z 785.1536     0   1E+4   12.5   13  RT (min)      lcqtof_t3_neg_4_001_49    m/z 785.1866     0   1E+4   12   12.5  RT (min)      lcqtof_t3_neg_4_064_17    m/z 785.5068     0   1E+5   12   12.5  RT (min)       lcqtof_t3_neg_1_019_23    m/z 785.5140     0   1   2E+2   10.5   11  RT (min)      lcqtof_t3_neg_3_038_06    m/z 785.5152     0   1   2E+2   11   11.5  RT (min)      lcqtof_t3_neg_3_038_06    m/z 785.5160     0   1   2E+2   10.5   11  RT (min)      lcqtof_t3_neg_3_038_06    m/z 785.6326     0   1   2E+3   11.5   12  RT (min)      lcqtof_t3_neg_3_032_24    m/z 785.6328     0   1   2E+3   11.5   12  RT (min)      lcqtof_t3_neg_3_032_24    m/z 785.6348     0   1   2E+2   10.5   11  RT (min)      lcqtof_t3_neg_3_038_06    m/z 785.6356     0   1   2E+2   10.5   11  RT (min)      lcqtof_t3_neg_3_038_06    m/z 785.6854     0   1   2   3E+2   12.5   13  RT (min)      lcqtof_t3_neg_3_032_24    m/z 785.6856     0   1   2   3E+2   12   12.5  RT (min)      lcqtof_t3_neg_3_032_24    m/z 785.6872     0   1   2   3E+4   13   13.5  RT (min)      lcqtof_t3_neg_2_016_05    m/z 785.6880     0   1   2E+2   11   11.5  RT (min)      lcqtof_t3_neg_3_038_06    m/z 785.6880     0   1   2E+5   13   13.5  RT (min)       lcqtof_t3_neg_2_016_05    m/z 785.6903     0   2   4E+2   12.5   13  RT (min)      lcqtof_t3_neg_3_032_24    m/z 785.6922     0   1   2E+2   11   11.5  RT (min)      lcqtof_t3_neg_3_038_06    m/z 785.6930     0   1   2E+2   11   11.5  RT (min)      lcqtof_t3_neg_3_038_06    m/z 785.7078     0   1E+4   14   14.5  RT (min)      lcqtof_t3_neg_4_018_23    m/z 785.9756     0   1E+4   13.5   14  RT (min)      lcqtof_t3_neg_4_040_24    m/z 786.1493     0   1E+4   12.5   13  RT (min)      lcqtof_t3_neg_4_057_49    m/z 786.1502     0   1E+4   12.5   13  RT (min)      lcqtof_t3_neg_4_001_49    m/z 786.1506     0   1E+4   12.5   13  RT (min)      lcqtof_t3_neg_4_001_49    m/z 786.1515     0   1E+4   12.5   13  RT (min)      lcqtof_t3_neg_4_001_49    m/z 786.3098     0   1   2E+3   15   15.5  RT (min)       lcqtof_t3_neg_4_000_dd    m/z 786.3104     0   1   2E+3   15   15.5  RT (min)       lcqtof_t3_neg_4_042_10    m/z 786.3108     0   1   2E+3   14.5   15  RT (min)       lcqtof_t3_neg_4_000_dd    m/z 786.3108     0   2   4   6E+3   16.5   17  RT (min)       lcqtof_t3_neg_4_002_17    m/z 786.3113     0   1   2E+3   15.5   16  RT (min)       lcqtof_t3_neg_4_000_cc    m/z 786.3113     0   2   4E+3   16   16.5  RT (min)       lcqtof_t3_neg_4_018_23    m/z 786.3115     0   1   2E+3   15.5   16  RT (min)      lcqtof_t3_neg_4_000_ff    m/z 786.3116     0   1   2E+3   15.5   16  RT (min)       lcqtof_t3_neg_4_000_bb    m/z 786.3120     0   2   4E+3   16.5   17  RT (min)      lcqtof_t3_neg_4_010_46    m/z 786.5127     0   2   4   6E+4   12   12.5  RT (min)       lcqtof_t3_neg_2_030_23    m/z 786.5225     0   2   4E+4   12   12.5  RT (min)      lcqtof_t3_neg_2_030_23    m/z 786.5243     0   1   2   3   4E+5   12.5   13  RT (min)      lcqtof_t3_neg_2_030_23    m/z 786.5276     0   1   2E+2   10.5   11  RT (min)      lcqtof_t3_neg_3_038_06    m/z 786.5326     0   1   2E+2   10.5   11  RT (min)      lcqtof_t3_neg_3_038_06    m/z 786.5642     0   1E+4   12   12.5  RT (min)       lcqtof_t3_neg_4_016_14    m/z 786.5654     0   1E+4   12   12.5  RT (min)      lcqtof_t3_neg_4_059_14    m/z 786.6285     0   1E+4   13.5   14  RT (min)      lcqtof_t3_neg_2_019_32    m/z 786.6348     0   1E+4   13.5   14  RT (min)      lcqtof_t3_neg_4_040_24    m/z 786.6434     0   1E+4   12.5   13  RT (min)      lcqtof_t3_neg_2_016_05    m/z 786.6900     0   1   2   3E+2   12.5   13  RT (min)      lcqtof_t3_neg_3_032_24    m/z 787.0730     0   1E+4   13.5   14  RT (min)      lcqtof_t3_neg_4_064_17    m/z 787.0735     0   1E+4   13.5   14  RT (min)      lcqtof_t3_neg_4_064_17    m/z 787.0748     0   1E+4   13.5   14  RT (min)      lcqtof_t3_neg_4_064_17    m/z 787.0750     0   1E+4   13   13.5  RT (min)      lcqtof_t3_neg_4_064_17    m/z 787.0751     0   1E+4   13.5   14  RT (min)      lcqtof_t3_neg_4_064_17    m/z 787.0764     0   1E+4   13   13.5  RT (min)      lcqtof_t3_neg_4_064_17    m/z 787.0767     0   1E+4   14   14.5  RT (min)      lcqtof_t3_neg_4_064_17    m/z 787.1435     0   1E+4   4.5   5  RT (min)      lcqtof_t3_neg_4_000_ee    m/z 787.1469     0   1E+4   12.5   13  RT (min)      lcqtof_t3_neg_4_057_49    m/z 787.1502     0   1E+4   12.5   13  RT (min)      lcqtof_t3_neg_4_057_49    m/z 787.1997     0   1   2E+4   12.5   13  RT (min)      lcqtof_t3_neg_4_064_17    m/z 787.2008     0   1   2E+4   12.5   13  RT (min)      lcqtof_t3_neg_4_064_17    m/z 787.3660     0   1E+4   4.5   5  RT (min)      lcqtof_t3_neg_4_000_ee    m/z 787.5334     0   1   2E+2   10.5   11  RT (min)      lcqtof_t3_neg_3_038_06    m/z 787.5350     0   1   2E+2   10.5   11  RT (min)      lcqtof_t3_neg_3_038_06    m/z 787.6361     0   1E+4   13.5   14  RT (min)      lcqtof_t3_neg_2_028_24    m/z 787.6419     0   1E+3   11.5   12  RT (min)      lcqtof_t3_neg_3_032_24    m/z 787.6502     0   1   2E+2   10.5   11  RT (min)      lcqtof_t3_neg_3_038_06    m/z 787.6507     0   1   2E+2   10.5   11  RT (min)      lcqtof_t3_neg_3_038_06    m/z 787.6508     0   1E+3   12   12.5  RT (min)      lcqtof_t3_neg_3_032_24    m/z 787.6510     0   2   4   6E+4   12.5   13  RT (min)      lcqtof_t3_neg_2_016_05    m/z 787.6510     0   1E+3   12   12.5  RT (min)      lcqtof_t3_neg_3_032_24    m/z 787.6576     0   1E+4   13.5   14  RT (min)      lcqtof_t3_neg_4_040_24    m/z 788.0753     0   1E+4   13.5   14  RT (min)      lcqtof_t3_neg_4_064_17    m/z 788.0768     0   1E+4   13.5   14  RT (min)      lcqtof_t3_neg_4_064_17    m/z 788.2026     0   1E+4   12.5   13  RT (min)      lcqtof_t3_neg_4_064_17    m/z 788.2969     0   1   2   3   4E+3   11   11.5  RT (min)       lcqtof_t3_neg_4_058_16    m/z 788.2969     0   1   2   3   4E+3   11   11.5  RT (min)       lcqtof_t3_neg_4_037_01    m/z 788.2981     0   1   2   3E+3   10.5   11  RT (min)       lcqtof_t3_neg_4_012_27    m/z 788.2981     0   1   2   3E+3   10.5   11  RT (min)      lcqtof_t3_neg_1_002_13    m/z 788.5393     0   1   2   3E+2   10.5   11  RT (min)      lcqtof_t3_neg_3_038_06    m/z 788.5398     0   1   2E+2   10.5   11  RT (min)      lcqtof_t3_neg_3_038_06    m/z 788.5538     0   1   2E+2   11   11.5  RT (min)      lcqtof_t3_neg_3_038_06    m/z 789.0622     0   1E+4   13   13.5  RT (min)      lcqtof_t3_neg_4_064_17    m/z 789.0644     0   1E+4   13   13.5  RT (min)      lcqtof_t3_neg_4_064_17    m/z 789.0646     0   1E+4   13   13.5  RT (min)      lcqtof_t3_neg_4_064_17    m/z 789.0646     0   1E+4   13   13.5  RT (min)      lcqtof_t3_neg_4_064_17    m/z 789.0666     0   1E+4   13   13.5  RT (min)      lcqtof_t3_neg_4_064_17    m/z 789.0705     0   1E+4   13.5   14  RT (min)      lcqtof_t3_neg_4_064_17    m/z 789.0711     0   1E+4   13.5   14  RT (min)      lcqtof_t3_neg_4_064_17    m/z 789.1445     0   1   2   3E+4   13   13.5  RT (min)      lcqtof_t3_neg_4_064_17    m/z 789.1448     0   1   2E+4   12.5   13  RT (min)      lcqtof_t3_neg_4_036_47    m/z 789.1449     0   1   2   3   4E+4   12.5   13  RT (min)      lcqtof_t3_neg_4_064_17    m/z 789.1452     0   1   2   3   4E+4   13   13.5  RT (min)      lcqtof_t3_neg_4_064_17    m/z 789.1455     0   1   2   3   4E+4   12.5   13  RT (min)      lcqtof_t3_neg_4_064_17    m/z 789.3354     0   1E+4   4.5   5  RT (min)      lcqtof_t3_neg_4_000_ee    m/z 789.3371     0   1E+4   4   4.5  RT (min)      lcqtof_t3_neg_4_000_ee    m/z 789.5297     0   1   2E+4   12.5   13  RT (min)      lcqtof_t3_neg_2_030_23    m/z 789.5383     0   1   2   3E+4   12.5   13  RT (min)      lcqtof_t3_neg_2_030_23    m/z 789.5446     0   1E+4   12.5   13  RT (min)      lcqtof_t3_neg_3_006_23    m/z 789.5540     0   2   4   6E+4   12.5   13  RT (min)      lcqtof_t3_neg_2_030_23    m/z 789.5664     0   1   2E+2   11   11.5  RT (min)      lcqtof_t3_neg_3_038_06    m/z 789.6116     0   2   4   6E+3   12.5   13  RT (min)      lcqtof_t3_neg_1_045_29    m/z 789.6128     0   2   4   6E+3   12.5   13  RT (min)      lcqtof_t3_neg_2_052_17    m/z 789.6156     0   2   4   6   8E+3   12.5   13  RT (min)      lcqtof_t3_neg_2_059_14    m/z 789.6162     0   1   2E+4   12   12.5  RT (min)      lcqtof_t3_neg_1_031_24    m/z 789.6169     0   2   4   6E+3   13   13.5  RT (min)      lcqtof_t3_neg_1_014_04    m/z 789.6173     0   2   4   6E+3   12   12.5  RT (min)      lcqtof_t3_neg_1_031_24    m/z 789.6174     0   1E+4   11   11.5  RT (min)      lcqtof_t3_neg_1_019_23    m/z 789.6177     0   1E+4   11   11.5  RT (min)      lcqtof_t3_neg_2_030_23    m/z 789.6177     0   1E+4   11   11.5  RT (min)      lcqtof_t3_neg_2_030_23    m/z 789.6178     0   1   2E+2   10   10.5  RT (min)      lcqtof_t3_neg_3_038_06    m/z 789.6186     0   1E+4   11   11.5  RT (min)      lcqtof_t3_neg_2_030_23    m/z 789.6191     0   1   2   3   4E+2   11   11.5  RT (min)      lcqtof_t3_neg_3_032_24    m/z 789.6200     0   2   4E+3   14   14.5  RT (min)      lcqtof_t3_neg_1_018_33    m/z 789.6206     0   1   2E+3   14.5   15  RT (min)      lcqtof_t3_neg_1_000_aa    m/z 789.6210     0   2   4E+3   14   14.5  RT (min)      lcqtof_t3_neg_1_015_34    m/z 790.1445     0   1   2   3E+4   12.5   13  RT (min)      lcqtof_t3_neg_4_064_17    m/z 790.1455     0   1   2   3E+4   12.5   13  RT (min)      lcqtof_t3_neg_4_064_17    m/z 790.1465     0   1E+4   12.5   13  RT (min)      lcqtof_t3_neg_4_036_47    m/z 790.1466     0   1   2   3E+4   13   13.5  RT (min)      lcqtof_t3_neg_4_064_17    m/z 790.2881     0   1   2E+3   10.5   11  RT (min)       lcqtof_t3_neg_4_012_27    m/z 790.5345     0   1E+4   12.5   13  RT (min)      lcqtof_t3_neg_2_030_23    m/z 790.5583     0   1E+5   12.5   13  RT (min)      lcqtof_t3_neg_2_030_23    m/z 790.5611     0   1   2E+2   11   11.5  RT (min)      lcqtof_t3_neg_3_038_06    m/z 790.5612     0   1   2E+2   11   11.5  RT (min)      lcqtof_t3_neg_3_038_06    m/z 790.5790     0   1E+4   12   12.5  RT (min)      lcqtof_t3_neg_4_040_24    m/z 790.5795     0   1E+4   12   12.5  RT (min)      lcqtof_t3_neg_4_040_24    m/z 790.5985     0   1   2   3E+4   14   14.5  RT (min)       lcqtof_t3_neg_2_026_36    m/z 790.5991     0   1   2E+4   14   14.5  RT (min)      lcqtof_t3_neg_4_015_36    m/z 790.6125     0   1   2E+4   14   14.5  RT (min)      lcqtof_t3_neg_4_015_36    m/z 790.6180     0   2   4   6   8E+3   13.5   14  RT (min)      lcqtof_t3_neg_3_017_51    m/z 790.6188     0   1E+4   13   13.5  RT (min)      lcqtof_t3_neg_1_008_02    m/z 790.6192     0   1E+4   14   14.5  RT (min)      lcqtof_t3_neg_1_044_37    m/z 790.6204     0   2   4   6E+3   14   14.5  RT (min)      lcqtof_t3_neg_1_017_51    m/z 790.6206     0   2   4   6E+3   13.5   14  RT (min)      lcqtof_t3_neg_2_004_48    m/z 790.6210     0   1E+4   11.5   12  RT (min)       lcqtof_t3_neg_1_001_49    m/z 790.6214     0   1E+4   11.5   12  RT (min)       lcqtof_t3_neg_1_002_13    m/z 790.6223     0   1E+4   11.5   12  RT (min)      lcqtof_t3_neg_1_027_21    m/z 790.6234     0   2   4   6E+3   14   14.5  RT (min)      lcqtof_t3_neg_1_009_50    m/z 790.6234     0   1   2   3   4E+3   14   14.5  RT (min)      lcqtof_t3_neg_2_004_48    m/z 790.6238     0   1   2E+4   12   12.5  RT (min)      lcqtof_t3_neg_4_040_24    m/z 790.6242     0   1   2E+4   12   12.5  RT (min)      lcqtof_t3_neg_4_040_24    m/z 790.6248     0   2   4E+3   13.5   14  RT (min)      lcqtof_t3_neg_1_006_05    m/z 790.6270     0   1   2E+4   12.5   13  RT (min)      lcqtof_t3_neg_4_040_24    m/z 790.6292     0   1   2E+3   14.5   15  RT (min)      lcqtof_t3_neg_1_008_02    m/z 790.6303     0   1E+4   12.5   13  RT (min)      lcqtof_t3_neg_4_011_32    m/z 790.6317     0   1   2E+3   16.5   17  RT (min)      lcqtof_t3_neg_1_000_bb    m/z 790.6465     0   1E+4   13.5   14  RT (min)      lcqtof_t3_neg_2_028_24    m/z 790.6610     0   1E+4   13.5   14  RT (min)       lcqtof_t3_neg_2_019_32    m/z 790.7927     0   1E+2   13   13.5  RT (min)      lcqtof_t3_neg_3_032_24    m/z 791.1413     0   1E+4   13   13.5  RT (min)      lcqtof_t3_neg_4_002_17    m/z 791.1422     0   1   2E+4   13   13.5  RT (min)      lcqtof_t3_neg_4_064_17    m/z 791.1429     0   1   2E+4   12.5   13  RT (min)      lcqtof_t3_neg_4_064_17    m/z 791.1450     0   1   2E+4   12.5   13  RT (min)      lcqtof_t3_neg_4_064_17    m/z 791.4665     0   1   2   3E+4   8.5   9  RT (min)      lcqtof_t3_neg_4_000_ee    m/z 791.4666     0   1   2   3E+4   8.5   9  RT (min)      lcqtof_t3_neg_4_000_ee    m/z 791.5620     0   1   2   3E+4   12.5   13  RT (min)      lcqtof_t3_neg_1_019_23    m/z 791.5628     0   1   2   3   4E+4   12.5   13  RT (min)      lcqtof_t3_neg_2_030_23    m/z 791.5634     0   1   2E+4   12.5   13  RT (min)      lcqtof_t3_neg_4_018_23    m/z 791.6261     0   2   4E+3   11.5   12  RT (min)      lcqtof_t3_neg_1_001_49    m/z 792.1437     0   1E+4   12.5   13  RT (min)      lcqtof_t3_neg_4_064_17    m/z 792.4686     0   1E+4   8.5   9  RT (min)      lcqtof_t3_neg_4_000_ee    m/z 792.4720     0   1E+4   8.5   9  RT (min)      lcqtof_t3_neg_4_000_ee    m/z 792.4929     0   1E+4   5.5   6  RT (min)      lcqtof_t3_neg_4_030_31    m/z 792.5520     0   1   2   3E+4   12.5   13  RT (min)      lcqtof_t3_neg_2_030_23    m/z 792.5534     0   1E+4   13.5   14  RT (min)       lcqtof_t3_neg_4_034_34    m/z 792.5544     0   1E+4   14   14.5  RT (min)      lcqtof_t3_neg_4_003_37    m/z 792.5918     0   2   4   6E+3   13.5   14  RT (min)      lcqtof_t3_neg_2_030_23    m/z 792.6472     0   1E+3   12   12.5  RT (min)      lcqtof_t3_neg_3_032_24    m/z 792.6488     0   1E+3   12   12.5  RT (min)      lcqtof_t3_neg_3_032_24    m/z 792.8597     0   2   4   6E+4   17   17.5  RT (min)      lcqtof_t3_neg_4_035_08    m/z 793.1674     0   1   2E+4   12   12.5  RT (min)      lcqtof_t3_neg_4_064_17    m/z 793.1679     0   1   2E+4   12   12.5  RT (min)      lcqtof_t3_neg_4_064_17    m/z 793.4520     0   1E+4   1   1.5  RT (min)      lcqtof_t3_neg_4_062_13    m/z 793.5367     0   1   2   3   4E+4   13   13.5  RT (min)      lcqtof_t3_neg_4_006_16    m/z 793.5560     0   2   4   6E+3   13.5   14  RT (min)      lcqtof_t3_neg_4_018_23    m/z 793.5594     0   1E+4   12.5   13  RT (min)      lcqtof_t3_neg_2_030_23    m/z 793.5972     0   1   2   3E+3   13.5   14  RT (min)      lcqtof_t3_neg_2_030_23    m/z 793.6026     0   2   4   6   8E+3   12   12.5  RT (min)      lcqtof_t3_neg_4_040_24    m/z 793.6282     0   1   2   3   4E+3   14.5   15  RT (min)      lcqtof_t3_neg_2_019_32    m/z 793.6286     0   1   2   3   4E+3   16   16.5  RT (min)      lcqtof_t3_neg_2_024_40    m/z 793.6296     0   2   4E+3   13   13.5  RT (min)      lcqtof_t3_neg_2_049_55    m/z 793.6374     0   2   4E+3   13   13.5  RT (min)      lcqtof_t3_neg_2_049_55    m/z 793.6492     0   1   2   3E+3   14   14.5  RT (min)       lcqtof_t3_neg_2_056_56    m/z 793.6572     0   1E+4   13   13.5  RT (min)      lcqtof_t3_neg_4_040_24    m/z 794.1662     0   1E+4   12   12.5  RT (min)      lcqtof_t3_neg_4_064_17    m/z 794.1664     0   1E+4   12   12.5  RT (min)      lcqtof_t3_neg_4_064_17    m/z 794.1667     0   1E+4   12   12.5  RT (min)      lcqtof_t3_neg_4_064_17    m/z 794.3783     0   1E+4   10   10.5  RT (min)      lcqtof_t3_neg_4_040_24    m/z 794.3789     0   1E+4   10   10.5  RT (min)       lcqtof_t3_neg_4_091_60    m/z 794.3799     0   1E+4   10   10.5  RT (min)      lcqtof_t3_neg_4_040_24    m/z 794.4991     0   1E+4   11.5   12  RT (min)      lcqtof_t3_neg_2_030_23    m/z 794.5029     0   1E+4   11.5   12  RT (min)      lcqtof_t3_neg_2_030_23    m/z 794.6008     0   2   4   6E+2   11.5   12  RT (min)      lcqtof_t3_neg_3_032_24    m/z 794.6012     0   1   2   3E+4   12   12.5  RT (min)       lcqtof_t3_neg_4_040_24    m/z 794.6015     0   1   2E+4   12   12.5  RT (min)      lcqtof_t3_neg_1_031_24    m/z 794.6610     0   1E+3   12   12.5  RT (min)      lcqtof_t3_neg_3_032_24    m/z 794.6625     0   1E+4   12.5   13  RT (min)      lcqtof_t3_neg_2_016_05    m/z 794.6627     0   2   4   6   8E+2   12   12.5  RT (min)      lcqtof_t3_neg_3_032_24    m/z 795.1640     0   1E+4   12   12.5  RT (min)      lcqtof_t3_neg_4_064_17    m/z 795.1664     0   1E+4   12   12.5  RT (min)      lcqtof_t3_neg_4_064_17    m/z 795.5451     0   2   4E+3   13.5   14  RT (min)      lcqtof_t3_neg_4_037_01    m/z 795.5886     0   1   2E+4   14.5   15  RT (min)      lcqtof_t3_neg_1_000_aa    m/z 795.5957     0   1   2E+4   14   14.5  RT (min)      lcqtof_t3_neg_1_000_aa    m/z 795.6032     0   2   4   6E+2   11.5   12  RT (min)      lcqtof_t3_neg_3_032_24    m/z 795.6093     0   2   4   6   8E+2   11.5   12  RT (min)      lcqtof_t3_neg_3_032_24    m/z 795.6106     0   1   2E+2   10.5   11  RT (min)      lcqtof_t3_neg_3_038_06    m/z 795.6108     0   2   4   6   8E+2   11.5   12  RT (min)      lcqtof_t3_neg_3_032_24    m/z 795.6117     0   1   2E+2   10.5   11  RT (min)      lcqtof_t3_neg_3_038_06    m/z 795.6120     0   2   4   6   8E+2   11.5   12  RT (min)      lcqtof_t3_neg_3_032_24    m/z 795.6136     0   1   2E+2   10   10.5  RT (min)      lcqtof_t3_neg_3_038_06    m/z 796.5118     0   1   2E+2   10   10.5  RT (min)      lcqtof_t3_neg_3_038_06    m/z 796.5130     0   1   2E+2   10.5   11  RT (min)      lcqtof_t3_neg_3_038_06    m/z 796.5132     0   1   2E+2   10.5   11  RT (min)      lcqtof_t3_neg_3_038_06    m/z 796.5942     0   1E+4   14.5   15  RT (min)      lcqtof_t3_neg_1_000_aa    m/z 796.6142     0   1E+2   11.5   12  RT (min)      lcqtof_t3_neg_3_032_24    m/z 796.6157     0   1   2   3   4E+2   10   10.5  RT (min)      lcqtof_t3_neg_3_038_06    m/z 796.6178     0   1   2   3   4E+2   10.5   11  RT (min)      lcqtof_t3_neg_3_038_06    m/z 796.6227     0   1   2E+2   11.5   12  RT (min)      lcqtof_t3_neg_3_032_24    m/z 796.6234     0   2   4   6E+4   11.5   12  RT (min)      lcqtof_t3_neg_3_032_24    m/z 796.6239     0   1   2   3   4E+2   10.5   11  RT (min)      lcqtof_t3_neg_3_038_06    m/z 796.6776     0   1   2   3E+2   12   12.5  RT (min)      lcqtof_t3_neg_3_032_24    m/z 796.6790     0   1   2E+2   11   11.5  RT (min)      lcqtof_t3_neg_3_038_06    m/z 796.6790     0   1   2   3E+2   12   12.5  RT (min)      lcqtof_t3_neg_3_032_24    m/z 796.6794     0   1   2E+2   11   11.5  RT (min)      lcqtof_t3_neg_3_038_06    m/z 796.6794     0   1   2E+4   12.5   13  RT (min)      lcqtof_t3_neg_2_016_05    m/z 796.6806     0   1   2E+2   11   11.5  RT (min)      lcqtof_t3_neg_3_038_06    m/z 797.0057     0   1E+4   1   1.5  RT (min)      lcqtof_t3_neg_4_059_14    m/z 797.1752     0   1E+4   12   12.5  RT (min)      lcqtof_t3_neg_4_064_17    m/z 797.5149     0   1   2   3   4E+4   12   12.5  RT (min)      lcqtof_t3_neg_1_019_23    m/z 797.5153     0   1   2   3   4E+4   12   12.5  RT (min)      lcqtof_t3_neg_2_030_23    m/z 797.6014     0   1E+4   16   16.5  RT (min)       lcqtof_t3_neg_2_053_38    m/z 797.6015     0   2   4   6   8E+3   16   16.5  RT (min)       lcqtof_t3_neg_2_059_14    m/z 797.6017     0   1E+4   12.5   13  RT (min)      lcqtof_t3_neg_2_030_23    m/z 797.6018     0   1E+4   14.5   15  RT (min)       lcqtof_t3_neg_2_016_05    m/z 797.6020     0   2   4   6   8E+3   15   15.5  RT (min)       lcqtof_t3_neg_2_000_aa    m/z 797.6020     0   2   4   6   8E+3   15.5   16  RT (min)       lcqtof_t3_neg_2_030_23    m/z 797.6031     0   1E+4   12   12.5  RT (min)      lcqtof_t3_neg_2_030_23    m/z 797.6048     0   2   4   6   8E+3   14   14.5  RT (min)      lcqtof_t3_neg_3_017_51    m/z 797.6163     0   2   4   6E+4   13.5   14  RT (min)      lcqtof_t3_neg_4_037_01    m/z 797.6208     0   1E+4   13.5   14  RT (min)      lcqtof_t3_neg_1_019_23    m/z 797.6237     0   1E+3   11.5   12  RT (min)      lcqtof_t3_neg_3_032_24    m/z 797.6263     0   1   2E+4   13.5   14  RT (min)      lcqtof_t3_neg_4_018_23    m/z 797.6272     0   1   2   3E+2   11.5   12  RT (min)      lcqtof_t3_neg_3_032_24    m/z 797.6280     0   1   2   3E+2   10.5   11  RT (min)      lcqtof_t3_neg_3_038_06    m/z 797.6281     0   1   2   3E+2   10.5   11  RT (min)      lcqtof_t3_neg_3_038_06    m/z 797.6282     0   1   2   3   4E+4   13.5   14  RT (min)      lcqtof_t3_neg_1_032_36    m/z 797.6287     0   1   2   3   4E+4   14   14.5  RT (min)      lcqtof_t3_neg_2_026_36    m/z 797.6288     0   1E+4   16.5   17  RT (min)      lcqtof_t3_neg_4_091_60    m/z 797.7229     0   1E+3   12   12.5  RT (min)      lcqtof_t3_neg_3_032_24    m/z 797.7255     0   1E+3   12   12.5  RT (min)      lcqtof_t3_neg_3_032_24    m/z 797.7674     0   1E+4   12.5   13  RT (min)      lcqtof_t3_neg_4_040_24    m/z 797.9126     0   2   4E+2   11.5   12  RT (min)      lcqtof_t3_neg_3_032_24    m/z 798.1352     0   1E+4   12.5   13  RT (min)      lcqtof_t3_neg_4_040_24    m/z 798.1761     0   1E+4   12   12.5  RT (min)      lcqtof_t3_neg_4_064_17    m/z 798.1779     0   1E+4   12   12.5  RT (min)      lcqtof_t3_neg_4_064_17    m/z 798.3115     0   1   2   3   4E+3   16.5   17  RT (min)      lcqtof_t3_neg_4_000_dd    m/z 798.5246     0   1   2   3E+5   13   13.5  RT (min)      lcqtof_t3_neg_4_018_23    m/z 798.5264     0   2   4E+5   12   12.5  RT (min)      lcqtof_t3_neg_2_030_23    m/z 798.5266     0   1E+4   13   13.5  RT (min)       lcqtof_t3_neg_4_085_59    m/z 798.5277     0   1   2   3   4E+2   10.5   11  RT (min)      lcqtof_t3_neg_3_038_06    m/z 798.5280     0   1   2   3   4E+2   10.5   11  RT (min)      lcqtof_t3_neg_3_038_06    m/z 798.6012     0   1E+4   13.5   14  RT (min)      lcqtof_t3_neg_4_002_17    m/z 798.6035     0   1   2   3E+4   13.5   14  RT (min)      lcqtof_t3_neg_4_037_01    m/z 798.6055     0   1   2   3   4E+3   15   15.5  RT (min)      lcqtof_t3_neg_2_020_46    m/z 798.6060     0   1   2   3E+3   15   15.5  RT (min)      lcqtof_t3_neg_2_020_46    m/z 798.6062     0   1   2   3   4E+4   13.5   14  RT (min)      lcqtof_t3_neg_4_037_01    m/z 798.6088     0   1   2E+4   13   13.5  RT (min)      lcqtof_t3_neg_2_047_28    m/z 798.6095     0   1   2E+4   13.5   14  RT (min)      lcqtof_t3_neg_4_051_15    m/z 798.6149     0   1E+5   13.5   14  RT (min)      lcqtof_t3_neg_4_011_32    m/z 798.6268     0   1   2E+5   13.5   14  RT (min)      lcqtof_t3_neg_4_011_32    m/z 798.6286     0   1   2E+2   11.5   12  RT (min)      lcqtof_t3_neg_3_032_24    m/z 798.6295     0   1E+3   11.5   12  RT (min)      lcqtof_t3_neg_3_032_24    m/z 798.6306     0   1   2   3E+2   10.5   11  RT (min)      lcqtof_t3_neg_3_038_06    m/z 798.6314     0   1   2   3   4E+2   10.5   11  RT (min)      lcqtof_t3_neg_3_038_06    m/z 798.6315     0   1   2E+4   16   16.5  RT (min)      lcqtof_t3_neg_4_041_54    m/z 798.6315     0   1   2   3   4E+4   16.5   17  RT (min)       lcqtof_t3_neg_4_091_60    m/z 798.6350     0   1   2E+4   13.5   14  RT (min)      lcqtof_t3_neg_2_015_33    m/z 798.6352     0   1   15   15.5  RT (min)       lcqtof_t3_neg_3_032_24    m/z 798.6913     0   1   2E+2   12   12.5  RT (min)      lcqtof_t3_neg_3_032_24    m/z 798.6933     0   1   2E+2   12.5   13  RT (min)      lcqtof_t3_neg_3_032_24    m/z 798.6944     0   1   2E+4   13   13.5  RT (min)      lcqtof_t3_neg_2_016_05    m/z 798.6956     0   1   2E+2   11   11.5  RT (min)      lcqtof_t3_neg_3_038_06    m/z 798.7708     0   1   2   3   4E+2   11.5   12  RT (min)      lcqtof_t3_neg_3_032_24    m/z 798.7710     0   1E+4   12   12.5  RT (min)      lcqtof_t3_neg_2_016_05    m/z 798.7712     0   1   2   3   4E+2   11.5   12  RT (min)      lcqtof_t3_neg_3_032_24    m/z 798.7714     0   1E+4   12   12.5  RT (min)      lcqtof_t3_neg_2_016_05    m/z 798.8113     0   1E+4   17   17.5  RT (min)      lcqtof_t3_neg_4_035_08    m/z 798.8122     0   1E+4   16   16.5  RT (min)      lcqtof_t3_neg_1_011_45    m/z 798.9160     0   1   2   3E+2   11.5   12  RT (min)      lcqtof_t3_neg_3_032_24    m/z 798.9164     0   2   4   6E+4   12   12.5  RT (min)      lcqtof_t3_neg_2_016_05    m/z 798.9167     0   2   4   6E+4   12   12.5  RT (min)      lcqtof_t3_neg_2_016_05    m/z 798.9170     0   1   2E+2   10.5   11  RT (min)      lcqtof_t3_neg_3_038_06    m/z 798.9172     0   1   2E+2   10.5   11  RT (min)      lcqtof_t3_neg_3_038_06    m/z 799.0494     0   1   2   3E+2   11.5   12  RT (min)      lcqtof_t3_neg_3_032_24    m/z 799.0504     0   1   2   3E+2   11.5   12  RT (min)      lcqtof_t3_neg_3_032_24    m/z 799.1374     0   2   4E+2   11.5   12  RT (min)      lcqtof_t3_neg_3_032_24    m/z 799.1375     0   2   4E+2   11.5   12  RT (min)      lcqtof_t3_neg_3_032_24    m/z 799.1380     0   1   2E+4   12   12.5  RT (min)      lcqtof_t3_neg_2_016_05    m/z 799.1479     0   2   4E+2   11.5   12  RT (min)      lcqtof_t3_neg_3_032_24    m/z 799.1616     0   1E+4   12   12.5  RT (min)      lcqtof_t3_neg_4_064_17    m/z 799.2970     0   1   2E+2   11.5   12  RT (min)      lcqtof_t3_neg_3_032_24    m/z 799.2974     0   1   2E+2   11.5   12  RT (min)      lcqtof_t3_neg_3_032_24    m/z 799.2979     0   1   2E+2   11.5   12  RT (min)      lcqtof_t3_neg_3_032_24    m/z 799.2984     0   1E+4   12   12.5  RT (min)      lcqtof_t3_neg_2_016_05    m/z 799.4264     0   1E+4   12.5   13  RT (min)      lcqtof_t3_neg_1_031_24    m/z 799.4810     0   1E+4   12.5   13  RT (min)      lcqtof_t3_neg_1_019_23    m/z 799.5308     0   1   2E+5   12   12.5  RT (min)      lcqtof_t3_neg_2_030_23    m/z 799.5314     0   1   2   3   4E+2   10.5   11  RT (min)      lcqtof_t3_neg_3_038_06    m/z 799.6344     0   1E+2   11.5   12  RT (min)      lcqtof_t3_neg_3_032_24    m/z 799.6356     0   1   2E+5   12.5   13  RT (min)      lcqtof_t3_neg_2_016_05    m/z 799.6361     0   1   2E+5   12.5   13  RT (min)      lcqtof_t3_neg_2_016_05    m/z 799.6370     0   1   2E+2   10.5   11  RT (min)      lcqtof_t3_neg_3_038_06    m/z 799.6411     0   1   2E+2   10.5   11  RT (min)      lcqtof_t3_neg_3_038_06    m/z 799.6422     0   1   2E+2   11.5   12  RT (min)      lcqtof_t3_neg_3_032_24    m/z 799.6430     0   1   2E+2   12   12.5  RT (min)      lcqtof_t3_neg_3_032_24    m/z 799.6433     0   1   2E+2   10.5   11  RT (min)      lcqtof_t3_neg_3_038_06    m/z 800.1638     0   1E+4   12   12.5  RT (min)      lcqtof_t3_neg_4_064_17    m/z 800.3306     0   1E+4   4   4.5  RT (min)      lcqtof_t3_neg_4_000_ee    m/z 800.3318     0   1E+4   12.5   13  RT (min)      lcqtof_t3_neg_4_040_24    m/z 800.4886     0   1   2E+4   13   13.5  RT (min)      lcqtof_t3_neg_1_019_23    m/z 800.4961     0   1E+4   12.5   13  RT (min)      lcqtof_t3_neg_1_019_23    m/z 800.5340     0   1   2E+2   10.5   11  RT (min)      lcqtof_t3_neg_3_038_06    m/z 800.5426     0   1   2E+2   10.5   11  RT (min)      lcqtof_t3_neg_3_038_06    m/z 800.5435     0   1E+6   12.5   13  RT (min)      lcqtof_t3_neg_2_030_23    m/z 800.5439     0   1   2E+2   10.5   11  RT (min)      lcqtof_t3_neg_3_038_06    m/z 800.5881     0   1   2   3E+5   12.5   13  RT (min)      lcqtof_t3_neg_2_030_23    m/z 800.5887     0   1   2E+5   12.5   13  RT (min)      lcqtof_t3_neg_1_019_23    m/z 800.5902     0   1   2E+2   11   11.5  RT (min)      lcqtof_t3_neg_3_038_06    m/z 800.6127     0   1   2E+5   13   13.5  RT (min)      lcqtof_t3_neg_4_019_05    m/z 800.6164     0   1E+5   13   13.5  RT (min)      lcqtof_t3_neg_4_010_46    m/z 800.6439     0   2   4E+4   13.5   14  RT (min)      lcqtof_t3_neg_4_040_24    m/z 800.6456     0   1   2   3E+2   10.5   11  RT (min)      lcqtof_t3_neg_3_038_06    m/z 800.6460     0   1   2E+2   11.5   12  RT (min)      lcqtof_t3_neg_3_032_24    m/z 800.6462     0   2   4   6E+2   15   15.5  RT (min)      lcqtof_t3_neg_3_032_24    m/z 800.6464     0   1   2   3E+2   10.5   11  RT (min)      lcqtof_t3_neg_3_038_06    m/z 800.6464     0   1   2E+2   12   12.5  RT (min)      lcqtof_t3_neg_3_032_24    m/z 800.8200     0   2   4   6E+2   11.5   12  RT (min)      lcqtof_t3_neg_3_038_06    m/z 800.8208     0   1   2E+4   13.5   14  RT (min)      lcqtof_t3_neg_2_016_05    m/z 800.8244     0   1   2   3   4E+2   13   13.5  RT (min)      lcqtof_t3_neg_3_032_24    m/z 800.8308     0   1   2E+4   12.5   13  RT (min)      lcqtof_t3_neg_2_030_23    m/z 800.9334     0   1   2   3   4E+2   11.5   12  RT (min)      lcqtof_t3_neg_3_032_24    m/z 800.9342     0   1   2   3   4E+2   12   12.5  RT (min)      lcqtof_t3_neg_3_032_24    m/z 801.1566     0   1   2E+4   12.5   13  RT (min)      lcqtof_t3_neg_4_040_24    m/z 801.3160     0   1E+4   12.5   13  RT (min)      lcqtof_t3_neg_4_040_24    m/z 801.3210     0   2   4E+4   4   4.5  RT (min)       lcqtof_t3_neg_4_000_ee    m/z 801.3235     0   2   4E+4   4.5   5  RT (min)      lcqtof_t3_neg_4_000_ee    m/z 801.3347     0   2   4E+4   4.5   5  RT (min)      lcqtof_t3_neg_4_000_ee    m/z 801.3760     0   1   2E+4   4.5   5  RT (min)      lcqtof_t3_neg_4_000_ee    m/z 801.4973     0   1E+4   12.5   13  RT (min)      lcqtof_t3_neg_1_019_23    m/z 801.4994     0   1E+4   12.5   13  RT (min)      lcqtof_t3_neg_1_019_23    m/z 801.5428     0   1   2   3   4E+5   12.5   13  RT (min)      lcqtof_t3_neg_2_030_23    m/z 801.5465     0   1   2E+2   10.5   11  RT (min)      lcqtof_t3_neg_3_038_06    m/z 801.5465     0   1   2E+2   10.5   11  RT (min)      lcqtof_t3_neg_3_038_06    m/z 801.5902     0   1E+5   12.5   13  RT (min)      lcqtof_t3_neg_2_030_23    m/z 801.5905     0   1E+5   12.5   13  RT (min)      lcqtof_t3_neg_1_019_23    m/z 801.5933     0   1   2E+2   11   11.5  RT (min)      lcqtof_t3_neg_3_038_06    m/z 801.6506     0   1E+4   12.5   13  RT (min)      lcqtof_t3_neg_2_016_05    m/z 801.6509     0   1E+4   12.5   13  RT (min)      lcqtof_t3_neg_2_016_05    m/z 801.6515     0   1   2   3E+2   12   12.5  RT (min)      lcqtof_t3_neg_3_032_24    m/z 801.6520     0   1E+2   12   12.5  RT (min)      lcqtof_t3_neg_3_032_24    m/z 802.2524     0   1   2   3E+3   10.5   11  RT (min)       lcqtof_t3_neg_4_036_47    m/z 802.4031     0   1E+4   1   1.5  RT (min)      lcqtof_t3_neg_4_037_01    m/z 802.4406     0   1E+4   1   1.5  RT (min)      lcqtof_t3_neg_4_014_13    m/z 802.4624     0   1   2E+4   11.5   12  RT (min)      lcqtof_t3_neg_1_019_23    m/z 802.4994     0   1E+4   13   13.5  RT (min)      lcqtof_t3_neg_1_019_23    m/z 802.5007     0   1E+4   12.5   13  RT (min)      lcqtof_t3_neg_1_019_23    m/z 802.5150     0   1   2E+4   13   13.5  RT (min)      lcqtof_t3_neg_4_018_23    m/z 802.5162     0   1   2E+4   13.5   14  RT (min)      lcqtof_t3_neg_4_089_59    m/z 802.5382     0   1   2   3E+5   13   13.5  RT (min)      lcqtof_t3_neg_4_018_23    m/z 802.5535     0   1E+5   12.5   13  RT (min)      lcqtof_t3_neg_2_030_23    m/z 802.5558     0   1   2E+5   12.5   13  RT (min)       lcqtof_t3_neg_1_019_23    m/z 802.5597     0   1E+6   12.5   13  RT (min)      lcqtof_t3_neg_2_030_23    m/z 802.5611     0   1   2   3E+2   10.5   11  RT (min)      lcqtof_t3_neg_3_038_06    m/z 802.5612     0   1   2   3E+2   11   11.5  RT (min)      lcqtof_t3_neg_3_038_06    m/z 802.5806     0   1E+4   14   14.5  RT (min)      lcqtof_t3_neg_4_038_29    m/z 802.6166     0   1E+4   13   13.5  RT (min)      lcqtof_t3_neg_1_036_14    m/z 802.6506     0   1   2E+2   12   12.5  RT (min)      lcqtof_t3_neg_3_032_24    m/z 802.6510     0   1   2E+2   12   12.5  RT (min)      lcqtof_t3_neg_3_032_24    m/z 802.6511     0   1E+4   12.5   13  RT (min)      lcqtof_t3_neg_2_016_05    m/z 802.6520     0   1   2E+2   11.5   12  RT (min)      lcqtof_t3_neg_3_032_24    m/z 802.6534     0   1E+4   12.5   13  RT (min)      lcqtof_t3_neg_2_016_05    m/z 802.8460     0   1E+4   12.5   13  RT (min)      lcqtof_t3_neg_2_030_23    m/z 803.1040     0   2   4   6   8E+5   12.5   13  RT (min)      lcqtof_t3_neg_4_064_17    m/z 803.1053     0   1   2   3E+5   14   14.5  RT (min)      lcqtof_t3_neg_4_002_17    m/z 803.1054     0   1E+5   12.5   13  RT (min)      lcqtof_t3_neg_4_064_17    m/z 803.1095     0   1   2   3E+5   12.5   13  RT (min)      lcqtof_t3_neg_4_036_47    m/z 803.2900     0   1   2   3E+3   10   10.5  RT (min)       lcqtof_t3_neg_4_023_45    m/z 803.2917     0   1   2E+3   15.5   16  RT (min)       lcqtof_t3_neg_4_022_28    m/z 803.2917     0   1   2E+3   16   16.5  RT (min)      lcqtof_t3_neg_4_011_32    m/z 803.2919     0   1   2E+3   14   14.5  RT (min)      lcqtof_t3_neg_4_000_cc    m/z 803.2919     0   1   2E+3   5   5.5  RT (min)      lcqtof_t3_neg_4_000_aa    m/z 803.2920     0   1   2   3E+3   16   16.5  RT (min)      lcqtof_t3_neg_4_004_20    m/z 803.2921     0   1   2E+3   6.5   7  RT (min)      lcqtof_t3_neg_4_000_cc    m/z 803.2926     0   1   2   3E+3   14.5   15  RT (min)      lcqtof_t3_neg_4_006_16    m/z 803.2928     0   1   2   3   4E+3   10.5   11  RT (min)      lcqtof_t3_neg_4_006_16    m/z 803.2928     0   1   2E+3   14.5   15  RT (min)       lcqtof_t3_neg_4_000_dd    m/z 803.2929     0   1   2E+3   15   15.5  RT (min)       lcqtof_t3_neg_4_000_dd    m/z 803.2930     0   1   2E+3   8   8.5  RT (min)      lcqtof_t3_neg_4_000_cc    m/z 803.2930     0   1   2E+3   16   16.5  RT (min)      lcqtof_t3_neg_4_011_32    m/z 803.2932     0   1   2   3   4E+3   15   16   17  RT (min)       lcqtof_t3_neg_4_001_49    m/z 803.2933     0   1   2E+3   8.5   9  RT (min)      lcqtof_t3_neg_4_000_bb    m/z 803.2937     0   1   2E+3   14   14.5  RT (min)      lcqtof_t3_neg_4_039_06    m/z 803.2938     0   1E+4   4.5   5  RT (min)      lcqtof_t3_neg_4_014_13    m/z 803.3473     0   1   2E+4   4.5   5  RT (min)      lcqtof_t3_neg_4_000_ee    m/z 803.3946     0   1E+4   13   13.5  RT (min)      lcqtof_t3_neg_4_064_17    m/z 803.4754     0   1E+4   12   12.5  RT (min)      lcqtof_t3_neg_4_018_23    m/z 803.5529     0   1   2E+4   12.5   13  RT (min)      lcqtof_t3_neg_2_030_23    m/z 803.5592     0   1   2   3E+4   12.5   13  RT (min)      lcqtof_t3_neg_3_006_23    m/z 803.5627     0   1   2   3E+2   10.5   11  RT (min)      lcqtof_t3_neg_3_038_06    m/z 803.5633     0   1   2   3   4E+5   12.5   13  RT (min)      lcqtof_t3_neg_2_030_23    m/z 803.5634     0   1   2E+2   11   11.5  RT (min)      lcqtof_t3_neg_3_038_06    m/z 803.5646     0   2   4   6   8E+3   14   14.5  RT (min)      lcqtof_t3_neg_4_003_37    m/z 803.5660     0   2   4   6   8E+3   14   14.5  RT (min)      lcqtof_t3_neg_4_059_14    m/z 804.0998     0   2   4   6   8E+4   12.5   13  RT (min)      lcqtof_t3_neg_4_064_17    m/z 804.1044     0   1   2   3E+4   14   14.5  RT (min)      lcqtof_t3_neg_4_004_20    m/z 804.1061     0   1   2   3E+4   14   14.5  RT (min)      lcqtof_t3_neg_4_004_20    m/z 804.1064     0   1   2E+5   14   14.5  RT (min)      lcqtof_t3_neg_4_002_17    m/z 804.1075     0   2   4   6E+5   12.5   13  RT (min)      lcqtof_t3_neg_4_064_17    m/z 804.1970     0   2   4   6E+4   12   12.5  RT (min)      lcqtof_t3_neg_4_064_17    m/z 804.1979     0   2   4   6E+4   12   12.5  RT (min)      lcqtof_t3_neg_4_064_17    m/z 804.1980     0   1   2   3E+4   12   12.5  RT (min)      lcqtof_t3_neg_4_036_47    m/z 804.1980     0   1   2E+4   12.5   13  RT (min)      lcqtof_t3_neg_4_002_17    m/z 804.4323     0   1E+4   13   13.5  RT (min)      lcqtof_t3_neg_4_037_01    m/z 804.4326     0   1E+4   13   13.5  RT (min)      lcqtof_t3_neg_2_042_13    m/z 804.4444     0   1E+4   1   1.5  RT (min)      lcqtof_t3_neg_4_014_13    m/z 804.4777     0   1   2   3E+2   10.5   11  RT (min)      lcqtof_t3_neg_3_038_06    m/z 804.4792     0   1   2   3E+2   10.5   11  RT (min)      lcqtof_t3_neg_3_038_06    m/z 804.4794     0   1   2   3   4E+4   12   12.5  RT (min)      lcqtof_t3_neg_1_019_23    m/z 804.5650     0   1   2E+2   11   11.5  RT (min)      lcqtof_t3_neg_3_038_06    m/z 804.5661     0   1E+5   12.5   13  RT (min)      lcqtof_t3_neg_2_030_23    m/z 804.5731     0   1   2E+4   16.5   17  RT (min)      lcqtof_t3_neg_4_037_01    m/z 804.5750     0   1E+6   13.5   14  RT (min)      lcqtof_t3_neg_2_026_36    m/z 804.5750     0   1   2   3   4E+2   11   11.5  RT (min)      lcqtof_t3_neg_3_038_06    m/z 804.5760     0   2   4   6E+2   11   11.5  RT (min)      lcqtof_t3_neg_3_038_06    m/z 804.5824     0   1   2   3   4E+3   11.5   12  RT (min)      lcqtof_t3_neg_1_019_23    m/z 804.5832     0   2   4   6E+3   12   12.5  RT (min)      lcqtof_t3_neg_4_034_34    m/z 804.6474     0   1E+4   12   12.5  RT (min)      lcqtof_t3_neg_2_016_05    m/z 804.6475     0   1E+2   11.5   12  RT (min)      lcqtof_t3_neg_3_032_24    m/z 804.6475     0   1E+2   11.5   12  RT (min)      lcqtof_t3_neg_3_032_24    m/z 804.6480     0   1E+2   11.5   12  RT (min)      lcqtof_t3_neg_3_032_24    m/z 804.6502     0   1E+4   12   12.5  RT (min)      lcqtof_t3_neg_2_016_05    m/z 804.6510     0   1E+4   13.5   14  RT (min)      lcqtof_t3_neg_4_018_23    m/z 804.6739     0   1E+4   13.5   14  RT (min)       lcqtof_t3_neg_4_011_32    m/z 805.1027     0   1E+5   14   14.5  RT (min)      lcqtof_t3_neg_4_036_47    m/z 805.1032     0   2   4E+5   12.5   13  RT (min)      lcqtof_t3_neg_4_064_17    m/z 805.1032     0   1   2   3E+4   14   14.5  RT (min)      lcqtof_t3_neg_4_004_20    m/z 805.1035     0   1   2   3E+4   14   14.5  RT (min)      lcqtof_t3_neg_4_004_20    m/z 805.1053     0   2   4   6E+4   12.5   13  RT (min)      lcqtof_t3_neg_4_064_17    m/z 805.1953     0   2   4   6E+4   12   12.5  RT (min)      lcqtof_t3_neg_4_064_17    m/z 805.1956     0   2   4   6E+4   12   12.5  RT (min)      lcqtof_t3_neg_4_064_17    m/z 805.1962     0   1   2E+4   12   12.5  RT (min)      lcqtof_t3_neg_4_002_17    m/z 805.5030     0   1   2E+4   12   12.5  RT (min)      lcqtof_t3_neg_4_018_23    m/z 805.5177     0   1   2E+4   12   12.5  RT (min)      lcqtof_t3_neg_1_019_23    m/z 805.5190     0   1E+4   12   12.5  RT (min)      lcqtof_t3_neg_3_006_23    m/z 805.5209     0   1E+4   12   12.5  RT (min)      lcqtof_t3_neg_3_006_23    m/z 805.5215     0   1   2E+4   12   12.5  RT (min)      lcqtof_t3_neg_2_030_23    m/z 805.5693     0   1   2E+4   12.5   13  RT (min)      lcqtof_t3_neg_2_030_23    m/z 805.5729     0   1   2E+4   12.5   13  RT (min)      lcqtof_t3_neg_2_030_23    m/z 805.5754     0   2   4   6E+4   12.5   13  RT (min)      lcqtof_t3_neg_2_030_23    m/z 805.5811     0   1E+3   11   11.5  RT (min)      lcqtof_t3_neg_3_038_06    m/z 805.5839     0   1E+6   13.5   14  RT (min)      lcqtof_t3_neg_4_062_13    m/z 805.5902     0   2   4E+5   13.5   14  RT (min)       lcqtof_t3_neg_4_041_54    m/z 805.5912     0   1   2E+4   14   14.5  RT (min)       lcqtof_t3_neg_4_002_17    m/z 805.5949     0   2   4E+3   13.5   14  RT (min)      lcqtof_t3_neg_1_000_aa    m/z 805.6124     0   2   4   6   8E+3   14   14.5  RT (min)       lcqtof_t3_neg_4_000_aa    m/z 805.6142     0   2   4E+3   14.5   15  RT (min)      lcqtof_t3_neg_4_016_14    m/z 805.6145     0   2   4   6   8E+4   14.5   15  RT (min)       lcqtof_t3_neg_1_000_aa    m/z 805.6250     0   1E+4   14.5   15  RT (min)       lcqtof_t3_neg_4_000_ee    m/z 805.6263     0   2   4   6   8E+3   14   14.5  RT (min)       lcqtof_t3_neg_4_008_35    m/z 805.6501     0   1E+2   11.5   12  RT (min)      lcqtof_t3_neg_3_032_24    m/z 805.6509     0   1   2E+2   10.5   11  RT (min)      lcqtof_t3_neg_3_038_06    m/z 805.6510     0   1E+2   11.5   12  RT (min)      lcqtof_t3_neg_3_032_24    m/z 805.6510     0   2   4   6E+4   12   12.5  RT (min)      lcqtof_t3_neg_2_016_05    m/z 805.6518     0   1   2E+2   10.5   11  RT (min)      lcqtof_t3_neg_3_038_06    m/z 805.6520     0   1   2E+2   10.5   11  RT (min)      lcqtof_t3_neg_3_038_06    m/z 806.1001     0   1   2   3   4E+4   13.5   14  RT (min)       lcqtof_t3_neg_3_073_57    m/z 806.1018     0   1   2E+5   13   13.5  RT (min)      lcqtof_t3_neg_4_064_17    m/z 806.1040     0   1   2   3   4E+4   12.5   13  RT (min)      lcqtof_t3_neg_4_064_17    m/z 806.1043     0   1   2E+5   12.5   13  RT (min)      lcqtof_t3_neg_4_064_17    m/z 806.1951     0   1   2E+4   12   12.5  RT (min)       lcqtof_t3_neg_4_064_17    m/z 806.1971     0   1   2E+4   12   12.5  RT (min)      lcqtof_t3_neg_4_064_17    m/z 806.2522     0   1E+4   13   13.5  RT (min)      lcqtof_t3_neg_2_042_13    m/z 806.2528     0   1E+4   13   13.5  RT (min)      lcqtof_t3_neg_4_037_01    m/z 806.4306     0   1E+4   1   1.5  RT (min)      lcqtof_t3_neg_4_014_13    m/z 806.4894     0   2   4E+4   12.5   13  RT (min)      lcqtof_t3_neg_2_030_23    m/z 806.4894     0   2   4   6E+4   12.5   13  RT (min)      lcqtof_t3_neg_1_019_23    m/z 806.4942     0   2   4   6E+4   12.5   13  RT (min)      lcqtof_t3_neg_1_019_23    m/z 806.4944     0   1   2   3E+2   11   11.5  RT (min)      lcqtof_t3_neg_3_038_06    m/z 806.4954     0   1   2   3E+2   10.5   11  RT (min)      lcqtof_t3_neg_3_038_06    m/z 806.4962     0   1E+4   12   12.5  RT (min)      lcqtof_t3_neg_1_019_23    m/z 806.5312     0   1   2E+4   12   12.5  RT (min)      lcqtof_t3_neg_1_019_23    m/z 806.5322     0   1E+4   12   12.5  RT (min)      lcqtof_t3_neg_1_019_23    m/z 806.5364     0   1   2E+4   12   12.5  RT (min)      lcqtof_t3_neg_1_019_23    m/z 806.5481     0   1E+4   12   12.5  RT (min)      lcqtof_t3_neg_2_030_23    m/z 806.5878     0   1E+4   12.5   13  RT (min)      lcqtof_t3_neg_1_019_23    m/z 806.6046     0   1E+4   13.5   14  RT (min)      lcqtof_t3_neg_1_015_34    m/z 806.6055     0   1   2E+4   12.5   13  RT (min)      lcqtof_t3_neg_2_030_23    m/z 806.6057     0   2   4   6E+3   12.5   13  RT (min)      lcqtof_t3_neg_1_019_23    m/z 806.6064     0   1E+2   9.5   10  RT (min)      lcqtof_t3_neg_3_038_06    m/z 806.6067     0   1E+4   14   14.5  RT (min)      lcqtof_t3_neg_2_026_36    m/z 806.6078     0   1   2E+4   12   12.5  RT (min)      lcqtof_t3_neg_2_026_36    m/z 806.6078     0   1E+4   11   11.5  RT (min)      lcqtof_t3_neg_2_030_23    m/z 806.6114     0   2   4E+4   14   14.5  RT (min)      lcqtof_t3_neg_1_000_aa    m/z 806.6117     0   2   4E+3   14   14.5  RT (min)      lcqtof_t3_neg_1_015_34    m/z 806.6145     0   2   4E+4   14.5   15  RT (min)       lcqtof_t3_neg_1_000_aa    m/z 806.6618     0   1E+2   12   12.5  RT (min)      lcqtof_t3_neg_3_032_24    m/z 806.6620     0   1E+2   11.5   12  RT (min)      lcqtof_t3_neg_3_032_24    m/z 806.6624     0   1E+2   11.5   12  RT (min)      lcqtof_t3_neg_3_032_24    m/z 806.6634     0   1   2E+2   10.5   11  RT (min)      lcqtof_t3_neg_3_038_06    m/z 806.6640     0   1   2   3E+4   12.5   13  RT (min)      lcqtof_t3_neg_2_016_05    m/z 806.6643     0   1   2E+2   10.5   11  RT (min)      lcqtof_t3_neg_3_038_06    m/z 807.0981     0   1E+5   12.5   13  RT (min)      lcqtof_t3_neg_4_064_17    m/z 807.0998     0   1E+4   12.5   13  RT (min)      lcqtof_t3_neg_4_064_17    m/z 807.1952     0   1E+4   12   12.5  RT (min)      lcqtof_t3_neg_4_064_17    m/z 807.1960     0   1E+4   12   12.5  RT (min)      lcqtof_t3_neg_4_064_17    m/z 807.3386     0   1   2E+4   4   4.5  RT (min)      lcqtof_t3_neg_4_000_ee    m/z 807.4968     0   1   2E+4   8.5   9  RT (min)      lcqtof_t3_neg_4_000_ee    m/z 807.4992     0   1   2E+4   8.5   9  RT (min)      lcqtof_t3_neg_4_000_ee    m/z 807.6089     0   2   4E+3   12   12.5  RT (min)      lcqtof_t3_neg_4_040_24    m/z 807.6110     0   2   4   6E+3   12   12.5  RT (min)      lcqtof_t3_neg_1_019_23    m/z 807.6134     0   1E+4   12.5   13  RT (min)      lcqtof_t3_neg_2_030_23    m/z 807.6162     0   1   2E+4   14   14.5  RT (min)      lcqtof_t3_neg_1_000_aa    m/z 807.6168     0   1   2E+4   14   14.5  RT (min)      lcqtof_t3_neg_1_000_aa    m/z 807.6191     0   1   2E+4   14   14.5  RT (min)      lcqtof_t3_neg_1_000_aa    m/z 807.6212     0   1   2E+4   14   14.5  RT (min)      lcqtof_t3_neg_1_000_aa    m/z 807.6295     0   1   2E+4   14.5   15  RT (min)      lcqtof_t3_neg_1_000_aa    m/z 807.6367     0   1   2E+4   14   14.5   15  RT (min)      lcqtof_t3_neg_1_000_aa    m/z 807.6650     0   1E+2   12   12.5  RT (min)      lcqtof_t3_neg_3_032_24    m/z 807.6654     0   1E+2   10.5   11  RT (min)      lcqtof_t3_neg_3_038_06    m/z 807.6657     0   1E+2   11.5   12  RT (min)      lcqtof_t3_neg_3_032_24    m/z 807.6659     0   1E+2   10.5   11  RT (min)      lcqtof_t3_neg_3_038_06    m/z 808.0912     0   1   2   3   4E+4   13.5   14  RT (min)      lcqtof_t3_neg_4_002_17    m/z 808.0955     0   2   4E+4   13   13.5  RT (min)      lcqtof_t3_neg_4_064_17    m/z 808.0978     0   1   2   3   4E+4   13.5   14  RT (min)      lcqtof_t3_neg_4_002_17    m/z 808.1004     0   1   2   3   4E+4   13.5   14  RT (min)      lcqtof_t3_neg_4_002_17    m/z 808.1007     0   2   4E+4   13.5   14  RT (min)      lcqtof_t3_neg_4_064_17    m/z 808.1019     0   1   2   3   4E+4   13   13.5  RT (min)      lcqtof_t3_neg_4_002_17    m/z 808.1019     0   2   4E+4   13   13.5  RT (min)      lcqtof_t3_neg_4_064_17    m/z 808.1020     0   1   2   3E+4   13   13.5  RT (min)      lcqtof_t3_neg_4_036_47    m/z 808.4606     0   1E+4   11   11.5  RT (min)      lcqtof_t3_neg_4_037_01    m/z 808.4693     0   1E+4   8.5   9  RT (min)      lcqtof_t3_neg_4_000_ee    m/z 808.5026     0   1E+4   8.5   9  RT (min)      lcqtof_t3_neg_4_000_ee    m/z 808.5108     0   1   2E+2   11   11.5  RT (min)      lcqtof_t3_neg_3_038_06    m/z 808.5114     0   1   2   3   4E+2   11   11.5  RT (min)      lcqtof_t3_neg_3_038_06    m/z 808.5128     0   1   2   3   4E+2   10   10.5  RT (min)      lcqtof_t3_neg_3_038_06    m/z 808.5129     0   2   4   6E+4   12   12.5  RT (min)      lcqtof_t3_neg_2_030_23    m/z 808.5362     0   1E+4   12   12.5  RT (min)      lcqtof_t3_neg_4_037_01    m/z 808.6200     0   1E+4   13.5   14  RT (min)       lcqtof_t3_neg_4_037_01    m/z 808.6206     0   2   4   6E+3   13.5   14  RT (min)      lcqtof_t3_neg_4_003_37    m/z 808.6715     0   1   2   3   4E+4   12.5   13  RT (min)      lcqtof_t3_neg_2_016_05    m/z 808.6762     0   1E+2   12   12.5  RT (min)      lcqtof_t3_neg_3_032_24    m/z 808.6770     0   1E+2   12   12.5  RT (min)      lcqtof_t3_neg_3_032_24    m/z 808.6771     0   1   2   3E+2   10.5   11  RT (min)      lcqtof_t3_neg_3_038_06    m/z 808.6786     0   1   2   3   4E+4   12.5   13  RT (min)      lcqtof_t3_neg_2_016_05    m/z 808.6788     0   1   2   3E+2   10.5   11  RT (min)      lcqtof_t3_neg_3_038_06    m/z 808.8316     0   1   2   3E+2   13.5   14  RT (min)      lcqtof_t3_neg_3_038_06    m/z 808.8322     0   1   2   3   4E+2   15   15.5  RT (min)      lcqtof_t3_neg_3_032_24    m/z 808.8324     0   1   2E+4   16   16.5  RT (min)      lcqtof_t3_neg_4_018_23    m/z 808.8329     0   2   4   6E+2   15.5   16  RT (min)      lcqtof_t3_neg_1_019_23    m/z 808.8338     0   1   2   3E+4   17   17.5  RT (min)      lcqtof_t3_neg_4_035_08    m/z 809.0989     0   1E+4   13   13.5  RT (min)      lcqtof_t3_neg_4_064_17    m/z 809.0998     0   1E+4   13   13.5  RT (min)      lcqtof_t3_neg_4_036_47    m/z 809.5122     0   1   2E+4   13   13.5  RT (min)      lcqtof_t3_neg_4_018_23    m/z 809.5161     0   1   2   3E+4   12   12.5  RT (min)      lcqtof_t3_neg_2_030_23    m/z 809.6806     0   1E+2   12   12.5  RT (min)      lcqtof_t3_neg_3_032_24    m/z 809.6811     0   1E+2   12   12.5  RT (min)      lcqtof_t3_neg_3_032_24    m/z 809.6815     0   1   2E+2   10.5   11  RT (min)      lcqtof_t3_neg_3_038_06    m/z 809.6823     0   1E+5   12.5   13  RT (min)      lcqtof_t3_neg_2_016_05    m/z 809.6834     0   1   2E+2   11   11.5  RT (min)      lcqtof_t3_neg_3_038_06    m/z 810.5268     0   1   2   3E+2   11   11.5  RT (min)      lcqtof_t3_neg_3_038_06    m/z 810.5270     0   1   2   3E+2   11   11.5  RT (min)      lcqtof_t3_neg_3_038_06    m/z 810.5273     0   1   2   3E+2   10.5   11  RT (min)      lcqtof_t3_neg_3_038_06    m/z 810.5280     0   1   2   3E+2   10.5   11  RT (min)      lcqtof_t3_neg_3_038_06    m/z 810.5282     0   1   2   3E+2   10.5   11  RT (min)      lcqtof_t3_neg_3_038_06    m/z 810.5294     0   1   2   3E+2   10.5   11  RT (min)      lcqtof_t3_neg_3_038_06    m/z 810.5558     0   1E+4   13   13.5  RT (min)       lcqtof_t3_neg_2_019_32    m/z 810.5613     0   1E+4   13   13.5  RT (min)      lcqtof_t3_neg_2_019_32    m/z 810.5639     0   1E+4   12.5   13  RT (min)      lcqtof_t3_neg_4_059_14    m/z 810.5639     0   1E+4   12.5   13  RT (min)      lcqtof_t3_neg_4_059_14    m/z 810.5648     0   1E+4   12.5   13  RT (min)      lcqtof_t3_neg_4_016_14    m/z 810.6308     0   1E+4   12   12.5  RT (min)       lcqtof_t3_neg_4_040_24    m/z 810.6325     0   1E+4   12   12.5  RT (min)      lcqtof_t3_neg_4_040_24    m/z 810.6386     0   1   2E+4   13.5   14  RT (min)      lcqtof_t3_neg_4_018_23    m/z 810.6438     0   1E+3   13.5   14  RT (min)      lcqtof_t3_neg_1_019_23    m/z 810.6930     0   1   2   3E+2   12   12.5  RT (min)      lcqtof_t3_neg_3_032_24    m/z 810.6931     0   1   2   3E+2   12   12.5  RT (min)      lcqtof_t3_neg_3_032_24    m/z 810.6957     0   1   2   3E+2   11   11.5  RT (min)      lcqtof_t3_neg_3_038_06    m/z 810.6958     0   1   2   3E+2   11   11.5  RT (min)      lcqtof_t3_neg_3_038_06    m/z 810.6982     0   1   2   3   4E+4   12.5   13  RT (min)      lcqtof_t3_neg_2_016_05    m/z 811.5304     0   1   2E+2   11   11.5  RT (min)      lcqtof_t3_neg_3_038_06    m/z 811.5315     0   1   2   3E+4   12   12.5  RT (min)      lcqtof_t3_neg_2_030_23    m/z 811.5976     0   1E+4   13   13.5  RT (min)      lcqtof_t3_neg_4_035_08    m/z 811.6371     0   2   4E+2   11.5   12  RT (min)      lcqtof_t3_neg_3_032_24    m/z 811.6372     0   1E+4   12   12.5  RT (min)      lcqtof_t3_neg_2_016_05    m/z 811.6379     0   2   4E+2   11.5   12  RT (min)      lcqtof_t3_neg_3_032_24    m/z 811.6477     0   1E+4   13.5   14  RT (min)      lcqtof_t3_neg_4_018_23    m/z 811.6959     0   1   2E+2   12   12.5  RT (min)      lcqtof_t3_neg_3_032_24    m/z 811.6978     0   1   2E+2   12   12.5  RT (min)      lcqtof_t3_neg_3_032_24    m/z 811.6980     0   1E+5   12.5   13  RT (min)      lcqtof_t3_neg_2_016_05    m/z 811.6981     0   1   2E+2   11   11.5  RT (min)      lcqtof_t3_neg_3_038_06    m/z 811.6989     0   1   2E+2   11   11.5  RT (min)      lcqtof_t3_neg_3_038_06    m/z 812.4989     0   1   2   3E+4   12   12.5  RT (min)      lcqtof_t3_neg_1_019_23    m/z 812.5426     0   1   2   3   4E+2   10.5   11  RT (min)      lcqtof_t3_neg_3_038_06    m/z 812.5429     0   1   2   3   4E+2   10.5   11  RT (min)      lcqtof_t3_neg_3_038_06    m/z 812.5431     0   1   2E+5   12   12.5  RT (min)      lcqtof_t3_neg_2_030_23    m/z 812.5434     0   1   2   3   4E+2   10.5   11  RT (min)      lcqtof_t3_neg_3_038_06    m/z 812.6316     0   1E+4   13.5   14  RT (min)      lcqtof_t3_neg_4_040_24    m/z 812.6457     0   2   4E+2   11.5   12  RT (min)      lcqtof_t3_neg_3_032_24    m/z 812.6462     0   2   4E+2   11.5   12  RT (min)      lcqtof_t3_neg_3_032_24    m/z 812.6479     0   1E+4   12   12.5  RT (min)      lcqtof_t3_neg_2_016_05    m/z 812.6484     0   1   2   3E+2   11.5   12  RT (min)      lcqtof_t3_neg_3_032_24    m/z 812.6496     0   2   4   6   8E+3   12   12.5  RT (min)      lcqtof_t3_neg_3_032_24    m/z 812.7037     0   1   2E+4   13   13.5  RT (min)      lcqtof_t3_neg_2_016_05    m/z 812.7038     0   1   2E+4   13   13.5  RT (min)      lcqtof_t3_neg_2_016_05    m/z 812.7082     0   2   4E+2   12.5   13  RT (min)      lcqtof_t3_neg_3_032_24    m/z 812.7106     0   1   2E+2   12.5   13  RT (min)      lcqtof_t3_neg_3_032_24    m/z 813.3142     0   2   4   6E+3   10.5   11  RT (min)       lcqtof_t3_neg_4_012_27    m/z 813.3204     0   2   4E+3   15   15.5  RT (min)       lcqtof_t3_neg_4_031_42    m/z 813.3208     0   2   4E+3   15.5   16  RT (min)       lcqtof_t3_neg_4_006_16    m/z 813.3210     0   1   2   3E+3   7   7.5  RT (min)       lcqtof_t3_neg_4_008_35    m/z 813.3211     0   1   2   3   4E+3   14   14.5  RT (min)       lcqtof_t3_neg_4_000_bb    m/z 813.3211     0   1   2   3E+3   12.5   13  RT (min)       lcqtof_t3_neg_4_005_18    m/z 813.3212     0   1   2   3   4E+3   12   12.5  RT (min)       lcqtof_t3_neg_4_016_14    m/z 813.3212     0   1   2E+3   5.5   6  RT (min)       lcqtof_t3_neg_4_002_17    m/z 813.3213     0   1   2   3E+3   11   11.5  RT (min)       lcqtof_t3_neg_4_000_cc    m/z 813.3214     0   1E+4   17   17.5  RT (min)       lcqtof_t3_neg_4_035_08    m/z 813.3215     0   1   2E+3   5.5   6  RT (min)      lcqtof_t3_neg_4_004_20    m/z 813.3216     0   2   4E+3   15.5   16  RT (min)       lcqtof_t3_neg_4_002_17    m/z 813.3216     0   1   2   3E+3   11.5   12  RT (min)       lcqtof_t3_neg_4_008_35    m/z 813.3217     0   1   2E+3   5   5.5  RT (min)       lcqtof_t3_neg_4_000_bb    m/z 813.3218     0   1   2   3   4E+3   5   5.5  RT (min)       lcqtof_t3_neg_4_000_cc    m/z 813.3218     0   1   2   3E+3   7.5   8  RT (min)       lcqtof_t3_neg_4_007_09    m/z 813.3218     0   1   2   3E+3   8   8.5  RT (min)       lcqtof_t3_neg_4_006_16    m/z 813.3218     0   1   2   3E+3   6   6.5  RT (min)       lcqtof_t3_neg_4_000_cc    m/z 813.3218     0   1   2   3E+3   6.5   7  RT (min)       lcqtof_t3_neg_4_014_13    m/z 813.3219     0   1   2   3   4E+3   13   13.5  RT (min)       lcqtof_t3_neg_4_013_39    m/z 813.3220     0   1   2   3E+3   11   11.5  RT (min)       lcqtof_t3_neg_4_000_aa    m/z 813.3220     0   1   2   3   4E+3   13.5   14   14.5  RT (min)       lcqtof_t3_neg_4_007_09    m/z 813.3220     0   1   2   3E+3   9  RT (min)       lcqtof_t3_neg_4_000_cc    m/z 813.3220     0   1   2   3E+3   5.5   6  RT (min)       lcqtof_t3_neg_4_007_09    m/z 813.3220     0   1   2   3E+3   11   11.5  RT (min)       lcqtof_t3_neg_4_010_46    m/z 813.3222     0   1   2   3E+3   7   7.5  RT (min)       lcqtof_t3_neg_4_000_aa    m/z 813.3222     0   1   2   3   4E+3   12.5   13  RT (min)       lcqtof_t3_neg_4_031_42    m/z 813.3224     0   1   2   3   4E+3   12   12.5  RT (min)       lcqtof_t3_neg_4_011_32    m/z 813.3224     0   1   2   3E+3   9   9.5  RT (min)       lcqtof_t3_neg_4_000_aa    m/z 813.3226     0   1   2   3E+3   6.5   7  RT (min)       lcqtof_t3_neg_4_000_bb    m/z 813.3227     0   1   2   3E+3   9.5   10  RT (min)       lcqtof_t3_neg_4_000_bb    m/z 813.3227     0   1   2   3E+3   7   7.5  RT (min)       lcqtof_t3_neg_4_033_53    m/z 813.3227     0   2   4   6E+3   9   9.5  RT (min)       lcqtof_t3_neg_4_000_ee    m/z 813.3228     0   1   2   3E+3   8.5   9  RT (min)       lcqtof_t3_neg_4_000_aa    m/z 813.3229     0   1   2   3E+3   8.5   9  RT (min)      lcqtof_t3_neg_4_013_39    m/z 813.3229     0   1   2   3E+3   12   12.5  RT (min)       lcqtof_t3_neg_4_002_17    m/z 813.3236     0   1   2   3E+3   7.5   8  RT (min)       lcqtof_t3_neg_4_000_aa    m/z 813.5003     0   1E+4   12   12.5  RT (min)      lcqtof_t3_neg_1_019_23    m/z 813.5444     0   2   4   6E+4   13   13.5  RT (min)      lcqtof_t3_neg_4_018_23    m/z 813.5472     0   1   2E+2   10.5   11  RT (min)      lcqtof_t3_neg_3_038_06    m/z 813.5480     0   1   2E+2   10.5   11  RT (min)      lcqtof_t3_neg_3_038_06    m/z 813.5959     0   1   2E+4   15.5   16  RT (min)       lcqtof_t3_neg_2_059_14    m/z 813.5960     0   1   2E+4   15   15.5  RT (min)       lcqtof_t3_neg_2_059_14    m/z 813.5961     0   1   2E+4   15   15.5  RT (min)      lcqtof_t3_neg_2_062_17    m/z 813.5962     0   2   4E+3   12.5   13  RT (min)      lcqtof_t3_neg_2_008_07    m/z 813.5963     0   1   2E+4   16   16.5  RT (min)       lcqtof_t3_neg_2_062_17    m/z 813.5963     0   1   2E+4   14.5   15  RT (min)       lcqtof_t3_neg_2_016_05    m/z 813.5964     0   1   2E+4   15   15.5  RT (min)       lcqtof_t3_neg_2_059_14    m/z 813.5964     0   1E+4   16   16.5  RT (min)      lcqtof_t3_neg_2_048_30    m/z 813.5964     0   1E+4   15.5   16  RT (min)      lcqtof_t3_neg_2_021_27    m/z 813.5965     0   2   4E+3   12.5   13  RT (min)      lcqtof_t3_neg_2_008_07    m/z 813.5965     0   1E+4   15   15.5  RT (min)       lcqtof_t3_neg_2_000_aa    m/z 813.5965     0   1E+4   16   16.5  RT (min)       lcqtof_t3_neg_2_048_30    m/z 813.5966     0   1   2E+4   15   15.5  RT (min)       lcqtof_t3_neg_2_002_35    m/z 813.5966     0   1   2E+4   15   15.5  RT (min)      lcqtof_t3_neg_2_062_17    m/z 813.5969     0   1E+4   16   16.5  RT (min)       lcqtof_t3_neg_2_047_28    m/z 813.5970     0   1E+4   16   16.5  RT (min)      lcqtof_t3_neg_2_047_28    m/z 813.5971     0   1   2E+4   16   16.5  RT (min)       lcqtof_t3_neg_2_058_15    m/z 813.5974     0   2   4   6E+3   12.5   13  RT (min)      lcqtof_t3_neg_2_041_54    m/z 813.5979     0   1   2E+4   11   11.5  RT (min)      lcqtof_t3_neg_2_016_05    m/z 813.5980     0   1E+4   16   16.5  RT (min)      lcqtof_t3_neg_2_021_27    m/z 813.5987     0   1   2E+2   10   10.5  RT (min)      lcqtof_t3_neg_3_038_06    m/z 813.5993     0   1   2   3E+2   11   11.5  RT (min)      lcqtof_t3_neg_3_032_24    m/z 813.6009     0   1   2E+4   13.5   14  RT (min)      lcqtof_t3_neg_1_031_24    m/z 813.6104     0   1E+4   13   13.5  RT (min)      lcqtof_t3_neg_2_028_24    m/z 813.6136     0   1E+4   13.5   14  RT (min)      lcqtof_t3_neg_2_028_24    m/z 813.6160     0   1E+4   13   13.5  RT (min)      lcqtof_t3_neg_4_047_07    m/z 813.6329     0   1E+4   13.5   14  RT (min)      lcqtof_t3_neg_2_028_24    m/z 813.6557     0   1   2E+2   11.5   12  RT (min)      lcqtof_t3_neg_3_032_24    m/z 813.6566     0   1   2   3E+2   10.5   11  RT (min)      lcqtof_t3_neg_3_038_06    m/z 813.6567     0   1   2E+2   12   12.5  RT (min)      lcqtof_t3_neg_3_032_24    m/z 813.6574     0   1   2   3E+2   10.5   11  RT (min)      lcqtof_t3_neg_3_038_06    m/z 813.7036     0   2   4   6E+4   13   13.5  RT (min)      lcqtof_t3_neg_2_016_05    m/z 813.7124     0   1   2   3   4E+2   12.5   13  RT (min)      lcqtof_t3_neg_3_032_24    m/z 813.7128     0   1   2   3   4E+2   12.5   13  RT (min)      lcqtof_t3_neg_3_032_24    m/z 813.7138     0   1   2E+2   11   11.5  RT (min)      lcqtof_t3_neg_3_038_06    m/z 813.7142     0   1   2E+2   11   11.5  RT (min)      lcqtof_t3_neg_3_038_06    m/z 814.3169     0   1   2   3E+3   10.5   11  RT (min)       lcqtof_t3_neg_4_014_13    m/z 814.3232     0   1   2   3   4E+3   16   16.5  RT (min)      lcqtof_t3_neg_4_018_23    m/z 814.3237     0   1   2E+3   15   15.5  RT (min)       lcqtof_t3_neg_4_000_dd    m/z 814.3238     0   1   2E+3   16   16.5  RT (min)       lcqtof_t3_neg_4_000_bb    m/z 814.4395     0   1E+4   12   12.5  RT (min)      lcqtof_t3_neg_1_019_23    m/z 814.4817     0   1E+4   1   1.5  RT (min)      lcqtof_t3_neg_4_037_01    m/z 814.4950     0   1E+4   12   12.5  RT (min)      lcqtof_t3_neg_4_018_23    m/z 814.5375     0   1   2   3E+4   12   12.5  RT (min)      lcqtof_t3_neg_2_030_23    m/z 814.5602     0   1E+4   10   10.5  RT (min)      lcqtof_t3_neg_4_059_14    m/z 814.5605     0   1   2   3E+2   10.5   11  RT (min)      lcqtof_t3_neg_3_038_06    m/z 814.5610     0   1   2   3E+2   11   11.5  RT (min)      lcqtof_t3_neg_3_038_06    m/z 814.5994     0   2   4   6   8E+3   16   16.5  RT (min)      lcqtof_t3_neg_2_051_14    m/z 814.6028     0   1E+4   11   11.5  RT (min)      lcqtof_t3_neg_2_016_05    m/z 814.6608     0   1   2E+2   11.5   12  RT (min)      lcqtof_t3_neg_3_032_24    m/z 814.6608     0   1   2E+2   10.5   11  RT (min)      lcqtof_t3_neg_3_038_06    m/z 814.6610     0   1   2E+2   12   12.5  RT (min)      lcqtof_t3_neg_3_032_24    m/z 814.6614     0   1   2E+4   12.5   13  RT (min)      lcqtof_t3_neg_2_016_05    m/z 814.6622     0   1   2E+2   11.5   12  RT (min)      lcqtof_t3_neg_3_032_24    m/z 814.7179     0   1E+4   13   13.5  RT (min)      lcqtof_t3_neg_4_040_24    m/z 815.1797     0   1E+4   13.5   14  RT (min)      lcqtof_t3_neg_4_064_17    m/z 815.3408     0   2   4E+4   4   4.5  RT (min)      lcqtof_t3_neg_4_000_ee    m/z 815.5237     0   1E+4   12   12.5  RT (min)      lcqtof_t3_neg_2_030_23    m/z 815.5238     0   1E+4   12   12.5  RT (min)      lcqtof_t3_neg_1_019_23    m/z 815.5636     0   1E+2   10.5   11  RT (min)      lcqtof_t3_neg_3_038_06    m/z 815.5637     0   1E+2   11   11.5  RT (min)      lcqtof_t3_neg_3_038_06    m/z 815.5816     0   1   2   3E+3   16   16.5  RT (min)      lcqtof_t3_neg_2_000_cc    m/z 815.5826     0   1   2   3   4E+3   16   16.5  RT (min)      lcqtof_t3_neg_1_030_42    m/z 815.5833     0   1   2   3   4E+3   15.5   16  RT (min)      lcqtof_t3_neg_1_021_39    m/z 815.5841     0   1   2   3E+3   16   16.5  RT (min)      lcqtof_t3_neg_2_024_40    m/z 815.5862     0   1   2   3E+3   15   15.5  RT (min)      lcqtof_t3_neg_1_007_03    m/z 815.5869     0   1   2   3E+3   14.5   15  RT (min)      lcqtof_t3_neg_1_019_23    m/z 815.5888     0   1   2   3   4E+3   16   16.5  RT (min)      lcqtof_t3_neg_1_009_50    m/z 815.5905     0   1   2   3   4E+3   15   15.5  RT (min)      lcqtof_t3_neg_2_000_aa    m/z 815.6659     0   1   2   3   4E+2   12   12.5  RT (min)      lcqtof_t3_neg_3_032_24    m/z 815.6664     0   1E+4   12.5   13  RT (min)      lcqtof_t3_neg_2_016_05    m/z 815.6668     0   1   2   3   4E+2   12   12.5  RT (min)      lcqtof_t3_neg_3_032_24    m/z 815.6675     0   1E+4   12.5   13  RT (min)      lcqtof_t3_neg_2_016_05    m/z 815.6676     0   1   2   3   4E+2   12   12.5  RT (min)      lcqtof_t3_neg_3_032_24    m/z 815.7288     0   2   4E+2   12.5   13  RT (min)      lcqtof_t3_neg_3_032_24    m/z 815.7298     0   2   4E+2   12.5   13  RT (min)      lcqtof_t3_neg_3_032_24    m/z 816.1800     0   1E+4   13.5   14  RT (min)      lcqtof_t3_neg_4_064_17    m/z 816.3418     0   1E+4   4   4.5  RT (min)      lcqtof_t3_neg_4_000_ee    m/z 816.5507     0   1   2   3   4E+4   12.5   13  RT (min)      lcqtof_t3_neg_2_030_23    m/z 816.5528     0   1   2   3E+4   12.5   13  RT (min)      lcqtof_t3_neg_1_019_23    m/z 816.5766     0   1   2E+2   11   11.5  RT (min)      lcqtof_t3_neg_3_038_06    m/z 816.6762     0   1E+3   12   12.5  RT (min)      lcqtof_t3_neg_3_032_24    m/z 816.6768     0   1E+3   12   12.5  RT (min)      lcqtof_t3_neg_3_032_24    m/z 816.6781     0   1E+4   12.5   13  RT (min)      lcqtof_t3_neg_2_016_05    m/z 817.1531     0   1E+4   13.5   14  RT (min)      lcqtof_t3_neg_4_064_17    m/z 817.1563     0   1E+4   13   13.5  RT (min)      lcqtof_t3_neg_4_064_17    m/z 817.1583     0   1E+4   12.5   13  RT (min)      lcqtof_t3_neg_4_036_47    m/z 817.1726     0   1E+4   13.5   14  RT (min)      lcqtof_t3_neg_4_064_17    m/z 817.5287     0   1E+4   12.5   13  RT (min)      lcqtof_t3_neg_2_030_23    m/z 817.5349     0   1E+4   12.5   13  RT (min)      lcqtof_t3_neg_2_030_23    m/z 817.5769     0   1   2E+4   12.5   13  RT (min)      lcqtof_t3_neg_2_016_05    m/z 817.5780     0   1   2E+4   12.5   13  RT (min)      lcqtof_t3_neg_2_016_05    m/z 817.5794     0   1E+2   11   11.5  RT (min)      lcqtof_t3_neg_3_038_06    m/z 818.5333     0   1E+4   13   13.5  RT (min)      lcqtof_t3_neg_3_030_03    m/z 818.5512     0   1E+4   12.5   13  RT (min)      lcqtof_t3_neg_4_064_17    m/z 818.5528     0   1E+4   12.5   13  RT (min)      lcqtof_t3_neg_4_051_15    m/z 818.5565     0   1   2   3E+4   12.5   13  RT (min)       lcqtof_t3_neg_2_030_23    m/z 818.5743     0   1   2   3   4E+4   12.5   13  RT (min)      lcqtof_t3_neg_1_019_23    m/z 818.5914     0   1   2   3E+4   13   13.5  RT (min)      lcqtof_t3_neg_2_030_23    m/z 818.5970     0   1   2   3E+2   11.5   12  RT (min)      lcqtof_t3_neg_3_032_24    m/z 818.5977     0   1E+2   11.5   12  RT (min)      lcqtof_t3_neg_3_032_24    m/z 818.5990     0   1   2E+2   10   10.5  RT (min)      lcqtof_t3_neg_3_038_06    m/z 818.5995     0   1E+4   12   12.5  RT (min)      lcqtof_t3_neg_2_016_05    m/z 818.6014     0   1E+4   12   12.5  RT (min)      lcqtof_t3_neg_2_016_05    m/z 819.0266     0   1E+5   14   14.5  RT (min)      lcqtof_t3_neg_4_064_17    m/z 819.0270     0   1E+5   14   14.5  RT (min)      lcqtof_t3_neg_4_036_47    m/z 819.0273     0   1   2E+5   13.5   14  RT (min)      lcqtof_t3_neg_4_064_17    m/z 819.0274     0   1   2E+5   13.5   14  RT (min)      lcqtof_t3_neg_4_064_17    m/z 819.0278     0   1   2E+5   12.5   13  RT (min)      lcqtof_t3_neg_4_064_17    m/z 819.0280     0   1E+5   14   14.5  RT (min)      lcqtof_t3_neg_4_064_17    m/z 819.0282     0   1   2E+5   13.5   14  RT (min)      lcqtof_t3_neg_4_002_17    m/z 819.0282     0   1   2E+5   13   13.5  RT (min)      lcqtof_t3_neg_4_064_17    m/z 819.0288     0   1E+4   13.5   14  RT (min)      lcqtof_t3_neg_4_003_37    m/z 819.4776     0   1E+4   9   9.5  RT (min)      lcqtof_t3_neg_4_000_ee    m/z 819.5158     0   1E+4   9   9.5  RT (min)      lcqtof_t3_neg_4_000_ee    m/z 819.5575     0   1   2E+4   12   12.5  RT (min)      lcqtof_t3_neg_4_016_14    m/z 819.5576     0   1   2E+4   12   12.5  RT (min)      lcqtof_t3_neg_4_059_14    m/z 819.5578     0   1   2E+4   12   12.5  RT (min)      lcqtof_t3_neg_4_016_14    m/z 819.5590     0   1   2E+4   12   12.5  RT (min)      lcqtof_t3_neg_4_059_14    m/z 819.6072     0   1   2E+2   10.5   11  RT (min)      lcqtof_t3_neg_3_038_06    m/z 819.6075     0   1E+4   12   12.5  RT (min)      lcqtof_t3_neg_2_016_05    m/z 819.6087     0   2   4   6   8E+2   11.5   12  RT (min)      lcqtof_t3_neg_3_032_24    m/z 819.6093     0   1   2E+2   10.5   11  RT (min)      lcqtof_t3_neg_3_038_06    m/z 819.6096     0   2   4   6   8E+2   11.5   12  RT (min)      lcqtof_t3_neg_3_032_24    m/z 820.0270     0   1   2   3   4E+4   14   14.5  RT (min)      lcqtof_t3_neg_4_036_47    m/z 820.0274     0   1   2   3   4E+4   14   14.5  RT (min)      lcqtof_t3_neg_4_036_47    m/z 820.0276     0   1E+5   14   14.5  RT (min)      lcqtof_t3_neg_4_002_17    m/z 820.0282     0   2   4   6   8E+4   13   13.5  RT (min)      lcqtof_t3_neg_4_036_47    m/z 820.0284     0   1   2E+4   13.5   14  RT (min)      lcqtof_t3_neg_4_022_28    m/z 820.0285     0   1E+5   13   13.5  RT (min)      lcqtof_t3_neg_4_064_17    m/z 820.0288     0   1E+5   13.5   14  RT (min)      lcqtof_t3_neg_4_002_17    m/z 820.0288     0   1E+5   13.5   14  RT (min)      lcqtof_t3_neg_4_064_17    m/z 820.0292     0   1E+5   13.5   14  RT (min)      lcqtof_t3_neg_4_064_17    m/z 820.0296     0   1   2E+4   13   13.5  RT (min)      lcqtof_t3_neg_4_022_28    m/z 820.0301     0   2   4   6   8E+4   13   13.5  RT (min)      lcqtof_t3_neg_4_036_47    m/z 820.0301     0   1E+5   12.5   13  RT (min)      lcqtof_t3_neg_4_064_17    m/z 820.5712     0   1   2   3   4E+4   12.5   13  RT (min)       lcqtof_t3_neg_4_037_01    m/z 820.6075     0   1   2   3   4E+4   12   12.5  RT (min)      lcqtof_t3_neg_2_016_05    m/z 820.6112     0   1   2   3   4E+3   11.5   12  RT (min)      lcqtof_t3_neg_3_032_24    m/z 820.6116     0   1   2E+3   11.5   12  RT (min)      lcqtof_t3_neg_3_032_24    m/z 820.6126     0   1   2E+2   10.5   11  RT (min)      lcqtof_t3_neg_3_038_06    m/z 820.6128     0   1   2E+2   10.5   11  RT (min)      lcqtof_t3_neg_3_038_06    m/z 820.6138     0   1   2E+2   10.5   11  RT (min)      lcqtof_t3_neg_3_038_06    m/z 821.0259     0   1   2   3   4E+4   14   14.5  RT (min)      lcqtof_t3_neg_4_036_47    m/z 821.0264     0   1E+5   13.5   14  RT (min)      lcqtof_t3_neg_4_064_17    m/z 821.0266     0   1E+5   13.5   14  RT (min)      lcqtof_t3_neg_4_064_17    m/z 821.0268     0   1E+5   12.5   13  RT (min)      lcqtof_t3_neg_4_064_17    m/z 821.0268     0   1E+5   13   13.5  RT (min)      lcqtof_t3_neg_4_064_17    m/z 821.0274     0   1E+5   14   14.5  RT (min)      lcqtof_t3_neg_4_064_17    m/z 821.6162     0   2   4   6E+2   11.5   12  RT (min)      lcqtof_t3_neg_3_032_24    m/z 821.6170     0   2   4   6E+2   11.5   12  RT (min)      lcqtof_t3_neg_3_032_24    m/z 821.6191     0   1E+4   12   12.5  RT (min)      lcqtof_t3_neg_3_032_24    m/z 821.6250     0   1   2   3E+4   12.5   13  RT (min)      lcqtof_t3_neg_2_016_05    m/z 821.6258     0   1   2E+2   11   11.5  RT (min)      lcqtof_t3_neg_3_038_06    m/z 821.6258     0   1   2   3E+4   12.5   13  RT (min)      lcqtof_t3_neg_2_016_05    m/z 821.6267     0   1   2E+2   11   11.5  RT (min)      lcqtof_t3_neg_3_038_06    m/z 822.0254     0   2   4   6E+4   12.5   13  RT (min)      lcqtof_t3_neg_4_064_17    m/z 822.0263     0   2   4   6E+4   13.5   14  RT (min)      lcqtof_t3_neg_4_064_17    m/z 822.0264     0   2   4   6E+4   13   13.5  RT (min)      lcqtof_t3_neg_4_064_17    m/z 822.0265     0   2   4   6E+4   13   13.5  RT (min)      lcqtof_t3_neg_4_064_17    m/z 822.0269     0   1   2   3   4E+4   14   14.5  RT (min)      lcqtof_t3_neg_4_002_17    m/z 822.0272     0   2   4   6E+4   14   14.5  RT (min)      lcqtof_t3_neg_4_002_17    m/z 822.0272     0   2   4   6E+4   13.5   14  RT (min)      lcqtof_t3_neg_4_002_17    m/z 822.0283     0   2   4   6E+4   13.5   14  RT (min)      lcqtof_t3_neg_4_064_17    m/z 822.0283     0   1   2   3E+4   14   14.5  RT (min)      lcqtof_t3_neg_4_002_17    m/z 822.3847     0   1E+4   1   1.5  RT (min)      lcqtof_t3_neg_4_051_15    m/z 822.5282     0   1E+6   12   12.5  RT (min)      lcqtof_t3_neg_2_030_23    m/z 822.5284     0   1   2   3E+2   10.5   11  RT (min)      lcqtof_t3_neg_3_038_06    m/z 822.5284     0   1E+6   12   12.5  RT (min)      lcqtof_t3_neg_1_019_23    m/z 822.5295     0   1   2   3E+2   10.5   11  RT (min)      lcqtof_t3_neg_3_038_06    m/z 822.6110     0   1E+4   14.5   15  RT (min)      lcqtof_t3_neg_1_000_aa    m/z 822.6210     0   1   2E+2   10   10.5  RT (min)      lcqtof_t3_neg_3_038_06    m/z 822.6212     0   1   2E+2   10   10.5  RT (min)      lcqtof_t3_neg_3_038_06    m/z 822.6220     0   1   2   3   4E+2   11.5   12  RT (min)      lcqtof_t3_neg_3_032_24    m/z 822.6223     0   1   2   3   4E+2   11.5   12  RT (min)      lcqtof_t3_neg_3_032_24    m/z 822.6224     0   1   2   3E+4   12   12.5  RT (min)      lcqtof_t3_neg_2_016_05    m/z 822.6242     0   1   2E+4   12   12.5  RT (min)      lcqtof_t3_neg_2_016_05    m/z 822.6331     0   2   4   6E+3   14   14.5  RT (min)       lcqtof_t3_neg_2_026_36    m/z 822.8171     0   1   2   3E+4   12   12.5  RT (min)      lcqtof_t3_neg_2_030_23    m/z 822.8178     0   1   2   3   4E+4   12   12.5  RT (min)      lcqtof_t3_neg_1_019_23    m/z 823.0230     0   1   2E+4   14   14.5  RT (min)      lcqtof_t3_neg_4_064_17    m/z 823.0250     0   1   2   3   4E+4   13   13.5  RT (min)      lcqtof_t3_neg_4_064_17    m/z 823.0252     0   1   2   3   4E+4   13   13.5  RT (min)      lcqtof_t3_neg_4_064_17    m/z 823.0256     0   1   2   3   4E+4   13.5   14  RT (min)      lcqtof_t3_neg_4_064_17    m/z 823.0259     0   1   2   3E+4   13.5   14  RT (min)      lcqtof_t3_neg_4_064_17    m/z 823.0261     0   1   2   3   4E+4   12.5   13  RT (min)      lcqtof_t3_neg_4_064_17    m/z 823.5308     0   1   2E+2   10.5   11  RT (min)      lcqtof_t3_neg_3_038_06    m/z 823.5310     0   1   2E+2   10.5   11  RT (min)      lcqtof_t3_neg_3_038_06    m/z 823.5314     0   2   4   6E+5   12   12.5  RT (min)      lcqtof_t3_neg_1_019_23    m/z 823.5316     0   2   4   6E+5   12   12.5  RT (min)      lcqtof_t3_neg_2_030_23    m/z 823.5323     0   1   2E+2   10.5   11  RT (min)      lcqtof_t3_neg_3_038_06    m/z 823.5398     0   1   2   3   4E+3   11.5   12  RT (min)      lcqtof_t3_neg_1_001_49    m/z 823.6270     0   2   4E+3   14   14.5  RT (min)      lcqtof_t3_neg_2_026_36    m/z 823.6275     0   2   4   6   8E+3   13.5   14  RT (min)      lcqtof_t3_neg_4_037_01    m/z 823.6322     0   1E+3   11.5   12  RT (min)      lcqtof_t3_neg_3_032_24    m/z 823.6322     0   1E+3   11.5   12  RT (min)      lcqtof_t3_neg_3_032_24    m/z 823.6323     0   1E+3   11.5   12  RT (min)      lcqtof_t3_neg_3_032_24    m/z 823.6339     0   1   2E+2   10.5   11  RT (min)      lcqtof_t3_neg_3_038_06    m/z 823.6340     0   1   2E+2   10.5   11  RT (min)      lcqtof_t3_neg_3_038_06    m/z 823.6342     0   1   2E+2   10.5   11  RT (min)      lcqtof_t3_neg_3_038_06    m/z 823.6407     0   1E+2   11   11.5  RT (min)      lcqtof_t3_neg_3_038_06    m/z 823.6410     0   1E+2   11   11.5  RT (min)      lcqtof_t3_neg_3_038_06    m/z 824.0229     0   1E+4   13   13.5  RT (min)      lcqtof_t3_neg_4_064_17    m/z 824.0250     0   1E+4   13.5   14  RT (min)      lcqtof_t3_neg_4_064_17    m/z 824.4053     0   1E+4   1   1.5  RT (min)      lcqtof_t3_neg_4_062_13    m/z 824.4674     0   1   2E+4   12   12.5  RT (min)      lcqtof_t3_neg_2_030_23    m/z 824.5354     0   1   2E+5   12   12.5  RT (min)      lcqtof_t3_neg_1_019_23    m/z 824.5355     0   1   2E+6   12   12.5  RT (min)      lcqtof_t3_neg_2_030_23    m/z 824.5406     0   1   2   3E+3   11.5   12  RT (min)      lcqtof_t3_neg_1_001_49    m/z 824.5432     0   1E+4   16   16.5  RT (min)      lcqtof_t3_neg_4_034_34    m/z 824.5436     0   1   2   3E+2   10.5   11  RT (min)      lcqtof_t3_neg_3_038_06    m/z 824.5440     0   1   2   3E+2   10.5   11  RT (min)      lcqtof_t3_neg_3_038_06    m/z 824.5448     0   1   2E+6   12   12.5  RT (min)      lcqtof_t3_neg_2_030_23    m/z 824.6362     0   1   2   3   4E+2   11.5   12  RT (min)      lcqtof_t3_neg_3_032_24    m/z 824.6368     0   1   2   3   4E+2   10.5   11  RT (min)      lcqtof_t3_neg_3_038_06    m/z 824.6370     0   1   2E+2   11.5   12  RT (min)      lcqtof_t3_neg_3_032_24    m/z 824.6370     0   1   2E+2   11.5   12  RT (min)      lcqtof_t3_neg_3_032_24    m/z 824.6375     0   1   2   3   4E+2   10.5   11  RT (min)      lcqtof_t3_neg_3_038_06    m/z 824.6380     0   1   2   3   4E+2   10.5   11  RT (min)      lcqtof_t3_neg_3_038_06    m/z 824.6386     0   1E+5   12   12.5  RT (min)      lcqtof_t3_neg_2_016_05    m/z 824.6591     0   1E+4   13.5   14  RT (min)      lcqtof_t3_neg_2_030_23    m/z 824.6622     0   1   2   3E+4   13.5   14  RT (min)      lcqtof_t3_neg_4_018_23    m/z 824.6857     0   1   2E+4   12   12.5  RT (min)      lcqtof_t3_neg_2_030_23    m/z 824.8338     0   2   4   6   8E+4   12   12.5  RT (min)      lcqtof_t3_neg_2_030_23    m/z 824.9702     0   1E+4   12.5   13  RT (min)      lcqtof_t3_neg_4_034_34    m/z 825.0593     0   1   2E+4   12   12.5  RT (min)      lcqtof_t3_neg_2_030_23    m/z 825.0600     0   1   2E+4   12   12.5  RT (min)      lcqtof_t3_neg_2_030_23    m/z 825.1817     0   1E+4   12   12.5  RT (min)      lcqtof_t3_neg_4_064_17    m/z 825.2205     0   1E+4   12   12.5  RT (min)      lcqtof_t3_neg_2_030_23    m/z 825.2208     0   1E+4   12.5   13  RT (min)      lcqtof_t3_neg_2_053_38    m/z 825.2221     0   1E+4   12.5   13  RT (min)      lcqtof_t3_neg_4_034_34    m/z 825.3200     0   1   2   3E+3   5   5.5  RT (min)       lcqtof_t3_neg_4_036_47    m/z 825.3202     0   1   2E+3   7   7.5  RT (min)       lcqtof_t3_neg_4_025_52    m/z 825.3205     0   1   2E+3   11.5   12  RT (min)       lcqtof_t3_neg_4_012_27    m/z 825.3209     0   1   2E+3   12   12.5  RT (min)       lcqtof_t3_neg_4_041_54    m/z 825.3211     0   1   2   3E+3   15   15.5  RT (min)       lcqtof_t3_neg_4_004_20    m/z 825.3212     0   1   2E+3   6   6.5  RT (min)       lcqtof_t3_neg_4_005_18    m/z 825.3213     0   1   2E+3   5.5   6  RT (min)       lcqtof_t3_neg_4_000_aa    m/z 825.3213     0   1E+4   16.5   17  RT (min)       lcqtof_t3_neg_4_000_cc    m/z 825.3214     0   1   2   3   4E+3   15.5   16  RT (min)       lcqtof_t3_neg_4_000_cc    m/z 825.3214     0   1   2   3E+3   15   15.5  RT (min)       lcqtof_t3_neg_4_018_23    m/z 825.3215     0   1   2   3E+3   15.5   16  RT (min)       lcqtof_t3_neg_4_000_bb    m/z 825.3216     0   1   2E+3   7   7.5  RT (min)       lcqtof_t3_neg_4_019_05    m/z 825.3216     0   1   2E+3   12   13  RT (min)       lcqtof_t3_neg_4_012_27    m/z 825.3216     0   1   2   3E+3   12.5   13  RT (min)      lcqtof_t3_neg_4_000_cc    m/z 825.3216     0   1   2E+3   14   14.5  RT (min)       lcqtof_t3_neg_4_004_20    m/z 825.3216     0   1   2   3E+3   7.5   8  RT (min)       lcqtof_t3_neg_4_000_cc    m/z 825.3216     0   1   2E+3   14.5   15  RT (min)       lcqtof_t3_neg_4_017_51    m/z 825.3216     0   1   2   3E+3   15   15.5  RT (min)       lcqtof_t3_neg_4_002_17    m/z 825.3217     0   1   2   3E+3   13.5   14  RT (min)       lcqtof_t3_neg_4_018_23    m/z 825.3217     0   1   2E+3   13.5   14  RT (min)      lcqtof_t3_neg_4_028_25    m/z 825.3219     0   1   2E+3   13.5   14  RT (min)      lcqtof_t3_neg_4_000_cc    m/z 825.3219     0   1   2E+3   10.5   11  RT (min)       lcqtof_t3_neg_4_022_28    m/z 825.3220     0   1   2E+3   8   8.5  RT (min)       lcqtof_t3_neg_4_029_40    m/z 825.3221     0   1   2E+3   9.5   10  RT (min)       lcqtof_t3_neg_4_019_05    m/z 825.3222     0   1   2E+3   10   10.5  RT (min)       lcqtof_t3_neg_4_026_30    m/z 825.3222     0   1   2E+3   11.5   12  RT (min)       lcqtof_t3_neg_4_017_51    m/z 825.3224     0   1   2E+3   6.5   7  RT (min)       lcqtof_t3_neg_4_031_42    m/z 825.3224     0   1   2E+3   14.5   15  RT (min)       lcqtof_t3_neg_4_040_24    m/z 825.3224     0   1   2E+3   11   11.5  RT (min)       lcqtof_t3_neg_4_014_13    m/z 825.3225     0   1   2E+3   14   14.5  RT (min)       lcqtof_t3_neg_4_000_ee    m/z 825.3226     0   1   2   3E+3   12.5   13  RT (min)       lcqtof_t3_neg_4_014_13    m/z 825.3227     0   1   2E+3   8.5   9  RT (min)      lcqtof_t3_neg_4_028_25    m/z 825.3228     0   1   2   3E+3   14.5   15  RT (min)       lcqtof_t3_neg_4_015_36    m/z 825.3230     0   1   2E+3   9   9.5  RT (min)       lcqtof_t3_neg_4_036_47    m/z 825.3230     0   1   2E+3   8   8.5  RT (min)       lcqtof_t3_neg_4_032_04    m/z 825.3232     0   1   2   3E+3   10   10.5  RT (min)       lcqtof_t3_neg_4_014_13    m/z 825.3233     0   1   2   3E+3   13.5   14  RT (min)       lcqtof_t3_neg_4_023_45    m/z 825.3236     0   1   2E+3   11   11.5  RT (min)       lcqtof_t3_neg_4_000_bb    m/z 825.4772     0   1E+4   12   12.5  RT (min)      lcqtof_t3_neg_3_006_23    m/z 825.4780     0   1E+4   12   12.5  RT (min)      lcqtof_t3_neg_2_030_23    m/z 825.5013     0   1E+4   12.5   13  RT (min)      lcqtof_t3_neg_1_019_23    m/z 825.5464     0   1E+6   12   12.5  RT (min)      lcqtof_t3_neg_2_030_23    m/z 825.5466     0   2   4   6   8E+5   12   12.5  RT (min)      lcqtof_t3_neg_1_019_23    m/z 825.5475     0   1   2E+2   10.5   11  RT (min)      lcqtof_t3_neg_3_038_06    m/z 825.5476     0   1   2E+2   10.5   11  RT (min)      lcqtof_t3_neg_3_038_06    m/z 825.6480     0   1   2E+2   11.5   12  RT (min)      lcqtof_t3_neg_3_032_24    m/z 825.6488     0   1   2E+2   11.5   12  RT (min)      lcqtof_t3_neg_3_032_24    m/z 825.6488     0   1   2E+3   12   12.5  RT (min)      lcqtof_t3_neg_3_032_24    m/z 825.6495     0   1   2E+2   10.5   11  RT (min)      lcqtof_t3_neg_3_038_06    m/z 825.6502     0   1   2E+2   10.5   11  RT (min)      lcqtof_t3_neg_3_038_06    m/z 825.6502     0   1   2E+2   10.5   11  RT (min)      lcqtof_t3_neg_3_038_06    m/z 825.6606     0   1   2   3E+3   13.5   14  RT (min)      lcqtof_t3_neg_2_030_23    m/z 826.1815     0   1E+4   12   12.5  RT (min)      lcqtof_t3_neg_4_064_17    m/z 826.4840     0   2   4   6E+4   12   12.5  RT (min)      lcqtof_t3_neg_1_019_23    m/z 826.4858     0   2   4   6E+4   12   12.5  RT (min)      lcqtof_t3_neg_1_019_23    m/z 826.4859     0   1   2   3E+2   10.5   11  RT (min)      lcqtof_t3_neg_3_038_06    m/z 826.5489     0   1   2   3E+5   12   12.5  RT (min)      lcqtof_t3_neg_2_030_23    m/z 826.5496     0   1E+6   12.5   13  RT (min)      lcqtof_t3_neg_2_030_23    m/z 826.5590     0   1   2   3   4E+2   10.5   11  RT (min)      lcqtof_t3_neg_3_038_06    m/z 826.5596     0   1   2   3   4E+2   10.5   11  RT (min)      lcqtof_t3_neg_3_038_06    m/z 826.6524     0   1   2   3   4E+2   11.5   12  RT (min)      lcqtof_t3_neg_3_032_24    m/z 826.6526     0   1   2   3   4E+2   12   12.5  RT (min)      lcqtof_t3_neg_3_032_24    m/z 826.6532     0   1   2E+2   10.5   11  RT (min)      lcqtof_t3_neg_3_038_06    m/z 826.6532     0   1   2E+2   10.5   11  RT (min)      lcqtof_t3_neg_3_038_06    m/z 826.6603     0   1E+4   12   12.5  RT (min)      lcqtof_t3_neg_3_032_24    m/z 826.6891     0   1E+4   1   1.5  RT (min)      lcqtof_t3_neg_4_037_01    m/z 826.7018     0   1E+4   12.5   13  RT (min)      lcqtof_t3_neg_2_030_23    m/z 826.7024     0   1E+4   12.5   13  RT (min)      lcqtof_t3_neg_2_030_23    m/z 826.8492     0   2   4   6E+4   12.5   13  RT (min)      lcqtof_t3_neg_2_030_23    m/z 826.8498     0   2   4   6E+4   12.5   13  RT (min)      lcqtof_t3_neg_2_030_23    m/z 827.0755     0   1E+4   12.5   13  RT (min)      lcqtof_t3_neg_2_030_23    m/z 827.0756     0   1E+4   12.5   13  RT (min)      lcqtof_t3_neg_2_030_23    m/z 827.1938     0   1E+4   12   12.5  RT (min)      lcqtof_t3_neg_4_064_17    m/z 827.1952     0   1E+4   12   12.5  RT (min)      lcqtof_t3_neg_4_002_17    m/z 827.4364     0   1E+4   8.5   9  RT (min)      lcqtof_t3_neg_4_000_ee    m/z 827.4883     0   1   2   3E+4   12   12.5  RT (min)      lcqtof_t3_neg_1_019_23    m/z 827.4922     0   1   2E+4   12   12.5  RT (min)      lcqtof_t3_neg_3_006_23    m/z 827.4925     0   1   2   3E+4   12   12.5  RT (min)       lcqtof_t3_neg_4_018_23    m/z 827.5069     0   1   2   3E+4   12   12.5  RT (min)      lcqtof_t3_neg_1_019_23    m/z 827.5155     0   1   2   3E+4   13   13.5  RT (min)      lcqtof_t3_neg_1_019_23    m/z 827.5158     0   1E+4   12.5   13  RT (min)      lcqtof_t3_neg_2_030_23    m/z 827.5510     0   2   4E+4   12   12.5  RT (min)      lcqtof_t3_neg_1_019_23    m/z 827.5608     0   2   4   6   8E+5   12.5   13  RT (min)      lcqtof_t3_neg_2_030_23    m/z 827.5620     0   2   4   6   8E+5   12.5   13  RT (min)      lcqtof_t3_neg_2_030_23    m/z 827.5626     0   1   2E+2   10.5   11  RT (min)      lcqtof_t3_neg_3_038_06    m/z 827.5634     0   1   2E+2   10.5   11  RT (min)      lcqtof_t3_neg_3_038_06    m/z 827.6640     0   1   2   3E+2   10.5   11  RT (min)      lcqtof_t3_neg_3_038_06    m/z 827.6668     0   1   2E+2   12   12.5  RT (min)      lcqtof_t3_neg_3_032_24    m/z 827.6684     0   1   2   3E+2   10.5   11  RT (min)      lcqtof_t3_neg_3_038_06    m/z 828.1958     0   1E+4   12   12.5  RT (min)      lcqtof_t3_neg_4_064_17    m/z 828.1965     0   1E+4   12   12.5  RT (min)      lcqtof_t3_neg_4_064_17    m/z 828.1972     0   1E+4   12   12.5  RT (min)      lcqtof_t3_neg_4_002_17    m/z 828.2569     0   1   2E+3   10.5   11  RT (min)       lcqtof_t3_neg_4_041_54    m/z 828.3437     0   1E+4   4   4.5  RT (min)      lcqtof_t3_neg_4_000_ee    m/z 828.3874     0   1E+4   4.5   5  RT (min)      lcqtof_t3_neg_4_000_ee    m/z 828.4596     0   1E+4   1   1.5  RT (min)      lcqtof_t3_neg_4_037_01    m/z 828.4946     0   1E+4   12   12.5  RT (min)      lcqtof_t3_neg_1_019_23    m/z 828.4959     0   1E+4   12   12.5  RT (min)      lcqtof_t3_neg_1_019_23    m/z 828.5493     0   1E+4   12   12.5  RT (min)      lcqtof_t3_neg_2_030_23    m/z 828.5518     0   2   4   6   8E+5   12.5   13  RT (min)      lcqtof_t3_neg_2_043_31    m/z 828.5647     0   1   2E+5   12.5   13  RT (min)      lcqtof_t3_neg_2_030_23    m/z 828.5650     0   1E+6   12.5   13  RT (min)      lcqtof_t3_neg_2_030_23    m/z 828.5658     0   1   2E+5   12.5   13  RT (min)      lcqtof_t3_neg_2_030_23    m/z 828.5750     0   1E+6   12.5   13  RT (min)      lcqtof_t3_neg_2_030_23    m/z 828.5754     0   1   2E+5   12.5   13  RT (min)       lcqtof_t3_neg_2_030_23    m/z 828.5755     0   1   2   3E+2   11   11.5  RT (min)      lcqtof_t3_neg_3_038_06    m/z 828.5756     0   1   2   3E+2   11   11.5  RT (min)      lcqtof_t3_neg_3_038_06    m/z 828.6690     0   1   2E+2   10.5   11  RT (min)      lcqtof_t3_neg_3_038_06    m/z 828.6704     0   2   4E+2   12   12.5  RT (min)      lcqtof_t3_neg_3_032_24    m/z 828.6720     0   1   2E+2   10.5   11  RT (min)      lcqtof_t3_neg_3_038_06    m/z 828.8669     0   1   2E+4   12.5   13  RT (min)      lcqtof_t3_neg_2_030_23    m/z 828.9620     0   1   2   3E+4   13   13.5  RT (min)      lcqtof_t3_neg_4_040_24    m/z 829.0030     0   1E+4   13   13.5  RT (min)      lcqtof_t3_neg_2_042_13    m/z 829.1925     0   1   2   3   4E+4   12   12.5  RT (min)      lcqtof_t3_neg_4_064_17    m/z 829.1932     0   1   2   3   4E+4   12   12.5  RT (min)       lcqtof_t3_neg_4_064_17    m/z 829.1939     0   1   2   3   4E+4   12   12.5  RT (min)      lcqtof_t3_neg_4_064_17    m/z 829.1964     0   1E+4   12.5   13  RT (min)      lcqtof_t3_neg_4_002_17    m/z 829.2540     0   1E+4   13   13.5  RT (min)      lcqtof_t3_neg_2_042_13    m/z 829.3814     0   1E+4   4.5   5  RT (min)      lcqtof_t3_neg_4_000_ee    m/z 829.5316     0   1E+4   13   13.5  RT (min)      lcqtof_t3_neg_1_019_23    m/z 829.5318     0   1E+4   13   13.5  RT (min)      lcqtof_t3_neg_1_019_23    m/z 829.5434     0   1E+4   13   13.5  RT (min)      lcqtof_t3_neg_4_018_23    m/z 829.5676     0   2   4E+5   12.5   13  RT (min)      lcqtof_t3_neg_2_030_23    m/z 829.5678     0   2   4E+4   12.5   13  RT (min)      lcqtof_t3_neg_2_030_23    m/z 829.5780     0   1   2   3   4E+5   12.5   13  RT (min)      lcqtof_t3_neg_2_030_23    m/z 829.5792     0   2   4E+2   11   11.5  RT (min)      lcqtof_t3_neg_3_038_06    m/z 829.5793     0   2   4E+2   11   11.5  RT (min)      lcqtof_t3_neg_3_038_06    m/z 829.6770     0   1   2   3   4E+2   12   12.5  RT (min)      lcqtof_t3_neg_3_032_24    m/z 829.6772     0   1   2   3   4E+2   12   12.5  RT (min)      lcqtof_t3_neg_3_032_24    m/z 829.6776     0   1   2E+4   12.5   13  RT (min)      lcqtof_t3_neg_2_016_05    m/z 829.6781     0   1   2E+4   12.5   13  RT (min)      lcqtof_t3_neg_2_016_05    m/z 829.6810     0   1   2   3E+2   12   12.5  RT (min)      lcqtof_t3_neg_3_032_24    m/z 829.6812     0   1   2   3   4E+2   12   12.5  RT (min)      lcqtof_t3_neg_3_032_24    m/z 829.6823     0   1   2   3E+2   10.5   11  RT (min)      lcqtof_t3_neg_3_038_06    m/z 829.6825     0   1   2   3E+4   12.5   13  RT (min)      lcqtof_t3_neg_2_016_05    m/z 829.6855     0   1   2   3E+2   11   11.5  RT (min)      lcqtof_t3_neg_3_038_06    m/z 830.1768     0   1E+4   13   13.5  RT (min)      lcqtof_t3_neg_4_057_49    m/z 830.1934     0   1   2E+4   12   12.5  RT (min)      lcqtof_t3_neg_4_064_17    m/z 830.1943     0   1   2E+4   12   12.5  RT (min)      lcqtof_t3_neg_4_064_17    m/z 830.1950     0   1E+4   12   12.5  RT (min)      lcqtof_t3_neg_4_002_17    m/z 830.1958     0   1E+4   12.5   13  RT (min)      lcqtof_t3_neg_4_002_17    m/z 830.2837     0   2   4E+3   11   11.5  RT (min)       lcqtof_t3_neg_4_051_15    m/z 830.4548     0   1E+4   1   1.5  RT (min)      lcqtof_t3_neg_4_037_01    m/z 830.5811     0   2   4   6   8E+4   12.5   13  RT (min)       lcqtof_t3_neg_3_006_23    m/z 830.5818     0   1   2   3E+2   11   11.5  RT (min)      lcqtof_t3_neg_3_038_06    m/z 830.5828     0   1   2   3E+2   11   11.5  RT (min)      lcqtof_t3_neg_3_038_06    m/z 830.5863     0   1E+5   12.5   13  RT (min)      lcqtof_t3_neg_2_030_23    m/z 830.5905     0   1   2   3E+5   12.5   13  RT (min)      lcqtof_t3_neg_2_030_23    m/z 830.5911     0   1E+4   16.5   17  RT (min)      lcqtof_t3_neg_4_060_15    m/z 830.5916     0   1E+3   11   11.5  RT (min)      lcqtof_t3_neg_3_038_06    m/z 830.5925     0   1E+6   14   14.5  RT (min)      lcqtof_t3_neg_1_022_28    m/z 830.6672     0   1E+5   13.5   14  RT (min)       lcqtof_t3_neg_4_040_24    m/z 830.6784     0   1E+5   12.5   13  RT (min)      lcqtof_t3_neg_2_016_05    m/z 830.6850     0   1E+3   12   12.5  RT (min)      lcqtof_t3_neg_3_032_24    m/z 830.6851     0   2   4E+2   12   12.5  RT (min)      lcqtof_t3_neg_3_032_24    m/z 830.6858     0   2   4E+2   12   12.5  RT (min)      lcqtof_t3_neg_3_032_24    m/z 830.6873     0   1   2   3E+2   11   11.5  RT (min)      lcqtof_t3_neg_3_038_06    m/z 831.1733     0   2   4E+4   13   13.5  RT (min)      lcqtof_t3_neg_4_057_49    m/z 831.1735     0   1   2   3E+4   13   13.5  RT (min)      lcqtof_t3_neg_4_001_49    m/z 831.1748     0   1   2   3E+4   13   13.5  RT (min)      lcqtof_t3_neg_4_001_49    m/z 831.1931     0   1E+4   12   12.5  RT (min)      lcqtof_t3_neg_4_064_17    m/z 831.3572     0   1E+4   5   5.5  RT (min)      lcqtof_t3_neg_4_000_ee    m/z 831.5016     0   1   2   3   4E+4   11.5   12  RT (min)      lcqtof_t3_neg_1_019_23    m/z 831.5247     0   2   4E+3   12.5   13  RT (min)      lcqtof_t3_neg_1_002_13    m/z 831.5301     0   2   4E+3   12.5   13  RT (min)       lcqtof_t3_neg_1_002_13    m/z 831.5572     0   1   2E+3   16   16.5  RT (min)      lcqtof_t3_neg_1_016_48    m/z 831.5812     0   1   2E+4   12.5   13  RT (min)      lcqtof_t3_neg_2_030_23    m/z 831.5938     0   1E+5   12.5   13  RT (min)      lcqtof_t3_neg_2_030_23    m/z 831.5944     0   2   4   6E+2   11   11.5  RT (min)      lcqtof_t3_neg_3_038_06    m/z 831.5944     0   2   4   6E+2   11   11.5  RT (min)      lcqtof_t3_neg_3_038_06    m/z 831.5965     0   1   2E+5   14   14.5  RT (min)      lcqtof_t3_neg_4_037_01    m/z 831.6928     0   2   4   6   8E+2   12   12.5  RT (min)      lcqtof_t3_neg_3_032_24    m/z 831.6932     0   2   4   6   8E+2   12   12.5  RT (min)      lcqtof_t3_neg_3_032_24    m/z 831.6949     0   1E+4   12.5   13  RT (min)      lcqtof_t3_neg_2_016_05    m/z 832.1735     0   1   2   3   4E+4   13   13.5  RT (min)      lcqtof_t3_neg_4_057_49    m/z 832.1750     0   1   2E+4   13   13.5  RT (min)      lcqtof_t3_neg_4_001_49    m/z 832.3513     0   1E+4   4   4.5  RT (min)      lcqtof_t3_neg_4_000_ee    m/z 832.5055     0   1   2E+4   11.5   12  RT (min)      lcqtof_t3_neg_1_019_23    m/z 832.5956     0   1   2   3   4E+4   12.5   13  RT (min)      lcqtof_t3_neg_2_030_23    m/z 832.6577     0   1E+4   12.5   13  RT (min)       lcqtof_t3_neg_1_003_07    m/z 832.6580     0   1E+4   12.5   13  RT (min)      lcqtof_t3_neg_1_031_24    m/z 832.6956     0   1   2E+2   12   12.5  RT (min)      lcqtof_t3_neg_3_032_24    m/z 832.6967     0   1   2E+2   12   12.5  RT (min)      lcqtof_t3_neg_3_032_24    m/z 832.6992     0   1E+4   12.5   13  RT (min)      lcqtof_t3_neg_2_016_05    m/z 833.0800     0   1   2   3E+4   14   14.5  RT (min)      lcqtof_t3_neg_4_064_17    m/z 833.0804     0   1   2   3E+4   13.5   14  RT (min)      lcqtof_t3_neg_4_064_17    m/z 833.0809     0   1   2   3E+4   13   13.5  RT (min)      lcqtof_t3_neg_4_064_17    m/z 833.0812     0   1   2   3E+4   14   14.5  RT (min)      lcqtof_t3_neg_4_064_17    m/z 833.0841     0   1   2   3E+4   13   13.5  RT (min)      lcqtof_t3_neg_4_064_17    m/z 833.0848     0   1   2   3E+4   13.5   14  RT (min)      lcqtof_t3_neg_4_064_17    m/z 833.1722     0   1   2   3E+4   13   13.5  RT (min)      lcqtof_t3_neg_4_057_49    m/z 833.1740     0   1   2E+4   13   13.5  RT (min)      lcqtof_t3_neg_4_001_49    m/z 833.5145     0   1   2   3E+5   12   12.5  RT (min)      lcqtof_t3_neg_2_030_23    m/z 833.5168     0   1   2   3   4E+5   12   12.5  RT (min)      lcqtof_t3_neg_1_019_23    m/z 833.5178     0   1   2   3E+2   10.5   11  RT (min)      lcqtof_t3_neg_3_038_06    m/z 833.5186     0   1   2   3E+2   10.5   11  RT (min)      lcqtof_t3_neg_3_038_06    m/z 833.6058     0   1E+4   13   13.5  RT (min)      lcqtof_t3_neg_1_019_23    m/z 833.6096     0   1   2E+4   13   13.5  RT (min)      lcqtof_t3_neg_2_030_23    m/z 833.7017     0   1   2   3E+2   12   12.5  RT (min)      lcqtof_t3_neg_3_032_24    m/z 833.7036     0   1   2   3E+2   11   11.5  RT (min)      lcqtof_t3_neg_3_038_06    m/z 833.7058     0   1   2   3E+2   11   11.5  RT (min)      lcqtof_t3_neg_3_038_06    m/z 834.0780     0   1   2E+4   13.5   14  RT (min)      lcqtof_t3_neg_4_064_17    m/z 834.0803     0   1   2E+4   13   13.5  RT (min)      lcqtof_t3_neg_4_064_17    m/z 834.0806     0   1   2E+4   13   13.5  RT (min)      lcqtof_t3_neg_4_064_17    m/z 834.0812     0   1   2E+4   13.5   14  RT (min)      lcqtof_t3_neg_4_064_17    m/z 834.1701     0   1E+4   13   13.5  RT (min)      lcqtof_t3_neg_4_057_49    m/z 834.3850     0   1E+4   1   1.5  RT (min)      lcqtof_t3_neg_4_037_01    m/z 834.5202     0   1E+5   12   12.5  RT (min)      lcqtof_t3_neg_1_019_23    m/z 834.5208     0   1E+5   12   12.5  RT (min)      lcqtof_t3_neg_3_006_23    m/z 835.0772     0   1   2E+4   13.5   14  RT (min)      lcqtof_t3_neg_4_064_17    m/z 835.0779     0   1   2E+4   13.5   14  RT (min)      lcqtof_t3_neg_4_064_17    m/z 835.0780     0   1   2E+4   14   14.5  RT (min)      lcqtof_t3_neg_4_064_17    m/z 835.0788     0   1   2E+4   14   14.5  RT (min)      lcqtof_t3_neg_4_064_17    m/z 835.0793     0   1   2E+4   13   13.5  RT (min)      lcqtof_t3_neg_4_064_17    m/z 835.0816     0   1   2E+4   13   13.5  RT (min)      lcqtof_t3_neg_4_064_17    m/z 835.0855     0   1   2E+4   13   13.5  RT (min)      lcqtof_t3_neg_4_064_17    m/z 835.0927     0   1E+4   13   13.5  RT (min)      lcqtof_t3_neg_4_064_17    m/z 835.4938     0   1E+4   8.5   9  RT (min)      lcqtof_t3_neg_4_000_ee    m/z 835.4946     0   1E+4   8.5   9  RT (min)      lcqtof_t3_neg_4_000_ee    m/z 835.4952     0   1E+4   8.5   9  RT (min)      lcqtof_t3_neg_4_000_ee    m/z 835.5218     0   1   2   3E+4   12   12.5  RT (min)      lcqtof_t3_neg_2_030_23    m/z 835.5229     0   2   4E+4   12   12.5  RT (min)      lcqtof_t3_neg_1_019_23    m/z 835.5240     0   1E+5   12   12.5  RT (min)      lcqtof_t3_neg_2_030_23    m/z 835.5324     0   1   2E+5   12   12.5  RT (min)      lcqtof_t3_neg_1_019_23    m/z 835.7024     0   1E+4   13   13.5  RT (min)      lcqtof_t3_neg_1_031_24    m/z 835.7029     0   1E+4   13   13.5  RT (min)      lcqtof_t3_neg_2_028_24    m/z 836.0788     0   1E+4   13.5   14  RT (min)      lcqtof_t3_neg_4_064_17    m/z 836.0798     0   1E+4   13   13.5  RT (min)      lcqtof_t3_neg_4_064_17    m/z 836.5326     0   2   4   6   8E+4   12   12.5  RT (min)      lcqtof_t3_neg_2_030_23    m/z 836.5342     0   1   2   3E+2   10.5   11  RT (min)      lcqtof_t3_neg_3_038_06    m/z 836.5371     0   1E+5   12   12.5  RT (min)      lcqtof_t3_neg_1_019_23    m/z 836.5390     0   1   2   3E+2   10.5   11  RT (min)      lcqtof_t3_neg_3_038_06    m/z 836.5393     0   2   4   6   8E+4   13   13.5  RT (min)       lcqtof_t3_neg_4_084_59    m/z 836.6374     0   1   2   3   4E+2   11.5   12  RT (min)      lcqtof_t3_neg_3_032_24    m/z 836.6399     0   1   2   3   4E+2   11.5   12  RT (min)      lcqtof_t3_neg_3_032_24    m/z 837.3378     0   1E+4   4   4.5  RT (min)      lcqtof_t3_neg_4_000_ee    m/z 837.5386     0   1   2E+4   12   12.5  RT (min)      lcqtof_t3_neg_1_019_23    m/z 837.6470     0   2   4   6E+2   11.5   12  RT (min)      lcqtof_t3_neg_3_032_24    m/z 837.6488     0   2   4   6E+2   11.5   12  RT (min)      lcqtof_t3_neg_3_032_24    m/z 837.6490     0   2   4   6E+2   11.5   12  RT (min)      lcqtof_t3_neg_3_032_24    m/z 837.6499     0   1E+4   12   12.5  RT (min)      lcqtof_t3_neg_2_016_05    m/z 838.5564     0   1E+4   13   13.5  RT (min)       lcqtof_t3_neg_4_084_59    m/z 838.6153     0   1E+4   13   13.5  RT (min)      lcqtof_t3_neg_1_003_07    m/z 838.6523     0   2   4E+4   12   12.5  RT (min)      lcqtof_t3_neg_2_016_05    m/z 838.6526     0   1   2E+2   10.5   11  RT (min)      lcqtof_t3_neg_3_038_06    m/z 838.6528     0   1   2E+2   10.5   11  RT (min)      lcqtof_t3_neg_3_038_06    m/z 838.6530     0   2   4E+2   11.5   12  RT (min)      lcqtof_t3_neg_3_032_24    m/z 838.6530     0   2   4E+2   11.5   12  RT (min)      lcqtof_t3_neg_3_032_24    m/z 838.6538     0   1   2E+2   10   10.5  RT (min)      lcqtof_t3_neg_3_038_06    m/z 838.6588     0   1E+4   13   13.5   14  RT (min)      lcqtof_t3_neg_4_040_24    m/z 838.6683     0   1E+4   1   1.5  RT (min)      lcqtof_t3_neg_4_059_14    m/z 839.5173     0   1   2   3   4E+4   12   12.5  RT (min)      lcqtof_t3_neg_1_019_23    m/z 839.5192     0   1   2   3   4E+4   12   12.5  RT (min)      lcqtof_t3_neg_1_019_23    m/z 839.6208     0   1   2   3   4E+3   14   14.5  RT (min)      lcqtof_t3_neg_2_015_33    m/z 839.6320     0   2   4E+3   12.5   13  RT (min)      lcqtof_t3_neg_2_004_48    m/z 839.6630     0   1   2   3E+2   11.5   12  RT (min)      lcqtof_t3_neg_3_032_24    m/z 839.6635     0   2   4   6   8E+2   11.5   12  RT (min)      lcqtof_t3_neg_3_032_24    m/z 839.6646     0   1   2E+2   10.5   11  RT (min)      lcqtof_t3_neg_3_038_06    m/z 839.6653     0   1   2E+2   10.5   11  RT (min)      lcqtof_t3_neg_3_038_06    m/z 840.3358     0   1E+4   4.5   5  RT (min)      lcqtof_t3_neg_4_000_ee    m/z 840.4340     0   1E+4   12   12.5  RT (min)      lcqtof_t3_neg_1_019_23    m/z 840.5177     0   1   2E+4   12   12.5  RT (min)      lcqtof_t3_neg_1_019_23    m/z 840.6227     0   1   2E+3   15.5   16  RT (min)      lcqtof_t3_neg_1_014_04    m/z 840.6665     0   1E+4   1   1.5  RT (min)      lcqtof_t3_neg_4_059_14    m/z 840.6682     0   1E+2   11.5   12  RT (min)      lcqtof_t3_neg_3_032_24    m/z 840.6684     0   1   2   3E+2   11.5   12  RT (min)      lcqtof_t3_neg_3_032_24    m/z 840.6685     0   1   2E+2   10.5   11  RT (min)      lcqtof_t3_neg_3_038_06    m/z 840.6685     0   1   2   3E+2   11.5   12  RT (min)      lcqtof_t3_neg_3_032_24    m/z 840.6690     0   1   2E+2   10.5   11  RT (min)      lcqtof_t3_neg_3_038_06    m/z 841.1761     0   1E+4   12   12.5  RT (min)      lcqtof_t3_neg_4_064_17    m/z 841.1771     0   1E+4   12   12.5  RT (min)      lcqtof_t3_neg_4_064_17    m/z 841.3785     0   1E+4   4.5   5  RT (min)      lcqtof_t3_neg_4_000_ee    m/z 841.4543     0   1   2E+4   12   12.5  RT (min)      lcqtof_t3_neg_1_019_23    m/z 841.5312     0   1   2E+4   12   12.5  RT (min)      lcqtof_t3_neg_2_030_23    m/z 841.5330     0   1   2E+4   12   12.5  RT (min)      lcqtof_t3_neg_1_019_23    m/z 841.5338     0   1   2E+4   12   12.5  RT (min)      lcqtof_t3_neg_2_030_23    m/z 841.6093     0   2   4   6E+3   12.5   13  RT (min)      lcqtof_t3_neg_2_059_14    m/z 841.6146     0   1   2   3   4E+3   11.5   12  RT (min)      lcqtof_t3_neg_1_001_49    m/z 841.6153     0   2   4E+3   11.5   12  RT (min)      lcqtof_t3_neg_2_000_aa    m/z 841.6343     0   2   4   6   8E+2   12.5   13  RT (min)      lcqtof_t3_neg_3_032_24    m/z 841.6344     0   2   4   6   8E+2   11   11.5  RT (min)      lcqtof_t3_neg_3_038_06    m/z 841.6797     0   1   2   3E+2   12   12.5  RT (min)      lcqtof_t3_neg_3_032_24    m/z 841.6799     0   1   2   3E+2   12   12.5  RT (min)      lcqtof_t3_neg_3_032_24    m/z 841.6806     0   2   4   6   8E+3   12   12.5  RT (min)      lcqtof_t3_neg_3_032_24    m/z 841.6809     0   1E+2   10.5   11  RT (min)      lcqtof_t3_neg_3_038_06    m/z 841.6816     0   1E+2   10.5   11  RT (min)      lcqtof_t3_neg_3_038_06    m/z 842.1784     0   1E+4   12   12.5  RT (min)      lcqtof_t3_neg_4_064_17    m/z 842.4483     0   1E+4   12   12.5  RT (min)      lcqtof_t3_neg_1_019_23    m/z 842.4521     0   1E+4   12   12.5  RT (min)      lcqtof_t3_neg_1_019_23    m/z 842.4603     0   1E+4   12   12.5  RT (min)      lcqtof_t3_neg_1_019_23    m/z 842.5335     0   1E+4   12   12.5  RT (min)      lcqtof_t3_neg_2_030_23    m/z 842.5559     0   1   2E+4   12   12.5  RT (min)      lcqtof_t3_neg_4_003_37    m/z 842.5562     0   1   2E+4   12   12.5  RT (min)      lcqtof_t3_neg_4_003_37    m/z 842.5738     0   1E+4   13   13.5  RT (min)      lcqtof_t3_neg_1_008_02    m/z 842.6504     0   1E+4   13.5   14  RT (min)      lcqtof_t3_neg_4_040_24    m/z 842.6516     0   1E+4   13.5   14  RT (min)      lcqtof_t3_neg_4_040_24    m/z 842.6522     0   1E+4   13.5   14  RT (min)       lcqtof_t3_neg_4_040_24    m/z 842.6575     0   1E+4   13.5   14  RT (min)      lcqtof_t3_neg_4_040_24    m/z 842.6630     0   1E+4   1   1.5  RT (min)      lcqtof_t3_neg_4_016_14    m/z 842.6826     0   1   2   3E+2   12   12.5  RT (min)      lcqtof_t3_neg_3_032_24    m/z 842.6827     0   1   2   3E+2   12   12.5  RT (min)      lcqtof_t3_neg_3_032_24    m/z 842.6848     0   1   2   3E+3   12   12.5  RT (min)      lcqtof_t3_neg_3_032_24    m/z 842.6849     0   1   2   3E+2   10.5   11  RT (min)      lcqtof_t3_neg_3_038_06    m/z 842.6850     0   1   2E+2   10.5   11  RT (min)      lcqtof_t3_neg_3_038_06    m/z 842.7218     0   2   4   6E+2   12   12.5  RT (min)      lcqtof_t3_neg_3_032_24    m/z 842.7248     0   1   2   3   4E+2   11   11.5  RT (min)      lcqtof_t3_neg_3_038_06    m/z 843.1740     0   1E+4   12   12.5  RT (min)      lcqtof_t3_neg_4_064_17    m/z 843.3467     0   1   2E+4   4   4.5  RT (min)      lcqtof_t3_neg_4_000_ee    m/z 843.3926     0   1E+4   4.5   5  RT (min)      lcqtof_t3_neg_4_000_ee    m/z 843.4659     0   1E+4   12   12.5  RT (min)      lcqtof_t3_neg_1_019_23    m/z 843.4667     0   1E+4   12   12.5  RT (min)      lcqtof_t3_neg_1_019_23    m/z 843.4691     0   1E+4   12   12.5  RT (min)      lcqtof_t3_neg_1_019_23    m/z 843.5551     0   1   2E+4   13   13.5  RT (min)      lcqtof_t3_neg_1_036_14    m/z 843.5579     0   1   2   3E+4   12   12.5  RT (min)      lcqtof_t3_neg_4_016_14    m/z 843.5583     0   1E+4   12   12.5  RT (min)      lcqtof_t3_neg_4_003_37    m/z 843.5588     0   1   2   3   4E+4   12   12.5  RT (min)      lcqtof_t3_neg_4_059_14    m/z 843.5588     0   1   2   3   4E+4   12   12.5  RT (min)      lcqtof_t3_neg_4_059_14    m/z 843.5602     0   1   2   3E+4   12.5   13  RT (min)      lcqtof_t3_neg_1_019_23    m/z 843.5738     0   1E+4   12   12.5  RT (min)      lcqtof_t3_neg_4_011_32    m/z 843.5743     0   1E+4   13   13.5  RT (min)      lcqtof_t3_neg_2_051_14    m/z 843.5751     0   1E+4   12.5   13  RT (min)      lcqtof_t3_neg_2_016_05    m/z 843.5790     0   1E+4   12   12.5  RT (min)      lcqtof_t3_neg_4_046_33    m/z 843.5920     0   1E+4   13   13.5  RT (min)      lcqtof_t3_neg_2_021_27    m/z 843.5929     0   1E+4   13   13.5  RT (min)      lcqtof_t3_neg_4_016_14    m/z 843.5933     0   2   4E+3   14.5   15  RT (min)      lcqtof_t3_neg_1_031_24    m/z 843.5947     0   1E+4   13   13.5  RT (min)      lcqtof_t3_neg_2_026_36    m/z 843.5994     0   1E+4   12   12.5  RT (min)      lcqtof_t3_neg_1_019_23    m/z 843.6000     0   1   2   3E+4   12.5   13  RT (min)      lcqtof_t3_neg_1_003_07    m/z 843.6132     0   1E+4   13   13.5  RT (min)      lcqtof_t3_neg_1_031_24    m/z 843.6490     0   2   4E+2   12.5   13  RT (min)      lcqtof_t3_neg_3_032_24    m/z 843.6525     0   1   2   3   4E+2   12.5   13  RT (min)      lcqtof_t3_neg_3_032_24    m/z 843.6885     0   1   2E+2   12   12.5  RT (min)      lcqtof_t3_neg_3_032_24    m/z 843.6890     0   1E+4   12.5   13  RT (min)      lcqtof_t3_neg_2_016_05    m/z 843.6930     0   1   2E+5   12.5   13  RT (min)      lcqtof_t3_neg_2_016_05    m/z 843.6959     0   1   2E+2   12   12.5  RT (min)      lcqtof_t3_neg_3_032_24    m/z 843.7192     0   1E+3   12   12.5  RT (min)      lcqtof_t3_neg_3_032_24    m/z 843.7201     0   1E+3   12   12.5  RT (min)      lcqtof_t3_neg_3_032_24    m/z 843.7238     0   1   2E+2   11   11.5  RT (min)      lcqtof_t3_neg_3_038_06    m/z 843.7242     0   1   2E+2   11   11.5  RT (min)      lcqtof_t3_neg_3_038_06    m/z 843.7341     0   1E+3   12   12.5  RT (min)      lcqtof_t3_neg_3_032_24    m/z 844.5525     0   1E+4   12.5   13  RT (min)      lcqtof_t3_neg_2_030_23    m/z 844.5674     0   1E+4   12.5   13  RT (min)      lcqtof_t3_neg_1_019_23    m/z 844.5691     0   1E+4   12   12.5  RT (min)      lcqtof_t3_neg_4_034_34    m/z 844.5691     0   1   2   3   4E+4   12   12.5  RT (min)      lcqtof_t3_neg_4_059_14    m/z 844.5693     0   1   2   3   4E+4   12   12.5  RT (min)      lcqtof_t3_neg_4_059_14    m/z 844.5715     0   1E+4   12   12.5  RT (min)      lcqtof_t3_neg_4_055_38    m/z 844.5852     0   1   2E+4   12   12.5  RT (min)      lcqtof_t3_neg_4_046_33    m/z 844.5936     0   1   2   3   4E+3   15   15.5  RT (min)      lcqtof_t3_neg_1_034_30    m/z 844.5942     0   1   2   3E+3   15   15.5  RT (min)      lcqtof_t3_neg_1_032_36    m/z 844.5944     0   1   2E+4   13.5   14  RT (min)      lcqtof_t3_neg_4_037_01    m/z 844.5983     0   1   2E+4   12   12.5  RT (min)       lcqtof_t3_neg_4_046_33    m/z 844.6022     0   1   2   3E+4   12   12.5  RT (min)       lcqtof_t3_neg_4_040_24    m/z 844.6043     0   2   4   6E+3   12   12.5  RT (min)      lcqtof_t3_neg_3_032_24    m/z 844.6046     0   1   2E+4   13.5   14  RT (min)       lcqtof_t3_neg_4_037_01    m/z 844.6051     0   1   2   3E+2   10.5   11  RT (min)      lcqtof_t3_neg_3_038_06    m/z 844.6060     0   1E+4   14   14.5  RT (min)      lcqtof_t3_neg_4_008_35    m/z 844.6087     0   2   4   6E+3   12   12.5  RT (min)      lcqtof_t3_neg_3_032_24    m/z 844.6989     0   2   4E+2   12   12.5  RT (min)      lcqtof_t3_neg_3_032_24    m/z 844.6990     0   2   4E+5   12.5   13  RT (min)      lcqtof_t3_neg_2_016_05    m/z 844.7000     0   2   4E+2   12   12.5  RT (min)      lcqtof_t3_neg_3_032_24    m/z 844.7012     0   1   2   3E+2   11   11.5  RT (min)      lcqtof_t3_neg_3_038_06    m/z 844.7013     0   1   2   3E+2   10.5   11  RT (min)      lcqtof_t3_neg_3_038_06    m/z 844.9954     0   1   2E+4   13   13.5  RT (min)      lcqtof_t3_neg_2_028_24    m/z 845.1529     0   1E+4   13.5   14  RT (min)      lcqtof_t3_neg_4_064_17    m/z 845.1979     0   1E+4   4   4.5  RT (min)      lcqtof_t3_neg_4_000_ee    m/z 845.2250     0   1E+4   13   13.5  RT (min)      lcqtof_t3_neg_4_036_47    m/z 845.2253     0   1E+4   13   13.5  RT (min)      lcqtof_t3_neg_4_064_17    m/z 845.2270     0   1E+4   13   13.5  RT (min)      lcqtof_t3_neg_4_002_17    m/z 845.5645     0   1E+4   12.5   13  RT (min)      lcqtof_t3_neg_1_019_23    m/z 845.5676     0   1E+4   12.5   13  RT (min)      lcqtof_t3_neg_1_019_23    m/z 845.5952     0   1   2E+3   16   16.5  RT (min)      lcqtof_t3_neg_1_016_48    m/z 845.6008     0   1   2   3E+4   12   12.5  RT (min)      lcqtof_t3_neg_4_011_32    m/z 845.6014     0   1E+4   12   12.5  RT (min)      lcqtof_t3_neg_4_046_33    m/z 845.6148     0   1   2   3E+3   12   12.5  RT (min)      lcqtof_t3_neg_3_032_24    m/z 845.6158     0   1   2   3E+2   10.5   11  RT (min)      lcqtof_t3_neg_3_038_06    m/z 845.6161     0   1   2   3E+2   10.5   11  RT (min)      lcqtof_t3_neg_3_038_06    m/z 845.6176     0   1   2   3E+4   12.5   13  RT (min)      lcqtof_t3_neg_2_016_05    m/z 845.6210     0   2   4E+3   12   12.5  RT (min)      lcqtof_t3_neg_3_032_24    m/z 845.6212     0   1   2   3E+4   12.5   13  RT (min)      lcqtof_t3_neg_2_016_05    m/z 845.6218     0   2   4   6E+4   12   12.5  RT (min)      lcqtof_t3_neg_4_040_24    m/z 845.6269     0   1E+4   13   13.5  RT (min)      lcqtof_t3_neg_4_034_34    m/z 845.7034     0   2   4   6   8E+2   12   12.5  RT (min)      lcqtof_t3_neg_3_032_24    m/z 845.7045     0   2   4   6   8E+2   12   12.5  RT (min)      lcqtof_t3_neg_3_032_24    m/z 845.7065     0   1   2E+4   12.5   13  RT (min)      lcqtof_t3_neg_2_016_05    m/z 845.7065     0   2   4   6   8E+2   12   12.5  RT (min)      lcqtof_t3_neg_3_032_24    m/z 845.7086     0   1   2   3   4E+2   11   11.5  RT (min)      lcqtof_t3_neg_3_038_06    m/z 845.7092     0   1   2   3   4E+2   11   11.5  RT (min)      lcqtof_t3_neg_3_038_06    m/z 845.7104     0   2   4   6   8E+2   12   12.5  RT (min)      lcqtof_t3_neg_3_032_24    m/z 845.7131     0   2   4   6   8E+2   12.5   13  RT (min)      lcqtof_t3_neg_3_032_24    m/z 846.1536     0   1E+4   13.5   14  RT (min)      lcqtof_t3_neg_4_064_17    m/z 846.2253     0   1E+4   13   13.5  RT (min)      lcqtof_t3_neg_4_064_17    m/z 846.2255     0   1E+4   13   13.5  RT (min)      lcqtof_t3_neg_4_036_47    m/z 846.4505     0   2   4E+4   12   12.5  RT (min)      lcqtof_t3_neg_1_019_23    m/z 846.5655     0   1   2E+4   13   13.5  RT (min)      lcqtof_t3_neg_1_001_49    m/z 846.5848     0   2   4E+4   12   12.5  RT (min)      lcqtof_t3_neg_4_059_14    m/z 846.6198     0   1   2   3E+2   11.5   12  RT (min)      lcqtof_t3_neg_3_032_24    m/z 846.6210     0   1   2E+2   11.5   12  RT (min)      lcqtof_t3_neg_3_032_24    m/z 846.6227     0   1   2E+4   12   12.5  RT (min)      lcqtof_t3_neg_4_041_54    m/z 846.6601     0   1E+4   12.5   13  RT (min)      lcqtof_t3_neg_4_046_33    m/z 846.7152     0   1   2E+2   11   11.5  RT (min)      lcqtof_t3_neg_3_038_06    m/z 846.7166     0   1   2E+2   11   11.5  RT (min)      lcqtof_t3_neg_3_038_06    m/z 846.7236     0   1   2E+3   12   12.5  RT (min)      lcqtof_t3_neg_3_032_24    m/z 846.7248     0   1   2E+3   12   12.5  RT (min)      lcqtof_t3_neg_3_032_24    m/z 847.1398     0   1E+4   13.5   14  RT (min)      lcqtof_t3_neg_4_064_17    m/z 847.2245     0   1E+4   13   13.5  RT (min)      lcqtof_t3_neg_4_036_47    m/z 847.4542     0   1   2E+4   12   12.5  RT (min)      lcqtof_t3_neg_1_019_23    m/z 847.5097     0   1E+4   9   9.5  RT (min)      lcqtof_t3_neg_4_000_ee    m/z 847.5346     0   1E+4   12   12.5  RT (min)      lcqtof_t3_neg_1_019_23    m/z 847.6154     0   1E+4   12   12.5  RT (min)      lcqtof_t3_neg_4_011_32    m/z 847.6310     0   1   2   3E+3   12   12.5  RT (min)      lcqtof_t3_neg_3_032_24    m/z 847.6312     0   1   2E+4   12.5   13  RT (min)      lcqtof_t3_neg_2_016_05    m/z 847.6320     0   1   2   3E+3   12   12.5  RT (min)      lcqtof_t3_neg_3_032_24    m/z 847.6323     0   1   2E+2   11   11.5  RT (min)      lcqtof_t3_neg_3_038_06    m/z 847.6327     0   1   2E+2   10.5   11  RT (min)      lcqtof_t3_neg_3_038_06    m/z 847.6329     0   1E+4   12.5   13  RT (min)      lcqtof_t3_neg_4_040_24    m/z 847.6578     0   1   2   3   4E+2   11.5   12  RT (min)      lcqtof_t3_neg_3_032_24    m/z 847.6584     0   1   2   3E+2   11.5   12  RT (min)      lcqtof_t3_neg_3_032_24    m/z 848.4673     0   1   2   3E+4   12   12.5  RT (min)      lcqtof_t3_neg_1_019_23    m/z 848.4676     0   1   2   3   4E+2   10.5   11  RT (min)      lcqtof_t3_neg_3_038_06    m/z 848.4686     0   1   2   3E+4   12   12.5  RT (min)      lcqtof_t3_neg_1_019_23    m/z 848.6391     0   1   2E+2   11.5   12  RT (min)      lcqtof_t3_neg_3_032_24    m/z 848.6406     0   1   2   3E+2   11.5   12  RT (min)      lcqtof_t3_neg_3_032_24    m/z 848.6621     0   1   2   3   4E+4   12.5   13  RT (min)      lcqtof_t3_neg_4_046_33    m/z 849.1140     0   1   2E+4   12.5   13  RT (min)      lcqtof_t3_neg_4_064_17    m/z 849.1174     0   1   2E+4   13   13.5  RT (min)      lcqtof_t3_neg_4_064_17    m/z 849.4686     0   1E+4   12   12.5  RT (min)      lcqtof_t3_neg_1_019_23    m/z 849.4687     0   1E+4   12   12.5  RT (min)      lcqtof_t3_neg_1_019_23    m/z 849.4697     0   1E+4   12   12.5  RT (min)      lcqtof_t3_neg_4_018_23    m/z 849.6469     0   1   2E+2   11   11.5  RT (min)      lcqtof_t3_neg_3_038_06    m/z 849.6478     0   1   2   3E+4   12.5   13  RT (min)      lcqtof_t3_neg_2_016_05    m/z 849.6495     0   1   2E+2   11   11.5  RT (min)      lcqtof_t3_neg_3_038_06    m/z 849.6558     0   1   2E+3   12   12.5  RT (min)      lcqtof_t3_neg_3_032_24    m/z 849.6615     0   1E+4   12   12.5  RT (min)      lcqtof_t3_neg_4_040_24    m/z 850.1122     0   1E+4   12.5   13  RT (min)      lcqtof_t3_neg_4_064_17    m/z 850.1123     0   1E+4   12.5   13  RT (min)      lcqtof_t3_neg_4_064_17    m/z 850.4725     0   1E+4   12   12.5  RT (min)      lcqtof_t3_neg_4_018_23    m/z 850.4830     0   1E+4   13   13.5  RT (min)      lcqtof_t3_neg_4_047_07    m/z 850.4832     0   1   2E+4   13   13.5  RT (min)      lcqtof_t3_neg_4_035_08    m/z 850.5566     0   1E+4   12.5   13  RT (min)      lcqtof_t3_neg_2_030_23    m/z 850.5566     0   1E+4   12   12.5  RT (min)      lcqtof_t3_neg_1_019_23    m/z 850.5571     0   1E+4   12.5   13  RT (min)      lcqtof_t3_neg_4_018_23    m/z 850.6494     0   1   2   3   4E+3   13.5   14  RT (min)      lcqtof_t3_neg_1_038_15    m/z 850.6602     0   1   2E+2   11   11.5  RT (min)      lcqtof_t3_neg_3_038_06    m/z 850.6608     0   1   2E+2   11   11.5  RT (min)      lcqtof_t3_neg_3_038_06    m/z 850.6673     0   2   4E+3   14   14.5  RT (min)      lcqtof_t3_neg_2_026_36    m/z 850.8162     0   1E+4   17   17.5  RT (min)      lcqtof_t3_neg_4_035_08    m/z 850.8182     0   1E+4   16   16.5  RT (min)      lcqtof_t3_neg_4_018_23    m/z 851.1088     0   1E+4   12.5   13  RT (min)      lcqtof_t3_neg_4_064_17    m/z 851.1096     0   1E+4   12.5   13  RT (min)      lcqtof_t3_neg_4_064_17    m/z 851.4857     0   1E+4   13   13.5  RT (min)      lcqtof_t3_neg_4_035_08    m/z 851.6547     0   1   2   3   4E+2   11.5   12  RT (min)      lcqtof_t3_neg_3_032_24    m/z 851.6560     0   1   2   3   4E+2   11.5   12  RT (min)      lcqtof_t3_neg_3_032_24    m/z 851.6567     0   1   2E+2   10   10.5  RT (min)      lcqtof_t3_neg_3_038_06    m/z 851.6644     0   1   2E+2   11   11.5  RT (min)      lcqtof_t3_neg_3_038_06    m/z 852.4829     0   1E+4   12   12.5  RT (min)      lcqtof_t3_neg_4_018_23    m/z 852.4834     0   1E+4   12   12.5  RT (min)      lcqtof_t3_neg_4_018_23    m/z 852.5734     0   1E+4   12.5   13  RT (min)      lcqtof_t3_neg_2_030_23    m/z 852.5749     0   1E+4   12.5   13  RT (min)      lcqtof_t3_neg_4_018_23    m/z 852.9665     0   1E+4   1   1.5  RT (min)      lcqtof_t3_neg_4_059_14    m/z 853.2712     0   1   2E+3   10   10.5  RT (min)      lcqtof_t3_neg_1_062_13    m/z 853.6696     0   1   2   3   4E+4   13   13.5  RT (min)      lcqtof_t3_neg_2_016_05    m/z 853.6708     0   1   2E+2   11.5   12  RT (min)      lcqtof_t3_neg_3_032_24    m/z 853.6715     0   1   2   3   4E+2   10.5   11  RT (min)      lcqtof_t3_neg_3_038_06    m/z 853.6716     0   1   2E+4   12   12.5  RT (min)      lcqtof_t3_neg_2_016_05    m/z 853.6722     0   1   2E+2   11.5   12  RT (min)      lcqtof_t3_neg_3_032_24    m/z 853.6727     0   1   2   3   4E+2   10.5   11  RT (min)      lcqtof_t3_neg_3_038_06    m/z 853.6729     0   1   2E+4   12   12.5  RT (min)      lcqtof_t3_neg_2_016_05    m/z 853.6735     0   1E+2   10.5   11  RT (min)      lcqtof_t3_neg_3_038_06    m/z 853.6738     0   1   2E+2   10.5   11  RT (min)      lcqtof_t3_neg_3_038_06    m/z 853.6780     0   1   2E+2   11   11.5  RT (min)      lcqtof_t3_neg_3_038_06    m/z 854.2985     0   2   4E+3   16.5   17  RT (min)       lcqtof_t3_neg_4_000_cc    m/z 854.5607     0   1E+4   13   13.5  RT (min)       lcqtof_t3_neg_4_084_59    m/z 854.5908     0   2   4E+4   12.5   13  RT (min)      lcqtof_t3_neg_2_030_23    m/z 854.6843     0   1   2   3E+2   12   12.5  RT (min)      lcqtof_t3_neg_3_032_24    m/z 854.6856     0   1E+4   12.5   13  RT (min)      lcqtof_t3_neg_2_016_05    m/z 854.6861     0   1   2   3E+2   11.5   12   12.5  RT (min)      lcqtof_t3_neg_3_032_24    m/z 854.9643     0   1E+4   1   1.5  RT (min)      lcqtof_t3_neg_4_059_14    m/z 855.5945     0   1   2E+4   12.5   13  RT (min)      lcqtof_t3_neg_2_030_23    m/z 855.6144     0   2   4   6E+5   14   14.5  RT (min)      lcqtof_t3_neg_4_037_01    m/z 855.6216     0   2   4   6E+5   13.5   14  RT (min)      lcqtof_t3_neg_4_016_14    m/z 855.6344     0   1   2   3E+5   13.5   14  RT (min)      lcqtof_t3_neg_4_046_33    m/z 855.6396     0   2   4   6E+3   12.5   13  RT (min)      lcqtof_t3_neg_1_031_24    m/z 855.6863     0   1   2E+2   12   12.5  RT (min)      lcqtof_t3_neg_3_032_24    m/z 855.6866     0   2   4E+4   12.5   13  RT (min)      lcqtof_t3_neg_2_016_05    m/z 855.6872     0   1   2E+2   10.5   11  RT (min)      lcqtof_t3_neg_3_038_06    m/z 855.6874     0   1   2E+2   11.5   12  RT (min)      lcqtof_t3_neg_3_032_24    m/z 855.6874     0   2   4E+4   12.5   13  RT (min)      lcqtof_t3_neg_2_016_05    m/z 855.6884     0   1   2E+2   10.5   11  RT (min)      lcqtof_t3_neg_3_038_06    m/z 856.5126     0   1E+4   12.5   13  RT (min)      lcqtof_t3_neg_4_037_01    m/z 856.5282     0   1E+4   13   13.5  RT (min)       lcqtof_t3_neg_4_038_29    m/z 856.5735     0   1E+4   13   13.5  RT (min)      lcqtof_t3_neg_4_018_23    m/z 856.6005     0   1E+4   12.5   13  RT (min)      lcqtof_t3_neg_1_003_07    m/z 856.6022     0   1   2   3E+4   13   13.5  RT (min)      lcqtof_t3_neg_2_030_23    m/z 856.6071     0   2   4   6E+5   14   14.5  RT (min)      lcqtof_t3_neg_4_002_17    m/z 856.6073     0   1E+4   13   13.5  RT (min)      lcqtof_t3_neg_2_016_05    m/z 856.6654     0   1E+4   13.5   14  RT (min)      lcqtof_t3_neg_4_038_29    m/z 856.6992     0   1   2   3   4E+2   12   12.5  RT (min)      lcqtof_t3_neg_3_032_24    m/z 856.6992     0   1   2   3   4E+2   12   12.5  RT (min)      lcqtof_t3_neg_3_032_24    m/z 856.6995     0   1E+4   12.5   13  RT (min)      lcqtof_t3_neg_2_016_05    m/z 857.2042     0   1E+4   12.5   13  RT (min)      lcqtof_t3_neg_4_002_17    m/z 857.2047     0   1E+4   12.5   13  RT (min)      lcqtof_t3_neg_4_002_17    m/z 857.2070     0   1   2   3E+4   12.5   13  RT (min)      lcqtof_t3_neg_4_064_17    m/z 857.3642     0   1E+4   4   4.5  RT (min)      lcqtof_t3_neg_4_000_ee    m/z 857.3978     0   1E+4   4.5   5  RT (min)      lcqtof_t3_neg_4_000_ee    m/z 857.5178     0   1E+4   12   12.5  RT (min)      lcqtof_t3_neg_2_030_23    m/z 857.5822     0   1   2E+3   16   16.5  RT (min)      lcqtof_t3_neg_1_025_52    m/z 857.5831     0   1   2E+3   15.5   16  RT (min)      lcqtof_t3_neg_1_022_28    m/z 857.5883     0   1   2E+3   14.5   15  RT (min)      lcqtof_t3_neg_1_012_16    m/z 857.6080     0   1E+4   13   13.5  RT (min)      lcqtof_t3_neg_2_030_23    m/z 857.6602     0   2   4   6   8E+3   12.5   13  RT (min)      lcqtof_t3_neg_1_003_07    m/z 857.6609     0   2   4   6   8E+3   12.5   13  RT (min)      lcqtof_t3_neg_1_003_07    m/z 857.7014     0   1   2   3E+2   12   12.5  RT (min)      lcqtof_t3_neg_3_032_24    m/z 857.7023     0   1   2   3E+2   12   12.5  RT (min)      lcqtof_t3_neg_3_032_24    m/z 857.7028     0   2   4E+4   12.5   13  RT (min)      lcqtof_t3_neg_2_016_05    m/z 857.7043     0   1   2   3   4E+2   10.5   11  RT (min)      lcqtof_t3_neg_3_038_06    m/z 857.7050     0   1   2   3   4E+2   11   11.5  RT (min)      lcqtof_t3_neg_3_038_06    m/z 857.9733     0   1E+4   13.5   14  RT (min)      lcqtof_t3_neg_4_037_01    m/z 858.2007     0   1E+4   12.5   13  RT (min)      lcqtof_t3_neg_4_002_17    m/z 858.2035     0   1E+4   12.5   13  RT (min)      lcqtof_t3_neg_4_002_17    m/z 858.2070     0   1   2E+4   12.5   13  RT (min)      lcqtof_t3_neg_4_064_17    m/z 858.3874     0   1   2E+4   4.5   5  RT (min)      lcqtof_t3_neg_4_000_ee    m/z 858.6485     0   2   4   6   8E+4   13.5   14  RT (min)       lcqtof_t3_neg_2_023_37    m/z 858.6631     0   1E+3   12   12.5  RT (min)      lcqtof_t3_neg_3_032_24    m/z 859.1691     0   1E+4   13   13.5  RT (min)      lcqtof_t3_neg_4_057_49    m/z 859.1715     0   1E+4   13   13.5  RT (min)      lcqtof_t3_neg_4_001_49    m/z 859.2023     0   1E+4   12.5   13  RT (min)      lcqtof_t3_neg_4_002_17    m/z 859.2029     0   1E+4   13   13.5  RT (min)      lcqtof_t3_neg_4_032_04    m/z 859.2047     0   1   2E+4   12.5   13  RT (min)      lcqtof_t3_neg_4_064_17    m/z 859.3890     0   1E+4   4.5   5  RT (min)      lcqtof_t3_neg_4_000_ee    m/z 859.5809     0   2   4   6   8E+3   12   12.5  RT (min)      lcqtof_t3_neg_1_015_34    m/z 859.5856     0   2   4   6   8E+3   13.5   14  RT (min)      lcqtof_t3_neg_1_020_47    m/z 859.5861     0   1E+4   15   15.5  RT (min)       lcqtof_t3_neg_2_000_aa    m/z 859.5868     0   1E+4   13   13.5  RT (min)      lcqtof_t3_neg_1_001_49    m/z 859.5886     0   2   4E+3   12.5   13  RT (min)      lcqtof_t3_neg_1_010_27    m/z 859.5898     0   1E+4   11   11.5  RT (min)      lcqtof_t3_neg_1_019_23    m/z 859.6500     0   2   4E+4   13.5   14  RT (min)      lcqtof_t3_neg_4_011_32    m/z 859.6758     0   2   4   6   8E+3   13   13.5  RT (min)      lcqtof_t3_neg_1_040_08    m/z 859.6761     0   1E+4   12.5   13  RT (min)      lcqtof_t3_neg_1_031_24    m/z 859.7168     0   1   2   3E+2   12   12.5  RT (min)      lcqtof_t3_neg_3_032_24    m/z 859.7175     0   1   2   3E+2   12   12.5  RT (min)      lcqtof_t3_neg_3_032_24    m/z 859.7183     0   1   2E+4   12.5   13  RT (min)      lcqtof_t3_neg_2_016_05    m/z 859.7214     0   2   4E+2   11   11.5  RT (min)      lcqtof_t3_neg_3_038_06    m/z 860.1695     0   1E+4   13   13.5  RT (min)      lcqtof_t3_neg_4_057_49    m/z 860.2034     0   1E+4   12.5   13  RT (min)      lcqtof_t3_neg_4_064_17    m/z 860.5345     0   1E+4   12   12.5  RT (min)      lcqtof_t3_neg_2_030_23    m/z 860.5371     0   1E+4   12   12.5  RT (min)      lcqtof_t3_neg_1_019_23    m/z 860.5383     0   1E+4   12   12.5  RT (min)      lcqtof_t3_neg_2_030_23    m/z 860.6760     0   1   2   3   4E+2   12   12.5  RT (min)      lcqtof_t3_neg_3_032_24    m/z 860.7021     0   1   2   3E+2   11   11.5  RT (min)      lcqtof_t3_neg_3_038_06    m/z 860.7077     0   2   4   6E+2   12   12.5  RT (min)      lcqtof_t3_neg_3_032_24    m/z 860.7085     0   1   2E+4   12.5   13  RT (min)      lcqtof_t3_neg_2_016_05    m/z 860.7198     0   1E+3   12   12.5  RT (min)      lcqtof_t3_neg_3_032_24    m/z 860.7198     0   1   2   3E+2   11   11.5  RT (min)      lcqtof_t3_neg_3_038_06    m/z 860.7218     0   1   2   3E+2   11   11.5  RT (min)      lcqtof_t3_neg_3_038_06    m/z 860.7219     0   1   2E+4   12.5   13  RT (min)      lcqtof_t3_neg_2_016_05    m/z 860.8469     0   2   4E+4   17   17.5  RT (min)      lcqtof_t3_neg_4_035_08    m/z 860.8483     0   2   4   6   8E+4   16   16.5  RT (min)      lcqtof_t3_neg_4_018_23    m/z 861.1708     0   1E+4   13   13.5  RT (min)      lcqtof_t3_neg_4_057_49    m/z 861.1855     0   1E+4   13.5   14  RT (min)      lcqtof_t3_neg_4_036_47    m/z 861.2142     0   1   2E+4   13   13.5  RT (min)      lcqtof_t3_neg_4_002_17    m/z 861.2194     0   1   2E+4   13   13.5  RT (min)      lcqtof_t3_neg_4_002_17    m/z 861.2196     0   1   2   3E+4   12.5   13  RT (min)      lcqtof_t3_neg_4_064_17    m/z 861.5204     0   1E+4   13   13.5  RT (min)      lcqtof_t3_neg_4_032_04    m/z 861.5483     0   2   4E+4   12   12.5  RT (min)      lcqtof_t3_neg_1_019_23    m/z 861.5935     0   1   2   3E+3   14.5   15  RT (min)      lcqtof_t3_neg_1_017_51    m/z 861.6916     0   1E+4   13   13.5  RT (min)      lcqtof_t3_neg_1_031_24    m/z 861.6918     0   1E+4   13   13.5  RT (min)      lcqtof_t3_neg_1_031_24    m/z 862.2205     0   1   2E+4   12.5   13  RT (min)      lcqtof_t3_neg_4_064_17    m/z 862.2212     0   1E+4   13   13.5  RT (min)      lcqtof_t3_neg_4_002_17    m/z 862.5521     0   1   2E+4   12   12.5  RT (min)      lcqtof_t3_neg_1_019_23    m/z 862.5996     0   1E+4   12.5   13  RT (min)      lcqtof_t3_neg_1_003_07    m/z 862.6694     0   2   4   6E+3   13   13.5  RT (min)      lcqtof_t3_neg_1_003_07    m/z 863.0732     0   2   4E+4   13.5   14  RT (min)      lcqtof_t3_neg_4_064_17    m/z 863.0736     0   2   4E+4   13   13.5  RT (min)      lcqtof_t3_neg_4_064_17    m/z 863.0747     0   1   2   3E+4   14   14.5  RT (min)      lcqtof_t3_neg_4_002_17    m/z 863.0752     0   1   2   3E+4   14   14.5  RT (min)      lcqtof_t3_neg_4_002_17    m/z 863.0756     0   2   4E+4   12.5   13  RT (min)      lcqtof_t3_neg_4_064_17    m/z 863.0769     0   2   4E+4   13   13.5  RT (min)      lcqtof_t3_neg_4_064_17    m/z 863.0774     0   1   2E+4   14   14.5  RT (min)      lcqtof_t3_neg_4_002_17    m/z 863.0776     0   2   4E+4   13.5   14  RT (min)      lcqtof_t3_neg_4_064_17    m/z 863.2120     0   1E+4   13   13.5  RT (min)      lcqtof_t3_neg_4_002_17    m/z 863.2140     0   1   2E+4   12.5   13   13.5  RT (min)      lcqtof_t3_neg_4_064_17    m/z 863.2214     0   1E+4   12.5   13  RT (min)      lcqtof_t3_neg_4_002_17    m/z 863.4380     0   1E+4   12   12.5  RT (min)      lcqtof_t3_neg_1_019_23    m/z 863.5408     0   1E+4   9   9.5  RT (min)      lcqtof_t3_neg_4_000_ee    m/z 863.5412     0   1E+4   9   9.5  RT (min)      lcqtof_t3_neg_4_000_ee    m/z 863.5569     0   1E+4   12.5   13  RT (min)      lcqtof_t3_neg_3_006_23    m/z 863.5642     0   1   2E+4   12.5   13  RT (min)      lcqtof_t3_neg_1_019_23    m/z 863.6674     0   1   2E+2   11.5   12  RT (min)      lcqtof_t3_neg_3_032_24    m/z 863.6702     0   1E+4   12   12.5  RT (min)      lcqtof_t3_neg_2_016_05    m/z 863.6722     0   1   2   3E+2   11.5   12  RT (min)      lcqtof_t3_neg_3_032_24    m/z 864.0706     0   1   2   3E+4   13.5   14  RT (min)      lcqtof_t3_neg_4_064_17    m/z 864.0713     0   1   2   3   4E+4   13.5   14  RT (min)      lcqtof_t3_neg_4_064_17    m/z 864.0721     0   1   2   3   4E+4   12.5   13  RT (min)      lcqtof_t3_neg_4_064_17    m/z 864.0725     0   1   2   3E+4   13.5   14  RT (min)      lcqtof_t3_neg_4_064_17    m/z 864.0739     0   1   2   3   4E+4   13   13.5  RT (min)      lcqtof_t3_neg_4_064_17    m/z 864.0744     0   1   2   3   4E+4   12.5   13  RT (min)      lcqtof_t3_neg_4_064_17    m/z 864.0745     0   1   2   3   4E+4   13   13.5  RT (min)      lcqtof_t3_neg_4_064_17    m/z 864.0752     0   1   2   3   4E+4   13   13.5  RT (min)      lcqtof_t3_neg_4_064_17    m/z 864.0773     0   1   2E+4   14   14.5  RT (min)      lcqtof_t3_neg_4_064_17    m/z 864.2097     0   1E+4   12.5   13  RT (min)      lcqtof_t3_neg_4_064_17    m/z 864.2164     0   1E+4   13   13.5  RT (min)      lcqtof_t3_neg_4_064_17    m/z 864.4911     0   1   2E+4   12   12.5  RT (min)      lcqtof_t3_neg_1_019_23    m/z 864.4947     0   1E+4   12   12.5  RT (min)      lcqtof_t3_neg_2_030_23    m/z 864.5620     0   1E+4   12.5   13  RT (min)      lcqtof_t3_neg_4_018_23    m/z 864.5694     0   1E+4   12.5   13  RT (min)      lcqtof_t3_neg_1_019_23    m/z 864.6031     0   2   4   6E+2   11.5   12  RT (min)      lcqtof_t3_neg_3_032_24    m/z 864.6224     0   1E+4   13   13.5  RT (min)      lcqtof_t3_neg_1_003_07    m/z 864.6738     0   1   2E+2   11.5   12  RT (min)      lcqtof_t3_neg_3_032_24    m/z 864.6745     0   1   2E+2   11.5   12  RT (min)      lcqtof_t3_neg_3_032_24    m/z 864.6746     0   2   4E+4   12   12.5  RT (min)      lcqtof_t3_neg_2_016_05    m/z 864.6753     0   1   2E+2   10.5   11  RT (min)      lcqtof_t3_neg_3_038_06    m/z 864.6768     0   1   2E+2   11.5   12  RT (min)      lcqtof_t3_neg_3_032_24    m/z 864.6801     0   1   2E+2   10.5   11  RT (min)      lcqtof_t3_neg_3_038_06    m/z 865.0680     0   1   2   3   4E+4   13.5   14  RT (min)      lcqtof_t3_neg_4_064_17    m/z 865.0709     0   1   2   3   4E+4   12.5   13  RT (min)      lcqtof_t3_neg_4_064_17    m/z 865.0710     0   1   2   3E+4   13.5   14  RT (min)      lcqtof_t3_neg_4_064_17    m/z 865.0716     0   1   2   3   4E+4   13   13.5  RT (min)      lcqtof_t3_neg_4_064_17    m/z 865.0723     0   1   2   3   4E+4   13.5   14  RT (min)      lcqtof_t3_neg_4_064_17    m/z 865.0726     0   1   2   3   4E+4   13   13.5  RT (min)      lcqtof_t3_neg_4_064_17    m/z 865.0728     0   1   2   3E+4   13.5   14  RT (min)      lcqtof_t3_neg_4_064_17    m/z 865.4985     0   1E+4   12   12.5  RT (min)      lcqtof_t3_neg_1_019_23    m/z 865.6697     0   1   2E+5   12   12.5  RT (min)      lcqtof_t3_neg_2_016_05    m/z 865.6706     0   1   2   3E+2   11.5   12  RT (min)      lcqtof_t3_neg_3_032_24    m/z 865.6719     0   1   2E+2   10.5   11  RT (min)      lcqtof_t3_neg_3_038_06    m/z 865.6723     0   1   2E+5   12   12.5  RT (min)      lcqtof_t3_neg_2_016_05    m/z 865.6724     0   1   2E+2   10.5   11  RT (min)      lcqtof_t3_neg_3_038_06    m/z 866.0705     0   1E+4   13.5   14  RT (min)      lcqtof_t3_neg_4_002_17    m/z 866.0709     0   1   2E+4   13.5   14  RT (min)      lcqtof_t3_neg_4_064_17    m/z 866.0713     0   1E+4   13   13.5  RT (min)      lcqtof_t3_neg_4_002_17    m/z 866.0721     0   1   2E+4   13   13.5  RT (min)      lcqtof_t3_neg_4_064_17    m/z 866.5161     0   1   2E+4   12   12.5  RT (min)      lcqtof_t3_neg_1_019_23    m/z 866.5547     0   1   2   3E+4   12   12.5  RT (min)      lcqtof_t3_neg_4_016_14    m/z 866.6227     0   1E+4   12.5   13  RT (min)      lcqtof_t3_neg_4_098_61    m/z 866.6413     0   1E+4   13   13.5  RT (min)      lcqtof_t3_neg_1_031_24    m/z 866.6430     0   1E+5   13   13.5  RT (min)      lcqtof_t3_neg_1_003_07    m/z 866.6755     0   1   2   3E+2   11.5   12  RT (min)      lcqtof_t3_neg_3_032_24    m/z 866.6756     0   2   4   6E+5   12   12.5  RT (min)      lcqtof_t3_neg_2_016_05    m/z 866.6756     0   2   4   6E+4   13   13.5  RT (min)      lcqtof_t3_neg_1_031_24    m/z 866.6757     0   1   2   3   4E+2   10.5   11  RT (min)      lcqtof_t3_neg_3_038_06    m/z 866.6761     0   1   2   3E+2   11.5   12  RT (min)      lcqtof_t3_neg_3_032_24    m/z 866.6777     0   1   2   3   4E+2   10.5   11  RT (min)      lcqtof_t3_neg_3_038_06    m/z 866.6810     0   1E+5   13   13.5  RT (min)      lcqtof_t3_neg_4_040_24    m/z 866.7976     0   1E+4   17   17.5  RT (min)      lcqtof_t3_neg_4_035_08    m/z 866.7993     0   1E+4   16   16.5  RT (min)      lcqtof_t3_neg_4_018_23    m/z 866.9730     0   1   2   3   4E+2   11.5   12  RT (min)      lcqtof_t3_neg_3_032_24    m/z 866.9738     0   1   2   3   4E+2   11.5   12  RT (min)      lcqtof_t3_neg_3_032_24    m/z 867.0675     0   1E+4   13   13.5  RT (min)      lcqtof_t3_neg_4_064_17    m/z 867.0688     0   1E+4   13   13.5  RT (min)      lcqtof_t3_neg_4_064_17    m/z 867.5217     0   1E+4   12   12.5  RT (min)      lcqtof_t3_neg_1_019_23    m/z 867.5497     0   2   4   6E+3   13   13.5  RT (min)      lcqtof_t3_neg_1_036_14    m/z 867.5567     0   1E+4   12   12.5  RT (min)      lcqtof_t3_neg_4_016_14    m/z 867.6428     0   2   4   6   8E+1   12.5   13  RT (min)      lcqtof_t3_neg_3_032_24    m/z 867.6865     0   1   2E+2   11.5   12  RT (min)      lcqtof_t3_neg_3_032_24    m/z 867.6869     0   1   2E+2   10.5   11  RT (min)      lcqtof_t3_neg_3_038_06    m/z 867.6872     0   1   2E+2   12   12.5  RT (min)      lcqtof_t3_neg_3_032_24    m/z 867.6876     0   1   2E+2   10.5   11  RT (min)      lcqtof_t3_neg_3_038_06    m/z 867.8079     0   1E+4   10.5   11  RT (min)      lcqtof_t3_neg_1_045_29    m/z 867.9868     0   1E+4   12.5   13  RT (min)      lcqtof_t3_neg_2_028_24    m/z 868.1862     0   1   2   3E+4   12.5   13  RT (min)      lcqtof_t3_neg_4_064_17    m/z 868.1887     0   1   2   3E+4   12.5   13  RT (min)      lcqtof_t3_neg_4_064_17    m/z 868.5309     0   1   2   3E+4   12.5   13  RT (min)      lcqtof_t3_neg_2_030_23    m/z 868.5324     0   1   2   3E+4   12   12.5   13  RT (min)      lcqtof_t3_neg_1_019_23    m/z 868.5690     0   1   2   3E+4   12   12.5  RT (min)      lcqtof_t3_neg_4_016_14    m/z 868.5714     0   1E+4   12.5   13  RT (min)       lcqtof_t3_neg_1_019_23    m/z 868.5789     0   1E+4   12.5   13  RT (min)      lcqtof_t3_neg_4_018_23    m/z 868.5806     0   1E+4   12.5   13  RT (min)      lcqtof_t3_neg_1_019_23    m/z 868.6423     0   1E+3   12.5   13  RT (min)      lcqtof_t3_neg_3_032_24    m/z 868.6512     0   1E+3   12   12.5  RT (min)      lcqtof_t3_neg_3_032_24    m/z 868.6526     0   1E+4   13   13.5  RT (min)      lcqtof_t3_neg_2_016_05    m/z 868.6532     0   1E+3   12.5   13  RT (min)      lcqtof_t3_neg_3_032_24    m/z 868.6534     0   1E+3   12.5   13  RT (min)      lcqtof_t3_neg_3_032_24    m/z 868.6555     0   1   2   3E+2   11   11.5  RT (min)      lcqtof_t3_neg_3_038_06    m/z 868.6639     0   1E+4   13   13.5  RT (min)      lcqtof_t3_neg_2_016_05    m/z 868.6896     0   1   2E+2   11.5   12  RT (min)      lcqtof_t3_neg_3_032_24    m/z 868.6899     0   1E+2   12   12.5  RT (min)      lcqtof_t3_neg_3_032_24    m/z 868.6915     0   1   2   3   4E+2   10.5   11  RT (min)      lcqtof_t3_neg_3_038_06    m/z 868.6915     0   1   2   3E+2   10.5   11  RT (min)      lcqtof_t3_neg_3_038_06    m/z 868.9385     0   1E+4   1   1.5  RT (min)      lcqtof_t3_neg_4_059_14    m/z 868.9880     0   1   2   3E+2   12   12.5  RT (min)      lcqtof_t3_neg_3_032_24    m/z 868.9888     0   1   2   3E+4   12.5   13  RT (min)      lcqtof_t3_neg_2_016_05    m/z 869.1299     0   1E+4   12.5   13  RT (min)      lcqtof_t3_neg_2_028_24    m/z 869.3877     0   1   2E+4   12.5   13  RT (min)      lcqtof_t3_neg_2_028_24    m/z 869.5311     0   1E+4   12.5   13  RT (min)      lcqtof_t3_neg_2_030_23    m/z 869.5323     0   1E+4   12.5   13  RT (min)      lcqtof_t3_neg_1_019_23    m/z 869.5350     0   1E+4   12   12.5  RT (min)      lcqtof_t3_neg_1_019_23    m/z 869.5730     0   1E+4   12   12.5  RT (min)      lcqtof_t3_neg_4_016_14    m/z 869.5740     0   1E+4   12   12.5  RT (min)      lcqtof_t3_neg_4_016_14    m/z 869.5740     0   1E+4   12   12.5  RT (min)       lcqtof_t3_neg_4_059_14    m/z 869.5748     0   1E+4   12   12.5  RT (min)      lcqtof_t3_neg_4_059_14    m/z 869.7017     0   1   2E+2   12   12.5  RT (min)      lcqtof_t3_neg_3_032_24    m/z 869.7025     0   1   2E+2   12   12.5  RT (min)      lcqtof_t3_neg_3_032_24    m/z 869.7032     0   1   2   3   4E+2   10.5   11  RT (min)      lcqtof_t3_neg_3_038_06    m/z 869.7038     0   1   2   3   4E+2   10.5   11  RT (min)      lcqtof_t3_neg_3_038_06    m/z 870.0012     0   1E+4   13   13.5  RT (min)      lcqtof_t3_neg_2_028_24    m/z 870.1840     0   1   2E+4   12.5   13  RT (min)      lcqtof_t3_neg_4_064_17    m/z 870.3947     0   1E+4   10   10.5  RT (min)       lcqtof_t3_neg_4_058_16    m/z 870.4519     0   1E+4   11.5   12  RT (min)      lcqtof_t3_neg_1_019_23    m/z 870.5442     0   1E+4   12.5   13  RT (min)      lcqtof_t3_neg_1_019_23    m/z 870.5477     0   1   2E+2   11   11.5  RT (min)      lcqtof_t3_neg_3_038_06    m/z 870.5482     0   1   2   3   4E+4   12.5   13  RT (min)      lcqtof_t3_neg_1_019_23    m/z 870.5490     0   1   2   3   4E+4   12.5   13  RT (min)      lcqtof_t3_neg_1_019_23    m/z 870.5506     0   1   2E+2   10.5   11  RT (min)      lcqtof_t3_neg_3_038_06    m/z 870.5841     0   1E+4   12   12.5  RT (min)      lcqtof_t3_neg_4_003_37    m/z 870.5843     0   1E+4   12   12.5  RT (min)      lcqtof_t3_neg_4_034_34    m/z 870.6037     0   1E+4   13.5   14  RT (min)      lcqtof_t3_neg_4_059_14    m/z 870.6187     0   1E+4   13.5   14  RT (min)      lcqtof_t3_neg_4_059_14    m/z 870.6676     0   2   4   6   8E+3   13   13.5  RT (min)      lcqtof_t3_neg_2_030_23    m/z 870.6685     0   2   4   6E+2   12.5   13  RT (min)      lcqtof_t3_neg_3_032_24    m/z 870.7056     0   1   2E+2   12   12.5  RT (min)      lcqtof_t3_neg_3_032_24    m/z 870.7062     0   1   2E+2   12   12.5  RT (min)      lcqtof_t3_neg_3_032_24    m/z 870.7072     0   1   2   3E+2   10.5   11  RT (min)      lcqtof_t3_neg_3_038_06    m/z 870.7076     0   1   2   3E+2   10.5   11  RT (min)      lcqtof_t3_neg_3_038_06    m/z 870.9354     0   1E+4   1   1.5  RT (min)      lcqtof_t3_neg_4_059_14    m/z 871.0047     0   1   2   3E+2   12   12.5  RT (min)      lcqtof_t3_neg_3_032_24    m/z 871.0051     0   1   2   3E+2   12   12.5  RT (min)      lcqtof_t3_neg_3_032_24    m/z 871.0052     0   1   2E+4   12.5   13  RT (min)      lcqtof_t3_neg_2_016_05    m/z 871.1467     0   1E+4   13   13.5  RT (min)      lcqtof_t3_neg_2_028_24    m/z 871.1835     0   1E+4   12.5   13  RT (min)      lcqtof_t3_neg_4_064_17    m/z 871.1929     0   1E+4   12   12.5  RT (min)      lcqtof_t3_neg_4_064_17    m/z 871.1943     0   1E+4   12   12.5  RT (min)      lcqtof_t3_neg_4_064_17    m/z 871.1960     0   1E+4   12   12.5  RT (min)      lcqtof_t3_neg_4_064_17    m/z 871.4051     0   1   2E+4   13   13.5  RT (min)      lcqtof_t3_neg_2_028_24    m/z 871.4052     0   1   2E+4   13   13.5  RT (min)      lcqtof_t3_neg_2_028_24    m/z 871.5383     0   1E+4   12.5   13  RT (min)      lcqtof_t3_neg_4_003_37    m/z 871.5420     0   1E+4   12.5   13  RT (min)       lcqtof_t3_neg_4_003_37    m/z 871.5489     0   1E+4   12.5   13  RT (min)       lcqtof_t3_neg_2_030_23    m/z 871.5651     0   1E+4   12.5   13  RT (min)      lcqtof_t3_neg_4_003_37    m/z 871.5740     0   1E+4   12.5   13  RT (min)      lcqtof_t3_neg_4_008_35    m/z 871.5861     0   1E+4   12.5   13  RT (min)      lcqtof_t3_neg_4_037_01    m/z 871.5881     0   1   2   3   4E+4   12.5   13  RT (min)       lcqtof_t3_neg_4_059_14    m/z 871.5891     0   1   2   3   4E+4   12.5   13  RT (min)      lcqtof_t3_neg_4_059_14    m/z 871.5894     0   1   2   3E+4   12   12.5  RT (min)      lcqtof_t3_neg_4_016_14    m/z 871.7103     0   1E+5   12.5   13  RT (min)      lcqtof_t3_neg_2_016_05    m/z 871.7104     0   1   2   3   4E+4   12.5   13  RT (min)      lcqtof_t3_neg_2_016_05    m/z 871.7170     0   1E+2   12   12.5  RT (min)      lcqtof_t3_neg_3_032_24    m/z 871.7173     0   1E+2   12   12.5  RT (min)      lcqtof_t3_neg_3_032_24    m/z 871.7191     0   1   2   3E+2   11   11.5  RT (min)      lcqtof_t3_neg_3_038_06    m/z 871.7192     0   1   2   3E+2   11   11.5  RT (min)      lcqtof_t3_neg_3_038_06    m/z 872.1953     0   1E+4   12   12.5  RT (min)      lcqtof_t3_neg_4_064_17    m/z 872.4671     0   1E+4   12   12.5  RT (min)      lcqtof_t3_neg_1_019_23    m/z 872.5261     0   1E+4   12.5   13  RT (min)      lcqtof_t3_neg_4_003_37    m/z 872.5350     0   1E+4   12.5   13  RT (min)       lcqtof_t3_neg_4_003_37    m/z 872.5430     0   1E+4   12.5   13  RT (min)      lcqtof_t3_neg_4_034_34    m/z 872.5541     0   1E+4   12.5   13  RT (min)      lcqtof_t3_neg_1_019_23    m/z 872.5693     0   1E+4   12.5   13  RT (min)      lcqtof_t3_neg_1_019_23    m/z 872.6010     0   1   2E+4   12   12.5  RT (min)      lcqtof_t3_neg_4_016_14    m/z 872.7211     0   1   2E+2   12   12.5  RT (min)      lcqtof_t3_neg_3_032_24    m/z 872.7217     0   1   2E+2   12   12.5  RT (min)      lcqtof_t3_neg_3_032_24    m/z 872.7232     0   1   2   3   4E+2   11   11.5  RT (min)      lcqtof_t3_neg_3_038_06    m/z 872.7234     0   1   2   3   4E+2   11   11.5  RT (min)      lcqtof_t3_neg_3_038_06    m/z 873.0197     0   1   2   3E+2   12   12.5  RT (min)      lcqtof_t3_neg_3_032_24    m/z 873.1752     0   1E+4   12.5   13  RT (min)      lcqtof_t3_neg_4_002_17    m/z 873.1831     0   1E+4   12   12.5  RT (min)      lcqtof_t3_neg_4_064_17    m/z 873.1837     0   1E+4   12   12.5  RT (min)      lcqtof_t3_neg_4_064_17    m/z 873.2541     0   1E+4   13   13.5  RT (min)      lcqtof_t3_neg_2_028_24    m/z 873.6163     0   1E+4   14.5   15  RT (min)      lcqtof_t3_neg_1_000_aa    m/z 873.6424     0   2   4E+2   11.5   12  RT (min)      lcqtof_t3_neg_3_032_24    m/z 873.6428     0   2   4E+2   11.5   12  RT (min)      lcqtof_t3_neg_3_032_24    m/z 873.7247     0   1   2E+2   12   12.5  RT (min)      lcqtof_t3_neg_3_032_24    m/z 873.7266     0   1E+2   12   12.5  RT (min)      lcqtof_t3_neg_3_032_24    m/z 873.7270     0   1   2   3E+4   12.5   13  RT (min)      lcqtof_t3_neg_2_016_05    m/z 873.7343     0   1   2   3   4E+2   12.5   13  RT (min)      lcqtof_t3_neg_3_032_24    m/z 873.7370     0   1   2   3E+2   11   11.5  RT (min)      lcqtof_t3_neg_3_038_06    m/z 873.7375     0   1   2   3E+2   11   11.5  RT (min)      lcqtof_t3_neg_3_038_06    m/z 874.1768     0   1E+4   12   12.5  RT (min)      lcqtof_t3_neg_4_064_17    m/z 874.1888     0   1E+4   12   12.5  RT (min)      lcqtof_t3_neg_4_064_17    m/z 874.6443     0   2   4   6E+2   11.5   12  RT (min)      lcqtof_t3_neg_3_032_24    m/z 874.6466     0   1   2E+2   10   10.5  RT (min)      lcqtof_t3_neg_3_038_06    m/z 874.6468     0   1   2E+2   10   10.5  RT (min)      lcqtof_t3_neg_3_038_06    m/z 874.6475     0   2   4   6E+2   11.5   12  RT (min)      lcqtof_t3_neg_3_032_24    m/z 874.6479     0   2   4   6E+2   11.5   12  RT (min)      lcqtof_t3_neg_3_032_24    m/z 874.6501     0   1   2E+2   10.5   11  RT (min)      lcqtof_t3_neg_3_038_06    m/z 874.6546     0   1   2   3   4E+4   11.5   12  RT (min)      lcqtof_t3_neg_3_032_24    m/z 874.6559     0   1   2E+2   10.5   11  RT (min)      lcqtof_t3_neg_3_038_06    m/z 874.7372     0   1   2   3E+2   12.5   13  RT (min)      lcqtof_t3_neg_3_032_24    m/z 874.7382     0   1   2   3E+2   11   11.5  RT (min)      lcqtof_t3_neg_3_038_06    m/z 875.1462     0   1E+5   13   13.5  RT (min)      lcqtof_t3_neg_4_064_17    m/z 875.6575     0   1   2E+2   10.5   11  RT (min)      lcqtof_t3_neg_3_038_06    m/z 875.6575     0   1   2   3E+2   11.5   12  RT (min)      lcqtof_t3_neg_3_032_24    m/z 875.6575     0   1E+5   12   12.5  RT (min)      lcqtof_t3_neg_2_016_05    m/z 875.6580     0   1   2E+2   10.5   11  RT (min)      lcqtof_t3_neg_3_038_06    m/z 875.6582     0   1   2   3E+2   11.5   12  RT (min)      lcqtof_t3_neg_3_032_24    m/z 875.6583     0   1E+3   11.5   12  RT (min)      lcqtof_t3_neg_3_032_24    m/z 875.6586     0   1   2E+2   10.5   11  RT (min)      lcqtof_t3_neg_3_038_06    m/z 875.7429     0   1   2   3E+2   12.5   13  RT (min)      lcqtof_t3_neg_3_032_24    m/z 875.7432     0   1   2   3E+2   12.5   13  RT (min)      lcqtof_t3_neg_3_032_24    m/z 875.7447     0   1   2   3E+4   13   13.5  RT (min)      lcqtof_t3_neg_4_040_24    m/z 875.7488     0   1   2   3E+4   13   13.5  RT (min)       lcqtof_t3_neg_4_040_24    m/z 875.7541     0   1   2   3   4E+2   12.5   13  RT (min)      lcqtof_t3_neg_3_032_24    m/z 875.7607     0   1   2   3   4E+2   12.5   13  RT (min)      lcqtof_t3_neg_3_032_24    m/z 875.7612     0   1   2   3   4E+2   12.5   13  RT (min)      lcqtof_t3_neg_3_032_24    m/z 875.7618     0   1E+4   12.5   13  RT (min)      lcqtof_t3_neg_4_019_05    m/z 876.1461     0   2   4   6   8E+4   13   13.5  RT (min)      lcqtof_t3_neg_4_064_17    m/z 876.4979     0   1E+4   13   13.5  RT (min)       lcqtof_t3_neg_4_035_08    m/z 876.4987     0   1E+4   12   12.5  RT (min)      lcqtof_t3_neg_1_019_23    m/z 876.5012     0   1E+4   12   12.5  RT (min)      lcqtof_t3_neg_1_019_23    m/z 876.6597     0   1   2   3E+2   11.5   12  RT (min)      lcqtof_t3_neg_3_032_24    m/z 876.6607     0   1   2   3E+2   11.5   12  RT (min)      lcqtof_t3_neg_3_032_24    m/z 876.6611     0   2   4   6   8E+2   11.5   12  RT (min)      lcqtof_t3_neg_3_032_24    m/z 876.6619     0   1   2E+2   10.5   11  RT (min)      lcqtof_t3_neg_3_038_06    m/z 876.6620     0   1   2E+2   10.5   11  RT (min)      lcqtof_t3_neg_3_038_06    m/z 876.6627     0   1   2E+2   10.5   11  RT (min)      lcqtof_t3_neg_3_038_06    m/z 876.7442     0   1   2   3E+4   13   13.5  RT (min)      lcqtof_t3_neg_2_016_05    m/z 876.7520     0   1E+2   12.5   13  RT (min)      lcqtof_t3_neg_3_032_24    m/z 876.7526     0   1E+2   12.5   13  RT (min)      lcqtof_t3_neg_3_032_24    m/z 876.7537     0   2   4E+2   11   11.5  RT (min)      lcqtof_t3_neg_3_038_06    m/z 876.7539     0   2   4E+2   11   11.5  RT (min)      lcqtof_t3_neg_3_038_06    m/z 876.8198     0   1   2E+4   16   16.5  RT (min)      lcqtof_t3_neg_4_018_23    m/z 876.8212     0   1   2E+4   17   17.5  RT (min)      lcqtof_t3_neg_4_035_08    m/z 876.9606     0   1   2   3E+2   11.5   12  RT (min)      lcqtof_t3_neg_3_032_24    m/z 876.9609     0   1   2   3E+2   11.5   12  RT (min)      lcqtof_t3_neg_3_032_24    m/z 877.1008     0   1E+4   12.5   13  RT (min)      lcqtof_t3_neg_4_040_24    m/z 877.1247     0   2   4E+4   13   13.5  RT (min)      lcqtof_t3_neg_4_002_17    m/z 877.1322     0   1E+5   13   13.5  RT (min)      lcqtof_t3_neg_4_064_17    m/z 877.3616     0   1E+4   12.5   13  RT (min)      lcqtof_t3_neg_4_040_24    m/z 877.5220     0   2   4E+3   12.5   13  RT (min)      lcqtof_t3_neg_1_002_13    m/z 877.6632     0   1   2   3E+4   14.5   15  RT (min)      lcqtof_t3_neg_1_000_aa    m/z 877.6640     0   1   2   3E+4   14.5   15  RT (min)      lcqtof_t3_neg_1_000_aa    m/z 877.6657     0   1   2E+2   11.5   12  RT (min)      lcqtof_t3_neg_3_032_24    m/z 877.6658     0   1E+4   12   12.5  RT (min)      lcqtof_t3_neg_2_016_05    m/z 877.6711     0   1   2E+2   11.5   12  RT (min)      lcqtof_t3_neg_3_032_24    m/z 877.6723     0   1E+3   12   12.5  RT (min)      lcqtof_t3_neg_3_032_24    m/z 877.6735     0   1   2   3E+2   10.5   11  RT (min)      lcqtof_t3_neg_3_038_06    m/z 878.1278     0   1   2   3E+4   13   13.5  RT (min)      lcqtof_t3_neg_4_002_17    m/z 878.1299     0   2   4   6E+4   13   13.5  RT (min)      lcqtof_t3_neg_4_064_17    m/z 878.5133     0   1   2   3   4E+4   13   13.5  RT (min)      lcqtof_t3_neg_1_019_23    m/z 878.5140     0   1   2E+4   12.5   13  RT (min)      lcqtof_t3_neg_1_019_23    m/z 878.5196     0   1E+4   12.5   13  RT (min)       lcqtof_t3_neg_4_030_31    m/z 878.6650     0   1   2E+4   14.5   15  RT (min)      lcqtof_t3_neg_1_000_aa    m/z 878.6655     0   1   2E+4   14.5   15  RT (min)      lcqtof_t3_neg_1_000_aa    m/z 878.6679     0   1   2E+5   12   12.5  RT (min)      lcqtof_t3_neg_2_016_05    m/z 878.6739     0   1   2E+5   12   12.5  RT (min)      lcqtof_t3_neg_2_016_05    m/z 878.6756     0   1   2   3E+2   12   12.5  RT (min)      lcqtof_t3_neg_3_032_24    m/z 878.6761     0   1   2   3E+2   11.5   12  RT (min)      lcqtof_t3_neg_3_032_24    m/z 878.6776     0   1   2E+2   10.5   11  RT (min)      lcqtof_t3_neg_3_038_06    m/z 878.6783     0   1   2E+2   10.5   11  RT (min)      lcqtof_t3_neg_3_038_06    m/z 878.9779     0   1E+4   12.5   13  RT (min)      lcqtof_t3_neg_4_040_24    m/z 879.1209     0   1   2   3E+4   13   13.5  RT (min)      lcqtof_t3_neg_4_002_17    m/z 879.1221     0   1   2E+4   13   13.5  RT (min)      lcqtof_t3_neg_4_002_17    m/z 879.1236     0   1   2   3   4E+4   13   13.5  RT (min)      lcqtof_t3_neg_4_064_17    m/z 879.1249     0   1   2   3   4E+4   13   13.5  RT (min)      lcqtof_t3_neg_4_002_17    m/z 879.1266     0   1   2   3   4E+4   13   13.5  RT (min)      lcqtof_t3_neg_4_064_17    m/z 879.5178     0   1E+4   13   13.5  RT (min)      lcqtof_t3_neg_1_019_23    m/z 879.5178     0   1E+4   12.5   13  RT (min)      lcqtof_t3_neg_1_019_23    m/z 879.5191     0   1E+4   8.5   9  RT (min)      lcqtof_t3_neg_4_000_ee    m/z 879.5267     0   1E+4   8.5   9  RT (min)      lcqtof_t3_neg_4_000_ee    m/z 879.6678     0   1E+4   14.5   15  RT (min)      lcqtof_t3_neg_1_000_aa    m/z 879.6743     0   2   4   6E+3   12   12.5  RT (min)      lcqtof_t3_neg_3_032_24    m/z 879.6851     0   1   2E+4   13   13.5  RT (min)      lcqtof_t3_neg_1_031_24    m/z 879.6856     0   1   2E+4   13   13.5  RT (min)       lcqtof_t3_neg_4_040_24    m/z 879.7076     0   1   2E+4   12.5   13  RT (min)      lcqtof_t3_neg_2_016_05    m/z 879.7092     0   1   2E+4   12.5   13  RT (min)      lcqtof_t3_neg_2_016_05    m/z 880.1198     0   1   2E+4   13   13.5  RT (min)      lcqtof_t3_neg_4_064_17    m/z 880.1229     0   1   2E+4   13   13.5  RT (min)      lcqtof_t3_neg_4_064_17    m/z 880.1237     0   1   2E+4   13.5   14  RT (min)      lcqtof_t3_neg_4_064_17    m/z 880.1238     0   1   2E+4   13.5   14  RT (min)      lcqtof_t3_neg_4_064_17    m/z 880.1255     0   1   2E+4   13   13.5  RT (min)      lcqtof_t3_neg_4_064_17    m/z 880.1280     0   1   2E+4   13.5   14  RT (min)      lcqtof_t3_neg_4_064_17    m/z 880.4862     0   1E+4   12   12.5  RT (min)      lcqtof_t3_neg_4_018_23    m/z 880.4874     0   1E+4   12   12.5  RT (min)      lcqtof_t3_neg_1_019_23    m/z 880.5273     0   1   2E+4   13   13.5  RT (min)       lcqtof_t3_neg_4_018_23    m/z 880.5291     0   1E+4   13.5   14  RT (min)       lcqtof_t3_neg_4_088_59    m/z 880.5318     0   1E+4   13   13.5  RT (min)      lcqtof_t3_neg_2_030_23    m/z 881.1047     0   1E+4   13   13.5  RT (min)      lcqtof_t3_neg_4_064_17    m/z 881.1078     0   1E+4   13   13.5  RT (min)      lcqtof_t3_neg_4_064_17    m/z 881.1134     0   1E+4   13   13.5  RT (min)      lcqtof_t3_neg_4_064_17    m/z 881.1216     0   1E+4   13.5   14  RT (min)      lcqtof_t3_neg_4_064_17    m/z 881.2859     0   1   2   3E+3   10.5   11  RT (min)       lcqtof_t3_neg_4_044_48    m/z 881.3084     0   1   2E+3   12   12.5  RT (min)       lcqtof_t3_neg_4_049_55    m/z 881.3087     0   1   2E+3   15   15.5  RT (min)       lcqtof_t3_neg_4_030_31    m/z 881.3091     0   1   2   3E+3   16   16.5  RT (min)       lcqtof_t3_neg_4_025_52    m/z 881.3094     0   2   4   6E+3   16   16.5  RT (min)      lcqtof_t3_neg_4_031_42    m/z 881.3096     0   1   2E+4   4.5   5  RT (min)       lcqtof_t3_neg_4_000_ee    m/z 881.3099     0   2   4   6E+3   16   16.5  RT (min)       lcqtof_t3_neg_4_018_23    m/z 881.3102     0   1   2E+3   15   15.5  RT (min)       lcqtof_t3_neg_4_038_29    m/z 881.3107     0   1   2E+3   15.5   16  RT (min)       lcqtof_t3_neg_4_018_23    m/z 881.3108     0   1   2E+3   15   15.5  RT (min)       lcqtof_t3_neg_4_034_34    m/z 881.3126     0   1   2E+3   14.5   15  RT (min)       lcqtof_t3_neg_4_032_04    m/z 882.2120     0   1E+4   12.5   13  RT (min)      lcqtof_t3_neg_4_036_47    m/z 882.5854     0   2   4   6E+2   11.5   12  RT (min)      lcqtof_t3_neg_3_032_24    m/z 883.6332     0   1E+4   12.5   13  RT (min)       lcqtof_t3_neg_1_040_08    m/z 883.6384     0   1   2E+4   12.5   13  RT (min)      lcqtof_t3_neg_1_056_56    m/z 883.6512     0   2   4   6E+2   11.5   12  RT (min)      lcqtof_t3_neg_3_032_24    m/z 883.6645     0   1   2E+2   10.5   11  RT (min)      lcqtof_t3_neg_3_038_06    m/z 883.6646     0   1E+4   12   12.5  RT (min)      lcqtof_t3_neg_2_016_05    m/z 884.3499     0   2   4   6   8E+3   7.5   8  RT (min)      lcqtof_t3_neg_4_033_53    m/z 885.6816     0   2   4E+2   12   12.5  RT (min)      lcqtof_t3_neg_3_032_24    m/z 885.8505     0   1   2   3E+4   12.5   13  RT (min)      lcqtof_t3_neg_4_037_01    m/z 886.5444     0   1E+4   12   12.5  RT (min)      lcqtof_t3_neg_4_016_14    m/z 886.5450     0   1E+4   11.5   12  RT (min)      lcqtof_t3_neg_4_016_14    m/z 886.5455     0   1E+4   12   12.5  RT (min)       lcqtof_t3_neg_4_059_14    m/z 886.5459     0   1E+4   12   12.5  RT (min)       lcqtof_t3_neg_4_059_14    m/z 886.5830     0   1   2   3   4E+2   11.5   12  RT (min)      lcqtof_t3_neg_3_032_24    m/z 887.5338     0   1E+4   13   13.5  RT (min)      lcqtof_t3_neg_1_007_03    m/z 887.5344     0   1E+4   13   13.5  RT (min)      lcqtof_t3_neg_1_007_03    m/z 887.6954     0   1   2E+2   12   12.5  RT (min)      lcqtof_t3_neg_3_032_24    m/z 887.6956     0   1   2E+2   12   12.5  RT (min)      lcqtof_t3_neg_3_032_24    m/z 887.6979     0   1E+4   12.5   13  RT (min)      lcqtof_t3_neg_2_016_05    m/z 887.6998     0   1E+4   12.5   13  RT (min)      lcqtof_t3_neg_2_016_05    m/z 887.7012     0   1   2   3E+2   10.5   11  RT (min)      lcqtof_t3_neg_3_038_06    m/z 888.5330     0   2   4   6   8E+3   13   13.5  RT (min)      lcqtof_t3_neg_1_001_49    m/z 888.5986     0   2   4   6E+2   11.5   12  RT (min)      lcqtof_t3_neg_3_032_24    m/z 888.5999     0   1E+3   11.5   12  RT (min)      lcqtof_t3_neg_3_032_24    m/z 888.6014     0   1   2   3E+2   10.5   11  RT (min)      lcqtof_t3_neg_3_038_06    m/z 889.2130     0   1E+4   12.5   13  RT (min)      lcqtof_t3_neg_4_064_17    m/z 889.7133     0   1   2E+2   12   12.5  RT (min)      lcqtof_t3_neg_3_032_24    m/z 890.5143     0   1   2E+2   10.5   11  RT (min)      lcqtof_t3_neg_3_038_06    m/z 890.5154     0   1E+5   12   12.5  RT (min)      lcqtof_t3_neg_1_019_23    m/z 890.5156     0   1   2E+2   10.5   11  RT (min)      lcqtof_t3_neg_3_038_06    m/z 890.5159     0   1E+5   12   12.5  RT (min)      lcqtof_t3_neg_1_019_23    m/z 891.2039     0   1E+4   4   4.5  RT (min)      lcqtof_t3_neg_4_000_ee    m/z 891.2391     0   1   2E+3   10.5   11  RT (min)      lcqtof_t3_neg_1_033_53    m/z 891.3694     0   1E+4   4   4.5  RT (min)      lcqtof_t3_neg_4_000_ee    m/z 891.5191     0   2   4   6   8E+4   12   12.5  RT (min)      lcqtof_t3_neg_1_019_23    m/z 891.5233     0   1   2   3E+2   10.5   11  RT (min)      lcqtof_t3_neg_3_038_06    m/z 891.5364     0   1E+4   9   9.5  RT (min)      lcqtof_t3_neg_4_000_ee    m/z 891.6108     0   1E+4   12   12.5  RT (min)       lcqtof_t3_neg_4_019_05    m/z 891.6782     0   1   2E+2   11.5   12  RT (min)      lcqtof_t3_neg_3_032_24    m/z 891.6786     0   1   2E+2   11.5   12  RT (min)      lcqtof_t3_neg_3_032_24    m/z 891.6814     0   1E+4   12   12.5  RT (min)      lcqtof_t3_neg_2_016_05    m/z 892.5220     0   2   4   6E+4   12   12.5  RT (min)      lcqtof_t3_neg_4_018_23    m/z 892.5226     0   2   4   6   8E+4   12   12.5  RT (min)      lcqtof_t3_neg_1_019_23    m/z 892.5321     0   1   2   3E+2   10.5   11  RT (min)      lcqtof_t3_neg_3_038_06    m/z 892.5322     0   1   2   3E+2   10.5   11  RT (min)      lcqtof_t3_neg_3_038_06    m/z 893.0473     0   1   2E+4   13.5   14  RT (min)      lcqtof_t3_neg_4_064_17    m/z 893.0475     0   1   2E+4   13   13.5  RT (min)      lcqtof_t3_neg_4_064_17    m/z 893.0482     0   1   2E+4   13.5   14  RT (min)      lcqtof_t3_neg_4_064_17    m/z 893.0482     0   1   2E+4   14   14.5  RT (min)      lcqtof_t3_neg_4_064_17    m/z 893.0485     0   1   2E+4   13   13.5  RT (min)      lcqtof_t3_neg_4_064_17    m/z 893.0508     0   1   2E+4   14   14.5  RT (min)      lcqtof_t3_neg_4_064_17    m/z 893.0525     0   1   2E+4   14   14.5  RT (min)      lcqtof_t3_neg_4_064_17    m/z 893.1555     0   1E+5   13   13.5  RT (min)      lcqtof_t3_neg_4_064_17    m/z 893.1556     0   1E+5   14   14.5  RT (min)      lcqtof_t3_neg_4_002_17    m/z 893.1565     0   1E+5   13   13.5  RT (min)      lcqtof_t3_neg_4_064_17    m/z 893.1566     0   1E+5   13   13.5  RT (min)      lcqtof_t3_neg_4_002_17    m/z 893.1568     0   1E+5   14   14.5  RT (min)      lcqtof_t3_neg_4_002_17    m/z 893.1577     0   1E+5   14   14.5  RT (min)      lcqtof_t3_neg_4_002_17    m/z 893.1581     0   2   4   6E+4   12.5   13  RT (min)      lcqtof_t3_neg_4_036_47    m/z 893.5226     0   1   2   3   4E+4   12   12.5  RT (min)      lcqtof_t3_neg_1_019_23    m/z 893.5351     0   1   2   3   4E+4   12   12.5  RT (min)       lcqtof_t3_neg_2_030_23    m/z 893.5357     0   1   2E+2   10.5   11  RT (min)      lcqtof_t3_neg_3_038_06    m/z 894.0468     0   1   2E+4   13.5   14  RT (min)      lcqtof_t3_neg_4_064_17    m/z 894.0472     0   1   2E+4   13.5   14  RT (min)      lcqtof_t3_neg_4_064_17    m/z 894.0474     0   1   2E+4   13.5   14  RT (min)      lcqtof_t3_neg_4_064_17    m/z 894.0475     0   1   2E+4   13   13.5  RT (min)      lcqtof_t3_neg_4_064_17    m/z 894.0477     0   1   2E+4   13   13.5  RT (min)      lcqtof_t3_neg_4_064_17    m/z 894.0494     0   1   2E+4   13   13.5  RT (min)      lcqtof_t3_neg_4_064_17    m/z 894.0497     0   1   2E+4   14   14.5  RT (min)      lcqtof_t3_neg_4_064_17    m/z 894.0506     0   1   2E+4   13   13.5  RT (min)      lcqtof_t3_neg_4_064_17    m/z 894.1556     0   1E+5   13   13.5  RT (min)      lcqtof_t3_neg_4_064_17    m/z 894.1557     0   1E+5   13   13.5  RT (min)      lcqtof_t3_neg_4_064_17    m/z 894.1559     0   2   4   6E+4   12.5   13  RT (min)      lcqtof_t3_neg_4_036_47    m/z 894.1566     0   2   4   6   8E+4   13   13.5  RT (min)      lcqtof_t3_neg_4_002_17    m/z 894.1573     0   2   4   6   8E+4   13   13.5  RT (min)      lcqtof_t3_neg_4_002_17    m/z 894.2056     0   1E+4   12.5   13  RT (min)       lcqtof_t3_neg_3_113_62    m/z 894.3903     0   2   4   6   8E+3   9.5   10  RT (min)      lcqtof_t3_neg_4_058_16    m/z 894.5376     0   1E+4   12   12.5  RT (min)      lcqtof_t3_neg_2_030_23    m/z 894.5381     0   1E+4   12   12.5  RT (min)      lcqtof_t3_neg_1_019_23    m/z 894.5382     0   2   4   6   8E+4   12.5   13  RT (min)      lcqtof_t3_neg_1_019_23    m/z 894.5391     0   2   4   6   8E+4   12.5   13  RT (min)      lcqtof_t3_neg_2_030_23    m/z 894.5469     0   1   2   3E+2   10.5   11  RT (min)      lcqtof_t3_neg_3_038_06    m/z 894.5504     0   1   2   3E+2   10.5   11  RT (min)      lcqtof_t3_neg_3_038_06    m/z 895.0427     0   1   2E+4   14   14.5  RT (min)      lcqtof_t3_neg_4_064_17    m/z 895.0444     0   1   2E+4   13.5   14  RT (min)      lcqtof_t3_neg_4_064_17    m/z 895.0455     0   1   2E+4   13.5   14  RT (min)      lcqtof_t3_neg_4_064_17    m/z 895.0460     0   1   2E+4   13   13.5  RT (min)      lcqtof_t3_neg_4_064_17    m/z 895.0474     0   1   2E+4   13.5   14  RT (min)      lcqtof_t3_neg_4_064_17    m/z 895.0482     0   1   2E+4   13   13.5  RT (min)      lcqtof_t3_neg_4_064_17    m/z 895.1541     0   2   4   6   8E+4   13   13.5  RT (min)      lcqtof_t3_neg_4_002_17    m/z 895.1547     0   1E+5   13   13.5  RT (min)      lcqtof_t3_neg_4_064_17    m/z 895.1552     0   2   4   6   8E+4   12.5   13  RT (min)      lcqtof_t3_neg_4_002_17    m/z 895.1554     0   2   4   6   8E+4   13   13.5  RT (min)      lcqtof_t3_neg_4_002_17    m/z 895.1555     0   2   4   6   8E+4   13   13.5  RT (min)      lcqtof_t3_neg_4_002_17    m/z 895.2045     0   1E+4   12.5   13  RT (min)      lcqtof_t3_neg_3_113_62    m/z 895.5506     0   1   2   3   4E+4   12.5   13  RT (min)      lcqtof_t3_neg_1_019_23    m/z 896.0429     0   1E+4   13.5   14  RT (min)      lcqtof_t3_neg_4_064_17    m/z 896.0467     0   1E+4   13.5   14  RT (min)      lcqtof_t3_neg_4_064_17    m/z 896.0474     0   1E+4   13   13.5  RT (min)      lcqtof_t3_neg_4_064_17    m/z 896.1535     0   1   2   3   4E+4   13   13.5  RT (min)      lcqtof_t3_neg_4_002_17    m/z 896.1543     0   2   4   6E+4   13   13.5  RT (min)      lcqtof_t3_neg_4_064_17    m/z 896.1552     0   1   2   3   4E+4   13   13.5  RT (min)      lcqtof_t3_neg_4_002_17    m/z 896.2049     0   1E+4   12.5   13  RT (min)      lcqtof_t3_neg_3_113_62    m/z 896.5517     0   1   2   3   4E+4   12.5   13  RT (min)      lcqtof_t3_neg_1_019_23    m/z 896.5521     0   1E+4   12.5   13  RT (min)      lcqtof_t3_neg_2_030_23    m/z 896.5525     0   2   4E+4   12.5   13  RT (min)      lcqtof_t3_neg_2_030_23    m/z 896.5534     0   2   4   6E+4   12.5   13  RT (min)       lcqtof_t3_neg_1_019_23    m/z 896.5638     0   1E+4   12.5   13  RT (min)      lcqtof_t3_neg_2_016_05    m/z 896.5645     0   1   2E+2   11   11.5  RT (min)      lcqtof_t3_neg_3_038_06    m/z 896.5961     0   1E+4   12.5   13  RT (min)      lcqtof_t3_neg_4_016_14    m/z 896.5997     0   1E+4   12.5   13  RT (min)      lcqtof_t3_neg_4_059_14    m/z 896.6002     0   1E+4   12.5   13  RT (min)       lcqtof_t3_neg_4_059_14    m/z 896.6879     0   2   4   6E+2   12   12.5  RT (min)      lcqtof_t3_neg_3_032_24    m/z 897.1526     0   1   2   3E+4   13.5   14  RT (min)      lcqtof_t3_neg_4_002_17    m/z 897.1526     0   1   2   3E+4   13   13.5  RT (min)      lcqtof_t3_neg_4_064_17    m/z 897.1527     0   1   2   3E+4   13.5   14  RT (min)      lcqtof_t3_neg_4_064_17    m/z 897.1541     0   1   2   3E+4   13   13.5  RT (min)      lcqtof_t3_neg_4_064_17    m/z 897.1544     0   1   2   3E+4   13.5   14  RT (min)      lcqtof_t3_neg_4_002_17    m/z 897.5569     0   1   2E+4   12.5   13  RT (min)      lcqtof_t3_neg_1_019_23    m/z 897.5656     0   1   2E+4   12.5   13  RT (min)      lcqtof_t3_neg_2_030_23    m/z 897.5678     0   1   2   3E+4   12.5   13  RT (min)      lcqtof_t3_neg_1_019_23    m/z 897.6763     0   1E+4   13.5   14  RT (min)      lcqtof_t3_neg_4_037_01    m/z 898.1489     0   1E+4   13   13.5  RT (min)      lcqtof_t3_neg_4_064_17    m/z 898.1518     0   1E+4   13.5   14  RT (min)      lcqtof_t3_neg_4_064_17    m/z 898.1528     0   1E+4   13   13.5  RT (min)      lcqtof_t3_neg_4_064_17    m/z 898.2510     0   1   2   3E+3   10   10.5  RT (min)       lcqtof_t3_neg_1_012_16    m/z 898.2515     0   1   2   3E+3   10   10.5  RT (min)      lcqtof_t3_neg_1_001_49    m/z 898.5661     0   1   2   3   4E+4   13   13.5  RT (min)       lcqtof_t3_neg_1_019_23    m/z 898.7032     0   1E+4   13   13.5  RT (min)      lcqtof_t3_neg_4_047_07    m/z 898.7260     0   1E+4   12.5   13  RT (min)      lcqtof_t3_neg_2_016_05    m/z 898.7265     0   2   4E+2   12   12.5  RT (min)      lcqtof_t3_neg_3_032_24    m/z 898.7269     0   2   4E+2   12   12.5  RT (min)      lcqtof_t3_neg_3_032_24    m/z 898.7285     0   1E+4   12.5   13  RT (min)      lcqtof_t3_neg_2_016_05    m/z 899.5815     0   1   2E+4   13   13.5  RT (min)      lcqtof_t3_neg_1_019_23    m/z 899.5842     0   2   4   6   8E+3   12.5   13  RT (min)      lcqtof_t3_neg_3_006_23    m/z 899.6980     0   2   4   6   8E+3   14   14.5  RT (min)      lcqtof_t3_neg_4_003_37    m/z 899.7405     0   1   2   3E+2   12   12.5  RT (min)      lcqtof_t3_neg_3_032_24    m/z 899.7410     0   1   2   3E+2   12.5   13  RT (min)      lcqtof_t3_neg_3_032_24    m/z 899.7432     0   1E+4   12.5   13  RT (min)      lcqtof_t3_neg_2_016_05    m/z 900.1986     0   1E+4   12   12.5  RT (min)      lcqtof_t3_neg_4_064_17    m/z 900.2004     0   1E+4   12   12.5  RT (min)      lcqtof_t3_neg_4_064_17    m/z 900.5836     0   2   4   6   8E+3   13   13.5  RT (min)      lcqtof_t3_neg_2_030_23    m/z 900.6506     0   2   4   6E+2   11.5   12  RT (min)      lcqtof_t3_neg_3_032_24    m/z 900.6526     0   1   2E+2   10   10.5  RT (min)      lcqtof_t3_neg_3_038_06    m/z 900.6528     0   1E+4   11.5   12  RT (min)      lcqtof_t3_neg_2_016_05    m/z 900.6530     0   1E+4   11.5   12  RT (min)      lcqtof_t3_neg_2_016_05    m/z 900.6531     0   2   4   6E+2   11.5   12  RT (min)      lcqtof_t3_neg_3_032_24    m/z 900.7438     0   1E+2   12   12.5  RT (min)      lcqtof_t3_neg_3_032_24    m/z 900.7452     0   1   2   3E+2   11   11.5  RT (min)      lcqtof_t3_neg_3_038_06    m/z 900.7454     0   1   2   3E+4   12.5   13  RT (min)      lcqtof_t3_neg_2_016_05    m/z 900.7469     0   1   2   3E+2   11   11.5  RT (min)      lcqtof_t3_neg_3_038_06    m/z 901.2047     0   1E+4   12.5   13  RT (min)      lcqtof_t3_neg_4_002_17    m/z 901.2080     0   1E+4   12.5   13  RT (min)      lcqtof_t3_neg_4_002_17    m/z 901.2131     0   1   2E+4   12.5   13  RT (min)      lcqtof_t3_neg_4_064_17    m/z 901.2133     0   1   2E+4   12.5   13  RT (min)      lcqtof_t3_neg_4_064_17    m/z 901.2138     0   1E+4   12.5   13  RT (min)      lcqtof_t3_neg_4_002_17    m/z 901.5045     0   1E+4   12   12.5  RT (min)      lcqtof_t3_neg_4_018_23    m/z 901.5054     0   1   2   3E+4   12   12.5  RT (min)      lcqtof_t3_neg_1_019_23    m/z 901.5417     0   1   2E+4   12   12.5  RT (min)      lcqtof_t3_neg_4_016_14    m/z 901.6635     0   1   2   3E+2   11.5   12  RT (min)      lcqtof_t3_neg_3_032_24    m/z 901.6641     0   1   2   3E+2   10.5   11  RT (min)      lcqtof_t3_neg_3_038_06    m/z 901.6642     0   2   4   6E+2   11.5   12  RT (min)      lcqtof_t3_neg_3_032_24    m/z 901.6646     0   1E+4   12   12.5  RT (min)      lcqtof_t3_neg_2_016_05    m/z 901.7568     0   1   2   3   4E+2   12.5   13  RT (min)      lcqtof_t3_neg_3_032_24    m/z 902.2154     0   1   2E+4   12.5   13  RT (min)      lcqtof_t3_neg_4_064_17    m/z 902.5095     0   1E+4   12   12.5  RT (min)      lcqtof_t3_neg_1_019_23    m/z 902.5460     0   1E+4   12   12.5  RT (min)      lcqtof_t3_neg_4_059_14    m/z 902.5669     0   1   2   3E+4   13   13.5  RT (min)      lcqtof_t3_neg_4_055_38    m/z 902.5685     0   1   2   3E+4   13   13.5  RT (min)      lcqtof_t3_neg_4_030_31    m/z 902.5688     0   1   2   3   4E+4   13   13.5  RT (min)       lcqtof_t3_neg_4_034_34    m/z 902.5914     0   1   2   3E+4   13   13.5  RT (min)      lcqtof_t3_neg_4_008_35    m/z 902.5965     0   1   2E+4   13   13.5  RT (min)       lcqtof_t3_neg_4_034_34    m/z 902.6672     0   1   2   3   4E+2   10.5   11  RT (min)      lcqtof_t3_neg_3_038_06    m/z 902.6675     0   1E+3   11.5   12  RT (min)      lcqtof_t3_neg_3_032_24    m/z 902.6685     0   1   2   3   4E+2   10   10.5  RT (min)      lcqtof_t3_neg_3_038_06    m/z 902.6686     0   1   2   3   4E+2   10   10.5  RT (min)      lcqtof_t3_neg_3_038_06    m/z 902.6686     0   1   2   3   4E+2   10.5   11  RT (min)      lcqtof_t3_neg_3_038_06    m/z 902.7598     0   1E+2   12.5   13  RT (min)      lcqtof_t3_neg_3_032_24    m/z 902.7634     0   2   4E+2   11   11.5  RT (min)      lcqtof_t3_neg_3_038_06    m/z 903.5693     0   1   2E+4   13   13.5  RT (min)      lcqtof_t3_neg_4_034_34    m/z 903.5700     0   1E+4   12.5   13  RT (min)      lcqtof_t3_neg_4_018_23    m/z 903.5708     0   1E+4   13   13.5  RT (min)      lcqtof_t3_neg_4_003_37    m/z 903.5712     0   1E+4   13   13.5  RT (min)      lcqtof_t3_neg_4_030_31    m/z 903.5721     0   1E+4   13   13.5  RT (min)      lcqtof_t3_neg_4_055_38    m/z 903.5751     0   1E+4   12.5   13  RT (min)      lcqtof_t3_neg_4_018_23    m/z 903.6791     0   2   4   6   8E+2   11.5   12  RT (min)      lcqtof_t3_neg_3_032_24    m/z 903.6791     0   1   2   3   4E+2   11.5   12  RT (min)      lcqtof_t3_neg_3_032_24    m/z 903.6805     0   1   2   3E+2   10.5   11  RT (min)      lcqtof_t3_neg_3_038_06    m/z 903.6813     0   1   2E+4   12   12.5  RT (min)      lcqtof_t3_neg_2_016_05    m/z 903.6827     0   1   2E+2   10.5   11  RT (min)      lcqtof_t3_neg_3_038_06    m/z 904.1967     0   1E+4   13   13.5  RT (min)      lcqtof_t3_neg_4_057_49    m/z 904.5806     0   1E+4   13   13.5  RT (min)      lcqtof_t3_neg_4_018_23    m/z 904.5830     0   1   2   3   4E+4   13   13.5  RT (min)       lcqtof_t3_neg_4_034_34    m/z 904.5835     0   1   2   3E+4   13   13.5  RT (min)      lcqtof_t3_neg_4_055_38    m/z 904.5841     0   1   2   3E+4   13   13.5  RT (min)       lcqtof_t3_neg_4_055_38    m/z 904.5848     0   1   2   3   4E+4   13   13.5  RT (min)      lcqtof_t3_neg_4_034_34    m/z 904.6833     0   1   2   3E+2   11.5   12  RT (min)      lcqtof_t3_neg_3_032_24    m/z 904.6833     0   1   2E+2   10.5   11  RT (min)      lcqtof_t3_neg_3_038_06    m/z 904.6838     0   1   2E+2   10.5   11  RT (min)      lcqtof_t3_neg_3_038_06    m/z 904.6841     0   1   2   3E+2   11.5   12  RT (min)      lcqtof_t3_neg_3_032_24    m/z 905.1918     0   2   4   6E+4   13   13.5  RT (min)      lcqtof_t3_neg_4_057_49    m/z 905.1942     0   1   2   3E+4   13   13.5  RT (min)      lcqtof_t3_neg_4_001_49    m/z 905.5846     0   1E+4   13   13.5  RT (min)       lcqtof_t3_neg_4_034_34    m/z 905.5850     0   1E+4   13   13.5  RT (min)      lcqtof_t3_neg_4_003_37    m/z 905.6243     0   1E+4   12   12.5  RT (min)      lcqtof_t3_neg_4_040_24    m/z 905.6951     0   1   2E+2   12   12.5  RT (min)      lcqtof_t3_neg_3_032_24    m/z 905.6958     0   1   2E+2   12   12.5  RT (min)      lcqtof_t3_neg_3_032_24    m/z 905.6958     0   1   2   3E+2   10.5   11  RT (min)      lcqtof_t3_neg_3_038_06    m/z 905.6959     0   1   2   3E+2   10.5   11  RT (min)      lcqtof_t3_neg_3_038_06    m/z 906.1922     0   1   2   3   4E+4   13   13.5  RT (min)      lcqtof_t3_neg_4_057_49    m/z 906.1924     0   1   2   3E+4   13   13.5  RT (min)      lcqtof_t3_neg_4_001_49    m/z 906.2121     0   1E+4   12.5   13  RT (min)      lcqtof_t3_neg_4_036_47    m/z 906.6403     0   1   2   3E+2   11.5   12  RT (min)      lcqtof_t3_neg_3_032_24    m/z 906.6981     0   1   2   3E+2   12   12.5  RT (min)      lcqtof_t3_neg_3_032_24    m/z 906.6985     0   1   2   3E+2   12   12.5  RT (min)      lcqtof_t3_neg_3_032_24    m/z 906.6995     0   1   2   3E+2   10.5   11  RT (min)      lcqtof_t3_neg_3_038_06    m/z 906.7000     0   1   2   3E+2   10.5   11  RT (min)      lcqtof_t3_neg_3_038_06    m/z 907.1684     0   1E+4   13.5   14  RT (min)      lcqtof_t3_neg_4_064_17    m/z 907.1760     0   1E+4   13   13.5  RT (min)      lcqtof_t3_neg_4_064_17    m/z 907.1802     0   1E+4   13.5   14  RT (min)      lcqtof_t3_neg_4_064_17    m/z 907.1889     0   1   2E+4   13   13.5  RT (min)      lcqtof_t3_neg_4_001_49    m/z 907.1893     0   1   2E+4   13   13.5  RT (min)      lcqtof_t3_neg_4_001_49    m/z 907.1898     0   1   2   3E+4   13   13.5  RT (min)      lcqtof_t3_neg_4_057_49    m/z 907.1906     0   1   2E+4   13   13.5  RT (min)      lcqtof_t3_neg_4_001_49    m/z 907.3138     0   1E+4   7.5   8  RT (min)      lcqtof_t3_neg_4_033_53    m/z 907.5034     0   1E+4   12   12.5  RT (min)      lcqtof_t3_neg_1_019_23    m/z 907.5690     0   1E+4   9   9.5  RT (min)      lcqtof_t3_neg_4_000_ee    m/z 907.5948     0   2   4E+3   12.5   13  RT (min)      lcqtof_t3_neg_1_003_07    m/z 907.7064     0   1   2E+4   12.5   13  RT (min)      lcqtof_t3_neg_2_016_05    m/z 907.7091     0   1   2E+2   12   12.5  RT (min)      lcqtof_t3_neg_3_032_24    m/z 907.7110     0   1   2E+2   10.5   11  RT (min)      lcqtof_t3_neg_3_038_06    m/z 907.7113     0   1   2E+2   12   12.5  RT (min)      lcqtof_t3_neg_3_032_24    m/z 907.7139     0   1   2E+2   11   11.5  RT (min)      lcqtof_t3_neg_3_038_06    m/z 908.1726     0   1E+4   13.5   14  RT (min)      lcqtof_t3_neg_4_064_17    m/z 908.1902     0   1E+4   13   13.5  RT (min)      lcqtof_t3_neg_4_001_49    m/z 908.1918     0   1   2E+4   13   13.5  RT (min)      lcqtof_t3_neg_4_057_49    m/z 908.3104     0   1E+4   7.5   8  RT (min)      lcqtof_t3_neg_4_041_54    m/z 908.6542     0   1   2E+2   11.5   12  RT (min)      lcqtof_t3_neg_3_032_24    m/z 908.6561     0   1   2E+2   11.5   12  RT (min)      lcqtof_t3_neg_3_032_24    m/z 908.6566     0   1   2   3E+2   10.5   11  RT (min)      lcqtof_t3_neg_3_038_06    m/z 908.6670     0   1   2E+2   11.5   12  RT (min)      lcqtof_t3_neg_3_032_24    m/z 908.7023     0   2   4   6   8E+4   12.5   13  RT (min)      lcqtof_t3_neg_2_016_05    m/z 908.7137     0   1E+2   12   12.5  RT (min)      lcqtof_t3_neg_3_032_24    m/z 908.7148     0   1   2   3E+2   10.5   11  RT (min)      lcqtof_t3_neg_3_038_06    m/z 908.7150     0   1E+2   12   12.5  RT (min)      lcqtof_t3_neg_3_032_24    m/z 909.1900     0   1E+4   13   13.5  RT (min)      lcqtof_t3_neg_4_057_49    m/z 909.6789     0   1E+4   15   15.5  RT (min)      lcqtof_t3_neg_1_000_aa    m/z 910.5093     0   2   4   6   8E+3   12.5   13  RT (min)      lcqtof_t3_neg_4_062_13    m/z 910.5211     0   1E+4   12.5   13  RT (min)       lcqtof_t3_neg_4_038_29    m/z 910.5436     0   1   2E+4   12   12.5  RT (min)       lcqtof_t3_neg_4_016_14    m/z 910.5442     0   1   2E+4   12   12.5  RT (min)      lcqtof_t3_neg_4_059_14    m/z 910.5443     0   1   2E+4   12   12.5  RT (min)      lcqtof_t3_neg_4_016_14    m/z 910.5443     0   1   2E+4   12   12.5  RT (min)       lcqtof_t3_neg_4_059_14    m/z 910.5445     0   1   2E+4   12   12.5  RT (min)      lcqtof_t3_neg_4_016_14    m/z 910.5466     0   1   2E+4   12   12.5  RT (min)      lcqtof_t3_neg_4_016_14    m/z 910.6708     0   1E+4   12.5   13  RT (min)      lcqtof_t3_neg_2_016_05    m/z 910.6710     0   1E+4   12.5   13  RT (min)      lcqtof_t3_neg_2_016_05    m/z 910.6714     0   1   2   3E+2   12   12.5  RT (min)      lcqtof_t3_neg_3_032_24    m/z 910.6718     0   1   2E+2   12   12.5  RT (min)      lcqtof_t3_neg_3_032_24    m/z 910.6727     0   1   2E+2   10.5   11  RT (min)      lcqtof_t3_neg_3_038_06    m/z 910.6747     0   1   2E+2   10.5   11  RT (min)      lcqtof_t3_neg_3_038_06    m/z 910.6752     0   1   2   3E+2   12   12.5  RT (min)      lcqtof_t3_neg_3_032_24    m/z 910.6839     0   1E+4   15   15.5  RT (min)      lcqtof_t3_neg_1_000_aa    m/z 911.5471     0   1E+4   12   12.5  RT (min)      lcqtof_t3_neg_4_059_14    m/z 911.5472     0   1E+4   12   12.5  RT (min)      lcqtof_t3_neg_4_059_14    m/z 911.6810     0   1E+4   15   15.5  RT (min)      lcqtof_t3_neg_1_000_aa    m/z 911.7041     0   1E+4   12.5   13  RT (min)      lcqtof_t3_neg_2_016_05    m/z 911.7074     0   1E+2   12   12.5  RT (min)      lcqtof_t3_neg_3_032_24    m/z 911.7079     0   1E+2   12   12.5  RT (min)      lcqtof_t3_neg_3_032_24    m/z 911.7095     0   1E+4   12.5   13  RT (min)      lcqtof_t3_neg_2_016_05    m/z 911.7104     0   1E+2   12   12.5  RT (min)      lcqtof_t3_neg_3_032_24    m/z 911.7158     0   1   2   3E+2   11   11.5  RT (min)      lcqtof_t3_neg_3_038_06    m/z 912.5580     0   1E+4   12   12.5  RT (min)      lcqtof_t3_neg_4_059_14    m/z 912.5585     0   1E+4   12   12.5  RT (min)      lcqtof_t3_neg_4_059_14    m/z 912.5589     0   1E+4   12   12.5  RT (min)       lcqtof_t3_neg_4_059_14    m/z 912.5598     0   1E+4   12   12.5  RT (min)      lcqtof_t3_neg_4_016_14    m/z 912.6618     0   1E+4   14   14.5  RT (min)       lcqtof_t3_neg_4_037_01    m/z 912.6873     0   1E+4   12.5   13  RT (min)      lcqtof_t3_neg_2_016_05    m/z 912.6876     0   1   2E+2   12   12.5  RT (min)      lcqtof_t3_neg_3_032_24    m/z 912.6879     0   1   2   3E+2   12   12.5  RT (min)      lcqtof_t3_neg_3_032_24    m/z 912.6880     0   1   2   3E+2   11   11.5  RT (min)      lcqtof_t3_neg_3_038_06    m/z 912.6907     0   1   2   3E+2   11   11.5  RT (min)      lcqtof_t3_neg_3_038_06    m/z 912.6909     0   1   2E+2   10.5   11  RT (min)      lcqtof_t3_neg_3_038_06    m/z 913.2980     0   1E+4   4   4.5  RT (min)      lcqtof_t3_neg_4_000_ee    m/z 913.5468     0   2   4   6   8E+3   13   13.5  RT (min)      lcqtof_t3_neg_1_002_13    m/z 913.6636     0   1   2E+4   14   14.5  RT (min)      lcqtof_t3_neg_4_037_01    m/z 913.6648     0   1E+4   14   14.5  RT (min)      lcqtof_t3_neg_4_014_13    m/z 914.4384     0   1E+4   12   12.5  RT (min)      lcqtof_t3_neg_2_030_23    m/z 914.4391     0   1   2E+4   12   12.5  RT (min)      lcqtof_t3_neg_4_018_23    m/z 914.4869     0   2   4   6   8E+3   12   12.5  RT (min)      lcqtof_t3_neg_4_018_23    m/z 914.5735     0   1E+4   12.5   13  RT (min)      lcqtof_t3_neg_4_059_14    m/z 914.5738     0   1E+4   12   12.5  RT (min)      lcqtof_t3_neg_4_059_14    m/z 914.5747     0   1E+4   12   12.5  RT (min)       lcqtof_t3_neg_4_059_14    m/z 915.1954     0   1E+4   12   12.5  RT (min)      lcqtof_t3_neg_4_064_17    m/z 915.4603     0   1E+4   12   12.5  RT (min)      lcqtof_t3_neg_1_019_23    m/z 915.4917     0   1E+4   12   12.5  RT (min)      lcqtof_t3_neg_4_046_33    m/z 916.1937     0   1E+4   12   12.5  RT (min)      lcqtof_t3_neg_4_064_17    m/z 916.6945     0   1E+4   12   12.5  RT (min)      lcqtof_t3_neg_2_016_05    m/z 916.6967     0   1   2   3E+2   11.5   12  RT (min)      lcqtof_t3_neg_3_032_24    m/z 917.1967     0   1E+4   12   12.5  RT (min)      lcqtof_t3_neg_4_064_17    m/z 917.2022     0   1E+4   12   12.5  RT (min)      lcqtof_t3_neg_4_064_17    m/z 917.2085     0   1E+4   12.5   13  RT (min)      lcqtof_t3_neg_4_064_17    m/z 917.2093     0   1E+4   12.5   13  RT (min)      lcqtof_t3_neg_4_064_17    m/z 917.2095     0   1E+4   12.5   13  RT (min)      lcqtof_t3_neg_4_064_17    m/z 918.2088     0   1E+4   12.5   13  RT (min)      lcqtof_t3_neg_4_064_17    m/z 918.5718     0   1E+4   13   13.5  RT (min)      lcqtof_t3_neg_4_055_38    m/z 918.6395     0   1E+4   12.5   13  RT (min)       lcqtof_t3_neg_1_003_07    m/z 918.6424     0   1E+4   13   13.5  RT (min)      lcqtof_t3_neg_4_035_08    m/z 918.6480     0   1E+4   13   13.5  RT (min)      lcqtof_t3_neg_4_040_24    m/z 918.6481     0   1E+4   13   13.5  RT (min)      lcqtof_t3_neg_4_095_60    m/z 918.8048     0   1E+4   17   17.5  RT (min)      lcqtof_t3_neg_4_035_08    m/z 918.8080     0   1E+4   16   16.5  RT (min)      lcqtof_t3_neg_4_018_23    m/z 919.2110     0   1E+4   12.5   13  RT (min)      lcqtof_t3_neg_4_064_17    m/z 919.2280     0   1E+4   1   1.5  RT (min)      lcqtof_t3_neg_2_051_14    m/z 919.2437     0   1E+4   13.5   14  RT (min)      lcqtof_t3_neg_4_064_17    m/z 919.6459     0   2   4   6E+3   12.5   13  RT (min)       lcqtof_t3_neg_3_032_24    m/z 919.6465     0   1E+4   12   12.5  RT (min)      lcqtof_t3_neg_4_011_32    m/z 919.6491     0   1   2   3E+2   11   11.5  RT (min)      lcqtof_t3_neg_3_038_06    m/z 919.6493     0   1E+4   13   13.5  RT (min)      lcqtof_t3_neg_2_016_05    m/z 919.7068     0   2   4E+4   14.5   15  RT (min)      lcqtof_t3_neg_1_000_aa    m/z 919.7088     0   2   4E+4   14.5   15  RT (min)      lcqtof_t3_neg_1_000_aa    m/z 920.2435     0   1E+4   13.5   14  RT (min)      lcqtof_t3_neg_4_057_49    m/z 920.2437     0   1E+4   13.5   14  RT (min)      lcqtof_t3_neg_4_002_17    m/z 920.6629     0   1E+4   13.5   14  RT (min)       lcqtof_t3_neg_4_040_24    m/z 920.7103     0   1   2   3E+4   15   15.5  RT (min)      lcqtof_t3_neg_1_000_aa    m/z 921.1490     0   1   2   3E+4   13   13.5  RT (min)      lcqtof_t3_neg_4_064_17    m/z 921.1500     0   1   2E+4   13.5   14  RT (min)      lcqtof_t3_neg_4_002_17    m/z 921.1500     0   1   2E+4   13   13.5  RT (min)      lcqtof_t3_neg_4_002_17    m/z 921.1500     0   1   2   3E+4   13.5   14  RT (min)      lcqtof_t3_neg_4_064_17    m/z 921.1505     0   1   2E+4   13   13.5  RT (min)      lcqtof_t3_neg_4_036_47    m/z 921.1506     0   1   2E+4   13.5   14  RT (min)      lcqtof_t3_neg_4_002_17    m/z 921.1527     0   1   2E+4   13.5   14  RT (min)      lcqtof_t3_neg_4_002_17    m/z 921.2428     0   1E+4   13.5   14  RT (min)      lcqtof_t3_neg_4_057_49    m/z 921.6639     0   1E+3   12.5   13  RT (min)      lcqtof_t3_neg_3_032_24    m/z 921.6639     0   1E+3   12.5   13  RT (min)      lcqtof_t3_neg_3_032_24    m/z 921.6650     0   1E+4   13   13.5  RT (min)      lcqtof_t3_neg_2_016_05    m/z 921.6661     0   1E+4   13   13.5  RT (min)      lcqtof_t3_neg_2_016_05    m/z 921.7104     0   1E+4   14.5   15  RT (min)      lcqtof_t3_neg_1_000_aa    m/z 922.1498     0   1   2E+4   13.5   14  RT (min)      lcqtof_t3_neg_4_064_17    m/z 922.1500     0   1   2E+4   13   13.5  RT (min)      lcqtof_t3_neg_4_064_17    m/z 922.1500     0   1   2E+4   13   13.5  RT (min)      lcqtof_t3_neg_4_002_17    m/z 922.1501     0   1   2E+4   13.5   14  RT (min)      lcqtof_t3_neg_4_002_17    m/z 922.1515     0   1   2E+4   13.5   14  RT (min)      lcqtof_t3_neg_4_002_17    m/z 922.1522     0   1   2E+4   13   13.5  RT (min)      lcqtof_t3_neg_4_036_47    m/z 923.1424     0   1   2E+4   13   13.5  RT (min)      lcqtof_t3_neg_4_064_17    m/z 923.1452     0   1   2E+4   13   13.5  RT (min)      lcqtof_t3_neg_4_064_17    m/z 923.1480     0   1   2E+4   13.5   14  RT (min)      lcqtof_t3_neg_4_064_17    m/z 923.1482     0   1   2E+4   13.5   14  RT (min)      lcqtof_t3_neg_4_064_17    m/z 923.6189     0   2   4E+4   13.5   14  RT (min)      lcqtof_t3_neg_4_012_27    m/z 924.1389     0   1E+4   13   13.5  RT (min)      lcqtof_t3_neg_4_064_17    m/z 924.1461     0   1E+4   13.5   14  RT (min)      lcqtof_t3_neg_4_002_17    m/z 924.1479     0   1E+4   13.5   14  RT (min)      lcqtof_t3_neg_4_002_17    m/z 924.3172     0   1E+4   1   1.5  RT (min)      lcqtof_t3_neg_2_042_13    m/z 924.5491     0   1E+4   12.5   13  RT (min)      lcqtof_t3_neg_4_018_23    m/z 924.5492     0   1E+4   12.5   13  RT (min)      lcqtof_t3_neg_4_018_23    m/z 924.6542     0   1   2   3E+2   11   11.5  RT (min)      lcqtof_t3_neg_3_038_06    m/z 924.6559     0   1   2   3E+2   11   11.5  RT (min)      lcqtof_t3_neg_3_038_06    m/z 924.7460     0   1E+4   1   1.5  RT (min)      lcqtof_t3_neg_3_012_14    m/z 925.0464     0   1E+4   13   13.5  RT (min)      lcqtof_t3_neg_4_064_17    m/z 925.5558     0   1E+4   12.5   13  RT (min)      lcqtof_t3_neg_4_018_23    m/z 925.6517     0   2   4   6E+4   12.5   13  RT (min)      lcqtof_t3_neg_1_019_23    m/z 925.6525     0   1E+4   13   13.5  RT (min)      lcqtof_t3_neg_2_016_05    m/z 925.6566     0   2   4   6E+4   12.5   13  RT (min)      lcqtof_t3_neg_2_030_23    m/z 926.5627     0   1E+4   12.5   13  RT (min)      lcqtof_t3_neg_4_018_23    m/z 926.6368     0   2   4E+4   13.5   14  RT (min)      lcqtof_t3_neg_4_014_13    m/z 926.6503     0   1E+4   12.5   13  RT (min)      lcqtof_t3_neg_1_003_07    m/z 926.6511     0   1E+4   12.5   13  RT (min)       lcqtof_t3_neg_2_030_23    m/z 926.6517     0   1E+4   12.5   13  RT (min)      lcqtof_t3_neg_1_019_23    m/z 927.5684     0   1E+4   12.5   13  RT (min)      lcqtof_t3_neg_4_034_34    m/z 927.5696     0   1E+4   13   13.5  RT (min)      lcqtof_t3_neg_4_055_38    m/z 927.6138     0   1E+4   13.5   14  RT (min)      lcqtof_t3_neg_4_034_34    m/z 928.5030     0   1E+4   13   13.5  RT (min)       lcqtof_t3_neg_4_032_04    m/z 928.5796     0   1E+4   13   13.5  RT (min)      lcqtof_t3_neg_4_003_37    m/z 928.5829     0   1   2E+4   13   13.5  RT (min)       lcqtof_t3_neg_4_030_31    m/z 928.5831     0   1E+4   13   13.5  RT (min)      lcqtof_t3_neg_4_055_38    m/z 928.8320     0   2   4E+2   15.5   16  RT (min)      lcqtof_t3_neg_1_019_23    m/z 928.8333     0   1   2   3E+4   17   17.5  RT (min)       lcqtof_t3_neg_4_035_08    m/z 929.5861     0   1E+4   13   13.5  RT (min)      lcqtof_t3_neg_4_034_34    m/z 930.3559     0   1E+4   7.5   8  RT (min)      lcqtof_t3_neg_4_033_53    m/z 930.5954     0   1   2   3E+4   13   13.5  RT (min)      lcqtof_t3_neg_4_030_31    m/z 931.5965     0   1E+4   13   13.5  RT (min)      lcqtof_t3_neg_4_055_38    m/z 931.5981     0   1E+4   13   13.5  RT (min)      lcqtof_t3_neg_4_003_37    m/z 931.7544     0   1E+4   15   15.5  RT (min)      lcqtof_t3_neg_1_051_26    m/z 932.6068     0   1   2E+4   13   13.5  RT (min)      lcqtof_t3_neg_4_034_34    m/z 932.6075     0   1E+4   13   13.5  RT (min)      lcqtof_t3_neg_4_055_38    m/z 933.1893     0   1E+4   13   13.5  RT (min)      lcqtof_t3_neg_4_057_49    m/z 933.2107     0   1E+4   13   13.5  RT (min)      lcqtof_t3_neg_4_032_04    m/z 933.2226     0   1   2   3E+4   12.5   13  RT (min)      lcqtof_t3_neg_4_064_17    m/z 933.2238     0   1E+4   12.5   13   13.5  RT (min)      lcqtof_t3_neg_4_002_17    m/z 933.3294     0   1E+4   7.5   8  RT (min)      lcqtof_t3_neg_1_020_47    m/z 933.4052     0   1E+4   5   5.5  RT (min)      lcqtof_t3_neg_4_000_ee    m/z 933.6587     0   1   2   3E+2   11.5   12  RT (min)      lcqtof_t3_neg_3_032_24    m/z 933.6594     0   1   2   3E+2   11.5   12  RT (min)      lcqtof_t3_neg_3_032_24    m/z 933.6597     0   1   2   3E+2   11.5   12  RT (min)      lcqtof_t3_neg_3_032_24    m/z 933.6600     0   1E+4   12   12.5  RT (min)      lcqtof_t3_neg_2_016_05    m/z 933.6609     0   1   2E+2   10.5   11  RT (min)      lcqtof_t3_neg_3_038_06    m/z 934.1875     0   1E+4   13   13.5  RT (min)      lcqtof_t3_neg_4_057_49    m/z 934.2224     0   1E+4   12.5   13  RT (min)      lcqtof_t3_neg_4_064_17    m/z 934.2244     0   1E+4   12.5   13  RT (min)      lcqtof_t3_neg_4_064_17    m/z 934.3317     0   2   4   6   8E+3   7.5   8  RT (min)      lcqtof_t3_neg_1_020_47    m/z 934.5429     0   2   4   6   8E+3   12   12.5  RT (min)       lcqtof_t3_neg_4_016_14    m/z 934.6613     0   2   4E+2   11.5   12  RT (min)      lcqtof_t3_neg_3_032_24    m/z 934.6630     0   2   4E+2   11.5   12  RT (min)      lcqtof_t3_neg_3_032_24    m/z 934.6630     0   2   4E+2   10.5   11  RT (min)      lcqtof_t3_neg_3_038_06    m/z 934.6631     0   2   4E+4   12   12.5  RT (min)      lcqtof_t3_neg_2_016_05    m/z 934.6636     0   1   2   3E+2   10.5   11  RT (min)      lcqtof_t3_neg_3_038_06    m/z 934.7867     0   1E+4   17   17.5  RT (min)      lcqtof_t3_neg_4_035_08    m/z 934.7871     0   1E+4   16   16.5  RT (min)      lcqtof_t3_neg_4_018_23    m/z 935.1118     0   2   4E+4   14   14.5  RT (min)      lcqtof_t3_neg_4_002_17    m/z 935.1122     0   2   4   6   8E+4   13.5   14  RT (min)      lcqtof_t3_neg_4_064_17    m/z 935.1124     0   2   4   6   8E+4   13.5   14  RT (min)      lcqtof_t3_neg_4_064_17    m/z 935.1124     0   2   4   6   8E+4   13   13.5  RT (min)      lcqtof_t3_neg_4_064_17    m/z 935.1128     0   2   4   6   8E+4   13.5   14  RT (min)      lcqtof_t3_neg_4_064_17    m/z 935.1130     0   2   4   6   8E+4   13   13.5  RT (min)      lcqtof_t3_neg_4_064_17    m/z 935.1137     0   1   2   3E+4   14   14.5  RT (min)      lcqtof_t3_neg_4_002_17    m/z 935.2021     0   1   2   3   4E+4   13.5   14  RT (min)      lcqtof_t3_neg_4_064_17    m/z 935.2025     0   1   2   3E+4   13.5   14  RT (min)      lcqtof_t3_neg_4_002_17    m/z 935.2025     0   1   2   3E+4   13.5   14  RT (min)      lcqtof_t3_neg_4_002_17    m/z 935.2032     0   1   2   3E+4   13.5   14  RT (min)      lcqtof_t3_neg_4_002_17    m/z 935.2046     0   1   2   3E+4   13.5   14  RT (min)      lcqtof_t3_neg_4_036_47    m/z 935.2063     0   1   2   3E+4   13   13.5  RT (min)      lcqtof_t3_neg_4_002_17    m/z 935.2330     0   1   2E+4   13   13.5  RT (min)      lcqtof_t3_neg_4_002_17    m/z 935.2369     0   1   2   3E+4   13   13.5  RT (min)      lcqtof_t3_neg_4_064_17    m/z 935.2402     0   1   2E+4   13   13.5  RT (min)      lcqtof_t3_neg_4_002_17    m/z 935.5654     0   1E+4   9   9.5  RT (min)      lcqtof_t3_neg_4_000_ee    m/z 935.6735     0   1E+4   12   12.5  RT (min)      lcqtof_t3_neg_2_016_05    m/z 935.6736     0   1E+3   12   12.5  RT (min)      lcqtof_t3_neg_3_032_24    m/z 935.6736     0   1   2   3E+2   11.5   12  RT (min)      lcqtof_t3_neg_3_032_24    m/z 935.6755     0   1   2   3E+2   12   12.5  RT (min)      lcqtof_t3_neg_3_032_24    m/z 935.6787     0   1   2   3E+2   10.5   11  RT (min)      lcqtof_t3_neg_3_038_06    m/z 936.1119     0   2   4   6E+4   13.5   14  RT (min)      lcqtof_t3_neg_4_064_17    m/z 936.1119     0   2   4   6E+4   13.5   14  RT (min)      lcqtof_t3_neg_4_064_17    m/z 936.1124     0   2   4   6E+4   13.5   14  RT (min)      lcqtof_t3_neg_4_064_17    m/z 936.1128     0   2   4   6E+4   13   13.5  RT (min)      lcqtof_t3_neg_4_064_17    m/z 936.1129     0   2   4   6E+4   13   13.5  RT (min)      lcqtof_t3_neg_4_064_17    m/z 936.1131     0   2   4E+4   14   14.5  RT (min)      lcqtof_t3_neg_4_002_17    m/z 936.1132     0   2   4E+4   13.5   14  RT (min)      lcqtof_t3_neg_4_002_17    m/z 936.1133     0   1   2   3   4E+4   14   14.5  RT (min)      lcqtof_t3_neg_4_002_17    m/z 936.1135     0   2   4E+4   14   14.5  RT (min)      lcqtof_t3_neg_4_002_17    m/z 936.2029     0   1   2   3E+4   13.5   14  RT (min)      lcqtof_t3_neg_4_064_17    m/z 936.2072     0   1   2   3E+4   13   13.5  RT (min)      lcqtof_t3_neg_4_064_17    m/z 936.2363     0   1   2E+4   13   13.5  RT (min)      lcqtof_t3_neg_4_064_17    m/z 936.5582     0   1E+4   12   12.5  RT (min)       lcqtof_t3_neg_4_059_14    m/z 936.6682     0   1   2   3   4E+4   12.5   13  RT (min)      lcqtof_t3_neg_2_016_05    m/z 936.6773     0   2   4   6E+2   11.5   12  RT (min)      lcqtof_t3_neg_3_032_24    m/z 936.6783     0   2   4   6E+2   12   12.5  RT (min)      lcqtof_t3_neg_3_032_24    m/z 936.6784     0   1   2   3   4E+2   10.5   11  RT (min)      lcqtof_t3_neg_3_038_06    m/z 936.6786     0   2   4E+2   10.5   11  RT (min)      lcqtof_t3_neg_3_038_06    m/z 936.6790     0   1   2   3   4E+2   10.5   11  RT (min)      lcqtof_t3_neg_3_038_06    m/z 937.0974     0   2   4   6E+4   13.5   14  RT (min)      lcqtof_t3_neg_4_002_17    m/z 937.1011     0   2   4   6E+4   13.5   14  RT (min)      lcqtof_t3_neg_4_002_17    m/z 937.1041     0   2   4   6   8E+4   13.5   14  RT (min)      lcqtof_t3_neg_4_064_17    m/z 937.1044     0   2   4   6   8E+4   13.5   14  RT (min)      lcqtof_t3_neg_4_064_17    m/z 937.1049     0   2   4   6E+4   13.5   14  RT (min)      lcqtof_t3_neg_4_002_17    m/z 937.1052     0   2   4   6E+4   14   14.5  RT (min)      lcqtof_t3_neg_4_002_17    m/z 937.1063     0   2   4   6   8E+4   13   13.5  RT (min)      lcqtof_t3_neg_4_064_17    m/z 937.1066     0   2   4   6E+4   14   14.5  RT (min)      lcqtof_t3_neg_4_002_17    m/z 937.1070     0   2   4   6   8E+4   13   13.5  RT (min)      lcqtof_t3_neg_4_064_17    m/z 937.2020     0   1   2   3E+4   13.5   14  RT (min)      lcqtof_t3_neg_4_064_17    m/z 937.2023     0   1   2E+4   13.5   14  RT (min)      lcqtof_t3_neg_4_002_17    m/z 937.2325     0   1   2E+4   13   13.5  RT (min)      lcqtof_t3_neg_4_064_17    m/z 937.2362     0   1   2E+4   13   13.5  RT (min)      lcqtof_t3_neg_4_002_17    m/z 937.2378     0   1   2E+4   13   13.5  RT (min)      lcqtof_t3_neg_4_002_17    m/z 937.6478     0   2   4   6E+3   13.5   14  RT (min)       lcqtof_t3_neg_4_037_01    m/z 937.6583     0   2   4   6E+3   14   14.5  RT (min)      lcqtof_t3_neg_4_037_01    m/z 937.6894     0   1   2   3E+2   12   12.5  RT (min)      lcqtof_t3_neg_3_032_24    m/z 937.6920     0   1   2   3E+2   10.5   11  RT (min)      lcqtof_t3_neg_3_038_06    m/z 937.6936     0   1   2   3E+2   12   12.5  RT (min)      lcqtof_t3_neg_3_032_24    m/z 937.6940     0   1   2   3E+2   12   12.5  RT (min)      lcqtof_t3_neg_3_032_24    m/z 937.6995     0   1   2E+4   12.5   13  RT (min)      lcqtof_t3_neg_2_016_05    m/z 938.0982     0   2   4E+4   13   13.5  RT (min)      lcqtof_t3_neg_4_064_17    m/z 938.1040     0   1   2   3   4E+4   14   14.5  RT (min)      lcqtof_t3_neg_4_064_17    m/z 938.1043     0   2   4E+4   13.5   14  RT (min)      lcqtof_t3_neg_4_064_17    m/z 938.1044     0   2   4E+4   13   13.5  RT (min)      lcqtof_t3_neg_4_064_17    m/z 938.1047     0   2   4E+4   13   13.5  RT (min)      lcqtof_t3_neg_4_064_17    m/z 938.1057     0   2   4E+4   13.5   14  RT (min)      lcqtof_t3_neg_4_064_17    m/z 938.2004     0   1   2E+4   13.5   14  RT (min)      lcqtof_t3_neg_4_064_17    m/z 938.2010     0   1E+4   13.5   14  RT (min)      lcqtof_t3_neg_4_036_47    m/z 938.5722     0   1E+4   12.5   13  RT (min)      lcqtof_t3_neg_4_016_14    m/z 938.5724     0   1E+4   12.5   13  RT (min)      lcqtof_t3_neg_4_016_14    m/z 938.5728     0   1E+4   12.5   13  RT (min)       lcqtof_t3_neg_4_059_14    m/z 938.5735     0   1E+4   12.5   13  RT (min)      lcqtof_t3_neg_4_059_14    m/z 938.5736     0   1E+4   12.5   13  RT (min)      lcqtof_t3_neg_4_016_14    m/z 938.6700     0   1   2   3E+4   12.5   13  RT (min)      lcqtof_t3_neg_2_030_23    m/z 938.6707     0   1   2E+4   12.5   13  RT (min)      lcqtof_t3_neg_2_030_23    m/z 938.6931     0   2   4E+2   12   12.5  RT (min)      lcqtof_t3_neg_3_032_24    m/z 938.6935     0   2   4E+2   12   12.5  RT (min)      lcqtof_t3_neg_3_032_24    m/z 938.6960     0   1   2   3E+2   10.5   11  RT (min)      lcqtof_t3_neg_3_038_06    m/z 939.0911     0   1   2   3E+4   13.5   14  RT (min)      lcqtof_t3_neg_4_064_17    m/z 939.0955     0   1   2   3E+4   14   14.5  RT (min)      lcqtof_t3_neg_4_064_17    m/z 939.1006     0   1   2   3E+4   13   13.5  RT (min)      lcqtof_t3_neg_4_064_17    m/z 939.1008     0   1   2   3E+4   13.5   14  RT (min)      lcqtof_t3_neg_4_064_17    m/z 939.1009     0   1   2   3E+4   13.5   14  RT (min)      lcqtof_t3_neg_4_064_17    m/z 939.1010     0   1   2   3E+4   13   13.5  RT (min)      lcqtof_t3_neg_4_064_17    m/z 939.1043     0   1   2   3E+4   13.5   14  RT (min)      lcqtof_t3_neg_4_064_17    m/z 939.1971     0   1E+4   13.5   14  RT (min)      lcqtof_t3_neg_4_064_17    m/z 939.6740     0   2   4   6E+3   13.5   14  RT (min)       lcqtof_t3_neg_4_037_01    m/z 939.6780     0   1E+4   12.5   13  RT (min)      lcqtof_t3_neg_1_019_23    m/z 939.6804     0   2   4E+3   14   14.5  RT (min)      lcqtof_t3_neg_4_016_14    m/z 939.6813     0   2   4   6E+3   14   14.5  RT (min)      lcqtof_t3_neg_4_037_01    m/z 939.6814     0   2   4   6E+3   14   14.5  RT (min)      lcqtof_t3_neg_4_037_01    m/z 939.7042     0   1   2   3   4E+2   12   12.5  RT (min)      lcqtof_t3_neg_3_032_24    m/z 939.7084     0   1   2   3   4E+2   12   12.5  RT (min)      lcqtof_t3_neg_3_032_24    m/z 940.0902     0   1E+4   13   13.5  RT (min)      lcqtof_t3_neg_4_064_17    m/z 940.0914     0   1E+4   13.5   14  RT (min)      lcqtof_t3_neg_4_064_17    m/z 940.0960     0   1E+4   13.5   14  RT (min)      lcqtof_t3_neg_4_064_17    m/z 940.0974     0   1E+4   13   13.5  RT (min)      lcqtof_t3_neg_4_064_17    m/z 940.0976     0   1E+4   13   13.5  RT (min)      lcqtof_t3_neg_4_064_17    m/z 940.0980     0   1E+4   13   13.5  RT (min)      lcqtof_t3_neg_4_064_17    m/z 940.6483     0   1E+4   12.5   13  RT (min)      lcqtof_t3_neg_2_030_23    m/z 940.6494     0   1   2E+4   12.5   13  RT (min)       lcqtof_t3_neg_4_018_23    m/z 940.6494     0   1   2E+4   12.5   13  RT (min)      lcqtof_t3_neg_4_018_23    m/z 940.6771     0   1   2E+4   13   13.5  RT (min)      lcqtof_t3_neg_4_046_33    m/z 940.7015     0   1   2   3   4E+4   12.5   13  RT (min)      lcqtof_t3_neg_2_016_05    m/z 940.7073     0   1   2   3E+2   12   12.5  RT (min)      lcqtof_t3_neg_3_032_24    m/z 940.7087     0   1   2   3   4E+4   12.5   13  RT (min)      lcqtof_t3_neg_2_016_05    m/z 940.7094     0   1   2   3E+2   12   12.5  RT (min)      lcqtof_t3_neg_3_032_24    m/z 940.7100     0   1   2E+2   11   11.5  RT (min)      lcqtof_t3_neg_3_038_06    m/z 940.7101     0   1   2   3   4E+4   12.5   13  RT (min)      lcqtof_t3_neg_2_016_05    m/z 940.7109     0   1   2   3E+2   11   11.5  RT (min)      lcqtof_t3_neg_3_038_06    m/z 941.0548     0   1   2E+4   13   13.5  RT (min)      lcqtof_t3_neg_4_064_17    m/z 941.0664     0   1   2E+4   13   13.5  RT (min)      lcqtof_t3_neg_4_064_17    m/z 941.0683     0   1   2E+4   13   13.5  RT (min)      lcqtof_t3_neg_4_064_17    m/z 941.0694     0   1   2E+4   13.5   14  RT (min)      lcqtof_t3_neg_4_064_17    m/z 941.0739     0   1E+4   13.5   14  RT (min)      lcqtof_t3_neg_4_002_17    m/z 941.0885     0   1   2E+4   13.5   14  RT (min)      lcqtof_t3_neg_4_064_17    m/z 941.6354     0   1   2   3   4E+3   13.5   14  RT (min)      lcqtof_t3_neg_4_037_01    m/z 941.6378     0   1   2   3E+3   13.5   14  RT (min)      lcqtof_t3_neg_1_038_15    m/z 941.6523     0   1E+4   12.5   13  RT (min)      lcqtof_t3_neg_4_018_23    m/z 941.6523     0   1E+4   12.5   13  RT (min)      lcqtof_t3_neg_4_018_23    m/z 941.6530     0   1E+4   12.5   13  RT (min)      lcqtof_t3_neg_4_018_23    m/z 941.6897     0   1   2E+4   13   13.5  RT (min)      lcqtof_t3_neg_4_028_25    m/z 941.6924     0   1   2E+4   13   13.5  RT (min)      lcqtof_t3_neg_4_046_33    m/z 941.6957     0   1   2   3   4E+4   13   13.5  RT (min)       lcqtof_t3_neg_4_040_24    m/z 942.0503     0   1E+4   13   13.5  RT (min)      lcqtof_t3_neg_4_064_17    m/z 942.0550     0   1E+4   13   13.5  RT (min)      lcqtof_t3_neg_4_064_17    m/z 942.0577     0   1E+4   13   13.5  RT (min)      lcqtof_t3_neg_4_064_17    m/z 942.6321     0   1   2   3   4E+2   11.5   12  RT (min)      lcqtof_t3_neg_3_032_24    m/z 942.6357     0   1   2   3   4E+2   11.5   12  RT (min)      lcqtof_t3_neg_3_032_24    m/z 942.7237     0   2   4E+2   12.5   13  RT (min)      lcqtof_t3_neg_3_032_24    m/z 942.7252     0   2   4E+2   11   11.5  RT (min)      lcqtof_t3_neg_3_038_06    m/z 942.7262     0   2   4E+2   11   11.5  RT (min)      lcqtof_t3_neg_3_038_06    m/z 942.7269     0   2   4E+2   11   11.5  RT (min)      lcqtof_t3_neg_3_038_06    m/z 943.0476     0   1E+4   13   13.5  RT (min)      lcqtof_t3_neg_4_064_17    m/z 943.0477     0   1E+4   13   13.5  RT (min)      lcqtof_t3_neg_4_064_17    m/z 943.0485     0   1E+4   13   13.5  RT (min)      lcqtof_t3_neg_4_064_17    m/z 943.6473     0   1   2E+2   11.5   12  RT (min)      lcqtof_t3_neg_3_032_24    m/z 943.7170     0   1   2   3E+4   13   13.5  RT (min)       lcqtof_t3_neg_4_033_53    m/z 943.7206     0   1   2   3E+4   13   13.5  RT (min)      lcqtof_t3_neg_4_040_24    m/z 943.7225     0   1   2   3E+4   13   13.5  RT (min)       lcqtof_t3_neg_4_011_32    m/z 943.7347     0   1E+4   13   13.5  RT (min)      lcqtof_t3_neg_4_028_25    m/z 944.2050     0   1   2E+4   12.5   13  RT (min)      lcqtof_t3_neg_4_064_17    m/z 944.6477     0   1   2   3E+2   11.5   12  RT (min)      lcqtof_t3_neg_3_032_24    m/z 944.6511     0   1   2   3E+2   10.5   11  RT (min)      lcqtof_t3_neg_3_038_06    m/z 944.6513     0   1   2E+4   12   12.5  RT (min)      lcqtof_t3_neg_2_016_05    m/z 944.6523     0   1   2   3E+2   10.5   11  RT (min)      lcqtof_t3_neg_3_038_06    m/z 944.6752     0   2   4   6E+4   12.5   13  RT (min)      lcqtof_t3_neg_4_040_24    m/z 944.8073     0   1   2E+4   17   17.5  RT (min)      lcqtof_t3_neg_4_035_08    m/z 944.8105     0   1E+4   16   16.5  RT (min)      lcqtof_t3_neg_4_018_23    m/z 945.2041     0   1   2E+4   12.5   13  RT (min)      lcqtof_t3_neg_4_064_17    m/z 945.2054     0   1   2E+4   12.5   13  RT (min)      lcqtof_t3_neg_4_064_17    m/z 945.2106     0   1   2E+4   12.5   13  RT (min)      lcqtof_t3_neg_4_064_17    m/z 945.2112     0   1   2E+4   12.5   13  RT (min)      lcqtof_t3_neg_4_064_17    m/z 945.4127     0   1E+4   7   7.5  RT (min)       lcqtof_t3_neg_4_004_20    m/z 945.6690     0   2   4   6E+3   12.5   13  RT (min)      lcqtof_t3_neg_1_003_07    m/z 945.6937     0   1E+4   12.5   13  RT (min)      lcqtof_t3_neg_4_040_24    m/z 945.6941     0   1E+4   12.5   13  RT (min)      lcqtof_t3_neg_4_040_24    m/z 945.6957     0   1E+4   12.5   13  RT (min)      lcqtof_t3_neg_4_040_24    m/z 945.7274     0   1E+4   13   13.5  RT (min)       lcqtof_t3_neg_4_011_32    m/z 946.2147     0   1E+4   12.5   13  RT (min)      lcqtof_t3_neg_4_002_17    m/z 946.2153     0   1E+4   12.5   13  RT (min)      lcqtof_t3_neg_4_064_17    m/z 946.6192     0   1E+4   13   13.5  RT (min)      lcqtof_t3_neg_4_030_31    m/z 946.6197     0   1E+4   13   13.5  RT (min)       lcqtof_t3_neg_4_034_34    m/z 946.6216     0   1E+4   13   13.5  RT (min)      lcqtof_t3_neg_4_034_34    m/z 946.6250     0   1E+4   13   13.5  RT (min)      lcqtof_t3_neg_4_034_34    m/z 947.1954     0   1E+4   12.5   13  RT (min)       lcqtof_t3_neg_4_002_17    m/z 947.2056     0   1   2E+4   12.5   13  RT (min)      lcqtof_t3_neg_4_064_17    m/z 947.2059     0   1E+4   12.5   13  RT (min)      lcqtof_t3_neg_4_002_17    m/z 947.2068     0   1   2E+4   12.5   13  RT (min)      lcqtof_t3_neg_4_064_17    m/z 947.6105     0   2   4E+3   10.5   11  RT (min)      lcqtof_t3_neg_4_037_01    m/z 948.2053     0   1E+4   12.5   13  RT (min)      lcqtof_t3_neg_4_064_17    m/z 948.2136     0   1E+4   12.5   13  RT (min)      lcqtof_t3_neg_4_064_17    m/z 948.2534     0   1   2   3   4E+3   10.5   11  RT (min)       lcqtof_t3_neg_4_079_58    m/z 948.6335     0   1E+4   13   13.5  RT (min)       lcqtof_t3_neg_4_034_34    m/z 949.1617     0   1E+4   13.5   14  RT (min)      lcqtof_t3_neg_4_064_17    m/z 949.2107     0   1E+4   12.5   13  RT (min)      lcqtof_t3_neg_4_064_17    m/z 949.2960     0   2   4E+3   16.5   17  RT (min)      lcqtof_t3_neg_4_000_bb    m/z 951.2334     0   1E+4   13.5   14  RT (min)      lcqtof_t3_neg_4_064_17    m/z 952.2332     0   1E+4   13.5   14  RT (min)      lcqtof_t3_neg_4_002_17    m/z 952.2356     0   1E+4   13.5   14  RT (min)      lcqtof_t3_neg_4_036_47    m/z 952.5793     0   1E+4   12.5   13  RT (min)       lcqtof_t3_neg_4_034_34    m/z 952.6839     0   1   2   3   4E+4   13   13.5  RT (min)      lcqtof_t3_neg_3_006_23    m/z 952.6849     0   2   4   6E+4   13   13.5  RT (min)      lcqtof_t3_neg_1_019_23    m/z 953.1213     0   1E+5   12.5   13  RT (min)      lcqtof_t3_neg_4_064_17    m/z 953.1222     0   1   2   3   4E+4   14   14.5  RT (min)      lcqtof_t3_neg_4_036_47    m/z 953.1232     0   1E+5   13.5   14  RT (min)       lcqtof_t3_neg_4_064_17    m/z 953.1233     0   2   4   6   8E+4   14   14.5  RT (min)      lcqtof_t3_neg_4_002_17    m/z 953.1235     0   2   4   6E+4   14   14.5  RT (min)      lcqtof_t3_neg_4_002_17    m/z 953.1236     0   1E+5   13   13.5  RT (min)      lcqtof_t3_neg_4_064_17    m/z 953.1237     0   1   2   3   4E+4   14   14.5  RT (min)      lcqtof_t3_neg_4_002_17    m/z 953.1239     0   2   4   6E+4   12.5   13  RT (min)      lcqtof_t3_neg_4_036_47    m/z 953.1246     0   1E+5   13.5   14  RT (min)      lcqtof_t3_neg_4_002_17    m/z 953.1249     0   1E+5   13.5   14  RT (min)      lcqtof_t3_neg_4_064_17    m/z 953.1250     0   2   4   6E+4   12.5   13  RT (min)      lcqtof_t3_neg_4_036_47    m/z 953.2320     0   1E+4   13.5   14  RT (min)      lcqtof_t3_neg_4_064_17    m/z 953.2358     0   1E+4   13.5   14  RT (min)      lcqtof_t3_neg_4_002_17    m/z 953.6886     0   1   2   3E+4   13   13.5  RT (min)      lcqtof_t3_neg_2_030_23    m/z 954.1216     0   2   4   6E+4   12.5   13  RT (min)      lcqtof_t3_neg_4_036_47    m/z 954.1222     0   1E+5   12.5   13  RT (min)      lcqtof_t3_neg_4_064_17    m/z 954.1223     0   1E+5   13.5   14  RT (min)      lcqtof_t3_neg_4_002_17    m/z 954.1225     0   1E+5   13.5   14  RT (min)      lcqtof_t3_neg_4_064_17    m/z 954.1228     0   1E+5   13   13.5  RT (min)      lcqtof_t3_neg_4_064_17    m/z 954.1237     0   1   2   3E+4   14   14.5  RT (min)      lcqtof_t3_neg_4_036_47    m/z 954.1239     0   1E+5   13.5   14  RT (min)      lcqtof_t3_neg_4_002_17    m/z 954.1249     0   2   4   6E+4   14   14.5  RT (min)      lcqtof_t3_neg_4_002_17    m/z 954.1270     0   1   2   3   4E+4   14   14.5  RT (min)      lcqtof_t3_neg_4_002_17    m/z 954.5946     0   1E+4   13   13.5  RT (min)       lcqtof_t3_neg_4_037_01    m/z 954.6724     0   2   4E+3   12.5   13  RT (min)      lcqtof_t3_neg_2_030_23    m/z 955.1191     0   2   4   6E+4   12.5   13  RT (min)      lcqtof_t3_neg_4_036_47    m/z 955.1196     0   1E+5   13   13.5  RT (min)      lcqtof_t3_neg_4_064_17    m/z 955.1204     0   2   4E+4   14   14.5  RT (min)      lcqtof_t3_neg_4_064_17    m/z 955.1208     0   2   4   6E+4   14   14.5  RT (min)      lcqtof_t3_neg_4_002_17    m/z 955.1209     0   1E+5   13.5   14  RT (min)      lcqtof_t3_neg_4_064_17    m/z 955.1209     0   1E+5   13.5   14  RT (min)      lcqtof_t3_neg_4_002_17    m/z 955.1213     0   1E+5   12.5   13  RT (min)      lcqtof_t3_neg_4_064_17    m/z 955.1219     0   1E+5   13.5   14  RT (min)      lcqtof_t3_neg_4_064_17    m/z 956.1165     0   2   4E+4   13.5   14  RT (min)      lcqtof_t3_neg_4_002_17    m/z 956.1168     0   2   4   6E+4   13.5   14  RT (min)      lcqtof_t3_neg_4_064_17    m/z 956.1194     0   2   4   6   8E+4   13.5   14  RT (min)      lcqtof_t3_neg_4_064_17    m/z 956.1196     0   2   4   6   8E+4   12.5   13  RT (min)      lcqtof_t3_neg_4_064_17    m/z 956.1213     0   2   4   6   8E+4   13   13.5  RT (min)      lcqtof_t3_neg_4_064_17    m/z 956.1221     0   2   4   6   8E+4   13   13.5  RT (min)      lcqtof_t3_neg_4_064_17    m/z 956.6058     0   1E+4   12.5   13  RT (min)      lcqtof_t3_neg_4_018_23    m/z 956.6094     0   1E+4   13   13.5  RT (min)      lcqtof_t3_neg_4_008_35    m/z 956.6096     0   1   2E+4   13   13.5  RT (min)      lcqtof_t3_neg_4_030_31    m/z 956.6100     0   1E+4   13   13.5  RT (min)      lcqtof_t3_neg_4_055_38    m/z 956.6117     0   1   2E+4   13   13.5  RT (min)      lcqtof_t3_neg_4_003_37    m/z 957.1076     0   1   2   3E+4   12.5   13  RT (min)      lcqtof_t3_neg_4_064_17    m/z 957.1179     0   1   2   3E+4   13   13.5  RT (min)      lcqtof_t3_neg_4_002_17    m/z 957.1181     0   1   2   3   4E+4   12.5   13  RT (min)      lcqtof_t3_neg_4_064_17    m/z 957.1184     0   1   2   3   4E+4   13.5   14  RT (min)      lcqtof_t3_neg_4_064_17    m/z 957.1184     0   1   2   3   4E+4   13   13.5  RT (min)      lcqtof_t3_neg_4_064_17    m/z 957.1196     0   1   2   3   4E+4   13   13.5  RT (min)      lcqtof_t3_neg_4_064_17    m/z 957.6066     0   1E+4   13   13.5  RT (min)      lcqtof_t3_neg_4_034_34    m/z 957.6124     0   1E+4   13   13.5  RT (min)      lcqtof_t3_neg_4_030_31    m/z 958.1148     0   1E+4   13   13.5  RT (min)      lcqtof_t3_neg_4_064_17    m/z 958.1150     0   1E+4   13   13.5  RT (min)      lcqtof_t3_neg_4_064_17    m/z 958.1172     0   1E+4   13   13.5  RT (min)      lcqtof_t3_neg_4_064_17    m/z 958.1188     0   1E+4   12.5   13  RT (min)      lcqtof_t3_neg_4_064_17    m/z 958.6230     0   1E+4   13   13.5  RT (min)       lcqtof_t3_neg_4_034_34    m/z 958.6250     0   1E+4   13   13.5  RT (min)      lcqtof_t3_neg_4_055_38    m/z 958.6263     0   1E+4   13   13.5  RT (min)      lcqtof_t3_neg_4_055_38    m/z 959.5062     0   1   2E+4   12   12.5  RT (min)      lcqtof_t3_neg_4_018_23    m/z 959.5062     0   1   2E+4   12   12.5  RT (min)      lcqtof_t3_neg_4_018_23    m/z 960.5104     0   1E+4   12   12.5  RT (min)      lcqtof_t3_neg_4_018_23    m/z 961.6404     0   1   2E+2   11.5   12  RT (min)      lcqtof_t3_neg_3_032_24    m/z 962.5287     0   1E+4   12.5   13  RT (min)       lcqtof_t3_neg_1_019_23    m/z 962.5298     0   1E+4   12.5   13  RT (min)      lcqtof_t3_neg_1_019_23    m/z 963.2361     0   1E+4   13   13.5  RT (min)      lcqtof_t3_neg_4_064_17    m/z 964.5455     0   1E+4   12.5   13  RT (min)      lcqtof_t3_neg_1_019_23    m/z 964.6646     0   2   4   6E+3   14   14.5  RT (min)      lcqtof_t3_neg_4_037_01    m/z 965.1983     0   1   2E+5   13   13.5  RT (min)      lcqtof_t3_neg_4_064_17    m/z 965.6837     0   1   2E+4   12.5   13  RT (min)       lcqtof_t3_neg_4_040_24    m/z 965.6839     0   1   2E+4   12.5   13  RT (min)      lcqtof_t3_neg_4_040_24    m/z 965.6846     0   1E+4   12.5   13  RT (min)      lcqtof_t3_neg_4_056_56    m/z 966.1961     0   1   2E+5   13   13.5  RT (min)      lcqtof_t3_neg_4_064_17    m/z 966.5643     0   1E+4   13   13.5  RT (min)      lcqtof_t3_neg_1_019_23    m/z 967.1714     0   1   2E+5   13   13.5  RT (min)      lcqtof_t3_neg_4_064_17    m/z 967.1763     0   1   2E+5   13   13.5  RT (min)      lcqtof_t3_neg_4_064_17    m/z 967.1905     0   1   2E+5   13   13.5  RT (min)      lcqtof_t3_neg_4_002_17    m/z 968.1405     0   1   2   3   4E+4   13   13.5  RT (min)      lcqtof_t3_neg_4_064_17    m/z 968.1678     0   1E+5   13   13.5  RT (min)      lcqtof_t3_neg_4_064_17    m/z 968.1884     0   1E+5   13   13.5  RT (min)      lcqtof_t3_neg_4_036_47    m/z 968.2237     0   1   2E+4   12.5   13  RT (min)       lcqtof_t3_neg_3_113_62    m/z 968.6578     0   1   2   3E+4   12   12.5  RT (min)      lcqtof_t3_neg_4_040_24    m/z 968.6797     0   1E+4   12.5   13  RT (min)      lcqtof_t3_neg_1_019_23    m/z 968.6807     0   1E+4   12.5   13  RT (min)      lcqtof_t3_neg_2_030_23    m/z 969.1534     0   1   2   3   4E+4   13   13.5  RT (min)      lcqtof_t3_neg_4_064_17    m/z 969.1624     0   2   4E+4   13   13.5  RT (min)      lcqtof_t3_neg_4_064_17    m/z 969.1734     0   2   4   6   8E+4   13   13.5  RT (min)      lcqtof_t3_neg_4_002_17    m/z 969.1735     0   2   4   6   8E+4   13   13.5  RT (min)      lcqtof_t3_neg_4_002_17    m/z 969.1753     0   2   4   6   8E+4   13   13.5  RT (min)      lcqtof_t3_neg_4_002_17    m/z 969.5144     0   1   2   3E+3   12.5   13  RT (min)      lcqtof_t3_neg_1_010_27    m/z 969.6832     0   1E+4   13   13.5  RT (min)      lcqtof_t3_neg_4_018_23    m/z 969.6837     0   2   4   6   8E+3   12.5   13  RT (min)      lcqtof_t3_neg_1_019_23    m/z 969.7151     0   1   2E+4   13   13.5  RT (min)      lcqtof_t3_neg_4_011_32    m/z 969.7165     0   1   2E+4   13   13.5  RT (min)       lcqtof_t3_neg_4_040_24    m/z 969.7176     0   1   2E+4   13   13.5  RT (min)      lcqtof_t3_neg_4_040_24    m/z 969.7237     0   1   2E+4   13   13.5  RT (min)       lcqtof_t3_neg_4_040_24    m/z 970.0475     0   1E+4   13   13.5  RT (min)      lcqtof_t3_neg_4_064_17    m/z 970.0487     0   1E+4   13   13.5  RT (min)      lcqtof_t3_neg_4_064_17    m/z 970.0498     0   1E+4   13   13.5  RT (min)      lcqtof_t3_neg_4_064_17    m/z 970.0555     0   1E+4   13   13.5  RT (min)      lcqtof_t3_neg_4_064_17    m/z 970.1568     0   1   2E+4   13   13.5  RT (min)      lcqtof_t3_neg_4_064_17    m/z 970.1645     0   1   2   3E+4   13   13.5  RT (min)      lcqtof_t3_neg_4_002_17    m/z 970.1651     0   1   2   3E+4   13   13.5  RT (min)      lcqtof_t3_neg_4_002_17    m/z 970.1828     0   1   2   3E+4   13.5   14  RT (min)      lcqtof_t3_neg_4_002_17    m/z 970.1842     0   1   2   3E+4   13.5   14  RT (min)      lcqtof_t3_neg_4_002_17    m/z 970.1921     0   1   2   3E+4   13.5   14  RT (min)      lcqtof_t3_neg_4_064_17    m/z 970.2709     0   1   2E+3   11   11.5  RT (min)       lcqtof_t3_neg_1_043_38    m/z 970.6547     0   2   4   6   8E+2   11.5   12  RT (min)      lcqtof_t3_neg_3_032_24    m/z 970.7297     0   1   2E+4   13   13.5  RT (min)      lcqtof_t3_neg_4_040_24    m/z 971.0421     0   1E+4   13   13.5  RT (min)      lcqtof_t3_neg_4_064_17    m/z 971.0439     0   1E+4   13   13.5  RT (min)      lcqtof_t3_neg_4_064_17    m/z 971.1476     0   1E+4   13   13.5  RT (min)      lcqtof_t3_neg_4_064_17    m/z 971.1852     0   1E+4   13.5   14  RT (min)      lcqtof_t3_neg_4_002_17    m/z 971.1888     0   1E+4   13.5   14  RT (min)      lcqtof_t3_neg_4_064_17    m/z 971.1907     0   1E+4   13.5   14  RT (min)      lcqtof_t3_neg_4_002_17    m/z 971.7318     0   2   4E+4   13   13.5  RT (min)      lcqtof_t3_neg_4_033_53    m/z 971.7333     0   1   2   3   4E+4   13   13.5  RT (min)      lcqtof_t3_neg_4_011_32    m/z 971.7333     0   1   2   3   4E+4   13   13.5  RT (min)      lcqtof_t3_neg_4_011_32    m/z 971.7334     0   1   2   3   4E+4   13   13.5  RT (min)      lcqtof_t3_neg_4_011_32    m/z 971.7360     0   1   2   3E+4   13   13.5  RT (min)       lcqtof_t3_neg_4_040_24    m/z 971.8094     0   1   2E+2   12   12.5  RT (min)      lcqtof_t3_neg_3_032_24    m/z 971.8099     0   1   2E+2   12   12.5  RT (min)      lcqtof_t3_neg_3_032_24    m/z 971.8107     0   1   2   3   4E+2   11   11.5  RT (min)      lcqtof_t3_neg_3_038_06    m/z 971.8114     0   1   2E+4   12.5   13  RT (min)      lcqtof_t3_neg_2_016_05    m/z 971.8133     0   1   2   3   4E+2   11   11.5  RT (min)      lcqtof_t3_neg_3_038_06    m/z 972.2207     0   1E+4   12.5   13  RT (min)      lcqtof_t3_neg_4_064_17    m/z 972.2214     0   1E+4   12.5   13  RT (min)      lcqtof_t3_neg_4_064_17    m/z 972.8127     0   1   2   3E+2   12   12.5  RT (min)      lcqtof_t3_neg_3_032_24    m/z 972.8129     0   1   2   3E+2   12   12.5  RT (min)      lcqtof_t3_neg_3_032_24    m/z 972.8145     0   1   2   3E+2   11   11.5  RT (min)      lcqtof_t3_neg_3_038_06    m/z 972.8149     0   1   2   3E+2   11   11.5  RT (min)      lcqtof_t3_neg_3_038_06    m/z 972.8153     0   2   4   6   8E+4   12.5   13  RT (min)      lcqtof_t3_neg_2_016_05    m/z 973.2147     0   1   2E+4   12.5   13  RT (min)      lcqtof_t3_neg_4_064_17    m/z 973.2162     0   1E+4   12.5   13  RT (min)      lcqtof_t3_neg_4_002_17    m/z 973.2175     0   1E+4   12.5   13  RT (min)      lcqtof_t3_neg_4_002_17    m/z 973.7478     0   1E+4   13   13.5  RT (min)      lcqtof_t3_neg_4_033_53    m/z 973.7483     0   1E+4   13.5   14  RT (min)       lcqtof_t3_neg_4_040_24    m/z 973.7558     0   1   2E+4   13   13.5  RT (min)      lcqtof_t3_neg_4_011_32    m/z 973.7652     0   1   2E+4   13   13.5  RT (min)      lcqtof_t3_neg_4_011_32    m/z 974.2162     0   1E+4   12.5   13  RT (min)      lcqtof_t3_neg_4_064_17    m/z 974.4881     0   1E+4   11.5   12  RT (min)      lcqtof_t3_neg_4_018_23    m/z 974.6403     0   1E+4   13   13.5  RT (min)       lcqtof_t3_neg_4_034_34    m/z 974.6486     0   1E+4   13   13.5  RT (min)      lcqtof_t3_neg_4_055_38    m/z 974.6507     0   1E+4   13   13.5  RT (min)       lcqtof_t3_neg_4_034_34    m/z 975.2177     0   1   2E+4   12.5   13  RT (min)      lcqtof_t3_neg_4_064_17    m/z 975.2302     0   1E+4   12.5   13  RT (min)      lcqtof_t3_neg_4_002_17    m/z 975.2333     0   1   2E+4   12.5   13  RT (min)      lcqtof_t3_neg_4_064_17    m/z 975.4916     0   2   4   6E+3   12   12.5  RT (min)      lcqtof_t3_neg_4_034_34    m/z 976.2326     0   1E+4   12.5   13  RT (min)      lcqtof_t3_neg_4_002_17    m/z 976.2336     0   1E+4   12.5   13  RT (min)      lcqtof_t3_neg_4_002_17    m/z 976.2343     0   1   2   3E+4   12.5   13  RT (min)      lcqtof_t3_neg_4_064_17    m/z 977.5100     0   1   2E+4   12   12.5  RT (min)      lcqtof_t3_neg_4_018_23    m/z 978.2089     0   1E+4   13   13.5  RT (min)      lcqtof_t3_neg_4_057_49    m/z 978.5192     0   1E+4   12   12.5  RT (min)      lcqtof_t3_neg_2_030_23    m/z 978.5194     0   1   2E+4   12   12.5  RT (min)      lcqtof_t3_neg_4_018_23    m/z 978.5206     0   1   2   3   4E+3   12.5   13  RT (min)      lcqtof_t3_neg_4_026_30    m/z 978.5212     0   1   2E+4   12   12.5  RT (min)      lcqtof_t3_neg_4_018_23    m/z 978.5286     0   1   2E+4   12   12.5  RT (min)      lcqtof_t3_neg_1_019_23    m/z 978.5336     0   1   2E+4   12   12.5  RT (min)      lcqtof_t3_neg_1_019_23    m/z 978.7135     0   1E+4   10.5   11  RT (min)      lcqtof_t3_neg_4_034_34    m/z 978.7523     0   2   4E+3   14   14.5  RT (min)      lcqtof_t3_neg_2_051_14    m/z 979.2107     0   2   4E+4   13   13.5  RT (min)      lcqtof_t3_neg_4_057_49    m/z 979.2129     0   1   2   3   4E+4   13.5   14  RT (min)      lcqtof_t3_neg_4_001_49    m/z 979.4235     0   1   2   3   4E+3   9.5   10  RT (min)       lcqtof_t3_neg_4_064_17    m/z 979.5260     0   1E+4   12   12.5  RT (min)      lcqtof_t3_neg_1_019_23    m/z 979.5262     0   1   2E+4   12   12.5  RT (min)      lcqtof_t3_neg_4_034_34    m/z 979.5350     0   1E+4   12   12.5  RT (min)      lcqtof_t3_neg_4_018_23    m/z 980.2095     0   1   2   3E+4   13.5   14  RT (min)      lcqtof_t3_neg_4_001_49    m/z 980.2106     0   2   4E+4   13   13.5  RT (min)      lcqtof_t3_neg_4_057_49    m/z 980.2336     0   1E+4   13   13.5  RT (min)      lcqtof_t3_neg_4_002_17    m/z 980.2346     0   1   2E+4   12.5   13  RT (min)      lcqtof_t3_neg_4_064_17    m/z 981.1153     0   2   4   6E+4   14   14.5  RT (min)      lcqtof_t3_neg_4_064_17    m/z 981.1174     0   2   4   6E+4   13   13.5  RT (min)      lcqtof_t3_neg_4_064_17    m/z 981.1175     0   2   4   6E+4   13   13.5  RT (min)      lcqtof_t3_neg_4_064_17    m/z 981.1186     0   2   4   6E+4   13.5   14  RT (min)      lcqtof_t3_neg_4_064_17    m/z 981.1200     0   1   2   3E+4   14   14.5  RT (min)      lcqtof_t3_neg_4_064_17    m/z 981.2081     0   1   2   3   4E+4   13   13.5  RT (min)      lcqtof_t3_neg_4_057_49    m/z 981.2097     0   1   2   3E+4   13.5   14  RT (min)      lcqtof_t3_neg_4_001_49    m/z 981.2113     0   1   2   3E+4   13   13.5  RT (min)      lcqtof_t3_neg_4_001_49    m/z 981.5929     0   1E+4   12.5   13  RT (min)      lcqtof_t3_neg_4_035_08    m/z 981.7708     0   1   2   3E+3   12.5   13  RT (min)      lcqtof_t3_neg_2_053_38    m/z 982.1161     0   1   2   3   4E+4   14   14.5  RT (min)      lcqtof_t3_neg_4_064_17    m/z 982.1168     0   2   4   6E+4   13.5   14  RT (min)      lcqtof_t3_neg_4_064_17    m/z 982.1171     0   2   4   6E+4   13.5   14  RT (min)      lcqtof_t3_neg_4_064_17    m/z 982.1171     0   2   4E+4   13   13.5  RT (min)      lcqtof_t3_neg_4_002_17    m/z 982.1172     0   2   4   6E+4   13   13.5  RT (min)      lcqtof_t3_neg_4_064_17    m/z 982.2091     0   1   2E+4   13   13.5  RT (min)      lcqtof_t3_neg_4_057_49    m/z 982.4282     0   1E+4   12   12.5  RT (min)      lcqtof_t3_neg_4_018_23    m/z 983.1112     0   1   2   3E+4   14   14.5  RT (min)      lcqtof_t3_neg_4_002_17    m/z 983.1118     0   2   4   6   8E+4   13   13.5  RT (min)      lcqtof_t3_neg_4_064_17    m/z 983.1119     0   2   4   6E+4   13.5   14  RT (min)      lcqtof_t3_neg_4_064_17    m/z 983.1121     0   2   4   6   8E+4   13.5   14  RT (min)      lcqtof_t3_neg_4_064_17    m/z 983.1134     0   2   4E+4   14   14.5  RT (min)      lcqtof_t3_neg_4_064_17    m/z 983.1135     0   2   4E+4   14   14.5  RT (min)      lcqtof_t3_neg_4_002_17    m/z 983.1136     0   2   4   6   8E+4   13   13.5  RT (min)      lcqtof_t3_neg_4_064_17    m/z 983.1147     0   1   2   3E+4   14   14.5  RT (min)      lcqtof_t3_neg_4_002_17    m/z 983.2069     0   1E+4   13   13.5  RT (min)      lcqtof_t3_neg_4_057_49    m/z 983.2084     0   1E+4   13   13.5  RT (min)      lcqtof_t3_neg_4_057_49    m/z 984.1035     0   2   4E+4   13   13.5  RT (min)      lcqtof_t3_neg_4_064_17    m/z 984.1092     0   2   4E+4   13   13.5  RT (min)      lcqtof_t3_neg_4_064_17    m/z 984.1099     0   1   2   3   4E+4   13.5   14  RT (min)      lcqtof_t3_neg_4_064_17    m/z 984.1109     0   2   4E+4   13   13.5  RT (min)      lcqtof_t3_neg_4_064_17    m/z 984.1121     0   2   4E+4   13.5   14  RT (min)      lcqtof_t3_neg_4_064_17    m/z 984.1132     0   1   2   3E+4   14   14.5  RT (min)      lcqtof_t3_neg_4_064_17    m/z 984.1162     0   1   2   3   4E+4   14   14.5  RT (min)      lcqtof_t3_neg_4_064_17    m/z 984.3255     0   1E+4   7.5   8  RT (min)      lcqtof_t3_neg_4_033_53    m/z 984.3260     0   1E+4   7.5   8  RT (min)      lcqtof_t3_neg_4_041_54    m/z 984.6007     0   1E+4   12   12.5  RT (min)      lcqtof_t3_neg_4_046_33    m/z 985.1033     0   1   2   3E+4   13   13.5  RT (min)      lcqtof_t3_neg_4_064_17    m/z 985.1037     0   1   2   3E+4   13.5   14  RT (min)      lcqtof_t3_neg_4_064_17    m/z 985.1060     0   1E+4   14   14.5  RT (min)      lcqtof_t3_neg_4_002_17    m/z 985.1063     0   1   2   3E+4   13   13.5  RT (min)      lcqtof_t3_neg_4_064_17    m/z 985.1068     0   1   2   3E+4   13   13.5  RT (min)      lcqtof_t3_neg_4_064_17    m/z 985.1078     0   1   2E+4   13.5   14  RT (min)      lcqtof_t3_neg_4_002_17    m/z 985.1088     0   1   2   3E+4   13.5   14  RT (min)      lcqtof_t3_neg_4_064_17    m/z 985.7510     0   1E+4   13   13.5  RT (min)      lcqtof_t3_neg_4_011_32    m/z 985.7523     0   1E+4   13   13.5  RT (min)      lcqtof_t3_neg_4_040_24    m/z 985.7527     0   1E+4   13   13.5  RT (min)       lcqtof_t3_neg_4_040_24    m/z 986.0874     0   1E+4   13   13.5  RT (min)      lcqtof_t3_neg_4_064_17    m/z 986.0945     0   1E+4   13.5   14  RT (min)      lcqtof_t3_neg_4_064_17    m/z 986.0955     0   1E+4   13   13.5  RT (min)      lcqtof_t3_neg_4_064_17    m/z 986.1005     0   1E+4   13.5   14  RT (min)      lcqtof_t3_neg_4_064_17    m/z 986.1016     0   1E+4   13   13.5  RT (min)      lcqtof_t3_neg_4_064_17    m/z 986.6783     0   1E+4   12.5   13  RT (min)       lcqtof_t3_neg_4_040_24    m/z 987.7775     0   1   2E+4   13   13.5  RT (min)      lcqtof_t3_neg_4_011_32    m/z 987.8322     0   1   2E+2   12.5   13  RT (min)      lcqtof_t3_neg_3_032_24    m/z 987.8331     0   1   2E+2   12.5   13  RT (min)      lcqtof_t3_neg_3_032_24    m/z 987.8354     0   1E+4   13   13.5  RT (min)      lcqtof_t3_neg_2_016_05    m/z 987.8356     0   1E+4   13   13.5  RT (min)      lcqtof_t3_neg_2_016_05    m/z 988.8076     0   1   2E+2   12   12.5  RT (min)      lcqtof_t3_neg_3_032_24    m/z 988.8093     0   1E+4   12.5   13  RT (min)      lcqtof_t3_neg_2_016_05    m/z 988.8112     0   1E+4   12.5   13  RT (min)      lcqtof_t3_neg_2_016_05    m/z 988.8114     0   2   4   6E+2   11   11.5  RT (min)      lcqtof_t3_neg_3_038_06    m/z 989.2066     0   1E+4   12.5   13  RT (min)      lcqtof_t3_neg_4_002_17    m/z 989.2140     0   1   2E+4   12.5   13  RT (min)      lcqtof_t3_neg_4_064_17    m/z 989.2141     0   1E+4   12.5   13  RT (min)      lcqtof_t3_neg_4_002_17    m/z 989.2150     0   1E+4   12.5   13  RT (min)      lcqtof_t3_neg_4_002_17    m/z 989.2155     0   1   2E+4   12.5   13  RT (min)      lcqtof_t3_neg_4_064_17    m/z 990.2149     0   1   2E+4   12.5   13  RT (min)      lcqtof_t3_neg_4_064_17    m/z 990.2167     0   1   2E+4   12.5   13  RT (min)      lcqtof_t3_neg_4_064_17    m/z 991.2132     0   1E+4   12.5   13  RT (min)      lcqtof_t3_neg_4_002_17    m/z 991.2253     0   1E+4   12.5   13  RT (min)      lcqtof_t3_neg_4_064_17    m/z 991.2344     0   1E+4   12.5   13  RT (min)      lcqtof_t3_neg_4_002_17    m/z 991.6312     0   1E+4   12.5   13  RT (min)      lcqtof_t3_neg_4_059_14    m/z 992.2311     0   1E+4   12.5   13  RT (min)      lcqtof_t3_neg_4_064_17    m/z 992.2332     0   1E+4   12.5   13  RT (min)      lcqtof_t3_neg_4_064_17    m/z 993.2254     0   1E+4   12.5   13  RT (min)      lcqtof_t3_neg_4_064_17    m/z 993.2287     0   1E+4   12.5   13  RT (min)      lcqtof_t3_neg_4_064_17    m/z 993.2599     0   1E+4   13.5   14  RT (min)      lcqtof_t3_neg_4_064_17    m/z 994.2622     0   1E+4   13.5   14  RT (min)      lcqtof_t3_neg_4_064_17    m/z 994.2631     0   1E+4   13.5   14  RT (min)      lcqtof_t3_neg_4_002_17    m/z 995.0774     0   2   4   6   8E+4   14   14.5  RT (min)      lcqtof_t3_neg_4_002_17    m/z 995.0780     0   1E+5   13.5   14  RT (min)      lcqtof_t3_neg_4_064_17    m/z 995.0782     0   2   4   6   8E+4   13.5   14  RT (min)      lcqtof_t3_neg_4_002_17    m/z 995.0786     0   1E+5   14   14.5  RT (min)      lcqtof_t3_neg_4_064_17    m/z 995.0788     0   1E+5   13.5   14  RT (min)      lcqtof_t3_neg_4_064_17    m/z 995.0792     0   2   4E+4   14   14.5  RT (min)      lcqtof_t3_neg_4_036_47    m/z 995.0792     0   2   4E+4   14   14.5  RT (min)      lcqtof_t3_neg_4_036_47    m/z 995.0793     0   2   4   6   8E+4   13   13.5  RT (min)      lcqtof_t3_neg_4_064_17    m/z 995.0793     0   1E+5   13   13.5  RT (min)      lcqtof_t3_neg_4_064_17    m/z 995.0795     0   2   4E+4   14   14.5  RT (min)      lcqtof_t3_neg_4_036_47    m/z 995.0798     0   2   4   6   8E+4   13.5   14  RT (min)      lcqtof_t3_neg_4_002_17    m/z 995.0802     0   2   4   6E+4   13   13.5  RT (min)      lcqtof_t3_neg_4_064_17    m/z 995.1661     0   1   2   3E+4   13   13.5  RT (min)      lcqtof_t3_neg_4_064_17    m/z 995.1678     0   1   2   3E+4   13   13.5  RT (min)      lcqtof_t3_neg_4_064_17    m/z 995.1680     0   1   2   3E+4   13.5   14  RT (min)      lcqtof_t3_neg_4_064_17    m/z 995.1714     0   1   2   3E+4   13   13.5  RT (min)      lcqtof_t3_neg_4_064_17    m/z 995.1764     0   1   2   3E+4   13.5   14  RT (min)      lcqtof_t3_neg_4_064_17    m/z 995.2614     0   1E+4   13.5   14  RT (min)      lcqtof_t3_neg_4_064_17    m/z 995.3284     0   2   4   6   8E+4   4   4.5  RT (min)      lcqtof_t3_neg_4_000_ee    m/z 995.7202     0   1E+4   12.5   13  RT (min)       lcqtof_t3_neg_4_011_32    m/z 995.7209     0   1E+4   12.5   13  RT (min)      lcqtof_t3_neg_4_040_24    m/z 995.8303     0   1   2   3E+4   4   4.5  RT (min)      lcqtof_t3_neg_4_000_ee    m/z 996.0773     0   2   4   6   8E+4   13   13.5  RT (min)      lcqtof_t3_neg_4_002_17    m/z 996.0783     0   1E+5   13.5   14  RT (min)      lcqtof_t3_neg_4_064_17    m/z 996.0793     0   2   4E+4   13   13.5  RT (min)      lcqtof_t3_neg_4_064_17    m/z 996.0793     0   1E+5   13.5   14  RT (min)      lcqtof_t3_neg_4_064_17    m/z 996.0807     0   1E+5   14   14.5  RT (min)      lcqtof_t3_neg_4_064_17    m/z 996.0815     0   2   4E+4   13   13.5  RT (min)      lcqtof_t3_neg_4_036_47    m/z 996.1690     0   1   2E+4   13   13.5  RT (min)      lcqtof_t3_neg_4_064_17    m/z 996.1690     0   1   2E+4   13.5   14  RT (min)      lcqtof_t3_neg_4_002_17    m/z 996.1698     0   1   2E+4   13.5   14  RT (min)      lcqtof_t3_neg_4_064_17    m/z 996.1700     0   1   2E+4   13   13.5  RT (min)      lcqtof_t3_neg_4_064_17    m/z 996.1705     0   1   2E+4   13.5   14  RT (min)      lcqtof_t3_neg_4_064_17    m/z 996.3298     0   1E+4   4   4.5  RT (min)      lcqtof_t3_neg_4_000_ee    m/z 996.8201     0   1   2E+4   17   17.5  RT (min)      lcqtof_t3_neg_4_035_08    m/z 997.0717     0   2   4   6E+4   14   14.5  RT (min)      lcqtof_t3_neg_4_002_17    m/z 997.0718     0   2   4   6   8E+4   14   14.5  RT (min)      lcqtof_t3_neg_4_002_17    m/z 997.0721     0   1E+5   13   13.5  RT (min)      lcqtof_t3_neg_4_064_17    m/z 997.0733     0   2   4   6E+4   13   13.5  RT (min)      lcqtof_t3_neg_4_064_17    m/z 997.0738     0   1E+5   13.5   14  RT (min)      lcqtof_t3_neg_4_064_17    m/z 997.0740     0   1E+5   13   13.5  RT (min)      lcqtof_t3_neg_4_064_17    m/z 997.0740     0   1   2E+4   14   14.5  RT (min)      lcqtof_t3_neg_4_004_20    m/z 997.0744     0   1E+5   14   14.5  RT (min)      lcqtof_t3_neg_4_064_17    m/z 997.0745     0   2   4   6E+4   13   13.5  RT (min)      lcqtof_t3_neg_4_036_47    m/z 997.0762     0   2   4   6   8E+4   13.5   14  RT (min)      lcqtof_t3_neg_4_002_17    m/z 997.0764     0   1E+5   13.5   14  RT (min)      lcqtof_t3_neg_4_064_17    m/z 997.1655     0   1   2E+4   13   13.5  RT (min)      lcqtof_t3_neg_4_064_17    m/z 997.1658     0   1   2E+4   13.5   14  RT (min)      lcqtof_t3_neg_4_064_17    m/z 997.1662     0   1   2E+4   13   13.5  RT (min)      lcqtof_t3_neg_4_064_17    m/z 997.1673     0   1   2E+4   13.5   14  RT (min)      lcqtof_t3_neg_4_002_17    m/z 997.1682     0   1   2E+4   13.5   14  RT (min)      lcqtof_t3_neg_4_064_17    m/z 997.7367     0   1   2E+4   13   13.5  RT (min)      lcqtof_t3_neg_4_040_24    m/z 997.7372     0   1   2E+4   13   13.5  RT (min)       lcqtof_t3_neg_4_040_24    m/z 997.7373     0   1E+4   13   13.5  RT (min)      lcqtof_t3_neg_4_011_32    m/z 997.8666     0   1E+4   12.5   13  RT (min)      lcqtof_t3_neg_2_030_23    m/z 997.8670     0   1E+4   12.5   13  RT (min)      lcqtof_t3_neg_2_030_23    m/z 997.8671     0   1E+4   12.5   13  RT (min)      lcqtof_t3_neg_2_016_05    m/z 997.8771     0   1   2E+3   14.5   15  RT (min)      lcqtof_t3_neg_1_035_31    m/z 998.0690     0   2   4   6E+4   13   13.5  RT (min)      lcqtof_t3_neg_4_064_17    m/z 998.0700     0   2   4E+4   13   13.5  RT (min)      lcqtof_t3_neg_4_064_17    m/z 998.0704     0   1   2   3   4E+4   13   13.5  RT (min)      lcqtof_t3_neg_4_064_17    m/z 998.0729     0   2   4   6   8E+4   13.5   14  RT (min)      lcqtof_t3_neg_4_064_17    m/z 998.0730     0   2   4   6   8E+4   14   14.5  RT (min)      lcqtof_t3_neg_4_064_17    m/z 998.0748     0   2   4   6   8E+4   13.5   14  RT (min)      lcqtof_t3_neg_4_064_17    m/z 998.0758     0   2   4   6   8E+4   14   14.5  RT (min)      lcqtof_t3_neg_4_064_17    m/z 998.0764     0   1E+4   13.5   14  RT (min)      lcqtof_t3_neg_4_063_39    m/z 998.1665     0   1E+4   13   13.5  RT (min)      lcqtof_t3_neg_4_064_17    m/z 998.1669     0   1E+4   13   13.5  RT (min)      lcqtof_t3_neg_4_002_17    m/z 998.1674     0   1E+4   13.5   14  RT (min)      lcqtof_t3_neg_4_064_17    m/z 998.3421     0   1E+4   8   8.5  RT (min)       lcqtof_t3_neg_4_004_20    m/z 998.4871     0   1E+4   12   12.5  RT (min)      lcqtof_t3_neg_4_018_23    m/z 998.4876     0   1E+4   11.5   12  RT (min)      lcqtof_t3_neg_1_019_23    m/z 998.7495     0   1E+4   13   13.5  RT (min)      lcqtof_t3_neg_4_040_24    m/z 998.8716     0   1E+4   12.5   13  RT (min)      lcqtof_t3_neg_2_030_23    m/z 999.0620     0   1   2   3E+4   13   13.5  RT (min)      lcqtof_t3_neg_4_064_17    m/z 999.0628     0   2   4E+4   13.5   14  RT (min)      lcqtof_t3_neg_4_064_17    m/z 999.0651     0   1   2   3   4E+4   13   13.5  RT (min)      lcqtof_t3_neg_4_064_17    m/z 999.0659     0   2   4E+4   14   14.5  RT (min)      lcqtof_t3_neg_4_064_17    m/z 999.0659     0   1   2   3E+4   13   13.5  RT (min)      lcqtof_t3_neg_4_064_17    m/z 999.0715     0   2   4E+4   14   14.5  RT (min)      lcqtof_t3_neg_4_064_17    m/z 999.0727     0   2   4E+4   13.5   14  RT (min)      lcqtof_t3_neg_4_064_17    m/z 999.0735     0   1   2   3E+4   13   13.5  RT (min)      lcqtof_t3_neg_4_064_17    m/z 999.7516     0   1   2E+4   13   13.5  RT (min)      lcqtof_t3_neg_4_046_33    m/z 999.7532     0   1   2   3E+4   13   13.5  RT (min)      lcqtof_t3_neg_4_011_32    m/z 999.7532     0   1   2   3E+4   13   13.5  RT (min)       lcqtof_t3_neg_4_011_32    m/z 999.7537     0   1   2   3E+4   13   13.5  RT (min)      lcqtof_t3_neg_4_011_32    m/z 1000.0594     0   1   2E+4   13   13.5  RT (min)      lcqtof_t3_neg_4_064_17    m/z 1000.0607     0   1   2E+4   13.5   14  RT (min)      lcqtof_t3_neg_4_064_17    m/z 1000.0626     0   1   2E+4   13.5   14  RT (min)      lcqtof_t3_neg_4_064_17    m/z 1000.0637     0   1   2E+4   13   13.5  RT (min)      lcqtof_t3_neg_4_064_17    m/z 1000.0649     0   1   2E+4   13.5   14  RT (min)      lcqtof_t3_neg_4_064_17    m/z 1000.0700     0   1   2E+4   14   14.5  RT (min)      lcqtof_t3_neg_4_064_17    m/z 1000.0715     0   1   2E+4   14   14.5  RT (min)      lcqtof_t3_neg_4_064_17    m/z 1000.0719     0   1   2E+4   13.5   14  RT (min)      lcqtof_t3_neg_4_064_17    m/z 1001.0526     0   1E+4   13   13.5  RT (min)      lcqtof_t3_neg_4_064_17    m/z 1001.0528     0   1E+4   13   13.5  RT (min)      lcqtof_t3_neg_4_064_17    m/z 1001.0636     0   1E+4   13.5   14  RT (min)      lcqtof_t3_neg_4_064_17    m/z 1001.1706     0   1E+4   13.5   14  RT (min)      lcqtof_t3_neg_4_036_47    m/z 1001.7683     0   1   2E+4   13   13.5  RT (min)       lcqtof_t3_neg_4_011_32    m/z 1001.7688     0   1E+4   13   13.5  RT (min)      lcqtof_t3_neg_4_033_53    m/z 1001.8402     0   1   2   3   4E+2   11.5   12  RT (min)      lcqtof_t3_neg_3_038_06    m/z 1001.8405     0   1E+3   13   13.5  RT (min)      lcqtof_t3_neg_3_032_24    m/z 1001.8408     0   1   2E+3   14.5   15  RT (min)      lcqtof_t3_neg_1_014_04    m/z 1001.8421     0   1   2E+3   16   16.5  RT (min)      lcqtof_t3_neg_2_059_14    m/z 1001.8474     0   1   2   3   4E+2   12.5   13  RT (min)      lcqtof_t3_neg_3_032_24    m/z 1001.8476     0   1   2   3   4E+2   12.5   13  RT (min)      lcqtof_t3_neg_3_032_24    m/z 1001.8488     0   1   2   3   4E+2   11   11.5  RT (min)      lcqtof_t3_neg_3_038_06    m/z 1001.8507     0   1E+4   13   13.5  RT (min)      lcqtof_t3_neg_2_016_05    m/z 1002.1708     0   1E+4   13.5   14  RT (min)      lcqtof_t3_neg_4_036_47    m/z 1002.6504     0   2   4   6E+2   10.5   11  RT (min)      lcqtof_t3_neg_3_038_06    m/z 1002.6505     0   1   2   3   4E+2   11.5   12  RT (min)      lcqtof_t3_neg_3_032_24    m/z 1002.6505     0   1E+4   12   12.5  RT (min)      lcqtof_t3_neg_2_016_05    m/z 1002.8458     0   1   2   3   4E+4   13   13.5  RT (min)      lcqtof_t3_neg_2_016_05    m/z 1002.8513     0   1   2E+2   12.5   13  RT (min)      lcqtof_t3_neg_3_032_24    m/z 1002.8531     0   2   4   6E+2   11   11.5  RT (min)      lcqtof_t3_neg_3_038_06    m/z 1002.8553     0   1   2   3   4E+4   13   13.5  RT (min)      lcqtof_t3_neg_2_016_05    m/z 1003.1691     0   1E+4   13.5   14  RT (min)      lcqtof_t3_neg_4_036_47    m/z 1004.1682     0   1E+4   13.5   14  RT (min)      lcqtof_t3_neg_4_036_47    m/z 1007.2054     0   1E+4   13   13.5  RT (min)      lcqtof_t3_neg_4_057_49    m/z 1007.8949     0   1   2   3   4E+4   12.5   13  RT (min)      lcqtof_t3_neg_2_030_23    m/z 1007.8950     0   1   2   3   4E+2   12   12.5  RT (min)      lcqtof_t3_neg_3_032_24    m/z 1007.8956     0   1   2   3   4E+2   12   12.5  RT (min)      lcqtof_t3_neg_3_032_24    m/z 1007.8962     0   1   2   3   4E+2   10.5   11  RT (min)      lcqtof_t3_neg_3_038_06    m/z 1007.8976     0   1   2   3   4E+2   10.5   11  RT (min)      lcqtof_t3_neg_3_038_06    m/z 1008.2061     0   1E+4   13   13.5  RT (min)      lcqtof_t3_neg_4_057_49    m/z 1008.2397     0   1E+4   13   13.5  RT (min)      lcqtof_t3_neg_4_002_17    m/z 1008.2457     0   1   2E+4   13   13.5  RT (min)      lcqtof_t3_neg_4_064_17    m/z 1008.8970     0   2   4E+3   13.5   14  RT (min)      lcqtof_t3_neg_2_046_45    m/z 1008.8972     0   1   2   3E+2   12   12.5  RT (min)      lcqtof_t3_neg_3_032_24    m/z 1008.8984     0   1   2   3E+3   13.5   14  RT (min)      lcqtof_t3_neg_1_055_32    m/z 1008.8986     0   1   2   3E+3   13.5   14  RT (min)      lcqtof_t3_neg_1_009_50    m/z 1008.8990     0   1   2E+4   12.5   13  RT (min)      lcqtof_t3_neg_2_030_23    m/z 1008.8992     0   1   2E+4   12.5   13  RT (min)      lcqtof_t3_neg_2_030_23    m/z 1008.9010     0   1   2   3E+2   12   12.5  RT (min)      lcqtof_t3_neg_3_032_24    m/z 1008.9038     0   1   2   3E+2   11   11.5  RT (min)      lcqtof_t3_neg_3_038_06    m/z 1009.1325     0   1   2E+4   13   13.5  RT (min)      lcqtof_t3_neg_4_064_17    m/z 1009.2066     0   1E+4   13   13.5  RT (min)      lcqtof_t3_neg_4_057_49    m/z 1009.2452     0   1   2E+4   13   13.5  RT (min)      lcqtof_t3_neg_4_064_17    m/z 1009.2563     0   1   2E+4   13   13.5  RT (min)      lcqtof_t3_neg_4_064_17    m/z 1009.5414     0   1E+4   13   13.5  RT (min)      lcqtof_t3_neg_4_032_04    m/z 1009.8942     0   1   2   3E+3   13.5   14  RT (min)      lcqtof_t3_neg_2_053_38    m/z 1010.2562     0   1   2E+4   13   13.5  RT (min)      lcqtof_t3_neg_4_064_17    m/z 1010.2568     0   1   2E+4   13   13.5  RT (min)      lcqtof_t3_neg_4_002_17    m/z 1010.7120     0   1   2E+2   12.5   13  RT (min)      lcqtof_t3_neg_3_032_24    m/z 1010.9084     0   1   2E+3   13   13.5  RT (min)      lcqtof_t3_neg_2_043_31    m/z 1011.1117     0   1   2   3E+4   13   13.5  RT (min)      lcqtof_t3_neg_4_002_17    m/z 1011.1216     0   1   2   3E+4   13   13.5  RT (min)      lcqtof_t3_neg_4_064_17    m/z 1011.1232     0   1   2   3E+4   14   14.5  RT (min)      lcqtof_t3_neg_4_002_17    m/z 1011.1234     0   1   2   3E+4   13   13.5  RT (min)      lcqtof_t3_neg_4_064_17    m/z 1011.1236     0   1   2   3E+4   13   13.5  RT (min)      lcqtof_t3_neg_4_064_17    m/z 1011.1989     0   1   2E+4   13   13.5  RT (min)      lcqtof_t3_neg_4_064_17    m/z 1011.2279     0   1   2   3E+4   13   13.5  RT (min)      lcqtof_t3_neg_4_002_17    m/z 1011.2559     0   1   2   3E+4   13   13.5  RT (min)      lcqtof_t3_neg_4_002_17    m/z 1011.7809     0   1   2   3   4E+3   12.5   13  RT (min)      lcqtof_t3_neg_2_065_51    m/z 1011.7815     0   1   2   3E+3   12   12.5  RT (min)      lcqtof_t3_neg_2_019_32    m/z 1011.7952     0   1   2   3   4E+3   12.5   13  RT (min)      lcqtof_t3_neg_2_057_49    m/z 1011.8682     0   2   4E+2   11.5   12  RT (min)      lcqtof_t3_neg_3_038_06    m/z 1011.8689     0   2   4E+2   11.5   12  RT (min)      lcqtof_t3_neg_3_038_06    m/z 1011.8689     0   1   2   3E+4   13   13.5  RT (min)      lcqtof_t3_neg_2_030_23    m/z 1011.8692     0   1   2E+4   13   13.5  RT (min)      lcqtof_t3_neg_2_030_23    m/z 1011.8704     0   1   2   3E+3   14.5   15  RT (min)      lcqtof_t3_neg_1_012_16    m/z 1011.8705     0   2   4E+4   14   14.5  RT (min)      lcqtof_t3_neg_1_000_dd    m/z 1011.8706     0   1E+2   12.5   13  RT (min)      lcqtof_t3_neg_3_032_24    m/z 1011.8708     0   1   2   3E+3   15   15.5  RT (min)      lcqtof_t3_neg_1_008_02    m/z 1011.8710     0   1   2E+3   15.5   16  RT (min)      lcqtof_t3_neg_1_014_04    m/z 1011.8713     0   1   2E+3   16   16.5  RT (min)      lcqtof_t3_neg_1_000_bb    m/z 1011.8715     0   1   2E+3   16   16.5  RT (min)      lcqtof_t3_neg_1_018_33    m/z 1011.8718     0   1   2E+3   16   16.5  RT (min)      lcqtof_t3_neg_1_008_02    m/z 1011.8720     0   1E+2   12.5   13  RT (min)      lcqtof_t3_neg_3_032_24    m/z 1011.8722     0   1E+2   13   13.5  RT (min)      lcqtof_t3_neg_3_032_24    m/z 1011.8723     0   1   2E+3   16.5   17  RT (min)      lcqtof_t3_neg_1_032_36    m/z 1012.1174     0   1   2E+4   14   14.5  RT (min)      lcqtof_t3_neg_4_002_17    m/z 1012.1203     0   1   2E+4   13.5   14  RT (min)      lcqtof_t3_neg_4_002_17    m/z 1012.1209     0   1   2E+4   13   13.5  RT (min)      lcqtof_t3_neg_4_064_17    m/z 1012.1216     0   1   2E+4   13   13.5  RT (min)      lcqtof_t3_neg_4_002_17    m/z 1012.1219     0   1   2E+4   13.5   14  RT (min)      lcqtof_t3_neg_4_002_17    m/z 1012.1243     0   1   2E+4   13   13.5  RT (min)      lcqtof_t3_neg_4_064_17    m/z 1012.1279     0   1   2E+4   13.5   14  RT (min)      lcqtof_t3_neg_4_002_17    m/z 1012.1280     0   1   2E+4   13.5   14  RT (min)      lcqtof_t3_neg_4_002_17    m/z 1012.1281     0   1   2E+4   13.5   14  RT (min)      lcqtof_t3_neg_4_002_17    m/z 1012.1970     0   1   2E+4   13.5   14  RT (min)      lcqtof_t3_neg_4_002_17    m/z 1012.1976     0   1   2E+4   13   13.5  RT (min)      lcqtof_t3_neg_4_064_17    m/z 1012.2097     0   1   2E+4   13   13.5  RT (min)      lcqtof_t3_neg_4_002_17    m/z 1012.2231     0   1   2E+4   13   13.5  RT (min)      lcqtof_t3_neg_4_002_17    m/z 1012.6348     0   2   4E+2   11.5   12  RT (min)      lcqtof_t3_neg_3_032_24    m/z 1012.7928     0   1E+4   17   17.5  RT (min)      lcqtof_t3_neg_4_035_08    m/z 1012.8692     0   1E+4   14.5   15  RT (min)      lcqtof_t3_neg_2_000_aa    m/z 1012.8717     0   1E+2   13   13.5  RT (min)      lcqtof_t3_neg_3_032_24    m/z 1012.8721     0   1E+4   13   13.5  RT (min)      lcqtof_t3_neg_2_030_23    m/z 1012.8726     0   1   2   3   4E+2   11.5   12  RT (min)      lcqtof_t3_neg_3_038_06    m/z 1012.8729     0   1   2   3E+4   14   14.5  RT (min)      lcqtof_t3_neg_1_033_53    m/z 1012.8734     0   1   2E+4   13   13.5  RT (min)      lcqtof_t3_neg_2_030_23    m/z 1012.8745     0   1   2   3   4E+2   11.5   12  RT (min)      lcqtof_t3_neg_3_038_06    m/z 1012.8745     0   1   2E+3   15   15.5  RT (min)      lcqtof_t3_neg_1_000_cc    m/z 1012.8746     0   1E+2   12.5   13  RT (min)      lcqtof_t3_neg_3_032_24    m/z 1012.8746     0   1   2   3E+4   14   14.5  RT (min)      lcqtof_t3_neg_1_011_45    m/z 1013.0894     0   2   4   6   8E+4   14   14.5  RT (min)      lcqtof_t3_neg_4_002_17    m/z 1013.0899     0   1E+5   12.5   13  RT (min)      lcqtof_t3_neg_4_064_17    m/z 1013.0899     0   1E+5   13.5   14  RT (min)      lcqtof_t3_neg_4_064_17    m/z 1013.0900     0   1E+5   14   14.5  RT (min)      lcqtof_t3_neg_4_064_17    m/z 1013.0902     0   1E+5   13   13.5  RT (min)      lcqtof_t3_neg_4_064_17    m/z 1013.0910     0   1E+5   14   14.5  RT (min)      lcqtof_t3_neg_4_002_17    m/z 1013.0919     0   1E+5   13   13.5  RT (min)      lcqtof_t3_neg_4_064_17    m/z 1013.0919     0   1E+4   13   13.5  RT (min)      lcqtof_t3_neg_4_022_28    m/z 1013.0919     0   1E+5   13.5   14  RT (min)      lcqtof_t3_neg_4_064_17    m/z 1013.0922     0   1   2E+4   13.5   14  RT (min)      lcqtof_t3_neg_4_004_20    m/z 1013.0924     0   1E+4   13   13.5  RT (min)      lcqtof_t3_neg_4_022_28    m/z 1013.0940     0   1E+5   13   13.5  RT (min)      lcqtof_t3_neg_4_064_17    m/z 1013.0941     0   1E+5   13.5   14  RT (min)      lcqtof_t3_neg_4_002_17    m/z 1013.0943     0   1E+5   13.5   14  RT (min)      lcqtof_t3_neg_4_002_17    m/z 1013.0997     0   1E+5   13.5   14  RT (min)      lcqtof_t3_neg_4_002_17    m/z 1013.2009     0   1   2E+4   13.5   14  RT (min)      lcqtof_t3_neg_4_002_17    m/z 1013.4979     0   1   2   3E+3   12.5   13  RT (min)       lcqtof_t3_neg_1_010_27    m/z 1013.8715     0   1   2   3   4E+3   14.5   15  RT (min)      lcqtof_t3_neg_2_000_aa    m/z 1013.8773     0   1E+4   13.5   14  RT (min)      lcqtof_t3_neg_2_016_05    m/z 1013.8786     0   2   4   6   8E+3   13   13.5  RT (min)      lcqtof_t3_neg_2_030_23    m/z 1014.0894     0   2   4   6E+4   14   14.5  RT (min)      lcqtof_t3_neg_4_036_47    m/z 1014.0895     0   1E+5   14   14.5  RT (min)      lcqtof_t3_neg_4_064_17    m/z 1014.0897     0   1E+5   12.5   13  RT (min)      lcqtof_t3_neg_4_064_17    m/z 1014.0899     0   2   4   6E+4   14   14.5  RT (min)      lcqtof_t3_neg_4_036_47    m/z 1014.0904     0   1E+5   13.5   14  RT (min)      lcqtof_t3_neg_4_064_17    m/z 1014.0906     0   1   2E+4   13.5   14  RT (min)      lcqtof_t3_neg_4_004_20    m/z 1014.0908     0   2   4   6   8E+4   13.5   14  RT (min)      lcqtof_t3_neg_4_002_17    m/z 1014.0908     0   1E+5   13.5   14  RT (min)      lcqtof_t3_neg_4_064_17    m/z 1014.0909     0   1E+5   13   13.5  RT (min)      lcqtof_t3_neg_4_064_17    m/z 1014.0909     0   1E+5   13   13.5  RT (min)      lcqtof_t3_neg_4_064_17    m/z 1014.0920     0   1E+5   13   13.5  RT (min)      lcqtof_t3_neg_4_064_17    m/z 1014.1976     0   1E+4   13   13.5  RT (min)      lcqtof_t3_neg_4_064_17    m/z 1015.0867     0   1   2E+4   14   14.5  RT (min)      lcqtof_t3_neg_4_004_20    m/z 1015.0875     0   1E+5   12.5   13  RT (min)      lcqtof_t3_neg_4_064_17    m/z 1015.0876     0   1E+5   14   14.5  RT (min)      lcqtof_t3_neg_4_064_17    m/z 1015.0882     0   1E+5   13.5   14  RT (min)      lcqtof_t3_neg_4_064_17    m/z 1015.0886     0   2   4   6E+4   14   14.5  RT (min)      lcqtof_t3_neg_4_036_47    m/z 1015.0888     0   1E+5   13   13.5  RT (min)      lcqtof_t3_neg_4_064_17    m/z 1015.0888     0   1E+5   13   13.5  RT (min)      lcqtof_t3_neg_4_064_17    m/z 1015.0891     0   1E+5   13.5   14  RT (min)      lcqtof_t3_neg_4_064_17    m/z 1015.0904     0   1   2E+4   13.5   14  RT (min)      lcqtof_t3_neg_4_004_20    m/z 1015.3294     0   1   2   3E+4   4   4.5  RT (min)      lcqtof_t3_neg_4_000_ee    m/z 1015.7871     0   1E+4   13   13.5  RT (min)       lcqtof_t3_neg_4_033_53    m/z 1015.7877     0   1E+4   13   13.5  RT (min)      lcqtof_t3_neg_4_011_32    m/z 1015.7883     0   1E+4   13   13.5  RT (min)       lcqtof_t3_neg_4_011_32    m/z 1015.8347     0   1   2E+4   4   4.5  RT (min)      lcqtof_t3_neg_4_000_ee    m/z 1016.0858     0   1   2   3E+4   14   14.5  RT (min)      lcqtof_t3_neg_4_036_47    m/z 1016.0863     0   1E+4   14   14.5  RT (min)      lcqtof_t3_neg_4_004_20    m/z 1016.0864     0   2   4   6E+4   13.5   14  RT (min)      lcqtof_t3_neg_4_064_17    m/z 1016.0877     0   2   4   6E+4   13.5   14  RT (min)      lcqtof_t3_neg_4_064_17    m/z 1016.0878     0   2   4   6E+4   14   14.5  RT (min)      lcqtof_t3_neg_4_064_17    m/z 1016.0881     0   2   4   6E+4   13   13.5  RT (min)      lcqtof_t3_neg_4_064_17    m/z 1016.0882     0   1   2   3E+4   13.5   14  RT (min)      lcqtof_t3_neg_4_036_47    m/z 1016.0886     0   2   4   6E+4   13   13.5  RT (min)      lcqtof_t3_neg_4_064_17    m/z 1016.0897     0   2   4   6E+4   12.5   13  RT (min)      lcqtof_t3_neg_4_064_17    m/z 1016.0928     0   1   2   3E+4   13   13.5  RT (min)      lcqtof_t3_neg_4_036_47    m/z 1016.3317     0   1E+4   4   4.5  RT (min)      lcqtof_t3_neg_4_000_ee    m/z 1017.0828     0   1   2   3   4E+4   13.5   14  RT (min)      lcqtof_t3_neg_4_064_17    m/z 1017.0838     0   1   2   3   4E+4   13   13.5  RT (min)      lcqtof_t3_neg_4_064_17    m/z 1017.0840     0   1   2   3E+4   14   14.5  RT (min)      lcqtof_t3_neg_4_064_17    m/z 1017.0845     0   1   2   3   4E+4   13   13.5  RT (min)      lcqtof_t3_neg_4_064_17    m/z 1017.0854     0   1   2   3E+4   13.5   14  RT (min)      lcqtof_t3_neg_4_064_17    m/z 1017.0883     0   1   2   3   4E+4   13   13.5  RT (min)      lcqtof_t3_neg_4_064_17    m/z 1017.3302     0   1E+5   4   4.5  RT (min)      lcqtof_t3_neg_4_000_ee    m/z 1017.8315     0   1E+5   4   4.5  RT (min)      lcqtof_t3_neg_4_000_ee    m/z 1017.8351     0   1   2E+4   13.5   14  RT (min)      lcqtof_t3_neg_1_031_24    m/z 1017.8352     0   2   4   6   8E+4   12.5   13  RT (min)      lcqtof_t3_neg_2_030_23    m/z 1017.8356     0   2   4   6   8E+4   12.5   13  RT (min)      lcqtof_t3_neg_2_016_05    m/z 1017.8357     0   2   4   6   8E+4   12.5   13  RT (min)      lcqtof_t3_neg_2_016_05    m/z 1017.8360     0   1E+4   16   16.5  RT (min)      lcqtof_t3_neg_1_000_cc    m/z 1017.8361     0   1E+4   15   15.5  RT (min)      lcqtof_t3_neg_2_065_51    m/z 1017.8363     0   1E+4   15   15.5  RT (min)      lcqtof_t3_neg_1_059_17    m/z 1017.8363     0   1E+4   14.5   15  RT (min)      lcqtof_t3_neg_2_059_14    m/z 1017.8363     0   1E+4   15   15.5  RT (min)      lcqtof_t3_neg_2_059_14    m/z 1017.8368     0   1E+4   15   15.5  RT (min)      lcqtof_t3_neg_1_007_03    m/z 1017.8368     0   2   4   6   8E+4   13.5   14  RT (min)      lcqtof_t3_neg_1_031_24    m/z 1017.8373     0   1E+4   16   16.5  RT (min)      lcqtof_t3_neg_1_000_cc    m/z 1017.8374     0   1E+4   16   16.5  RT (min)      lcqtof_t3_neg_1_000_cc    m/z 1017.8376     0   1E+4   14.5   15  RT (min)      lcqtof_t3_neg_1_057_49    m/z 1017.8380     0   1E+4   13   13.5  RT (min)      lcqtof_t3_neg_4_011_32    m/z 1017.8680     0   1E+4   13.5   14  RT (min)       lcqtof_t3_neg_4_011_32    m/z 1018.0829     0   1E+4   13   13.5  RT (min)      lcqtof_t3_neg_4_064_17    m/z 1018.0830     0   1E+4   13.5   14  RT (min)      lcqtof_t3_neg_4_064_17    m/z 1018.2199     0   1E+4   13   13.5  RT (min)      lcqtof_t3_neg_4_064_17    m/z 1018.2199     0   1E+4   13   13.5  RT (min)      lcqtof_t3_neg_4_036_47    m/z 1018.3322     0   2   4   6E+4   4   4.5  RT (min)      lcqtof_t3_neg_4_000_ee    m/z 1018.8327     0   1   2E+4   4   4.5  RT (min)      lcqtof_t3_neg_4_000_ee    m/z 1018.8378     0   1   2   3E+2   12   12.5  RT (min)      lcqtof_t3_neg_3_032_24    m/z 1018.8384     0   1   2   3E+2   12   12.5  RT (min)      lcqtof_t3_neg_3_032_24    m/z 1018.8390     0   2   4   6   8E+3   13.5   14  RT (min)      lcqtof_t3_neg_1_001_49    m/z 1018.8391     0   2   4   6E+4   12.5   13  RT (min)      lcqtof_t3_neg_2_030_23    m/z 1018.8395     0   2   4   6E+3   14   14.5  RT (min)      lcqtof_t3_neg_1_056_56    m/z 1018.8397     0   1   2E+4   13.5   14  RT (min)      lcqtof_t3_neg_1_011_45    m/z 1018.8398     0   2   4   6E+2   11   11.5  RT (min)      lcqtof_t3_neg_3_038_06    m/z 1018.8401     0   1   2   3   4E+4   12.5   13  RT (min)      lcqtof_t3_neg_1_019_23    m/z 1018.8405     0   2   4   6   8E+3   16   16.5  RT (min)      lcqtof_t3_neg_2_059_14    m/z 1018.8413     0   2   4   6E+2   10.5   11  RT (min)      lcqtof_t3_neg_3_038_06    m/z 1018.8444     0   1   2E+2   12   12.5  RT (min)      lcqtof_t3_neg_3_032_24    m/z 1018.8452     0   1   2   3E+2   11   11.5  RT (min)      lcqtof_t3_neg_3_038_06    m/z 1018.8481     0   1   2   3E+2   11   11.5  RT (min)      lcqtof_t3_neg_3_038_06    m/z 1019.2218     0   1E+4   12.5   13  RT (min)      lcqtof_t3_neg_4_002_17    m/z 1019.2292     0   1   2E+4   12.5   13  RT (min)      lcqtof_t3_neg_4_064_17    m/z 1019.2316     0   1E+4   12.5   13  RT (min)      lcqtof_t3_neg_4_002_17    m/z 1019.2333     0   1E+4   12.5   13  RT (min)      lcqtof_t3_neg_4_002_17    m/z 1019.4506     0   1E+4   9.5   10  RT (min)      lcqtof_t3_neg_4_016_14    m/z 1019.8137     0   1E+4   1   1.5  RT (min)      lcqtof_t3_neg_2_042_13    m/z 1019.8347     0   1   2   3E+2   12   12.5  RT (min)      lcqtof_t3_neg_3_032_24    m/z 1019.8359     0   2   4E+4   12.5   13  RT (min)      lcqtof_t3_neg_2_030_23    m/z 1019.8364     0   1   2   3E+2   12   12.5  RT (min)      lcqtof_t3_neg_3_032_24    m/z 1019.8378     0   2   4   6   8E+3   13.5   14  RT (min)      lcqtof_t3_neg_1_044_37    m/z 1019.8381     0   2   4E+3   14   14.5  RT (min)      lcqtof_t3_neg_1_061_50    m/z 1019.8383     0   1   2   3   4E+2   11   11.5  RT (min)      lcqtof_t3_neg_3_038_06    m/z 1019.8385     0   2   4E+3   14   14.5  RT (min)      lcqtof_t3_neg_1_061_50    m/z 1019.8387     0   1   2   3   4E+2   10.5   11  RT (min)      lcqtof_t3_neg_3_038_06    m/z 1019.8388     0   1   2   3   4E+4   12.5   13  RT (min)      lcqtof_t3_neg_2_016_05    m/z 1019.8390     0   1   2   3E+4   13.5   14  RT (min)      lcqtof_t3_neg_2_020_46    m/z 1020.2361     0   1   2E+4   12.5   13  RT (min)      lcqtof_t3_neg_4_064_17    m/z 1020.6697     0   1E+4   13   13.5  RT (min)      lcqtof_t3_neg_4_018_23    m/z 1020.8374     0   1   2E+4   12.5   13  RT (min)      lcqtof_t3_neg_2_030_23    m/z 1020.8374     0   1E+4   12.5   13  RT (min)      lcqtof_t3_neg_2_030_23    m/z 1020.8380     0   1   2   3E+2   12   12.5  RT (min)      lcqtof_t3_neg_3_032_24    m/z 1020.8380     0   1   2E+3   16   16.5  RT (min)      lcqtof_t3_neg_1_000_cc    m/z 1020.8386     0   1   2   3   4E+2   11   11.5  RT (min)      lcqtof_t3_neg_3_038_06    m/z 1020.8390     0   1   2E+3   14.5   15  RT (min)      lcqtof_t3_neg_2_051_14    m/z 1020.8390     0   1E+4   12.5   13  RT (min)      lcqtof_t3_neg_2_016_05    m/z 1020.8391     0   1   2E+3   16   16.5  RT (min)      lcqtof_t3_neg_2_048_30    m/z 1020.8391     0   1   2E+3   15   15.5  RT (min)      lcqtof_t3_neg_1_022_28    m/z 1021.2114     0   1E+4   12.5   13  RT (min)       lcqtof_t3_neg_3_113_62    m/z 1021.2274     0   1E+4   12.5   13  RT (min)      lcqtof_t3_neg_4_002_17    m/z 1021.2276     0   1E+4   12.5   13  RT (min)      lcqtof_t3_neg_4_002_17    m/z 1021.2279     0   1E+4   12.5   13  RT (min)      lcqtof_t3_neg_4_002_17    m/z 1021.2294     0   1   2   3E+4   12.5   13  RT (min)      lcqtof_t3_neg_4_064_17    m/z 1021.6993     0   1E+4   10   10.5  RT (min)       lcqtof_t3_neg_4_019_05    m/z 1022.2202     0   1E+4   12.5   13  RT (min)      lcqtof_t3_neg_4_002_17    m/z 1022.2310     0   1   2E+4   12.5   13  RT (min)      lcqtof_t3_neg_4_064_17    m/z 1022.6586     0   1   2   3   4E+2   11.5   12  RT (min)      lcqtof_t3_neg_3_032_24    m/z 1023.2032     0   1E+4   12.5   13  RT (min)      lcqtof_t3_neg_3_113_62    m/z 1023.2185     0   1E+4   12.5   13  RT (min)      lcqtof_t3_neg_4_064_17    m/z 1024.8855     0   1   2   3E+4   12.5   13  RT (min)      lcqtof_t3_neg_2_030_23    m/z 1024.8855     0   1   2E+2   12   12.5  RT (min)      lcqtof_t3_neg_3_032_24    m/z 1024.8856     0   1   2   3E+4   12.5   13  RT (min)      lcqtof_t3_neg_2_016_05    m/z 1024.8862     0   1   2   3E+4   12.5   13  RT (min)      lcqtof_t3_neg_2_030_23    m/z 1024.8866     0   1   2E+2   12   12.5  RT (min)      lcqtof_t3_neg_3_032_24    m/z 1024.8873     0   1   2   3   4E+2   11   11.5  RT (min)      lcqtof_t3_neg_3_038_06    m/z 1024.8887     0   1   2   3   4E+2   10.5   11  RT (min)      lcqtof_t3_neg_3_038_06    m/z 1025.1607     0   2   4E+4   14   14.5  RT (min)      lcqtof_t3_neg_4_004_20    m/z 1025.1610     0   2   4   6E+5   13   13.5  RT (min)      lcqtof_t3_neg_4_064_17    m/z 1025.1614     0   1   2E+5   14   14.5  RT (min)      lcqtof_t3_neg_4_064_17    m/z 1025.1615     0   2   4   6E+5   13.5   14  RT (min)       lcqtof_t3_neg_4_064_17    m/z 1025.1616     0   2   4   6E+5   13   13.5  RT (min)       lcqtof_t3_neg_4_064_17    m/z 1025.1622     0   1   2   3E+5   14   14.5  RT (min)      lcqtof_t3_neg_4_002_17    m/z 1025.7963     0   1   2   3   4E+3   12.5   13  RT (min)      lcqtof_t3_neg_2_023_37    m/z 1025.7965     0   1   2   3   4E+3   12.5   13  RT (min)      lcqtof_t3_neg_2_023_37    m/z 1025.8833     0   1E+4   12.5   13  RT (min)      lcqtof_t3_neg_2_016_05    m/z 1025.8851     0   1   2   3E+2   12   12.5  RT (min)      lcqtof_t3_neg_3_032_24    m/z 1025.8873     0   1   2E+4   12.5   13  RT (min)      lcqtof_t3_neg_2_030_23    m/z 1025.8887     0   1   2   3E+2   12   12.5  RT (min)      lcqtof_t3_neg_3_032_24    m/z 1025.8890     0   1   2   3   4E+2   11   11.5  RT (min)      lcqtof_t3_neg_3_038_06    m/z 1025.8894     0   1   2E+4   12.5   13  RT (min)      lcqtof_t3_neg_2_030_23    m/z 1026.1617     0   1   2   3E+5   14   14.5  RT (min)      lcqtof_t3_neg_4_002_17    m/z 1026.1619     0   2   4E+4   13.5   14  RT (min)      lcqtof_t3_neg_4_004_20    m/z 1026.1619     0   2   4E+5   13.5   14  RT (min)       lcqtof_t3_neg_4_064_17    m/z 1026.1621     0   2   4E+5   13   13.5  RT (min)       lcqtof_t3_neg_4_064_17    m/z 1026.1622     0   1   2   3   4E+5   13   13.5  RT (min)       lcqtof_t3_neg_4_002_17    m/z 1026.1625     0   2   4E+5   13   13.5  RT (min)      lcqtof_t3_neg_4_064_17    m/z 1026.1635     0   1   2E+4   13   13.5  RT (min)      lcqtof_t3_neg_4_003_37    m/z 1026.4922     0   1E+4   12   12.5  RT (min)      lcqtof_t3_neg_4_018_23    m/z 1026.8418     0   2   4E+3   13.5   14  RT (min)      lcqtof_t3_neg_4_011_32    m/z 1026.8489     0   2   4   6E+3   13.5   14  RT (min)      lcqtof_t3_neg_4_040_24    m/z 1026.8508     0   2   4   6E+3   13.5   14  RT (min)      lcqtof_t3_neg_4_040_24    m/z 1027.1578     0   2   4E+4   14   14.5  RT (min)      lcqtof_t3_neg_4_004_20    m/z 1027.1594     0   1   2   3   4E+5   13   13.5  RT (min)       lcqtof_t3_neg_4_002_17    m/z 1027.1595     0   2   4E+4   14   14.5  RT (min)       lcqtof_t3_neg_4_004_20    m/z 1027.1597     0   1   2   3E+5   14   14.5  RT (min)      lcqtof_t3_neg_4_036_47    m/z 1027.1598     0   2   4E+5   13.5   14  RT (min)       lcqtof_t3_neg_4_064_17    m/z 1027.1599     0   2   4E+5   13   13.5  RT (min)       lcqtof_t3_neg_4_064_17    m/z 1027.1610     0   2   4E+5   13   13.5  RT (min)      lcqtof_t3_neg_4_064_17    m/z 1027.8640     0   1   2   3   4E+2   12   12.5  RT (min)      lcqtof_t3_neg_3_032_24    m/z 1027.8643     0   2   4   6E+2   12   12.5  RT (min)      lcqtof_t3_neg_3_032_24    m/z 1027.8646     0   1E+4   13.5   14  RT (min)       lcqtof_t3_neg_1_006_05    m/z 1027.8650     0   1E+4   14   14.5  RT (min)      lcqtof_t3_neg_1_015_34    m/z 1027.8651     0   1E+4   14.5   15  RT (min)       lcqtof_t3_neg_1_012_16    m/z 1027.8652     0   1   2E+5   12.5   13  RT (min)      lcqtof_t3_neg_2_016_05    m/z 1027.8652     0   1E+4   14   14.5  RT (min)       lcqtof_t3_neg_1_015_34    m/z 1027.8654     0   1E+4   16   16.5  RT (min)      lcqtof_t3_neg_1_000_cc    m/z 1027.8655     0   1E+4   16   16.5  RT (min)      lcqtof_t3_neg_1_030_42    m/z 1027.8655     0   1   2E+5   12.5   13  RT (min)      lcqtof_t3_neg_2_030_23    m/z 1027.8655     0   1   2E+5   12.5   13  RT (min)      lcqtof_t3_neg_2_016_05    m/z 1027.8656     0   1E+4   16   16.5  RT (min)      lcqtof_t3_neg_1_000_cc    m/z 1027.8656     0   1E+4   13.5   14  RT (min)       lcqtof_t3_neg_1_006_05    m/z 1027.8657     0   2   4E+2   11   11.5  RT (min)      lcqtof_t3_neg_3_038_06    m/z 1027.8657     0   1E+4   14.5   15  RT (min)       lcqtof_t3_neg_1_007_03    m/z 1027.8658     0   1E+4   14   14.5  RT (min)       lcqtof_t3_neg_1_061_50    m/z 1027.8662     0   1E+5   13.5   14  RT (min)      lcqtof_t3_neg_1_042_46    m/z 1027.8675     0   1E+5   13.5   14  RT (min)      lcqtof_t3_neg_1_042_46    m/z 1027.8679     0   2   4E+2   10.5   11  RT (min)      lcqtof_t3_neg_3_038_06    m/z 1028.1540     0   1   2   3E+4   14   14.5  RT (min)      lcqtof_t3_neg_4_004_20    m/z 1028.1574     0   1   2E+5   14   14.5  RT (min)       lcqtof_t3_neg_4_036_47    m/z 1028.1587     0   1   2   3E+4   13.5   14  RT (min)      lcqtof_t3_neg_4_063_39    m/z 1028.1593     0   1   2   3E+5   13   13.5  RT (min)      lcqtof_t3_neg_4_064_17    m/z 1028.1593     0   1   2   3E+5   13   13.5  RT (min)       lcqtof_t3_neg_4_064_17    m/z 1028.1596     0   1   2   3E+5   13.5   14  RT (min)       lcqtof_t3_neg_4_064_17    m/z 1028.1596     0   1   2   3E+5   13   13.5  RT (min)      lcqtof_t3_neg_4_064_17    m/z 1028.1600     0   1   2   3E+4   13.5   14  RT (min)      lcqtof_t3_neg_4_022_28    m/z 1028.1630     0   1E+4   14   14.5  RT (min)      lcqtof_t3_neg_4_013_39    m/z 1028.8617     0   2   4E+4   13.5   14  RT (min)       lcqtof_t3_neg_2_000_aa    m/z 1028.8630     0   2   4E+4   13.5   14  RT (min)       lcqtof_t3_neg_2_051_14    m/z 1028.8639     0   1   2   3   4E+4   13.5   14  RT (min)      lcqtof_t3_neg_1_005_18    m/z 1028.8652     0   2   4E+4   13.5   14  RT (min)      lcqtof_t3_neg_2_036_06    m/z 1028.8665     0   1E+4   16.5   17  RT (min)      lcqtof_t3_neg_1_035_31    m/z 1028.8670     0   1   2   3   4E+4   14   14.5  RT (min)       lcqtof_t3_neg_1_009_50    m/z 1028.8673     0   1E+4   16   16.5  RT (min)      lcqtof_t3_neg_1_000_cc    m/z 1028.8673     0   1E+4   16   16.5  RT (min)      lcqtof_t3_neg_1_007_03    m/z 1028.8676     0   1   2   3   4E+4   14   14.5  RT (min)       lcqtof_t3_neg_2_053_38    m/z 1028.8677     0   1E+5   12.5   13  RT (min)      lcqtof_t3_neg_2_030_23    m/z 1028.8680     0   1E+4   16   16.5  RT (min)      lcqtof_t3_neg_1_013_09    m/z 1028.8680     0   1E+5   12.5   13  RT (min)      lcqtof_t3_neg_2_030_23    m/z 1028.8684     0   1E+5   13.5   14  RT (min)      lcqtof_t3_neg_2_016_05    m/z 1028.8699     0   1E+5   13.5   14  RT (min)      lcqtof_t3_neg_2_055_16    m/z 1028.8701     0   1E+4   15.5   16  RT (min)      lcqtof_t3_neg_1_063_39    m/z 1029.0151     0   1E+4   13   13.5  RT (min)      lcqtof_t3_neg_4_064_17    m/z 1029.0171     0   1E+4   13   13.5  RT (min)      lcqtof_t3_neg_4_064_17    m/z 1029.0197     0   1E+4   13   13.5  RT (min)      lcqtof_t3_neg_4_064_17    m/z 1029.1386     0   1E+4   1   1.5  RT (min)      lcqtof_t3_neg_4_059_14    m/z 1029.1491     0   1E+5   13   13.5  RT (min)      lcqtof_t3_neg_4_002_17    m/z 1029.1562     0   1E+5   14   14.5  RT (min)      lcqtof_t3_neg_4_064_17    m/z 1029.1574     0   1   2E+4   13   13.5  RT (min)      lcqtof_t3_neg_4_022_28    m/z 1029.1577     0   1   2E+5   13   13.5  RT (min)      lcqtof_t3_neg_4_064_17    m/z 1029.8674     0   1   2E+2   11.5   12  RT (min)      lcqtof_t3_neg_3_038_06    m/z 1029.8687     0   2   4E+3   16   16.5  RT (min)      lcqtof_t3_neg_2_047_28    m/z 1029.8688     0   1   2E+2   12   12.5  RT (min)      lcqtof_t3_neg_3_032_24    m/z 1029.8690     0   2   4   6   8E+3   14.5   15  RT (min)      lcqtof_t3_neg_1_004_11    m/z 1029.8691     0   1   2   3   4E+2   12   12.5  RT (min)      lcqtof_t3_neg_3_032_24    m/z 1029.8696     0   1   2E+4   14   14.5  RT (min)      lcqtof_t3_neg_1_016_48    m/z 1029.8697     0   2   4E+3   14.5   15  RT (min)      lcqtof_t3_neg_1_043_38    m/z 1029.8699     0   2   4E+4   12.5   13  RT (min)      lcqtof_t3_neg_2_030_23    m/z 1029.8704     0   1   2E+4   14   14.5  RT (min)      lcqtof_t3_neg_1_024_10    m/z 1029.8705     0   2   4E+3   15.5   16  RT (min)      lcqtof_t3_neg_1_006_05    m/z 1029.8706     0   2   4   6E+3   14.5   15  RT (min)      lcqtof_t3_neg_1_005_18    m/z 1029.8711     0   1   2   3   4E+4   12.5   13  RT (min)      lcqtof_t3_neg_2_016_05    m/z 1029.8713     0   1   2   3   4E+4   12.5   13  RT (min)      lcqtof_t3_neg_2_016_05    m/z 1029.8725     0   1   2   3E+2   12   12.5  RT (min)      lcqtof_t3_neg_3_032_24    m/z 1029.8727     0   1   2   3   4E+4   13.5   14  RT (min)      lcqtof_t3_neg_2_016_05    m/z 1029.8732     0   1   2   3   4E+2   11   11.5  RT (min)      lcqtof_t3_neg_3_038_06    m/z 1029.8737     0   1   2   3   4E+2   10.5   11  RT (min)      lcqtof_t3_neg_3_038_06    m/z 1030.0160     0   1E+4   13   13.5  RT (min)      lcqtof_t3_neg_4_064_17    m/z 1030.1558     0   1E+5   13   13.5  RT (min)      lcqtof_t3_neg_4_064_17    m/z 1030.8724     0   1E+4   12.5   13  RT (min)      lcqtof_t3_neg_2_030_23    m/z 1031.1475     0   1   2   3E+4   13.5   14  RT (min)      lcqtof_t3_neg_4_002_17    m/z 1031.1489     0   1   2   3   4E+4   13.5   14  RT (min)      lcqtof_t3_neg_4_064_17    m/z 1031.1492     0   1   2   3   4E+4   13.5   14  RT (min)      lcqtof_t3_neg_4_064_17    m/z 1031.1533     0   1   2   3E+4   13.5   14  RT (min)      lcqtof_t3_neg_4_002_17    m/z 1031.1540     0   1   2   3E+4   13   13.5  RT (min)      lcqtof_t3_neg_4_002_17    m/z 1031.1543     0   1   2   3   4E+4   13   13.5  RT (min)      lcqtof_t3_neg_4_064_17    m/z 1031.1549     0   1   2   3   4E+4   13.5   14  RT (min)      lcqtof_t3_neg_4_064_17    m/z 1031.1571     0   1   2   3E+4   13.5   14  RT (min)      lcqtof_t3_neg_4_002_17    m/z 1031.2501     0   1E+4   13   13.5  RT (min)      lcqtof_t3_neg_4_036_47    m/z 1031.2504     0   1   2E+4   13   13.5  RT (min)      lcqtof_t3_neg_4_064_17    m/z 1032.1533     0   1E+4   13   13.5  RT (min)      lcqtof_t3_neg_4_064_17    m/z 1032.1566     0   1E+4   13.5   14  RT (min)      lcqtof_t3_neg_4_064_17    m/z 1037.2025     0   1E+4   13.5   14  RT (min)      lcqtof_t3_neg_4_036_47    m/z 1038.1986     0   1E+4   13.5   14  RT (min)      lcqtof_t3_neg_4_064_17    m/z 1038.3330     0   1E+4   4   4.5  RT (min)      lcqtof_t3_neg_4_000_ee    m/z 1038.8351     0   1E+4   4   4.5  RT (min)      lcqtof_t3_neg_4_000_ee    m/z 1039.2081     0   1E+4   13.5   14  RT (min)      lcqtof_t3_neg_4_002_17    m/z 1039.2145     0   1E+4   13.5   14  RT (min)      lcqtof_t3_neg_4_036_47    m/z 1039.3379     0   1E+4   4   4.5  RT (min)      lcqtof_t3_neg_4_000_ee    m/z 1041.0827     0   1   2   3E+5   14   14.5  RT (min)      lcqtof_t3_neg_4_064_17    m/z 1041.0828     0   1   2   3E+5   14   14.5  RT (min)      lcqtof_t3_neg_4_064_17    m/z 1041.0838     0   1   2E+4   13.5   14  RT (min)      lcqtof_t3_neg_4_013_39    m/z 1041.0838     0   1   2E+4   13.5   14  RT (min)      lcqtof_t3_neg_4_013_39    m/z 1041.0840     0   1   2E+5   13.5   14  RT (min)      lcqtof_t3_neg_4_002_17    m/z 1041.0840     0   1   2E+5   13   13.5  RT (min)      lcqtof_t3_neg_4_064_17    m/z 1041.0841     0   1   2   3E+5   13.5   14  RT (min)      lcqtof_t3_neg_4_064_17    m/z 1041.0841     0   1   2E+5   13.5   14  RT (min)      lcqtof_t3_neg_4_002_17    m/z 1041.0845     0   1E+5   14   14.5  RT (min)       lcqtof_t3_neg_4_036_47    m/z 1041.0856     0   1E+5   13   13.5  RT (min)      lcqtof_t3_neg_4_064_17    m/z 1042.0830     0   1   2   3E+5   14   14.5  RT (min)      lcqtof_t3_neg_4_064_17    m/z 1042.0831     0   1   2   3E+5   14   14.5  RT (min)      lcqtof_t3_neg_4_064_17    m/z 1042.0831     0   1   2E+4   14   14.5  RT (min)      lcqtof_t3_neg_4_013_39    m/z 1042.0833     0   2   4E+4   13   13.5  RT (min)      lcqtof_t3_neg_4_036_47    m/z 1042.0835     0   2   4   6E+4   13.5   14  RT (min)      lcqtof_t3_neg_4_004_20    m/z 1042.0838     0   1   2   3E+5   13.5   14  RT (min)       lcqtof_t3_neg_4_064_17    m/z 1042.0839     0   1   2E+5   13.5   14  RT (min)      lcqtof_t3_neg_4_002_17    m/z 1042.0844     0   1   2   3E+5   13.5   14  RT (min)      lcqtof_t3_neg_4_064_17    m/z 1042.0846     0   2   4   6E+4   13.5   14  RT (min)      lcqtof_t3_neg_4_004_20    m/z 1042.0846     0   1E+5   14   14.5  RT (min)       lcqtof_t3_neg_4_036_47    m/z 1042.0850     0   2   4   6E+4   14.5   15  RT (min)      lcqtof_t3_neg_4_004_20    m/z 1042.0855     0   1E+5   13   13.5  RT (min)      lcqtof_t3_neg_4_064_17    m/z 1043.0819     0   1   2   3E+5   14   14.5  RT (min)      lcqtof_t3_neg_4_064_17    m/z 1043.0819     0   1   2   3E+5   13.5   14  RT (min)      lcqtof_t3_neg_4_064_17    m/z 1043.0821     0   1   2   3E+5   13.5   14  RT (min)      lcqtof_t3_neg_4_064_17    m/z 1043.0822     0   1   2   3E+5   14   14.5  RT (min)      lcqtof_t3_neg_4_064_17    m/z 1043.0828     0   1   2E+5   14   14.5  RT (min)       lcqtof_t3_neg_4_002_17    m/z 1043.0829     0   1E+5   13   13.5  RT (min)      lcqtof_t3_neg_4_064_17    m/z 1043.1371     0   1E+4   13   13.5  RT (min)      lcqtof_t3_neg_4_008_35    m/z 1043.1376     0   1E+4   13   13.5  RT (min)      lcqtof_t3_neg_4_008_35    m/z 1043.2434     0   1   2E+4   13   13.5  RT (min)      lcqtof_t3_neg_3_113_62    m/z 1044.0806     0   1E+5   14   14.5  RT (min)      lcqtof_t3_neg_4_002_17    m/z 1044.0806     0   1E+5   13.5   14  RT (min)      lcqtof_t3_neg_4_064_17    m/z 1044.0808     0   1E+5   14   14.5  RT (min)      lcqtof_t3_neg_4_002_17    m/z 1044.0809     0   2   4   6   8E+4   14   14.5  RT (min)      lcqtof_t3_neg_4_036_47    m/z 1044.0813     0   1E+5   14   14.5  RT (min)      lcqtof_t3_neg_4_064_17    m/z 1044.0814     0   2   4   6   8E+4   13   13.5  RT (min)      lcqtof_t3_neg_4_064_17    m/z 1044.0818     0   1E+5   13   13.5  RT (min)      lcqtof_t3_neg_4_064_17    m/z 1044.0820     0   1E+5   13.5   14  RT (min)      lcqtof_t3_neg_4_064_17    m/z 1044.0831     0   1E+5   14   14.5  RT (min)      lcqtof_t3_neg_4_002_17    m/z 1044.0843     0   1E+5   13   13.5  RT (min)      lcqtof_t3_neg_4_064_17    m/z 1044.7441     0   1E+4   1   1.5  RT (min)      lcqtof_t3_neg_1_036_14    m/z 1044.7745     0   1E+4   10   10.5  RT (min)       lcqtof_t3_neg_4_019_05    m/z 1044.8529     0   1E+5   12.5   13  RT (min)      lcqtof_t3_neg_3_006_23    m/z 1044.8546     0   2   4E+2   12   12.5  RT (min)      lcqtof_t3_neg_3_032_24    m/z 1044.8547     0   1   2   3   4E+2   12   12.5  RT (min)      lcqtof_t3_neg_3_032_24    m/z 1044.8551     0   1   2E+4   16.5   17  RT (min)      lcqtof_t3_neg_2_053_38    m/z 1044.8551     0   1   2E+4   16   16.5  RT (min)      lcqtof_t3_neg_1_006_05    m/z 1044.8553     0   1   2E+5   12.5   13  RT (min)      lcqtof_t3_neg_2_030_23    m/z 1044.8553     0   1E+4   16.5   17  RT (min)      lcqtof_t3_neg_2_065_51    m/z 1044.8553     0   1   2E+4   16   16.5  RT (min)      lcqtof_t3_neg_1_009_50    m/z 1044.8556     0   1E+4   13.5   14  RT (min)      lcqtof_t3_neg_1_055_32    m/z 1044.8558     0   1   2E+4   13.5   14  RT (min)      lcqtof_t3_neg_1_055_32    m/z 1044.8566     0   1   2   3   4E+2   11   11.5  RT (min)      lcqtof_t3_neg_3_038_06    m/z 1044.8575     0   1   2   3   4E+2   10.5   11  RT (min)      lcqtof_t3_neg_3_038_06    m/z 1045.0775     0   2   4   6   8E+4   13   13.5  RT (min)      lcqtof_t3_neg_4_064_17    m/z 1045.0784     0   1   2E+4   14   14.5  RT (min)      lcqtof_t3_neg_4_004_20    m/z 1045.0795     0   1E+5   13.5   14  RT (min)      lcqtof_t3_neg_4_064_17    m/z 1045.0797     0   1   2E+4   13.5   14  RT (min)      lcqtof_t3_neg_4_004_20    m/z 1045.0799     0   1E+5   14   14.5  RT (min)      lcqtof_t3_neg_4_064_17    m/z 1045.0799     0   2   4E+4   13   13.5  RT (min)      lcqtof_t3_neg_4_064_17    m/z 1045.0799     0   2   4   6   8E+4   14   14.5  RT (min)      lcqtof_t3_neg_4_002_17    m/z 1045.0801     0   1E+5   13   13.5  RT (min)      lcqtof_t3_neg_4_064_17    m/z 1045.0806     0   1E+5   13.5   14  RT (min)      lcqtof_t3_neg_4_064_17    m/z 1045.0816     0   2   4   6   8E+4   14   14.5  RT (min)      lcqtof_t3_neg_4_002_17    m/z 1045.8220     0   1E+4   13   13.5  RT (min)      lcqtof_t3_neg_1_019_23    m/z 1045.8501     0   1E+4   13.5   14  RT (min)      lcqtof_t3_neg_1_056_56    m/z 1045.8573     0   1E+5   12.5   13  RT (min)      lcqtof_t3_neg_2_030_23    m/z 1045.8574     0   1E+4   16   16.5  RT (min)      lcqtof_t3_neg_2_009_50    m/z 1045.8575     0   2   4   6E+3   14   14.5  RT (min)      lcqtof_t3_neg_1_023_06    m/z 1045.8575     0   1E+5   12.5   13  RT (min)      lcqtof_t3_neg_2_016_05    m/z 1045.8578     0   1   2   3E+3   15   15.5  RT (min)      lcqtof_t3_neg_3_032_24    m/z 1045.8580     0   1   2E+4   13.5   14  RT (min)      lcqtof_t3_neg_1_017_51    m/z 1045.8582     0   1E+5   12.5   13  RT (min)      lcqtof_t3_neg_2_016_05    m/z 1045.8583     0   2   4E+2   11   11.5  RT (min)      lcqtof_t3_neg_3_038_06    m/z 1045.8584     0   1E+4   16.5   17  RT (min)      lcqtof_t3_neg_2_009_50    m/z 1045.8592     0   1   2   3   4E+2   12   12.5  RT (min)      lcqtof_t3_neg_3_032_24    m/z 1045.8593     0   2   4E+2   10.5   11  RT (min)      lcqtof_t3_neg_3_038_06    m/z 1045.8595     0   1   2   3E+2   12   12.5  RT (min)      lcqtof_t3_neg_3_032_24    m/z 1046.0778     0   1   2E+4   13   13.5  RT (min)      lcqtof_t3_neg_4_064_17    m/z 1046.0782     0   1   2   3   4E+4   13   13.5  RT (min)      lcqtof_t3_neg_4_064_17    m/z 1046.0782     0   2   4E+4   14   14.5  RT (min)      lcqtof_t3_neg_4_064_17    m/z 1046.0787     0   2   4E+4   14   14.5  RT (min)      lcqtof_t3_neg_4_064_17    m/z 1046.0800     0   2   4E+4   13.5   14  RT (min)      lcqtof_t3_neg_4_064_17    m/z 1046.0803     0   1   2E+4   14   14.5  RT (min)      lcqtof_t3_neg_4_036_47    m/z 1046.0807     0   2   4E+4   13.5   14  RT (min)      lcqtof_t3_neg_4_064_17    m/z 1046.8566     0   1   2   3E+3   14   14.5  RT (min)      lcqtof_t3_neg_1_015_34    m/z 1046.8586     0   2   4E+3   16   16.5  RT (min)      lcqtof_t3_neg_1_011_45    m/z 1046.8591     0   1   2E+3   15   15.5  RT (min)      lcqtof_t3_neg_1_000_dd    m/z 1046.8591     0   1   2E+3   15.5   16  RT (min)      lcqtof_t3_neg_1_019_23    m/z 1046.8591     0   1   2   3E+3   15.5   16  RT (min)      lcqtof_t3_neg_1_040_08    m/z 1046.8601     0   1   2   3   4E+3   16   16.5  RT (min)      lcqtof_t3_neg_1_041_54    m/z 1046.8605     0   1   2E+4   12.5   13  RT (min)      lcqtof_t3_neg_3_006_23    m/z 1046.8605     0   2   4   6E+4   12.5   13  RT (min)      lcqtof_t3_neg_2_030_23    m/z 1046.8606     0   1   2E+3   14.5   15  RT (min)      lcqtof_t3_neg_1_013_09    m/z 1046.8608     0   2   4E+3   16   16.5  RT (min)      lcqtof_t3_neg_1_006_05    m/z 1046.8612     0   1   2   3E+2   12   12.5  RT (min)      lcqtof_t3_neg_3_032_24    m/z 1046.8614     0   2   4E+4   12.5   13  RT (min)      lcqtof_t3_neg_2_016_05    m/z 1046.8620     0   1   2   3E+2   12   12.5  RT (min)      lcqtof_t3_neg_3_032_24    m/z 1046.8639     0   1   2   3   4E+2   11   11.5  RT (min)      lcqtof_t3_neg_3_038_06    m/z 1047.0772     0   1   2E+4   13.5   14  RT (min)      lcqtof_t3_neg_4_064_17    m/z 1047.0796     0   1   2E+4   14   14.5  RT (min)      lcqtof_t3_neg_4_064_17    m/z 1047.0802     0   1   2E+4   14   14.5  RT (min)      lcqtof_t3_neg_4_064_17    m/z 1047.2343     0   1E+4   12.5   13  RT (min)      lcqtof_t3_neg_4_002_17    m/z 1047.2347     0   1   2E+4   12.5   13  RT (min)      lcqtof_t3_neg_4_064_17    m/z 1047.7910     0   2   4   6E+3   13   13.5  RT (min)      lcqtof_t3_neg_2_030_23    m/z 1047.8644     0   1E+4   12.5   13  RT (min)      lcqtof_t3_neg_2_016_05    m/z 1047.8654     0   1E+4   12.5   13  RT (min)      lcqtof_t3_neg_2_030_23    m/z 1048.2336     0   1E+4   12.5   13  RT (min)      lcqtof_t3_neg_4_064_17    m/z 1048.2340     0   1E+4   12.5   13  RT (min)      lcqtof_t3_neg_4_002_17    m/z 1048.2341     0   1E+4   12.5   13  RT (min)      lcqtof_t3_neg_4_002_17    m/z 1049.2355     0   1   2E+4   12.5   13  RT (min)      lcqtof_t3_neg_4_064_17    m/z 1049.2373     0   1E+4   12.5   13  RT (min)      lcqtof_t3_neg_4_002_17    m/z 1049.2446     0   1   2E+4   13   13.5  RT (min)      lcqtof_t3_neg_4_064_17    m/z 1049.2518     0   1E+4   13   13.5  RT (min)      lcqtof_t3_neg_4_002_17    m/z 1049.2524     0   1E+4   13   13.5  RT (min)      lcqtof_t3_neg_4_002_17    m/z 1049.2535     0   1E+4   13   13.5  RT (min)      lcqtof_t3_neg_4_002_17    m/z 1050.2360     0   1   2   3   4E+4   12.5   13  RT (min)      lcqtof_t3_neg_4_064_17    m/z 1050.2485     0   1   2E+4   13   13.5  RT (min)      lcqtof_t3_neg_4_002_17    m/z 1050.2523     0   1   2   3   4E+4   13   13.5  RT (min)      lcqtof_t3_neg_4_064_17    m/z 1051.2411     0   1E+4   4   4.5  RT (min)      lcqtof_t3_neg_4_000_ee    m/z 1052.2299     0   1E+4   13.5   14  RT (min)      lcqtof_t3_neg_4_057_49    m/z 1053.2290     0   1   2   3   4E+4   13.5   14  RT (min)      lcqtof_t3_neg_4_057_49    m/z 1053.8856     0   1E+4   14.5   15  RT (min)      lcqtof_t3_neg_4_057_49    m/z 1054.2289     0   1   2   3   4E+4   13.5   14  RT (min)      lcqtof_t3_neg_4_057_49    m/z 1054.6605     0   1   2   3E+3   10.5   11  RT (min)      lcqtof_t3_neg_1_002_13    m/z 1055.0434     0   1E+5   13.5   14  RT (min)      lcqtof_t3_neg_4_002_17    m/z 1055.0441     0   2   4   6   8E+4   13   13.5  RT (min)      lcqtof_t3_neg_4_064_17    m/z 1055.0445     0   1E+5   13   13.5  RT (min)      lcqtof_t3_neg_4_064_17    m/z 1055.0446     0   1E+5   13   13.5  RT (min)      lcqtof_t3_neg_4_064_17    m/z 1055.0447     0   1E+5   13.5   14  RT (min)      lcqtof_t3_neg_4_064_17    m/z 1055.0449     0   2   4   6E+4   13   13.5  RT (min)      lcqtof_t3_neg_4_002_17    m/z 1055.0450     0   1E+5   13.5   14  RT (min)      lcqtof_t3_neg_4_002_17    m/z 1055.0453     0   1E+5   14   14.5  RT (min)      lcqtof_t3_neg_4_064_17    m/z 1055.0467     0   2   4E+4   14   14.5  RT (min)      lcqtof_t3_neg_4_036_47    m/z 1055.0470     0   2   4   6E+4   14   14.5  RT (min)      lcqtof_t3_neg_4_036_47    m/z 1055.0471     0   1E+5   13.5   14  RT (min)      lcqtof_t3_neg_4_002_17    m/z 1055.0473     0   1E+5   13.5   14  RT (min)      lcqtof_t3_neg_4_002_17    m/z 1055.1336     0   1   2   3E+4   14   14.5  RT (min)      lcqtof_t3_neg_4_036_47    m/z 1055.1341     0   1   2   3E+4   13.5   14  RT (min)      lcqtof_t3_neg_4_036_47    m/z 1055.1341     0   2   4E+4   13   13.5  RT (min)      lcqtof_t3_neg_4_064_17    m/z 1055.1357     0   2   4E+4   13.5   14  RT (min)      lcqtof_t3_neg_4_064_17    m/z 1055.1362     0   1   2   3E+4   14   14.5  RT (min)      lcqtof_t3_neg_4_036_47    m/z 1055.1363     0   2   4E+4   14   14.5  RT (min)      lcqtof_t3_neg_4_064_17    m/z 1055.1400     0   2   4E+4   13.5   14  RT (min)      lcqtof_t3_neg_4_064_17    m/z 1055.2280     0   1   2   3   4E+4   13.5   14  RT (min)      lcqtof_t3_neg_4_057_49    m/z 1055.2282     0   1   2E+4   13.5   14  RT (min)      lcqtof_t3_neg_4_001_49    m/z 1055.2461     0   1E+4   13   13.5  RT (min)      lcqtof_t3_neg_4_002_17    m/z 1055.2461     0   1   2E+4   13   13.5  RT (min)      lcqtof_t3_neg_4_064_17    m/z 1056.0442     0   1   2E+4   13.5   14  RT (min)      lcqtof_t3_neg_4_004_20    m/z 1056.0448     0   1E+5   13.5   14  RT (min)      lcqtof_t3_neg_4_064_17    m/z 1056.0449     0   1E+5   14   14.5  RT (min)      lcqtof_t3_neg_4_064_17    m/z 1056.0451     0   1E+5   13   13.5  RT (min)      lcqtof_t3_neg_4_064_17    m/z 1056.0458     0   1E+5   13.5   14  RT (min)      lcqtof_t3_neg_4_064_17    m/z 1056.0459     0   1E+5   13.5   14  RT (min)      lcqtof_t3_neg_4_064_17    m/z 1056.0460     0   1E+5   13   13.5  RT (min)      lcqtof_t3_neg_4_064_17    m/z 1056.0463     0   2   4   6   8E+4   13   13.5  RT (min)      lcqtof_t3_neg_4_064_17    m/z 1056.0471     0   1E+5   13.5   14  RT (min)      lcqtof_t3_neg_4_002_17    m/z 1056.0476     0   2   4   6E+4   14   14.5  RT (min)      lcqtof_t3_neg_4_036_47    m/z 1056.0486     0   1   2E+4   14   14.5  RT (min)      lcqtof_t3_neg_4_004_20    m/z 1056.0498     0   2   4   6E+4   14   14.5  RT (min)      lcqtof_t3_neg_4_036_47    m/z 1056.1329     0   2   4E+4   13.5   14  RT (min)      lcqtof_t3_neg_4_064_17    m/z 1056.1349     0   2   4E+4   14   14.5  RT (min)      lcqtof_t3_neg_4_064_17    m/z 1056.1356     0   2   4E+4   13.5   14  RT (min)      lcqtof_t3_neg_4_064_17    m/z 1056.1366     0   2   4E+4   13   13.5  RT (min)      lcqtof_t3_neg_4_064_17    m/z 1056.1433     0   1   2   3E+4   13   13.5  RT (min)      lcqtof_t3_neg_4_064_17    m/z 1056.2269     0   1   2E+4   13.5   14  RT (min)      lcqtof_t3_neg_4_057_49    m/z 1056.8892     0   1   2   3E+3   13.5   14  RT (min)      lcqtof_t3_neg_2_000_aa    m/z 1056.9016     0   1   2   3E+3   13   13.5  RT (min)      lcqtof_t3_neg_2_000_aa    m/z 1056.9030     0   1   2   3E+3   13   13.5  RT (min)      lcqtof_t3_neg_2_000_aa    m/z 1057.0429     0   1   2   3E+4   13   13.5  RT (min)      lcqtof_t3_neg_4_002_17    m/z 1057.0432     0   1E+5   13.5   14  RT (min)      lcqtof_t3_neg_4_002_17    m/z 1057.0436     0   1E+5   14   14.5  RT (min)      lcqtof_t3_neg_4_002_17    m/z 1057.0437     0   1E+5   13.5   14  RT (min)      lcqtof_t3_neg_4_002_17    m/z 1057.0439     0   1E+5   13   13.5  RT (min)      lcqtof_t3_neg_4_064_17    m/z 1057.0440     0   1E+5   13   13.5  RT (min)      lcqtof_t3_neg_4_064_17    m/z 1057.0441     0   2   4   6E+4   14   14.5  RT (min)      lcqtof_t3_neg_4_036_47    m/z 1057.0446     0   1E+5   13.5   14  RT (min)      lcqtof_t3_neg_4_064_17    m/z 1057.0446     0   1E+5   14   14.5  RT (min)      lcqtof_t3_neg_4_002_17    m/z 1057.0447     0   2   4   6   8E+4   13   13.5  RT (min)      lcqtof_t3_neg_4_064_17    m/z 1057.0450     0   1E+5   14   14.5  RT (min)      lcqtof_t3_neg_4_064_17    m/z 1057.0457     0   1E+5   13.5   14  RT (min)      lcqtof_t3_neg_4_002_17    m/z 1057.0457     0   1E+5   13.5   14  RT (min)      lcqtof_t3_neg_4_002_17    m/z 1057.1319     0   2   4E+4   13   13.5  RT (min)      lcqtof_t3_neg_4_064_17    m/z 1057.1333     0   2   4E+4   14   14.5  RT (min)      lcqtof_t3_neg_4_064_17    m/z 1057.1334     0   1   2   3   4E+4   14   14.5  RT (min)      lcqtof_t3_neg_4_002_17    m/z 1057.1337     0   1   2   3E+4   13   13.5  RT (min)      lcqtof_t3_neg_4_064_17    m/z 1057.1341     0   2   4E+4   13.5   14  RT (min)      lcqtof_t3_neg_4_064_17    m/z 1057.1343     0   1   2   3E+4   14   14.5  RT (min)      lcqtof_t3_neg_4_036_47    m/z 1057.1350     0   1   2   3   4E+4   13.5   14  RT (min)      lcqtof_t3_neg_4_002_17    m/z 1057.1356     0   1   2   3   4E+4   13.5   14  RT (min)      lcqtof_t3_neg_4_002_17    m/z 1057.1367     0   1   2   3   4E+4   13.5   14  RT (min)      lcqtof_t3_neg_4_002_17    m/z 1057.1409     0   1   2   3   4E+4   13.5   14  RT (min)      lcqtof_t3_neg_4_002_17    m/z 1057.2257     0   1E+4   13.5   14  RT (min)      lcqtof_t3_neg_4_057_49    m/z 1058.0424     0   2   4   6E+4   13   13.5  RT (min)      lcqtof_t3_neg_4_064_17    m/z 1058.0425     0   1E+5   13   13.5  RT (min)      lcqtof_t3_neg_4_064_17    m/z 1058.0430     0   1E+5   14   14.5  RT (min)      lcqtof_t3_neg_4_064_17    m/z 1058.0435     0   2   4   6E+4   13.5   14  RT (min)      lcqtof_t3_neg_4_002_17    m/z 1058.0438     0   2   4   6E+4   13.5   14  RT (min)      lcqtof_t3_neg_4_002_17    m/z 1058.0438     0   1E+5   13   13.5  RT (min)      lcqtof_t3_neg_4_064_17    m/z 1058.0440     0   2   4   6E+4   13.5   14  RT (min)      lcqtof_t3_neg_4_002_17    m/z 1058.0443     0   2   4   6E+4   14   14.5  RT (min)      lcqtof_t3_neg_4_002_17    m/z 1058.0444     0   2   4   6E+4   13   13.5  RT (min)      lcqtof_t3_neg_4_064_17    m/z 1058.0447     0   1E+5   13.5   14  RT (min)      lcqtof_t3_neg_4_064_17    m/z 1058.0451     0   2   4   6E+4   13.5   14  RT (min)      lcqtof_t3_neg_4_002_17    m/z 1058.0469     0   2   4   6E+4   13.5   14  RT (min)      lcqtof_t3_neg_4_002_17    m/z 1058.0472     0   1   2   3   4E+4   14   14.5  RT (min)      lcqtof_t3_neg_4_036_47    m/z 1058.1290     0   1   2E+4   13   13.5  RT (min)      lcqtof_t3_neg_4_064_17    m/z 1058.1343     0   1   2   3E+4   14   14.5  RT (min)      lcqtof_t3_neg_4_064_17    m/z 1058.1344     0   1E+4   13.5   14  RT (min)      lcqtof_t3_neg_4_036_47    m/z 1058.1349     0   1   2   3E+4   13.5   14  RT (min)      lcqtof_t3_neg_4_064_17    m/z 1058.1354     0   1   2E+4   13   13.5  RT (min)      lcqtof_t3_neg_4_064_17    m/z 1058.1366     0   1   2   3E+4   13.5   14  RT (min)      lcqtof_t3_neg_4_064_17    m/z 1059.0385     0   2   4   6E+4   13   13.5  RT (min)      lcqtof_t3_neg_4_064_17    m/z 1059.0426     0   1   2   3   4E+4   13   13.5  RT (min)      lcqtof_t3_neg_4_064_17    m/z 1059.0435     0   2   4   6E+4   13   13.5  RT (min)      lcqtof_t3_neg_4_064_17    m/z 1059.0442     0   2   4   6E+4   14   14.5  RT (min)      lcqtof_t3_neg_4_064_17    m/z 1059.0448     0   1   2E+4   13.5   14  RT (min)      lcqtof_t3_neg_4_036_47    m/z 1059.0454     0   1   2E+4   13.5   14  RT (min)      lcqtof_t3_neg_4_036_47    m/z 1059.0460     0   1   2E+4   14   14.5  RT (min)      lcqtof_t3_neg_4_036_47    m/z 1059.0464     0   1   2E+4   13.5   14  RT (min)      lcqtof_t3_neg_4_036_47    m/z 1059.0467     0   1   2   3   4E+4   13   13.5  RT (min)      lcqtof_t3_neg_4_064_17    m/z 1059.1314     0   1   2E+4   13.5   14  RT (min)      lcqtof_t3_neg_4_064_17    m/z 1059.1321     0   1   2E+4   13.5   14  RT (min)      lcqtof_t3_neg_4_064_17    m/z 1059.1327     0   1E+4   13   13.5  RT (min)      lcqtof_t3_neg_4_064_17    m/z 1059.1471     0   1   2E+4   13   13.5  RT (min)      lcqtof_t3_neg_4_064_17    m/z 1060.0394     0   1   2   3E+4   13   13.5  RT (min)      lcqtof_t3_neg_4_064_17    m/z 1060.0414     0   1   2   3E+4   13.5   14  RT (min)      lcqtof_t3_neg_4_064_17    m/z 1060.0416     0   1   2   3E+4   14   14.5  RT (min)      lcqtof_t3_neg_4_064_17    m/z 1060.0422     0   1   2   3E+4   13.5   14  RT (min)      lcqtof_t3_neg_4_064_17    m/z 1060.0431     0   1   2   3E+4   13   13.5  RT (min)      lcqtof_t3_neg_4_064_17    m/z 1060.0432     0   1E+4   13.5   14  RT (min)      lcqtof_t3_neg_4_036_47    m/z 1060.0446     0   1   2   3E+4   13   13.5  RT (min)      lcqtof_t3_neg_4_064_17    m/z 1061.0380     0   1E+4   13.5   14  RT (min)      lcqtof_t3_neg_4_064_17    m/z 1061.0421     0   1E+4   13.5   14  RT (min)      lcqtof_t3_neg_4_064_17    m/z 1061.7863     0   2   4E+3   16.5   17  RT (min)      lcqtof_t3_neg_1_008_02    m/z 1061.7866     0   2   4E+3   16   16.5  RT (min)      lcqtof_t3_neg_2_051_14    m/z 1061.7870     0   1   2   3   4E+3   16   16.5  RT (min)      lcqtof_t3_neg_2_028_24    m/z 1061.7874     0   1   2   3   4E+3   15   15.5  RT (min)      lcqtof_t3_neg_1_000_aa    m/z 1061.7885     0   1E+4   12.5   13  RT (min)      lcqtof_t3_neg_2_016_05    m/z 1061.7915     0   1   2   3   4E+3   14.5   15  RT (min)      lcqtof_t3_neg_2_051_14    m/z 1061.7919     0   2   4   6E+2   11   11.5  RT (min)      lcqtof_t3_neg_3_038_06    m/z 1061.7937     0   1   2   3E+3   14   14.5  RT (min)      lcqtof_t3_neg_2_061_50    m/z 1061.7940     0   1E+4   12.5   13  RT (min)      lcqtof_t3_neg_2_030_23    m/z 1061.7943     0   1E+4   12.5   13  RT (min)      lcqtof_t3_neg_2_030_23    m/z 1061.7963     0   1E+4   12.5   13  RT (min)      lcqtof_t3_neg_2_016_05    m/z 1062.2383     0   1E+4   12.5   13  RT (min)      lcqtof_t3_neg_4_064_17    m/z 1062.7903     0   1   2   3E+3   15   15.5  RT (min)      lcqtof_t3_neg_1_000_aa    m/z 1062.7903     0   1   2   3E+3   16   16.5  RT (min)      lcqtof_t3_neg_2_058_15    m/z 1062.7906     0   1   2   3E+3   15.5   16  RT (min)      lcqtof_t3_neg_2_051_14    m/z 1062.7922     0   1   2E+3   14.5   15  RT (min)      lcqtof_t3_neg_2_032_39    m/z 1062.7955     0   1   2E+3   14   14.5  RT (min)      lcqtof_t3_neg_1_018_33    m/z 1062.7969     0   1E+4   12.5   13  RT (min)      lcqtof_t3_neg_2_030_23    m/z 1062.7971     0   1E+4   12.5   13  RT (min)      lcqtof_t3_neg_2_016_05    m/z 1063.2322     0   1E+4   12.5   13  RT (min)      lcqtof_t3_neg_4_002_17    m/z 1063.2328     0   1   2E+4   12.5   13  RT (min)      lcqtof_t3_neg_4_064_17    m/z 1063.2332     0   1E+4   12.5   13  RT (min)      lcqtof_t3_neg_4_002_17    m/z 1063.6709     0   1E+4   9.5   10  RT (min)      lcqtof_t3_neg_4_019_05    m/z 1063.7865     0   2   4E+3   16   16.5  RT (min)      lcqtof_t3_neg_2_063_16    m/z 1063.7870     0   1   2   3E+3   14.5   15  RT (min)      lcqtof_t3_neg_1_024_10    m/z 1063.7874     0   1E+4   12.5   13  RT (min)      lcqtof_t3_neg_2_030_23    m/z 1063.7876     0   1   2   3   4E+3   14.5   15  RT (min)      lcqtof_t3_neg_1_011_45    m/z 1063.7882     0   1E+4   12.5   13  RT (min)      lcqtof_t3_neg_2_016_05    m/z 1063.7883     0   2   4E+2   11   11.5  RT (min)      lcqtof_t3_neg_3_038_06    m/z 1063.7896     0   1E+4   13   13.5  RT (min)      lcqtof_t3_neg_2_046_45    m/z 1063.7897     0   1   2   3E+3   14   14.5  RT (min)      lcqtof_t3_neg_1_018_33    m/z 1063.7899     0   1E+4   13   13.5  RT (min)      lcqtof_t3_neg_1_001_49    m/z 1063.7905     0   2   4   6E+3   13.5   14  RT (min)      lcqtof_t3_neg_2_016_05    m/z 1063.7908     0   1E+4   13.5   14  RT (min)      lcqtof_t3_neg_3_038_06    m/z 1064.2279     0   1E+4   12.5   13  RT (min)      lcqtof_t3_neg_4_002_17    m/z 1064.2327     0   1   2E+4   12.5   13  RT (min)      lcqtof_t3_neg_4_064_17    m/z 1064.2339     0   1   2E+4   12.5   13  RT (min)      lcqtof_t3_neg_4_064_17    m/z 1064.7891     0   1   2   3E+3   14.5   15  RT (min)      lcqtof_t3_neg_2_045_04    m/z 1064.7892     0   1   2   3E+3   16   16.5  RT (min)      lcqtof_t3_neg_2_062_17    m/z 1064.7896     0   1E+4   16   16.5  RT (min)      lcqtof_t3_neg_1_011_45    m/z 1064.7897     0   1   2E+3   14.5   15  RT (min)      lcqtof_t3_neg_2_041_54    m/z 1064.7902     0   1   2E+3   14.5   15  RT (min)      lcqtof_t3_neg_1_019_23    m/z 1064.7904     0   1   2   3E+3   15.5   16  RT (min)      lcqtof_t3_neg_1_009_50    m/z 1064.7906     0   1E+4   12.5   13  RT (min)      lcqtof_t3_neg_2_030_23    m/z 1064.7906     0   1   2E+3   14.5   15  RT (min)      lcqtof_t3_neg_2_030_23    m/z 1064.8090     0   1E+4   17   17.5  RT (min)      lcqtof_t3_neg_4_035_08    m/z 1065.2473     0   1   2   3E+4   13   13.5  RT (min)      lcqtof_t3_neg_4_064_17    m/z 1066.2465     0   1   2E+4   13   13.5  RT (min)      lcqtof_t3_neg_4_064_17    m/z 1066.2500     0   1E+4   13   13.5  RT (min)      lcqtof_t3_neg_4_002_17    m/z 1067.2482     0   1   2E+4   13   13.5  RT (min)      lcqtof_t3_neg_4_064_17    m/z 1067.2811     0   1E+4   13.5   14  RT (min)      lcqtof_t3_neg_4_064_17    m/z 1068.2476     0   1E+4   13   13.5  RT (min)      lcqtof_t3_neg_4_064_17    m/z 1068.2809     0   1E+4   13.5   14  RT (min)      lcqtof_t3_neg_4_064_17    m/z 1069.0955     0   1   2   3E+4   14   14.5  RT (min)      lcqtof_t3_neg_4_064_17    m/z 1069.0961     0   1   2   3E+4   13   13.5  RT (min)      lcqtof_t3_neg_4_064_17    m/z 1069.0964     0   1   2   3E+4   14   14.5  RT (min)      lcqtof_t3_neg_4_064_17    m/z 1069.0969     0   1   2   3E+4   14   14.5  RT (min)      lcqtof_t3_neg_4_064_17    m/z 1069.0977     0   1   2   3E+4   13.5   14  RT (min)      lcqtof_t3_neg_4_064_17    m/z 1069.0980     0   1   2   3E+4   13.5   14  RT (min)      lcqtof_t3_neg_4_064_17    m/z 1069.0980     0   1   2   3E+4   13.5   14  RT (min)      lcqtof_t3_neg_4_064_17    m/z 1069.0981     0   1   2   3E+4   14   14.5  RT (min)      lcqtof_t3_neg_4_064_17    m/z 1069.0984     0   1   2   3E+4   14   14.5  RT (min)      lcqtof_t3_neg_4_064_17    m/z 1069.0988     0   1   2   3E+4   13   13.5  RT (min)      lcqtof_t3_neg_4_064_17    m/z 1069.1010     0   1   2   3E+4   13   13.5  RT (min)      lcqtof_t3_neg_4_064_17    m/z 1069.2793     0   1E+4   13.5   14  RT (min)      lcqtof_t3_neg_4_064_17    m/z 1070.0953     0   1   2   3E+4   13.5   14  RT (min)      lcqtof_t3_neg_4_064_17    m/z 1070.0956     0   1   2   3E+4   14   14.5  RT (min)      lcqtof_t3_neg_4_064_17    m/z 1070.0960     0   1   2   3E+4   13.5   14  RT (min)      lcqtof_t3_neg_4_064_17    m/z 1070.0977     0   1   2   3E+4   13.5   14  RT (min)      lcqtof_t3_neg_4_064_17    m/z 1070.0983     0   1   2   3E+4   14   14.5  RT (min)      lcqtof_t3_neg_4_064_17    m/z 1070.0991     0   1   2   3E+4   14   14.5  RT (min)      lcqtof_t3_neg_4_064_17    m/z 1070.0992     0   1   2   3E+4   13   13.5  RT (min)      lcqtof_t3_neg_4_064_17    m/z 1070.1005     0   1   2   3E+4   13   13.5  RT (min)      lcqtof_t3_neg_4_064_17    m/z 1070.1010     0   1   2   3E+4   13.5   14  RT (min)      lcqtof_t3_neg_4_064_17    m/z 1070.6892     0   1   2E+4   13   13.5  RT (min)      lcqtof_t3_neg_4_034_34    m/z 1070.7106     0   1E+4   13   13.5  RT (min)      lcqtof_t3_neg_4_018_23    m/z 1070.7108     0   2   4   6   8E+3   12.5   13  RT (min)      lcqtof_t3_neg_2_030_23    m/z 1070.7118     0   1E+4   13   13.5  RT (min)      lcqtof_t3_neg_4_018_23    m/z 1071.0957     0   1   2   3E+4   14   14.5  RT (min)      lcqtof_t3_neg_4_064_17    m/z 1071.0965     0   2   4E+4   13.5   14  RT (min)      lcqtof_t3_neg_4_064_17    m/z 1071.0965     0   2   4E+4   14   14.5  RT (min)      lcqtof_t3_neg_4_064_17    m/z 1071.0985     0   2   4E+4   13   13.5  RT (min)      lcqtof_t3_neg_4_064_17    m/z 1071.0986     0   1   2   3   4E+4   13   13.5  RT (min)      lcqtof_t3_neg_4_002_17    m/z 1071.1020     0   2   4E+4   13.5   14  RT (min)      lcqtof_t3_neg_4_064_17    m/z 1071.1231     0   2   4E+4   13   13.5  RT (min)      lcqtof_t3_neg_4_064_17    m/z 1071.1665     0   1E+4   13   13.5  RT (min)      lcqtof_t3_neg_4_036_47    m/z 1072.0900     0   1   2   3E+4   14   14.5  RT (min)      lcqtof_t3_neg_4_064_17    m/z 1072.0907     0   1   2   3E+4   13.5   14  RT (min)      lcqtof_t3_neg_4_064_17    m/z 1072.0908     0   1   2   3E+4   13   13.5  RT (min)      lcqtof_t3_neg_4_064_17    m/z 1072.0932     0   1   2E+4   14   14.5  RT (min)      lcqtof_t3_neg_4_064_17    m/z 1072.0937     0   1   2   3E+4   13.5   14  RT (min)      lcqtof_t3_neg_4_064_17    m/z 1072.0946     0   1   2E+4   14   14.5  RT (min)      lcqtof_t3_neg_4_064_17    m/z 1072.1031     0   1   2E+4   13   13.5  RT (min)      lcqtof_t3_neg_4_002_17    m/z 1072.1648     0   1E+4   13   13.5  RT (min)      lcqtof_t3_neg_4_002_17    m/z 1072.1695     0   1E+4   13   13.5  RT (min)      lcqtof_t3_neg_4_036_47    m/z 1072.6042     0   1E+4   12.5   13  RT (min)      lcqtof_t3_neg_4_058_16    m/z 1072.6865     0   1E+4   12.5   13  RT (min)      lcqtof_t3_neg_4_018_23    m/z 1072.6890     0   1E+4   12.5   13  RT (min)       lcqtof_t3_neg_4_018_23    m/z 1073.1553     0   1   2E+4   13   13.5  RT (min)      lcqtof_t3_neg_4_064_17    m/z 1073.1565     0   1   2E+4   13.5   14  RT (min)      lcqtof_t3_neg_4_064_17    m/z 1073.1646     0   1   2E+4   13.5   14  RT (min)      lcqtof_t3_neg_4_064_17    m/z 1074.0707     0   1E+4   13   13.5  RT (min)      lcqtof_t3_neg_4_064_17    m/z 1074.0763     0   1E+4   13.5   14  RT (min)      lcqtof_t3_neg_4_064_17    m/z 1074.0776     0   1E+4   13   13.5  RT (min)      lcqtof_t3_neg_4_064_17    m/z 1074.0779     0   1E+4   13.5   14  RT (min)      lcqtof_t3_neg_4_064_17    m/z 1074.0823     0   1E+4   13   13.5  RT (min)      lcqtof_t3_neg_4_064_17    m/z 1074.0841     0   1E+4   13.5   14  RT (min)      lcqtof_t3_neg_4_064_17    m/z 1074.0886     0   1E+4   13.5   14  RT (min)      lcqtof_t3_neg_4_064_17    m/z 1074.1595     0   1E+4   13   13.5  RT (min)      lcqtof_t3_neg_4_064_17    m/z 1074.1601     0   1E+4   13.5   14  RT (min)      lcqtof_t3_neg_4_064_17    m/z 1074.6719     0   1E+4   13   13.5  RT (min)       lcqtof_t3_neg_4_035_08    m/z 1078.5447     0   1E+4   6   6.5  RT (min)      lcqtof_t3_neg_4_005_18    m/z 1079.1877     0   1E+4   13.5   14  RT (min)      lcqtof_t3_neg_4_002_17    m/z 1079.1919     0   1   2E+4   13.5   14  RT (min)      lcqtof_t3_neg_4_036_47    m/z 1079.5471     0   1E+4   6   6.5  RT (min)      lcqtof_t3_neg_4_005_18    m/z 1080.1914     0   1   2E+4   13.5   14  RT (min)      lcqtof_t3_neg_4_036_47    m/z 1080.1916     0   1E+4   13.5   14  RT (min)      lcqtof_t3_neg_4_002_17    m/z 1080.7811     0   1E+4   17   17.5  RT (min)      lcqtof_t3_neg_4_035_08    m/z 1081.1904     0   1E+4   13.5   14  RT (min)      lcqtof_t3_neg_4_002_17    m/z 1081.1912     0   1   2E+4   13.5   14  RT (min)      lcqtof_t3_neg_4_036_47    m/z 1081.1949     0   1E+4   13.5   14  RT (min)      lcqtof_t3_neg_4_002_17    m/z 1081.2011     0   1E+4   13.5   14  RT (min)      lcqtof_t3_neg_4_002_17    m/z 1081.2246     0   1E+4   13.5   14  RT (min)      lcqtof_t3_neg_4_057_49    m/z 1081.6980     0   2   4   6   8E+3   11   11.5  RT (min)      lcqtof_t3_neg_4_037_01    m/z 1081.8687     0   1E+4   4   4.5  RT (min)      lcqtof_t3_neg_4_000_ee    m/z 1082.1926     0   1E+4   13.5   14  RT (min)      lcqtof_t3_neg_4_036_47    m/z 1082.2279     0   1E+4   13.5   14  RT (min)      lcqtof_t3_neg_4_057_49    m/z 1082.2611     0   1E+4   13   13.5  RT (min)      lcqtof_t3_neg_4_002_17    m/z 1082.2615     0   1   2E+4   13   13.5  RT (min)      lcqtof_t3_neg_4_064_17    m/z 1082.2747     0   1   2E+4   13.5   14  RT (min)      lcqtof_t3_neg_4_064_17    m/z 1082.3680     0   1E+4   4   4.5  RT (min)      lcqtof_t3_neg_4_000_ee    m/z 1082.7117     0   2   4   6E+3   13.5   14  RT (min)      lcqtof_t3_neg_1_019_23    m/z 1083.1837     0   1E+4   13.5   14  RT (min)      lcqtof_t3_neg_4_036_47    m/z 1083.2263     0   1E+4   13.5   14  RT (min)      lcqtof_t3_neg_4_057_49    m/z 1083.2543     0   1   2   3   4E+4   13   13.5  RT (min)      lcqtof_t3_neg_4_064_17    m/z 1083.2756     0   1   2   3   4E+4   13.5   14  RT (min)      lcqtof_t3_neg_4_064_17    m/z 1084.2750     0   1   2   3   4E+4   13.5   14  RT (min)      lcqtof_t3_neg_4_064_17    m/z 1085.1283     0   1   2   3E+5   13   13.5  RT (min)      lcqtof_t3_neg_4_064_17    m/z 1085.1293     0   1   2E+5   14   14.5  RT (min)      lcqtof_t3_neg_4_002_17    m/z 1085.1296     0   1   2E+5   13   13.5  RT (min)      lcqtof_t3_neg_4_064_17    m/z 1085.1301     0   1   2   3E+5   13.5   14  RT (min)      lcqtof_t3_neg_4_064_17    m/z 1085.1301     0   1   2E+5   13   13.5  RT (min)      lcqtof_t3_neg_4_064_17    m/z 1085.2732     0   1   2   3   4E+4   13.5   14  RT (min)      lcqtof_t3_neg_4_064_17    m/z 1086.1279     0   1   2   3E+5   13   13.5  RT (min)      lcqtof_t3_neg_4_064_17    m/z 1086.1294     0   1   2   3E+5   14   14.5  RT (min)      lcqtof_t3_neg_4_064_17    m/z 1086.1295     0   1   2   3E+5   13.5   14  RT (min)      lcqtof_t3_neg_4_064_17    m/z 1086.1298     0   1   2E+5   13   13.5  RT (min)      lcqtof_t3_neg_4_064_17    m/z 1086.1301     0   1   2E+5   13   13.5  RT (min)      lcqtof_t3_neg_4_064_17    m/z 1086.2726     0   1   2E+4   13.5   14  RT (min)      lcqtof_t3_neg_4_064_17    m/z 1086.2737     0   1   2E+4   13.5   14  RT (min)      lcqtof_t3_neg_4_002_17    m/z 1086.6802     0   1E+4   10   10.5  RT (min)      lcqtof_t3_neg_4_062_13    m/z 1086.6814     0   1E+4   10.5   11  RT (min)      lcqtof_t3_neg_4_062_13    m/z 1086.7067     0   2   4   6E+5   12.5   13  RT (min)      lcqtof_t3_neg_1_019_23    m/z 1086.7070     0   2   4E+5   12.5   13  RT (min)      lcqtof_t3_neg_2_030_23    m/z 1086.7075     0   2   4E+2   16   16.5  RT (min)      lcqtof_t3_neg_2_030_23    m/z 1086.7078     0   1   2E+4   16   16.5  RT (min)      lcqtof_t3_neg_4_018_23    m/z 1086.7090     0   2   4E+2   11   11.5  RT (min)      lcqtof_t3_neg_3_038_06    m/z 1086.7091     0   2   4E+2   11   11.5  RT (min)      lcqtof_t3_neg_3_038_06    m/z 1087.0397     0   1   2E+4   13   13.5  RT (min)      lcqtof_t3_neg_4_034_34    m/z 1087.0405     0   1   2E+4   13   13.5  RT (min)      lcqtof_t3_neg_4_018_23    m/z 1087.0410     0   1E+4   13   13.5  RT (min)      lcqtof_t3_neg_1_032_36    m/z 1087.1222     0   1   2   3E+5   14   14.5  RT (min)      lcqtof_t3_neg_4_002_17    m/z 1087.1251     0   1   2   3E+5   13.5   14  RT (min)      lcqtof_t3_neg_4_064_17    m/z 1087.1269     0   1   2E+5   13   13.5  RT (min)      lcqtof_t3_neg_4_064_17    m/z 1087.1273     0   1   2E+5   13   13.5  RT (min)      lcqtof_t3_neg_4_064_17    m/z 1087.1275     0   1   2   3E+5   13   13.5  RT (min)      lcqtof_t3_neg_4_064_17    m/z 1087.2666     0   1E+4   13.5   14  RT (min)      lcqtof_t3_neg_4_002_17    m/z 1087.2745     0   1E+4   13.5   14  RT (min)      lcqtof_t3_neg_4_064_17    m/z 1087.7095     0   1E+4   16   16.5  RT (min)      lcqtof_t3_neg_4_018_23    m/z 1087.7099     0   1   2   3E+5   12.5   13  RT (min)      lcqtof_t3_neg_1_019_23    m/z 1087.7101     0   1   2   3E+5   12.5   13  RT (min)      lcqtof_t3_neg_2_030_23    m/z 1087.7120     0   1   2   3   4E+2   11   11.5  RT (min)      lcqtof_t3_neg_3_038_06    m/z 1088.1224     0   1   2E+5   14   14.5  RT (min)      lcqtof_t3_neg_4_064_17    m/z 1088.1247     0   1   2E+5   14   14.5  RT (min)      lcqtof_t3_neg_4_064_17    m/z 1088.1262     0   1E+5   13.5   14  RT (min)      lcqtof_t3_neg_4_002_17    m/z 1088.1266     0   1E+5   13   13.5  RT (min)      lcqtof_t3_neg_4_064_17    m/z 1088.1270     0   1   2E+5   13   13.5  RT (min)      lcqtof_t3_neg_4_064_17    m/z 1088.1274     0   1   2E+5   13.5   14  RT (min)      lcqtof_t3_neg_4_064_17    m/z 1088.1282     0   1E+5   14   14.5  RT (min)       lcqtof_t3_neg_4_036_47    m/z 1088.7126     0   1E+5   12.5   13  RT (min)      lcqtof_t3_neg_2_030_23    m/z 1088.7139     0   1E+5   12.5   13  RT (min)      lcqtof_t3_neg_1_019_23    m/z 1088.7162     0   1   2   3E+2   11   11.5  RT (min)      lcqtof_t3_neg_3_038_06    m/z 1089.1172     0   2   4   6   8E+4   13.5   14  RT (min)      lcqtof_t3_neg_4_036_47    m/z 1089.1229     0   1E+5   13.5   14  RT (min)      lcqtof_t3_neg_4_064_17    m/z 1089.1238     0   2   4   6   8E+4   13   13.5  RT (min)      lcqtof_t3_neg_4_036_47    m/z 1089.1239     0   2   4   6   8E+4   13   13.5  RT (min)      lcqtof_t3_neg_4_064_17    m/z 1089.1244     0   2   4   6   8E+4   13   13.5  RT (min)      lcqtof_t3_neg_4_064_17    m/z 1089.1253     0   1E+5   14   14.5  RT (min)      lcqtof_t3_neg_4_064_17    m/z 1089.1259     0   1E+5   13   13.5  RT (min)      lcqtof_t3_neg_4_064_17    m/z 1089.7166     0   1   2E+4   12.5   13  RT (min)      lcqtof_t3_neg_1_019_23    m/z 1090.1200     0   2   4E+4   13.5   14  RT (min)      lcqtof_t3_neg_4_002_17    m/z 1090.1217     0   2   4E+4   14   14.5  RT (min)      lcqtof_t3_neg_4_002_17    m/z 1090.1222     0   2   4   6E+4   13   13.5  RT (min)      lcqtof_t3_neg_4_064_17    m/z 1090.1228     0   2   4   6E+4   13.5   14  RT (min)      lcqtof_t3_neg_4_064_17    m/z 1090.1244     0   1   2   3   4E+4   13   13.5  RT (min)      lcqtof_t3_neg_4_064_17    m/z 1090.1247     0   1   2   3   4E+4   13   13.5  RT (min)      lcqtof_t3_neg_4_064_17    m/z 1090.1254     0   2   4   6E+4   13.5   14  RT (min)      lcqtof_t3_neg_4_064_17    m/z 1090.7206     0   1E+4   13   13.5  RT (min)      lcqtof_t3_neg_4_018_23    m/z 1091.1173     0   1   2   3E+4   13.5   14  RT (min)      lcqtof_t3_neg_4_064_17    m/z 1091.1198     0   1E+4   13   13.5  RT (min)      lcqtof_t3_neg_4_064_17    m/z 1091.1207     0   1   2   3E+4   13.5   14  RT (min)      lcqtof_t3_neg_4_064_17    m/z 1091.1217     0   1E+4   13   13.5  RT (min)      lcqtof_t3_neg_4_064_17    m/z 1093.2408     0   1E+4   13   13.5  RT (min)      lcqtof_t3_neg_4_036_47    m/z 1093.2416     0   1E+4   13   13.5  RT (min)      lcqtof_t3_neg_4_002_17    m/z 1093.2422     0   1E+4   13   13.5  RT (min)      lcqtof_t3_neg_4_063_39    m/z 1093.2487     0   1E+4   13   13.5  RT (min)      lcqtof_t3_neg_4_002_17    m/z 1093.2511     0   1   2   3E+4   13   13.5  RT (min)      lcqtof_t3_neg_4_064_17    m/z 1093.2567     0   1E+4   13   13.5  RT (min)      lcqtof_t3_neg_4_002_17    m/z 1094.2423     0   1E+4   13   13.5  RT (min)      lcqtof_t3_neg_4_002_17    m/z 1094.2509     0   1   2   3   4E+4   13   13.5  RT (min)      lcqtof_t3_neg_4_064_17    m/z 1094.2550     0   1E+4   13   13.5  RT (min)      lcqtof_t3_neg_4_002_17    m/z 1094.6949     0   1   2E+4   13   13.5  RT (min)      lcqtof_t3_neg_4_018_23    m/z 1095.6942     0   1E+4   13   13.5  RT (min)      lcqtof_t3_neg_4_018_23    m/z 1095.8521     0   1E+4   12.5   13  RT (min)      lcqtof_t3_neg_2_030_23    m/z 1095.8533     0   1E+4   12.5   13  RT (min)      lcqtof_t3_neg_2_016_05    m/z 1095.8542     0   1E+4   12.5   13  RT (min)      lcqtof_t3_neg_2_030_23    m/z 1095.8551     0   1   2E+3   16   16.5  RT (min)      lcqtof_t3_neg_2_052_17    m/z 1095.8555     0   1   2E+3   16   16.5  RT (min)      lcqtof_t3_neg_2_045_04    m/z 1095.8556     0   1   2E+3   14.5   15  RT (min)      lcqtof_t3_neg_1_010_27    m/z 1095.8557     0   1   2E+3   15.5   16  RT (min)      lcqtof_t3_neg_2_030_23    m/z 1095.8560     0   1   2E+3   15   15.5  RT (min)      lcqtof_t3_neg_2_030_23    m/z 1096.2418     0   1E+4   13   13.5  RT (min)      lcqtof_t3_neg_4_002_17    m/z 1096.2552     0   1   2   3E+4   13   13.5  RT (min)      lcqtof_t3_neg_4_064_17    m/z 1097.2017     0   1E+4   13.5   14  RT (min)      lcqtof_t3_neg_4_064_17    m/z 1097.2024     0   1E+4   14   14.5  RT (min)      lcqtof_t3_neg_4_064_17    m/z 1097.2032     0   1E+4   14   14.5  RT (min)      lcqtof_t3_neg_4_002_17    m/z 1097.2033     0   1E+4   14   14.5  RT (min)      lcqtof_t3_neg_4_036_47    m/z 1097.2034     0   1E+4   14   14.5  RT (min)      lcqtof_t3_neg_4_036_47    m/z 1097.2270     0   1E+4   13.5   14  RT (min)      lcqtof_t3_neg_4_057_49    m/z 1097.2513     0   1   2E+4   13   13.5  RT (min)      lcqtof_t3_neg_4_064_17    m/z 1098.2015     0   1   2E+4   13.5   14  RT (min)      lcqtof_t3_neg_4_064_17    m/z 1098.2036     0   1E+4   13.5   14  RT (min)      lcqtof_t3_neg_4_002_17    m/z 1098.2040     0   1   2E+4   14   14.5  RT (min)      lcqtof_t3_neg_4_064_17    m/z 1098.7445     0   1E+4   13   13.5  RT (min)      lcqtof_t3_neg_4_018_23    m/z 1099.1796     0   1   2E+4   13.5   14  RT (min)      lcqtof_t3_neg_4_036_47    m/z 1099.1812     0   1   2E+4   13.5   14  RT (min)      lcqtof_t3_neg_4_036_47    m/z 1099.1820     0   1   2   3E+4   13.5   14  RT (min)      lcqtof_t3_neg_4_064_17    m/z 1099.1838     0   1   2   3E+4   13.5   14  RT (min)      lcqtof_t3_neg_4_064_17    m/z 1099.1887     0   1   2   3E+4   13.5   14  RT (min)      lcqtof_t3_neg_4_064_17    m/z 1099.1941     0   1   2E+4   13.5   14  RT (min)      lcqtof_t3_neg_4_036_47    m/z 1099.1956     0   1   2   3E+4   13.5   14  RT (min)      lcqtof_t3_neg_4_064_17    m/z 1099.1968     0   1   2   3E+4   14   14.5  RT (min)      lcqtof_t3_neg_4_064_17    m/z 1099.1971     0   1   2   3E+4   14   14.5  RT (min)      lcqtof_t3_neg_4_064_17    m/z 1099.1991     0   1E+4   14   14.5  RT (min)      lcqtof_t3_neg_4_004_20    m/z 1100.1823     0   1   2E+4   13.5   14  RT (min)      lcqtof_t3_neg_4_064_17    m/z 1100.1876     0   1   2E+4   13.5   14  RT (min)      lcqtof_t3_neg_4_064_17    m/z 1100.1941     0   1   2E+4   14   14.5  RT (min)      lcqtof_t3_neg_4_064_17    m/z 1100.1943     0   1   2E+4   13.5   14  RT (min)      lcqtof_t3_neg_4_064_17    m/z 1100.7240     0   1   2E+5   12.5   13  RT (min)      lcqtof_t3_neg_1_019_23    m/z 1100.7240     0   1   2   3   4E+2   11   11.5  RT (min)      lcqtof_t3_neg_3_038_06    m/z 1100.7245     0   1   2   3   4E+2   11   11.5  RT (min)      lcqtof_t3_neg_3_038_06    m/z 1101.0477     0   1   2   3   4E+4   13   13.5  RT (min)      lcqtof_t3_neg_4_002_17    m/z 1101.0486     0   1   2   3   4E+4   14   14.5  RT (min)      lcqtof_t3_neg_4_036_47    m/z 1101.0490     0   1E+5   13   13.5  RT (min)      lcqtof_t3_neg_4_064_17    m/z 1101.0498     0   2   4E+4   13.5   14  RT (min)      lcqtof_t3_neg_4_036_47    m/z 1101.0498     0   2   4E+4   13.5   14  RT (min)      lcqtof_t3_neg_4_036_47    m/z 1101.0502     0   1E+5   13   13.5  RT (min)      lcqtof_t3_neg_4_064_17    m/z 1101.0507     0   1E+5   14   14.5  RT (min)      lcqtof_t3_neg_4_064_17    m/z 1101.0508     0   1E+5   13   13.5  RT (min)      lcqtof_t3_neg_4_064_17    m/z 1101.0525     0   1E+5   13.5   14  RT (min)      lcqtof_t3_neg_4_064_17    m/z 1101.7267     0   1   2   3   4E+2   11   11.5  RT (min)      lcqtof_t3_neg_3_038_06    m/z 1101.7271     0   1E+5   12.5   13  RT (min)      lcqtof_t3_neg_1_019_23    m/z 1102.0489     0   1E+5   13.5   14  RT (min)      lcqtof_t3_neg_4_002_17    m/z 1102.0497     0   1E+5   13   13.5  RT (min)      lcqtof_t3_neg_4_064_17    m/z 1102.0498     0   1E+5   13.5   14  RT (min)      lcqtof_t3_neg_4_002_17    m/z 1102.0500     0   1E+5   13.5   14  RT (min)      lcqtof_t3_neg_4_064_17    m/z 1102.0501     0   1E+5   13   13.5  RT (min)      lcqtof_t3_neg_4_064_17    m/z 1102.0503     0   1   2   3   4E+4   13   13.5  RT (min)      lcqtof_t3_neg_4_002_17    m/z 1102.0504     0   1E+5   14   14.5  RT (min)      lcqtof_t3_neg_4_064_17    m/z 1102.0505     0   1E+5   14   14.5  RT (min)      lcqtof_t3_neg_4_064_17    m/z 1102.0510     0   2   4   6E+4   13.5   14  RT (min)      lcqtof_t3_neg_4_036_47    m/z 1102.0514     0   2   4   6E+4   13   13.5  RT (min)      lcqtof_t3_neg_4_002_17    m/z 1102.0539     0   1E+5   13   13.5  RT (min)      lcqtof_t3_neg_4_064_17    m/z 1102.7035     0   1   2   3   4E+4   12.5   13  RT (min)      lcqtof_t3_neg_2_030_23    m/z 1102.7288     0   1   2   3   4E+4   12.5   13  RT (min)      lcqtof_t3_neg_1_019_23    m/z 1103.0475     0   1   2   3   4E+4   13   13.5  RT (min)      lcqtof_t3_neg_4_002_17    m/z 1103.0478     0   1E+5   13.5   14  RT (min)      lcqtof_t3_neg_4_002_17    m/z 1103.0481     0   1E+5   13   13.5  RT (min)      lcqtof_t3_neg_4_064_17    m/z 1103.0482     0   1   2E+4   13.5   14  RT (min)      lcqtof_t3_neg_4_004_20    m/z 1103.0482     0   1E+5   13   13.5  RT (min)      lcqtof_t3_neg_4_064_17    m/z 1103.0482     0   2   4   6E+4   13.5   14  RT (min)      lcqtof_t3_neg_4_036_47    m/z 1103.0487     0   1E+5   13   13.5  RT (min)      lcqtof_t3_neg_4_064_17    m/z 1103.0493     0   2   4   6E+4   14   14.5  RT (min)      lcqtof_t3_neg_4_036_47    m/z 1103.0493     0   1E+5   14   14.5  RT (min)      lcqtof_t3_neg_4_064_17    m/z 1103.0507     0   1E+5   13.5   14  RT (min)      lcqtof_t3_neg_4_064_17    m/z 1103.0515     0   2   4   6E+4   14   14.5  RT (min)      lcqtof_t3_neg_4_036_47    m/z 1103.0516     0   2   4E+4   13   13.5  RT (min)      lcqtof_t3_neg_4_002_17    m/z 1103.7053     0   1   2E+4   12.5   13  RT (min)      lcqtof_t3_neg_1_019_23    m/z 1103.7069     0   1   2E+4   12.5   13  RT (min)      lcqtof_t3_neg_2_030_23    m/z 1104.0467     0   2   4   6   8E+4   13.5   14  RT (min)      lcqtof_t3_neg_4_002_17    m/z 1104.0470     0   1E+5   13   13.5  RT (min)      lcqtof_t3_neg_4_064_17    m/z 1104.0478     0   1E+5   13   13.5  RT (min)      lcqtof_t3_neg_4_064_17    m/z 1104.0487     0   1E+5   13   13.5  RT (min)      lcqtof_t3_neg_4_064_17    m/z 1104.0489     0   1E+5   13.5   14  RT (min)      lcqtof_t3_neg_4_064_17    m/z 1104.0489     0   2   4   6   8E+4   13.5   14  RT (min)      lcqtof_t3_neg_4_002_17    m/z 1104.0491     0   2   4   6   8E+4   13.5   14  RT (min)      lcqtof_t3_neg_4_002_17    m/z 1104.0494     0   2   4   6   8E+4   14   14.5  RT (min)      lcqtof_t3_neg_4_002_17    m/z 1104.0505     0   2   4   6E+4   13   13.5  RT (min)      lcqtof_t3_neg_4_064_17    m/z 1104.0529     0   2   4   6   8E+4   13.5   14  RT (min)      lcqtof_t3_neg_4_002_17    m/z 1104.8722     0   1   2E+4   4   4.5  RT (min)      lcqtof_t3_neg_4_000_ee    m/z 1105.0463     0   1   2   3   4E+4   13.5   14  RT (min)      lcqtof_t3_neg_4_002_17    m/z 1105.0464     0   2   4   6E+4   14   14.5  RT (min)      lcqtof_t3_neg_4_064_17    m/z 1105.0468     0   1   2   3   4E+4   13.5   14  RT (min)      lcqtof_t3_neg_4_002_17    m/z 1105.0471     0   1   2   3   4E+4   13   13.5  RT (min)      lcqtof_t3_neg_4_064_17    m/z 1105.0474     0   1   2   3   4E+4   13   13.5  RT (min)      lcqtof_t3_neg_4_064_17    m/z 1105.0476     0   2   4   6E+4   13   13.5  RT (min)      lcqtof_t3_neg_4_064_17    m/z 1105.0481     0   2   4   6E+4   13   13.5  RT (min)      lcqtof_t3_neg_4_064_17    m/z 1105.0486     0   2   4   6E+4   13.5   14  RT (min)      lcqtof_t3_neg_4_064_17    m/z 1105.0492     0   2   4   6E+4   13   13.5  RT (min)      lcqtof_t3_neg_4_064_17    m/z 1105.1363     0   1E+4   13   13.5  RT (min)      lcqtof_t3_neg_4_064_17    m/z 1105.2666     0   1E+4   13   13.5  RT (min)      lcqtof_t3_neg_4_002_17    m/z 1105.2668     0   1   2E+4   13   13.5  RT (min)      lcqtof_t3_neg_4_064_17    m/z 1105.3724     0   1   2E+4   4   4.5  RT (min)      lcqtof_t3_neg_4_000_ee    m/z 1105.8767     0   1E+4   4   4.5  RT (min)      lcqtof_t3_neg_4_000_ee    m/z 1106.0471     0   1   2   3E+4   13.5   14  RT (min)      lcqtof_t3_neg_4_064_17    m/z 1106.0480     0   1   2   3E+4   13.5   14  RT (min)      lcqtof_t3_neg_4_064_17    m/z 1106.0483     0   1   2   3E+4   14   14.5  RT (min)      lcqtof_t3_neg_4_064_17    m/z 1106.0491     0   1   2   3E+4   13   13.5  RT (min)      lcqtof_t3_neg_4_064_17    m/z 1106.0494     0   1   2   3E+4   13   13.5  RT (min)      lcqtof_t3_neg_4_064_17    m/z 1106.0495     0   1   2E+4   13   13.5  RT (min)      lcqtof_t3_neg_4_064_17    m/z 1106.1140     0   1E+4   13   13.5  RT (min)      lcqtof_t3_neg_4_063_39    m/z 1107.0469     0   1E+4   13   13.5  RT (min)      lcqtof_t3_neg_4_064_17    m/z 1107.0497     0   1E+4   13.5   14  RT (min)      lcqtof_t3_neg_4_064_17    m/z 1109.6696     0   1E+4   10   10.5  RT (min)      lcqtof_t3_neg_4_062_13    m/z 1109.6748     0   1E+4   10   10.5  RT (min)      lcqtof_t3_neg_4_062_13    m/z 1109.7304     0   1E+4   13.5   14  RT (min)      lcqtof_t3_neg_4_018_23    m/z 1110.4660     0   1E+4   1   1.5  RT (min)      lcqtof_t3_neg_2_042_13    m/z 1112.9010     0   2   4   6E+3   15   15.5  RT (min)      lcqtof_t3_neg_1_036_14    m/z 1113.7143     0   1E+4   13   13.5  RT (min)      lcqtof_t3_neg_4_018_23    m/z 1114.7376     0   1   2E+4   16   16.5  RT (min)      lcqtof_t3_neg_4_018_23    m/z 1114.7385     0   2   4E+2   11   11.5  RT (min)      lcqtof_t3_neg_3_038_06    m/z 1114.7393     0   2   4   6E+2   16   16.5  RT (min)      lcqtof_t3_neg_2_030_23    m/z 1114.7394     0   1   2   3   4E+5   13   13.5  RT (min)      lcqtof_t3_neg_2_030_23    m/z 1114.7405     0   1   2   3   4E+5   13   13.5  RT (min)      lcqtof_t3_neg_1_019_23    m/z 1114.7418     0   2   4E+2   11   11.5  RT (min)      lcqtof_t3_neg_3_038_06    m/z 1115.0767     0   1E+4   13   13.5  RT (min)      lcqtof_t3_neg_4_018_23    m/z 1115.7405     0   1E+4   16   16.5  RT (min)      lcqtof_t3_neg_4_018_23    m/z 1115.7410     0   1   2   3   4E+2   11   11.5  RT (min)      lcqtof_t3_neg_3_038_06    m/z 1115.7429     0   1   2E+5   13   13.5  RT (min)      lcqtof_t3_neg_2_030_23    m/z 1115.7432     0   1   2E+5   13   13.5  RT (min)      lcqtof_t3_neg_1_019_23    m/z 1115.7434     0   1   2   3   4E+2   11   11.5  RT (min)      lcqtof_t3_neg_3_038_06    m/z 1116.2104     0   2   4   6E+4   14   14.5  RT (min)      lcqtof_t3_neg_4_002_17    m/z 1116.2105     0   2   4   6E+4   13.5   14  RT (min)      lcqtof_t3_neg_4_002_17    m/z 1116.7160     0   1E+5   13   13.5  RT (min)      lcqtof_t3_neg_4_018_23    m/z 1116.7187     0   1E+4   12.5   13  RT (min)      lcqtof_t3_neg_2_030_23    m/z 1116.7192     0   2   4   6   8E+4   12.5   13  RT (min)      lcqtof_t3_neg_4_018_23    m/z 1116.7454     0   1E+5   13   13.5  RT (min)      lcqtof_t3_neg_1_019_23    m/z 1116.7458     0   2   4E+2   11   11.5  RT (min)      lcqtof_t3_neg_3_038_06    m/z 1116.7459     0   2   4   6   8E+4   13   13.5  RT (min)      lcqtof_t3_neg_2_030_23    m/z 1116.7487     0   2   4E+2   11   11.5  RT (min)      lcqtof_t3_neg_3_038_06    m/z 1117.7492     0   1   2E+4   13   13.5  RT (min)      lcqtof_t3_neg_2_030_23    m/z 1118.2097     0   2   4E+4   13.5   14  RT (min)      lcqtof_t3_neg_4_002_17    m/z 1118.2107     0   2   4E+4   13.5   14  RT (min)      lcqtof_t3_neg_4_002_17    m/z 1118.2123     0   2   4E+4   13.5   14  RT (min)      lcqtof_t3_neg_4_002_17    m/z 1119.2048     0   1   2   3E+4   13.5   14  RT (min)      lcqtof_t3_neg_4_002_17    m/z 1119.2072     0   1   2   3E+4   13.5   14  RT (min)      lcqtof_t3_neg_4_002_17    m/z 1119.2084     0   1   2   3E+4   13.5   14  RT (min)      lcqtof_t3_neg_4_002_17    m/z 1119.2118     0   1   2   3E+4   13.5   14  RT (min)      lcqtof_t3_neg_4_002_17    m/z 1119.2622     0   1E+4   13   13.5  RT (min)      lcqtof_t3_neg_3_089_59    m/z 1120.2070     0   1E+4   13.5   14  RT (min)      lcqtof_t3_neg_4_002_17    m/z 1120.2078     0   1E+4   13.5   14  RT (min)      lcqtof_t3_neg_4_002_17    m/z 1121.2539     0   1   2E+4   13   13.5  RT (min)      lcqtof_t3_neg_4_064_17    m/z 1122.2573     0   1E+4   13   13.5  RT (min)      lcqtof_t3_neg_4_064_17    m/z 1122.7214     0   1   2   3E+4   13   13.5  RT (min)      lcqtof_t3_neg_4_018_23    m/z 1122.8792     0   1   2   3E+3   13   13.5  RT (min)      lcqtof_t3_neg_1_001_49    m/z 1123.2290     0   1E+4   13.5   14  RT (min)      lcqtof_t3_neg_4_057_49    m/z 1123.2551     0   1E+4   13   13.5  RT (min)      lcqtof_t3_neg_4_064_17    m/z 1123.2676     0   1E+4   13   13.5  RT (min)      lcqtof_t3_neg_4_064_17    m/z 1123.7244     0   1E+4   13   13.5  RT (min)      lcqtof_t3_neg_4_018_23    m/z 1123.7267     0   1E+4   13   13.5  RT (min)      lcqtof_t3_neg_4_018_23    m/z 1124.2338     0   1E+4   13.5   14  RT (min)      lcqtof_t3_neg_4_057_49    m/z 1124.2345     0   1E+4   13.5   14  RT (min)      lcqtof_t3_neg_4_057_49    m/z 1124.2481     0   1   2   3E+4   13   13.5  RT (min)      lcqtof_t3_neg_4_064_17    m/z 1124.2676     0   1E+4   13   13.5  RT (min)      lcqtof_t3_neg_4_002_17    m/z 1124.2706     0   1   2   3E+4   13   13.5  RT (min)      lcqtof_t3_neg_4_064_17    m/z 1125.2260     0   1E+4   13.5   14  RT (min)      lcqtof_t3_neg_4_057_49    m/z 1125.2260     0   1E+4   13.5   14  RT (min)      lcqtof_t3_neg_4_001_49    m/z 1126.2256     0   1E+4   13.5   14  RT (min)      lcqtof_t3_neg_4_057_49    m/z 1127.2476     0   1   2   3E+4   13.5   14  RT (min)      lcqtof_t3_neg_4_057_49    m/z 1128.2486     0   1   2   3E+4   13.5   14  RT (min)      lcqtof_t3_neg_4_057_49    m/z 1128.7517     0   1   2   3   4E+4   13   13.5  RT (min)      lcqtof_t3_neg_4_018_23    m/z 1128.7543     0   1E+4   13   13.5  RT (min)      lcqtof_t3_neg_2_030_23    m/z 1129.0588     0   1   2E+4   13.5   14  RT (min)      lcqtof_t3_neg_4_064_17    m/z 1129.0618     0   1E+4   14   14.5  RT (min)      lcqtof_t3_neg_4_064_17    m/z 1129.0623     0   1   2E+4   13.5   14  RT (min)      lcqtof_t3_neg_4_064_17    m/z 1129.0627     0   1   2E+4   13   13.5  RT (min)      lcqtof_t3_neg_4_064_17    m/z 1129.0642     0   1   2E+4   13.5   14  RT (min)      lcqtof_t3_neg_4_064_17    m/z 1129.0660     0   1E+4   14   14.5  RT (min)      lcqtof_t3_neg_4_064_17    m/z 1129.0672     0   1   2E+4   13.5   14  RT (min)      lcqtof_t3_neg_4_064_17    m/z 1129.0691     0   1   2E+4   13   13.5  RT (min)      lcqtof_t3_neg_4_064_17    m/z 1129.0700     0   1   2E+4   13   13.5  RT (min)      lcqtof_t3_neg_4_064_17    m/z 1129.2488     0   1   2   3E+4   13.5   14  RT (min)      lcqtof_t3_neg_4_057_49    m/z 1129.2662     0   1E+4   13   13.5  RT (min)      lcqtof_t3_neg_4_002_17    m/z 1129.2680     0   1   2E+4   13   13.5  RT (min)      lcqtof_t3_neg_4_064_17    m/z 1129.7583     0   1E+4   13   13.5  RT (min)      lcqtof_t3_neg_2_030_23    m/z 1129.7587     0   1E+4   13   13.5  RT (min)      lcqtof_t3_neg_1_019_23    m/z 1130.0629     0   1   2E+4   14   14.5  RT (min)      lcqtof_t3_neg_4_064_17    m/z 1130.0631     0   1   2E+4   14   14.5  RT (min)      lcqtof_t3_neg_4_064_17    m/z 1130.0647     0   1   2E+4   13.5   14  RT (min)      lcqtof_t3_neg_4_064_17    m/z 1130.0647     0   1   2E+4   13   13.5  RT (min)      lcqtof_t3_neg_4_064_17    m/z 1130.0651     0   1   2E+4   13.5   14  RT (min)      lcqtof_t3_neg_4_064_17    m/z 1130.0659     0   1   2E+4   13.5   14  RT (min)      lcqtof_t3_neg_4_064_17    m/z 1130.0661     0   1   2E+4   13.5   14  RT (min)      lcqtof_t3_neg_4_064_17    m/z 1130.0674     0   1   2E+4   13   13.5  RT (min)      lcqtof_t3_neg_4_064_17    m/z 1130.0675     0   1   2E+4   13.5   14  RT (min)      lcqtof_t3_neg_4_064_17    m/z 1130.2429     0   1E+4   13.5   14  RT (min)      lcqtof_t3_neg_4_001_49    m/z 1130.2476     0   1E+4   13.5   14  RT (min)      lcqtof_t3_neg_4_057_49    m/z 1130.7332     0   2   4E+4   12.5   13  RT (min)      lcqtof_t3_neg_2_030_23    m/z 1131.0614     0   1   2E+4   13.5   14  RT (min)      lcqtof_t3_neg_4_064_17    m/z 1131.0630     0   1   2E+4   14   14.5  RT (min)      lcqtof_t3_neg_4_064_17    m/z 1131.0642     0   1   2   3E+4   13.5   14  RT (min)      lcqtof_t3_neg_4_064_17    m/z 1131.0788     0   1   2   3   4E+4   13.5   14  RT (min)      lcqtof_t3_neg_4_064_17    m/z 1131.0963     0   2   4   6   8E+4   13   13.5  RT (min)      lcqtof_t3_neg_4_064_17    m/z 1131.1047     0   1   2   3   4E+4   13   13.5  RT (min)      lcqtof_t3_neg_4_002_17    m/z 1131.1145     0   1   2   3   4E+4   13   13.5  RT (min)      lcqtof_t3_neg_4_002_17    m/z 1131.1152     0   1   2   3   4E+4   13.5   14  RT (min)      lcqtof_t3_neg_4_002_17    m/z 1131.1271     0   2   4   6   8E+4   13.5   14  RT (min)      lcqtof_t3_neg_4_064_17    m/z 1131.2460     0   1E+4   13.5   14  RT (min)      lcqtof_t3_neg_4_057_49    m/z 1131.6450     0   1E+4   10   10.5  RT (min)       lcqtof_t3_neg_4_051_15    m/z 1131.6540     0   1E+4   10   10.5  RT (min)      lcqtof_t3_neg_4_014_13    m/z 1131.6622     0   1   2E+4   10   10.5  RT (min)      lcqtof_t3_neg_4_062_13    m/z 1131.6631     0   1E+4   10   10.5  RT (min)      lcqtof_t3_neg_4_014_13    m/z 1131.7376     0   1   2   3E+4   12.5   13  RT (min)      lcqtof_t3_neg_2_030_23    m/z 1131.8767     0   1E+4   4   4.5  RT (min)      lcqtof_t3_neg_4_000_ee    m/z 1132.0614     0   1E+4   13   13.5  RT (min)      lcqtof_t3_neg_4_064_17    m/z 1132.0634     0   1E+4   13.5   14  RT (min)      lcqtof_t3_neg_4_064_17    m/z 1132.0638     0   1E+4   14   14.5  RT (min)      lcqtof_t3_neg_4_064_17    m/z 1132.0655     0   1E+4   13.5   14  RT (min)      lcqtof_t3_neg_4_064_17    m/z 1132.0730     0   1   2E+4   13.5   14  RT (min)      lcqtof_t3_neg_4_064_17    m/z 1132.0754     0   1   2E+4   13.5   14  RT (min)      lcqtof_t3_neg_4_064_17    m/z 1132.1046     0   1   2   3E+4   13.5   14  RT (min)      lcqtof_t3_neg_4_002_17    m/z 1132.1174     0   2   4   6   8E+4   13   13.5  RT (min)      lcqtof_t3_neg_4_064_17    m/z 1132.1198     0   1   2   3E+4   13   13.5  RT (min)      lcqtof_t3_neg_4_002_17    m/z 1132.1249     0   2   4   6   8E+4   13   13.5  RT (min)      lcqtof_t3_neg_4_064_17    m/z 1132.3776     0   1E+4   4   4.5  RT (min)      lcqtof_t3_neg_4_000_ee    m/z 1132.6631     0   1E+4   10   10.5  RT (min)      lcqtof_t3_neg_4_062_13    m/z 1132.6655     0   1E+4   10   10.5  RT (min)      lcqtof_t3_neg_4_058_16    m/z 1132.7417     0   1E+4   12.5   13  RT (min)      lcqtof_t3_neg_2_030_23    m/z 1132.7905     0   1E+4   17   17.5  RT (min)      lcqtof_t3_neg_4_035_08    m/z 1132.8828     0   1E+4   4   4.5  RT (min)      lcqtof_t3_neg_4_000_ee    m/z 1133.0569     0   1E+4   13   13.5  RT (min)      lcqtof_t3_neg_4_064_17    m/z 1133.0614     0   1E+4   14   14.5  RT (min)      lcqtof_t3_neg_4_064_17    m/z 1133.0617     0   1E+4   14   14.5  RT (min)      lcqtof_t3_neg_4_064_17    m/z 1133.0696     0   1   2E+4   13.5   14  RT (min)      lcqtof_t3_neg_4_064_17    m/z 1133.0715     0   1   2E+4   13.5   14  RT (min)      lcqtof_t3_neg_4_064_17    m/z 1133.1191     0   1   2   3E+4   13.5   14  RT (min)      lcqtof_t3_neg_4_002_17    m/z 1133.1255     0   1   2   3E+4   13   13.5  RT (min)      lcqtof_t3_neg_4_002_17    m/z 1133.1268     0   2   4   6   8E+4   13   13.5  RT (min)      lcqtof_t3_neg_4_064_17    m/z 1134.1275     0   2   4E+4   13   13.5  RT (min)      lcqtof_t3_neg_4_064_17    m/z 1134.6310     0   1E+4   12.5   13  RT (min)      lcqtof_t3_neg_4_006_16    m/z 1134.6376     0   1   2E+4   13   13.5  RT (min)      lcqtof_t3_neg_4_058_16    m/z 1135.1250     0   1   2   3E+4   13   13.5  RT (min)      lcqtof_t3_neg_4_064_17    m/z 1135.1279     0   1   2   3E+4   13   13.5  RT (min)      lcqtof_t3_neg_4_064_17    m/z 1135.5594     0   1E+4   9.5   10  RT (min)      lcqtof_t3_neg_2_051_14    m/z 1135.6429     0   1E+4   13   13.5  RT (min)      lcqtof_t3_neg_4_058_16    m/z 1136.1273     0   1E+4   13   13.5  RT (min)      lcqtof_t3_neg_4_064_17    m/z 1137.2509     0   1   2E+4   13   13.5  RT (min)      lcqtof_t3_neg_4_064_17    m/z 1137.2565     0   1E+4   13   13.5  RT (min)      lcqtof_t3_neg_4_002_17    m/z 1138.2517     0   1E+4   13   13.5  RT (min)      lcqtof_t3_neg_4_064_17    m/z 1138.2569     0   1E+4   13   13.5  RT (min)      lcqtof_t3_neg_4_064_17    m/z 1138.8818     0   2   4E+2   12   12.5  RT (min)      lcqtof_t3_neg_3_032_24    m/z 1138.8840     0   2   4   6E+2   11   11.5  RT (min)      lcqtof_t3_neg_3_038_06    m/z 1138.8844     0   2   4E+2   12   12.5  RT (min)      lcqtof_t3_neg_3_032_24    m/z 1138.8846     0   1E+4   12.5   13  RT (min)      lcqtof_t3_neg_2_016_05    m/z 1139.2491     0   1   2   3E+4   13   13.5  RT (min)      lcqtof_t3_neg_4_064_17    m/z 1139.2676     0   1E+4   13   13.5  RT (min)      lcqtof_t3_neg_4_002_17    m/z 1139.2694     0   1   2   3E+4   13   13.5  RT (min)      lcqtof_t3_neg_4_064_17    m/z 1139.8837     0   1   2   3   4E+2   12   12.5  RT (min)      lcqtof_t3_neg_3_032_24    m/z 1139.8857     0   2   4   6   8E+4   12.5   13  RT (min)      lcqtof_t3_neg_2_016_05    m/z 1139.8859     0   1   2   3   4E+2   12   12.5  RT (min)      lcqtof_t3_neg_3_032_24    m/z 1139.8875     0   2   4E+2   11   11.5  RT (min)      lcqtof_t3_neg_3_038_06    m/z 1139.8876     0   2   4   6   8E+4   12.5   13  RT (min)      lcqtof_t3_neg_2_016_05    m/z 1139.8885     0   2   4E+2   11   11.5  RT (min)      lcqtof_t3_neg_3_038_06    m/z 1140.1589     0   1E+4   13.5   14  RT (min)      lcqtof_t3_neg_4_036_47    m/z 1140.2494     0   1   2E+4   13   13.5  RT (min)      lcqtof_t3_neg_4_064_17    m/z 1140.2704     0   1   2E+4   13   13.5  RT (min)      lcqtof_t3_neg_4_064_17    m/z 1140.7350     0   1   2E+4   13   13.5  RT (min)       lcqtof_t3_neg_4_018_23    m/z 1140.7372     0   1   2E+4   13   13.5  RT (min)       lcqtof_t3_neg_4_018_23    m/z 1140.8881     0   1   2   3E+2   12   12.5  RT (min)      lcqtof_t3_neg_3_032_24    m/z 1140.8889     0   1   2   3E+2   12   12.5  RT (min)      lcqtof_t3_neg_3_032_24    m/z 1140.8903     0   1   2   3E+2   11   11.5  RT (min)      lcqtof_t3_neg_3_038_06    m/z 1140.8906     0   1   2E+5   12.5   13  RT (min)      lcqtof_t3_neg_2_016_05    m/z 1140.8917     0   1   2   3E+2   11   11.5  RT (min)      lcqtof_t3_neg_3_038_06    m/z 1141.1581     0   1E+4   13.5   14  RT (min)      lcqtof_t3_neg_4_036_47    m/z 1141.1766     0   1E+4   13.5   14  RT (min)      lcqtof_t3_neg_4_036_47    m/z 1141.1771     0   1E+4   13.5   14  RT (min)      lcqtof_t3_neg_4_002_17    m/z 1141.2294     0   1E+4   13   13.5  RT (min)      lcqtof_t3_neg_4_040_24    m/z 1141.2670     0   1   2E+4   13   13.5  RT (min)      lcqtof_t3_neg_4_064_17    m/z 1141.2971     0   1E+4   13.5   14  RT (min)      lcqtof_t3_neg_4_064_17    m/z 1141.7378     0   1E+4   13   13.5  RT (min)      lcqtof_t3_neg_4_018_23    m/z 1141.7400     0   1E+4   13   13.5  RT (min)      lcqtof_t3_neg_4_018_23    m/z 1141.8446     0   1E+4   12.5   13  RT (min)      lcqtof_t3_neg_1_019_23    m/z 1141.8922     0   1   2   3E+2   12   12.5  RT (min)      lcqtof_t3_neg_3_032_24    m/z 1142.2704     0   1E+4   13   13.5  RT (min)      lcqtof_t3_neg_4_064_17    m/z 1142.2963     0   1E+4   13.5   14  RT (min)      lcqtof_t3_neg_4_064_17    m/z 1142.7698     0   2   4   6   8E+3   13   13.5  RT (min)      lcqtof_t3_neg_1_019_23    m/z 1142.7735     0   1E+4   13   13.5  RT (min)      lcqtof_t3_neg_1_019_23    m/z 1143.1302     0   1E+4   13.5   14  RT (min)      lcqtof_t3_neg_4_064_17    m/z 1143.1552     0   1E+4   13.5   14  RT (min)      lcqtof_t3_neg_4_036_47    m/z 1143.2643     0   1E+4   13   13.5  RT (min)      lcqtof_t3_neg_4_064_17    m/z 1143.2976     0   1E+4   13.5   14  RT (min)      lcqtof_t3_neg_4_064_17    m/z 1143.7751     0   1   2E+4   13   13.5  RT (min)      lcqtof_t3_neg_4_018_23    m/z 1144.7073     0   1E+4   13   13.5  RT (min)      lcqtof_t3_neg_4_018_23    m/z 1144.7179     0   1E+4   13   13.5  RT (min)      lcqtof_t3_neg_4_018_23    m/z 1144.7192     0   1E+4   13   13.5  RT (min)      lcqtof_t3_neg_4_018_23    m/z 1145.0961     0   2   4   6E+4   13.5   14  RT (min)      lcqtof_t3_neg_4_002_17    m/z 1145.0972     0   2   4   6   8E+4   13   13.5  RT (min)      lcqtof_t3_neg_4_064_17    m/z 1145.0983     0   2   4   6   8E+4   13   13.5  RT (min)      lcqtof_t3_neg_4_064_17    m/z 1145.0997     0   2   4   6   8E+4   14   14.5  RT (min)      lcqtof_t3_neg_4_064_17    m/z 1145.1073     0   2   4   6E+4   13   13.5  RT (min)      lcqtof_t3_neg_4_002_17    m/z 1146.0974     0   1   2   3E+4   13.5   14  RT (min)      lcqtof_t3_neg_4_036_47    m/z 1146.0975     0   1E+5   13   13.5  RT (min)      lcqtof_t3_neg_4_064_17    m/z 1146.0979     0   1E+5   13   13.5  RT (min)      lcqtof_t3_neg_4_064_17    m/z 1146.0984     0   1E+5   14   14.5  RT (min)      lcqtof_t3_neg_4_064_17    m/z 1146.0989     0   1E+5   13.5   14  RT (min)      lcqtof_t3_neg_4_064_17    m/z 1146.0990     0   2   4   6E+4   13.5   14  RT (min)      lcqtof_t3_neg_4_002_17    m/z 1146.0992     0   2   4   6E+4   13.5   14  RT (min)      lcqtof_t3_neg_4_002_17    m/z 1146.0992     0   2   4   6E+4   14   14.5  RT (min)      lcqtof_t3_neg_4_002_17    m/z 1146.1001     0   2   4   6E+4   13   13.5  RT (min)      lcqtof_t3_neg_4_002_17    m/z 1146.6583     0   1   2   3E+3   10.5   11  RT (min)      lcqtof_t3_neg_1_018_33    m/z 1147.0928     0   1E+4   14   14.5  RT (min)      lcqtof_t3_neg_4_004_20    m/z 1147.0934     0   1E+5   13   13.5  RT (min)      lcqtof_t3_neg_4_064_17    m/z 1147.0945     0   1E+5   13   13.5  RT (min)      lcqtof_t3_neg_4_064_17    m/z 1147.0953     0   2   4   6E+4   13   13.5  RT (min)      lcqtof_t3_neg_4_002_17    m/z 1147.0973     0   1E+5   14   14.5  RT (min)      lcqtof_t3_neg_4_064_17    m/z 1148.0926     0   2   4   6E+4   13   13.5  RT (min)      lcqtof_t3_neg_4_064_17    m/z 1148.0929     0   2   4   6E+4   13.5   14  RT (min)      lcqtof_t3_neg_4_064_17    m/z 1148.0934     0   2   4   6E+4   13   13.5  RT (min)      lcqtof_t3_neg_4_064_17    m/z 1148.0936     0   2   4   6E+4   13.5   14  RT (min)      lcqtof_t3_neg_4_064_17    m/z 1148.0941     0   2   4E+4   13.5   14  RT (min)      lcqtof_t3_neg_4_002_17    m/z 1148.0943     0   2   4E+4   14   14.5  RT (min)      lcqtof_t3_neg_4_002_17    m/z 1148.0953     0   1   2   3   4E+4   13   13.5  RT (min)      lcqtof_t3_neg_4_002_17    m/z 1148.0956     0   2   4E+4   13.5   14  RT (min)      lcqtof_t3_neg_4_002_17    m/z 1148.0967     0   2   4E+4   13.5   14  RT (min)      lcqtof_t3_neg_4_002_17    m/z 1148.0969     0   2   4E+4   14   14.5  RT (min)      lcqtof_t3_neg_4_002_17    m/z 1149.0860     0   1   2   3   4E+4   13   13.5  RT (min)      lcqtof_t3_neg_4_064_17    m/z 1149.0905     0   1E+4   13.5   14  RT (min)      lcqtof_t3_neg_4_036_47    m/z 1149.0906     0   1   2   3   4E+4   13.5   14  RT (min)      lcqtof_t3_neg_4_064_17    m/z 1149.0911     0   1   2   3   4E+4   13.5   14  RT (min)      lcqtof_t3_neg_4_064_17    m/z 1149.0911     0   1   2   3E+4   13   13.5  RT (min)      lcqtof_t3_neg_4_064_17    m/z 1149.0911     0   1   2   3   4E+4   13   13.5  RT (min)      lcqtof_t3_neg_4_064_17    m/z 1149.0912     0   1   2   3   4E+4   13.5   14  RT (min)      lcqtof_t3_neg_4_064_17    m/z 1149.0974     0   1   2   3   4E+4   14   14.5  RT (min)      lcqtof_t3_neg_4_064_17    m/z 1149.6863     0   2   4   6   8E+3   11   11.5  RT (min)      lcqtof_t3_neg_4_037_01    m/z 1150.0850     0   1   2E+4   13   13.5  RT (min)      lcqtof_t3_neg_4_064_17    m/z 1150.0871     0   1   2E+4   13.5   14  RT (min)      lcqtof_t3_neg_4_064_17    m/z 1150.0910     0   1   2E+4   13.5   14  RT (min)      lcqtof_t3_neg_4_064_17    m/z 1150.0916     0   1   2E+4   13   13.5  RT (min)      lcqtof_t3_neg_4_064_17    m/z 1153.2815     0   1   2E+4   13   13.5  RT (min)      lcqtof_t3_neg_4_064_17    m/z 1153.6613     0   1E+4   10   10.5  RT (min)      lcqtof_t3_neg_4_062_13    m/z 1153.6631     0   1E+4   10   10.5  RT (min)      lcqtof_t3_neg_4_062_13    m/z 1154.2823     0   1E+4   13   13.5  RT (min)      lcqtof_t3_neg_4_002_17    m/z 1154.2826     0   1   2E+4   13   13.5  RT (min)      lcqtof_t3_neg_4_064_17    m/z 1154.6957     0   1   2E+4   12.5   13  RT (min)      lcqtof_t3_neg_2_030_23    m/z 1154.6969     0   1   2   3E+4   12.5   13  RT (min)      lcqtof_t3_neg_1_019_23    m/z 1154.9050     0   1   2E+2   12   12.5  RT (min)      lcqtof_t3_neg_3_032_24    m/z 1154.9053     0   1   2E+2   12.5   13  RT (min)      lcqtof_t3_neg_3_032_24    m/z 1154.9058     0   1E+4   12.5   13  RT (min)      lcqtof_t3_neg_2_016_05    m/z 1154.9060     0   1   2   3   4E+2   11   11.5  RT (min)      lcqtof_t3_neg_3_038_06    m/z 1154.9083     0   1E+4   13   13.5  RT (min)      lcqtof_t3_neg_2_016_05    m/z 1155.2755     0   1E+4   13.5   14  RT (min)      lcqtof_t3_neg_4_002_17    m/z 1155.2800     0   1E+4   13   13.5  RT (min)      lcqtof_t3_neg_4_036_47    m/z 1155.2817     0   1   2E+4   13   13.5  RT (min)      lcqtof_t3_neg_4_064_17    m/z 1155.7001     0   1E+4   12.5   13  RT (min)      lcqtof_t3_neg_1_019_23    m/z 1155.7015     0   1E+4   12.5   13  RT (min)      lcqtof_t3_neg_1_019_23    m/z 1155.9083     0   1   2E+2   12   12.5  RT (min)      lcqtof_t3_neg_3_032_24    m/z 1155.9106     0   2   4E+2   11   11.5  RT (min)      lcqtof_t3_neg_3_038_06    m/z 1155.9115     0   2   4E+2   11   11.5  RT (min)      lcqtof_t3_neg_3_038_06    m/z 1156.2788     0   1E+4   13   13.5  RT (min)      lcqtof_t3_neg_4_036_47    m/z 1156.2793     0   1E+4   13   13.5  RT (min)      lcqtof_t3_neg_4_002_17    m/z 1156.2794     0   1E+4   13   13.5  RT (min)      lcqtof_t3_neg_4_064_17    m/z 1156.2941     0   1E+4   13.5   14  RT (min)      lcqtof_t3_neg_4_064_17    m/z 1156.6211     0   1E+4   12.5   13  RT (min)      lcqtof_t3_neg_4_002_17    m/z 1156.6233     0   1   2E+4   12.5   13  RT (min)       lcqtof_t3_neg_4_058_16    m/z 1156.6249     0   1E+4   12.5   13  RT (min)      lcqtof_t3_neg_4_037_01    m/z 1156.6875     0   2   4   6   8E+3   10.5   11  RT (min)      lcqtof_t3_neg_4_037_01    m/z 1156.8817     0   1   2   3E+2   12   12.5  RT (min)      lcqtof_t3_neg_3_032_24    m/z 1156.8835     0   1   2   3E+2   12   12.5  RT (min)      lcqtof_t3_neg_3_032_24    m/z 1156.8843     0   1   2E+4   12.5   13  RT (min)      lcqtof_t3_neg_2_016_05    m/z 1156.8866     0   1   2   3   4E+2   10.5   11  RT (min)      lcqtof_t3_neg_3_038_06    m/z 1156.8884     0   1   2   3   4E+2   10.5   11  RT (min)      lcqtof_t3_neg_3_038_06    m/z 1157.2933     0   2   4E+4   13.5   14  RT (min)      lcqtof_t3_neg_4_064_17    m/z 1157.6253     0   1E+4   12.5   13  RT (min)      lcqtof_t3_neg_4_006_16    m/z 1157.6271     0   1E+4   12.5   13  RT (min)      lcqtof_t3_neg_4_058_16    m/z 1158.2934     0   1   2   3E+4   13.5   14  RT (min)      lcqtof_t3_neg_4_002_17    m/z 1158.2935     0   2   4E+4   13.5   14  RT (min)      lcqtof_t3_neg_4_064_17    m/z 1158.6352     0   1E+4   12.5   13  RT (min)      lcqtof_t3_neg_4_006_16    m/z 1158.6366     0   1   2E+4   12.5   13  RT (min)      lcqtof_t3_neg_4_058_16    m/z 1158.6392     0   1E+4   12.5   13  RT (min)      lcqtof_t3_neg_4_022_28    m/z 1158.7405     0   1E+4   13   13.5  RT (min)       lcqtof_t3_neg_4_018_23    m/z 1158.7407     0   1E+4   13   13.5  RT (min)      lcqtof_t3_neg_4_018_23    m/z 1158.7513     0   1E+4   13   13.5  RT (min)      lcqtof_t3_neg_4_018_23    m/z 1159.2932     0   1   2   3E+4   13.5   14  RT (min)      lcqtof_t3_neg_4_002_17    m/z 1159.2935     0   2   4E+4   13.5   14  RT (min)      lcqtof_t3_neg_4_064_17    m/z 1159.6399     0   1E+4   12.5   13  RT (min)      lcqtof_t3_neg_4_058_16    m/z 1159.6423     0   1E+4   12.5   13  RT (min)      lcqtof_t3_neg_4_058_16    m/z 1160.2905     0   1   2   3E+4   13.5   14  RT (min)      lcqtof_t3_neg_4_064_17    m/z 1160.2906     0   1   2E+4   13.5   14  RT (min)      lcqtof_t3_neg_4_002_17    m/z 1160.6558     0   1E+4   13   13.5  RT (min)      lcqtof_t3_neg_4_058_16    m/z 1160.9016     0   2   4   6E+3   14.5   15  RT (min)      lcqtof_t3_neg_4_003_37    m/z 1161.2880     0   1E+4   13.5   14  RT (min)      lcqtof_t3_neg_4_064_17    m/z 1161.2933     0   1E+4   13.5   14  RT (min)      lcqtof_t3_neg_4_002_17    m/z 1161.7756     0   1E+4   1   1.5  RT (min)      lcqtof_t3_neg_3_028_13    m/z 1162.1520     0   2   4   6E+4   13.5   14  RT (min)      lcqtof_t3_neg_4_002_17    m/z 1162.1615     0   2   4   6   8E+4   13.5   14  RT (min)      lcqtof_t3_neg_4_064_17    m/z 1162.1622     0   2   4   6E+4   14   14.5  RT (min)      lcqtof_t3_neg_4_002_17    m/z 1162.1642     0   2   4   6   8E+4   14   14.5  RT (min)      lcqtof_t3_neg_4_064_17    m/z 1162.1667     0   2   4   6E+4   14   14.5  RT (min)       lcqtof_t3_neg_4_036_47    m/z 1163.1584     0   1   2   3   4E+4   13.5   14  RT (min)      lcqtof_t3_neg_4_064_17    m/z 1163.1590     0   1   2   3E+4   13.5   14  RT (min)      lcqtof_t3_neg_4_002_17    m/z 1163.1592     0   1   2   3E+4   13.5   14  RT (min)      lcqtof_t3_neg_4_002_17    m/z 1163.1609     0   1   2   3E+4   14   14.5  RT (min)      lcqtof_t3_neg_4_002_17    m/z 1163.1620     0   1   2   3   4E+4   14   14.5  RT (min)      lcqtof_t3_neg_4_064_17    m/z 1163.1666     0   1   2   3E+4   14   14.5  RT (min)      lcqtof_t3_neg_4_036_47    m/z 1164.1534     0   1E+4   13.5   14  RT (min)      lcqtof_t3_neg_4_002_17    m/z 1164.1554     0   1E+4   13.5   14  RT (min)      lcqtof_t3_neg_4_064_17    m/z 1164.1613     0   1E+4   14   14.5  RT (min)      lcqtof_t3_neg_4_064_17    m/z 1164.1621     0   1E+4   14   14.5  RT (min)      lcqtof_t3_neg_4_002_17    m/z 1164.1622     0   1E+4   14   14.5  RT (min)      lcqtof_t3_neg_4_036_47    m/z 1164.1654     0   1E+4   14   14.5  RT (min)      lcqtof_t3_neg_4_036_47    m/z 1165.2524     0   2   4E+4   13   13.5  RT (min)      lcqtof_t3_neg_4_064_17    m/z 1168.7124     0   1   2E+4   13   13.5  RT (min)      lcqtof_t3_neg_4_018_23    m/z 1168.7132     0   1E+4   12.5   13  RT (min)      lcqtof_t3_neg_1_019_23    m/z 1168.9116     0   1E+4   13   13.5  RT (min)      lcqtof_t3_neg_2_016_05    m/z 1168.9227     0   1   2   3E+2   12.5   13  RT (min)      lcqtof_t3_neg_3_032_24    m/z 1168.9229     0   1   2   3E+2   12.5   13  RT (min)      lcqtof_t3_neg_3_032_24    m/z 1169.7170     0   1E+4   13   13.5  RT (min)      lcqtof_t3_neg_4_018_23    m/z 1169.9237     0   1   2   3E+2   12.5   13  RT (min)      lcqtof_t3_neg_3_032_24    m/z 1169.9244     0   1   2   3E+2   12.5   13  RT (min)      lcqtof_t3_neg_3_032_24    m/z 1169.9263     0   2   4E+2   11   11.5  RT (min)      lcqtof_t3_neg_3_038_06    m/z 1169.9265     0   2   4E+2   11   11.5  RT (min)      lcqtof_t3_neg_3_038_06    m/z 1169.9270     0   2   4E+4   13   13.5  RT (min)      lcqtof_t3_neg_2_016_05    m/z 1169.9274     0   2   4E+4   13   13.5  RT (min)      lcqtof_t3_neg_2_016_05    m/z 1169.9280     0   2   4E+2   11   11.5  RT (min)      lcqtof_t3_neg_3_038_06    m/z 1170.9272     0   1   2   3E+2   12.5   13  RT (min)      lcqtof_t3_neg_3_032_24    m/z 1170.9279     0   1E+5   13   13.5  RT (min)      lcqtof_t3_neg_2_016_05    m/z 1170.9280     0   1   2   3E+2   12.5   13  RT (min)      lcqtof_t3_neg_3_032_24    m/z 1170.9283     0   1   2   3E+2   11   11.5  RT (min)      lcqtof_t3_neg_3_038_06    m/z 1170.9285     0   1   2   3E+2   11   11.5  RT (min)      lcqtof_t3_neg_3_038_06    m/z 1172.2689     0   1E+4   13   13.5  RT (min)      lcqtof_t3_neg_4_064_17    m/z 1172.6937     0   1E+4   13   13.5  RT (min)      lcqtof_t3_neg_4_018_23    m/z 1175.1762     0   1   2E+4   14   14.5  RT (min)      lcqtof_t3_neg_4_004_20    m/z 1175.1773     0   1   2E+4   13.5   14  RT (min)      lcqtof_t3_neg_4_004_20    m/z 1175.1780     0   2   4   6E+4   13   13.5  RT (min)      lcqtof_t3_neg_4_036_47    m/z 1175.1797     0   1E+5   13.5   14  RT (min)      lcqtof_t3_neg_4_002_17    m/z 1175.1803     0   1E+5   14   14.5  RT (min)      lcqtof_t3_neg_4_002_17    m/z 1175.1806     0   1E+5   13.5   14  RT (min)      lcqtof_t3_neg_4_064_17    m/z 1175.1816     0   1E+5   13.5   14  RT (min)      lcqtof_t3_neg_4_064_17    m/z 1175.1818     0   1E+5   13.5   14  RT (min)      lcqtof_t3_neg_4_064_17    m/z 1176.1784     0   1E+5   13.5   14  RT (min)      lcqtof_t3_neg_4_064_17    m/z 1176.1795     0   1E+5   13.5   14  RT (min)      lcqtof_t3_neg_4_064_17    m/z 1176.1808     0   2   4   6E+4   13   13.5  RT (min)      lcqtof_t3_neg_4_036_47    m/z 1176.1810     0   1E+4   13.5   14  RT (min)      lcqtof_t3_neg_4_013_39    m/z 1176.1813     0   2   4   6E+4   14   14.5  RT (min)       lcqtof_t3_neg_4_036_47    m/z 1176.1816     0   1E+5   14   14.5  RT (min)       lcqtof_t3_neg_4_064_17    m/z 1176.1840     0   1E+5   13.5   14  RT (min)      lcqtof_t3_neg_4_002_17    m/z 1177.1760     0   1   2E+4   14   14.5  RT (min)      lcqtof_t3_neg_4_004_20    m/z 1177.1761     0   1   2E+4   14   14.5  RT (min)      lcqtof_t3_neg_4_004_20    m/z 1177.1767     0   1   2E+4   14   14.5  RT (min)      lcqtof_t3_neg_4_004_20    m/z 1177.1771     0   1   2E+4   13.5   14  RT (min)      lcqtof_t3_neg_4_004_20    m/z 1177.1775     0   1E+5   13.5   14  RT (min)      lcqtof_t3_neg_4_064_17    m/z 1177.1775     0   2   4   6E+4   13.5   14  RT (min)      lcqtof_t3_neg_4_036_47    m/z 1177.1786     0   2   4   6E+4   13   13.5  RT (min)      lcqtof_t3_neg_4_036_47    m/z 1177.1798     0   1E+5   13.5   14  RT (min)      lcqtof_t3_neg_4_064_17    m/z 1177.1803     0   1E+5   14   14.5  RT (min)       lcqtof_t3_neg_4_064_17    m/z 1177.6573     0   1E+4   10   10.5  RT (min)      lcqtof_t3_neg_4_062_13    m/z 1178.1758     0   2   4E+4   13   13.5  RT (min)      lcqtof_t3_neg_4_036_47    m/z 1178.1761     0   1E+4   14   14.5  RT (min)      lcqtof_t3_neg_4_004_20    m/z 1178.1765     0   2   4E+4   13   13.5  RT (min)      lcqtof_t3_neg_4_036_47    m/z 1178.1772     0   1E+4   13.5   14  RT (min)      lcqtof_t3_neg_4_004_20    m/z 1178.1789     0   1E+4   13.5   14  RT (min)      lcqtof_t3_neg_4_004_20    m/z 1178.1793     0   1E+5   13.5   14  RT (min)      lcqtof_t3_neg_4_064_17    m/z 1178.1807     0   1E+5   14   14.5  RT (min)      lcqtof_t3_neg_4_064_17    m/z 1178.1814     0   1E+5   13.5   14  RT (min)      lcqtof_t3_neg_4_064_17    m/z 1178.1821     0   2   4   6   8E+4   13.5   14  RT (min)      lcqtof_t3_neg_4_002_17    m/z 1179.1747     0   2   4   6E+4   14   14.5  RT (min)      lcqtof_t3_neg_4_064_17    m/z 1179.1750     0   2   4   6E+4   13.5   14  RT (min)      lcqtof_t3_neg_4_064_17    m/z 1179.1758     0   2   4E+4   13   13.5  RT (min)      lcqtof_t3_neg_4_002_17    m/z 1179.1776     0   2   4E+4   13.5   14  RT (min)      lcqtof_t3_neg_4_002_17    m/z 1179.1786     0   2   4   6E+4   13.5   14  RT (min)      lcqtof_t3_neg_4_064_17    m/z 1179.1788     0   1   2   3E+4   14   14.5  RT (min)      lcqtof_t3_neg_4_036_47    m/z 1179.1832     0   2   4   6E+4   13.5   14  RT (min)      lcqtof_t3_neg_4_064_17    m/z 1179.2838     0   1E+4   13   13.5  RT (min)      lcqtof_t3_neg_4_002_17    m/z 1179.2881     0   1   2   3E+4   13   13.5  RT (min)      lcqtof_t3_neg_4_064_17    m/z 1179.7042     0   1E+4   7   7.5  RT (min)      lcqtof_t3_neg_4_048_19    m/z 1180.1265     0   1E+4   13   13.5  RT (min)      lcqtof_t3_neg_4_063_39    m/z 1180.1306     0   1E+4   13   13.5  RT (min)      lcqtof_t3_neg_4_063_39    m/z 1180.1726     0   1   2E+4   13.5   14  RT (min)      lcqtof_t3_neg_4_002_17    m/z 1180.1746     0   1   2   3E+4   13.5   14  RT (min)      lcqtof_t3_neg_4_064_17    m/z 1180.1760     0   1   2   3E+4   13.5   14  RT (min)      lcqtof_t3_neg_4_064_17    m/z 1180.1771     0   1   2E+4   13.5   14  RT (min)      lcqtof_t3_neg_4_002_17    m/z 1180.2877     0   1E+4   13   13.5  RT (min)      lcqtof_t3_neg_4_064_17    m/z 1181.1258     0   1E+4   13   13.5  RT (min)      lcqtof_t3_neg_4_063_39    m/z 1181.1699     0   1E+4   13.5   14  RT (min)      lcqtof_t3_neg_4_064_17    m/z 1181.1729     0   1E+4   13.5   14  RT (min)      lcqtof_t3_neg_4_064_17    m/z 1181.1771     0   1E+4   13.5   14  RT (min)      lcqtof_t3_neg_4_064_17    m/z 1182.7291     0   1   2   3   4E+4   13   13.5  RT (min)      lcqtof_t3_neg_1_019_23    m/z 1183.7212     0   2   4   6E+4   13   13.5  RT (min)      lcqtof_t3_neg_4_018_23    m/z 1183.7216     0   2   4   6E+4   13   13.5  RT (min)      lcqtof_t3_neg_4_018_23    m/z 1183.7313     0   1   2   3E+4   13   13.5  RT (min)      lcqtof_t3_neg_1_019_23    m/z 1183.7331     0   1   2E+4   13   13.5  RT (min)      lcqtof_t3_neg_1_019_23    m/z 1184.7188     0   1   2E+4   13   13.5  RT (min)      lcqtof_t3_neg_4_018_23    m/z 1184.7222     0   1   2E+4   13   13.5  RT (min)      lcqtof_t3_neg_4_018_23    m/z 1184.7245     0   1   2E+4   13   13.5  RT (min)      lcqtof_t3_neg_4_018_23    m/z 1184.7271     0   1E+4   13   13.5  RT (min)      lcqtof_t3_neg_4_089_59    m/z 1184.7345     0   1E+4   13   13.5  RT (min)      lcqtof_t3_neg_4_019_05    m/z 1184.7352     0   2   4   6   8E+3   13   13.5  RT (min)      lcqtof_t3_neg_2_030_23    m/z 1184.9026     0   1E+4   13   13.5  RT (min)       lcqtof_t3_neg_4_040_24    m/z 1184.9375     0   1E+4   13.5   14  RT (min)      lcqtof_t3_neg_4_040_24    m/z 1184.9390     0   1E+4   13.5   14  RT (min)      lcqtof_t3_neg_4_011_32    m/z 1185.7334     0   1   2E+4   13   13.5  RT (min)      lcqtof_t3_neg_4_018_23    m/z 1185.7336     0   1   2E+4   13   13.5  RT (min)      lcqtof_t3_neg_4_018_23    m/z 1186.7373     0   1E+4   13   13.5  RT (min)      lcqtof_t3_neg_4_018_23    m/z 1186.7389     0   1E+4   13   13.5  RT (min)      lcqtof_t3_neg_4_018_23    m/z 1186.9229     0   1   2   3   4E+2   12   12.5  RT (min)      lcqtof_t3_neg_3_032_24    m/z 1186.9249     0   1   2E+4   12.5   13  RT (min)      lcqtof_t3_neg_2_016_05    m/z 1186.9285     0   1   2   3   4E+2   11   11.5  RT (min)      lcqtof_t3_neg_3_038_06    m/z 1187.7500     0   1   2E+4   13   13.5  RT (min)      lcqtof_t3_neg_4_018_23    m/z 1187.7513     0   1   2E+4   13   13.5  RT (min)      lcqtof_t3_neg_4_018_23    m/z 1187.7532     0   1   2E+4   13   13.5  RT (min)      lcqtof_t3_neg_4_018_23    m/z 1188.7535     0   1E+4   13   13.5  RT (min)      lcqtof_t3_neg_4_018_23    m/z 1190.2367     0   1   2E+4   13.5   14  RT (min)      lcqtof_t3_neg_4_002_17    m/z 1190.2392     0   1   2E+4   14   14.5  RT (min)      lcqtof_t3_neg_4_036_47    m/z 1190.2498     0   1   2E+4   14   14.5  RT (min)      lcqtof_t3_neg_4_064_17    m/z 1190.2546     0   1   2E+4   14   14.5  RT (min)      lcqtof_t3_neg_4_036_47    m/z 1190.7163     0   1E+4   13   13.5  RT (min)      lcqtof_t3_neg_4_018_23    m/z 1190.7239     0   1E+4   13   13.5  RT (min)      lcqtof_t3_neg_4_018_23    m/z 1191.0991     0   2   4   6E+4   13   13.5  RT (min)      lcqtof_t3_neg_4_064_17    m/z 1191.1006     0   1   2E+4   13.5   14  RT (min)      lcqtof_t3_neg_4_002_17    m/z 1191.1013     0   1   2E+4   13.5   14  RT (min)      lcqtof_t3_neg_4_002_17    m/z 1191.1018     0   2   4   6E+4   13   13.5  RT (min)      lcqtof_t3_neg_4_064_17    m/z 1191.1025     0   2   4   6E+4   13.5   14  RT (min)      lcqtof_t3_neg_4_064_17    m/z 1191.1037     0   2   4   6E+4   13.5   14  RT (min)      lcqtof_t3_neg_4_064_17    m/z 1191.1102     0   1   2E+4   13.5   14  RT (min)      lcqtof_t3_neg_4_002_17    m/z 1191.1160     0   1   2E+4   14   14.5  RT (min)      lcqtof_t3_neg_4_002_17    m/z 1191.2437     0   1E+4   14   14.5  RT (min)      lcqtof_t3_neg_4_064_17    m/z 1191.2467     0   1E+4   14   14.5  RT (min)      lcqtof_t3_neg_4_002_17    m/z 1191.2473     0   1E+4   14   14.5  RT (min)      lcqtof_t3_neg_4_036_47    m/z 1192.0977     0   2   4   6E+4   13   13.5  RT (min)      lcqtof_t3_neg_4_064_17    m/z 1192.0984     0   2   4   6E+4   13   13.5  RT (min)      lcqtof_t3_neg_4_064_17    m/z 1192.0985     0   2   4   6E+4   13.5   14  RT (min)      lcqtof_t3_neg_4_064_17    m/z 1192.1008     0   1   2E+4   13.5   14  RT (min)      lcqtof_t3_neg_4_064_17    m/z 1192.1029     0   2   4   6E+4   13.5   14  RT (min)      lcqtof_t3_neg_4_064_17    m/z 1193.0958     0   2   4   6E+4   13   13.5  RT (min)      lcqtof_t3_neg_4_064_17    m/z 1193.0963     0   2   4   6E+4   13   13.5  RT (min)      lcqtof_t3_neg_4_064_17    m/z 1193.0970     0   2   4   6E+4   13   13.5  RT (min)      lcqtof_t3_neg_4_064_17    m/z 1193.0991     0   1   2E+4   14   14.5  RT (min)      lcqtof_t3_neg_4_064_17    m/z 1193.1039     0   2   4   6E+4   13.5   14  RT (min)      lcqtof_t3_neg_4_064_17    m/z 1193.1063     0   2   4   6E+4   13.5   14  RT (min)      lcqtof_t3_neg_4_064_17    m/z 1193.5201     0   1E+4   9.5   10  RT (min)       lcqtof_t3_neg_4_059_14    m/z 1194.0954     0   2   4E+4   13   13.5  RT (min)      lcqtof_t3_neg_4_064_17    m/z 1194.0973     0   2   4E+4   13   13.5  RT (min)      lcqtof_t3_neg_4_064_17    m/z 1194.0977     0   2   4E+4   13.5   14  RT (min)      lcqtof_t3_neg_4_064_17    m/z 1194.0986     0   2   4E+4   13.5   14  RT (min)      lcqtof_t3_neg_4_064_17    m/z 1195.0946     0   1   2   3E+4   13   13.5  RT (min)      lcqtof_t3_neg_4_064_17    m/z 1195.0967     0   1   2   3E+4   13.5   14  RT (min)      lcqtof_t3_neg_4_064_17    m/z 1195.0972     0   1   2   3E+4   13   13.5  RT (min)      lcqtof_t3_neg_4_064_17    m/z 1195.2722     0   1E+4   13   13.5  RT (min)      lcqtof_t3_neg_4_064_17    m/z 1195.9207     0   1E+4   13.5   14  RT (min)      lcqtof_t3_neg_4_040_24    m/z 1196.0974     0   1E+4   13   13.5  RT (min)      lcqtof_t3_neg_4_064_17    m/z 1196.2715     0   1E+4   13   13.5  RT (min)      lcqtof_t3_neg_4_064_17    m/z 1196.9167     0   1E+4   13.5   14  RT (min)      lcqtof_t3_neg_4_019_05    m/z 1196.9230     0   1E+4   13.5   14  RT (min)       lcqtof_t3_neg_4_040_24    m/z 1197.2426     0   1E+4   13.5   14  RT (min)      lcqtof_t3_neg_4_057_49    m/z 1197.2721     0   1E+4   13   13.5  RT (min)      lcqtof_t3_neg_4_064_17    m/z 1198.2447     0   1E+4   13.5   14  RT (min)      lcqtof_t3_neg_4_057_49    m/z 1198.2546     0   1E+4   13   13.5  RT (min)      lcqtof_t3_neg_4_064_17    m/z 1198.2913     0   1E+4   13   13.5  RT (min)      lcqtof_t3_neg_4_064_17    m/z 1198.2917     0   1E+4   13   13.5  RT (min)      lcqtof_t3_neg_4_036_47    m/z 1199.2399     0   1E+4   13.5   14  RT (min)      lcqtof_t3_neg_4_057_49    m/z 1199.6481     0   1E+4   10   10.5  RT (min)      lcqtof_t3_neg_4_058_16   
