## Supplementary material for "The Everything Bagel Feature Finder: Ultra-fast automated feature finding for untargeted metabolomics": SI Data File: tool_unique_xic_st_Masscube_ST004651.html

 
 
 Masscube-unique features (not detected by the other two tools) - ST004651 
 
 Masscube-unique features (not detected by the other two tools) - ST004651 
 13746 features · qtof · XIC window m/z ± 25 ppm (≥ 0.002 Da) · ± 0.5 min 
 Every panel is a feature  Masscube  detected that neither of the other two tools detected — three-way unique across EB (standard mode (min_detection_rate=0.1)), Masscube and XCMS, under the standard matching rule (m/z ± 25 ppm; apex within the reference RT window ± 0.2 min). It shows the MS1 extracted-ion chromatogram at the feature's m/z, drawn from the raw mzML named in  alignment_reference_file , zoomed to ± 0.5 min around the feature's reported apex RT.  Gray solid line  = the feature's reported apex RT;  red dashed line  = masscube's assigned MS/MS —  MS2_scan_id  in  MS2_source_file ; the panel widens to keep an off-apex MS/MS on-screen. Features with no assigned MS/MS show only the gray line. y is per-cell autoscaled. Rows keep the table order (m/z ascending). 
   m/z 61.9876     0   1   2   3E+3   5.5   6  RT (min)      lcqtof_hilic_neg_1_008_13    m/z 61.9877     0   1   2   3   4E+3   1.5   2  RT (min)      lcqtof_hilic_neg_4_061_50    m/z 61.9877     0   1   2   3   4E+3   3.5   4  RT (min)      lcqtof_hilic_neg_4_045_26    m/z 61.9878     0   1   2   3   4E+3   2   2.5  RT (min)      lcqtof_hilic_neg_4_061_50    m/z 73.0475     0   1E+4   2   2.5  RT (min)      lcqtof_hilic_neg_4_023_15    m/z 78.9580     0   1   2   3   4E+4   5   5.5  RT (min)      lcqtof_hilic_neg_1_000_aa    m/z 78.9582     0   1   2   3   4E+4   5.5   6  RT (min)      lcqtof_hilic_neg_1_000_aa    m/z 78.9583     0   1   2   3E+4   7.5   8  RT (min)      lcqtof_hilic_neg_1_000_aa    m/z 78.9583     0   1   2   3E+4   8   8.5  RT (min)      lcqtof_hilic_neg_1_000_aa    m/z 78.9583     0   1   2   3E+5   7   7.5  RT (min)      lcqtof_hilic_neg_4_040_05    m/z 78.9583     0   1E+4   3.5   4  RT (min)      lcqtof_hilic_neg_3_048_05    m/z 78.9583     0   1   2   3   4E+4   5.5   6  RT (min)      lcqtof_hilic_neg_1_000_aa    m/z 78.9584     0   1E+4   9   9.5  RT (min)      lcqtof_hilic_neg_1_000_aa    m/z 78.9584     0   1   2   3E+4   8   8.5  RT (min)      lcqtof_hilic_neg_1_000_aa    m/z 78.9584     0   1E+4   4.5   5  RT (min)      lcqtof_hilic_neg_4_002_25    m/z 78.9584     0   1   2   3E+4   8   8.5  RT (min)      lcqtof_hilic_neg_1_000_aa    m/z 78.9584     0   1E+4   3.5   4  RT (min)      lcqtof_hilic_neg_3_048_05    m/z 78.9585     0   1   2   3E+4   7.5   8  RT (min)      lcqtof_hilic_neg_1_000_aa    m/z 78.9586     0   1   2   3E+4   8   8.5  RT (min)      lcqtof_hilic_neg_1_000_aa    m/z 78.9587     0   1   2   3   4E+4   5.5   6  RT (min)      lcqtof_hilic_neg_1_000_aa    m/z 79.9568     0   1E+4   7   7.5  RT (min)      lcqtof_hilic_neg_4_047_06    m/z 80.9163     0   1   2E+3   4   4.5  RT (min)      lcqtof_hilic_neg_4_001_49    m/z 80.9166     0   1   2   3   4E+4   2.5   3  RT (min)      lcqtof_hilic_neg_4_019_03    m/z 80.9646     0   2   4   6   8E+3   1.5   2  RT (min)      lcqtof_hilic_neg_4_047_06    m/z 83.0138     0   1E+4   1.5   2  RT (min)      lcqtof_hilic_neg_4_023_15    m/z 83.0139     0   1E+4   1.5   2  RT (min)      lcqtof_hilic_neg_4_010_28    m/z 83.0139     0   1E+4   1.5   2  RT (min)      lcqtof_hilic_neg_4_064_15    m/z 83.9303     0   1   2   3E+3   1   1.5  RT (min)      lcqtof_hilic_neg_2_012_43    m/z 83.9303     0   1   2   3E+3   2   2.5  RT (min)      lcqtof_hilic_neg_3_068_57    m/z 84.9380     0   2   4   6E+3   5   5.5  RT (min)      lcqtof_hilic_neg_2_047_32    m/z 84.9380     0   2   4   6E+3   5   5.5  RT (min)      lcqtof_hilic_neg_2_044_37    m/z 84.9381     0   2   4   6E+3   5   5.5  RT (min)      lcqtof_hilic_neg_1_025_52    m/z 84.9381     0   2   4   6E+3   5   5.5  RT (min)      lcqtof_hilic_neg_2_047_32    m/z 85.0288     0   1   2   3E+4   3   3.5  RT (min)      lcqtof_hilic_neg_4_039_18    m/z 85.0289     0   1E+4   3.5   4  RT (min)      lcqtof_hilic_neg_1_032_18    m/z 85.0290     0   2   4   6E+3   4   4.5  RT (min)      lcqtof_hilic_neg_4_048_23    m/z 85.9398     0   1   2E+3   2   2.5  RT (min)      lcqtof_hilic_neg_3_062_14    m/z 86.0312     0   1E+3   1   1.5  RT (min)      lcqtof_hilic_neg_4_050_02    m/z 86.0318     0   2   4   6E+2   1.5   2  RT (min)      lcqtof_hilic_neg_4_019_03    m/z 87.0027     0   1   2E+4   0.5   1  RT (min)      lcqtof_hilic_neg_2_024_10    m/z 87.0084     0   1E+4   4.5   5  RT (min)      lcqtof_hilic_neg_1_004_14    m/z 87.9483     0   1E+4   1   1.5  RT (min)      lcqtof_hilic_neg_2_045_11    m/z 88.0395     0   2   4   6   8E+4   6   6.5  RT (min)      lcqtof_hilic_neg_4_082_59    m/z 89.0237     0   2   4E+3   2.5   3  RT (min)      lcqtof_hilic_neg_4_019_03    m/z 89.0237     0   2   4   6E+3   5.5   6  RT (min)      lcqtof_hilic_neg_4_032_01    m/z 89.0237     0   1   2   3   4E+4   3   3.5  RT (min)      lcqtof_hilic_neg_4_049_55    m/z 89.0237     0   2   4   6   8E+4   1.5   2  RT (min)      lcqtof_hilic_neg_4_004_47    m/z 89.0237     0   2   4E+4   3   3.5  RT (min)      lcqtof_hilic_neg_4_014_46    m/z 89.0238     0   1E+4   0.5   1  RT (min)      lcqtof_hilic_neg_1_040_20    m/z 89.0238     0   2   4   6E+3   3.5   4  RT (min)      lcqtof_hilic_neg_4_050_02    m/z 89.0238     0   1E+4   5.5   6  RT (min)      lcqtof_hilic_neg_1_063_13    m/z 89.0238     0   1   2   3E+3   3.5   4  RT (min)      lcqtof_hilic_neg_4_062_17    m/z 89.0238     0   2   4   6E+3   3.5   4  RT (min)      lcqtof_hilic_neg_4_050_02    m/z 89.0238     0   2   4E+3   2.5   3  RT (min)      lcqtof_hilic_neg_4_019_03    m/z 89.0239     0   1   2   3   4E+3   5.5   6  RT (min)      lcqtof_hilic_neg_4_045_26    m/z 89.0239     0   1   2   3E+4   2   2.5  RT (min)      lcqtof_hilic_neg_4_017_51    m/z 89.0239     0   1   2E+4   2   2.5  RT (min)      lcqtof_hilic_neg_4_014_46    m/z 89.0240     0   1E+4   5.5   6  RT (min)      lcqtof_hilic_neg_1_025_52    m/z 90.0181     0   1E+4   2   2.5  RT (min)      lcqtof_hilic_neg_4_087_59    m/z 91.0499     0   1E+4   2   2.5  RT (min)      lcqtof_hilic_neg_4_055_32    m/z 94.9802     0   2   4E+3   3.5   4  RT (min)       lcqtof_hilic_neg_4_014_46    m/z 94.9804     0   1E+4   5.5   6  RT (min)      lcqtof_hilic_neg_1_000_aa    m/z 96.9593     0   1E+5   1.5   2  RT (min)      lcqtof_hilic_neg_4_032_01    m/z 96.9594     0   1E+5   8   8.5  RT (min)      lcqtof_hilic_neg_1_000_aa    m/z 96.9594     0   1   2   3E+4   2   2.5  RT (min)       lcqtof_hilic_neg_3_068_57    m/z 96.9594     0   1E+4   2   2.5  RT (min)       lcqtof_hilic_neg_2_021_17    m/z 96.9594     0   1   2E+4   2   2.5  RT (min)       lcqtof_hilic_neg_4_009_50    m/z 96.9594     0   1E+4   0   0.5  RT (min)      lcqtof_hilic_neg_4_000_aa    m/z 96.9595     0   2   4   6E+4   1.5   2  RT (min)       lcqtof_hilic_neg_4_004_47    m/z 96.9595     0   1   2   3   4E+5   7   7.5  RT (min)       lcqtof_hilic_neg_4_092_60    m/z 96.9595     0   2   4   6E+3   2.5   3  RT (min)       lcqtof_hilic_neg_4_085_59    m/z 96.9595     0   1E+4   0   0.5  RT (min)       lcqtof_hilic_neg_4_000_aa    m/z 96.9596     0   1   2   3   4E+3   2.5   3  RT (min)      lcqtof_hilic_neg_2_052_24    m/z 96.9596     0   1E+4   0   0.5  RT (min)       lcqtof_hilic_neg_4_000_aa    m/z 96.9596     0   1   2   3   4E+3   3   3.5  RT (min)      lcqtof_hilic_neg_4_034_31    m/z 96.9596     0   1E+4   2   2.5  RT (min)      lcqtof_hilic_neg_4_061_50    m/z 96.9597     0   1E+4   0   0.5  RT (min)      lcqtof_hilic_neg_4_000_aa    m/z 96.9604     0   2   4   6E+4   8.5   9  RT (min)       lcqtof_hilic_neg_1_000_aa    m/z 96.9617     0   1   2   3E+5   7   7.5  RT (min)      lcqtof_hilic_neg_4_061_50    m/z 96.9621     0   2   4   6   8E+3   8.5   9  RT (min)       lcqtof_hilic_neg_1_000_aa    m/z 96.9627     0   2   4   6E+3   7.5   8  RT (min)      lcqtof_hilic_neg_2_000_aa    m/z 96.9629     0   1   2E+3   7.5   8   8.5  RT (min)      lcqtof_hilic_neg_1_000_aa    m/z 96.9631     0   1   2E+3   8   8.5  RT (min)       lcqtof_hilic_neg_1_000_aa    m/z 96.9632     0   1E+4   4   4.5  RT (min)       lcqtof_hilic_neg_4_008_24    m/z 96.9686     0   2   4   6   8E+4   7   7.5  RT (min)      lcqtof_hilic_neg_1_000_aa    m/z 97.0101     0   1E+4   7.5   8  RT (min)      lcqtof_hilic_neg_4_008_24    m/z 97.9597     0   2   4E+3   7   7.5  RT (min)      lcqtof_hilic_neg_4_040_05    m/z 98.9389     0   1   2E+3   4   4.5  RT (min)      lcqtof_hilic_neg_2_048_35    m/z 98.9394     0   1   2   3E+3   1   1.5  RT (min)      lcqtof_hilic_neg_4_019_03    m/z 98.9556     0   1   2   3E+4   7   7.5  RT (min)      lcqtof_hilic_neg_1_000_aa    m/z 98.9560     0   1   2E+4   7   7.5  RT (min)      lcqtof_hilic_neg_4_047_06    m/z 99.9250     0   1   2E+4   2   4  RT (min)       lcqtof_hilic_neg_2_049_55    m/z 99.9250     0   1   2E+4   2  RT (min)       lcqtof_hilic_neg_2_031_01    m/z 99.9250     0   1   2E+5   1   1.5  RT (min)       lcqtof_hilic_neg_4_050_02    m/z 99.9250     0   2   4E+3   18   18.5  RT (min)      lcqtof_hilic_neg_3_067_57    m/z 99.9251     0   1E+4   5   5.5  RT (min)      lcqtof_hilic_neg_2_016_33    m/z 99.9251     0   1E+4   5   5.5  RT (min)      lcqtof_hilic_neg_2_013_29    m/z 99.9251     0   2   4E+3   18.5   19  RT (min)      lcqtof_hilic_neg_3_111_62    m/z 99.9251     0   1E+4   4.5   5  RT (min)      lcqtof_hilic_neg_2_031_01    m/z 99.9251     0   1E+4   5   5.5  RT (min)      lcqtof_hilic_neg_2_018_16    m/z 99.9251     0   1E+4   5   5.5  RT (min)      lcqtof_hilic_neg_2_031_01    m/z 99.9252     0   1   2   3E+3   2.5   3  RT (min)      lcqtof_hilic_neg_4_074_58    m/z 99.9253     0   1E+4   6   6.5  RT (min)      lcqtof_hilic_neg_2_036_34    m/z 100.9324     0   2   4E+4   1.5   2  RT (min)       lcqtof_hilic_neg_4_063_16    m/z 100.9325     0   2   4   6   8E+3   4.5   5  RT (min)      lcqtof_hilic_neg_3_112_62    m/z 100.9325     0   1E+4   1.5   2  RT (min)       lcqtof_hilic_neg_4_060_13    m/z 100.9325     0   2   4E+4   19.5   20  RT (min)       lcqtof_hilic_neg_1_036_03    m/z 100.9325     0   2   4   6E+3   5   5.5  RT (min)      lcqtof_hilic_neg_3_011_34    m/z 100.9325     0   1   2   3E+4   19   19.5  RT (min)       lcqtof_hilic_neg_2_060_14    m/z 100.9326     0   2   4E+4   19.5   20  RT (min)       lcqtof_hilic_neg_2_000_bb    m/z 100.9326     0   2   4   6E+3   3   3.5  RT (min)       lcqtof_hilic_neg_3_104_61    m/z 100.9326     0   1   2   3   4E+4   20   20.5  RT (min)       lcqtof_hilic_neg_1_055_25    m/z 100.9326     0   1   2   3E+4   17.5   18  RT (min)      lcqtof_hilic_neg_1_059_15    m/z 100.9326     0   1   2   3   4E+4   18.5   19  RT (min)       lcqtof_hilic_neg_1_060_14    m/z 100.9326     0   1   2   3   4E+4   18.5   19  RT (min)       lcqtof_hilic_neg_1_058_39    m/z 100.9326     0   1   2   3   4E+4   18   18.5  RT (min)       lcqtof_hilic_neg_1_058_39    m/z 100.9326     0   1   2   3E+4   17.5   18  RT (min)      lcqtof_hilic_neg_1_057_49    m/z 100.9326     0   1   2   3E+4   17.5   18  RT (min)      lcqtof_hilic_neg_1_059_15    m/z 100.9327     0   1   2   3E+4   17   18  RT (min)       lcqtof_hilic_neg_1_061_50    m/z 100.9327     0   1E+4   0   0.5  RT (min)      lcqtof_hilic_neg_4_065_51    m/z 100.9327     0   1   2   3E+4   17.5   18  RT (min)      lcqtof_hilic_neg_1_060_14    m/z 100.9327     0   1   2   3E+4   17.5   18  RT (min)      lcqtof_hilic_neg_1_057_49    m/z 100.9327     0   1E+4   2   2.5  RT (min)       lcqtof_hilic_neg_4_059_14    m/z 100.9327     0   1   2   3   4E+4   18.5   19  RT (min)       lcqtof_hilic_neg_1_065_51    m/z 100.9327     0   1   2   3   4E+4   18.5   19  RT (min)       lcqtof_hilic_neg_1_057_49    m/z 100.9327     0   2   4   6E+4   1.5   2  RT (min)       lcqtof_hilic_neg_2_027_07    m/z 100.9327     0   1   2   3   4E+4   18.5   19  RT (min)       lcqtof_hilic_neg_1_057_49    m/z 100.9327     0   1   2E+4   17   17.5  RT (min)      lcqtof_hilic_neg_1_057_49    m/z 100.9327     0   1   2E+4   17   17.5  RT (min)      lcqtof_hilic_neg_1_060_14    m/z 100.9327     0   1E+5   2   2.5  RT (min)       lcqtof_hilic_neg_1_054_01    m/z 100.9327     0   1E+4   0   0.5  RT (min)       lcqtof_hilic_neg_4_059_14    m/z 100.9327     0   1E+4   0.5   1  RT (min)       lcqtof_hilic_neg_4_057_49    m/z 100.9327     0   2   4   6   8E+3   2.5   3  RT (min)       lcqtof_hilic_neg_4_082_59    m/z 100.9328     0   1E+4   2   2.5  RT (min)       lcqtof_hilic_neg_4_056_56    m/z 100.9328     0   1   2E+4   16.5   17  RT (min)       lcqtof_hilic_neg_4_000_cc    m/z 100.9328     0   1   2E+4   17   17.5  RT (min)       lcqtof_hilic_neg_1_057_49    m/z 100.9328     0   1E+4   0   0.5  RT (min)       lcqtof_hilic_neg_4_061_50    m/z 100.9328     0   2   4E+4   1.5   2  RT (min)       lcqtof_hilic_neg_4_078_58    m/z 100.9328     0   1E+4   0.5   1  RT (min)      lcqtof_hilic_neg_4_061_50    m/z 100.9328     0   2   4E+4   0.5   1  RT (min)      lcqtof_hilic_neg_4_061_50    m/z 100.9328     0   1E+4   0.5   1  RT (min)       lcqtof_hilic_neg_4_060_13    m/z 100.9328     0   1   2E+4   17   17.5  RT (min)      lcqtof_hilic_neg_1_057_49    m/z 100.9328     0   1E+4   0   0.5  RT (min)       lcqtof_hilic_neg_4_056_56    m/z 100.9329     0   1   2E+4   17   17.5  RT (min)      lcqtof_hilic_neg_1_057_49    m/z 100.9329     0   1E+4   0   0.5  RT (min)       lcqtof_hilic_neg_4_060_13    m/z 100.9329     0   2   4E+3   3   3.5  RT (min)      lcqtof_hilic_neg_3_107_62    m/z 100.9329     0   1E+4   2.5   3  RT (min)       lcqtof_hilic_neg_4_060_13    m/z 100.9329     0   1E+4   2   2.5  RT (min)       lcqtof_hilic_neg_4_076_58    m/z 101.0239     0   1   2   3E+3   5   5.5  RT (min)      lcqtof_hilic_neg_4_055_32    m/z 101.0240     0   1   2   3E+3   3.5   4  RT (min)      lcqtof_hilic_neg_1_017_51    m/z 101.0241     0   2   4   6   8E+3   5   5.5  RT (min)      lcqtof_hilic_neg_4_016_37    m/z 101.9340     0   1   2   3   4E+3   5   5.5  RT (min)      lcqtof_hilic_neg_2_043_15    m/z 101.9341     0   2   4E+3   5   5.5  RT (min)      lcqtof_hilic_neg_2_048_35    m/z 101.9342     0   2   4E+3   5   5.5  RT (min)      lcqtof_hilic_neg_2_049_55    m/z 102.9330     0   1   2E+3   4   4.5  RT (min)      lcqtof_hilic_neg_2_029_31    m/z 102.9474     0   2   4   6   8E+3   5   5.5  RT (min)      lcqtof_hilic_neg_4_044_08    m/z 102.9489     0   1E+4   10   10.5  RT (min)      lcqtof_hilic_neg_4_014_46    m/z 105.9689     0   1E+4   5.5   6  RT (min)      lcqtof_hilic_neg_1_000_aa    m/z 107.0470     0   1   2E+4   4   4.5  RT (min)      lcqtof_hilic_neg_4_055_32    m/z 108.9014     0   2   4E+4   4.5   5  RT (min)      lcqtof_hilic_neg_4_050_02    m/z 109.0357     0   1   2   3E+3   1   1.5  RT (min)      lcqtof_hilic_neg_4_052_13    m/z 110.8984     0   1   2   3   4E+4   4.5   5  RT (min)      lcqtof_hilic_neg_4_050_02    m/z 110.9844     0   1E+4   3.5   4  RT (min)       lcqtof_hilic_neg_4_091_60    m/z 110.9846     0   1E+4   4   4.5  RT (min)      lcqtof_hilic_neg_4_065_51    m/z 111.0070     0   2   4   6E+4   0   0.5  RT (min)      lcqtof_hilic_neg_4_000_aa    m/z 111.0074     0   2   4   6E+4   0   0.5  RT (min)      lcqtof_hilic_neg_4_000_aa    m/z 111.0075     0   2   4   6E+4   0   0.5  RT (min)      lcqtof_hilic_neg_4_000_aa    m/z 111.0078     0   1   2   3   4E+4   1   1.5  RT (min)      lcqtof_hilic_neg_4_050_02    m/z 111.0078     0   2   4   6E+4   0.5   1  RT (min)      lcqtof_hilic_neg_4_000_aa    m/z 111.0079     0   1   2E+4   7.5   8  RT (min)      lcqtof_hilic_neg_4_016_37    m/z 111.0080     0   2   4E+4   8   8.5  RT (min)      lcqtof_hilic_neg_4_035_33    m/z 111.0081     0   1E+5   7.5   8  RT (min)      lcqtof_hilic_neg_4_053_14    m/z 111.0081     0   1E+5   3   3.5  RT (min)      lcqtof_hilic_neg_4_050_02    m/z 111.0081     0   1E+5   2.5   3  RT (min)      lcqtof_hilic_neg_4_050_02    m/z 111.0082     0   1   2   3E+4   3.5   4  RT (min)      lcqtof_hilic_neg_4_064_15    m/z 111.0082     0   1   2   3E+4   8.5   9  RT (min)      lcqtof_hilic_neg_4_055_32    m/z 111.0082     0   1E+5   3   3.5  RT (min)      lcqtof_hilic_neg_4_050_02    m/z 111.0082     0   1E+5   3   3.5  RT (min)      lcqtof_hilic_neg_4_050_02    m/z 111.0082     0   1   2E+4   3.5   4  RT (min)      lcqtof_hilic_neg_4_063_16    m/z 111.0083     0   2   4   6E+4   0.5   1  RT (min)      lcqtof_hilic_neg_4_000_aa    m/z 111.0084     0   2   4   6E+4   4   4.5  RT (min)      lcqtof_hilic_neg_4_050_02    m/z 111.0085     0   1   2   3E+4   4   4.5  RT (min)      lcqtof_hilic_neg_4_050_02    m/z 111.0085     0   2   4   6   8E+4   4   4.5  RT (min)      lcqtof_hilic_neg_4_050_02    m/z 111.0089     0   1E+5   3.5   4  RT (min)      lcqtof_hilic_neg_4_050_02    m/z 111.0096     0   2   4   6   8E+4   3.5   4  RT (min)      lcqtof_hilic_neg_4_050_02    m/z 111.0116     0   1E+4   4   4.5  RT (min)      lcqtof_hilic_neg_4_050_02    m/z 111.0292     0   1   2   3E+4   1   1.5  RT (min)      lcqtof_hilic_neg_4_050_02    m/z 111.1637     0   1   2E+4   1   1.5  RT (min)      lcqtof_hilic_neg_4_050_02    m/z 112.0112     0   1E+4   2   2.5  RT (min)      lcqtof_hilic_neg_4_053_14    m/z 112.0118     0   1   2E+4   4.5   5  RT (min)      lcqtof_hilic_neg_1_004_14    m/z 112.8961     0   1E+4   4.5   5  RT (min)      lcqtof_hilic_neg_4_050_02    m/z 112.9727     0   2   4   6E+4   1   1.5  RT (min)      lcqtof_hilic_neg_4_050_02    m/z 112.9850     0   1E+4   6.5   7  RT (min)      lcqtof_hilic_neg_1_007_05    m/z 112.9850     0   1E+4   6   6.5  RT (min)      lcqtof_hilic_neg_1_011_46    m/z 112.9850     0   1   2   3   4E+3   6   6.5  RT (min)      lcqtof_hilic_neg_2_005_13    m/z 112.9850     0   2   4   6   8E+3   2   2.5  RT (min)      lcqtof_hilic_neg_1_003_37    m/z 112.9850     0   2   4   6   8E+4   0.5   1  RT (min)      lcqtof_hilic_neg_1_028_47    m/z 112.9851     0   1E+4   3.5   4  RT (min)      lcqtof_hilic_neg_1_007_05    m/z 112.9851     0   1E+4   3.5   4  RT (min)      lcqtof_hilic_neg_2_028_05    m/z 112.9851     0   2   4   6   8E+4   0.5   1  RT (min)      lcqtof_hilic_neg_1_028_47    m/z 112.9851     0   1   2   3   4E+3   0   0.5  RT (min)       lcqtof_hilic_neg_4_072_57    m/z 112.9851     0   2   4   6E+3   6.5   7  RT (min)      lcqtof_hilic_neg_1_009_50    m/z 112.9851     0   1E+4   6   6.5  RT (min)      lcqtof_hilic_neg_1_011_46    m/z 112.9851     0   1   2E+4   1.5   2  RT (min)      lcqtof_hilic_neg_1_007_05    m/z 112.9851     0   2   4   6   8E+3   6   6.5  RT (min)      lcqtof_hilic_neg_2_022_46    m/z 112.9851     0   1   2   3   4E+3   0   0.5  RT (min)      lcqtof_hilic_neg_4_061_50    m/z 112.9851     0   1   2   3   4E+3   7.5   8  RT (min)      lcqtof_hilic_neg_4_025_52    m/z 112.9852     0   1   2   3   4E+3   0.5   1  RT (min)      lcqtof_hilic_neg_4_061_50    m/z 112.9852     0   1E+4   3   3.5  RT (min)      lcqtof_hilic_neg_1_011_46    m/z 112.9852     0   1   2   3E+3   7   7.5  RT (min)      lcqtof_hilic_neg_4_025_52    m/z 112.9852     0   2   4E+3   2.5   3  RT (min)      lcqtof_hilic_neg_1_007_05    m/z 112.9852     0   1   2   3E+3   7   7.5  RT (min)      lcqtof_hilic_neg_4_025_52    m/z 112.9852     0   2   4   6   8E+3   3.5   4  RT (min)      lcqtof_hilic_neg_1_055_25    m/z 112.9852     0   2   4   6E+3   2   2.5  RT (min)      lcqtof_hilic_neg_1_007_05    m/z 112.9852     0   1   2   3E+3   6.5   7  RT (min)      lcqtof_hilic_neg_1_034_08    m/z 112.9853     0   1   2   3E+3   7.5   8  RT (min)      lcqtof_hilic_neg_1_009_50    m/z 112.9853     0   2   4   6E+3   0.5   1  RT (min)      lcqtof_hilic_neg_4_033_53    m/z 112.9853     0   1E+4   3.5   4  RT (min)      lcqtof_hilic_neg_1_007_05    m/z 113.0238     0   1E+4   3   3.5  RT (min)      lcqtof_hilic_neg_4_103_61    m/z 114.9484     0   1   2   3E+4   1   1.5  RT (min)       lcqtof_hilic_neg_3_103_61    m/z 114.9484     0   1E+4   2   4  RT (min)       lcqtof_hilic_neg_3_068_57    m/z 115.0032     0   1E+4   4   4.5  RT (min)      lcqtof_hilic_neg_4_016_37    m/z 115.0033     0   2   4   6   8E+3   2   2.5  RT (min)      lcqtof_hilic_neg_4_048_23    m/z 115.0034     0   1E+4   3.5   4  RT (min)      lcqtof_hilic_neg_4_088_59    m/z 115.0203     0   1E+4   2.5   3  RT (min)      lcqtof_hilic_neg_4_007_45    m/z 115.0203     0   1E+4   2.5   3  RT (min)      lcqtof_hilic_neg_1_028_47    m/z 115.0204     0   1   2   3   4E+3   2.5   3  RT (min)      lcqtof_hilic_neg_1_028_47    m/z 115.0204     0   2   4   6E+3   3   3.5  RT (min)      lcqtof_hilic_neg_4_076_58    m/z 116.0248     0   2   4   6E+4   7.5   8  RT (min)       lcqtof_hilic_neg_4_095_60    m/z 116.0249     0   1   2   3E+4   7   7.5  RT (min)       lcqtof_hilic_neg_4_055_32    m/z 116.0249     0   2   4   6E+4   7   7.5  RT (min)       lcqtof_hilic_neg_4_095_60    m/z 116.0250     0   1   2   3   4E+4   7   7.5  RT (min)       lcqtof_hilic_neg_4_091_60    m/z 116.0250     0   1   2E+4   7   7.5  RT (min)      lcqtof_hilic_neg_4_047_06    m/z 116.0250     0   1   2E+4   7   7.5  RT (min)      lcqtof_hilic_neg_4_047_06    m/z 116.0250     0   1E+4   7   7.5  RT (min)      lcqtof_hilic_neg_4_065_51    m/z 116.0251     0   1E+5   4.5   5  RT (min)      lcqtof_hilic_neg_1_005_32    m/z 116.0251     0   1E+4   4.5   5  RT (min)      lcqtof_hilic_neg_1_037_24    m/z 116.0252     0   1E+5   4.5   5  RT (min)      lcqtof_hilic_neg_1_005_32    m/z 116.0253     0   1   2   3E+4   4   4.5  RT (min)      lcqtof_hilic_neg_1_015_33    m/z 116.0253     0   1   2E+4   7   7.5  RT (min)      lcqtof_hilic_neg_1_037_24    m/z 116.0284     0   1   2E+3   2   2.5  RT (min)      lcqtof_hilic_neg_4_032_01    m/z 116.0313     0   1   2   3   4E+3   2.5   3  RT (min)      lcqtof_hilic_neg_1_005_32    m/z 116.9275     0   1E+4   6.5   7  RT (min)      lcqtof_hilic_neg_3_011_34    m/z 116.9276     0   2   4   6E+4   2   4  RT (min)       lcqtof_hilic_neg_2_000_cc    m/z 116.9277     0   1E+5   4   4.5  RT (min)       lcqtof_hilic_neg_2_028_05    m/z 116.9277     0   1   2E+4   6   6.5  RT (min)      lcqtof_hilic_neg_1_030_17    m/z 116.9277     0   1   2   3E+4   6   6.5  RT (min)       lcqtof_hilic_neg_2_044_37    m/z 116.9277     0   2   4   6E+4   4   4.5  RT (min)      lcqtof_hilic_neg_1_013_38    m/z 116.9277     0   1E+5   0   0.5  RT (min)       lcqtof_hilic_neg_4_058_39    m/z 116.9277     0   2   4   6   8E+4   4   4.5  RT (min)       lcqtof_hilic_neg_3_094_60    m/z 116.9277     0   1   2   3E+4   6.5   7  RT (min)       lcqtof_hilic_neg_2_018_16    m/z 116.9278     0   1   2   3E+4   7   7.5  RT (min)       lcqtof_hilic_neg_4_058_39    m/z 116.9278     0   1   2E+4   6.5   7  RT (min)       lcqtof_hilic_neg_2_016_33    m/z 116.9278     0   1   2E+4   17.5   18  RT (min)       lcqtof_hilic_neg_4_058_39    m/z 116.9278     0   1E+4   18.5   19  RT (min)       lcqtof_hilic_neg_3_067_57    m/z 116.9278     0   1   2E+4   7.5   8  RT (min)       lcqtof_hilic_neg_1_023_34    m/z 116.9278     0   2   4   6E+4   4   4.5  RT (min)      lcqtof_hilic_neg_1_014_31    m/z 116.9278     0   1   2E+4   6   6.5  RT (min)      lcqtof_hilic_neg_2_016_33    m/z 116.9278     0   1   2E+4   7   7.5  RT (min)       lcqtof_hilic_neg_3_088_59    m/z 116.9278     0   1   2E+4   7.5   8  RT (min)       lcqtof_hilic_neg_3_027_47    m/z 116.9278     0   2   4   6E+4   0   0.5  RT (min)       lcqtof_hilic_neg_4_058_39    m/z 116.9278     0   1E+4   19.5   20  RT (min)       lcqtof_hilic_neg_3_097_60    m/z 116.9278     0   1E+4   19.5   20  RT (min)       lcqtof_hilic_neg_3_102_61    m/z 116.9278     0   1E+4   18   18.5  RT (min)       lcqtof_hilic_neg_3_067_57    m/z 116.9278     0   2   4E+4   4.5   5  RT (min)       lcqtof_hilic_neg_2_011_39    m/z 116.9278     0   1E+4   18   18.5  RT (min)       lcqtof_hilic_neg_3_067_57    m/z 116.9278     0   1E+4   3.5   4  RT (min)      lcqtof_hilic_neg_2_029_31    m/z 116.9278     0   2   4   6E+4   3   3.5  RT (min)       lcqtof_hilic_neg_4_058_39    m/z 116.9278     0   2   4   6   8E+4   2   4  RT (min)       lcqtof_hilic_neg_4_058_39    m/z 116.9279     0   2   4   6E+4   2.5   3  RT (min)       lcqtof_hilic_neg_4_058_39    m/z 116.9279     0   2   4   6   8E+4   3.5   4  RT (min)       lcqtof_hilic_neg_2_039_38    m/z 116.9279     0   1   2   3   4E+4   5   5.5  RT (min)       lcqtof_hilic_neg_1_030_17    m/z 116.9279     0   1   2E+4   5.5   6  RT (min)      lcqtof_hilic_neg_2_001_49    m/z 116.9279     0   1E+4   19   19.5  RT (min)       lcqtof_hilic_neg_3_067_57    m/z 116.9279     0   1E+4   19   19.5   20  RT (min)       lcqtof_hilic_neg_3_068_57    m/z 116.9279     0   1E+5   1.5   2  RT (min)       lcqtof_hilic_neg_4_058_39    m/z 116.9279     0   1   2E+4   5.5   6  RT (min)       lcqtof_hilic_neg_1_027_27    m/z 116.9279     0   1E+5   1.5   2  RT (min)       lcqtof_hilic_neg_4_058_39    m/z 116.9279     0   1E+4   16   16.5  RT (min)       lcqtof_hilic_neg_4_001_49    m/z 116.9279     0   1   2E+4   6.5   7  RT (min)      lcqtof_hilic_neg_2_016_33    m/z 116.9279     0   1   2   3E+3   15.5   16  RT (min)       lcqtof_hilic_neg_3_067_57    m/z 116.9279     0   1E+4   18.5   19  RT (min)       lcqtof_hilic_neg_3_067_57    m/z 116.9279     0   1   2E+4   5.5   6  RT (min)       lcqtof_hilic_neg_1_027_27    m/z 116.9279     0   2   4   6E+4   0   0.5  RT (min)       lcqtof_hilic_neg_4_058_39    m/z 116.9279     0   2   4   6E+4   0   0.5  RT (min)       lcqtof_hilic_neg_4_058_39    m/z 116.9279     0   2   4   6   8E+3   17   17.5  RT (min)       lcqtof_hilic_neg_3_067_57    m/z 116.9279     0   1   2   3   4E+5   1.5   2  RT (min)       lcqtof_hilic_neg_4_058_39    m/z 116.9279     0   2   4   6E+4   2.5   3  RT (min)       lcqtof_hilic_neg_4_058_39    m/z 116.9279     0   1E+5   0   0.5  RT (min)      lcqtof_hilic_neg_4_058_39    m/z 116.9279     0   2   4   6E+4   0   0.5  RT (min)       lcqtof_hilic_neg_4_058_39    m/z 116.9280     0   2   4   6E+4   2.5   3  RT (min)       lcqtof_hilic_neg_4_058_39    m/z 116.9280     0   1E+4   16   16.5  RT (min)       lcqtof_hilic_neg_4_000_bb    m/z 116.9280     0   1   2E+4   5   5.5  RT (min)      lcqtof_hilic_neg_1_023_34    m/z 116.9280     0   1   2E+4   5   5.5  RT (min)       lcqtof_hilic_neg_1_023_34    m/z 116.9280     0   1   2E+4   7.5   8  RT (min)       lcqtof_hilic_neg_3_096_60    m/z 116.9280     0   2   4   6   8E+4   2.5   3  RT (min)       lcqtof_hilic_neg_4_058_39    m/z 116.9280     0   1E+4   20   20.5  RT (min)       lcqtof_hilic_neg_2_003_25    m/z 116.9280     0   2   4   6E+4   4   4.5  RT (min)       lcqtof_hilic_neg_1_013_38    m/z 116.9280     0   1   2E+4   5.5   6  RT (min)      lcqtof_hilic_neg_1_027_27    m/z 116.9280     0   1E+4   8   8.5  RT (min)       lcqtof_hilic_neg_3_093_60    m/z 116.9281     0   2   4   6E+4   3   3.5  RT (min)       lcqtof_hilic_neg_4_058_39    m/z 116.9283     0   1   2   3E+4   5   5.5  RT (min)      lcqtof_hilic_neg_1_027_27    m/z 117.0385     0   1E+4   1   1.5  RT (min)      lcqtof_hilic_neg_4_050_02    m/z 117.9282     0   2   4   6   8E+3   4.5   5  RT (min)      lcqtof_hilic_neg_2_020_23    m/z 117.9283     0   2   4E+3   2   2.5  RT (min)      lcqtof_hilic_neg_3_068_57    m/z 117.9283     0   2   4   6   8E+3   4.5   5  RT (min)      lcqtof_hilic_neg_2_015_06    m/z 117.9283     0   1   2E+3   7   7.5  RT (min)      lcqtof_hilic_neg_2_047_32    m/z 117.9284     0   1   2E+3   7   7.5  RT (min)      lcqtof_hilic_neg_1_045_07    m/z 117.9284     0   1   2   3E+3   6   6.5  RT (min)      lcqtof_hilic_neg_2_028_05    m/z 117.9284     0   1   2   3E+3   6   6.5  RT (min)      lcqtof_hilic_neg_2_028_05    m/z 117.9284     0   1   2   3E+3   2   2.5  RT (min)      lcqtof_hilic_neg_1_030_17    m/z 117.9284     0   1   2E+3   7.5   8  RT (min)      lcqtof_hilic_neg_3_085_59    m/z 117.9284     0   1   2E+3   3.5   4  RT (min)      lcqtof_hilic_neg_3_108_62    m/z 117.9284     0   1   2E+3   6.5   7  RT (min)      lcqtof_hilic_neg_1_035_23    m/z 117.9284     0   1   2E+3   6   6.5  RT (min)      lcqtof_hilic_neg_1_027_27    m/z 117.9284     0   1   2E+3   5.5   6  RT (min)      lcqtof_hilic_neg_1_007_05    m/z 117.9286     0   1E+3   3.5   4  RT (min)      lcqtof_hilic_neg_1_063_13    m/z 118.0408     0   2   4E+3   2.5   3  RT (min)      lcqtof_hilic_neg_1_036_03    m/z 118.0504     0   1E+4   5   5.5  RT (min)      lcqtof_hilic_neg_1_000_aa    m/z 118.8990     0   1   2E+4   4.5   5  RT (min)      lcqtof_hilic_neg_4_050_02    m/z 118.8991     0   1   2E+4   4   4.5  RT (min)      lcqtof_hilic_neg_4_050_02    m/z 118.8996     0   1   2E+4   4   4.5  RT (min)      lcqtof_hilic_neg_4_050_02    m/z 118.9269     0   1E+4   4.5   5  RT (min)      lcqtof_hilic_neg_4_050_02    m/z 119.0164     0   2   4   6   8E+4   6   6.5  RT (min)      lcqtof_hilic_neg_4_008_24    m/z 119.0165     0   1   2   3   4E+4   3.5   4  RT (min)      lcqtof_hilic_neg_4_090_60    m/z 119.0165     0   2   4   6   8E+4   3.5   4  RT (min)      lcqtof_hilic_neg_1_005_32    m/z 119.0165     0   2   4   6   8E+4   6   6.5  RT (min)      lcqtof_hilic_neg_4_008_24    m/z 119.0166     0   1   2   3   4E+4   3.5   4  RT (min)      lcqtof_hilic_neg_4_055_32    m/z 119.0166     0   1   2   3   4E+4   4   4.5  RT (min)      lcqtof_hilic_neg_4_055_32    m/z 119.0167     0   1   2   3   4E+4   4   4.5  RT (min)      lcqtof_hilic_neg_4_040_05    m/z 119.0167     0   1   2   3   4E+4   3.5   4  RT (min)      lcqtof_hilic_neg_4_055_32    m/z 119.0167     0   1   2   3   4E+4   4   4.5  RT (min)      lcqtof_hilic_neg_4_055_32    m/z 119.0338     0   1E+4   2   2.5  RT (min)      lcqtof_hilic_neg_4_050_02    m/z 119.9466     0   1   2E+3   4.5   5  RT (min)      lcqtof_hilic_neg_4_012_07    m/z 120.0547     0   1   2   3E+3   1   1.5  RT (min)       lcqtof_hilic_neg_4_107_62    m/z 123.9014     0   1E+4   4.5   5  RT (min)      lcqtof_hilic_neg_4_080_58    m/z 123.9017     0   2   4   6   8E+3   0.5   1  RT (min)      lcqtof_hilic_neg_2_000_aa    m/z 124.0065     0   1   2E+4   6.5   7  RT (min)      lcqtof_hilic_neg_1_000_aa    m/z 124.0481     0   1   2E+4   0.5   1  RT (min)      lcqtof_hilic_neg_4_039_18    m/z 124.0491     0   1   2   3E+3   0.5   1  RT (min)       lcqtof_hilic_neg_4_059_14    m/z 125.0342     0   1E+3   1   1.5  RT (min)      lcqtof_hilic_neg_4_039_18    m/z 125.8992     0   2   4   6E+3   0.5   1  RT (min)      lcqtof_hilic_neg_2_000_aa    m/z 125.8993     0   2   4   6E+3   4.5   5  RT (min)      lcqtof_hilic_neg_4_064_15    m/z 127.0027     0   1E+4   7   7.5  RT (min)      lcqtof_hilic_neg_4_022_34    m/z 127.0029     0   1E+4   6.5   7  RT (min)      lcqtof_hilic_neg_4_022_34    m/z 127.0030     0   1E+4   7   7.5  RT (min)      lcqtof_hilic_neg_4_022_34    m/z 127.0031     0   1E+4   6.5   7  RT (min)      lcqtof_hilic_neg_4_022_34    m/z 127.0032     0   1   2E+4   1.5   2  RT (min)      lcqtof_hilic_neg_4_064_15    m/z 128.0342     0   2   4E+3   1.5   2  RT (min)      lcqtof_hilic_neg_4_038_10    m/z 128.0346     0   2   4   6E+4   6.5   7  RT (min)      lcqtof_hilic_neg_4_002_25    m/z 128.0347     0   1   2   3   4E+3   4.5   5  RT (min)      lcqtof_hilic_neg_2_002_36    m/z 128.0348     0   1   2E+4   2   2.5  RT (min)      lcqtof_hilic_neg_4_052_13    m/z 128.0348     0   1   2   3E+4   5.5   6  RT (min)      lcqtof_hilic_neg_4_053_14    m/z 128.0348     0   1   2E+4   0.5   1  RT (min)      lcqtof_hilic_neg_4_052_13    m/z 128.0348     0   1   2E+4   6.5   7  RT (min)      lcqtof_hilic_neg_2_028_05    m/z 128.0348     0   1   2   3E+4   4   4.5  RT (min)      lcqtof_hilic_neg_4_040_05    m/z 128.0348     0   1   2   3E+4   4   4.5  RT (min)      lcqtof_hilic_neg_4_040_05    m/z 128.0349     0   2   4E+3   1.5   2  RT (min)      lcqtof_hilic_neg_4_060_13    m/z 128.0349     0   2   4   6E+3   1   1.5  RT (min)       lcqtof_hilic_neg_4_110_62    m/z 128.0355     0   2   4   6E+3   2   2.5  RT (min)      lcqtof_hilic_neg_4_026_04    m/z 128.0927     0   1E+4   6   6.5  RT (min)      lcqtof_hilic_neg_4_048_23    m/z 128.0928     0   1E+4   2.5   3  RT (min)      lcqtof_hilic_neg_4_047_06    m/z 128.1246     0   1E+4   2.5   3  RT (min)      lcqtof_hilic_neg_4_048_23    m/z 128.1499     0   2   4   6   8E+4   2.5   3  RT (min)      lcqtof_hilic_neg_4_048_23    m/z 128.1500     0   1   2   3E+4   6   6.5  RT (min)      lcqtof_hilic_neg_4_048_23    m/z 128.1501     0   1   2   3E+4   2.5   3  RT (min)      lcqtof_hilic_neg_4_047_06    m/z 128.1728     0   1   2E+4   2.5   3  RT (min)      lcqtof_hilic_neg_4_048_23    m/z 128.2021     0   1   2E+4   2.5   3  RT (min)      lcqtof_hilic_neg_4_048_23    m/z 128.2371     0   1E+4   2.5   3  RT (min)      lcqtof_hilic_neg_4_047_06    m/z 128.2372     0   1   2   3   4E+4   3   3.5  RT (min)      lcqtof_hilic_neg_4_016_37    m/z 128.2373     0   1   2   3   4E+4   2.5   3  RT (min)      lcqtof_hilic_neg_4_048_23    m/z 128.3022     0   1   2   3E+4   2.5   3  RT (min)      lcqtof_hilic_neg_4_048_23    m/z 128.3022     0   1E+4   3   3.5  RT (min)      lcqtof_hilic_neg_4_031_38    m/z 128.8779     0   1   2E+3   10   10.5  RT (min)      lcqtof_hilic_neg_3_013_48    m/z 128.9270     0   1E+4   7.5   8  RT (min)      lcqtof_hilic_neg_3_067_57    m/z 128.9273     0   1   2E+4   1.5   2  RT (min)       lcqtof_hilic_neg_3_067_57    m/z 128.9274     0   1E+4   6.5   7  RT (min)      lcqtof_hilic_neg_3_027_47    m/z 128.9274     0   1E+4   19   19.5  RT (min)       lcqtof_hilic_neg_3_066_57    m/z 128.9274     0   1   2   3E+4   2.5   3  RT (min)      lcqtof_hilic_neg_3_067_57    m/z 128.9274     0   1   2E+4   5.5   6  RT (min)      lcqtof_hilic_neg_3_011_34    m/z 128.9275     0   1   2E+4   6   6.5  RT (min)      lcqtof_hilic_neg_3_024_35    m/z 128.9275     0   1E+4   19   19.5  RT (min)       lcqtof_hilic_neg_3_112_62    m/z 128.9275     0   1E+4   18.5   19  RT (min)       lcqtof_hilic_neg_3_112_62    m/z 128.9275     0   1E+4   6.5   7  RT (min)      lcqtof_hilic_neg_3_027_47    m/z 128.9275     0   1E+4   19.5   20  RT (min)       lcqtof_hilic_neg_3_104_61    m/z 128.9275     0   1   2E+4   5.5   6  RT (min)      lcqtof_hilic_neg_3_040_14    m/z 128.9276     0   1E+4   6.5   7  RT (min)      lcqtof_hilic_neg_3_067_57    m/z 128.9276     0   1E+4   18.5   19  RT (min)       lcqtof_hilic_neg_3_111_62    m/z 128.9276     0   1   2E+4   19.5   20  RT (min)       lcqtof_hilic_neg_3_107_62    m/z 128.9276     0   1   2E+4   5.5   6  RT (min)      lcqtof_hilic_neg_3_053_26    m/z 128.9276     0   1E+4   18   18.5  RT (min)       lcqtof_hilic_neg_3_111_62    m/z 128.9276     0   1E+4   17.5   18  RT (min)       lcqtof_hilic_neg_3_111_62    m/z 128.9276     0   1E+4   19   19.5  RT (min)       lcqtof_hilic_neg_3_111_62    m/z 128.9276     0   1E+4   19   19.5  RT (min)       lcqtof_hilic_neg_3_113_62    m/z 128.9276     0   1E+4   17.5   18  RT (min)      lcqtof_hilic_neg_3_090_60    m/z 128.9276     0   1   2   3   4E+4   2   2.5  RT (min)       lcqtof_hilic_neg_3_068_57    m/z 128.9276     0   1E+4   6   6.5  RT (min)      lcqtof_hilic_neg_3_011_34    m/z 128.9277     0   1E+4   6.5   7  RT (min)      lcqtof_hilic_neg_3_010_45    m/z 128.9277     0   1   2   3E+4   1   1.5  RT (min)       lcqtof_hilic_neg_3_044_09    m/z 128.9277     0   1   2E+4   5.5   6  RT (min)      lcqtof_hilic_neg_3_024_35    m/z 128.9277     0   1   2E+4   5.5   6  RT (min)      lcqtof_hilic_neg_3_040_14    m/z 128.9277     0   1E+4   6.5   7  RT (min)      lcqtof_hilic_neg_3_027_47    m/z 128.9277     0   1   2E+4   5   5.5  RT (min)      lcqtof_hilic_neg_3_054_37    m/z 128.9277     0   1E+4   18   18.5  RT (min)      lcqtof_hilic_neg_3_105_61    m/z 128.9277     0   1   2E+4   2   4   6  RT (min)       lcqtof_hilic_neg_3_089_59    m/z 128.9277     0   1   2   3E+4   2   4   6  RT (min)       lcqtof_hilic_neg_3_054_37    m/z 128.9278     0   1E+4   5   5.5  RT (min)      lcqtof_hilic_neg_3_053_26    m/z 128.9278     0   1E+4   5   5.5  RT (min)      lcqtof_hilic_neg_3_055_01    m/z 128.9278     0   1   2E+4   5   5.5  RT (min)      lcqtof_hilic_neg_3_054_37    m/z 128.9278     0   1E+4   5   5.5  RT (min)      lcqtof_hilic_neg_3_024_35    m/z 128.9278     0   1E+4   5   5.5  RT (min)      lcqtof_hilic_neg_3_031_36    m/z 128.9278     0   1   2E+4   5.5   6  RT (min)      lcqtof_hilic_neg_3_040_14    m/z 128.9278     0   1E+4   6.5   7  RT (min)      lcqtof_hilic_neg_3_009_50    m/z 128.9280     0   1   2   3E+4   1.5   2  RT (min)       lcqtof_hilic_neg_3_067_57    m/z 129.0181     0   1   2   3   4E+4   3   3.5  RT (min)      lcqtof_hilic_neg_4_039_18    m/z 129.0187     0   1   2E+4   4.5   5  RT (min)      lcqtof_hilic_neg_1_004_14    m/z 129.0187     0   1   2E+4   3.5   4  RT (min)      lcqtof_hilic_neg_1_032_18    m/z 129.0188     0   1   2E+4   4.5   5  RT (min)      lcqtof_hilic_neg_1_004_14    m/z 129.0194     0   2   4   6E+3   2   2.5  RT (min)      lcqtof_hilic_neg_4_004_47    m/z 129.0195     0   1E+4   2   2.5  RT (min)      lcqtof_hilic_neg_4_053_14    m/z 129.9314     0   2   4   6   8E+3   2   2.5  RT (min)      lcqtof_hilic_neg_3_067_57    m/z 129.9940     0   1E+4   4.5   5  RT (min)      lcqtof_hilic_neg_4_022_34    m/z 130.0136     0   1   2   3E+3   1   1.5  RT (min)      lcqtof_hilic_neg_4_008_24    m/z 130.0392     0   1E+4   2.5   3  RT (min)      lcqtof_hilic_neg_4_086_59    m/z 130.0434     0   1   2   3E+3   1.5   2  RT (min)      lcqtof_hilic_neg_4_014_46    m/z 130.0604     0   1   2E+4   4   4.5  RT (min)      lcqtof_hilic_neg_4_055_32    m/z 130.9655     0   1   2   3E+4   8.5   9  RT (min)      lcqtof_hilic_neg_4_074_58    m/z 130.9655     0   1   2   3E+4   8.5   9  RT (min)      lcqtof_hilic_neg_4_032_01    m/z 130.9656     0   2   4   6E+4   8.5   9  RT (min)      lcqtof_hilic_neg_4_053_14    m/z 131.0165     0   1   2   3E+4   1   1.5  RT (min)      lcqtof_hilic_neg_4_050_02    m/z 131.9411     0   1E+4   1   1.5  RT (min)      lcqtof_hilic_neg_2_045_11    m/z 132.0292     0   1E+4   5   5.5  RT (min)      lcqtof_hilic_neg_1_000_aa    m/z 132.0294     0   2   4   6E+4   7   7.5  RT (min)      lcqtof_hilic_neg_4_016_37    m/z 132.0295     0   1E+5   7   7.5  RT (min)      lcqtof_hilic_neg_4_016_37    m/z 132.0479     0   1E+5   2.5   3  RT (min)      lcqtof_hilic_neg_4_055_32    m/z 132.0480     0   2   4   6   8E+4   2.5   3  RT (min)      lcqtof_hilic_neg_4_097_60    m/z 132.0650     0   1E+4   1   1.5  RT (min)      lcqtof_hilic_neg_4_050_02    m/z 132.1462     0   1E+4   6.5   7  RT (min)      lcqtof_hilic_neg_4_048_23    m/z 132.8669     0   1   2   3   4E+4   5   5.5  RT (min)      lcqtof_hilic_neg_4_033_53    m/z 132.8671     0   2   4E+4   3.5   4  RT (min)      lcqtof_hilic_neg_4_063_16    m/z 132.8671     0   1E+4   2   2.5  RT (min)      lcqtof_hilic_neg_4_065_51    m/z 132.8671     0   1E+4   3   3.5  RT (min)      lcqtof_hilic_neg_2_040_20    m/z 132.8671     0   1   2   3   4E+4   5   5.5  RT (min)      lcqtof_hilic_neg_4_054_48    m/z 132.8672     0   1E+4   3   3.5  RT (min)      lcqtof_hilic_neg_2_040_20    m/z 132.8672     0   1E+5   0.5   1  RT (min)      lcqtof_hilic_neg_2_000_aa    m/z 132.8673     0   2   4   6   8E+3   1.5   2  RT (min)      lcqtof_hilic_neg_4_064_15    m/z 132.8673     0   1   2   3E+4   3   3.5  RT (min)      lcqtof_hilic_neg_4_064_15    m/z 132.8673     0   1   2   3   4E+4   5   5.5  RT (min)      lcqtof_hilic_neg_4_035_33    m/z 132.8673     0   2   4E+4   4   4.5  RT (min)      lcqtof_hilic_neg_4_063_16    m/z 132.8673     0   1   2   3   4E+4   3.5   4  RT (min)      lcqtof_hilic_neg_4_063_16    m/z 132.8674     0   2   4E+4   4   4.5  RT (min)      lcqtof_hilic_neg_4_064_15    m/z 132.8674     0   1E+4   3.5   4  RT (min)      lcqtof_hilic_neg_2_040_20    m/z 132.8998     0   1E+4   4   4.5  RT (min)      lcqtof_hilic_neg_2_034_47    m/z 132.8998     0   1   2   3   4E+3   3.5   4  RT (min)      lcqtof_hilic_neg_1_040_20    m/z 132.8999     0   1   2   3E+3   3.5   4  RT (min)      lcqtof_hilic_neg_2_022_46    m/z 132.8999     0   1   2E+4   4   4.5  RT (min)      lcqtof_hilic_neg_2_051_14    m/z 132.9000     0   1   2E+3   3.5   4  RT (min)      lcqtof_hilic_neg_2_028_05    m/z 132.9000     0   2   4E+3   3.5   4  RT (min)      lcqtof_hilic_neg_2_010_45    m/z 133.0130     0   1E+4   6   6.5  RT (min)      lcqtof_hilic_neg_4_048_23    m/z 133.0136     0   2   4E+4   4   4.5  RT (min)      lcqtof_hilic_neg_4_048_23    m/z 133.0136     0   1E+4   6   6.5  RT (min)      lcqtof_hilic_neg_4_048_23    m/z 133.0136     0   2   4E+4   4   4.5  RT (min)      lcqtof_hilic_neg_4_048_23    m/z 133.0138     0   1   2   3   4E+4   3.5   4  RT (min)      lcqtof_hilic_neg_4_034_31    m/z 133.0138     0   2   4E+4   4   4.5  RT (min)      lcqtof_hilic_neg_4_048_23    m/z 133.0143     0   1   2   3   4E+4   4.5   5  RT (min)      lcqtof_hilic_neg_2_020_23    m/z 133.0514     0   1   2   3E+6   2.5   3  RT (min)       lcqtof_hilic_neg_4_055_32    m/z 133.0515     0   2   4   6E+5   5.5   6  RT (min)      lcqtof_hilic_neg_4_008_24    m/z 133.0518     0   1   2E+6   2.5   3  RT (min)      lcqtof_hilic_neg_4_092_60    m/z 133.0519     0   2   4   6E+5   5.5   6  RT (min)      lcqtof_hilic_neg_4_008_24    m/z 133.1106     0   1   2E+4   2.5   3  RT (min)      lcqtof_hilic_neg_4_047_06    m/z 133.1107     0   2   4E+4   3   3.5  RT (min)      lcqtof_hilic_neg_4_008_24    m/z 133.1108     0   2   4   6E+4   2.5   3  RT (min)      lcqtof_hilic_neg_4_055_32    m/z 133.1435     0   1E+4   3   3.5  RT (min)      lcqtof_hilic_neg_4_008_24    m/z 133.1436     0   1   2E+4   2.5   3  RT (min)      lcqtof_hilic_neg_4_055_32    m/z 133.1688     0   2   4   6   8E+4   3   3.5  RT (min)      lcqtof_hilic_neg_4_008_24    m/z 133.1689     0   1E+5   2.5   3  RT (min)      lcqtof_hilic_neg_4_055_32    m/z 133.1690     0   1   2   3   4E+4   2.5   3  RT (min)      lcqtof_hilic_neg_4_047_06    m/z 133.1921     0   1   2E+4   3   3.5  RT (min)      lcqtof_hilic_neg_4_008_24    m/z 133.1922     0   1   2   3E+4   2.5   3  RT (min)      lcqtof_hilic_neg_4_055_32    m/z 133.2217     0   1   2E+4   3   3.5  RT (min)      lcqtof_hilic_neg_4_008_24    m/z 133.2221     0   1   2   3E+4   2.5   3  RT (min)      lcqtof_hilic_neg_4_055_32    m/z 133.2579     0   1   2   3   4E+4   3   3.5  RT (min)      lcqtof_hilic_neg_4_008_24    m/z 133.2580     0   1E+4   2.5   3  RT (min)      lcqtof_hilic_neg_4_047_06    m/z 133.2583     0   2   4   6E+4   2.5   3  RT (min)      lcqtof_hilic_neg_4_055_32    m/z 133.3242     0   1   2   3   4E+4   2.5   3  RT (min)      lcqtof_hilic_neg_4_055_32    m/z 133.3245     0   1   2   3E+4   3   3.5  RT (min)      lcqtof_hilic_neg_4_008_24    m/z 133.9910     0   1   2   3   4E+3   1   1.5  RT (min)      lcqtof_hilic_neg_4_102_61    m/z 134.0174     0   1   2   3   4E+3   3   3.5  RT (min)      lcqtof_hilic_neg_1_014_31    m/z 134.0463     0   1E+4   2.5   3  RT (min)      lcqtof_hilic_neg_4_092_60    m/z 134.0494     0   1E+4   3   3.5  RT (min)      lcqtof_hilic_neg_4_008_24    m/z 134.1646     0   1   2E+4   3.5   4  RT (min)      lcqtof_hilic_neg_4_048_23    m/z 134.8644     0   2   4   6   8E+4   3.5   4  RT (min)       lcqtof_hilic_neg_1_000_aa    m/z 134.8644     0   1   2   3   4E+4   4.5   5  RT (min)      lcqtof_hilic_neg_4_063_16    m/z 134.8645     0   1   2   3   4E+4   5   5.5  RT (min)      lcqtof_hilic_neg_4_063_16    m/z 134.8645     0   2   4   6E+4   3.5   4  RT (min)      lcqtof_hilic_neg_4_063_16    m/z 134.8646     0   2   4   6E+4   2   4  RT (min)       lcqtof_hilic_neg_1_000_bb    m/z 134.8646     0   1E+4   3   3.5  RT (min)      lcqtof_hilic_neg_2_040_20    m/z 134.8646     0   1E+4   2   2.5  RT (min)      lcqtof_hilic_neg_4_033_53    m/z 134.8646     0   2   4E+4   2  RT (min)       lcqtof_hilic_neg_1_032_18    m/z 134.8646     0   1E+4   0   0.5  RT (min)      lcqtof_hilic_neg_4_000_aa    m/z 134.8646     0   2   4E+4   1   1.5  RT (min)      lcqtof_hilic_neg_4_033_53    m/z 134.8646     0   2   4E+4   3.5   4  RT (min)      lcqtof_hilic_neg_4_063_16    m/z 134.8647     0   1E+5   0.5   1  RT (min)      lcqtof_hilic_neg_2_000_aa    m/z 134.8647     0   1E+4   17   18  RT (min)       lcqtof_hilic_neg_1_000_cc    m/z 134.8647     0   1E+4   3   4  RT (min)       lcqtof_hilic_neg_1_025_52    m/z 134.8647     0   1   2   3E+4   16.5   17   17.5  RT (min)       lcqtof_hilic_neg_1_000_aa    m/z 134.8648     0   1E+4   2.5   3  RT (min)      lcqtof_hilic_neg_4_033_53    m/z 134.8648     0   1E+4   2   2.5  RT (min)      lcqtof_hilic_neg_4_033_53    m/z 134.8648     0   1E+4   2   2.5  RT (min)      lcqtof_hilic_neg_4_033_53    m/z 134.8649     0   1E+4   3   3.5  RT (min)      lcqtof_hilic_neg_2_040_20    m/z 134.8649     0   2   4   6   8E+3   1.5   2  RT (min)      lcqtof_hilic_neg_4_063_16    m/z 134.8650     0   1E+4   17.5   18  RT (min)       lcqtof_hilic_neg_1_007_05    m/z 134.8938     0   1   2E+4   0.5   1  RT (min)       lcqtof_hilic_neg_3_068_57    m/z 134.8938     0   2   4   6   8E+4   3.5   4  RT (min)       lcqtof_hilic_neg_2_028_05    m/z 134.8938     0   1E+5   3.5   4  RT (min)       lcqtof_hilic_neg_2_060_14    m/z 134.8938     0   1   2   3   4E+4   1.5   2  RT (min)       lcqtof_hilic_neg_2_014_08    m/z 134.8938     0   2   4   6E+4   1   1.5  RT (min)       lcqtof_hilic_neg_2_011_39    m/z 134.8938     0   2   4E+4   0.5   1  RT (min)       lcqtof_hilic_neg_1_060_14    m/z 134.8938     0   1E+4   19   19.5  RT (min)       lcqtof_hilic_neg_3_067_57    m/z 134.8938     0   1E+4   18.5   19  RT (min)       lcqtof_hilic_neg_3_067_57    m/z 134.8938     0   1E+5   3.5   4  RT (min)       lcqtof_hilic_neg_2_032_02    m/z 134.8938     0   1   2   3   4E+4   3   3.5  RT (min)       lcqtof_hilic_neg_2_027_07    m/z 134.8938     0   2   4E+4   3   3.5  RT (min)       lcqtof_hilic_neg_2_053_18    m/z 134.8938     0   1E+4   0   0.5  RT (min)       lcqtof_hilic_neg_2_042_27    m/z 134.8938     0   1   2   3   4E+4   1.5   2  RT (min)      lcqtof_hilic_neg_2_037_21    m/z 134.8938     0   1   2   3E+4   0.5   1  RT (min)      lcqtof_hilic_neg_2_034_47    m/z 134.8938     0   1E+5   3.5   4  RT (min)       lcqtof_hilic_neg_2_000_dd    m/z 134.8939     0   1   2   3   4E+4   2   2.5  RT (min)       lcqtof_hilic_neg_2_051_14    m/z 134.8939     0   1   2   3   4E+4   1.5   2  RT (min)       lcqtof_hilic_neg_2_037_21    m/z 134.8939     0   1   2   3   4E+4   2   2.5  RT (min)      lcqtof_hilic_neg_2_051_14    m/z 134.8939     0   1E+4   0   0.5  RT (min)       lcqtof_hilic_neg_1_055_25    m/z 134.8939     0   1   2   3   4E+4   1   1.5  RT (min)       lcqtof_hilic_neg_2_024_10    m/z 134.8939     0   2   4   6E+3   17.5   18  RT (min)       lcqtof_hilic_neg_3_081_58    m/z 134.8939     0   1E+4   0   0.5  RT (min)      lcqtof_hilic_neg_1_061_50    m/z 134.8939     0   1   2   3E+4   2   2.5  RT (min)      lcqtof_hilic_neg_2_053_18    m/z 134.8939     0   2   4E+4   3   3.5  RT (min)      lcqtof_hilic_neg_2_037_21    m/z 134.8939     0   2   4   6   8E+3   17.5   18  RT (min)       lcqtof_hilic_neg_4_058_39    m/z 134.8939     0   1E+4   0   0.5  RT (min)      lcqtof_hilic_neg_1_055_25    m/z 134.8939     0   2   4E+4   3   3.5  RT (min)      lcqtof_hilic_neg_2_053_18    m/z 134.8939     0   2   4E+4   1   1.5  RT (min)      lcqtof_hilic_neg_2_045_11    m/z 134.8939     0   1E+4   19   19.5  RT (min)       lcqtof_hilic_neg_3_067_57    m/z 134.8939     0   1   2E+4   7   7.5  RT (min)       lcqtof_hilic_neg_2_034_47    m/z 134.8939     0   1   2   3   4E+4   1.5   2  RT (min)      lcqtof_hilic_neg_2_037_21    m/z 134.8939     0   1   2   3   4E+4   1.5   2  RT (min)       lcqtof_hilic_neg_2_037_21    m/z 134.8939     0   1E+4   19.5   20  RT (min)       lcqtof_hilic_neg_3_068_57    m/z 134.8939     0   1   2E+4   1   1.5  RT (min)       lcqtof_hilic_neg_4_058_39    m/z 134.8940     0   1   2   3   4E+4   2.5   3  RT (min)       lcqtof_hilic_neg_2_035_09    m/z 134.8940     0   1   2E+4   2.5   3  RT (min)      lcqtof_hilic_neg_3_000_bb    m/z 134.8940     0   1E+4   2   2.5  RT (min)      lcqtof_hilic_neg_3_000_bb    m/z 134.8940     0   1   2   3E+4   1.5   2  RT (min)      lcqtof_hilic_neg_2_030_40    m/z 134.8940     0   1   2   3E+4   6   6.5  RT (min)       lcqtof_hilic_neg_1_023_34    m/z 134.8940     0   1E+4   7.5   8  RT (min)       lcqtof_hilic_neg_2_044_37    m/z 134.8940     0   1E+4   7   7.5  RT (min)       lcqtof_hilic_neg_2_049_55    m/z 134.8940     0   2   4E+4   1   1.5  RT (min)      lcqtof_hilic_neg_2_045_11    m/z 134.8941     0   1   2   3E+4   2.5   3  RT (min)      lcqtof_hilic_neg_2_000_cc    m/z 134.8942     0   1E+4   7   7.5  RT (min)      lcqtof_hilic_neg_1_025_52    m/z 134.8942     0   1   2   3E+4   5.5   6  RT (min)       lcqtof_hilic_neg_1_023_34    m/z 134.8942     0   1E+4   6   6.5  RT (min)      lcqtof_hilic_neg_3_040_14    m/z 134.8942     0   1E+4   5.5   6  RT (min)      lcqtof_hilic_neg_2_031_01    m/z 134.8942     0   1   2E+4   2.5   3  RT (min)      lcqtof_hilic_neg_3_000_bb    m/z 134.8943     0   2   4   6   8E+3   5.5   6  RT (min)      lcqtof_hilic_neg_2_031_01    m/z 135.0290     0   1E+4   6   6.5  RT (min)      lcqtof_hilic_neg_4_016_37    m/z 135.0290     0   1E+4   5.5   6  RT (min)      lcqtof_hilic_neg_4_032_01    m/z 135.0290     0   1   2E+4   5.5   6  RT (min)      lcqtof_hilic_neg_4_053_14    m/z 135.0291     0   1   2E+4   5.5   6  RT (min)      lcqtof_hilic_neg_4_053_14    m/z 135.0291     0   1   2E+4   5.5   6  RT (min)      lcqtof_hilic_neg_4_053_14    m/z 135.0292     0   1   2E+4   5.5   6  RT (min)      lcqtof_hilic_neg_4_053_14    m/z 135.0297     0   1   2E+4   5.5   6  RT (min)      lcqtof_hilic_neg_4_053_14    m/z 135.0345     0   1E+3   6   6.5  RT (min)      lcqtof_hilic_neg_4_047_06    m/z 135.0396     0   1   2E+4   1   1.5  RT (min)      lcqtof_hilic_neg_4_053_14    m/z 135.0444     0   1   2E+6   0.5   1  RT (min)      lcqtof_hilic_neg_4_018_27    m/z 135.0448     0   1   2   3E+3   1.5   2  RT (min)      lcqtof_hilic_neg_1_041_54    m/z 135.0557     0   1E+4   2.5   3  RT (min)      lcqtof_hilic_neg_4_055_32    m/z 135.1367     0   1   2E+4   0.5   1  RT (min)      lcqtof_hilic_neg_4_027_19    m/z 135.1368     0   1E+4   1   1.5  RT (min)      lcqtof_hilic_neg_4_013_20    m/z 135.1864     0   1   2E+4   1   1.5  RT (min)      lcqtof_hilic_neg_4_013_20    m/z 135.1866     0   1   2   3   4E+4   0.5   1  RT (min)      lcqtof_hilic_neg_4_027_19    m/z 135.2165     0   1   2   3E+4   1   1.5  RT (min)      lcqtof_hilic_neg_4_013_20    m/z 135.2166     0   1   2   3   4E+4   0.5   1  RT (min)      lcqtof_hilic_neg_4_027_19    m/z 135.3192     0   1   2   3   4E+4   1   1.5  RT (min)      lcqtof_hilic_neg_4_013_20    m/z 135.3193     0   2   4E+4   0.5   1  RT (min)      lcqtof_hilic_neg_4_027_19    m/z 135.8942     0   1E+4   3.5   4  RT (min)      lcqtof_hilic_neg_2_053_18    m/z 135.8944     0   1   2   3   4E+4   4   4.5  RT (min)      lcqtof_hilic_neg_2_009_50    m/z 135.8944     0   2   4E+4   4   4.5  RT (min)      lcqtof_hilic_neg_2_051_14    m/z 135.8944     0   1   2E+3   6   6.5  RT (min)      lcqtof_hilic_neg_3_043_13    m/z 135.8944     0   2   4E+3   1   1.5  RT (min)      lcqtof_hilic_neg_2_000_cc    m/z 135.8944     0   1   2   3   4E+3   0.5   1  RT (min)      lcqtof_hilic_neg_2_051_14    m/z 135.8944     0   1   2   3E+3   0.5   1  RT (min)      lcqtof_hilic_neg_2_036_34    m/z 135.8944     0   2   4   6E+4   4   4.5  RT (min)      lcqtof_hilic_neg_2_052_24    m/z 135.8945     0   2   4   6E+3   3.5   4  RT (min)      lcqtof_hilic_neg_2_028_05    m/z 135.8945     0   1E+4   3.5   4  RT (min)      lcqtof_hilic_neg_2_032_02    m/z 135.8945     0   1   2   3E+3   1.5   2  RT (min)      lcqtof_hilic_neg_3_014_44    m/z 135.8945     0   1   2   3E+3   2   2.5  RT (min)      lcqtof_hilic_neg_2_051_14    m/z 135.8945     0   1   2   3E+3   2   2.5  RT (min)      lcqtof_hilic_neg_1_042_26    m/z 135.8946     0   1   2E+3   6.5   7  RT (min)      lcqtof_hilic_neg_2_003_25    m/z 135.8946     0   2   4   6   8E+3   3.5   4  RT (min)      lcqtof_hilic_neg_1_026_10    m/z 135.8947     0   2   4   6   8E+3   3.5   4  RT (min)      lcqtof_hilic_neg_2_032_02    m/z 135.9706     0   1   2E+4   1.5   2   2.5  RT (min)       lcqtof_hilic_neg_4_050_02    m/z 135.9706     0   1   2   3E+3   2   2.5  RT (min)      lcqtof_hilic_neg_2_043_15    m/z 136.0240     0   2   4   6E+3   3   3.5  RT (min)      lcqtof_hilic_neg_1_005_32    m/z 136.0428     0   1   2E+4   4.5   5  RT (min)      lcqtof_hilic_neg_4_055_32    m/z 136.0429     0   1E+4   4   4.5  RT (min)      lcqtof_hilic_neg_4_095_60    m/z 136.0451     0   2   4   6   8E+4   7   7.5  RT (min)      lcqtof_hilic_neg_4_008_24    m/z 136.0464     0   1   2E+5   7   7.5  RT (min)      lcqtof_hilic_neg_4_008_24    m/z 136.0490     0   1E+3   7   7.5  RT (min)      lcqtof_hilic_neg_4_008_24    m/z 136.0508     0   1E+4   1.5   2  RT (min)      lcqtof_hilic_neg_4_008_24    m/z 136.0599     0   1E+4   1   1.5  RT (min)      lcqtof_hilic_neg_4_055_32    m/z 136.8619     0   1   2E+4   5   5.5  RT (min)      lcqtof_hilic_neg_4_017_51    m/z 136.8619     0   1   2E+4   5   5.5  RT (min)      lcqtof_hilic_neg_4_016_37    m/z 136.8621     0   1   2E+4   5   5.5  RT (min)      lcqtof_hilic_neg_4_017_51    m/z 136.8622     0   1   2E+4   5   5.5  RT (min)      lcqtof_hilic_neg_4_016_37    m/z 136.8623     0   1   2   3E+4   4.5   5  RT (min)       lcqtof_hilic_neg_4_093_60    m/z 136.8625     0   1E+4   0   0.5  RT (min)      lcqtof_hilic_neg_4_018_27    m/z 136.8910     0   1   2   3   4E+4   3.5   4  RT (min)      lcqtof_hilic_neg_2_053_18    m/z 136.8911     0   2   4E+4   3.5   4  RT (min)      lcqtof_hilic_neg_2_065_51    m/z 136.8911     0   1   2   3   4E+4   3   3.5  RT (min)      lcqtof_hilic_neg_2_053_18    m/z 136.8912     0   1   2E+4   3.5   4  RT (min)      lcqtof_hilic_neg_2_028_05    m/z 136.8912     0   1   2   3E+4   3.5   4  RT (min)      lcqtof_hilic_neg_2_008_44    m/z 136.8912     0   1   2   3E+4   3.5   4  RT (min)      lcqtof_hilic_neg_2_000_cc    m/z 136.8912     0   1E+4   1   1.5  RT (min)      lcqtof_hilic_neg_1_026_10    m/z 136.8912     0   1E+4   1.5   2  RT (min)      lcqtof_hilic_neg_2_035_09    m/z 136.8912     0   1E+4   1.5   2  RT (min)      lcqtof_hilic_neg_2_014_08    m/z 136.8912     0   1E+4   0.5   1  RT (min)      lcqtof_hilic_neg_1_060_14    m/z 136.8912     0   2   4   6   8E+3   6   6.5  RT (min)      lcqtof_hilic_neg_1_022_36    m/z 136.8912     0   1   2E+4   3.5   4  RT (min)      lcqtof_hilic_neg_1_007_05    m/z 136.8913     0   1E+4   2   2.5  RT (min)      lcqtof_hilic_neg_2_051_14    m/z 136.8913     0   1   2   3E+3   7   7.5  RT (min)      lcqtof_hilic_neg_3_082_59    m/z 136.8913     0   2   4E+4   3.5   4  RT (min)      lcqtof_hilic_neg_1_034_08    m/z 136.8913     0   1   2E+4   3.5   4  RT (min)      lcqtof_hilic_neg_2_012_43    m/z 136.8913     0   2   4   6E+3   0.5   1  RT (min)      lcqtof_hilic_neg_3_072_57    m/z 136.8913     0   2   4   6E+3   7   7.5  RT (min)      lcqtof_hilic_neg_1_034_08    m/z 136.8913     0   1E+4   1.5   2  RT (min)      lcqtof_hilic_neg_2_053_18    m/z 136.8913     0   1E+4   3   3.5  RT (min)      lcqtof_hilic_neg_2_035_09    m/z 136.8913     0   2   4   6   8E+3   6   6.5  RT (min)      lcqtof_hilic_neg_3_064_13    m/z 136.8913     0   1E+4   1   1.5  RT (min)      lcqtof_hilic_neg_2_030_40    m/z 136.8913     0   1E+4   1.5   2  RT (min)      lcqtof_hilic_neg_1_024_21    m/z 136.8913     0   2   4   6   8E+3   1   1.5  RT (min)      lcqtof_hilic_neg_4_058_39    m/z 136.8914     0   1E+4   2   2.5  RT (min)      lcqtof_hilic_neg_2_037_21    m/z 136.8914     0   2   4E+3   6   6.5  RT (min)      lcqtof_hilic_neg_1_025_52    m/z 136.8914     0   1E+5   5   5.5  RT (min)      lcqtof_hilic_neg_3_055_01    m/z 136.8914     0   1E+4   3   3.5  RT (min)      lcqtof_hilic_neg_2_037_21    m/z 136.8915     0   2   4   6E+3   3   3.5  RT (min)      lcqtof_hilic_neg_1_010_15    m/z 136.8915     0   1E+4   3   3.5  RT (min)      lcqtof_hilic_neg_2_037_21    m/z 136.8915     0   2   4   6E+3   7   7.5  RT (min)      lcqtof_hilic_neg_2_034_47    m/z 136.8915     0   1E+4   5.5   6  RT (min)      lcqtof_hilic_neg_1_023_34    m/z 136.8916     0   2   4   6E+3   2.5   3  RT (min)      lcqtof_hilic_neg_3_059_39    m/z 136.8916     0   2   4   6E+3   2.5   3  RT (min)      lcqtof_hilic_neg_3_059_39    m/z 136.8916     0   1   2   3   4E+3   7   7.5  RT (min)      lcqtof_hilic_neg_2_049_55    m/z 136.9098     0   1E+4   10   10.5  RT (min)      lcqtof_hilic_neg_4_014_46    m/z 137.0271     0   1   2   3E+4   1.5   2  RT (min)       lcqtof_hilic_neg_4_104_61    m/z 137.0271     0   2   4E+4   6   6.5  RT (min)       lcqtof_hilic_neg_4_008_24    m/z 137.0274     0   1E+5   4   4.5  RT (min)      lcqtof_hilic_neg_4_055_32    m/z 137.0275     0   1E+5   4   4.5  RT (min)       lcqtof_hilic_neg_4_055_32    m/z 137.0276     0   1   2   3   4E+4   3.5   4  RT (min)      lcqtof_hilic_neg_4_054_48    m/z 137.0276     0   1E+5   3.5   4  RT (min)      lcqtof_hilic_neg_4_055_32    m/z 137.0279     0   1   2   3E+4   6   6.5  RT (min)      lcqtof_hilic_neg_4_002_25    m/z 137.0280     0   1E+5   4   4.5  RT (min)      lcqtof_hilic_neg_4_055_32    m/z 137.0281     0   1E+5   4   4.5  RT (min)      lcqtof_hilic_neg_4_055_32    m/z 137.0284     0   1   2   3E+4   6   6.5  RT (min)      lcqtof_hilic_neg_4_002_25    m/z 137.0292     0   1E+5   4   4.5  RT (min)      lcqtof_hilic_neg_4_055_32    m/z 137.0302     0   1   2   3E+4   6.5   7  RT (min)      lcqtof_hilic_neg_4_002_25    m/z 137.0303     0   1   2   3E+4   6.5   7  RT (min)      lcqtof_hilic_neg_4_002_25    m/z 137.0344     0   1E+4   2   2.5  RT (min)      lcqtof_hilic_neg_4_032_01    m/z 137.0344     0   1E+3   5.5   6  RT (min)      lcqtof_hilic_neg_4_002_25    m/z 137.0907     0   1E+4   1   1.5  RT (min)      lcqtof_hilic_neg_4_055_32    m/z 137.8915     0   1E+4   4   4.5  RT (min)      lcqtof_hilic_neg_2_034_47    m/z 137.8915     0   1   2   3E+3   3.5   4  RT (min)       lcqtof_hilic_neg_3_100_61    m/z 137.8915     0   1   2E+3   3.5   4  RT (min)      lcqtof_hilic_neg_2_028_05    m/z 137.8917     0   1E+4   4   4.5  RT (min)      lcqtof_hilic_neg_2_051_14    m/z 137.8917     0   1E+4   4   4.5  RT (min)      lcqtof_hilic_neg_2_032_02    m/z 138.0190     0   1   2E+3   1.5   2  RT (min)       lcqtof_hilic_neg_4_102_61    m/z 138.0191     0   1   2E+3   0   0.5  RT (min)       lcqtof_hilic_neg_4_012_07    m/z 138.0192     0   1   2E+3   2.5   3  RT (min)       lcqtof_hilic_neg_4_090_60    m/z 138.0195     0   1   2E+3   1.5   2  RT (min)       lcqtof_hilic_neg_4_058_39    m/z 138.8594     0   1   2   3E+3   4   4.5  RT (min)      lcqtof_hilic_neg_4_097_60    m/z 138.8596     0   1   2   3E+3   5   5.5  RT (min)      lcqtof_hilic_neg_4_007_45    m/z 138.9697     0   1   2E+4   5.5   6  RT (min)      lcqtof_hilic_neg_1_000_aa    m/z 139.0034     0   1E+4   2   2.5  RT (min)      lcqtof_hilic_neg_4_052_13    m/z 139.0064     0   1E+4   2.5   3  RT (min)      lcqtof_hilic_neg_4_038_10    m/z 139.1239     0   1   2   3   4E+4   3.5   4  RT (min)      lcqtof_hilic_neg_4_008_24    m/z 139.1833     0   2   4   6E+4   3.5   4  RT (min)      lcqtof_hilic_neg_4_008_24    m/z 139.2744     0   1   2   3   4E+4   3.5   4  RT (min)      lcqtof_hilic_neg_4_008_24    m/z 139.3424     0   1E+4   3.5   4  RT (min)      lcqtof_hilic_neg_4_008_24    m/z 139.9982     0   1   2E+3   1   1.5  RT (min)      lcqtof_hilic_neg_4_002_25    m/z 139.9982     0   2   4   6   8E+3   1   1.5  RT (min)       lcqtof_hilic_neg_4_058_39    m/z 139.9984     0   1   2   3E+3   0.5   1  RT (min)      lcqtof_hilic_neg_4_009_50    m/z 140.0344     0   1E+4   1.5   2  RT (min)      lcqtof_hilic_neg_4_031_38    m/z 140.9817     0   1E+4   6.5   7  RT (min)      lcqtof_hilic_neg_1_000_aa    m/z 140.9856     0   1   2   3E+3   1   1.5  RT (min)       lcqtof_hilic_neg_4_012_07    m/z 141.9223     0   1E+4   17.5   18  RT (min)       lcqtof_hilic_neg_3_067_57    m/z 141.9226     0   1E+4   19   19.5  RT (min)       lcqtof_hilic_neg_3_066_57    m/z 141.9226     0   1E+4   18   18.5  RT (min)       lcqtof_hilic_neg_3_066_57    m/z 141.9226     0   1E+4   19   19.5  RT (min)      lcqtof_hilic_neg_3_067_57    m/z 141.9227     0   1E+4   19   19.5  RT (min)       lcqtof_hilic_neg_3_066_57    m/z 141.9228     0   1E+4   19.5   20  RT (min)       lcqtof_hilic_neg_3_067_57    m/z 141.9228     0   1E+4   17.5   18  RT (min)       lcqtof_hilic_neg_3_066_57    m/z 141.9229     0   1E+4   17   17.5  RT (min)       lcqtof_hilic_neg_3_066_57    m/z 141.9229     0   1E+4   19.5   20  RT (min)       lcqtof_hilic_neg_3_066_57    m/z 141.9230     0   1E+4   18   18.5  RT (min)       lcqtof_hilic_neg_3_066_57    m/z 141.9230     0   1E+4   17   17.5  RT (min)       lcqtof_hilic_neg_3_067_57    m/z 141.9230     0   1E+4   18.5   19  RT (min)       lcqtof_hilic_neg_3_067_57    m/z 141.9232     0   1E+4   17.5   18  RT (min)       lcqtof_hilic_neg_3_066_57    m/z 141.9248     0   2   4   6   8E+3   1   1.5  RT (min)       lcqtof_hilic_neg_4_050_02    m/z 141.9274     0   1   2   3   4E+3   2   2.5  RT (min)      lcqtof_hilic_neg_2_031_01    m/z 142.0502     0   2   4   6   8E+5   2.5   3  RT (min)      lcqtof_hilic_neg_4_048_23    m/z 142.0503     0   2   4   6   8E+5   2.5   3  RT (min)      lcqtof_hilic_neg_4_089_59    m/z 142.0503     0   1E+4   4   4.5  RT (min)      lcqtof_hilic_neg_4_054_48    m/z 142.0503     0   1E+4   4   4.5  RT (min)      lcqtof_hilic_neg_4_031_38    m/z 142.1117     0   1E+4   2.5   3  RT (min)      lcqtof_hilic_neg_4_048_23    m/z 142.1719     0   1   2E+4   2.5   3  RT (min)      lcqtof_hilic_neg_4_048_23    m/z 142.9248     0   2   4E+3   3   3.5  RT (min)      lcqtof_hilic_neg_3_102_61    m/z 142.9249     0   1   2E+3   3   3.5  RT (min)      lcqtof_hilic_neg_1_030_17    m/z 142.9250     0   1   2E+3   3.5   4  RT (min)      lcqtof_hilic_neg_2_018_16    m/z 142.9250     0   2   4E+3   3   3.5  RT (min)      lcqtof_hilic_neg_3_099_61    m/z 142.9432     0   1E+4   0.5   1  RT (min)      lcqtof_hilic_neg_3_051_12    m/z 142.9434     0   1E+4   1   1.5  RT (min)      lcqtof_hilic_neg_3_113_62    m/z 142.9831     0   1E+4   2   2.5  RT (min)      lcqtof_hilic_neg_4_009_50    m/z 142.9980     0   2   4   6   8E+3   1.5   2  RT (min)      lcqtof_hilic_neg_1_042_26    m/z 142.9982     0   1E+4   3   3.5  RT (min)      lcqtof_hilic_neg_4_011_35    m/z 143.0344     0   1E+4   2   2.5  RT (min)      lcqtof_hilic_neg_1_016_28    m/z 143.0534     0   2   4   6E+4   2.5   3  RT (min)      lcqtof_hilic_neg_4_048_23    m/z 143.0534     0   2   4E+4   2.5   3  RT (min)      lcqtof_hilic_neg_4_089_59    m/z 143.0628     0   1   2E+4   5.5   6  RT (min)      lcqtof_hilic_neg_4_055_32    m/z 143.9274     0   1   2E+3   2.5   3  RT (min)      lcqtof_hilic_neg_3_066_57    m/z 143.9275     0   1   2E+3   2   2.5  RT (min)      lcqtof_hilic_neg_3_066_57    m/z 143.9506     0   1E+4   5   5.5  RT (min)      lcqtof_hilic_neg_1_000_aa    m/z 144.0659     0   1E+4   3   3.5  RT (min)      lcqtof_hilic_neg_4_047_06    m/z 144.9036     0   1   2E+4   7.5   8  RT (min)      lcqtof_hilic_neg_1_000_aa    m/z 144.9230     0   1   2   3E+3   1   1.5  RT (min)      lcqtof_hilic_neg_1_004_14    m/z 144.9230     0   1   2E+3   4   4.5  RT (min)      lcqtof_hilic_neg_1_048_45    m/z 144.9230     0   2   4   6   8E+3   1   1.5  RT (min)       lcqtof_hilic_neg_4_058_39    m/z 144.9232     0   1   2E+3   4   4.5  RT (min)      lcqtof_hilic_neg_2_043_15    m/z 144.9395     0   1E+4   1   1.5  RT (min)      lcqtof_hilic_neg_4_058_39    m/z 145.0499     0   2   4   6   8E+3   0.5   1  RT (min)      lcqtof_hilic_neg_4_050_02    m/z 145.0606     0   2   4   6E+4   2   2.5  RT (min)      lcqtof_hilic_neg_4_050_02    m/z 145.9188     0   1   2E+4   1   1.5  RT (min)      lcqtof_hilic_neg_4_058_39    m/z 145.9303     0   1   2   3E+3   5   5.5  RT (min)      lcqtof_hilic_neg_3_054_37    m/z 146.0456     0   1E+4   2.5   3  RT (min)      lcqtof_hilic_neg_4_050_02    m/z 146.0457     0   1E+4   2   2.5  RT (min)      lcqtof_hilic_neg_4_032_01    m/z 146.0630     0   2   4   6E+3   2   2.5  RT (min)      lcqtof_hilic_neg_4_064_15    m/z 146.0630     0   2   4   6   8E+3   2   2.5  RT (min)      lcqtof_hilic_neg_4_003_16    m/z 146.0818     0   1   2E+4   4.5   5  RT (min)      lcqtof_hilic_neg_4_050_02    m/z 146.1072     0   2   4E+4   6   6.5  RT (min)      lcqtof_hilic_neg_4_048_23    m/z 146.1417     0   1E+4   6   6.5  RT (min)      lcqtof_hilic_neg_4_048_23    m/z 146.1931     0   1   2E+4   6   6.5  RT (min)      lcqtof_hilic_neg_4_048_23    m/z 146.2242     0   1   2E+4   6   6.5  RT (min)      lcqtof_hilic_neg_4_048_23    m/z 146.2615     0   2   4E+4   6   6.5  RT (min)      lcqtof_hilic_neg_4_048_23    m/z 146.3310     0   1   2   3E+4   6   6.5  RT (min)      lcqtof_hilic_neg_4_048_23    m/z 146.9354     0   2   4E+4   10   10.5  RT (min)      lcqtof_hilic_neg_4_014_46    m/z 146.9358     0   1   2E+4   8   9  RT (min)       lcqtof_hilic_neg_4_107_62    m/z 146.9359     0   1   2E+4   7.5   8   8.5  RT (min)       lcqtof_hilic_neg_4_042_22    m/z 146.9359     0   1   2E+4   9   9.5   10  RT (min)       lcqtof_hilic_neg_4_019_03    m/z 146.9361     0   1   2E+4   5   5.5  RT (min)      lcqtof_hilic_neg_4_026_04    m/z 146.9367     0   1   2E+4   11.5   12  RT (min)      lcqtof_hilic_neg_4_014_46    m/z 146.9368     0   2   4E+4   9.5   10  RT (min)      lcqtof_hilic_neg_4_014_46    m/z 146.9368     0   1   2E+4   6   6.5  RT (min)      lcqtof_hilic_neg_4_044_08    m/z 146.9369     0   1   2E+4   7   7.5  RT (min)      lcqtof_hilic_neg_4_030_44    m/z 146.9370     0   1   2E+4   14  RT (min)       lcqtof_hilic_neg_4_100_61    m/z 146.9372     0   1   2E+4   7   7.5  RT (min)      lcqtof_hilic_neg_4_039_18    m/z 146.9373     0   1   2E+4   6   6.5  RT (min)       lcqtof_hilic_neg_4_047_06    m/z 146.9373     0   1   2E+4   6   6.5  RT (min)      lcqtof_hilic_neg_4_039_18    m/z 146.9374     0   1   2E+4   7   7.5  RT (min)      lcqtof_hilic_neg_4_019_03    m/z 146.9375     0   1   2E+4   6.5   7   7.5  RT (min)       lcqtof_hilic_neg_4_105_61    m/z 146.9376     0   2   4E+4   9.5   10  RT (min)      lcqtof_hilic_neg_4_014_46    m/z 146.9377     0   1   2   3E+4   4   4.5  RT (min)      lcqtof_hilic_neg_4_001_49    m/z 146.9379     0   1   2E+4   4   5   6  RT (min)       lcqtof_hilic_neg_4_104_61    m/z 146.9379     0   1   2E+4   4.5   5  RT (min)      lcqtof_hilic_neg_4_046_41    m/z 146.9379     0   1   2E+4   5   5.5  RT (min)      lcqtof_hilic_neg_4_019_03    m/z 146.9379     0   1E+4   4.5   5  RT (min)      lcqtof_hilic_neg_4_014_46    m/z 146.9379     0   1E+4   4.5   5  RT (min)      lcqtof_hilic_neg_4_020_42    m/z 146.9380     0   1   2   3E+4   4   4.5  RT (min)      lcqtof_hilic_neg_4_014_46    m/z 146.9380     0   1   2E+4   0.5   1  RT (min)       lcqtof_hilic_neg_4_095_60    m/z 146.9381     0   2   4E+4   10   10.5  RT (min)      lcqtof_hilic_neg_4_014_46    m/z 146.9384     0   1   2E+4   5.5   6  RT (min)      lcqtof_hilic_neg_4_014_46    m/z 146.9399     0   1E+4   0   0.5  RT (min)      lcqtof_hilic_neg_1_000_aa    m/z 146.9622     0   1   2   3E+4   8   9  RT (min)       lcqtof_hilic_neg_4_053_14    m/z 146.9647     0   1E+4   15   15.5   16  RT (min)       lcqtof_hilic_neg_4_082_59    m/z 146.9650     0   1E+4   15.5   16  RT (min)       lcqtof_hilic_neg_4_099_61    m/z 146.9651     0   2   4E+3   8   8.5  RT (min)      lcqtof_hilic_neg_1_029_06    m/z 146.9651     0   1E+4   15.5   16   16.5  RT (min)       lcqtof_hilic_neg_4_112_62    m/z 146.9651     0   1E+4   16   16.5  RT (min)       lcqtof_hilic_neg_4_012_07    m/z 146.9652     0   2   4   6   8E+3   0   0.5  RT (min)       lcqtof_hilic_neg_4_113_62    m/z 146.9652     0   1   2   3E+3   6   6.5  RT (min)      lcqtof_hilic_neg_1_008_13    m/z 146.9653     0   1E+4   3   3.5  RT (min)      lcqtof_hilic_neg_1_000_aa    m/z 146.9654     0   2   4E+3   2   2.5  RT (min)      lcqtof_hilic_neg_4_113_62    m/z 146.9654     0   2   4E+3   8   8.5  RT (min)      lcqtof_hilic_neg_1_033_53    m/z 146.9654     0   2   4   6   8E+3   0   0.5  RT (min)       lcqtof_hilic_neg_4_047_06    m/z 146.9654     0   1   2   3E+3   5.5   6  RT (min)       lcqtof_hilic_neg_1_000_bb    m/z 146.9654     0   2   4E+3   8   8.5  RT (min)      lcqtof_hilic_neg_2_015_06    m/z 146.9654     0   1   2   3E+3   5.5   6  RT (min)       lcqtof_hilic_neg_2_015_06    m/z 146.9654     0   2   4   6E+3   6   6.5  RT (min)       lcqtof_hilic_neg_4_012_07    m/z 146.9655     0   2   4   6E+3   6   6.5  RT (min)       lcqtof_hilic_neg_4_012_07    m/z 146.9655     0   1   2   3E+3   5.5   6  RT (min)      lcqtof_hilic_neg_1_008_13    m/z 147.0288     0   1   2   3E+4   5.5   6  RT (min)      lcqtof_hilic_neg_4_048_23    m/z 147.0289     0   1   2   3E+4   6   6.5  RT (min)      lcqtof_hilic_neg_4_016_37    m/z 147.0290     0   1   2   3E+4   6   6.5  RT (min)      lcqtof_hilic_neg_4_016_37    m/z 147.0293     0   1E+4   4   4.5  RT (min)       lcqtof_hilic_neg_4_088_59    m/z 147.0294     0   1   2E+4   3.5   4  RT (min)       lcqtof_hilic_neg_4_088_59    m/z 147.0294     0   1   2   3E+4   6.5   7  RT (min)      lcqtof_hilic_neg_4_016_37    m/z 147.0294     0   1   2   3E+4   6   6.5  RT (min)      lcqtof_hilic_neg_4_048_23    m/z 147.0294     0   1   2   3E+4   6   6.5  RT (min)      lcqtof_hilic_neg_4_048_23    m/z 147.0294     0   1   2E+4   3   3.5  RT (min)       lcqtof_hilic_neg_4_089_59    m/z 147.0294     0   2   4E+3   4   4.5  RT (min)      lcqtof_hilic_neg_1_013_38    m/z 147.0296     0   1   2E+4   3.5   4  RT (min)       lcqtof_hilic_neg_4_082_59    m/z 147.0297     0   1   2   3E+4   6   6.5  RT (min)      lcqtof_hilic_neg_4_016_37    m/z 147.0299     0   1   2E+4   3   3.5  RT (min)      lcqtof_hilic_neg_4_086_59    m/z 147.0301     0   1   2   3E+4   6   6.5  RT (min)      lcqtof_hilic_neg_4_048_23    m/z 147.0656     0   1E+4   1.5   2  RT (min)      lcqtof_hilic_neg_4_028_36    m/z 147.0669     0   1E+4   2.5   3  RT (min)      lcqtof_hilic_neg_4_055_32    m/z 148.0149     0   1   2E+4   3   3.5  RT (min)      lcqtof_hilic_neg_4_008_24    m/z 148.0150     0   1E+4   3   3.5  RT (min)      lcqtof_hilic_neg_4_002_25    m/z 148.0330     0   1E+4   2   2.5  RT (min)      lcqtof_hilic_neg_1_003_37    m/z 148.0704     0   1   2   3E+5   2.5   3  RT (min)      lcqtof_hilic_neg_4_055_32    m/z 148.0706     0   1   2E+5   2.5   3  RT (min)      lcqtof_hilic_neg_4_097_60    m/z 149.0083     0   1   2E+4   7   7.5  RT (min)      lcqtof_hilic_neg_4_022_34    m/z 149.0084     0   1   2E+4   7   7.5  RT (min)      lcqtof_hilic_neg_4_010_28    m/z 149.0084     0   1   2E+4   6.5   7  RT (min)      lcqtof_hilic_neg_4_006_30    m/z 150.0116     0   1E+4   6   6.5  RT (min)      lcqtof_hilic_neg_4_022_34    m/z 150.0856     0   1E+4   3   3.5  RT (min)       lcqtof_hilic_neg_4_091_60    m/z 150.1406     0   1E+4   6   6.5  RT (min)      lcqtof_hilic_neg_4_008_24    m/z 150.2027     0   1   2   3E+4   6   6.5  RT (min)      lcqtof_hilic_neg_4_008_24    m/z 150.8888     0   1   2E+4   4   4.5  RT (min)      lcqtof_hilic_neg_4_050_02    m/z 151.0436     0   1E+4   4.5   5  RT (min)      lcqtof_hilic_neg_2_020_23    m/z 151.1234     0   1   2   3   4E+4   4   4.5  RT (min)      lcqtof_hilic_neg_4_048_23    m/z 151.1250     0   1   2   3   4E+4   6   6.5  RT (min)      lcqtof_hilic_neg_4_008_24    m/z 151.1855     0   2   4   6E+4   4   4.5  RT (min)      lcqtof_hilic_neg_4_048_23    m/z 151.1871     0   2   4   6E+4   6   6.5  RT (min)      lcqtof_hilic_neg_4_008_24    m/z 151.2806     0   1   2   3E+4   4   4.5  RT (min)      lcqtof_hilic_neg_4_048_23    m/z 151.2820     0   1   2   3E+4   6   6.5  RT (min)      lcqtof_hilic_neg_4_008_24    m/z 151.3525     0   1E+4   6   6.5  RT (min)      lcqtof_hilic_neg_4_008_24    m/z 152.0015     0   1E+4   6.5   7  RT (min)      lcqtof_hilic_neg_1_000_aa    m/z 152.0457     0   1E+4   4   4.5  RT (min)       lcqtof_hilic_neg_4_096_60    m/z 152.0457     0   1   2E+4   3.5   4  RT (min)       lcqtof_hilic_neg_4_090_60    m/z 152.0458     0   1   2   3   4E+4   6   6.5  RT (min)      lcqtof_hilic_neg_4_008_24    m/z 152.0459     0   1   2   3   4E+4   6.5   7  RT (min)      lcqtof_hilic_neg_4_008_24    m/z 152.0459     0   1   2E+5   2   2.5  RT (min)       lcqtof_hilic_neg_1_005_32    m/z 152.0459     0   2   4E+4   3   3.5  RT (min)       lcqtof_hilic_neg_1_005_32    m/z 152.0459     0   1   2   3   4E+4   6   6.5  RT (min)      lcqtof_hilic_neg_4_008_24    m/z 152.0459     0   1   2E+5   2   2.5  RT (min)      lcqtof_hilic_neg_1_005_32    m/z 152.0459     0   1   2   3   4E+4   5.5   6  RT (min)      lcqtof_hilic_neg_4_008_24    m/z 152.0459     0   2   4E+4   3   3.5  RT (min)       lcqtof_hilic_neg_2_047_32    m/z 152.0460     0   1   2E+4   2.5   3  RT (min)       lcqtof_hilic_neg_4_091_60    m/z 152.0460     0   1   2   3   4E+4   6   6.5  RT (min)      lcqtof_hilic_neg_4_008_24    m/z 152.0460     0   1   2E+4   3.5   4  RT (min)       lcqtof_hilic_neg_4_091_60    m/z 152.0460     0   1   2   3   4E+4   6   6.5  RT (min)      lcqtof_hilic_neg_4_008_24    m/z 152.0460     0   1   2E+4   2   2.5  RT (min)      lcqtof_hilic_neg_2_003_25    m/z 152.0461     0   1   2   3   4E+4   6.5   7  RT (min)      lcqtof_hilic_neg_4_008_24    m/z 152.0461     0   1   2E+4   3   3.5  RT (min)       lcqtof_hilic_neg_4_093_60    m/z 152.0464     0   1   2   3   4E+4   6   6.5  RT (min)      lcqtof_hilic_neg_4_008_24    m/z 152.9857     0   1   2E+4   5.5   6  RT (min)      lcqtof_hilic_neg_1_000_aa    m/z 152.9953     0   1E+4   3.5   4  RT (min)      lcqtof_hilic_neg_4_040_05    m/z 153.0209     0   2   4   6E+3   1.5   2  RT (min)       lcqtof_hilic_neg_4_060_13    m/z 153.0282     0   2   4   6E+3   0.5   1  RT (min)      lcqtof_hilic_neg_4_034_31    m/z 153.0460     0   1E+4   0.5   1  RT (min)      lcqtof_hilic_neg_4_036_11    m/z 153.8679     0   1   2   3E+4   4   4.5  RT (min)      lcqtof_hilic_neg_1_018_44    m/z 153.8680     0   1E+4   4   4.5  RT (min)      lcqtof_hilic_neg_1_000_dd    m/z 153.8680     0   1E+4   3.5   4  RT (min)      lcqtof_hilic_neg_1_006_22    m/z 153.8680     0   1E+4   3.5   4  RT (min)      lcqtof_hilic_neg_1_006_22    m/z 153.8681     0   1E+4   3.5   4  RT (min)      lcqtof_hilic_neg_1_024_21    m/z 153.8681     0   1E+4   3   3.5  RT (min)      lcqtof_hilic_neg_1_006_22    m/z 153.8682     0   1   2   3E+3   2   2.5  RT (min)      lcqtof_hilic_neg_1_004_14    m/z 153.8682     0   1E+4   3.5   4  RT (min)      lcqtof_hilic_neg_1_006_22    m/z 153.8682     0   1E+4   3   3.5  RT (min)      lcqtof_hilic_neg_1_006_22    m/z 153.8683     0   1E+4   3   3.5  RT (min)      lcqtof_hilic_neg_1_006_22    m/z 153.8684     0   1   2   3E+3   2   2.5  RT (min)      lcqtof_hilic_neg_1_004_14    m/z 154.0553     0   1E+4   4.5   5  RT (min)      lcqtof_hilic_neg_2_052_24    m/z 154.0613     0   1E+5   7.5   8  RT (min)      lcqtof_hilic_neg_4_063_16    m/z 154.9468     0   2   4E+3   5.5   6  RT (min)      lcqtof_hilic_neg_1_004_14    m/z 154.9470     0   1   2   3E+3   6   6.5  RT (min)      lcqtof_hilic_neg_1_010_15    m/z 154.9472     0   2   4E+3   5   5.5  RT (min)      lcqtof_hilic_neg_4_026_04    m/z 154.9992     0   1E+4   2   2.5  RT (min)      lcqtof_hilic_neg_4_053_14    m/z 155.0018     0   1E+4   3.5   4  RT (min)      lcqtof_hilic_neg_4_091_60    m/z 155.0021     0   1E+4   4   4.5  RT (min)      lcqtof_hilic_neg_4_055_32    m/z 155.0039     0   1E+4   1   1.5  RT (min)       lcqtof_hilic_neg_4_050_02    m/z 155.0277     0   1E+4   2   2.5  RT (min)      lcqtof_hilic_neg_4_052_13    m/z 155.8652     0   2   4   6   8E+3   3.5   4  RT (min)      lcqtof_hilic_neg_1_006_22    m/z 155.8652     0   2   4E+4   4   4.5  RT (min)      lcqtof_hilic_neg_1_002_39    m/z 155.8653     0   2   4   6   8E+3   3.5   4  RT (min)      lcqtof_hilic_neg_1_006_22    m/z 155.8653     0   2   4E+4   4   4.5  RT (min)      lcqtof_hilic_neg_1_002_39    m/z 155.8654     0   2   4   6   8E+3   3.5   4  RT (min)      lcqtof_hilic_neg_1_006_22    m/z 155.8654     0   2   4   6E+3   3.5   4  RT (min)      lcqtof_hilic_neg_1_024_21    m/z 155.8655     0   1E+4   4   4.5  RT (min)      lcqtof_hilic_neg_1_006_22    m/z 155.8657     0   1   2E+3   2   2.5  RT (min)      lcqtof_hilic_neg_1_004_14    m/z 156.0052     0   1E+4   3.5   4  RT (min)      lcqtof_hilic_neg_4_040_05    m/z 156.0054     0   1E+4   6   6.5  RT (min)      lcqtof_hilic_neg_4_008_24    m/z 156.0293     0   1E+4   2.5   3  RT (min)      lcqtof_hilic_neg_4_085_59    m/z 156.0294     0   1   2   3   4E+4   3   3.5  RT (min)      lcqtof_hilic_neg_4_040_05    m/z 156.0295     0   2   4   6E+4   2.5   3  RT (min)       lcqtof_hilic_neg_4_048_23    m/z 156.0296     0   1E+4   3   3.5  RT (min)      lcqtof_hilic_neg_4_033_53    m/z 156.0770     0   2   4   6E+4   6.5   7  RT (min)      lcqtof_hilic_neg_4_008_24    m/z 156.0773     0   2   4   6E+3   4.5   5  RT (min)      lcqtof_hilic_neg_4_055_32    m/z 156.0774     0   1E+6   3.5   4  RT (min)      lcqtof_hilic_neg_4_008_24    m/z 156.0774     0   2   4   6   8E+5   3.5   4  RT (min)      lcqtof_hilic_neg_1_037_24    m/z 156.0775     0   1E+4   6.5   7  RT (min)      lcqtof_hilic_neg_4_002_25    m/z 156.0775     0   1   2   3E+6   3.5   4  RT (min)      lcqtof_hilic_neg_4_008_24    m/z 156.0776     0   1   2E+4   6.5   7  RT (min)      lcqtof_hilic_neg_1_037_24    m/z 156.1298     0   2   4E+4   1.5   2  RT (min)      lcqtof_hilic_neg_4_048_23    m/z 156.1652     0   1E+4   1.5   2  RT (min)      lcqtof_hilic_neg_4_048_23    m/z 156.1769     0   1   2E+4   4   4.5  RT (min)      lcqtof_hilic_neg_4_055_32    m/z 156.1932     0   2   4   6   8E+4   1.5   2  RT (min)      lcqtof_hilic_neg_4_048_23    m/z 156.2185     0   1E+4   1.5   2  RT (min)      lcqtof_hilic_neg_4_048_23    m/z 156.2301     0   1   2   3E+4   4   4.5  RT (min)      lcqtof_hilic_neg_4_055_32    m/z 156.2509     0   1   2E+4   1.5   2  RT (min)      lcqtof_hilic_neg_4_048_23    m/z 156.2623     0   1   2   3E+4   4   4.5  RT (min)      lcqtof_hilic_neg_4_055_32    m/z 156.2895     0   2   4E+4   1.5   2  RT (min)      lcqtof_hilic_neg_4_048_23    m/z 156.3614     0   1   2E+4   1.5   2  RT (min)      lcqtof_hilic_neg_4_048_23    m/z 156.3728     0   2   4E+4   4   4.5  RT (min)      lcqtof_hilic_neg_4_055_32    m/z 156.9446     0   2   4E+4   0   0.5  RT (min)      lcqtof_hilic_neg_4_000_aa    m/z 156.9446     0   1E+5   9   9.5  RT (min)      lcqtof_hilic_neg_1_000_aa    m/z 156.9447     0   2   4E+4   10.5   11  RT (min)      lcqtof_hilic_neg_1_000_aa    m/z 156.9448     0   1   2   3   4E+4   11   11.5  RT (min)      lcqtof_hilic_neg_1_000_aa    m/z 156.9448     0   1E+5   9   9.5  RT (min)      lcqtof_hilic_neg_1_000_aa    m/z 156.9449     0   2   4E+4   0.5   1  RT (min)      lcqtof_hilic_neg_4_000_aa    m/z 156.9450     0   2   4E+4   0.5   1  RT (min)      lcqtof_hilic_neg_4_000_aa    m/z 156.9450     0   2   4E+4   10.5   11  RT (min)      lcqtof_hilic_neg_1_000_aa    m/z 156.9450     0   1E+5   12.5   13  RT (min)      lcqtof_hilic_neg_1_000_aa    m/z 156.9451     0   1   2   3   4E+5   8   8.5  RT (min)      lcqtof_hilic_neg_1_000_aa    m/z 156.9451     0   1   2   3   4E+5   7.5   8  RT (min)      lcqtof_hilic_neg_1_000_aa    m/z 156.9452     0   1   2   3   4E+5   7.5   8  RT (min)      lcqtof_hilic_neg_1_000_aa    m/z 156.9452     0   1   2   3   4E+5   8   8.5  RT (min)      lcqtof_hilic_neg_1_000_aa    m/z 156.9452     0   1   2   3   4E+5   7.5   8  RT (min)      lcqtof_hilic_neg_1_000_aa    m/z 156.9452     0   2   4E+4   10.5   11  RT (min)      lcqtof_hilic_neg_1_000_aa    m/z 156.9452     0   1E+5   12.5   13  RT (min)      lcqtof_hilic_neg_1_000_aa    m/z 156.9452     0   2   4E+4   0   0.5  RT (min)      lcqtof_hilic_neg_4_000_aa    m/z 156.9455     0   2   4E+4   0   0.5  RT (min)      lcqtof_hilic_neg_4_000_aa    m/z 157.0504     0   1   2   3   4E+3   1.5   2  RT (min)      lcqtof_hilic_neg_4_053_14    m/z 157.9307     0   1   2E+4   5   5.5  RT (min)      lcqtof_hilic_neg_1_000_aa    m/z 157.9307     0   2   4E+3   4.5   5  RT (min)      lcqtof_hilic_neg_1_000_bb    m/z 158.9242     0   1   2   3   4E+4   9   9.5  RT (min)      lcqtof_hilic_neg_1_000_aa    m/z 158.9243     0   1E+5   8   8.5  RT (min)      lcqtof_hilic_neg_1_000_aa    m/z 158.9244     0   2   4   6E+4   8.5   9  RT (min)      lcqtof_hilic_neg_1_000_aa    m/z 158.9247     0   1E+5   8.5   9  RT (min)      lcqtof_hilic_neg_1_000_aa    m/z 158.9247     0   1E+5   8.5   9  RT (min)      lcqtof_hilic_neg_1_000_aa    m/z 158.9248     0   1E+5   8   8.5  RT (min)      lcqtof_hilic_neg_1_000_aa    m/z 158.9248     0   1E+4   9.5   10  RT (min)      lcqtof_hilic_neg_2_000_aa    m/z 158.9380     0   2   4   6E+3   19.5   20  RT (min)       lcqtof_hilic_neg_3_068_57    m/z 158.9380     0   1   2   3   4E+4   1   1.5  RT (min)      lcqtof_hilic_neg_3_112_62    m/z 158.9380     0   1E+4   4  RT (min)       lcqtof_hilic_neg_3_086_59    m/z 158.9382     0   1E+4   4.5   5  RT (min)       lcqtof_hilic_neg_3_107_62    m/z 158.9382     0   2   4E+3   18   18.5  RT (min)      lcqtof_hilic_neg_3_111_62    m/z 158.9382     0   2   4   6E+3   19   19.5  RT (min)       lcqtof_hilic_neg_3_081_58    m/z 158.9385     0   1   2   3   4E+3   17.5   18  RT (min)      lcqtof_hilic_neg_3_103_61    m/z 158.9386     0   2   4   6   8E+3   4   4.5  RT (min)       lcqtof_hilic_neg_3_054_37    m/z 159.0102     0   2   4   6   8E+3   3   3.5  RT (min)       lcqtof_hilic_neg_4_084_59    m/z 159.0102     0   2   4   6   8E+3   3   3.5  RT (min)       lcqtof_hilic_neg_4_084_59    m/z 159.0287     0   2   4   6   8E+3   2   2.5  RT (min)      lcqtof_hilic_neg_1_016_28    m/z 159.0292     0   2   4   6E+4   1.5   2  RT (min)      lcqtof_hilic_neg_4_064_15    m/z 159.0292     0   1   2E+4   1   1.5  RT (min)      lcqtof_hilic_neg_4_050_02    m/z 159.0293     0   1E+4   1.5   2  RT (min)      lcqtof_hilic_neg_4_022_34    m/z 159.0293     0   1E+4   2   2.5  RT (min)      lcqtof_hilic_neg_1_042_26    m/z 159.0654     0   1   2   3   4E+3   1   1.5  RT (min)       lcqtof_hilic_neg_4_050_02    m/z 159.8861     0   1   2   3E+3   0   0.5  RT (min)       lcqtof_hilic_neg_1_000_cc    m/z 159.8862     0   1   2E+4   0.5   1  RT (min)      lcqtof_hilic_neg_1_000_cc    m/z 159.8863     0   1E+4   0.5   1  RT (min)      lcqtof_hilic_neg_4_020_42    m/z 159.8863     0   1E+3   0   0.5  RT (min)       lcqtof_hilic_neg_4_000_aa    m/z 159.9388     0   2   4   6   8E+3   2   2.5  RT (min)      lcqtof_hilic_neg_1_054_01    m/z 159.9392     0   1   2   3E+3   3   3.5  RT (min)      lcqtof_hilic_neg_3_067_57    m/z 159.9392     0   1   2E+3   3.5   4  RT (min)      lcqtof_hilic_neg_3_053_26    m/z 159.9393     0   1   2   3   4E+3   1   1.5  RT (min)      lcqtof_hilic_neg_4_050_02    m/z 160.0432     0   1E+4   2   2.5  RT (min)      lcqtof_hilic_neg_4_050_02    m/z 160.0612     0   1E+4   2   2.5  RT (min)      lcqtof_hilic_neg_4_050_02    m/z 160.8382     0   1   2   3E+5   5   5.5  RT (min)      lcqtof_hilic_neg_4_032_01    m/z 160.8413     0   2   4   6E+5   5   5.5  RT (min)       lcqtof_hilic_neg_3_066_57    m/z 160.8413     0   1E+5   3.5   4  RT (min)       lcqtof_hilic_neg_3_066_57    m/z 160.8414     0   2   4   6E+4   3.5   4  RT (min)      lcqtof_hilic_neg_4_013_20    m/z 160.8414     0   1E+5   3.5   4  RT (min)      lcqtof_hilic_neg_4_013_20    m/z 160.8414     0   2   4   6E+3   1.5   2  RT (min)       lcqtof_hilic_neg_3_066_57    m/z 160.8414     0   1E+4   16   16.5  RT (min)       lcqtof_hilic_neg_3_066_57    m/z 160.8414     0   2   4   6   8E+3   1   1.5  RT (min)       lcqtof_hilic_neg_4_106_62    m/z 160.8415     0   1E+4   6.5   7  RT (min)      lcqtof_hilic_neg_4_032_01    m/z 160.8415     0   1   2   3   4E+4   3   3.5  RT (min)       lcqtof_hilic_neg_3_066_57    m/z 160.8415     0   2   4   6   8E+4   3.5   4  RT (min)       lcqtof_hilic_neg_3_066_57    m/z 160.8415     0   1   2E+3   0   0.5  RT (min)       lcqtof_hilic_neg_3_000_aa    m/z 160.8416     0   2   4   6   8E+4   3.5   4  RT (min)      lcqtof_hilic_neg_4_013_20    m/z 160.8416     0   1E+5   3.5   4  RT (min)      lcqtof_hilic_neg_4_051_43    m/z 160.8416     0   1   2   3   4E+5   3.5   4  RT (min)      lcqtof_hilic_neg_4_038_10    m/z 160.8417     0   2   4   6   8E+3   0.5   1  RT (min)       lcqtof_hilic_neg_3_066_57    m/z 160.8417     0   1   2E+5   3.5   4  RT (min)      lcqtof_hilic_neg_4_020_42    m/z 160.8417     0   1E+4   6.5   7  RT (min)      lcqtof_hilic_neg_4_005_29    m/z 160.8418     0   1E+5   3.5   4  RT (min)      lcqtof_hilic_neg_4_013_20    m/z 160.9349     0   1   2   3   4E+4   2   2.5  RT (min)       lcqtof_hilic_neg_1_046_30    m/z 160.9349     0   2   4   6   8E+3   0.5   1  RT (min)      lcqtof_hilic_neg_2_018_16    m/z 160.9350     0   2   4   6   8E+3   2   2.5  RT (min)      lcqtof_hilic_neg_4_058_39    m/z 160.9350     0   1   2   3   4E+4   2   2.5  RT (min)      lcqtof_hilic_neg_2_051_14    m/z 160.9351     0   1E+4   3.5   4  RT (min)       lcqtof_hilic_neg_2_045_11    m/z 160.9351     0   1   2   3E+4   2.5   3  RT (min)       lcqtof_hilic_neg_1_041_54    m/z 160.9351     0   2   4   6E+3   0   0.5  RT (min)       lcqtof_hilic_neg_2_062_17    m/z 160.9351     0   1   2   3E+4   1   1.5  RT (min)       lcqtof_hilic_neg_4_058_39    m/z 160.9352     0   1   2   3E+4   2.5   3  RT (min)      lcqtof_hilic_neg_1_011_46    m/z 160.9352     0   2   4   6   8E+3   4   4.5  RT (min)       lcqtof_hilic_neg_2_055_30    m/z 160.9352     0   1   2   3   4E+3   19.5   20  RT (min)       lcqtof_hilic_neg_2_012_43    m/z 160.9352     0   1   2E+4   2.5   3  RT (min)      lcqtof_hilic_neg_1_048_45    m/z 160.9352     0   1   2   3E+4   2.5   3  RT (min)      lcqtof_hilic_neg_1_011_46    m/z 160.9353     0   2   4   6E+3   4   4.5  RT (min)       lcqtof_hilic_neg_3_076_58    m/z 160.9353     0   1   2   3E+4   2.5   3  RT (min)       lcqtof_hilic_neg_1_028_47    m/z 160.9354     0   1E+4   2.5   3  RT (min)      lcqtof_hilic_neg_1_028_47    m/z 160.9355     0   1   2E+3   6.5   7  RT (min)      lcqtof_hilic_neg_2_021_17    m/z 160.9355     0   1   2   3E+3   6.5   7  RT (min)      lcqtof_hilic_neg_2_058_15    m/z 160.9355     0   2   4   6   8E+3   1.5   2  RT (min)      lcqtof_hilic_neg_4_058_39    m/z 160.9357     0   1   2   3E+4   1   1.5  RT (min)      lcqtof_hilic_neg_4_058_39    m/z 161.0234     0   1   2E+3   2   2.5  RT (min)      lcqtof_hilic_neg_4_017_51    m/z 161.0451     0   1   2E+4   1   1.5  RT (min)      lcqtof_hilic_neg_4_050_02    m/z 161.8838     0   1E+4   0.5   1  RT (min)      lcqtof_hilic_neg_1_000_cc    m/z 161.9130     0   2   4   6   8E+3   0.5   1  RT (min)       lcqtof_hilic_neg_2_015_06    m/z 161.9130     0   1   2E+3   0   0.5  RT (min)       lcqtof_hilic_neg_1_065_51    m/z 162.0495     0   1E+4   2   2.5  RT (min)      lcqtof_hilic_neg_4_073_57    m/z 162.0496     0   1   2E+4   3   3.5  RT (min)      lcqtof_hilic_neg_4_040_05    m/z 162.0499     0   2   4   6   8E+4   2.5   3  RT (min)       lcqtof_hilic_neg_4_091_60    m/z 162.0499     0   1E+4   2   2.5  RT (min)      lcqtof_hilic_neg_4_050_02    m/z 162.0500     0   1E+4   3   3.5  RT (min)      lcqtof_hilic_neg_4_033_53    m/z 162.1416     0   1E+4   1   1.5  RT (min)      lcqtof_hilic_neg_4_039_18    m/z 162.1748     0   1E+4   0.5   1   1.5  RT (min)      lcqtof_hilic_neg_4_039_18    m/z 162.2139     0   2   4E+4   0.5   1   1.5  RT (min)      lcqtof_hilic_neg_4_039_18    m/z 162.2857     0   1   2E+4   1   1.5  RT (min)      lcqtof_hilic_neg_4_039_18    m/z 162.8383     0   1E+5   3.5   4  RT (min)      lcqtof_hilic_neg_3_066_57    m/z 162.8384     0   2   4   6   8E+4   3.5   4  RT (min)      lcqtof_hilic_neg_4_013_20    m/z 162.8384     0   1E+4   6.5   7  RT (min)      lcqtof_hilic_neg_4_005_29    m/z 162.8385     0   2   4E+4   5   5.5  RT (min)      lcqtof_hilic_neg_3_000_aa    m/z 162.8385     0   2   4   6E+4   3.5   4  RT (min)      lcqtof_hilic_neg_4_015_09    m/z 162.8385     0   1E+4   6.5   7  RT (min)      lcqtof_hilic_neg_4_032_01    m/z 162.8386     0   1   2   3   4E+4   3   3.5  RT (min)      lcqtof_hilic_neg_3_066_57    m/z 162.8386     0   1E+4   6.5   7  RT (min)      lcqtof_hilic_neg_4_032_01    m/z 162.8386     0   1E+4   6.5   7  RT (min)      lcqtof_hilic_neg_4_005_29    m/z 162.8388     0   2   4   6   8E+3   0.5   1  RT (min)      lcqtof_hilic_neg_3_066_57    m/z 162.8388     0   1   2   3   4E+5   3.5   4  RT (min)      lcqtof_hilic_neg_4_039_18    m/z 162.8389     0   1   2   3   4E+3   8   8.5  RT (min)      lcqtof_hilic_neg_4_081_58    m/z 162.8928     0   1E+4   5   5.5  RT (min)      lcqtof_hilic_neg_4_050_02    m/z 163.0029     0   1E+4   2   2.5  RT (min)      lcqtof_hilic_neg_4_058_39    m/z 163.0029     0   1E+4   2   2.5  RT (min)      lcqtof_hilic_neg_4_058_39    m/z 163.0031     0   1E+4   2   2.5  RT (min)      lcqtof_hilic_neg_4_058_39    m/z 163.0032     0   1E+4   2   2.5  RT (min)      lcqtof_hilic_neg_4_058_39    m/z 163.0236     0   1E+4   3   3.5  RT (min)       lcqtof_hilic_neg_4_093_60    m/z 163.0241     0   1E+4   3   3.5  RT (min)      lcqtof_hilic_neg_4_045_26    m/z 163.0334     0   2   4   6   8E+4   2   2.5  RT (min)      lcqtof_hilic_neg_4_023_15    m/z 163.0336     0   1   2E+4   2.5   3  RT (min)      lcqtof_hilic_neg_4_001_49    m/z 163.0337     0   1E+4   2   2.5  RT (min)      lcqtof_hilic_neg_4_013_20    m/z 163.0339     0   1E+4   2   2.5  RT (min)      lcqtof_hilic_neg_4_013_20    m/z 163.0340     0   2   4   6E+3   2.5   3  RT (min)      lcqtof_hilic_neg_1_029_06    m/z 163.0386     0   2   4   6E+4   2   2.5  RT (min)      lcqtof_hilic_neg_4_032_01    m/z 163.0421     0   2   4   6   8E+4   1   1.5  RT (min)      lcqtof_hilic_neg_4_050_02    m/z 164.0457     0   1E+4   1.5   2  RT (min)      lcqtof_hilic_neg_4_097_60    m/z 164.8352     0   1   2   3   4E+4   3.5   4  RT (min)      lcqtof_hilic_neg_4_000_bb    m/z 164.8352     0   2   4   6E+4   4   4.5  RT (min)      lcqtof_hilic_neg_4_000_dd    m/z 164.8354     0   2   4   6E+4   4   4.5  RT (min)      lcqtof_hilic_neg_4_000_dd    m/z 164.8354     0   1   2   3E+4   3.5   4  RT (min)      lcqtof_hilic_neg_3_066_57    m/z 164.8358     0   2   4E+4   3.5   4  RT (min)      lcqtof_hilic_neg_4_000_bb    m/z 165.0202     0   1E+5   1.5   2  RT (min)      lcqtof_hilic_neg_1_064_16    m/z 165.0306     0   1   2   3   4E+2   1   1.5  RT (min)      lcqtof_hilic_neg_4_000_aa    m/z 165.0311     0   1   2E+2   1   1.5  RT (min)      lcqtof_hilic_neg_4_000_aa    m/z 165.0494     0   2   4   6   8E+3   2.5   3  RT (min)      lcqtof_hilic_neg_4_028_36    m/z 165.0499     0   2   4   6E+4   2   2.5  RT (min)      lcqtof_hilic_neg_4_053_14    m/z 165.0661     0   1E+3   1   1.5  RT (min)      lcqtof_hilic_neg_4_005_29    m/z 165.8376     0   2   4   6   8E+3   4.5   5  RT (min)      lcqtof_hilic_neg_4_014_46    m/z 166.0221     0   1E+4   1.5   2  RT (min)      lcqtof_hilic_neg_1_064_16    m/z 166.0812     0   1E+4   3   3.5  RT (min)      lcqtof_hilic_neg_2_047_32    m/z 166.0822     0   1   2   3   4E+4   2.5   3  RT (min)      lcqtof_hilic_neg_4_055_32    m/z 167.0036     0   1E+4   2   2.5  RT (min)      lcqtof_hilic_neg_4_062_17    m/z 167.0652     0   2   4   6E+4   2.5   3  RT (min)       lcqtof_hilic_neg_1_005_32    m/z 167.1372     0   1   2   3E+4   1   1.5  RT (min)      lcqtof_hilic_neg_4_039_18    m/z 167.1660     0   1E+5   0.5   1  RT (min)      lcqtof_hilic_neg_4_039_18    m/z 167.1921     0   2   4E+4   1   1.5  RT (min)      lcqtof_hilic_neg_4_039_18    m/z 167.2257     0   2   4   6E+4   1   1.5  RT (min)      lcqtof_hilic_neg_4_039_18    m/z 167.2645     0   2   4   6   8E+4   0.5   1  RT (min)      lcqtof_hilic_neg_4_039_18    m/z 167.8361     0   1E+4   4   4.5  RT (min)      lcqtof_hilic_neg_4_050_02    m/z 167.8361     0   1E+4   3.5   4  RT (min)       lcqtof_hilic_neg_1_001_49    m/z 167.8362     0   1   2   3   4E+3   0   0.5  RT (min)       lcqtof_hilic_neg_2_008_44    m/z 167.8362     0   2   4E+3   16   16.5  RT (min)       lcqtof_hilic_neg_4_000_cc    m/z 167.8362     0   1E+4   4   4.5  RT (min)      lcqtof_hilic_neg_4_050_02    m/z 167.8363     0   1   2   3E+3   0   0.5  RT (min)       lcqtof_hilic_neg_2_000_dd    m/z 167.8364     0   1E+4   4   4.5  RT (min)      lcqtof_hilic_neg_1_001_49    m/z 167.8364     0   2   4E+3   15.5   16  RT (min)       lcqtof_hilic_neg_4_006_30    m/z 167.8365     0   2   4   6E+3   1   1.5  RT (min)      lcqtof_hilic_neg_1_020_09    m/z 167.8995     0   1E+4   1   1.5  RT (min)      lcqtof_hilic_neg_4_039_18    m/z 168.0276     0   1E+3   0.5   1  RT (min)      lcqtof_hilic_neg_1_062_17    m/z 168.8668     0   1E+4   4   4.5  RT (min)      lcqtof_hilic_neg_4_050_02    m/z 168.8862     0   1E+4   1   1.5  RT (min)      lcqtof_hilic_neg_4_051_43    m/z 168.9803     0   1   2   3   4E+4   5.5   6  RT (min)      lcqtof_hilic_neg_1_000_aa    m/z 168.9813     0   1   2E+4   2   2.5  RT (min)       lcqtof_hilic_neg_3_063_17    m/z 168.9814     0   2   4E+4   2   2.5  RT (min)       lcqtof_hilic_neg_2_021_17    m/z 168.9885     0   2   4E+3   1   1.5  RT (min)       lcqtof_hilic_neg_4_071_57    m/z 168.9885     0   1E+4   0   0.5  RT (min)       lcqtof_hilic_neg_4_091_60    m/z 168.9886     0   1   2   3   4E+3   2   2.5  RT (min)       lcqtof_hilic_neg_4_099_61    m/z 168.9886     0   1   2E+4   0.5   1  RT (min)       lcqtof_hilic_neg_4_098_61    m/z 168.9886     0   1   2   3   4E+3   0   0.5  RT (min)       lcqtof_hilic_neg_4_080_58    m/z 168.9886     0   2   4   6E+3   2   2.5  RT (min)       lcqtof_hilic_neg_4_050_02    m/z 168.9886     0   1E+4   0.5   1  RT (min)       lcqtof_hilic_neg_4_107_62    m/z 168.9887     0   2   4E+3   3   3.5  RT (min)       lcqtof_hilic_neg_4_050_02    m/z 168.9887     0   1E+4   1   1.5  RT (min)       lcqtof_hilic_neg_4_096_60    m/z 168.9888     0   1E+4   1   1.5  RT (min)      lcqtof_hilic_neg_4_075_58    m/z 168.9888     0   1   2   3   4E+3   2.5   3  RT (min)       lcqtof_hilic_neg_4_098_61    m/z 168.9889     0   2   4E+3   1.5   2  RT (min)      lcqtof_hilic_neg_4_057_49    m/z 168.9889     0   1E+4   16   18  RT (min)       lcqtof_hilic_neg_4_050_02    m/z 168.9890     0   1   2   3   4E+3   0   0.5  RT (min)       lcqtof_hilic_neg_4_098_61    m/z 168.9890     0   2   4   6   8E+3   0.5   1  RT (min)       lcqtof_hilic_neg_4_062_17    m/z 168.9891     0   1E+4   0.5   1  RT (min)       lcqtof_hilic_neg_4_099_61    m/z 169.0237     0   1E+4   0.5   1  RT (min)      lcqtof_hilic_neg_4_039_18    m/z 169.0245     0   2   4   6   8E+3   1   2   3  RT (min)       lcqtof_hilic_neg_4_039_18    m/z 169.0248     0   2   4   6   8E+3   1   2   3  RT (min)       lcqtof_hilic_neg_4_039_18    m/z 169.8334     0   2   4   6E+3   0   0.5  RT (min)       lcqtof_hilic_neg_1_063_13    m/z 169.8334     0   2   4   6E+4   3.5   4  RT (min)       lcqtof_hilic_neg_1_000_bb    m/z 169.8334     0   2   4   6E+3   15.5   16  RT (min)       lcqtof_hilic_neg_4_080_58    m/z 169.8334     0   2   4E+3   0   0.5  RT (min)       lcqtof_hilic_neg_1_063_13    m/z 169.8334     0   2   4E+3   0   0.5  RT (min)       lcqtof_hilic_neg_1_064_16    m/z 169.8335     0   2   4   6E+3   0   0.5  RT (min)       lcqtof_hilic_neg_1_064_16    m/z 169.8335     0   1   2E+4   3   3.5  RT (min)      lcqtof_hilic_neg_1_008_13    m/z 169.8336     0   1   2   3E+3   1.5   2  RT (min)      lcqtof_hilic_neg_4_000_aa    m/z 169.8336     0   2   4   6   8E+3   2.5   3  RT (min)       lcqtof_hilic_neg_1_004_14    m/z 169.8336     0   1E+4   18   19  RT (min)       lcqtof_hilic_neg_1_000_bb    m/z 169.8336     0   2   4   6E+3   1   1.5  RT (min)      lcqtof_hilic_neg_4_023_15    m/z 169.8337     0   2   4   6E+3   1.5   2  RT (min)       lcqtof_hilic_neg_2_005_13    m/z 169.8338     0   1   2   3E+3   2   2.5  RT (min)       lcqtof_hilic_neg_4_012_07    m/z 169.8338     0   1   2   3E+3   1   1.5  RT (min)      lcqtof_hilic_neg_4_000_aa    m/z 169.8338     0   2   4   6E+3   2.5   3  RT (min)       lcqtof_hilic_neg_1_007_05    m/z 169.9728     0   1E+4   6   6.5  RT (min)      lcqtof_hilic_neg_2_052_24    m/z 170.0314     0   1E+4   4   4.5  RT (min)       lcqtof_hilic_neg_4_092_60    m/z 170.0438     0   1E+3   4   4.5  RT (min)       lcqtof_hilic_neg_4_004_47    m/z 170.0455     0   1   2E+4   3   3.5  RT (min)      lcqtof_hilic_neg_4_047_06    m/z 170.0562     0   1E+4   5.5   6  RT (min)      lcqtof_hilic_neg_1_005_32    m/z 170.8324     0   1E+4   7.5   8  RT (min)      lcqtof_hilic_neg_4_010_28    m/z 170.9963     0   1   2E+4   3.5   4  RT (min)       lcqtof_hilic_neg_4_052_13    m/z 171.8309     0   1   2   3   4E+3   16   16.5  RT (min)       lcqtof_hilic_neg_4_000_cc    m/z 171.8310     0   1E+4   3.5   4  RT (min)       lcqtof_hilic_neg_1_001_49    m/z 171.8310     0   2   4   6   8E+3   4   4.5  RT (min)      lcqtof_hilic_neg_1_001_49    m/z 171.8311     0   2   4   6   8E+3   3.5   4  RT (min)      lcqtof_hilic_neg_1_004_14    m/z 171.8311     0   1   2   3E+3   1.5   2  RT (min)       lcqtof_hilic_neg_2_005_13    m/z 171.8311     0   1E+4   4   4.5  RT (min)      lcqtof_hilic_neg_4_050_02    m/z 171.8311     0   1   2   3E+3   0   0.5  RT (min)      lcqtof_hilic_neg_1_043_41    m/z 171.8311     0   2   4   6   8E+3   4   4.5  RT (min)      lcqtof_hilic_neg_1_001_49    m/z 171.9462     0   2   4   6E+3   6.5   7  RT (min)       lcqtof_hilic_neg_2_060_14    m/z 171.9462     0   2   4   6E+3   6.5   7  RT (min)      lcqtof_hilic_neg_2_060_14    m/z 171.9463     0   2   4E+3   7   7.5  RT (min)      lcqtof_hilic_neg_1_016_28    m/z 171.9463     0   2   4   6E+3   6.5   7  RT (min)      lcqtof_hilic_neg_2_060_14    m/z 171.9463     0   2   4   6   8E+3   6   6.5  RT (min)      lcqtof_hilic_neg_2_038_04    m/z 171.9463     0   2   4   6   8E+3   6   6.5  RT (min)       lcqtof_hilic_neg_2_038_04    m/z 171.9463     0   1   2   3E+3   1   1.5  RT (min)      lcqtof_hilic_neg_1_002_39    m/z 171.9464     0   2   4   6E+3   2   2.5  RT (min)      lcqtof_hilic_neg_1_060_14    m/z 171.9464     0   1E+4   1.5   2  RT (min)       lcqtof_hilic_neg_3_066_57    m/z 171.9464     0   2   4   6E+3   6.5   7  RT (min)       lcqtof_hilic_neg_2_060_14    m/z 171.9464     0   1E+4   1.5   2  RT (min)       lcqtof_hilic_neg_3_066_57    m/z 171.9465     0   1   2   3E+3   0   0.5  RT (min)       lcqtof_hilic_neg_2_035_09    m/z 171.9465     0   1   2   3E+3   8   8.5  RT (min)       lcqtof_hilic_neg_2_000_bb    m/z 171.9465     0   1   2   3E+3   1.5   2  RT (min)      lcqtof_hilic_neg_1_002_39    m/z 171.9468     0   1   2E+3   8.5   9  RT (min)      lcqtof_hilic_neg_2_044_37    m/z 171.9886     0   1E+4   6   6.5  RT (min)      lcqtof_hilic_neg_3_034_24    m/z 172.0413     0   1   2E+4   2.5   3  RT (min)       lcqtof_hilic_neg_4_088_59    m/z 172.1946     0   1E+4   1.5   2  RT (min)      lcqtof_hilic_neg_4_031_38    m/z 172.8302     0   1E+4   7.5   8  RT (min)      lcqtof_hilic_neg_4_010_28    m/z 172.9542     0   1E+4   1   1.5  RT (min)      lcqtof_hilic_neg_4_050_02    m/z 172.9544     0   1   2   3   4E+2   6.5   7  RT (min)      lcqtof_hilic_neg_1_000_aa    m/z 172.9572     0   1E+4   5   5.5  RT (min)      lcqtof_hilic_neg_1_001_49    m/z 172.9574     0   2   4   6   8E+3   4.5   5  RT (min)       lcqtof_hilic_neg_1_000_bb    m/z 172.9574     0   1E+4   5   5.5  RT (min)      lcqtof_hilic_neg_1_008_13    m/z 172.9574     0   1E+4   5.5   6  RT (min)       lcqtof_hilic_neg_1_001_49    m/z 172.9574     0   1E+4   5.5   6  RT (min)      lcqtof_hilic_neg_1_004_14    m/z 172.9574     0   1E+4   5   5.5  RT (min)       lcqtof_hilic_neg_1_000_aa    m/z 172.9574     0   1E+4   5   5.5  RT (min)      lcqtof_hilic_neg_1_012_04    m/z 173.0081     0   1E+4   0.5   1  RT (min)      lcqtof_hilic_neg_4_000_aa    m/z 173.0084     0   1E+4   7   7.5  RT (min)      lcqtof_hilic_neg_4_062_17    m/z 173.0084     0   1   2E+4   7   7.5  RT (min)       lcqtof_hilic_neg_4_059_14    m/z 173.0085     0   1E+4   3.5   4  RT (min)      lcqtof_hilic_neg_4_064_15    m/z 173.0085     0   1   2   3E+4   3.5   4  RT (min)      lcqtof_hilic_neg_4_050_02    m/z 173.0085     0   1   2E+4   7   7.5  RT (min)      lcqtof_hilic_neg_4_059_14    m/z 173.0085     0   1   2E+4   7.5   8  RT (min)      lcqtof_hilic_neg_4_059_14    m/z 173.0086     0   1   2   3E+4   3   3.5  RT (min)      lcqtof_hilic_neg_4_050_02    m/z 173.0086     0   1   2E+4   6.5   7  RT (min)      lcqtof_hilic_neg_4_059_14    m/z 173.0086     0   1   2E+4   7   7.5  RT (min)      lcqtof_hilic_neg_4_059_14    m/z 173.0087     0   1   2E+4   2   2.5  RT (min)      lcqtof_hilic_neg_1_016_28    m/z 173.0087     0   1   2E+4   7.5   8  RT (min)      lcqtof_hilic_neg_4_059_14    m/z 173.0087     0   1   2E+4   3   3.5  RT (min)      lcqtof_hilic_neg_4_039_18    m/z 173.0087     0   1   2E+4   4   4.5  RT (min)      lcqtof_hilic_neg_4_050_02    m/z 173.0088     0   1   2   3E+4   3   3.5  RT (min)      lcqtof_hilic_neg_4_050_02    m/z 173.0097     0   1   2E+4   2   2.5  RT (min)      lcqtof_hilic_neg_1_016_28    m/z 173.0202     0   1E+4   1.5   2  RT (min)       lcqtof_hilic_neg_4_059_14    m/z 173.0257     0   1E+4   2.5   3  RT (min)      lcqtof_hilic_neg_4_082_59    m/z 173.0449     0   1   2E+4   1   1.5  RT (min)      lcqtof_hilic_neg_4_050_02    m/z 173.0460     0   2   4   6   8E+3   1.5   2  RT (min)      lcqtof_hilic_neg_1_051_35    m/z 173.0817     0   2   4E+3   1   1.5  RT (min)      lcqtof_hilic_neg_4_050_02    m/z 173.8286     0   2   4   6E+3   3.5   4  RT (min)       lcqtof_hilic_neg_1_000_bb    m/z 173.9484     0   1E+4   1.5   2  RT (min)      lcqtof_hilic_neg_4_050_02    m/z 173.9856     0   1E+4   1   1.5  RT (min)      lcqtof_hilic_neg_4_011_35    m/z 174.0016     0   1   2E+3   6   6.5  RT (min)       lcqtof_hilic_neg_2_038_04    m/z 174.0018     0   1   2E+3   5.5   6  RT (min)      lcqtof_hilic_neg_4_019_03    m/z 174.0018     0   1   2E+3   6.5   7  RT (min)       lcqtof_hilic_neg_4_019_03    m/z 174.0161     0   1E+5   6.5   7  RT (min)      lcqtof_hilic_neg_1_007_05    m/z 174.8604     0   1   2   3   4E+4   4.5   5  RT (min)      lcqtof_hilic_neg_4_050_02    m/z 174.9334     0   1   2   3E+3   4   4.5  RT (min)      lcqtof_hilic_neg_3_028_31    m/z 174.9336     0   1   2   3E+3   5   5.5  RT (min)      lcqtof_hilic_neg_3_031_36    m/z 174.9365     0   1E+4   7.5   8  RT (min)       lcqtof_hilic_neg_1_000_aa    m/z 174.9529     0   2   4   6   8E+5   7.5   8  RT (min)      lcqtof_hilic_neg_1_000_aa    m/z 174.9531     0   2   4   6   8E+5   7.5   8  RT (min)      lcqtof_hilic_neg_1_000_aa    m/z 174.9535     0   2   4   6   8E+5   7.5   8  RT (min)      lcqtof_hilic_neg_1_000_aa    m/z 174.9549     0   1E+5   9   9.5  RT (min)      lcqtof_hilic_neg_4_018_27    m/z 174.9551     0   1   2   3   4E+4   9.5   10  RT (min)      lcqtof_hilic_neg_4_050_02    m/z 174.9551     0   2   4   6   8E+5   8   8.5  RT (min)       lcqtof_hilic_neg_1_000_aa    m/z 174.9552     0   1E+5   13.5   14  RT (min)       lcqtof_hilic_neg_1_000_aa    m/z 174.9552     0   1   2   3   4E+4   9.5   10  RT (min)      lcqtof_hilic_neg_4_050_02    m/z 174.9552     0   2   4   6E+5   8.5   9  RT (min)       lcqtof_hilic_neg_1_000_aa    m/z 174.9552     0   1   2E+5   9   9.5  RT (min)      lcqtof_hilic_neg_1_000_aa    m/z 174.9553     0   2   4   6   8E+4   9   9.5  RT (min)      lcqtof_hilic_neg_4_018_27    m/z 174.9553     0   1E+5   10   10.5  RT (min)       lcqtof_hilic_neg_1_000_aa    m/z 174.9553     0   2   4   6E+4   0.5   1  RT (min)      lcqtof_hilic_neg_4_000_aa    m/z 174.9553     0   1   2   3E+5   8.5   9  RT (min)       lcqtof_hilic_neg_4_053_14    m/z 174.9553     0   1E+5   10   10.5  RT (min)       lcqtof_hilic_neg_1_000_aa    m/z 174.9554     0   1E+5   9.5   10  RT (min)       lcqtof_hilic_neg_1_000_aa    m/z 174.9554     0   1E+5   10   10.5  RT (min)       lcqtof_hilic_neg_1_000_aa    m/z 174.9554     0   1E+5   9.5   10  RT (min)       lcqtof_hilic_neg_1_000_aa    m/z 174.9554     0   2   4   6   8E+4   11   11.5  RT (min)       lcqtof_hilic_neg_1_000_aa    m/z 174.9554     0   1E+4   13   13.5  RT (min)       lcqtof_hilic_neg_1_000_bb    m/z 174.9554     0   2   4   6   8E+5   8   8.5  RT (min)       lcqtof_hilic_neg_1_000_aa    m/z 174.9554     0   2   4   6E+4   11.5   12  RT (min)       lcqtof_hilic_neg_1_000_aa    m/z 174.9555     0   1E+5   9.5   10  RT (min)       lcqtof_hilic_neg_1_000_aa    m/z 174.9555     0   1E+5   9.5   10  RT (min)       lcqtof_hilic_neg_1_000_aa    m/z 174.9555     0   2   4   6   8E+4   11.5   12  RT (min)       lcqtof_hilic_neg_1_000_aa    m/z 174.9555     0   1E+5   9.5   10  RT (min)       lcqtof_hilic_neg_1_000_aa    m/z 174.9555     0   1E+5   10   10.5  RT (min)       lcqtof_hilic_neg_1_000_aa    m/z 174.9555     0   1E+5   9   9.5  RT (min)       lcqtof_hilic_neg_1_000_aa    m/z 174.9555     0   2   4   6   8E+4   0   0.5  RT (min)      lcqtof_hilic_neg_4_000_aa    m/z 174.9555     0   1   2   3E+5   9   9.5  RT (min)       lcqtof_hilic_neg_1_000_aa    m/z 174.9555     0   1   2   3E+5   9   9.5  RT (min)       lcqtof_hilic_neg_1_000_aa    m/z 174.9555     0   1E+5   12.5   13  RT (min)       lcqtof_hilic_neg_1_000_aa    m/z 174.9555     0   2   4   6   8E+4   11   11.5  RT (min)       lcqtof_hilic_neg_1_000_aa    m/z 174.9555     0   1E+5   12.5   13  RT (min)       lcqtof_hilic_neg_1_000_aa    m/z 174.9556     0   2   4   6   8E+4   11   11.5  RT (min)       lcqtof_hilic_neg_1_000_aa    m/z 174.9556     0   1   2E+5   9   9.5  RT (min)      lcqtof_hilic_neg_1_000_aa    m/z 174.9556     0   1E+5   10   10.5   11  RT (min)       lcqtof_hilic_neg_1_000_aa    m/z 174.9557     0   2   4   6   8E+4   0.5   1  RT (min)      lcqtof_hilic_neg_4_000_aa    m/z 174.9557     0   2   4   6   8E+4   11   11.5  RT (min)       lcqtof_hilic_neg_1_000_aa    m/z 174.9559     0   2   4   6   8E+5   7.5   8  RT (min)      lcqtof_hilic_neg_1_000_aa    m/z 174.9588     0   2   4   6   8E+4   0   0.5  RT (min)      lcqtof_hilic_neg_4_000_aa    m/z 175.0211     0   1   2   3E+4   2.5   3  RT (min)      lcqtof_hilic_neg_4_004_47    m/z 175.0240     0   1E+4   4   4.5  RT (min)      lcqtof_hilic_neg_4_050_02    m/z 175.0387     0   1E+4   1   1.5  RT (min)      lcqtof_hilic_neg_4_005_29    m/z 175.0602     0   1   2   3E+4   3.5   4  RT (min)      lcqtof_hilic_neg_4_048_23    m/z 175.0602     0   1   2   3E+4   3   3.5  RT (min)      lcqtof_hilic_neg_4_028_36    m/z 175.0605     0   1E+5   1   1.5  RT (min)       lcqtof_hilic_neg_1_022_36    m/z 175.0605     0   2   4E+4   2.5   3  RT (min)       lcqtof_hilic_neg_4_048_23    m/z 175.0605     0   1   2   3E+4   3.5   4  RT (min)       lcqtof_hilic_neg_4_048_23    m/z 175.0606     0   2   4E+4   2.5   3  RT (min)       lcqtof_hilic_neg_4_048_23    m/z 175.0607     0   1E+4   3.5   4   4.5  RT (min)       lcqtof_hilic_neg_4_028_36    m/z 175.0607     0   1   2   3E+4   3   3.5  RT (min)      lcqtof_hilic_neg_4_028_36    m/z 175.0607     0   1   2   3E+4   3   3.5  RT (min)      lcqtof_hilic_neg_4_028_36    m/z 175.0607     0   1   2   3E+4   3   3.5  RT (min)      lcqtof_hilic_neg_4_028_36    m/z 175.0608     0   1E+4   3.5   4  RT (min)      lcqtof_hilic_neg_4_028_36    m/z 175.0609     0   1E+5   2   2.5  RT (min)       lcqtof_hilic_neg_4_034_31    m/z 175.0609     0   1   2   3   4E+4   2   2.5  RT (min)      lcqtof_hilic_neg_4_047_06    m/z 175.0616     0   2   4E+4   2   2.5  RT (min)      lcqtof_hilic_neg_4_047_06    m/z 175.0900     0   1E+4   7.5   8  RT (min)      lcqtof_hilic_neg_1_000_aa    m/z 175.0905     0   1E+4   7.5   8  RT (min)      lcqtof_hilic_neg_1_000_aa    m/z 175.9590     0   1E+4   8.5   9  RT (min)      lcqtof_hilic_neg_4_053_14    m/z 175.9592     0   1   2   3   4E+3   8.5   9  RT (min)      lcqtof_hilic_neg_4_019_03    m/z 176.0069     0   1E+4   4   4.5  RT (min)      lcqtof_hilic_neg_4_048_23    m/z 176.0640     0   1E+4   1.5   2  RT (min)      lcqtof_hilic_neg_4_085_59    m/z 176.9350     0   1   2E+5   8   8.5  RT (min)      lcqtof_hilic_neg_1_000_aa    m/z 177.0501     0   1E+4   1   1.5  RT (min)      lcqtof_hilic_neg_4_050_02    m/z 177.0503     0   1E+4   1.5   2  RT (min)      lcqtof_hilic_neg_4_005_29    m/z 177.0588     0   1E+4   2.5   3  RT (min)       lcqtof_hilic_neg_4_092_60    m/z 177.0682     0   1   2   3   4E+4   3   3.5  RT (min)       lcqtof_hilic_neg_4_090_60    m/z 177.8447     0   1   2E+3   4   4.5  RT (min)      lcqtof_hilic_neg_4_003_16    m/z 177.8590     0   1   2   3   4E+4   4.5   5  RT (min)      lcqtof_hilic_neg_4_050_02    m/z 177.9024     0   1E+4   6.5   7  RT (min)       lcqtof_hilic_neg_1_003_37    m/z 177.9025     0   1E+4   6.5   7  RT (min)       lcqtof_hilic_neg_1_003_37    m/z 177.9026     0   2   4   6E+2   5.5   6  RT (min)       lcqtof_hilic_neg_1_000_aa    m/z 177.9026     0   1E+4   6.5   7  RT (min)       lcqtof_hilic_neg_1_001_49    m/z 177.9026     0   2   4   6E+3   7.5   8  RT (min)       lcqtof_hilic_neg_2_025_52    m/z 177.9026     0   2   4   6E+2   5.5   6  RT (min)      lcqtof_hilic_neg_1_000_aa    m/z 177.9028     0   1E+4   7.5   8  RT (min)       lcqtof_hilic_neg_1_001_49    m/z 177.9029     0   2   4E+3   7   7.5  RT (min)       lcqtof_hilic_neg_3_066_57    m/z 177.9029     0   1   2E+4   6.5   7   7.5  RT (min)       lcqtof_hilic_neg_1_004_14    m/z 177.9038     0   2   4   6E+3   0   0.5  RT (min)       lcqtof_hilic_neg_3_068_57    m/z 177.9038     0   1   2   3E+3   2   2.5  RT (min)      lcqtof_hilic_neg_1_000_dd    m/z 177.9038     0   2   4   6   8E+3   0   0.5  RT (min)      lcqtof_hilic_neg_3_068_57    m/z 177.9040     0   2   4   6   8E+3   0.5   1  RT (min)      lcqtof_hilic_neg_1_004_14    m/z 177.9040     0   2   4   6E+3   0   0.5  RT (min)      lcqtof_hilic_neg_3_068_57    m/z 177.9042     0   1   2   3E+4   3.5   4  RT (min)       lcqtof_hilic_neg_1_000_dd    m/z 177.9042     0   1   2   3E+4   3.5   4  RT (min)       lcqtof_hilic_neg_1_000_dd    m/z 177.9042     0   1   2E+4   3.5   4  RT (min)       lcqtof_hilic_neg_1_000_dd    m/z 177.9059     0   1   2E+4   1   1.5  RT (min)      lcqtof_hilic_neg_4_058_39    m/z 178.0808     0   1E+4   1.5   2  RT (min)      lcqtof_hilic_neg_4_055_32    m/z 178.8140     0   1E+4   5   10  RT (min)       lcqtof_hilic_neg_4_112_62    m/z 178.8140     0   1E+4   13  RT (min)       lcqtof_hilic_neg_4_084_59    m/z 178.8141     0   1   2   3E+4   13   13.5  RT (min)       lcqtof_hilic_neg_4_000_bb    m/z 178.8142     0   1   2   3E+4   10   12.5  RT (min)       lcqtof_hilic_neg_4_006_30    m/z 178.8143     0   1   2E+4   12.5   13  RT (min)       lcqtof_hilic_neg_4_027_19    m/z 178.8143     0   1   2   3E+4   12   12.5  RT (min)       lcqtof_hilic_neg_4_000_cc    m/z 178.8144     0   1   2   3E+4   12   13  RT (min)       lcqtof_hilic_neg_4_015_09    m/z 178.8144     0   1   2E+4   14.5   15  RT (min)       lcqtof_hilic_neg_4_033_53    m/z 178.8144     0   1   2   3E+4   14.5   15  RT (min)       lcqtof_hilic_neg_4_005_29    m/z 178.8145     0   1E+4   5   5.5  RT (min)      lcqtof_hilic_neg_4_009_50    m/z 178.8146     0   2   4   6   8E+3   5   5.5  RT (min)      lcqtof_hilic_neg_4_026_04    m/z 178.8147     0   1E+4   4.5   5  RT (min)      lcqtof_hilic_neg_4_008_24    m/z 178.8147     0   1   2E+4   12   13  RT (min)       lcqtof_hilic_neg_4_068_57    m/z 178.8147     0   1E+4   0.5   1  RT (min)      lcqtof_hilic_neg_2_000_aa    m/z 178.8149     0   1   2E+4   14   15  RT (min)       lcqtof_hilic_neg_4_032_01    m/z 178.8162     0   1E+4   10   12  RT (min)       lcqtof_hilic_neg_4_085_59    m/z 178.8437     0   1   2   3   4E+3   6.5   7  RT (min)      lcqtof_hilic_neg_2_014_08    m/z 178.8440     0   1   2   3E+3   1.5   2  RT (min)      lcqtof_hilic_neg_3_023_08    m/z 178.9026     0   1   2   3E+3   7.5   8  RT (min)      lcqtof_hilic_neg_1_004_14    m/z 178.9027     0   1   2E+3   8   8.5  RT (min)      lcqtof_hilic_neg_2_018_16    m/z 178.9031     0   2   4   6E+3   4.5   5  RT (min)      lcqtof_hilic_neg_1_008_13    m/z 178.9483     0   2   4   6E+3   8.5   9  RT (min)      lcqtof_hilic_neg_4_053_14    m/z 178.9484     0   2   4   6E+3   8.5   9  RT (min)      lcqtof_hilic_neg_4_053_14    m/z 178.9770     0   2   4   6   8E+3   14   15  RT (min)       lcqtof_hilic_neg_4_003_16    m/z 179.0160     0   1   2E+4   0.5   1  RT (min)      lcqtof_hilic_neg_4_016_37    m/z 179.0287     0   1E+4   4.5   5  RT (min)      lcqtof_hilic_neg_1_005_32    m/z 179.0349     0   1E+4   1.5   2  RT (min)       lcqtof_hilic_neg_4_055_32    m/z 179.0552     0   1E+5   4.5   5  RT (min)      lcqtof_hilic_neg_4_053_14    m/z 179.0553     0   1   2   3   4E+4   3.5   4  RT (min)      lcqtof_hilic_neg_4_039_18    m/z 179.0554     0   1E+5   4.5   5  RT (min)      lcqtof_hilic_neg_1_023_34    m/z 179.0554     0   1   2   3E+4   4   4.5  RT (min)       lcqtof_hilic_neg_4_053_14    m/z 179.0555     0   1E+4   6   6.5  RT (min)      lcqtof_hilic_neg_4_013_20    m/z 179.0556     0   1   2   3E+4   4.5   5  RT (min)      lcqtof_hilic_neg_4_053_14    m/z 179.0765     0   1E+4   4   4.5  RT (min)      lcqtof_hilic_neg_4_055_32    m/z 179.0767     0   1E+4   4   4.5  RT (min)       lcqtof_hilic_neg_4_097_60    m/z 179.1776     0   1   2   3E+4   0.5   1  RT (min)      lcqtof_hilic_neg_4_027_19    m/z 179.2341     0   1   2   3   4E+4   0.5   1  RT (min)      lcqtof_hilic_neg_4_027_19    m/z 179.2601     0   1E+4   7   7.5  RT (min)      lcqtof_hilic_neg_4_008_24    m/z 179.2687     0   2   4E+4   0.5   1  RT (min)      lcqtof_hilic_neg_4_027_19    m/z 179.3871     0   2   4   6E+4   0.5   1  RT (min)      lcqtof_hilic_neg_4_027_19    m/z 179.8407     0   2   4E+3   4.5   5  RT (min)       lcqtof_hilic_neg_4_014_46    m/z 179.8578     0   1   2   3   4E+4   4.5   5  RT (min)      lcqtof_hilic_neg_4_050_02    m/z 179.8581     0   1   2   3   4E+4   4.5   5  RT (min)      lcqtof_hilic_neg_4_050_02    m/z 180.8122     0   1E+4   7.5   10   12.5  RT (min)       lcqtof_hilic_neg_4_008_24    m/z 180.8418     0   1   2   3   4E+3   7   7.5  RT (min)      lcqtof_hilic_neg_3_023_08    m/z 180.8418     0   1   2   3E+3   6.5   7  RT (min)      lcqtof_hilic_neg_2_020_23    m/z 180.8419     0   1   2   3   4E+3   1.5   2  RT (min)      lcqtof_hilic_neg_2_045_11    m/z 180.8420     0   1   2   3   4E+3   6.5   7  RT (min)      lcqtof_hilic_neg_3_023_08    m/z 180.9224     0   1   2E+4   8   10  RT (min)       lcqtof_hilic_neg_4_104_61    m/z 180.9227     0   1   2E+4   14   14.5  RT (min)       lcqtof_hilic_neg_4_078_58    m/z 180.9227     0   1   2E+4   14  RT (min)       lcqtof_hilic_neg_4_075_58    m/z 180.9227     0   1E+4   4   4.5  RT (min)      lcqtof_hilic_neg_4_040_05    m/z 180.9228     0   1   2E+4   14  RT (min)       lcqtof_hilic_neg_4_006_30    m/z 180.9228     0   1   2   3E+4   5   7.5   10  RT (min)       lcqtof_hilic_neg_4_084_59    m/z 180.9228     0   1   2E+4   4.5   5  RT (min)      lcqtof_hilic_neg_4_017_51    m/z 180.9228     0   1   2E+4   5   7.5   10  RT (min)       lcqtof_hilic_neg_4_007_45    m/z 180.9228     0   1   2   3E+4   5   5.5  RT (min)       lcqtof_hilic_neg_4_020_42    m/z 180.9228     0   1   2E+4   3.5   4  RT (min)       lcqtof_hilic_neg_4_015_09    m/z 180.9228     0   1E+4   4   4.5  RT (min)      lcqtof_hilic_neg_4_047_06    m/z 180.9229     0   1   2E+4   12   14  RT (min)       lcqtof_hilic_neg_4_026_04    m/z 180.9229     0   1   2E+4   5   5.5  RT (min)      lcqtof_hilic_neg_4_017_51    m/z 180.9229     0   1E+4   4   4.5  RT (min)      lcqtof_hilic_neg_4_033_53    m/z 180.9231     0   1E+4   3.5   4  RT (min)      lcqtof_hilic_neg_4_040_05    m/z 180.9795     0   1E+4   2   2.5  RT (min)      lcqtof_hilic_neg_4_058_39    m/z 180.9803     0   1E+4   2   2.5  RT (min)      lcqtof_hilic_neg_4_058_39    m/z 180.9806     0   1E+4   2   2.5  RT (min)      lcqtof_hilic_neg_4_058_39    m/z 180.9900     0   1E+4   4   4.5  RT (min)      lcqtof_hilic_neg_4_048_23    m/z 180.9999     0   1   2E+4   2.5   3  RT (min)      lcqtof_hilic_neg_4_032_01    m/z 181.0454     0   1E+4   1.5   2  RT (min)      lcqtof_hilic_neg_4_019_03    m/z 181.0465     0   1E+4   2   2.5  RT (min)      lcqtof_hilic_neg_4_019_03    m/z 182.0455     0   1   2   3   4E+3   2   2.5  RT (min)      lcqtof_hilic_neg_1_054_01    m/z 182.0604     0   2   4   6   8E+4   1   1.5  RT (min)      lcqtof_hilic_neg_4_027_19    m/z 182.0760     0   1E+4   4   4.5  RT (min)       lcqtof_hilic_neg_4_093_60    m/z 182.0837     0   2   4E+4   3   3.5  RT (min)      lcqtof_hilic_neg_4_055_32    m/z 182.0837     0   2   4E+4   2.5   3  RT (min)       lcqtof_hilic_neg_4_093_60    m/z 182.0838     0   1E+5   2   2.5  RT (min)       lcqtof_hilic_neg_4_055_32    m/z 182.0838     0   1   2E+4   3.5   4  RT (min)      lcqtof_hilic_neg_4_055_32    m/z 182.0838     0   1   2   3   4E+4   3   3.5  RT (min)       lcqtof_hilic_neg_4_040_05    m/z 182.0838     0   2   4E+4   2.5   3  RT (min)       lcqtof_hilic_neg_4_055_32    m/z 182.0838     0   2   4E+4   3   3.5  RT (min)      lcqtof_hilic_neg_4_055_32    m/z 182.0839     0   1   2   3   4E+4   3   3.5  RT (min)      lcqtof_hilic_neg_4_040_05    m/z 182.0840     0   2   4E+4   2.5   3  RT (min)      lcqtof_hilic_neg_4_040_05    m/z 182.0840     0   2   4   6   8E+4   2   2.5   3  RT (min)      lcqtof_hilic_neg_4_040_05    m/z 182.0840     0   2   4E+4   3   3.5  RT (min)       lcqtof_hilic_neg_4_055_32    m/z 182.0841     0   1E+4   3.5   4  RT (min)       lcqtof_hilic_neg_4_055_32    m/z 182.0843     0   2   4   6   8E+4   2   2.5  RT (min)      lcqtof_hilic_neg_4_040_05    m/z 182.0990     0   1   2   3E+4   1   1.5  RT (min)      lcqtof_hilic_neg_4_027_19    m/z 182.1283     0   1E+5   1   1.5  RT (min)      lcqtof_hilic_neg_4_027_19    m/z 182.1557     0   2   4E+4   1   1.5  RT (min)      lcqtof_hilic_neg_4_027_19    m/z 182.1909     0   2   4E+4   1   1.5  RT (min)      lcqtof_hilic_neg_4_027_19    m/z 182.2326     0   2   4   6   8E+4   1   1.5  RT (min)      lcqtof_hilic_neg_4_027_19    m/z 182.3101     0   2   4   6E+4   1   1.5  RT (min)      lcqtof_hilic_neg_4_027_19    m/z 182.9203     0   1E+4   5   5.5  RT (min)      lcqtof_hilic_neg_4_026_04    m/z 182.9203     0   1E+4   5   7.5   10  RT (min)       lcqtof_hilic_neg_4_069_57    m/z 182.9203     0   1   2E+4   7.5   10  RT (min)       lcqtof_hilic_neg_4_017_51    m/z 182.9203     0   1   2E+4   4.5   5  RT (min)      lcqtof_hilic_neg_4_014_46    m/z 182.9204     0   1   2   3E+4   5   10  RT (min)       lcqtof_hilic_neg_4_015_09    m/z 182.9204     0   1   2E+4   10   12   14  RT (min)       lcqtof_hilic_neg_4_104_61    m/z 182.9205     0   1E+4   10   11   12  RT (min)       lcqtof_hilic_neg_4_112_62    m/z 182.9205     0   1   2E+4   3   4   5  RT (min)       lcqtof_hilic_neg_4_083_59    m/z 182.9205     0   1   2E+4   4   4.5  RT (min)      lcqtof_hilic_neg_4_008_24    m/z 182.9205     0   1   2E+4   5   5.5  RT (min)      lcqtof_hilic_neg_4_018_27    m/z 182.9206     0   1E+4   3   4  RT (min)       lcqtof_hilic_neg_4_015_09    m/z 182.9206     0   1   2E+4   5   5.5  RT (min)      lcqtof_hilic_neg_4_018_27    m/z 182.9206     0   1E+4   14  RT (min)       lcqtof_hilic_neg_4_078_58    m/z 182.9206     0   1   2E+4   10   15  RT (min)       lcqtof_hilic_neg_4_005_29    m/z 182.9207     0   1E+4   4   4.5  RT (min)      lcqtof_hilic_neg_4_047_06    m/z 182.9207     0   1E+4   4   4.5  RT (min)      lcqtof_hilic_neg_4_031_38    m/z 182.9207     0   1   2E+4   4   4.5  RT (min)      lcqtof_hilic_neg_4_008_24    m/z 182.9209     0   1E+4   3.5   4  RT (min)      lcqtof_hilic_neg_4_008_24    m/z 182.9966     0   1   2   3E+4   2   2.5  RT (min)       lcqtof_hilic_neg_4_070_57    m/z 182.9980     0   1   2E+4   2   2.5  RT (min)      lcqtof_hilic_neg_4_050_02    m/z 183.0084     0   1   2E+4   2   2.5  RT (min)      lcqtof_hilic_neg_4_050_02    m/z 183.0098     0   1E+4   2   2.5  RT (min)      lcqtof_hilic_neg_4_003_16    m/z 183.0104     0   2   4   6E+3   3   3.5  RT (min)      lcqtof_hilic_neg_4_002_25    m/z 183.0106     0   2   4   6E+3   1.5   2  RT (min)      lcqtof_hilic_neg_4_019_03    m/z 183.0106     0   2   4   6   8E+3   2   2.5  RT (min)      lcqtof_hilic_neg_4_050_02    m/z 183.0107     0   1   2E+2   2   2.5  RT (min)       lcqtof_hilic_neg_3_067_57    m/z 183.0110     0   1E+4   2.5   3  RT (min)      lcqtof_hilic_neg_4_088_59    m/z 183.0111     0   2   4E+4   15   15.5  RT (min)      lcqtof_hilic_neg_4_000_aa    m/z 183.0113     0   2   4E+4   15   15.5  RT (min)      lcqtof_hilic_neg_4_000_aa    m/z 183.0116     0   2   4E+3   0   0.5  RT (min)      lcqtof_hilic_neg_4_024_40    m/z 184.0000     0   2   4E+4   3.5   4  RT (min)       lcqtof_hilic_neg_4_091_60    m/z 184.0004     0   1   2   3E+4   4   4.5  RT (min)      lcqtof_hilic_neg_4_055_32    m/z 184.0007     0   1   2   3E+4   4   4.5  RT (min)      lcqtof_hilic_neg_4_055_32    m/z 184.8630     0   1E+4   1   1.5  RT (min)      lcqtof_hilic_neg_4_058_39    m/z 184.8757     0   1   2   3E+4   4   4.5  RT (min)      lcqtof_hilic_neg_3_067_57    m/z 184.8758     0   1E+4   4   4.5  RT (min)      lcqtof_hilic_neg_4_027_19    m/z 184.9182     0   2   4   6E+3   5   5.5  RT (min)       lcqtof_hilic_neg_4_015_09    m/z 184.9183     0   1   2   3   4E+3   4   4.5  RT (min)      lcqtof_hilic_neg_4_008_24    m/z 184.9183     0   1   2   3E+3   4   4.5  RT (min)      lcqtof_hilic_neg_4_017_51    m/z 184.9711     0   1E+4   6   6.5  RT (min)      lcqtof_hilic_neg_2_020_23    m/z 184.9749     0   1   2E+4   5.5   6  RT (min)      lcqtof_hilic_neg_1_000_aa    m/z 185.0140     0   1E+4   2   2.5  RT (min)      lcqtof_hilic_neg_4_047_06    m/z 185.0168     0   2   4E+3   1   1.5  RT (min)      lcqtof_hilic_neg_4_010_28    m/z 185.0252     0   1   2E+4   1   1.5  RT (min)      lcqtof_hilic_neg_4_010_28    m/z 185.0269     0   2   4E+3   4   4.5  RT (min)      lcqtof_hilic_neg_4_035_33    m/z 185.0741     0   1   2E+4   4   4.5  RT (min)      lcqtof_hilic_neg_4_061_50    m/z 185.9615     0   1E+4   1   1.5  RT (min)       lcqtof_hilic_neg_3_066_57    m/z 185.9618     0   1E+4   1   1.5  RT (min)      lcqtof_hilic_neg_3_066_57    m/z 186.0400     0   2   4   6E+4   4   4.5  RT (min)      lcqtof_hilic_neg_4_031_38    m/z 186.0401     0   1E+4   4   4.5  RT (min)      lcqtof_hilic_neg_4_084_59    m/z 186.0402     0   1E+4   4   4.5  RT (min)       lcqtof_hilic_neg_4_048_23    m/z 186.0403     0   1   2E+4   4.5   5  RT (min)      lcqtof_hilic_neg_4_016_37    m/z 186.0426     0   2   4   6E+3   2   2.5  RT (min)      lcqtof_hilic_neg_4_052_13    m/z 186.0441     0   1E+4   2   2.5  RT (min)      lcqtof_hilic_neg_4_059_14    m/z 187.0235     0   1E+4   1   1.5  RT (min)      lcqtof_hilic_neg_4_023_15    m/z 187.0238     0   1E+4   1   1.5  RT (min)      lcqtof_hilic_neg_4_064_15    m/z 187.0414     0   1   2   3E+4   2.5   3  RT (min)      lcqtof_hilic_neg_4_034_31    m/z 187.0416     0   2   4   6   8E+3   2.5   3  RT (min)       lcqtof_hilic_neg_4_048_23    m/z 187.0417     0   2   4E+3   2.5   3  RT (min)       lcqtof_hilic_neg_4_012_07    m/z 187.0417     0   2   4   6   8E+3   2.5   3  RT (min)      lcqtof_hilic_neg_4_048_23    m/z 187.0419     0   1   2   3   4E+3   3.5   4  RT (min)       lcqtof_hilic_neg_4_113_62    m/z 187.0421     0   2   4   6   8E+3   2.5   3  RT (min)       lcqtof_hilic_neg_4_084_59    m/z 187.0421     0   2   4   6   8E+3   3   3.5  RT (min)       lcqtof_hilic_neg_4_002_25    m/z 187.0431     0   1E+5   1.5   2  RT (min)      lcqtof_hilic_neg_4_050_02    m/z 187.0457     0   1E+5   0.5   1  RT (min)      lcqtof_hilic_neg_4_050_02    m/z 187.0458     0   1   2E+5   1.5   2  RT (min)       lcqtof_hilic_neg_4_022_34    m/z 187.0477     0   1   2E+4   1   1.5  RT (min)      lcqtof_hilic_neg_4_050_02    m/z 187.0481     0   1   2E+4   1   1.5  RT (min)      lcqtof_hilic_neg_4_050_02    m/z 187.0701     0   1E+3   0.5   1  RT (min)      lcqtof_hilic_neg_4_053_14    m/z 187.0969     0   2   4   6E+3   0   0.5  RT (min)       lcqtof_hilic_neg_4_050_02    m/z 187.0970     0   1   2   3   4E+3   0   0.5  RT (min)       lcqtof_hilic_neg_4_050_02    m/z 187.0972     0   2   4   6   8E+4   1   1.5  RT (min)       lcqtof_hilic_neg_4_039_18    m/z 187.0972     0   1   2E+3   2.5   3  RT (min)       lcqtof_hilic_neg_4_012_07    m/z 187.0973     0   1   2E+3   2   2.5  RT (min)      lcqtof_hilic_neg_4_109_62    m/z 187.0974     0   1   2   3E+3   1.5   2  RT (min)      lcqtof_hilic_neg_4_055_32    m/z 187.0974     0   1   2E+3   2.5   3  RT (min)      lcqtof_hilic_neg_4_111_62    m/z 187.0974     0   1   2   3E+3   0   0.5  RT (min)       lcqtof_hilic_neg_4_050_02    m/z 187.1153     0   1E+4   0.5   1  RT (min)      lcqtof_hilic_neg_4_053_14    m/z 187.1736     0   1   2E+4   0.5   1  RT (min)      lcqtof_hilic_neg_4_059_14    m/z 187.2086     0   1   2   3E+4   0.5   1  RT (min)      lcqtof_hilic_neg_4_053_14    m/z 187.2514     0   2   4E+4   0.5   1  RT (min)      lcqtof_hilic_neg_4_053_14    m/z 187.2515     0   2   4E+4   0.5   1  RT (min)      lcqtof_hilic_neg_4_053_14    m/z 187.2516     0   2   4E+4   0.5   1  RT (min)      lcqtof_hilic_neg_4_053_14    m/z 187.2516     0   2   4E+4   0.5   1  RT (min)      lcqtof_hilic_neg_4_059_14    m/z 187.3299     0   1   2   3   4E+4   0.5   1  RT (min)      lcqtof_hilic_neg_4_053_14    m/z 187.8604     0   1E+4   3.5   4  RT (min)       lcqtof_hilic_neg_3_066_57    m/z 187.8608     0   1E+4   16   16.5  RT (min)       lcqtof_hilic_neg_3_066_57    m/z 187.8612     0   1E+4   15.5   16  RT (min)       lcqtof_hilic_neg_1_000_aa    m/z 187.9385     0   1   2   3E+4   10   10.5  RT (min)      lcqtof_hilic_neg_4_014_46    m/z 187.9408     0   1   2E+3   0   0.5  RT (min)       lcqtof_hilic_neg_4_000_dd    m/z 187.9409     0   2   4   6   8E+3   1   1.5  RT (min)      lcqtof_hilic_neg_3_000_cc    m/z 187.9410     0   2   4E+4   5   5.5  RT (min)      lcqtof_hilic_neg_1_000_bb    m/z 187.9411     0   1E+4   3.5   4  RT (min)       lcqtof_hilic_neg_3_066_57    m/z 187.9412     0   1E+4   7   7.5  RT (min)       lcqtof_hilic_neg_1_005_32    m/z 187.9412     0   1E+4   2  RT (min)       lcqtof_hilic_neg_1_020_09    m/z 187.9412     0   2   4E+3   0   0.5  RT (min)      lcqtof_hilic_neg_1_000_bb    m/z 187.9413     0   1   2E+3   0   0.5  RT (min)      lcqtof_hilic_neg_1_020_09    m/z 187.9413     0   1   2   3   4E+3   8   8.5  RT (min)      lcqtof_hilic_neg_1_000_dd    m/z 187.9414     0   1   2E+3   3.5   4  RT (min)       lcqtof_hilic_neg_1_036_03    m/z 187.9415     0   2   4   6   8E+3   7.5   8  RT (min)      lcqtof_hilic_neg_4_003_16    m/z 187.9415     0   1E+4   7   7.5  RT (min)      lcqtof_hilic_neg_1_005_32    m/z 187.9868     0   1E+4   4   4.5  RT (min)      lcqtof_hilic_neg_1_005_32    m/z 187.9951     0   1   2E+4   6.5   7  RT (min)       lcqtof_hilic_neg_4_055_32    m/z 187.9952     0   1E+4   6.5   7  RT (min)      lcqtof_hilic_neg_4_008_24    m/z 188.0398     0   2   4   6   8E+2   2   2.5  RT (min)      lcqtof_hilic_neg_3_025_52    m/z 188.0553     0   1   2   3E+4   2.5   3  RT (min)      lcqtof_hilic_neg_4_055_32    m/z 188.0555     0   1E+4   2.5   3  RT (min)      lcqtof_hilic_neg_1_005_32    m/z 188.0557     0   1   2E+5   3   3.5  RT (min)      lcqtof_hilic_neg_4_047_06    m/z 188.9422     0   1   2E+4   5   5.5  RT (min)      lcqtof_hilic_neg_1_000_aa    m/z 188.9425     0   1   2   3E+3   6   6.5  RT (min)      lcqtof_hilic_neg_1_001_49    m/z 188.9837     0   1E+4   6   6.5  RT (min)      lcqtof_hilic_neg_1_037_24    m/z 189.0030     0   2   4E+4   7   7.5  RT (min)      lcqtof_hilic_neg_4_033_53    m/z 189.0031     0   2   4E+4   7.5   8  RT (min)      lcqtof_hilic_neg_4_006_30    m/z 189.0032     0   1   2   3E+4   7   7.5  RT (min)      lcqtof_hilic_neg_4_049_55    m/z 189.0034     0   1   2   3E+4   7.5   8  RT (min)      lcqtof_hilic_neg_4_025_52    m/z 189.0387     0   1E+4   2.5   3  RT (min)      lcqtof_hilic_neg_4_003_16    m/z 189.0394     0   1   2E+4   2   2.5  RT (min)      lcqtof_hilic_neg_4_050_02    m/z 189.0420     0   1   2E+4   2   2.5  RT (min)      lcqtof_hilic_neg_4_050_02    m/z 189.0591     0   1   2E+4   3   3.5  RT (min)      lcqtof_hilic_neg_4_047_06    m/z 189.9977     0   1E+4   0.5   1  RT (min)      lcqtof_hilic_neg_4_053_14    m/z 190.0029     0   1E+4   18.5   19  RT (min)      lcqtof_hilic_neg_3_066_57    m/z 190.0031     0   1E+4   17.5   18  RT (min)      lcqtof_hilic_neg_3_066_57    m/z 190.0042     0   1E+4   19.5   20  RT (min)      lcqtof_hilic_neg_3_066_57    m/z 190.0045     0   1E+4   19   19.5  RT (min)      lcqtof_hilic_neg_3_066_57    m/z 190.0145     0   1   16   16.5  RT (min)      lcqtof_hilic_neg_3_066_57    m/z 190.0534     0   1E+4   2   2.5  RT (min)      lcqtof_hilic_neg_4_031_38    m/z 190.1637     0   1   2   3E+4   1   1.5  RT (min)      lcqtof_hilic_neg_4_027_19    m/z 190.2224     0   2   4E+4   1   1.5  RT (min)      lcqtof_hilic_neg_4_027_19    m/z 190.2576     0   2   4   6E+4   1   1.5  RT (min)      lcqtof_hilic_neg_4_027_19    m/z 190.3798     0   2   4   6E+4   1   1.5  RT (min)      lcqtof_hilic_neg_4_027_19    m/z 190.9246     0   1E+4   5   5.5  RT (min)      lcqtof_hilic_neg_4_044_08    m/z 190.9246     0   1E+4   14  RT (min)       lcqtof_hilic_neg_4_033_53    m/z 190.9246     0   1E+4   6   6.5  RT (min)      lcqtof_hilic_neg_4_044_08    m/z 190.9254     0   1E+4   12.5   13  RT (min)       lcqtof_hilic_neg_4_102_61    m/z 190.9258     0   1E+4   13   13.5  RT (min)       lcqtof_hilic_neg_4_012_07    m/z 190.9258     0   1E+4   14   14.5  RT (min)       lcqtof_hilic_neg_4_046_41    m/z 190.9262     0   1E+4   12   12.5  RT (min)       lcqtof_hilic_neg_4_030_44    m/z 190.9262     0   1E+4   14   16  RT (min)       lcqtof_hilic_neg_4_105_61    m/z 190.9264     0   1E+4   6   6.5  RT (min)      lcqtof_hilic_neg_4_012_07    m/z 190.9266     0   1E+4   11.5   12  RT (min)       lcqtof_hilic_neg_4_014_46    m/z 190.9268     0   1E+4   6.5   7  RT (min)      lcqtof_hilic_neg_4_109_62    m/z 190.9273     0   1E+4   14.5   15  RT (min)       lcqtof_hilic_neg_4_027_19    m/z 190.9274     0   1E+4   10.5   11   11.5  RT (min)       lcqtof_hilic_neg_4_101_61    m/z 190.9280     0   1E+4   10   10.5  RT (min)       lcqtof_hilic_neg_4_100_61    m/z 190.9281     0   1   2   3E+4   10   11  RT (min)       lcqtof_hilic_neg_4_014_46    m/z 190.9282     0   1E+4   6   6.5  RT (min)      lcqtof_hilic_neg_4_039_18    m/z 190.9284     0   1   2   3E+4   10   11  RT (min)       lcqtof_hilic_neg_4_014_46    m/z 190.9761     0   1E+4   1   1.5  RT (min)      lcqtof_hilic_neg_4_027_19    m/z 190.9997     0   2   4   6E+4   0.5   1  RT (min)       lcqtof_hilic_neg_3_066_57    m/z 191.0017     0   2   4   6E+4   18   20  RT (min)       lcqtof_hilic_neg_3_067_57    m/z 191.0044     0   2   4   6E+4   10  RT (min)       lcqtof_hilic_neg_3_068_57    m/z 191.0111     0   1E+4   15.5   16  RT (min)       lcqtof_hilic_neg_1_025_52    m/z 191.0151     0   2   4   6E+3   16   16.5  RT (min)       lcqtof_hilic_neg_1_025_52    m/z 191.0164     0   2   4E+3   15.5   16  RT (min)       lcqtof_hilic_neg_1_030_17    m/z 191.0172     0   1   2E+5   5.5   6  RT (min)      lcqtof_hilic_neg_1_003_37    m/z 191.0174     0   2   4   6E+3   15.5   16  RT (min)      lcqtof_hilic_neg_1_032_18    m/z 191.0175     0   1   2   3E+5   5   5.5  RT (min)      lcqtof_hilic_neg_1_022_36    m/z 191.0181     0   1   2E+5   5.5   6  RT (min)      lcqtof_hilic_neg_1_023_34    m/z 191.0182     0   1   2E+5   0.5   1  RT (min)      lcqtof_hilic_neg_4_000_aa    m/z 191.0183     0   1   2E+4   8.5   9  RT (min)      lcqtof_hilic_neg_4_049_55    m/z 191.0185     0   1   2   3E+5   0   0.5  RT (min)      lcqtof_hilic_neg_4_000_aa    m/z 191.0185     0   1   2E+5   7.5   8  RT (min)      lcqtof_hilic_neg_4_034_31    m/z 191.0187     0   1   2   3E+5   0.5   1  RT (min)      lcqtof_hilic_neg_4_000_aa    m/z 191.0187     0   1E+5   8   8.5  RT (min)       lcqtof_hilic_neg_4_028_36    m/z 191.0187     0   1   2   3E+5   0   0.5  RT (min)      lcqtof_hilic_neg_4_000_aa    m/z 191.0187     0   1   2E+5   7.5   8  RT (min)       lcqtof_hilic_neg_4_034_31    m/z 191.0189     0   1   2   3E+5   0.5   1  RT (min)      lcqtof_hilic_neg_4_000_aa    m/z 191.0190     0   2   4   6E+4   8.5   9  RT (min)      lcqtof_hilic_neg_4_053_14    m/z 191.0558     0   2   4E+3   6   6.5  RT (min)      lcqtof_hilic_neg_4_016_37    m/z 191.0558     0   1   2   3E+4   3.5   4  RT (min)      lcqtof_hilic_neg_1_032_18    m/z 191.0561     0   2   4   6E+3   2   2.5  RT (min)      lcqtof_hilic_neg_4_005_29    m/z 191.0896     0   1   2   3   4E+4   4.5   5  RT (min)      lcqtof_hilic_neg_1_004_14    m/z 191.1268     0   1E+4   1   1.5  RT (min)      lcqtof_hilic_neg_4_027_19    m/z 191.1595     0   2   4   6E+4   4.5   5  RT (min)      lcqtof_hilic_neg_1_004_14    m/z 191.1597     0   2   4   6E+4   4.5   5  RT (min)      lcqtof_hilic_neg_1_004_14    m/z 191.2668     0   1   2   3E+4   4.5   5  RT (min)      lcqtof_hilic_neg_1_004_14    m/z 191.8545     0   2   4E+3   4.5   5  RT (min)       lcqtof_hilic_neg_4_000_bb    m/z 192.0224     0   1   2   3E+4   7.5   8  RT (min)      lcqtof_hilic_neg_4_059_14    m/z 192.0224     0   1   2   3E+4   6.5   7  RT (min)      lcqtof_hilic_neg_4_059_14    m/z 192.0224     0   1   2   3E+4   7   7.5  RT (min)      lcqtof_hilic_neg_4_059_14    m/z 192.0225     0   1   2   3E+4   7.5   8  RT (min)      lcqtof_hilic_neg_4_059_14    m/z 192.0229     0   1   2   3E+4   7.5   8  RT (min)      lcqtof_hilic_neg_4_059_14    m/z 192.0823     0   1   2   3E+4   4   4.5  RT (min)      lcqtof_hilic_neg_4_009_50    m/z 192.9279     0   1   2   3   4E+3   8   8.5  RT (min)      lcqtof_hilic_neg_4_058_39    m/z 192.9282     0   1   2E+3   7.5   8  RT (min)       lcqtof_hilic_neg_1_052_40    m/z 192.9981     0   1   2   3   4E+3   1   1.5  RT (min)      lcqtof_hilic_neg_4_050_02    m/z 193.0235     0   1   2   3E+4   4.5   5  RT (min)      lcqtof_hilic_neg_1_004_14    m/z 193.0237     0   2   4   6   8E+3   7   7.5  RT (min)      lcqtof_hilic_neg_4_059_14    m/z 193.0238     0   1   2   3E+4   4.5   5  RT (min)      lcqtof_hilic_neg_1_004_14    m/z 193.0635     0   1E+4   4   4.5  RT (min)       lcqtof_hilic_neg_4_092_60    m/z 193.0635     0   1   2   3E+4   4   4.5  RT (min)      lcqtof_hilic_neg_4_040_05    m/z 193.0640     0   1E+4   4.5   5  RT (min)      lcqtof_hilic_neg_4_017_51    m/z 193.0641     0   1   2E+5   4   4.5  RT (min)      lcqtof_hilic_neg_4_055_32    m/z 193.0768     0   1E+4   3.5   4  RT (min)      lcqtof_hilic_neg_4_038_10    m/z 193.0862     0   1E+4   15   15.5  RT (min)      lcqtof_hilic_neg_4_000_aa    m/z 193.0899     0   1E+3   1.5   2  RT (min)      lcqtof_hilic_neg_4_002_25    m/z 193.8970     0   1E+4   7   7.5  RT (min)      lcqtof_hilic_neg_1_000_aa    m/z 193.8981     0   1E+4   7   7.5  RT (min)      lcqtof_hilic_neg_1_000_aa    m/z 194.0292     0   2   4   6E+3   7   7.5  RT (min)       lcqtof_hilic_neg_4_047_06    m/z 194.0475     0   1E+4   6   6.5  RT (min)      lcqtof_hilic_neg_4_055_32    m/z 194.0478     0   1E+4   6   6.5  RT (min)      lcqtof_hilic_neg_4_055_32    m/z 194.0479     0   1E+4   6   6.5  RT (min)      lcqtof_hilic_neg_4_055_32    m/z 194.0480     0   1E+4   6.5   7  RT (min)      lcqtof_hilic_neg_4_055_32    m/z 194.0481     0   1   2E+4   3.5   4  RT (min)      lcqtof_hilic_neg_1_005_32    m/z 194.0526     0   2   4E+3   2   2.5  RT (min)      lcqtof_hilic_neg_4_032_01    m/z 194.0758     0   1E+4   3   3.5  RT (min)      lcqtof_hilic_neg_4_047_06    m/z 194.8383     0   1   2E+4   2.5   3  RT (min)      lcqtof_hilic_neg_4_050_02    m/z 194.9051     0   1   2   3E+4   6.5   7  RT (min)       lcqtof_hilic_neg_1_012_04    m/z 194.9051     0   1   2E+4   6.5   7  RT (min)       lcqtof_hilic_neg_1_031_16    m/z 194.9053     0   2   4E+4   4   4.5  RT (min)       lcqtof_hilic_neg_1_012_04    m/z 194.9053     0   1   2   3E+4   4   4.5  RT (min)       lcqtof_hilic_neg_1_002_39    m/z 194.9053     0   2   4E+4   4   4.5  RT (min)       lcqtof_hilic_neg_1_019_02    m/z 194.9053     0   1E+4   6.5   7  RT (min)       lcqtof_hilic_neg_1_004_14    m/z 194.9053     0   1   2   3E+3   0   0.5  RT (min)      lcqtof_hilic_neg_4_050_02    m/z 194.9053     0   1   2   3E+3   0   0.5  RT (min)      lcqtof_hilic_neg_4_050_02    m/z 194.9054     0   2   4   6   8E+3   8   8.5  RT (min)       lcqtof_hilic_neg_1_014_31    m/z 194.9054     0   2   4   6   8E+3   3.5   4  RT (min)       lcqtof_hilic_neg_2_050_19    m/z 194.9054     0   1E+4   8   8.5  RT (min)       lcqtof_hilic_neg_1_005_32    m/z 194.9054     0   1E+4   8   8.5  RT (min)       lcqtof_hilic_neg_1_015_33    m/z 194.9055     0   1E+4   0   0.5  RT (min)      lcqtof_hilic_neg_4_050_02    m/z 194.9055     0   1E+4   8   8.5  RT (min)       lcqtof_hilic_neg_1_005_32    m/z 194.9056     0   1E+4   8   8.5  RT (min)       lcqtof_hilic_neg_1_014_31    m/z 194.9056     0   2   4E+3   0   0.5  RT (min)       lcqtof_hilic_neg_3_068_57    m/z 194.9057     0   1E+4   7.5   8  RT (min)       lcqtof_hilic_neg_1_004_14    m/z 194.9058     0   1E+4   3.5   4  RT (min)      lcqtof_hilic_neg_2_040_20    m/z 194.9058     0   1   2   3E+3   0   0.5  RT (min)       lcqtof_hilic_neg_4_050_02    m/z 194.9058     0   1   2E+3   19   19.5  RT (min)       lcqtof_hilic_neg_2_000_cc    m/z 194.9268     0   2   4   6E+4   7   7.5  RT (min)      lcqtof_hilic_neg_4_061_50    m/z 194.9269     0   2   4   6   8E+4   7   7.5  RT (min)       lcqtof_hilic_neg_4_047_06    m/z 195.0406     0   2   4   6   8E+3   0.5   1  RT (min)      lcqtof_hilic_neg_4_050_02    m/z 195.0500     0   2   4   6   8E+3   7   7.5  RT (min)      lcqtof_hilic_neg_4_002_25    m/z 195.0508     0   1E+4   6   6.5  RT (min)      lcqtof_hilic_neg_4_055_32    m/z 195.0522     0   1E+3   2   2.5  RT (min)      lcqtof_hilic_neg_4_012_07    m/z 195.0725     0   1E+4   0.5   1  RT (min)      lcqtof_hilic_neg_4_050_02    m/z 195.0770     0   1E+4   4   4.5  RT (min)      lcqtof_hilic_neg_4_022_34    m/z 195.0791     0   1   2   3   4E+5   3   3.5  RT (min)       lcqtof_hilic_neg_4_090_60    m/z 195.8102     0   1E+4   1   1.5  RT (min)      lcqtof_hilic_neg_3_066_57    m/z 195.8102     0   2   4   6   8E+3   1.5   2  RT (min)       lcqtof_hilic_neg_4_101_61    m/z 195.8103     0   2   4   6E+3   0   0.5  RT (min)       lcqtof_hilic_neg_4_072_57    m/z 195.8104     0   1   2   3   4E+4   3   3.5  RT (min)       lcqtof_hilic_neg_4_043_12    m/z 195.8104     0   2   4   6E+4   3.5   4  RT (min)      lcqtof_hilic_neg_4_051_43    m/z 195.8104     0   2   4   6E+4   3.5   4  RT (min)       lcqtof_hilic_neg_3_067_57    m/z 195.8104     0   2   4   6E+3   0.5   1  RT (min)       lcqtof_hilic_neg_4_072_57    m/z 195.8104     0   1   2   3E+4   3   3.5  RT (min)       lcqtof_hilic_neg_4_043_12    m/z 195.8104     0   1   2   3E+4   3   3.5  RT (min)       lcqtof_hilic_neg_4_044_08    m/z 195.8104     0   2   4   6   8E+4   3.5   4  RT (min)       lcqtof_hilic_neg_3_067_57    m/z 195.8104     0   2   4   6   8E+3   2   2.5  RT (min)       lcqtof_hilic_neg_4_108_62    m/z 195.8105     0   1   2   3E+4   3   3.5  RT (min)       lcqtof_hilic_neg_4_044_08    m/z 195.8105     0   1   2E+4   3.5   4  RT (min)      lcqtof_hilic_neg_1_000_bb    m/z 195.8105     0   2   4   6E+4   3.5   4  RT (min)       lcqtof_hilic_neg_4_051_43    m/z 195.8105     0   1   2E+4   2.5   3  RT (min)       lcqtof_hilic_neg_4_024_40    m/z 195.8105     0   1   2   3E+4   3   3.5  RT (min)       lcqtof_hilic_neg_4_044_08    m/z 195.8106     0   1   2E+4   1   1.5  RT (min)      lcqtof_hilic_neg_4_046_41    m/z 195.8107     0   2   4   6E+4   3.5   4  RT (min)      lcqtof_hilic_neg_4_012_07    m/z 195.8108     0   1   2   3E+4   3.5   4  RT (min)      lcqtof_hilic_neg_4_048_23    m/z 195.8816     0   1E+4   4.5   5  RT (min)      lcqtof_hilic_neg_4_014_46    m/z 195.9051     0   1   2   3   4E+3   7   7.5  RT (min)      lcqtof_hilic_neg_1_055_25    m/z 195.9053     0   2   4E+3   7   7.5  RT (min)      lcqtof_hilic_neg_2_052_24    m/z 195.9054     0   1   2   3   4E+3   6   6.5  RT (min)      lcqtof_hilic_neg_2_028_05    m/z 195.9055     0   2   4   6   8E+3   5.5   6  RT (min)      lcqtof_hilic_neg_1_022_36    m/z 195.9055     0   2   4   6E+3   5.5   6  RT (min)      lcqtof_hilic_neg_1_005_32    m/z 195.9058     0   1   2   3   4E+3   7   7.5  RT (min)      lcqtof_hilic_neg_1_034_08    m/z 195.9533     0   1   2   3E+4   4.5   5  RT (min)      lcqtof_hilic_neg_4_014_46    m/z 195.9910     0   1   2E+4   6   6.5  RT (min)      lcqtof_hilic_neg_1_000_aa    m/z 196.0246     0   1   2   3   4E+4   0.5   1  RT (min)      lcqtof_hilic_neg_2_045_11    m/z 196.0364     0   1E+4   4.5   5  RT (min)      lcqtof_hilic_neg_1_005_32    m/z 196.8369     0   1   2E+4   2.5   3  RT (min)      lcqtof_hilic_neg_4_050_02    m/z 196.8770     0   1   2   3E+3   8   8.5  RT (min)      lcqtof_hilic_neg_1_010_15    m/z 196.8937     0   1   2E+3   7.5   8  RT (min)      lcqtof_hilic_neg_1_003_37    m/z 196.9008     0   1   2   3E+3   7   7.5  RT (min)      lcqtof_hilic_neg_1_037_24    m/z 196.9037     0   1   2   3E+3   5.5   6  RT (min)      lcqtof_hilic_neg_1_007_05    m/z 196.9038     0   2   4   6E+3   6   6.5  RT (min)      lcqtof_hilic_neg_2_031_01    m/z 196.9232     0   1E+4   7   7.5  RT (min)      lcqtof_hilic_neg_4_047_06    m/z 196.9753     0   1E+5   5.5   6  RT (min)      lcqtof_hilic_neg_1_000_aa    m/z 197.0225     0   1E+4   2   2.5  RT (min)       lcqtof_hilic_neg_4_003_16    m/z 197.0240     0   2   4   6   8E+2   2   2.5  RT (min)      lcqtof_hilic_neg_1_051_35    m/z 197.0249     0   1   2   3   4E+3   4   4.5  RT (min)      lcqtof_hilic_neg_4_096_60    m/z 197.0260     0   1   2   3E+4   0.5   1  RT (min)       lcqtof_hilic_neg_4_045_26    m/z 197.0387     0   2   4   6E+4   7   7.5  RT (min)      lcqtof_hilic_neg_1_005_32    m/z 197.0389     0   1E+5   7   7.5  RT (min)       lcqtof_hilic_neg_4_091_60    m/z 197.0389     0   1   2   3   4E+4   7.5   8  RT (min)      lcqtof_hilic_neg_4_008_24    m/z 197.0389     0   1   2E+5   7   7.5  RT (min)       lcqtof_hilic_neg_4_095_60    m/z 197.0389     0   2   4   6E+4   7   7.5  RT (min)      lcqtof_hilic_neg_1_005_32    m/z 197.0390     0   1   2   3   4E+4   7.5   8  RT (min)      lcqtof_hilic_neg_4_008_24    m/z 197.0390     0   1   2   3   4E+4   8   8.5  RT (min)       lcqtof_hilic_neg_4_008_24    m/z 197.0390     0   1   2E+5   7.5   8  RT (min)       lcqtof_hilic_neg_4_094_60    m/z 197.0390     0   1   2   3   4E+4   7   7.5  RT (min)      lcqtof_hilic_neg_4_065_51    m/z 197.0392     0   1E+5   4   4.5  RT (min)      lcqtof_hilic_neg_1_015_33    m/z 197.0392     0   1E+5   7   7.5  RT (min)       lcqtof_hilic_neg_4_055_32    m/z 197.0668     0   2   4   6   8E+3   3   3.5  RT (min)      lcqtof_hilic_neg_4_052_13    m/z 197.0668     0   2   4E+3   2.5   3  RT (min)      lcqtof_hilic_neg_4_050_02    m/z 197.0672     0   1E+3   3   3.5  RT (min)      lcqtof_hilic_neg_4_008_24    m/z 197.0672     0   2   4   6   8E+3   2.5   3  RT (min)       lcqtof_hilic_neg_4_052_13    m/z 197.1028     0   1E+4   1.5   2  RT (min)       lcqtof_hilic_neg_4_095_60    m/z 197.1028     0   1   2   3E+4   2   2.5  RT (min)       lcqtof_hilic_neg_4_052_13    m/z 197.1739     0   2   4   6E+4   2   2.5  RT (min)      lcqtof_hilic_neg_4_052_13    m/z 197.2823     0   1   2E+4   2   2.5  RT (min)      lcqtof_hilic_neg_4_052_13    m/z 197.8070     0   2   4   6E+3   0   0.5  RT (min)       lcqtof_hilic_neg_4_070_57    m/z 197.8071     0   1E+4   1.5   2  RT (min)       lcqtof_hilic_neg_4_106_62    m/z 197.8071     0   1E+4   1.5   2  RT (min)       lcqtof_hilic_neg_4_108_62    m/z 197.8071     0   1E+4   2   2.5  RT (min)       lcqtof_hilic_neg_4_099_61    m/z 197.8071     0   1   2   3E+4   1   1.5  RT (min)      lcqtof_hilic_neg_4_046_41    m/z 197.8071     0   2   4   6   8E+3   0.5   1  RT (min)       lcqtof_hilic_neg_4_073_57    m/z 197.8072     0   1E+4   1   1.5  RT (min)       lcqtof_hilic_neg_3_066_57    m/z 197.8073     0   2   4   6   8E+3   16.5   17  RT (min)       lcqtof_hilic_neg_3_066_57    m/z 197.8073     0   1   2E+4   2.5   3  RT (min)       lcqtof_hilic_neg_4_013_20    m/z 197.8073     0   1   2   3E+4   0.5   1  RT (min)      lcqtof_hilic_neg_4_037_39    m/z 197.8074     0   1E+4   15   15.5   16  RT (min)       lcqtof_hilic_neg_4_000_aa    m/z 197.8075     0   2   4   6E+3   0.5   1  RT (min)       lcqtof_hilic_neg_4_061_50    m/z 197.8781     0   1   2E+4   4.5   5  RT (min)      lcqtof_hilic_neg_4_014_46    m/z 197.9857     0   1E+4   6   6.5  RT (min)      lcqtof_hilic_neg_1_022_36    m/z 197.9869     0   1E+4   2   2.5  RT (min)      lcqtof_hilic_neg_4_013_20    m/z 197.9912     0   1E+4   2   2.5  RT (min)      lcqtof_hilic_neg_4_027_19    m/z 197.9915     0   2   4E+3   2   2.5  RT (min)       lcqtof_hilic_neg_4_031_38    m/z 197.9934     0   1   2E+4   0.5   1  RT (min)      lcqtof_hilic_neg_4_004_47    m/z 198.0528     0   1E+4   1.5   2  RT (min)      lcqtof_hilic_neg_4_055_32    m/z 198.0530     0   1E+4   2   2.5  RT (min)      lcqtof_hilic_neg_4_055_32    m/z 198.0768     0   1E+4   1   1.5  RT (min)       lcqtof_hilic_neg_4_048_23    m/z 198.9925     0   1E+4   0.5   1  RT (min)      lcqtof_hilic_neg_4_004_47    m/z 199.0050     0   1E+4   2   2.5  RT (min)      lcqtof_hilic_neg_4_045_26    m/z 199.0056     0   1E+4   5.5   6  RT (min)      lcqtof_hilic_neg_1_000_aa    m/z 199.0074     0   1   2   3   4E+3   4   4.5  RT (min)      lcqtof_hilic_neg_4_034_31    m/z 199.0242     0   1E+4   1.5   2  RT (min)       lcqtof_hilic_neg_4_028_36    m/z 199.0349     0   1   2E+3   1   1.5  RT (min)       lcqtof_hilic_neg_4_042_22    m/z 199.0353     0   1E+4   2   2.5  RT (min)      lcqtof_hilic_neg_4_052_13    m/z 199.0715     0   1E+4   3   3.5  RT (min)      lcqtof_hilic_neg_4_050_02    m/z 199.0720     0   1   2   3E+4   4   4.5  RT (min)      lcqtof_hilic_neg_4_011_35    m/z 199.1697     0   2   4E+4   1   1.5  RT (min)      lcqtof_hilic_neg_4_038_10    m/z 199.1698     0   2   4E+4   0.5   1  RT (min)      lcqtof_hilic_neg_4_038_10    m/z 199.8045     0   1E+4   3   3.5  RT (min)       lcqtof_hilic_neg_4_051_43    m/z 199.8046     0   1E+4   1   1.5  RT (min)      lcqtof_hilic_neg_4_046_41    m/z 199.8046     0   1   2   3E+3   0   0.5  RT (min)       lcqtof_hilic_neg_4_073_57    m/z 199.8046     0   1   2   3   4E+3   0.5   1  RT (min)       lcqtof_hilic_neg_4_065_51    m/z 199.8046     0   2   4E+3   1.5   2  RT (min)       lcqtof_hilic_neg_4_106_62    m/z 199.8048     0   1   2   3   4E+3   0.5   1  RT (min)       lcqtof_hilic_neg_4_061_50    m/z 199.8048     0   1E+4   3   3.5  RT (min)      lcqtof_hilic_neg_4_039_18    m/z 199.8506     0   1   2E+3   8   8.5  RT (min)      lcqtof_hilic_neg_2_043_15    m/z 199.8508     0   1   2E+3   4.5   5  RT (min)      lcqtof_hilic_neg_3_016_39    m/z 199.8510     0   1E+4   1   1.5  RT (min)      lcqtof_hilic_neg_4_058_39    m/z 200.0557     0   1E+4   2.5   3  RT (min)      lcqtof_hilic_neg_4_048_23    m/z 200.0557     0   1E+4   2   2.5  RT (min)      lcqtof_hilic_neg_4_031_38    m/z 200.0560     0   1E+4   2.5   3  RT (min)      lcqtof_hilic_neg_4_034_31    m/z 200.0562     0   1E+4   4   4.5  RT (min)      lcqtof_hilic_neg_4_085_59    m/z 200.8579     0   1   2   3   4E+3   0   0.5  RT (min)       lcqtof_hilic_neg_1_045_07    m/z 200.8579     0   1   2   3E+4   0.5   1  RT (min)       lcqtof_hilic_neg_1_053_19    m/z 200.8580     0   1   2   3E+3   19.5   20  RT (min)       lcqtof_hilic_neg_2_000_bb    m/z 200.8581     0   1E+4   0.5   1  RT (min)      lcqtof_hilic_neg_1_045_07    m/z 200.8587     0   1E+4   1.5   2  RT (min)      lcqtof_hilic_neg_1_050_12    m/z 200.9461     0   1E+4   7.5   8  RT (min)      lcqtof_hilic_neg_1_000_aa    m/z 200.9857     0   2   4   6E+3   6.5   7  RT (min)       lcqtof_hilic_neg_4_012_07    m/z 200.9858     0   1   2E+3   0   0.5  RT (min)       lcqtof_hilic_neg_4_047_06    m/z 200.9866     0   1   2   3E+3   2   2.5  RT (min)      lcqtof_hilic_neg_4_079_58    m/z 201.8014     0   1   2   3   4E+4   3.5   4  RT (min)       lcqtof_hilic_neg_4_024_40    m/z 202.0275     0   1   2E+4   4.5   5  RT (min)      lcqtof_hilic_neg_4_000_aa    m/z 202.0281     0   1   2E+4   4.5   5  RT (min)      lcqtof_hilic_neg_4_000_aa    m/z 202.0285     0   1   2E+4   4.5   5  RT (min)      lcqtof_hilic_neg_4_000_aa    m/z 202.0287     0   1   2E+4   4.5   5  RT (min)      lcqtof_hilic_neg_4_000_aa    m/z 202.0292     0   1   2E+4   4.5   5  RT (min)      lcqtof_hilic_neg_4_000_aa    m/z 202.0702     0   1E+4   2.5   3  RT (min)      lcqtof_hilic_neg_4_047_06    m/z 202.9282     0   1E+4   2   4  RT (min)       lcqtof_hilic_neg_3_059_39    m/z 202.9283     0   1   2   3E+3   0   0.5  RT (min)       lcqtof_hilic_neg_3_069_57    m/z 202.9283     0   2   4E+3   5   5.5  RT (min)       lcqtof_hilic_neg_3_089_59    m/z 203.0332     0   1   2   3E+4   2.5   3  RT (min)       lcqtof_hilic_neg_3_112_62    m/z 203.0496     0   2   4   6E+3   1.5   2  RT (min)      lcqtof_hilic_neg_4_048_23    m/z 203.9323     0   1E+4   0.5   1  RT (min)      lcqtof_hilic_neg_4_023_15    m/z 203.9341     0   2   4   6   8E+3   1   1.5  RT (min)      lcqtof_hilic_neg_4_018_27    m/z 203.9355     0   1   2E+3   7.5   8  RT (min)       lcqtof_hilic_neg_4_008_24    m/z 203.9366     0   1   2   3E+3   7   7.5  RT (min)       lcqtof_hilic_neg_4_060_13    m/z 203.9376     0   2   4E+3   2.5   3  RT (min)      lcqtof_hilic_neg_3_079_58    m/z 203.9377     0   1   2   3   4E+3   4   4.5  RT (min)      lcqtof_hilic_neg_4_012_07    m/z 203.9381     0   1   2   3   4E+3   4   4.5  RT (min)      lcqtof_hilic_neg_4_008_24    m/z 203.9382     0   2   4E+3   4   4.5  RT (min)       lcqtof_hilic_neg_4_012_07    m/z 203.9388     0   2   4   6E+3   4.5   5  RT (min)       lcqtof_hilic_neg_4_019_03    m/z 203.9389     0   1   2   3   4E+3   4.5   5  RT (min)       lcqtof_hilic_neg_4_006_30    m/z 203.9398     0   1   2   3E+3   2   2.5  RT (min)      lcqtof_hilic_neg_2_041_54    m/z 204.0509     0   1E+4   4   4.5  RT (min)      lcqtof_hilic_neg_4_034_31    m/z 204.9319     0   1   2E+3   1.5   2  RT (min)      lcqtof_hilic_neg_2_045_11    m/z 204.9659     0   1   2E+4   6.5   7  RT (min)      lcqtof_hilic_neg_1_000_aa    m/z 204.9659     0   1   2E+4   6.5   7  RT (min)      lcqtof_hilic_neg_1_000_aa    m/z 204.9762     0   1E+3   1   1.5  RT (min)      lcqtof_hilic_neg_2_000_aa    m/z 204.9958     0   1E+4   0.5   1  RT (min)      lcqtof_hilic_neg_3_012_27    m/z 205.0343     0   1   2E+4   6.5   7  RT (min)       lcqtof_hilic_neg_4_010_28    m/z 205.0344     0   1   2E+5   2.5   3  RT (min)      lcqtof_hilic_neg_4_050_02    m/z 205.0344     0   1   2   3E+4   3   3.5  RT (min)      lcqtof_hilic_neg_2_026_28    m/z 205.0344     0   1   2E+4   7   7.5  RT (min)       lcqtof_hilic_neg_4_010_28    m/z 205.0345     0   1E+4   7   7.5  RT (min)      lcqtof_hilic_neg_4_023_15    m/z 205.0346     0   2   4E+4   6.5   7  RT (min)       lcqtof_hilic_neg_4_034_31    m/z 205.0346     0   1E+4   7   7.5  RT (min)       lcqtof_hilic_neg_4_034_31    m/z 205.0346     0   1   2E+5   3   3.5  RT (min)      lcqtof_hilic_neg_4_050_02    m/z 205.0347     0   2   4   6   8E+4   3.5   4  RT (min)       lcqtof_hilic_neg_4_064_15    m/z 205.0347     0   2   4   6E+4   3.5   4  RT (min)      lcqtof_hilic_neg_4_062_17    m/z 205.0348     0   1   2E+5   3   3.5  RT (min)       lcqtof_hilic_neg_4_050_02    m/z 205.0348     0   1   2E+5   3.5   4  RT (min)       lcqtof_hilic_neg_4_050_02    m/z 205.0348     0   1   2E+5   3   3.5  RT (min)      lcqtof_hilic_neg_4_050_02    m/z 205.0348     0   1   2E+5   3.5   4  RT (min)       lcqtof_hilic_neg_4_050_02    m/z 205.0349     0   1   2E+4   3   3.5  RT (min)      lcqtof_hilic_neg_2_018_16    m/z 205.0349     0   1   2   3   4E+4   4   4.5  RT (min)      lcqtof_hilic_neg_2_002_36    m/z 205.0353     0   1   2   3   4E+4   4.5   5  RT (min)      lcqtof_hilic_neg_1_051_35    m/z 205.8394     0   1   2E+4   4.5   5  RT (min)       lcqtof_hilic_neg_4_014_46    m/z 205.9366     0   2   4E+3   4.5   5  RT (min)       lcqtof_hilic_neg_4_083_59    m/z 205.9368     0   1   2   3   4E+3   4   4.5  RT (min)       lcqtof_hilic_neg_4_086_59    m/z 205.9368     0   1   2   3   4E+3   5   10  RT (min)       lcqtof_hilic_neg_4_080_58    m/z 205.9370     0   1   2E+3   5.5   6  RT (min)       lcqtof_hilic_neg_4_007_45    m/z 206.0382     0   1E+4   2   2.5  RT (min)      lcqtof_hilic_neg_1_016_28    m/z 206.0392     0   2   4   6E+2   4   4.5  RT (min)      lcqtof_hilic_neg_4_054_48    m/z 206.0430     0   1   2E+4   3   3.5  RT (min)       lcqtof_hilic_neg_4_050_02    m/z 206.0486     0   2   4E+4   3.5   4  RT (min)       lcqtof_hilic_neg_4_110_62    m/z 206.0487     0   1E+5   4.5   5  RT (min)      lcqtof_hilic_neg_4_016_37    m/z 206.7878     0   1   2E+4   3   3.5  RT (min)       lcqtof_hilic_neg_3_066_57    m/z 206.7896     0   1E+3   9   9.5  RT (min)       lcqtof_hilic_neg_2_031_01    m/z 206.9686     0   1E+4   14.5   15   15.5  RT (min)       lcqtof_hilic_neg_4_000_bb    m/z 206.9691     0   1E+4   14   15  RT (min)       lcqtof_hilic_neg_4_050_02    m/z 206.9696     0   2   4   6   8E+3   11   12  RT (min)       lcqtof_hilic_neg_4_000_cc    m/z 206.9717     0   2   4E+4   0.5   1  RT (min)      lcqtof_hilic_neg_4_000_aa    m/z 206.9717     0   2   4E+4   0   0.5  RT (min)      lcqtof_hilic_neg_4_000_aa    m/z 206.9718     0   2   4E+4   0.5   1  RT (min)      lcqtof_hilic_neg_4_000_aa    m/z 206.9718     0   2   4E+4   0   0.5  RT (min)      lcqtof_hilic_neg_4_000_aa    m/z 206.9721     0   1E+4   8.5   9  RT (min)       lcqtof_hilic_neg_4_053_14    m/z 206.9788     0   1   2   3E+4   1   1.5  RT (min)      lcqtof_hilic_neg_4_000_aa    m/z 207.0104     0   2   4   6E+3   2   2.5  RT (min)       lcqtof_hilic_neg_4_075_58    m/z 207.0115     0   1E+5   1   1.5  RT (min)      lcqtof_hilic_neg_4_038_10    m/z 207.0116     0   1E+5   1   1.5  RT (min)      lcqtof_hilic_neg_4_038_10    m/z 207.0518     0   1   2E+4   4   4.5  RT (min)      lcqtof_hilic_neg_4_054_48    m/z 207.0756     0   1   2E+3   1   1.5  RT (min)       lcqtof_hilic_neg_4_011_35    m/z 207.0791     0   1   2E+3   1.5   2  RT (min)       lcqtof_hilic_neg_4_064_15    m/z 207.0811     0   1E+4   2   2.5  RT (min)      lcqtof_hilic_neg_4_055_32    m/z 207.8362     0   1   2E+4   4.5   5  RT (min)       lcqtof_hilic_neg_4_014_46    m/z 207.8779     0   1E+4   1   1.5  RT (min)      lcqtof_hilic_neg_4_050_02    m/z 208.0459     0   1E+4   4   4.5  RT (min)      lcqtof_hilic_neg_4_054_48    m/z 208.0799     0   1E+4   1   1.5  RT (min)      lcqtof_hilic_neg_4_050_02    m/z 208.0800     0   1E+4   2   2.5  RT (min)       lcqtof_hilic_neg_4_095_60    m/z 208.0817     0   1E+4   2.5   3  RT (min)       lcqtof_hilic_neg_4_008_24    m/z 208.0827     0   1   2   3   4E+4   2.5   3  RT (min)      lcqtof_hilic_neg_4_055_32    m/z 208.0967     0   1E+4   15   15.5  RT (min)      lcqtof_hilic_neg_4_000_aa    m/z 208.8444     0   1E+4   4   4.5  RT (min)      lcqtof_hilic_neg_4_050_02    m/z 208.9343     0   2   4   6   8E+3   4.5   5  RT (min)      lcqtof_hilic_neg_1_000_bb    m/z 209.0121     0   2   4   6   8E+3   1   1.5  RT (min)       lcqtof_hilic_neg_4_047_06    m/z 209.0939     0   1   2   3E+4   3   3.5  RT (min)       lcqtof_hilic_neg_4_071_57    m/z 209.8487     0   1E+4   4   4.5  RT (min)      lcqtof_hilic_neg_4_050_02    m/z 210.0903     0   1E+4   4   4.5  RT (min)       lcqtof_hilic_neg_4_055_32    m/z 210.1127     0   1E+4   2   2.5  RT (min)      lcqtof_hilic_neg_4_055_32    m/z 210.1132     0   1E+4   2   2.5  RT (min)      lcqtof_hilic_neg_4_055_32    m/z 210.8430     0   1E+4   4   4.5  RT (min)      lcqtof_hilic_neg_4_050_02    m/z 210.8880     0   1   2E+4   0.5   1  RT (min)      lcqtof_hilic_neg_2_000_aa    m/z 210.9336     0   1E+4   3.5   4  RT (min)      lcqtof_hilic_neg_1_032_18    m/z 211.0004     0   2   4   6   8E+2   5.5   6  RT (min)      lcqtof_hilic_neg_1_000_aa    m/z 211.0553     0   2   4E+3   2   2.5  RT (min)      lcqtof_hilic_neg_1_005_32    m/z 211.0743     0   2   4E+4   4   4.5  RT (min)       lcqtof_hilic_neg_4_055_32    m/z 211.0824     0   2   4   6E+3   1.5   2  RT (min)      lcqtof_hilic_neg_4_052_13    m/z 211.8480     0   1E+4   4   4.5  RT (min)      lcqtof_hilic_neg_4_050_02    m/z 212.0582     0   2   4E+4   4.5   5  RT (min)      lcqtof_hilic_neg_1_005_32    m/z 212.0582     0   1E+5   3.5   4  RT (min)       lcqtof_hilic_neg_1_005_32    m/z 212.0582     0   1   2   3   4E+4   6.5   7  RT (min)      lcqtof_hilic_neg_4_040_05    m/z 212.0582     0   2   4E+4   6.5   7  RT (min)      lcqtof_hilic_neg_4_035_33    m/z 212.0583     0   2   4E+4   6.5   7  RT (min)      lcqtof_hilic_neg_4_035_33    m/z 212.0584     0   1   2E+4   7   7.5  RT (min)       lcqtof_hilic_neg_4_035_33    m/z 212.0584     0   2   4   6E+4   3.5   4  RT (min)      lcqtof_hilic_neg_2_047_32    m/z 212.0584     0   1E+4   7.5   8  RT (min)       lcqtof_hilic_neg_4_002_25    m/z 212.0584     0   1E+5   4   4.5  RT (min)      lcqtof_hilic_neg_1_005_32    m/z 212.0584     0   2   4E+4   6.5   7  RT (min)      lcqtof_hilic_neg_4_035_33    m/z 212.0587     0   2   4   6E+4   4.5   5  RT (min)      lcqtof_hilic_neg_2_047_32    m/z 212.8476     0   1   2E+4   4.5   5  RT (min)      lcqtof_hilic_neg_4_050_02    m/z 212.8476     0   1   2E+4   4   4.5  RT (min)      lcqtof_hilic_neg_4_050_02    m/z 212.9103     0   1   2   3E+3   0   0.5  RT (min)       lcqtof_hilic_neg_3_068_57    m/z 212.9106     0   1   2   3E+3   0.5   1  RT (min)       lcqtof_hilic_neg_3_068_57    m/z 212.9107     0   1   2   3E+3   0   0.5  RT (min)       lcqtof_hilic_neg_3_068_57    m/z 212.9700     0   1   2   3E+4   5.5   6  RT (min)      lcqtof_hilic_neg_1_000_aa    m/z 213.0143     0   1E+4   17   17.5  RT (min)       lcqtof_hilic_neg_4_078_58    m/z 213.0143     0   1E+4   19.5   20  RT (min)      lcqtof_hilic_neg_4_070_57    m/z 213.0144     0   1E+4   2.5   3  RT (min)       lcqtof_hilic_neg_4_086_59    m/z 213.0144     0   1E+4   19   19.5  RT (min)       lcqtof_hilic_neg_4_102_61    m/z 213.0144     0   1E+4   15   15.5   16  RT (min)       lcqtof_hilic_neg_4_083_59    m/z 213.0144     0   1E+4   15.5   16  RT (min)       lcqtof_hilic_neg_4_082_59    m/z 213.0145     0   1E+4   18.5   19  RT (min)       lcqtof_hilic_neg_4_101_61    m/z 213.0145     0   1E+4   19   20  RT (min)       lcqtof_hilic_neg_4_080_58    m/z 213.0145     0   1   2E+4   2.5   3  RT (min)       lcqtof_hilic_neg_4_059_14    m/z 213.0145     0   1E+4   15.5   16   16.5  RT (min)       lcqtof_hilic_neg_4_083_59    m/z 213.0146     0   1   2E+4   2   2.5  RT (min)       lcqtof_hilic_neg_4_059_14    m/z 213.0146     0   1E+4   17   18  RT (min)       lcqtof_hilic_neg_4_107_62    m/z 213.0146     0   1E+4   3.5   4  RT (min)       lcqtof_hilic_neg_4_016_37    m/z 213.0146     0   1E+4   15.5   16   16.5  RT (min)       lcqtof_hilic_neg_4_082_59    m/z 213.0146     0   1E+4   16.5   17  RT (min)       lcqtof_hilic_neg_4_083_59    m/z 213.0146     0   1E+4   0   0.5  RT (min)       lcqtof_hilic_neg_4_031_38    m/z 213.0146     0   1E+4   17  RT (min)       lcqtof_hilic_neg_4_111_62    m/z 213.0146     0   1E+4   18.5   19  RT (min)       lcqtof_hilic_neg_4_098_61    m/z 213.0146     0   1E+4   18   18.5  RT (min)      lcqtof_hilic_neg_4_101_61    m/z 213.0146     0   1E+4   17   17.5  RT (min)       lcqtof_hilic_neg_4_000_cc    m/z 213.0146     0   1E+4   16   16.5  RT (min)       lcqtof_hilic_neg_4_108_62    m/z 213.0147     0   1E+4   18   19  RT (min)       lcqtof_hilic_neg_4_097_60    m/z 213.0147     0   1E+4   19   19.5  RT (min)       lcqtof_hilic_neg_4_098_61    m/z 213.0147     0   1E+4   0.5   1  RT (min)       lcqtof_hilic_neg_4_072_57    m/z 213.0147     0   1E+4   1.5   2  RT (min)       lcqtof_hilic_neg_4_072_57    m/z 213.0147     0   1E+4   16   16.5  RT (min)       lcqtof_hilic_neg_4_098_61    m/z 213.0148     0   1E+4   0   0.5  RT (min)       lcqtof_hilic_neg_4_103_61    m/z 213.0148     0   1   2E+4   2   2.5  RT (min)      lcqtof_hilic_neg_4_035_33    m/z 213.0148     0   1   2E+4   2   2.5  RT (min)       lcqtof_hilic_neg_4_053_14    m/z 213.0148     0   1E+4   18   19  RT (min)       lcqtof_hilic_neg_4_112_62    m/z 213.0148     0   1   2E+4   1.5   2  RT (min)       lcqtof_hilic_neg_4_074_58    m/z 213.0148     0   1   2   3E+4   0   0.5  RT (min)      lcqtof_hilic_neg_4_081_58    m/z 213.0149     0   1E+4   17.5   18  RT (min)       lcqtof_hilic_neg_4_110_62    m/z 213.0150     0   1   2E+4   2   2.5  RT (min)      lcqtof_hilic_neg_4_055_32    m/z 213.0150     0   1   2E+4   2   4   6  RT (min)       lcqtof_hilic_neg_4_012_07    m/z 213.0150     0   1   2E+4   2.5   3  RT (min)      lcqtof_hilic_neg_4_028_36    m/z 213.0152     0   1   2E+4   2   2.5  RT (min)       lcqtof_hilic_neg_4_016_37    m/z 213.0152     0   1   2E+4   2.5   3  RT (min)      lcqtof_hilic_neg_4_075_58    m/z 213.7837     0   2   4E+3   3   3.5  RT (min)      lcqtof_hilic_neg_1_006_22    m/z 213.8792     0   2   4   6E+3   4   4.5  RT (min)      lcqtof_hilic_neg_1_004_14    m/z 213.8792     0   2   4E+3   7.5   8  RT (min)       lcqtof_hilic_neg_1_004_14    m/z 213.8793     0   2   4   6E+3   6.5   7  RT (min)      lcqtof_hilic_neg_1_004_14    m/z 213.8793     0   1E+4   4.5   5  RT (min)       lcqtof_hilic_neg_1_008_13    m/z 213.8795     0   1E+4   4   4.5  RT (min)      lcqtof_hilic_neg_1_008_13    m/z 214.0176     0   1   2   3   4E+3   3   3.5  RT (min)      lcqtof_hilic_neg_4_053_14    m/z 214.0232     0   2   4   6E+3   2   2.5  RT (min)      lcqtof_hilic_neg_4_018_27    m/z 214.0263     0   1   2E+3   2.5   3  RT (min)       lcqtof_hilic_neg_4_086_59    m/z 214.0270     0   1   2   3   4E+3   2   2.5  RT (min)      lcqtof_hilic_neg_4_053_14    m/z 214.1446     0   2   4   6E+4   1   1.5  RT (min)      lcqtof_hilic_neg_4_026_04    m/z 214.8488     0   1E+4   4   4.5  RT (min)      lcqtof_hilic_neg_4_050_02    m/z 214.8798     0   1E+4   1   1.5  RT (min)      lcqtof_hilic_neg_4_051_43    m/z 214.9458     0   2   4   6E+3   2   2.5  RT (min)      lcqtof_hilic_neg_1_060_14    m/z 214.9460     0   1   2   3E+3   1   1.5  RT (min)      lcqtof_hilic_neg_2_045_11    m/z 214.9662     0   1E+4   3   3.5  RT (min)      lcqtof_hilic_neg_2_003_25    m/z 215.0296     0   2   4   6   8E+3   5.5   6  RT (min)      lcqtof_hilic_neg_4_053_14    m/z 215.0322     0   1E+5   5.5   6  RT (min)      lcqtof_hilic_neg_1_025_52    m/z 215.0322     0   2   4   6E+3   6   6.5  RT (min)      lcqtof_hilic_neg_1_054_01    m/z 215.0324     0   1E+4   3.5   4  RT (min)      lcqtof_hilic_neg_1_004_14    m/z 215.0329     0   1   2   3   4E+3   5.5   6  RT (min)      lcqtof_hilic_neg_4_023_15    m/z 215.0666     0   2   4   6   8E+3   1   1.5  RT (min)      lcqtof_hilic_neg_4_050_02    m/z 215.0669     0   1   2E+4   1   1.5  RT (min)      lcqtof_hilic_neg_3_004_32    m/z 215.8768     0   2   4E+3   4.5   5  RT (min)      lcqtof_hilic_neg_1_012_04    m/z 216.0185     0   1E+5   0   0.5  RT (min)      lcqtof_hilic_neg_4_000_aa    m/z 216.0188     0   1E+5   0.5   1  RT (min)      lcqtof_hilic_neg_4_000_aa    m/z 216.0333     0   2   4   6E+3   4.5   5  RT (min)      lcqtof_hilic_neg_4_053_14    m/z 216.0535     0   2   4   6E+3   2   2.5  RT (min)      lcqtof_hilic_neg_2_031_01    m/z 216.8532     0   1E+4   1   1.5  RT (min)      lcqtof_hilic_neg_4_058_39    m/z 216.8534     0   1   2   3   4E+3   4   4.5  RT (min)      lcqtof_hilic_neg_2_038_04    m/z 216.8535     0   1   2   3   4E+3   7.5   8  RT (min)      lcqtof_hilic_neg_3_065_51    m/z 216.8538     0   2   4E+3   4   4.5  RT (min)      lcqtof_hilic_neg_2_050_19    m/z 216.9090     0   1   2E+4   7   7.5  RT (min)       lcqtof_hilic_neg_4_047_06    m/z 217.0034     0   2   4E+3   1   1.5  RT (min)      lcqtof_hilic_neg_4_033_53    m/z 217.0132     0   1   2   3E+3   6.5   7  RT (min)      lcqtof_hilic_neg_2_025_52    m/z 217.0154     0   1E+4   7   7.5  RT (min)       lcqtof_hilic_neg_4_081_58    m/z 217.0155     0   1E+4   7.5   8  RT (min)       lcqtof_hilic_neg_4_059_14    m/z 217.0158     0   2   4   6   8E+4   0   0.5  RT (min)      lcqtof_hilic_neg_4_000_aa    m/z 217.0160     0   1E+4   7   7.5  RT (min)      lcqtof_hilic_neg_4_064_15    m/z 217.0170     0   2   4   6   8E+4   0   0.5  RT (min)      lcqtof_hilic_neg_4_000_aa    m/z 217.0182     0   1   2   3   4E+5   0   0.5  RT (min)      lcqtof_hilic_neg_4_000_aa    m/z 217.0283     0   1E+5   4.5   5  RT (min)      lcqtof_hilic_neg_4_053_14    m/z 217.0283     0   1   2E+4   4.5   5  RT (min)      lcqtof_hilic_neg_4_053_14    m/z 217.0289     0   1E+4   4.5   5  RT (min)      lcqtof_hilic_neg_1_014_31    m/z 217.0290     0   1   2E+4   4   4.5  RT (min)      lcqtof_hilic_neg_4_053_14    m/z 217.0292     0   2   4E+4   4.5   5  RT (min)      lcqtof_hilic_neg_4_035_33    m/z 217.0293     0   1   2E+4   4   4.5  RT (min)      lcqtof_hilic_neg_4_059_14    m/z 217.0489     0   1   2   3E+4   2   2.5  RT (min)      lcqtof_hilic_neg_1_054_01    m/z 217.0501     0   2   4E+5   1   1.5  RT (min)       lcqtof_hilic_neg_4_108_62    m/z 218.0183     0   1   2   3E+4   0.5   1  RT (min)      lcqtof_hilic_neg_4_000_aa    m/z 218.0204     0   1   2   3E+4   0.5   1  RT (min)      lcqtof_hilic_neg_4_000_aa    m/z 218.9416     0   1E+4   2   2.5  RT (min)      lcqtof_hilic_neg_4_046_41    m/z 218.9416     0   1   2   3   4E+4   1.5   2  RT (min)       lcqtof_hilic_neg_4_058_39    m/z 219.0142     0   1E+4   0.5   1  RT (min)      lcqtof_hilic_neg_4_000_aa    m/z 219.0506     0   2   4E+4   1   1.5  RT (min)      lcqtof_hilic_neg_4_050_02    m/z 219.0583     0   2   4   6   8E+2   1   1.5  RT (min)      lcqtof_hilic_neg_4_045_26    m/z 219.0680     0   2   4   6E+4   2   2.5  RT (min)      lcqtof_hilic_neg_4_010_28    m/z 219.0769     0   1E+4   1   1.5  RT (min)      lcqtof_hilic_neg_4_050_02    m/z 219.0798     0   1E+4   1   1.5  RT (min)      lcqtof_hilic_neg_4_050_02    m/z 219.0799     0   1   2E+4   1   1.5  RT (min)      lcqtof_hilic_neg_4_003_16    m/z 219.0801     0   1E+4   1   1.5  RT (min)      lcqtof_hilic_neg_4_050_02    m/z 219.1835     0   1E+4   1   1.5  RT (min)      lcqtof_hilic_neg_4_013_20    m/z 219.1836     0   1   2   3E+4   0.5   1  RT (min)      lcqtof_hilic_neg_4_027_19    m/z 219.2462     0   1   2E+4   1   1.5  RT (min)      lcqtof_hilic_neg_4_013_20    m/z 219.2464     0   2   4E+4   0.5   1   1.5  RT (min)      lcqtof_hilic_neg_4_027_19    m/z 219.2848     0   1   2   3E+4   1   1.5  RT (min)      lcqtof_hilic_neg_4_013_20    m/z 219.2850     0   2   4   6E+4   0.5   1   1.5  RT (min)      lcqtof_hilic_neg_4_027_19    m/z 219.4156     0   1   2   3   4E+4   1   1.5  RT (min)      lcqtof_hilic_neg_4_013_20    m/z 219.4156     0   2   4   6E+4   0.5   1   1.5  RT (min)      lcqtof_hilic_neg_4_027_19    m/z 219.8446     0   1E+4   6.5   7  RT (min)       lcqtof_hilic_neg_4_032_01    m/z 219.8447     0   1E+5   5   5.5  RT (min)      lcqtof_hilic_neg_3_000_aa    m/z 219.8448     0   2   4   6E+3   0.5   1  RT (min)       lcqtof_hilic_neg_3_067_57    m/z 219.8448     0   1   2E+4   16   16.5  RT (min)       lcqtof_hilic_neg_3_068_57    m/z 219.8449     0   1   2E+4   16.5   17  RT (min)       lcqtof_hilic_neg_3_067_57    m/z 219.8450     0   1   2   3E+3   9   9.5  RT (min)      lcqtof_hilic_neg_1_060_14    m/z 219.8450     0   1E+4   1.5   2  RT (min)       lcqtof_hilic_neg_3_066_57    m/z 219.8450     0   2   4   6E+4   3.5   4  RT (min)       lcqtof_hilic_neg_3_067_57    m/z 219.8450     0   1   2E+4   3   3.5  RT (min)       lcqtof_hilic_neg_4_024_40    m/z 219.8450     0   1   2   3E+4   3.5   4  RT (min)      lcqtof_hilic_neg_4_024_40    m/z 219.8450     0   1E+5   3.5   4  RT (min)       lcqtof_hilic_neg_3_067_57    m/z 219.8450     0   1E+4   0   0.5  RT (min)       lcqtof_hilic_neg_3_066_57    m/z 219.8451     0   1   2E+4   0.5   1  RT (min)       lcqtof_hilic_neg_3_066_57    m/z 219.8451     0   1E+4   1   1.5  RT (min)      lcqtof_hilic_neg_1_000_bb    m/z 219.8452     0   1E+4   1.5   2  RT (min)       lcqtof_hilic_neg_3_066_57    m/z 219.8452     0   1   2   3E+4   3.5   4  RT (min)      lcqtof_hilic_neg_4_024_40    m/z 219.8452     0   1   2   3E+4   3.5   4  RT (min)      lcqtof_hilic_neg_4_024_40    m/z 219.8453     0   1   2   3   4E+3   9   9.5  RT (min)      lcqtof_hilic_neg_3_077_58    m/z 219.8453     0   1   2   3   4E+3   9   9.5  RT (min)      lcqtof_hilic_neg_3_078_58    m/z 219.8453     0   1   2   3E+4   3.5   4  RT (min)       lcqtof_hilic_neg_4_020_42    m/z 219.8960     0   1   2E+4   7.5   8  RT (min)      lcqtof_hilic_neg_1_000_aa    m/z 220.0621     0   1E+4   0.5   1  RT (min)      lcqtof_hilic_neg_4_055_32    m/z 220.0699     0   1   2E+4   2   2.5  RT (min)      lcqtof_hilic_neg_4_050_02    m/z 220.2197     0   1   2E+4   0.5   1  RT (min)      lcqtof_hilic_neg_4_027_19    m/z 220.8082     0   1E+4   2.5   3  RT (min)      lcqtof_hilic_neg_4_050_02    m/z 220.8456     0   1   2E+4   4   4.5  RT (min)      lcqtof_hilic_neg_3_067_57    m/z 220.8798     0   2   4   6   8E+4   8   8.5  RT (min)      lcqtof_hilic_neg_1_000_aa    m/z 220.8801     0   2   4   6   8E+4   7.5   8  RT (min)      lcqtof_hilic_neg_1_000_aa    m/z 221.0654     0   2   4   6   8E+3   2   2.5  RT (min)      lcqtof_hilic_neg_4_052_13    m/z 221.0655     0   1   2E+4   3.5   4  RT (min)      lcqtof_hilic_neg_4_050_02    m/z 221.0656     0   1   2   3E+4   4.5   5  RT (min)      lcqtof_hilic_neg_4_035_33    m/z 221.0659     0   1E+4   4.5   5  RT (min)      lcqtof_hilic_neg_4_028_36    m/z 221.0660     0   1   2E+4   4.5   5  RT (min)      lcqtof_hilic_neg_4_034_31    m/z 221.0661     0   2   4   6   8E+4   4   4.5  RT (min)       lcqtof_hilic_neg_4_055_32    m/z 221.1068     0   1E+4   4   4.5  RT (min)      lcqtof_hilic_neg_4_004_47    m/z 221.8418     0   1   2E+4   3.5   4  RT (min)      lcqtof_hilic_neg_4_030_44    m/z 221.8418     0   1   0   0.5  RT (min)       lcqtof_hilic_neg_3_066_57    m/z 221.8420     0   1E+4   1.5   2  RT (min)      lcqtof_hilic_neg_3_066_57    m/z 221.8420     0   1E+4   16.5   17  RT (min)       lcqtof_hilic_neg_3_066_57    m/z 221.8421     0   1E+4   1   1.5  RT (min)      lcqtof_hilic_neg_1_000_bb    m/z 221.8422     0   1E+4   16   17  RT (min)       lcqtof_hilic_neg_3_068_57    m/z 221.8422     0   1E+5   3.5   4  RT (min)       lcqtof_hilic_neg_3_067_57    m/z 221.8422     0   1   2   3   4E+4   3.5   4  RT (min)      lcqtof_hilic_neg_1_000_cc    m/z 221.8423     0   1E+4   0.5   1  RT (min)       lcqtof_hilic_neg_3_066_57    m/z 221.8426     0   1   2   3E+3   9   9.5  RT (min)      lcqtof_hilic_neg_3_075_58    m/z 221.9024     0   1   2E+3   4.5   5  RT (min)      lcqtof_hilic_neg_1_004_14    m/z 222.8074     0   1E+4   2.5   3  RT (min)      lcqtof_hilic_neg_4_050_02    m/z 222.8421     0   1E+4   4   4.5  RT (min)      lcqtof_hilic_neg_3_067_57    m/z 223.0264     0   2   4   6E+4   1.5   2  RT (min)      lcqtof_hilic_neg_4_000_aa    m/z 223.0268     0   2   4   6   8E+4   1.5   2  RT (min)      lcqtof_hilic_neg_4_000_aa    m/z 223.0272     0   2   4   6   8E+4   1   1.5  RT (min)      lcqtof_hilic_neg_4_000_aa    m/z 223.0274     0   2   4   6   8E+4   1.5   2  RT (min)      lcqtof_hilic_neg_4_000_aa    m/z 223.0274     0   2   4   6   8E+4   1.5   2  RT (min)      lcqtof_hilic_neg_4_000_aa    m/z 223.0280     0   2   4   6   8E+4   1   1.5  RT (min)      lcqtof_hilic_neg_4_000_aa    m/z 223.0282     0   2   4   6   8E+4   1   1.5  RT (min)      lcqtof_hilic_neg_4_000_aa    m/z 223.0282     0   2   4   6   8E+4   1.5   2  RT (min)      lcqtof_hilic_neg_4_000_aa    m/z 223.0723     0   1E+4   2   2.5  RT (min)      lcqtof_hilic_neg_4_053_14    m/z 223.0818     0   1E+4   3.5   4  RT (min)      lcqtof_hilic_neg_4_057_49    m/z 223.0836     0   1E+4   6   6.5  RT (min)      lcqtof_hilic_neg_4_008_24    m/z 223.1069     0   1E+4   15   15.5  RT (min)      lcqtof_hilic_neg_4_000_aa    m/z 223.1106     0   1E+4   1   1.5  RT (min)      lcqtof_hilic_neg_4_008_24    m/z 223.7978     0   2   4   6E+3   4.5   5  RT (min)       lcqtof_hilic_neg_4_018_27    m/z 223.7980     0   1   2E+4   1   1.5  RT (min)      lcqtof_hilic_neg_1_000_aa    m/z 223.7980     0   1   2   3E+4   3   3.5  RT (min)       lcqtof_hilic_neg_1_000_aa    m/z 223.7982     0   1   2   3   4E+3   4   4.5  RT (min)       lcqtof_hilic_neg_4_040_05    m/z 223.7982     0   1   2E+2   2.5   3  RT (min)      lcqtof_hilic_neg_4_050_02    m/z 223.9848     0   1   2   3E+3   2   2.5  RT (min)       lcqtof_hilic_neg_4_077_58    m/z 224.0284     0   1E+4   1   1.5  RT (min)      lcqtof_hilic_neg_4_000_aa    m/z 224.0289     0   1E+4   1   1.5  RT (min)      lcqtof_hilic_neg_4_000_aa    m/z 224.0613     0   2   4   6E+4   2   2.5  RT (min)       lcqtof_hilic_neg_4_060_13    m/z 224.0639     0   2   4   6E+3   0.5   1  RT (min)      lcqtof_hilic_neg_4_013_20    m/z 224.0917     0   1E+4   15   15.5  RT (min)      lcqtof_hilic_neg_4_000_aa    m/z 224.1255     0   1E+4   4   4.5  RT (min)      lcqtof_hilic_neg_4_035_33    m/z 224.1656     0   1E+4   0.5   1  RT (min)      lcqtof_hilic_neg_2_000_dd    m/z 224.8073     0   1   2E+4   2.5   3  RT (min)      lcqtof_hilic_neg_4_050_02    m/z 225.0076     0   1   2E+4   1   1.5  RT (min)      lcqtof_hilic_neg_4_040_05    m/z 225.0076     0   2   4   6E+3   1   1.5  RT (min)       lcqtof_hilic_neg_4_000_aa    m/z 225.0573     0   2   4   6   8E+3   0.5   1  RT (min)      lcqtof_hilic_neg_4_047_06    m/z 225.0575     0   2   4   6E+3   0.5   1  RT (min)      lcqtof_hilic_neg_4_036_11    m/z 225.0676     0   1   2E+4   3   3.5  RT (min)       lcqtof_hilic_neg_4_050_02    m/z 225.0870     0   1E+4   1.5   2  RT (min)       lcqtof_hilic_neg_4_048_23    m/z 226.0181     0   1E+4   0.5   1  RT (min)       lcqtof_hilic_neg_1_062_17    m/z 226.0724     0   2   4   6E+3   3   3.5  RT (min)       lcqtof_hilic_neg_4_008_24    m/z 226.0736     0   1E+4   2.5   3  RT (min)       lcqtof_hilic_neg_4_055_32    m/z 226.8075     0   1E+4   2.5   3  RT (min)      lcqtof_hilic_neg_4_050_02    m/z 226.8702     0   1E+4   8   10  RT (min)       lcqtof_hilic_neg_4_015_09    m/z 226.8706     0   1E+4   10   12.5   15  RT (min)       lcqtof_hilic_neg_4_013_20    m/z 226.9968     0   2   4E+3   0.5   1  RT (min)      lcqtof_hilic_neg_4_040_05    m/z 227.1286     0   1E+4   0.5   1  RT (min)      lcqtof_hilic_neg_4_050_02    m/z 227.1404     0   1E+4   4   4.5  RT (min)      lcqtof_hilic_neg_4_048_23    m/z 227.9597     0   1E+4   2.5   3  RT (min)      lcqtof_hilic_neg_4_050_02    m/z 227.9601     0   1   2   3E+4   2.5   3  RT (min)       lcqtof_hilic_neg_4_088_59    m/z 227.9662     0   1E+4   1   1.5  RT (min)      lcqtof_hilic_neg_4_022_34    m/z 228.0510     0   1E+4   4   4.5  RT (min)      lcqtof_hilic_neg_4_028_36    m/z 228.0630     0   2   4   6E+3   0.5   1  RT (min)      lcqtof_hilic_neg_4_050_02    m/z 228.1589     0   1E+4   15   15.5  RT (min)      lcqtof_hilic_neg_4_000_aa    m/z 228.1602     0   1E+4   1.5   2  RT (min)       lcqtof_hilic_neg_1_000_aa    m/z 228.9500     0   1   2E+4   1.5   2  RT (min)      lcqtof_hilic_neg_4_022_34    m/z 228.9501     0   1   2E+4   1.5   2  RT (min)      lcqtof_hilic_neg_4_022_34    m/z 228.9502     0   1   2   3E+4   1   1.5  RT (min)      lcqtof_hilic_neg_4_022_34    m/z 228.9502     0   1   2   3E+4   1   1.5  RT (min)      lcqtof_hilic_neg_4_022_34    m/z 228.9504     0   1   2   3E+4   1   1.5  RT (min)      lcqtof_hilic_neg_4_022_34    m/z 228.9596     0   1E+3   9   9.5  RT (min)       lcqtof_hilic_neg_3_029_29    m/z 228.9613     0   1E+4   3   3.5  RT (min)      lcqtof_hilic_neg_2_020_23    m/z 228.9613     0   1   2   3   4E+3   13.5   14   14.5  RT (min)       lcqtof_hilic_neg_4_038_10    m/z 228.9680     0   2   4   6E+3   14   14.5  RT (min)       lcqtof_hilic_neg_4_013_20    m/z 228.9685     0   2   4   6E+3   13   14  RT (min)       lcqtof_hilic_neg_4_013_20    m/z 228.9687     0   2   4   6E+3   14.5   15  RT (min)       lcqtof_hilic_neg_4_015_09    m/z 228.9688     0   2   4E+3   13.5   14  RT (min)       lcqtof_hilic_neg_4_014_46    m/z 228.9688     0   2   4   6E+3   14.5   15  RT (min)       lcqtof_hilic_neg_4_021_21    m/z 228.9690     0   2   4E+3   13.5   14  RT (min)       lcqtof_hilic_neg_4_015_09    m/z 228.9691     0   2   4   6E+3   13.5   14   14.5  RT (min)       lcqtof_hilic_neg_4_013_20    m/z 228.9692     0   2   4   6E+3   13   13.5   14  RT (min)       lcqtof_hilic_neg_4_013_20    m/z 228.9693     0   2   4   6E+3   15   15.5  RT (min)       lcqtof_hilic_neg_4_008_24    m/z 228.9697     0   2   4E+3   12.5   13  RT (min)       lcqtof_hilic_neg_4_021_21    m/z 228.9701     0   2   4E+3   12.5   13  RT (min)       lcqtof_hilic_neg_4_021_21    m/z 228.9706     0   1   2   3   4E+3   12  RT (min)       lcqtof_hilic_neg_4_009_50    m/z 229.0172     0   1   2E+4   0.5   1  RT (min)      lcqtof_hilic_neg_4_053_14    m/z 229.0387     0   1E+4   2   2.5  RT (min)       lcqtof_hilic_neg_4_040_05    m/z 229.1486     0   1E+4   0.5   1  RT (min)      lcqtof_hilic_neg_3_000_aa    m/z 229.1565     0   1   2E+4   1   1.5  RT (min)      lcqtof_hilic_neg_1_000_dd    m/z 229.8605     0   1   2E+4   0.5   1  RT (min)      lcqtof_hilic_neg_4_055_32    m/z 229.9519     0   1   2E+4   1   1.5  RT (min)       lcqtof_hilic_neg_3_066_57    m/z 229.9525     0   1   2   3   4E+3   1   1.5  RT (min)      lcqtof_hilic_neg_1_023_34    m/z 229.9636     0   1E+4   1   1.5  RT (min)      lcqtof_hilic_neg_4_022_34    m/z 230.1249     0   1E+4   6   6.5  RT (min)      lcqtof_hilic_neg_4_008_24    m/z 230.8815     0   1   2   3E+4   4   4.5  RT (min)       lcqtof_hilic_neg_3_067_57    m/z 230.8817     0   1   2   3E+4   4   4.5  RT (min)       lcqtof_hilic_neg_3_067_57    m/z 230.9481     0   1   2   3E+4   1   1.5  RT (min)      lcqtof_hilic_neg_4_022_34    m/z 230.9483     0   1   2E+4   1.5   2  RT (min)      lcqtof_hilic_neg_4_022_34    m/z 230.9484     0   1   2   3E+4   1   1.5  RT (min)      lcqtof_hilic_neg_4_022_34    m/z 230.9492     0   1   2E+4   1.5   2  RT (min)      lcqtof_hilic_neg_4_022_34    m/z 230.9493     0   1   2E+4   1.5   2  RT (min)      lcqtof_hilic_neg_4_022_34    m/z 230.9567     0   1E+3   13   14  RT (min)       lcqtof_hilic_neg_4_009_50    m/z 230.9615     0   1   2   3E+3   14.5   15  RT (min)       lcqtof_hilic_neg_4_000_dd    m/z 230.9620     0   1   2   3E+3   14.5   15  RT (min)       lcqtof_hilic_neg_4_003_16    m/z 230.9625     0   1   2   3E+3   14   14.5  RT (min)       lcqtof_hilic_neg_4_015_09    m/z 230.9631     0   1   2   3E+3   15   15.5  RT (min)       lcqtof_hilic_neg_4_008_24    m/z 230.9968     0   1   2   3E+4   1   1.5  RT (min)      lcqtof_hilic_neg_4_039_18    m/z 231.0175     0   2   4   6E+4   5   5.5  RT (min)       lcqtof_hilic_neg_4_048_23    m/z 231.0177     0   2   4E+4   4   4.5  RT (min)       lcqtof_hilic_neg_4_048_23    m/z 231.0227     0   1E+4   6   6.5  RT (min)      lcqtof_hilic_neg_4_012_07    m/z 231.0232     0   1E+4   6   6.5  RT (min)       lcqtof_hilic_neg_4_012_07    m/z 231.0441     0   1E+4   2   2.5  RT (min)      lcqtof_hilic_neg_4_060_13    m/z 231.0662     0   2   4   6E+3   2   2.5  RT (min)      lcqtof_hilic_neg_4_016_37    m/z 231.9307     0   1   2E+4   10   10.5  RT (min)      lcqtof_hilic_neg_4_014_46    m/z 231.9312     0   1   2   3E+4   4.5   5  RT (min)      lcqtof_hilic_neg_1_000_bb    m/z 231.9312     0   1E+4   7   7.5  RT (min)      lcqtof_hilic_neg_1_003_37    m/z 231.9312     0   1E+4   7   7.5  RT (min)       lcqtof_hilic_neg_1_003_37    m/z 231.9313     0   2   4   6   8E+3   1   1.5  RT (min)      lcqtof_hilic_neg_3_109_62    m/z 231.9313     0   1   2   3E+3   8   8.5  RT (min)       lcqtof_hilic_neg_1_003_37    m/z 231.9314     0   1E+4   7   7.5  RT (min)      lcqtof_hilic_neg_1_003_37    m/z 231.9314     0   1   2   3E+4   4   4.5  RT (min)      lcqtof_hilic_neg_1_000_bb    m/z 231.9314     0   2   4E+3   0.5   1  RT (min)      lcqtof_hilic_neg_4_017_51    m/z 231.9315     0   1E+4   2   3   4  RT (min)       lcqtof_hilic_neg_3_066_57    m/z 231.9316     0   1E+4   6.5   7  RT (min)      lcqtof_hilic_neg_1_014_31    m/z 231.9317     0   1E+4   6.5   7  RT (min)      lcqtof_hilic_neg_1_009_50    m/z 231.9317     0   1   2E+4   6.5   7  RT (min)      lcqtof_hilic_neg_3_066_57    m/z 231.9317     0   1   2   3E+4   4   4.5  RT (min)      lcqtof_hilic_neg_1_000_bb    m/z 231.9318     0   1   2   3E+4   4.5   5  RT (min)      lcqtof_hilic_neg_1_000_bb    m/z 231.9320     0   1   2   3   4E+3   1   1.5  RT (min)      lcqtof_hilic_neg_1_013_38    m/z 231.9321     0   2   4E+3   4   4.5  RT (min)      lcqtof_hilic_neg_4_014_46    m/z 231.9323     0   2   4E+3   0.5   1  RT (min)      lcqtof_hilic_neg_2_024_10    m/z 231.9330     0   1   2   3E+3   3.5   4  RT (min)      lcqtof_hilic_neg_4_014_46    m/z 231.9338     0   1E+3   17   17.5  RT (min)       lcqtof_hilic_neg_4_037_39    m/z 231.9342     0   1E+3   16.5   17  RT (min)      lcqtof_hilic_neg_4_038_10    m/z 231.9443     0   1E+4   3.5   4  RT (min)       lcqtof_hilic_neg_4_048_23    m/z 231.9511     0   2   4E+4   0.5   1  RT (min)       lcqtof_hilic_neg_4_100_61    m/z 232.0209     0   1E+4   4   4.5  RT (min)      lcqtof_hilic_neg_4_002_25    m/z 232.0694     0   2   4E+3   0.5   1  RT (min)      lcqtof_hilic_neg_3_003_20    m/z 232.9246     0   1   2   3E+4   4.5   5  RT (min)       lcqtof_hilic_neg_1_000_bb    m/z 232.9247     0   1   2E+4   5   5.5  RT (min)      lcqtof_hilic_neg_1_000_dd    m/z 232.9250     0   1   2   3E+3   6   6.5  RT (min)      lcqtof_hilic_neg_1_010_15    m/z 232.9258     0   1   2   3E+3   6   6.5  RT (min)      lcqtof_hilic_neg_4_001_49    m/z 232.9763     0   1E+4   2.5   3  RT (min)      lcqtof_hilic_neg_4_047_06    m/z 232.9765     0   1   2   3E+4   2.5   3  RT (min)       lcqtof_hilic_neg_4_090_60    m/z 232.9767     0   1E+4   6   6.5  RT (min)      lcqtof_hilic_neg_1_037_24    m/z 233.0147     0   1E+4   4   4.5  RT (min)      lcqtof_hilic_neg_4_002_25    m/z 233.0570     0   2   4E+4   4   4.5  RT (min)      lcqtof_hilic_neg_4_055_32    m/z 233.0660     0   1E+4   1.5   2  RT (min)       lcqtof_hilic_neg_4_050_02    m/z 233.0665     0   1   2E+4   1   1.5  RT (min)      lcqtof_hilic_neg_4_050_02    m/z 233.0668     0   1E+4   1.5   2  RT (min)      lcqtof_hilic_neg_4_052_13    m/z 233.0961     0   1E+4   0.5   1  RT (min)      lcqtof_hilic_neg_4_008_24    m/z 234.0040     0   1E+4   5.5   6  RT (min)      lcqtof_hilic_neg_1_000_aa    m/z 234.0040     0   1E+4   0   0.5  RT (min)      lcqtof_hilic_neg_1_000_bb    m/z 234.0254     0   1   2   3E+4   2.5   3  RT (min)      lcqtof_hilic_neg_4_004_47    m/z 234.0254     0   1E+4   1   1.5  RT (min)      lcqtof_hilic_neg_4_034_31    m/z 234.0671     0   1   2E+4   2   2.5  RT (min)      lcqtof_hilic_neg_4_050_02    m/z 234.8196     0   1   2   3E+3   0   0.5  RT (min)       lcqtof_hilic_neg_1_063_13    m/z 234.8196     0   2   4E+3   0.5   1  RT (min)       lcqtof_hilic_neg_2_024_10    m/z 234.8198     0   1   2   3E+3   0   0.5  RT (min)       lcqtof_hilic_neg_1_053_19    m/z 235.0300     0   1E+4   1   1.5  RT (min)      lcqtof_hilic_neg_4_055_32    m/z 235.0310     0   2   4   6E+4   4   4.5  RT (min)      lcqtof_hilic_neg_4_008_24    m/z 235.0310     0   2   4   6E+4   4   4.5  RT (min)      lcqtof_hilic_neg_4_008_24    m/z 235.0311     0   1E+4   5   5.5  RT (min)      lcqtof_hilic_neg_4_008_24    m/z 235.0631     0   2   4   6   8E+4   2   2.5  RT (min)      lcqtof_hilic_neg_4_050_02    m/z 235.1210     0   1   2E+4   1.5   2  RT (min)       lcqtof_hilic_neg_4_055_32    m/z 235.1214     0   2   4   6   8E+3   1.5   2  RT (min)      lcqtof_hilic_neg_1_005_32    m/z 235.1215     0   1E+4   3.5   4  RT (min)      lcqtof_hilic_neg_4_008_24    m/z 236.0194     0   1E+4   0   0.5  RT (min)      lcqtof_hilic_neg_1_000_bb    m/z 236.0342     0   1   2   3E+5   4   4.5  RT (min)       lcqtof_hilic_neg_4_090_60    m/z 236.0394     0   1E+4   6   6.5  RT (min)       lcqtof_hilic_neg_4_090_60    m/z 236.0437     0   1   2E+5   6.5   7  RT (min)       lcqtof_hilic_neg_4_008_24    m/z 236.0439     0   1   2E+5   6.5   7  RT (min)       lcqtof_hilic_neg_4_008_24    m/z 236.0439     0   1   2E+5   6.5   7  RT (min)      lcqtof_hilic_neg_4_008_24    m/z 236.0542     0   1   2   3E+4   2   2.5  RT (min)      lcqtof_hilic_neg_4_055_32    m/z 236.0869     0   1   2   3E+3   2   2.5  RT (min)      lcqtof_hilic_neg_4_060_13    m/z 236.1042     0   1   2E+4   15   15.5  RT (min)      lcqtof_hilic_neg_4_000_aa    m/z 236.1130     0   1   2E+4   4   4.5  RT (min)      lcqtof_hilic_neg_4_008_24    m/z 236.1909     0   2   4   6E+4   4   4.5  RT (min)      lcqtof_hilic_neg_4_008_24    m/z 236.3095     0   1E+4   4   4.5  RT (min)      lcqtof_hilic_neg_4_008_24    m/z 236.9113     0   1E+5   8   8.5  RT (min)      lcqtof_hilic_neg_1_000_aa    m/z 236.9530     0   1   2   3   4E+3   5   5.5  RT (min)       lcqtof_hilic_neg_2_000_bb    m/z 236.9532     0   1   2E+3   6   6.5  RT (min)       lcqtof_hilic_neg_1_054_01    m/z 236.9880     0   1E+4   3   3.5  RT (min)       lcqtof_hilic_neg_1_032_18    m/z 236.9881     0   1E+4   3   3.5  RT (min)      lcqtof_hilic_neg_1_032_18    m/z 237.0249     0   1E+4   4   4.5  RT (min)      lcqtof_hilic_neg_4_086_59    m/z 237.0364     0   1   2   3E+4   4   4.5  RT (min)      lcqtof_hilic_neg_4_008_24    m/z 237.0610     0   1E+4   6.5   7  RT (min)      lcqtof_hilic_neg_4_031_38    m/z 237.0780     0   1   2E+4   2   2.5  RT (min)      lcqtof_hilic_neg_4_019_03    m/z 237.0787     0   1E+4   2   2.5  RT (min)      lcqtof_hilic_neg_4_003_16    m/z 237.0817     0   2   4E+4   2   2.5  RT (min)       lcqtof_hilic_neg_4_060_13    m/z 237.0910     0   2   4   6   8E+2   2   2.5  RT (min)      lcqtof_hilic_neg_4_050_02    m/z 237.1028     0   1E+4   2   2.5  RT (min)      lcqtof_hilic_neg_2_047_32    m/z 237.9064     0   2   4   6   8E+4   8.5   9  RT (min)      lcqtof_hilic_neg_1_000_aa    m/z 238.0315     0   1   2E+4   4   4.5  RT (min)      lcqtof_hilic_neg_4_090_60    m/z 238.0322     0   1   2   3E+4   5   5.5  RT (min)      lcqtof_hilic_neg_4_008_24    m/z 238.0388     0   2   4   6E+3   3.5   4  RT (min)      lcqtof_hilic_neg_2_005_13    m/z 238.0391     0   1E+4   3   3.5  RT (min)      lcqtof_hilic_neg_4_050_02    m/z 238.0464     0   1E+4   0.5   1  RT (min)      lcqtof_hilic_neg_4_002_25    m/z 238.0768     0   1E+4   1.5   2  RT (min)      lcqtof_hilic_neg_4_019_03    m/z 238.0793     0   1E+4   1.5   2  RT (min)      lcqtof_hilic_neg_4_019_03    m/z 238.2338     0   1E+4   2   2.5  RT (min)      lcqtof_hilic_neg_4_023_15    m/z 238.5746     0   1   2E+4   1   1.5  RT (min)      lcqtof_hilic_neg_4_050_02    m/z 238.8908     0   2   4E+6   8   8.5  RT (min)       lcqtof_hilic_neg_1_000_aa    m/z 238.8909     0   1E+4   12   12.5  RT (min)       lcqtof_hilic_neg_4_110_62    m/z 238.8909     0   1   2   3   4E+6   8.5   9  RT (min)       lcqtof_hilic_neg_1_000_aa    m/z 238.8909     0   1E+4   12   12.5  RT (min)       lcqtof_hilic_neg_4_110_62    m/z 238.8910     0   1   2   3   4E+5   9   9.5  RT (min)       lcqtof_hilic_neg_1_000_aa    m/z 238.8910     0   2   4E+5   9   9.5  RT (min)       lcqtof_hilic_neg_1_000_aa    m/z 238.8910     0   1   2   3E+6   8.5   9  RT (min)       lcqtof_hilic_neg_1_000_aa    m/z 238.8910     0   1   2   3   4E+5   9   9.5  RT (min)       lcqtof_hilic_neg_1_000_aa    m/z 238.8910     0   2   4E+3   10   10.5  RT (min)       lcqtof_hilic_neg_4_095_60    m/z 238.8910     0   1E+4   13   13.5  RT (min)       lcqtof_hilic_neg_4_109_62    m/z 238.8910     0   2   4   6E+4   9.5   10  RT (min)       lcqtof_hilic_neg_2_000_aa    m/z 238.8911     0   1   2   3E+5   9.5   10  RT (min)       lcqtof_hilic_neg_1_000_aa    m/z 238.8911     0   1   2   3   4E+3   7   7.5  RT (min)      lcqtof_hilic_neg_4_044_08    m/z 238.8911     0   1E+4   13.5   14  RT (min)       lcqtof_hilic_neg_4_100_61    m/z 238.8911     0   1E+4   13   13.5  RT (min)       lcqtof_hilic_neg_4_012_07    m/z 238.8911     0   2   4E+3   10   10.5  RT (min)       lcqtof_hilic_neg_4_105_61    m/z 238.8911     0   2   4E+3   10.5   11  RT (min)       lcqtof_hilic_neg_4_113_62    m/z 238.8912     0   1E+4   13   13.5  RT (min)       lcqtof_hilic_neg_4_019_03    m/z 238.8912     0   2   4   6E+3   13.5   14  RT (min)       lcqtof_hilic_neg_4_100_61    m/z 238.8920     0   2   4   6E+3   6.5   7  RT (min)      lcqtof_hilic_neg_4_012_07    m/z 238.8920     0   2   4   6E+3   7   7.5  RT (min)       lcqtof_hilic_neg_4_001_49    m/z 238.9113     0   1   2   3E+3   7.5   8  RT (min)      lcqtof_hilic_neg_2_062_17    m/z 238.9128     0   2   4   6   8E+3   1   1.5  RT (min)      lcqtof_hilic_neg_4_058_39    m/z 238.9132     0   1   2   3E+3   3.5   4  RT (min)       lcqtof_hilic_neg_3_104_61    m/z 238.9132     0   1   2   3E+3   2   2.5  RT (min)      lcqtof_hilic_neg_1_004_14    m/z 238.9312     0   1   2   3E+3   5   5.5  RT (min)      lcqtof_hilic_neg_4_060_13    m/z 238.9315     0   1   2   3E+3   5.5   6  RT (min)      lcqtof_hilic_neg_4_005_29    m/z 238.9316     0   1   2   3E+3   5.5   6  RT (min)      lcqtof_hilic_neg_4_029_17    m/z 238.9693     0   2   4   6   8E+4   8   8.5  RT (min)      lcqtof_hilic_neg_1_000_aa    m/z 238.9702     0   1   2   3   4E+4   8   8.5  RT (min)      lcqtof_hilic_neg_2_000_aa    m/z 239.0144     0   1   2   3E+4   8   8.5  RT (min)      lcqtof_hilic_neg_1_000_aa    m/z 239.0476     0   1E+5   8   8.5  RT (min)      lcqtof_hilic_neg_1_000_aa    m/z 239.0482     0   2   4   6E+4   8   8.5  RT (min)      lcqtof_hilic_neg_2_000_aa    m/z 239.0556     0   1E+4   4   4.5  RT (min)      lcqtof_hilic_neg_4_050_02    m/z 239.0620     0   1   2E+5   1   1.5  RT (min)      lcqtof_hilic_neg_4_000_aa    m/z 239.0651     0   1   2E+5   1   1.5  RT (min)      lcqtof_hilic_neg_4_000_aa    m/z 239.0670     0   2   4E+1   1.5   2  RT (min)       lcqtof_hilic_neg_4_000_aa    m/z 239.0709     0   2   4   6   8E+3   1   1.5  RT (min)      lcqtof_hilic_neg_4_000_aa    m/z 239.0748     0   2   4E+4   2   2.5  RT (min)      lcqtof_hilic_neg_4_062_17    m/z 239.0802     0   2   4E+4   8   8.5  RT (min)      lcqtof_hilic_neg_1_000_aa    m/z 239.0806     0   2   4   6E+3   2.5   3  RT (min)      lcqtof_hilic_neg_1_054_01    m/z 239.0829     0   2   4E+3   3   3.5  RT (min)      lcqtof_hilic_neg_1_004_14    m/z 239.0834     0   2   4E+3   2.5   3  RT (min)      lcqtof_hilic_neg_2_031_01    m/z 239.0994     0   1E+4   4   4.5  RT (min)      lcqtof_hilic_neg_4_008_24    m/z 239.1214     0   2   4   6E+4   8   8.5  RT (min)      lcqtof_hilic_neg_1_000_aa    m/z 239.1680     0   1   2   3E+4   8   8.5  RT (min)      lcqtof_hilic_neg_2_000_aa    m/z 239.1698     0   2   4   6   8E+4   8   8.5  RT (min)      lcqtof_hilic_neg_1_000_aa    m/z 239.2560     0   2   4   6E+4   8   8.5  RT (min)      lcqtof_hilic_neg_1_000_aa    m/z 239.2565     0   1E+4   8   8.5  RT (min)      lcqtof_hilic_neg_2_000_aa    m/z 239.5841     0   1E+4   1   1.5  RT (min)      lcqtof_hilic_neg_4_050_02    m/z 239.9007     0   1   2   3   4E+4   8   8.5  RT (min)       lcqtof_hilic_neg_1_000_aa    m/z 239.9515     0   1E+4   8   8.5  RT (min)      lcqtof_hilic_neg_1_000_aa    m/z 240.0104     0   1   2   3E+4   8   8.5  RT (min)      lcqtof_hilic_neg_1_000_aa    m/z 240.0551     0   2   4E+4   1   1.5  RT (min)      lcqtof_hilic_neg_4_000_aa    m/z 240.0608     0   2   4E+4   1   1.5  RT (min)      lcqtof_hilic_neg_4_000_aa    m/z 240.0610     0   2   4E+4   1   1.5  RT (min)      lcqtof_hilic_neg_4_000_aa    m/z 240.0893     0   1E+4   1.5   2  RT (min)      lcqtof_hilic_neg_1_005_32    m/z 240.0974     0   1E+4   2   2.5  RT (min)      lcqtof_hilic_neg_4_054_48    m/z 240.0983     0   1   2   3   4E+3   2.5   3  RT (min)      lcqtof_hilic_neg_1_033_53    m/z 240.0985     0   1E+4   2.5   3  RT (min)      lcqtof_hilic_neg_4_054_48    m/z 240.0986     0   1E+4   2.5   3  RT (min)      lcqtof_hilic_neg_4_054_48    m/z 240.1007     0   1E+4   4   4.5  RT (min)      lcqtof_hilic_neg_4_055_32    m/z 240.8666     0   1E+4   0.5   1  RT (min)      lcqtof_hilic_neg_4_050_02    m/z 240.8702     0   1E+4   4.5   5  RT (min)      lcqtof_hilic_neg_4_050_02    m/z 240.8704     0   1   2E+3   15   15.5  RT (min)       lcqtof_hilic_neg_1_000_aa    m/z 240.8706     0   1E+3   15.5   16  RT (min)       lcqtof_hilic_neg_3_066_57    m/z 240.8955     0   1E+5   8   8.5  RT (min)      lcqtof_hilic_neg_1_000_aa    m/z 241.0010     0   2   4   6E+3   3.5   4  RT (min)       lcqtof_hilic_neg_4_091_60    m/z 241.0496     0   1E+4   2.5   3  RT (min)      lcqtof_hilic_neg_4_082_59    m/z 241.0917     0   1E+4   6   6.5  RT (min)       lcqtof_hilic_neg_4_034_31    m/z 241.8987     0   1   2E+4   1   1.5  RT (min)      lcqtof_hilic_neg_4_051_43    m/z 241.9916     0   1   2E+5   0.5   1  RT (min)      lcqtof_hilic_neg_4_050_02    m/z 241.9918     0   1   2E+5   1   1.5  RT (min)      lcqtof_hilic_neg_4_050_02    m/z 241.9955     0   1   2E+4   2   2.5  RT (min)      lcqtof_hilic_neg_4_050_02    m/z 242.0461     0   1   2E+4   2.5   3  RT (min)       lcqtof_hilic_neg_4_089_59    m/z 242.0596     0   1E+4   0.5   1  RT (min)      lcqtof_hilic_neg_1_062_17    m/z 242.0674     0   1E+4   2.5   3  RT (min)      lcqtof_hilic_neg_4_047_06    m/z 242.0740     0   1   2   3   4E+3   4   4.5  RT (min)      lcqtof_hilic_neg_4_010_28    m/z 242.0783     0   1   2   3E+3   3.5   4  RT (min)      lcqtof_hilic_neg_4_011_35    m/z 242.0808     0   2   4E+3   1   1.5  RT (min)      lcqtof_hilic_neg_4_018_27    m/z 242.0968     0   1E+4   2   2.5  RT (min)       lcqtof_hilic_neg_4_008_24    m/z 242.1752     0   1   2   3   4E+4   15   15.5  RT (min)      lcqtof_hilic_neg_4_000_aa    m/z 242.1760     0   1   2   3   4E+3   0   0.5  RT (min)      lcqtof_hilic_neg_1_010_15    m/z 242.1760     0   1   2   3   4E+3   0   0.5  RT (min)       lcqtof_hilic_neg_1_010_15    m/z 242.1761     0   2   4   6   8E+3   4   4.5  RT (min)       lcqtof_hilic_neg_4_047_06    m/z 242.1761     0   2   4   6   8E+3   0   0.5  RT (min)       lcqtof_hilic_neg_1_002_39    m/z 242.1761     0   1E+4   0   0.5  RT (min)      lcqtof_hilic_neg_1_008_13    m/z 242.1762     0   1   2   3E+3   2   2.5  RT (min)       lcqtof_hilic_neg_1_002_39    m/z 242.1762     0   1E+4   0.5   1  RT (min)       lcqtof_hilic_neg_1_010_15    m/z 242.1762     0   1   2   3E+3   2   3  RT (min)       lcqtof_hilic_neg_1_006_22    m/z 242.1762     0   1   2   3E+3   2.5   3  RT (min)       lcqtof_hilic_neg_1_004_14    m/z 242.1762     0   1   2E+4   0.5   1  RT (min)      lcqtof_hilic_neg_1_002_39    m/z 242.1762     0   1   2   3   4E+3   0   0.5  RT (min)       lcqtof_hilic_neg_1_002_39    m/z 242.1762     0   1E+4   2   2.5  RT (min)       lcqtof_hilic_neg_1_004_14    m/z 242.1763     0   1   2E+3   2.5   3  RT (min)       lcqtof_hilic_neg_4_105_61    m/z 242.1763     0   1   2   3   4E+3   0   0.5  RT (min)       lcqtof_hilic_neg_1_002_39    m/z 242.1764     0   1   2   3E+4   1.5   2  RT (min)       lcqtof_hilic_neg_1_000_aa    m/z 242.2550     0   1   2   3   4E+4   1   1.5  RT (min)      lcqtof_hilic_neg_4_026_04    m/z 242.3342     0   2   4   6E+4   1   1.5  RT (min)      lcqtof_hilic_neg_4_026_04    m/z 242.4970     0   1   2   3   4E+4   1   1.5  RT (min)      lcqtof_hilic_neg_4_050_02    m/z 242.9429     0   2   4   6E+3   8.5   9  RT (min)       lcqtof_hilic_neg_4_046_41    m/z 242.9430     0   2   4   6E+3   8.5   9  RT (min)       lcqtof_hilic_neg_4_021_21    m/z 242.9902     0   2   4   6   8E+4   1   1.5  RT (min)      lcqtof_hilic_neg_4_050_02    m/z 243.0175     0   1   2E+4   2   2.5  RT (min)       lcqtof_hilic_neg_4_052_13    m/z 243.8626     0   1   2E+4   6.5   7  RT (min)      lcqtof_hilic_neg_1_000_aa    m/z 243.8628     0   1   2E+4   6.5   7  RT (min)       lcqtof_hilic_neg_1_000_aa    m/z 243.8630     0   1   2E+4   6   6.5  RT (min)      lcqtof_hilic_neg_1_000_aa    m/z 243.8636     0   1   2E+4   6.5   7  RT (min)      lcqtof_hilic_neg_1_000_aa    m/z 243.8985     0   1   2E+4   0.5   1  RT (min)      lcqtof_hilic_neg_4_020_42    m/z 243.8986     0   1   2E+4   1   1.5  RT (min)      lcqtof_hilic_neg_4_030_44    m/z 245.0474     0   1E+4   3   3.5  RT (min)       lcqtof_hilic_neg_2_036_34    m/z 245.0634     0   1   2   3E+4   2   2.5  RT (min)       lcqtof_hilic_neg_4_005_29    m/z 245.0926     0   1E+4   1   1.5  RT (min)      lcqtof_hilic_neg_4_052_13    m/z 245.2018     0   1   2E+4   6   6.5  RT (min)      lcqtof_hilic_neg_4_048_23    m/z 245.8786     0   1   2   3   4E+3   4.5   5  RT (min)      lcqtof_hilic_neg_4_001_49    m/z 246.0554     0   1E+4   2   2.5  RT (min)      lcqtof_hilic_neg_4_031_38    m/z 246.0670     0   1E+4   2   2.5  RT (min)      lcqtof_hilic_neg_4_050_02    m/z 247.0467     0   1   2E+4   0.5   1  RT (min)      lcqtof_hilic_neg_4_059_14    m/z 247.0554     0   2   4   6E+3   2   2.5  RT (min)      lcqtof_hilic_neg_4_052_13    m/z 247.0592     0   1E+4   1   1.5  RT (min)       lcqtof_hilic_neg_4_094_60    m/z 247.0926     0   2   4E+4   4   4.5  RT (min)      lcqtof_hilic_neg_4_047_06    m/z 248.9211     0   1   2   3E+3   1.5   2  RT (min)      lcqtof_hilic_neg_2_045_11    m/z 248.9213     0   1   2E+3   0.5   1  RT (min)      lcqtof_hilic_neg_2_040_20    m/z 248.9214     0   1   2E+3   0.5   1  RT (min)      lcqtof_hilic_neg_1_037_24    m/z 248.9600     0   1   2   3E+4   5.5   6  RT (min)       lcqtof_hilic_neg_4_012_07    m/z 248.9601     0   2   4   6E+3   7.5   8  RT (min)       lcqtof_hilic_neg_4_112_62    m/z 248.9602     0   1E+4   5.5   6  RT (min)      lcqtof_hilic_neg_4_012_07    m/z 248.9603     0   2   4   6   8E+3   7   7.5  RT (min)      lcqtof_hilic_neg_4_104_61    m/z 248.9603     0   1   2   3E+4   5.5   6  RT (min)      lcqtof_hilic_neg_4_012_07    m/z 248.9604     0   1E+4   6   6.5  RT (min)       lcqtof_hilic_neg_4_099_61    m/z 248.9605     0   2   4   6   8E+3   6.5   7  RT (min)       lcqtof_hilic_neg_4_111_62    m/z 248.9606     0   1E+4   6   6.5  RT (min)      lcqtof_hilic_neg_4_052_13    m/z 248.9606     0   1E+4   5.5   6  RT (min)      lcqtof_hilic_neg_4_008_24    m/z 248.9607     0   2   4   6E+3   7.5   8  RT (min)       lcqtof_hilic_neg_4_104_61    m/z 248.9726     0   2   4   6   8E+4   6.5   7  RT (min)      lcqtof_hilic_neg_1_000_aa    m/z 248.9731     0   2   4   6   8E+4   6.5   7  RT (min)      lcqtof_hilic_neg_1_000_aa    m/z 249.0068     0   1E+4   1   1.5  RT (min)      lcqtof_hilic_neg_4_059_14    m/z 249.0159     0   1E+4   0.5   1  RT (min)      lcqtof_hilic_neg_4_024_40    m/z 249.0283     0   1   2E+4   4   4.5  RT (min)      lcqtof_hilic_neg_4_055_32    m/z 249.0613     0   1E+4   3   3.5  RT (min)      lcqtof_hilic_neg_4_050_02    m/z 249.1109     0   1   2   3   4E+4   6   6.5  RT (min)       lcqtof_hilic_neg_4_055_32    m/z 249.1526     0   2   4   6E+3   2.5   3  RT (min)      lcqtof_hilic_neg_4_085_59    m/z 250.0729     0   1   2   3E+3   1.5   2  RT (min)      lcqtof_hilic_neg_4_032_01    m/z 250.7384     0   2   4E+4   3   3.5  RT (min)       lcqtof_hilic_neg_3_066_57    m/z 251.0591     0   1E+4   2   2.5  RT (min)      lcqtof_hilic_neg_2_045_11    m/z 251.0761     0   2   4   6E+4   3.5   4  RT (min)      lcqtof_hilic_neg_4_050_02    m/z 251.0767     0   1   2   3E+4   5   5.5  RT (min)      lcqtof_hilic_neg_4_019_03    m/z 251.0768     0   2   4   6   8E+4   5   5.5  RT (min)       lcqtof_hilic_neg_4_066_57    m/z 251.0771     0   1   2E+4   4   4.5  RT (min)      lcqtof_hilic_neg_4_026_04    m/z 251.0772     0   1   2   3E+4   4.5   5  RT (min)      lcqtof_hilic_neg_4_019_03    m/z 251.0773     0   1E+5   4   4.5  RT (min)       lcqtof_hilic_neg_4_019_03    m/z 251.0774     0   1   2   3E+4   4   4.5  RT (min)      lcqtof_hilic_neg_4_014_46    m/z 251.0781     0   1E+5   3.5   4  RT (min)       lcqtof_hilic_neg_4_019_03    m/z 251.0785     0   2   4   6E+4   3.5   4  RT (min)      lcqtof_hilic_neg_4_050_02    m/z 251.0815     0   2   4   6E+4   3.5   4  RT (min)      lcqtof_hilic_neg_4_050_02    m/z 251.0947     0   2   4   6   8E+3   3   3.5  RT (min)       lcqtof_hilic_neg_4_008_24    m/z 251.0950     0   1   2E+4   2   2.5  RT (min)       lcqtof_hilic_neg_4_045_26    m/z 251.0951     0   1E+4   2.5   3  RT (min)       lcqtof_hilic_neg_4_091_60    m/z 251.0952     0   2   4   6   8E+3   0.5   1  RT (min)       lcqtof_hilic_neg_4_091_60    m/z 251.0952     0   1   2   3   4E+3   0   0.5  RT (min)       lcqtof_hilic_neg_4_095_60    m/z 251.0952     0   1E+4   2   2.5  RT (min)      lcqtof_hilic_neg_4_032_01    m/z 251.0952     0   2   4   6   8E+3   3   3.5  RT (min)      lcqtof_hilic_neg_4_004_47    m/z 251.0953     0   2   4   6E+3   4   4.5  RT (min)       lcqtof_hilic_neg_4_009_50    m/z 251.0953     0   1   2E+4   2   2.5  RT (min)       lcqtof_hilic_neg_4_045_26    m/z 251.0953     0   2   4   6E+3   2.5   3  RT (min)       lcqtof_hilic_neg_4_049_55    m/z 251.0953     0   1   2   3   4E+3   0   0.5  RT (min)       lcqtof_hilic_neg_4_096_60    m/z 251.0954     0   1E+4   2   2.5  RT (min)       lcqtof_hilic_neg_4_041_54    m/z 251.0955     0   1E+4   2   2.5  RT (min)       lcqtof_hilic_neg_4_057_49    m/z 251.0956     0   2   4   6E+3   1.5   2  RT (min)       lcqtof_hilic_neg_4_097_60    m/z 251.1029     0   1E+5   1   1.5  RT (min)      lcqtof_hilic_neg_4_048_23    m/z 251.1032     0   1E+5   1   1.5  RT (min)      lcqtof_hilic_neg_4_034_31    m/z 251.1129     0   1E+4   4.5   5  RT (min)      lcqtof_hilic_neg_4_047_06    m/z 251.2243     0   1E+4   6   6.5  RT (min)      lcqtof_hilic_neg_4_008_24    m/z 252.0575     0   1E+4   1.5   2  RT (min)      lcqtof_hilic_neg_4_008_24    m/z 252.0819     0   1E+4   3.5   4  RT (min)      lcqtof_hilic_neg_4_019_03    m/z 252.0834     0   1   2E+4   2.5   3  RT (min)      lcqtof_hilic_neg_4_055_32    m/z 252.0869     0   1E+4   15   15.5  RT (min)      lcqtof_hilic_neg_4_000_aa    m/z 252.0987     0   2   4   6   8E+3   0.5   1  RT (min)      lcqtof_hilic_neg_4_058_39    m/z 252.0996     0   1E+4   2.5   3  RT (min)      lcqtof_hilic_neg_4_047_06    m/z 252.0998     0   1   2E+4   3   3.5  RT (min)      lcqtof_hilic_neg_4_055_32    m/z 252.9984     0   1E+4   0   0.5  RT (min)      lcqtof_hilic_neg_1_000_cc    m/z 253.0763     0   1E+4   4.5   5  RT (min)      lcqtof_hilic_neg_4_050_02    m/z 253.0763     0   1   2E+4   4   4.5  RT (min)      lcqtof_hilic_neg_4_040_05    m/z 253.0894     0   2   4   6E+3   0.5   1  RT (min)      lcqtof_hilic_neg_4_040_05    m/z 254.1076     0   1E+4   3.5   4  RT (min)      lcqtof_hilic_neg_4_008_24    m/z 254.7972     0   1E+4   2.5   3  RT (min)      lcqtof_hilic_neg_4_050_02    m/z 254.9194     0   2   4   6E+3   7   7.5  RT (min)       lcqtof_hilic_neg_4_058_39    m/z 254.9222     0   2   4E+3   7.5   8  RT (min)       lcqtof_hilic_neg_4_046_41    m/z 254.9240     0   1E+4   8.5   9  RT (min)      lcqtof_hilic_neg_1_000_aa    m/z 254.9243     0   1E+4   8   8.5  RT (min)       lcqtof_hilic_neg_1_000_aa    m/z 254.9260     0   1   2   3   4E+3   6.5   7  RT (min)       lcqtof_hilic_neg_4_098_61    m/z 254.9264     0   1   2   3   4E+3   7   7.5  RT (min)      lcqtof_hilic_neg_4_012_07    m/z 254.9810     0   1E+4   2.5   3  RT (min)       lcqtof_hilic_neg_4_077_58    m/z 254.9813     0   1E+4   3   3.5  RT (min)       lcqtof_hilic_neg_4_059_14    m/z 255.0203     0   1E+4   3   3.5  RT (min)      lcqtof_hilic_neg_1_042_26    m/z 255.1090     0   1E+4   0.5   1  RT (min)      lcqtof_hilic_neg_4_050_02    m/z 255.1130     0   1E+4   1.5   2  RT (min)       lcqtof_hilic_neg_4_032_01    m/z 255.2328     0   2   4   6E+3   3   3.5  RT (min)      lcqtof_hilic_neg_4_005_29    m/z 255.7966     0   1E+4   2.5   3  RT (min)      lcqtof_hilic_neg_4_050_02    m/z 255.8214     0   1   0   0.5  RT (min)       lcqtof_hilic_neg_3_066_57    m/z 255.8216     0   1   2   3E+4   3.5   4  RT (min)      lcqtof_hilic_neg_4_020_42    m/z 255.8216     0   1E+4   16.5   17  RT (min)       lcqtof_hilic_neg_3_066_57    m/z 255.8216     0   1E+4   1.5   2  RT (min)       lcqtof_hilic_neg_3_066_57    m/z 255.8217     0   2   4   6E+4   3.5   4  RT (min)       lcqtof_hilic_neg_3_067_57    m/z 255.8218     0   1E+5   3.5   4  RT (min)       lcqtof_hilic_neg_3_067_57    m/z 255.8218     0   2   4   6E+4   3.5   4  RT (min)      lcqtof_hilic_neg_1_000_dd    m/z 255.8218     0   1E+4   1.5   2  RT (min)       lcqtof_hilic_neg_3_066_57    m/z 255.8219     0   1E+4   0.5   1  RT (min)       lcqtof_hilic_neg_3_066_57    m/z 255.8220     0   1E+4   16   16.5  RT (min)       lcqtof_hilic_neg_3_068_57    m/z 255.8220     0   2   4   6E+4   3.5   4  RT (min)      lcqtof_hilic_neg_1_000_dd    m/z 255.8220     0   1   2   3E+4   3.5   4  RT (min)       lcqtof_hilic_neg_4_020_42    m/z 255.8220     0   1E+4   16.5   17  RT (min)       lcqtof_hilic_neg_3_067_57    m/z 255.8221     0   1E+4   1   1.5  RT (min)      lcqtof_hilic_neg_1_000_bb    m/z 256.0127     0   1   2   3E+4   6.5   7  RT (min)      lcqtof_hilic_neg_4_053_14    m/z 256.0144     0   1   2   3E+4   6.5   7  RT (min)      lcqtof_hilic_neg_4_053_14    m/z 256.0146     0   1   2   3E+4   6.5   7  RT (min)      lcqtof_hilic_neg_4_053_14    m/z 256.0147     0   1   2   3E+4   6.5   7  RT (min)      lcqtof_hilic_neg_4_053_14    m/z 256.0148     0   1   2   3E+4   6.5   7  RT (min)       lcqtof_hilic_neg_4_053_14    m/z 256.0826     0   1E+4   2.5   3  RT (min)      lcqtof_hilic_neg_4_048_23    m/z 256.0846     0   1E+4   2.5   3  RT (min)       lcqtof_hilic_neg_4_048_23    m/z 256.0846     0   1E+4   2   2.5  RT (min)       lcqtof_hilic_neg_4_048_23    m/z 256.0901     0   2   4   6E+3   1   1.5  RT (min)      lcqtof_hilic_neg_4_052_13    m/z 256.1231     0   1E+5   4   4.5  RT (min)       lcqtof_hilic_neg_4_055_32    m/z 256.1235     0   1E+5   4   4.5  RT (min)      lcqtof_hilic_neg_4_097_60    m/z 256.1237     0   1   2E+4   4.5   5  RT (min)      lcqtof_hilic_neg_4_008_24    m/z 256.2217     0   1E+4   4.5   5  RT (min)      lcqtof_hilic_neg_4_054_48    m/z 256.7659     0   1   2E+4   4.5   5  RT (min)      lcqtof_hilic_neg_4_050_02    m/z 256.7975     0   1E+4   2.5   3  RT (min)      lcqtof_hilic_neg_4_050_02    m/z 256.8224     0   1E+4   4   4.5  RT (min)      lcqtof_hilic_neg_3_067_57    m/z 256.9004     0   1E+5   9   9.5  RT (min)      lcqtof_hilic_neg_1_000_aa    m/z 256.9012     0   1E+5   8.5   9  RT (min)      lcqtof_hilic_neg_1_000_aa    m/z 256.9013     0   1E+5   8.5   9  RT (min)      lcqtof_hilic_neg_1_000_aa    m/z 256.9017     0   1E+5   8.5   9  RT (min)      lcqtof_hilic_neg_1_000_aa    m/z 256.9020     0   1E+5   8.5   9  RT (min)      lcqtof_hilic_neg_1_000_aa    m/z 256.9032     0   1E+4   8   8.5  RT (min)      lcqtof_hilic_neg_1_000_aa    m/z 256.9545     0   1E+4   14   15  RT (min)       lcqtof_hilic_neg_4_000_dd    m/z 256.9546     0   1E+4   14  RT (min)       lcqtof_hilic_neg_3_067_57    m/z 256.9548     0   1E+4   14   15  RT (min)       lcqtof_hilic_neg_4_001_49    m/z 256.9550     0   1E+4   13   13.5  RT (min)       lcqtof_hilic_neg_3_068_57    m/z 256.9576     0   1   2E+4   5.5   6  RT (min)      lcqtof_hilic_neg_4_016_37    m/z 257.0512     0   1E+4   0.5   1  RT (min)      lcqtof_hilic_neg_2_045_11    m/z 257.0754     0   1E+4   6   6.5  RT (min)      lcqtof_hilic_neg_4_053_14    m/z 257.1134     0   1   2   3E+5   6   6.5  RT (min)       lcqtof_hilic_neg_4_089_59    m/z 257.8184     0   1E+4   0.5   1  RT (min)       lcqtof_hilic_neg_3_066_57    m/z 257.8186     0   1   2   3   4E+3   0.5   1  RT (min)       lcqtof_hilic_neg_3_067_57    m/z 257.8187     0   1E+4   16   16.5  RT (min)       lcqtof_hilic_neg_3_066_57    m/z 257.8188     0   2   4   6   8E+3   0   0.5  RT (min)       lcqtof_hilic_neg_3_066_57    m/z 257.8188     0   2   4E+4   3.5   4  RT (min)       lcqtof_hilic_neg_3_067_57    m/z 257.8188     0   2   4   6   8E+4   3.5   4  RT (min)      lcqtof_hilic_neg_1_000_cc    m/z 257.8188     0   1E+4   1   1.5  RT (min)      lcqtof_hilic_neg_1_000_bb    m/z 257.8188     0   1E+4   16   16.5  RT (min)       lcqtof_hilic_neg_3_068_57    m/z 257.8189     0   1   2   3E+4   3.5   4  RT (min)      lcqtof_hilic_neg_4_030_44    m/z 257.8190     0   1E+5   3.5   4  RT (min)       lcqtof_hilic_neg_3_067_57    m/z 257.8190     0   1E+4   1.5   2  RT (min)       lcqtof_hilic_neg_3_066_57    m/z 257.8190     0   1   2E+4   3.5   4  RT (min)      lcqtof_hilic_neg_4_046_41    m/z 257.8191     0   1E+4   1.5   2  RT (min)       lcqtof_hilic_neg_3_066_57    m/z 257.8191     0   1   2   3E+4   3.5   4  RT (min)      lcqtof_hilic_neg_4_030_44    m/z 257.9373     0   1   2E+4   2.5   3  RT (min)       lcqtof_hilic_neg_4_000_aa    m/z 257.9376     0   1   2E+4   2.5   3  RT (min)      lcqtof_hilic_neg_4_000_aa    m/z 257.9380     0   1   2   3   4E+4   2   2.5  RT (min)      lcqtof_hilic_neg_4_000_aa    m/z 257.9381     0   1   2   3   4E+4   2   2.5  RT (min)       lcqtof_hilic_neg_4_000_aa    m/z 257.9382     0   1   2   3E+4   2.5   3  RT (min)      lcqtof_hilic_neg_4_000_aa    m/z 257.9382     0   1   2   3   4E+4   2   2.5  RT (min)      lcqtof_hilic_neg_4_000_aa    m/z 257.9384     0   1E+5   1   1.5  RT (min)      lcqtof_hilic_neg_4_000_aa    m/z 257.9385     0   1E+5   1   1.5   2  RT (min)       lcqtof_hilic_neg_4_000_aa    m/z 257.9386     0   2   4E+4   2   2.5  RT (min)       lcqtof_hilic_neg_4_000_aa    m/z 257.9387     0   1E+5   1   1.5  RT (min)      lcqtof_hilic_neg_4_000_aa    m/z 257.9388     0   1   2   3   4E+4   2   2.5  RT (min)       lcqtof_hilic_neg_4_000_aa    m/z 257.9393     0   1E+5   1   1.5  RT (min)      lcqtof_hilic_neg_4_000_aa    m/z 257.9405     0   1E+5   1   1.5   2  RT (min)       lcqtof_hilic_neg_4_000_aa    m/z 257.9798     0   1   2E+4   7   7.5  RT (min)      lcqtof_hilic_neg_4_008_24    m/z 257.9801     0   1   2E+4   7   7.5  RT (min)       lcqtof_hilic_neg_4_008_24    m/z 257.9801     0   1   2E+4   7   7.5  RT (min)      lcqtof_hilic_neg_4_008_24    m/z 258.0000     0   1   2   3E+4   0.5   1  RT (min)      lcqtof_hilic_neg_2_062_17    m/z 258.0910     0   1   2   3E+4   3.5   4  RT (min)       lcqtof_hilic_neg_4_092_60    m/z 258.0914     0   2   4   6   8E+4   4   4.5  RT (min)      lcqtof_hilic_neg_4_040_05    m/z 258.0916     0   2   4   6E+4   4.5   5  RT (min)      lcqtof_hilic_neg_4_008_24    m/z 258.0949     0   2   4   6E+4   4.5   5  RT (min)      lcqtof_hilic_neg_4_008_24    m/z 258.1082     0   1E+4   4   4.5  RT (min)       lcqtof_hilic_neg_4_085_59    m/z 258.1086     0   1E+4   4   4.5  RT (min)      lcqtof_hilic_neg_4_048_23    m/z 258.1466     0   1E+4   5.5   6  RT (min)      lcqtof_hilic_neg_4_008_24    m/z 258.7634     0   1   2   3E+4   4.5   5  RT (min)      lcqtof_hilic_neg_4_050_02    m/z 258.7975     0   1E+4   2.5   3  RT (min)      lcqtof_hilic_neg_4_050_02    m/z 258.9099     0   1E+4   1.5   2  RT (min)      lcqtof_hilic_neg_4_050_02    m/z 259.0126     0   2   4   6E+4   4   4.5  RT (min)       lcqtof_hilic_neg_4_060_13    m/z 259.0226     0   1   2E+4   7.5   8  RT (min)      lcqtof_hilic_neg_4_040_05    m/z 259.0590     0   1   2E+4   3.5   4  RT (min)      lcqtof_hilic_neg_1_049_55    m/z 259.0928     0   1   2E+4   3.5   4  RT (min)       lcqtof_hilic_neg_4_048_23    m/z 259.0931     0   1   2E+4   3.5   4  RT (min)      lcqtof_hilic_neg_4_048_23    m/z 259.0934     0   1E+4   3.5   4  RT (min)      lcqtof_hilic_neg_4_047_06    m/z 259.8158     0   1   2   3E+4   4   4.5  RT (min)      lcqtof_hilic_neg_4_030_44    m/z 259.8159     0   1   2   3E+4   3.5   4  RT (min)      lcqtof_hilic_neg_1_000_cc    m/z 259.8160     0   1E+4   3.5   4  RT (min)       lcqtof_hilic_neg_3_067_57    m/z 259.8160     0   1   2   3   4E+4   3.5   4  RT (min)       lcqtof_hilic_neg_3_067_57    m/z 259.8388     0   1E+4   7   7.5  RT (min)      lcqtof_hilic_neg_4_050_02    m/z 259.8742     0   1E+4   9   9.5  RT (min)      lcqtof_hilic_neg_1_000_aa    m/z 259.9779     0   1E+4   7   7.5  RT (min)      lcqtof_hilic_neg_4_008_24    m/z 259.9782     0   1E+4   7   7.5  RT (min)      lcqtof_hilic_neg_4_008_24    m/z 259.9784     0   1E+4   7   7.5  RT (min)      lcqtof_hilic_neg_4_008_24    m/z 260.0254     0   2   4E+4   0.5   1  RT (min)      lcqtof_hilic_neg_4_000_aa    m/z 260.0262     0   2   4E+4   0.5   1  RT (min)      lcqtof_hilic_neg_4_000_aa    m/z 260.0515     0   1   2   3   4E+4   4   4.5  RT (min)       lcqtof_hilic_neg_4_095_60    m/z 260.0525     0   1   2E+4   4   4.5  RT (min)      lcqtof_hilic_neg_4_055_32    m/z 260.0768     0   2   4   6   8E+4   4   4.5  RT (min)       lcqtof_hilic_neg_4_085_59    m/z 260.1068     0   1E+4   3.5   4  RT (min)       lcqtof_hilic_neg_4_008_24    m/z 260.7605     0   1   2E+4   4.5   5  RT (min)      lcqtof_hilic_neg_4_050_02    m/z 260.7613     0   1   2E+4   4.5   5  RT (min)      lcqtof_hilic_neg_4_050_02    m/z 260.8731     0   1E+4   7   7.5  RT (min)       lcqtof_hilic_neg_4_102_61    m/z 260.8732     0   2   4   6   8E+3   9   9.5  RT (min)       lcqtof_hilic_neg_4_098_61    m/z 260.8732     0   1E+4   8.5   9  RT (min)       lcqtof_hilic_neg_4_110_62    m/z 260.8735     0   1   2E+3   12   12.5  RT (min)       lcqtof_hilic_neg_4_112_62    m/z 260.8735     0   1   2E+3   12.5   13  RT (min)       lcqtof_hilic_neg_4_106_62    m/z 260.8736     0   1   2E+3   9.5   10  RT (min)       lcqtof_hilic_neg_4_093_60    m/z 260.8736     0   1   2E+3   9   9.5  RT (min)       lcqtof_hilic_neg_4_089_59    m/z 260.8736     0   1E+3   9.5   10  RT (min)       lcqtof_hilic_neg_4_055_32    m/z 260.8739     0   1E+4   7   7.5  RT (min)       lcqtof_hilic_neg_4_052_13    m/z 260.8740     0   2   4   6E+3   7.5   8  RT (min)      lcqtof_hilic_neg_4_003_16    m/z 260.8740     0   1E+4   7.5   8  RT (min)       lcqtof_hilic_neg_4_052_13    m/z 261.0228     0   1   2   3   4E+4   1.5   2  RT (min)       lcqtof_hilic_neg_4_058_39    m/z 261.0229     0   1   2   3   4E+4   2   2.5  RT (min)      lcqtof_hilic_neg_4_046_41    m/z 261.0409     0   1E+4   2.5   3  RT (min)      lcqtof_hilic_neg_3_053_26    m/z 261.0420     0   1E+4   3   3.5  RT (min)       lcqtof_hilic_neg_2_004_26    m/z 261.0562     0   2   4   6E+4   4.5   5  RT (min)      lcqtof_hilic_neg_4_061_50    m/z 261.0601     0   2   4   6E+3   3.5   4  RT (min)      lcqtof_hilic_neg_2_001_49    m/z 261.0764     0   1E+5   1   1.5  RT (min)       lcqtof_hilic_neg_4_098_61    m/z 261.1252     0   1E+4   4   4.5  RT (min)      lcqtof_hilic_neg_4_008_24    m/z 261.1359     0   2   4   6   8E+4   1   1.5  RT (min)       lcqtof_hilic_neg_4_096_60    m/z 261.1372     0   2   4   6E+3   1.5   2  RT (min)      lcqtof_hilic_neg_4_082_59    m/z 261.8920     0   1   2E+4   4   4.5  RT (min)      lcqtof_hilic_neg_4_003_16    m/z 261.8921     0   1E+4   4   4.5  RT (min)      lcqtof_hilic_neg_4_026_04    m/z 261.8927     0   1   2E+4   3.5   4  RT (min)      lcqtof_hilic_neg_1_000_bb    m/z 261.9061     0   1E+4   1.5   2  RT (min)      lcqtof_hilic_neg_4_050_02    m/z 261.9784     0   2   4   6E+3   5   5.5  RT (min)       lcqtof_hilic_neg_3_067_57    m/z 261.9785     0   2   4   6E+3   5   5.5  RT (min)      lcqtof_hilic_neg_1_004_14    m/z 261.9787     0   2   4   6E+3   5.5   6  RT (min)      lcqtof_hilic_neg_1_004_14    m/z 261.9787     0   2   4   6E+3   5   5.5  RT (min)      lcqtof_hilic_neg_1_012_04    m/z 262.0260     0   1E+4   0.5   1  RT (min)      lcqtof_hilic_neg_4_000_aa    m/z 262.0796     0   1   2E+4   1   1.5  RT (min)      lcqtof_hilic_neg_4_098_61    m/z 262.0798     0   1   2   3E+4   1   1.5  RT (min)      lcqtof_hilic_neg_4_104_61    m/z 262.9059     0   1E+4   1.5   2  RT (min)      lcqtof_hilic_neg_4_050_02    m/z 262.9603     0   1   2   3   4E+3   2   2.5  RT (min)      lcqtof_hilic_neg_4_026_04    m/z 262.9608     0   1   2   3   4E+3   2.5   3  RT (min)      lcqtof_hilic_neg_2_007_03    m/z 262.9777     0   2   4E+4   5.5   6  RT (min)       lcqtof_hilic_neg_4_008_24    m/z 263.0592     0   1E+4   2   2.5  RT (min)      lcqtof_hilic_neg_4_050_02    m/z 263.8893     0   1   2E+4   4   4.5  RT (min)       lcqtof_hilic_neg_3_067_57    m/z 264.0752     0   2   4   6   8E+4   3.5   4  RT (min)      lcqtof_hilic_neg_1_049_55    m/z 264.0950     0   1   2   3E+3   0.5   1  RT (min)      lcqtof_hilic_neg_3_003_20    m/z 264.1088     0   1E+4   4   4.5  RT (min)      lcqtof_hilic_neg_4_048_23    m/z 264.9074     0   1E+4   1.5   2  RT (min)      lcqtof_hilic_neg_4_050_02    m/z 264.9436     0   1   2E+4   1.5   2  RT (min)      lcqtof_hilic_neg_1_064_16    m/z 265.0704     0   1E+4   1.5   2  RT (min)      lcqtof_hilic_neg_4_095_60    m/z 265.0774     0   1E+4   3   3.5  RT (min)      lcqtof_hilic_neg_4_014_46    m/z 265.0925     0   1E+4   3.5   4  RT (min)       lcqtof_hilic_neg_4_072_57    m/z 265.0925     0   1   2   3E+4   1.5   2  RT (min)      lcqtof_hilic_neg_4_052_13    m/z 265.0925     0   1E+4   1.5   2  RT (min)      lcqtof_hilic_neg_4_050_02    m/z 265.1078     0   1E+4   2   2.5  RT (min)      lcqtof_hilic_neg_4_050_02    m/z 265.1355     0   2   4E+5   0.5   1  RT (min)      lcqtof_hilic_neg_2_044_37    m/z 265.1418     0   1E+4   13   13.5  RT (min)      lcqtof_hilic_neg_4_111_62    m/z 265.1426     0   1   2E+4   4   4.5  RT (min)      lcqtof_hilic_neg_4_030_44    m/z 265.1438     0   1E+4   9.5   10  RT (min)      lcqtof_hilic_neg_4_113_62    m/z 265.1448     0   1E+4   7   7.5  RT (min)      lcqtof_hilic_neg_4_092_60    m/z 265.1449     0   1E+4   8.5   9  RT (min)      lcqtof_hilic_neg_4_102_61    m/z 265.1449     0   1E+4   12.5   13  RT (min)      lcqtof_hilic_neg_4_085_59    m/z 265.1451     0   1E+4   8   8.5  RT (min)      lcqtof_hilic_neg_4_113_62    m/z 265.1454     0   1E+4   10.5   11  RT (min)      lcqtof_hilic_neg_4_105_61    m/z 265.1454     0   1E+4   11.5   12  RT (min)      lcqtof_hilic_neg_4_092_60    m/z 265.1456     0   1E+4   9   9.5  RT (min)      lcqtof_hilic_neg_4_078_58    m/z 265.1456     0   1E+4   11   11.5  RT (min)      lcqtof_hilic_neg_4_092_60    m/z 265.1456     0   1E+4   11   11.5  RT (min)      lcqtof_hilic_neg_4_083_59    m/z 265.1457     0   1E+4   14.5   15  RT (min)      lcqtof_hilic_neg_4_110_62    m/z 265.1458     0   1E+4   10   10.5  RT (min)      lcqtof_hilic_neg_4_096_60    m/z 265.1459     0   1E+4   10   10.5  RT (min)      lcqtof_hilic_neg_4_104_61    m/z 265.1463     0   1   2E+5   15   15.5  RT (min)      lcqtof_hilic_neg_4_000_aa    m/z 265.1465     0   1E+4   5.5   6  RT (min)      lcqtof_hilic_neg_4_087_59    m/z 265.1466     0   1E+4   5   5.5  RT (min)       lcqtof_hilic_neg_4_002_25    m/z 265.1466     0   1E+4   8   8.5  RT (min)       lcqtof_hilic_neg_4_110_62    m/z 265.1467     0   1   2E+5   15   15.5  RT (min)      lcqtof_hilic_neg_4_000_aa    m/z 265.1468     0   1   2   3E+4   19.5   20  RT (min)       lcqtof_hilic_neg_4_097_60    m/z 265.1470     0   1E+4   8.5   9  RT (min)      lcqtof_hilic_neg_4_073_57    m/z 265.1470     0   1E+4   8.5   9  RT (min)      lcqtof_hilic_neg_4_078_58    m/z 265.1471     0   1   2E+4   6   6.5  RT (min)       lcqtof_hilic_neg_4_048_23    m/z 265.1471     0   1   2   3E+4   19   19.5  RT (min)       lcqtof_hilic_neg_4_097_60    m/z 265.1471     0   1   2E+4   3.5   4  RT (min)       lcqtof_hilic_neg_4_112_62    m/z 265.1472     0   1   2   3E+4   0   0.5  RT (min)       lcqtof_hilic_neg_4_055_32    m/z 265.1472     0   1E+4   7.5   8  RT (min)      lcqtof_hilic_neg_4_105_61    m/z 265.1472     0   1E+4   5   5.5  RT (min)      lcqtof_hilic_neg_4_005_29    m/z 265.1472     0   1E+5   2   2.5  RT (min)       lcqtof_hilic_neg_4_018_27    m/z 265.1472     0   1   2   3E+4   17   17.5  RT (min)       lcqtof_hilic_neg_4_111_62    m/z 265.1472     0   1   2   3E+5   0   0.5  RT (min)       lcqtof_hilic_neg_4_053_14    m/z 265.1472     0   1   2   3E+4   0   0.5  RT (min)       lcqtof_hilic_neg_4_053_14    m/z 265.1473     0   2   4   6E+4   1.5   2  RT (min)       lcqtof_hilic_neg_4_019_03    m/z 265.1473     0   1   2E+4   16   16.5  RT (min)       lcqtof_hilic_neg_4_111_62    m/z 265.1473     0   1E+5   2   2.5  RT (min)       lcqtof_hilic_neg_4_003_16    m/z 265.1473     0   1   2E+4   15.5   16  RT (min)       lcqtof_hilic_neg_4_083_59    m/z 265.1473     0   1   2   3E+4   2.5   3  RT (min)       lcqtof_hilic_neg_4_113_62    m/z 265.1474     0   1   2E+4   16.5   17  RT (min)       lcqtof_hilic_neg_4_083_59    m/z 265.1474     0   1   2   3E+4   17.5   18  RT (min)       lcqtof_hilic_neg_4_112_62    m/z 265.1474     0   1   2   3E+4   0   0.5  RT (min)       lcqtof_hilic_neg_4_101_61    m/z 265.1474     0   1   2E+4   15.5   16  RT (min)       lcqtof_hilic_neg_4_083_59    m/z 265.1474     0   2   4E+4   2.5   3   3.5  RT (min)       lcqtof_hilic_neg_4_055_32    m/z 265.1474     0   1   2   3E+4   16.5   17  RT (min)       lcqtof_hilic_neg_4_080_58    m/z 265.1474     0   1   2   3E+5   0.5   1  RT (min)       lcqtof_hilic_neg_4_072_57    m/z 265.1474     0   1E+5   1.5   2  RT (min)       lcqtof_hilic_neg_4_019_03    m/z 265.1474     0   1   2   3E+4   17   18  RT (min)       lcqtof_hilic_neg_4_083_59    m/z 265.1474     0   1   2   3E+4   16.5   17   17.5  RT (min)       lcqtof_hilic_neg_4_014_46    m/z 265.1474     0   1   2   3E+4   18   18.5   19  RT (min)       lcqtof_hilic_neg_4_013_20    m/z 265.1474     0   1   2E+5   0   0.5  RT (min)       lcqtof_hilic_neg_4_108_62    m/z 265.1474     0   1   19   20  RT (min)       lcqtof_hilic_neg_4_000_dd    m/z 265.1474     0   1   2   3E+4   18   18.5   19  RT (min)       lcqtof_hilic_neg_4_072_57    m/z 265.1474     0   1   2   3E+4   17.5   18  RT (min)       lcqtof_hilic_neg_4_110_62    m/z 265.1474     0   1E+5   1.5   2  RT (min)       lcqtof_hilic_neg_4_006_30    m/z 265.1474     0   1E+4   7   7.5  RT (min)       lcqtof_hilic_neg_4_058_39    m/z 265.1474     0   1E+4   5   5.5  RT (min)      lcqtof_hilic_neg_4_081_58    m/z 265.1474     0   1E+4   6.5   7  RT (min)       lcqtof_hilic_neg_4_107_62    m/z 265.1474     0   1E+5   2.5   3  RT (min)       lcqtof_hilic_neg_4_088_59    m/z 265.1474     0   1E+5   2   2.5  RT (min)       lcqtof_hilic_neg_4_032_01    m/z 265.1474     0   1   2E+4   16   16.5  RT (min)       lcqtof_hilic_neg_4_110_62    m/z 265.1474     0   1   2   3E+4   18.5   19  RT (min)       lcqtof_hilic_neg_4_097_60    m/z 265.1475     0   1E+5   2   2.5  RT (min)       lcqtof_hilic_neg_4_053_14    m/z 265.1475     0   2   4   6E+4   1.5   2  RT (min)       lcqtof_hilic_neg_4_106_62    m/z 265.1475     0   1   2   3E+4   0   0.5  RT (min)      lcqtof_hilic_neg_4_053_14    m/z 265.1475     0   1E+5   2   2.5  RT (min)      lcqtof_hilic_neg_4_018_27    m/z 265.1475     0   2   4E+4   3   3.5  RT (min)       lcqtof_hilic_neg_4_008_24    m/z 265.1475     0   1   2   3E+4   18   19   20  RT (min)       lcqtof_hilic_neg_4_019_03    m/z 265.1476     0   1E+4   6.5   7  RT (min)      lcqtof_hilic_neg_4_087_59    m/z 265.1476     0   1E+4   6   6.5  RT (min)       lcqtof_hilic_neg_4_088_59    m/z 265.1476     0   1   2   3   4E+4   4.5   5  RT (min)       lcqtof_hilic_neg_4_016_37    m/z 265.1476     0   1E+5   1.5   2  RT (min)      lcqtof_hilic_neg_4_019_03    m/z 265.1476     0   1   2   3E+4   18   19  RT (min)       lcqtof_hilic_neg_4_038_10    m/z 265.1476     0   1E+4   3.5   4  RT (min)      lcqtof_hilic_neg_4_074_58    m/z 265.1476     0   1   2   3E+4   19   19.5  RT (min)       lcqtof_hilic_neg_4_109_62    m/z 265.1476     0   2   4E+4   3.5   4  RT (min)       lcqtof_hilic_neg_4_008_24    m/z 265.1477     0   1E+4   6.5   7  RT (min)      lcqtof_hilic_neg_4_113_62    m/z 265.1477     0   1E+4   4.5   5  RT (min)       lcqtof_hilic_neg_4_019_03    m/z 265.1477     0   1   2   3E+4   0   0.5  RT (min)      lcqtof_hilic_neg_4_055_32    m/z 265.1477     0   1   2E+4   3   3.5  RT (min)       lcqtof_hilic_neg_2_003_25    m/z 265.1477     0   1   2E+4   3   3.5  RT (min)      lcqtof_hilic_neg_4_006_30    m/z 265.1478     0   1E+4   5   5.5  RT (min)      lcqtof_hilic_neg_4_031_38    m/z 265.1478     0   1   2E+4   3.5   4  RT (min)      lcqtof_hilic_neg_4_010_28    m/z 265.1478     0   1E+4   3.5   4  RT (min)       lcqtof_hilic_neg_4_074_58    m/z 265.1480     0   1E+5   1   1.5  RT (min)      lcqtof_hilic_neg_2_038_04    m/z 265.1583     0   1E+5   0.5   1  RT (min)      lcqtof_hilic_neg_3_000_bb    m/z 265.1601     0   1E+5   0.5   1  RT (min)      lcqtof_hilic_neg_2_045_11    m/z 265.2058     0   2   4E+4   0.5   1  RT (min)      lcqtof_hilic_neg_1_032_18    m/z 265.2736     0   1   2E+4   1   1.5  RT (min)      lcqtof_hilic_neg_4_005_29    m/z 265.3467     0   2   4   6E+4   0.5   1  RT (min)      lcqtof_hilic_neg_2_016_33    m/z 265.8504     0   1   2   3E+4   4.5   5  RT (min)       lcqtof_hilic_neg_3_067_57    m/z 265.8505     0   1   2   3E+4   4   4.5  RT (min)       lcqtof_hilic_neg_3_067_57    m/z 265.9016     0   1E+4   8   8.5  RT (min)      lcqtof_hilic_neg_1_000_aa    m/z 266.0429     0   1E+4   1.5   2  RT (min)      lcqtof_hilic_neg_4_034_31    m/z 266.0568     0   1   2   3E+4   2   2.5  RT (min)      lcqtof_hilic_neg_4_055_32    m/z 266.0578     0   1   2   3E+4   2   2.5  RT (min)      lcqtof_hilic_neg_4_055_32    m/z 266.0730     0   1   2E+4   3.5   4  RT (min)      lcqtof_hilic_neg_1_049_55    m/z 266.1014     0   1   2   3E+4   6   6.5  RT (min)       lcqtof_hilic_neg_4_008_24    m/z 266.1014     0   1   2E+4   6   6.5  RT (min)      lcqtof_hilic_neg_1_037_24    m/z 266.1117     0   1   2   3   4E+4   0.5   1  RT (min)      lcqtof_hilic_neg_1_047_29    m/z 266.1141     0   1E+4   2   2.5  RT (min)      lcqtof_hilic_neg_4_014_46    m/z 266.1142     0   1E+4   2   2.5  RT (min)      lcqtof_hilic_neg_4_014_46    m/z 266.1503     0   1E+4   2.5   3  RT (min)      lcqtof_hilic_neg_4_086_59    m/z 266.1507     0   1E+5   1   1.5  RT (min)       lcqtof_hilic_neg_4_026_04    m/z 266.1508     0   1E+4   1   1.5  RT (min)      lcqtof_hilic_neg_4_008_24    m/z 266.1509     0   1   2   3E+4   15   15.5  RT (min)      lcqtof_hilic_neg_4_000_aa    m/z 266.1609     0   1E+4   0.5   1  RT (min)      lcqtof_hilic_neg_3_051_12    m/z 266.2332     0   2   4E+4   0.5   1  RT (min)      lcqtof_hilic_neg_2_016_33    m/z 267.0502     0   1   2   3   4E+4   2.5   3  RT (min)      lcqtof_hilic_neg_1_032_18    m/z 267.0505     0   1E+4   3   3.5  RT (min)      lcqtof_hilic_neg_4_039_18    m/z 267.0506     0   1E+4   3   3.5  RT (min)      lcqtof_hilic_neg_4_039_18    m/z 267.1046     0   2   4   6E+3   2   2.5  RT (min)      lcqtof_hilic_neg_4_008_24    m/z 267.1047     0   2   4   6E+3   0.5   1  RT (min)      lcqtof_hilic_neg_4_040_05    m/z 267.1448     0   1E+4   15   15.5  RT (min)      lcqtof_hilic_neg_4_000_aa    m/z 267.1456     0   2   4   6E+3   2   2.5  RT (min)      lcqtof_hilic_neg_4_062_17    m/z 267.1461     0   2   4   6E+3   2.5   3  RT (min)      lcqtof_hilic_neg_4_088_59    m/z 267.8479     0   1   2E+4   4   4.5  RT (min)       lcqtof_hilic_neg_3_067_57    m/z 268.0374     0   1   2E+4   3   3.5  RT (min)       lcqtof_hilic_neg_4_108_62    m/z 268.0493     0   1   2   3E+3   3.5   4  RT (min)      lcqtof_hilic_neg_1_008_13    m/z 268.0548     0   1   2   3E+3   0.5   1  RT (min)      lcqtof_hilic_neg_4_002_25    m/z 268.0754     0   2   4   6E+4   1   1.5  RT (min)      lcqtof_hilic_neg_4_050_02    m/z 268.0761     0   1   2   3   4E+3   0.5   1  RT (min)      lcqtof_hilic_neg_4_053_14    m/z 268.0761     0   2   4   6E+4   1   1.5  RT (min)      lcqtof_hilic_neg_4_050_02    m/z 268.0767     0   2   4   6E+4   1.5   2  RT (min)      lcqtof_hilic_neg_4_050_02    m/z 268.0785     0   2   4E+3   2   2.5  RT (min)      lcqtof_hilic_neg_4_010_28    m/z 268.0863     0   1   2   3E+3   1   1.5  RT (min)      lcqtof_hilic_neg_4_055_32    m/z 268.0933     0   1   2   3   4E+3   0.5   1  RT (min)      lcqtof_hilic_neg_4_050_02    m/z 268.0946     0   1E+4   4.5   5  RT (min)      lcqtof_hilic_neg_1_005_32    m/z 268.0998     0   1E+4   6   6.5  RT (min)      lcqtof_hilic_neg_4_008_24    m/z 268.1132     0   2   4   6E+3   2   2.5  RT (min)      lcqtof_hilic_neg_4_055_32    m/z 268.1427     0   1   2   3E+4   4   4.5  RT (min)       lcqtof_hilic_neg_4_092_60    m/z 268.1430     0   1   2E+4   4   4.5  RT (min)      lcqtof_hilic_neg_4_040_05    m/z 268.1430     0   1   2E+4   4   4.5  RT (min)      lcqtof_hilic_neg_4_040_05    m/z 268.1503     0   1E+5   6   6.5  RT (min)       lcqtof_hilic_neg_4_092_60    m/z 269.0874     0   2   4   6   8E+3   5.5   6  RT (min)      lcqtof_hilic_neg_4_063_16    m/z 269.1241     0   1   2   3E+3   2   2.5  RT (min)      lcqtof_hilic_neg_4_055_32    m/z 269.1264     0   1   2   3E+4   3.5   4  RT (min)       lcqtof_hilic_neg_4_047_06    m/z 269.1302     0   1E+4   1.5   2  RT (min)      lcqtof_hilic_neg_4_093_60    m/z 269.8109     0   1E+4   0.5   1  RT (min)      lcqtof_hilic_neg_4_016_37    m/z 269.8778     0   1   2E+4   7.5   8  RT (min)      lcqtof_hilic_neg_1_000_aa    m/z 269.8779     0   1   2E+4   7.5   8  RT (min)      lcqtof_hilic_neg_1_000_aa    m/z 269.8780     0   1   2E+4   8   8.5  RT (min)      lcqtof_hilic_neg_1_000_aa    m/z 269.8782     0   1   2E+4   7.5   8  RT (min)      lcqtof_hilic_neg_1_000_aa    m/z 269.8782     0   1   2E+4   7.5   8  RT (min)      lcqtof_hilic_neg_1_000_aa    m/z 269.8787     0   1   2E+4   7.5   8  RT (min)      lcqtof_hilic_neg_1_000_aa    m/z 270.0730     0   1E+4   1   1.5  RT (min)      lcqtof_hilic_neg_4_048_23    m/z 270.1018     0   1E+4   5   5.5  RT (min)      lcqtof_hilic_neg_2_047_32    m/z 270.1020     0   1E+4   6.5   7  RT (min)      lcqtof_hilic_neg_2_028_05    m/z 270.1023     0   1E+4   6.5   7  RT (min)      lcqtof_hilic_neg_4_040_05    m/z 270.1032     0   1E+4   6.5   7  RT (min)      lcqtof_hilic_neg_4_008_24    m/z 270.1101     0   1   2   3   4E+4   4   4.5  RT (min)       lcqtof_hilic_neg_4_090_60    m/z 270.1101     0   1   2   3E+4   4.5   5  RT (min)       lcqtof_hilic_neg_4_055_32    m/z 270.7964     0   1   2E+3   4   4.5  RT (min)      lcqtof_hilic_neg_2_049_55    m/z 270.8280     0   1   2   3E+3   7   7.5  RT (min)       lcqtof_hilic_neg_4_069_57    m/z 270.9170     0   1E+4   8   8.5  RT (min)      lcqtof_hilic_neg_1_000_aa    m/z 270.9314     0   1   2   3E+4   6.5   7  RT (min)      lcqtof_hilic_neg_2_028_05    m/z 270.9752     0   2   4   6   8E+3   2   2.5  RT (min)      lcqtof_hilic_neg_1_060_14    m/z 271.0512     0   1   2   3   4E+3   6   6.5   7  RT (min)       lcqtof_hilic_neg_2_031_01    m/z 271.0513     0   1   2   3E+3   5.5   6  RT (min)      lcqtof_hilic_neg_1_051_35    m/z 271.0689     0   2   4E+4   4   4.5  RT (min)      lcqtof_hilic_neg_4_060_13    m/z 271.0699     0   2   4E+4   4   4.5  RT (min)      lcqtof_hilic_neg_4_019_03    m/z 271.1043     0   1   2E+4   6.5   7  RT (min)      lcqtof_hilic_neg_2_028_05    m/z 271.1045     0   1   2E+4   6.5   7  RT (min)      lcqtof_hilic_neg_4_040_05    m/z 271.1247     0   1E+4   2   2.5  RT (min)      lcqtof_hilic_neg_4_046_41    m/z 271.1291     0   1   2E+4   1.5   2  RT (min)       lcqtof_hilic_neg_4_046_41    m/z 271.2859     0   1   2E+4   1   1.5  RT (min)      lcqtof_hilic_neg_2_028_05    m/z 272.0547     0   1E+4   4   4.5  RT (min)       lcqtof_hilic_neg_4_086_59    m/z 272.0773     0   1E+4   3.5   4  RT (min)      lcqtof_hilic_neg_4_034_31    m/z 272.0922     0   1E+4   1.5   2  RT (min)      lcqtof_hilic_neg_4_018_27    m/z 272.0953     0   1E+4   4   4.5  RT (min)      lcqtof_hilic_neg_4_022_34    m/z 272.1260     0   1E+4   6   6.5  RT (min)       lcqtof_hilic_neg_4_008_24    m/z 272.1857     0   1E+4   15   15.5  RT (min)      lcqtof_hilic_neg_4_000_aa    m/z 272.1862     0   2   4   6   8E+3   1.5   2  RT (min)      lcqtof_hilic_neg_4_032_01    m/z 272.2493     0   1E+4   0.5   1  RT (min)      lcqtof_hilic_neg_4_018_27    m/z 272.2661     0   1   2E+4   0.5   1  RT (min)      lcqtof_hilic_neg_4_050_02    m/z 272.2977     0   1E+4   0.5   1  RT (min)      lcqtof_hilic_neg_4_035_33    m/z 272.8828     0   2   4   6   8E+3   6   6.5  RT (min)      lcqtof_hilic_neg_1_027_27    m/z 272.8829     0   2   4   6E+3   5.5   6  RT (min)       lcqtof_hilic_neg_1_029_06    m/z 272.8830     0   1   2   3E+3   6.5   7  RT (min)       lcqtof_hilic_neg_1_029_06    m/z 272.8831     0   2   4   6   8E+3   5.5   6  RT (min)      lcqtof_hilic_neg_2_004_26    m/z 272.8831     0   2   4E+3   6.5   7  RT (min)       lcqtof_hilic_neg_1_045_07    m/z 272.9224     0   1E+5   8.5   9  RT (min)      lcqtof_hilic_neg_1_000_aa    m/z 272.9224     0   1E+5   8.5   9  RT (min)      lcqtof_hilic_neg_1_000_aa    m/z 272.9226     0   1E+5   8.5   9  RT (min)      lcqtof_hilic_neg_1_000_aa    m/z 272.9227     0   1E+5   9   9.5  RT (min)      lcqtof_hilic_neg_1_000_aa    m/z 272.9228     0   1   2E+5   8.5   9  RT (min)      lcqtof_hilic_neg_1_000_aa    m/z 272.9228     0   1   2E+5   8.5   9  RT (min)      lcqtof_hilic_neg_1_000_aa    m/z 272.9231     0   1E+5   8.5   9  RT (min)      lcqtof_hilic_neg_1_000_aa    m/z 272.9232     0   1   2E+5   7.5   8  RT (min)      lcqtof_hilic_neg_1_000_aa    m/z 272.9334     0   1   2E+4   3.5   4  RT (min)      lcqtof_hilic_neg_1_032_18    m/z 273.0281     0   1   2E+4   1.5   2  RT (min)      lcqtof_hilic_neg_4_050_02    m/z 273.0286     0   1   2   3   4E+4   2   2.5  RT (min)       lcqtof_hilic_neg_4_052_13    m/z 273.0288     0   1   2E+4   1.5   2  RT (min)      lcqtof_hilic_neg_4_050_02    m/z 273.0288     0   1   2E+4   2   2.5  RT (min)      lcqtof_hilic_neg_4_050_02    m/z 273.0333     0   1E+4   4   4.5  RT (min)      lcqtof_hilic_neg_4_050_02    m/z 273.0351     0   1   2   3E+4   2   2.5  RT (min)       lcqtof_hilic_neg_4_060_13    m/z 273.0672     0   1E+4   3.5   4  RT (min)      lcqtof_hilic_neg_1_063_13    m/z 273.0673     0   1E+4   4   4.5  RT (min)      lcqtof_hilic_neg_4_019_03    m/z 273.1052     0   1E+4   4   4.5  RT (min)      lcqtof_hilic_neg_4_055_32    m/z 273.3609     0   1   2E+4   9   9.5  RT (min)      lcqtof_hilic_neg_1_000_aa    m/z 273.8725     0   1   2E+4   0   0.5  RT (min)      lcqtof_hilic_neg_4_000_aa    m/z 273.8729     0   1   2E+4   0.5   1  RT (min)      lcqtof_hilic_neg_4_000_aa    m/z 273.8729     0   1   2E+4   0   0.5  RT (min)      lcqtof_hilic_neg_4_000_aa    m/z 273.8729     0   1   2E+4   0.5   1  RT (min)      lcqtof_hilic_neg_4_000_aa    m/z 273.8730     0   1   2E+4   0   0.5  RT (min)      lcqtof_hilic_neg_4_000_aa    m/z 273.8741     0   1   2E+4   0   0.5  RT (min)      lcqtof_hilic_neg_4_000_aa    m/z 273.8856     0   1   2   3   4E+3   4.5   5  RT (min)      lcqtof_hilic_neg_1_002_39    m/z 273.8858     0   1   2   3   4E+3   6   6.5  RT (min)      lcqtof_hilic_neg_1_004_14    m/z 273.9245     0   1E+4   2.5   3  RT (min)      lcqtof_hilic_neg_4_050_02    m/z 273.9421     0   1   2   3   4E+4   6.5   7  RT (min)      lcqtof_hilic_neg_2_028_05    m/z 274.0358     0   1E+4   3.5   4  RT (min)      lcqtof_hilic_neg_4_038_10    m/z 274.9035     0   1   2E+4   8   8.5  RT (min)      lcqtof_hilic_neg_1_000_aa    m/z 274.9209     0   1E+4   7.5   8  RT (min)      lcqtof_hilic_neg_1_000_aa    m/z 274.9219     0   1E+4   7.5   8  RT (min)      lcqtof_hilic_neg_1_000_aa    m/z 274.9220     0   1E+4   7.5   8  RT (min)      lcqtof_hilic_neg_1_000_aa    m/z 275.0224     0   1E+4   1   1.5  RT (min)      lcqtof_hilic_neg_4_016_37    m/z 275.0885     0   1E+4   4   4.5  RT (min)       lcqtof_hilic_neg_4_085_59    m/z 276.1016     0   1   2E+4   6.5   7  RT (min)       lcqtof_hilic_neg_4_008_24    m/z 276.1016     0   2   4   6   8E+4   5   5.5  RT (min)      lcqtof_hilic_neg_4_008_24    m/z 276.1017     0   1E+5   4.5   5  RT (min)       lcqtof_hilic_neg_4_097_60    m/z 276.1017     0   1E+4   6.5   7  RT (min)      lcqtof_hilic_neg_4_047_06    m/z 276.1020     0   2   4   6   8E+4   5   5.5  RT (min)       lcqtof_hilic_neg_4_008_24    m/z 276.1307     0   1E+4   6   6.5  RT (min)      lcqtof_hilic_neg_4_008_24    m/z 276.8232     0   1E+4   4.5   5  RT (min)      lcqtof_hilic_neg_4_050_02    m/z 277.0158     0   1E+4   0.5   1  RT (min)      lcqtof_hilic_neg_4_050_02    m/z 277.1450     0   1E+4   0.5   1  RT (min)      lcqtof_hilic_neg_2_000_aa    m/z 277.2164     0   1E+5   1   1.5  RT (min)      lcqtof_hilic_neg_2_028_05    m/z 277.2166     0   1E+4   4   4.5  RT (min)      lcqtof_hilic_neg_4_040_05    m/z 277.3528     0   1E+5   1   1.5  RT (min)      lcqtof_hilic_neg_4_061_50    m/z 277.4218     0   2   4   6E+5   0.5   1  RT (min)      lcqtof_hilic_neg_4_050_02    m/z 277.6476     0   1E+4   0.5   1  RT (min)      lcqtof_hilic_neg_4_050_02    m/z 278.0451     0   2   4   6   8E+3   2   2.5  RT (min)      lcqtof_hilic_neg_4_004_47    m/z 278.0734     0   2   4   6   8E+4   2   2.5  RT (min)      lcqtof_hilic_neg_4_050_02    m/z 278.0736     0   2   4   6   8E+4   2   2.5  RT (min)      lcqtof_hilic_neg_4_050_02    m/z 278.1583     0   1E+4   4   4.5  RT (min)      lcqtof_hilic_neg_4_055_32    m/z 278.8213     0   1E+4   4.5   5  RT (min)      lcqtof_hilic_neg_4_050_02    m/z 278.9300     0   2   4   6E+3   4.5   5  RT (min)       lcqtof_hilic_neg_1_000_bb    m/z 278.9765     0   1E+4   5.5   6  RT (min)      lcqtof_hilic_neg_2_004_26    m/z 278.9767     0   1   2   3E+3   5.5   6  RT (min)       lcqtof_hilic_neg_2_053_18    m/z 278.9768     0   1   2   3   4E+3   5.5   6  RT (min)       lcqtof_hilic_neg_1_032_18    m/z 279.1620     0   1   2   3E+4   1   1.5  RT (min)       lcqtof_hilic_neg_4_053_14    m/z 279.1621     0   1   2   3E+4   15   15.5  RT (min)      lcqtof_hilic_neg_4_000_aa    m/z 279.1623     0   1E+4   18   18.5  RT (min)       lcqtof_hilic_neg_4_107_62    m/z 279.1625     0   1E+4   17   17.5  RT (min)       lcqtof_hilic_neg_4_083_59    m/z 279.1626     0   2   4   6   8E+3   15.5   16  RT (min)       lcqtof_hilic_neg_4_083_59    m/z 279.1626     0   1E+4   19   19.5  RT (min)       lcqtof_hilic_neg_4_098_61    m/z 279.1628     0   1   2   3E+3   5   5.5  RT (min)      lcqtof_hilic_neg_4_011_35    m/z 279.1628     0   1E+4   17.5   18  RT (min)       lcqtof_hilic_neg_4_066_57    m/z 279.1628     0   1E+4   0   0.5  RT (min)       lcqtof_hilic_neg_4_019_03    m/z 279.1629     0   1E+4   16   16.5  RT (min)       lcqtof_hilic_neg_4_083_59    m/z 279.1629     0   1   2E+4   1.5   2  RT (min)       lcqtof_hilic_neg_4_047_06    m/z 279.1629     0   1E+4   3   3.5  RT (min)      lcqtof_hilic_neg_4_017_51    m/z 279.1629     0   1E+4   2.5   3  RT (min)      lcqtof_hilic_neg_4_113_62    m/z 279.1630     0   1   2E+4   2   2.5  RT (min)       lcqtof_hilic_neg_4_026_04    m/z 279.1630     0   1   2   3   4E+4   2   3  RT (min)       lcqtof_hilic_neg_4_088_59    m/z 279.1630     0   1   2   3   4E+4   1.5   2  RT (min)       lcqtof_hilic_neg_4_019_03    m/z 279.1630     0   1   2E+4   2.5   3  RT (min)       lcqtof_hilic_neg_4_055_32    m/z 279.1630     0   1E+4   2   2.5  RT (min)      lcqtof_hilic_neg_4_021_21    m/z 279.1630     0   1   2E+4   3   3.5  RT (min)       lcqtof_hilic_neg_4_004_47    m/z 279.1630     0   2   4   6E+4   1   1.5  RT (min)      lcqtof_hilic_neg_4_026_04    m/z 279.1630     0   2   4E+3   6.5   7  RT (min)       lcqtof_hilic_neg_4_012_07    m/z 279.1630     0   1   2   3   4E+4   0   0.5  RT (min)       lcqtof_hilic_neg_4_019_03    m/z 279.1630     0   2   4   6   8E+4   1   1.5  RT (min)      lcqtof_hilic_neg_4_020_42    m/z 279.1630     0   1E+4   17.5   18  RT (min)       lcqtof_hilic_neg_4_108_62    m/z 279.1630     0   1E+4   16.5   17  RT (min)       lcqtof_hilic_neg_4_083_59    m/z 279.1630     0   1   2E+4   2   2.5  RT (min)      lcqtof_hilic_neg_4_039_18    m/z 279.1630     0   1   2   3   4E+4   0   0.5  RT (min)       lcqtof_hilic_neg_4_019_03    m/z 279.1630     0   1   2   3E+4   1.5   2  RT (min)      lcqtof_hilic_neg_4_047_06    m/z 279.1631     0   1E+4   0   0.5  RT (min)      lcqtof_hilic_neg_4_055_32    m/z 279.1631     0   2   4E+4   2   2.5  RT (min)       lcqtof_hilic_neg_4_003_16    m/z 279.1631     0   2   4   6E+4   1   1.5  RT (min)       lcqtof_hilic_neg_4_026_04    m/z 279.1631     0   1E+4   20   20.5  RT (min)       lcqtof_hilic_neg_4_107_62    m/z 279.1631     0   1   2   3E+4   0   0.5  RT (min)      lcqtof_hilic_neg_4_006_30    m/z 279.1631     0   1E+4   17   18   19  RT (min)       lcqtof_hilic_neg_4_073_57    m/z 279.1632     0   1   2   3   4E+3   3.5   4  RT (min)       lcqtof_hilic_neg_4_059_14    m/z 279.1632     0   1E+4   18   18.5   19  RT (min)       lcqtof_hilic_neg_4_029_17    m/z 279.1632     0   1E+4   19   19.5  RT (min)       lcqtof_hilic_neg_4_012_07    m/z 279.1632     0   1   2   3   4E+4   2.5   3  RT (min)       lcqtof_hilic_neg_4_045_26    m/z 279.1632     0   2   4   6E+4   1   1.5  RT (min)       lcqtof_hilic_neg_4_056_56    m/z 279.1632     0   1E+4   1.5   2  RT (min)       lcqtof_hilic_neg_4_021_21    m/z 279.1632     0   1E+4   18   18.5  RT (min)       lcqtof_hilic_neg_4_113_62    m/z 279.1632     0   1E+4   19   19.5   20  RT (min)       lcqtof_hilic_neg_4_007_45    m/z 279.1632     0   1E+4   18  RT (min)       lcqtof_hilic_neg_4_097_60    m/z 279.1633     0   1E+4   18.5   19  RT (min)       lcqtof_hilic_neg_4_009_50    m/z 279.1633     0   2   4   6E+3   3.5   4  RT (min)       lcqtof_hilic_neg_4_104_61    m/z 279.1634     0   1E+4   19.5   20  RT (min)       lcqtof_hilic_neg_4_047_06    m/z 279.1636     0   1E+4   16   16.5  RT (min)       lcqtof_hilic_neg_4_102_61    m/z 279.1636     0   1E+5   1   1.5  RT (min)      lcqtof_hilic_neg_4_032_01    m/z 279.2325     0   2   4   6E+4   1   1.5  RT (min)      lcqtof_hilic_neg_4_002_25    m/z 279.2326     0   1E+5   1   1.5  RT (min)      lcqtof_hilic_neg_2_028_05    m/z 279.2326     0   1   2E+4   3   3.5  RT (min)      lcqtof_hilic_neg_4_028_36    m/z 279.3719     0   1   2   3   4E+2   0.5   1  RT (min)      lcqtof_hilic_neg_4_050_02    m/z 279.5254     0   2   4E+4   0.5   1  RT (min)      lcqtof_hilic_neg_4_050_02    m/z 280.0614     0   1E+4   2   2.5  RT (min)      lcqtof_hilic_neg_4_055_32    m/z 280.0698     0   1E+4   2   2.5  RT (min)      lcqtof_hilic_neg_4_050_02    m/z 280.1298     0   1E+4   1   1.5  RT (min)      lcqtof_hilic_neg_4_015_09    m/z 280.1302     0   1E+4   0.5   1  RT (min)      lcqtof_hilic_neg_4_038_10    m/z 280.1661     0   1   2E+3   0   0.5  RT (min)      lcqtof_hilic_neg_4_002_25    m/z 280.1662     0   1E+4   1   1.5  RT (min)      lcqtof_hilic_neg_4_019_03    m/z 280.2592     0   2   4   6E+4   0.5   1  RT (min)      lcqtof_hilic_neg_4_050_02    m/z 280.8596     0   1E+4   8.5   9  RT (min)      lcqtof_hilic_neg_1_000_aa    m/z 280.8597     0   1E+4   8.5   9  RT (min)      lcqtof_hilic_neg_1_000_aa    m/z 280.8598     0   1E+4   8.5   9  RT (min)      lcqtof_hilic_neg_1_000_aa    m/z 280.9858     0   1   2   3E+3   5   5.5  RT (min)      lcqtof_hilic_neg_1_060_14    m/z 281.0266     0   1E+4   3.5   4  RT (min)      lcqtof_hilic_neg_1_032_18    m/z 281.0282     0   1   2   3E+4   3   3.5  RT (min)      lcqtof_hilic_neg_4_039_18    m/z 281.0591     0   1   2E+4   2   2.5  RT (min)      lcqtof_hilic_neg_4_055_32    m/z 281.0640     0   1E+4   4   4.5  RT (min)      lcqtof_hilic_neg_4_050_02    m/z 281.0985     0   1E+4   3.5   4  RT (min)       lcqtof_hilic_neg_4_052_13    m/z 281.1382     0   1E+5   2   2.5  RT (min)      lcqtof_hilic_neg_4_045_26    m/z 281.1386     0   1   2E+4   2.5   3  RT (min)      lcqtof_hilic_neg_4_105_61    m/z 281.1387     0   2   4   6E+5   1.5   2  RT (min)      lcqtof_hilic_neg_4_045_26    m/z 281.1391     0   1   2E+4   3   3.5  RT (min)       lcqtof_hilic_neg_4_104_61    m/z 281.1924     0   2   4   6E+4   0.5   1  RT (min)      lcqtof_hilic_neg_4_050_02    m/z 281.2479     0   2   4   6E+3   3   3.5  RT (min)      lcqtof_hilic_neg_4_028_36    m/z 281.2480     0   1E+4   3   3.5  RT (min)      lcqtof_hilic_neg_4_048_23    m/z 281.2734     0   2   4E+4   0.5   1  RT (min)      lcqtof_hilic_neg_4_039_18    m/z 281.3900     0   1E+5   0.5   1  RT (min)      lcqtof_hilic_neg_4_039_18    m/z 281.3961     0   1E+5   0.5   1  RT (min)      lcqtof_hilic_neg_4_006_30    m/z 281.4156     0   1   2E+5   0.5   1  RT (min)      lcqtof_hilic_neg_4_050_02    m/z 281.4312     0   1E+5   0.5   1  RT (min)      lcqtof_hilic_neg_4_010_28    m/z 281.4992     0   2   4E+4   0.5   1  RT (min)      lcqtof_hilic_neg_4_039_18    m/z 281.6112     0   2   4E+4   0.5   1  RT (min)      lcqtof_hilic_neg_4_039_18    m/z 281.8517     0   1E+4   7.5   8  RT (min)      lcqtof_hilic_neg_1_000_aa    m/z 281.9163     0   1E+5   1   1.5  RT (min)      lcqtof_hilic_neg_2_050_19    m/z 282.0743     0   1E+4   3   3.5  RT (min)      lcqtof_hilic_neg_4_008_24    m/z 282.0880     0   1E+4   1   1.5  RT (min)      lcqtof_hilic_neg_4_003_16    m/z 282.1422     0   1E+5   1.5   2  RT (min)      lcqtof_hilic_neg_4_045_26    m/z 282.1923     0   1E+5   0.5   1  RT (min)      lcqtof_hilic_neg_4_050_02    m/z 282.2070     0   2   4E+4   0.5   1  RT (min)      lcqtof_hilic_neg_4_023_15    m/z 282.2264     0   1E+5   0.5   1  RT (min)      lcqtof_hilic_neg_4_039_18    m/z 282.2744     0   1   2   3E+4   1   1.5  RT (min)      lcqtof_hilic_neg_4_039_18    m/z 282.3098     0   1   2E+5   0.5   1  RT (min)      lcqtof_hilic_neg_4_039_18    m/z 282.3494     0   2   4   6E+4   0.5   1  RT (min)      lcqtof_hilic_neg_4_039_18    m/z 282.3881     0   2   4   6   8E+4   0.5   1  RT (min)      lcqtof_hilic_neg_4_039_18    m/z 282.4130     0   2   4   6E+4   0.5   1  RT (min)      lcqtof_hilic_neg_4_050_02    m/z 282.4435     0   2   4   6   8E+4   0.5   1  RT (min)      lcqtof_hilic_neg_4_039_18    m/z 283.0729     0   1E+4   2.5   3  RT (min)       lcqtof_hilic_neg_4_085_59    m/z 283.1007     0   1   2   3E+4   5   5.5  RT (min)       lcqtof_hilic_neg_4_048_23    m/z 283.1189     0   2   4E+3   1   1.5  RT (min)      lcqtof_hilic_neg_4_055_32    m/z 283.1348     0   1   2   3   4E+4   0.5   1  RT (min)      lcqtof_hilic_neg_4_050_02    m/z 283.1378     0   2   4E+3   2   2.5  RT (min)      lcqtof_hilic_neg_4_005_29    m/z 283.2639     0   1E+4   14.5   15  RT (min)      lcqtof_hilic_neg_4_081_58    m/z 283.2716     0   1   2E+4   1   1.5  RT (min)      lcqtof_hilic_neg_4_045_26    m/z 283.2780     0   2   4   6E+4   0.5   1  RT (min)      lcqtof_hilic_neg_4_050_02    m/z 283.3102     0   2   4   6E+4   1   1.5  RT (min)      lcqtof_hilic_neg_4_039_18    m/z 283.3117     0   2   4   6E+4   0.5   1  RT (min)      lcqtof_hilic_neg_4_039_18    m/z 283.8850     0   1E+4   0.5   1   1.5  RT (min)      lcqtof_hilic_neg_4_027_19    m/z 284.0346     0   1E+4   1.5   2  RT (min)      lcqtof_hilic_neg_4_064_15    m/z 284.0348     0   1E+4   1.5   2  RT (min)      lcqtof_hilic_neg_4_023_15    m/z 284.0758     0   1E+4   2.5   3  RT (min)      lcqtof_hilic_neg_4_054_48    m/z 284.0762     0   1   2   3   4E+4   2.5   3  RT (min)      lcqtof_hilic_neg_4_047_06    m/z 284.0765     0   1   2E+4   2.5   3  RT (min)      lcqtof_hilic_neg_4_044_08    m/z 285.0380     0   2   4E+4   1.5   2  RT (min)      lcqtof_hilic_neg_4_039_18    m/z 285.0396     0   2   4   6   8E+4   1.5   2  RT (min)      lcqtof_hilic_neg_1_032_18    m/z 285.0401     0   2   4E+4   1   1.5  RT (min)      lcqtof_hilic_neg_4_039_18    m/z 285.1204     0   1   2E+4   4   4.5  RT (min)       lcqtof_hilic_neg_4_095_60    m/z 285.1209     0   1   2   3E+4   4.5   5  RT (min)       lcqtof_hilic_neg_4_047_06    m/z 285.2070     0   1E+4   1   1.5  RT (min)      lcqtof_hilic_neg_4_053_14    m/z 285.2641     0   1   2E+4   1   1.5  RT (min)      lcqtof_hilic_neg_4_008_24    m/z 285.9164     0   1E+4   4.5   5  RT (min)      lcqtof_hilic_neg_4_039_18    m/z 285.9200     0   1   2   3E+3   7   7.5  RT (min)      lcqtof_hilic_neg_1_023_34    m/z 286.0429     0   1E+4   1.5   2  RT (min)      lcqtof_hilic_neg_1_032_18    m/z 286.1037     0   1E+4   4   4.5  RT (min)       lcqtof_hilic_neg_4_040_05    m/z 286.9116     0   2   4   6   8E+3   4.5   5  RT (min)      lcqtof_hilic_neg_1_000_bb    m/z 287.0020     0   1   2   3   4E+4   1   1.5  RT (min)      lcqtof_hilic_neg_4_058_39    m/z 287.0472     0   1   2   3E+4   4.5   5  RT (min)      lcqtof_hilic_neg_4_045_26    m/z 287.0537     0   1E+5   3.5   4  RT (min)      lcqtof_hilic_neg_4_050_02    m/z 287.0579     0   1E+4   2   2.5  RT (min)      lcqtof_hilic_neg_4_053_14    m/z 287.1228     0   1E+4   5.5   6  RT (min)      lcqtof_hilic_neg_4_028_36    m/z 287.6110     0   1E+4   4   4.5  RT (min)      lcqtof_hilic_neg_4_055_32    m/z 287.8360     0   1   2E+4   0.5   1  RT (min)      lcqtof_hilic_neg_4_055_32    m/z 288.0044     0   1E+4   1   1.5  RT (min)      lcqtof_hilic_neg_4_019_03    m/z 288.1195     0   1E+4   5   5.5  RT (min)      lcqtof_hilic_neg_4_040_05    m/z 288.1294     0   1E+4   3.5   4  RT (min)      lcqtof_hilic_neg_4_008_24    m/z 288.9357     0   2   4   6E+3   8.5   9  RT (min)       lcqtof_hilic_neg_4_053_14    m/z 288.9359     0   2   4   6E+3   8.5   9  RT (min)      lcqtof_hilic_neg_4_053_14    m/z 288.9985     0   2   4E+4   1   1.5  RT (min)      lcqtof_hilic_neg_4_019_03    m/z 289.0372     0   1   2   3E+4   0.5   1  RT (min)      lcqtof_hilic_neg_4_000_aa    m/z 289.0649     0   1E+3   2   2.5  RT (min)       lcqtof_hilic_neg_4_074_58    m/z 289.0926     0   2   4   6E+4   2.5   3  RT (min)       lcqtof_hilic_neg_4_091_60    m/z 289.1185     0   1   2E+4   1.5   2  RT (min)       lcqtof_hilic_neg_4_053_14    m/z 289.1363     0   1E+4   6.5   7  RT (min)      lcqtof_hilic_neg_4_008_24    m/z 289.9872     0   1   2   3E+4   1   1.5  RT (min)      lcqtof_hilic_neg_4_004_47    m/z 290.0062     0   1E+4   1   1.5  RT (min)      lcqtof_hilic_neg_4_019_03    m/z 290.0879     0   1   2E+4   5   5.5  RT (min)       lcqtof_hilic_neg_4_034_31    m/z 290.1330     0   1E+4   5.5   6  RT (min)      lcqtof_hilic_neg_4_055_32    m/z 290.9240     0   1   2   3E+4   0.5   1  RT (min)      lcqtof_hilic_neg_4_000_aa    m/z 290.9244     0   1   2   3E+4   0   0.5  RT (min)      lcqtof_hilic_neg_4_000_aa    m/z 290.9245     0   1   2   3E+4   0   0.5  RT (min)      lcqtof_hilic_neg_4_000_aa    m/z 290.9245     0   1   2   3E+4   0   0.5  RT (min)      lcqtof_hilic_neg_4_000_aa    m/z 290.9249     0   1   2   3E+4   0.5   1  RT (min)      lcqtof_hilic_neg_4_000_aa    m/z 290.9250     0   1   2   3E+4   0.5   1  RT (min)      lcqtof_hilic_neg_4_000_aa    m/z 290.9251     0   1   2   3E+4   0.5   1  RT (min)      lcqtof_hilic_neg_4_000_aa    m/z 290.9851     0   1   2E+4   1   1.5  RT (min)      lcqtof_hilic_neg_4_004_47    m/z 291.0075     0   1E+4   1   1.5  RT (min)      lcqtof_hilic_neg_4_019_03    m/z 291.0708     0   2   4   6E+3   3.5   4  RT (min)      lcqtof_hilic_neg_1_003_37    m/z 291.1207     0   1   2E+4   4   4.5  RT (min)       lcqtof_hilic_neg_4_090_60    m/z 291.2190     0   1   2E+4   7   7.5  RT (min)      lcqtof_hilic_neg_1_060_14    m/z 291.2912     0   2   4   6E+4   7   7.5  RT (min)      lcqtof_hilic_neg_1_060_14    m/z 291.3352     0   2   4   6E+4   7   7.5  RT (min)      lcqtof_hilic_neg_1_060_14    m/z 291.4860     0   2   4   6E+4   7   7.5  RT (min)      lcqtof_hilic_neg_1_060_14    m/z 291.9710     0   2   4   6   8E+3   3   3.5  RT (min)      lcqtof_hilic_neg_1_037_24    m/z 291.9711     0   2   4   6   8E+3   3   3.5  RT (min)      lcqtof_hilic_neg_1_037_24    m/z 291.9711     0   2   4   6   8E+3   3   3.5  RT (min)      lcqtof_hilic_neg_1_037_24    m/z 291.9899     0   1E+4   1   1.5  RT (min)      lcqtof_hilic_neg_4_038_10    m/z 292.1740     0   1E+4   7   7.5  RT (min)      lcqtof_hilic_neg_2_051_14    m/z 292.2208     0   1E+4   7   7.5  RT (min)      lcqtof_hilic_neg_2_051_14    m/z 292.2572     0   2   4   6E+4   7   7.5  RT (min)      lcqtof_hilic_neg_1_060_14    m/z 292.2993     0   1E+4   7   7.5  RT (min)      lcqtof_hilic_neg_2_051_14    m/z 292.3353     0   1E+4   7   7.5  RT (min)      lcqtof_hilic_neg_1_004_14    m/z 292.8100     0   1E+4   8   8.5  RT (min)      lcqtof_hilic_neg_1_000_aa    m/z 292.9972     0   2   4   6   8E+3   2   2.5  RT (min)      lcqtof_hilic_neg_4_050_02    m/z 293.0751     0   1   2E+4   2.5   3  RT (min)      lcqtof_hilic_neg_4_086_59    m/z 293.0756     0   1E+4   2.5   3  RT (min)      lcqtof_hilic_neg_4_047_06    m/z 293.0773     0   1E+4   2   2.5  RT (min)      lcqtof_hilic_neg_4_053_14    m/z 293.0876     0   1E+4   0.5   1  RT (min)      lcqtof_hilic_neg_4_004_47    m/z 293.1363     0   1   2   3   4E+4   5   5.5  RT (min)      lcqtof_hilic_neg_1_037_24    m/z 293.1414     0   1   2   3E+4   5   5.5  RT (min)      lcqtof_hilic_neg_4_008_24    m/z 293.1418     0   2   4E+4   15   15.5  RT (min)      lcqtof_hilic_neg_4_000_aa    m/z 293.1424     0   2   4   6   8E+3   1.5   2  RT (min)      lcqtof_hilic_neg_4_002_25    m/z 293.1424     0   2   4   6   8E+4   1.5   2  RT (min)      lcqtof_hilic_neg_4_002_25    m/z 293.1426     0   1E+4   3   3.5  RT (min)      lcqtof_hilic_neg_4_008_24    m/z 293.1426     0   1E+4   3.5   4  RT (min)      lcqtof_hilic_neg_4_002_25    m/z 293.1426     0   1E+4   1.5   2  RT (min)      lcqtof_hilic_neg_4_039_18    m/z 293.1426     0   1E+4   3   3.5  RT (min)      lcqtof_hilic_neg_4_034_31    m/z 293.1427     0   1   2   3   4E+3   3.5   4  RT (min)      lcqtof_hilic_neg_4_075_58    m/z 293.1427     0   1   2E+4   2   2.5  RT (min)      lcqtof_hilic_neg_4_029_17    m/z 293.1427     0   1   2E+4   2.5   3  RT (min)      lcqtof_hilic_neg_4_048_23    m/z 293.1427     0   1   2E+4   2   2.5  RT (min)       lcqtof_hilic_neg_4_052_13    m/z 293.1427     0   1E+4   3   3.5  RT (min)      lcqtof_hilic_neg_4_002_25    m/z 293.1427     0   1E+4   2   2.5  RT (min)      lcqtof_hilic_neg_4_026_04    m/z 293.1427     0   1E+4   1.5   2  RT (min)      lcqtof_hilic_neg_4_039_18    m/z 293.1427     0   2   4   6   8E+4   1.5   2  RT (min)      lcqtof_hilic_neg_4_008_24    m/z 293.1427     0   1   2E+4   2   2.5  RT (min)      lcqtof_hilic_neg_4_032_01    m/z 293.1428     0   1   2   3E+4   4  RT (min)       lcqtof_hilic_neg_4_089_59    m/z 293.1428     0   1   2E+4   2.5   3  RT (min)      lcqtof_hilic_neg_4_005_29    m/z 293.1429     0   1   2E+4   2.5   3  RT (min)      lcqtof_hilic_neg_4_016_37    m/z 293.1431     0   1E+4   4   4.5  RT (min)      lcqtof_hilic_neg_4_031_38    m/z 293.1431     0   1E+4   3.5   4  RT (min)      lcqtof_hilic_neg_4_008_24    m/z 293.1755     0   1E+5   15   15.5  RT (min)      lcqtof_hilic_neg_4_000_aa    m/z 293.1764     0   1E+4   6   7   8  RT (min)       lcqtof_hilic_neg_4_040_05    m/z 293.1768     0   1E+5   15   15.5  RT (min)      lcqtof_hilic_neg_4_000_aa    m/z 293.1770     0   1E+4   5   5.5  RT (min)       lcqtof_hilic_neg_4_051_43    m/z 293.1771     0   1E+4   6   6.5  RT (min)      lcqtof_hilic_neg_4_000_cc    m/z 293.1777     0   1E+4   3.5   4  RT (min)      lcqtof_hilic_neg_4_084_59    m/z 293.1778     0   2   4   6   8E+3   5   5.5  RT (min)      lcqtof_hilic_neg_4_047_06    m/z 293.1780     0   2   4   6E+3   5   5.5  RT (min)      lcqtof_hilic_neg_4_005_29    m/z 293.1780     0   1E+5   0   0.5  RT (min)      lcqtof_hilic_neg_4_081_58    m/z 293.1781     0   1E+6   1   1.5  RT (min)      lcqtof_hilic_neg_4_004_47    m/z 293.1782     0   2   4   6E+4   2   2.5  RT (min)       lcqtof_hilic_neg_4_003_16    m/z 293.1782     0   1E+4   4   4.5  RT (min)      lcqtof_hilic_neg_4_019_03    m/z 293.1782     0   1E+4   3   3.5  RT (min)       lcqtof_hilic_neg_4_053_14    m/z 293.1782     0   1E+4   18   19  RT (min)       lcqtof_hilic_neg_4_083_59    m/z 293.1782     0   2   4   6E+4   2   2.5  RT (min)      lcqtof_hilic_neg_4_053_14    m/z 293.1782     0   2   4   6E+4   2.5   3  RT (min)       lcqtof_hilic_neg_4_086_59    m/z 293.1782     0   1E+5   1   1.5  RT (min)       lcqtof_hilic_neg_4_027_19    m/z 293.1783     0   1E+4   0   0.5  RT (min)       lcqtof_hilic_neg_4_100_61    m/z 293.1783     0   1   2   3E+4   1.5   2  RT (min)       lcqtof_hilic_neg_4_019_03    m/z 293.1783     0   1E+4   3   3.5  RT (min)      lcqtof_hilic_neg_4_100_61    m/z 293.1783     0   1E+4   17.5   18  RT (min)       lcqtof_hilic_neg_4_110_62    m/z 293.1784     0   1E+4   4   4.5  RT (min)      lcqtof_hilic_neg_4_051_43    m/z 293.1784     0   1   2E+4   4.5   5  RT (min)      lcqtof_hilic_neg_4_017_51    m/z 293.1784     0   1E+4   16.5   17   17.5  RT (min)       lcqtof_hilic_neg_4_081_58    m/z 293.1784     0   1   2E+4   3.5   4  RT (min)      lcqtof_hilic_neg_4_008_24    m/z 293.1784     0   2   4   6E+4   0   0.5  RT (min)       lcqtof_hilic_neg_4_101_61    m/z 293.1784     0   1E+4   16.5   17  RT (min)       lcqtof_hilic_neg_4_111_62    m/z 293.1784     0   1E+4   19   19.5  RT (min)       lcqtof_hilic_neg_4_012_07    m/z 293.1784     0   2   4   6E+4   2   2.5  RT (min)      lcqtof_hilic_neg_4_053_14    m/z 293.1784     0   1E+4   17   18  RT (min)       lcqtof_hilic_neg_4_052_13    m/z 293.1784     0   1E+4   19   20  RT (min)       lcqtof_hilic_neg_4_100_61    m/z 293.1785     0   2   4E+3   15   15.5  RT (min)      lcqtof_hilic_neg_2_055_30    m/z 293.1785     0   1   2E+4   19   19.5  RT (min)       lcqtof_hilic_neg_4_019_03    m/z 293.1785     0   2   4   6E+4   2   2.5  RT (min)       lcqtof_hilic_neg_4_032_01    m/z 293.1786     0   1E+5   1.5   2  RT (min)      lcqtof_hilic_neg_4_012_07    m/z 293.1786     0   2   4   6   8E+3   4.5   5  RT (min)      lcqtof_hilic_neg_4_019_03    m/z 293.1787     0   1E+5   1   1.5  RT (min)       lcqtof_hilic_neg_4_001_49    m/z 293.1787     0   1E+4   16   17   18  RT (min)       lcqtof_hilic_neg_4_005_29    m/z 293.1788     0   2   4E+4   1.5   2  RT (min)      lcqtof_hilic_neg_4_019_03    m/z 293.1788     0   2   4   6   8E+3   6   6.5  RT (min)      lcqtof_hilic_neg_4_095_60    m/z 293.1788     0   1   2E+4   3.5   4  RT (min)      lcqtof_hilic_neg_4_056_56    m/z 293.1788     0   1E+4   3.5   4  RT (min)      lcqtof_hilic_neg_4_098_61    m/z 293.1789     0   2   4   6E+3   6.5   7  RT (min)      lcqtof_hilic_neg_4_089_59    m/z 293.1789     0   2   4   6E+4   2   2.5  RT (min)      lcqtof_hilic_neg_4_023_15    m/z 293.1889     0   1   2E+5   0.5   1  RT (min)      lcqtof_hilic_neg_3_003_20    m/z 293.1894     0   1   2E+4   0.5   1  RT (min)      lcqtof_hilic_neg_3_000_bb    m/z 293.1898     0   1E+4   0.5   1  RT (min)      lcqtof_hilic_neg_3_000_aa    m/z 293.1920     0   1   2   3   4E+4   1   1.5  RT (min)      lcqtof_hilic_neg_4_003_16    m/z 293.2003     0   2   4   6E+4   4   4.5  RT (min)      lcqtof_hilic_neg_2_036_34    m/z 293.3666     0   1   2   3E+4   4   4.5  RT (min)      lcqtof_hilic_neg_2_036_34    m/z 293.4202     0   2   4E+4   4   4.5  RT (min)      lcqtof_hilic_neg_2_036_34    m/z 293.4861     0   2   4   6E+4   0.5   1  RT (min)      lcqtof_hilic_neg_2_016_33    m/z 293.5182     0   1   2   3E+4   4   4.5  RT (min)      lcqtof_hilic_neg_2_036_34    m/z 294.0666     0   1   2   3E+4   4.5   5  RT (min)       lcqtof_hilic_neg_4_008_24    m/z 294.0670     0   1   2   3E+4   4.5   5  RT (min)      lcqtof_hilic_neg_4_008_24    m/z 294.0671     0   1E+4   3.5   4  RT (min)       lcqtof_hilic_neg_4_090_60    m/z 294.0897     0   1E+4   0.5   1  RT (min)      lcqtof_hilic_neg_4_055_32    m/z 294.1455     0   1   2   3   4E+3   2   2.5  RT (min)      lcqtof_hilic_neg_4_003_16    m/z 294.1456     0   1   2   3   4E+3   2   2.5  RT (min)      lcqtof_hilic_neg_4_033_53    m/z 294.1810     0   1   2E+4   15   15.5  RT (min)      lcqtof_hilic_neg_4_000_aa    m/z 294.1815     0   2   4   6   8E+3   2.5   3  RT (min)      lcqtof_hilic_neg_4_086_59    m/z 294.1816     0   1   2E+4   1.5   2  RT (min)      lcqtof_hilic_neg_4_013_20    m/z 294.1819     0   1E+4   2   2.5  RT (min)      lcqtof_hilic_neg_4_010_28    m/z 294.1826     0   2   4   6E+4   1   1.5  RT (min)      lcqtof_hilic_neg_4_003_16    m/z 294.1924     0   2   4E+4   0.5   1  RT (min)      lcqtof_hilic_neg_3_003_20    m/z 294.2904     0   1   2E+4   4   4.5  RT (min)      lcqtof_hilic_neg_2_036_34    m/z 294.9524     0   1   2E+3   8   8.5  RT (min)       lcqtof_hilic_neg_4_030_44    m/z 294.9528     0   1   2E+3   8   8.5  RT (min)       lcqtof_hilic_neg_4_019_03    m/z 294.9928     0   1E+4   0   0.5  RT (min)      lcqtof_hilic_neg_4_000_aa    m/z 294.9929     0   1   2   3E+4   7.5   8  RT (min)      lcqtof_hilic_neg_4_053_14    m/z 294.9938     0   1E+4   0.5   1  RT (min)      lcqtof_hilic_neg_4_000_aa    m/z 295.0044     0   1E+4   6   6.5  RT (min)       lcqtof_hilic_neg_4_064_15    m/z 295.0048     0   1E+4   6   6.5  RT (min)       lcqtof_hilic_neg_4_032_01    m/z 295.0342     0   1E+4   2.5   3  RT (min)      lcqtof_hilic_neg_4_082_59    m/z 295.0344     0   1E+4   2.5   3  RT (min)      lcqtof_hilic_neg_4_050_02    m/z 295.0451     0   1E+4   4   4.5  RT (min)      lcqtof_hilic_neg_4_050_02    m/z 295.0526     0   1   2E+4   2   2.5  RT (min)      lcqtof_hilic_neg_4_039_18    m/z 295.0589     0   1   2   3E+4   4   4.5  RT (min)      lcqtof_hilic_neg_4_048_23    m/z 295.0650     0   1   2   3E+4   4.5   5  RT (min)      lcqtof_hilic_neg_4_048_23    m/z 295.0662     0   1   2   3E+4   4   4.5  RT (min)       lcqtof_hilic_neg_4_048_23    m/z 295.0666     0   2   4   6   8E+3   4   4.5  RT (min)      lcqtof_hilic_neg_1_035_23    m/z 295.0670     0   2   4   6E+3   2   2.5  RT (min)      lcqtof_hilic_neg_4_036_11    m/z 295.0675     0   1   2   3E+4   2   2.5  RT (min)      lcqtof_hilic_neg_4_050_02    m/z 295.0818     0   1E+4   0.5   1  RT (min)      lcqtof_hilic_neg_2_005_13    m/z 295.0951     0   1E+4   2   2.5  RT (min)      lcqtof_hilic_neg_4_053_14    m/z 295.1038     0   2   4   6E+4   5.5   6  RT (min)      lcqtof_hilic_neg_1_005_32    m/z 295.1375     0   1E+4   0.5   1  RT (min)      lcqtof_hilic_neg_4_013_20    m/z 295.1398     0   1E+4   0.5   1   1.5  RT (min)      lcqtof_hilic_neg_4_038_10    m/z 295.1398     0   2   4E+3   2   2.5  RT (min)      lcqtof_hilic_neg_4_006_30    m/z 295.1402     0   2   4   6   8E+3   0.5   1  RT (min)      lcqtof_hilic_neg_4_007_45    m/z 295.1408     0   1   2E+3   17.5   18  RT (min)      lcqtof_hilic_neg_4_008_24    m/z 295.1421     0   1E+4   0.5   1  RT (min)      lcqtof_hilic_neg_4_040_05    m/z 295.2771     0   1   2E+5   1   1.5  RT (min)      lcqtof_hilic_neg_2_028_05    m/z 295.2772     0   1E+4   4   4.5  RT (min)      lcqtof_hilic_neg_2_028_05    m/z 295.2774     0   2   4   6   8E+4   1   1.5  RT (min)      lcqtof_hilic_neg_4_002_25    m/z 295.2776     0   1   2E+4   4   4.5  RT (min)      lcqtof_hilic_neg_4_008_24    m/z 295.2779     0   1   2E+4   4   4.5  RT (min)      lcqtof_hilic_neg_4_008_24    m/z 296.0621     0   1E+4   4.5   5  RT (min)      lcqtof_hilic_neg_4_008_24    m/z 297.0820     0   1E+4   3.5   4  RT (min)       lcqtof_hilic_neg_4_050_02    m/z 297.0825     0   1E+4   3.5   4  RT (min)      lcqtof_hilic_neg_4_052_13    m/z 297.1519     0   2   4E+5   15   15.5  RT (min)      lcqtof_hilic_neg_4_000_aa    m/z 297.1520     0   1   2E+4   19.5   20  RT (min)       lcqtof_hilic_neg_4_090_60    m/z 297.1521     0   1   2E+4   16   16.5  RT (min)       lcqtof_hilic_neg_4_104_61    m/z 297.1521     0   1E+3   14   14.5  RT (min)      lcqtof_hilic_neg_4_083_59    m/z 297.1521     0   1E+5   1.5   2  RT (min)      lcqtof_hilic_neg_4_016_37    m/z 297.1522     0   1   2   3E+4   17.5   18  RT (min)       lcqtof_hilic_neg_4_111_62    m/z 297.1522     0   1   2   3E+4   19.5   20  RT (min)       lcqtof_hilic_neg_4_106_62    m/z 297.1522     0   1   2   3E+4   16   16.5   17  RT (min)       lcqtof_hilic_neg_4_103_61    m/z 297.1522     0   1   2   3E+5   1.5   2  RT (min)       lcqtof_hilic_neg_4_050_02    m/z 297.1522     0   1   2   3E+4   16.5   17  RT (min)       lcqtof_hilic_neg_4_107_62    m/z 297.1522     0   1   2   3E+4   19   19.5  RT (min)       lcqtof_hilic_neg_4_105_61    m/z 297.1522     0   1   2E+4   15.5   16  RT (min)       lcqtof_hilic_neg_4_110_62    m/z 297.1522     0   1   2E+4   15.5   16   16.5  RT (min)       lcqtof_hilic_neg_4_082_59    m/z 297.1522     0   1   2   3E+4   18  RT (min)       lcqtof_hilic_neg_4_083_59    m/z 297.1523     0   2   4E+5   15   15.5   16  RT (min)       lcqtof_hilic_neg_4_000_aa    m/z 297.1523     0   1   2   3E+4   19   19.5  RT (min)       lcqtof_hilic_neg_4_073_57    m/z 297.1523     0   1   2   3E+4   17.5   18  RT (min)       lcqtof_hilic_neg_4_012_07    m/z 297.1523     0   1   2   3E+5   0   0.5  RT (min)       lcqtof_hilic_neg_4_019_03    m/z 297.1523     0   1   2   3E+4   17   17.5  RT (min)       lcqtof_hilic_neg_4_106_62    m/z 297.1523     0   1   2   3E+4   19   19.5  RT (min)       lcqtof_hilic_neg_4_109_62    m/z 297.1523     0   1   2   3E+4   16   16.5  RT (min)       lcqtof_hilic_neg_4_103_61    m/z 297.1523     0   1   2E+4   15.5   16   16.5  RT (min)       lcqtof_hilic_neg_4_082_59    m/z 297.1523     0   2   4   6E+4   1.5   2  RT (min)       lcqtof_hilic_neg_4_019_03    m/z 297.1523     0   1   2   3E+4   16.5   17  RT (min)       lcqtof_hilic_neg_4_111_62    m/z 297.1523     0   1   2E+4   4.5   5  RT (min)      lcqtof_hilic_neg_4_040_05    m/z 297.1523     0   1   2   3E+4   0   0.5  RT (min)       lcqtof_hilic_neg_4_012_07    m/z 297.1524     0   1   2   3E+4   0   0.5  RT (min)       lcqtof_hilic_neg_4_019_03    m/z 297.1524     0   1   2   3E+4   17   18  RT (min)       lcqtof_hilic_neg_4_105_61    m/z 297.1524     0   1   2   3E+4   17.5   18  RT (min)       lcqtof_hilic_neg_4_112_62    m/z 297.1524     0   1E+5   2.5   3  RT (min)       lcqtof_hilic_neg_4_086_59    m/z 297.1524     0   2   4   6E+4   3   4  RT (min)       lcqtof_hilic_neg_4_002_25    m/z 297.1524     0   1   2E+4   16   16.5  RT (min)       lcqtof_hilic_neg_4_073_57    m/z 297.1524     0   1   2   3   4E+4   2   2.5  RT (min)      lcqtof_hilic_neg_4_039_18    m/z 297.1524     0   1   2   3E+4   0   0.5  RT (min)       lcqtof_hilic_neg_4_019_03    m/z 297.1524     0   1   2   3E+4   0   0.5  RT (min)       lcqtof_hilic_neg_4_019_03    m/z 297.1524     0   1E+4   14.5   15  RT (min)      lcqtof_hilic_neg_4_083_59    m/z 297.1524     0   2   4E+3   5   5.5  RT (min)      lcqtof_hilic_neg_4_053_14    m/z 297.1524     0   2   4   6E+4   3   4  RT (min)       lcqtof_hilic_neg_4_048_23    m/z 297.1524     0   1   2   3E+4   18   18.5  RT (min)       lcqtof_hilic_neg_4_022_34    m/z 297.1524     0   1   2   3E+4   0   0.5  RT (min)      lcqtof_hilic_neg_4_019_03    m/z 297.1524     0   1   2E+4   0   0.5  RT (min)      lcqtof_hilic_neg_4_055_32    m/z 297.1524     0   1   2   3E+4   19.5   20  RT (min)       lcqtof_hilic_neg_4_019_03    m/z 297.1525     0   2   4   6E+4   3.5   4   4.5  RT (min)       lcqtof_hilic_neg_4_008_24    m/z 297.1526     0   1E+4   6.5   7  RT (min)       lcqtof_hilic_neg_4_047_06    m/z 297.1526     0   1E+4   6.5   7  RT (min)       lcqtof_hilic_neg_4_012_07    m/z 297.1526     0   2   4   6E+4   4.5   5  RT (min)       lcqtof_hilic_neg_4_083_59    m/z 297.1526     0   1   2   3E+4   3   3.5  RT (min)      lcqtof_hilic_neg_4_034_31    m/z 297.1526     0   1E+4   15   15.5  RT (min)      lcqtof_hilic_neg_4_110_62    m/z 297.1527     0   2   4   6E+4   3.5   4  RT (min)       lcqtof_hilic_neg_4_008_24    m/z 297.1527     0   1E+4   3.5   4  RT (min)      lcqtof_hilic_neg_4_061_50    m/z 297.1527     0   1E+4   3   3.5  RT (min)      lcqtof_hilic_neg_4_109_62    m/z 297.1527     0   1E+4   6   6.5  RT (min)       lcqtof_hilic_neg_4_047_06    m/z 297.1528     0   1   2   3E+3   6.5   7  RT (min)      lcqtof_hilic_neg_1_003_37    m/z 297.1528     0   1   2E+4   3.5   4  RT (min)      lcqtof_hilic_neg_1_015_33    m/z 297.2428     0   2   4   6E+4   1   1.5  RT (min)      lcqtof_hilic_neg_4_002_25    m/z 297.2918     0   1E+5   1   1.5  RT (min)      lcqtof_hilic_neg_4_002_25    m/z 297.2926     0   1   2E+5   1   1.5  RT (min)      lcqtof_hilic_neg_2_028_05    m/z 297.2928     0   1   2   3E+4   4   4.5  RT (min)      lcqtof_hilic_neg_4_040_05    m/z 297.2929     0   1E+4   3.5   4  RT (min)      lcqtof_hilic_neg_2_028_05    m/z 297.2930     0   1   2   3   4E+4   4   4.5  RT (min)      lcqtof_hilic_neg_4_008_24    m/z 297.2930     0   1   2   3   4E+4   4   4.5  RT (min)      lcqtof_hilic_neg_4_008_24    m/z 297.3280     0   1E+4   0.5   1  RT (min)      lcqtof_hilic_neg_4_047_06    m/z 297.8224     0   1   2E+4   4.5   5  RT (min)       lcqtof_hilic_neg_1_000_bb    m/z 297.8680     0   1   2E+3   4.5   5  RT (min)      lcqtof_hilic_neg_4_001_49    m/z 297.8686     0   1   2E+3   4   4.5  RT (min)       lcqtof_hilic_neg_4_001_49    m/z 298.0670     0   1E+4   3   3.5  RT (min)      lcqtof_hilic_neg_3_091_60    m/z 298.0709     0   1E+4   0.5   1  RT (min)      lcqtof_hilic_neg_4_045_26    m/z 298.0888     0   1E+4   3.5   4  RT (min)      lcqtof_hilic_neg_4_052_13    m/z 298.0928     0   1E+4   2.5   3  RT (min)      lcqtof_hilic_neg_4_047_06    m/z 298.1526     0   1   2E+3   3.5   4  RT (min)       lcqtof_hilic_neg_4_110_62    m/z 298.1551     0   1E+5   15   15.5  RT (min)      lcqtof_hilic_neg_4_000_aa    m/z 298.1557     0   2   4   6E+3   2   2.5  RT (min)      lcqtof_hilic_neg_4_039_18    m/z 298.1557     0   2   4   6   8E+3   3   3.5  RT (min)      lcqtof_hilic_neg_4_002_25    m/z 298.1558     0   1E+4   2.5   3  RT (min)      lcqtof_hilic_neg_4_047_06    m/z 298.1558     0   2   4   6E+3   3.5   4  RT (min)       lcqtof_hilic_neg_4_008_24    m/z 298.1558     0   2   4   6E+3   1.5   2  RT (min)      lcqtof_hilic_neg_4_047_06    m/z 298.1559     0   2   4E+3   0   0.5  RT (min)      lcqtof_hilic_neg_4_022_34    m/z 298.1559     0   1E+5   15   15.5  RT (min)      lcqtof_hilic_neg_4_000_aa    m/z 298.1560     0   1   2E+4   2   2.5  RT (min)      lcqtof_hilic_neg_4_003_16    m/z 298.1560     0   1E+4   0   0.5  RT (min)      lcqtof_hilic_neg_4_017_51    m/z 298.1560     0   1E+4   2   2.5  RT (min)      lcqtof_hilic_neg_4_026_04    m/z 298.1560     0   1   2E+4   2.5   3  RT (min)      lcqtof_hilic_neg_4_088_59    m/z 298.1560     0   1   2   3   4E+4   0.5   1  RT (min)      lcqtof_hilic_neg_4_060_13    m/z 298.1560     0   1E+4   2.5   3  RT (min)      lcqtof_hilic_neg_4_048_23    m/z 298.1562     0   2   4   6   8E+3   2.5   3  RT (min)      lcqtof_hilic_neg_4_055_32    m/z 298.6838     0   1   2   3E+4   3   3.5  RT (min)       lcqtof_hilic_neg_3_066_57    m/z 298.8702     0   2   4E+4   9   9.5  RT (min)      lcqtof_hilic_neg_2_000_aa    m/z 298.8709     0   1E+5   8.5   9  RT (min)      lcqtof_hilic_neg_1_000_aa    m/z 298.8709     0   1E+5   8.5   9  RT (min)       lcqtof_hilic_neg_1_000_aa    m/z 298.8710     0   2   4E+4   9   9.5  RT (min)      lcqtof_hilic_neg_2_000_aa    m/z 298.8712     0   1E+5   8.5   9  RT (min)      lcqtof_hilic_neg_1_000_aa    m/z 299.1039     0   1   2   3   4E+3   1   1.5  RT (min)      lcqtof_hilic_neg_4_004_47    m/z 299.1464     0   1E+3   3   3.5  RT (min)      lcqtof_hilic_neg_4_008_24    m/z 299.1485     0   2   4   6E+3   2   2.5  RT (min)      lcqtof_hilic_neg_4_035_33    m/z 299.1490     0   1   2   3E+3   1.5   2  RT (min)      lcqtof_hilic_neg_4_103_61    m/z 299.1494     0   2   4   6E+3   2.5   3  RT (min)      lcqtof_hilic_neg_4_047_06    m/z 299.1500     0   1   2   3E+4   15   15.5  RT (min)      lcqtof_hilic_neg_4_000_aa    m/z 299.1502     0   1   2   3E+4   15   15.5  RT (min)      lcqtof_hilic_neg_4_000_aa    m/z 299.1506     0   2   4   6E+3   2.5   3  RT (min)      lcqtof_hilic_neg_4_086_59    m/z 299.1513     0   2   4   6E+3   2   2.5  RT (min)      lcqtof_hilic_neg_4_052_13    m/z 299.1868     0   2   4E+3   1   1.5  RT (min)      lcqtof_hilic_neg_4_020_42    m/z 299.1872     0   1E+3   0.5   1  RT (min)      lcqtof_hilic_neg_3_003_20    m/z 299.2585     0   1   2E+5   1   1.5  RT (min)      lcqtof_hilic_neg_4_034_31    m/z 299.2587     0   1   2E+5   1   1.5  RT (min)      lcqtof_hilic_neg_4_002_25    m/z 299.3083     0   1E+5   1   1.5  RT (min)      lcqtof_hilic_neg_2_028_05    m/z 299.3086     0   1E+4   3   3.5  RT (min)      lcqtof_hilic_neg_4_008_24    m/z 299.8198     0   1E+4   4.5   5  RT (min)      lcqtof_hilic_neg_1_000_bb    m/z 299.8657     0   1   2   3E+3   4.5   5  RT (min)      lcqtof_hilic_neg_4_001_49    m/z 299.9505     0   2   4   6E+4   7.5   8  RT (min)      lcqtof_hilic_neg_1_000_aa    m/z 300.0042     0   1E+4   7.5   8  RT (min)       lcqtof_hilic_neg_1_000_aa    m/z 300.0043     0   2   4   6E+4   7   7.5  RT (min)      lcqtof_hilic_neg_4_032_01    m/z 300.0043     0   1   2   3E+4   7   7.5  RT (min)       lcqtof_hilic_neg_4_032_01    m/z 300.0044     0   1E+5   6.5   7  RT (min)      lcqtof_hilic_neg_4_023_15    m/z 300.0044     0   1   2E+4   7.5   8  RT (min)      lcqtof_hilic_neg_4_032_01    m/z 300.0045     0   2   4E+4   7   7.5  RT (min)      lcqtof_hilic_neg_4_032_01    m/z 300.0045     0   1E+5   6.5   7  RT (min)       lcqtof_hilic_neg_4_053_14    m/z 300.0046     0   2   4   6   8E+4   6.5   7  RT (min)      lcqtof_hilic_neg_4_064_15    m/z 300.0046     0   1E+4   7.5   8  RT (min)      lcqtof_hilic_neg_4_059_14    m/z 300.0381     0   1E+5   7.5   8  RT (min)      lcqtof_hilic_neg_1_000_aa    m/z 300.0738     0   1   2   3E+4   7.5   8  RT (min)      lcqtof_hilic_neg_1_000_aa    m/z 300.1146     0   1   2E+4   3.5   4  RT (min)       lcqtof_hilic_neg_4_055_32    m/z 300.1186     0   1   2   3   4E+4   7.5   8  RT (min)      lcqtof_hilic_neg_1_000_aa    m/z 300.1734     0   2   4   6E+4   7.5   8  RT (min)      lcqtof_hilic_neg_1_000_aa    m/z 300.2717     0   2   4E+4   7.5   8  RT (min)      lcqtof_hilic_neg_1_000_aa    m/z 300.8650     0   1   2E+4   7.5   8  RT (min)      lcqtof_hilic_neg_1_000_aa    m/z 300.9986     0   1E+4   1.5   2  RT (min)      lcqtof_hilic_neg_4_098_61    m/z 301.0077     0   1E+4   6.5   7  RT (min)      lcqtof_hilic_neg_4_059_14    m/z 301.0392     0   1E+4   0.5   1  RT (min)      lcqtof_hilic_neg_4_000_aa    m/z 301.0687     0   1   2E+4   1.5   2  RT (min)      lcqtof_hilic_neg_4_060_13    m/z 301.0692     0   1   2E+4   1.5   2  RT (min)      lcqtof_hilic_neg_4_060_13    m/z 301.0865     0   1   2E+4   4   4.5  RT (min)      lcqtof_hilic_neg_4_048_23    m/z 301.1245     0   2   4E+4   5   5.5  RT (min)       lcqtof_hilic_neg_4_055_32    m/z 301.1506     0   1E+4   5   5.5  RT (min)      lcqtof_hilic_neg_4_055_32    m/z 301.1796     0   1E+4   0.5   1  RT (min)      lcqtof_hilic_neg_4_038_10    m/z 301.8666     0   2   4   6E+4   7.5   8  RT (min)      lcqtof_hilic_neg_1_000_aa    m/z 302.0981     0   1E+4   4   4.5  RT (min)      lcqtof_hilic_neg_4_040_05    m/z 302.0983     0   1   2E+4   4   4.5  RT (min)       lcqtof_hilic_neg_4_055_32    m/z 302.0986     0   1   2E+4   4   4.5  RT (min)       lcqtof_hilic_neg_4_008_24    m/z 302.1082     0   1E+4   2.5   3  RT (min)       lcqtof_hilic_neg_4_008_24    m/z 302.1083     0   1E+4   2.5   3  RT (min)       lcqtof_hilic_neg_4_040_05    m/z 302.1084     0   1E+4   2.5   3  RT (min)       lcqtof_hilic_neg_4_008_24    m/z 302.1090     0   1E+4   2.5   3  RT (min)      lcqtof_hilic_neg_4_008_24    m/z 302.8960     0   1E+4   0.5   1  RT (min)      lcqtof_hilic_neg_4_018_27    m/z 303.0136     0   1E+4   0.5   1  RT (min)      lcqtof_hilic_neg_3_003_20    m/z 303.0384     0   1E+4   4   4.5  RT (min)      lcqtof_hilic_neg_4_014_46    m/z 303.1796     0   1   2   3   4E+4   0.5   1  RT (min)      lcqtof_hilic_neg_4_004_47    m/z 303.1998     0   1E+4   0.5   1  RT (min)      lcqtof_hilic_neg_4_046_41    m/z 303.4092     0   2   4E+4   0.5   1  RT (min)      lcqtof_hilic_neg_1_004_14    m/z 303.6653     0   1E+4   4   4.5  RT (min)      lcqtof_hilic_neg_4_016_37    m/z 303.9736     0   1E+4   0.5   1  RT (min)      lcqtof_hilic_neg_4_031_38    m/z 303.9903     0   1   2   3   4E+4   1   1.5  RT (min)      lcqtof_hilic_neg_2_045_11    m/z 304.0336     0   1E+4   0.5   1  RT (min)      lcqtof_hilic_neg_4_000_aa    m/z 304.0386     0   1E+4   2   2.5  RT (min)       lcqtof_hilic_neg_4_004_47    m/z 304.0969     0   1E+4   3.5   4  RT (min)       lcqtof_hilic_neg_4_034_31    m/z 304.1120     0   1E+4   2.5   3  RT (min)       lcqtof_hilic_neg_4_097_60    m/z 304.1523     0   1   2E+4   5.5   6  RT (min)       lcqtof_hilic_neg_4_008_24    m/z 304.9130     0   1   2   3   4E+3   10   10.5  RT (min)       lcqtof_hilic_neg_4_050_02    m/z 304.9130     0   1   2E+3   10   10.5  RT (min)       lcqtof_hilic_neg_4_000_aa    m/z 304.9132     0   1   2E+3   10   10.5  RT (min)       lcqtof_hilic_neg_4_000_aa    m/z 304.9134     0   1   2   3E+3   9.5   10  RT (min)       lcqtof_hilic_neg_2_000_aa    m/z 304.9138     0   1   2E+3   9.5   10  RT (min)       lcqtof_hilic_neg_4_066_57    m/z 304.9618     0   1E+4   2   2.5  RT (min)      lcqtof_hilic_neg_4_050_02    m/z 305.0286     0   1E+4   0.5   1  RT (min)      lcqtof_hilic_neg_4_000_aa    m/z 305.0298     0   1E+4   0.5   1  RT (min)      lcqtof_hilic_neg_4_000_aa    m/z 305.0449     0   1E+4   4.5   5  RT (min)      lcqtof_hilic_neg_1_022_36    m/z 305.0680     0   1E+4   2.5   3  RT (min)      lcqtof_hilic_neg_4_090_60    m/z 305.0687     0   1E+4   3.5   4  RT (min)      lcqtof_hilic_neg_4_034_31    m/z 305.0934     0   1E+4   0.5   1  RT (min)      lcqtof_hilic_neg_3_000_aa    m/z 305.0950     0   1E+4   4   4.5  RT (min)      lcqtof_hilic_neg_2_028_05    m/z 305.1824     0   1   2   3   4E+4   0.5   1  RT (min)      lcqtof_hilic_neg_4_000_aa    m/z 305.1857     0   1   2   3   4E+4   1   1.5  RT (min)       lcqtof_hilic_neg_4_000_aa    m/z 305.1857     0   1   2   3   4E+4   1   1.5  RT (min)      lcqtof_hilic_neg_4_000_aa    m/z 305.1858     0   2   4E+4   15   15.5   16  RT (min)      lcqtof_hilic_neg_4_000_aa    m/z 305.2255     0   1   2E+4   0.5   1  RT (min)      lcqtof_hilic_neg_3_003_20    m/z 305.9171     0   2   4E+3   8.5   9  RT (min)      lcqtof_hilic_neg_4_050_02    m/z 305.9176     0   2   4   6   8E+3   8.5   9  RT (min)      lcqtof_hilic_neg_4_053_14    m/z 306.1184     0   1   2   3E+4   4.5   5  RT (min)      lcqtof_hilic_neg_4_048_23    m/z 306.1193     0   1   2   3E+4   4   4.5  RT (min)       lcqtof_hilic_neg_4_048_23    m/z 306.2536     0   1   2   3E+4   6   6.5  RT (min)      lcqtof_hilic_neg_4_048_23    m/z 306.9189     0   1E+4   5.5   6  RT (min)       lcqtof_hilic_neg_4_008_24    m/z 306.9190     0   1   2   3E+3   5   5.5  RT (min)       lcqtof_hilic_neg_4_060_13    m/z 306.9192     0   1   2   3E+3   5.5   6  RT (min)      lcqtof_hilic_neg_4_032_01    m/z 306.9369     0   1   2   3E+3   8   8.5  RT (min)       lcqtof_hilic_neg_4_064_15    m/z 306.9369     0   1   2E+3   14   14.5  RT (min)       lcqtof_hilic_neg_3_068_57    m/z 307.0994     0   1E+4   4   4.5  RT (min)      lcqtof_hilic_neg_4_093_60    m/z 307.1140     0   1E+4   6.5   7  RT (min)      lcqtof_hilic_neg_4_008_24    m/z 307.1563     0   1E+5   15   15.5  RT (min)      lcqtof_hilic_neg_4_000_aa    m/z 307.1574     0   1E+5   15   15.5  RT (min)      lcqtof_hilic_neg_4_000_aa    m/z 307.1578     0   1   2   3E+4   3   3.5  RT (min)       lcqtof_hilic_neg_4_002_25    m/z 307.1578     0   1E+4   19.5   20  RT (min)       lcqtof_hilic_neg_4_026_04    m/z 307.1578     0   1E+4   0   0.5  RT (min)       lcqtof_hilic_neg_4_012_07    m/z 307.1578     0   1E+4   0   0.5  RT (min)       lcqtof_hilic_neg_4_047_06    m/z 307.1578     0   2   4   6   8E+3   6.5   7  RT (min)       lcqtof_hilic_neg_4_012_07    m/z 307.1578     0   1E+4   18   19  RT (min)       lcqtof_hilic_neg_4_047_06    m/z 307.1578     0   1   2E+4   1.5   2  RT (min)       lcqtof_hilic_neg_4_101_61    m/z 307.1579     0   1E+4   0   0.5  RT (min)       lcqtof_hilic_neg_4_106_62    m/z 307.1579     0   2   4   6E+4   2.5   3  RT (min)       lcqtof_hilic_neg_4_086_59    m/z 307.1579     0   1   2   3E+4   0.5   1  RT (min)      lcqtof_hilic_neg_4_048_23    m/z 307.1579     0   1E+4   16   16.5  RT (min)       lcqtof_hilic_neg_4_103_61    m/z 307.1579     0   1   2   3E+4   0   0.5  RT (min)       lcqtof_hilic_neg_4_047_06    m/z 307.1579     0   1E+4   18   18.5  RT (min)       lcqtof_hilic_neg_4_012_07    m/z 307.1580     0   1   2   3E+4   3.5   4  RT (min)       lcqtof_hilic_neg_4_090_60    m/z 307.1580     0   1   2   3   4E+4   2   2.5  RT (min)      lcqtof_hilic_neg_4_018_27    m/z 307.1580     0   2   4   6E+3   6   6.5  RT (min)       lcqtof_hilic_neg_4_082_59    m/z 307.1580     0   1E+4   18   19  RT (min)       lcqtof_hilic_neg_4_019_03    m/z 307.1581     0   2   4   6   8E+3   3.5   4  RT (min)       lcqtof_hilic_neg_4_105_61    m/z 307.1581     0   1E+4   17.5   18  RT (min)       lcqtof_hilic_neg_4_110_62    m/z 307.1581     0   1E+4   16.5   17  RT (min)       lcqtof_hilic_neg_4_107_62    m/z 307.1582     0   1   2   3   4E+4   2.5   3  RT (min)       lcqtof_hilic_neg_4_048_23    m/z 307.1582     0   1E+4   3   3.5  RT (min)      lcqtof_hilic_neg_4_017_51    m/z 307.1582     0   1E+4   17   18  RT (min)       lcqtof_hilic_neg_4_002_25    m/z 307.1582     0   1   2   3   4E+4   2.5   3  RT (min)      lcqtof_hilic_neg_4_048_23    m/z 307.1582     0   1E+4   18   20  RT (min)       lcqtof_hilic_neg_4_098_61    m/z 307.1583     0   1E+4   4.5   5  RT (min)      lcqtof_hilic_neg_4_040_05    m/z 307.1583     0   1   2E+4   3.5   4  RT (min)      lcqtof_hilic_neg_4_002_25    m/z 307.1583     0   2   4E+3   6.5   7  RT (min)      lcqtof_hilic_neg_4_049_55    m/z 307.1583     0   1E+4   3   3.5  RT (min)      lcqtof_hilic_neg_4_033_53    m/z 307.1583     0   1   2E+3   5.5   6  RT (min)       lcqtof_hilic_neg_4_053_14    m/z 307.1584     0   1   2E+4   3   3.5  RT (min)      lcqtof_hilic_neg_4_034_31    m/z 307.1584     0   2   4E+4   2   2.5  RT (min)      lcqtof_hilic_neg_4_053_14    m/z 307.1585     0   1   2   3E+3   5   5.5  RT (min)      lcqtof_hilic_neg_4_005_29    m/z 307.1687     0   1E+4   0.5   1  RT (min)      lcqtof_hilic_neg_3_003_20    m/z 307.1936     0   1   2   3   4E+4   1   1.5  RT (min)      lcqtof_hilic_neg_4_026_04    m/z 307.1949     0   1E+4   15   15.5  RT (min)      lcqtof_hilic_neg_4_000_aa    m/z 308.0252     0   1E+4   3   3.5  RT (min)      lcqtof_hilic_neg_1_011_46    m/z 308.0551     0   1   2   3   4E+4   4   4.5  RT (min)      lcqtof_hilic_neg_4_014_46    m/z 308.0554     0   1   2   3   4E+4   4   4.5  RT (min)      lcqtof_hilic_neg_4_014_46    m/z 308.0846     0   1E+4   3.5   4  RT (min)      lcqtof_hilic_neg_4_055_32    m/z 308.0969     0   2   4E+3   5   5.5  RT (min)      lcqtof_hilic_neg_4_048_23    m/z 308.0971     0   1   2   3   4E+3   5   5.5  RT (min)      lcqtof_hilic_neg_1_000_aa    m/z 308.1036     0   1E+5   4   4.5  RT (min)       lcqtof_hilic_neg_4_008_24    m/z 308.1565     0   2   4E+3   0.5   1  RT (min)      lcqtof_hilic_neg_2_015_06    m/z 308.1611     0   1   2E+4   15   15.5  RT (min)      lcqtof_hilic_neg_4_000_aa    m/z 308.1611     0   1   2E+3   0   0.5  RT (min)      lcqtof_hilic_neg_4_019_03    m/z 308.1612     0   1E+4   2.5   3  RT (min)      lcqtof_hilic_neg_4_086_59    m/z 308.1613     0   2   4   6   8E+3   2   2.5  RT (min)      lcqtof_hilic_neg_4_023_15    m/z 308.9706     0   1E+4   4.5   5  RT (min)      lcqtof_hilic_neg_4_013_20    m/z 308.9966     0   1E+4   3   3.5  RT (min)      lcqtof_hilic_neg_4_050_02    m/z 309.0590     0   1E+4   6   6.5  RT (min)       lcqtof_hilic_neg_4_040_05    m/z 309.0835     0   1   2   3   4E+4   1   1.5  RT (min)      lcqtof_hilic_neg_4_003_16    m/z 309.0836     0   1E+5   0.5   1  RT (min)      lcqtof_hilic_neg_4_050_02    m/z 309.0874     0   1E+4   2   2.5  RT (min)      lcqtof_hilic_neg_4_008_24    m/z 309.0981     0   1   2   3   4E+4   0.5   1  RT (min)      lcqtof_hilic_neg_2_005_13    m/z 309.1001     0   1E+4   2   2.5  RT (min)      lcqtof_hilic_neg_4_052_13    m/z 309.1113     0   1E+4   6   6.5  RT (min)       lcqtof_hilic_neg_4_094_60    m/z 309.1514     0   2   4   6E+4   0.5   1  RT (min)      lcqtof_hilic_neg_4_052_13    m/z 309.1599     0   2   4E+3   16.5   17  RT (min)      lcqtof_hilic_neg_4_005_29    m/z 309.1600     0   1   2   3   4E+4   15   15.5  RT (min)      lcqtof_hilic_neg_4_000_aa    m/z 309.1601     0   2   4E+3   16   16.5  RT (min)      lcqtof_hilic_neg_4_039_18    m/z 309.1602     0   2   4E+3   16   16.5  RT (min)      lcqtof_hilic_neg_4_103_61    m/z 309.1604     0   2   4E+3   19   19.5  RT (min)      lcqtof_hilic_neg_4_005_29    m/z 309.1604     0   1E+4   2.5   3  RT (min)      lcqtof_hilic_neg_4_047_06    m/z 309.1606     0   2   4E+3   0   0.5  RT (min)      lcqtof_hilic_neg_4_010_28    m/z 309.1607     0   2   4E+3   17   17.5  RT (min)      lcqtof_hilic_neg_4_002_25    m/z 309.1611     0   2   4   6   8E+3   2.5   3  RT (min)      lcqtof_hilic_neg_4_048_23    m/z 309.1612     0   1   2   3   4E+4   15   15.5  RT (min)      lcqtof_hilic_neg_4_000_aa    m/z 309.1614     0   2   4   6E+3   3   3.5  RT (min)      lcqtof_hilic_neg_4_008_24    m/z 309.1622     0   1E+4   2.5   3  RT (min)      lcqtof_hilic_neg_4_086_59    m/z 309.1624     0   2   4   6   8E+3   3   3.5  RT (min)      lcqtof_hilic_neg_4_002_25    m/z 309.1634     0   1E+4   2   2.5  RT (min)      lcqtof_hilic_neg_4_045_26    m/z 309.1646     0   2   4E+3   0   0.5  RT (min)      lcqtof_hilic_neg_4_009_50    m/z 309.1651     0   2   4E+3   18.5   19  RT (min)      lcqtof_hilic_neg_4_007_45    m/z 309.1654     0   2   4   6E+3   19   19.5  RT (min)      lcqtof_hilic_neg_4_054_48    m/z 309.1664     0   2   4E+3   16.5   17  RT (min)      lcqtof_hilic_neg_4_008_24    m/z 309.1673     0   1E+4   2   2.5  RT (min)      lcqtof_hilic_neg_4_003_16    m/z 309.4889     0   1   2   3   4E+4   0.5   1  RT (min)      lcqtof_hilic_neg_4_048_23    m/z 310.0232     0   1E+4   3   3.5  RT (min)      lcqtof_hilic_neg_1_011_46    m/z 310.0833     0   1E+4   5.5   6  RT (min)      lcqtof_hilic_neg_4_008_24    m/z 310.0936     0   1E+4   3   3.5  RT (min)      lcqtof_hilic_neg_4_039_18    m/z 310.1098     0   1E+4   6.5   7  RT (min)      lcqtof_hilic_neg_4_097_60    m/z 310.1160     0   1E+4   6   6.5  RT (min)      lcqtof_hilic_neg_3_008_25    m/z 310.1186     0   1   2   3E+4   6   6.5  RT (min)      lcqtof_hilic_neg_4_002_25    m/z 310.1488     0   1E+4   0.5   1  RT (min)      lcqtof_hilic_neg_4_060_13    m/z 310.1764     0   1   2   3   4E+3   1.5   2  RT (min)      lcqtof_hilic_neg_4_090_60    m/z 310.2300     0   1   2   3   4E+4   3.5   4  RT (min)      lcqtof_hilic_neg_4_013_20    m/z 310.4557     0   1   2   3E+4   3.5   4  RT (min)      lcqtof_hilic_neg_4_013_20    m/z 310.5562     0   1E+4   3.5   4  RT (min)      lcqtof_hilic_neg_4_013_20    m/z 310.8760     0   1   2E+3   8.5   9  RT (min)      lcqtof_hilic_neg_4_028_36    m/z 311.0811     0   1   2E+4   4   4.5  RT (min)      lcqtof_hilic_neg_4_019_03    m/z 311.0939     0   1E+4   4   4.5  RT (min)      lcqtof_hilic_neg_4_008_24    m/z 311.0974     0   1   2E+4   1.5   2  RT (min)      lcqtof_hilic_neg_1_008_13    m/z 311.0981     0   1E+4   2.5   3  RT (min)      lcqtof_hilic_neg_1_019_02    m/z 311.1014     0   1   2E+4   0.5   1  RT (min)      lcqtof_hilic_neg_2_045_11    m/z 311.1563     0   1   2   3   4E+4   3.5   4  RT (min)      lcqtof_hilic_neg_4_113_62    m/z 311.1675     0   1   2   3E+4   3   3.5  RT (min)      lcqtof_hilic_neg_3_016_39    m/z 311.1676     0   1E+4   5   5.5  RT (min)      lcqtof_hilic_neg_4_016_37    m/z 311.1677     0   2   4   6   8E+4   20   20.5  RT (min)       lcqtof_hilic_neg_4_075_58    m/z 311.1677     0   1E+5   19.5   20  RT (min)       lcqtof_hilic_neg_4_096_60    m/z 311.1678     0   2   4   6   8E+4   0   0.5  RT (min)      lcqtof_hilic_neg_4_031_38    m/z 311.1678     0   2   4   6   8E+4   0   0.5  RT (min)       lcqtof_hilic_neg_4_004_47    m/z 311.1679     0   2   4   6E+4   3   3.5  RT (min)      lcqtof_hilic_neg_4_109_62    m/z 311.1679     0   2   4   6   8E+4   3   3.5   4  RT (min)       lcqtof_hilic_neg_4_107_62    m/z 311.1679     0   1   2   3E+4   15.5   16  RT (min)      lcqtof_hilic_neg_3_034_24    m/z 311.1679     0   1E+6   0.5   1  RT (min)       lcqtof_hilic_neg_4_004_47    m/z 311.1679     0   1E+5   0   0.5  RT (min)       lcqtof_hilic_neg_4_113_62    m/z 311.1679     0   1E+5   0   0.5  RT (min)       lcqtof_hilic_neg_4_113_62    m/z 311.1679     0   2   4   6   8E+5   0   0.5  RT (min)       lcqtof_hilic_neg_4_100_61    m/z 311.1680     0   1E+5   16   17  RT (min)       lcqtof_hilic_neg_4_019_03    m/z 311.1680     0   1E+5   19   19.5  RT (min)       lcqtof_hilic_neg_4_109_62    m/z 311.1680     0   1   2   3E+4   2.5   3  RT (min)      lcqtof_hilic_neg_3_000_cc    m/z 311.1680     0   2   4   6   8E+4   14.5   15   15.5  RT (min)       lcqtof_hilic_neg_4_083_59    m/z 311.1680     0   1E+5   17   17.5  RT (min)       lcqtof_hilic_neg_4_112_62    m/z 311.1680     0   1   2E+5   4.5   5  RT (min)       lcqtof_hilic_neg_4_055_32    m/z 311.1680     0   1E+5   18.5   19  RT (min)       lcqtof_hilic_neg_4_105_61    m/z 311.1680     0   1E+5   17   17.5  RT (min)       lcqtof_hilic_neg_4_112_62    m/z 311.1680     0   1E+5   3   3.5  RT (min)       lcqtof_hilic_neg_4_002_25    m/z 311.1680     0   1   2E+4   5.5   6  RT (min)       lcqtof_hilic_neg_4_096_60    m/z 311.1680     0   2   4   6E+5   1.5   2  RT (min)       lcqtof_hilic_neg_4_050_02    m/z 311.1680     0   1E+5   16   16.5   17  RT (min)       lcqtof_hilic_neg_4_019_03    m/z 311.1680     0   1   2E+6   15   16  RT (min)       lcqtof_hilic_neg_4_000_aa    m/z 311.1680     0   1E+5   0   0.5  RT (min)       lcqtof_hilic_neg_4_011_35    m/z 311.1680     0   2   4   6   8E+4   15   15.5  RT (min)       lcqtof_hilic_neg_4_110_62    m/z 311.1680     0   1E+5   17   18  RT (min)       lcqtof_hilic_neg_4_046_41    m/z 311.1680     0   1E+5   19   19.5  RT (min)       lcqtof_hilic_neg_4_097_60    m/z 311.1680     0   1E+5   17  RT (min)       lcqtof_hilic_neg_4_107_62    m/z 311.1680     0   1E+5   2.5   3  RT (min)       lcqtof_hilic_neg_4_048_23    m/z 311.1680     0   1E+5   19.5   20  RT (min)       lcqtof_hilic_neg_4_019_03    m/z 311.1680     0   1   2E+5   4   4.5  RT (min)       lcqtof_hilic_neg_4_096_60    m/z 311.1680     0   1E+5   17.5   18  RT (min)       lcqtof_hilic_neg_4_113_62    m/z 311.1680     0   1   2   3   4E+4   3.5   4  RT (min)       lcqtof_hilic_neg_4_107_62    m/z 311.1680     0   2   4   6E+4   15   15.5  RT (min)       lcqtof_hilic_neg_4_021_21    m/z 311.1680     0   1E+5   4   4.5  RT (min)      lcqtof_hilic_neg_4_001_49    m/z 311.1680     0   1E+4   5   5.5  RT (min)      lcqtof_hilic_neg_4_010_28    m/z 311.1680     0   1E+5   1.5   2  RT (min)       lcqtof_hilic_neg_4_019_03    m/z 311.1680     0   1   2E+4   7   8   9  RT (min)       lcqtof_hilic_neg_4_087_59    m/z 311.1680     0   1E+4   8   9  RT (min)       lcqtof_hilic_neg_4_044_08    m/z 311.1680     0   1E+5   15.5   16  RT (min)       lcqtof_hilic_neg_4_058_39    m/z 311.1680     0   1E+5   16   17  RT (min)       lcqtof_hilic_neg_4_111_62    m/z 311.1681     0   1E+5   18   19  RT (min)       lcqtof_hilic_neg_4_087_59    m/z 311.1681     0   1E+5   18   18.5  RT (min)      lcqtof_hilic_neg_4_030_44    m/z 311.1681     0   1E+5   19   20  RT (min)       lcqtof_hilic_neg_4_021_21    m/z 311.1681     0   1   2   3E+5   2.5   3  RT (min)       lcqtof_hilic_neg_4_086_59    m/z 311.1681     0   1E+5   16.5   17  RT (min)       lcqtof_hilic_neg_4_113_62    m/z 311.1681     0   1   2E+4   6   6.5  RT (min)       lcqtof_hilic_neg_4_090_60    m/z 311.1681     0   1   2E+6   15   15.5  RT (min)       lcqtof_hilic_neg_4_000_aa    m/z 311.1681     0   1   2   3E+4   3   3.5  RT (min)      lcqtof_hilic_neg_2_035_09    m/z 311.1681     0   1   2E+4   5   5.5  RT (min)      lcqtof_hilic_neg_4_011_35    m/z 311.1682     0   1E+5   17.5   18  RT (min)       lcqtof_hilic_neg_4_110_62    m/z 311.1682     0   1E+5   16   16.5  RT (min)       lcqtof_hilic_neg_4_019_03    m/z 311.1682     0   2   4   6   8E+4   15.5   16  RT (min)      lcqtof_hilic_neg_4_051_43    m/z 311.1682     0   2   4   6   8E+4   16   16.5  RT (min)      lcqtof_hilic_neg_4_021_21    m/z 311.1682     0   1E+5   17.5   18   18.5  RT (min)       lcqtof_hilic_neg_4_088_59    m/z 311.1682     0   1E+5   18.5   19  RT (min)       lcqtof_hilic_neg_4_113_62    m/z 311.1682     0   2   4E+4   14   14.5   15  RT (min)       lcqtof_hilic_neg_4_083_59    m/z 311.1682     0   1E+4   5   5.5  RT (min)      lcqtof_hilic_neg_4_053_14    m/z 311.1682     0   1E+5   17   17.5  RT (min)       lcqtof_hilic_neg_4_097_60    m/z 311.1682     0   1E+5   15.5   16   16.5  RT (min)       lcqtof_hilic_neg_4_058_39    m/z 311.1682     0   1E+5   16   16.5  RT (min)       lcqtof_hilic_neg_4_112_62    m/z 311.1682     0   2   4   6   8E+4   14.5   15   15.5  RT (min)       lcqtof_hilic_neg_4_099_61    m/z 311.1683     0   1   2E+4   6.5   7  RT (min)      lcqtof_hilic_neg_4_048_23    m/z 311.1683     0   1   2E+4   3.5   4  RT (min)      lcqtof_hilic_neg_3_016_39    m/z 311.1683     0   1   2E+4   7   8   9  RT (min)       lcqtof_hilic_neg_4_002_25    m/z 311.1683     0   2   4   6   8E+4   16.5   17  RT (min)      lcqtof_hilic_neg_4_051_43    m/z 311.1683     0   1E+5   17   17.5  RT (min)       lcqtof_hilic_neg_4_104_61    m/z 311.1683     0   1E+5   19.5   20  RT (min)       lcqtof_hilic_neg_4_106_62    m/z 311.1684     0   2   4   6   8E+4   15.5   16  RT (min)      lcqtof_hilic_neg_4_051_43    m/z 311.1684     0   1E+5   18.5   19   19.5  RT (min)       lcqtof_hilic_neg_4_097_60    m/z 311.1684     0   2   4   6   8E+3   8   8.5   9  RT (min)       lcqtof_hilic_neg_4_094_60    m/z 311.1684     0   1E+5   17.5   18  RT (min)      lcqtof_hilic_neg_4_046_41    m/z 311.1685     0   2   4   6E+4   15   15.5  RT (min)      lcqtof_hilic_neg_4_021_21    m/z 311.1686     0   1E+5   18.5   19  RT (min)       lcqtof_hilic_neg_4_113_62    m/z 311.1794     0   2   4E+4   0.5   1  RT (min)      lcqtof_hilic_neg_3_000_bb    m/z 311.1796     0   2   4   6E+4   0   0.5  RT (min)      lcqtof_hilic_neg_3_003_20    m/z 311.1796     0   1   2   3E+4   0   0.5  RT (min)      lcqtof_hilic_neg_3_000_bb    m/z 311.1797     0   1   2E+5   0.5   1  RT (min)      lcqtof_hilic_neg_3_003_20    m/z 311.1798     0   1   2E+5   0.5   1  RT (min)      lcqtof_hilic_neg_3_003_20    m/z 311.1804     0   2   4E+4   0.5   1  RT (min)      lcqtof_hilic_neg_3_000_bb    m/z 311.2052     0   1   2E+4   4   4.5  RT (min)      lcqtof_hilic_neg_4_011_35    m/z 311.2217     0   1   2   3   4E+4   1   1.5  RT (min)      lcqtof_hilic_neg_4_053_14    m/z 311.2570     0   1   2   3   4E+4   15   15.5  RT (min)      lcqtof_hilic_neg_4_000_aa    m/z 311.2702     0   1   2E+4   0.5   1  RT (min)      lcqtof_hilic_neg_2_028_05    m/z 311.2956     0   2   4E+4   4   4.5  RT (min)      lcqtof_hilic_neg_4_016_37    m/z 311.3092     0   1   2   3E+4   0.5   1  RT (min)      lcqtof_hilic_neg_4_047_06    m/z 311.3216     0   1E+4   3.5   4  RT (min)      lcqtof_hilic_neg_4_013_20    m/z 311.3463     0   2   4   6   8E+4   15   15.5  RT (min)      lcqtof_hilic_neg_4_000_aa    m/z 311.3833     0   1E+4   15   15.5  RT (min)      lcqtof_hilic_neg_4_000_aa    m/z 311.3843     0   2   4   6E+4   0.5   1  RT (min)      lcqtof_hilic_neg_4_047_06    m/z 311.4294     0   1E+4   15   15.5  RT (min)      lcqtof_hilic_neg_4_000_aa    m/z 311.4303     0   1E+5   0.5   1  RT (min)      lcqtof_hilic_neg_4_047_06    m/z 311.4314     0   1E+4   4   4.5  RT (min)      lcqtof_hilic_neg_4_016_37    m/z 311.4839     0   1   2   3E+4   15   15.5  RT (min)      lcqtof_hilic_neg_4_000_aa    m/z 311.5448     0   1   2E+4   0.5   1  RT (min)      lcqtof_hilic_neg_4_047_06    m/z 311.5842     0   1E+4   15   15.5  RT (min)      lcqtof_hilic_neg_4_000_aa    m/z 311.5856     0   2   4   6   8E+4   0.5   1  RT (min)      lcqtof_hilic_neg_4_047_06    m/z 311.6643     0   1   2E+4   0.5   1  RT (min)      lcqtof_hilic_neg_4_047_06    m/z 312.0988     0   1E+4   2   2.5  RT (min)      lcqtof_hilic_neg_4_050_02    m/z 312.1320     0   1E+4   6   6.5  RT (min)      lcqtof_hilic_neg_4_055_32    m/z 312.1706     0   1   2   3   4E+5   15   15.5  RT (min)      lcqtof_hilic_neg_4_000_aa    m/z 312.1709     0   1   2   3   4E+4   3.5   4  RT (min)      lcqtof_hilic_neg_4_008_24    m/z 312.1709     0   1   2   3E+4   3.5   4  RT (min)      lcqtof_hilic_neg_4_014_46    m/z 312.1709     0   1   2   3E+4   3.5   4  RT (min)      lcqtof_hilic_neg_4_092_60    m/z 312.1710     0   1   2   3E+4   4.5   5  RT (min)      lcqtof_hilic_neg_4_014_46    m/z 312.1711     0   1   2   3   4E+3   5   5.5  RT (min)      lcqtof_hilic_neg_4_011_35    m/z 312.1713     0   1   2   3E+4   4.5   5  RT (min)      lcqtof_hilic_neg_4_017_51    m/z 312.1713     0   1   2E+5   0.5   1  RT (min)      lcqtof_hilic_neg_4_050_02    m/z 312.1713     0   1   2E+4   0   0.5  RT (min)      lcqtof_hilic_neg_4_050_02    m/z 312.1713     0   1   2   3E+4   2.5   3  RT (min)      lcqtof_hilic_neg_4_048_23    m/z 312.1713     0   1   2E+4   0   0.5  RT (min)      lcqtof_hilic_neg_4_103_61    m/z 312.1714     0   1E+4   3   3.5  RT (min)      lcqtof_hilic_neg_4_107_62    m/z 312.1714     0   1   2   3   4E+5   15   15.5  RT (min)      lcqtof_hilic_neg_4_000_aa    m/z 312.1714     0   1   2   3E+4   1.5   2  RT (min)      lcqtof_hilic_neg_4_019_03    m/z 312.1714     0   1   2   3E+4   3   3.5  RT (min)      lcqtof_hilic_neg_4_002_25    m/z 312.1714     0   1   2E+4   3.5   4  RT (min)      lcqtof_hilic_neg_4_001_49    m/z 312.1714     0   2   4   6E+4   2.5   3  RT (min)      lcqtof_hilic_neg_4_086_59    m/z 312.1714     0   1   2E+4   1.5   2  RT (min)      lcqtof_hilic_neg_4_042_22    m/z 312.1715     0   1E+4   3   3.5  RT (min)      lcqtof_hilic_neg_4_049_55    m/z 312.1716     0   1E+4   15   15.5  RT (min)      lcqtof_hilic_neg_4_049_55    m/z 312.1716     0   1E+5   1.5   2  RT (min)      lcqtof_hilic_neg_4_050_02    m/z 312.1716     0   1   2E+5   1.5   2  RT (min)      lcqtof_hilic_neg_4_046_41    m/z 312.1717     0   1E+4   3   3.5  RT (min)      lcqtof_hilic_neg_4_011_35    m/z 312.1822     0   1   2   3E+4   0.5   1  RT (min)      lcqtof_hilic_neg_3_003_20    m/z 312.1827     0   1   2   3E+4   0.5   1  RT (min)      lcqtof_hilic_neg_3_003_20    m/z 312.2622     0   2   4E+4   0.5   1  RT (min)      lcqtof_hilic_neg_4_047_06    m/z 312.3109     0   1   2   3E+4   0.5   1  RT (min)      lcqtof_hilic_neg_4_047_06    m/z 312.3495     0   1   2E+4   15   15.5  RT (min)      lcqtof_hilic_neg_4_000_aa    m/z 312.3505     0   1E+5   0.5   1  RT (min)      lcqtof_hilic_neg_4_047_06    m/z 312.3865     0   1   2   3E+4   0.5   1  RT (min)      lcqtof_hilic_neg_4_047_06    m/z 312.4344     0   2   4   6E+4   0.5   1  RT (min)      lcqtof_hilic_neg_4_047_06    m/z 312.4885     0   1   2   3   4E+4   0.5   1  RT (min)      lcqtof_hilic_neg_4_047_06    m/z 312.5413     0   1   2E+4   0.5   1  RT (min)      lcqtof_hilic_neg_4_047_06    m/z 312.5856     0   1   2E+4   0.5   1  RT (min)      lcqtof_hilic_neg_4_047_06    m/z 313.0781     0   1   2   3E+4   2   2.5  RT (min)      lcqtof_hilic_neg_4_050_02    m/z 313.0782     0   1E+5   1.5   2  RT (min)      lcqtof_hilic_neg_4_000_aa    m/z 313.0784     0   2   4   6E+4   1.5   2  RT (min)      lcqtof_hilic_neg_4_000_aa    m/z 313.0784     0   2   4E+4   2   2.5  RT (min)      lcqtof_hilic_neg_4_000_aa    m/z 313.0898     0   1E+4   0.5   1  RT (min)      lcqtof_hilic_neg_3_000_aa    m/z 313.0939     0   1E+4   0.5   1  RT (min)      lcqtof_hilic_neg_4_009_50    m/z 313.0950     0   1E+4   3.5   4  RT (min)      lcqtof_hilic_neg_4_039_18    m/z 313.0955     0   1E+4   3.5   4  RT (min)      lcqtof_hilic_neg_4_048_23    m/z 313.1013     0   1E+4   3.5   4  RT (min)      lcqtof_hilic_neg_4_048_23    m/z 313.1167     0   1   2   3   4E+3   3   3.5  RT (min)      lcqtof_hilic_neg_4_049_55    m/z 313.1172     0   1   2   3E+3   3   3.5  RT (min)      lcqtof_hilic_neg_4_109_62    m/z 313.1179     0   1E+4   2   2.5  RT (min)      lcqtof_hilic_neg_4_070_57    m/z 313.1180     0   1E+4   1.5   2  RT (min)       lcqtof_hilic_neg_4_053_14    m/z 313.1182     0   1E+4   1.5   2  RT (min)      lcqtof_hilic_neg_4_053_14    m/z 313.1521     0   2   4   6E+4   4   4.5  RT (min)      lcqtof_hilic_neg_4_008_24    m/z 313.1655     0   1E+4   3.5   4  RT (min)       lcqtof_hilic_neg_4_016_37    m/z 313.1664     0   2   4   6   8E+3   3.5   4  RT (min)      lcqtof_hilic_neg_4_004_47    m/z 313.1666     0   1E+4   2.5   3  RT (min)      lcqtof_hilic_neg_4_048_23    m/z 313.1668     0   1E+4   3   3.5  RT (min)      lcqtof_hilic_neg_4_002_25    m/z 313.1669     0   2   4   6   8E+3   3   3.5  RT (min)      lcqtof_hilic_neg_4_008_24    m/z 313.1669     0   2   4   6E+4   1.5   2  RT (min)      lcqtof_hilic_neg_4_048_23    m/z 313.1669     0   1E+5   15   15.5  RT (min)      lcqtof_hilic_neg_4_000_aa    m/z 313.1670     0   1E+4   2   2.5  RT (min)      lcqtof_hilic_neg_4_026_04    m/z 313.1671     0   2   4   6E+4   1.5   2  RT (min)      lcqtof_hilic_neg_4_013_20    m/z 313.1671     0   1   2E+4   2   2.5  RT (min)      lcqtof_hilic_neg_4_032_01    m/z 313.1671     0   1   2E+4   0   0.5  RT (min)      lcqtof_hilic_neg_4_051_43    m/z 313.1672     0   1   2E+4   2.5   3  RT (min)      lcqtof_hilic_neg_4_086_59    m/z 313.1672     0   1   2E+4   2   2.5  RT (min)      lcqtof_hilic_neg_4_052_13    m/z 313.1672     0   1   2E+4   2   2.5  RT (min)      lcqtof_hilic_neg_4_053_14    m/z 313.1672     0   2   4   6   8E+3   3   3.5  RT (min)      lcqtof_hilic_neg_4_040_05    m/z 313.1672     0   2   4   6   8E+3   1.5   2  RT (min)      lcqtof_hilic_neg_4_039_18    m/z 313.1673     0   2   4   6E+3   0   0.5  RT (min)      lcqtof_hilic_neg_4_046_41    m/z 313.1673     0   2   4   6E+3   2.5   3  RT (min)      lcqtof_hilic_neg_4_020_42    m/z 313.1679     0   1E+5   15   15.5  RT (min)      lcqtof_hilic_neg_4_000_aa    m/z 313.2255     0   1   2E+3   16.5   17  RT (min)      lcqtof_hilic_neg_4_003_16    m/z 313.2256     0   1   2E+3   0   0.5  RT (min)      lcqtof_hilic_neg_4_006_30    m/z 313.2260     0   1   2   3E+4   0.5   1  RT (min)      lcqtof_hilic_neg_4_050_02    m/z 313.2260     0   1E+4   0.5   1  RT (min)      lcqtof_hilic_neg_4_017_51    m/z 313.2262     0   1   2E+3   18   18.5  RT (min)      lcqtof_hilic_neg_4_006_30    m/z 313.2265     0   1   2E+3   18.5   19  RT (min)      lcqtof_hilic_neg_4_001_49    m/z 313.2266     0   1   2E+3   16   16.5  RT (min)      lcqtof_hilic_neg_4_006_30    m/z 313.2269     0   1   2E+3   16   16.5  RT (min)      lcqtof_hilic_neg_4_000_dd    m/z 313.2283     0   2   4E+3   2   2.5  RT (min)      lcqtof_hilic_neg_4_064_15    m/z 313.2765     0   1E+4   1   1.5  RT (min)      lcqtof_hilic_neg_4_000_aa    m/z 313.3489     0   1   2   3E+4   0.5   1  RT (min)      lcqtof_hilic_neg_4_047_06    m/z 314.0766     0   2   4   6E+4   1   1.5  RT (min)      lcqtof_hilic_neg_4_000_aa    m/z 314.0772     0   2   4   6E+4   1   1.5  RT (min)      lcqtof_hilic_neg_4_000_aa    m/z 314.0778     0   2   4   6E+4   1   1.5  RT (min)      lcqtof_hilic_neg_4_000_aa    m/z 314.0780     0   2   4   6E+4   1   1.5  RT (min)      lcqtof_hilic_neg_4_000_aa    m/z 314.0782     0   2   4   6E+4   1   1.5  RT (min)      lcqtof_hilic_neg_4_000_aa    m/z 314.0785     0   2   4   6E+4   1   1.5  RT (min)      lcqtof_hilic_neg_4_000_aa    m/z 314.0786     0   1   2   3   4E+4   1.5   2  RT (min)      lcqtof_hilic_neg_4_000_aa    m/z 314.0787     0   2   4   6E+4   1.5   2  RT (min)      lcqtof_hilic_neg_4_000_aa    m/z 314.1692     0   1E+4   15   15.5  RT (min)      lcqtof_hilic_neg_4_000_aa    m/z 314.3335     0   1E+4   9.5   10  RT (min)      lcqtof_hilic_neg_1_000_aa    m/z 315.0407     0   1   2   3   4E+4   6.5   7  RT (min)       lcqtof_hilic_neg_4_059_14    m/z 315.0407     0   2   4   6E+4   5.5   6  RT (min)      lcqtof_hilic_neg_4_026_04    m/z 315.0408     0   1   2   3E+4   6.5   7  RT (min)       lcqtof_hilic_neg_4_059_14    m/z 315.0408     0   1E+5   5   5.5  RT (min)       lcqtof_hilic_neg_4_019_03    m/z 315.0408     0   1E+4   8   8.5  RT (min)       lcqtof_hilic_neg_4_059_14    m/z 315.0410     0   1   2   3   4E+4   5.5   6  RT (min)      lcqtof_hilic_neg_4_026_04    m/z 315.0410     0   1E+5   5.5   6  RT (min)      lcqtof_hilic_neg_4_053_14    m/z 315.0410     0   1   2E+4   7.5   8  RT (min)      lcqtof_hilic_neg_4_059_14    m/z 315.0412     0   1   2E+4   7.5   8  RT (min)      lcqtof_hilic_neg_4_059_14    m/z 315.0412     0   1   2E+4   6.5   7  RT (min)      lcqtof_hilic_neg_4_053_14    m/z 315.0412     0   2   4   6   8E+4   5   5.5  RT (min)       lcqtof_hilic_neg_4_026_04    m/z 315.0412     0   1E+5   5   5.5  RT (min)       lcqtof_hilic_neg_4_019_03    m/z 315.0414     0   1   2E+4   7.5   8  RT (min)       lcqtof_hilic_neg_4_059_14    m/z 315.0414     0   2   4E+3   0.5   1  RT (min)      lcqtof_hilic_neg_4_000_aa    m/z 315.0414     0   2   4   6E+3   4   4.5  RT (min)      lcqtof_hilic_neg_4_054_48    m/z 315.0415     0   1E+5   4.5   5  RT (min)       lcqtof_hilic_neg_4_019_03    m/z 315.0416     0   2   4   6   8E+3   4   4.5  RT (min)      lcqtof_hilic_neg_2_052_24    m/z 315.0422     0   2   4   6E+4   4.5   5  RT (min)      lcqtof_hilic_neg_4_050_02    m/z 315.0588     0   1   2   3   4E+3   1   1.5  RT (min)      lcqtof_hilic_neg_4_000_aa    m/z 315.0744     0   1   2   3   4E+4   1   1.5  RT (min)      lcqtof_hilic_neg_4_000_aa    m/z 315.0746     0   1   2   3   4E+4   1   1.5  RT (min)      lcqtof_hilic_neg_4_000_aa    m/z 315.0747     0   1   2   3   4E+4   1   1.5  RT (min)      lcqtof_hilic_neg_4_000_aa    m/z 315.0752     0   1   2   3   4E+4   1   1.5  RT (min)      lcqtof_hilic_neg_4_000_aa    m/z 315.0753     0   1   2   3   4E+4   1   1.5   2  RT (min)      lcqtof_hilic_neg_4_000_aa    m/z 315.0778     0   1E+4   4.5   5  RT (min)      lcqtof_hilic_neg_1_005_32    m/z 315.1182     0   1   2   3E+3   3   3.5  RT (min)       lcqtof_hilic_neg_4_100_61    m/z 315.1209     0   1   2   3E+3   2   2.5  RT (min)      lcqtof_hilic_neg_4_011_35    m/z 315.1222     0   2   4   6E+3   0.5   1  RT (min)      lcqtof_hilic_neg_4_054_48    m/z 315.1336     0   1E+4   3.5   4  RT (min)       lcqtof_hilic_neg_4_092_60    m/z 315.1802     0   1E+4   2   2.5  RT (min)       lcqtof_hilic_neg_4_052_13    m/z 315.1805     0   1   2E+3   3.5   4  RT (min)      lcqtof_hilic_neg_4_032_01    m/z 315.1806     0   1   2   3   4E+3   4.5   5  RT (min)      lcqtof_hilic_neg_4_019_03    m/z 315.1806     0   2   4   6E+3   4   4.5  RT (min)       lcqtof_hilic_neg_4_008_24    m/z 315.1810     0   1E+4   1   2  RT (min)       lcqtof_hilic_neg_4_026_04    m/z 315.1810     0   1E+4   16   18  RT (min)       lcqtof_hilic_neg_4_000_aa    m/z 315.8333     0   2   4   6   8E+4   4   4.5  RT (min)       lcqtof_hilic_neg_1_000_bb    m/z 315.8337     0   1   2   3E+4   4   4.5  RT (min)      lcqtof_hilic_neg_3_067_57    m/z 316.0437     0   2   4   6E+3   5   5.5  RT (min)      lcqtof_hilic_neg_4_034_31    m/z 316.0442     0   1   2   3   4E+3   7.5   8  RT (min)      lcqtof_hilic_neg_1_065_51    m/z 316.0453     0   1   2   3E+4   0.5   1  RT (min)      lcqtof_hilic_neg_4_000_aa    m/z 316.2129     0   1   2   3E+4   1.5   2  RT (min)      lcqtof_hilic_neg_4_032_01    m/z 316.3130     0   1   2   3E+4   4   4.5  RT (min)      lcqtof_hilic_neg_4_004_47    m/z 316.3156     0   1   2   3   4E+4   1   1.5  RT (min)      lcqtof_hilic_neg_4_002_25    m/z 316.8898     0   1   2   3E+3   4   4.5  RT (min)       lcqtof_hilic_neg_1_031_16    m/z 316.8900     0   2   4E+3   4.5   5  RT (min)       lcqtof_hilic_neg_2_000_dd    m/z 316.8901     0   1   2   3E+3   1   1.5  RT (min)      lcqtof_hilic_neg_4_058_39    m/z 316.8901     0   1   2   3E+3   4.5   5  RT (min)      lcqtof_hilic_neg_2_018_16    m/z 316.8901     0   1   2E+3   1   1.5  RT (min)      lcqtof_hilic_neg_1_026_10    m/z 316.9466     0   2   4   6E+3   6.5   7  RT (min)       lcqtof_hilic_neg_4_058_39    m/z 316.9470     0   2   4   6   8E+3   6   6.5  RT (min)       lcqtof_hilic_neg_4_058_39    m/z 316.9470     0   2   4   6   8E+3   6   6.5  RT (min)       lcqtof_hilic_neg_4_058_39    m/z 316.9471     0   1E+4   6   6.5  RT (min)       lcqtof_hilic_neg_4_112_62    m/z 316.9472     0   1E+4   6   6.5  RT (min)       lcqtof_hilic_neg_4_058_39    m/z 316.9473     0   2   4E+3   7.5   8  RT (min)       lcqtof_hilic_neg_4_000_aa    m/z 316.9474     0   2   4   6E+3   6.5   7  RT (min)       lcqtof_hilic_neg_4_058_39    m/z 317.0315     0   1   2E+3   0.5   1  RT (min)      lcqtof_hilic_neg_4_000_aa    m/z 317.0381     0   1E+5   0.5   1  RT (min)      lcqtof_hilic_neg_4_000_aa    m/z 317.0466     0   1E+4   4.5   5  RT (min)      lcqtof_hilic_neg_4_050_02    m/z 317.0479     0   1E+4   4   4.5  RT (min)      lcqtof_hilic_neg_4_050_02    m/z 317.0508     0   2   4   6E+3   6   6.5  RT (min)      lcqtof_hilic_neg_4_059_14    m/z 317.0515     0   2   4E+3   5.5   6  RT (min)      lcqtof_hilic_neg_3_004_32    m/z 317.0521     0   2   4   6   8E+3   5.5   6  RT (min)      lcqtof_hilic_neg_1_023_34    m/z 317.0530     0   1   2   3   4E+3   5   5.5  RT (min)      lcqtof_hilic_neg_3_055_01    m/z 317.0552     0   2   4E+3   2   2.5  RT (min)      lcqtof_hilic_neg_4_029_17    m/z 317.1236     0   1E+4   1   1.5  RT (min)      lcqtof_hilic_neg_4_019_03    m/z 317.1544     0   2   4   6E+4   0.5   1  RT (min)      lcqtof_hilic_neg_4_039_18    m/z 317.2089     0   1E+4   1   1.5  RT (min)      lcqtof_hilic_neg_4_039_18    m/z 317.2473     0   1E+5   0.5   1  RT (min)      lcqtof_hilic_neg_4_039_18    m/z 317.2837     0   1   2   3E+4   1   1.5  RT (min)      lcqtof_hilic_neg_4_039_18    m/z 317.3190     0   1   2   3E+5   1   1.5  RT (min)      lcqtof_hilic_neg_4_002_25    m/z 317.3298     0   1   2   3   4E+4   1   1.5  RT (min)      lcqtof_hilic_neg_4_039_18    m/z 317.4870     0   1   2   3   4E+4   1   1.5  RT (min)      lcqtof_hilic_neg_4_039_18    m/z 317.8304     0   1   2E+4   4   4.5  RT (min)      lcqtof_hilic_neg_3_067_57    m/z 317.8304     0   2   4E+4   4   4.5  RT (min)      lcqtof_hilic_neg_1_000_bb    m/z 318.0468     0   1   2E+4   0.5   1  RT (min)      lcqtof_hilic_neg_4_000_aa    m/z 318.1184     0   1   2   3E+4   6   6.5  RT (min)      lcqtof_hilic_neg_4_055_32    m/z 318.1300     0   2   4   6E+3   3   3.5  RT (min)      lcqtof_hilic_neg_2_022_46    m/z 318.1555     0   2   4E+4   4   4.5  RT (min)       lcqtof_hilic_neg_4_040_05    m/z 318.1581     0   1   2   3E+4   4   4.5  RT (min)      lcqtof_hilic_neg_4_008_24    m/z 318.1598     0   1E+5   4   4.5  RT (min)       lcqtof_hilic_neg_4_091_60    m/z 318.2514     0   1   2   3   4E+4   1   1.5  RT (min)      lcqtof_hilic_neg_4_039_18    m/z 318.2576     0   1E+4   6.5   7  RT (min)      lcqtof_hilic_neg_4_048_23    m/z 318.8580     0   1E+6   9   9.5  RT (min)      lcqtof_hilic_neg_1_000_aa    m/z 318.8583     0   1   2E+5   9.5   10  RT (min)      lcqtof_hilic_neg_2_000_aa    m/z 318.9001     0   1E+4   0.5   1  RT (min)      lcqtof_hilic_neg_4_005_29    m/z 318.9472     0   1   2E+4   9   9.5  RT (min)      lcqtof_hilic_neg_1_000_aa    m/z 319.0376     0   2   4   6E+4   9   9.5  RT (min)      lcqtof_hilic_neg_1_000_aa    m/z 319.0475     0   1E+4   0.5   1  RT (min)      lcqtof_hilic_neg_4_000_aa    m/z 319.1767     0   1E+4   9   9.5  RT (min)      lcqtof_hilic_neg_1_000_aa    m/z 319.2418     0   1E+4   0.5   1  RT (min)      lcqtof_hilic_neg_3_003_20    m/z 320.1017     0   2   4E+3   2.5   3  RT (min)      lcqtof_hilic_neg_4_033_53    m/z 320.1876     0   1   2   3E+4   1   1.5  RT (min)      lcqtof_hilic_neg_4_038_10    m/z 320.1892     0   1   2   3E+4   1.5   2  RT (min)      lcqtof_hilic_neg_4_038_10    m/z 320.8609     0   1   2   3E+4   9   9.5  RT (min)      lcqtof_hilic_neg_1_000_aa    m/z 320.9762     0   1   2   3   4E+4   0.5   1  RT (min)      lcqtof_hilic_neg_4_006_30    m/z 320.9764     0   1   2   3E+4   1.5   2  RT (min)       lcqtof_hilic_neg_4_039_18    m/z 320.9765     0   1   2   3E+4   1.5   2  RT (min)      lcqtof_hilic_neg_4_039_18    m/z 321.0222     0   1E+4   0.5   1  RT (min)      lcqtof_hilic_neg_4_031_38    m/z 321.0852     0   1E+4   3.5   4  RT (min)       lcqtof_hilic_neg_4_090_60    m/z 321.0996     0   1   2E+4   3   3.5  RT (min)      lcqtof_hilic_neg_4_004_47    m/z 321.0999     0   1E+4   3   3.5  RT (min)      lcqtof_hilic_neg_4_014_46    m/z 321.1510     0   1E+4   4.5   5  RT (min)      lcqtof_hilic_neg_4_008_24    m/z 321.1732     0   2   4   6   8E+3   3   3.5  RT (min)      lcqtof_hilic_neg_4_002_25    m/z 321.1733     0   2   4E+3   1.5   2  RT (min)       lcqtof_hilic_neg_4_021_21    m/z 321.1734     0   1E+4   2.5   3  RT (min)       lcqtof_hilic_neg_4_088_59    m/z 321.1734     0   2   4   6   8E+3   2.5   3  RT (min)      lcqtof_hilic_neg_4_048_23    m/z 321.1737     0   1E+4   2   2.5  RT (min)      lcqtof_hilic_neg_4_035_33    m/z 321.1738     0   1E+4   2   2.5  RT (min)       lcqtof_hilic_neg_4_003_16    m/z 321.1738     0   2   4   6E+3   0   0.5  RT (min)      lcqtof_hilic_neg_4_006_30    m/z 321.1738     0   1   2   3   4E+3   0   0.5  RT (min)      lcqtof_hilic_neg_4_012_07    m/z 321.1740     0   1E+4   2.5   3  RT (min)      lcqtof_hilic_neg_4_047_06    m/z 321.1740     0   2   4   6   8E+3   2   2.5  RT (min)      lcqtof_hilic_neg_4_033_53    m/z 321.1944     0   1E+4   1.5   2  RT (min)      lcqtof_hilic_neg_4_038_10    m/z 321.2103     0   2   4   6   8E+4   1   1.5  RT (min)       lcqtof_hilic_neg_4_000_aa    m/z 321.2104     0   1   2   3E+3   0   0.5  RT (min)      lcqtof_hilic_neg_4_006_30    m/z 321.2105     0   2   4   6   8E+3   2.5   3  RT (min)      lcqtof_hilic_neg_4_089_59    m/z 321.2106     0   2   4E+3   2   2.5  RT (min)       lcqtof_hilic_neg_4_038_10    m/z 321.2107     0   1E+4   15   15.5  RT (min)      lcqtof_hilic_neg_4_000_aa    m/z 321.2107     0   2   4E+4   1   1.5  RT (min)      lcqtof_hilic_neg_4_019_03    m/z 321.2110     0   1   2E+4   1   1.5  RT (min)       lcqtof_hilic_neg_4_009_50    m/z 321.2244     0   1   2   3E+4   1   1.5  RT (min)      lcqtof_hilic_neg_4_061_50    m/z 321.2398     0   1   2E+4   4   4.5  RT (min)      lcqtof_hilic_neg_4_055_32    m/z 321.2398     0   1   2   3E+4   4   4.5  RT (min)      lcqtof_hilic_neg_4_004_47    m/z 321.3311     0   2   4   6E+4   4   4.5  RT (min)      lcqtof_hilic_neg_4_004_47    m/z 321.3312     0   2   4   6E+4   4   4.5  RT (min)      lcqtof_hilic_neg_4_008_24    m/z 321.4697     0   1   2E+4   4   4.5  RT (min)      lcqtof_hilic_neg_4_004_47    m/z 321.9028     0   1   2   3E+4   4   4.5  RT (min)      lcqtof_hilic_neg_1_000_bb    m/z 321.9031     0   1   2   3E+4   4   4.5  RT (min)       lcqtof_hilic_neg_1_000_bb    m/z 321.9033     0   1   2   3E+4   4.5   5  RT (min)      lcqtof_hilic_neg_1_000_bb    m/z 321.9033     0   1   2   3E+4   4.5   5  RT (min)       lcqtof_hilic_neg_1_000_bb    m/z 321.9033     0   1   2   3E+4   4   4.5  RT (min)      lcqtof_hilic_neg_1_000_bb    m/z 321.9804     0   1E+4   1   1.5  RT (min)      lcqtof_hilic_neg_4_045_26    m/z 322.0575     0   1E+4   1   1.5  RT (min)      lcqtof_hilic_neg_4_055_32    m/z 322.2139     0   1E+4   1   1.5  RT (min)      lcqtof_hilic_neg_4_000_aa    m/z 322.8438     0   1   2   3   4E+3   8.5   9  RT (min)      lcqtof_hilic_neg_4_053_14    m/z 322.8441     0   1   2   3E+3   8.5   9  RT (min)      lcqtof_hilic_neg_4_059_14    m/z 322.9134     0   1   2   3E+3   6.5   7  RT (min)      lcqtof_hilic_neg_4_098_61    m/z 322.9742     0   2   4   6E+4   1   1.5  RT (min)      lcqtof_hilic_neg_4_045_26    m/z 322.9746     0   2   4   6E+4   0.5   1  RT (min)      lcqtof_hilic_neg_4_045_26    m/z 323.0919     0   1   2E+4   2   2.5  RT (min)      lcqtof_hilic_neg_4_039_18    m/z 323.0930     0   1   2E+4   1   1.5  RT (min)      lcqtof_hilic_neg_3_045_18    m/z 323.0934     0   1   2   3   4E+4   1   1.5  RT (min)      lcqtof_hilic_neg_4_039_18    m/z 323.0974     0   1   2E+4   3.5   4  RT (min)      lcqtof_hilic_neg_4_050_02    m/z 323.0979     0   1   2E+4   3.5   4  RT (min)      lcqtof_hilic_neg_4_050_02    m/z 323.0980     0   1E+4   4   4.5  RT (min)      lcqtof_hilic_neg_4_014_46    m/z 323.1748     0   1   2   3   4E+3   19   19.5  RT (min)      lcqtof_hilic_neg_4_093_60    m/z 323.1749     0   1   2   3E+3   16   16.5  RT (min)      lcqtof_hilic_neg_4_103_61    m/z 323.1749     0   1   2   3E+3   19.5   20  RT (min)      lcqtof_hilic_neg_4_000_cc    m/z 323.1750     0   1   2   3E+3   16   16.5  RT (min)      lcqtof_hilic_neg_4_012_07    m/z 323.1751     0   1   2E+4   15   15.5  RT (min)      lcqtof_hilic_neg_4_000_aa    m/z 323.1756     0   1   2   3E+3   0   0.5  RT (min)      lcqtof_hilic_neg_4_008_24    m/z 323.1757     0   1   2   3E+3   16.5   17  RT (min)      lcqtof_hilic_neg_4_019_03    m/z 323.1758     0   1   2   3E+3   19   19.5  RT (min)      lcqtof_hilic_neg_4_019_03    m/z 323.1758     0   1E+4   2.5   3  RT (min)      lcqtof_hilic_neg_4_089_59    m/z 323.1758     0   1   2   3E+3   0   0.5  RT (min)      lcqtof_hilic_neg_4_007_45    m/z 323.1773     0   2   4   6E+3   3   3.5  RT (min)      lcqtof_hilic_neg_4_002_25    m/z 323.9781     0   1E+4   1   1.5  RT (min)      lcqtof_hilic_neg_4_045_26    m/z 324.1329     0   1   2   3   4E+4   6   6.5  RT (min)      lcqtof_hilic_neg_4_008_24    m/z 324.1335     0   1   2   3   4E+4   6   6.5  RT (min)      lcqtof_hilic_neg_4_008_24    m/z 325.0171     0   1E+4   3   3.5  RT (min)      lcqtof_hilic_neg_4_039_18    m/z 325.0691     0   1E+4   1   1.5  RT (min)      lcqtof_hilic_neg_4_055_32    m/z 325.1135     0   1E+4   0.5   1  RT (min)      lcqtof_hilic_neg_4_033_53    m/z 325.1136     0   1   2E+4   5.5   6  RT (min)      lcqtof_hilic_neg_4_050_02    m/z 325.1136     0   1E+4   5.5   6  RT (min)      lcqtof_hilic_neg_4_052_13    m/z 325.1136     0   1   2E+4   5.5   6  RT (min)      lcqtof_hilic_neg_4_050_02    m/z 325.1150     0   1E+4   1.5   2  RT (min)      lcqtof_hilic_neg_4_060_13    m/z 325.1832     0   1E+5   0   0.5  RT (min)      lcqtof_hilic_neg_4_020_42    m/z 325.1833     0   1   2   3E+5   1.5   2  RT (min)      lcqtof_hilic_neg_2_040_20    m/z 325.1833     0   1E+5   0   0.5  RT (min)       lcqtof_hilic_neg_4_011_35    m/z 325.1833     0   1E+6   0.5   1  RT (min)       lcqtof_hilic_neg_4_072_57    m/z 325.1833     0   1E+5   0   0.5  RT (min)       lcqtof_hilic_neg_4_081_58    m/z 325.1833     0   1E+5   16.5   17  RT (min)       lcqtof_hilic_neg_4_019_03    m/z 325.1834     0   1   2   3   4E+5   2.5   3  RT (min)       lcqtof_hilic_neg_4_086_59    m/z 325.1834     0   1   2E+6   10   15  RT (min)       lcqtof_hilic_neg_4_000_aa    m/z 325.1834     0   1E+5   18.5   19  RT (min)       lcqtof_hilic_neg_4_104_61    m/z 325.1834     0   1E+5   16.5   17  RT (min)       lcqtof_hilic_neg_4_019_03    m/z 325.1834     0   1E+5   15.5   16  RT (min)       lcqtof_hilic_neg_4_113_62    m/z 325.1834     0   1   2E+5   16   17  RT (min)       lcqtof_hilic_neg_4_050_02    m/z 325.1834     0   2   4   6E+5   0   0.5  RT (min)       lcqtof_hilic_neg_4_100_61    m/z 325.1834     0   1E+5   17   17.5  RT (min)       lcqtof_hilic_neg_4_104_61    m/z 325.1834     0   1E+5   0   0.5  RT (min)       lcqtof_hilic_neg_4_112_62    m/z 325.1834     0   1E+5   15   15.5  RT (min)      lcqtof_hilic_neg_4_110_62    m/z 325.1834     0   1E+5   18.5   19  RT (min)       lcqtof_hilic_neg_4_105_61    m/z 325.1834     0   2   4E+4   3   3.5  RT (min)      lcqtof_hilic_neg_1_024_21    m/z 325.1834     0   2   4   6   8E+4   15.5   16  RT (min)      lcqtof_hilic_neg_4_030_44    m/z 325.1834     0   1E+5   1.5   2  RT (min)      lcqtof_hilic_neg_3_047_16    m/z 325.1834     0   2   4   6E+4   15   15.5  RT (min)      lcqtof_hilic_neg_4_027_19    m/z 325.1834     0   1E+5   19   19.5   20  RT (min)       lcqtof_hilic_neg_4_109_62    m/z 325.1834     0   1E+5   19.5   20  RT (min)       lcqtof_hilic_neg_4_070_57    m/z 325.1834     0   2   4   6E+4   15   15.5  RT (min)      lcqtof_hilic_neg_4_027_19    m/z 325.1834     0   1   2   3   4E+4   3   3.5  RT (min)      lcqtof_hilic_neg_1_026_10    m/z 325.1834     0   1E+5   1.5   2  RT (min)       lcqtof_hilic_neg_4_019_03    m/z 325.1835     0   1E+5   18.5   19  RT (min)       lcqtof_hilic_neg_4_063_16    m/z 325.1835     0   1E+5   17   18  RT (min)       lcqtof_hilic_neg_4_107_62    m/z 325.1835     0   1E+5   3.5   4  RT (min)       lcqtof_hilic_neg_4_008_24    m/z 325.1835     0   1E+5   15.5   16  RT (min)       lcqtof_hilic_neg_4_019_03    m/z 325.1835     0   1E+5   18.5   19  RT (min)      lcqtof_hilic_neg_4_062_17    m/z 325.1835     0   1E+5   17.5   18  RT (min)       lcqtof_hilic_neg_4_110_62    m/z 325.1835     0   1E+5   17   17.5  RT (min)       lcqtof_hilic_neg_4_019_03    m/z 325.1835     0   1E+5   17.5   18   18.5  RT (min)       lcqtof_hilic_neg_4_080_58    m/z 325.1835     0   1   2   3   4E+5   1.5   2  RT (min)       lcqtof_hilic_neg_4_006_30    m/z 325.1835     0   2   4   6E+4   14.5   15  RT (min)       lcqtof_hilic_neg_4_078_58    m/z 325.1835     0   1E+5   3   3.5  RT (min)       lcqtof_hilic_neg_4_002_25    m/z 325.1835     0   1E+5   14.5   15   15.5  RT (min)       lcqtof_hilic_neg_4_083_59    m/z 325.1835     0   1E+5   16   16.5  RT (min)       lcqtof_hilic_neg_4_019_03    m/z 325.1835     0   1E+5   17.5   18  RT (min)       lcqtof_hilic_neg_4_047_06    m/z 325.1835     0   2   4   6E+4   2.5   3  RT (min)      lcqtof_hilic_neg_1_039_43    m/z 325.1835     0   1   2E+5   2.5   3  RT (min)       lcqtof_hilic_neg_4_048_23    m/z 325.1835     0   2   4   6   8E+4   14.5   15  RT (min)       lcqtof_hilic_neg_4_099_61    m/z 325.1835     0   1   2   3   4E+4   3   3.5  RT (min)      lcqtof_hilic_neg_1_026_10    m/z 325.1836     0   1   2E+6   15   15.5  RT (min)       lcqtof_hilic_neg_4_000_aa    m/z 325.1836     0   1E+5   19   19.5  RT (min)       lcqtof_hilic_neg_4_101_61    m/z 325.1836     0   1E+5   16.5   17  RT (min)       lcqtof_hilic_neg_4_019_03    m/z 325.1836     0   1E+5   19   19.5  RT (min)       lcqtof_hilic_neg_4_021_21    m/z 325.1836     0   1E+5   3.5   4  RT (min)       lcqtof_hilic_neg_4_091_60    m/z 325.1836     0   1   2E+4   6.5   7  RT (min)       lcqtof_hilic_neg_4_044_08    m/z 325.1836     0   2   4   6E+4   3.5   4  RT (min)      lcqtof_hilic_neg_2_003_25    m/z 325.1836     0   1E+5   16   16.5  RT (min)      lcqtof_hilic_neg_4_019_03    m/z 325.1836     0   2   4   6   8E+4   15.5   16  RT (min)      lcqtof_hilic_neg_4_021_21    m/z 325.1836     0   1   2E+4   6   6.5  RT (min)       lcqtof_hilic_neg_4_085_59    m/z 325.1836     0   2   4E+4   6   8  RT (min)       lcqtof_hilic_neg_4_019_03    m/z 325.1837     0   2   4   6   8E+4   4   4.5  RT (min)      lcqtof_hilic_neg_4_044_08    m/z 325.1837     0   1E+5   18  RT (min)       lcqtof_hilic_neg_4_019_03    m/z 325.1838     0   1E+4   5   5.5  RT (min)      lcqtof_hilic_neg_4_059_14    m/z 325.1838     0   1E+5   19  RT (min)       lcqtof_hilic_neg_4_062_17    m/z 325.1838     0   1   2E+4   3.5   4  RT (min)      lcqtof_hilic_neg_3_000_aa    m/z 325.1838     0   2   4   6E+4   4.5   5  RT (min)      lcqtof_hilic_neg_4_008_24    m/z 325.1841     0   1   2E+4   5   5.5  RT (min)      lcqtof_hilic_neg_4_022_34    m/z 325.1842     0   1   2E+4   7   8  RT (min)       lcqtof_hilic_neg_4_048_23    m/z 325.1949     0   1   2E+5   0.5   1  RT (min)      lcqtof_hilic_neg_3_035_46    m/z 325.1956     0   1   2E+5   0.5   1  RT (min)      lcqtof_hilic_neg_3_003_20    m/z 325.1957     0   1   2E+5   0.5   1  RT (min)      lcqtof_hilic_neg_3_003_20    m/z 325.1961     0   2   4E+4   0.5   1  RT (min)      lcqtof_hilic_neg_3_000_bb    m/z 325.1961     0   1   2E+5   0.5   1  RT (min)      lcqtof_hilic_neg_3_003_20    m/z 325.2745     0   1   2   3   4E+4   15   15.5  RT (min)      lcqtof_hilic_neg_4_000_aa    m/z 325.3659     0   2   4   6   8E+4   15   15.5  RT (min)      lcqtof_hilic_neg_4_000_aa    m/z 325.4036     0   1E+4   15   15.5  RT (min)      lcqtof_hilic_neg_4_000_aa    m/z 325.4501     0   1E+4   15   15.5  RT (min)      lcqtof_hilic_neg_4_000_aa    m/z 325.5066     0   1   2   3E+4   15   15.5  RT (min)      lcqtof_hilic_neg_4_000_aa    m/z 325.5697     0   1   2   3   4E+4   0.5   1  RT (min)      lcqtof_hilic_neg_4_047_06    m/z 325.6094     0   1E+4   15   15.5  RT (min)      lcqtof_hilic_neg_4_000_aa    m/z 325.6909     0   2   4E+4   0.5   1  RT (min)      lcqtof_hilic_neg_4_047_06    m/z 326.0874     0   1   2   3   4E+5   4   4.5  RT (min)      lcqtof_hilic_neg_4_050_02    m/z 326.1077     0   1E+4   5   5.5  RT (min)      lcqtof_hilic_neg_4_034_31    m/z 326.1866     0   1   2   3E+4   15.5   16  RT (min)      lcqtof_hilic_neg_4_109_62    m/z 326.1866     0   1   2   3   4E+5   15   15.5  RT (min)      lcqtof_hilic_neg_4_000_aa    m/z 326.1866     0   1   2   3   4E+5   15   15.5  RT (min)      lcqtof_hilic_neg_4_000_aa    m/z 326.1868     0   1E+4   0   0.5  RT (min)      lcqtof_hilic_neg_1_011_46    m/z 326.1868     0   1E+4   0   0.5  RT (min)      lcqtof_hilic_neg_1_003_37    m/z 326.1868     0   1   2   3E+4   1.5   2  RT (min)      lcqtof_hilic_neg_4_042_22    m/z 326.1868     0   1   2E+4   0   0.5  RT (min)      lcqtof_hilic_neg_4_089_59    m/z 326.1868     0   2   4   6   8E+4   1.5   2  RT (min)      lcqtof_hilic_neg_4_029_17    m/z 326.1868     0   1E+5   0   0.5  RT (min)      lcqtof_hilic_neg_4_100_61    m/z 326.1868     0   2   4E+3   6   6.5  RT (min)      lcqtof_hilic_neg_4_092_60    m/z 326.1868     0   1E+4   0   0.5  RT (min)      lcqtof_hilic_neg_1_049_55    m/z 326.1869     0   1   2E+5   1.5   2  RT (min)      lcqtof_hilic_neg_4_013_20    m/z 326.1869     0   2   4   6   8E+4   2.5   3  RT (min)      lcqtof_hilic_neg_4_086_59    m/z 326.1869     0   1   2E+5   0.5   1  RT (min)      lcqtof_hilic_neg_4_089_59    m/z 326.1869     0   1   2E+4   0   0.5  RT (min)      lcqtof_hilic_neg_4_108_62    m/z 326.1869     0   1   2E+5   0.5   1  RT (min)      lcqtof_hilic_neg_4_083_59    m/z 326.1870     0   1E+4   3   3.5  RT (min)      lcqtof_hilic_neg_4_017_51    m/z 326.1870     0   1   2   3   4E+4   2.5   3  RT (min)      lcqtof_hilic_neg_4_048_23    m/z 326.1870     0   1   2   3   4E+3   5   5.5  RT (min)      lcqtof_hilic_neg_4_059_14    m/z 326.1871     0   1   2E+4   1.5   2  RT (min)      lcqtof_hilic_neg_4_021_21    m/z 326.1872     0   1E+4   3   3.5  RT (min)      lcqtof_hilic_neg_4_098_61    m/z 326.1872     0   1   2   3E+4   3   3.5  RT (min)      lcqtof_hilic_neg_4_002_25    m/z 326.1872     0   1   2   3   4E+4   3.5   4  RT (min)      lcqtof_hilic_neg_4_008_24    m/z 326.1872     0   1   2   3   4E+4   4.5   5  RT (min)      lcqtof_hilic_neg_4_055_32    m/z 326.1873     0   1   2   3   4E+4   3.5   4  RT (min)      lcqtof_hilic_neg_4_004_47    m/z 326.1874     0   1E+4   4   4.5  RT (min)      lcqtof_hilic_neg_4_044_08    m/z 326.1874     0   2   4   6   8E+3   6.5   7  RT (min)       lcqtof_hilic_neg_4_012_07    m/z 326.1874     0   2   4   6E+3   6.5   7  RT (min)      lcqtof_hilic_neg_4_088_59    m/z 326.1875     0   1   2E+3   6.5   7  RT (min)      lcqtof_hilic_neg_1_009_50    m/z 326.1980     0   1   2   3E+4   0.5   1  RT (min)      lcqtof_hilic_neg_3_003_20    m/z 326.1997     0   1   2   3E+4   0.5   1  RT (min)      lcqtof_hilic_neg_3_003_20    m/z 326.3329     0   1   2   3E+4   0.5   1  RT (min)      lcqtof_hilic_neg_4_047_06    m/z 326.3714     0   1   2E+4   15   15.5  RT (min)      lcqtof_hilic_neg_4_000_aa    m/z 326.4140     0   2   4   6E+4   0.5   1  RT (min)      lcqtof_hilic_neg_4_047_06    m/z 326.4647     0   1E+5   0.5   1  RT (min)      lcqtof_hilic_neg_4_047_06    m/z 326.5720     0   1   2   3   4E+4   0.5   1  RT (min)      lcqtof_hilic_neg_4_047_06    m/z 326.6130     0   2   4E+4   0.5   1  RT (min)      lcqtof_hilic_neg_4_047_06    m/z 326.8645     0   1E+4   8.5   9  RT (min)      lcqtof_hilic_neg_1_000_aa    m/z 326.8653     0   1E+4   8.5   9  RT (min)      lcqtof_hilic_neg_1_000_aa    m/z 327.0589     0   1E+4   4   4.5  RT (min)      lcqtof_hilic_neg_4_050_02    m/z 327.0698     0   1   2E+4   6.5   7  RT (min)       lcqtof_hilic_neg_4_097_60    m/z 327.0915     0   2   4   6E+4   4   4.5  RT (min)      lcqtof_hilic_neg_4_050_02    m/z 327.1056     0   1   2E+4   6.5   7  RT (min)      lcqtof_hilic_neg_4_055_32    m/z 327.1257     0   2   4   6   8E+3   3   3.5  RT (min)      lcqtof_hilic_neg_4_004_47    m/z 327.1300     0   1E+4   4   4.5  RT (min)      lcqtof_hilic_neg_4_040_05    m/z 327.1315     0   1   2   3E+4   1   1.5  RT (min)      lcqtof_hilic_neg_4_052_13    m/z 327.1820     0   1E+4   2.5   3  RT (min)      lcqtof_hilic_neg_4_048_23    m/z 327.1820     0   2   4   6   8E+3   3   3.5  RT (min)      lcqtof_hilic_neg_4_007_45    m/z 327.1820     0   2   4   6E+4   1.5   2  RT (min)      lcqtof_hilic_neg_4_013_20    m/z 327.1820     0   1E+5   15   15.5  RT (min)      lcqtof_hilic_neg_4_000_aa    m/z 327.1821     0   1E+4   3   3.5  RT (min)      lcqtof_hilic_neg_4_031_38    m/z 327.1824     0   1   2   3E+4   0.5   1  RT (min)      lcqtof_hilic_neg_4_020_42    m/z 327.1825     0   1E+4   3   3.5  RT (min)      lcqtof_hilic_neg_4_002_25    m/z 327.1826     0   2   4   6   8E+3   0   0.5  RT (min)      lcqtof_hilic_neg_4_051_43    m/z 327.1827     0   1   2   3E+4   2   2.5  RT (min)      lcqtof_hilic_neg_4_005_29    m/z 327.1827     0   1E+4   4   4.5  RT (min)      lcqtof_hilic_neg_4_012_07    m/z 327.1827     0   2   4   6   8E+3   0   0.5  RT (min)      lcqtof_hilic_neg_4_010_28    m/z 327.1828     0   1   2   3E+4   2.5   3  RT (min)      lcqtof_hilic_neg_4_006_30    m/z 327.1828     0   2   4   6E+4   1.5   2  RT (min)      lcqtof_hilic_neg_4_021_21    m/z 327.1828     0   1E+4   4   4.5  RT (min)      lcqtof_hilic_neg_4_004_47    m/z 327.1828     0   2   4   6   8E+3   0   0.5  RT (min)      lcqtof_hilic_neg_4_024_40    m/z 327.1828     0   1   2E+4   2   2.5  RT (min)      lcqtof_hilic_neg_4_053_14    m/z 327.1829     0   1   2E+4   2.5   3  RT (min)      lcqtof_hilic_neg_4_088_59    m/z 327.1829     0   1E+4   2   2.5  RT (min)      lcqtof_hilic_neg_4_026_04    m/z 327.1829     0   1E+4   3   3.5  RT (min)      lcqtof_hilic_neg_4_040_05    m/z 327.1830     0   1E+5   15   15.5  RT (min)      lcqtof_hilic_neg_4_000_aa    m/z 327.1831     0   1E+4   3.5   4  RT (min)      lcqtof_hilic_neg_4_008_24    m/z 327.1831     0   1E+4   4   4.5  RT (min)      lcqtof_hilic_neg_4_002_25    m/z 327.1832     0   2   4   6   8E+3   1.5   2  RT (min)      lcqtof_hilic_neg_4_021_21    m/z 327.1832     0   2   4E+4   1.5   2  RT (min)      lcqtof_hilic_neg_4_050_02    m/z 327.1834     0   2   4   6E+3   3.5   4  RT (min)      lcqtof_hilic_neg_4_055_32    m/z 327.1962     0   1E+4   0.5   1  RT (min)      lcqtof_hilic_neg_3_003_20    m/z 327.1975     0   1E+4   0.5   1  RT (min)      lcqtof_hilic_neg_4_048_23    m/z 327.2528     0   1   2E+4   1   1.5  RT (min)      lcqtof_hilic_neg_4_000_aa    m/z 327.2671     0   1E+4   0.5   1  RT (min)      lcqtof_hilic_neg_4_047_06    m/z 327.3724     0   2   4   6E+4   0.5   1  RT (min)      lcqtof_hilic_neg_4_047_06    m/z 327.4641     0   1   2   3   4E+4   0.5   1  RT (min)      lcqtof_hilic_neg_4_047_06    m/z 327.5122     0   2   4   6E+4   0.5   1  RT (min)      lcqtof_hilic_neg_4_047_06    m/z 327.6038     0   1   2   3   4E+4   0.5   1  RT (min)      lcqtof_hilic_neg_4_047_06    m/z 328.0768     0   1E+4   1   1.5  RT (min)      lcqtof_hilic_neg_4_055_32    m/z 328.0912     0   1E+4   4   4.5  RT (min)      lcqtof_hilic_neg_4_050_02    m/z 328.1087     0   1   2E+4   6   6.5  RT (min)      lcqtof_hilic_neg_4_047_06    m/z 328.1512     0   2   4E+3   3.5   4  RT (min)      lcqtof_hilic_neg_4_059_14    m/z 328.1840     0   1E+4   15   15.5  RT (min)      lcqtof_hilic_neg_4_000_aa    m/z 328.2475     0   1E+4   15   15.5  RT (min)      lcqtof_hilic_neg_4_000_aa    m/z 328.2491     0   1   2E+4   0.5   1  RT (min)      lcqtof_hilic_neg_3_035_46    m/z 328.6109     0   1   2   3E+4   0.5   1  RT (min)      lcqtof_hilic_neg_4_047_06    m/z 328.7019     0   1   2   3E+4   0.5   1  RT (min)      lcqtof_hilic_neg_4_047_06    m/z 328.9033     0   1   2   3E+3   8   8.5  RT (min)       lcqtof_hilic_neg_4_058_39    m/z 329.0490     0   1E+4   2   2.5  RT (min)      lcqtof_hilic_neg_4_053_14    m/z 329.0566     0   1   2   3E+4   3.5   4  RT (min)      lcqtof_hilic_neg_4_026_04    m/z 329.0567     0   1   2   3E+4   3.5   4  RT (min)      lcqtof_hilic_neg_4_026_04    m/z 329.0570     0   2   4   6   8E+4   4   4.5  RT (min)      lcqtof_hilic_neg_4_026_04    m/z 329.0570     0   1   2   3E+4   3.5   4  RT (min)      lcqtof_hilic_neg_4_026_04    m/z 329.0577     0   2   4   6   8E+4   4   4.5  RT (min)      lcqtof_hilic_neg_4_026_04    m/z 329.0844     0   1   2E+4   0.5   1  RT (min)      lcqtof_hilic_neg_4_020_42    m/z 329.0852     0   1   2E+4   1   1.5  RT (min)      lcqtof_hilic_neg_4_020_42    m/z 329.1388     0   1E+4   4   4.5  RT (min)       lcqtof_hilic_neg_4_047_06    m/z 330.1061     0   1   2E+4   6.5   7  RT (min)      lcqtof_hilic_neg_4_055_32    m/z 330.6988     0   1E+4   4.5   5  RT (min)      lcqtof_hilic_neg_4_050_02    m/z 331.0554     0   1E+4   6.5   7  RT (min)      lcqtof_hilic_neg_4_008_24    m/z 331.0728     0   1E+4   5   5.5  RT (min)      lcqtof_hilic_neg_4_052_13    m/z 331.0800     0   1E+4   3.5   4  RT (min)      lcqtof_hilic_neg_1_009_50    m/z 331.1002     0   1E+4   3.5   4  RT (min)      lcqtof_hilic_neg_4_004_47    m/z 331.1363     0   1E+4   3   3.5  RT (min)       lcqtof_hilic_neg_4_008_24    m/z 331.1746     0   1E+4   2   2.5  RT (min)      lcqtof_hilic_neg_4_050_02    m/z 331.1767     0   1E+4   1   1.5  RT (min)      lcqtof_hilic_neg_4_060_13    m/z 331.2396     0   1E+4   0.5   1  RT (min)      lcqtof_hilic_neg_3_035_46    m/z 332.6964     0   1   2   3E+4   4.5   5  RT (min)      lcqtof_hilic_neg_4_050_02    m/z 332.9166     0   1   2E+3   4.5   5  RT (min)      lcqtof_hilic_neg_4_004_47    m/z 333.0983     0   1E+4   0.5   1  RT (min)       lcqtof_hilic_neg_4_100_61    m/z 333.1293     0   1E+4   1   1.5  RT (min)      lcqtof_hilic_neg_4_050_02    m/z 333.2306     0   1E+4   0.5   1  RT (min)      lcqtof_hilic_neg_4_006_30    m/z 334.6929     0   1   2E+4   4.5   5  RT (min)      lcqtof_hilic_neg_4_050_02    m/z 334.6936     0   1   2E+4   4.5   5  RT (min)      lcqtof_hilic_neg_4_050_02    m/z 334.9376     0   1E+5   1   1.5  RT (min)      lcqtof_hilic_neg_4_004_47    m/z 334.9377     0   1E+5   1   1.5  RT (min)      lcqtof_hilic_neg_4_004_47    m/z 334.9379     0   2   4   6   8E+4   1.5   2  RT (min)      lcqtof_hilic_neg_4_004_47    m/z 334.9380     0   2   4   6   8E+4   1.5   2  RT (min)      lcqtof_hilic_neg_4_004_47    m/z 335.0786     0   1E+4   3   3.5  RT (min)      lcqtof_hilic_neg_4_050_02    m/z 335.1218     0   1   2E+4   4   4.5  RT (min)       lcqtof_hilic_neg_4_090_60    m/z 335.1226     0   1E+4   4   4.5  RT (min)      lcqtof_hilic_neg_1_037_24    m/z 335.1887     0   1   2   3   4E+4   15   15.5  RT (min)      lcqtof_hilic_neg_4_000_aa    m/z 335.1889     0   2   4E+3   0   0.5  RT (min)      lcqtof_hilic_neg_4_005_29    m/z 335.1889     0   1E+4   2.5   3  RT (min)      lcqtof_hilic_neg_4_048_23    m/z 335.1889     0   2   4   6E+3   1.5   2  RT (min)      lcqtof_hilic_neg_4_047_06    m/z 335.1890     0   1E+4   3   3.5  RT (min)      lcqtof_hilic_neg_4_002_25    m/z 335.1890     0   1E+4   2   2.5  RT (min)       lcqtof_hilic_neg_4_005_29    m/z 335.1890     0   1E+4   2.5   3  RT (min)       lcqtof_hilic_neg_4_088_59    m/z 335.1891     0   2   4   6E+4   1.5   2  RT (min)      lcqtof_hilic_neg_4_013_20    m/z 335.1891     0   1E+4   2   2.5  RT (min)      lcqtof_hilic_neg_4_041_54    m/z 335.1892     0   2   4   6E+3   1.5   2  RT (min)       lcqtof_hilic_neg_4_039_18    m/z 335.1892     0   1E+4   2   2.5  RT (min)      lcqtof_hilic_neg_4_033_53    m/z 335.1892     0   1   2E+4   0.5   1  RT (min)      lcqtof_hilic_neg_4_044_08    m/z 335.1893     0   1E+4   2   2.5  RT (min)      lcqtof_hilic_neg_4_003_16    m/z 335.1893     0   1   2   3E+4   0   0.5  RT (min)      lcqtof_hilic_neg_4_013_20    m/z 335.1894     0   1   2   3   4E+3   0   0.5  RT (min)      lcqtof_hilic_neg_4_025_52    m/z 335.1895     0   2   4E+3   2.5   3  RT (min)      lcqtof_hilic_neg_4_106_62    m/z 335.2221     0   1   2   3E+4   1   1.5  RT (min)      lcqtof_hilic_neg_4_053_14    m/z 335.9382     0   1   2E+4   1   1.5  RT (min)      lcqtof_hilic_neg_4_004_47    m/z 335.9881     0   2   4   6   8E+3   2.5   3  RT (min)      lcqtof_hilic_neg_1_012_04    m/z 336.0960     0   1E+4   1.5   2  RT (min)      lcqtof_hilic_neg_4_014_46    m/z 336.1062     0   1E+4   2   2.5  RT (min)      lcqtof_hilic_neg_4_008_24    m/z 336.1137     0   2   4E+4   6   6.5  RT (min)       lcqtof_hilic_neg_4_090_60    m/z 336.1141     0   1   2E+4   7   7.5  RT (min)       lcqtof_hilic_neg_4_008_24    m/z 336.1144     0   1E+4   6.5   7  RT (min)       lcqtof_hilic_neg_4_040_05    m/z 336.1457     0   1E+4   0.5   1  RT (min)      lcqtof_hilic_neg_2_000_aa    m/z 336.1574     0   1E+4   4   4.5  RT (min)      lcqtof_hilic_neg_4_022_34    m/z 336.1926     0   1   2   3   4E+4   0.5   1  RT (min)      lcqtof_hilic_neg_4_038_10    m/z 336.6908     0   1E+4   4.5   5  RT (min)      lcqtof_hilic_neg_4_050_02    m/z 336.8585     0   1E+4   10   10.5  RT (min)      lcqtof_hilic_neg_4_014_46    m/z 336.8587     0   1E+4   7   7.5  RT (min)      lcqtof_hilic_neg_4_056_56    m/z 336.8604     0   1E+4   7   7.5  RT (min)       lcqtof_hilic_neg_4_047_06    m/z 337.0043     0   2   4   6E+3   2   2.5  RT (min)      lcqtof_hilic_neg_4_050_02    m/z 337.1671     0   1   2E+4   4   4.5  RT (min)      lcqtof_hilic_neg_4_016_37    m/z 337.1864     0   1   2   3E+4   0.5   1  RT (min)      lcqtof_hilic_neg_4_038_10    m/z 337.1876     0   1   2   3E+3   16   16.5  RT (min)      lcqtof_hilic_neg_4_058_39    m/z 337.1897     0   1   2E+3   19.5   20  RT (min)      lcqtof_hilic_neg_4_000_cc    m/z 337.1899     0   1   2   3E+3   0   0.5  RT (min)      lcqtof_hilic_neg_4_000_bb    m/z 337.1899     0   1   2   3E+3   17.5   18  RT (min)      lcqtof_hilic_neg_4_042_22    m/z 337.1902     0   1   2E+4   15   15.5  RT (min)      lcqtof_hilic_neg_4_000_aa    m/z 337.1902     0   1   2   3E+3   18   18.5  RT (min)      lcqtof_hilic_neg_4_045_26    m/z 337.1904     0   1   2   3E+3   18.5   19  RT (min)      lcqtof_hilic_neg_4_015_09    m/z 337.1909     0   1   2   3E+3   19   19.5  RT (min)      lcqtof_hilic_neg_4_005_29    m/z 337.1915     0   2   4   6   8E+3   2   2.5  RT (min)      lcqtof_hilic_neg_4_035_33    m/z 337.1915     0   2   4E+3   1.5   2  RT (min)      lcqtof_hilic_neg_4_000_bb    m/z 337.1916     0   2   4   6E+3   2.5   3  RT (min)      lcqtof_hilic_neg_4_089_59    m/z 337.1918     0   2   4   6E+3   2.5   3  RT (min)      lcqtof_hilic_neg_4_047_06    m/z 337.1920     0   1   2   3   4E+3   2   2.5  RT (min)      lcqtof_hilic_neg_4_039_18    m/z 337.1931     0   1   2   3E+3   16.5   17  RT (min)      lcqtof_hilic_neg_4_000_cc    m/z 337.1939     0   1   2   3E+3   17   17.5  RT (min)      lcqtof_hilic_neg_4_004_47    m/z 337.1942     0   1   2   3E+3   17   17.5  RT (min)      lcqtof_hilic_neg_4_019_03    m/z 337.1942     0   1   2   3   4E+3   2   2.5  RT (min)      lcqtof_hilic_neg_3_055_01    m/z 337.1979     0   2   4   6   8E+3   2   2.5  RT (min)      lcqtof_hilic_neg_4_005_29    m/z 337.2028     0   2   4E+4   1   1.5  RT (min)      lcqtof_hilic_neg_4_000_aa    m/z 337.2169     0   1E+4   0.5   1  RT (min)      lcqtof_hilic_neg_3_003_20    m/z 337.3918     0   1   2   3   4E+4   0.5   1  RT (min)      lcqtof_hilic_neg_4_048_23    m/z 338.0411     0   1E+4   4.5   5  RT (min)      lcqtof_hilic_neg_1_004_14    m/z 338.0752     0   1E+4   2   2.5  RT (min)      lcqtof_hilic_neg_4_050_02    m/z 338.0760     0   1E+4   1   1.5  RT (min)      lcqtof_hilic_neg_1_043_41    m/z 338.0940     0   1E+4   2   2.5  RT (min)      lcqtof_hilic_neg_4_061_50    m/z 338.1021     0   1E+4   1.5   2  RT (min)      lcqtof_hilic_neg_4_005_29    m/z 338.1562     0   1E+4   5   5.5  RT (min)      lcqtof_hilic_neg_4_022_34    m/z 339.0720     0   1   2E+4   2   2.5  RT (min)      lcqtof_hilic_neg_4_050_02    m/z 339.0734     0   1   2   3E+4   1   1.5  RT (min)      lcqtof_hilic_neg_2_046_41    m/z 339.0738     0   1   2   3E+4   1   1.5  RT (min)      lcqtof_hilic_neg_2_046_41    m/z 339.0784     0   1E+4   4   4.5  RT (min)      lcqtof_hilic_neg_4_040_05    m/z 339.0951     0   1   2   3   4E+3   0.5   1  RT (min)      lcqtof_hilic_neg_4_031_38    m/z 339.0974     0   1E+4   1.5   2  RT (min)      lcqtof_hilic_neg_4_091_60    m/z 339.1361     0   1E+4   5.5   6  RT (min)      lcqtof_hilic_neg_4_028_36    m/z 339.1417     0   1E+4   5   5.5  RT (min)       lcqtof_hilic_neg_1_037_24    m/z 339.1988     0   1E+6   15   15.5  RT (min)      lcqtof_hilic_neg_4_000_aa    m/z 339.1989     0   2   4   6E+4   17   18  RT (min)       lcqtof_hilic_neg_4_111_62    m/z 339.1989     0   1E+4   3   3.5  RT (min)      lcqtof_hilic_neg_1_055_25    m/z 339.1990     0   1E+6   0   0.5  RT (min)      lcqtof_hilic_neg_4_019_03    m/z 339.1990     0   2   4   6E+4   16.5   17  RT (min)       lcqtof_hilic_neg_4_107_62    m/z 339.1990     0   2   4   6E+4   17.5   18  RT (min)       lcqtof_hilic_neg_4_110_62    m/z 339.1990     0   2   4   6E+4   0   0.5  RT (min)      lcqtof_hilic_neg_4_019_03    m/z 339.1991     0   2   4   6E+5   1.5   2  RT (min)      lcqtof_hilic_neg_4_041_54    m/z 339.1991     0   1E+6   0.5   1  RT (min)      lcqtof_hilic_neg_4_019_03    m/z 339.1991     0   2   4   6E+4   0   0.5  RT (min)      lcqtof_hilic_neg_4_019_03    m/z 339.1991     0   2   4   6   8E+4   0   0.5  RT (min)      lcqtof_hilic_neg_4_025_52    m/z 339.1991     0   2   4   6E+4   0   0.5  RT (min)       lcqtof_hilic_neg_4_113_62    m/z 339.1991     0   2   4   6E+4   17   18   19  RT (min)       lcqtof_hilic_neg_4_012_07    m/z 339.1991     0   1   2E+4   18.5   19  RT (min)      lcqtof_hilic_neg_3_109_62    m/z 339.1991     0   2   4   6   8E+5   0   0.5  RT (min)       lcqtof_hilic_neg_4_019_03    m/z 339.1992     0   2   4   6E+5   1.5   2  RT (min)      lcqtof_hilic_neg_4_021_21    m/z 339.1992     0   2   4   6   8E+4   1.5   2  RT (min)       lcqtof_hilic_neg_4_019_03    m/z 339.1992     0   2   4   6E+5   1.5   2  RT (min)      lcqtof_hilic_neg_4_003_16    m/z 339.1992     0   1E+5   5   6  RT (min)       lcqtof_hilic_neg_4_055_32    m/z 339.1992     0   2   4   6E+4   16   18  RT (min)       lcqtof_hilic_neg_4_002_25    m/z 339.1992     0   1E+5   2.5   3  RT (min)       lcqtof_hilic_neg_4_048_23    m/z 339.1992     0   2   4   6E+4   1.5   2  RT (min)      lcqtof_hilic_neg_4_021_21    m/z 339.1992     0   1E+5   3   3.5  RT (min)       lcqtof_hilic_neg_4_002_25    m/z 339.1992     0   2   4   6   8E+4   2   2.5  RT (min)       lcqtof_hilic_neg_4_039_18    m/z 339.1992     0   2   4   6E+4   18   19   20  RT (min)       lcqtof_hilic_neg_4_019_03    m/z 339.1993     0   1   2   3   4E+4   3   3.5  RT (min)       lcqtof_hilic_neg_4_098_61    m/z 339.1993     0   1   2E+4   19   19.5  RT (min)      lcqtof_hilic_neg_3_113_62    m/z 339.1993     0   1   2E+5   2.5   3  RT (min)       lcqtof_hilic_neg_4_086_59    m/z 339.1994     0   1   2E+4   3.5   4  RT (min)      lcqtof_hilic_neg_4_098_61    m/z 339.1994     0   1E+4   6   6.5  RT (min)       lcqtof_hilic_neg_4_085_59    m/z 339.1995     0   1E+4   6.5   7   7.5  RT (min)       lcqtof_hilic_neg_4_001_49    m/z 339.1995     0   1E+4   5   5.5  RT (min)       lcqtof_hilic_neg_4_035_33    m/z 339.1995     0   1E+4   7   7.5  RT (min)       lcqtof_hilic_neg_4_048_23    m/z 339.1995     0   1   2E+4   15   15.5  RT (min)      lcqtof_hilic_neg_4_048_23    m/z 339.1995     0   1E+4   5   5.5  RT (min)      lcqtof_hilic_neg_4_016_37    m/z 339.1996     0   1   2   3   4E+3   7   8  RT (min)       lcqtof_hilic_neg_4_019_03    m/z 339.1996     0   2   4   6E+3   5   5.5  RT (min)      lcqtof_hilic_neg_4_045_26    m/z 339.2110     0   1   2E+5   0.5   1  RT (min)      lcqtof_hilic_neg_3_003_20    m/z 339.2115     0   2   4E+4   0.5   1  RT (min)      lcqtof_hilic_neg_3_000_bb    m/z 339.2120     0   1   2E+4   0   0.5  RT (min)      lcqtof_hilic_neg_3_000_bb    m/z 339.2128     0   1   2E+5   0   0.5  RT (min)      lcqtof_hilic_neg_3_035_46    m/z 339.2129     0   2   4E+4   0.5   1  RT (min)      lcqtof_hilic_neg_3_000_bb    m/z 339.2131     0   1   2E+5   0.5   1  RT (min)      lcqtof_hilic_neg_2_005_13    m/z 339.2522     0   1   2   3E+4   0.5   1  RT (min)      lcqtof_hilic_neg_4_038_10    m/z 339.2923     0   1   2E+4   15   15.5  RT (min)      lcqtof_hilic_neg_4_000_aa    m/z 339.3468     0   1   2   3   4E+4   0.5   1  RT (min)      lcqtof_hilic_neg_4_040_05    m/z 339.3856     0   2   4E+4   15   15.5  RT (min)      lcqtof_hilic_neg_4_000_aa    m/z 339.5290     0   1E+4   15   15.5  RT (min)      lcqtof_hilic_neg_4_000_aa    m/z 339.5907     0   1   2   3E+4   0.5   1  RT (min)      lcqtof_hilic_neg_4_038_10    m/z 339.7172     0   1   2   3E+4   0.5   1  RT (min)      lcqtof_hilic_neg_4_038_10    m/z 340.0002     0   1   2E+3   6   6.5  RT (min)      lcqtof_hilic_neg_1_054_01    m/z 340.0005     0   2   4   6E+3   6   6.5  RT (min)      lcqtof_hilic_neg_1_004_14    m/z 340.0576     0   1   2   3   4E+4   6   6.5  RT (min)       lcqtof_hilic_neg_4_083_59    m/z 340.1134     0   1E+4   6   6.5  RT (min)       lcqtof_hilic_neg_4_008_24    m/z 340.1355     0   1E+4   6   6.5  RT (min)       lcqtof_hilic_neg_4_060_13    m/z 340.2022     0   1E+4   0   0.5  RT (min)      lcqtof_hilic_neg_4_045_26    m/z 340.2025     0   1   2   3E+5   15   15.5  RT (min)      lcqtof_hilic_neg_4_000_aa    m/z 340.2026     0   2   4E+4   2   2.5  RT (min)      lcqtof_hilic_neg_4_005_29    m/z 340.2026     0   1E+4   3   3.5  RT (min)      lcqtof_hilic_neg_4_007_45    m/z 340.2026     0   2   4E+4   2   2.5  RT (min)      lcqtof_hilic_neg_4_032_01    m/z 340.2027     0   1   2E+4   3   3.5  RT (min)      lcqtof_hilic_neg_4_002_25    m/z 340.2027     0   1E+4   3   3.5  RT (min)      lcqtof_hilic_neg_4_034_31    m/z 340.2027     0   1   2E+4   2.5   3  RT (min)      lcqtof_hilic_neg_4_048_23    m/z 340.2027     0   1E+4   0   0.5  RT (min)      lcqtof_hilic_neg_4_010_28    m/z 340.2028     0   1   2   3   4E+4   2.5   3  RT (min)      lcqtof_hilic_neg_4_086_59    m/z 340.2028     0   2   4   6E+3   3   3.5  RT (min)      lcqtof_hilic_neg_4_104_61    m/z 340.2028     0   2   4   6E+3   3.5   4  RT (min)      lcqtof_hilic_neg_4_065_51    m/z 340.2028     0   1   2E+4   2   2.5  RT (min)      lcqtof_hilic_neg_4_026_04    m/z 340.2028     0   1   2E+3   15   15.5  RT (min)      lcqtof_hilic_neg_2_012_43    m/z 340.2029     0   1E+4   2.5   3  RT (min)      lcqtof_hilic_neg_4_113_62    m/z 340.2029     0   1   2   3   4E+4   2.5   3  RT (min)      lcqtof_hilic_neg_4_014_46    m/z 340.2029     0   1E+4   0   0.5  RT (min)      lcqtof_hilic_neg_4_010_28    m/z 340.2029     0   1   2   3E+5   0.5   1  RT (min)      lcqtof_hilic_neg_4_010_28    m/z 340.2029     0   1   2   3   4E+5   0.5   1  RT (min)      lcqtof_hilic_neg_4_052_13    m/z 340.2029     0   2   4   6E+3   15   15.5  RT (min)      lcqtof_hilic_neg_4_083_59    m/z 340.2030     0   1E+5   1.5   2  RT (min)      lcqtof_hilic_neg_4_027_19    m/z 340.2030     0   2   4E+4   2   2.5  RT (min)      lcqtof_hilic_neg_4_010_28    m/z 340.2030     0   1   2E+4   2   2.5  RT (min)      lcqtof_hilic_neg_4_026_04    m/z 340.2031     0   2   4E+3   3   3.5  RT (min)      lcqtof_hilic_neg_2_003_25    m/z 340.2033     0   1   2   3   4E+4   2.5   3  RT (min)      lcqtof_hilic_neg_4_014_46    m/z 340.2142     0   2   4E+4   0.5   1  RT (min)      lcqtof_hilic_neg_3_003_20    m/z 340.2146     0   2   4E+4   0.5   1  RT (min)      lcqtof_hilic_neg_3_003_20    m/z 340.2459     0   1   2   3E+5   0.5   1  RT (min)      lcqtof_hilic_neg_3_035_46    m/z 340.2978     0   2   4   6E+4   0.5   1  RT (min)      lcqtof_hilic_neg_4_038_10    m/z 340.3490     0   1   2   3   4E+4   0.5   1  RT (min)      lcqtof_hilic_neg_4_038_10    m/z 340.4723     0   2   4   6E+4   0.5   1  RT (min)      lcqtof_hilic_neg_4_038_10    m/z 340.5057     0   1   2   3   4E+4   0.5   1  RT (min)      lcqtof_hilic_neg_4_052_13    m/z 340.5359     0   2   4   6   8E+4   0.5   1  RT (min)      lcqtof_hilic_neg_4_038_10    m/z 340.5923     0   1   2E+4   0.5   1  RT (min)      lcqtof_hilic_neg_4_038_10    m/z 340.6361     0   2   4E+4   0.5   1  RT (min)      lcqtof_hilic_neg_4_038_10    m/z 340.6692     0   1   2   3E+4   0.5   1  RT (min)      lcqtof_hilic_neg_4_052_13    m/z 340.9682     0   1E+4   6.5   7  RT (min)       lcqtof_hilic_neg_4_048_23    m/z 341.0567     0   1E+4   3.5   4  RT (min)      lcqtof_hilic_neg_4_008_24    m/z 341.0766     0   1E+4   3.5   4  RT (min)      lcqtof_hilic_neg_1_037_24    m/z 341.0768     0   1E+4   3.5   4  RT (min)      lcqtof_hilic_neg_4_008_24    m/z 341.0816     0   1E+4   0.5   1  RT (min)      lcqtof_hilic_neg_4_023_15    m/z 341.1031     0   2   4   6E+3   1.5   2  RT (min)      lcqtof_hilic_neg_4_010_28    m/z 341.1078     0   1   2E+4   5   5.5  RT (min)      lcqtof_hilic_neg_2_000_aa    m/z 341.1078     0   2   4   6   8E+4   5.5   6  RT (min)      lcqtof_hilic_neg_2_000_aa    m/z 341.1079     0   2   4   6   8E+4   5.5   6  RT (min)      lcqtof_hilic_neg_2_000_aa    m/z 341.1080     0   2   4   6   8E+4   5.5   6  RT (min)      lcqtof_hilic_neg_2_000_aa    m/z 341.1080     0   1   2E+4   5   5.5  RT (min)      lcqtof_hilic_neg_2_000_aa    m/z 341.1080     0   1E+5   5   5.5  RT (min)      lcqtof_hilic_neg_1_000_aa    m/z 341.1083     0   1E+4   3.5   4  RT (min)      lcqtof_hilic_neg_4_014_46    m/z 341.1083     0   2   4   6   8E+3   1   1.5  RT (min)      lcqtof_hilic_neg_4_027_19    m/z 341.1084     0   1   2E+4   5   5.5  RT (min)      lcqtof_hilic_neg_2_000_aa    m/z 341.1087     0   1E+4   0.5   1  RT (min)      lcqtof_hilic_neg_4_050_02    m/z 341.1089     0   1   2E+4   5.5   6  RT (min)      lcqtof_hilic_neg_2_000_aa    m/z 341.1956     0   2   4   6   8E+3   3   3.5  RT (min)      lcqtof_hilic_neg_4_002_25    m/z 341.1960     0   2   4   6   8E+3   2.5   3  RT (min)      lcqtof_hilic_neg_4_048_23    m/z 341.1962     0   1E+5   15   15.5  RT (min)      lcqtof_hilic_neg_4_000_aa    m/z 341.1967     0   2   4   6E+3   1.5   2  RT (min)      lcqtof_hilic_neg_4_021_21    m/z 341.1970     0   2   4   6E+3   2   2.5  RT (min)      lcqtof_hilic_neg_4_038_10    m/z 341.1972     0   2   4   6E+3   2.5   3  RT (min)      lcqtof_hilic_neg_4_013_20    m/z 341.1973     0   1E+4   2   2.5  RT (min)      lcqtof_hilic_neg_4_026_04    m/z 341.1973     0   2   4E+3   0   0.5  RT (min)      lcqtof_hilic_neg_4_051_43    m/z 341.1973     0   1   2   3E+4   0   0.5  RT (min)      lcqtof_hilic_neg_4_037_39    m/z 341.1974     0   1   2   3   4E+3   15   15.5  RT (min)      lcqtof_hilic_neg_4_047_06    m/z 341.1974     0   1E+4   2.5   3  RT (min)      lcqtof_hilic_neg_4_088_59    m/z 341.1976     0   2   4   6E+3   0   0.5  RT (min)      lcqtof_hilic_neg_4_107_62    m/z 341.1976     0   1E+4   4   4.5  RT (min)      lcqtof_hilic_neg_4_028_36    m/z 341.1976     0   2   4   6   8E+3   2   2.5  RT (min)      lcqtof_hilic_neg_4_019_03    m/z 341.1977     0   1E+4   4   4.5  RT (min)      lcqtof_hilic_neg_4_096_60    m/z 341.1978     0   1   2E+4   2   2.5  RT (min)      lcqtof_hilic_neg_4_003_16    m/z 341.2012     0   1   2E+4   6   6.5  RT (min)      lcqtof_hilic_neg_3_011_34    m/z 341.2492     0   1   2   3   4E+4   0.5   1  RT (min)      lcqtof_hilic_neg_3_035_46    m/z 341.2694     0   1   2   3   4E+4   1   1.5  RT (min)      lcqtof_hilic_neg_4_000_aa    m/z 341.2952     0   2   4   6E+4   6   6.5  RT (min)      lcqtof_hilic_neg_3_011_34    m/z 341.3083     0   1E+4   1   1.5  RT (min)      lcqtof_hilic_neg_4_000_aa    m/z 341.3893     0   1E+4   0.5   1  RT (min)      lcqtof_hilic_neg_4_038_10    m/z 341.4392     0   1E+4   6   6.5  RT (min)      lcqtof_hilic_neg_3_011_34    m/z 341.5340     0   1   2E+4   0.5   1  RT (min)      lcqtof_hilic_neg_4_038_10    m/z 341.9986     0   1   2E+4   7.5   8  RT (min)      lcqtof_hilic_neg_4_064_15    m/z 341.9988     0   1E+4   7.5   8  RT (min)      lcqtof_hilic_neg_4_059_14    m/z 341.9988     0   1E+5   6.5   7  RT (min)      lcqtof_hilic_neg_4_023_15    m/z 341.9988     0   1   2   3E+4   7   7.5  RT (min)      lcqtof_hilic_neg_1_003_37    m/z 341.9989     0   1   2   3E+4   7.5   8  RT (min)      lcqtof_hilic_neg_4_032_01    m/z 341.9992     0   2   4E+4   7   7.5  RT (min)      lcqtof_hilic_neg_4_032_01    m/z 341.9992     0   1E+4   7.5   8  RT (min)      lcqtof_hilic_neg_4_059_14    m/z 341.9992     0   1E+5   6.5   7  RT (min)      lcqtof_hilic_neg_4_032_01    m/z 341.9992     0   1E+4   8   8.5  RT (min)      lcqtof_hilic_neg_4_059_14    m/z 342.1113     0   1   2E+4   5.5   6  RT (min)      lcqtof_hilic_neg_1_000_aa    m/z 342.1456     0   1E+4   5.5   6  RT (min)      lcqtof_hilic_neg_4_055_32    m/z 342.1993     0   1E+4   15   15.5  RT (min)      lcqtof_hilic_neg_4_000_aa    m/z 342.1998     0   1   2E+3   2   2.5  RT (min)      lcqtof_hilic_neg_4_004_47    m/z 342.2276     0   1E+4   1.5   2  RT (min)      lcqtof_hilic_neg_4_032_01    m/z 342.8549     0   1   2   3E+3   7.5   8  RT (min)      lcqtof_hilic_neg_2_059_39    m/z 343.0020     0   1   2E+4   6.5   7  RT (min)      lcqtof_hilic_neg_4_053_14    m/z 343.0026     0   1   2E+4   6.5   7  RT (min)      lcqtof_hilic_neg_4_053_14    m/z 343.0028     0   1   2E+4   6.5   7  RT (min)      lcqtof_hilic_neg_4_053_14    m/z 343.0611     0   1E+4   4.5   5  RT (min)      lcqtof_hilic_neg_1_005_32    m/z 343.1120     0   1   2   3   4E+4   6   6.5  RT (min)      lcqtof_hilic_neg_3_011_34    m/z 343.1199     0   1   2   3E+3   1.5   2  RT (min)       lcqtof_hilic_neg_4_006_30    m/z 343.1200     0   2   4   6E+3   0.5   1  RT (min)       lcqtof_hilic_neg_4_005_29    m/z 343.1201     0   1   2   3E+3   18   18.5  RT (min)      lcqtof_hilic_neg_4_058_39    m/z 343.2121     0   2   4   6E+3   15   15.5  RT (min)      lcqtof_hilic_neg_4_000_aa    m/z 343.2124     0   2   4   6   8E+3   2   2.5  RT (min)       lcqtof_hilic_neg_4_052_13    m/z 343.2126     0   1E+4   0.5   1   1.5  RT (min)       lcqtof_hilic_neg_4_050_02    m/z 343.9282     0   2   4   6E+4   1   1.5  RT (min)      lcqtof_hilic_neg_4_027_19    m/z 343.9289     0   2   4   6E+4   1   1.5  RT (min)      lcqtof_hilic_neg_4_027_19    m/z 343.9303     0   2   4   6E+4   1   1.5  RT (min)      lcqtof_hilic_neg_4_027_19    m/z 343.9305     0   2   4   6E+4   0.5   1  RT (min)      lcqtof_hilic_neg_4_027_19    m/z 343.9305     0   2   4   6E+4   1   1.5  RT (min)       lcqtof_hilic_neg_4_027_19    m/z 343.9935     0   1   2E+5   7.5   8  RT (min)       lcqtof_hilic_neg_1_004_14    m/z 343.9936     0   1   2   3E+6   6.5   7  RT (min)      lcqtof_hilic_neg_4_053_14    m/z 343.9937     0   1E+6   7   7.5  RT (min)       lcqtof_hilic_neg_4_032_01    m/z 343.9937     0   1E+5   7.5   8  RT (min)      lcqtof_hilic_neg_3_054_37    m/z 343.9937     0   2   4E+5   7.5   8  RT (min)       lcqtof_hilic_neg_4_032_01    m/z 343.9938     0   1E+5   7.5   8  RT (min)       lcqtof_hilic_neg_2_016_33    m/z 343.9938     0   1   2   3E+5   7.5   8  RT (min)      lcqtof_hilic_neg_4_023_15    m/z 343.9938     0   1   2   3   4E+5   7.5   8  RT (min)       lcqtof_hilic_neg_4_023_15    m/z 343.9939     0   1   2E+5   8.5   9  RT (min)       lcqtof_hilic_neg_4_059_14    m/z 343.9939     0   1   2   3E+6   6.5   7  RT (min)      lcqtof_hilic_neg_4_053_14    m/z 343.9939     0   1E+6   7   7.5  RT (min)       lcqtof_hilic_neg_4_032_01    m/z 343.9940     0   1   2E+5   8   8.5  RT (min)       lcqtof_hilic_neg_4_059_14    m/z 343.9940     0   1E+5   8.5   9  RT (min)       lcqtof_hilic_neg_4_064_15    m/z 343.9940     0   1   2E+5   7.5   8  RT (min)      lcqtof_hilic_neg_4_078_58    m/z 343.9940     0   1   2E+4   8   8.5  RT (min)       lcqtof_hilic_neg_2_051_14    m/z 343.9940     0   1   2   3E+6   6.5   7  RT (min)       lcqtof_hilic_neg_4_053_14    m/z 343.9940     0   2   4E+5   7   7.5  RT (min)       lcqtof_hilic_neg_1_003_37    m/z 343.9942     0   1E+4   10   10.5  RT (min)       lcqtof_hilic_neg_4_059_14    m/z 343.9942     0   1   2E+5   8   8.5  RT (min)       lcqtof_hilic_neg_4_059_14    m/z 343.9943     0   1E+4   9.5   10  RT (min)       lcqtof_hilic_neg_4_080_58    m/z 343.9943     0   2   4E+4   9   9.5  RT (min)      lcqtof_hilic_neg_4_059_14    m/z 343.9943     0   2   4   6   8E+5   7   7.5  RT (min)       lcqtof_hilic_neg_4_032_01    m/z 343.9956     0   1E+5   8   8.5  RT (min)      lcqtof_hilic_neg_4_081_58    m/z 344.0876     0   2   4E+4   6.5   7  RT (min)      lcqtof_hilic_neg_4_032_01    m/z 344.0880     0   1   2   3   4E+4   6.5   7  RT (min)      lcqtof_hilic_neg_4_023_15    m/z 344.0881     0   2   4E+4   6.5   7  RT (min)      lcqtof_hilic_neg_4_023_15    m/z 344.1038     0   1E+4   6.5   7  RT (min)      lcqtof_hilic_neg_1_003_37    m/z 344.1825     0   1E+5   6.5   7  RT (min)      lcqtof_hilic_neg_4_059_14    m/z 344.1826     0   1   2E+5   6.5   7  RT (min)      lcqtof_hilic_neg_4_035_33    m/z 344.2206     0   1   2   3E+4   6.5   7  RT (min)      lcqtof_hilic_neg_4_053_14    m/z 344.2212     0   1   2   3E+4   6.5   7  RT (min)      lcqtof_hilic_neg_4_053_14    m/z 344.2439     0   1   2E+4   1.5   2  RT (min)      lcqtof_hilic_neg_4_032_01    m/z 344.2442     0   1   2E+4   1.5   2  RT (min)      lcqtof_hilic_neg_4_032_01    m/z 344.2688     0   1   2   3   4E+4   6.5   7  RT (min)      lcqtof_hilic_neg_4_053_14    m/z 344.2690     0   1   2   3   4E+4   6.5   7  RT (min)      lcqtof_hilic_neg_4_053_14    m/z 344.3267     0   2   4E+4   6.5   7  RT (min)      lcqtof_hilic_neg_4_032_01    m/z 344.3267     0   1   2   3   4E+4   6.5   7  RT (min)      lcqtof_hilic_neg_4_023_15    m/z 344.3270     0   2   4   6E+4   6.5   7  RT (min)      lcqtof_hilic_neg_4_053_14    m/z 344.4324     0   1   2   3   4E+4   6.5   7  RT (min)      lcqtof_hilic_neg_4_053_14    m/z 344.4325     0   1   2   3   4E+4   6.5   7  RT (min)      lcqtof_hilic_neg_4_053_14    m/z 344.8255     0   1   2   3E+3   8.5   9  RT (min)       lcqtof_hilic_neg_4_019_03    m/z 344.8261     0   1   2E+3   9   9.5  RT (min)      lcqtof_hilic_neg_4_100_61    m/z 344.8444     0   1   2E+3   7.5   8  RT (min)      lcqtof_hilic_neg_2_046_41    m/z 344.9332     0   1E+4   1   1.5  RT (min)      lcqtof_hilic_neg_4_027_19    m/z 344.9679     0   2   4   6E+4   6   6.5  RT (min)       lcqtof_hilic_neg_2_020_23    m/z 344.9968     0   1   2   3E+4   8   8.5  RT (min)      lcqtof_hilic_neg_4_059_14    m/z 344.9969     0   1   2   3E+4   6.5   7  RT (min)      lcqtof_hilic_neg_3_040_14    m/z 344.9971     0   1   2   3   4E+4   7.5   8  RT (min)      lcqtof_hilic_neg_4_023_15    m/z 344.9972     0   1   2E+5   7   7.5  RT (min)      lcqtof_hilic_neg_4_032_01    m/z 344.9972     0   2   4   6E+4   7.5   8  RT (min)      lcqtof_hilic_neg_4_023_15    m/z 344.9972     0   2   4E+4   7.5   8  RT (min)      lcqtof_hilic_neg_4_064_15    m/z 344.9973     0   1   2   3E+4   8   8.5  RT (min)      lcqtof_hilic_neg_4_059_14    m/z 344.9973     0   1E+5   7   7.5  RT (min)      lcqtof_hilic_neg_4_032_01    m/z 344.9974     0   1E+5   7   7.5  RT (min)      lcqtof_hilic_neg_1_004_14    m/z 344.9974     0   1E+5   7   7.5  RT (min)      lcqtof_hilic_neg_1_004_14    m/z 344.9981     0   1E+4   8   8.5  RT (min)      lcqtof_hilic_neg_4_079_58    m/z 345.0019     0   1   2   3E+4   8.5   9  RT (min)       lcqtof_hilic_neg_4_059_14    m/z 345.0535     0   1E+4   6   6.5  RT (min)      lcqtof_hilic_neg_4_048_23    m/z 345.0567     0   1E+4   6.5   7  RT (min)      lcqtof_hilic_neg_4_048_23    m/z 345.0578     0   1E+4   6.5   7  RT (min)      lcqtof_hilic_neg_4_048_23    m/z 345.0578     0   1E+4   6.5   7  RT (min)      lcqtof_hilic_neg_4_048_23    m/z 345.0618     0   1E+4   1.5   2  RT (min)      lcqtof_hilic_neg_4_010_28    m/z 345.0933     0   1E+4   2   2.5  RT (min)      lcqtof_hilic_neg_4_055_32    m/z 345.1369     0   1E+4   1   1.5  RT (min)       lcqtof_hilic_neg_4_039_18    m/z 345.1549     0   1E+4   5.5   6  RT (min)      lcqtof_hilic_neg_4_055_32    m/z 345.1871     0   1   2   3E+4   6.5   7  RT (min)      lcqtof_hilic_neg_4_053_14    m/z 345.1876     0   1   2   3E+4   6.5   7  RT (min)      lcqtof_hilic_neg_4_053_14    m/z 345.1878     0   1   2   3E+4   6.5   7  RT (min)      lcqtof_hilic_neg_4_053_14    m/z 345.3901     0   1E+4   6   6.5  RT (min)      lcqtof_hilic_neg_1_003_37    m/z 345.9276     0   1   2   3   4E+4   1   1.5  RT (min)      lcqtof_hilic_neg_4_027_19    m/z 345.9276     0   1   2   3   4E+4   1   1.5  RT (min)      lcqtof_hilic_neg_4_027_19    m/z 345.9985     0   2   4   6   8E+4   6.5   7  RT (min)      lcqtof_hilic_neg_4_023_15    m/z 345.9986     0   1   2E+4   7   7.5  RT (min)      lcqtof_hilic_neg_4_032_01    m/z 345.9986     0   2   4   6E+4   6.5   7  RT (min)      lcqtof_hilic_neg_4_032_01    m/z 345.9989     0   1   2   3   4E+4   7   7.5  RT (min)      lcqtof_hilic_neg_4_032_01    m/z 345.9992     0   1E+4   7   7.5  RT (min)      lcqtof_hilic_neg_1_004_14    m/z 346.0691     0   1   2   3E+5   3   3.5  RT (min)      lcqtof_hilic_neg_4_039_18    m/z 346.1116     0   1E+4   6   6.5  RT (min)      lcqtof_hilic_neg_4_008_24    m/z 346.1116     0   1E+4   6   6.5  RT (min)       lcqtof_hilic_neg_4_008_24    m/z 346.9439     0   1   2E+4   4.5   5  RT (min)      lcqtof_hilic_neg_4_013_20    m/z 347.0014     0   2   4   6E+4   7   7.5  RT (min)      lcqtof_hilic_neg_4_078_58    m/z 347.0020     0   1   2   3   4E+4   7.5   8  RT (min)      lcqtof_hilic_neg_4_055_32    m/z 347.0022     0   1   2   3   4E+4   7.5   8  RT (min)      lcqtof_hilic_neg_4_055_32    m/z 347.0028     0   1   2   3E+4   7.5   8  RT (min)       lcqtof_hilic_neg_4_053_14    m/z 347.0180     0   1E+4   6   6.5  RT (min)       lcqtof_hilic_neg_4_048_23    m/z 347.0668     0   1   2   3   4E+4   4   4.5  RT (min)      lcqtof_hilic_neg_4_026_04    m/z 347.0670     0   1   2   3   4E+4   3.5   4  RT (min)      lcqtof_hilic_neg_4_026_04    m/z 347.0671     0   1   2E+4   3.5   4  RT (min)      lcqtof_hilic_neg_4_026_04    m/z 347.0671     0   1   2   3   4E+4   4   4.5  RT (min)      lcqtof_hilic_neg_4_026_04    m/z 347.0721     0   2   4E+4   3   3.5  RT (min)      lcqtof_hilic_neg_4_039_18    m/z 347.1330     0   1   2   3E+3   1   1.5  RT (min)       lcqtof_hilic_neg_4_109_62    m/z 347.2429     0   1   2E+4   1   1.5  RT (min)      lcqtof_hilic_neg_4_053_14    m/z 347.2430     0   1   2E+4   1   1.5  RT (min)      lcqtof_hilic_neg_4_053_14    m/z 347.6076     0   1E+4   6   6.5  RT (min)      lcqtof_hilic_neg_1_003_37    m/z 348.0666     0   1E+5   3   3.5  RT (min)      lcqtof_hilic_neg_4_039_18    m/z 348.2202     0   1   2   3E+4   1.5   2  RT (min)      lcqtof_hilic_neg_4_038_10    m/z 348.2208     0   1   2   3E+4   0.5   1   1.5  RT (min)      lcqtof_hilic_neg_4_038_10    m/z 348.7134     0   1E+4   6   6.5  RT (min)      lcqtof_hilic_neg_1_003_37    m/z 348.9646     0   1E+4   0.5   1  RT (min)      lcqtof_hilic_neg_3_003_20    m/z 349.0611     0   1   2   3E+3   3   3.5  RT (min)      lcqtof_hilic_neg_4_039_18    m/z 349.0720     0   1   2E+4   3.5   4  RT (min)      lcqtof_hilic_neg_1_035_23    m/z 349.0726     0   1E+4   3.5   4  RT (min)      lcqtof_hilic_neg_1_032_18    m/z 349.0753     0   1E+4   1.5   2  RT (min)       lcqtof_hilic_neg_4_052_13    m/z 349.0771     0   1   2E+4   3   3.5  RT (min)       lcqtof_hilic_neg_4_039_18    m/z 349.0919     0   2   4   6   8E+3   4   4.5  RT (min)       lcqtof_hilic_neg_4_050_02    m/z 349.2050     0   1   2   3   4E+3   2   2.5  RT (min)      lcqtof_hilic_neg_4_033_53    m/z 349.2052     0   1   2E+3   0   0.5  RT (min)       lcqtof_hilic_neg_4_013_20    m/z 349.2052     0   1E+4   15   15.5  RT (min)      lcqtof_hilic_neg_4_000_aa    m/z 349.2055     0   2   4   6   8E+3   2   2.5  RT (min)       lcqtof_hilic_neg_4_032_01    m/z 349.9540     0   2   4   6   8E+4   1   1.5  RT (min)      lcqtof_hilic_neg_4_004_47    m/z 350.0505     0   1E+4   4   4.5  RT (min)      lcqtof_hilic_neg_4_047_06    m/z 350.0756     0   1E+4   6.5   7  RT (min)      lcqtof_hilic_neg_4_002_25    m/z 350.0911     0   1E+4   6.5   7  RT (min)      lcqtof_hilic_neg_4_055_32    m/z 350.0916     0   1E+4   2   2.5  RT (min)      lcqtof_hilic_neg_4_050_02    m/z 350.1247     0   2   4E+3   16.5   17  RT (min)       lcqtof_hilic_neg_4_012_07    m/z 350.1248     0   1   2   3E+3   16   16.5  RT (min)       lcqtof_hilic_neg_4_043_12    m/z 350.1249     0   1   2E+4   0   0.5  RT (min)       lcqtof_hilic_neg_4_012_07    m/z 350.1250     0   1   2E+4   0   0.5  RT (min)      lcqtof_hilic_neg_4_012_07    m/z 350.1250     0   1   2   3   4E+3   0   0.5  RT (min)       lcqtof_hilic_neg_4_047_06    m/z 350.1254     0   1   2   3   4E+3   0   0.5  RT (min)       lcqtof_hilic_neg_4_047_06    m/z 350.8953     0   1E+4   1   1.5  RT (min)       lcqtof_hilic_neg_4_042_22    m/z 350.9188     0   1   2   3   4E+5   8.5   9  RT (min)      lcqtof_hilic_neg_1_000_aa    m/z 350.9190     0   1   2   3E+5   8.5   9  RT (min)      lcqtof_hilic_neg_1_000_aa    m/z 350.9190     0   1E+5   9   9.5  RT (min)      lcqtof_hilic_neg_1_000_aa    m/z 350.9190     0   1E+5   9   9.5  RT (min)      lcqtof_hilic_neg_1_000_aa    m/z 350.9190     0   1   2   3   4E+5   7.5   8   8.5  RT (min)      lcqtof_hilic_neg_1_000_aa    m/z 350.9190     0   1   2   3   4E+5   8   8.5  RT (min)      lcqtof_hilic_neg_1_000_aa    m/z 350.9191     0   1   2   3   4E+5   7.5   8  RT (min)      lcqtof_hilic_neg_1_000_aa    m/z 350.9192     0   1   2   3   4E+5   7.5   8  RT (min)      lcqtof_hilic_neg_1_000_aa    m/z 350.9193     0   1E+5   12.5   13  RT (min)      lcqtof_hilic_neg_1_000_aa    m/z 350.9194     0   1E+5   9   9.5  RT (min)      lcqtof_hilic_neg_1_000_aa    m/z 350.9196     0   1E+5   12.5   13  RT (min)      lcqtof_hilic_neg_1_000_aa    m/z 350.9198     0   2   4   6   8E+4   9   9.5  RT (min)      lcqtof_hilic_neg_1_000_aa    m/z 350.9522     0   2   4   6E+4   1   1.5  RT (min)      lcqtof_hilic_neg_4_004_47    m/z 351.0557     0   2   4   6E+3   2   2.5  RT (min)      lcqtof_hilic_neg_4_053_14    m/z 351.0941     0   1   2E+4   2   2.5  RT (min)      lcqtof_hilic_neg_4_050_02    m/z 351.1634     0   1E+4   3   3.5  RT (min)      lcqtof_hilic_neg_4_045_26    m/z 351.2021     0   1E+4   0.5   1  RT (min)      lcqtof_hilic_neg_4_032_01    m/z 351.2170     0   1   2E+4   1   1.5  RT (min)      lcqtof_hilic_neg_4_000_aa    m/z 351.2179     0   1E+4   1   1.5  RT (min)      lcqtof_hilic_neg_4_053_14    m/z 351.2193     0   1   2E+4   1   1.5  RT (min)      lcqtof_hilic_neg_4_000_aa    m/z 351.2646     0   1E+4   1   1.5  RT (min)       lcqtof_hilic_neg_4_063_16    m/z 351.3037     0   1E+5   0.5   1  RT (min)      lcqtof_hilic_neg_3_000_aa    m/z 352.0346     0   1E+4   6   6.5  RT (min)      lcqtof_hilic_neg_4_001_49    m/z 352.0752     0   1E+5   1   1.5  RT (min)      lcqtof_hilic_neg_3_027_47    m/z 352.0753     0   1E+5   1   1.5  RT (min)      lcqtof_hilic_neg_3_027_47    m/z 352.0894     0   1E+4   4   4.5  RT (min)      lcqtof_hilic_neg_4_008_24    m/z 352.3084     0   1   2   3E+4   0.5   1  RT (min)      lcqtof_hilic_neg_3_035_46    m/z 352.8918     0   1E+4   1   1.5  RT (min)       lcqtof_hilic_neg_4_107_62    m/z 352.8990     0   1E+5   8   8.5  RT (min)      lcqtof_hilic_neg_1_000_aa    m/z 352.9201     0   1E+4   7.5   8  RT (min)      lcqtof_hilic_neg_1_000_aa    m/z 353.0099     0   1   2   3   4E+3   6   6.5  RT (min)       lcqtof_hilic_neg_4_059_14    m/z 353.0270     0   1E+4   1.5   2  RT (min)      lcqtof_hilic_neg_1_031_16    m/z 353.0505     0   1E+4   0.5   1  RT (min)      lcqtof_hilic_neg_4_058_39    m/z 353.0805     0   1   2E+4   1   1.5  RT (min)      lcqtof_hilic_neg_3_027_47    m/z 353.0869     0   2   4   6E+4   3.5   4  RT (min)      lcqtof_hilic_neg_2_053_18    m/z 353.0871     0   1E+5   3.5   4  RT (min)      lcqtof_hilic_neg_1_032_18    m/z 353.0881     0   1   2E+4   4   4.5  RT (min)      lcqtof_hilic_neg_4_039_18    m/z 353.0881     0   1E+5   4   4.5  RT (min)      lcqtof_hilic_neg_1_032_18    m/z 353.0882     0   1   2E+4   4   4.5  RT (min)      lcqtof_hilic_neg_4_039_18    m/z 353.1031     0   1   2   3   4E+3   5.5   6  RT (min)      lcqtof_hilic_neg_1_001_49    m/z 353.1245     0   1E+4   3   3.5  RT (min)      lcqtof_hilic_neg_4_004_47    m/z 353.1254     0   1E+4   3   3.5  RT (min)      lcqtof_hilic_neg_4_014_46    m/z 353.1322     0   1E+4   6.5   7  RT (min)      lcqtof_hilic_neg_4_008_24    m/z 353.1954     0   1   2   3E+4   6   6.5  RT (min)      lcqtof_hilic_neg_4_008_24    m/z 353.1958     0   2   4   6E+4   6   6.5  RT (min)       lcqtof_hilic_neg_1_037_24    m/z 353.2067     0   1   2   3E+4   15   15.5  RT (min)      lcqtof_hilic_neg_4_000_aa    m/z 353.2078     0   2   4   6   8E+4   1   1.5  RT (min)      lcqtof_hilic_neg_4_000_aa    m/z 353.2088     0   1E+4   2   2.5  RT (min)       lcqtof_hilic_neg_4_032_01    m/z 353.2091     0   1   2   3   4E+3   2   2.5  RT (min)       lcqtof_hilic_neg_4_042_22    m/z 353.2091     0   1E+4   1.5   2  RT (min)      lcqtof_hilic_neg_4_060_13    m/z 353.2096     0   1   2   3   4E+3   2.5   3  RT (min)      lcqtof_hilic_neg_4_055_32    m/z 353.2097     0   2   4E+3   3   3.5  RT (min)      lcqtof_hilic_neg_4_004_47    m/z 353.2097     0   2   4E+4   1   1.5  RT (min)      lcqtof_hilic_neg_4_026_04    m/z 353.2099     0   2   4   6   8E+3   2.5   3  RT (min)      lcqtof_hilic_neg_4_089_59    m/z 353.2134     0   2   4E+4   1   1.5  RT (min)       lcqtof_hilic_neg_4_026_04    m/z 353.2216     0   2   4   6E+4   1   1.5  RT (min)      lcqtof_hilic_neg_4_059_14    m/z 353.2245     0   1E+4   1.5   2  RT (min)      lcqtof_hilic_neg_4_059_14    m/z 353.3188     0   1   2E+4   0.5   1  RT (min)      lcqtof_hilic_neg_3_000_aa    m/z 353.4779     0   1E+4   0.5   1  RT (min)      lcqtof_hilic_neg_1_014_31    m/z 353.6439     0   1   2E+4   0.5   1  RT (min)      lcqtof_hilic_neg_1_003_37    m/z 354.1755     0   1E+4   7.5   8  RT (min)      lcqtof_hilic_neg_1_037_24    m/z 354.3168     0   1E+4   10   10.5  RT (min)      lcqtof_hilic_neg_1_000_aa    m/z 354.8782     0   2   4   6   8E+4   8   8.5  RT (min)      lcqtof_hilic_neg_1_000_aa    m/z 355.0887     0   1E+4   0.5   1  RT (min)      lcqtof_hilic_neg_4_010_28    m/z 355.0908     0   1   2E+4   6   6.5  RT (min)       lcqtof_hilic_neg_4_090_60    m/z 355.1205     0   2   4E+3   0.5   1  RT (min)      lcqtof_hilic_neg_4_004_47    m/z 355.1207     0   1   2   3E+3   18   18.5  RT (min)      lcqtof_hilic_neg_4_050_02    m/z 355.1216     0   1   2E+4   5.5   6  RT (min)      lcqtof_hilic_neg_2_032_02    m/z 355.1224     0   2   4   6E+3   2   2.5  RT (min)      lcqtof_hilic_neg_4_055_32    m/z 355.1232     0   2   4   6   8E+3   0.5   1  RT (min)      lcqtof_hilic_neg_4_062_17    m/z 355.1233     0   1E+4   4.5   5  RT (min)      lcqtof_hilic_neg_4_060_13    m/z 355.1236     0   1   2   3E+4   5.5   6  RT (min)      lcqtof_hilic_neg_4_050_02    m/z 355.1590     0   1E+4   6   6.5  RT (min)      lcqtof_hilic_neg_4_008_24    m/z 355.1705     0   2   4   6   8E+4   0.5   1  RT (min)      lcqtof_hilic_neg_3_035_46    m/z 355.2160     0   1   2E+3   1   1.5  RT (min)      lcqtof_hilic_neg_4_000_aa    m/z 355.2492     0   1   2E+3   15   15.5  RT (min)      lcqtof_hilic_neg_4_012_07    m/z 355.2858     0   1   2E+4   1   1.5  RT (min)      lcqtof_hilic_neg_4_000_aa    m/z 355.2860     0   1   2E+4   1   1.5  RT (min)      lcqtof_hilic_neg_4_000_aa    m/z 356.0416     0   1   2   3E+4   4   4.5  RT (min)       lcqtof_hilic_neg_4_016_37    m/z 356.1287     0   1E+4   3   3.5  RT (min)      lcqtof_hilic_neg_4_014_46    m/z 356.1288     0   1E+4   3   3.5  RT (min)      lcqtof_hilic_neg_4_004_47    m/z 356.1445     0   1E+4   6   6.5  RT (min)      lcqtof_hilic_neg_4_048_23    m/z 356.1714     0   1   2E+4   0.5   1  RT (min)      lcqtof_hilic_neg_3_035_46    m/z 356.8312     0   1E+4   8   8.5  RT (min)       lcqtof_hilic_neg_4_100_61    m/z 357.0512     0   1E+4   6   6.5  RT (min)       lcqtof_hilic_neg_4_008_24    m/z 357.1078     0   1   2   3   4E+4   4.5   5  RT (min)      lcqtof_hilic_neg_4_055_32    m/z 357.1394     0   1   2E+4   4   4.5  RT (min)      lcqtof_hilic_neg_4_061_50    m/z 357.1399     0   1   2E+4   4   4.5  RT (min)      lcqtof_hilic_neg_4_061_50    m/z 357.1403     0   1E+4   4   4.5  RT (min)      lcqtof_hilic_neg_4_009_50    m/z 357.1405     0   1   2E+4   4   4.5  RT (min)      lcqtof_hilic_neg_4_061_50    m/z 357.2057     0   1   2E+4   19.5   20  RT (min)       lcqtof_hilic_neg_4_070_57    m/z 357.2058     0   1   2E+4   19   19.5  RT (min)       lcqtof_hilic_neg_4_076_58    m/z 357.2060     0   1E+4   19.5   20  RT (min)       lcqtof_hilic_neg_4_090_60    m/z 357.2060     0   1   2E+4   16   18  RT (min)       lcqtof_hilic_neg_4_068_57    m/z 357.2060     0   1E+4   6   6.5  RT (min)      lcqtof_hilic_neg_4_016_37    m/z 357.2061     0   1   2   3E+4   17   17.5   18  RT (min)       lcqtof_hilic_neg_4_000_cc    m/z 357.2061     0   1E+4   14.5   15  RT (min)       lcqtof_hilic_neg_4_078_58    m/z 357.2061     0   1   2E+4   19   19.5  RT (min)       lcqtof_hilic_neg_4_070_57    m/z 357.2061     0   1E+4   19   19.5  RT (min)       lcqtof_hilic_neg_4_090_60    m/z 357.2062     0   1   2E+4   17.5   18  RT (min)       lcqtof_hilic_neg_4_071_57    m/z 357.2063     0   1   2   3E+4   16   18  RT (min)       lcqtof_hilic_neg_4_072_57    m/z 357.2063     0   1   2E+4   17   18  RT (min)       lcqtof_hilic_neg_4_073_57    m/z 357.2063     0   1E+4   5.5   6  RT (min)      lcqtof_hilic_neg_4_059_14    m/z 357.2063     0   1   2   3E+4   16   16.5  RT (min)       lcqtof_hilic_neg_4_021_21    m/z 357.2063     0   1   2   3E+4   17   17.5  RT (min)       lcqtof_hilic_neg_4_002_25    m/z 357.2064     0   1E+4   5   5.5  RT (min)      lcqtof_hilic_neg_4_006_30    m/z 357.2064     0   1E+4   15   15.5  RT (min)       lcqtof_hilic_neg_4_070_57    m/z 357.2064     0   1   2E+5   16   17  RT (min)       lcqtof_hilic_neg_4_000_aa    m/z 357.2065     0   1E+4   5   5.5  RT (min)      lcqtof_hilic_neg_4_001_49    m/z 357.2066     0   1   2   3E+4   17   18  RT (min)       lcqtof_hilic_neg_4_000_bb    m/z 357.2066     0   1E+4   5   5.5  RT (min)      lcqtof_hilic_neg_4_011_35    m/z 357.2066     0   1   2   3E+4   17   18   19  RT (min)       lcqtof_hilic_neg_4_000_cc    m/z 357.2066     0   2   4   6E+4   1   2  RT (min)       lcqtof_hilic_neg_4_028_36    m/z 357.2067     0   1E+4   5   5.5  RT (min)      lcqtof_hilic_neg_4_010_28    m/z 357.2071     0   1   2E+4   19  RT (min)       lcqtof_hilic_neg_4_068_57    m/z 357.2071     0   1   2E+4   15.5   16  RT (min)      lcqtof_hilic_neg_4_078_58    m/z 357.2081     0   1E+4   16.5   17  RT (min)       lcqtof_hilic_neg_4_090_60    m/z 357.2082     0   1   2   3E+5   1   1.5  RT (min)      lcqtof_hilic_neg_4_000_aa    m/z 358.0100     0   2   4   6E+4   11   11.5  RT (min)       lcqtof_hilic_neg_4_050_02    m/z 358.0461     0   1   2E+4   2   2.5  RT (min)      lcqtof_hilic_neg_4_039_18    m/z 358.0462     0   1   2E+4   2   2.5  RT (min)      lcqtof_hilic_neg_4_039_18    m/z 358.0684     0   1   2E+4   0.5   1  RT (min)      lcqtof_hilic_neg_4_000_aa    m/z 358.1371     0   1E+4   3   3.5  RT (min)      lcqtof_hilic_neg_4_025_52    m/z 358.1439     0   1   2E+4   3   3.5  RT (min)      lcqtof_hilic_neg_4_004_47    m/z 358.2093     0   2   4   6E+4   15.5   16  RT (min)      lcqtof_hilic_neg_4_000_aa    m/z 358.2111     0   2   4   6   8E+4   1   1.5  RT (min)      lcqtof_hilic_neg_4_000_aa    m/z 359.0751     0   1E+4   4.5   5  RT (min)      lcqtof_hilic_neg_4_019_03    m/z 359.1050     0   1E+4   4.5   5  RT (min)      lcqtof_hilic_neg_4_055_32    m/z 359.1441     0   2   4   6E+3   2.5   3  RT (min)      lcqtof_hilic_neg_4_057_49    m/z 359.1441     0   2   4   6E+3   2.5   3  RT (min)      lcqtof_hilic_neg_1_001_49    m/z 359.1444     0   2   4E+3   2   2.5  RT (min)      lcqtof_hilic_neg_2_057_49    m/z 359.1449     0   2   4   6   8E+3   0.5   1  RT (min)      lcqtof_hilic_neg_4_032_01    m/z 359.2159     0   1E+4   15.5   16  RT (min)      lcqtof_hilic_neg_4_000_aa    m/z 359.2177     0   1   2E+4   1   1.5  RT (min)      lcqtof_hilic_neg_4_053_14    m/z 359.9790     0   2   4   6   8E+3   2   2.5  RT (min)      lcqtof_hilic_neg_1_016_28    m/z 359.9892     0   1   2E+4   6   6.5  RT (min)       lcqtof_hilic_neg_4_050_02    m/z 359.9894     0   1   2E+4   6   6.5  RT (min)      lcqtof_hilic_neg_4_050_02    m/z 360.0455     0   1E+4   2   2.5  RT (min)      lcqtof_hilic_neg_4_039_18    m/z 360.0532     0   1   2E+4   0.5   1  RT (min)      lcqtof_hilic_neg_4_000_aa    m/z 360.1594     0   1E+4   5   5.5  RT (min)      lcqtof_hilic_neg_4_055_32    m/z 360.2380     0   1   2E+5   2   2.5  RT (min)      lcqtof_hilic_neg_4_032_01    m/z 360.2386     0   1   2E+5   1.5   2  RT (min)      lcqtof_hilic_neg_4_032_01    m/z 360.7847     0   1E+4   4.5   5  RT (min)      lcqtof_hilic_neg_4_050_02    m/z 360.8265     0   1   2E+3   8.5   9  RT (min)       lcqtof_hilic_neg_3_099_61    m/z 360.9901     0   1E+4   4.5   5  RT (min)      lcqtof_hilic_neg_2_004_26    m/z 361.0493     0   1   2   3   4E+4   0.5   1  RT (min)      lcqtof_hilic_neg_4_000_aa    m/z 361.0583     0   1   2E+4   4   4.5  RT (min)      lcqtof_hilic_neg_4_002_25    m/z 361.0621     0   1E+3   0.5   1  RT (min)      lcqtof_hilic_neg_4_000_aa    m/z 361.0898     0   1E+4   6   6.5  RT (min)      lcqtof_hilic_neg_4_050_02    m/z 361.1372     0   1E+4   6   6.5  RT (min)       lcqtof_hilic_neg_4_060_13    m/z 361.1543     0   1E+4   3   3.5  RT (min)      lcqtof_hilic_neg_4_014_46    m/z 361.2228     0   1E+4   2.5   3  RT (min)      lcqtof_hilic_neg_4_053_14    m/z 361.2229     0   1   2E+4   1.5   2  RT (min)      lcqtof_hilic_neg_4_053_14    m/z 361.2231     0   1   2E+4   1.5   2  RT (min)      lcqtof_hilic_neg_4_053_14    m/z 361.2234     0   1E+4   1.5   2  RT (min)      lcqtof_hilic_neg_4_032_01    m/z 361.2369     0   1E+4   1   1.5  RT (min)      lcqtof_hilic_neg_4_053_14    m/z 361.2417     0   1   2   3   4E+4   2   2.5  RT (min)      lcqtof_hilic_neg_4_032_01    m/z 361.9941     0   1E+4   4   4.5  RT (min)      lcqtof_hilic_neg_1_014_31    m/z 362.0583     0   1E+4   0.5   1  RT (min)      lcqtof_hilic_neg_4_000_aa    m/z 362.1240     0   1   2E+4   5.5   6  RT (min)      lcqtof_hilic_neg_4_008_24    m/z 362.1241     0   1E+4   2   2.5  RT (min)      lcqtof_hilic_neg_4_050_02    m/z 362.7826     0   1E+4   4.5   5  RT (min)      lcqtof_hilic_neg_4_050_02    m/z 362.9402     0   1   2   3E+3   8   8.5  RT (min)       lcqtof_hilic_neg_4_038_10    m/z 362.9794     0   2   4   6   8E+4   0.5   1  RT (min)      lcqtof_hilic_neg_3_035_46    m/z 362.9806     0   2   4   6   8E+4   0.5   1  RT (min)      lcqtof_hilic_neg_3_035_46    m/z 363.0094     0   1E+4   7.5   8  RT (min)      lcqtof_hilic_neg_4_008_24    m/z 363.0684     0   1E+4   7   7.5  RT (min)      lcqtof_hilic_neg_4_048_23    m/z 363.0694     0   1E+4   7.5   8  RT (min)      lcqtof_hilic_neg_4_048_23    m/z 363.0889     0   1E+4   5.5   6  RT (min)      lcqtof_hilic_neg_4_055_32    m/z 363.0945     0   1E+4   4   4.5  RT (min)      lcqtof_hilic_neg_4_050_02    m/z 363.2192     0   1E+4   15   15.5  RT (min)      lcqtof_hilic_neg_4_000_aa    m/z 363.2206     0   1   2   3   4E+3   2   2.5  RT (min)      lcqtof_hilic_neg_4_023_15    m/z 363.9817     0   1E+4   0.5   1  RT (min)      lcqtof_hilic_neg_3_003_20    m/z 364.0800     0   1E+4   4   4.5   5  RT (min)      lcqtof_hilic_neg_4_039_18    m/z 364.9800     0   1E+4   0.5   1  RT (min)      lcqtof_hilic_neg_3_035_46    m/z 365.0897     0   1E+4   2   2.5  RT (min)      lcqtof_hilic_neg_4_050_02    m/z 365.1398     0   1   2E+4   15   15.5  RT (min)       lcqtof_hilic_neg_4_000_aa    m/z 365.2838     0   1E+4   0.5   1  RT (min)      lcqtof_hilic_neg_3_000_bb    m/z 366.0315     0   1E+4   4.5   5  RT (min)      lcqtof_hilic_neg_4_064_15    m/z 366.1105     0   1   2   3E+4   1   1.5  RT (min)       lcqtof_hilic_neg_4_039_18    m/z 366.1338     0   1E+4   1   1.5  RT (min)      lcqtof_hilic_neg_4_005_29    m/z 366.9643     0   2   4E+4   1   1.5  RT (min)      lcqtof_hilic_neg_4_004_47    m/z 366.9644     0   2   4E+4   1   1.5  RT (min)      lcqtof_hilic_neg_4_004_47    m/z 366.9644     0   1   2E+4   1.5   2  RT (min)      lcqtof_hilic_neg_4_004_47    m/z 367.1043     0   1E+4   3   3.5  RT (min)      lcqtof_hilic_neg_4_039_18    m/z 367.1055     0   1E+4   4   4.5  RT (min)      lcqtof_hilic_neg_4_050_02    m/z 367.1576     0   1E+4   7.5   8  RT (min)       lcqtof_hilic_neg_4_008_24    m/z 367.1834     0   2   4E+5   5.5   6  RT (min)      lcqtof_hilic_neg_4_097_60    m/z 367.2244     0   1E+4   1   1.5  RT (min)      lcqtof_hilic_neg_4_002_25    m/z 367.2283     0   1E+4   1   1.5  RT (min)      lcqtof_hilic_neg_4_026_04    m/z 367.2291     0   1   2E+4   0.5   1  RT (min)      lcqtof_hilic_neg_4_047_06    m/z 367.2304     0   1   2E+4   0.5   1  RT (min)       lcqtof_hilic_neg_4_038_10    m/z 367.2305     0   1E+4   15   15.5  RT (min)      lcqtof_hilic_neg_4_000_aa    m/z 367.2550     0   1   2   3E+4   1   1.5  RT (min)      lcqtof_hilic_neg_4_053_14    m/z 367.2853     0   1   2   3   4E+4   1   1.5  RT (min)      lcqtof_hilic_neg_4_038_10    m/z 367.2856     0   1   2E+4   1   1.5  RT (min)      lcqtof_hilic_neg_4_000_aa    m/z 367.2991     0   1E+4   0.5   1  RT (min)      lcqtof_hilic_neg_3_000_bb    m/z 368.0278     0   1E+4   4.5   5  RT (min)      lcqtof_hilic_neg_4_064_15    m/z 368.0858     0   1E+4   7   7.5  RT (min)      lcqtof_hilic_neg_3_048_05    m/z 368.0862     0   1E+4   7.5   8  RT (min)      lcqtof_hilic_neg_1_007_05    m/z 368.0862     0   1   2E+4   7   7.5  RT (min)      lcqtof_hilic_neg_4_040_05    m/z 368.0944     0   1E+4   1   1.5  RT (min)      lcqtof_hilic_neg_4_004_47    m/z 368.1491     0   1E+4   4   4.5  RT (min)      lcqtof_hilic_neg_4_040_05    m/z 368.1824     0   1   2E+3   1   1.5  RT (min)      lcqtof_hilic_neg_4_003_16    m/z 368.2258     0   1E+4   2   2.5  RT (min)      lcqtof_hilic_neg_4_032_01    m/z 368.9958     0   1E+4   7.5   8  RT (min)      lcqtof_hilic_neg_4_059_14    m/z 368.9959     0   1   2E+4   7   7.5  RT (min)      lcqtof_hilic_neg_4_080_58    m/z 369.0038     0   1   2   3   4E+4   7   7.5  RT (min)       lcqtof_hilic_neg_4_064_15    m/z 369.0042     0   1   2   3   4E+4   7   7.5  RT (min)      lcqtof_hilic_neg_4_059_14    m/z 369.0044     0   1   2   3   4E+4   6.5   7  RT (min)       lcqtof_hilic_neg_4_064_15    m/z 369.0045     0   2   4E+4   5.5   6  RT (min)       lcqtof_hilic_neg_4_050_02    m/z 369.0046     0   1   2   3   4E+4   6.5   7  RT (min)      lcqtof_hilic_neg_4_059_14    m/z 369.0048     0   1   2   3   4E+4   6.5   7  RT (min)      lcqtof_hilic_neg_4_064_15    m/z 369.0048     0   2   4E+4   5.5   6  RT (min)      lcqtof_hilic_neg_4_050_02    m/z 369.0050     0   1   2E+4   6   6.5  RT (min)      lcqtof_hilic_neg_4_064_15    m/z 369.0050     0   2   4E+4   5.5   6  RT (min)      lcqtof_hilic_neg_4_050_02    m/z 369.0051     0   1   2   3   4E+4   6.5   7  RT (min)       lcqtof_hilic_neg_4_059_14    m/z 369.0057     0   1   2   3   4E+4   6.5   7  RT (min)      lcqtof_hilic_neg_4_059_14    m/z 369.0065     0   1   2   3E+4   3.5   4  RT (min)      lcqtof_hilic_neg_1_032_18    m/z 369.0065     0   1E+4   1.5   2  RT (min)      lcqtof_hilic_neg_4_059_14    m/z 369.0066     0   2   4   6E+4   3   3.5  RT (min)      lcqtof_hilic_neg_4_039_18    m/z 369.0582     0   1   2   3E+4   6   6.5  RT (min)      lcqtof_hilic_neg_4_048_23    m/z 369.1084     0   1E+4   6   6.5  RT (min)      lcqtof_hilic_neg_4_008_24    m/z 369.1226     0   1   2   3E+3   0.5   1  RT (min)       lcqtof_hilic_neg_4_013_20    m/z 369.1254     0   1   2   3E+3   0   0.5  RT (min)       lcqtof_hilic_neg_4_000_bb    m/z 369.1255     0   1   2   3   4E+3   19.5   20  RT (min)       lcqtof_hilic_neg_4_000_aa    m/z 369.1262     0   2   4   6E+3   1   1.5  RT (min)      lcqtof_hilic_neg_1_002_39    m/z 369.1262     0   1   2   3E+3   2.5   3  RT (min)       lcqtof_hilic_neg_4_015_09    m/z 369.1263     0   1E+4   15   15.5  RT (min)       lcqtof_hilic_neg_4_000_aa    m/z 369.1264     0   1   2   3   4E+3   19   19.5  RT (min)       lcqtof_hilic_neg_4_000_aa    m/z 369.1264     0   1   2E+3   0   0.5  RT (min)       lcqtof_hilic_neg_4_004_47    m/z 369.1267     0   1   2   3E+3   15.5   16  RT (min)       lcqtof_hilic_neg_4_058_39    m/z 369.1277     0   1   2   3   4E+3   19  RT (min)       lcqtof_hilic_neg_4_000_aa    m/z 369.1278     0   1   2   3E+3   18   18.5   19  RT (min)       lcqtof_hilic_neg_4_000_aa    m/z 369.1278     0   1   2   3   4E+3   18.5   19  RT (min)       lcqtof_hilic_neg_4_000_aa    m/z 369.1292     0   1   2   3E+3   17.5   18   18.5  RT (min)       lcqtof_hilic_neg_4_000_aa    m/z 369.1293     0   1   2   3E+3   18   18.5  RT (min)       lcqtof_hilic_neg_4_000_aa    m/z 369.1298     0   1   2   3E+3   17   17.5   18  RT (min)       lcqtof_hilic_neg_4_000_aa    m/z 369.1308     0   1   2   3E+3   17   17.5  RT (min)       lcqtof_hilic_neg_4_000_aa    m/z 369.1901     0   1E+4   6   6.5  RT (min)      lcqtof_hilic_neg_4_008_24    m/z 369.2281     0   1   2E+4   1   1.5  RT (min)      lcqtof_hilic_neg_4_053_14    m/z 369.2282     0   1   2E+4   1   1.5  RT (min)      lcqtof_hilic_neg_4_053_14    m/z 369.3008     0   1   2   3   4E+4   1   1.5  RT (min)      lcqtof_hilic_neg_4_000_aa    m/z 369.3135     0   1E+4   0.5   1  RT (min)      lcqtof_hilic_neg_3_000_aa    m/z 369.3686     0   1   2   3E+4   1   1.5  RT (min)      lcqtof_hilic_neg_4_032_01    m/z 369.9878     0   1   2   3E+3   7.5   8  RT (min)      lcqtof_hilic_neg_4_050_02    m/z 370.0761     0   1E+4   5.5   6  RT (min)       lcqtof_hilic_neg_1_005_32    m/z 370.0906     0   1E+4   4   4.5  RT (min)      lcqtof_hilic_neg_4_061_50    m/z 370.1017     0   1E+5   1   1.5  RT (min)      lcqtof_hilic_neg_1_043_41    m/z 370.1020     0   2   4   6   8E+4   1   1.5  RT (min)      lcqtof_hilic_neg_2_046_41    m/z 370.1073     0   1E+4   4   4.5  RT (min)       lcqtof_hilic_neg_4_093_60    m/z 370.1102     0   1E+4   4   4.5  RT (min)      lcqtof_hilic_neg_4_054_48    m/z 370.1392     0   1E+4   3   3.5  RT (min)       lcqtof_hilic_neg_4_004_47    m/z 371.0292     0   1E+4   6.5   7  RT (min)      lcqtof_hilic_neg_2_028_05    m/z 371.1050     0   2   4   6   8E+3   4   4.5  RT (min)      lcqtof_hilic_neg_4_016_37    m/z 371.1051     0   2   4   6E+3   4   4.5  RT (min)      lcqtof_hilic_neg_1_008_13    m/z 371.1052     0   1   2E+3   3.5   4  RT (min)       lcqtof_hilic_neg_1_037_24    m/z 371.1073     0   1   2E+3   3.5   4  RT (min)      lcqtof_hilic_neg_4_046_41    m/z 371.1179     0   1E+4   5.5   6  RT (min)      lcqtof_hilic_neg_4_050_02    m/z 371.1182     0   1E+4   6   6.5  RT (min)       lcqtof_hilic_neg_4_050_02    m/z 371.1183     0   1E+4   6   6.5  RT (min)      lcqtof_hilic_neg_4_050_02    m/z 371.1240     0   1E+4   0.5   1  RT (min)      lcqtof_hilic_neg_4_013_20    m/z 371.1557     0   1   2   3   4E+4   4   4.5  RT (min)      lcqtof_hilic_neg_4_061_50    m/z 371.1560     0   1   2   3   4E+4   4   4.5  RT (min)      lcqtof_hilic_neg_4_061_50    m/z 371.1567     0   1   2   3   4E+4   4   4.5  RT (min)      lcqtof_hilic_neg_4_061_50    m/z 371.2274     0   1E+4   1   2  RT (min)       lcqtof_hilic_neg_2_004_26    m/z 371.2416     0   1   2E+3   2.5   3  RT (min)       lcqtof_hilic_neg_4_002_25    m/z 371.2433     0   1   2   3E+3   16  RT (min)       lcqtof_hilic_neg_4_041_54    m/z 371.2436     0   2   4   6   8E+3   1   1.5  RT (min)       lcqtof_hilic_neg_4_048_23    m/z 371.2436     0   2   4   6E+3   2   2.5  RT (min)       lcqtof_hilic_neg_4_052_13    m/z 371.2438     0   2   4   6   8E+3   0   0.5  RT (min)       lcqtof_hilic_neg_4_055_32    m/z 371.2441     0   1   2E+3   0   0.5  RT (min)       lcqtof_hilic_neg_4_031_38    m/z 371.2442     0   1   2   3E+3   15   15.5  RT (min)      lcqtof_hilic_neg_4_047_06    m/z 371.3036     0   1E+4   0.5   1  RT (min)      lcqtof_hilic_neg_1_003_37    m/z 371.3156     0   1E+4   1   1.5  RT (min)      lcqtof_hilic_neg_2_002_36    m/z 371.3304     0   1E+4   0.5   1  RT (min)      lcqtof_hilic_neg_3_000_aa    m/z 372.0516     0   1E+4   4   4.5  RT (min)       lcqtof_hilic_neg_3_092_60    m/z 372.1036     0   1E+4   1   1.5  RT (min)      lcqtof_hilic_neg_2_046_41    m/z 372.1039     0   1   2   3E+4   1   1.5  RT (min)      lcqtof_hilic_neg_4_004_47    m/z 372.7760     0   1E+4   9   9.5  RT (min)      lcqtof_hilic_neg_1_000_aa    m/z 372.8119     0   1   2E+4   4.5   5  RT (min)      lcqtof_hilic_neg_4_050_02    m/z 372.8120     0   1E+4   4.5   5  RT (min)      lcqtof_hilic_neg_4_053_14    m/z 373.0082     0   2   4   6   8E+3   7   7.5  RT (min)       lcqtof_hilic_neg_3_078_58    m/z 373.0877     0   1E+5   3   3.5  RT (min)      lcqtof_hilic_neg_4_039_18    m/z 373.0945     0   2   4   6   8E+4   6.5   7  RT (min)      lcqtof_hilic_neg_4_008_24    m/z 373.0947     0   1   2   3E+4   6.5   7  RT (min)       lcqtof_hilic_neg_1_037_24    m/z 373.1020     0   2   4   6   8E+4   7   7.5  RT (min)      lcqtof_hilic_neg_4_008_24    m/z 373.1032     0   2   4   6   8E+4   7   7.5  RT (min)      lcqtof_hilic_neg_4_008_24    m/z 373.1036     0   1E+4   0.5   1  RT (min)      lcqtof_hilic_neg_3_003_20    m/z 373.1832     0   1E+4   0.5   1  RT (min)      lcqtof_hilic_neg_3_003_20    m/z 373.2217     0   1E+4   1   1.5  RT (min)      lcqtof_hilic_neg_4_000_aa    m/z 374.0913     0   1   2E+4   3   3.5  RT (min)      lcqtof_hilic_neg_4_039_18    m/z 374.1396     0   1E+4   5   5.5  RT (min)      lcqtof_hilic_neg_4_055_32    m/z 374.2565     0   1   2   3E+4   0.5   1  RT (min)      lcqtof_hilic_neg_3_035_46    m/z 374.8764     0   1E+5   8.5   9  RT (min)      lcqtof_hilic_neg_1_000_aa    m/z 374.8764     0   2   4   6E+4   8.5   9  RT (min)      lcqtof_hilic_neg_1_000_aa    m/z 374.8768     0   2   4   6E+4   9   9.5  RT (min)      lcqtof_hilic_neg_1_000_aa    m/z 374.8768     0   2   4   6E+4   8.5   9  RT (min)      lcqtof_hilic_neg_1_000_aa    m/z 374.8768     0   2   4   6E+4   9   9.5  RT (min)      lcqtof_hilic_neg_1_000_aa    m/z 374.8768     0   1E+5   8   8.5  RT (min)      lcqtof_hilic_neg_1_000_aa    m/z 374.8769     0   1   2E+4   12.5   13  RT (min)      lcqtof_hilic_neg_1_000_aa    m/z 374.8769     0   1   2   3E+4   0   0.5  RT (min)      lcqtof_hilic_neg_4_000_aa    m/z 374.8769     0   1   2   3E+4   0.5   1  RT (min)      lcqtof_hilic_neg_4_000_aa    m/z 374.8770     0   1E+5   7.5   8  RT (min)      lcqtof_hilic_neg_1_000_aa    m/z 374.8770     0   1   2E+4   12.5   13  RT (min)      lcqtof_hilic_neg_1_000_aa    m/z 374.8770     0   1E+5   8   8.5  RT (min)      lcqtof_hilic_neg_1_000_aa    m/z 374.8770     0   1   2   3E+4   0   0.5  RT (min)      lcqtof_hilic_neg_4_000_aa    m/z 374.8772     0   1E+5   7.5   8  RT (min)      lcqtof_hilic_neg_1_000_aa    m/z 374.8772     0   1   2   3E+4   0   0.5  RT (min)      lcqtof_hilic_neg_4_000_aa    m/z 374.8772     0   1E+4   11.5   12  RT (min)      lcqtof_hilic_neg_1_000_aa    m/z 374.8774     0   1   2E+4   12.5   13  RT (min)      lcqtof_hilic_neg_1_000_aa    m/z 374.8775     0   1   2E+4   12.5   13  RT (min)      lcqtof_hilic_neg_1_000_aa    m/z 374.8776     0   1   2   3E+4   0   0.5  RT (min)      lcqtof_hilic_neg_4_000_aa    m/z 374.8778     0   1   2   3E+4   0   0.5  RT (min)      lcqtof_hilic_neg_4_000_aa    m/z 374.8782     0   1   2   3E+4   0   0.5  RT (min)      lcqtof_hilic_neg_4_000_aa    m/z 374.9058     0   1   2E+3   5.5   6  RT (min)       lcqtof_hilic_neg_4_059_14    m/z 375.0424     0   1E+5   6.5   7  RT (min)       lcqtof_hilic_neg_1_037_24    m/z 375.0590     0   1   2E+4   7   7.5  RT (min)       lcqtof_hilic_neg_4_090_60    m/z 375.0618     0   2   4   6   8E+4   4.5   5  RT (min)       lcqtof_hilic_neg_4_019_03    m/z 375.0619     0   1E+5   4   4.5  RT (min)       lcqtof_hilic_neg_4_026_04    m/z 375.0620     0   1   2E+4   3.5   4  RT (min)       lcqtof_hilic_neg_4_019_03    m/z 375.0621     0   1   2E+4   3.5   4  RT (min)       lcqtof_hilic_neg_4_019_03    m/z 375.0621     0   2   4   6   8E+4   4.5   5  RT (min)       lcqtof_hilic_neg_4_019_03    m/z 375.0622     0   1   2E+4   3.5   4  RT (min)      lcqtof_hilic_neg_4_019_03    m/z 375.0624     0   1E+4   3.5   4  RT (min)      lcqtof_hilic_neg_4_062_17    m/z 375.0632     0   1   2E+4   3   3.5  RT (min)      lcqtof_hilic_neg_4_019_03    m/z 375.0892     0   1E+4   0.5   1  RT (min)      lcqtof_hilic_neg_3_035_46    m/z 375.1450     0   1   2E+4   5   5.5  RT (min)      lcqtof_hilic_neg_4_055_32    m/z 375.1477     0   1E+4   4   4.5  RT (min)      lcqtof_hilic_neg_4_055_32    m/z 375.1596     0   1E+4   3   3.5  RT (min)      lcqtof_hilic_neg_4_004_47    m/z 375.2740     0   1   2   3   4E+4   1   1.5  RT (min)      lcqtof_hilic_neg_4_000_aa    m/z 375.9415     0   1E+4   4   4.5  RT (min)      lcqtof_hilic_neg_3_066_57    m/z 375.9416     0   1   2E+4   4   4.5  RT (min)       lcqtof_hilic_neg_4_003_16    m/z 376.0802     0   1E+4   2   2.5  RT (min)      lcqtof_hilic_neg_4_060_13    m/z 376.0923     0   1E+4   3.5   4  RT (min)      lcqtof_hilic_neg_4_050_02    m/z 376.1088     0   1E+4   5.5   6  RT (min)      lcqtof_hilic_neg_4_057_49    m/z 376.1090     0   1   2   3   4E+4   5.5   6  RT (min)       lcqtof_hilic_neg_4_048_23    m/z 376.1196     0   1E+4   6   6.5  RT (min)      lcqtof_hilic_neg_4_035_33    m/z 376.1214     0   1   2E+4   6   6.5  RT (min)       lcqtof_hilic_neg_4_008_24    m/z 376.1463     0   1E+4   4   4.5  RT (min)      lcqtof_hilic_neg_4_055_32    m/z 376.1714     0   1E+4   6   6.5  RT (min)      lcqtof_hilic_neg_4_019_03    m/z 376.2512     0   1   2E+4   1   1.5  RT (min)      lcqtof_hilic_neg_4_038_10    m/z 376.8503     0   1E+4   8.5   9  RT (min)      lcqtof_hilic_neg_1_000_aa    m/z 376.8514     0   1E+4   8.5   9  RT (min)      lcqtof_hilic_neg_1_000_aa    m/z 376.8521     0   1   2E+4   8.5   9  RT (min)      lcqtof_hilic_neg_1_000_aa    m/z 376.8563     0   1   2E+4   8   8.5  RT (min)      lcqtof_hilic_neg_1_000_aa    m/z 376.9980     0   1   2E+4   7   7.5  RT (min)      lcqtof_hilic_neg_1_005_32    m/z 377.0523     0   1   2   3E+4   5.5   6  RT (min)      lcqtof_hilic_neg_4_011_35    m/z 377.0528     0   1E+4   3   3.5  RT (min)      lcqtof_hilic_neg_4_004_47    m/z 377.0532     0   1E+4   3   3.5  RT (min)      lcqtof_hilic_neg_4_004_47    m/z 377.0692     0   1   7   7.5  RT (min)       lcqtof_hilic_neg_2_000_aa    m/z 377.0799     0   1   2   3E+5   0.5   1  RT (min)      lcqtof_hilic_neg_3_035_46    m/z 377.0836     0   1E+4   4.5   5  RT (min)      lcqtof_hilic_neg_2_000_aa    m/z 377.0837     0   1E+5   5.5   6  RT (min)      lcqtof_hilic_neg_1_000_aa    m/z 377.0840     0   1E+5   5   5.5  RT (min)      lcqtof_hilic_neg_1_000_aa    m/z 377.0841     0   1E+5   5   5.5  RT (min)      lcqtof_hilic_neg_1_000_aa    m/z 377.0845     0   1E+5   5.5   6  RT (min)       lcqtof_hilic_neg_1_000_aa    m/z 377.0848     0   2   4E+4   6.5   7  RT (min)       lcqtof_hilic_neg_4_050_02    m/z 377.0848     0   2   4E+4   6.5   7  RT (min)       lcqtof_hilic_neg_4_050_02    m/z 377.0849     0   1   2   3   4E+4   6.5   7  RT (min)      lcqtof_hilic_neg_4_052_13    m/z 377.0849     0   2   4E+4   6.5   7  RT (min)      lcqtof_hilic_neg_4_050_02    m/z 377.0850     0   1   2   3E+4   5.5   6  RT (min)      lcqtof_hilic_neg_2_007_03    m/z 377.0854     0   1   2   3E+4   5   5.5  RT (min)      lcqtof_hilic_neg_1_000_aa    m/z 377.0855     0   1   2   3E+4   4.5   5  RT (min)      lcqtof_hilic_neg_1_000_aa    m/z 377.1085     0   1   2E+4   1   1.5  RT (min)      lcqtof_hilic_neg_3_002_10    m/z 377.1086     0   1   2E+4   1   1.5  RT (min)      lcqtof_hilic_neg_3_002_10    m/z 377.1789     0   1E+4   3   3.5  RT (min)      lcqtof_hilic_neg_4_009_50    m/z 377.2076     0   1E+6   2.5   3  RT (min)       lcqtof_hilic_neg_2_050_19    m/z 377.2076     0   2   4   6   8E+5   2.5   3  RT (min)      lcqtof_hilic_neg_3_022_38    m/z 377.2081     0   1   2   3E+5   2.5   3  RT (min)      lcqtof_hilic_neg_1_023_34    m/z 377.2236     0   1E+4   2.5   3  RT (min)      lcqtof_hilic_neg_4_034_31    m/z 377.2245     0   1E+4   2   2.5  RT (min)      lcqtof_hilic_neg_4_034_31    m/z 377.2354     0   1   2E+3   2   2.5  RT (min)       lcqtof_hilic_neg_4_054_48    m/z 377.2358     0   1E+4   2   2.5  RT (min)       lcqtof_hilic_neg_4_034_31    m/z 377.2360     0   1   2E+3   0   0.5  RT (min)       lcqtof_hilic_neg_4_047_06    m/z 377.2363     0   1E+4   2   2.5  RT (min)      lcqtof_hilic_neg_4_034_31    m/z 377.2371     0   1   2E+4   15   15.5  RT (min)      lcqtof_hilic_neg_4_000_aa    m/z 377.2494     0   1E+4   0.5   1  RT (min)      lcqtof_hilic_neg_1_060_14    m/z 377.3623     0   1E+4   3   3.5  RT (min)      lcqtof_hilic_neg_4_027_19    m/z 377.4454     0   1   2E+4   3   3.5  RT (min)      lcqtof_hilic_neg_4_027_19    m/z 377.4951     0   1   2   3E+4   3   3.5  RT (min)      lcqtof_hilic_neg_4_027_19    m/z 377.5560     0   2   4   6E+4   3   3.5  RT (min)      lcqtof_hilic_neg_4_027_19    m/z 377.6664     0   1   2   3   4E+4   3   3.5  RT (min)      lcqtof_hilic_neg_4_027_19    m/z 377.9393     0   1E+4   4   4.5  RT (min)      lcqtof_hilic_neg_4_003_16    m/z 377.9878     0   1E+4   6   6.5  RT (min)      lcqtof_hilic_neg_2_007_03    m/z 378.0870     0   2   4   6   8E+4   0.5   1  RT (min)      lcqtof_hilic_neg_3_035_46    m/z 378.0880     0   1E+4   5.5   6  RT (min)      lcqtof_hilic_neg_1_000_aa    m/z 378.0881     0   1E+4   6.5   7  RT (min)       lcqtof_hilic_neg_4_040_05    m/z 378.1091     0   1E+4   1.5   2  RT (min)      lcqtof_hilic_neg_4_027_19    m/z 378.1550     0   1   2   3E+3   16   16.5  RT (min)       lcqtof_hilic_neg_4_019_03    m/z 378.2113     0   1   2   3   4E+5   2.5   3  RT (min)      lcqtof_hilic_neg_2_050_19    m/z 378.2114     0   1   2   3   4E+5   2.5   3  RT (min)      lcqtof_hilic_neg_2_050_19    m/z 378.2116     0   1   2   3   4E+4   2   2.5  RT (min)      lcqtof_hilic_neg_4_005_29    m/z 378.3067     0   1E+4   1   1.5  RT (min)      lcqtof_hilic_neg_4_019_03    m/z 378.4091     0   1   2   3   4E+4   3   3.5  RT (min)      lcqtof_hilic_neg_4_027_19    m/z 379.0330     0   1E+4   6   6.5  RT (min)       lcqtof_hilic_neg_4_008_24    m/z 379.0820     0   1   2   3E+4   5.5   6  RT (min)      lcqtof_hilic_neg_1_000_aa    m/z 379.0820     0   1   2   3E+4   5   5.5  RT (min)      lcqtof_hilic_neg_1_000_aa    m/z 379.0824     0   1E+4   6.5   7  RT (min)      lcqtof_hilic_neg_4_050_02    m/z 379.0825     0   1E+4   6.5   7  RT (min)      lcqtof_hilic_neg_4_050_02    m/z 379.0826     0   1E+4   6.5   7  RT (min)      lcqtof_hilic_neg_4_060_13    m/z 379.0828     0   1E+4   6.5   7  RT (min)      lcqtof_hilic_neg_4_050_02    m/z 379.1553     0   2   4   6E+3   15   15.5  RT (min)       lcqtof_hilic_neg_4_112_62    m/z 379.1555     0   2   4E+4   15   15.5  RT (min)       lcqtof_hilic_neg_4_000_aa    m/z 379.1560     0   2   4   6E+3   15.5   16  RT (min)       lcqtof_hilic_neg_4_113_62    m/z 379.1561     0   2   4   6   8E+3   15.5   16  RT (min)       lcqtof_hilic_neg_4_058_39    m/z 379.1844     0   1E+4   6.5   7  RT (min)      lcqtof_hilic_neg_1_037_24    m/z 379.1868     0   1   2E+4   0.5   1  RT (min)      lcqtof_hilic_neg_4_050_02    m/z 379.1944     0   2   4E+4   2   2.5  RT (min)      lcqtof_hilic_neg_4_023_15    m/z 379.2710     0   1   2E+4   1   1.5  RT (min)      lcqtof_hilic_neg_4_000_aa    m/z 379.2719     0   1   2E+4   1   1.5  RT (min)      lcqtof_hilic_neg_4_000_aa    m/z 379.8281     0   1E+5   9   9.5  RT (min)      lcqtof_hilic_neg_1_000_aa    m/z 379.9069     0   1E+4   1   1.5  RT (min)       lcqtof_hilic_neg_4_111_62    m/z 380.0690     0   2   4E+3   0   0.5  RT (min)      lcqtof_hilic_neg_4_058_39    m/z 380.0815     0   1E+4   2.5   3  RT (min)      lcqtof_hilic_neg_4_054_48    m/z 380.1500     0   1E+4   1.5   2  RT (min)      lcqtof_hilic_neg_4_005_29    m/z 380.1572     0   1E+4   15   15.5  RT (min)      lcqtof_hilic_neg_4_000_aa    m/z 380.1716     0   1E+4   0.5   1  RT (min)      lcqtof_hilic_neg_3_035_46    m/z 380.1974     0   1E+4   2   2.5  RT (min)      lcqtof_hilic_neg_4_023_15    m/z 380.8895     0   1   2   3   4E+4   1   1.5  RT (min)      lcqtof_hilic_neg_4_004_47    m/z 380.9236     0   1   2E+4   4.5   5  RT (min)       lcqtof_hilic_neg_4_013_20    m/z 380.9904     0   1   2   3E+4   0.5   1  RT (min)      lcqtof_hilic_neg_3_035_46    m/z 380.9906     0   1   2   3E+4   0.5   1  RT (min)      lcqtof_hilic_neg_3_035_46    m/z 381.0496     0   1   2E+4   0.5   1  RT (min)      lcqtof_hilic_neg_4_021_21    m/z 381.0643     0   1E+4   0.5   1  RT (min)      lcqtof_hilic_neg_4_039_18    m/z 381.0664     0   1   2   3E+4   3   3.5  RT (min)       lcqtof_hilic_neg_4_004_47    m/z 381.0678     0   1   2   3E+4   3   3.5  RT (min)      lcqtof_hilic_neg_4_004_47    m/z 381.1057     0   1E+4   2   2.5  RT (min)      lcqtof_hilic_neg_4_050_02    m/z 381.1256     0   1   2   3   4E+4   5.5   6  RT (min)      lcqtof_hilic_neg_4_004_47    m/z 381.1846     0   2   4   6E+4   0.5   1  RT (min)      lcqtof_hilic_neg_3_035_46    m/z 381.2036     0   1E+4   0.5   1  RT (min)      lcqtof_hilic_neg_4_050_02    m/z 381.2136     0   1   2E+4   2   2.5  RT (min)      lcqtof_hilic_neg_4_032_01    m/z 381.2342     0   1   2   3   4E+3   2   2.5  RT (min)      lcqtof_hilic_neg_4_033_53    m/z 381.2361     0   1E+4   1   1.5  RT (min)      lcqtof_hilic_neg_4_053_14    m/z 381.2408     0   1E+4   15   15.5  RT (min)      lcqtof_hilic_neg_4_000_aa    m/z 381.3353     0   1   2   3   4E+4   1   1.5  RT (min)      lcqtof_hilic_neg_4_000_aa    m/z 381.3387     0   1   2   3   4E+4   1   1.5  RT (min)      lcqtof_hilic_neg_4_000_aa    m/z 381.3395     0   1   2   3   4E+4   1   1.5  RT (min)      lcqtof_hilic_neg_4_000_aa    m/z 382.0360     0   1E+4   2.5   3  RT (min)      lcqtof_hilic_neg_4_014_46    m/z 382.0706     0   1E+4   3   3.5  RT (min)      lcqtof_hilic_neg_4_004_47    m/z 382.0798     0   1E+4   2.5   3  RT (min)      lcqtof_hilic_neg_4_054_48    m/z 382.0920     0   1   2E+4   4   4.5  RT (min)      lcqtof_hilic_neg_4_040_05    m/z 382.1246     0   1   2   3E+3   0.5   1  RT (min)      lcqtof_hilic_neg_2_053_18    m/z 382.1308     0   1   2   3E+3   0   0.5  RT (min)       lcqtof_hilic_neg_4_110_62    m/z 382.1308     0   1   2   3   4E+3   0   1  RT (min)       lcqtof_hilic_neg_4_105_61    m/z 382.1313     0   1   2   3E+3   0   0.5  RT (min)       lcqtof_hilic_neg_4_081_58    m/z 382.1775     0   1E+4   6.5   7  RT (min)      lcqtof_hilic_neg_1_037_24    m/z 382.1785     0   1E+4   5.5   6  RT (min)      lcqtof_hilic_neg_4_028_36    m/z 382.1872     0   1E+4   0.5   1  RT (min)      lcqtof_hilic_neg_3_035_46    m/z 382.1964     0   1E+4   4   4.5  RT (min)       lcqtof_hilic_neg_4_040_05    m/z 382.2501     0   1   2   3E+4   0.5   1  RT (min)      lcqtof_hilic_neg_3_035_46    m/z 382.9206     0   1   2E+4   4.5   5  RT (min)      lcqtof_hilic_neg_4_013_20    m/z 383.0112     0   1E+5   3.5   4  RT (min)      lcqtof_hilic_neg_4_050_02    m/z 383.0122     0   1E+5   3.5   4  RT (min)      lcqtof_hilic_neg_4_050_02    m/z 383.0196     0   1E+5   3.5   4  RT (min)      lcqtof_hilic_neg_4_050_02    m/z 383.0202     0   1   2   3E+4   4   4.5  RT (min)      lcqtof_hilic_neg_4_050_02    m/z 383.0206     0   2   4   6E+4   4   4.5  RT (min)      lcqtof_hilic_neg_4_050_02    m/z 383.0206     0   2   4E+4   4   4.5  RT (min)      lcqtof_hilic_neg_4_050_02    m/z 383.0208     0   1E+5   3   3.5  RT (min)      lcqtof_hilic_neg_4_050_02    m/z 383.1077     0   1E+4   3   3.5  RT (min)      lcqtof_hilic_neg_4_050_02    m/z 383.1207     0   1E+4   2   2.5  RT (min)      lcqtof_hilic_neg_4_052_13    m/z 383.1245     0   1E+4   2   2.5  RT (min)      lcqtof_hilic_neg_4_050_02    m/z 383.1292     0   1E+4   7   7.5  RT (min)      lcqtof_hilic_neg_4_052_13    m/z 383.1296     0   1E+4   7   7.5  RT (min)      lcqtof_hilic_neg_4_052_13    m/z 383.1402     0   1   2E+3   18.5   19  RT (min)       lcqtof_hilic_neg_4_021_21    m/z 383.1413     0   1   2   3E+3   16   16.5  RT (min)       lcqtof_hilic_neg_4_005_29    m/z 383.1413     0   1   2   3   4E+3   1.5   2  RT (min)       lcqtof_hilic_neg_4_021_21    m/z 383.1415     0   1   2   3E+3   16   16.5  RT (min)       lcqtof_hilic_neg_4_070_57    m/z 383.1417     0   1   2   3   4E+3   15.5   16  RT (min)       lcqtof_hilic_neg_4_058_39    m/z 383.1417     0   1E+4   15   15.5  RT (min)       lcqtof_hilic_neg_4_000_aa    m/z 383.1418     0   1   2   3E+3   2.5   3  RT (min)       lcqtof_hilic_neg_4_042_22    m/z 383.1420     0   1   2   3E+3   16.5   17  RT (min)       lcqtof_hilic_neg_2_002_36    m/z 383.1421     0   1   2   3   4E+3   2   4  RT (min)       lcqtof_hilic_neg_4_109_62    m/z 383.1547     0   2   4   6   8E+4   1   1.5  RT (min)      lcqtof_hilic_neg_4_050_02    m/z 383.2025     0   1   2   3E+4   0.5   1  RT (min)      lcqtof_hilic_neg_3_035_46    m/z 383.2806     0   1E+4   1   1.5  RT (min)      lcqtof_hilic_neg_4_003_16    m/z 383.3161     0   1   2   3E+4   1   1.5  RT (min)      lcqtof_hilic_neg_4_000_aa    m/z 383.3166     0   1   2   3E+4   1   1.5  RT (min)      lcqtof_hilic_neg_4_000_aa    m/z 383.3508     0   1   2E+4   15.5   16  RT (min)      lcqtof_hilic_neg_4_000_aa    m/z 383.3647     0   1E+4   0.5   1  RT (min)      lcqtof_hilic_neg_3_035_46    m/z 384.1474     0   1   2   3E+3   0   0.5  RT (min)       lcqtof_hilic_neg_4_002_25    m/z 384.1476     0   1   2   3E+3   0   0.5  RT (min)       lcqtof_hilic_neg_4_000_cc    m/z 384.1485     0   1   2   3E+3   0.5   1  RT (min)       lcqtof_hilic_neg_4_027_19    m/z 384.1491     0   1E+4   1   1.5  RT (min)       lcqtof_hilic_neg_4_050_02    m/z 384.1509     0   2   4E+3   4   4.5  RT (min)       lcqtof_hilic_neg_4_002_25    m/z 384.2499     0   1E+5   2   2.5  RT (min)      lcqtof_hilic_neg_4_016_37    m/z 384.9344     0   2   4   6   8E+3   6   6.5  RT (min)       lcqtof_hilic_neg_4_058_39    m/z 384.9348     0   1   2E+4   5.5   6  RT (min)       lcqtof_hilic_neg_4_012_07    m/z 384.9806     0   1   2E+4   3.5   4  RT (min)      lcqtof_hilic_neg_1_032_18    m/z 384.9814     0   1E+4   3   3.5  RT (min)      lcqtof_hilic_neg_4_039_18    m/z 385.0437     0   1E+4   0.5   1  RT (min)      lcqtof_hilic_neg_4_050_02    m/z 385.0514     0   1E+4   6   6.5  RT (min)       lcqtof_hilic_neg_4_093_60    m/z 385.1391     0   2   4E+3   0.5   1  RT (min)      lcqtof_hilic_neg_4_029_17    m/z 385.1678     0   2   4   6E+4   1   1.5  RT (min)      lcqtof_hilic_neg_4_032_01    m/z 385.1680     0   2   4E+4   1   1.5  RT (min)       lcqtof_hilic_neg_4_032_01    m/z 385.1710     0   1E+4   4   4.5  RT (min)      lcqtof_hilic_neg_4_061_50    m/z 385.1714     0   1E+4   4   4.5  RT (min)      lcqtof_hilic_neg_4_009_50    m/z 385.1715     0   1E+4   4   4.5  RT (min)      lcqtof_hilic_neg_4_009_50    m/z 385.1718     0   1E+4   4   4.5  RT (min)      lcqtof_hilic_neg_4_009_50    m/z 386.1714     0   1E+4   2   2.5  RT (min)      lcqtof_hilic_neg_4_053_14    m/z 386.2559     0   2   4   6   8E+3   2   2.5  RT (min)      lcqtof_hilic_neg_4_048_23    m/z 386.2562     0   1   2   3   4E+3   2   2.5  RT (min)      lcqtof_hilic_neg_1_007_05    m/z 386.8308     0   1E+4   9   9.5  RT (min)      lcqtof_hilic_neg_1_000_aa    m/z 386.8312     0   1E+4   9   9.5  RT (min)      lcqtof_hilic_neg_1_000_aa    m/z 387.0946     0   1   2   3   4E+4   1   1.5  RT (min)      lcqtof_hilic_neg_4_000_aa    m/z 387.0952     0   1   2   3   4E+4   1   1.5  RT (min)      lcqtof_hilic_neg_4_000_aa    m/z 387.0962     0   1   2   3   4E+4   1   1.5  RT (min)      lcqtof_hilic_neg_4_000_aa    m/z 387.1120     0   1E+4   2   2.5  RT (min)      lcqtof_hilic_neg_4_052_13    m/z 387.1131     0   1   2E+5   5.5   6  RT (min)      lcqtof_hilic_neg_1_000_aa    m/z 387.1133     0   1E+5   5.5   6  RT (min)      lcqtof_hilic_neg_2_000_aa    m/z 387.1135     0   1   2E+4   5   5.5  RT (min)      lcqtof_hilic_neg_2_000_aa    m/z 387.1136     0   1   2E+4   5   5.5  RT (min)      lcqtof_hilic_neg_2_000_aa    m/z 387.1136     0   1   2E+4   5   5.5  RT (min)      lcqtof_hilic_neg_2_000_aa    m/z 387.1453     0   1   2   3E+4   5.5   6  RT (min)      lcqtof_hilic_neg_4_040_05    m/z 387.1455     0   1   2   3   4E+4   5.5   6  RT (min)      lcqtof_hilic_neg_4_008_24    m/z 387.1808     0   1   2E+4   1.5   2  RT (min)      lcqtof_hilic_neg_4_039_18    m/z 387.1829     0   1   2E+4   2   2.5  RT (min)      lcqtof_hilic_neg_4_053_14    m/z 387.1945     0   1E+4   1   1.5  RT (min)      lcqtof_hilic_neg_4_053_14    m/z 387.2132     0   1   2   3   4E+4   6   6.5  RT (min)      lcqtof_hilic_neg_3_011_34    m/z 387.3132     0   2   4   6   8E+4   6   6.5  RT (min)      lcqtof_hilic_neg_3_011_34    m/z 387.4046     0   1E+4   6   6.5  RT (min)      lcqtof_hilic_neg_3_011_34    m/z 387.4660     0   1   2   3   4E+4   6   6.5  RT (min)      lcqtof_hilic_neg_3_011_34    m/z 387.5780     0   1   2E+4   6   6.5  RT (min)      lcqtof_hilic_neg_3_011_34    m/z 388.0797     0   1E+4   0.5   1  RT (min)      lcqtof_hilic_neg_4_000_aa    m/z 388.0950     0   1E+4   1   1.5  RT (min)      lcqtof_hilic_neg_4_000_aa    m/z 388.0972     0   1E+4   1   1.5  RT (min)      lcqtof_hilic_neg_4_000_aa    m/z 388.1099     0   1   2   3E+4   5.5   6  RT (min)      lcqtof_hilic_neg_1_000_aa    m/z 388.1171     0   1   2   3E+5   6   6.5  RT (min)      lcqtof_hilic_neg_3_011_34    m/z 388.1664     0   1E+4   5   5.5  RT (min)      lcqtof_hilic_neg_1_005_32    m/z 388.2697     0   1E+4   2   2.5  RT (min)      lcqtof_hilic_neg_4_032_01    m/z 388.3250     0   1E+4   0.5   1  RT (min)      lcqtof_hilic_neg_1_008_13    m/z 388.9000     0   1E+4   0.5   1  RT (min)      lcqtof_hilic_neg_4_021_21    m/z 389.0592     0   2   4   6   8E+3   6.5   7  RT (min)      lcqtof_hilic_neg_4_035_33    m/z 389.0780     0   1E+4   6.5   7  RT (min)      lcqtof_hilic_neg_3_034_24    m/z 389.0830     0   1E+4   3.5   4  RT (min)      lcqtof_hilic_neg_4_055_32    m/z 389.0924     0   1   2   3E+4   5.5   6  RT (min)      lcqtof_hilic_neg_4_008_24    m/z 389.1956     0   1   2E+4   2   2.5  RT (min)       lcqtof_hilic_neg_4_052_13    m/z 389.1957     0   1   2E+4   0.5   1  RT (min)       lcqtof_hilic_neg_4_106_62    m/z 389.1958     0   1E+4   17   18  RT (min)       lcqtof_hilic_neg_4_110_62    m/z 389.1959     0   1   2E+4   16   18  RT (min)       lcqtof_hilic_neg_4_112_62    m/z 389.1959     0   1E+4   17   18  RT (min)       lcqtof_hilic_neg_4_110_62    m/z 389.1959     0   1E+4   0   0.5  RT (min)       lcqtof_hilic_neg_4_106_62    m/z 389.1960     0   1E+4   4   4.5  RT (min)       lcqtof_hilic_neg_4_100_61    m/z 389.1960     0   1   2E+4   1.5   2  RT (min)       lcqtof_hilic_neg_4_101_61    m/z 389.1960     0   1E+4   16   17  RT (min)       lcqtof_hilic_neg_4_110_62    m/z 389.1960     0   1   2E+4   0   0.5  RT (min)       lcqtof_hilic_neg_4_112_62    m/z 389.1960     0   1   2E+4   0   0.5  RT (min)       lcqtof_hilic_neg_4_112_62    m/z 389.1961     0   1E+4   19  RT (min)       lcqtof_hilic_neg_4_107_62    m/z 389.1961     0   1   2E+4   19   19.5  RT (min)       lcqtof_hilic_neg_4_111_62    m/z 389.1962     0   1E+4   3   3.5  RT (min)       lcqtof_hilic_neg_4_103_61    m/z 389.1962     0   1   2E+4   16.5   17  RT (min)       lcqtof_hilic_neg_4_107_62    m/z 389.1962     0   1   2   3E+4   1.5   2  RT (min)       lcqtof_hilic_neg_4_106_62    m/z 389.1962     0   1E+4   2.5   3  RT (min)       lcqtof_hilic_neg_4_108_62    m/z 389.1962     0   2   4   6   8E+3   6   7  RT (min)       lcqtof_hilic_neg_4_019_03    m/z 389.1962     0   1E+4   1.5   2  RT (min)      lcqtof_hilic_neg_4_111_62    m/z 389.1963     0   1E+4   3.5   4  RT (min)       lcqtof_hilic_neg_4_112_62    m/z 389.1963     0   1E+4   16.5   17  RT (min)       lcqtof_hilic_neg_4_105_61    m/z 389.1964     0   1   2E+4   16   16.5  RT (min)       lcqtof_hilic_neg_4_012_07    m/z 389.1964     0   1E+4   17.5   18  RT (min)      lcqtof_hilic_neg_4_093_60    m/z 389.1965     0   1E+4   16   16.5  RT (min)       lcqtof_hilic_neg_4_107_62    m/z 389.1965     0   2   4   6   8E+3   5   5.5  RT (min)       lcqtof_hilic_neg_4_096_60    m/z 389.1965     0   2   4   6E+3   5   5.5  RT (min)       lcqtof_hilic_neg_4_101_61    m/z 389.1965     0   1   2E+4   4   4.5  RT (min)       lcqtof_hilic_neg_4_096_60    m/z 389.1966     0   1E+4   19.5   20  RT (min)       lcqtof_hilic_neg_4_109_62    m/z 389.1966     0   1E+4   19   19.5  RT (min)       lcqtof_hilic_neg_4_086_59    m/z 389.1967     0   1E+4   17   18  RT (min)       lcqtof_hilic_neg_4_109_62    m/z 389.1970     0   1E+4   2.5   3  RT (min)       lcqtof_hilic_neg_4_106_62    m/z 389.2700     0   1E+4   1   1.5  RT (min)      lcqtof_hilic_neg_4_000_aa    m/z 390.0177     0   2   4E+4   3   3.5  RT (min)      lcqtof_hilic_neg_1_032_18    m/z 390.1175     0   1   2   3   4E+5   5.5   6  RT (min)      lcqtof_hilic_neg_4_048_23    m/z 390.1512     0   1E+4   5.5   6  RT (min)      lcqtof_hilic_neg_2_061_50    m/z 390.1515     0   1   2   3   4E+4   5.5   6  RT (min)      lcqtof_hilic_neg_4_014_46    m/z 390.1996     0   1   2   3   4E+3   2   2.5  RT (min)      lcqtof_hilic_neg_4_060_13    m/z 390.1996     0   1E+4   0.5   1  RT (min)      lcqtof_hilic_neg_4_111_62    m/z 390.1998     0   1E+4   1   1.5  RT (min)      lcqtof_hilic_neg_4_110_62    m/z 390.9863     0   1E+4   5.5   6  RT (min)      lcqtof_hilic_neg_4_050_02    m/z 390.9864     0   1E+4   5.5   6  RT (min)      lcqtof_hilic_neg_4_050_02    m/z 390.9867     0   1E+4   5.5   6  RT (min)      lcqtof_hilic_neg_4_050_02    m/z 390.9877     0   1E+4   5.5   6  RT (min)      lcqtof_hilic_neg_4_050_02    m/z 390.9894     0   1E+4   3   3.5  RT (min)      lcqtof_hilic_neg_4_039_18    m/z 391.0695     0   1   2E+4   1   1.5  RT (min)      lcqtof_hilic_neg_4_004_47    m/z 391.0822     0   1E+4   1   1.5  RT (min)      lcqtof_hilic_neg_4_013_20    m/z 391.0903     0   1E+4   5.5   6  RT (min)      lcqtof_hilic_neg_4_008_24    m/z 391.1000     0   2   4E+4   5   5.5  RT (min)      lcqtof_hilic_neg_4_050_02    m/z 391.1004     0   1   2E+4   5.5   6  RT (min)      lcqtof_hilic_neg_4_050_02    m/z 391.1010     0   1E+4   5   5.5  RT (min)      lcqtof_hilic_neg_4_050_02    m/z 391.1039     0   1   2E+4   2   2.5  RT (min)      lcqtof_hilic_neg_4_060_13    m/z 391.1239     0   1   2E+4   1   1.5  RT (min)      lcqtof_hilic_neg_3_002_10    m/z 391.1908     0   1E+4   0.5   1  RT (min)      lcqtof_hilic_neg_4_013_20    m/z 391.2149     0   1   2   3E+4   0.5   1  RT (min)      lcqtof_hilic_neg_2_062_17    m/z 392.0164     0   2   4E+4   3   3.5  RT (min)      lcqtof_hilic_neg_1_032_18    m/z 392.1046     0   1E+4   4.5   5  RT (min)      lcqtof_hilic_neg_4_052_13    m/z 392.1482     0   1E+4   4.5   5  RT (min)       lcqtof_hilic_neg_4_091_60    m/z 392.2553     0   1   2E+4   1   1.5  RT (min)      lcqtof_hilic_neg_4_040_05    m/z 393.0802     0   1   2E+4   7.5   8  RT (min)      lcqtof_hilic_neg_4_048_23    m/z 393.0943     0   1   2E+4   4   4.5  RT (min)      lcqtof_hilic_neg_4_048_23    m/z 393.0984     0   1   2   3E+4   4.5   5  RT (min)      lcqtof_hilic_neg_4_052_13    m/z 393.1165     0   1E+4   6   6.5  RT (min)       lcqtof_hilic_neg_4_048_23    m/z 393.1572     0   1   2   3E+3   1.5   2  RT (min)       lcqtof_hilic_neg_1_063_13    m/z 393.1708     0   1E+4   15.5   16  RT (min)       lcqtof_hilic_neg_4_113_62    m/z 393.1710     0   2   4   6E+3   15   15.5  RT (min)       lcqtof_hilic_neg_4_110_62    m/z 393.1710     0   2   4E+4   15   15.5  RT (min)       lcqtof_hilic_neg_4_000_aa    m/z 393.1712     0   1E+4   15.5   16  RT (min)       lcqtof_hilic_neg_4_058_39    m/z 393.1713     0   1E+4   0.5   1  RT (min)      lcqtof_hilic_neg_4_038_10    m/z 393.1714     0   1E+4   0.5   1  RT (min)      lcqtof_hilic_neg_4_000_aa    m/z 393.2490     0   1E+4   0.5   1  RT (min)      lcqtof_hilic_neg_4_038_10    m/z 393.2968     0   1E+4   2.5   3  RT (min)      lcqtof_hilic_neg_4_027_19    m/z 393.3897     0   1   2   3   4E+4   1   1.5  RT (min)      lcqtof_hilic_neg_1_005_32    m/z 393.3900     0   1   2E+4   0.5   1  RT (min)      lcqtof_hilic_neg_4_055_32    m/z 394.0232     0   1   2E+4   7   7.5  RT (min)       lcqtof_hilic_neg_4_055_32    m/z 394.0756     0   1E+4   1   1.5  RT (min)      lcqtof_hilic_neg_4_004_47    m/z 394.1103     0   1   2   3E+4   4   4.5  RT (min)      lcqtof_hilic_neg_4_048_23    m/z 394.1390     0   1E+4   3   3.5  RT (min)      lcqtof_hilic_neg_4_050_02    m/z 394.1744     0   1E+4   15   15.5  RT (min)      lcqtof_hilic_neg_4_000_aa    m/z 394.1986     0   1E+4   0.5   1  RT (min)      lcqtof_hilic_neg_1_037_24    m/z 394.2961     0   1E+4   1.5   2  RT (min)      lcqtof_hilic_neg_3_025_52    m/z 394.8902     0   2   4   6E+3   5.5   6  RT (min)       lcqtof_hilic_neg_4_000_bb    m/z 394.8904     0   1   2   3E+3   5   5.5  RT (min)       lcqtof_hilic_neg_4_036_11    m/z 394.9046     0   2   4   6   8E+4   1   1.5  RT (min)      lcqtof_hilic_neg_2_034_47    m/z 394.9046     0   1E+5   1   1.5  RT (min)       lcqtof_hilic_neg_4_004_47    m/z 394.9048     0   1E+5   1   1.5  RT (min)       lcqtof_hilic_neg_4_004_47    m/z 394.9166     0   1   2E+4   0.5   1  RT (min)      lcqtof_hilic_neg_3_035_46    m/z 395.0076     0   1E+5   0.5   1  RT (min)      lcqtof_hilic_neg_3_035_46    m/z 395.0737     0   1   2   3   4E+4   1   1.5  RT (min)      lcqtof_hilic_neg_4_004_47    m/z 395.0812     0   1   2   3   4E+4   1   1.5  RT (min)      lcqtof_hilic_neg_4_004_47    m/z 395.1322     0   1   2E+4   4   4.5  RT (min)      lcqtof_hilic_neg_4_008_24    m/z 395.1365     0   2   4   6E+4   3   3.5  RT (min)      lcqtof_hilic_neg_4_050_02    m/z 395.2647     0   1   2   3E+4   1   1.5  RT (min)      lcqtof_hilic_neg_4_000_aa    m/z 395.3381     0   1E+4   2   2.5  RT (min)      lcqtof_hilic_neg_4_052_13    m/z 395.3522     0   1E+4   1   1.5  RT (min)      lcqtof_hilic_neg_4_000_aa    m/z 396.0078     0   1   2E+4   0.5   1  RT (min)      lcqtof_hilic_neg_3_035_46    m/z 396.0086     0   1   2E+4   0.5   1  RT (min)      lcqtof_hilic_neg_3_035_46    m/z 396.0858     0   1   2E+4   1   1.5  RT (min)      lcqtof_hilic_neg_4_004_47    m/z 396.1293     0   1   2   3   4E+4   4   4.5  RT (min)      lcqtof_hilic_neg_4_008_24    m/z 396.1402     0   2   4   6   8E+4   2   2.5  RT (min)      lcqtof_hilic_neg_4_052_13    m/z 396.1409     0   2   4   6E+4   2   2.5  RT (min)      lcqtof_hilic_neg_4_060_13    m/z 396.1426     0   1E+4   0.5   1  RT (min)       lcqtof_hilic_neg_4_107_62    m/z 396.1450     0   1E+4   19   19.5  RT (min)      lcqtof_hilic_neg_4_107_62    m/z 396.1452     0   1E+4   16   16.5  RT (min)       lcqtof_hilic_neg_4_112_62    m/z 396.1454     0   1E+4   1.5   2  RT (min)       lcqtof_hilic_neg_4_108_62    m/z 396.1454     0   1E+4   1.5   2  RT (min)       lcqtof_hilic_neg_4_104_61    m/z 396.1454     0   1E+4   0   0.5  RT (min)       lcqtof_hilic_neg_4_104_61    m/z 396.1455     0   1E+4   19   19.5  RT (min)      lcqtof_hilic_neg_4_107_62    m/z 396.1457     0   1E+4   17   17.5  RT (min)       lcqtof_hilic_neg_4_107_62    m/z 396.1458     0   1E+4   1   2   3  RT (min)       lcqtof_hilic_neg_4_051_43    m/z 396.1458     0   1E+4   18   18.5  RT (min)       lcqtof_hilic_neg_4_107_62    m/z 396.1458     0   1E+4   16.5   17  RT (min)       lcqtof_hilic_neg_4_047_06    m/z 396.1458     0   1E+4   0   0.5  RT (min)       lcqtof_hilic_neg_4_047_06    m/z 396.1458     0   2   4   6   8E+3   0   0.5  RT (min)       lcqtof_hilic_neg_4_062_17    m/z 396.1459     0   1E+4   15.5   16  RT (min)       lcqtof_hilic_neg_4_107_62    m/z 396.1459     0   1E+4   16   17  RT (min)       lcqtof_hilic_neg_4_019_03    m/z 396.1462     0   2   4   6E+3   1.5   2  RT (min)       lcqtof_hilic_neg_4_102_61    m/z 396.1463     0   1E+4   17   17.5  RT (min)       lcqtof_hilic_neg_4_022_34    m/z 396.1465     0   1E+4   15   16  RT (min)       lcqtof_hilic_neg_4_079_58    m/z 396.1465     0   1E+4   16   18  RT (min)       lcqtof_hilic_neg_4_097_60    m/z 396.1465     0   1E+4   17.5   18  RT (min)       lcqtof_hilic_neg_4_012_07    m/z 396.1470     0   1E+4   19.5   20  RT (min)      lcqtof_hilic_neg_4_107_62    m/z 396.2310     0   1E+4   0.5   1  RT (min)      lcqtof_hilic_neg_3_003_20    m/z 396.8909     0   1   2   3E+3   8   8.5  RT (min)       lcqtof_hilic_neg_4_027_19    m/z 396.8910     0   1   2   3E+3   8   8.5  RT (min)      lcqtof_hilic_neg_4_109_62    m/z 396.8912     0   1   2E+3   7.5   8  RT (min)       lcqtof_hilic_neg_2_024_10    m/z 396.8912     0   1   2E+3   7.5   8  RT (min)       lcqtof_hilic_neg_1_050_12    m/z 397.1383     0   1E+4   3   3.5  RT (min)       lcqtof_hilic_neg_4_099_61    m/z 397.1402     0   1   2E+4   2   2.5  RT (min)      lcqtof_hilic_neg_4_052_13    m/z 397.1540     0   1E+4   15   15.5  RT (min)      lcqtof_hilic_neg_4_000_aa    m/z 397.1566     0   1E+4   1   1.5  RT (min)      lcqtof_hilic_neg_4_053_14    m/z 397.2430     0   1   2   3   4E+5   0.5   1  RT (min)      lcqtof_hilic_neg_3_035_46    m/z 397.3314     0   1E+4   14.5   15  RT (min)       lcqtof_hilic_neg_4_103_61    m/z 397.3667     0   1E+4   15.5   16  RT (min)      lcqtof_hilic_neg_4_000_aa    m/z 397.3821     0   1   2   3E+4   0.5   1  RT (min)      lcqtof_hilic_neg_3_035_46    m/z 397.5829     0   1   2   3   4E+4   0.5   1  RT (min)      lcqtof_hilic_neg_2_016_33    m/z 398.0810     0   1   2E+4   6   6.5  RT (min)      lcqtof_hilic_neg_4_014_46    m/z 398.0870     0   1   2E+4   6.5   7  RT (min)      lcqtof_hilic_neg_4_014_46    m/z 398.1068     0   2   4E+3   0.5   1  RT (min)      lcqtof_hilic_neg_2_019_12    m/z 398.1120     0   1   2   3E+3   0.5   1  RT (min)      lcqtof_hilic_neg_4_058_39    m/z 398.1282     0   1E+4   4.5   5  RT (min)       lcqtof_hilic_neg_4_090_60    m/z 398.2428     0   2   4   6   8E+4   0.5   1  RT (min)      lcqtof_hilic_neg_3_035_46    m/z 398.3347     0   1   2E+4   1   1.5  RT (min)      lcqtof_hilic_neg_4_000_aa    m/z 398.3700     0   1E+4   1   1.5  RT (min)      lcqtof_hilic_neg_4_000_aa    m/z 398.8226     0   1   2E+4   10   10.5  RT (min)      lcqtof_hilic_neg_1_000_aa    m/z 398.9482     0   1   2E+4   0.5   1  RT (min)      lcqtof_hilic_neg_3_035_46    m/z 399.0522     0   1E+4   0.5   1  RT (min)      lcqtof_hilic_neg_4_016_37    m/z 399.0638     0   1E+4   0.5   1  RT (min)      lcqtof_hilic_neg_3_003_20    m/z 399.1653     0   1E+4   3.5   4  RT (min)      lcqtof_hilic_neg_4_039_18    m/z 399.2436     0   2   4   6E+4   0.5   1  RT (min)      lcqtof_hilic_neg_3_003_20    m/z 400.0472     0   1   2   3E+4   0.5   1  RT (min)      lcqtof_hilic_neg_4_016_37    m/z 400.0856     0   1E+4   6   6.5  RT (min)      lcqtof_hilic_neg_4_048_23    m/z 400.1239     0   1E+4   3   3.5  RT (min)      lcqtof_hilic_neg_1_032_18    m/z 400.1295     0   1E+4   4   4.5  RT (min)      lcqtof_hilic_neg_4_040_05    m/z 400.2169     0   1   2E+4   1   1.5  RT (min)      lcqtof_hilic_neg_4_050_02    m/z 401.0929     0   1E+4   1   1.5  RT (min)      lcqtof_hilic_neg_4_004_47    m/z 401.1225     0   1   2   3E+4   1   1.5  RT (min)      lcqtof_hilic_neg_4_000_aa    m/z 401.1300     0   1   2   3E+4   1   1.5  RT (min)      lcqtof_hilic_neg_4_000_aa    m/z 401.1310     0   1   2   3E+4   1   1.5  RT (min)      lcqtof_hilic_neg_4_000_aa    m/z 401.1314     0   1E+4   4.5   5  RT (min)      lcqtof_hilic_neg_4_052_13    m/z 401.2400     0   1   2E+5   2   2.5  RT (min)       lcqtof_hilic_neg_4_048_23    m/z 402.0436     0   1   2E+4   1   1.5  RT (min)       lcqtof_hilic_neg_4_098_61    m/z 402.1192     0   1   2   3   4E+4   1   1.5  RT (min)      lcqtof_hilic_neg_1_052_40    m/z 402.1193     0   1   2E+4   0.5   1  RT (min)      lcqtof_hilic_neg_4_058_39    m/z 402.1322     0   1E+4   1   1.5  RT (min)      lcqtof_hilic_neg_4_000_aa    m/z 402.1338     0   1E+4   5.5   6  RT (min)      lcqtof_hilic_neg_4_050_02    m/z 402.2430     0   2   4E+4   2   2.5  RT (min)      lcqtof_hilic_neg_4_048_23    m/z 402.2430     0   2   4E+4   2   2.5  RT (min)      lcqtof_hilic_neg_4_048_23    m/z 402.2670     0   1E+4   0.5   1   1.5  RT (min)      lcqtof_hilic_neg_4_038_10    m/z 402.8096     0   1E+4   9.5   10  RT (min)      lcqtof_hilic_neg_2_000_aa    m/z 402.8290     0   1E+4   8   8.5  RT (min)       lcqtof_hilic_neg_4_099_61    m/z 402.9623     0   2   4E+4   1   1.5  RT (min)      lcqtof_hilic_neg_4_005_29    m/z 403.0648     0   1   2E+4   4   4.5  RT (min)       lcqtof_hilic_neg_4_048_23    m/z 403.1172     0   1E+5   1   1.5  RT (min)      lcqtof_hilic_neg_1_052_40    m/z 403.1174     0   2   4   6   8E+4   0.5   1  RT (min)      lcqtof_hilic_neg_4_058_39    m/z 403.1323     0   1   2E+4   1   1.5  RT (min)      lcqtof_hilic_neg_4_050_02    m/z 403.1486     0   1E+4   5.5   6  RT (min)      lcqtof_hilic_neg_4_054_48    m/z 403.1614     0   1   2   3   4E+5   5.5   6  RT (min)      lcqtof_hilic_neg_4_008_24    m/z 403.1924     0   1E+4   15   15.5  RT (min)      lcqtof_hilic_neg_4_000_aa    m/z 403.1941     0   2   4   6E+3   2   2.5  RT (min)      lcqtof_hilic_neg_4_005_29    m/z 403.2250     0   1   2   3   4E+4   2.5   3  RT (min)      lcqtof_hilic_neg_4_013_20    m/z 403.2630     0   1E+4   15.5   16  RT (min)      lcqtof_hilic_neg_4_000_aa    m/z 403.2633     0   1   2   3E+4   1   1.5  RT (min)      lcqtof_hilic_neg_4_000_aa    m/z 403.2693     0   1   2   3E+4   1   1.5  RT (min)      lcqtof_hilic_neg_4_000_aa    m/z 403.3690     0   1E+4   2.5   3  RT (min)      lcqtof_hilic_neg_4_013_20    m/z 403.4211     0   1E+4   2.5   3  RT (min)      lcqtof_hilic_neg_4_013_20    m/z 403.4833     0   1   2   3   4E+4   2.5   3  RT (min)      lcqtof_hilic_neg_4_013_20    m/z 403.5979     0   1   2E+4   2.5   3  RT (min)      lcqtof_hilic_neg_4_013_20    m/z 403.8306     0   1   2   3E+4   0.5   1  RT (min)      lcqtof_hilic_neg_4_000_aa    m/z 403.8309     0   1   2   3E+4   0   0.5  RT (min)      lcqtof_hilic_neg_4_000_aa    m/z 403.8310     0   1   2   3E+4   0   0.5  RT (min)      lcqtof_hilic_neg_4_000_aa    m/z 403.8310     0   1   2   3E+4   0.5   1  RT (min)      lcqtof_hilic_neg_4_000_aa    m/z 403.8312     0   1   2   3E+4   0   0.5  RT (min)      lcqtof_hilic_neg_4_000_aa    m/z 404.0144     0   1   2   3E+4   10.5   11  RT (min)       lcqtof_hilic_neg_4_078_58    m/z 404.0149     0   1E+5   11.5   12  RT (min)       lcqtof_hilic_neg_4_064_15    m/z 404.0150     0   1   2   3E+4   11   11.5  RT (min)       lcqtof_hilic_neg_4_080_58    m/z 404.0150     0   1E+4   8.5   9  RT (min)       lcqtof_hilic_neg_4_064_15    m/z 404.0152     0   1   2E+4   10.5   11  RT (min)      lcqtof_hilic_neg_4_078_58    m/z 404.0152     0   1   2   3E+4   10.5   11  RT (min)       lcqtof_hilic_neg_4_078_58    m/z 404.0152     0   1   2   3   4E+4   12.5   13  RT (min)       lcqtof_hilic_neg_4_064_15    m/z 404.0153     0   1E+4   8   8.5  RT (min)       lcqtof_hilic_neg_4_064_15    m/z 404.0154     0   1E+4   8.5   9  RT (min)       lcqtof_hilic_neg_4_064_15    m/z 404.0154     0   2   4   6   8E+4   12   12.5  RT (min)       lcqtof_hilic_neg_4_050_02    m/z 404.0155     0   1E+4   13   13.5  RT (min)       lcqtof_hilic_neg_4_064_15    m/z 404.0156     0   1E+5   11.5   12  RT (min)       lcqtof_hilic_neg_4_064_15    m/z 404.0156     0   1E+4   12.5   13  RT (min)      lcqtof_hilic_neg_4_059_14    m/z 404.0156     0   1   2E+4   5.5   6  RT (min)      lcqtof_hilic_neg_1_016_28    m/z 404.0156     0   2   4   6E+4   12   12.5  RT (min)      lcqtof_hilic_neg_4_064_15    m/z 404.0158     0   1E+4   9   9.5  RT (min)       lcqtof_hilic_neg_4_064_15    m/z 404.0159     0   1E+4   8   8.5  RT (min)      lcqtof_hilic_neg_4_064_15    m/z 404.0160     0   2   4E+3   8.5   9  RT (min)       lcqtof_hilic_neg_2_043_15    m/z 404.0780     0   1E+4   6   6.5  RT (min)      lcqtof_hilic_neg_4_008_24    m/z 404.1026     0   1E+5   5.5   6  RT (min)      lcqtof_hilic_neg_1_000_aa    m/z 404.1030     0   1E+5   5.5   6  RT (min)      lcqtof_hilic_neg_1_000_aa    m/z 404.1033     0   1E+5   5   5.5  RT (min)      lcqtof_hilic_neg_1_000_aa    m/z 404.1047     0   1E+4   8   8.5  RT (min)      lcqtof_hilic_neg_2_000_aa    m/z 404.1051     0   1E+4   7.5   8  RT (min)      lcqtof_hilic_neg_2_000_aa    m/z 404.1055     0   1   2E+4   7.5   8  RT (min)      lcqtof_hilic_neg_2_000_aa    m/z 404.1078     0   1E+4   8   8.5  RT (min)       lcqtof_hilic_neg_2_000_aa    m/z 404.1124     0   1E+4   7.5   8  RT (min)      lcqtof_hilic_neg_4_008_24    m/z 404.1135     0   1   2E+4   7   7.5  RT (min)      lcqtof_hilic_neg_2_000_aa    m/z 404.1140     0   1E+4   7.5   8  RT (min)      lcqtof_hilic_neg_4_008_24    m/z 404.1151     0   2   4   6   8E+3   7   7.5  RT (min)       lcqtof_hilic_neg_2_000_aa    m/z 404.1449     0   1   2E+5   4.5   5  RT (min)      lcqtof_hilic_neg_1_037_24    m/z 404.3321     0   1E+4   2.5   3  RT (min)      lcqtof_hilic_neg_4_013_20    m/z 404.8252     0   1E+4   8   8.5  RT (min)       lcqtof_hilic_neg_4_099_61    m/z 404.8260     0   1E+4   8   8.5  RT (min)       lcqtof_hilic_neg_4_100_61    m/z 405.0281     0   1E+4   4   4.5  RT (min)      lcqtof_hilic_neg_1_060_14    m/z 405.0563     0   1E+4   0.5   1  RT (min)      lcqtof_hilic_neg_4_015_09    m/z 405.0674     0   1E+4   0.5   1  RT (min)      lcqtof_hilic_neg_4_050_02    m/z 405.0796     0   1E+4   6.5   7  RT (min)      lcqtof_hilic_neg_4_002_25    m/z 405.1411     0   1E+4   4   4.5  RT (min)      lcqtof_hilic_neg_4_090_60    m/z 405.2646     0   1E+4   1.5   2  RT (min)      lcqtof_hilic_neg_4_039_18    m/z 405.2649     0   2   4   6   8E+4   1   1.5  RT (min)      lcqtof_hilic_neg_4_000_aa    m/z 406.0036     0   1   2   3E+4   0.5   1  RT (min)      lcqtof_hilic_neg_4_038_10    m/z 406.0037     0   1   2   3E+4   1   1.5  RT (min)      lcqtof_hilic_neg_4_038_10    m/z 406.1038     0   1E+4   3   3.5  RT (min)      lcqtof_hilic_neg_4_050_02    m/z 406.1380     0   1E+4   2   2.5  RT (min)      lcqtof_hilic_neg_4_055_32    m/z 406.2677     0   1E+4   1   1.5  RT (min)      lcqtof_hilic_neg_4_000_aa    m/z 406.2681     0   1E+4   1   1.5  RT (min)      lcqtof_hilic_neg_4_000_aa    m/z 406.2699     0   1E+4   1   1.5  RT (min)      lcqtof_hilic_neg_4_000_aa    m/z 406.4291     0   1E+4   0.5   1  RT (min)      lcqtof_hilic_neg_2_028_05    m/z 406.8247     0   1E+4   8   8.5  RT (min)       lcqtof_hilic_neg_4_099_61    m/z 406.9632     0   1E+4   3.5   4  RT (min)      lcqtof_hilic_neg_1_032_18    m/z 407.0452     0   1E+4   2.5   3  RT (min)      lcqtof_hilic_neg_4_045_26    m/z 407.0544     0   1E+4   0.5   1  RT (min)      lcqtof_hilic_neg_4_015_09    m/z 407.0991     0   2   4   6   8E+4   3   3.5  RT (min)      lcqtof_hilic_neg_4_050_02    m/z 407.4331     0   2   4   6   8E+4   1   1.5  RT (min)      lcqtof_hilic_neg_4_040_05    m/z 407.7024     0   1E+4   2   2.5  RT (min)      lcqtof_hilic_neg_4_016_37    m/z 408.0006     0   1E+4   1   1.5  RT (min)      lcqtof_hilic_neg_4_039_18    m/z 408.0009     0   1E+4   0.5   1  RT (min)      lcqtof_hilic_neg_4_038_10    m/z 408.0013     0   1E+4   1   1.5  RT (min)      lcqtof_hilic_neg_4_038_10    m/z 408.1038     0   1E+4   3   3.5  RT (min)      lcqtof_hilic_neg_4_050_02    m/z 408.1178     0   1   2E+4   4.5   5  RT (min)      lcqtof_hilic_neg_4_009_50    m/z 408.1619     0   1E+4   6   6.5  RT (min)      lcqtof_hilic_neg_2_022_46    m/z 408.1878     0   1E+4   4.5   5  RT (min)      lcqtof_hilic_neg_4_008_24    m/z 408.1906     0   1E+4   15   15.5  RT (min)      lcqtof_hilic_neg_4_000_aa    m/z 408.2016     0   1   2E+4   3   3.5  RT (min)      lcqtof_hilic_neg_4_008_24    m/z 408.2843     0   1E+4   2   2.5  RT (min)      lcqtof_hilic_neg_4_039_18    m/z 408.9077     0   1E+4   0.5   1  RT (min)      lcqtof_hilic_neg_4_021_21    m/z 408.9204     0   1   2   3E+5   1   1.5  RT (min)      lcqtof_hilic_neg_4_004_47    m/z 408.9206     0   1   2   3E+5   1   1.5  RT (min)      lcqtof_hilic_neg_4_004_47    m/z 408.9881     0   1E+4   0.5   1  RT (min)      lcqtof_hilic_neg_3_035_46    m/z 409.0227     0   2   4   6E+4   0.5   1  RT (min)      lcqtof_hilic_neg_3_035_46    m/z 409.0954     0   1E+4   6.5   7  RT (min)      lcqtof_hilic_neg_4_052_13    m/z 409.0958     0   1E+4   6.5   7  RT (min)      lcqtof_hilic_neg_4_052_13    m/z 409.0965     0   1E+4   6.5   7  RT (min)      lcqtof_hilic_neg_4_052_13    m/z 409.1050     0   1E+4   2   2.5  RT (min)      lcqtof_hilic_neg_4_004_47    m/z 409.2360     0   1E+5   4   4.5  RT (min)      lcqtof_hilic_neg_4_040_05    m/z 409.2708     0   1   2   3E+4   1   1.5  RT (min)      lcqtof_hilic_neg_4_000_aa    m/z 409.2774     0   1E+4   0.5   1  RT (min)      lcqtof_hilic_neg_3_035_46    m/z 409.3121     0   1   2   3E+4   1   1.5  RT (min)      lcqtof_hilic_neg_4_000_aa    m/z 409.3124     0   1   2   3E+4   1   1.5  RT (min)      lcqtof_hilic_neg_4_000_aa    m/z 409.3680     0   1E+4   1   1.5  RT (min)      lcqtof_hilic_neg_4_000_aa    m/z 409.3807     0   1   2E+4   0.5   1  RT (min)      lcqtof_hilic_neg_3_035_46    m/z 409.9336     0   1E+4   0.5   1  RT (min)      lcqtof_hilic_neg_3_035_46    m/z 410.0174     0   1   2E+4   3   3.5  RT (min)      lcqtof_hilic_neg_2_053_18    m/z 410.0238     0   1E+4   0.5   1  RT (min)      lcqtof_hilic_neg_3_035_46    m/z 410.1238     0   1E+4   1   1.5  RT (min)      lcqtof_hilic_neg_4_005_29    m/z 410.1604     0   1E+4   1   1.5  RT (min)       lcqtof_hilic_neg_4_013_20    m/z 410.1609     0   1E+4   19.5   20  RT (min)       lcqtof_hilic_neg_4_106_62    m/z 410.1609     0   1E+4   18.5   19  RT (min)       lcqtof_hilic_neg_4_106_62    m/z 410.1610     0   1   2E+4   16   16.5  RT (min)       lcqtof_hilic_neg_4_012_07    m/z 410.1610     0   1E+4   16   18  RT (min)       lcqtof_hilic_neg_4_104_61    m/z 410.1611     0   1E+4   17.5   18  RT (min)       lcqtof_hilic_neg_4_096_60    m/z 410.1611     0   1E+4   18   18.5  RT (min)       lcqtof_hilic_neg_4_093_60    m/z 410.1611     0   1E+4   0   0.5  RT (min)       lcqtof_hilic_neg_4_106_62    m/z 410.1612     0   1E+4   19   19.5  RT (min)       lcqtof_hilic_neg_4_106_62    m/z 410.1613     0   1E+4   15.5   16  RT (min)       lcqtof_hilic_neg_4_106_62    m/z 410.1613     0   1E+4   16   16.5  RT (min)       lcqtof_hilic_neg_4_109_62    m/z 410.1613     0   1E+4   17.5   18  RT (min)       lcqtof_hilic_neg_4_106_62    m/z 410.1614     0   1E+4   16.5   17  RT (min)       lcqtof_hilic_neg_4_106_62    m/z 410.1614     0   1E+4   16.5   17  RT (min)       lcqtof_hilic_neg_4_111_62    m/z 410.1614     0   1E+4   1   1.5  RT (min)       lcqtof_hilic_neg_4_051_43    m/z 410.1615     0   1E+4   0.5   1  RT (min)       lcqtof_hilic_neg_4_113_62    m/z 410.1615     0   1E+4   1   1.5  RT (min)       lcqtof_hilic_neg_4_000_cc    m/z 410.1615     0   1E+4   15   16   17  RT (min)       lcqtof_hilic_neg_4_110_62    m/z 410.1615     0   1E+4   16   17  RT (min)       lcqtof_hilic_neg_4_107_62    m/z 410.1615     0   1E+4   0   0.5  RT (min)       lcqtof_hilic_neg_4_012_07    m/z 410.1616     0   1E+4   0   0.5  RT (min)       lcqtof_hilic_neg_4_065_51    m/z 410.1616     0   1E+4   17.5   18  RT (min)       lcqtof_hilic_neg_4_106_62    m/z 410.1618     0   1E+4   16.5   17  RT (min)       lcqtof_hilic_neg_4_017_51    m/z 410.1619     0   1E+4   17   17.5  RT (min)       lcqtof_hilic_neg_4_106_62    m/z 410.1621     0   1E+4   18.5   19  RT (min)       lcqtof_hilic_neg_4_106_62    m/z 410.1622     0   1E+4   19.5   20  RT (min)       lcqtof_hilic_neg_4_106_62    m/z 410.2892     0   1E+4   2   2.5  RT (min)       lcqtof_hilic_neg_4_008_24    m/z 410.3215     0   1E+4   1   1.5  RT (min)      lcqtof_hilic_neg_4_000_aa    m/z 410.3260     0   1E+4   1   1.5  RT (min)      lcqtof_hilic_neg_4_000_aa    m/z 410.9896     0   1   2E+4   4.5   5  RT (min)       lcqtof_hilic_neg_4_039_18    m/z 411.0652     0   1E+4   0.5   1  RT (min)      lcqtof_hilic_neg_4_038_10    m/z 411.0929     0   2   4   6E+3   1.5   2  RT (min)       lcqtof_hilic_neg_4_008_24    m/z 411.0949     0   1E+4   0.5   1  RT (min)      lcqtof_hilic_neg_4_047_06    m/z 411.0962     0   2   4   6E+3   1   1.5  RT (min)      lcqtof_hilic_neg_2_008_44    m/z 411.0967     0   1E+4   0.5   1  RT (min)      lcqtof_hilic_neg_4_050_02    m/z 411.2033     0   1E+4   6   6.5  RT (min)      lcqtof_hilic_neg_4_008_24    m/z 411.2424     0   1   2   3   4E+4   1   1.5  RT (min)      lcqtof_hilic_neg_4_000_aa    m/z 411.3469     0   2   4E+3   15.5   16  RT (min)      lcqtof_hilic_neg_1_007_05    m/z 411.3474     0   1   2E+3   19.5   20  RT (min)      lcqtof_hilic_neg_4_022_34    m/z 411.3476     0   2   4   6E+4   1   1.5  RT (min)      lcqtof_hilic_neg_4_000_aa    m/z 411.3477     0   1E+4   14.5   15  RT (min)      lcqtof_hilic_neg_4_081_58    m/z 411.3479     0   2   4   6E+4   1   1.5  RT (min)      lcqtof_hilic_neg_4_000_aa    m/z 411.3482     0   2   4E+3   2   2.5  RT (min)      lcqtof_hilic_neg_4_032_01    m/z 411.3822     0   1E+4   15.5   16  RT (min)      lcqtof_hilic_neg_4_000_aa    m/z 411.3825     0   2   4   6E+4   1   1.5  RT (min)      lcqtof_hilic_neg_4_000_aa    m/z 411.3964     0   1   2   3   4E+4   0.5   1  RT (min)      lcqtof_hilic_neg_3_035_46    m/z 411.9177     0   1E+4   4   4.5  RT (min)       lcqtof_hilic_neg_4_003_16    m/z 412.1496     0   1   2   3E+3   18.5   19  RT (min)      lcqtof_hilic_neg_4_020_42    m/z 412.2238     0   1   2E+4   6   6.5  RT (min)      lcqtof_hilic_neg_4_008_24    m/z 412.3497     0   1E+4   1   1.5  RT (min)      lcqtof_hilic_neg_4_000_aa    m/z 412.3502     0   1E+4   1   1.5  RT (min)      lcqtof_hilic_neg_4_000_aa    m/z 412.3504     0   1E+4   1   1.5  RT (min)      lcqtof_hilic_neg_4_000_aa    m/z 412.3855     0   1   2E+4   1   1.5  RT (min)      lcqtof_hilic_neg_4_000_aa    m/z 412.4016     0   1E+4   0.5   1  RT (min)      lcqtof_hilic_neg_3_035_46    m/z 412.8124     0   1   2   3E+3   8.5   9  RT (min)       lcqtof_hilic_neg_4_019_03    m/z 412.9278     0   1   2E+4   0.5   1  RT (min)      lcqtof_hilic_neg_3_035_46    m/z 412.9873     0   1E+4   4.5   5  RT (min)      lcqtof_hilic_neg_4_039_18    m/z 413.0280     0   2   4   6E+4   0.5   1  RT (min)      lcqtof_hilic_neg_2_046_41    m/z 413.1426     0   1E+4   0.5   1  RT (min)      lcqtof_hilic_neg_4_050_02    m/z 413.1438     0   1E+4   3.5   4  RT (min)      lcqtof_hilic_neg_1_032_18    m/z 413.2092     0   1E+4   2.5   3  RT (min)      lcqtof_hilic_neg_4_002_25    m/z 413.9152     0   1E+4   4   4.5  RT (min)       lcqtof_hilic_neg_4_003_16    m/z 414.1839     0   1E+4   7   7.5  RT (min)       lcqtof_hilic_neg_4_092_60    m/z 414.2530     0   1E+4   1   1.5  RT (min)      lcqtof_hilic_neg_4_000_aa    m/z 414.8544     0   1   2E+4   8   8.5  RT (min)      lcqtof_hilic_neg_1_000_aa    m/z 414.8552     0   1   2E+4   8   8.5  RT (min)      lcqtof_hilic_neg_1_000_aa    m/z 415.1651     0   1E+4   0.5   1  RT (min)      lcqtof_hilic_neg_4_050_02    m/z 415.1771     0   1E+4   5.5   6  RT (min)       lcqtof_hilic_neg_4_008_24    m/z 415.2153     0   1E+4   1.5   2  RT (min)      lcqtof_hilic_neg_4_053_14    m/z 415.2160     0   1E+4   1.5   2  RT (min)      lcqtof_hilic_neg_4_053_14    m/z 415.2160     0   2   4E+3   1.5   2  RT (min)      lcqtof_hilic_neg_4_003_16    m/z 415.2264     0   1E+4   1   1.5  RT (min)      lcqtof_hilic_neg_4_002_25    m/z 415.2264     0   1E+4   1   1.5  RT (min)      lcqtof_hilic_neg_4_040_05    m/z 415.2561     0   1   2   3E+4   1   1.5  RT (min)      lcqtof_hilic_neg_4_000_aa    m/z 415.9683     0   1   2E+4   0.5   1  RT (min)      lcqtof_hilic_neg_4_021_21    m/z 416.1123     0   1E+4   3   3.5  RT (min)      lcqtof_hilic_neg_4_004_47    m/z 416.8323     0   2   4   6E+4   8   8.5  RT (min)      lcqtof_hilic_neg_1_000_aa    m/z 416.8332     0   2   4   6E+4   8   8.5  RT (min)      lcqtof_hilic_neg_1_000_aa    m/z 416.9632     0   1E+5   0.5   1  RT (min)      lcqtof_hilic_neg_4_021_21    m/z 416.9747     0   1E+4   0.5   1  RT (min)      lcqtof_hilic_neg_4_005_29    m/z 416.9795     0   1E+4   1   1.5  RT (min)      lcqtof_hilic_neg_4_017_51    m/z 417.0299     0   1   2E+4   0.5   1  RT (min)      lcqtof_hilic_neg_4_050_02    m/z 417.1333     0   1E+4   0.5   1  RT (min)      lcqtof_hilic_neg_4_004_47    m/z 417.2087     0   2   4   6E+4   15   15.5  RT (min)      lcqtof_hilic_neg_4_000_aa    m/z 417.2104     0   2   4   6E+3   3   3.5  RT (min)      lcqtof_hilic_neg_4_002_25    m/z 417.2106     0   2   4   6   8E+3   3   3.5  RT (min)       lcqtof_hilic_neg_1_000_aa    m/z 417.2108     0   2   4E+3   2   2.5  RT (min)      lcqtof_hilic_neg_4_043_12    m/z 417.2108     0   2   4   6   8E+3   2.5   3  RT (min)      lcqtof_hilic_neg_4_047_06    m/z 417.2108     0   1   2   3E+4   0.5   1  RT (min)      lcqtof_hilic_neg_4_007_45    m/z 417.2108     0   1E+4   2.5   3  RT (min)       lcqtof_hilic_neg_4_088_59    m/z 417.2113     0   2   4   6E+3   2   2.5  RT (min)      lcqtof_hilic_neg_4_026_04    m/z 417.2114     0   2   4   6E+3   2   2.5  RT (min)      lcqtof_hilic_neg_4_039_18    m/z 417.2118     0   1E+4   2   2.5  RT (min)       lcqtof_hilic_neg_4_004_47    m/z 417.2119     0   2   4   6E+3   1.5   2  RT (min)      lcqtof_hilic_neg_4_039_18    m/z 417.2120     0   2   4   6E+3   1.5   2  RT (min)      lcqtof_hilic_neg_2_053_18    m/z 417.2126     0   1   2   3E+3   3.5   4  RT (min)      lcqtof_hilic_neg_1_027_27    m/z 417.2426     0   1E+4   0.5   1  RT (min)      lcqtof_hilic_neg_2_047_32    m/z 417.3182     0   1E+4   1   1.5  RT (min)      lcqtof_hilic_neg_4_000_aa    m/z 417.3533     0   1E+4   0.5   1  RT (min)      lcqtof_hilic_neg_4_008_24    m/z 417.9660     0   1E+4   0.5   1  RT (min)      lcqtof_hilic_neg_4_021_21    m/z 418.1187     0   1E+4   5.5   6  RT (min)      lcqtof_hilic_neg_1_019_02    m/z 418.1472     0   1E+4   4   4.5  RT (min)       lcqtof_hilic_neg_4_092_60    m/z 418.1703     0   1E+4   2.5   3  RT (min)       lcqtof_hilic_neg_4_034_31    m/z 418.2116     0   1E+4   15   15.5  RT (min)      lcqtof_hilic_neg_4_000_aa    m/z 418.2199     0   1E+4   0.5   1  RT (min)      lcqtof_hilic_neg_4_046_41    m/z 418.9608     0   1   2E+4   0.5   1  RT (min)      lcqtof_hilic_neg_4_021_21    m/z 419.0286     0   1E+4   0.5   1  RT (min)      lcqtof_hilic_neg_4_050_02    m/z 419.0698     0   1   2   3E+4   0.5   1  RT (min)      lcqtof_hilic_neg_4_010_28    m/z 419.1069     0   1   2E+4   6.5   7  RT (min)       lcqtof_hilic_neg_4_052_13    m/z 419.1400     0   1E+4   6   6.5  RT (min)      lcqtof_hilic_neg_4_048_23    m/z 420.0289     0   1   2   3E+4   0.5   1  RT (min)      lcqtof_hilic_neg_4_050_02    m/z 420.0445     0   1   2E+4   4.5   5  RT (min)       lcqtof_hilic_neg_4_027_19    m/z 420.1440     0   1E+4   2   2.5  RT (min)      lcqtof_hilic_neg_3_035_46    m/z 420.1687     0   1E+4   2.5   3  RT (min)       lcqtof_hilic_neg_4_034_31    m/z 420.1689     0   1E+4   2.5   3  RT (min)      lcqtof_hilic_neg_4_034_31    m/z 420.2651     0   1E+5   0.5   1  RT (min)      lcqtof_hilic_neg_3_035_46    m/z 420.2862     0   1   2E+4   1   1.5  RT (min)      lcqtof_hilic_neg_4_000_aa    m/z 420.3594     0   1   2   3E+4   3.5   4  RT (min)      lcqtof_hilic_neg_4_039_18    m/z 420.7818     0   1E+4   9   9.5  RT (min)      lcqtof_hilic_neg_1_000_aa    m/z 421.0043     0   1E+4   6.5   7  RT (min)       lcqtof_hilic_neg_4_085_59    m/z 421.0189     0   1E+4   4.5   5  RT (min)      lcqtof_hilic_neg_4_039_18    m/z 421.0283     0   1   2   3E+4   0.5   1  RT (min)      lcqtof_hilic_neg_4_050_02    m/z 421.0696     0   1E+4   0.5   1  RT (min)      lcqtof_hilic_neg_4_038_10    m/z 421.1289     0   1E+4   3   3.5  RT (min)      lcqtof_hilic_neg_4_004_47    m/z 421.1495     0   1E+4   1   1.5  RT (min)      lcqtof_hilic_neg_4_059_14    m/z 421.2260     0   1E+4   1   1.5  RT (min)      lcqtof_hilic_neg_4_024_40    m/z 421.2265     0   1E+5   1   1.5  RT (min)      lcqtof_hilic_neg_4_000_aa    m/z 421.2276     0   1E+5   1   1.5  RT (min)      lcqtof_hilic_neg_4_000_aa    m/z 421.2348     0   1E+4   3.5   4  RT (min)      lcqtof_hilic_neg_1_007_05    m/z 421.2353     0   1E+4   3.5   4  RT (min)      lcqtof_hilic_neg_1_007_05    m/z 421.2632     0   1   2E+4   0.5   1  RT (min)      lcqtof_hilic_neg_3_035_46    m/z 421.3802     0   1E+4   0.5   1  RT (min)      lcqtof_hilic_neg_3_035_46    m/z 422.0290     0   1   2E+4   0.5   1  RT (min)      lcqtof_hilic_neg_4_050_02    m/z 422.2826     0   1   2E+4   1   1.5  RT (min)      lcqtof_hilic_neg_4_000_aa    m/z 422.2828     0   1   2E+4   1   1.5  RT (min)      lcqtof_hilic_neg_4_000_aa    m/z 422.3270     0   1E+4   1.5   2  RT (min)      lcqtof_hilic_neg_3_025_52    m/z 422.4519     0   1   2E+4   0.5   1  RT (min)      lcqtof_hilic_neg_2_028_05    m/z 422.9363     0   1   2   3E+5   1   1.5  RT (min)      lcqtof_hilic_neg_4_004_47    m/z 422.9525     0   2   4   6E+4   0.5   1  RT (min)      lcqtof_hilic_neg_3_035_46    m/z 423.0282     0   1   2   3   4E+4   0.5   1  RT (min)      lcqtof_hilic_neg_4_050_02    m/z 423.0892     0   1E+5   6   6.5  RT (min)      lcqtof_hilic_neg_1_000_aa    m/z 423.0892     0   1E+5   5.5   6  RT (min)      lcqtof_hilic_neg_1_000_aa    m/z 423.0896     0   1E+5   5.5   6  RT (min)      lcqtof_hilic_neg_1_000_aa    m/z 423.0954     0   1   2   3E+4   3   3.5  RT (min)       lcqtof_hilic_neg_4_098_61    m/z 423.0971     0   1E+5   5.5   6  RT (min)      lcqtof_hilic_neg_1_000_aa    m/z 423.2256     0   1E+4   1   1.5  RT (min)      lcqtof_hilic_neg_4_000_aa    m/z 423.2506     0   1E+4   1   1.5  RT (min)      lcqtof_hilic_neg_4_052_13    m/z 423.2872     0   1E+4   1   1.5  RT (min)      lcqtof_hilic_neg_4_000_aa    m/z 423.2971     0   1E+4   4   4.5  RT (min)      lcqtof_hilic_neg_4_047_06    m/z 423.3799     0   2   4E+4   1   1.5  RT (min)      lcqtof_hilic_neg_4_000_aa    m/z 423.3818     0   2   4E+4   1   1.5  RT (min)      lcqtof_hilic_neg_4_000_aa    m/z 423.3991     0   1E+4   0.5   1  RT (min)      lcqtof_hilic_neg_3_035_46    m/z 423.4199     0   1E+4   1.5   2  RT (min)      lcqtof_hilic_neg_4_000_aa    m/z 423.9491     0   1E+4   0.5   1  RT (min)      lcqtof_hilic_neg_3_035_46    m/z 423.9636     0   1   2   3E+4   7   7.5  RT (min)      lcqtof_hilic_neg_4_050_02    m/z 424.0754     0   1E+4   6   6.5  RT (min)      lcqtof_hilic_neg_2_052_24    m/z 424.0858     0   1E+4   3   3.5  RT (min)      lcqtof_hilic_neg_1_032_18    m/z 424.0928     0   1E+4   5.5   6  RT (min)      lcqtof_hilic_neg_1_000_aa    m/z 424.0932     0   1E+4   5.5   6  RT (min)      lcqtof_hilic_neg_1_000_aa    m/z 424.0937     0   1E+4   7.5   8  RT (min)      lcqtof_hilic_neg_4_048_23    m/z 424.0938     0   1E+4   7.5   8  RT (min)      lcqtof_hilic_neg_4_048_23    m/z 424.1766     0   2   4   6   8E+3   0   0.5  RT (min)       lcqtof_hilic_neg_4_012_07    m/z 424.1769     0   1   2E+4   0.5   1  RT (min)       lcqtof_hilic_neg_4_012_07    m/z 424.1772     0   1   2   3E+3   2.5   3  RT (min)       lcqtof_hilic_neg_4_112_62    m/z 424.1772     0   2   4   6   8E+3   17   18  RT (min)       lcqtof_hilic_neg_4_047_06    m/z 424.1772     0   1E+4   0   0.5  RT (min)       lcqtof_hilic_neg_4_019_03    m/z 424.1774     0   2   4   6   8E+3   1   1.5  RT (min)       lcqtof_hilic_neg_4_037_39    m/z 424.1774     0   1E+4   0.5   1  RT (min)      lcqtof_hilic_neg_2_014_08    m/z 424.3312     0   1E+4   0.5   1  RT (min)      lcqtof_hilic_neg_3_035_46    m/z 424.3873     0   1E+4   1   1.5  RT (min)      lcqtof_hilic_neg_4_000_aa    m/z 424.3886     0   1E+4   1   1.5  RT (min)      lcqtof_hilic_neg_4_000_aa    m/z 424.9475     0   1E+4   0.5   1  RT (min)      lcqtof_hilic_neg_3_035_46    m/z 424.9629     0   1   2   3E+4   7   7.5  RT (min)      lcqtof_hilic_neg_4_050_02    m/z 425.0241     0   1E+4   0.5   1  RT (min)      lcqtof_hilic_neg_4_050_02    m/z 425.0924     0   1   2E+4   0.5   1  RT (min)       lcqtof_hilic_neg_4_038_10    m/z 425.1103     0   1   2E+3   0   0.5  RT (min)       lcqtof_hilic_neg_2_016_33    m/z 425.1108     0   1   2E+3   19.5   20  RT (min)       lcqtof_hilic_neg_3_044_09    m/z 425.1143     0   1E+4   6   6.5  RT (min)      lcqtof_hilic_neg_4_008_24    m/z 425.1234     0   2   4   6   8E+4   0.5   1  RT (min)      lcqtof_hilic_neg_2_015_06    m/z 425.1404     0   2   4E+4   6   6.5  RT (min)       lcqtof_hilic_neg_4_036_11    m/z 425.1410     0   1   2   3E+4   6   6.5  RT (min)       lcqtof_hilic_neg_4_027_19    m/z 425.1600     0   1E+4   0.5   1  RT (min)      lcqtof_hilic_neg_3_003_20    m/z 425.2188     0   1   2   3E+4   6.5   7  RT (min)      lcqtof_hilic_neg_4_008_24    m/z 425.2844     0   1E+4   3.5   4  RT (min)      lcqtof_hilic_neg_4_040_05    m/z 425.3308     0   1E+4   3   3.5  RT (min)      lcqtof_hilic_neg_4_013_20    m/z 425.3626     0   1E+5   1   1.5  RT (min)      lcqtof_hilic_neg_4_000_aa    m/z 425.3626     0   1   2E+4   14.5   15  RT (min)      lcqtof_hilic_neg_4_112_62    m/z 425.3626     0   1E+2   14.5   15  RT (min)      lcqtof_hilic_neg_4_083_59    m/z 425.3630     0   1E+4   15.5   16  RT (min)      lcqtof_hilic_neg_4_000_aa    m/z 425.3634     0   1E+4   15.5   16  RT (min)      lcqtof_hilic_neg_1_007_05    m/z 425.3636     0   1E+5   1   1.5  RT (min)      lcqtof_hilic_neg_4_000_aa    m/z 425.3986     0   1E+4   1   1.5  RT (min)      lcqtof_hilic_neg_4_000_aa    m/z 425.4666     0   1   2   3   4E+4   0.5   1  RT (min)      lcqtof_hilic_neg_2_016_33    m/z 425.6098     0   1E+4   5   5.5  RT (min)      lcqtof_hilic_neg_3_003_20    m/z 426.0218     0   2   4   6   8E+3   8   8.5  RT (min)      lcqtof_hilic_neg_2_029_31    m/z 426.0219     0   1E+4   7.5   8  RT (min)      lcqtof_hilic_neg_1_003_37    m/z 426.1255     0   1   2E+4   0.5   1  RT (min)      lcqtof_hilic_neg_4_040_05    m/z 426.1266     0   2   4   6E+3   0.5   1  RT (min)      lcqtof_hilic_neg_4_047_06    m/z 426.1282     0   1   2E+3   19.5   20  RT (min)      lcqtof_hilic_neg_4_033_53    m/z 426.1282     0   1E+4   5   5.5  RT (min)      lcqtof_hilic_neg_4_014_46    m/z 426.1303     0   1   2   3E+3   1   1.5  RT (min)      lcqtof_hilic_neg_4_030_44    m/z 426.1326     0   1   2E+3   2.5   3  RT (min)      lcqtof_hilic_neg_4_055_32    m/z 426.1341     0   1E+4   6   6.5  RT (min)      lcqtof_hilic_neg_4_036_11    m/z 426.1351     0   2   4   6E+3   0   0.5  RT (min)      lcqtof_hilic_neg_4_012_07    m/z 426.2036     0   1   2E+4   6.5   7  RT (min)       lcqtof_hilic_neg_4_008_24    m/z 426.2837     0   2   4E+4   3.5   4  RT (min)      lcqtof_hilic_neg_1_007_05    m/z 426.2841     0   2   4E+4   3.5   4  RT (min)      lcqtof_hilic_neg_1_007_05    m/z 426.2844     0   1E+4   3.5   4  RT (min)      lcqtof_hilic_neg_4_002_25    m/z 426.2854     0   1   2   3   4E+4   4   4.5  RT (min)      lcqtof_hilic_neg_4_040_05    m/z 426.3673     0   1   2   3E+4   1   1.5  RT (min)      lcqtof_hilic_neg_4_000_aa    m/z 426.9125     0   1   2   3E+4   0.5   1   1.5  RT (min)      lcqtof_hilic_neg_4_027_19    m/z 426.9310     0   2   4   6   8E+4   1   1.5  RT (min)      lcqtof_hilic_neg_4_004_47    m/z 426.9435     0   1   2   3E+4   0.5   1  RT (min)      lcqtof_hilic_neg_3_035_46    m/z 426.9440     0   1   2   3E+4   0.5   1  RT (min)      lcqtof_hilic_neg_3_035_46    m/z 426.9640     0   1   2   3E+4   7   7.5  RT (min)       lcqtof_hilic_neg_4_050_02    m/z 427.0836     0   1E+4   0.5   1  RT (min)      lcqtof_hilic_neg_4_050_02    m/z 427.1333     0   1E+4   0.5   1  RT (min)      lcqtof_hilic_neg_4_050_02    m/z 427.2455     0   1E+4   0.5   1  RT (min)      lcqtof_hilic_neg_4_016_37    m/z 428.0086     0   1E+4   2.5   3  RT (min)      lcqtof_hilic_neg_1_012_04    m/z 428.0092     0   1E+4   2.5   3  RT (min)      lcqtof_hilic_neg_1_004_14    m/z 428.0399     0   1E+4   6   6.5  RT (min)       lcqtof_hilic_neg_4_093_60    m/z 428.0411     0   1   2E+4   1   1.5  RT (min)      lcqtof_hilic_neg_2_045_11    m/z 428.2762     0   1E+4   2   2.5  RT (min)       lcqtof_hilic_neg_4_016_37    m/z 428.2771     0   2   4   6   8E+3   2   2.5  RT (min)      lcqtof_hilic_neg_4_011_35    m/z 428.2988     0   2   4E+4   4   4.5  RT (min)      lcqtof_hilic_neg_4_040_05    m/z 428.9101     0   1E+4   0.5   1   1.5  RT (min)      lcqtof_hilic_neg_4_027_19    m/z 428.9450     0   1E+4   3.5   4  RT (min)      lcqtof_hilic_neg_1_032_18    m/z 429.0370     0   1   2E+4   0.5   1  RT (min)      lcqtof_hilic_neg_4_058_39    m/z 429.0382     0   1   2E+4   1   1.5  RT (min)      lcqtof_hilic_neg_4_058_39    m/z 429.0564     0   1E+4   1   1.5  RT (min)      lcqtof_hilic_neg_4_050_02    m/z 429.1271     0   1   2   3   4E+4   1   1.5  RT (min)      lcqtof_hilic_neg_4_000_aa    m/z 429.1271     0   1   2   3   4E+4   1   1.5  RT (min)      lcqtof_hilic_neg_4_000_aa    m/z 429.1277     0   1   2   3   4E+4   1   1.5  RT (min)      lcqtof_hilic_neg_4_000_aa    m/z 429.1344     0   1   2   3E+4   7   7.5  RT (min)      lcqtof_hilic_neg_4_052_13    m/z 429.1354     0   1   2   3E+4   6.5   7  RT (min)       lcqtof_hilic_neg_4_052_13    m/z 429.1356     0   1   2   3E+4   7   7.5  RT (min)       lcqtof_hilic_neg_4_050_02    m/z 429.1357     0   1   2   3E+4   7   7.5  RT (min)       lcqtof_hilic_neg_4_052_13    m/z 429.1358     0   1E+4   6.5   7  RT (min)      lcqtof_hilic_neg_4_071_57    m/z 429.1924     0   2   4E+3   2   2.5  RT (min)      lcqtof_hilic_neg_4_050_02    m/z 429.1944     0   2   4   6   8E+3   2   2.5  RT (min)      lcqtof_hilic_neg_4_005_29    m/z 429.2670     0   1   2E+4   1   1.5  RT (min)      lcqtof_hilic_neg_4_000_aa    m/z 429.2692     0   1   2E+4   1   1.5  RT (min)      lcqtof_hilic_neg_4_000_aa    m/z 430.1268     0   1E+4   1   1.5  RT (min)      lcqtof_hilic_neg_4_000_aa    m/z 430.1270     0   1E+4   1   1.5  RT (min)      lcqtof_hilic_neg_4_000_aa    m/z 430.2078     0   1E+4   1   1.5  RT (min)      lcqtof_hilic_neg_4_039_18    m/z 430.2623     0   1E+4   1   1.5  RT (min)      lcqtof_hilic_neg_2_021_17    m/z 430.2830     0   1   2E+4   1   1.5  RT (min)      lcqtof_hilic_neg_1_007_05    m/z 431.1266     0   1E+4   1   1.5  RT (min)      lcqtof_hilic_neg_4_000_aa    m/z 431.1281     0   1E+4   1   1.5  RT (min)      lcqtof_hilic_neg_4_000_aa    m/z 431.1415     0   1E+4   5.5   6  RT (min)      lcqtof_hilic_neg_1_000_aa    m/z 431.1629     0   1   2E+4   4   4.5  RT (min)      lcqtof_hilic_neg_4_008_24    m/z 431.2079     0   1   2   3   4E+4   1   1.5  RT (min)      lcqtof_hilic_neg_4_000_aa    m/z 431.2238     0   1E+4   15   15.5  RT (min)      lcqtof_hilic_neg_4_000_aa    m/z 431.2240     0   2   4   6E+3   2   2.5  RT (min)      lcqtof_hilic_neg_4_032_01    m/z 431.2245     0   2   4E+4   4   4.5  RT (min)      lcqtof_hilic_neg_4_040_05    m/z 431.2251     0   1   2E+5   4   4.5  RT (min)      lcqtof_hilic_neg_4_040_05    m/z 431.2271     0   2   4   6   8E+3   1   1.5  RT (min)       lcqtof_hilic_neg_4_039_18    m/z 431.2273     0   1E+4   15   15.5  RT (min)      lcqtof_hilic_neg_4_000_aa    m/z 431.9830     0   1E+4   6.5   7  RT (min)       lcqtof_hilic_neg_1_037_24    m/z 432.1067     0   1E+4   0.5   1  RT (min)      lcqtof_hilic_neg_4_005_29    m/z 432.1232     0   1   2E+4   5.5   6  RT (min)      lcqtof_hilic_neg_4_004_47    m/z 432.1439     0   1E+4   6   6.5  RT (min)      lcqtof_hilic_neg_4_036_11    m/z 432.2025     0   1   2   3E+4   6   6.5  RT (min)      lcqtof_hilic_neg_1_037_24    m/z 432.2120     0   1E+4   0.5   1  RT (min)      lcqtof_hilic_neg_2_026_28    m/z 432.2228     0   1E+4   3.5   4  RT (min)      lcqtof_hilic_neg_4_002_25    m/z 432.2233     0   2   4   6E+4   3.5   4  RT (min)      lcqtof_hilic_neg_1_007_05    m/z 432.2241     0   2   4   6E+4   3.5   4  RT (min)      lcqtof_hilic_neg_1_007_05    m/z 432.2244     0   2   4E+4   4   4.5  RT (min)      lcqtof_hilic_neg_4_040_05    m/z 432.2791     0   1E+5   1   1.5  RT (min)      lcqtof_hilic_neg_2_021_17    m/z 433.0975     0   1E+4   2   2.5  RT (min)      lcqtof_hilic_neg_4_045_26    m/z 433.1194     0   1E+4   1   1.5  RT (min)      lcqtof_hilic_neg_4_055_32    m/z 433.1276     0   1E+4   1.5   2  RT (min)      lcqtof_hilic_neg_4_039_18    m/z 433.1426     0   1E+4   4.5   5  RT (min)       lcqtof_hilic_neg_4_090_60    m/z 433.1433     0   1E+4   4.5   5  RT (min)       lcqtof_hilic_neg_4_008_24    m/z 433.2350     0   1   2   3   4E+5   4   4.5  RT (min)      lcqtof_hilic_neg_4_040_05    m/z 433.2355     0   1E+5   4   4.5  RT (min)      lcqtof_hilic_neg_4_040_05    m/z 433.2836     0   1   2E+4   1   1.5  RT (min)      lcqtof_hilic_neg_2_021_17    m/z 434.1508     0   1   2E+4   6.5   7  RT (min)      lcqtof_hilic_neg_4_008_24    m/z 434.1882     0   1E+4   1.5   2  RT (min)      lcqtof_hilic_neg_4_008_24    m/z 434.2390     0   1E+5   4   4.5  RT (min)      lcqtof_hilic_neg_4_040_05    m/z 434.3513     0   1E+4   1   1.5  RT (min)      lcqtof_hilic_neg_2_028_05    m/z 434.9752     0   1E+4   1   1.5  RT (min)      lcqtof_hilic_neg_4_000_aa    m/z 434.9762     0   1E+4   1   1.5  RT (min)      lcqtof_hilic_neg_4_000_aa    m/z 435.1204     0   1E+4   4   4.5  RT (min)      lcqtof_hilic_neg_4_052_13    m/z 435.1912     0   1E+4   1.5   2  RT (min)      lcqtof_hilic_neg_4_008_24    m/z 435.2511     0   1E+5   4   4.5  RT (min)      lcqtof_hilic_neg_4_040_05    m/z 435.2553     0   1   2E+4   3   3.5  RT (min)      lcqtof_hilic_neg_4_013_20    m/z 435.3453     0   1   2E+4   1   1.5  RT (min)       lcqtof_hilic_neg_4_027_19    m/z 435.3615     0   2   4   6E+4   3   3.5  RT (min)      lcqtof_hilic_neg_4_027_19    m/z 435.5241     0   1E+4   3   3.5  RT (min)      lcqtof_hilic_neg_4_027_19    m/z 435.6126     0   1E+4   1   1.5  RT (min)      lcqtof_hilic_neg_4_027_19    m/z 436.0552     0   1E+4   7.5   8  RT (min)      lcqtof_hilic_neg_2_047_32    m/z 436.0552     0   1   2E+4   7.5   8  RT (min)       lcqtof_hilic_neg_1_005_32    m/z 436.0554     0   1E+4   8   8.5  RT (min)      lcqtof_hilic_neg_2_016_33    m/z 436.0555     0   2   4   6   8E+3   8   8.5  RT (min)      lcqtof_hilic_neg_3_004_32    m/z 436.0640     0   1   2   3E+4   6   6.5  RT (min)       lcqtof_hilic_neg_4_048_23    m/z 436.1569     0   1   2E+4   5.5   6  RT (min)      lcqtof_hilic_neg_4_014_46    m/z 436.1571     0   1E+4   5.5   6  RT (min)      lcqtof_hilic_neg_2_022_46    m/z 436.1986     0   1E+4   4   4.5  RT (min)      lcqtof_hilic_neg_4_091_60    m/z 436.2518     0   1   2   3   4E+4   4   4.5  RT (min)      lcqtof_hilic_neg_4_040_05    m/z 436.3675     0   1E+4   1   1.5  RT (min)      lcqtof_hilic_neg_2_028_05    m/z 436.4672     0   1E+4   0.5   1  RT (min)      lcqtof_hilic_neg_3_048_05    m/z 437.1604     0   1E+4   2   2.5  RT (min)       lcqtof_hilic_neg_4_008_24    m/z 437.1968     0   1   2   3   4E+4   4   4.5  RT (min)       lcqtof_hilic_neg_4_091_60    m/z 437.2666     0   1   2E+4   1   1.5  RT (min)      lcqtof_hilic_neg_1_031_16    m/z 437.2680     0   1   2E+4   1   1.5  RT (min)      lcqtof_hilic_neg_1_031_16    m/z 437.3791     0   2   4   6E+4   0.5   1  RT (min)      lcqtof_hilic_neg_4_053_14    m/z 437.3911     0   1   2E+4   1   1.5  RT (min)      lcqtof_hilic_neg_4_000_aa    m/z 437.3964     0   1   2E+4   1   1.5  RT (min)      lcqtof_hilic_neg_4_000_aa    m/z 437.3968     0   1   2E+4   1   1.5  RT (min)      lcqtof_hilic_neg_4_000_aa    m/z 437.4706     0   2   4E+4   1   1.5  RT (min)      lcqtof_hilic_neg_4_040_05    m/z 438.1466     0   1   2E+4   2   2.5  RT (min)      lcqtof_hilic_neg_4_003_16    m/z 438.1648     0   1E+4   2   2.5  RT (min)       lcqtof_hilic_neg_4_008_24    m/z 438.3986     0   1E+4   1   1.5  RT (min)      lcqtof_hilic_neg_4_000_aa    m/z 439.0997     0   1E+4   0.5   1  RT (min)      lcqtof_hilic_neg_4_015_09    m/z 439.1263     0   2   4   6   8E+4   3   3.5  RT (min)      lcqtof_hilic_neg_4_039_18    m/z 439.1360     0   2   4   6   8E+4   3   3.5  RT (min)      lcqtof_hilic_neg_4_039_18    m/z 439.2852     0   1E+4   0.5   1  RT (min)      lcqtof_hilic_neg_3_035_46    m/z 439.3780     0   2   4   6E+3   16   17   18  RT (min)       lcqtof_hilic_neg_4_000_aa    m/z 439.3781     0   1   2   3   4E+3   2   2.5  RT (min)      lcqtof_hilic_neg_4_010_28    m/z 439.3782     0   1   2E+3   0   0.5  RT (min)       lcqtof_hilic_neg_4_042_22    m/z 439.3784     0   1   2   3E+4   0.5   1  RT (min)       lcqtof_hilic_neg_4_006_30    m/z 439.3786     0   2   4   6E+4   1   1.5  RT (min)      lcqtof_hilic_neg_4_000_aa    m/z 439.3786     0   1   2E+3   16.5   17  RT (min)       lcqtof_hilic_neg_4_000_dd    m/z 439.3787     0   1   2E+1   14.5   15  RT (min)      lcqtof_hilic_neg_4_082_59    m/z 439.3787     0   1E+4   14.5   15  RT (min)      lcqtof_hilic_neg_4_081_58    m/z 439.3787     0   1   2E+3   19.5   20  RT (min)       lcqtof_hilic_neg_4_026_04    m/z 439.3790     0   1   2E+3   17   17.5   18  RT (min)       lcqtof_hilic_neg_4_026_04    m/z 439.4116     0   1E+4   1   1.5  RT (min)      lcqtof_hilic_neg_4_000_aa    m/z 439.4138     0   1E+4   0.5   1  RT (min)      lcqtof_hilic_neg_3_007_04    m/z 439.4146     0   1E+4   1.5   2  RT (min)      lcqtof_hilic_neg_4_000_aa    m/z 440.0867     0   1E+4   6   6.5  RT (min)      lcqtof_hilic_neg_1_000_aa    m/z 440.1387     0   1E+4   3   3.5  RT (min)      lcqtof_hilic_neg_4_039_18    m/z 440.3122     0   1E+5   1   1.5  RT (min)      lcqtof_hilic_neg_4_040_05    m/z 440.3272     0   1E+4   4   4.5  RT (min)      lcqtof_hilic_neg_4_008_24    m/z 440.3818     0   1   2E+4   1   1.5  RT (min)      lcqtof_hilic_neg_4_000_aa    m/z 440.3821     0   1   2E+4   1   1.5  RT (min)      lcqtof_hilic_neg_4_000_aa    m/z 440.9464     0   1E+5   1   1.5  RT (min)       lcqtof_hilic_neg_4_004_47    m/z 440.9468     0   2   4   6E+4   1   1.5  RT (min)      lcqtof_hilic_neg_1_028_47    m/z 440.9468     0   1E+5   1   1.5  RT (min)      lcqtof_hilic_neg_4_004_47    m/z 440.9616     0   1   2   3   4E+4   0.5   1  RT (min)      lcqtof_hilic_neg_3_035_46    m/z 441.0484     0   1E+4   2.5   3  RT (min)      lcqtof_hilic_neg_4_018_27    m/z 441.0986     0   1E+4   0.5   1  RT (min)      lcqtof_hilic_neg_4_050_02    m/z 441.1154     0   1E+4   4.5   5  RT (min)      lcqtof_hilic_neg_4_046_41    m/z 441.1204     0   1E+4   3   3.5  RT (min)      lcqtof_hilic_neg_2_053_18    m/z 441.2377     0   1E+5   0.5   1  RT (min)      lcqtof_hilic_neg_4_052_13    m/z 441.2522     0   1E+6   1   1.5  RT (min)      lcqtof_hilic_neg_4_016_37    m/z 441.2703     0   1   2E+5   0.5   1  RT (min)      lcqtof_hilic_neg_3_035_46    m/z 441.2820     0   1E+4   1   1.5  RT (min)      lcqtof_hilic_neg_4_000_aa    m/z 441.2840     0   1E+4   1   1.5  RT (min)      lcqtof_hilic_neg_4_000_aa    m/z 441.3026     0   1E+4   3.5   4  RT (min)      lcqtof_hilic_neg_4_040_05    m/z 441.3158     0   1   2E+4   1   1.5  RT (min)      lcqtof_hilic_neg_4_040_05    m/z 441.3822     0   1E+4   0.5   1  RT (min)      lcqtof_hilic_neg_2_063_13    m/z 441.4088     0   1E+4   0.5   1  RT (min)      lcqtof_hilic_neg_3_035_46    m/z 441.6285     0   1   2E+4   0.5   1  RT (min)      lcqtof_hilic_neg_4_016_37    m/z 441.9602     0   1E+4   0.5   1  RT (min)      lcqtof_hilic_neg_3_035_46    m/z 442.2158     0   1   2E+4   6   6.5  RT (min)       lcqtof_hilic_neg_4_090_60    m/z 442.2440     0   1   2E+4   0.5   1  RT (min)      lcqtof_hilic_neg_4_010_28    m/z 442.2558     0   1   2   3   4E+5   1   1.5  RT (min)      lcqtof_hilic_neg_4_016_37    m/z 442.2692     0   1   2   3   4E+4   0.5   1  RT (min)      lcqtof_hilic_neg_3_035_46    m/z 442.2825     0   1   2   3   4E+4   1   1.5  RT (min)      lcqtof_hilic_neg_4_000_aa    m/z 442.3044     0   1E+4   1   1.5  RT (min)      lcqtof_hilic_neg_1_042_26    m/z 442.3340     0   1   2   3   4E+4   1   1.5  RT (min)      lcqtof_hilic_neg_3_003_20    m/z 442.5354     0   1   2   3E+4   1   1.5  RT (min)      lcqtof_hilic_neg_4_026_04    m/z 442.8934     0   1   2   3E+3   5   5.5  RT (min)       lcqtof_hilic_neg_4_032_01    m/z 442.8936     0   1   2   3E+3   5.5   6  RT (min)       lcqtof_hilic_neg_4_060_13    m/z 442.8938     0   1   2   3E+3   5.5   6  RT (min)       lcqtof_hilic_neg_4_052_13    m/z 443.0374     0   1E+4   1.5   2  RT (min)      lcqtof_hilic_neg_4_013_20    m/z 443.1770     0   1   2   3E+4   1   1.5  RT (min)      lcqtof_hilic_neg_4_000_aa    m/z 443.1772     0   1   2   3E+4   1   1.5  RT (min)      lcqtof_hilic_neg_4_000_aa    m/z 443.1775     0   1   2   3E+4   1   1.5  RT (min)      lcqtof_hilic_neg_4_000_aa    m/z 443.2562     0   1E+4   2   2.5  RT (min)      lcqtof_hilic_neg_4_039_18    m/z 443.2666     0   1E+4   0.5   1  RT (min)      lcqtof_hilic_neg_3_035_46    m/z 443.2790     0   2   4   6E+4   1   1.5  RT (min)      lcqtof_hilic_neg_4_000_aa    m/z 443.2823     0   2   4   6E+4   1   1.5  RT (min)      lcqtof_hilic_neg_4_000_aa    m/z 443.2971     0   2   4E+3   2   2.5  RT (min)       lcqtof_hilic_neg_4_077_58    m/z 444.0322     0   1   2E+4   6.5   7  RT (min)       lcqtof_hilic_neg_4_088_59    m/z 444.0946     0   1E+4   4.5   5  RT (min)      lcqtof_hilic_neg_4_009_50    m/z 444.1430     0   1E+4   3   3.5  RT (min)      lcqtof_hilic_neg_4_004_47    m/z 444.1778     0   1E+4   1   1.5  RT (min)      lcqtof_hilic_neg_4_000_aa    m/z 444.2814     0   1E+4   1   1.5  RT (min)      lcqtof_hilic_neg_4_000_aa    m/z 444.2825     0   1E+4   1   1.5  RT (min)      lcqtof_hilic_neg_4_000_aa    m/z 445.1435     0   1E+4   3.5   4  RT (min)      lcqtof_hilic_neg_1_007_05    m/z 445.1749     0   1E+4   6   6.5  RT (min)      lcqtof_hilic_neg_4_050_02    m/z 445.2352     0   1E+4   3   3.5  RT (min)      lcqtof_hilic_neg_4_040_05    m/z 445.3704     0   1E+4   0.5   1  RT (min)      lcqtof_hilic_neg_4_008_24    m/z 446.1513     0   1E+4   6   6.5  RT (min)      lcqtof_hilic_neg_1_000_aa    m/z 446.1775     0   1E+4   6   6.5  RT (min)       lcqtof_hilic_neg_4_090_60    m/z 446.9050     0   1   2E+3   9   9.5  RT (min)       lcqtof_hilic_neg_4_038_10    m/z 447.0579     0   1E+4   5.5   6  RT (min)      lcqtof_hilic_neg_1_010_15    m/z 447.0926     0   1E+5   3.5   4  RT (min)      lcqtof_hilic_neg_2_053_18    m/z 447.0965     0   1E+4   6   6.5  RT (min)      lcqtof_hilic_neg_4_008_24    m/z 447.1248     0   1E+5   1   1.5  RT (min)      lcqtof_hilic_neg_4_027_19    m/z 447.1440     0   1E+4   15   15.5  RT (min)       lcqtof_hilic_neg_4_000_aa    m/z 447.1444     0   1   2   3E+3   15.5   16  RT (min)       lcqtof_hilic_neg_4_058_39    m/z 447.1770     0   1   2   3   4E+4   6   6.5  RT (min)       lcqtof_hilic_neg_4_008_24    m/z 447.1774     0   1   2   3   4E+4   6   6.5  RT (min)      lcqtof_hilic_neg_4_008_24    m/z 447.2509     0   1E+4   1   1.5  RT (min)      lcqtof_hilic_neg_1_007_05    m/z 447.2510     0   1E+4   3   3.5  RT (min)      lcqtof_hilic_neg_4_052_13    m/z 447.2512     0   1   2E+4   2.5   3  RT (min)      lcqtof_hilic_neg_4_040_05    m/z 447.9584     0   1E+4   4.5   5  RT (min)      lcqtof_hilic_neg_1_004_14    m/z 448.0845     0   1E+5   5.5   6  RT (min)      lcqtof_hilic_neg_1_000_aa    m/z 448.0962     0   1   2   3E+4   3.5   4  RT (min)      lcqtof_hilic_neg_1_032_18    m/z 448.1406     0   1E+4   0.5   1  RT (min)      lcqtof_hilic_neg_3_003_20    m/z 448.1607     0   1   2E+4   4.5   5  RT (min)       lcqtof_hilic_neg_4_008_24    m/z 448.1850     0   1E+4   6.5   7  RT (min)      lcqtof_hilic_neg_4_008_24    m/z 448.8861     0   1E+4   7.5   8  RT (min)      lcqtof_hilic_neg_1_000_aa    m/z 448.8862     0   1E+4   7.5   8  RT (min)      lcqtof_hilic_neg_1_000_aa    m/z 448.8867     0   1E+4   7.5   8  RT (min)      lcqtof_hilic_neg_1_000_aa    m/z 448.8880     0   1E+4   7.5   8  RT (min)      lcqtof_hilic_neg_1_000_aa    m/z 449.0188     0   2   4E+4   0.5   1  RT (min)      lcqtof_hilic_neg_4_050_02    m/z 449.0880     0   1E+4   5.5   6  RT (min)      lcqtof_hilic_neg_1_000_aa    m/z 449.1302     0   1E+4   2.5   3  RT (min)       lcqtof_hilic_neg_4_073_57    m/z 449.1304     0   1E+4   2.5   3  RT (min)      lcqtof_hilic_neg_4_019_03    m/z 449.1311     0   1E+4   2.5   3  RT (min)      lcqtof_hilic_neg_4_052_13    m/z 449.1704     0   1   2E+4   6.5   7  RT (min)      lcqtof_hilic_neg_4_008_24    m/z 449.2540     0   1E+5   2   2.5  RT (min)      lcqtof_hilic_neg_4_050_02    m/z 449.2646     0   1E+4   1   1.5  RT (min)      lcqtof_hilic_neg_1_007_05    m/z 449.3044     0   1   2   3E+5   1   1.5  RT (min)      lcqtof_hilic_neg_4_000_aa    m/z 449.3050     0   1   2   3E+5   1   1.5  RT (min)      lcqtof_hilic_neg_4_000_aa    m/z 449.3096     0   2   4   6E+4   2   2.5  RT (min)      lcqtof_hilic_neg_4_050_02    m/z 449.3104     0   1   2   3E+5   1   1.5  RT (min)      lcqtof_hilic_neg_4_000_aa    m/z 449.3114     0   1   2   3E+5   1   1.5  RT (min)      lcqtof_hilic_neg_4_000_aa    m/z 449.3165     0   1   2   3E+5   1   1.5  RT (min)      lcqtof_hilic_neg_4_000_aa    m/z 450.1524     0   1E+4   3   3.5  RT (min)      lcqtof_hilic_neg_4_008_24    m/z 450.1687     0   1E+4   7   7.5  RT (min)       lcqtof_hilic_neg_4_093_60    m/z 450.2930     0   1E+4   4   4.5  RT (min)      lcqtof_hilic_neg_4_008_24    m/z 450.3132     0   1E+5   1   1.5  RT (min)      lcqtof_hilic_neg_4_000_aa    m/z 450.3144     0   1E+5   1   1.5  RT (min)      lcqtof_hilic_neg_4_000_aa    m/z 450.7998     0   1E+4   4   4.5  RT (min)      lcqtof_hilic_neg_4_008_24    m/z 450.8673     0   1E+4   8   8.5  RT (min)      lcqtof_hilic_neg_1_000_aa    m/z 451.0178     0   1   2   3E+4   0.5   1  RT (min)      lcqtof_hilic_neg_4_050_02    m/z 451.0623     0   1E+4   0.5   1  RT (min)      lcqtof_hilic_neg_4_018_27    m/z 451.1527     0   1E+4   5.5   6  RT (min)       lcqtof_hilic_neg_4_008_24    m/z 451.1562     0   1E+4   2   2.5  RT (min)      lcqtof_hilic_neg_4_050_02    m/z 451.1574     0   1E+4   6   6.5  RT (min)      lcqtof_hilic_neg_4_008_24    m/z 451.2154     0   1E+4   6   6.5  RT (min)       lcqtof_hilic_neg_4_091_60    m/z 451.2162     0   1E+4   5.5   6  RT (min)      lcqtof_hilic_neg_4_008_24    m/z 451.2824     0   1E+4   2.5   3  RT (min)      lcqtof_hilic_neg_4_052_13    m/z 451.2831     0   1E+4   3   3.5  RT (min)      lcqtof_hilic_neg_4_050_02    m/z 451.2858     0   2   4E+4   4   4.5  RT (min)      lcqtof_hilic_neg_4_040_05    m/z 451.2874     0   2   4E+4   3.5   4  RT (min)      lcqtof_hilic_neg_1_007_05    m/z 451.2876     0   1E+4   3.5   4  RT (min)      lcqtof_hilic_neg_4_002_25    m/z 451.2878     0   2   4E+4   3.5   4  RT (min)      lcqtof_hilic_neg_1_007_05    m/z 451.2977     0   1   2   3   4E+4   4   4.5  RT (min)      lcqtof_hilic_neg_4_008_24    m/z 451.3066     0   2   4E+4   1   1.5  RT (min)      lcqtof_hilic_neg_4_000_aa    m/z 451.3094     0   2   4E+4   1   1.5  RT (min)      lcqtof_hilic_neg_4_000_aa    m/z 451.3108     0   2   4E+4   1   1.5  RT (min)      lcqtof_hilic_neg_4_000_aa    m/z 451.3492     0   1   2   3   4E+4   1   1.5  RT (min)      lcqtof_hilic_neg_1_007_05    m/z 451.3611     0   1   2E+4   0.5   1  RT (min)      lcqtof_hilic_neg_4_035_33    m/z 451.4144     0   1E+4   1   1.5  RT (min)      lcqtof_hilic_neg_4_000_aa    m/z 451.4520     0   1E+4   1.5   2  RT (min)      lcqtof_hilic_neg_4_000_aa    m/z 451.8048     0   1E+4   4   4.5  RT (min)      lcqtof_hilic_neg_4_008_24    m/z 452.0181     0   2   4   6E+4   0.5   1  RT (min)      lcqtof_hilic_neg_4_050_02    m/z 452.1007     0   1E+4   3   3.5  RT (min)      lcqtof_hilic_neg_1_032_18    m/z 452.1880     0   1E+4   0.5   1  RT (min)      lcqtof_hilic_neg_4_045_26    m/z 452.2677     0   2   4E+4   1   1.5  RT (min)      lcqtof_hilic_neg_1_007_05    m/z 452.2901     0   1E+5   4   4.5  RT (min)      lcqtof_hilic_neg_4_040_05    m/z 452.2902     0   1   2   3E+5   3.5   4  RT (min)       lcqtof_hilic_neg_1_007_05    m/z 452.2902     0   1   2   3E+5   3.5   4  RT (min)      lcqtof_hilic_neg_2_028_05    m/z 452.2910     0   1   2E+4   3.5   4  RT (min)      lcqtof_hilic_neg_4_008_24    m/z 452.3062     0   2   4E+4   4   4.5  RT (min)      lcqtof_hilic_neg_4_008_24    m/z 452.3196     0   1E+4   1   1.5  RT (min)      lcqtof_hilic_neg_4_040_05    m/z 452.9222     0   1E+4   6   6.5  RT (min)       lcqtof_hilic_neg_4_044_08    m/z 452.9222     0   1   2   3E+4   5.5   6  RT (min)      lcqtof_hilic_neg_4_012_07    m/z 452.9224     0   1E+4   6   6.5  RT (min)       lcqtof_hilic_neg_4_112_62    m/z 452.9226     0   1   2   3E+4   5.5   6  RT (min)       lcqtof_hilic_neg_4_012_07    m/z 452.9228     0   2   4E+3   6.5   7  RT (min)       lcqtof_hilic_neg_4_112_62    m/z 453.0180     0   2   4   6E+4   0.5   1  RT (min)      lcqtof_hilic_neg_4_050_02    m/z 453.0993     0   2   4   6E+4   5.5   6  RT (min)      lcqtof_hilic_neg_1_000_aa    m/z 453.0994     0   2   4   6E+4   5   5.5  RT (min)      lcqtof_hilic_neg_1_000_aa    m/z 453.0995     0   2   4   6E+4   5.5   6  RT (min)      lcqtof_hilic_neg_1_000_aa    m/z 453.0998     0   2   4   6E+4   5.5   6  RT (min)      lcqtof_hilic_neg_1_000_aa    m/z 453.1013     0   2   4   6E+4   5.5   6  RT (min)      lcqtof_hilic_neg_1_000_aa    m/z 453.2405     0   2   4   6E+3   2   2.5  RT (min)      lcqtof_hilic_neg_1_007_05    m/z 453.2409     0   2   4   6E+3   2   2.5  RT (min)      lcqtof_hilic_neg_1_003_37    m/z 453.2956     0   2   4   6   8E+4   4   4.5  RT (min)      lcqtof_hilic_neg_4_040_05    m/z 453.3033     0   1E+4   4.5   5  RT (min)      lcqtof_hilic_neg_4_040_05    m/z 453.3942     0   1E+5   1   1.5  RT (min)      lcqtof_hilic_neg_4_000_aa    m/z 453.3943     0   1E+4   15   15.5  RT (min)      lcqtof_hilic_neg_4_112_62    m/z 453.3944     0   1E+5   1   1.5  RT (min)      lcqtof_hilic_neg_4_000_aa    m/z 453.3944     0   1   2E+3   16   16.5  RT (min)      lcqtof_hilic_neg_4_003_16    m/z 453.3945     0   1E+5   1   1.5  RT (min)      lcqtof_hilic_neg_4_000_aa    m/z 453.3946     0   1   2E+3   0   0.5  RT (min)       lcqtof_hilic_neg_4_007_45    m/z 453.9796     0   1E+4   7   7.5  RT (min)       lcqtof_hilic_neg_4_008_24    m/z 454.0186     0   2   4E+4   0.5   1  RT (min)      lcqtof_hilic_neg_4_050_02    m/z 454.0654     0   2   4   6E+4   6.5   7  RT (min)       lcqtof_hilic_neg_4_093_60    m/z 454.1396     0   1   2E+4   2   2.5  RT (min)      lcqtof_hilic_neg_4_050_02    m/z 454.1512     0   1E+4   2   2.5  RT (min)       lcqtof_hilic_neg_4_008_24    m/z 454.1518     0   1E+4   2   2.5  RT (min)      lcqtof_hilic_neg_4_008_24    m/z 454.1612     0   1E+4   3   3.5  RT (min)      lcqtof_hilic_neg_4_014_46    m/z 454.2267     0   1E+4   7   7.5  RT (min)      lcqtof_hilic_neg_2_052_24    m/z 454.2886     0   1E+4   2   2.5  RT (min)      lcqtof_hilic_neg_4_039_18    m/z 454.3064     0   1E+5   4   4.5  RT (min)      lcqtof_hilic_neg_4_040_05    m/z 454.3496     0   1   2   3   4E+4   1   1.5  RT (min)      lcqtof_hilic_neg_4_040_05    m/z 454.3506     0   1   2   3   4E+4   1   1.5  RT (min)      lcqtof_hilic_neg_1_007_05    m/z 454.3693     0   1E+4   0.5   1  RT (min)      lcqtof_hilic_neg_4_006_30    m/z 454.3900     0   1   2   3E+4   1   1.5  RT (min)      lcqtof_hilic_neg_4_000_aa    m/z 454.3968     0   1   2   3E+4   1   1.5  RT (min)      lcqtof_hilic_neg_4_000_aa    m/z 454.3979     0   1   2   3E+4   1   1.5  RT (min)      lcqtof_hilic_neg_4_000_aa    m/z 454.5225     0   1E+4   3.5   4  RT (min)      lcqtof_hilic_neg_1_007_05    m/z 454.9768     0   2   4   6E+4   0.5   1  RT (min)      lcqtof_hilic_neg_3_035_46    m/z 455.1002     0   1   2   3   4E+4   5.5   6  RT (min)      lcqtof_hilic_neg_1_000_aa    m/z 455.1009     0   2   4   6   8E+4   5.5   6  RT (min)      lcqtof_hilic_neg_2_000_aa    m/z 455.1012     0   1   2   3E+4   6.5   7  RT (min)       lcqtof_hilic_neg_1_008_13    m/z 455.1014     0   1E+4   6.5   7  RT (min)      lcqtof_hilic_neg_4_050_02    m/z 455.1388     0   2   4E+4   2   2.5  RT (min)      lcqtof_hilic_neg_4_050_02    m/z 455.1540     0   1E+4   2   2.5  RT (min)      lcqtof_hilic_neg_4_008_24    m/z 455.2492     0   1   2   3   4E+4   1   1.5  RT (min)      lcqtof_hilic_neg_4_000_aa    m/z 455.2750     0   1E+4   1.5   2  RT (min)      lcqtof_hilic_neg_4_055_32    m/z 455.3218     0   1   2E+4   1   1.5  RT (min)      lcqtof_hilic_neg_4_000_aa    m/z 455.3244     0   1   2   3   4E+4   4   4.5  RT (min)      lcqtof_hilic_neg_4_040_05    m/z 455.3854     0   1   2E+4   1   1.5  RT (min)      lcqtof_hilic_neg_4_000_aa    m/z 455.4605     0   1E+4   1   1.5  RT (min)      lcqtof_hilic_neg_4_013_20    m/z 455.5692     0   1   2   3   4E+4   1   1.5  RT (min)      lcqtof_hilic_neg_4_013_20    m/z 455.7348     0   1E+4   1   1.5  RT (min)      lcqtof_hilic_neg_4_013_20    m/z 455.9762     0   1E+4   0.5   1  RT (min)      lcqtof_hilic_neg_3_035_46    m/z 456.0184     0   1E+4   0.5   1  RT (min)      lcqtof_hilic_neg_4_050_02    m/z 456.0213     0   1   2   3E+4   4.5   5  RT (min)       lcqtof_hilic_neg_4_027_19    m/z 456.1340     0   1E+4   6.5   7  RT (min)      lcqtof_hilic_neg_4_008_24    m/z 456.1571     0   1   2   3E+3   2   2.5  RT (min)      lcqtof_hilic_neg_1_049_55    m/z 456.2332     0   1   2E+4   6.5   7  RT (min)      lcqtof_hilic_neg_4_008_24    m/z 456.2916     0   1   2E+4   1   1.5  RT (min)      lcqtof_hilic_neg_2_028_05    m/z 456.3211     0   2   4   6E+4   4   4.5  RT (min)      lcqtof_hilic_neg_4_040_05    m/z 456.3224     0   1   2E+5   4   4.5  RT (min)      lcqtof_hilic_neg_4_040_05    m/z 456.3683     0   2   4   6   8E+4   0.5   1  RT (min)       lcqtof_hilic_neg_3_003_20    m/z 456.3687     0   1   2   3   4E+4   1   1.5  RT (min)      lcqtof_hilic_neg_4_002_25    m/z 456.3914     0   1E+4   1   1.5  RT (min)      lcqtof_hilic_neg_4_008_24    m/z 456.7735     0   1E+4   4.5   5  RT (min)      lcqtof_hilic_neg_4_050_02    m/z 456.8084     0   1   2E+4   7   7.5  RT (min)       lcqtof_hilic_neg_4_047_06    m/z 456.8230     0   1   2E+4   8.5   9  RT (min)      lcqtof_hilic_neg_1_000_aa    m/z 456.8232     0   1   2E+4   8.5   9  RT (min)      lcqtof_hilic_neg_1_000_aa    m/z 456.9733     0   1E+4   0.5   1  RT (min)      lcqtof_hilic_neg_3_035_46    m/z 457.0200     0   1   2   3   4E+4   0.5   1  RT (min)      lcqtof_hilic_neg_4_050_02    m/z 457.1146     0   1E+4   0.5   1  RT (min)      lcqtof_hilic_neg_4_050_02    m/z 457.1711     0   1E+4   2   2.5  RT (min)      lcqtof_hilic_neg_4_052_13    m/z 457.2157     0   1E+4   6   6.5  RT (min)      lcqtof_hilic_neg_1_037_24    m/z 457.2342     0   1   2E+4   6   6.5  RT (min)       lcqtof_hilic_neg_4_090_60    m/z 457.2487     0   2   4   6E+4   1   1.5  RT (min)      lcqtof_hilic_neg_4_000_aa    m/z 457.2504     0   2   4   6E+4   1   1.5  RT (min)      lcqtof_hilic_neg_4_000_aa    m/z 457.2697     0   1E+4   0.5   1  RT (min)      lcqtof_hilic_neg_4_005_29    m/z 457.2803     0   1   2   3   4E+4   1   1.5  RT (min)      lcqtof_hilic_neg_4_040_05    m/z 457.3124     0   2   4   6E+3   2   2.5  RT (min)      lcqtof_hilic_neg_4_052_13    m/z 457.3144     0   1   2   3E+3   2.5   3  RT (min)      lcqtof_hilic_neg_4_007_45    m/z 457.3158     0   2   4   6E+4   1   1.5  RT (min)      lcqtof_hilic_neg_4_000_aa    m/z 457.3762     0   1E+4   0.5   1  RT (min)      lcqtof_hilic_neg_3_003_20    m/z 458.0180     0   1   2E+4   4.5   5  RT (min)       lcqtof_hilic_neg_4_027_19    m/z 458.2206     0   1   2E+4   5   5.5  RT (min)      lcqtof_hilic_neg_4_008_24    m/z 458.2221     0   1E+4   0.5   1  RT (min)      lcqtof_hilic_neg_3_003_20    m/z 458.2529     0   1   2E+4   1   1.5  RT (min)      lcqtof_hilic_neg_4_000_aa    m/z 458.3469     0   1E+4   15.5   16  RT (min)      lcqtof_hilic_neg_4_000_aa    m/z 458.3619     0   1E+4   1   1.5  RT (min)      lcqtof_hilic_neg_4_013_20    m/z 458.9743     0   1E+4   5.5   6  RT (min)       lcqtof_hilic_neg_4_050_02    m/z 458.9746     0   1E+4   5   5.5  RT (min)      lcqtof_hilic_neg_4_050_02    m/z 458.9746     0   1E+4   5.5   6  RT (min)       lcqtof_hilic_neg_4_050_02    m/z 458.9747     0   1E+4   5.5   6  RT (min)      lcqtof_hilic_neg_4_050_02    m/z 459.0575     0   1E+4   0.5   1  RT (min)      lcqtof_hilic_neg_4_024_40    m/z 459.1194     0   1E+4   0.5   1  RT (min)      lcqtof_hilic_neg_4_050_02    m/z 459.2167     0   1E+4   6   6.5  RT (min)      lcqtof_hilic_neg_4_008_24    m/z 459.2230     0   1E+4   5   5.5  RT (min)      lcqtof_hilic_neg_4_008_24    m/z 459.2313     0   1E+4   1   1.5  RT (min)      lcqtof_hilic_neg_4_000_aa    m/z 459.2930     0   1E+4   14.5   15  RT (min)      lcqtof_hilic_neg_4_039_18    m/z 459.2934     0   1E+4   2.5   3  RT (min)       lcqtof_hilic_neg_4_047_06    m/z 459.2934     0   2   4   6   8E+3   3.5   4  RT (min)      lcqtof_hilic_neg_4_055_32    m/z 459.2936     0   1E+4   4   4.5  RT (min)      lcqtof_hilic_neg_4_016_37    m/z 459.3229     0   1   2   3E+5   1   1.5  RT (min)      lcqtof_hilic_neg_4_000_aa    m/z 459.3242     0   1   2   3E+4   15.5   16  RT (min)      lcqtof_hilic_neg_4_000_aa    m/z 459.3257     0   1   2   3E+5   1   1.5  RT (min)      lcqtof_hilic_neg_4_000_aa    m/z 459.3258     0   1   2   3E+4   15.5   16  RT (min)      lcqtof_hilic_neg_4_000_aa    m/z 459.3266     0   1E+4   15.5   16  RT (min)      lcqtof_hilic_neg_1_007_05    m/z 459.3278     0   1   2   3E+5   1   1.5  RT (min)      lcqtof_hilic_neg_4_000_aa    m/z 459.9550     0   1E+4   3   3.5  RT (min)       lcqtof_hilic_neg_4_039_18    m/z 460.0414     0   1   2E+4   1   1.5  RT (min)      lcqtof_hilic_neg_4_039_18    m/z 460.0957     0   1   2   3   4E+4   3   3.5  RT (min)      lcqtof_hilic_neg_4_108_62    m/z 460.1656     0   1E+5   5.5   6  RT (min)      lcqtof_hilic_neg_1_000_aa    m/z 460.1658     0   1E+5   5   5.5  RT (min)      lcqtof_hilic_neg_1_000_aa    m/z 460.1660     0   1E+5   5   5.5  RT (min)      lcqtof_hilic_neg_1_000_aa    m/z 460.2958     0   2   4   6E+3   1.5   2  RT (min)      lcqtof_hilic_neg_4_035_33    m/z 460.2972     0   1E+4   1   1.5  RT (min)      lcqtof_hilic_neg_4_040_05    m/z 460.3236     0   1E+5   1   1.5  RT (min)      lcqtof_hilic_neg_4_000_aa    m/z 460.3292     0   1E+4   15.5   16  RT (min)      lcqtof_hilic_neg_4_000_aa    m/z 460.3294     0   1E+5   1   1.5  RT (min)      lcqtof_hilic_neg_4_000_aa    m/z 460.3296     0   1E+5   1   1.5  RT (min)      lcqtof_hilic_neg_4_000_aa    m/z 461.0980     0   1   2E+4   1   1.5  RT (min)      lcqtof_hilic_neg_4_000_aa    m/z 461.0992     0   1   2E+4   1   1.5  RT (min)      lcqtof_hilic_neg_4_000_aa    m/z 461.1000     0   1   2E+4   1   1.5  RT (min)      lcqtof_hilic_neg_4_000_aa    m/z 461.1322     0   1E+4   2.5   3  RT (min)      lcqtof_hilic_neg_4_027_19    m/z 461.1574     0   1   2E+3   18.5   19  RT (min)      lcqtof_hilic_neg_4_004_47    m/z 461.1588     0   2   4   6E+3   15.5   16  RT (min)       lcqtof_hilic_neg_4_058_39    m/z 461.1588     0   1E+4   15   15.5  RT (min)       lcqtof_hilic_neg_4_000_aa    m/z 461.1601     0   1   2E+3   18   18.5  RT (min)      lcqtof_hilic_neg_4_037_39    m/z 461.1679     0   1   2E+4   5.5   6  RT (min)      lcqtof_hilic_neg_1_000_aa    m/z 461.1689     0   1   2E+4   5   5.5  RT (min)      lcqtof_hilic_neg_1_000_aa    m/z 461.1782     0   1   2   3E+4   3   3.5  RT (min)      lcqtof_hilic_neg_4_004_47    m/z 461.2831     0   1E+4   0.5   1  RT (min)      lcqtof_hilic_neg_4_000_aa    m/z 461.2867     0   1E+4   1   1.5  RT (min)      lcqtof_hilic_neg_4_000_aa    m/z 461.3088     0   1   2E+4   1   1.5  RT (min)      lcqtof_hilic_neg_4_040_05    m/z 461.3455     0   2   4E+4   1   1.5  RT (min)      lcqtof_hilic_neg_4_000_aa    m/z 462.2684     0   1   2E+4   4   4.5  RT (min)      lcqtof_hilic_neg_4_055_32    m/z 462.2852     0   2   4   6   8E+4   3   3.5  RT (min)      lcqtof_hilic_neg_2_053_18    m/z 462.2861     0   1E+5   3   3.5  RT (min)      lcqtof_hilic_neg_4_039_18    m/z 462.2971     0   1E+5   1   1.5  RT (min)      lcqtof_hilic_neg_1_007_05    m/z 462.2972     0   1E+5   1   1.5  RT (min)      lcqtof_hilic_neg_4_040_05    m/z 462.3502     0   1E+4   1   1.5  RT (min)      lcqtof_hilic_neg_4_000_aa    m/z 463.0396     0   1E+4   0.5   1  RT (min)      lcqtof_hilic_neg_4_039_18    m/z 463.0399     0   1E+4   1   1.5  RT (min)      lcqtof_hilic_neg_4_039_18    m/z 463.1216     0   2   4   6E+4   4.5   5  RT (min)       lcqtof_hilic_neg_4_052_13    m/z 463.1220     0   1   2E+4   4.5   5  RT (min)      lcqtof_hilic_neg_4_019_03    m/z 463.1220     0   1   2E+4   4.5   5  RT (min)      lcqtof_hilic_neg_4_050_02    m/z 463.1223     0   1   2E+4   4   4.5  RT (min)      lcqtof_hilic_neg_4_050_02    m/z 463.1224     0   1   2E+4   4   4.5  RT (min)      lcqtof_hilic_neg_4_019_03    m/z 463.2835     0   1E+4   1   1.5  RT (min)      lcqtof_hilic_neg_1_031_16    m/z 463.2891     0   1   2E+4   3   3.5  RT (min)      lcqtof_hilic_neg_2_053_18    m/z 463.2896     0   1   2   3   4E+4   3   3.5  RT (min)      lcqtof_hilic_neg_4_039_18    m/z 463.2914     0   2   4   6   8E+3   1   1.5  RT (min)      lcqtof_hilic_neg_4_063_16    m/z 463.2964     0   1   2   3E+4   1   1.5  RT (min)      lcqtof_hilic_neg_1_007_05    m/z 463.3012     0   1   2   3E+4   1   1.5  RT (min)      lcqtof_hilic_neg_4_040_05    m/z 463.9439     0   1E+4   7   7.5  RT (min)       lcqtof_hilic_neg_4_059_14    m/z 463.9536     0   2   4   6   8E+3   7   7.5  RT (min)      lcqtof_hilic_neg_4_034_31    m/z 464.0176     0   1E+4   1   1.5  RT (min)      lcqtof_hilic_neg_2_045_11    m/z 464.0498     0   1E+4   6   6.5  RT (min)       lcqtof_hilic_neg_4_084_59    m/z 464.0912     0   1   2E+4   0.5   1  RT (min)      lcqtof_hilic_neg_3_035_46    m/z 464.1151     0   1E+4   0.5   1  RT (min)      lcqtof_hilic_neg_1_062_17    m/z 464.1251     0   1E+4   4.5   5  RT (min)      lcqtof_hilic_neg_4_052_13    m/z 464.1674     0   1E+4   2   2.5  RT (min)      lcqtof_hilic_neg_4_014_46    m/z 464.1969     0   1E+4   4.5   5  RT (min)      lcqtof_hilic_neg_4_019_03    m/z 464.2738     0   1E+4   0.5   1  RT (min)      lcqtof_hilic_neg_4_005_29    m/z 464.2844     0   1   2E+4   0.5   1  RT (min)      lcqtof_hilic_neg_2_062_17    m/z 464.3091     0   1   2E+5   1   1.5  RT (min)      lcqtof_hilic_neg_4_040_05    m/z 464.4106     0   1   2E+4   3.5   4  RT (min)      lcqtof_hilic_neg_4_039_18    m/z 464.5197     0   2   4   6E+4   3.5   4  RT (min)      lcqtof_hilic_neg_4_039_18    m/z 464.6881     0   1E+4   3.5   4  RT (min)      lcqtof_hilic_neg_4_039_18    m/z 465.0384     0   1E+4   0.5   1  RT (min)      lcqtof_hilic_neg_4_039_18    m/z 465.1199     0   1   2E+4   4.5   5  RT (min)      lcqtof_hilic_neg_4_052_13    m/z 465.1269     0   1   2E+4   2.5   3  RT (min)       lcqtof_hilic_neg_4_091_60    m/z 465.1271     0   1E+4   1   1.5  RT (min)       lcqtof_hilic_neg_4_097_60    m/z 465.3036     0   1E+4   15.5   16  RT (min)      lcqtof_hilic_neg_4_000_aa    m/z 465.3048     0   1   2   3   4E+3   2   2.5  RT (min)      lcqtof_hilic_neg_4_003_16    m/z 465.3055     0   1   2   3   4E+3   4   4.5  RT (min)      lcqtof_hilic_neg_4_016_37    m/z 465.3066     0   1   2   3E+4   1.5   2  RT (min)      lcqtof_hilic_neg_4_034_31    m/z 465.3196     0   1E+4   0.5   1  RT (min)      lcqtof_hilic_neg_3_003_20    m/z 465.3396     0   2   4   6E+4   1   1.5  RT (min)      lcqtof_hilic_neg_4_000_aa    m/z 465.3953     0   1E+4   1   1.5  RT (min)      lcqtof_hilic_neg_4_000_aa    m/z 465.3988     0   2   4   6E+4   0.5   1  RT (min)      lcqtof_hilic_neg_4_050_02    m/z 465.4298     0   1E+5   1   1.5  RT (min)      lcqtof_hilic_neg_4_000_aa    m/z 465.4650     0   1E+4   1.5   2  RT (min)      lcqtof_hilic_neg_4_000_aa    m/z 465.5698     0   1E+5   0.5   1  RT (min)      lcqtof_hilic_neg_1_031_16    m/z 465.7664     0   1   2   3   4E+4   0.5   1  RT (min)      lcqtof_hilic_neg_1_031_16    m/z 465.9108     0   1   2   3   4E+4   0.5   1  RT (min)      lcqtof_hilic_neg_2_021_17    m/z 466.1266     0   1E+4   6   6.5  RT (min)      lcqtof_hilic_neg_4_048_23    m/z 466.1602     0   1E+4   2   2.5  RT (min)      lcqtof_hilic_neg_4_050_02    m/z 466.3064     0   1   2   3E+4   3   3.5  RT (min)       lcqtof_hilic_neg_4_040_05    m/z 466.3070     0   1E+4   3   3.5  RT (min)      lcqtof_hilic_neg_4_008_24    m/z 466.3272     0   1   2E+5   0.5   1  RT (min)      lcqtof_hilic_neg_3_035_46    m/z 466.3426     0   1   2E+4   1   1.5  RT (min)      lcqtof_hilic_neg_4_000_aa    m/z 466.4347     0   1   2   3E+4   1   1.5  RT (min)      lcqtof_hilic_neg_4_000_aa    m/z 466.6312     0   2   4   6   8E+4   0.5   1  RT (min)      lcqtof_hilic_neg_1_031_16    m/z 466.7557     0   2   4   6   8E+4   0.5   1  RT (min)      lcqtof_hilic_neg_1_031_16    m/z 466.9011     0   2   4   6E+4   0.5   1  RT (min)      lcqtof_hilic_neg_1_031_16    m/z 467.1584     0   2   4E+4   2   2.5  RT (min)      lcqtof_hilic_neg_4_050_02    m/z 467.1632     0   2   4   6E+4   0.5   1  RT (min)      lcqtof_hilic_neg_3_003_20    m/z 467.1843     0   1E+4   6.5   7  RT (min)      lcqtof_hilic_neg_4_008_24    m/z 467.2068     0   1   2   3   4E+4   0.5   1  RT (min)      lcqtof_hilic_neg_4_050_02    m/z 467.2080     0   2   4   6E+3   0.5   1  RT (min)      lcqtof_hilic_neg_4_023_15    m/z 467.2206     0   1   2   3E+4   0.5   1  RT (min)       lcqtof_hilic_neg_4_027_19    m/z 467.2468     0   1   2E+4   0.5   1  RT (min)      lcqtof_hilic_neg_4_052_13    m/z 467.3120     0   1E+4   3.5   4  RT (min)      lcqtof_hilic_neg_4_039_18    m/z 467.3292     0   1   2   3   4E+4   1   1.5  RT (min)      lcqtof_hilic_neg_2_028_05    m/z 467.3352     0   1   2   3E+4   0.5   1  RT (min)      lcqtof_hilic_neg_4_023_15    m/z 467.4072     0   2   4   6E+4   1   1.5  RT (min)      lcqtof_hilic_neg_4_000_aa    m/z 467.4092     0   2   4   6E+4   1   1.5  RT (min)      lcqtof_hilic_neg_4_000_aa    m/z 467.4098     0   2   4E+1   14.5   15  RT (min)      lcqtof_hilic_neg_4_083_59    m/z 467.4099     0   2   4   6E+4   1   1.5  RT (min)      lcqtof_hilic_neg_4_000_aa    m/z 467.4444     0   1   2   3   4E+4   1   1.5  RT (min)      lcqtof_hilic_neg_4_000_aa    m/z 468.1591     0   1E+4   6   6.5  RT (min)       lcqtof_hilic_neg_4_091_60    m/z 468.1662     0   1E+4   0.5   1  RT (min)      lcqtof_hilic_neg_3_003_20    m/z 468.3214     0   1   2   3   4E+4   3   3.5  RT (min)      lcqtof_hilic_neg_4_040_05    m/z 468.3221     0   1   2   3   4E+4   2.5   3  RT (min)       lcqtof_hilic_neg_4_040_05    m/z 468.3257     0   1E+4   1   1.5  RT (min)       lcqtof_hilic_neg_4_040_05    m/z 468.3426     0   1   2   3E+4   1   1.5  RT (min)      lcqtof_hilic_neg_1_007_05    m/z 468.4120     0   1   2E+4   1   1.5  RT (min)      lcqtof_hilic_neg_4_000_aa    m/z 468.4125     0   1   2E+4   1   1.5  RT (min)      lcqtof_hilic_neg_4_000_aa    m/z 468.4126     0   1   2E+4   1   1.5  RT (min)      lcqtof_hilic_neg_4_000_aa    m/z 468.4494     0   1E+4   1   1.5  RT (min)      lcqtof_hilic_neg_4_000_aa    m/z 468.8872     0   1   2   3E+4   1   1.5  RT (min)      lcqtof_hilic_neg_4_007_45    m/z 468.8872     0   1E+5   1   1.5  RT (min)       lcqtof_hilic_neg_4_004_47    m/z 468.8956     0   2   4E+3   5   5.5  RT (min)       lcqtof_hilic_neg_4_058_39    m/z 468.9501     0   2   4   6E+3   7.5   8  RT (min)      lcqtof_hilic_neg_4_045_26    m/z 468.9904     0   2   4   6E+4   0.5   1  RT (min)      lcqtof_hilic_neg_3_035_46    m/z 468.9926     0   2   4   6E+4   0.5   1  RT (min)      lcqtof_hilic_neg_3_035_46    m/z 469.1178     0   1E+4   5.5   6  RT (min)      lcqtof_hilic_neg_4_050_02    m/z 469.1178     0   1E+4   1.5   2  RT (min)      lcqtof_hilic_neg_1_008_13    m/z 469.1198     0   1   2   3E+4   4.5   5  RT (min)       lcqtof_hilic_neg_1_037_24    m/z 469.1357     0   1E+4   6   6.5  RT (min)      lcqtof_hilic_neg_1_035_23    m/z 469.1398     0   2   4   6E+3   4.5   5  RT (min)      lcqtof_hilic_neg_1_007_05    m/z 469.1401     0   1   2E+3   4.5   5  RT (min)      lcqtof_hilic_neg_1_037_24    m/z 469.1417     0   1   2   3   4E+3   3.5   4  RT (min)      lcqtof_hilic_neg_4_038_10    m/z 469.1716     0   1   2   3E+4   0.5   1  RT (min)      lcqtof_hilic_neg_4_021_21    m/z 469.1998     0   2   4   6E+4   0.5   1  RT (min)      lcqtof_hilic_neg_3_003_20    m/z 469.2016     0   2   4   6E+4   0.5   1  RT (min)      lcqtof_hilic_neg_3_003_20    m/z 469.2332     0   1E+4   0.5   1  RT (min)      lcqtof_hilic_neg_2_015_06    m/z 469.2420     0   1E+4   6   6.5  RT (min)      lcqtof_hilic_neg_4_008_24    m/z 469.2628     0   2   4   6   8E+4   0.5   1  RT (min)      lcqtof_hilic_neg_4_060_13    m/z 469.2980     0   1E+5   0.5   1  RT (min)      lcqtof_hilic_neg_3_035_46    m/z 469.3102     0   1   2   3E+4   0.5   1  RT (min)      lcqtof_hilic_neg_1_031_16    m/z 469.4000     0   1   2E+4   1   1.5  RT (min)      lcqtof_hilic_neg_4_000_aa    m/z 469.8874     0   1   2   3E+4   1   1.5  RT (min)      lcqtof_hilic_neg_4_004_47    m/z 469.9455     0   1E+4   0.5   1  RT (min)      lcqtof_hilic_neg_4_004_47    m/z 469.9918     0   1E+4   0.5   1  RT (min)      lcqtof_hilic_neg_3_035_46    m/z 470.0470     0   1E+4   5.5   6  RT (min)      lcqtof_hilic_neg_1_037_24    m/z 470.1501     0   1E+4   5   5.5  RT (min)      lcqtof_hilic_neg_1_000_aa    m/z 470.2000     0   2   4   6   8E+4   6   6.5  RT (min)      lcqtof_hilic_neg_4_014_46    m/z 470.2070     0   1E+4   0.5   1  RT (min)      lcqtof_hilic_neg_3_003_20    m/z 470.2083     0   1E+4   0.5   1  RT (min)      lcqtof_hilic_neg_3_003_20    m/z 470.2392     0   1E+4   5   5.5  RT (min)      lcqtof_hilic_neg_4_008_24    m/z 470.2655     0   1   2E+4   0.5   1  RT (min)      lcqtof_hilic_neg_4_060_13    m/z 470.2684     0   2   4   6E+4   6.5   7  RT (min)       lcqtof_hilic_neg_1_037_24    m/z 470.3014     0   1   2   3E+4   0.5   1  RT (min)      lcqtof_hilic_neg_3_035_46    m/z 470.3844     0   1E+4   1   1.5  RT (min)      lcqtof_hilic_neg_4_000_aa    m/z 470.9977     0   1E+4   0.5   1  RT (min)      lcqtof_hilic_neg_3_035_46    m/z 471.0699     0   1E+4   7   7.5  RT (min)       lcqtof_hilic_neg_4_091_60    m/z 471.0702     0   1E+4   7.5   8  RT (min)      lcqtof_hilic_neg_4_040_05    m/z 471.1168     0   1E+4   4.5   5  RT (min)      lcqtof_hilic_neg_2_052_24    m/z 471.1502     0   1E+4   6   6.5  RT (min)      lcqtof_hilic_neg_4_040_05    m/z 471.1515     0   1E+4   6.5   7  RT (min)      lcqtof_hilic_neg_4_008_24    m/z 471.1704     0   1E+4   4   4.5  RT (min)       lcqtof_hilic_neg_4_048_23    m/z 471.1713     0   1E+4   4   4.5  RT (min)      lcqtof_hilic_neg_4_011_35    m/z 471.2207     0   1   2   3E+3   2   2.5  RT (min)      lcqtof_hilic_neg_4_004_47    m/z 471.2410     0   1E+4   5   5.5  RT (min)      lcqtof_hilic_neg_4_008_24    m/z 471.2676     0   1E+4   1   1.5  RT (min)      lcqtof_hilic_neg_4_000_aa    m/z 471.3449     0   1E+4   1   1.5  RT (min)      lcqtof_hilic_neg_2_028_05    m/z 471.4097     0   1E+4   0.5   1  RT (min)      lcqtof_hilic_neg_4_008_24    m/z 472.0086     0   1E+4   15   15.5  RT (min)      lcqtof_hilic_neg_4_000_aa    m/z 472.2478     0   1E+4   0.5   1  RT (min)      lcqtof_hilic_neg_2_028_05    m/z 472.2842     0   1   2   3E+4   1   1.5  RT (min)      lcqtof_hilic_neg_4_000_aa    m/z 472.2868     0   1   2   3E+4   1   1.5  RT (min)      lcqtof_hilic_neg_4_000_aa    m/z 472.2878     0   1   2   3E+4   1   1.5  RT (min)      lcqtof_hilic_neg_4_000_aa    m/z 472.9439     0   1E+4   7   7.5  RT (min)      lcqtof_hilic_neg_4_050_02    m/z 472.9519     0   1E+4   7   7.5  RT (min)      lcqtof_hilic_neg_4_050_02    m/z 473.0780     0   1E+4   6   6.5  RT (min)      lcqtof_hilic_neg_1_035_23    m/z 473.0952     0   2   4E+4   7   7.5  RT (min)       lcqtof_hilic_neg_4_008_24    m/z 473.1213     0   1E+4   5.5   6  RT (min)      lcqtof_hilic_neg_4_048_23    m/z 473.1495     0   1E+4   5   5.5  RT (min)      lcqtof_hilic_neg_4_050_02    m/z 473.1517     0   1E+4   4.5   5  RT (min)      lcqtof_hilic_neg_4_027_19    m/z 473.1960     0   1   2   3   4E+5   6   6.5  RT (min)       lcqtof_hilic_neg_4_048_23    m/z 473.2901     0   2   4E+3   3   3.5  RT (min)      lcqtof_hilic_neg_4_011_35    m/z 473.2902     0   2   4E+3   3.5   4  RT (min)      lcqtof_hilic_neg_4_002_25    m/z 473.2930     0   2   4E+4   15.5   16  RT (min)       lcqtof_hilic_neg_4_000_aa    m/z 473.2936     0   2   4E+4   15.5   16  RT (min)       lcqtof_hilic_neg_4_000_aa    m/z 473.2936     0   2   4E+4   15.5   16  RT (min)      lcqtof_hilic_neg_4_000_aa    m/z 473.2943     0   1E+6   1   1.5  RT (min)      lcqtof_hilic_neg_4_000_aa    m/z 473.2944     0   2   4   6E+3   1.5   2  RT (min)       lcqtof_hilic_neg_4_034_31    m/z 473.2944     0   2   4   6E+3   2.5   3  RT (min)       lcqtof_hilic_neg_4_047_06    m/z 473.2944     0   1E+6   1   1.5  RT (min)      lcqtof_hilic_neg_4_000_aa    m/z 473.2946     0   2   4E+3   1.5   2  RT (min)       lcqtof_hilic_neg_4_019_03    m/z 473.2948     0   2   4   6E+3   1.5   2  RT (min)      lcqtof_hilic_neg_4_034_31    m/z 473.2969     0   1   2   3   4E+3   0   0.5  RT (min)       lcqtof_hilic_neg_4_005_29    m/z 473.2970     0   1   2   3   4E+3   19   19.5  RT (min)       lcqtof_hilic_neg_4_008_24    m/z 473.2970     0   1   2   3   4E+3   19.5   20  RT (min)       lcqtof_hilic_neg_4_019_03    m/z 473.2978     0   1   2   3   4E+3   17   18   19  RT (min)       lcqtof_hilic_neg_4_000_dd    m/z 473.2978     0   1   2   3   4E+3   19  RT (min)       lcqtof_hilic_neg_4_013_20    m/z 473.2984     0   1   2   3   4E+3   19   19.5  RT (min)       lcqtof_hilic_neg_4_002_25    m/z 473.2988     0   1   2   3   4E+3   17.5   18  RT (min)       lcqtof_hilic_neg_4_006_30    m/z 473.2989     0   1   2   3   4E+3   17   17.5  RT (min)       lcqtof_hilic_neg_4_013_20    m/z 473.2991     0   1   2   3E+3   16   16.5  RT (min)       lcqtof_hilic_neg_4_016_37    m/z 473.2992     0   1   2   3   4E+3   16.5   17   17.5  RT (min)       lcqtof_hilic_neg_4_000_bb    m/z 473.3001     0   1   2   3   4E+3   16   16.5  RT (min)       lcqtof_hilic_neg_4_004_47    m/z 473.3002     0   1   2   3   4E+3   16   16.5  RT (min)      lcqtof_hilic_neg_4_008_24    m/z 473.3002     0   1   2   3   4E+3   16.5   17  RT (min)       lcqtof_hilic_neg_4_009_50    m/z 473.3008     0   1   2   3E+3   16   16.5  RT (min)       lcqtof_hilic_neg_4_025_52    m/z 473.3010     0   1   2   3   4E+3   16.5   17  RT (min)       lcqtof_hilic_neg_4_000_dd    m/z 473.3102     0   2   4E+4   0.5   1  RT (min)      lcqtof_hilic_neg_4_003_16    m/z 473.3188     0   1   2E+4   0.5   1  RT (min)      lcqtof_hilic_neg_4_032_01    m/z 473.3767     0   1   2   3   4E+4   0.5   1  RT (min)      lcqtof_hilic_neg_3_035_46    m/z 473.9601     0   1   2   3E+4   1   1.5  RT (min)       lcqtof_hilic_neg_4_046_41    m/z 474.0056     0   1E+4   15   15.5  RT (min)      lcqtof_hilic_neg_4_000_aa    m/z 474.0379     0   1E+4   2.5   3  RT (min)      lcqtof_hilic_neg_4_045_26    m/z 474.0458     0   1   2E+4   1   1.5  RT (min)      lcqtof_hilic_neg_2_045_11    m/z 474.1453     0   1E+4   7   7.5  RT (min)      lcqtof_hilic_neg_4_048_23    m/z 474.1676     0   2   4E+4   1   1.5  RT (min)      lcqtof_hilic_neg_4_000_aa    m/z 474.1962     0   1   2   3   4E+4   6   6.5  RT (min)       lcqtof_hilic_neg_4_008_24    m/z 474.1994     0   2   4   6E+4   6   6.5  RT (min)      lcqtof_hilic_neg_4_048_23    m/z 474.2335     0   1E+4   6   6.5  RT (min)      lcqtof_hilic_neg_1_037_24    m/z 474.2628     0   1E+4   0.5   1  RT (min)      lcqtof_hilic_neg_2_028_05    m/z 474.2837     0   1E+4   15.5   16  RT (min)      lcqtof_hilic_neg_4_000_aa    m/z 474.2960     0   1   2   3   4E+5   1   1.5  RT (min)      lcqtof_hilic_neg_4_000_aa    m/z 474.2971     0   1   2   3   4E+5   1   1.5  RT (min)      lcqtof_hilic_neg_4_000_aa    m/z 474.2982     0   1   2   3   4E+3   2   2.5  RT (min)      lcqtof_hilic_neg_4_005_29    m/z 474.3806     0   1E+4   0.5   1  RT (min)      lcqtof_hilic_neg_3_035_46    m/z 474.9351     0   1E+4   7   7.5  RT (min)      lcqtof_hilic_neg_4_050_02    m/z 474.9351     0   1E+4   7   7.5  RT (min)      lcqtof_hilic_neg_4_050_02    m/z 475.1243     0   1   2   3   4E+4   3.5   4  RT (min)      lcqtof_hilic_neg_1_032_18    m/z 475.1739     0   1E+4   15   15.5  RT (min)       lcqtof_hilic_neg_4_000_aa    m/z 475.2044     0   1E+4   6   6.5  RT (min)      lcqtof_hilic_neg_4_048_23    m/z 475.2164     0   1   2   3E+4   6   6.5  RT (min)      lcqtof_hilic_neg_4_008_24    m/z 475.2403     0   1   2   3E+4   0.5   1  RT (min)      lcqtof_hilic_neg_2_026_28    m/z 475.2758     0   1E+5   1   1.5  RT (min)      lcqtof_hilic_neg_4_000_aa    m/z 475.3103     0   1   2   3E+4   0.5   1  RT (min)      lcqtof_hilic_neg_3_035_46    m/z 475.3262     0   2   4E+3   2   2.5  RT (min)      lcqtof_hilic_neg_4_003_16    m/z 475.3401     0   2   4   6   8E+4   1   1.5  RT (min)      lcqtof_hilic_neg_2_028_05    m/z 475.9571     0   1   2   3E+4   1   1.5  RT (min)       lcqtof_hilic_neg_4_046_41    m/z 476.0475     0   1   2E+4   6.5   7  RT (min)       lcqtof_hilic_neg_4_090_60    m/z 476.1632     0   1   2E+4   1   1.5  RT (min)      lcqtof_hilic_neg_4_000_aa    m/z 476.1645     0   1   2E+4   1   1.5  RT (min)      lcqtof_hilic_neg_4_000_aa    m/z 476.2738     0   1   2E+4   1   1.5  RT (min)      lcqtof_hilic_neg_4_000_aa    m/z 476.2808     0   1   2E+4   1   1.5  RT (min)      lcqtof_hilic_neg_4_000_aa    m/z 476.3568     0   1   2E+4   1   1.5  RT (min)      lcqtof_hilic_neg_2_028_05    m/z 476.4994     0   1   2E+4   4   4.5  RT (min)      lcqtof_hilic_neg_4_040_05    m/z 477.1437     0   1   2E+4   5   5.5  RT (min)      lcqtof_hilic_neg_1_000_aa    m/z 477.1563     0   1   2E+4   1   1.5  RT (min)      lcqtof_hilic_neg_4_000_aa    m/z 477.1615     0   1   2E+3   2.5   3  RT (min)      lcqtof_hilic_neg_1_016_28    m/z 477.2646     0   1E+4   1   1.5  RT (min)      lcqtof_hilic_neg_4_000_aa    m/z 477.3555     0   1E+5   1   1.5  RT (min)      lcqtof_hilic_neg_1_007_05    m/z 478.3166     0   1E+4   3   3.5  RT (min)      lcqtof_hilic_neg_4_039_18    m/z 478.3602     0   1E+4   1   1.5  RT (min)      lcqtof_hilic_neg_4_040_05    m/z 478.7900     0   2   4   6E+5   8   8.5  RT (min)      lcqtof_hilic_neg_1_000_aa    m/z 479.1367     0   1   2   3E+4   6   6.5  RT (min)      lcqtof_hilic_neg_4_048_23    m/z 479.1597     0   1E+4   0.5   1  RT (min)      lcqtof_hilic_neg_4_058_39    m/z 479.1768     0   1   2   3E+4   6   6.5  RT (min)      lcqtof_hilic_neg_4_008_24    m/z 479.2828     0   1E+4   1   1.5  RT (min)      lcqtof_hilic_neg_4_000_aa    m/z 479.2846     0   1E+4   1   1.5  RT (min)      lcqtof_hilic_neg_4_000_aa    m/z 479.3529     0   2   4   6   8E+4   1   1.5  RT (min)      lcqtof_hilic_neg_4_000_aa    m/z 479.3673     0   2   4   6   8E+4   1   1.5  RT (min)      lcqtof_hilic_neg_4_040_05    m/z 479.3906     0   1   2E+4   0.5   1  RT (min)      lcqtof_hilic_neg_4_005_29    m/z 479.4434     0   1   2   3E+4   1   1.5  RT (min)      lcqtof_hilic_neg_4_000_aa    m/z 479.4486     0   1   2   3E+4   1   1.5  RT (min)      lcqtof_hilic_neg_4_000_aa    m/z 480.1832     0   1E+4   7   7.5  RT (min)      lcqtof_hilic_neg_2_022_46    m/z 480.1833     0   1E+4   7   7.5  RT (min)      lcqtof_hilic_neg_4_014_46    m/z 480.2384     0   1E+4   2   2.5  RT (min)      lcqtof_hilic_neg_4_002_25    m/z 480.3084     0   1E+6   4   4.5  RT (min)      lcqtof_hilic_neg_4_055_32    m/z 480.3574     0   1   2E+5   1   1.5  RT (min)      lcqtof_hilic_neg_4_040_05    m/z 480.4468     0   1E+4   1   1.5  RT (min)      lcqtof_hilic_neg_4_000_aa    m/z 480.7940     0   1E+4   8   8.5  RT (min)      lcqtof_hilic_neg_1_000_aa    m/z 481.1372     0   1E+4   2   2.5  RT (min)      lcqtof_hilic_neg_4_050_02    m/z 481.2178     0   1E+4   3   3.5  RT (min)      lcqtof_hilic_neg_4_005_29    m/z 481.3282     0   2   4   6E+4   1   1.5  RT (min)      lcqtof_hilic_neg_4_000_aa    m/z 481.3286     0   2   4   6E+4   1   1.5  RT (min)      lcqtof_hilic_neg_4_000_aa    m/z 481.3298     0   1E+4   15.5   16  RT (min)      lcqtof_hilic_neg_4_000_aa    m/z 481.3310     0   2   4   6E+4   1   1.5  RT (min)      lcqtof_hilic_neg_4_000_aa    m/z 481.3325     0   2   4   6E+4   1   1.5  RT (min)      lcqtof_hilic_neg_4_000_aa    m/z 481.3655     0   2   4   6E+4   1   1.5  RT (min)      lcqtof_hilic_neg_1_007_05    m/z 481.4253     0   1E+5   1   1.5  RT (min)      lcqtof_hilic_neg_4_000_aa    m/z 481.4254     0   1E+4   15   15.5  RT (min)      lcqtof_hilic_neg_4_103_61    m/z 481.4254     0   1E+5   1   1.5  RT (min)      lcqtof_hilic_neg_4_000_aa    m/z 481.4256     0   1E+5   1.5   2  RT (min)      lcqtof_hilic_neg_4_000_aa    m/z 481.4256     0   1   2   3   4E+3   2   2.5  RT (min)      lcqtof_hilic_neg_4_052_13    m/z 481.4257     0   1   2E+3   16   16.5  RT (min)       lcqtof_hilic_neg_4_012_07    m/z 481.4258     0   2   4   6   8E+3   15.5   16  RT (min)      lcqtof_hilic_neg_1_007_05    m/z 481.4266     0   1E+4   15.5   16  RT (min)       lcqtof_hilic_neg_4_000_aa    m/z 481.8082     0   1   2E+4   4   4.5  RT (min)      lcqtof_hilic_neg_4_008_24    m/z 482.2777     0   1E+4   0.5   1  RT (min)      lcqtof_hilic_neg_1_007_05    m/z 482.2839     0   1E+4   2   2.5  RT (min)      lcqtof_hilic_neg_4_053_14    m/z 482.3336     0   1E+4   1   1.5  RT (min)      lcqtof_hilic_neg_4_000_aa    m/z 482.3360     0   1E+4   1   1.5  RT (min)      lcqtof_hilic_neg_4_000_aa    m/z 482.3724     0   1   2   3E+5   1   1.5  RT (min)       lcqtof_hilic_neg_4_040_05    m/z 482.4252     0   2   4E+4   1   1.5  RT (min)      lcqtof_hilic_neg_4_000_aa    m/z 482.4258     0   2   4E+4   1   1.5  RT (min)      lcqtof_hilic_neg_4_000_aa    m/z 482.4280     0   2   4E+4   1   1.5  RT (min)      lcqtof_hilic_neg_4_000_aa    m/z 482.4284     0   2   4E+4   1.5   2  RT (min)      lcqtof_hilic_neg_4_000_aa    m/z 482.5580     0   1E+4   4.5   5  RT (min)      lcqtof_hilic_neg_4_050_02    m/z 482.8152     0   1   2E+4   4   4.5  RT (min)      lcqtof_hilic_neg_4_008_24    m/z 482.9028     0   1   2   3E+5   1   1.5  RT (min)      lcqtof_hilic_neg_4_004_47    m/z 482.9183     0   2   4   6E+4   0.5   1  RT (min)      lcqtof_hilic_neg_3_035_46    m/z 483.0700     0   1   2E+4   3.5   4  RT (min)      lcqtof_hilic_neg_1_032_18    m/z 483.1540     0   1   2   3E+4   6   6.5  RT (min)       lcqtof_hilic_neg_4_008_24    m/z 483.1542     0   1   2   3E+4   6   6.5  RT (min)      lcqtof_hilic_neg_4_008_24    m/z 483.3480     0   1E+4   0.5   1  RT (min)      lcqtof_hilic_neg_4_039_18    m/z 483.3707     0   2   4   6   8E+3   1   1.5  RT (min)      lcqtof_hilic_neg_1_007_05    m/z 483.4053     0   1   2   3E+4   1   1.5  RT (min)      lcqtof_hilic_neg_4_000_aa    m/z 483.8220     0   1   2E+4   4   4.5  RT (min)      lcqtof_hilic_neg_4_008_24    m/z 483.9180     0   1   2E+4   0.5   1  RT (min)      lcqtof_hilic_neg_3_035_46    m/z 484.1958     0   1E+4   0.5   1  RT (min)      lcqtof_hilic_neg_4_064_15    m/z 484.4014     0   1E+4   1   1.5  RT (min)      lcqtof_hilic_neg_4_000_aa    m/z 484.5569     0   1E+4   4.5   5  RT (min)      lcqtof_hilic_neg_4_050_02    m/z 484.9191     0   1E+4   0.5   1  RT (min)      lcqtof_hilic_neg_3_035_46    m/z 485.1517     0   1   2E+4   6   6.5  RT (min)      lcqtof_hilic_neg_4_008_24    m/z 485.1685     0   1E+4   2   2.5  RT (min)      lcqtof_hilic_neg_4_052_13    m/z 485.1685     0   1E+4   2   2.5  RT (min)      lcqtof_hilic_neg_4_052_13    m/z 485.2046     0   1E+4   0.5   1  RT (min)      lcqtof_hilic_neg_4_039_18    m/z 485.2079     0   1   2   3E+4   0.5   1  RT (min)      lcqtof_hilic_neg_3_003_20    m/z 485.2201     0   2   4E+3   2   2.5  RT (min)      lcqtof_hilic_neg_4_032_01    m/z 485.2558     0   1   2   3E+4   0.5   1  RT (min)      lcqtof_hilic_neg_3_003_20    m/z 485.3350     0   2   4E+3   2.5   3  RT (min)       lcqtof_hilic_neg_4_048_23    m/z 485.3842     0   1E+4   1   1.5  RT (min)      lcqtof_hilic_neg_4_053_14    m/z 485.5023     0   1   2   3   4E+4   0.5   1  RT (min)      lcqtof_hilic_neg_4_034_31    m/z 486.0244     0   1E+4   15   15.5  RT (min)      lcqtof_hilic_neg_4_000_aa    m/z 486.1442     0   1   2E+4   5.5   6  RT (min)      lcqtof_hilic_neg_1_000_aa    m/z 486.1444     0   1   2E+4   5   5.5  RT (min)      lcqtof_hilic_neg_1_000_aa    m/z 486.1446     0   1   2E+4   5   5.5  RT (min)      lcqtof_hilic_neg_1_000_aa    m/z 486.2197     0   1E+4   4   4.5  RT (min)       lcqtof_hilic_neg_4_008_24    m/z 486.2541     0   1E+4   0.5   1  RT (min)      lcqtof_hilic_neg_3_003_20    m/z 486.2609     0   1E+4   1   1.5  RT (min)      lcqtof_hilic_neg_4_052_13    m/z 486.2822     0   1   2   3E+5   1   1.5  RT (min)      lcqtof_hilic_neg_4_016_37    m/z 486.2963     0   1E+4   0.5   1  RT (min)      lcqtof_hilic_neg_3_035_46    m/z 486.4032     0   1   2   3E+4   1   1.5  RT (min)       lcqtof_hilic_neg_1_007_05    m/z 486.4039     0   1   2   3E+4   1   1.5  RT (min)      lcqtof_hilic_neg_4_040_05    m/z 487.1646     0   2   4   6E+4   2.5   3  RT (min)      lcqtof_hilic_neg_4_054_48    m/z 487.1794     0   1E+4   1   1.5  RT (min)      lcqtof_hilic_neg_4_000_aa    m/z 487.2130     0   1E+4   7   7.5  RT (min)      lcqtof_hilic_neg_4_048_23    m/z 487.2480     0   2   4   6   8E+4   0.5   1  RT (min)      lcqtof_hilic_neg_4_013_20    m/z 487.2658     0   1E+4   0.5   1  RT (min)      lcqtof_hilic_neg_4_005_29    m/z 487.3247     0   1E+4   2   2.5  RT (min)      lcqtof_hilic_neg_4_050_02    m/z 487.5294     0   1E+4   3   3.5  RT (min)      lcqtof_hilic_neg_4_034_31    m/z 487.9986     0   1E+4   1   1.5  RT (min)      lcqtof_hilic_neg_4_050_02    m/z 488.0212     0   1E+4   15   15.5  RT (min)      lcqtof_hilic_neg_4_000_aa    m/z 488.0779     0   1E+4   4.5   5  RT (min)      lcqtof_hilic_neg_4_013_20    m/z 488.1670     0   1E+4   2.5   3  RT (min)      lcqtof_hilic_neg_4_054_48    m/z 488.2518     0   1   2E+4   0.5   1  RT (min)      lcqtof_hilic_neg_4_027_19    m/z 488.9927     0   1E+4   0.5   1  RT (min)      lcqtof_hilic_neg_4_050_02    m/z 488.9969     0   1E+4   1   1.5  RT (min)      lcqtof_hilic_neg_4_050_02    m/z 489.2522     0   1   2E+4   6   6.5  RT (min)      lcqtof_hilic_neg_4_008_24    m/z 489.2548     0   1   2   3E+4   0.5   1  RT (min)      lcqtof_hilic_neg_4_005_29    m/z 489.2815     0   1E+5   1   1.5  RT (min)      lcqtof_hilic_neg_4_000_aa    m/z 489.3083     0   1E+4   1   1.5  RT (min)      lcqtof_hilic_neg_4_052_13    m/z 489.3154     0   1   2   3   4E+4   1.5   2  RT (min)      lcqtof_hilic_neg_4_003_16    m/z 490.0754     0   1E+4   4.5   5  RT (min)      lcqtof_hilic_neg_4_027_19    m/z 490.1680     0   1E+4   6   6.5  RT (min)      lcqtof_hilic_neg_4_047_06    m/z 490.1844     0   1E+4   1   1.5  RT (min)      lcqtof_hilic_neg_4_000_aa    m/z 490.1852     0   1E+4   1   1.5  RT (min)      lcqtof_hilic_neg_4_000_aa    m/z 490.2571     0   1E+4   0.5   1  RT (min)      lcqtof_hilic_neg_4_005_29    m/z 490.2577     0   1E+4   4   4.5  RT (min)      lcqtof_hilic_neg_4_040_05    m/z 490.2732     0   2   4   6   8E+4   1   1.5  RT (min)      lcqtof_hilic_neg_4_000_aa    m/z 490.2746     0   2   4   6   8E+4   1   1.5  RT (min)      lcqtof_hilic_neg_4_000_aa    m/z 490.2896     0   2   4   6E+4   1   1.5  RT (min)      lcqtof_hilic_neg_4_000_aa    m/z 490.3007     0   1E+4   4.5   5  RT (min)       lcqtof_hilic_neg_4_008_24    m/z 490.3067     0   1E+4   0.5   1  RT (min)      lcqtof_hilic_neg_2_028_05    m/z 490.3206     0   1E+4   2   2.5  RT (min)      lcqtof_hilic_neg_4_003_16    m/z 490.4250     0   1E+4   0.5   1  RT (min)      lcqtof_hilic_neg_3_036_17    m/z 490.9864     0   1E+4   0.5   1  RT (min)      lcqtof_hilic_neg_4_050_02    m/z 490.9968     0   1E+4   0.5   1  RT (min)      lcqtof_hilic_neg_4_058_39    m/z 490.9984     0   1E+4   1   1.5  RT (min)      lcqtof_hilic_neg_4_050_02    m/z 491.0258     0   1E+4   1   1.5  RT (min)      lcqtof_hilic_neg_4_000_aa    m/z 491.0315     0   1E+4   1   1.5  RT (min)      lcqtof_hilic_neg_4_000_aa    m/z 491.0901     0   1E+4   6.5   7  RT (min)      lcqtof_hilic_neg_4_008_24    m/z 491.1778     0   1   2E+5   2.5   3  RT (min)      lcqtof_hilic_neg_4_057_49    m/z 491.1792     0   1E+4   1   1.5  RT (min)      lcqtof_hilic_neg_4_000_aa    m/z 491.2670     0   1E+5   1   1.5  RT (min)      lcqtof_hilic_neg_4_000_aa    m/z 491.2684     0   1E+5   1   1.5  RT (min)      lcqtof_hilic_neg_4_000_aa    m/z 491.2688     0   1E+5   1   1.5  RT (min)      lcqtof_hilic_neg_4_000_aa    m/z 491.3356     0   1E+4   0.5   1  RT (min)      lcqtof_hilic_neg_4_005_29    m/z 491.3582     0   1E+6   1   1.5  RT (min)      lcqtof_hilic_neg_4_000_aa    m/z 491.3588     0   1E+6   1   1.5  RT (min)      lcqtof_hilic_neg_4_000_aa    m/z 491.3588     0   1E+6   1   1.5  RT (min)      lcqtof_hilic_neg_4_000_aa    m/z 491.3590     0   1E+6   1   1.5  RT (min)      lcqtof_hilic_neg_4_000_aa    m/z 491.4712     0   1E+4   1   1.5  RT (min)      lcqtof_hilic_neg_4_000_aa    m/z 491.5834     0   1   2   3   4E+4   1   1.5  RT (min)      lcqtof_hilic_neg_4_000_aa    m/z 491.5840     0   1   2   3   4E+4   1   1.5  RT (min)      lcqtof_hilic_neg_4_000_aa    m/z 492.0490     0   1E+4   0.5   1   1.5  RT (min)      lcqtof_hilic_neg_4_027_19    m/z 492.1062     0   1   2   3E+4   1   1.5  RT (min)      lcqtof_hilic_neg_4_000_aa    m/z 492.1066     0   1   2   3E+4   1   1.5  RT (min)      lcqtof_hilic_neg_4_000_aa    m/z 492.1071     0   1   2   3E+4   0.5   1  RT (min)      lcqtof_hilic_neg_4_000_aa    m/z 492.1084     0   1   2   3E+4   1   1.5  RT (min)      lcqtof_hilic_neg_4_000_aa    m/z 492.1807     0   1   2   3   4E+4   2.5   3  RT (min)      lcqtof_hilic_neg_4_054_48    m/z 492.2150     0   1   2E+4   6   6.5  RT (min)       lcqtof_hilic_neg_4_092_60    m/z 492.2696     0   1   2   3E+4   1   1.5  RT (min)      lcqtof_hilic_neg_4_000_aa    m/z 492.2702     0   1   2   3E+4   1   1.5  RT (min)      lcqtof_hilic_neg_4_000_aa    m/z 492.2704     0   1   2   3E+4   1   1.5  RT (min)      lcqtof_hilic_neg_4_000_aa    m/z 492.3608     0   2   4   6E+5   1   1.5  RT (min)      lcqtof_hilic_neg_4_000_aa    m/z 492.3608     0   2   4   6E+5   1   1.5  RT (min)      lcqtof_hilic_neg_4_000_aa    m/z 492.3608     0   2   4   6E+5   1   1.5  RT (min)      lcqtof_hilic_neg_4_000_aa    m/z 493.0226     0   1   2   3E+4   1   1.5  RT (min)      lcqtof_hilic_neg_4_000_aa    m/z 493.0232     0   1   2   3E+4   1   1.5  RT (min)      lcqtof_hilic_neg_4_000_aa    m/z 493.0292     0   1E+4   0.5   1  RT (min)      lcqtof_hilic_neg_4_046_41    m/z 493.0607     0   1E+4   7.5   8  RT (min)       lcqtof_hilic_neg_4_008_24    m/z 493.1102     0   1E+4   1   1.5  RT (min)      lcqtof_hilic_neg_4_000_aa    m/z 493.1553     0   1E+4   1   1.5  RT (min)      lcqtof_hilic_neg_4_004_47    m/z 493.2668     0   1   2   3E+4   1   1.5  RT (min)      lcqtof_hilic_neg_4_000_aa    m/z 493.3542     0   1   2E+5   1   1.5  RT (min)      lcqtof_hilic_neg_4_000_aa    m/z 493.3554     0   1   2E+5   1   1.5  RT (min)      lcqtof_hilic_neg_4_000_aa    m/z 493.3592     0   1   2E+5   1   1.5  RT (min)      lcqtof_hilic_neg_4_000_aa    m/z 493.3618     0   1   2E+5   1   1.5  RT (min)      lcqtof_hilic_neg_4_000_aa    m/z 493.3890     0   1   2E+5   1   1.5  RT (min)      lcqtof_hilic_neg_4_053_14    m/z 493.4032     0   1   2E+4   0.5   1  RT (min)      lcqtof_hilic_neg_3_035_46    m/z 493.4252     0   1E+4   1   1.5  RT (min)      lcqtof_hilic_neg_4_000_aa    m/z 493.4633     0   1E+4   1   1.5  RT (min)      lcqtof_hilic_neg_4_000_aa    m/z 493.4649     0   1E+4   1   1.5  RT (min)      lcqtof_hilic_neg_4_000_aa    m/z 493.4974     0   1E+4   1.5   2  RT (min)      lcqtof_hilic_neg_4_000_aa    m/z 494.0990     0   1   2   3E+4   1   1.5  RT (min)      lcqtof_hilic_neg_4_000_aa    m/z 494.1028     0   1   2   3E+4   1   1.5  RT (min)      lcqtof_hilic_neg_4_000_aa    m/z 494.1048     0   1   2   3E+4   0.5   1  RT (min)      lcqtof_hilic_neg_4_000_aa    m/z 494.1063     0   1   2   3E+4   1   1.5  RT (min)      lcqtof_hilic_neg_4_000_aa    m/z 494.2690     0   1E+5   1   1.5  RT (min)      lcqtof_hilic_neg_4_032_01    m/z 494.3560     0   2   4   6E+4   1   1.5  RT (min)      lcqtof_hilic_neg_4_000_aa    m/z 494.3600     0   2   4   6E+4   1   1.5  RT (min)      lcqtof_hilic_neg_4_000_aa    m/z 494.3602     0   2   4   6E+4   1   1.5  RT (min)      lcqtof_hilic_neg_4_000_aa    m/z 494.3619     0   2   4   6E+4   1   1.5  RT (min)      lcqtof_hilic_neg_4_000_aa    m/z 494.3666     0   2   4   6E+4   1   1.5  RT (min)      lcqtof_hilic_neg_4_000_aa    m/z 495.0144     0   1   2E+4   1   1.5  RT (min)      lcqtof_hilic_neg_4_000_aa    m/z 495.0201     0   1   2E+4   1   1.5  RT (min)      lcqtof_hilic_neg_4_000_aa    m/z 495.0770     0   1   2E+4   6   6.5  RT (min)       lcqtof_hilic_neg_4_082_59    m/z 495.1643     0   1E+4   6.5   7  RT (min)       lcqtof_hilic_neg_4_008_24    m/z 495.1906     0   1   2E+4   6   6.5  RT (min)       lcqtof_hilic_neg_4_091_60    m/z 495.2210     0   1E+4   4   4.5  RT (min)       lcqtof_hilic_neg_4_055_32    m/z 495.2958     0   1   2   3   4E+3   1.5   2  RT (min)      lcqtof_hilic_neg_4_063_16    m/z 495.3603     0   1E+4   1   1.5  RT (min)      lcqtof_hilic_neg_4_000_aa    m/z 495.3622     0   1E+4   1   1.5  RT (min)      lcqtof_hilic_neg_4_000_aa    m/z 495.3859     0   1   2E+4   0.5   1  RT (min)      lcqtof_hilic_neg_2_051_14    m/z 495.4404     0   1E+5   1   1.5  RT (min)      lcqtof_hilic_neg_4_000_aa    m/z 495.4420     0   2   4   6   8E+3   15   15.5  RT (min)      lcqtof_hilic_neg_4_038_10    m/z 495.4430     0   1E+5   1.5   2  RT (min)      lcqtof_hilic_neg_4_000_aa    m/z 495.4622     0   1   2E+4   1   1.5  RT (min)      lcqtof_hilic_neg_4_000_aa    m/z 495.4770     0   1   2E+4   1.5   2  RT (min)      lcqtof_hilic_neg_4_000_aa    m/z 496.1080     0   1   2E+4   6   6.5  RT (min)       lcqtof_hilic_neg_4_048_23    m/z 496.2736     0   1E+4   3   3.5  RT (min)      lcqtof_hilic_neg_4_039_18    m/z 496.4395     0   2   4E+4   1   1.5  RT (min)      lcqtof_hilic_neg_4_000_aa    m/z 496.4461     0   2   4E+4   1   1.5  RT (min)      lcqtof_hilic_neg_4_000_aa    m/z 496.4462     0   2   4E+4   1.5   2  RT (min)      lcqtof_hilic_neg_4_000_aa    m/z 496.9145     0   1E+4   7   7.5  RT (min)      lcqtof_hilic_neg_4_050_02    m/z 497.0088     0   1E+4   1   1.5  RT (min)      lcqtof_hilic_neg_4_000_aa    m/z 497.1656     0   1   2E+4   1   1.5  RT (min)       lcqtof_hilic_neg_4_058_39    m/z 497.1660     0   2   4   6E+3   3   3.5  RT (min)      lcqtof_hilic_neg_4_039_18    m/z 497.1666     0   1E+4   0.5   1  RT (min)       lcqtof_hilic_neg_4_000_aa    m/z 497.1668     0   1   2E+4   1   1.5  RT (min)       lcqtof_hilic_neg_4_058_39    m/z 497.1670     0   1   2   3E+3   15.5   16  RT (min)       lcqtof_hilic_neg_4_113_62    m/z 497.1670     0   1   2E+4   1   1.5  RT (min)      lcqtof_hilic_neg_4_058_39    m/z 497.1672     0   1E+5   3   3.5  RT (min)       lcqtof_hilic_neg_4_038_10    m/z 497.1673     0   2   4   6   8E+3   3   3.5  RT (min)      lcqtof_hilic_neg_3_000_aa    m/z 497.1673     0   2   4   6   8E+3   3   3.5  RT (min)      lcqtof_hilic_neg_3_000_aa    m/z 497.1674     0   1E+5   3   3.5  RT (min)      lcqtof_hilic_neg_4_038_10    m/z 497.1712     0   1   2E+4   4   4.5  RT (min)      lcqtof_hilic_neg_1_000_aa    m/z 497.1760     0   1   2   3E+4   15   15.5  RT (min)       lcqtof_hilic_neg_4_000_aa    m/z 497.2592     0   2   4E+4   0.5   1  RT (min)      lcqtof_hilic_neg_4_013_20    m/z 497.3344     0   1   2   3E+5   3   3.5  RT (min)      lcqtof_hilic_neg_4_011_35    m/z 497.3975     0   1E+4   0.5   1  RT (min)      lcqtof_hilic_neg_4_020_42    m/z 497.4328     0   1   2E+4   1   1.5  RT (min)      lcqtof_hilic_neg_4_000_aa    m/z 497.5896     0   2   4E+4   0.5   1  RT (min)      lcqtof_hilic_neg_1_032_18    m/z 498.0464     0   1E+4   4.5   5  RT (min)      lcqtof_hilic_neg_4_027_19    m/z 498.1429     0   1   2   3E+4   3   3.5  RT (min)      lcqtof_hilic_neg_4_039_18    m/z 498.1674     0   1E+4   15   15.5  RT (min)      lcqtof_hilic_neg_4_000_aa    m/z 498.1856     0   1   2E+4   3.5   4  RT (min)      lcqtof_hilic_neg_4_055_32    m/z 498.1984     0   1E+4   1.5   2  RT (min)      lcqtof_hilic_neg_4_040_05    m/z 498.2620     0   1E+4   0.5   1  RT (min)      lcqtof_hilic_neg_4_013_20    m/z 498.2972     0   1   2   3E+4   1   1.5  RT (min)      lcqtof_hilic_neg_4_000_aa    m/z 498.2976     0   1   2   3E+4   1   1.5  RT (min)      lcqtof_hilic_neg_4_000_aa    m/z 498.2980     0   1   2   3E+4   1   1.5  RT (min)      lcqtof_hilic_neg_4_000_aa    m/z 498.4157     0   1   2E+4   1   1.5  RT (min)      lcqtof_hilic_neg_4_000_aa    m/z 498.9446     0   2   4   6   8E+4   0.5   1  RT (min)      lcqtof_hilic_neg_3_035_46    m/z 499.1112     0   1E+4   2   2.5  RT (min)      lcqtof_hilic_neg_4_003_16    m/z 499.1818     0   1E+4   3.5   4  RT (min)      lcqtof_hilic_neg_4_050_02    m/z 499.1874     0   1E+4   6.5   7  RT (min)      lcqtof_hilic_neg_1_007_05    m/z 499.1973     0   1   2E+4   1   1.5  RT (min)       lcqtof_hilic_neg_4_100_61    m/z 499.2224     0   2   4E+3   2   2.5  RT (min)       lcqtof_hilic_neg_4_050_02    m/z 499.2224     0   1   2   3E+4   15   15.5  RT (min)      lcqtof_hilic_neg_4_000_aa    m/z 499.2224     0   2   4E+4   1   1.5  RT (min)      lcqtof_hilic_neg_4_050_02    m/z 499.3621     0   1   2E+3   16   16.5  RT (min)       lcqtof_hilic_neg_4_002_25    m/z 499.3624     0   1E+3   0   0.5  RT (min)      lcqtof_hilic_neg_4_000_dd    m/z 499.5821     0   1   2   3E+4   4   4.5  RT (min)      lcqtof_hilic_neg_4_040_05    m/z 500.1769     0   2   4E+4   2.5   3  RT (min)      lcqtof_hilic_neg_4_054_48    m/z 500.2228     0   1E+4   15   15.5  RT (min)      lcqtof_hilic_neg_4_000_aa    m/z 500.2519     0   1   2E+4   0.5   1  RT (min)      lcqtof_hilic_neg_4_027_19    m/z 500.3383     0   1   2   3E+4   0.5   1  RT (min)      lcqtof_hilic_neg_1_007_05    m/z 500.6097     0   1   2E+4   4   4.5  RT (min)      lcqtof_hilic_neg_4_032_01    m/z 500.8063     0   1   2E+4   4   4.5  RT (min)      lcqtof_hilic_neg_4_032_01    m/z 500.8780     0   2   4E+3   1.5   2  RT (min)       lcqtof_hilic_neg_3_078_58    m/z 500.8830     0   2   4   6   8E+3   1   1.5  RT (min)      lcqtof_hilic_neg_3_081_58    m/z 500.8956     0   1   2   3E+4   1   1.5  RT (min)      lcqtof_hilic_neg_3_075_58    m/z 500.9088     0   2   4   6E+4   1   1.5  RT (min)      lcqtof_hilic_neg_4_004_47    m/z 500.9239     0   1E+4   0.5   1  RT (min)      lcqtof_hilic_neg_3_035_46    m/z 501.1184     0   1   2E+4   6   6.5  RT (min)       lcqtof_hilic_neg_4_048_23    m/z 501.1721     0   1E+5   0.5   1  RT (min)      lcqtof_hilic_neg_4_058_39    m/z 501.1816     0   1E+4   2.5   3  RT (min)      lcqtof_hilic_neg_4_054_48    m/z 501.2550     0   1   2   3E+4   0.5   1  RT (min)      lcqtof_hilic_neg_4_050_02    m/z 501.2916     0   1E+4   3.5   4  RT (min)      lcqtof_hilic_neg_4_039_18    m/z 501.3710     0   1   2E+5   3.5   4  RT (min)      lcqtof_hilic_neg_4_008_24    m/z 501.3783     0   1E+5   0.5   1  RT (min)      lcqtof_hilic_neg_3_003_20    m/z 501.3787     0   2   4   6   8E+4   1   1.5  RT (min)      lcqtof_hilic_neg_4_001_49    m/z 501.5102     0   1   2E+4   4   4.5  RT (min)      lcqtof_hilic_neg_4_032_01    m/z 502.1716     0   1   2E+4   0.5   1  RT (min)      lcqtof_hilic_neg_4_050_02    m/z 502.1794     0   1E+4   6   6.5  RT (min)      lcqtof_hilic_neg_4_048_23    m/z 502.2626     0   1E+4   0.5   1  RT (min)      lcqtof_hilic_neg_4_050_02    m/z 502.2890     0   1E+4   2   2.5  RT (min)      lcqtof_hilic_neg_4_016_37    m/z 502.3196     0   2   4   6   8E+3   2.5   3  RT (min)      lcqtof_hilic_neg_4_016_37    m/z 502.4243     0   1E+4   1   1.5  RT (min)      lcqtof_hilic_neg_4_000_aa    m/z 502.9171     0   1E+4   8   8.5  RT (min)       lcqtof_hilic_neg_4_059_14    m/z 503.0600     0   1   2E+4   1   1.5  RT (min)      lcqtof_hilic_neg_4_000_aa    m/z 503.0613     0   1   2E+4   1   1.5  RT (min)      lcqtof_hilic_neg_4_000_aa    m/z 503.1266     0   1E+4   1   1.5  RT (min)      lcqtof_hilic_neg_4_024_40    m/z 503.2977     0   2   4   6E+4   3.5   4  RT (min)      lcqtof_hilic_neg_4_040_05    m/z 503.3044     0   1   2E+4   1   1.5  RT (min)      lcqtof_hilic_neg_4_000_aa    m/z 503.3048     0   1   2E+4   1   1.5  RT (min)      lcqtof_hilic_neg_4_000_aa    m/z 503.3121     0   1   2E+4   1   1.5  RT (min)      lcqtof_hilic_neg_4_000_aa    m/z 503.3180     0   1   2E+4   1   1.5  RT (min)      lcqtof_hilic_neg_4_000_aa    m/z 503.4050     0   1   2   3E+4   1   1.5  RT (min)      lcqtof_hilic_neg_4_000_aa    m/z 503.4289     0   1   2   3E+4   0.5   1  RT (min)      lcqtof_hilic_neg_3_035_46    m/z 503.4358     0   1E+4   0.5   1  RT (min)      lcqtof_hilic_neg_4_008_24    m/z 504.1295     0   1E+4   6   6.5  RT (min)       lcqtof_hilic_neg_4_048_23    m/z 504.1438     0   1E+4   5.5   6  RT (min)      lcqtof_hilic_neg_2_022_46    m/z 504.2083     0   1   2   3E+4   6.5   7  RT (min)      lcqtof_hilic_neg_1_007_05    m/z 504.2869     0   1E+4   1   1.5  RT (min)      lcqtof_hilic_neg_4_000_aa    m/z 504.2912     0   1E+4   1   1.5  RT (min)      lcqtof_hilic_neg_4_000_aa    m/z 504.4130     0   1E+4   1   1.5  RT (min)      lcqtof_hilic_neg_4_000_aa    m/z 504.4277     0   1E+4   0.5   1  RT (min)      lcqtof_hilic_neg_3_035_46    m/z 505.0559     0   2   4   6E+4   1   1.5  RT (min)      lcqtof_hilic_neg_4_000_aa    m/z 505.0570     0   2   4   6E+4   1   1.5  RT (min)      lcqtof_hilic_neg_4_000_aa    m/z 505.1806     0   1E+5   0.5   1  RT (min)      lcqtof_hilic_neg_3_003_20    m/z 505.1926     0   1E+4   6.5   7  RT (min)      lcqtof_hilic_neg_4_002_25    m/z 505.2172     0   1   2E+4   6   6.5  RT (min)      lcqtof_hilic_neg_4_008_24    m/z 505.2176     0   1   2E+4   6   6.5  RT (min)       lcqtof_hilic_neg_4_094_60    m/z 505.2571     0   1E+4   3.5   4  RT (min)      lcqtof_hilic_neg_4_040_05    m/z 505.2744     0   1E+4   7   7.5  RT (min)       lcqtof_hilic_neg_4_008_24    m/z 505.3315     0   2   4E+4   15.5   16  RT (min)      lcqtof_hilic_neg_4_000_aa    m/z 505.3331     0   1   2E+3   16   16.5  RT (min)       lcqtof_hilic_neg_1_006_22    m/z 505.3396     0   1E+6   1   1.5  RT (min)      lcqtof_hilic_neg_4_000_aa    m/z 505.3524     0   1E+4   1   1.5  RT (min)      lcqtof_hilic_neg_4_053_14    m/z 505.4090     0   2   4E+4   0.5   1  RT (min)      lcqtof_hilic_neg_4_053_14    m/z 505.4399     0   1E+4   0.5   1  RT (min)      lcqtof_hilic_neg_3_035_46    m/z 505.5599     0   1   2E+4   1   1.5  RT (min)      lcqtof_hilic_neg_4_000_aa    m/z 506.0973     0   1E+4   6   6.5  RT (min)       lcqtof_hilic_neg_4_048_23    m/z 506.1675     0   1E+4   6   6.5  RT (min)       lcqtof_hilic_neg_4_008_24    m/z 506.1830     0   1   2E+4   0.5   1  RT (min)      lcqtof_hilic_neg_3_003_20    m/z 506.1978     0   1E+4   2.5   3  RT (min)      lcqtof_hilic_neg_4_049_55    m/z 506.2324     0   1   2   3E+4   2   2.5  RT (min)      lcqtof_hilic_neg_4_057_49    m/z 506.2330     0   1   2   3E+4   2   2.5  RT (min)      lcqtof_hilic_neg_4_057_49    m/z 506.2491     0   1E+4   6   6.5  RT (min)      lcqtof_hilic_neg_4_008_24    m/z 506.2504     0   1   2   3E+5   3   3.5  RT (min)      lcqtof_hilic_neg_4_031_38    m/z 506.3076     0   1E+4   2   2.5  RT (min)      lcqtof_hilic_neg_4_063_16    m/z 506.3254     0   2   4E+5   1   1.5  RT (min)      lcqtof_hilic_neg_4_000_aa    m/z 506.3353     0   1E+4   15.5   16  RT (min)      lcqtof_hilic_neg_4_000_aa    m/z 506.8432     0   1   2E+3   8.5   9  RT (min)      lcqtof_hilic_neg_4_003_16    m/z 507.0530     0   2   4   6E+4   1   1.5  RT (min)      lcqtof_hilic_neg_4_000_aa    m/z 507.0533     0   2   4   6E+4   1   1.5  RT (min)      lcqtof_hilic_neg_4_000_aa    m/z 507.1603     0   1E+4   7   7.5  RT (min)      lcqtof_hilic_neg_4_002_25    m/z 507.2544     0   1E+5   3   3.5  RT (min)      lcqtof_hilic_neg_4_031_38    m/z 507.3382     0   1E+5   1   1.5  RT (min)      lcqtof_hilic_neg_4_000_aa    m/z 507.3868     0   2   4E+4   1   1.5  RT (min)      lcqtof_hilic_neg_4_000_aa    m/z 507.4748     0   1E+5   1   1.5  RT (min)      lcqtof_hilic_neg_4_000_aa    m/z 507.4772     0   1E+5   1   1.5  RT (min)      lcqtof_hilic_neg_4_000_aa    m/z 507.5116     0   1E+4   1.5   2  RT (min)      lcqtof_hilic_neg_4_000_aa    m/z 507.5145     0   1E+4   1.5   2  RT (min)      lcqtof_hilic_neg_4_000_aa    m/z 508.0134     0   1E+4   6   6.5  RT (min)       lcqtof_hilic_neg_4_084_59    m/z 508.0151     0   1E+4   6.5   7  RT (min)      lcqtof_hilic_neg_3_030_23    m/z 508.0520     0   1E+4   1   1.5  RT (min)      lcqtof_hilic_neg_4_000_aa    m/z 508.0559     0   1E+4   1   1.5  RT (min)      lcqtof_hilic_neg_4_000_aa    m/z 508.0936     0   1   2   3E+4   1   1.5  RT (min)      lcqtof_hilic_neg_4_000_aa    m/z 508.1033     0   1   2   3E+4   1   1.5  RT (min)      lcqtof_hilic_neg_4_000_aa    m/z 508.1044     0   1   2   3E+4   1   1.5  RT (min)      lcqtof_hilic_neg_4_000_aa    m/z 508.1227     0   1   2E+5   4   4.5   5  RT (min)      lcqtof_hilic_neg_4_039_18    m/z 508.2440     0   1E+4   2   2.5  RT (min)      lcqtof_hilic_neg_4_001_49    m/z 508.2560     0   1E+4   3   3.5  RT (min)      lcqtof_hilic_neg_4_031_38    m/z 508.4781     0   1   2   3   4E+4   1   1.5  RT (min)      lcqtof_hilic_neg_4_000_aa    m/z 508.4796     0   1   2   3   4E+4   1   1.5  RT (min)      lcqtof_hilic_neg_4_000_aa    m/z 509.0476     0   1   2E+4   1   1.5  RT (min)      lcqtof_hilic_neg_4_000_aa    m/z 509.0484     0   1   2E+4   1   1.5  RT (min)      lcqtof_hilic_neg_4_000_aa    m/z 509.1019     0   1   2E+4   1   1.5  RT (min)      lcqtof_hilic_neg_4_000_aa    m/z 509.1197     0   1   2   3   4E+4   0.5   1  RT (min)      lcqtof_hilic_neg_1_062_17    m/z 509.1256     0   1   2   3   4E+4   4   4.5   5  RT (min)      lcqtof_hilic_neg_4_039_18    m/z 509.1630     0   1E+4   3.5   4  RT (min)      lcqtof_hilic_neg_2_028_05    m/z 509.2181     0   2   4   6E+4   0.5   1  RT (min)      lcqtof_hilic_neg_3_003_20    m/z 509.2249     0   1   2   3E+4   6.5   7  RT (min)       lcqtof_hilic_neg_1_037_24    m/z 509.2529     0   2   4E+4   0.5   1  RT (min)      lcqtof_hilic_neg_4_050_02    m/z 509.2626     0   1   2E+4   0.5   1  RT (min)      lcqtof_hilic_neg_4_000_aa    m/z 509.2772     0   2   4   6E+4   1   1.5  RT (min)      lcqtof_hilic_neg_4_000_aa    m/z 509.4210     0   1   2   3   4E+4   1   1.5  RT (min)      lcqtof_hilic_neg_4_000_aa    m/z 509.4531     0   2   4   6   8E+4   1   1.5  RT (min)      lcqtof_hilic_neg_4_000_aa    m/z 509.4568     0   2   4   6   8E+4   1.5   2  RT (min)      lcqtof_hilic_neg_4_000_aa    m/z 509.4576     0   2   4   6   8E+4   1   1.5  RT (min)      lcqtof_hilic_neg_4_000_aa    m/z 509.4708     0   2   4   6E+4   1.5   2  RT (min)      lcqtof_hilic_neg_4_000_aa    m/z 510.0886     0   1E+4   3.5   4  RT (min)      lcqtof_hilic_neg_1_032_18    m/z 510.1110     0   1E+4   2   2.5  RT (min)      lcqtof_hilic_neg_4_034_31    m/z 510.1206     0   2   4   6E+4   4   4.5   5  RT (min)      lcqtof_hilic_neg_4_039_18    m/z 510.1442     0   1   2E+4   0.5   1  RT (min)      lcqtof_hilic_neg_4_010_28    m/z 510.1506     0   1   2E+4   3.5   4  RT (min)      lcqtof_hilic_neg_4_050_02    m/z 510.1732     0   1E+4   6   6.5  RT (min)      lcqtof_hilic_neg_4_008_24    m/z 510.2070     0   1   2E+4   6.5   7  RT (min)       lcqtof_hilic_neg_4_008_24    m/z 510.2462     0   1E+4   1   1.5  RT (min)      lcqtof_hilic_neg_4_050_02    m/z 510.2532     0   1   2   3E+4   1.5   2  RT (min)      lcqtof_hilic_neg_4_019_03    m/z 510.2570     0   1E+4   0.5   1  RT (min)      lcqtof_hilic_neg_4_050_02    m/z 510.2704     0   1E+4   1.5   2  RT (min)      lcqtof_hilic_neg_4_063_16    m/z 510.2786     0   1   2E+4   1   1.5  RT (min)      lcqtof_hilic_neg_4_000_aa    m/z 510.2788     0   1   2E+4   1   1.5  RT (min)      lcqtof_hilic_neg_4_000_aa    m/z 510.2796     0   1   2E+4   1   1.5  RT (min)      lcqtof_hilic_neg_4_000_aa    m/z 510.4214     0   1E+4   1   1.5  RT (min)      lcqtof_hilic_neg_4_000_aa    m/z 510.4510     0   1   2   3E+4   1   1.5  RT (min)      lcqtof_hilic_neg_4_000_aa    m/z 510.4596     0   1   2   3E+4   1.5   2  RT (min)      lcqtof_hilic_neg_4_000_aa    m/z 510.8813     0   1   2   3   4E+3   5   5.5  RT (min)       lcqtof_hilic_neg_4_032_01    m/z 511.1234     0   1E+4   4   4.5   5  RT (min)      lcqtof_hilic_neg_4_039_18    m/z 511.1431     0   1E+4   7   7.5  RT (min)      lcqtof_hilic_neg_1_011_46    m/z 511.2555     0   1E+4   1.5   2  RT (min)      lcqtof_hilic_neg_4_019_03    m/z 511.2670     0   2   4E+4   3   3.5  RT (min)      lcqtof_hilic_neg_4_033_53    m/z 511.2735     0   1   2E+4   1   1.5  RT (min)      lcqtof_hilic_neg_4_000_aa    m/z 511.2907     0   1E+4   4   4.5  RT (min)      lcqtof_hilic_neg_4_053_14    m/z 511.3142     0   2   4   6E+3   1.5   2  RT (min)      lcqtof_hilic_neg_4_063_16    m/z 511.4326     0   2   4E+4   1   1.5  RT (min)      lcqtof_hilic_neg_4_000_aa    m/z 511.4414     0   2   4E+4   1   1.5  RT (min)      lcqtof_hilic_neg_4_000_aa    m/z 512.1514     0   1   2E+4   3.5   4  RT (min)      lcqtof_hilic_neg_4_050_02    m/z 512.1711     0   2   4   6E+4   7   7.5  RT (min)      lcqtof_hilic_neg_4_008_24    m/z 512.1772     0   2   4   6E+4   7.5   8  RT (min)      lcqtof_hilic_neg_4_008_24    m/z 512.1773     0   2   4   6E+4   7   7.5  RT (min)      lcqtof_hilic_neg_4_008_24    m/z 512.1775     0   2   4   6E+4   7   7.5  RT (min)      lcqtof_hilic_neg_4_008_24    m/z 512.1778     0   2   4   6E+4   7   7.5  RT (min)       lcqtof_hilic_neg_4_008_24    m/z 512.1984     0   1E+4   1   1.5   2  RT (min)      lcqtof_hilic_neg_4_000_aa    m/z 512.2504     0   1   2   3E+3   2.5   3  RT (min)      lcqtof_hilic_neg_1_004_14    m/z 512.2506     0   2   4   6E+3   2   2.5  RT (min)      lcqtof_hilic_neg_4_059_14    m/z 512.2677     0   1E+5   4   4.5  RT (min)      lcqtof_hilic_neg_3_045_18    m/z 512.2688     0   1E+5   4   4.5  RT (min)      lcqtof_hilic_neg_4_039_18    m/z 512.2688     0   1E+5   3.5   4  RT (min)      lcqtof_hilic_neg_4_039_18    m/z 512.2700     0   1E+4   3   3.5  RT (min)      lcqtof_hilic_neg_4_033_53    m/z 512.2747     0   1E+4   0.5   1  RT (min)      lcqtof_hilic_neg_4_013_20    m/z 512.3032     0   1E+4   2   2.5  RT (min)      lcqtof_hilic_neg_4_059_14    m/z 512.3692     0   1   2E+4   15.5   16  RT (min)      lcqtof_hilic_neg_4_000_aa    m/z 512.3700     0   1E+4   1   1.5  RT (min)      lcqtof_hilic_neg_4_000_aa    m/z 512.4316     0   1E+4   1   1.5  RT (min)      lcqtof_hilic_neg_4_000_aa    m/z 512.4368     0   1E+4   1   1.5  RT (min)      lcqtof_hilic_neg_4_000_aa    m/z 512.8284     0   1E+4   8   8.5  RT (min)      lcqtof_hilic_neg_1_000_aa    m/z 512.9729     0   1E+4   1   1.5  RT (min)      lcqtof_hilic_neg_4_000_aa    m/z 512.9784     0   1E+4   1   1.5  RT (min)      lcqtof_hilic_neg_4_000_aa    m/z 513.2345     0   1   2   3   4E+4   0   0.5  RT (min)       lcqtof_hilic_neg_4_058_39    m/z 513.2362     0   1   2E+3   19   19.5  RT (min)       lcqtof_hilic_neg_4_006_30    m/z 513.2368     0   1   2   3E+3   18.5   19   19.5  RT (min)       lcqtof_hilic_neg_4_058_39    m/z 513.2368     0   1   2   3E+3   0   0.5  RT (min)       lcqtof_hilic_neg_4_058_39    m/z 513.2371     0   1   2E+3   19   19.5  RT (min)       lcqtof_hilic_neg_4_016_37    m/z 513.2374     0   1   2E+3   19.5   20  RT (min)       lcqtof_hilic_neg_4_004_47    m/z 513.2376     0   1   2   3E+4   15   15.5  RT (min)      lcqtof_hilic_neg_4_000_aa    m/z 513.2378     0   1   2E+3   18.5   19  RT (min)       lcqtof_hilic_neg_4_006_30    m/z 513.2380     0   1   2E+3   18   18.5  RT (min)       lcqtof_hilic_neg_4_012_07    m/z 513.2382     0   1   2   3   4E+3   2.5   3  RT (min)       lcqtof_hilic_neg_4_047_06    m/z 513.2387     0   1   2E+3   18   18.5  RT (min)       lcqtof_hilic_neg_4_035_33    m/z 513.2716     0   1   2E+4   4   4.5  RT (min)      lcqtof_hilic_neg_2_053_18    m/z 513.2720     0   2   4E+4   4   4.5  RT (min)      lcqtof_hilic_neg_4_039_18    m/z 513.3080     0   1   2   3   4E+5   1   1.5  RT (min)      lcqtof_hilic_neg_4_016_37    m/z 513.3250     0   2   4   6E+4   0.5   1  RT (min)      lcqtof_hilic_neg_3_035_46    m/z 513.3337     0   1   2   3E+4   4   4.5  RT (min)       lcqtof_hilic_neg_4_008_24    m/z 513.4143     0   1E+4   1   1.5  RT (min)      lcqtof_hilic_neg_4_053_14    m/z 513.4544     0   1   2   3   4E+4   1.5   2  RT (min)      lcqtof_hilic_neg_4_039_18    m/z 514.0092     0   1E+4   3   3.5  RT (min)      lcqtof_hilic_neg_4_027_19    m/z 514.2640     0   1E+4   4   4.5  RT (min)      lcqtof_hilic_neg_4_039_18    m/z 514.2701     0   1E+4   2   2.5  RT (min)      lcqtof_hilic_neg_4_053_14    m/z 514.2702     0   1E+4   2   2.5  RT (min)      lcqtof_hilic_neg_4_032_01    m/z 514.3288     0   1E+4   0.5   1  RT (min)      lcqtof_hilic_neg_3_035_46    m/z 514.3980     0   1   2E+4   4   4.5  RT (min)      lcqtof_hilic_neg_4_039_18    m/z 514.5136     0   2   4   6E+4   4   4.5  RT (min)      lcqtof_hilic_neg_4_039_18    m/z 514.5279     0   1E+4   1.5   2  RT (min)      lcqtof_hilic_neg_4_039_18    m/z 514.5548     0   1   2   3E+4   3   3.5  RT (min)      lcqtof_hilic_neg_4_048_23    m/z 514.5860     0   1   2   3E+4   1.5   2  RT (min)      lcqtof_hilic_neg_4_039_18    m/z 514.6908     0   1   2E+4   4   4.5  RT (min)      lcqtof_hilic_neg_4_039_18    m/z 514.8158     0   1   2E+4   1.5   2  RT (min)      lcqtof_hilic_neg_4_039_18    m/z 514.8554     0   1E+4   0.5   1  RT (min)      lcqtof_hilic_neg_4_021_21    m/z 514.8667     0   1E+4   0.5   1  RT (min)      lcqtof_hilic_neg_4_015_09    m/z 514.9405     0   1   2   3   4E+4   0.5   1  RT (min)      lcqtof_hilic_neg_3_035_46    m/z 515.0016     0   1E+4   0.5   1  RT (min)      lcqtof_hilic_neg_2_049_55    m/z 515.1824     0   1   2E+5   1   1.5  RT (min)      lcqtof_hilic_neg_2_037_21    m/z 515.1826     0   1   2E+5   1   1.5  RT (min)      lcqtof_hilic_neg_4_039_18    m/z 515.2062     0   1   2E+5   1   1.5  RT (min)      lcqtof_hilic_neg_4_000_aa    m/z 515.2086     0   2   4   6E+4   6   6.5  RT (min)      lcqtof_hilic_neg_4_048_23    m/z 515.2104     0   1   2E+5   0.5   1  RT (min)      lcqtof_hilic_neg_3_003_20    m/z 515.2200     0   1E+4   7   7.5  RT (min)      lcqtof_hilic_neg_4_048_23    m/z 515.2408     0   1E+4   0.5   1  RT (min)      lcqtof_hilic_neg_4_050_02    m/z 515.2497     0   1E+4   15   15.5  RT (min)      lcqtof_hilic_neg_4_000_aa    m/z 515.3267     0   2   4   6   8E+4   3   3.5  RT (min)      lcqtof_hilic_neg_4_054_48    m/z 515.3318     0   1E+4   2   2.5  RT (min)       lcqtof_hilic_neg_4_003_16    m/z 515.3491     0   1   2E+4   4   4.5  RT (min)       lcqtof_hilic_neg_4_094_60    m/z 515.3554     0   1   2   3E+4   1   1.5  RT (min)      lcqtof_hilic_neg_1_007_05    m/z 515.3559     0   2   4   6E+4   1.5   2  RT (min)      lcqtof_hilic_neg_4_040_05    m/z 515.3692     0   2   4E+4   1   1.5  RT (min)      lcqtof_hilic_neg_4_053_14    m/z 515.3900     0   1   2   3E+4   1   1.5  RT (min)      lcqtof_hilic_neg_4_000_aa    m/z 515.4234     0   1E+4   0.5   1  RT (min)      lcqtof_hilic_neg_4_027_19    m/z 516.1590     0   2   4E+4   7   7.5  RT (min)      lcqtof_hilic_neg_1_011_46    m/z 516.1603     0   2   4E+4   7   7.5  RT (min)      lcqtof_hilic_neg_1_011_46    m/z 516.1849     0   1   2   3   4E+4   1   1.5  RT (min)      lcqtof_hilic_neg_4_039_18    m/z 516.2124     0   1E+4   6   6.5  RT (min)      lcqtof_hilic_neg_4_048_23    m/z 516.2134     0   2   4   6   8E+4   0.5   1  RT (min)      lcqtof_hilic_neg_3_003_20    m/z 516.2151     0   1E+5   1   1.5  RT (min)      lcqtof_hilic_neg_4_000_aa    m/z 516.2176     0   1E+5   1   1.5  RT (min)      lcqtof_hilic_neg_4_000_aa    m/z 516.3049     0   1   2E+4   1   1.5  RT (min)      lcqtof_hilic_neg_4_000_aa    m/z 516.3298     0   1   2E+4   3   3.5  RT (min)      lcqtof_hilic_neg_4_031_38    m/z 516.3612     0   1   2E+4   1.5   2  RT (min)      lcqtof_hilic_neg_4_040_05    m/z 516.3929     0   1E+4   1   1.5  RT (min)      lcqtof_hilic_neg_4_000_aa    m/z 516.8507     0   1E+4   0.5   1  RT (min)      lcqtof_hilic_neg_4_021_21    m/z 517.0762     0   1E+4   0.5   1  RT (min)      lcqtof_hilic_neg_3_003_20    m/z 517.0774     0   1E+4   1   1.5  RT (min)      lcqtof_hilic_neg_4_000_aa    m/z 517.0798     0   1   2E+4   1   1.5  RT (min)      lcqtof_hilic_neg_4_004_47    m/z 517.1742     0   1   2E+4   6   6.5  RT (min)       lcqtof_hilic_neg_4_090_60    m/z 517.2824     0   1E+4   3   3.5  RT (min)      lcqtof_hilic_neg_4_039_18    m/z 517.2878     0   2   4E+4   1   1.5  RT (min)      lcqtof_hilic_neg_4_000_aa    m/z 517.2998     0   2   4E+4   1   1.5  RT (min)      lcqtof_hilic_neg_4_000_aa    m/z 517.3484     0   1E+4   1.5   2  RT (min)       lcqtof_hilic_neg_4_003_16    m/z 517.3674     0   1E+4   0.5   1  RT (min)      lcqtof_hilic_neg_1_030_17    m/z 517.3705     0   1E+4   1   1.5  RT (min)      lcqtof_hilic_neg_4_000_aa    m/z 517.3899     0   1   2E+5   1   1.5  RT (min)      lcqtof_hilic_neg_4_053_14    m/z 517.4161     0   1   2E+4   1   1.5  RT (min)       lcqtof_hilic_neg_4_008_24    m/z 517.8464     0   1   2E+4   4   4.5  RT (min)      lcqtof_hilic_neg_4_008_24    m/z 517.9603     0   1E+4   2.5   3  RT (min)       lcqtof_hilic_neg_4_027_19    m/z 518.0810     0   1E+4   0.5   1  RT (min)      lcqtof_hilic_neg_3_003_20    m/z 518.1001     0   1   2E+4   0.5   1   1.5  RT (min)      lcqtof_hilic_neg_4_027_19    m/z 518.1576     0   1E+4   7   7.5  RT (min)      lcqtof_hilic_neg_1_011_46    m/z 518.1720     0   1   2E+4   3   3.5  RT (min)      lcqtof_hilic_neg_4_013_20    m/z 518.2126     0   1   2E+4   1   1.5  RT (min)      lcqtof_hilic_neg_4_000_aa    m/z 518.2150     0   1   2E+4   1   1.5  RT (min)      lcqtof_hilic_neg_4_000_aa    m/z 518.2158     0   1   2E+4   1   1.5  RT (min)      lcqtof_hilic_neg_4_000_aa    m/z 518.2781     0   2   4   6E+4   1   1.5  RT (min)      lcqtof_hilic_neg_4_000_aa    m/z 518.2827     0   2   4   6E+4   1   1.5  RT (min)      lcqtof_hilic_neg_4_000_aa    m/z 518.2838     0   2   4   6E+4   1   1.5  RT (min)      lcqtof_hilic_neg_4_000_aa    m/z 518.2838     0   2   4   6E+4   1   1.5  RT (min)      lcqtof_hilic_neg_4_000_aa    m/z 518.3925     0   2   4   6E+4   1   1.5  RT (min)      lcqtof_hilic_neg_4_059_14    m/z 518.8548     0   1E+4   4   4.5  RT (min)      lcqtof_hilic_neg_4_008_24    m/z 519.0036     0   1   2   3E+3   5.5   6  RT (min)       lcqtof_hilic_neg_2_051_14    m/z 519.0036     0   1   2E+3   5.5   6  RT (min)      lcqtof_hilic_neg_1_060_14    m/z 519.0041     0   1   2   3E+3   5   5.5  RT (min)      lcqtof_hilic_neg_4_019_03    m/z 519.0734     0   1   2E+4   1   1.5  RT (min)      lcqtof_hilic_neg_4_000_aa    m/z 519.0756     0   1E+4   6   6.5  RT (min)       lcqtof_hilic_neg_4_083_59    m/z 519.0877     0   1   2   3E+4   0.5   1  RT (min)      lcqtof_hilic_neg_3_003_20    m/z 519.0900     0   1   2   3E+4   0.5   1  RT (min)      lcqtof_hilic_neg_3_003_20    m/z 519.1137     0   1   2   3E+4   1.5   2  RT (min)      lcqtof_hilic_neg_4_010_28    m/z 519.1755     0   1   2E+5   1   1.5  RT (min)      lcqtof_hilic_neg_4_000_aa    m/z 519.2531     0   1   2E+4   6   6.5  RT (min)      lcqtof_hilic_neg_4_008_24    m/z 519.2762     0   1E+5   1   1.5  RT (min)      lcqtof_hilic_neg_4_000_aa    m/z 519.2802     0   1E+5   1   1.5  RT (min)      lcqtof_hilic_neg_4_000_aa    m/z 519.2817     0   1E+5   1   1.5  RT (min)      lcqtof_hilic_neg_4_000_aa    m/z 519.2847     0   1E+5   1   1.5  RT (min)      lcqtof_hilic_neg_4_000_aa    m/z 519.2961     0   1E+4   1   1.5  RT (min)      lcqtof_hilic_neg_4_031_38    m/z 519.4301     0   1   2   3   4E+4   0.5   1  RT (min)      lcqtof_hilic_neg_3_035_46    m/z 519.4311     0   1E+4   1   1.5  RT (min)      lcqtof_hilic_neg_4_008_24    m/z 520.0937     0   1E+4   0.5   1  RT (min)      lcqtof_hilic_neg_3_003_20    m/z 520.1506     0   1E+4   15   15.5  RT (min)      lcqtof_hilic_neg_4_000_aa    m/z 520.1522     0   1E+4   3.5   4  RT (min)      lcqtof_hilic_neg_4_038_10    m/z 520.1758     0   1E+5   1   1.5  RT (min)      lcqtof_hilic_neg_4_000_aa    m/z 520.1758     0   1E+5   1   1.5  RT (min)      lcqtof_hilic_neg_4_000_aa    m/z 520.1927     0   1E+4   4   4.5  RT (min)       lcqtof_hilic_neg_4_092_60    m/z 520.1930     0   1E+4   3   3.5  RT (min)      lcqtof_hilic_neg_4_005_29    m/z 520.2253     0   1E+4   6   6.5  RT (min)      lcqtof_hilic_neg_4_040_05    m/z 520.2660     0   2   4   6   8E+4   3   3.5  RT (min)      lcqtof_hilic_neg_4_028_36    m/z 520.2816     0   1   2E+4   1   1.5  RT (min)      lcqtof_hilic_neg_4_000_aa    m/z 520.4990     0   1E+4   4.5   5  RT (min)      lcqtof_hilic_neg_4_040_05    m/z 520.6771     0   1E+4   4.5   5  RT (min)      lcqtof_hilic_neg_4_050_02    m/z 520.9096     0   1   2   3   4E+4   5.5   6  RT (min)       lcqtof_hilic_neg_4_012_07    m/z 520.9097     0   1E+4   5.5   6  RT (min)       lcqtof_hilic_neg_4_112_62    m/z 520.9098     0   1   2   3   4E+3   6.5   7  RT (min)       lcqtof_hilic_neg_4_102_61    m/z 520.9100     0   1E+4   5.5   6  RT (min)       lcqtof_hilic_neg_3_050_41    m/z 520.9101     0   1   2E+4   5.5   6  RT (min)       lcqtof_hilic_neg_1_034_08    m/z 520.9104     0   1E+4   6   6.5  RT (min)       lcqtof_hilic_neg_4_021_21    m/z 521.0697     0   1   2E+4   1   1.5  RT (min)      lcqtof_hilic_neg_4_000_aa    m/z 521.1520     0   1E+4   2.5   3  RT (min)      lcqtof_hilic_neg_1_016_28    m/z 521.1696     0   2   4   6E+4   1   1.5  RT (min)      lcqtof_hilic_neg_4_000_aa    m/z 521.1726     0   2   4   6E+4   1   1.5  RT (min)      lcqtof_hilic_neg_4_000_aa    m/z 521.2349     0   1E+4   0.5   1  RT (min)      lcqtof_hilic_neg_2_004_26    m/z 521.2390     0   1E+4   2   2.5  RT (min)      lcqtof_hilic_neg_4_052_13    m/z 521.2696     0   1   2E+4   3   3.5  RT (min)       lcqtof_hilic_neg_4_028_36    m/z 521.3252     0   1   2   3E+4   1   1.5  RT (min)      lcqtof_hilic_neg_4_000_aa    m/z 521.3257     0   1   2   3E+4   0.5   1  RT (min)      lcqtof_hilic_neg_4_000_aa    m/z 521.3413     0   1   2E+4   1   1.5  RT (min)      lcqtof_hilic_neg_4_000_aa    m/z 521.3433     0   1E+4   1.5   2  RT (min)      lcqtof_hilic_neg_4_003_16    m/z 521.3948     0   1   2   3   4E+4   1   1.5  RT (min)      lcqtof_hilic_neg_4_000_aa    m/z 521.4028     0   1E+5   0.5   1  RT (min)      lcqtof_hilic_neg_4_005_29    m/z 521.4193     0   1   2E+5   1   1.5  RT (min)      lcqtof_hilic_neg_4_053_14    m/z 521.4352     0   1   2   3E+4   0.5   1  RT (min)      lcqtof_hilic_neg_3_035_46    m/z 521.4607     0   1   2   3E+4   1   1.5  RT (min)      lcqtof_hilic_neg_4_000_aa    m/z 521.4657     0   1   2   3E+4   1   1.5  RT (min)      lcqtof_hilic_neg_4_000_aa    m/z 521.4916     0   1   2   3E+4   1   1.5  RT (min)      lcqtof_hilic_neg_4_000_aa    m/z 521.4919     0   1   2   3E+4   1   1.5  RT (min)      lcqtof_hilic_neg_4_000_aa    m/z 522.1716     0   1E+4   7   7.5  RT (min)      lcqtof_hilic_neg_4_040_05    m/z 522.1728     0   1E+4   1   1.5  RT (min)      lcqtof_hilic_neg_4_000_aa    m/z 522.1754     0   1E+4   1   1.5  RT (min)      lcqtof_hilic_neg_4_000_aa    m/z 522.3294     0   1E+4   7   7.5  RT (min)      lcqtof_hilic_neg_1_035_23    m/z 522.4057     0   1E+4   1   1.5  RT (min)      lcqtof_hilic_neg_4_000_aa    m/z 522.4101     0   1E+4   0.5   1  RT (min)      lcqtof_hilic_neg_4_029_17    m/z 522.4990     0   1E+4   1   1.5  RT (min)      lcqtof_hilic_neg_4_000_aa    m/z 523.0555     0   1E+4   6.5   7  RT (min)      lcqtof_hilic_neg_4_048_23    m/z 523.0664     0   1E+4   1   1.5  RT (min)      lcqtof_hilic_neg_4_000_aa    m/z 523.0884     0   1   2   3E+4   6   6.5  RT (min)       lcqtof_hilic_neg_4_036_11    m/z 523.2291     0   1E+4   6   6.5  RT (min)       lcqtof_hilic_neg_4_091_60    m/z 523.2640     0   1E+4   3.5   4  RT (min)      lcqtof_hilic_neg_4_039_18    m/z 523.3509     0   1E+4   4   4.5  RT (min)      lcqtof_hilic_neg_4_040_05    m/z 523.4340     0   1E+4   1   1.5  RT (min)      lcqtof_hilic_neg_4_008_24    m/z 523.4556     0   1E+4   0.5   1  RT (min)      lcqtof_hilic_neg_2_052_24    m/z 523.4712     0   2   4   6E+4   1   1.5  RT (min)      lcqtof_hilic_neg_4_000_aa    m/z 523.4732     0   2   4   6E+4   1.5   2  RT (min)      lcqtof_hilic_neg_4_000_aa    m/z 523.4744     0   2   4   6E+4   1   1.5  RT (min)      lcqtof_hilic_neg_4_000_aa    m/z 523.7916     0   1E+4   10   10.5  RT (min)      lcqtof_hilic_neg_4_014_46    m/z 524.2586     0   1E+4   3.5   4  RT (min)      lcqtof_hilic_neg_4_039_18    m/z 524.2688     0   1   2E+4   1   1.5  RT (min)      lcqtof_hilic_neg_4_050_02    m/z 524.3278     0   1   2E+4   1   1.5  RT (min)      lcqtof_hilic_neg_4_000_aa    m/z 524.3370     0   1   2E+4   1   1.5  RT (min)      lcqtof_hilic_neg_4_000_aa    m/z 524.3442     0   1   2E+4   1   1.5  RT (min)      lcqtof_hilic_neg_4_000_aa    m/z 524.3444     0   1   2E+4   1   1.5  RT (min)      lcqtof_hilic_neg_4_000_aa    m/z 524.3692     0   1E+4   15.5   16  RT (min)      lcqtof_hilic_neg_4_000_aa    m/z 524.3800     0   1E+4   1   1.5  RT (min)      lcqtof_hilic_neg_2_028_05    m/z 524.4750     0   1   2E+4   1   1.5  RT (min)      lcqtof_hilic_neg_4_000_aa    m/z 524.5108     0   1E+4   4   4.5  RT (min)      lcqtof_hilic_neg_4_032_01    m/z 525.0014     0   1   2E+4   1   1.5  RT (min)      lcqtof_hilic_neg_4_000_aa    m/z 525.0030     0   1   2E+4   1   1.5  RT (min)      lcqtof_hilic_neg_4_000_aa    m/z 525.3291     0   1   2E+4   1   1.5  RT (min)      lcqtof_hilic_neg_4_000_aa    m/z 525.3324     0   1   2E+4   1   1.5  RT (min)      lcqtof_hilic_neg_4_000_aa    m/z 525.4547     0   1   2   3E+4   1   1.5  RT (min)      lcqtof_hilic_neg_4_000_aa    m/z 525.4628     0   1   2E+4   1   1.5  RT (min)      lcqtof_hilic_neg_4_000_aa    m/z 526.1676     0   1E+4   0.5   1  RT (min)      lcqtof_hilic_neg_4_013_20    m/z 526.1882     0   1   2E+4   7   7.5  RT (min)      lcqtof_hilic_neg_4_014_46    m/z 526.1885     0   1E+4   7   7.5  RT (min)      lcqtof_hilic_neg_1_011_46    m/z 526.1890     0   1   2E+4   7   7.5  RT (min)      lcqtof_hilic_neg_4_014_46    m/z 526.3738     0   2   4   6E+4   4   4.5  RT (min)      lcqtof_hilic_neg_4_008_24    m/z 526.3997     0   1E+4   1   1.5  RT (min)      lcqtof_hilic_neg_2_028_05    m/z 526.4520     0   1E+4   1   1.5  RT (min)      lcqtof_hilic_neg_4_000_aa    m/z 526.8824     0   2   4   6   8E+4   8.5   9  RT (min)      lcqtof_hilic_neg_1_000_aa    m/z 526.8824     0   1E+5   7.5   8  RT (min)      lcqtof_hilic_neg_1_000_aa    m/z 526.8828     0   1E+5   7.5   8  RT (min)      lcqtof_hilic_neg_1_000_aa    m/z 526.8836     0   1E+5   8.5   9  RT (min)       lcqtof_hilic_neg_1_000_aa    m/z 526.8842     0   1E+5   8.5   9  RT (min)      lcqtof_hilic_neg_1_000_aa    m/z 526.8858     0   1E+5   7.5   8   8.5  RT (min)      lcqtof_hilic_neg_1_000_aa    m/z 527.1812     0   1E+4   0.5   1  RT (min)      lcqtof_hilic_neg_3_003_20    m/z 527.1890     0   1   2E+4   7   7.5  RT (min)       lcqtof_hilic_neg_4_008_24    m/z 527.1950     0   1   2E+4   7   7.5  RT (min)      lcqtof_hilic_neg_4_008_24    m/z 527.2528     0   1   2E+4   15   15.5  RT (min)      lcqtof_hilic_neg_4_000_aa    m/z 527.2730     0   2   4   6   8E+4   1   1.5  RT (min)      lcqtof_hilic_neg_4_000_aa    m/z 527.2764     0   1E+5   1   1.5  RT (min)      lcqtof_hilic_neg_4_000_aa    m/z 527.2838     0   1E+5   1   1.5  RT (min)      lcqtof_hilic_neg_4_000_aa    m/z 527.2853     0   1E+5   1   1.5  RT (min)      lcqtof_hilic_neg_4_000_aa    m/z 527.2867     0   1E+5   1   1.5  RT (min)      lcqtof_hilic_neg_4_000_aa    m/z 527.2969     0   1   2   3E+4   3.5   4  RT (min)      lcqtof_hilic_neg_4_039_18    m/z 527.3028     0   1E+4   2   2.5  RT (min)      lcqtof_hilic_neg_4_039_18    m/z 527.3166     0   1E+4   4   4.5  RT (min)      lcqtof_hilic_neg_4_031_38    m/z 527.4322     0   1E+4   1.5   2  RT (min)      lcqtof_hilic_neg_4_000_aa    m/z 527.4469     0   1E+4   1.5   2  RT (min)      lcqtof_hilic_neg_4_000_aa    m/z 528.1655     0   1E+4   0.5   1  RT (min)      lcqtof_hilic_neg_4_013_20    m/z 528.2282     0   1E+4   3.5   4  RT (min)      lcqtof_hilic_neg_1_007_05    m/z 528.2303     0   1E+4   3.5   4  RT (min)      lcqtof_hilic_neg_1_007_05    m/z 528.2324     0   1E+4   3   3.5  RT (min)      lcqtof_hilic_neg_4_038_10    m/z 528.2530     0   1E+4   6   6.5  RT (min)      lcqtof_hilic_neg_4_040_05    m/z 528.2636     0   1   2E+4   3   3.5  RT (min)      lcqtof_hilic_neg_4_019_03    m/z 528.2870     0   1E+4   4   4.5  RT (min)      lcqtof_hilic_neg_1_015_33    m/z 528.3350     0   1E+4   4   4.5  RT (min)      lcqtof_hilic_neg_4_008_24    m/z 528.3590     0   1E+4   4   4.5  RT (min)      lcqtof_hilic_neg_4_053_14    m/z 528.5427     0   1E+4   3.5   4  RT (min)      lcqtof_hilic_neg_4_052_13    m/z 528.7078     0   1E+4   4.5   5  RT (min)      lcqtof_hilic_neg_4_050_02    m/z 528.8346     0   1   2   3E+4   4   4.5  RT (min)      lcqtof_hilic_neg_4_008_24    m/z 528.8616     0   2   4   6E+4   8   8.5  RT (min)      lcqtof_hilic_neg_1_000_aa    m/z 528.9588     0   1E+5   0.5   1  RT (min)      lcqtof_hilic_neg_3_035_46    m/z 528.9842     0   1   2   3E+4   1   1.5  RT (min)      lcqtof_hilic_neg_4_000_aa    m/z 529.1597     0   1E+4   0.5   1  RT (min)      lcqtof_hilic_neg_4_039_18    m/z 529.2332     0   1E+4   1   1.5  RT (min)      lcqtof_hilic_neg_4_000_aa    m/z 529.2904     0   1   2   3E+4   1   1.5  RT (min)      lcqtof_hilic_neg_4_000_aa    m/z 529.3038     0   1   2   3E+4   1   1.5  RT (min)       lcqtof_hilic_neg_4_000_aa    m/z 529.3379     0   1E+4   3.5   4  RT (min)      lcqtof_hilic_neg_4_040_05    m/z 529.3384     0   1   2   3   4E+4   4   4.5  RT (min)      lcqtof_hilic_neg_4_008_24    m/z 529.3657     0   2   4   6   8E+4   1   1.5  RT (min)      lcqtof_hilic_neg_4_000_aa    m/z 529.3756     0   1E+4   0.5   1  RT (min)      lcqtof_hilic_neg_4_050_02    m/z 529.4414     0   1   2   3E+4   0.5   1  RT (min)      lcqtof_hilic_neg_3_035_46    m/z 529.5384     0   1   2E+4   0.5   1  RT (min)      lcqtof_hilic_neg_4_048_23    m/z 529.8418     0   2   4   6   8E+4   4   4.5  RT (min)      lcqtof_hilic_neg_4_008_24    m/z 529.9524     0   1E+4   6.5   7  RT (min)       lcqtof_hilic_neg_4_029_17    m/z 529.9525     0   1   2   3   4E+3   6.5   7  RT (min)      lcqtof_hilic_neg_4_053_14    m/z 529.9592     0   1   2   3   4E+4   0.5   1  RT (min)      lcqtof_hilic_neg_3_035_46    m/z 530.3022     0   2   4   6E+4   4   4.5  RT (min)       lcqtof_hilic_neg_4_059_14    m/z 530.3083     0   1   2E+5   1   1.5  RT (min)      lcqtof_hilic_neg_4_016_37    m/z 530.3492     0   1   2   3   4E+4   4   4.5  RT (min)      lcqtof_hilic_neg_4_008_24    m/z 530.3498     0   1E+4   3.5   4  RT (min)      lcqtof_hilic_neg_4_040_05    m/z 530.3722     0   1   2   3E+4   1   1.5  RT (min)      lcqtof_hilic_neg_4_000_aa    m/z 530.8422     0   1   2   3   4E+4   8   8.5  RT (min)      lcqtof_hilic_neg_1_000_aa    m/z 530.8500     0   1E+5   4   4.5  RT (min)      lcqtof_hilic_neg_4_008_24    m/z 530.9561     0   1   2   3E+4   0.5   1  RT (min)      lcqtof_hilic_neg_3_035_46    m/z 530.9798     0   1   2E+4   1   1.5  RT (min)      lcqtof_hilic_neg_4_000_aa    m/z 530.9815     0   1   2E+4   1   1.5  RT (min)      lcqtof_hilic_neg_4_000_aa    m/z 531.1455     0   1E+4   3.5   4  RT (min)      lcqtof_hilic_neg_1_007_05    m/z 531.1579     0   1E+4   0.5   1  RT (min)      lcqtof_hilic_neg_4_039_18    m/z 531.2561     0   1E+4   2.5   3  RT (min)      lcqtof_hilic_neg_4_031_38    m/z 531.2844     0   1   2   3E+5   0.5   1  RT (min)      lcqtof_hilic_neg_4_000_aa    m/z 531.3451     0   1   2E+4   4   4.5  RT (min)       lcqtof_hilic_neg_4_008_24    m/z 531.4058     0   1E+4   1   1.5  RT (min)      lcqtof_hilic_neg_4_000_aa    m/z 531.4066     0   1E+4   1   1.5  RT (min)      lcqtof_hilic_neg_4_000_aa    m/z 531.4570     0   1   2E+4   0.5   1  RT (min)      lcqtof_hilic_neg_3_035_46    m/z 531.8574     0   1   2   3E+4   4   4.5  RT (min)      lcqtof_hilic_neg_4_008_24    m/z 531.9253     0   1E+4   0.5   1  RT (min)      lcqtof_hilic_neg_4_004_47    m/z 532.1867     0   1E+4   6.5   7  RT (min)      lcqtof_hilic_neg_4_008_24    m/z 532.1869     0   1E+4   6   6.5  RT (min)       lcqtof_hilic_neg_4_052_13    m/z 532.2426     0   1E+4   1   1.5  RT (min)      lcqtof_hilic_neg_4_027_19    m/z 532.2826     0   1E+5   0.5   1  RT (min)      lcqtof_hilic_neg_4_000_aa    m/z 532.2882     0   1E+5   1   1.5  RT (min)      lcqtof_hilic_neg_4_000_aa    m/z 532.2896     0   1   2   3E+4   3.5   4  RT (min)      lcqtof_hilic_neg_4_039_18    m/z 532.3654     0   1E+4   3.5   4  RT (min)      lcqtof_hilic_neg_4_008_24    m/z 532.8210     0   1E+4   8   8.5  RT (min)      lcqtof_hilic_neg_1_000_aa    m/z 532.8494     0   2   4E+4   1   1.5  RT (min)      lcqtof_hilic_neg_4_004_47    m/z 532.8667     0   1E+4   4   4.5  RT (min)      lcqtof_hilic_neg_4_008_24    m/z 532.8674     0   1E+4   0.5   1  RT (min)      lcqtof_hilic_neg_3_035_46    m/z 532.9803     0   1E+4   1   1.5  RT (min)      lcqtof_hilic_neg_4_000_aa    m/z 533.0983     0   1E+4   7.5   8  RT (min)      lcqtof_hilic_neg_4_008_24    m/z 533.1388     0   1E+4   1   1.5  RT (min)      lcqtof_hilic_neg_4_000_aa    m/z 533.1410     0   1E+4   1   1.5  RT (min)      lcqtof_hilic_neg_4_000_aa    m/z 533.1606     0   1   2   3   4E+4   3.5   4  RT (min)      lcqtof_hilic_neg_2_028_05    m/z 533.1607     0   1   2   3   4E+4   3.5   4  RT (min)      lcqtof_hilic_neg_2_028_05    m/z 533.2803     0   1   2E+4   7   7.5  RT (min)       lcqtof_hilic_neg_4_008_24    m/z 533.2924     0   1E+4   3.5   4  RT (min)      lcqtof_hilic_neg_4_039_18    m/z 533.3014     0   1   2E+4   1   1.5  RT (min)      lcqtof_hilic_neg_4_000_aa    m/z 533.3079     0   1E+4   1   1.5  RT (min)      lcqtof_hilic_neg_4_000_aa    m/z 533.3512     0   1E+4   0.5   1  RT (min)      lcqtof_hilic_neg_1_046_30    m/z 533.3841     0   1   2   3   4E+4   1   1.5  RT (min)      lcqtof_hilic_neg_4_053_14    m/z 533.4386     0   1E+4   0.5   1  RT (min)      lcqtof_hilic_neg_3_035_46    m/z 533.4833     0   1E+4   0.5   1  RT (min)      lcqtof_hilic_neg_3_035_46    m/z 534.1654     0   1E+4   3.5   4  RT (min)      lcqtof_hilic_neg_2_028_05    m/z 534.1655     0   1E+4   3.5   4  RT (min)      lcqtof_hilic_neg_2_028_05    m/z 534.2720     0   1   2E+4   6   6.5  RT (min)      lcqtof_hilic_neg_4_008_24    m/z 534.3122     0   1   2E+4   2   2.5  RT (min)      lcqtof_hilic_neg_4_028_36    m/z 534.3131     0   1   2E+4   2   2.5  RT (min)      lcqtof_hilic_neg_4_034_31    m/z 535.0756     0   1E+4   2   2.5  RT (min)      lcqtof_hilic_neg_4_039_18    m/z 535.1746     0   1E+4   3.5   4  RT (min)      lcqtof_hilic_neg_2_028_05    m/z 535.1750     0   1E+4   3.5   4  RT (min)      lcqtof_hilic_neg_2_028_05    m/z 535.3608     0   1E+4   3   3.5  RT (min)      lcqtof_hilic_neg_4_031_38    m/z 535.3643     0   2   4E+4   0.5   1  RT (min)      lcqtof_hilic_neg_4_000_aa    m/z 535.3650     0   2   4E+4   1   1.5  RT (min)      lcqtof_hilic_neg_4_000_aa    m/z 535.3666     0   2   4E+4   1   1.5  RT (min)      lcqtof_hilic_neg_4_000_aa    m/z 535.3673     0   2   4E+4   1   1.5  RT (min)      lcqtof_hilic_neg_4_000_aa    m/z 535.4530     0   1   2E+4   0.5   1  RT (min)      lcqtof_hilic_neg_3_035_46    m/z 535.4724     0   1   2   3E+4   1   1.5  RT (min)      lcqtof_hilic_neg_4_000_aa    m/z 535.5432     0   1E+4   1.5   2  RT (min)      lcqtof_hilic_neg_4_000_aa    m/z 535.5434     0   1E+4   1.5   2  RT (min)      lcqtof_hilic_neg_4_000_aa    m/z 535.9682     0   1E+4   1   1.5  RT (min)      lcqtof_hilic_neg_4_054_48    m/z 536.0646     0   1E+4   4.5   5  RT (min)      lcqtof_hilic_neg_4_039_18    m/z 536.3575     0   1E+4   1   1.5  RT (min)      lcqtof_hilic_neg_4_000_aa    m/z 536.3660     0   1E+4   1   1.5  RT (min)      lcqtof_hilic_neg_4_000_aa    m/z 536.3670     0   1E+4   1   1.5  RT (min)      lcqtof_hilic_neg_4_000_aa    m/z 536.4786     0   1E+4   1   1.5  RT (min)      lcqtof_hilic_neg_4_000_aa    m/z 536.8830     0   1   2   3E+3   5.5   6  RT (min)       lcqtof_hilic_neg_4_110_62    m/z 536.8833     0   2   4   6   8E+3   5   5.5  RT (min)       lcqtof_hilic_neg_4_058_39    m/z 536.8841     0   1   2   3E+3   5   5.5  RT (min)       lcqtof_hilic_neg_4_108_62    m/z 537.0054     0   1E+4   0.5   1  RT (min)      lcqtof_hilic_neg_3_035_46    m/z 537.0140     0   1   2   3   4E+4   1   1.5  RT (min)      lcqtof_hilic_neg_4_000_aa    m/z 537.0931     0   1E+4   1   1.5  RT (min)      lcqtof_hilic_neg_4_000_aa    m/z 537.2082     0   1   2   3   4E+4   0.5   1  RT (min)      lcqtof_hilic_neg_3_003_20    m/z 537.2167     0   1E+4   6   6.5  RT (min)      lcqtof_hilic_neg_4_048_23    m/z 537.2204     0   1   2   3   4E+4   0.5   1  RT (min)      lcqtof_hilic_neg_3_003_20    m/z 537.2657     0   1E+4   6   6.5  RT (min)      lcqtof_hilic_neg_4_008_24    m/z 537.3482     0   1   2E+4   1   1.5  RT (min)      lcqtof_hilic_neg_4_000_aa    m/z 537.3958     0   1E+4   4   4.5  RT (min)      lcqtof_hilic_neg_4_008_24    m/z 537.4303     0   1E+4   0.5   1  RT (min)      lcqtof_hilic_neg_3_035_46    m/z 537.4544     0   1   2   3E+4   1   1.5  RT (min)      lcqtof_hilic_neg_4_000_aa    m/z 537.4876     0   1   2E+5   1   1.5  RT (min)      lcqtof_hilic_neg_4_000_aa    m/z 537.4882     0   1   2E+5   1.5   2  RT (min)      lcqtof_hilic_neg_4_000_aa    m/z 537.4884     0   1   2E+5   1.5   2  RT (min)      lcqtof_hilic_neg_4_000_aa    m/z 537.4905     0   1   2E+5   1   1.5  RT (min)      lcqtof_hilic_neg_4_000_aa    m/z 538.1126     0   1E+4   6   6.5  RT (min)      lcqtof_hilic_neg_4_002_25    m/z 538.1131     0   1E+4   6   6.5  RT (min)      lcqtof_hilic_neg_4_012_07    m/z 538.1619     0   1E+4   7   7.5  RT (min)      lcqtof_hilic_neg_2_022_46    m/z 538.1994     0   1   2   3E+4   7   7.5  RT (min)       lcqtof_hilic_neg_4_052_13    m/z 538.2331     0   1   2   3E+4   3.5   4  RT (min)      lcqtof_hilic_neg_1_032_18    m/z 538.3334     0   2   4   6E+4   1   1.5  RT (min)      lcqtof_hilic_neg_4_000_aa    m/z 538.3372     0   1   2E+4   0.5   1  RT (min)      lcqtof_hilic_neg_1_032_18    m/z 538.4583     0   1E+4   1   1.5  RT (min)      lcqtof_hilic_neg_4_000_aa    m/z 538.4910     0   2   4   6   8E+4   1   1.5  RT (min)      lcqtof_hilic_neg_4_000_aa    m/z 539.1104     0   1E+4   0.5   1  RT (min)       lcqtof_hilic_neg_4_073_57    m/z 539.1383     0   1E+4   7   7.5  RT (min)      lcqtof_hilic_neg_1_000_aa    m/z 539.2070     0   1E+4   0.5   1  RT (min)      lcqtof_hilic_neg_3_003_20    m/z 539.2488     0   1E+5   4   4.5  RT (min)      lcqtof_hilic_neg_4_055_32    m/z 539.2950     0   1E+4   1   1.5  RT (min)      lcqtof_hilic_neg_4_000_aa    m/z 539.3334     0   1   2E+4   0.5   1  RT (min)      lcqtof_hilic_neg_1_032_18    m/z 539.3388     0   1   2   3E+3   1.5   2  RT (min)      lcqtof_hilic_neg_1_064_16    m/z 539.4038     0   1E+4   4   4.5  RT (min)      lcqtof_hilic_neg_4_040_05    m/z 539.4304     0   1   2   3E+5   1   1.5  RT (min)      lcqtof_hilic_neg_4_053_14    m/z 539.4362     0   1E+4   1.5   2  RT (min)      lcqtof_hilic_neg_4_040_05    m/z 539.4654     0   1   2   3   4E+4   1   1.5  RT (min)      lcqtof_hilic_neg_4_000_aa    m/z 539.4848     0   1   2   3   4E+4   1   1.5  RT (min)      lcqtof_hilic_neg_4_000_aa    m/z 540.0205     0   1   2E+4   1   1.5  RT (min)      lcqtof_hilic_neg_4_000_aa    m/z 540.0214     0   1   2E+4   1   1.5  RT (min)      lcqtof_hilic_neg_4_000_aa    m/z 540.0226     0   1   2E+4   1   1.5  RT (min)      lcqtof_hilic_neg_4_000_aa    m/z 540.2320     0   1E+4   3.5   4  RT (min)      lcqtof_hilic_neg_1_032_18    m/z 540.4340     0   1E+5   1   1.5  RT (min)      lcqtof_hilic_neg_4_053_14    m/z 540.4515     0   1E+4   0.5   1  RT (min)      lcqtof_hilic_neg_3_035_46    m/z 540.4656     0   1E+4   1   1.5  RT (min)      lcqtof_hilic_neg_4_000_aa    m/z 540.6170     0   2   4   6   8E+4   4   4.5  RT (min)      lcqtof_hilic_neg_4_053_14    m/z 541.0155     0   1E+5   1   1.5  RT (min)      lcqtof_hilic_neg_4_000_aa    m/z 541.2569     0   1E+4   0.5   1  RT (min)      lcqtof_hilic_neg_4_038_10    m/z 541.3017     0   1E+4   1   1.5  RT (min)      lcqtof_hilic_neg_4_000_aa    m/z 541.3189     0   1E+4   1   1.5  RT (min)      lcqtof_hilic_neg_4_000_aa    m/z 541.3568     0   1E+4   3   3.5  RT (min)       lcqtof_hilic_neg_4_031_38    m/z 541.9720     0   1   2   3E+4   0.5   1  RT (min)      lcqtof_hilic_neg_4_021_21    m/z 541.9825     0   1   2   3E+4   0.5   1  RT (min)      lcqtof_hilic_neg_1_024_21    m/z 542.0182     0   1   2E+4   1   1.5  RT (min)      lcqtof_hilic_neg_4_000_aa    m/z 542.0186     0   1   2E+4   1   1.5  RT (min)      lcqtof_hilic_neg_4_000_aa    m/z 542.0192     0   1   2E+4   1   1.5  RT (min)      lcqtof_hilic_neg_4_000_aa    m/z 542.0204     0   1   2E+4   1   1.5  RT (min)      lcqtof_hilic_neg_4_000_aa    m/z 542.4462     0   1E+4   1.5   2  RT (min)      lcqtof_hilic_neg_4_040_05    m/z 543.0110     0   2   4   6E+4   1   1.5  RT (min)      lcqtof_hilic_neg_4_000_aa    m/z 543.1130     0   1E+4   1   1.5  RT (min)      lcqtof_hilic_neg_4_013_20    m/z 543.1147     0   1E+4   0.5   1  RT (min)      lcqtof_hilic_neg_4_027_19    m/z 543.1284     0   1   2   3E+4   6   6.5  RT (min)      lcqtof_hilic_neg_4_008_24    m/z 543.1290     0   1   2   3   4E+4   6   6.5  RT (min)       lcqtof_hilic_neg_1_037_24    m/z 543.1762     0   1   2   3E+4   7   7.5  RT (min)       lcqtof_hilic_neg_1_011_46    m/z 543.1778     0   1   2   3   4E+4   7   7.5  RT (min)      lcqtof_hilic_neg_2_022_46    m/z 543.2827     0   1E+4   1.5   2  RT (min)      lcqtof_hilic_neg_4_052_13    m/z 543.4276     0   1E+4   0.5   1  RT (min)      lcqtof_hilic_neg_2_054_42    m/z 543.4280     0   1   2E+4   1   1.5  RT (min)      lcqtof_hilic_neg_4_046_41    m/z 543.9801     0   1   2E+4   0.5   1  RT (min)      lcqtof_hilic_neg_4_021_21    m/z 544.0143     0   1E+4   1   1.5  RT (min)      lcqtof_hilic_neg_4_000_aa    m/z 544.0156     0   1E+4   1   1.5  RT (min)      lcqtof_hilic_neg_4_000_aa    m/z 544.1723     0   1E+4   4   4.5  RT (min)       lcqtof_hilic_neg_4_085_59    m/z 544.2855     0   1E+4   4   4.5  RT (min)      lcqtof_hilic_neg_4_032_01    m/z 544.5852     0   1   2   3E+4   4.5   5  RT (min)      lcqtof_hilic_neg_4_040_05    m/z 545.0052     0   1   2E+4   1   1.5  RT (min)      lcqtof_hilic_neg_4_000_aa    m/z 545.0088     0   1   2E+4   1   1.5  RT (min)      lcqtof_hilic_neg_4_000_aa    m/z 545.0103     0   1   2E+4   1   1.5  RT (min)      lcqtof_hilic_neg_4_000_aa    m/z 545.0628     0   1   2E+4   0.5   1  RT (min)      lcqtof_hilic_neg_3_003_20    m/z 545.2068     0   1E+4   1   1.5  RT (min)      lcqtof_hilic_neg_4_000_aa    m/z 545.2295     0   1   2E+4   0.5   1  RT (min)      lcqtof_hilic_neg_4_050_02    m/z 545.2343     0   1   2E+4   1   1.5  RT (min)      lcqtof_hilic_neg_4_050_02    m/z 545.2874     0   1E+4   1   1.5  RT (min)      lcqtof_hilic_neg_4_000_aa    m/z 545.2913     0   1E+4   0.5   1  RT (min)      lcqtof_hilic_neg_4_000_aa    m/z 545.3527     0   1E+4   0.5   1  RT (min)      lcqtof_hilic_neg_4_019_03    m/z 545.4202     0   2   4   6E+4   1   1.5  RT (min)       lcqtof_hilic_neg_4_000_aa    m/z 546.0482     0   2   4E+4   0.5   1  RT (min)      lcqtof_hilic_neg_2_053_18    m/z 546.2258     0   1E+4   3.5   4  RT (min)      lcqtof_hilic_neg_1_032_18    m/z 546.4220     0   1   2E+4   1   1.5  RT (min)      lcqtof_hilic_neg_4_000_aa    m/z 546.8652     0   2   4   6   8E+4   1   1.5  RT (min)      lcqtof_hilic_neg_4_004_47    m/z 546.8658     0   2   4   6   8E+4   1   1.5  RT (min)      lcqtof_hilic_neg_4_004_47    m/z 546.8833     0   1E+4   0.5   1  RT (min)      lcqtof_hilic_neg_3_035_46    m/z 546.9421     0   1E+4   1   1.5  RT (min)      lcqtof_hilic_neg_4_000_aa    m/z 547.0594     0   1E+4   0.5   1  RT (min)      lcqtof_hilic_neg_3_003_20    m/z 547.2224     0   1E+4   4   4.5  RT (min)      lcqtof_hilic_neg_4_055_32    m/z 547.2342     0   1   2   3E+4   0.5   1  RT (min)      lcqtof_hilic_neg_3_003_20    m/z 547.2644     0   1E+4   3   3.5  RT (min)      lcqtof_hilic_neg_4_039_18    m/z 547.2685     0   1E+4   6   6.5  RT (min)      lcqtof_hilic_neg_4_008_24    m/z 547.2932     0   1E+4   0.5   1  RT (min)      lcqtof_hilic_neg_4_039_18    m/z 547.3130     0   1E+4   1   1.5  RT (min)      lcqtof_hilic_neg_4_000_aa    m/z 547.3218     0   1E+4   1   1.5  RT (min)      lcqtof_hilic_neg_4_000_aa    m/z 547.3501     0   1   2E+4   4   4.5  RT (min)      lcqtof_hilic_neg_4_040_05    m/z 547.3994     0   1E+4   1   1.5  RT (min)      lcqtof_hilic_neg_4_000_aa    m/z 547.4355     0   2   4   6   8E+4   1   1.5  RT (min)      lcqtof_hilic_neg_4_053_14    m/z 547.5268     0   1E+4   1   1.5  RT (min)      lcqtof_hilic_neg_4_000_aa    m/z 548.0462     0   2   4E+4   0.5   1  RT (min)       lcqtof_hilic_neg_3_045_18    m/z 548.2295     0   1E+4   4   4.5  RT (min)      lcqtof_hilic_neg_2_028_05    m/z 548.2515     0   1   2E+4   6   6.5  RT (min)       lcqtof_hilic_neg_4_093_60    m/z 548.3078     0   1   2   3E+4   6   6.5  RT (min)      lcqtof_hilic_neg_4_008_24    m/z 548.3436     0   1E+4   1   1.5  RT (min)      lcqtof_hilic_neg_1_007_05    m/z 548.3808     0   1   2   3E+4   0.5   1  RT (min)      lcqtof_hilic_neg_1_032_18    m/z 549.0511     0   1E+4   0.5   1  RT (min)      lcqtof_hilic_neg_4_039_18    m/z 549.1527     0   1   2   3E+4   1   1.5  RT (min)      lcqtof_hilic_neg_1_032_18    m/z 549.1534     0   1   2E+4   1   1.5  RT (min)      lcqtof_hilic_neg_4_000_aa    m/z 549.1540     0   1   2E+4   1   1.5  RT (min)      lcqtof_hilic_neg_4_000_aa    m/z 549.1661     0   1   2E+4   7   7.5  RT (min)       lcqtof_hilic_neg_1_000_aa    m/z 549.1668     0   1E+4   7.5   8  RT (min)       lcqtof_hilic_neg_4_050_02    m/z 549.1672     0   1   2E+4   7   7.5  RT (min)       lcqtof_hilic_neg_4_039_18    m/z 549.2760     0   1   2E+4   3.5   4  RT (min)      lcqtof_hilic_neg_4_039_18    m/z 549.3704     0   1E+4   4   4.5  RT (min)      lcqtof_hilic_neg_4_040_05    m/z 549.4683     0   2   4   6E+4   0.5   1  RT (min)      lcqtof_hilic_neg_4_053_14    m/z 549.4861     0   1E+4   0.5   1  RT (min)      lcqtof_hilic_neg_1_004_14    m/z 549.5212     0   2   4   6   8E+4   1   1.5  RT (min)      lcqtof_hilic_neg_4_000_aa    m/z 549.5233     0   2   4   6   8E+4   1   1.5  RT (min)      lcqtof_hilic_neg_4_000_aa    m/z 549.5598     0   1E+4   1.5   2  RT (min)      lcqtof_hilic_neg_4_000_aa    m/z 550.1879     0   1E+4   1   1.5  RT (min)      lcqtof_hilic_neg_3_103_61    m/z 550.3147     0   1E+4   4   4.5  RT (min)      lcqtof_hilic_neg_4_002_25    m/z 550.3149     0   1   2E+4   3.5   4  RT (min)      lcqtof_hilic_neg_1_007_05    m/z 550.3598     0   1E+4   1   1.5  RT (min)      lcqtof_hilic_neg_1_007_05    m/z 550.4839     0   1E+4   0.5   1  RT (min)      lcqtof_hilic_neg_4_048_23    m/z 550.5206     0   1   2   3E+4   1   1.5  RT (min)      lcqtof_hilic_neg_4_000_aa    m/z 550.5274     0   1   2   3E+4   1   1.5  RT (min)      lcqtof_hilic_neg_4_000_aa    m/z 551.0382     0   1   2E+4   1   1.5  RT (min)      lcqtof_hilic_neg_4_000_aa    m/z 551.1848     0   1   2   3   4E+4   0.5   1  RT (min)       lcqtof_hilic_neg_4_105_61    m/z 551.2443     0   2   4E+4   6.5   7  RT (min)      lcqtof_hilic_neg_4_008_24    m/z 551.2672     0   1E+4   1   1.5  RT (min)      lcqtof_hilic_neg_4_052_13    m/z 551.3246     0   1E+5   1   1.5  RT (min)      lcqtof_hilic_neg_4_000_aa    m/z 551.3252     0   1E+5   1   1.5  RT (min)      lcqtof_hilic_neg_4_000_aa    m/z 551.3260     0   1E+5   1   1.5  RT (min)      lcqtof_hilic_neg_4_000_aa    m/z 551.4688     0   1   2   3   4E+4   1   1.5  RT (min)       lcqtof_hilic_neg_4_000_aa    m/z 551.4694     0   1   2   3   4E+4   1   1.5  RT (min)      lcqtof_hilic_neg_4_000_aa    m/z 551.5017     0   1E+5   1   1.5  RT (min)      lcqtof_hilic_neg_4_000_aa    m/z 551.5036     0   1E+5   1.5   2  RT (min)      lcqtof_hilic_neg_4_000_aa    m/z 551.5037     0   1E+5   1   1.5  RT (min)      lcqtof_hilic_neg_4_000_aa    m/z 552.1869     0   1E+4   1   1.5  RT (min)      lcqtof_hilic_neg_3_103_61    m/z 552.2478     0   1E+4   4   4.5  RT (min)      lcqtof_hilic_neg_4_008_24    m/z 552.3258     0   2   4E+4   1   1.5  RT (min)      lcqtof_hilic_neg_4_000_aa    m/z 552.3270     0   2   4E+4   1   1.5  RT (min)      lcqtof_hilic_neg_4_000_aa    m/z 552.3271     0   2   4   6E+4   3.5   4  RT (min)      lcqtof_hilic_neg_1_007_05    m/z 552.3276     0   2   4E+4   1   1.5  RT (min)      lcqtof_hilic_neg_4_000_aa    m/z 552.3276     0   2   4E+4   1   1.5  RT (min)      lcqtof_hilic_neg_4_000_aa    m/z 552.3278     0   2   4E+4   1   1.5  RT (min)      lcqtof_hilic_neg_4_000_aa    m/z 552.3279     0   2   4E+4   1   1.5  RT (min)      lcqtof_hilic_neg_4_000_aa    m/z 552.3284     0   2   4E+4   1   1.5  RT (min)      lcqtof_hilic_neg_4_000_aa    m/z 552.3285     0   2   4E+4   1   1.5  RT (min)      lcqtof_hilic_neg_4_000_aa    m/z 552.3293     0   2   4   6E+4   4   4.5  RT (min)      lcqtof_hilic_neg_4_040_05    m/z 552.3693     0   1E+4   4   4.5  RT (min)      lcqtof_hilic_neg_4_040_05    m/z 552.3746     0   1E+4   1   1.5  RT (min)      lcqtof_hilic_neg_1_007_05    m/z 552.4600     0   1E+5   0.5   1  RT (min)      lcqtof_hilic_neg_3_035_46    m/z 552.5064     0   1   2   3   4E+4   1   1.5  RT (min)      lcqtof_hilic_neg_4_000_aa    m/z 552.6756     0   1   2E+4   0.5   1  RT (min)      lcqtof_hilic_neg_4_053_14    m/z 553.0334     0   1   2   3   4E+4   1   1.5  RT (min)      lcqtof_hilic_neg_4_000_aa    m/z 553.1872     0   1E+4   2.5   3  RT (min)      lcqtof_hilic_neg_4_027_19    m/z 553.1988     0   1E+4   1   1.5  RT (min)      lcqtof_hilic_neg_4_000_aa    m/z 553.2458     0   2   4   6E+4   4   4.5  RT (min)       lcqtof_hilic_neg_4_090_60    m/z 553.2576     0   1E+4   0.5   1  RT (min)      lcqtof_hilic_neg_4_045_26    m/z 553.3184     0   1   2   3   4E+4   1   1.5  RT (min)      lcqtof_hilic_neg_4_000_aa    m/z 553.3255     0   1   2   3   4E+4   1   1.5  RT (min)      lcqtof_hilic_neg_4_000_aa    m/z 553.3263     0   1   2   3   4E+4   1   1.5  RT (min)      lcqtof_hilic_neg_4_000_aa    m/z 553.3334     0   1E+4   4   4.5  RT (min)      lcqtof_hilic_neg_4_040_05    m/z 553.3450     0   2   4   6E+4   1   1.5  RT (min)      lcqtof_hilic_neg_4_000_aa    m/z 553.3657     0   1E+4   1   1.5  RT (min)      lcqtof_hilic_neg_4_053_14    m/z 553.4583     0   2   4   6E+4   1   1.5  RT (min)      lcqtof_hilic_neg_4_046_41    m/z 553.4660     0   2   4   6E+4   1   1.5  RT (min)      lcqtof_hilic_neg_4_000_aa    m/z 553.4815     0   2   4   6E+4   1   1.5  RT (min)      lcqtof_hilic_neg_4_000_aa    m/z 553.4886     0   2   4   6E+4   1   1.5  RT (min)      lcqtof_hilic_neg_4_000_aa    m/z 554.0256     0   1E+4   1   1.5  RT (min)      lcqtof_hilic_neg_4_000_aa    m/z 554.0340     0   1E+4   1   1.5  RT (min)      lcqtof_hilic_neg_4_000_aa    m/z 554.2315     0   2   4   6E+4   3.5   4  RT (min)      lcqtof_hilic_neg_2_028_05    m/z 554.2542     0   1   2   3   4E+5   2   2.5  RT (min)      lcqtof_hilic_neg_4_010_28    m/z 554.2568     0   2   4E+5   3   3.5  RT (min)      lcqtof_hilic_neg_4_000_bb    m/z 554.2601     0   1   2   3   4E+5   1   1.5  RT (min)      lcqtof_hilic_neg_4_060_13    m/z 554.2648     0   1   2   3E+5   0.5   1  RT (min)      lcqtof_hilic_neg_4_082_59    m/z 554.3247     0   1E+4   1   1.5  RT (min)      lcqtof_hilic_neg_4_000_aa    m/z 554.3354     0   1E+5   4   4.5  RT (min)      lcqtof_hilic_neg_4_055_32    m/z 554.4651     0   1   2   3   4E+4   0.5   1  RT (min)      lcqtof_hilic_neg_3_035_46    m/z 554.4837     0   1   2E+4   1   1.5  RT (min)      lcqtof_hilic_neg_4_000_aa    m/z 555.0314     0   1   2   3   4E+4   1   1.5  RT (min)      lcqtof_hilic_neg_4_000_aa    m/z 555.0317     0   1   2   3   4E+4   1   1.5  RT (min)      lcqtof_hilic_neg_4_000_aa    m/z 555.0776     0   1E+4   6   6.5  RT (min)      lcqtof_hilic_neg_2_015_06    m/z 555.0780     0   1E+4   6   6.5  RT (min)      lcqtof_hilic_neg_4_002_25    m/z 555.0782     0   1   2E+4   6   6.5  RT (min)      lcqtof_hilic_neg_4_012_07    m/z 555.1928     0   1E+4   7   7.5  RT (min)       lcqtof_hilic_neg_4_091_60    m/z 555.2097     0   1E+4   7   7.5  RT (min)      lcqtof_hilic_neg_4_008_24    m/z 555.2097     0   1E+4   7   7.5  RT (min)      lcqtof_hilic_neg_4_008_24    m/z 555.2145     0   2   4   6E+4   1   1.5  RT (min)      lcqtof_hilic_neg_4_050_02    m/z 555.2626     0   1E+5   1   1.5  RT (min)      lcqtof_hilic_neg_4_058_39    m/z 555.3403     0   1E+4   1   1.5  RT (min)      lcqtof_hilic_neg_4_000_aa    m/z 555.3532     0   1   2   3   4E+3   15.5   16  RT (min)       lcqtof_hilic_neg_4_019_03    m/z 555.3544     0   1E+4   1   1.5  RT (min)      lcqtof_hilic_neg_4_000_aa    m/z 555.4260     0   1E+4   0.5   1  RT (min)      lcqtof_hilic_neg_2_015_06    m/z 556.0194     0   1E+4   1   1.5  RT (min)      lcqtof_hilic_neg_4_000_aa    m/z 556.0294     0   1E+4   1   1.5  RT (min)      lcqtof_hilic_neg_4_000_aa    m/z 556.2460     0   1E+4   3.5   4  RT (min)      lcqtof_hilic_neg_2_028_05    m/z 556.2474     0   1   2E+4   3.5   4  RT (min)      lcqtof_hilic_neg_4_039_18    m/z 556.4915     0   1E+4   4.5   5  RT (min)      lcqtof_hilic_neg_4_050_02    m/z 557.0256     0   1   2E+4   1   1.5  RT (min)      lcqtof_hilic_neg_4_000_aa    m/z 557.0258     0   1   2E+4   1   1.5  RT (min)      lcqtof_hilic_neg_4_000_aa    m/z 557.0301     0   1   2E+4   1   1.5  RT (min)      lcqtof_hilic_neg_4_000_aa    m/z 557.1622     0   1E+4   1   1.5  RT (min)      lcqtof_hilic_neg_4_000_aa    m/z 557.2704     0   1   2   3E+4   1   1.5  RT (min)      lcqtof_hilic_neg_4_000_aa    m/z 557.2895     0   1E+4   0.5   1  RT (min)      lcqtof_hilic_neg_4_050_02    m/z 557.2899     0   1E+4   15   15.5  RT (min)      lcqtof_hilic_neg_4_000_aa    m/z 557.3320     0   1   2E+5   1   1.5  RT (min)      lcqtof_hilic_neg_4_048_23    m/z 557.3339     0   2   4   6E+3   2   2.5  RT (min)      lcqtof_hilic_neg_4_029_17    m/z 557.3345     0   1   2   3E+5   1   1.5  RT (min)      lcqtof_hilic_neg_4_016_37    m/z 557.3346     0   1   2   3E+3   15.5   16  RT (min)      lcqtof_hilic_neg_4_019_03    m/z 557.3349     0   1   2   3E+3   15   15.5  RT (min)      lcqtof_hilic_neg_4_004_47    m/z 557.3553     0   1E+4   3   3.5  RT (min)      lcqtof_hilic_neg_1_003_37    m/z 557.4736     0   1   2   3   4E+5   0.5   1  RT (min)      lcqtof_hilic_neg_3_035_46    m/z 557.8815     0   1   2E+4   0.5   1  RT (min)      lcqtof_hilic_neg_4_053_14    m/z 558.0030     0   1E+4   7   7.5  RT (min)       lcqtof_hilic_neg_4_059_14    m/z 558.0056     0   1E+4   1   1.5  RT (min)      lcqtof_hilic_neg_4_000_aa    m/z 558.0396     0   1E+4   0.5   1  RT (min)      lcqtof_hilic_neg_4_027_19    m/z 558.0628     0   1   2E+4   7.5   8  RT (min)      lcqtof_hilic_neg_4_016_37    m/z 558.2619     0   1E+4   0.5   1  RT (min)      lcqtof_hilic_neg_4_045_26    m/z 558.2627     0   1E+4   1   1.5  RT (min)      lcqtof_hilic_neg_4_000_aa    m/z 558.4446     0   1E+4   0.5   1  RT (min)      lcqtof_hilic_neg_2_052_24    m/z 558.4770     0   1E+5   0.5   1  RT (min)      lcqtof_hilic_neg_3_035_46    m/z 558.4875     0   1E+4   4.5   5  RT (min)      lcqtof_hilic_neg_4_050_02    m/z 558.7044     0   1E+4   0.5   1  RT (min)      lcqtof_hilic_neg_4_053_14    m/z 558.9634     0   1E+4   1   1.5  RT (min)      lcqtof_hilic_neg_4_000_aa    m/z 558.9674     0   1E+4   1   1.5  RT (min)      lcqtof_hilic_neg_4_000_aa    m/z 558.9681     0   1E+4   1   1.5  RT (min)      lcqtof_hilic_neg_4_000_aa    m/z 559.2571     0   1   2   3E+4   0.5   1  RT (min)      lcqtof_hilic_neg_2_004_26    m/z 559.2720     0   1E+4   2   2.5  RT (min)      lcqtof_hilic_neg_4_003_16    m/z 559.3072     0   1   2E+5   1   1.5  RT (min)      lcqtof_hilic_neg_4_000_aa    m/z 559.3302     0   2   4   6E+4   1   1.5  RT (min)      lcqtof_hilic_neg_4_061_50    m/z 559.4876     0   1E+5   0.5   1  RT (min)      lcqtof_hilic_neg_3_035_46    m/z 559.7138     0   1E+4   0.5   1  RT (min)      lcqtof_hilic_neg_4_053_14    m/z 560.0948     0   1E+4   6   6.5  RT (min)      lcqtof_hilic_neg_4_012_07    m/z 560.2190     0   1   2E+4   0.5   1  RT (min)      lcqtof_hilic_neg_4_050_02    m/z 560.2834     0   1   2   3   4E+4   1   1.5  RT (min)      lcqtof_hilic_neg_4_039_18    m/z 560.3097     0   2   4   6   8E+4   1   1.5  RT (min)      lcqtof_hilic_neg_4_000_aa    m/z 560.3106     0   2   4   6   8E+4   1   1.5  RT (min)      lcqtof_hilic_neg_4_000_aa    m/z 560.3139     0   2   4   6   8E+4   1   1.5  RT (min)      lcqtof_hilic_neg_4_000_aa    m/z 560.3334     0   1E+4   1   1.5  RT (min)      lcqtof_hilic_neg_4_061_50    m/z 560.3512     0   1E+4   4   4.5  RT (min)      lcqtof_hilic_neg_4_008_24    m/z 560.4922     0   2   4   6E+4   0.5   1  RT (min)      lcqtof_hilic_neg_3_035_46    m/z 560.8811     0   1E+5   1   1.5  RT (min)      lcqtof_hilic_neg_4_004_47    m/z 560.8989     0   1E+4   0.5   1  RT (min)      lcqtof_hilic_neg_3_035_46    m/z 560.8992     0   1E+4   0.5   1  RT (min)      lcqtof_hilic_neg_3_035_46    m/z 560.9558     0   1   2E+4   1   1.5  RT (min)      lcqtof_hilic_neg_4_000_aa    m/z 560.9594     0   1   2E+4   1   1.5  RT (min)      lcqtof_hilic_neg_4_000_aa    m/z 561.1386     0   1   2E+4   0.5   1  RT (min)      lcqtof_hilic_neg_4_021_21    m/z 561.1444     0   1E+4   2.5   3  RT (min)      lcqtof_hilic_neg_4_010_28    m/z 561.1490     0   1   2   3E+4   0.5   1  RT (min)      lcqtof_hilic_neg_2_053_18    m/z 561.2181     0   2   4   6E+5   1   1.5  RT (min)      lcqtof_hilic_neg_4_000_aa    m/z 561.2187     0   2   4   6E+5   1   1.5  RT (min)      lcqtof_hilic_neg_4_000_aa    m/z 561.2697     0   1   2E+4   1   1.5  RT (min)       lcqtof_hilic_neg_4_100_61    m/z 561.2699     0   1   2   3E+5   1.5   2  RT (min)       lcqtof_hilic_neg_4_099_61    m/z 561.2705     0   1   2E+5   1   1.5  RT (min)       lcqtof_hilic_neg_4_098_61    m/z 561.2826     0   1E+4   6   6.5  RT (min)      lcqtof_hilic_neg_4_008_24    m/z 561.2866     0   1E+4   1   1.5  RT (min)      lcqtof_hilic_neg_4_039_18    m/z 561.3042     0   1E+5   1   1.5  RT (min)      lcqtof_hilic_neg_4_000_aa    m/z 561.3084     0   1E+5   1   1.5  RT (min)      lcqtof_hilic_neg_4_000_aa    m/z 561.3840     0   1E+5   1   1.5  RT (min)      lcqtof_hilic_neg_4_000_aa    m/z 561.3930     0   1E+5   1   1.5  RT (min)      lcqtof_hilic_neg_4_000_aa    m/z 561.3938     0   1E+5   1   1.5  RT (min)      lcqtof_hilic_neg_4_000_aa    m/z 561.3946     0   1E+5   1   1.5  RT (min)      lcqtof_hilic_neg_4_000_aa    m/z 561.4872     0   1   2   3E+4   1   1.5  RT (min)      lcqtof_hilic_neg_4_000_aa    m/z 561.5028     0   2   4   6E+4   0.5   1  RT (min)      lcqtof_hilic_neg_3_035_46    m/z 561.9859     0   1   2   3E+4   0.5   1  RT (min)      lcqtof_hilic_neg_4_015_09    m/z 562.1928     0   1   2   3   4E+4   1   1.5  RT (min)       lcqtof_hilic_neg_4_100_61    m/z 562.2730     0   2   4   6   8E+4   1.5   2  RT (min)      lcqtof_hilic_neg_4_099_61    m/z 562.2860     0   2   4   6   8E+4   6   6.5  RT (min)       lcqtof_hilic_neg_4_008_24    m/z 562.3072     0   1   2   3E+4   1   1.5  RT (min)      lcqtof_hilic_neg_4_000_aa    m/z 562.3078     0   1   2   3E+4   1   1.5  RT (min)      lcqtof_hilic_neg_4_000_aa    m/z 562.3092     0   1   2   3E+4   1   1.5  RT (min)      lcqtof_hilic_neg_4_000_aa    m/z 562.3292     0   1   2E+4   0.5   1  RT (min)      lcqtof_hilic_neg_3_003_20    m/z 562.3642     0   1E+4   4   4.5  RT (min)      lcqtof_hilic_neg_4_008_24    m/z 562.3660     0   1E+4   1   1.5  RT (min)      lcqtof_hilic_neg_4_040_05    m/z 562.3837     0   1E+4   2   2.5  RT (min)      lcqtof_hilic_neg_4_050_02    m/z 562.3954     0   2   4E+4   1   1.5  RT (min)      lcqtof_hilic_neg_4_000_aa    m/z 562.3972     0   2   4E+4   1   1.5  RT (min)      lcqtof_hilic_neg_4_000_aa    m/z 562.4912     0   1E+4   1   1.5  RT (min)      lcqtof_hilic_neg_4_000_aa    m/z 562.5072     0   1E+4   0.5   1  RT (min)      lcqtof_hilic_neg_3_035_46    m/z 562.5568     0   1   2   3E+4   3.5   4  RT (min)      lcqtof_hilic_neg_1_007_05    m/z 562.9450     0   1   2E+4   1   1.5  RT (min)      lcqtof_hilic_neg_4_000_aa    m/z 562.9493     0   1   2E+4   1   1.5  RT (min)      lcqtof_hilic_neg_4_000_aa    m/z 562.9886     0   1E+4   0.5   1  RT (min)      lcqtof_hilic_neg_4_015_09    m/z 563.1470     0   1   2   3E+4   0.5   1  RT (min)       lcqtof_hilic_neg_4_102_61    m/z 563.1822     0   1E+4   6.5   7  RT (min)       lcqtof_hilic_neg_4_050_02    m/z 563.2133     0   1E+5   1   1.5  RT (min)      lcqtof_hilic_neg_4_000_aa    m/z 563.2203     0   1E+5   1   1.5  RT (min)      lcqtof_hilic_neg_4_000_aa    m/z 563.2210     0   1E+5   1   1.5  RT (min)      lcqtof_hilic_neg_4_000_aa    m/z 563.2714     0   1   2E+4   6   6.5  RT (min)      lcqtof_hilic_neg_4_008_24    m/z 563.2751     0   1   2E+4   1.5   2  RT (min)      lcqtof_hilic_neg_4_099_61    m/z 563.2752     0   1   2E+4   6   6.5  RT (min)      lcqtof_hilic_neg_4_008_24    m/z 563.3050     0   1   2E+4   1   1.5  RT (min)      lcqtof_hilic_neg_4_000_aa    m/z 563.3056     0   1   2E+4   1   1.5  RT (min)      lcqtof_hilic_neg_4_000_aa    m/z 563.3080     0   1   2E+4   1   1.5  RT (min)      lcqtof_hilic_neg_4_000_aa    m/z 563.3328     0   1E+4   3   3.5  RT (min)      lcqtof_hilic_neg_4_011_35    m/z 563.3369     0   1E+4   2   2.5  RT (min)       lcqtof_hilic_neg_4_052_13    m/z 563.3449     0   1E+4   0.5   1  RT (min)      lcqtof_hilic_neg_1_003_37    m/z 563.3696     0   1E+4   1   1.5  RT (min)      lcqtof_hilic_neg_4_000_aa    m/z 563.3750     0   1E+4   1   1.5  RT (min)      lcqtof_hilic_neg_4_000_aa    m/z 563.3938     0   1   2E+4   4   4.5  RT (min)      lcqtof_hilic_neg_4_040_05    m/z 563.3987     0   1E+4   1   1.5  RT (min)      lcqtof_hilic_neg_4_000_aa    m/z 563.5016     0   2   4E+4   1   1.5  RT (min)      lcqtof_hilic_neg_4_000_aa    m/z 563.5122     0   2   4E+4   1   1.5  RT (min)      lcqtof_hilic_neg_4_000_aa    m/z 563.5150     0   2   4E+4   1   1.5  RT (min)      lcqtof_hilic_neg_4_000_aa    m/z 563.5152     0   2   4E+4   1.5   2  RT (min)      lcqtof_hilic_neg_4_000_aa    m/z 563.5169     0   1E+4   0.5   1  RT (min)      lcqtof_hilic_neg_3_035_46    m/z 563.9848     0   1   2E+4   0.5   1  RT (min)      lcqtof_hilic_neg_4_015_09    m/z 564.2250     0   1   2   3   4E+4   0.5   1  RT (min)      lcqtof_hilic_neg_3_003_20    m/z 564.2261     0   2   4   6E+4   1   1.5  RT (min)      lcqtof_hilic_neg_4_000_aa    m/z 564.2265     0   2   4   6   8E+4   3.5   4  RT (min)      lcqtof_hilic_neg_2_053_18    m/z 564.2278     0   2   4   6E+4   1   1.5  RT (min)      lcqtof_hilic_neg_4_000_aa    m/z 564.4296     0   1   2   3   4E+4   0.5   1  RT (min)      lcqtof_hilic_neg_3_035_46    m/z 564.5150     0   1   2E+4   1   1.5  RT (min)      lcqtof_hilic_neg_4_000_aa    m/z 564.5208     0   1   2E+4   1   1.5  RT (min)      lcqtof_hilic_neg_4_000_aa    m/z 564.9446     0   1E+4   1   1.5  RT (min)      lcqtof_hilic_neg_4_000_aa    m/z 564.9453     0   1E+4   1   1.5  RT (min)      lcqtof_hilic_neg_4_000_aa    m/z 565.0474     0   1   2E+4   7.5   8  RT (min)       lcqtof_hilic_neg_4_082_59    m/z 565.1112     0   1   2E+4   6   6.5  RT (min)       lcqtof_hilic_neg_4_090_60    m/z 565.1123     0   1   2E+4   6   6.5  RT (min)      lcqtof_hilic_neg_4_008_24    m/z 565.2188     0   1   2E+4   1   1.5  RT (min)      lcqtof_hilic_neg_4_000_aa    m/z 565.2327     0   1   2   3   4E+4   3.5   4  RT (min)      lcqtof_hilic_neg_2_053_18    m/z 565.2615     0   1E+4   3   3.5  RT (min)       lcqtof_hilic_neg_4_027_19    m/z 565.2810     0   1E+4   4   4.5  RT (min)      lcqtof_hilic_neg_4_002_25    m/z 565.3354     0   1   2   3   4E+4   0.5   1  RT (min)      lcqtof_hilic_neg_4_021_21    m/z 565.3573     0   1E+4   4   4.5  RT (min)      lcqtof_hilic_neg_4_008_24    m/z 565.3678     0   1   2   3E+4   1   1.5  RT (min)      lcqtof_hilic_neg_4_000_aa    m/z 565.3744     0   1   2   3E+4   1   1.5  RT (min)      lcqtof_hilic_neg_4_000_aa    m/z 565.3762     0   1   2   3E+4   1   1.5  RT (min)      lcqtof_hilic_neg_4_000_aa    m/z 565.3776     0   1   2   3E+4   1   1.5  RT (min)      lcqtof_hilic_neg_4_000_aa    m/z 565.3794     0   1   2   3E+4   1   1.5  RT (min)      lcqtof_hilic_neg_4_000_aa    m/z 565.3804     0   1   2   3E+4   1   1.5  RT (min)      lcqtof_hilic_neg_4_000_aa    m/z 565.3810     0   1   2   3E+4   1   1.5  RT (min)      lcqtof_hilic_neg_4_000_aa    m/z 565.4099     0   1   2E+4   4   4.5  RT (min)      lcqtof_hilic_neg_4_040_05    m/z 565.4464     0   2   4E+4   1   1.5  RT (min)      lcqtof_hilic_neg_4_059_14    m/z 565.4645     0   1E+4   0.5   1  RT (min)      lcqtof_hilic_neg_3_035_46    m/z 565.4994     0   2   4   6E+4   1   1.5  RT (min)      lcqtof_hilic_neg_4_000_aa    m/z 565.5147     0   2   4   6   8E+4   1   1.5  RT (min)      lcqtof_hilic_neg_4_000_aa    m/z 565.5189     0   2   4   6   8E+4   1   1.5  RT (min)      lcqtof_hilic_neg_4_000_aa    m/z 565.5198     0   2   4   6   8E+4   1.5   2  RT (min)      lcqtof_hilic_neg_4_000_aa    m/z 565.5198     0   2   4   6   8E+4   1.5   2  RT (min)      lcqtof_hilic_neg_4_000_aa    m/z 565.5790     0   1   2   3E+4   3.5   4  RT (min)      lcqtof_hilic_neg_4_052_13    m/z 565.7600     0   1   2   3   4E+4   3.5   4  RT (min)      lcqtof_hilic_neg_4_052_13    m/z 565.8994     0   2   4E+4   3.5   4  RT (min)      lcqtof_hilic_neg_4_052_13    m/z 566.0158     0   1   2E+4   3.5   4  RT (min)      lcqtof_hilic_neg_4_052_13    m/z 566.1556     0   1E+4   7   7.5  RT (min)      lcqtof_hilic_neg_1_000_aa    m/z 566.2622     0   1E+4   6.5   7  RT (min)      lcqtof_hilic_neg_4_008_24    m/z 566.2809     0   1E+4   3.5   4  RT (min)      lcqtof_hilic_neg_4_039_18    m/z 566.3691     0   1E+4   4   4.5  RT (min)      lcqtof_hilic_neg_4_008_24    m/z 566.3747     0   1E+4   1   1.5  RT (min)      lcqtof_hilic_neg_4_000_aa    m/z 566.5131     0   1   2   3E+4   1   1.5  RT (min)      lcqtof_hilic_neg_4_000_aa    m/z 566.5205     0   1   2   3E+4   1   1.5  RT (min)      lcqtof_hilic_neg_4_000_aa    m/z 566.5252     0   1   2   3E+4   1.5   2  RT (min)      lcqtof_hilic_neg_4_000_aa    m/z 566.9448     0   1E+4   1   1.5  RT (min)      lcqtof_hilic_neg_4_000_aa    m/z 567.0502     0   1E+4   1   1.5  RT (min)      lcqtof_hilic_neg_4_000_aa    m/z 567.1140     0   1E+4   6.5   7  RT (min)      lcqtof_hilic_neg_4_008_24    m/z 567.2437     0   1E+4   6   6.5  RT (min)      lcqtof_hilic_neg_4_008_24    m/z 567.2705     0   1E+4   0.5   1  RT (min)      lcqtof_hilic_neg_4_045_26    m/z 567.2924     0   1E+4   4   4.5  RT (min)      lcqtof_hilic_neg_4_002_25    m/z 567.2935     0   1E+4   4   4.5  RT (min)      lcqtof_hilic_neg_4_040_05    m/z 567.4264     0   1E+4   4   4.5  RT (min)      lcqtof_hilic_neg_4_040_05    m/z 567.4620     0   1E+5   1   1.5  RT (min)       lcqtof_hilic_neg_4_053_14    m/z 567.4906     0   2   4   6E+4   1   1.5  RT (min)      lcqtof_hilic_neg_4_000_aa    m/z 568.2677     0   1E+4   0.5   1  RT (min)      lcqtof_hilic_neg_4_002_25    m/z 568.4334     0   1E+4   0.5   1  RT (min)      lcqtof_hilic_neg_4_053_14    m/z 568.4812     0   1   2E+4   1   1.5  RT (min)      lcqtof_hilic_neg_4_000_aa    m/z 568.6550     0   1   2   3   4E+4   3.5   4  RT (min)      lcqtof_hilic_neg_4_053_14    m/z 569.0493     0   1E+4   1   1.5  RT (min)      lcqtof_hilic_neg_4_000_aa    m/z 569.1092     0   1E+4   6.5   7  RT (min)      lcqtof_hilic_neg_4_008_24    m/z 569.2018     0   1   2   3E+4   4.5   5  RT (min)      lcqtof_hilic_neg_4_013_20    m/z 569.3344     0   2   4   6E+5   1   1.5  RT (min)      lcqtof_hilic_neg_4_000_aa    m/z 569.3413     0   2   4   6E+5   1   1.5  RT (min)      lcqtof_hilic_neg_4_000_aa    m/z 569.3452     0   2   4   6E+5   1   1.5  RT (min)      lcqtof_hilic_neg_4_000_aa    m/z 569.3656     0   2   4E+3   0   0.5  RT (min)      lcqtof_hilic_neg_4_009_50    m/z 569.3677     0   1   2E+3   19   19.5  RT (min)      lcqtof_hilic_neg_4_002_25    m/z 569.3679     0   1   2E+3   0   0.5  RT (min)      lcqtof_hilic_neg_4_018_27    m/z 569.4368     0   1E+4   0.5   1  RT (min)      lcqtof_hilic_neg_4_032_01    m/z 569.4417     0   1   2E+4   1   1.5  RT (min)      lcqtof_hilic_neg_4_000_aa    m/z 570.2282     0   1E+4   4   4.5  RT (min)      lcqtof_hilic_neg_4_039_18    m/z 570.2754     0   1   2E+4   0.5   1  RT (min)      lcqtof_hilic_neg_4_039_18    m/z 570.3377     0   1   2E+5   1   1.5  RT (min)      lcqtof_hilic_neg_4_000_aa    m/z 570.3384     0   1   2E+5   1   1.5  RT (min)      lcqtof_hilic_neg_4_000_aa    m/z 570.3386     0   1   2E+5   1   1.5  RT (min)      lcqtof_hilic_neg_4_000_aa    m/z 570.4281     0   1E+4   4   4.5  RT (min)      lcqtof_hilic_neg_4_040_05    m/z 570.4483     0   2   4E+4   0.5   1  RT (min)      lcqtof_hilic_neg_4_020_42    m/z 570.9822     0   1E+4   1   1.5  RT (min)      lcqtof_hilic_neg_4_000_aa    m/z 570.9843     0   1E+4   1   1.5  RT (min)      lcqtof_hilic_neg_4_000_aa    m/z 570.9890     0   1E+4   1   1.5  RT (min)      lcqtof_hilic_neg_4_000_aa    m/z 571.2339     0   1E+4   0.5   1  RT (min)      lcqtof_hilic_neg_4_050_02    m/z 571.2718     0   1E+5   0.5   1  RT (min)      lcqtof_hilic_neg_2_053_18    m/z 571.3170     0   1E+4   0.5   1  RT (min)      lcqtof_hilic_neg_4_058_39    m/z 571.3308     0   2   4   6   8E+3   2   2.5  RT (min)      lcqtof_hilic_neg_4_052_13    m/z 571.3364     0   1   2   3   4E+3   16   16.5  RT (min)      lcqtof_hilic_neg_1_002_39    m/z 571.3372     0   1   2   3   4E+3   16.5   17  RT (min)      lcqtof_hilic_neg_1_007_05    m/z 571.3379     0   1   2   3   4E+3   16   16.5  RT (min)      lcqtof_hilic_neg_1_006_22    m/z 571.3388     0   1E+5   1   1.5  RT (min)      lcqtof_hilic_neg_4_000_aa    m/z 571.3390     0   1   2   3   4E+3   16.5   17  RT (min)      lcqtof_hilic_neg_1_007_05    m/z 571.3397     0   2   4   6   8E+3   2   2.5  RT (min)      lcqtof_hilic_neg_4_010_28    m/z 571.3402     0   1   2   3   4E+3   17   17.5  RT (min)      lcqtof_hilic_neg_1_010_15    m/z 571.3416     0   1   2   3E+3   18.5   19  RT (min)       lcqtof_hilic_neg_4_013_20    m/z 571.3421     0   1   2   3   4E+3   0.5   1  RT (min)      lcqtof_hilic_neg_4_000_cc    m/z 571.3468     0   2   4E+3   1   1.5  RT (min)      lcqtof_hilic_neg_4_000_dd    m/z 571.3470     0   1   2   3   4E+3   1.5   2  RT (min)      lcqtof_hilic_neg_4_021_21    m/z 571.3487     0   1   2   3E+3   2.5   3  RT (min)      lcqtof_hilic_neg_4_000_cc    m/z 571.3491     0   2   4E+3   15   15.5  RT (min)      lcqtof_hilic_neg_4_034_31    m/z 571.3799     0   1E+4   0.5   1  RT (min)      lcqtof_hilic_neg_4_021_21    m/z 571.4543     0   1   2E+4   0.5   1  RT (min)      lcqtof_hilic_neg_4_046_41    m/z 571.4900     0   1   2E+4   0.5   1  RT (min)      lcqtof_hilic_neg_3_035_46    m/z 571.4944     0   1   2   3E+4   1   1.5  RT (min)      lcqtof_hilic_neg_4_000_aa    m/z 572.2216     0   1E+4   3.5   4  RT (min)      lcqtof_hilic_neg_1_032_18    m/z 572.2425     0   1   2   3   4E+4   4   4.5  RT (min)      lcqtof_hilic_neg_4_039_18    m/z 572.2473     0   1E+4   0.5   1  RT (min)      lcqtof_hilic_neg_3_003_20    m/z 572.2745     0   1   2   3E+4   0.5   1  RT (min)      lcqtof_hilic_neg_4_039_18    m/z 572.3103     0   1E+4   4   4.5  RT (min)      lcqtof_hilic_neg_4_050_02    m/z 572.3351     0   1   2   3E+4   1   1.5  RT (min)      lcqtof_hilic_neg_4_000_aa    m/z 572.3361     0   1   2   3E+4   1   1.5  RT (min)      lcqtof_hilic_neg_4_000_aa    m/z 572.3432     0   1   2   3E+4   1   1.5  RT (min)      lcqtof_hilic_neg_4_000_aa    m/z 572.4974     0   1   2   3   4E+4   0.5   1  RT (min)      lcqtof_hilic_neg_3_035_46    m/z 572.5004     0   1E+4   1   1.5  RT (min)      lcqtof_hilic_neg_4_000_aa    m/z 572.9792     0   2   4   6E+4   1   1.5  RT (min)      lcqtof_hilic_neg_4_000_aa    m/z 573.2240     0   1   2E+4   6.5   7  RT (min)      lcqtof_hilic_neg_4_008_24    m/z 573.2461     0   1E+4   4   4.5  RT (min)      lcqtof_hilic_neg_4_039_18    m/z 573.3153     0   1E+4   0.5   1  RT (min)      lcqtof_hilic_neg_4_000_aa    m/z 573.3250     0   1E+4   1   1.5  RT (min)      lcqtof_hilic_neg_4_000_aa    m/z 573.3312     0   1   2   3   4E+5   1   1.5  RT (min)      lcqtof_hilic_neg_4_034_31    m/z 573.3452     0   1E+5   1   1.5  RT (min)      lcqtof_hilic_neg_4_061_50    m/z 573.4235     0   1   2E+4   0.5   1  RT (min)      lcqtof_hilic_neg_4_023_15    m/z 573.4248     0   1   2   3   4E+5   1   1.5  RT (min)      lcqtof_hilic_neg_4_000_aa    m/z 573.4258     0   1E+4   14.5   15   15.5  RT (min)       lcqtof_hilic_neg_4_082_59    m/z 573.4258     0   1E+4   14.5   15  RT (min)      lcqtof_hilic_neg_4_013_20    m/z 573.4261     0   1   2E+4   15.5   16  RT (min)      lcqtof_hilic_neg_4_000_aa    m/z 573.4510     0   2   4   6E+5   1   1.5  RT (min)      lcqtof_hilic_neg_4_053_14    m/z 573.4940     0   1E+4   1   1.5  RT (min)      lcqtof_hilic_neg_4_000_aa    m/z 573.9864     0   1E+4   1   1.5  RT (min)      lcqtof_hilic_neg_4_000_aa    m/z 573.9867     0   1E+4   1   1.5  RT (min)      lcqtof_hilic_neg_4_000_aa    m/z 574.0240     0   1E+4   6.5   7  RT (min)       lcqtof_hilic_neg_4_093_60    m/z 574.2402     0   1E+4   4   4.5  RT (min)      lcqtof_hilic_neg_4_039_18    m/z 574.2911     0   1   2E+4   0.5   1  RT (min)      lcqtof_hilic_neg_4_000_aa    m/z 574.3500     0   1   2   3   4E+4   0.5   1  RT (min)      lcqtof_hilic_neg_4_038_10    m/z 574.4289     0   1E+4   15.5   16  RT (min)      lcqtof_hilic_neg_4_000_aa    m/z 574.4303     0   1E+5   1   1.5  RT (min)      lcqtof_hilic_neg_4_000_aa    m/z 574.4551     0   1   2E+5   1   1.5  RT (min)      lcqtof_hilic_neg_4_059_14    m/z 574.9766     0   2   4   6E+4   1   1.5  RT (min)      lcqtof_hilic_neg_4_000_aa    m/z 575.0672     0   1E+4   6   6.5  RT (min)       lcqtof_hilic_neg_4_008_24    m/z 575.0678     0   1E+4   6   6.5  RT (min)      lcqtof_hilic_neg_4_008_24    m/z 575.1800     0   1   2E+4   1   1.5  RT (min)      lcqtof_hilic_neg_4_000_aa    m/z 575.1814     0   1   2E+4   1   1.5  RT (min)      lcqtof_hilic_neg_4_000_aa    m/z 575.1834     0   1   2E+4   1   1.5  RT (min)      lcqtof_hilic_neg_4_000_aa    m/z 575.1836     0   1   2E+4   1   1.5  RT (min)      lcqtof_hilic_neg_4_000_aa    m/z 575.2358     0   1E+4   1   1.5  RT (min)      lcqtof_hilic_neg_4_000_aa    m/z 575.2386     0   1E+4   0.5   1  RT (min)      lcqtof_hilic_neg_4_045_26    m/z 575.2820     0   1   2   3E+4   7   7.5  RT (min)       lcqtof_hilic_neg_4_091_60    m/z 575.2876     0   1   2   3   4E+4   0.5   1  RT (min)      lcqtof_hilic_neg_4_000_aa    m/z 575.3972     0   1E+4   4   4.5  RT (min)      lcqtof_hilic_neg_4_008_24    m/z 575.3989     0   1E+4   3.5   4  RT (min)      lcqtof_hilic_neg_1_007_05    m/z 575.4100     0   1E+4   0.5   1  RT (min)      lcqtof_hilic_neg_4_013_20    m/z 575.4316     0   1   2   3E+4   1   1.5  RT (min)      lcqtof_hilic_neg_4_000_aa    m/z 575.9766     0   1E+4   1   1.5  RT (min)      lcqtof_hilic_neg_4_000_aa    m/z 576.2210     0   1   2   3   4E+4   6   6.5  RT (min)      lcqtof_hilic_neg_4_008_24    m/z 576.2885     0   1E+4   0.5   1  RT (min)      lcqtof_hilic_neg_4_000_aa    m/z 576.4704     0   1   2E+5   1   1.5  RT (min)      lcqtof_hilic_neg_4_059_14    m/z 576.7640     0   1E+4   8   8.5  RT (min)      lcqtof_hilic_neg_1_000_aa    m/z 576.8617     0   1   2   3E+3   8.5   9  RT (min)       lcqtof_hilic_neg_4_020_42    m/z 576.8620     0   1   2   3E+3   8.5   9  RT (min)       lcqtof_hilic_neg_4_051_43    m/z 576.9734     0   2   4E+4   1   1.5  RT (min)      lcqtof_hilic_neg_4_000_aa    m/z 576.9740     0   1   2E+4   0.5   1  RT (min)      lcqtof_hilic_neg_1_028_47    m/z 576.9778     0   1E+4   0.5   1  RT (min)      lcqtof_hilic_neg_4_004_47    m/z 577.0924     0   1E+4   6.5   7  RT (min)      lcqtof_hilic_neg_4_048_23    m/z 577.1909     0   1E+4   7   7.5  RT (min)       lcqtof_hilic_neg_4_008_24    m/z 577.2365     0   1E+4   0.5   1  RT (min)      lcqtof_hilic_neg_4_045_26    m/z 577.3978     0   1E+4   1   1.5  RT (min)      lcqtof_hilic_neg_4_000_aa    m/z 577.4992     0   2   4   6E+4   0.5   1  RT (min)      lcqtof_hilic_neg_3_035_46    m/z 577.5028     0   1   2   3   4E+4   1   1.5  RT (min)      lcqtof_hilic_neg_4_000_aa    m/z 577.5124     0   1   2   3   4E+4   1   1.5  RT (min)      lcqtof_hilic_neg_4_000_aa    m/z 577.5882     0   1E+4   1.5   2  RT (min)      lcqtof_hilic_neg_4_000_aa    m/z 577.5910     0   1E+4   1.5   2  RT (min)      lcqtof_hilic_neg_4_000_aa    m/z 577.9704     0   1E+4   1   1.5  RT (min)      lcqtof_hilic_neg_4_000_aa    m/z 578.1592     0   1   2E+4   7   7.5  RT (min)      lcqtof_hilic_neg_4_014_46    m/z 578.2467     0   1E+4   4.5   5  RT (min)      lcqtof_hilic_neg_4_013_20    m/z 578.5146     0   2   4E+4   0.5   1  RT (min)      lcqtof_hilic_neg_4_048_23    m/z 578.5219     0   1E+4   1   1.5  RT (min)      lcqtof_hilic_neg_4_000_aa    m/z 578.8684     0   1   2   3   4E+3   5   5.5  RT (min)       lcqtof_hilic_neg_4_053_14    m/z 578.9664     0   1   2E+4   1   1.5  RT (min)      lcqtof_hilic_neg_4_000_aa    m/z 578.9696     0   1   2E+4   1   1.5  RT (min)      lcqtof_hilic_neg_4_000_aa    m/z 579.1413     0   2   4   6E+4   1   1.5  RT (min)      lcqtof_hilic_neg_4_000_aa    m/z 579.1417     0   2   4   6E+4   1   1.5  RT (min)      lcqtof_hilic_neg_4_000_aa    m/z 579.1422     0   2   4   6E+4   1   1.5  RT (min)      lcqtof_hilic_neg_4_000_aa    m/z 579.3856     0   1E+4   3.5   4  RT (min)      lcqtof_hilic_neg_1_007_05    m/z 579.3857     0   1   2E+4   4   4.5  RT (min)      lcqtof_hilic_neg_1_037_24    m/z 579.3880     0   1E+4   1   1.5  RT (min)      lcqtof_hilic_neg_4_000_aa    m/z 579.4814     0   2   4   6E+4   0.5   1  RT (min)      lcqtof_hilic_neg_4_053_14    m/z 579.4978     0   2   4E+4   1   1.5  RT (min)       lcqtof_hilic_neg_4_000_aa    m/z 579.5258     0   1E+5   1   1.5  RT (min)      lcqtof_hilic_neg_4_000_aa    m/z 579.5336     0   1E+5   1   1.5  RT (min)      lcqtof_hilic_neg_4_000_aa    m/z 579.5342     0   1E+5   1   1.5  RT (min)      lcqtof_hilic_neg_4_000_aa    m/z 580.1419     0   1   2   3E+4   1   1.5  RT (min)      lcqtof_hilic_neg_4_000_aa    m/z 580.1433     0   1   2   3E+4   1   1.5  RT (min)      lcqtof_hilic_neg_4_000_aa    m/z 580.1439     0   1   2   3E+4   1   1.5  RT (min)      lcqtof_hilic_neg_4_000_aa    m/z 580.2524     0   1   2   3   4E+4   4.5   5  RT (min)      lcqtof_hilic_neg_4_013_20    m/z 580.2809     0   1E+4   3.5   4  RT (min)      lcqtof_hilic_neg_4_040_05    m/z 580.3026     0   1E+4   0.5   1  RT (min)      lcqtof_hilic_neg_4_015_09    m/z 580.3708     0   1   2   3E+4   3   3.5  RT (min)      lcqtof_hilic_neg_4_034_31    m/z 580.4800     0   1   2E+4   0.5   1  RT (min)      lcqtof_hilic_neg_4_032_01    m/z 580.5023     0   1E+4   1   1.5  RT (min)      lcqtof_hilic_neg_4_000_aa    m/z 580.5286     0   2   4   6E+4   1   1.5  RT (min)      lcqtof_hilic_neg_4_000_aa    m/z 581.1227     0   1   2   3E+4   7   7.5  RT (min)      lcqtof_hilic_neg_1_022_36    m/z 581.1394     0   1   2E+4   1   1.5  RT (min)      lcqtof_hilic_neg_4_000_aa    m/z 581.1406     0   1   2E+4   1   1.5  RT (min)      lcqtof_hilic_neg_4_000_aa    m/z 581.1502     0   1   2E+4   1   1.5  RT (min)      lcqtof_hilic_neg_4_000_aa    m/z 581.2120     0   1   2E+4   0.5   1  RT (min)      lcqtof_hilic_neg_4_050_02    m/z 581.2806     0   1E+4   6.5   7  RT (min)      lcqtof_hilic_neg_4_008_24    m/z 581.3600     0   1E+4   1   1.5  RT (min)      lcqtof_hilic_neg_4_000_aa    m/z 581.3608     0   1E+4   1   1.5  RT (min)      lcqtof_hilic_neg_4_000_aa    m/z 581.3664     0   1E+4   1   1.5  RT (min)      lcqtof_hilic_neg_4_000_aa    m/z 581.3717     0   1E+4   1   1.5  RT (min)      lcqtof_hilic_neg_4_000_aa    m/z 581.3750     0   1E+4   3   3.5  RT (min)      lcqtof_hilic_neg_4_034_31    m/z 581.5008     0   2   4E+4   1   1.5  RT (min)      lcqtof_hilic_neg_4_000_aa    m/z 581.5108     0   1   2   3E+5   1   1.5  RT (min)      lcqtof_hilic_neg_4_000_aa    m/z 581.5237     0   2   4   6E+4   15.5   16  RT (min)      lcqtof_hilic_neg_4_000_aa    m/z 581.5252     0   1   2   3E+5   1   1.5  RT (min)      lcqtof_hilic_neg_4_000_aa    m/z 582.2713     0   1E+4   4   4.5  RT (min)      lcqtof_hilic_neg_4_039_18    m/z 582.5156     0   1E+5   1   1.5  RT (min)      lcqtof_hilic_neg_4_000_aa    m/z 582.5263     0   1   2E+4   15.5   16  RT (min)      lcqtof_hilic_neg_4_000_aa    m/z 582.5274     0   1E+5   1   1.5  RT (min)      lcqtof_hilic_neg_4_000_aa    m/z 583.2497     0   1E+4   2   2.5  RT (min)      lcqtof_hilic_neg_4_052_13    m/z 583.2740     0   1   2E+5   0.5   1  RT (min)      lcqtof_hilic_neg_3_035_46    m/z 583.3528     0   2   4   6E+4   1   1.5  RT (min)      lcqtof_hilic_neg_4_000_aa    m/z 583.3542     0   2   4   6E+4   1   1.5  RT (min)      lcqtof_hilic_neg_4_000_aa    m/z 583.3600     0   2   4   6E+4   1   1.5  RT (min)      lcqtof_hilic_neg_4_000_aa    m/z 583.3770     0   1E+4   1.5   2  RT (min)      lcqtof_hilic_neg_1_064_16    m/z 583.3840     0   2   4E+3   15.5   16  RT (min)       lcqtof_hilic_neg_4_012_07    m/z 583.4758     0   1   2E+4   1   1.5  RT (min)      lcqtof_hilic_neg_4_000_aa    m/z 583.4854     0   1   2E+4   1   1.5  RT (min)      lcqtof_hilic_neg_4_000_aa    m/z 583.5286     0   1   2   3   4E+4   0.5   1  RT (min)      lcqtof_hilic_neg_3_035_46    m/z 583.5308     0   1   2E+4   1   1.5  RT (min)      lcqtof_hilic_neg_4_000_aa    m/z 583.5439     0   1E+4   0.5   1  RT (min)      lcqtof_hilic_neg_4_048_23    m/z 584.2738     0   2   4   6   8E+4   0.5   1  RT (min)      lcqtof_hilic_neg_3_035_46    m/z 584.3013     0   1E+4   4   4.5  RT (min)       lcqtof_hilic_neg_4_008_24    m/z 584.3516     0   1   2E+4   1   1.5  RT (min)      lcqtof_hilic_neg_4_000_aa    m/z 584.3542     0   1   2E+4   1   1.5  RT (min)      lcqtof_hilic_neg_4_000_aa    m/z 584.3610     0   1   2E+4   1   1.5  RT (min)      lcqtof_hilic_neg_4_000_aa    m/z 584.3614     0   1   2E+4   1   1.5  RT (min)      lcqtof_hilic_neg_4_000_aa    m/z 584.3616     0   1   2E+4   1   1.5  RT (min)      lcqtof_hilic_neg_4_000_aa    m/z 584.4035     0   1E+4   3.5   4  RT (min)      lcqtof_hilic_neg_1_007_05    m/z 584.4040     0   1   2E+4   3.5   4  RT (min)       lcqtof_hilic_neg_1_037_24    m/z 584.5146     0   1E+4   4   4.5  RT (min)      lcqtof_hilic_neg_4_008_24    m/z 584.5430     0   1   2E+4   0.5   1  RT (min)      lcqtof_hilic_neg_3_035_46    m/z 584.9555     0   1E+4   1   1.5  RT (min)      lcqtof_hilic_neg_4_054_48    m/z 585.1490     0   1   2E+4   1   1.5  RT (min)       lcqtof_hilic_neg_4_004_47    m/z 585.2810     0   1   2E+4   0.5   1  RT (min)      lcqtof_hilic_neg_4_045_26    m/z 585.2840     0   1E+4   15   15.5  RT (min)      lcqtof_hilic_neg_4_000_aa    m/z 585.3128     0   1E+4   3.5   4  RT (min)      lcqtof_hilic_neg_4_003_16    m/z 585.3185     0   1   2   3   4E+3   3.5   4  RT (min)      lcqtof_hilic_neg_4_033_53    m/z 585.3340     0   1E+4   15.5   16  RT (min)      lcqtof_hilic_neg_1_007_05    m/z 585.3343     0   2   4   6E+3   15.5   16  RT (min)      lcqtof_hilic_neg_2_003_25    m/z 585.3355     0   2   4   6E+3   15.5   16  RT (min)      lcqtof_hilic_neg_1_055_25    m/z 585.3363     0   2   4   6E+3   16   16.5  RT (min)      lcqtof_hilic_neg_1_003_37    m/z 585.3369     0   1   2   3   4E+3   16   16.5  RT (min)      lcqtof_hilic_neg_4_004_47    m/z 585.3380     0   2   4   6E+3   16   16.5  RT (min)      lcqtof_hilic_neg_1_003_37    m/z 585.3380     0   1   2   3   4E+3   16   16.5  RT (min)      lcqtof_hilic_neg_4_049_55    m/z 585.3392     0   2   4   6E+3   16.5   17  RT (min)      lcqtof_hilic_neg_1_015_33    m/z 585.3396     0   1E+4   4   4.5  RT (min)      lcqtof_hilic_neg_1_005_32    m/z 585.3396     0   1   2E+4   1   1.5  RT (min)      lcqtof_hilic_neg_4_000_aa    m/z 585.3397     0   2   4   6E+3   17.5   18  RT (min)      lcqtof_hilic_neg_1_000_bb    m/z 585.3397     0   1E+4   0   0.5  RT (min)      lcqtof_hilic_neg_1_012_04    m/z 585.3398     0   2   4   6E+3   1.5   2  RT (min)      lcqtof_hilic_neg_1_006_22    m/z 585.3398     0   2   4   6E+3   19   19.5  RT (min)      lcqtof_hilic_neg_1_003_37    m/z 585.3399     0   2   4   6E+3   19   19.5  RT (min)      lcqtof_hilic_neg_1_003_37    m/z 585.3400     0   2   4   6E+3   19   19.5  RT (min)      lcqtof_hilic_neg_1_003_37    m/z 585.3400     0   1E+4   0.5   1  RT (min)      lcqtof_hilic_neg_1_008_13    m/z 585.3400     0   1E+4   1.5   2  RT (min)      lcqtof_hilic_neg_1_006_22    m/z 585.3400     0   2   4   6E+3   2   2.5  RT (min)      lcqtof_hilic_neg_1_006_22    m/z 585.3401     0   2   4   6E+3   17   17.5  RT (min)      lcqtof_hilic_neg_1_012_04    m/z 585.3401     0   2   4   6E+3   19.5   20  RT (min)      lcqtof_hilic_neg_1_008_13    m/z 585.3405     0   2   4   6E+3   19   19.5  RT (min)      lcqtof_hilic_neg_1_003_37    m/z 585.3406     0   2   4   6E+3   1.5   2  RT (min)      lcqtof_hilic_neg_1_006_22    m/z 585.3407     0   2   4   6E+3   19.5   20  RT (min)      lcqtof_hilic_neg_1_000_dd    m/z 585.3408     0   1E+4   0   0.5  RT (min)      lcqtof_hilic_neg_1_003_37    m/z 585.3410     0   2   4E+3   2.5   3  RT (min)      lcqtof_hilic_neg_1_000_dd    m/z 585.3410     0   1E+4   1   1.5  RT (min)      lcqtof_hilic_neg_1_006_22    m/z 585.3410     0   2   4   6E+3   2   2.5  RT (min)      lcqtof_hilic_neg_1_006_22    m/z 585.3416     0   2   4E+3   2.5   3  RT (min)      lcqtof_hilic_neg_1_006_22    m/z 585.3420     0   2   4E+3   2.5   3  RT (min)      lcqtof_hilic_neg_1_000_dd    m/z 585.3423     0   2   4   6   8E+3   4.5   5  RT (min)      lcqtof_hilic_neg_1_007_05    m/z 585.3430     0   1E+4   3   3.5  RT (min)      lcqtof_hilic_neg_1_035_23    m/z 585.3442     0   1   2   3   4E+3   3   3.5  RT (min)      lcqtof_hilic_neg_1_022_36    m/z 585.3470     0   1E+4   3   3.5  RT (min)      lcqtof_hilic_neg_2_020_23    m/z 585.3550     0   2   4   6   8E+3   15.5   16  RT (min)      lcqtof_hilic_neg_4_027_19    m/z 585.3551     0   2   4   6   8E+3   15.5   16  RT (min)      lcqtof_hilic_neg_4_000_aa    m/z 585.3562     0   2   4E+3   19.5   20  RT (min)      lcqtof_hilic_neg_1_000_dd    m/z 585.3565     0   1E+4   15.5   16  RT (min)      lcqtof_hilic_neg_4_000_aa    m/z 585.3568     0   1E+4   4.5   5  RT (min)      lcqtof_hilic_neg_4_019_03    m/z 585.3570     0   1E+4   4   4.5  RT (min)      lcqtof_hilic_neg_4_002_25    m/z 585.3575     0   1E+4   4   4.5  RT (min)      lcqtof_hilic_neg_4_012_07    m/z 585.3578     0   1   2   3E+4   4   4.5  RT (min)      lcqtof_hilic_neg_4_019_03    m/z 585.3580     0   1E+4   15   15.5  RT (min)      lcqtof_hilic_neg_4_012_07    m/z 585.3583     0   2   4   6E+3   19   19.5  RT (min)      lcqtof_hilic_neg_4_007_45    m/z 585.3584     0   2   4E+3   2.5   3  RT (min)      lcqtof_hilic_neg_4_000_dd    m/z 585.3584     0   1E+4   2   2.5  RT (min)      lcqtof_hilic_neg_4_052_13    m/z 585.3585     0   2   4   6   8E+3   1.5   2  RT (min)       lcqtof_hilic_neg_4_003_16    m/z 585.3587     0   2   4E+3   0   0.5  RT (min)      lcqtof_hilic_neg_4_025_52    m/z 585.3618     0   1   2   3   4E+3   3   3.5  RT (min)      lcqtof_hilic_neg_4_000_dd    m/z 585.3874     0   1E+4   3   3.5  RT (min)      lcqtof_hilic_neg_4_040_05    m/z 585.3948     0   1E+5   1   1.5  RT (min)      lcqtof_hilic_neg_4_000_aa    m/z 585.3972     0   1E+4   15.5   16  RT (min)      lcqtof_hilic_neg_4_000_aa    m/z 585.4278     0   1   2E+4   15.5   16  RT (min)      lcqtof_hilic_neg_4_000_aa    m/z 585.4286     0   1   2E+4   1   1.5  RT (min)      lcqtof_hilic_neg_4_000_aa    m/z 585.4298     0   1   2   3   4E+3   2   2.5  RT (min)      lcqtof_hilic_neg_4_032_01    m/z 586.0982     0   1E+4   7   7.5  RT (min)      lcqtof_hilic_neg_4_035_33    m/z 586.3234     0   2   4   6E+4   1   1.5  RT (min)      lcqtof_hilic_neg_4_000_aa    m/z 586.3358     0   1   2   3E+3   15.5   16  RT (min)      lcqtof_hilic_neg_1_003_37    m/z 586.3424     0   1   2E+3   16   16.5  RT (min)      lcqtof_hilic_neg_1_010_15    m/z 586.3437     0   1   2E+3   16.5   17  RT (min)      lcqtof_hilic_neg_1_011_46    m/z 586.3446     0   1   2E+3   16.5   17  RT (min)      lcqtof_hilic_neg_1_008_13    m/z 586.3448     0   2   4   6E+4   1   1.5  RT (min)      lcqtof_hilic_neg_4_000_aa    m/z 586.3618     0   1   2   3   4E+3   15   15.5  RT (min)      lcqtof_hilic_neg_4_046_41    m/z 586.3630     0   2   4   6E+3   15.5   16  RT (min)      lcqtof_hilic_neg_4_000_aa    m/z 586.3640     0   1   2E+3   16.5   17  RT (min)      lcqtof_hilic_neg_1_006_22    m/z 586.3670     0   2   4E+3   1   1.5  RT (min)      lcqtof_hilic_neg_4_030_44    m/z 586.4006     0   2   4   6E+4   1   1.5  RT (min)      lcqtof_hilic_neg_4_000_aa    m/z 586.7604     0   1   2   3   4E+3   6.5   7  RT (min)       lcqtof_hilic_neg_4_012_07    m/z 586.9916     0   1   2E+3   5.5   6  RT (min)      lcqtof_hilic_neg_4_004_47    m/z 586.9917     0   1   2E+3   5.5   6  RT (min)      lcqtof_hilic_neg_1_010_15    m/z 586.9921     0   1   2E+3   5.5   6  RT (min)       lcqtof_hilic_neg_1_010_15    m/z 586.9958     0   1   2E+4   1   1.5  RT (min)      lcqtof_hilic_neg_4_000_aa    m/z 587.2537     0   1   2   3E+4   1   1.5  RT (min)      lcqtof_hilic_neg_4_000_aa    m/z 587.2560     0   1   2   3E+4   1   1.5  RT (min)      lcqtof_hilic_neg_4_000_aa    m/z 587.2604     0   1   2   3E+4   1   1.5  RT (min)      lcqtof_hilic_neg_4_000_aa    m/z 587.2632     0   1   2   3E+4   1   1.5  RT (min)      lcqtof_hilic_neg_4_000_aa    m/z 587.3117     0   1   2   3   4E+3   0.5   1  RT (min)      lcqtof_hilic_neg_1_005_32    m/z 587.3252     0   1   2   3E+4   1   1.5  RT (min)      lcqtof_hilic_neg_4_000_aa    m/z 587.3305     0   1   2   3E+4   1   1.5  RT (min)      lcqtof_hilic_neg_4_000_aa    m/z 587.3320     0   1   2   3E+4   1   1.5  RT (min)      lcqtof_hilic_neg_4_000_aa    m/z 587.3934     0   1E+4   1   1.5  RT (min)      lcqtof_hilic_neg_4_000_aa    m/z 588.2336     0   1E+4   1   1.5  RT (min)      lcqtof_hilic_neg_4_005_29    m/z 588.2437     0   1   2E+4   6.5   7  RT (min)      lcqtof_hilic_neg_4_008_24    m/z 588.2554     0   1E+4   1   1.5  RT (min)      lcqtof_hilic_neg_4_000_aa    m/z 588.2569     0   1E+4   1   1.5  RT (min)      lcqtof_hilic_neg_4_000_aa    m/z 588.2572     0   1E+4   1   1.5  RT (min)      lcqtof_hilic_neg_4_000_aa    m/z 588.3646     0   1E+4   1   1.5  RT (min)      lcqtof_hilic_neg_4_061_50    m/z 588.4313     0   1   2E+4   0.5   1  RT (min)      lcqtof_hilic_neg_3_035_46    m/z 588.5242     0   1   2E+4   4   4.5  RT (min)      lcqtof_hilic_neg_4_008_24    m/z 588.6473     0   2   4   6E+4   3.5   4  RT (min)      lcqtof_hilic_neg_1_007_05    m/z 588.6901     0   1   2   3E+4   3.5   4  RT (min)      lcqtof_hilic_neg_4_052_13    m/z 588.8370     0   1   2E+4   4   4.5  RT (min)      lcqtof_hilic_neg_4_008_24    m/z 588.8974     0   2   4E+4   5.5   6  RT (min)       lcqtof_hilic_neg_4_012_07    m/z 588.8977     0   2   4E+4   5.5   6  RT (min)      lcqtof_hilic_neg_4_012_07    m/z 588.8982     0   2   4   6   8E+3   6   6.5  RT (min)       lcqtof_hilic_neg_4_107_62    m/z 588.9031     0   1   2   3E+4   3.5   4  RT (min)      lcqtof_hilic_neg_4_052_13    m/z 588.9284     0   1   2   3   4E+4   0.5   1  RT (min)      lcqtof_hilic_neg_3_035_46    m/z 588.9299     0   1   2   3   4E+4   0.5   1  RT (min)      lcqtof_hilic_neg_3_035_46    m/z 588.9746     0   1E+4   4   4.5  RT (min)      lcqtof_hilic_neg_4_008_24    m/z 588.9920     0   1   2   3E+4   1   1.5  RT (min)      lcqtof_hilic_neg_4_000_aa    m/z 588.9922     0   1   2   3E+4   1   1.5  RT (min)      lcqtof_hilic_neg_4_000_aa    m/z 589.1999     0   1E+4   3   3.5  RT (min)      lcqtof_hilic_neg_4_048_23    m/z 589.2322     0   1E+4   4.5   5  RT (min)      lcqtof_hilic_neg_4_055_32    m/z 589.2832     0   1E+4   1   1.5  RT (min)      lcqtof_hilic_neg_4_042_22    m/z 589.3045     0   2   4   6E+4   0.5   1  RT (min)      lcqtof_hilic_neg_4_038_10    m/z 589.3326     0   1E+4   5   5.5  RT (min)      lcqtof_hilic_neg_4_032_01    m/z 589.3610     0   1   2E+4   1   1.5  RT (min)      lcqtof_hilic_neg_4_000_aa    m/z 589.3611     0   1   2E+4   1   1.5  RT (min)      lcqtof_hilic_neg_4_000_aa    m/z 589.3630     0   1E+4   3   3.5  RT (min)      lcqtof_hilic_neg_2_028_05    m/z 589.7729     0   1E+4   3.5   4  RT (min)      lcqtof_hilic_neg_4_052_13    m/z 589.9072     0   1E+4   3.5   4  RT (min)      lcqtof_hilic_neg_4_052_13    m/z 589.9320     0   1E+4   0.5   1  RT (min)      lcqtof_hilic_neg_3_035_46    m/z 589.9824     0   1E+4   1   1.5  RT (min)      lcqtof_hilic_neg_4_000_aa    m/z 590.2140     0   1E+4   4   4.5  RT (min)      lcqtof_hilic_neg_1_032_18    m/z 590.2495     0   1E+4   1   1.5  RT (min)      lcqtof_hilic_neg_4_005_29    m/z 590.3077     0   1   2E+4   0.5   1  RT (min)      lcqtof_hilic_neg_4_015_09    m/z 590.3084     0   1   2E+4   1   1.5  RT (min)      lcqtof_hilic_neg_4_015_09    m/z 590.5396     0   1   2   3E+4   4   4.5  RT (min)      lcqtof_hilic_neg_4_008_24    m/z 590.6634     0   2   4   6   8E+4   4   4.5  RT (min)      lcqtof_hilic_neg_4_008_24    m/z 590.7772     0   1E+4   4   4.5  RT (min)      lcqtof_hilic_neg_4_008_24    m/z 590.8172     0   2   4   6E+4   8   8.5  RT (min)      lcqtof_hilic_neg_1_000_aa    m/z 590.8176     0   2   4   6E+4   8   8.5  RT (min)      lcqtof_hilic_neg_1_000_aa    m/z 590.8536     0   1   2   3   4E+4   4   4.5  RT (min)      lcqtof_hilic_neg_4_008_24    m/z 590.9291     0   1E+4   0.5   1  RT (min)      lcqtof_hilic_neg_3_035_46    m/z 590.9864     0   1   2E+4   1   1.5  RT (min)      lcqtof_hilic_neg_4_000_aa    m/z 590.9878     0   1   2E+4   1   1.5  RT (min)      lcqtof_hilic_neg_4_000_aa    m/z 590.9886     0   1   2E+4   1   1.5  RT (min)      lcqtof_hilic_neg_4_000_aa    m/z 590.9916     0   1   2E+4   4   4.5  RT (min)      lcqtof_hilic_neg_4_008_24    m/z 591.2115     0   1   2E+4   1   1.5  RT (min)      lcqtof_hilic_neg_4_000_aa    m/z 591.2130     0   1   2E+4   1   1.5  RT (min)      lcqtof_hilic_neg_4_000_aa    m/z 591.2140     0   1   2E+4   1   1.5  RT (min)      lcqtof_hilic_neg_4_000_aa    m/z 591.2165     0   1   2E+4   1   1.5  RT (min)      lcqtof_hilic_neg_4_000_aa    m/z 591.2908     0   1E+5   0.5   1  RT (min)      lcqtof_hilic_neg_3_003_20    m/z 591.3641     0   2   4E+4   4   4.5  RT (min)      lcqtof_hilic_neg_4_008_24    m/z 591.3686     0   1E+4   0.5   1  RT (min)      lcqtof_hilic_neg_4_010_28    m/z 591.4284     0   1E+4   3   3.5  RT (min)      lcqtof_hilic_neg_4_008_24    m/z 591.4347     0   1E+4   1   1.5  RT (min)      lcqtof_hilic_neg_4_000_aa    m/z 591.5127     0   1E+4   0.5   1  RT (min)      lcqtof_hilic_neg_3_035_46    m/z 591.5588     0   2   4   6E+4   1   1.5  RT (min)      lcqtof_hilic_neg_4_000_aa    m/z 591.5674     0   2   4   6E+4   1   1.5  RT (min)      lcqtof_hilic_neg_4_000_aa    m/z 591.5710     0   2   4   6E+4   1   1.5  RT (min)      lcqtof_hilic_neg_4_000_aa    m/z 591.6092     0   1E+4   1.5   2  RT (min)      lcqtof_hilic_neg_4_000_aa    m/z 591.9750     0   1E+4   1   1.5  RT (min)      lcqtof_hilic_neg_4_000_aa    m/z 592.2178     0   1E+4   1   1.5  RT (min)      lcqtof_hilic_neg_4_000_aa    m/z 592.2942     0   1   2   3   4E+4   0.5   1  RT (min)      lcqtof_hilic_neg_3_003_20    m/z 592.3388     0   1E+4   3.5   4  RT (min)       lcqtof_hilic_neg_4_040_05    m/z 592.3756     0   1E+4   4   4.5  RT (min)      lcqtof_hilic_neg_4_008_24    m/z 592.5296     0   1E+4   0.5   1  RT (min)      lcqtof_hilic_neg_4_040_05    m/z 592.5744     0   1   2   3E+4   1   1.5  RT (min)      lcqtof_hilic_neg_4_000_aa    m/z 592.6798     0   1E+4   3.5   4  RT (min)      lcqtof_hilic_neg_4_008_24    m/z 592.7976     0   1E+5   8   8.5  RT (min)      lcqtof_hilic_neg_1_000_aa    m/z 592.7978     0   1E+5   8   8.5  RT (min)      lcqtof_hilic_neg_1_000_aa    m/z 592.9833     0   1E+4   1   1.5  RT (min)      lcqtof_hilic_neg_4_000_aa    m/z 593.1200     0   1E+4   7   7.5  RT (min)      lcqtof_hilic_neg_1_005_32    m/z 593.1498     0   1   2E+4   4.5   5  RT (min)      lcqtof_hilic_neg_1_032_18    m/z 593.1560     0   1E+4   7   7.5  RT (min)      lcqtof_hilic_neg_1_000_aa    m/z 593.2141     0   1E+4   1   1.5  RT (min)      lcqtof_hilic_neg_4_000_aa    m/z 593.2313     0   1E+4   6.5   7  RT (min)      lcqtof_hilic_neg_4_002_25    m/z 593.2922     0   2   4E+4   0.5   1  RT (min)      lcqtof_hilic_neg_3_003_20    m/z 593.3806     0   2   4   6E+4   4   4.5  RT (min)       lcqtof_hilic_neg_4_008_24    m/z 593.3811     0   2   4   6E+4   4   4.5  RT (min)      lcqtof_hilic_neg_4_008_24    m/z 593.5160     0   2   4   6E+4   1   1.5  RT (min)      lcqtof_hilic_neg_4_000_aa    m/z 593.5494     0   2   4   6E+4   1   1.5  RT (min)      lcqtof_hilic_neg_4_000_aa    m/z 593.5528     0   2   4   6E+4   1.5   2  RT (min)      lcqtof_hilic_neg_4_000_aa    m/z 593.5627     0   2   4   6E+4   1   1.5  RT (min)      lcqtof_hilic_neg_4_000_aa    m/z 593.5667     0   1   2E+4   1   1.5  RT (min)      lcqtof_hilic_neg_4_000_aa    m/z 593.6023     0   1   2   3E+4   0.5   1  RT (min)      lcqtof_hilic_neg_4_053_14    m/z 594.1761     0   1E+4   7   7.5  RT (min)      lcqtof_hilic_neg_2_022_46    m/z 594.2948     0   1E+4   0.5   1  RT (min)      lcqtof_hilic_neg_3_003_20    m/z 594.3189     0   1   2   3E+4   0.5   1  RT (min)      lcqtof_hilic_neg_4_015_09    m/z 594.3264     0   1   2E+4   0.5   1  RT (min)      lcqtof_hilic_neg_1_032_18    m/z 594.5180     0   1   2   3E+4   1   1.5  RT (min)      lcqtof_hilic_neg_4_000_aa    m/z 594.5506     0   1   2E+4   1   1.5  RT (min)      lcqtof_hilic_neg_4_000_aa    m/z 594.7774     0   1   2   3E+4   8   8.5  RT (min)      lcqtof_hilic_neg_1_000_aa    m/z 595.2351     0   2   4   6E+4   0.5   1  RT (min)      lcqtof_hilic_neg_1_042_26    m/z 595.3683     0   2   4E+4   1   1.5  RT (min)      lcqtof_hilic_neg_4_000_aa    m/z 595.3686     0   2   4E+4   1   1.5  RT (min)      lcqtof_hilic_neg_4_000_aa    m/z 595.3696     0   2   4E+4   1   1.5  RT (min)      lcqtof_hilic_neg_4_000_aa    m/z 595.4936     0   2   4   6E+4   1   1.5  RT (min)      lcqtof_hilic_neg_4_000_aa    m/z 595.5222     0   1E+5   1   1.5  RT (min)      lcqtof_hilic_neg_4_000_aa    m/z 596.2043     0   1E+4   2   2.5  RT (min)      lcqtof_hilic_neg_4_003_16    m/z 596.2589     0   1   2E+4   1   1.5  RT (min)      lcqtof_hilic_neg_4_052_13    m/z 596.3080     0   1E+4   7   7.5  RT (min)      lcqtof_hilic_neg_1_037_24    m/z 596.3084     0   1E+4   1   1.5  RT (min)      lcqtof_hilic_neg_4_005_29    m/z 596.3709     0   1   2E+4   1   1.5  RT (min)      lcqtof_hilic_neg_4_000_aa    m/z 596.3714     0   1   2E+4   1   1.5  RT (min)      lcqtof_hilic_neg_4_000_aa    m/z 596.3725     0   1   2E+4   1   1.5  RT (min)      lcqtof_hilic_neg_4_000_aa    m/z 596.3736     0   1   2E+4   1   1.5  RT (min)      lcqtof_hilic_neg_4_000_aa    m/z 596.5148     0   1   2   3E+4   1   1.5  RT (min)      lcqtof_hilic_neg_4_000_aa    m/z 596.5241     0   1   2   3E+4   1   1.5  RT (min)      lcqtof_hilic_neg_4_000_aa    m/z 596.5424     0   1E+4   0.5   1  RT (min)      lcqtof_hilic_neg_3_035_46    m/z 596.9672     0   1E+4   0.5   1  RT (min)      lcqtof_hilic_neg_3_035_46    m/z 597.1474     0   1E+4   0.5   1  RT (min)      lcqtof_hilic_neg_3_003_20    m/z 597.1958     0   1E+4   4.5   5  RT (min)      lcqtof_hilic_neg_2_028_05    m/z 597.2032     0   1   2E+4   2   2.5  RT (min)      lcqtof_hilic_neg_4_003_16    m/z 597.2112     0   1E+4   0.5   1  RT (min)      lcqtof_hilic_neg_4_050_02    m/z 597.2268     0   1   2E+4   6   6.5  RT (min)      lcqtof_hilic_neg_4_048_23    m/z 597.2602     0   1   2E+4   0.5   1  RT (min)      lcqtof_hilic_neg_4_013_20    m/z 597.2850     0   1   2   3E+4   6   6.5  RT (min)      lcqtof_hilic_neg_4_008_24    m/z 597.3314     0   2   4E+4   1   1.5  RT (min)      lcqtof_hilic_neg_4_000_aa    m/z 597.3344     0   2   4E+4   1   1.5  RT (min)      lcqtof_hilic_neg_4_000_aa    m/z 597.3363     0   2   4E+4   1   1.5  RT (min)      lcqtof_hilic_neg_4_000_aa    m/z 597.3386     0   2   4E+4   1   1.5  RT (min)      lcqtof_hilic_neg_4_000_aa    m/z 597.3405     0   2   4E+4   1   1.5  RT (min)      lcqtof_hilic_neg_4_000_aa    m/z 597.4026     0   1   2   3E+4   1   1.5  RT (min)      lcqtof_hilic_neg_4_000_aa    m/z 597.4029     0   1   2   3E+4   1   1.5  RT (min)      lcqtof_hilic_neg_4_000_aa    m/z 597.4408     0   1   2E+4   1   1.5  RT (min)      lcqtof_hilic_neg_4_000_aa    m/z 597.4736     0   1E+4   1   1.5  RT (min)      lcqtof_hilic_neg_4_000_aa    m/z 598.0971     0   1E+4   0.5   1  RT (min)      lcqtof_hilic_neg_1_062_17    m/z 598.1291     0   1   2   3E+4   0.5   1  RT (min)      lcqtof_hilic_neg_1_032_18    m/z 598.2748     0   2   4   6   8E+3   1   1.5  RT (min)      lcqtof_hilic_neg_2_018_16    m/z 598.4051     0   1E+4   1   1.5  RT (min)      lcqtof_hilic_neg_4_000_aa    m/z 598.4792     0   1E+4   0.5   1  RT (min)      lcqtof_hilic_neg_4_046_41    m/z 598.5142     0   1E+4   1   1.5  RT (min)      lcqtof_hilic_neg_4_000_aa    m/z 599.1434     0   1E+4   0.5   1  RT (min)      lcqtof_hilic_neg_3_003_20    m/z 599.2295     0   1E+4   6   6.5  RT (min)       lcqtof_hilic_neg_4_048_23    m/z 599.2518     0   1   2E+4   6   6.5  RT (min)       lcqtof_hilic_neg_4_090_60    m/z 599.2600     0   1E+4   4   4.5  RT (min)      lcqtof_hilic_neg_4_039_18    m/z 599.3056     0   1E+4   15   15.5  RT (min)      lcqtof_hilic_neg_4_000_aa    m/z 599.3284     0   1E+5   1   1.5  RT (min)      lcqtof_hilic_neg_4_000_aa    m/z 599.3297     0   1E+5   1   1.5  RT (min)      lcqtof_hilic_neg_4_000_aa    m/z 599.3301     0   1E+5   1   1.5  RT (min)      lcqtof_hilic_neg_4_000_aa    m/z 599.3796     0   2   4   6   8E+3   1   1.5  RT (min)      lcqtof_hilic_neg_4_026_04    m/z 599.3799     0   2   4   6   8E+3   1   1.5  RT (min)      lcqtof_hilic_neg_4_002_25    m/z 599.3802     0   1   2   3E+3   16.5   17  RT (min)      lcqtof_hilic_neg_4_048_23    m/z 599.3807     0   2   4E+3   4   4.5  RT (min)      lcqtof_hilic_neg_4_001_49    m/z 599.3809     0   2   4   6E+3   2   2.5  RT (min)      lcqtof_hilic_neg_4_032_01    m/z 599.3811     0   1   2E+3   0   0.5  RT (min)      lcqtof_hilic_neg_4_001_49    m/z 599.3812     0   2   4E+3   4.5   5  RT (min)      lcqtof_hilic_neg_4_047_06    m/z 599.3816     0   2   4   6E+3   15.5   16  RT (min)      lcqtof_hilic_neg_4_012_07    m/z 599.3817     0   2   4   6E+3   15.5   16  RT (min)      lcqtof_hilic_neg_4_019_03    m/z 599.4225     0   1   2E+4   1   1.5  RT (min)      lcqtof_hilic_neg_4_000_aa    m/z 599.5284     0   2   4E+4   1.5   2  RT (min)      lcqtof_hilic_neg_4_000_aa    m/z 600.1158     0   1E+4   0.5   1  RT (min)      lcqtof_hilic_neg_4_021_21    m/z 600.1266     0   2   4E+4   0.5   1  RT (min)      lcqtof_hilic_neg_1_032_18    m/z 600.2055     0   1   2   3   4E+4   0.5   1  RT (min)      lcqtof_hilic_neg_4_021_21    m/z 600.2169     0   1   2   3   4E+4   0.5   1  RT (min)      lcqtof_hilic_neg_4_039_18    m/z 600.3293     0   2   4E+4   1   1.5  RT (min)      lcqtof_hilic_neg_4_000_aa    m/z 600.3312     0   2   4E+4   1   1.5  RT (min)      lcqtof_hilic_neg_4_000_aa    m/z 600.3318     0   2   4E+4   1   1.5  RT (min)      lcqtof_hilic_neg_4_000_aa    m/z 600.3354     0   2   4E+4   1   1.5  RT (min)      lcqtof_hilic_neg_4_000_aa    m/z 600.3809     0   1   2E+4   1   1.5  RT (min)      lcqtof_hilic_neg_4_000_aa    m/z 600.5313     0   1E+4   1.5   2  RT (min)      lcqtof_hilic_neg_4_000_aa    m/z 601.0120     0   1E+4   1   1.5  RT (min)      lcqtof_hilic_neg_4_000_aa    m/z 601.1219     0   1   2   3   4E+4   0.5   1  RT (min)      lcqtof_hilic_neg_1_032_18    m/z 601.1382     0   1E+4   6.5   7  RT (min)      lcqtof_hilic_neg_1_000_aa    m/z 601.2986     0   1E+4   4   4.5  RT (min)      lcqtof_hilic_neg_1_037_24    m/z 601.2992     0   1   2E+4   4   4.5  RT (min)      lcqtof_hilic_neg_1_007_05    m/z 601.3116     0   1   2   3E+3   3.5   4  RT (min)      lcqtof_hilic_neg_4_060_13    m/z 601.3224     0   1E+4   1   1.5  RT (min)      lcqtof_hilic_neg_3_011_34    m/z 601.3230     0   1   2   3   4E+3   0   0.5  RT (min)      lcqtof_hilic_neg_1_004_14    m/z 601.3230     0   1E+4   0.5   1  RT (min)      lcqtof_hilic_neg_1_010_15    m/z 601.3230     0   2   4   6E+3   1.5   2  RT (min)      lcqtof_hilic_neg_1_032_18    m/z 601.3232     0   1   2   3   4E+3   2.5   3  RT (min)      lcqtof_hilic_neg_4_101_61    m/z 601.3238     0   2   4E+4   1   1.5  RT (min)      lcqtof_hilic_neg_4_000_aa    m/z 601.3238     0   1   2E+5   0.5   1  RT (min)      lcqtof_hilic_neg_3_003_20    m/z 601.3243     0   2   4   6E+3   0.5   1  RT (min)      lcqtof_hilic_neg_4_019_03    m/z 601.3250     0   2   4E+3   2   2.5  RT (min)      lcqtof_hilic_neg_4_023_15    m/z 601.3378     0   2   4   6E+4   1   1.5  RT (min)      lcqtof_hilic_neg_4_000_aa    m/z 601.3422     0   2   4   6E+4   1   1.5  RT (min)      lcqtof_hilic_neg_4_000_aa    m/z 601.3578     0   2   4   6E+4   1   1.5  RT (min)      lcqtof_hilic_neg_4_000_aa    m/z 601.4260     0   1E+4   0.5   1  RT (min)      lcqtof_hilic_neg_4_010_28    m/z 601.5326     0   1E+4   1.5   2  RT (min)      lcqtof_hilic_neg_4_034_31    m/z 601.5329     0   1   2   3E+4   1   1.5  RT (min)      lcqtof_hilic_neg_4_034_31    m/z 601.5330     0   1   2   3E+4   1   1.5  RT (min)      lcqtof_hilic_neg_4_034_31    m/z 601.5332     0   1   2E+4   1.5   2  RT (min)      lcqtof_hilic_neg_4_000_aa    m/z 602.2416     0   1E+4   6   6.5  RT (min)      lcqtof_hilic_neg_4_040_05    m/z 602.3287     0   2   4   6   8E+4   0.5   1  RT (min)      lcqtof_hilic_neg_3_003_20    m/z 602.3449     0   1   2E+4   1   1.5  RT (min)      lcqtof_hilic_neg_4_000_aa    m/z 602.3491     0   1   2E+4   1   1.5  RT (min)      lcqtof_hilic_neg_4_000_aa    m/z 602.3811     0   1E+4   0.5   1  RT (min)      lcqtof_hilic_neg_4_038_10    m/z 602.8408     0   1E+4   2.5   3  RT (min)      lcqtof_hilic_neg_4_045_26    m/z 603.0076     0   1E+4   1   1.5  RT (min)      lcqtof_hilic_neg_4_000_aa    m/z 603.0076     0   1E+4   1   1.5  RT (min)      lcqtof_hilic_neg_4_000_aa    m/z 603.2485     0   1E+4   3   3.5  RT (min)      lcqtof_hilic_neg_4_003_16    m/z 603.2630     0   1   2   3   4E+4   1   1.5  RT (min)      lcqtof_hilic_neg_4_000_aa    m/z 603.2632     0   1E+4   6.5   7  RT (min)      lcqtof_hilic_neg_4_008_24    m/z 603.2638     0   1   2   3   4E+4   1   1.5  RT (min)      lcqtof_hilic_neg_4_000_aa    m/z 603.2644     0   1E+4   6.5   7  RT (min)      lcqtof_hilic_neg_4_008_24    m/z 603.2650     0   1   2   3   4E+4   1   1.5  RT (min)      lcqtof_hilic_neg_4_000_aa    m/z 603.2668     0   1   2   3   4E+4   1   1.5  RT (min)      lcqtof_hilic_neg_4_000_aa    m/z 603.3183     0   2   4   6   8E+4   1   1.5  RT (min)      lcqtof_hilic_neg_4_038_10    m/z 603.3496     0   1   2E+4   1   1.5  RT (min)      lcqtof_hilic_neg_4_000_aa    m/z 603.3500     0   1   2E+4   1   1.5  RT (min)      lcqtof_hilic_neg_4_000_aa    m/z 603.3535     0   1   2E+4   1   1.5  RT (min)      lcqtof_hilic_neg_4_000_aa    m/z 603.4239     0   1E+4   1   1.5  RT (min)      lcqtof_hilic_neg_4_000_aa    m/z 603.4288     0   1E+4   1   1.5  RT (min)      lcqtof_hilic_neg_4_000_aa    m/z 603.5767     0   1E+4   0.5   1  RT (min)      lcqtof_hilic_neg_1_037_24    m/z 604.0702     0   1E+4   8   8.5  RT (min)       lcqtof_hilic_neg_4_082_59    m/z 604.2652     0   1E+4   1   1.5  RT (min)      lcqtof_hilic_neg_4_000_aa    m/z 604.2657     0   1E+4   1   1.5  RT (min)      lcqtof_hilic_neg_4_000_aa    m/z 604.3233     0   1   2E+4   1   1.5  RT (min)      lcqtof_hilic_neg_4_038_10    m/z 604.3560     0   1E+4   1   1.5  RT (min)      lcqtof_hilic_neg_4_000_aa    m/z 604.3566     0   1E+4   1   1.5  RT (min)      lcqtof_hilic_neg_4_000_aa    m/z 604.8693     0   1   2   3E+3   5   5.5  RT (min)       lcqtof_hilic_neg_4_000_dd    m/z 604.8705     0   2   4   6   8E+3   5   5.5  RT (min)       lcqtof_hilic_neg_4_058_39    m/z 604.8706     0   2   4   6   8E+3   5   5.5  RT (min)       lcqtof_hilic_neg_4_058_39    m/z 604.8707     0   2   4   6   8E+3   5   5.5  RT (min)       lcqtof_hilic_neg_4_058_39    m/z 604.8713     0   2   4   6   8E+3   5.5   6  RT (min)       lcqtof_hilic_neg_4_058_39    m/z 605.0953     0   1E+4   6   6.5  RT (min)      lcqtof_hilic_neg_4_048_23    m/z 605.2546     0   1   2   3E+4   1   1.5  RT (min)      lcqtof_hilic_neg_4_000_aa    m/z 605.2554     0   1   2   3E+4   1   1.5  RT (min)      lcqtof_hilic_neg_4_000_aa    m/z 605.2578     0   1   2   3E+4   1   1.5  RT (min)      lcqtof_hilic_neg_4_000_aa    m/z 605.2578     0   1   2   3E+4   1   1.5  RT (min)      lcqtof_hilic_neg_4_000_aa    m/z 605.2602     0   1E+4   0.5   1  RT (min)      lcqtof_hilic_neg_3_035_46    m/z 605.2898     0   1E+4   0.5   1  RT (min)      lcqtof_hilic_neg_4_021_21    m/z 605.3210     0   1E+4   0.5   1  RT (min)      lcqtof_hilic_neg_4_038_10    m/z 605.3522     0   2   4E+4   1   1.5  RT (min)      lcqtof_hilic_neg_4_000_aa    m/z 605.4043     0   2   4E+3   2   2.5  RT (min)      lcqtof_hilic_neg_4_011_35    m/z 605.4774     0   1   2E+5   1   1.5  RT (min)      lcqtof_hilic_neg_4_053_14    m/z 605.4951     0   2   4E+4   0.5   1  RT (min)      lcqtof_hilic_neg_3_035_46    m/z 605.4956     0   2   4E+4   0.5   1  RT (min)      lcqtof_hilic_neg_3_035_46    m/z 605.5536     0   1   2   3E+4   1   1.5  RT (min)      lcqtof_hilic_neg_4_000_aa    m/z 605.5866     0   1   2E+4   1   1.5  RT (min)      lcqtof_hilic_neg_4_000_aa    m/z 605.8937     0   2   4E+3   7   7.5  RT (min)       lcqtof_hilic_neg_4_032_01    m/z 606.0735     0   1   2   3   4E+5   7.5   8  RT (min)       lcqtof_hilic_neg_4_083_59    m/z 606.1970     0   1E+4   7.5   8  RT (min)      lcqtof_hilic_neg_1_022_36    m/z 606.3015     0   1   2E+4   6   6.5  RT (min)      lcqtof_hilic_neg_4_008_24    m/z 606.3146     0   1   2E+4   3   3.5  RT (min)      lcqtof_hilic_neg_4_061_50    m/z 606.3222     0   2   4   6E+4   7.5   8  RT (min)      lcqtof_hilic_neg_1_022_36    m/z 606.3510     0   1E+4   1   1.5  RT (min)      lcqtof_hilic_neg_4_000_aa    m/z 606.3526     0   1E+4   1   1.5  RT (min)      lcqtof_hilic_neg_4_000_aa    m/z 606.4590     0   1   2   3E+4   0.5   1  RT (min)      lcqtof_hilic_neg_4_023_15    m/z 606.4990     0   1E+4   0.5   1  RT (min)      lcqtof_hilic_neg_3_035_46    m/z 606.5576     0   1E+4   1   1.5  RT (min)      lcqtof_hilic_neg_4_000_aa    m/z 606.6126     0   1   2E+4   0.5   1  RT (min)      lcqtof_hilic_neg_3_035_46    m/z 606.9370     0   1E+4   1   1.5  RT (min)      lcqtof_hilic_neg_4_000_aa    m/z 606.9386     0   1E+4   1   1.5  RT (min)      lcqtof_hilic_neg_4_000_aa    m/z 606.9974     0   1   2   3E+4   0.5   1  RT (min)      lcqtof_hilic_neg_3_035_46    m/z 607.0754     0   2   4   6   8E+4   7.5   8  RT (min)      lcqtof_hilic_neg_4_083_59    m/z 607.1694     0   1E+5   1   1.5  RT (min)      lcqtof_hilic_neg_4_039_18    m/z 607.3300     0   2   4   6   8E+4   0.5   1  RT (min)      lcqtof_hilic_neg_4_038_10    m/z 607.3403     0   1   2   3E+3   15.5   16  RT (min)      lcqtof_hilic_neg_4_000_aa    m/z 607.3724     0   1E+4   4   4.5  RT (min)      lcqtof_hilic_neg_4_008_24    m/z 607.3844     0   1   2E+4   1.5   2  RT (min)      lcqtof_hilic_neg_4_053_14    m/z 607.4580     0   1E+4   0.5   1  RT (min)      lcqtof_hilic_neg_1_005_32    m/z 607.5380     0   2   4E+4   1   1.5  RT (min)      lcqtof_hilic_neg_4_000_aa    m/z 607.5493     0   2   4   6E+4   1   1.5  RT (min)      lcqtof_hilic_neg_4_000_aa    m/z 607.5622     0   2   4   6E+4   1   1.5  RT (min)      lcqtof_hilic_neg_4_000_aa    m/z 607.5675     0   2   4   6E+4   1.5   2  RT (min)      lcqtof_hilic_neg_4_000_aa    m/z 607.5681     0   2   4   6E+4   1.5   2  RT (min)      lcqtof_hilic_neg_4_000_aa    m/z 607.5768     0   2   4   6E+4   1.5   2  RT (min)      lcqtof_hilic_neg_4_000_aa    m/z 607.5836     0   1   2E+4   1   1.5  RT (min)      lcqtof_hilic_neg_4_000_aa    m/z 607.6174     0   1   2E+4   0.5   1  RT (min)      lcqtof_hilic_neg_4_023_15    m/z 607.8840     0   1E+4   4   4.5  RT (min)      lcqtof_hilic_neg_4_008_24    m/z 607.9968     0   1E+4   0.5   1  RT (min)      lcqtof_hilic_neg_3_035_46    m/z 608.1834     0   1   2E+4   6.5   7  RT (min)      lcqtof_hilic_neg_4_008_24    m/z 608.2024     0   1   2E+4   1   1.5  RT (min)      lcqtof_hilic_neg_4_039_18    m/z 608.3330     0   1   2   3E+4   0.5   1  RT (min)      lcqtof_hilic_neg_1_032_18    m/z 608.3848     0   1   2E+4   4   4.5  RT (min)      lcqtof_hilic_neg_4_008_24    m/z 608.5511     0   1   2E+4   1   1.5  RT (min)      lcqtof_hilic_neg_4_000_aa    m/z 608.5631     0   1   2E+4   1   1.5  RT (min)      lcqtof_hilic_neg_4_000_aa    m/z 608.8917     0   1E+4   4   4.5  RT (min)      lcqtof_hilic_neg_4_008_24    m/z 608.9356     0   1   2E+4   1   1.5  RT (min)      lcqtof_hilic_neg_4_000_aa    m/z 608.9364     0   1   2E+4   1   1.5  RT (min)      lcqtof_hilic_neg_4_000_aa    m/z 609.1461     0   1E+4   4.5   5  RT (min)      lcqtof_hilic_neg_2_053_18    m/z 609.1690     0   1E+5   1   1.5  RT (min)      lcqtof_hilic_neg_4_039_18    m/z 609.1718     0   1   2   3E+4   0.5   1  RT (min)      lcqtof_hilic_neg_3_003_20    m/z 609.2660     0   1E+4   6   6.5  RT (min)      lcqtof_hilic_neg_4_040_05    m/z 609.3925     0   1   2   3E+4   4   4.5  RT (min)      lcqtof_hilic_neg_4_008_24    m/z 609.4030     0   2   4E+4   1   1.5  RT (min)      lcqtof_hilic_neg_4_000_aa    m/z 609.5416     0   1   2   3E+5   1   1.5  RT (min)      lcqtof_hilic_neg_4_000_aa    m/z 609.5573     0   1   2   3   4E+5   1.5   2  RT (min)      lcqtof_hilic_neg_4_000_aa    m/z 609.5581     0   2   4E+4   15.5   16  RT (min)      lcqtof_hilic_neg_4_000_aa    m/z 609.7584     0   2   4E+4   0.5   1  RT (min)      lcqtof_hilic_neg_4_005_29    m/z 610.1554     0   2   4   6E+4   0.5   1  RT (min)      lcqtof_hilic_neg_1_032_18    m/z 610.2655     0   1   2   3   4E+4   2.5   3  RT (min)      lcqtof_hilic_neg_4_039_18    m/z 610.2702     0   1E+4   6   6.5  RT (min)      lcqtof_hilic_neg_4_040_05    m/z 610.2755     0   1   2E+4   0.5   1  RT (min)      lcqtof_hilic_neg_4_052_13    m/z 610.4014     0   1E+4   4   4.5  RT (min)      lcqtof_hilic_neg_4_008_24    m/z 610.4022     0   1E+4   1   1.5  RT (min)      lcqtof_hilic_neg_4_000_aa    m/z 610.4126     0   1E+4   0.5   1  RT (min)      lcqtof_hilic_neg_1_042_26    m/z 610.4746     0   1E+4   0.5   1  RT (min)      lcqtof_hilic_neg_4_053_14    m/z 610.5236     0   1   2   3E+4   1   1.5  RT (min)      lcqtof_hilic_neg_4_000_aa    m/z 610.5462     0   1E+5   1   1.5  RT (min)      lcqtof_hilic_neg_4_000_aa    m/z 610.5590     0   1   2E+4   15.5   16  RT (min)      lcqtof_hilic_neg_4_000_aa    m/z 610.5600     0   1E+5   1.5   2  RT (min)      lcqtof_hilic_neg_4_000_aa    m/z 610.9308     0   1E+4   1   1.5  RT (min)      lcqtof_hilic_neg_4_000_aa    m/z 610.9314     0   1E+4   1   1.5  RT (min)      lcqtof_hilic_neg_4_000_aa    m/z 610.9319     0   1E+4   1   1.5  RT (min)      lcqtof_hilic_neg_4_000_aa    m/z 611.1442     0   1E+5   7.5   8  RT (min)       lcqtof_hilic_neg_4_083_59    m/z 611.1570     0   1   2E+4   1   1.5  RT (min)      lcqtof_hilic_neg_4_000_aa    m/z 611.1585     0   1   2E+4   1   1.5  RT (min)      lcqtof_hilic_neg_4_000_aa    m/z 611.2508     0   1   2   3E+3   4.5   5  RT (min)      lcqtof_hilic_neg_4_002_25    m/z 611.2514     0   1   2   3E+3   5   5.5  RT (min)       lcqtof_hilic_neg_4_058_39    m/z 611.2517     0   1   2   3   4E+3   4.5   5  RT (min)      lcqtof_hilic_neg_4_032_01    m/z 611.2536     0   2   4E+3   1   1.5  RT (min)      lcqtof_hilic_neg_4_024_40    m/z 611.2540     0   1   2   3E+3   0.5   1  RT (min)      lcqtof_hilic_neg_4_036_11    m/z 611.2553     0   1   2E+3   0   0.5  RT (min)      lcqtof_hilic_neg_4_000_bb    m/z 611.2555     0   1E+4   2   2.5  RT (min)      lcqtof_hilic_neg_4_029_17    m/z 611.2565     0   1   2   3E+3   0.5   1  RT (min)      lcqtof_hilic_neg_4_005_29    m/z 611.2566     0   1E+4   2.5   3  RT (min)      lcqtof_hilic_neg_4_039_18    m/z 611.2571     0   1E+4   2   2.5  RT (min)       lcqtof_hilic_neg_4_027_19    m/z 611.2816     0   1E+4   2   2.5  RT (min)      lcqtof_hilic_neg_4_045_26    m/z 611.2865     0   1   2E+4   2   2.5  RT (min)      lcqtof_hilic_neg_4_010_28    m/z 611.3818     0   1   2   3E+5   1   1.5  RT (min)      lcqtof_hilic_neg_4_000_aa    m/z 611.3824     0   1   2   3E+5   1   1.5  RT (min)      lcqtof_hilic_neg_4_000_aa    m/z 611.3912     0   1   2   3E+5   1   1.5  RT (min)      lcqtof_hilic_neg_4_000_aa    m/z 611.3912     0   1E+4   4.5   5  RT (min)      lcqtof_hilic_neg_4_008_24    m/z 611.4132     0   1   2   3   4E+3   15   15.5  RT (min)       lcqtof_hilic_neg_4_006_30    m/z 611.4159     0   1   2   3E+3   15.5   16  RT (min)      lcqtof_hilic_neg_4_010_28    m/z 611.5136     0   1   2E+4   1   1.5  RT (min)      lcqtof_hilic_neg_4_000_aa    m/z 611.5617     0   1   2   3   4E+4   0.5   1  RT (min)      lcqtof_hilic_neg_3_035_46    m/z 611.5631     0   1   2   3E+4   1.5   2  RT (min)      lcqtof_hilic_neg_4_000_aa    m/z 612.1457     0   1   2   3E+4   7.5   8  RT (min)      lcqtof_hilic_neg_4_083_59    m/z 612.1590     0   1E+4   1   1.5  RT (min)      lcqtof_hilic_neg_4_000_aa    m/z 612.2655     0   1E+4   0.5   1  RT (min)      lcqtof_hilic_neg_4_050_02    m/z 612.2805     0   1E+4   0.5   1  RT (min)      lcqtof_hilic_neg_4_002_25    m/z 612.2942     0   1E+4   2   2.5  RT (min)      lcqtof_hilic_neg_4_010_28    m/z 612.3228     0   2   4   6E+4   0.5   1  RT (min)      lcqtof_hilic_neg_3_003_20    m/z 612.3842     0   1E+5   1   1.5  RT (min)      lcqtof_hilic_neg_4_000_aa    m/z 612.3854     0   1E+5   1   1.5  RT (min)      lcqtof_hilic_neg_4_000_aa    m/z 612.3928     0   1E+5   1   1.5  RT (min)      lcqtof_hilic_neg_4_000_aa    m/z 612.4720     0   1E+4   1   1.5  RT (min)      lcqtof_hilic_neg_4_046_41    m/z 612.5774     0   1   2   3E+4   0.5   1  RT (min)      lcqtof_hilic_neg_3_035_46    m/z 612.6101     0   1E+4   0.5   1  RT (min)      lcqtof_hilic_neg_4_008_24    m/z 612.6689     0   1E+4   4.5   5  RT (min)      lcqtof_hilic_neg_4_050_02    m/z 612.9170     0   1E+4   1   1.5  RT (min)      lcqtof_hilic_neg_4_000_aa    m/z 613.1322     0   1E+4   0.5   1  RT (min)       lcqtof_hilic_neg_4_072_57    m/z 613.1556     0   1E+4   1   1.5  RT (min)      lcqtof_hilic_neg_4_000_aa    m/z 613.1568     0   1E+4   1   1.5  RT (min)      lcqtof_hilic_neg_4_000_aa    m/z 613.1572     0   1E+4   1   1.5  RT (min)      lcqtof_hilic_neg_4_000_aa    m/z 613.1594     0   1E+4   6   6.5  RT (min)      lcqtof_hilic_neg_1_014_31    m/z 613.1988     0   1E+4   0.5   1  RT (min)      lcqtof_hilic_neg_3_035_46    m/z 613.3136     0   1E+4   15   15.5  RT (min)      lcqtof_hilic_neg_4_000_aa    m/z 613.3792     0   1E+4   0.5   1  RT (min)      lcqtof_hilic_neg_4_053_14    m/z 613.3944     0   2   4   6   8E+3   2   2.5  RT (min)      lcqtof_hilic_neg_4_032_01    m/z 613.3946     0   1   2   3   4E+3   15.5   16  RT (min)      lcqtof_hilic_neg_1_007_05    m/z 613.3948     0   2   4   6   8E+3   15.5   16  RT (min)      lcqtof_hilic_neg_4_012_07    m/z 613.3952     0   2   4   6   8E+3   15.5   16  RT (min)      lcqtof_hilic_neg_4_019_03    m/z 613.3954     0   1E+5   1   1.5  RT (min)       lcqtof_hilic_neg_4_000_aa    m/z 613.3976     0   1E+4   0.5   1  RT (min)      lcqtof_hilic_neg_4_018_27    m/z 613.4406     0   1   2E+4   1   1.5  RT (min)      lcqtof_hilic_neg_4_053_14    m/z 613.5812     0   1E+4   0.5   1  RT (min)      lcqtof_hilic_neg_3_035_46    m/z 614.2090     0   1E+4   4   4.5  RT (min)      lcqtof_hilic_neg_4_039_18    m/z 614.3809     0   2   4E+4   1   1.5  RT (min)      lcqtof_hilic_neg_4_000_aa    m/z 614.3920     0   2   4E+4   1   1.5  RT (min)      lcqtof_hilic_neg_4_000_aa    m/z 614.3967     0   2   4E+4   1   1.5  RT (min)      lcqtof_hilic_neg_4_000_aa    m/z 614.3988     0   2   4E+4   1   1.5  RT (min)      lcqtof_hilic_neg_4_000_aa    m/z 614.3992     0   1   2   3E+3   15.5   16  RT (min)      lcqtof_hilic_neg_4_019_03    m/z 614.4013     0   2   4E+4   1   1.5  RT (min)      lcqtof_hilic_neg_4_000_aa    m/z 615.3995     0   1   2E+4   0.5   1  RT (min)      lcqtof_hilic_neg_4_032_01    m/z 615.4085     0   1E+4   1   1.5  RT (min)      lcqtof_hilic_neg_4_000_aa    m/z 615.4110     0   1E+4   1   1.5  RT (min)      lcqtof_hilic_neg_4_000_aa    m/z 615.8704     0   1E+4   2   2.5  RT (min)      lcqtof_hilic_neg_4_016_37    m/z 616.0623     0   1   2   3   4E+4   0   0.5  RT (min)      lcqtof_hilic_neg_1_000_bb    m/z 616.0636     0   2   4   6E+3   2   2.5  RT (min)      lcqtof_hilic_neg_4_005_29    m/z 616.1409     0   1E+4   1   1.5  RT (min)      lcqtof_hilic_neg_4_027_19    m/z 616.3802     0   1   2E+4   0.5   1  RT (min)      lcqtof_hilic_neg_4_053_14    m/z 617.1544     0   1E+4   7   7.5  RT (min)       lcqtof_hilic_neg_4_039_18    m/z 617.2930     0   1E+4   15   15.5  RT (min)      lcqtof_hilic_neg_4_000_aa    m/z 617.2932     0   1   2   3E+4   6   6.5  RT (min)       lcqtof_hilic_neg_4_008_24    m/z 617.3343     0   1   2E+4   0.5   1  RT (min)       lcqtof_hilic_neg_4_038_10    m/z 617.3356     0   1   2E+4   1   1.5  RT (min)      lcqtof_hilic_neg_4_015_09    m/z 617.4206     0   1E+4   1   1.5  RT (min)      lcqtof_hilic_neg_4_000_aa    m/z 617.4289     0   1E+4   1   1.5  RT (min)      lcqtof_hilic_neg_4_000_aa    m/z 617.4488     0   1E+4   1   1.5  RT (min)      lcqtof_hilic_neg_4_000_aa    m/z 618.3114     0   1E+5   0.5   1  RT (min)      lcqtof_hilic_neg_3_003_20    m/z 618.5272     0   1   2   3E+4   0.5   1  RT (min)      lcqtof_hilic_neg_3_035_46    m/z 619.2704     0   1E+4   1   1.5  RT (min)      lcqtof_hilic_neg_4_000_aa    m/z 619.2718     0   1E+4   1   1.5  RT (min)      lcqtof_hilic_neg_4_000_aa    m/z 619.3160     0   1   2   3E+4   0.5   1  RT (min)      lcqtof_hilic_neg_3_003_20    m/z 619.4136     0   1E+4   4.5   5  RT (min)      lcqtof_hilic_neg_4_003_16    m/z 619.4315     0   2   4   6E+4   1   1.5  RT (min)      lcqtof_hilic_neg_4_000_aa    m/z 619.5401     0   2   4E+4   4.5   5  RT (min)      lcqtof_hilic_neg_4_003_16    m/z 619.5634     0   1   2E+4   1   1.5  RT (min)      lcqtof_hilic_neg_4_000_aa    m/z 620.2552     0   1   2   3E+4   0.5   1  RT (min)      lcqtof_hilic_neg_4_010_28    m/z 620.3062     0   1E+4   7   7.5  RT (min)      lcqtof_hilic_neg_4_048_23    m/z 620.3814     0   1E+4   4   4.5  RT (min)      lcqtof_hilic_neg_4_008_24    m/z 620.4348     0   1   2E+4   1   1.5  RT (min)      lcqtof_hilic_neg_4_000_aa    m/z 620.5726     0   1E+4   1   1.5  RT (min)      lcqtof_hilic_neg_4_000_aa    m/z 620.8485     0   1   2E+4   1   1.5  RT (min)      lcqtof_hilic_neg_4_004_47    m/z 620.9570     0   1E+4   1   1.5  RT (min)      lcqtof_hilic_neg_4_000_aa    m/z 621.1489     0   1E+4   6   6.5  RT (min)      lcqtof_hilic_neg_4_008_24    m/z 621.1798     0   1E+4   0.5   1  RT (min)      lcqtof_hilic_neg_1_004_14    m/z 621.1873     0   1E+5   1   1.5  RT (min)      lcqtof_hilic_neg_4_000_aa    m/z 621.1960     0   1   2E+4   4.5   5  RT (min)      lcqtof_hilic_neg_2_028_05    m/z 621.2111     0   1E+4   6   6.5  RT (min)      lcqtof_hilic_neg_4_048_23    m/z 621.2203     0   1E+4   3   3.5  RT (min)      lcqtof_hilic_neg_4_004_47    m/z 621.2721     0   1E+4   1   1.5  RT (min)      lcqtof_hilic_neg_4_000_aa    m/z 621.2756     0   1E+4   1   1.5  RT (min)      lcqtof_hilic_neg_4_000_aa    m/z 621.3205     0   1E+5   1   1.5  RT (min)      lcqtof_hilic_neg_4_045_26    m/z 621.5452     0   2   4   6E+4   1   1.5  RT (min)      lcqtof_hilic_neg_4_000_aa    m/z 621.5739     0   2   4   6   8E+4   1   1.5  RT (min)      lcqtof_hilic_neg_4_000_aa    m/z 621.5808     0   2   4   6   8E+4   1   1.5  RT (min)      lcqtof_hilic_neg_4_000_aa    m/z 621.5823     0   2   4   6   8E+4   1.5   2  RT (min)      lcqtof_hilic_neg_4_000_aa    m/z 621.5829     0   2   4   6   8E+4   1.5   2  RT (min)      lcqtof_hilic_neg_4_000_aa    m/z 622.1880     0   2   4   6   8E+4   1   1.5  RT (min)      lcqtof_hilic_neg_4_000_aa    m/z 622.1894     0   2   4   6   8E+4   1   1.5  RT (min)      lcqtof_hilic_neg_4_000_aa    m/z 622.2536     0   1   2   3E+4   0.5   1  RT (min)      lcqtof_hilic_neg_2_004_26    m/z 622.2688     0   1   2E+4   3   3.5  RT (min)      lcqtof_hilic_neg_4_039_18    m/z 622.3519     0   1E+5   0.5   1  RT (min)      lcqtof_hilic_neg_3_003_20    m/z 622.5545     0   1   2E+4   1   1.5  RT (min)      lcqtof_hilic_neg_4_000_aa    m/z 622.5844     0   1   2   3   4E+4   1   1.5  RT (min)      lcqtof_hilic_neg_4_000_aa    m/z 622.6056     0   1E+4   0.5   1  RT (min)      lcqtof_hilic_neg_3_035_46    m/z 622.9533     0   1E+4   1   1.5  RT (min)      lcqtof_hilic_neg_4_000_aa    m/z 622.9552     0   1E+4   1   1.5  RT (min)      lcqtof_hilic_neg_4_000_aa    m/z 622.9596     0   1E+4   4   4.5  RT (min)      lcqtof_hilic_neg_4_008_24    m/z 623.1880     0   2   4E+4   1   1.5  RT (min)      lcqtof_hilic_neg_4_000_aa    m/z 623.2159     0   1E+4   4   4.5  RT (min)      lcqtof_hilic_neg_4_050_02    m/z 623.2618     0   1   2E+4   0.5   1  RT (min)      lcqtof_hilic_neg_1_042_26    m/z 623.2796     0   1   2   3E+4   1   1.5  RT (min)      lcqtof_hilic_neg_4_039_18    m/z 623.3000     0   1   2   3E+4   1   1.5  RT (min)      lcqtof_hilic_neg_4_000_aa    m/z 623.3438     0   1E+4   15   15.5  RT (min)      lcqtof_hilic_neg_4_000_aa    m/z 623.3528     0   1   2E+4   0.5   1  RT (min)      lcqtof_hilic_neg_3_003_20    m/z 623.3834     0   1   2   3E+4   7.5   8  RT (min)      lcqtof_hilic_neg_1_005_32    m/z 623.5212     0   2   4   6E+4   1   1.5  RT (min)      lcqtof_hilic_neg_4_000_aa    m/z 623.5404     0   2   4   6E+4   1   1.5  RT (min)      lcqtof_hilic_neg_4_000_aa    m/z 623.5572     0   2   4   6E+4   1   1.5  RT (min)      lcqtof_hilic_neg_4_000_aa    m/z 623.5838     0   1E+4   1   1.5  RT (min)      lcqtof_hilic_neg_4_000_aa    m/z 623.9678     0   1E+4   4   4.5  RT (min)      lcqtof_hilic_neg_4_008_24    m/z 624.1803     0   1   2E+4   1   1.5  RT (min)      lcqtof_hilic_neg_4_000_aa    m/z 624.1864     0   1   2E+4   1   1.5  RT (min)      lcqtof_hilic_neg_4_000_aa    m/z 624.1864     0   1   2E+4   1   1.5  RT (min)      lcqtof_hilic_neg_4_000_aa    m/z 624.1873     0   1   2E+4   1   1.5  RT (min)      lcqtof_hilic_neg_4_000_aa    m/z 624.1882     0   1   2E+4   1   1.5  RT (min)      lcqtof_hilic_neg_4_000_aa    m/z 624.2828     0   1E+4   1   1.5  RT (min)      lcqtof_hilic_neg_4_039_18    m/z 624.3368     0   1E+4   4.5   5  RT (min)      lcqtof_hilic_neg_4_022_34    m/z 624.3380     0   1E+4   4   4.5  RT (min)      lcqtof_hilic_neg_4_019_03    m/z 624.3383     0   1E+4   4.5   5  RT (min)      lcqtof_hilic_neg_4_050_02    m/z 624.4819     0   1   2   3   4E+4   0.5   1  RT (min)      lcqtof_hilic_neg_4_032_01    m/z 624.4890     0   1E+4   0.5   1  RT (min)      lcqtof_hilic_neg_2_021_17    m/z 624.4942     0   1   2E+4   0.5   1  RT (min)      lcqtof_hilic_neg_4_005_29    m/z 624.5447     0   1   2E+4   1   1.5  RT (min)      lcqtof_hilic_neg_4_000_aa    m/z 624.5559     0   1   2E+4   0.5   1  RT (min)      lcqtof_hilic_neg_4_016_37    m/z 624.5595     0   1   2E+4   1   1.5  RT (min)      lcqtof_hilic_neg_4_000_aa    m/z 624.5646     0   1   2E+4   1   1.5  RT (min)      lcqtof_hilic_neg_4_000_aa    m/z 624.9480     0   1E+4   1   1.5  RT (min)      lcqtof_hilic_neg_4_000_aa    m/z 624.9753     0   1E+4   4   4.5  RT (min)       lcqtof_hilic_neg_4_008_24    m/z 625.1366     0   1   2E+4   0.5   1  RT (min)      lcqtof_hilic_neg_4_021_21    m/z 625.1476     0   1   2E+4   0.5   1  RT (min)      lcqtof_hilic_neg_1_032_18    m/z 625.1616     0   1E+4   1.5   2  RT (min)      lcqtof_hilic_neg_1_042_26    m/z 625.4254     0   1E+4   1   1.5  RT (min)      lcqtof_hilic_neg_4_000_aa    m/z 625.4337     0   1   2   3   4E+3   15.5   16  RT (min)      lcqtof_hilic_neg_4_012_07    m/z 625.4343     0   1   2   3E+3   15   15.5  RT (min)      lcqtof_hilic_neg_4_006_30    m/z 625.4668     0   1   2   3   4E+4   1   1.5  RT (min)      lcqtof_hilic_neg_4_000_aa    m/z 625.4701     0   1   2   3   4E+4   1   1.5  RT (min)      lcqtof_hilic_neg_4_000_aa    m/z 625.5126     0   1   2E+4   1   1.5  RT (min)      lcqtof_hilic_neg_4_000_aa    m/z 626.0953     0   1E+4   1   1.5  RT (min)      lcqtof_hilic_neg_4_027_19    m/z 626.2374     0   1   2   3E+4   3   3.5  RT (min)      lcqtof_hilic_neg_4_004_47    m/z 626.3625     0   1E+4   4   4.5  RT (min)      lcqtof_hilic_neg_4_055_32    m/z 626.4680     0   1E+4   1   1.5  RT (min)      lcqtof_hilic_neg_4_000_aa    m/z 626.5468     0   1   2   3E+4   1.5   2  RT (min)      lcqtof_hilic_neg_4_000_aa    m/z 626.6621     0   1E+4   0.5   1  RT (min)      lcqtof_hilic_neg_4_048_23    m/z 626.7900     0   2   4   6E+4   0.5   1  RT (min)      lcqtof_hilic_neg_4_048_23    m/z 626.9856     0   1E+4   0.5   1  RT (min)      lcqtof_hilic_neg_4_048_23    m/z 627.1456     0   1   2E+4   0.5   1  RT (min)      lcqtof_hilic_neg_1_032_18    m/z 627.3208     0   1E+4   15   15.5  RT (min)      lcqtof_hilic_neg_4_000_aa    m/z 627.3703     0   1E+4   1   1.5  RT (min)      lcqtof_hilic_neg_4_026_04    m/z 627.3937     0   1   2E+4   1   1.5  RT (min)      lcqtof_hilic_neg_4_038_10    m/z 627.4120     0   2   4   6E+4   1   1.5  RT (min)      lcqtof_hilic_neg_4_000_aa    m/z 627.4122     0   1   2   3   4E+3   15.5   16  RT (min)      lcqtof_hilic_neg_4_019_03    m/z 627.4450     0   1E+4   1   1.5  RT (min)      lcqtof_hilic_neg_4_000_aa    m/z 627.4974     0   2   4   6E+4   1   1.5  RT (min)      lcqtof_hilic_neg_4_026_04    m/z 627.5507     0   1   2E+4   1.5   2  RT (min)      lcqtof_hilic_neg_4_000_aa    m/z 627.5582     0   1   2E+4   1.5   2  RT (min)      lcqtof_hilic_neg_4_000_aa    m/z 628.0558     0   2   4   6E+4   7.5   8  RT (min)       lcqtof_hilic_neg_4_082_59    m/z 628.3488     0   1E+4   0.5   1  RT (min)      lcqtof_hilic_neg_4_010_28    m/z 628.4124     0   1   2E+4   1   1.5  RT (min)      lcqtof_hilic_neg_4_000_aa    m/z 628.6506     0   1E+4   0.5   1  RT (min)      lcqtof_hilic_neg_4_008_24    m/z 629.1922     0   1   2E+4   1   1.5  RT (min)      lcqtof_hilic_neg_4_000_aa    m/z 629.2033     0   1   2E+4   1   1.5  RT (min)      lcqtof_hilic_neg_4_000_aa    m/z 629.3020     0   1E+5   1   1.5  RT (min)      lcqtof_hilic_neg_4_000_aa    m/z 629.3026     0   1E+5   1   1.5  RT (min)      lcqtof_hilic_neg_4_000_aa    m/z 629.3036     0   1E+5   1   1.5  RT (min)      lcqtof_hilic_neg_4_000_aa    m/z 629.3043     0   1E+5   1   1.5  RT (min)      lcqtof_hilic_neg_4_000_aa    m/z 629.3389     0   1E+4   0.5   1  RT (min)      lcqtof_hilic_neg_4_038_10    m/z 629.4778     0   1E+4   0.5   1  RT (min)      lcqtof_hilic_neg_1_063_13    m/z 629.4798     0   1E+4   1   1.5  RT (min)      lcqtof_hilic_neg_4_000_aa    m/z 630.2046     0   1E+4   1   1.5  RT (min)      lcqtof_hilic_neg_4_000_aa    m/z 630.2988     0   2   4E+4   1   1.5  RT (min)      lcqtof_hilic_neg_4_000_aa    m/z 630.3024     0   2   4E+4   1   1.5  RT (min)      lcqtof_hilic_neg_4_000_aa    m/z 630.3025     0   2   4E+4   1   1.5  RT (min)      lcqtof_hilic_neg_4_000_aa    m/z 630.3036     0   2   4E+4   1   1.5  RT (min)      lcqtof_hilic_neg_4_000_aa    m/z 630.3044     0   2   4E+4   1   1.5  RT (min)      lcqtof_hilic_neg_4_000_aa    m/z 630.6209     0   1   2E+4   0.5   1  RT (min)      lcqtof_hilic_neg_3_035_46    m/z 631.1621     0   1E+4   0.5   1  RT (min)      lcqtof_hilic_neg_4_026_04    m/z 631.2112     0   2   4   6   8E+4   8.5   9  RT (min)      lcqtof_hilic_neg_4_097_60    m/z 631.2613     0   1   2E+4   0.5   1  RT (min)      lcqtof_hilic_neg_3_003_20    m/z 631.3086     0   1   2   3   4E+4   15   15.5  RT (min)      lcqtof_hilic_neg_4_000_aa    m/z 631.3408     0   1E+5   1   1.5  RT (min)      lcqtof_hilic_neg_4_000_aa    m/z 631.3478     0   1E+5   1   1.5  RT (min)      lcqtof_hilic_neg_4_000_aa    m/z 631.4651     0   1   2   3   4E+4   8   8.5  RT (min)      lcqtof_hilic_neg_2_052_24    m/z 631.5408     0   1E+4   1.5   2  RT (min)      lcqtof_hilic_neg_4_000_aa    m/z 632.2154     0   1E+4   6.5   7  RT (min)      lcqtof_hilic_neg_4_008_24    m/z 632.3114     0   1E+4   15   15.5  RT (min)      lcqtof_hilic_neg_4_000_aa    m/z 632.3462     0   2   4   6E+4   1   1.5  RT (min)      lcqtof_hilic_neg_4_000_aa    m/z 632.3488     0   2   4   6E+4   1   1.5  RT (min)      lcqtof_hilic_neg_4_000_aa    m/z 632.3496     0   2   4   6E+4   1   1.5  RT (min)      lcqtof_hilic_neg_4_000_aa    m/z 632.3501     0   2   4   6E+4   1   1.5  RT (min)      lcqtof_hilic_neg_4_000_aa    m/z 632.3898     0   1E+4   3.5   4  RT (min)      lcqtof_hilic_neg_4_031_38    m/z 632.3905     0   1E+4   3   3.5  RT (min)       lcqtof_hilic_neg_4_048_23    m/z 632.3908     0   1E+4   3   3.5  RT (min)      lcqtof_hilic_neg_4_031_38    m/z 632.4204     0   1E+4   4.5   5  RT (min)      lcqtof_hilic_neg_4_050_02    m/z 632.5428     0   1E+4   0.5   1  RT (min)      lcqtof_hilic_neg_3_035_46    m/z 632.6386     0   1   2E+4   0.5   1  RT (min)      lcqtof_hilic_neg_3_035_46    m/z 633.1259     0   2   4   6E+4   7.5   8  RT (min)       lcqtof_hilic_neg_4_083_59    m/z 633.3480     0   2   4   6E+4   1   1.5  RT (min)      lcqtof_hilic_neg_4_000_aa    m/z 633.3495     0   2   4   6E+4   1   1.5  RT (min)      lcqtof_hilic_neg_4_000_aa    m/z 633.6034     0   2   4   6E+4   1   1.5  RT (min)      lcqtof_hilic_neg_4_000_aa    m/z 633.6179     0   2   4   6E+4   1   1.5  RT (min)      lcqtof_hilic_neg_4_000_aa    m/z 633.6498     0   1E+4   1.5   2  RT (min)      lcqtof_hilic_neg_4_000_aa    m/z 633.6510     0   1E+4   2   2.5  RT (min)      lcqtof_hilic_neg_4_000_aa    m/z 633.6530     0   1E+4   2   2.5  RT (min)      lcqtof_hilic_neg_4_000_aa    m/z 634.1279     0   1E+4   7.5   8  RT (min)      lcqtof_hilic_neg_4_083_59    m/z 634.2442     0   1E+4   4.5   5  RT (min)      lcqtof_hilic_neg_4_008_24    m/z 634.2449     0   1E+4   4.5   5  RT (min)      lcqtof_hilic_neg_4_008_24    m/z 634.2630     0   1E+4   1   1.5  RT (min)      lcqtof_hilic_neg_4_000_aa    m/z 634.3504     0   1   2E+4   1   1.5  RT (min)      lcqtof_hilic_neg_4_000_aa    m/z 634.3538     0   1   2E+4   1   1.5  RT (min)      lcqtof_hilic_neg_4_000_aa    m/z 634.3603     0   1   2E+4   1   1.5  RT (min)      lcqtof_hilic_neg_4_000_aa    m/z 634.4037     0   2   4   6E+3   3.5   4  RT (min)       lcqtof_hilic_neg_3_068_57    m/z 634.4659     0   1   2E+4   0.5   1  RT (min)      lcqtof_hilic_neg_2_028_05    m/z 634.6230     0   1   2   3E+4   1   1.5  RT (min)      lcqtof_hilic_neg_4_000_aa    m/z 634.8843     0   1E+4   0.5   1  RT (min)      lcqtof_hilic_neg_3_035_46    m/z 635.1424     0   1   2   3E+4   6   6.5  RT (min)       lcqtof_hilic_neg_4_082_59    m/z 635.2257     0   1   2   3   4E+3   6.5   7  RT (min)       lcqtof_hilic_neg_4_003_16    m/z 635.2276     0   1   2   3E+3   7   7.5  RT (min)       lcqtof_hilic_neg_4_005_29    m/z 635.2280     0   2   4E+3   4   4.5  RT (min)       lcqtof_hilic_neg_4_012_07    m/z 635.2281     0   1   2   3E+3   8   8.5  RT (min)       lcqtof_hilic_neg_4_010_28    m/z 635.2283     0   1   2E+3   9   9.5  RT (min)       lcqtof_hilic_neg_4_005_29    m/z 635.2284     0   1   2   3E+3   7.5   8  RT (min)       lcqtof_hilic_neg_4_006_30    m/z 635.2440     0   1E+4   1   1.5  RT (min)      lcqtof_hilic_neg_4_000_aa    m/z 635.2522     0   1E+4   7   7.5  RT (min)      lcqtof_hilic_neg_4_014_46    m/z 635.2550     0   1E+4   1   1.5  RT (min)      lcqtof_hilic_neg_4_000_aa    m/z 635.3474     0   1E+4   1   1.5  RT (min)      lcqtof_hilic_neg_4_000_aa    m/z 635.5442     0   2   4   6   8E+4   1   1.5  RT (min)      lcqtof_hilic_neg_4_000_aa    m/z 635.5600     0   2   4   6   8E+4   1   1.5  RT (min)      lcqtof_hilic_neg_4_000_aa    m/z 635.5956     0   1   2   3   4E+4   1   1.5  RT (min)      lcqtof_hilic_neg_4_000_aa    m/z 635.5974     0   1   2   3   4E+4   1.5   2  RT (min)      lcqtof_hilic_neg_4_000_aa    m/z 635.5996     0   1   2   3   4E+4   1.5   2  RT (min)      lcqtof_hilic_neg_4_000_aa    m/z 635.6097     0   1   2   3   4E+4   1.5   2  RT (min)      lcqtof_hilic_neg_4_000_aa    m/z 636.2834     0   1   2E+4   3   3.5  RT (min)      lcqtof_hilic_neg_4_039_18    m/z 636.3473     0   1E+4   4   4.5  RT (min)       lcqtof_hilic_neg_4_048_23    m/z 636.4144     0   1   2E+4   4   4.5  RT (min)      lcqtof_hilic_neg_4_008_24    m/z 636.4202     0   1E+4   14.5   15   15.5  RT (min)       lcqtof_hilic_neg_4_082_59    m/z 636.4227     0   1E+4   14.5   15  RT (min)      lcqtof_hilic_neg_4_013_20    m/z 636.5608     0   1   2   3E+4   1   1.5  RT (min)      lcqtof_hilic_neg_4_000_aa    m/z 636.5620     0   1   2   3E+4   1   1.5  RT (min)      lcqtof_hilic_neg_4_000_aa    m/z 636.5991     0   1   2E+4   1   1.5  RT (min)      lcqtof_hilic_neg_4_000_aa    m/z 636.6510     0   1   2E+4   0.5   1  RT (min)      lcqtof_hilic_neg_3_035_46    m/z 636.9688     0   1E+4   0.5   1  RT (min)      lcqtof_hilic_neg_3_035_46    m/z 637.1413     0   1E+4   1   1.5  RT (min)      lcqtof_hilic_neg_4_004_47    m/z 637.3229     0   1E+4   15   15.5   16  RT (min)      lcqtof_hilic_neg_4_000_aa    m/z 637.3584     0   1E+4   15   15.5  RT (min)      lcqtof_hilic_neg_4_000_aa    m/z 637.4128     0   2   4   6E+4   1   1.5  RT (min)      lcqtof_hilic_neg_4_000_aa    m/z 637.4172     0   2   4   6E+4   1   1.5  RT (min)      lcqtof_hilic_neg_4_000_aa    m/z 637.4173     0   2   4   6E+4   1   1.5  RT (min)      lcqtof_hilic_neg_4_000_aa    m/z 637.4548     0   1E+4   4   4.5  RT (min)      lcqtof_hilic_neg_4_008_24    m/z 637.5224     0   1   2   3E+4   1   1.5  RT (min)      lcqtof_hilic_neg_4_000_aa    m/z 637.5490     0   2   4   6   8E+4   1   1.5  RT (min)      lcqtof_hilic_neg_4_000_aa    m/z 637.5555     0   2   4   6   8E+4   1   1.5  RT (min)      lcqtof_hilic_neg_4_000_aa    m/z 637.5650     0   2   4   6   8E+4   1   1.5  RT (min)      lcqtof_hilic_neg_4_000_aa    m/z 637.5861     0   1E+4   16   16.5  RT (min)      lcqtof_hilic_neg_4_000_aa    m/z 637.5880     0   1E+5   1.5   2  RT (min)      lcqtof_hilic_neg_4_000_aa    m/z 638.2374     0   1E+4   0.5   1  RT (min)      lcqtof_hilic_neg_3_035_46    m/z 638.3252     0   2   4E+4   1   1.5  RT (min)      lcqtof_hilic_neg_4_000_aa    m/z 638.3285     0   1E+4   4   4.5  RT (min)      lcqtof_hilic_neg_4_040_05    m/z 638.3400     0   2   4E+4   0.5   1  RT (min)      lcqtof_hilic_neg_3_003_20    m/z 638.3621     0   1   2E+4   0.5   1  RT (min)      lcqtof_hilic_neg_4_047_06    m/z 638.4179     0   1   2   3E+4   1   1.5  RT (min)      lcqtof_hilic_neg_4_000_aa    m/z 638.4190     0   1   2   3E+4   1   1.5  RT (min)      lcqtof_hilic_neg_4_000_aa    m/z 638.4192     0   1   2   3E+4   1   1.5  RT (min)      lcqtof_hilic_neg_4_000_aa    m/z 638.4198     0   1   2   3E+4   1   1.5  RT (min)      lcqtof_hilic_neg_4_000_aa    m/z 638.4204     0   1   2   3E+4   1   1.5  RT (min)      lcqtof_hilic_neg_4_000_aa    m/z 638.4214     0   1   2   3E+4   1   1.5  RT (min)      lcqtof_hilic_neg_4_000_aa    m/z 638.4231     0   1   2   3E+4   1   1.5  RT (min)      lcqtof_hilic_neg_4_000_aa    m/z 638.4310     0   1E+4   4   4.5  RT (min)      lcqtof_hilic_neg_4_008_24    m/z 638.5445     0   1   2   3E+4   1   1.5  RT (min)      lcqtof_hilic_neg_4_000_aa    m/z 638.5476     0   1   2   3E+4   1   1.5  RT (min)      lcqtof_hilic_neg_4_000_aa    m/z 638.5560     0   1   2   3E+4   1   1.5  RT (min)      lcqtof_hilic_neg_4_000_aa    m/z 638.5714     0   1   2   3E+4   1   1.5  RT (min)      lcqtof_hilic_neg_4_000_aa    m/z 638.5914     0   2   4   6E+4   1.5   2  RT (min)      lcqtof_hilic_neg_4_000_aa    m/z 638.6618     0   1E+4   0.5   1  RT (min)      lcqtof_hilic_neg_3_035_46    m/z 638.7208     0   2   4   6E+4   9   9.5  RT (min)      lcqtof_hilic_neg_1_000_aa    m/z 638.8324     0   1   2   3   4E+3   8.5   9  RT (min)       lcqtof_hilic_neg_4_038_10    m/z 638.8326     0   1   2   3   4E+3   8.5   9  RT (min)       lcqtof_hilic_neg_4_019_03    m/z 638.8332     0   1   2   3   4E+3   8.5   9  RT (min)      lcqtof_hilic_neg_4_019_03    m/z 639.2506     0   1   2E+4   1   1.5  RT (min)      lcqtof_hilic_neg_4_047_06    m/z 639.3218     0   1   2E+4   1   1.5  RT (min)      lcqtof_hilic_neg_4_000_aa    m/z 639.4240     0   1   2   3E+4   1   1.5  RT (min)      lcqtof_hilic_neg_4_000_aa    m/z 639.4248     0   1   2   3E+4   1   1.5  RT (min)      lcqtof_hilic_neg_4_000_aa    m/z 639.4790     0   2   4   6   8E+4   1   1.5  RT (min)      lcqtof_hilic_neg_4_000_aa    m/z 639.4792     0   2   4   6   8E+4   1   1.5  RT (min)      lcqtof_hilic_neg_4_000_aa    m/z 639.5368     0   1   2   3E+4   1   1.5  RT (min)      lcqtof_hilic_neg_4_000_aa    m/z 639.5432     0   1   2   3E+4   1   1.5  RT (min)      lcqtof_hilic_neg_4_000_aa    m/z 639.5930     0   1E+4   1.5   2  RT (min)      lcqtof_hilic_neg_4_000_aa    m/z 640.2562     0   2   4   6E+3   4   4.5  RT (min)      lcqtof_hilic_neg_4_014_46    m/z 640.3454     0   1E+4   4   4.5  RT (min)      lcqtof_hilic_neg_4_008_24    m/z 640.3456     0   1   2E+4   4   4.5  RT (min)       lcqtof_hilic_neg_4_059_14    m/z 640.3476     0   1   2E+4   4   4.5  RT (min)      lcqtof_hilic_neg_4_059_14    m/z 640.3923     0   1E+4   3   3.5  RT (min)      lcqtof_hilic_neg_4_016_37    m/z 640.4515     0   1E+4   4   4.5  RT (min)       lcqtof_hilic_neg_4_097_60    m/z 640.4861     0   1   2   3E+4   1   1.5  RT (min)      lcqtof_hilic_neg_4_000_aa    m/z 640.6251     0   1   2E+4   1.5   2  RT (min)      lcqtof_hilic_neg_4_005_29    m/z 640.6264     0   1   2E+4   1   1.5  RT (min)      lcqtof_hilic_neg_4_000_aa    m/z 641.1554     0   1E+4   4   4.5  RT (min)      lcqtof_hilic_neg_4_021_21    m/z 641.2624     0   1   2   3E+3   5.5   6  RT (min)      lcqtof_hilic_neg_4_018_27    m/z 641.2626     0   2   4E+3   5.5   6  RT (min)       lcqtof_hilic_neg_4_008_24    m/z 641.2717     0   1E+4   3.5   4  RT (min)       lcqtof_hilic_neg_1_063_13    m/z 641.2832     0   1E+4   7   7.5  RT (min)      lcqtof_hilic_neg_4_048_23    m/z 641.3731     0   2   4   6E+5   1   1.5  RT (min)      lcqtof_hilic_neg_4_000_aa    m/z 641.3750     0   2   4   6E+5   1   1.5  RT (min)      lcqtof_hilic_neg_4_000_aa    m/z 641.3760     0   2   4   6E+5   1   1.5  RT (min)      lcqtof_hilic_neg_4_000_aa    m/z 641.4104     0   1E+4   1   1.5  RT (min)      lcqtof_hilic_neg_4_061_50    m/z 641.4278     0   1   2   3   4E+3   15.5   16  RT (min)      lcqtof_hilic_neg_4_001_49    m/z 641.4924     0   1E+4   1   1.5  RT (min)      lcqtof_hilic_neg_4_000_aa    m/z 641.5540     0   1E+4   0.5   1  RT (min)      lcqtof_hilic_neg_4_040_05    m/z 641.6275     0   1E+4   1   1.5  RT (min)      lcqtof_hilic_neg_4_000_aa    m/z 641.6292     0   1E+4   1.5   2  RT (min)      lcqtof_hilic_neg_4_005_29    m/z 642.1250     0   1E+4   0.5   1  RT (min)      lcqtof_hilic_neg_1_062_17    m/z 642.2635     0   2   4   6E+3   5   5.5  RT (min)      lcqtof_hilic_neg_4_050_02    m/z 642.2636     0   2   4   6E+3   5   5.5  RT (min)      lcqtof_hilic_neg_4_058_39    m/z 642.3766     0   1   2   3E+5   1   1.5  RT (min)      lcqtof_hilic_neg_4_000_aa    m/z 642.3772     0   1   2   3E+5   1   1.5  RT (min)      lcqtof_hilic_neg_4_000_aa    m/z 642.3778     0   1   2   3E+5   1   1.5  RT (min)      lcqtof_hilic_neg_4_000_aa    m/z 642.4959     0   1E+4   0.5   1  RT (min)      lcqtof_hilic_neg_4_040_05    m/z 643.2704     0   1E+4   4.5   5  RT (min)      lcqtof_hilic_neg_2_028_05    m/z 643.3194     0   1E+4   15   15.5  RT (min)      lcqtof_hilic_neg_4_000_aa    m/z 643.3574     0   1E+4   0.5   1  RT (min)      lcqtof_hilic_neg_4_038_10    m/z 643.3758     0   1   2E+5   1   1.5  RT (min)      lcqtof_hilic_neg_4_000_aa    m/z 643.3758     0   1   2E+5   1   1.5  RT (min)      lcqtof_hilic_neg_4_000_aa    m/z 643.3759     0   1   2E+5   1   1.5  RT (min)      lcqtof_hilic_neg_4_000_aa    m/z 643.3761     0   1   2E+5   1   1.5  RT (min)      lcqtof_hilic_neg_4_000_aa    m/z 643.3778     0   1   2E+5   1   1.5  RT (min)      lcqtof_hilic_neg_4_000_aa    m/z 643.3960     0   2   4E+4   0.5   1  RT (min)       lcqtof_hilic_neg_4_013_20    m/z 643.4082     0   2   4E+4   1   1.5  RT (min)       lcqtof_hilic_neg_4_013_20    m/z 643.4287     0   1E+4   0.5   1  RT (min)      lcqtof_hilic_neg_2_016_33    m/z 643.4832     0   1   2E+4   1.5   2  RT (min)      lcqtof_hilic_neg_4_000_aa    m/z 643.4870     0   1E+4   0.5   1  RT (min)      lcqtof_hilic_neg_3_035_46    m/z 643.5300     0   1E+4   1.5   2  RT (min)      lcqtof_hilic_neg_4_000_aa    m/z 644.2727     0   2   4   6   8E+4   4.5   5  RT (min)       lcqtof_hilic_neg_2_028_05    m/z 644.3577     0   1E+4   4   4.5  RT (min)      lcqtof_hilic_neg_4_053_14    m/z 644.3599     0   1E+4   4   4.5  RT (min)       lcqtof_hilic_neg_4_059_14    m/z 644.3736     0   2   4   6E+4   1   1.5  RT (min)      lcqtof_hilic_neg_4_000_aa    m/z 644.3772     0   2   4   6E+4   1   1.5  RT (min)      lcqtof_hilic_neg_4_000_aa    m/z 644.3777     0   2   4   6E+4   1   1.5  RT (min)      lcqtof_hilic_neg_4_000_aa    m/z 644.3785     0   2   4   6E+4   1   1.5  RT (min)      lcqtof_hilic_neg_4_000_aa    m/z 644.3829     0   2   4   6E+4   1   1.5  RT (min)      lcqtof_hilic_neg_4_000_aa    m/z 644.3990     0   1E+4   0.5   1  RT (min)      lcqtof_hilic_neg_4_039_18    m/z 644.5034     0   1   2E+4   0.5   1  RT (min)      lcqtof_hilic_neg_1_007_05    m/z 644.6221     0   1E+4   0.5   1  RT (min)      lcqtof_hilic_neg_4_008_24    m/z 645.1163     0   1E+4   1   1.5  RT (min)      lcqtof_hilic_neg_4_004_47    m/z 645.2559     0   1   2E+3   8   8.5  RT (min)       lcqtof_hilic_neg_4_027_19    m/z 645.2568     0   1   2E+3   8.5   9  RT (min)       lcqtof_hilic_neg_4_044_08    m/z 645.2573     0   1   2E+3   8.5   9  RT (min)       lcqtof_hilic_neg_4_006_30    m/z 645.2776     0   1E+4   4.5   5  RT (min)      lcqtof_hilic_neg_3_048_05    m/z 645.3265     0   1E+5   15   15.5  RT (min)      lcqtof_hilic_neg_4_000_aa    m/z 645.3484     0   1E+4   1   1.5  RT (min)      lcqtof_hilic_neg_4_015_09    m/z 645.3687     0   1   2E+4   0.5   1  RT (min)      lcqtof_hilic_neg_4_005_29    m/z 645.4765     0   1E+4   1.5   2  RT (min)      lcqtof_hilic_neg_4_000_aa    m/z 645.5039     0   1E+4   0.5   1  RT (min)      lcqtof_hilic_neg_3_035_46    m/z 646.2892     0   1E+4   4.5   5  RT (min)      lcqtof_hilic_neg_3_048_05    m/z 646.3272     0   1   2   3   4E+4   15   15.5  RT (min)      lcqtof_hilic_neg_4_000_aa    m/z 646.3997     0   1E+4   3   3.5  RT (min)      lcqtof_hilic_neg_4_048_23    m/z 646.4111     0   2   4   6E+3   2   2.5  RT (min)       lcqtof_hilic_neg_4_016_37    m/z 646.6810     0   1   2   3   4E+4   0.5   1  RT (min)      lcqtof_hilic_neg_3_035_46    m/z 647.2284     0   1E+4   4.5   5  RT (min)       lcqtof_hilic_neg_4_027_19    m/z 647.2284     0   1   2   3   4E+3   3.5   4  RT (min)       lcqtof_hilic_neg_4_100_61    m/z 647.2776     0   1E+4   0.5   1  RT (min)      lcqtof_hilic_neg_2_046_41    m/z 647.3252     0   1E+4   15   15.5  RT (min)      lcqtof_hilic_neg_4_000_aa    m/z 647.4066     0   1E+4   3   3.5  RT (min)      lcqtof_hilic_neg_4_002_25    m/z 647.6014     0   1   2E+4   1   1.5  RT (min)      lcqtof_hilic_neg_4_000_aa    m/z 647.6029     0   1   2E+4   1   1.5  RT (min)      lcqtof_hilic_neg_4_000_aa    m/z 647.6355     0   1   2E+4   1.5   2  RT (min)      lcqtof_hilic_neg_4_000_aa    m/z 648.2745     0   1E+4   0.5   1  RT (min)      lcqtof_hilic_neg_4_006_30    m/z 648.4864     0   1E+4   0.5   1  RT (min)      lcqtof_hilic_neg_1_010_15    m/z 648.6080     0   1E+4   1   1.5  RT (min)      lcqtof_hilic_neg_4_000_aa    m/z 648.6972     0   1E+4   0.5   1  RT (min)      lcqtof_hilic_neg_3_035_46    m/z 648.9009     0   2   4E+4   0.5   1  RT (min)      lcqtof_hilic_neg_3_035_46    m/z 648.9022     0   2   4E+4   0.5   1  RT (min)      lcqtof_hilic_neg_3_035_46    m/z 649.3234     0   1E+4   1   1.5  RT (min)      lcqtof_hilic_neg_4_000_aa    m/z 649.5377     0   1   2   3E+4   1   1.5  RT (min)      lcqtof_hilic_neg_4_000_aa    m/z 649.5818     0   2   4E+4   1   1.5  RT (min)      lcqtof_hilic_neg_4_000_aa    m/z 649.6034     0   1   2   3   4E+4   1   1.5  RT (min)      lcqtof_hilic_neg_4_000_aa    m/z 649.6230     0   1   2   3E+4   1.5   2  RT (min)      lcqtof_hilic_neg_4_000_aa    m/z 649.8989     0   1   2E+4   0.5   1  RT (min)      lcqtof_hilic_neg_3_035_46    m/z 650.1912     0   1E+4   0.5   1  RT (min)      lcqtof_hilic_neg_4_052_13    m/z 650.2117     0   1E+4   7   7.5  RT (min)      lcqtof_hilic_neg_4_055_32    m/z 650.2995     0   1E+4   3   3.5  RT (min)      lcqtof_hilic_neg_4_039_18    m/z 650.3716     0   1E+4   2   2.5  RT (min)      lcqtof_hilic_neg_4_003_16    m/z 650.4614     0   1E+4   4   4.5  RT (min)      lcqtof_hilic_neg_4_008_24    m/z 650.5892     0   1   2E+4   1   1.5  RT (min)      lcqtof_hilic_neg_4_000_aa    m/z 650.6042     0   1   2E+4   1   1.5  RT (min)      lcqtof_hilic_neg_4_000_aa    m/z 650.6162     0   1   2E+4   1   1.5  RT (min)      lcqtof_hilic_neg_4_000_aa    m/z 650.6323     0   1   2E+4   0.5   1  RT (min)      lcqtof_hilic_neg_3_035_46    m/z 650.8970     0   1   2E+4   0.5   1  RT (min)      lcqtof_hilic_neg_3_035_46    m/z 651.3388     0   1   2   3E+4   15.5   16  RT (min)      lcqtof_hilic_neg_4_000_aa    m/z 651.3734     0   1E+4   15   15.5  RT (min)      lcqtof_hilic_neg_4_000_aa    m/z 651.4055     0   1E+4   3.5   4  RT (min)      lcqtof_hilic_neg_4_052_13    m/z 651.4388     0   1   2   3E+4   4   4.5  RT (min)      lcqtof_hilic_neg_4_008_24    m/z 651.5556     0   2   4   6E+4   1   1.5  RT (min)      lcqtof_hilic_neg_4_000_aa    m/z 651.5627     0   2   4   6   8E+4   1   1.5  RT (min)      lcqtof_hilic_neg_4_000_aa    m/z 651.5786     0   2   4   6   8E+4   1   1.5  RT (min)      lcqtof_hilic_neg_4_000_aa    m/z 651.6415     0   1E+4   0.5   1  RT (min)      lcqtof_hilic_neg_3_035_46    m/z 652.3416     0   1E+4   15.5   16  RT (min)      lcqtof_hilic_neg_4_000_aa    m/z 652.3538     0   1E+4   4   4.5  RT (min)      lcqtof_hilic_neg_4_048_23    m/z 652.3661     0   1   2E+4   4   4.5  RT (min)       lcqtof_hilic_neg_4_059_14    m/z 652.4738     0   2   4   6E+4   4   4.5  RT (min)       lcqtof_hilic_neg_4_008_24    m/z 652.5522     0   1E+4   0.5   1  RT (min)      lcqtof_hilic_neg_4_040_05    m/z 652.5596     0   1   2   3E+4   1   1.5  RT (min)      lcqtof_hilic_neg_4_000_aa    m/z 652.5837     0   1   2   3E+4   1   1.5  RT (min)      lcqtof_hilic_neg_4_000_aa    m/z 652.6448     0   2   4E+4   0.5   1  RT (min)      lcqtof_hilic_neg_3_035_46    m/z 653.0888     0   1E+4   0.5   1  RT (min)      lcqtof_hilic_neg_2_063_13    m/z 653.1590     0   1   2E+4   7   7.5  RT (min)       lcqtof_hilic_neg_4_090_60    m/z 653.3208     0   1E+4   0.5   1  RT (min)      lcqtof_hilic_neg_4_002_25    m/z 653.3384     0   1E+5   1   1.5  RT (min)      lcqtof_hilic_neg_4_000_aa    m/z 653.3402     0   1E+5   1   1.5  RT (min)      lcqtof_hilic_neg_4_000_aa    m/z 653.3554     0   1E+5   1   1.5  RT (min)      lcqtof_hilic_neg_4_000_aa    m/z 653.3611     0   1   2E+4   1   1.5  RT (min)      lcqtof_hilic_neg_4_000_aa    m/z 653.3968     0   1E+4   0.5   1  RT (min)      lcqtof_hilic_neg_4_053_14    m/z 653.4600     0   1   2   3   4E+4   1   1.5  RT (min)      lcqtof_hilic_neg_4_000_aa    m/z 653.4738     0   1E+4   0.5   1  RT (min)      lcqtof_hilic_neg_1_045_07    m/z 653.4868     0   1   2   3E+3   16   16.5  RT (min)      lcqtof_hilic_neg_4_000_dd    m/z 653.4872     0   1   2E+3   19.5   20  RT (min)      lcqtof_hilic_neg_4_008_24    m/z 653.4876     0   1   2E+3   17   17.5  RT (min)      lcqtof_hilic_neg_4_006_30    m/z 653.4879     0   1   2   3E+3   18   18.5  RT (min)      lcqtof_hilic_neg_4_000_cc    m/z 653.4880     0   1   2E+3   18  RT (min)       lcqtof_hilic_neg_4_017_51    m/z 653.4880     0   1   2E+3   17.5   18  RT (min)      lcqtof_hilic_neg_4_009_50    m/z 653.4889     0   2   4E+3   2   2.5  RT (min)      lcqtof_hilic_neg_4_010_28    m/z 653.4890     0   1   2E+3   19   19.5  RT (min)      lcqtof_hilic_neg_4_006_30    m/z 653.4890     0   1   2E+3   18.5   19  RT (min)      lcqtof_hilic_neg_4_022_34    m/z 653.4890     0   1   2E+3   18   18.5  RT (min)       lcqtof_hilic_neg_4_005_29    m/z 653.4890     0   1   2   3   4E+3   1   1.5  RT (min)      lcqtof_hilic_neg_4_024_40    m/z 653.4891     0   1   2E+3   0   0.5  RT (min)      lcqtof_hilic_neg_4_000_bb    m/z 653.4892     0   1   2   3E+3   19   19.5  RT (min)      lcqtof_hilic_neg_4_011_35    m/z 653.4893     0   1   2E+3   0   0.5  RT (min)      lcqtof_hilic_neg_4_004_47    m/z 653.4894     0   1   2E+3   0   0.5  RT (min)      lcqtof_hilic_neg_3_000_aa    m/z 653.4904     0   1E+5   1   1.5  RT (min)      lcqtof_hilic_neg_4_000_aa    m/z 653.4973     0   1E+4   14.5   15  RT (min)      lcqtof_hilic_neg_4_013_20    m/z 653.4987     0   1E+4   15   15.5  RT (min)      lcqtof_hilic_neg_4_038_10    m/z 653.5483     0   1   2   3E+4   1   1.5  RT (min)      lcqtof_hilic_neg_4_000_aa    m/z 653.5633     0   1   2   3E+4   1   1.5  RT (min)      lcqtof_hilic_neg_4_000_aa    m/z 653.5834     0   1   2E+4   1   1.5  RT (min)      lcqtof_hilic_neg_4_000_aa    m/z 653.5842     0   1   2E+4   1   1.5  RT (min)      lcqtof_hilic_neg_4_000_aa    m/z 654.0318     0   1E+4   6.5   7  RT (min)      lcqtof_hilic_neg_4_011_35    m/z 654.0328     0   1E+4   6.5   7  RT (min)      lcqtof_hilic_neg_4_034_31    m/z 654.1737     0   1E+4   7   7.5  RT (min)      lcqtof_hilic_neg_4_035_33    m/z 654.1936     0   1E+4   6   6.5  RT (min)      lcqtof_hilic_neg_1_037_24    m/z 654.1966     0   1E+4   6.5   7  RT (min)      lcqtof_hilic_neg_4_008_24    m/z 654.3395     0   1   2E+4   1   1.5  RT (min)      lcqtof_hilic_neg_4_000_aa    m/z 654.3615     0   1   2E+4   0.5   1  RT (min)      lcqtof_hilic_neg_4_050_02    m/z 654.4612     0   1   2E+4   1   1.5  RT (min)      lcqtof_hilic_neg_4_000_aa    m/z 654.5003     0   2   4   6E+4   1   1.5  RT (min)      lcqtof_hilic_neg_4_000_aa    m/z 654.5884     0   1E+4   1.5   2  RT (min)      lcqtof_hilic_neg_4_000_aa    m/z 654.6452     0   1E+4   0.5   1  RT (min)      lcqtof_hilic_neg_3_035_46    m/z 654.7514     0   1E+5   8   8.5  RT (min)      lcqtof_hilic_neg_1_000_aa    m/z 654.7515     0   1E+5   8   8.5  RT (min)      lcqtof_hilic_neg_1_000_aa    m/z 654.9782     0   1   2   3E+3   5.5   6  RT (min)       lcqtof_hilic_neg_1_010_15    m/z 654.9785     0   1   2E+3   5.5   6  RT (min)      lcqtof_hilic_neg_1_060_14    m/z 654.9787     0   1   2   3E+3   5.5   6  RT (min)      lcqtof_hilic_neg_1_010_15    m/z 655.1073     0   1   2   3E+4   7.5   8  RT (min)       lcqtof_hilic_neg_4_083_59    m/z 655.1501     0   1E+4   1   1.5  RT (min)      lcqtof_hilic_neg_4_000_aa    m/z 655.3282     0   1E+4   15   15.5  RT (min)      lcqtof_hilic_neg_4_000_aa    m/z 655.4042     0   2   4E+4   1   1.5  RT (min)      lcqtof_hilic_neg_4_000_aa    m/z 655.4054     0   2   4E+4   1   1.5  RT (min)      lcqtof_hilic_neg_4_000_aa    m/z 655.4408     0   2   4   6   8E+4   1   1.5  RT (min)      lcqtof_hilic_neg_4_000_aa    m/z 655.4460     0   1   2   3   4E+3   15.5   16  RT (min)      lcqtof_hilic_neg_4_000_aa    m/z 655.4954     0   1E+4   1.5   2  RT (min)      lcqtof_hilic_neg_4_000_aa    m/z 655.5056     0   1E+4   1   1.5  RT (min)      lcqtof_hilic_neg_4_000_aa    m/z 656.0106     0   1E+4   0.5   1  RT (min)      lcqtof_hilic_neg_3_035_46    m/z 656.1693     0   1E+4   1   1.5  RT (min)      lcqtof_hilic_neg_4_000_aa    m/z 656.4484     0   1   2   3E+4   1   1.5  RT (min)      lcqtof_hilic_neg_4_000_aa    m/z 656.7332     0   2   4   6E+4   8   8.5  RT (min)      lcqtof_hilic_neg_1_000_aa    m/z 656.7350     0   2   4   6E+4   8   8.5  RT (min)      lcqtof_hilic_neg_1_000_aa    m/z 657.1226     0   1   2   3E+4   6   6.5  RT (min)       lcqtof_hilic_neg_4_083_59    m/z 657.2566     0   1   2   3   4E+3   4.5   5  RT (min)       lcqtof_hilic_neg_4_003_16    m/z 657.2582     0   1   2E+3   0   0.5  RT (min)       lcqtof_hilic_neg_4_027_19    m/z 657.2583     0   2   4   6E+3   3.5   4  RT (min)       lcqtof_hilic_neg_4_027_19    m/z 657.2588     0   1   2E+3   0   0.5  RT (min)       lcqtof_hilic_neg_4_007_45    m/z 657.2719     0   1E+4   0.5   1  RT (min)      lcqtof_hilic_neg_4_050_02    m/z 657.5122     0   1   2E+4   1.5   2  RT (min)      lcqtof_hilic_neg_4_000_aa    m/z 657.5450     0   1E+4   1.5   2  RT (min)      lcqtof_hilic_neg_4_000_aa    m/z 658.1142     0   2   4   6   8E+4   0.5   1  RT (min)      lcqtof_hilic_neg_4_021_21    m/z 658.1251     0   1E+5   0.5   1  RT (min)      lcqtof_hilic_neg_1_024_21    m/z 658.2049     0   1E+4   0.5   1  RT (min)      lcqtof_hilic_neg_4_039_18    m/z 658.3653     0   1   2   3E+4   1   1.5  RT (min)      lcqtof_hilic_neg_4_000_aa    m/z 658.3690     0   1   2   3E+4   1   1.5  RT (min)      lcqtof_hilic_neg_4_000_aa    m/z 658.3734     0   1   2   3E+4   1   1.5  RT (min)      lcqtof_hilic_neg_4_000_aa    m/z 658.4049     0   1   2E+4   4   4.5  RT (min)       lcqtof_hilic_neg_4_008_24    m/z 658.6320     0   1   2E+4   0.5   1  RT (min)      lcqtof_hilic_neg_4_008_24    m/z 659.2936     0   1E+4   1   1.5  RT (min)      lcqtof_hilic_neg_4_000_aa    m/z 659.2942     0   1E+4   1   1.5  RT (min)      lcqtof_hilic_neg_4_000_aa    m/z 659.2960     0   1E+4   1   1.5  RT (min)      lcqtof_hilic_neg_4_000_aa    m/z 659.2966     0   1E+4   1   1.5  RT (min)      lcqtof_hilic_neg_4_000_aa    m/z 659.3406     0   1E+5   15   15.5  RT (min)      lcqtof_hilic_neg_4_000_aa    m/z 659.3422     0   1E+4   4.5   5  RT (min)      lcqtof_hilic_neg_4_040_05    m/z 659.3628     0   1   2E+4   1   1.5  RT (min)      lcqtof_hilic_neg_4_000_aa    m/z 659.3722     0   1   2E+4   1   1.5  RT (min)      lcqtof_hilic_neg_4_000_aa    m/z 659.3762     0   1   2E+4   1   1.5  RT (min)      lcqtof_hilic_neg_4_000_aa    m/z 660.0042     0   1E+4   7.5   8  RT (min)      lcqtof_hilic_neg_4_016_37    m/z 660.1914     0   1E+4   0.5   1  RT (min)      lcqtof_hilic_neg_4_021_21    m/z 660.2032     0   1E+4   0.5   1  RT (min)      lcqtof_hilic_neg_4_039_18    m/z 660.3432     0   2   4   6E+4   15   15.5  RT (min)      lcqtof_hilic_neg_4_000_aa    m/z 660.4008     0   1E+4   0.5   1  RT (min)      lcqtof_hilic_neg_4_005_29    m/z 660.4203     0   1E+4   3.5   4  RT (min)      lcqtof_hilic_neg_4_008_24    m/z 660.5090     0   1E+4   0.5   1  RT (min)      lcqtof_hilic_neg_4_040_05    m/z 660.5306     0   1E+4   0.5   1  RT (min)      lcqtof_hilic_neg_2_028_05    m/z 660.6606     0   1   2   3   4E+4   0.5   1  RT (min)      lcqtof_hilic_neg_3_035_46    m/z 661.2918     0   1E+4   0.5   1  RT (min)      lcqtof_hilic_neg_2_046_41    m/z 661.3357     0   1E+4   0.5   1  RT (min)      lcqtof_hilic_neg_4_000_aa    m/z 661.3390     0   1   2E+4   15   15.5  RT (min)      lcqtof_hilic_neg_4_000_aa    m/z 661.4039     0   1E+4   3.5   4  RT (min)      lcqtof_hilic_neg_4_047_06    m/z 661.4179     0   1E+4   0.5   1  RT (min)      lcqtof_hilic_neg_4_026_04    m/z 661.5805     0   1E+4   1   1.5  RT (min)      lcqtof_hilic_neg_4_000_aa    m/z 661.6110     0   1   2E+4   1   1.5  RT (min)      lcqtof_hilic_neg_4_000_aa    m/z 661.6142     0   1   2E+4   1   1.5  RT (min)      lcqtof_hilic_neg_4_000_aa    m/z 661.6848     0   1E+4   0.5   1  RT (min)      lcqtof_hilic_neg_3_035_46    m/z 662.3478     0   1E+4   4   4.5  RT (min)      lcqtof_hilic_neg_1_037_24    m/z 662.3678     0   1E+4   4   4.5  RT (min)       lcqtof_hilic_neg_4_048_23    m/z 662.4873     0   1E+4   4   4.5  RT (min)      lcqtof_hilic_neg_4_055_32    m/z 662.5121     0   1   2E+4   0.5   1  RT (min)      lcqtof_hilic_neg_2_038_04    m/z 662.6182     0   1E+4   1   1.5  RT (min)      lcqtof_hilic_neg_4_000_aa    m/z 662.6757     0   2   4   6   8E+4   0.5   1  RT (min)      lcqtof_hilic_neg_3_035_46    m/z 663.2204     0   1E+4   4.5   5  RT (min)      lcqtof_hilic_neg_4_013_20    m/z 663.3599     0   1E+4   4.5   5  RT (min)      lcqtof_hilic_neg_4_035_33    m/z 663.5957     0   2   4   6E+4   1   1.5  RT (min)      lcqtof_hilic_neg_4_000_aa    m/z 663.6144     0   2   4   6E+4   1.5   2  RT (min)      lcqtof_hilic_neg_4_000_aa    m/z 663.6197     0   2   4   6E+4   1   1.5  RT (min)      lcqtof_hilic_neg_4_000_aa    m/z 663.6280     0   2   4   6E+4   1   1.5  RT (min)      lcqtof_hilic_neg_4_000_aa    m/z 664.1560     0   1E+4   6   6.5  RT (min)      lcqtof_hilic_neg_4_048_23    m/z 664.1887     0   1E+4   6   6.5  RT (min)       lcqtof_hilic_neg_4_096_60    m/z 664.3331     0   1E+4   4   4.5  RT (min)      lcqtof_hilic_neg_4_050_02    m/z 664.3362     0   1E+4   4   4.5  RT (min)      lcqtof_hilic_neg_4_050_02    m/z 664.3585     0   1   2E+4   4   4.5  RT (min)      lcqtof_hilic_neg_4_048_23    m/z 664.4992     0   1E+4   4   4.5  RT (min)      lcqtof_hilic_neg_4_055_32    m/z 664.5977     0   1   2   3E+4   1   1.5  RT (min)      lcqtof_hilic_neg_4_000_aa    m/z 664.6314     0   1   2   3E+4   1   1.5  RT (min)      lcqtof_hilic_neg_4_000_aa    m/z 665.1231     0   1   2E+4   6.5   7  RT (min)      lcqtof_hilic_neg_4_008_24    m/z 665.2395     0   1E+4   0.5   1  RT (min)      lcqtof_hilic_neg_1_032_18    m/z 665.3552     0   2   4E+3   3   3.5  RT (min)       lcqtof_hilic_neg_4_042_22    m/z 665.3553     0   2   4E+3   3   3.5  RT (min)       lcqtof_hilic_neg_4_042_22    m/z 665.3668     0   1E+5   15.5   16  RT (min)      lcqtof_hilic_neg_4_000_aa    m/z 665.3730     0   1E+4   0.5   1  RT (min)      lcqtof_hilic_neg_3_003_20    m/z 665.3748     0   1E+4   4   4.5  RT (min)       lcqtof_hilic_neg_4_075_58    m/z 665.3908     0   1E+5   0.5   1  RT (min)      lcqtof_hilic_neg_4_047_06    m/z 665.4251     0   1   2E+4   4   4.5  RT (min)      lcqtof_hilic_neg_4_008_24    m/z 665.4859     0   1   2   3   4E+4   0.5   1  RT (min)      lcqtof_hilic_neg_2_028_05    m/z 665.5786     0   2   4   6E+4   1   1.5  RT (min)      lcqtof_hilic_neg_4_000_aa    m/z 665.5915     0   2   4   6E+4   1   1.5  RT (min)      lcqtof_hilic_neg_4_000_aa    m/z 665.6098     0   2   4E+4   1   1.5  RT (min)      lcqtof_hilic_neg_4_000_aa    m/z 665.6142     0   1   2   3   4E+4   1   1.5  RT (min)      lcqtof_hilic_neg_4_000_aa    m/z 666.1417     0   1E+4   6   6.5  RT (min)      lcqtof_hilic_neg_4_055_32    m/z 666.3560     0   1   2   3   4E+4   15.5   16  RT (min)      lcqtof_hilic_neg_4_000_aa    m/z 666.3575     0   1   2E+4   4   4.5  RT (min)       lcqtof_hilic_neg_4_048_23    m/z 666.3949     0   1   2   3   4E+4   0.5   1  RT (min)      lcqtof_hilic_neg_4_047_06    m/z 666.4495     0   1   2E+4   0.5   1  RT (min)      lcqtof_hilic_neg_4_032_01    m/z 666.4514     0   1   2   3E+4   4   4.5  RT (min)       lcqtof_hilic_neg_4_008_24    m/z 666.4812     0   2   4   6   8E+4   0.5   1  RT (min)      lcqtof_hilic_neg_2_028_05    m/z 666.5765     0   1   2E+4   1   1.5  RT (min)      lcqtof_hilic_neg_4_000_aa    m/z 666.5855     0   1   2E+4   1   1.5  RT (min)      lcqtof_hilic_neg_4_000_aa    m/z 666.6020     0   1   2E+4   1.5   2  RT (min)      lcqtof_hilic_neg_4_000_aa    m/z 666.6092     0   1   2E+4   1   1.5  RT (min)      lcqtof_hilic_neg_4_000_aa    m/z 666.6150     0   1E+4   1   1.5  RT (min)      lcqtof_hilic_neg_4_000_aa    m/z 666.9641     0   1E+4   0.5   1  RT (min)      lcqtof_hilic_neg_3_035_46    m/z 667.2718     0   1   2   3E+4   0.5   1   1.5  RT (min)       lcqtof_hilic_neg_4_039_18    m/z 667.2736     0   1E+4   6   6.5  RT (min)      lcqtof_hilic_neg_1_037_24    m/z 667.3548     0   1   2E+4   15.5   16  RT (min)      lcqtof_hilic_neg_4_000_aa    m/z 667.4254     0   1E+4   4   4.5  RT (min)      lcqtof_hilic_neg_4_008_24    m/z 667.4298     0   1E+4   4   4.5  RT (min)      lcqtof_hilic_neg_4_040_05    m/z 667.4699     0   1E+4   0.5   1  RT (min)      lcqtof_hilic_neg_2_051_14    m/z 667.4842     0   1   2E+4   0.5   1  RT (min)      lcqtof_hilic_neg_1_007_05    m/z 667.5137     0   1   2E+4   15.5   16  RT (min)      lcqtof_hilic_neg_1_007_05    m/z 667.5140     0   1   2   3E+4   14.5   15   15.5  RT (min)       lcqtof_hilic_neg_4_083_59    m/z 667.5140     0   1   2   3E+5   1   1.5  RT (min)       lcqtof_hilic_neg_4_000_aa    m/z 667.5146     0   1   2   3E+3   1.5   2  RT (min)       lcqtof_hilic_neg_4_013_20    m/z 667.5146     0   1   2   3E+4   15   15.5  RT (min)       lcqtof_hilic_neg_4_081_58    m/z 667.5146     0   2   4   6E+3   2   2.5  RT (min)      lcqtof_hilic_neg_4_032_01    m/z 667.5148     0   1   2   3E+5   1   1.5  RT (min)      lcqtof_hilic_neg_4_000_aa    m/z 668.2752     0   1E+4   0.5   1   1.5  RT (min)      lcqtof_hilic_neg_4_039_18    m/z 668.3576     0   1E+5   1   1.5  RT (min)      lcqtof_hilic_neg_4_000_aa    m/z 668.3588     0   1E+5   1   1.5  RT (min)      lcqtof_hilic_neg_4_000_aa    m/z 668.4668     0   1E+4   4   4.5  RT (min)       lcqtof_hilic_neg_4_008_24    m/z 668.5171     0   1E+4   15   15.5  RT (min)      lcqtof_hilic_neg_4_013_20    m/z 668.5174     0   1E+5   1   1.5  RT (min)      lcqtof_hilic_neg_4_000_aa    m/z 668.5175     0   1E+5   1   1.5  RT (min)      lcqtof_hilic_neg_4_000_aa    m/z 668.5178     0   1E+4   15   15.5  RT (min)      lcqtof_hilic_neg_4_081_58    m/z 668.6186     0   1   2E+4   1.5   2  RT (min)      lcqtof_hilic_neg_4_000_aa    m/z 669.1267     0   1E+4   0.5   1  RT (min)      lcqtof_hilic_neg_1_062_17    m/z 669.2816     0   1E+4   6   6.5  RT (min)      lcqtof_hilic_neg_1_037_24    m/z 669.3608     0   1   2E+4   1   1.5  RT (min)      lcqtof_hilic_neg_4_000_aa    m/z 669.3912     0   1   2E+4   1   1.5  RT (min)      lcqtof_hilic_neg_4_005_29    m/z 669.4556     0   1   2E+5   1   1.5  RT (min)      lcqtof_hilic_neg_4_000_aa    m/z 669.4558     0   1   2   3E+4   1   1.5  RT (min)      lcqtof_hilic_neg_4_002_25    m/z 669.4562     0   1   2   3E+4   1   1.5  RT (min)      lcqtof_hilic_neg_4_002_25    m/z 669.4565     0   2   4   6E+3   1.5   2  RT (min)      lcqtof_hilic_neg_4_011_35    m/z 669.4566     0   1   2E+4   4   4.5  RT (min)      lcqtof_hilic_neg_4_040_05    m/z 669.4568     0   1E+5   2.5   3  RT (min)       lcqtof_hilic_neg_1_007_05    m/z 669.4574     0   2   4E+3   1.5   2  RT (min)      lcqtof_hilic_neg_4_011_35    m/z 669.4577     0   2   4   6   8E+3   2   2.5  RT (min)      lcqtof_hilic_neg_4_010_28    m/z 669.4580     0   1E+4   15.5   16  RT (min)      lcqtof_hilic_neg_4_000_aa    m/z 669.4584     0   1E+4   15.5   16  RT (min)      lcqtof_hilic_neg_4_047_06    m/z 669.4584     0   2   4   6   8E+3   2   2.5  RT (min)      lcqtof_hilic_neg_4_010_28    m/z 669.4588     0   1   2   3E+3   0   0.5  RT (min)      lcqtof_hilic_neg_1_004_14    m/z 669.4588     0   1E+4   0   0.5  RT (min)      lcqtof_hilic_neg_1_009_50    m/z 669.4588     0   1E+4   15.5   16  RT (min)      lcqtof_hilic_neg_1_007_05    m/z 669.4597     0   1   2   3E+3   19.5   20  RT (min)      lcqtof_hilic_neg_4_009_50    m/z 669.4886     0   1E+4   1   1.5  RT (min)      lcqtof_hilic_neg_4_000_aa    m/z 669.5196     0   1   2   3E+4   1   1.5  RT (min)      lcqtof_hilic_neg_4_000_aa    m/z 669.5448     0   1E+4   0.5   1  RT (min)      lcqtof_hilic_neg_4_048_23    m/z 669.6618     0   1E+4   0.5   1  RT (min)      lcqtof_hilic_neg_3_035_46    m/z 670.1528     0   1E+4   6   6.5  RT (min)       lcqtof_hilic_neg_4_048_23    m/z 670.4579     0   1E+4   1   1.5  RT (min)      lcqtof_hilic_neg_4_002_25    m/z 670.4618     0   1   2   3   4E+3   15.5   16  RT (min)      lcqtof_hilic_neg_4_000_aa    m/z 670.4620     0   1   2   3   4E+3   15.5   16  RT (min)      lcqtof_hilic_neg_4_052_13    m/z 670.5106     0   1E+4   0.5   1  RT (min)      lcqtof_hilic_neg_1_035_23    m/z 671.2882     0   1E+4   0.5   1  RT (min)      lcqtof_hilic_neg_3_035_46    m/z 671.3092     0   1E+4   0.5   1  RT (min)      lcqtof_hilic_neg_4_053_14    m/z 671.3503     0   1   2   3   4E+4   1   1.5  RT (min)      lcqtof_hilic_neg_4_000_aa    m/z 671.3516     0   1   2   3   4E+4   1   1.5  RT (min)      lcqtof_hilic_neg_4_000_aa    m/z 671.3538     0   1   2   3   4E+4   1   1.5  RT (min)      lcqtof_hilic_neg_4_000_aa    m/z 671.3578     0   1   2   3   4E+4   1   1.5  RT (min)      lcqtof_hilic_neg_4_000_aa    m/z 671.3596     0   1   2   3   4E+4   1   1.5  RT (min)      lcqtof_hilic_neg_4_000_aa    m/z 671.3614     0   1   2   3   4E+4   1   1.5  RT (min)      lcqtof_hilic_neg_4_000_aa    m/z 671.3855     0   1   2   3E+4   0.5   1   1.5  RT (min)      lcqtof_hilic_neg_4_038_10    m/z 671.4643     0   1   2   3E+5   4.5   5  RT (min)      lcqtof_hilic_neg_4_048_23    m/z 671.4647     0   1   2   3E+4   1.5   2  RT (min)      lcqtof_hilic_neg_4_048_23    m/z 671.4649     0   1E+5   4.5   5  RT (min)      lcqtof_hilic_neg_4_016_37    m/z 671.4650     0   1   2   3E+5   4   4.5  RT (min)       lcqtof_hilic_neg_4_048_23    m/z 671.4652     0   1E+4   1.5   2  RT (min)       lcqtof_hilic_neg_4_048_23    m/z 671.4652     0   1E+4   1.5   2  RT (min)      lcqtof_hilic_neg_4_048_23    m/z 671.4660     0   1E+4   1.5   2  RT (min)      lcqtof_hilic_neg_4_048_23    m/z 671.5194     0   1E+4   1.5   2  RT (min)      lcqtof_hilic_neg_4_000_aa    m/z 671.5591     0   1   2E+4   1.5   2  RT (min)      lcqtof_hilic_neg_4_000_aa    m/z 671.5836     0   1   2   3E+4   1   1.5  RT (min)      lcqtof_hilic_neg_4_000_aa    m/z 671.6773     0   1   2E+4   0.5   1  RT (min)      lcqtof_hilic_neg_3_035_46    m/z 671.7272     0   1   2E+4   3   3.5  RT (min)      lcqtof_hilic_neg_4_022_34    m/z 671.7278     0   1E+4   3   3.5  RT (min)      lcqtof_hilic_neg_4_048_23    m/z 672.0009     0   1E+4   6.5   7  RT (min)       lcqtof_hilic_neg_4_096_60    m/z 672.1020     0   1   2E+4   0.5   1  RT (min)      lcqtof_hilic_neg_1_062_17    m/z 672.2550     0   1E+4   6   6.5  RT (min)      lcqtof_hilic_neg_4_008_24    m/z 672.3551     0   1E+4   1   1.5  RT (min)      lcqtof_hilic_neg_4_000_aa    m/z 672.3557     0   1E+4   1   1.5  RT (min)      lcqtof_hilic_neg_4_000_aa    m/z 672.3632     0   1E+4   4   4.5  RT (min)      lcqtof_hilic_neg_1_037_24    m/z 672.3886     0   1E+4   0.5   1  RT (min)      lcqtof_hilic_neg_4_038_10    m/z 672.3899     0   1E+4   3.5   4  RT (min)      lcqtof_hilic_neg_4_019_03    m/z 672.4693     0   1E+5   4   4.5  RT (min)      lcqtof_hilic_neg_4_031_38    m/z 672.5656     0   1E+4   1.5   2  RT (min)      lcqtof_hilic_neg_4_000_aa    m/z 672.8566     0   2   4   6E+3   5   5.5  RT (min)       lcqtof_hilic_neg_4_058_39    m/z 672.8578     0   2   4   6E+3   5.5   6  RT (min)      lcqtof_hilic_neg_4_058_39    m/z 672.8582     0   2   4   6E+3   5   5.5  RT (min)      lcqtof_hilic_neg_4_058_39    m/z 673.0986     0   1E+4   0.5   1  RT (min)      lcqtof_hilic_neg_1_062_17    m/z 673.1671     0   1   2   3   4E+4   0.5   1  RT (min)      lcqtof_hilic_neg_3_035_46    m/z 673.3247     0   1E+4   4   4.5  RT (min)      lcqtof_hilic_neg_4_035_33    m/z 673.3561     0   1E+5   15   15.5  RT (min)      lcqtof_hilic_neg_4_000_aa    m/z 673.3572     0   1   2E+3   18.5   19  RT (min)      lcqtof_hilic_neg_4_012_07    m/z 673.3930     0   1E+5   1   1.5  RT (min)      lcqtof_hilic_neg_4_000_aa    m/z 673.3936     0   1E+5   1   1.5  RT (min)      lcqtof_hilic_neg_4_000_aa    m/z 673.4798     0   1E+5   4   4.5  RT (min)      lcqtof_hilic_neg_4_016_37    m/z 673.5067     0   1   2E+4   1   1.5  RT (min)      lcqtof_hilic_neg_4_000_aa    m/z 673.5447     0   1   2E+4   0.5   1  RT (min)      lcqtof_hilic_neg_4_026_04    m/z 673.6932     0   1E+4   0.5   1  RT (min)      lcqtof_hilic_neg_3_035_46    m/z 674.1685     0   1E+4   0.5   1  RT (min)      lcqtof_hilic_neg_3_035_46    m/z 674.2469     0   1   2   3   4E+3   1   1.5  RT (min)      lcqtof_hilic_neg_4_039_18    m/z 674.3578     0   2   4E+4   15   15.5  RT (min)      lcqtof_hilic_neg_4_000_aa    m/z 674.3776     0   1E+4   4   4.5  RT (min)      lcqtof_hilic_neg_1_037_24    m/z 674.3930     0   2   4   6E+4   1   1.5  RT (min)      lcqtof_hilic_neg_4_000_aa    m/z 674.3932     0   2   4   6E+4   1   1.5  RT (min)      lcqtof_hilic_neg_4_000_aa    m/z 674.3962     0   2   4   6E+4   1.5   2  RT (min)      lcqtof_hilic_neg_4_000_aa    m/z 675.1721     0   1E+4   6.5   7  RT (min)      lcqtof_hilic_neg_1_007_05    m/z 675.2994     0   1E+4   0.5   1  RT (min)      lcqtof_hilic_neg_4_053_14    m/z 675.3548     0   1   2E+4   15   15.5  RT (min)      lcqtof_hilic_neg_4_000_aa    m/z 675.4931     0   1E+4   15.5   16  RT (min)      lcqtof_hilic_neg_4_000_aa    m/z 675.5572     0   1E+4   1   1.5  RT (min)      lcqtof_hilic_neg_4_000_aa    m/z 675.6616     0   2   4E+4   1   1.5  RT (min)      lcqtof_hilic_neg_4_000_aa    m/z 675.6644     0   2   4E+4   1   1.5  RT (min)      lcqtof_hilic_neg_4_000_aa    m/z 675.6649     0   2   4E+4   1.5   2  RT (min)      lcqtof_hilic_neg_4_000_aa    m/z 675.6716     0   2   4E+4   1   1.5  RT (min)      lcqtof_hilic_neg_4_000_aa    m/z 675.7040     0   1E+4   2   2.5  RT (min)      lcqtof_hilic_neg_4_000_aa    m/z 676.0150     0   1E+4   6.5   7  RT (min)      lcqtof_hilic_neg_4_034_31    m/z 676.0153     0   1E+4   6.5   7  RT (min)      lcqtof_hilic_neg_4_011_35    m/z 676.1292     0   1E+4   0.5   1  RT (min)      lcqtof_hilic_neg_4_017_51    m/z 676.3916     0   1   2E+4   1.5   2  RT (min)      lcqtof_hilic_neg_4_000_aa    m/z 676.3962     0   1E+4   7   7.5  RT (min)       lcqtof_hilic_neg_4_031_38    m/z 676.5095     0   2   4   6E+4   0.5   1  RT (min)      lcqtof_hilic_neg_1_007_05    m/z 676.6672     0   1   2E+4   1.5   2  RT (min)      lcqtof_hilic_neg_4_000_aa    m/z 676.6749     0   1   2   3E+4   0.5   1  RT (min)      lcqtof_hilic_neg_4_040_05    m/z 677.0901     0   1   2E+4   7.5   8  RT (min)       lcqtof_hilic_neg_4_083_59    m/z 677.2376     0   1   2   3E+4   5.5   6  RT (min)      lcqtof_hilic_neg_4_003_16    m/z 677.2377     0   2   4   6E+3   7   8  RT (min)       lcqtof_hilic_neg_4_003_16    m/z 677.2383     0   2   4   6   8E+4   5.5   6  RT (min)       lcqtof_hilic_neg_4_002_25    m/z 677.2384     0   2   4E+3   8.5   9  RT (min)       lcqtof_hilic_neg_4_011_35    m/z 677.2386     0   2   4E+3   8  RT (min)       lcqtof_hilic_neg_4_029_17    m/z 677.2914     0   1E+4   1   1.5  RT (min)      lcqtof_hilic_neg_4_000_aa    m/z 677.2926     0   1E+4   1   1.5  RT (min)      lcqtof_hilic_neg_4_000_aa    m/z 677.2990     0   1E+4   1   1.5  RT (min)      lcqtof_hilic_neg_4_000_aa    m/z 677.3539     0   1   2E+4   1   1.5  RT (min)      lcqtof_hilic_neg_4_000_aa    m/z 677.4310     0   2   4   6   8E+4   1   1.5  RT (min)      lcqtof_hilic_neg_4_000_aa    m/z 677.6015     0   2   4   6E+4   1   1.5  RT (min)      lcqtof_hilic_neg_4_000_aa    m/z 677.6064     0   2   4   6E+4   1   1.5  RT (min)      lcqtof_hilic_neg_4_000_aa    m/z 677.6435     0   1   2   3E+4   1   1.5  RT (min)      lcqtof_hilic_neg_4_000_aa    m/z 678.2401     0   1   2   3E+3   5.5   6  RT (min)      lcqtof_hilic_neg_4_005_29    m/z 678.3522     0   1E+4   4   4.5  RT (min)      lcqtof_hilic_neg_4_028_36    m/z 678.3828     0   1E+4   4   4.5  RT (min)       lcqtof_hilic_neg_4_048_23    m/z 678.4340     0   1   2   3E+4   1   1.5  RT (min)      lcqtof_hilic_neg_4_000_aa    m/z 678.4755     0   1E+4   4   4.5  RT (min)      lcqtof_hilic_neg_4_008_24    m/z 678.6065     0   1   2   3E+4   1   1.5  RT (min)      lcqtof_hilic_neg_4_000_aa    m/z 678.6488     0   1E+4   1   1.5  RT (min)      lcqtof_hilic_neg_4_000_aa    m/z 679.1046     0   1   2   3E+4   6   6.5  RT (min)       lcqtof_hilic_neg_4_082_59    m/z 679.2347     0   1   2   3   4E+3   5.5   6  RT (min)      lcqtof_hilic_neg_4_053_14    m/z 679.2359     0   1E+4   5.5   6  RT (min)      lcqtof_hilic_neg_4_006_30    m/z 679.3683     0   1   2E+4   15.5   16  RT (min)      lcqtof_hilic_neg_4_000_aa    m/z 679.4034     0   2   4E+3   3   3.5  RT (min)       lcqtof_hilic_neg_4_051_43    m/z 679.4048     0   1   2E+3   5.5   6  RT (min)      lcqtof_hilic_neg_4_001_49    m/z 679.4054     0   1E+4   4   4.5  RT (min)       lcqtof_hilic_neg_4_002_25    m/z 679.4063     0   1   2   3   4E+3   5   5.5  RT (min)      lcqtof_hilic_neg_4_055_32    m/z 679.4558     0   1   2E+4   4   4.5  RT (min)      lcqtof_hilic_neg_4_008_24    m/z 679.4589     0   1E+4   1   1.5  RT (min)      lcqtof_hilic_neg_4_000_aa    m/z 679.5898     0   2   4   6E+4   1   1.5  RT (min)      lcqtof_hilic_neg_4_000_aa    m/z 679.6025     0   2   4   6E+4   1   1.5  RT (min)      lcqtof_hilic_neg_4_000_aa    m/z 679.6671     0   1E+4   0.5   1  RT (min)      lcqtof_hilic_neg_3_035_46    m/z 680.3437     0   1   2E+4   4   4.5  RT (min)       lcqtof_hilic_neg_4_048_23    m/z 680.3546     0   1   2E+4   4   4.5  RT (min)      lcqtof_hilic_neg_4_048_23    m/z 680.3676     0   1E+4   4   4.5  RT (min)       lcqtof_hilic_neg_4_089_59    m/z 680.3759     0   1   2E+5   1   1.5  RT (min)      lcqtof_hilic_neg_4_000_aa    m/z 680.3788     0   1   2E+4   4   4.5  RT (min)      lcqtof_hilic_neg_4_047_06    m/z 680.4082     0   1   2E+4   0.5   1  RT (min)      lcqtof_hilic_neg_4_047_06    m/z 680.4090     0   1E+4   3.5   4  RT (min)      lcqtof_hilic_neg_4_032_01    m/z 680.4657     0   1E+4   4   4.5  RT (min)      lcqtof_hilic_neg_4_047_06    m/z 680.4669     0   1   2   3   4E+4   4   4.5  RT (min)      lcqtof_hilic_neg_4_055_32    m/z 680.4671     0   2   4   6E+4   4   4.5  RT (min)      lcqtof_hilic_neg_4_008_24    m/z 680.4950     0   1E+4   0.5   1  RT (min)      lcqtof_hilic_neg_2_028_05    m/z 680.5695     0   1   2   3E+4   0.5   1  RT (min)      lcqtof_hilic_neg_2_005_13    m/z 680.5989     0   1   2   3E+4   1   1.5  RT (min)      lcqtof_hilic_neg_4_000_aa    m/z 680.6005     0   1   2   3E+4   1   1.5  RT (min)      lcqtof_hilic_neg_4_000_aa    m/z 680.6071     0   1   2   3E+4   1   1.5  RT (min)      lcqtof_hilic_neg_4_000_aa    m/z 680.6172     0   1   2   3E+4   1.5   2  RT (min)      lcqtof_hilic_neg_4_000_aa    m/z 681.0070     0   1E+4   0.5   1  RT (min)      lcqtof_hilic_neg_4_024_40    m/z 681.2764     0   1   2   3E+4   0.5   1  RT (min)      lcqtof_hilic_neg_4_050_02    m/z 681.3152     0   1   2E+4   0.5   1  RT (min)      lcqtof_hilic_neg_3_035_46    m/z 681.5081     0   2   4E+3   15.5   16  RT (min)      lcqtof_hilic_neg_2_056_56    m/z 681.5258     0   1   2   3   4E+4   15.5   16  RT (min)      lcqtof_hilic_neg_4_027_19    m/z 681.5258     0   1   2   3E+3   17.5   18  RT (min)      lcqtof_hilic_neg_4_009_50    m/z 681.5258     0   1   2   3E+4   15.5   16  RT (min)      lcqtof_hilic_neg_1_007_05    m/z 681.5268     0   2   4   6E+3   1   1.5  RT (min)      lcqtof_hilic_neg_4_043_12    m/z 681.5268     0   1   2   3   4E+3   16.5   17  RT (min)      lcqtof_hilic_neg_4_017_51    m/z 681.5273     0   1   2   3   4E+5   1   1.5  RT (min)      lcqtof_hilic_neg_4_000_aa    m/z 681.5274     0   1   2E+3   3   3.5  RT (min)       lcqtof_hilic_neg_4_008_24    m/z 681.5275     0   1   2   3   4E+3   1.5   2  RT (min)      lcqtof_hilic_neg_4_055_32    m/z 681.5276     0   1   2   3   4E+3   18   18.5  RT (min)       lcqtof_hilic_neg_4_007_45    m/z 681.5277     0   1   2E+3   2.5   3  RT (min)      lcqtof_hilic_neg_4_008_24    m/z 681.5280     0   1   2   3   4E+5   1   1.5  RT (min)      lcqtof_hilic_neg_4_000_aa    m/z 681.5280     0   1   2   3   4E+5   1   1.5  RT (min)      lcqtof_hilic_neg_4_000_aa    m/z 681.5283     0   2   4   6   8E+3   2   2.5  RT (min)      lcqtof_hilic_neg_4_032_01    m/z 681.5290     0   1   2   3E+3   2   2.5  RT (min)      lcqtof_hilic_neg_4_000_dd    m/z 681.5296     0   1   2   3   4E+4   15   15.5  RT (min)      lcqtof_hilic_neg_4_015_09    m/z 681.5297     0   1   2   3E+4   14.5   15   15.5  RT (min)       lcqtof_hilic_neg_4_083_59    m/z 681.5979     0   1   2   3E+4   0.5   1  RT (min)      lcqtof_hilic_neg_2_028_05    m/z 681.6106     0   1E+5   1   1.5  RT (min)      lcqtof_hilic_neg_4_000_aa    m/z 682.3704     0   1E+4   1   1.5  RT (min)      lcqtof_hilic_neg_4_000_aa    m/z 682.3762     0   1E+4   1   1.5  RT (min)      lcqtof_hilic_neg_4_000_aa    m/z 682.4190     0   1   2E+4   0.5   1  RT (min)      lcqtof_hilic_neg_4_059_14    m/z 682.4445     0   1E+4   1   1.5  RT (min)      lcqtof_hilic_neg_4_013_20    m/z 682.4504     0   1   2E+4   4   4.5  RT (min)      lcqtof_hilic_neg_4_008_24    m/z 682.4829     0   1   2E+4   1   1.5  RT (min)      lcqtof_hilic_neg_4_000_aa    m/z 682.5110     0   1   2   3E+4   1   1.5  RT (min)      lcqtof_hilic_neg_4_000_aa    m/z 682.5121     0   1   2   3E+4   1   1.5  RT (min)      lcqtof_hilic_neg_4_000_aa    m/z 682.5121     0   1   2   3E+4   1   1.5  RT (min)      lcqtof_hilic_neg_4_000_aa    m/z 682.5304     0   1E+5   1   1.5  RT (min)      lcqtof_hilic_neg_4_000_aa    m/z 682.5314     0   1E+4   15.5   16  RT (min)      lcqtof_hilic_neg_1_007_05    m/z 682.5321     0   1E+5   1   1.5  RT (min)      lcqtof_hilic_neg_4_000_aa    m/z 682.5321     0   2   4E+1   14.5   15  RT (min)      lcqtof_hilic_neg_4_083_59    m/z 682.5325     0   1E+4   15   15.5  RT (min)      lcqtof_hilic_neg_4_013_20    m/z 682.6009     0   1   2   3   4E+4   1.5   2  RT (min)      lcqtof_hilic_neg_4_000_aa    m/z 682.6098     0   1   2   3   4E+4   1   1.5  RT (min)      lcqtof_hilic_neg_4_000_aa    m/z 682.6150     0   1   2   3   4E+4   1   1.5  RT (min)      lcqtof_hilic_neg_4_000_aa    m/z 682.6344     0   1   2E+4   1.5   2  RT (min)      lcqtof_hilic_neg_4_000_aa    m/z 682.9782     0   1   2E+4   1   1.5  RT (min)      lcqtof_hilic_neg_4_000_aa    m/z 683.2252     0   1   2   3E+5   6   6.5  RT (min)       lcqtof_hilic_neg_3_011_34    m/z 683.4194     0   2   4   6   8E+5   1   1.5  RT (min)      lcqtof_hilic_neg_4_000_aa    m/z 683.4208     0   2   4   6   8E+5   1   1.5  RT (min)      lcqtof_hilic_neg_4_000_aa    m/z 683.4218     0   1E+4   4.5   5  RT (min)      lcqtof_hilic_neg_4_040_05    m/z 683.4220     0   2   4   6   8E+5   1   1.5  RT (min)      lcqtof_hilic_neg_4_000_aa    m/z 683.5160     0   1   2   3E+4   1.5   2  RT (min)      lcqtof_hilic_neg_4_000_aa    m/z 683.5287     0   2   4E+4   1   1.5  RT (min)      lcqtof_hilic_neg_4_000_aa    m/z 683.5346     0   2   4E+4   1   1.5  RT (min)      lcqtof_hilic_neg_4_000_aa    m/z 684.1834     0   1E+4   7   7.5  RT (min)      lcqtof_hilic_neg_1_005_32    m/z 684.2276     0   2   4   6   8E+4   6   6.5  RT (min)      lcqtof_hilic_neg_3_011_34    m/z 684.3166     0   1E+4   0.5   1  RT (min)      lcqtof_hilic_neg_3_003_20    m/z 684.4238     0   2   4E+5   1   1.5  RT (min)      lcqtof_hilic_neg_4_000_aa    m/z 684.4260     0   2   4E+5   1   1.5  RT (min)      lcqtof_hilic_neg_4_000_aa    m/z 684.6126     0   1E+4   1.5   2  RT (min)      lcqtof_hilic_neg_4_000_aa    m/z 685.0595     0   1E+4   6.5   7  RT (min)      lcqtof_hilic_neg_4_034_31    m/z 685.2308     0   1   2E+4   6   6.5  RT (min)      lcqtof_hilic_neg_3_011_34    m/z 685.3250     0   1E+4   1   1.5  RT (min)      lcqtof_hilic_neg_4_000_aa    m/z 685.3980     0   1E+4   0.5   1  RT (min)      lcqtof_hilic_neg_4_021_21    m/z 685.4119     0   1E+4   3.5   4  RT (min)      lcqtof_hilic_neg_4_032_01    m/z 685.5410     0   1E+4   1.5   2  RT (min)      lcqtof_hilic_neg_4_000_aa    m/z 685.5892     0   1E+5   1   1.5  RT (min)      lcqtof_hilic_neg_4_000_aa    m/z 686.3006     0   1E+4   4   4.5  RT (min)      lcqtof_hilic_neg_4_039_18    m/z 686.4208     0   2   4   6   8E+4   1   1.5  RT (min)      lcqtof_hilic_neg_4_000_aa    m/z 686.4230     0   2   4   6   8E+4   1   1.5  RT (min)      lcqtof_hilic_neg_4_000_aa    m/z 686.4235     0   2   4   6   8E+4   1   1.5  RT (min)      lcqtof_hilic_neg_4_000_aa    m/z 686.4239     0   2   4   6   8E+4   1   1.5  RT (min)      lcqtof_hilic_neg_4_000_aa    m/z 686.4689     0   2   4E+4   3   3.5  RT (min)      lcqtof_hilic_neg_4_016_37    m/z 686.4769     0   1   2   3   4E+4   3   3.5  RT (min)      lcqtof_hilic_neg_4_028_36    m/z 686.4835     0   1   2   3   4E+4   3   3.5  RT (min)      lcqtof_hilic_neg_4_034_31    m/z 686.4838     0   2   4   6   8E+4   3   3.5  RT (min)      lcqtof_hilic_neg_1_014_31    m/z 686.5920     0   1   2   3   4E+4   1   1.5  RT (min)      lcqtof_hilic_neg_4_000_aa    m/z 687.0004     0   2   4E+4   6.5   7  RT (min)      lcqtof_hilic_neg_4_053_14    m/z 687.0010     0   2   4E+4   6.5   7  RT (min)      lcqtof_hilic_neg_4_053_14    m/z 687.0014     0   1E+4   6.5   7  RT (min)      lcqtof_hilic_neg_4_032_01    m/z 687.2658     0   1   2E+3   10   10.5  RT (min)       lcqtof_hilic_neg_4_024_40    m/z 687.2660     0   1E+4   5.5   6  RT (min)       lcqtof_hilic_neg_4_092_60    m/z 687.2660     0   1   2   3E+4   0   0.5  RT (min)      lcqtof_hilic_neg_4_000_aa    m/z 687.2662     0   1   2E+4   5.5   6  RT (min)       lcqtof_hilic_neg_4_053_14    m/z 687.2663     0   1   2E+3   10.5   11  RT (min)       lcqtof_hilic_neg_4_003_16    m/z 687.2666     0   2   4   6E+4   5   5.5  RT (min)      lcqtof_hilic_neg_4_000_bb    m/z 687.2669     0   1E+4   6   6.5  RT (min)       lcqtof_hilic_neg_4_052_13    m/z 687.2669     0   1   2   3   4E+3   9   9.5  RT (min)       lcqtof_hilic_neg_4_000_dd    m/z 687.2669     0   2   4E+3   8.5   9  RT (min)       lcqtof_hilic_neg_4_000_dd    m/z 687.2670     0   1   2E+3   10.5   11  RT (min)       lcqtof_hilic_neg_4_000_bb    m/z 687.2671     0   2   4E+3   8   8.5  RT (min)       lcqtof_hilic_neg_4_000_dd    m/z 687.2672     0   1   2E+3   11   11.5  RT (min)       lcqtof_hilic_neg_4_003_16    m/z 687.2673     0   1   2   3   4E+3   8   8.5  RT (min)       lcqtof_hilic_neg_4_039_18    m/z 687.2674     0   1   2   3E+3   9.5   10  RT (min)       lcqtof_hilic_neg_4_051_43    m/z 687.2677     0   1   2E+4   15   15.5  RT (min)       lcqtof_hilic_neg_4_000_aa    m/z 687.3364     0   2   4   6   8E+4   15.5   16  RT (min)      lcqtof_hilic_neg_4_000_aa    m/z 687.3656     0   1E+5   15   15.5  RT (min)       lcqtof_hilic_neg_4_000_aa    m/z 687.4195     0   1   2   3E+4   1   1.5  RT (min)      lcqtof_hilic_neg_4_000_aa    m/z 687.4206     0   1   2   3E+4   1   1.5  RT (min)      lcqtof_hilic_neg_4_000_aa    m/z 687.4221     0   1   2   3E+4   1.5   2  RT (min)      lcqtof_hilic_neg_4_000_aa    m/z 687.4240     0   1   2   3E+4   1   1.5  RT (min)      lcqtof_hilic_neg_4_000_aa    m/z 687.4637     0   1   2E+4   2.5   3  RT (min)      lcqtof_hilic_neg_4_048_23    m/z 687.5515     0   1E+4   0.5   1  RT (min)      lcqtof_hilic_neg_4_040_05    m/z 687.5920     0   1E+4   1   1.5  RT (min)      lcqtof_hilic_neg_4_000_aa    m/z 688.0047     0   1E+4   6.5   7  RT (min)      lcqtof_hilic_neg_4_053_14    m/z 688.0050     0   1E+4   6.5   7  RT (min)      lcqtof_hilic_neg_4_053_14    m/z 688.2694     0   1   2   3   4E+4   5   5.5  RT (min)      lcqtof_hilic_neg_4_002_25    m/z 688.2704     0   2   4   6E+3   5.5   6  RT (min)      lcqtof_hilic_neg_4_059_14    m/z 688.3076     0   1E+5   3   3.5  RT (min)      lcqtof_hilic_neg_4_039_18    m/z 688.3384     0   1   2   3E+4   15.5   16  RT (min)      lcqtof_hilic_neg_4_000_aa    m/z 688.3391     0   2   4E+4   1   1.5  RT (min)      lcqtof_hilic_neg_4_000_aa    m/z 688.3393     0   2   4E+4   1   1.5  RT (min)      lcqtof_hilic_neg_4_000_aa    m/z 688.3746     0   1   2   3   4E+4   15   15.5  RT (min)      lcqtof_hilic_neg_4_000_aa    m/z 688.4220     0   1   2   3E+5   1.5   2  RT (min)      lcqtof_hilic_neg_4_000_aa    m/z 688.5280     0   1   2E+4   0.5   1  RT (min)      lcqtof_hilic_neg_4_040_05    m/z 688.5994     0   1E+4   4.5   5  RT (min)      lcqtof_hilic_neg_4_050_02    m/z 688.9958     0   1   2E+5   7   7.5  RT (min)       lcqtof_hilic_neg_4_059_14    m/z 688.9960     0   1   2   3   4E+5   6.5   7  RT (min)       lcqtof_hilic_neg_4_053_14    m/z 688.9961     0   1   2   3   4E+5   6.5   7  RT (min)       lcqtof_hilic_neg_4_053_14    m/z 688.9964     0   1   2   3E+5   6.5   7  RT (min)      lcqtof_hilic_neg_4_035_33    m/z 688.9967     0   1E+5   7   7.5  RT (min)      lcqtof_hilic_neg_4_064_15    m/z 689.1403     0   1E+4   1   1.5  RT (min)      lcqtof_hilic_neg_4_000_aa    m/z 689.1456     0   1E+4   1   1.5  RT (min)      lcqtof_hilic_neg_4_000_aa    m/z 689.1554     0   1   2   3E+4   6.5   7  RT (min)       lcqtof_hilic_neg_4_008_24    m/z 689.1658     0   1   2   3E+4   0.5   1  RT (min)      lcqtof_hilic_neg_4_024_40    m/z 689.3103     0   1   2   3   4E+4   3   3.5  RT (min)      lcqtof_hilic_neg_4_039_18    m/z 689.3362     0   1E+4   15.5   16  RT (min)      lcqtof_hilic_neg_4_000_aa    m/z 689.3649     0   1E+4   0.5   1  RT (min)      lcqtof_hilic_neg_4_038_10    m/z 689.3718     0   1   2E+4   15   15.5  RT (min)      lcqtof_hilic_neg_4_000_aa    m/z 689.4254     0   1E+5   1.5   2  RT (min)      lcqtof_hilic_neg_4_000_aa    m/z 689.4985     0   1E+4   0.5   1  RT (min)      lcqtof_hilic_neg_4_018_27    m/z 689.5315     0   1E+4   0.5   1  RT (min)      lcqtof_hilic_neg_4_040_05    m/z 689.6495     0   1   2E+4   1   1.5  RT (min)      lcqtof_hilic_neg_4_000_aa    m/z 689.6584     0   1   2E+4   1   1.5  RT (min)      lcqtof_hilic_neg_4_000_aa    m/z 689.6748     0   1   2   3E+4   0.5   1  RT (min)      lcqtof_hilic_neg_4_040_05    m/z 689.6806     0   1E+4   1.5   2  RT (min)      lcqtof_hilic_neg_4_000_aa    m/z 689.9982     0   1   2   3E+4   6.5   7  RT (min)      lcqtof_hilic_neg_4_032_01    m/z 689.9986     0   1   2E+4   7.5   8  RT (min)      lcqtof_hilic_neg_1_014_31    m/z 689.9994     0   1E+5   6.5   7  RT (min)      lcqtof_hilic_neg_4_053_14    m/z 689.9994     0   1   2   3E+4   6.5   7  RT (min)      lcqtof_hilic_neg_4_032_01    m/z 690.3147     0   1E+4   3   3.5  RT (min)      lcqtof_hilic_neg_4_039_18    m/z 690.3660     0   1E+4   4.5   5  RT (min)       lcqtof_hilic_neg_4_008_24    m/z 690.4282     0   1   2   3E+4   1.5   2  RT (min)      lcqtof_hilic_neg_4_000_aa    m/z 690.5267     0   1E+4   0.5   1  RT (min)      lcqtof_hilic_neg_4_040_05    m/z 690.6572     0   1E+4   1   1.5  RT (min)      lcqtof_hilic_neg_4_000_aa    m/z 691.0014     0   1   2E+4   6.5   7  RT (min)      lcqtof_hilic_neg_4_059_14    m/z 691.0014     0   1   2   3E+4   6.5   7  RT (min)      lcqtof_hilic_neg_4_053_14    m/z 691.1656     0   1E+4   0.5   1  RT (min)      lcqtof_hilic_neg_4_024_40    m/z 691.2250     0   1E+4   6   6.5  RT (min)       lcqtof_hilic_neg_4_091_60    m/z 691.3091     0   1   2E+4   1   1.5  RT (min)      lcqtof_hilic_neg_4_000_aa    m/z 691.3106     0   1   2E+4   1   1.5  RT (min)      lcqtof_hilic_neg_4_000_aa    m/z 691.3113     0   1   2E+4   1   1.5  RT (min)      lcqtof_hilic_neg_4_000_aa    m/z 691.3133     0   1   2E+4   1   1.5  RT (min)      lcqtof_hilic_neg_4_000_aa    m/z 691.3152     0   1   2E+4   1   1.5  RT (min)      lcqtof_hilic_neg_4_000_aa    m/z 691.3679     0   1E+4   1   1.5  RT (min)      lcqtof_hilic_neg_4_000_aa    m/z 691.5014     0   2   4   6   8E+4   0.5   1  RT (min)      lcqtof_hilic_neg_1_007_05    m/z 691.5331     0   1   2   3   4E+5   1   1.5  RT (min)      lcqtof_hilic_neg_4_000_aa    m/z 691.5359     0   1   2   3   4E+5   1   1.5  RT (min)      lcqtof_hilic_neg_4_000_aa    m/z 691.5363     0   1   2   3   4E+5   1.5   2  RT (min)      lcqtof_hilic_neg_4_000_aa    m/z 691.5391     0   1   2   3   4E+5   1   1.5  RT (min)      lcqtof_hilic_neg_4_000_aa    m/z 691.5444     0   1   2   3   4E+5   1   1.5  RT (min)      lcqtof_hilic_neg_4_000_aa    m/z 691.6412     0   1   2   3E+4   1   1.5  RT (min)      lcqtof_hilic_neg_4_000_aa    m/z 691.6461     0   1   2   3E+4   1   1.5  RT (min)      lcqtof_hilic_neg_4_000_aa    m/z 691.6548     0   1   2   3E+4   1   1.5  RT (min)      lcqtof_hilic_neg_4_000_aa    m/z 691.6590     0   1   2   3E+4   1   1.5  RT (min)      lcqtof_hilic_neg_4_000_aa    m/z 691.6610     0   1   2E+4   1   1.5  RT (min)      lcqtof_hilic_neg_4_000_aa    m/z 692.0750     0   1E+4   6.5   7  RT (min)      lcqtof_hilic_neg_4_048_23    m/z 692.1358     0   1E+4   6   6.5  RT (min)       lcqtof_hilic_neg_4_048_23    m/z 692.3088     0   1E+4   1   1.5  RT (min)      lcqtof_hilic_neg_4_000_aa    m/z 692.3144     0   1E+4   1   1.5  RT (min)      lcqtof_hilic_neg_4_000_aa    m/z 692.3596     0   1E+4   3.5   4  RT (min)       lcqtof_hilic_neg_4_019_03    m/z 692.3722     0   1E+4   4   4.5  RT (min)      lcqtof_hilic_neg_4_048_23    m/z 692.3984     0   1E+4   4   4.5  RT (min)      lcqtof_hilic_neg_4_031_38    m/z 692.4133     0   1E+4   4   4.5  RT (min)      lcqtof_hilic_neg_4_016_37    m/z 692.4812     0   1   2E+4   0.5   1  RT (min)      lcqtof_hilic_neg_1_007_05    m/z 692.5053     0   1   2   3E+4   0.5   1  RT (min)      lcqtof_hilic_neg_1_007_05    m/z 692.5344     0   1   2E+5   1.5   2  RT (min)      lcqtof_hilic_neg_4_000_aa    m/z 692.5378     0   1   2E+5   1   1.5  RT (min)      lcqtof_hilic_neg_4_000_aa    m/z 692.5380     0   1   2E+5   1   1.5  RT (min)      lcqtof_hilic_neg_4_000_aa    m/z 692.5406     0   1   2E+5   1   1.5  RT (min)      lcqtof_hilic_neg_4_000_aa    m/z 692.5451     0   1   2E+5   1   1.5  RT (min)      lcqtof_hilic_neg_4_000_aa    m/z 692.5460     0   1   2E+5   1   1.5  RT (min)      lcqtof_hilic_neg_4_000_aa    m/z 692.6388     0   1E+4   1   1.5  RT (min)      lcqtof_hilic_neg_4_000_aa    m/z 692.6568     0   1E+4   1   1.5  RT (min)      lcqtof_hilic_neg_4_000_aa    m/z 692.6580     0   1E+4   1   1.5  RT (min)      lcqtof_hilic_neg_4_000_aa    m/z 693.2256     0   1   2E+4   1   1.5  RT (min)      lcqtof_hilic_neg_4_000_aa    m/z 693.2266     0   1   2E+4   1   1.5  RT (min)      lcqtof_hilic_neg_4_000_aa    m/z 693.3857     0   1E+5   1   1.5  RT (min)      lcqtof_hilic_neg_4_000_aa    m/z 693.5376     0   1E+5   1   1.5  RT (min)      lcqtof_hilic_neg_4_000_aa    m/z 693.5384     0   1E+5   1.5   2  RT (min)      lcqtof_hilic_neg_4_000_aa    m/z 693.5548     0   1E+5   1   1.5  RT (min)      lcqtof_hilic_neg_4_000_aa    m/z 693.6266     0   2   4E+4   1   1.5  RT (min)      lcqtof_hilic_neg_4_000_aa    m/z 693.6496     0   2   4   6E+4   1   1.5  RT (min)      lcqtof_hilic_neg_4_000_aa    m/z 693.9575     0   1E+4   5   5.5  RT (min)       lcqtof_hilic_neg_4_008_24    m/z 693.9720     0   1E+5   6.5   7  RT (min)       lcqtof_hilic_neg_4_090_60    m/z 694.2287     0   1E+4   1   1.5  RT (min)      lcqtof_hilic_neg_4_000_aa    m/z 694.2326     0   1E+4   1   1.5  RT (min)      lcqtof_hilic_neg_4_000_aa    m/z 694.2362     0   1E+4   6   6.5  RT (min)      lcqtof_hilic_neg_4_008_24    m/z 694.3868     0   2   4   6E+4   1   1.5  RT (min)      lcqtof_hilic_neg_4_000_aa    m/z 694.4315     0   1E+4   3   3.5  RT (min)       lcqtof_hilic_neg_4_001_49    m/z 694.4712     0   1   2   3   4E+4   4   4.5  RT (min)       lcqtof_hilic_neg_4_008_24    m/z 694.4718     0   1E+4   0.5   1  RT (min)      lcqtof_hilic_neg_4_003_16    m/z 694.4734     0   1E+4   0.5   1  RT (min)      lcqtof_hilic_neg_2_028_05    m/z 694.4886     0   1E+4   0.5   1  RT (min)      lcqtof_hilic_neg_1_007_05    m/z 694.5356     0   1   2   3E+4   1   1.5  RT (min)      lcqtof_hilic_neg_4_000_aa    m/z 694.5388     0   1   2   3E+4   1.5   2  RT (min)      lcqtof_hilic_neg_4_000_aa    m/z 694.5406     0   1   2   3   4E+4   1   1.5  RT (min)      lcqtof_hilic_neg_4_000_aa    m/z 694.5594     0   2   4E+4   1   1.5  RT (min)      lcqtof_hilic_neg_4_000_aa    m/z 694.6040     0   1E+4   0.5   1  RT (min)      lcqtof_hilic_neg_4_040_05    m/z 694.6270     0   1   2E+4   1   1.5  RT (min)      lcqtof_hilic_neg_4_000_aa    m/z 694.6330     0   1   2E+4   1.5   2  RT (min)      lcqtof_hilic_neg_4_000_aa    m/z 694.6363     0   1   2   3E+4   1   1.5  RT (min)      lcqtof_hilic_neg_4_000_aa    m/z 694.6530     0   1   2   3E+4   1   1.5  RT (min)      lcqtof_hilic_neg_4_000_aa    m/z 694.7864     0   1   2E+4   8.5   9  RT (min)       lcqtof_hilic_neg_4_053_14    m/z 694.9348     0   2   4   6E+4   0.5   1  RT (min)      lcqtof_hilic_neg_3_035_46    m/z 694.9380     0   2   4   6E+4   0.5   1  RT (min)      lcqtof_hilic_neg_3_035_46    m/z 695.0226     0   1E+4   0.5   1  RT (min)      lcqtof_hilic_neg_4_024_40    m/z 695.2212     0   1E+4   1   1.5  RT (min)      lcqtof_hilic_neg_4_000_aa    m/z 695.3895     0   1   2E+4   1   1.5  RT (min)      lcqtof_hilic_neg_4_000_aa    m/z 695.4353     0   2   4E+3   3.5   4  RT (min)      lcqtof_hilic_neg_4_063_16    m/z 695.4635     0   1E+4   1   1.5  RT (min)      lcqtof_hilic_neg_1_053_19    m/z 695.5444     0   1   2   3   4E+5   1.5   2  RT (min)      lcqtof_hilic_neg_4_000_aa    m/z 695.5453     0   2   4   6E+4   15   15.5  RT (min)       lcqtof_hilic_neg_4_039_18    m/z 695.5454     0   2   4E+4   14.5   15   15.5  RT (min)       lcqtof_hilic_neg_4_083_59    m/z 695.5456     0   1   2   3   4E+5   1   1.5  RT (min)      lcqtof_hilic_neg_4_000_aa    m/z 695.5456     0   1   2   3   4E+3   2.5   3  RT (min)      lcqtof_hilic_neg_4_008_24    m/z 695.5458     0   1   2   3   4E+3   2.5   3  RT (min)       lcqtof_hilic_neg_4_008_24    m/z 695.5458     0   1   2   3   4E+4   15.5   16  RT (min)      lcqtof_hilic_neg_1_007_05    m/z 695.5460     0   2   4   6   8E+3   2   2.5  RT (min)      lcqtof_hilic_neg_4_052_13    m/z 695.5460     0   1   2   3   4E+5   1   1.5  RT (min)      lcqtof_hilic_neg_4_000_aa    m/z 695.5462     0   1   2   3   4E+5   1   1.5  RT (min)      lcqtof_hilic_neg_4_000_aa    m/z 695.5464     0   1   2   3   4E+5   1   1.5  RT (min)      lcqtof_hilic_neg_4_000_aa    m/z 695.5471     0   1E+4   4   4.5  RT (min)      lcqtof_hilic_neg_4_008_24    m/z 695.5703     0   2   4   6E+4   1   1.5  RT (min)      lcqtof_hilic_neg_4_000_aa    m/z 695.6023     0   1   2E+4   0.5   1  RT (min)      lcqtof_hilic_neg_2_028_05    m/z 695.6145     0   1   2E+4   1   1.5  RT (min)      lcqtof_hilic_neg_4_000_aa    m/z 695.9350     0   1   2   3E+4   0.5   1  RT (min)      lcqtof_hilic_neg_3_035_46    m/z 695.9384     0   1   2   3E+4   0.5   1  RT (min)      lcqtof_hilic_neg_3_035_46    m/z 696.1356     0   1E+4   6.5   7  RT (min)      lcqtof_hilic_neg_4_008_24    m/z 696.4109     0   1E+4   3.5   4  RT (min)      lcqtof_hilic_neg_4_032_01    m/z 696.4407     0   1E+4   7   7.5  RT (min)      lcqtof_hilic_neg_1_035_23    m/z 696.5010     0   1E+4   4   4.5  RT (min)      lcqtof_hilic_neg_4_055_32    m/z 696.5484     0   1   2   3E+2   14.5   15  RT (min)      lcqtof_hilic_neg_4_083_59    m/z 696.5490     0   1E+4   15.5   16  RT (min)      lcqtof_hilic_neg_1_007_05    m/z 696.5492     0   1   2E+4   15   15.5  RT (min)      lcqtof_hilic_neg_4_013_20    m/z 696.5496     0   1   2E+5   1   1.5  RT (min)      lcqtof_hilic_neg_4_000_aa    m/z 696.6174     0   1E+4   1.5   2  RT (min)      lcqtof_hilic_neg_4_000_aa    m/z 696.6490     0   1   2   3E+4   1.5   2  RT (min)      lcqtof_hilic_neg_4_000_aa    m/z 696.9272     0   1   2   3   4E+4   0.5   1  RT (min)      lcqtof_hilic_neg_4_098_61    m/z 696.9380     0   1   2E+4   0.5   1  RT (min)      lcqtof_hilic_neg_3_035_46    m/z 697.0502     0   1E+5   0.5   1  RT (min)      lcqtof_hilic_neg_4_021_21    m/z 697.0622     0   2   4   6E+4   0.5   1  RT (min)      lcqtof_hilic_neg_4_039_18    m/z 697.1134     0   1E+4   6   6.5  RT (min)      lcqtof_hilic_neg_1_035_23    m/z 697.1144     0   1E+4   6   6.5  RT (min)       lcqtof_hilic_neg_4_048_23    m/z 697.4432     0   1   2   3E+4   1.5   2  RT (min)      lcqtof_hilic_neg_4_000_aa    m/z 697.4440     0   1   2   3E+4   1   1.5  RT (min)      lcqtof_hilic_neg_4_000_aa    m/z 697.4591     0   2   4   6   8E+4   1   1.5  RT (min)      lcqtof_hilic_neg_4_000_aa    m/z 697.4833     0   2   4   6E+4   1   1.5  RT (min)      lcqtof_hilic_neg_2_020_23    m/z 697.4846     0   1E+5   1   1.5  RT (min)      lcqtof_hilic_neg_4_000_aa    m/z 697.4846     0   1   2   3E+4   1   1.5  RT (min)      lcqtof_hilic_neg_4_011_35    m/z 697.4874     0   2   4E+3   2   2.5  RT (min)      lcqtof_hilic_neg_4_029_17    m/z 697.4896     0   2   4   6E+3   15.5   16  RT (min)      lcqtof_hilic_neg_1_009_50    m/z 697.4906     0   2   4   6E+3   15.5   16  RT (min)      lcqtof_hilic_neg_1_010_15    m/z 697.4907     0   1   2E+3   18   18.5  RT (min)      lcqtof_hilic_neg_4_005_29    m/z 697.4916     0   1   2E+3   16.5   17  RT (min)      lcqtof_hilic_neg_1_009_50    m/z 697.4917     0   2   4E+3   15.5   16  RT (min)      lcqtof_hilic_neg_1_007_05    m/z 697.5236     0   2   4   6   8E+4   1   1.5  RT (min)      lcqtof_hilic_neg_4_000_aa    m/z 697.5511     0   2   4E+4   1   1.5  RT (min)      lcqtof_hilic_neg_4_000_aa    m/z 697.6540     0   1E+4   1.5   2  RT (min)      lcqtof_hilic_neg_4_000_aa    m/z 697.6853     0   1E+4   0.5   1  RT (min)      lcqtof_hilic_neg_3_035_46    m/z 698.0537     0   1   2   3E+4   0.5   1  RT (min)      lcqtof_hilic_neg_4_021_21    m/z 698.0663     0   1   2E+4   0.5   1  RT (min)      lcqtof_hilic_neg_4_039_18    m/z 698.4418     0   1E+4   1   1.5  RT (min)      lcqtof_hilic_neg_4_000_aa    m/z 698.4541     0   1   2   3E+4   1   1.5  RT (min)      lcqtof_hilic_neg_4_000_aa    m/z 698.4647     0   1   2   3E+4   1   1.5  RT (min)      lcqtof_hilic_neg_4_000_aa    m/z 698.4676     0   1   2   3   4E+4   1   1.5  RT (min)      lcqtof_hilic_neg_4_000_aa    m/z 698.4755     0   1E+4   0.5   1  RT (min)      lcqtof_hilic_neg_4_029_17    m/z 698.4932     0   1   2E+3   15.5   16  RT (min)      lcqtof_hilic_neg_4_012_07    m/z 698.5269     0   1   2   3   4E+4   1   1.5  RT (min)      lcqtof_hilic_neg_4_000_aa    m/z 698.6074     0   1E+4   1   1.5  RT (min)      lcqtof_hilic_neg_4_000_aa    m/z 698.6276     0   1   2E+4   1.5   2  RT (min)      lcqtof_hilic_neg_4_000_aa    m/z 699.0498     0   2   4E+4   0.5   1  RT (min)      lcqtof_hilic_neg_4_021_21    m/z 699.0616     0   1   2   3E+4   0.5   1  RT (min)      lcqtof_hilic_neg_4_039_18    m/z 699.4472     0   1E+4   1   1.5  RT (min)      lcqtof_hilic_neg_4_000_aa    m/z 699.4780     0   1E+5   1   1.5  RT (min)      lcqtof_hilic_neg_4_000_aa    m/z 699.5439     0   1E+4   0.5   1  RT (min)      lcqtof_hilic_neg_4_047_06    m/z 699.6162     0   1   2E+5   1   1.5  RT (min)      lcqtof_hilic_neg_4_000_aa    m/z 699.7003     0   1   2E+4   0.5   1  RT (min)      lcqtof_hilic_neg_3_035_46    m/z 700.0518     0   1E+4   0.5   1  RT (min)      lcqtof_hilic_neg_4_021_21    m/z 700.0952     0   1E+4   0.5   1  RT (min)      lcqtof_hilic_neg_3_007_04    m/z 700.4120     0   2   4E+4   1   1.5  RT (min)      lcqtof_hilic_neg_4_000_aa    m/z 700.4122     0   2   4E+4   1   1.5  RT (min)      lcqtof_hilic_neg_4_000_aa    m/z 700.4175     0   2   4E+4   1   1.5  RT (min)      lcqtof_hilic_neg_4_000_aa    m/z 700.4800     0   1E+5   1   1.5  RT (min)      lcqtof_hilic_neg_4_000_aa    m/z 700.5249     0   1E+4   0.5   1  RT (min)      lcqtof_hilic_neg_4_040_05    m/z 700.5423     0   1E+4   0.5   1  RT (min)      lcqtof_hilic_neg_1_031_16    m/z 700.6075     0   2   4   6   8E+4   1.5   2  RT (min)      lcqtof_hilic_neg_4_000_aa    m/z 700.6144     0   2   4   6   8E+4   1   1.5  RT (min)      lcqtof_hilic_neg_4_000_aa    m/z 701.3368     0   1E+5   1   1.5  RT (min)      lcqtof_hilic_neg_4_000_aa    m/z 701.3412     0   1E+5   1   1.5  RT (min)      lcqtof_hilic_neg_4_000_aa    m/z 701.3413     0   1E+5   1   1.5  RT (min)      lcqtof_hilic_neg_4_000_aa    m/z 701.3420     0   1E+5   1   1.5  RT (min)      lcqtof_hilic_neg_4_000_aa    m/z 701.3534     0   1E+4   15.5   16  RT (min)      lcqtof_hilic_neg_4_000_aa    m/z 701.3860     0   2   4E+4   15   15.5  RT (min)      lcqtof_hilic_neg_4_000_aa    m/z 701.4644     0   1   2   3E+4   1   1.5  RT (min)      lcqtof_hilic_neg_4_000_aa    m/z 701.4825     0   1   2   3E+4   1   1.5  RT (min)      lcqtof_hilic_neg_4_000_aa    m/z 701.5832     0   2   4   6E+5   1   1.5  RT (min)      lcqtof_hilic_neg_4_000_aa    m/z 701.7154     0   1   2E+4   0.5   1  RT (min)      lcqtof_hilic_neg_3_035_46    m/z 702.2835     0   1E+4   3   3.5  RT (min)      lcqtof_hilic_neg_4_004_47    m/z 702.3424     0   2   4E+4   1   1.5  RT (min)      lcqtof_hilic_neg_4_000_aa    m/z 702.3427     0   2   4E+4   1   1.5  RT (min)      lcqtof_hilic_neg_4_000_aa    m/z 702.3435     0   2   4E+4   1   1.5  RT (min)      lcqtof_hilic_neg_4_000_aa    m/z 702.3440     0   2   4E+4   1   1.5  RT (min)      lcqtof_hilic_neg_4_000_aa    m/z 702.3444     0   2   4E+4   1   1.5  RT (min)      lcqtof_hilic_neg_4_000_aa    m/z 702.3896     0   1   2E+4   15   15.5  RT (min)      lcqtof_hilic_neg_4_000_aa    m/z 702.3910     0   1E+4   0.5   1  RT (min)      lcqtof_hilic_neg_4_047_06    m/z 702.4055     0   1E+4   0.5   1  RT (min)      lcqtof_hilic_neg_4_038_10    m/z 702.4394     0   1   2   3E+4   1.5   2  RT (min)      lcqtof_hilic_neg_4_000_aa    m/z 702.4467     0   1E+4   3.5   4  RT (min)      lcqtof_hilic_neg_4_040_05    m/z 702.5076     0   2   4   6E+3   2   2.5  RT (min)      lcqtof_hilic_neg_4_028_36    m/z 702.5400     0   1   2E+4   0.5   1  RT (min)      lcqtof_hilic_neg_2_028_05    m/z 702.5862     0   1   2   3E+5   1   1.5  RT (min)      lcqtof_hilic_neg_4_000_aa    m/z 702.8466     0   1E+4   7.5   8  RT (min)      lcqtof_hilic_neg_1_000_aa    m/z 702.8468     0   1E+4   7.5   8  RT (min)      lcqtof_hilic_neg_1_000_aa    m/z 702.8476     0   1E+4   7.5   8  RT (min)      lcqtof_hilic_neg_1_000_aa    m/z 703.0853     0   1   2   3E+4   1   1.5  RT (min)       lcqtof_hilic_neg_4_103_61    m/z 703.2976     0   1E+4   0.5   1  RT (min)      lcqtof_hilic_neg_4_052_13    m/z 703.3380     0   1   2   3   4E+4   1   1.5  RT (min)      lcqtof_hilic_neg_4_000_aa    m/z 703.3874     0   1E+4   15   15.5  RT (min)      lcqtof_hilic_neg_4_000_aa    m/z 703.4288     0   1E+4   0.5   1  RT (min)      lcqtof_hilic_neg_4_005_29    m/z 703.4397     0   1E+4   1.5   2  RT (min)      lcqtof_hilic_neg_4_000_aa    m/z 703.5182     0   1   2   3E+4   3   3.5  RT (min)       lcqtof_hilic_neg_1_014_31    m/z 703.5187     0   1E+4   4   4.5  RT (min)      lcqtof_hilic_neg_4_008_24    m/z 703.5888     0   2   4   6E+4   1   1.5  RT (min)      lcqtof_hilic_neg_4_000_aa    m/z 703.5901     0   2   4   6E+4   1.5   2  RT (min)      lcqtof_hilic_neg_4_000_aa    m/z 703.6617     0   1E+4   1   1.5  RT (min)      lcqtof_hilic_neg_4_000_aa    m/z 704.2558     0   1   2E+3   8.5   9  RT (min)       lcqtof_hilic_neg_4_000_bb    m/z 704.2561     0   1   2   3E+4   5   5.5  RT (min)      lcqtof_hilic_neg_4_000_cc    m/z 704.2562     0   2   4   6   8E+3   5.5   6  RT (min)       lcqtof_hilic_neg_4_012_07    m/z 704.2568     0   1   2   3E+3   7.5   8  RT (min)      lcqtof_hilic_neg_4_004_47    m/z 704.2569     0   1   2   3   4E+4   5.5   6  RT (min)      lcqtof_hilic_neg_4_000_bb    m/z 704.2571     0   1   2   3   4E+3   6   6.5  RT (min)       lcqtof_hilic_neg_4_068_57    m/z 704.2578     0   1   2   3E+3   8   8.5  RT (min)       lcqtof_hilic_neg_4_040_05    m/z 704.2579     0   1   2   3E+4   5   5.5  RT (min)      lcqtof_hilic_neg_4_043_12    m/z 704.3356     0   1E+4   1   1.5  RT (min)      lcqtof_hilic_neg_4_000_aa    m/z 704.3561     0   1E+4   4.5   5  RT (min)      lcqtof_hilic_neg_4_050_02    m/z 704.5170     0   1E+4   0.5   1  RT (min)      lcqtof_hilic_neg_4_053_14    m/z 704.5930     0   1E+4   1.5   2  RT (min)      lcqtof_hilic_neg_4_000_aa    m/z 704.6674     0   1E+4   0.5   1  RT (min)      lcqtof_hilic_neg_1_037_24    m/z 704.8894     0   1E+4   1   1.5  RT (min)       lcqtof_hilic_neg_4_054_48    m/z 705.3894     0   1E+4   0.5   1  RT (min)      lcqtof_hilic_neg_4_023_15    m/z 705.5256     0   1E+4   3   3.5  RT (min)      lcqtof_hilic_neg_4_008_24    m/z 705.5263     0   1E+4   3   3.5  RT (min)      lcqtof_hilic_neg_4_008_24    m/z 705.5506     0   1   2E+4   1.5   2  RT (min)      lcqtof_hilic_neg_4_000_aa    m/z 705.5512     0   1   2E+4   1.5   2  RT (min)      lcqtof_hilic_neg_4_000_aa    m/z 705.5516     0   1   2E+4   1   1.5  RT (min)      lcqtof_hilic_neg_4_000_aa    m/z 705.5731     0   1   2E+4   1.5   2  RT (min)      lcqtof_hilic_neg_4_000_aa    m/z 705.5992     0   1E+4   0.5   1  RT (min)      lcqtof_hilic_neg_2_028_05    m/z 705.6538     0   1   2   3   4E+4   1   1.5  RT (min)      lcqtof_hilic_neg_4_000_aa    m/z 705.6695     0   2   4E+4   1   1.5  RT (min)      lcqtof_hilic_neg_4_000_aa    m/z 705.6740     0   2   4E+4   1   1.5  RT (min)      lcqtof_hilic_neg_4_000_aa    m/z 705.6803     0   2   4E+4   1   1.5  RT (min)      lcqtof_hilic_neg_4_000_aa    m/z 706.0517     0   2   4   6   8E+4   7.5   8  RT (min)      lcqtof_hilic_neg_1_005_32    m/z 706.3407     0   1E+4   4.5   5  RT (min)       lcqtof_hilic_neg_4_008_24    m/z 706.3534     0   1E+4   4.5   5  RT (min)      lcqtof_hilic_neg_4_050_02    m/z 706.3934     0   1   2   3   4E+4   4   4.5  RT (min)       lcqtof_hilic_neg_4_048_23    m/z 706.5572     0   1E+4   1   1.5  RT (min)      lcqtof_hilic_neg_4_000_aa    m/z 706.6632     0   1   2E+4   1   1.5  RT (min)      lcqtof_hilic_neg_4_000_aa    m/z 706.6790     0   1   2E+4   1   1.5  RT (min)      lcqtof_hilic_neg_4_000_aa    m/z 706.6795     0   1   2E+4   1   1.5   2  RT (min)      lcqtof_hilic_neg_4_000_aa    m/z 706.6821     0   1   2   3E+4   0.5   1  RT (min)      lcqtof_hilic_neg_4_040_05    m/z 707.0016     0   1E+4   7   7.5  RT (min)      lcqtof_hilic_neg_1_054_01    m/z 707.3960     0   1E+4   4   4.5  RT (min)      lcqtof_hilic_neg_4_048_23    m/z 707.4058     0   1   2   3E+4   1   1.5  RT (min)      lcqtof_hilic_neg_4_000_aa    m/z 707.4774     0   1   2   3E+4   4   4.5  RT (min)      lcqtof_hilic_neg_4_008_24    m/z 707.5861     0   1E+5   4   4.5  RT (min)      lcqtof_hilic_neg_4_008_24    m/z 707.6146     0   2   4   6E+4   1   1.5  RT (min)      lcqtof_hilic_neg_4_000_aa    m/z 707.6194     0   2   4   6E+4   1   1.5  RT (min)      lcqtof_hilic_neg_4_000_aa    m/z 707.6281     0   2   4   6E+4   1   1.5  RT (min)      lcqtof_hilic_neg_4_000_aa    m/z 707.6564     0   1   2   3E+4   1   1.5  RT (min)      lcqtof_hilic_neg_4_000_aa    m/z 707.6791     0   1E+4   1.5   2  RT (min)      lcqtof_hilic_neg_4_000_aa    m/z 707.6851     0   1E+5   0.5   1  RT (min)      lcqtof_hilic_neg_4_040_05    m/z 708.4030     0   1E+4   1   1.5  RT (min)      lcqtof_hilic_neg_4_000_aa    m/z 708.4763     0   1   2E+4   4   4.5  RT (min)      lcqtof_hilic_neg_4_008_24    m/z 708.5896     0   2   4E+5   4.5   5  RT (min)      lcqtof_hilic_neg_4_008_24    m/z 708.5896     0   1   2   3   4E+4   1.5   2  RT (min)      lcqtof_hilic_neg_4_008_24    m/z 708.5899     0   1   2   3E+5   4   4.5  RT (min)       lcqtof_hilic_neg_4_055_32    m/z 708.5906     0   1E+4   1.5   2  RT (min)      lcqtof_hilic_neg_4_008_24    m/z 708.6070     0   1   2   3E+4   1   1.5  RT (min)      lcqtof_hilic_neg_4_000_aa    m/z 708.6505     0   1E+4   1.5   2  RT (min)      lcqtof_hilic_neg_4_000_aa    m/z 708.6548     0   1E+4   1   1.5  RT (min)      lcqtof_hilic_neg_4_000_aa    m/z 708.8516     0   1E+4   0.5   1  RT (min)      lcqtof_hilic_neg_4_004_47    m/z 708.8585     0   1E+4   3   3.5  RT (min)      lcqtof_hilic_neg_1_037_24    m/z 708.9967     0   1E+4   7   7.5  RT (min)      lcqtof_hilic_neg_1_054_01    m/z 709.3476     0   2   4E+4   1   1.5  RT (min)      lcqtof_hilic_neg_4_000_aa    m/z 709.5604     0   1   2   3E+4   15   15.5  RT (min)      lcqtof_hilic_neg_4_013_20    m/z 709.5605     0   1   2   3E+3   16.5   17  RT (min)      lcqtof_hilic_neg_4_001_49    m/z 709.5608     0   1   2   3E+4   14.5   15   15.5  RT (min)       lcqtof_hilic_neg_4_083_59    m/z 709.5608     0   1   2   3E+4   15.5   16  RT (min)      lcqtof_hilic_neg_1_007_05    m/z 709.5612     0   1   2   3E+4   15   15.5  RT (min)      lcqtof_hilic_neg_4_039_18    m/z 709.5612     0   1   2   3E+3   17   17.5  RT (min)      lcqtof_hilic_neg_4_008_24    m/z 709.5612     0   1   2E+3   19.5   20  RT (min)      lcqtof_hilic_neg_4_006_30    m/z 709.5613     0   1   2E+3   19   19.5  RT (min)      lcqtof_hilic_neg_4_009_50    m/z 709.5614     0   1   2E+5   1   1.5  RT (min)      lcqtof_hilic_neg_4_000_aa    m/z 709.5615     0   1   2   3E+3   18.5   19  RT (min)      lcqtof_hilic_neg_4_020_42    m/z 709.5628     0   2   4E+3   2   2.5  RT (min)      lcqtof_hilic_neg_4_032_01    m/z 709.5632     0   1   2   3   4E+3   2   2.5  RT (min)      lcqtof_hilic_neg_4_008_24    m/z 709.5633     0   1   2E+5   1.5   2  RT (min)      lcqtof_hilic_neg_4_000_aa    m/z 709.6446     0   2   4   6E+5   1   1.5  RT (min)      lcqtof_hilic_neg_4_000_aa    m/z 710.2606     0   1E+4   3.5   4  RT (min)      lcqtof_hilic_neg_4_014_46    m/z 710.3604     0   1   2E+4   1   1.5  RT (min)      lcqtof_hilic_neg_4_000_aa    m/z 710.3656     0   1   2E+4   1   1.5  RT (min)      lcqtof_hilic_neg_4_000_aa    m/z 710.4602     0   1   2   3E+4   4   4.5  RT (min)      lcqtof_hilic_neg_4_008_24    m/z 710.4685     0   1   2   3E+4   4   4.5  RT (min)      lcqtof_hilic_neg_4_008_24    m/z 710.4709     0   2   4   6   8E+4   3.5   4  RT (min)      lcqtof_hilic_neg_4_048_23    m/z 710.4726     0   1E+4   1.5   2  RT (min)      lcqtof_hilic_neg_4_082_59    m/z 710.4726     0   1E+4   0.5   1  RT (min)      lcqtof_hilic_neg_4_040_05    m/z 710.4748     0   1E+4   2   2.5  RT (min)      lcqtof_hilic_neg_4_082_59    m/z 710.4751     0   2   4   6E+4   2.5   3  RT (min)       lcqtof_hilic_neg_4_088_59    m/z 710.4760     0   1E+4   1   1.5  RT (min)      lcqtof_hilic_neg_4_011_35    m/z 710.4762     0   1E+4   2.5   3  RT (min)      lcqtof_hilic_neg_4_002_25    m/z 710.5629     0   1   2E+2   14.5   15  RT (min)      lcqtof_hilic_neg_4_083_59    m/z 710.5638     0   1E+4   15.5   16  RT (min)      lcqtof_hilic_neg_1_007_05    m/z 710.5644     0   1E+5   1.5   2  RT (min)      lcqtof_hilic_neg_4_000_aa    m/z 710.5650     0   1E+4   15   15.5  RT (min)      lcqtof_hilic_neg_4_015_09    m/z 710.6043     0   1   2E+5   4.5   5  RT (min)      lcqtof_hilic_neg_4_008_24    m/z 710.6045     0   2   4   6E+4   4   4.5  RT (min)      lcqtof_hilic_neg_4_040_05    m/z 710.6046     0   1   2E+5   4   4.5  RT (min)      lcqtof_hilic_neg_4_008_24    m/z 710.6278     0   1   2   3   4E+4   1.5   2  RT (min)      lcqtof_hilic_neg_4_000_aa    m/z 710.6322     0   1   2   3E+5   1.5   2  RT (min)      lcqtof_hilic_neg_4_000_aa    m/z 710.6637     0   2   4   6E+4   1.5   2  RT (min)      lcqtof_hilic_neg_4_000_aa    m/z 710.9783     0   2   4E+4   7   7.5  RT (min)       lcqtof_hilic_neg_4_059_14    m/z 710.9784     0   2   4   6E+4   6.5   7  RT (min)      lcqtof_hilic_neg_4_023_15    m/z 710.9784     0   2   4   6E+4   6.5   7  RT (min)       lcqtof_hilic_neg_4_023_15    m/z 710.9784     0   2   4E+4   7   7.5  RT (min)       lcqtof_hilic_neg_4_059_14    m/z 710.9785     0   2   4   6E+4   7   7.5  RT (min)       lcqtof_hilic_neg_4_059_14    m/z 710.9786     0   2   4   6E+4   6.5   7  RT (min)       lcqtof_hilic_neg_4_075_58    m/z 710.9790     0   2   4   6E+4   7   7.5  RT (min)       lcqtof_hilic_neg_4_059_14    m/z 710.9792     0   2   4   6E+4   6.5   7  RT (min)      lcqtof_hilic_neg_4_053_14    m/z 710.9969     0   1E+4   7   7.5  RT (min)      lcqtof_hilic_neg_1_054_01    m/z 711.1370     0   1   2E+4   6.5   7  RT (min)       lcqtof_hilic_neg_4_008_24    m/z 711.3673     0   1   2E+4   1   1.5  RT (min)      lcqtof_hilic_neg_4_000_aa    m/z 711.4521     0   1   2   3   4E+4   4   4.5  RT (min)      lcqtof_hilic_neg_4_008_24    m/z 711.4624     0   1   2E+4   1   1.5  RT (min)      lcqtof_hilic_neg_4_000_aa    m/z 711.4633     0   2   4   6E+4   3   3.5  RT (min)      lcqtof_hilic_neg_4_034_31    m/z 711.4640     0   2   4   6   8E+4   2.5   3  RT (min)      lcqtof_hilic_neg_4_048_23    m/z 711.4679     0   1   2E+4   0.5   1  RT (min)      lcqtof_hilic_neg_4_040_05    m/z 711.4719     0   1   2   3   4E+4   2.5   3  RT (min)      lcqtof_hilic_neg_4_088_59    m/z 711.4766     0   1E+4   2   2.5  RT (min)      lcqtof_hilic_neg_4_034_31    m/z 711.4784     0   1E+4   2.5   3  RT (min)      lcqtof_hilic_neg_4_047_06    m/z 711.4789     0   1   2   3E+4   2.5   3  RT (min)      lcqtof_hilic_neg_4_031_38    m/z 711.4804     0   1E+4   1   1.5  RT (min)      lcqtof_hilic_neg_4_048_23    m/z 711.4808     0   1   2   3   4E+4   1   1.5  RT (min)      lcqtof_hilic_neg_4_000_aa    m/z 711.5084     0   1   2E+4   4   4.5  RT (min)      lcqtof_hilic_neg_4_092_60    m/z 711.5199     0   1   2   3E+4   1   1.5  RT (min)      lcqtof_hilic_neg_4_000_aa    m/z 711.5456     0   1E+4   1   1.5  RT (min)      lcqtof_hilic_neg_4_000_aa    m/z 711.5816     0   1   2   3   4E+4   1   1.5  RT (min)      lcqtof_hilic_neg_4_000_aa    m/z 711.6316     0   1   2   3E+4   1.5   2  RT (min)      lcqtof_hilic_neg_4_000_aa    m/z 711.6318     0   1   2   3   4E+4   1.5   2  RT (min)      lcqtof_hilic_neg_4_000_aa    m/z 711.9814     0   1   2E+4   6.5   7  RT (min)      lcqtof_hilic_neg_4_064_15    m/z 711.9818     0   1   2E+4   6.5   7  RT (min)      lcqtof_hilic_neg_4_064_15    m/z 711.9819     0   1   2E+4   6.5   7  RT (min)      lcqtof_hilic_neg_4_059_14    m/z 711.9822     0   1   2E+4   6.5   7  RT (min)      lcqtof_hilic_neg_4_064_15    m/z 712.2730     0   1E+4   3.5   4  RT (min)      lcqtof_hilic_neg_4_014_46    m/z 712.4881     0   1   2   3E+5   3   3.5  RT (min)      lcqtof_hilic_neg_4_034_31    m/z 712.4902     0   1   2   3E+4   1.5   2  RT (min)      lcqtof_hilic_neg_4_022_34    m/z 712.4908     0   1   2   3E+4   1.5   2  RT (min)      lcqtof_hilic_neg_4_022_34    m/z 712.4910     0   1   2   3   4E+4   2   2.5  RT (min)      lcqtof_hilic_neg_4_048_23    m/z 712.4911     0   1   2   3   4E+4   2   2.5  RT (min)      lcqtof_hilic_neg_4_048_23    m/z 712.4911     0   1   2   3E+4   1.5   2  RT (min)      lcqtof_hilic_neg_4_022_34    m/z 712.4914     0   1   2   3E+4   2   2.5  RT (min)      lcqtof_hilic_neg_2_029_31    m/z 712.4923     0   2   4   6E+4   2.5   3  RT (min)      lcqtof_hilic_neg_4_048_23    m/z 712.4945     0   1   2   3E+4   1.5   2  RT (min)      lcqtof_hilic_neg_4_022_34    m/z 712.4976     0   2   4   6E+4   2   2.5  RT (min)      lcqtof_hilic_neg_4_048_23    m/z 712.5026     0   1E+4   0.5   1  RT (min)      lcqtof_hilic_neg_4_038_10    m/z 712.6046     0   1E+4   1   1.5  RT (min)      lcqtof_hilic_neg_4_000_aa    m/z 712.6446     0   2   4E+4   1.5   2  RT (min)      lcqtof_hilic_neg_4_000_aa    m/z 712.8397     0   1   2E+4   1   1.5  RT (min)      lcqtof_hilic_neg_4_004_47    m/z 713.2051     0   1E+4   6   6.5  RT (min)       lcqtof_hilic_neg_4_092_60    m/z 713.4161     0   2   4   6   8E+4   1   1.5  RT (min)      lcqtof_hilic_neg_4_000_aa    m/z 713.4437     0   1E+4   3.5   4  RT (min)      lcqtof_hilic_neg_4_032_01    m/z 713.4533     0   1   2E+5   1   1.5  RT (min)      lcqtof_hilic_neg_4_000_aa    m/z 713.4776     0   2   4   6E+5   2.5   3  RT (min)      lcqtof_hilic_neg_4_048_23    m/z 713.4784     0   2   4E+5   3   3.5  RT (min)       lcqtof_hilic_neg_4_034_31    m/z 713.4867     0   1   2E+4   1.5   2  RT (min)      lcqtof_hilic_neg_4_011_35    m/z 713.4904     0   2   4   6   8E+4   3   3.5  RT (min)      lcqtof_hilic_neg_4_033_53    m/z 713.4908     0   1   2E+4   2   2.5  RT (min)      lcqtof_hilic_neg_4_016_37    m/z 713.4914     0   1   2E+5   3   3.5  RT (min)      lcqtof_hilic_neg_1_035_23    m/z 713.4916     0   1E+4   1.5   2  RT (min)      lcqtof_hilic_neg_4_002_25    m/z 713.4919     0   1   2E+5   2.5   3  RT (min)      lcqtof_hilic_neg_2_002_36    m/z 713.4936     0   1   2E+4   2   2.5  RT (min)      lcqtof_hilic_neg_4_016_37    m/z 713.5182     0   1   2E+4   1   1.5  RT (min)      lcqtof_hilic_neg_4_000_aa    m/z 713.5654     0   1E+4   0.5   1  RT (min)      lcqtof_hilic_neg_4_040_05    m/z 713.5752     0   1E+4   1.5   2  RT (min)      lcqtof_hilic_neg_4_000_aa    m/z 713.5892     0   1E+4   1   1.5  RT (min)      lcqtof_hilic_neg_4_000_aa    m/z 713.6196     0   1   2   3   4E+5   1.5   2  RT (min)      lcqtof_hilic_neg_4_000_aa    m/z 713.6441     0   1   2E+4   1.5   2  RT (min)      lcqtof_hilic_neg_4_000_aa    m/z 713.7469     0   1E+4   2.5   3  RT (min)      lcqtof_hilic_neg_4_048_23    m/z 714.0031     0   1E+4   6.5   7  RT (min)       lcqtof_hilic_neg_4_059_14    m/z 714.2911     0   1E+4   3.5   4  RT (min)      lcqtof_hilic_neg_4_014_46    m/z 714.4530     0   1E+5   1   1.5  RT (min)      lcqtof_hilic_neg_4_000_aa    m/z 714.4748     0   1   2E+4   0.5   1  RT (min)      lcqtof_hilic_neg_4_045_26    m/z 714.5064     0   1   2E+5   1.5   2  RT (min)      lcqtof_hilic_neg_4_048_23    m/z 714.5064     0   1   2E+5   2   2.5  RT (min)       lcqtof_hilic_neg_4_048_23    m/z 714.5064     0   1   2E+5   1.5   2  RT (min)      lcqtof_hilic_neg_4_048_23    m/z 714.5064     0   2   4   6   8E+4   1.5   2  RT (min)      lcqtof_hilic_neg_4_040_05    m/z 714.5065     0   1   2E+5   2   2.5  RT (min)       lcqtof_hilic_neg_2_029_31    m/z 714.5066     0   1   2E+5   2   2.5  RT (min)       lcqtof_hilic_neg_4_048_23    m/z 714.5066     0   1   2E+5   1.5   2  RT (min)      lcqtof_hilic_neg_4_048_23    m/z 714.5067     0   1E+6   3.5   4   4.5  RT (min)       lcqtof_hilic_neg_4_089_59    m/z 714.5068     0   1   2E+5   1   1.5   2  RT (min)       lcqtof_hilic_neg_4_011_35    m/z 714.5070     0   1   2E+5   2   2.5  RT (min)      lcqtof_hilic_neg_4_048_23    m/z 714.5070     0   1E+5   2.5   3  RT (min)       lcqtof_hilic_neg_2_020_23    m/z 714.5071     0   2   4   6E+4   4   4.5  RT (min)      lcqtof_hilic_neg_4_048_23    m/z 714.5095     0   2   4   6E+4   1   1.5  RT (min)      lcqtof_hilic_neg_4_054_48    m/z 714.6219     0   1   2E+5   1.5   2  RT (min)      lcqtof_hilic_neg_4_000_aa    m/z 714.6511     0   1E+4   0.5   1  RT (min)      lcqtof_hilic_neg_3_007_04    m/z 715.0502     0   1   2E+4   0.5   1  RT (min)      lcqtof_hilic_neg_3_007_04    m/z 715.4065     0   1   2   3E+4   1   1.5  RT (min)      lcqtof_hilic_neg_4_000_aa    m/z 715.4647     0   1E+5   1   1.5  RT (min)      lcqtof_hilic_neg_4_000_aa    m/z 715.5086     0   1   2E+4   2   2.5  RT (min)      lcqtof_hilic_neg_4_001_49    m/z 715.5092     0   1E+6   3   3.5  RT (min)      lcqtof_hilic_neg_1_014_31    m/z 715.5094     0   1   2   3E+5   3   3.5  RT (min)      lcqtof_hilic_neg_3_048_05    m/z 715.5096     0   2   4   6   8E+4   2   2.5  RT (min)      lcqtof_hilic_neg_4_048_23    m/z 715.5098     0   1E+5   2   2.5  RT (min)      lcqtof_hilic_neg_2_029_31    m/z 715.5099     0   2   4   6   8E+4   1.5   2  RT (min)      lcqtof_hilic_neg_4_011_35    m/z 715.5100     0   2   4   6   8E+4   1.5   2  RT (min)      lcqtof_hilic_neg_4_048_23    m/z 715.5100     0   2   4   6   8E+4   2   2.5  RT (min)      lcqtof_hilic_neg_4_048_23    m/z 715.5100     0   2   4   6E+4   2.5   3  RT (min)      lcqtof_hilic_neg_2_020_23    m/z 715.5102     0   1   2   3E+4   4   4.5  RT (min)      lcqtof_hilic_neg_4_048_23    m/z 715.5237     0   1E+4   0.5   1  RT (min)      lcqtof_hilic_neg_4_000_aa    m/z 715.5964     0   1E+4   0.5   1  RT (min)      lcqtof_hilic_neg_4_040_05    m/z 715.6250     0   1   2E+5   1.5   2  RT (min)      lcqtof_hilic_neg_4_000_aa    m/z 716.0520     0   1E+4   0.5   1  RT (min)      lcqtof_hilic_neg_3_007_04    m/z 716.3424     0   1E+4   0.5   1  RT (min)      lcqtof_hilic_neg_4_023_15    m/z 716.4680     0   1   2   3E+4   1   1.5  RT (min)      lcqtof_hilic_neg_4_000_aa    m/z 716.4696     0   1   2   3E+4   1   1.5  RT (min)      lcqtof_hilic_neg_4_000_aa    m/z 716.5021     0   1   2E+4   3.5   4  RT (min)      lcqtof_hilic_neg_4_034_31    m/z 716.5182     0   1   2   3   4E+5   3.5   4  RT (min)      lcqtof_hilic_neg_4_089_59    m/z 716.5194     0   2   4   6   8E+4   1.5   2  RT (min)      lcqtof_hilic_neg_4_011_35    m/z 716.5194     0   2   4   6   8E+4   1.5   2  RT (min)      lcqtof_hilic_neg_4_011_35    m/z 716.5196     0   2   4   6   8E+4   1.5   2  RT (min)      lcqtof_hilic_neg_4_011_35    m/z 716.5196     0   2   4   6   8E+4   1.5   2  RT (min)      lcqtof_hilic_neg_4_011_35    m/z 716.5196     0   2   4   6   8E+4   1.5   2  RT (min)      lcqtof_hilic_neg_4_011_35    m/z 716.5198     0   2   4   6   8E+4   1.5   2  RT (min)      lcqtof_hilic_neg_4_048_23    m/z 716.5200     0   1E+5   2   2.5  RT (min)      lcqtof_hilic_neg_4_028_36    m/z 716.5200     0   2   4   6   8E+4   2   2.5  RT (min)      lcqtof_hilic_neg_4_048_23    m/z 716.5204     0   2   4   6E+5   3   3.5  RT (min)      lcqtof_hilic_neg_4_034_31    m/z 716.5204     0   2   4   6   8E+4   2   2.5  RT (min)      lcqtof_hilic_neg_4_048_23    m/z 716.5208     0   1E+5   2.5   3  RT (min)      lcqtof_hilic_neg_4_048_23    m/z 716.5210     0   1   2E+4   4   4.5  RT (min)      lcqtof_hilic_neg_4_048_23    m/z 717.1802     0   1E+4   6   6.5  RT (min)       lcqtof_hilic_neg_1_037_24    m/z 717.2723     0   1   2E+4   6   6.5  RT (min)      lcqtof_hilic_neg_4_048_23    m/z 717.4814     0   1   2E+4   3   3.5  RT (min)      lcqtof_hilic_neg_2_028_05    m/z 717.5017     0   1   2   3E+4   3   3.5  RT (min)       lcqtof_hilic_neg_4_082_59    m/z 717.5098     0   1   2E+5   3   3.5  RT (min)      lcqtof_hilic_neg_4_048_23    m/z 717.5142     0   1   2E+5   3   3.5  RT (min)      lcqtof_hilic_neg_4_028_36    m/z 717.5240     0   1   2E+5   2.5   3  RT (min)       lcqtof_hilic_neg_4_048_23    m/z 717.5240     0   1   2E+4   1.5   2  RT (min)      lcqtof_hilic_neg_4_048_23    m/z 717.5240     0   1   2E+4   1.5   2  RT (min)      lcqtof_hilic_neg_4_048_23    m/z 717.5242     0   1   2E+4   2   2.5  RT (min)       lcqtof_hilic_neg_4_048_23    m/z 717.5246     0   1   2   3E+4   2   2.5  RT (min)       lcqtof_hilic_neg_4_028_36    m/z 717.5247     0   1   2E+4   2   2.5  RT (min)      lcqtof_hilic_neg_2_039_38    m/z 717.5250     0   1   2   3E+4   2   2.5  RT (min)       lcqtof_hilic_neg_4_016_37    m/z 717.5251     0   1   2   3E+4   1.5   2  RT (min)       lcqtof_hilic_neg_4_011_35    m/z 717.5251     0   1   2E+4   1   2  RT (min)       lcqtof_hilic_neg_4_048_23    m/z 717.5252     0   1E+4   4   4.5  RT (min)      lcqtof_hilic_neg_4_048_23    m/z 717.5252     0   1   2E+4   2   2.5  RT (min)       lcqtof_hilic_neg_4_048_23    m/z 717.5614     0   1   2   3E+4   1   1.5  RT (min)      lcqtof_hilic_neg_4_000_aa    m/z 717.5888     0   1E+4   0.5   1  RT (min)      lcqtof_hilic_neg_2_028_05    m/z 717.6114     0   1E+5   1   1.5  RT (min)      lcqtof_hilic_neg_4_000_aa    m/z 717.6154     0   1E+4   0.5   1  RT (min)      lcqtof_hilic_neg_2_028_05    m/z 717.7043     0   1   2   3   4E+4   1   1.5  RT (min)      lcqtof_hilic_neg_4_000_aa    m/z 717.7114     0   1   2   3   4E+4   1.5   2  RT (min)      lcqtof_hilic_neg_4_000_aa    m/z 717.7118     0   1   2   3   4E+4   1.5   2  RT (min)      lcqtof_hilic_neg_4_000_aa    m/z 717.7130     0   1   2   3   4E+4   1   1.5  RT (min)      lcqtof_hilic_neg_4_000_aa    m/z 718.3938     0   1   2E+4   4   4.5  RT (min)       lcqtof_hilic_neg_4_048_23    m/z 718.4889     0   1E+4   3   3.5  RT (min)      lcqtof_hilic_neg_4_002_25    m/z 718.5000     0   1E+4   3   3.5  RT (min)      lcqtof_hilic_neg_4_048_23    m/z 718.5716     0   1E+4   0.5   1  RT (min)      lcqtof_hilic_neg_4_008_24    m/z 718.5722     0   2   4   6   8E+4   1   1.5  RT (min)      lcqtof_hilic_neg_4_000_aa    m/z 718.5909     0   1   2   3   4E+4   0.5   1  RT (min)      lcqtof_hilic_neg_2_028_05    m/z 718.6884     0   1   2   3E+5   8   8.5  RT (min)      lcqtof_hilic_neg_1_000_aa    m/z 718.7159     0   1   2E+4   1.5   2  RT (min)      lcqtof_hilic_neg_4_000_aa    m/z 719.0171     0   1E+4   6   6.5  RT (min)      lcqtof_hilic_neg_4_001_49    m/z 719.1488     0   1E+4   0.5   1  RT (min)      lcqtof_hilic_neg_1_062_17    m/z 719.2007     0   1E+4   6   6.5  RT (min)      lcqtof_hilic_neg_3_011_34    m/z 719.2086     0   1E+4   0.5   1  RT (min)      lcqtof_hilic_neg_3_035_46    m/z 719.3401     0   1   2E+4   1   1.5  RT (min)      lcqtof_hilic_neg_4_000_aa    m/z 719.3424     0   1   2E+4   1   1.5  RT (min)      lcqtof_hilic_neg_4_000_aa    m/z 719.3426     0   1   2E+4   1   1.5  RT (min)      lcqtof_hilic_neg_4_000_aa    m/z 719.3446     0   1   2E+4   1   1.5  RT (min)      lcqtof_hilic_neg_4_000_aa    m/z 719.5309     0   2   4   6E+4   1.5   2  RT (min)      lcqtof_hilic_neg_4_000_aa    m/z 719.5321     0   2   4   6   8E+4   1   1.5  RT (min)      lcqtof_hilic_neg_4_000_aa    m/z 719.5322     0   2   4   6   8E+4   1.5   2  RT (min)      lcqtof_hilic_neg_4_000_aa    m/z 719.5334     0   1E+5   1   1.5  RT (min)      lcqtof_hilic_neg_4_000_aa    m/z 719.5664     0   1E+6   1   1.5  RT (min)      lcqtof_hilic_neg_4_000_aa    m/z 719.5739     0   1   2   3   4E+4   0.5   1  RT (min)      lcqtof_hilic_neg_2_028_05    m/z 719.5843     0   1E+6   1   1.5  RT (min)      lcqtof_hilic_neg_4_000_aa    m/z 719.5884     0   1   2   3   4E+4   0.5   1  RT (min)      lcqtof_hilic_neg_2_028_05    m/z 719.6556     0   1   2   3E+4   1   1.5  RT (min)      lcqtof_hilic_neg_4_000_aa    m/z 719.6891     0   1   2   3E+4   1.5   2  RT (min)      lcqtof_hilic_neg_4_000_aa    m/z 719.6950     0   1   2   3E+4   1.5   2  RT (min)      lcqtof_hilic_neg_4_000_aa    m/z 719.7160     0   1E+4   1.5   2  RT (min)      lcqtof_hilic_neg_4_000_aa    m/z 720.5331     0   1   2   3E+4   1.5   2  RT (min)      lcqtof_hilic_neg_4_000_aa    m/z 720.5336     0   1   2   3E+4   1.5   2  RT (min)      lcqtof_hilic_neg_4_000_aa    m/z 720.5337     0   1   2   3   4E+4   1   1.5  RT (min)      lcqtof_hilic_neg_4_000_aa    m/z 720.5338     0   1   2   3   4E+4   1.5   2  RT (min)      lcqtof_hilic_neg_4_000_aa    m/z 720.5346     0   1   2   3   4E+4   1.5   2  RT (min)      lcqtof_hilic_neg_4_000_aa    m/z 720.5352     0   1   2   3   4E+4   1   1.5  RT (min)      lcqtof_hilic_neg_4_000_aa    m/z 720.5489     0   1   2   3   4E+4   1   1.5  RT (min)      lcqtof_hilic_neg_4_000_aa    m/z 720.6584     0   1E+4   1   1.5  RT (min)      lcqtof_hilic_neg_4_000_aa    m/z 720.6916     0   1E+4   8   8.5  RT (min)      lcqtof_hilic_neg_2_000_aa    m/z 720.6926     0   1E+4   8   8.5  RT (min)      lcqtof_hilic_neg_1_000_aa    m/z 721.0122     0   1E+4   7   7.5  RT (min)      lcqtof_hilic_neg_4_016_37    m/z 721.2138     0   1E+4   3.5   4  RT (min)      lcqtof_hilic_neg_4_013_20    m/z 721.3802     0   1E+4   1   1.5  RT (min)      lcqtof_hilic_neg_4_013_20    m/z 721.3806     0   1E+4   1   1.5  RT (min)      lcqtof_hilic_neg_4_006_30    m/z 721.4154     0   1E+4   1   1.5  RT (min)      lcqtof_hilic_neg_4_000_aa    m/z 721.5170     0   1E+4   4   4.5  RT (min)      lcqtof_hilic_neg_4_055_32    m/z 721.5338     0   1   2E+4   1.5   2  RT (min)      lcqtof_hilic_neg_4_000_aa    m/z 721.5482     0   1   2E+4   1   1.5  RT (min)      lcqtof_hilic_neg_4_000_aa    m/z 721.5497     0   1   2E+4   1.5   2  RT (min)      lcqtof_hilic_neg_4_000_aa    m/z 721.5848     0   1   2   3   4E+5   1   1.5  RT (min)      lcqtof_hilic_neg_4_000_aa    m/z 721.5930     0   1E+4   3   3.5  RT (min)      lcqtof_hilic_neg_4_008_24    m/z 721.5935     0   1E+4   3   3.5  RT (min)      lcqtof_hilic_neg_4_008_24    m/z 721.6559     0   1   2   3E+4   1   1.5  RT (min)      lcqtof_hilic_neg_4_000_aa    m/z 721.6618     0   1   2   3E+4   1   1.5  RT (min)      lcqtof_hilic_neg_4_000_aa    m/z 721.6628     0   1   2   3E+4   1   1.5  RT (min)      lcqtof_hilic_neg_4_000_aa    m/z 721.7021     0   1   2   3E+4   0.5   1  RT (min)      lcqtof_hilic_neg_4_040_05    m/z 722.3046     0   1E+4   4.5   5  RT (min)      lcqtof_hilic_neg_3_048_05    m/z 722.3742     0   1E+4   3.5   4  RT (min)      lcqtof_hilic_neg_4_050_02    m/z 722.4751     0   1   2   3   4E+3   1.5   2  RT (min)       lcqtof_hilic_neg_4_034_31    m/z 722.4962     0   1   2   3E+4   4   4.5  RT (min)      lcqtof_hilic_neg_4_008_24    m/z 722.5094     0   2   4E+4   4   4.5  RT (min)      lcqtof_hilic_neg_4_055_32    m/z 722.5746     0   1E+4   4   4.5  RT (min)      lcqtof_hilic_neg_4_008_24    m/z 722.5859     0   1E+4   3   3.5  RT (min)      lcqtof_hilic_neg_4_008_24    m/z 722.6640     0   1   2E+4   1   1.5  RT (min)      lcqtof_hilic_neg_4_000_aa    m/z 722.6666     0   1   2E+4   1.5   2  RT (min)      lcqtof_hilic_neg_4_000_aa    m/z 722.7605     0   1E+4   0.5   1  RT (min)      lcqtof_hilic_neg_4_040_05    m/z 722.9666     0   1   2E+3   5.5   6  RT (min)      lcqtof_hilic_neg_2_043_15    m/z 722.9668     0   1   2E+3   5   5.5  RT (min)      lcqtof_hilic_neg_2_051_14    m/z 722.9694     0   2   4E+3   3.5   4  RT (min)      lcqtof_hilic_neg_1_003_37    m/z 722.9694     0   1   2   3   4E+3   3.5   4  RT (min)      lcqtof_hilic_neg_4_011_35    m/z 723.0554     0   1   2   3E+4   7.5   8  RT (min)      lcqtof_hilic_neg_2_047_32    m/z 723.4986     0   1   2   3E+4   4   4.5  RT (min)      lcqtof_hilic_neg_4_008_24    m/z 723.5762     0   1   2   3E+4   15   15.5  RT (min)      lcqtof_hilic_neg_4_013_20    m/z 723.5765     0   1   2E+3   18.5   19  RT (min)      lcqtof_hilic_neg_4_006_30    m/z 723.5766     0   1   2E+4   15.5   16  RT (min)      lcqtof_hilic_neg_1_007_05    m/z 723.5766     0   1   2   3E+4   14.5   15   15.5  RT (min)       lcqtof_hilic_neg_4_083_59    m/z 723.5772     0   1E+4   0   0.5  RT (min)      lcqtof_hilic_neg_4_002_25    m/z 723.5774     0   1   2E+3   19   19.5  RT (min)      lcqtof_hilic_neg_4_017_51    m/z 723.5782     0   1   2E+5   1   1.5  RT (min)      lcqtof_hilic_neg_4_000_aa    m/z 723.5792     0   1   2E+5   1.5   2  RT (min)      lcqtof_hilic_neg_4_000_aa    m/z 723.6562     0   1E+4   1   1.5  RT (min)      lcqtof_hilic_neg_4_000_aa    m/z 724.1129     0   1E+4   6   6.5  RT (min)      lcqtof_hilic_neg_4_012_07    m/z 724.3891     0   1E+4   3.5   4  RT (min)      lcqtof_hilic_neg_4_005_29    m/z 724.5256     0   1   2E+4   0.5   1  RT (min)      lcqtof_hilic_neg_4_040_05    m/z 724.5798     0   1E+4   15   15.5  RT (min)      lcqtof_hilic_neg_4_112_62    m/z 724.5800     0   1E+5   1.5   2  RT (min)      lcqtof_hilic_neg_4_000_aa    m/z 724.5802     0   1E+4   16   16.5  RT (min)      lcqtof_hilic_neg_1_007_05    m/z 724.5802     0   1E+2   14.5   15  RT (min)      lcqtof_hilic_neg_4_082_59    m/z 724.5811     0   1E+4   15   15.5  RT (min)      lcqtof_hilic_neg_4_013_20    m/z 724.5851     0   1E+4   4   4.5  RT (min)      lcqtof_hilic_neg_4_008_24    m/z 724.5926     0   1E+4   3   3.5  RT (min)      lcqtof_hilic_neg_4_008_24    m/z 724.5977     0   1   2E+4   3   3.5  RT (min)      lcqtof_hilic_neg_4_008_24    m/z 724.6255     0   1   2   3E+4   1   1.5  RT (min)      lcqtof_hilic_neg_4_000_aa    m/z 724.6432     0   1   2E+4   1.5   2  RT (min)      lcqtof_hilic_neg_4_000_aa    m/z 724.6484     0   1   2E+4   1.5   2  RT (min)      lcqtof_hilic_neg_4_000_aa    m/z 724.6744     0   1   2E+4   0.5   1  RT (min)      lcqtof_hilic_neg_4_040_05    m/z 724.6800     0   1   2   3E+4   1.5   2  RT (min)      lcqtof_hilic_neg_4_000_aa    m/z 724.8724     0   1   2   3   4E+4   5.5   6  RT (min)      lcqtof_hilic_neg_4_012_07    m/z 725.1966     0   1E+4   1   1.5  RT (min)      lcqtof_hilic_neg_4_000_aa    m/z 725.2000     0   1E+4   1   1.5  RT (min)      lcqtof_hilic_neg_4_000_aa    m/z 725.2936     0   2   4E+3   5   5.5  RT (min)       lcqtof_hilic_neg_4_012_07    m/z 725.3812     0   1E+4   0.5   1  RT (min)      lcqtof_hilic_neg_4_050_02    m/z 725.4698     0   2   4   6   8E+4   1.5   2  RT (min)      lcqtof_hilic_neg_4_000_aa    m/z 725.4700     0   2   4   6   8E+4   1   1.5  RT (min)      lcqtof_hilic_neg_4_000_aa    m/z 725.4702     0   2   4   6   8E+4   1   1.5  RT (min)      lcqtof_hilic_neg_4_000_aa    m/z 725.4705     0   2   4   6   8E+4   1.5   2  RT (min)      lcqtof_hilic_neg_4_000_aa    m/z 725.4720     0   2   4   6   8E+4   1   1.5  RT (min)      lcqtof_hilic_neg_4_000_aa    m/z 725.4724     0   2   4   6   8E+4   1   1.5  RT (min)      lcqtof_hilic_neg_4_000_aa    m/z 725.4774     0   2   4   6   8E+4   1   1.5  RT (min)      lcqtof_hilic_neg_4_000_aa    m/z 725.5224     0   1   2   3E+4   1   1.5  RT (min)      lcqtof_hilic_neg_4_000_aa    m/z 725.5226     0   1   2   3E+3   15.5   16  RT (min)       lcqtof_hilic_neg_4_000_dd    m/z 725.5231     0   1   2   3E+4   1   1.5  RT (min)      lcqtof_hilic_neg_4_000_aa    m/z 725.5495     0   1   2   3E+4   1   1.5  RT (min)      lcqtof_hilic_neg_4_000_aa    m/z 725.5825     0   1   2   3E+4   1.5   2  RT (min)      lcqtof_hilic_neg_4_000_aa    m/z 725.6178     0   1E+5   3   3.5  RT (min)      lcqtof_hilic_neg_4_008_24    m/z 725.6200     0   1E+5   3   3.5  RT (min)       lcqtof_hilic_neg_4_008_24    m/z 725.6201     0   1E+5   3   3.5  RT (min)      lcqtof_hilic_neg_4_008_24    m/z 725.6215     0   1E+5   3   3.5  RT (min)      lcqtof_hilic_neg_1_005_32    m/z 725.6461     0   1E+5   1.5   2  RT (min)      lcqtof_hilic_neg_4_000_aa    m/z 725.6502     0   1   2E+6   1   1.5  RT (min)      lcqtof_hilic_neg_4_000_aa    m/z 725.6530     0   1   2   3   4E+4   0.5   1  RT (min)      lcqtof_hilic_neg_1_007_05    m/z 725.6837     0   1E+4   1.5   2  RT (min)      lcqtof_hilic_neg_4_000_aa    m/z 725.6858     0   1E+4   0.5   1  RT (min)      lcqtof_hilic_neg_3_007_04    m/z 725.9138     0   1   2E+4   1   1.5  RT (min)      lcqtof_hilic_neg_4_000_aa    m/z 726.0312     0   1E+4   7.5   8  RT (min)       lcqtof_hilic_neg_4_082_59    m/z 726.1968     0   1E+4   1   1.5  RT (min)      lcqtof_hilic_neg_4_000_aa    m/z 726.3716     0   1E+4   0.5   1  RT (min)      lcqtof_hilic_neg_4_050_02    m/z 726.4547     0   1E+4   3.5   4  RT (min)      lcqtof_hilic_neg_4_032_01    m/z 726.4588     0   1E+4   4   4.5  RT (min)      lcqtof_hilic_neg_4_008_24    m/z 726.4940     0   1   2   3E+4   1   1.5  RT (min)      lcqtof_hilic_neg_4_000_aa    m/z 726.5086     0   2   4   6   8E+3   1.5   2  RT (min)      lcqtof_hilic_neg_4_082_59    m/z 726.5550     0   1E+4   1   1.5  RT (min)      lcqtof_hilic_neg_4_000_aa    m/z 726.6240     0   1E+5   1   1.5  RT (min)      lcqtof_hilic_neg_4_000_aa    m/z 726.6275     0   1E+6   1.5   2  RT (min)      lcqtof_hilic_neg_4_000_aa    m/z 726.6335     0   1E+6   1   1.5  RT (min)      lcqtof_hilic_neg_4_000_aa    m/z 726.6601     0   1   2   3E+4   1.5   2  RT (min)      lcqtof_hilic_neg_4_000_aa    m/z 726.7994     0   1E+4   7.5   8  RT (min)      lcqtof_hilic_neg_1_000_aa    m/z 726.8010     0   1E+4   7.5   8  RT (min)      lcqtof_hilic_neg_1_000_aa    m/z 726.8014     0   1E+4   7.5   8  RT (min)      lcqtof_hilic_neg_1_000_aa    m/z 726.8026     0   1E+4   7.5   8  RT (min)      lcqtof_hilic_neg_1_000_aa    m/z 726.8552     0   2   4   6E+4   1   1.5  RT (min)      lcqtof_hilic_neg_4_004_47    m/z 726.8784     0   1E+4   0.5   1  RT (min)      lcqtof_hilic_neg_3_035_46    m/z 726.9579     0   1E+4   0.5   1  RT (min)       lcqtof_hilic_neg_4_099_61    m/z 727.1692     0   1E+4   0.5   1  RT (min)      lcqtof_hilic_neg_4_024_40    m/z 727.3272     0   1E+4   15   15.5  RT (min)      lcqtof_hilic_neg_4_000_aa    m/z 727.4563     0   1   2   3   4E+4   1   1.5  RT (min)      lcqtof_hilic_neg_4_000_aa    m/z 727.4650     0   1   2   3   4E+4   1   1.5  RT (min)      lcqtof_hilic_neg_4_000_aa    m/z 727.4654     0   1   2   3   4E+4   1   1.5  RT (min)      lcqtof_hilic_neg_4_000_aa    m/z 727.4672     0   1   2   3   4E+4   1   1.5  RT (min)      lcqtof_hilic_neg_4_000_aa    m/z 727.4684     0   1   2   3   4E+4   1   1.5  RT (min)      lcqtof_hilic_neg_4_000_aa    m/z 727.4707     0   1   2   3   4E+4   1.5   2  RT (min)      lcqtof_hilic_neg_4_000_aa    m/z 727.4916     0   1   2   3   4E+4   2.5   3  RT (min)      lcqtof_hilic_neg_4_048_23    m/z 727.5121     0   1   2   3E+4   3   3.5  RT (min)      lcqtof_hilic_neg_4_028_36    m/z 727.5279     0   1   2   3E+4   1   1.5  RT (min)      lcqtof_hilic_neg_4_000_aa    m/z 727.5388     0   1   2   3E+4   1   1.5  RT (min)      lcqtof_hilic_neg_4_000_aa    m/z 727.5492     0   1E+4   2   2.5  RT (min)      lcqtof_hilic_neg_4_016_37    m/z 727.5989     0   1E+5   1   1.5  RT (min)      lcqtof_hilic_neg_4_000_aa    m/z 727.5992     0   1E+5   1   1.5  RT (min)      lcqtof_hilic_neg_4_000_aa    m/z 727.6282     0   1   2   3E+5   1   1.5  RT (min)      lcqtof_hilic_neg_4_000_aa    m/z 727.6635     0   1E+4   1.5   2  RT (min)      lcqtof_hilic_neg_4_000_aa    m/z 728.4462     0   1E+4   1   1.5  RT (min)      lcqtof_hilic_neg_4_000_aa    m/z 728.4521     0   1E+4   1   1.5  RT (min)      lcqtof_hilic_neg_4_000_aa    m/z 728.5097     0   1   2E+5   2.5   3  RT (min)      lcqtof_hilic_neg_4_048_23    m/z 728.5217     0   1   2E+4   1.5   2  RT (min)      lcqtof_hilic_neg_4_011_35    m/z 728.5221     0   1   2E+4   2   2.5  RT (min)       lcqtof_hilic_neg_4_048_23    m/z 728.5222     0   1   2E+4   1.5   2  RT (min)      lcqtof_hilic_neg_4_011_35    m/z 728.5223     0   1   2E+5   3   3.5  RT (min)       lcqtof_hilic_neg_4_048_23    m/z 728.5224     0   1   2E+4   2   2.5  RT (min)      lcqtof_hilic_neg_4_048_23    m/z 728.5225     0   2   4E+4   3   3.5  RT (min)      lcqtof_hilic_neg_4_033_53    m/z 728.5227     0   1   2E+4   1.5   2  RT (min)       lcqtof_hilic_neg_4_011_35    m/z 728.5229     0   1   2   3E+4   2   2.5  RT (min)      lcqtof_hilic_neg_4_028_36    m/z 728.5229     0   1   2E+4   2   2.5  RT (min)      lcqtof_hilic_neg_2_020_23    m/z 728.5231     0   1   2E+5   2.5   3  RT (min)      lcqtof_hilic_neg_4_048_23    m/z 728.5232     0   1E+4   1.5   2  RT (min)       lcqtof_hilic_neg_2_020_23    m/z 728.5235     0   1E+4   1.5   2  RT (min)      lcqtof_hilic_neg_2_020_23    m/z 728.5246     0   1   2E+4   2   2.5  RT (min)      lcqtof_hilic_neg_4_048_23    m/z 728.5266     0   1E+4   1   1.5  RT (min)      lcqtof_hilic_neg_4_000_aa    m/z 728.6031     0   1   2   3   4E+4   1   1.5  RT (min)      lcqtof_hilic_neg_4_000_aa    m/z 728.6381     0   2   4   6E+4   1.5   2  RT (min)      lcqtof_hilic_neg_4_000_aa    m/z 728.6400     0   1E+5   1   1.5  RT (min)      lcqtof_hilic_neg_4_002_25    m/z 728.6661     0   1E+5   0.5   1  RT (min)      lcqtof_hilic_neg_3_035_46    m/z 728.8772     0   1E+4   0.5   1  RT (min)      lcqtof_hilic_neg_4_007_45    m/z 729.0516     0   1E+4   6   6.5  RT (min)       lcqtof_hilic_neg_4_048_23    m/z 729.0524     0   1E+4   6   6.5  RT (min)      lcqtof_hilic_neg_4_048_23    m/z 729.0904     0   1   2   3E+4   0.5   1  RT (min)      lcqtof_hilic_neg_1_062_17    m/z 729.1296     0   1E+4   6   6.5  RT (min)      lcqtof_hilic_neg_4_012_07    m/z 729.1771     0   1   2   3E+4   1   1.5  RT (min)      lcqtof_hilic_neg_4_000_aa    m/z 729.1788     0   1   2   3E+4   1   1.5  RT (min)      lcqtof_hilic_neg_4_000_aa    m/z 729.2298     0   1E+5   6   6.5  RT (min)      lcqtof_hilic_neg_3_011_34    m/z 729.4757     0   1   2E+4   3   3.5  RT (min)      lcqtof_hilic_neg_4_011_35    m/z 729.4776     0   1   2E+4   1   1.5  RT (min)      lcqtof_hilic_neg_4_000_aa    m/z 729.5262     0   1E+4   2   2.5  RT (min)      lcqtof_hilic_neg_4_028_36    m/z 729.6020     0   1   2   3   4E+4   0.5   1  RT (min)      lcqtof_hilic_neg_1_007_05    m/z 729.6145     0   2   4E+6   1   1.5  RT (min)      lcqtof_hilic_neg_4_000_aa    m/z 729.6172     0   2   4E+6   1   1.5  RT (min)      lcqtof_hilic_neg_4_000_aa    m/z 729.6413     0   1   2   3   4E+4   1.5   2  RT (min)      lcqtof_hilic_neg_4_000_aa    m/z 729.6656     0   1   2   3   4E+4   0.5   1  RT (min)      lcqtof_hilic_neg_3_035_46    m/z 729.6748     0   1E+4   2   2.5  RT (min)      lcqtof_hilic_neg_4_000_aa    m/z 729.6760     0   1E+4   2   2.5  RT (min)      lcqtof_hilic_neg_4_000_aa    m/z 729.7497     0   1   2   3E+4   1   1.5  RT (min)      lcqtof_hilic_neg_4_000_aa    m/z 729.8866     0   1E+5   1   1.5  RT (min)      lcqtof_hilic_neg_4_000_aa    m/z 730.0142     0   1E+4   1   1.5  RT (min)      lcqtof_hilic_neg_4_000_aa    m/z 730.0880     0   1E+4   0.5   1  RT (min)      lcqtof_hilic_neg_1_062_17    m/z 730.0984     0   1   2E+4   1   1.5  RT (min)      lcqtof_hilic_neg_4_000_aa    m/z 730.1766     0   1   2E+4   1   1.5  RT (min)      lcqtof_hilic_neg_4_000_aa    m/z 730.2334     0   1   2   3   4E+4   6   6.5  RT (min)      lcqtof_hilic_neg_3_011_34    m/z 730.3194     0   1E+4   5.5   6  RT (min)      lcqtof_hilic_neg_4_062_17    m/z 730.4797     0   1E+4   1   1.5  RT (min)      lcqtof_hilic_neg_4_040_05    m/z 730.4867     0   1   2E+4   4   4.5  RT (min)      lcqtof_hilic_neg_4_040_05    m/z 730.5032     0   1   2   3E+4   4   4.5  RT (min)      lcqtof_hilic_neg_4_008_24    m/z 730.5067     0   1E+4   3.5   4  RT (min)      lcqtof_hilic_neg_4_016_37    m/z 730.5348     0   2   4   6E+3   1.5   2  RT (min)      lcqtof_hilic_neg_4_016_37    m/z 730.5355     0   1E+4   2   2.5  RT (min)      lcqtof_hilic_neg_4_034_31    m/z 730.5367     0   2   4   6   8E+3   2   2.5  RT (min)      lcqtof_hilic_neg_1_035_23    m/z 730.5635     0   1E+4   3   3.5  RT (min)      lcqtof_hilic_neg_1_005_32    m/z 730.6170     0   1   2E+6   1   1.5  RT (min)      lcqtof_hilic_neg_4_000_aa    m/z 730.6263     0   2   4E+4   0.5   1  RT (min)      lcqtof_hilic_neg_1_007_05    m/z 730.6693     0   1E+4   0.5   1  RT (min)      lcqtof_hilic_neg_3_035_46    m/z 730.8914     0   1E+4   1   1.5  RT (min)      lcqtof_hilic_neg_4_000_aa    m/z 730.9783     0   1E+4   7   7.5  RT (min)      lcqtof_hilic_neg_2_007_03    m/z 731.1766     0   1   2E+4   1   1.5  RT (min)      lcqtof_hilic_neg_4_000_aa    m/z 731.2344     0   1E+4   6   6.5  RT (min)      lcqtof_hilic_neg_3_011_34    m/z 731.4634     0   1   2   3E+4   1   1.5  RT (min)      lcqtof_hilic_neg_4_000_aa    m/z 731.4635     0   1   2   3E+4   1   1.5  RT (min)      lcqtof_hilic_neg_4_000_aa    m/z 731.4640     0   1   2   3E+4   1   1.5  RT (min)      lcqtof_hilic_neg_4_000_aa    m/z 731.4750     0   1   2   3E+4   1   1.5  RT (min)      lcqtof_hilic_neg_4_000_aa    m/z 731.4918     0   1   2   3   4E+4   3   3.5  RT (min)      lcqtof_hilic_neg_4_031_38    m/z 731.4922     0   1E+4   3   3.5  RT (min)      lcqtof_hilic_neg_4_002_25    m/z 731.5664     0   1E+4   1   1.5  RT (min)      lcqtof_hilic_neg_1_005_32    m/z 731.5946     0   1E+4   1   1.5  RT (min)      lcqtof_hilic_neg_4_000_aa    m/z 731.6000     0   1   2   3E+4   0.5   1  RT (min)      lcqtof_hilic_neg_2_028_05    m/z 731.6192     0   2   4   6E+5   1   1.5  RT (min)      lcqtof_hilic_neg_4_000_aa    m/z 731.6979     0   1E+4   1   1.5   2  RT (min)      lcqtof_hilic_neg_4_000_aa    m/z 732.1781     0   1E+4   1   1.5  RT (min)      lcqtof_hilic_neg_4_000_aa    m/z 732.2964     0   1   2   3   4E+3   5   5.5  RT (min)       lcqtof_hilic_neg_4_047_06    m/z 732.4328     0   1E+4   1   1.5  RT (min)      lcqtof_hilic_neg_4_040_05    m/z 732.4345     0   1E+4   1   1.5  RT (min)      lcqtof_hilic_neg_4_040_05    m/z 732.4892     0   1   2E+4   2.5   3  RT (min)       lcqtof_hilic_neg_4_048_23    m/z 732.4912     0   1E+4   3   3.5  RT (min)      lcqtof_hilic_neg_4_034_31    m/z 732.9762     0   1E+4   7   7.5  RT (min)      lcqtof_hilic_neg_2_007_03    m/z 733.3571     0   1   2   3   4E+4   1   1.5  RT (min)      lcqtof_hilic_neg_4_000_aa    m/z 733.3579     0   1   2   3   4E+4   1   1.5  RT (min)      lcqtof_hilic_neg_4_000_aa    m/z 733.3582     0   1   2   3   4E+4   1   1.5  RT (min)      lcqtof_hilic_neg_4_000_aa    m/z 733.3598     0   1   2   3   4E+4   1   1.5  RT (min)      lcqtof_hilic_neg_4_000_aa    m/z 733.3618     0   1   2   3   4E+4   1   1.5  RT (min)      lcqtof_hilic_neg_4_000_aa    m/z 733.4067     0   1E+4   1.5   2  RT (min)      lcqtof_hilic_neg_4_000_aa    m/z 733.5740     0   1E+5   1   1.5  RT (min)      lcqtof_hilic_neg_4_000_aa    m/z 733.5808     0   1E+5   1   1.5  RT (min)      lcqtof_hilic_neg_4_000_aa    m/z 733.5820     0   1E+5   1.5   2  RT (min)      lcqtof_hilic_neg_4_000_aa    m/z 733.5824     0   1E+5   1   1.5  RT (min)      lcqtof_hilic_neg_4_000_aa    m/z 733.5825     0   1E+5   1.5   2  RT (min)      lcqtof_hilic_neg_4_000_aa    m/z 733.5895     0   1E+5   1   1.5  RT (min)      lcqtof_hilic_neg_4_000_aa    m/z 733.6878     0   1   2E+4   1   1.5  RT (min)      lcqtof_hilic_neg_4_000_aa    m/z 733.6894     0   1   2E+4   1   1.5  RT (min)      lcqtof_hilic_neg_4_000_aa    m/z 733.7005     0   1   2E+4   1   1.5  RT (min)      lcqtof_hilic_neg_4_000_aa    m/z 733.7036     0   1   2E+4   1.5   2  RT (min)      lcqtof_hilic_neg_4_000_aa    m/z 734.2594     0   1E+4   3.5   4  RT (min)      lcqtof_hilic_neg_4_014_46    m/z 734.3616     0   1   2E+4   1   1.5  RT (min)      lcqtof_hilic_neg_4_000_aa    m/z 734.3625     0   1   2E+4   1   1.5  RT (min)      lcqtof_hilic_neg_4_000_aa    m/z 734.3648     0   1   2E+4   1   1.5  RT (min)      lcqtof_hilic_neg_4_000_aa    m/z 734.3706     0   1   2E+4   1   1.5  RT (min)      lcqtof_hilic_neg_4_000_aa    m/z 734.4734     0   1E+4   1.5   2  RT (min)      lcqtof_hilic_neg_2_020_23    m/z 734.4744     0   1E+4   1.5   2  RT (min)      lcqtof_hilic_neg_2_020_23    m/z 734.4750     0   1E+4   1.5   2  RT (min)      lcqtof_hilic_neg_2_020_23    m/z 734.4751     0   1E+4   1.5   2  RT (min)      lcqtof_hilic_neg_2_020_23    m/z 734.4922     0   1E+4   2.5   3  RT (min)      lcqtof_hilic_neg_4_016_37    m/z 734.4932     0   1E+4   2.5   3  RT (min)      lcqtof_hilic_neg_4_016_37    m/z 734.5097     0   1E+4   0.5   1  RT (min)      lcqtof_hilic_neg_4_022_34    m/z 734.5257     0   1E+4   1   1.5  RT (min)      lcqtof_hilic_neg_1_007_05    m/z 734.6060     0   1E+4   1   1.5  RT (min)      lcqtof_hilic_neg_1_035_23    m/z 734.6249     0   2   4   6   8E+3   1   1.5  RT (min)      lcqtof_hilic_neg_4_031_38    m/z 734.6916     0   1E+4   1   1.5  RT (min)      lcqtof_hilic_neg_4_000_aa    m/z 735.2774     0   1E+4   1   1.5  RT (min)      lcqtof_hilic_neg_4_000_aa    m/z 735.2774     0   1E+4   1   1.5  RT (min)      lcqtof_hilic_neg_4_000_aa    m/z 735.3426     0   1E+4   7   7.5  RT (min)      lcqtof_hilic_neg_4_008_24    m/z 735.3578     0   1   2E+4   1   1.5  RT (min)      lcqtof_hilic_neg_4_000_aa    m/z 735.3594     0   1   2E+4   1   1.5  RT (min)      lcqtof_hilic_neg_4_000_aa    m/z 735.3596     0   1   2E+4   1   1.5  RT (min)      lcqtof_hilic_neg_4_000_aa    m/z 735.3604     0   1   2E+4   1   1.5  RT (min)      lcqtof_hilic_neg_4_000_aa    m/z 735.3612     0   1   2E+4   1   1.5  RT (min)      lcqtof_hilic_neg_4_000_aa    m/z 735.4317     0   1E+4   1   1.5  RT (min)      lcqtof_hilic_neg_4_000_aa    m/z 735.4350     0   1E+4   1.5   2  RT (min)      lcqtof_hilic_neg_4_000_aa    m/z 735.4387     0   1E+4   1   1.5  RT (min)      lcqtof_hilic_neg_4_000_aa    m/z 735.4452     0   1E+4   1   1.5  RT (min)      lcqtof_hilic_neg_4_000_aa    m/z 735.4464     0   1E+4   1.5   2  RT (min)      lcqtof_hilic_neg_4_000_aa    m/z 735.4980     0   1   2E+4   1   1.5  RT (min)      lcqtof_hilic_neg_4_000_aa    m/z 735.5008     0   1   2E+4   1   1.5  RT (min)      lcqtof_hilic_neg_4_000_aa    m/z 735.5808     0   2   4   6   8E+5   1   1.5  RT (min)      lcqtof_hilic_neg_4_000_aa    m/z 735.6743     0   1   2   3E+4   1   1.5  RT (min)      lcqtof_hilic_neg_4_000_aa    m/z 735.9853     0   1E+4   6   6.5  RT (min)       lcqtof_hilic_neg_4_011_35    m/z 736.0614     0   1E+4   6   6.5  RT (min)      lcqtof_hilic_neg_4_055_32    m/z 736.1354     0   1E+4   6   6.5  RT (min)      lcqtof_hilic_neg_4_036_11    m/z 736.3538     0   1   2E+4   1   1.5  RT (min)      lcqtof_hilic_neg_4_000_aa    m/z 736.3539     0   1   2E+4   1   1.5  RT (min)      lcqtof_hilic_neg_4_000_aa    m/z 736.3548     0   1   2E+4   1   1.5  RT (min)      lcqtof_hilic_neg_4_000_aa    m/z 736.4906     0   1   2E+6   3   3.5  RT (min)       lcqtof_hilic_neg_4_011_35    m/z 736.4908     0   1   2E+5   1.5   2  RT (min)       lcqtof_hilic_neg_2_020_23    m/z 736.4908     0   1   2E+6   3   3.5  RT (min)       lcqtof_hilic_neg_4_048_23    m/z 736.4909     0   1   2E+5   1.5   2  RT (min)       lcqtof_hilic_neg_2_020_23    m/z 736.4910     0   1   2E+5   2   2.5  RT (min)      lcqtof_hilic_neg_2_020_23    m/z 736.4910     0   1   2E+5   2   2.5  RT (min)       lcqtof_hilic_neg_2_002_36    m/z 736.4910     0   1   2E+5   2   2.5  RT (min)      lcqtof_hilic_neg_4_028_36    m/z 736.4910     0   1   2E+5   2   2.5  RT (min)       lcqtof_hilic_neg_4_011_35    m/z 736.4910     0   1   2E+5   1.5   2  RT (min)       lcqtof_hilic_neg_4_011_35    m/z 736.4911     0   1   2E+5   2   2.5  RT (min)      lcqtof_hilic_neg_2_029_31    m/z 736.4911     0   1   2E+5   1.5   2  RT (min)      lcqtof_hilic_neg_4_016_37    m/z 736.4911     0   1   2E+5   2   2.5  RT (min)       lcqtof_hilic_neg_2_020_23    m/z 736.4912     0   1   2   3   4E+4   4   4.5  RT (min)      lcqtof_hilic_neg_4_048_23    m/z 736.4913     0   1   2E+5   1.5   2  RT (min)      lcqtof_hilic_neg_4_011_35    m/z 736.4914     0   2   4   6E+5   2.5   3  RT (min)      lcqtof_hilic_neg_3_027_47    m/z 736.5273     0   1E+4   3   3.5  RT (min)      lcqtof_hilic_neg_4_055_32    m/z 736.5835     0   1   2   3E+5   1   1.5  RT (min)      lcqtof_hilic_neg_4_000_aa    m/z 736.6755     0   1E+4   1   1.5  RT (min)      lcqtof_hilic_neg_4_000_aa    m/z 737.4250     0   1E+4   1   1.5  RT (min)      lcqtof_hilic_neg_4_000_aa    m/z 737.4300     0   1E+4   1   1.5  RT (min)      lcqtof_hilic_neg_4_000_aa    m/z 737.4938     0   1   2   3E+5   2.5   3  RT (min)      lcqtof_hilic_neg_4_047_06    m/z 737.4939     0   1E+6   3   3.5  RT (min)      lcqtof_hilic_neg_4_048_23    m/z 737.4942     0   2   4   6   8E+4   1   1.5  RT (min)      lcqtof_hilic_neg_4_011_35    m/z 737.4943     0   1E+5   2   2.5  RT (min)      lcqtof_hilic_neg_2_020_23    m/z 737.4943     0   1   2E+5   2.5   3  RT (min)      lcqtof_hilic_neg_4_017_51    m/z 737.4944     0   1E+5   2   2.5  RT (min)      lcqtof_hilic_neg_2_002_36    m/z 737.4944     0   1E+5   2   2.5  RT (min)      lcqtof_hilic_neg_2_020_23    m/z 737.4944     0   2   4   6   8E+4   1.5   2  RT (min)      lcqtof_hilic_neg_4_022_34    m/z 737.4944     0   2   4   6   8E+4   1.5   2  RT (min)      lcqtof_hilic_neg_4_016_37    m/z 737.4944     0   1E+5   1.5   2  RT (min)      lcqtof_hilic_neg_2_020_23    m/z 737.4944     0   2   4   6   8E+4   1.5   2  RT (min)      lcqtof_hilic_neg_4_016_37    m/z 737.4946     0   2   4   6   8E+4   2   2.5  RT (min)      lcqtof_hilic_neg_4_011_35    m/z 737.4948     0   1E+5   2   2.5  RT (min)      lcqtof_hilic_neg_2_020_23    m/z 737.5010     0   1E+4   4   4.5  RT (min)      lcqtof_hilic_neg_4_048_23    m/z 737.5146     0   1E+4   4   4.5  RT (min)      lcqtof_hilic_neg_4_008_24    m/z 737.5879     0   1   2E+3   0   0.5  RT (min)      lcqtof_hilic_neg_4_000_cc    m/z 737.5895     0   1   2E+4   16   16.5  RT (min)      lcqtof_hilic_neg_1_007_05    m/z 737.5896     0   2   4   6   8E+3   17   17.5  RT (min)      lcqtof_hilic_neg_4_001_49    m/z 737.5896     0   2   4E+3   16   16.5  RT (min)      lcqtof_hilic_neg_4_000_cc    m/z 737.5897     0   2   4   6   8E+3   16   16.5  RT (min)      lcqtof_hilic_neg_4_017_51    m/z 737.5898     0   1   2E+3   18.5   19  RT (min)      lcqtof_hilic_neg_4_010_28    m/z 737.5899     0   2   4   6   8E+3   17   17.5  RT (min)      lcqtof_hilic_neg_4_001_49    m/z 737.5901     0   1   2   3E+5   1   1.5  RT (min)      lcqtof_hilic_neg_4_000_aa    m/z 737.5904     0   1   2E+3   17.5   18  RT (min)      lcqtof_hilic_neg_4_018_27    m/z 737.5914     0   1E+4   2   2.5  RT (min)      lcqtof_hilic_neg_4_055_32    m/z 737.5915     0   1   2E+3   19.5   20  RT (min)      lcqtof_hilic_neg_4_002_25    m/z 737.5916     0   1   2   3E+5   1.5   2  RT (min)      lcqtof_hilic_neg_4_000_aa    m/z 737.5917     0   1   2   3E+5   1   1.5  RT (min)      lcqtof_hilic_neg_4_000_aa    m/z 737.5920     0   1   2   3E+4   14.5   15   15.5  RT (min)       lcqtof_hilic_neg_4_083_59    m/z 737.5932     0   1   2   3E+4   15   15.5  RT (min)      lcqtof_hilic_neg_4_027_19    m/z 737.5932     0   1   2   3E+4   15   15.5  RT (min)      lcqtof_hilic_neg_4_039_18    m/z 737.6711     0   1   2   3   4E+4   1   1.5  RT (min)      lcqtof_hilic_neg_4_000_aa    m/z 738.4917     0   1   2E+4   4   4.5  RT (min)      lcqtof_hilic_neg_4_097_60    m/z 738.5052     0   1E+5   2   2.5  RT (min)      lcqtof_hilic_neg_2_020_23    m/z 738.5052     0   1E+5   1.5   2  RT (min)      lcqtof_hilic_neg_4_048_23    m/z 738.5052     0   1E+5   1.5   2  RT (min)       lcqtof_hilic_neg_2_020_23    m/z 738.5053     0   1E+5   2   2.5  RT (min)      lcqtof_hilic_neg_2_002_36    m/z 738.5053     0   1E+5   2   2.5  RT (min)      lcqtof_hilic_neg_4_048_23    m/z 738.5054     0   1E+5   1.5   2  RT (min)      lcqtof_hilic_neg_4_048_23    m/z 738.5055     0   1E+5   2   2.5  RT (min)      lcqtof_hilic_neg_2_020_23    m/z 738.5055     0   2   4   6E+4   2   2.5  RT (min)      lcqtof_hilic_neg_4_035_33    m/z 738.5058     0   1E+6   3   3.5  RT (min)      lcqtof_hilic_neg_4_048_23    m/z 738.5060     0   1E+5   1   1.5  RT (min)      lcqtof_hilic_neg_4_048_23    m/z 738.5061     0   1E+5   2   2.5  RT (min)      lcqtof_hilic_neg_2_002_36    m/z 738.5066     0   1E+5   1   1.5  RT (min)      lcqtof_hilic_neg_4_048_23    m/z 738.5066     0   2   4E+5   2.5   3  RT (min)      lcqtof_hilic_neg_4_035_33    m/z 738.5066     0   1E+5   1   1.5  RT (min)      lcqtof_hilic_neg_4_016_37    m/z 738.5067     0   2   4E+5   3   3.5  RT (min)      lcqtof_hilic_neg_4_047_06    m/z 738.5071     0   1E+5   1.5   2  RT (min)      lcqtof_hilic_neg_4_016_37    m/z 738.5072     0   1   2E+4   4   4.5  RT (min)      lcqtof_hilic_neg_4_048_23    m/z 738.5796     0   1E+5   1   1.5  RT (min)      lcqtof_hilic_neg_4_000_aa    m/z 738.5954     0   1   2   3E+2   14.5   15  RT (min)      lcqtof_hilic_neg_4_083_59    m/z 738.5955     0   1E+5   1.5   2  RT (min)      lcqtof_hilic_neg_4_000_aa    m/z 738.5968     0   1E+4   15   15.5  RT (min)      lcqtof_hilic_neg_4_013_20    m/z 738.5971     0   1E+4   16   16.5  RT (min)      lcqtof_hilic_neg_1_007_05    m/z 738.6243     0   1E+4   1.5   2  RT (min)      lcqtof_hilic_neg_4_008_24    m/z 738.6252     0   1E+5   4.5   5  RT (min)      lcqtof_hilic_neg_4_008_24    m/z 738.6267     0   1E+5   3.5   4  RT (min)      lcqtof_hilic_neg_4_008_24    m/z 738.6270     0   2   4   6   8E+4   4   4.5  RT (min)      lcqtof_hilic_neg_4_055_32    m/z 738.6272     0   1E+5   4   4.5  RT (min)      lcqtof_hilic_neg_4_008_24    m/z 738.6588     0   1   2   3   4E+4   1.5   2  RT (min)      lcqtof_hilic_neg_4_000_aa    m/z 738.6605     0   1   2   3   4E+4   1.5   2  RT (min)      lcqtof_hilic_neg_4_000_aa    m/z 738.6726     0   1   2   3   4E+4   1   1.5  RT (min)      lcqtof_hilic_neg_4_000_aa    m/z 738.6961     0   1E+4   1.5   2  RT (min)      lcqtof_hilic_neg_4_000_aa    m/z 739.1634     0   1   2E+4   6   6.5  RT (min)       lcqtof_hilic_neg_2_052_24    m/z 739.5064     0   1   2   3E+4   4   4.5  RT (min)      lcqtof_hilic_neg_4_008_24    m/z 739.5086     0   1   2E+4   2.5   3  RT (min)      lcqtof_hilic_neg_2_003_25    m/z 739.5094     0   1   2   3   4E+4   2.5   3  RT (min)      lcqtof_hilic_neg_3_030_23    m/z 739.5095     0   1   2E+4   2.5   3  RT (min)      lcqtof_hilic_neg_2_003_25    m/z 739.5095     0   2   4   6E+4   1.5   2  RT (min)      lcqtof_hilic_neg_4_048_23    m/z 739.5096     0   2   4   6E+4   1.5   2  RT (min)      lcqtof_hilic_neg_4_048_23    m/z 739.5096     0   2   4   6E+4   1.5   2  RT (min)      lcqtof_hilic_neg_4_048_23    m/z 739.5098     0   2   4   6E+4   2   2.5  RT (min)      lcqtof_hilic_neg_2_020_23    m/z 739.5100     0   2   4   6E+4   2   2.5  RT (min)      lcqtof_hilic_neg_2_020_23    m/z 739.5100     0   2   4   6E+4   2.5   3  RT (min)      lcqtof_hilic_neg_2_020_23    m/z 739.5102     0   2   4   6E+4   2   2.5  RT (min)      lcqtof_hilic_neg_4_048_23    m/z 739.5102     0   2   4E+4   2   2.5  RT (min)      lcqtof_hilic_neg_1_022_36    m/z 739.5102     0   2   4   6E+4   1   1.5  RT (min)      lcqtof_hilic_neg_4_048_23    m/z 739.5102     0   2   4E+4   2   2.5  RT (min)      lcqtof_hilic_neg_2_002_36    m/z 739.5102     0   2   4   6E+4   1.5   2  RT (min)      lcqtof_hilic_neg_4_048_23    m/z 739.5123     0   2   4   6E+4   2   2.5  RT (min)      lcqtof_hilic_neg_4_048_23    m/z 739.5133     0   2   4E+4   1.5   2  RT (min)      lcqtof_hilic_neg_4_011_35    m/z 739.5382     0   1   2E+4   1   1.5  RT (min)      lcqtof_hilic_neg_4_000_aa    m/z 739.5597     0   1   2E+4   3.5   4  RT (min)      lcqtof_hilic_neg_4_003_16    m/z 739.5700     0   2   4   6E+4   1   1.5  RT (min)      lcqtof_hilic_neg_4_000_aa    m/z 739.5700     0   1E+4   3.5   4  RT (min)      lcqtof_hilic_neg_4_018_27    m/z 739.5813     0   1   2E+4   0.5   1  RT (min)      lcqtof_hilic_neg_2_028_05    m/z 739.5990     0   1   2   3E+4   1.5   2  RT (min)      lcqtof_hilic_neg_4_000_aa    m/z 739.6378     0   1   2   3   4E+4   3   3.5  RT (min)      lcqtof_hilic_neg_1_005_32    m/z 739.6378     0   1   2   3E+4   3   3.5  RT (min)      lcqtof_hilic_neg_4_008_24    m/z 739.6614     0   1   2E+4   1.5   2  RT (min)      lcqtof_hilic_neg_4_000_aa    m/z 739.6644     0   1   2E+4   1.5   2  RT (min)      lcqtof_hilic_neg_4_000_aa    m/z 740.5072     0   1   2   3   4E+4   4   4.5  RT (min)       lcqtof_hilic_neg_4_092_60    m/z 740.5156     0   2   4   6   8E+4   4   4.5  RT (min)       lcqtof_hilic_neg_4_008_24    m/z 740.5211     0   1E+5   1.5   2  RT (min)       lcqtof_hilic_neg_4_011_35    m/z 740.5211     0   1E+5   2   2.5  RT (min)       lcqtof_hilic_neg_2_020_23    m/z 740.5212     0   1E+5   1.5   2  RT (min)       lcqtof_hilic_neg_4_011_35    m/z 740.5214     0   1E+5   1.5   2  RT (min)       lcqtof_hilic_neg_4_011_35    m/z 740.5214     0   1E+5   1.5   2  RT (min)      lcqtof_hilic_neg_2_020_23    m/z 740.5214     0   1   2   3   4E+5   2.5   3  RT (min)      lcqtof_hilic_neg_4_004_47    m/z 740.5214     0   1E+5   2   2.5  RT (min)      lcqtof_hilic_neg_4_011_35    m/z 740.5215     0   1E+5   1.5   2  RT (min)       lcqtof_hilic_neg_4_048_23    m/z 740.5216     0   2   4   6E+4   2   2.5  RT (min)      lcqtof_hilic_neg_4_002_25    m/z 740.5216     0   1E+6   3   3.5  RT (min)       lcqtof_hilic_neg_4_048_23    m/z 740.5216     0   1E+5   1.5   2  RT (min)       lcqtof_hilic_neg_4_011_35    m/z 740.5216     0   1E+6   2.5   3  RT (min)       lcqtof_hilic_neg_4_048_23    m/z 740.5218     0   1E+5   2   2.5  RT (min)       lcqtof_hilic_neg_2_020_23    m/z 740.5218     0   1   2   3E+5   2.5   3  RT (min)      lcqtof_hilic_neg_1_017_51    m/z 740.5220     0   2   4   6E+4   2   2.5  RT (min)      lcqtof_hilic_neg_4_002_25    m/z 740.5224     0   1   2E+5   2   2.5  RT (min)       lcqtof_hilic_neg_4_034_31    m/z 740.5248     0   1E+4   4   4.5  RT (min)      lcqtof_hilic_neg_4_083_59    m/z 740.5475     0   1   2   3E+4   0.5   1  RT (min)      lcqtof_hilic_neg_3_035_46    m/z 740.5723     0   1   2E+4   0.5   1  RT (min)      lcqtof_hilic_neg_2_028_05    m/z 740.5724     0   1   2E+4   1   1.5  RT (min)      lcqtof_hilic_neg_4_000_aa    m/z 740.6425     0   1   2   3E+4   1.5   2  RT (min)      lcqtof_hilic_neg_4_000_aa    m/z 740.6754     0   2   4E+4   1.5   2  RT (min)      lcqtof_hilic_neg_4_000_aa    m/z 740.7033     0   1E+4   0.5   1  RT (min)      lcqtof_hilic_neg_1_029_06    m/z 740.7246     0   1   2E+4   0.5   1  RT (min)      lcqtof_hilic_neg_2_052_24    m/z 740.8445     0   2   4   6   8E+3   5   5.5  RT (min)       lcqtof_hilic_neg_4_058_39    m/z 740.8451     0   2   4   6   8E+3   5   5.5  RT (min)      lcqtof_hilic_neg_4_058_39    m/z 740.8961     0   1   2E+4   0.5   1  RT (min)      lcqtof_hilic_neg_3_003_20    m/z 740.9737     0   2   4E+4   0.5   1  RT (min)       lcqtof_hilic_neg_4_098_61    m/z 741.0801     0   1   2E+4   6   6.5  RT (min)       lcqtof_hilic_neg_2_020_23    m/z 741.3430     0   1E+4   15   15.5  RT (min)      lcqtof_hilic_neg_4_000_aa    m/z 741.4831     0   1E+4   4   4.5  RT (min)      lcqtof_hilic_neg_4_055_32    m/z 741.4982     0   1E+4   4   4.5  RT (min)      lcqtof_hilic_neg_4_008_24    m/z 741.5248     0   1   2   3   4E+5   3   3.5  RT (min)      lcqtof_hilic_neg_4_034_31    m/z 741.5250     0   2   4E+4   1.5   2  RT (min)      lcqtof_hilic_neg_1_003_37    m/z 741.5251     0   2   4E+5   2.5   3  RT (min)      lcqtof_hilic_neg_4_048_23    m/z 741.5251     0   2   4   6E+4   2   2.5  RT (min)      lcqtof_hilic_neg_2_020_23    m/z 741.5252     0   2   4   6E+4   1.5   2  RT (min)      lcqtof_hilic_neg_2_020_23    m/z 741.5252     0   2   4   6E+4   1.5   2  RT (min)      lcqtof_hilic_neg_2_020_23    m/z 741.5254     0   2   4   6E+4   1.5   2  RT (min)      lcqtof_hilic_neg_2_020_23    m/z 741.5254     0   2   4   6E+4   1.5   2  RT (min)      lcqtof_hilic_neg_2_020_23    m/z 741.5256     0   1   2E+5   2.5   3  RT (min)      lcqtof_hilic_neg_4_047_06    m/z 741.5256     0   2   4   6E+4   2   2.5  RT (min)      lcqtof_hilic_neg_2_020_23    m/z 741.5263     0   1   2   3E+4   1   1.5  RT (min)      lcqtof_hilic_neg_4_088_59    m/z 741.5265     0   2   4   6   8E+4   2   2.5  RT (min)      lcqtof_hilic_neg_4_028_36    m/z 741.5281     0   1E+5   1   1.5  RT (min)      lcqtof_hilic_neg_4_048_23    m/z 741.5509     0   1   2   3   4E+4   1   1.5  RT (min)      lcqtof_hilic_neg_4_000_aa    m/z 741.5512     0   1E+4   0.5   1  RT (min)      lcqtof_hilic_neg_3_035_46    m/z 741.6430     0   1   2   3E+4   1.5   2  RT (min)      lcqtof_hilic_neg_4_000_aa    m/z 741.6496     0   1   2   3E+4   1.5   2  RT (min)      lcqtof_hilic_neg_4_000_aa    m/z 741.6784     0   1   2E+4   1.5   2  RT (min)      lcqtof_hilic_neg_4_000_aa    m/z 741.8955     0   1E+4   0.5   1  RT (min)      lcqtof_hilic_neg_3_003_20    m/z 742.3413     0   1   2   3   4E+4   3.5   4  RT (min)      lcqtof_hilic_neg_4_050_02    m/z 742.5346     0   1E+6   3.5   4  RT (min)      lcqtof_hilic_neg_4_031_38    m/z 742.5362     0   2   4   6   8E+5   3.5   4  RT (min)      lcqtof_hilic_neg_4_016_37    m/z 742.5372     0   1   2   3   4E+5   1.5   2  RT (min)       lcqtof_hilic_neg_4_011_35    m/z 742.5376     0   1   2   3   4E+5   1.5   2  RT (min)       lcqtof_hilic_neg_4_011_35    m/z 742.5378     0   2   4E+5   1   1.5  RT (min)      lcqtof_hilic_neg_4_052_13    m/z 742.5378     0   1   2   3   4E+5   2.5   3  RT (min)       lcqtof_hilic_neg_2_002_36    m/z 742.5378     0   1   2   3   4E+5   2.5   3  RT (min)      lcqtof_hilic_neg_4_031_38    m/z 742.5383     0   1   2E+4   4   4.5  RT (min)      lcqtof_hilic_neg_4_048_23    m/z 742.5404     0   1E+4   4   4.5  RT (min)      lcqtof_hilic_neg_4_035_33    m/z 742.5615     0   1   2E+5   0.5   1  RT (min)      lcqtof_hilic_neg_3_035_46    m/z 742.6302     0   2   4   6   8E+4   1   1.5  RT (min)      lcqtof_hilic_neg_4_000_aa    m/z 742.6532     0   1   2   3   4E+4   1   1.5  RT (min)      lcqtof_hilic_neg_4_000_aa    m/z 742.6557     0   1   2   3   4E+4   1.5   2  RT (min)      lcqtof_hilic_neg_4_000_aa    m/z 742.6804     0   1   2   3   4E+4   0.5   1  RT (min)      lcqtof_hilic_neg_3_035_46    m/z 742.8956     0   1E+4   0.5   1  RT (min)      lcqtof_hilic_neg_3_003_20    m/z 743.1689     0   1E+4   0.5   1  RT (min)      lcqtof_hilic_neg_3_000_aa    m/z 743.3446     0   1E+4   3.5   4  RT (min)      lcqtof_hilic_neg_4_050_02    m/z 743.3595     0   1E+4   6   6.5  RT (min)      lcqtof_hilic_neg_4_008_24    m/z 743.3888     0   1E+5   1   1.5  RT (min)      lcqtof_hilic_neg_4_000_aa    m/z 743.3890     0   1E+5   1   1.5  RT (min)      lcqtof_hilic_neg_4_000_aa    m/z 743.3974     0   1E+5   1   1.5  RT (min)      lcqtof_hilic_neg_4_000_aa    m/z 743.4545     0   1E+4   3.5   4  RT (min)      lcqtof_hilic_neg_2_031_01    m/z 743.4588     0   1   2   3   4E+3   0.5   1  RT (min)      lcqtof_hilic_neg_1_003_37    m/z 743.4756     0   1   2E+4   1   1.5  RT (min)      lcqtof_hilic_neg_4_000_aa    m/z 743.4856     0   1E+4   1   1.5  RT (min)      lcqtof_hilic_neg_1_035_23    m/z 743.5114     0   1E+4   0.5   1  RT (min)      lcqtof_hilic_neg_4_038_10    m/z 743.5403     0   1E+5   2.5   3  RT (min)      lcqtof_hilic_neg_4_031_38    m/z 743.5408     0   1E+5   2.5   3  RT (min)      lcqtof_hilic_neg_2_002_36    m/z 743.5410     0   2   4E+5   3   3.5  RT (min)       lcqtof_hilic_neg_4_016_37    m/z 743.5411     0   1E+5   1.5   2  RT (min)       lcqtof_hilic_neg_4_011_35    m/z 743.5412     0   1E+5   1.5   2  RT (min)       lcqtof_hilic_neg_4_028_36    m/z 743.5415     0   2   4E+4   1.5   2  RT (min)      lcqtof_hilic_neg_4_028_36    m/z 743.5417     0   2   4   6E+4   1.5   2  RT (min)      lcqtof_hilic_neg_4_028_36    m/z 743.5418     0   1   2E+5   1   1.5  RT (min)      lcqtof_hilic_neg_4_028_36    m/z 743.5421     0   1   2E+5   1   1.5  RT (min)      lcqtof_hilic_neg_4_028_36    m/z 743.5572     0   2   4E+4   1   1.5  RT (min)      lcqtof_hilic_neg_4_000_aa    m/z 743.5962     0   1   2   3E+5   1   1.5  RT (min)      lcqtof_hilic_neg_4_000_aa    m/z 743.6250     0   1   2E+4   1   1.5  RT (min)      lcqtof_hilic_neg_4_000_aa    m/z 743.6574     0   1   2E+4   1   1.5  RT (min)      lcqtof_hilic_neg_4_000_aa    m/z 743.6584     0   1   2E+4   1.5   2  RT (min)      lcqtof_hilic_neg_4_000_aa    m/z 743.6820     0   1E+4   0.5   1  RT (min)      lcqtof_hilic_neg_3_035_46    m/z 743.9152     0   1E+4   0.5   1  RT (min)      lcqtof_hilic_neg_3_003_20    m/z 744.3568     0   1E+4   3.5   4  RT (min)      lcqtof_hilic_neg_4_050_02    m/z 744.3896     0   1E+5   1   1.5  RT (min)      lcqtof_hilic_neg_4_000_aa    m/z 744.3896     0   1E+5   1   1.5  RT (min)      lcqtof_hilic_neg_4_000_aa    m/z 744.3900     0   1E+5   1   1.5  RT (min)      lcqtof_hilic_neg_4_000_aa    m/z 744.3900     0   1E+5   1   1.5  RT (min)      lcqtof_hilic_neg_4_000_aa    m/z 744.3904     0   1E+5   1   1.5  RT (min)      lcqtof_hilic_neg_4_000_aa    m/z 744.3906     0   1E+5   1   1.5  RT (min)      lcqtof_hilic_neg_4_000_aa    m/z 744.4740     0   1E+4   3   3.5  RT (min)      lcqtof_hilic_neg_4_016_37    m/z 744.5171     0   1E+4   1   1.5  RT (min)      lcqtof_hilic_neg_4_040_05    m/z 744.5497     0   1   2   3   4E+4   2   2.5  RT (min)      lcqtof_hilic_neg_4_002_25    m/z 744.5498     0   2   4   6   8E+4   2   2.5  RT (min)      lcqtof_hilic_neg_4_034_31    m/z 744.5500     0   1E+5   2   2.5  RT (min)      lcqtof_hilic_neg_1_035_23    m/z 744.5507     0   1   2   3   4E+5   2.5   3  RT (min)      lcqtof_hilic_neg_4_029_17    m/z 744.5508     0   2   4   6E+4   2.5   3  RT (min)      lcqtof_hilic_neg_4_029_17    m/z 744.5511     0   2   4E+5   2.5   3  RT (min)      lcqtof_hilic_neg_4_032_01    m/z 744.5516     0   1   2E+4   1.5   2  RT (min)      lcqtof_hilic_neg_4_003_16    m/z 744.5526     0   1   2   3   4E+4   1.5   2  RT (min)      lcqtof_hilic_neg_4_003_16    m/z 744.5532     0   2   4   6   8E+4   1   1.5  RT (min)      lcqtof_hilic_neg_3_066_57    m/z 744.5700     0   2   4E+4   1   1.5  RT (min)      lcqtof_hilic_neg_4_000_aa    m/z 744.5746     0   1   2   3   4E+4   1   1.5  RT (min)      lcqtof_hilic_neg_1_005_32    m/z 744.5748     0   1E+4   1.5   2  RT (min)      lcqtof_hilic_neg_4_008_24    m/z 744.6016     0   2   4   6   8E+4   0.5   1  RT (min)      lcqtof_hilic_neg_2_028_05    m/z 744.6026     0   1E+5   1   1.5  RT (min)      lcqtof_hilic_neg_4_000_aa    m/z 745.3868     0   2   4   6E+4   1   1.5  RT (min)      lcqtof_hilic_neg_4_000_aa    m/z 745.3879     0   2   4   6E+4   1   1.5  RT (min)      lcqtof_hilic_neg_4_000_aa    m/z 745.3886     0   2   4   6E+4   1   1.5  RT (min)      lcqtof_hilic_neg_4_000_aa    m/z 745.3886     0   2   4   6E+4   1   1.5  RT (min)      lcqtof_hilic_neg_4_000_aa    m/z 745.3890     0   2   4   6E+4   1   1.5  RT (min)      lcqtof_hilic_neg_4_000_aa    m/z 745.3943     0   2   4   6E+4   1   1.5  RT (min)      lcqtof_hilic_neg_4_000_aa    m/z 745.3987     0   2   4   6E+4   1   1.5  RT (min)      lcqtof_hilic_neg_4_000_aa    m/z 745.4802     0   1E+4   15.5   16  RT (min)      lcqtof_hilic_neg_4_000_aa    m/z 745.4821     0   1   2   3   4E+5   1   1.5  RT (min)      lcqtof_hilic_neg_4_000_aa    m/z 745.4843     0   1   2E+5   3   3.5  RT (min)      lcqtof_hilic_neg_4_011_35    m/z 745.5017     0   1   2E+5   1   1.5  RT (min)      lcqtof_hilic_neg_2_020_23    m/z 745.5050     0   1   2E+5   3   3.5  RT (min)      lcqtof_hilic_neg_1_035_23    m/z 745.5142     0   1E+4   3   3.5  RT (min)      lcqtof_hilic_neg_4_002_25    m/z 745.5258     0   1   2E+4   3   3.5  RT (min)      lcqtof_hilic_neg_4_028_36    m/z 745.5443     0   1E+5   2.5   3  RT (min)      lcqtof_hilic_neg_4_034_31    m/z 745.5603     0   1   2E+4   1.5   2  RT (min)      lcqtof_hilic_neg_4_034_31    m/z 745.5634     0   1E+4   1.5   2  RT (min)       lcqtof_hilic_neg_4_003_16    m/z 745.5762     0   1   2E+4   0.5   1  RT (min)      lcqtof_hilic_neg_3_035_46    m/z 745.5972     0   1   2E+4   0.5   1  RT (min)      lcqtof_hilic_neg_4_040_05    m/z 745.6090     0   2   4   6   8E+6   1   1.5  RT (min)      lcqtof_hilic_neg_4_000_aa    m/z 745.6097     0   1   2E+4   0.5   1  RT (min)      lcqtof_hilic_neg_2_028_05    m/z 745.7070     0   1E+4   1.5   2  RT (min)      lcqtof_hilic_neg_4_000_aa    m/z 745.7464     0   1E+5   1   1.5  RT (min)      lcqtof_hilic_neg_4_000_aa    m/z 746.0170     0   1   2   3   4E+4   1   1.5  RT (min)      lcqtof_hilic_neg_4_000_aa    m/z 746.0962     0   1E+4   6   6.5  RT (min)      lcqtof_hilic_neg_4_012_07    m/z 746.0998     0   1E+5   1   1.5  RT (min)      lcqtof_hilic_neg_4_000_aa    m/z 746.2530     0   1   2   3   4E+4   1   1.5  RT (min)      lcqtof_hilic_neg_4_000_aa    m/z 746.3123     0   1E+4   4.5   5  RT (min)      lcqtof_hilic_neg_4_016_37    m/z 746.4854     0   1   2E+5   1   1.5  RT (min)      lcqtof_hilic_neg_4_000_aa    m/z 746.5056     0   1E+5   1   1.5  RT (min)      lcqtof_hilic_neg_2_020_23    m/z 746.5240     0   2   4   6E+4   1   1.5  RT (min)      lcqtof_hilic_neg_4_040_05    m/z 746.5308     0   1   2   3E+4   1   1.5  RT (min)      lcqtof_hilic_neg_4_000_aa    m/z 746.5310     0   1   2   3E+4   0.5   1  RT (min)      lcqtof_hilic_neg_4_000_aa    m/z 746.5948     0   2   4   6E+4   0.5   1  RT (min)       lcqtof_hilic_neg_4_040_05    m/z 746.5961     0   2   4E+4   0.5   1  RT (min)      lcqtof_hilic_neg_4_008_24    m/z 746.6138     0   1   2   3   4E+6   1   1.5  RT (min)      lcqtof_hilic_neg_4_000_aa    m/z 746.7446     0   1E+4   1   1.5  RT (min)      lcqtof_hilic_neg_4_000_aa    m/z 746.8945     0   2   4E+4   1   1.5  RT (min)      lcqtof_hilic_neg_4_000_aa    m/z 746.9850     0   1E+4   1   1.5  RT (min)      lcqtof_hilic_neg_4_000_aa    m/z 747.1054     0   1   2E+4   1   1.5  RT (min)      lcqtof_hilic_neg_4_000_aa    m/z 747.2578     0   1   2   3E+4   1   1.5  RT (min)      lcqtof_hilic_neg_4_000_aa    m/z 747.3612     0   1E+4   0.5   1  RT (min)      lcqtof_hilic_neg_3_035_46    m/z 747.3884     0   1   2E+4   1   1.5  RT (min)      lcqtof_hilic_neg_4_000_aa    m/z 747.4460     0   1E+4   1   1.5  RT (min)      lcqtof_hilic_neg_4_000_aa    m/z 747.4527     0   1E+4   1   1.5  RT (min)      lcqtof_hilic_neg_4_000_aa    m/z 747.4900     0   2   4   6E+4   1   1.5  RT (min)      lcqtof_hilic_neg_4_000_aa    m/z 747.4942     0   2   4   6E+4   1.5   2  RT (min)      lcqtof_hilic_neg_4_000_aa    m/z 747.5166     0   1   2   3E+5   1   1.5  RT (min)       lcqtof_hilic_neg_2_020_23    m/z 747.5603     0   1E+4   1.5   2  RT (min)      lcqtof_hilic_neg_4_000_aa    m/z 747.5684     0   1E+4   1   1.5  RT (min)      lcqtof_hilic_neg_4_000_aa    m/z 747.6074     0   1E+6   1   1.5  RT (min)      lcqtof_hilic_neg_4_000_aa    m/z 747.6197     0   1E+6   1   1.5  RT (min)      lcqtof_hilic_neg_4_000_aa    m/z 747.6231     0   1E+4   0.5   1  RT (min)      lcqtof_hilic_neg_2_028_05    m/z 747.6976     0   1   2E+4   1   1.5  RT (min)      lcqtof_hilic_neg_4_000_aa    m/z 747.7050     0   2   4E+4   1   1.5  RT (min)      lcqtof_hilic_neg_4_000_aa    m/z 747.7214     0   2   4E+4   1.5   2  RT (min)      lcqtof_hilic_neg_4_000_aa    m/z 747.8942     0   1E+4   1   1.5  RT (min)       lcqtof_hilic_neg_4_054_48    m/z 748.0113     0   1   2E+4   7.5   8  RT (min)       lcqtof_hilic_neg_4_082_59    m/z 748.3064     0   1E+4   0.5   1  RT (min)      lcqtof_hilic_neg_4_023_15    m/z 748.5211     0   1E+5   1   1.5  RT (min)      lcqtof_hilic_neg_2_020_23    m/z 748.5215     0   1   2   3   4E+4   1   1.5  RT (min)      lcqtof_hilic_neg_1_007_05    m/z 748.5218     0   2   4E+4   1   1.5  RT (min)      lcqtof_hilic_neg_1_013_38    m/z 748.5413     0   1   2E+4   1   1.5  RT (min)      lcqtof_hilic_neg_4_040_05    m/z 748.5432     0   1E+4   1   1.5  RT (min)      lcqtof_hilic_neg_3_035_46    m/z 748.6038     0   1   2E+4   2.5   3  RT (min)      lcqtof_hilic_neg_4_097_60    m/z 748.6114     0   1   2E+5   1   1.5  RT (min)      lcqtof_hilic_neg_4_000_aa    m/z 748.6204     0   1E+4   0.5   1  RT (min)      lcqtof_hilic_neg_4_040_05    m/z 748.6967     0   1E+4   1   1.5  RT (min)      lcqtof_hilic_neg_4_000_aa    m/z 748.7019     0   1E+4   1   1.5  RT (min)      lcqtof_hilic_neg_4_000_aa    m/z 748.7260     0   1   2E+4   1.5   2  RT (min)      lcqtof_hilic_neg_4_000_aa    m/z 749.4876     0   1E+4   3   3.5  RT (min)      lcqtof_hilic_neg_4_002_25    m/z 749.4878     0   1E+4   3   3.5  RT (min)       lcqtof_hilic_neg_2_020_23    m/z 749.5244     0   1E+4   1   1.5  RT (min)      lcqtof_hilic_neg_4_000_aa    m/z 749.5247     0   1E+4   1.5   2  RT (min)      lcqtof_hilic_neg_4_000_aa    m/z 749.6070     0   2   4E+4   2   2.5  RT (min)      lcqtof_hilic_neg_4_055_32    m/z 749.6070     0   1   2   3E+4   1   1.5  RT (min)       lcqtof_hilic_neg_4_008_24    m/z 749.6074     0   2   4   6   8E+4   2.5   3  RT (min)       lcqtof_hilic_neg_4_097_60    m/z 749.6078     0   2   4   6E+4   2.5   3  RT (min)      lcqtof_hilic_neg_4_055_32    m/z 749.6082     0   1E+5   3   3.5  RT (min)      lcqtof_hilic_neg_4_008_24    m/z 749.6086     0   1   2E+4   1.5   2  RT (min)      lcqtof_hilic_neg_4_091_60    m/z 749.6087     0   1   2E+4   2   2.5  RT (min)      lcqtof_hilic_neg_4_095_60    m/z 749.6398     0   2   4   6E+4   1   1.5  RT (min)      lcqtof_hilic_neg_4_000_aa    m/z 749.6448     0   2   4   6E+4   1   1.5  RT (min)      lcqtof_hilic_neg_4_000_aa    m/z 749.7046     0   1   2E+4   1   1.5  RT (min)      lcqtof_hilic_neg_4_000_aa    m/z 749.7218     0   1   2E+4   1.5   2  RT (min)      lcqtof_hilic_neg_4_000_aa    m/z 750.3070     0   1E+4   0.5   1  RT (min)      lcqtof_hilic_neg_4_010_28    m/z 750.4572     0   1E+4   5.5   6  RT (min)      lcqtof_hilic_neg_4_040_05    m/z 750.4912     0   1   2   3E+4   1   1.5  RT (min)      lcqtof_hilic_neg_4_040_05    m/z 750.5040     0   2   4   6E+4   3   3.5  RT (min)      lcqtof_hilic_neg_4_028_36    m/z 750.5299     0   1E+5   2.5   3  RT (min)      lcqtof_hilic_neg_1_022_36    m/z 750.5387     0   1E+5   2.5   3  RT (min)      lcqtof_hilic_neg_1_060_14    m/z 750.5610     0   1   2E+4   1   1.5  RT (min)      lcqtof_hilic_neg_1_007_05    m/z 750.6176     0   1   2E+4   2   2.5   3  RT (min)      lcqtof_hilic_neg_4_008_24    m/z 750.6198     0   1E+4   2   2.5  RT (min)      lcqtof_hilic_neg_4_008_24    m/z 750.6203     0   1E+4   2   2.5  RT (min)      lcqtof_hilic_neg_4_055_32    m/z 750.6326     0   1E+4   1   1.5  RT (min)      lcqtof_hilic_neg_4_000_aa    m/z 750.6944     0   1E+4   1.5   2  RT (min)      lcqtof_hilic_neg_4_000_aa    m/z 750.7034     0   1E+4   1   1.5  RT (min)      lcqtof_hilic_neg_4_000_aa    m/z 751.1123     0   1E+4   6   6.5  RT (min)      lcqtof_hilic_neg_4_012_07    m/z 751.1262     0   1E+4   7   7.5  RT (min)       lcqtof_hilic_neg_4_092_60    m/z 751.2015     0   1   2E+4   6   6.5  RT (min)      lcqtof_hilic_neg_3_011_34    m/z 751.2119     0   1   2E+4   6   6.5  RT (min)      lcqtof_hilic_neg_3_011_34    m/z 751.3866     0   2   4   6E+4   1   1.5  RT (min)      lcqtof_hilic_neg_4_000_aa    m/z 751.3958     0   2   4   6E+4   1   1.5  RT (min)      lcqtof_hilic_neg_4_000_aa    m/z 751.4019     0   2   4   6E+4   1   1.5  RT (min)      lcqtof_hilic_neg_4_000_aa    m/z 751.4026     0   2   4   6E+4   1   1.5  RT (min)      lcqtof_hilic_neg_4_000_aa    m/z 751.4040     0   2   4   6E+4   1   1.5  RT (min)      lcqtof_hilic_neg_4_000_aa    m/z 751.4115     0   2   4   6E+4   1   1.5  RT (min)      lcqtof_hilic_neg_4_000_aa    m/z 751.5016     0   2   4   6   8E+4   1   1.5  RT (min)      lcqtof_hilic_neg_4_000_aa    m/z 751.5024     0   2   4   6   8E+4   1   1.5  RT (min)      lcqtof_hilic_neg_4_000_aa    m/z 751.5030     0   2   4   6   8E+4   1.5   2  RT (min)      lcqtof_hilic_neg_4_000_aa    m/z 751.5051     0   2   4   6   8E+4   1   1.5  RT (min)      lcqtof_hilic_neg_4_000_aa    m/z 751.6072     0   1E+4   15   15.5  RT (min)       lcqtof_hilic_neg_4_039_18    m/z 751.6074     0   1E+4   15   15.5  RT (min)      lcqtof_hilic_neg_4_013_20    m/z 751.6075     0   1E+4   15   15.5  RT (min)      lcqtof_hilic_neg_4_015_09    m/z 751.6150     0   1   2   3E+5   2.5   3  RT (min)      lcqtof_hilic_neg_4_055_32    m/z 751.6178     0   1   2E+5   3   3.5  RT (min)      lcqtof_hilic_neg_4_035_33    m/z 751.6187     0   1   2E+4   4   4.5  RT (min)      lcqtof_hilic_neg_4_008_24    m/z 751.6222     0   2   4   6   8E+4   1.5   2  RT (min)      lcqtof_hilic_neg_4_000_aa    m/z 751.6226     0   1   2   3   4E+4   1.5   2  RT (min)      lcqtof_hilic_neg_4_055_32    m/z 751.6226     0   1   2E+4   1.5   2  RT (min)      lcqtof_hilic_neg_4_040_05    m/z 751.6227     0   2   4   6E+4   2   2.5  RT (min)      lcqtof_hilic_neg_4_055_32    m/z 751.6236     0   1   2   3   4E+4   2   2.5  RT (min)      lcqtof_hilic_neg_2_047_32    m/z 751.6236     0   2   4E+4   2   2.5  RT (min)      lcqtof_hilic_neg_4_008_24    m/z 751.6236     0   1   2   3   4E+5   2.5   3   3.5  RT (min)      lcqtof_hilic_neg_3_004_32    m/z 751.6237     0   2   4   6E+4   1.5   2  RT (min)      lcqtof_hilic_neg_4_055_32    m/z 751.6239     0   2   4   6E+4   2   2.5  RT (min)      lcqtof_hilic_neg_4_055_32    m/z 751.6246     0   1   2E+4   1.5   2  RT (min)      lcqtof_hilic_neg_4_035_33    m/z 751.6258     0   2   4   6   8E+4   1   1.5  RT (min)      lcqtof_hilic_neg_4_000_aa    m/z 751.6269     0   2   4   6   8E+4   1   1.5  RT (min)      lcqtof_hilic_neg_4_000_aa    m/z 751.6427     0   1   2   3   4E+4   1   1.5  RT (min)      lcqtof_hilic_neg_4_000_aa    m/z 752.3804     0   1   2E+4   0.5   1  RT (min)      lcqtof_hilic_neg_4_050_02    m/z 752.3881     0   1   2   3E+4   1   1.5  RT (min)      lcqtof_hilic_neg_4_000_aa    m/z 752.4005     0   1   2   3E+4   1   1.5  RT (min)      lcqtof_hilic_neg_4_000_aa    m/z 752.4106     0   1   2E+4   1   1.5  RT (min)      lcqtof_hilic_neg_4_000_aa    m/z 752.4114     0   1   2E+4   1   1.5  RT (min)      lcqtof_hilic_neg_4_000_aa    m/z 752.5225     0   1E+4   2   2.5  RT (min)       lcqtof_hilic_neg_4_011_35    m/z 752.5242     0   1E+4   4   4.5  RT (min)      lcqtof_hilic_neg_4_091_60    m/z 752.5285     0   1   2E+4   4   4.5  RT (min)      lcqtof_hilic_neg_4_008_24    m/z 752.5851     0   1E+4   0.5   1  RT (min)      lcqtof_hilic_neg_4_005_29    m/z 752.6080     0   2   4   6   8E+4   2.5   3  RT (min)      lcqtof_hilic_neg_4_047_06    m/z 752.6081     0   1   2E+5   2.5   3  RT (min)       lcqtof_hilic_neg_4_093_60    m/z 752.6124     0   1   2E+5   3   3.5  RT (min)      lcqtof_hilic_neg_4_008_24    m/z 752.6134     0   1   2   3   4E+4   1.5   2  RT (min)      lcqtof_hilic_neg_4_000_aa    m/z 752.6157     0   1   2   3   4E+4   1.5   2  RT (min)      lcqtof_hilic_neg_4_000_aa    m/z 752.6172     0   1   2   3E+5   3   3.5  RT (min)      lcqtof_hilic_neg_4_035_33    m/z 752.6313     0   1   2E+4   4   4.5  RT (min)      lcqtof_hilic_neg_4_008_24    m/z 752.6327     0   2   4   6E+5   3   3.5  RT (min)      lcqtof_hilic_neg_1_005_32    m/z 752.6329     0   2   4   6E+5   3   3.5  RT (min)      lcqtof_hilic_neg_2_047_32    m/z 752.6330     0   2   4   6E+4   2   2.5  RT (min)      lcqtof_hilic_neg_4_055_32    m/z 752.6332     0   2   4   6E+4   1.5   2  RT (min)      lcqtof_hilic_neg_4_055_32    m/z 752.6332     0   2   4   6E+4   2   2.5  RT (min)      lcqtof_hilic_neg_4_008_24    m/z 752.6334     0   2   4   6E+4   1.5   2  RT (min)      lcqtof_hilic_neg_4_055_32    m/z 752.6338     0   2   4   6E+4   1.5   2  RT (min)      lcqtof_hilic_neg_4_055_32    m/z 752.6342     0   2   4   6E+4   2.5   3  RT (min)      lcqtof_hilic_neg_2_047_32    m/z 752.6385     0   2   4   6E+4   2   2.5  RT (min)      lcqtof_hilic_neg_4_008_24    m/z 752.6451     0   1   2E+4   2   2.5  RT (min)      lcqtof_hilic_neg_4_054_48    m/z 752.6480     0   1E+4   2.5   3  RT (min)      lcqtof_hilic_neg_4_033_53    m/z 752.6745     0   1   2E+4   1.5   2  RT (min)      lcqtof_hilic_neg_4_000_aa    m/z 752.6750     0   1   2E+4   1.5   2  RT (min)      lcqtof_hilic_neg_4_000_aa    m/z 752.7102     0   1   2E+4   1.5   2  RT (min)      lcqtof_hilic_neg_4_000_aa    m/z 752.7126     0   1E+4   0.5   1  RT (min)      lcqtof_hilic_neg_3_007_04    m/z 752.8848     0   1E+4   2.5   3  RT (min)      lcqtof_hilic_neg_4_055_32    m/z 753.3770     0   1   2   3E+4   1   1.5  RT (min)      lcqtof_hilic_neg_4_000_aa    m/z 753.3814     0   1E+4   0.5   1  RT (min)      lcqtof_hilic_neg_4_050_02    m/z 753.3844     0   1   2   3E+4   1   1.5  RT (min)      lcqtof_hilic_neg_4_000_aa    m/z 753.3890     0   1   2   3E+4   1   1.5  RT (min)      lcqtof_hilic_neg_4_000_aa    m/z 753.3898     0   1   2   3E+4   1   1.5  RT (min)      lcqtof_hilic_neg_4_000_aa    m/z 753.4080     0   1   2E+4   1   1.5  RT (min)      lcqtof_hilic_neg_4_000_aa    m/z 753.5116     0   1   2   3E+4   4   4.5  RT (min)       lcqtof_hilic_neg_4_097_60    m/z 753.5244     0   1   2   3E+3   2   2.5  RT (min)      lcqtof_hilic_neg_1_035_23    m/z 753.5248     0   2   4E+4   1.5   2  RT (min)      lcqtof_hilic_neg_4_000_aa    m/z 753.5381     0   2   4   6   8E+4   1.5   2  RT (min)      lcqtof_hilic_neg_4_000_aa    m/z 753.5387     0   2   4   6   8E+4   1.5   2  RT (min)      lcqtof_hilic_neg_4_000_aa    m/z 753.5459     0   1E+4   3.5   4  RT (min)      lcqtof_hilic_neg_4_094_60    m/z 753.5485     0   2   4   6   8E+4   1   1.5  RT (min)      lcqtof_hilic_neg_4_000_aa    m/z 753.5492     0   2   4   6   8E+4   1   1.5  RT (min)      lcqtof_hilic_neg_4_000_aa    m/z 753.5534     0   1   2E+3   15.5   16  RT (min)       lcqtof_hilic_neg_1_009_50    m/z 753.5842     0   1E+5   1   1.5  RT (min)      lcqtof_hilic_neg_4_000_aa    m/z 753.6360     0   1E+6   3.5   4  RT (min)       lcqtof_hilic_neg_4_092_60    m/z 753.6372     0   1E+6   2.5   3  RT (min)      lcqtof_hilic_neg_4_055_32    m/z 753.6375     0   1E+6   3   3.5  RT (min)      lcqtof_hilic_neg_4_055_32    m/z 753.6375     0   1   2E+5   2   2.5  RT (min)       lcqtof_hilic_neg_2_047_32    m/z 753.6375     0   1   2E+6   2.5   3  RT (min)       lcqtof_hilic_neg_4_008_24    m/z 753.6376     0   1   2E+5   1.5   2  RT (min)      lcqtof_hilic_neg_4_008_24    m/z 753.6376     0   1   2E+6   3   3.5  RT (min)       lcqtof_hilic_neg_4_008_24    m/z 753.6377     0   1   2E+5   1.5   2  RT (min)      lcqtof_hilic_neg_4_008_24    m/z 753.6377     0   1   2E+5   2   2.5  RT (min)       lcqtof_hilic_neg_4_008_24    m/z 753.6378     0   1   2E+5   1.5   2  RT (min)       lcqtof_hilic_neg_4_008_24    m/z 753.6378     0   1   2E+5   2   2.5  RT (min)      lcqtof_hilic_neg_4_055_32    m/z 753.6378     0   1   2E+5   2   2.5  RT (min)       lcqtof_hilic_neg_4_008_24    m/z 753.6379     0   1   2E+5   1.5   2  RT (min)       lcqtof_hilic_neg_4_008_24    m/z 753.6386     0   1   2E+5   1.5   2  RT (min)      lcqtof_hilic_neg_4_008_24    m/z 753.6387     0   2   4   6E+4   4   4.5  RT (min)       lcqtof_hilic_neg_4_092_60    m/z 753.6390     0   1   2E+4   1   1.5  RT (min)      lcqtof_hilic_neg_4_000_aa    m/z 753.6712     0   1   2E+4   1   1.5  RT (min)      lcqtof_hilic_neg_4_000_aa    m/z 753.7124     0   1E+4   1.5   2  RT (min)      lcqtof_hilic_neg_4_000_aa    m/z 754.0444     0   1E+4   3   3.5  RT (min)      lcqtof_hilic_neg_1_005_32    m/z 754.2859     0   1   2E+4   3   3.5  RT (min)      lcqtof_hilic_neg_1_005_32    m/z 754.4857     0   1   2E+4   4   4.5  RT (min)      lcqtof_hilic_neg_4_008_24    m/z 754.5475     0   1   2E+4   1   1.5  RT (min)      lcqtof_hilic_neg_4_000_aa    m/z 754.5496     0   1   2E+4   4   4.5  RT (min)      lcqtof_hilic_neg_4_008_24    m/z 754.5680     0   2   4E+4   1   1.5  RT (min)      lcqtof_hilic_neg_4_000_aa    m/z 754.5874     0   2   4E+4   1   1.5  RT (min)      lcqtof_hilic_neg_4_000_aa    m/z 754.6238     0   1E+5   2.5   3  RT (min)      lcqtof_hilic_neg_4_093_60    m/z 754.6368     0   1E+5   2.5   3  RT (min)      lcqtof_hilic_neg_4_091_60    m/z 754.6466     0   2   4   6E+4   1.5   2  RT (min)      lcqtof_hilic_neg_4_000_aa    m/z 754.6472     0   1   2   3E+4   2   2.5  RT (min)      lcqtof_hilic_neg_4_055_32    m/z 754.6476     0   1E+5   2.5   3  RT (min)      lcqtof_hilic_neg_4_008_24    m/z 754.6480     0   1E+4   4   4.5  RT (min)      lcqtof_hilic_neg_4_008_24    m/z 754.6484     0   2   4   6E+4   2.5   3  RT (min)      lcqtof_hilic_neg_4_055_32    m/z 754.6484     0   1E+4   1.5   2  RT (min)      lcqtof_hilic_neg_1_037_24    m/z 754.6486     0   1   2   3E+4   2   2.5  RT (min)      lcqtof_hilic_neg_4_008_24    m/z 754.6488     0   1   2   3E+4   2   2.5  RT (min)      lcqtof_hilic_neg_4_055_32    m/z 754.6900     0   1   2   3E+4   1.5   2  RT (min)      lcqtof_hilic_neg_4_000_aa    m/z 754.9106     0   2   4   6   8E+4   0.5   1  RT (min)      lcqtof_hilic_neg_3_035_46    m/z 754.9156     0   1E+4   0.5   1  RT (min)      lcqtof_hilic_neg_3_003_20    m/z 754.9884     0   2   4   6E+4   0.5   1  RT (min)       lcqtof_hilic_neg_4_098_61    m/z 755.2545     0   1   2   3   4E+3   5.5   6  RT (min)       lcqtof_hilic_neg_4_003_16    m/z 755.4599     0   1E+5   1.5   2  RT (min)      lcqtof_hilic_neg_4_000_aa    m/z 755.4604     0   1E+5   1.5   2  RT (min)      lcqtof_hilic_neg_4_000_aa    m/z 755.4610     0   1E+5   1   1.5  RT (min)      lcqtof_hilic_neg_4_000_aa    m/z 755.4611     0   1E+5   1   1.5  RT (min)      lcqtof_hilic_neg_4_000_aa    m/z 755.4612     0   1E+5   1   1.5  RT (min)      lcqtof_hilic_neg_4_000_aa    m/z 755.4618     0   1E+5   1   1.5  RT (min)      lcqtof_hilic_neg_4_000_aa    m/z 755.4730     0   1E+4   0.5   1  RT (min)      lcqtof_hilic_neg_4_026_04    m/z 755.5304     0   1E+4   4   4.5  RT (min)      lcqtof_hilic_neg_4_008_24    m/z 755.5465     0   1   2   3E+4   1   1.5  RT (min)      lcqtof_hilic_neg_4_000_aa    m/z 755.5476     0   1   2   3E+4   1.5   2  RT (min)      lcqtof_hilic_neg_4_000_aa    m/z 755.5540     0   1   2   3E+4   1   1.5  RT (min)      lcqtof_hilic_neg_4_000_aa    m/z 755.5645     0   1   2   3E+4   1.5   2  RT (min)      lcqtof_hilic_neg_4_000_aa    m/z 755.5789     0   1   2   3E+4   1   1.5  RT (min)      lcqtof_hilic_neg_4_000_aa    m/z 755.5868     0   1   2E+4   1   1.5  RT (min)      lcqtof_hilic_neg_4_048_23    m/z 755.6521     0   1   2   3   4E+5   3.5   4  RT (min)      lcqtof_hilic_neg_4_096_60    m/z 755.6528     0   2   4   6   8E+5   2.5   3  RT (min)      lcqtof_hilic_neg_4_008_24    m/z 755.6528     0   1   2E+4   2   2.5  RT (min)      lcqtof_hilic_neg_2_001_49    m/z 755.6528     0   2   4   6   8E+4   2   2.5  RT (min)      lcqtof_hilic_neg_4_008_24    m/z 755.6529     0   1E+5   2   2.5  RT (min)      lcqtof_hilic_neg_4_055_32    m/z 755.6530     0   2   4   6   8E+4   2   2.5  RT (min)      lcqtof_hilic_neg_4_008_24    m/z 755.6530     0   1E+5   2   2.5  RT (min)      lcqtof_hilic_neg_4_055_32    m/z 755.6530     0   2   4   6   8E+5   3   3.5  RT (min)       lcqtof_hilic_neg_4_008_24    m/z 755.6531     0   2   4   6   8E+4   1.5   2  RT (min)      lcqtof_hilic_neg_4_008_24    m/z 755.6532     0   2   4   6   8E+5   3   3.5  RT (min)      lcqtof_hilic_neg_4_008_24    m/z 755.6533     0   2   4   6   8E+4   1.5   2  RT (min)      lcqtof_hilic_neg_4_008_24    m/z 755.6536     0   1   2   3E+4   4   4.5  RT (min)      lcqtof_hilic_neg_4_055_32    m/z 755.6537     0   1   2   3E+4   1.5   2  RT (min)      lcqtof_hilic_neg_4_040_05    m/z 755.6928     0   1E+4   1.5   2  RT (min)      lcqtof_hilic_neg_4_000_aa    m/z 755.9113     0   1   2   3   4E+4   0.5   1  RT (min)      lcqtof_hilic_neg_3_035_46    m/z 755.9128     0   1   2   3   4E+4   0.5   1  RT (min)      lcqtof_hilic_neg_3_035_46    m/z 755.9314     0   1   2E+4   3   3.5  RT (min)      lcqtof_hilic_neg_1_005_32    m/z 756.2570     0   1E+4   5   5.5  RT (min)      lcqtof_hilic_neg_4_058_39    m/z 756.4624     0   2   4   6E+4   1.5   2  RT (min)      lcqtof_hilic_neg_4_000_aa    m/z 756.4630     0   2   4   6E+4   1   1.5  RT (min)      lcqtof_hilic_neg_4_000_aa    m/z 756.4636     0   2   4   6E+4   1   1.5  RT (min)      lcqtof_hilic_neg_4_000_aa    m/z 756.4637     0   2   4   6E+4   1.5   2  RT (min)      lcqtof_hilic_neg_4_000_aa    m/z 756.4644     0   2   4   6E+4   1   1.5  RT (min)      lcqtof_hilic_neg_4_000_aa    m/z 756.4646     0   2   4   6E+4   1.5   2  RT (min)      lcqtof_hilic_neg_4_000_aa    m/z 756.4804     0   2   4   6E+5   4   4.5  RT (min)      lcqtof_hilic_neg_4_048_23    m/z 756.4813     0   2   4   6E+5   4   4.5  RT (min)      lcqtof_hilic_neg_4_028_36    m/z 756.4824     0   1   2E+5   4.5   5  RT (min)      lcqtof_hilic_neg_4_040_05    m/z 756.5112     0   1E+4   4   4.5  RT (min)      lcqtof_hilic_neg_4_008_24    m/z 756.5242     0   1E+4   4   4.5  RT (min)      lcqtof_hilic_neg_4_008_24    m/z 756.6680     0   2   4   6E+4   1.5   2  RT (min)      lcqtof_hilic_neg_4_000_aa    m/z 756.6706     0   2   4   6E+4   1.5   2  RT (min)      lcqtof_hilic_neg_4_000_aa    m/z 756.6717     0   2   4   6   8E+4   1   1.5  RT (min)      lcqtof_hilic_neg_4_002_25    m/z 756.6938     0   2   4   6E+4   0.5   1  RT (min)      lcqtof_hilic_neg_3_035_46    m/z 756.8952     0   1   2   3E+4   0.5   1  RT (min)      lcqtof_hilic_neg_4_098_61    m/z 756.9093     0   1   2   3E+4   0.5   1  RT (min)      lcqtof_hilic_neg_3_035_46    m/z 757.3998     0   1E+4   1   1.5  RT (min)      lcqtof_hilic_neg_4_005_29    m/z 757.4818     0   1   2E+5   4   4.5  RT (min)      lcqtof_hilic_neg_4_016_37    m/z 757.5478     0   1E+4   4   4.5  RT (min)      lcqtof_hilic_neg_4_008_24    m/z 757.6431     0   1E+4   0.5   1  RT (min)      lcqtof_hilic_neg_2_028_05    m/z 757.6440     0   1   2E+5   1   1.5  RT (min)      lcqtof_hilic_neg_4_000_aa    m/z 757.6746     0   1   2   3E+4   1.5   2  RT (min)      lcqtof_hilic_neg_4_000_aa    m/z 757.6754     0   1   2   3E+4   1.5   2  RT (min)      lcqtof_hilic_neg_4_000_aa    m/z 757.9112     0   1E+4   0.5   1  RT (min)      lcqtof_hilic_neg_3_035_46    m/z 758.4615     0   1E+4   1.5   2  RT (min)      lcqtof_hilic_neg_4_000_aa    m/z 758.4636     0   1E+4   1.5   2  RT (min)      lcqtof_hilic_neg_4_000_aa    m/z 758.4637     0   1E+4   1.5   2  RT (min)      lcqtof_hilic_neg_4_000_aa    m/z 758.4706     0   1E+4   0.5   1  RT (min)      lcqtof_hilic_neg_4_021_21    m/z 758.4922     0   2   4   6E+4   1   1.5  RT (min)      lcqtof_hilic_neg_4_000_aa    m/z 758.4961     0   1E+4   14.5   15  RT (min)       lcqtof_hilic_neg_4_083_59    m/z 758.4968     0   1   2   3E+6   4.5   5  RT (min)       lcqtof_hilic_neg_4_048_23    m/z 758.4970     0   1   2E+6   4.5   5  RT (min)      lcqtof_hilic_neg_4_016_37    m/z 758.4970     0   1   2   3E+6   4   4.5  RT (min)       lcqtof_hilic_neg_4_048_23    m/z 758.5418     0   1   2   3   4E+3   1.5   2  RT (min)       lcqtof_hilic_neg_2_020_23    m/z 758.6324     0   1E+4   0.5   1  RT (min)      lcqtof_hilic_neg_2_028_05    m/z 758.6531     0   1E+5   1   1.5  RT (min)      lcqtof_hilic_neg_4_000_aa    m/z 758.7336     0   1E+4   0.5   1  RT (min)      lcqtof_hilic_neg_1_037_24    m/z 759.3239     0   1   2E+4   0.5   1  RT (min)      lcqtof_hilic_neg_4_045_26    m/z 759.4618     0   2   4E+4   1   1.5  RT (min)      lcqtof_hilic_neg_4_000_aa    m/z 759.4714     0   2   4E+4   1   1.5  RT (min)      lcqtof_hilic_neg_4_000_aa    m/z 759.4892     0   2   4E+4   1   1.5  RT (min)      lcqtof_hilic_neg_4_000_aa    m/z 759.4931     0   1E+4   0.5   1  RT (min)      lcqtof_hilic_neg_4_039_18    m/z 759.5004     0   1E+6   4.5   5  RT (min)      lcqtof_hilic_neg_4_016_37    m/z 759.5009     0   1E+6   4   4.5  RT (min)      lcqtof_hilic_neg_4_028_36    m/z 759.5367     0   1   2E+4   1.5   2  RT (min)      lcqtof_hilic_neg_4_000_aa    m/z 759.5748     0   2   4   6   8E+4   1   1.5  RT (min)      lcqtof_hilic_neg_4_000_aa    m/z 759.5797     0   2   4   6   8E+4   1   1.5  RT (min)      lcqtof_hilic_neg_4_000_aa    m/z 759.6448     0   1   2   3   4E+4   1.5   2  RT (min)      lcqtof_hilic_neg_4_000_aa    m/z 759.6578     0   1   2   3   4E+4   1.5   2  RT (min)      lcqtof_hilic_neg_4_000_aa    m/z 759.7581     0   1   2   3E+4   1.5   2  RT (min)      lcqtof_hilic_neg_4_000_aa    m/z 759.7582     0   1   2   3E+4   1.5   2  RT (min)      lcqtof_hilic_neg_4_000_aa    m/z 759.7634     0   1   2   3E+4   1.5   2  RT (min)      lcqtof_hilic_neg_4_000_aa    m/z 759.7636     0   1   2   3E+4   1.5   2  RT (min)      lcqtof_hilic_neg_4_000_aa    m/z 760.2773     0   1E+4   4   4.5  RT (min)      lcqtof_hilic_neg_4_048_23    m/z 760.2799     0   1E+4   4   4.5  RT (min)      lcqtof_hilic_neg_4_016_37    m/z 760.4180     0   1E+4   1   1.5  RT (min)      lcqtof_hilic_neg_4_005_29    m/z 760.4748     0   1   2   3E+4   1   1.5  RT (min)      lcqtof_hilic_neg_4_000_aa    m/z 760.4952     0   1   2   3E+4   1   1.5  RT (min)      lcqtof_hilic_neg_4_000_aa    m/z 760.5095     0   1E+6   4.5   5  RT (min)      lcqtof_hilic_neg_4_048_23    m/z 760.5100     0   2   4   6   8E+5   4.5   5  RT (min)      lcqtof_hilic_neg_4_016_37    m/z 760.5103     0   1E+6   4   4.5  RT (min)      lcqtof_hilic_neg_4_048_23    m/z 760.5136     0   1E+5   4.5   5  RT (min)      lcqtof_hilic_neg_4_009_50    m/z 760.5338     0   1E+4   4   4.5  RT (min)      lcqtof_hilic_neg_4_008_24    m/z 760.5650     0   1   2   3   4E+4   1   1.5  RT (min)      lcqtof_hilic_neg_4_000_aa    m/z 760.5765     0   1   2   3   4E+4   1   1.5  RT (min)      lcqtof_hilic_neg_4_000_aa    m/z 760.7614     0   1E+4   1.5   2  RT (min)      lcqtof_hilic_neg_4_000_aa    m/z 761.0888     0   1E+4   4   4.5  RT (min)      lcqtof_hilic_neg_4_008_24    m/z 761.1448     0   1E+4   6   6.5  RT (min)       lcqtof_hilic_neg_3_034_24    m/z 761.1677     0   1E+4   0.5   1  RT (min)       lcqtof_hilic_neg_4_072_57    m/z 761.3437     0   1E+4   6   6.5  RT (min)      lcqtof_hilic_neg_4_008_24    m/z 761.5156     0   1   2   3   4E+5   4   4.5  RT (min)       lcqtof_hilic_neg_4_028_36    m/z 761.5166     0   1   2E+5   4.5   5  RT (min)      lcqtof_hilic_neg_4_016_37    m/z 761.5860     0   1   2E+5   1   1.5  RT (min)      lcqtof_hilic_neg_4_000_aa    m/z 761.5932     0   1   2E+5   1   1.5  RT (min)      lcqtof_hilic_neg_4_000_aa    m/z 761.6009     0   1   2E+5   1   1.5  RT (min)      lcqtof_hilic_neg_4_000_aa    m/z 761.6613     0   1   2E+4   0.5   1  RT (min)      lcqtof_hilic_neg_2_028_05    m/z 761.7025     0   1   2E+4   1   1.5  RT (min)      lcqtof_hilic_neg_4_000_aa    m/z 761.7172     0   1   2E+4   1.5   2  RT (min)      lcqtof_hilic_neg_4_000_aa    m/z 761.7402     0   1   2E+4   1.5   2  RT (min)      lcqtof_hilic_neg_4_000_aa    m/z 761.7456     0   1   2E+4   1.5   2  RT (min)      lcqtof_hilic_neg_4_000_aa    m/z 761.9454     0   1E+4   5   5.5  RT (min)       lcqtof_hilic_neg_4_008_24    m/z 762.0267     0   1E+4   0.5   1  RT (min)      lcqtof_hilic_neg_3_035_46    m/z 762.4808     0   2   4E+4   1   1.5  RT (min)      lcqtof_hilic_neg_4_000_aa    m/z 762.4810     0   2   4E+4   1   1.5  RT (min)      lcqtof_hilic_neg_4_000_aa    m/z 762.5058     0   1E+5   2.5   3  RT (min)      lcqtof_hilic_neg_4_011_35    m/z 762.5063     0   2   4   6E+3   2   2.5  RT (min)      lcqtof_hilic_neg_4_032_01    m/z 762.5064     0   2   4   6E+4   2.5   3  RT (min)      lcqtof_hilic_neg_4_011_35    m/z 762.5067     0   1   2   3E+4   2   2.5  RT (min)      lcqtof_hilic_neg_1_003_37    m/z 762.5068     0   2   4   6   8E+4   1   1.5  RT (min)      lcqtof_hilic_neg_4_011_35    m/z 762.5073     0   2   4   6E+3   2   2.5  RT (min)      lcqtof_hilic_neg_4_032_01    m/z 762.5086     0   2   4   6E+3   1.5   2  RT (min)      lcqtof_hilic_neg_1_051_35    m/z 762.5092     0   2   4   6E+3   1.5   2  RT (min)      lcqtof_hilic_neg_4_032_01    m/z 762.5097     0   1E+4   1.5   2  RT (min)      lcqtof_hilic_neg_4_011_35    m/z 762.5110     0   2   4E+3   1.5   2  RT (min)      lcqtof_hilic_neg_4_060_13    m/z 762.5127     0   1   2E+4   1.5   2  RT (min)      lcqtof_hilic_neg_1_051_35    m/z 762.5191     0   1E+5   4   4.5  RT (min)      lcqtof_hilic_neg_4_048_23    m/z 762.5754     0   2   4   6E+4   1   1.5  RT (min)      lcqtof_hilic_neg_4_000_aa    m/z 762.5986     0   1   2   3E+5   1   1.5  RT (min)      lcqtof_hilic_neg_4_000_aa    m/z 762.6003     0   1E+5   3.5   4  RT (min)      lcqtof_hilic_neg_2_047_32    m/z 762.6064     0   1   2   3E+5   1   1.5  RT (min)      lcqtof_hilic_neg_4_000_aa    m/z 762.7094     0   1E+4   1   1.5  RT (min)      lcqtof_hilic_neg_4_000_aa    m/z 762.7456     0   1E+4   1.5   2  RT (min)      lcqtof_hilic_neg_4_000_aa    m/z 762.7737     0   1E+4   8.5   9  RT (min)       lcqtof_hilic_neg_4_059_14    m/z 763.0580     0   1E+4   0.5   1  RT (min)      lcqtof_hilic_neg_3_007_04    m/z 763.1035     0   1E+4   3.5   4  RT (min)      lcqtof_hilic_neg_4_055_32    m/z 763.1043     0   1   2E+4   3.5   4  RT (min)      lcqtof_hilic_neg_4_008_24    m/z 763.1046     0   1   2E+4   3.5   4  RT (min)       lcqtof_hilic_neg_1_005_32    m/z 763.1047     0   2   4   6E+4   3.5   4  RT (min)      lcqtof_hilic_neg_4_008_24    m/z 763.1054     0   2   4   6E+4   3.5   4  RT (min)      lcqtof_hilic_neg_4_008_24    m/z 763.5082     0   1E+5   3   3.5  RT (min)      lcqtof_hilic_neg_4_048_23    m/z 763.5110     0   1E+4   2   2.5  RT (min)      lcqtof_hilic_neg_1_003_37    m/z 763.5650     0   1   2   3E+5   1   1.5  RT (min)      lcqtof_hilic_neg_4_000_aa    m/z 763.5758     0   1   2   3E+5   1.5   2  RT (min)      lcqtof_hilic_neg_4_000_aa    m/z 763.5765     0   1   2   3E+5   1   1.5  RT (min)      lcqtof_hilic_neg_4_000_aa    m/z 763.5776     0   1   2   3E+5   1.5   2  RT (min)      lcqtof_hilic_neg_4_000_aa    m/z 763.5894     0   1   2   3E+5   1   1.5  RT (min)      lcqtof_hilic_neg_4_000_aa    m/z 763.6760     0   1E+4   0.5   1  RT (min)      lcqtof_hilic_neg_1_007_05    m/z 763.6938     0   1   2   3   4E+4   1   1.5  RT (min)      lcqtof_hilic_neg_4_000_aa    m/z 763.7007     0   1   2   3   4E+4   1   1.5  RT (min)      lcqtof_hilic_neg_4_000_aa    m/z 764.1117     0   2   4   6   8E+4   3.5   4  RT (min)      lcqtof_hilic_neg_4_008_24    m/z 764.1120     0   1E+4   3.5   4  RT (min)      lcqtof_hilic_neg_4_008_24    m/z 764.1120     0   1E+4   3.5   4  RT (min)      lcqtof_hilic_neg_4_008_24    m/z 764.1120     0   1E+4   3.5   4  RT (min)      lcqtof_hilic_neg_1_005_32    m/z 764.1120     0   2   4   6   8E+4   3.5   4  RT (min)      lcqtof_hilic_neg_4_008_24    m/z 764.1121     0   1E+4   3.5   4  RT (min)      lcqtof_hilic_neg_4_055_32    m/z 764.1128     0   1   2   3   4E+4   3.5   4  RT (min)      lcqtof_hilic_neg_4_008_24    m/z 764.4636     0   1   2E+4   4   4.5  RT (min)      lcqtof_hilic_neg_4_008_24    m/z 764.4750     0   1E+4   3.5   4  RT (min)      lcqtof_hilic_neg_4_032_01    m/z 764.5209     0   1   2E+4   2   2.5  RT (min)      lcqtof_hilic_neg_4_012_07    m/z 764.5218     0   1   2E+5   2.5   3  RT (min)      lcqtof_hilic_neg_4_011_35    m/z 764.5218     0   2   4   6E+3   2   2.5  RT (min)      lcqtof_hilic_neg_2_062_17    m/z 764.5221     0   2   4   6E+4   2.5   3  RT (min)      lcqtof_hilic_neg_4_011_35    m/z 764.5222     0   2   4   6E+3   2   2.5  RT (min)      lcqtof_hilic_neg_4_005_29    m/z 764.5222     0   1   2E+5   2.5   3  RT (min)      lcqtof_hilic_neg_4_011_35    m/z 764.5222     0   1   2   3E+5   2.5   3  RT (min)       lcqtof_hilic_neg_4_016_37    m/z 764.5223     0   1E+5   2   2.5  RT (min)       lcqtof_hilic_neg_4_034_31    m/z 764.5223     0   1E+5   2   2.5  RT (min)       lcqtof_hilic_neg_1_035_23    m/z 764.5224     0   1E+5   1   1.5  RT (min)      lcqtof_hilic_neg_2_044_37    m/z 764.5227     0   2   4   6E+5   1   1.5  RT (min)       lcqtof_hilic_neg_4_048_23    m/z 764.5230     0   1   2E+4   1.5   2  RT (min)       lcqtof_hilic_neg_4_086_59    m/z 764.5243     0   1E+5   2.5   3  RT (min)      lcqtof_hilic_neg_2_020_23    m/z 764.5526     0   1   2   3   4E+4   3   3.5  RT (min)      lcqtof_hilic_neg_1_010_15    m/z 764.5738     0   1E+5   1   1.5  RT (min)      lcqtof_hilic_neg_4_000_aa    m/z 764.5780     0   1E+5   1.5   2  RT (min)      lcqtof_hilic_neg_4_000_aa    m/z 764.5782     0   1E+5   1.5   2  RT (min)      lcqtof_hilic_neg_4_000_aa    m/z 764.5786     0   1E+5   1.5   2  RT (min)      lcqtof_hilic_neg_4_000_aa    m/z 764.5857     0   1E+5   1   1.5  RT (min)      lcqtof_hilic_neg_4_000_aa    m/z 764.6286     0   2   4E+4   3   3.5  RT (min)      lcqtof_hilic_neg_4_008_24    m/z 764.6996     0   1E+4   1   1.5  RT (min)      lcqtof_hilic_neg_4_000_aa    m/z 765.5256     0   1E+5   2.5   3  RT (min)      lcqtof_hilic_neg_4_011_35    m/z 765.5260     0   1   2   3E+4   2.5   3  RT (min)      lcqtof_hilic_neg_4_011_35    m/z 765.5260     0   1   2E+5   1   1.5  RT (min)      lcqtof_hilic_neg_4_048_23    m/z 765.5457     0   1   2E+5   1   1.5  RT (min)      lcqtof_hilic_neg_4_000_aa    m/z 765.5836     0   1E+5   1   1.5  RT (min)      lcqtof_hilic_neg_4_000_aa    m/z 765.5870     0   1E+5   1   1.5  RT (min)      lcqtof_hilic_neg_4_000_aa    m/z 765.5950     0   1E+5   1   1.5  RT (min)      lcqtof_hilic_neg_4_000_aa    m/z 765.6226     0   1   2E+5   1   1.5  RT (min)      lcqtof_hilic_neg_4_000_aa    m/z 765.6232     0   1E+4   15   15.5  RT (min)      lcqtof_hilic_neg_4_013_20    m/z 765.6234     0   1E+4   14.5   15   15.5  RT (min)       lcqtof_hilic_neg_4_082_59    m/z 765.6237     0   1E+4   16   16.5  RT (min)      lcqtof_hilic_neg_1_007_05    m/z 765.6260     0   1   2E+5   1.5   2  RT (min)      lcqtof_hilic_neg_4_000_aa    m/z 765.6400     0   1E+4   3   3.5  RT (min)      lcqtof_hilic_neg_4_008_24    m/z 766.3432     0   1E+4   3.5   4  RT (min)      lcqtof_hilic_neg_4_050_02    m/z 766.4933     0   1E+4   3.5   4  RT (min)      lcqtof_hilic_neg_2_031_01    m/z 766.5072     0   1E+4   0.5   1  RT (min)      lcqtof_hilic_neg_4_040_05    m/z 766.5374     0   1E+5   1   1.5  RT (min)      lcqtof_hilic_neg_2_021_17    m/z 766.5376     0   2   4   6   8E+4   1   1.5  RT (min)      lcqtof_hilic_neg_4_000_aa    m/z 766.5376     0   1E+4   1.5   2  RT (min)       lcqtof_hilic_neg_4_053_14    m/z 766.5376     0   1   2E+5   2   2.5  RT (min)       lcqtof_hilic_neg_1_003_37    m/z 766.5376     0   1   2E+5   2   2.5  RT (min)       lcqtof_hilic_neg_4_034_31    m/z 766.5376     0   1E+5   1.5   2  RT (min)      lcqtof_hilic_neg_4_059_14    m/z 766.5378     0   1E+4   1.5   2  RT (min)       lcqtof_hilic_neg_3_006_30    m/z 766.5378     0   1E+5   2   2.5  RT (min)       lcqtof_hilic_neg_3_084_59    m/z 766.5379     0   1   2E+4   2   2.5  RT (min)      lcqtof_hilic_neg_2_031_01    m/z 766.5379     0   1E+4   1.5   2  RT (min)       lcqtof_hilic_neg_4_053_14    m/z 766.5380     0   1   2E+5   2.5   3  RT (min)      lcqtof_hilic_neg_2_048_35    m/z 766.5381     0   1   2E+4   1.5   2  RT (min)      lcqtof_hilic_neg_2_031_01    m/z 766.5384     0   2   4   6   8E+4   2.5   3  RT (min)      lcqtof_hilic_neg_4_011_35    m/z 766.5823     0   1   2E+4   1   1.5  RT (min)      lcqtof_hilic_neg_4_000_aa    m/z 766.5852     0   1   2E+4   1.5   2  RT (min)      lcqtof_hilic_neg_4_000_aa    m/z 766.6085     0   2   4   6   8E+4   1   1.5  RT (min)      lcqtof_hilic_neg_4_000_aa    m/z 766.6273     0   1E+4   15   15.5  RT (min)      lcqtof_hilic_neg_4_015_09    m/z 766.6407     0   2   4   6   8E+4   1.5   2  RT (min)      lcqtof_hilic_neg_4_000_aa    m/z 766.6910     0   1   2   3E+4   1.5   2  RT (min)      lcqtof_hilic_neg_4_000_aa    m/z 766.6923     0   1   2   3E+4   1.5   2  RT (min)      lcqtof_hilic_neg_4_000_aa    m/z 766.6923     0   1E+4   0.5   1  RT (min)      lcqtof_hilic_neg_4_040_05    m/z 766.7254     0   1E+4   2   2.5  RT (min)      lcqtof_hilic_neg_4_000_aa    m/z 766.7501     0   2   4E+3   6.5   7  RT (min)       lcqtof_hilic_neg_4_012_07    m/z 767.2456     0   1E+4   1   1.5  RT (min)      lcqtof_hilic_neg_4_000_aa    m/z 767.2476     0   1E+4   1   1.5  RT (min)      lcqtof_hilic_neg_4_000_aa    m/z 767.2504     0   1E+4   1   1.5  RT (min)      lcqtof_hilic_neg_4_000_aa    m/z 767.3596     0   1E+4   3   3.5  RT (min)       lcqtof_hilic_neg_4_108_62    m/z 767.5407     0   2   4   6E+4   1.5   2  RT (min)      lcqtof_hilic_neg_4_059_14    m/z 767.5408     0   1   2   3E+5   3   3.5  RT (min)       lcqtof_hilic_neg_4_062_17    m/z 767.5410     0   1   2   3   4E+4   2   2.5  RT (min)      lcqtof_hilic_neg_4_035_33    m/z 767.5412     0   1E+5   2   2.5  RT (min)      lcqtof_hilic_neg_1_003_37    m/z 767.5412     0   1   2E+4   2   2.5  RT (min)      lcqtof_hilic_neg_3_009_50    m/z 767.5414     0   1   2   3   4E+4   2   2.5  RT (min)      lcqtof_hilic_neg_4_035_33    m/z 767.5624     0   1   2   3E+4   1   1.5  RT (min)      lcqtof_hilic_neg_4_000_aa    m/z 767.5670     0   1   2   3E+4   1   1.5  RT (min)      lcqtof_hilic_neg_4_000_aa    m/z 767.5746     0   1   2   3E+4   1   1.5  RT (min)      lcqtof_hilic_neg_4_000_aa    m/z 767.5992     0   1   2   3E+4   1   1.5  RT (min)      lcqtof_hilic_neg_4_000_aa    m/z 767.6172     0   1   2   3E+4   2.5   3  RT (min)      lcqtof_hilic_neg_4_055_32    m/z 767.6178     0   1E+4   2.5   3  RT (min)      lcqtof_hilic_neg_4_097_60    m/z 767.6219     0   1   2E+4   3   3.5  RT (min)      lcqtof_hilic_neg_4_008_24    m/z 767.6465     0   1   2E+4   1.5   2  RT (min)      lcqtof_hilic_neg_4_000_aa    m/z 767.6563     0   1E+4   1   1.5  RT (min)      lcqtof_hilic_neg_4_000_aa    m/z 767.6830     0   1E+4   1   1.5  RT (min)      lcqtof_hilic_neg_4_008_24    m/z 767.6918     0   1E+4   1.5   2  RT (min)      lcqtof_hilic_neg_4_000_aa    m/z 767.6938     0   1E+4   1.5   2  RT (min)      lcqtof_hilic_neg_4_000_aa    m/z 767.7677     0   1E+4   0.5   1  RT (min)      lcqtof_hilic_neg_4_040_05    m/z 767.7899     0   1E+4   0.5   1  RT (min)      lcqtof_hilic_neg_4_008_24    m/z 768.5048     0   2   4E+4   3.5   4  RT (min)      lcqtof_hilic_neg_4_032_01    m/z 768.5214     0   1E+4   0.5   1  RT (min)      lcqtof_hilic_neg_4_040_05    m/z 768.5296     0   1E+5   3.5   4  RT (min)      lcqtof_hilic_neg_4_050_02    m/z 768.5514     0   2   4   6E+4   2.5   3  RT (min)      lcqtof_hilic_neg_4_032_01    m/z 768.5518     0   2   4E+5   1   1.5  RT (min)       lcqtof_hilic_neg_4_048_23    m/z 768.5520     0   2   4   6E+4   2.5   3  RT (min)      lcqtof_hilic_neg_4_016_37    m/z 768.5520     0   2   4   6E+4   2.5   3  RT (min)      lcqtof_hilic_neg_4_032_01    m/z 768.5521     0   2   4   6   8E+4   2.5   3  RT (min)       lcqtof_hilic_neg_4_032_01    m/z 768.5524     0   1   2   3E+5   2.5   3  RT (min)      lcqtof_hilic_neg_4_032_01    m/z 768.5860     0   1E+5   0.5   1  RT (min)      lcqtof_hilic_neg_3_035_46    m/z 768.6015     0   1E+4   1   1.5  RT (min)      lcqtof_hilic_neg_4_000_aa    m/z 768.6018     0   1E+4   2.5   3  RT (min)      lcqtof_hilic_neg_4_047_06    m/z 768.6023     0   2   4   6E+4   2.5   3  RT (min)      lcqtof_hilic_neg_4_055_32    m/z 768.6098     0   1E+4   3   3.5  RT (min)      lcqtof_hilic_neg_4_002_25    m/z 768.6296     0   1   2   3   4E+4   3   3.5  RT (min)      lcqtof_hilic_neg_4_008_24    m/z 768.6558     0   2   4   6   8E+4   1   1.5  RT (min)      lcqtof_hilic_neg_4_008_24    m/z 768.6559     0   1   2E+4   2   2.5  RT (min)      lcqtof_hilic_neg_4_008_24    m/z 768.6560     0   1   2E+4   2   2.5  RT (min)      lcqtof_hilic_neg_4_008_24    m/z 768.6570     0   1E+4   2   2.5  RT (min)       lcqtof_hilic_neg_2_047_32    m/z 768.6571     0   1   2E+4   1.5   2  RT (min)      lcqtof_hilic_neg_4_008_24    m/z 768.6574     0   1   2E+4   2   2.5  RT (min)       lcqtof_hilic_neg_4_008_24    m/z 768.6575     0   1   2E+4   1.5   2  RT (min)       lcqtof_hilic_neg_4_008_24    m/z 768.6577     0   1   2E+4   1.5   2  RT (min)       lcqtof_hilic_neg_4_008_24    m/z 768.6580     0   1   2E+4   1.5   2  RT (min)      lcqtof_hilic_neg_4_008_24    m/z 768.6584     0   1   2E+4   1.5   2  RT (min)      lcqtof_hilic_neg_4_008_24    m/z 768.6590     0   1   2   3E+4   1.5   2  RT (min)      lcqtof_hilic_neg_4_008_24    m/z 768.6732     0   1   2   3   4E+4   1.5   2  RT (min)      lcqtof_hilic_neg_4_000_aa    m/z 768.7074     0   1   2   3   4E+4   1.5   2  RT (min)      lcqtof_hilic_neg_4_000_aa    m/z 769.5056     0   1E+4   3.5   4  RT (min)      lcqtof_hilic_neg_4_032_01    m/z 769.5243     0   1E+4   1   1.5  RT (min)      lcqtof_hilic_neg_4_000_aa    m/z 769.5298     0   1E+4   1   1.5  RT (min)      lcqtof_hilic_neg_4_000_aa    m/z 769.5570     0   1E+5   2.5   3  RT (min)      lcqtof_hilic_neg_4_032_01    m/z 769.5808     0   2   4E+4   0.5   1  RT (min)      lcqtof_hilic_neg_3_035_46    m/z 769.5812     0   1   2   3E+4   1.5   2  RT (min)      lcqtof_hilic_neg_4_000_aa    m/z 769.6215     0   1   2E+4   2.5   3  RT (min)      lcqtof_hilic_neg_4_055_32    m/z 769.6296     0   1E+4   3   3.5  RT (min)      lcqtof_hilic_neg_4_008_24    m/z 769.6784     0   1   2E+4   1.5   2  RT (min)      lcqtof_hilic_neg_4_000_aa    m/z 769.7096     0   1   2E+4   1.5   2  RT (min)      lcqtof_hilic_neg_4_000_aa    m/z 769.8058     0   1E+4   0.5   1  RT (min)      lcqtof_hilic_neg_3_035_46    m/z 770.3935     0   1E+4   3.5   4  RT (min)      lcqtof_hilic_neg_4_005_29    m/z 770.4740     0   1E+4   4   4.5  RT (min)      lcqtof_hilic_neg_4_093_60    m/z 770.5692     0   2   4E+5   3   3.5  RT (min)      lcqtof_hilic_neg_4_032_01    m/z 770.5694     0   2   4   6   8E+4   2.5   3  RT (min)      lcqtof_hilic_neg_4_032_01    m/z 770.5694     0   1   2E+4   1.5   2  RT (min)      lcqtof_hilic_neg_4_052_13    m/z 770.5694     0   2   4   6E+5   2.5   3  RT (min)       lcqtof_hilic_neg_4_010_28    m/z 770.5696     0   2   4   6   8E+4   2.5   3  RT (min)       lcqtof_hilic_neg_4_032_01    m/z 770.5711     0   2   4   6   8E+3   1.5   2  RT (min)      lcqtof_hilic_neg_4_052_13    m/z 770.5847     0   1   2E+5   2.5   3  RT (min)      lcqtof_hilic_neg_4_049_55    m/z 770.5924     0   1E+5   0.5   1  RT (min)      lcqtof_hilic_neg_3_035_46    m/z 770.6015     0   2   4   6   8E+4   3   3.5  RT (min)      lcqtof_hilic_neg_4_055_32    m/z 770.6169     0   1   2   3E+4   2.5   3  RT (min)      lcqtof_hilic_neg_4_097_60    m/z 770.6170     0   1E+4   2.5   3  RT (min)      lcqtof_hilic_neg_4_047_06    m/z 770.6175     0   2   4   6   8E+4   2.5   3  RT (min)       lcqtof_hilic_neg_4_055_32    m/z 770.6282     0   1   2   3   4E+4   3   3.5  RT (min)      lcqtof_hilic_neg_4_008_24    m/z 770.6762     0   1   2   3   4E+4   1.5   2  RT (min)      lcqtof_hilic_neg_4_000_aa    m/z 770.6858     0   1   2   3   4E+4   1.5   2  RT (min)      lcqtof_hilic_neg_4_000_aa    m/z 770.7820     0   2   4   6E+4   1   1.5  RT (min)      lcqtof_hilic_neg_4_002_25    m/z 770.8055     0   2   4E+4   0.5   1  RT (min)      lcqtof_hilic_neg_3_035_46    m/z 770.8501     0   1E+4   3   3.5  RT (min)      lcqtof_hilic_neg_4_005_29    m/z 771.4325     0   1E+4   4   4.5  RT (min)      lcqtof_hilic_neg_4_047_06    m/z 771.5736     0   1   2   3E+4   2.5   3  RT (min)      lcqtof_hilic_neg_4_032_01    m/z 771.5738     0   1   2E+4   2.5   3  RT (min)      lcqtof_hilic_neg_4_060_13    m/z 771.5768     0   1   2   3   4E+3   1   1.5  RT (min)      lcqtof_hilic_neg_4_008_24    m/z 771.6058     0   1   2   3E+4   1   1.5  RT (min)      lcqtof_hilic_neg_4_000_aa    m/z 771.6436     0   1E+4   0.5   1  RT (min)      lcqtof_hilic_neg_4_053_14    m/z 771.6892     0   1E+4   1.5   2  RT (min)      lcqtof_hilic_neg_4_000_aa    m/z 772.4894     0   2   4E+4   4   4.5  RT (min)       lcqtof_hilic_neg_4_095_60    m/z 772.5012     0   1   2   3E+5   4   4.5  RT (min)       lcqtof_hilic_neg_4_048_23    m/z 772.5028     0   1   2E+5   4.5   5  RT (min)      lcqtof_hilic_neg_4_048_23    m/z 772.5571     0   1E+4   1.5   2  RT (min)       lcqtof_hilic_neg_2_020_23    m/z 772.5576     0   1   2E+4   1   1.5  RT (min)       lcqtof_hilic_neg_4_048_23    m/z 772.5578     0   1   2E+4   1   1.5  RT (min)      lcqtof_hilic_neg_4_048_23    m/z 772.5586     0   1E+4   1.5   2  RT (min)      lcqtof_hilic_neg_1_035_23    m/z 772.5587     0   1E+4   1.5   2  RT (min)      lcqtof_hilic_neg_2_020_23    m/z 772.5599     0   1E+4   4.5   5  RT (min)      lcqtof_hilic_neg_4_050_02    m/z 772.5784     0   1   2E+4   2   2.5  RT (min)      lcqtof_hilic_neg_4_016_37    m/z 772.5786     0   2   4E+3   2   2.5  RT (min)      lcqtof_hilic_neg_2_033_53    m/z 772.8016     0   1E+4   0.5   1  RT (min)      lcqtof_hilic_neg_4_004_47    m/z 772.9097     0   1E+4   0.5   1  RT (min)      lcqtof_hilic_neg_4_104_61    m/z 772.9818     0   1E+4   1   1.5  RT (min)      lcqtof_hilic_neg_4_000_aa    m/z 773.1115     0   1E+4   7   7.5  RT (min)       lcqtof_hilic_neg_4_091_60    m/z 773.3386     0   1   2E+4   0.5   1  RT (min)      lcqtof_hilic_neg_4_045_26    m/z 773.3814     0   1E+4   0.5   1  RT (min)      lcqtof_hilic_neg_4_038_10    m/z 773.5041     0   1E+5   4   4.5  RT (min)      lcqtof_hilic_neg_4_048_23    m/z 773.5252     0   1E+4   1   1.5  RT (min)      lcqtof_hilic_neg_4_048_23    m/z 773.5943     0   1E+4   3   3.5  RT (min)      lcqtof_hilic_neg_1_005_32    m/z 773.6245     0   1E+5   1   1.5  RT (min)      lcqtof_hilic_neg_4_000_aa    m/z 773.7144     0   1E+4   0.5   1  RT (min)      lcqtof_hilic_neg_4_008_24    m/z 773.7442     0   1E+4   1.5   2  RT (min)      lcqtof_hilic_neg_4_000_aa    m/z 773.7760     0   1E+4   1.5   2  RT (min)      lcqtof_hilic_neg_4_000_aa    m/z 774.0238     0   1E+4   0.5   1  RT (min)      lcqtof_hilic_neg_3_007_04    m/z 774.5012     0   2   4   6E+3   2   2.5  RT (min)       lcqtof_hilic_neg_4_053_14    m/z 774.5108     0   1E+5   4.5   5  RT (min)      lcqtof_hilic_neg_4_048_23    m/z 774.5316     0   2   4E+3   2.5   3  RT (min)      lcqtof_hilic_neg_4_048_23    m/z 774.5470     0   1   2E+4   4   4.5  RT (min)      lcqtof_hilic_neg_4_008_24    m/z 774.5486     0   1   2E+4   4   4.5  RT (min)      lcqtof_hilic_neg_4_008_24    m/z 774.6276     0   1E+4   0.5   1  RT (min)      lcqtof_hilic_neg_4_055_32    m/z 774.6290     0   2   4   6E+4   1   1.5  RT (min)      lcqtof_hilic_neg_4_000_aa    m/z 774.6303     0   2   4   6E+4   1   1.5  RT (min)      lcqtof_hilic_neg_4_000_aa    m/z 775.0218     0   1E+4   0.5   1  RT (min)      lcqtof_hilic_neg_3_007_04    m/z 775.3374     0   1   2E+4   0.5   1  RT (min)      lcqtof_hilic_neg_4_045_26    m/z 775.3654     0   1E+4   1   1.5  RT (min)      lcqtof_hilic_neg_4_000_aa    m/z 775.3690     0   1E+4   1   1.5  RT (min)      lcqtof_hilic_neg_4_000_aa    m/z 775.3794     0   1E+4   1   1.5  RT (min)      lcqtof_hilic_neg_4_000_aa    m/z 775.4872     0   1   2   3E+4   1   1.5  RT (min)      lcqtof_hilic_neg_4_000_aa    m/z 775.4904     0   1   2   3E+4   1   1.5  RT (min)      lcqtof_hilic_neg_4_000_aa    m/z 775.5310     0   1E+4   1   1.5  RT (min)       lcqtof_hilic_neg_4_048_23    m/z 775.5426     0   1   2E+4   2   2.5  RT (min)      lcqtof_hilic_neg_4_034_31    m/z 775.5480     0   1E+4   2   2.5  RT (min)       lcqtof_hilic_neg_4_086_59    m/z 775.5666     0   1   2   3E+4   2   2.5  RT (min)      lcqtof_hilic_neg_4_048_23    m/z 775.5946     0   1E+4   2   2.5  RT (min)      lcqtof_hilic_neg_4_040_05    m/z 775.6166     0   1   2   3E+4   1.5   2  RT (min)      lcqtof_hilic_neg_2_052_24    m/z 775.6168     0   1   2   3E+4   2   2.5  RT (min)      lcqtof_hilic_neg_2_047_32    m/z 775.6168     0   1   2   3E+4   1   1.5  RT (min)      lcqtof_hilic_neg_4_008_24    m/z 775.6168     0   1   2   3E+4   1   1.5  RT (min)      lcqtof_hilic_neg_4_008_24    m/z 775.6172     0   1   2   3E+4   1.5   2  RT (min)      lcqtof_hilic_neg_1_037_24    m/z 775.6172     0   1   2   3   4E+4   2   2.5  RT (min)      lcqtof_hilic_neg_4_008_24    m/z 775.6173     0   1   2E+4   1.5   2  RT (min)      lcqtof_hilic_neg_4_091_60    m/z 775.6173     0   1   2   3   4E+5   2.5   3  RT (min)      lcqtof_hilic_neg_4_008_24    m/z 775.6175     0   1   2   3   4E+4   2   2.5  RT (min)      lcqtof_hilic_neg_4_008_24    m/z 775.6176     0   1   2   3E+4   1.5   2  RT (min)      lcqtof_hilic_neg_2_052_24    m/z 775.6178     0   2   4E+5   2.5   3  RT (min)      lcqtof_hilic_neg_1_037_24    m/z 775.6178     0   1   2   3E+4   2   2.5  RT (min)      lcqtof_hilic_neg_2_052_24    m/z 775.6180     0   1   2   3   4E+4   2   2.5  RT (min)      lcqtof_hilic_neg_4_008_24    m/z 775.6426     0   1   2   3   4E+4   1   1.5  RT (min)      lcqtof_hilic_neg_4_000_aa    m/z 775.6442     0   1   2   3   4E+4   1   1.5  RT (min)      lcqtof_hilic_neg_4_000_aa    m/z 775.6479     0   1   2   3   4E+4   1   1.5  RT (min)      lcqtof_hilic_neg_4_000_aa    m/z 775.6542     0   1   2   3   4E+4   1   1.5  RT (min)      lcqtof_hilic_neg_4_000_aa    m/z 775.7350     0   1E+4   1   1.5  RT (min)      lcqtof_hilic_neg_4_000_aa    m/z 775.7489     0   1E+4   1.5   2  RT (min)      lcqtof_hilic_neg_4_000_aa    m/z 775.7562     0   1E+4   1.5   2  RT (min)      lcqtof_hilic_neg_4_000_aa    m/z 776.0798     0   1E+4   6   6.5  RT (min)      lcqtof_hilic_neg_4_036_11    m/z 776.4765     0   1E+4   3.5   4  RT (min)      lcqtof_hilic_neg_4_039_18    m/z 776.4906     0   1E+4   1   1.5  RT (min)      lcqtof_hilic_neg_4_000_aa    m/z 776.4958     0   1E+4   1   1.5  RT (min)      lcqtof_hilic_neg_4_000_aa    m/z 776.5424     0   1E+4   1.5   2  RT (min)       lcqtof_hilic_neg_2_002_36    m/z 776.5432     0   2   4   6   8E+3   1.5   2  RT (min)      lcqtof_hilic_neg_2_020_23    m/z 776.5433     0   1E+4   1.5   2  RT (min)       lcqtof_hilic_neg_2_002_36    m/z 776.5435     0   1   2E+5   3   3.5  RT (min)       lcqtof_hilic_neg_4_052_13    m/z 776.5447     0   1   2E+4   1   1.5  RT (min)      lcqtof_hilic_neg_4_048_23    m/z 776.5453     0   2   4   6E+3   2.5   3  RT (min)      lcqtof_hilic_neg_1_008_13    m/z 776.5676     0   1E+4   1   1.5  RT (min)      lcqtof_hilic_neg_3_035_46    m/z 776.6245     0   1   2   3   4E+4   1   1.5  RT (min)      lcqtof_hilic_neg_4_040_05    m/z 776.6247     0   2   4   6   8E+4   1.5   2  RT (min)      lcqtof_hilic_neg_4_055_32    m/z 776.6250     0   1E+5   1   1.5  RT (min)      lcqtof_hilic_neg_4_008_24    m/z 776.6250     0   1E+6   2.5   3  RT (min)      lcqtof_hilic_neg_4_008_24    m/z 776.6250     0   1   2   3E+4   2   2.5  RT (min)      lcqtof_hilic_neg_1_007_05    m/z 776.6252     0   1E+5   1.5   2  RT (min)      lcqtof_hilic_neg_4_008_24    m/z 776.6252     0   2   4   6   8E+4   2   2.5  RT (min)      lcqtof_hilic_neg_2_047_32    m/z 776.6253     0   2   4   6E+4   2   2.5  RT (min)      lcqtof_hilic_neg_1_005_32    m/z 776.6254     0   1E+5   1.5   2  RT (min)      lcqtof_hilic_neg_4_008_24    m/z 776.6254     0   1   2   3   4E+5   2.5   3  RT (min)      lcqtof_hilic_neg_3_048_05    m/z 776.6256     0   1E+5   2   2.5  RT (min)      lcqtof_hilic_neg_4_008_24    m/z 776.6262     0   2   4E+4   2   2.5  RT (min)      lcqtof_hilic_neg_3_034_24    m/z 776.6264     0   1E+5   1.5   2  RT (min)      lcqtof_hilic_neg_4_008_24    m/z 776.6276     0   1E+4   4   4.5  RT (min)      lcqtof_hilic_neg_4_055_32    m/z 776.6298     0   1E+4   4   4.5  RT (min)      lcqtof_hilic_neg_4_055_32    m/z 777.3365     0   1E+4   5   5.5  RT (min)      lcqtof_hilic_neg_4_047_06    m/z 777.3399     0   1E+4   0.5   1  RT (min)      lcqtof_hilic_neg_4_045_26    m/z 777.4748     0   2   4   6E+4   1   1.5  RT (min)      lcqtof_hilic_neg_4_000_aa    m/z 777.5168     0   1E+4   0.5   1  RT (min)      lcqtof_hilic_neg_4_000_aa    m/z 777.5198     0   1E+4   1   1.5  RT (min)      lcqtof_hilic_neg_4_016_37    m/z 777.5202     0   1   2   3E+4   2.5   3  RT (min)      lcqtof_hilic_neg_4_048_23    m/z 777.5218     0   1   2E+4   3   3.5  RT (min)      lcqtof_hilic_neg_4_034_31    m/z 777.5872     0   1E+4   1.5   2  RT (min)      lcqtof_hilic_neg_4_000_aa    m/z 777.5892     0   1E+4   1.5   2  RT (min)      lcqtof_hilic_neg_4_000_aa    m/z 777.5898     0   1E+4   1.5   2  RT (min)      lcqtof_hilic_neg_4_000_aa    m/z 777.5948     0   1E+4   1.5   2  RT (min)      lcqtof_hilic_neg_4_000_aa    m/z 777.6283     0   2   4   6   8E+4   1   1.5  RT (min)      lcqtof_hilic_neg_4_007_45    m/z 777.6284     0   1   2   3E+5   1.5   2  RT (min)      lcqtof_hilic_neg_4_055_32    m/z 777.6286     0   1   2   3E+5   1.5   2  RT (min)       lcqtof_hilic_neg_1_037_24    m/z 777.6286     0   1   2   3E+5   1.5   2  RT (min)      lcqtof_hilic_neg_4_055_32    m/z 777.6286     0   1   2   3E+6   3   3.5  RT (min)       lcqtof_hilic_neg_2_052_24    m/z 777.6286     0   1   2   3E+5   2   2.5  RT (min)       lcqtof_hilic_neg_4_055_32    m/z 777.6286     0   1   2   3E+5   2   2.5  RT (min)       lcqtof_hilic_neg_2_047_32    m/z 777.6286     0   1   2   3E+5   1.5   2  RT (min)       lcqtof_hilic_neg_4_055_32    m/z 777.6286     0   1   2   3   4E+5   1.5   2  RT (min)       lcqtof_hilic_neg_4_008_24    m/z 777.6286     0   1   2   3E+5   1.5   2  RT (min)      lcqtof_hilic_neg_4_055_32    m/z 777.6286     0   1   2   3E+5   2   2.5  RT (min)       lcqtof_hilic_neg_2_047_32    m/z 777.6286     0   1   2   3E+5   2   2.5  RT (min)      lcqtof_hilic_neg_4_055_32    m/z 777.6287     0   1   2E+5   1   1.5  RT (min)       lcqtof_hilic_neg_4_002_25    m/z 777.6287     0   1E+5   1.5   2  RT (min)      lcqtof_hilic_neg_4_040_05    m/z 777.6287     0   1   2   3E+6   2.5   3  RT (min)      lcqtof_hilic_neg_4_055_32    m/z 777.6287     0   1   2   3E+6   3   3.5  RT (min)       lcqtof_hilic_neg_1_005_32    m/z 777.6289     0   1   2   3E+5   1   1.5  RT (min)      lcqtof_hilic_neg_4_055_32    m/z 777.6290     0   1   2   3E+5   2   2.5  RT (min)      lcqtof_hilic_neg_4_055_32    m/z 777.6290     0   1   2   3E+5   1.5   2  RT (min)       lcqtof_hilic_neg_4_055_32    m/z 777.6302     0   2   4   6   8E+4   4   4.5  RT (min)       lcqtof_hilic_neg_4_055_32    m/z 777.6638     0   2   4   6   8E+4   1   1.5  RT (min)      lcqtof_hilic_neg_4_000_aa    m/z 777.6700     0   2   4   6   8E+4   1   1.5  RT (min)      lcqtof_hilic_neg_4_000_aa    m/z 777.7087     0   1   2   3E+4   1   1.5  RT (min)      lcqtof_hilic_neg_4_000_aa    m/z 778.4710     0   1   2   3E+4   1   1.5  RT (min)      lcqtof_hilic_neg_4_000_aa    m/z 778.4758     0   1   2   3E+4   1   1.5  RT (min)      lcqtof_hilic_neg_4_000_aa    m/z 778.5234     0   1   2E+4   3   3.5  RT (min)      lcqtof_hilic_neg_2_020_23    m/z 778.5250     0   1E+4   3   3.5  RT (min)      lcqtof_hilic_neg_4_034_31    m/z 778.5254     0   1E+4   3   3.5  RT (min)      lcqtof_hilic_neg_4_028_36    m/z 778.5522     0   2   4   6E+3   1.5   2  RT (min)      lcqtof_hilic_neg_2_020_23    m/z 778.5523     0   1E+4   1.5   2  RT (min)       lcqtof_hilic_neg_2_002_36    m/z 778.5523     0   1E+4   1.5   2  RT (min)      lcqtof_hilic_neg_1_051_35    m/z 778.6379     0   1   2   3   4E+5   2.5   3  RT (min)      lcqtof_hilic_neg_4_055_32    m/z 778.6386     0   1   2   3E+4   1   1.5  RT (min)      lcqtof_hilic_neg_4_092_60    m/z 778.6388     0   2   4E+4   1   1.5  RT (min)      lcqtof_hilic_neg_4_008_24    m/z 778.6388     0   2   4   6E+4   1   1.5  RT (min)      lcqtof_hilic_neg_4_008_24    m/z 778.6389     0   2   4   6E+4   1   1.5  RT (min)      lcqtof_hilic_neg_4_008_24    m/z 778.6392     0   2   4E+5   2.5   3  RT (min)      lcqtof_hilic_neg_4_008_24    m/z 778.6392     0   2   4   6E+4   2   2.5  RT (min)      lcqtof_hilic_neg_4_008_24    m/z 778.6394     0   2   4   6E+4   2   2.5  RT (min)      lcqtof_hilic_neg_4_008_24    m/z 778.6396     0   1   2   3   4E+4   1.5   2  RT (min)      lcqtof_hilic_neg_1_037_24    m/z 778.6396     0   1   2   3   4E+4   1.5   2  RT (min)      lcqtof_hilic_neg_1_037_24    m/z 778.6396     0   2   4   6E+4   1   1.5  RT (min)      lcqtof_hilic_neg_4_008_24    m/z 778.6397     0   1   2E+4   1   1.5  RT (min)      lcqtof_hilic_neg_4_040_05    m/z 778.6398     0   1   2   3   4E+4   1.5   2  RT (min)      lcqtof_hilic_neg_2_052_24    m/z 778.6398     0   1   2   3   4E+4   1.5   2  RT (min)      lcqtof_hilic_neg_2_052_24    m/z 778.6400     0   1   2   3   4E+4   2   2.5  RT (min)      lcqtof_hilic_neg_2_052_24    m/z 778.6402     0   2   4E+4   2   2.5  RT (min)      lcqtof_hilic_neg_4_055_32    m/z 778.6402     0   1   2   3   4E+4   2   2.5  RT (min)      lcqtof_hilic_neg_2_052_24    m/z 778.6660     0   1   2   3   4E+4   1   1.5  RT (min)      lcqtof_hilic_neg_4_000_aa    m/z 778.6816     0   1   2   3   4E+4   1   1.5  RT (min)      lcqtof_hilic_neg_4_000_aa    m/z 778.7200     0   1E+4   1   1.5  RT (min)      lcqtof_hilic_neg_4_000_aa    m/z 778.8732     0   1E+4   1   1.5  RT (min)      lcqtof_hilic_neg_4_004_47    m/z 778.9651     0   1E+4   6.5   7  RT (min)       lcqtof_hilic_neg_4_023_15    m/z 778.9659     0   1E+4   6.5   7  RT (min)       lcqtof_hilic_neg_4_023_15    m/z 778.9664     0   1E+4   6.5   7  RT (min)      lcqtof_hilic_neg_4_023_15    m/z 779.1791     0   1   2   3   4E+4   0.5   1  RT (min)      lcqtof_hilic_neg_3_035_46    m/z 779.5284     0   1   2E+5   1   1.5  RT (min)      lcqtof_hilic_neg_4_000_aa    m/z 779.6338     0   1   2   3   4E+4   1   1.5  RT (min)      lcqtof_hilic_neg_4_000_aa    m/z 779.6441     0   1   2E+5   2   2.5  RT (min)      lcqtof_hilic_neg_4_008_24    m/z 779.6441     0   1E+6   2.5   3  RT (min)      lcqtof_hilic_neg_4_008_24    m/z 779.6443     0   1   2E+5   1.5   2  RT (min)      lcqtof_hilic_neg_4_008_24    m/z 779.6444     0   1   2E+5   1.5   2  RT (min)      lcqtof_hilic_neg_4_008_24    m/z 779.6444     0   1   2E+5   1   1.5  RT (min)      lcqtof_hilic_neg_4_008_24    m/z 779.6444     0   1E+5   2   2.5  RT (min)      lcqtof_hilic_neg_4_055_32    m/z 779.6445     0   1   2E+5   1.5   2  RT (min)      lcqtof_hilic_neg_4_008_24    m/z 779.6446     0   1E+5   2   2.5  RT (min)      lcqtof_hilic_neg_2_047_32    m/z 779.6447     0   1   2E+5   1.5   2  RT (min)      lcqtof_hilic_neg_4_008_24    m/z 779.6449     0   1   2   3E+4   1   1.5  RT (min)      lcqtof_hilic_neg_4_041_54    m/z 779.6457     0   1   2   3   4E+4   4   4.5  RT (min)      lcqtof_hilic_neg_4_055_32    m/z 779.6699     0   1E+4   0.5   1  RT (min)      lcqtof_hilic_neg_3_035_46    m/z 780.1474     0   1E+4   3   3.5  RT (min)      lcqtof_hilic_neg_4_008_24    m/z 780.1782     0   1   2E+4   0.5   1  RT (min)      lcqtof_hilic_neg_3_035_46    m/z 780.2844     0   1E+4   4.5   5  RT (min)      lcqtof_hilic_neg_4_050_02    m/z 780.5113     0   2   4   6E+4   0.5   1  RT (min)      lcqtof_hilic_neg_4_000_aa    m/z 780.5135     0   2   4   6   8E+4   1   1.5  RT (min)      lcqtof_hilic_neg_4_000_aa    m/z 780.5246     0   2   4   6   8E+4   1   1.5  RT (min)      lcqtof_hilic_neg_4_000_aa    m/z 780.6398     0   1   2   3E+5   2.5   3  RT (min)      lcqtof_hilic_neg_4_055_32    m/z 780.6422     0   1   2E+4   1.5   2  RT (min)      lcqtof_hilic_neg_4_000_aa    m/z 780.6430     0   1   2E+4   1.5   2  RT (min)      lcqtof_hilic_neg_4_000_aa    m/z 780.6451     0   1E+5   2.5   3  RT (min)      lcqtof_hilic_neg_4_049_55    m/z 780.6548     0   1   2E+4   1.5   2  RT (min)      lcqtof_hilic_neg_2_028_05    m/z 780.6550     0   1   2   3E+4   1.5   2  RT (min)      lcqtof_hilic_neg_1_037_24    m/z 780.6551     0   1E+5   1   1.5  RT (min)      lcqtof_hilic_neg_4_008_24    m/z 780.6554     0   2   4E+4   1   1.5  RT (min)      lcqtof_hilic_neg_4_092_60    m/z 780.6554     0   1   2   3   4E+5   2.5   3  RT (min)      lcqtof_hilic_neg_4_008_24    m/z 780.6555     0   1   2   3   4E+4   1.5   2  RT (min)      lcqtof_hilic_neg_1_037_24    m/z 780.6556     0   1   2   3   4E+5   3   3.5  RT (min)      lcqtof_hilic_neg_4_008_24    m/z 780.6558     0   1   2E+4   2.5   3  RT (min)      lcqtof_hilic_neg_2_003_25    m/z 780.6560     0   1E+5   1   1.5  RT (min)      lcqtof_hilic_neg_4_008_24    m/z 780.6562     0   2   4E+4   1.5   2  RT (min)      lcqtof_hilic_neg_1_037_24    m/z 780.6562     0   2   4E+4   1.5   2  RT (min)      lcqtof_hilic_neg_2_047_32    m/z 780.6567     0   2   4   6E+4   2   2.5  RT (min)      lcqtof_hilic_neg_4_055_32    m/z 780.6567     0   1E+5   0.5   1  RT (min)      lcqtof_hilic_neg_4_008_24    m/z 780.6568     0   1   2E+4   2   2.5  RT (min)      lcqtof_hilic_neg_4_002_25    m/z 780.6568     0   1   2   3   4E+4   2   2.5  RT (min)      lcqtof_hilic_neg_3_034_24    m/z 780.6819     0   1E+4   0.5   1  RT (min)      lcqtof_hilic_neg_3_035_46    m/z 780.7448     0   1E+4   2   2.5  RT (min)      lcqtof_hilic_neg_4_000_aa    m/z 781.1809     0   1E+4   0.5   1  RT (min)      lcqtof_hilic_neg_3_035_46    m/z 781.3965     0   1E+4   4   4.5  RT (min)      lcqtof_hilic_neg_4_008_24    m/z 781.5053     0   1E+4   3.5   4  RT (min)      lcqtof_hilic_neg_4_032_01    m/z 781.5142     0   1E+5   0.5   1  RT (min)      lcqtof_hilic_neg_4_000_aa    m/z 781.5309     0   1E+5   1   1.5  RT (min)      lcqtof_hilic_neg_4_000_aa    m/z 781.5826     0   1   2E+4   1   1.5  RT (min)      lcqtof_hilic_neg_4_000_aa    m/z 781.5874     0   1   2E+4   1   1.5  RT (min)      lcqtof_hilic_neg_4_000_aa    m/z 781.6598     0   2   4   6E+5   2.5   3  RT (min)      lcqtof_hilic_neg_2_003_25    m/z 781.6599     0   1E+5   1.5   2  RT (min)      lcqtof_hilic_neg_4_008_24    m/z 781.6600     0   1   2   3   4E+5   1   1.5  RT (min)      lcqtof_hilic_neg_4_008_24    m/z 781.6600     0   1   2E+5   1.5   2  RT (min)       lcqtof_hilic_neg_4_008_24    m/z 781.6600     0   1E+6   2.5   3  RT (min)      lcqtof_hilic_neg_4_008_24    m/z 781.6602     0   1   2E+5   2.5   3  RT (min)      lcqtof_hilic_neg_4_047_06    m/z 781.6602     0   1   2E+5   1.5   2  RT (min)       lcqtof_hilic_neg_4_008_24    m/z 781.6602     0   1E+6   3   3.5  RT (min)      lcqtof_hilic_neg_4_008_24    m/z 781.6602     0   1E+5   1.5   2  RT (min)       lcqtof_hilic_neg_4_008_24    m/z 781.6603     0   1   2   3   4E+4   4   4.5  RT (min)       lcqtof_hilic_neg_4_091_60    m/z 781.6603     0   1   2E+5   1.5   2  RT (min)       lcqtof_hilic_neg_4_008_24    m/z 781.6604     0   1   2   3   4E+5   1   1.5  RT (min)      lcqtof_hilic_neg_4_008_24    m/z 781.6604     0   1   2E+5   2   2.5  RT (min)       lcqtof_hilic_neg_4_008_24    m/z 781.6604     0   1E+5   1.5   2  RT (min)       lcqtof_hilic_neg_4_008_24    m/z 781.6604     0   1E+5   1.5   2  RT (min)       lcqtof_hilic_neg_4_008_24    m/z 781.6608     0   1   2   3   4E+5   0.5   1  RT (min)       lcqtof_hilic_neg_4_008_24    m/z 781.6608     0   1   2E+5   2   2.5  RT (min)      lcqtof_hilic_neg_4_008_24    m/z 781.6612     0   1   2E+5   2   2.5  RT (min)      lcqtof_hilic_neg_4_055_32    m/z 781.6612     0   1   2   3   4E+5   1   1.5  RT (min)      lcqtof_hilic_neg_4_008_24    m/z 781.6848     0   2   4E+4   0.5   1  RT (min)      lcqtof_hilic_neg_3_035_46    m/z 782.1634     0   1E+4   3   3.5  RT (min)      lcqtof_hilic_neg_4_008_24    m/z 782.4818     0   1E+4   3.5   4  RT (min)      lcqtof_hilic_neg_2_031_01    m/z 782.4913     0   1   2   3E+5   4   4.5  RT (min)      lcqtof_hilic_neg_4_034_31    m/z 782.4993     0   1   2   3E+5   4   4.5  RT (min)      lcqtof_hilic_neg_4_016_37    m/z 782.5362     0   1E+4   0.5   1  RT (min)      lcqtof_hilic_neg_4_008_24    m/z 782.5370     0   2   4   6E+4   1   1.5  RT (min)      lcqtof_hilic_neg_4_000_aa    m/z 782.5424     0   1E+4   4   4.5  RT (min)      lcqtof_hilic_neg_4_035_33    m/z 782.6638     0   1   2E+5   3   3.5  RT (min)      lcqtof_hilic_neg_4_035_33    m/z 782.6698     0   2   4E+5   3   3.5  RT (min)      lcqtof_hilic_neg_4_008_24    m/z 782.6708     0   2   4E+5   3   3.5  RT (min)      lcqtof_hilic_neg_4_008_24    m/z 782.6716     0   1E+4   4   4.5  RT (min)      lcqtof_hilic_neg_4_008_24    m/z 782.6718     0   2   4E+4   1.5   2  RT (min)      lcqtof_hilic_neg_4_000_aa    m/z 782.6720     0   1   2E+5   0.5   1  RT (min)      lcqtof_hilic_neg_4_008_24    m/z 782.6724     0   1E+5   1.5   2  RT (min)      lcqtof_hilic_neg_1_005_32    m/z 782.6726     0   1   2E+4   1.5   2  RT (min)      lcqtof_hilic_neg_3_034_24    m/z 782.6728     0   2   4   6   8E+4   1.5   2  RT (min)      lcqtof_hilic_neg_3_034_24    m/z 782.6730     0   2   4   6   8E+4   1   1.5  RT (min)      lcqtof_hilic_neg_4_092_60    m/z 782.6731     0   1   2E+5   1   1.5  RT (min)      lcqtof_hilic_neg_4_008_24    m/z 782.6738     0   2   4   6   8E+4   1.5   2  RT (min)      lcqtof_hilic_neg_3_034_24    m/z 782.6742     0   1   2E+5   1   1.5  RT (min)      lcqtof_hilic_neg_4_008_24    m/z 782.6974     0   1   2E+4   0.5   1  RT (min)      lcqtof_hilic_neg_3_035_46    m/z 782.7232     0   1   2E+4   1.5   2  RT (min)      lcqtof_hilic_neg_4_000_aa    m/z 783.3260     0   1   2   3   4E+4   0.5   1  RT (min)      lcqtof_hilic_neg_4_039_18    m/z 783.5377     0   1   2E+4   1   1.5  RT (min)      lcqtof_hilic_neg_4_000_aa    m/z 783.5977     0   1   2   3E+4   1.5   2  RT (min)      lcqtof_hilic_neg_4_000_aa    m/z 783.6700     0   2   4   6E+5   3.5   4  RT (min)      lcqtof_hilic_neg_4_049_55    m/z 783.6728     0   1   2   3E+5   3.5   4  RT (min)      lcqtof_hilic_neg_4_035_33    m/z 783.6733     0   2   4   6E+4   4   4.5  RT (min)       lcqtof_hilic_neg_4_092_60    m/z 783.6751     0   2   4   6E+4   1.5   2  RT (min)       lcqtof_hilic_neg_4_040_05    m/z 783.6753     0   2   4E+5   2   2.5  RT (min)       lcqtof_hilic_neg_4_008_24    m/z 783.6754     0   1E+5   1   1.5  RT (min)      lcqtof_hilic_neg_4_009_50    m/z 783.6754     0   1   2E+6   3   3.5  RT (min)       lcqtof_hilic_neg_4_008_24    m/z 783.6754     0   2   4E+5   1.5   2  RT (min)      lcqtof_hilic_neg_4_055_32    m/z 783.6754     0   2   4E+5   1.5   2  RT (min)      lcqtof_hilic_neg_4_055_32    m/z 783.6754     0   2   4E+4   4   4.5  RT (min)      lcqtof_hilic_neg_4_055_32    m/z 783.6755     0   1   2   3E+5   1.5   2  RT (min)       lcqtof_hilic_neg_4_055_32    m/z 783.6756     0   1E+5   1.5   2  RT (min)       lcqtof_hilic_neg_4_055_32    m/z 783.6756     0   2   4E+5   1.5   2  RT (min)       lcqtof_hilic_neg_4_008_24    m/z 783.6756     0   1   2E+6   3   3.5  RT (min)       lcqtof_hilic_neg_4_008_24    m/z 783.6756     0   1   2E+5   1.5   2  RT (min)       lcqtof_hilic_neg_4_055_32    m/z 783.6756     0   1E+5   1.5   2  RT (min)       lcqtof_hilic_neg_4_055_32    m/z 783.6756     0   2   4E+5   2   2.5  RT (min)       lcqtof_hilic_neg_4_008_24    m/z 783.6898     0   1   2E+4   1.5   2  RT (min)      lcqtof_hilic_neg_4_000_aa    m/z 783.6996     0   2   4   6   8E+4   0.5   1  RT (min)      lcqtof_hilic_neg_3_035_46    m/z 783.7242     0   1E+4   1.5   2  RT (min)      lcqtof_hilic_neg_4_000_aa    m/z 783.7585     0   1   2E+4   0.5   1  RT (min)      lcqtof_hilic_neg_4_055_32    m/z 783.9596     0   1E+4   0.5   1  RT (min)      lcqtof_hilic_neg_4_008_24    m/z 784.1784     0   1   2E+4   3   3.5  RT (min)      lcqtof_hilic_neg_4_008_24    m/z 784.3288     0   1   2E+4   0.5   1  RT (min)      lcqtof_hilic_neg_4_039_18    m/z 784.5106     0   2   4   6   8E+5   4   4.5  RT (min)      lcqtof_hilic_neg_4_048_23    m/z 784.5457     0   1E+4   3.5   4  RT (min)      lcqtof_hilic_neg_4_053_14    m/z 784.6003     0   1E+4   1.5   2  RT (min)      lcqtof_hilic_neg_4_000_aa    m/z 784.6594     0   1   2E+4   2.5   3  RT (min)      lcqtof_hilic_neg_4_008_24    m/z 784.6604     0   2   4E+4   2.5   3  RT (min)      lcqtof_hilic_neg_4_055_32    m/z 784.6867     0   1   2   3   4E+4   2   2.5  RT (min)      lcqtof_hilic_neg_4_055_32    m/z 784.6886     0   2   4E+4   1.5   2  RT (min)      lcqtof_hilic_neg_4_000_aa    m/z 784.6896     0   2   4E+4   1.5   2  RT (min)       lcqtof_hilic_neg_4_000_aa    m/z 784.6908     0   1   2   3E+4   2   2.5  RT (min)      lcqtof_hilic_neg_4_008_24    m/z 784.7061     0   1   2E+4   2   2.5  RT (min)      lcqtof_hilic_neg_4_035_33    m/z 784.7112     0   1   2E+4   2   2.5  RT (min)      lcqtof_hilic_neg_4_008_24    m/z 784.9986     0   1   2   3E+3   4.5   5  RT (min)      lcqtof_hilic_neg_4_052_13    m/z 785.2514     0   1   2   3E+4   5.5   6  RT (min)      lcqtof_hilic_neg_4_013_20    m/z 785.5922     0   1E+4   1   1.5  RT (min)      lcqtof_hilic_neg_4_000_aa    m/z 785.6374     0   1   2E+5   1   1.5  RT (min)      lcqtof_hilic_neg_2_052_24    m/z 785.6639     0   1   2   3E+4   1   1.5  RT (min)      lcqtof_hilic_neg_3_035_46    m/z 785.6737     0   1E+5   2.5   3  RT (min)      lcqtof_hilic_neg_4_040_05    m/z 785.6907     0   1E+5   3   3.5  RT (min)      lcqtof_hilic_neg_4_055_32    m/z 785.6919     0   1   2E+4   1.5   2  RT (min)       lcqtof_hilic_neg_4_008_24    m/z 785.6937     0   1   2E+4   1.5   2  RT (min)       lcqtof_hilic_neg_4_000_aa    m/z 785.6940     0   2   4   6E+4   1   1.5  RT (min)      lcqtof_hilic_neg_4_000_aa    m/z 786.2874     0   1E+4   3   3.5  RT (min)      lcqtof_hilic_neg_4_013_20    m/z 786.3101     0   1E+4   5.5   6  RT (min)      lcqtof_hilic_neg_4_058_39    m/z 786.3266     0   1E+4   0.5   1  RT (min)      lcqtof_hilic_neg_4_039_18    m/z 786.5269     0   1   2E+5   4.5   5  RT (min)      lcqtof_hilic_neg_4_048_23    m/z 786.5277     0   1E+6   4   4.5  RT (min)       lcqtof_hilic_neg_4_048_23    m/z 786.5566     0   1   2E+4   1   1.5  RT (min)      lcqtof_hilic_neg_4_027_19    m/z 786.5725     0   1   2   3   4E+4   1   1.5  RT (min)       lcqtof_hilic_neg_4_048_23    m/z 786.5726     0   1   2   3   4E+4   1   1.5  RT (min)      lcqtof_hilic_neg_4_048_23    m/z 786.5726     0   1   2E+4   1   1.5  RT (min)      lcqtof_hilic_neg_4_034_31    m/z 786.5728     0   1   2   3   4E+4   1   1.5  RT (min)       lcqtof_hilic_neg_4_048_23    m/z 786.5731     0   1   2E+4   1.5   2  RT (min)      lcqtof_hilic_neg_2_020_23    m/z 786.5733     0   1   2E+4   1.5   2  RT (min)       lcqtof_hilic_neg_2_020_23    m/z 786.6472     0   2   4   6E+4   1   1.5  RT (min)      lcqtof_hilic_neg_4_000_aa    m/z 786.7765     0   1E+4   5   5.5  RT (min)      lcqtof_hilic_neg_4_055_32    m/z 786.8116     0   1   2   3E+4   4   4.5  RT (min)      lcqtof_hilic_neg_1_035_23    m/z 786.9732     0   2   4   6E+4   6.5   7  RT (min)       lcqtof_hilic_neg_4_059_14    m/z 787.3348     0   1E+4   1   1.5  RT (min)      lcqtof_hilic_neg_4_000_aa    m/z 787.3467     0   1E+4   1   1.5  RT (min)      lcqtof_hilic_neg_4_000_aa    m/z 787.5327     0   1   2   3   4E+5   4   4.5  RT (min)      lcqtof_hilic_neg_4_048_23    m/z 787.5623     0   1E+4   1   1.5  RT (min)      lcqtof_hilic_neg_4_027_19    m/z 787.5774     0   1E+4   1   1.5  RT (min)      lcqtof_hilic_neg_4_048_23    m/z 787.5776     0   1E+4   1   1.5  RT (min)      lcqtof_hilic_neg_4_048_23    m/z 787.5782     0   1E+4   1.5   2  RT (min)      lcqtof_hilic_neg_2_020_23    m/z 787.6221     0   1   2   3E+4   0.5   1  RT (min)      lcqtof_hilic_neg_2_016_33    m/z 787.6512     0   1E+5   1   1.5  RT (min)      lcqtof_hilic_neg_3_034_24    m/z 787.6513     0   1   2E+5   1   1.5  RT (min)      lcqtof_hilic_neg_2_052_24    m/z 787.6789     0   1   2   3E+4   1   1.5  RT (min)      lcqtof_hilic_neg_3_035_46    m/z 787.7591     0   1E+4   1.5   2  RT (min)      lcqtof_hilic_neg_4_000_aa    m/z 787.9776     0   1E+4   6.5   7  RT (min)      lcqtof_hilic_neg_4_059_14    m/z 788.2885     0   1E+4   3   3.5  RT (min)      lcqtof_hilic_neg_4_013_20    m/z 788.3142     0   1E+4   6   6.5  RT (min)       lcqtof_hilic_neg_4_048_23    m/z 788.3156     0   1E+4   6   6.5  RT (min)      lcqtof_hilic_neg_4_048_23    m/z 788.4035     0   1   2E+4   1   1.5  RT (min)      lcqtof_hilic_neg_4_000_aa    m/z 788.4062     0   1   2E+4   1   1.5  RT (min)      lcqtof_hilic_neg_4_000_aa    m/z 788.4090     0   1   2E+4   1   1.5  RT (min)      lcqtof_hilic_neg_4_000_aa    m/z 788.4722     0   1E+5   4.5   5  RT (min)      lcqtof_hilic_neg_4_048_23    m/z 788.4726     0   1E+5   4   4.5  RT (min)      lcqtof_hilic_neg_4_048_23    m/z 788.5630     0   1   2   3   4E+4   3   3.5  RT (min)      lcqtof_hilic_neg_4_032_01    m/z 788.5872     0   1E+4   1.5   2  RT (min)       lcqtof_hilic_neg_2_020_23    m/z 788.5893     0   1E+5   1.5   2  RT (min)      lcqtof_hilic_neg_1_035_23    m/z 788.6131     0   1   2   3   4E+4   2.5   3  RT (min)      lcqtof_hilic_neg_4_035_33    m/z 788.6223     0   1   2   3E+4   1   1.5  RT (min)      lcqtof_hilic_neg_4_000_aa    m/z 788.9212     0   1E+4   6   6.5  RT (min)      lcqtof_hilic_neg_4_001_49    m/z 789.0668     0   1E+4   0.5   1  RT (min)      lcqtof_hilic_neg_1_062_17    m/z 789.4112     0   1E+4   1   1.5  RT (min)      lcqtof_hilic_neg_4_000_aa    m/z 789.5189     0   1   2   3E+4   4   4.5  RT (min)      lcqtof_hilic_neg_4_047_06    m/z 789.5750     0   1E+4   1.5   2  RT (min)      lcqtof_hilic_neg_2_020_23    m/z 789.5802     0   1E+4   1.5   2  RT (min)      lcqtof_hilic_neg_2_002_36    m/z 789.5820     0   1E+4   1.5   2  RT (min)      lcqtof_hilic_neg_2_020_23    m/z 789.5882     0   1E+4   1.5   2  RT (min)      lcqtof_hilic_neg_1_035_23    m/z 789.6230     0   1   2E+5   1   1.5  RT (min)      lcqtof_hilic_neg_4_000_aa    m/z 789.7286     0   1   2E+4   1   1.5  RT (min)      lcqtof_hilic_neg_4_000_aa    m/z 789.7481     0   1   2   3E+4   1   1.5  RT (min)      lcqtof_hilic_neg_4_000_aa    m/z 789.7516     0   1   2   3E+4   1   1.5  RT (min)      lcqtof_hilic_neg_4_000_aa    m/z 789.7632     0   1   2   3E+4   1.5   2  RT (min)      lcqtof_hilic_neg_4_000_aa    m/z 789.7692     0   1   2   3E+4   1.5   2  RT (min)      lcqtof_hilic_neg_4_000_aa    m/z 790.1448     0   1E+4   0.5   1  RT (min)      lcqtof_hilic_neg_1_062_17    m/z 790.4031     0   1E+4   0.5   1  RT (min)      lcqtof_hilic_neg_4_000_aa    m/z 790.4960     0   1E+4   3.5   4  RT (min)      lcqtof_hilic_neg_4_039_18    m/z 790.5149     0   1E+4   3   3.5  RT (min)       lcqtof_hilic_neg_1_034_08    m/z 790.5397     0   1E+4   2.5   3  RT (min)      lcqtof_hilic_neg_4_062_17    m/z 790.5401     0   1   2E+4   1   1.5  RT (min)      lcqtof_hilic_neg_1_048_45    m/z 790.5411     0   1   2E+5   1.5   2  RT (min)      lcqtof_hilic_neg_1_013_38    m/z 790.5416     0   1   2E+5   2   2.5  RT (min)      lcqtof_hilic_neg_1_014_31    m/z 790.5572     0   1   2   3E+4   1   1.5  RT (min)       lcqtof_hilic_neg_4_048_23    m/z 790.5574     0   1   2   3E+4   1.5   2  RT (min)      lcqtof_hilic_neg_4_048_23    m/z 790.5578     0   1E+5   1.5   2  RT (min)       lcqtof_hilic_neg_1_051_35    m/z 790.5578     0   1   2   3E+4   1.5   2  RT (min)       lcqtof_hilic_neg_2_002_36    m/z 790.5583     0   1   2   3E+4   1   1.5  RT (min)       lcqtof_hilic_neg_1_035_23    m/z 790.5604     0   2   4   6   8E+4   3   3.5  RT (min)       lcqtof_hilic_neg_4_005_29    m/z 790.5644     0   1   2E+4   3.5   4  RT (min)      lcqtof_hilic_neg_4_014_46    m/z 790.5837     0   1   2   3   4E+4   3   3.5  RT (min)      lcqtof_hilic_neg_2_062_17    m/z 790.5883     0   1E+4   2   2.5  RT (min)      lcqtof_hilic_neg_4_004_47    m/z 790.6326     0   2   4   6   8E+4   1   1.5  RT (min)      lcqtof_hilic_neg_4_000_aa    m/z 790.6522     0   1E+4   0.5   1  RT (min)      lcqtof_hilic_neg_4_055_32    m/z 790.7732     0   1E+4   1.5   2  RT (min)      lcqtof_hilic_neg_4_000_aa    m/z 790.7912     0   1E+4   1   1.5  RT (min)      lcqtof_hilic_neg_4_055_32    m/z 790.9530     0   1   2E+3   5.5   6  RT (min)      lcqtof_hilic_neg_2_026_28    m/z 790.9536     0   1   2E+3   5   5.5  RT (min)      lcqtof_hilic_neg_2_013_29    m/z 791.1438     0   1E+4   0.5   1  RT (min)      lcqtof_hilic_neg_1_062_17    m/z 791.4874     0   1E+4   1   1.5  RT (min)      lcqtof_hilic_neg_4_000_aa    m/z 791.4896     0   1   2E+4   4   4.5  RT (min)      lcqtof_hilic_neg_4_048_23    m/z 791.5408     0   2   4   6E+4   3   3.5  RT (min)      lcqtof_hilic_neg_4_034_31    m/z 791.5417     0   1E+4   3   3.5  RT (min)      lcqtof_hilic_neg_4_048_23    m/z 791.5595     0   1E+4   1.5   2  RT (min)      lcqtof_hilic_neg_2_020_23    m/z 791.5604     0   1E+4   1.5   2  RT (min)      lcqtof_hilic_neg_4_000_aa    m/z 791.5606     0   2   4   6E+4   1.5   2  RT (min)      lcqtof_hilic_neg_1_051_35    m/z 791.5606     0   1E+4   1.5   2  RT (min)      lcqtof_hilic_neg_1_035_23    m/z 791.5606     0   1E+4   1.5   2  RT (min)      lcqtof_hilic_neg_2_002_36    m/z 791.5610     0   1E+4   1.5   2  RT (min)      lcqtof_hilic_neg_4_000_aa    m/z 791.5620     0   1E+4   1   1.5  RT (min)      lcqtof_hilic_neg_4_087_59    m/z 791.5634     0   1E+4   1   1.5  RT (min)      lcqtof_hilic_neg_4_000_aa    m/z 791.6354     0   1   2   3E+4   1   1.5  RT (min)      lcqtof_hilic_neg_4_000_aa    m/z 791.6694     0   1   2   3E+4   1   1.5  RT (min)      lcqtof_hilic_neg_4_000_aa    m/z 791.7032     0   1   2   3E+4   1   1.5  RT (min)      lcqtof_hilic_neg_4_000_aa    m/z 791.7514     0   1   2E+4   1.5   2  RT (min)      lcqtof_hilic_neg_4_000_aa    m/z 791.9617     0   1E+4   6.5   7  RT (min)       lcqtof_hilic_neg_4_093_60    m/z 792.3296     0   1   2E+4   0.5   1  RT (min)      lcqtof_hilic_neg_3_003_20    m/z 792.5340     0   1   2   3E+4   2.5   3  RT (min)       lcqtof_hilic_neg_1_016_28    m/z 792.5345     0   1   2   3   4E+4   1   1.5  RT (min)       lcqtof_hilic_neg_2_063_13    m/z 792.5348     0   2   4E+4   2.5   3  RT (min)      lcqtof_hilic_neg_1_016_28    m/z 792.5350     0   2   4   6E+4   2.5   3  RT (min)       lcqtof_hilic_neg_1_030_17    m/z 792.5362     0   1   2   3   4E+4   2.5   3  RT (min)       lcqtof_hilic_neg_4_053_14    m/z 792.5371     0   1   2   3   4E+4   1   1.5  RT (min)      lcqtof_hilic_neg_2_063_13    m/z 792.5372     0   1   2E+4   2.5   3  RT (min)      lcqtof_hilic_neg_1_016_28    m/z 792.5378     0   1   2   3   4E+4   1   1.5  RT (min)      lcqtof_hilic_neg_2_005_13    m/z 792.5395     0   2   4   6   8E+4   1.5   2  RT (min)       lcqtof_hilic_neg_1_051_35    m/z 792.5490     0   1   2   3   4E+4   2.5   3  RT (min)      lcqtof_hilic_neg_4_045_26    m/z 792.5494     0   1E+4   1.5   2  RT (min)      lcqtof_hilic_neg_2_002_36    m/z 792.5499     0   1E+4   1   1.5  RT (min)      lcqtof_hilic_neg_1_003_37    m/z 792.5562     0   1E+4   1.5   2  RT (min)      lcqtof_hilic_neg_2_020_23    m/z 792.5590     0   1E+4   1.5   2  RT (min)      lcqtof_hilic_neg_2_020_23    m/z 792.5852     0   1   2E+4   1   1.5  RT (min)      lcqtof_hilic_neg_3_035_46    m/z 792.6458     0   2   4   6   8E+3   1.5   2  RT (min)       lcqtof_hilic_neg_4_008_24    m/z 792.6883     0   1E+4   1   1.5  RT (min)      lcqtof_hilic_neg_4_000_aa    m/z 792.7525     0   1E+4   1.5   2  RT (min)      lcqtof_hilic_neg_4_000_aa    m/z 792.9438     0   1E+4   0.5   1  RT (min)      lcqtof_hilic_neg_4_004_47    m/z 793.3317     0   1E+4   6   6.5  RT (min)       lcqtof_hilic_neg_4_093_60    m/z 793.3335     0   1E+4   0.5   1  RT (min)      lcqtof_hilic_neg_3_003_20    m/z 793.5377     0   1E+4   1   1.5  RT (min)      lcqtof_hilic_neg_2_005_13    m/z 793.5391     0   1E+4   2.5   3  RT (min)      lcqtof_hilic_neg_4_060_13    m/z 793.5405     0   2   4   6   8E+3   1.5   2  RT (min)      lcqtof_hilic_neg_2_064_16    m/z 793.5548     0   1E+4   2   2.5  RT (min)      lcqtof_hilic_neg_4_048_23    m/z 793.5584     0   1   2   3E+4   2   2.5  RT (min)       lcqtof_hilic_neg_1_014_31    m/z 793.6314     0   1   2E+4   1   1.5  RT (min)      lcqtof_hilic_neg_4_000_aa    m/z 793.6432     0   1   2   3   4E+4   1   1.5  RT (min)      lcqtof_hilic_neg_4_000_aa    m/z 793.6544     0   2   4   6E+3   15   15.5  RT (min)      lcqtof_hilic_neg_4_013_20    m/z 793.6546     0   2   4   6   8E+4   1.5   2  RT (min)      lcqtof_hilic_neg_4_000_aa    m/z 793.6548     0   1E+4   15   15.5  RT (min)      lcqtof_hilic_neg_4_039_18    m/z 793.6552     0   2   4   6   8E+4   1.5   2  RT (min)      lcqtof_hilic_neg_4_000_aa    m/z 793.6573     0   2   4   6   8E+4   1.5   2  RT (min)      lcqtof_hilic_neg_4_000_aa    m/z 793.6829     0   1   2   3   4E+4   1   1.5  RT (min)      lcqtof_hilic_neg_4_000_aa    m/z 794.1686     0   1E+4   0.5   1  RT (min)      lcqtof_hilic_neg_1_062_17    m/z 794.5426     0   1E+4   1   1.5  RT (min)      lcqtof_hilic_neg_1_004_14    m/z 794.5434     0   1   2E+4   2.5   3  RT (min)      lcqtof_hilic_neg_1_008_13    m/z 794.5470     0   2   4   6   8E+4   1   1.5  RT (min)       lcqtof_hilic_neg_2_063_13    m/z 794.5506     0   1E+4   2   2.5  RT (min)       lcqtof_hilic_neg_1_008_13    m/z 794.5513     0   1E+4   2   2.5  RT (min)      lcqtof_hilic_neg_1_008_13    m/z 794.5514     0   1E+4   2   2.5  RT (min)       lcqtof_hilic_neg_1_008_13    m/z 794.5520     0   1E+4   1.5   2  RT (min)       lcqtof_hilic_neg_3_006_30    m/z 794.5541     0   1E+4   1.5   2  RT (min)       lcqtof_hilic_neg_3_067_57    m/z 794.5578     0   1E+4   2   2.5  RT (min)       lcqtof_hilic_neg_3_006_30    m/z 794.5596     0   1   2   3E+4   2   2.5  RT (min)      lcqtof_hilic_neg_4_060_13    m/z 794.5606     0   2   4E+4   2.5   3  RT (min)       lcqtof_hilic_neg_4_059_14    m/z 794.5608     0   1   2   3E+4   2   2.5  RT (min)      lcqtof_hilic_neg_4_059_14    m/z 794.5612     0   2   4E+4   2.5   3  RT (min)       lcqtof_hilic_neg_4_053_14    m/z 794.5636     0   1   2   3E+4   2   2.5  RT (min)       lcqtof_hilic_neg_4_059_14    m/z 794.6004     0   1   2   3   4E+4   0.5   1  RT (min)      lcqtof_hilic_neg_3_035_46    m/z 794.6617     0   1   2   3   4E+4   1.5   2  RT (min)      lcqtof_hilic_neg_4_000_aa    m/z 794.6621     0   1   2   3   4E+4   1.5   2  RT (min)      lcqtof_hilic_neg_4_000_aa    m/z 794.6626     0   1   2   3   4E+4   1.5   2  RT (min)      lcqtof_hilic_neg_4_000_aa    m/z 794.6648     0   2   4   6E+3   2   2.5  RT (min)      lcqtof_hilic_neg_1_005_32    m/z 794.7201     0   1E+4   1.5   2  RT (min)      lcqtof_hilic_neg_4_000_aa    m/z 794.8561     0   1E+4   3   3.5  RT (min)      lcqtof_hilic_neg_4_005_29    m/z 795.1644     0   1E+4   0.5   1  RT (min)      lcqtof_hilic_neg_1_062_17    m/z 795.5463     0   1   2E+4   2.5   3  RT (min)       lcqtof_hilic_neg_2_063_13    m/z 795.5484     0   1   2E+4   2.5   3  RT (min)       lcqtof_hilic_neg_2_063_13    m/z 795.5494     0   2   4   6   8E+3   1   1.5  RT (min)      lcqtof_hilic_neg_1_064_16    m/z 795.5550     0   2   4E+3   1.5   2  RT (min)       lcqtof_hilic_neg_3_068_57    m/z 795.5599     0   2   4E+3   1.5   2  RT (min)       lcqtof_hilic_neg_3_067_57    m/z 795.5636     0   2   4E+3   2   2.5  RT (min)       lcqtof_hilic_neg_3_068_57    m/z 795.5641     0   1E+4   2.5   3  RT (min)      lcqtof_hilic_neg_4_060_13    m/z 795.5649     0   1E+4   2.5   3  RT (min)      lcqtof_hilic_neg_4_023_15    m/z 795.5678     0   1   2E+5   2.5   3  RT (min)       lcqtof_hilic_neg_4_023_15    m/z 795.5829     0   1E+4   4   4.5  RT (min)      lcqtof_hilic_neg_4_047_06    m/z 795.5912     0   2   4   6E+4   1.5   2  RT (min)      lcqtof_hilic_neg_4_000_aa    m/z 795.5924     0   2   4   6E+4   1.5   2  RT (min)      lcqtof_hilic_neg_4_000_aa    m/z 795.5926     0   2   4   6E+4   1.5   2  RT (min)      lcqtof_hilic_neg_4_000_aa    m/z 795.5943     0   2   4   6E+4   1   1.5  RT (min)      lcqtof_hilic_neg_4_000_aa    m/z 795.5949     0   2   4   6E+4   1   1.5  RT (min)      lcqtof_hilic_neg_4_000_aa    m/z 795.5952     0   2   4   6E+4   1   1.5  RT (min)      lcqtof_hilic_neg_4_000_aa    m/z 795.5966     0   1E+4   0.5   1  RT (min)      lcqtof_hilic_neg_3_035_46    m/z 795.7194     0   1E+4   1.5   2  RT (min)      lcqtof_hilic_neg_4_000_aa    m/z 795.9006     0   1E+4   5   5.5  RT (min)      lcqtof_hilic_neg_4_055_32    m/z 796.5738     0   1   2   3   4E+3   2   2.5  RT (min)      lcqtof_hilic_neg_3_055_01    m/z 796.5908     0   1   2   3   4E+4   1   1.5  RT (min)      lcqtof_hilic_neg_4_000_aa    m/z 796.5926     0   1   2   3   4E+4   1.5   2  RT (min)      lcqtof_hilic_neg_4_000_aa    m/z 796.5943     0   1   2E+4   1.5   2  RT (min)      lcqtof_hilic_neg_4_000_aa    m/z 796.5944     0   1   2E+4   1.5   2  RT (min)      lcqtof_hilic_neg_4_000_aa    m/z 796.5955     0   1   2   3   4E+4   1.5   2  RT (min)      lcqtof_hilic_neg_4_000_aa    m/z 796.5958     0   1   2E+4   1.5   2  RT (min)      lcqtof_hilic_neg_4_000_aa    m/z 796.5984     0   1   2   3   4E+4   1   1.5  RT (min)      lcqtof_hilic_neg_4_000_aa    m/z 796.6126     0   1E+4   0.5   1  RT (min)      lcqtof_hilic_neg_3_035_46    m/z 796.6183     0   1   2   3E+5   4.5   5  RT (min)      lcqtof_hilic_neg_4_002_25    m/z 796.6190     0   1   2   3   4E+5   4   4.5  RT (min)      lcqtof_hilic_neg_4_090_60    m/z 796.6237     0   1E+4   2.5   3  RT (min)      lcqtof_hilic_neg_4_055_32    m/z 796.6252     0   1E+4   0.5   1  RT (min)      lcqtof_hilic_neg_2_020_23    m/z 796.6784     0   1   2E+5   3   3.5  RT (min)       lcqtof_hilic_neg_4_008_24    m/z 796.6795     0   1   2E+4   1.5   2  RT (min)       lcqtof_hilic_neg_4_008_24    m/z 796.6798     0   1   2E+4   1.5   2  RT (min)       lcqtof_hilic_neg_4_008_24    m/z 796.6806     0   1   2E+4   2   2.5  RT (min)       lcqtof_hilic_neg_4_008_24    m/z 796.6880     0   1E+4   1.5   2  RT (min)       lcqtof_hilic_neg_4_000_aa    m/z 796.7062     0   1E+4   0.5   1  RT (min)      lcqtof_hilic_neg_3_035_46    m/z 796.7186     0   1   2   3E+4   1   1.5  RT (min)      lcqtof_hilic_neg_4_008_24    m/z 796.7368     0   1   2E+4   1.5   2  RT (min)      lcqtof_hilic_neg_4_000_aa    m/z 796.8822     0   1E+4   1   1.5  RT (min)       lcqtof_hilic_neg_4_054_48    m/z 797.2164     0   1E+4   6   6.5  RT (min)      lcqtof_hilic_neg_4_036_11    m/z 797.5060     0   2   4E+4   1   1.5  RT (min)      lcqtof_hilic_neg_4_000_aa    m/z 797.5067     0   2   4E+4   1.5   2  RT (min)      lcqtof_hilic_neg_4_000_aa    m/z 797.5070     0   2   4E+4   1.5   2  RT (min)      lcqtof_hilic_neg_4_000_aa    m/z 797.5078     0   1   2   3   4E+4   1.5   2  RT (min)      lcqtof_hilic_neg_4_000_aa    m/z 797.5078     0   2   4E+4   1   1.5  RT (min)      lcqtof_hilic_neg_4_000_aa    m/z 797.5084     0   2   4E+4   1   1.5  RT (min)      lcqtof_hilic_neg_4_000_aa    m/z 797.5086     0   2   4E+4   1.5   2  RT (min)      lcqtof_hilic_neg_4_000_aa    m/z 797.5100     0   2   4E+4   1   1.5  RT (min)      lcqtof_hilic_neg_4_000_aa    m/z 797.5104     0   2   4E+4   1   1.5  RT (min)      lcqtof_hilic_neg_4_000_aa    m/z 797.5912     0   1   2E+4   1.5   2  RT (min)      lcqtof_hilic_neg_4_000_aa    m/z 797.5921     0   1   2E+4   1.5   2  RT (min)      lcqtof_hilic_neg_4_000_aa    m/z 797.5950     0   2   4   6   8E+4   1   1.5  RT (min)      lcqtof_hilic_neg_4_000_aa    m/z 797.5960     0   2   4   6   8E+4   1.5   2  RT (min)      lcqtof_hilic_neg_4_000_aa    m/z 797.6012     0   2   4   6   8E+4   1.5   2  RT (min)      lcqtof_hilic_neg_4_000_aa    m/z 797.6034     0   2   4   6   8E+4   1   1.5  RT (min)      lcqtof_hilic_neg_4_000_aa    m/z 797.6273     0   1E+6   4.5   5  RT (min)      lcqtof_hilic_neg_4_055_32    m/z 797.6280     0   2   4   6   8E+5   4   4.5  RT (min)      lcqtof_hilic_neg_4_090_60    m/z 797.6283     0   1   2   3E+5   4.5   5  RT (min)      lcqtof_hilic_neg_4_004_47    m/z 797.6422     0   1E+4   0.5   1  RT (min)      lcqtof_hilic_neg_4_010_28    m/z 797.6944     0   1   2E+4   2   2.5  RT (min)      lcqtof_hilic_neg_4_008_24    m/z 797.7195     0   1E+5   1.5   2  RT (min)      lcqtof_hilic_neg_4_008_24    m/z 797.7216     0   1E+5   1.5   2  RT (min)       lcqtof_hilic_neg_4_008_24    m/z 797.7250     0   1   2E+4   1   1.5  RT (min)      lcqtof_hilic_neg_4_009_50    m/z 797.7268     0   1E+5   1   1.5  RT (min)       lcqtof_hilic_neg_4_008_24    m/z 797.7394     0   1E+4   1.5   2  RT (min)      lcqtof_hilic_neg_4_000_aa    m/z 798.1358     0   1E+4   4   4.5  RT (min)      lcqtof_hilic_neg_4_008_24    m/z 798.3835     0   1E+4   0.5   1  RT (min)      lcqtof_hilic_neg_4_015_09    m/z 798.5005     0   2   4   6E+4   3   3.5  RT (min)      lcqtof_hilic_neg_4_016_37    m/z 798.5146     0   1   2   3E+5   3   3.5  RT (min)      lcqtof_hilic_neg_4_034_31    m/z 798.5276     0   1   2   3   4E+5   3   3.5  RT (min)      lcqtof_hilic_neg_4_011_35    m/z 798.5280     0   1   2   3   4E+5   3   3.5  RT (min)      lcqtof_hilic_neg_4_011_35    m/z 798.5282     0   2   4E+5   2.5   3  RT (min)      lcqtof_hilic_neg_2_002_36    m/z 798.5282     0   1E+4   2.5   3  RT (min)      lcqtof_hilic_neg_1_014_31    m/z 798.5520     0   1   2   3   4E+4   1   1.5  RT (min)      lcqtof_hilic_neg_3_035_46    m/z 798.6070     0   1E+5   1   1.5  RT (min)      lcqtof_hilic_neg_4_000_aa    m/z 798.6274     0   1E+5   1   1.5  RT (min)      lcqtof_hilic_neg_4_000_aa    m/z 798.6315     0   1   2E+6   4.5   5  RT (min)      lcqtof_hilic_neg_4_002_25    m/z 798.6318     0   1   2E+6   4   4.5  RT (min)       lcqtof_hilic_neg_4_090_60    m/z 798.6394     0   1E+4   2.5   3  RT (min)       lcqtof_hilic_neg_4_055_32    m/z 798.6779     0   2   4   6E+4   1   1.5  RT (min)       lcqtof_hilic_neg_4_008_24    m/z 798.6834     0   2   4   6   8E+3   1.5   2  RT (min)       lcqtof_hilic_neg_3_094_60    m/z 798.6847     0   2   4   6   8E+3   1.5   2  RT (min)       lcqtof_hilic_neg_3_096_60    m/z 798.6948     0   1   2E+5   2   2.5  RT (min)       lcqtof_hilic_neg_4_008_24    m/z 798.6952     0   2   4   6E+4   2   2.5  RT (min)      lcqtof_hilic_neg_3_004_32    m/z 798.7176     0   1   2E+4   1.5   2  RT (min)      lcqtof_hilic_neg_4_000_aa    m/z 799.0040     0   1E+4   4   4.5  RT (min)      lcqtof_hilic_neg_4_048_23    m/z 799.4853     0   1E+4   1   1.5  RT (min)      lcqtof_hilic_neg_4_016_37    m/z 799.5050     0   1   2E+4   3   3.5  RT (min)      lcqtof_hilic_neg_4_022_34    m/z 799.5059     0   1E+4   4   4.5  RT (min)       lcqtof_hilic_neg_4_048_23    m/z 799.5316     0   1   2E+5   3   3.5  RT (min)      lcqtof_hilic_neg_4_048_23    m/z 799.5317     0   1   2E+5   3   3.5  RT (min)      lcqtof_hilic_neg_4_048_23    m/z 799.5435     0   1   2   3E+4   4   4.5  RT (min)      lcqtof_hilic_neg_4_008_24    m/z 799.5531     0   1E+4   1   1.5  RT (min)      lcqtof_hilic_neg_3_035_46    m/z 799.6421     0   2   4   6   8E+5   4.5   5  RT (min)      lcqtof_hilic_neg_4_008_24    m/z 799.6423     0   2   4E+5   4   4.5  RT (min)      lcqtof_hilic_neg_4_055_32    m/z 800.3225     0   1E+4   4   4.5  RT (min)      lcqtof_hilic_neg_4_008_24    m/z 800.4790     0   1   2   3E+4   0.5   1  RT (min)      lcqtof_hilic_neg_4_021_21    m/z 800.5141     0   1   2E+5   3   3.5  RT (min)      lcqtof_hilic_neg_4_022_34    m/z 800.5220     0   1E+5   3   3.5  RT (min)      lcqtof_hilic_neg_4_034_31    m/z 800.5428     0   2   4   6   8E+5   3   3.5  RT (min)      lcqtof_hilic_neg_4_048_23    m/z 800.5428     0   1   2E+5   2.5   3  RT (min)       lcqtof_hilic_neg_4_014_46    m/z 800.5429     0   2   4   6E+5   2.5   3  RT (min)       lcqtof_hilic_neg_4_011_35    m/z 800.5641     0   1   2   3E+4   2   2.5  RT (min)       lcqtof_hilic_neg_4_086_59    m/z 800.5850     0   1   2   3   4E+4   1   1.5  RT (min)      lcqtof_hilic_neg_4_009_50    m/z 800.5858     0   2   4   6E+4   1.5   2  RT (min)      lcqtof_hilic_neg_3_030_23    m/z 800.5860     0   1E+5   1   1.5  RT (min)       lcqtof_hilic_neg_4_048_23    m/z 800.6464     0   2   4E+4   1   1.5  RT (min)      lcqtof_hilic_neg_4_000_aa    m/z 800.6546     0   1E+4   2.5   3  RT (min)       lcqtof_hilic_neg_4_055_32    m/z 800.7292     0   1E+4   2   2.5  RT (min)      lcqtof_hilic_neg_4_016_37    m/z 800.8186     0   1E+4   1.5   2  RT (min)      lcqtof_hilic_neg_4_000_aa    m/z 800.8304     0   1   2   3E+4   3   3.5  RT (min)      lcqtof_hilic_neg_1_035_23    m/z 800.9440     0   1E+4   0.5   1  RT (min)      lcqtof_hilic_neg_3_035_46    m/z 800.9450     0   1E+4   0.5   1  RT (min)      lcqtof_hilic_neg_3_035_46    m/z 801.0962     0   1E+4   2   2.5  RT (min)      lcqtof_hilic_neg_4_016_37    m/z 801.1138     0   2   4   6E+4   1   1.5  RT (min)      lcqtof_hilic_neg_4_004_47    m/z 801.5182     0   1E+5   3   3.5  RT (min)      lcqtof_hilic_neg_4_022_34    m/z 801.5222     0   2   4   6E+4   3   3.5  RT (min)      lcqtof_hilic_neg_4_034_31    m/z 801.5462     0   1   2   3   4E+5   3   3.5  RT (min)      lcqtof_hilic_neg_4_048_23    m/z 801.5470     0   1   2   3   4E+5   3   3.5  RT (min)      lcqtof_hilic_neg_4_048_23    m/z 801.5474     0   1   2   3E+5   2.5   3  RT (min)      lcqtof_hilic_neg_4_016_37    m/z 801.5476     0   1   2   3E+5   2.5   3  RT (min)      lcqtof_hilic_neg_4_011_35    m/z 801.5477     0   2   4   6E+4   2.5   3  RT (min)      lcqtof_hilic_neg_4_016_37    m/z 801.5505     0   1E+4   1   1.5  RT (min)      lcqtof_hilic_neg_4_000_aa    m/z 801.5597     0   2   4   6   8E+4   4   4.5  RT (min)      lcqtof_hilic_neg_4_008_24    m/z 801.5743     0   1   2   3   4E+4   1   1.5  RT (min)      lcqtof_hilic_neg_3_035_46    m/z 801.5896     0   2   4   6E+4   1   1.5  RT (min)      lcqtof_hilic_neg_4_048_23    m/z 801.5911     0   1   2   3   4E+5   1.5   2  RT (min)      lcqtof_hilic_neg_1_022_36    m/z 801.5912     0   1   2E+5   1.5   2  RT (min)      lcqtof_hilic_neg_1_035_23    m/z 801.5912     0   1   2   3E+4   1.5   2  RT (min)      lcqtof_hilic_neg_1_035_23    m/z 801.5912     0   1   2   3   4E+4   1.5   2  RT (min)      lcqtof_hilic_neg_2_002_36    m/z 801.5914     0   1   2   3   4E+4   1.5   2  RT (min)      lcqtof_hilic_neg_2_020_23    m/z 801.5919     0   2   4   6E+4   1.5   2  RT (min)      lcqtof_hilic_neg_1_035_23    m/z 801.8055     0   1   2E+4   1.5   2  RT (min)      lcqtof_hilic_neg_4_000_aa    m/z 802.3126     0   1E+4   4.5   5  RT (min)       lcqtof_hilic_neg_4_039_18    m/z 802.4795     0   1E+4   3.5   4  RT (min)      lcqtof_hilic_neg_1_037_24    m/z 802.5272     0   1   2E+4   4   4.5  RT (min)      lcqtof_hilic_neg_4_040_05    m/z 802.5590     0   1   2E+5   2   2.5  RT (min)       lcqtof_hilic_neg_1_008_13    m/z 802.5592     0   1E+5   1.5   2  RT (min)       lcqtof_hilic_neg_4_050_02    m/z 802.5592     0   1   2E+5   2.5   3  RT (min)      lcqtof_hilic_neg_1_008_13    m/z 802.5593     0   1E+5   2.5   3  RT (min)      lcqtof_hilic_neg_1_016_28    m/z 802.5594     0   1E+5   1.5   2  RT (min)      lcqtof_hilic_neg_4_050_02    m/z 802.5594     0   1   2   3   4E+6   3   3.5  RT (min)      lcqtof_hilic_neg_4_032_01    m/z 802.5594     0   1   2E+5   2.5   3  RT (min)       lcqtof_hilic_neg_2_031_01    m/z 802.5594     0   1E+6   1   1.5  RT (min)       lcqtof_hilic_neg_4_050_02    m/z 802.5595     0   2   4E+6   2.5   3  RT (min)       lcqtof_hilic_neg_4_005_29    m/z 802.5597     0   1E+5   1.5   2  RT (min)      lcqtof_hilic_neg_4_050_02    m/z 802.5602     0   1E+5   1.5   2  RT (min)       lcqtof_hilic_neg_4_050_02    m/z 802.5631     0   1   2   3   4E+6   3.5   4  RT (min)      lcqtof_hilic_neg_4_053_14    m/z 802.5667     0   1E+5   1.5   2  RT (min)      lcqtof_hilic_neg_4_050_02    m/z 802.5742     0   2   4   6E+5   1   1.5  RT (min)      lcqtof_hilic_neg_4_048_23    m/z 802.5806     0   2   4   6   8E+4   1.5   2  RT (min)      lcqtof_hilic_neg_1_035_23    m/z 802.5818     0   1   2   3E+4   1.5   2  RT (min)      lcqtof_hilic_neg_4_048_23    m/z 802.5883     0   2   4   6   8E+5   1   1.5  RT (min)      lcqtof_hilic_neg_3_035_46    m/z 802.5918     0   2   4   6E+4   3.5   4  RT (min)      lcqtof_hilic_neg_4_034_31    m/z 802.8114     0   1E+4   1.5   2  RT (min)      lcqtof_hilic_neg_4_000_aa    m/z 802.8452     0   1E+5   3   3.5  RT (min)      lcqtof_hilic_neg_4_032_01    m/z 803.1378     0   1   2E+4   0.5   1  RT (min)      lcqtof_hilic_neg_3_000_aa    m/z 803.1946     0   2   4   6E+4   1   1.5  RT (min)      lcqtof_hilic_neg_4_000_aa    m/z 803.5066     0   1E+4   3   3.5  RT (min)      lcqtof_hilic_neg_1_035_23    m/z 803.5136     0   1E+4   3.5   4  RT (min)      lcqtof_hilic_neg_1_035_23    m/z 803.5235     0   1E+4   3   3.5  RT (min)      lcqtof_hilic_neg_4_002_25    m/z 803.5384     0   1   2E+4   0.5   1  RT (min)      lcqtof_hilic_neg_4_000_aa    m/z 803.5619     0   1E+5   3   3.5  RT (min)      lcqtof_hilic_neg_4_002_25    m/z 803.5621     0   1E+5   2.5   3  RT (min)      lcqtof_hilic_neg_2_031_01    m/z 803.5622     0   1   2E+6   3   3.5  RT (min)      lcqtof_hilic_neg_4_032_01    m/z 803.5622     0   1   2E+6   2.5   3  RT (min)      lcqtof_hilic_neg_4_006_30    m/z 803.5623     0   1   2   3   4E+5   2.5   3  RT (min)      lcqtof_hilic_neg_1_030_17    m/z 803.5624     0   1E+5   2   2.5  RT (min)      lcqtof_hilic_neg_1_008_13    m/z 803.5628     0   1E+5   1.5   2  RT (min)      lcqtof_hilic_neg_4_050_02    m/z 803.5628     0   2   4   6E+4   1.5   2  RT (min)      lcqtof_hilic_neg_4_050_02    m/z 803.5630     0   2   4   6   8E+4   2.5   3  RT (min)      lcqtof_hilic_neg_1_016_28    m/z 803.5631     0   1   2   3   4E+4   2   2.5  RT (min)      lcqtof_hilic_neg_1_005_32    m/z 803.5631     0   2   4   6E+4   1.5   2  RT (min)      lcqtof_hilic_neg_4_050_02    m/z 803.5634     0   1E+5   2   2.5  RT (min)       lcqtof_hilic_neg_1_008_13    m/z 803.5634     0   1E+5   2   2.5  RT (min)      lcqtof_hilic_neg_1_008_13    m/z 803.5638     0   2   4   6E+4   1.5   2  RT (min)      lcqtof_hilic_neg_4_050_02    m/z 803.5641     0   2   4   6   8E+4   1.5   2  RT (min)       lcqtof_hilic_neg_4_050_02    m/z 803.5644     0   1   2E+5   3.5   4  RT (min)      lcqtof_hilic_neg_4_014_46    m/z 803.5647     0   1   2E+6   3.5   4  RT (min)      lcqtof_hilic_neg_4_052_13    m/z 803.5659     0   1   2   3E+4   4   4.5  RT (min)      lcqtof_hilic_neg_4_008_24    m/z 803.5798     0   2   4   6   8E+4   2   2.5  RT (min)      lcqtof_hilic_neg_4_016_37    m/z 803.5876     0   1   2   3E+5   1   1.5  RT (min)      lcqtof_hilic_neg_3_035_46    m/z 803.6778     0   1   2E+4   1   1.5  RT (min)      lcqtof_hilic_neg_4_000_aa    m/z 803.6975     0   1   2E+4   1.5   2  RT (min)      lcqtof_hilic_neg_4_000_aa    m/z 803.7475     0   1   2E+4   1   1.5  RT (min)      lcqtof_hilic_neg_4_000_aa    m/z 803.7508     0   1   2E+4   1.5   2  RT (min)      lcqtof_hilic_neg_4_000_aa    m/z 803.7740     0   1E+4   1.5   2  RT (min)      lcqtof_hilic_neg_4_000_aa    m/z 803.7820     0   1E+4   1.5   2  RT (min)      lcqtof_hilic_neg_4_000_aa    m/z 804.1381     0   1E+4   0.5   1  RT (min)      lcqtof_hilic_neg_3_000_bb    m/z 804.1956     0   2   4E+4   1   1.5  RT (min)      lcqtof_hilic_neg_4_000_aa    m/z 804.5404     0   1E+4   4   4.5  RT (min)      lcqtof_hilic_neg_4_047_06    m/z 804.5710     0   2   4   6   8E+4   1.5   2  RT (min)       lcqtof_hilic_neg_4_050_02    m/z 804.5716     0   2   4   6E+4   1.5   2  RT (min)      lcqtof_hilic_neg_4_050_02    m/z 804.5721     0   1E+5   2.5   3  RT (min)      lcqtof_hilic_neg_1_008_13    m/z 804.5722     0   1E+6   2.5   3  RT (min)      lcqtof_hilic_neg_2_004_26    m/z 804.5724     0   2   4E+5   1   1.5  RT (min)      lcqtof_hilic_neg_4_050_02    m/z 804.5724     0   2   4   6E+4   1.5   2  RT (min)      lcqtof_hilic_neg_4_050_02    m/z 804.5724     0   1   2E+6   2.5   3  RT (min)      lcqtof_hilic_neg_4_006_30    m/z 804.5724     0   1E+5   2   2.5  RT (min)       lcqtof_hilic_neg_1_008_13    m/z 804.5724     0   1   2E+6   3   3.5  RT (min)      lcqtof_hilic_neg_4_032_01    m/z 804.5726     0   1E+5   2   2.5  RT (min)      lcqtof_hilic_neg_1_008_13    m/z 804.5728     0   1E+5   2   2.5  RT (min)      lcqtof_hilic_neg_1_008_13    m/z 804.5729     0   1   2E+4   2   2.5  RT (min)      lcqtof_hilic_neg_3_001_49    m/z 804.5730     0   2   4   6E+5   1   1.5  RT (min)      lcqtof_hilic_neg_2_029_31    m/z 804.5732     0   1E+5   1.5   2  RT (min)      lcqtof_hilic_neg_4_050_02    m/z 804.5734     0   2   4   6E+4   1.5   2  RT (min)      lcqtof_hilic_neg_4_050_02    m/z 804.5760     0   1E+6   3.5   4  RT (min)      lcqtof_hilic_neg_4_052_13    m/z 804.5765     0   1E+6   3.5   4  RT (min)      lcqtof_hilic_neg_4_005_29    m/z 804.5974     0   1   2   3E+5   1   1.5  RT (min)      lcqtof_hilic_neg_3_035_46    m/z 804.6947     0   1E+4   1   1.5  RT (min)      lcqtof_hilic_neg_4_000_aa    m/z 804.7019     0   1E+4   1.5   2  RT (min)      lcqtof_hilic_neg_4_000_aa    m/z 804.7561     0   1E+4   1   1.5  RT (min)      lcqtof_hilic_neg_4_000_aa    m/z 804.9540     0   1E+4   1   1.5  RT (min)      lcqtof_hilic_neg_4_000_aa    m/z 805.1336     0   1E+4   6   6.5  RT (min)      lcqtof_hilic_neg_4_036_11    m/z 805.1340     0   1E+4   0.5   1  RT (min)      lcqtof_hilic_neg_3_000_aa    m/z 805.1944     0   1   2   3   4E+4   1   1.5  RT (min)      lcqtof_hilic_neg_4_000_aa    m/z 805.5066     0   1E+4   1   1.5  RT (min)      lcqtof_hilic_neg_4_000_aa    m/z 805.5113     0   1E+4   1   1.5  RT (min)      lcqtof_hilic_neg_4_000_aa    m/z 805.5224     0   2   4   6   8E+4   3.5   4  RT (min)      lcqtof_hilic_neg_1_035_23    m/z 805.5229     0   1   2   3E+4   2.5   3  RT (min)      lcqtof_hilic_neg_1_003_37    m/z 805.5230     0   1   2E+4   3   3.5  RT (min)      lcqtof_hilic_neg_4_002_25    m/z 805.5230     0   1E+5   3   3.5  RT (min)       lcqtof_hilic_neg_2_002_36    m/z 805.5238     0   1E+4   1   1.5  RT (min)      lcqtof_hilic_neg_2_020_23    m/z 805.5252     0   1E+4   1   1.5  RT (min)      lcqtof_hilic_neg_1_035_23    m/z 805.5273     0   1   2E+4   3   3.5  RT (min)      lcqtof_hilic_neg_4_040_05    m/z 805.5276     0   1E+5   2   2.5  RT (min)       lcqtof_hilic_neg_4_028_36    m/z 805.5390     0   1   2   3E+4   2.5   3  RT (min)      lcqtof_hilic_neg_2_002_36    m/z 805.5401     0   1E+5   3   3.5  RT (min)      lcqtof_hilic_neg_2_039_38    m/z 805.5453     0   2   4   6   8E+4   2   2.5  RT (min)      lcqtof_hilic_neg_4_016_37    m/z 805.5639     0   1   2   3E+4   2.5   3  RT (min)      lcqtof_hilic_neg_2_048_35    m/z 805.5657     0   1   2   3E+4   2.5   3  RT (min)      lcqtof_hilic_neg_1_014_31    m/z 805.5704     0   1   2   3E+4   1.5   2  RT (min)      lcqtof_hilic_neg_4_028_36    m/z 805.5756     0   1   2   3E+4   1.5   2  RT (min)      lcqtof_hilic_neg_4_031_38    m/z 805.5761     0   2   4   6E+4   2.5   3  RT (min)      lcqtof_hilic_neg_2_031_01    m/z 805.5768     0   2   4   6   8E+5   2.5   3  RT (min)       lcqtof_hilic_neg_4_005_29    m/z 805.5768     0   1   2   3   4E+4   1.5   2  RT (min)       lcqtof_hilic_neg_1_003_37    m/z 805.5770     0   1   2   3   4E+4   2   2.5  RT (min)       lcqtof_hilic_neg_1_008_13    m/z 805.5770     0   1   2   3E+4   2   2.5  RT (min)      lcqtof_hilic_neg_1_003_37    m/z 805.5770     0   1   2   3E+4   1   1.5  RT (min)       lcqtof_hilic_neg_4_048_23    m/z 805.5770     0   2   4   6   8E+5   3   3.5  RT (min)       lcqtof_hilic_neg_4_032_01    m/z 805.5770     0   2   4   6   8E+5   2.5   3  RT (min)       lcqtof_hilic_neg_4_005_29    m/z 805.5770     0   2   4E+4   2   2.5  RT (min)       lcqtof_hilic_neg_4_048_23    m/z 805.5772     0   1   2   3   4E+4   2   2.5  RT (min)       lcqtof_hilic_neg_1_008_13    m/z 805.5772     0   1   2E+4   1.5   2  RT (min)      lcqtof_hilic_neg_2_002_36    m/z 805.5774     0   1   2E+5   1   1.5  RT (min)      lcqtof_hilic_neg_4_034_31    m/z 805.5774     0   1E+4   1.5   2  RT (min)       lcqtof_hilic_neg_4_003_16    m/z 805.5777     0   1   2   3   4E+4   2   2.5  RT (min)       lcqtof_hilic_neg_1_008_13    m/z 805.5778     0   1   2   3   4E+4   2   2.5  RT (min)      lcqtof_hilic_neg_1_008_13    m/z 805.5780     0   1E+5   2.5   3  RT (min)       lcqtof_hilic_neg_3_055_01    m/z 805.5783     0   1   2E+4   1.5   2  RT (min)      lcqtof_hilic_neg_4_035_33    m/z 805.6142     0   2   4E+5   1   1.5  RT (min)      lcqtof_hilic_neg_4_000_aa    m/z 805.6218     0   2   4E+5   1   1.5  RT (min)      lcqtof_hilic_neg_4_000_aa    m/z 805.6229     0   1   2   3   4E+5   1.5   2  RT (min)      lcqtof_hilic_neg_4_000_aa    m/z 805.6232     0   2   4E+5   1   1.5  RT (min)      lcqtof_hilic_neg_4_000_aa    m/z 805.6522     0   2   4E+4   2.5   3  RT (min)      lcqtof_hilic_neg_4_008_24    m/z 805.6524     0   1   2   3E+5   2.5   3  RT (min)      lcqtof_hilic_neg_4_008_24    m/z 805.6529     0   1   2E+5   2.5   3  RT (min)      lcqtof_hilic_neg_4_008_24    m/z 805.7310     0   1   2   3E+4   1   1.5  RT (min)      lcqtof_hilic_neg_4_000_aa    m/z 805.7310     0   1   2   3E+4   1   1.5  RT (min)      lcqtof_hilic_neg_4_000_aa    m/z 805.7602     0   1   2E+4   1.5   2  RT (min)      lcqtof_hilic_neg_4_000_aa    m/z 806.1920     0   1   2E+4   1   1.5  RT (min)      lcqtof_hilic_neg_4_000_aa    m/z 806.3466     0   1   2E+4   0.5   1  RT (min)      lcqtof_hilic_neg_3_003_20    m/z 806.5282     0   1   2   3   4E+4   2   2.5  RT (min)      lcqtof_hilic_neg_4_028_36    m/z 806.5336     0   1E+4   1   1.5  RT (min)      lcqtof_hilic_neg_2_002_36    m/z 806.5357     0   1   2   3E+4   2   2.5  RT (min)      lcqtof_hilic_neg_4_011_35    m/z 806.5539     0   1E+4   3   3.5  RT (min)      lcqtof_hilic_neg_3_018_33    m/z 806.5808     0   1E+4   1.5   2  RT (min)      lcqtof_hilic_neg_1_023_34    m/z 806.5808     0   2   4   6   8E+3   2   2.5  RT (min)      lcqtof_hilic_neg_1_013_38    m/z 806.5811     0   2   4   6E+4   1   1.5  RT (min)       lcqtof_hilic_neg_4_028_36    m/z 806.5812     0   1E+4   2.5   3  RT (min)      lcqtof_hilic_neg_2_005_13    m/z 806.5818     0   1   2E+4   2   2.5  RT (min)      lcqtof_hilic_neg_4_048_23    m/z 806.5819     0   1   2E+4   2.5   3  RT (min)      lcqtof_hilic_neg_2_005_13    m/z 806.5822     0   1   2E+5   3   3.5  RT (min)      lcqtof_hilic_neg_4_005_29    m/z 806.5824     0   1   2E+5   2.5   3  RT (min)      lcqtof_hilic_neg_4_005_29    m/z 806.5824     0   1   2E+5   3   3.5  RT (min)      lcqtof_hilic_neg_4_006_30    m/z 806.6078     0   1   2E+4   1   1.5  RT (min)      lcqtof_hilic_neg_3_035_46    m/z 806.6238     0   1   2   3E+5   1.5   2  RT (min)      lcqtof_hilic_neg_4_000_aa    m/z 806.6242     0   1   2   3E+5   1   1.5  RT (min)      lcqtof_hilic_neg_4_000_aa    m/z 806.6245     0   1   2   3E+5   1   1.5  RT (min)      lcqtof_hilic_neg_4_000_aa    m/z 806.6248     0   1   2E+5   1.5   2  RT (min)      lcqtof_hilic_neg_4_000_aa    m/z 806.6253     0   1   2   3E+5   1   1.5  RT (min)      lcqtof_hilic_neg_4_000_aa    m/z 806.6260     0   1   2E+5   1.5   2  RT (min)      lcqtof_hilic_neg_4_000_aa    m/z 806.6625     0   1E+5   3   3.5  RT (min)      lcqtof_hilic_neg_4_008_24    m/z 806.6635     0   1E+5   2.5   3  RT (min)      lcqtof_hilic_neg_4_008_24    m/z 806.6637     0   1   2   3E+4   2.5   3  RT (min)      lcqtof_hilic_neg_4_008_24    m/z 806.6637     0   1   2E+4   2.5   3  RT (min)      lcqtof_hilic_neg_4_008_24    m/z 806.6899     0   1   2   3E+4   0.5   1  RT (min)      lcqtof_hilic_neg_3_035_46    m/z 806.7324     0   1E+4   1   1.5  RT (min)      lcqtof_hilic_neg_4_000_aa    m/z 806.7394     0   1E+4   1   1.5  RT (min)      lcqtof_hilic_neg_4_000_aa    m/z 807.3476     0   1E+4   5   5.5  RT (min)       lcqtof_hilic_neg_4_008_24    m/z 807.5348     0   1   2E+4   2   2.5  RT (min)      lcqtof_hilic_neg_4_034_31    m/z 807.6244     0   1E+5   1   1.5  RT (min)      lcqtof_hilic_neg_4_000_aa    m/z 807.6244     0   1E+5   1   1.5  RT (min)      lcqtof_hilic_neg_4_000_aa    m/z 807.6253     0   1E+5   1   1.5  RT (min)      lcqtof_hilic_neg_4_000_aa    m/z 807.6384     0   1   2E+5   1.5   2  RT (min)      lcqtof_hilic_neg_4_000_aa    m/z 807.6671     0   1E+5   2.5   3  RT (min)      lcqtof_hilic_neg_4_008_24    m/z 807.6673     0   1   2E+5   2.5   3  RT (min)      lcqtof_hilic_neg_4_008_24    m/z 807.6677     0   1   2   3E+4   1.5   2  RT (min)       lcqtof_hilic_neg_4_097_60    m/z 807.6704     0   2   4   6   8E+3   15   15.5  RT (min)      lcqtof_hilic_neg_4_013_20    m/z 807.6706     0   1E+4   15   15.5  RT (min)      lcqtof_hilic_neg_4_015_09    m/z 807.6926     0   1E+5   0.5   1  RT (min)      lcqtof_hilic_neg_3_035_46    m/z 807.7306     0   1   2E+4   1   1.5  RT (min)      lcqtof_hilic_neg_4_000_aa    m/z 808.3508     0   1E+4   5   5.5  RT (min)       lcqtof_hilic_neg_4_091_60    m/z 808.5481     0   1E+4   1   1.5  RT (min)      lcqtof_hilic_neg_4_000_aa    m/z 808.5863     0   1E+4   3   3.5  RT (min)      lcqtof_hilic_neg_4_053_14    m/z 808.6240     0   2   4   6E+4   1   1.5  RT (min)      lcqtof_hilic_neg_4_000_aa    m/z 808.6252     0   2   4   6E+4   1   1.5  RT (min)      lcqtof_hilic_neg_4_000_aa    m/z 808.6266     0   2   4   6   8E+4   1.5   2  RT (min)      lcqtof_hilic_neg_4_000_aa    m/z 808.6271     0   2   4   6E+4   1   1.5  RT (min)      lcqtof_hilic_neg_4_000_aa    m/z 808.6276     0   2   4   6E+4   1   1.5  RT (min)      lcqtof_hilic_neg_4_000_aa    m/z 808.6297     0   2   4   6E+4   1   1.5  RT (min)      lcqtof_hilic_neg_4_000_aa    m/z 808.6302     0   2   4   6   8E+4   1   1.5  RT (min)      lcqtof_hilic_neg_4_000_aa    m/z 808.6471     0   2   4   6   8E+4   1.5   2  RT (min)      lcqtof_hilic_neg_4_000_aa    m/z 808.6750     0   2   4   6E+3   15.5   16  RT (min)      lcqtof_hilic_neg_4_015_09    m/z 808.7039     0   1   2   3E+4   0.5   1  RT (min)      lcqtof_hilic_neg_3_035_46    m/z 808.9548     0   1   2E+4   7   7.5  RT (min)      lcqtof_hilic_neg_4_053_14    m/z 808.9548     0   1   2E+4   7   7.5  RT (min)      lcqtof_hilic_neg_4_053_14    m/z 808.9550     0   1   2E+4   6.5   7  RT (min)       lcqtof_hilic_neg_4_059_14    m/z 809.6214     0   1   2E+4   1.5   2  RT (min)      lcqtof_hilic_neg_4_000_aa    m/z 809.6247     0   1   2   3   4E+4   1.5   2  RT (min)      lcqtof_hilic_neg_4_000_aa    m/z 809.6252     0   1   2   3   4E+4   1.5   2  RT (min)      lcqtof_hilic_neg_4_000_aa    m/z 809.6255     0   1   2   3   4E+4   1   1.5  RT (min)      lcqtof_hilic_neg_4_000_aa    m/z 809.6271     0   1   2   3E+4   1.5   2  RT (min)      lcqtof_hilic_neg_4_000_aa    m/z 809.6283     0   1   2   3   4E+4   1.5   2  RT (min)      lcqtof_hilic_neg_4_000_aa    m/z 809.6290     0   1   2   3   4E+4   1   1.5  RT (min)      lcqtof_hilic_neg_4_000_aa    m/z 809.6291     0   1   2   3   4E+4   1   1.5  RT (min)      lcqtof_hilic_neg_4_000_aa    m/z 809.6400     0   1   2E+4   1.5   2  RT (min)      lcqtof_hilic_neg_4_000_aa    m/z 809.6414     0   1   2E+4   1.5   2  RT (min)      lcqtof_hilic_neg_4_000_aa    m/z 809.6499     0   1   2   3   4E+4   1   1.5  RT (min)      lcqtof_hilic_neg_4_000_aa    m/z 809.6818     0   1   2E+5   2.5   3  RT (min)      lcqtof_hilic_neg_4_055_32    m/z 809.6828     0   2   4E+4   2.5   3  RT (min)      lcqtof_hilic_neg_4_096_60    m/z 809.6828     0   1   2E+5   2.5   3  RT (min)      lcqtof_hilic_neg_4_055_32    m/z 809.6840     0   1   2   3E+4   2.5   3  RT (min)      lcqtof_hilic_neg_4_004_47    m/z 809.7082     0   1E+5   0.5   1  RT (min)      lcqtof_hilic_neg_3_035_46    m/z 810.6022     0   1E+4   4   4.5  RT (min)      lcqtof_hilic_neg_4_008_24    m/z 810.7160     0   1E+4   1.5   2  RT (min)      lcqtof_hilic_neg_4_000_aa    m/z 810.7204     0   1   2   3E+4   0.5   1  RT (min)      lcqtof_hilic_neg_3_035_46    m/z 811.1294     0   1E+4   1   1.5  RT (min)      lcqtof_hilic_neg_4_061_50    m/z 811.3394     0   1   2E+4   1   1.5  RT (min)      lcqtof_hilic_neg_4_000_aa    m/z 811.3514     0   1   2E+4   1   1.5  RT (min)      lcqtof_hilic_neg_4_000_aa    m/z 811.3520     0   1   2E+4   1   1.5  RT (min)      lcqtof_hilic_neg_4_000_aa    m/z 811.3700     0   1   2E+4   1   1.5  RT (min)      lcqtof_hilic_neg_4_000_aa    m/z 811.3719     0   1E+4   1   1.5  RT (min)      lcqtof_hilic_neg_4_000_aa    m/z 811.5180     0   1E+4   3.5   4  RT (min)      lcqtof_hilic_neg_4_048_23    m/z 811.6323     0   1   2E+4   1.5   2  RT (min)      lcqtof_hilic_neg_4_000_aa    m/z 811.6410     0   1   2E+4   3   3.5  RT (min)      lcqtof_hilic_neg_4_008_24    m/z 811.6424     0   1E+4   2.5   3  RT (min)      lcqtof_hilic_neg_4_040_05    m/z 811.6446     0   1E+4   0.5   1  RT (min)      lcqtof_hilic_neg_4_005_29    m/z 811.6562     0   1E+4   0.5   1  RT (min)      lcqtof_hilic_neg_1_042_26    m/z 811.6983     0   1   2   3   4E+4   2.5   3  RT (min)      lcqtof_hilic_neg_4_035_33    m/z 811.6984     0   1   2   3   4E+5   3   3.5  RT (min)       lcqtof_hilic_neg_4_008_24    m/z 811.6986     0   1   2   3   4E+5   2.5   3  RT (min)      lcqtof_hilic_neg_4_055_32    m/z 811.6992     0   2   4   6   8E+4   2.5   3  RT (min)      lcqtof_hilic_neg_4_096_60    m/z 811.7234     0   1E+5   0.5   1  RT (min)      lcqtof_hilic_neg_3_035_46    m/z 811.7738     0   1   2   3E+4   4   4.5  RT (min)      lcqtof_hilic_neg_4_013_20    m/z 811.9866     0   1E+4   0.5   1  RT (min)      lcqtof_hilic_neg_4_008_24    m/z 812.2253     0   1E+4   6   6.5  RT (min)      lcqtof_hilic_neg_4_048_23    m/z 812.5172     0   1E+4   3   3.5  RT (min)      lcqtof_hilic_neg_4_022_34    m/z 812.5316     0   2   4   6   8E+4   3   3.5  RT (min)      lcqtof_hilic_neg_4_002_25    m/z 812.5398     0   1E+5   3   3.5  RT (min)       lcqtof_hilic_neg_4_048_23    m/z 812.5477     0   1E+5   2.5   3  RT (min)      lcqtof_hilic_neg_4_016_37    m/z 812.5558     0   2   4   6   8E+4   2.5   3  RT (min)      lcqtof_hilic_neg_4_034_31    m/z 812.5592     0   2   4   6   8E+4   2.5   3  RT (min)      lcqtof_hilic_neg_4_011_35    m/z 812.5799     0   2   4   6E+3   2   2.5  RT (min)      lcqtof_hilic_neg_4_004_47    m/z 812.5802     0   1   2   3E+4   2.5   3  RT (min)       lcqtof_hilic_neg_4_005_29    m/z 812.6135     0   1E+5   4   4.5  RT (min)      lcqtof_hilic_neg_4_055_32    m/z 812.6456     0   2   4   6E+4   3   3.5  RT (min)       lcqtof_hilic_neg_4_008_24    m/z 812.7088     0   1E+5   1   1.5  RT (min)      lcqtof_hilic_neg_4_008_24    m/z 812.7336     0   1E+4   1.5   2  RT (min)      lcqtof_hilic_neg_4_000_aa    m/z 812.7351     0   1E+4   0.5   1  RT (min)      lcqtof_hilic_neg_3_035_46    m/z 813.3200     0   1E+4   6   6.5  RT (min)       lcqtof_hilic_neg_4_058_39    m/z 813.3334     0   1E+4   1   1.5  RT (min)      lcqtof_hilic_neg_4_000_aa    m/z 813.3366     0   1E+4   1   1.5  RT (min)      lcqtof_hilic_neg_4_000_aa    m/z 813.3698     0   1E+4   4   4.5  RT (min)      lcqtof_hilic_neg_4_011_35    m/z 813.3702     0   1   2   3   4E+4   3.5   4  RT (min)       lcqtof_hilic_neg_4_048_23    m/z 813.5421     0   2   4   6E+4   3   3.5  RT (min)      lcqtof_hilic_neg_4_048_23    m/z 813.6474     0   1E+5   4   4.5  RT (min)       lcqtof_hilic_neg_4_055_32    m/z 813.6562     0   1E+4   2.5   3  RT (min)       lcqtof_hilic_neg_4_008_24    m/z 813.6974     0   1E+5   1.5   2  RT (min)      lcqtof_hilic_neg_1_037_24    m/z 813.6979     0   1E+5   1.5   2  RT (min)      lcqtof_hilic_neg_1_037_24    m/z 813.6980     0   1   2E+5   1.5   2  RT (min)       lcqtof_hilic_neg_4_008_24    m/z 813.7008     0   1   2   3E+5   1   1.5  RT (min)      lcqtof_hilic_neg_4_008_24    m/z 813.7008     0   1   2   3E+5   1   1.5  RT (min)       lcqtof_hilic_neg_4_008_24    m/z 813.7014     0   1E+5   1   1.5  RT (min)      lcqtof_hilic_neg_4_002_25    m/z 813.7389     0   1   2   3E+4   0.5   1  RT (min)      lcqtof_hilic_neg_3_035_46    m/z 814.3728     0   1E+4   3.5   4  RT (min)       lcqtof_hilic_neg_4_048_23    m/z 814.5157     0   1E+4   1   1.5  RT (min)      lcqtof_hilic_neg_2_020_23    m/z 814.5187     0   1E+4   1   1.5  RT (min)      lcqtof_hilic_neg_1_035_23    m/z 814.5238     0   1   2E+4   1   1.5  RT (min)      lcqtof_hilic_neg_1_035_23    m/z 814.5246     0   1   2   3E+4   3   3.5  RT (min)       lcqtof_hilic_neg_2_020_23    m/z 814.5258     0   1   2   3E+4   2.5   3  RT (min)      lcqtof_hilic_neg_1_035_23    m/z 814.5328     0   1   2   3E+4   2.5   3  RT (min)      lcqtof_hilic_neg_1_035_23    m/z 814.5616     0   1E+4   2   2.5  RT (min)       lcqtof_hilic_neg_4_016_37    m/z 814.5647     0   1E+4   2   2.5  RT (min)      lcqtof_hilic_neg_1_003_37    m/z 814.6610     0   2   4   6   8E+4   3   3.5  RT (min)      lcqtof_hilic_neg_4_008_24    m/z 814.6852     0   1E+4   9   9.5  RT (min)      lcqtof_hilic_neg_1_000_aa    m/z 814.8759     0   1   2   3E+4   0.5   1  RT (min)      lcqtof_hilic_neg_3_035_46    m/z 814.8789     0   1   2   3E+4   0.5   1  RT (min)      lcqtof_hilic_neg_3_035_46    m/z 815.3690     0   1E+4   3.5   4  RT (min)      lcqtof_hilic_neg_4_048_23    m/z 815.4203     0   1   2   3E+4   0.5   1  RT (min)      lcqtof_hilic_neg_4_000_aa    m/z 815.5104     0   1   2   3E+4   1   1.5  RT (min)      lcqtof_hilic_neg_4_013_20    m/z 815.5197     0   1E+4   3   3.5  RT (min)      lcqtof_hilic_neg_4_016_37    m/z 815.5230     0   1E+4   1   1.5  RT (min)      lcqtof_hilic_neg_2_020_23    m/z 815.5311     0   1E+4   3.5   4  RT (min)      lcqtof_hilic_neg_4_016_37    m/z 815.5342     0   1   2E+4   4   4.5  RT (min)      lcqtof_hilic_neg_4_008_24    m/z 815.8802     0   1E+4   0.5   1  RT (min)      lcqtof_hilic_neg_3_035_46    m/z 816.4558     0   1E+4   4   4.5  RT (min)      lcqtof_hilic_neg_4_016_37    m/z 816.5141     0   1   2E+4   1   1.5  RT (min)      lcqtof_hilic_neg_4_013_20    m/z 816.5215     0   1   2   3   4E+4   3   3.5  RT (min)      lcqtof_hilic_neg_4_019_03    m/z 816.5310     0   1E+4   1   1.5  RT (min)      lcqtof_hilic_neg_1_014_31    m/z 816.5328     0   1E+4   2.5   3  RT (min)       lcqtof_hilic_neg_2_062_17    m/z 816.5330     0   1E+4   2   2.5  RT (min)       lcqtof_hilic_neg_1_016_28    m/z 816.5332     0   1E+4   2   2.5  RT (min)      lcqtof_hilic_neg_1_016_28    m/z 816.5336     0   1E+4   1.5   2  RT (min)       lcqtof_hilic_neg_1_016_28    m/z 816.5337     0   1E+4   2   2.5  RT (min)       lcqtof_hilic_neg_2_062_17    m/z 816.5340     0   2   4   6   8E+3   1.5   2  RT (min)       lcqtof_hilic_neg_1_010_15    m/z 816.5340     0   2   4   6   8E+3   1   1.5  RT (min)      lcqtof_hilic_neg_2_002_36    m/z 816.5344     0   1   2E+4   3.5   4  RT (min)      lcqtof_hilic_neg_4_016_37    m/z 816.5360     0   1E+4   2   2.5  RT (min)      lcqtof_hilic_neg_1_027_27    m/z 816.5502     0   1   2   3E+4   3   3.5  RT (min)      lcqtof_hilic_neg_4_052_13    m/z 816.5776     0   1E+4   2   2.5  RT (min)      lcqtof_hilic_neg_4_016_37    m/z 816.6003     0   1E+4   1   1.5  RT (min)      lcqtof_hilic_neg_3_035_46    m/z 816.6510     0   1   2   3E+4   4   4.5  RT (min)      lcqtof_hilic_neg_4_008_24    m/z 816.6816     0   1   2   3   4E+4   3   3.5  RT (min)       lcqtof_hilic_neg_4_008_24    m/z 816.8160     0   1E+4   5   5.5  RT (min)      lcqtof_hilic_neg_4_055_32    m/z 816.8786     0   1E+4   0.5   1  RT (min)      lcqtof_hilic_neg_3_035_46    m/z 816.9886     0   1E+4   0.5   1  RT (min)      lcqtof_hilic_neg_4_000_aa    m/z 817.1581     0   1   2E+4   0.5   1  RT (min)      lcqtof_hilic_neg_1_062_17    m/z 817.4182     0   1   2   3E+4   1   1.5  RT (min)      lcqtof_hilic_neg_4_000_aa    m/z 817.4237     0   1   2   3E+4   1   1.5  RT (min)      lcqtof_hilic_neg_4_000_aa    m/z 817.5358     0   2   4   6   8E+3   2.5   3  RT (min)      lcqtof_hilic_neg_4_062_17    m/z 817.5360     0   2   4   6   8E+3   3   3.5  RT (min)       lcqtof_hilic_neg_4_059_14    m/z 817.5364     0   2   4   6   8E+3   2.5   3  RT (min)       lcqtof_hilic_neg_4_005_29    m/z 817.5367     0   2   4   6   8E+3   2.5   3  RT (min)      lcqtof_hilic_neg_4_059_14    m/z 817.5657     0   1E+5   1.5   2  RT (min)      lcqtof_hilic_neg_2_048_35    m/z 817.5766     0   1E+5   1.5   2  RT (min)      lcqtof_hilic_neg_1_051_35    m/z 817.5771     0   1E+5   1.5   2  RT (min)      lcqtof_hilic_neg_1_035_23    m/z 817.7332     0   1E+4   1   1.5  RT (min)      lcqtof_hilic_neg_4_000_aa    m/z 817.7728     0   1E+4   1   1.5  RT (min)      lcqtof_hilic_neg_4_000_aa    m/z 817.7973     0   1E+4   1.5   2  RT (min)      lcqtof_hilic_neg_4_000_aa    m/z 817.7984     0   1E+4   1.5   2  RT (min)      lcqtof_hilic_neg_4_000_aa    m/z 818.1596     0   1   2E+4   0.5   1  RT (min)      lcqtof_hilic_neg_1_062_17    m/z 818.4194     0   1E+4   1   1.5  RT (min)      lcqtof_hilic_neg_4_000_aa    m/z 818.5444     0   1E+4   2   2.5  RT (min)      lcqtof_hilic_neg_2_063_13    m/z 818.5445     0   2   4   6   8E+3   1   1.5  RT (min)      lcqtof_hilic_neg_1_054_01    m/z 818.5445     0   2   4   6   8E+3   1   1.5  RT (min)      lcqtof_hilic_neg_1_054_01    m/z 818.5447     0   1E+4   1   1.5  RT (min)      lcqtof_hilic_neg_2_005_13    m/z 818.5452     0   2   4   6   8E+3   2   2.5  RT (min)      lcqtof_hilic_neg_2_064_16    m/z 818.5454     0   1E+4   1   1.5  RT (min)      lcqtof_hilic_neg_1_063_13    m/z 818.5465     0   1E+4   2   2.5  RT (min)      lcqtof_hilic_neg_2_063_13    m/z 818.5470     0   2   4   6   8E+3   1   1.5  RT (min)      lcqtof_hilic_neg_1_008_13    m/z 818.5512     0   1E+4   1.5   2  RT (min)      lcqtof_hilic_neg_1_023_34    m/z 818.5528     0   1E+4   2.5   3  RT (min)      lcqtof_hilic_neg_1_054_01    m/z 818.5609     0   1   2   3E+4   2   2.5  RT (min)      lcqtof_hilic_neg_1_014_31    m/z 818.5746     0   2   4   6E+4   1.5   2  RT (min)       lcqtof_hilic_neg_1_051_35    m/z 818.5775     0   2   4   6E+4   2.5   3  RT (min)       lcqtof_hilic_neg_4_034_31    m/z 818.5974     0   1   2E+4   2.5   3  RT (min)      lcqtof_hilic_neg_4_022_34    m/z 818.6850     0   1E+4   1   1.5  RT (min)       lcqtof_hilic_neg_4_008_24    m/z 818.6852     0   2   4   6   8E+3   1.5   2  RT (min)      lcqtof_hilic_neg_1_037_24    m/z 819.1555     0   1   2E+4   0.5   1  RT (min)      lcqtof_hilic_neg_1_062_17    m/z 819.5254     0   1   2   3E+4   1   1.5  RT (min)      lcqtof_hilic_neg_4_000_aa    m/z 819.5485     0   1   2   3E+4   2.5   3  RT (min)      lcqtof_hilic_neg_2_031_01    m/z 819.5494     0   1   2   3   4E+4   2.5   3  RT (min)       lcqtof_hilic_neg_4_032_01    m/z 819.5494     0   1   2   3E+4   2.5   3  RT (min)       lcqtof_hilic_neg_1_008_13    m/z 819.5494     0   1   2   3E+4   2.5   3  RT (min)       lcqtof_hilic_neg_2_005_13    m/z 819.5496     0   1   2   3E+4   2.5   3  RT (min)       lcqtof_hilic_neg_2_064_16    m/z 819.5496     0   1   2   3E+4   2.5   3  RT (min)      lcqtof_hilic_neg_2_055_30    m/z 819.5497     0   1   2E+4   2   2.5  RT (min)       lcqtof_hilic_neg_1_008_13    m/z 819.5502     0   1   2E+4   2   2.5  RT (min)      lcqtof_hilic_neg_1_008_13    m/z 819.5505     0   1   2E+4   2   2.5  RT (min)       lcqtof_hilic_neg_1_008_13    m/z 819.5506     0   2   4   6E+4   1.5   2  RT (min)       lcqtof_hilic_neg_4_063_16    m/z 819.5739     0   1   2E+4   1.5   2  RT (min)      lcqtof_hilic_neg_2_020_23    m/z 819.5824     0   1   2E+4   2   2.5  RT (min)      lcqtof_hilic_neg_4_048_23    m/z 819.6393     0   1   2   3E+4   1   1.5  RT (min)      lcqtof_hilic_neg_4_000_aa    m/z 819.6590     0   1E+4   0.5   1  RT (min)      lcqtof_hilic_neg_2_016_33    m/z 819.7308     0   1   2   3E+4   1   1.5  RT (min)      lcqtof_hilic_neg_4_000_aa    m/z 819.7445     0   1   2   3E+4   1   1.5  RT (min)      lcqtof_hilic_neg_4_000_aa    m/z 819.7594     0   1   2E+4   1.5   2  RT (min)      lcqtof_hilic_neg_4_000_aa    m/z 820.1568     0   1E+4   0.5   1  RT (min)      lcqtof_hilic_neg_1_062_17    m/z 820.4351     0   1E+4   4   4.5  RT (min)      lcqtof_hilic_neg_4_016_37    m/z 820.4762     0   1E+4   3   3.5  RT (min)      lcqtof_hilic_neg_4_022_34    m/z 820.5277     0   1E+4   1   1.5  RT (min)      lcqtof_hilic_neg_4_000_aa    m/z 820.5575     0   1   2   3E+4   2.5   3  RT (min)       lcqtof_hilic_neg_1_031_16    m/z 820.5575     0   1E+4   2   2.5  RT (min)      lcqtof_hilic_neg_1_031_16    m/z 820.5581     0   1   2   3E+4   1.5   2  RT (min)       lcqtof_hilic_neg_4_005_29    m/z 820.5584     0   1   2   3E+4   2.5   3  RT (min)       lcqtof_hilic_neg_1_031_16    m/z 820.5584     0   1   2E+4   2   2.5  RT (min)       lcqtof_hilic_neg_1_063_13    m/z 820.5584     0   1   2   3   4E+4   2.5   3  RT (min)       lcqtof_hilic_neg_1_063_13    m/z 820.5586     0   1   2   3E+4   2.5   3  RT (min)       lcqtof_hilic_neg_1_031_16    m/z 820.5587     0   1   2   3E+4   2.5   3  RT (min)      lcqtof_hilic_neg_1_031_16    m/z 820.5590     0   1   2E+4   2.5   3  RT (min)      lcqtof_hilic_neg_1_031_16    m/z 820.5592     0   1E+4   2   2.5  RT (min)       lcqtof_hilic_neg_1_063_13    m/z 820.5592     0   1E+4   2   2.5  RT (min)       lcqtof_hilic_neg_1_031_16    m/z 820.6383     0   1E+4   1.5   2  RT (min)      lcqtof_hilic_neg_4_000_aa    m/z 820.6386     0   1E+4   2   2.5  RT (min)      lcqtof_hilic_neg_4_000_aa    m/z 820.6398     0   1E+4   1.5   2  RT (min)      lcqtof_hilic_neg_4_000_aa    m/z 820.6926     0   1   2   3   4E+3   1.5   2  RT (min)      lcqtof_hilic_neg_1_037_24    m/z 820.7302     0   1E+4   1   1.5  RT (min)      lcqtof_hilic_neg_4_000_aa    m/z 820.7607     0   1E+4   1.5   2  RT (min)      lcqtof_hilic_neg_4_000_aa    m/z 821.5625     0   1E+4   2.5   3  RT (min)      lcqtof_hilic_neg_2_005_13    m/z 821.5630     0   1E+4   2.5   3  RT (min)      lcqtof_hilic_neg_2_005_13    m/z 821.5634     0   1   2E+4   2.5   3  RT (min)      lcqtof_hilic_neg_2_063_13    m/z 821.5636     0   2   4   6   8E+3   2   2.5  RT (min)      lcqtof_hilic_neg_2_013_29    m/z 821.5636     0   1E+4   2   2.5  RT (min)      lcqtof_hilic_neg_1_027_27    m/z 821.5638     0   1   2E+4   2.5   3  RT (min)      lcqtof_hilic_neg_2_063_13    m/z 821.5638     0   1   2   3   4E+4   1.5   2  RT (min)      lcqtof_hilic_neg_4_052_13    m/z 821.5640     0   1E+4   1.5   2  RT (min)      lcqtof_hilic_neg_4_032_01    m/z 821.5642     0   1E+4   1.5   2  RT (min)      lcqtof_hilic_neg_4_052_13    m/z 821.5644     0   1   2E+4   3   3.5  RT (min)      lcqtof_hilic_neg_1_019_02    m/z 821.5645     0   1   2E+4   2.5   3  RT (min)      lcqtof_hilic_neg_2_005_13    m/z 821.6148     0   1E+5   1   1.5  RT (min)      lcqtof_hilic_neg_4_000_aa    m/z 821.6842     0   2   4E+4   1.5   2  RT (min)      lcqtof_hilic_neg_4_000_aa    m/z 821.6850     0   2   4E+4   1.5   2  RT (min)      lcqtof_hilic_neg_4_000_aa    m/z 821.6864     0   2   4E+4   1.5   2  RT (min)      lcqtof_hilic_neg_4_000_aa    m/z 821.9933     0   1E+4   0.5   1  RT (min)      lcqtof_hilic_neg_3_035_46    m/z 822.5183     0   2   4   6E+5   3   3.5  RT (min)      lcqtof_hilic_neg_4_022_34    m/z 822.5286     0   1E+6   3   3.5  RT (min)      lcqtof_hilic_neg_4_048_23    m/z 822.5294     0   2   4   6   8E+5   3   3.5  RT (min)      lcqtof_hilic_neg_4_011_35    m/z 822.5294     0   2   4   6E+5   3   3.5  RT (min)      lcqtof_hilic_neg_4_022_34    m/z 822.5304     0   2   4   6   8E+5   2.5   3  RT (min)      lcqtof_hilic_neg_2_048_35    m/z 822.5304     0   1E+5   2.5   3  RT (min)      lcqtof_hilic_neg_2_048_35    m/z 822.5311     0   1   2   3   4E+4   2.5   3  RT (min)      lcqtof_hilic_neg_2_036_34    m/z 822.5318     0   1   2   3   4E+4   2.5   3  RT (min)      lcqtof_hilic_neg_3_011_34    m/z 822.5318     0   2   4E+4   2.5   3  RT (min)      lcqtof_hilic_neg_1_013_38    m/z 822.5322     0   1   2   3E+4   2.5   3  RT (min)      lcqtof_hilic_neg_2_036_34    m/z 822.5328     0   1   2   3   4E+4   2.5   3  RT (min)      lcqtof_hilic_neg_2_029_31    m/z 822.5634     0   1E+4   3   3.5  RT (min)       lcqtof_hilic_neg_4_019_03    m/z 822.5641     0   1E+4   3   3.5  RT (min)       lcqtof_hilic_neg_2_005_13    m/z 822.5642     0   1E+4   3   3.5  RT (min)       lcqtof_hilic_neg_4_019_03    m/z 822.5644     0   1E+4   3   3.5  RT (min)       lcqtof_hilic_neg_4_019_03    m/z 822.5645     0   1E+4   3   3.5  RT (min)       lcqtof_hilic_neg_4_019_03    m/z 822.5660     0   1E+4   1   1.5  RT (min)      lcqtof_hilic_neg_4_006_30    m/z 822.5668     0   1E+4   1.5   2  RT (min)       lcqtof_hilic_neg_3_068_57    m/z 822.5670     0   1E+4   1   2  RT (min)       lcqtof_hilic_neg_4_053_14    m/z 822.6166     0   2   4   6E+4   1   1.5  RT (min)      lcqtof_hilic_neg_4_000_aa    m/z 822.6865     0   1   2   3E+4   1.5   2  RT (min)      lcqtof_hilic_neg_4_000_aa    m/z 822.6904     0   1   2   3E+4   1.5   2  RT (min)      lcqtof_hilic_neg_4_000_aa    m/z 822.6960     0   1   2   3E+4   1.5   2  RT (min)      lcqtof_hilic_neg_4_000_aa    m/z 822.8194     0   1   2   3   4E+4   3   3.5  RT (min)      lcqtof_hilic_neg_1_035_23    m/z 822.8208     0   1E+4   1   1.5  RT (min)      lcqtof_hilic_neg_4_048_23    m/z 823.5304     0   1   2   3E+5   3   3.5  RT (min)      lcqtof_hilic_neg_4_011_35    m/z 823.5305     0   1   2   3   4E+4   2.5   3  RT (min)      lcqtof_hilic_neg_2_048_35    m/z 823.5306     0   1   2   3E+5   2.5   3  RT (min)      lcqtof_hilic_neg_4_011_35    m/z 823.5306     0   1   2   3   4E+5   2.5   3  RT (min)      lcqtof_hilic_neg_2_048_35    m/z 823.5309     0   1E+4   2.5   3  RT (min)      lcqtof_hilic_neg_1_014_31    m/z 823.5310     0   1E+4   2   2.5  RT (min)      lcqtof_hilic_neg_1_014_31    m/z 823.5312     0   1E+5   3   3.5  RT (min)      lcqtof_hilic_neg_4_047_06    m/z 823.5550     0   1   2   3   4E+4   1   1.5  RT (min)      lcqtof_hilic_neg_3_035_46    m/z 823.6222     0   1   2   3E+4   1   1.5  RT (min)      lcqtof_hilic_neg_4_000_aa    m/z 823.6268     0   1   2   3E+4   1   1.5  RT (min)      lcqtof_hilic_neg_4_000_aa    m/z 823.6272     0   1   2   3E+4   1   1.5  RT (min)      lcqtof_hilic_neg_4_000_aa    m/z 823.7182     0   1E+4   1   1.5  RT (min)      lcqtof_hilic_neg_4_000_aa    m/z 824.5425     0   2   4   6   8E+4   2.5   3  RT (min)      lcqtof_hilic_neg_1_022_36    m/z 824.5430     0   1   2E+6   1   1.5  RT (min)       lcqtof_hilic_neg_1_035_23    m/z 824.5431     0   1E+6   2.5   3  RT (min)       lcqtof_hilic_neg_4_011_35    m/z 824.5431     0   1E+6   2.5   3  RT (min)       lcqtof_hilic_neg_4_011_35    m/z 824.5432     0   2   4   6   8E+4   2.5   3  RT (min)      lcqtof_hilic_neg_1_022_36    m/z 824.5432     0   1   2E+5   2.5   3  RT (min)       lcqtof_hilic_neg_2_044_37    m/z 824.5432     0   1E+5   1.5   2  RT (min)       lcqtof_hilic_neg_4_082_59    m/z 824.5432     0   1E+4   1.5   2  RT (min)      lcqtof_hilic_neg_3_001_49    m/z 824.5433     0   1E+6   3   3.5  RT (min)      lcqtof_hilic_neg_4_011_35    m/z 824.5435     0   1   2E+5   2.5   3  RT (min)      lcqtof_hilic_neg_2_044_37    m/z 824.5439     0   2   4   6E+4   2   2.5  RT (min)       lcqtof_hilic_neg_1_014_31    m/z 824.5445     0   2   4   6E+4   2   2.5  RT (min)       lcqtof_hilic_neg_1_013_38    m/z 824.5448     0   1   2E+4   2   2.5  RT (min)      lcqtof_hilic_neg_4_011_35    m/z 824.5460     0   1E+4   2   2.5  RT (min)      lcqtof_hilic_neg_3_009_50    m/z 824.5667     0   1   2E+5   1   1.5  RT (min)      lcqtof_hilic_neg_3_035_46    m/z 824.6284     0   1   2E+4   1   1.5  RT (min)      lcqtof_hilic_neg_4_000_aa    m/z 824.7440     0   1E+4   8.5   9  RT (min)      lcqtof_hilic_neg_4_053_14    m/z 824.9130     0   1E+4   1   1.5  RT (min)      lcqtof_hilic_neg_4_004_47    m/z 824.9696     0   1E+4   3   3.5  RT (min)      lcqtof_hilic_neg_1_035_23    m/z 825.0617     0   1E+4   1   1.5  RT (min)      lcqtof_hilic_neg_4_048_23    m/z 825.2218     0   1E+4   3   3.5  RT (min)      lcqtof_hilic_neg_1_035_23    m/z 825.5470     0   2   4   6E+5   2.5   3  RT (min)      lcqtof_hilic_neg_4_011_35    m/z 825.5470     0   2   4   6   8E+5   3   3.5  RT (min)      lcqtof_hilic_neg_4_048_23    m/z 825.5472     0   2   4   6   8E+5   3   3.5  RT (min)      lcqtof_hilic_neg_4_048_23    m/z 825.5473     0   2   4   6E+5   2.5   3  RT (min)      lcqtof_hilic_neg_4_011_35    m/z 825.5474     0   2   4   6E+5   2.5   3  RT (min)      lcqtof_hilic_neg_2_048_35    m/z 825.5474     0   2   4   6E+4   2.5   3  RT (min)      lcqtof_hilic_neg_2_048_35    m/z 825.5476     0   1E+6   1   1.5  RT (min)      lcqtof_hilic_neg_1_035_23    m/z 825.5478     0   1E+4   2   2.5  RT (min)      lcqtof_hilic_neg_1_007_05    m/z 825.5481     0   1   2   3E+4   2.5   3  RT (min)      lcqtof_hilic_neg_2_048_35    m/z 825.6246     0   1E+4   1.5   2  RT (min)      lcqtof_hilic_neg_4_000_aa    m/z 826.5220     0   1E+4   6   6.5  RT (min)      lcqtof_hilic_neg_4_008_24    m/z 826.5480     0   1   2E+4   2   2.5  RT (min)      lcqtof_hilic_neg_4_040_05    m/z 826.5567     0   1E+5   4   4.5  RT (min)       lcqtof_hilic_neg_1_035_23    m/z 826.5572     0   1   2E+5   4   4.5  RT (min)       lcqtof_hilic_neg_2_020_23    m/z 826.5583     0   1E+5   4   4.5  RT (min)      lcqtof_hilic_neg_2_002_36    m/z 826.5584     0   2   4   6   8E+4   1.5   2  RT (min)      lcqtof_hilic_neg_4_050_02    m/z 826.5588     0   1E+5   2   2.5  RT (min)      lcqtof_hilic_neg_1_014_31    m/z 826.5588     0   1E+5   2   2.5  RT (min)       lcqtof_hilic_neg_1_014_31    m/z 826.5588     0   1   2E+5   2.5   3  RT (min)      lcqtof_hilic_neg_4_005_29    m/z 826.5589     0   1   2E+6   3   3.5  RT (min)      lcqtof_hilic_neg_1_041_54    m/z 826.5589     0   1E+6   2.5   3  RT (min)       lcqtof_hilic_neg_4_011_35    m/z 826.5589     0   1   2   3E+5   2.5   3  RT (min)       lcqtof_hilic_neg_2_048_35    m/z 826.5590     0   1   2E+6   2.5   3  RT (min)       lcqtof_hilic_neg_4_011_35    m/z 826.5590     0   1   2E+6   2.5   3  RT (min)       lcqtof_hilic_neg_4_016_37    m/z 826.5592     0   1E+6   3.5   4  RT (min)      lcqtof_hilic_neg_4_086_59    m/z 826.5592     0   1E+5   2   2.5  RT (min)       lcqtof_hilic_neg_1_014_31    m/z 826.5593     0   1E+5   2.5   3  RT (min)      lcqtof_hilic_neg_2_048_35    m/z 826.5594     0   1E+5   2   2.5  RT (min)       lcqtof_hilic_neg_2_062_17    m/z 826.5713     0   1   2E+4   2   2.5  RT (min)      lcqtof_hilic_neg_4_086_59    m/z 826.5846     0   2   4   6E+5   0.5   1  RT (min)      lcqtof_hilic_neg_3_035_46    m/z 826.6543     0   2   4   6   8E+5   4   4.5  RT (min)      lcqtof_hilic_neg_4_055_32    m/z 826.6560     0   2   4E+5   4   4.5  RT (min)      lcqtof_hilic_neg_4_047_06    m/z 827.5377     0   1   2E+4   3   3.5  RT (min)      lcqtof_hilic_neg_1_035_23    m/z 827.5592     0   1   2   3   4E+4   1.5   2  RT (min)      lcqtof_hilic_neg_4_050_02    m/z 827.5618     0   1   2   3   4E+4   1.5   2  RT (min)      lcqtof_hilic_neg_4_050_02    m/z 827.5620     0   2   4E+4   2   2.5  RT (min)       lcqtof_hilic_neg_1_013_38    m/z 827.5621     0   1   2   3   4E+5   1   1.5  RT (min)      lcqtof_hilic_neg_4_050_02    m/z 827.5622     0   2   4E+4   1.5   2  RT (min)      lcqtof_hilic_neg_1_013_38    m/z 827.5623     0   1E+6   3   3.5  RT (min)      lcqtof_hilic_neg_4_016_37    m/z 827.5623     0   2   4   6E+5   2.5   3  RT (min)       lcqtof_hilic_neg_4_011_35    m/z 827.5624     0   2   4   6E+4   2.5   3  RT (min)      lcqtof_hilic_neg_2_062_17    m/z 827.5625     0   1E+6   2.5   3  RT (min)      lcqtof_hilic_neg_4_022_34    m/z 827.5628     0   1   2E+4   1.5   2  RT (min)      lcqtof_hilic_neg_4_050_02    m/z 827.5628     0   1E+5   2.5   3  RT (min)      lcqtof_hilic_neg_4_005_29    m/z 827.5630     0   2   4   6E+4   2.5   3  RT (min)      lcqtof_hilic_neg_2_002_36    m/z 827.5632     0   1   2   3E+4   2   2.5  RT (min)      lcqtof_hilic_neg_1_003_37    m/z 827.5642     0   1   2   3E+4   2.5   3  RT (min)      lcqtof_hilic_neg_3_021_28    m/z 827.6646     0   1   2E+5   4   4.5  RT (min)      lcqtof_hilic_neg_4_047_06    m/z 827.6648     0   1   2   3E+5   4   4.5  RT (min)      lcqtof_hilic_neg_4_055_32    m/z 827.7092     0   1E+4   0.5   1  RT (min)      lcqtof_hilic_neg_4_053_14    m/z 827.7112     0   1E+4   1   1.5  RT (min)      lcqtof_hilic_neg_4_008_24    m/z 827.7114     0   1E+4   1   1.5  RT (min)      lcqtof_hilic_neg_4_008_24    m/z 827.7122     0   1E+4   1   1.5  RT (min)      lcqtof_hilic_neg_4_008_24    m/z 827.7123     0   1E+4   1   1.5  RT (min)      lcqtof_hilic_neg_4_055_32    m/z 827.7133     0   1E+4   1.5   2  RT (min)      lcqtof_hilic_neg_4_008_24    m/z 827.7136     0   2   4   6E+3   1.5   2  RT (min)      lcqtof_hilic_neg_1_037_24    m/z 828.5712     0   1   2   3   4E+4   1.5   2  RT (min)      lcqtof_hilic_neg_4_050_02    m/z 828.5722     0   2   4   6E+4   1.5   2  RT (min)      lcqtof_hilic_neg_4_050_02    m/z 828.5725     0   2   4   6   8E+4   2   2.5  RT (min)       lcqtof_hilic_neg_1_014_31    m/z 828.5728     0   1   2E+6   2.5   3  RT (min)       lcqtof_hilic_neg_4_011_35    m/z 828.5729     0   2   4   6E+4   2   2.5  RT (min)      lcqtof_hilic_neg_1_008_13    m/z 828.5729     0   2   4   6   8E+4   2   2.5  RT (min)       lcqtof_hilic_neg_1_014_31    m/z 828.5730     0   1   2   3   4E+5   2.5   3  RT (min)       lcqtof_hilic_neg_4_011_35    m/z 828.5731     0   1E+5   2.5   3  RT (min)      lcqtof_hilic_neg_2_005_13    m/z 828.5732     0   2   4   6E+4   2.5   3  RT (min)      lcqtof_hilic_neg_2_001_49    m/z 828.5732     0   2   4   6E+4   1.5   2  RT (min)       lcqtof_hilic_neg_4_050_02    m/z 828.5733     0   1   2   3   4E+4   2.5   3  RT (min)      lcqtof_hilic_neg_3_012_27    m/z 828.5734     0   1   2E+5   2.5   3  RT (min)      lcqtof_hilic_neg_2_048_35    m/z 828.5734     0   2   4   6E+4   1.5   2  RT (min)      lcqtof_hilic_neg_4_050_02    m/z 828.5734     0   2   4   6   8E+4   2   2.5  RT (min)      lcqtof_hilic_neg_1_014_31    m/z 828.5734     0   2   4   6   8E+4   2   2.5  RT (min)      lcqtof_hilic_neg_2_062_17    m/z 828.5734     0   1E+5   2.5   3  RT (min)      lcqtof_hilic_neg_2_005_13    m/z 828.5734     0   1E+6   3   3.5  RT (min)       lcqtof_hilic_neg_4_032_01    m/z 828.5735     0   2   4   6E+5   1   1.5  RT (min)      lcqtof_hilic_neg_4_050_02    m/z 828.5976     0   1E+4   2   2.5  RT (min)      lcqtof_hilic_neg_4_048_23    m/z 828.5994     0   1   2   3   4E+5   0.5   1  RT (min)      lcqtof_hilic_neg_3_035_46    m/z 828.6691     0   1E+6   4.5   5  RT (min)      lcqtof_hilic_neg_4_008_24    m/z 828.6693     0   1E+6   4   4.5  RT (min)       lcqtof_hilic_neg_4_055_32    m/z 828.6702     0   1E+4   1   1.5  RT (min)      lcqtof_hilic_neg_4_000_aa    m/z 828.7140     0   2   4   6E+4   1   1.5  RT (min)       lcqtof_hilic_neg_4_008_24    m/z 828.7142     0   1   2   3E+4   1.5   2  RT (min)       lcqtof_hilic_neg_1_037_24    m/z 828.7143     0   1E+4   1.5   2  RT (min)      lcqtof_hilic_neg_2_028_05    m/z 829.0944     0   1E+4   4   4.5  RT (min)      lcqtof_hilic_neg_4_008_24    m/z 829.3892     0   1E+4   1   1.5  RT (min)      lcqtof_hilic_neg_4_000_aa    m/z 829.3927     0   1E+4   1   1.5  RT (min)      lcqtof_hilic_neg_4_000_aa    m/z 829.5619     0   1   2E+5   3   3.5  RT (min)      lcqtof_hilic_neg_4_040_05    m/z 829.5758     0   1E+4   1.5   2  RT (min)      lcqtof_hilic_neg_4_050_02    m/z 829.5762     0   1E+4   1.5   2  RT (min)      lcqtof_hilic_neg_4_050_02    m/z 829.5766     0   1   2   3E+4   1.5   2  RT (min)      lcqtof_hilic_neg_4_050_02    m/z 829.5766     0   2   4   6E+5   2.5   3  RT (min)      lcqtof_hilic_neg_4_032_01    m/z 829.5769     0   1   2   3   4E+4   2.5   3  RT (min)      lcqtof_hilic_neg_2_005_13    m/z 829.5770     0   1E+5   2.5   3  RT (min)       lcqtof_hilic_neg_4_005_29    m/z 829.5771     0   2   4   6   8E+5   2.5   3  RT (min)       lcqtof_hilic_neg_4_011_35    m/z 829.5772     0   2   4E+4   2.5   3  RT (min)      lcqtof_hilic_neg_2_005_13    m/z 829.5772     0   1   2   3E+4   1.5   2  RT (min)      lcqtof_hilic_neg_1_008_13    m/z 829.5772     0   1E+4   1.5   2  RT (min)      lcqtof_hilic_neg_4_050_02    m/z 829.5772     0   1   2E+4   2   2.5  RT (min)      lcqtof_hilic_neg_1_013_38    m/z 829.5773     0   2   4   6E+5   2.5   3  RT (min)       lcqtof_hilic_neg_4_032_01    m/z 829.5776     0   1   2   3E+4   2   2.5  RT (min)       lcqtof_hilic_neg_2_062_17    m/z 829.5778     0   1   2E+5   1   1.5  RT (min)      lcqtof_hilic_neg_4_050_02    m/z 829.5788     0   1   2E+4   2   2.5  RT (min)      lcqtof_hilic_neg_1_051_35    m/z 829.6040     0   1E+5   0.5   1  RT (min)      lcqtof_hilic_neg_3_035_46    m/z 829.6844     0   1E+4   1   1.5  RT (min)      lcqtof_hilic_neg_4_055_32    m/z 829.9316     0   1E+4   5   5.5  RT (min)      lcqtof_hilic_neg_4_008_24    m/z 830.5694     0   1   2E+5   3.5   4  RT (min)      lcqtof_hilic_neg_4_040_05    m/z 830.5886     0   2   4   6E+4   1.5   2  RT (min)      lcqtof_hilic_neg_4_050_02    m/z 830.5890     0   2   4   6E+4   1.5   2  RT (min)      lcqtof_hilic_neg_4_050_02    m/z 830.5891     0   1E+5   2.5   3  RT (min)      lcqtof_hilic_neg_2_031_01    m/z 830.5893     0   2   4   6   8E+4   1.5   2  RT (min)      lcqtof_hilic_neg_4_050_02    m/z 830.5894     0   1E+5   2   2.5  RT (min)       lcqtof_hilic_neg_1_008_13    m/z 830.5895     0   1   2E+6   2.5   3  RT (min)       lcqtof_hilic_neg_4_005_29    m/z 830.5895     0   1   2E+6   2.5   3  RT (min)       lcqtof_hilic_neg_4_032_01    m/z 830.5895     0   1E+5   2.5   3  RT (min)      lcqtof_hilic_neg_2_005_13    m/z 830.5896     0   1E+5   2   2.5  RT (min)       lcqtof_hilic_neg_2_062_17    m/z 830.5897     0   1E+5   2   2.5  RT (min)      lcqtof_hilic_neg_1_008_13    m/z 830.5897     0   1E+5   2.5   3  RT (min)      lcqtof_hilic_neg_3_043_13    m/z 830.5898     0   2   4   6   8E+5   1   1.5  RT (min)      lcqtof_hilic_neg_4_050_02    m/z 830.5901     0   1E+5   2   2.5  RT (min)      lcqtof_hilic_neg_1_008_13    m/z 830.5908     0   2   4   6E+4   1.5   2  RT (min)      lcqtof_hilic_neg_4_050_02    m/z 830.5912     0   2   4   6   8E+4   1.5   2  RT (min)      lcqtof_hilic_neg_4_050_02    m/z 830.5985     0   2   4   6   8E+5   3.5   4  RT (min)      lcqtof_hilic_neg_4_053_14    m/z 830.6162     0   1   2   3E+4   4   4.5  RT (min)      lcqtof_hilic_neg_4_035_33    m/z 830.6162     0   1   2   3E+4   4   4.5  RT (min)      lcqtof_hilic_neg_4_035_33    m/z 830.6162     0   1   2   3E+4   4   4.5  RT (min)      lcqtof_hilic_neg_4_035_33    m/z 830.6167     0   1   2   3   4E+4   4   4.5  RT (min)       lcqtof_hilic_neg_4_055_32    m/z 830.6231     0   2   4E+4   3.5   4  RT (min)      lcqtof_hilic_neg_4_045_26    m/z 830.6787     0   1   2   3E+5   4   4.5  RT (min)      lcqtof_hilic_neg_4_055_32    m/z 830.6808     0   1E+5   4.5   5  RT (min)      lcqtof_hilic_neg_4_040_05    m/z 830.6844     0   1   2E+5   4.5   5  RT (min)      lcqtof_hilic_neg_4_004_47    m/z 830.6853     0   2   4E+5   4.5   5  RT (min)      lcqtof_hilic_neg_4_008_24    m/z 830.7166     0   1E+4   1.5   2  RT (min)      lcqtof_hilic_neg_3_093_60    m/z 830.7168     0   1   2E+4   1   1.5  RT (min)      lcqtof_hilic_neg_4_008_24    m/z 830.7288     0   1E+4   1   1.5  RT (min)      lcqtof_hilic_neg_4_008_24    m/z 830.9366     0   1E+4   7   7.5  RT (min)      lcqtof_hilic_neg_4_059_14    m/z 830.9368     0   1E+4   7   7.5  RT (min)      lcqtof_hilic_neg_4_059_14    m/z 830.9370     0   1E+4   6.5   7  RT (min)       lcqtof_hilic_neg_4_064_15    m/z 831.5269     0   1E+4   1   1.5  RT (min)      lcqtof_hilic_neg_4_000_aa    m/z 831.5556     0   1   2E+4   1   1.5  RT (min)      lcqtof_hilic_neg_4_000_aa    m/z 831.5932     0   1E+5   2.5   3  RT (min)      lcqtof_hilic_neg_4_005_29    m/z 831.5933     0   2   4   6E+4   2.5   3  RT (min)      lcqtof_hilic_neg_2_005_13    m/z 831.5934     0   1E+6   2.5   3  RT (min)      lcqtof_hilic_neg_4_032_01    m/z 831.5934     0   2   4   6   8E+4   2.5   3  RT (min)      lcqtof_hilic_neg_2_005_13    m/z 831.5936     0   1E+6   2.5   3  RT (min)      lcqtof_hilic_neg_4_005_29    m/z 831.5936     0   1E+6   3   3.5  RT (min)      lcqtof_hilic_neg_4_045_26    m/z 831.5939     0   1   2   3E+4   2   2.5  RT (min)      lcqtof_hilic_neg_1_013_38    m/z 831.5940     0   2   4E+4   2   2.5  RT (min)      lcqtof_hilic_neg_2_062_17    m/z 831.5941     0   1   2   3   4E+4   2   2.5  RT (min)      lcqtof_hilic_neg_1_008_13    m/z 831.5948     0   1E+4   1.5   2  RT (min)      lcqtof_hilic_neg_3_001_49    m/z 831.7150     0   1   2E+4   1   1.5  RT (min)      lcqtof_hilic_neg_4_000_aa    m/z 831.7970     0   1   2E+4   1   1.5  RT (min)      lcqtof_hilic_neg_4_000_aa    m/z 831.7990     0   1   2E+4   1.5   2  RT (min)      lcqtof_hilic_neg_4_000_aa    m/z 831.8053     0   1   2E+4   1   1.5  RT (min)      lcqtof_hilic_neg_4_000_aa    m/z 831.8150     0   1   2E+4   1.5   2  RT (min)      lcqtof_hilic_neg_4_000_aa    m/z 832.6016     0   2   4E+5   3   3.5  RT (min)      lcqtof_hilic_neg_4_032_01    m/z 832.6021     0   1   2E+4   2   2.5  RT (min)      lcqtof_hilic_neg_2_062_17    m/z 832.6024     0   1   2E+4   2   2.5  RT (min)      lcqtof_hilic_neg_2_031_01    m/z 832.6037     0   1E+4   1.5   2  RT (min)      lcqtof_hilic_neg_1_008_13    m/z 832.6042     0   1   2   3   4E+4   2   2.5  RT (min)      lcqtof_hilic_neg_4_040_05    m/z 832.6048     0   1E+4   1.5   2  RT (min)      lcqtof_hilic_neg_1_008_13    m/z 832.6169     0   1   2   3   4E+5   3.5   4  RT (min)      lcqtof_hilic_neg_4_011_35    m/z 832.6171     0   1E+4   2   2.5  RT (min)       lcqtof_hilic_neg_3_048_05    m/z 832.6544     0   1E+4   3   3.5  RT (min)       lcqtof_hilic_neg_1_034_08    m/z 832.6845     0   1E+4   1   1.5  RT (min)      lcqtof_hilic_neg_2_052_24    m/z 832.6986     0   1E+4   1.5   2  RT (min)      lcqtof_hilic_neg_1_037_24    m/z 832.6993     0   1E+4   1   1.5  RT (min)      lcqtof_hilic_neg_4_008_24    m/z 832.6996     0   1E+4   1   1.5  RT (min)      lcqtof_hilic_neg_1_037_24    m/z 832.7010     0   1E+4   1.5   2  RT (min)      lcqtof_hilic_neg_1_037_24    m/z 832.7013     0   1E+4   1.5   2  RT (min)      lcqtof_hilic_neg_2_052_24    m/z 832.7019     0   1E+4   1   1.5  RT (min)      lcqtof_hilic_neg_4_008_24    m/z 832.7028     0   1E+4   1   1.5  RT (min)      lcqtof_hilic_neg_4_008_24    m/z 832.8172     0   1E+4   1.5   2  RT (min)      lcqtof_hilic_neg_4_000_aa    m/z 832.8762     0   1E+4   0.5   1  RT (min)      lcqtof_hilic_neg_4_098_61    m/z 833.4444     0   1E+4   4   4.5  RT (min)       lcqtof_hilic_neg_4_022_34    m/z 833.4718     0   2   4E+4   0.5   1  RT (min)      lcqtof_hilic_neg_3_035_46    m/z 833.4829     0   1E+4   4   4.5  RT (min)      lcqtof_hilic_neg_4_039_18    m/z 833.5285     0   1E+4   3   3.5  RT (min)      lcqtof_hilic_neg_1_004_14    m/z 833.5406     0   1   2   3   4E+4   1   1.5  RT (min)      lcqtof_hilic_neg_4_000_aa    m/z 833.5422     0   1   2   3   4E+4   1   1.5  RT (min)      lcqtof_hilic_neg_4_000_aa    m/z 833.5564     0   1E+4   3   3.5  RT (min)      lcqtof_hilic_neg_4_040_05    m/z 833.6078     0   1E+5   3   3.5  RT (min)       lcqtof_hilic_neg_4_005_29    m/z 833.6092     0   1E+5   2.5   3  RT (min)       lcqtof_hilic_neg_4_005_29    m/z 833.6096     0   1E+4   2.5   3  RT (min)      lcqtof_hilic_neg_4_003_16    m/z 833.6102     0   1E+4   2   2.5  RT (min)       lcqtof_hilic_neg_4_040_05    m/z 833.6378     0   1   2   3E+4   1   1.5  RT (min)      lcqtof_hilic_neg_4_000_aa    m/z 833.7036     0   1E+4   1   1.5  RT (min)      lcqtof_hilic_neg_4_035_33    m/z 833.7038     0   1   2   3   4E+4   1   1.5  RT (min)      lcqtof_hilic_neg_4_008_24    m/z 833.7038     0   1   2   3   4E+4   1   1.5  RT (min)      lcqtof_hilic_neg_4_008_24    m/z 833.7042     0   1   2   3   4E+4   1   1.5  RT (min)       lcqtof_hilic_neg_4_008_24    m/z 833.7323     0   1   2   3E+4   1   1.5  RT (min)      lcqtof_hilic_neg_4_000_aa    m/z 833.7821     0   1   2E+4   1.5   2  RT (min)      lcqtof_hilic_neg_4_000_aa    m/z 833.7862     0   1   2E+4   1.5   2  RT (min)      lcqtof_hilic_neg_4_000_aa    m/z 833.7970     0   1E+4   1.5   2  RT (min)      lcqtof_hilic_neg_4_000_aa    m/z 834.3728     0   1E+4   3.5   4  RT (min)      lcqtof_hilic_neg_4_050_02    m/z 834.4764     0   1   2E+4   0.5   1  RT (min)      lcqtof_hilic_neg_3_035_46    m/z 834.5304     0   1E+4   2   2.5  RT (min)       lcqtof_hilic_neg_4_048_23    m/z 834.5435     0   1   2E+4   4   4.5  RT (min)      lcqtof_hilic_neg_4_040_05    m/z 834.5469     0   1E+4   1   1.5  RT (min)      lcqtof_hilic_neg_4_000_aa    m/z 834.5662     0   1E+4   4   4.5  RT (min)      lcqtof_hilic_neg_4_008_24    m/z 834.6206     0   1E+4   1.5   2  RT (min)      lcqtof_hilic_neg_4_000_aa    m/z 834.6216     0   1E+4   1.5   2  RT (min)      lcqtof_hilic_neg_4_000_aa    m/z 834.6261     0   1E+4   1.5   2  RT (min)      lcqtof_hilic_neg_4_000_aa    m/z 834.6642     0   2   4   6E+3   3   3.5  RT (min)       lcqtof_hilic_neg_1_037_24    m/z 834.6677     0   2   4   6E+3   3   3.5  RT (min)       lcqtof_hilic_neg_1_034_08    m/z 834.9794     0   1E+4   1   1.5  RT (min)      lcqtof_hilic_neg_4_000_aa    m/z 835.2495     0   1   2E+4   3.5   4  RT (min)      lcqtof_hilic_neg_4_013_20    m/z 835.4592     0   2   4   6E+5   4.5   5  RT (min)      lcqtof_hilic_neg_4_013_20    m/z 835.4594     0   2   4E+5   4   4.5  RT (min)      lcqtof_hilic_neg_4_109_62    m/z 835.4597     0   2   4E+5   4   4.5  RT (min)      lcqtof_hilic_neg_4_042_22    m/z 835.4960     0   1   2E+4   0.5   1  RT (min)      lcqtof_hilic_neg_1_007_05    m/z 835.5063     0   1   2E+4   1   1.5  RT (min)      lcqtof_hilic_neg_4_000_aa    m/z 835.5164     0   1   2E+4   1   1.5  RT (min)      lcqtof_hilic_neg_4_000_aa    m/z 835.5421     0   1E+4   3   3.5  RT (min)       lcqtof_hilic_neg_1_060_14    m/z 835.5618     0   1   2E+4   4   4.5  RT (min)      lcqtof_hilic_neg_4_047_06    m/z 835.5820     0   1   2E+4   4   4.5  RT (min)      lcqtof_hilic_neg_4_008_24    m/z 835.7021     0   1   2   3E+4   1.5   2  RT (min)      lcqtof_hilic_neg_4_000_aa    m/z 835.7029     0   1   2   3E+4   1.5   2  RT (min)      lcqtof_hilic_neg_4_000_aa    m/z 835.7029     0   1E+4   1   1.5  RT (min)      lcqtof_hilic_neg_4_008_24    m/z 835.7322     0   1   2   3E+4   1   1.5  RT (min)      lcqtof_hilic_neg_4_000_aa    m/z 836.4311     0   1   2   3   4E+4   0.5   1  RT (min)      lcqtof_hilic_neg_2_004_26    m/z 836.4630     0   1   2   3   4E+4   4   4.5  RT (min)      lcqtof_hilic_neg_4_028_36    m/z 836.4970     0   1E+4   0.5   1  RT (min)      lcqtof_hilic_neg_1_007_05    m/z 836.5531     0   1E+4   3   3.5  RT (min)      lcqtof_hilic_neg_1_004_14    m/z 836.5584     0   1E+5   4   4.5  RT (min)       lcqtof_hilic_neg_2_039_38    m/z 836.5644     0   1   2   3   4E+4   2   2.5  RT (min)      lcqtof_hilic_neg_1_007_05    m/z 836.6465     0   1E+4   0.5   1  RT (min)      lcqtof_hilic_neg_4_008_24    m/z 836.7053     0   1   2E+4   1.5   2  RT (min)      lcqtof_hilic_neg_4_000_aa    m/z 836.7060     0   1   2E+4   1.5   2  RT (min)      lcqtof_hilic_neg_4_000_aa    m/z 836.7146     0   1   2E+4   1.5   2  RT (min)      lcqtof_hilic_neg_4_000_aa    m/z 836.7210     0   1   2E+4   1.5   2  RT (min)      lcqtof_hilic_neg_4_000_aa    m/z 837.3978     0   1   2   3E+4   0.5   1  RT (min)      lcqtof_hilic_neg_4_005_29    m/z 837.4105     0   2   4   6E+4   0.5   1  RT (min)      lcqtof_hilic_neg_1_042_26    m/z 837.4767     0   1   2   3E+5   4.5   5  RT (min)      lcqtof_hilic_neg_4_013_20    m/z 837.4818     0   1   2   3E+4   2.5   3  RT (min)      lcqtof_hilic_neg_4_032_01    m/z 837.4930     0   1   2E+4   0.5   1  RT (min)      lcqtof_hilic_neg_1_007_05    m/z 837.5199     0   1E+4   1   1.5  RT (min)      lcqtof_hilic_neg_1_013_38    m/z 837.5686     0   1   2E+4   2   2.5  RT (min)      lcqtof_hilic_neg_4_040_05    m/z 837.5804     0   1E+4   3   3.5  RT (min)      lcqtof_hilic_neg_1_005_32    m/z 837.6164     0   1   2E+4   1   1.5  RT (min)      lcqtof_hilic_neg_4_000_aa    m/z 837.6236     0   1   2E+4   1.5   2  RT (min)      lcqtof_hilic_neg_4_000_aa    m/z 837.6252     0   1E+4   2   2.5  RT (min)      lcqtof_hilic_neg_4_000_aa    m/z 837.6254     0   1E+4   1.5   2  RT (min)      lcqtof_hilic_neg_4_000_aa    m/z 837.6258     0   1E+4   1.5   2  RT (min)      lcqtof_hilic_neg_4_000_aa    m/z 837.6262     0   1E+4   2   2.5  RT (min)      lcqtof_hilic_neg_4_000_aa    m/z 837.6286     0   1   2E+4   1   1.5  RT (min)      lcqtof_hilic_neg_4_000_aa    m/z 837.6298     0   1E+4   2   2.5  RT (min)      lcqtof_hilic_neg_4_000_aa    m/z 837.6330     0   1   2E+4   1   1.5  RT (min)      lcqtof_hilic_neg_4_000_aa    m/z 837.6334     0   1   2E+4   1   1.5  RT (min)      lcqtof_hilic_neg_4_000_aa    m/z 837.6342     0   1E+4   3   3.5  RT (min)      lcqtof_hilic_neg_4_035_33    m/z 838.3080     0   1   2   3   4E+4   1   1.5  RT (min)      lcqtof_hilic_neg_4_027_19    m/z 838.3081     0   2   4E+4   1   1.5  RT (min)       lcqtof_hilic_neg_4_013_20    m/z 838.3368     0   1   2E+4   0.5   1  RT (min)      lcqtof_hilic_neg_3_003_20    m/z 838.4014     0   1E+4   0.5   1  RT (min)      lcqtof_hilic_neg_4_005_29    m/z 838.4041     0   1E+4   3.5   4  RT (min)      lcqtof_hilic_neg_4_019_03    m/z 838.4134     0   1   2E+4   0.5   1  RT (min)      lcqtof_hilic_neg_1_042_26    m/z 838.4519     0   1E+4   4   4.5  RT (min)      lcqtof_hilic_neg_4_016_37    m/z 838.4961     0   1E+4   0.5   1  RT (min)      lcqtof_hilic_neg_1_007_05    m/z 838.5593     0   1E+4   3.5   4  RT (min)      lcqtof_hilic_neg_1_027_27    m/z 838.5664     0   1   2E+4   2   2.5  RT (min)      lcqtof_hilic_neg_4_040_05    m/z 838.5945     0   1   2E+4   2   2.5  RT (min)       lcqtof_hilic_neg_4_016_37    m/z 838.5992     0   1   2   3   4E+5   3   3.5  RT (min)      lcqtof_hilic_neg_1_054_01    m/z 838.6324     0   2   4   6   8E+4   0.5   1  RT (min)      lcqtof_hilic_neg_1_005_32    m/z 838.6557     0   1   2E+5   3   3.5  RT (min)      lcqtof_hilic_neg_4_008_24    m/z 838.6566     0   1   2E+5   3   3.5  RT (min)      lcqtof_hilic_neg_4_008_24    m/z 839.1490     0   1   2E+4   0.5   1  RT (min)      lcqtof_hilic_neg_3_035_46    m/z 839.3116     0   1   2E+4   1   1.5  RT (min)      lcqtof_hilic_neg_4_013_20    m/z 839.4751     0   1   2E+4   4.5   5  RT (min)      lcqtof_hilic_neg_4_013_20    m/z 839.5176     0   2   4   6   8E+3   2.5   3  RT (min)      lcqtof_hilic_neg_4_082_59    m/z 839.5929     0   1E+4   2   2.5  RT (min)      lcqtof_hilic_neg_4_016_37    m/z 839.5944     0   2   4   6   8E+4   3.5   4  RT (min)      lcqtof_hilic_neg_4_055_32    m/z 839.6580     0   1E+4   1.5   2  RT (min)      lcqtof_hilic_neg_4_000_aa    m/z 839.6608     0   1E+4   1.5   2  RT (min)      lcqtof_hilic_neg_4_000_aa    m/z 839.6649     0   1   2E+4   2.5   3  RT (min)      lcqtof_hilic_neg_4_008_24    m/z 839.6659     0   1   2E+5   3   3.5  RT (min)      lcqtof_hilic_neg_4_008_24    m/z 839.6668     0   1   2E+5   3   3.5  RT (min)      lcqtof_hilic_neg_4_008_24    m/z 839.6675     0   1   2E+5   2.5   3  RT (min)      lcqtof_hilic_neg_4_008_24    m/z 840.1493     0   1E+4   0.5   1  RT (min)      lcqtof_hilic_neg_3_035_46    m/z 840.4803     0   1E+4   4.5   5  RT (min)      lcqtof_hilic_neg_4_013_20    m/z 840.5190     0   1   2E+4   1   1.5  RT (min)      lcqtof_hilic_neg_4_048_23    m/z 840.5199     0   2   4   6E+3   1.5   2  RT (min)      lcqtof_hilic_neg_2_036_34    m/z 840.5336     0   2   4   6E+3   2   2.5  RT (min)      lcqtof_hilic_neg_4_016_37    m/z 840.5381     0   1E+4   3.5   4  RT (min)      lcqtof_hilic_neg_4_028_36    m/z 840.5776     0   1E+4   1   1.5  RT (min)      lcqtof_hilic_neg_4_055_32    m/z 840.6042     0   1E+4   2   2.5  RT (min)      lcqtof_hilic_neg_4_016_37    m/z 840.6575     0   1E+4   1.5   2  RT (min)      lcqtof_hilic_neg_1_037_24    m/z 840.6640     0   1E+4   1.5   2  RT (min)      lcqtof_hilic_neg_4_008_24    m/z 840.6660     0   1   2   3E+5   1   1.5  RT (min)      lcqtof_hilic_neg_2_047_32    m/z 840.6672     0   1   2E+4   2   2.5  RT (min)      lcqtof_hilic_neg_1_005_32    m/z 840.6689     0   2   4   6E+5   3   3.5  RT (min)      lcqtof_hilic_neg_4_055_32    m/z 840.6692     0   2   4   6   8E+5   2.5   3  RT (min)      lcqtof_hilic_neg_4_008_24    m/z 840.6693     0   2   4   6   8E+5   3   3.5  RT (min)      lcqtof_hilic_neg_4_008_24    m/z 840.6694     0   1   2E+4   2.5   3  RT (min)      lcqtof_hilic_neg_1_005_32    m/z 840.6695     0   2   4   6   8E+5   2.5   3  RT (min)      lcqtof_hilic_neg_2_052_24    m/z 840.6696     0   1   2   3   4E+4   2.5   3  RT (min)      lcqtof_hilic_neg_3_034_24    m/z 840.6696     0   1E+4   2.5   3  RT (min)      lcqtof_hilic_neg_1_015_33    m/z 840.6697     0   2   4   6   8E+4   2.5   3  RT (min)      lcqtof_hilic_neg_2_047_32    m/z 840.6703     0   1   2E+4   2   2.5  RT (min)      lcqtof_hilic_neg_1_005_32    m/z 840.6705     0   1   2E+4   2.5   3  RT (min)      lcqtof_hilic_neg_1_005_32    m/z 840.6911     0   2   4   6   8E+4   1   1.5  RT (min)      lcqtof_hilic_neg_3_035_46    m/z 840.7238     0   1E+4   2   2.5  RT (min)      lcqtof_hilic_neg_4_008_24    m/z 841.4980     0   1E+4   2.5   3  RT (min)      lcqtof_hilic_neg_4_052_13    m/z 841.5331     0   1E+4   3   3.5  RT (min)      lcqtof_hilic_neg_4_047_06    m/z 841.5336     0   1   2   3E+4   1.5   2  RT (min)      lcqtof_hilic_neg_4_048_23    m/z 841.5339     0   1E+4   3   3.5  RT (min)       lcqtof_hilic_neg_4_087_59    m/z 841.6582     0   2   4   6E+4   3   3.5  RT (min)      lcqtof_hilic_neg_4_008_24    m/z 841.6810     0   1E+5   3   3.5  RT (min)      lcqtof_hilic_neg_4_002_25    m/z 841.6813     0   1   2   3E+5   2.5   3  RT (min)      lcqtof_hilic_neg_4_008_24    m/z 841.6816     0   1   2   3E+5   2.5   3  RT (min)      lcqtof_hilic_neg_2_052_24    m/z 841.6821     0   1   2   3   4E+4   2.5   3  RT (min)      lcqtof_hilic_neg_2_047_32    m/z 841.7110     0   1E+4   1   1.5  RT (min)      lcqtof_hilic_neg_4_055_32    m/z 841.7131     0   1   2E+4   1   1.5  RT (min)      lcqtof_hilic_neg_4_008_24    m/z 842.5377     0   1   2   3   4E+4   1   1.5  RT (min)      lcqtof_hilic_neg_4_048_23    m/z 842.5379     0   1E+4   1.5   2  RT (min)      lcqtof_hilic_neg_2_002_36    m/z 842.5382     0   1E+4   1   1.5  RT (min)      lcqtof_hilic_neg_4_011_35    m/z 842.5429     0   1   2   3E+3   1.5   2  RT (min)      lcqtof_hilic_neg_1_042_26    m/z 842.5437     0   2   4   6   8E+3   1.5   2   2.5  RT (min)      lcqtof_hilic_neg_1_014_31    m/z 842.5439     0   2   4   6   8E+3   2.5   3  RT (min)      lcqtof_hilic_neg_4_016_37    m/z 842.5441     0   1E+4   2.5   3  RT (min)      lcqtof_hilic_neg_4_085_59    m/z 842.5444     0   1E+4   3   3.5  RT (min)      lcqtof_hilic_neg_4_088_59    m/z 842.5499     0   2   4   6E+4   3   3.5  RT (min)      lcqtof_hilic_neg_4_028_36    m/z 842.5506     0   1E+4   1   1.5  RT (min)      lcqtof_hilic_neg_2_044_37    m/z 842.5607     0   1E+4   2   2.5  RT (min)      lcqtof_hilic_neg_2_039_38    m/z 842.5790     0   1E+4   4   4.5  RT (min)      lcqtof_hilic_neg_4_091_60    m/z 842.5966     0   1E+5   2   2.5  RT (min)      lcqtof_hilic_neg_4_016_37    m/z 842.5980     0   1E+4   0.5   1   1.5  RT (min)      lcqtof_hilic_neg_4_032_01    m/z 842.6074     0   1E+4   0.5   1  RT (min)      lcqtof_hilic_neg_1_008_13    m/z 842.6283     0   1   2E+4   4   4.5  RT (min)      lcqtof_hilic_neg_4_008_24    m/z 842.6590     0   1   2E+5   3   3.5  RT (min)      lcqtof_hilic_neg_4_008_24    m/z 842.6840     0   2   4E+4   2.5   3  RT (min)       lcqtof_hilic_neg_3_096_60    m/z 842.6852     0   1   2E+4   2.5   3  RT (min)      lcqtof_hilic_neg_3_041_54    m/z 842.6866     0   1   2E+4   1.5   2  RT (min)       lcqtof_hilic_neg_4_091_60    m/z 842.6893     0   1   2   3E+5   1   1.5  RT (min)      lcqtof_hilic_neg_3_004_32    m/z 842.7056     0   1E+5   1   1.5  RT (min)      lcqtof_hilic_neg_2_052_24    m/z 842.7096     0   1E+5   1   1.5  RT (min)      lcqtof_hilic_neg_3_035_46    m/z 842.7193     0   1   2   3E+4   1.5   2  RT (min)      lcqtof_hilic_neg_4_047_06    m/z 842.7197     0   1E+5   1   1.5  RT (min)      lcqtof_hilic_neg_4_008_24    m/z 842.7209     0   1E+4   1.5   2  RT (min)      lcqtof_hilic_neg_2_001_49    m/z 842.7214     0   2   4E+4   1.5   2  RT (min)      lcqtof_hilic_neg_1_037_24    m/z 842.7215     0   2   4E+4   1.5   2  RT (min)      lcqtof_hilic_neg_1_037_24    m/z 842.7217     0   1E+5   1.5   2  RT (min)      lcqtof_hilic_neg_4_008_24    m/z 842.7224     0   1   2   3E+5   1.5   2  RT (min)      lcqtof_hilic_neg_2_016_33    m/z 842.7230     0   1E+5   1   1.5  RT (min)      lcqtof_hilic_neg_4_008_24    m/z 842.7241     0   1   2   3E+4   1.5   2  RT (min)      lcqtof_hilic_neg_2_047_32    m/z 843.4784     0   1E+4   3.5   4  RT (min)      lcqtof_hilic_neg_1_007_05    m/z 843.4791     0   1E+4   3.5   4  RT (min)      lcqtof_hilic_neg_1_007_05    m/z 843.5480     0   1E+4   2.5   3  RT (min)      lcqtof_hilic_neg_4_060_13    m/z 843.5491     0   1E+4   2.5   3  RT (min)       lcqtof_hilic_neg_4_059_14    m/z 843.5492     0   1   2E+4   2.5   3  RT (min)       lcqtof_hilic_neg_4_006_30    m/z 843.5492     0   1E+4   2.5   3  RT (min)      lcqtof_hilic_neg_4_060_13    m/z 843.5496     0   1   2   3   4E+4   1.5   2  RT (min)       lcqtof_hilic_neg_1_008_13    m/z 843.5497     0   1   2   3   4E+4   1.5   2  RT (min)      lcqtof_hilic_neg_1_008_13    m/z 843.5501     0   1E+4   2   2.5  RT (min)       lcqtof_hilic_neg_2_031_01    m/z 843.5501     0   1   2   3   4E+4   2.5   3  RT (min)       lcqtof_hilic_neg_4_074_58    m/z 843.5501     0   1E+4   2   2.5  RT (min)       lcqtof_hilic_neg_2_060_14    m/z 843.5502     0   1E+4   2   2.5  RT (min)      lcqtof_hilic_neg_2_031_01    m/z 843.5503     0   1   2   3   4E+4   3   3.5  RT (min)       lcqtof_hilic_neg_4_081_58    m/z 843.5505     0   1E+5   3   3.5  RT (min)      lcqtof_hilic_neg_1_060_14    m/z 843.5901     0   1E+4   3   3.5  RT (min)      lcqtof_hilic_neg_4_053_14    m/z 843.5922     0   2   4E+4   1   1.5  RT (min)      lcqtof_hilic_neg_4_000_aa    m/z 843.6815     0   1   2E+5   3   3.5  RT (min)      lcqtof_hilic_neg_4_008_24    m/z 843.6968     0   1E+5   3   3.5  RT (min)      lcqtof_hilic_neg_1_055_25    m/z 843.6972     0   1E+5   2.5   3  RT (min)      lcqtof_hilic_neg_4_002_25    m/z 843.7318     0   1   2   3E+5   1   1.5  RT (min)       lcqtof_hilic_neg_4_008_24    m/z 843.7318     0   1   2   3E+5   1   1.5  RT (min)       lcqtof_hilic_neg_4_008_24    m/z 843.7319     0   1E+5   1.5   2  RT (min)       lcqtof_hilic_neg_1_037_24    m/z 843.7320     0   1E+5   1   1.5  RT (min)      lcqtof_hilic_neg_4_002_25    m/z 843.7321     0   2   4   6E+5   1.5   2  RT (min)      lcqtof_hilic_neg_1_029_06    m/z 843.7322     0   1   2   3E+5   1   1.5  RT (min)      lcqtof_hilic_neg_4_008_24    m/z 843.7324     0   1   2   3   4E+4   1   1.5  RT (min)      lcqtof_hilic_neg_1_049_55    m/z 843.8563     0   1E+4   1.5   2   2.5  RT (min)      lcqtof_hilic_neg_4_000_aa    m/z 843.8574     0   1E+4   1.5   2  RT (min)      lcqtof_hilic_neg_4_000_aa    m/z 843.8771     0   1E+4   2   2.5  RT (min)      lcqtof_hilic_neg_1_005_32    m/z 844.2550     0   1   2E+4   2   2.5  RT (min)      lcqtof_hilic_neg_1_005_32    m/z 844.4185     0   1E+4   2   2.5  RT (min)      lcqtof_hilic_neg_1_005_32    m/z 844.5129     0   1E+4   4   4.5  RT (min)      lcqtof_hilic_neg_4_039_18    m/z 844.5540     0   1   2E+4   1   2   3  RT (min)       lcqtof_hilic_neg_4_034_31    m/z 844.5563     0   1E+4   2.5   3  RT (min)       lcqtof_hilic_neg_4_059_14    m/z 844.5582     0   1E+4   2   2.5  RT (min)      lcqtof_hilic_neg_2_062_17    m/z 844.5584     0   1E+4   2   2.5  RT (min)      lcqtof_hilic_neg_2_060_14    m/z 844.5584     0   1   2   3   4E+4   2.5   3  RT (min)       lcqtof_hilic_neg_1_054_01    m/z 844.5584     0   1E+4   2   2.5  RT (min)       lcqtof_hilic_neg_2_060_14    m/z 844.5587     0   1E+4   2.5   3  RT (min)      lcqtof_hilic_neg_4_059_14    m/z 844.5588     0   1E+4   1.5   2  RT (min)       lcqtof_hilic_neg_1_016_28    m/z 844.5594     0   2   4E+4   2.5   3  RT (min)      lcqtof_hilic_neg_4_019_03    m/z 844.5596     0   1   2E+4   2.5   3  RT (min)       lcqtof_hilic_neg_4_011_35    m/z 844.5605     0   1E+4   2.5   3  RT (min)       lcqtof_hilic_neg_2_060_14    m/z 844.5749     0   1   2E+4   1   1.5   2  RT (min)      lcqtof_hilic_neg_4_034_31    m/z 844.6736     0   1E+5   3   3.5  RT (min)      lcqtof_hilic_neg_4_008_24    m/z 844.7004     0   2   4   6   8E+5   2.5   3  RT (min)      lcqtof_hilic_neg_4_008_24    m/z 844.7006     0   2   4E+4   2.5   3  RT (min)      lcqtof_hilic_neg_2_028_05    m/z 844.7014     0   1   2   3E+4   2.5   3  RT (min)      lcqtof_hilic_neg_3_034_24    m/z 844.7014     0   2   4   6E+4   2   2.5  RT (min)      lcqtof_hilic_neg_1_005_32    m/z 844.7055     0   1   2   3E+4   2   2.5  RT (min)      lcqtof_hilic_neg_4_091_60    m/z 844.7068     0   1   2E+4   1.5   2  RT (min)      lcqtof_hilic_neg_4_091_60    m/z 844.7096     0   1   2E+4   1.5   2  RT (min)      lcqtof_hilic_neg_4_055_32    m/z 844.7164     0   1   2   3E+5   1   1.5  RT (min)      lcqtof_hilic_neg_4_055_32    m/z 844.7376     0   2   4   6   8E+4   1   1.5  RT (min)      lcqtof_hilic_neg_3_035_46    m/z 844.9775     0   1E+4   0.5   1  RT (min)      lcqtof_hilic_neg_4_000_aa    m/z 844.9954     0   1E+4   3   3.5  RT (min)      lcqtof_hilic_neg_1_037_24    m/z 844.9958     0   1E+4   0.5   1  RT (min)      lcqtof_hilic_neg_4_008_24    m/z 845.5646     0   1E+4   2.5   3  RT (min)       lcqtof_hilic_neg_4_045_26    m/z 845.5646     0   1E+4   2.5   3  RT (min)       lcqtof_hilic_neg_4_006_30    m/z 845.5654     0   1   2   3   4E+4   3   3.5  RT (min)      lcqtof_hilic_neg_4_059_14    m/z 845.5656     0   1E+4   2.5   3  RT (min)      lcqtof_hilic_neg_2_051_14    m/z 845.5657     0   1   2E+4   1.5   2  RT (min)       lcqtof_hilic_neg_1_008_13    m/z 845.5660     0   1   2E+4   2   2.5  RT (min)      lcqtof_hilic_neg_1_008_13    m/z 845.5662     0   1   2E+4   1.5   2  RT (min)      lcqtof_hilic_neg_1_008_13    m/z 845.5668     0   2   4   6   8E+3   2.5   3  RT (min)      lcqtof_hilic_neg_2_060_14    m/z 845.5668     0   2   4   6   8E+3   2   2.5  RT (min)      lcqtof_hilic_neg_1_008_13    m/z 845.5673     0   2   4   6E+3   2   2.5  RT (min)      lcqtof_hilic_neg_1_004_14    m/z 845.5889     0   1   2E+4   1.5   2  RT (min)      lcqtof_hilic_neg_2_029_31    m/z 845.5991     0   1   2E+4   1   1.5  RT (min)      lcqtof_hilic_neg_2_039_38    m/z 845.6078     0   1   2E+4   1   1.5  RT (min)      lcqtof_hilic_neg_1_003_37    m/z 845.6086     0   1E+4   1.5   2  RT (min)      lcqtof_hilic_neg_4_011_35    m/z 845.6208     0   1E+4   3   3.5  RT (min)      lcqtof_hilic_neg_2_052_24    m/z 845.6235     0   2   4E+4   1   1.5  RT (min)      lcqtof_hilic_neg_4_011_35    m/z 845.6295     0   2   4   6E+4   0.5   1  RT (min)      lcqtof_hilic_neg_4_022_34    m/z 845.6559     0   1   2E+4   2   2.5  RT (min)      lcqtof_hilic_neg_4_055_32    m/z 845.6780     0   1E+4   4   4.5  RT (min)      lcqtof_hilic_neg_4_008_24    m/z 845.7123     0   1   2   3   4E+4   3   3.5  RT (min)      lcqtof_hilic_neg_4_008_24    m/z 845.7135     0   1   2   3   4E+4   3   3.5  RT (min)      lcqtof_hilic_neg_4_008_24    m/z 845.7999     0   1E+4   1.5   2  RT (min)      lcqtof_hilic_neg_4_000_aa    m/z 845.8113     0   1E+4   1.5   2  RT (min)      lcqtof_hilic_neg_4_000_aa    m/z 845.8339     0   1E+4   1.5   2  RT (min)      lcqtof_hilic_neg_4_000_aa    m/z 845.9221     0   1   2E+4   4   4.5  RT (min)      lcqtof_hilic_neg_4_013_20    m/z 846.0122     0   1   2   3   4E+4   4   4.5  RT (min)      lcqtof_hilic_neg_4_013_20    m/z 846.1765     0   1   2E+4   4   4.5  RT (min)      lcqtof_hilic_neg_4_013_20    m/z 846.5742     0   2   4   6   8E+3   2   2.5  RT (min)      lcqtof_hilic_neg_1_027_27    m/z 846.5746     0   2   4   6   8E+3   2   2.5  RT (min)      lcqtof_hilic_neg_1_027_27    m/z 846.5764     0   2   4E+4   1.5   2  RT (min)      lcqtof_hilic_neg_4_059_14    m/z 846.5765     0   1   2   3E+4   2   2.5  RT (min)       lcqtof_hilic_neg_1_035_23    m/z 846.5769     0   1E+4   2.5   3  RT (min)      lcqtof_hilic_neg_2_031_01    m/z 846.5772     0   1E+4   2.5   3  RT (min)      lcqtof_hilic_neg_2_062_17    m/z 846.5773     0   1   2E+4   1.5   2  RT (min)      lcqtof_hilic_neg_4_006_30    m/z 846.5784     0   1E+5   3   3.5  RT (min)       lcqtof_hilic_neg_4_059_14    m/z 846.6237     0   1   2   3   4E+4   3   3.5  RT (min)       lcqtof_hilic_neg_2_052_24    m/z 846.6276     0   1   2   3E+4   0.5   1  RT (min)      lcqtof_hilic_neg_4_022_34    m/z 846.6312     0   1   2E+4   1   1.5  RT (min)      lcqtof_hilic_neg_4_011_35    m/z 846.6632     0   1   2   3E+4   2   2.5  RT (min)      lcqtof_hilic_neg_4_055_32    m/z 846.6958     0   2   4   6   8E+4   1   1.5  RT (min)      lcqtof_hilic_neg_4_008_24    m/z 846.7046     0   1   2   3E+4   1   1.5  RT (min)      lcqtof_hilic_neg_4_055_32    m/z 846.7127     0   2   4   6E+3   2   2.5  RT (min)      lcqtof_hilic_neg_1_005_32    m/z 846.7155     0   1E+4   2   2.5  RT (min)      lcqtof_hilic_neg_4_008_24    m/z 846.7166     0   1E+5   2.5   3  RT (min)      lcqtof_hilic_neg_4_008_24    m/z 846.9741     0   1E+4   0.5   1  RT (min)      lcqtof_hilic_neg_4_000_aa    m/z 847.4407     0   1   2   3   4E+4   3.5   4  RT (min)      lcqtof_hilic_neg_4_039_18    m/z 847.4520     0   1   2E+4   4.5   5  RT (min)      lcqtof_hilic_neg_1_007_05    m/z 847.4989     0   1E+4   0.5   1  RT (min)      lcqtof_hilic_neg_4_000_aa    m/z 847.5810     0   1E+4   2   2.5  RT (min)      lcqtof_hilic_neg_1_004_14    m/z 847.5811     0   1   2E+4   2   2.5  RT (min)      lcqtof_hilic_neg_1_004_14    m/z 847.5811     0   1E+4   2   2.5  RT (min)      lcqtof_hilic_neg_1_004_14    m/z 847.5812     0   1E+4   2.5   3  RT (min)      lcqtof_hilic_neg_4_060_13    m/z 847.5813     0   1   2E+4   3   3.5  RT (min)       lcqtof_hilic_neg_4_019_03    m/z 847.5814     0   1E+4   2   2.5  RT (min)       lcqtof_hilic_neg_1_035_23    m/z 847.5817     0   1   2   3E+4   1.5   2  RT (min)      lcqtof_hilic_neg_4_053_14    m/z 847.5819     0   2   4E+4   1.5   2  RT (min)       lcqtof_hilic_neg_4_053_14    m/z 847.5819     0   1E+4   2.5   3  RT (min)       lcqtof_hilic_neg_2_063_13    m/z 847.5820     0   1E+4   3   3.5  RT (min)       lcqtof_hilic_neg_4_060_13    m/z 847.5820     0   2   4   6E+4   3   3.5  RT (min)      lcqtof_hilic_neg_4_059_14    m/z 847.5824     0   2   4   6E+4   3   3.5  RT (min)       lcqtof_hilic_neg_4_059_14    m/z 847.5824     0   2   4E+4   1.5   2  RT (min)      lcqtof_hilic_neg_4_053_14    m/z 847.6587     0   1E+4   4.5   5  RT (min)      lcqtof_hilic_neg_4_008_24    m/z 847.7928     0   1   2E+4   1.5   2  RT (min)      lcqtof_hilic_neg_4_000_aa    m/z 847.8044     0   1   2E+4   1.5   2  RT (min)      lcqtof_hilic_neg_4_000_aa    m/z 848.3937     0   1E+4   4   4.5  RT (min)      lcqtof_hilic_neg_4_011_35    m/z 848.4668     0   1E+4   4   4.5  RT (min)       lcqtof_hilic_neg_4_048_23    m/z 848.5875     0   1E+4   1.5   2  RT (min)       lcqtof_hilic_neg_4_018_27    m/z 848.5876     0   2   4   6E+3   2   2.5  RT (min)      lcqtof_hilic_neg_1_035_23    m/z 848.5877     0   2   4   6   8E+3   3   3.5  RT (min)      lcqtof_hilic_neg_2_018_16    m/z 848.5879     0   2   4   6   8E+3   2.5   3  RT (min)      lcqtof_hilic_neg_4_006_30    m/z 848.6345     0   1E+5   4   4.5  RT (min)      lcqtof_hilic_neg_4_008_24    m/z 848.6385     0   1E+4   3   3.5  RT (min)      lcqtof_hilic_neg_2_052_24    m/z 848.6713     0   1   2   3   4E+4   4.5   5  RT (min)      lcqtof_hilic_neg_4_008_24    m/z 848.7836     0   1E+4   1.5   2  RT (min)      lcqtof_hilic_neg_4_000_aa    m/z 849.4670     0   1   2   3E+4   4   4.5  RT (min)      lcqtof_hilic_neg_2_028_05    m/z 849.4680     0   2   4E+4   4.5   5  RT (min)      lcqtof_hilic_neg_1_007_05    m/z 849.5166     0   1   2   3   4E+4   1   1.5  RT (min)      lcqtof_hilic_neg_4_000_aa    m/z 849.5206     0   1   2   3   4E+4   1   1.5  RT (min)      lcqtof_hilic_neg_4_000_aa    m/z 849.6717     0   1   2   3   4E+4   2   2.5  RT (min)      lcqtof_hilic_neg_4_000_aa    m/z 849.6729     0   2   4   6E+4   1.5   2  RT (min)      lcqtof_hilic_neg_4_000_aa    m/z 849.6731     0   1   2   3   4E+4   2   2.5  RT (min)      lcqtof_hilic_neg_4_000_aa    m/z 849.6751     0   2   4   6E+4   1   1.5  RT (min)      lcqtof_hilic_neg_4_000_aa    m/z 849.6751     0   2   4   6E+4   1.5   2  RT (min)      lcqtof_hilic_neg_4_000_aa    m/z 849.6880     0   2   4   6E+4   1.5   2  RT (min)      lcqtof_hilic_neg_4_000_aa    m/z 849.6884     0   2   4   6E+4   1   1.5  RT (min)      lcqtof_hilic_neg_4_000_aa    m/z 849.6913     0   1   2   3   4E+4   2   2.5  RT (min)      lcqtof_hilic_neg_4_000_aa    m/z 849.6972     0   1   2   3   4E+4   1.5   2  RT (min)      lcqtof_hilic_neg_4_000_aa    m/z 850.4717     0   1E+4   4.5   5  RT (min)      lcqtof_hilic_neg_1_007_05    m/z 850.4757     0   1E+4   4   4.5  RT (min)      lcqtof_hilic_neg_4_048_23    m/z 850.5240     0   1   2E+4   1   1.5  RT (min)      lcqtof_hilic_neg_4_000_aa    m/z 850.5256     0   1   2E+4   1   1.5  RT (min)      lcqtof_hilic_neg_4_000_aa    m/z 850.5298     0   1E+4   2   2.5  RT (min)      lcqtof_hilic_neg_4_031_38    m/z 850.5371     0   1   2E+4   2   2.5  RT (min)      lcqtof_hilic_neg_4_031_38    m/z 850.5592     0   1E+4   1.5   2  RT (min)      lcqtof_hilic_neg_1_015_33    m/z 850.5595     0   1   2E+4   2   2.5  RT (min)       lcqtof_hilic_neg_2_062_17    m/z 850.5597     0   2   4E+4   2.5   3  RT (min)      lcqtof_hilic_neg_4_032_01    m/z 850.5597     0   1E+4   1.5   2  RT (min)      lcqtof_hilic_neg_1_010_15    m/z 850.5598     0   1   2E+4   2   2.5  RT (min)       lcqtof_hilic_neg_2_062_17    m/z 850.5599     0   1   2   3   4E+4   2.5   3  RT (min)      lcqtof_hilic_neg_4_053_14    m/z 850.5602     0   2   4E+4   2.5   3  RT (min)      lcqtof_hilic_neg_4_032_01    m/z 850.5603     0   1   2   3   4E+4   2.5   3  RT (min)       lcqtof_hilic_neg_4_053_14    m/z 850.5604     0   1   2E+4   2   2.5  RT (min)      lcqtof_hilic_neg_2_031_01    m/z 850.5614     0   1   2E+5   1   1.5  RT (min)      lcqtof_hilic_neg_4_050_02    m/z 850.5857     0   1E+5   0.5   1  RT (min)      lcqtof_hilic_neg_3_035_46    m/z 850.6609     0   1E+4   0.5   1  RT (min)      lcqtof_hilic_neg_4_047_06    m/z 850.6964     0   1   2E+4   2   2.5  RT (min)      lcqtof_hilic_neg_4_000_aa    m/z 850.7012     0   1   2E+4   1.5   2  RT (min)      lcqtof_hilic_neg_4_000_aa    m/z 850.7074     0   1   2E+4   1.5   2  RT (min)      lcqtof_hilic_neg_4_000_aa    m/z 850.7939     0   1E+4   0.5   1  RT (min)       lcqtof_hilic_neg_4_103_61    m/z 850.9291     0   1E+4   6.5   7  RT (min)      lcqtof_hilic_neg_4_005_29    m/z 850.9760     0   1E+4   0.5   1  RT (min)      lcqtof_hilic_neg_4_000_aa    m/z 851.4818     0   1E+4   4.5   5  RT (min)      lcqtof_hilic_neg_1_007_05    m/z 851.5626     0   1E+4   2   2.5  RT (min)      lcqtof_hilic_neg_2_062_17    m/z 851.5628     0   2   4   6E+3   2   2.5  RT (min)      lcqtof_hilic_neg_1_030_17    m/z 851.5630     0   1E+4   2.5   3  RT (min)      lcqtof_hilic_neg_4_059_14    m/z 851.5922     0   1   2E+4   3.5   4  RT (min)       lcqtof_hilic_neg_4_019_03    m/z 851.5970     0   1E+4   2   2.5  RT (min)      lcqtof_hilic_neg_4_016_37    m/z 851.6267     0   2   4   6E+4   1   1.5  RT (min)      lcqtof_hilic_neg_4_000_aa    m/z 851.6946     0   1E+4   0.5   1  RT (min)      lcqtof_hilic_neg_4_047_06    m/z 851.6980     0   1   2   3E+4   1   1.5  RT (min)      lcqtof_hilic_neg_4_000_aa    m/z 851.7175     0   1E+4   4   4.5  RT (min)      lcqtof_hilic_neg_4_055_32    m/z 852.2269     0   1E+4   6.5   7  RT (min)      lcqtof_hilic_neg_4_008_24    m/z 852.3508     0   2   4E+4   0.5   1  RT (min)      lcqtof_hilic_neg_3_003_20    m/z 852.5725     0   1   2   3E+4   1.5   2  RT (min)      lcqtof_hilic_neg_1_014_31    m/z 852.5726     0   1E+4   1.5   2  RT (min)      lcqtof_hilic_neg_4_050_02    m/z 852.5728     0   1   2E+4   2   2.5  RT (min)      lcqtof_hilic_neg_2_031_01    m/z 852.5731     0   1   2E+4   1.5   2  RT (min)      lcqtof_hilic_neg_4_050_02    m/z 852.5741     0   2   4   6E+4   2.5   3  RT (min)      lcqtof_hilic_neg_4_032_01    m/z 852.6008     0   1E+5   0.5   1  RT (min)      lcqtof_hilic_neg_3_035_46    m/z 852.6268     0   1   2E+4   1   1.5  RT (min)      lcqtof_hilic_neg_4_000_aa    m/z 852.6660     0   1E+4   0.5   1  RT (min)      lcqtof_hilic_neg_4_008_24    m/z 852.9045     0   1E+4   0.5   1  RT (min)      lcqtof_hilic_neg_4_004_47    m/z 853.3269     0   2   4   6E+4   1   1.5  RT (min)      lcqtof_hilic_neg_4_013_20    m/z 853.3550     0   1   2E+4   0.5   1  RT (min)      lcqtof_hilic_neg_3_003_20    m/z 853.4716     0   1E+4   3.5   4  RT (min)      lcqtof_hilic_neg_4_005_29    m/z 853.5020     0   1   2E+4   0.5   1  RT (min)      lcqtof_hilic_neg_3_035_46    m/z 853.5756     0   2   4   6E+4   2.5   3  RT (min)      lcqtof_hilic_neg_4_028_36    m/z 853.5759     0   1   2   3E+5   2.5   3  RT (min)      lcqtof_hilic_neg_1_051_35    m/z 853.5761     0   1   2   3E+5   3   3.5  RT (min)      lcqtof_hilic_neg_4_059_14    m/z 853.5777     0   1E+4   2   2.5  RT (min)      lcqtof_hilic_neg_2_062_17    m/z 853.5995     0   1   2   3E+4   2   2.5  RT (min)      lcqtof_hilic_neg_4_016_37    m/z 853.6314     0   1E+4   1   1.5  RT (min)      lcqtof_hilic_neg_4_000_aa    m/z 853.6777     0   1E+4   1.5   2  RT (min)      lcqtof_hilic_neg_4_000_aa    m/z 854.5734     0   2   4   6E+5   1   1.5  RT (min)      lcqtof_hilic_neg_4_028_36    m/z 854.5884     0   2   4E+4   1.5   2  RT (min)      lcqtof_hilic_neg_1_008_13    m/z 854.5887     0   2   4   6E+5   1   1.5  RT (min)      lcqtof_hilic_neg_4_050_02    m/z 854.5892     0   2   4   6E+4   1.5   2  RT (min)      lcqtof_hilic_neg_2_031_01    m/z 854.5893     0   2   4   6   8E+4   2   2.5  RT (min)       lcqtof_hilic_neg_2_062_17    m/z 854.5895     0   2   4   6   8E+4   2   2.5  RT (min)       lcqtof_hilic_neg_2_062_17    m/z 854.5895     0   2   4E+4   1.5   2  RT (min)      lcqtof_hilic_neg_1_008_13    m/z 854.5897     0   1   2   3   4E+4   1.5   2  RT (min)      lcqtof_hilic_neg_4_050_02    m/z 854.5897     0   2   4   6E+5   2.5   3  RT (min)       lcqtof_hilic_neg_4_032_01    m/z 854.5899     0   1   2E+6   2.5   3  RT (min)      lcqtof_hilic_neg_4_005_29    m/z 854.5909     0   1E+4   1.5   2  RT (min)      lcqtof_hilic_neg_3_049_55    m/z 855.5629     0   1   2E+4   1   1.5  RT (min)      lcqtof_hilic_neg_4_000_aa    m/z 855.5920     0   1E+4   1.5   2  RT (min)      lcqtof_hilic_neg_4_000_aa    m/z 855.5923     0   1E+4   1.5   2  RT (min)      lcqtof_hilic_neg_1_013_38    m/z 855.5930     0   1   2   3E+4   1.5   2  RT (min)      lcqtof_hilic_neg_2_031_01    m/z 855.5932     0   1E+4   1.5   2  RT (min)      lcqtof_hilic_neg_3_024_35    m/z 855.5932     0   1   2   3   4E+4   2.5   3  RT (min)      lcqtof_hilic_neg_4_045_26    m/z 855.5935     0   1E+5   1   1.5  RT (min)      lcqtof_hilic_neg_3_001_49    m/z 855.5935     0   1   2E+4   2.5   3  RT (min)      lcqtof_hilic_neg_3_055_01    m/z 855.5936     0   1   2   3E+4   2   2.5  RT (min)      lcqtof_hilic_neg_2_062_17    m/z 855.5939     0   2   4   6E+4   2.5   3  RT (min)      lcqtof_hilic_neg_4_032_01    m/z 855.5952     0   1E+4   1.5   2  RT (min)      lcqtof_hilic_neg_4_050_02    m/z 855.6799     0   1E+5   2.5   3  RT (min)       lcqtof_hilic_neg_4_008_24    m/z 855.6847     0   1E+5   3   3.5  RT (min)      lcqtof_hilic_neg_4_008_24    m/z 855.6861     0   1E+5   2.5   3  RT (min)      lcqtof_hilic_neg_4_008_24    m/z 855.6867     0   1E+4   2.5   3  RT (min)      lcqtof_hilic_neg_4_008_24    m/z 855.7178     0   1E+4   0.5   1  RT (min)      lcqtof_hilic_neg_3_035_46    m/z 856.5179     0   1E+4   2   2.5  RT (min)       lcqtof_hilic_neg_4_048_23    m/z 856.5677     0   1E+4   1   1.5  RT (min)      lcqtof_hilic_neg_4_000_aa    m/z 856.5812     0   1   2E+4   2   2.5  RT (min)      lcqtof_hilic_neg_1_035_23    m/z 856.5903     0   1E+4   4   4.5  RT (min)       lcqtof_hilic_neg_4_055_32    m/z 856.6029     0   1   2E+4   2   2.5  RT (min)      lcqtof_hilic_neg_2_062_17    m/z 856.6036     0   1   2E+4   2   2.5  RT (min)      lcqtof_hilic_neg_2_062_17    m/z 856.6301     0   2   4   6   8E+4   0.5   1  RT (min)      lcqtof_hilic_neg_3_035_46    m/z 856.8494     0   1E+4   1   1.5  RT (min)       lcqtof_hilic_neg_4_054_48    m/z 857.4159     0   1   2E+4   0.5   1  RT (min)      lcqtof_hilic_neg_4_045_26    m/z 857.4711     0   1   2E+4   1   1.5  RT (min)      lcqtof_hilic_neg_4_000_aa    m/z 857.4728     0   1   2E+4   1   1.5  RT (min)      lcqtof_hilic_neg_4_000_aa    m/z 857.4767     0   1   2E+4   1   1.5  RT (min)      lcqtof_hilic_neg_4_000_aa    m/z 857.4791     0   1   2E+4   1   1.5  RT (min)      lcqtof_hilic_neg_4_000_aa    m/z 857.5223     0   1E+4   0.5   1  RT (min)      lcqtof_hilic_neg_4_039_18    m/z 857.5727     0   1E+4   3   3.5  RT (min)      lcqtof_hilic_neg_1_003_37    m/z 857.5839     0   1E+4   2   2.5  RT (min)      lcqtof_hilic_neg_1_035_23    m/z 857.5876     0   1   2E+5   3   3.5  RT (min)      lcqtof_hilic_neg_1_005_32    m/z 857.6082     0   2   4   6   8E+3   2   2.5  RT (min)      lcqtof_hilic_neg_2_062_17    m/z 857.6089     0   2   4   6   8E+3   2   2.5  RT (min)      lcqtof_hilic_neg_2_062_17    m/z 857.6101     0   1E+5   2.5   3  RT (min)       lcqtof_hilic_neg_4_053_14    m/z 857.6108     0   1E+4   2.5   3  RT (min)       lcqtof_hilic_neg_4_053_14    m/z 857.6109     0   1   2   3   4E+3   1.5   2  RT (min)      lcqtof_hilic_neg_1_004_14    m/z 857.6109     0   1   2E+5   2.5   3  RT (min)       lcqtof_hilic_neg_4_053_14    m/z 857.6117     0   1   2E+5   2.5   3  RT (min)      lcqtof_hilic_neg_4_053_14    m/z 857.6351     0   1E+4   1   1.5  RT (min)      lcqtof_hilic_neg_4_000_aa    m/z 857.6368     0   1   2   3E+4   0.5   1  RT (min)      lcqtof_hilic_neg_3_035_46    m/z 857.6611     0   1   2   3   4E+4   1   1.5  RT (min)      lcqtof_hilic_neg_4_008_24    m/z 857.6620     0   1E+4   1.5   2  RT (min)      lcqtof_hilic_neg_3_004_32    m/z 857.6627     0   1E+4   1   1.5  RT (min)      lcqtof_hilic_neg_4_055_32    m/z 857.6645     0   1   2E+4   3   3.5  RT (min)      lcqtof_hilic_neg_4_008_24    m/z 857.6732     0   1   2E+4   2.5   3  RT (min)      lcqtof_hilic_neg_4_008_24    m/z 857.6799     0   1   2E+4   3   3.5  RT (min)      lcqtof_hilic_neg_4_008_24    m/z 857.6801     0   1   2E+4   2.5   3  RT (min)      lcqtof_hilic_neg_4_008_24    m/z 857.6900     0   1E+5   2.5   3  RT (min)      lcqtof_hilic_neg_4_008_24    m/z 857.7189     0   1   2   3E+4   1   1.5  RT (min)      lcqtof_hilic_neg_4_000_aa    m/z 857.7320     0   1E+4   1   1.5  RT (min)      lcqtof_hilic_neg_3_035_46    m/z 858.0344     0   1   2E+4   7.5   8  RT (min)      lcqtof_hilic_neg_4_016_37    m/z 858.4745     0   1E+4   1   1.5  RT (min)      lcqtof_hilic_neg_4_000_aa    m/z 858.5455     0   1E+4   2   2.5  RT (min)      lcqtof_hilic_neg_4_016_37    m/z 858.5847     0   1E+4   3.5   4  RT (min)       lcqtof_hilic_neg_4_040_05    m/z 858.6168     0   2   4   6E+4   3   3.5  RT (min)      lcqtof_hilic_neg_4_059_14    m/z 858.6257     0   1E+4   1   1.5  RT (min)      lcqtof_hilic_neg_4_031_38    m/z 858.6413     0   1   2   3   4E+4   4   4.5  RT (min)       lcqtof_hilic_neg_4_008_24    m/z 858.6520     0   1E+4   3   3.5  RT (min)      lcqtof_hilic_neg_1_025_52    m/z 858.6584     0   1   2E+4   3   3.5  RT (min)       lcqtof_hilic_neg_4_096_60    m/z 858.6590     0   1E+4   2.5   3  RT (min)       lcqtof_hilic_neg_3_034_24    m/z 858.6594     0   2   4   6   8E+3   2   2.5  RT (min)       lcqtof_hilic_neg_1_005_32    m/z 858.6595     0   1   2E+4   2.5   3  RT (min)       lcqtof_hilic_neg_4_008_24    m/z 858.6596     0   2   4   6   8E+3   2.5   3  RT (min)      lcqtof_hilic_neg_3_034_24    m/z 858.6597     0   1E+4   2.5   3  RT (min)       lcqtof_hilic_neg_3_091_60    m/z 858.6608     0   1   2E+4   1   1.5  RT (min)      lcqtof_hilic_neg_2_052_24    m/z 858.6966     0   1   2E+4   3   3.5  RT (min)      lcqtof_hilic_neg_4_008_24    m/z 858.7107     0   1E+5   0.5   1  RT (min)      lcqtof_hilic_neg_3_035_46    m/z 858.7304     0   1E+4   1   1.5  RT (min)      lcqtof_hilic_neg_4_000_aa    m/z 859.5814     0   1E+4   3   3.5  RT (min)      lcqtof_hilic_neg_1_003_37    m/z 859.5883     0   2   4   6E+4   1   1.5  RT (min)      lcqtof_hilic_neg_4_000_aa    m/z 859.6762     0   1E+4   2.5   3  RT (min)      lcqtof_hilic_neg_4_008_24    m/z 859.6765     0   1E+4   3   3.5  RT (min)      lcqtof_hilic_neg_1_034_08    m/z 859.6771     0   1E+4   2.5   3  RT (min)       lcqtof_hilic_neg_4_008_24    m/z 859.6795     0   1E+4   3   3.5  RT (min)      lcqtof_hilic_neg_4_008_24    m/z 859.7112     0   2   4   6E+4   0.5   1  RT (min)      lcqtof_hilic_neg_3_035_46    m/z 859.7195     0   1E+4   0.5   1   1.5  RT (min)      lcqtof_hilic_neg_1_037_24    m/z 859.7881     0   1E+4   1   1.5  RT (min)      lcqtof_hilic_neg_4_000_aa    m/z 860.5319     0   1E+4   3   3.5  RT (min)       lcqtof_hilic_neg_4_048_23    m/z 860.5555     0   1E+4   2   2.5  RT (min)      lcqtof_hilic_neg_4_011_35    m/z 860.5755     0   1   2E+4   4   4.5  RT (min)      lcqtof_hilic_neg_4_055_32    m/z 860.5892     0   1   2E+4   4   4.5  RT (min)       lcqtof_hilic_neg_4_055_32    m/z 860.6490     0   1E+4   0.5   1  RT (min)      lcqtof_hilic_neg_4_022_34    m/z 860.6680     0   1E+4   4.5   5  RT (min)      lcqtof_hilic_neg_4_008_24    m/z 860.6744     0   1   2   3E+4   2.5   3  RT (min)      lcqtof_hilic_neg_2_052_24    m/z 860.6747     0   1   2   3E+4   2.5   3  RT (min)       lcqtof_hilic_neg_2_052_24    m/z 860.6754     0   1E+4   2.5   3  RT (min)      lcqtof_hilic_neg_2_052_24    m/z 860.7168     0   1E+5   1   1.5  RT (min)      lcqtof_hilic_neg_4_008_24    m/z 860.7219     0   2   4E+4   1   1.5  RT (min)      lcqtof_hilic_neg_4_002_25    m/z 860.9165     0   2   4   6E+4   0.5   1  RT (min)      lcqtof_hilic_neg_3_035_46    m/z 860.9226     0   2   4   6E+4   0.5   1  RT (min)      lcqtof_hilic_neg_3_035_46    m/z 861.0954     0   1E+4   0.5   1  RT (min)      lcqtof_hilic_neg_1_062_17    m/z 861.4246     0   1   2E+4   0.5   1  RT (min)      lcqtof_hilic_neg_2_004_26    m/z 861.6426     0   1E+4   3   3.5  RT (min)      lcqtof_hilic_neg_1_005_32    m/z 861.6864     0   1E+4   3   3.5  RT (min)       lcqtof_hilic_neg_1_034_08    m/z 861.7303     0   1E+4   0.5   1  RT (min)      lcqtof_hilic_neg_3_035_46    m/z 861.7309     0   1E+4   2   2.5  RT (min)      lcqtof_hilic_neg_4_008_24    m/z 861.7467     0   1   2E+4   0.5   1  RT (min)      lcqtof_hilic_neg_1_005_32    m/z 861.7709     0   1   2E+4   1   1.5  RT (min)      lcqtof_hilic_neg_4_000_aa    m/z 861.7770     0   1   2E+4   1   1.5  RT (min)      lcqtof_hilic_neg_4_000_aa    m/z 861.8030     0   1   2E+4   1.5   2  RT (min)      lcqtof_hilic_neg_4_000_aa    m/z 861.8129     0   1E+4   1.5   2  RT (min)      lcqtof_hilic_neg_4_000_aa    m/z 861.9235     0   1   2   3E+4   0.5   1  RT (min)      lcqtof_hilic_neg_3_035_46    m/z 862.0832     0   1E+4   0.5   1  RT (min)      lcqtof_hilic_neg_4_026_04    m/z 862.5670     0   1   2E+4   2   2.5  RT (min)      lcqtof_hilic_neg_4_011_35    m/z 862.6150     0   1E+4   4   4.5  RT (min)       lcqtof_hilic_neg_4_040_05    m/z 862.6608     0   1E+4   0.5   1  RT (min)      lcqtof_hilic_neg_4_040_05    m/z 862.7750     0   1E+4   0.5   1  RT (min)      lcqtof_hilic_neg_3_035_46    m/z 862.8130     0   1E+4   1.5   2  RT (min)      lcqtof_hilic_neg_4_000_aa    m/z 862.9231     0   1   2   3E+4   0.5   1  RT (min)      lcqtof_hilic_neg_3_035_46    m/z 863.4225     0   1   2E+4   0.5   1  RT (min)      lcqtof_hilic_neg_4_005_29    m/z 863.4333     0   2   4E+4   0.5   1  RT (min)      lcqtof_hilic_neg_1_042_26    m/z 863.4877     0   1   2   3E+4   2.5   3  RT (min)      lcqtof_hilic_neg_4_052_13    m/z 863.5328     0   1   2   3E+4   1   1.5  RT (min)      lcqtof_hilic_neg_4_000_aa    m/z 863.5386     0   1   2   3E+4   1   1.5  RT (min)      lcqtof_hilic_neg_4_000_aa    m/z 863.7361     0   1E+4   1.5   2  RT (min)      lcqtof_hilic_neg_4_000_aa    m/z 863.7392     0   1   2   3E+4   1.5   2  RT (min)      lcqtof_hilic_neg_4_000_aa    m/z 863.7510     0   1   2E+4   1.5   2  RT (min)      lcqtof_hilic_neg_4_000_aa    m/z 863.7550     0   1   2   3E+4   1   1.5  RT (min)      lcqtof_hilic_neg_4_000_aa    m/z 863.7703     0   1   2   3E+4   1   1.5  RT (min)      lcqtof_hilic_neg_4_000_aa    m/z 863.8020     0   1E+4   0.5   1  RT (min)      lcqtof_hilic_neg_4_004_47    m/z 863.9202     0   1E+4   0.5   1  RT (min)      lcqtof_hilic_neg_3_035_46    m/z 864.4362     0   1   2E+4   0.5   1  RT (min)      lcqtof_hilic_neg_2_004_26    m/z 864.5370     0   1   2E+4   1   1.5  RT (min)      lcqtof_hilic_neg_4_000_aa    m/z 864.5515     0   1   2   3E+4   3.5   4  RT (min)      lcqtof_hilic_neg_1_007_05    m/z 864.5989     0   1   2E+4   4   4.5  RT (min)      lcqtof_hilic_neg_4_008_24    m/z 864.7000     0   1   2E+4   3   3.5  RT (min)      lcqtof_hilic_neg_1_037_24    m/z 864.7575     0   1E+4   1   1.5  RT (min)      lcqtof_hilic_neg_4_000_aa    m/z 864.7760     0   1E+4   1   1.5  RT (min)      lcqtof_hilic_neg_4_000_aa    m/z 864.9965     0   1E+4   4.5   5  RT (min)      lcqtof_hilic_neg_4_008_24    m/z 865.4418     0   1   2   3E+4   0.5   1  RT (min)      lcqtof_hilic_neg_2_004_26    m/z 865.4615     0   1E+4   3.5   4  RT (min)      lcqtof_hilic_neg_2_028_05    m/z 865.4629     0   1E+4   3.5   4  RT (min)      lcqtof_hilic_neg_4_040_05    m/z 865.5875     0   1E+5   1.5   2  RT (min)      lcqtof_hilic_neg_4_000_aa    m/z 865.5877     0   1E+5   1   1.5  RT (min)      lcqtof_hilic_neg_4_000_aa    m/z 865.5880     0   1E+5   1.5   2  RT (min)      lcqtof_hilic_neg_4_000_aa    m/z 865.5880     0   2   4   6E+4   1.5   2  RT (min)      lcqtof_hilic_neg_4_000_aa    m/z 865.5883     0   1E+5   1   1.5  RT (min)      lcqtof_hilic_neg_4_000_aa    m/z 865.5884     0   1E+5   1   1.5  RT (min)      lcqtof_hilic_neg_4_000_aa    m/z 865.5900     0   1E+5   1.5   2  RT (min)      lcqtof_hilic_neg_4_000_aa    m/z 865.6718     0   1   2   3   4E+5   2.5   3  RT (min)      lcqtof_hilic_neg_4_008_24    m/z 865.6722     0   1   2   3   4E+5   3   3.5  RT (min)      lcqtof_hilic_neg_4_008_24    m/z 865.6723     0   1E+5   2.5   3  RT (min)      lcqtof_hilic_neg_2_052_24    m/z 865.6724     0   1   2   3   4E+5   2.5   3  RT (min)      lcqtof_hilic_neg_4_008_24    m/z 865.6726     0   1E+4   2.5   3  RT (min)      lcqtof_hilic_neg_2_047_32    m/z 865.6734     0   1E+4   2.5   3  RT (min)      lcqtof_hilic_neg_2_047_32    m/z 865.7056     0   1   2E+4   1   1.5  RT (min)      lcqtof_hilic_neg_4_000_aa    m/z 865.7296     0   1E+4   2   2.5  RT (min)      lcqtof_hilic_neg_4_040_05    m/z 865.9960     0   1   2E+4   6.5   7  RT (min)      lcqtof_hilic_neg_4_034_31    m/z 865.9975     0   1E+4   6.5   7  RT (min)      lcqtof_hilic_neg_4_011_35    m/z 865.9980     0   1   2E+4   6.5   7  RT (min)      lcqtof_hilic_neg_4_034_31    m/z 866.0697     0   1E+4   0.5   1  RT (min)      lcqtof_hilic_neg_1_062_17    m/z 866.4444     0   1E+4   0.5   1  RT (min)      lcqtof_hilic_neg_2_004_26    m/z 866.5898     0   2   4   6   8E+4   1   1.5   2  RT (min)      lcqtof_hilic_neg_4_000_aa    m/z 866.5904     0   2   4   6   8E+4   1   1.5  RT (min)      lcqtof_hilic_neg_4_000_aa    m/z 866.5906     0   2   4   6   8E+4   1.5   2  RT (min)      lcqtof_hilic_neg_4_000_aa    m/z 866.5906     0   1   2   3   4E+4   1.5   2  RT (min)      lcqtof_hilic_neg_4_000_aa    m/z 866.5907     0   2   4   6E+4   1.5   2  RT (min)      lcqtof_hilic_neg_4_000_aa    m/z 866.5920     0   2   4   6   8E+4   1   1.5  RT (min)      lcqtof_hilic_neg_4_000_aa    m/z 866.5922     0   2   4   6   8E+4   1   1.5  RT (min)      lcqtof_hilic_neg_4_000_aa    m/z 866.5923     0   2   4   6   8E+4   1   1.5  RT (min)      lcqtof_hilic_neg_4_000_aa    m/z 866.5936     0   2   4   6   8E+4   1   1.5  RT (min)      lcqtof_hilic_neg_4_000_aa    m/z 866.6490     0   2   4   6E+4   3   3.5  RT (min)      lcqtof_hilic_neg_4_035_33    m/z 866.6711     0   1E+6   3   3.5  RT (min)      lcqtof_hilic_neg_4_055_32    m/z 866.6748     0   1E+4   1.5   2  RT (min)      lcqtof_hilic_neg_3_001_49    m/z 866.6753     0   2   4   6   8E+5   1   1.5  RT (min)      lcqtof_hilic_neg_2_015_06    m/z 866.6754     0   1   2   3E+4   2   2.5  RT (min)      lcqtof_hilic_neg_3_004_32    m/z 866.6759     0   2   4   6   8E+4   2.5   3  RT (min)      lcqtof_hilic_neg_2_047_32    m/z 866.6760     0   2   4E+4   2   2.5  RT (min)      lcqtof_hilic_neg_1_005_32    m/z 866.6760     0   1E+6   2.5   3  RT (min)      lcqtof_hilic_neg_2_052_24    m/z 866.6761     0   1   2   3E+4   2.5   3  RT (min)      lcqtof_hilic_neg_3_048_05    m/z 866.6761     0   1E+6   2.5   3  RT (min)      lcqtof_hilic_neg_4_008_24    m/z 866.6763     0   1   2E+5   2.5   3  RT (min)      lcqtof_hilic_neg_2_052_24    m/z 866.6763     0   1E+6   1   1.5  RT (min)      lcqtof_hilic_neg_2_052_24    m/z 866.6764     0   1   2   3E+5   2.5   3  RT (min)      lcqtof_hilic_neg_2_052_24    m/z 866.6766     0   1   2E+4   2   2.5  RT (min)      lcqtof_hilic_neg_4_055_32    m/z 866.7136     0   1E+4   0.5   1  RT (min)      lcqtof_hilic_neg_4_047_06    m/z 866.7687     0   1E+4   1   1.5  RT (min)      lcqtof_hilic_neg_4_008_24    m/z 866.9761     0   2   4E+4   0.5   1  RT (min)      lcqtof_hilic_neg_4_008_24    m/z 867.2065     0   1E+4   3   3.5  RT (min)      lcqtof_hilic_neg_1_037_24    m/z 867.4778     0   1   2   3E+4   3.5   4  RT (min)      lcqtof_hilic_neg_1_007_05    m/z 867.4779     0   1   2   3E+4   3.5   4  RT (min)      lcqtof_hilic_neg_1_007_05    m/z 867.5490     0   2   4   6   8E+3   2.5   3  RT (min)       lcqtof_hilic_neg_4_019_03    m/z 867.5502     0   1   2   3   4E+3   2.5   3  RT (min)      lcqtof_hilic_neg_4_059_14    m/z 867.5903     0   2   4   6E+4   1   1.5  RT (min)      lcqtof_hilic_neg_4_000_aa    m/z 867.5919     0   2   4   6E+4   1   1.5  RT (min)      lcqtof_hilic_neg_4_000_aa    m/z 867.5926     0   2   4   6E+4   1   1.5  RT (min)      lcqtof_hilic_neg_4_000_aa    m/z 867.5933     0   2   4E+4   1.5   2  RT (min)      lcqtof_hilic_neg_4_000_aa    m/z 867.5938     0   2   4   6E+4   1   1.5  RT (min)      lcqtof_hilic_neg_4_000_aa    m/z 867.5947     0   2   4   6E+4   1.5   2  RT (min)      lcqtof_hilic_neg_4_000_aa    m/z 867.5953     0   2   4   6E+4   1   1.5  RT (min)      lcqtof_hilic_neg_4_000_aa    m/z 867.5965     0   1   2E+4   1.5   2  RT (min)      lcqtof_hilic_neg_4_000_aa    m/z 867.5986     0   2   4   6E+4   1   1.5  RT (min)      lcqtof_hilic_neg_4_000_aa    m/z 867.6001     0   2   4   6E+4   1   1.5  RT (min)      lcqtof_hilic_neg_4_000_aa    m/z 867.6046     0   1   2E+4   1.5   2  RT (min)      lcqtof_hilic_neg_4_000_aa    m/z 867.6176     0   1   2E+4   2   2.5  RT (min)      lcqtof_hilic_neg_4_000_aa    m/z 867.6653     0   1   2   3   4E+4   3   3.5  RT (min)      lcqtof_hilic_neg_4_035_33    m/z 867.6860     0   1   2E+4   1.5   2  RT (min)      lcqtof_hilic_neg_3_004_32    m/z 867.6868     0   1E+6   2.5   3  RT (min)      lcqtof_hilic_neg_4_008_24    m/z 867.6870     0   1   2E+4   2   2.5  RT (min)      lcqtof_hilic_neg_3_004_32    m/z 867.6870     0   1   2   3   4E+4   2.5   3  RT (min)      lcqtof_hilic_neg_2_047_32    m/z 867.6870     0   1E+6   3   3.5  RT (min)      lcqtof_hilic_neg_4_008_24    m/z 867.6871     0   1   2   3   4E+5   1   1.5  RT (min)      lcqtof_hilic_neg_2_047_32    m/z 867.6872     0   1E+6   2.5   3  RT (min)      lcqtof_hilic_neg_4_008_24    m/z 867.6872     0   1   2   3   4E+4   2.5   3  RT (min)      lcqtof_hilic_neg_2_047_32    m/z 867.6873     0   1   2E+4   2   2.5  RT (min)      lcqtof_hilic_neg_1_005_32    m/z 867.6874     0   2   4   6   8E+4   2.5   3  RT (min)      lcqtof_hilic_neg_2_052_24    m/z 867.6878     0   2   4   6   8E+5   1   1.5  RT (min)      lcqtof_hilic_neg_2_052_24    m/z 867.6880     0   1   2E+4   2   2.5  RT (min)      lcqtof_hilic_neg_3_004_32    m/z 867.7139     0   1E+5   1   1.5  RT (min)      lcqtof_hilic_neg_3_035_46    m/z 868.0359     0   1E+4   0.5   1  RT (min)      lcqtof_hilic_neg_3_035_46    m/z 868.1876     0   1E+4   0.5   1  RT (min)      lcqtof_hilic_neg_1_062_17    m/z 868.4783     0   1E+4   3.5   4  RT (min)      lcqtof_hilic_neg_1_007_05    m/z 868.4803     0   1E+4   3.5   4  RT (min)      lcqtof_hilic_neg_1_007_05    m/z 868.5252     0   1E+4   2.5   3  RT (min)      lcqtof_hilic_neg_4_032_01    m/z 868.5589     0   1E+4   1.5   2  RT (min)      lcqtof_hilic_neg_4_016_37    m/z 868.5620     0   1   2   3   4E+3   2   2.5  RT (min)      lcqtof_hilic_neg_2_060_14    m/z 868.5640     0   1E+4   2.5   3  RT (min)      lcqtof_hilic_neg_1_051_35    m/z 868.5682     0   1   2   3E+4   1   1.5  RT (min)       lcqtof_hilic_neg_2_002_36    m/z 868.5685     0   1E+4   1.5   2  RT (min)      lcqtof_hilic_neg_4_022_34    m/z 868.6024     0   1   2E+4   1   1.5  RT (min)      lcqtof_hilic_neg_4_000_aa    m/z 868.6101     0   1   2   3   4E+4   3   3.5  RT (min)      lcqtof_hilic_neg_1_004_14    m/z 868.6898     0   1   2E+5   1.5   2  RT (min)       lcqtof_hilic_neg_2_052_24    m/z 868.6898     0   1E+4   1.5   2  RT (min)      lcqtof_hilic_neg_4_047_06    m/z 868.6901     0   1E+6   1   1.5  RT (min)      lcqtof_hilic_neg_2_015_06    m/z 868.6908     0   1E+6   2.5   3  RT (min)      lcqtof_hilic_neg_3_034_24    m/z 868.6908     0   1E+5   1.5   2   2.5  RT (min)       lcqtof_hilic_neg_1_005_32    m/z 868.6909     0   1E+5   2.5   3  RT (min)      lcqtof_hilic_neg_2_047_32    m/z 868.6911     0   1   2   3E+6   2.5   3  RT (min)       lcqtof_hilic_neg_4_008_24    m/z 868.6914     0   1   2   3E+6   3   3.5  RT (min)       lcqtof_hilic_neg_4_008_24    m/z 868.6914     0   2   4E+5   2.5   3  RT (min)       lcqtof_hilic_neg_2_052_24    m/z 868.6915     0   2   4E+4   2   2.5  RT (min)      lcqtof_hilic_neg_2_016_33    m/z 868.6915     0   1   2   3E+6   2.5   3  RT (min)      lcqtof_hilic_neg_4_008_24    m/z 868.6916     0   1   2E+6   1   1.5  RT (min)       lcqtof_hilic_neg_2_052_24    m/z 868.6916     0   1E+5   2.5   3  RT (min)      lcqtof_hilic_neg_1_037_24    m/z 868.6918     0   2   4   6   8E+4   2   2.5  RT (min)       lcqtof_hilic_neg_3_004_32    m/z 868.6918     0   1E+5   2.5   3  RT (min)      lcqtof_hilic_neg_3_034_24    m/z 869.1290     0   1   2   3E+4   3   3.5  RT (min)      lcqtof_hilic_neg_1_037_24    m/z 869.1304     0   1E+4   0.5   1  RT (min)      lcqtof_hilic_neg_4_008_24    m/z 869.1831     0   1E+4   1   1.5  RT (min)      lcqtof_hilic_neg_4_000_aa    m/z 869.1840     0   1   2E+4   0.5   1  RT (min)      lcqtof_hilic_neg_1_062_17    m/z 869.2218     0   2   4E+4   3   3.5  RT (min)      lcqtof_hilic_neg_1_037_24    m/z 869.2219     0   1   2   3   4E+4   0.5   1  RT (min)      lcqtof_hilic_neg_4_008_24    m/z 869.3876     0   1   2   3E+4   3   3.5  RT (min)      lcqtof_hilic_neg_1_037_24    m/z 869.3891     0   1   2E+4   0.5   1  RT (min)      lcqtof_hilic_neg_4_008_24    m/z 869.4914     0   1E+4   3.5   4  RT (min)      lcqtof_hilic_neg_2_028_05    m/z 869.4915     0   1E+4   3.5   4  RT (min)      lcqtof_hilic_neg_1_007_05    m/z 869.5658     0   1   2   3   4E+3   2   2.5  RT (min)      lcqtof_hilic_neg_2_062_17    m/z 869.5663     0   2   4   6E+3   3   3.5  RT (min)      lcqtof_hilic_neg_4_069_57    m/z 869.5666     0   1   2E+4   1.5   2  RT (min)      lcqtof_hilic_neg_1_008_13    m/z 869.5668     0   1E+4   2   2.5  RT (min)      lcqtof_hilic_neg_2_062_17    m/z 869.5669     0   2   4   6E+3   2.5   3  RT (min)       lcqtof_hilic_neg_4_023_15    m/z 869.5670     0   2   4   6E+3   2.5   3  RT (min)      lcqtof_hilic_neg_4_029_17    m/z 869.5670     0   2   4   6   8E+3   2.5   3  RT (min)      lcqtof_hilic_neg_4_075_58    m/z 869.6886     0   1   2   3E+4   2   2.5  RT (min)      lcqtof_hilic_neg_1_007_05    m/z 869.7014     0   1   2E+4   2   2.5  RT (min)      lcqtof_hilic_neg_1_005_32    m/z 869.7019     0   1E+6   2.5   3  RT (min)      lcqtof_hilic_neg_4_008_24    m/z 869.7020     0   2   4   6E+4   2.5   3  RT (min)      lcqtof_hilic_neg_4_001_49    m/z 869.7020     0   1   2E+5   3   3.5  RT (min)      lcqtof_hilic_neg_4_014_46    m/z 869.7021     0   1   2E+4   2   2.5  RT (min)      lcqtof_hilic_neg_1_005_32    m/z 869.7023     0   1   2   3E+4   2.5   3  RT (min)      lcqtof_hilic_neg_1_005_32    m/z 869.7025     0   1   2   3E+5   2.5   3  RT (min)      lcqtof_hilic_neg_4_001_49    m/z 869.7025     0   2   4   6E+5   1   1.5  RT (min)      lcqtof_hilic_neg_2_052_24    m/z 869.7030     0   1   2E+4   1.5   2  RT (min)      lcqtof_hilic_neg_1_005_32    m/z 869.7031     0   1   2E+4   2   2.5  RT (min)      lcqtof_hilic_neg_1_005_32    m/z 869.7034     0   1   2E+4   2   2.5  RT (min)      lcqtof_hilic_neg_1_005_32    m/z 869.7128     0   1   2E+4   2   2.5  RT (min)      lcqtof_hilic_neg_1_005_32    m/z 869.7286     0   2   4   6   8E+4   1   1.5  RT (min)      lcqtof_hilic_neg_3_035_46    m/z 870.0106     0   1   2E+4   6.5   7  RT (min)      lcqtof_hilic_neg_4_055_32    m/z 870.1860     0   1E+4   0.5   1  RT (min)      lcqtof_hilic_neg_1_062_17    m/z 870.3618     0   2   4   6E+4   0.5   1  RT (min)      lcqtof_hilic_neg_3_003_20    m/z 870.4775     0   1E+4   0.5   1  RT (min)      lcqtof_hilic_neg_4_015_09    m/z 870.5576     0   1E+4   1   1.5  RT (min)      lcqtof_hilic_neg_1_063_13    m/z 870.5703     0   1E+4   2.5   3  RT (min)      lcqtof_hilic_neg_4_006_30    m/z 870.5717     0   1   2E+4   2.5   3  RT (min)      lcqtof_hilic_neg_1_030_17    m/z 870.5757     0   1   2   3E+4   1.5   2  RT (min)      lcqtof_hilic_neg_1_054_01    m/z 870.5765     0   1   2E+4   1.5   2  RT (min)      lcqtof_hilic_neg_4_053_14    m/z 870.6485     0   1E+4   0.5   1  RT (min)      lcqtof_hilic_neg_2_039_38    m/z 870.6947     0   1   2   3   4E+4   2   2.5  RT (min)      lcqtof_hilic_neg_4_035_33    m/z 870.7064     0   1   2   3E+6   3   3.5  RT (min)       lcqtof_hilic_neg_4_008_24    m/z 870.7065     0   1E+5   2.5   3  RT (min)       lcqtof_hilic_neg_2_052_24    m/z 870.7065     0   1E+4   1.5   2  RT (min)      lcqtof_hilic_neg_3_001_49    m/z 870.7065     0   1   2   3E+6   2.5   3  RT (min)       lcqtof_hilic_neg_4_008_24    m/z 870.7066     0   1E+5   2.5   3  RT (min)      lcqtof_hilic_neg_2_052_24    m/z 870.7066     0   2   4   6   8E+4   2   2.5  RT (min)       lcqtof_hilic_neg_1_005_32    m/z 870.7067     0   1   2   3E+6   2.5   3  RT (min)      lcqtof_hilic_neg_4_008_24    m/z 870.7072     0   2   4   6   8E+4   1.5   2   2.5  RT (min)      lcqtof_hilic_neg_1_005_32    m/z 870.7074     0   1   2E+5   2.5   3  RT (min)      lcqtof_hilic_neg_3_034_24    m/z 871.1451     0   1   2E+4   3   3.5  RT (min)      lcqtof_hilic_neg_1_037_24    m/z 871.2374     0   1E+4   0.5   1  RT (min)      lcqtof_hilic_neg_4_008_24    m/z 871.2386     0   2   4E+4   3   3.5  RT (min)      lcqtof_hilic_neg_1_037_24    m/z 871.3680     0   1   2E+4   0.5   1  RT (min)      lcqtof_hilic_neg_3_003_20    m/z 871.4058     0   1   2   3E+4   3   3.5  RT (min)      lcqtof_hilic_neg_1_037_24    m/z 871.5463     0   1E+4   3   3.5  RT (min)      lcqtof_hilic_neg_4_048_23    m/z 871.5528     0   1   2E+4   3   3.5  RT (min)      lcqtof_hilic_neg_4_011_35    m/z 871.5798     0   1   2   3   4E+4   3   3.5  RT (min)       lcqtof_hilic_neg_4_019_03    m/z 871.5798     0   1   2   3   4E+4   3   3.5  RT (min)       lcqtof_hilic_neg_4_019_03    m/z 871.5800     0   1   2   3   4E+4   2.5   3  RT (min)      lcqtof_hilic_neg_4_019_03    m/z 871.5800     0   1   2   3   4E+4   2.5   3  RT (min)       lcqtof_hilic_neg_4_019_03    m/z 871.5801     0   1E+4   2.5   3  RT (min)       lcqtof_hilic_neg_4_006_30    m/z 871.5806     0   1E+4   2.5   3  RT (min)       lcqtof_hilic_neg_4_053_14    m/z 871.5815     0   1E+4   2   2.5  RT (min)       lcqtof_hilic_neg_2_051_14    m/z 871.5816     0   1E+4   2.5   3  RT (min)       lcqtof_hilic_neg_4_059_14    m/z 871.5817     0   1E+4   2   2.5  RT (min)      lcqtof_hilic_neg_2_051_14    m/z 871.5818     0   1E+4   2   2.5  RT (min)      lcqtof_hilic_neg_1_030_17    m/z 871.5818     0   1E+4   2   2.5  RT (min)       lcqtof_hilic_neg_2_018_16    m/z 871.5819     0   1E+4   2   2.5  RT (min)      lcqtof_hilic_neg_2_051_14    m/z 871.5823     0   1   2   3E+4   1.5   2  RT (min)      lcqtof_hilic_neg_1_004_14    m/z 871.5824     0   1   2E+4   1.5   2  RT (min)      lcqtof_hilic_neg_1_030_17    m/z 871.5826     0   1E+4   2   2.5  RT (min)      lcqtof_hilic_neg_4_053_14    m/z 871.5970     0   1   2   3E+4   2   2.5  RT (min)      lcqtof_hilic_neg_4_011_35    m/z 871.6461     0   1E+4   0.5   1  RT (min)      lcqtof_hilic_neg_4_053_14    m/z 871.6587     0   1E+4   3.5   4  RT (min)       lcqtof_hilic_neg_4_059_14    m/z 871.6594     0   1E+4   3.5   4  RT (min)      lcqtof_hilic_neg_4_053_14    m/z 871.7160     0   1E+4   2.5   3  RT (min)      lcqtof_hilic_neg_2_047_32    m/z 871.7164     0   1   2E+4   2.5   3  RT (min)      lcqtof_hilic_neg_2_052_24    m/z 871.7178     0   1E+4   2   2.5  RT (min)      lcqtof_hilic_neg_1_005_32    m/z 871.7180     0   1E+4   2   2.5  RT (min)      lcqtof_hilic_neg_1_005_32    m/z 871.7186     0   2   4   6   8E+5   2.5   3  RT (min)      lcqtof_hilic_neg_4_008_24    m/z 871.7190     0   1E+4   2.5   3  RT (min)      lcqtof_hilic_neg_4_004_47    m/z 871.7443     0   2   4E+4   0.5   1  RT (min)      lcqtof_hilic_neg_3_035_46    m/z 872.1042     0   1E+4   7.5   8  RT (min)       lcqtof_hilic_neg_4_048_23    m/z 872.4330     0   1   2   3E+4   0.5   1  RT (min)      lcqtof_hilic_neg_4_045_26    m/z 872.4729     0   1E+4   0.5   1  RT (min)      lcqtof_hilic_neg_1_007_05    m/z 872.5481     0   1E+4   3   3.5  RT (min)      lcqtof_hilic_neg_4_011_35    m/z 872.5656     0   1E+4   2.5   3  RT (min)      lcqtof_hilic_neg_1_014_31    m/z 872.5858     0   1E+4   2   2.5  RT (min)      lcqtof_hilic_neg_1_022_36    m/z 872.5875     0   1E+4   2   2.5  RT (min)      lcqtof_hilic_neg_4_053_14    m/z 872.5884     0   1E+4   2.5   3  RT (min)      lcqtof_hilic_neg_4_059_14    m/z 872.5909     0   1E+4   2   2.5  RT (min)      lcqtof_hilic_neg_4_011_35    m/z 872.7127     0   1   2   3E+4   2   2.5  RT (min)      lcqtof_hilic_neg_4_035_33    m/z 872.7188     0   2   4   6E+4   2.5   3  RT (min)       lcqtof_hilic_neg_2_052_24    m/z 872.7220     0   2   4E+4   2   2.5  RT (min)       lcqtof_hilic_neg_1_005_32    m/z 872.7222     0   1   2E+6   2.5   3  RT (min)       lcqtof_hilic_neg_4_008_24    m/z 872.7228     0   1   2   3   4E+4   1.5   2   2.5  RT (min)      lcqtof_hilic_neg_1_005_32    m/z 872.7228     0   1E+6   2.5   3  RT (min)       lcqtof_hilic_neg_4_008_24    m/z 872.7233     0   1E+4   2   2.5  RT (min)      lcqtof_hilic_neg_3_008_25    m/z 872.7233     0   1E+4   2   2.5  RT (min)      lcqtof_hilic_neg_3_008_25    m/z 872.7234     0   2   4E+4   2   2.5  RT (min)       lcqtof_hilic_neg_1_005_32    m/z 872.7242     0   1E+5   1   1.5  RT (min)      lcqtof_hilic_neg_3_001_49    m/z 872.7254     0   1E+6   1   1.5  RT (min)       lcqtof_hilic_neg_2_052_24    m/z 872.7275     0   1   2   3   4E+4   1.5   2  RT (min)       lcqtof_hilic_neg_1_005_32    m/z 872.7490     0   1E+5   1   1.5  RT (min)      lcqtof_hilic_neg_3_035_46    m/z 873.0214     0   1   2   3E+4   0.5   1  RT (min)      lcqtof_hilic_neg_4_008_24    m/z 873.2537     0   1   2E+4   3   3.5  RT (min)      lcqtof_hilic_neg_4_008_24    m/z 873.3110     0   1   2E+4   0.5   1  RT (min)      lcqtof_hilic_neg_4_021_21    m/z 873.3238     0   2   4E+4   0.5   1  RT (min)      lcqtof_hilic_neg_4_015_09    m/z 873.5718     0   1E+4   3   3.5  RT (min)      lcqtof_hilic_neg_4_016_37    m/z 873.5921     0   2   4   6   8E+3   3   3.5  RT (min)       lcqtof_hilic_neg_4_022_34    m/z 873.5927     0   2   4E+3   2.5   3  RT (min)      lcqtof_hilic_neg_1_013_38    m/z 873.5936     0   1   2   3   4E+3   2.5   3  RT (min)      lcqtof_hilic_neg_4_059_14    m/z 873.5939     0   2   4   6E+3   2   2.5  RT (min)      lcqtof_hilic_neg_4_011_35    m/z 873.5941     0   1   2   3   4E+3   2.5   3  RT (min)      lcqtof_hilic_neg_4_064_15    m/z 873.5949     0   1E+4   3   3.5  RT (min)      lcqtof_hilic_neg_4_059_14    m/z 873.5958     0   2   4   6   8E+3   2   2.5  RT (min)      lcqtof_hilic_neg_4_032_01    m/z 873.5961     0   2   4   6   8E+3   2   2.5  RT (min)       lcqtof_hilic_neg_1_054_01    m/z 873.5990     0   1E+4   1.5   2  RT (min)      lcqtof_hilic_neg_4_000_aa    m/z 873.6000     0   1E+4   1.5   2  RT (min)      lcqtof_hilic_neg_4_000_aa    m/z 873.6225     0   1E+4   1.5   2  RT (min)      lcqtof_hilic_neg_4_000_aa    m/z 873.6598     0   1   2   3   4E+4   1   1.5  RT (min)      lcqtof_hilic_neg_4_022_34    m/z 873.6605     0   1   2   3   4E+4   0.5   1  RT (min)      lcqtof_hilic_neg_4_022_34    m/z 873.6630     0   1E+4   2   2.5  RT (min)       lcqtof_hilic_neg_4_055_32    m/z 873.6652     0   1E+4   2.5   3  RT (min)      lcqtof_hilic_neg_1_005_32    m/z 873.7375     0   1E+4   2   2.5  RT (min)      lcqtof_hilic_neg_1_005_32    m/z 873.8317     0   1E+4   1.5   2  RT (min)      lcqtof_hilic_neg_4_000_aa    m/z 873.8595     0   1E+4   1.5   2  RT (min)      lcqtof_hilic_neg_4_000_aa    m/z 874.0013     0   1   2   3E+3   4.5   5  RT (min)      lcqtof_hilic_neg_4_022_34    m/z 874.3136     0   1E+4   0.5   1  RT (min)      lcqtof_hilic_neg_4_021_21    m/z 874.3281     0   1   2E+4   0.5   1  RT (min)      lcqtof_hilic_neg_4_015_09    m/z 874.4701     0   1E+4   0.5   1  RT (min)      lcqtof_hilic_neg_1_007_05    m/z 874.4906     0   1   2   3E+3   3   3.5  RT (min)      lcqtof_hilic_neg_2_058_15    m/z 874.4907     0   1   2   3E+3   3   3.5  RT (min)      lcqtof_hilic_neg_2_032_02    m/z 874.4908     0   2   4E+3   3   3.5  RT (min)      lcqtof_hilic_neg_2_038_04    m/z 874.5715     0   2   4   6E+3   4   4.5  RT (min)      lcqtof_hilic_neg_4_028_36    m/z 874.6055     0   2   4   6E+3   1   1.5  RT (min)       lcqtof_hilic_neg_4_019_03    m/z 874.6104     0   1   2E+4   1.5   2  RT (min)      lcqtof_hilic_neg_4_053_14    m/z 874.6115     0   2   4   6E+3   3   3.5  RT (min)       lcqtof_hilic_neg_4_006_30    m/z 874.6118     0   2   4   6E+3   2.5   3  RT (min)       lcqtof_hilic_neg_4_006_30    m/z 874.6644     0   1   2E+4   1   1.5  RT (min)      lcqtof_hilic_neg_4_022_34    m/z 874.6647     0   1   2E+4   0.5   1  RT (min)      lcqtof_hilic_neg_4_022_34    m/z 874.7384     0   2   4E+5   2.5   3  RT (min)       lcqtof_hilic_neg_4_008_24    m/z 874.7385     0   2   4   6E+5   2.5   3  RT (min)      lcqtof_hilic_neg_4_008_24    m/z 874.7386     0   2   4   6E+5   2.5   3  RT (min)       lcqtof_hilic_neg_4_008_24    m/z 874.7390     0   1   2   3E+4   2.5   3  RT (min)      lcqtof_hilic_neg_4_008_24    m/z 874.7490     0   1   2E+4   2   2.5  RT (min)      lcqtof_hilic_neg_4_040_05    m/z 874.7649     0   1   2E+4   0.5   1  RT (min)      lcqtof_hilic_neg_3_035_46    m/z 874.8636     0   1E+4   0.5   1  RT (min)      lcqtof_hilic_neg_3_035_46    m/z 875.0349     0   1E+4   3   3.5  RT (min)      lcqtof_hilic_neg_4_008_24    m/z 875.3102     0   1   2E+4   0.5   1  RT (min)      lcqtof_hilic_neg_4_021_21    m/z 875.3240     0   2   4   6E+4   0.5   1  RT (min)      lcqtof_hilic_neg_4_015_09    m/z 875.4689     0   1E+4   0.5   1  RT (min)      lcqtof_hilic_neg_2_028_05    m/z 875.7593     0   2   4   6E+4   2   2.5  RT (min)       lcqtof_hilic_neg_4_040_05    m/z 875.7605     0   1   2   3E+4   4   4.5  RT (min)      lcqtof_hilic_neg_4_040_05    m/z 875.8075     0   1   2E+4   1   1.5  RT (min)      lcqtof_hilic_neg_4_000_aa    m/z 875.8213     0   1E+4   1.5   2  RT (min)      lcqtof_hilic_neg_4_000_aa    m/z 875.8437     0   1E+4   1.5   2  RT (min)      lcqtof_hilic_neg_4_000_aa    m/z 876.1453     0   1   2   3E+4   0.5   1  RT (min)      lcqtof_hilic_neg_1_062_17    m/z 876.3117     0   1E+4   0.5   1  RT (min)      lcqtof_hilic_neg_4_021_21    m/z 876.3272     0   1   2E+4   0.5   1  RT (min)      lcqtof_hilic_neg_4_015_09    m/z 876.5031     0   1   2   3E+3   1   1.5  RT (min)      lcqtof_hilic_neg_1_063_13    m/z 876.5787     0   1   2   3E+4   2   2.5  RT (min)       lcqtof_hilic_neg_4_053_14    m/z 876.5961     0   1   2   3   4E+4   2.5   3  RT (min)      lcqtof_hilic_neg_4_055_32    m/z 876.6617     0   1   2E+6   4.5   5  RT (min)       lcqtof_hilic_neg_4_092_60    m/z 876.6663     0   1E+4   1   1.5  RT (min)      lcqtof_hilic_neg_4_000_aa    m/z 877.2126     0   1E+5   1   1.5  RT (min)      lcqtof_hilic_neg_4_000_aa    m/z 877.5823     0   1E+4   2   2.5  RT (min)      lcqtof_hilic_neg_4_053_14    m/z 877.6596     0   1E+5   1   1.5  RT (min)      lcqtof_hilic_neg_4_000_aa    m/z 877.6599     0   1E+5   1.5   2   2.5  RT (min)      lcqtof_hilic_neg_4_000_aa    m/z 877.6603     0   1E+5   2   2.5  RT (min)      lcqtof_hilic_neg_4_000_aa    m/z 877.6605     0   1E+5   1.5   2  RT (min)      lcqtof_hilic_neg_4_000_aa    m/z 877.6613     0   1E+5   1   1.5  RT (min)      lcqtof_hilic_neg_4_000_aa    m/z 877.6633     0   1E+5   1   1.5  RT (min)      lcqtof_hilic_neg_4_000_aa    m/z 877.6651     0   1E+5   2   2.5  RT (min)      lcqtof_hilic_neg_4_000_aa    m/z 877.6941     0   1   2E+5   4.5   5  RT (min)      lcqtof_hilic_neg_4_008_24    m/z 877.7882     0   1   2E+4   1   1.5  RT (min)      lcqtof_hilic_neg_4_000_aa    m/z 878.2136     0   2   4   6   8E+4   1   1.5  RT (min)      lcqtof_hilic_neg_4_000_aa    m/z 878.5897     0   1E+4   2   2.5  RT (min)      lcqtof_hilic_neg_4_052_13    m/z 878.5914     0   1   2   3E+4   2   2.5  RT (min)       lcqtof_hilic_neg_2_031_01    m/z 878.5917     0   1E+4   2.5   3  RT (min)       lcqtof_hilic_neg_1_010_15    m/z 878.5919     0   1   2   3E+4   1.5   2  RT (min)      lcqtof_hilic_neg_4_050_02    m/z 878.5919     0   1   2E+4   2   2.5  RT (min)      lcqtof_hilic_neg_4_003_16    m/z 878.6185     0   1   2   3E+4   0.5   1  RT (min)      lcqtof_hilic_neg_3_035_46    m/z 878.6622     0   1E+5   1   1.5  RT (min)      lcqtof_hilic_neg_4_000_aa    m/z 878.6630     0   1E+5   1   1.5   2  RT (min)      lcqtof_hilic_neg_4_000_aa    m/z 878.6632     0   2   4   6E+4   2   2.5  RT (min)      lcqtof_hilic_neg_4_000_aa    m/z 878.6635     0   1E+5   1   1.5  RT (min)      lcqtof_hilic_neg_4_000_aa    m/z 878.6636     0   2   4   6E+4   2   2.5  RT (min)      lcqtof_hilic_neg_4_000_aa    m/z 878.6637     0   2   4   6E+4   2   2.5  RT (min)      lcqtof_hilic_neg_4_000_aa    m/z 878.6646     0   2   4   6E+4   2   2.5  RT (min)      lcqtof_hilic_neg_4_000_aa    m/z 878.6652     0   1E+5   1   1.5  RT (min)      lcqtof_hilic_neg_4_000_aa    m/z 878.6784     0   1E+6   4.5   5  RT (min)      lcqtof_hilic_neg_4_092_60    m/z 878.7060     0   1   2E+4   2   2.5  RT (min)      lcqtof_hilic_neg_4_040_05    m/z 878.7499     0   1E+4   0.5   1  RT (min)      lcqtof_hilic_neg_1_022_36    m/z 878.7803     0   1E+4   1   1.5  RT (min)      lcqtof_hilic_neg_4_000_aa    m/z 879.2087     0   1E+4   4   4.5  RT (min)      lcqtof_hilic_neg_4_008_24    m/z 879.2112     0   2   4   6E+4   1   1.5  RT (min)      lcqtof_hilic_neg_4_000_aa    m/z 879.5893     0   1   2   3   4E+3   2   2.5  RT (min)      lcqtof_hilic_neg_4_053_14    m/z 879.5934     0   1E+4   1.5   2  RT (min)      lcqtof_hilic_neg_4_050_02    m/z 879.5952     0   1   2   3E+5   3   3.5  RT (min)      lcqtof_hilic_neg_1_054_01    m/z 879.5956     0   2   4   6   8E+3   2   2.5  RT (min)      lcqtof_hilic_neg_2_031_01    m/z 879.5961     0   2   4   6   8E+3   2   2.5  RT (min)       lcqtof_hilic_neg_4_005_29    m/z 879.7072     0   1   2E+4   4   4.5  RT (min)      lcqtof_hilic_neg_4_040_05    m/z 879.7080     0   1   2   3E+4   1   1.5  RT (min)      lcqtof_hilic_neg_4_040_05    m/z 879.7094     0   1E+4   4.5   5  RT (min)      lcqtof_hilic_neg_4_008_24    m/z 879.7633     0   1E+4   1   1.5  RT (min)      lcqtof_hilic_neg_4_000_aa    m/z 880.0181     0   1   2E+4   7.5   8  RT (min)      lcqtof_hilic_neg_4_016_37    m/z 880.1232     0   1   2   3E+4   0.5   1  RT (min)      lcqtof_hilic_neg_1_062_17    m/z 880.2109     0   1   2   3   4E+4   1   1.5  RT (min)      lcqtof_hilic_neg_4_000_aa    m/z 880.5077     0   1   2   3E+4   4.5   5  RT (min)       lcqtof_hilic_neg_4_048_23    m/z 880.6031     0   2   4   6E+3   1.5   2  RT (min)      lcqtof_hilic_neg_4_055_32    m/z 880.6157     0   1E+5   2.5   3  RT (min)      lcqtof_hilic_neg_4_055_32    m/z 880.6307     0   1E+4   0.5   1  RT (min)      lcqtof_hilic_neg_3_035_46    m/z 880.6651     0   1   2E+4   1   1.5  RT (min)      lcqtof_hilic_neg_4_000_aa    m/z 880.6660     0   1   2E+4   1   1.5  RT (min)      lcqtof_hilic_neg_4_000_aa    m/z 880.6688     0   1   2E+4   1   1.5  RT (min)      lcqtof_hilic_neg_4_000_aa    m/z 880.7067     0   1E+4   2   2.5  RT (min)      lcqtof_hilic_neg_4_040_05    m/z 880.7277     0   1   2E+4   2   2.5  RT (min)      lcqtof_hilic_neg_4_055_32    m/z 880.7337     0   1E+4   0.5   1  RT (min)      lcqtof_hilic_neg_4_034_31    m/z 881.1198     0   1   2E+4   0.5   1  RT (min)      lcqtof_hilic_neg_1_062_17    m/z 881.2109     0   1   2E+4   1   1.5  RT (min)      lcqtof_hilic_neg_4_000_aa    m/z 881.5113     0   1E+4   4.5   5  RT (min)      lcqtof_hilic_neg_4_048_23    m/z 881.5185     0   1E+4   0.5   1  RT (min)      lcqtof_hilic_neg_3_003_20    m/z 881.5648     0   1   2E+4   2.5   3  RT (min)      lcqtof_hilic_neg_4_048_23    m/z 881.5973     0   1E+4   3.5   4  RT (min)      lcqtof_hilic_neg_1_007_05    m/z 881.5992     0   1E+4   3.5   4  RT (min)      lcqtof_hilic_neg_4_040_05    m/z 881.6615     0   1   2   3E+4   1   1.5  RT (min)      lcqtof_hilic_neg_2_047_32    m/z 881.6746     0   1   2E+4   1   1.5  RT (min)      lcqtof_hilic_neg_4_000_aa    m/z 881.6854     0   1   2E+4   1   1.5  RT (min)      lcqtof_hilic_neg_4_000_aa    m/z 881.6895     0   1E+4   1.5   2  RT (min)      lcqtof_hilic_neg_4_000_aa    m/z 881.7001     0   1E+4   1.5   2  RT (min)      lcqtof_hilic_neg_4_000_aa    m/z 881.7051     0   1E+4   0.5   1  RT (min)      lcqtof_hilic_neg_4_047_06    m/z 881.7065     0   1   2   3E+4   2   2.5  RT (min)      lcqtof_hilic_neg_4_040_05    m/z 882.2091     0   1E+4   1   1.5  RT (min)      lcqtof_hilic_neg_4_000_aa    m/z 882.5069     0   1E+4   4.5   5  RT (min)      lcqtof_hilic_neg_4_048_23    m/z 882.5686     0   1E+4   2.5   3  RT (min)      lcqtof_hilic_neg_4_048_23    m/z 882.5687     0   1   2E+4   3   3.5  RT (min)      lcqtof_hilic_neg_1_003_37    m/z 882.6129     0   1E+4   2   2.5  RT (min)      lcqtof_hilic_neg_1_022_36    m/z 882.6523     0   1E+4   4   4.5  RT (min)      lcqtof_hilic_neg_4_040_05    m/z 882.6611     0   2   4   6   8E+3   1.5   2  RT (min)      lcqtof_hilic_neg_2_052_24    m/z 882.6699     0   1E+4   3   3.5  RT (min)       lcqtof_hilic_neg_4_008_24    m/z 882.6894     0   1E+4   0.5   1  RT (min)      lcqtof_hilic_neg_4_040_05    m/z 882.9714     0   1E+4   0.5   1  RT (min)      lcqtof_hilic_neg_4_000_aa    m/z 883.4549     0   1   2E+4   0.5   1  RT (min)      lcqtof_hilic_neg_3_053_26    m/z 883.4628     0   1E+5   0.5   1  RT (min)      lcqtof_hilic_neg_4_045_26    m/z 883.5787     0   1E+4   2.5   3  RT (min)      lcqtof_hilic_neg_4_087_59    m/z 883.5802     0   2   4E+4   3   3.5  RT (min)      lcqtof_hilic_neg_4_034_31    m/z 883.5806     0   2   4E+4   2.5   3  RT (min)       lcqtof_hilic_neg_4_048_23    m/z 883.6129     0   1   2E+4   3.5   4  RT (min)      lcqtof_hilic_neg_1_007_05    m/z 883.6140     0   1   2E+4   3.5   4  RT (min)      lcqtof_hilic_neg_1_007_05    m/z 883.6649     0   1E+4   2.5   3  RT (min)      lcqtof_hilic_neg_2_052_24    m/z 883.6649     0   1E+4   2.5   3  RT (min)      lcqtof_hilic_neg_2_052_24    m/z 883.6661     0   1E+4   2.5   3  RT (min)      lcqtof_hilic_neg_2_052_24    m/z 883.6661     0   1E+4   3   3.5  RT (min)      lcqtof_hilic_neg_4_095_60    m/z 883.6693     0   1E+4   3   3.5  RT (min)       lcqtof_hilic_neg_4_012_07    m/z 884.3773     0   1   2E+4   0.5   1  RT (min)      lcqtof_hilic_neg_3_003_20    m/z 884.4576     0   1   2   3   4E+4   0.5   1  RT (min)      lcqtof_hilic_neg_4_045_26    m/z 884.4929     0   1E+4   0.5   1  RT (min)      lcqtof_hilic_neg_4_015_09    m/z 884.5501     0   1E+4   2   2.5  RT (min)      lcqtof_hilic_neg_4_016_37    m/z 884.5830     0   1E+4   2.5   3  RT (min)      lcqtof_hilic_neg_4_082_59    m/z 884.6791     0   1E+4   1.5   2  RT (min)      lcqtof_hilic_neg_2_047_32    m/z 884.6798     0   1   2   3E+4   1   1.5  RT (min)      lcqtof_hilic_neg_4_008_24    m/z 884.6799     0   1E+4   1   1.5  RT (min)      lcqtof_hilic_neg_4_055_32    m/z 884.6816     0   1   2   3   4E+3   2.5   3  RT (min)      lcqtof_hilic_neg_1_037_24    m/z 884.6820     0   1E+4   3   3.5  RT (min)      lcqtof_hilic_neg_4_091_60    m/z 884.6920     0   1E+4   1   1.5  RT (min)      lcqtof_hilic_neg_2_047_32    m/z 884.7391     0   1E+4   2   2.5  RT (min)      lcqtof_hilic_neg_4_035_33    m/z 884.9724     0   1E+4   1   1.5  RT (min)      lcqtof_hilic_neg_4_000_aa    m/z 885.4389     0   1E+4   0.5   1  RT (min)      lcqtof_hilic_neg_4_005_29    m/z 885.4494     0   2   4   6   8E+4   0.5   1  RT (min)      lcqtof_hilic_neg_4_045_26    m/z 885.5943     0   1   2E+4   2.5   3  RT (min)      lcqtof_hilic_neg_4_050_02    m/z 885.6821     0   1   2   3E+4   2.5   3  RT (min)       lcqtof_hilic_neg_4_008_24    m/z 885.6823     0   1   2   3E+4   2.5   3  RT (min)      lcqtof_hilic_neg_4_008_24    m/z 885.6829     0   1   2   3E+4   2.5   3  RT (min)       lcqtof_hilic_neg_4_008_24    m/z 885.6830     0   1   2   3E+4   2.5   3  RT (min)      lcqtof_hilic_neg_4_008_24    m/z 885.6832     0   1   2   3E+4   2.5   3  RT (min)       lcqtof_hilic_neg_4_008_24    m/z 885.6833     0   1   2   3E+4   3   3.5  RT (min)       lcqtof_hilic_neg_4_008_24    m/z 885.6833     0   1   2   3E+4   2.5   3  RT (min)      lcqtof_hilic_neg_4_008_24    m/z 885.6837     0   2   4   6   8E+3   2   2.5  RT (min)       lcqtof_hilic_neg_4_097_60    m/z 885.6843     0   1   2   3E+4   3   3.5  RT (min)      lcqtof_hilic_neg_4_008_24    m/z 885.6848     0   2   4   6E+3   2   2.5  RT (min)      lcqtof_hilic_neg_4_047_06    m/z 885.6849     0   1   2   3E+4   3   3.5  RT (min)       lcqtof_hilic_neg_4_008_24    m/z 885.6853     0   1   2   3E+4   3   3.5  RT (min)      lcqtof_hilic_neg_4_008_24    m/z 885.7095     0   1E+4   3   3.5  RT (min)      lcqtof_hilic_neg_4_008_24    m/z 886.4512     0   1   2   3E+4   0.5   1  RT (min)      lcqtof_hilic_neg_4_045_26    m/z 886.5231     0   1E+4   3   3.5  RT (min)       lcqtof_hilic_neg_1_012_04    m/z 886.5251     0   2   4   6E+3   3.5   4  RT (min)      lcqtof_hilic_neg_4_050_02    m/z 886.5653     0   1E+4   2   2.5  RT (min)      lcqtof_hilic_neg_4_016_37    m/z 886.5952     0   1E+4   2.5   3  RT (min)      lcqtof_hilic_neg_4_050_02    m/z 886.6010     0   2   4   6   8E+4   2.5   3  RT (min)       lcqtof_hilic_neg_4_053_14    m/z 886.6565     0   1   2   3   4E+4   0.5   1  RT (min)      lcqtof_hilic_neg_1_007_05    m/z 886.6974     0   1   2E+4   1   1.5  RT (min)      lcqtof_hilic_neg_4_008_24    m/z 886.6994     0   2   4   6   8E+3   3   3.5  RT (min)      lcqtof_hilic_neg_4_094_60    m/z 886.7021     0   2   4   6E+3   3.5   4  RT (min)       lcqtof_hilic_neg_4_055_32    m/z 886.7106     0   1   2E+4   2   2.5  RT (min)      lcqtof_hilic_neg_1_037_24    m/z 886.7245     0   1E+4   2   2.5  RT (min)      lcqtof_hilic_neg_4_008_24    m/z 886.7394     0   1   2E+4   2   2.5  RT (min)      lcqtof_hilic_neg_4_055_32    m/z 886.7432     0   1E+4   2   2.5  RT (min)      lcqtof_hilic_neg_1_037_24    m/z 886.9703     0   1E+4   0.5   1  RT (min)      lcqtof_hilic_neg_4_000_aa    m/z 887.3258     0   1E+4   0.5   1  RT (min)      lcqtof_hilic_neg_4_021_21    m/z 887.3404     0   2   4E+4   0.5   1  RT (min)      lcqtof_hilic_neg_4_015_09    m/z 887.3915     0   1E+4   4   4.5  RT (min)      lcqtof_hilic_neg_4_032_01    m/z 887.5301     0   1E+4   0.5   1  RT (min)      lcqtof_hilic_neg_1_032_18    m/z 887.5801     0   1E+4   4.5   5  RT (min)      lcqtof_hilic_neg_2_028_05    m/z 887.5802     0   1E+4   4   4.5  RT (min)      lcqtof_hilic_neg_4_040_05    m/z 887.6082     0   1E+4   2.5   3  RT (min)      lcqtof_hilic_neg_4_076_58    m/z 887.6083     0   1E+4   2.5   3  RT (min)      lcqtof_hilic_neg_4_050_02    m/z 887.6085     0   2   4   6   8E+4   3   3.5  RT (min)      lcqtof_hilic_neg_4_034_31    m/z 887.6967     0   1E+4   3   3.5  RT (min)      lcqtof_hilic_neg_4_002_25    m/z 887.6990     0   1   2E+4   2.5   3  RT (min)       lcqtof_hilic_neg_4_008_24    m/z 887.6992     0   1   2E+4   3   3.5  RT (min)       lcqtof_hilic_neg_4_095_60    m/z 887.6995     0   1E+4   2.5   3  RT (min)      lcqtof_hilic_neg_4_002_25    m/z 887.7003     0   1E+4   3   3.5  RT (min)       lcqtof_hilic_neg_4_044_08    m/z 887.7004     0   2   4   6E+3   2   2.5  RT (min)      lcqtof_hilic_neg_4_092_60    m/z 887.7149     0   1E+4   3   3.5  RT (min)      lcqtof_hilic_neg_4_002_25    m/z 887.7368     0   1   2   3E+4   2   2.5  RT (min)      lcqtof_hilic_neg_4_008_24    m/z 887.7580     0   1E+4   2   2.5  RT (min)      lcqtof_hilic_neg_3_034_24    m/z 887.7582     0   1   2   3   4E+4   2   2.5  RT (min)       lcqtof_hilic_neg_4_055_32    m/z 887.8376     0   1E+4   1.5   2  RT (min)      lcqtof_hilic_neg_4_000_aa    m/z 887.8421     0   1E+4   1.5   2  RT (min)      lcqtof_hilic_neg_4_000_aa    m/z 887.8480     0   1E+4   1.5   2  RT (min)      lcqtof_hilic_neg_4_000_aa    m/z 887.9794     0   1E+4   6.5   7  RT (min)      lcqtof_hilic_neg_4_034_31    m/z 888.3267     0   1E+4   1   1.5  RT (min)      lcqtof_hilic_neg_4_027_19    m/z 888.3317     0   1E+5   0.5   1  RT (min)      lcqtof_hilic_neg_3_003_20    m/z 888.3433     0   1   2E+4   0.5   1  RT (min)      lcqtof_hilic_neg_4_015_09    m/z 888.5306     0   1E+4   4   4.5  RT (min)      lcqtof_hilic_neg_4_014_46    m/z 888.5350     0   2   4E+3   3   3.5  RT (min)      lcqtof_hilic_neg_1_012_04    m/z 888.5494     0   1E+5   3.5   4  RT (min)      lcqtof_hilic_neg_1_007_05    m/z 888.5798     0   1   2E+4   2   2.5  RT (min)      lcqtof_hilic_neg_4_016_37    m/z 888.5942     0   1E+4   4.5   5  RT (min)      lcqtof_hilic_neg_1_007_05    m/z 888.5974     0   1   2E+4   4   4.5  RT (min)      lcqtof_hilic_neg_4_008_24    m/z 888.5992     0   1E+4   2.5   3  RT (min)      lcqtof_hilic_neg_4_054_48    m/z 888.6015     0   1   2E+4   4   4.5  RT (min)      lcqtof_hilic_neg_4_008_24    m/z 888.7543     0   1   2E+4   2   2.5  RT (min)      lcqtof_hilic_neg_4_008_24    m/z 888.7549     0   1E+4   2   2.5  RT (min)      lcqtof_hilic_neg_2_047_32    m/z 889.3253     0   1   2E+4   0.5   1  RT (min)      lcqtof_hilic_neg_4_021_21    m/z 889.3391     0   2   4   6E+4   0.5   1  RT (min)      lcqtof_hilic_neg_4_015_09    m/z 889.5538     0   1   2E+4   3.5   4  RT (min)      lcqtof_hilic_neg_1_007_05    m/z 889.5546     0   1E+4   0.5   1  RT (min)      lcqtof_hilic_neg_4_000_aa    m/z 889.6121     0   1E+4   1   1.5  RT (min)      lcqtof_hilic_neg_4_000_aa    m/z 889.7118     0   1E+4   2.5   3  RT (min)      lcqtof_hilic_neg_4_008_24    m/z 889.7131     0   1E+4   2.5   3  RT (min)       lcqtof_hilic_neg_4_008_24    m/z 889.7136     0   1   2   3   4E+4   1   1.5  RT (min)       lcqtof_hilic_neg_4_008_24    m/z 889.7139     0   1   2   3E+4   1.5   2  RT (min)       lcqtof_hilic_neg_2_052_24    m/z 889.7148     0   1   2E+4   2.5   3  RT (min)       lcqtof_hilic_neg_4_008_24    m/z 889.7156     0   1E+4   1   1.5  RT (min)      lcqtof_hilic_neg_4_055_32    m/z 889.8262     0   1E+4   1   1.5  RT (min)      lcqtof_hilic_neg_4_000_aa    m/z 889.8350     0   1E+4   1.5   2  RT (min)      lcqtof_hilic_neg_4_000_aa    m/z 889.8378     0   1E+4   1.5   2  RT (min)      lcqtof_hilic_neg_4_000_aa    m/z 890.3294     0   2   4E+4   0.5   1  RT (min)      lcqtof_hilic_neg_3_003_20    m/z 890.3423     0   1   2E+4   0.5   1  RT (min)      lcqtof_hilic_neg_4_015_09    m/z 890.4506     0   1   2   3   4E+4   0.5   1  RT (min)      lcqtof_hilic_neg_4_027_19    m/z 890.5741     0   1   2E+4   3   3.5  RT (min)      lcqtof_hilic_neg_1_003_37    m/z 890.5766     0   1E+4   3   3.5  RT (min)      lcqtof_hilic_neg_2_029_31    m/z 890.5945     0   1E+4   1   1.5  RT (min)      lcqtof_hilic_neg_4_052_13    m/z 890.6057     0   1   2   3   4E+4   4.5   5  RT (min)      lcqtof_hilic_neg_1_007_05    m/z 890.6062     0   1   2E+4   4   4.5  RT (min)      lcqtof_hilic_neg_2_028_05    m/z 890.6131     0   1E+5   2.5   3  RT (min)      lcqtof_hilic_neg_4_055_32    m/z 890.7775     0   1E+4   0.5   1  RT (min)      lcqtof_hilic_neg_1_005_32    m/z 891.3363     0   1E+4   0.5   1  RT (min)      lcqtof_hilic_neg_3_003_20    m/z 891.3655     0   1E+4   0.5   1  RT (min)      lcqtof_hilic_neg_4_023_15    m/z 891.6093     0   1E+5   4.5   5  RT (min)       lcqtof_hilic_neg_1_007_05    m/z 891.6095     0   1E+5   4   4.5  RT (min)       lcqtof_hilic_neg_2_028_05    m/z 891.6173     0   1   2   3   4E+4   2.5   3  RT (min)      lcqtof_hilic_neg_4_055_32    m/z 891.7626     0   1   2   3E+4   1   1.5  RT (min)      lcqtof_hilic_neg_4_000_aa    m/z 891.7632     0   1   2   3E+4   1   1.5  RT (min)      lcqtof_hilic_neg_4_000_aa    m/z 891.7726     0   1E+4   1.5   2  RT (min)      lcqtof_hilic_neg_4_000_aa    m/z 891.9929     0   1E+4   6.5   7  RT (min)      lcqtof_hilic_neg_4_055_32    m/z 892.5524     0   1E+4   4   4.5  RT (min)      lcqtof_hilic_neg_1_007_05    m/z 892.5799     0   1E+4   3.5   4  RT (min)      lcqtof_hilic_neg_1_007_05    m/z 892.5888     0   1E+4   3.5   4  RT (min)      lcqtof_hilic_neg_2_028_05    m/z 892.6183     0   1E+4   4.5   5  RT (min)      lcqtof_hilic_neg_1_007_05    m/z 892.6259     0   2   4E+4   2.5   3  RT (min)      lcqtof_hilic_neg_4_055_32    m/z 892.7134     0   1E+4   4.5   5  RT (min)      lcqtof_hilic_neg_4_008_24    m/z 892.7509     0   1E+4   2   2.5  RT (min)      lcqtof_hilic_neg_4_040_05    m/z 892.7607     0   2   4   6E+4   0.5   1  RT (min)      lcqtof_hilic_neg_4_034_31    m/z 892.7735     0   1E+4   1   1.5  RT (min)      lcqtof_hilic_neg_4_000_aa    m/z 892.7858     0   1E+4   1   1.5  RT (min)      lcqtof_hilic_neg_4_000_aa    m/z 892.9758     0   1E+4   0.5   1  RT (min)      lcqtof_hilic_neg_4_000_aa    m/z 892.9764     0   1E+4   1   1.5  RT (min)      lcqtof_hilic_neg_4_000_aa    m/z 893.4954     0   1   2E+4   3.5   4  RT (min)      lcqtof_hilic_neg_3_066_57    m/z 893.6242     0   1   2   3   4E+4   4.5   5  RT (min)      lcqtof_hilic_neg_1_007_05    m/z 893.6246     0   1   2   3   4E+4   4   4.5  RT (min)      lcqtof_hilic_neg_2_028_05    m/z 893.6344     0   1E+4   2.5   3  RT (min)      lcqtof_hilic_neg_4_055_32    m/z 893.7443     0   1   2   3   4E+4   0.5   1  RT (min)      lcqtof_hilic_neg_4_047_06    m/z 893.8242     0   1   2E+4   0.5   1  RT (min)      lcqtof_hilic_neg_3_035_46    m/z 894.2015     0   1E+4   6.5   7  RT (min)      lcqtof_hilic_neg_4_008_24    m/z 894.5546     0   2   4   6E+3   2.5   3  RT (min)       lcqtof_hilic_neg_4_006_30    m/z 894.5568     0   1   2   3   4E+3   2.5   3  RT (min)      lcqtof_hilic_neg_1_014_31    m/z 894.5574     0   1   2   3E+3   2   2.5  RT (min)      lcqtof_hilic_neg_1_022_36    m/z 894.5584     0   1   2   3E+3   1.5   2  RT (min)      lcqtof_hilic_neg_4_011_35    m/z 894.5592     0   1   2   3E+3   3   3.5  RT (min)      lcqtof_hilic_neg_4_059_14    m/z 894.5782     0   2   4   6E+3   1   1.5  RT (min)      lcqtof_hilic_neg_4_052_13    m/z 894.6508     0   1   2E+4   8   8.5  RT (min)      lcqtof_hilic_neg_1_000_aa    m/z 894.7745     0   1   2   3E+4   0.5   1  RT (min)      lcqtof_hilic_neg_4_047_06    m/z 894.8294     0   2   4   6   8E+4   0.5   1  RT (min)      lcqtof_hilic_neg_3_035_46    m/z 894.9703     0   1E+4   1   1.5  RT (min)      lcqtof_hilic_neg_4_000_aa    m/z 895.5347     0   1E+4   0.5   1  RT (min)      lcqtof_hilic_neg_3_003_20    m/z 895.5646     0   2   4   6E+3   1   1.5  RT (min)      lcqtof_hilic_neg_4_050_02    m/z 895.5778     0   1E+4   2.5   3  RT (min)       lcqtof_hilic_neg_4_048_23    m/z 895.5792     0   1E+4   1   1.5  RT (min)      lcqtof_hilic_neg_4_000_aa    m/z 895.5800     0   2   4   6E+3   1.5   2  RT (min)      lcqtof_hilic_neg_1_008_13    m/z 895.5811     0   1E+4   1   1.5  RT (min)      lcqtof_hilic_neg_4_000_aa    m/z 895.6409     0   1E+4   4.5   5  RT (min)      lcqtof_hilic_neg_2_028_05    m/z 895.7710     0   1E+4   0.5   1  RT (min)      lcqtof_hilic_neg_4_008_24    m/z 896.5765     0   1E+4   1   1.5  RT (min)       lcqtof_hilic_neg_4_064_15    m/z 896.5809     0   2   4   6E+3   2.5   3  RT (min)       lcqtof_hilic_neg_4_048_23    m/z 896.5837     0   1   2E+4   2   2.5  RT (min)      lcqtof_hilic_neg_4_002_25    m/z 896.5897     0   2   4   6   8E+3   1   1.5  RT (min)      lcqtof_hilic_neg_2_063_13    m/z 896.6343     0   1E+4   8   8.5  RT (min)      lcqtof_hilic_neg_1_000_aa    m/z 896.6999     0   1E+4   2   2.5  RT (min)       lcqtof_hilic_neg_4_040_05    m/z 896.7028     0   1E+4   2   2.5  RT (min)      lcqtof_hilic_neg_4_055_32    m/z 896.7728     0   2   4E+4   0.5   1  RT (min)      lcqtof_hilic_neg_4_008_24    m/z 896.9739     0   1E+4   0.5   1  RT (min)      lcqtof_hilic_neg_4_000_aa    m/z 897.1545     0   1   2   3E+4   0.5   1  RT (min)      lcqtof_hilic_neg_1_062_17    m/z 897.3737     0   1E+4   1   1.5  RT (min)      lcqtof_hilic_neg_4_000_aa    m/z 897.4127     0   1E+4   3.5   4  RT (min)      lcqtof_hilic_neg_4_050_02    m/z 897.5449     0   1E+4   1   1.5  RT (min)      lcqtof_hilic_neg_4_000_aa    m/z 897.5792     0   1E+4   1   1.5  RT (min)      lcqtof_hilic_neg_4_050_02    m/z 897.5957     0   1   2E+4   2.5   3  RT (min)       lcqtof_hilic_neg_4_048_23    m/z 897.5959     0   2   4   6   8E+3   2   2.5  RT (min)       lcqtof_hilic_neg_4_016_37    m/z 897.5969     0   2   4E+3   1   1.5  RT (min)      lcqtof_hilic_neg_1_010_15    m/z 897.5978     0   1   2   3   4E+3   1   1.5  RT (min)      lcqtof_hilic_neg_4_026_04    m/z 897.5982     0   2   4E+3   2   2.5  RT (min)      lcqtof_hilic_neg_2_042_27    m/z 897.6171     0   1E+4   2   2.5  RT (min)      lcqtof_hilic_neg_4_053_14    m/z 897.6548     0   1   2E+4   1.5   2  RT (min)       lcqtof_hilic_neg_4_091_60    m/z 897.6923     0   1E+4   0.5   1  RT (min)      lcqtof_hilic_neg_4_013_20    m/z 897.7824     0   1   2E+4   0.5   1  RT (min)      lcqtof_hilic_neg_4_008_24    m/z 897.8155     0   1   2E+4   0.5   1  RT (min)      lcqtof_hilic_neg_4_055_32    m/z 897.9504     0   1E+4   0.5   1  RT (min)      lcqtof_hilic_neg_4_004_47    m/z 898.3575     0   1   2E+5   0.5   1  RT (min)      lcqtof_hilic_neg_3_003_20    m/z 898.5700     0   1E+4   3   3.5  RT (min)       lcqtof_hilic_neg_4_032_01    m/z 898.5838     0   1E+4   3   3.5  RT (min)      lcqtof_hilic_neg_4_006_30    m/z 898.5903     0   2   4   6E+3   1   1.5  RT (min)      lcqtof_hilic_neg_4_005_29    m/z 898.5983     0   1E+4   3   3.5  RT (min)      lcqtof_hilic_neg_4_034_31    m/z 898.5992     0   2   4E+3   2   2.5  RT (min)      lcqtof_hilic_neg_2_055_30    m/z 898.6029     0   2   4   6   8E+3   2   2.5  RT (min)      lcqtof_hilic_neg_4_053_14    m/z 898.6033     0   2   4E+3   2   2.5  RT (min)      lcqtof_hilic_neg_2_060_14    m/z 898.6357     0   1E+4   2   2.5  RT (min)      lcqtof_hilic_neg_4_016_37    m/z 898.6813     0   1   2E+4   0.5   1  RT (min)      lcqtof_hilic_neg_4_016_37    m/z 898.7896     0   1   2   3   4E+4   0.5   1  RT (min)      lcqtof_hilic_neg_4_008_24    m/z 899.0468     0   1E+4   1   1.5  RT (min)      lcqtof_hilic_neg_4_000_aa    m/z 899.3610     0   1E+5   0.5   1  RT (min)      lcqtof_hilic_neg_3_003_20    m/z 899.4532     0   1E+4   0.5   1  RT (min)      lcqtof_hilic_neg_4_005_29    m/z 899.5897     0   1E+4   1.5   2  RT (min)      lcqtof_hilic_neg_1_014_31    m/z 899.6064     0   2   4   6E+3   2.5   3  RT (min)      lcqtof_hilic_neg_4_054_48    m/z 899.6107     0   1   2   3E+3   2   2.5  RT (min)      lcqtof_hilic_neg_1_027_27    m/z 899.6146     0   2   4   6E+3   1.5   2  RT (min)      lcqtof_hilic_neg_1_054_01    m/z 899.6157     0   2   4   6E+3   1.5   2  RT (min)      lcqtof_hilic_neg_1_022_36    m/z 899.6699     0   1E+4   2   2.5  RT (min)      lcqtof_hilic_neg_4_055_32    m/z 899.7933     0   1E+4   0.5   1  RT (min)      lcqtof_hilic_neg_4_008_24    m/z 900.3662     0   1   2   3   4E+4   0.5   1  RT (min)      lcqtof_hilic_neg_3_003_20    m/z 900.4670     0   1   2E+4   0.5   1  RT (min)      lcqtof_hilic_neg_4_045_26    m/z 900.5035     0   1   2E+3   1   1.5  RT (min)      lcqtof_hilic_neg_1_062_17    m/z 900.5763     0   1E+4   3   3.5  RT (min)      lcqtof_hilic_neg_4_014_46    m/z 900.5974     0   1E+4   2.5   3  RT (min)      lcqtof_hilic_neg_4_055_32    m/z 900.6133     0   1   2   3E+4   3   3.5  RT (min)      lcqtof_hilic_neg_4_028_36    m/z 900.9819     0   1E+4   0.5   1  RT (min)      lcqtof_hilic_neg_4_000_aa    m/z 901.6277     0   2   4E+3   2   2.5  RT (min)      lcqtof_hilic_neg_2_031_01    m/z 901.6995     0   1   2E+4   2   2.5  RT (min)      lcqtof_hilic_neg_4_008_24    m/z 901.7737     0   1E+4   1   1.5  RT (min)      lcqtof_hilic_neg_4_055_32    m/z 902.0037     0   1E+4   7.5   8  RT (min)      lcqtof_hilic_neg_4_022_34    m/z 902.4916     0   2   4   6E+3   1   1.5  RT (min)      lcqtof_hilic_neg_2_064_16    m/z 902.4934     0   1   2   3   4E+3   2.5   3  RT (min)       lcqtof_hilic_neg_2_062_17    m/z 902.4946     0   1   2   3   4E+3   2.5   3  RT (min)       lcqtof_hilic_neg_2_064_16    m/z 902.4946     0   2   4   6E+3   1   1.5  RT (min)      lcqtof_hilic_neg_1_054_01    m/z 902.4960     0   1   2   3E+3   2   2.5  RT (min)      lcqtof_hilic_neg_1_008_13    m/z 902.4986     0   1   2   3E+3   1.5   2  RT (min)      lcqtof_hilic_neg_3_068_57    m/z 902.5023     0   1   2   3   4E+3   3   3.5  RT (min)      lcqtof_hilic_neg_1_063_13    m/z 902.5926     0   1E+4   2   2.5  RT (min)       lcqtof_hilic_neg_4_052_13    m/z 902.7679     0   1E+4   7.5   8  RT (min)      lcqtof_hilic_neg_1_000_aa    m/z 902.9254     0   1E+4   0.5   1  RT (min)      lcqtof_hilic_neg_4_008_24    m/z 903.0910     0   1E+4   0.5   1  RT (min)      lcqtof_hilic_neg_4_004_47    m/z 903.3553     0   1   2   3E+4   0.5   1  RT (min)      lcqtof_hilic_neg_1_032_18    m/z 903.4931     0   1   2   3E+3   1   1.5  RT (min)      lcqtof_hilic_neg_1_064_16    m/z 903.6135     0   1E+4   2.5   3  RT (min)      lcqtof_hilic_neg_4_055_32    m/z 903.7006     0   2   4   6E+4   2   2.5  RT (min)      lcqtof_hilic_neg_4_008_24    m/z 903.7047     0   1E+4   2   2.5  RT (min)      lcqtof_hilic_neg_4_002_25    m/z 903.8279     0   1E+4   1.5   2  RT (min)      lcqtof_hilic_neg_4_000_aa    m/z 903.8460     0   1E+4   1.5   2  RT (min)      lcqtof_hilic_neg_4_000_aa    m/z 904.3519     0   1   2E+4   0.5   1  RT (min)      lcqtof_hilic_neg_3_003_20    m/z 904.3581     0   1E+4   0.5   1  RT (min)      lcqtof_hilic_neg_1_032_18    m/z 904.5007     0   1   2   3E+3   1   1.5  RT (min)      lcqtof_hilic_neg_2_064_16    m/z 904.6079     0   1E+4   2   2.5  RT (min)      lcqtof_hilic_neg_4_032_01    m/z 904.6096     0   1E+4   2   2.5  RT (min)      lcqtof_hilic_neg_4_053_14    m/z 904.8333     0   1E+4   4   4.5  RT (min)      lcqtof_hilic_neg_4_008_24    m/z 904.9717     0   1E+4   0.5   1  RT (min)      lcqtof_hilic_neg_4_000_aa    m/z 904.9903     0   2   4   6E+4   4   4.5  RT (min)      lcqtof_hilic_neg_4_008_24    m/z 905.2255     0   1E+4   4   4.5  RT (min)      lcqtof_hilic_neg_4_055_32    m/z 905.5623     0   1E+4   2.5   3  RT (min)      lcqtof_hilic_neg_4_088_59    m/z 905.5643     0   2   4   6E+4   2.5   3  RT (min)      lcqtof_hilic_neg_4_048_23    m/z 905.6135     0   1E+4   1.5   2  RT (min)      lcqtof_hilic_neg_4_050_02    m/z 905.6290     0   1E+4   2.5   3  RT (min)      lcqtof_hilic_neg_4_055_32    m/z 905.6333     0   1E+4   3   3.5  RT (min)      lcqtof_hilic_neg_2_047_32    m/z 905.7981     0   1   2E+4   1   1.5  RT (min)      lcqtof_hilic_neg_4_000_aa    m/z 905.8122     0   1E+4   1   1.5  RT (min)      lcqtof_hilic_neg_4_000_aa    m/z 906.3045     0   1   2E+4   0.5   1  RT (min)       lcqtof_hilic_neg_4_106_62    m/z 906.3422     0   1E+4   0.5   1  RT (min)      lcqtof_hilic_neg_3_003_20    m/z 906.5671     0   1   2   3E+4   3   3.5  RT (min)      lcqtof_hilic_neg_4_048_23    m/z 906.6601     0   1E+4   0.5   1  RT (min)      lcqtof_hilic_neg_4_053_14    m/z 906.6987     0   1   2E+6   4   4.5  RT (min)      lcqtof_hilic_neg_4_055_32    m/z 906.8496     0   1   2   3E+4   4   4.5  RT (min)      lcqtof_hilic_neg_4_008_24    m/z 906.9496     0   1   2E+4   0.5   1  RT (min)       lcqtof_hilic_neg_4_099_61    m/z 906.9750     0   1E+4   1   1.5  RT (min)      lcqtof_hilic_neg_4_000_aa    m/z 907.1486     0   1   2E+4   4   4.5  RT (min)      lcqtof_hilic_neg_4_008_24    m/z 907.2426     0   1   2   3   4E+4   4   4.5  RT (min)      lcqtof_hilic_neg_4_008_24    m/z 907.4570     0   1E+4   0.5   1  RT (min)      lcqtof_hilic_neg_1_007_05    m/z 907.5472     0   1E+4   1   1.5  RT (min)      lcqtof_hilic_neg_4_000_aa    m/z 907.5781     0   2   4E+4   2.5   3  RT (min)       lcqtof_hilic_neg_4_087_59    m/z 907.5797     0   1E+5   2.5   3  RT (min)       lcqtof_hilic_neg_4_048_23    m/z 907.6279     0   1E+4   4   4.5  RT (min)      lcqtof_hilic_neg_4_035_33    m/z 907.6361     0   1   2E+4   1   1.5  RT (min)      lcqtof_hilic_neg_4_000_aa    m/z 907.6419     0   2   4E+4   4   4.5  RT (min)      lcqtof_hilic_neg_4_008_24    m/z 907.7258     0   1   2E+4   1   1.5  RT (min)      lcqtof_hilic_neg_4_000_aa    m/z 907.7507     0   1   2E+4   1   1.5  RT (min)      lcqtof_hilic_neg_4_000_aa    m/z 908.5833     0   2   4   6   8E+4   2.5   3  RT (min)      lcqtof_hilic_neg_4_048_23    m/z 908.5836     0   1   2E+4   2.5   3  RT (min)      lcqtof_hilic_neg_4_087_59    m/z 908.6450     0   1E+4   2.5   3  RT (min)      lcqtof_hilic_neg_4_055_32    m/z 908.6576     0   1   2E+4   1   1.5  RT (min)      lcqtof_hilic_neg_4_000_aa    m/z 908.6650     0   1   2E+4   1   1.5  RT (min)      lcqtof_hilic_neg_4_000_aa    m/z 908.7324     0   1E+4   1   1.5  RT (min)      lcqtof_hilic_neg_4_000_aa    m/z 909.4465     0   1E+4   0.5   1  RT (min)      lcqtof_hilic_neg_1_007_05    m/z 909.4657     0   1   2E+4   0.5   1  RT (min)      lcqtof_hilic_neg_4_045_26    m/z 909.5945     0   2   4E+4   2.5   3  RT (min)       lcqtof_hilic_neg_4_083_59    m/z 909.5948     0   1E+4   2.5   3  RT (min)      lcqtof_hilic_neg_4_050_02    m/z 909.6193     0   2   4   6E+4   3.5   4  RT (min)      lcqtof_hilic_neg_1_007_05    m/z 909.6197     0   2   4   6E+4   3.5   4  RT (min)      lcqtof_hilic_neg_1_007_05    m/z 909.6562     0   1E+4   4   4.5  RT (min)      lcqtof_hilic_neg_4_008_24    m/z 909.6779     0   1   2   3E+4   1.5   2  RT (min)      lcqtof_hilic_neg_4_000_aa    m/z 909.6784     0   1E+4   2   2.5  RT (min)      lcqtof_hilic_neg_4_000_aa    m/z 909.6787     0   1   2   3E+4   1.5   2  RT (min)      lcqtof_hilic_neg_4_000_aa    m/z 909.6807     0   1E+4   1.5   2  RT (min)      lcqtof_hilic_neg_4_000_aa    m/z 909.6822     0   1E+4   2   2.5  RT (min)      lcqtof_hilic_neg_4_008_24    m/z 909.6841     0   1E+4   1.5   2  RT (min)      lcqtof_hilic_neg_4_000_aa    m/z 909.6891     0   1E+4   1.5   2  RT (min)      lcqtof_hilic_neg_4_000_aa    m/z 910.1143     0   1E+4   6   6.5  RT (min)      lcqtof_hilic_neg_2_020_23    m/z 910.5978     0   1   2E+4   2.5   3  RT (min)       lcqtof_hilic_neg_4_084_59    m/z 910.6925     0   1E+4   1   1.5  RT (min)      lcqtof_hilic_neg_4_000_aa    m/z 910.7407     0   1E+4   0.5   1  RT (min)      lcqtof_hilic_neg_4_016_37    m/z 910.9791     0   1E+4   0.5   1  RT (min)      lcqtof_hilic_neg_4_000_aa    m/z 911.3069     0   1   2E+4   1   1.5  RT (min)      lcqtof_hilic_neg_4_031_38    m/z 911.4431     0   1E+4   0.5   1  RT (min)      lcqtof_hilic_neg_1_007_05    m/z 911.4637     0   1E+4   0.5   1  RT (min)      lcqtof_hilic_neg_4_005_29    m/z 911.6091     0   1   2   3E+4   2.5   3  RT (min)       lcqtof_hilic_neg_4_084_59    m/z 911.6340     0   1   2E+4   3.5   4  RT (min)      lcqtof_hilic_neg_2_028_05    m/z 911.6343     0   1   2E+4   3.5   4  RT (min)      lcqtof_hilic_neg_2_028_05    m/z 911.8303     0   1   2E+4   0.5   1  RT (min)      lcqtof_hilic_neg_4_008_24    m/z 912.3736     0   1   2   3E+5   0.5   1  RT (min)      lcqtof_hilic_neg_3_003_20    m/z 912.4774     0   1   2   3E+4   0.5   1  RT (min)      lcqtof_hilic_neg_4_045_26    m/z 912.5583     0   1   2E+4   3   3.5  RT (min)       lcqtof_hilic_neg_1_035_23    m/z 912.6144     0   1E+4   2.5   3  RT (min)       lcqtof_hilic_neg_4_084_59    m/z 912.6280     0   1E+4   0.5   1  RT (min)      lcqtof_hilic_neg_1_007_05    m/z 912.6477     0   1E+4   0.5   1  RT (min)      lcqtof_hilic_neg_4_045_26    m/z 912.7131     0   2   4E+3   3   3.5  RT (min)       lcqtof_hilic_neg_2_014_08    m/z 912.7160     0   2   4   6E+3   2.5   3  RT (min)       lcqtof_hilic_neg_4_008_24    m/z 912.7161     0   2   4   6E+3   2.5   3  RT (min)       lcqtof_hilic_neg_4_008_24    m/z 912.7163     0   2   4E+3   3   3.5  RT (min)      lcqtof_hilic_neg_4_047_06    m/z 912.7165     0   2   4   6E+3   2.5   3  RT (min)      lcqtof_hilic_neg_2_052_24    m/z 912.7471     0   1E+4   2   2.5  RT (min)      lcqtof_hilic_neg_1_005_32    m/z 912.8378     0   1   2   3E+4   0.5   1  RT (min)      lcqtof_hilic_neg_4_047_06    m/z 912.8603     0   1E+4   0.5   1  RT (min)      lcqtof_hilic_neg_4_004_47    m/z 912.9843     0   1E+4   0.5   1  RT (min)      lcqtof_hilic_neg_4_000_aa    m/z 913.3484     0   1   2   3E+5   1   1.5  RT (min)      lcqtof_hilic_neg_4_013_20    m/z 913.3773     0   1E+5   0.5   1  RT (min)      lcqtof_hilic_neg_3_003_20    m/z 913.4797     0   1   2   3   4E+4   0.5   1  RT (min)      lcqtof_hilic_neg_4_045_26    m/z 913.5799     0   1E+4   1   1.5  RT (min)      lcqtof_hilic_neg_4_000_aa    m/z 913.6255     0   2   4   6   8E+4   2.5   3  RT (min)      lcqtof_hilic_neg_4_053_14    m/z 913.6295     0   1E+4   2.5   3  RT (min)      lcqtof_hilic_neg_4_050_02    m/z 913.6359     0   1E+4   1   1.5  RT (min)      lcqtof_hilic_neg_4_000_aa    m/z 913.6497     0   1E+4   3.5   4  RT (min)      lcqtof_hilic_neg_1_007_05    m/z 913.6514     0   1E+4   3.5   4  RT (min)      lcqtof_hilic_neg_1_007_05    m/z 913.8075     0   1   2   3E+4   0.5   1  RT (min)      lcqtof_hilic_neg_4_055_32    m/z 914.3828     0   1   2   3   4E+4   0.5   1  RT (min)      lcqtof_hilic_neg_3_003_20    m/z 914.4832     0   1E+4   0.5   1  RT (min)      lcqtof_hilic_neg_4_045_26    m/z 914.5953     0   1E+4   1   1.5  RT (min)      lcqtof_hilic_neg_4_050_02    m/z 914.6254     0   1E+4   0.5   1  RT (min)      lcqtof_hilic_neg_1_007_05    m/z 914.6845     0   1   2E+4   3   3.5  RT (min)      lcqtof_hilic_neg_4_008_24    m/z 914.8117     0   2   4E+4   0.5   1  RT (min)      lcqtof_hilic_neg_4_055_32    m/z 915.3473     0   1   2E+4   1   1.5  RT (min)      lcqtof_hilic_neg_4_018_27    m/z 915.3511     0   1E+5   0.5   1  RT (min)      lcqtof_hilic_neg_3_003_20    m/z 915.4601     0   1E+4   7   7.5  RT (min)       lcqtof_hilic_neg_4_094_60    m/z 915.6674     0   1E+4   3   3.5  RT (min)      lcqtof_hilic_neg_1_037_24    m/z 915.6805     0   1E+4   0.5   1  RT (min)      lcqtof_hilic_neg_3_040_14    m/z 915.6902     0   1   2   3   4E+4   3   3.5  RT (min)      lcqtof_hilic_neg_4_008_24    m/z 915.6918     0   1   2   3   4E+4   3   3.5  RT (min)      lcqtof_hilic_neg_4_008_24    m/z 915.8862     0   1E+4   1.5   2  RT (min)      lcqtof_hilic_neg_4_000_aa    m/z 916.1828     0   1E+4   6.5   7  RT (min)      lcqtof_hilic_neg_4_008_24    m/z 916.6855     0   1E+4   0.5   1  RT (min)      lcqtof_hilic_neg_1_007_05    m/z 916.7029     0   1E+5   3.5   4  RT (min)      lcqtof_hilic_neg_4_008_24    m/z 916.7178     0   1E+4   2   2.5  RT (min)      lcqtof_hilic_neg_4_008_24    m/z 916.7213     0   1E+4   3   3.5  RT (min)      lcqtof_hilic_neg_3_034_24    m/z 916.7500     0   1E+4   1   1.5  RT (min)       lcqtof_hilic_neg_2_047_32    m/z 916.9787     0   1E+4   0.5   1  RT (min)      lcqtof_hilic_neg_4_000_aa    m/z 917.3588     0   1E+4   0.5   1  RT (min)      lcqtof_hilic_neg_3_003_20    m/z 917.5395     0   1E+4   4   4.5  RT (min)      lcqtof_hilic_neg_4_035_33    m/z 917.5765     0   1E+4   1   1.5  RT (min)      lcqtof_hilic_neg_4_011_35    m/z 917.6316     0   1E+4   2.5   3  RT (min)      lcqtof_hilic_neg_4_055_32    m/z 917.6330     0   1E+4   2.5   3  RT (min)      lcqtof_hilic_neg_4_055_32    m/z 917.7070     0   1E+4   3   3.5  RT (min)      lcqtof_hilic_neg_4_008_24    m/z 917.8072     0   1E+4   1   1.5  RT (min)      lcqtof_hilic_neg_4_000_aa    m/z 918.5431     0   1E+4   4   4.5  RT (min)      lcqtof_hilic_neg_4_016_37    m/z 918.5539     0   2   4E+3   1.5   2  RT (min)      lcqtof_hilic_neg_1_008_13    m/z 918.6216     0   1E+4   1   1.5  RT (min)      lcqtof_hilic_neg_4_052_13    m/z 918.6449     0   2   4E+4   2.5   3  RT (min)      lcqtof_hilic_neg_4_055_32    m/z 918.6794     0   1E+4   0.5   1  RT (min)      lcqtof_hilic_neg_2_028_05    m/z 918.7094     0   1   2   3E+4   3   3.5  RT (min)      lcqtof_hilic_neg_4_008_24    m/z 918.7110     0   1   2   3E+4   3   3.5  RT (min)      lcqtof_hilic_neg_4_008_24    m/z 918.7378     0   1E+4   1.5   2  RT (min)      lcqtof_hilic_neg_4_008_24    m/z 919.3644     0   1   2   3   4E+4   0.5   1  RT (min)      lcqtof_hilic_neg_3_035_46    m/z 919.3772     0   1E+4   1   1.5  RT (min)      lcqtof_hilic_neg_4_000_aa    m/z 919.3974     0   1E+4   3.5   4  RT (min)      lcqtof_hilic_neg_4_050_02    m/z 919.5769     0   1E+4   1   1.5  RT (min)      lcqtof_hilic_neg_3_035_46    m/z 919.6328     0   1E+4   0.5   1  RT (min)      lcqtof_hilic_neg_2_028_05    m/z 919.6506     0   1   2E+4   2.5   3  RT (min)      lcqtof_hilic_neg_4_055_32    m/z 919.7067     0   1   2   3E+4   15.5   16  RT (min)      lcqtof_hilic_neg_4_000_aa    m/z 919.8242     0   1E+4   1   1.5  RT (min)      lcqtof_hilic_neg_4_000_aa    m/z 920.3681     0   1   2E+4   0.5   1  RT (min)      lcqtof_hilic_neg_3_035_46    m/z 920.4068     0   1E+4   3.5   4  RT (min)      lcqtof_hilic_neg_4_050_02    m/z 920.5958     0   2   4E+3   2.5   3  RT (min)      lcqtof_hilic_neg_2_062_17    m/z 920.6005     0   1E+4   2   2.5  RT (min)      lcqtof_hilic_neg_1_008_13    m/z 920.6042     0   1E+4   1.5   2  RT (min)      lcqtof_hilic_neg_4_052_13    m/z 920.6047     0   1E+4   1.5   2  RT (min)      lcqtof_hilic_neg_4_052_13    m/z 920.6088     0   1E+4   1   1.5  RT (min)      lcqtof_hilic_neg_4_034_31    m/z 920.6348     0   1   2   3   4E+4   0.5   1  RT (min)      lcqtof_hilic_neg_1_007_05    m/z 920.6826     0   1E+4   4   4.5  RT (min)      lcqtof_hilic_neg_4_008_24    m/z 920.7085     0   1   2E+4   15.5   16  RT (min)      lcqtof_hilic_neg_4_000_aa    m/z 920.7086     0   1E+4   3.5   4  RT (min)      lcqtof_hilic_neg_4_008_24    m/z 920.7087     0   1E+4   3.5   4  RT (min)      lcqtof_hilic_neg_4_008_24    m/z 920.7100     0   1   2E+4   15.5   16  RT (min)      lcqtof_hilic_neg_4_000_aa    m/z 920.8859     0   1   2   3E+4   0.5   1  RT (min)      lcqtof_hilic_neg_3_035_46    m/z 920.9645     0   1E+4   0.5   1  RT (min)       lcqtof_hilic_neg_4_099_61    m/z 920.9741     0   1E+4   1   1.5  RT (min)      lcqtof_hilic_neg_4_000_aa    m/z 921.1974     0   1E+4   4   4.5  RT (min)      lcqtof_hilic_neg_4_008_24    m/z 921.3660     0   1E+4   0.5   1  RT (min)      lcqtof_hilic_neg_3_035_46    m/z 921.5941     0   1   2   3   4E+4   2.5   3  RT (min)       lcqtof_hilic_neg_4_048_23    m/z 921.6013     0   1E+4   2.5   3  RT (min)      lcqtof_hilic_neg_4_089_59    m/z 921.6290     0   1E+4   0.5   1  RT (min)      lcqtof_hilic_neg_2_028_05    m/z 921.7073     0   1E+4   15.5   16  RT (min)      lcqtof_hilic_neg_4_000_aa    m/z 921.7085     0   1E+4   15.5   16  RT (min)      lcqtof_hilic_neg_4_000_aa    m/z 921.8898     0   1   2E+4   0.5   1  RT (min)      lcqtof_hilic_neg_3_035_46    m/z 922.2955     0   1   2E+4   1   1.5  RT (min)      lcqtof_hilic_neg_4_010_28    m/z 922.5826     0   1E+4   1.5   2  RT (min)      lcqtof_hilic_neg_1_064_16    m/z 922.5981     0   1   2E+4   2.5   3  RT (min)      lcqtof_hilic_neg_4_048_23    m/z 922.6019     0   1   2   3E+3   2.5   3  RT (min)      lcqtof_hilic_neg_2_062_17    m/z 922.6318     0   2   4E+4   0.5   1  RT (min)      lcqtof_hilic_neg_1_007_05    m/z 922.6467     0   1   2E+4   4.5   5  RT (min)       lcqtof_hilic_neg_4_008_24    m/z 922.6790     0   1E+4   5   5.5  RT (min)      lcqtof_hilic_neg_4_048_23    m/z 922.6873     0   1   2E+4   4   4.5  RT (min)      lcqtof_hilic_neg_4_008_24    m/z 922.8246     0   1E+4   0.5   1  RT (min)      lcqtof_hilic_neg_1_005_32    m/z 922.8860     0   1E+4   0.5   1  RT (min)      lcqtof_hilic_neg_3_035_46    m/z 922.8873     0   1E+4   0.5   1  RT (min)      lcqtof_hilic_neg_3_035_46    m/z 922.9778     0   1E+4   1   1.5  RT (min)      lcqtof_hilic_neg_4_000_aa    m/z 923.2991     0   1E+4   1   1.5  RT (min)      lcqtof_hilic_neg_4_005_29    m/z 923.4811     0   1E+4   0.5   1  RT (min)      lcqtof_hilic_neg_4_045_26    m/z 923.6093     0   1E+4   2.5   3  RT (min)      lcqtof_hilic_neg_4_084_59    m/z 923.6952     0   1E+4   2   2.5  RT (min)      lcqtof_hilic_neg_4_008_24    m/z 923.7266     0   1E+4   3.5   4  RT (min)      lcqtof_hilic_neg_4_008_24    m/z 923.7278     0   1   2E+4   1   1.5  RT (min)      lcqtof_hilic_neg_4_000_aa    m/z 924.6131     0   1   2E+4   2.5   3  RT (min)      lcqtof_hilic_neg_4_048_23    m/z 924.6133     0   1   2E+4   3   3.5  RT (min)      lcqtof_hilic_neg_4_028_36    m/z 924.6294     0   1E+4   0.5   1  RT (min)      lcqtof_hilic_neg_1_007_05    m/z 924.6523     0   1   2   3E+4   1   1.5  RT (min)      lcqtof_hilic_neg_4_000_aa    m/z 924.7026     0   1   2   3   4E+4   2   2.5  RT (min)      lcqtof_hilic_neg_4_008_24    m/z 924.9756     0   1E+4   0.5   1  RT (min)      lcqtof_hilic_neg_4_000_aa    m/z 924.9770     0   1E+4   1   1.5  RT (min)      lcqtof_hilic_neg_4_000_aa    m/z 925.3229     0   1   2E+4   1   1.5  RT (min)      lcqtof_hilic_neg_4_031_38    m/z 925.4837     0   2   4   6E+4   4   4.5  RT (min)      lcqtof_hilic_neg_4_005_29    m/z 925.4891     0   1   2E+4   0.5   1  RT (min)      lcqtof_hilic_neg_2_004_26    m/z 925.5708     0   1   2E+4   1   1.5  RT (min)      lcqtof_hilic_neg_4_000_aa    m/z 925.5827     0   1E+4   3   3.5  RT (min)      lcqtof_hilic_neg_1_003_37    m/z 925.5927     0   1   2E+4   3   3.5  RT (min)      lcqtof_hilic_neg_1_004_14    m/z 925.7216     0   1   2E+4   3   3.5  RT (min)      lcqtof_hilic_neg_4_040_05    m/z 925.7264     0   1E+4   1   1.5  RT (min)      lcqtof_hilic_neg_4_000_aa    m/z 926.0331     0   1E+4   6.5   7  RT (min)      lcqtof_hilic_neg_4_034_31    m/z 926.3267     0   1E+4   0.5   1   1.5  RT (min)      lcqtof_hilic_neg_4_031_38    m/z 926.4921     0   1   2E+3   1   1.5  RT (min)      lcqtof_hilic_neg_1_031_16    m/z 926.4941     0   1E+4   0.5   1  RT (min)      lcqtof_hilic_neg_2_004_26    m/z 926.4950     0   1   2   3E+3   2   2.5  RT (min)      lcqtof_hilic_neg_2_062_17    m/z 926.5942     0   1E+4   3   3.5  RT (min)      lcqtof_hilic_neg_1_004_14    m/z 926.6271     0   1E+4   2.5   3  RT (min)      lcqtof_hilic_neg_4_048_23    m/z 926.6297     0   1E+4   3   3.5  RT (min)      lcqtof_hilic_neg_4_028_36    m/z 926.6637     0   1E+4   0.5   1  RT (min)      lcqtof_hilic_neg_4_040_05    m/z 926.7120     0   1E+4   2   2.5  RT (min)      lcqtof_hilic_neg_4_008_24    m/z 926.7172     0   1   2E+4   3   3.5  RT (min)      lcqtof_hilic_neg_1_005_32    m/z 926.9707     0   1E+4   0.5   1  RT (min)      lcqtof_hilic_neg_4_000_aa    m/z 927.4968     0   1E+4   0.5   1  RT (min)      lcqtof_hilic_neg_2_004_26    m/z 927.6948     0   1   2E+4   2   2.5  RT (min)      lcqtof_hilic_neg_4_008_24    m/z 927.7186     0   1E+4   2.5   3  RT (min)      lcqtof_hilic_neg_4_055_32    m/z 927.7337     0   1E+4   3   3.5  RT (min)      lcqtof_hilic_neg_4_040_05    m/z 927.7346     0   1E+4   3   3.5  RT (min)      lcqtof_hilic_neg_4_035_33    m/z 927.8554     0   1E+4   0.5   1  RT (min)      lcqtof_hilic_neg_4_047_06    m/z 928.5964     0   1   2   3E+3   3.5   4  RT (min)      lcqtof_hilic_neg_1_051_35    m/z 928.6090     0   1E+4   2   2.5  RT (min)      lcqtof_hilic_neg_4_053_14    m/z 928.6103     0   1E+4   2   2.5  RT (min)      lcqtof_hilic_neg_4_032_01    m/z 928.6470     0   1E+4   3   3.5  RT (min)      lcqtof_hilic_neg_4_028_36    m/z 928.6675     0   1E+4   0.5   1  RT (min)      lcqtof_hilic_neg_4_040_05    m/z 928.7072     0   2   4   6E+4   3   3.5  RT (min)      lcqtof_hilic_neg_4_008_24    m/z 928.7244     0   1   2E+4   2.5   3  RT (min)      lcqtof_hilic_neg_4_055_32    m/z 928.8608     0   1   2E+4   0.5   1  RT (min)      lcqtof_hilic_neg_4_047_06    m/z 928.8721     0   1   2E+4   0.5   1  RT (min)      lcqtof_hilic_neg_4_055_32    m/z 928.9119     0   1E+4   7   7.5  RT (min)      lcqtof_hilic_neg_4_053_14    m/z 928.9133     0   1E+4   6.5   7  RT (min)       lcqtof_hilic_neg_4_053_14    m/z 929.3925     0   2   4   6   8E+4   0.5   1  RT (min)      lcqtof_hilic_neg_3_035_46    m/z 929.6367     0   1   2E+4   3   3.5  RT (min)      lcqtof_hilic_neg_1_005_32    m/z 929.6819     0   1E+4   3   3.5  RT (min)      lcqtof_hilic_neg_2_047_32    m/z 929.7094     0   1   2E+4   2   2.5  RT (min)      lcqtof_hilic_neg_4_008_24    m/z 929.7334     0   1E+4   2.5   3  RT (min)      lcqtof_hilic_neg_4_055_32    m/z 929.7417     0   1E+4   2.5   3  RT (min)      lcqtof_hilic_neg_4_055_32    m/z 929.8306     0   2   4E+4   0.5   1  RT (min)      lcqtof_hilic_neg_4_055_32    m/z 929.9005     0   1E+4   5   5.5  RT (min)      lcqtof_hilic_neg_4_048_23    m/z 930.3956     0   2   4E+4   0.5   1  RT (min)      lcqtof_hilic_neg_3_035_46    m/z 930.6048     0   2   4   6   8E+3   3.5   4  RT (min)      lcqtof_hilic_neg_2_051_14    m/z 930.6070     0   1E+5   3.5   4  RT (min)      lcqtof_hilic_neg_4_039_18    m/z 930.6300     0   1E+4   3   3.5  RT (min)      lcqtof_hilic_neg_1_005_32    m/z 930.6680     0   1E+4   0.5   1  RT (min)      lcqtof_hilic_neg_1_007_05    m/z 930.7393     0   1   2   3   4E+4   2.5   3  RT (min)       lcqtof_hilic_neg_4_055_32    m/z 930.7394     0   1E+4   3   3.5  RT (min)      lcqtof_hilic_neg_4_035_33    m/z 930.8345     0   1   2E+4   0.5   1  RT (min)      lcqtof_hilic_neg_4_055_32    m/z 931.1449     0   1E+4   0.5   1  RT (min)      lcqtof_hilic_neg_3_035_46    m/z 931.2589     0   1E+4   7   7.5  RT (min)       lcqtof_hilic_neg_4_091_60    m/z 931.3931     0   1E+4   0.5   1  RT (min)      lcqtof_hilic_neg_3_035_46    m/z 931.6066     0   1   2   3   4E+4   3.5   4  RT (min)      lcqtof_hilic_neg_1_032_18    m/z 931.6068     0   1   2   3E+3   3.5   4  RT (min)      lcqtof_hilic_neg_1_010_15    m/z 931.6173     0   2   4E+4   3.5   4  RT (min)      lcqtof_hilic_neg_4_039_18    m/z 931.7511     0   1   2E+4   3   3.5  RT (min)      lcqtof_hilic_neg_4_008_24    m/z 931.7858     0   1E+4   0.5   1  RT (min)      lcqtof_hilic_neg_2_028_05    m/z 931.8767     0   1E+4   1.5   2  RT (min)      lcqtof_hilic_neg_4_000_aa    m/z 931.8838     0   1E+4   1.5   2  RT (min)      lcqtof_hilic_neg_4_000_aa    m/z 931.8866     0   1E+4   1.5   2  RT (min)      lcqtof_hilic_neg_4_000_aa    m/z 932.5480     0   1   2   3   4E+4   4   4.5  RT (min)       lcqtof_hilic_neg_4_048_23    m/z 932.6199     0   1E+4   3.5   4  RT (min)      lcqtof_hilic_neg_4_039_18    m/z 932.6656     0   1E+4   0.5   1  RT (min)      lcqtof_hilic_neg_1_007_05    m/z 932.6927     0   1E+4   4   4.5  RT (min)      lcqtof_hilic_neg_1_007_05    m/z 932.7565     0   1   2   3E+4   2.5   3  RT (min)      lcqtof_hilic_neg_4_055_32    m/z 932.7650     0   1   2   3E+4   4   4.5  RT (min)      lcqtof_hilic_neg_4_055_32    m/z 932.9639     0   1   2E+4   0.5   1  RT (min)      lcqtof_hilic_neg_4_008_24    m/z 932.9753     0   1E+4   0.5   1  RT (min)      lcqtof_hilic_neg_4_000_aa    m/z 933.3983     0   1E+4   1   1.5  RT (min)      lcqtof_hilic_neg_4_000_aa    m/z 933.5513     0   1E+4   4   4.5  RT (min)      lcqtof_hilic_neg_4_048_23    m/z 933.5956     0   1   2E+4   2.5   3  RT (min)       lcqtof_hilic_neg_4_053_14    m/z 933.7408     0   1   2E+4   1   1.5  RT (min)      lcqtof_hilic_neg_4_000_aa    m/z 933.7888     0   1   2E+4   1   1.5  RT (min)      lcqtof_hilic_neg_4_000_aa    m/z 933.8145     0   1   2E+4   1   1.5  RT (min)      lcqtof_hilic_neg_4_000_aa    m/z 934.5602     0   1E+4   4   4.5  RT (min)      lcqtof_hilic_neg_4_048_23    m/z 934.5999     0   1E+4   2.5   3  RT (min)      lcqtof_hilic_neg_4_053_14    m/z 934.6195     0   1E+4   0.5   1  RT (min)      lcqtof_hilic_neg_4_003_16    m/z 934.7163     0   1E+4   3   3.5  RT (min)      lcqtof_hilic_neg_1_037_24    m/z 934.9207     0   1E+4   5   5.5  RT (min)       lcqtof_hilic_neg_4_092_60    m/z 935.1972     0   1E+4   6   6.5  RT (min)      lcqtof_hilic_neg_2_052_24    m/z 935.5766     0   1   2E+4   1   1.5  RT (min)      lcqtof_hilic_neg_4_000_aa    m/z 935.5819     0   1   2E+4   1   1.5  RT (min)      lcqtof_hilic_neg_4_000_aa    m/z 935.6096     0   1E+4   3   3.5  RT (min)      lcqtof_hilic_neg_4_032_01    m/z 935.6102     0   1   2E+4   2.5   3  RT (min)      lcqtof_hilic_neg_4_053_14    m/z 935.6432     0   1   2   3   4E+4   3   3.5  RT (min)      lcqtof_hilic_neg_1_051_35    m/z 935.7568     0   1   2E+5   1   1.5  RT (min)      lcqtof_hilic_neg_4_000_aa    m/z 935.7621     0   1   2E+5   1   1.5  RT (min)      lcqtof_hilic_neg_4_000_aa    m/z 936.5020     0   1   2E+4   4   4.5  RT (min)      lcqtof_hilic_neg_2_029_31    m/z 936.6153     0   1E+4   2.5   3  RT (min)      lcqtof_hilic_neg_4_053_14    m/z 936.6352     0   1E+4   2   2.5  RT (min)      lcqtof_hilic_neg_4_048_23    m/z 936.6361     0   1E+4   2   2.5  RT (min)      lcqtof_hilic_neg_4_034_31    m/z 936.7051     0   1E+4   4   4.5  RT (min)      lcqtof_hilic_neg_1_037_24    m/z 936.7065     0   1E+4   4   4.5  RT (min)      lcqtof_hilic_neg_1_007_05    m/z 936.7574     0   1E+5   1   1.5  RT (min)      lcqtof_hilic_neg_4_000_aa    m/z 936.9772     0   1E+4   1   1.5  RT (min)      lcqtof_hilic_neg_4_000_aa    m/z 937.3137     0   1   2E+4   1   1.5  RT (min)      lcqtof_hilic_neg_4_010_28    m/z 937.5662     0   1E+4   4   4.5  RT (min)      lcqtof_hilic_neg_4_002_25    m/z 937.5817     0   1E+4   1   1.5  RT (min)      lcqtof_hilic_neg_4_031_38    m/z 937.6256     0   1   2E+4   1.5   2  RT (min)      lcqtof_hilic_neg_4_000_aa    m/z 937.6262     0   1   2E+4   2   2.5  RT (min)      lcqtof_hilic_neg_4_000_aa    m/z 937.6262     0   2   4   6E+4   1   1.5  RT (min)      lcqtof_hilic_neg_4_000_aa    m/z 937.6265     0   2   4   6E+4   2.5   3  RT (min)       lcqtof_hilic_neg_4_053_14    m/z 937.6267     0   2   4   6E+4   1.5   2  RT (min)      lcqtof_hilic_neg_4_000_aa    m/z 937.6271     0   1   2E+4   2   2.5  RT (min)      lcqtof_hilic_neg_4_000_aa    m/z 937.6272     0   2   4E+4   1.5   2  RT (min)      lcqtof_hilic_neg_4_000_aa    m/z 937.6273     0   1   2   3E+4   1.5   2  RT (min)      lcqtof_hilic_neg_4_000_aa    m/z 937.6279     0   1   2E+4   2   2.5  RT (min)      lcqtof_hilic_neg_4_000_aa    m/z 937.6281     0   1   2E+4   2   2.5  RT (min)      lcqtof_hilic_neg_4_000_aa    m/z 937.6305     0   1   2E+4   2   2.5  RT (min)      lcqtof_hilic_neg_4_000_aa    m/z 937.6346     0   1   2E+4   2   2.5  RT (min)      lcqtof_hilic_neg_4_000_aa    m/z 937.6352     0   2   4   6E+4   1   1.5  RT (min)      lcqtof_hilic_neg_4_000_aa    m/z 937.6359     0   2   4   6E+4   1   1.5  RT (min)      lcqtof_hilic_neg_4_000_aa    m/z 937.6964     0   1E+4   3   3.5  RT (min)      lcqtof_hilic_neg_4_014_46    m/z 937.7593     0   1E+4   2   2.5  RT (min)      lcqtof_hilic_neg_1_005_32    m/z 937.7608     0   1   2   3   4E+4   1   1.5  RT (min)      lcqtof_hilic_neg_4_000_aa    m/z 938.6065     0   2   4   6E+3   2.5   3  RT (min)      lcqtof_hilic_neg_1_004_14    m/z 938.6212     0   1E+4   2.5   3  RT (min)      lcqtof_hilic_neg_4_055_32    m/z 938.6243     0   1E+4   3   3.5  RT (min)      lcqtof_hilic_neg_4_031_38    m/z 938.6276     0   1   2   3E+4   2.5   3  RT (min)      lcqtof_hilic_neg_4_053_14    m/z 938.6675     0   1   2   3E+4   4   4.5  RT (min)       lcqtof_hilic_neg_4_008_24    m/z 938.7113     0   1E+4   2   2.5  RT (min)      lcqtof_hilic_neg_4_002_25    m/z 938.7144     0   1   2   3E+4   2   2.5  RT (min)      lcqtof_hilic_neg_4_008_24    m/z 938.9280     0   1E+4   0.5   1  RT (min)      lcqtof_hilic_neg_4_004_47    m/z 939.5166     0   1E+4   1   1.5  RT (min)      lcqtof_hilic_neg_4_022_34    m/z 939.6115     0   1E+4   2   2.5  RT (min)      lcqtof_hilic_neg_4_034_31    m/z 939.6351     0   1E+4   2   2.5  RT (min)      lcqtof_hilic_neg_4_034_31    m/z 939.6375     0   1   2E+4   2.5   3  RT (min)      lcqtof_hilic_neg_4_053_14    m/z 939.7219     0   1E+4   4   4.5  RT (min)      lcqtof_hilic_neg_1_007_05    m/z 939.7223     0   1E+5   2   2.5  RT (min)      lcqtof_hilic_neg_4_008_24    m/z 939.7237     0   1E+4   4   4.5  RT (min)      lcqtof_hilic_neg_1_037_24    m/z 939.8568     0   1E+4   0.5   1  RT (min)      lcqtof_hilic_neg_4_008_24    m/z 940.0939     0   1E+4   0.5   1  RT (min)      lcqtof_hilic_neg_1_062_17    m/z 940.6299     0   1E+4   2.5   3  RT (min)      lcqtof_hilic_neg_4_055_32    m/z 940.6399     0   1E+4   2.5   3  RT (min)      lcqtof_hilic_neg_4_055_32    m/z 940.6418     0   1E+4   3   3.5  RT (min)      lcqtof_hilic_neg_4_035_33    m/z 940.6821     0   2   4   6   8E+4   0.5   1  RT (min)      lcqtof_hilic_neg_4_040_05    m/z 940.8523     0   2   4E+4   0.5   1  RT (min)      lcqtof_hilic_neg_4_004_47    m/z 940.8637     0   2   4   6E+4   0.5   1  RT (min)      lcqtof_hilic_neg_4_008_24    m/z 940.9753     0   1E+4   0.5   1  RT (min)      lcqtof_hilic_neg_4_000_aa    m/z 941.1751     0   1E+4   3.5   4  RT (min)      lcqtof_hilic_neg_4_013_20    m/z 941.2038     0   1   2E+4   1   1.5  RT (min)      lcqtof_hilic_neg_4_000_aa    m/z 941.6859     0   2   4E+4   0.5   1  RT (min)      lcqtof_hilic_neg_4_040_05    m/z 941.8704     0   1   2E+4   0.5   1  RT (min)      lcqtof_hilic_neg_4_008_24    m/z 942.2039     0   1   2E+4   1   1.5  RT (min)      lcqtof_hilic_neg_4_000_aa    m/z 942.2043     0   1E+4   0.5   1  RT (min)      lcqtof_hilic_neg_1_062_17    m/z 942.6081     0   1   2   3   4E+3   2   2.5  RT (min)      lcqtof_hilic_neg_4_011_35    m/z 942.6182     0   1E+4   4   4.5  RT (min)      lcqtof_hilic_neg_4_008_24    m/z 942.6454     0   1   2   3E+4   2.5   3  RT (min)      lcqtof_hilic_neg_4_055_32    m/z 942.6556     0   1E+4   3.5   4  RT (min)      lcqtof_hilic_neg_4_052_13    m/z 942.6830     0   2   4E+4   0.5   1  RT (min)      lcqtof_hilic_neg_4_040_05    m/z 942.8773     0   1   2E+4   0.5   1  RT (min)      lcqtof_hilic_neg_4_055_32    m/z 943.2030     0   1   2E+4   1   1.5  RT (min)      lcqtof_hilic_neg_4_000_aa    m/z 943.6110     0   1E+4   4   4.5  RT (min)      lcqtof_hilic_neg_4_008_24    m/z 943.6478     0   1E+4   2.5   3  RT (min)      lcqtof_hilic_neg_4_055_32    m/z 943.6829     0   1   2E+4   0.5   1  RT (min)      lcqtof_hilic_neg_4_040_05    m/z 943.7420     0   1E+4   3   3.5  RT (min)      lcqtof_hilic_neg_4_008_24    m/z 944.1992     0   1E+4   1   1.5  RT (min)      lcqtof_hilic_neg_4_000_aa    m/z 944.2040     0   1E+4   0.5   1  RT (min)      lcqtof_hilic_neg_1_062_17    m/z 944.3584     0   1E+4   0.5   1  RT (min)      lcqtof_hilic_neg_3_003_20    m/z 944.7269     0   1E+4   2   2.5  RT (min)      lcqtof_hilic_neg_4_055_32    m/z 944.9727     0   1E+4   0.5   1  RT (min)      lcqtof_hilic_neg_4_000_aa    m/z 945.7575     0   1E+4   3   3.5  RT (min)      lcqtof_hilic_neg_4_008_24    m/z 945.8798     0   1E+4   1.5   2  RT (min)      lcqtof_hilic_neg_4_000_aa    m/z 946.3837     0   1E+4   0.5   1  RT (min)      lcqtof_hilic_neg_3_003_20    m/z 946.4220     0   1E+4   4   4.5  RT (min)      lcqtof_hilic_neg_4_013_20    m/z 946.6260     0   1E+4   4   4.5  RT (min)      lcqtof_hilic_neg_4_008_24    m/z 946.7116     0   1E+4   3.5   4  RT (min)      lcqtof_hilic_neg_4_008_24    m/z 947.6099     0   1E+4   4   4.5  RT (min)      lcqtof_hilic_neg_4_053_14    m/z 947.7008     0   1E+4   0.5   1  RT (min)      lcqtof_hilic_neg_4_045_26    m/z 947.7789     0   1   2E+4   1   1.5  RT (min)      lcqtof_hilic_neg_4_000_aa    m/z 948.0163     0   1E+4   6.5   7  RT (min)      lcqtof_hilic_neg_4_034_31    m/z 948.4915     0   1E+4   3   3.5  RT (min)       lcqtof_hilic_neg_3_066_57    m/z 948.5224     0   1   2E+4   4   4.5  RT (min)       lcqtof_hilic_neg_4_048_23    m/z 948.6274     0   1   2   3E+3   2.5   3  RT (min)      lcqtof_hilic_neg_2_062_17    m/z 948.6346     0   2   4   6E+3   2   2.5  RT (min)      lcqtof_hilic_neg_1_008_13    m/z 948.9725     0   1E+4   1   1.5  RT (min)      lcqtof_hilic_neg_4_000_aa    m/z 949.7264     0   2   4E+4   1   1.5  RT (min)      lcqtof_hilic_neg_4_000_aa    m/z 949.7339     0   1   2E+4   3.5   4  RT (min)      lcqtof_hilic_neg_4_008_24    m/z 949.7524     0   1   2   3E+4   0.5   1  RT (min)      lcqtof_hilic_neg_1_007_05    m/z 949.7789     0   1E+4   1   1.5  RT (min)      lcqtof_hilic_neg_4_008_24    m/z 949.7979     0   1E+4   2.5   3  RT (min)      lcqtof_hilic_neg_4_055_32    m/z 950.5299     0   1E+4   4   4.5  RT (min)      lcqtof_hilic_neg_4_048_23    m/z 950.6321     0   1E+4   3   3.5  RT (min)      lcqtof_hilic_neg_1_014_31    m/z 950.6492     0   1   2   3E+4   2   2.5  RT (min)      lcqtof_hilic_neg_4_048_23    m/z 950.7280     0   1   2   3E+4   1   1.5  RT (min)      lcqtof_hilic_neg_4_000_aa    m/z 950.7377     0   1   2   3E+4   1   1.5  RT (min)      lcqtof_hilic_neg_4_000_aa    m/z 950.8940     0   1E+4   6.5   7  RT (min)       lcqtof_hilic_neg_4_032_01    m/z 950.9704     0   1E+4   1   1.5  RT (min)      lcqtof_hilic_neg_4_000_aa    m/z 951.2318     0   1E+5   1   1.5  RT (min)      lcqtof_hilic_neg_4_000_aa    m/z 951.5919     0   1E+5   3.5   4  RT (min)      lcqtof_hilic_neg_4_039_18    m/z 951.6429     0   1E+4   3   3.5  RT (min)      lcqtof_hilic_neg_4_034_31    m/z 951.6548     0   1E+4   2   2.5  RT (min)      lcqtof_hilic_neg_4_048_23    m/z 951.7283     0   1E+4   2.5   3  RT (min)      lcqtof_hilic_neg_4_047_06    m/z 951.7388     0   2   4   6E+4   1   1.5  RT (min)      lcqtof_hilic_neg_4_000_aa    m/z 951.7491     0   1   2E+4   0.5   1  RT (min)      lcqtof_hilic_neg_1_007_05    m/z 952.5960     0   2   4   6   8E+4   3.5   4  RT (min)      lcqtof_hilic_neg_4_039_18    m/z 952.6829     0   1E+4   1   1.5  RT (min)      lcqtof_hilic_neg_4_000_aa    m/z 952.7346     0   1E+4   3   3.5  RT (min)      lcqtof_hilic_neg_4_040_05    m/z 952.7410     0   1   2   3E+4   1   1.5  RT (min)      lcqtof_hilic_neg_4_000_aa    m/z 952.8160     0   1   2   3   4E+4   0.5   1  RT (min)      lcqtof_hilic_neg_4_004_47    m/z 952.9753     0   1E+4   1   1.5  RT (min)      lcqtof_hilic_neg_4_000_aa    m/z 953.2301     0   1E+5   1   1.5  RT (min)      lcqtof_hilic_neg_4_000_aa    m/z 953.5985     0   1E+4   1   1.5  RT (min)      lcqtof_hilic_neg_4_000_aa    m/z 953.5991     0   1   2E+4   3.5   4  RT (min)      lcqtof_hilic_neg_4_039_18    m/z 953.6054     0   1E+4   1   1.5  RT (min)      lcqtof_hilic_neg_4_000_aa    m/z 953.6218     0   1E+4   3   3.5  RT (min)      lcqtof_hilic_neg_1_004_14    m/z 953.6532     0   1   2E+4   3   3.5  RT (min)      lcqtof_hilic_neg_1_014_31    m/z 953.7389     0   1E+4   2.5   3  RT (min)      lcqtof_hilic_neg_4_047_06    m/z 953.7396     0   2   4E+4   3   3.5  RT (min)      lcqtof_hilic_neg_4_040_05    m/z 953.7415     0   1   2   3E+4   1   1.5  RT (min)      lcqtof_hilic_neg_4_000_aa    m/z 953.7505     0   1   2   3E+4   1   1.5  RT (min)      lcqtof_hilic_neg_4_000_aa    m/z 954.2316     0   2   4   6E+4   1   1.5  RT (min)      lcqtof_hilic_neg_4_000_aa    m/z 954.7308     0   2   4   6E+4   2.5   3  RT (min)      lcqtof_hilic_neg_4_055_32    m/z 954.7549     0   1E+4   1   1.5  RT (min)      lcqtof_hilic_neg_4_000_aa    m/z 954.9745     0   1E+4   0.5   1  RT (min)      lcqtof_hilic_neg_4_000_aa    m/z 954.9757     0   1E+4   1   1.5  RT (min)      lcqtof_hilic_neg_4_000_aa    m/z 954.9811     0   1E+4   1   1.5  RT (min)      lcqtof_hilic_neg_4_000_aa    m/z 955.0920     0   1E+4   0.5   1  RT (min)      lcqtof_hilic_neg_3_036_17    m/z 955.2287     0   1   2   3E+4   1   1.5  RT (min)      lcqtof_hilic_neg_4_000_aa    m/z 955.6215     0   1E+4   1   1.5  RT (min)      lcqtof_hilic_neg_4_000_aa    m/z 955.6226     0   1E+4   1   1.5  RT (min)      lcqtof_hilic_neg_4_000_aa    m/z 955.6301     0   1E+4   1   1.5  RT (min)      lcqtof_hilic_neg_4_000_aa    m/z 955.6309     0   1E+4   1   1.5  RT (min)      lcqtof_hilic_neg_4_000_aa    m/z 955.6327     0   1E+4   1   1.5  RT (min)      lcqtof_hilic_neg_4_000_aa    m/z 955.7439     0   1   2   3   4E+4   2.5   3  RT (min)      lcqtof_hilic_neg_4_055_32    m/z 955.7448     0   1   2   3E+4   2.5   3  RT (min)      lcqtof_hilic_neg_4_091_60    m/z 955.7529     0   1   2   3E+4   3   3.5  RT (min)      lcqtof_hilic_neg_4_040_05    m/z 955.7563     0   1E+4   2.5   3  RT (min)      lcqtof_hilic_neg_4_047_06    m/z 956.2303     0   1E+4   1   1.5  RT (min)      lcqtof_hilic_neg_4_000_aa    m/z 956.6996     0   2   4E+4   0.5   1  RT (min)      lcqtof_hilic_neg_4_040_05    m/z 956.7454     0   1E+5   2.5   3  RT (min)       lcqtof_hilic_neg_4_055_32    m/z 956.7476     0   1E+4   2.5   3  RT (min)      lcqtof_hilic_neg_4_047_06    m/z 956.7648     0   1E+4   2   2.5  RT (min)      lcqtof_hilic_neg_1_005_32    m/z 956.8865     0   1E+4   0.5   1  RT (min)      lcqtof_hilic_neg_3_035_46    m/z 956.9770     0   1E+4   0.5   1  RT (min)      lcqtof_hilic_neg_4_000_aa    m/z 957.1174     0   1   2E+4   0.5   1  RT (min)      lcqtof_hilic_neg_1_062_17    m/z 957.6074     0   1E+4   4   4.5  RT (min)      lcqtof_hilic_neg_1_007_05    m/z 957.6671     0   1E+4   3   3.5  RT (min)      lcqtof_hilic_neg_1_005_32    m/z 957.6985     0   1   2E+4   0.5   1  RT (min)      lcqtof_hilic_neg_4_040_05    m/z 957.7369     0   2   4   6E+4   1   1.5  RT (min)      lcqtof_hilic_neg_4_000_aa    m/z 957.7563     0   1   2   3E+4   2.5   3  RT (min)      lcqtof_hilic_neg_4_055_32    m/z 957.7629     0   1E+4   3   3.5  RT (min)      lcqtof_hilic_neg_4_035_33    m/z 957.7631     0   1   2   3E+4   2.5   3  RT (min)      lcqtof_hilic_neg_4_055_32    m/z 958.5864     0   1E+5   8   8.5  RT (min)      lcqtof_hilic_neg_2_000_aa    m/z 958.5878     0   1E+5   8   8.5  RT (min)      lcqtof_hilic_neg_1_000_aa    m/z 958.6095     0   1   2   3E+4   0.5   1  RT (min)      lcqtof_hilic_neg_1_007_05    m/z 958.7400     0   1   2   3E+4   1   1.5  RT (min)      lcqtof_hilic_neg_4_000_aa    m/z 958.7621     0   2   4   6   8E+4   2.5   3  RT (min)       lcqtof_hilic_neg_4_055_32    m/z 958.7644     0   1E+4   2.5   3  RT (min)      lcqtof_hilic_neg_4_047_06    m/z 958.9164     0   1E+4   0.5   1  RT (min)      lcqtof_hilic_neg_4_004_47    m/z 958.9714     0   1E+4   1   1.5  RT (min)      lcqtof_hilic_neg_4_000_aa    m/z 958.9729     0   1E+4   1   1.5  RT (min)      lcqtof_hilic_neg_4_000_aa    m/z 958.9739     0   1E+4   1   1.5  RT (min)      lcqtof_hilic_neg_4_000_aa    m/z 959.6880     0   1E+4   3.5   4  RT (min)      lcqtof_hilic_neg_4_019_03    m/z 959.7504     0   2   4   6E+5   1   1.5  RT (min)      lcqtof_hilic_neg_4_000_aa    m/z 959.7535     0   2   4   6E+5   1   1.5  RT (min)      lcqtof_hilic_neg_4_000_aa    m/z 959.7721     0   1E+4   2.5   3  RT (min)      lcqtof_hilic_neg_4_091_60    m/z 959.7722     0   1E+4   2.5   3  RT (min)      lcqtof_hilic_neg_4_055_32    m/z 959.8980     0   1E+4   1   1.5  RT (min)      lcqtof_hilic_neg_4_000_aa    m/z 959.9062     0   1E+4   1.5   2  RT (min)      lcqtof_hilic_neg_4_000_aa    m/z 960.6076     0   1E+4   0.5   1  RT (min)      lcqtof_hilic_neg_1_007_05    m/z 960.6270     0   1E+4   2   2.5  RT (min)      lcqtof_hilic_neg_4_034_31    m/z 960.7563     0   1   2   3   4E+5   1   1.5  RT (min)      lcqtof_hilic_neg_4_000_aa    m/z 960.7578     0   1   2   3   4E+5   1   1.5  RT (min)      lcqtof_hilic_neg_4_000_aa    m/z 960.7764     0   1   2   3E+4   2.5   3  RT (min)      lcqtof_hilic_neg_4_097_60    m/z 960.7765     0   1   2   3   4E+4   2.5   3  RT (min)       lcqtof_hilic_neg_4_055_32    m/z 960.7788     0   1E+4   3   3.5  RT (min)      lcqtof_hilic_neg_4_035_33    m/z 961.1453     0   1E+4   0.5   1  RT (min)      lcqtof_hilic_neg_4_004_47    m/z 961.7692     0   2   4   6E+5   1   1.5  RT (min)      lcqtof_hilic_neg_4_000_aa    m/z 961.7750     0   2   4   6E+5   1   1.5  RT (min)      lcqtof_hilic_neg_4_000_aa    m/z 961.7890     0   2   4E+4   0.5   1  RT (min)      lcqtof_hilic_neg_4_016_37    m/z 961.7902     0   2   4   6E+5   1   1.5  RT (min)      lcqtof_hilic_neg_4_000_aa    m/z 961.8240     0   1E+4   1.5   2  RT (min)      lcqtof_hilic_neg_4_008_24    m/z 962.0553     0   1E+4   0.5   1  RT (min)      lcqtof_hilic_neg_4_004_47    m/z 962.3215     0   1E+4   4   4.5  RT (min)      lcqtof_hilic_neg_4_027_19    m/z 962.7715     0   1   2   3E+5   1   1.5  RT (min)      lcqtof_hilic_neg_4_000_aa    m/z 962.7808     0   1   2   3E+5   1   1.5  RT (min)      lcqtof_hilic_neg_4_000_aa    m/z 962.9757     0   1E+4   0.5   1  RT (min)      lcqtof_hilic_neg_4_000_aa    m/z 963.5271     0   1E+4   4   4.5  RT (min)      lcqtof_hilic_neg_4_013_20    m/z 963.6133     0   1   2   3E+4   1   1.5  RT (min)      lcqtof_hilic_neg_4_000_aa    m/z 963.6137     0   1   2   3E+4   1   1.5  RT (min)      lcqtof_hilic_neg_4_000_aa    m/z 963.7546     0   1E+4   1   1.5  RT (min)      lcqtof_hilic_neg_4_055_32    m/z 963.7776     0   1   2E+5   1   1.5  RT (min)      lcqtof_hilic_neg_4_000_aa    m/z 963.7780     0   1   2E+5   1   1.5  RT (min)      lcqtof_hilic_neg_4_000_aa    m/z 963.7835     0   1   2E+5   1   1.5  RT (min)      lcqtof_hilic_neg_4_000_aa    m/z 963.7846     0   1   2E+5   1   1.5  RT (min)      lcqtof_hilic_neg_4_000_aa    m/z 963.7875     0   1   2E+4   0.5   1  RT (min)      lcqtof_hilic_neg_4_040_05    m/z 963.7973     0   1E+4   3   3.5  RT (min)      lcqtof_hilic_neg_2_052_24    m/z 964.2078     0   1E+4   6   6.5  RT (min)      lcqtof_hilic_neg_1_014_31    m/z 964.6143     0   1   2   3E+3   2.5   3  RT (min)      lcqtof_hilic_neg_2_060_14    m/z 964.6262     0   1   2E+4   4   4.5  RT (min)      lcqtof_hilic_neg_4_008_24    m/z 964.6916     0   1   2E+4   4   4.5  RT (min)      lcqtof_hilic_neg_4_008_24    m/z 964.7520     0   1E+4   3   3.5  RT (min)       lcqtof_hilic_neg_2_052_24    m/z 964.7526     0   1E+4   3   3.5  RT (min)       lcqtof_hilic_neg_1_037_24    m/z 964.7811     0   1   2   3E+4   3   3.5  RT (min)      lcqtof_hilic_neg_1_037_24    m/z 964.7873     0   1E+5   1   1.5  RT (min)      lcqtof_hilic_neg_4_000_aa    m/z 964.7929     0   1E+5   1   1.5  RT (min)      lcqtof_hilic_neg_4_000_aa    m/z 964.9725     0   1E+4   0.5   1  RT (min)      lcqtof_hilic_neg_4_000_aa    m/z 965.3089     0   1E+4   1   1.5  RT (min)      lcqtof_hilic_neg_2_026_28    m/z 965.7252     0   1E+4   2   2.5  RT (min)      lcqtof_hilic_neg_4_055_32    m/z 965.7438     0   1E+4   2.5   3  RT (min)      lcqtof_hilic_neg_4_040_05    m/z 965.7496     0   1   2E+4   1   1.5  RT (min)      lcqtof_hilic_neg_4_000_aa    m/z 965.7550     0   1   2E+4   1   1.5  RT (min)      lcqtof_hilic_neg_4_000_aa    m/z 965.7867     0   2   4E+4   1   1.5  RT (min)      lcqtof_hilic_neg_4_000_aa    m/z 965.7904     0   2   4E+4   1   1.5  RT (min)      lcqtof_hilic_neg_4_000_aa    m/z 965.8697     0   1E+4   5   5.5  RT (min)      lcqtof_hilic_neg_4_055_32    m/z 966.7003     0   1E+4   4.5   5  RT (min)      lcqtof_hilic_neg_4_048_23    m/z 966.7596     0   1E+4   1   1.5  RT (min)      lcqtof_hilic_neg_4_000_aa    m/z 966.9315     0   1   2E+4   0.5   1  RT (min)      lcqtof_hilic_neg_3_035_46    m/z 966.9698     0   1E+4   0.5   1  RT (min)      lcqtof_hilic_neg_4_000_aa    m/z 966.9803     0   1E+4   1   1.5  RT (min)      lcqtof_hilic_neg_4_000_aa    m/z 967.5670     0   2   4   6E+4   3.5   4  RT (min)      lcqtof_hilic_neg_2_053_18    m/z 967.5989     0   1   2   3E+4   3   3.5  RT (min)       lcqtof_hilic_neg_4_048_23    m/z 967.6695     0   1   2   3E+4   0.5   1  RT (min)      lcqtof_hilic_neg_1_042_26    m/z 967.7171     0   1E+4   4   4.5  RT (min)      lcqtof_hilic_neg_4_008_24    m/z 967.7386     0   1   2E+5   1   1.5  RT (min)      lcqtof_hilic_neg_4_000_aa    m/z 967.7597     0   1E+4   2.5   3  RT (min)      lcqtof_hilic_neg_4_040_05    m/z 967.8164     0   2   4   6E+4   1.5   2  RT (min)      lcqtof_hilic_neg_4_000_aa    m/z 967.8402     0   2   4   6E+4   1.5   2  RT (min)      lcqtof_hilic_neg_4_000_aa    m/z 967.9327     0   1E+4   0.5   1  RT (min)      lcqtof_hilic_neg_3_035_46    m/z 968.5693     0   1   2   3E+4   3.5   4  RT (min)      lcqtof_hilic_neg_2_053_18    m/z 968.6057     0   1E+4   3   3.5  RT (min)      lcqtof_hilic_neg_4_048_23    m/z 968.6144     0   1E+4   2   2.5  RT (min)      lcqtof_hilic_neg_4_034_31    m/z 968.6551     0   1   2E+4   3.5   4  RT (min)      lcqtof_hilic_neg_4_055_32    m/z 968.6744     0   1E+4   0.5   1  RT (min)      lcqtof_hilic_neg_1_042_26    m/z 968.7126     0   1   2   3E+4   0.5   1  RT (min)      lcqtof_hilic_neg_3_048_05    m/z 968.7411     0   1E+5   1   1.5  RT (min)      lcqtof_hilic_neg_4_000_aa    m/z 968.7677     0   1E+4   3   3.5  RT (min)      lcqtof_hilic_neg_1_007_05    m/z 968.8187     0   1   2   3   4E+4   1.5   2  RT (min)      lcqtof_hilic_neg_4_000_aa    m/z 968.8566     0   1E+4   0.5   1  RT (min)      lcqtof_hilic_neg_4_008_24    m/z 968.9315     0   1E+4   0.5   1  RT (min)      lcqtof_hilic_neg_3_035_46    m/z 968.9758     0   1E+4   0.5   1  RT (min)      lcqtof_hilic_neg_4_000_aa    m/z 969.5714     0   1E+4   3.5   4  RT (min)      lcqtof_hilic_neg_2_053_18    m/z 969.5915     0   1E+4   1   1.5  RT (min)      lcqtof_hilic_neg_4_000_aa    m/z 969.5975     0   1E+4   1   1.5  RT (min)      lcqtof_hilic_neg_4_000_aa    m/z 969.5985     0   1E+4   0.5   1  RT (min)      lcqtof_hilic_neg_3_003_20    m/z 969.6633     0   1E+4   3.5   4  RT (min)      lcqtof_hilic_neg_4_055_32    m/z 969.6703     0   1E+4   4.5   5  RT (min)      lcqtof_hilic_neg_4_008_24    m/z 969.7332     0   2   4   6   8E+4   1   1.5  RT (min)      lcqtof_hilic_neg_4_000_aa    m/z 969.7739     0   1E+4   2.5   3  RT (min)      lcqtof_hilic_neg_4_040_05    m/z 969.7760     0   2   4E+4   3   3.5  RT (min)      lcqtof_hilic_neg_1_007_05    m/z 969.8202     0   1E+4   1.5   2  RT (min)      lcqtof_hilic_neg_4_000_aa    m/z 970.2933     0   1E+4   1   1.5  RT (min)      lcqtof_hilic_neg_3_003_20    m/z 970.7139     0   1E+4   0.5   1  RT (min)      lcqtof_hilic_neg_3_048_05    m/z 970.7352     0   1   2   3E+4   1   1.5  RT (min)      lcqtof_hilic_neg_4_000_aa    m/z 970.9776     0   1E+4   1   1.5  RT (min)      lcqtof_hilic_neg_4_000_aa    m/z 971.6302     0   1E+4   3   3.5  RT (min)       lcqtof_hilic_neg_4_048_23    m/z 971.6849     0   1E+4   4.5   5  RT (min)      lcqtof_hilic_neg_4_008_24    m/z 971.7636     0   1E+4   2.5   3  RT (min)       lcqtof_hilic_neg_4_055_32    m/z 971.7914     0   1E+4   3   3.5  RT (min)      lcqtof_hilic_neg_1_007_05    m/z 972.6371     0   2   4E+3   2   2.5  RT (min)      lcqtof_hilic_neg_2_063_13    m/z 972.6852     0   1   2   3E+4   4   4.5  RT (min)      lcqtof_hilic_neg_4_008_24    m/z 972.9676     0   1E+4   4.5   5  RT (min)      lcqtof_hilic_neg_4_055_32    m/z 972.9720     0   1E+4   1   1.5  RT (min)      lcqtof_hilic_neg_4_000_aa    m/z 972.9821     0   1E+4   0.5   1  RT (min)      lcqtof_hilic_neg_4_000_aa    m/z 973.1294     0   2   4   6E+4   4.5   5  RT (min)      lcqtof_hilic_neg_4_055_32    m/z 973.3755     0   1E+4   4.5   5  RT (min)      lcqtof_hilic_neg_4_055_32    m/z 973.6467     0   1E+4   3   3.5  RT (min)      lcqtof_hilic_neg_4_034_31    m/z 973.7792     0   1E+4   3   3.5  RT (min)      lcqtof_hilic_neg_4_008_24    m/z 973.8133     0   1E+4   1   1.5  RT (min)      lcqtof_hilic_neg_4_000_aa    m/z 973.8559     0   1   2E+4   0.5   1  RT (min)      lcqtof_hilic_neg_4_040_05    m/z 974.3589     0   1E+4   1   1.5  RT (min)      lcqtof_hilic_neg_2_045_11    m/z 974.6651     0   1E+4   4   4.5  RT (min)      lcqtof_hilic_neg_4_008_24    m/z 974.9742     0   1E+4   0.5   1  RT (min)      lcqtof_hilic_neg_4_000_aa    m/z 974.9789     0   1E+4   1   1.5  RT (min)      lcqtof_hilic_neg_4_000_aa    m/z 975.4959     0   1   2E+4   0.5   1  RT (min)      lcqtof_hilic_neg_4_047_06    m/z 975.7487     0   1   2E+4   1   1.5  RT (min)      lcqtof_hilic_neg_4_000_aa    m/z 975.7947     0   1E+4   0.5   1  RT (min)      lcqtof_hilic_neg_4_040_05    m/z 975.7947     0   1E+4   3   3.5  RT (min)      lcqtof_hilic_neg_1_007_05    m/z 975.8477     0   1E+4   0.5   1  RT (min)      lcqtof_hilic_neg_4_009_50    m/z 976.4995     0   1E+4   0.5   1  RT (min)      lcqtof_hilic_neg_4_047_06    m/z 976.6963     0   1E+4   4   4.5  RT (min)       lcqtof_hilic_neg_4_008_24    m/z 976.9758     0   1E+4   0.5   1  RT (min)      lcqtof_hilic_neg_4_000_aa    m/z 977.4824     0   1E+4   0.5   1  RT (min)      lcqtof_hilic_neg_4_047_06    m/z 977.6993     0   1   2   3   4E+4   4   4.5  RT (min)       lcqtof_hilic_neg_4_008_24    m/z 977.7640     0   1   2   3   4E+6   1   1.5  RT (min)      lcqtof_hilic_neg_4_000_aa    m/z 977.7740     0   1   2   3   4E+6   1   1.5  RT (min)      lcqtof_hilic_neg_4_000_aa    m/z 977.8044     0   1E+5   0.5   1  RT (min)      lcqtof_hilic_neg_4_040_05    m/z 977.9841     0   1E+4   7.5   8  RT (min)      lcqtof_hilic_neg_4_016_37    m/z 978.0808     0   1E+5   1   1.5  RT (min)      lcqtof_hilic_neg_4_000_aa    m/z 978.6552     0   1E+4   4   4.5  RT (min)      lcqtof_hilic_neg_1_007_05    m/z 978.7711     0   1   2E+6   1   1.5  RT (min)      lcqtof_hilic_neg_4_000_aa    m/z 978.7749     0   1E+4   2   2.5  RT (min)      lcqtof_hilic_neg_4_055_32    m/z 978.8102     0   2   4E+4   0.5   1  RT (min)      lcqtof_hilic_neg_4_040_05    m/z 978.9800     0   1E+4   0.5   1  RT (min)      lcqtof_hilic_neg_4_000_aa    m/z 979.0640     0   1E+4   0.5   1  RT (min)      lcqtof_hilic_neg_4_004_47    m/z 979.0829     0   1E+4   1   1.5  RT (min)      lcqtof_hilic_neg_4_000_aa    m/z 979.4783     0   1E+4   15   15.5  RT (min)      lcqtof_hilic_neg_4_000_aa    m/z 979.7643     0   1E+4   3   3.5  RT (min)      lcqtof_hilic_neg_4_008_24    m/z 979.7690     0   1E+4   0.5   1  RT (min)      lcqtof_hilic_neg_1_014_31    m/z 979.7714     0   2   4   6   8E+5   1   1.5  RT (min)      lcqtof_hilic_neg_4_000_aa    m/z 979.8041     0   2   4   6E+4   0.5   1  RT (min)      lcqtof_hilic_neg_4_040_05    m/z 980.6711     0   2   4E+4   4   4.5  RT (min)      lcqtof_hilic_neg_1_007_05    m/z 980.6801     0   1   2   3E+4   1   1.5  RT (min)      lcqtof_hilic_neg_4_000_aa    m/z 980.7746     0   1   2E+5   1   1.5  RT (min)      lcqtof_hilic_neg_4_000_aa    m/z 980.7801     0   1   2E+5   1.5   2  RT (min)      lcqtof_hilic_neg_4_000_aa    m/z 980.7935     0   1E+4   1   1.5  RT (min)      lcqtof_hilic_neg_4_047_06    m/z 980.8710     0   1E+4   0.5   1  RT (min)      lcqtof_hilic_neg_3_035_46    m/z 980.9645     0   1E+4   1   1.5  RT (min)      lcqtof_hilic_neg_4_000_aa    m/z 980.9686     0   1E+4   0.5   1  RT (min)      lcqtof_hilic_neg_4_000_aa    m/z 980.9765     0   1E+4   1   1.5  RT (min)      lcqtof_hilic_neg_4_000_aa    m/z 981.6747     0   1   2E+4   1   1.5  RT (min)      lcqtof_hilic_neg_4_000_aa    m/z 981.6753     0   1E+4   4   4.5  RT (min)      lcqtof_hilic_neg_1_007_05    m/z 981.7736     0   1E+5   1   1.5  RT (min)      lcqtof_hilic_neg_4_000_aa    m/z 981.7785     0   1E+5   1   1.5  RT (min)      lcqtof_hilic_neg_4_000_aa    m/z 981.8629     0   1E+4   0.5   1  RT (min)      lcqtof_hilic_neg_3_035_46    m/z 982.0052     0   1   2   3E+3   4.5   5  RT (min)      lcqtof_hilic_neg_4_032_01    m/z 982.3178     0   1E+4   1   1.5  RT (min)      lcqtof_hilic_neg_4_045_26    m/z 982.5214     0   1E+4   4   4.5  RT (min)      lcqtof_hilic_neg_4_018_27    m/z 982.7740     0   1   2   3E+4   1   1.5  RT (min)      lcqtof_hilic_neg_4_000_aa    m/z 982.8638     0   1E+4   0.5   1  RT (min)      lcqtof_hilic_neg_3_035_46    m/z 982.9025     0   1   2E+4   0.5   1  RT (min)      lcqtof_hilic_neg_1_005_32    m/z 982.9737     0   1E+4   0.5   1  RT (min)      lcqtof_hilic_neg_4_000_aa    m/z 983.0432     0   1E+4   0.5   1  RT (min)      lcqtof_hilic_neg_3_036_17    m/z 983.8102     0   1   2   3E+5   1   1.5  RT (min)      lcqtof_hilic_neg_4_000_aa    m/z 983.8329     0   1   2   3E+5   1   1.5  RT (min)      lcqtof_hilic_neg_4_000_aa    m/z 983.8368     0   2   4   6E+4   1   1.5  RT (min)      lcqtof_hilic_neg_4_000_aa    m/z 984.0383     0   1E+4   0.5   1  RT (min)      lcqtof_hilic_neg_3_036_17    m/z 984.1155     0   1E+4   0.5   1  RT (min)      lcqtof_hilic_neg_3_007_04    m/z 984.3101     0   1   2   3   4E+4   1   1.5  RT (min)      lcqtof_hilic_neg_3_003_20    m/z 984.8143     0   1   2E+5   1   1.5  RT (min)      lcqtof_hilic_neg_4_000_aa    m/z 984.8387     0   1   2E+5   1   1.5  RT (min)      lcqtof_hilic_neg_4_000_aa    m/z 984.8401     0   1   2   3   4E+4   1   1.5  RT (min)      lcqtof_hilic_neg_4_000_aa    m/z 984.8979     0   1E+4   0.5   1  RT (min)      lcqtof_hilic_neg_3_004_32    m/z 984.9085     0   1   2   3   4E+4   1   1.5  RT (min)      lcqtof_hilic_neg_4_000_aa    m/z 984.9766     0   1   2E+4   0.5   1  RT (min)      lcqtof_hilic_neg_4_000_aa    m/z 985.6287     0   1   2   3E+4   1   1.5  RT (min)      lcqtof_hilic_neg_4_000_aa    m/z 985.6301     0   1   2   3E+4   1   1.5  RT (min)      lcqtof_hilic_neg_4_000_aa    m/z 985.8199     0   2   4   6E+4   1   1.5  RT (min)      lcqtof_hilic_neg_4_000_aa    m/z 985.8401     0   2   4   6E+4   1   1.5  RT (min)      lcqtof_hilic_neg_4_000_aa    m/z 986.0213     0   1E+4   0.5   1  RT (min)      lcqtof_hilic_neg_3_036_17    m/z 986.1891     0   1E+4   6   6.5  RT (min)      lcqtof_hilic_neg_2_020_23    m/z 986.3618     0   1   2E+4   1   1.5  RT (min)      lcqtof_hilic_neg_4_010_28    m/z 986.5974     0   1   2   3E+3   2.5   3  RT (min)      lcqtof_hilic_neg_4_005_29    m/z 986.6327     0   1   2E+4   1   1.5   2  RT (min)      lcqtof_hilic_neg_4_000_aa    m/z 986.6335     0   1   2E+4   1   1.5  RT (min)      lcqtof_hilic_neg_4_000_aa    m/z 986.8177     0   1E+4   0.5   1  RT (min)      lcqtof_hilic_neg_4_052_13    m/z 986.9718     0   1E+4   1   1.5  RT (min)      lcqtof_hilic_neg_4_000_aa    m/z 986.9758     0   1E+4   0.5   1  RT (min)      lcqtof_hilic_neg_4_000_aa    m/z 987.3676     0   1E+4   1   1.5  RT (min)      lcqtof_hilic_neg_4_010_28    m/z 987.6436     0   1E+4   1   1.5  RT (min)      lcqtof_hilic_neg_4_000_aa    m/z 987.7857     0   1   2E+4   15.5   16  RT (min)      lcqtof_hilic_neg_4_000_aa    m/z 987.7864     0   1   2E+4   15.5   16  RT (min)      lcqtof_hilic_neg_4_000_aa    m/z 987.7867     0   1   2E+4   15.5   16  RT (min)      lcqtof_hilic_neg_4_000_aa    m/z 987.7886     0   1   2E+4   15.5   16  RT (min)      lcqtof_hilic_neg_4_000_aa    m/z 987.7912     0   2   4   6E+4   1   1.5  RT (min)      lcqtof_hilic_neg_4_000_aa    m/z 987.7994     0   1E+4   3   3.5  RT (min)      lcqtof_hilic_neg_1_037_24    m/z 988.6241     0   1E+4   4.5   5  RT (min)      lcqtof_hilic_neg_4_008_24    m/z 988.6491     0   1E+4   3.5   4  RT (min)      lcqtof_hilic_neg_4_028_36    m/z 988.7504     0   1E+4   2   2.5  RT (min)      lcqtof_hilic_neg_4_055_32    m/z 988.7877     0   1E+4   15.5   16  RT (min)      lcqtof_hilic_neg_4_000_aa    m/z 988.7888     0   1E+4   15.5   16  RT (min)      lcqtof_hilic_neg_4_000_aa    m/z 988.7915     0   1E+4   15.5   16  RT (min)      lcqtof_hilic_neg_4_000_aa    m/z 988.7917     0   1E+4   15.5   16  RT (min)      lcqtof_hilic_neg_4_000_aa    m/z 988.7941     0   1   2   3E+4   1   1.5  RT (min)      lcqtof_hilic_neg_4_000_aa    m/z 988.8005     0   1   2   3E+4   0.5   1  RT (min)      lcqtof_hilic_neg_4_034_31    m/z 988.8184     0   2   4   6E+4   0.5   1  RT (min)      lcqtof_hilic_neg_4_016_37    m/z 988.8346     0   1E+4   0.5   1  RT (min)      lcqtof_hilic_neg_4_054_48    m/z 988.8517     0   1   2   3E+4   0.5   1  RT (min)      lcqtof_hilic_neg_4_009_50    m/z 988.9757     0   1E+4   0.5   1  RT (min)      lcqtof_hilic_neg_4_000_aa    m/z 989.7894     0   1   2   3E+4   2   2.5  RT (min)      lcqtof_hilic_neg_4_000_aa    m/z 989.7899     0   2   4   6E+4   1.5   2  RT (min)      lcqtof_hilic_neg_4_000_aa    m/z 989.7900     0   2   4   6E+4   1.5   2  RT (min)      lcqtof_hilic_neg_4_000_aa    m/z 989.7913     0   1   2E+4   2   2.5  RT (min)      lcqtof_hilic_neg_4_000_aa    m/z 989.7913     0   2   4   6E+4   1.5   2  RT (min)      lcqtof_hilic_neg_4_000_aa    m/z 989.7940     0   1   2E+4   2   2.5  RT (min)      lcqtof_hilic_neg_4_000_aa    m/z 989.7961     0   2   4   6E+4   1   1.5  RT (min)      lcqtof_hilic_neg_4_000_aa    m/z 989.7999     0   2   4   6E+4   1   1.5  RT (min)      lcqtof_hilic_neg_4_000_aa    m/z 989.8043     0   2   4   6E+4   1   1.5  RT (min)      lcqtof_hilic_neg_4_000_aa    m/z 989.8061     0   1E+4   0.5   1  RT (min)      lcqtof_hilic_neg_4_040_05    m/z 989.8111     0   1E+4   3   3.5  RT (min)      lcqtof_hilic_neg_1_037_24    m/z 989.8228     0   1   2   3E+4   0.5   1  RT (min)      lcqtof_hilic_neg_4_011_35    m/z 990.3553     0   1E+4   1   1.5  RT (min)      lcqtof_hilic_neg_4_016_37    m/z 990.6288     0   1   2   3E+3   2.5   3  RT (min)      lcqtof_hilic_neg_2_051_14    m/z 990.6363     0   1E+4   4.5   5  RT (min)      lcqtof_hilic_neg_4_008_24    m/z 990.8007     0   1   2   3E+4   1   1.5  RT (min)      lcqtof_hilic_neg_4_000_aa    m/z 990.8151     0   2   4E+4   3   3.5  RT (min)      lcqtof_hilic_neg_1_037_24    m/z 990.8328     0   1E+4   1   1.5  RT (min)      lcqtof_hilic_neg_4_000_aa    m/z 990.9734     0   1E+4   0.5   1  RT (min)      lcqtof_hilic_neg_4_000_aa    m/z 991.0815     0   1E+4   6   6.5  RT (min)      lcqtof_hilic_neg_4_034_31    m/z 991.3578     0   1E+4   1   1.5  RT (min)      lcqtof_hilic_neg_4_011_35    m/z 991.8043     0   1E+5   1   1.5  RT (min)      lcqtof_hilic_neg_4_000_aa    m/z 991.8987     0   2   4   6E+4   1   1.5  RT (min)      lcqtof_hilic_neg_4_000_aa    m/z 991.9027     0   2   4   6E+4   1   1.5  RT (min)      lcqtof_hilic_neg_4_000_aa    m/z 992.6844     0   1E+4   4   4.5  RT (min)      lcqtof_hilic_neg_4_008_24    m/z 992.7935     0   1   2E+4   2   2.5  RT (min)      lcqtof_hilic_neg_4_055_32    m/z 992.9028     0   1   2   3E+4   1   1.5  RT (min)      lcqtof_hilic_neg_4_000_aa    m/z 992.9035     0   1   2   3E+4   1   1.5  RT (min)      lcqtof_hilic_neg_4_000_aa    m/z 992.9695     0   1E+4   1   1.5  RT (min)      lcqtof_hilic_neg_4_000_aa    m/z 992.9748     0   1E+4   0.5   1  RT (min)      lcqtof_hilic_neg_4_000_aa    m/z 993.4979     0   1E+4   15   15.5  RT (min)      lcqtof_hilic_neg_4_000_aa    m/z 993.6661     0   1E+4   3.5   4  RT (min)      lcqtof_hilic_neg_4_035_33    m/z 993.7604     0   1   2E+4   1   1.5  RT (min)      lcqtof_hilic_neg_4_000_aa    m/z 993.7947     0   2   4   6   8E+4   2   2.5  RT (min)      lcqtof_hilic_neg_4_055_32    m/z 993.7974     0   2   4E+4   1.5   2  RT (min)      lcqtof_hilic_neg_4_000_aa    m/z 993.8187     0   2   4   6E+4   0.5   1  RT (min)      lcqtof_hilic_neg_4_040_05    m/z 994.6469     0   1E+4   3   3.5  RT (min)      lcqtof_hilic_neg_4_059_14    m/z 994.7149     0   1E+4   4   4.5  RT (min)      lcqtof_hilic_neg_4_008_24    m/z 994.7565     0   1   2   3   4E+4   1   1.5  RT (min)      lcqtof_hilic_neg_4_000_aa    m/z 994.7631     0   1E+4   2   2.5  RT (min)      lcqtof_hilic_neg_4_055_32    m/z 994.7784     0   1E+4   2   2.5  RT (min)      lcqtof_hilic_neg_4_055_32    m/z 994.8247     0   1E+4   0.5   1  RT (min)      lcqtof_hilic_neg_4_040_05    m/z 994.9525     0   1E+4   0.5   1  RT (min)      lcqtof_hilic_neg_4_000_aa    m/z 995.6800     0   1E+4   3.5   4  RT (min)      lcqtof_hilic_neg_4_055_32    m/z 995.7766     0   1E+5   1   1.5  RT (min)      lcqtof_hilic_neg_4_000_aa    m/z 995.7809     0   1E+5   1   1.5  RT (min)      lcqtof_hilic_neg_4_000_aa    m/z 995.8739     0   1   2   3E+4   1.5   2  RT (min)      lcqtof_hilic_neg_4_000_aa    m/z 996.3317     0   1   2E+4   1   1.5  RT (min)      lcqtof_hilic_neg_4_007_45    m/z 996.7410     0   1E+4   2   2.5  RT (min)      lcqtof_hilic_neg_4_055_32    m/z 996.7440     0   1   2E+4   4   4.5  RT (min)      lcqtof_hilic_neg_4_008_24    m/z 996.8760     0   1   2E+4   1.5   2  RT (min)      lcqtof_hilic_neg_4_000_aa    m/z 996.9713     0   1E+4   1   1.5  RT (min)      lcqtof_hilic_neg_4_000_aa    m/z 996.9792     0   1E+4   0.5   1  RT (min)      lcqtof_hilic_neg_4_000_aa    m/z 997.3344     0   1E+4   1   1.5  RT (min)      lcqtof_hilic_neg_4_007_45    m/z 997.5870     0   1E+4   3.5   4  RT (min)      lcqtof_hilic_neg_4_039_18    m/z 997.6492     0   1   2   3E+4   3   3.5  RT (min)      lcqtof_hilic_neg_4_011_35    m/z 997.7030     0   1E+4   4   4.5  RT (min)       lcqtof_hilic_neg_4_040_05    m/z 997.7101     0   1E+4   3.5   4  RT (min)      lcqtof_hilic_neg_2_028_05    m/z 997.8471     0   1E+4   0.5   1  RT (min)      lcqtof_hilic_neg_4_047_06    m/z 997.8644     0   1   2   3   4E+4   1   1.5  RT (min)      lcqtof_hilic_neg_4_000_aa    m/z 997.8806     0   1E+4   5   5.5  RT (min)      lcqtof_hilic_neg_4_048_23    m/z 998.3265     0   1E+4   1   1.5  RT (min)      lcqtof_hilic_neg_3_003_20    m/z 998.6516     0   1   2E+4   3   3.5  RT (min)      lcqtof_hilic_neg_4_011_35    m/z 998.6571     0   1E+4   3.5   4  RT (min)      lcqtof_hilic_neg_4_040_05    m/z 998.8693     0   1   2E+4   1   1.5  RT (min)      lcqtof_hilic_neg_4_000_aa    m/z 999.0532     0   1E+4   1   1.5  RT (min)      lcqtof_hilic_neg_4_000_aa    m/z 999.0647     0   1   2E+4   0.5   1  RT (min)      lcqtof_hilic_neg_1_062_17    m/z 999.0745     0   1E+4   0.5   1  RT (min)      lcqtof_hilic_neg_3_007_04    m/z 999.3320     0   1E+4   1   1.5  RT (min)      lcqtof_hilic_neg_3_003_20    m/z 999.5881     0   1E+4   1   1.5  RT (min)      lcqtof_hilic_neg_4_000_aa    m/z 999.7188     0   1   2E+4   4   4.5  RT (min)       lcqtof_hilic_neg_4_092_60    m/z 999.7418     0   1E+4   0.5   1  RT (min)      lcqtof_hilic_neg_4_026_04    m/z 999.8597     0   1   2E+4   1   1.5  RT (min)      lcqtof_hilic_neg_4_000_aa    m/z 999.8607     0   1E+4   0.5   1  RT (min)      lcqtof_hilic_neg_4_047_06    m/z 999.9681     0   1E+4   7.5   8  RT (min)      lcqtof_hilic_neg_4_016_37    m/z 1000.0634     0   1E+4   0.5   1  RT (min)      lcqtof_hilic_neg_1_062_17    m/z 1000.0847     0   1E+4   4.5   5  RT (min)      lcqtof_hilic_neg_4_050_02    m/z 1000.9700     0   1E+4   0.5   1  RT (min)      lcqtof_hilic_neg_4_000_aa    m/z 1000.9753     0   1E+4   1   1.5  RT (min)      lcqtof_hilic_neg_4_000_aa    m/z 1001.5924     0   1E+4   3.5   4  RT (min)      lcqtof_hilic_neg_3_066_57    m/z 1001.6151     0   1   2E+4   1   1.5  RT (min)      lcqtof_hilic_neg_4_000_aa    m/z 1001.6221     0   1   2E+4   1   1.5  RT (min)      lcqtof_hilic_neg_4_000_aa    m/z 1001.6234     0   1   2E+4   1   1.5  RT (min)      lcqtof_hilic_neg_4_000_aa    m/z 1001.6252     0   1   2E+4   1   1.5  RT (min)      lcqtof_hilic_neg_4_000_aa    m/z 1001.6256     0   1   2E+4   1   1.5  RT (min)      lcqtof_hilic_neg_4_000_aa    m/z 1001.7117     0   1E+4   1   1.5  RT (min)      lcqtof_hilic_neg_4_000_aa    m/z 1001.7367     0   1E+4   4   4.5  RT (min)      lcqtof_hilic_neg_1_007_05    m/z 1001.8395     0   2   4   6E+4   1   1.5  RT (min)      lcqtof_hilic_neg_4_000_aa    m/z 1001.8406     0   2   4   6E+4   1   1.5  RT (min)      lcqtof_hilic_neg_4_000_aa    m/z 1001.8452     0   2   4   6E+4   1   1.5  RT (min)      lcqtof_hilic_neg_4_000_aa    m/z 1001.8480     0   2   4   6E+4   1   1.5  RT (min)      lcqtof_hilic_neg_4_000_aa    m/z 1002.0807     0   1   2   3E+4   4.5   5  RT (min)      lcqtof_hilic_neg_4_050_02    m/z 1002.1321     0   1E+4   6   6.5  RT (min)      lcqtof_hilic_neg_4_011_35    m/z 1002.6256     0   1E+4   1   1.5  RT (min)      lcqtof_hilic_neg_4_000_aa    m/z 1002.7626     0   1E+4   2   2.5  RT (min)      lcqtof_hilic_neg_4_055_32    m/z 1002.8439     0   2   4E+4   1   1.5  RT (min)      lcqtof_hilic_neg_4_000_aa    m/z 1002.8466     0   2   4E+4   1   1.5  RT (min)      lcqtof_hilic_neg_4_000_aa    m/z 1002.8474     0   2   4E+4   1   1.5  RT (min)      lcqtof_hilic_neg_4_000_aa    m/z 1002.8526     0   2   4E+4   1   1.5  RT (min)      lcqtof_hilic_neg_4_000_aa    m/z 1002.9061     0   1E+4   5   5.5  RT (min)       lcqtof_hilic_neg_4_092_60    m/z 1003.6034     0   1E+4   3.5   4  RT (min)      lcqtof_hilic_neg_3_066_57    m/z 1003.7500     0   1   2   3E+4   4   4.5  RT (min)      lcqtof_hilic_neg_1_007_05    m/z 1003.7596     0   1   2   3E+4   1   1.5  RT (min)      lcqtof_hilic_neg_4_000_aa    m/z 1003.7705     0   2   4E+4   2   2.5  RT (min)      lcqtof_hilic_neg_4_055_32    m/z 1003.8419     0   1   2   3   4E+4   1   1.5  RT (min)      lcqtof_hilic_neg_4_000_aa    m/z 1003.8442     0   1   2   3   4E+4   1   1.5  RT (min)      lcqtof_hilic_neg_4_000_aa    m/z 1004.0781     0   1   2   3   4E+4   4.5   5  RT (min)      lcqtof_hilic_neg_4_050_02    m/z 1004.3708     0   1E+4   1   1.5  RT (min)      lcqtof_hilic_neg_4_016_37    m/z 1004.6710     0   1E+4   3   3.5  RT (min)      lcqtof_hilic_neg_4_004_47    m/z 1004.7684     0   1   2E+4   1   1.5  RT (min)      lcqtof_hilic_neg_4_000_aa    m/z 1004.8444     0   1E+4   1   1.5  RT (min)      lcqtof_hilic_neg_4_000_aa    m/z 1004.9791     0   1E+4   0.5   1  RT (min)      lcqtof_hilic_neg_4_000_aa    m/z 1005.7775     0   1E+5   1   1.5  RT (min)      lcqtof_hilic_neg_4_000_aa    m/z 1005.7962     0   1E+5   1   1.5  RT (min)      lcqtof_hilic_neg_4_000_aa    m/z 1006.0756     0   1   2   3E+4   4.5   5  RT (min)      lcqtof_hilic_neg_4_050_02    m/z 1006.7783     0   2   4E+4   1   1.5  RT (min)      lcqtof_hilic_neg_4_000_aa    m/z 1006.7785     0   2   4E+4   1   1.5  RT (min)      lcqtof_hilic_neg_4_000_aa    m/z 1006.7970     0   2   4E+4   1   1.5  RT (min)      lcqtof_hilic_neg_4_000_aa    m/z 1007.5130     0   1E+4   15   15.5  RT (min)      lcqtof_hilic_neg_4_000_aa    m/z 1007.7347     0   1E+4   4   4.5  RT (min)      lcqtof_hilic_neg_4_008_24    m/z 1007.7751     0   1   2   3E+4   1   1.5  RT (min)      lcqtof_hilic_neg_4_000_aa    m/z 1007.7787     0   1   2   3E+4   1   1.5  RT (min)      lcqtof_hilic_neg_4_000_aa    m/z 1007.7933     0   1   2   3E+4   1   1.5  RT (min)      lcqtof_hilic_neg_4_000_aa    m/z 1007.7937     0   1   2   3E+4   1   1.5  RT (min)      lcqtof_hilic_neg_4_000_aa    m/z 1007.8046     0   2   4   6   8E+4   1   1.5  RT (min)      lcqtof_hilic_neg_4_000_aa    m/z 1007.8937     0   1   2   3E+5   1   1.5  RT (min)      lcqtof_hilic_neg_4_000_aa    m/z 1008.0723     0   1   2E+4   4.5   5  RT (min)      lcqtof_hilic_neg_4_050_02    m/z 1008.1716     0   1E+4   6   6.5  RT (min)      lcqtof_hilic_neg_2_020_23    m/z 1008.5140     0   1E+4   15   15.5  RT (min)      lcqtof_hilic_neg_4_000_aa    m/z 1008.7960     0   1E+4   1   1.5  RT (min)      lcqtof_hilic_neg_4_000_aa    m/z 1008.8273     0   1   2E+4   1   1.5  RT (min)      lcqtof_hilic_neg_4_047_06    m/z 1008.8972     0   1   2E+5   1   1.5  RT (min)      lcqtof_hilic_neg_4_000_aa    m/z 1009.7861     0   1   2   3E+4   2   2.5  RT (min)      lcqtof_hilic_neg_4_055_32    m/z 1009.7862     0   1   2E+5   1   1.5  RT (min)      lcqtof_hilic_neg_4_000_aa    m/z 1009.8032     0   1   2E+5   1   1.5  RT (min)      lcqtof_hilic_neg_4_000_aa    m/z 1009.8305     0   1E+4   1   1.5  RT (min)      lcqtof_hilic_neg_4_012_07    m/z 1009.8528     0   1E+4   1   1.5  RT (min)      lcqtof_hilic_neg_4_000_aa    m/z 1009.9025     0   2   4   6E+4   1   1.5  RT (min)      lcqtof_hilic_neg_4_000_aa    m/z 1010.0700     0   1E+4   4.5   5  RT (min)      lcqtof_hilic_neg_4_050_02    m/z 1010.6436     0   1E+4   4   4.5  RT (min)      lcqtof_hilic_neg_4_008_24    m/z 1010.7357     0   1   2E+4   1   1.5  RT (min)      lcqtof_hilic_neg_4_055_32    m/z 1010.7563     0   1   2   3E+4   2   2.5  RT (min)      lcqtof_hilic_neg_4_055_32    m/z 1010.7997     0   1E+5   1   1.5  RT (min)      lcqtof_hilic_neg_4_000_aa    m/z 1010.9109     0   1E+4   1   1.5  RT (min)      lcqtof_hilic_neg_4_000_aa    m/z 1010.9675     0   1E+4   1   1.5  RT (min)      lcqtof_hilic_neg_4_000_aa    m/z 1011.6750     0   1E+4   1   1.5  RT (min)      lcqtof_hilic_neg_4_000_aa    m/z 1011.7559     0   2   4   6E+4   2   2.5  RT (min)      lcqtof_hilic_neg_4_055_32    m/z 1011.8609     0   1E+4   0.5   1  RT (min)      lcqtof_hilic_neg_4_008_24    m/z 1011.8663     0   1   2   3   4E+5   1   1.5  RT (min)      lcqtof_hilic_neg_4_000_aa    m/z 1011.8678     0   1   2   3   4E+5   1   1.5  RT (min)      lcqtof_hilic_neg_4_000_aa    m/z 1011.8688     0   1   2   3   4E+5   1   1.5  RT (min)      lcqtof_hilic_neg_4_000_aa    m/z 1012.8702     0   1   2   3E+5   1   1.5  RT (min)      lcqtof_hilic_neg_4_000_aa    m/z 1012.8716     0   1   2   3E+5   1   1.5  RT (min)      lcqtof_hilic_neg_4_000_aa    m/z 1012.8716     0   1   2   3E+5   1   1.5  RT (min)      lcqtof_hilic_neg_4_000_aa    m/z 1012.9639     0   1E+4   1   1.5  RT (min)      lcqtof_hilic_neg_4_000_aa    m/z 1012.9792     0   1E+4   0.5   1  RT (min)      lcqtof_hilic_neg_4_000_aa    m/z 1013.0869     0   1E+4   1   1.5  RT (min)      lcqtof_hilic_neg_4_000_aa    m/z 1013.4228     0   1E+4   7   7.5  RT (min)       lcqtof_hilic_neg_4_095_60    m/z 1013.7314     0   1E+4   0.5   1  RT (min)      lcqtof_hilic_neg_4_022_34    m/z 1013.7710     0   2   4   6   8E+4   1   1.5  RT (min)      lcqtof_hilic_neg_4_000_aa    m/z 1013.8736     0   1E+5   1   1.5  RT (min)      lcqtof_hilic_neg_4_000_aa    m/z 1013.8752     0   1E+5   1   1.5  RT (min)      lcqtof_hilic_neg_4_000_aa    m/z 1013.8755     0   1E+5   1   1.5  RT (min)      lcqtof_hilic_neg_4_000_aa    m/z 1014.0799     0   1E+4   1   1.5  RT (min)      lcqtof_hilic_neg_4_000_aa    m/z 1014.7580     0   1E+4   0.5   1  RT (min)      lcqtof_hilic_neg_4_004_47    m/z 1014.8756     0   1   2E+4   1   1.5  RT (min)      lcqtof_hilic_neg_4_000_aa    m/z 1014.8789     0   1   2E+4   1   1.5  RT (min)      lcqtof_hilic_neg_4_000_aa    m/z 1014.9693     0   1E+4   0.5   1  RT (min)      lcqtof_hilic_neg_4_000_aa    m/z 1014.9705     0   1E+4   1   1.5  RT (min)      lcqtof_hilic_neg_4_000_aa    m/z 1014.9711     0   1E+4   1   1.5  RT (min)      lcqtof_hilic_neg_4_000_aa    m/z 1015.0784     0   1E+4   1   1.5  RT (min)      lcqtof_hilic_neg_4_000_aa    m/z 1015.0800     0   1E+4   1   1.5  RT (min)      lcqtof_hilic_neg_4_000_aa    m/z 1015.2219     0   1   2   3E+4   1   1.5  RT (min)      lcqtof_hilic_neg_4_000_aa    m/z 1015.3517     0   1E+5   1   1.5  RT (min)      lcqtof_hilic_neg_3_003_20    m/z 1015.6173     0   1E+4   4.5   5  RT (min)      lcqtof_hilic_neg_2_028_05    m/z 1015.7759     0   1E+4   3   3.5  RT (min)      lcqtof_hilic_neg_4_002_25    m/z 1016.2219     0   1   2E+4   1   1.5  RT (min)      lcqtof_hilic_neg_4_000_aa    m/z 1016.2242     0   1   2E+4   0.5   1  RT (min)      lcqtof_hilic_neg_1_062_17    m/z 1016.3564     0   2   4   6E+4   1   1.5  RT (min)      lcqtof_hilic_neg_3_003_20    m/z 1016.6739     0   1E+4   4   4.5  RT (min)      lcqtof_hilic_neg_2_028_05    m/z 1016.7609     0   1E+4   3   3.5  RT (min)      lcqtof_hilic_neg_4_008_24    m/z 1016.7712     0   1E+4   1   1.5  RT (min)      lcqtof_hilic_neg_4_104_61    m/z 1016.7723     0   1E+4   2   2.5  RT (min)      lcqtof_hilic_neg_4_055_32    m/z 1016.9265     0   1E+4   0.5   1  RT (min)      lcqtof_hilic_neg_4_055_32    m/z 1016.9711     0   1E+4   1   1.5  RT (min)      lcqtof_hilic_neg_4_000_aa    m/z 1016.9780     0   1E+4   0.5   1  RT (min)      lcqtof_hilic_neg_4_009_50    m/z 1017.2183     0   1   2   3   4E+4   1   1.5  RT (min)      lcqtof_hilic_neg_4_000_aa    m/z 1017.3213     0   1   2E+4   1   1.5  RT (min)      lcqtof_hilic_neg_4_006_30    m/z 1017.3573     0   1   2   3E+4   1   1.5  RT (min)      lcqtof_hilic_neg_3_003_20    m/z 1017.5437     0   1E+4   4.5   5  RT (min)      lcqtof_hilic_neg_2_028_05    m/z 1017.6104     0   1E+4   1   1.5  RT (min)      lcqtof_hilic_neg_4_000_aa    m/z 1017.6217     0   1E+4   1   1.5  RT (min)      lcqtof_hilic_neg_4_000_aa    m/z 1017.7757     0   1   2   3E+4   2   2.5  RT (min)      lcqtof_hilic_neg_4_055_32    m/z 1017.8331     0   1   2E+5   1   1.5  RT (min)      lcqtof_hilic_neg_4_000_aa    m/z 1018.0619     0   1E+4   6   6.5  RT (min)      lcqtof_hilic_neg_4_016_37    m/z 1018.2202     0   1   2E+4   1   1.5  RT (min)      lcqtof_hilic_neg_4_000_aa    m/z 1018.2220     0   1E+4   0.5   1  RT (min)      lcqtof_hilic_neg_1_062_17    m/z 1018.7490     0   1   2E+4   0.5   1  RT (min)      lcqtof_hilic_neg_3_003_20    m/z 1018.7694     0   1E+4   3   3.5  RT (min)       lcqtof_hilic_neg_4_091_60    m/z 1018.7736     0   1   2E+4   3   3.5  RT (min)      lcqtof_hilic_neg_4_008_24    m/z 1018.8358     0   1E+5   1   1.5  RT (min)      lcqtof_hilic_neg_4_000_aa    m/z 1018.8804     0   1E+4   0.5   1  RT (min)      lcqtof_hilic_neg_4_004_47    m/z 1018.9715     0   1E+4   0.5   1  RT (min)      lcqtof_hilic_neg_4_000_aa    m/z 1019.2175     0   1E+4   1   1.5  RT (min)      lcqtof_hilic_neg_4_000_aa    m/z 1019.5683     0   1E+4   3.5   4  RT (min)      lcqtof_hilic_neg_4_039_18    m/z 1019.6091     0   1E+4   7   7.5  RT (min)      lcqtof_hilic_neg_4_008_24    m/z 1019.7269     0   1E+4   3.5   4  RT (min)      lcqtof_hilic_neg_4_040_05    m/z 1019.8331     0   1   2E+5   1   1.5  RT (min)      lcqtof_hilic_neg_4_000_aa    m/z 1020.0253     0   1E+4   0.5   1  RT (min)      lcqtof_hilic_neg_4_004_47    m/z 1020.5695     0   1E+4   3.5   4  RT (min)      lcqtof_hilic_neg_4_039_18    m/z 1020.6139     0   1E+4   3   3.5  RT (min)      lcqtof_hilic_neg_1_042_26    m/z 1020.6210     0   1E+4   4   4.5  RT (min)      lcqtof_hilic_neg_4_008_24    m/z 1020.7032     0   1E+4   3.5   4  RT (min)      lcqtof_hilic_neg_2_028_05    m/z 1020.7861     0   1E+4   3   3.5  RT (min)       lcqtof_hilic_neg_4_095_60    m/z 1020.7876     0   1   2E+4   3   3.5  RT (min)      lcqtof_hilic_neg_4_008_24    m/z 1020.8314     0   1E+5   1   1.5  RT (min)      lcqtof_hilic_neg_4_000_aa    m/z 1020.9692     0   1E+4   0.5   1  RT (min)      lcqtof_hilic_neg_4_000_aa    m/z 1021.0306     0   1E+4   0.5   1  RT (min)      lcqtof_hilic_neg_4_004_47    m/z 1021.5291     0   1E+4   15   15.5  RT (min)      lcqtof_hilic_neg_4_000_aa    m/z 1021.6480     0   1E+4   3   3.5  RT (min)      lcqtof_hilic_neg_4_016_37    m/z 1021.6990     0   1E+4   4   4.5  RT (min)      lcqtof_hilic_neg_4_040_05    m/z 1021.7419     0   1E+4   3.5   4  RT (min)      lcqtof_hilic_neg_4_040_05    m/z 1021.7793     0   2   4   6   8E+4   1   1.5  RT (min)      lcqtof_hilic_neg_4_000_aa    m/z 1021.7797     0   2   4   6   8E+4   1   1.5  RT (min)      lcqtof_hilic_neg_4_000_aa    m/z 1021.8412     0   2   4   6   8E+4   1   1.5  RT (min)      lcqtof_hilic_neg_4_000_aa    m/z 1022.0861     0   1E+4   6   6.5  RT (min)      lcqtof_hilic_neg_4_031_38    m/z 1022.5298     0   1E+4   15   15.5  RT (min)      lcqtof_hilic_neg_4_000_aa    m/z 1022.6790     0   1   2E+3   2.5   3  RT (min)      lcqtof_hilic_neg_2_062_17    m/z 1022.7780     0   1   2   3   4E+4   1   1.5  RT (min)      lcqtof_hilic_neg_4_000_aa    m/z 1022.8029     0   1   2E+4   3   3.5  RT (min)      lcqtof_hilic_neg_4_008_24    m/z 1022.9794     0   1E+4   0.5   1  RT (min)      lcqtof_hilic_neg_4_000_aa    m/z 1023.3024     0   1E+4   1   1.5  RT (min)      lcqtof_hilic_neg_4_000_aa    m/z 1023.7248     0   1E+4   4   4.5  RT (min)      lcqtof_hilic_neg_4_055_32    m/z 1023.7789     0   1   2   3E+4   1   1.5  RT (min)      lcqtof_hilic_neg_4_000_aa    m/z 1023.7966     0   1E+4   0.5   1  RT (min)      lcqtof_hilic_neg_4_040_05    m/z 1023.8106     0   1E+5   1   1.5  RT (min)      lcqtof_hilic_neg_4_000_aa    m/z 1024.8189     0   1E+4   3   3.5  RT (min)      lcqtof_hilic_neg_4_008_24    m/z 1024.9722     0   1E+4   0.5   1  RT (min)      lcqtof_hilic_neg_4_000_aa    m/z 1025.2489     0   1E+5   1   1.5  RT (min)      lcqtof_hilic_neg_4_000_aa    m/z 1025.6516     0   1E+4   1   1.5  RT (min)      lcqtof_hilic_neg_4_000_aa    m/z 1025.8322     0   1E+5   1   1.5  RT (min)      lcqtof_hilic_neg_4_000_aa    m/z 1025.8680     0   1E+4   0.5   1  RT (min)      lcqtof_hilic_neg_4_047_06    m/z 1026.1078     0   1E+4   0.5   1  RT (min)      lcqtof_hilic_neg_4_004_47    m/z 1026.2504     0   1E+5   1   1.5  RT (min)      lcqtof_hilic_neg_4_000_aa    m/z 1026.6438     0   1E+4   4.5   5  RT (min)      lcqtof_hilic_neg_4_028_36    m/z 1026.8445     0   1   2   3   4E+4   1   1.5  RT (min)      lcqtof_hilic_neg_4_000_aa    m/z 1026.8811     0   1E+4   0.5   1  RT (min)       lcqtof_hilic_neg_4_105_61    m/z 1026.9014     0   1E+4   0.5   1  RT (min)      lcqtof_hilic_neg_3_035_46    m/z 1026.9688     0   1   2E+4   0.5   1  RT (min)      lcqtof_hilic_neg_4_000_aa    m/z 1026.9691     0   1   2E+4   1   1.5  RT (min)      lcqtof_hilic_neg_4_000_aa    m/z 1026.9717     0   1   2E+4   0.5   1  RT (min)      lcqtof_hilic_neg_4_000_aa    m/z 1027.1078     0   1E+4   0.5   1  RT (min)      lcqtof_hilic_neg_4_004_47    m/z 1027.1964     0   1E+4   0.5   1  RT (min)      lcqtof_hilic_neg_3_000_bb    m/z 1027.2491     0   1E+5   1   1.5  RT (min)      lcqtof_hilic_neg_4_000_aa    m/z 1027.7467     0   1E+4   0.5   1  RT (min)      lcqtof_hilic_neg_4_004_47    m/z 1027.8636     0   1   2E+6   1   1.5  RT (min)      lcqtof_hilic_neg_4_000_aa    m/z 1027.9016     0   1E+4   0.5   1  RT (min)      lcqtof_hilic_neg_3_035_46    m/z 1028.0165     0   1E+4   1   1.5  RT (min)      lcqtof_hilic_neg_4_000_aa    m/z 1028.2324     0   1E+5   1   1.5  RT (min)      lcqtof_hilic_neg_4_000_aa    m/z 1028.8650     0   1E+6   1   1.5  RT (min)      lcqtof_hilic_neg_4_000_aa    m/z 1028.9010     0   1E+4   0.5   1  RT (min)      lcqtof_hilic_neg_3_035_46    m/z 1028.9685     0   1   2E+4   1   1.5  RT (min)      lcqtof_hilic_neg_4_000_aa    m/z 1028.9697     0   1   2E+4   1   1.5  RT (min)      lcqtof_hilic_neg_4_000_aa    m/z 1028.9730     0   1   2E+4   1   1.5  RT (min)      lcqtof_hilic_neg_4_000_aa    m/z 1029.0065     0   1E+4   1   1.5  RT (min)      lcqtof_hilic_neg_4_000_aa    m/z 1029.2443     0   2   4E+4   1   1.5  RT (min)      lcqtof_hilic_neg_4_000_aa    m/z 1029.3326     0   2   4E+4   1   1.5  RT (min)      lcqtof_hilic_neg_4_006_30    m/z 1029.3682     0   2   4   6   8E+4   1   1.5  RT (min)      lcqtof_hilic_neg_3_003_20    m/z 1029.5751     0   2   4   6   8E+4   4   4.5  RT (min)      lcqtof_hilic_neg_4_039_18    m/z 1029.7368     0   1   2   3   4E+4   1   1.5  RT (min)      lcqtof_hilic_neg_4_000_aa    m/z 1029.8620     0   2   4   6E+5   1   1.5   2  RT (min)      lcqtof_hilic_neg_4_000_aa    m/z 1029.8656     0   2   4   6E+5   1   1.5  RT (min)      lcqtof_hilic_neg_4_000_aa    m/z 1029.8680     0   2   4   6E+5   1   1.5  RT (min)      lcqtof_hilic_neg_4_000_aa    m/z 1030.0131     0   1E+4   1   1.5  RT (min)      lcqtof_hilic_neg_4_000_aa    m/z 1030.1157     0   1   2E+4   6   6.5  RT (min)      lcqtof_hilic_neg_4_014_46    m/z 1030.1550     0   1E+4   6   6.5  RT (min)      lcqtof_hilic_neg_2_020_23    m/z 1030.2374     0   1   2E+4   1   1.5  RT (min)      lcqtof_hilic_neg_4_000_aa    m/z 1030.3359     0   1   2   3E+4   1   1.5  RT (min)      lcqtof_hilic_neg_4_006_30    m/z 1030.3723     0   1   2   3   4E+4   0.5   1  RT (min)      lcqtof_hilic_neg_3_003_20    m/z 1030.5789     0   2   4E+4   4   4.5  RT (min)      lcqtof_hilic_neg_4_039_18    m/z 1030.8722     0   1E+5   1   1.5  RT (min)      lcqtof_hilic_neg_4_000_aa    m/z 1030.9422     0   1   2E+4   0.5   1  RT (min)      lcqtof_hilic_neg_4_008_24    m/z 1030.9740     0   1   2E+4   0.5   1  RT (min)      lcqtof_hilic_neg_4_000_aa    m/z 1031.3372     0   1E+4   1   1.5  RT (min)      lcqtof_hilic_neg_4_006_30    m/z 1031.3759     0   1E+4   1   1.5  RT (min)      lcqtof_hilic_neg_3_003_20    m/z 1031.5797     0   1   2E+4   4   4.5  RT (min)      lcqtof_hilic_neg_4_039_18    m/z 1031.7191     0   1E+5   0.5   1  RT (min)      lcqtof_hilic_neg_3_003_20    m/z 1031.8727     0   1   2   3E+4   1   1.5  RT (min)      lcqtof_hilic_neg_4_000_aa    m/z 1031.8771     0   1   2   3E+4   1   1.5  RT (min)      lcqtof_hilic_neg_4_000_aa    m/z 1032.0009     0   1   2E+4   6.5   7  RT (min)      lcqtof_hilic_neg_4_053_14    m/z 1032.0020     0   1   2   3E+4   6.5   7  RT (min)      lcqtof_hilic_neg_4_053_14    m/z 1032.0032     0   1   2E+4   6.5   7  RT (min)      lcqtof_hilic_neg_4_053_14    m/z 1032.7133     0   2   4   6   8E+4   0.5   1  RT (min)      lcqtof_hilic_neg_3_003_20    m/z 1033.1239     0   1E+4   6   6.5  RT (min)      lcqtof_hilic_neg_4_055_32    m/z 1033.2804     0   1   2E+4   1   1.5  RT (min)      lcqtof_hilic_neg_4_000_aa    m/z 1033.3671     0   1E+4   1   1.5  RT (min)      lcqtof_hilic_neg_3_003_20    m/z 1033.7216     0   1E+4   1   1.5  RT (min)      lcqtof_hilic_neg_4_011_35    m/z 1033.9965     0   1E+5   6.5   7  RT (min)      lcqtof_hilic_neg_4_053_14    m/z 1033.9967     0   1E+5   6.5   7  RT (min)      lcqtof_hilic_neg_4_053_14    m/z 1033.9970     0   1E+5   6.5   7  RT (min)       lcqtof_hilic_neg_4_053_14    m/z 1034.5615     0   1   2   3   4E+3   1   1.5  RT (min)      lcqtof_hilic_neg_4_032_01    m/z 1034.7681     0   1E+4   0.5   1  RT (min)      lcqtof_hilic_neg_1_047_29    m/z 1034.9556     0   1E+4   1   1.5  RT (min)      lcqtof_hilic_neg_4_000_aa    m/z 1034.9995     0   2   4   6E+4   6.5   7  RT (min)      lcqtof_hilic_neg_4_053_14    m/z 1034.9997     0   2   4   6E+4   6.5   7  RT (min)      lcqtof_hilic_neg_4_053_14    m/z 1035.0006     0   2   4   6   8E+4   6.5   7  RT (min)      lcqtof_hilic_neg_4_053_14    m/z 1035.3607     0   2   4E+4   1   1.5  RT (min)      lcqtof_hilic_neg_3_003_20    m/z 1035.5417     0   1E+4   15   15.5  RT (min)      lcqtof_hilic_neg_4_000_aa    m/z 1036.0026     0   1   2E+4   6.5   7  RT (min)      lcqtof_hilic_neg_4_053_14    m/z 1036.0032     0   1   2   3E+4   6.5   7  RT (min)      lcqtof_hilic_neg_4_053_14    m/z 1036.0035     0   1   2E+4   6.5   7  RT (min)      lcqtof_hilic_neg_4_053_14    m/z 1036.0040     0   1   2E+4   6.5   7  RT (min)      lcqtof_hilic_neg_4_053_14    m/z 1036.3608     0   1   2   3E+4   1   1.5  RT (min)      lcqtof_hilic_neg_3_003_20    m/z 1036.9660     0   1E+4   1   1.5  RT (min)      lcqtof_hilic_neg_4_000_aa    m/z 1036.9806     0   1E+4   1   1.5  RT (min)      lcqtof_hilic_neg_4_000_aa    m/z 1037.3660     0   1E+4   1   1.5  RT (min)      lcqtof_hilic_neg_3_003_20    m/z 1037.7977     0   1   2   3   4E+4   1   1.5  RT (min)      lcqtof_hilic_neg_4_000_aa    m/z 1037.8019     0   1E+4   0.5   1  RT (min)      lcqtof_hilic_neg_1_007_05    m/z 1037.8266     0   1   2E+4   1   1.5  RT (min)      lcqtof_hilic_neg_4_000_aa    m/z 1038.3207     0   1   2   3E+4   0.5   1   1.5  RT (min)      lcqtof_hilic_neg_4_027_19    m/z 1038.3269     0   1   2   3E+4   1   1.5  RT (min)      lcqtof_hilic_neg_4_027_19    m/z 1038.7013     0   1E+4   7   7.5  RT (min)      lcqtof_hilic_neg_4_008_24    m/z 1038.7965     0   1   2E+4   1   1.5  RT (min)      lcqtof_hilic_neg_4_000_aa    m/z 1038.9802     0   1E+4   0.5   1  RT (min)      lcqtof_hilic_neg_4_000_aa    m/z 1039.6169     0   1   2E+4   4.5   5  RT (min)      lcqtof_hilic_neg_2_028_05    m/z 1039.8100     0   2   4   6   8E+5   1   1.5  RT (min)      lcqtof_hilic_neg_4_000_aa    m/z 1039.8992     0   1E+4   1.5   2  RT (min)      lcqtof_hilic_neg_4_000_aa    m/z 1040.0436     0   1E+4   6   6.5  RT (min)      lcqtof_hilic_neg_4_011_35    m/z 1040.2978     0   1E+4   1   1.5  RT (min)      lcqtof_hilic_neg_4_000_aa    m/z 1040.8021     0   1E+4   4.5   5  RT (min)       lcqtof_hilic_neg_4_012_07    m/z 1040.9712     0   1   2E+4   1   1.5  RT (min)      lcqtof_hilic_neg_4_000_aa    m/z 1040.9742     0   1   2E+4   0.5   1  RT (min)      lcqtof_hilic_neg_4_000_aa    m/z 1041.0812     0   1E+4   1   1.5  RT (min)      lcqtof_hilic_neg_4_000_aa    m/z 1041.5430     0   1   2E+4   4.5   5  RT (min)      lcqtof_hilic_neg_2_028_05    m/z 1041.6208     0   1   2E+4   4.5   5  RT (min)      lcqtof_hilic_neg_2_028_05    m/z 1041.7565     0   1E+4   4   4.5  RT (min)      lcqtof_hilic_neg_1_037_24    m/z 1041.8271     0   1   2   3E+5   1   1.5  RT (min)      lcqtof_hilic_neg_4_000_aa    m/z 1042.5497     0   1E+4   4.5   5  RT (min)      lcqtof_hilic_neg_2_028_05    m/z 1042.7594     0   1   2   3   4E+4   4   4.5  RT (min)       lcqtof_hilic_neg_1_037_24    m/z 1042.9674     0   1   2E+4   1   1.5  RT (min)      lcqtof_hilic_neg_4_000_aa    m/z 1042.9763     0   1   2E+4   0.5   1  RT (min)      lcqtof_hilic_neg_4_000_aa    m/z 1043.0514     0   1E+4   1   1.5  RT (min)      lcqtof_hilic_neg_4_000_aa    m/z 1043.7716     0   1E+4   4   4.5  RT (min)      lcqtof_hilic_neg_4_008_24    m/z 1044.7766     0   1   2   3E+4   4   4.5  RT (min)       lcqtof_hilic_neg_4_008_24    m/z 1044.7833     0   1E+4   1   1.5  RT (min)      lcqtof_hilic_neg_4_000_aa    m/z 1044.8508     0   2   4   6   8E+4   1   1.5  RT (min)      lcqtof_hilic_neg_4_000_aa    m/z 1044.8585     0   2   4   6   8E+4   1   1.5  RT (min)      lcqtof_hilic_neg_4_000_aa    m/z 1045.0552     0   1   2E+4   0.5   1  RT (min)      lcqtof_hilic_neg_3_063_17    m/z 1045.0673     0   1E+4   1   1.5  RT (min)      lcqtof_hilic_neg_4_000_aa    m/z 1045.7931     0   1E+4   1   1.5  RT (min)      lcqtof_hilic_neg_4_000_aa    m/z 1045.8551     0   2   4E+4   1   1.5  RT (min)      lcqtof_hilic_neg_4_000_aa    m/z 1046.0433     0   1E+4   0.5   1  RT (min)      lcqtof_hilic_neg_3_063_17    m/z 1046.0808     0   1E+4   0.5   1  RT (min)      lcqtof_hilic_neg_3_007_04    m/z 1046.7916     0   1   2E+4   4   4.5  RT (min)      lcqtof_hilic_neg_4_008_24    m/z 1046.8613     0   1E+4   1   1.5  RT (min)      lcqtof_hilic_neg_4_000_aa    m/z 1046.9670     0   1E+4   0.5   1  RT (min)      lcqtof_hilic_neg_4_000_aa    m/z 1046.9739     0   1E+4   1   1.5  RT (min)      lcqtof_hilic_neg_4_000_aa    m/z 1047.7460     0   1E+4   1   1.5  RT (min)      lcqtof_hilic_neg_4_000_aa    m/z 1047.7635     0   1E+4   1   1.5  RT (min)      lcqtof_hilic_neg_4_000_aa    m/z 1047.7786     0   1E+4   1   1.5   2  RT (min)      lcqtof_hilic_neg_4_000_aa    m/z 1047.8666     0   1E+4   1   1.5  RT (min)      lcqtof_hilic_neg_4_000_aa    m/z 1048.9666     0   1E+4   1   1.5  RT (min)      lcqtof_hilic_neg_4_000_aa    m/z 1049.3765     0   1   2   3E+4   1   1.5  RT (min)      lcqtof_hilic_neg_3_003_20    m/z 1049.7428     0   1E+4   4   4.5  RT (min)      lcqtof_hilic_neg_4_048_23    m/z 1050.3809     0   1E+4   1   1.5  RT (min)      lcqtof_hilic_neg_3_003_20    m/z 1050.6832     0   1E+4   3   3.5  RT (min)      lcqtof_hilic_neg_4_031_38    m/z 1050.8663     0   1   2E+4   0.5   1  RT (min)      lcqtof_hilic_neg_4_011_35    m/z 1050.9670     0   1E+4   0.5   1  RT (min)      lcqtof_hilic_neg_4_000_aa    m/z 1051.0284     0   1E+4   6.5   7  RT (min)      lcqtof_hilic_neg_4_055_32    m/z 1051.3403     0   2   4   6   8E+4   1   1.5  RT (min)      lcqtof_hilic_neg_4_027_19    m/z 1051.5590     0   2   4   6   8E+4   4   4.5  RT (min)      lcqtof_hilic_neg_4_039_18    m/z 1051.8734     0   1E+4   0.5   1  RT (min)      lcqtof_hilic_neg_4_011_35    m/z 1052.3423     0   1   2E+4   0.5   1   1.5  RT (min)      lcqtof_hilic_neg_4_027_19    m/z 1052.3685     0   1   2   3E+4   0.5   1   1.5  RT (min)      lcqtof_hilic_neg_4_042_22    m/z 1052.5624     0   1   2   3   4E+4   4   4.5  RT (min)      lcqtof_hilic_neg_4_039_18    m/z 1052.7381     0   1E+4   1   1.5  RT (min)      lcqtof_hilic_neg_4_048_23    m/z 1052.9740     0   1E+4   0.5   1  RT (min)      lcqtof_hilic_neg_4_000_aa    m/z 1053.3595     0   1   2E+4   0.5   1  RT (min)      lcqtof_hilic_neg_4_042_22    m/z 1053.4846     0   1E+4   0.5   1  RT (min)      lcqtof_hilic_neg_3_035_46    m/z 1053.5622     0   1   2E+4   4   4.5  RT (min)      lcqtof_hilic_neg_4_039_18    m/z 1053.7045     0   2   4   6E+4   0.5   1  RT (min)      lcqtof_hilic_neg_3_003_20    m/z 1053.8197     0   1   2   3E+4   1   1.5  RT (min)      lcqtof_hilic_neg_4_000_aa    m/z 1053.9833     0   1E+4   6.5   7  RT (min)      lcqtof_hilic_neg_4_023_15    m/z 1053.9833     0   1E+4   6.5   7  RT (min)      lcqtof_hilic_neg_4_059_14    m/z 1053.9843     0   1E+4   6.5   7  RT (min)      lcqtof_hilic_neg_4_059_14    m/z 1053.9849     0   1E+4   6.5   7  RT (min)      lcqtof_hilic_neg_4_023_15    m/z 1054.6915     0   1   2E+4   1   1.5  RT (min)      lcqtof_hilic_neg_4_048_23    m/z 1054.7064     0   2   4E+4   0.5   1  RT (min)      lcqtof_hilic_neg_3_003_20    m/z 1054.7129     0   1   2E+4   1   1.5  RT (min)      lcqtof_hilic_neg_4_048_23    m/z 1054.8167     0   1   2E+4   1   1.5  RT (min)      lcqtof_hilic_neg_4_000_aa    m/z 1054.8224     0   1   2E+4   1   1.5  RT (min)      lcqtof_hilic_neg_4_000_aa    m/z 1054.9767     0   1E+4   0.5   1  RT (min)      lcqtof_hilic_neg_4_000_aa    m/z 1054.9781     0   1E+4   1   1.5  RT (min)      lcqtof_hilic_neg_4_000_aa    m/z 1055.6334     0   1E+4   3.5   4  RT (min)      lcqtof_hilic_neg_1_007_05    m/z 1055.6953     0   1E+4   1   1.5  RT (min)      lcqtof_hilic_neg_4_048_23    m/z 1055.7123     0   1E+4   0.5   1  RT (min)      lcqtof_hilic_neg_3_003_20    m/z 1055.7913     0   1E+5   1   1.5  RT (min)      lcqtof_hilic_neg_4_000_aa    m/z 1055.7950     0   1E+5   1   1.5  RT (min)      lcqtof_hilic_neg_4_000_aa    m/z 1055.8036     0   1E+5   1   1.5  RT (min)      lcqtof_hilic_neg_4_000_aa    m/z 1055.8945     0   1   2   3E+4   1   1.5  RT (min)      lcqtof_hilic_neg_4_000_aa    m/z 1055.9781     0   2   4   6E+4   6.5   7  RT (min)      lcqtof_hilic_neg_4_059_14    m/z 1055.9788     0   2   4   6E+4   6.5   7  RT (min)      lcqtof_hilic_neg_4_053_14    m/z 1055.9789     0   2   4   6E+4   6.5   7  RT (min)       lcqtof_hilic_neg_4_053_14    m/z 1055.9794     0   2   4   6E+4   6.5   7  RT (min)       lcqtof_hilic_neg_4_023_15    m/z 1056.0491     0   1E+4   1   1.5  RT (min)      lcqtof_hilic_neg_4_000_aa    m/z 1056.8079     0   2   4   6   8E+4   1   1.5  RT (min)      lcqtof_hilic_neg_4_000_aa    m/z 1056.9023     0   1   2E+4   1   1.5  RT (min)      lcqtof_hilic_neg_4_000_aa    m/z 1056.9647     0   1   2E+4   1   1.5  RT (min)      lcqtof_hilic_neg_4_000_aa    m/z 1056.9796     0   1   2E+4   0.5   1  RT (min)      lcqtof_hilic_neg_4_000_aa    m/z 1056.9819     0   1   2   3E+4   6.5   7  RT (min)      lcqtof_hilic_neg_4_032_01    m/z 1056.9823     0   1   2   3E+4   6.5   7  RT (min)      lcqtof_hilic_neg_4_059_14    m/z 1056.9832     0   1   2   3E+4   6.5   7  RT (min)      lcqtof_hilic_neg_4_023_15    m/z 1056.9835     0   1   2   3E+4   6.5   7  RT (min)      lcqtof_hilic_neg_4_059_14    m/z 1056.9836     0   1   2   3E+4   6.5   7  RT (min)      lcqtof_hilic_neg_4_023_15    m/z 1057.0466     0   1E+4   1   1.5  RT (min)      lcqtof_hilic_neg_4_000_aa    m/z 1057.6326     0   1E+4   0.5   1  RT (min)      lcqtof_hilic_neg_2_004_26    m/z 1057.6503     0   1E+4   1   1.5  RT (min)      lcqtof_hilic_neg_4_000_aa    m/z 1057.6599     0   1E+4   0.5   1  RT (min)      lcqtof_hilic_neg_4_013_20    m/z 1057.8107     0   2   4E+4   1   1.5  RT (min)      lcqtof_hilic_neg_4_000_aa    m/z 1057.9869     0   1E+4   6.5   7  RT (min)      lcqtof_hilic_neg_4_023_15    m/z 1057.9879     0   1E+4   6.5   7  RT (min)      lcqtof_hilic_neg_4_059_14    m/z 1058.0408     0   1E+4   1   1.5  RT (min)      lcqtof_hilic_neg_4_000_aa    m/z 1058.6359     0   1E+4   1   1.5  RT (min)      lcqtof_hilic_neg_4_022_34    m/z 1058.9769     0   1   2E+4   1   1.5  RT (min)      lcqtof_hilic_neg_4_000_aa    m/z 1058.9783     0   1   2E+4   0.5   1  RT (min)      lcqtof_hilic_neg_4_000_aa    m/z 1059.5018     0   1   2E+4   0.5   1  RT (min)      lcqtof_hilic_neg_4_021_21    m/z 1059.5201     0   2   4   6E+4   0.5   1  RT (min)      lcqtof_hilic_neg_2_053_18    m/z 1059.6352     0   1   2E+5   4   4.5  RT (min)      lcqtof_hilic_neg_4_004_47    m/z 1059.6451     0   1E+4   1   1.5  RT (min)      lcqtof_hilic_neg_4_000_aa    m/z 1060.3414     0   1E+4   1   1.5  RT (min)      lcqtof_hilic_neg_4_038_10    m/z 1060.5051     0   1E+4   0.5   1  RT (min)      lcqtof_hilic_neg_4_021_21    m/z 1060.5236     0   1   2   3   4E+4   0.5   1  RT (min)      lcqtof_hilic_neg_1_032_18    m/z 1060.6863     0   1E+4   0.5   1  RT (min)      lcqtof_hilic_neg_4_022_34    m/z 1060.8372     0   1E+4   1   1.5  RT (min)      lcqtof_hilic_neg_4_000_aa    m/z 1060.9636     0   1   2E+4   1   1.5  RT (min)      lcqtof_hilic_neg_4_000_aa    m/z 1060.9695     0   1   2E+4   1   1.5  RT (min)      lcqtof_hilic_neg_4_000_aa    m/z 1060.9755     0   1   2E+4   1   1.5  RT (min)      lcqtof_hilic_neg_4_000_aa    m/z 1061.3469     0   1E+4   1   1.5  RT (min)      lcqtof_hilic_neg_4_038_10    m/z 1061.5000     0   1   2   3E+4   0.5   1  RT (min)      lcqtof_hilic_neg_4_021_21    m/z 1061.5195     0   2   4   6   8E+4   0.5   1  RT (min)      lcqtof_hilic_neg_2_053_18    m/z 1062.5032     0   1   2E+4   0.5   1  RT (min)      lcqtof_hilic_neg_4_021_21    m/z 1062.5224     0   1   2   3   4E+4   0.5   1  RT (min)      lcqtof_hilic_neg_2_053_18    m/z 1062.7689     0   1E+4   0.5   1  RT (min)      lcqtof_hilic_neg_4_009_50    m/z 1062.9764     0   1   2E+4   0.5   1  RT (min)      lcqtof_hilic_neg_4_000_aa    m/z 1062.9769     0   1   2E+4   1   1.5  RT (min)      lcqtof_hilic_neg_4_000_aa    m/z 1063.5263     0   1E+4   0.5   1  RT (min)      lcqtof_hilic_neg_2_053_18    m/z 1063.6025     0   1E+4   4.5   5  RT (min)      lcqtof_hilic_neg_4_040_05    m/z 1063.6984     0   1   2E+4   4.5   5  RT (min)      lcqtof_hilic_neg_2_028_05    m/z 1063.7088     0   1E+4   0.5   1  RT (min)      lcqtof_hilic_neg_2_004_26    m/z 1063.7919     0   1   2   3   4E+4   1   1.5  RT (min)      lcqtof_hilic_neg_4_000_aa    m/z 1063.9228     0   1E+4   0.5   1  RT (min)      lcqtof_hilic_neg_4_004_47    m/z 1064.6207     0   1E+4   4.5   5  RT (min)      lcqtof_hilic_neg_4_040_05    m/z 1064.9614     0   1E+4   0.5   1  RT (min)      lcqtof_hilic_neg_4_000_aa    m/z 1064.9705     0   1E+4   1   1.5  RT (min)      lcqtof_hilic_neg_4_000_aa    m/z 1065.6244     0   2   4   6E+4   4.5   5  RT (min)      lcqtof_hilic_neg_2_028_05    m/z 1066.0635     0   1E+4   0.5   1  RT (min)      lcqtof_hilic_neg_4_004_47    m/z 1066.3855     0   1   2E+4   0.5   1   1.5  RT (min)      lcqtof_hilic_neg_4_042_22    m/z 1066.6278     0   1E+4   4.5   5  RT (min)      lcqtof_hilic_neg_4_040_05    m/z 1066.7596     0   1E+4   1   1.5  RT (min)      lcqtof_hilic_neg_4_002_25    m/z 1066.7604     0   1   2E+4   1   1.5  RT (min)       lcqtof_hilic_neg_4_048_23    m/z 1066.7659     0   1E+4   0.5   1  RT (min)      lcqtof_hilic_neg_4_005_29    m/z 1066.7804     0   1E+4   0.5   1  RT (min)      lcqtof_hilic_neg_4_010_28    m/z 1067.3475     0   1   2E+4   0.5   1   1.5  RT (min)      lcqtof_hilic_neg_4_027_19    m/z 1067.3823     0   1E+4   0.5   1   1.5  RT (min)      lcqtof_hilic_neg_4_042_22    m/z 1067.5327     0   1   2   3E+4   4   4.5  RT (min)      lcqtof_hilic_neg_4_039_18    m/z 1067.6377     0   1E+4   4.5   5  RT (min)      lcqtof_hilic_neg_4_040_05    m/z 1067.7598     0   1E+4   1   1.5  RT (min)      lcqtof_hilic_neg_4_048_23    m/z 1067.8403     0   1E+4   1   1.5  RT (min)      lcqtof_hilic_neg_4_000_aa    m/z 1068.3530     0   1E+4   0.5   1   1.5  RT (min)      lcqtof_hilic_neg_4_027_19    m/z 1068.5358     0   1E+4   4   4.5  RT (min)      lcqtof_hilic_neg_4_039_18    m/z 1068.7665     0   1E+4   1   1.5  RT (min)      lcqtof_hilic_neg_4_000_aa    m/z 1068.9717     0   1   2E+4   0.5   1  RT (min)      lcqtof_hilic_neg_4_000_aa    m/z 1068.9718     0   1   2E+4   1   1.5  RT (min)      lcqtof_hilic_neg_4_000_aa    m/z 1068.9816     0   1   2E+4   1   1.5  RT (min)      lcqtof_hilic_neg_4_000_aa    m/z 1069.5345     0   1E+4   4   4.5  RT (min)      lcqtof_hilic_neg_4_039_18    m/z 1069.6566     0   1E+4   4.5   5  RT (min)      lcqtof_hilic_neg_2_028_05    m/z 1069.8043     0   1   2E+4   1   1.5  RT (min)      lcqtof_hilic_neg_4_000_aa    m/z 1070.7087     0   1E+4   1   1.5  RT (min)      lcqtof_hilic_neg_4_048_23    m/z 1070.8019     0   1E+4   1   1.5  RT (min)      lcqtof_hilic_neg_4_000_aa    m/z 1070.9645     0   1E+4   1   1.5  RT (min)      lcqtof_hilic_neg_4_000_aa    m/z 1070.9736     0   1E+4   1   1.5  RT (min)      lcqtof_hilic_neg_4_000_aa    m/z 1071.1476     0   1E+4   0.5   1  RT (min)      lcqtof_hilic_neg_4_057_49    m/z 1071.7649     0   1   2   3E+4   4   4.5  RT (min)       lcqtof_hilic_neg_4_008_24    m/z 1071.7656     0   1   2   3E+4   4   4.5  RT (min)      lcqtof_hilic_neg_4_008_24    m/z 1071.7657     0   1   2   3E+4   4   4.5  RT (min)      lcqtof_hilic_neg_4_008_24    m/z 1071.8757     0   1   2   3   4E+4   1   1.5  RT (min)      lcqtof_hilic_neg_4_000_aa    m/z 1072.0977     0   1   2E+4   0.5   1  RT (min)      lcqtof_hilic_neg_1_062_17    m/z 1072.6542     0   1   2E+4   1   1.5  RT (min)      lcqtof_hilic_neg_4_022_34    m/z 1072.7464     0   1E+4   4.5   5  RT (min)      lcqtof_hilic_neg_3_034_24    m/z 1072.8783     0   1   2E+4   1   1.5  RT (min)      lcqtof_hilic_neg_4_000_aa    m/z 1072.9727     0   1   2E+4   1   1.5  RT (min)      lcqtof_hilic_neg_4_000_aa    m/z 1072.9758     0   1   2E+4   1   1.5  RT (min)      lcqtof_hilic_neg_4_000_aa    m/z 1072.9799     0   1   2E+4   0.5   1  RT (min)      lcqtof_hilic_neg_4_000_aa    m/z 1073.0983     0   1E+4   0.5   1  RT (min)      lcqtof_hilic_neg_1_062_17    m/z 1073.6578     0   1E+4   1   1.5  RT (min)      lcqtof_hilic_neg_4_022_34    m/z 1073.7330     0   1E+4   0.5   1  RT (min)      lcqtof_hilic_neg_3_003_20    m/z 1074.3585     0   1E+4   1   1.5  RT (min)      lcqtof_hilic_neg_4_038_10    m/z 1074.8419     0   1E+4   1   1.5  RT (min)      lcqtof_hilic_neg_4_055_32    m/z 1074.9646     0   1   2E+4   1   1.5  RT (min)      lcqtof_hilic_neg_4_000_aa    m/z 1074.9684     0   1   2E+4   1   1.5  RT (min)      lcqtof_hilic_neg_4_000_aa    m/z 1074.9694     0   1   2E+4   1   1.5  RT (min)      lcqtof_hilic_neg_4_000_aa    m/z 1074.9819     0   1   2E+4   0.5   1  RT (min)      lcqtof_hilic_neg_4_000_aa    m/z 1075.8733     0   1   2E+4   1   1.5  RT (min)      lcqtof_hilic_neg_4_004_47    m/z 1076.9735     0   1   2E+4   1   1.5  RT (min)      lcqtof_hilic_neg_4_000_aa    m/z 1076.9747     0   1   2E+4   0.5   1  RT (min)      lcqtof_hilic_neg_4_000_aa    m/z 1077.9606     0   1   2   3E+4   6.5   7  RT (min)      lcqtof_hilic_neg_4_032_01    m/z 1077.9608     0   1   2   3E+4   6.5   7  RT (min)      lcqtof_hilic_neg_4_064_15    m/z 1077.9608     0   1   2   3E+4   6.5   7  RT (min)      lcqtof_hilic_neg_4_032_01    m/z 1077.9610     0   1   2   3E+4   6.5   7  RT (min)      lcqtof_hilic_neg_4_059_14    m/z 1077.9614     0   1   2   3E+4   6.5   7  RT (min)       lcqtof_hilic_neg_4_064_15    m/z 1078.5447     0   1   2   3E+4   5.5   6  RT (min)      lcqtof_hilic_neg_4_039_18    m/z 1078.9650     0   1   2E+4   6.5   7  RT (min)      lcqtof_hilic_neg_4_075_58    m/z 1078.9665     0   1E+4   6.5   7  RT (min)      lcqtof_hilic_neg_4_064_15    m/z 1078.9672     0   1E+4   6.5   7  RT (min)      lcqtof_hilic_neg_4_059_14    m/z 1078.9788     0   1   2E+4   1   1.5  RT (min)      lcqtof_hilic_neg_4_000_aa    m/z 1079.5488     0   1E+4   5.5   6  RT (min)      lcqtof_hilic_neg_4_039_18    m/z 1079.6350     0   1   2   3   4E+4   3.5   4  RT (min)      lcqtof_hilic_neg_1_007_05    m/z 1080.6355     0   1   2E+4   3.5   4  RT (min)      lcqtof_hilic_neg_1_007_05    m/z 1080.7732     0   1E+4   1   1.5  RT (min)      lcqtof_hilic_neg_4_002_25    m/z 1080.7733     0   1   2E+4   1   1.5  RT (min)      lcqtof_hilic_neg_4_048_23    m/z 1080.9716     0   1   2E+4   0.5   1  RT (min)      lcqtof_hilic_neg_4_000_aa    m/z 1080.9731     0   1   2E+4   1   1.5  RT (min)      lcqtof_hilic_neg_4_000_aa    m/z 1081.0605     0   1E+4   6   6.5  RT (min)      lcqtof_hilic_neg_4_055_32    m/z 1081.3515     0   1   2E+4   1   1.5  RT (min)      lcqtof_hilic_neg_4_018_27    m/z 1081.6444     0   1E+4   0.5   1  RT (min)      lcqtof_hilic_neg_2_004_26    m/z 1081.7732     0   1E+4   1   1.5  RT (min)      lcqtof_hilic_neg_4_048_23    m/z 1081.8127     0   1E+4   1   1.5  RT (min)      lcqtof_hilic_neg_4_008_24    m/z 1082.3610     0   1E+4   2.5   3  RT (min)      lcqtof_hilic_neg_4_027_19    m/z 1082.6529     0   1E+4   4   4.5  RT (min)      lcqtof_hilic_neg_4_040_05    m/z 1083.6554     0   1   2   3E+4   0.5   1  RT (min)      lcqtof_hilic_neg_2_004_26    m/z 1083.8175     0   1   2   3E+4   1   1.5  RT (min)      lcqtof_hilic_neg_4_000_aa    m/z 1084.6583     0   1E+4   0.5   1  RT (min)      lcqtof_hilic_neg_2_004_26    m/z 1084.9819     0   1E+4   0.5   1  RT (min)      lcqtof_hilic_neg_4_000_aa    m/z 1085.1235     0   1E+4   1   1.5  RT (min)      lcqtof_hilic_neg_4_000_aa    m/z 1085.6613     0   1   2E+4   0.5   1  RT (min)      lcqtof_hilic_neg_2_004_26    m/z 1085.6832     0   1E+4   4.5   5  RT (min)      lcqtof_hilic_neg_4_008_24    m/z 1085.8009     0   1   2E+4   1   1.5  RT (min)      lcqtof_hilic_neg_4_000_aa    m/z 1086.0742     0   1   2E+4   0.5   1  RT (min)      lcqtof_hilic_neg_4_004_47    m/z 1086.1207     0   1E+4   1   1.5  RT (min)      lcqtof_hilic_neg_4_000_aa    m/z 1086.1266     0   1E+4   1   1.5  RT (min)      lcqtof_hilic_neg_4_000_aa    m/z 1086.4620     0   1E+4   0.5   1  RT (min)      lcqtof_hilic_neg_4_005_29    m/z 1086.6654     0   1E+4   0.5   1  RT (min)      lcqtof_hilic_neg_2_004_26    m/z 1086.7029     0   1   2   3E+4   0.5   1  RT (min)      lcqtof_hilic_neg_4_000_aa    m/z 1086.8674     0   1E+4   5   5.5  RT (min)      lcqtof_hilic_neg_4_048_23    m/z 1086.8757     0   1E+4   0.5   1  RT (min)      lcqtof_hilic_neg_3_035_46    m/z 1086.9695     0   1   2E+4   1   1.5  RT (min)      lcqtof_hilic_neg_4_000_aa    m/z 1087.0731     0   1E+4   0.5   1  RT (min)      lcqtof_hilic_neg_4_004_47    m/z 1087.2974     0   1   2E+4   5   5.5  RT (min)      lcqtof_hilic_neg_4_048_23    m/z 1087.4813     0   1E+4   5   5.5  RT (min)      lcqtof_hilic_neg_4_048_23    m/z 1087.6069     0   1   2E+4   4.5   5  RT (min)      lcqtof_hilic_neg_4_040_05    m/z 1087.6923     0   1E+4   7   7.5  RT (min)      lcqtof_hilic_neg_4_008_24    m/z 1087.7094     0   1   2E+4   0.5   1  RT (min)      lcqtof_hilic_neg_4_000_aa    m/z 1087.8771     0   1E+4   0.5   1  RT (min)      lcqtof_hilic_neg_3_035_46    m/z 1088.0443     0   1E+4   5   5.5  RT (min)      lcqtof_hilic_neg_4_048_23    m/z 1088.1217     0   1E+4   1   1.5  RT (min)      lcqtof_hilic_neg_4_000_aa    m/z 1088.8756     0   1E+4   0.5   1  RT (min)      lcqtof_hilic_neg_3_035_46    m/z 1088.9762     0   1   2E+4   1   1.5  RT (min)      lcqtof_hilic_neg_4_000_aa    m/z 1088.9777     0   1   2E+4   1   1.5  RT (min)      lcqtof_hilic_neg_4_000_aa    m/z 1089.2386     0   1   2   3E+4   1   1.5  RT (min)      lcqtof_hilic_neg_4_000_aa    m/z 1089.6593     0   1   2E+4   3.5   4  RT (min)      lcqtof_hilic_neg_4_022_34    m/z 1089.6988     0   1   2E+4   4.5   5  RT (min)      lcqtof_hilic_neg_4_002_25    m/z 1089.7051     0   1   2   3E+4   4.5   5  RT (min)      lcqtof_hilic_neg_2_028_05    m/z 1089.7599     0   1   2E+4   0.5   1  RT (min)      lcqtof_hilic_neg_1_031_16    m/z 1090.2174     0   1E+4   6.5   7  RT (min)      lcqtof_hilic_neg_4_008_24    m/z 1090.2396     0   1   2   3E+4   1   1.5  RT (min)      lcqtof_hilic_neg_4_000_aa    m/z 1090.6642     0   1E+4   3.5   4  RT (min)      lcqtof_hilic_neg_4_022_34    m/z 1090.9015     0   1E+4   0.5   1  RT (min)      lcqtof_hilic_neg_4_011_35    m/z 1091.1211     0   1E+4   0.5   1  RT (min)      lcqtof_hilic_neg_1_062_17    m/z 1091.2379     0   1   2   3   4E+4   1   1.5  RT (min)      lcqtof_hilic_neg_4_000_aa    m/z 1091.6693     0   1E+4   3.5   4  RT (min)      lcqtof_hilic_neg_4_022_34    m/z 1091.7164     0   1E+4   4.5   5  RT (min)      lcqtof_hilic_neg_4_040_05    m/z 1092.2423     0   1   2E+4   0.5   1  RT (min)      lcqtof_hilic_neg_1_062_17    m/z 1092.2426     0   1   2   3E+4   1   1.5  RT (min)      lcqtof_hilic_neg_4_000_aa    m/z 1092.2654     0   1E+4   6.5   7  RT (min)      lcqtof_hilic_neg_4_008_24    m/z 1092.7432     0   1E+4   4   4.5  RT (min)      lcqtof_hilic_neg_4_008_24    m/z 1092.9691     0   1   2E+4   1   1.5  RT (min)      lcqtof_hilic_neg_4_000_aa    m/z 1092.9698     0   1   2E+4   0.5   1  RT (min)      lcqtof_hilic_neg_4_000_aa    m/z 1092.9711     0   1   2E+4   1   1.5  RT (min)      lcqtof_hilic_neg_4_000_aa    m/z 1093.2341     0   1   2E+4   1   1.5  RT (min)      lcqtof_hilic_neg_4_000_aa    m/z 1093.2425     0   1E+4   0.5   1  RT (min)      lcqtof_hilic_neg_1_062_17    m/z 1093.7456     0   2   4E+4   4   4.5  RT (min)       lcqtof_hilic_neg_4_008_24    m/z 1094.7892     0   1E+4   1   1.5  RT (min)      lcqtof_hilic_neg_4_048_23    m/z 1094.8807     0   1E+4   1   1.5  RT (min)      lcqtof_hilic_neg_4_008_24    m/z 1095.6040     0   1E+4   7   7.5  RT (min)      lcqtof_hilic_neg_4_008_24    m/z 1095.6942     0   1   2E+4   3.5   4  RT (min)      lcqtof_hilic_neg_1_007_05    m/z 1095.7607     0   1E+4   4   4.5  RT (min)      lcqtof_hilic_neg_4_008_24    m/z 1095.8191     0   1E+4   1   1.5  RT (min)      lcqtof_hilic_neg_4_008_24    m/z 1095.8819     0   1E+4   1   1.5  RT (min)      lcqtof_hilic_neg_4_008_24    m/z 1096.8307     0   1   2   3E+4   1   1.5  RT (min)      lcqtof_hilic_neg_4_008_24    m/z 1096.8608     0   1E+4   1   1.5  RT (min)      lcqtof_hilic_neg_4_008_24    m/z 1096.9695     0   1   2E+4   0.5   1  RT (min)      lcqtof_hilic_neg_4_000_aa    m/z 1096.9783     0   1   2E+4   1   1.5  RT (min)      lcqtof_hilic_neg_4_000_aa    m/z 1097.7042     0   1   2E+4   3.5   4  RT (min)      lcqtof_hilic_neg_1_007_05    m/z 1097.8344     0   1E+4   1   1.5  RT (min)      lcqtof_hilic_neg_4_008_24    m/z 1098.8208     0   1E+4   1   1.5  RT (min)      lcqtof_hilic_neg_4_055_32    m/z 1099.1852     0   1E+4   0.5   1  RT (min)      lcqtof_hilic_neg_2_063_13    m/z 1099.2695     0   1E+5   1   1.5  RT (min)      lcqtof_hilic_neg_4_000_aa    m/z 1099.7219     0   1   2   3E+4   3.5   4  RT (min)      lcqtof_hilic_neg_1_007_05    m/z 1099.7708     0   1E+4   4   4.5  RT (min)      lcqtof_hilic_neg_4_004_47    m/z 1099.8042     0   1E+4   4   4.5  RT (min)      lcqtof_hilic_neg_1_037_24    m/z 1099.8050     0   1   2E+4   1   1.5  RT (min)      lcqtof_hilic_neg_4_000_aa    m/z 1099.8196     0   1   2E+4   1   1.5  RT (min)      lcqtof_hilic_neg_4_000_aa    m/z 1100.1858     0   1E+4   0.5   1  RT (min)      lcqtof_hilic_neg_1_059_15    m/z 1100.2681     0   1E+5   1   1.5  RT (min)      lcqtof_hilic_neg_4_000_aa    m/z 1100.9731     0   1   2E+4   0.5   1  RT (min)      lcqtof_hilic_neg_4_000_aa    m/z 1101.0483     0   1E+4   1   1.5  RT (min)      lcqtof_hilic_neg_4_000_aa    m/z 1101.1781     0   1   2E+4   0.5   1  RT (min)      lcqtof_hilic_neg_1_062_17    m/z 1101.2677     0   1E+5   1   1.5  RT (min)      lcqtof_hilic_neg_4_000_aa    m/z 1101.7229     0   1   2   3   4E+4   1   1.5  RT (min)      lcqtof_hilic_neg_4_000_aa    m/z 1101.7324     0   1E+4   3.5   4  RT (min)      lcqtof_hilic_neg_1_007_05    m/z 1101.8116     0   1   2E+4   1   1.5  RT (min)      lcqtof_hilic_neg_4_000_aa    m/z 1101.8445     0   1   2   3E+4   1   1.5  RT (min)      lcqtof_hilic_neg_4_000_aa    m/z 1101.8549     0   1   2   3E+4   1   1.5  RT (min)      lcqtof_hilic_neg_4_000_aa    m/z 1102.0385     0   1E+4   1   1.5  RT (min)      lcqtof_hilic_neg_4_000_aa    m/z 1102.0869     0   1E+4   0.5   1  RT (min)      lcqtof_hilic_neg_4_004_47    m/z 1102.2661     0   2   4   6   8E+4   1   1.5  RT (min)      lcqtof_hilic_neg_4_000_aa    m/z 1102.7238     0   1E+4   1   1.5  RT (min)      lcqtof_hilic_neg_4_000_aa    m/z 1103.1721     0   1E+4   0.5   1  RT (min)      lcqtof_hilic_neg_1_062_17    m/z 1103.2678     0   2   4   6E+4   1   1.5  RT (min)      lcqtof_hilic_neg_4_000_aa    m/z 1103.4491     0   1E+4   0.5   1  RT (min)      lcqtof_hilic_neg_4_042_22    m/z 1103.5134     0   1   2E+3   4.5   5  RT (min)       lcqtof_hilic_neg_4_063_16    m/z 1104.2711     0   1   2E+4   1   1.5  RT (min)      lcqtof_hilic_neg_4_000_aa    m/z 1104.9639     0   1   2E+4   1   1.5  RT (min)      lcqtof_hilic_neg_4_000_aa    m/z 1104.9761     0   1   2E+4   0.5   1  RT (min)      lcqtof_hilic_neg_4_000_aa    m/z 1105.2656     0   1E+4   1   1.5  RT (min)      lcqtof_hilic_neg_4_000_aa    m/z 1105.7200     0   1E+5   3.5   4  RT (min)      lcqtof_hilic_neg_1_007_05    m/z 1105.8010     0   1   2E+4   4   4.5  RT (min)      lcqtof_hilic_neg_1_037_24    m/z 1106.7253     0   2   4E+4   3.5   4  RT (min)      lcqtof_hilic_neg_1_007_05    m/z 1106.9760     0   1   2E+4   1   1.5  RT (min)      lcqtof_hilic_neg_4_000_aa    m/z 1107.8163     0   1   2E+4   4   4.5  RT (min)      lcqtof_hilic_neg_4_008_24    m/z 1108.7433     0   1   2   3E+4   3.5   4  RT (min)      lcqtof_hilic_neg_1_007_05    m/z 1108.8210     0   1E+4   4   4.5  RT (min)      lcqtof_hilic_neg_4_008_24    m/z 1108.8992     0   1E+4   1   1.5  RT (min)      lcqtof_hilic_neg_4_008_24    m/z 1108.9657     0   1   2E+4   0.5   1  RT (min)      lcqtof_hilic_neg_4_000_aa    m/z 1108.9689     0   1   2E+4   1   1.5  RT (min)      lcqtof_hilic_neg_4_000_aa    m/z 1109.5169     0   1E+4   4   4.5  RT (min)      lcqtof_hilic_neg_4_039_18    m/z 1109.7208     0   1   2E+4   4   4.5  RT (min)       lcqtof_hilic_neg_4_008_24    m/z 1109.8252     0   1E+4   1   1.5  RT (min)      lcqtof_hilic_neg_4_000_aa    m/z 1109.8310     0   1   2   3E+4   4   4.5  RT (min)      lcqtof_hilic_neg_4_008_24    m/z 1109.9003     0   1E+4   1   1.5  RT (min)      lcqtof_hilic_neg_4_002_25    m/z 1109.9010     0   1   2   3E+4   1   1.5  RT (min)       lcqtof_hilic_neg_4_008_24    m/z 1110.8450     0   1   2   3   4E+4   1   1.5  RT (min)      lcqtof_hilic_neg_4_008_24    m/z 1110.8777     0   1E+4   0.5   1  RT (min)      lcqtof_hilic_neg_4_004_47    m/z 1110.8807     0   1   2E+4   1   1.5  RT (min)      lcqtof_hilic_neg_4_008_24    m/z 1110.9711     0   1   2E+4   1   1.5  RT (min)      lcqtof_hilic_neg_4_000_aa    m/z 1111.6432     0   1E+4   3.5   4  RT (min)      lcqtof_hilic_neg_4_022_34    m/z 1111.6880     0   1E+4   4.5   5  RT (min)      lcqtof_hilic_neg_4_008_24    m/z 1111.7656     0   2   4E+4   4   4.5  RT (min)      lcqtof_hilic_neg_4_040_05    m/z 1111.8502     0   1E+5   1   1.5  RT (min)       lcqtof_hilic_neg_4_008_24    m/z 1112.7722     0   1E+4   4   4.5  RT (min)      lcqtof_hilic_neg_4_040_05    m/z 1112.9718     0   1   2E+4   1   1.5  RT (min)      lcqtof_hilic_neg_4_000_aa    m/z 1113.1041     0   1E+4   0.5   1  RT (min)      lcqtof_hilic_neg_4_004_47    m/z 1113.6543     0   1E+4   3.5   4  RT (min)      lcqtof_hilic_neg_4_022_34    m/z 1114.1025     0   1E+4   0.5   1  RT (min)      lcqtof_hilic_neg_4_004_47    m/z 1114.2467     0   1E+4   6.5   7  RT (min)      lcqtof_hilic_neg_4_008_24    m/z 1114.7336     0   2   4   6   8E+4   1   1.5  RT (min)      lcqtof_hilic_neg_4_000_aa    m/z 1114.9810     0   1   2E+4   1   1.5  RT (min)      lcqtof_hilic_neg_4_000_aa    m/z 1115.0565     0   1E+4   0.5   1  RT (min)      lcqtof_hilic_neg_3_035_46    m/z 1115.7356     0   2   4E+4   1   1.5  RT (min)      lcqtof_hilic_neg_4_000_aa    m/z 1115.7382     0   2   4E+4   1   1.5  RT (min)      lcqtof_hilic_neg_4_000_aa    m/z 1115.7998     0   1E+4   1   1.5  RT (min)      lcqtof_hilic_neg_4_004_47    m/z 1116.2607     0   1E+4   1   1.5  RT (min)      lcqtof_hilic_neg_4_000_aa    m/z 1116.7430     0   1E+4   1   1.5  RT (min)      lcqtof_hilic_neg_4_000_aa    m/z 1117.2628     0   1E+4   1   1.5  RT (min)      lcqtof_hilic_neg_4_000_aa    m/z 1117.8142     0   1E+4   1   1.5  RT (min)      lcqtof_hilic_neg_4_000_aa    m/z 1118.2593     0   1E+4   1   1.5  RT (min)      lcqtof_hilic_neg_4_000_aa    m/z 1118.8213     0   1E+4   1   1.5  RT (min)      lcqtof_hilic_neg_4_000_aa    m/z 1119.5416     0   1E+4   4.5   5  RT (min)      lcqtof_hilic_neg_4_040_05    m/z 1120.7815     0   1E+4   4   4.5  RT (min)      lcqtof_hilic_neg_1_037_24    m/z 1120.9753     0   1E+4   0.5   1  RT (min)      lcqtof_hilic_neg_4_000_aa    m/z 1121.7787     0   1   2   3   4E+4   4   4.5  RT (min)      lcqtof_hilic_neg_1_037_24    m/z 1121.7814     0   1E+4   3.5   4  RT (min)      lcqtof_hilic_neg_1_007_05    m/z 1122.7927     0   1E+4   4   4.5  RT (min)      lcqtof_hilic_neg_2_052_24    m/z 1122.9120     0   1E+4   1   1.5  RT (min)      lcqtof_hilic_neg_4_008_24    m/z 1122.9673     0   1   2E+4   1   1.5  RT (min)      lcqtof_hilic_neg_4_000_aa    m/z 1122.9760     0   1   2E+4   0.5   1  RT (min)      lcqtof_hilic_neg_4_000_aa    m/z 1123.7910     0   1   2   3E+4   4   4.5  RT (min)      lcqtof_hilic_neg_1_037_24    m/z 1123.7949     0   1E+4   3.5   4  RT (min)      lcqtof_hilic_neg_1_007_05    m/z 1123.8230     0   1E+4   1   1.5  RT (min)      lcqtof_hilic_neg_4_000_aa    m/z 1123.8882     0   1E+4   0.5   1  RT (min)      lcqtof_hilic_neg_4_004_47    m/z 1123.9180     0   1   2   3E+4   1   1.5  RT (min)       lcqtof_hilic_neg_4_008_24    m/z 1123.9180     0   1E+4   1   1.5  RT (min)      lcqtof_hilic_neg_4_002_25    m/z 1124.8086     0   1E+4   4   4.5  RT (min)      lcqtof_hilic_neg_1_037_24    m/z 1124.9193     0   1E+4   1   1.5  RT (min)      lcqtof_hilic_neg_4_008_24    m/z 1125.8091     0   2   4E+4   4   4.5  RT (min)      lcqtof_hilic_neg_1_037_24    m/z 1125.8099     0   1   2E+4   3.5   4  RT (min)      lcqtof_hilic_neg_1_007_05    m/z 1126.9578     0   1   2E+4   1   1.5  RT (min)      lcqtof_hilic_neg_4_000_aa    m/z 1127.8164     0   1   2E+4   3.5   4  RT (min)      lcqtof_hilic_neg_1_037_24    m/z 1128.7686     0   1E+4   7   7.5  RT (min)      lcqtof_hilic_neg_4_008_24    m/z 1129.0661     0   1E+4   0.5   1  RT (min)      lcqtof_hilic_neg_1_062_17    m/z 1129.8375     0   1E+4   3.5   4  RT (min)      lcqtof_hilic_neg_1_037_24    m/z 1130.0668     0   1E+4   0.5   1  RT (min)      lcqtof_hilic_neg_1_062_17    m/z 1130.9749     0   1   2E+4   0.5   1  RT (min)      lcqtof_hilic_neg_4_000_aa    m/z 1131.0609     0   1E+4   1   1.5  RT (min)      lcqtof_hilic_neg_4_000_aa    m/z 1131.0992     0   1E+4   0.5   1  RT (min)      lcqtof_hilic_neg_4_026_04    m/z 1131.1135     0   1E+4   0.5   1  RT (min)      lcqtof_hilic_neg_4_004_47    m/z 1132.1175     0   1E+4   0.5   1  RT (min)      lcqtof_hilic_neg_4_004_47    m/z 1132.5117     0   1   2   3E+3   4.5   5  RT (min)       lcqtof_hilic_neg_4_054_48    m/z 1132.8189     0   1   2   3E+4   4   4.5  RT (min)      lcqtof_hilic_neg_4_008_24    m/z 1132.9664     0   1   2E+4   0.5   1  RT (min)      lcqtof_hilic_neg_4_000_aa    m/z 1133.6257     0   1E+4   0.5   1   1.5  RT (min)      lcqtof_hilic_neg_4_019_03    m/z 1133.8226     0   1E+5   4   4.5  RT (min)       lcqtof_hilic_neg_4_008_24    m/z 1133.8748     0   1E+4   1   1.5  RT (min)      lcqtof_hilic_neg_4_000_aa    m/z 1133.8784     0   1E+4   1   1.5  RT (min)      lcqtof_hilic_neg_4_000_aa    m/z 1134.6353     0   1E+4   1   1.5  RT (min)      lcqtof_hilic_neg_4_019_03    m/z 1134.8349     0   2   4E+4   4   4.5  RT (min)      lcqtof_hilic_neg_4_008_24    m/z 1134.9839     0   1E+4   0.5   1  RT (min)      lcqtof_hilic_neg_4_000_aa    m/z 1135.8392     0   1E+5   4   4.5  RT (min)       lcqtof_hilic_neg_4_008_24    m/z 1136.8484     0   1   2E+4   4   4.5  RT (min)      lcqtof_hilic_neg_4_008_24    m/z 1137.8540     0   2   4E+4   4   4.5  RT (min)      lcqtof_hilic_neg_4_008_24    m/z 1137.9318     0   1   2E+4   1   1.5  RT (min)       lcqtof_hilic_neg_4_008_24    m/z 1138.9305     0   1E+4   1   1.5  RT (min)      lcqtof_hilic_neg_4_000_aa    m/z 1139.8724     0   1E+4   3.5   4  RT (min)      lcqtof_hilic_neg_4_008_24    m/z 1139.9982     0   1E+4   1   1.5  RT (min)      lcqtof_hilic_neg_4_022_34    m/z 1142.9645     0   1   2E+4   1   1.5  RT (min)      lcqtof_hilic_neg_4_000_aa    m/z 1142.9836     0   1   2E+4   1   1.5  RT (min)      lcqtof_hilic_neg_4_000_aa    m/z 1144.9573     0   1   2E+4   1   1.5  RT (min)      lcqtof_hilic_neg_4_000_aa    m/z 1146.0874     0   1E+4   1   1.5  RT (min)      lcqtof_hilic_neg_4_000_aa    m/z 1146.4823     0   1   2   3E+4   1   1.5  RT (min)      lcqtof_hilic_neg_4_027_19    m/z 1146.9734     0   1   2E+4   0.5   1  RT (min)      lcqtof_hilic_neg_4_000_aa    m/z 1146.9758     0   1   2E+4   1   1.5  RT (min)      lcqtof_hilic_neg_4_000_aa    m/z 1146.9783     0   1   2E+4   1   1.5  RT (min)      lcqtof_hilic_neg_4_000_aa    m/z 1147.1280     0   1E+4   0.5   1  RT (min)      lcqtof_hilic_neg_4_004_47    m/z 1147.4729     0   1E+4   0.5   1  RT (min)      lcqtof_hilic_neg_4_021_21    m/z 1147.5525     0   1   2E+4   0.5   1  RT (min)      lcqtof_hilic_neg_3_003_20    m/z 1148.5224     0   1E+4   1   1.5  RT (min)      lcqtof_hilic_neg_4_061_50    m/z 1148.9717     0   1   2E+4   1   1.5  RT (min)      lcqtof_hilic_neg_4_000_aa    m/z 1148.9784     0   1   2E+4   0.5   1  RT (min)      lcqtof_hilic_neg_4_000_aa    m/z 1149.0574     0   1E+4   0.5   1  RT (min)      lcqtof_hilic_neg_3_036_17    m/z 1149.4692     0   1E+4   0.5   1  RT (min)      lcqtof_hilic_neg_4_021_21    m/z 1149.5250     0   1   2E+4   1   1.5  RT (min)      lcqtof_hilic_neg_4_009_50    m/z 1149.5491     0   1E+4   0.5   1  RT (min)      lcqtof_hilic_neg_3_035_46    m/z 1149.6150     0   1E+4   2.5   3  RT (min)      lcqtof_hilic_neg_4_019_03    m/z 1149.7998     0   1E+4   4   4.5  RT (min)      lcqtof_hilic_neg_1_037_24    m/z 1150.5504     0   1E+4   0.5   1  RT (min)      lcqtof_hilic_neg_3_035_46    m/z 1150.6159     0   1E+4   2.5   3  RT (min)      lcqtof_hilic_neg_4_019_03    m/z 1150.9708     0   1   2E+4   1   1.5  RT (min)      lcqtof_hilic_neg_4_000_aa    m/z 1150.9749     0   1   2E+4   0.5   1  RT (min)      lcqtof_hilic_neg_4_000_aa    m/z 1151.6265     0   1E+4   2.5   3  RT (min)      lcqtof_hilic_neg_4_019_03    m/z 1152.9710     0   1E+4   1   1.5  RT (min)      lcqtof_hilic_neg_4_000_aa    m/z 1153.9546     0   1   2   3E+4   6.5   7  RT (min)       lcqtof_hilic_neg_4_059_14    m/z 1153.9563     0   1   2E+4   7   7.5  RT (min)      lcqtof_hilic_neg_4_053_14    m/z 1156.6219     0   1E+4   1   1.5  RT (min)      lcqtof_hilic_neg_4_019_03    m/z 1156.7522     0   1E+4   3   3.5  RT (min)      lcqtof_hilic_neg_4_016_37    m/z 1156.9674     0   1   2E+4   1   1.5  RT (min)      lcqtof_hilic_neg_4_000_aa    m/z 1158.6334     0   1E+4   0.5   1   1.5  RT (min)      lcqtof_hilic_neg_4_019_03    m/z 1158.9674     0   1   2E+4   1   1.5  RT (min)      lcqtof_hilic_neg_4_000_aa    m/z 1160.4800     0   1   2E+4   0.5   1  RT (min)      lcqtof_hilic_neg_4_021_21    m/z 1160.4928     0   1   2E+5   0.5   1  RT (min)       lcqtof_hilic_neg_4_038_10    m/z 1160.5037     0   1   2   3E+4   0.5   1  RT (min)      lcqtof_hilic_neg_4_042_22    m/z 1160.6530     0   1E+4   1   1.5  RT (min)      lcqtof_hilic_neg_4_019_03    m/z 1161.0600     0   1   2E+4   0.5   1  RT (min)      lcqtof_hilic_neg_4_004_47    m/z 1161.5448     0   1E+4   0.5   1  RT (min)      lcqtof_hilic_neg_1_053_19    m/z 1161.7845     0   1E+4   3   3.5  RT (min)      lcqtof_hilic_neg_4_005_29    m/z 1162.0557     0   1E+4   0.5   1  RT (min)      lcqtof_hilic_neg_4_004_47    m/z 1163.1562     0   1E+4   0.5   1  RT (min)      lcqtof_hilic_neg_1_062_17    m/z 1163.2561     0   1   2   3E+4   1   1.5  RT (min)      lcqtof_hilic_neg_4_000_aa    m/z 1163.5441     0   1E+4   1   1.5  RT (min)      lcqtof_hilic_neg_4_009_50    m/z 1163.8056     0   1E+4   3   3.5  RT (min)      lcqtof_hilic_neg_4_018_27    m/z 1163.8451     0   1E+4   1   1.5  RT (min)      lcqtof_hilic_neg_4_000_aa    m/z 1164.2620     0   1   2   3E+4   1   1.5  RT (min)      lcqtof_hilic_neg_4_000_aa    m/z 1164.2650     0   1   2E+4   0.5   1  RT (min)      lcqtof_hilic_neg_1_062_17    m/z 1164.5444     0   2   4   6   8E+4   1   1.5  RT (min)      lcqtof_hilic_neg_3_003_20    m/z 1164.9778     0   1   2E+4   0.5   1  RT (min)      lcqtof_hilic_neg_4_000_aa    m/z 1165.2502     0   2   4E+4   1   1.5  RT (min)      lcqtof_hilic_neg_4_000_aa    m/z 1165.2559     0   2   4E+4   1   1.5  RT (min)      lcqtof_hilic_neg_4_000_aa    m/z 1165.4550     0   1E+4   1   1.5  RT (min)      lcqtof_hilic_neg_4_038_10    m/z 1165.4653     0   1E+4   1   1.5  RT (min)      lcqtof_hilic_neg_4_042_22    m/z 1165.5475     0   2   4E+4   1   1.5  RT (min)      lcqtof_hilic_neg_3_003_20    m/z 1165.8912     0   1E+4   1   1.5  RT (min)      lcqtof_hilic_neg_4_055_32    m/z 1166.2582     0   1   2   3   4E+4   1   1.5  RT (min)      lcqtof_hilic_neg_4_000_aa    m/z 1166.4961     0   1   2E+4   1   1.5  RT (min)      lcqtof_hilic_neg_4_045_26    m/z 1166.5430     0   1E+4   2.5   3  RT (min)      lcqtof_hilic_neg_4_063_16    m/z 1166.7938     0   1   2   3E+4   1   1.5  RT (min)      lcqtof_hilic_neg_4_013_20    m/z 1166.9768     0   1   2E+4   1   1.5  RT (min)      lcqtof_hilic_neg_4_000_aa    m/z 1167.2473     0   1   2E+4   1   1.5  RT (min)      lcqtof_hilic_neg_4_000_aa    m/z 1167.2644     0   1E+4   0.5   1  RT (min)      lcqtof_hilic_neg_1_062_17    m/z 1167.6824     0   1E+4   3.5   4  RT (min)      lcqtof_hilic_neg_4_045_26    m/z 1168.5455     0   1E+4   2.5   3  RT (min)      lcqtof_hilic_neg_4_063_16    m/z 1168.9384     0   1   2E+4   1   1.5  RT (min)      lcqtof_hilic_neg_4_000_aa    m/z 1168.9600     0   1   2E+4   1   1.5  RT (min)      lcqtof_hilic_neg_4_000_aa    m/z 1168.9606     0   1   2E+4   1   1.5  RT (min)      lcqtof_hilic_neg_4_000_aa    m/z 1168.9716     0   1   2E+4   0.5   1  RT (min)      lcqtof_hilic_neg_4_000_aa    m/z 1169.9227     0   1   2   3   4E+4   1   1.5  RT (min)      lcqtof_hilic_neg_4_000_aa    m/z 1170.4443     0   1E+5   0.5   1  RT (min)      lcqtof_hilic_neg_3_003_20    m/z 1170.4584     0   1E+4   1   1.5  RT (min)      lcqtof_hilic_neg_4_013_20    m/z 1170.8435     0   1E+4   0.5   1  RT (min)      lcqtof_hilic_neg_4_004_47    m/z 1170.9233     0   1E+5   1   1.5  RT (min)      lcqtof_hilic_neg_4_000_aa    m/z 1170.9716     0   1   2E+4   1   1.5  RT (min)      lcqtof_hilic_neg_4_000_aa    m/z 1171.4459     0   1E+5   0.5   1  RT (min)      lcqtof_hilic_neg_3_003_20    m/z 1171.7363     0   1   2E+4   1   1.5  RT (min)      lcqtof_hilic_neg_4_000_aa    m/z 1172.4009     0   1E+5   1   1.5  RT (min)      lcqtof_hilic_neg_4_013_20    m/z 1172.4473     0   1   2   3E+4   0.5   1  RT (min)      lcqtof_hilic_neg_3_003_20    m/z 1172.7351     0   1E+4   1   1.5  RT (min)      lcqtof_hilic_neg_4_000_aa    m/z 1172.9550     0   1   2E+4   1   1.5  RT (min)      lcqtof_hilic_neg_4_000_aa    m/z 1172.9755     0   1   2E+4   0.5   1  RT (min)      lcqtof_hilic_neg_4_000_aa    m/z 1173.2876     0   1E+5   1   1.5  RT (min)      lcqtof_hilic_neg_4_000_aa    m/z 1173.3849     0   1   2E+4   0.5   1  RT (min)      lcqtof_hilic_neg_4_021_21    m/z 1173.4070     0   1   2   3E+4   0.5   1  RT (min)       lcqtof_hilic_neg_2_037_21    m/z 1173.7554     0   2   4E+4   1   1.5  RT (min)      lcqtof_hilic_neg_4_000_aa    m/z 1173.8930     0   2   4   6   8E+4   1   1.5  RT (min)      lcqtof_hilic_neg_4_000_aa    m/z 1174.2887     0   1E+5   1   1.5  RT (min)      lcqtof_hilic_neg_4_000_aa    m/z 1174.4927     0   1E+4   0.5   1  RT (min)      lcqtof_hilic_neg_4_021_21    m/z 1174.5029     0   1   2   3E+5   1   1.5  RT (min)      lcqtof_hilic_neg_4_038_10    m/z 1174.5044     0   1   2   3E+5   0.5   1  RT (min)      lcqtof_hilic_neg_4_038_10    m/z 1174.5345     0   1E+4   0.5   1  RT (min)      lcqtof_hilic_neg_4_042_22    m/z 1174.5630     0   1   2E+4   0.5   1  RT (min)      lcqtof_hilic_neg_3_002_10    m/z 1174.7607     0   1   2   3   4E+4   1   1.5  RT (min)      lcqtof_hilic_neg_4_000_aa    m/z 1174.8967     0   2   4   6E+4   1   1.5  RT (min)      lcqtof_hilic_neg_4_000_aa    m/z 1174.9705     0   1   2E+4   1   1.5  RT (min)      lcqtof_hilic_neg_4_000_aa    m/z 1175.2798     0   1E+5   1   1.5  RT (min)      lcqtof_hilic_neg_4_000_aa    m/z 1175.2833     0   1E+5   1   1.5  RT (min)      lcqtof_hilic_neg_4_000_aa    m/z 1175.2852     0   1E+5   1   1.5  RT (min)      lcqtof_hilic_neg_4_000_aa    m/z 1175.2914     0   1E+5   1   1.5  RT (min)      lcqtof_hilic_neg_4_000_aa    m/z 1175.5046     0   1   2E+5   1   1.5  RT (min)      lcqtof_hilic_neg_4_038_10    m/z 1175.5090     0   1   2E+5   0.5   1  RT (min)      lcqtof_hilic_neg_4_038_10    m/z 1175.7676     0   1   2E+4   1   1.5  RT (min)      lcqtof_hilic_neg_4_000_aa    m/z 1175.8955     0   1   2E+4   1   1.5  RT (min)      lcqtof_hilic_neg_4_000_aa    m/z 1175.9364     0   1E+4   7   7.5  RT (min)      lcqtof_hilic_neg_4_059_14    m/z 1175.9388     0   1E+4   7   7.5  RT (min)      lcqtof_hilic_neg_4_059_14    m/z 1175.9388     0   1E+4   6.5   7  RT (min)       lcqtof_hilic_neg_4_059_14    m/z 1176.2878     0   2   4   6   8E+4   1   1.5  RT (min)      lcqtof_hilic_neg_4_000_aa    m/z 1176.5104     0   1E+5   0.5   1  RT (min)      lcqtof_hilic_neg_4_038_10    m/z 1176.5534     0   1E+4   1   1.5  RT (min)      lcqtof_hilic_neg_4_009_50    m/z 1176.5550     0   1   2   3E+4   0.5   1  RT (min)      lcqtof_hilic_neg_4_013_20    m/z 1176.7773     0   1E+4   1   1.5  RT (min)      lcqtof_hilic_neg_4_000_aa    m/z 1176.9677     0   1   2E+4   1   1.5  RT (min)      lcqtof_hilic_neg_4_000_aa    m/z 1177.2864     0   2   4E+4   1   1.5  RT (min)      lcqtof_hilic_neg_4_000_aa    m/z 1177.5590     0   1E+4   1   1.5  RT (min)      lcqtof_hilic_neg_4_009_50    m/z 1177.7097     0   1   2E+4   3.5   4  RT (min)      lcqtof_hilic_neg_4_045_26    m/z 1177.7623     0   1   2E+4   7   7.5  RT (min)      lcqtof_hilic_neg_4_008_24    m/z 1177.9791     0   1E+4   1   1.5  RT (min)      lcqtof_hilic_neg_4_000_aa    m/z 1178.2854     0   1   2   3E+4   1   1.5  RT (min)      lcqtof_hilic_neg_4_000_aa    m/z 1178.7092     0   1E+4   3   3.5  RT (min)      lcqtof_hilic_neg_4_006_30    m/z 1178.7145     0   1E+4   3.5   4  RT (min)      lcqtof_hilic_neg_4_045_26    m/z 1178.9742     0   1   2E+4   1   1.5  RT (min)      lcqtof_hilic_neg_4_000_aa    m/z 1179.2833     0   1E+4   1   1.5  RT (min)      lcqtof_hilic_neg_4_000_aa    m/z 1179.7241     0   1E+4   3.5   4  RT (min)      lcqtof_hilic_neg_4_045_26    m/z 1179.8477     0   1E+4   1   1.5  RT (min)      lcqtof_hilic_neg_4_000_aa    m/z 1180.5139     0   1E+5   0.5   1  RT (min)      lcqtof_hilic_neg_1_026_10    m/z 1180.8113     0   1E+4   1   1.5  RT (min)      lcqtof_hilic_neg_4_013_20    m/z 1180.9767     0   1   2E+4   0.5   1  RT (min)      lcqtof_hilic_neg_4_000_aa    m/z 1181.8582     0   1E+4   0.5   1  RT (min)      lcqtof_hilic_neg_4_004_47    m/z 1182.5130     0   2   4   6E+4   1   1.5  RT (min)      lcqtof_hilic_neg_2_053_18    m/z 1182.8585     0   1E+4   0.5   1  RT (min)      lcqtof_hilic_neg_4_004_47    m/z 1182.9752     0   1   2E+4   0.5   1  RT (min)      lcqtof_hilic_neg_4_000_aa    m/z 1183.5167     0   1   2   3E+4   1   1.5  RT (min)      lcqtof_hilic_neg_2_053_18    m/z 1183.8576     0   1E+4   0.5   1  RT (min)      lcqtof_hilic_neg_4_004_47    m/z 1184.6521     0   1E+4   0.5   1   1.5  RT (min)      lcqtof_hilic_neg_4_019_03    m/z 1184.9701     0   1   2E+4   0.5   1  RT (min)      lcqtof_hilic_neg_4_000_aa    m/z 1184.9733     0   1   2E+4   0.5   1  RT (min)      lcqtof_hilic_neg_4_000_aa    m/z 1185.4165     0   1   2E+5   1   1.5  RT (min)      lcqtof_hilic_neg_4_013_20    m/z 1186.3990     0   2   4   6E+4   0.5   1  RT (min)      lcqtof_hilic_neg_4_021_21    m/z 1186.4248     0   2   4   6   8E+4   0.5   1  RT (min)      lcqtof_hilic_neg_1_024_21    m/z 1186.4407     0   1   2   3E+4   0.5   1  RT (min)      lcqtof_hilic_neg_4_027_19    m/z 1186.4640     0   1E+4   1   1.5  RT (min)      lcqtof_hilic_neg_4_004_47    m/z 1186.5820     0   1   2E+4   0.5   1  RT (min)      lcqtof_hilic_neg_4_038_10    m/z 1186.6523     0   1E+4   5   5.5  RT (min)       lcqtof_hilic_neg_4_048_23    m/z 1187.3998     0   1E+4   0.5   1  RT (min)      lcqtof_hilic_neg_4_021_21    m/z 1187.4214     0   1E+4   0.5   1  RT (min)      lcqtof_hilic_neg_4_042_22    m/z 1187.5803     0   1E+4   0.5   1  RT (min)      lcqtof_hilic_neg_4_038_10    m/z 1188.3882     0   2   4   6E+4   0.5   1  RT (min)      lcqtof_hilic_neg_4_021_21    m/z 1188.4132     0   1E+5   0.5   1  RT (min)       lcqtof_hilic_neg_1_032_18    m/z 1188.4540     0   1   2E+4   0.5   1  RT (min)      lcqtof_hilic_neg_3_003_20    m/z 1188.5219     0   1E+5   0.5   1  RT (min)      lcqtof_hilic_neg_4_038_10    m/z 1188.9701     0   1   2E+4   1   1.5  RT (min)      lcqtof_hilic_neg_4_000_aa    m/z 1189.3896     0   1   2   3   4E+4   0.5   1  RT (min)      lcqtof_hilic_neg_4_021_21    m/z 1189.4148     0   1E+5   0.5   1  RT (min)      lcqtof_hilic_neg_1_032_18    m/z 1190.2751     0   1E+4   1   1.5  RT (min)      lcqtof_hilic_neg_4_000_aa    m/z 1190.3898     0   1E+4   0.5   1  RT (min)      lcqtof_hilic_neg_4_021_21    m/z 1190.4123     0   1   2   3E+4   0.5   1  RT (min)      lcqtof_hilic_neg_4_039_18    m/z 1190.5735     0   1E+4   0.5   1  RT (min)      lcqtof_hilic_neg_4_013_20    m/z 1190.9696     0   1   2E+4   1   1.5  RT (min)      lcqtof_hilic_neg_4_000_aa    m/z 1191.2749     0   1E+4   1   1.5  RT (min)      lcqtof_hilic_neg_4_000_aa    m/z 1192.5066     0   1E+5   0.5   1  RT (min)      lcqtof_hilic_neg_4_021_21    m/z 1192.5770     0   1   2   3   4E+4   0.5   1  RT (min)      lcqtof_hilic_neg_3_003_20    m/z 1192.7522     0   1E+4   0.5   1  RT (min)      lcqtof_hilic_neg_4_004_47    m/z 1192.9731     0   1   2E+4   0.5   1  RT (min)      lcqtof_hilic_neg_4_000_aa    m/z 1193.2764     0   1E+4   1   1.5  RT (min)      lcqtof_hilic_neg_4_000_aa    m/z 1193.5793     0   1   2   3E+4   0.5   1  RT (min)      lcqtof_hilic_neg_3_003_20    m/z 1194.5823     0   1E+4   1   1.5  RT (min)      lcqtof_hilic_neg_3_003_20    m/z 1194.7532     0   1   2E+4   0.5   1  RT (min)      lcqtof_hilic_neg_4_004_47    m/z 1194.9613     0   1   2E+4   1   1.5  RT (min)      lcqtof_hilic_neg_4_000_aa    m/z 1194.9681     0   1   2E+4   1   1.5  RT (min)      lcqtof_hilic_neg_4_000_aa    m/z 1195.5349     0   1E+4   1   1.5  RT (min)      lcqtof_hilic_neg_4_038_10    m/z 1195.7495     0   1E+4   0.5   1  RT (min)      lcqtof_hilic_neg_4_004_47    m/z 1196.5735     0   1E+4   1   1.5  RT (min)      lcqtof_hilic_neg_3_003_20    m/z 1196.9675     0   1   2E+4   1   1.5  RT (min)      lcqtof_hilic_neg_4_000_aa    m/z 1196.9761     0   1   2E+4   1   1.5  RT (min)      lcqtof_hilic_neg_4_000_aa    m/z 1198.4880     0   1   2E+4   8   8.5  RT (min)      lcqtof_hilic_neg_2_000_aa    m/z 1198.4902     0   1   2E+4   8   8.5  RT (min)      lcqtof_hilic_neg_1_000_aa    m/z 1198.9744     0   1   2E+4   1   1.5  RT (min)      lcqtof_hilic_neg_4_000_aa    m/z 1199.7096     0   1E+4   5   5.5  RT (min)      lcqtof_hilic_neg_4_048_23    m/z 1199.7267     0   1E+4   5   5.5  RT (min)      lcqtof_hilic_neg_4_048_23    m/z 1199.7437     0   1E+4   7   7.5  RT (min)      lcqtof_hilic_neg_4_008_24   
