## Supplementary material for "The Everything Bagel Feature Finder: Ultra-fast automated feature finding for untargeted metabolomics": SI Data File: tool_unique_xic_st_XCMS_ST004500.html

 
 
 XCMS-unique features (not detected by the other two tools) - ST004500 
 
 XCMS-unique features (not detected by the other two tools) - ST004500 
 11462 features · orbitrap · XIC window m/z ± 10 ppm (≥ 0.002 Da) · ± 0.5 min 
 Every panel is a feature  XCMS  detected that neither of the other two tools detected — three-way unique across EB (standard mode (min_detection_rate=0.1)), Masscube and XCMS, under the standard matching rule (m/z ± 10 ppm; apex within the reference RT window ± 0.2 min). It shows the MS1 extracted-ion chromatogram at the feature's m/z, drawn from the raw mzML named in  alignment_reference_file , zoomed to ± 0.5 min around the feature's reported apex RT.  Gray solid line  = the feature's reported apex RT;  red dashed line  = XCMS's assigned MS/MS —  RTINSECONDS  in its  aligned_features.mgf ; the panel widens to keep an off-apex MS/MS on-screen. Features with no assigned MS/MS show only the gray line. y is per-cell autoscaled. Rows keep the table order (m/z ascending). 
   m/z 61.0102     0   2   4E+3   18   18.5  RT (min)      lcorbi_t3_neg_4_095_60    m/z 74.0239     0   1   2   3E+4   1.5   2   2.5  RT (min)       lcorbi_t3_neg_4_097_60    m/z 78.9189     0   1   1   1.5  RT (min)       lcorbi_t3_neg_4_042_14    m/z 78.9590     0   2   4   6E+3   18   18.5  RT (min)      lcorbi_t3_neg_4_097_60    m/z 80.9168     0   1   2E+4   1   1.5  RT (min)       lcorbi_t3_neg_4_079_58    m/z 84.9907     0   1E+4   17.5   18  RT (min)      lcorbi_t3_neg_4_084_59    m/z 88.0087     0   2   4   6   8E+3   18   18.5  RT (min)      lcorbi_t3_neg_4_095_60    m/z 89.0051     0   1   2   3E+4   18   18.5  RT (min)       lcorbi_t3_neg_4_095_60    m/z 89.9410     0   2   4   6E+3   1.5   2  RT (min)      lcorbi_t3_neg_4_091_60    m/z 90.0188     0   2   4   6E+4   1   1.5  RT (min)       lcorbi_t3_neg_4_076_58    m/z 90.0471     0   1E+4   9.5   10  RT (min)      lcorbi_t3_neg_4_100_61    m/z 91.0522     0   1   2   3   4E+5   9.5   10  RT (min)       lcorbi_t3_neg_4_102_61    m/z 92.9280     0   1   2   3   4E+4   17.5   18  RT (min)       lcorbi_t3_neg_4_068_57    m/z 94.9250     0   1   2   3E+4   17.5   18  RT (min)       lcorbi_t3_neg_4_068_57    m/z 96.9220     0   1   2   3   4E+4   1   1.5  RT (min)       lcorbi_t3_neg_4_076_58    m/z 96.9600     0   1   2   3E+4   18   18.5  RT (min)       lcorbi_t3_neg_4_095_60    m/z 96.9695     0   1E+4   18   18.5  RT (min)      lcorbi_t3_neg_4_094_60    m/z 98.9067     0   1E+4   0.5   1  RT (min)       lcorbi_t3_neg_4_069_57    m/z 98.9068     0   1   2   3E+4   18.5   19  RT (min)       lcorbi_t3_neg_4_111_62    m/z 100.0379     0   1   2   3E+4   9   9.5  RT (min)      lcorbi_t3_neg_2_048_27    m/z 100.0667     0   1E+3   14.5   15  RT (min)      lcorbi_t3_neg_4_039_32    m/z 100.0835     0   1E+4   14   14.5  RT (min)      lcorbi_t3_neg_3_002_24    m/z 100.1857     0   1E+4   13.5   14  RT (min)      lcorbi_t3_neg_2_052_32    m/z 100.2580     0   1   2   3E+4   9   9.5  RT (min)      lcorbi_t3_neg_2_048_27    m/z 100.2591     0   1   2   3E+4   9   9.5  RT (min)      lcorbi_t3_neg_2_048_27    m/z 100.3695     0   1E+4   9   9.5  RT (min)      lcorbi_t3_neg_2_048_27    m/z 100.4097     0   1E+4   13.5   14  RT (min)      lcorbi_t3_neg_2_022_24    m/z 100.6337     0   1E+4   13.5   14  RT (min)      lcorbi_t3_neg_2_052_32    m/z 100.7812     0   1E+4   5   5.5  RT (min)      lcorbi_t3_neg_2_048_27    m/z 100.9050     0   1E+3   18.5   19  RT (min)      lcorbi_t3_neg_1_046_11    m/z 100.9335     0   1   2E+5   7   7.5  RT (min)      lcorbi_t3_neg_4_007_13    m/z 100.9335     0   1   2   3E+5   12   13  RT (min)       lcorbi_t3_neg_2_001_49    m/z 100.9335     0   1E+5   16.5   17  RT (min)       lcorbi_t3_neg_3_035_03    m/z 100.9337     0   1E+6   0   0.5  RT (min)       lcorbi_t3_neg_3_017_51    m/z 100.9691     0   1   2E+4   14   14.5  RT (min)      lcorbi_t3_neg_2_052_32    m/z 101.5952     0   1   2E+4   9   9.5  RT (min)      lcorbi_t3_neg_2_048_27    m/z 101.9336     0   2   4E+4   13   13.5  RT (min)      lcorbi_t3_neg_1_047_02    m/z 101.9336     0   2   4E+4   18   18.5  RT (min)      lcorbi_t3_neg_1_044_23    m/z 101.9336     0   1   2   3E+4   14.5   15  RT (min)      lcorbi_t3_neg_3_022_27    m/z 101.9336     0   1   2   3E+4   8   8.5  RT (min)      lcorbi_t3_neg_4_007_13    m/z 101.9336     0   1E+4   10   10.5  RT (min)      lcorbi_t3_neg_3_011_04    m/z 102.2978     0   2   4   6E+3   14.5   15  RT (min)      lcorbi_t3_neg_4_007_13    m/z 102.5218     0   1   2   3E+3   14.5   15  RT (min)      lcorbi_t3_neg_3_018_13    m/z 102.8520     0   1   14.5   15  RT (min)      lcorbi_t3_neg_1_048_12    m/z 102.9318     0   1E+4   14   14.5  RT (min)      lcorbi_t3_neg_2_023_39    m/z 102.9335     0   1E+4   18   18.5  RT (min)      lcorbi_t3_neg_4_100_61    m/z 102.9487     0   2   4   6E+3   15.5   16  RT (min)      lcorbi_t3_neg_4_112_62    m/z 102.9487     0   1E+4   18.5   19  RT (min)      lcorbi_t3_neg_4_113_62    m/z 102.9567     0   2   4   6   8E+4   13   13.5  RT (min)      lcorbi_t3_neg_4_104_61    m/z 102.9567     0   2   4   6   8E+4   18   18.5  RT (min)      lcorbi_t3_neg_4_077_58    m/z 102.9635     0   1   14.5   15  RT (min)      lcorbi_t3_neg_4_104_61    m/z 103.0037     0   2   4E+3   0   0.5  RT (min)      lcorbi_t3_neg_1_005_07    m/z 103.0220     0   1E+2   1   1.5  RT (min)       lcorbi_t3_neg_3_022_27    m/z 103.0244     0   1E+4   18   18.5  RT (min)      lcorbi_t3_neg_4_093_60    m/z 103.0400     0   1   2   3E+4   4.5   5  RT (min)      lcorbi_t3_neg_1_003_41    m/z 103.0400     0   1   2E+5   2.5   3  RT (min)      lcorbi_t3_neg_4_007_13    m/z 103.0593     0   2   4   6E+3   1   1.5  RT (min)      lcorbi_t3_neg_3_005_23    m/z 103.2627     0   2   4E+3   13   13.5  RT (min)      lcorbi_t3_neg_4_034_01    m/z 103.8522     0   2   4   6   8E+3   13.5   14  RT (min)      lcorbi_t3_neg_2_052_32    m/z 104.0208     0   2   4   6   8E+4   13   13.5  RT (min)      lcorbi_t3_neg_4_098_61    m/z 104.0208     0   2   4   6E+4   10   10.5  RT (min)      lcorbi_t3_neg_4_106_62    m/z 104.0762     0   2   4   6   8E+3   13.5   14  RT (min)      lcorbi_t3_neg_4_039_32    m/z 104.3002     0   2   4   6E+3   14   14.5  RT (min)      lcorbi_t3_neg_3_002_24    m/z 104.9538     0   1   2   3E+4   18   18.5  RT (min)       lcorbi_t3_neg_4_076_58    m/z 104.9538     0   1   2E+4   10   10.5  RT (min)       lcorbi_t3_neg_4_079_58    m/z 104.9538     0   1   2E+4   13   13.5   14  RT (min)       lcorbi_t3_neg_4_067_57    m/z 105.0171     0   1   2E+5   13   13.5  RT (min)      lcorbi_t3_neg_4_100_61    m/z 105.0171     0   1E+5   10   10.5  RT (min)      lcorbi_t3_neg_4_085_59    m/z 106.9119     0   1E+4   1   1.5  RT (min)      lcorbi_t3_neg_4_050_02    m/z 107.0501     0   2   4   6E+3   10.5   11  RT (min)      lcorbi_t3_neg_2_007_14    m/z 108.0216     0   1E+4   7.5   8  RT (min)      lcorbi_t3_neg_4_089_59    m/z 108.8098     0   2   4   6   8E+3   6   6.5  RT (min)      lcorbi_t3_neg_2_048_27    m/z 109.0294     0   2   4   6   8E+3   4.5   5  RT (min)      lcorbi_t3_neg_2_048_27    m/z 109.0380     0   1   2   3   4E+4   9   9.5  RT (min)      lcorbi_t3_neg_2_048_27    m/z 110.9429     0   2   4   6E+3   18   18.5  RT (min)      lcorbi_t3_neg_4_107_62    m/z 111.0451     0   1   2E+2   0   0.5  RT (min)      lcorbi_t3_neg_3_002_24    m/z 111.9789     0   1   1   1.5  RT (min)      lcorbi_t3_neg_2_039_31    m/z 112.9830     0   1E+4   1   1.5  RT (min)      lcorbi_t3_neg_4_036_16    m/z 112.9877     0   2   4E+4   7   7.5  RT (min)      lcorbi_t3_neg_4_080_58    m/z 112.9877     0   1   2   3   4E+4   4.5   5  RT (min)      lcorbi_t3_neg_4_102_61    m/z 112.9878     0   1   2   3E+4   1   1.5  RT (min)      lcorbi_t3_neg_4_098_61    m/z 112.9878     0   2   4   6E+4   18   18.5  RT (min)       lcorbi_t3_neg_4_087_59    m/z 113.0155     0   1   1   1.5  RT (min)      lcorbi_t3_neg_4_113_62    m/z 113.0356     0   1   2E+3   1   1.5  RT (min)      lcorbi_t3_neg_3_039_29    m/z 113.0608     0   2   4E+3   0   0.5  RT (min)      lcorbi_t3_neg_1_011_48    m/z 113.9888     0   1   2   3E+4   7   7.5  RT (min)      lcorbi_t3_neg_4_080_58    m/z 113.9889     0   1   2E+4   4.5   5  RT (min)      lcorbi_t3_neg_4_109_62    m/z 113.9889     0   2   4   6E+4   18   18.5  RT (min)      lcorbi_t3_neg_4_066_57    m/z 114.0244     0   1E+4   1.5   2  RT (min)      lcorbi_t3_neg_4_075_58    m/z 114.0575     0   1   2   3   4E+4   1   1.5  RT (min)      lcorbi_t3_neg_3_002_24    m/z 114.9516     0   1E+4   18.5   19  RT (min)      lcorbi_t3_neg_4_113_62    m/z 114.9516     0   2   4   6   8E+3   0.5   1  RT (min)      lcorbi_t3_neg_4_070_57    m/z 114.9897     0   1   2E+4   18   18.5  RT (min)       lcorbi_t3_neg_4_078_58    m/z 115.0207     0   2   4   6E+3   14   14.5  RT (min)      lcorbi_t3_neg_4_111_62    m/z 115.0207     0   2   4E+3   11   11.5  RT (min)      lcorbi_t3_neg_4_083_59    m/z 115.0400     0   1E+3   2.5   3  RT (min)      lcorbi_t3_neg_4_113_62    m/z 115.2975     0   2   4   6   8E+3   11   11.5  RT (min)      lcorbi_t3_neg_4_034_01    m/z 115.9206     0   1   2   3E+4   18   18.5  RT (min)      lcorbi_t3_neg_1_046_11    m/z 115.9206     0   2   4   6E+4   10.5   11  RT (min)      lcorbi_t3_neg_4_007_13    m/z 115.9206     0   1   2E+4   8   8.5  RT (min)      lcorbi_t3_neg_3_021_38    m/z 115.9206     0   2   4   6   8E+4   5.5   6  RT (min)      lcorbi_t3_neg_4_007_13    m/z 116.0036     0   1E+4   18   18.5  RT (min)      lcorbi_t3_neg_4_097_60    m/z 116.0071     0   1   2   3   4E+2   1   1.5  RT (min)      lcorbi_t3_neg_4_039_32    m/z 116.0263     0   1E+3   9   9.5  RT (min)      lcorbi_t3_neg_4_076_58    m/z 116.3262     0   1   2   3   4E+3   12.5   13  RT (min)      lcorbi_t3_neg_4_007_13    m/z 116.9284     0   1   2   3   4E+4   14   14.5  RT (min)      lcorbi_t3_neg_2_038_22    m/z 116.9284     0   1   2E+4   18   18.5  RT (min)      lcorbi_t3_neg_4_066_57    m/z 116.9285     0   2   4E+4   8   8.5  RT (min)      lcorbi_t3_neg_2_048_27    m/z 116.9285     0   1E+4   10   10.5  RT (min)      lcorbi_t3_neg_4_012_17    m/z 116.9285     0   1   2   3E+4   4.5   5  RT (min)      lcorbi_t3_neg_3_054_18    m/z 117.0000     0   2   4E+4   18   18.5  RT (min)       lcorbi_t3_neg_4_095_60    m/z 117.0192     0   1   2E+4   14   14.5  RT (min)      lcorbi_t3_neg_4_073_57    m/z 117.0193     0   2   4   6E+4   2.5   3  RT (min)       lcorbi_t3_neg_4_044_38    m/z 117.0557     0   1E+3   1   1.5  RT (min)      lcorbi_t3_neg_4_110_62    m/z 117.9290     0   1   2E+5   0   0.5  RT (min)      lcorbi_t3_neg_3_001_49    m/z 118.0137     0   2   4   6E+2   1   1.5  RT (min)      lcorbi_t3_neg_4_034_01    m/z 118.0364     0   1E+4   14   14.5  RT (min)      lcorbi_t3_neg_4_090_60    m/z 118.0364     0   1   2E+4   18   18.5  RT (min)       lcorbi_t3_neg_4_093_60    m/z 118.9241     0   2   4   6E+4   15   16  RT (min)       lcorbi_t3_neg_2_011_04    m/z 118.9241     0   1   2E+4   17.5   18  RT (min)       lcorbi_t3_neg_3_039_29    m/z 119.0327     0   1   2   3E+4   13.5   14   14.5  RT (min)       lcorbi_t3_neg_4_088_59    m/z 119.0328     0   1   2   3E+4   18   18.5  RT (min)       lcorbi_t3_neg_4_096_60    m/z 119.0349     0   1   2   3E+4   2.5   3  RT (min)      lcorbi_t3_neg_1_039_38    m/z 119.0502     0   1   2E+4   6   6.5  RT (min)      lcorbi_t3_neg_2_048_27    m/z 119.0698     0   2   4   6   8E+3   11   11.5  RT (min)      lcorbi_t3_neg_1_055_05    m/z 119.9469     0   2   4   6E+3   17.5   18  RT (min)      lcorbi_t3_neg_4_042_14    m/z 120.0124     0   1   2   3E+4   11.5   12  RT (min)      lcorbi_t3_neg_4_007_13    m/z 120.0293     0   1   2   3   4E+4   2.5   3  RT (min)       lcorbi_t3_neg_4_093_60    m/z 120.0383     0   1E+4   0   0.5  RT (min)      lcorbi_t3_neg_1_029_21    m/z 120.0777     0   2   4   6E+3   11.5   12  RT (min)      lcorbi_t3_neg_4_007_13    m/z 120.7462     0   2   4E+4   14   14.5  RT (min)      lcorbi_t3_neg_2_053_23    m/z 120.8577     0   1   2E+4   14   14.5  RT (min)      lcorbi_t3_neg_3_005_23    m/z 120.9964     0   1E+3   14   14.5  RT (min)      lcorbi_t3_neg_1_003_41    m/z 121.0272     0   1E+4   10.5   11  RT (min)      lcorbi_t3_neg_4_042_14    m/z 121.0294     0   1   2   3   4E+4   5   6  RT (min)       lcorbi_t3_neg_1_001_49    m/z 121.4153     0   1E+4   12   12.5  RT (min)      lcorbi_t3_neg_4_005_27    m/z 121.8266     0   1   2   3   4E+3   5.5   6  RT (min)      lcorbi_t3_neg_4_032_23    m/z 121.9440     0   2   4   6E+4   1   1.5  RT (min)      lcorbi_t3_neg_4_007_13    m/z 121.9440     0   1   2E+3   17.5   18  RT (min)      lcorbi_t3_neg_3_005_23    m/z 122.0328     0   1E+3   5   5.5  RT (min)      lcorbi_t3_neg_4_023_18    m/z 122.0328     0   1   10.5   11  RT (min)      lcorbi_t3_neg_4_067_57    m/z 122.6289     0   2   4   6   8E+3   11   11.5  RT (min)      lcorbi_t3_neg_3_053_14    m/z 122.9757     0   1   2   3E+4   1   1.5  RT (min)      lcorbi_t3_neg_2_048_27    m/z 123.0120     0   2   4   6   8E+3   14   14.5  RT (min)      lcorbi_t3_neg_4_030_04    m/z 123.0451     0   1E+4   5   5.5  RT (min)      lcorbi_t3_neg_4_005_27    m/z 123.8609     0   1   2   3E+4   14   14.5  RT (min)      lcorbi_t3_neg_3_005_23    m/z 124.0156     0   1   2E+4   10   10.5  RT (min)      lcorbi_t3_neg_4_023_18    m/z 124.0484     0   1   8   8.5  RT (min)      lcorbi_t3_neg_1_002_16    m/z 125.0244     0   2   4E+2   1   1.5  RT (min)      lcorbi_t3_neg_3_002_24    m/z 125.0710     0   1E+4   11   11.5  RT (min)      lcorbi_t3_neg_4_034_01    m/z 125.0971     0   1   2   3   4E+3   7   7.5  RT (min)      lcorbi_t3_neg_4_023_18    m/z 125.0971     0   1E+4   10.5   11  RT (min)      lcorbi_t3_neg_3_053_14    m/z 125.3206     0   2   4E+3   11   11.5  RT (min)      lcorbi_t3_neg_4_007_13    m/z 125.8731     0   2   4   6E+4   4.5   5  RT (min)      lcorbi_t3_neg_1_047_02    m/z 126.0197     0   2   4   6E+2   1   1.5  RT (min)      lcorbi_t3_neg_3_002_24    m/z 126.0308     0   1   2E+4   3   3.5  RT (min)      lcorbi_t3_neg_2_048_27    m/z 126.7665     0   1   2E+4   14   14.5  RT (min)      lcorbi_t3_neg_4_011_24    m/z 126.8810     0   1   2E+4   5   5.5  RT (min)      lcorbi_t3_neg_1_010_17    m/z 126.8810     0   1   2E+4   3   3.5  RT (min)      lcorbi_t3_neg_1_033_53    m/z 126.9049     0   1   2   3   4E+3   14   14.5  RT (min)      lcorbi_t3_neg_1_007_26    m/z 126.9049     0   2   4   6E+3   18   18.5  RT (min)      lcorbi_t3_neg_1_003_41    m/z 127.0297     0   1   2E+4   2.5   3  RT (min)      lcorbi_t3_neg_3_019_12    m/z 127.0400     0   1E+4   0   0.5  RT (min)      lcorbi_t3_neg_1_011_48    m/z 127.0764     0   2   4   6E+3   6.5   7  RT (min)      lcorbi_t3_neg_4_006_47    m/z 127.8702     0   2   4E+4   4.5   5  RT (min)      lcorbi_t3_neg_1_045_44    m/z 127.8972     0   1   1.5   2  RT (min)      lcorbi_t3_neg_1_045_44    m/z 128.0716     0   1   2E+4   5   5.5  RT (min)      lcorbi_t3_neg_4_007_13    m/z 128.2943     0   1E+4   9   9.5  RT (min)      lcorbi_t3_neg_4_023_18    m/z 128.5451     0   1   2   3E+3   9   9.5  RT (min)      lcorbi_t3_neg_4_023_18    m/z 128.8581     0   2   4   6E+3   12   12.5  RT (min)      lcorbi_t3_neg_4_034_01    m/z 128.8780     0   1   2E+4   5   5.5  RT (min)      lcorbi_t3_neg_1_043_14    m/z 128.8780     0   1E+4   3   3.5  RT (min)      lcorbi_t3_neg_2_029_12    m/z 128.8781     0   1   2E+4   1  RT (min)       lcorbi_t3_neg_4_069_57    m/z 129.0193     0   1E+5   1.5   2  RT (min)       lcorbi_t3_neg_4_042_14    m/z 129.0208     0   1   2   3   4E+3   18   18.5  RT (min)      lcorbi_t3_neg_4_108_62    m/z 129.9325     0   1E+3   11   11.5  RT (min)      lcorbi_t3_neg_1_047_02    m/z 129.9363     0   1E+4   18   18.5  RT (min)      lcorbi_t3_neg_4_066_57    m/z 129.9757     0   1E+4   18   18.5  RT (min)       lcorbi_t3_neg_4_051_40    m/z 129.9757     0   1E+4   10   10.5  RT (min)       lcorbi_t3_neg_4_038_15    m/z 129.9758     0   1   2E+4   0.5   1  RT (min)       lcorbi_t3_neg_4_082_59    m/z 130.0000     0   1E+3   18   18.5  RT (min)      lcorbi_t3_neg_4_110_62    m/z 130.0171     0   1   2   3   4E+4   14   14.5  RT (min)      lcorbi_t3_neg_4_030_04    m/z 130.0227     0   1   1   1.5  RT (min)      lcorbi_t3_neg_4_111_62    m/z 130.0395     0   2   4   6   8E+2   1   1.5  RT (min)      lcorbi_t3_neg_4_111_62    m/z 130.0622     0   1E+3   1   1.5  RT (min)      lcorbi_t3_neg_4_054_36    m/z 130.0873     0   2   4   6E+4   2.5   3  RT (min)      lcorbi_t3_neg_2_048_27    m/z 130.2278     0   1E+3   18   18.5  RT (min)      lcorbi_t3_neg_4_030_04    m/z 130.9926     0   1   2E+4   0   0.5  RT (min)      lcorbi_t3_neg_1_001_49    m/z 130.9964     0   1E+3   18   18.5  RT (min)      lcorbi_t3_neg_1_003_41    m/z 131.0135     0   1E+4   14   14.5  RT (min)      lcorbi_t3_neg_2_040_43    m/z 131.0164     0   1   2   3   4E+3   14   14.5  RT (min)      lcorbi_t3_neg_2_023_39    m/z 131.0350     0   1   2E+4   3   4  RT (min)       lcorbi_t3_neg_2_051_34    m/z 131.0350     0   1   2   3E+4   1   1.5  RT (min)      lcorbi_t3_neg_4_007_13    m/z 131.0698     0   2   4   6   8E+3   11   11.5  RT (min)      lcorbi_t3_neg_1_002_16    m/z 131.0713     0   2   4   6   8E+3   6   6.5  RT (min)      lcorbi_t3_neg_4_042_14    m/z 132.0128     0   2   4   6   8E+3   13.5   14  RT (min)      lcorbi_t3_neg_4_046_37    m/z 132.0128     0   2   4   6   8E+3   12   12.5  RT (min)      lcorbi_t3_neg_4_029_31    m/z 132.0128     0   1   2E+3   18   18.5  RT (min)      lcorbi_t3_neg_4_093_60    m/z 132.0217     0   1E+4   7   7.5  RT (min)      lcorbi_t3_neg_2_048_27    m/z 132.0270     0   1E+4   1   1.5  RT (min)      lcorbi_t3_neg_4_029_31    m/z 132.7156     0   2   4E+3   11   11.5  RT (min)      lcorbi_t3_neg_4_034_01    m/z 132.8678     0   1   2   3E+4   15.5   16  RT (min)      lcorbi_t3_neg_4_018_42    m/z 132.8678     0   1   2   3E+4   10   10.5  RT (min)      lcorbi_t3_neg_4_105_61    m/z 132.8678     0   1   2E+5   5   5.5  RT (min)      lcorbi_t3_neg_4_089_59    m/z 132.8679     0   1E+5   2.5   3  RT (min)      lcorbi_t3_neg_2_042_44    m/z 132.8706     0   2   4   6   8E+3   0.5   1  RT (min)      lcorbi_t3_neg_4_070_57    m/z 132.9234     0   1   18   18.5  RT (min)      lcorbi_t3_neg_4_086_59    m/z 133.0092     0   1E+4   13.5   14  RT (min)      lcorbi_t3_neg_1_055_05    m/z 133.0092     0   1E+4   12   12.5  RT (min)      lcorbi_t3_neg_4_017_51    m/z 133.0092     0   1   2   3E+3   18   18.5  RT (min)       lcorbi_t3_neg_4_096_60    m/z 133.0272     0   1   2   3E+2   1   1.5  RT (min)      lcorbi_t3_neg_4_023_18    m/z 133.0484     0   2   4   6E+3   18   18.5  RT (min)      lcorbi_t3_neg_4_071_57    m/z 133.0506     0   2   4   6E+3   4.5   5  RT (min)      lcorbi_t3_neg_1_021_31    m/z 133.0548     0   1E+4   1   1.5  RT (min)      lcorbi_t3_neg_4_093_60    m/z 133.0658     0   1E+3   9.5   10  RT (min)      lcorbi_t3_neg_4_039_32    m/z 133.0658     0   1   2   3   4E+2   7   7.5  RT (min)      lcorbi_t3_neg_4_017_51    m/z 133.9917     0   1E+5   1   1.5  RT (min)      lcorbi_t3_neg_3_043_25    m/z 134.0540     0   1   2   3   4E+3   0   0.5  RT (min)      lcorbi_t3_neg_1_029_21    m/z 134.8653     0   1E+5   5   5.5  RT (min)      lcorbi_t3_neg_1_052_24    m/z 134.8654     0   1E+5   3   3.5  RT (min)      lcorbi_t3_neg_2_042_44    m/z 134.8946     0   1   2   3   4E+4   10.5   11  RT (min)      lcorbi_t3_neg_2_002_26    m/z 134.8946     0   1   2   3E+4   17.5   18  RT (min)      lcorbi_t3_neg_3_038_39    m/z 135.0106     0   1E+4   18   18.5  RT (min)      lcorbi_t3_neg_4_084_59    m/z 135.0270     0   1E+4   1.5   2  RT (min)      lcorbi_t3_neg_4_076_58    m/z 135.0562     0   2   4E+3   1   1.5  RT (min)      lcorbi_t3_neg_1_055_05    m/z 135.5844     0   1   2   3   4E+3   11.5   12  RT (min)      lcorbi_t3_neg_4_042_14    m/z 135.8947     0   1   2E+4   14.5   15  RT (min)      lcorbi_t3_neg_2_042_44    m/z 136.0459     0   1   2   3   4E+4   1   1.5  RT (min)       lcorbi_t3_neg_3_052_32    m/z 136.0513     0   1   2E+4   1   1.5  RT (min)      lcorbi_t3_neg_2_022_24    m/z 136.8628     0   2   4   6E+4   3   3.5  RT (min)      lcorbi_t3_neg_4_028_39    m/z 136.8917     0   1   2E+4   14.5   15  RT (min)       lcorbi_t3_neg_4_004_34    m/z 136.8945     0   1   2E+3   17.5   18  RT (min)      lcorbi_t3_neg_3_012_47    m/z 136.9097     0   1E+4   18.5   19  RT (min)      lcorbi_t3_neg_4_113_62    m/z 137.0243     0   2   4   6E+4   5   5.5  RT (min)       lcorbi_t3_neg_4_023_18    m/z 137.0245     0   1   2E+4   1   1.5  RT (min)       lcorbi_t3_neg_1_052_24    m/z 137.0328     0   1   2   3E+3   1   1.5  RT (min)       lcorbi_t3_neg_4_019_08    m/z 137.0406     0   1E+3   1   1.5  RT (min)      lcorbi_t3_neg_2_012_36    m/z 138.0348     0   1   11   11.5  RT (min)      lcorbi_t3_neg_1_005_07    m/z 138.0673     0   1   2   3E+4   1   1.5  RT (min)      lcorbi_t3_neg_4_011_24    m/z 138.9068     0   2   4   6E+3   0   0.5  RT (min)      lcorbi_t3_neg_4_093_60    m/z 138.9706     0   1   2   3E+3   1   1.5  RT (min)      lcorbi_t3_neg_4_096_60    m/z 138.9875     0   1   2E+4   13   13.5  RT (min)       lcorbi_t3_neg_4_083_59    m/z 138.9875     0   1E+4   18   18.5  RT (min)      lcorbi_t3_neg_4_076_58    m/z 139.0230     0   1   2   3E+3   7   7.5  RT (min)      lcorbi_t3_neg_4_098_61    m/z 139.0432     0   1E+4   1   1.5  RT (min)      lcorbi_t3_neg_3_052_32    m/z 139.0639     0   1   2   3E+4   4.5   5  RT (min)      lcorbi_t3_neg_3_002_24    m/z 140.8679     0   2   4   6E+3   18   18.5  RT (min)      lcorbi_t3_neg_1_044_23    m/z 140.9957     0   1E+3   1   1.5  RT (min)      lcorbi_t3_neg_2_009_50    m/z 141.2090     0   2   4   6E+3   11   11.5  RT (min)      lcorbi_t3_neg_4_007_13    m/z 141.3337     0   1E+4   11   11.5  RT (min)      lcorbi_t3_neg_4_007_13    m/z 141.8375     0   2   4   6E+3   11.5   12  RT (min)      lcorbi_t3_neg_4_007_13    m/z 141.8392     0   2   4   6E+3   11.5   12  RT (min)      lcorbi_t3_neg_4_007_13    m/z 141.8683     0   1E+3   1   1.5  RT (min)      lcorbi_t3_neg_1_045_44    m/z 142.0510     0   1   2   3   4E+3   4.5   5  RT (min)       lcorbi_t3_neg_4_082_59    m/z 142.2168     0   2   4   6   8E+3   12   12.5  RT (min)      lcorbi_t3_neg_4_034_01    m/z 142.3414     0   1E+4   12   12.5  RT (min)      lcorbi_t3_neg_4_034_01    m/z 142.5923     0   1   2E+3   12   12.5  RT (min)      lcorbi_t3_neg_4_005_27    m/z 143.0463     0   1   2E+3   1   1.5  RT (min)      lcorbi_t3_neg_4_007_13    m/z 143.0543     0   2   4E+4   2.5   3  RT (min)      lcorbi_t3_neg_4_029_31    m/z 143.8653     0   1   2   3   4E+3   1   1.5  RT (min)      lcorbi_t3_neg_4_102_61    m/z 143.9327     0   2   4E+3   18   18.5  RT (min)      lcorbi_t3_neg_1_046_11    m/z 144.0368     0   1E+3   11   11.5  RT (min)      lcorbi_t3_neg_1_009_50    m/z 144.0489     0   1   2E+3   6.5   7  RT (min)      lcorbi_t3_neg_2_053_23    m/z 144.3199     0   1E+4   10.5   11  RT (min)      lcorbi_t3_neg_2_048_27    m/z 144.8697     0   2   4   6E+3   1   1.5  RT (min)      lcorbi_t3_neg_1_010_17    m/z 144.8716     0   1   2E+3   17   17.5  RT (min)      lcorbi_t3_neg_4_066_57    m/z 144.9230     0   1E+5   0   0.5  RT (min)      lcorbi_t3_neg_3_036_41    m/z 144.9234     0   2   4   6E+4   13   13.5  RT (min)      lcorbi_t3_neg_4_066_57    m/z 144.9234     0   1   2   3   4E+4   18   18.5  RT (min)      lcorbi_t3_neg_4_066_57    m/z 144.9234     0   1   2   3   4E+4   10.5   11  RT (min)      lcorbi_t3_neg_4_066_57    m/z 144.9234     0   1E+4   9   9.5  RT (min)      lcorbi_t3_neg_1_047_02    m/z 144.9235     0   1   2E+4   15.5   16  RT (min)       lcorbi_t3_neg_4_068_57    m/z 144.9617     0   1E+4   8.5   9  RT (min)      lcorbi_t3_neg_2_048_27    m/z 144.9648     0   1E+3   1   1.5  RT (min)       lcorbi_t3_neg_2_039_31    m/z 145.0488     0   1E+4   6.5   7  RT (min)      lcorbi_t3_neg_2_051_34    m/z 145.0506     0   1E+4   4.5   5  RT (min)      lcorbi_t3_neg_2_018_40    m/z 145.0506     0   1   2E+4   2.5   3  RT (min)       lcorbi_t3_neg_4_005_27    m/z 145.0652     0   2   4   6   8E+3   1   1.5  RT (min)      lcorbi_t3_neg_4_089_59    m/z 145.0870     0   1   2E+4   7.5   8  RT (min)      lcorbi_t3_neg_2_048_27    m/z 145.9308     0   1   2   3   4E+4   4.5   5  RT (min)      lcorbi_t3_neg_1_044_23    m/z 145.9308     0   1E+4   0.5   1  RT (min)      lcorbi_t3_neg_1_045_44    m/z 145.9417     0   1E+4   18   18.5  RT (min)      lcorbi_t3_neg_4_085_59    m/z 146.0177     0   1E+3   1   1.5  RT (min)      lcorbi_t3_neg_2_031_25    m/z 146.0314     0   1E+4   18   18.5  RT (min)      lcorbi_t3_neg_4_095_60    m/z 146.0491     0   1   2   3   4E+4   1   1.5  RT (min)      lcorbi_t3_neg_4_009_50    m/z 146.0677     0   1   2E+4   13.5   14   14.5  RT (min)       lcorbi_t3_neg_4_073_57    m/z 146.0677     0   1   2E+4   18   18.5  RT (min)       lcorbi_t3_neg_4_093_60    m/z 146.8668     0   2   4   6   8E+2   1   1.5  RT (min)      lcorbi_t3_neg_1_003_41    m/z 146.9233     0   2   4E+3   18   18.5  RT (min)       lcorbi_t3_neg_1_001_49    m/z 146.9234     0   2   4   6E+4   0   0.5  RT (min)      lcorbi_t3_neg_4_052_26    m/z 146.9386     0   1E+5   15.5   16  RT (min)      lcorbi_t3_neg_4_042_14    m/z 146.9588     0   1E+4   8.5   9  RT (min)      lcorbi_t3_neg_2_048_27    m/z 146.9612     0   1   2E+4   18   18.5  RT (min)       lcorbi_t3_neg_1_003_41    m/z 146.9659     0   2   4   6E+3   18   18.5  RT (min)      lcorbi_t3_neg_2_049_55    m/z 146.9831     0   1   2   3   4E+3   1.5   2  RT (min)      lcorbi_t3_neg_4_046_37    m/z 147.0084     0   1E+3   18   18.5  RT (min)      lcorbi_t3_neg_1_001_49    m/z 147.0277     0   1   2E+4   18   18.5  RT (min)       lcorbi_t3_neg_4_092_60    m/z 147.0299     0   1   2   3E+5   2   3  RT (min)       lcorbi_t3_neg_4_054_36    m/z 147.0429     0   1E+2   1   1.5  RT (min)      lcorbi_t3_neg_3_035_03    m/z 147.0597     0   2   4   6   8E+3   1   1.5  RT (min)      lcorbi_t3_neg_2_007_14    m/z 147.0641     0   2   4E+4   14   14.5  RT (min)      lcorbi_t3_neg_4_097_60    m/z 147.0829     0   1   11   11.5  RT (min)      lcorbi_t3_neg_4_080_58    m/z 147.3338     0   1   2E+3   11   11.5  RT (min)      lcorbi_t3_neg_4_036_16    m/z 147.9392     0   1   2E+4   0   0.5  RT (min)      lcorbi_t3_neg_4_050_02    m/z 148.0332     0   2   4   6E+2   1   1.5  RT (min)      lcorbi_t3_neg_3_022_27    m/z 148.0530     0   2   4   6E+3   9.5   10  RT (min)      lcorbi_t3_neg_2_048_27    m/z 148.0696     0   2   4E+3   0   0.5  RT (min)      lcorbi_t3_neg_1_007_26    m/z 148.9188     0   1E+4   18   18.5  RT (min)      lcorbi_t3_neg_4_077_58    m/z 149.0055     0   1E+4   1   1.5  RT (min)      lcorbi_t3_neg_4_010_28    m/z 149.0095     0   2   4   6   8E+4   13   13.5  RT (min)      lcorbi_t3_neg_2_036_35    m/z 149.0306     0   1E+3   1   1.5  RT (min)      lcorbi_t3_neg_2_002_26    m/z 149.0971     0   1E+4   7.5   8  RT (min)      lcorbi_t3_neg_2_048_27    m/z 149.0971     0   2   4   6E+3   9.5   10  RT (min)      lcorbi_t3_neg_2_048_27    m/z 150.0263     0   1E+4   18   18.5  RT (min)       lcorbi_t3_neg_4_110_62    m/z 150.0339     0   1   2   3E+3   1   1.5  RT (min)      lcorbi_t3_neg_4_006_47    m/z 150.8893     0   1E+4   1   1.5  RT (min)      lcorbi_t3_neg_4_007_13    m/z 150.9829     0   1   2   3   4E+3   18.5   19  RT (min)      lcorbi_t3_neg_4_077_58    m/z 150.9829     0   1E+4   0.5   1  RT (min)      lcorbi_t3_neg_1_044_23    m/z 151.0132     0   1E+4   1   1.5  RT (min)      lcorbi_t3_neg_2_048_27    m/z 151.0226     0   1   2   3   4E+4   13   13.5  RT (min)       lcorbi_t3_neg_4_103_61    m/z 151.0261     0   1   2E+4   2   2.5  RT (min)      lcorbi_t3_neg_2_048_27    m/z 151.0652     0   1   2   3E+4   1   1.5  RT (min)      lcorbi_t3_neg_1_053_32    m/z 151.1128     0   1   2   3E+2   8.5   9  RT (min)      lcorbi_t3_neg_4_032_23    m/z 152.0434     0   1E+4   5   5.5  RT (min)      lcorbi_t3_neg_4_101_61    m/z 152.0466     0   2   4   6E+5   2   3  RT (min)       lcorbi_t3_neg_4_090_60    m/z 152.0723     0   1   2E+4   6.5   7  RT (min)      lcorbi_t3_neg_4_039_32    m/z 152.0799     0   1   2E+3   11.5   12  RT (min)      lcorbi_t3_neg_4_046_37    m/z 152.9785     0   1E+4   18.5   19  RT (min)      lcorbi_t3_neg_4_039_32    m/z 152.9785     0   1E+4   0.5   1  RT (min)      lcorbi_t3_neg_1_045_44    m/z 152.9863     0   1   2   3E+4   1   1.5  RT (min)      lcorbi_t3_neg_3_043_25    m/z 153.0154     0   1   2   3E+3   13   13.5  RT (min)      lcorbi_t3_neg_2_023_39    m/z 153.0386     0   1   2   3E+3   8   8.5  RT (min)      lcorbi_t3_neg_4_097_60    m/z 153.0921     0   1   2   3   4E+3   8   8.5  RT (min)      lcorbi_t3_neg_4_082_59    m/z 153.8686     0   1E+4   11   11.5  RT (min)      lcorbi_t3_neg_4_016_05    m/z 153.8686     0   1E+4   13   13.5  RT (min)      lcorbi_t3_neg_4_041_54    m/z 153.8686     0   1   2E+4   15   15.5  RT (min)      lcorbi_t3_neg_3_048_08    m/z 153.8686     0   1   2   3E+3   17.5   18  RT (min)      lcorbi_t3_neg_4_041_54    m/z 153.9000     0   2   4   6   8E+3   0   0.5  RT (min)      lcorbi_t3_neg_4_090_60    m/z 154.8986     0   1   2   3   4E+3   1   1.5  RT (min)      lcorbi_t3_neg_4_039_32    m/z 154.9328     0   1E+4   18   18.5  RT (min)      lcorbi_t3_neg_4_106_62    m/z 154.9598     0   2   4   6   8E+3   18.5   19  RT (min)      lcorbi_t3_neg_4_079_58    m/z 154.9598     0   2   4   6   8E+3   0.5   1  RT (min)      lcorbi_t3_neg_2_015_21    m/z 154.9742     0   2   4   6   8E+3   18   18.5  RT (min)      lcorbi_t3_neg_1_043_14    m/z 154.9742     0   1E+4   0   0.5  RT (min)      lcorbi_t3_neg_1_044_23    m/z 155.3322     0   1E+4   8   8.5  RT (min)      lcorbi_t3_neg_2_048_27    m/z 155.3343     0   2   4   6E+3   8   8.5  RT (min)      lcorbi_t3_neg_2_048_27    m/z 155.8656     0   1E+4   13   13.5  RT (min)      lcorbi_t3_neg_3_019_12    m/z 155.8741     0   2   4   6E+3   18.5   19  RT (min)      lcorbi_t3_neg_4_106_62    m/z 156.0812     0   2   4   6E+4   1   1.5  RT (min)       lcorbi_t3_neg_2_052_32    m/z 156.0853     0   1   2   3   4E+3   1   1.5  RT (min)      lcorbi_t3_neg_4_006_47    m/z 156.6345     0   1   2   3   4E+3   14.5   15  RT (min)      lcorbi_t3_neg_3_005_23    m/z 156.8948     0   1   2E+3   18   18.5  RT (min)      lcorbi_t3_neg_4_072_57    m/z 156.8956     0   1   2   3   4E+3   1   1.5  RT (min)      lcorbi_t3_neg_4_039_32    m/z 156.9297     0   1   2E+3   18   18.5  RT (min)      lcorbi_t3_neg_1_001_49    m/z 156.9555     0   2   4   6   8E+3   18.5   19  RT (min)      lcorbi_t3_neg_4_080_58    m/z 156.9810     0   2   4   6   8E+3   14   14.5  RT (min)      lcorbi_t3_neg_1_055_05    m/z 157.0157     0   2   4   6E+4   14   14.5  RT (min)      lcorbi_t3_neg_4_030_04    m/z 157.0368     0   1E+3   1   1.5  RT (min)      lcorbi_t3_neg_1_021_31    m/z 157.0506     0   1E+4   5   5.5  RT (min)      lcorbi_t3_neg_2_036_35    m/z 157.0700     0   1   2   3E+4   3   3.5  RT (min)      lcorbi_t3_neg_4_029_31    m/z 157.0837     0   2   4   6   8E+3   1   1.5  RT (min)      lcorbi_t3_neg_4_011_24    m/z 157.0980     0   2   4   6E+3   1   1.5  RT (min)      lcorbi_t3_neg_4_038_15    m/z 157.8633     0   1   2E+4   1   1.5  RT (min)      lcorbi_t3_neg_2_027_13    m/z 157.9313     0   1   2E+3   16.5   17  RT (min)      lcorbi_t3_neg_3_029_02    m/z 158.0121     0   1   2   3   4E+4   12   12.5  RT (min)      lcorbi_t3_neg_4_046_37    m/z 158.0313     0   1   2E+3   18   18.5  RT (min)      lcorbi_t3_neg_4_039_32    m/z 158.8931     0   1   2E+3   18   18.5  RT (min)      lcorbi_t3_neg_1_045_44    m/z 158.9254     0   1   2E+4   1   1.5  RT (min)      lcorbi_t3_neg_1_055_05    m/z 158.9391     0   2   4   6E+3   18   18.5  RT (min)      lcorbi_t3_neg_1_045_44    m/z 158.9755     0   1E+5   10   10.5   11  RT (min)       lcorbi_t3_neg_3_007_16    m/z 158.9755     0   2   4   6E+4   7   7.5  RT (min)      lcorbi_t3_neg_4_017_51    m/z 158.9822     0   2   4   6E+3   1   1.5  RT (min)      lcorbi_t3_neg_2_032_45    m/z 159.0107     0   1   2   3E+2   1   1.5  RT (min)      lcorbi_t3_neg_4_053_43    m/z 159.0277     0   1E+3   18   18.5  RT (min)      lcorbi_t3_neg_1_003_41    m/z 159.0279     0   2   4   6   8E+2   1.5   2  RT (min)      lcorbi_t3_neg_1_012_20    m/z 159.0379     0   1   9   9.5  RT (min)      lcorbi_t3_neg_1_045_44    m/z 159.0878     0   1E+3   12   12.5  RT (min)      lcorbi_t3_neg_1_009_50    m/z 159.1027     0   1   2E+4   8.5   9  RT (min)      lcorbi_t3_neg_2_048_27    m/z 159.1137     0   2   4   6E+3   1   1.5  RT (min)      lcorbi_t3_neg_2_007_14    m/z 159.8789     0   2   4   6   8E+3   1   1.5  RT (min)      lcorbi_t3_neg_1_042_18    m/z 159.8938     0   2   4   6   8E+3   18   18.5  RT (min)      lcorbi_t3_neg_1_056_56    m/z 159.9756     0   1   2   3E+4   8   8.5  RT (min)      lcorbi_t3_neg_2_048_27    m/z 159.9756     0   1E+4   9.5   10  RT (min)      lcorbi_t3_neg_2_028_38    m/z 160.0107     0   2   4   6   8E+2   18   18.5  RT (min)      lcorbi_t3_neg_1_001_49    m/z 160.0277     0   1   2   3   4E+3   14   14.5  RT (min)      lcorbi_t3_neg_1_046_11    m/z 160.0277     0   1   2E+3   18   18.5  RT (min)      lcorbi_t3_neg_1_003_41    m/z 160.0452     0   2   4   6   8E+3   7   7.5  RT (min)      lcorbi_t3_neg_4_005_27    m/z 160.8421     0   2   4   6E+4   17   17.5  RT (min)      lcorbi_t3_neg_4_037_29    m/z 160.8466     0   1E+4   1   1.5  RT (min)      lcorbi_t3_neg_4_112_62    m/z 160.8927     0   1   2   3   4E+3   18   18.5  RT (min)      lcorbi_t3_neg_4_066_57    m/z 160.9355     0   1   2   3   4E+4   13   13.5   14  RT (min)       lcorbi_t3_neg_1_048_12    m/z 160.9355     0   1   2   3E+4   12  RT (min)       lcorbi_t3_neg_3_035_03    m/z 160.9355     0   1E+4   18   18.5  RT (min)      lcorbi_t3_neg_4_066_57    m/z 160.9355     0   1   2   3   4E+4   6   6.5  RT (min)      lcorbi_t3_neg_3_039_29    m/z 160.9355     0   1   2E+4   9   9.5  RT (min)      lcorbi_t3_neg_4_010_28    m/z 160.9566     0   1   6.5   7  RT (min)      lcorbi_t3_neg_4_040_19    m/z 160.9738     0   1E+4   14   14.5  RT (min)      lcorbi_t3_neg_2_037_09    m/z 160.9825     0   2   4   6   8E+3   1   1.5  RT (min)      lcorbi_t3_neg_4_044_38    m/z 161.0241     0   1E+4   14   14.5  RT (min)      lcorbi_t3_neg_1_040_42    m/z 161.0366     0   1   2E+3   9   9.5  RT (min)      lcorbi_t3_neg_2_049_55    m/z 161.0456     0   2   4   6E+4   3   4   5  RT (min)       lcorbi_t3_neg_4_046_37    m/z 161.0627     0   1   2   3   4E+3   18   18.5  RT (min)      lcorbi_t3_neg_4_090_60    m/z 161.0798     0   2   4   6E+3   18   18.5  RT (min)      lcorbi_t3_neg_4_091_60    m/z 161.0820     0   1E+4   2.5   3  RT (min)      lcorbi_t3_neg_1_019_29    m/z 161.8498     0   2   4   6E+3   13.5   14  RT (min)      lcorbi_t3_neg_4_040_19    m/z 161.8499     0   1E+4   4.5   5  RT (min)      lcorbi_t3_neg_3_040_05    m/z 161.8500     0   1E+4   1   1.5  RT (min)      lcorbi_t3_neg_2_027_13    m/z 161.8759     0   1   2   3E+3   1   1.5  RT (min)      lcorbi_t3_neg_3_015_11    m/z 161.8940     0   2   4   6E+3   18   18.5  RT (min)      lcorbi_t3_neg_1_053_32    m/z 161.9904     0   1   2E+4   2.5   3  RT (min)      lcorbi_t3_neg_2_051_34    m/z 162.0277     0   1E+4   7   7.5  RT (min)      lcorbi_t3_neg_2_048_27    m/z 162.0409     0   1   2   3   4E+3   1   1.5  RT (min)      lcorbi_t3_neg_4_046_37    m/z 162.0489     0   1   2   3E+3   1   1.5  RT (min)      lcorbi_t3_neg_4_046_37    m/z 162.0772     0   1E+4   1   1.5  RT (min)      lcorbi_t3_neg_4_053_43    m/z 162.0853     0   1   2   3   4E+3   0   0.5  RT (min)      lcorbi_t3_neg_1_017_51    m/z 162.8391     0   2   4   6E+4   17   17.5  RT (min)      lcorbi_t3_neg_4_042_14    m/z 162.8454     0   2   4   6E+3   1   1.5  RT (min)      lcorbi_t3_neg_3_004_30    m/z 162.9827     0   1   2E+4   0.5   1  RT (min)      lcorbi_t3_neg_1_001_49    m/z 163.0245     0   2   4   6   8E+2   1   1.5  RT (min)      lcorbi_t3_neg_1_004_22    m/z 163.0431     0   2   4   6E+3   2.5   3   3.5  RT (min)       lcorbi_t3_neg_4_052_26    m/z 163.0612     0   2   4E+4   0   0.5  RT (min)      lcorbi_t3_neg_1_005_07    m/z 163.0820     0   2   4   6E+4   1   1.5  RT (min)      lcorbi_t3_neg_3_002_24    m/z 163.0976     0   2   4E+3   0   0.5  RT (min)      lcorbi_t3_neg_2_022_24    m/z 163.5761     0   2   4E+3   13.5   14  RT (min)      lcorbi_t3_neg_1_023_03    m/z 163.8467     0   2   4   6   8E+3   1   1.5  RT (min)       lcorbi_t3_neg_4_067_57    m/z 163.8469     0   2   4   6   8E+3   4.5   5  RT (min)      lcorbi_t3_neg_4_036_16    m/z 163.9905     0   1E+4   2.5   3  RT (min)      lcorbi_t3_neg_2_048_27    m/z 164.0049     0   1   2E+4   17.5   18  RT (min)      lcorbi_t3_neg_1_006_40    m/z 164.0635     0   2   4   6E+3   1   1.5  RT (min)      lcorbi_t3_neg_1_055_05    m/z 164.0842     0   2   4   6E+3   8   8.5  RT (min)      lcorbi_t3_neg_4_007_13    m/z 164.8361     0   1   2   3E+4   4.5   5  RT (min)       lcorbi_t3_neg_1_052_24    m/z 164.8361     0   1   2E+4   17   17.5  RT (min)      lcorbi_t3_neg_4_070_57    m/z 164.8362     0   1   2   3E+4   14.5   15  RT (min)      lcorbi_t3_neg_4_038_15    m/z 164.8401     0   1E+4   1   1.5  RT (min)      lcorbi_t3_neg_1_047_02    m/z 164.8954     0   1   2E+3   18   18.5  RT (min)      lcorbi_t3_neg_4_072_57    m/z 164.9273     0   1   2E+4   18   18.5  RT (min)       lcorbi_t3_neg_4_068_57    m/z 164.9570     0   1   2E+3   18   18.5  RT (min)      lcorbi_t3_neg_1_015_27    m/z 165.0012     0   2   4   6E+4   13.5   14   14.5  RT (min)       lcorbi_t3_neg_1_046_11    m/z 165.0012     0   2   4   6   8E+4   17.5   18  RT (min)       lcorbi_t3_neg_1_006_40    m/z 165.0054     0   1   2   3E+4   1.5   2  RT (min)      lcorbi_t3_neg_1_047_02    m/z 165.0193     0   2   4   6   8E+3   14  RT (min)       lcorbi_t3_neg_2_036_35    m/z 165.0193     0   1E+3   10.5   11  RT (min)       lcorbi_t3_neg_1_005_07    m/z 165.0198     0   2   4E+2   1   1.5  RT (min)      lcorbi_t3_neg_3_022_27    m/z 165.0417     0   2   4   6E+4   3   3.5  RT (min)      lcorbi_t3_neg_2_048_27    m/z 165.0500     0   1   1   1.5  RT (min)      lcorbi_t3_neg_4_046_37    m/z 165.0557     0   2   4   6E+4   6   6.5  RT (min)      lcorbi_t3_neg_2_048_27    m/z 165.0669     0   2   4E+2   6   6.5  RT (min)      lcorbi_t3_neg_4_023_18    m/z 165.0709     0   2   4   6E+3   9   9.5  RT (min)      lcorbi_t3_neg_3_019_12    m/z 165.0750     0   1E+4   3   3.5  RT (min)      lcorbi_t3_neg_2_048_27    m/z 165.0768     0   1   2E+4   0   0.5  RT (min)      lcorbi_t3_neg_2_022_24    m/z 165.8441     0   1   2   3   4E+3   1   1.5  RT (min)      lcorbi_t3_neg_4_007_13    m/z 165.9790     0   2   4   6   8E+2   7.5   8  RT (min)      lcorbi_t3_neg_1_026_19    m/z 166.0134     0   1   2E+3   1.5   2  RT (min)      lcorbi_t3_neg_2_052_32    m/z 166.0404     0   1   2   3   4E+4   13   13.5  RT (min)      lcorbi_t3_neg_4_045_35    m/z 166.0404     0   1   2E+4   11   11.5  RT (min)      lcorbi_t3_neg_4_007_13    m/z 166.0873     0   1   2   3E+4   3.5   4  RT (min)      lcorbi_t3_neg_4_007_13    m/z 166.9244     0   1E+4   18   18.5  RT (min)      lcorbi_t3_neg_4_066_57    m/z 166.9266     0   1   2E+4   0.5   1  RT (min)      lcorbi_t3_neg_1_006_40    m/z 166.9751     0   1   1   1.5  RT (min)      lcorbi_t3_neg_4_007_13    m/z 167.0250     0   1E+4   1   1.5  RT (min)      lcorbi_t3_neg_2_045_30    m/z 167.0304     0   2   4   6E+3   8   8.5  RT (min)      lcorbi_t3_neg_4_023_18    m/z 167.0377     0   1E+4   11   11.5  RT (min)      lcorbi_t3_neg_4_007_13    m/z 167.0383     0   1   2E+3   1   1.5  RT (min)      lcorbi_t3_neg_2_054_15    m/z 167.0713     0   1   2   3E+3   7.5   8  RT (min)      lcorbi_t3_neg_1_044_23    m/z 167.8669     0   2   4   6E+3   13.5   14  RT (min)      lcorbi_t3_neg_4_046_37    m/z 167.9091     0   1E+4   1   1.5  RT (min)      lcorbi_t3_neg_3_019_12    m/z 168.0302     0   2   4   6E+3   7   7.5  RT (min)      lcorbi_t3_neg_4_050_02    m/z 168.0666     0   2   4   6E+3   4   5  RT (min)       lcorbi_t3_neg_4_034_01    m/z 168.6684     0   1   2   3   4E+4   13.5   14  RT (min)      lcorbi_t3_neg_3_005_23    m/z 168.8363     0   1   2E+4   7.5   8  RT (min)      lcorbi_t3_neg_1_047_02    m/z 168.8363     0   1   2E+4   5   5.5  RT (min)      lcorbi_t3_neg_3_006_48    m/z 168.8726     0   1E+3   1   1.5  RT (min)      lcorbi_t3_neg_4_067_57    m/z 168.9438     0   1   2E+2   1   1.5  RT (min)      lcorbi_t3_neg_4_048_03    m/z 169.0253     0   2   4E+3   1   1.5  RT (min)      lcorbi_t3_neg_2_032_45    m/z 169.0392     0   2   4   6   8E+3   8   8.5  RT (min)      lcorbi_t3_neg_4_023_18    m/z 169.0659     0   1E+4   9   9.5  RT (min)      lcorbi_t3_neg_2_048_27    m/z 169.8996     0   2   4   6   8E+3   1.5   2  RT (min)      lcorbi_t3_neg_4_007_13    m/z 170.0458     0   1   1   1.5  RT (min)      lcorbi_t3_neg_4_091_60    m/z 170.0573     0   1   2E+4   1   1.5  RT (min)      lcorbi_t3_neg_1_053_32    m/z 170.6714     0   1   2E+4   13.5   14  RT (min)      lcorbi_t3_neg_4_044_38    m/z 170.8333     0   1   2   3   4E+4   8   8.5  RT (min)      lcorbi_t3_neg_2_006_10    m/z 170.8333     0   1   2   3E+4   5   5.5  RT (min)      lcorbi_t3_neg_2_050_17    m/z 170.8547     0   1   1   1.5  RT (min)      lcorbi_t3_neg_3_002_24    m/z 170.8708     0   1   2E+4   5   6  RT (min)       lcorbi_t3_neg_1_045_44    m/z 170.8709     0   1   2   3E+4   3  RT (min)       lcorbi_t3_neg_1_032_13    m/z 170.9415     0   1   2E+3   0.5   1  RT (min)      lcorbi_t3_neg_4_069_57    m/z 170.9442     0   1E+4   9.5   10  RT (min)      lcorbi_t3_neg_1_038_43    m/z 170.9443     0   1   2   3   4E+4   15   15.5  RT (min)      lcorbi_t3_neg_4_069_57    m/z 170.9443     0   2   4E+4   17.5   18  RT (min)      lcorbi_t3_neg_4_033_53    m/z 171.0122     0   1E+4   4.5   5  RT (min)      lcorbi_t3_neg_2_048_27    m/z 171.0275     0   1E+4   4.5   5  RT (min)      lcorbi_t3_neg_4_050_02    m/z 171.1027     0   1E+4   7   7.5  RT (min)      lcorbi_t3_neg_4_021_25    m/z 171.8319     0   1E+4   1   1.5  RT (min)      lcorbi_t3_neg_4_098_61    m/z 171.8343     0   2   4   6   8E+3   18.5   19  RT (min)      lcorbi_t3_neg_1_053_32    m/z 171.9470     0   1   2   3   4E+4   1.5   2  RT (min)      lcorbi_t3_neg_1_047_02    m/z 172.0278     0   2   4   6E+3   18   18.5  RT (min)      lcorbi_t3_neg_1_053_32    m/z 172.0696     0   1E+3   6.5   7  RT (min)      lcorbi_t3_neg_1_007_26    m/z 172.0702     0   1   2E+4   8   8.5  RT (min)      lcorbi_t3_neg_2_052_32    m/z 172.8303     0   1   2E+4   7   7.5  RT (min)      lcorbi_t3_neg_1_008_08    m/z 172.8304     0   1   2   3E+4   0.5   1  RT (min)      lcorbi_t3_neg_3_011_04    m/z 172.8426     0   1E+3   1   1.5  RT (min)      lcorbi_t3_neg_2_046_20    m/z 172.8679     0   1   2   3E+4   5   5.5  RT (min)      lcorbi_t3_neg_1_043_14    m/z 172.8679     0   1E+4   3   3.5  RT (min)      lcorbi_t3_neg_1_033_53    m/z 172.9413     0   1E+4   17.5   18  RT (min)       lcorbi_t3_neg_2_043_33    m/z 172.9413     0   1E+4   15   15.5  RT (min)      lcorbi_t3_neg_4_067_57    m/z 172.9954     0   1   2   3E+3   4.5   5  RT (min)      lcorbi_t3_neg_3_004_30    m/z 173.0242     0   2   4   6   8E+3   18   18.5  RT (min)      lcorbi_t3_neg_1_053_32    m/z 173.0263     0   2   4E+3   1.5   2  RT (min)      lcorbi_t3_neg_4_042_14    m/z 173.0456     0   2   4   6E+3   2.5   3  RT (min)      lcorbi_t3_neg_4_081_58    m/z 173.0819     0   2   4E+4   6   6.5  RT (min)      lcorbi_t3_neg_3_043_25    m/z 173.0933     0   1   2   3   4E+3   1   1.5  RT (min)      lcorbi_t3_neg_3_021_38    m/z 173.4206     0   2   4   6   8E+3   18   18.5  RT (min)      lcorbi_t3_neg_4_020_44    m/z 173.4206     0   2   4   6E+4   11   11.5  RT (min)      lcorbi_t3_neg_4_007_13    m/z 173.8313     0   2   4   6   8E+3   18.5   19  RT (min)      lcorbi_t3_neg_1_054_37    m/z 173.8314     0   2   4E+3   1   1.5  RT (min)      lcorbi_t3_neg_2_023_39    m/z 173.9262     0   1E+4   18   18.5  RT (min)      lcorbi_t3_neg_4_066_57    m/z 173.9729     0   2   4E+3   9   9.5  RT (min)      lcorbi_t3_neg_4_007_13    m/z 173.9909     0   1   2   3   4E+3   4.5   5  RT (min)      lcorbi_t3_neg_1_031_33    m/z 174.0070     0   1   2E+3   18   18.5  RT (min)      lcorbi_t3_neg_4_054_36    m/z 174.0241     0   2   4   6E+3   14   14.5  RT (min)      lcorbi_t3_neg_1_005_07    m/z 174.0242     0   2   4E+3   18   18.5  RT (min)      lcorbi_t3_neg_1_021_31    m/z 174.0322     0   1E+4   8   8.5  RT (min)      lcorbi_t3_neg_4_040_19    m/z 174.0409     0   1   2E+3   1   1.5  RT (min)      lcorbi_t3_neg_4_021_25    m/z 174.0560     0   1E+4   6   6.5  RT (min)      lcorbi_t3_neg_2_048_27    m/z 174.8397     0   2   4   6E+4   1   1.5  RT (min)      lcorbi_t3_neg_1_047_02    m/z 174.8650     0   1   2   3E+3   18.5   19  RT (min)      lcorbi_t3_neg_4_081_58    m/z 174.8650     0   2   4   6   8E+3   0.5   1  RT (min)      lcorbi_t3_neg_4_068_57    m/z 174.9561     0   1   2   3E+4   13   13.5  RT (min)      lcorbi_t3_neg_1_002_16    m/z 174.9561     0   1E+5   9.5   10  RT (min)      lcorbi_t3_neg_4_099_61    m/z 174.9561     0   2   4   6E+4   5   5.5  RT (min)      lcorbi_t3_neg_4_030_04    m/z 174.9598     0   1E+4   18   18.5  RT (min)      lcorbi_t3_neg_4_072_57    m/z 174.9602     0   1E+4   1   1.5  RT (min)      lcorbi_t3_neg_3_032_17    m/z 174.9954     0   1   2E+2   4.5   5  RT (min)      lcorbi_t3_neg_1_001_49    m/z 175.0033     0   1E+3   18   18.5  RT (min)      lcorbi_t3_neg_4_072_57    m/z 175.0205     0   2   4   6E+3   13.5   14  RT (min)      lcorbi_t3_neg_1_002_16    m/z 175.0205     0   2   4   6E+3   14.5   15  RT (min)      lcorbi_t3_neg_1_003_41    m/z 175.0226     0   1   2E+3   18   18.5  RT (min)      lcorbi_t3_neg_4_013_09    m/z 175.0400     0   1E+3   7   7.5  RT (min)      lcorbi_t3_neg_4_103_61    m/z 175.0591     0   2   4   6   8E+3   18   18.5  RT (min)      lcorbi_t3_neg_4_092_60    m/z 175.0612     0   1   2E+4   0   0.5  RT (min)      lcorbi_t3_neg_1_007_26    m/z 175.0918     0   1E+3   1   1.5  RT (min)      lcorbi_t3_neg_1_017_51    m/z 175.8283     0   1   2E+3   18.5   19  RT (min)      lcorbi_t3_neg_1_054_37    m/z 175.9594     0   1E+4   18   18.5  RT (min)      lcorbi_t3_neg_4_070_57    m/z 176.0055     0   1   2E+3   18   18.5  RT (min)      lcorbi_t3_neg_1_002_16    m/z 176.0198     0   1E+3   18   18.5  RT (min)      lcorbi_t3_neg_4_070_57    m/z 176.0198     0   2   4   6   8E+3   14   14.5  RT (min)      lcorbi_t3_neg_2_038_22    m/z 176.0566     0   1E+4   1   1.5  RT (min)      lcorbi_t3_neg_4_044_38    m/z 176.0843     0   2   4E+3   9.5   10  RT (min)      lcorbi_t3_neg_2_048_27    m/z 176.8173     0   1E+4   1   1.5  RT (min)      lcorbi_t3_neg_4_101_61    m/z 176.8262     0   1   2E+3   1   1.5  RT (min)      lcorbi_t3_neg_4_103_61    m/z 176.8369     0   1E+3   1   1.5  RT (min)      lcorbi_t3_neg_4_101_61    m/z 176.8942     0   1E+3   1   1.5  RT (min)      lcorbi_t3_neg_1_055_05    m/z 176.9399     0   1E+4   1   1.5  RT (min)      lcorbi_t3_neg_4_092_60    m/z 176.9516     0   2   4   6E+3   7.5   8  RT (min)      lcorbi_t3_neg_4_075_58    m/z 176.9520     0   1E+4   1   1.5  RT (min)      lcorbi_t3_neg_2_048_27    m/z 176.9603     0   2   4   6   8E+3   18   18.5  RT (min)      lcorbi_t3_neg_4_069_57    m/z 177.0019     0   1E+4   18   18.5  RT (min)      lcorbi_t3_neg_4_096_60    m/z 177.0019     0   1E+4   14   14.5  RT (min)      lcorbi_t3_neg_4_071_57    m/z 177.0212     0   2   4   6E+3   18   18.5  RT (min)      lcorbi_t3_neg_1_053_32    m/z 177.0219     0   1E+4   13.5   14  RT (min)      lcorbi_t3_neg_4_078_58    m/z 177.0595     0   2   4E+3   6   6.5  RT (min)      lcorbi_t3_neg_4_041_54    m/z 177.0769     0   1E+4   2.5   3  RT (min)      lcorbi_t3_neg_1_019_29    m/z 177.8451     0   1   2   3E+4   1   1.5  RT (min)      lcorbi_t3_neg_3_019_12    m/z 177.9132     0   1E+3   1   1.5  RT (min)      lcorbi_t3_neg_4_045_35    m/z 178.0067     0   1   1   1.5  RT (min)      lcorbi_t3_neg_3_026_44    m/z 178.0184     0   2   4   6E+4   13   13.5   14  RT (min)       lcorbi_t3_neg_4_079_58    m/z 178.0184     0   1   2   3E+4   11   12  RT (min)       lcorbi_t3_neg_2_051_34    m/z 178.0377     0   1   2   3E+3   1.5   2  RT (min)      lcorbi_t3_neg_3_044_01    m/z 178.0591     0   1   2E+4   7   7.5  RT (min)      lcorbi_t3_neg_2_048_27    m/z 178.0802     0   2   4   6   8E+3   0   0.5  RT (min)      lcorbi_t3_neg_1_035_25    m/z 178.0954     0   2   4   6E+3   9.5   10  RT (min)      lcorbi_t3_neg_3_019_12    m/z 178.1172     0   2   4   6E+4   8.5   9  RT (min)       lcorbi_t3_neg_4_011_24    m/z 178.1236     0   1E+4   14   14.5  RT (min)      lcorbi_t3_neg_4_029_31    m/z 178.8152     0   1   2E+4   1   1.5  RT (min)       lcorbi_t3_neg_4_102_61    m/z 178.9204     0   1E+4   1.5   2  RT (min)      lcorbi_t3_neg_1_032_13    m/z 178.9402     0   1   2E+4   1   1.5  RT (min)      lcorbi_t3_neg_1_055_05    m/z 178.9490     0   1E+4   18   18.5  RT (min)      lcorbi_t3_neg_4_067_57    m/z 178.9778     0   1   2   3E+4   4.5   5  RT (min)      lcorbi_t3_neg_1_014_28    m/z 178.9778     0   2   4   6E+4   3   3.5  RT (min)      lcorbi_t3_neg_1_046_11    m/z 178.9778     0   1   2E+4   7   7.5  RT (min)      lcorbi_t3_neg_1_046_11    m/z 179.0147     0   2   4   6E+4   13.5   14  RT (min)       lcorbi_t3_neg_4_085_59    m/z 179.0147     0   2   4E+4   11   11.5  RT (min)       lcorbi_t3_neg_2_002_26    m/z 179.0147     0   1E+4   8   8.5  RT (min)      lcorbi_t3_neg_3_046_10    m/z 179.0197     0   1   2   3   4E+2   0   0.5  RT (min)      lcorbi_t3_neg_3_026_44    m/z 179.0340     0   1   2   3   4E+2   2  RT (min)       lcorbi_t3_neg_1_046_11    m/z 179.0349     0   2   4   6   8E+2   5   5.5  RT (min)      lcorbi_t3_neg_1_044_23    m/z 179.0350     0   1E+3   10.5   11  RT (min)      lcorbi_t3_neg_3_043_25    m/z 179.0604     0   1   2   3   4E+3   1   1.5  RT (min)      lcorbi_t3_neg_4_039_32    m/z 179.0755     0   1   2   3E+3   8.5   9  RT (min)      lcorbi_t3_neg_4_109_62    m/z 179.0925     0   2   4   6   8E+3   0   0.5  RT (min)      lcorbi_t3_neg_2_022_24    m/z 179.1053     0   1   1   1.5  RT (min)      lcorbi_t3_neg_2_048_27    m/z 179.1288     0   1   2   3   4E+4   1   1.5  RT (min)      lcorbi_t3_neg_4_011_24    m/z 179.1441     0   1E+4   10   10.5  RT (min)      lcorbi_t3_neg_2_048_27    m/z 179.3822     0   1E+4   14   14.5  RT (min)      lcorbi_t3_neg_3_005_23    m/z 179.8405     0   2   4   6   8E+3   17.5   18  RT (min)      lcorbi_t3_neg_1_036_09    m/z 179.8406     0   2   4   6   8E+3   15.5   16  RT (min)      lcorbi_t3_neg_1_039_38    m/z 179.8841     0   1   2   3   4E+3   13.5   14  RT (min)      lcorbi_t3_neg_4_007_13    m/z 180.0417     0   1E+4   1   1.5  RT (min)      lcorbi_t3_neg_2_048_27    m/z 180.1030     0   1E+4   9   9.5  RT (min)      lcorbi_t3_neg_2_048_27    m/z 180.1167     0   1   1   1.5  RT (min)      lcorbi_t3_neg_2_048_27    m/z 180.8598     0   1   2   3   4E+2   1   1.5  RT (min)      lcorbi_t3_neg_3_002_24    m/z 180.8997     0   1E+4   15.5   16  RT (min)      lcorbi_t3_neg_4_112_62    m/z 180.9731     0   1E+5   10.5   11  RT (min)      lcorbi_t3_neg_4_079_58    m/z 180.9731     0   1   2   3   4E+4   15.5   16  RT (min)      lcorbi_t3_neg_1_018_46    m/z 180.9776     0   1   2E+4   1   1.5  RT (min)      lcorbi_t3_neg_4_036_16    m/z 180.9894     0   1   2E+4   0   0.5  RT (min)      lcorbi_t3_neg_1_001_49    m/z 181.0367     0   1   2.5   3  RT (min)       lcorbi_t3_neg_1_044_23    m/z 181.0450     0   1   2   3   4E+5   1.5   2  RT (min)       lcorbi_t3_neg_2_048_27    m/z 181.0495     0   1   2   3   4E+2   1   1.5  RT (min)      lcorbi_t3_neg_4_030_04    m/z 181.0540     0   1E+4   14   14.5  RT (min)      lcorbi_t3_neg_4_030_04    m/z 181.0719     0   2   4   6E+2   1   1.5  RT (min)      lcorbi_t3_neg_3_002_24    m/z 181.0813     0   1   2E+4   5   5.5  RT (min)      lcorbi_t3_neg_4_039_32    m/z 181.8375     0   2   4   6   8E+3   17.5   18  RT (min)      lcorbi_t3_neg_1_043_14    m/z 181.8376     0   2   4   6E+3   15.5   16  RT (min)      lcorbi_t3_neg_1_021_31    m/z 181.9960     0   2   4   6E+4   4.5   5  RT (min)      lcorbi_t3_neg_4_110_62    m/z 182.0436     0   1   1   1.5  RT (min)      lcorbi_t3_neg_3_035_03    m/z 182.0450     0   1E+3   5   5.5  RT (min)      lcorbi_t3_neg_4_105_61    m/z 182.0890     0   2   4   6   8E+3   5   5.5  RT (min)      lcorbi_t3_neg_4_039_32    m/z 182.1314     0   2   4   6E+3   14   14.5  RT (min)      lcorbi_t3_neg_3_005_23    m/z 182.1604     0   1   2   3   4E+3   14.5   15  RT (min)      lcorbi_t3_neg_2_039_31    m/z 182.8597     0   2   4   6   8E+3   18   18.5  RT (min)      lcorbi_t3_neg_4_078_58    m/z 182.8966     0   2   4   6   8E+3   18.5   19  RT (min)      lcorbi_t3_neg_4_111_62    m/z 182.9886     0   1   2E+4   13.5   14  RT (min)       lcorbi_t3_neg_4_101_61    m/z 182.9886     0   2   4   6E+3   18   18.5  RT (min)      lcorbi_t3_neg_4_079_58    m/z 182.9886     0   1   2   3E+4   10   11  RT (min)       lcorbi_t3_neg_4_071_57    m/z 183.0044     0   1   2E+3   7   7.5  RT (min)      lcorbi_t3_neg_1_029_21    m/z 183.0067     0   2   4   6   8E+3   1   1.5  RT (min)      lcorbi_t3_neg_2_050_17    m/z 183.0161     0   2   4E+3   1.5   2  RT (min)      lcorbi_t3_neg_2_048_27    m/z 183.1390     0   1E+4   9   9.5  RT (min)      lcorbi_t3_neg_2_048_27    m/z 183.9956     0   1E+4   4.5   5  RT (min)      lcorbi_t3_neg_4_107_62    m/z 184.0981     0   2   4   6   8E+3   1   1.5  RT (min)      lcorbi_t3_neg_4_007_13    m/z 184.8578     0   1   2   3   4E+3   18   18.5  RT (min)      lcorbi_t3_neg_1_053_32    m/z 184.8765     0   2   4   6   8E+3   4.5   5  RT (min)      lcorbi_t3_neg_1_047_02    m/z 184.8766     0   1E+4   0.5   1  RT (min)      lcorbi_t3_neg_1_047_02    m/z 184.8783     0   2   4   6   8E+3   14   14.5  RT (min)      lcorbi_t3_neg_1_044_23    m/z 184.9928     0   2   4E+3   7   7.5  RT (min)      lcorbi_t3_neg_4_080_58    m/z 185.0223     0   1   2E+3   1   1.5  RT (min)      lcorbi_t3_neg_1_044_23    m/z 185.0569     0   1   2   3   4E+3   1   1.5  RT (min)      lcorbi_t3_neg_1_053_32    m/z 185.3822     0   2   4   6   8E+3   14   14.5  RT (min)      lcorbi_t3_neg_3_005_23    m/z 185.3844     0   2   4   6E+3   14   14.5  RT (min)      lcorbi_t3_neg_3_020_31    m/z 185.8861     0   1E+4   14   14.5  RT (min)      lcorbi_t3_neg_3_005_23    m/z 185.9625     0   1   2   3E+4   4.5   5  RT (min)      lcorbi_t3_neg_3_032_17    m/z 185.9940     0   1E+3   1   1.5  RT (min)      lcorbi_t3_neg_2_007_14    m/z 186.0070     0   1   2E+3   18   18.5  RT (min)      lcorbi_t3_neg_4_056_56    m/z 186.0384     0   1E+4   1   1.5  RT (min)      lcorbi_t3_neg_4_005_27    m/z 186.0521     0   1E+3   1   1.5  RT (min)      lcorbi_t3_neg_2_052_32    m/z 186.0867     0   2   4E+4   1.5   2  RT (min)      lcorbi_t3_neg_3_019_12    m/z 186.1212     0   2   4E+3   13.5   14  RT (min)      lcorbi_t3_neg_4_046_37    m/z 186.1392     0   1   2   3   4E+3   14.5   15  RT (min)      lcorbi_t3_neg_2_053_23    m/z 186.3897     0   1E+4   13.5   14  RT (min)      lcorbi_t3_neg_4_007_13    m/z 186.6385     0   2   4   6E+3   13.5   14  RT (min)      lcorbi_t3_neg_4_007_13    m/z 186.6411     0   1   2   3   4E+3   13.5   14  RT (min)      lcorbi_t3_neg_4_007_13    m/z 186.8760     0   1   2   3E+2   1   1.5  RT (min)      lcorbi_t3_neg_4_039_32    m/z 186.9294     0   1E+4   18   18.5  RT (min)      lcorbi_t3_neg_4_092_60    m/z 187.0034     0   1   2   3E+3   18   18.5  RT (min)      lcorbi_t3_neg_4_031_48    m/z 187.0045     0   1   2E+3   2   2.5  RT (min)      lcorbi_t3_neg_4_081_58    m/z 187.0113     0   1   2   3E+3   5.5   6  RT (min)      lcorbi_t3_neg_4_036_16    m/z 187.0464     0   1   2   3E+4   1.5   2  RT (min)      lcorbi_t3_neg_4_077_58    m/z 187.0611     0   1E+3   4.5   5   5.5  RT (min)       lcorbi_t3_neg_1_003_41    m/z 187.1022     0   1   2E+3   7   7.5  RT (min)      lcorbi_t3_neg_4_046_37    m/z 187.1168     0   1E+3   6   6.5  RT (min)      lcorbi_t3_neg_1_045_44    m/z 187.1424     0   1   13.5   14  RT (min)      lcorbi_t3_neg_1_006_40    m/z 187.1451     0   1   13.5   14  RT (min)      lcorbi_t3_neg_1_004_22    m/z 187.3935     0   2   4E+3   13.5   14  RT (min)      lcorbi_t3_neg_4_034_01    m/z 187.6365     0   1   2   3E+3   14.5   15  RT (min)      lcorbi_t3_neg_2_052_32    m/z 187.8610     0   1   2   3E+3   18.5   19  RT (min)      lcorbi_t3_neg_1_001_49    m/z 187.8610     0   2   4E+4   0.5   1   1.5  RT (min)       lcorbi_t3_neg_4_090_60    m/z 187.8611     0   1   2   3   4E+3   15.5   16  RT (min)      lcorbi_t3_neg_2_024_37    m/z 187.9063     0   1   14   14.5  RT (min)      lcorbi_t3_neg_1_047_02    m/z 187.9418     0   1   2E+4   13.5   14  RT (min)       lcorbi_t3_neg_3_011_04    m/z 187.9418     0   1   2   3   4E+4   18   18.5  RT (min)       lcorbi_t3_neg_3_050_40    m/z 187.9421     0   1E+3   0   0.5  RT (min)      lcorbi_t3_neg_4_082_59    m/z 187.9782     0   1   2E+3   18   18.5  RT (min)      lcorbi_t3_neg_4_046_37    m/z 188.0065     0   1   2E+3   5.5   6  RT (min)      lcorbi_t3_neg_3_032_17    m/z 188.0282     0   1   1.5   2  RT (min)      lcorbi_t3_neg_4_081_58    m/z 188.0609     0   1   2   3E+3   1   1.5  RT (min)      lcorbi_t3_neg_4_086_59    m/z 188.1009     0   1   2   3E+3   7   7.5  RT (min)      lcorbi_t3_neg_3_019_12    m/z 188.1147     0   1   2E+3   18   18.5  RT (min)      lcorbi_t3_neg_4_035_33    m/z 188.1594     0   1E+4   14   14.5  RT (min)      lcorbi_t3_neg_3_002_24    m/z 188.6632     0   2   4   6E+3   14   14.5  RT (min)      lcorbi_t3_neg_3_002_24    m/z 188.8938     0   2   4E+3   18   18.5  RT (min)      lcorbi_t3_neg_4_104_61    m/z 188.9128     0   1E+4   0   0.5  RT (min)      lcorbi_t3_neg_1_004_22    m/z 188.9492     0   1E+4   1.5   2  RT (min)      lcorbi_t3_neg_4_067_57    m/z 189.0110     0   1   2E+3   5.5   6  RT (min)      lcorbi_t3_neg_1_014_28    m/z 189.0190     0   1E+3   18   18.5  RT (min)      lcorbi_t3_neg_1_004_22    m/z 189.0227     0   1E+4   14   14.5  RT (min)      lcorbi_t3_neg_4_008_22    m/z 189.0557     0   1E+4   7   7.5  RT (min)      lcorbi_t3_neg_2_048_27    m/z 189.0598     0   1E+4   1   1.5  RT (min)      lcorbi_t3_neg_3_005_23    m/z 189.0769     0   1E+4   0   0.5  RT (min)      lcorbi_t3_neg_1_023_03    m/z 189.1111     0   2   4E+3   18   18.5  RT (min)      lcorbi_t3_neg_3_002_24    m/z 189.1132     0   1E+4   7   7.5  RT (min)      lcorbi_t3_neg_2_048_27    m/z 189.1133     0   1E+4   0   0.5  RT (min)      lcorbi_t3_neg_1_017_51    m/z 189.1620     0   1   13.5   14  RT (min)      lcorbi_t3_neg_4_029_31    m/z 189.3717     0   2   4   6   8E+3   13.5   14  RT (min)      lcorbi_t3_neg_4_046_37    m/z 189.3742     0   2   4   6E+3   13.5   14  RT (min)      lcorbi_t3_neg_2_039_31    m/z 189.8581     0   1   2   3   4E+4   0.5   1  RT (min)       lcorbi_t3_neg_4_082_59    m/z 189.8706     0   2   4   6E+2   1   1.5  RT (min)      lcorbi_t3_neg_1_038_43    m/z 189.9315     0   1   2E+4   18   18.5  RT (min)       lcorbi_t3_neg_4_084_59    m/z 189.9549     0   2   4   6E+3   8.5   9  RT (min)      lcorbi_t3_neg_2_048_27    m/z 189.9896     0   1E+4   4.5   5  RT (min)      lcorbi_t3_neg_2_048_27    m/z 190.0041     0   1   2E+4   14   14.5  RT (min)      lcorbi_t3_neg_1_056_56    m/z 190.0041     0   1   2E+4   17.5   18  RT (min)      lcorbi_t3_neg_1_006_40    m/z 190.0060     0   1   2   3   4E+3   5.5   6  RT (min)      lcorbi_t3_neg_2_048_27    m/z 190.0191     0   2   4E+3   14   14.5  RT (min)      lcorbi_t3_neg_1_047_02    m/z 190.0212     0   1   2E+3   18   18.5  RT (min)      lcorbi_t3_neg_1_047_02    m/z 190.0494     0   1E+4   4.5   5  RT (min)      lcorbi_t3_neg_4_044_38    m/z 190.0999     0   2   4   6   8E+3   9.5   10  RT (min)      lcorbi_t3_neg_2_048_27    m/z 190.8908     0   2   4E+3   18   18.5  RT (min)      lcorbi_t3_neg_4_111_62    m/z 190.9284     0   2   4   6E+4   4.5   5  RT (min)      lcorbi_t3_neg_1_044_23    m/z 190.9285     0   1   2E+5   15.5   16  RT (min)      lcorbi_t3_neg_4_023_18    m/z 190.9820     0   2   4   6   8E+3   4.5   5  RT (min)      lcorbi_t3_neg_2_048_27    m/z 190.9922     0   2   4   6   8E+2   1   1.5  RT (min)      lcorbi_t3_neg_2_007_14    m/z 191.0005     0   2   4   6E+4   13.5   14  RT (min)       lcorbi_t3_neg_3_023_07    m/z 191.0005     0   2   4   6E+4   17.5   18  RT (min)       lcorbi_t3_neg_2_014_29    m/z 191.0111     0   1   2E+4   1   1.5  RT (min)      lcorbi_t3_neg_4_077_58    m/z 191.0154     0   2   4   6E+3   14   14.5  RT (min)      lcorbi_t3_neg_1_025_52    m/z 191.0176     0   1   2   3E+3   18   18.5  RT (min)      lcorbi_t3_neg_2_038_22    m/z 191.0391     0   1E+3   1   1.5  RT (min)      lcorbi_t3_neg_2_054_15    m/z 191.0769     0   1   2   3E+3   9   9.5  RT (min)      lcorbi_t3_neg_2_048_27    m/z 191.0925     0   1   2E+4   0   0.5  RT (min)      lcorbi_t3_neg_1_023_03    m/z 191.1289     0   2   4   6E+4   0   0.5  RT (min)      lcorbi_t3_neg_2_022_24    m/z 191.6353     0   1E+4   14   14.5  RT (min)      lcorbi_t3_neg_4_007_13    m/z 191.8552     0   1E+3   1   1.5  RT (min)      lcorbi_t3_neg_4_082_59    m/z 191.8604     0   1   2   3   4E+2   13.5   14  RT (min)      lcorbi_t3_neg_3_002_24    m/z 191.8810     0   2   4   6E+3   13.5   14  RT (min)      lcorbi_t3_neg_2_039_31    m/z 191.9318     0   2   4   6E+4   0   0.5  RT (min)      lcorbi_t3_neg_4_051_40    m/z 191.9519     0   1E+3   8.5   9  RT (min)      lcorbi_t3_neg_4_023_18    m/z 192.0271     0   2   4E+3   1   1.5  RT (min)      lcorbi_t3_neg_4_038_15    m/z 192.0349     0   1E+4   8   8.5  RT (min)      lcorbi_t3_neg_3_024_26    m/z 192.0480     0   1   1   1.5  RT (min)      lcorbi_t3_neg_2_024_37    m/z 192.0649     0   2   4E+3   1   1.5  RT (min)      lcorbi_t3_neg_4_033_53    m/z 192.0732     0   2   4   6E+3   18   18.5  RT (min)      lcorbi_t3_neg_4_097_60    m/z 192.0958     0   1   14   14.5  RT (min)      lcorbi_t3_neg_4_110_62    m/z 192.1110     0   2   4   6E+3   7.5   8  RT (min)      lcorbi_t3_neg_2_048_27    m/z 192.1196     0   2   4E+3   9.5   10  RT (min)      lcorbi_t3_neg_2_006_10    m/z 192.9288     0   1   2   3   4E+4   18   18.5  RT (min)       lcorbi_t3_neg_4_079_58    m/z 192.9288     0   2   4   6E+3   0   0.5  RT (min)      lcorbi_t3_neg_4_011_24    m/z 192.9288     0   1E+4   14.5   15  RT (min)      lcorbi_t3_neg_4_089_59    m/z 192.9456     0   1E+3   8.5   9  RT (min)      lcorbi_t3_neg_4_104_61    m/z 192.9889     0   1   2E+3   1   1.5  RT (min)      lcorbi_t3_neg_4_039_32    m/z 192.9988     0   2   4   6E+2   1   1.5  RT (min)      lcorbi_t3_neg_2_007_14    m/z 193.0182     0   1   2   3E+2   1   1.5  RT (min)      lcorbi_t3_neg_2_007_14    m/z 193.0303     0   1   2E+3   18   18.5  RT (min)      lcorbi_t3_neg_1_001_49    m/z 193.0505     0   2   4   6E+3   7   7.5  RT (min)      lcorbi_t3_neg_4_001_49    m/z 193.0605     0   1E+4   1   1.5  RT (min)      lcorbi_t3_neg_3_052_32    m/z 193.0696     0   1E+4   18   18.5  RT (min)      lcorbi_t3_neg_4_092_60    m/z 193.0716     0   1   2E+3   1   1.5  RT (min)      lcorbi_t3_neg_1_029_21    m/z 193.0903     0   1E+4   8   8.5  RT (min)      lcorbi_t3_neg_2_048_27    m/z 193.0982     0   2   4   6   8E+3   5   5.5  RT (min)      lcorbi_t3_neg_4_029_31    m/z 193.1081     0   2   4   6   8E+3   0   0.5  RT (min)      lcorbi_t3_neg_2_022_24    m/z 193.3957     0   1   2   3E+3   14   14.5  RT (min)      lcorbi_t3_neg_4_047_30    m/z 193.6467     0   1   2E+3   14   14.5  RT (min)      lcorbi_t3_neg_4_007_13    m/z 193.8156     0   1E+4   17.5   18  RT (min)      lcorbi_t3_neg_4_066_57    m/z 193.8156     0   1E+4   15.5   16  RT (min)      lcorbi_t3_neg_1_035_25    m/z 193.8574     0   2   4   6E+3   13   13.5  RT (min)      lcorbi_t3_neg_4_040_19    m/z 193.9267     0   1E+4   1   1.5  RT (min)      lcorbi_t3_neg_4_072_57    m/z 194.0303     0   2   4   6E+3   13.5   14  RT (min)      lcorbi_t3_neg_4_016_05    m/z 194.0347     0   1   2   3E+3   8   8.5  RT (min)      lcorbi_t3_neg_4_023_18    m/z 194.0458     0   1   2   3E+3   8   8.5  RT (min)      lcorbi_t3_neg_4_098_61    m/z 194.0764     0   1   2   3   4E+3   1   1.5  RT (min)      lcorbi_t3_neg_4_091_60    m/z 194.0863     0   2   4E+3   10   10.5  RT (min)      lcorbi_t3_neg_4_049_55    m/z 194.0957     0   1   2E+3   1   1.5  RT (min)      lcorbi_t3_neg_3_001_49    m/z 194.1368     0   1E+4   14   14.5  RT (min)      lcorbi_t3_neg_4_007_13    m/z 194.1570     0   1E+4   13.5   14  RT (min)      lcorbi_t3_neg_4_039_32    m/z 194.1596     0   1E+4   13.5   14  RT (min)      lcorbi_t3_neg_4_039_32    m/z 194.3873     0   1   14   14.5  RT (min)      lcorbi_t3_neg_4_011_24    m/z 194.6611     0   1E+4   14   14.5  RT (min)      lcorbi_t3_neg_4_039_32    m/z 194.6760     0   2   4   6   8E+3   13.5   14  RT (min)      lcorbi_t3_neg_1_055_05    m/z 194.9243     0   1E+4   18   18.5  RT (min)      lcorbi_t3_neg_4_101_61    m/z 194.9830     0   2   4   6E+3   7   7.5  RT (min)      lcorbi_t3_neg_4_039_32    m/z 194.9886     0   2   4E+4   0   0.5  RT (min)      lcorbi_t3_neg_1_001_49    m/z 194.9897     0   1   2   3   4E+3   18.5   19  RT (min)      lcorbi_t3_neg_1_044_23    m/z 195.0267     0   2   4   6   8E+3   13.5   14  RT (min)      lcorbi_t3_neg_2_020_05    m/z 195.0330     0   2   4   6E+3   1   1.5  RT (min)      lcorbi_t3_neg_1_055_05    m/z 195.0605     0   1   2E+3   1   1.5  RT (min)      lcorbi_t3_neg_4_011_24    m/z 195.0639     0   1E+4   6   6.5  RT (min)      lcorbi_t3_neg_4_087_59    m/z 195.0873     0   1E+4   0   0.5  RT (min)      lcorbi_t3_neg_2_022_24    m/z 195.1040     0   1   2E+2   4.5   5  RT (min)      lcorbi_t3_neg_4_007_13    m/z 195.1207     0   1   2E+3   13   13.5  RT (min)      lcorbi_t3_neg_4_054_36    m/z 195.1651     0   2   4   6   8E+3   14   14.5  RT (min)      lcorbi_t3_neg_3_002_24    m/z 195.6249     0   2   4   6E+3   13.5   14  RT (min)      lcorbi_t3_neg_4_044_38    m/z 195.6689     0   1E+4   14   14.5  RT (min)      lcorbi_t3_neg_4_039_32    m/z 195.8007     0   1   2   3   4E+3   1   1.5  RT (min)      lcorbi_t3_neg_3_018_13    m/z 195.8109     0   2   4   6E+4   4.5   5  RT (min)      lcorbi_t3_neg_2_048_27    m/z 195.8110     0   1   2E+5   15.5   16  RT (min)      lcorbi_t3_neg_1_035_25    m/z 195.8164     0   1   2   3E+4   1   1.5  RT (min)      lcorbi_t3_neg_2_010_02    m/z 195.8756     0   1   2   3   4E+3   13.5   14  RT (min)      lcorbi_t3_neg_3_047_37    m/z 196.0226     0   1   1.5   2  RT (min)       lcorbi_t3_neg_4_011_24    m/z 196.0251     0   2   4   6   8E+3   6   6.5  RT (min)      lcorbi_t3_neg_2_048_27    m/z 196.1277     0   2   4   6   8E+4   7   7.5  RT (min)       lcorbi_t3_neg_4_021_25    m/z 196.1289     0   2   4   6   8E+3   13.5   14  RT (min)      lcorbi_t3_neg_3_005_23    m/z 196.1423     0   1E+4   9.5   10  RT (min)      lcorbi_t3_neg_2_048_27    m/z 196.4233     0   2   4   6   8E+3   14.5   15  RT (min)      lcorbi_t3_neg_4_011_24    m/z 196.4261     0   2   4   6E+3   14.5   15  RT (min)      lcorbi_t3_neg_2_022_24    m/z 196.6328     0   2   4   6   8E+3   14   14.5  RT (min)      lcorbi_t3_neg_3_020_31    m/z 196.8946     0   2   4   6E+3   1   1.5  RT (min)      lcorbi_t3_neg_1_044_23    m/z 196.9410     0   1   2   3E+3   0.5   1  RT (min)      lcorbi_t3_neg_4_001_49    m/z 196.9507     0   1E+3   1   1.5  RT (min)      lcorbi_t3_neg_3_006_48    m/z 196.9854     0   1   2   3E+3   18   18.5  RT (min)      lcorbi_t3_neg_1_043_14    m/z 197.0204     0   1E+4   8.5   9  RT (min)      lcorbi_t3_neg_4_082_59    m/z 197.0317     0   1   2   3E+3   2   2.5  RT (min)      lcorbi_t3_neg_4_095_60    m/z 197.0429     0   2   4   6   8E+2   5   5.5  RT (min)      lcorbi_t3_neg_4_042_14    m/z 197.0819     0   1E+3   6.5   7  RT (min)      lcorbi_t3_neg_4_088_59    m/z 197.3868     0   1E+4   13.5   14  RT (min)      lcorbi_t3_neg_1_013_35    m/z 197.8012     0   1E+4   1   1.5  RT (min)      lcorbi_t3_neg_1_019_29    m/z 197.8079     0   1E+5   12.5   13  RT (min)      lcorbi_t3_neg_4_030_04    m/z 197.8080     0   1   2   3E+5   15.5   16  RT (min)      lcorbi_t3_neg_1_026_19    m/z 197.8155     0   1   2   3   4E+4   1   1.5  RT (min)      lcorbi_t3_neg_2_027_13    m/z 197.9401     0   1   2E+3   0.5   1  RT (min)      lcorbi_t3_neg_1_045_44    m/z 197.9631     0   1   2E+4   10   10.5  RT (min)      lcorbi_t3_neg_4_011_24    m/z 197.9631     0   1   2E+4   0.5   1  RT (min)      lcorbi_t3_neg_4_082_59    m/z 198.9037     0   1   13.5   14  RT (min)      lcorbi_t3_neg_4_104_61    m/z 198.9226     0   1   2   3E+3   18   18.5  RT (min)      lcorbi_t3_neg_4_020_44    m/z 198.9363     0   1   2   3E+4   4.5   5  RT (min)      lcorbi_t3_neg_4_003_21    m/z 198.9829     0   1E+3   1   1.5  RT (min)      lcorbi_t3_neg_2_048_27    m/z 199.0014     0   1   2   3E+2   1   1.5  RT (min)      lcorbi_t3_neg_2_048_27    m/z 199.0400     0   1E+3   8   8.5  RT (min)      lcorbi_t3_neg_2_048_27    m/z 199.0433     0   1   6.5   7  RT (min)      lcorbi_t3_neg_4_075_58    m/z 199.0764     0   2   4   6   8E+3   9   9.5  RT (min)      lcorbi_t3_neg_2_048_27    m/z 199.1286     0   1   2   3E+3   13.5   14  RT (min)      lcorbi_t3_neg_4_044_38    m/z 199.1542     0   1E+4   13.5   14  RT (min)      lcorbi_t3_neg_4_039_32    m/z 199.5341     0   1E+4   13.5   14  RT (min)      lcorbi_t3_neg_4_039_32    m/z 199.7995     0   1   2E+4   1   1.5  RT (min)      lcorbi_t3_neg_3_018_13    m/z 199.8050     0   1E+5   15.5   16  RT (min)      lcorbi_t3_neg_1_035_25    m/z 199.8509     0   1E+4   15.5   16  RT (min)      lcorbi_t3_neg_1_044_23    m/z 199.8834     0   1E+4   13.5   14  RT (min)      lcorbi_t3_neg_2_039_31    m/z 199.8868     0   1   2E+3   0   0.5  RT (min)      lcorbi_t3_neg_3_003_09    m/z 199.9116     0   1E+4   13.5   14  RT (min)      lcorbi_t3_neg_3_002_24    m/z 199.9362     0   1   2E+3   0.5   1  RT (min)      lcorbi_t3_neg_4_021_25    m/z 200.0103     0   1E+4   6   6.5  RT (min)      lcorbi_t3_neg_2_048_27    m/z 200.0591     0   1   2   3   4E+3   18   18.5  RT (min)      lcorbi_t3_neg_1_021_31    m/z 200.0611     0   1   2   3E+3   1.5   2  RT (min)      lcorbi_t3_neg_2_050_17    m/z 200.1292     0   1E+4   6.5   7  RT (min)      lcorbi_t3_neg_4_002_45    m/z 200.3875     0   1E+4   13.5   14  RT (min)      lcorbi_t3_neg_4_029_31    m/z 200.8588     0   2   4   6E+4   15.5   16  RT (min)      lcorbi_t3_neg_1_045_44    m/z 200.9196     0   1E+3   18   18.5  RT (min)      lcorbi_t3_neg_1_001_49    m/z 200.9332     0   1   2E+3   0.5   1  RT (min)      lcorbi_t3_neg_4_016_05    m/z 201.0226     0   2   4   6E+4   6.5   7  RT (min)      lcorbi_t3_neg_2_048_27    m/z 201.0227     0   1   2   3   4E+3   18   18.5  RT (min)      lcorbi_t3_neg_1_053_32    m/z 201.0380     0   1   2   3   4E+3   1   1.5  RT (min)       lcorbi_t3_neg_4_038_15    m/z 201.0554     0   2   4E+3   18   18.5  RT (min)      lcorbi_t3_neg_4_046_37    m/z 201.0557     0   2   4   6   8E+3   7   7.5  RT (min)      lcorbi_t3_neg_4_037_29    m/z 201.0750     0   1E+4   5.5   6  RT (min)      lcorbi_t3_neg_4_023_18    m/z 201.0773     0   1E+4   7.5   8  RT (min)      lcorbi_t3_neg_4_011_24    m/z 201.0877     0   1   1   1.5  RT (min)      lcorbi_t3_neg_4_011_24    m/z 201.1132     0   2   4E+4   7.5   8  RT (min)      lcorbi_t3_neg_4_023_18    m/z 201.2711     0   1E+4   14   14.5  RT (min)      lcorbi_t3_neg_4_007_13    m/z 201.6461     0   1E+4   14   14.5  RT (min)      lcorbi_t3_neg_4_007_13    m/z 201.6574     0   1E+4   14   14.5  RT (min)      lcorbi_t3_neg_2_052_32    m/z 201.8020     0   1   2E+4   13   13.5  RT (min)      lcorbi_t3_neg_4_043_11    m/z 201.8020     0   1   2E+4   17   17.5   18  RT (min)       lcorbi_t3_neg_1_036_09    m/z 201.8020     0   1   2   3E+4   15.5   16  RT (min)       lcorbi_t3_neg_1_049_55    m/z 202.0191     0   1   2E+4   14   14.5  RT (min)      lcorbi_t3_neg_1_040_42    m/z 202.0383     0   2   4   6   8E+4   1.5   2  RT (min)      lcorbi_t3_neg_2_048_27    m/z 202.0384     0   1   2   3E+4   11.5   12  RT (min)      lcorbi_t3_neg_1_021_31    m/z 202.0384     0   2   4   6E+4   14   14.5  RT (min)      lcorbi_t3_neg_1_034_04    m/z 202.0510     0   2   4E+2   7   7.5  RT (min)      lcorbi_t3_neg_1_055_05    m/z 202.0576     0   1E+3   18   18.5  RT (min)      lcorbi_t3_neg_4_111_62    m/z 202.0788     0   1   2E+5   15.5   16  RT (min)      lcorbi_t3_neg_4_032_23    m/z 202.0788     0   2   4E+5   0   0.5  RT (min)      lcorbi_t3_neg_1_043_14    m/z 202.0873     0   1   2E+3   7.5   8  RT (min)      lcorbi_t3_neg_4_017_51    m/z 202.1165     0   1E+3   7.5   8  RT (min)      lcorbi_t3_neg_4_039_32    m/z 202.4729     0   1   13.5   14  RT (min)      lcorbi_t3_neg_2_022_24    m/z 202.8024     0   1E+4   1   1.5  RT (min)      lcorbi_t3_neg_2_027_13    m/z 202.9289     0   1E+4   18   18.5  RT (min)      lcorbi_t3_neg_1_044_23    m/z 202.9323     0   1   1   1.5  RT (min)      lcorbi_t3_neg_3_019_12    m/z 202.9648     0   1   2E+4   0   0.5  RT (min)      lcorbi_t3_neg_1_017_51    m/z 203.0346     0   2   4   6E+3   18   18.5  RT (min)      lcorbi_t3_neg_1_010_17    m/z 203.0346     0   2   4   6E+3   1.5   2  RT (min)      lcorbi_t3_neg_4_075_58    m/z 203.0346     0   1   2E+4   14   14.5  RT (min)      lcorbi_t3_neg_1_029_21    m/z 203.0346     0   1E+4   12   12.5  RT (min)      lcorbi_t3_neg_4_032_23    m/z 203.0468     0   1E+3   13.5   14  RT (min)      lcorbi_t3_neg_3_029_02    m/z 203.0520     0   1   2   3E+4   6   6.5  RT (min)      lcorbi_t3_neg_4_001_49    m/z 203.0600     0   2   4   6E+3   18   18.5  RT (min)      lcorbi_t3_neg_2_006_10    m/z 203.0673     0   1E+3   1   1.5  RT (min)      lcorbi_t3_neg_4_039_32    m/z 203.0867     0   1E+4   1   1.5  RT (min)      lcorbi_t3_neg_4_011_24    m/z 203.1034     0   2   4   6   8E+3   1   1.5  RT (min)      lcorbi_t3_neg_4_038_15    m/z 203.1456     0   1   2   3   4E+3   14   14.5  RT (min)      lcorbi_t3_neg_4_044_38    m/z 203.7312     0   1E+4   14   14.5  RT (min)      lcorbi_t3_neg_4_007_13    m/z 203.8443     0   1E+4   1   1.5  RT (min)      lcorbi_t3_neg_1_047_02    m/z 203.8547     0   1E+4   14   14.5  RT (min)      lcorbi_t3_neg_4_007_13    m/z 204.0052     0   1   2E+4   5   5.5  RT (min)      lcorbi_t3_neg_2_048_27    m/z 204.0175     0   1   2E+4   8   8.5  RT (min)      lcorbi_t3_neg_4_011_24    m/z 204.0175     0   1E+5   10   10.5  RT (min)      lcorbi_t3_neg_4_032_23    m/z 204.0229     0   1E+4   1   1.5  RT (min)      lcorbi_t3_neg_4_054_36    m/z 204.0236     0   1   2   3E+3   18   18.5  RT (min)      lcorbi_t3_neg_1_055_05    m/z 204.0560     0   1E+4   14.5   15  RT (min)      lcorbi_t3_neg_2_011_04    m/z 204.0561     0   1   2   3   4E+4   1   1.5  RT (min)       lcorbi_t3_neg_2_048_27    m/z 204.1126     0   1   14   14.5  RT (min)      lcorbi_t3_neg_1_036_09    m/z 204.1474     0   1E+4   10.5   11  RT (min)      lcorbi_t3_neg_4_042_14    m/z 204.7915     0   1E+4   1   1.5  RT (min)      lcorbi_t3_neg_4_067_57    m/z 205.0083     0   2   4E+3   18   18.5  RT (min)      lcorbi_t3_neg_1_013_35    m/z 205.0139     0   1E+5   10   10.5  RT (min)      lcorbi_t3_neg_2_043_33    m/z 205.0139     0   1   2E+4   7.5   8  RT (min)      lcorbi_t3_neg_2_025_52    m/z 205.0202     0   1E+4   14   14.5  RT (min)      lcorbi_t3_neg_4_011_24    m/z 205.0329     0   1   2   3E+3   18   18.5  RT (min)      lcorbi_t3_neg_4_021_25    m/z 205.0523     0   2   4   6E+4   14   14.5  RT (min)       lcorbi_t3_neg_1_036_09    m/z 205.0524     0   1   2   3E+4   17.5   18  RT (min)       lcorbi_t3_neg_1_032_13    m/z 205.0699     0   1E+4   6   6.5  RT (min)      lcorbi_t3_neg_2_048_27    m/z 205.0717     0   1E+4   0   0.5  RT (min)      lcorbi_t3_neg_1_009_50    m/z 205.0869     0   1   2E+4   6.5   7  RT (min)      lcorbi_t3_neg_2_048_27    m/z 205.1080     0   1   2E+4   0   0.5  RT (min)      lcorbi_t3_neg_1_017_51    m/z 205.8395     0   1   2   3E+4   15.5   16  RT (min)      lcorbi_t3_neg_1_042_18    m/z 205.8397     0   1E+4   17.5   18  RT (min)      lcorbi_t3_neg_1_033_53    m/z 205.8397     0   1E+5   4.5   5  RT (min)      lcorbi_t3_neg_3_032_17    m/z 205.8397     0   1   2E+5   0.5   1  RT (min)       lcorbi_t3_neg_2_016_11    m/z 205.9417     0   2   4   6E+3   8.5   9  RT (min)      lcorbi_t3_neg_2_048_27    m/z 205.9628     0   1   2   3E+3   1   1.5  RT (min)      lcorbi_t3_neg_4_017_51    m/z 206.0173     0   1E+4   14   14.5  RT (min)      lcorbi_t3_neg_1_029_21    m/z 206.0217     0   2   4   6E+3   18   18.5  RT (min)      lcorbi_t3_neg_4_009_50    m/z 206.0386     0   1   2   3E+4   2   2.5  RT (min)      lcorbi_t3_neg_4_034_01    m/z 206.1034     0   1   2E+4   1   1.5  RT (min)      lcorbi_t3_neg_2_048_27    m/z 206.1600     0   1E+4   13.5   14  RT (min)      lcorbi_t3_neg_2_052_32    m/z 206.2632     0   1E+4   13.5   14  RT (min)      lcorbi_t3_neg_4_029_31    m/z 206.6382     0   2   4E+4   13.5   14  RT (min)      lcorbi_t3_neg_4_029_31    m/z 206.6638     0   1E+4   13.5   14  RT (min)      lcorbi_t3_neg_2_052_32    m/z 206.7671     0   1E+4   14   14.5  RT (min)      lcorbi_t3_neg_1_039_38    m/z 206.7891     0   1   2E+4   1   1.5  RT (min)       lcorbi_t3_neg_4_067_57    m/z 206.9726     0   1   2   3   4E+4   4.5   5  RT (min)      lcorbi_t3_neg_2_027_13    m/z 206.9726     0   1E+5   3   3.5  RT (min)      lcorbi_t3_neg_1_044_23    m/z 206.9726     0   1   2E+4   7   7.5  RT (min)      lcorbi_t3_neg_2_052_32    m/z 207.0049     0   1   1   1.5  RT (min)      lcorbi_t3_neg_2_026_03    m/z 207.0180     0   2   4   6   8E+3   14   14.5  RT (min)      lcorbi_t3_neg_1_007_26    m/z 207.0298     0   1E+3   6.5   7  RT (min)      lcorbi_t3_neg_1_005_07    m/z 207.0492     0   1E+3   18   18.5  RT (min)      lcorbi_t3_neg_1_002_16    m/z 207.0509     0   2   4E+4   1   1.5  RT (min)      lcorbi_t3_neg_2_048_27    m/z 207.0873     0   1E+4   0   0.5  RT (min)      lcorbi_t3_neg_1_029_21    m/z 207.1138     0   1E+4   6   6.5  RT (min)      lcorbi_t3_neg_4_029_31    m/z 207.1237     0   1   2   3   4E+4   0   0.5  RT (min)      lcorbi_t3_neg_2_022_24    m/z 207.1677     0   1E+4   14   14.5  RT (min)      lcorbi_t3_neg_4_011_24    m/z 207.1706     0   1E+4   14   14.5  RT (min)      lcorbi_t3_neg_3_002_24    m/z 207.2709     0   1E+4   14   14.5  RT (min)      lcorbi_t3_neg_4_007_13    m/z 207.4115     0   1   2   3   4E+3   14.5   15  RT (min)      lcorbi_t3_neg_4_007_13    m/z 207.8366     0   1   2   3E+4   15.5   16  RT (min)      lcorbi_t3_neg_1_042_18    m/z 207.8367     0   1E+4   17.5   18  RT (min)      lcorbi_t3_neg_1_023_03    m/z 207.8368     0   1E+5   0.5   1  RT (min)       lcorbi_t3_neg_2_016_11    m/z 207.8839     0   1   1   1.5  RT (min)      lcorbi_t3_neg_1_045_44    m/z 207.9153     0   1   2   3   4E+3   14.5   15  RT (min)      lcorbi_t3_neg_4_034_01    m/z 207.9185     0   1   2E+3   0   0.5  RT (min)      lcorbi_t3_neg_1_045_44    m/z 207.9310     0   1E+5   4   4.5  RT (min)      lcorbi_t3_neg_1_046_11    m/z 207.9387     0   1   2   3E+2   8.5   9  RT (min)      lcorbi_t3_neg_1_044_23    m/z 207.9760     0   1   2   3E+3   18   18.5  RT (min)      lcorbi_t3_neg_1_044_23    m/z 208.0153     0   2   4E+3   6   6.5  RT (min)      lcorbi_t3_neg_2_048_27    m/z 208.0178     0   2   4   6E+3   1   1.5  RT (min)      lcorbi_t3_neg_2_035_46    m/z 208.0695     0   1E+4   7.5   8  RT (min)      lcorbi_t3_neg_2_048_27    m/z 208.0695     0   1   10.5   11  RT (min)      lcorbi_t3_neg_4_083_59    m/z 208.1035     0   1E+3   10   10.5  RT (min)      lcorbi_t3_neg_3_023_07    m/z 208.3799     0   1E+4   13.5   14  RT (min)      lcorbi_t3_neg_4_046_37    m/z 208.4029     0   2   4   6   8E+3   14.5   15  RT (min)      lcorbi_t3_neg_4_007_13    m/z 208.4090     0   1E+4   13.5   14  RT (min)      lcorbi_t3_neg_2_052_32    m/z 208.6307     0   2   4   6E+3   13.5   14  RT (min)      lcorbi_t3_neg_3_045_34    m/z 208.7865     0   1E+3   1   1.5  RT (min)      lcorbi_t3_neg_3_032_17    m/z 208.9842     0   1   2   3E+4   0   0.5  RT (min)      lcorbi_t3_neg_1_001_49    m/z 209.0852     0   1E+4   8   8.5  RT (min)      lcorbi_t3_neg_2_037_09    m/z 209.1029     0   2   4   6E+3   0   0.5  RT (min)      lcorbi_t3_neg_2_022_24    m/z 209.6639     0   2   4E+3   13.5   14  RT (min)      lcorbi_t3_neg_2_052_32    m/z 209.8336     0   1   2E+4   4.5   5  RT (min)      lcorbi_t3_neg_3_010_15    m/z 209.8337     0   1E+4   13   13.5  RT (min)      lcorbi_t3_neg_4_007_13    m/z 209.9171     0   2   4   6   8E+3   13.5   14  RT (min)      lcorbi_t3_neg_4_039_32    m/z 209.9763     0   1E+5   1   1.5  RT (min)      lcorbi_t3_neg_3_002_24    m/z 210.0692     0   2   4   6E+3   1   1.5  RT (min)      lcorbi_t3_neg_4_046_37    m/z 210.0771     0   2   4E+3   8   8.5  RT (min)      lcorbi_t3_neg_3_019_12    m/z 210.0907     0   1   1   1.5  RT (min)      lcorbi_t3_neg_2_048_27    m/z 210.1045     0   1E+3   10   10.5  RT (min)      lcorbi_t3_neg_4_088_59    m/z 210.1134     0   2   4   6E+3   14   14.5  RT (min)      lcorbi_t3_neg_4_034_01    m/z 210.1535     0   2   4E+3   14.5   15  RT (min)      lcorbi_t3_neg_4_007_13    m/z 210.1579     0   2   4   6E+3   9   9.5  RT (min)      lcorbi_t3_neg_2_048_27    m/z 210.4210     0   2   4   6   8E+3   13.5   14  RT (min)      lcorbi_t3_neg_1_053_32    m/z 210.7403     0   2   4   6   8E+3   14   14.5  RT (min)      lcorbi_t3_neg_4_034_01    m/z 210.8839     0   1E+4   18.5   19  RT (min)      lcorbi_t3_neg_4_103_61    m/z 210.8839     0   1E+4   0.5   1  RT (min)      lcorbi_t3_neg_1_003_41    m/z 210.9154     0   1   2   3E+3   14.5   15  RT (min)      lcorbi_t3_neg_1_053_32    m/z 210.9338     0   1E+4   14   14.5  RT (min)      lcorbi_t3_neg_3_002_24    m/z 210.9470     0   1   2E+2   7   7.5  RT (min)      lcorbi_t3_neg_1_006_40    m/z 210.9495     0   2   4   6   8E+3   18.5   19  RT (min)      lcorbi_t3_neg_4_109_62    m/z 210.9624     0   1   1   1.5  RT (min)      lcorbi_t3_neg_4_052_26    m/z 210.9943     0   1E+4   4.5   5  RT (min)      lcorbi_t3_neg_4_034_01    m/z 211.0167     0   1E+4   6.5   7  RT (min)      lcorbi_t3_neg_4_040_19    m/z 211.0247     0   1   6.5   7  RT (min)      lcorbi_t3_neg_4_108_62    m/z 211.0360     0   1   2   3E+3   8.5   9  RT (min)      lcorbi_t3_neg_4_044_38    m/z 211.3727     0   1E+4   7.5   8  RT (min)      lcorbi_t3_neg_2_048_27    m/z 211.5484     0   1E+4   13.5   14  RT (min)      lcorbi_t3_neg_1_055_05    m/z 211.6787     0   2   4E+3   13.5   14  RT (min)      lcorbi_t3_neg_1_052_24    m/z 211.8310     0   2   4   6   8E+3   1   1.5  RT (min)      lcorbi_t3_neg_1_046_11    m/z 211.8940     0   2   4   6   8E+3   1.5   2  RT (min)      lcorbi_t3_neg_4_007_13    m/z 212.0200     0   1E+4   6.5   7  RT (min)      lcorbi_t3_neg_2_022_24    m/z 212.0526     0   1E+4   14   14.5  RT (min)      lcorbi_t3_neg_1_055_05    m/z 212.0927     0   1   2   3E+4   5.5   6  RT (min)      lcorbi_t3_neg_2_048_27    m/z 212.1023     0   1   2E+4   6   6.5  RT (min)      lcorbi_t3_neg_3_019_12    m/z 212.1372     0   2   4E+3   9.5   10  RT (min)      lcorbi_t3_neg_4_013_09    m/z 212.8817     0   1   2E+3   0.5   1  RT (min)      lcorbi_t3_neg_2_052_32    m/z 212.8818     0   2   4   6E+3   18.5   19  RT (min)      lcorbi_t3_neg_4_109_62    m/z 212.8913     0   1   14   14.5  RT (min)      lcorbi_t3_neg_1_046_11    m/z 212.9107     0   1   2   3   4E+4   12   12.5  RT (min)      lcorbi_t3_neg_1_051_15    m/z 213.0055     0   1   2   3E+4   4.5   5  RT (min)      lcorbi_t3_neg_2_048_27    m/z 213.0152     0   1E+4   13.5   14  RT (min)       lcorbi_t3_neg_4_024_07    m/z 213.0152     0   2   4E+4   11   11.5  RT (min)      lcorbi_t3_neg_2_054_15    m/z 213.0152     0   2   4E+4   7.5   8  RT (min)       lcorbi_t3_neg_4_021_25    m/z 213.0152     0   1   2E+4   18   18.5  RT (min)      lcorbi_t3_neg_2_022_24    m/z 213.0556     0   2   4E+3   8.5   9  RT (min)      lcorbi_t3_neg_4_007_13    m/z 213.1160     0   1   2E+4   4.5   5  RT (min)      lcorbi_t3_neg_3_019_12    m/z 213.1495     0   1E+4   10   10.5  RT (min)      lcorbi_t3_neg_2_048_27    m/z 213.3952     0   1E+4   14   14.5  RT (min)      lcorbi_t3_neg_4_007_13    m/z 213.9109     0   2   4   6E+3   17.5   18  RT (min)      lcorbi_t3_neg_3_018_13    m/z 213.9110     0   1E+4   14.5   15  RT (min)      lcorbi_t3_neg_2_038_22    m/z 213.9637     0   2   4   6E+3   5   5.5  RT (min)      lcorbi_t3_neg_2_016_11    m/z 213.9979     0   1E+4   4.5   5  RT (min)      lcorbi_t3_neg_2_048_27    m/z 214.0178     0   2   4E+3   1   1.5  RT (min)      lcorbi_t3_neg_4_032_23    m/z 214.0260     0   1   7   7.5  RT (min)      lcorbi_t3_neg_4_033_53    m/z 214.0509     0   1E+4   9.5   10  RT (min)      lcorbi_t3_neg_4_029_31    m/z 214.0745     0   1   2E+4   1   1.5  RT (min)      lcorbi_t3_neg_2_022_24    m/z 214.0768     0   2   4   6E+4   2.5   3  RT (min)       lcorbi_t3_neg_4_079_58    m/z 214.1549     0   1E+4   14   14.5   15  RT (min)      lcorbi_t3_neg_4_007_13    m/z 214.1658     0   2   4   6E+3   14.5   15  RT (min)      lcorbi_t3_neg_4_034_01    m/z 214.4028     0   2   4   6E+3   14.5   15  RT (min)      lcorbi_t3_neg_4_034_01    m/z 214.4058     0   2   4E+3   14.5   15  RT (min)      lcorbi_t3_neg_4_034_01    m/z 214.4189     0   1E+4   14.5   15  RT (min)      lcorbi_t3_neg_4_034_01    m/z 214.6698     0   2   4E+3   14.5   15  RT (min)      lcorbi_t3_neg_4_038_15    m/z 214.8970     0   1E+4   1.5   2  RT (min)      lcorbi_t3_neg_1_032_13    m/z 214.9304     0   1   2E+4   7.5   8  RT (min)      lcorbi_t3_neg_4_023_18    m/z 214.9349     0   1E+4   7   7.5  RT (min)      lcorbi_t3_neg_4_101_61    m/z 215.0018     0   1   2   3   4E+4   5.5   6  RT (min)      lcorbi_t3_neg_2_048_27    m/z 215.0157     0   1   2E+4   1   1.5  RT (min)      lcorbi_t3_neg_2_048_27    m/z 215.0194     0   1   2E+3   6.5   7  RT (min)      lcorbi_t3_neg_4_034_01    m/z 215.0382     0   1   2   3   4E+3   7   7.5  RT (min)      lcorbi_t3_neg_4_034_01    m/z 215.0418     0   1   2   3   4E+3   14   14.5  RT (min)      lcorbi_t3_neg_2_023_39    m/z 215.0427     0   1   2   3   4E+4   1   1.5  RT (min)      lcorbi_t3_neg_4_046_37    m/z 215.0511     0   2   4   6   8E+3   1   1.5  RT (min)      lcorbi_t3_neg_4_005_27    m/z 215.1287     0   2   4   6   8E+4   8   8.5  RT (min)      lcorbi_t3_neg_4_023_18    m/z 215.1651     0   2   4   6E+3   10.5   11  RT (min)      lcorbi_t3_neg_2_028_38    m/z 215.3879     0   2   4   6E+3   13.5   14  RT (min)      lcorbi_t3_neg_4_034_01    m/z 215.9997     0   1   1   1.5  RT (min)      lcorbi_t3_neg_3_032_17    m/z 216.0382     0   1   2E+3   18   18.5  RT (min)      lcorbi_t3_neg_4_039_32    m/z 216.0383     0   1   2E+4   13.5   14  RT (min)      lcorbi_t3_neg_4_021_25    m/z 216.0539     0   1E+4   14   14.5  RT (min)      lcorbi_t3_neg_4_030_04    m/z 216.0539     0   2   4   6E+3   12   12.5  RT (min)      lcorbi_t3_neg_1_023_03    m/z 216.0560     0   1E+4   1   1.5  RT (min)      lcorbi_t3_neg_4_077_58    m/z 216.0848     0   1   2   3E+4   1   1.5  RT (min)      lcorbi_t3_neg_4_039_32    m/z 216.1321     0   2   4   6   8E+3   8   8.5  RT (min)      lcorbi_t3_neg_4_023_18    m/z 216.4187     0   1E+4   13.5   14  RT (min)      lcorbi_t3_neg_1_053_32    m/z 216.8940     0   1   2   3E+2   1.5   2  RT (min)      lcorbi_t3_neg_1_003_41    m/z 216.9252     0   1   2   3   4E+4   1   1.5  RT (min)      lcorbi_t3_neg_1_004_22    m/z 216.9285     0   2   4   6E+2   7.5   8  RT (min)      lcorbi_t3_neg_1_001_49    m/z 216.9340     0   1   2   3   4E+3   17.5   18  RT (min)      lcorbi_t3_neg_3_013_28    m/z 216.9340     0   1E+4   15.5   16  RT (min)      lcorbi_t3_neg_1_055_05    m/z 216.9809     0   1E+3   18   18.5  RT (min)      lcorbi_t3_neg_4_039_32    m/z 217.0175     0   2   4   6E+4   6   6.5  RT (min)      lcorbi_t3_neg_2_048_27    m/z 217.0502     0   1   2   3   4E+3   18   18.5  RT (min)      lcorbi_t3_neg_4_054_36    m/z 217.0502     0   1   2E+4   13.5   14  RT (min)      lcorbi_t3_neg_4_030_04    m/z 217.0502     0   1E+4   12   12.5  RT (min)      lcorbi_t3_neg_4_044_38    m/z 217.0716     0   1   2   3E+3   0   0.5  RT (min)      lcorbi_t3_neg_1_029_21    m/z 217.0828     0   1   1   1.5  RT (min)      lcorbi_t3_neg_4_075_58    m/z 217.0869     0   1E+4   9.5   10  RT (min)      lcorbi_t3_neg_2_048_27    m/z 217.1080     0   2   4E+3   0   0.5  RT (min)      lcorbi_t3_neg_1_017_51    m/z 217.1444     0   1   2E+4   8   8.5  RT (min)      lcorbi_t3_neg_2_048_27    m/z 217.4264     0   1E+4   14   14.5  RT (min)      lcorbi_t3_neg_2_052_32    m/z 217.4295     0   1E+4   14   14.5  RT (min)      lcorbi_t3_neg_2_052_32    m/z 217.8654     0   1E+5   4.5   5  RT (min)      lcorbi_t3_neg_1_046_11    m/z 217.8833     0   1   2   3E+2   1.5   2  RT (min)      lcorbi_t3_neg_1_045_44    m/z 217.9280     0   1E+3   8.5   9  RT (min)      lcorbi_t3_neg_1_044_23    m/z 217.9305     0   2   4   6   8E+3   14   14.5  RT (min)      lcorbi_t3_neg_1_052_24    m/z 217.9311     0   1   2   2.5  RT (min)      lcorbi_t3_neg_1_045_44    m/z 218.0136     0   2   4   6   8E+2   18   18.5  RT (min)      lcorbi_t3_neg_1_053_32    m/z 218.0136     0   1E+3   13   13.5  RT (min)       lcorbi_t3_neg_4_034_01    m/z 218.0193     0   2   4   6E+3   1.5   2  RT (min)      lcorbi_t3_neg_2_007_14    m/z 218.0311     0   1   2   3E+5   14.5   15  RT (min)       lcorbi_t3_neg_2_037_09    m/z 218.0505     0   1   2   3   4E+3   1   1.5  RT (min)      lcorbi_t3_neg_2_052_32    m/z 218.0695     0   1   2   3E+3   18   18.5  RT (min)      lcorbi_t3_neg_4_045_35    m/z 218.0822     0   1   6.5   7  RT (min)      lcorbi_t3_neg_4_102_61    m/z 218.1185     0   1E+4   5   5.5  RT (min)      lcorbi_t3_neg_2_048_27    m/z 218.6852     0   1E+4   14   14.5  RT (min)      lcorbi_t3_neg_3_002_24    m/z 218.9151     0   1E+4   13.5   14  RT (min)      lcorbi_t3_neg_1_053_32    m/z 218.9652     0   2   4E+3   7   7.5  RT (min)      lcorbi_t3_neg_4_080_58    m/z 218.9822     0   1   2   3E+3   1.5   2  RT (min)      lcorbi_t3_neg_4_075_58    m/z 218.9861     0   1   2   3E+3   7.5   8  RT (min)      lcorbi_t3_neg_4_100_61    m/z 218.9941     0   1E+2   1   1.5  RT (min)      lcorbi_t3_neg_3_019_12    m/z 219.0089     0   1E+3   4.5   5  RT (min)      lcorbi_t3_neg_2_048_27    m/z 219.0100     0   1E+3   18   18.5  RT (min)      lcorbi_t3_neg_1_053_32    m/z 219.0100     0   2   4E+4   8   10   12  RT (min)       lcorbi_t3_neg_4_084_59    m/z 219.0451     0   1E+3   1   1.5  RT (min)      lcorbi_t3_neg_4_074_58    m/z 219.0486     0   2   4   6   8E+3   14   14.5  RT (min)      lcorbi_t3_neg_3_023_07    m/z 219.0659     0   2   4E+3   18   18.5  RT (min)      lcorbi_t3_neg_2_048_27    m/z 219.0659     0   1E+4   14.5   15  RT (min)      lcorbi_t3_neg_2_048_27    m/z 219.0720     0   1   2E+4   7   7.5  RT (min)      lcorbi_t3_neg_4_040_19    m/z 219.0873     0   1E+4   0   0.5  RT (min)      lcorbi_t3_neg_1_029_21    m/z 219.0982     0   1E+3   9.5   10  RT (min)      lcorbi_t3_neg_4_014_20    m/z 219.1236     0   1E+4   0   0.5  RT (min)      lcorbi_t3_neg_1_017_51    m/z 219.1389     0   1   11   11.5  RT (min)      lcorbi_t3_neg_4_014_20    m/z 219.6483     0   1E+4   14   14.5  RT (min)      lcorbi_t3_neg_2_050_17    m/z 219.6697     0   1E+4   13.5   14  RT (min)      lcorbi_t3_neg_4_011_24    m/z 219.8453     0   2   4E+4   4.5   5  RT (min)      lcorbi_t3_neg_1_047_02    m/z 219.8453     0   1E+5   3   3.5  RT (min)      lcorbi_t3_neg_3_052_32    m/z 219.8624     0   1E+4   1   1.5  RT (min)      lcorbi_t3_neg_1_045_44    m/z 219.9282     0   1   2   2.5  RT (min)      lcorbi_t3_neg_1_045_44    m/z 219.9466     0   1   2E+4   7   7.5  RT (min)      lcorbi_t3_neg_4_007_13    m/z 220.0266     0   1   2   3E+4   13   13.5  RT (min)       lcorbi_t3_neg_2_031_25    m/z 220.0295     0   1E+4   14   14.5  RT (min)      lcorbi_t3_neg_4_043_11    m/z 220.0651     0   1   2   3E+3   18   18.5  RT (min)      lcorbi_t3_neg_4_046_37    m/z 220.0827     0   2   4   6E+4   1   1.5  RT (min)       lcorbi_t3_neg_2_048_27    m/z 220.1521     0   1   2   3E+3   14   14.5   15  RT (min)      lcorbi_t3_neg_4_034_01    m/z 220.6560     0   1   2   3   4E+3   14.5   15  RT (min)      lcorbi_t3_neg_4_007_13    m/z 220.7664     0   1   1   1.5  RT (min)      lcorbi_t3_neg_4_005_27    m/z 220.9861     0   1   2   3E+3   18   18.5  RT (min)      lcorbi_t3_neg_1_016_39    m/z 221.0231     0   1   2E+4   13.5   14  RT (min)      lcorbi_t3_neg_3_043_25    m/z 221.0280     0   1   2E+3   18   18.5  RT (min)      lcorbi_t3_neg_1_008_08    m/z 221.0302     0   1   2E+4   1   1.5  RT (min)      lcorbi_t3_neg_4_039_32    m/z 221.0315     0   1E+3   14   14.5  RT (min)      lcorbi_t3_neg_4_019_08    m/z 221.0615     0   2   4   6E+3   18   18.5  RT (min)      lcorbi_t3_neg_2_043_33    m/z 221.0665     0   1E+3   1   1.5  RT (min)       lcorbi_t3_neg_1_055_05    m/z 221.0716     0   1E+4   7   7.5  RT (min)      lcorbi_t3_neg_4_040_19    m/z 221.0816     0   1   2   3E+3   14   14.5  RT (min)      lcorbi_t3_neg_3_035_03    m/z 221.1029     0   1E+4   0   0.5  RT (min)      lcorbi_t3_neg_1_023_03    m/z 221.1393     0   1E+4   0   0.5  RT (min)      lcorbi_t3_neg_2_022_24    m/z 221.1481     0   1   2   3   4E+5   9.5   10  RT (min)      lcorbi_t3_neg_4_019_08    m/z 221.1546     0   1   2E+4   11.5   12  RT (min)      lcorbi_t3_neg_2_039_31    m/z 221.3876     0   1E+4   13.5   14  RT (min)      lcorbi_t3_neg_4_034_01    m/z 221.3909     0   2   4   6   8E+3   13.5   14  RT (min)      lcorbi_t3_neg_4_034_01    m/z 221.6386     0   2   4   6   8E+3   13.5   14  RT (min)      lcorbi_t3_neg_4_034_01    m/z 221.8423     0   1   2   3E+4   4.5   5  RT (min)      lcorbi_t3_neg_1_045_44    m/z 221.8516     0   1E+4   14.5   15  RT (min)      lcorbi_t3_neg_1_040_42    m/z 221.9026     0   1   2   3E+3   1   1.5  RT (min)      lcorbi_t3_neg_4_045_35    m/z 221.9027     0   2   4   6   8E+3   14.5   15  RT (min)      lcorbi_t3_neg_3_049_55    m/z 221.9028     0   2   4   6E+3   17.5   18  RT (min)      lcorbi_t3_neg_3_020_31    m/z 221.9753     0   1   1   1.5  RT (min)      lcorbi_t3_neg_3_022_27    m/z 221.9994     0   1   1   1.5  RT (min)      lcorbi_t3_neg_2_048_27    m/z 222.0017     0   1   2E+4   7   7.5  RT (min)      lcorbi_t3_neg_4_039_32    m/z 222.0251     0   2   4E+3   13.5   14  RT (min)      lcorbi_t3_neg_4_027_12    m/z 222.0252     0   2   4E+3   18   18.5  RT (min)      lcorbi_t3_neg_1_013_35    m/z 222.0397     0   1E+4   1.5   2  RT (min)      lcorbi_t3_neg_4_011_24    m/z 222.1134     0   2   4   6E+3   8.5   9  RT (min)      lcorbi_t3_neg_2_048_27    m/z 222.1134     0   1   2   3E+3   10.5   11  RT (min)      lcorbi_t3_neg_4_029_31    m/z 222.8987     0   1   2   3E+3   8   8.5  RT (min)      lcorbi_t3_neg_4_080_58    m/z 222.9479     0   1E+4   2   2.5  RT (min)      lcorbi_t3_neg_2_022_24    m/z 222.9802     0   2   4   6   8E+3   6   6.5  RT (min)      lcorbi_t3_neg_2_048_27    m/z 222.9846     0   1   2   3E+3   0.5   1  RT (min)      lcorbi_t3_neg_1_045_44    m/z 222.9846     0   2   4   6   8E+3   18   18.5  RT (min)      lcorbi_t3_neg_1_044_23    m/z 223.0068     0   1E+4   6.5   7  RT (min)      lcorbi_t3_neg_4_042_14    m/z 223.0216     0   1   2   3E+3   18   18.5  RT (min)      lcorbi_t3_neg_4_086_59    m/z 223.0282     0   1E+4   9.5   10  RT (min)      lcorbi_t3_neg_4_007_13    m/z 223.0282     0   1E+4   7.5   8  RT (min)      lcorbi_t3_neg_2_048_27    m/z 223.0822     0   2   4   6E+3   0   0.5  RT (min)      lcorbi_t3_neg_1_035_25    m/z 223.1186     0   1E+4   0   0.5  RT (min)      lcorbi_t3_neg_2_022_24    m/z 223.1702     0   2   4   6E+3   9   9.5  RT (min)      lcorbi_t3_neg_2_048_27    m/z 223.8393     0   1   2   3   4E+3   18.5   19  RT (min)      lcorbi_t3_neg_2_048_27    m/z 223.8393     0   1E+4   1   1.5  RT (min)      lcorbi_t3_neg_3_019_12    m/z 223.8486     0   1E+4   14.5   15  RT (min)      lcorbi_t3_neg_1_042_18    m/z 223.8999     0   1E+3   1   1.5  RT (min)      lcorbi_t3_neg_4_045_35    m/z 223.9538     0   1E+4   8.5   9  RT (min)      lcorbi_t3_neg_4_007_13    m/z 224.0278     0   1   2   3E+4   11.5   12  RT (min)      lcorbi_t3_neg_4_007_13    m/z 224.0539     0   1E+4   3   3.5  RT (min)      lcorbi_t3_neg_4_032_23    m/z 224.0616     0   1   2   3E+4   14   14.5  RT (min)      lcorbi_t3_neg_4_008_22    m/z 224.0618     0   1E+4   18   18.5  RT (min)      lcorbi_t3_neg_2_048_27    m/z 224.0927     0   2   4   6E+3   10   10.5  RT (min)      lcorbi_t3_neg_4_047_30    m/z 224.1008     0   1E+3   8.5   9  RT (min)      lcorbi_t3_neg_4_007_13    m/z 224.1372     0   1E+4   10   10.5  RT (min)      lcorbi_t3_neg_2_048_27    m/z 224.5824     0   1   2E+4   9   9.5  RT (min)      lcorbi_t3_neg_2_048_27    m/z 224.5860     0   1E+4   9   9.5  RT (min)      lcorbi_t3_neg_2_048_27    m/z 224.7624     0   1   1   1.5  RT (min)      lcorbi_t3_neg_3_022_27    m/z 224.8034     0   1   2E+3   0.5   1  RT (min)      lcorbi_t3_neg_2_007_14    m/z 224.8332     0   1E+4   9   9.5  RT (min)      lcorbi_t3_neg_2_048_27    m/z 224.8495     0   2   4E+2   1   1.5  RT (min)      lcorbi_t3_neg_4_042_14    m/z 224.9950     0   1   2   3E+4   1   1.5  RT (min)      lcorbi_t3_neg_3_002_24    m/z 225.0075     0   2   4   6E+3   12   12.5  RT (min)      lcorbi_t3_neg_2_022_24    m/z 225.0075     0   1E+4   6.5   7  RT (min)      lcorbi_t3_neg_3_019_12    m/z 225.0250     0   1   2E+4   11.5   12  RT (min)      lcorbi_t3_neg_4_007_13    m/z 225.0318     0   1E+4   1.5   2  RT (min)      lcorbi_t3_neg_2_052_32    m/z 225.0597     0   2   4   6   8E+2   6   6.5  RT (min)      lcorbi_t3_neg_4_023_18    m/z 225.0626     0   1   2   3E+4   2.5   3  RT (min)      lcorbi_t3_neg_2_048_27    m/z 225.0643     0   1   4.5   5  RT (min)      lcorbi_t3_neg_4_023_18    m/z 225.0855     0   2   4   6   8E+3   1   1.5  RT (min)      lcorbi_t3_neg_4_047_30    m/z 225.1096     0   1   1   1.5  RT (min)      lcorbi_t3_neg_3_022_27    m/z 225.1495     0   1E+4   8.5   9  RT (min)      lcorbi_t3_neg_4_002_45    m/z 225.1607     0   1E+4   4.5   5  RT (min)      lcorbi_t3_neg_4_054_36    m/z 225.1858     0   2   4E+3   9.5   10  RT (min)      lcorbi_t3_neg_4_007_13    m/z 225.2222     0   1   2   3E+3   12   12.5  RT (min)      lcorbi_t3_neg_4_005_27    m/z 225.5712     0   2   4   6E+3   6   6.5  RT (min)      lcorbi_t3_neg_4_007_13    m/z 225.6677     0   2   4   6E+3   13.5   14  RT (min)      lcorbi_t3_neg_4_039_32    m/z 225.8971     0   1E+5   4   4.5  RT (min)      lcorbi_t3_neg_1_044_23    m/z 226.0178     0   1   2E+4   5.5   6  RT (min)      lcorbi_t3_neg_2_048_27    m/z 226.0648     0   1   2   3   4E+4   1   1.5  RT (min)      lcorbi_t3_neg_2_048_27    m/z 226.0833     0   1   2E+4   1   1.5  RT (min)      lcorbi_t3_neg_2_035_46    m/z 226.9202     0   2   4   6   8E+3   7   7.5  RT (min)      lcorbi_t3_neg_1_021_31    m/z 226.9629     0   1E+3   18   18.5  RT (min)      lcorbi_t3_neg_4_104_61    m/z 226.9629     0   1E+3   15.5   16  RT (min)      lcorbi_t3_neg_4_080_58    m/z 226.9659     0   1E+4   10   10.5  RT (min)      lcorbi_t3_neg_4_075_58    m/z 226.9783     0   1E+3   1   1.5  RT (min)      lcorbi_t3_neg_4_045_35    m/z 226.9784     0   1   2   3E+4   9.5   10  RT (min)      lcorbi_t3_neg_4_102_61    m/z 226.9784     0   2   4   6E+4   15   15.5  RT (min)      lcorbi_t3_neg_1_014_28    m/z 226.9784     0   2   4   6   8E+4   17.5   18  RT (min)      lcorbi_t3_neg_4_069_57    m/z 227.0252     0   1E+3   9   9.5  RT (min)      lcorbi_t3_neg_4_101_61    m/z 227.0382     0   2   4   6   8E+2   7   7.5  RT (min)      lcorbi_t3_neg_4_103_61    m/z 227.0923     0   1   2   3   4E+3   6.5   7  RT (min)      lcorbi_t3_neg_4_017_51    m/z 227.1036     0   1   2   3E+2   4.5   5  RT (min)      lcorbi_t3_neg_2_022_24    m/z 227.2015     0   1E+3   12.5   13  RT (min)      lcorbi_t3_neg_1_055_05    m/z 227.8942     0   2   4E+4   4   4.5  RT (min)      lcorbi_t3_neg_1_044_23    m/z 227.9666     0   1E+4   18   18.5  RT (min)      lcorbi_t3_neg_1_001_49    m/z 227.9693     0   1   2   3   4E+3   18   18.5  RT (min)      lcorbi_t3_neg_1_006_40    m/z 228.0863     0   1E+4   9   9.5  RT (min)      lcorbi_t3_neg_2_048_27    m/z 228.9027     0   1E+4   17.5   18  RT (min)      lcorbi_t3_neg_2_043_33    m/z 228.9503     0   1E+4   5.5   6  RT (min)      lcorbi_t3_neg_4_017_51    m/z 228.9569     0   2   4   6   8E+3   7.5   8  RT (min)      lcorbi_t3_neg_3_034_21    m/z 228.9600     0   2   4E+3   0.5   1  RT (min)      lcorbi_t3_neg_1_029_21    m/z 228.9865     0   1   1   1.5  RT (min)      lcorbi_t3_neg_1_002_16    m/z 229.0030     0   1E+3   14   14.5  RT (min)      lcorbi_t3_neg_4_069_57    m/z 229.0118     0   2   4   6   8E+2   7   7.5  RT (min)      lcorbi_t3_neg_1_051_15    m/z 229.0828     0   2   4   6   8E+2   1   1.5  RT (min)      lcorbi_t3_neg_4_011_24    m/z 229.0933     0   1E+4   1   1.5  RT (min)      lcorbi_t3_neg_1_052_24    m/z 229.1556     0   1E+4   4.5   5  RT (min)      lcorbi_t3_neg_4_034_01    m/z 229.1637     0   1E+4   7.5   8  RT (min)      lcorbi_t3_neg_2_048_27    m/z 229.1807     0   2   4   6   8E+3   11   11.5  RT (min)      lcorbi_t3_neg_2_035_46    m/z 229.8741     0   1   2   3   4E+3   1   1.5  RT (min)       lcorbi_t3_neg_3_019_12    m/z 229.8818     0   1E+4   13.5   14  RT (min)      lcorbi_t3_neg_4_053_43    m/z 229.9018     0   1   2E+4   2.5   3  RT (min)      lcorbi_t3_neg_4_007_13    m/z 229.9371     0   2   4   6E+3   17.5   18  RT (min)      lcorbi_t3_neg_1_040_42    m/z 229.9973     0   1E+3   7   7.5  RT (min)      lcorbi_t3_neg_4_023_18    m/z 230.0069     0   1E+5   1   1.5  RT (min)       lcorbi_t3_neg_3_005_23    m/z 230.0127     0   1E+5   2.5   3  RT (min)      lcorbi_t3_neg_2_048_27    m/z 230.0572     0   1   2E+4   7.5   8  RT (min)      lcorbi_t3_neg_2_048_27    m/z 230.1477     0   2   4E+3   8.5   9  RT (min)      lcorbi_t3_neg_2_048_27    m/z 230.8998     0   2   4   6E+3   17.5   18  RT (min)      lcorbi_t3_neg_2_028_38    m/z 230.9418     0   2   4   6   8E+3   7.5   8  RT (min)      lcorbi_t3_neg_4_008_22    m/z 230.9483     0   1E+4   5.5   6  RT (min)      lcorbi_t3_neg_4_017_51    m/z 230.9544     0   1   2E+4   7.5   8  RT (min)      lcorbi_t3_neg_4_023_18    m/z 230.9557     0   2   4   6   8E+3   18.5   19  RT (min)      lcorbi_t3_neg_4_081_58    m/z 230.9557     0   1   2   3   4E+3   0.5   1  RT (min)      lcorbi_t3_neg_1_011_48    m/z 230.9869     0   2   4   6   8E+2   1   1.5  RT (min)      lcorbi_t3_neg_4_021_25    m/z 231.0160     0   2   4   6   8E+3   18   18.5  RT (min)      lcorbi_t3_neg_4_045_35    m/z 231.0331     0   2   4E+3   7   7.5  RT (min)      lcorbi_t3_neg_4_034_01    m/z 231.0496     0   1E+4   7.5   8  RT (min)      lcorbi_t3_neg_2_048_27    m/z 231.0502     0   2   4   6   8E+3   14   14.5  RT (min)      lcorbi_t3_neg_3_050_40    m/z 231.0524     0   1   2   3E+3   18   18.5  RT (min)      lcorbi_t3_neg_4_046_37    m/z 231.0872     0   1   2   3E+3   0   0.5  RT (min)      lcorbi_t3_neg_1_023_03    m/z 231.1137     0   1   2   3   4E+3   6   6.5  RT (min)      lcorbi_t3_neg_2_048_27    m/z 231.1753     0   2   4   6E+3   9.5   10  RT (min)      lcorbi_t3_neg_2_048_27    m/z 231.1792     0   2   4   6E+3   7.5   8  RT (min)      lcorbi_t3_neg_3_019_12    m/z 231.8988     0   1   2E+3   4.5   5  RT (min)      lcorbi_t3_neg_1_050_45    m/z 231.9316     0   2   4   6E+5   0   0.5  RT (min)      lcorbi_t3_neg_4_056_56    m/z 231.9316     0   2   4   6E+4   18   18.5  RT (min)      lcorbi_t3_neg_4_110_62    m/z 231.9454     0   1E+2   1.5   2  RT (min)      lcorbi_t3_neg_1_052_24    m/z 231.9674     0   1E+4   1.5   2  RT (min)      lcorbi_t3_neg_4_007_13    m/z 232.0124     0   1   2E+4   14   14.5  RT (min)      lcorbi_t3_neg_4_042_14    m/z 232.0350     0   1E+4   1   1.5  RT (min)      lcorbi_t3_neg_4_044_38    m/z 232.0466     0   1E+4   13.5   14  RT (min)      lcorbi_t3_neg_3_053_14    m/z 232.0589     0   1E+4   3   3.5  RT (min)      lcorbi_t3_neg_2_048_27    m/z 232.1424     0   1E+4   10.5   11  RT (min)      lcorbi_t3_neg_2_048_27    m/z 232.9147     0   1E+4   18   18.5  RT (min)      lcorbi_t3_neg_4_066_57    m/z 232.9316     0   1E+4   13.5   14  RT (min)      lcorbi_t3_neg_4_005_27    m/z 232.9316     0   1E+4   17.5   18  RT (min)      lcorbi_t3_neg_3_015_11    m/z 232.9317     0   1E+4   0   0.5  RT (min)      lcorbi_t3_neg_1_044_23    m/z 232.9523     0   1E+4   7.5   8  RT (min)      lcorbi_t3_neg_4_023_18    m/z 233.0088     0   1E+4   14   14.5  RT (min)      lcorbi_t3_neg_3_041_54    m/z 233.0099     0   1E+4   2   2.5  RT (min)      lcorbi_t3_neg_4_077_58    m/z 233.0429     0   1   2   3E+3   14   14.5  RT (min)      lcorbi_t3_neg_4_012_17    m/z 233.0644     0   1   18   18.5  RT (min)      lcorbi_t3_neg_4_019_08    m/z 233.0777     0   2   4   6E+3   1   1.5  RT (min)      lcorbi_t3_neg_3_020_31    m/z 233.0815     0   1E+3   18   18.5  RT (min)      lcorbi_t3_neg_2_012_36    m/z 233.0930     0   1E+4   10   10.5  RT (min)      lcorbi_t3_neg_3_023_07    m/z 233.0961     0   1E+4   5.5   6  RT (min)      lcorbi_t3_neg_4_011_24    m/z 233.1029     0   2   4   6   8E+3   0   0.5  RT (min)      lcorbi_t3_neg_1_023_03    m/z 233.1757     0   1E+4   0   0.5  RT (min)      lcorbi_t3_neg_2_022_24    m/z 233.9305     0   2   4   6E+3   0.5   1  RT (min)      lcorbi_t3_neg_1_044_23    m/z 233.9466     0   1E+3   1.5   2  RT (min)      lcorbi_t3_neg_1_045_44    m/z 233.9551     0   2   4   6   8E+3   1   1.5  RT (min)      lcorbi_t3_neg_1_052_24    m/z 234.0258     0   1E+4   14   14.5  RT (min)      lcorbi_t3_neg_3_050_40    m/z 234.0451     0   1   2   3E+3   18   18.5  RT (min)      lcorbi_t3_neg_4_046_37    m/z 234.0486     0   2   4   6E+4   1   1.5  RT (min)      lcorbi_t3_neg_4_039_32    m/z 234.0837     0   1   2   3   4E+3   18   18.5  RT (min)      lcorbi_t3_neg_4_021_25    m/z 234.0981     0   2   4   6E+3   1   1.5  RT (min)      lcorbi_t3_neg_3_005_23    m/z 234.1134     0   1E+4   6.5   7  RT (min)      lcorbi_t3_neg_2_048_27    m/z 234.9357     0   1   2E+4   18.5   19  RT (min)      lcorbi_t3_neg_1_001_49    m/z 234.9357     0   1   2   3E+4   0.5   1  RT (min)      lcorbi_t3_neg_1_001_49    m/z 235.0083     0   2   4   6E+2   1.5   2  RT (min)      lcorbi_t3_neg_4_032_23    m/z 235.0222     0   1E+4   14   14.5  RT (min)      lcorbi_t3_neg_4_023_18    m/z 235.0414     0   1   2E+3   18   18.5  RT (min)      lcorbi_t3_neg_4_044_38    m/z 235.0722     0   1   2   3E+3   9.5   10  RT (min)      lcorbi_t3_neg_4_053_43    m/z 235.0800     0   2   4   6E+3   18   18.5  RT (min)      lcorbi_t3_neg_4_039_32    m/z 235.0822     0   2   4   6E+3   0   0.5  RT (min)      lcorbi_t3_neg_1_035_25    m/z 235.1185     0   1   2E+4   0   0.5  RT (min)      lcorbi_t3_neg_1_023_03    m/z 235.1531     0   1E+4   11.5   12  RT (min)      lcorbi_t3_neg_4_034_01    m/z 235.1613     0   1E+4   10.5   11  RT (min)      lcorbi_t3_neg_3_053_14    m/z 235.9259     0   1E+5   18.5   19  RT (min)      lcorbi_t3_neg_1_044_23    m/z 235.9259     0   1E+5   0   0.5   1  RT (min)       lcorbi_t3_neg_1_047_02    m/z 235.9950     0   1   2   3   4E+3   7.5   8  RT (min)      lcorbi_t3_neg_4_106_62    m/z 236.0716     0   2   4E+3   1   1.5  RT (min)      lcorbi_t3_neg_4_039_32    m/z 236.0959     0   1E+3   7.5   8  RT (min)      lcorbi_t3_neg_2_048_27    m/z 236.0984     0   1E+4   9.5   10  RT (min)      lcorbi_t3_neg_2_048_27    m/z 236.1114     0   1E+4   9.5   10  RT (min)      lcorbi_t3_neg_2_028_38    m/z 236.1290     0   1   2E+3   9   9.5  RT (min)      lcorbi_t3_neg_1_013_35    m/z 236.7588     0   2   4   6   8E+3   0   0.5  RT (min)      lcorbi_t3_neg_4_113_62    m/z 236.9026     0   1   1   1.5  RT (min)      lcorbi_t3_neg_4_046_37    m/z 236.9805     0   2   4   6   8E+3   7.5   8  RT (min)      lcorbi_t3_neg_2_048_27    m/z 237.0049     0   1   1.5   2  RT (min)      lcorbi_t3_neg_4_090_60    m/z 237.0225     0   2   4   6   8E+3   7   7.5  RT (min)      lcorbi_t3_neg_4_042_14    m/z 237.0491     0   1E+3   1   1.5  RT (min)      lcorbi_t3_neg_3_002_24    m/z 237.0767     0   1E+4   8.5   9  RT (min)      lcorbi_t3_neg_2_048_27    m/z 237.0879     0   2   4   6   8E+3   9.5   10  RT (min)      lcorbi_t3_neg_2_035_46    m/z 237.0978     0   2   4   6E+3   0   0.5  RT (min)      lcorbi_t3_neg_1_035_25    m/z 237.1107     0   1E+4   8   8.5  RT (min)      lcorbi_t3_neg_4_023_18    m/z 237.1341     0   1   2E+4   0   0.5  RT (min)      lcorbi_t3_neg_2_022_24    m/z 237.1494     0   1   10   10.5  RT (min)      lcorbi_t3_neg_4_073_57    m/z 238.0304     0   1E+5   1   1.5  RT (min)       lcorbi_t3_neg_4_011_24    m/z 238.0746     0   1   2E+3   10   10.5  RT (min)      lcorbi_t3_neg_2_048_27    m/z 238.0790     0   2   4   6   8E+3   3   3.5  RT (min)      lcorbi_t3_neg_3_019_12    m/z 238.0912     0   1E+4   5   5.5  RT (min)      lcorbi_t3_neg_4_029_31    m/z 238.1082     0   1E+4   7   7.5  RT (min)      lcorbi_t3_neg_2_048_27    m/z 238.1161     0   1   2E+3   9.5   10  RT (min)      lcorbi_t3_neg_4_056_56    m/z 238.5790     0   2   4   6   8E+3   6.5   7  RT (min)      lcorbi_t3_neg_4_034_01    m/z 238.7563     0   2   4E+3   18.5   19  RT (min)      lcorbi_t3_neg_4_111_62    m/z 238.8580     0   1E+4   3.5   4  RT (min)      lcorbi_t3_neg_3_044_01    m/z 238.8580     0   1   2E+4   15.5   16  RT (min)      lcorbi_t3_neg_4_028_39    m/z 238.8581     0   1   2   3   4E+3   0.5   1  RT (min)      lcorbi_t3_neg_1_002_16    m/z 238.9315     0   1   2E+4   13   13.5  RT (min)      lcorbi_t3_neg_1_048_12    m/z 238.9315     0   1E+4   9.5   10  RT (min)       lcorbi_t3_neg_4_036_16    m/z 238.9315     0   2   4E+4   18   18.5  RT (min)       lcorbi_t3_neg_2_037_09    m/z 238.9966     0   1   2E+3   1   1.5  RT (min)      lcorbi_t3_neg_3_019_12    m/z 239.0727     0   2   4   6E+3   11   11.5  RT (min)      lcorbi_t3_neg_2_035_46    m/z 239.0922     0   1E+3   7   7.5  RT (min)      lcorbi_t3_neg_4_034_01    m/z 239.1130     0   1E+4   4.5   5  RT (min)      lcorbi_t3_neg_4_001_49    m/z 239.1651     0   1E+4   10.5   11  RT (min)      lcorbi_t3_neg_2_048_27    m/z 239.1803     0   1E+4   7   7.5  RT (min)      lcorbi_t3_neg_4_023_18    m/z 239.7605     0   2   4   6   8E+3   1   1.5  RT (min)      lcorbi_t3_neg_2_027_13    m/z 240.0590     0   1E+4   11.5   12  RT (min)      lcorbi_t3_neg_4_036_16    m/z 240.0591     0   1E+5   13.5   14  RT (min)      lcorbi_t3_neg_4_045_35    m/z 240.0705     0   1   2   3E+3   9.5   10  RT (min)      lcorbi_t3_neg_4_039_32    m/z 240.0795     0   1   2   3   4E+3   1   1.5  RT (min)      lcorbi_t3_neg_4_001_49    m/z 240.1321     0   1E+4   9.5   10  RT (min)      lcorbi_t3_neg_2_035_46    m/z 240.8551     0   2   4   6E+3   3.5   4  RT (min)      lcorbi_t3_neg_3_004_30    m/z 240.8551     0   2   4   6E+3   0.5   1  RT (min)      lcorbi_t3_neg_4_068_57    m/z 240.9286     0   1E+4   18   18.5  RT (min)      lcorbi_t3_neg_4_067_57    m/z 241.0011     0   1   1.5   2  RT (min)      lcorbi_t3_neg_4_030_04    m/z 241.0563     0   1E+4   11.5   12  RT (min)      lcorbi_t3_neg_4_007_13    m/z 241.0713     0   1   2   3   4E+3   11   11.5  RT (min)      lcorbi_t3_neg_2_035_46    m/z 241.0869     0   1E+4   9   9.5  RT (min)      lcorbi_t3_neg_2_048_27    m/z 241.1192     0   1E+4   5.5   6  RT (min)      lcorbi_t3_neg_2_048_27    m/z 241.1807     0   1E+4   11   11.5  RT (min)      lcorbi_t3_neg_4_034_01    m/z 241.5793     0   1E+4   9   9.5  RT (min)      lcorbi_t3_neg_2_048_27    m/z 241.7580     0   1E+4   1   1.5  RT (min)      lcorbi_t3_neg_2_027_13    m/z 241.8839     0   1   2E+4   2.5   3  RT (min)      lcorbi_t3_neg_2_048_27    m/z 241.9203     0   1E+4   5   5.5  RT (min)      lcorbi_t3_neg_4_007_13    m/z 241.9203     0   1   2   3   4E+3   3.5   4  RT (min)      lcorbi_t3_neg_4_009_50    m/z 242.0282     0   1E+4   1.5   2  RT (min)      lcorbi_t3_neg_4_089_59    m/z 242.0951     0   1   1   1.5  RT (min)      lcorbi_t3_neg_1_043_14    m/z 242.1477     0   2   4   6E+3   8.5   9  RT (min)      lcorbi_t3_neg_2_048_27    m/z 242.7901     0   1E+4   1   1.5  RT (min)      lcorbi_t3_neg_4_042_14    m/z 242.9433     0   1   2   3E+4   10   10.5  RT (min)       lcorbi_t3_neg_4_105_61    m/z 242.9492     0   1   2E+3   18   18.5  RT (min)      lcorbi_t3_neg_4_072_57    m/z 243.0175     0   1E+3   1   1.5  RT (min)      lcorbi_t3_neg_2_048_27    m/z 243.0468     0   1   2E+4   6   6.5  RT (min)      lcorbi_t3_neg_3_019_12    m/z 243.0773     0   1   2E+2   5   5.5  RT (min)      lcorbi_t3_neg_1_055_05    m/z 243.0830     0   1E+3   1   1.5  RT (min)      lcorbi_t3_neg_1_002_16    m/z 243.1963     0   2   4   6E+3   11   11.5  RT (min)      lcorbi_t3_neg_4_021_25    m/z 243.7552     0   1E+4   1   1.5  RT (min)      lcorbi_t3_neg_2_027_13    m/z 243.8809     0   1E+4   2.5   3  RT (min)      lcorbi_t3_neg_2_048_27    m/z 243.9467     0   2   4   6E+3   18   18.5  RT (min)      lcorbi_t3_neg_4_080_58    m/z 244.0072     0   2   4E+4   9   9.5  RT (min)      lcorbi_t3_neg_2_048_27    m/z 244.0272     0   2   4   6E+3   14   14.5  RT (min)      lcorbi_t3_neg_4_028_39    m/z 244.0298     0   1   2E+3   1   1.5  RT (min)      lcorbi_t3_neg_4_089_59    m/z 244.0660     0   1E+4   7   7.5  RT (min)      lcorbi_t3_neg_4_044_38    m/z 244.1633     0   1   2   3   4E+3   9   9.5  RT (min)      lcorbi_t3_neg_2_048_27    m/z 244.1745     0   1E+4   5.5   6  RT (min)      lcorbi_t3_neg_4_042_14    m/z 244.1915     0   2   4   6E+3   7   7.5  RT (min)      lcorbi_t3_neg_3_019_12    m/z 244.8896     0   1E+3   2   2.5  RT (min)      lcorbi_t3_neg_1_003_41    m/z 244.9140     0   1   2E+3   7.5   8  RT (min)      lcorbi_t3_neg_4_004_34    m/z 244.9200     0   2   4   6E+4   0.5   1  RT (min)      lcorbi_t3_neg_1_001_49    m/z 244.9654     0   2   4   6   8E+3   5.5   6  RT (min)      lcorbi_t3_neg_4_034_01    m/z 244.9680     0   1   2   3   4E+3   1   1.5  RT (min)      lcorbi_t3_neg_4_054_36    m/z 245.0149     0   1   7   7.5  RT (min)      lcorbi_t3_neg_4_080_58    m/z 245.0295     0   1   2   3E+3   18   18.5  RT (min)      lcorbi_t3_neg_1_050_45    m/z 245.0643     0   1   2E+3   5   5.5  RT (min)      lcorbi_t3_neg_2_048_27    m/z 245.0679     0   2   4   6   8E+3   14   14.5  RT (min)      lcorbi_t3_neg_1_004_22    m/z 245.0814     0   1   2E+3   18   18.5  RT (min)      lcorbi_t3_neg_1_022_34    m/z 245.1140     0   2   4   6E+3   4.5   5  RT (min)      lcorbi_t3_neg_4_029_31    m/z 245.1141     0   2   4   6   8E+3   3   3.5  RT (min)      lcorbi_t3_neg_2_048_27    m/z 245.1157     0   2   4   6E+3   10   10.5  RT (min)      lcorbi_t3_neg_2_048_27    m/z 245.8973     0   1E+4   2.5   3  RT (min)      lcorbi_t3_neg_3_052_32    m/z 245.9466     0   1   2   3E+3   1.5   2  RT (min)      lcorbi_t3_neg_1_047_02    m/z 245.9890     0   2   4E+3   4.5   5  RT (min)      lcorbi_t3_neg_1_026_19    m/z 246.0227     0   1   2   3   4E+4   7   7.5  RT (min)      lcorbi_t3_neg_2_048_27    m/z 246.0258     0   2   4E+3   13.5   14  RT (min)      lcorbi_t3_neg_2_037_09    m/z 246.0259     0   2   4   6E+3   18   18.5  RT (min)      lcorbi_t3_neg_1_018_46    m/z 246.0301     0   1E+4   1.5   2  RT (min)      lcorbi_t3_neg_2_051_34    m/z 246.0317     0   1   2E+3   18   18.5  RT (min)      lcorbi_t3_neg_2_022_24    m/z 246.0464     0   1   2   3E+4   1   1.5  RT (min)      lcorbi_t3_neg_2_048_27    m/z 246.0644     0   1   2   3E+4   14   14.5  RT (min)       lcorbi_t3_neg_1_005_07    m/z 246.0657     0   1E+4   7   7.5  RT (min)      lcorbi_t3_neg_4_044_38    m/z 246.0706     0   1   2E+3   18   18.5  RT (min)      lcorbi_t3_neg_4_009_50    m/z 246.0747     0   1E+4   6   6.5  RT (min)      lcorbi_t3_neg_4_034_01    m/z 246.1004     0   2   4   6E+3   1   1.5  RT (min)      lcorbi_t3_neg_4_039_32    m/z 246.8522     0   1   2   3E+3   18   18.5  RT (min)      lcorbi_t3_neg_1_053_32    m/z 246.8865     0   2   4   6E+3   2   2.5  RT (min)      lcorbi_t3_neg_1_023_03    m/z 246.9100     0   1   2   3E+3   7.5   8  RT (min)       lcorbi_t3_neg_4_042_14    m/z 246.9362     0   1   2   3E+3   18   18.5  RT (min)      lcorbi_t3_neg_4_072_57    m/z 246.9642     0   2   4   6E+3   2.5   3  RT (min)      lcorbi_t3_neg_1_044_23    m/z 246.9809     0   2   4E+3   8   8.5  RT (min)      lcorbi_t3_neg_2_048_27    m/z 247.0244     0   2   4   6E+3   18   18.5  RT (min)      lcorbi_t3_neg_4_046_37    m/z 247.0280     0   1E+4   14   14.5  RT (min)      lcorbi_t3_neg_3_034_21    m/z 247.0444     0   2   4   6   8E+3   6.5   7  RT (min)      lcorbi_t3_neg_4_034_01    m/z 247.0674     0   1   18   18.5  RT (min)      lcorbi_t3_neg_4_019_08    m/z 247.0933     0   2   4E+4   1   1.5  RT (min)       lcorbi_t3_neg_4_042_14    m/z 247.1549     0   2   4E+3   0   0.5  RT (min)      lcorbi_t3_neg_2_022_24    m/z 247.1701     0   1   2E+3   10.5   11  RT (min)      lcorbi_t3_neg_1_003_41    m/z 247.8943     0   1E+4   2.5   3  RT (min)      lcorbi_t3_neg_3_052_32    m/z 247.9084     0   1E+4   7.5   8  RT (min)      lcorbi_t3_neg_4_034_01    m/z 247.9622     0   1   2   3E+4   4.5   5  RT (min)      lcorbi_t3_neg_1_044_23    m/z 247.9622     0   1E+4   0   0.5  RT (min)      lcorbi_t3_neg_1_017_51    m/z 247.9630     0   1E+4   7   7.5  RT (min)      lcorbi_t3_neg_4_007_13    m/z 247.9938     0   2   4E+3   1   1.5  RT (min)      lcorbi_t3_neg_2_053_23    m/z 248.0127     0   1E+4   4.5   5  RT (min)      lcorbi_t3_neg_4_040_19    m/z 248.0244     0   1   2   3E+4   13.5   14  RT (min)       lcorbi_t3_neg_2_013_08    m/z 248.0298     0   1   2   3E+3   1   1.5  RT (min)      lcorbi_t3_neg_3_019_12    m/z 248.0439     0   1E+3   1   1.5   2  RT (min)       lcorbi_t3_neg_4_088_59    m/z 248.0540     0   1   2   2.5  RT (min)      lcorbi_t3_neg_3_002_24    m/z 248.0640     0   2   4   6   8E+3   18   18.5  RT (min)      lcorbi_t3_neg_1_053_32    m/z 248.1007     0   1   2   3   4E+4   9   9.5  RT (min)      lcorbi_t3_neg_2_048_27    m/z 248.1247     0   1   2E+4   5   5.5  RT (min)      lcorbi_t3_neg_4_039_32    m/z 248.8492     0   1   2   3   4E+3   18   18.5  RT (min)      lcorbi_t3_neg_1_054_37    m/z 248.8868     0   2   4   6   8E+3   0   0.5  RT (min)      lcorbi_t3_neg_4_072_57    m/z 248.9070     0   1   2E+3   8.5   9  RT (min)       lcorbi_t3_neg_4_081_58    m/z 248.9602     0   2   4   6E+4   5   5.5  RT (min)      lcorbi_t3_neg_4_010_28    m/z 248.9602     0   2   4   6E+4   14.5   15  RT (min)      lcorbi_t3_neg_3_016_42    m/z 248.9602     0   1E+5   9   9.5  RT (min)      lcorbi_t3_neg_4_021_25    m/z 248.9740     0   2   4   6E+3   1   1.5  RT (min)      lcorbi_t3_neg_3_039_29    m/z 249.0207     0   1E+4   13.5   14  RT (min)       lcorbi_t3_neg_4_104_61    m/z 249.0319     0   1   2   3   4E+2   6.5   7  RT (min)      lcorbi_t3_neg_4_021_25    m/z 249.0592     0   1   2   3   4E+3   18   18.5  RT (min)      lcorbi_t3_neg_1_013_35    m/z 249.0613     0   2   4   6E+2   1   1.5  RT (min)      lcorbi_t3_neg_4_044_38    m/z 249.0649     0   1   2E+3   18   18.5  RT (min)      lcorbi_t3_neg_3_037_36    m/z 249.1242     0   1E+4   5.5   6  RT (min)      lcorbi_t3_neg_4_023_18    m/z 249.1341     0   2   4   6E+3   0   0.5  RT (min)      lcorbi_t3_neg_2_022_24    m/z 249.1705     0   1   2E+4   0   0.5  RT (min)      lcorbi_t3_neg_2_022_24    m/z 249.9054     0   1   2   3   4E+3   7.5   8  RT (min)      lcorbi_t3_neg_4_034_01    m/z 249.9573     0   1   8   8.5  RT (min)      lcorbi_t3_neg_1_044_23    m/z 249.9636     0   1   2E+4   18   18.5  RT (min)      lcorbi_t3_neg_3_004_30    m/z 249.9779     0   1E+4   5   5.5  RT (min)       lcorbi_t3_neg_3_035_03    m/z 249.9779     0   1   2E+4   14   14.5  RT (min)      lcorbi_t3_neg_2_018_40    m/z 249.9779     0   1E+4   17.5   18  RT (min)      lcorbi_t3_neg_1_035_25    m/z 249.9779     0   2   4   6   8E+4   0   0.5  RT (min)      lcorbi_t3_neg_2_030_47    m/z 250.1446     0   1   2E+3   9   9.5  RT (min)      lcorbi_t3_neg_1_001_49    m/z 250.1527     0   2   4   6   8E+3   11.5   12  RT (min)      lcorbi_t3_neg_4_046_37    m/z 250.8838     0   1   2   3E+3   0   0.5  RT (min)      lcorbi_t3_neg_4_068_57    m/z 250.8990     0   2   4E+3   7.5   8  RT (min)      lcorbi_t3_neg_4_034_01    m/z 250.9645     0   2   4   6E+3   18   18.5  RT (min)      lcorbi_t3_neg_3_027_43    m/z 251.0943     0   1E+2   1   1.5  RT (min)      lcorbi_t3_neg_1_052_24    m/z 251.1134     0   1E+4   0   0.5  RT (min)      lcorbi_t3_neg_1_035_25    m/z 251.1263     0   1   2E+3   8.5   9  RT (min)      lcorbi_t3_neg_1_044_23    m/z 251.1497     0   2   4   6   8E+3   0   0.5  RT (min)      lcorbi_t3_neg_2_022_24    m/z 251.1963     0   2   4   6   8E+3   10.5   11  RT (min)      lcorbi_t3_neg_1_052_24    m/z 252.0012     0   2   4   6E+3   14   14.5  RT (min)      lcorbi_t3_neg_1_012_20    m/z 252.0921     0   1E+4   1   1.5  RT (min)      lcorbi_t3_neg_3_002_24    m/z 252.0957     0   1   2E+3   8.5   9  RT (min)      lcorbi_t3_neg_4_090_60    m/z 252.1364     0   2   4E+3   6.5   7  RT (min)      lcorbi_t3_neg_2_054_15    m/z 252.8573     0   2   4E+3   1   1.5  RT (min)      lcorbi_t3_neg_1_053_32    m/z 252.9046     0   1   2   3E+3   1   1.5  RT (min)      lcorbi_t3_neg_1_044_23    m/z 252.9976     0   1E+3   14   14.5  RT (min)      lcorbi_t3_neg_1_021_31    m/z 253.0271     0   1E+4   8.5   9  RT (min)      lcorbi_t3_neg_4_001_49    m/z 253.0504     0   1   7   7.5  RT (min)      lcorbi_t3_neg_3_002_24    m/z 253.0568     0   1   2E+5   1   1.5  RT (min)       lcorbi_t3_neg_2_022_24    m/z 253.0755     0   1   2   3E+3   10.5   11  RT (min)      lcorbi_t3_neg_4_005_27    m/z 253.0760     0   1E+4   1   1.5   2  RT (min)       lcorbi_t3_neg_4_093_60    m/z 253.0828     0   1E+4   4.5   5  RT (min)      lcorbi_t3_neg_4_044_38    m/z 253.0868     0   1E+4   8   8.5  RT (min)      lcorbi_t3_neg_4_023_18    m/z 253.0927     0   1   2E+2   1   1.5  RT (min)      lcorbi_t3_neg_1_052_24    m/z 253.0988     0   1   2   3E+2   1.5   2  RT (min)      lcorbi_t3_neg_3_002_24    m/z 253.1290     0   2   4   6   8E+3   0   0.5  RT (min)      lcorbi_t3_neg_2_033_53    m/z 253.1443     0   1   2E+3   6.5   7  RT (min)      lcorbi_t3_neg_2_052_32    m/z 253.8268     0   2   4E+3   1.5   2  RT (min)      lcorbi_t3_neg_4_048_03    m/z 253.9078     0   1E+4   2   2.5  RT (min)      lcorbi_t3_neg_4_032_23    m/z 254.0076     0   1   2   3   4E+4   1   1.5  RT (min)      lcorbi_t3_neg_4_052_26    m/z 254.0740     0   1E+4   2   2.5  RT (min)      lcorbi_t3_neg_3_019_12    m/z 254.1014     0   1   10   10.5  RT (min)      lcorbi_t3_neg_4_102_61    m/z 254.1031     0   1   2E+2   4.5   5  RT (min)      lcorbi_t3_neg_4_026_46    m/z 254.1477     0   1E+4   9   9.5  RT (min)      lcorbi_t3_neg_2_048_27    m/z 254.2487     0   1   2E+4   13.5   14  RT (min)      lcorbi_t3_neg_1_023_03    m/z 254.8970     0   1E+4   2   2.5  RT (min)      lcorbi_t3_neg_4_032_23    m/z 254.9366     0   2   4   6   8E+3   7   7.5  RT (min)      lcorbi_t3_neg_4_007_13    m/z 254.9564     0   2   4E+3   18   18.5  RT (min)      lcorbi_t3_neg_1_043_14    m/z 254.9885     0   1E+4   5   5.5  RT (min)      lcorbi_t3_neg_4_081_58    m/z 254.9918     0   2   4E+3   2   2.5  RT (min)      lcorbi_t3_neg_3_019_12    m/z 255.0242     0   1   2   3   4E+3   8.5   9  RT (min)      lcorbi_t3_neg_4_001_49    m/z 255.0427     0   2   4E+3   7   7.5  RT (min)      lcorbi_t3_neg_4_023_18    m/z 255.0870     0   1   2   3E+3   10.5   11  RT (min)      lcorbi_t3_neg_4_046_37    m/z 255.0870     0   1E+4   6.5   7  RT (min)      lcorbi_t3_neg_4_039_32    m/z 255.0876     0   1   2   3   4E+4   4.5   5   5.5  RT (min)       lcorbi_t3_neg_4_039_32    m/z 255.1136     0   1   2E+4   6.5   7  RT (min)      lcorbi_t3_neg_4_007_13    m/z 255.8219     0   1   2E+4   4.5   5  RT (min)      lcorbi_t3_neg_1_053_32    m/z 255.8219     0   2   4   6   8E+3   18.5   19  RT (min)      lcorbi_t3_neg_1_051_15    m/z 255.9048     0   1E+4   2   2.5  RT (min)      lcorbi_t3_neg_4_032_23    m/z 255.9290     0   2   4   6E+3   14.5   15  RT (min)      lcorbi_t3_neg_4_046_37    m/z 255.9290     0   2   4   6   8E+3   17.5   18  RT (min)      lcorbi_t3_neg_3_042_22    m/z 255.9598     0   2   4   6E+2   2.5   3  RT (min)      lcorbi_t3_neg_4_042_14    m/z 256.0654     0   1   2   3E+4   1   1.5  RT (min)      lcorbi_t3_neg_4_005_27    m/z 256.8219     0   1E+4   1   1.5  RT (min)      lcorbi_t3_neg_2_008_16    m/z 256.8762     0   1E+4   1   1.5  RT (min)      lcorbi_t3_neg_2_052_32    m/z 256.9549     0   2   4   6   8E+3   3   3.5  RT (min)      lcorbi_t3_neg_1_022_34    m/z 256.9549     0   1E+4   4.5   5  RT (min)      lcorbi_t3_neg_1_004_22    m/z 257.0162     0   1   1.5   2  RT (min)      lcorbi_t3_neg_4_024_07    m/z 257.0431     0   1   2E+3   1   1.5  RT (min)      lcorbi_t3_neg_4_089_59    m/z 257.0484     0   1E+4   10.5   11  RT (min)      lcorbi_t3_neg_3_019_12    m/z 257.0778     0   2   4E+2   1   1.5  RT (min)      lcorbi_t3_neg_2_050_17    m/z 257.2022     0   1E+4   8   8.5  RT (min)      lcorbi_t3_neg_2_048_27    m/z 257.2120     0   2   4   6E+3   12   12.5  RT (min)      lcorbi_t3_neg_3_021_38    m/z 257.2433     0   1E+4   11   11.5  RT (min)      lcorbi_t3_neg_3_002_24    m/z 257.8188     0   1   2E+4   4.5   5  RT (min)      lcorbi_t3_neg_1_053_32    m/z 257.8188     0   2   4   6   8E+3   18.5   19  RT (min)      lcorbi_t3_neg_1_051_15    m/z 258.0382     0   1E+4   1   1.5  RT (min)      lcorbi_t3_neg_1_055_05    m/z 258.0513     0   1   2   3   4E+3   1   1.5  RT (min)      lcorbi_t3_neg_4_010_28    m/z 258.0745     0   2   4   6E+3   1   1.5  RT (min)      lcorbi_t3_neg_4_005_27    m/z 258.1174     0   1   1   1.5  RT (min)      lcorbi_t3_neg_3_022_27    m/z 258.1789     0   1   2   3   4E+3   9.5   10  RT (min)      lcorbi_t3_neg_2_048_27    m/z 258.2072     0   1E+4   8.5   9  RT (min)      lcorbi_t3_neg_3_019_12    m/z 258.8188     0   2   4   6E+3   1.5   2  RT (min)      lcorbi_t3_neg_3_007_16    m/z 258.9005     0   1   15.5   16  RT (min)      lcorbi_t3_neg_4_112_62    m/z 258.9155     0   2   4E+4   15.5   16  RT (min)      lcorbi_t3_neg_4_030_04    m/z 258.9157     0   1   2E+4   18.5   19  RT (min)       lcorbi_t3_neg_4_108_62    m/z 258.9244     0   1E+4   2   2.5  RT (min)      lcorbi_t3_neg_2_022_24    m/z 259.0069     0   1   1   1.5  RT (min)      lcorbi_t3_neg_4_007_13    m/z 259.0278     0   2   4   6E+3   7   7.5  RT (min)      lcorbi_t3_neg_2_048_27    m/z 259.0449     0   2   4   6E+3   4.5   5  RT (min)      lcorbi_t3_neg_4_112_62    m/z 259.0755     0   1E+5   2.5   3  RT (min)      lcorbi_t3_neg_1_055_05    m/z 259.0779     0   2   4   6E+3   7   7.5  RT (min)      lcorbi_t3_neg_4_040_19    m/z 259.0835     0   2   4   6E+3   18   18.5  RT (min)      lcorbi_t3_neg_4_011_24    m/z 259.0973     0   1   2E+3   9.5   10  RT (min)      lcorbi_t3_neg_4_005_27    m/z 259.1660     0   1E+4   6   6.5  RT (min)      lcorbi_t3_neg_4_054_36    m/z 259.1701     0   2   4   6E+3   11   11.5  RT (min)      lcorbi_t3_neg_3_007_16    m/z 259.1912     0   1   2   3   4E+3   11.5   12  RT (min)      lcorbi_t3_neg_4_042_14    m/z 259.8158     0   1E+4   3   3.5  RT (min)      lcorbi_t3_neg_3_030_33    m/z 259.8848     0   1   2E+4   2.5   3   3.5  RT (min)       lcorbi_t3_neg_4_102_61    m/z 260.0235     0   2   4   6   8E+2   1.5   2  RT (min)      lcorbi_t3_neg_4_032_23    m/z 260.0451     0   2   4   6   8E+3   13.5   14  RT (min)      lcorbi_t3_neg_1_044_23    m/z 260.0509     0   1   2   3E+2   1   1.5  RT (min)      lcorbi_t3_neg_4_053_43    m/z 260.1163     0   1   2E+3   18   18.5  RT (min)      lcorbi_t3_neg_1_053_32    m/z 260.9214     0   2   4   6   8E+3   2   2.5  RT (min)      lcorbi_t3_neg_2_022_24    m/z 261.0224     0   1E+3   7.5   8  RT (min)      lcorbi_t3_neg_2_048_27    m/z 261.0228     0   2   4   6E+3   1.5   2  RT (min)      lcorbi_t3_neg_4_052_26    m/z 261.0354     0   1E+4   6.5   7  RT (min)      lcorbi_t3_neg_4_098_61    m/z 261.0369     0   1   1   1.5  RT (min)      lcorbi_t3_neg_2_050_17    m/z 261.0415     0   1   2   3E+4   13.5   14  RT (min)       lcorbi_t3_neg_1_044_23    m/z 261.0415     0   2   4E+3   18   18.5  RT (min)      lcorbi_t3_neg_4_046_37    m/z 261.0726     0   1   2E+3   1   1.5  RT (min)      lcorbi_t3_neg_1_018_46    m/z 261.0829     0   1   2E+3   18   18.5  RT (min)      lcorbi_t3_neg_2_024_37    m/z 261.1089     0   2   4   6   8E+3   2   2.5  RT (min)      lcorbi_t3_neg_4_038_15    m/z 261.1127     0   2   4   6E+3   14.5   15  RT (min)      lcorbi_t3_neg_4_040_19    m/z 261.1127     0   1   2   3E+3   18   18.5  RT (min)      lcorbi_t3_neg_1_037_06    m/z 261.1341     0   1E+3   6   7  RT (min)       lcorbi_t3_neg_4_088_59    m/z 261.8482     0   1E+4   1   1.5  RT (min)      lcorbi_t3_neg_3_032_17    m/z 261.8922     0   2   4   6   8E+3   1   1.5  RT (min)      lcorbi_t3_neg_1_050_45    m/z 261.9828     0   1   1   1.5  RT (min)      lcorbi_t3_neg_1_055_05    m/z 262.0378     0   2   4E+4   13.5   14  RT (min)       lcorbi_t3_neg_4_016_05    m/z 262.0384     0   1E+4   18   18.5  RT (min)      lcorbi_t3_neg_4_046_37    m/z 262.0597     0   1E+4   1   1.5  RT (min)      lcorbi_t3_neg_4_017_51    m/z 262.0746     0   1E+5   9   9.5  RT (min)      lcorbi_t3_neg_3_041_54    m/z 262.0763     0   1   2   3E+3   18   18.5  RT (min)      lcorbi_t3_neg_2_012_36    m/z 262.0954     0   2   4   6E+3   1.5   2  RT (min)      lcorbi_t3_neg_4_021_25    m/z 262.1149     0   1   2E+3   18   18.5  RT (min)      lcorbi_t3_neg_4_105_61    m/z 262.7838     0   1E+4   1   1.5  RT (min)      lcorbi_t3_neg_2_007_14    m/z 262.8394     0   1E+4   11.5   12  RT (min)      lcorbi_t3_neg_4_007_13    m/z 262.8559     0   1E+3   1   1.5  RT (min)      lcorbi_t3_neg_1_003_41    m/z 262.9305     0   1E+5   0.5   1  RT (min)      lcorbi_t3_neg_1_001_49    m/z 262.9757     0   1E+4   7.5   8  RT (min)      lcorbi_t3_neg_2_048_27    m/z 263.0342     0   1   2   3E+4   13   13.5  RT (min)      lcorbi_t3_neg_3_040_05    m/z 263.0347     0   2   4E+3   8   8.5  RT (min)      lcorbi_t3_neg_4_013_09    m/z 263.0363     0   2   4   6E+3   18   18.5  RT (min)      lcorbi_t3_neg_4_039_32    m/z 263.0722     0   1E+4   1   1.5  RT (min)      lcorbi_t3_neg_2_035_46    m/z 263.0727     0   1   2   3   4E+3   18   18.5  RT (min)      lcorbi_t3_neg_4_046_37    m/z 263.0729     0   1   2E+4   7   7.5  RT (min)      lcorbi_t3_neg_2_048_27    m/z 263.0915     0   2   4   6   8E+3   1   1.5  RT (min)      lcorbi_t3_neg_3_002_24    m/z 263.1113     0   2   4   6   8E+3   18   18.5  RT (min)      lcorbi_t3_neg_4_011_24    m/z 263.1133     0   1   2   3   4E+3   0   0.5  RT (min)      lcorbi_t3_neg_1_023_03    m/z 263.1497     0   2   4   6E+3   0   0.5  RT (min)      lcorbi_t3_neg_2_022_24    m/z 263.1650     0   1   2   3E+3   9.5   10  RT (min)      lcorbi_t3_neg_2_048_27    m/z 263.8521     0   1E+4   3   3.5  RT (min)      lcorbi_t3_neg_2_007_14    m/z 263.8859     0   2   4   6E+4   7   7.5  RT (min)      lcorbi_t3_neg_4_007_13    m/z 263.9598     0   1   2   3E+4   1   1.5  RT (min)      lcorbi_t3_neg_4_054_36    m/z 264.0093     0   1E+4   8.5   9  RT (min)      lcorbi_t3_neg_2_048_27    m/z 264.0236     0   1E+4   1   1.5  RT (min)      lcorbi_t3_neg_3_040_05    m/z 264.0335     0   1   2   3   4E+3   13   13.5  RT (min)      lcorbi_t3_neg_2_048_27    m/z 264.0922     0   1E+3   6.5   7  RT (min)      lcorbi_t3_neg_4_001_49    m/z 264.1320     0   1   10   10.5  RT (min)      lcorbi_t3_neg_3_042_22    m/z 264.1376     0   1E+4   4.5   5  RT (min)      lcorbi_t3_neg_4_032_23    m/z 264.1578     0   1E+4   8   8.5  RT (min)      lcorbi_t3_neg_4_002_45    m/z 264.8365     0   1E+4   11.5   12  RT (min)      lcorbi_t3_neg_4_007_13    m/z 264.8528     0   2   4E+3   1   1.5  RT (min)      lcorbi_t3_neg_3_051_46    m/z 264.9730     0   1   1   1.5  RT (min)      lcorbi_t3_neg_1_055_05    m/z 264.9882     0   1E+3   1.5   2  RT (min)      lcorbi_t3_neg_4_007_13    m/z 264.9977     0   2   4   6   8E+2   1   1.5  RT (min)      lcorbi_t3_neg_1_055_05    m/z 265.0303     0   2   4   6E+3   5   5.5  RT (min)      lcorbi_t3_neg_4_044_38    m/z 265.0554     0   2   4   6   8E+2   1   1.5  RT (min)      lcorbi_t3_neg_4_045_35    m/z 265.0685     0   2   4   6   8E+3   7   7.5  RT (min)      lcorbi_t3_neg_4_005_27    m/z 265.0790     0   2   4   6E+3   1   1.5  RT (min)      lcorbi_t3_neg_2_039_31    m/z 265.0926     0   1   0   0.5  RT (min)      lcorbi_t3_neg_1_055_05    m/z 265.0926     0   2   4   6E+4   2.5   3  RT (min)      lcorbi_t3_neg_4_007_13    m/z 265.1078     0   1E+3   7.5   8  RT (min)      lcorbi_t3_neg_1_019_29    m/z 265.1290     0   2   4E+3   0   0.5  RT (min)      lcorbi_t3_neg_2_020_05    m/z 265.1476     0   2   4   6   8E+4   13.5   14   14.5  RT (min)       lcorbi_t3_neg_4_021_25    m/z 265.1477     0   2   4   6   8E+3   17.5   18  RT (min)      lcorbi_t3_neg_4_044_38    m/z 265.1555     0   1E+4   10   10.5  RT (min)      lcorbi_t3_neg_4_044_38    m/z 265.1635     0   2   4   6E+3   11.5   12  RT (min)      lcorbi_t3_neg_4_029_31    m/z 265.1653     0   1E+4   0   0.5  RT (min)      lcorbi_t3_neg_2_022_24    m/z 265.1806     0   2   4   6   8E+3   11.5   12  RT (min)      lcorbi_t3_neg_4_032_23    m/z 265.8424     0   1   2   3   4E+2   1   1.5  RT (min)      lcorbi_t3_neg_3_014_19    m/z 265.8499     0   1E+4   4.5   5  RT (min)      lcorbi_t3_neg_1_047_02    m/z 265.8503     0   2   4   6E+3   18.5   19  RT (min)      lcorbi_t3_neg_1_047_02    m/z 265.9504     0   2   4E+3   18   18.5  RT (min)      lcorbi_t3_neg_4_050_02    m/z 266.1031     0   1   2   3E+3   10.5   11  RT (min)      lcorbi_t3_neg_4_049_55    m/z 266.1395     0   1   2E+4   11   11.5  RT (min)      lcorbi_t3_neg_4_043_11    m/z 266.1510     0   1   2E+3   13.5   14  RT (min)      lcorbi_t3_neg_3_023_07    m/z 266.8335     0   1E+4   11.5   12  RT (min)      lcorbi_t3_neg_4_007_13    m/z 266.9460     0   2   4   6   8E+3   1.5   2  RT (min)      lcorbi_t3_neg_1_047_02    m/z 266.9881     0   2   4   6E+2   0   0.5  RT (min)      lcorbi_t3_neg_2_052_32    m/z 267.0077     0   1E+3   1   1.5  RT (min)      lcorbi_t3_neg_2_052_32    m/z 267.1082     0   2   4E+3   0   0.5  RT (min)      lcorbi_t3_neg_1_035_25    m/z 267.1109     0   1   1   1.5  RT (min)      lcorbi_t3_neg_2_026_03    m/z 267.1300     0   1   10   10.5  RT (min)      lcorbi_t3_neg_2_053_23    m/z 267.1433     0   2   4   6E+3   14   14.5  RT (min)      lcorbi_t3_neg_3_023_07    m/z 267.1599     0   1E+4   9.5   10  RT (min)      lcorbi_t3_neg_2_048_27    m/z 267.1963     0   1   2E+3   11   11.5  RT (min)      lcorbi_t3_neg_4_021_25    m/z 267.2690     0   2   4E+3   12   12.5  RT (min)      lcorbi_t3_neg_4_029_31    m/z 267.8475     0   1   2   3   4E+3   18.5   19  RT (min)      lcorbi_t3_neg_1_043_14    m/z 267.8476     0   2   4   6E+3   0.5   1  RT (min)      lcorbi_t3_neg_4_023_18    m/z 268.0373     0   2   4   6E+3   7   7.5  RT (min)      lcorbi_t3_neg_2_048_27    m/z 268.0375     0   2   4   6   8E+3   8.5   9  RT (min)      lcorbi_t3_neg_4_040_19    m/z 268.1468     0   2   4   6E+2   10   10.5  RT (min)      lcorbi_t3_neg_2_048_27    m/z 268.8856     0   2   4E+3   8   8.5  RT (min)      lcorbi_t3_neg_4_005_27    m/z 269.0453     0   1E+4   11.5   12  RT (min)      lcorbi_t3_neg_1_007_26    m/z 269.0542     0   1   1   1.5  RT (min)      lcorbi_t3_neg_4_039_32    m/z 269.0752     0   1   2   3   4E+4   1   1.5  RT (min)      lcorbi_t3_neg_2_048_27    m/z 269.1028     0   1   2E+4   7.5   8  RT (min)      lcorbi_t3_neg_2_048_27    m/z 269.1309     0   2   4   6   8E+3   13.5   14  RT (min)      lcorbi_t3_neg_4_005_27    m/z 269.1504     0   1E+4   7   7.5  RT (min)      lcorbi_t3_neg_2_048_27    m/z 269.8789     0   1E+4   18   18.5  RT (min)      lcorbi_t3_neg_1_002_16    m/z 269.9790     0   1   2   3E+2   1   1.5  RT (min)      lcorbi_t3_neg_1_053_32    m/z 270.0371     0   1E+4   1   1.5  RT (min)      lcorbi_t3_neg_4_039_32    m/z 270.0487     0   1E+4   8.5   9  RT (min)      lcorbi_t3_neg_2_048_27    m/z 270.0707     0   1   2E+3   1   1.5  RT (min)      lcorbi_t3_neg_4_035_33    m/z 270.1026     0   1   2   3E+3   1   1.5  RT (min)      lcorbi_t3_neg_4_009_50    m/z 270.1094     0   1E+4   2   2.5  RT (min)      lcorbi_t3_neg_4_032_23    m/z 270.1279     0   1E+4   6   6.5  RT (min)      lcorbi_t3_neg_2_048_27    m/z 270.1343     0   1   9   9.5  RT (min)      lcorbi_t3_neg_4_112_62    m/z 270.2072     0   1E+4   9   9.5  RT (min)      lcorbi_t3_neg_4_002_45    m/z 271.0514     0   1E+3   1   1.5  RT (min)      lcorbi_t3_neg_1_055_05    m/z 271.0609     0   1   2E+4   8.5   9  RT (min)      lcorbi_t3_neg_4_040_19    m/z 271.9762     0   1   2E+5   1   1.5  RT (min)      lcorbi_t3_neg_3_002_24    m/z 272.0047     0   1   2E+4   14.5   15  RT (min)      lcorbi_t3_neg_2_018_40    m/z 272.0048     0   2   4E+3   18   18.5  RT (min)      lcorbi_t3_neg_2_052_32    m/z 272.0538     0   1   6   6.5  RT (min)      lcorbi_t3_neg_4_092_60    m/z 272.0650     0   1   2E+4   1   1.5  RT (min)      lcorbi_t3_neg_4_032_23    m/z 272.0776     0   1E+4   4.5   5  RT (min)      lcorbi_t3_neg_3_019_12    m/z 272.1289     0   2   4   6E+3   7.5   8  RT (min)      lcorbi_t3_neg_4_007_13    m/z 272.8985     0   2   4   6   8E+3   18.5   19  RT (min)      lcorbi_t3_neg_1_002_16    m/z 272.9587     0   1E+4   18   18.5  RT (min)      lcorbi_t3_neg_1_001_49    m/z 273.0011     0   1   2   3E+4   14.5   15  RT (min)      lcorbi_t3_neg_1_040_42    m/z 273.0011     0   2   4E+3   18   18.5  RT (min)      lcorbi_t3_neg_1_001_49    m/z 273.0596     0   1   1   1.5  RT (min)      lcorbi_t3_neg_4_042_14    m/z 273.0726     0   1   2   3   4E+4   1.5   2  RT (min)       lcorbi_t3_neg_4_042_14    m/z 273.1493     0   1E+4   9.5   10  RT (min)      lcorbi_t3_neg_4_013_09    m/z 273.2069     0   2   4   6E+3   11   11.5  RT (min)      lcorbi_t3_neg_4_034_01    m/z 273.2342     0   1   2E+3   12   12.5  RT (min)      lcorbi_t3_neg_4_005_27    m/z 273.8270     0   2   4E+4   16  RT (min)       lcorbi_t3_neg_1_036_09    m/z 273.8270     0   2   4E+4   17.5   18  RT (min)       lcorbi_t3_neg_1_049_55    m/z 273.8734     0   1   2E+4   0   0.5  RT (min)      lcorbi_t3_neg_1_010_17    m/z 273.8734     0   1   2E+4   18.5   19  RT (min)      lcorbi_t3_neg_1_002_16    m/z 273.8921     0   1   1.5   2  RT (min)      lcorbi_t3_neg_1_003_41    m/z 273.9232     0   1   1   1.5  RT (min)      lcorbi_t3_neg_4_039_32    m/z 273.9289     0   2   4E+3   8.5   9  RT (min)      lcorbi_t3_neg_4_034_01    m/z 274.0207     0   1   2   3   4E+3   18   18.5  RT (min)      lcorbi_t3_neg_1_010_17    m/z 274.0592     0   2   4   6E+3   18   18.5  RT (min)      lcorbi_t3_neg_4_011_24    m/z 274.0935     0   1   2E+3   18   18.5  RT (min)      lcorbi_t3_neg_1_052_24    m/z 275.0094     0   1E+4   2.5   3  RT (min)      lcorbi_t3_neg_4_007_13    m/z 275.0170     0   2   4   6   8E+2   18   18.5  RT (min)      lcorbi_t3_neg_4_019_08    m/z 275.0315     0   1E+4   4.5   5  RT (min)      lcorbi_t3_neg_4_040_19    m/z 275.0534     0   1E+4   7.5   8  RT (min)      lcorbi_t3_neg_4_042_14    m/z 275.0535     0   1E+3   1   1.5  RT (min)      lcorbi_t3_neg_4_011_24    m/z 275.0882     0   1   1   1.5  RT (min)      lcorbi_t3_neg_1_052_24    m/z 275.0985     0   1   2   3   4E+2   1   1.5  RT (min)      lcorbi_t3_neg_4_046_37    m/z 275.8239     0   2   4E+4   15.5   16  RT (min)       lcorbi_t3_neg_1_040_42    m/z 275.8239     0   1   2   3   4E+4   17.5   18  RT (min)       lcorbi_t3_neg_3_040_05    m/z 275.9057     0   2   4E+3   1   1.5  RT (min)      lcorbi_t3_neg_1_044_23    m/z 275.9259     0   1   2   3E+3   8.5   9  RT (min)      lcorbi_t3_neg_4_038_15    m/z 276.0173     0   1E+4   2.5   3  RT (min)      lcorbi_t3_neg_4_007_13    m/z 276.0191     0   1   2E+3   18   18.5  RT (min)      lcorbi_t3_neg_1_018_46    m/z 276.0222     0   2   4   6E+3   10.5   11  RT (min)      lcorbi_t3_neg_2_035_46    m/z 276.0247     0   1   2   3E+4   1   1.5  RT (min)      lcorbi_t3_neg_4_083_59    m/z 276.0533     0   2   4   6   8E+3   14   14.5  RT (min)      lcorbi_t3_neg_3_042_22    m/z 276.0770     0   1   1   1.5  RT (min)       lcorbi_t3_neg_3_002_24    m/z 276.1068     0   1E+4   5.5   6  RT (min)      lcorbi_t3_neg_4_023_18    m/z 276.1601     0   1   2E+3   11.5   12  RT (min)      lcorbi_t3_neg_4_011_24    m/z 276.1683     0   1E+3   9.5   10  RT (min)      lcorbi_t3_neg_4_102_61    m/z 276.9129     0   1E+3   1.5   2  RT (min)      lcorbi_t3_neg_1_002_16    m/z 276.9527     0   2   4E+3   1   1.5  RT (min)      lcorbi_t3_neg_3_006_48    m/z 277.0155     0   1   2E+3   18   18.5  RT (min)      lcorbi_t3_neg_1_010_17    m/z 277.0666     0   1   2E+3   7   7.5  RT (min)      lcorbi_t3_neg_4_050_02    m/z 277.0808     0   1   2E+3   1   1.5  RT (min)      lcorbi_t3_neg_4_032_23    m/z 277.1012     0   1E+5   1   1.5  RT (min)      lcorbi_t3_neg_3_002_24    m/z 277.1160     0   1   2   3E+4   10.5   11  RT (min)      lcorbi_t3_neg_2_048_27    m/z 277.1251     0   1   2   3E+4   10.5   11  RT (min)      lcorbi_t3_neg_2_036_35    m/z 277.1289     0   1   2   3E+3   0   0.5  RT (min)      lcorbi_t3_neg_1_023_03    m/z 277.1442     0   1   2   3   4E+4   18   18.5  RT (min)      lcorbi_t3_neg_4_042_14    m/z 277.1442     0   2   4   6E+7   11   11.5  RT (min)       lcorbi_t3_neg_2_048_27    m/z 277.1443     0   1   2   3E+4   14   14.5  RT (min)      lcorbi_t3_neg_2_048_27    m/z 277.1840     0   1E+4   13.5   14  RT (min)      lcorbi_t3_neg_4_034_01    m/z 277.2170     0   1E+5   11   11.5  RT (min)      lcorbi_t3_neg_1_055_05    m/z 277.8209     0   1E+4   15.5   16  RT (min)      lcorbi_t3_neg_1_036_09    m/z 277.8209     0   1E+4   17.5   18  RT (min)      lcorbi_t3_neg_3_001_49    m/z 277.8211     0   2   4E+3   1   1.5  RT (min)      lcorbi_t3_neg_1_046_11    m/z 277.9365     0   1   1   1.5  RT (min)      lcorbi_t3_neg_4_013_09    m/z 278.0554     0   1   2E+4   1   1.5  RT (min)      lcorbi_t3_neg_2_048_27    m/z 278.0850     0   2   4   6E+4   1   1.5  RT (min)      lcorbi_t3_neg_3_002_24    m/z 278.1030     0   1   2   3E+4   5   5.5  RT (min)      lcorbi_t3_neg_2_048_27    m/z 278.1873     0   1E+4   10   10.5  RT (min)      lcorbi_t3_neg_2_012_36    m/z 278.2123     0   2   4   6E+4   10   10.5  RT (min)      lcorbi_t3_neg_3_024_26    m/z 278.2203     0   1   2E+4   12.5   13  RT (min)      lcorbi_t3_neg_1_055_05    m/z 278.8482     0   2   4E+3   8   8.5  RT (min)      lcorbi_t3_neg_4_023_18    m/z 278.9099     0   2   4E+3   1.5   2  RT (min)      lcorbi_t3_neg_3_007_16    m/z 278.9155     0   1   2   3   4E+3   7   7.5  RT (min)      lcorbi_t3_neg_4_034_01    m/z 279.0620     0   1E+4   1.5   2  RT (min)       lcorbi_t3_neg_4_075_58    m/z 279.0718     0   1   2E+2   1   1.5  RT (min)      lcorbi_t3_neg_2_048_27    m/z 279.1024     0   2   4   6   8E+3   8   8.5  RT (min)      lcorbi_t3_neg_4_023_18    m/z 279.1446     0   2   4E+3   0   0.5  RT (min)      lcorbi_t3_neg_2_033_53    m/z 279.1632     0   1E+4   10   10.5  RT (min)      lcorbi_t3_neg_4_043_11    m/z 279.1632     0   1E+3   14   14.5  RT (min)      lcorbi_t3_neg_4_034_01    m/z 279.1809     0   1E+4   0   0.5  RT (min)      lcorbi_t3_neg_2_022_24    m/z 279.1962     0   2   4   6E+2   11.5   12  RT (min)      lcorbi_t3_neg_1_055_05    m/z 279.6139     0   1E+4   7   7.5  RT (min)      lcorbi_t3_neg_4_034_01    m/z 279.9076     0   2   4   6   8E+2   1.5   2  RT (min)      lcorbi_t3_neg_4_030_04    m/z 279.9579     0   1E+4   1   1.5  RT (min)      lcorbi_t3_neg_2_039_31    m/z 279.9893     0   1   4.5   5  RT (min)      lcorbi_t3_neg_1_026_19    m/z 280.0202     0   1   2   3E+3   1   1.5  RT (min)      lcorbi_t3_neg_2_022_24    m/z 280.0404     0   2   4   6   8E+3   3   3.5  RT (min)      lcorbi_t3_neg_2_048_27    m/z 280.0703     0   1   1   1.5  RT (min)      lcorbi_t3_neg_3_005_23    m/z 280.1034     0   1   2   3   4E+4   1   1.5  RT (min)      lcorbi_t3_neg_3_005_23    m/z 280.1317     0   2   4   6   8E+3   10   10.5  RT (min)      lcorbi_t3_neg_4_023_18    m/z 280.6218     0   2   4   6   8E+3   7.5   8  RT (min)      lcorbi_t3_neg_4_007_13    m/z 280.8550     0   2   4   6E+3   1   1.5  RT (min)      lcorbi_t3_neg_4_046_37    m/z 280.9421     0   1   2E+4   1   1.5  RT (min)      lcorbi_t3_neg_3_048_08    m/z 280.9521     0   2   4   6   8E+3   7   7.5  RT (min)      lcorbi_t3_neg_2_035_46    m/z 280.9826     0   1   2   3E+4   0   0.5  RT (min)      lcorbi_t3_neg_1_001_49    m/z 280.9852     0   1E+4   14   14.5  RT (min)      lcorbi_t3_neg_4_030_04    m/z 280.9890     0   1E+3   1.5   2  RT (min)      lcorbi_t3_neg_3_007_16    m/z 280.9928     0   1E+4   6   6.5  RT (min)      lcorbi_t3_neg_2_048_27    m/z 281.0095     0   2   4   6   8E+3   7   7.5  RT (min)      lcorbi_t3_neg_1_050_45    m/z 281.0357     0   2   4E+4   2   2.5  RT (min)      lcorbi_t3_neg_2_048_27    m/z 281.0868     0   2   4E+4   1   1.5  RT (min)      lcorbi_t3_neg_1_055_05    m/z 281.1028     0   1E+4   7.5   8  RT (min)      lcorbi_t3_neg_2_048_27    m/z 281.1196     0   1E+4   10   10.5  RT (min)      lcorbi_t3_neg_3_004_30    m/z 281.1238     0   1E+4   5   5.5  RT (min)      lcorbi_t3_neg_4_023_18    m/z 281.1238     0   1   2   3E+3   0   0.5  RT (min)      lcorbi_t3_neg_2_020_05    m/z 281.1380     0   1E+4   1   1.5  RT (min)      lcorbi_t3_neg_3_002_24    m/z 281.1628     0   1E+4   10   10.5  RT (min)      lcorbi_t3_neg_2_002_26    m/z 281.1755     0   1E+4   9.5   10  RT (min)      lcorbi_t3_neg_4_023_18    m/z 281.2342     0   1E+4   11.5   12  RT (min)      lcorbi_t3_neg_4_011_24    m/z 281.2482     0   1   2   3   4E+3   0   0.5  RT (min)      lcorbi_t3_neg_1_002_16    m/z 281.2483     0   1E+4   11   11.5  RT (min)      lcorbi_t3_neg_4_007_13    m/z 281.9047     0   1   2E+2   1.5   2  RT (min)      lcorbi_t3_neg_4_030_04    m/z 281.9389     0   2   4   6E+4   3   3.5  RT (min)      lcorbi_t3_neg_4_045_35    m/z 281.9469     0   1E+4   4.5   5  RT (min)      lcorbi_t3_neg_1_026_19    m/z 281.9782     0   2   4   6E+4   1   1.5  RT (min)      lcorbi_t3_neg_3_005_23    m/z 282.0170     0   1   2   3E+3   1.5   2  RT (min)      lcorbi_t3_neg_2_008_16    m/z 282.0198     0   2   4   6E+3   8.5   9  RT (min)      lcorbi_t3_neg_2_048_27    m/z 282.0446     0   1   2E+4   1   1.5  RT (min)      lcorbi_t3_neg_4_011_24    m/z 282.1219     0   1E+3   1   1.5  RT (min)      lcorbi_t3_neg_4_092_60    m/z 282.9807     0   1E+4   14   14.5  RT (min)      lcorbi_t3_neg_4_030_04    m/z 282.9907     0   1E+4   6   6.5  RT (min)      lcorbi_t3_neg_2_048_27    m/z 283.0608     0   1E+4   7   7.5  RT (min)      lcorbi_t3_neg_2_048_27    m/z 283.0676     0   2   4   6E+2   8.5   9  RT (min)      lcorbi_t3_neg_1_012_20    m/z 283.0700     0   2   4   6E+3   12.5   13  RT (min)      lcorbi_t3_neg_4_007_13    m/z 283.0782     0   2   4   6E+4   1   1.5  RT (min)       lcorbi_t3_neg_2_048_27    m/z 283.0820     0   1E+4   5.5   6  RT (min)      lcorbi_t3_neg_2_048_27    m/z 283.0853     0   1   1   1.5  RT (min)      lcorbi_t3_neg_2_048_27    m/z 283.1013     0   1E+4   10.5   11  RT (min)      lcorbi_t3_neg_4_029_31    m/z 283.1184     0   1   2   3E+2   7   7.5  RT (min)      lcorbi_t3_neg_1_036_09    m/z 283.1188     0   1   2   3   4E+4   6   6.5  RT (min)       lcorbi_t3_neg_4_095_60    m/z 283.1371     0   1   2E+3   9.5   10  RT (min)      lcorbi_t3_neg_4_010_28    m/z 283.1700     0   1E+4   12   12.5  RT (min)      lcorbi_t3_neg_4_007_13    m/z 283.1911     0   1   2E+4   10   10.5  RT (min)      lcorbi_t3_neg_2_048_27    m/z 283.2275     0   1   2   3   4E+3   12   12.5  RT (min)      lcorbi_t3_neg_4_007_13    m/z 283.2428     0   2   4   6E+3   13   13.5  RT (min)      lcorbi_t3_neg_4_042_14    m/z 283.2499     0   2   4E+3   11   11.5  RT (min)      lcorbi_t3_neg_3_043_25    m/z 283.2549     0   2   4   6E+4   13   13.5  RT (min)      lcorbi_t3_neg_4_019_08    m/z 283.2833     0   1   2E+3   14   14.5  RT (min)      lcorbi_t3_neg_1_001_49    m/z 283.8524     0   1   2   2.5  RT (min)      lcorbi_t3_neg_1_045_44    m/z 283.8555     0   1E+4   4.5   5  RT (min)      lcorbi_t3_neg_3_038_39    m/z 283.9360     0   1   2   3E+4   3   3.5  RT (min)      lcorbi_t3_neg_4_045_35    m/z 283.9714     0   1   2E+4   1   1.5  RT (min)      lcorbi_t3_neg_4_039_32    m/z 284.0643     0   2   4   6   8E+3   9.5   10  RT (min)      lcorbi_t3_neg_4_010_28    m/z 284.0691     0   1   1   1.5  RT (min)      lcorbi_t3_neg_4_007_13    m/z 284.0924     0   1E+4   6.5   7  RT (min)      lcorbi_t3_neg_2_035_46    m/z 284.1054     0   1E+3   1   1.5  RT (min)      lcorbi_t3_neg_4_011_24    m/z 284.1374     0   1   1   1.5  RT (min)      lcorbi_t3_neg_4_007_13    m/z 284.2672     0   1   2   3   4E+3   0   0.5  RT (min)      lcorbi_t3_neg_1_002_16    m/z 285.0184     0   2   4   6   8E+3   1   1.5  RT (min)      lcorbi_t3_neg_1_050_45    m/z 285.0720     0   1E+3   11.5   12  RT (min)      lcorbi_t3_neg_4_007_13    m/z 285.1163     0   1   2E+3   9.5   10  RT (min)      lcorbi_t3_neg_2_048_27    m/z 285.1255     0   1   2E+4   6   6.5  RT (min)      lcorbi_t3_neg_4_039_32    m/z 285.1606     0   1   2E+4   8   8.5  RT (min)      lcorbi_t3_neg_2_048_27    m/z 285.2432     0   1E+4   12.5   13  RT (min)      lcorbi_t3_neg_4_029_31    m/z 285.2655     0   1   2E+4   11.5   12  RT (min)      lcorbi_t3_neg_3_002_24    m/z 285.8526     0   1E+4   4.5   5  RT (min)      lcorbi_t3_neg_1_045_44    m/z 285.9162     0   1   2.5   3  RT (min)      lcorbi_t3_neg_1_045_44    m/z 285.9331     0   1   3   3.5  RT (min)      lcorbi_t3_neg_1_045_44    m/z 285.9395     0   2   4   6   8E+3   17.5   18  RT (min)      lcorbi_t3_neg_2_015_21    m/z 285.9917     0   1E+3   8.5   9  RT (min)      lcorbi_t3_neg_2_048_27    m/z 286.0510     0   2   4   6   8E+2   10.5   11  RT (min)      lcorbi_t3_neg_4_113_62    m/z 286.0606     0   1E+4   3   3.5  RT (min)      lcorbi_t3_neg_3_019_12    m/z 286.1036     0   1E+4   1   1.5  RT (min)      lcorbi_t3_neg_1_055_05    m/z 286.1292     0   2   4   6E+3   9   9.5  RT (min)      lcorbi_t3_neg_2_048_27    m/z 286.1515     0   1E+5   4.5   5  RT (min)      lcorbi_t3_neg_4_028_39    m/z 286.1615     0   1   2   3E+4   1   1.5  RT (min)      lcorbi_t3_neg_3_002_24    m/z 286.1891     0   2   4   6E+3   11   11.5  RT (min)      lcorbi_t3_neg_2_048_27    m/z 286.2102     0   1   2   3   4E+3   10.5   11  RT (min)      lcorbi_t3_neg_4_034_01    m/z 286.2385     0   2   4   6   8E+3   9.5   10  RT (min)      lcorbi_t3_neg_2_048_27    m/z 286.8612     0   2   4   6   8E+3   17.5   18  RT (min)      lcorbi_t3_neg_2_007_14    m/z 286.8813     0   1   2   3   4E+2   1.5   2  RT (min)      lcorbi_t3_neg_4_030_04    m/z 286.9088     0   1E+4   7.5   8  RT (min)      lcorbi_t3_neg_4_023_18    m/z 287.0227     0   2   4E+4   4.5   5  RT (min)      lcorbi_t3_neg_2_048_27    m/z 287.1149     0   2   4   6   8E+3   18   18.5  RT (min)      lcorbi_t3_neg_4_011_24    m/z 287.1497     0   1   2   3E+3   8   8.5  RT (min)      lcorbi_t3_neg_2_048_27    m/z 287.2812     0   1E+4   12   12.5  RT (min)      lcorbi_t3_neg_2_022_24    m/z 287.9260     0   1   2E+4   1   1.5  RT (min)      lcorbi_t3_neg_2_048_27    m/z 288.0658     0   1   2E+3   1   1.5  RT (min)      lcorbi_t3_neg_2_053_23    m/z 288.0727     0   1   2E+3   18   18.5  RT (min)      lcorbi_t3_neg_1_053_32    m/z 288.1561     0   2   4   6   8E+3   4.5   5  RT (min)      lcorbi_t3_neg_4_032_23    m/z 288.8583     0   2   4   6E+3   17.5   18  RT (min)      lcorbi_t3_neg_2_028_38    m/z 288.8765     0   1E+4   8.5   9  RT (min)      lcorbi_t3_neg_2_048_27    m/z 288.9065     0   1E+4   7.5   8  RT (min)      lcorbi_t3_neg_4_023_18    m/z 288.9363     0   1E+4   3   3.5  RT (min)      lcorbi_t3_neg_4_007_13    m/z 288.9624     0   1   2E+4   3   3.5  RT (min)      lcorbi_t3_neg_2_052_32    m/z 288.9898     0   1   1   1.5  RT (min)      lcorbi_t3_neg_4_032_23    m/z 289.0508     0   2   4   6   8E+4   1   1.5  RT (min)       lcorbi_t3_neg_4_007_13    m/z 289.0681     0   1E+3   1   1.5  RT (min)      lcorbi_t3_neg_2_048_27    m/z 289.0749     0   1   2E+3   18   18.5  RT (min)      lcorbi_t3_neg_1_037_06    m/z 289.1053     0   1   1   1.5  RT (min)      lcorbi_t3_neg_4_007_13    m/z 289.1146     0   1   2E+3   18   18.5  RT (min)      lcorbi_t3_neg_3_028_35    m/z 289.1232     0   1   2   3E+4   2   2.5  RT (min)      lcorbi_t3_neg_4_032_23    m/z 289.8839     0   2   4E+4   5   5.5  RT (min)      lcorbi_t3_neg_2_048_27    m/z 289.8870     0   1E+3   2   2.5  RT (min)      lcorbi_t3_neg_3_039_29    m/z 289.8900     0   2   4   6E+3   17.5   18  RT (min)      lcorbi_t3_neg_3_047_37    m/z 289.9378     0   2   4E+3   18   18.5  RT (min)      lcorbi_t3_neg_4_072_57    m/z 290.0053     0   1   1   1.5  RT (min)      lcorbi_t3_neg_1_044_23    m/z 290.0380     0   1   2   3E+3   10.5   11  RT (min)      lcorbi_t3_neg_4_007_13    m/z 290.0459     0   1E+4   3   3.5  RT (min)      lcorbi_t3_neg_1_054_37    m/z 290.0490     0   1   2E+4   7.5   8  RT (min)      lcorbi_t3_neg_2_048_27    m/z 290.0715     0   2   4   6   8E+3   7   7.5  RT (min)      lcorbi_t3_neg_4_040_19    m/z 290.1089     0   2   4E+3   10   10.5  RT (min)      lcorbi_t3_neg_2_048_27    m/z 290.1109     0   1   2   3E+3   18   18.5  RT (min)      lcorbi_t3_neg_1_047_02    m/z 290.1759     0   2   4   6   8E+3   10   10.5  RT (min)      lcorbi_t3_neg_2_048_27    m/z 290.8735     0   2   4   6   8E+2   8.5   9  RT (min)      lcorbi_t3_neg_1_001_49    m/z 290.9254     0   2   4   6E+4   0.5   1  RT (min)      lcorbi_t3_neg_1_001_49    m/z 290.9333     0   1   2   3   4E+2   3   3.5  RT (min)      lcorbi_t3_neg_1_001_49    m/z 290.9338     0   2   4   6E+3   18   18.5  RT (min)      lcorbi_t3_neg_1_001_49    m/z 290.9443     0   1E+2   1.5   2  RT (min)      lcorbi_t3_neg_1_003_41    m/z 290.9475     0   2   4E+2   3   3.5  RT (min)      lcorbi_t3_neg_1_001_49    m/z 290.9595     0   1E+4   3   3.5  RT (min)      lcorbi_t3_neg_2_052_32    m/z 291.0391     0   2   4   6   8E+3   1.5   2  RT (min)      lcorbi_t3_neg_4_081_58    m/z 291.0482     0   1   2   3E+4   6   6.5  RT (min)      lcorbi_t3_neg_4_001_49    m/z 291.0527     0   1   2   3   4E+3   1   1.5  RT (min)      lcorbi_t3_neg_2_048_27    m/z 291.0678     0   1   2   3   4E+4   7   7.5  RT (min)       lcorbi_t3_neg_2_048_27    m/z 291.0882     0   1E+4   1.5   2  RT (min)      lcorbi_t3_neg_4_042_14    m/z 291.1235     0   1E+4   8.5   9  RT (min)      lcorbi_t3_neg_4_042_14    m/z 291.1598     0   1   2   3E+3   10   10.5  RT (min)      lcorbi_t3_neg_4_102_61    m/z 291.1997     0   2   4E+3   10.5   11  RT (min)       lcorbi_t3_neg_1_055_05    m/z 291.8809     0   1   2E+4   5   5.5  RT (min)      lcorbi_t3_neg_2_048_27    m/z 291.9024     0   1   2   3E+4   7   7.5  RT (min)      lcorbi_t3_neg_4_007_13    m/z 291.9301     0   1   2E+2   3   3.5  RT (min)      lcorbi_t3_neg_4_005_27    m/z 291.9677     0   1   2   3   4E+3   3   3.5  RT (min)      lcorbi_t3_neg_3_020_31    m/z 292.0413     0   2   4   6   8E+3   3   3.5  RT (min)      lcorbi_t3_neg_4_044_38    m/z 292.0694     0   1   2   3E+3   8   8.5  RT (min)      lcorbi_t3_neg_4_040_19    m/z 292.0864     0   2   4   6E+4   1.5   2  RT (min)       lcorbi_t3_neg_4_047_30    m/z 292.8934     0   1   1.5   2  RT (min)      lcorbi_t3_neg_4_032_23    m/z 292.9256     0   2   4E+3   2.5   3  RT (min)      lcorbi_t3_neg_3_007_16    m/z 293.0138     0   2   4E+4   1   1.5  RT (min)      lcorbi_t3_neg_4_038_15    m/z 293.0487     0   1   2   3   4E+3   1   1.5  RT (min)      lcorbi_t3_neg_2_052_32    m/z 293.0873     0   1E+3   1   1.5  RT (min)      lcorbi_t3_neg_2_053_23    m/z 293.1044     0   1E+4   5.5   6  RT (min)      lcorbi_t3_neg_4_052_26    m/z 293.1392     0   1E+4   9   9.5  RT (min)      lcorbi_t3_neg_3_053_14    m/z 293.1602     0   2   4   6E+3   0   0.5  RT (min)      lcorbi_t3_neg_2_033_53    m/z 293.1660     0   2   4   6   8E+3   9.5   10  RT (min)      lcorbi_t3_neg_4_035_33    m/z 293.1755     0   1E+4   14   14.5  RT (min)      lcorbi_t3_neg_2_026_03    m/z 293.1756     0   1   2   3E+6   10   10.5   11  RT (min)       lcorbi_t3_neg_4_013_09    m/z 293.1756     0   2   4E+3   17   17.5  RT (min)      lcorbi_t3_neg_1_003_41    m/z 293.1790     0   1   2E+4   14   14.5  RT (min)      lcorbi_t3_neg_3_023_07    m/z 293.1909     0   1E+4   9.5   10  RT (min)      lcorbi_t3_neg_2_048_27    m/z 293.2483     0   1   11.5   12  RT (min)      lcorbi_t3_neg_3_019_12    m/z 293.8404     0   1   9   9.5  RT (min)      lcorbi_t3_neg_4_023_18    m/z 293.9382     0   1   3   3.5  RT (min)      lcorbi_t3_neg_1_045_44    m/z 294.0256     0   1   2E+3   1   1.5  RT (min)      lcorbi_t3_neg_4_046_37    m/z 294.0292     0   2   4   6   8E+3   3   3.5  RT (min)      lcorbi_t3_neg_2_048_27    m/z 294.0467     0   1E+3   1   1.5  RT (min)      lcorbi_t3_neg_1_055_05    m/z 294.1424     0   2   4   6   8E+3   8   8.5  RT (min)      lcorbi_t3_neg_2_048_27    m/z 294.1692     0   2   4E+3   9.5   10  RT (min)      lcorbi_t3_neg_2_048_27    m/z 294.1883     0   2   4   6E+3   9.5   10  RT (min)      lcorbi_t3_neg_2_048_27    m/z 294.2434     0   1   10   10.5  RT (min)      lcorbi_t3_neg_4_073_57    m/z 294.8848     0   1   18   18.5  RT (min)      lcorbi_t3_neg_4_019_08    m/z 294.8904     0   2   4   6E+3   1.5   2  RT (min)      lcorbi_t3_neg_1_032_13    m/z 294.9500     0   1   2E+4   15.5   16  RT (min)      lcorbi_t3_neg_1_045_44    m/z 294.9500     0   1   2E+4   17.5   18  RT (min)       lcorbi_t3_neg_3_037_36    m/z 294.9562     0   2   4   6   8E+2   3   3.5  RT (min)      lcorbi_t3_neg_4_054_36    m/z 294.9818     0   1   2E+4   0   0.5  RT (min)      lcorbi_t3_neg_1_001_49    m/z 295.0047     0   1   2   3   4E+3   2   2.5  RT (min)      lcorbi_t3_neg_4_081_58    m/z 295.0295     0   1E+3   1   1.5  RT (min)      lcorbi_t3_neg_4_042_14    m/z 295.0543     0   1   2E+4   4.5   5  RT (min)      lcorbi_t3_neg_4_001_49    m/z 295.0569     0   1   2   3   4E+4   1.5   2  RT (min)      lcorbi_t3_neg_2_052_32    m/z 295.0668     0   1   2   3   4E+3   1   1.5  RT (min)      lcorbi_t3_neg_2_043_33    m/z 295.0819     0   1   5   5.5  RT (min)      lcorbi_t3_neg_2_048_27    m/z 295.1296     0   2   4E+3   11   11.5  RT (min)      lcorbi_t3_neg_4_007_13    m/z 295.1854     0   1   2   3E+3   10.5   11  RT (min)      lcorbi_t3_neg_1_055_05    m/z 295.2638     0   2   4   6E+3   12   12.5  RT (min)      lcorbi_t3_neg_4_035_33    m/z 295.8383     0   2   4   6   8E+3   9   9.5  RT (min)      lcorbi_t3_neg_4_005_27    m/z 295.9539     0   2   4   6   8E+3   18   18.5  RT (min)      lcorbi_t3_neg_1_005_07    m/z 295.9962     0   1   1.5   2  RT (min)      lcorbi_t3_neg_1_052_24    m/z 296.0562     0   1E+5   1   1.5  RT (min)       lcorbi_t3_neg_3_002_24    m/z 296.0818     0   1   2E+4   4.5   5  RT (min)      lcorbi_t3_neg_4_029_31    m/z 296.1041     0   1   1   1.5  RT (min)      lcorbi_t3_neg_3_002_24    m/z 296.1136     0   2   4   6E+3   11   11.5  RT (min)      lcorbi_t3_neg_4_017_51    m/z 296.1204     0   1   2   3E+4   5   5.5  RT (min)      lcorbi_t3_neg_2_048_27    m/z 296.1581     0   2   4E+3   10.5   11  RT (min)      lcorbi_t3_neg_4_023_18    m/z 296.1612     0   1E+4   6   6.5  RT (min)      lcorbi_t3_neg_4_046_37    m/z 296.1826     0   2   4   6E+3   9.5   10  RT (min)      lcorbi_t3_neg_2_048_27    m/z 296.8900     0   1   2E+4   18   18.5  RT (min)      lcorbi_t3_neg_2_035_46    m/z 296.9506     0   1E+4   18   18.5  RT (min)      lcorbi_t3_neg_1_001_49    m/z 296.9533     0   2   4   6E+3   3   3.5  RT (min)      lcorbi_t3_neg_3_023_07    m/z 296.9786     0   1E+3   1.5   2  RT (min)      lcorbi_t3_neg_4_078_58    m/z 296.9895     0   1   2   3   4E+3   1.5   2  RT (min)      lcorbi_t3_neg_2_007_14    m/z 297.0376     0   1   7   7.5  RT (min)      lcorbi_t3_neg_1_043_14    m/z 297.0469     0   2   4E+4   9   9.5  RT (min)      lcorbi_t3_neg_2_048_27    m/z 297.0850     0   2   4   6E+4   1   1.5  RT (min)       lcorbi_t3_neg_2_022_24    m/z 297.1088     0   1   2E+4   6   6.5  RT (min)      lcorbi_t3_neg_2_048_27    m/z 297.1526     0   2   4E+3   13.5   14  RT (min)      lcorbi_t3_neg_3_023_07    m/z 297.2431     0   2   4   6E+4   11   11.5  RT (min)      lcorbi_t3_neg_4_044_38    m/z 297.2795     0   2   4   6   8E+3   12.5   13  RT (min)      lcorbi_t3_neg_4_035_33    m/z 297.7329     0   2   4   6E+3   15.5   16  RT (min)      lcorbi_t3_neg_1_055_05    m/z 297.8363     0   1   2   3   4E+3   9   9.5  RT (min)      lcorbi_t3_neg_4_044_38    m/z 297.9242     0   2   4   6E+3   14.5   15  RT (min)      lcorbi_t3_neg_4_046_37    m/z 297.9243     0   2   4   6   8E+3   17.5   18  RT (min)      lcorbi_t3_neg_3_052_32    m/z 297.9382     0   1E+4   3   3.5  RT (min)      lcorbi_t3_neg_4_007_13    m/z 297.9409     0   2   4   6   8E+3   7.5   8  RT (min)      lcorbi_t3_neg_4_023_18    m/z 297.9844     0   2   4   6E+3   7   7.5  RT (min)      lcorbi_t3_neg_1_024_01    m/z 298.0289     0   1   1.5   2  RT (min)      lcorbi_t3_neg_1_043_14    m/z 298.0694     0   2   4E+4   2.5   3  RT (min)      lcorbi_t3_neg_4_032_23    m/z 298.0828     0   2   4E+3   9.5   10  RT (min)      lcorbi_t3_neg_4_021_25    m/z 298.1080     0   1E+4   7   7.5  RT (min)      lcorbi_t3_neg_2_035_46    m/z 298.1139     0   1E+3   1   1.5  RT (min)      lcorbi_t3_neg_4_031_48    m/z 298.1532     0   1E+4   2   2.5  RT (min)      lcorbi_t3_neg_2_022_24    m/z 298.1560     0   1E+4   14   14.5  RT (min)      lcorbi_t3_neg_3_023_07    m/z 298.1632     0   2   4   6   8E+3   7.5   8  RT (min)      lcorbi_t3_neg_3_019_12    m/z 298.2384     0   1E+4   13.5   14  RT (min)      lcorbi_t3_neg_1_023_03    m/z 298.3052     0   1E+4   13   13.5  RT (min)      lcorbi_t3_neg_1_055_05    m/z 298.8347     0   2   4   6   8E+3   0   0.5  RT (min)      lcorbi_t3_neg_4_067_57    m/z 298.8870     0   1E+4   18   18.5  RT (min)      lcorbi_t3_neg_2_031_25    m/z 298.9332     0   1E+4   7.5   8  RT (min)      lcorbi_t3_neg_2_030_47    m/z 298.9418     0   2   4E+3   3   3.5  RT (min)      lcorbi_t3_neg_3_037_36    m/z 298.9520     0   2   4   6   8E+3   7.5   8  RT (min)      lcorbi_t3_neg_2_048_27    m/z 298.9649     0   1E+4   3   3.5  RT (min)      lcorbi_t3_neg_4_007_13    m/z 299.0260     0   2   4   6   8E+3   12.5   13  RT (min)      lcorbi_t3_neg_1_003_41    m/z 299.0556     0   2   4   6   8E+3   7   7.5  RT (min)      lcorbi_t3_neg_3_019_12    m/z 299.0730     0   1E+4   8   8.5  RT (min)      lcorbi_t3_neg_4_023_18    m/z 299.0921     0   2   4   6   8E+3   10   10.5  RT (min)      lcorbi_t3_neg_2_048_27    m/z 299.0952     0   1E+4   11.5   12  RT (min)      lcorbi_t3_neg_3_022_27    m/z 299.0971     0   1E+4   1   1.5  RT (min)      lcorbi_t3_neg_4_035_33    m/z 299.1484     0   1   2E+4   10   10.5  RT (min)      lcorbi_t3_neg_4_007_13    m/z 299.1649     0   2   4E+3   11.5   12  RT (min)      lcorbi_t3_neg_3_054_18    m/z 299.1649     0   1   2E+4   8.5   9  RT (min)      lcorbi_t3_neg_4_023_18    m/z 299.2012     0   1   12.5   13  RT (min)      lcorbi_t3_neg_4_019_08    m/z 299.8290     0   1   2   3   4E+3   1   1.5  RT (min)      lcorbi_t3_neg_3_029_02    m/z 299.9187     0   1E+5   14.5   15  RT (min)      lcorbi_t3_neg_4_011_24    m/z 299.9187     0   1E+5   17.5   18  RT (min)       lcorbi_t3_neg_2_024_37    m/z 299.9341     0   1   2   3E+4   7   7.5  RT (min)      lcorbi_t3_neg_4_007_13    m/z 299.9352     0   1E+4   3   3.5  RT (min)      lcorbi_t3_neg_4_007_13    m/z 299.9673     0   1E+4   8.5   9  RT (min)      lcorbi_t3_neg_2_048_27    m/z 299.9728     0   2   4   6E+2   3   3.5  RT (min)      lcorbi_t3_neg_4_042_14    m/z 300.0046     0   2   4   6E+4   1.5   2  RT (min)      lcorbi_t3_neg_1_043_14    m/z 300.0166     0   1   2   3   4E+2   1.5   2  RT (min)      lcorbi_t3_neg_1_043_14    m/z 300.0389     0   2   4   6E+3   1   1.5  RT (min)      lcorbi_t3_neg_4_034_01    m/z 300.0487     0   1   1   1.5  RT (min)      lcorbi_t3_neg_1_043_14    m/z 300.0493     0   1E+3   3   3.5  RT (min)      lcorbi_t3_neg_2_007_14    m/z 300.0667     0   1   2   3   4E+3   1   1.5  RT (min)      lcorbi_t3_neg_1_050_45    m/z 300.0833     0   2   4   6   8E+3   1   1.5  RT (min)      lcorbi_t3_neg_2_052_32    m/z 300.8657     0   1   1   1.5  RT (min)      lcorbi_t3_neg_3_022_27    m/z 300.9018     0   2   4   6   8E+3   18   18.5  RT (min)      lcorbi_t3_neg_4_066_57    m/z 300.9189     0   1E+4   14.5   15  RT (min)      lcorbi_t3_neg_4_004_34    m/z 300.9189     0   1E+4   17.5   18  RT (min)      lcorbi_t3_neg_3_037_36    m/z 300.9467     0   1   2E+3   7.5   8  RT (min)      lcorbi_t3_neg_1_031_33    m/z 300.9959     0   2   4   6E+3   14.5   15  RT (min)      lcorbi_t3_neg_4_030_04    m/z 301.0004     0   1E+4   1   1.5  RT (min)      lcorbi_t3_neg_4_054_36    m/z 301.0132     0   1E+4   1.5   2  RT (min)      lcorbi_t3_neg_2_052_32    m/z 301.0405     0   1   2   3   4E+3   17.5   18  RT (min)      lcorbi_t3_neg_2_003_42    m/z 301.1190     0   1E+4   8.5   9  RT (min)      lcorbi_t3_neg_2_048_27    m/z 301.1918     0   2   4E+4   8   8.5  RT (min)      lcorbi_t3_neg_2_048_27    m/z 301.2381     0   1   2E+3   13   13.5  RT (min)      lcorbi_t3_neg_3_032_17    m/z 301.2604     0   1   2   3E+3   9.5   10  RT (min)      lcorbi_t3_neg_4_021_25    m/z 301.2653     0   2   4   6E+3   12.5   13  RT (min)      lcorbi_t3_neg_3_005_23    m/z 301.8260     0   2   4E+3   1   1.5  RT (min)      lcorbi_t3_neg_4_030_04    m/z 301.9171     0   1   2   3   4E+3   17.5   18  RT (min)      lcorbi_t3_neg_3_025_52    m/z 301.9311     0   1E+4   7   7.5  RT (min)      lcorbi_t3_neg_4_007_13    m/z 301.9328     0   2   4   6E+3   3   3.5  RT (min)      lcorbi_t3_neg_4_007_13    m/z 301.9883     0   1   2   3E+4   7   7.5  RT (min)      lcorbi_t3_neg_3_022_27    m/z 302.0747     0   1E+4   6.5   7  RT (min)      lcorbi_t3_neg_4_023_18    m/z 302.1047     0   1E+3   1   1.5  RT (min)      lcorbi_t3_neg_3_055_06    m/z 302.1247     0   1   2E+4   13.5   14  RT (min)      lcorbi_t3_neg_4_040_19    m/z 302.2203     0   2   4E+3   12.5   13  RT (min)      lcorbi_t3_neg_4_042_14    m/z 302.9846     0   1E+4   7.5   8  RT (min)      lcorbi_t3_neg_2_048_27    m/z 303.0015     0   2   4   6   8E+2   1   1.5  RT (min)      lcorbi_t3_neg_3_005_23    m/z 303.0070     0   1E+4   14   14.5  RT (min)      lcorbi_t3_neg_4_030_04    m/z 303.0218     0   2   4   6   8E+3   14   14.5  RT (min)      lcorbi_t3_neg_3_035_03    m/z 303.0483     0   1   2E+4   1   1.5  RT (min)      lcorbi_t3_neg_3_005_23    m/z 303.0863     0   1   2   3E+4   1   1.5  RT (min)      lcorbi_t3_neg_1_055_05    m/z 303.1175     0   1E+4   1   1.5  RT (min)      lcorbi_t3_neg_4_039_32    m/z 303.1210     0   1   2E+4   13.5   14  RT (min)      lcorbi_t3_neg_4_040_19    m/z 303.1446     0   2   4   6E+3   9   9.5  RT (min)      lcorbi_t3_neg_2_048_27    m/z 303.1996     0   2   4   6   8E+4   11   11.5  RT (min)       lcorbi_t3_neg_4_102_61    m/z 303.2761     0   1E+4   10   10.5  RT (min)      lcorbi_t3_neg_4_039_32    m/z 303.9171     0   1   2   3E+2   6   6.5  RT (min)      lcorbi_t3_neg_1_044_23    m/z 303.9583     0   1   2E+4   3   3.5  RT (min)      lcorbi_t3_neg_3_019_12    m/z 303.9601     0   1E+4   1   1.5  RT (min)      lcorbi_t3_neg_4_029_31    m/z 304.0305     0   2   4E+2   8   8.5  RT (min)      lcorbi_t3_neg_4_044_38    m/z 304.0847     0   2   4   6   8E+3   13.5   14  RT (min)      lcorbi_t3_neg_1_055_05    m/z 304.0866     0   1   2   3E+3   18   18.5  RT (min)      lcorbi_t3_neg_4_039_32    m/z 304.1301     0   1   2E+4   5   5.5  RT (min)      lcorbi_t3_neg_4_001_49    m/z 304.1914     0   1   2   3   4E+3   10   10.5  RT (min)      lcorbi_t3_neg_4_023_18    m/z 304.2360     0   1   11   11.5  RT (min)      lcorbi_t3_neg_4_102_61    m/z 304.8992     0   1E+3   18   18.5  RT (min)      lcorbi_t3_neg_4_067_57    m/z 304.9353     0   1   1   1.5  RT (min)      lcorbi_t3_neg_3_035_03    m/z 304.9850     0   1E+4   3   3.5  RT (min)      lcorbi_t3_neg_3_019_12    m/z 305.0483     0   2   4   6E+3   13.5   14  RT (min)      lcorbi_t3_neg_3_026_44    m/z 305.0559     0   1   9   9.5  RT (min)      lcorbi_t3_neg_4_032_23    m/z 305.0645     0   1E+3   10   10.5  RT (min)      lcorbi_t3_neg_1_009_50    m/z 305.0831     0   1   2   3E+3   18   18.5  RT (min)      lcorbi_t3_neg_4_044_38    m/z 305.2152     0   1   2   3E+4   11   11.5  RT (min)      lcorbi_t3_neg_2_012_36    m/z 305.9057     0   2   4   6E+2   1   1.5  RT (min)      lcorbi_t3_neg_1_032_13    m/z 305.9171     0   2   4   6E+3   18   18.5  RT (min)      lcorbi_t3_neg_4_070_57    m/z 305.9207     0   1   2E+4   4.5   5  RT (min)      lcorbi_t3_neg_1_046_11    m/z 305.9552     0   1   2E+4   3   3.5  RT (min)      lcorbi_t3_neg_3_019_12    m/z 305.9929     0   1   2E+4   3   3.5  RT (min)      lcorbi_t3_neg_3_019_12    m/z 306.0447     0   1E+4   13.5   14  RT (min)      lcorbi_t3_neg_2_053_23    m/z 306.0817     0   1E+3   1   1.5  RT (min)      lcorbi_t3_neg_1_022_34    m/z 306.0834     0   1E+4   7   7.5  RT (min)      lcorbi_t3_neg_2_048_27    m/z 306.2516     0   2   4   6E+3   13   13.5  RT (min)      lcorbi_t3_neg_4_034_01    m/z 306.7720     0   2   4   6E+3   0   0.5  RT (min)      lcorbi_t3_neg_4_067_57    m/z 306.9092     0   2   4   6E+4   18   18.5  RT (min)      lcorbi_t3_neg_4_019_08    m/z 306.9187     0   1   2E+4   13   13.5  RT (min)      lcorbi_t3_neg_1_012_20    m/z 306.9188     0   1   2E+4   10   10.5  RT (min)       lcorbi_t3_neg_4_036_16    m/z 306.9323     0   1E+4   1   1.5  RT (min)      lcorbi_t3_neg_3_039_29    m/z 306.9371     0   2   4E+3   18.5   19  RT (min)      lcorbi_t3_neg_4_076_58    m/z 307.0411     0   1   2   3   4E+3   13.5   14  RT (min)      lcorbi_t3_neg_2_023_39    m/z 307.0796     0   1E+4   9   9.5  RT (min)      lcorbi_t3_neg_2_048_27    m/z 307.0798     0   1   2E+4   7   7.5  RT (min)      lcorbi_t3_neg_2_048_27    m/z 307.0995     0   1   2   3   4  RT (min)       lcorbi_t3_neg_4_082_59    m/z 307.1160     0   1E+4   9.5   10  RT (min)      lcorbi_t3_neg_4_043_11    m/z 307.1187     0   1   2   3E+4   4.5   5  RT (min)      lcorbi_t3_neg_4_039_32    m/z 307.1945     0   1   2   3E+3   11   11.5  RT (min)      lcorbi_t3_neg_3_043_25    m/z 307.2275     0   1   2   3E+4   11   11.5  RT (min)      lcorbi_t3_neg_1_044_23    m/z 307.2638     0   1   2E+4   13.5   14  RT (min)      lcorbi_t3_neg_4_042_14    m/z 307.3004     0   2   4E+3   13   13.5  RT (min)      lcorbi_t3_neg_4_044_38    m/z 307.8556     0   2   4E+3   0   0.5  RT (min)      lcorbi_t3_neg_1_049_55    m/z 307.9027     0   1   2   3E+2   1   1.5  RT (min)      lcorbi_t3_neg_2_050_17    m/z 307.9529     0   1E+4   3   3.5  RT (min)      lcorbi_t3_neg_3_019_12    m/z 308.0748     0   2   4   6E+3   10   10.5  RT (min)      lcorbi_t3_neg_4_032_23    m/z 308.0759     0   1   2   3E+3   1   1.5  RT (min)      lcorbi_t3_neg_2_051_34    m/z 308.1028     0   1   2   3   4E+3   1   2  RT (min)       lcorbi_t3_neg_3_045_34    m/z 308.7690     0   2   4E+3   0   0.5  RT (min)      lcorbi_t3_neg_4_067_57    m/z 308.8400     0   1E+4   8   8.5  RT (min)      lcorbi_t3_neg_4_034_01    m/z 308.8635     0   1   2   3E+5   0   0.5  RT (min)      lcorbi_t3_neg_4_056_56    m/z 308.9065     0   2   4   6E+3   18   18.5  RT (min)      lcorbi_t3_neg_4_104_61    m/z 308.9158     0   1   2E+4   18   18.5  RT (min)      lcorbi_t3_neg_2_034_19    m/z 308.9618     0   1E+4   7   7.5  RT (min)      lcorbi_t3_neg_1_039_38    m/z 308.9770     0   1E+4   1   1.5  RT (min)      lcorbi_t3_neg_1_055_05    m/z 308.9937     0   1   2   3E+4   3   3.5  RT (min)      lcorbi_t3_neg_4_007_13    m/z 309.0363     0   2   4   6E+4   1   1.5  RT (min)      lcorbi_t3_neg_4_032_23    m/z 309.0741     0   1   9   9.5  RT (min)      lcorbi_t3_neg_4_040_19    m/z 309.1408     0   2   4   6E+3   10   10.5  RT (min)      lcorbi_t3_neg_4_047_30    m/z 309.1526     0   1   2E+3   10   10.5  RT (min)      lcorbi_t3_neg_1_039_38    m/z 309.1551     0   2   4E+3   0   0.5  RT (min)      lcorbi_t3_neg_2_020_05    m/z 309.1704     0   1   2   3   4E+3   10   10.5  RT (min)       lcorbi_t3_neg_4_032_23    m/z 309.1840     0   2   4E+3   10   10.5  RT (min)      lcorbi_t3_neg_4_032_23    m/z 309.2432     0   2   4   6E+3   12   12.5  RT (min)      lcorbi_t3_neg_4_034_01    m/z 309.2795     0   1   2   3E+3   13.5   14  RT (min)      lcorbi_t3_neg_4_007_13    m/z 309.8622     0   2   4   6E+3   8.5   9  RT (min)      lcorbi_t3_neg_4_023_18    m/z 309.9472     0   1   1   1.5  RT (min)      lcorbi_t3_neg_3_052_32    m/z 309.9629     0   2   4   6   8E+3   7   7.5  RT (min)      lcorbi_t3_neg_4_007_13    m/z 310.0016     0   1E+4   3   3.5  RT (min)      lcorbi_t3_neg_4_007_13    m/z 310.0715     0   1   1   1.5  RT (min)      lcorbi_t3_neg_1_044_23    m/z 310.0716     0   2   4   6E+3   6.5   7  RT (min)      lcorbi_t3_neg_4_017_51    m/z 310.0847     0   2   4   6E+3   1   1.5  RT (min)      lcorbi_t3_neg_2_036_35    m/z 310.1007     0   2   4   6E+2   1   1.5  RT (min)      lcorbi_t3_neg_4_039_32    m/z 310.1292     0   1E+4   11.5   12  RT (min)      lcorbi_t3_neg_4_029_31    m/z 310.1634     0   1E+4   8   8.5  RT (min)      lcorbi_t3_neg_4_002_45    m/z 310.2101     0   1   2E+3   10.5   11  RT (min)      lcorbi_t3_neg_4_032_23    m/z 310.2690     0   1E+5   11   11.5  RT (min)       lcorbi_t3_neg_4_011_24    m/z 310.7855     0   1   2E+3   1   1.5  RT (min)      lcorbi_t3_neg_2_052_32    m/z 310.8813     0   1E+3   1   1.5  RT (min)      lcorbi_t3_neg_3_035_03    m/z 310.9306     0   1E+4   8   9   10  RT (min)       lcorbi_t3_neg_2_041_54    m/z 311.0673     0   2   4   6E+2   1   1.5  RT (min)      lcorbi_t3_neg_2_053_23    m/z 311.0980     0   2   4E+4   3   3.5  RT (min)      lcorbi_t3_neg_4_007_13    m/z 311.1285     0   1   2E+4   6   6.5  RT (min)      lcorbi_t3_neg_2_048_27    m/z 311.1385     0   2   4E+4   5   5.5  RT (min)      lcorbi_t3_neg_2_048_27    m/z 311.1497     0   1E+4   9.5   10  RT (min)      lcorbi_t3_neg_2_048_27    m/z 311.1683     0   1E+4   13   13.5   14  RT (min)       lcorbi_t3_neg_4_032_23    m/z 311.1683     0   1E+4   17.5   18  RT (min)      lcorbi_t3_neg_2_028_38    m/z 311.1785     0   2   4E+3   10.5   11  RT (min)      lcorbi_t3_neg_4_043_11    m/z 311.2011     0   1   2   3   4E+3   14   14.5  RT (min)      lcorbi_t3_neg_4_032_23    m/z 311.8827     0   1   2   3E+4   1   1.5  RT (min)      lcorbi_t3_neg_1_007_26    m/z 311.9162     0   1   1   1.5  RT (min)      lcorbi_t3_neg_1_010_17    m/z 311.9163     0   1   2   3   4E+3   17.5   18  RT (min)      lcorbi_t3_neg_2_016_11    m/z 311.9340     0   2   4E+3   18   18.5  RT (min)      lcorbi_t3_neg_3_019_12    m/z 311.9742     0   1E+4   9.5   10  RT (min)      lcorbi_t3_neg_2_048_27    m/z 312.0800     0   1   1   1.5  RT (min)      lcorbi_t3_neg_2_036_35    m/z 312.1716     0   1   2   3E+3   13.5   14  RT (min)      lcorbi_t3_neg_3_023_07    m/z 312.2046     0   1   2E+4   11.5   12  RT (min)      lcorbi_t3_neg_1_029_21    m/z 312.2257     0   2   4E+2   11   11.5  RT (min)      lcorbi_t3_neg_4_043_11    m/z 312.2541     0   1E+4   12   12.5  RT (min)      lcorbi_t3_neg_4_029_31    m/z 312.7825     0   1E+4   1   1.5  RT (min)      lcorbi_t3_neg_4_007_13    m/z 312.7953     0   1   1   1.5  RT (min)      lcorbi_t3_neg_4_007_13    m/z 312.8359     0   1E+4   8   8.5  RT (min)      lcorbi_t3_neg_4_034_01    m/z 313.0359     0   1E+4   6.5   7  RT (min)      lcorbi_t3_neg_4_034_01    m/z 313.0444     0   1E+4   1   1.5  RT (min)      lcorbi_t3_neg_4_032_23    m/z 313.0650     0   1E+6   1.5   2  RT (min)       lcorbi_t3_neg_2_048_27    m/z 313.0714     0   1E+4   7.5   8  RT (min)      lcorbi_t3_neg_2_048_27    m/z 313.0781     0   1   10.5   11  RT (min)      lcorbi_t3_neg_4_098_61    m/z 313.0968     0   1E+4   10   10.5  RT (min)      lcorbi_t3_neg_3_031_45    m/z 313.1141     0   1   2   3E+4   13.5   14  RT (min)      lcorbi_t3_neg_1_044_23    m/z 313.1141     0   1E+4   17.5   18  RT (min)      lcorbi_t3_neg_1_018_46    m/z 313.1183     0   1   2E+4   9   9.5  RT (min)      lcorbi_t3_neg_3_019_12    m/z 313.1190     0   1   2E+4   8   8.5  RT (min)      lcorbi_t3_neg_4_007_13    m/z 313.1368     0   2   4E+3   6   6.5  RT (min)      lcorbi_t3_neg_2_028_38    m/z 313.1642     0   1E+4   14   14.5  RT (min)      lcorbi_t3_neg_3_023_07    m/z 313.1748     0   1E+4   10.5   11  RT (min)      lcorbi_t3_neg_4_007_13    m/z 313.1912     0   2   4   6   8E+3   13   13.5  RT (min)      lcorbi_t3_neg_2_001_49    m/z 313.1977     0   2   4E+3   7   7.5  RT (min)      lcorbi_t3_neg_4_023_18    m/z 313.2939     0   1E+4   14   14.5  RT (min)      lcorbi_t3_neg_3_023_07    m/z 313.8798     0   1   2E+4   1   1.5  RT (min)      lcorbi_t3_neg_1_007_26    m/z 313.9277     0   1   2   3E+4   7.5   8  RT (min)      lcorbi_t3_neg_4_023_18    m/z 313.9606     0   1   2   3E+4   1   1.5  RT (min)      lcorbi_t3_neg_4_039_32    m/z 313.9708     0   1   2E+4   17   17.5  RT (min)      lcorbi_t3_neg_1_036_09    m/z 313.9758     0   2   4E+3   8   8.5  RT (min)      lcorbi_t3_neg_2_048_27    m/z 314.0253     0   1E+4   5   5.5  RT (min)      lcorbi_t3_neg_4_034_01    m/z 314.0370     0   1   2   3   4E+3   1.5   2  RT (min)      lcorbi_t3_neg_4_042_14    m/z 314.0556     0   1E+4   8.5   9  RT (min)      lcorbi_t3_neg_2_048_27    m/z 314.0777     0   2   4E+4   13   13.5  RT (min)      lcorbi_t3_neg_4_045_35    m/z 314.1162     0   1   2   3E+3   1   1.5  RT (min)      lcorbi_t3_neg_1_033_53    m/z 314.1319     0   1   2E+3   1   1.5  RT (min)      lcorbi_t3_neg_4_001_49    m/z 314.1448     0   1   2   3E+4   6   6.5  RT (min)      lcorbi_t3_neg_2_048_27    m/z 314.1673     0   2   4   6   8E+3   10.5   11  RT (min)      lcorbi_t3_neg_4_007_13    m/z 314.2122     0   2   4   6E+3   11   11.5  RT (min)      lcorbi_t3_neg_4_017_51    m/z 314.2203     0   1E+4   12   12.5  RT (min)      lcorbi_t3_neg_4_005_27    m/z 314.2782     0   1   2E+4   12.5   13  RT (min)      lcorbi_t3_neg_4_097_60    m/z 314.3002     0   2   4   6   8E+3   11   11.5  RT (min)      lcorbi_t3_neg_3_002_24    m/z 315.0138     0   1E+3   3   3.5  RT (min)      lcorbi_t3_neg_2_007_14    m/z 315.0332     0   1E+3   1.5   2  RT (min)      lcorbi_t3_neg_1_001_49    m/z 315.0479     0   1   2E+3   18   18.5  RT (min)      lcorbi_t3_neg_3_032_17    m/z 315.0570     0   2   4E+3   1   1.5  RT (min)      lcorbi_t3_neg_4_024_07    m/z 315.0928     0   1   2E+3   1   1.5  RT (min)      lcorbi_t3_neg_2_044_06    m/z 315.1375     0   1   2E+4   6   6.5  RT (min)      lcorbi_t3_neg_2_048_27    m/z 315.1512     0   2   4   6   8E+3   1   1.5  RT (min)      lcorbi_t3_neg_4_011_24    m/z 315.1882     0   1   2   3E+3   13   13.5  RT (min)      lcorbi_t3_neg_4_040_19    m/z 315.2343     0   1   2   3E+3   12   12.5  RT (min)      lcorbi_t3_neg_4_005_27    m/z 315.3124     0   2   4   6   8E+3   12.5   13  RT (min)      lcorbi_t3_neg_4_011_24    m/z 315.8841     0   2   4   6   8E+3   1.5   2  RT (min)      lcorbi_t3_neg_3_039_29    m/z 315.9256     0   1   2   3E+4   7.5   8  RT (min)      lcorbi_t3_neg_4_023_18    m/z 316.0115     0   2   4   6   8E+3   14.5   15  RT (min)      lcorbi_t3_neg_2_011_04    m/z 316.0216     0   1E+4   3   3.5  RT (min)      lcorbi_t3_neg_3_019_12    m/z 316.0280     0   1E+4   6.5   7  RT (min)      lcorbi_t3_neg_2_048_27    m/z 316.0444     0   1   2E+4   1.5   2  RT (min)      lcorbi_t3_neg_1_039_38    m/z 316.0839     0   1   2   3E+2   1   1.5  RT (min)      lcorbi_t3_neg_4_024_07    m/z 316.0879     0   2   4E+3   9   9.5  RT (min)      lcorbi_t3_neg_4_042_14    m/z 316.1216     0   2   4   6   8E+3   1   1.5  RT (min)      lcorbi_t3_neg_4_009_50    m/z 316.1843     0   1   2   3   4E+3   8.5   9  RT (min)      lcorbi_t3_neg_2_039_31    m/z 316.8775     0   1   2E+3   1   1.5  RT (min)      lcorbi_t3_neg_3_035_03    m/z 316.9077     0   2   4   6   8E+4   3   3.5  RT (min)      lcorbi_t3_neg_3_045_34    m/z 316.9474     0   1   2   3E+4   14.5   15  RT (min)      lcorbi_t3_neg_4_014_20    m/z 316.9475     0   2   4   6   8E+4   17.5   18  RT (min)       lcorbi_t3_neg_3_014_19    m/z 316.9644     0   1   2   3E+3   5.5   6  RT (min)      lcorbi_t3_neg_2_048_27    m/z 316.9805     0   1E+4   1   1.5  RT (min)      lcorbi_t3_neg_3_047_37    m/z 317.0078     0   2   4   6   8E+3   14.5   15  RT (min)      lcorbi_t3_neg_2_048_27    m/z 317.0097     0   1   2E+4   1   1.5  RT (min)      lcorbi_t3_neg_4_034_01    m/z 317.0882     0   1E+3   9.5   10  RT (min)      lcorbi_t3_neg_4_023_18    m/z 317.1390     0   1   2E+3   9.5   10  RT (min)      lcorbi_t3_neg_2_015_21    m/z 317.1868     0   1E+4   8   8.5  RT (min)      lcorbi_t3_neg_2_048_27    m/z 317.2118     0   1E+4   11   11.5  RT (min)      lcorbi_t3_neg_4_044_38    m/z 317.2361     0   1E+4   12.5   13  RT (min)      lcorbi_t3_neg_4_034_01    m/z 317.8811     0   1   1.5   2  RT (min)      lcorbi_t3_neg_4_007_13    m/z 317.9360     0   1   2   3   4E+4   1   1.5  RT (min)      lcorbi_t3_neg_4_050_02    m/z 318.0406     0   2   4   6E+3   3   3.5  RT (min)      lcorbi_t3_neg_3_020_31    m/z 318.0590     0   1   2   3   4E+3   8   8.5  RT (min)      lcorbi_t3_neg_4_023_18    m/z 318.0691     0   2   4   6   8E+3   11.5   12  RT (min)      lcorbi_t3_neg_3_022_27    m/z 318.0696     0   2   4   6   8E+5   8.5   9  RT (min)       lcorbi_t3_neg_4_023_18    m/z 318.1454     0   1E+4   5   5.5  RT (min)      lcorbi_t3_neg_4_032_23    m/z 318.2089     0   2   4   6   8E+3   10   10.5  RT (min)      lcorbi_t3_neg_2_048_27    m/z 318.7977     0   2   4   6   8E+3   0   0.5  RT (min)      lcorbi_t3_neg_4_067_57    m/z 318.8505     0   1E+4   1   1.5  RT (min)      lcorbi_t3_neg_3_028_35    m/z 318.8745     0   1   2   3   4E+3   17.5   18  RT (min)      lcorbi_t3_neg_1_021_31    m/z 318.9047     0   2   4   6   8E+4   3   3.5  RT (min)      lcorbi_t3_neg_3_045_34    m/z 319.0326     0   2   4   6E+3   8.5   9  RT (min)      lcorbi_t3_neg_2_048_27    m/z 319.0794     0   1   2E+4   1   1.5  RT (min)      lcorbi_t3_neg_3_005_23    m/z 319.1009     0   1   1   1.5  RT (min)       lcorbi_t3_neg_1_052_24    m/z 319.1142     0   2   4   6   8E+4   2.5   3  RT (min)      lcorbi_t3_neg_4_050_02    m/z 319.1334     0   2   4E+3   1   1.5  RT (min)      lcorbi_t3_neg_1_052_24    m/z 319.1910     0   1   2E+3   10   10.5  RT (min)      lcorbi_t3_neg_4_102_61    m/z 319.8891     0   2   4   6   8E+3   18   18.5  RT (min)      lcorbi_t3_neg_4_080_58    m/z 319.9250     0   1   2E+3   3   3.5  RT (min)      lcorbi_t3_neg_3_018_13    m/z 319.9924     0   1E+4   11   11.5  RT (min)      lcorbi_t3_neg_4_029_31    m/z 320.0118     0   1E+4   4.5   5  RT (min)      lcorbi_t3_neg_4_109_62    m/z 320.0560     0   2   4   6E+3   7.5   8  RT (min)      lcorbi_t3_neg_4_017_51    m/z 320.0666     0   2   4   6   8E+2   1   1.5  RT (min)      lcorbi_t3_neg_3_048_08    m/z 320.0716     0   1E+4   9.5   10  RT (min)      lcorbi_t3_neg_4_007_13    m/z 320.0737     0   1E+4   8   8.5  RT (min)      lcorbi_t3_neg_4_023_18    m/z 320.0788     0   1   1   1.5  RT (min)      lcorbi_t3_neg_3_048_08    m/z 320.1029     0   1   2   3   4E+3   1.5   2  RT (min)      lcorbi_t3_neg_4_026_46    m/z 320.1129     0   1   2   3E+5   1   1.5  RT (min)      lcorbi_t3_neg_1_055_05    m/z 320.1161     0   1   2E+4   10   10.5  RT (min)      lcorbi_t3_neg_4_023_18    m/z 320.8859     0   1E+4   0.5   1  RT (min)       lcorbi_t3_neg_1_001_49    m/z 320.8859     0   1   2E+4   18.5   19  RT (min)       lcorbi_t3_neg_4_112_62    m/z 320.9017     0   1   2E+4   3   3.5  RT (min)      lcorbi_t3_neg_3_045_34    m/z 320.9763     0   1   2E+4   5.5   6  RT (min)      lcorbi_t3_neg_4_052_26    m/z 321.0994     0   1E+4   1.5   2  RT (min)      lcorbi_t3_neg_2_035_46    m/z 321.1315     0   2   4E+3   9   9.5  RT (min)      lcorbi_t3_neg_4_029_31    m/z 321.1468     0   2   4   6   8E+3   8.5   9  RT (min)      lcorbi_t3_neg_2_048_27    m/z 321.2043     0   1   2   3   4E+2   12.5   13  RT (min)      lcorbi_t3_neg_4_019_08    m/z 321.2196     0   1E+4   11   11.5  RT (min)      lcorbi_t3_neg_4_007_13    m/z 321.9220     0   2   4   6E+3   3   3.5  RT (min)      lcorbi_t3_neg_4_017_51    m/z 321.9817     0   1   2E+4   9   9.5  RT (min)      lcorbi_t3_neg_4_007_13    m/z 321.9903     0   1E+4   11   11.5  RT (min)      lcorbi_t3_neg_4_029_31    m/z 322.0009     0   2   4   6   8E+3   14   14.5  RT (min)      lcorbi_t3_neg_4_048_03    m/z 322.0664     0   2   4   6   8E+2   1   1.5  RT (min)      lcorbi_t3_neg_2_048_27    m/z 322.0903     0   2   4   6   8E+3   10   10.5  RT (min)      lcorbi_t3_neg_2_053_23    m/z 322.2134     0   1E+3   11.5   12  RT (min)      lcorbi_t3_neg_4_042_14    m/z 322.8864     0   1E+4   18   18.5  RT (min)      lcorbi_t3_neg_4_080_58    m/z 322.9083     0   1   2E+4   17.5   18  RT (min)      lcorbi_t3_neg_2_020_05    m/z 322.9087     0   2   4   6   8E+3   17.5   18  RT (min)      lcorbi_t3_neg_4_073_57    m/z 322.9787     0   1   1   1.5  RT (min)      lcorbi_t3_neg_4_032_23    m/z 323.0615     0   1   2   3   4E+3   1   1.5  RT (min)      lcorbi_t3_neg_2_048_27    m/z 323.1285     0   1   2E+4   9.5   10  RT (min)      lcorbi_t3_neg_4_040_19    m/z 323.1496     0   1E+4   9   9.5  RT (min)      lcorbi_t3_neg_4_044_38    m/z 323.1625     0   1   9   9.5  RT (min)      lcorbi_t3_neg_4_011_24    m/z 323.1687     0   2   4   6E+3   10.5   11  RT (min)      lcorbi_t3_neg_4_102_61    m/z 323.1894     0   1   2E+4   10.5   11  RT (min)      lcorbi_t3_neg_4_038_15    m/z 323.2058     0   1   11.5   12  RT (min)      lcorbi_t3_neg_4_089_59    m/z 323.9162     0   1E+4   17.5   18  RT (min)      lcorbi_t3_neg_2_020_05    m/z 323.9208     0   1E+4   4.5   5  RT (min)      lcorbi_t3_neg_4_040_19    m/z 323.9311     0   1   2   3E+4   3   3.5  RT (min)      lcorbi_t3_neg_3_052_32    m/z 323.9847     0   2   4E+3   5.5   6  RT (min)      lcorbi_t3_neg_4_034_01    m/z 324.0109     0   2   4   6   8E+3   8.5   9  RT (min)      lcorbi_t3_neg_2_048_27    m/z 324.0225     0   1   2E+3   4.5   5  RT (min)      lcorbi_t3_neg_2_048_27    m/z 324.0295     0   1E+4   3   3.5  RT (min)      lcorbi_t3_neg_3_019_12    m/z 324.0873     0   1E+4   7   7.5  RT (min)      lcorbi_t3_neg_2_048_27    m/z 324.0979     0   1E+3   1   1.5  RT (min)      lcorbi_t3_neg_1_055_05    m/z 324.1062     0   2   4E+3   9.5   10  RT (min)      lcorbi_t3_neg_4_029_31    m/z 324.1173     0   2   4   6E+4   1   1.5  RT (min)      lcorbi_t3_neg_1_055_05    m/z 324.1495     0   1E+4   10   10.5  RT (min)      lcorbi_t3_neg_2_055_18    m/z 324.1501     0   1   2   3   4E+4   1   1.5  RT (min)      lcorbi_t3_neg_2_022_24    m/z 324.2087     0   1   2E+4   13   13.5  RT (min)      lcorbi_t3_neg_4_007_13    m/z 324.2233     0   1   2   3   4E+3   13   13.5  RT (min)      lcorbi_t3_neg_3_018_13    m/z 324.2258     0   1   2   3E+4   11   11.5  RT (min)      lcorbi_t3_neg_4_032_23    m/z 324.2307     0   2   4E+3   9.5   10  RT (min)      lcorbi_t3_neg_4_017_51    m/z 324.2620     0   2   4   6   8E+3   12.5   13  RT (min)      lcorbi_t3_neg_3_007_16    m/z 324.8342     0   1   2   3E+3   0   0.5  RT (min)      lcorbi_t3_neg_4_067_57    m/z 324.8819     0   2   4E+3   18   18.5  RT (min)      lcorbi_t3_neg_4_087_59    m/z 324.9128     0   1E+4   3   3.5  RT (min)      lcorbi_t3_neg_3_022_27    m/z 325.0665     0   2   4   6E+3   7.5   8  RT (min)      lcorbi_t3_neg_4_034_01    m/z 325.0765     0   1E+3   1   1.5  RT (min)      lcorbi_t3_neg_2_036_35    m/z 325.0820     0   2   4E+3   10   10.5  RT (min)      lcorbi_t3_neg_2_048_27    m/z 325.1137     0   2   4E+2   1   1.5  RT (min)      lcorbi_t3_neg_2_036_35    m/z 325.1199     0   1E+4   1.5   2  RT (min)      lcorbi_t3_neg_2_035_46    m/z 325.1686     0   1   2   3   4E+3   9.5   10  RT (min)      lcorbi_t3_neg_4_032_23    m/z 325.1729     0   1   11   11.5  RT (min)      lcorbi_t3_neg_4_102_61    m/z 325.1839     0   1E+4   13.5   14  RT (min)       lcorbi_t3_neg_4_032_23    m/z 325.1839     0   1E+4   17.5   18  RT (min)      lcorbi_t3_neg_4_008_22    m/z 325.2016     0   1   2E+4   9   9.5  RT (min)      lcorbi_t3_neg_2_048_27    m/z 325.2493     0   1E+4   9   9.5  RT (min)      lcorbi_t3_neg_3_002_24    m/z 325.2551     0   1E+4   13   13.5  RT (min)      lcorbi_t3_neg_4_046_37    m/z 325.2743     0   2   4   6E+3   12.5   13  RT (min)      lcorbi_t3_neg_4_044_38    m/z 325.8606     0   1E+4   5   5.5  RT (min)      lcorbi_t3_neg_2_048_27    m/z 325.9177     0   1   2E+4   4.5   5  RT (min)      lcorbi_t3_neg_4_040_19    m/z 325.9282     0   1   2   3E+4   3   3.5  RT (min)      lcorbi_t3_neg_3_052_32    m/z 326.0875     0   1   2   3   4E+3   1   1.5  RT (min)      lcorbi_t3_neg_1_055_05    m/z 326.0971     0   2   4   6   8E+2   1   1.5  RT (min)      lcorbi_t3_neg_1_055_05    m/z 326.1451     0   1E+4   10   10.5  RT (min)      lcorbi_t3_neg_4_023_18    m/z 326.1480     0   1   2.5   3  RT (min)      lcorbi_t3_neg_2_048_27    m/z 326.1873     0   1   2   3E+4   14   14.5  RT (min)      lcorbi_t3_neg_3_023_07    m/z 326.2912     0   2   4E+4   11   11.5  RT (min)      lcorbi_t3_neg_1_052_24    m/z 326.9029     0   1E+4   0.5   1  RT (min)      lcorbi_t3_neg_4_080_58    m/z 326.9029     0   1   2E+4   9.5   10  RT (min)       lcorbi_t3_neg_4_105_61    m/z 326.9033     0   1E+4   18   18.5  RT (min)       lcorbi_t3_neg_4_101_61    m/z 326.9099     0   1E+4   3   3.5  RT (min)      lcorbi_t3_neg_3_022_27    m/z 326.9230     0   1E+3   8.5   9  RT (min)      lcorbi_t3_neg_4_104_61    m/z 326.9365     0   1   2   3E+4   3   3.5  RT (min)      lcorbi_t3_neg_3_037_36    m/z 327.0702     0   1   2   3   4E+4   1   1.5  RT (min)      lcorbi_t3_neg_3_002_24    m/z 327.0920     0   1   1   1.5  RT (min)      lcorbi_t3_neg_3_002_24    m/z 327.1797     0   2   4   6   8E+3   14   14.5  RT (min)      lcorbi_t3_neg_3_023_07    m/z 327.1903     0   1E+4   11   11.5  RT (min)      lcorbi_t3_neg_4_007_13    m/z 327.1961     0   2   4   6E+3   12.5   13  RT (min)      lcorbi_t3_neg_4_005_27    m/z 327.2284     0   2   4   6   8E+3   7   7.5  RT (min)      lcorbi_t3_neg_4_032_23    m/z 327.2537     0   1E+3   12.5   13  RT (min)      lcorbi_t3_neg_4_032_23    m/z 327.2649     0   1   2E+4   12.5   13  RT (min)      lcorbi_t3_neg_4_034_01    m/z 327.8699     0   2   4   6E+3   1.5   2  RT (min)      lcorbi_t3_neg_4_050_02    m/z 328.0949     0   1   2E+3   1   1.5  RT (min)      lcorbi_t3_neg_2_025_52    m/z 328.0979     0   1   2   3E+3   11.5   12  RT (min)      lcorbi_t3_neg_4_044_38    m/z 328.1095     0   1   2   2.5  RT (min)      lcorbi_t3_neg_4_082_59    m/z 328.1295     0   1   2   3E+3   12.5   13  RT (min)      lcorbi_t3_neg_4_044_38    m/z 328.1334     0   1   2E+4   6   6.5  RT (min)      lcorbi_t3_neg_4_046_37    m/z 328.1343     0   2   4   6   8E+3   12.5   13  RT (min)      lcorbi_t3_neg_4_007_13    m/z 328.1451     0   1E+4   10   10.5  RT (min)      lcorbi_t3_neg_4_023_18    m/z 328.1617     0   1E+4   8.5   9  RT (min)      lcorbi_t3_neg_4_034_01    m/z 328.1829     0   2   4   6   8E+3   11   11.5  RT (min)      lcorbi_t3_neg_4_007_13    m/z 328.1962     0   2   4   6E+3   11   11.5  RT (min)      lcorbi_t3_neg_2_048_27    m/z 328.2040     0   2   4E+3   12.5   13  RT (min)      lcorbi_t3_neg_1_036_09    m/z 328.2489     0   1E+4   9.5   10  RT (min)      lcorbi_t3_neg_1_023_03    m/z 328.2935     0   1E+4   13   13.5  RT (min)      lcorbi_t3_neg_1_050_45    m/z 328.8266     0   2   4   6E+3   0   0.5  RT (min)      lcorbi_t3_neg_4_068_57    m/z 328.9033     0   1E+4   18   18.5  RT (min)      lcorbi_t3_neg_1_023_03    m/z 328.9069     0   1E+3   3   3.5  RT (min)      lcorbi_t3_neg_4_045_35    m/z 328.9335     0   1   2E+4   3   3.5  RT (min)      lcorbi_t3_neg_3_037_36    m/z 328.9548     0   1E+4   4.5   5  RT (min)      lcorbi_t3_neg_1_010_17    m/z 329.0119     0   1   2E+4   9   9.5  RT (min)      lcorbi_t3_neg_2_048_27    m/z 329.0290     0   1   2   3   4E+3   9   9.5  RT (min)      lcorbi_t3_neg_1_050_45    m/z 329.1142     0   2   4   6E+3   1   1.5  RT (min)      lcorbi_t3_neg_1_055_05    m/z 329.1237     0   1E+3   6   6.5  RT (min)      lcorbi_t3_neg_2_048_27    m/z 329.1754     0   2   4E+3   12.5   13  RT (min)      lcorbi_t3_neg_4_042_14    m/z 329.1754     0   1   7   7.5  RT (min)      lcorbi_t3_neg_3_048_08    m/z 329.2393     0   2   4   6   8E+3   13   13.5  RT (min)      lcorbi_t3_neg_4_034_01    m/z 329.2693     0   2   4E+3   13.5   14  RT (min)      lcorbi_t3_neg_4_040_19    m/z 329.8454     0   1   2   3   4E+3   13   13.5  RT (min)      lcorbi_t3_neg_1_047_02    m/z 329.9313     0   1   2   3   4E+3   1   1.5  RT (min)      lcorbi_t3_neg_4_017_51    m/z 330.0933     0   1   2E+4   1   1.5  RT (min)      lcorbi_t3_neg_1_055_05    m/z 330.1603     0   1   2   3   4E+4   6   6.5  RT (min)      lcorbi_t3_neg_2_048_27    m/z 330.1960     0   2   4   6E+3   11   11.5  RT (min)      lcorbi_t3_neg_2_048_27    m/z 330.2516     0   1E+4   13   13.5  RT (min)      lcorbi_t3_neg_4_034_01    m/z 330.8791     0   1E+5   1   1.5  RT (min)      lcorbi_t3_neg_1_055_05    m/z 330.9329     0   1E+4   3   3.5  RT (min)      lcorbi_t3_neg_1_055_05    m/z 330.9794     0   1E+4   0   0.5  RT (min)      lcorbi_t3_neg_1_001_49    m/z 331.0431     0   1   2   3E+4   6   6.5  RT (min)       lcorbi_t3_neg_4_040_19    m/z 331.0623     0   1   1.5   2  RT (min)      lcorbi_t3_neg_1_055_05    m/z 331.0740     0   1E+4   1   1.5  RT (min)      lcorbi_t3_neg_4_039_32    m/z 331.0907     0   1   2E+4   5   5.5  RT (min)      lcorbi_t3_neg_2_048_27    m/z 331.0990     0   2   4   6   8E+3   7   7.5  RT (min)      lcorbi_t3_neg_4_046_37    m/z 331.1159     0   1E+4   10   10.5  RT (min)      lcorbi_t3_neg_2_048_27    m/z 331.1471     0   1E+4   6   6.5  RT (min)      lcorbi_t3_neg_4_044_38    m/z 331.1890     0   1E+3   10.5   11  RT (min)      lcorbi_t3_neg_1_008_08    m/z 331.2274     0   1E+4   12   12.5  RT (min)      lcorbi_t3_neg_4_032_23    m/z 331.2392     0   2   4   6   8E+3   9.5   10  RT (min)      lcorbi_t3_neg_4_019_08    m/z 331.2984     0   1E+4   12.5   13  RT (min)      lcorbi_t3_neg_2_022_24    m/z 331.2984     0   1   2E+3   10   10.5  RT (min)      lcorbi_t3_neg_3_023_07    m/z 332.0649     0   1   2E+2   1   1.5  RT (min)      lcorbi_t3_neg_3_002_24    m/z 332.0768     0   1   2   3   4E+4   1   1.5  RT (min)      lcorbi_t3_neg_4_039_32    m/z 332.0797     0   1E+4   7   7.5  RT (min)      lcorbi_t3_neg_4_005_27    m/z 332.0853     0   1E+4   9.5   10  RT (min)      lcorbi_t3_neg_4_023_18    m/z 332.1525     0   1E+4   9.5   10  RT (min)      lcorbi_t3_neg_2_048_27    m/z 332.1792     0   2   4   6E+2   8.5   9  RT (min)      lcorbi_t3_neg_4_019_08    m/z 332.2268     0   1   7.5   8  RT (min)      lcorbi_t3_neg_4_103_61    m/z 332.2519     0   1   2E+3   9.5   10  RT (min)      lcorbi_t3_neg_4_023_18    m/z 332.2672     0   1   2   3   4E+3   13   13.5  RT (min)      lcorbi_t3_neg_4_010_28    m/z 332.2967     0   1E+4   14   14.5  RT (min)      lcorbi_t3_neg_1_023_03    m/z 332.9300     0   2   4   6   8E+3   3   3.5  RT (min)      lcorbi_t3_neg_1_055_05    m/z 332.9373     0   1E+4   3   3.5  RT (min)      lcorbi_t3_neg_1_049_55    m/z 333.0401     0   2   4   6E+3   4.5   5  RT (min)      lcorbi_t3_neg_4_038_15    m/z 333.0604     0   1   2   3E+4   1   1.5  RT (min)      lcorbi_t3_neg_3_005_23    m/z 333.0760     0   2   4   6E+3   7   7.5  RT (min)      lcorbi_t3_neg_4_110_62    m/z 333.1700     0   2   4E+3   10   10.5  RT (min)      lcorbi_t3_neg_4_090_60    m/z 333.1875     0   2   4   6   8E+3   11.5   12  RT (min)      lcorbi_t3_neg_1_032_13    m/z 333.1974     0   1E+4   7   7.5  RT (min)      lcorbi_t3_neg_4_023_18    m/z 333.2045     0   2   4   6E+3   11   11.5  RT (min)      lcorbi_t3_neg_4_032_23    m/z 333.2180     0   1   2   3E+4   8   8.5  RT (min)      lcorbi_t3_neg_2_048_27    m/z 333.2431     0   2   4E+3   12   12.5  RT (min)      lcorbi_t3_neg_4_038_15    m/z 333.3141     0   1E+4   10.5   11  RT (min)      lcorbi_t3_neg_4_011_24    m/z 333.8943     0   1E+3   18   18.5  RT (min)      lcorbi_t3_neg_1_001_49    m/z 333.9377     0   1E+4   18   18.5  RT (min)      lcorbi_t3_neg_2_015_21    m/z 333.9599     0   1E+4   3   3.5  RT (min)      lcorbi_t3_neg_3_052_32    m/z 334.0985     0   1   1   1.5  RT (min)      lcorbi_t3_neg_4_040_19    m/z 334.1704     0   2   4   6E+3   11.5   12  RT (min)      lcorbi_t3_neg_4_032_23    m/z 334.1990     0   1E+4   5   5.5  RT (min)      lcorbi_t3_neg_3_002_24    m/z 334.2231     0   1E+4   12   12.5  RT (min)      lcorbi_t3_neg_4_042_14    m/z 334.7474     0   1E+4   1   1.5  RT (min)      lcorbi_t3_neg_4_036_16    m/z 334.8629     0   2   4   6   8E+4   0   0.5  RT (min)      lcorbi_t3_neg_4_046_37    m/z 334.9112     0   2   4   6   8E+2   1   1.5  RT (min)      lcorbi_t3_neg_1_055_05    m/z 334.9343     0   1   3   3.5  RT (min)      lcorbi_t3_neg_4_066_57    m/z 334.9764     0   1   2E+4   4.5   5  RT (min)      lcorbi_t3_neg_3_022_27    m/z 334.9910     0   2   4   6E+3   1   1.5  RT (min)      lcorbi_t3_neg_2_048_27    m/z 335.0264     0   1E+3   1   1.5  RT (min)      lcorbi_t3_neg_2_048_27    m/z 335.0596     0   1   2E+3   1   1.5  RT (min)      lcorbi_t3_neg_4_024_07    m/z 335.1108     0   1E+4   10   10.5  RT (min)      lcorbi_t3_neg_2_048_27    m/z 335.1307     0   2   4E+3   10   10.5  RT (min)      lcorbi_t3_neg_4_044_38    m/z 335.1894     0   1   2E+4   10.5   11  RT (min)      lcorbi_t3_neg_2_022_24    m/z 335.2467     0   1E+4   12.5   13  RT (min)      lcorbi_t3_neg_4_007_13    m/z 335.8614     0   1E+3   0   0.5  RT (min)      lcorbi_t3_neg_3_034_21    m/z 335.9256     0   1   2E+4   6   6.5  RT (min)      lcorbi_t3_neg_3_054_18    m/z 335.9570     0   1E+4   3   3.5  RT (min)      lcorbi_t3_neg_3_052_32    m/z 336.0523     0   2   4   6   8E+4   1   1.5  RT (min)      lcorbi_t3_neg_4_042_14    m/z 336.0783     0   1   1   1.5  RT (min)      lcorbi_t3_neg_2_048_27    m/z 336.1139     0   2   4   6   8E+4   1   1.5  RT (min)      lcorbi_t3_neg_3_002_24    m/z 336.1701     0   1   2   3   4E+3   11.5   12  RT (min)      lcorbi_t3_neg_2_053_23    m/z 336.1859     0   1E+3   9.5   10  RT (min)      lcorbi_t3_neg_4_039_32    m/z 336.2115     0   1   2   3E+2   11   11.5  RT (min)      lcorbi_t3_neg_4_019_08    m/z 336.2201     0   1   2   3   4E+3   12   12.5  RT (min)      lcorbi_t3_neg_4_042_14    m/z 336.2257     0   1E+4   10   10.5  RT (min)      lcorbi_t3_neg_2_048_27    m/z 336.8166     0   1   1   1.5  RT (min)      lcorbi_t3_neg_2_048_27    m/z 336.9083     0   1   2   3E+3   1   1.5  RT (min)      lcorbi_t3_neg_1_053_32    m/z 336.9653     0   1E+4   3   3.5  RT (min)      lcorbi_t3_neg_3_037_36    m/z 336.9943     0   1   2E+4   2   2.5  RT (min)      lcorbi_t3_neg_2_048_27    m/z 337.0326     0   2   4   6   8E+3   8.5   9  RT (min)      lcorbi_t3_neg_4_040_19    m/z 337.0400     0   1   2   3   4E+4   1   1.5  RT (min)      lcorbi_t3_neg_2_052_32    m/z 337.0709     0   2   4   6E+3   8.5   9  RT (min)      lcorbi_t3_neg_2_048_27    m/z 337.0944     0   1E+3   1   1.5  RT (min)      lcorbi_t3_neg_2_030_47    m/z 337.1116     0   1E+3   1   1.5  RT (min)      lcorbi_t3_neg_3_045_34    m/z 337.1418     0   1E+4   8   8.5  RT (min)      lcorbi_t3_neg_2_048_27    m/z 337.1824     0   1E+4   9.5   10  RT (min)      lcorbi_t3_neg_2_048_27    m/z 337.1914     0   2   4   6   8E+3   13   13.5  RT (min)      lcorbi_t3_neg_2_024_37    m/z 337.2050     0   2   4   6   8E+3   14   14.5  RT (min)      lcorbi_t3_neg_3_023_07    m/z 337.9395     0   1   1   1.5  RT (min)      lcorbi_t3_neg_3_034_21    m/z 338.0029     0   1   2   3E+4   5.5   6  RT (min)      lcorbi_t3_neg_4_047_30    m/z 338.0612     0   1   2E+3   1   1.5  RT (min)      lcorbi_t3_neg_4_044_38    m/z 338.0773     0   1   2   3   4E+3   1   1.5  RT (min)      lcorbi_t3_neg_1_021_31    m/z 338.0917     0   1E+4   1.5   2  RT (min)      lcorbi_t3_neg_4_039_32    m/z 338.1606     0   1E+4   7.5   8  RT (min)      lcorbi_t3_neg_2_048_27    m/z 338.2381     0   1E+3   11   11.5  RT (min)      lcorbi_t3_neg_1_055_05    m/z 338.2527     0   1   2E+4   10   10.5  RT (min)      lcorbi_t3_neg_4_034_01    m/z 338.2695     0   2   4E+3   12   12.5  RT (min)      lcorbi_t3_neg_4_007_13    m/z 338.8135     0   2   4   6E+2   1   1.5  RT (min)      lcorbi_t3_neg_2_048_27    m/z 338.8875     0   1   2   3   4E+3   6.5   7  RT (min)      lcorbi_t3_neg_1_009_50    m/z 338.8920     0   1   2E+4   5   5.5  RT (min)      lcorbi_t3_neg_3_022_27    m/z 339.0398     0   1   2   3E+4   1   1.5  RT (min)      lcorbi_t3_neg_4_039_32    m/z 339.0560     0   1E+3   9   9.5  RT (min)      lcorbi_t3_neg_4_023_18    m/z 339.0737     0   1E+3   1   1.5  RT (min)       lcorbi_t3_neg_1_050_45    m/z 339.0921     0   2   4   6   8E+2   1   1.5  RT (min)      lcorbi_t3_neg_4_045_35    m/z 339.1095     0   1   2E+4   9   9.5  RT (min)      lcorbi_t3_neg_4_023_18    m/z 339.1597     0   1E+4   8   8.5  RT (min)      lcorbi_t3_neg_2_048_27    m/z 339.1615     0   1E+4   10   10.5  RT (min)      lcorbi_t3_neg_4_023_18    m/z 339.1781     0   2   4   6E+3   9.5   10  RT (min)      lcorbi_t3_neg_2_048_27    m/z 339.1995     0   1E+4   13   13.5  RT (min)       lcorbi_t3_neg_3_023_07    m/z 339.1995     0   2   4   6   8E+3   17.5   18  RT (min)      lcorbi_t3_neg_1_016_39    m/z 339.2147     0   2   4   6E+4   13   13.5  RT (min)      lcorbi_t3_neg_4_007_13    m/z 339.2570     0   1E+4   13   13.5  RT (min)      lcorbi_t3_neg_4_073_57    m/z 339.3012     0   1E+4   13.5   14  RT (min)      lcorbi_t3_neg_1_023_03    m/z 339.8696     0   2   4   6E+3   2   2.5  RT (min)      lcorbi_t3_neg_4_036_16    m/z 339.8972     0   2   4E+4   1   1.5  RT (min)      lcorbi_t3_neg_2_027_13    m/z 340.0008     0   1   2   3E+4   5.5   6  RT (min)      lcorbi_t3_neg_4_047_30    m/z 340.0821     0   1   1   1.5  RT (min)      lcorbi_t3_neg_4_044_38    m/z 340.1382     0   1   1   1.5  RT (min)      lcorbi_t3_neg_2_036_35    m/z 340.2028     0   1   2E+4   14   14.5  RT (min)      lcorbi_t3_neg_3_023_07    m/z 340.2031     0   2   4E+3   13   13.5  RT (min)      lcorbi_t3_neg_3_007_16    m/z 340.2126     0   1   2E+3   13   13.5  RT (min)      lcorbi_t3_neg_2_027_13    m/z 340.2240     0   1E+4   13   13.5  RT (min)      lcorbi_t3_neg_4_012_17    m/z 340.2377     0   1   11   11.5  RT (min)      lcorbi_t3_neg_3_002_24    m/z 340.8350     0   2   4E+3   0   0.5  RT (min)      lcorbi_t3_neg_4_066_57    m/z 340.8890     0   1   2E+4   5   5.5  RT (min)      lcorbi_t3_neg_3_022_27    m/z 340.9082     0   1   2   3E+3   2   2.5  RT (min)      lcorbi_t3_neg_2_048_27    m/z 340.9157     0   2   4   6E+3   8.5   9  RT (min)      lcorbi_t3_neg_1_054_37    m/z 340.9457     0   1E+4   1   1.5  RT (min)      lcorbi_t3_neg_2_048_27    m/z 340.9783     0   1E+4   1   1.5  RT (min)      lcorbi_t3_neg_4_034_01    m/z 341.0036     0   1E+4   13   13.5  RT (min)      lcorbi_t3_neg_1_023_03    m/z 341.0612     0   1E+4   7   7.5  RT (min)      lcorbi_t3_neg_4_007_13    m/z 341.0726     0   1E+4   6   6.5  RT (min)      lcorbi_t3_neg_4_087_59    m/z 341.0946     0   1   2E+4   10   10.5  RT (min)      lcorbi_t3_neg_4_023_18    m/z 341.0985     0   1E+3   4.5   5  RT (min)      lcorbi_t3_neg_4_039_32    m/z 341.0987     0   1   2E+4   1   1.5  RT (min)      lcorbi_t3_neg_1_055_05    m/z 341.1025     0   1   2   3   4E+3   6.5   7  RT (min)      lcorbi_t3_neg_4_023_18    m/z 341.1066     0   2   4   6E+3   9   9.5  RT (min)      lcorbi_t3_neg_4_023_18    m/z 341.1085     0   1   2E+5   1   1.5  RT (min)      lcorbi_t3_neg_4_023_18    m/z 341.1144     0   2   4E+2   1   1.5  RT (min)      lcorbi_t3_neg_2_048_27    m/z 341.1390     0   1   2E+4   6   6.5  RT (min)      lcorbi_t3_neg_2_048_27    m/z 341.1452     0   1   2   3   4E+3   10.5   11  RT (min)      lcorbi_t3_neg_4_023_18    m/z 341.1452     0   1   2E+4   13.5   14  RT (min)      lcorbi_t3_neg_2_044_06    m/z 341.1453     0   2   4   6   8E+3   17.5   18  RT (min)      lcorbi_t3_neg_3_052_32    m/z 341.2060     0   2   4   6   8E+3   11.5   12  RT (min)      lcorbi_t3_neg_4_043_11    m/z 341.2224     0   1E+4   13.5   14  RT (min)      lcorbi_t3_neg_2_023_39    m/z 341.2500     0   1E+3   11   11.5  RT (min)      lcorbi_t3_neg_2_022_24    m/z 341.2805     0   2   4E+3   9.5   10  RT (min)      lcorbi_t3_neg_1_055_05    m/z 341.2827     0   1   2   3E+4   11   11.5  RT (min)      lcorbi_t3_neg_2_022_24    m/z 341.8666     0   1   2   3   4E+3   2   2.5  RT (min)      lcorbi_t3_neg_3_025_52    m/z 341.9149     0   1   2E+4   6   6.5  RT (min)      lcorbi_t3_neg_3_019_12    m/z 341.9848     0   1E+4   9   9.5  RT (min)      lcorbi_t3_neg_4_053_43    m/z 342.1107     0   1   2E+3   1   1.5  RT (min)      lcorbi_t3_neg_1_055_05    m/z 342.1275     0   1E+4   1   1.5  RT (min)      lcorbi_t3_neg_1_055_05    m/z 342.1528     0   1   2E+3   10.5   11  RT (min)      lcorbi_t3_neg_4_033_53    m/z 342.1986     0   2   4   6   8E+3   11   11.5  RT (min)      lcorbi_t3_neg_4_102_61    m/z 342.2645     0   2   4E+3   11.5   12  RT (min)      lcorbi_t3_neg_4_034_01    m/z 342.3091     0   1E+4   13   13.5  RT (min)      lcorbi_t3_neg_4_097_60    m/z 342.9050     0   1   2   3E+3   2   2.5  RT (min)      lcorbi_t3_neg_2_048_27    m/z 342.9128     0   2   4E+3   8.5   9  RT (min)      lcorbi_t3_neg_1_054_37    m/z 343.0406     0   2   4   6   8E+3   7.5   8  RT (min)      lcorbi_t3_neg_4_042_14    m/z 343.0434     0   2   4   6E+2   1.5   2  RT (min)      lcorbi_t3_neg_1_043_14    m/z 343.0599     0   1   2E+4   1   1.5  RT (min)      lcorbi_t3_neg_4_011_24    m/z 343.0626     0   2   4   6E+4   8.5   9  RT (min)       lcorbi_t3_neg_2_015_21    m/z 343.0688     0   2   4   6E+3   8   8.5  RT (min)      lcorbi_t3_neg_4_023_18    m/z 343.0785     0   1   2   3E+3   1   1.5  RT (min)      lcorbi_t3_neg_1_050_45    m/z 343.0941     0   1   2E+3   1   1.5  RT (min)      lcorbi_t3_neg_4_039_32    m/z 343.1560     0   1   2E+4   10.5   11  RT (min)       lcorbi_t3_neg_4_102_61    m/z 343.1729     0   1   2E+3   10   10.5  RT (min)      lcorbi_t3_neg_1_013_35    m/z 343.2121     0   1E+4   9.5   10  RT (min)      lcorbi_t3_neg_4_029_31    m/z 343.2194     0   1   2   3E+3   13.5   14  RT (min)      lcorbi_t3_neg_2_037_09    m/z 343.2500     0   2   4   6E+3   12   12.5  RT (min)      lcorbi_t3_neg_4_046_37    m/z 343.2656     0   1E+4   11.5   12  RT (min)      lcorbi_t3_neg_4_032_23    m/z 343.9065     0   1E+4   8.5   9  RT (min)      lcorbi_t3_neg_1_054_37    m/z 343.9934     0   2   4   6E+4   8.5   9  RT (min)       lcorbi_t3_neg_1_015_27    m/z 343.9943     0   1   2   3   4E+5   4.5   5  RT (min)       lcorbi_t3_neg_2_007_14    m/z 344.0852     0   2   4   6   8E+3   8   8.5  RT (min)      lcorbi_t3_neg_4_023_18    m/z 344.0898     0   1   2E+3   1   1.5  RT (min)      lcorbi_t3_neg_1_025_52    m/z 344.0997     0   2   4   6E+3   1.5   2  RT (min)      lcorbi_t3_neg_4_011_24    m/z 344.1403     0   1   2E+4   12   12.5  RT (min)      lcorbi_t3_neg_3_045_34    m/z 344.1944     0   1   2   3E+3   11   11.5  RT (min)      lcorbi_t3_neg_1_053_32    m/z 344.2417     0   1E+3   11   11.5  RT (min)      lcorbi_t3_neg_4_071_57    m/z 344.2773     0   2   4   6   8E+3   10.5   11  RT (min)      lcorbi_t3_neg_4_032_23    m/z 344.2803     0   2   4   6   8E+3   14   14.5  RT (min)      lcorbi_t3_neg_1_023_03    m/z 344.9338     0   1E+4   7   7.5  RT (min)      lcorbi_t3_neg_1_054_37    m/z 344.9475     0   1   2   3E+4   1   1.5  RT (min)      lcorbi_t3_neg_3_014_19    m/z 344.9785     0   1   2E+4   0   0.5  RT (min)      lcorbi_t3_neg_1_001_49    m/z 344.9959     0   1E+4   8.5   9  RT (min)      lcorbi_t3_neg_2_004_28    m/z 345.0024     0   1   2E+4   1   1.5  RT (min)      lcorbi_t3_neg_1_043_14    m/z 345.0396     0   1   1   1.5  RT (min)       lcorbi_t3_neg_2_007_14    m/z 345.0539     0   2   4   6   8E+3   7   7.5  RT (min)      lcorbi_t3_neg_4_023_18    m/z 345.0589     0   2   4   6E+4   1   1.5  RT (min)      lcorbi_t3_neg_3_005_23    m/z 345.0687     0   1   2E+4   10   10.5  RT (min)      lcorbi_t3_neg_4_002_45    m/z 345.1451     0   2   4   6E+3   8   8.5  RT (min)      lcorbi_t3_neg_4_038_15    m/z 345.1549     0   2   4   6E+3   7   7.5  RT (min)      lcorbi_t3_neg_2_048_27    m/z 345.1712     0   1E+4   10   10.5  RT (min)      lcorbi_t3_neg_2_012_36    m/z 345.1774     0   1E+4   4.5   5  RT (min)      lcorbi_t3_neg_4_032_23    m/z 345.1840     0   2   4   6E+3   10   10.5  RT (min)      lcorbi_t3_neg_1_055_05    m/z 345.2043     0   2   4   6   8E+3   12.5   13  RT (min)      lcorbi_t3_neg_1_055_05    m/z 345.2066     0   1   2E+3   9.5   10  RT (min)      lcorbi_t3_neg_2_053_23    m/z 345.2673     0   1   2E+4   13   13.5  RT (min)      lcorbi_t3_neg_4_034_01    m/z 345.3413     0   1E+4   12.5   13  RT (min)      lcorbi_t3_neg_3_002_24    m/z 345.9276     0   1E+5   7   7.5  RT (min)       lcorbi_t3_neg_2_048_27    m/z 345.9628     0   1E+5   1.5   2  RT (min)      lcorbi_t3_neg_4_034_01    m/z 345.9987     0   1   2E+4   2   2.5  RT (min)      lcorbi_t3_neg_1_051_15    m/z 346.0478     0   2   4   6   8E+3   10   10.5  RT (min)      lcorbi_t3_neg_4_040_19    m/z 346.1305     0   1   1   1.5  RT (min)      lcorbi_t3_neg_4_032_23    m/z 346.1482     0   1   1   1.5  RT (min)      lcorbi_t3_neg_4_032_23    m/z 346.2788     0   2   4   6   8E+3   12.5   13  RT (min)      lcorbi_t3_neg_4_034_01    m/z 346.3537     0   1E+4   13   13.5  RT (min)      lcorbi_t3_neg_4_039_32    m/z 346.8167     0   1   2   3   4E+3   17.5   18  RT (min)      lcorbi_t3_neg_2_007_14    m/z 346.9308     0   1E+4   7   7.5  RT (min)      lcorbi_t3_neg_2_048_27    m/z 346.9446     0   1   1   1.5  RT (min)      lcorbi_t3_neg_4_039_32    m/z 346.9778     0   1E+3   7   7.5  RT (min)      lcorbi_t3_neg_2_007_14    m/z 346.9936     0   1E+4   8.5   9  RT (min)      lcorbi_t3_neg_4_052_26    m/z 347.0024     0   1   2E+4   1.5   2  RT (min)      lcorbi_t3_neg_1_010_17    m/z 347.0026     0   1E+4   18   18.5  RT (min)      lcorbi_t3_neg_1_043_14    m/z 347.0224     0   1E+4   9   9.5  RT (min)      lcorbi_t3_neg_2_048_27    m/z 347.0585     0   1   2   3   4E+4   1.5   2  RT (min)      lcorbi_t3_neg_3_005_23    m/z 347.0639     0   2   4   6   8E+3   10   10.5  RT (min)      lcorbi_t3_neg_2_048_27    m/z 347.0918     0   1E+3   1   1.5  RT (min)      lcorbi_t3_neg_3_048_08    m/z 347.1131     0   2   4   6E+3   7.5   8  RT (min)      lcorbi_t3_neg_4_047_30    m/z 347.1471     0   2   4   6   8E+3   6   6.5  RT (min)      lcorbi_t3_neg_4_046_37    m/z 347.1624     0   2   4E+3   9.5   10  RT (min)      lcorbi_t3_neg_4_042_14    m/z 347.1873     0   1   2E+3   11.5   12  RT (min)      lcorbi_t3_neg_2_048_27    m/z 347.1970     0   1E+4   6.5   7  RT (min)      lcorbi_t3_neg_4_029_31    m/z 347.2435     0   1E+3   11   11.5  RT (min)      lcorbi_t3_neg_1_055_05    m/z 347.2644     0   2   4   6   8E+3   13   13.5  RT (min)      lcorbi_t3_neg_4_007_13    m/z 347.9246     0   1E+4   7   7.5  RT (min)      lcorbi_t3_neg_1_054_37    m/z 347.9459     0   1   2   3   4E+3   8.5   9  RT (min)      lcorbi_t3_neg_2_048_27    m/z 347.9858     0   1E+4   9   9.5  RT (min)      lcorbi_t3_neg_2_048_27    m/z 348.0002     0   1E+4   1.5   2  RT (min)      lcorbi_t3_neg_2_054_15    m/z 348.0005     0   1E+4   18   18.5  RT (min)      lcorbi_t3_neg_1_043_14    m/z 348.0611     0   2   4   6   8E+4   1   1.5  RT (min)      lcorbi_t3_neg_2_051_34    m/z 348.0953     0   1   1   1.5  RT (min)      lcorbi_t3_neg_2_012_36    m/z 348.0974     0   2   4   6   8E+3   12.5   13  RT (min)      lcorbi_t3_neg_4_007_13    m/z 348.1404     0   1   1   1.5  RT (min)      lcorbi_t3_neg_2_051_34    m/z 348.2233     0   1   2E+3   13   13.5  RT (min)      lcorbi_t3_neg_2_007_14    m/z 348.2468     0   2   4E+3   8.5   9  RT (min)      lcorbi_t3_neg_2_048_27    m/z 348.2620     0   1   2   3   4E+3   11.5   12  RT (min)      lcorbi_t3_neg_4_042_14    m/z 348.8874     0   2   4   6   8E+3   2   2.5  RT (min)      lcorbi_t3_neg_4_032_23    m/z 348.8972     0   1   2   2.5  RT (min)      lcorbi_t3_neg_3_053_14    m/z 348.9001     0   1   2E+4   4.5   5  RT (min)      lcorbi_t3_neg_3_022_27    m/z 348.9393     0   2   4   6E+3   8   8.5  RT (min)      lcorbi_t3_neg_2_048_27    m/z 348.9881     0   2   4   6E+3   1   1.5  RT (min)      lcorbi_t3_neg_2_043_33    m/z 348.9896     0   2   4E+3   6.5   7  RT (min)      lcorbi_t3_neg_4_021_25    m/z 349.0122     0   1E+4   4.5   5  RT (min)      lcorbi_t3_neg_1_043_14    m/z 349.0325     0   2   4   6E+3   8   8.5  RT (min)      lcorbi_t3_neg_4_013_09    m/z 349.0609     0   1   10   10.5  RT (min)      lcorbi_t3_neg_4_023_18    m/z 349.0728     0   1E+4   1.5   2  RT (min)      lcorbi_t3_neg_2_048_27    m/z 349.1138     0   1   2   3   4E+3   10   10.5  RT (min)      lcorbi_t3_neg_2_006_10    m/z 349.2622     0   1   2E+4   12.5   13  RT (min)      lcorbi_t3_neg_4_034_01    m/z 349.2744     0   1   2   3E+4   12.5   13  RT (min)      lcorbi_t3_neg_4_010_28    m/z 349.2937     0   1   2   3E+4   12   12.5  RT (min)      lcorbi_t3_neg_3_024_26    m/z 349.7927     0   2   4   6E+3   8.5   9  RT (min)      lcorbi_t3_neg_2_003_42    m/z 350.0608     0   1E+3   1   1.5  RT (min)      lcorbi_t3_neg_1_021_31    m/z 350.0931     0   1E+3   1   1.5  RT (min)      lcorbi_t3_neg_2_024_37    m/z 350.0981     0   1E+4   9   9.5  RT (min)      lcorbi_t3_neg_3_019_12    m/z 350.1001     0   1E+4   6.5   7  RT (min)      lcorbi_t3_neg_2_051_34    m/z 350.1354     0   1   2E+4   5   5.5  RT (min)      lcorbi_t3_neg_4_001_49    m/z 350.1458     0   2   4   6   8E+3   10   10.5  RT (min)      lcorbi_t3_neg_4_023_18    m/z 350.1463     0   1   1   1.5  RT (min)      lcorbi_t3_neg_2_024_37    m/z 350.1831     0   1   2   3E+3   11.5   12  RT (min)      lcorbi_t3_neg_4_011_24    m/z 350.2049     0   2   4   6E+3   7   7.5  RT (min)      lcorbi_t3_neg_4_023_18    m/z 350.2389     0   1   2   3   4E+3   13   13.5  RT (min)      lcorbi_t3_neg_4_034_01    m/z 350.2542     0   1E+4   12   12.5  RT (min)      lcorbi_t3_neg_4_029_31    m/z 350.2777     0   2   4   6E+3   12.5   13  RT (min)      lcorbi_t3_neg_4_010_28    m/z 350.8845     0   2   4E+3   2   2.5  RT (min)      lcorbi_t3_neg_4_032_23    m/z 350.9139     0   1   2E+4   1   1.5  RT (min)      lcorbi_t3_neg_4_039_32    m/z 350.9445     0   2   4   6   8E+3   8.5   9  RT (min)      lcorbi_t3_neg_2_048_27    m/z 351.0081     0   1E+4   14   14.5  RT (min)      lcorbi_t3_neg_1_023_03    m/z 351.0148     0   2   4   6E+3   1   1.5  RT (min)      lcorbi_t3_neg_3_035_03    m/z 351.0767     0   2   4E+3   10.5   11  RT (min)      lcorbi_t3_neg_2_028_38    m/z 351.0919     0   1   2E+4   9   9.5  RT (min)      lcorbi_t3_neg_3_019_12    m/z 351.1423     0   1   10   10.5  RT (min)      lcorbi_t3_neg_4_067_57    m/z 351.1476     0   1   2   3   4E+3   12.5   13  RT (min)      lcorbi_t3_neg_4_007_13    m/z 351.1785     0   1   2   3E+3   11.5   12  RT (min)      lcorbi_t3_neg_3_002_24    m/z 351.1841     0   1E+4   10.5   11  RT (min)      lcorbi_t3_neg_4_039_32    m/z 351.2648     0   1   2   3   4E+4   12   12.5  RT (min)      lcorbi_t3_neg_4_007_13    m/z 351.2801     0   1E+4   12   12.5  RT (min)      lcorbi_t3_neg_4_011_24    m/z 351.2899     0   1   2E+4   13   13.5  RT (min)      lcorbi_t3_neg_3_007_16    m/z 351.7897     0   2   4   6   8E+3   8.5   9  RT (min)      lcorbi_t3_neg_4_007_13    m/z 351.8429     0   1   2   3E+3   17.5   18  RT (min)      lcorbi_t3_neg_3_030_33    m/z 351.8985     0   1   2   3E+4   4.5   5  RT (min)      lcorbi_t3_neg_3_022_27    m/z 351.9968     0   1E+4   18.5   19  RT (min)      lcorbi_t3_neg_4_042_14    m/z 352.0138     0   1E+4   1.5   2  RT (min)      lcorbi_t3_neg_3_041_54    m/z 352.0283     0   1E+4   4.5   5  RT (min)      lcorbi_t3_neg_2_034_19    m/z 352.0765     0   1   2E+2   1   1.5  RT (min)      lcorbi_t3_neg_4_011_24    m/z 352.0855     0   1E+4   6.5   7  RT (min)      lcorbi_t3_neg_4_034_01    m/z 352.0918     0   1   2E+3   1   1.5  RT (min)      lcorbi_t3_neg_4_011_24    m/z 352.1124     0   1E+4   1   1.5  RT (min)      lcorbi_t3_neg_3_002_24    m/z 352.1343     0   1   2   3E+3   12.5   13  RT (min)      lcorbi_t3_neg_4_007_13    m/z 352.1664     0   2   4   6E+3   6   6.5  RT (min)      lcorbi_t3_neg_4_007_13    m/z 352.2125     0   1E+4   10   10.5  RT (min)      lcorbi_t3_neg_4_023_18    m/z 352.2206     0   1E+4   8   8.5  RT (min)      lcorbi_t3_neg_4_023_18    m/z 352.2546     0   2   4   6E+3   13.5   14  RT (min)      lcorbi_t3_neg_4_040_19    m/z 352.2851     0   1E+4   14   14.5  RT (min)      lcorbi_t3_neg_1_023_03    m/z 352.2852     0   1   2E+3   12.5   13  RT (min)      lcorbi_t3_neg_4_034_01    m/z 352.2977     0   1   2E+4   12   12.5  RT (min)      lcorbi_t3_neg_3_002_24    m/z 352.8607     0   2   4   6   8E+4   1   1.5  RT (min)      lcorbi_t3_neg_1_055_05    m/z 352.9416     0   2   4E+3   8.5   9  RT (min)      lcorbi_t3_neg_2_048_27    m/z 352.9584     0   1   1   1.5  RT (min)      lcorbi_t3_neg_2_054_15    m/z 352.9967     0   2   4   6E+3   1.5   2  RT (min)      lcorbi_t3_neg_3_044_01    m/z 353.0360     0   1   2   3   4E+2   1   1.5  RT (min)      lcorbi_t3_neg_2_054_15    m/z 353.0694     0   1   2   3E+3   5.5   6  RT (min)      lcorbi_t3_neg_4_031_48    m/z 353.0696     0   2   4E+3   9   9.5  RT (min)      lcorbi_t3_neg_4_034_01    m/z 353.1032     0   1   1   1.5  RT (min)      lcorbi_t3_neg_4_011_24    m/z 353.1477     0   1   2E+4   10   10.5  RT (min)      lcorbi_t3_neg_4_023_18    m/z 353.1862     0   1   11   11.5  RT (min)      lcorbi_t3_neg_4_019_08    m/z 353.1999     0   1   2E+4   14   14.5  RT (min)      lcorbi_t3_neg_4_032_23    m/z 353.2328     0   1E+3   10   10.5  RT (min)      lcorbi_t3_neg_4_032_23    m/z 353.2653     0   1   2E+4   10.5   11  RT (min)      lcorbi_t3_neg_2_053_23    m/z 353.2693     0   1E+4   13   13.5  RT (min)      lcorbi_t3_neg_4_007_13    m/z 353.2805     0   1E+4   13   13.5  RT (min)      lcorbi_t3_neg_4_034_01    m/z 353.3056     0   1E+3   13   13.5  RT (min)      lcorbi_t3_neg_4_038_15    m/z 353.7867     0   2   4   6E+3   8.5   9  RT (min)      lcorbi_t3_neg_4_007_13    m/z 353.8109     0   1   2   3   4E+3   0.5   1  RT (min)      lcorbi_t3_neg_1_013_35    m/z 353.9268     0   1   2   3   4E+3   17.5   18  RT (min)      lcorbi_t3_neg_2_020_05    m/z 353.9354     0   2   4   6E+3   8.5   9  RT (min)      lcorbi_t3_neg_2_048_27    m/z 353.9593     0   1   2   3   4E+3   7   7.5  RT (min)      lcorbi_t3_neg_2_038_22    m/z 353.9950     0   1   2   3   4E+3   18.5   19  RT (min)      lcorbi_t3_neg_3_010_15    m/z 354.0131     0   1E+4   1.5   2  RT (min)      lcorbi_t3_neg_4_034_01    m/z 354.0587     0   1   2   3   4E+3   1   1.5  RT (min)      lcorbi_t3_neg_2_009_50    m/z 354.0801     0   2   4   6E+3   1.5   2  RT (min)      lcorbi_t3_neg_1_052_24    m/z 354.1270     0   2   4   6   8E+3   5.5   6  RT (min)      lcorbi_t3_neg_4_001_49    m/z 354.1344     0   2   4   6   8E+3   9.5   10  RT (min)      lcorbi_t3_neg_3_019_12    m/z 354.1446     0   2   4   6   8E+3   12.5   13  RT (min)      lcorbi_t3_neg_4_007_13    m/z 354.1667     0   2   4   6E+3   13   13.5  RT (min)      lcorbi_t3_neg_3_018_13    m/z 354.2151     0   2   4   6E+3   12   12.5  RT (min)      lcorbi_t3_neg_4_044_38    m/z 354.2151     0   2   4   6   8E+3   7.5   8  RT (min)      lcorbi_t3_neg_4_017_51    m/z 354.2393     0   1E+4   6.5   7  RT (min)      lcorbi_t3_neg_4_029_31    m/z 354.8485     0   2   4E+3   17.5   18  RT (min)      lcorbi_t3_neg_2_003_42    m/z 354.9932     0   2   4   6E+3   1.5   2  RT (min)      lcorbi_t3_neg_4_038_15    m/z 354.9991     0   2   4   6E+2   3.5   4  RT (min)      lcorbi_t3_neg_4_038_15    m/z 355.0067     0   1E+4   1.5   2  RT (min)      lcorbi_t3_neg_4_034_01    m/z 355.0282     0   1E+4   9   9.5  RT (min)      lcorbi_t3_neg_1_001_49    m/z 355.0698     0   1   1.5   2  RT (min)      lcorbi_t3_neg_2_026_03    m/z 355.0753     0   2   4   6   8E+3   4.5   5  RT (min)      lcorbi_t3_neg_3_007_16    m/z 355.1111     0   1E+4   10   10.5  RT (min)      lcorbi_t3_neg_4_007_13    m/z 355.1453     0   1   13   13.5  RT (min)      lcorbi_t3_neg_4_019_08    m/z 355.1674     0   1E+4   13   13.5  RT (min)      lcorbi_t3_neg_3_018_13    m/z 355.2053     0   1E+4   10   10.5  RT (min)      lcorbi_t3_neg_4_032_23    m/z 355.2274     0   1   2   3E+3   12   12.5  RT (min)      lcorbi_t3_neg_4_007_13    m/z 355.2293     0   1   2   3   4E+3   9.5   10  RT (min)      lcorbi_t3_neg_2_048_27    m/z 355.2623     0   2   4   6   8E+3   10.5   11  RT (min)      lcorbi_t3_neg_2_053_23    m/z 355.9078     0   2   4   6E+2   1   1.5  RT (min)      lcorbi_t3_neg_3_040_05    m/z 356.0746     0   2   4   6   8E+3   10   10.5  RT (min)      lcorbi_t3_neg_4_029_31    m/z 356.1287     0   1E+4   9   9.5  RT (min)      lcorbi_t3_neg_3_019_12    m/z 356.1291     0   1E+4   12.5   13  RT (min)      lcorbi_t3_neg_4_007_13    m/z 356.1434     0   1   2   3   4E+3   5   5.5  RT (min)      lcorbi_t3_neg_4_044_38    m/z 356.1710     0   1E+4   9.5   10  RT (min)      lcorbi_t3_neg_2_048_27    m/z 356.1910     0   2   4E+3   7   7.5  RT (min)      lcorbi_t3_neg_4_023_18    m/z 356.1989     0   1   2   3   4E+3   10   10.5  RT (min)      lcorbi_t3_neg_4_032_23    m/z 356.2261     0   1E+4   11   11.5  RT (min)      lcorbi_t3_neg_4_042_14    m/z 356.2883     0   2   4   6   8E+3   13.5   14  RT (min)      lcorbi_t3_neg_4_034_01    m/z 356.2996     0   2   4   6   8E+3   13   13.5  RT (min)      lcorbi_t3_neg_4_034_01    m/z 356.3247     0   1   2   3E+3   12.5   13  RT (min)      lcorbi_t3_neg_4_032_23    m/z 356.8241     0   1   1   1.5  RT (min)      lcorbi_t3_neg_4_036_16    m/z 356.8455     0   2   4E+3   17.5   18  RT (min)      lcorbi_t3_neg_4_049_55    m/z 356.9235     0   1E+4   18   18.5  RT (min)      lcorbi_t3_neg_1_008_08    m/z 356.9921     0   1E+4   18   18.5  RT (min)      lcorbi_t3_neg_2_007_14    m/z 357.0611     0   1   2E+3   8   8.5  RT (min)      lcorbi_t3_neg_3_024_26    m/z 357.0856     0   1E+4   1.5   2  RT (min)      lcorbi_t3_neg_2_022_24    m/z 357.0935     0   2   4   6E+4   1   1.5  RT (min)      lcorbi_t3_neg_1_055_05    m/z 357.1075     0   1   2E+4   1   1.5  RT (min)      lcorbi_t3_neg_2_035_46    m/z 357.1411     0   1E+3   1   1.5  RT (min)      lcorbi_t3_neg_1_055_05    m/z 357.1689     0   1E+3   13   13.5  RT (min)      lcorbi_t3_neg_1_054_37    m/z 357.1874     0   1E+4   7   7.5  RT (min)      lcorbi_t3_neg_4_023_18    m/z 357.1875     0   2   4   6E+3   11   11.5  RT (min)      lcorbi_t3_neg_4_023_18    m/z 357.2066     0   1E+3   13   13.5  RT (min)      lcorbi_t3_neg_2_051_34    m/z 357.2173     0   2   4E+3   12.5   13  RT (min)      lcorbi_t3_neg_4_029_31    m/z 357.2179     0   1   2   3E+3   10   10.5  RT (min)      lcorbi_t3_neg_3_012_47    m/z 357.2671     0   2   4E+2   13   13.5  RT (min)      lcorbi_t3_neg_4_005_27    m/z 357.2794     0   2   4   6E+3   13.5   14  RT (min)      lcorbi_t3_neg_4_010_28    m/z 357.8234     0   2   4   6E+3   7.5   8  RT (min)      lcorbi_t3_neg_1_022_34    m/z 357.8383     0   2   4   6   8E+3   1   1.5  RT (min)      lcorbi_t3_neg_4_007_13    m/z 357.8773     0   2   4   6   8E+3   17.5   18  RT (min)      lcorbi_t3_neg_2_043_33    m/z 357.8779     0   2   4   6   8E+3   7.5   8  RT (min)      lcorbi_t3_neg_3_054_18    m/z 358.0099     0   1   2E+4   2   2.5  RT (min)      lcorbi_t3_neg_3_007_16    m/z 358.0873     0   2   4   6E+3   9.5   10  RT (min)      lcorbi_t3_neg_2_048_27    m/z 358.0928     0   1E+4   1.5   2  RT (min)      lcorbi_t3_neg_3_002_24    m/z 358.0961     0   2   4E+3   10   10.5  RT (min)      lcorbi_t3_neg_2_048_27    m/z 358.1056     0   1   2E+4   1   1.5  RT (min)      lcorbi_t3_neg_3_043_25    m/z 358.1221     0   1   2   3   4E+4   5   5.5  RT (min)      lcorbi_t3_neg_2_048_27    m/z 358.1285     0   1E+4   9   9.5  RT (min)      lcorbi_t3_neg_3_019_12    m/z 358.1593     0   1   2   3E+3   13   13.5  RT (min)      lcorbi_t3_neg_3_001_49    m/z 358.1650     0   1E+4   9   9.5  RT (min)      lcorbi_t3_neg_2_048_27    m/z 358.2015     0   2   4E+3   7   7.5  RT (min)      lcorbi_t3_neg_4_034_01    m/z 358.2173     0   2   4   6   8E+3   5.5   6  RT (min)      lcorbi_t3_neg_4_032_23    m/z 358.2272     0   2   4   6   8E+3   12   12.5  RT (min)      lcorbi_t3_neg_4_005_27    m/z 358.2401     0   2   4E+3   10.5   11  RT (min)      lcorbi_t3_neg_2_006_10    m/z 358.9078     0   1E+4   7.5   8  RT (min)      lcorbi_t3_neg_4_023_18    m/z 359.0143     0   1E+4   13.5   14  RT (min)      lcorbi_t3_neg_1_023_03    m/z 359.0177     0   1   2E+4   2   2.5  RT (min)      lcorbi_t3_neg_1_010_17    m/z 359.1184     0   1E+4   1   1.5  RT (min)      lcorbi_t3_neg_2_035_46    m/z 359.1208     0   2   4   6E+3   9   9.5  RT (min)      lcorbi_t3_neg_3_019_12    m/z 359.1494     0   1   2E+3   5.5   6  RT (min)      lcorbi_t3_neg_4_056_56    m/z 359.1563     0   1E+4   1   1.5  RT (min)      lcorbi_t3_neg_1_055_05    m/z 359.2223     0   1E+4   9.5   10  RT (min)      lcorbi_t3_neg_2_048_27    m/z 359.2952     0   1   2   3E+3   13   13.5  RT (min)      lcorbi_t3_neg_2_054_15    m/z 359.3104     0   1E+4   11   11.5  RT (min)      lcorbi_t3_neg_4_011_24    m/z 359.8759     0   1E+4   7.5   8  RT (min)      lcorbi_t3_neg_3_054_18    m/z 359.9701     0   1   2   3E+4   8.5   9  RT (min)      lcorbi_t3_neg_2_048_27    m/z 360.0002     0   1E+4   5   5.5  RT (min)      lcorbi_t3_neg_3_019_12    m/z 360.0492     0   1E+4   4.5   5  RT (min)      lcorbi_t3_neg_3_019_12    m/z 360.0509     0   1E+3   1.5   2  RT (min)      lcorbi_t3_neg_4_045_35    m/z 360.1160     0   2   4E+3   12.5   13  RT (min)      lcorbi_t3_neg_4_017_51    m/z 360.1813     0   1   2E+4   9   9.5  RT (min)      lcorbi_t3_neg_2_048_27    m/z 360.2008     0   1   2E+4   8   8.5  RT (min)      lcorbi_t3_neg_2_048_27    m/z 360.2386     0   1E+5   7   7.5  RT (min)       lcorbi_t3_neg_4_071_57    m/z 360.2559     0   1   2   3   4E+3   9.5   10  RT (min)      lcorbi_t3_neg_2_006_10    m/z 360.2751     0   1   2E+3   10   10.5  RT (min)      lcorbi_t3_neg_4_053_43    m/z 360.9058     0   1E+4   7.5   8  RT (min)      lcorbi_t3_neg_4_023_18    m/z 361.0191     0   2   4   6E+3   3   3.5  RT (min)      lcorbi_t3_neg_3_019_12    m/z 361.0732     0   2   4   6   8E+3   8   8.5  RT (min)      lcorbi_t3_neg_4_023_18    m/z 361.0906     0   1E+3   1   1.5  RT (min)      lcorbi_t3_neg_4_011_24    m/z 361.1139     0   2   4   6E+4   1   1.5  RT (min)      lcorbi_t3_neg_4_011_24    m/z 361.1509     0   1   2   3   4E+3   9.5   10  RT (min)      lcorbi_t3_neg_2_028_38    m/z 361.1911     0   2   4E+3   13   13.5  RT (min)      lcorbi_t3_neg_2_007_14    m/z 361.2165     0   1E+3   11   11.5  RT (min)      lcorbi_t3_neg_4_034_01    m/z 361.2226     0   1E+4   6.5   7  RT (min)      lcorbi_t3_neg_4_034_01    m/z 361.2493     0   2   4E+4   10.5   11  RT (min)      lcorbi_t3_neg_4_034_01    m/z 361.2571     0   1   12   12.5  RT (min)      lcorbi_t3_neg_4_007_13    m/z 361.3260     0   1E+4   12.5   13  RT (min)      lcorbi_t3_neg_4_011_24    m/z 361.8713     0   1E+4   1   1.5  RT (min)      lcorbi_t3_neg_2_024_37    m/z 361.8892     0   2   4E+3   18   18.5  RT (min)      lcorbi_t3_neg_4_074_58    m/z 361.9185     0   1E+4   1   1.5  RT (min)      lcorbi_t3_neg_4_046_37    m/z 361.9671     0   1E+4   8.5   9  RT (min)      lcorbi_t3_neg_2_048_27    m/z 361.9972     0   2   4   6   8E+3   5   5.5  RT (min)      lcorbi_t3_neg_3_019_12    m/z 362.0774     0   1   1   1.5  RT (min)       lcorbi_t3_neg_4_011_24    m/z 362.1237     0   1E+4   1   1.5  RT (min)      lcorbi_t3_neg_1_052_24    m/z 362.1578     0   1   1   1.5  RT (min)      lcorbi_t3_neg_4_039_32    m/z 362.1695     0   2   4   6   8E+3   10.5   11  RT (min)      lcorbi_t3_neg_4_029_31    m/z 362.2366     0   1E+4   11   11.5  RT (min)      lcorbi_t3_neg_2_048_27    m/z 362.7487     0   1   1   1.5  RT (min)      lcorbi_t3_neg_3_053_14    m/z 362.8168     0   1E+4   7.5   8  RT (min)      lcorbi_t3_neg_4_023_18    m/z 362.8748     0   2   4   6E+3   0.5   1  RT (min)      lcorbi_t3_neg_1_010_17    m/z 362.8749     0   1E+4   18   18.5  RT (min)      lcorbi_t3_neg_1_001_49    m/z 363.0079     0   2   4   6E+3   5.5   6  RT (min)      lcorbi_t3_neg_4_017_51    m/z 363.0158     0   2   4   6   8E+3   9   9.5  RT (min)      lcorbi_t3_neg_4_017_51    m/z 363.1687     0   2   4   6   8E+3   9.5   10  RT (min)      lcorbi_t3_neg_3_019_12    m/z 363.2245     0   1E+4   6   6.5  RT (min)      lcorbi_t3_neg_4_044_38    m/z 363.2345     0   2   4E+4   11   11.5  RT (min)      lcorbi_t3_neg_2_007_14    m/z 363.2536     0   1   2E+3   11.5   12  RT (min)      lcorbi_t3_neg_1_055_05    m/z 363.2646     0   1E+4   12.5   13  RT (min)      lcorbi_t3_neg_1_055_05    m/z 363.3521     0   1E+4   11   11.5  RT (min)      lcorbi_t3_neg_4_039_32    m/z 363.9809     0   1   2E+4   1   1.5  RT (min)      lcorbi_t3_neg_3_037_36    m/z 363.9933     0   2   4   6   8E+3   8.5   9  RT (min)      lcorbi_t3_neg_2_048_27    m/z 364.1956     0   2   4   6   8E+3   7.5   8  RT (min)      lcorbi_t3_neg_2_035_46    m/z 364.2115     0   1   2   3E+3   13   13.5  RT (min)      lcorbi_t3_neg_4_007_13    m/z 364.2182     0   1   2E+4   6   6.5  RT (min)      lcorbi_t3_neg_4_044_38    m/z 364.2182     0   1E+4   11   11.5  RT (min)      lcorbi_t3_neg_4_042_14    m/z 364.2254     0   1   10.5   11  RT (min)      lcorbi_t3_neg_2_022_24    m/z 364.2620     0   1E+4   11   11.5  RT (min)      lcorbi_t3_neg_2_053_23    m/z 364.8142     0   1   2E+4   7.5   8  RT (min)      lcorbi_t3_neg_4_023_18    m/z 364.8514     0   1   2E+4   7.5   8  RT (min)      lcorbi_t3_neg_1_042_18    m/z 364.8773     0   1E+4   18   18.5  RT (min)      lcorbi_t3_neg_3_019_12    m/z 364.9237     0   2   4   6E+3   9   9.5  RT (min)      lcorbi_t3_neg_4_007_13    m/z 364.9445     0   1   2   3E+3   18   18.5  RT (min)      lcorbi_t3_neg_1_014_28    m/z 365.0058     0   2   4   6E+3   5.5   6  RT (min)      lcorbi_t3_neg_4_017_51    m/z 365.0225     0   2   4   6E+3   9   9.5  RT (min)      lcorbi_t3_neg_4_017_51    m/z 365.0304     0   2   4   6   8E+3   2   2.5  RT (min)      lcorbi_t3_neg_4_050_02    m/z 365.0481     0   2   4   6   8E+3   10   10.5  RT (min)      lcorbi_t3_neg_3_004_30    m/z 365.0645     0   1   10   10.5  RT (min)      lcorbi_t3_neg_4_007_13    m/z 365.0682     0   1E+4   1.5   2  RT (min)      lcorbi_t3_neg_4_007_13    m/z 365.1422     0   1   2   3   4E+3   7   7.5  RT (min)      lcorbi_t3_neg_4_034_01    m/z 365.1771     0   1E+4   9.5   10  RT (min)      lcorbi_t3_neg_2_048_27    m/z 365.2062     0   2   4E+3   13   13.5  RT (min)      lcorbi_t3_neg_4_007_13    m/z 365.2095     0   2   4   6E+3   9.5   10  RT (min)      lcorbi_t3_neg_4_034_01    m/z 365.2305     0   1E+3   12   12.5  RT (min)      lcorbi_t3_neg_4_032_23    m/z 365.2501     0   1E+4   11.5   12  RT (min)      lcorbi_t3_neg_4_042_14    m/z 365.2750     0   1   13   13.5  RT (min)      lcorbi_t3_neg_1_055_05    m/z 365.9116     0   2   4   6E+3   17.5   18  RT (min)      lcorbi_t3_neg_3_042_22    m/z 365.9210     0   1   2   3E+4   9   9.5   10  RT (min)      lcorbi_t3_neg_4_007_13    m/z 365.9533     0   2   4   6E+3   8.5   9  RT (min)      lcorbi_t3_neg_1_054_37    m/z 365.9903     0   1   2   3   4E+3   8.5   9  RT (min)      lcorbi_t3_neg_2_048_27    m/z 366.0125     0   2   4E+3   2   2.5  RT (min)      lcorbi_t3_neg_1_002_16    m/z 366.0295     0   2   4   6E+3   2   2.5  RT (min)      lcorbi_t3_neg_4_050_02    m/z 366.0720     0   1   1   1.5  RT (min)      lcorbi_t3_neg_2_048_27    m/z 366.1166     0   1E+4   10   10.5  RT (min)      lcorbi_t3_neg_4_023_18    m/z 366.1416     0   1   1   1.5  RT (min)      lcorbi_t3_neg_2_048_27    m/z 366.1433     0   2   4   6E+3   10   10.5  RT (min)      lcorbi_t3_neg_4_007_13    m/z 366.1602     0   1E+4   10.5   11  RT (min)      lcorbi_t3_neg_3_032_17    m/z 366.1665     0   2   4   6E+3   5   5.5  RT (min)      lcorbi_t3_neg_2_035_46    m/z 366.1965     0   2   4   6   8E+3   10   10.5  RT (min)      lcorbi_t3_neg_2_002_26    m/z 366.2396     0   2   4   6E+3   11.5   12  RT (min)      lcorbi_t3_neg_4_032_23    m/z 366.2644     0   1   2   3E+3   11.5   12  RT (min)      lcorbi_t3_neg_4_029_31    m/z 366.2726     0   1   11.5   12  RT (min)      lcorbi_t3_neg_1_052_24    m/z 366.8116     0   1   2E+4   7.5   8  RT (min)      lcorbi_t3_neg_4_023_18    m/z 366.8743     0   2   4   6   8E+3   18   18.5  RT (min)      lcorbi_t3_neg_4_025_52    m/z 366.9840     0   2   4   6   8E+2   7   7.5  RT (min)      lcorbi_t3_neg_4_042_14    m/z 367.0096     0   1E+4   1.5   2  RT (min)      lcorbi_t3_neg_4_036_16    m/z 367.1006     0   1E+4   9   9.5  RT (min)      lcorbi_t3_neg_4_023_18    m/z 367.1289     0   1E+5   6   6.5  RT (min)      lcorbi_t3_neg_2_048_27    m/z 367.1444     0   1E+4   7.5   8  RT (min)      lcorbi_t3_neg_2_048_27    m/z 367.1576     0   1   2   3   4E+4   13   13.5  RT (min)      lcorbi_t3_neg_1_055_05    m/z 367.1580     0   1E+6   8   8.5  RT (min)       lcorbi_t3_neg_4_034_01    m/z 367.1687     0   1E+4   7.5   8  RT (min)      lcorbi_t3_neg_4_005_27    m/z 367.1835     0   1   2   3E+4   1   1.5  RT (min)      lcorbi_t3_neg_2_022_24    m/z 367.2010     0   1E+4   4.5   5  RT (min)      lcorbi_t3_neg_4_011_24    m/z 367.2155     0   1E+3   10.5   11  RT (min)      lcorbi_t3_neg_2_007_14    m/z 367.2959     0   1   2   3E+3   13   13.5  RT (min)      lcorbi_t3_neg_4_005_27    m/z 367.3577     0   2   4   6E+3   14.5   15  RT (min)      lcorbi_t3_neg_2_056_56    m/z 367.9060     0   1E+5   16   16.5  RT (min)       lcorbi_t3_neg_2_026_03    m/z 367.9181     0   2   4   6E+3   9   9.5  RT (min)      lcorbi_t3_neg_4_007_13    m/z 367.9600     0   1   2   3   4E+2   3   3.5  RT (min)      lcorbi_t3_neg_4_042_14    m/z 368.0088     0   1E+4   1.5   2  RT (min)      lcorbi_t3_neg_3_044_01    m/z 368.0289     0   1   2E+2   2   2.5  RT (min)      lcorbi_t3_neg_4_042_14    m/z 368.0664     0   1E+4   8.5   9  RT (min)      lcorbi_t3_neg_4_052_26    m/z 368.0861     0   1   1   1.5  RT (min)      lcorbi_t3_neg_2_048_27    m/z 368.0990     0   1   5   5.5  RT (min)      lcorbi_t3_neg_4_039_32    m/z 368.1356     0   1   2   3E+3   10   10.5  RT (min)      lcorbi_t3_neg_4_044_38    m/z 368.1718     0   1   2   3   4E+3   7.5   8  RT (min)      lcorbi_t3_neg_4_042_14    m/z 368.2494     0   2   4   6   8E+3   12.5   13  RT (min)      lcorbi_t3_neg_2_039_31    m/z 368.2802     0   1E+4   12   12.5  RT (min)      lcorbi_t3_neg_4_046_37    m/z 368.3165     0   2   4   6E+3   13   13.5  RT (min)      lcorbi_t3_neg_4_034_01    m/z 368.8094     0   1E+4   7.5   8  RT (min)      lcorbi_t3_neg_4_023_18    m/z 368.8465     0   2   4   6   8E+3   7.5   8  RT (min)      lcorbi_t3_neg_1_042_18    m/z 368.8891     0   1E+3   18   18.5  RT (min)      lcorbi_t3_neg_4_019_08    m/z 368.9066     0   1E+4   17.5   18  RT (min)      lcorbi_t3_neg_3_032_17    m/z 369.0049     0   2   4E+4   5   5.5  RT (min)       lcorbi_t3_neg_3_025_52    m/z 369.0050     0   1   2   3   4E+4   10.5   11  RT (min)      lcorbi_t3_neg_2_048_27    m/z 369.0223     0   1   2E+3   2   2.5  RT (min)      lcorbi_t3_neg_4_037_29    m/z 369.0734     0   1   1   1.5  RT (min)      lcorbi_t3_neg_2_048_27    m/z 369.1203     0   1E+4   1   1.5  RT (min)      lcorbi_t3_neg_2_048_27    m/z 369.1291     0   2   4   6   8E+3   10   10.5  RT (min)      lcorbi_t3_neg_4_029_31    m/z 369.1599     0   1E+4   10   10.5  RT (min)      lcorbi_t3_neg_2_002_26    m/z 369.1957     0   2   4   6   8E+4   12.5   13  RT (min)      lcorbi_t3_neg_1_055_05    m/z 369.2025     0   2   4E+3   9.5   10  RT (min)      lcorbi_t3_neg_4_023_18    m/z 369.2278     0   2   4   6E+2   8.5   9  RT (min)      lcorbi_t3_neg_4_007_13    m/z 369.2386     0   1E+4   13.5   14  RT (min)      lcorbi_t3_neg_2_035_46    m/z 369.2446     0   1E+4   12   12.5  RT (min)      lcorbi_t3_neg_4_007_13    m/z 369.2754     0   1   2E+4   11.5   12  RT (min)      lcorbi_t3_neg_4_007_13    m/z 369.3370     0   1E+4   13   13.5  RT (min)      lcorbi_t3_neg_4_034_01    m/z 369.9043     0   2   4   6E+3   7.5   8  RT (min)      lcorbi_t3_neg_2_007_14    m/z 369.9235     0   1E+4   1   1.5  RT (min)      lcorbi_t3_neg_4_045_35    m/z 369.9824     0   2   4   6   8E+3   18   18.5  RT (min)      lcorbi_t3_neg_1_043_14    m/z 370.0078     0   1E+4   1.5   2  RT (min)      lcorbi_t3_neg_3_017_51    m/z 370.0735     0   1   2E+3   9   9.5  RT (min)      lcorbi_t3_neg_3_019_12    m/z 370.1301     0   2   4E+3   10.5   11  RT (min)      lcorbi_t3_neg_4_007_13    m/z 370.1660     0   1   2   3E+3   7.5   8  RT (min)      lcorbi_t3_neg_2_054_15    m/z 370.1841     0   1   12.5   13  RT (min)      lcorbi_t3_neg_4_005_27    m/z 370.2779     0   1   2   3E+4   13   13.5  RT (min)      lcorbi_t3_neg_4_029_31    m/z 370.8009     0   2   4   6   8E+3   1   1.5  RT (min)      lcorbi_t3_neg_4_046_37    m/z 370.8943     0   2   4   6E+3   9   9.5  RT (min)      lcorbi_t3_neg_4_005_27    m/z 371.0017     0   2   4   6   8E+3   1.5   2  RT (min)      lcorbi_t3_neg_3_044_01    m/z 371.0145     0   2   4   6E+3   8   8.5  RT (min)      lcorbi_t3_neg_4_013_09    m/z 371.0235     0   2   4E+4   1   1.5  RT (min)      lcorbi_t3_neg_2_048_27    m/z 371.0654     0   1   2E+3   10   10.5  RT (min)      lcorbi_t3_neg_4_019_08    m/z 371.0883     0   1E+4   1   1.5  RT (min)      lcorbi_t3_neg_2_035_46    m/z 371.1237     0   2   4   6E+3   10.5   11  RT (min)      lcorbi_t3_neg_4_102_61    m/z 371.1319     0   1   2E+3   9   9.5  RT (min)      lcorbi_t3_neg_4_056_56    m/z 371.1485     0   1E+4   10   10.5  RT (min)      lcorbi_t3_neg_4_023_18    m/z 371.1797     0   1E+4   8   8.5  RT (min)      lcorbi_t3_neg_2_048_27    m/z 371.1874     0   1   2E+4   11.5   12  RT (min)       lcorbi_t3_neg_4_102_61    m/z 371.2434     0   2   4   6   8E+3   10.5   11  RT (min)      lcorbi_t3_neg_4_029_31    m/z 371.2605     0   2   4   6   8E+3   13   13.5  RT (min)      lcorbi_t3_neg_4_054_36    m/z 371.2814     0   1   2E+3   13   13.5  RT (min)      lcorbi_t3_neg_2_043_33    m/z 371.3274     0   1   2   3   4E+3   10.5   11  RT (min)      lcorbi_t3_neg_4_029_31    m/z 371.3482     0   1E+4   13.5   14  RT (min)      lcorbi_t3_neg_3_002_24    m/z 371.9005     0   1E+4   18   18.5  RT (min)      lcorbi_t3_neg_2_048_27    m/z 371.9471     0   1   2E+4   1   1.5  RT (min)      lcorbi_t3_neg_2_048_27    m/z 371.9819     0   1   0   0.5  RT (min)      lcorbi_t3_neg_4_011_24    m/z 372.1033     0   1E+4   1   1.5  RT (min)      lcorbi_t3_neg_3_002_24    m/z 372.1241     0   2   4   6   8E+3   12.5   13  RT (min)      lcorbi_t3_neg_4_007_13    m/z 372.1372     0   1E+4   13   13.5  RT (min)      lcorbi_t3_neg_4_038_15    m/z 372.1715     0   1   2E+4   12.5   13  RT (min)      lcorbi_t3_neg_4_005_27    m/z 372.1728     0   1   2   3   4E+3   8   8.5  RT (min)      lcorbi_t3_neg_4_034_01    m/z 372.1882     0   1   2E+2   4.5   5  RT (min)      lcorbi_t3_neg_4_011_24    m/z 372.1926     0   2   4E+3   8   8.5  RT (min)      lcorbi_t3_neg_2_048_27    m/z 372.2233     0   1   13   13.5  RT (min)      lcorbi_t3_neg_4_016_05    m/z 372.2256     0   1E+4   7.5   8  RT (min)      lcorbi_t3_neg_4_017_51    m/z 372.2257     0   1E+4   11   11.5  RT (min)      lcorbi_t3_neg_3_024_26    m/z 372.2436     0   1E+3   13   13.5  RT (min)      lcorbi_t3_neg_1_055_05    m/z 372.2945     0   2   4   6E+3   12.5   13  RT (min)      lcorbi_t3_neg_4_007_13    m/z 372.3196     0   1E+4   13   13.5  RT (min)      lcorbi_t3_neg_4_054_36    m/z 372.7510     0   1E+4   1   1.5  RT (min)      lcorbi_t3_neg_4_046_37    m/z 372.9432     0   1   2   3   4E+4   1   1.5  RT (min)      lcorbi_t3_neg_4_039_32    m/z 372.9967     0   1E+4   8   8.5  RT (min)      lcorbi_t3_neg_3_004_30    m/z 373.0322     0   1E+4   10   10.5  RT (min)      lcorbi_t3_neg_1_042_18    m/z 373.0448     0   1   2E+4   13   13.5  RT (min)      lcorbi_t3_neg_1_003_41    m/z 373.0448     0   1E+4   11   11.5  RT (min)      lcorbi_t3_neg_2_048_27    m/z 373.1190     0   2   4   6   8E+3   12.5   13  RT (min)      lcorbi_t3_neg_4_007_13    m/z 373.1653     0   1E+4   9   9.5  RT (min)      lcorbi_t3_neg_2_048_27    m/z 373.2089     0   1E+4   2.5   3  RT (min)      lcorbi_t3_neg_4_032_23    m/z 373.2146     0   1   2   3   4E+4   7.5   8  RT (min)      lcorbi_t3_neg_4_023_18    m/z 373.2674     0   1   2E+4   5.5   6  RT (min)      lcorbi_t3_neg_4_039_32    m/z 373.9798     0   2   4   6   8E+3   3   3.5  RT (min)      lcorbi_t3_neg_1_023_03    m/z 374.0140     0   1   2   3   4E+3   11.5   12  RT (min)      lcorbi_t3_neg_4_040_19    m/z 374.0221     0   2   4   6   8E+3   8.5   9  RT (min)      lcorbi_t3_neg_2_048_27    m/z 374.0772     0   2   4   6E+3   9   9.5  RT (min)      lcorbi_t3_neg_3_019_12    m/z 374.0782     0   1E+3   1   1.5  RT (min)       lcorbi_t3_neg_1_018_46    m/z 374.0961     0   2   4   6E+3   7.5   8  RT (min)      lcorbi_t3_neg_4_023_18    m/z 374.1264     0   1E+4   10   10.5  RT (min)      lcorbi_t3_neg_4_023_18    m/z 374.1363     0   1   1   1.5  RT (min)      lcorbi_t3_neg_2_022_24    m/z 374.2033     0   2   4   6E+3   12.5   13  RT (min)      lcorbi_t3_neg_1_043_14    m/z 374.2071     0   1E+4   5   5.5  RT (min)      lcorbi_t3_neg_4_054_36    m/z 374.3059     0   1   2   3E+4   12   12.5  RT (min)      lcorbi_t3_neg_3_024_26    m/z 374.8802     0   1   7.5   8  RT (min)      lcorbi_t3_neg_4_028_39    m/z 374.9060     0   1E+4   12.5   13  RT (min)      lcorbi_t3_neg_4_024_07    m/z 374.9061     0   1E+4   10   10.5  RT (min)      lcorbi_t3_neg_3_018_13    m/z 374.9395     0   1   1   1.5  RT (min)      lcorbi_t3_neg_3_039_29    m/z 374.9526     0   1E+4   9   9.5  RT (min)      lcorbi_t3_neg_4_007_13    m/z 374.9784     0   1E+4   18   18.5  RT (min)      lcorbi_t3_neg_4_034_01    m/z 375.0869     0   2   4   6E+3   7.5   8  RT (min)      lcorbi_t3_neg_4_013_09    m/z 375.1032     0   1E+4   9.5   10  RT (min)      lcorbi_t3_neg_2_035_46    m/z 375.1979     0   1E+4   11   11.5  RT (min)      lcorbi_t3_neg_2_048_27    m/z 375.1981     0   2   4   6E+3   7   7.5  RT (min)      lcorbi_t3_neg_4_023_18    m/z 375.2383     0   2   4   6E+3   7   7.5  RT (min)      lcorbi_t3_neg_2_048_27    m/z 375.2537     0   1   2E+3   12.5   13   13.5  RT (min)      lcorbi_t3_neg_2_054_15    m/z 375.2779     0   1   2E+4   12.5   13  RT (min)      lcorbi_t3_neg_4_007_13    m/z 375.2942     0   2   4   6   8E+3   11   11.5  RT (min)      lcorbi_t3_neg_4_044_38    m/z 375.9605     0   1   2E+4   1   1.5  RT (min)      lcorbi_t3_neg_4_039_32    m/z 376.0222     0   1   2   3   4E+4   1.5   2  RT (min)      lcorbi_t3_neg_3_002_24    m/z 376.0282     0   1E+4   8   8.5  RT (min)      lcorbi_t3_neg_4_023_18    m/z 376.0615     0   2   4   6E+3   10.5   11  RT (min)      lcorbi_t3_neg_3_010_15    m/z 376.0673     0   1   2E+5   9   9.5  RT (min)      lcorbi_t3_neg_1_036_09    m/z 376.1138     0   1E+4   9   9.5  RT (min)      lcorbi_t3_neg_3_019_12    m/z 376.1360     0   1   2   3E+3   13   13.5  RT (min)      lcorbi_t3_neg_4_039_32    m/z 376.3064     0   2   4   6   8E+3   11   11.5  RT (min)      lcorbi_t3_neg_4_046_37    m/z 376.3216     0   1   2   3   4E+4   12.5   13  RT (min)      lcorbi_t3_neg_3_024_26    m/z 376.8091     0   1   2E+4   1   1.5  RT (min)      lcorbi_t3_neg_4_039_32    m/z 376.8782     0   1E+4   7.5   8  RT (min)      lcorbi_t3_neg_4_023_18    m/z 376.9032     0   1   2E+4   18   18.5  RT (min)      lcorbi_t3_neg_2_026_03    m/z 376.9187     0   1   2   3E+4   3   3.5  RT (min)      lcorbi_t3_neg_4_046_37    m/z 376.9753     0   2   4   6E+3   18   18.5  RT (min)      lcorbi_t3_neg_4_042_14    m/z 376.9848     0   1E+4   10   10.5  RT (min)      lcorbi_t3_neg_1_023_03    m/z 377.0379     0   1E+4   7   7.5  RT (min)      lcorbi_t3_neg_4_007_13    m/z 377.0662     0   2   4   6E+3   8.5   9  RT (min)      lcorbi_t3_neg_2_048_27    m/z 377.0849     0   1   2E+4   6   6.5  RT (min)      lcorbi_t3_neg_4_017_51    m/z 377.1008     0   1   2E+3   9.5   10  RT (min)      lcorbi_t3_neg_2_056_56    m/z 377.1318     0   1E+4   13   13.5  RT (min)      lcorbi_t3_neg_4_038_15    m/z 377.1367     0   1E+4   9.5   10  RT (min)      lcorbi_t3_neg_2_048_27    m/z 377.1600     0   1   7   7.5  RT (min)      lcorbi_t3_neg_4_023_18    m/z 377.1855     0   1E+4   8   8.5  RT (min)      lcorbi_t3_neg_3_022_27    m/z 377.2267     0   2   4   6E+3   12   12.5  RT (min)      lcorbi_t3_neg_4_042_14    m/z 377.2287     0   1   10.5   11  RT (min)      lcorbi_t3_neg_2_053_23    m/z 377.2572     0   1E+4   9.5   10  RT (min)      lcorbi_t3_neg_2_053_23    m/z 377.2724     0   1   2E+4   13   13.5  RT (min)      lcorbi_t3_neg_1_055_05    m/z 377.2805     0   1E+3   11.5   12  RT (min)      lcorbi_t3_neg_4_007_13    m/z 377.3249     0   1   12.5   13  RT (min)      lcorbi_t3_neg_4_034_01    m/z 377.3951     0   1   2   3   4E+4   13   13.5  RT (min)      lcorbi_t3_neg_4_039_32    m/z 378.0252     0   1   2   3   4E+3   8   8.5  RT (min)      lcorbi_t3_neg_4_023_18    m/z 378.0619     0   1   2   3   4E+3   10   10.5  RT (min)      lcorbi_t3_neg_3_004_30    m/z 378.0722     0   1   2E+3   1   1.5  RT (min)      lcorbi_t3_neg_3_039_29    m/z 378.0741     0   1E+4   8.5   9  RT (min)      lcorbi_t3_neg_4_023_18    m/z 378.0913     0   1   2   3E+4   5.5   6  RT (min)      lcorbi_t3_neg_4_023_18    m/z 378.1011     0   2   4   6E+3   7   7.5  RT (min)      lcorbi_t3_neg_4_034_01    m/z 378.1081     0   1   2   3   4E+2   1   1.5  RT (min)      lcorbi_t3_neg_3_022_27    m/z 378.1335     0   2   4   6E+3   3   3.5  RT (min)      lcorbi_t3_neg_3_005_23    m/z 378.1499     0   1   2   3E+3   12.5   13  RT (min)      lcorbi_t3_neg_1_055_05    m/z 378.1562     0   1E+3   10.5   11  RT (min)      lcorbi_t3_neg_4_034_01    m/z 378.1644     0   1E+4   10   10.5  RT (min)      lcorbi_t3_neg_4_029_31    m/z 378.1667     0   2   4E+3   13   13.5  RT (min)      lcorbi_t3_neg_4_007_13    m/z 378.1999     0   2   4   6   8E+2   7   7.5  RT (min)      lcorbi_t3_neg_2_048_27    m/z 378.2242     0   1E+4   5   5.5  RT (min)      lcorbi_t3_neg_4_054_36    m/z 378.2663     0   1   2   3E+3   9.5   10  RT (min)      lcorbi_t3_neg_4_002_45    m/z 378.2774     0   1E+3   11   11.5  RT (min)      lcorbi_t3_neg_1_006_40    m/z 378.2970     0   1E+4   13   13.5  RT (min)      lcorbi_t3_neg_4_038_15    m/z 378.3121     0   1   12.5   13  RT (min)      lcorbi_t3_neg_1_023_03    m/z 378.9161     0   1E+4   3   3.5  RT (min)      lcorbi_t3_neg_4_046_37    m/z 378.9179     0   1E+4   9.5   10  RT (min)      lcorbi_t3_neg_3_049_55    m/z 378.9741     0   2   4E+3   18   18.5  RT (min)      lcorbi_t3_neg_4_034_01    m/z 378.9806     0   1   2   3   4E+3   1   1.5  RT (min)      lcorbi_t3_neg_4_045_35    m/z 378.9818     0   1   2E+4   7.5   8  RT (min)      lcorbi_t3_neg_4_023_18    m/z 379.0349     0   1E+4   7   7.5  RT (min)      lcorbi_t3_neg_4_007_13    m/z 379.0456     0   1   2   3   4E+4   1   1.5  RT (min)      lcorbi_t3_neg_4_054_36    m/z 379.0675     0   1   10.5   11  RT (min)      lcorbi_t3_neg_4_019_08    m/z 379.0739     0   1   2   3   4E+4   10.5   11  RT (min)      lcorbi_t3_neg_2_007_14    m/z 379.0854     0   1   2   3E+2   3   3.5  RT (min)      lcorbi_t3_neg_3_018_13    m/z 379.1055     0   1   2   3   4E+3   1   1.5  RT (min)      lcorbi_t3_neg_2_048_27    m/z 379.1314     0   2   4   6   8E+3   5   5.5  RT (min)      lcorbi_t3_neg_2_048_27    m/z 379.1452     0   1   1   1.5  RT (min)      lcorbi_t3_neg_4_007_13    m/z 379.1943     0   2   4E+3   6.5   7  RT (min)      lcorbi_t3_neg_4_007_13    m/z 379.2543     0   2   4   6E+2   9.5   10  RT (min)      lcorbi_t3_neg_4_011_24    m/z 379.9073     0   2   4E+3   7   7.5  RT (min)      lcorbi_t3_neg_2_028_38    m/z 379.9214     0   2   4   6E+3   18   18.5  RT (min)      lcorbi_t3_neg_4_034_01    m/z 380.1824     0   1   2   3E+3   13   13.5  RT (min)      lcorbi_t3_neg_4_046_37    m/z 380.1928     0   1   2   3E+3   9.5   10  RT (min)      lcorbi_t3_neg_2_048_27    m/z 380.2158     0   2   4E+4   8   8.5  RT (min)      lcorbi_t3_neg_2_048_27    m/z 380.2940     0   2   4   6E+3   13   13.5  RT (min)      lcorbi_t3_neg_4_034_01    m/z 380.9223     0   1   2   3E+3   18   18.5  RT (min)      lcorbi_t3_neg_4_028_39    m/z 380.9243     0   1   2   3E+3   1   1.5  RT (min)      lcorbi_t3_neg_4_046_37    m/z 380.9796     0   1   2E+4   7.5   8  RT (min)      lcorbi_t3_neg_4_023_18    m/z 381.0086     0   2   4   6E+4   5.5   6  RT (min)      lcorbi_t3_neg_4_047_30    m/z 381.0231     0   2   4   6   8E+3   6.5   7  RT (min)      lcorbi_t3_neg_4_034_01    m/z 381.0295     0   1   2E+4   1.5   2  RT (min)      lcorbi_t3_neg_2_048_27    m/z 381.0575     0   2   4   6E+3   10   10.5  RT (min)      lcorbi_t3_neg_4_023_18    m/z 381.0841     0   1   2E+3   1   1.5  RT (min)      lcorbi_t3_neg_3_028_35    m/z 381.0913     0   1   9   9.5  RT (min)      lcorbi_t3_neg_4_019_08    m/z 381.1017     0   1E+4   2   2.5  RT (min)      lcorbi_t3_neg_3_002_24    m/z 381.1052     0   1E+3   13   13.5  RT (min)      lcorbi_t3_neg_3_024_26    m/z 381.1412     0   1   2E+4   2.5   3  RT (min)      lcorbi_t3_neg_4_032_23    m/z 381.1513     0   1   2   3   4E+3   10.5   11  RT (min)      lcorbi_t3_neg_4_007_13    m/z 381.1551     0   1   2E+4   9   9.5  RT (min)      lcorbi_t3_neg_2_048_27    m/z 381.1558     0   1E+4   13   13.5  RT (min)      lcorbi_t3_neg_2_007_14    m/z 381.1893     0   2   4   6   8E+3   9.5   10  RT (min)      lcorbi_t3_neg_2_048_27    m/z 381.2598     0   2   4   6   8E+2   11   11.5  RT (min)      lcorbi_t3_neg_4_021_25    m/z 381.3371     0   1   2E+4   13.5   14  RT (min)      lcorbi_t3_neg_4_034_01    m/z 381.8058     0   2   4E+3   0.5   1  RT (min)      lcorbi_t3_neg_1_003_41    m/z 381.9043     0   2   4   6E+3   7   7.5  RT (min)      lcorbi_t3_neg_2_028_38    m/z 381.9581     0   1E+4   17.5   18  RT (min)      lcorbi_t3_neg_2_020_05    m/z 381.9594     0   2   4   6E+3   7.5   8  RT (min)      lcorbi_t3_neg_4_007_13    m/z 382.0673     0   1E+4   10   10.5  RT (min)      lcorbi_t3_neg_4_023_18    m/z 382.1898     0   1   6   6.5  RT (min)      lcorbi_t3_neg_4_011_24    m/z 382.1980     0   1   2E+3   13.5   14  RT (min)      lcorbi_t3_neg_4_030_04    m/z 382.2201     0   2   4E+3   5   5.5  RT (min)      lcorbi_t3_neg_4_001_49    m/z 382.3322     0   2   4   6E+3   13   13.5  RT (min)      lcorbi_t3_neg_4_034_01    m/z 382.7333     0   2   4E+3   1   1.5  RT (min)      lcorbi_t3_neg_4_007_13    m/z 382.9109     0   1   2   3E+3   18   18.5  RT (min)      lcorbi_t3_neg_4_051_40    m/z 382.9213     0   1   1   1.5  RT (min)      lcorbi_t3_neg_4_045_35    m/z 382.9344     0   2   4   6   8E+3   1   1.5  RT (min)      lcorbi_t3_neg_4_046_37    m/z 383.0067     0   2   4   6E+4   5.5   6  RT (min)      lcorbi_t3_neg_4_047_30    m/z 383.0556     0   2   4   6E+3   7   7.5  RT (min)      lcorbi_t3_neg_1_013_35    m/z 383.0713     0   1E+3   9.5   10  RT (min)      lcorbi_t3_neg_4_042_14    m/z 383.0873     0   1   2   3E+4   10.5   11  RT (min)      lcorbi_t3_neg_2_048_27    m/z 383.0996     0   2   4   6E+4   1   1.5  RT (min)      lcorbi_t3_neg_1_055_05    m/z 383.1112     0   1E+4   9   9.5  RT (min)      lcorbi_t3_neg_3_019_12    m/z 383.1138     0   2   4   6   8E+3   1   1.5  RT (min)      lcorbi_t3_neg_4_029_31    m/z 383.1243     0   1E+4   6   6.5  RT (min)      lcorbi_t3_neg_2_035_46    m/z 383.1849     0   1   2   3   4E+3   9.5   10  RT (min)      lcorbi_t3_neg_2_048_27    m/z 383.2294     0   2   4   6E+3   5.5   6  RT (min)      lcorbi_t3_neg_4_042_14    m/z 383.2410     0   1   2   3   4E+3   10.5   11  RT (min)      lcorbi_t3_neg_2_007_14    m/z 383.2448     0   1   2   3   4E+3   12   12.5  RT (min)      lcorbi_t3_neg_4_034_01    m/z 383.2799     0   1E+4   12.5   13  RT (min)      lcorbi_t3_neg_3_007_16    m/z 383.2911     0   2   4   6   8E+3   11.5   12  RT (min)      lcorbi_t3_neg_4_007_13    m/z 383.9097     0   1   2E+4   1   1.5  RT (min)      lcorbi_t3_neg_2_052_32    m/z 383.9563     0   2   4   6E+3   7.5   8  RT (min)      lcorbi_t3_neg_4_007_13    m/z 383.9816     0   1   2E+4   9   9.5  RT (min)      lcorbi_t3_neg_4_007_13    m/z 384.0222     0   1E+4   10   10.5  RT (min)      lcorbi_t3_neg_1_054_37    m/z 384.1170     0   2   4   6   8E+3   1   1.5  RT (min)      lcorbi_t3_neg_2_035_46    m/z 384.1236     0   2   4E+3   9   9.5  RT (min)      lcorbi_t3_neg_3_019_12    m/z 384.1458     0   1   2   3   4E+3   11   11.5  RT (min)      lcorbi_t3_neg_4_043_11    m/z 384.2235     0   1   10.5   11  RT (min)      lcorbi_t3_neg_4_042_14    m/z 384.2303     0   1   11   11.5  RT (min)      lcorbi_t3_neg_4_034_01    m/z 384.2468     0   1   2E+4   7.5   8  RT (min)      lcorbi_t3_neg_4_023_18    m/z 384.2665     0   1E+4   14.5   15  RT (min)      lcorbi_t3_neg_4_017_51    m/z 384.3031     0   2   4   6   8E+3   12.5   13  RT (min)      lcorbi_t3_neg_2_052_32    m/z 384.3114     0   2   4   6E+3   12   12.5  RT (min)      lcorbi_t3_neg_4_054_36    m/z 384.3560     0   1   2E+4   14   14.5  RT (min)      lcorbi_t3_neg_1_055_05    m/z 384.8744     0   1   2E+3   18   18.5  RT (min)      lcorbi_t3_neg_4_053_43    m/z 384.9349     0   1   2   3E+4   12.5   13  RT (min)      lcorbi_t3_neg_4_024_07    m/z 384.9349     0   1   2   3   4E+4   10   10.5  RT (min)       lcorbi_t3_neg_4_036_16    m/z 385.0682     0   1E+4   7   7.5  RT (min)      lcorbi_t3_neg_4_007_13    m/z 385.1096     0   2   4E+3   8   8.5  RT (min)      lcorbi_t3_neg_2_055_18    m/z 385.1108     0   1E+4   11   11.5  RT (min)      lcorbi_t3_neg_3_034_21    m/z 385.1288     0   1E+3   9   9.5  RT (min)      lcorbi_t3_neg_4_013_09    m/z 385.1393     0   2   4   6E+3   11   11.5  RT (min)      lcorbi_t3_neg_4_043_11    m/z 385.1486     0   2   4   6E+3   6.5   7  RT (min)      lcorbi_t3_neg_2_048_27    m/z 385.1527     0   1E+4   1   1.5  RT (min)      lcorbi_t3_neg_4_002_45    m/z 385.1634     0   2   4E+3   13   13.5  RT (min)      lcorbi_t3_neg_4_054_36    m/z 385.1873     0   1E+4   13.5   14  RT (min)      lcorbi_t3_neg_3_018_13    m/z 385.2014     0   1E+3   11   11.5  RT (min)      lcorbi_t3_neg_4_081_58    m/z 385.2087     0   1E+4   5   5.5  RT (min)      lcorbi_t3_neg_4_032_23    m/z 385.2173     0   1   2E+3   11   11.5  RT (min)      lcorbi_t3_neg_4_042_14    m/z 385.2380     0   1   2E+4   9   9.5  RT (min)      lcorbi_t3_neg_2_048_27    m/z 385.2492     0   1   2   3E+3   10.5   11  RT (min)      lcorbi_t3_neg_2_007_14    m/z 385.3067     0   1E+3   10.5   11  RT (min)      lcorbi_t3_neg_4_007_13    m/z 385.3318     0   1   12.5   13  RT (min)      lcorbi_t3_neg_1_055_05    m/z 386.0193     0   1   2   3E+3   10   10.5  RT (min)      lcorbi_t3_neg_1_054_37    m/z 386.1008     0   2   4   6E+3   1   1.5  RT (min)      lcorbi_t3_neg_2_035_46    m/z 386.1187     0   1   2   3   4E+3   11   11.5  RT (min)      lcorbi_t3_neg_4_008_22    m/z 386.1191     0   2   4E+4   2   2.5  RT (min)       lcorbi_t3_neg_2_022_24    m/z 386.1233     0   1   2   3   4E+3   9   9.5  RT (min)      lcorbi_t3_neg_3_019_12    m/z 386.1604     0   1   9.5   10  RT (min)      lcorbi_t3_neg_1_023_03    m/z 386.1719     0   1   2E+4   6   6.5  RT (min)      lcorbi_t3_neg_2_048_27    m/z 386.2206     0   2   4   6E+3   12.5   13  RT (min)      lcorbi_t3_neg_4_034_01    m/z 386.2365     0   1E+3   10.5   11  RT (min)      lcorbi_t3_neg_4_075_58    m/z 386.2403     0   1   12   12.5  RT (min)      lcorbi_t3_neg_4_090_60    m/z 386.8407     0   2   4E+3   0   0.5  RT (min)      lcorbi_t3_neg_4_067_57    m/z 386.8618     0   2   4E+3   18   18.5  RT (min)      lcorbi_t3_neg_3_056_56    m/z 387.0098     0   1E+4   1   1.5  RT (min)      lcorbi_t3_neg_3_005_23    m/z 387.0201     0   1   2   3   4E+3   8.5   9  RT (min)      lcorbi_t3_neg_4_034_01    m/z 387.0505     0   1   2E+3   9   9.5  RT (min)      lcorbi_t3_neg_3_019_12    m/z 387.0589     0   2   4   6E+3   8   8.5  RT (min)      lcorbi_t3_neg_4_023_18    m/z 387.0652     0   2   4   6   8E+3   7   7.5  RT (min)      lcorbi_t3_neg_4_007_13    m/z 387.0969     0   2   4E+4   13   13.5  RT (min)      lcorbi_t3_neg_3_028_35    m/z 387.1556     0   2   4   6   8E+2   12   12.5  RT (min)      lcorbi_t3_neg_4_024_07    m/z 387.1917     0   2   4   6E+3   9.5   10  RT (min)      lcorbi_t3_neg_2_048_27    m/z 387.2169     0   1   2E+3   12.5   13  RT (min)      lcorbi_t3_neg_2_048_27    m/z 387.2362     0   2   4   6E+2   13   13.5  RT (min)      lcorbi_t3_neg_2_048_27    m/z 387.2514     0   2   4   6E+3   11.5   12  RT (min)      lcorbi_t3_neg_4_005_27    m/z 387.2538     0   1E+4   11.5   12  RT (min)      lcorbi_t3_neg_3_024_26    m/z 387.2972     0   1E+4   11.5   12  RT (min)      lcorbi_t3_neg_1_039_38    m/z 387.3224     0   1   2   3E+4   12.5   13  RT (min)      lcorbi_t3_neg_4_034_01    m/z 388.0428     0   2   4E+3   12.5   13  RT (min)      lcorbi_t3_neg_1_043_14    m/z 388.0718     0   2   4E+3   8   8.5  RT (min)      lcorbi_t3_neg_4_023_18    m/z 388.0854     0   1E+5   1   1.5  RT (min)      lcorbi_t3_neg_4_039_32    m/z 388.1156     0   2   4   6   8E+4   1   1.5  RT (min)      lcorbi_t3_neg_3_002_24    m/z 388.1473     0   1   2E+3   13.5   14  RT (min)      lcorbi_t3_neg_4_045_35    m/z 388.1697     0   1E+4   12.5   13  RT (min)      lcorbi_t3_neg_4_016_05    m/z 388.1732     0   1   2E+4   10.5   11  RT (min)      lcorbi_t3_neg_2_048_27    m/z 388.2523     0   1E+4   11   11.5  RT (min)      lcorbi_t3_neg_4_034_01    m/z 388.2620     0   1   2E+4   12   12.5  RT (min)      lcorbi_t3_neg_2_053_23    m/z 388.3065     0   1E+4   11   11.5  RT (min)      lcorbi_t3_neg_3_020_31    m/z 388.8053     0   2   4E+3   0   0.5  RT (min)      lcorbi_t3_neg_1_031_33    m/z 388.8735     0   1E+4   18.5   19  RT (min)      lcorbi_t3_neg_4_111_62    m/z 388.9081     0   1   2E+4   1   1.5  RT (min)      lcorbi_t3_neg_2_048_27    m/z 388.9848     0   1   0   0.5  RT (min)      lcorbi_t3_neg_3_002_24    m/z 389.1576     0   2   4E+3   10.5   11  RT (min)      lcorbi_t3_neg_2_036_35    m/z 389.1816     0   2   4   6E+3   7   7.5  RT (min)      lcorbi_t3_neg_4_023_18    m/z 389.1968     0   1   2E+3   11   11.5  RT (min)      lcorbi_t3_neg_4_102_61    m/z 389.2307     0   2   4   6   8E+3   11.5   12  RT (min)      lcorbi_t3_neg_4_042_14    m/z 389.2402     0   1   6   6.5  RT (min)      lcorbi_t3_neg_2_048_27    m/z 389.2545     0   1   2   3   4E+3   11   11.5  RT (min)      lcorbi_t3_neg_4_029_31    m/z 389.2562     0   1E+4   13   13.5  RT (min)      lcorbi_t3_neg_4_034_01    m/z 389.2692     0   2   4   6   8E+3   12   12.5  RT (min)      lcorbi_t3_neg_4_042_14    m/z 389.2694     0   2   4   6   8E+3   10   10.5  RT (min)      lcorbi_t3_neg_2_048_27    m/z 389.2884     0   1   2E+3   12.5   13  RT (min)      lcorbi_t3_neg_2_052_32    m/z 389.2921     0   1E+4   9.5   10  RT (min)      lcorbi_t3_neg_4_029_31    m/z 389.3075     0   2   4   6   8E+3   13.5   14  RT (min)      lcorbi_t3_neg_4_007_13    m/z 389.3209     0   1   2E+4   12.5   13  RT (min)      lcorbi_t3_neg_3_002_24    m/z 389.3506     0   1E+4   12   12.5  RT (min)      lcorbi_t3_neg_4_039_32    m/z 389.4226     0   1   2   3E+3   14.5   15  RT (min)      lcorbi_t3_neg_4_048_03    m/z 389.7916     0   2   4   6E+3   7.5   8  RT (min)      lcorbi_t3_neg_1_042_18    m/z 389.9282     0   1   2E+4   9   9.5  RT (min)      lcorbi_t3_neg_4_007_13    m/z 390.0398     0   2   4E+3   12.5   13  RT (min)      lcorbi_t3_neg_1_043_14    m/z 390.0715     0   1   2   3   4E+3   8   8.5  RT (min)      lcorbi_t3_neg_3_024_26    m/z 390.1044     0   1E+4   6   6.5  RT (min)      lcorbi_t3_neg_2_048_27    m/z 390.1171     0   1   2   3E+4   1   1.5  RT (min)      lcorbi_t3_neg_4_032_23    m/z 390.1266     0   2   4   6   8E+3   13   13.5  RT (min)      lcorbi_t3_neg_4_047_30    m/z 390.1854     0   2   4   6E+4   13   13.5  RT (min)      lcorbi_t3_neg_4_016_05    m/z 390.2160     0   2   4   6E+3   6.5   7  RT (min)      lcorbi_t3_neg_4_029_31    m/z 390.2160     0   1E+4   5   5.5  RT (min)      lcorbi_t3_neg_4_032_23    m/z 390.2364     0   1E+4   6   6.5  RT (min)      lcorbi_t3_neg_2_048_27    m/z 390.2591     0   2   4   6E+3   12   12.5  RT (min)      lcorbi_t3_neg_2_053_23    m/z 390.3222     0   1E+4   11   11.5  RT (min)      lcorbi_t3_neg_4_054_36    m/z 390.8007     0   1   2E+4   4   4.5  RT (min)      lcorbi_t3_neg_1_046_11    m/z 390.8737     0   2   4   6E+3   18   18.5  RT (min)      lcorbi_t3_neg_4_079_58    m/z 390.9063     0   1   2E+4   14   14.5  RT (min)      lcorbi_t3_neg_3_035_03    m/z 390.9688     0   1   2   3E+3   5.5   6  RT (min)      lcorbi_t3_neg_4_077_58    m/z 390.9801     0   1   1   1.5  RT (min)      lcorbi_t3_neg_2_050_17    m/z 391.0820     0   2   4   6E+4   9   9.5  RT (min)      lcorbi_t3_neg_4_040_19    m/z 391.0935     0   2   4   6E+3   10   10.5  RT (min)      lcorbi_t3_neg_4_029_31    m/z 391.1009     0   2   4E+4   1   1.5  RT (min)      lcorbi_t3_neg_4_007_13    m/z 391.1058     0   2   4   6E+3   9.5   10  RT (min)      lcorbi_t3_neg_4_007_13    m/z 391.1757     0   1E+4   7.5   8  RT (min)      lcorbi_t3_neg_2_048_27    m/z 391.1784     0   1   2   3   4E+3   11.5   12  RT (min)      lcorbi_t3_neg_2_007_14    m/z 391.1928     0   1   2   3E+3   9.5   10  RT (min)      lcorbi_t3_neg_2_048_27    m/z 391.2073     0   1   2   3   4E+3   13   13.5  RT (min)      lcorbi_t3_neg_4_020_44    m/z 391.2519     0   1   2E+3   12   12.5  RT (min)      lcorbi_t3_neg_3_024_26    m/z 391.2849     0   1   2E+4   12   12.5  RT (min)      lcorbi_t3_neg_4_042_14    m/z 391.3092     0   1E+4   13   13.5  RT (min)      lcorbi_t3_neg_4_034_01    m/z 391.3579     0   1E+4   14   14.5  RT (min)      lcorbi_t3_neg_4_034_01    m/z 391.3662     0   1E+5   12   12.5  RT (min)      lcorbi_t3_neg_4_039_32    m/z 391.3744     0   1E+4   13   13.5  RT (min)      lcorbi_t3_neg_4_039_32    m/z 391.7890     0   1E+4   7.5   8  RT (min)      lcorbi_t3_neg_1_042_18    m/z 391.8963     0   1   2   3   4E+3   18   18.5  RT (min)      lcorbi_t3_neg_2_041_54    m/z 391.9252     0   1   2E+4   9   9.5  RT (min)      lcorbi_t3_neg_4_007_13    m/z 391.9849     0   1E+4   10.5   11  RT (min)      lcorbi_t3_neg_4_023_18    m/z 391.9911     0   2   4   6E+3   2   2.5  RT (min)      lcorbi_t3_neg_3_007_16    m/z 392.0878     0   1E+4   1   1.5  RT (min)      lcorbi_t3_neg_4_031_48    m/z 392.1238     0   1   2   3E+3   13   13.5  RT (min)      lcorbi_t3_neg_3_018_13    m/z 392.1909     0   2   4   6E+3   11.5   12  RT (min)      lcorbi_t3_neg_2_054_15    m/z 392.2028     0   1   2   3   4E+3   11   11.5  RT (min)      lcorbi_t3_neg_1_043_14    m/z 392.2285     0   2   4   6   8E+3   11   11.5  RT (min)      lcorbi_t3_neg_4_017_51    m/z 392.9133     0   2   4   6E+3   8.5   9  RT (min)      lcorbi_t3_neg_4_053_43    m/z 392.9386     0   1E+4   9.5   10  RT (min)      lcorbi_t3_neg_1_039_38    m/z 393.0106     0   1   2   3E+3   1.5   2  RT (min)      lcorbi_t3_neg_3_053_14    m/z 393.0539     0   2   4   6E+3   7.5   8  RT (min)      lcorbi_t3_neg_4_034_01    m/z 393.0634     0   1   2E+4   10.5   11  RT (min)      lcorbi_t3_neg_3_010_15    m/z 393.0979     0   1   2E+4   7   7.5  RT (min)      lcorbi_t3_neg_2_048_27    m/z 393.1013     0   2   4   6   8E+4   1   1.5  RT (min)      lcorbi_t3_neg_2_048_27    m/z 393.1243     0   1   2   3E+4   5   5.5  RT (min)      lcorbi_t3_neg_2_048_27    m/z 393.1446     0   2   4   6   8E+4   6   6.5  RT (min)      lcorbi_t3_neg_2_048_27    m/z 393.1550     0   2   4   6E+3   6   6.5  RT (min)      lcorbi_t3_neg_4_007_13    m/z 393.1872     0   1   11.5   12  RT (min)      lcorbi_t3_neg_4_105_61    m/z 393.2079     0   1   2   3E+4   12.5   13  RT (min)      lcorbi_t3_neg_1_055_05    m/z 393.2956     0   2   4   6E+3   10   10.5  RT (min)      lcorbi_t3_neg_3_019_12    m/z 393.3006     0   1E+4   12   12.5  RT (min)      lcorbi_t3_neg_4_034_01    m/z 393.3734     0   2   4E+3   14.5   15  RT (min)      lcorbi_t3_neg_4_046_37    m/z 393.3900     0   1E+5   13   13.5  RT (min)      lcorbi_t3_neg_4_039_32    m/z 393.7862     0   2   4   6   8E+3   7.5   8  RT (min)      lcorbi_t3_neg_1_042_18    m/z 393.9293     0   1   1   1.5  RT (min)      lcorbi_t3_neg_2_052_32    m/z 394.0160     0   1   4.5   5  RT (min)      lcorbi_t3_neg_1_043_14    m/z 394.0368     0   1E+4   1   1.5  RT (min)      lcorbi_t3_neg_4_011_24    m/z 394.0489     0   1   2E+4   10   10.5  RT (min)      lcorbi_t3_neg_3_010_15    m/z 394.1030     0   1E+4   9.5   10  RT (min)      lcorbi_t3_neg_4_007_13    m/z 394.1171     0   1   5.5   6  RT (min)      lcorbi_t3_neg_4_097_60    m/z 394.1746     0   1   2E+4   11   11.5  RT (min)      lcorbi_t3_neg_4_102_61    m/z 394.1991     0   2   4   6   8E+4   12.5   13  RT (min)      lcorbi_t3_neg_1_055_05    m/z 394.2675     0   1E+4   12.5   13   13.5  RT (min)      lcorbi_t3_neg_4_023_18    m/z 394.2957     0   2   4   6   8E+3   12   12.5  RT (min)      lcorbi_t3_neg_2_053_23    m/z 394.2958     0   1E+5   9.5   10  RT (min)      lcorbi_t3_neg_2_016_11    m/z 394.3320     0   2   4   6   8E+3   13   13.5  RT (min)      lcorbi_t3_neg_4_034_01    m/z 394.7978     0   1E+4   1   1.5  RT (min)      lcorbi_t3_neg_4_039_32    m/z 394.8668     0   1E+3   1   1.5  RT (min)      lcorbi_t3_neg_3_029_02    m/z 394.8902     0   1E+4   0.5   1  RT (min)      lcorbi_t3_neg_3_027_43    m/z 394.8904     0   1E+4   9   10  RT (min)       lcorbi_t3_neg_3_023_07    m/z 394.8909     0   1E+4   18   18.5  RT (min)       lcorbi_t3_neg_4_041_54    m/z 394.9177     0   1   8.5   9  RT (min)      lcorbi_t3_neg_4_104_61    m/z 394.9355     0   1   9.5   10  RT (min)      lcorbi_t3_neg_4_104_61    m/z 394.9898     0   1E+4   2   2.5  RT (min)      lcorbi_t3_neg_3_007_16    m/z 395.0532     0   1E+4   6   6.5  RT (min)      lcorbi_t3_neg_4_045_35    m/z 395.0632     0   1   1   1.5  RT (min)      lcorbi_t3_neg_1_055_05    m/z 395.0976     0   2   4   6E+3   6.5   7  RT (min)      lcorbi_t3_neg_4_007_13    m/z 395.1001     0   2   4E+3   5   5.5  RT (min)      lcorbi_t3_neg_2_036_35    m/z 395.1053     0   1E+3   10   10.5  RT (min)      lcorbi_t3_neg_4_023_18    m/z 395.1166     0   1   2E+4   10   10.5  RT (min)      lcorbi_t3_neg_4_023_18    m/z 395.1331     0   1   2   3   4E+3   1   1.5  RT (min)      lcorbi_t3_neg_2_048_27    m/z 395.1558     0   1E+4   10   10.5  RT (min)      lcorbi_t3_neg_4_045_35    m/z 395.1626     0   1E+3   7   7.5  RT (min)      lcorbi_t3_neg_4_023_18    m/z 395.1667     0   1   2E+3   11   11.5  RT (min)      lcorbi_t3_neg_1_055_05    m/z 395.1873     0   2   4E+3   10   10.5  RT (min)      lcorbi_t3_neg_2_045_30    m/z 395.2434     0   2   4   6E+3   9.5   10  RT (min)      lcorbi_t3_neg_4_038_15    m/z 395.2546     0   1   2   3E+4   10.5   11  RT (min)      lcorbi_t3_neg_4_011_24    m/z 395.2775     0   2   4   6E+3   13   13.5  RT (min)      lcorbi_t3_neg_1_050_45    m/z 395.2952     0   1E+4   13.5   14  RT (min)      lcorbi_t3_neg_4_007_13    m/z 395.3526     0   2   4E+3   13   13.5  RT (min)      lcorbi_t3_neg_4_034_01    m/z 395.3890     0   1E+4   14.5   15  RT (min)      lcorbi_t3_neg_2_056_56    m/z 395.8578     0   2   4E+3   18.5   19  RT (min)      lcorbi_t3_neg_1_006_40    m/z 395.8981     0   1   2   3E+4   1   1.5  RT (min)      lcorbi_t3_neg_4_039_32    m/z 395.9768     0   2   4   6E+3   5   5.5  RT (min)      lcorbi_t3_neg_3_019_12    m/z 396.0922     0   2   4   6E+3   12   12.5  RT (min)      lcorbi_t3_neg_4_035_33    m/z 396.1190     0   1   2E+4   1   1.5  RT (min)      lcorbi_t3_neg_4_032_23    m/z 396.1372     0   2   4   6   8E+3   13   13.5  RT (min)      lcorbi_t3_neg_4_042_14    m/z 396.1379     0   1   2E+3   7   7.5  RT (min)      lcorbi_t3_neg_4_044_38    m/z 396.1821     0   2   4   6E+3   6   6.5  RT (min)      lcorbi_t3_neg_4_044_38    m/z 396.1927     0   1E+4   8   8.5  RT (min)      lcorbi_t3_neg_4_034_01    m/z 396.2025     0   2   4   6E+4   12.5   13  RT (min)      lcorbi_t3_neg_1_055_05    m/z 396.2233     0   2   4E+2   13   13.5  RT (min)      lcorbi_t3_neg_1_055_05    m/z 396.3010     0   1   10   10.5  RT (min)      lcorbi_t3_neg_4_019_08    m/z 396.3317     0   1   2   3E+3   10   10.5  RT (min)      lcorbi_t3_neg_2_048_27    m/z 396.8907     0   1   2E+4   18   18.5  RT (min)      lcorbi_t3_neg_2_048_27    m/z 396.9149     0   1E+4   8.5   9  RT (min)      lcorbi_t3_neg_1_026_19    m/z 396.9909     0   1   2   3E+2   1   1.5  RT (min)      lcorbi_t3_neg_1_052_24    m/z 397.0068     0   2   4   6   8E+3   8   8.5  RT (min)      lcorbi_t3_neg_4_023_18    m/z 397.0433     0   1E+4   9   9.5  RT (min)      lcorbi_t3_neg_3_019_12    m/z 397.0489     0   1E+4   8.5   9  RT (min)      lcorbi_t3_neg_4_034_01    m/z 397.0794     0   1E+4   9   9.5  RT (min)      lcorbi_t3_neg_3_019_12    m/z 397.0980     0   2   4E+3   5   5.5  RT (min)      lcorbi_t3_neg_2_035_46    m/z 397.1004     0   1E+4   7.5   8  RT (min)      lcorbi_t3_neg_3_023_07    m/z 397.1189     0   2   4   6   8E+3   12.5   13  RT (min)      lcorbi_t3_neg_4_016_05    m/z 397.1463     0   1E+4   10   10.5  RT (min)      lcorbi_t3_neg_1_007_26    m/z 397.1499     0   1   2E+4   6   6.5  RT (min)      lcorbi_t3_neg_2_048_27    m/z 397.1550     0   1   2   3   4E+4   9   9.5  RT (min)      lcorbi_t3_neg_2_048_27    m/z 397.1581     0   1E+4   11   11.5  RT (min)      lcorbi_t3_neg_4_102_61    m/z 397.1685     0   2   4   6E+3   7.5   8  RT (min)      lcorbi_t3_neg_4_034_01    m/z 397.1841     0   1   2   3   4E+2   10.5   11  RT (min)      lcorbi_t3_neg_2_053_23    m/z 397.2268     0   1   2   3E+3   13   13.5  RT (min)      lcorbi_t3_neg_4_034_01    m/z 397.2392     0   1   2   3   4E+4   13   13.5  RT (min)      lcorbi_t3_neg_1_055_05    m/z 397.2701     0   1E+5   11   11.5  RT (min)       lcorbi_t3_neg_4_011_24    m/z 397.3107     0   1E+3   13   13.5  RT (min)      lcorbi_t3_neg_4_013_09    m/z 397.8884     0   1   2E+3   18   18.5  RT (min)      lcorbi_t3_neg_1_040_42    m/z 397.9426     0   1   2E+4   1   1.5  RT (min)      lcorbi_t3_neg_4_039_32    m/z 397.9943     0   1   2   3   4E+3   8   8.5  RT (min)      lcorbi_t3_neg_3_024_26    m/z 398.0060     0   1   2E+4   5.5   6  RT (min)      lcorbi_t3_neg_2_048_27    m/z 398.0479     0   2   4   6   8E+3   10.5   11  RT (min)      lcorbi_t3_neg_3_010_15    m/z 398.0892     0   1   2   3   4E+3   12   12.5  RT (min)      lcorbi_t3_neg_4_042_14    m/z 398.1131     0   1   2E+4   12.5   13  RT (min)      lcorbi_t3_neg_4_016_05    m/z 398.1175     0   1E+4   2.5   3  RT (min)      lcorbi_t3_neg_4_001_49    m/z 398.1396     0   1E+4   10   10.5  RT (min)      lcorbi_t3_neg_2_055_18    m/z 398.1529     0   1E+3   13   13.5  RT (min)      lcorbi_t3_neg_2_020_05    m/z 398.1815     0   2   4   6   8E+3   7   7.5  RT (min)      lcorbi_t3_neg_4_034_01    m/z 398.2034     0   2   4   6   8E+3   9.5   10  RT (min)      lcorbi_t3_neg_2_048_27    m/z 398.2151     0   1E+4   7   7.5  RT (min)      lcorbi_t3_neg_4_023_18    m/z 398.2181     0   2   4   6   8E+4   13   13.5  RT (min)      lcorbi_t3_neg_1_055_05    m/z 398.2236     0   1   9.5   10  RT (min)      lcorbi_t3_neg_4_019_08    m/z 398.2592     0   2   4E+3   13   13.5  RT (min)      lcorbi_t3_neg_4_034_01    m/z 398.8861     0   1E+4   18   18.5  RT (min)      lcorbi_t3_neg_2_048_27    m/z 399.0038     0   1   2   3E+3   8   8.5  RT (min)      lcorbi_t3_neg_4_023_18    m/z 399.0990     0   1   2E+4   4.5   5  RT (min)      lcorbi_t3_neg_4_032_23    m/z 399.1041     0   1E+4   1   1.5  RT (min)      lcorbi_t3_neg_4_001_49    m/z 399.1520     0   1   2E+4   9   9.5  RT (min)      lcorbi_t3_neg_2_048_27    m/z 399.1542     0   1   2E+4   1   1.5  RT (min)      lcorbi_t3_neg_4_011_24    m/z 399.1551     0   2   4E+3   11   11.5  RT (min)      lcorbi_t3_neg_4_043_11    m/z 399.1632     0   1   2   3   4E+3   8.5   9  RT (min)      lcorbi_t3_neg_2_048_27    m/z 399.1752     0   1E+4   7   7.5  RT (min)      lcorbi_t3_neg_4_034_01    m/z 399.2007     0   1E+4   8   8.5  RT (min)      lcorbi_t3_neg_4_034_01    m/z 399.2110     0   1E+3   8   8.5  RT (min)      lcorbi_t3_neg_4_041_54    m/z 399.2171     0   1E+3   11.5   12  RT (min)      lcorbi_t3_neg_4_016_05    m/z 399.2315     0   2   4   6   8E+3   10   10.5  RT (min)      lcorbi_t3_neg_4_032_23    m/z 399.2682     0   1E+4   12.5   13  RT (min)      lcorbi_t3_neg_4_029_31    m/z 399.2858     0   1E+4   11   11.5  RT (min)      lcorbi_t3_neg_4_011_24    m/z 399.2941     0   1E+4   12   12.5  RT (min)      lcorbi_t3_neg_4_032_23    m/z 399.3476     0   1E+4   13.5   14  RT (min)      lcorbi_t3_neg_4_054_36    m/z 399.8203     0   1E+4   7.5   8  RT (min)      lcorbi_t3_neg_1_042_18    m/z 399.9237     0   2   4   6   8E+3   8.5   9  RT (min)      lcorbi_t3_neg_2_048_27    m/z 399.9395     0   2   4   6   8E+3   8.5   9  RT (min)      lcorbi_t3_neg_2_048_27    m/z 400.1077     0   1   2E+4   1   1.5  RT (min)      lcorbi_t3_neg_1_052_24    m/z 400.1320     0   1E+4   6   6.5  RT (min)      lcorbi_t3_neg_2_048_27    m/z 400.1599     0   1   2   3E+3   11   11.5  RT (min)      lcorbi_t3_neg_4_056_56    m/z 400.1686     0   2   4   6   8E+3   13.5   14  RT (min)      lcorbi_t3_neg_4_040_19    m/z 400.1868     0   1   2   3E+3   13.5   14  RT (min)      lcorbi_t3_neg_4_038_15    m/z 400.2251     0   1   2   3E+3   10   10.5  RT (min)      lcorbi_t3_neg_4_032_23    m/z 400.2749     0   1   2   3E+3   13.5   14  RT (min)      lcorbi_t3_neg_3_018_13    m/z 400.3063     0   1   2   3E+3   11   11.5  RT (min)      lcorbi_t3_neg_1_013_35    m/z 400.3298     0   1   2E+3   12.5   13  RT (min)      lcorbi_t3_neg_4_036_16    m/z 400.9319     0   2   4   6E+3   8.5   9  RT (min)      lcorbi_t3_neg_2_048_27    m/z 400.9914     0   1E+4   8   8.5  RT (min)      lcorbi_t3_neg_3_024_26    m/z 401.0214     0   1E+4   10.5   11  RT (min)      lcorbi_t3_neg_1_039_38    m/z 401.1315     0   1E+4   13   13.5  RT (min)      lcorbi_t3_neg_4_042_14    m/z 401.1420     0   2   4   6   8E+3   12.5   13  RT (min)      lcorbi_t3_neg_4_042_14    m/z 401.1576     0   1   2   3E+3   13   13.5  RT (min)      lcorbi_t3_neg_2_005_01    m/z 401.1632     0   2   4   6   8E+3   13   13.5  RT (min)      lcorbi_t3_neg_2_007_14    m/z 401.1965     0   1E+4   8   8.5  RT (min)      lcorbi_t3_neg_4_013_09    m/z 401.2321     0   1   2E+3   13   13.5  RT (min)      lcorbi_t3_neg_4_038_15    m/z 401.2518     0   1E+4   13.5   14  RT (min)      lcorbi_t3_neg_4_034_01    m/z 401.2712     0   1E+4   13.5   14  RT (min)      lcorbi_t3_neg_4_040_19    m/z 401.2905     0   1E+4   12.5   13  RT (min)      lcorbi_t3_neg_3_007_16    m/z 401.8179     0   1   2E+4   7.5   8  RT (min)      lcorbi_t3_neg_1_042_18    m/z 401.8817     0   2   4   6E+3   18   18.5  RT (min)      lcorbi_t3_neg_3_016_42    m/z 401.9251     0   2   4   6   8E+3   18   18.5  RT (min)      lcorbi_t3_neg_3_026_44    m/z 401.9370     0   2   4E+3   1   1.5  RT (min)      lcorbi_t3_neg_4_045_35    m/z 402.0201     0   1E+4   2   2.5  RT (min)      lcorbi_t3_neg_3_007_16    m/z 402.1257     0   2   4   6   8E+3   10   10.5  RT (min)      lcorbi_t3_neg_4_032_23    m/z 402.1319     0   1   2   3E+4   1   1.5  RT (min)      lcorbi_t3_neg_2_044_06    m/z 402.1354     0   2   4   6E+3   9   9.5  RT (min)      lcorbi_t3_neg_3_019_12    m/z 402.1522     0   1   2   3   4E+4   13   13.5  RT (min)      lcorbi_t3_neg_4_019_08    m/z 402.2105     0   1   2   3   4E+3   12   12.5  RT (min)      lcorbi_t3_neg_4_042_14    m/z 402.8844     0   2   4   6E+3   18   18.5  RT (min)      lcorbi_t3_neg_4_052_26    m/z 403.0141     0   2   4   6E+3   9   9.5  RT (min)      lcorbi_t3_neg_1_023_03    m/z 403.0281     0   1   2   3E+4   1   1.5  RT (min)      lcorbi_t3_neg_3_005_23    m/z 403.0815     0   1E+5   1   1.5  RT (min)      lcorbi_t3_neg_2_048_27    m/z 403.1232     0   1E+4   10   10.5  RT (min)      lcorbi_t3_neg_4_023_18    m/z 403.1346     0   1E+4   7.5   8  RT (min)      lcorbi_t3_neg_4_050_02    m/z 403.1473     0   1E+4   13   13.5  RT (min)      lcorbi_t3_neg_4_038_15    m/z 403.1601     0   1   2E+4   1   1.5  RT (min)      lcorbi_t3_neg_4_011_24    m/z 403.1931     0   1   2E+3   11   11.5  RT (min)      lcorbi_t3_neg_4_043_11    m/z 403.2042     0   2   4   6E+3   12   12.5  RT (min)      lcorbi_t3_neg_4_042_14    m/z 403.2042     0   1E+4   8   8.5  RT (min)      lcorbi_t3_neg_2_048_27    m/z 403.2292     0   1E+4   12   12.5  RT (min)      lcorbi_t3_neg_4_005_27    m/z 403.2485     0   2   4   6E+3   11.5   12  RT (min)      lcorbi_t3_neg_3_023_07    m/z 403.2640     0   1   2E+4   13.5   14  RT (min)      lcorbi_t3_neg_4_039_32    m/z 403.2828     0   1   2E+4   11.5   12  RT (min)      lcorbi_t3_neg_4_007_13    m/z 403.2849     0   1   2   3E+3   10   10.5  RT (min)      lcorbi_t3_neg_4_042_14    m/z 403.3577     0   2   4E+3   13   13.5  RT (min)      lcorbi_t3_neg_4_042_14    m/z 403.5551     0   1E+4   4.5   5  RT (min)      lcorbi_t3_neg_2_048_27    m/z 403.8151     0   2   4   6   8E+3   7.5   8  RT (min)      lcorbi_t3_neg_1_042_18    m/z 403.9500     0   2   4E+3   1   1.5  RT (min)      lcorbi_t3_neg_4_054_36    m/z 404.0084     0   1E+4   10   10.5  RT (min)      lcorbi_t3_neg_3_045_34    m/z 404.0983     0   1E+4   11   11.5  RT (min)      lcorbi_t3_neg_2_035_46    m/z 404.1451     0   1   1   1.5  RT (min)      lcorbi_t3_neg_2_053_23    m/z 404.1668     0   1E+3   13   13.5  RT (min)      lcorbi_t3_neg_2_007_14    m/z 404.2673     0   2   4   6E+3   13.5   14  RT (min)      lcorbi_t3_neg_1_055_05    m/z 404.2978     0   2   4   6   8E+3   13   13.5  RT (min)      lcorbi_t3_neg_1_023_03    m/z 404.3126     0   2   4   6   8E+3   13   13.5  RT (min)      lcorbi_t3_neg_2_050_17    m/z 404.3391     0   1E+4   11.5   12  RT (min)      lcorbi_t3_neg_2_035_46    m/z 404.8456     0   2   4   6E+3   0   0.5  RT (min)      lcorbi_t3_neg_4_067_57    m/z 404.8814     0   1   2   3   4E+3   18   18.5  RT (min)      lcorbi_t3_neg_4_051_40    m/z 405.0186     0   1   2   3E+4   2   2.5  RT (min)      lcorbi_t3_neg_3_007_16    m/z 405.0282     0   1E+4   1.5   2  RT (min)      lcorbi_t3_neg_2_007_14    m/z 405.0801     0   1   2   3E+3   8   8.5  RT (min)      lcorbi_t3_neg_2_051_34    m/z 405.0876     0   2   4E+3   11   11.5  RT (min)      lcorbi_t3_neg_2_035_46    m/z 405.1629     0   1E+4   13.5   14  RT (min)      lcorbi_t3_neg_4_040_19    m/z 405.1735     0   1E+4   13   13.5  RT (min)      lcorbi_t3_neg_2_007_14    m/z 405.1913     0   1E+4   7   7.5  RT (min)      lcorbi_t3_neg_4_034_01    m/z 405.2085     0   1   2E+3   9.5   10  RT (min)      lcorbi_t3_neg_4_016_05    m/z 405.2278     0   2   4   6E+2   11   11.5  RT (min)      lcorbi_t3_neg_4_032_23    m/z 405.2673     0   1E+3   12   12.5  RT (min)      lcorbi_t3_neg_3_024_26    m/z 405.2883     0   2   4   6E+3   10.5   11  RT (min)      lcorbi_t3_neg_4_032_23    m/z 405.3006     0   1   2   3E+3   12   12.5  RT (min)      lcorbi_t3_neg_1_007_26    m/z 405.3117     0   1   2E+4   12   12.5  RT (min)      lcorbi_t3_neg_3_024_26    m/z 405.8143     0   1   1   1.5  RT (min)      lcorbi_t3_neg_1_053_32    m/z 405.9544     0   1   2   3   4E+3   8   8.5  RT (min)      lcorbi_t3_neg_3_037_36    m/z 406.0819     0   2   4   6E+3   11   11.5  RT (min)      lcorbi_t3_neg_2_037_09    m/z 406.0953     0   2   4E+3   11   11.5  RT (min)      lcorbi_t3_neg_2_035_46    m/z 406.1323     0   1E+4   10   10.5  RT (min)      lcorbi_t3_neg_3_004_30    m/z 406.1448     0   1E+4   12.5   13  RT (min)      lcorbi_t3_neg_4_054_36    m/z 406.1538     0   2   4E+3   7.5   8  RT (min)      lcorbi_t3_neg_4_034_01    m/z 406.1820     0   1   2   3   4E+4   10   10.5  RT (min)      lcorbi_t3_neg_2_048_27    m/z 406.1896     0   2   4   6   8E+3   9.5   10  RT (min)      lcorbi_t3_neg_2_048_27    m/z 406.2770     0   1E+4   13   13.5  RT (min)      lcorbi_t3_neg_1_023_03    m/z 406.3095     0   1   2   3   4E+2   13   13.5  RT (min)      lcorbi_t3_neg_4_017_51    m/z 406.3283     0   1   2E+4   13   13.5  RT (min)      lcorbi_t3_neg_4_034_01    m/z 406.9474     0   2   4   6E+3   1   1.5  RT (min)      lcorbi_t3_neg_4_046_37    m/z 407.0443     0   2   4E+3   4.5   5  RT (min)      lcorbi_t3_neg_4_036_16    m/z 407.0748     0   2   4E+3   10.5   11  RT (min)      lcorbi_t3_neg_1_043_14    m/z 407.1472     0   2   4   6   8E+3   7.5   8  RT (min)      lcorbi_t3_neg_4_005_27    m/z 407.1556     0   1   2   3E+3   9.5   10  RT (min)      lcorbi_t3_neg_4_017_51    m/z 407.1566     0   2   4   6   8E+3   4.5   5  RT (min)      lcorbi_t3_neg_4_032_23    m/z 407.1682     0   1   2   3   4E+3   13.5   14  RT (min)      lcorbi_t3_neg_4_034_01    m/z 407.1850     0   1E+4   9.5   10  RT (min)      lcorbi_t3_neg_4_034_01    m/z 407.2022     0   2   4   6   8E+3   12   12.5  RT (min)      lcorbi_t3_neg_4_034_01    m/z 407.2202     0   1E+5   10.5   11  RT (min)      lcorbi_t3_neg_1_055_05    m/z 407.2622     0   2   4   6E+3   12   12.5  RT (min)      lcorbi_t3_neg_4_029_31    m/z 407.2805     0   2   4   6   8E+3   12   12.5  RT (min)      lcorbi_t3_neg_4_007_13    m/z 407.3526     0   2   4   6   8E+3   13   13.5  RT (min)      lcorbi_t3_neg_4_034_01    m/z 407.9386     0   2   4   6   8E+3   9   9.5  RT (min)      lcorbi_t3_neg_1_043_14    m/z 408.0389     0   2   4   6E+3   8   8.5  RT (min)      lcorbi_t3_neg_4_023_18    m/z 408.1053     0   2   4E+3   10.5   11  RT (min)      lcorbi_t3_neg_2_024_37    m/z 408.1311     0   2   4E+4   1   1.5  RT (min)      lcorbi_t3_neg_2_048_27    m/z 408.1903     0   1   2E+4   11   11.5  RT (min)      lcorbi_t3_neg_4_102_61    m/z 408.2136     0   1E+4   9   9.5  RT (min)      lcorbi_t3_neg_2_048_27    m/z 408.2256     0   1   2   3E+4   10   10.5  RT (min)      lcorbi_t3_neg_1_055_05    m/z 408.2516     0   2   4   6E+3   11   11.5  RT (min)      lcorbi_t3_neg_3_005_23    m/z 408.2832     0   2   4   6   8E+4   9.5   10  RT (min)      lcorbi_t3_neg_4_023_18    m/z 408.3254     0   2   4   6   8E+3   13   13.5  RT (min)      lcorbi_t3_neg_4_034_01    m/z 408.7747     0   1   1   1.5  RT (min)      lcorbi_t3_neg_4_007_13    m/z 408.8226     0   1   2   3E+3   0   0.5  RT (min)      lcorbi_t3_neg_4_067_57    m/z 408.9594     0   1E+4   8   8.5  RT (min)      lcorbi_t3_neg_1_007_26    m/z 408.9795     0   1   2E+4   1   1.5  RT (min)      lcorbi_t3_neg_4_042_14    m/z 408.9916     0   1   2E+2   1   1.5  RT (min)      lcorbi_t3_neg_4_042_14    m/z 409.0039     0   1E+4   10   10.5  RT (min)      lcorbi_t3_neg_3_045_34    m/z 409.0600     0   2   4   6   8E+3   6   6.5  RT (min)      lcorbi_t3_neg_4_029_31    m/z 409.0928     0   2   4   6   8E+3   9.5   10  RT (min)      lcorbi_t3_neg_3_019_12    m/z 409.1443     0   1   2E+3   9.5   10  RT (min)      lcorbi_t3_neg_2_048_27    m/z 409.1541     0   2   4   6E+4   2.5   3  RT (min)      lcorbi_t3_neg_2_048_27    m/z 409.1584     0   1   2E+3   1   1.5  RT (min)      lcorbi_t3_neg_4_040_19    m/z 409.1633     0   1   2   3   4E+3   13   13.5  RT (min)      lcorbi_t3_neg_4_042_14    m/z 409.1740     0   2   4   6   8E+2   9.5   10  RT (min)      lcorbi_t3_neg_4_042_14    m/z 409.2047     0   1   2   3E+3   13.5   14  RT (min)      lcorbi_t3_neg_3_022_27    m/z 409.2264     0   1   2   3   4E+3   11   11.5  RT (min)      lcorbi_t3_neg_1_055_05    m/z 409.2722     0   2   4   6E+3   12.5   13  RT (min)      lcorbi_t3_neg_4_017_51    m/z 409.3324     0   1   2   3   4E+4   12   12.5  RT (min)      lcorbi_t3_neg_2_022_24    m/z 409.4047     0   1   2   3E+3   14.5   15  RT (min)      lcorbi_t3_neg_4_017_51    m/z 409.8750     0   1   2E+4   1   1.5  RT (min)      lcorbi_t3_neg_2_052_32    m/z 409.9977     0   2   4   6   8E+3   10   10.5  RT (min)      lcorbi_t3_neg_3_045_34    m/z 410.1638     0   1   2   3E+3   8   8.5  RT (min)      lcorbi_t3_neg_4_034_01    m/z 410.1782     0   1E+4   7   7.5  RT (min)      lcorbi_t3_neg_4_023_18    m/z 410.1994     0   2   4   6E+3   8.5   9  RT (min)      lcorbi_t3_neg_4_017_51    m/z 410.2212     0   1   2E+4   9   9.5  RT (min)      lcorbi_t3_neg_2_048_27    m/z 410.2311     0   2   4E+3   10   10.5  RT (min)      lcorbi_t3_neg_3_040_05    m/z 410.2381     0   1   2E+4   13   13.5  RT (min)      lcorbi_t3_neg_1_055_05    m/z 410.2659     0   1E+3   11.5   12  RT (min)      lcorbi_t3_neg_2_022_24    m/z 410.2890     0   1   2   3E+4   11   11.5  RT (min)      lcorbi_t3_neg_4_011_24    m/z 410.3717     0   1E+4   13   13.5  RT (min)      lcorbi_t3_neg_4_034_01    m/z 410.9066     0   1   2E+4   4.5   5  RT (min)      lcorbi_t3_neg_3_022_27    m/z 410.9171     0   1   2   3E+4   1   1.5  RT (min)      lcorbi_t3_neg_2_052_32    m/z 411.0280     0   2   4   6E+3   7.5   8  RT (min)      lcorbi_t3_neg_4_042_14    m/z 411.0415     0   1E+4   1   1.5  RT (min)      lcorbi_t3_neg_3_030_33    m/z 411.0488     0   1   2E+4   9   9.5  RT (min)      lcorbi_t3_neg_4_007_13    m/z 411.0898     0   1E+3   9.5   10  RT (min)      lcorbi_t3_neg_2_055_18    m/z 411.1292     0   1   2E+4   6.5   7  RT (min)      lcorbi_t3_neg_2_048_27    m/z 411.1684     0   1   2   3E+4   10.5   11  RT (min)      lcorbi_t3_neg_2_048_27    m/z 411.1790     0   2   4E+3   13   13.5  RT (min)      lcorbi_t3_neg_4_054_36    m/z 411.1842     0   2   4   6   8E+3   8.5   9  RT (min)      lcorbi_t3_neg_4_001_49    m/z 411.2020     0   2   4   6   8E+3   9.5   10  RT (min)      lcorbi_t3_neg_4_034_01    m/z 411.2203     0   1E+4   14   14.5  RT (min)      lcorbi_t3_neg_4_040_19    m/z 411.2384     0   2   4   6   8E+3   8   8.5  RT (min)      lcorbi_t3_neg_2_048_27    m/z 411.2400     0   1   12.5   13  RT (min)      lcorbi_t3_neg_3_002_24    m/z 411.2517     0   1E+3   11.5   12  RT (min)      lcorbi_t3_neg_4_007_13    m/z 411.2563     0   1E+4   13   13.5  RT (min)      lcorbi_t3_neg_1_055_05    m/z 411.2607     0   2   4   6   8E+3   5.5   6  RT (min)      lcorbi_t3_neg_2_048_27    m/z 411.2901     0   2   4   6   8E+3   12   12.5  RT (min)      lcorbi_t3_neg_4_007_13    m/z 411.3112     0   2   4   6   8E+3   13   13.5  RT (min)      lcorbi_t3_neg_3_007_16    m/z 411.3294     0   1E+4   12   12.5  RT (min)      lcorbi_t3_neg_2_022_24    m/z 411.3475     0   1E+4   13   13.5  RT (min)      lcorbi_t3_neg_4_034_01    m/z 411.8710     0   1E+4   1   1.5  RT (min)      lcorbi_t3_neg_2_052_32    m/z 412.0651     0   1   2   3E+4   1   1.5  RT (min)      lcorbi_t3_neg_2_035_46    m/z 412.1329     0   2   4   6   8E+3   6   6.5  RT (min)      lcorbi_t3_neg_4_007_13    m/z 412.1343     0   1   2   3E+3   12   12.5  RT (min)      lcorbi_t3_neg_3_014_19    m/z 412.1721     0   1   2E+4   5.5   6  RT (min)      lcorbi_t3_neg_2_035_46    m/z 412.1775     0   1   2E+3   7   7.5  RT (min)      lcorbi_t3_neg_1_025_52    m/z 412.2125     0   1   2E+4   13   13.5  RT (min)      lcorbi_t3_neg_4_034_01    m/z 412.2593     0   2   4   6E+3   13   13.5  RT (min)      lcorbi_t3_neg_4_034_01    m/z 412.2727     0   1   2   3E+4   13   13.5  RT (min)      lcorbi_t3_neg_2_053_23    m/z 412.2852     0   1E+4   12.5   13  RT (min)      lcorbi_t3_neg_4_007_13    m/z 412.3063     0   1   11.5   12  RT (min)      lcorbi_t3_neg_4_007_13    m/z 412.3874     0   1E+4   13.5   14  RT (min)      lcorbi_t3_neg_4_034_01    m/z 412.8518     0   2   4E+3   18   18.5  RT (min)      lcorbi_t3_neg_3_003_09    m/z 412.9661     0   1   2   3   4E+2   9   9.5  RT (min)      lcorbi_t3_neg_2_053_23    m/z 413.0412     0   2   4   6   8E+3   7   7.5  RT (min)      lcorbi_t3_neg_4_023_18    m/z 413.0714     0   2   4E+4   6   6.5  RT (min)      lcorbi_t3_neg_2_048_27    m/z 413.0772     0   1E+4   9.5   10  RT (min)      lcorbi_t3_neg_2_048_27    m/z 413.0933     0   1E+4   11   11.5  RT (min)      lcorbi_t3_neg_4_007_13    m/z 413.0954     0   1E+4   7.5   8  RT (min)      lcorbi_t3_neg_3_023_07    m/z 413.1300     0   1   2E+4   1   1.5  RT (min)      lcorbi_t3_neg_1_055_05    m/z 413.1342     0   1   2E+3   9.5   10  RT (min)      lcorbi_t3_neg_3_022_27    m/z 413.1421     0   1E+4   12   12.5  RT (min)      lcorbi_t3_neg_4_040_19    m/z 413.1446     0   2   4   6E+3   9   9.5  RT (min)      lcorbi_t3_neg_3_019_12    m/z 413.1556     0   1E+4   10   10.5  RT (min)      lcorbi_t3_neg_1_043_14    m/z 413.1731     0   1E+4   12.5   13  RT (min)      lcorbi_t3_neg_4_016_05    m/z 413.1946     0   2   4   6E+3   13.5   14  RT (min)      lcorbi_t3_neg_4_047_30    m/z 413.2036     0   1   2   3E+3   5   5.5  RT (min)      lcorbi_t3_neg_4_032_23    m/z 413.2158     0   1   2   3E+4   13   13.5  RT (min)      lcorbi_t3_neg_4_016_05    m/z 413.2176     0   1   2E+4   5.5   6  RT (min)      lcorbi_t3_neg_4_023_18    m/z 413.2313     0   1E+4   11   11.5  RT (min)      lcorbi_t3_neg_2_005_01    m/z 413.3075     0   1   2E+3   12.5   13  RT (min)      lcorbi_t3_neg_3_040_05    m/z 413.3632     0   2   4   6E+3   13   13.5  RT (min)      lcorbi_t3_neg_4_034_01    m/z 414.0428     0   1E+4   1.5   2  RT (min)      lcorbi_t3_neg_4_032_23    m/z 414.0584     0   1E+4   12.5   13  RT (min)      lcorbi_t3_neg_4_007_13    m/z 414.0642     0   2   4   6   8E+4   1   1.5  RT (min)      lcorbi_t3_neg_2_022_24    m/z 414.0716     0   1E+4   9.5   10  RT (min)      lcorbi_t3_neg_2_048_27    m/z 414.0788     0   2   4   6E+3   13   13.5  RT (min)      lcorbi_t3_neg_3_018_13    m/z 414.1247     0   1E+4   8   8.5  RT (min)      lcorbi_t3_neg_2_048_27    m/z 414.1266     0   1E+4   13   13.5  RT (min)      lcorbi_t3_neg_4_007_13    m/z 414.1267     0   1   2   3E+4   11   11.5  RT (min)      lcorbi_t3_neg_2_007_14    m/z 414.1750     0   1   7   7.5  RT (min)      lcorbi_t3_neg_1_051_15    m/z 414.2360     0   2   4   6E+2   12.5   13  RT (min)      lcorbi_t3_neg_4_019_08    m/z 414.2890     0   1   2E+4   12.5   13  RT (min)      lcorbi_t3_neg_4_017_51    m/z 414.3120     0   1   2E+4   11   11.5  RT (min)      lcorbi_t3_neg_4_039_32    m/z 414.3415     0   1   13   13.5  RT (min)      lcorbi_t3_neg_4_040_19    m/z 414.8893     0   1   2   3E+3   18   18.5  RT (min)      lcorbi_t3_neg_2_008_16    m/z 415.0057     0   1E+4   10.5   11  RT (min)      lcorbi_t3_neg_1_007_26    m/z 415.0428     0   2   4   6   8E+3   9   9.5  RT (min)      lcorbi_t3_neg_3_019_12    m/z 415.0514     0   2   4E+3   10   10.5  RT (min)      lcorbi_t3_neg_4_034_01    m/z 415.0820     0   1E+4   4.5   5  RT (min)      lcorbi_t3_neg_2_015_21    m/z 415.0937     0   1E+3   4.5   5  RT (min)      lcorbi_t3_neg_2_015_21    m/z 415.1163     0   1E+4   11   11.5  RT (min)      lcorbi_t3_neg_4_013_09    m/z 415.1247     0   2   4   6E+3   10.5   11  RT (min)      lcorbi_t3_neg_3_020_31    m/z 415.1265     0   1   2E+4   6   6.5  RT (min)      lcorbi_t3_neg_2_048_27    m/z 415.1555     0   2   4E+3   10   10.5  RT (min)      lcorbi_t3_neg_1_022_34    m/z 415.1766     0   2   4   6   8E+3   12.5   13  RT (min)      lcorbi_t3_neg_4_016_05    m/z 415.1864     0   2   4   6E+3   10.5   11  RT (min)      lcorbi_t3_neg_4_042_14    m/z 415.1956     0   1   2E+2   7   7.5  RT (min)      lcorbi_t3_neg_4_021_25    m/z 415.2061     0   1   2   3E+3   7   7.5  RT (min)      lcorbi_t3_neg_4_023_18    m/z 415.2074     0   1E+3   13   13.5  RT (min)      lcorbi_t3_neg_4_016_05    m/z 415.2194     0   1   6   6.5  RT (min)      lcorbi_t3_neg_4_005_27    m/z 415.2287     0   1   13   13.5  RT (min)      lcorbi_t3_neg_4_090_60    m/z 415.2819     0   2   4   6E+3   12   12.5  RT (min)      lcorbi_t3_neg_3_005_23    m/z 415.2876     0   1E+3   13.5   14  RT (min)      lcorbi_t3_neg_1_055_05    m/z 415.2962     0   1E+4   12   12.5  RT (min)      lcorbi_t3_neg_3_024_26    m/z 415.3002     0   1   2E+4   14.5   15  RT (min)      lcorbi_t3_neg_4_005_27    m/z 415.3233     0   2   4E+2   13.5   14  RT (min)      lcorbi_t3_neg_4_040_19    m/z 415.3445     0   1E+3   13   13.5  RT (min)      lcorbi_t3_neg_4_005_27    m/z 415.8831     0   1   2E+4   1   1.5  RT (min)      lcorbi_t3_neg_4_039_32    m/z 416.0327     0   1   2   3   4E+3   10   10.5  RT (min)      lcorbi_t3_neg_3_037_36    m/z 416.0448     0   2   4   6E+3   9.5   10  RT (min)      lcorbi_t3_neg_4_007_13    m/z 416.0554     0   1E+4   12.5   13  RT (min)      lcorbi_t3_neg_4_007_13    m/z 416.1224     0   2   4   6   8E+3   9   9.5  RT (min)      lcorbi_t3_neg_4_023_18    m/z 416.1237     0   1   2   3E+3   13   13.5  RT (min)      lcorbi_t3_neg_4_010_28    m/z 416.1423     0   2   4   6E+2   13   13.5  RT (min)      lcorbi_t3_neg_4_053_43    m/z 416.1425     0   1   2   3   4E+3   10.5   11  RT (min)      lcorbi_t3_neg_1_031_33    m/z 416.1735     0   1   2   3E+4   12.5   13  RT (min)      lcorbi_t3_neg_4_016_05    m/z 416.1850     0   2   4   6E+4   8   8.5  RT (min)      lcorbi_t3_neg_2_048_27    m/z 416.2133     0   1E+4   13   13.5  RT (min)      lcorbi_t3_neg_4_016_05    m/z 416.2283     0   1E+4   11   11.5  RT (min)      lcorbi_t3_neg_1_055_05    m/z 416.2746     0   1   12   12.5  RT (min)      lcorbi_t3_neg_4_019_08    m/z 416.3741     0   2   4   6   8E+3   10.5   11  RT (min)      lcorbi_t3_neg_3_009_50    m/z 416.7797     0   1   1   1.5  RT (min)      lcorbi_t3_neg_3_032_17    m/z 416.8272     0   1   2   3E+2   1   1.5  RT (min)      lcorbi_t3_neg_2_050_17    m/z 417.0388     0   1E+3   9   9.5  RT (min)      lcorbi_t3_neg_4_023_18    m/z 417.1011     0   1E+4   10   10.5  RT (min)      lcorbi_t3_neg_4_023_18    m/z 417.1104     0   1E+4   1   1.5  RT (min)      lcorbi_t3_neg_2_035_46    m/z 417.1500     0   2   4   6E+3   9   9.5  RT (min)      lcorbi_t3_neg_3_019_12    m/z 417.1551     0   1   2   3   4E+3   8   8.5  RT (min)      lcorbi_t3_neg_4_026_46    m/z 417.2101     0   1E+4   11   11.5  RT (min)      lcorbi_t3_neg_4_043_11    m/z 417.2126     0   2   4   6   8E+3   8.5   9  RT (min)      lcorbi_t3_neg_2_048_27    m/z 417.2231     0   1   2   3E+2   13   13.5  RT (min)      lcorbi_t3_neg_4_040_19    m/z 417.2407     0   1   2   3E+4   11.5   12  RT (min)      lcorbi_t3_neg_1_055_05    m/z 417.2537     0   2   4   6   8E+3   13   13.5  RT (min)      lcorbi_t3_neg_4_034_01    m/z 417.2854     0   2   4E+3   12   12.5  RT (min)      lcorbi_t3_neg_4_005_27    m/z 417.3114     0   1   2E+4   14.5   15  RT (min)      lcorbi_t3_neg_4_044_38    m/z 417.3219     0   1E+4   10   10.5  RT (min)      lcorbi_t3_neg_4_023_18    m/z 417.3390     0   2   4   6   8E+3   14   14.5  RT (min)      lcorbi_t3_neg_4_034_01    m/z 417.3729     0   1E+4   12   12.5  RT (min)      lcorbi_t3_neg_3_002_24    m/z 417.9287     0   2   4   6   8E+3   8.5   9  RT (min)      lcorbi_t3_neg_1_054_37    m/z 417.9786     0   1   2E+4   5.5   6  RT (min)      lcorbi_t3_neg_3_024_26    m/z 417.9941     0   1E+4   8   8.5  RT (min)      lcorbi_t3_neg_4_004_34    m/z 418.0525     0   1   2   3   4E+3   12.5   13  RT (min)      lcorbi_t3_neg_4_007_13    m/z 418.0812     0   2   4   6   8E+3   4.5   5  RT (min)      lcorbi_t3_neg_4_023_18    m/z 418.1004     0   1   12.5   13  RT (min)      lcorbi_t3_neg_4_019_08    m/z 418.1110     0   1   1   1.5  RT (min)      lcorbi_t3_neg_4_011_24    m/z 418.1172     0   1   2   3E+4   10   10.5  RT (min)      lcorbi_t3_neg_4_023_18    m/z 418.1394     0   1   13   13.5  RT (min)      lcorbi_t3_neg_4_053_43    m/z 418.1892     0   1   13   13.5  RT (min)      lcorbi_t3_neg_4_019_08    m/z 418.2058     0   2   4   6E+2   11   11.5  RT (min)      lcorbi_t3_neg_4_043_11    m/z 418.2396     0   1   11   11.5  RT (min)      lcorbi_t3_neg_1_055_05    m/z 418.2838     0   1   2E+4   13.5   14  RT (min)      lcorbi_t3_neg_3_018_13    m/z 418.3039     0   1E+4   12.5   13  RT (min)      lcorbi_t3_neg_1_055_05    m/z 418.3577     0   1   2E+3   12   12.5  RT (min)      lcorbi_t3_neg_1_053_32    m/z 418.8940     0   2   4   6E+3   18   18.5  RT (min)      lcorbi_t3_neg_1_001_49    m/z 418.9730     0   2   4   6E+3   9.5   10  RT (min)      lcorbi_t3_neg_2_048_27    m/z 418.9832     0   1   2   3E+4   5.5   6  RT (min)      lcorbi_t3_neg_4_047_30    m/z 419.0435     0   1E+4   9.5   10  RT (min)      lcorbi_t3_neg_4_007_13    m/z 419.0597     0   2   4E+3   5.5   6  RT (min)      lcorbi_t3_neg_4_023_18    m/z 419.0617     0   2   4E+2   9   9.5  RT (min)      lcorbi_t3_neg_1_023_03    m/z 419.1071     0   2   4   6E+4   1   1.5  RT (min)      lcorbi_t3_neg_4_007_13    m/z 419.1920     0   1   2   3E+3   10   10.5  RT (min)      lcorbi_t3_neg_3_045_34    m/z 419.2023     0   2   4   6E+3   11   11.5  RT (min)      lcorbi_t3_neg_1_055_05    m/z 419.2079     0   2   4   6   8E+3   13   13.5  RT (min)      lcorbi_t3_neg_3_040_05    m/z 419.2341     0   2   4   6E+3   8.5   9  RT (min)      lcorbi_t3_neg_4_017_51    m/z 419.2387     0   2   4   6E+3   13.5   14  RT (min)      lcorbi_t3_neg_4_020_44    m/z 419.3526     0   2   4E+3   13   13.5  RT (min)      lcorbi_t3_neg_4_038_15    m/z 419.3886     0   1E+5   12   12.5  RT (min)      lcorbi_t3_neg_4_039_32    m/z 419.9257     0   1   2   3   4E+3   8.5   9  RT (min)      lcorbi_t3_neg_1_054_37    m/z 420.0451     0   1E+4   6   6.5  RT (min)      lcorbi_t3_neg_2_048_27    m/z 420.0924     0   1E+4   11   11.5  RT (min)      lcorbi_t3_neg_1_043_14    m/z 420.1078     0   1   2.5   3  RT (min)      lcorbi_t3_neg_2_048_27    m/z 420.1161     0   1E+4   12.5   13  RT (min)      lcorbi_t3_neg_3_048_08    m/z 420.1248     0   2   4   6E+3   9.5   10  RT (min)      lcorbi_t3_neg_3_019_12    m/z 420.1461     0   1E+4   9   9.5  RT (min)      lcorbi_t3_neg_3_019_12    m/z 420.1551     0   2   4E+3   13.5   14  RT (min)      lcorbi_t3_neg_4_040_19    m/z 420.1998     0   2   4   6E+3   10   10.5  RT (min)      lcorbi_t3_neg_4_023_18    m/z 420.2070     0   1E+4   7   7.5  RT (min)      lcorbi_t3_neg_4_023_18    m/z 420.2598     0   1   2   3   4E+3   12   12.5  RT (min)      lcorbi_t3_neg_4_017_51    m/z 420.2599     0   2   4   6E+3   10   10.5  RT (min)      lcorbi_t3_neg_1_001_49    m/z 420.3044     0   2   4   6   8E+3   12.5   13  RT (min)      lcorbi_t3_neg_4_017_51    m/z 420.3197     0   1   2   3E+3   12.5   13  RT (min)      lcorbi_t3_neg_3_024_26    m/z 420.7073     0   1   1   1.5  RT (min)      lcorbi_t3_neg_4_034_01    m/z 421.0405     0   2   4E+3   9.5   10  RT (min)      lcorbi_t3_neg_1_039_38    m/z 421.0449     0   2   4   6E+3   7   7.5  RT (min)      lcorbi_t3_neg_4_007_13    m/z 421.0713     0   2   4E+3   13   13.5  RT (min)      lcorbi_t3_neg_3_018_13    m/z 421.1175     0   1   5   5.5  RT (min)      lcorbi_t3_neg_4_039_32    m/z 421.1185     0   2   4   6   8E+3   9.5   10  RT (min)      lcorbi_t3_neg_3_019_12    m/z 421.1735     0   1   11   11.5  RT (min)      lcorbi_t3_neg_2_020_05    m/z 421.2262     0   2   4   6E+3   14   14.5  RT (min)      lcorbi_t3_neg_1_039_38    m/z 421.2534     0   1   2E+4   9   9.5  RT (min)      lcorbi_t3_neg_2_048_27    m/z 421.4043     0   2   4   6   8E+4   12.5   13  RT (min)      lcorbi_t3_neg_4_039_32    m/z 421.8744     0   1E+4   1   1.5  RT (min)      lcorbi_t3_neg_4_054_36    m/z 422.0343     0   1E+4   9.5   10  RT (min)      lcorbi_t3_neg_4_007_13    m/z 422.0894     0   2   4   6E+3   11   11.5  RT (min)      lcorbi_t3_neg_1_043_14    m/z 422.0944     0   1   2   3E+3   8   8.5  RT (min)      lcorbi_t3_neg_3_045_34    m/z 422.0983     0   1E+4   1   1.5  RT (min)      lcorbi_t3_neg_2_035_46    m/z 422.1217     0   1E+3   9.5   10  RT (min)      lcorbi_t3_neg_4_007_13    m/z 422.1398     0   1E+4   12.5   13  RT (min)      lcorbi_t3_neg_4_016_05    m/z 422.1581     0   1E+3   10   10.5  RT (min)      lcorbi_t3_neg_4_042_14    m/z 422.1827     0   1   2E+3   11   11.5  RT (min)      lcorbi_t3_neg_1_055_05    m/z 422.1907     0   2   4   6   8E+3   10   10.5  RT (min)      lcorbi_t3_neg_4_029_31    m/z 422.2067     0   1E+4   7   7.5  RT (min)      lcorbi_t3_neg_4_023_18    m/z 422.2295     0   2   4   6E+4   10.5   11  RT (min)      lcorbi_t3_neg_3_039_29    m/z 422.2674     0   2   4   6   8E+3   11.5   12  RT (min)      lcorbi_t3_neg_2_039_31    m/z 422.3272     0   1E+5   10.5   11  RT (min)      lcorbi_t3_neg_2_016_11    m/z 422.3353     0   1E+4   12.5   13  RT (min)      lcorbi_t3_neg_4_034_01    m/z 422.3717     0   1E+4   13   13.5  RT (min)      lcorbi_t3_neg_4_034_01    m/z 422.8359     0   2   4   6E+3   17.5   18  RT (min)      lcorbi_t3_neg_2_003_42    m/z 423.0096     0   1E+4   8   8.5  RT (min)      lcorbi_t3_neg_2_051_34    m/z 423.0419     0   2   4   6E+3   7   7.5  RT (min)      lcorbi_t3_neg_4_007_13    m/z 423.0716     0   1E+4   5.5   6  RT (min)      lcorbi_t3_neg_4_023_18    m/z 423.0908     0   2   4   6E+3   8   8.5  RT (min)      lcorbi_t3_neg_4_023_18    m/z 423.1227     0   2   4   6E+3   10   10.5  RT (min)      lcorbi_t3_neg_4_019_08    m/z 423.1344     0   1E+4   13   13.5  RT (min)      lcorbi_t3_neg_4_042_14    m/z 423.1517     0   1   2E+3   10   10.5  RT (min)      lcorbi_t3_neg_1_043_14    m/z 423.1633     0   1E+4   9   9.5  RT (min)      lcorbi_t3_neg_4_017_51    m/z 423.1943     0   2   4   6   8E+3   6   6.5  RT (min)      lcorbi_t3_neg_4_037_29    m/z 423.2725     0   2   4E+3   13.5   14  RT (min)      lcorbi_t3_neg_4_034_01    m/z 423.2865     0   1E+3   11.5   12  RT (min)      lcorbi_t3_neg_3_040_05    m/z 423.3143     0   1   12.5   13  RT (min)      lcorbi_t3_neg_4_019_08    m/z 423.3841     0   1E+4   13.5   14  RT (min)      lcorbi_t3_neg_2_007_14    m/z 423.9189     0   2   4   6   8E+3   1   1.5  RT (min)      lcorbi_t3_neg_4_029_31    m/z 423.9692     0   1   2   3   4E+3   10   10.5  RT (min)      lcorbi_t3_neg_2_048_27    m/z 424.0762     0   1   2E+4   1.5   2  RT (min)      lcorbi_t3_neg_2_022_24    m/z 424.0776     0   2   4   6   8E+3   10.5   11  RT (min)      lcorbi_t3_neg_3_037_36    m/z 424.0980     0   1   2E+4   1   1.5  RT (min)      lcorbi_t3_neg_3_005_23    m/z 424.1167     0   2   4   6   8E+2   12.5   13  RT (min)      lcorbi_t3_neg_1_055_05    m/z 424.1332     0   2   4   6E+3   5   5.5  RT (min)      lcorbi_t3_neg_4_040_19    m/z 424.1366     0   1   2   3E+3   9.5   10  RT (min)      lcorbi_t3_neg_3_019_12    m/z 424.1474     0   1E+4   13   13.5  RT (min)      lcorbi_t3_neg_4_042_14    m/z 424.1666     0   1   2   3   4E+3   11   11.5  RT (min)      lcorbi_t3_neg_1_022_34    m/z 424.2052     0   1E+4   12.5   13  RT (min)      lcorbi_t3_neg_4_016_05    m/z 424.2191     0   1E+3   10.5   11  RT (min)      lcorbi_t3_neg_1_055_05    m/z 424.2465     0   1E+4   10.5   11  RT (min)      lcorbi_t3_neg_1_055_05    m/z 424.2548     0   2   4   6   8E+3   11   11.5  RT (min)      lcorbi_t3_neg_1_055_05    m/z 424.2830     0   1   2   3   4E+3   11.5   12  RT (min)      lcorbi_t3_neg_1_055_05    m/z 424.3510     0   1E+4   13   13.5  RT (min)      lcorbi_t3_neg_4_034_01    m/z 424.8330     0   2   4E+3   17.5   18  RT (min)      lcorbi_t3_neg_2_049_55    m/z 424.9811     0   1E+4   4.5   5  RT (min)      lcorbi_t3_neg_3_034_21    m/z 425.0801     0   1   1   1.5  RT (min)      lcorbi_t3_neg_2_048_27    m/z 425.0802     0   1E+3   7.5   8  RT (min)      lcorbi_t3_neg_4_042_14    m/z 425.1316     0   2   4E+3   13   13.5  RT (min)      lcorbi_t3_neg_4_042_14    m/z 425.1338     0   2   4   6E+3   9   9.5  RT (min)      lcorbi_t3_neg_3_019_12    m/z 425.1580     0   1E+4   13   13.5  RT (min)      lcorbi_t3_neg_4_054_36    m/z 425.1929     0   2   4   6E+3   6   6.5  RT (min)      lcorbi_t3_neg_3_022_27    m/z 425.2000     0   2   4E+3   8.5   9  RT (min)      lcorbi_t3_neg_4_007_13    m/z 425.2271     0   1   2   3   4E+3   6   6.5  RT (min)      lcorbi_t3_neg_1_026_19    m/z 425.2671     0   1   2E+4   12   12.5  RT (min)      lcorbi_t3_neg_1_055_05    m/z 425.3098     0   1E+4   12   12.5  RT (min)      lcorbi_t3_neg_2_053_23    m/z 425.3382     0   2   4   6E+4   12   12.5  RT (min)      lcorbi_t3_neg_4_007_13    m/z 425.3633     0   1E+4   14   14.5  RT (min)      lcorbi_t3_neg_4_023_18    m/z 425.3997     0   1E+4   13.5   14  RT (min)      lcorbi_t3_neg_4_034_01    m/z 425.8648     0   2   4   6   8E+3   18   18.5  RT (min)      lcorbi_t3_neg_4_006_47    m/z 426.0487     0   2   4   6   8E+3   8.5   9  RT (min)      lcorbi_t3_neg_2_048_27    m/z 426.0836     0   1   2   3E+3   10   10.5  RT (min)      lcorbi_t3_neg_4_005_27    m/z 426.1029     0   1E+4   13   13.5  RT (min)      lcorbi_t3_neg_3_018_13    m/z 426.1087     0   1   2   3   4E+3   9.5   10  RT (min)      lcorbi_t3_neg_2_048_27    m/z 426.1200     0   1   7.5   8  RT (min)      lcorbi_t3_neg_4_019_08    m/z 426.1206     0   2   4E+3   12.5   13  RT (min)      lcorbi_t3_neg_4_054_36    m/z 426.1296     0   1   2   3   4E+4   1   1.5  RT (min)      lcorbi_t3_neg_2_035_46    m/z 426.1445     0   1E+3   13   13.5  RT (min)      lcorbi_t3_neg_3_018_13    m/z 426.1629     0   2   4   6E+3   13.5   14  RT (min)      lcorbi_t3_neg_4_034_01    m/z 426.2208     0   1E+4   13   13.5  RT (min)      lcorbi_t3_neg_4_016_05    m/z 426.2493     0   2   4   6   8E+3   9   9.5  RT (min)      lcorbi_t3_neg_2_048_27    m/z 426.2506     0   1   13   13.5  RT (min)      lcorbi_t3_neg_4_110_62    m/z 426.2541     0   1E+4   7.5   8  RT (min)      lcorbi_t3_neg_4_023_18    m/z 426.3030     0   1   11   11.5  RT (min)      lcorbi_t3_neg_4_019_08    m/z 426.3220     0   1   2   3E+4   12.5   13  RT (min)      lcorbi_t3_neg_4_032_23    m/z 426.4030     0   2   4E+3   13.5   14  RT (min)      lcorbi_t3_neg_4_034_01    m/z 426.9635     0   1   2   3   4E+4   1   1.5  RT (min)      lcorbi_t3_neg_3_002_24    m/z 426.9677     0   1E+4   10   10.5  RT (min)      lcorbi_t3_neg_1_046_11    m/z 427.0101     0   1E+3   1.5   2  RT (min)      lcorbi_t3_neg_2_030_47    m/z 427.0984     0   2   4   6   8E+3   9   9.5  RT (min)      lcorbi_t3_neg_3_019_12    m/z 427.1033     0   2   4   6   8E+2   10   10.5  RT (min)      lcorbi_t3_neg_4_019_08    m/z 427.1135     0   1E+4   7.5   8  RT (min)      lcorbi_t3_neg_4_034_01    m/z 427.1737     0   2   4   6E+3   13   13.5  RT (min)      lcorbi_t3_neg_4_007_13    m/z 427.1790     0   2   4E+3   7   7.5  RT (min)      lcorbi_t3_neg_4_034_01    m/z 427.2155     0   1E+4   7.5   8  RT (min)      lcorbi_t3_neg_4_007_13    m/z 427.2336     0   1E+4   13   13.5  RT (min)      lcorbi_t3_neg_4_044_38    m/z 427.2469     0   1E+4   13   13.5  RT (min)      lcorbi_t3_neg_4_007_13    m/z 427.2531     0   1   11   11.5  RT (min)      lcorbi_t3_neg_1_032_13    m/z 427.2632     0   1   11   11.5  RT (min)      lcorbi_t3_neg_4_007_13    m/z 427.2728     0   1E+4   12   12.5  RT (min)      lcorbi_t3_neg_1_007_26    m/z 427.2865     0   1E+4   12.5   13  RT (min)      lcorbi_t3_neg_4_007_13    m/z 427.2962     0   1E+5   11   11.5  RT (min)      lcorbi_t3_neg_1_055_05    m/z 427.2996     0   1   13   13.5  RT (min)      lcorbi_t3_neg_4_032_23    m/z 427.3538     0   2   4   6E+4   12.5   13  RT (min)      lcorbi_t3_neg_4_007_13    m/z 427.3696     0   2   4   6   8E+3   13   13.5  RT (min)      lcorbi_t3_neg_4_034_01    m/z 427.3789     0   1E+4   13   13.5  RT (min)      lcorbi_t3_neg_4_034_01    m/z 427.8619     0   1   2   3E+3   18   18.5  RT (min)      lcorbi_t3_neg_3_012_47    m/z 427.8822     0   1E+4   1   1.5  RT (min)      lcorbi_t3_neg_4_045_35    m/z 428.0671     0   1   2   3   4E+4   3   3.5  RT (min)      lcorbi_t3_neg_4_007_13    m/z 428.0998     0   1E+4   13   13.5  RT (min)      lcorbi_t3_neg_3_018_13    m/z 428.1008     0   1E+3   9   9.5  RT (min)      lcorbi_t3_neg_2_048_27    m/z 428.1315     0   1   9   9.5  RT (min)      lcorbi_t3_neg_4_019_08    m/z 428.1601     0   2   4E+3   13   13.5  RT (min)      lcorbi_t3_neg_2_007_14    m/z 428.1888     0   2   4   6   8E+3   7   7.5  RT (min)      lcorbi_t3_neg_4_023_18    m/z 428.2075     0   1E+4   10   10.5  RT (min)      lcorbi_t3_neg_2_026_03    m/z 428.2538     0   1E+4   7.5   8  RT (min)      lcorbi_t3_neg_4_023_18    m/z 428.2695     0   1   2   3E+4   12.5   13  RT (min)      lcorbi_t3_neg_4_047_30    m/z 428.2880     0   1E+4   13   13.5  RT (min)      lcorbi_t3_neg_1_055_05    m/z 428.3571     0   1E+4   12.5   13  RT (min)      lcorbi_t3_neg_4_007_13    m/z 428.7596     0   1   1   1.5  RT (min)      lcorbi_t3_neg_4_034_01    m/z 428.9115     0   1E+4   4.5   5  RT (min)      lcorbi_t3_neg_3_042_22    m/z 428.9613     0   1   2   3E+3   18   18.5  RT (min)      lcorbi_t3_neg_2_053_23    m/z 428.9646     0   1E+4   10   10.5  RT (min)      lcorbi_t3_neg_1_046_11    m/z 429.0359     0   1E+4   9   9.5  RT (min)      lcorbi_t3_neg_4_011_24    m/z 429.0471     0   2   4   6E+3   10.5   11  RT (min)      lcorbi_t3_neg_1_007_26    m/z 429.0658     0   1   2   3   4E+4   6   6.5  RT (min)      lcorbi_t3_neg_2_048_27    m/z 429.0806     0   1   2   3E+3   8   8.5  RT (min)      lcorbi_t3_neg_4_023_18    m/z 429.0928     0   2   4   6E+3   10.5   11  RT (min)      lcorbi_t3_neg_4_005_27    m/z 429.0995     0   1   2   3   4E+3   8   8.5  RT (min)      lcorbi_t3_neg_4_052_26    m/z 429.1051     0   2   4E+3   13   13.5  RT (min)      lcorbi_t3_neg_4_042_14    m/z 429.1195     0   1   2E+3   10   10.5  RT (min)      lcorbi_t3_neg_2_048_27    m/z 429.1206     0   1   2E+4   5   5.5  RT (min)      lcorbi_t3_neg_4_040_19    m/z 429.1292     0   1   2   3   4E+3   8   8.5  RT (min)      lcorbi_t3_neg_4_034_01    m/z 429.1368     0   1   2E+3   9.5   10  RT (min)      lcorbi_t3_neg_4_017_51    m/z 429.1734     0   2   4   6E+3   13   13.5  RT (min)      lcorbi_t3_neg_2_007_14    m/z 429.1788     0   1E+4   13   13.5  RT (min)      lcorbi_t3_neg_4_007_13    m/z 429.1895     0   2   4   6E+3   13.5   14  RT (min)      lcorbi_t3_neg_4_053_43    m/z 429.2131     0   1E+4   6   6.5  RT (min)      lcorbi_t3_neg_2_048_27    m/z 429.2446     0   1E+3   12   12.5  RT (min)      lcorbi_t3_neg_4_006_47    m/z 429.2826     0   1   2E+4   12.5   13  RT (min)      lcorbi_t3_neg_1_055_05    m/z 429.3167     0   1   13   13.5  RT (min)      lcorbi_t3_neg_3_018_13    m/z 429.3218     0   2   4E+3   9.5   10  RT (min)      lcorbi_t3_neg_4_023_18    m/z 429.3350     0   1E+4   11   11.5  RT (min)      lcorbi_t3_neg_4_007_13    m/z 429.8767     0   1   2   3E+3   18   18.5  RT (min)      lcorbi_t3_neg_3_015_11    m/z 429.8867     0   1E+4   8.5   9  RT (min)      lcorbi_t3_neg_2_048_27    m/z 430.0229     0   1E+4   9   9.5  RT (min)      lcorbi_t3_neg_3_019_12    m/z 430.0357     0   2   4   6   8E+3   10.5   11  RT (min)      lcorbi_t3_neg_1_024_01    m/z 430.0663     0   1   9.5   10  RT (min)      lcorbi_t3_neg_4_019_08    m/z 430.0803     0   1E+4   8   8.5  RT (min)      lcorbi_t3_neg_4_052_26    m/z 430.0919     0   1E+4   5.5   6  RT (min)      lcorbi_t3_neg_4_023_18    m/z 430.1086     0   1   2   3   4E+3   11   11.5  RT (min)      lcorbi_t3_neg_4_008_22    m/z 430.1131     0   1E+4   1   1.5  RT (min)      lcorbi_t3_neg_1_055_05    m/z 430.1256     0   1   2E+3   11   11.5  RT (min)      lcorbi_t3_neg_4_004_34    m/z 430.1570     0   1   2E+4   1   1.5  RT (min)      lcorbi_t3_neg_4_026_46    m/z 430.1881     0   2   4   6E+3   7   7.5  RT (min)      lcorbi_t3_neg_2_051_34    m/z 430.2276     0   1E+3   12.5   13  RT (min)      lcorbi_t3_neg_2_048_27    m/z 430.2664     0   1E+3   13   13.5  RT (min)      lcorbi_t3_neg_4_011_24    m/z 430.2694     0   1   2   3E+4   12.5   13  RT (min)      lcorbi_t3_neg_4_047_30    m/z 430.2838     0   2   4   6E+4   12   12.5  RT (min)      lcorbi_t3_neg_1_055_05    m/z 430.2956     0   2   4   6E+2   10.5   11  RT (min)      lcorbi_t3_neg_2_048_27    m/z 430.2959     0   2   4E+3   9   9.5  RT (min)      lcorbi_t3_neg_2_048_27    m/z 430.2997     0   2   4   6   8E+3   13   13.5  RT (min)      lcorbi_t3_neg_1_055_05    m/z 430.3039     0   1   2E+4   10.5   11  RT (min)      lcorbi_t3_neg_2_048_27    m/z 430.3331     0   2   4   6E+4   12.5   13  RT (min)      lcorbi_t3_neg_4_011_24    m/z 430.7568     0   1   1   1.5  RT (min)      lcorbi_t3_neg_4_037_29    m/z 430.8601     0   1   2   3E+3   18   18.5  RT (min)      lcorbi_t3_neg_4_053_43    m/z 430.9618     0   1   2   3E+3   10   10.5  RT (min)      lcorbi_t3_neg_1_054_37    m/z 431.0325     0   1   2   3   4E+3   9   9.5  RT (min)      lcorbi_t3_neg_2_013_08    m/z 431.1164     0   2   4   6E+3   5   5.5  RT (min)      lcorbi_t3_neg_2_048_27    m/z 431.1559     0   1E+3   9.5   10  RT (min)      lcorbi_t3_neg_3_043_25    m/z 431.1707     0   1   8   8.5  RT (min)      lcorbi_t3_neg_4_019_08    m/z 431.1743     0   2   4   6E+3   13   13.5  RT (min)      lcorbi_t3_neg_4_007_13    m/z 431.1821     0   1E+4   7   7.5  RT (min)      lcorbi_t3_neg_4_023_18    m/z 431.1917     0   1   5.5   6  RT (min)      lcorbi_t3_neg_4_096_60    m/z 431.1944     0   1E+4   13.5   14  RT (min)      lcorbi_t3_neg_4_040_19    m/z 431.2241     0   1E+4   12   12.5  RT (min)      lcorbi_t3_neg_2_048_27    m/z 431.2282     0   2   4   6E+3   8   8.5  RT (min)      lcorbi_t3_neg_4_023_18    m/z 431.2628     0   1   2   3E+3   13   13.5  RT (min)      lcorbi_t3_neg_3_043_25    m/z 431.2980     0   1E+5   13   13.5  RT (min)      lcorbi_t3_neg_1_055_05    m/z 431.3374     0   2   4   6E+3   10   10.5  RT (min)      lcorbi_t3_neg_4_023_18    m/z 431.3679     0   1E+4   12.5   13  RT (min)      lcorbi_t3_neg_4_011_24    m/z 431.8837     0   1E+4   8.5   9  RT (min)      lcorbi_t3_neg_2_048_27    m/z 431.9295     0   2   4   6   8E+3   18   18.5  RT (min)      lcorbi_t3_neg_1_004_22    m/z 432.0165     0   1   2   3   4E+4   6.5   7  RT (min)      lcorbi_t3_neg_4_023_18    m/z 432.0625     0   1E+4   8   8.5  RT (min)      lcorbi_t3_neg_4_023_18    m/z 432.0787     0   1   2   3E+3   8   8.5  RT (min)      lcorbi_t3_neg_4_023_18    m/z 432.0962     0   2   4   6   8E+3   1.5   2  RT (min)      lcorbi_t3_neg_1_035_25    m/z 432.1423     0   1   2E+4   1   1.5  RT (min)      lcorbi_t3_neg_2_043_33    m/z 432.1605     0   2   4   6   8E+3   13   13.5  RT (min)      lcorbi_t3_neg_4_042_14    m/z 432.1943     0   2   4   6E+3   13   13.5  RT (min)      lcorbi_t3_neg_4_034_01    m/z 432.2057     0   2   4   6   8E+2   11   11.5  RT (min)      lcorbi_t3_neg_1_055_05    m/z 432.2233     0   1   2E+5   10.5   11  RT (min)      lcorbi_t3_neg_1_055_05    m/z 432.2468     0   2   4   6E+3   7.5   8  RT (min)      lcorbi_t3_neg_4_017_51    m/z 432.2753     0   1E+3   11.5   12  RT (min)      lcorbi_t3_neg_1_055_05    m/z 432.2931     0   1   2   3E+4   10   10.5  RT (min)      lcorbi_t3_neg_3_019_12    m/z 432.3035     0   1E+4   12.5   13  RT (min)      lcorbi_t3_neg_4_017_51    m/z 432.3115     0   1E+4   10   10.5  RT (min)      lcorbi_t3_neg_4_007_13    m/z 432.3196     0   2   4   6E+3   10.5   11  RT (min)      lcorbi_t3_neg_2_048_27    m/z 432.3196     0   2   4   6   8E+3   12.5   13  RT (min)      lcorbi_t3_neg_4_042_14    m/z 432.3288     0   1   2   3E+4   13   13.5  RT (min)      lcorbi_t3_neg_1_055_05    m/z 432.3440     0   2   4E+3   13   13.5  RT (min)      lcorbi_t3_neg_4_038_15    m/z 432.8648     0   1E+4   18   18.5  RT (min)      lcorbi_t3_neg_2_027_13    m/z 432.9122     0   1E+4   7.5   8  RT (min)      lcorbi_t3_neg_4_023_18    m/z 432.9253     0   2   4   6E+3   18   18.5  RT (min)      lcorbi_t3_neg_1_002_16    m/z 432.9506     0   2   4   6   8E+3   1   1.5  RT (min)      lcorbi_t3_neg_4_036_16    m/z 432.9762     0   1   2   3   4E+3   1   1.5  RT (min)      lcorbi_t3_neg_4_035_33    m/z 433.0236     0   1   2   3E+4   1   1.5  RT (min)      lcorbi_t3_neg_4_039_32    m/z 433.0473     0   1   9   9.5  RT (min)      lcorbi_t3_neg_4_019_08    m/z 433.0674     0   1E+4   9   9.5  RT (min)      lcorbi_t3_neg_1_055_05    m/z 433.0770     0   1   2E+3   9   9.5  RT (min)      lcorbi_t3_neg_3_022_27    m/z 433.1021     0   1E+4   11   11.5  RT (min)      lcorbi_t3_neg_4_007_13    m/z 433.1188     0   2   4E+2   12   12.5  RT (min)      lcorbi_t3_neg_4_019_08    m/z 433.1278     0   1   2E+4   10   10.5  RT (min)      lcorbi_t3_neg_4_007_13    m/z 433.1351     0   1E+4   1   1.5  RT (min)      lcorbi_t3_neg_2_035_46    m/z 433.1500     0   2   4   6E+3   9.5   10  RT (min)      lcorbi_t3_neg_4_023_18    m/z 433.1630     0   1E+4   7   7.5  RT (min)      lcorbi_t3_neg_2_036_35    m/z 433.1644     0   2   4   6E+3   9.5   10  RT (min)      lcorbi_t3_neg_2_048_27    m/z 433.1724     0   2   4   6E+3   5   5.5  RT (min)      lcorbi_t3_neg_4_032_23    m/z 433.1900     0   2   4   6E+3   13.5   14  RT (min)      lcorbi_t3_neg_4_040_19    m/z 433.2261     0   1E+3   11   11.5  RT (min)      lcorbi_t3_neg_4_017_51    m/z 433.2803     0   1E+4   9   9.5  RT (min)      lcorbi_t3_neg_2_048_27    m/z 433.3267     0   1   2E+4   13   13.5  RT (min)      lcorbi_t3_neg_4_034_01    m/z 433.3452     0   2   4E+4   13   13.5  RT (min)      lcorbi_t3_neg_1_055_05    m/z 433.3487     0   1E+3   11   11.5  RT (min)      lcorbi_t3_neg_1_055_05    m/z 433.3683     0   1E+4   13.5   14  RT (min)      lcorbi_t3_neg_4_005_27    m/z 434.0066     0   1   2E+2   4.5   5  RT (min)      lcorbi_t3_neg_3_052_32    m/z 434.0144     0   1   2   3   4E+4   6.5   7  RT (min)      lcorbi_t3_neg_4_023_18    m/z 434.0352     0   2   4E+3   11   11.5  RT (min)      lcorbi_t3_neg_4_023_18    m/z 434.0872     0   2   4E+4   3   3.5  RT (min)      lcorbi_t3_neg_4_007_13    m/z 434.1033     0   2   4   6   8E+3   9.5   10  RT (min)      lcorbi_t3_neg_3_019_12    m/z 434.1132     0   2   4   6E+3   12   12.5  RT (min)      lcorbi_t3_neg_2_048_27    m/z 434.1351     0   1   2E+4   9   9.5  RT (min)      lcorbi_t3_neg_2_048_27    m/z 434.2027     0   2   4   6   8E+3   13   13.5  RT (min)      lcorbi_t3_neg_4_016_05    m/z 434.2180     0   1E+4   8.5   9  RT (min)      lcorbi_t3_neg_4_017_51    m/z 434.2594     0   1   2   3   4E+3   13   13.5  RT (min)      lcorbi_t3_neg_4_042_14    m/z 434.3083     0   1   2E+4   12.5   13  RT (min)      lcorbi_t3_neg_1_023_03    m/z 434.8618     0   1E+4   18   18.5  RT (min)      lcorbi_t3_neg_2_031_25    m/z 434.8715     0   2   4   6   8E+3   18   18.5  RT (min)      lcorbi_t3_neg_2_014_29    m/z 434.9172     0   1   2E+3   7.5   8  RT (min)      lcorbi_t3_neg_3_054_18    m/z 435.0170     0   1E+4   9   9.5  RT (min)      lcorbi_t3_neg_3_019_12    m/z 435.1198     0   1   10   10.5  RT (min)      lcorbi_t3_neg_3_053_14    m/z 435.1310     0   2   4   6E+4   1   1.5  RT (min)      lcorbi_t3_neg_3_002_24    m/z 435.2415     0   2   4   6E+4   11   11.5  RT (min)      lcorbi_t3_neg_1_055_05    m/z 435.2561     0   1   2E+4   6   6.5  RT (min)      lcorbi_t3_neg_4_040_19    m/z 435.2834     0   1E+3   13   13.5  RT (min)      lcorbi_t3_neg_4_005_27    m/z 435.8936     0   1   2   3E+4   18   18.5  RT (min)      lcorbi_t3_neg_4_052_26    m/z 435.9656     0   1   2E+3   8   8.5  RT (min)      lcorbi_t3_neg_2_048_27    m/z 436.0550     0   2   4   6E+3   1.5   2  RT (min)      lcorbi_t3_neg_1_011_48    m/z 436.0661     0   1E+4   11.5   12  RT (min)      lcorbi_t3_neg_1_029_21    m/z 436.0827     0   1   2   3   4E+3   10   10.5  RT (min)      lcorbi_t3_neg_3_013_28    m/z 436.1322     0   1   2E+4   9   9.5  RT (min)      lcorbi_t3_neg_2_048_27    m/z 436.1339     0   2   4E+3   1.5   2  RT (min)      lcorbi_t3_neg_2_052_32    m/z 436.1997     0   1   2   3   4E+3   10   10.5  RT (min)      lcorbi_t3_neg_3_028_35    m/z 436.2364     0   2   4   6E+4   12.5   13  RT (min)      lcorbi_t3_neg_1_055_05    m/z 436.2527     0   1   2E+3   13   13.5  RT (min)      lcorbi_t3_neg_1_055_05    m/z 436.2957     0   1E+5   11   11.5  RT (min)      lcorbi_t3_neg_1_055_05    m/z 436.3511     0   1E+4   13   13.5  RT (min)      lcorbi_t3_neg_4_034_01    m/z 436.8766     0   2   4E+3   18   18.5  RT (min)      lcorbi_t3_neg_4_052_26    m/z 436.9487     0   1   2   3E+3   18   18.5  RT (min)      lcorbi_t3_neg_2_007_14    m/z 436.9576     0   2   4   6E+3   8.5   9  RT (min)      lcorbi_t3_neg_4_040_19    m/z 436.9957     0   1E+4   9   9.5  RT (min)      lcorbi_t3_neg_1_023_03    m/z 437.1143     0   2   4   6E+3   10.5   11  RT (min)      lcorbi_t3_neg_4_043_11    m/z 437.1258     0   1   2   3E+4   9   9.5  RT (min)      lcorbi_t3_neg_2_048_27    m/z 437.1319     0   2   4   6E+3   10.5   11  RT (min)      lcorbi_t3_neg_3_002_24    m/z 437.1411     0   2   4   6E+3   7.5   8  RT (min)      lcorbi_t3_neg_4_034_01    m/z 437.1512     0   1   2   3   4E+3   7.5   8  RT (min)      lcorbi_t3_neg_4_034_01    m/z 437.1790     0   2   4   6E+3   10.5   11  RT (min)      lcorbi_t3_neg_4_029_31    m/z 437.1962     0   2   4   6E+3   10   10.5  RT (min)      lcorbi_t3_neg_3_045_34    m/z 437.2327     0   2   4   6E+2   11   11.5  RT (min)      lcorbi_t3_neg_4_032_23    m/z 437.2401     0   1E+4   9   9.5  RT (min)      lcorbi_t3_neg_2_048_27    m/z 437.2481     0   1E+4   13   13.5  RT (min)      lcorbi_t3_neg_4_007_13    m/z 437.2863     0   1   11.5   12  RT (min)      lcorbi_t3_neg_4_016_05    m/z 437.3019     0   1   2   3   4E+4   12   12.5  RT (min)      lcorbi_t3_neg_3_024_26    m/z 437.3422     0   1   2E+4   13.5   14  RT (min)      lcorbi_t3_neg_2_035_46    m/z 437.3632     0   1E+3   13   13.5  RT (min)      lcorbi_t3_neg_2_022_24    m/z 437.9298     0   1   2E+4   1   1.5  RT (min)      lcorbi_t3_neg_1_043_14    m/z 437.9429     0   1E+4   10.5   11  RT (min)      lcorbi_t3_neg_4_007_13    m/z 437.9699     0   1   2   3E+3   18   18.5  RT (min)      lcorbi_t3_neg_2_007_14    m/z 438.0788     0   2   4   6   8E+3   13   13.5  RT (min)      lcorbi_t3_neg_4_007_13    m/z 438.1268     0   1   2   3   4E+3   13   13.5  RT (min)      lcorbi_t3_neg_4_007_13    m/z 438.1580     0   2   4E+4   5   5.5  RT (min)      lcorbi_t3_neg_2_048_27    m/z 438.1646     0   1E+4   8   8.5  RT (min)      lcorbi_t3_neg_4_034_01    m/z 438.1776     0   2   4   6E+3   10   10.5  RT (min)      lcorbi_t3_neg_4_032_23    m/z 438.1935     0   1E+3   7   7.5  RT (min)      lcorbi_t3_neg_4_102_61    m/z 438.2360     0   1E+4   12.5   13  RT (min)      lcorbi_t3_neg_1_055_05    m/z 438.2468     0   1   2E+4   6   6.5  RT (min)      lcorbi_t3_neg_2_048_27    m/z 438.2622     0   1   2   3E+4   11   11.5  RT (min)      lcorbi_t3_neg_1_055_05    m/z 438.2659     0   1   2E+3   13   13.5  RT (min)      lcorbi_t3_neg_4_035_33    m/z 438.2704     0   1   2E+4   12   12.5  RT (min)      lcorbi_t3_neg_1_055_05    m/z 438.2833     0   1   12   12.5  RT (min)      lcorbi_t3_neg_4_019_08    m/z 438.3111     0   2   4E+4   11.5   12  RT (min)      lcorbi_t3_neg_1_055_05    m/z 438.3333     0   2   4   6   8E+3   12.5   13  RT (min)      lcorbi_t3_neg_4_017_51    m/z 438.3456     0   2   4   6   8E+3   13.5   14  RT (min)      lcorbi_t3_neg_2_035_46    m/z 438.3666     0   2   4   6E+3   13   13.5  RT (min)      lcorbi_t3_neg_4_034_01    m/z 438.7208     0   1   1   1.5  RT (min)      lcorbi_t3_neg_1_055_05    m/z 438.8960     0   1E+4   1   1.5  RT (min)      lcorbi_t3_neg_4_045_35    m/z 439.0617     0   1   2   3   4E+3   12.5   13  RT (min)      lcorbi_t3_neg_1_043_14    m/z 439.0961     0   1E+4   10   10.5  RT (min)      lcorbi_t3_neg_4_023_18    m/z 439.1316     0   1   2   3E+3   10   10.5  RT (min)      lcorbi_t3_neg_4_029_31    m/z 439.1393     0   1   2E+4   7   7.5  RT (min)      lcorbi_t3_neg_4_040_19    m/z 439.1475     0   1E+3   9   9.5  RT (min)      lcorbi_t3_neg_2_046_20    m/z 439.1603     0   1   2   3   4E+3   6.5   7  RT (min)      lcorbi_t3_neg_2_048_27    m/z 439.1899     0   1   2E+3   11   11.5  RT (min)      lcorbi_t3_neg_4_017_51    m/z 439.2075     0   2   4   6E+2   13   13.5  RT (min)      lcorbi_t3_neg_1_055_05    m/z 439.2251     0   1E+4   11   11.5  RT (min)      lcorbi_t3_neg_4_017_51    m/z 439.2330     0   1   2E+3   7   7.5  RT (min)      lcorbi_t3_neg_4_008_22    m/z 439.2827     0   1   2   3E+3   12.5   13  RT (min)      lcorbi_t3_neg_4_007_13    m/z 439.3010     0   1E+4   10   10.5  RT (min)      lcorbi_t3_neg_3_019_12    m/z 439.3020     0   1E+4   11.5   12  RT (min)      lcorbi_t3_neg_1_055_05    m/z 439.3216     0   1   2E+3   13   13.5  RT (min)      lcorbi_t3_neg_4_034_01    m/z 439.3286     0   1E+4   12   12.5  RT (min)      lcorbi_t3_neg_4_007_13    m/z 439.3425     0   1   2E+4   12.5   13  RT (min)      lcorbi_t3_neg_4_034_01    m/z 439.3519     0   1   2   3E+4   12   12.5  RT (min)      lcorbi_t3_neg_2_022_24    m/z 439.3703     0   1   2E+4   12.5   13  RT (min)      lcorbi_t3_neg_2_022_24    m/z 439.8713     0   1   2E+4   1   1.5  RT (min)      lcorbi_t3_neg_2_052_32    m/z 439.8881     0   1E+4   18   18.5  RT (min)      lcorbi_t3_neg_2_048_27    m/z 439.9267     0   1E+4   1   1.5  RT (min)      lcorbi_t3_neg_1_043_14    m/z 439.9357     0   1   8   8.5  RT (min)      lcorbi_t3_neg_1_045_44    m/z 439.9399     0   1E+4   10.5   11  RT (min)      lcorbi_t3_neg_4_007_13    m/z 440.0677     0   1   2   3   4E+3   9   9.5  RT (min)      lcorbi_t3_neg_3_019_12    m/z 440.0885     0   2   4   6   8E+3   1   1.5  RT (min)      lcorbi_t3_neg_3_043_25    m/z 440.0945     0   1   2   3E+3   13   13.5  RT (min)      lcorbi_t3_neg_4_007_13    m/z 440.0981     0   1E+4   5   5.5  RT (min)      lcorbi_t3_neg_2_048_27    m/z 440.1549     0   1E+4   13   13.5  RT (min)      lcorbi_t3_neg_3_018_13    m/z 440.2002     0   1   2E+4   12.5   13  RT (min)      lcorbi_t3_neg_4_016_05    m/z 440.2285     0   1   2   3E+3   11   11.5  RT (min)      lcorbi_t3_neg_4_017_51    m/z 440.2865     0   1   2   3E+3   13   13.5  RT (min)      lcorbi_t3_neg_3_037_36    m/z 440.3047     0   1E+3   12.5   13  RT (min)      lcorbi_t3_neg_1_055_05    m/z 440.3124     0   2   4   6   8E+3   7   7.5  RT (min)      lcorbi_t3_neg_2_039_31    m/z 440.3234     0   1E+4   10   10.5  RT (min)      lcorbi_t3_neg_3_019_12    m/z 440.3821     0   2   4   6E+3   13   13.5  RT (min)      lcorbi_t3_neg_4_034_01    m/z 440.6923     0   1   1   1.5  RT (min)      lcorbi_t3_neg_4_007_13    m/z 440.8889     0   2   4   6   8E+4   1   1.5  RT (min)      lcorbi_t3_neg_2_048_27    m/z 441.0587     0   1   2   3E+3   12.5   13  RT (min)      lcorbi_t3_neg_1_043_14    m/z 441.0613     0   2   4   6   8E+3   9   9.5  RT (min)      lcorbi_t3_neg_3_019_12    m/z 441.1217     0   1E+4   10   10.5  RT (min)      lcorbi_t3_neg_2_002_26    m/z 441.1300     0   1   2E+4   6   6.5  RT (min)      lcorbi_t3_neg_2_048_27    m/z 441.1344     0   1E+4   10.5   11  RT (min)      lcorbi_t3_neg_1_043_14    m/z 441.1501     0   1E+4   10.5   11  RT (min)      lcorbi_t3_neg_1_044_23    m/z 441.2070     0   1   2   3E+4   7.5   8  RT (min)      lcorbi_t3_neg_2_048_27    m/z 441.2408     0   1   6   6.5  RT (min)      lcorbi_t3_neg_4_051_40    m/z 441.2537     0   2   4E+3   13   13.5  RT (min)      lcorbi_t3_neg_4_034_01    m/z 441.2869     0   2   4   6   8E+4   13   13.5  RT (min)      lcorbi_t3_neg_4_038_15    m/z 441.3161     0   2   4   6E+4   12   12.5  RT (min)      lcorbi_t3_neg_1_055_05    m/z 441.3445     0   1E+4   11.5   12  RT (min)      lcorbi_t3_neg_3_018_13    m/z 441.3581     0   1   2   3   4E+3   12   12.5  RT (min)      lcorbi_t3_neg_4_042_14    m/z 441.3753     0   1E+3   13   13.5  RT (min)      lcorbi_t3_neg_4_007_13    m/z 441.5492     0   1E+4   4.5   5  RT (min)      lcorbi_t3_neg_2_048_27    m/z 441.8867     0   1E+4   8   8.5  RT (min)      lcorbi_t3_neg_2_037_09    m/z 441.8920     0   1   2   3E+3   18   18.5  RT (min)      lcorbi_t3_neg_2_011_04    m/z 441.9312     0   1   2   3E+4   8.5   9  RT (min)      lcorbi_t3_neg_2_015_21    m/z 442.0526     0   2   4   6E+3   12.5   13  RT (min)      lcorbi_t3_neg_1_043_14    m/z 442.0721     0   2   4   6   8E+4   8.5   9  RT (min)      lcorbi_t3_neg_2_048_27    m/z 442.1101     0   1E+4   13.5   14  RT (min)      lcorbi_t3_neg_4_040_19    m/z 442.1840     0   1E+4   12.5   13  RT (min)      lcorbi_t3_neg_4_039_32    m/z 442.1892     0   1   2   3   4E+3   11   11.5  RT (min)      lcorbi_t3_neg_4_004_34    m/z 442.2158     0   1   2E+4   13   13.5  RT (min)      lcorbi_t3_neg_4_016_05    m/z 442.2665     0   1E+4   6   6.5  RT (min)      lcorbi_t3_neg_2_048_27    m/z 442.3037     0   1   10   10.5  RT (min)      lcorbi_t3_neg_4_019_08    m/z 442.3399     0   1   2   3   4E+3   10   10.5  RT (min)      lcorbi_t3_neg_2_048_27    m/z 442.3404     0   1E+4   13.5   14  RT (min)      lcorbi_t3_neg_4_046_37    m/z 442.3979     0   1E+4   13   13.5  RT (min)      lcorbi_t3_neg_4_034_01    m/z 442.8936     0   1E+4   10   10.5  RT (min)      lcorbi_t3_neg_4_053_43    m/z 442.9659     0   2   4   6   8E+3   18   18.5  RT (min)      lcorbi_t3_neg_4_042_14    m/z 442.9851     0   1   2E+4   9   9.5  RT (min)      lcorbi_t3_neg_1_023_03    m/z 443.0123     0   2   4E+3   8   8.5  RT (min)      lcorbi_t3_neg_4_023_18    m/z 443.0487     0   1   2   3   4E+2   9   9.5  RT (min)      lcorbi_t3_neg_1_023_03    m/z 443.0930     0   1E+4   13   13.5  RT (min)      lcorbi_t3_neg_4_007_13    m/z 443.1102     0   2   4   6E+3   10   10.5  RT (min)      lcorbi_t3_neg_3_054_18    m/z 443.1476     0   2   4   6   8E+3   12   12.5  RT (min)      lcorbi_t3_neg_4_040_19    m/z 443.1687     0   1   2   3   4E+3   13   13.5  RT (min)      lcorbi_t3_neg_4_044_38    m/z 443.1971     0   1   2E+4   11   11.5  RT (min)      lcorbi_t3_neg_2_050_17    m/z 443.2089     0   1E+4   13   13.5  RT (min)      lcorbi_t3_neg_2_035_46    m/z 443.2244     0   1   2E+3   11   11.5  RT (min)      lcorbi_t3_neg_4_010_28    m/z 443.2281     0   1E+3   13   13.5  RT (min)      lcorbi_t3_neg_4_034_01    m/z 443.2423     0   1E+4   11   11.5  RT (min)      lcorbi_t3_neg_4_017_51    m/z 443.2473     0   2   4   6E+3   13.5   14  RT (min)      lcorbi_t3_neg_4_017_51    m/z 443.2580     0   1   2   3   4E+2   10   10.5  RT (min)      lcorbi_t3_neg_3_019_12    m/z 443.2605     0   1   2E+3   13   13.5  RT (min)      lcorbi_t3_neg_4_044_38    m/z 443.2695     0   1E+5   13   13.5  RT (min)      lcorbi_t3_neg_4_019_08    m/z 443.2987     0   1   2E+3   13   13.5  RT (min)      lcorbi_t3_neg_3_048_08    m/z 443.3069     0   2   4   6E+2   12.5   13  RT (min)      lcorbi_t3_neg_4_034_01    m/z 443.3239     0   1E+3   13   13.5  RT (min)      lcorbi_t3_neg_3_024_26    m/z 443.8654     0   1E+3   1   1.5  RT (min)      lcorbi_t3_neg_3_022_27    m/z 443.8837     0   1E+4   8   8.5  RT (min)      lcorbi_t3_neg_2_037_09    m/z 443.9422     0   1   2   3E+3   8   8.5  RT (min)      lcorbi_t3_neg_3_054_18    m/z 444.0795     0   2   4E+3   9   9.5  RT (min)      lcorbi_t3_neg_3_019_12    m/z 444.0924     0   1E+4   11   11.5  RT (min)      lcorbi_t3_neg_2_048_27    m/z 444.1274     0   1   2E+5   2.5   3   3.5  RT (min)       lcorbi_t3_neg_2_048_27    m/z 444.1589     0   2   4   6   8E+3   8.5   9  RT (min)      lcorbi_t3_neg_4_007_13    m/z 444.1810     0   2   4   6E+3   12.5   13  RT (min)      lcorbi_t3_neg_4_039_32    m/z 444.1888     0   1   2   3   4E+3   11   11.5  RT (min)      lcorbi_t3_neg_2_048_27    m/z 444.2127     0   2   4E+3   10.5   11  RT (min)      lcorbi_t3_neg_4_034_01    m/z 444.2458     0   2   4   6   8E+3   5.5   6  RT (min)      lcorbi_t3_neg_4_042_14    m/z 444.2625     0   1   12.5   13  RT (min)      lcorbi_t3_neg_3_053_14    m/z 444.2906     0   1   14   14.5  RT (min)      lcorbi_t3_neg_4_042_14    m/z 444.3195     0   1E+4   12.5   13  RT (min)      lcorbi_t3_neg_4_034_01    m/z 444.3327     0   1   2   3   4E+3   11   11.5  RT (min)      lcorbi_t3_neg_4_034_01    m/z 444.6870     0   1   2   3E+3   1   1.5  RT (min)      lcorbi_t3_neg_4_007_13    m/z 444.8907     0   1   2E+4   18   18.5  RT (min)      lcorbi_t3_neg_2_021_41    m/z 444.9628     0   1   2   3   4E+3   18   18.5  RT (min)      lcorbi_t3_neg_4_042_14    m/z 445.0165     0   2   4E+3   9.5   10  RT (min)      lcorbi_t3_neg_4_020_44    m/z 445.0713     0   2   4   6E+3   13   13.5  RT (min)      lcorbi_t3_neg_4_007_13    m/z 445.0900     0   1   13   13.5  RT (min)      lcorbi_t3_neg_4_045_35    m/z 445.0927     0   1E+4   5   5.5  RT (min)      lcorbi_t3_neg_2_048_27    m/z 445.1192     0   1   2   3E+4   3   3.5  RT (min)      lcorbi_t3_neg_2_048_27    m/z 445.1223     0   1E+4   7   7.5  RT (min)      lcorbi_t3_neg_2_007_14    m/z 445.1242     0   2   4E+4   9.5   10  RT (min)      lcorbi_t3_neg_1_007_26    m/z 445.1346     0   1   2E+4   5   5.5  RT (min)      lcorbi_t3_neg_2_048_27    m/z 445.2419     0   1   11   11.5  RT (min)      lcorbi_t3_neg_1_055_05    m/z 445.2446     0   1   2E+3   9.5   10  RT (min)      lcorbi_t3_neg_4_029_31    m/z 445.2590     0   2   4E+2   12.5   13  RT (min)      lcorbi_t3_neg_2_045_30    m/z 445.2764     0   1E+3   12.5   13  RT (min)      lcorbi_t3_neg_4_042_14    m/z 445.2853     0   1E+4   13.5   14  RT (min)      lcorbi_t3_neg_3_018_13    m/z 445.2955     0   2   4E+2   9.5   10  RT (min)      lcorbi_t3_neg_4_032_23    m/z 445.3028     0   2   4   6E+3   12   12.5  RT (min)      lcorbi_t3_neg_4_042_14    m/z 445.3304     0   2   4   6E+3   10   10.5  RT (min)      lcorbi_t3_neg_3_019_12    m/z 445.3449     0   1E+4   13   13.5  RT (min)      lcorbi_t3_neg_3_024_26    m/z 445.3679     0   1E+4   12   12.5  RT (min)      lcorbi_t3_neg_4_011_24    m/z 445.9252     0   1E+3   8   8.5  RT (min)      lcorbi_t3_neg_1_007_26    m/z 445.9391     0   1E+4   8   8.5  RT (min)      lcorbi_t3_neg_1_007_26    m/z 445.9746     0   2   4   6E+3   10   10.5  RT (min)      lcorbi_t3_neg_2_036_35    m/z 445.9943     0   1   2   3   4E+3   8   8.5  RT (min)      lcorbi_t3_neg_3_045_34    m/z 446.0129     0   1   2E+4   9.5   10  RT (min)      lcorbi_t3_neg_4_020_44    m/z 446.0220     0   1   2E+3   10   10.5  RT (min)      lcorbi_t3_neg_2_051_34    m/z 446.1152     0   2   4   6E+3   5   5.5  RT (min)      lcorbi_t3_neg_4_040_19    m/z 446.1319     0   1E+4   10   10.5  RT (min)      lcorbi_t3_neg_1_007_26    m/z 446.1374     0   2   4E+4   6   6.5  RT (min)      lcorbi_t3_neg_2_048_27    m/z 446.1385     0   2   4   6   8E+3   9   9.5  RT (min)      lcorbi_t3_neg_3_019_12    m/z 446.1519     0   2   4   6E+3   10   10.5  RT (min)      lcorbi_t3_neg_4_044_38    m/z 446.1713     0   1E+4   10.5   11  RT (min)      lcorbi_t3_neg_2_048_27    m/z 446.2285     0   1E+4   11   11.5  RT (min)      lcorbi_t3_neg_4_042_14    m/z 446.2371     0   1E+3   12   12.5  RT (min)      lcorbi_t3_neg_1_055_05    m/z 446.3200     0   2   4   6   8E+2   12.5   13  RT (min)      lcorbi_t3_neg_4_034_01    m/z 446.3802     0   1   2   3E+4   12   12.5  RT (min)      lcorbi_t3_neg_4_011_24    m/z 446.9054     0   1E+4   9.5   10  RT (min)      lcorbi_t3_neg_2_002_26    m/z 446.9331     0   1   8   8.5  RT (min)      lcorbi_t3_neg_4_028_39    m/z 447.0242     0   2   4   6E+3   8   8.5  RT (min)      lcorbi_t3_neg_4_023_18    m/z 447.0305     0   2   4   6E+3   11   11.5  RT (min)      lcorbi_t3_neg_1_042_18    m/z 447.0636     0   1E+4   12   12.5  RT (min)      lcorbi_t3_neg_3_002_24    m/z 447.0870     0   1E+3   13   13.5  RT (min)      lcorbi_t3_neg_4_035_33    m/z 447.0884     0   1E+3   5   5.5  RT (min)      lcorbi_t3_neg_4_023_18    m/z 447.0977     0   1E+3   9   9.5  RT (min)      lcorbi_t3_neg_2_048_27    m/z 447.1657     0   1   2   3   4E+3   9   9.5  RT (min)      lcorbi_t3_neg_4_044_38    m/z 447.1975     0   1E+4   11   11.5  RT (min)      lcorbi_t3_neg_4_021_25    m/z 447.2012     0   1E+4   8   8.5  RT (min)      lcorbi_t3_neg_2_048_27    m/z 447.2335     0   2   4   6   8E+3   12   12.5  RT (min)      lcorbi_t3_neg_4_005_27    m/z 447.2392     0   2   4E+3   13   13.5  RT (min)      lcorbi_t3_neg_4_016_05    m/z 447.2514     0   1   2   3   4E+4   11   11.5  RT (min)      lcorbi_t3_neg_1_055_05    m/z 447.2556     0   1   2E+3   13.5   14  RT (min)      lcorbi_t3_neg_4_042_14    m/z 447.2942     0   2   4   6E+3   8.5   9  RT (min)      lcorbi_t3_neg_2_048_27    m/z 447.3802     0   2   4   6E+4   14   14.5  RT (min)      lcorbi_t3_neg_4_007_13    m/z 447.8467     0   1   2   3E+3   1   1.5  RT (min)      lcorbi_t3_neg_4_039_32    m/z 447.9089     0   2   4   6   8E+3   18   18.5  RT (min)      lcorbi_t3_neg_4_051_40    m/z 447.9716     0   1   2   3   4E+4   10.5   11  RT (min)      lcorbi_t3_neg_4_007_13    m/z 447.9884     0   1E+3   9   9.5  RT (min)      lcorbi_t3_neg_1_047_02    m/z 448.0635     0   2   4E+3   12   12.5  RT (min)      lcorbi_t3_neg_4_011_24    m/z 448.0897     0   1   2   3   4E+3   9   9.5  RT (min)      lcorbi_t3_neg_3_019_12    m/z 448.1288     0   1   2   3   4E+3   13   13.5  RT (min)      lcorbi_t3_neg_4_042_14    m/z 448.1464     0   2   4   6   8E+3   10.5   11  RT (min)      lcorbi_t3_neg_4_102_61    m/z 448.1474     0   2   4   6   8E+3   13   13.5  RT (min)      lcorbi_t3_neg_4_042_14    m/z 448.1554     0   1E+4   13   13.5  RT (min)      lcorbi_t3_neg_4_042_14    m/z 448.1641     0   2   4   6E+3   10.5   11  RT (min)      lcorbi_t3_neg_3_002_24    m/z 448.1656     0   2   4E+4   6   6.5  RT (min)      lcorbi_t3_neg_2_048_27    m/z 448.2159     0   1E+4   13   13.5  RT (min)      lcorbi_t3_neg_4_005_27    m/z 448.2782     0   2   4   6   8E+3   10   10.5  RT (min)      lcorbi_t3_neg_2_007_14    m/z 448.2876     0   1E+3   12   12.5  RT (min)      lcorbi_t3_neg_1_055_05    m/z 448.2881     0   1   2E+3   9   9.5  RT (min)      lcorbi_t3_neg_1_043_14    m/z 448.3167     0   1   2   3   4E+2   12   12.5  RT (min)      lcorbi_t3_neg_3_007_16    m/z 448.3249     0   2   4E+3   9   9.5  RT (min)      lcorbi_t3_neg_2_048_27    m/z 448.3357     0   1   2   3   4E+3   13   13.5  RT (min)      lcorbi_t3_neg_1_055_05    m/z 448.7624     0   1E+3   1   1.5  RT (min)      lcorbi_t3_neg_3_022_27    m/z 448.9099     0   1   2   3   4E+3   18   18.5  RT (min)      lcorbi_t3_neg_4_053_43    m/z 448.9331     0   1E+4   9   9.5  RT (min)      lcorbi_t3_neg_2_048_27    m/z 448.9456     0   1   2   3E+3   1   1.5  RT (min)      lcorbi_t3_neg_4_046_37    m/z 448.9472     0   2   4   6E+3   8   8.5  RT (min)      lcorbi_t3_neg_2_048_27    m/z 449.0176     0   1   2E+4   10.5   11  RT (min)      lcorbi_t3_neg_1_043_14    m/z 449.0197     0   1E+3   9   9.5  RT (min)      lcorbi_t3_neg_4_023_18    m/z 449.0275     0   2   4E+3   11   11.5  RT (min)      lcorbi_t3_neg_1_042_18    m/z 449.0512     0   1   2E+4   10   10.5  RT (min)      lcorbi_t3_neg_4_023_18    m/z 449.1026     0   1E+4   13.5   14  RT (min)      lcorbi_t3_neg_4_040_19    m/z 449.1267     0   2   4E+2   7   7.5  RT (min)      lcorbi_t3_neg_4_007_13    m/z 449.1404     0   1E+3   9   9.5  RT (min)      lcorbi_t3_neg_3_052_32    m/z 449.1496     0   1   2E+3   9   9.5  RT (min)      lcorbi_t3_neg_2_053_23    m/z 449.1529     0   1E+4   5   5.5  RT (min)      lcorbi_t3_neg_4_010_28    m/z 449.1625     0   1E+4   13   13.5  RT (min)      lcorbi_t3_neg_4_042_14    m/z 449.2023     0   1   2   3   4E+4   5   5.5  RT (min)      lcorbi_t3_neg_2_048_27    m/z 449.2208     0   1E+4   8   8.5  RT (min)      lcorbi_t3_neg_2_035_46    m/z 449.2672     0   1   2E+4   11.5   12  RT (min)      lcorbi_t3_neg_2_020_05    m/z 449.3019     0   1E+3   12.5   13  RT (min)      lcorbi_t3_neg_4_019_08    m/z 449.3122     0   1   2E+4   13.5   14  RT (min)      lcorbi_t3_neg_1_009_50    m/z 449.3540     0   2   4E+3   13   13.5  RT (min)      lcorbi_t3_neg_1_020_36    m/z 449.9686     0   1   2E+4   10.5   11  RT (min)      lcorbi_t3_neg_4_007_13    m/z 449.9846     0   2   4   6E+3   9   9.5  RT (min)      lcorbi_t3_neg_3_019_12    m/z 450.0091     0   2   4   6   8E+3   9   9.5  RT (min)      lcorbi_t3_neg_4_007_13    m/z 450.0213     0   1E+4   11   11.5  RT (min)      lcorbi_t3_neg_3_034_21    m/z 450.1028     0   1   2E+4   13   13.5  RT (min)      lcorbi_t3_neg_4_007_13    m/z 450.1324     0   2   4E+3   12.5   13  RT (min)      lcorbi_t3_neg_3_018_13    m/z 450.1631     0   2   4   6E+3   13   13.5  RT (min)      lcorbi_t3_neg_4_042_14    m/z 450.1653     0   2   4E+4   6   6.5  RT (min)      lcorbi_t3_neg_2_048_27    m/z 450.2221     0   1   2   3E+3   11   11.5  RT (min)      lcorbi_t3_neg_4_032_23    m/z 450.7595     0   1   1   1.5  RT (min)      lcorbi_t3_neg_3_039_29    m/z 450.8441     0   1   2   3E+3   18.5   19  RT (min)      lcorbi_t3_neg_1_001_49    m/z 450.8720     0   1E+3   8.5   9  RT (min)      lcorbi_t3_neg_4_023_18    m/z 450.8985     0   1   2   3E+3   18   18.5  RT (min)      lcorbi_t3_neg_2_003_42    m/z 451.0595     0   1E+4   9   9.5  RT (min)      lcorbi_t3_neg_4_042_14    m/z 451.0896     0   1   2   3E+4   1   1.5  RT (min)      lcorbi_t3_neg_4_036_16    m/z 451.0985     0   2   4   6E+3   9   9.5  RT (min)      lcorbi_t3_neg_3_019_12    m/z 451.1299     0   2   4   6   8E+3   11   11.5  RT (min)      lcorbi_t3_neg_4_043_11    m/z 451.1393     0   1   2E+4   3   3.5  RT (min)      lcorbi_t3_neg_2_048_27    m/z 451.1400     0   1E+4   6.5   7  RT (min)      lcorbi_t3_neg_2_048_27    m/z 451.1631     0   1E+4   10.5   11  RT (min)      lcorbi_t3_neg_2_048_27    m/z 451.1632     0   2   4   6   8E+3   13   13.5  RT (min)      lcorbi_t3_neg_1_043_14    m/z 451.1946     0   1   11.5   12  RT (min)      lcorbi_t3_neg_1_055_05    m/z 451.2310     0   1E+4   7.5   8  RT (min)      lcorbi_t3_neg_4_023_18    m/z 451.2656     0   2   4E+4   11   11.5  RT (min)      lcorbi_t3_neg_1_055_05    m/z 451.2827     0   1   2E+4   12   12.5  RT (min)      lcorbi_t3_neg_1_055_05    m/z 451.3079     0   1E+4   11.5   12  RT (min)      lcorbi_t3_neg_4_029_31    m/z 451.3174     0   2   4   6   8E+2   12   12.5  RT (min)      lcorbi_t3_neg_1_055_05    m/z 451.3272     0   1E+3   12   12.5  RT (min)      lcorbi_t3_neg_1_055_05    m/z 451.3423     0   1   2   3E+3   13   13.5  RT (min)      lcorbi_t3_neg_4_017_51    m/z 451.8598     0   1   2E+4   1   1.5  RT (min)      lcorbi_t3_neg_2_052_32    m/z 452.0054     0   1E+4   10   10.5  RT (min)      lcorbi_t3_neg_1_043_14    m/z 452.0215     0   2   4E+3   9.5   10  RT (min)      lcorbi_t3_neg_1_046_11    m/z 452.0998     0   1E+4   13   13.5  RT (min)      lcorbi_t3_neg_1_043_14    m/z 452.1186     0   1E+4   13   13.5  RT (min)      lcorbi_t3_neg_4_007_13    m/z 452.1479     0   2   4   6E+3   13   13.5  RT (min)      lcorbi_t3_neg_4_053_43    m/z 452.1877     0   1E+4   13.5   14  RT (min)      lcorbi_t3_neg_4_047_30    m/z 452.2677     0   1E+3   13   13.5  RT (min)      lcorbi_t3_neg_2_030_47    m/z 452.3822     0   1   2E+4   13   13.5  RT (min)      lcorbi_t3_neg_4_034_01    m/z 452.7565     0   2   4   6   8E+3   1   1.5  RT (min)      lcorbi_t3_neg_3_021_38    m/z 452.8619     0   1   2E+3   18   18.5  RT (min)      lcorbi_t3_neg_4_019_08    m/z 452.8693     0   2   4   6E+2   8.5   9  RT (min)      lcorbi_t3_neg_4_022_41    m/z 452.8715     0   1   2E+3   18   18.5  RT (min)      lcorbi_t3_neg_4_053_43    m/z 452.9223     0   1   2E+4   12.5   13  RT (min)      lcorbi_t3_neg_1_037_06    m/z 452.9224     0   1   2   3   4E+4   9.5   10  RT (min)      lcorbi_t3_neg_4_018_42    m/z 452.9774     0   1   2E+3   9   9.5  RT (min)      lcorbi_t3_neg_1_042_18    m/z 453.0411     0   1   2   3   4E+5   8.5   9  RT (min)       lcorbi_t3_neg_4_023_18    m/z 453.1129     0   1E+4   9   9.5  RT (min)      lcorbi_t3_neg_3_019_12    m/z 453.1318     0   1   2   3E+3   8   8.5  RT (min)      lcorbi_t3_neg_4_023_18    m/z 453.1360     0   2   4   6   8E+3   1   1.5  RT (min)      lcorbi_t3_neg_3_002_24    m/z 453.1389     0   1   2   3   4E+3   7.5   8  RT (min)      lcorbi_t3_neg_4_034_01    m/z 453.1559     0   1   6   6.5  RT (min)      lcorbi_t3_neg_2_046_20    m/z 453.1601     0   1E+4   10.5   11  RT (min)      lcorbi_t3_neg_4_042_14    m/z 453.1656     0   2   4   6   8E+3   7   7.5  RT (min)      lcorbi_t3_neg_4_023_18    m/z 453.1742     0   1E+4   10.5   11  RT (min)      lcorbi_t3_neg_2_048_27    m/z 453.2275     0   2   4   6   8E+3   11   11.5  RT (min)      lcorbi_t3_neg_4_042_14    m/z 453.2312     0   1   2   3E+3   8.5   9  RT (min)      lcorbi_t3_neg_4_034_01    m/z 453.2465     0   2   4   6E+3   7.5   8  RT (min)      lcorbi_t3_neg_4_023_18    m/z 453.2621     0   1E+4   12   12.5  RT (min)      lcorbi_t3_neg_4_005_27    m/z 453.2854     0   1E+5   9.5   10  RT (min)      lcorbi_t3_neg_4_023_18    m/z 453.3008     0   1   2E+4   12.5   13  RT (min)      lcorbi_t3_neg_1_055_05    m/z 453.3027     0   1   2   3E+5   11   11.5  RT (min)      lcorbi_t3_neg_1_055_05    m/z 453.3333     0   1   12.5   13  RT (min)      lcorbi_t3_neg_4_039_32    m/z 453.3946     0   1   2   3E+4   13.5   14  RT (min)      lcorbi_t3_neg_4_034_01    m/z 453.8568     0   1   2E+3   1   1.5  RT (min)      lcorbi_t3_neg_3_039_29    m/z 453.9330     0   2   4E+3   8   8.5  RT (min)      lcorbi_t3_neg_4_023_18    m/z 453.9553     0   1E+4   5.5   6  RT (min)      lcorbi_t3_neg_3_024_26    m/z 454.1015     0   1   2E+4   4.5   5  RT (min)      lcorbi_t3_neg_4_029_31    m/z 454.1156     0   2   4   6   8E+3   13   13.5  RT (min)      lcorbi_t3_neg_4_007_13    m/z 454.1342     0   1   2   3E+4   13.5   14  RT (min)      lcorbi_t3_neg_4_040_19    m/z 454.1432     0   1   2E+4   4.5   5  RT (min)      lcorbi_t3_neg_4_011_24    m/z 454.1513     0   2   4   6   8E+3   8   8.5  RT (min)      lcorbi_t3_neg_4_034_01    m/z 454.1863     0   1E+4   13   13.5  RT (min)      lcorbi_t3_neg_3_024_26    m/z 454.2695     0   1   12   12.5  RT (min)      lcorbi_t3_neg_4_019_08    m/z 454.2833     0   2   4   6   8E+3   13   13.5  RT (min)      lcorbi_t3_neg_4_034_01    m/z 454.2855     0   1   2   3E+3   12.5   13  RT (min)      lcorbi_t3_neg_4_021_25    m/z 454.2888     0   1   2E+4   9.5   10  RT (min)      lcorbi_t3_neg_4_023_18    m/z 454.2969     0   2   4   6   8E+4   12.5   13  RT (min)      lcorbi_t3_neg_1_055_05    m/z 454.3253     0   2   4   6E+2   11   11.5  RT (min)      lcorbi_t3_neg_2_022_24    m/z 454.3478     0   1E+5   11   11.5  RT (min)      lcorbi_t3_neg_1_055_05    m/z 454.3978     0   1E+4   13.5   14  RT (min)      lcorbi_t3_neg_4_034_01    m/z 454.4119     0   2   4   6   8E+3   13   13.5  RT (min)      lcorbi_t3_neg_4_039_32    m/z 454.8493     0   1   2   3   4E+3   18   18.5  RT (min)      lcorbi_t3_neg_3_028_35    m/z 455.0057     0   1   2   3   4E+4   1   1.5  RT (min)      lcorbi_t3_neg_4_039_32    m/z 455.0075     0   1   8.5   9  RT (min)      lcorbi_t3_neg_4_019_08    m/z 455.0482     0   1   2E+3   9   9.5  RT (min)      lcorbi_t3_neg_2_051_34    m/z 455.1211     0   1E+4   1   1.5  RT (min)      lcorbi_t3_neg_1_044_23    m/z 455.1528     0   2   4   6E+3   9   9.5  RT (min)      lcorbi_t3_neg_3_019_12    m/z 455.1635     0   2   4E+3   8   8.5  RT (min)      lcorbi_t3_neg_4_034_01    m/z 455.1846     0   2   4   6E+3   10   10.5  RT (min)      lcorbi_t3_neg_4_029_31    m/z 455.2283     0   2   4   6E+3   8.5   9  RT (min)      lcorbi_t3_neg_2_048_27    m/z 455.2386     0   2   4   6E+2   11.5   12  RT (min)      lcorbi_t3_neg_4_011_24    m/z 455.2470     0   1E+4   9   9.5  RT (min)      lcorbi_t3_neg_4_034_01    m/z 455.2472     0   1E+3   12.5   13  RT (min)      lcorbi_t3_neg_4_032_23    m/z 455.2629     0   1   2   3   4E+3   13   13.5  RT (min)      lcorbi_t3_neg_4_007_13    m/z 455.2784     0   1E+4   10   10.5  RT (min)      lcorbi_t3_neg_3_019_12    m/z 455.2930     0   1E+5   12.5   13  RT (min)      lcorbi_t3_neg_1_055_05    m/z 455.3027     0   2   4E+4   13   13.5  RT (min)      lcorbi_t3_neg_3_024_26    m/z 455.3711     0   1E+4   10   10.5  RT (min)      lcorbi_t3_neg_4_029_31    m/z 455.3732     0   1E+3   12   12.5  RT (min)      lcorbi_t3_neg_4_023_18    m/z 455.9013     0   1   2E+4   1   1.5  RT (min)      lcorbi_t3_neg_4_039_32    m/z 455.9523     0   1E+4   5.5   6  RT (min)      lcorbi_t3_neg_2_002_26    m/z 456.0215     0   2   4   6   8E+3   6   6.5  RT (min)      lcorbi_t3_neg_1_039_38    m/z 456.0691     0   1   11   11.5  RT (min)      lcorbi_t3_neg_4_007_13    m/z 456.0949     0   1E+4   6   6.5  RT (min)      lcorbi_t3_neg_4_040_19    m/z 456.1040     0   1E+4   11   11.5  RT (min)      lcorbi_t3_neg_4_007_13    m/z 456.1312     0   1   2E+4   13.5   14  RT (min)      lcorbi_t3_neg_4_040_19    m/z 456.2042     0   2   4   6E+3   7   7.5  RT (min)      lcorbi_t3_neg_4_023_18    m/z 456.2431     0   2   4   6E+3   12.5   13  RT (min)      lcorbi_t3_neg_4_007_13    m/z 456.2503     0   1   2   3E+3   9   9.5  RT (min)      lcorbi_t3_neg_4_007_13    m/z 456.2573     0   1E+4   1   1.5  RT (min)      lcorbi_t3_neg_4_042_14    m/z 456.2597     0   1   13   13.5  RT (min)      lcorbi_t3_neg_4_011_24    m/z 456.2739     0   1E+3   11.5   12  RT (min)      lcorbi_t3_neg_2_020_05    m/z 456.2767     0   1   2   3   4E+3   9.5   10  RT (min)      lcorbi_t3_neg_4_029_31    m/z 456.2853     0   1E+4   12.5   13  RT (min)      lcorbi_t3_neg_4_007_13    m/z 456.2942     0   2   4   6   8E+2   11   11.5  RT (min)      lcorbi_t3_neg_1_052_24    m/z 456.3122     0   1E+5   13   13.5  RT (min)      lcorbi_t3_neg_1_055_05    m/z 456.3194     0   2   4   6E+3   13   13.5  RT (min)      lcorbi_t3_neg_1_055_05    m/z 456.3439     0   2   4   6   8E+2   12   12.5  RT (min)      lcorbi_t3_neg_2_031_25    m/z 456.3912     0   1   12   12.5  RT (min)      lcorbi_t3_neg_4_007_13    m/z 456.8611     0   1E+4   18   18.5  RT (min)      lcorbi_t3_neg_4_113_62    m/z 456.9233     0   1   2   3   4E+4   4.5   5  RT (min)      lcorbi_t3_neg_3_037_36    m/z 456.9256     0   2   4   6E+3   8   8.5  RT (min)      lcorbi_t3_neg_4_023_18    m/z 457.0171     0   1E+4   9.5   10  RT (min)      lcorbi_t3_neg_4_007_13    m/z 457.0435     0   1   2E+4   5   5.5  RT (min)      lcorbi_t3_neg_2_048_27    m/z 457.0559     0   1   2   3   4E+3   8.5   9  RT (min)      lcorbi_t3_neg_4_040_19    m/z 457.0593     0   2   4   6   8E+3   11   11.5  RT (min)      lcorbi_t3_neg_3_034_21    m/z 457.1006     0   1   2E+4   11   11.5  RT (min)      lcorbi_t3_neg_4_007_13    m/z 457.1196     0   1   2   3E+4   1   1.5  RT (min)      lcorbi_t3_neg_3_005_23    m/z 457.1651     0   2   4   6E+3   9.5   10  RT (min)      lcorbi_t3_neg_1_013_35    m/z 457.2017     0   1   2E+4   12   12.5  RT (min)      lcorbi_t3_neg_4_040_19    m/z 457.2183     0   1   2   3   4E+3   11.5   12  RT (min)      lcorbi_t3_neg_4_042_14    m/z 457.2369     0   2   4E+3   13   13.5  RT (min)      lcorbi_t3_neg_3_018_13    m/z 457.2558     0   2   4   6E+2   11   11.5  RT (min)      lcorbi_t3_neg_1_044_23    m/z 457.2560     0   2   4   6   8E+2   13   13.5  RT (min)      lcorbi_t3_neg_4_011_24    m/z 457.2754     0   2   4E+3   10   10.5  RT (min)      lcorbi_t3_neg_3_019_12    m/z 457.2778     0   1E+4   13   13.5  RT (min)      lcorbi_t3_neg_4_034_01    m/z 457.2804     0   1E+5   11.5   12  RT (min)      lcorbi_t3_neg_1_055_05    m/z 457.3146     0   1E+3   13   13.5  RT (min)      lcorbi_t3_neg_2_020_05    m/z 457.3317     0   1   2E+4   13   13.5  RT (min)      lcorbi_t3_neg_1_055_05    m/z 457.3433     0   1E+4   12.5   13  RT (min)      lcorbi_t3_neg_4_007_13    m/z 457.3534     0   1   2E+3   12   12.5  RT (min)      lcorbi_t3_neg_3_024_26    m/z 457.9852     0   1E+4   8.5   9  RT (min)      lcorbi_t3_neg_4_052_26    m/z 457.9897     0   1   2E+3   9   9.5  RT (min)      lcorbi_t3_neg_3_010_15    m/z 458.0004     0   1E+4   10.5   11  RT (min)      lcorbi_t3_neg_4_007_13    m/z 458.0661     0   1   11   11.5  RT (min)      lcorbi_t3_neg_4_007_13    m/z 458.0880     0   1E+3   12   12.5  RT (min)      lcorbi_t3_neg_4_023_18    m/z 458.0918     0   1   2   3   4E+3   6   6.5  RT (min)      lcorbi_t3_neg_4_040_19    m/z 458.1012     0   1E+4   11   11.5  RT (min)      lcorbi_t3_neg_4_007_13    m/z 458.1345     0   1   2   3   4E+4   13   13.5  RT (min)      lcorbi_t3_neg_3_040_05    m/z 458.1548     0   1   2   3E+3   13   13.5  RT (min)      lcorbi_t3_neg_4_040_19    m/z 458.2540     0   1E+4   11   11.5  RT (min)      lcorbi_t3_neg_4_017_51    m/z 458.3192     0   1   2E+4   12.5   13  RT (min)      lcorbi_t3_neg_1_055_05    m/z 458.3374     0   1   2E+5   12   12.5  RT (min)      lcorbi_t3_neg_1_055_05    m/z 458.3612     0   1E+4   11   11.5  RT (min)      lcorbi_t3_neg_2_035_46    m/z 458.4068     0   1E+4   12   12.5  RT (min)      lcorbi_t3_neg_4_039_32    m/z 458.8613     0   1   2   3E+3   18   18.5  RT (min)      lcorbi_t3_neg_4_019_08    m/z 458.9202     0   1   2   3E+4   4.5   5  RT (min)      lcorbi_t3_neg_3_037_36    m/z 458.9561     0   1E+3   5.5   6  RT (min)      lcorbi_t3_neg_4_012_17    m/z 459.0142     0   1E+3   9.5   10  RT (min)      lcorbi_t3_neg_4_042_14    m/z 459.0478     0   1E+4   9   9.5  RT (min)      lcorbi_t3_neg_4_007_13    m/z 459.0563     0   2   4   6E+3   11   11.5  RT (min)      lcorbi_t3_neg_3_034_21    m/z 459.0809     0   1E+4   10   10.5  RT (min)      lcorbi_t3_neg_4_029_31    m/z 459.0986     0   1   2   3E+4   11   11.5  RT (min)      lcorbi_t3_neg_4_007_13    m/z 459.1114     0   1   2   3   4E+3   1.5   2  RT (min)      lcorbi_t3_neg_4_038_15    m/z 459.1468     0   2   4   6   8E+3   13   13.5  RT (min)      lcorbi_t3_neg_4_040_19    m/z 459.1948     0   1   13   13.5  RT (min)      lcorbi_t3_neg_4_019_08    m/z 459.2129     0   1E+4   11   11.5  RT (min)      lcorbi_t3_neg_4_023_18    m/z 459.2389     0   1E+3   11   11.5  RT (min)      lcorbi_t3_neg_1_055_05    m/z 459.2515     0   1E+4   11   11.5  RT (min)      lcorbi_t3_neg_4_034_01    m/z 459.2837     0   1   12   12.5  RT (min)      lcorbi_t3_neg_2_053_23    m/z 459.2960     0   1   2E+5   12   12.5  RT (min)      lcorbi_t3_neg_1_055_05    m/z 459.3262     0   1E+5   13.5   14  RT (min)      lcorbi_t3_neg_1_055_05    m/z 459.3357     0   1   2E+3   11.5   12  RT (min)      lcorbi_t3_neg_4_021_25    m/z 459.8839     0   1   2   3   4E+3   18   18.5  RT (min)      lcorbi_t3_neg_4_004_34    m/z 459.9118     0   1   2   3E+4   1   1.5  RT (min)      lcorbi_t3_neg_1_043_14    m/z 459.9592     0   1   2E+4   6.5   7  RT (min)      lcorbi_t3_neg_3_045_34    m/z 459.9831     0   1E+4   8.5   9  RT (min)      lcorbi_t3_neg_4_052_26    m/z 460.0428     0   1E+4   12.5   13  RT (min)      lcorbi_t3_neg_3_048_08    m/z 460.0500     0   1   2   3   4E+3   11   11.5  RT (min)      lcorbi_t3_neg_4_008_22    m/z 460.0984     0   1   2E+4   11   11.5  RT (min)      lcorbi_t3_neg_4_007_13    m/z 460.1096     0   2   4   6   8E+3   5   5.5  RT (min)      lcorbi_t3_neg_2_012_36    m/z 460.1122     0   1E+4   10.5   11  RT (min)      lcorbi_t3_neg_3_010_15    m/z 460.1130     0   2   4   6E+3   12.5   13  RT (min)      lcorbi_t3_neg_1_055_05    m/z 460.1314     0   1   2   3E+4   13   13.5  RT (min)      lcorbi_t3_neg_3_040_05    m/z 460.1372     0   1   11.5   12  RT (min)      lcorbi_t3_neg_4_019_08    m/z 460.2173     0   2   4E+3   11   11.5  RT (min)      lcorbi_t3_neg_4_023_18    m/z 460.2384     0   2   4   6E+3   8.5   9  RT (min)      lcorbi_t3_neg_4_017_51    m/z 460.2465     0   1   2E+4   10.5   11  RT (min)      lcorbi_t3_neg_1_055_05    m/z 460.2530     0   1   2   3   4E+3   8   8.5  RT (min)      lcorbi_t3_neg_2_035_46    m/z 460.2587     0   2   4   6E+3   13   13.5  RT (min)      lcorbi_t3_neg_4_038_15    m/z 460.2751     0   1   2E+3   13   13.5  RT (min)      lcorbi_t3_neg_1_055_05    m/z 460.2778     0   2   4   6E+3   10   10.5  RT (min)      lcorbi_t3_neg_3_019_12    m/z 460.2811     0   1E+4   7   7.5  RT (min)      lcorbi_t3_neg_4_032_23    m/z 460.2993     0   2   4E+4   12   12.5  RT (min)      lcorbi_t3_neg_1_055_05    m/z 460.3509     0   1E+3   13   13.5  RT (min)      lcorbi_t3_neg_1_055_05    m/z 460.8567     0   1   2   3E+3   18   18.5  RT (min)      lcorbi_t3_neg_4_031_48    m/z 460.9008     0   2   4   6E+3   8.5   9  RT (min)      lcorbi_t3_neg_4_053_43    m/z 460.9331     0   2   4   6   8E+3   9   9.5  RT (min)      lcorbi_t3_neg_2_048_27    m/z 461.0259     0   2   4   6E+3   5.5   6  RT (min)      lcorbi_t3_neg_4_023_18    m/z 461.0414     0   2   4E+3   7.5   8  RT (min)      lcorbi_t3_neg_4_034_01    m/z 461.0487     0   2   4E+3   10.5   11  RT (min)      lcorbi_t3_neg_2_011_04    m/z 461.0765     0   1E+3   11   11.5  RT (min)      lcorbi_t3_neg_4_007_13    m/z 461.0829     0   2   4   6   8E+3   5   5.5  RT (min)      lcorbi_t3_neg_4_040_19    m/z 461.0984     0   1E+4   7   7.5  RT (min)      lcorbi_t3_neg_2_048_27    m/z 461.1438     0   2   4   6   8E+3   13   13.5  RT (min)      lcorbi_t3_neg_4_040_19    m/z 461.1467     0   1   2E+4   5   5.5  RT (min)      lcorbi_t3_neg_2_048_27    m/z 461.1692     0   2   4E+4   8   8.5  RT (min)      lcorbi_t3_neg_2_048_27    m/z 461.2138     0   1   11   11.5  RT (min)      lcorbi_t3_neg_4_112_62    m/z 461.2363     0   1   2   3E+4   10.5   11  RT (min)      lcorbi_t3_neg_4_032_23    m/z 461.2479     0   1   2E+4   13   13.5  RT (min)      lcorbi_t3_neg_4_007_13    m/z 461.2905     0   1   2E+4   11   11.5  RT (min)      lcorbi_t3_neg_4_032_23    m/z 461.3116     0   1E+5   12   12.5  RT (min)      lcorbi_t3_neg_1_055_05    m/z 461.3400     0   2   4E+3   13   13.5  RT (min)      lcorbi_t3_neg_4_034_01    m/z 461.3801     0   1   11.5   12  RT (min)      lcorbi_t3_neg_4_039_32    m/z 461.3912     0   2   4   6   8E+3   10   10.5  RT (min)      lcorbi_t3_neg_4_001_49    m/z 461.9088     0   2   4   6E+3   1   1.5  RT (min)      lcorbi_t3_neg_4_054_36    m/z 462.0036     0   1   2E+3   4.5   5  RT (min)      lcorbi_t3_neg_3_004_30    m/z 462.0398     0   1E+4   12.5   13  RT (min)      lcorbi_t3_neg_3_048_08    m/z 462.0586     0   1E+4   12.5   13  RT (min)      lcorbi_t3_neg_3_048_08    m/z 462.0754     0   1   2E+4   5.5   6  RT (min)      lcorbi_t3_neg_2_048_27    m/z 462.0787     0   1   13   13.5  RT (min)      lcorbi_t3_neg_3_040_05    m/z 462.0908     0   1   2E+3   9.5   10  RT (min)      lcorbi_t3_neg_3_019_12    m/z 462.0951     0   1   2E+4   11   11.5  RT (min)      lcorbi_t3_neg_4_007_13    m/z 462.1029     0   1E+4   7.5   8  RT (min)      lcorbi_t3_neg_3_054_18    m/z 462.1281     0   1E+4   13   13.5  RT (min)      lcorbi_t3_neg_3_040_05    m/z 462.1621     0   1E+4   11   11.5  RT (min)      lcorbi_t3_neg_4_102_61    m/z 462.2382     0   2   4   6E+3   8.5   9  RT (min)      lcorbi_t3_neg_4_017_51    m/z 462.2622     0   2   4E+4   11   11.5  RT (min)      lcorbi_t3_neg_1_055_05    m/z 462.2906     0   1   2   3   4E+3   13.5   14  RT (min)      lcorbi_t3_neg_4_040_19    m/z 462.3666     0   2   4   6   8E+2   13   13.5  RT (min)      lcorbi_t3_neg_1_055_05    m/z 462.8043     0   1   2   3E+3   0   0.5  RT (min)      lcorbi_t3_neg_4_067_57    m/z 462.8779     0   2   4   6E+3   9.5   10  RT (min)      lcorbi_t3_neg_2_054_15    m/z 462.8790     0   1E+4   18   18.5  RT (min)      lcorbi_t3_neg_4_037_29    m/z 462.8980     0   1   2   3   4E+2   8.5   9  RT (min)      lcorbi_t3_neg_3_036_41    m/z 463.0394     0   2   4E+4   7.5   8  RT (min)      lcorbi_t3_neg_4_023_18    m/z 463.0733     0   1E+4   11   11.5  RT (min)      lcorbi_t3_neg_1_043_14    m/z 463.0881     0   2   4   6E+2   9.5   10  RT (min)      lcorbi_t3_neg_3_007_16    m/z 463.1606     0   2   4   6E+3   8   8.5  RT (min)      lcorbi_t3_neg_4_044_38    m/z 463.1650     0   1   2   3   4E+3   9.5   10  RT (min)      lcorbi_t3_neg_4_039_32    m/z 463.1768     0   2   4   6E+3   8   8.5  RT (min)      lcorbi_t3_neg_4_034_01    m/z 463.1799     0   2   4   6   8E+3   6.5   7  RT (min)      lcorbi_t3_neg_2_048_27    m/z 463.2075     0   1E+4   13   13.5  RT (min)      lcorbi_t3_neg_4_034_01    m/z 463.2635     0   2   4E+4   13   13.5  RT (min)      lcorbi_t3_neg_4_007_13    m/z 463.2828     0   2   4   6E+3   12   12.5  RT (min)      lcorbi_t3_neg_4_007_13    m/z 463.2974     0   1E+4   10   10.5  RT (min)      lcorbi_t3_neg_3_019_12    m/z 463.3019     0   1E+5   12   12.5  RT (min)      lcorbi_t3_neg_1_055_05    m/z 463.3060     0   1E+4   10   10.5  RT (min)      lcorbi_t3_neg_4_034_01    m/z 463.3172     0   2   4   6   8E+2   12   12.5  RT (min)      lcorbi_t3_neg_4_034_01    m/z 463.3425     0   1   2E+4   11.5   12  RT (min)      lcorbi_t3_neg_4_034_01    m/z 463.3789     0   1   2E+4   13   13.5  RT (min)      lcorbi_t3_neg_4_034_01    m/z 463.9531     0   2   4   6   8E+3   1   1.5  RT (min)      lcorbi_t3_neg_1_031_33    m/z 464.0555     0   2   4   6   8E+3   12.5   13  RT (min)      lcorbi_t3_neg_4_038_15    m/z 464.0999     0   2   4   6   8E+3   7.5   8  RT (min)      lcorbi_t3_neg_3_054_18    m/z 464.1441     0   1E+4   13   13.5  RT (min)      lcorbi_t3_neg_4_040_19    m/z 464.1506     0   2   4E+4   1   1.5  RT (min)      lcorbi_t3_neg_3_052_32    m/z 464.1548     0   1E+4   13   13.5  RT (min)      lcorbi_t3_neg_4_007_13    m/z 464.2068     0   1   2   3E+3   10   10.5  RT (min)      lcorbi_t3_neg_4_023_18    m/z 464.2390     0   2   4   6E+3   11   11.5  RT (min)      lcorbi_t3_neg_4_054_36    m/z 464.2516     0   1E+4   13   13.5  RT (min)      lcorbi_t3_neg_4_034_01    m/z 464.2676     0   1   2   3   4E+3   13   13.5  RT (min)      lcorbi_t3_neg_4_044_38    m/z 464.2943     0   1   2   3E+3   12   12.5  RT (min)      lcorbi_t3_neg_1_055_05    m/z 464.2990     0   1E+4   12   12.5  RT (min)      lcorbi_t3_neg_1_055_05    m/z 464.3095     0   1   10   10.5  RT (min)      lcorbi_t3_neg_4_023_18    m/z 464.3736     0   1E+4   12.5   13  RT (min)      lcorbi_t3_neg_4_011_24    m/z 464.8782     0   1   2E+4   18   18.5  RT (min)      lcorbi_t3_neg_2_048_27    m/z 464.9943     0   2   4   6E+3   8   8.5  RT (min)      lcorbi_t3_neg_4_023_18    m/z 465.0363     0   2   4   6E+2   8.5   9  RT (min)      lcorbi_t3_neg_4_023_18    m/z 465.0532     0   2   4   6E+3   10   10.5  RT (min)      lcorbi_t3_neg_2_048_27    m/z 465.0669     0   1   2   3   4E+3   9   9.5  RT (min)      lcorbi_t3_neg_3_019_12    m/z 465.1266     0   1E+4   9.5   10  RT (min)      lcorbi_t3_neg_2_022_24    m/z 465.2079     0   1   2   3   4E+2   9.5   10  RT (min)      lcorbi_t3_neg_4_032_23    m/z 465.2295     0   1   2E+3   7   7.5  RT (min)      lcorbi_t3_neg_2_015_21    m/z 465.2576     0   1   2   3   4E+4   11   11.5  RT (min)      lcorbi_t3_neg_4_011_24    m/z 465.2639     0   2   4   6E+3   13   13.5  RT (min)      lcorbi_t3_neg_4_042_14    m/z 465.2979     0   1E+4   13   13.5  RT (min)      lcorbi_t3_neg_3_007_16    m/z 465.2984     0   1   9.5   10  RT (min)      lcorbi_t3_neg_4_019_08    m/z 465.3310     0   2   4   6E+4   14   14.5  RT (min)      lcorbi_t3_neg_2_048_27    m/z 465.3323     0   1   2   3   4E+3   12.5   13  RT (min)      lcorbi_t3_neg_3_032_17    m/z 465.3581     0   1E+4   12.5   13  RT (min)      lcorbi_t3_neg_2_035_46    m/z 465.3920     0   1   2E+4   13   13.5  RT (min)      lcorbi_t3_neg_3_002_24    m/z 465.9476     0   2   4   6E+3   1   1.5  RT (min)      lcorbi_t3_neg_4_054_36    m/z 466.0832     0   1   2   3E+3   9   9.5  RT (min)      lcorbi_t3_neg_3_019_12    m/z 466.0848     0   1E+4   11   11.5  RT (min)      lcorbi_t3_neg_3_034_21    m/z 466.1517     0   1   2   3E+3   13   13.5  RT (min)      lcorbi_t3_neg_3_004_30    m/z 466.1660     0   1E+4   13   13.5  RT (min)      lcorbi_t3_neg_4_042_14    m/z 466.1706     0   1E+4   13   13.5  RT (min)      lcorbi_t3_neg_4_007_13    m/z 466.1958     0   2   4E+3   8   8.5  RT (min)      lcorbi_t3_neg_4_034_01    m/z 466.2168     0   2   4E+3   10   10.5  RT (min)      lcorbi_t3_neg_4_029_31    m/z 466.2523     0   1   8.5   9  RT (min)      lcorbi_t3_neg_4_023_18    m/z 466.2830     0   2   4E+3   12   12.5  RT (min)      lcorbi_t3_neg_4_032_23    m/z 466.3076     0   1E+5   8.5   9  RT (min)      lcorbi_t3_neg_4_023_18    m/z 466.3100     0   1E+4   10   10.5  RT (min)      lcorbi_t3_neg_3_019_12    m/z 466.8161     0   2   4   6   8E+2   0   0.5  RT (min)      lcorbi_t3_neg_1_055_05    m/z 466.8736     0   2   4E+3   18   18.5  RT (min)      lcorbi_t3_neg_4_040_19    m/z 466.9382     0   1   2E+4   1   1.5  RT (min)      lcorbi_t3_neg_4_042_14    m/z 467.0807     0   1E+4   11   11.5  RT (min)      lcorbi_t3_neg_3_007_16    m/z 467.0976     0   1   2   3E+4   13   13.5  RT (min)      lcorbi_t3_neg_4_007_13    m/z 467.0979     0   2   4   6   8E+3   9.5   10  RT (min)      lcorbi_t3_neg_4_023_18    m/z 467.1270     0   1E+4   6   6.5  RT (min)      lcorbi_t3_neg_4_040_19    m/z 467.1686     0   1   2   3E+3   13   13.5  RT (min)      lcorbi_t3_neg_4_042_14    m/z 467.1918     0   2   4   6   8E+2   7.5   8  RT (min)      lcorbi_t3_neg_2_047_48    m/z 467.2646     0   2   4E+3   8   8.5  RT (min)      lcorbi_t3_neg_2_048_27    m/z 467.2795     0   1   2E+4   13.5   14  RT (min)      lcorbi_t3_neg_4_034_01    m/z 467.2797     0   1   2E+3   12   12.5  RT (min)      lcorbi_t3_neg_3_040_05    m/z 467.3334     0   2   4   6E+4   12   12.5  RT (min)      lcorbi_t3_neg_1_055_05    m/z 467.4103     0   2   4   6E+3   14.5   15  RT (min)      lcorbi_t3_neg_4_029_31    m/z 468.0617     0   1E+4   9   9.5  RT (min)      lcorbi_t3_neg_4_007_13    m/z 468.0720     0   1E+4   10.5   11  RT (min)      lcorbi_t3_neg_1_043_14    m/z 468.0813     0   2   4   6E+3   11   11.5  RT (min)      lcorbi_t3_neg_4_007_13    m/z 468.0868     0   1   2E+4   13   13.5  RT (min)      lcorbi_t3_neg_4_007_13    m/z 468.0913     0   2   4   6E+3   13   13.5  RT (min)      lcorbi_t3_neg_4_042_14    m/z 468.1053     0   1   2   3   4E+3   13.5   14  RT (min)      lcorbi_t3_neg_4_038_15    m/z 468.1068     0   2   4   6E+3   8   8.5  RT (min)      lcorbi_t3_neg_1_026_19    m/z 468.1428     0   1   2E+3   13   13.5  RT (min)      lcorbi_t3_neg_4_007_13    m/z 468.1674     0   1   2E+3   13   13.5  RT (min)      lcorbi_t3_neg_4_035_33    m/z 468.2727     0   2   4   6   8E+2   11   11.5  RT (min)      lcorbi_t3_neg_2_031_25    m/z 468.3326     0   2   4   6E+2   10.5   11  RT (min)      lcorbi_t3_neg_1_055_05    m/z 468.3439     0   1E+5   12.5   13  RT (min)      lcorbi_t3_neg_1_055_05    m/z 468.3522     0   2   4E+3   11   11.5  RT (min)      lcorbi_t3_neg_4_054_36    m/z 469.0060     0   2   4   6   8E+3   8   8.5  RT (min)      lcorbi_t3_neg_4_023_18    m/z 469.0713     0   2   4E+3   13   13.5  RT (min)      lcorbi_t3_neg_1_043_14    m/z 469.0778     0   2   4   6   8E+3   11   11.5  RT (min)      lcorbi_t3_neg_3_007_16    m/z 469.1021     0   2   4   6   8E+3   8   8.5  RT (min)      lcorbi_t3_neg_1_026_19    m/z 469.1529     0   1E+4   8   8.5  RT (min)      lcorbi_t3_neg_4_023_18    m/z 469.1843     0   2   4   6E+3   13   13.5  RT (min)      lcorbi_t3_neg_4_007_13    m/z 469.1946     0   1E+4   12   12.5  RT (min)      lcorbi_t3_neg_4_042_14    m/z 469.2262     0   1   2   3E+3   7.5   8  RT (min)      lcorbi_t3_neg_1_050_45    m/z 469.2298     0   1E+4   6   6.5  RT (min)      lcorbi_t3_neg_4_032_23    m/z 469.2569     0   2   4E+3   11   11.5  RT (min)      lcorbi_t3_neg_1_055_05    m/z 469.2953     0   1E+4   12.5   13  RT (min)      lcorbi_t3_neg_1_055_05    m/z 469.3168     0   1E+4   10.5   11  RT (min)      lcorbi_t3_neg_1_055_05    m/z 469.3200     0   1   2E+3   12.5   13  RT (min)      lcorbi_t3_neg_4_007_13    m/z 469.3319     0   1   2E+3   11   11.5  RT (min)      lcorbi_t3_neg_2_020_05    m/z 469.3530     0   1E+3   12.5   13  RT (min)      lcorbi_t3_neg_1_055_05    m/z 469.3802     0   2   4   6   8E+2   12.5   13  RT (min)      lcorbi_t3_neg_4_011_24    m/z 469.4009     0   1E+4   13.5   14  RT (min)      lcorbi_t3_neg_4_007_13    m/z 469.8295     0   1E+4   1   1.5  RT (min)      lcorbi_t3_neg_2_052_32    m/z 469.8692     0   2   4   6   8E+3   18   18.5  RT (min)      lcorbi_t3_neg_3_011_04    m/z 469.8999     0   1E+4   18   18.5  RT (min)      lcorbi_t3_neg_4_019_08    m/z 469.9126     0   1E+4   18   18.5  RT (min)      lcorbi_t3_neg_2_055_18    m/z 469.9411     0   2   4E+3   1   1.5  RT (min)      lcorbi_t3_neg_3_029_02    m/z 470.0556     0   2   4   6E+3   8.5   9  RT (min)      lcorbi_t3_neg_2_048_27    m/z 470.1130     0   2   4   6E+3   10   10.5  RT (min)      lcorbi_t3_neg_4_029_31    m/z 470.1392     0   2   4   6   8E+3   9   9.5  RT (min)      lcorbi_t3_neg_3_019_12    m/z 470.1603     0   2   4   6   8E+3   13   13.5  RT (min)      lcorbi_t3_neg_1_055_05    m/z 470.1832     0   1   2   3E+3   13.5   14  RT (min)      lcorbi_t3_neg_4_034_01    m/z 470.1896     0   1E+4   7   7.5  RT (min)      lcorbi_t3_neg_2_048_27    m/z 470.2442     0   2   4   6E+3   12   12.5  RT (min)      lcorbi_t3_neg_1_055_05    m/z 470.2670     0   1   11.5   12  RT (min)      lcorbi_t3_neg_2_020_05    m/z 470.2781     0   2   4   6E+3   12.5   13  RT (min)      lcorbi_t3_neg_4_047_30    m/z 470.2952     0   2   4   6   8E+3   13.5   14  RT (min)      lcorbi_t3_neg_3_018_13    m/z 470.3293     0   1   2   3   4E+4   11   11.5  RT (min)      lcorbi_t3_neg_1_055_05    m/z 470.3465     0   2   4   6E+3   12.5   13  RT (min)      lcorbi_t3_neg_4_044_38    m/z 470.3487     0   2   4   6   8E+3   11   11.5  RT (min)      lcorbi_t3_neg_4_007_13    m/z 470.3849     0   1E+4   11   11.5  RT (min)      lcorbi_t3_neg_2_039_31    m/z 470.4343     0   1   2E+4   13.5   14  RT (min)      lcorbi_t3_neg_4_039_32    m/z 470.8716     0   1E+3   18   18.5  RT (min)      lcorbi_t3_neg_1_003_41    m/z 470.8943     0   1   2E+3   18   18.5  RT (min)      lcorbi_t3_neg_1_013_35    m/z 471.0197     0   1   2E+4   11   11.5  RT (min)      lcorbi_t3_neg_4_007_13    m/z 471.0519     0   1E+4   8.5   9  RT (min)      lcorbi_t3_neg_2_048_27    m/z 471.1355     0   2   4   6   8E+3   9   9.5  RT (min)      lcorbi_t3_neg_3_019_12    m/z 471.1503     0   2   4   6   8E+3   8   8.5  RT (min)      lcorbi_t3_neg_4_023_18    m/z 471.2000     0   1E+4   13.5   14  RT (min)      lcorbi_t3_neg_4_038_15    m/z 471.2110     0   1E+4   11   11.5  RT (min)      lcorbi_t3_neg_4_017_51    m/z 471.2134     0   2   4   6   8E+3   6   6.5  RT (min)      lcorbi_t3_neg_2_048_27    m/z 471.2174     0   1   2E+4   12   12.5  RT (min)      lcorbi_t3_neg_4_010_28    m/z 471.2417     0   1E+4   8.5   9  RT (min)      lcorbi_t3_neg_4_007_13    m/z 471.2543     0   1E+3   11   11.5  RT (min)      lcorbi_t3_neg_2_053_23    m/z 471.2665     0   2   4   6   8E+3   11.5   12  RT (min)      lcorbi_t3_neg_4_007_13    m/z 471.2745     0   1   2   3   4E+3   12.5   13  RT (min)      lcorbi_t3_neg_3_005_23    m/z 471.2898     0   1   11   11.5  RT (min)      lcorbi_t3_neg_4_007_13    m/z 471.2914     0   1E+4   13.5   14  RT (min)      lcorbi_t3_neg_3_018_13    m/z 471.2961     0   2   4   6E+3   12   12.5  RT (min)      lcorbi_t3_neg_3_007_16    m/z 471.3325     0   1   2   3   4E+3   12.5   13  RT (min)      lcorbi_t3_neg_4_029_31    m/z 471.3460     0   1   2   3   4E+4   13   13.5  RT (min)      lcorbi_t3_neg_4_034_01    m/z 471.3708     0   2   4   6E+3   11.5   12  RT (min)      lcorbi_t3_neg_4_019_08    m/z 471.9644     0   1E+4   9   9.5  RT (min)      lcorbi_t3_neg_2_048_27    m/z 472.0170     0   1E+4   11   11.5  RT (min)      lcorbi_t3_neg_4_007_13    m/z 472.0458     0   2   4E+3   10   10.5  RT (min)      lcorbi_t3_neg_2_048_27    m/z 472.0640     0   1E+3   11   11.5  RT (min)      lcorbi_t3_neg_2_048_27    m/z 472.0809     0   1   2   3E+3   10.5   11  RT (min)      lcorbi_t3_neg_2_048_27    m/z 472.1004     0   2   4   6E+3   11   11.5  RT (min)      lcorbi_t3_neg_4_007_13    m/z 472.1332     0   1   2E+2   5.5   6  RT (min)      lcorbi_t3_neg_2_048_27    m/z 472.1367     0   2   4   6   8E+3   14   14.5  RT (min)      lcorbi_t3_neg_4_040_19    m/z 472.1731     0   1   2   3   4E+3   1   1.5  RT (min)      lcorbi_t3_neg_1_052_24    m/z 472.1761     0   1E+4   13   13.5  RT (min)      lcorbi_t3_neg_4_007_13    m/z 472.1868     0   1   2   3E+4   1   1.5  RT (min)      lcorbi_t3_neg_1_052_24    m/z 472.2002     0   2   4   6E+3   11   11.5  RT (min)      lcorbi_t3_neg_4_023_18    m/z 472.2106     0   1   2E+3   13   13.5  RT (min)      lcorbi_t3_neg_1_018_46    m/z 472.2202     0   2   4   6E+3   12   12.5  RT (min)      lcorbi_t3_neg_4_010_28    m/z 472.2364     0   2   4E+3   12   12.5  RT (min)      lcorbi_t3_neg_4_005_27    m/z 472.2506     0   2   4E+2   11.5   12  RT (min)      lcorbi_t3_neg_4_011_24    m/z 472.2522     0   1   6.5   7  RT (min)      lcorbi_t3_neg_4_011_24    m/z 472.2746     0   1   2E+3   13.5   14  RT (min)      lcorbi_t3_neg_4_016_05    m/z 472.2948     0   1   13.5   14  RT (min)      lcorbi_t3_neg_1_055_05    m/z 472.3129     0   1   2   3E+3   13   13.5  RT (min)      lcorbi_t3_neg_4_042_14    m/z 472.3451     0   1E+3   12.5   13  RT (min)      lcorbi_t3_neg_2_020_05    m/z 472.8332     0   1E+4   0   0.5  RT (min)      lcorbi_t3_neg_4_067_57    m/z 472.9564     0   2   4E+3   9   9.5  RT (min)      lcorbi_t3_neg_4_007_13    m/z 473.0397     0   1   2   3E+3   10   10.5  RT (min)      lcorbi_t3_neg_2_048_27    m/z 473.0745     0   2   4   6E+3   10.5   11  RT (min)      lcorbi_t3_neg_2_011_04    m/z 473.0875     0   1E+4   5   5.5  RT (min)      lcorbi_t3_neg_2_048_27    m/z 473.1424     0   1E+4   6   6.5  RT (min)      lcorbi_t3_neg_4_005_27    m/z 473.1787     0   1   2   3E+4   7   7.5  RT (min)      lcorbi_t3_neg_4_034_01    m/z 473.1944     0   1   2E+4   6   6.5  RT (min)      lcorbi_t3_neg_2_048_27    m/z 473.1965     0   2   4   6E+3   11   11.5  RT (min)      lcorbi_t3_neg_4_023_18    m/z 473.2070     0   1   2   3E+3   13   13.5  RT (min)      lcorbi_t3_neg_1_030_47    m/z 473.2328     0   2   4   6E+3   12   12.5  RT (min)      lcorbi_t3_neg_4_005_27    m/z 473.2710     0   2   4   6   8E+3   13.5   14  RT (min)      lcorbi_t3_neg_4_019_08    m/z 473.2821     0   1E+4   14   14.5  RT (min)      lcorbi_t3_neg_4_024_07    m/z 473.3074     0   1E+5   13   13.5  RT (min)      lcorbi_t3_neg_1_029_21    m/z 473.3343     0   1E+3   12.5   13  RT (min)      lcorbi_t3_neg_4_007_13    m/z 473.3743     0   2   4   6E+2   13   13.5  RT (min)      lcorbi_t3_neg_4_034_01    m/z 473.3998     0   1   2   3   4E+3   14.5   15  RT (min)      lcorbi_t3_neg_1_050_45    m/z 474.0383     0   1   2E+4   6   6.5  RT (min)      lcorbi_t3_neg_4_047_30    m/z 474.0600     0   1E+4   10.5   11  RT (min)      lcorbi_t3_neg_1_043_14    m/z 474.0655     0   1   2E+4   1   1.5  RT (min)      lcorbi_t3_neg_2_035_46    m/z 474.0663     0   2   4   6   8E+3   7   7.5  RT (min)      lcorbi_t3_neg_3_045_34    m/z 474.0726     0   1   2   3E+4   3   3.5  RT (min)      lcorbi_t3_neg_4_007_13    m/z 474.1193     0   2   4   6   8E+3   5   5.5  RT (min)      lcorbi_t3_neg_4_032_23    m/z 474.1336     0   2   4   6E+3   1   1.5  RT (min)      lcorbi_t3_neg_1_050_45    m/z 474.1833     0   1   2   3   4E+3   11   11.5  RT (min)      lcorbi_t3_neg_4_032_23    m/z 474.1918     0   2   4E+3   13.5   14  RT (min)      lcorbi_t3_neg_4_040_19    m/z 474.2535     0   1   12   12.5  RT (min)      lcorbi_t3_neg_4_019_08    m/z 474.2732     0   2   4   6   8E+3   12   12.5  RT (min)      lcorbi_t3_neg_1_021_31    m/z 474.2855     0   2   4E+3   14   14.5  RT (min)      lcorbi_t3_neg_4_024_07    m/z 474.3148     0   1   12.5   13  RT (min)      lcorbi_t3_neg_1_029_21    m/z 474.3513     0   1E+3   13   13.5  RT (min)      lcorbi_t3_neg_1_055_05    m/z 474.7858     0   1   2E+2   1   1.5  RT (min)      lcorbi_t3_neg_3_022_27    m/z 475.0316     0   1   2   3E+3   4.5   5  RT (min)      lcorbi_t3_neg_4_036_16    m/z 475.0617     0   1E+4   12   12.5  RT (min)      lcorbi_t3_neg_4_040_19    m/z 475.0723     0   1E+4   10   10.5  RT (min)      lcorbi_t3_neg_4_023_18    m/z 475.0895     0   1E+4   10.5   11  RT (min)      lcorbi_t3_neg_1_007_26    m/z 475.1294     0   1   2   3E+4   13   13.5  RT (min)      lcorbi_t3_neg_4_007_13    m/z 475.1411     0   1E+4   9.5   10  RT (min)      lcorbi_t3_neg_3_019_12    m/z 475.1454     0   1E+4   5.5   6  RT (min)      lcorbi_t3_neg_2_035_46    m/z 475.1892     0   1   2   3E+4   1   1.5  RT (min)      lcorbi_t3_neg_1_055_05    m/z 475.1897     0   2   4   6E+3   12   12.5  RT (min)      lcorbi_t3_neg_4_034_01    m/z 475.2674     0   2   4E+4   9.5   10  RT (min)      lcorbi_t3_neg_4_023_18    m/z 475.3407     0   1E+4   13   13.5  RT (min)      lcorbi_t3_neg_4_011_24    m/z 475.3524     0   1E+4   11.5   12  RT (min)      lcorbi_t3_neg_1_055_05    m/z 475.3691     0   1   2   3   4E+3   13   13.5  RT (min)      lcorbi_t3_neg_3_037_36    m/z 475.8862     0   1E+4   18   18.5  RT (min)      lcorbi_t3_neg_2_014_29    m/z 476.0264     0   1   2   3   4E+3   8   8.5  RT (min)      lcorbi_t3_neg_4_023_18    m/z 476.0363     0   1   2E+4   6   6.5  RT (min)      lcorbi_t3_neg_4_047_30    m/z 476.0633     0   2   4   6E+3   7   7.5  RT (min)      lcorbi_t3_neg_2_002_26    m/z 476.0911     0   1   2E+4   6   6.5  RT (min)      lcorbi_t3_neg_2_048_27    m/z 476.0928     0   1   2   3E+3   10.5   11  RT (min)      lcorbi_t3_neg_1_001_49    m/z 476.0998     0   1E+4   13   13.5  RT (min)      lcorbi_t3_neg_1_043_14    m/z 476.1105     0   1E+4   11   11.5  RT (min)      lcorbi_t3_neg_4_007_13    m/z 476.1230     0   1   2   3   4E+4   13   13.5  RT (min)      lcorbi_t3_neg_4_007_13    m/z 476.1321     0   1E+4   1   1.5  RT (min)      lcorbi_t3_neg_4_026_46    m/z 476.1479     0   1   2   3E+3   13   13.5  RT (min)      lcorbi_t3_neg_1_043_14    m/z 476.1781     0   2   4   6   8E+3   11   11.5  RT (min)      lcorbi_t3_neg_4_102_61    m/z 476.1977     0   1   2E+4   7   7.5  RT (min)      lcorbi_t3_neg_4_034_01    m/z 476.2577     0   2   4   6   8E+3   11   11.5  RT (min)      lcorbi_t3_neg_2_027_13    m/z 476.2683     0   2   4   6E+4   11   11.5  RT (min)      lcorbi_t3_neg_1_055_05    m/z 476.2732     0   1   8.5   9  RT (min)      lcorbi_t3_neg_4_023_18    m/z 476.2980     0   1E+4   11   11.5  RT (min)      lcorbi_t3_neg_4_007_13    m/z 476.3040     0   2   4E+3   11   11.5  RT (min)      lcorbi_t3_neg_4_007_13    m/z 476.3085     0   1E+4   6.5   7  RT (min)      lcorbi_t3_neg_4_032_23    m/z 476.3307     0   1E+4   9.5   10  RT (min)      lcorbi_t3_neg_4_023_18    m/z 476.3356     0   1   2   3E+4   13   13.5  RT (min)      lcorbi_t3_neg_3_007_16    m/z 476.3488     0   1   2   3E+4   12.5   13  RT (min)      lcorbi_t3_neg_1_055_05    m/z 476.7755     0   2   4   6   8E+3   0   0.5  RT (min)      lcorbi_t3_neg_4_067_57    m/z 476.9208     0   1E+4   18   18.5  RT (min)      lcorbi_t3_neg_1_001_49    m/z 477.0587     0   1E+4   12   12.5  RT (min)      lcorbi_t3_neg_4_040_19    m/z 477.0668     0   1E+4   11   11.5  RT (min)      lcorbi_t3_neg_4_007_13    m/z 477.1069     0   1E+4   11   11.5  RT (min)      lcorbi_t3_neg_4_007_13    m/z 477.1884     0   1E+4   1   1.5  RT (min)      lcorbi_t3_neg_2_035_46    m/z 477.2006     0   1   2   3   4E+4   13.5   14  RT (min)      lcorbi_t3_neg_4_047_30    m/z 477.2235     0   1   2   3E+4   11   11.5  RT (min)      lcorbi_t3_neg_4_016_05    m/z 477.2621     0   1   11.5   12  RT (min)      lcorbi_t3_neg_4_084_59    m/z 477.3022     0   1E+4   6.5   7  RT (min)      lcorbi_t3_neg_4_032_23    m/z 477.3321     0   1E+4   13   13.5  RT (min)      lcorbi_t3_neg_1_055_05    m/z 477.3334     0   1   11.5   12  RT (min)      lcorbi_t3_neg_1_055_05    m/z 477.3563     0   2   4   6E+3   13   13.5  RT (min)      lcorbi_t3_neg_4_034_01    m/z 477.3712     0   1   2E+3   13   13.5  RT (min)      lcorbi_t3_neg_2_054_15    m/z 477.8358     0   1   2E+4   1   1.5  RT (min)      lcorbi_t3_neg_2_039_31    m/z 477.9079     0   1E+4   8.5   9  RT (min)      lcorbi_t3_neg_2_015_21    m/z 478.0526     0   1E+4   12   12.5  RT (min)      lcorbi_t3_neg_4_040_19    m/z 478.0807     0   1   2   3E+3   10   10.5  RT (min)      lcorbi_t3_neg_3_037_36    m/z 478.0917     0   1   12   12.5  RT (min)      lcorbi_t3_neg_3_008_20    m/z 478.1357     0   2   4   6E+4   8   8.5  RT (min)      lcorbi_t3_neg_2_048_27    m/z 478.1589     0   1E+5   1   1.5  RT (min)       lcorbi_t3_neg_4_011_24    m/z 478.1636     0   1   2   3E+3   13   13.5  RT (min)      lcorbi_t3_neg_4_050_02    m/z 478.1796     0   2   4   6   8E+3   8   8.5  RT (min)      lcorbi_t3_neg_4_023_18    m/z 478.1841     0   1   2   3   4E+4   1   1.5  RT (min)      lcorbi_t3_neg_2_035_46    m/z 478.2217     0   1E+3   12   12.5  RT (min)      lcorbi_t3_neg_1_055_05    m/z 478.2353     0   1E+4   12.5   13  RT (min)      lcorbi_t3_neg_4_034_01    m/z 478.2671     0   1   2E+3   11.5   12  RT (min)      lcorbi_t3_neg_3_043_25    m/z 478.3647     0   2   4   6   8E+3   13   13.5  RT (min)      lcorbi_t3_neg_4_017_51    m/z 478.3980     0   2   4   6   8E+3   13.5   14  RT (min)      lcorbi_t3_neg_4_054_36    m/z 478.9219     0   2   4E+3   18   18.5  RT (min)      lcorbi_t3_neg_1_001_49    m/z 478.9568     0   1E+4   9   9.5  RT (min)      lcorbi_t3_neg_4_007_13    m/z 479.0155     0   2   4   6E+3   7.5   8  RT (min)      lcorbi_t3_neg_4_042_14    m/z 479.0557     0   1   2   3E+3   12   12.5  RT (min)      lcorbi_t3_neg_4_040_19    m/z 479.0753     0   1E+4   9.5   10  RT (min)      lcorbi_t3_neg_3_037_36    m/z 479.0844     0   1   2E+4   10   10.5  RT (min)      lcorbi_t3_neg_4_023_18    m/z 479.1331     0   1E+4   8   8.5  RT (min)      lcorbi_t3_neg_2_048_27    m/z 479.1459     0   1   10   10.5  RT (min)      lcorbi_t3_neg_4_102_61    m/z 479.1506     0   1E+4   1   1.5  RT (min)      lcorbi_t3_neg_2_053_23    m/z 479.1746     0   1   2E+4   10.5   11  RT (min)      lcorbi_t3_neg_4_042_14    m/z 479.1765     0   1   2   3E+4   5   5.5  RT (min)      lcorbi_t3_neg_2_048_27    m/z 479.1985     0   1   2   3   4E+4   13.5   14  RT (min)      lcorbi_t3_neg_3_024_26    m/z 479.2550     0   2   4   6   8E+3   7.5   8  RT (min)      lcorbi_t3_neg_4_017_51    m/z 479.2647     0   2   4   6E+3   9.5   10  RT (min)      lcorbi_t3_neg_2_048_27    m/z 479.2649     0   1E+4   6   6.5  RT (min)      lcorbi_t3_neg_2_048_27    m/z 479.2778     0   1E+3   12   12.5  RT (min)      lcorbi_t3_neg_4_034_01    m/z 479.2867     0   1   2E+4   11   11.5  RT (min)      lcorbi_t3_neg_1_055_05    m/z 479.3011     0   1   2   3   4E+3   9   9.5  RT (min)      lcorbi_t3_neg_2_048_27    m/z 479.3338     0   1E+4   11.5   12  RT (min)      lcorbi_t3_neg_4_034_01    m/z 479.3716     0   2   4   6   8E+2   13.5   14  RT (min)      lcorbi_t3_neg_1_055_05    m/z 479.3985     0   1E+4   14   14.5  RT (min)      lcorbi_t3_neg_4_040_19    m/z 479.4103     0   1E+4   13.5   14  RT (min)      lcorbi_t3_neg_4_005_27    m/z 479.8330     0   1E+4   1   1.5  RT (min)      lcorbi_t3_neg_4_054_36    m/z 479.9049     0   1E+4   8   8.5   9  RT (min)      lcorbi_t3_neg_2_015_21    m/z 479.9508     0   2   4   6   8E+3   9   9.5  RT (min)      lcorbi_t3_neg_4_007_13    m/z 480.0691     0   2   4   6   8E+3   11.5   12  RT (min)      lcorbi_t3_neg_1_043_14    m/z 480.0878     0   1   2E+4   6   6.5  RT (min)      lcorbi_t3_neg_4_044_38    m/z 480.0927     0   1   2   3   4E+4   3.5   4  RT (min)      lcorbi_t3_neg_4_007_13    m/z 480.1793     0   2   4   6E+3   13.5   14  RT (min)      lcorbi_t3_neg_4_051_40    m/z 480.1827     0   2   4   6E+2   1   1.5  RT (min)      lcorbi_t3_neg_1_055_05    m/z 480.2431     0   2   4   6   8E+3   12.5   13  RT (min)      lcorbi_t3_neg_1_055_05    m/z 480.2509     0   1E+4   13   13.5  RT (min)      lcorbi_t3_neg_4_042_14    m/z 480.2598     0   1   2E+3   13   13.5  RT (min)      lcorbi_t3_neg_4_016_05    m/z 480.2777     0   2   4   6   8E+3   10   10.5  RT (min)      lcorbi_t3_neg_3_019_12    m/z 480.2866     0   2   4   6E+3   10   10.5  RT (min)      lcorbi_t3_neg_4_034_01    m/z 480.2989     0   1E+4   13   13.5  RT (min)      lcorbi_t3_neg_4_034_01    m/z 480.3229     0   1E+4   12.5   13  RT (min)      lcorbi_t3_neg_4_007_13    m/z 480.3806     0   1   13.5   14  RT (min)      lcorbi_t3_neg_1_055_05    m/z 481.0304     0   2   4   6E+3   9   9.5  RT (min)      lcorbi_t3_neg_3_019_12    m/z 481.0417     0   1   2   3E+3   8   8.5  RT (min)      lcorbi_t3_neg_4_028_39    m/z 481.0919     0   1   2   3E+3   9.5   10  RT (min)      lcorbi_t3_neg_2_006_10    m/z 481.0933     0   1   2   3E+3   1.5   2  RT (min)      lcorbi_t3_neg_2_048_27    m/z 481.1669     0   1E+4   10.5   11  RT (min)      lcorbi_t3_neg_3_053_14    m/z 481.1872     0   1E+4   7   7.5  RT (min)      lcorbi_t3_neg_4_034_01    m/z 481.2075     0   1   2   3   4E+3   8   8.5  RT (min)      lcorbi_t3_neg_2_048_27    m/z 481.2144     0   1   2   3   4E+4   13.5   14  RT (min)      lcorbi_t3_neg_3_024_26    m/z 481.2394     0   2   4E+3   12.5   13  RT (min)      lcorbi_t3_neg_1_055_05    m/z 481.2438     0   1E+4   7   7.5  RT (min)      lcorbi_t3_neg_4_034_01    m/z 481.2514     0   2   4E+3   6   6.5  RT (min)      lcorbi_t3_neg_4_040_19    m/z 481.2626     0   1   13   13.5  RT (min)      lcorbi_t3_neg_3_046_10    m/z 481.2952     0   1E+4   13   13.5  RT (min)      lcorbi_t3_neg_4_034_01    m/z 481.3021     0   1E+3   12.5   13  RT (min)      lcorbi_t3_neg_1_055_05    m/z 481.3258     0   1   2   3E+4   12.5   13  RT (min)      lcorbi_t3_neg_4_019_08    m/z 481.3756     0   1   2   3E+3   12   12.5  RT (min)      lcorbi_t3_neg_1_055_05    m/z 481.4259     0   1E+4   14.5   15  RT (min)      lcorbi_t3_neg_4_029_31    m/z 481.9019     0   1   2   3   4E+3   8   8.5  RT (min)      lcorbi_t3_neg_2_002_26    m/z 482.1239     0   2   4   6   8E+2   10.5   11  RT (min)      lcorbi_t3_neg_4_030_04    m/z 482.1605     0   2   4   6   8E+3   7.5   8  RT (min)      lcorbi_t3_neg_3_045_34    m/z 482.2054     0   1E+4   1   1.5  RT (min)      lcorbi_t3_neg_2_035_46    m/z 482.2153     0   1   13   13.5  RT (min)      lcorbi_t3_neg_3_024_26    m/z 482.2245     0   2   4E+3   7   7.5  RT (min)      lcorbi_t3_neg_4_023_18    m/z 482.2391     0   2   4   6E+3   11   11.5  RT (min)      lcorbi_t3_neg_4_038_15    m/z 482.2394     0   1E+4   13   13.5  RT (min)      lcorbi_t3_neg_3_018_13    m/z 482.2472     0   1   2   3   4E+3   7   7.5  RT (min)      lcorbi_t3_neg_4_034_01    m/z 482.2519     0   1E+4   10.5   11  RT (min)      lcorbi_t3_neg_1_055_05    m/z 482.2561     0   1E+4   13   13.5  RT (min)      lcorbi_t3_neg_4_017_51    m/z 482.2837     0   2   4   6   8E+3   9.5   10  RT (min)      lcorbi_t3_neg_2_048_27    m/z 482.8768     0   1   2   3E+3   18   18.5  RT (min)      lcorbi_t3_neg_2_027_13    m/z 482.8877     0   2   4   6E+4   7.5   8  RT (min)      lcorbi_t3_neg_4_023_18    m/z 482.9866     0   2   4   6   8E+3   5   5.5  RT (min)      lcorbi_t3_neg_1_054_37    m/z 483.0308     0   2   4E+3   9   9.5  RT (min)      lcorbi_t3_neg_3_019_12    m/z 483.0388     0   1   2   3   4E+3   10   10.5  RT (min)      lcorbi_t3_neg_4_034_01    m/z 483.0540     0   1E+4   5.5   6  RT (min)      lcorbi_t3_neg_2_048_27    m/z 483.1010     0   1E+4   10.5   11  RT (min)      lcorbi_t3_neg_2_007_14    m/z 483.1119     0   1   2E+3   10.5   11  RT (min)      lcorbi_t3_neg_3_010_15    m/z 483.2208     0   1E+4   7   7.5  RT (min)      lcorbi_t3_neg_4_023_18    m/z 483.2525     0   1E+4   13   13.5  RT (min)      lcorbi_t3_neg_4_035_33    m/z 483.2782     0   2   4   6   8E+3   13   13.5  RT (min)      lcorbi_t3_neg_4_046_37    m/z 483.3048     0   2   4   6   8E+3   10.5   11  RT (min)      lcorbi_t3_neg_4_032_23    m/z 483.3182     0   1E+4   5   5.5  RT (min)      lcorbi_t3_neg_4_032_23    m/z 483.3184     0   1E+4   11.5   12  RT (min)      lcorbi_t3_neg_4_034_01    m/z 483.3384     0   2   4   6   8E+3   13   13.5  RT (min)      lcorbi_t3_neg_3_032_17    m/z 483.3588     0   2   4   6E+3   12.5   13  RT (min)      lcorbi_t3_neg_4_007_13    m/z 483.3689     0   1E+4   12   12.5  RT (min)      lcorbi_t3_neg_2_007_14    m/z 483.3911     0   2   4E+2   12.5   13  RT (min)      lcorbi_t3_neg_1_055_05    m/z 483.8235     0   1   2   3   4E+3   17.5   18  RT (min)      lcorbi_t3_neg_2_010_02    m/z 484.0144     0   1E+4   5.5   6  RT (min)      lcorbi_t3_neg_3_014_19    m/z 484.1181     0   1E+4   11   11.5  RT (min)      lcorbi_t3_neg_4_007_13    m/z 484.2586     0   1   2   3E+3   13   13.5  RT (min)      lcorbi_t3_neg_4_042_14    m/z 484.2677     0   2   4   6E+3   9.5   10  RT (min)      lcorbi_t3_neg_4_007_13    m/z 484.2759     0   1   2E+4   11   11.5  RT (min)      lcorbi_t3_neg_1_055_05    m/z 484.2891     0   1   12   12.5  RT (min)      lcorbi_t3_neg_4_019_08    m/z 484.3143     0   1   2E+3   13   13.5  RT (min)      lcorbi_t3_neg_1_013_35    m/z 484.3361     0   1   2   3   4E+4   11   11.5  RT (min)      lcorbi_t3_neg_1_055_05    m/z 484.3751     0   2   4   6E+2   13   13.5  RT (min)      lcorbi_t3_neg_1_055_05    m/z 484.8703     0   1   2E+3   18   18.5  RT (min)      lcorbi_t3_neg_3_015_11    m/z 484.9786     0   2   4   6   8E+3   8   8.5  RT (min)      lcorbi_t3_neg_3_045_34    m/z 484.9836     0   2   4E+3   5   5.5  RT (min)      lcorbi_t3_neg_1_054_37    m/z 484.9916     0   1   2   3   4E+3   9   9.5  RT (min)      lcorbi_t3_neg_1_044_23    m/z 485.0178     0   1   2E+4   8.5   9  RT (min)      lcorbi_t3_neg_2_048_27    m/z 485.0262     0   1E+3   9   9.5  RT (min)      lcorbi_t3_neg_1_055_05    m/z 485.0571     0   1   2E+3   9   9.5  RT (min)      lcorbi_t3_neg_1_008_08    m/z 485.0890     0   1   2E+4   10   10.5  RT (min)      lcorbi_t3_neg_4_023_18    m/z 485.0905     0   2   4   6   8E+3   12   12.5  RT (min)      lcorbi_t3_neg_4_040_19    m/z 485.1146     0   1   2   3E+4   11   11.5  RT (min)      lcorbi_t3_neg_4_007_13    m/z 485.1660     0   1E+4   6   6.5  RT (min)      lcorbi_t3_neg_2_048_27    m/z 485.1812     0   2   4   6E+3   7   7.5  RT (min)      lcorbi_t3_neg_4_023_18    m/z 485.2024     0   2   4   6E+3   9   9.5  RT (min)      lcorbi_t3_neg_4_050_02    m/z 485.2106     0   1   2   3E+3   13   13.5  RT (min)      lcorbi_t3_neg_4_034_01    m/z 485.2329     0   1   2E+4   12.5   13  RT (min)      lcorbi_t3_neg_4_040_19    m/z 485.2937     0   2   4   6E+3   13.5   14  RT (min)      lcorbi_t3_neg_4_032_23    m/z 485.3263     0   1   2E+3   12   12.5  RT (min)      lcorbi_t3_neg_4_007_13    m/z 485.3745     0   1E+4   13   13.5  RT (min)      lcorbi_t3_neg_4_007_13    m/z 485.8410     0   1   2E+4   1   1.5  RT (min)      lcorbi_t3_neg_4_045_35    m/z 485.9774     0   1   2   3   4E+3   5   5.5  RT (min)      lcorbi_t3_neg_1_054_37    m/z 485.9855     0   1E+4   9   9.5  RT (min)      lcorbi_t3_neg_1_044_23    m/z 486.0145     0   2   4   6   8E+3   11   11.5  RT (min)      lcorbi_t3_neg_4_007_13    m/z 486.1047     0   1E+4   10   10.5  RT (min)      lcorbi_t3_neg_4_023_18    m/z 486.1155     0   1E+4   11.5   12  RT (min)      lcorbi_t3_neg_4_007_13    m/z 486.1962     0   2   4   6E+3   11   11.5  RT (min)      lcorbi_t3_neg_2_048_27    m/z 486.2136     0   1   2   3E+3   11   11.5  RT (min)      lcorbi_t3_neg_1_022_34    m/z 486.2187     0   1E+4   8.5   9  RT (min)      lcorbi_t3_neg_4_017_51    m/z 486.2282     0   1E+3   13   13.5  RT (min)      lcorbi_t3_neg_4_016_05    m/z 486.2420     0   1E+4   13   13.5  RT (min)      lcorbi_t3_neg_1_055_05    m/z 486.2520     0   2   4   6E+3   13   13.5  RT (min)      lcorbi_t3_neg_4_034_01    m/z 486.2743     0   1E+3   13   13.5  RT (min)      lcorbi_t3_neg_2_048_27    m/z 486.2804     0   1   2E+3   14   14.5  RT (min)      lcorbi_t3_neg_4_019_08    m/z 486.2856     0   1   2E+3   11   11.5  RT (min)      lcorbi_t3_neg_3_040_05    m/z 486.3114     0   1   13.5   14  RT (min)      lcorbi_t3_neg_4_019_08    m/z 486.3288     0   1   2E+3   10   10.5  RT (min)      lcorbi_t3_neg_4_032_23    m/z 486.3300     0   1   13.5   14  RT (min)      lcorbi_t3_neg_4_019_08    m/z 486.3374     0   2   4   6E+3   6.5   7  RT (min)      lcorbi_t3_neg_4_032_23    m/z 486.3885     0   1E+4   12   12.5  RT (min)      lcorbi_t3_neg_1_055_05    m/z 486.4043     0   1E+5   12.5   13  RT (min)      lcorbi_t3_neg_1_055_05    m/z 486.7184     0   1E+4   1   1.5  RT (min)      lcorbi_t3_neg_2_035_46    m/z 486.8813     0   1   2   3E+4   1   1.5  RT (min)      lcorbi_t3_neg_4_042_14    m/z 486.8815     0   2   4   6E+3   18   18.5  RT (min)      lcorbi_t3_neg_1_008_08    m/z 487.0148     0   1E+4   8.5   9  RT (min)      lcorbi_t3_neg_2_048_27    m/z 487.0222     0   1E+4   11   11.5  RT (min)      lcorbi_t3_neg_4_007_13    m/z 487.0472     0   1   2   3E+3   5.5   6  RT (min)      lcorbi_t3_neg_4_023_18    m/z 487.0680     0   2   4   6E+3   1   1.5  RT (min)      lcorbi_t3_neg_3_020_31    m/z 487.0875     0   2   4E+3   12   12.5  RT (min)      lcorbi_t3_neg_4_040_19    m/z 487.0939     0   1   2E+4   11   11.5  RT (min)      lcorbi_t3_neg_4_007_13    m/z 487.1288     0   2   4E+3   5   5.5  RT (min)      lcorbi_t3_neg_4_009_50    m/z 487.1790     0   2   4   6   8E+3   1   1.5  RT (min)      lcorbi_t3_neg_3_005_23    m/z 487.1926     0   1E+4   11   11.5  RT (min)      lcorbi_t3_neg_2_048_27    m/z 487.2262     0   1   2   3   4E+3   13.5   14  RT (min)      lcorbi_t3_neg_4_023_18    m/z 487.2483     0   2   4   6   8E+2   13   13.5  RT (min)      lcorbi_t3_neg_4_040_19    m/z 487.2740     0   1   2   3   4E+3   10   10.5  RT (min)      lcorbi_t3_neg_4_032_23    m/z 487.2845     0   1   2   3   4E+3   10   10.5  RT (min)      lcorbi_t3_neg_4_032_23    m/z 487.2845     0   2   4   6   8E+3   5.5   6  RT (min)      lcorbi_t3_neg_4_004_34    m/z 487.3062     0   1   2   3E+3   13.5   14  RT (min)      lcorbi_t3_neg_4_032_23    m/z 487.3150     0   1E+4   12.5   13  RT (min)      lcorbi_t3_neg_3_024_26    m/z 487.3264     0   2   4   6E+3   5.5   6  RT (min)      lcorbi_t3_neg_4_017_51    m/z 487.3272     0   1E+4   10   10.5  RT (min)      lcorbi_t3_neg_2_048_27    m/z 487.3325     0   1   2   3   4E+3   5.5   6  RT (min)      lcorbi_t3_neg_4_046_37    m/z 487.3789     0   1E+4   13   13.5  RT (min)      lcorbi_t3_neg_4_034_01    m/z 487.4052     0   1   2   3   4E+4   12   12.5  RT (min)      lcorbi_t3_neg_1_052_24    m/z 488.0086     0   2   4   6   8E+3   8.5   9  RT (min)      lcorbi_t3_neg_2_048_27    m/z 488.0777     0   1   2E+4   6   6.5  RT (min)      lcorbi_t3_neg_2_048_27    m/z 488.0942     0   2   4   6E+3   11   11.5  RT (min)      lcorbi_t3_neg_4_007_13    m/z 488.1122     0   1   2E+3   9.5   10  RT (min)      lcorbi_t3_neg_1_052_24    m/z 488.1124     0   1   2   3E+4   11.5   12  RT (min)      lcorbi_t3_neg_4_007_13    m/z 488.1549     0   1E+4   13   13.5  RT (min)      lcorbi_t3_neg_1_043_14    m/z 488.1918     0   1   2E+4   1   1.5  RT (min)      lcorbi_t3_neg_1_055_05    m/z 488.1943     0   1E+4   7   7.5  RT (min)      lcorbi_t3_neg_4_052_26    m/z 488.1960     0   1   2   3E+3   11   11.5  RT (min)      lcorbi_t3_neg_2_002_26    m/z 488.2485     0   1E+4   12   12.5  RT (min)      lcorbi_t3_neg_4_005_27    m/z 488.2695     0   1E+4   12   12.5  RT (min)      lcorbi_t3_neg_4_029_31    m/z 488.2896     0   1   2E+4   12.5   13  RT (min)      lcorbi_t3_neg_4_029_31    m/z 488.2943     0   1E+4   11.5   12  RT (min)      lcorbi_t3_neg_4_005_27    m/z 488.3127     0   1E+3   12   12.5  RT (min)      lcorbi_t3_neg_4_029_31    m/z 488.3306     0   1   10   10.5  RT (min)      lcorbi_t3_neg_1_055_05    m/z 488.3673     0   1   2   3   4E+4   11.5   12  RT (min)      lcorbi_t3_neg_1_055_05    m/z 488.8865     0   1   2E+3   18   18.5  RT (min)      lcorbi_t3_neg_3_014_19    m/z 488.9642     0   1   2   3E+4   1   1.5  RT (min)      lcorbi_t3_neg_3_002_24    m/z 489.0773     0   2   4   6   8E+3   13   13.5  RT (min)      lcorbi_t3_neg_4_007_13    m/z 489.1059     0   2   4   6   8E+3   9.5   10  RT (min)      lcorbi_t3_neg_2_023_39    m/z 489.1289     0   2   4   6E+3   6   6.5  RT (min)      lcorbi_t3_neg_4_006_47    m/z 489.1582     0   1E+4   6   6.5  RT (min)      lcorbi_t3_neg_3_039_29    m/z 489.1883     0   1   2E+4   1   1.5  RT (min)      lcorbi_t3_neg_1_055_05    m/z 489.2312     0   1E+4   1   1.5  RT (min)      lcorbi_t3_neg_3_005_23    m/z 489.2450     0   1E+4   12   12.5  RT (min)      lcorbi_t3_neg_4_005_27    m/z 489.2851     0   1   2E+4   12.5   13  RT (min)      lcorbi_t3_neg_4_029_31    m/z 489.3947     0   1   14.5   15  RT (min)      lcorbi_t3_neg_4_034_01    m/z 489.4207     0   2   4E+4   12.5   13  RT (min)      lcorbi_t3_neg_4_011_24    m/z 490.0543     0   2   4   6   8E+3   10   10.5  RT (min)      lcorbi_t3_neg_3_054_18    m/z 490.0899     0   1E+4   13   13.5  RT (min)      lcorbi_t3_neg_4_007_13    m/z 490.1109     0   1E+4   12.5   13  RT (min)      lcorbi_t3_neg_4_047_30    m/z 490.1453     0   1   2   3   4E+3   9   9.5  RT (min)      lcorbi_t3_neg_3_019_12    m/z 490.1519     0   1E+4   13   13.5  RT (min)      lcorbi_t3_neg_1_043_14    m/z 490.1660     0   2   4   6E+3   13   13.5  RT (min)      lcorbi_t3_neg_4_042_14    m/z 490.1780     0   2   4   6E+3   10   10.5  RT (min)      lcorbi_t3_neg_4_032_23    m/z 490.1942     0   1E+4   7   7.5  RT (min)      lcorbi_t3_neg_2_051_34    m/z 490.2223     0   2   4E+3   8   8.5  RT (min)      lcorbi_t3_neg_4_040_19    m/z 490.2706     0   1E+3   12   12.5  RT (min)      lcorbi_t3_neg_4_007_13    m/z 490.2883     0   2   4   6E+3   12.5   13  RT (min)      lcorbi_t3_neg_4_029_31    m/z 490.3102     0   1E+4   12   12.5  RT (min)      lcorbi_t3_neg_4_007_13    m/z 490.3219     0   1   2   3   4E+4   12   12.5  RT (min)      lcorbi_t3_neg_4_007_13    m/z 490.3273     0   1E+3   11   11.5  RT (min)      lcorbi_t3_neg_4_029_31    m/z 490.3596     0   1E+4   12.5   13  RT (min)      lcorbi_t3_neg_2_039_31    m/z 490.3746     0   2   4E+3   11.5   12  RT (min)      lcorbi_t3_neg_3_033_53    m/z 490.7128     0   1   1   1.5  RT (min)      lcorbi_t3_neg_4_037_29    m/z 490.8235     0   2   4   6E+3   18   18.5  RT (min)      lcorbi_t3_neg_2_006_10    m/z 490.9905     0   2   4   6   8E+3   6   6.5  RT (min)      lcorbi_t3_neg_2_048_27    m/z 491.0591     0   2   4   6E+3   5.5   6  RT (min)      lcorbi_t3_neg_4_023_18    m/z 491.0618     0   1E+4   10.5   11  RT (min)      lcorbi_t3_neg_3_037_36    m/z 491.0743     0   2   4   6   8E+3   13   13.5  RT (min)      lcorbi_t3_neg_4_007_13    m/z 491.1416     0   1   2   3   4E+3   9   9.5  RT (min)      lcorbi_t3_neg_3_019_12    m/z 491.2387     0   1   2   3   4E+3   10   10.5  RT (min)      lcorbi_t3_neg_4_042_14    m/z 491.2405     0   1   2   3E+3   12   12.5  RT (min)      lcorbi_t3_neg_1_055_05    m/z 491.2834     0   1   2   3   4E+3   9.5   10  RT (min)      lcorbi_t3_neg_4_034_01    m/z 491.2978     0   1   2   3E+4   11   11.5  RT (min)      lcorbi_t3_neg_1_055_05    m/z 491.3591     0   2   4   6E+3   17.5   18  RT (min)      lcorbi_t3_neg_1_004_22    m/z 492.0681     0   1E+4   13   13.5  RT (min)      lcorbi_t3_neg_4_007_13    m/z 492.0845     0   1   2E+4   1   1.5  RT (min)      lcorbi_t3_neg_3_052_32    m/z 492.0869     0   1E+4   13   13.5  RT (min)      lcorbi_t3_neg_4_007_13    m/z 492.1057     0   2   4E+3   13   13.5  RT (min)      lcorbi_t3_neg_4_052_26    m/z 492.1241     0   1E+3   9.5   10  RT (min)      lcorbi_t3_neg_1_052_24    m/z 492.1428     0   1   2   3   4E+3   13   13.5  RT (min)      lcorbi_t3_neg_4_007_13    m/z 492.1926     0   1E+3   12.5   13  RT (min)      lcorbi_t3_neg_1_055_05    m/z 492.2938     0   1   2E+4   12   12.5  RT (min)      lcorbi_t3_neg_4_032_23    m/z 492.3090     0   2   4   6E+4   11.5   12  RT (min)      lcorbi_t3_neg_1_055_05    m/z 492.3251     0   1E+4   12.5   13  RT (min)      lcorbi_t3_neg_4_035_33    m/z 492.3539     0   2   4E+4   11   11.5  RT (min)      lcorbi_t3_neg_1_006_40    m/z 492.3621     0   1   2E+4   14   14.5  RT (min)      lcorbi_t3_neg_2_020_05    m/z 492.3773     0   1   2E+4   13   13.5  RT (min)      lcorbi_t3_neg_4_034_01    m/z 492.8205     0   2   4   6E+3   18   18.5  RT (min)      lcorbi_t3_neg_4_033_53    m/z 492.9151     0   1E+3   18   18.5  RT (min)      lcorbi_t3_neg_4_039_32    m/z 492.9875     0   1E+4   6   6.5  RT (min)      lcorbi_t3_neg_2_048_27    m/z 493.0675     0   1E+4   10   10.5  RT (min)      lcorbi_t3_neg_4_034_01    m/z 493.1041     0   1E+4   7.5   8  RT (min)      lcorbi_t3_neg_4_034_01    m/z 493.1252     0   1   2   3   4E+3   8   8.5  RT (min)      lcorbi_t3_neg_2_034_19    m/z 493.2067     0   1E+4   12.5   13  RT (min)      lcorbi_t3_neg_3_024_26    m/z 493.2089     0   1   2E+4   1   1.5  RT (min)      lcorbi_t3_neg_1_055_05    m/z 493.2440     0   2   4   6E+3   9   9.5  RT (min)      lcorbi_t3_neg_2_048_27    m/z 493.2451     0   2   4   6   8E+3   11   11.5  RT (min)      lcorbi_t3_neg_4_032_23    m/z 493.2625     0   2   4   6E+3   10.5   11  RT (min)      lcorbi_t3_neg_4_010_28    m/z 493.2804     0   1E+4   6   6.5  RT (min)      lcorbi_t3_neg_4_005_27    m/z 493.3492     0   2   4   6   8E+2   12   12.5  RT (min)      lcorbi_t3_neg_2_053_23    m/z 493.8523     0   1E+4   18   18.5  RT (min)      lcorbi_t3_neg_2_011_04    m/z 493.9003     0   2   4E+3   18   18.5  RT (min)      lcorbi_t3_neg_1_006_40    m/z 494.0092     0   2   4E+3   5   5.5  RT (min)      lcorbi_t3_neg_1_054_37    m/z 494.1027     0   2   4   6E+3   13   13.5  RT (min)      lcorbi_t3_neg_4_007_13    m/z 494.1161     0   2   4   6E+3   12   12.5  RT (min)      lcorbi_t3_neg_4_040_19    m/z 494.1586     0   1   2   3E+3   13   13.5  RT (min)      lcorbi_t3_neg_4_007_13    m/z 494.2051     0   1   2   3E+4   1   1.5  RT (min)      lcorbi_t3_neg_1_055_05    m/z 494.2520     0   2   4   6E+4   10.5   11  RT (min)      lcorbi_t3_neg_1_055_05    m/z 494.2695     0   1   2   3E+4   11   11.5  RT (min)      lcorbi_t3_neg_4_007_13    m/z 494.3611     0   2   4   6   8E+3   12.5   13  RT (min)      lcorbi_t3_neg_4_034_01    m/z 494.3929     0   1   2   3E+4   13   13.5  RT (min)      lcorbi_t3_neg_4_034_01    m/z 494.8958     0   1   2   3E+3   18   18.5  RT (min)      lcorbi_t3_neg_1_006_40    m/z 494.9421     0   2   4E+3   6.5   7  RT (min)      lcorbi_t3_neg_3_045_34    m/z 494.9997     0   1   2   3E+4   1   1.5  RT (min)      lcorbi_t3_neg_4_007_13    m/z 495.0465     0   1   2E+4   8.5   9  RT (min)      lcorbi_t3_neg_2_048_27    m/z 495.1010     0   1   2   3E+3   7.5   8  RT (min)      lcorbi_t3_neg_4_034_01    m/z 495.1218     0   2   4E+3   8   8.5  RT (min)      lcorbi_t3_neg_4_034_01    m/z 495.1318     0   2   4   6E+3   7   7.5  RT (min)      lcorbi_t3_neg_4_023_18    m/z 495.1698     0   1E+4   8   8.5  RT (min)      lcorbi_t3_neg_4_023_18    m/z 495.2047     0   1E+4   12.5   13  RT (min)      lcorbi_t3_neg_3_024_26    m/z 495.2231     0   2   4   6   8E+3   7   7.5  RT (min)      lcorbi_t3_neg_2_048_27    m/z 495.2474     0   1E+4   11.5   12  RT (min)      lcorbi_t3_neg_1_055_05    m/z 495.2739     0   1E+4   6   6.5  RT (min)      lcorbi_t3_neg_4_044_38    m/z 495.2781     0   1   2   3E+3   9   9.5  RT (min)      lcorbi_t3_neg_4_034_01    m/z 495.2963     0   2   4   6E+2   12   12.5  RT (min)      lcorbi_t3_neg_1_044_23    m/z 495.3030     0   1   2E+3   14   14.5  RT (min)      lcorbi_t3_neg_4_007_13    m/z 495.3285     0   1E+3   11.5   12  RT (min)      lcorbi_t3_neg_1_055_05    m/z 495.3299     0   1   2   3E+3   13.5   14  RT (min)      lcorbi_t3_neg_2_053_23    m/z 495.3956     0   1   2E+3   13   13.5  RT (min)      lcorbi_t3_neg_4_007_13    m/z 495.4167     0   1E+4   13.5   14  RT (min)      lcorbi_t3_neg_4_007_13    m/z 495.9917     0   2   4   6   8E+3   11   11.5  RT (min)      lcorbi_t3_neg_4_007_13    m/z 496.0110     0   2   4E+3   17.5   18  RT (min)      lcorbi_t3_neg_2_053_23    m/z 496.0403     0   1   2E+4   8.5   9  RT (min)      lcorbi_t3_neg_2_048_27    m/z 496.0873     0   1   2   3   4E+3   9   9.5  RT (min)      lcorbi_t3_neg_3_019_12    m/z 496.1742     0   1   2   3   4E+3   13.5   14  RT (min)      lcorbi_t3_neg_4_051_40    m/z 496.2239     0   2   4E+3   13   13.5  RT (min)      lcorbi_t3_neg_4_014_20    m/z 496.2410     0   1E+4   11.5   12  RT (min)      lcorbi_t3_neg_1_055_05    m/z 496.2413     0   1E+4   7.5   8  RT (min)      lcorbi_t3_neg_4_023_18    m/z 496.2416     0   1E+4   10   10.5  RT (min)      lcorbi_t3_neg_3_019_12    m/z 496.2596     0   1   12   12.5  RT (min)      lcorbi_t3_neg_4_019_08    m/z 496.2849     0   1   2   3E+4   11.5   12  RT (min)      lcorbi_t3_neg_4_016_05    m/z 496.2996     0   1E+4   9   9.5  RT (min)      lcorbi_t3_neg_2_048_27    m/z 496.3041     0   1E+4   11.5   12  RT (min)      lcorbi_t3_neg_4_007_13    m/z 496.3179     0   1   2   3   4E+4   13   13.5  RT (min)      lcorbi_t3_neg_4_034_01    m/z 496.3197     0   1E+3   11   11.5  RT (min)      lcorbi_t3_neg_3_043_25    m/z 496.3404     0   1E+4   12.5   13  RT (min)      lcorbi_t3_neg_4_054_36    m/z 496.3487     0   1E+4   13   13.5  RT (min)      lcorbi_t3_neg_4_034_01    m/z 496.7157     0   1E+4   1   1.5  RT (min)      lcorbi_t3_neg_4_038_15    m/z 496.9489     0   1   2   3   4E+3   18   18.5  RT (min)      lcorbi_t3_neg_4_027_12    m/z 497.0229     0   1   2   3   4E+3   9   9.5  RT (min)      lcorbi_t3_neg_3_019_12    m/z 497.0533     0   2   4   6E+3   6   6.5  RT (min)      lcorbi_t3_neg_2_048_27    m/z 497.0803     0   1   2   3   4E+3   10.5   11  RT (min)      lcorbi_t3_neg_4_029_31    m/z 497.0925     0   1E+4   11.5   12  RT (min)      lcorbi_t3_neg_4_007_13    m/z 497.1079     0   2   4E+3   5   5.5  RT (min)      lcorbi_t3_neg_3_022_27    m/z 497.1159     0   2   4   6E+3   7.5   8  RT (min)      lcorbi_t3_neg_4_034_01    m/z 497.1258     0   1   2   3E+3   9.5   10  RT (min)      lcorbi_t3_neg_3_019_12    m/z 497.1411     0   1E+4   10.5   11  RT (min)      lcorbi_t3_neg_4_032_23    m/z 497.1570     0   2   4E+3   8   8.5  RT (min)      lcorbi_t3_neg_4_044_38    m/z 497.1670     0   1   2E+4   8.5   9  RT (min)      lcorbi_t3_neg_4_007_13    m/z 497.2753     0   1E+4   8.5   9  RT (min)      lcorbi_t3_neg_2_048_27    m/z 497.2901     0   1E+3   12.5   13  RT (min)      lcorbi_t3_neg_3_005_23    m/z 497.3077     0   1   2   3   4E+3   13.5   14  RT (min)      lcorbi_t3_neg_4_016_05    m/z 497.3150     0   2   4   6E+3   11.5   12  RT (min)      lcorbi_t3_neg_2_053_23    m/z 497.3229     0   1   13   13.5  RT (min)      lcorbi_t3_neg_4_019_08    m/z 497.3488     0   1E+5   11   11.5  RT (min)      lcorbi_t3_neg_1_055_05    m/z 497.4116     0   1E+4   13   13.5  RT (min)      lcorbi_t3_neg_4_034_01    m/z 497.4210     0   2   4E+3   13.5   14  RT (min)      lcorbi_t3_neg_4_005_27    m/z 497.8643     0   1   2   3   4E+3   18   18.5  RT (min)      lcorbi_t3_neg_3_027_43    m/z 497.9330     0   2   4   6   8E+3   6.5   7  RT (min)      lcorbi_t3_neg_3_045_34    m/z 497.9969     0   2   4   6E+3   10.5   11  RT (min)      lcorbi_t3_neg_1_047_02    m/z 498.0406     0   2   4   6   8E+3   9   9.5  RT (min)      lcorbi_t3_neg_4_007_13    m/z 498.0611     0   1E+4   6   6.5  RT (min)      lcorbi_t3_neg_2_048_27    m/z 498.1028     0   2   4   6   8E+3   13   13.5  RT (min)      lcorbi_t3_neg_1_043_14    m/z 498.1042     0   1E+4   8   8.5  RT (min)      lcorbi_t3_neg_2_048_27    m/z 498.1340     0   1E+4   11   11.5  RT (min)      lcorbi_t3_neg_4_007_13    m/z 498.1379     0   1E+3   13   13.5  RT (min)      lcorbi_t3_neg_3_014_19    m/z 498.1720     0   2   4E+3   10.5   11  RT (min)      lcorbi_t3_neg_2_048_27    m/z 498.1869     0   1   2E+4   5.5   6  RT (min)      lcorbi_t3_neg_4_039_32    m/z 498.1984     0   2   4   6   8E+4   8.5   9  RT (min)      lcorbi_t3_neg_1_055_05    m/z 498.2736     0   1   11   11.5  RT (min)      lcorbi_t3_neg_4_012_17    m/z 498.2785     0   2   4   6   8E+2   8.5   9  RT (min)      lcorbi_t3_neg_4_023_18    m/z 498.2981     0   1E+4   12   12.5  RT (min)      lcorbi_t3_neg_4_007_13    m/z 498.3095     0   1   2E+3   13   13.5  RT (min)      lcorbi_t3_neg_2_020_05    m/z 498.3198     0   1E+3   11   11.5  RT (min)      lcorbi_t3_neg_3_031_45    m/z 498.3271     0   1E+4   13.5   14  RT (min)      lcorbi_t3_neg_4_040_19    m/z 498.8478     0   1   2   3   4E+3   18   18.5  RT (min)      lcorbi_t3_neg_2_021_41    m/z 499.0006     0   1   2E+4   6.5   7  RT (min)      lcorbi_t3_neg_4_023_18    m/z 499.0199     0   1   2   3E+3   9   9.5  RT (min)      lcorbi_t3_neg_2_013_08    m/z 499.0311     0   1E+4   8   8.5  RT (min)      lcorbi_t3_neg_3_024_26    m/z 499.0417     0   2   4   6   8E+3   6   6.5  RT (min)      lcorbi_t3_neg_4_040_19    m/z 499.1433     0   1   2   3E+3   9.5   10  RT (min)      lcorbi_t3_neg_2_023_39    m/z 499.1710     0   1E+4   10   10.5  RT (min)      lcorbi_t3_neg_3_002_24    m/z 499.1899     0   1E+4   11   11.5  RT (min)      lcorbi_t3_neg_4_042_14    m/z 499.2110     0   1   2   3   4E+2   10   10.5  RT (min)      lcorbi_t3_neg_3_002_24    m/z 499.2296     0   1   2   3E+4   5   5.5  RT (min)      lcorbi_t3_neg_3_037_36    m/z 499.2592     0   2   4   6   8E+2   12.5   13  RT (min)      lcorbi_t3_neg_4_016_05    m/z 499.2748     0   1E+4   13   13.5  RT (min)      lcorbi_t3_neg_4_046_37    m/z 499.2885     0   1E+3   12.5   13  RT (min)      lcorbi_t3_neg_4_021_25    m/z 499.3061     0   1   2   3E+3   13   13.5  RT (min)      lcorbi_t3_neg_4_044_38    m/z 499.3191     0   1E+4   12   12.5  RT (min)      lcorbi_t3_neg_4_007_13    m/z 499.3436     0   2   4   6   8E+4   10.5   11  RT (min)      lcorbi_t3_neg_4_019_08    m/z 499.3640     0   1   2E+3   11.5   12  RT (min)      lcorbi_t3_neg_4_090_60    m/z 499.9172     0   1E+4   18   18.5  RT (min)      lcorbi_t3_neg_1_004_22    m/z 500.0006     0   2   4E+3   9   9.5  RT (min)      lcorbi_t3_neg_1_044_23    m/z 500.0132     0   1E+4   4.5   5  RT (min)      lcorbi_t3_neg_3_024_26    m/z 500.0376     0   2   4   6E+3   9   9.5  RT (min)      lcorbi_t3_neg_1_044_23    m/z 500.0767     0   1   2E+4   7.5   8  RT (min)      lcorbi_t3_neg_2_002_26    m/z 500.0994     0   2   4   6E+3   12   12.5  RT (min)      lcorbi_t3_neg_4_040_19    m/z 500.1311     0   1E+4   11   11.5  RT (min)      lcorbi_t3_neg_4_007_13    m/z 500.2313     0   2   4   6E+3   12   12.5  RT (min)      lcorbi_t3_neg_4_005_27    m/z 500.2419     0   1   2   3E+3   13.5   14  RT (min)      lcorbi_t3_neg_4_023_18    m/z 500.2684     0   1E+4   10.5   11  RT (min)      lcorbi_t3_neg_4_007_13    m/z 500.2944     0   1E+4   8.5   9  RT (min)      lcorbi_t3_neg_4_023_18    m/z 500.3081     0   1   2   3   4E+3   13   13.5  RT (min)      lcorbi_t3_neg_4_047_30    m/z 500.3123     0   2   4   6E+3   11   11.5  RT (min)      lcorbi_t3_neg_3_007_16    m/z 500.3264     0   1   13.5   14  RT (min)      lcorbi_t3_neg_1_055_05    m/z 500.3329     0   1   2   3   4E+3   11   11.5  RT (min)      lcorbi_t3_neg_4_034_01    m/z 500.3601     0   2   4   6   8E+3   14   14.5  RT (min)      lcorbi_t3_neg_4_037_29    m/z 500.3607     0   2   4   6   8E+4   11   11.5  RT (min)      lcorbi_t3_neg_1_055_05    m/z 500.3727     0   1E+4   13.5   14  RT (min)      lcorbi_t3_neg_4_023_18    m/z 500.3821     0   1E+4   13   13.5  RT (min)      lcorbi_t3_neg_4_042_14    m/z 500.6476     0   1   1   1.5  RT (min)      lcorbi_t3_neg_4_007_13    m/z 500.8523     0   1   2E+4   18   18.5  RT (min)      lcorbi_t3_neg_2_009_50    m/z 500.9129     0   1   2   3E+3   18   18.5  RT (min)      lcorbi_t3_neg_3_052_32    m/z 500.9255     0   1E+4   9.5   10  RT (min)      lcorbi_t3_neg_4_007_13    m/z 500.9986     0   1   2E+4   6.5   7  RT (min)      lcorbi_t3_neg_4_023_18    m/z 501.0070     0   1   2   3   4E+4   4.5   5  RT (min)      lcorbi_t3_neg_3_024_26    m/z 501.0348     0   1   2   3E+3   9   9.5  RT (min)      lcorbi_t3_neg_3_019_12    m/z 501.0548     0   1   2E+3   9   9.5  RT (min)      lcorbi_t3_neg_1_055_05    m/z 501.0665     0   1   2E+4   10   10.5  RT (min)      lcorbi_t3_neg_4_023_18    m/z 501.0683     0   1E+4   5   5.5  RT (min)      lcorbi_t3_neg_2_048_27    m/z 501.0799     0   2   4E+3   7.5   8  RT (min)      lcorbi_t3_neg_2_002_26    m/z 501.0902     0   1   2E+3   11   11.5  RT (min)      lcorbi_t3_neg_1_010_17    m/z 501.1149     0   1E+4   10   10.5  RT (min)      lcorbi_t3_neg_4_007_13    m/z 501.1269     0   1   2   3E+4   1   1.5  RT (min)      lcorbi_t3_neg_4_032_23    m/z 501.1275     0   1   2E+3   11   11.5  RT (min)      lcorbi_t3_neg_2_054_15    m/z 501.1584     0   1E+4   9   9.5  RT (min)      lcorbi_t3_neg_2_048_27    m/z 501.2233     0   2   4   6   8E+4   11   11.5  RT (min)      lcorbi_t3_neg_1_055_05    m/z 501.2384     0   1   2   3E+3   13.5   14  RT (min)      lcorbi_t3_neg_4_023_18    m/z 501.2677     0   1E+4   12.5   13  RT (min)      lcorbi_t3_neg_4_047_30    m/z 501.2717     0   2   4   6E+2   13   13.5  RT (min)      lcorbi_t3_neg_3_007_16    m/z 501.3032     0   2   4   6   8E+3   11   11.5  RT (min)      lcorbi_t3_neg_3_007_16    m/z 501.3042     0   1E+4   13   13.5  RT (min)      lcorbi_t3_neg_4_007_13    m/z 501.3065     0   1   2   3   4E+3   9.5   10  RT (min)      lcorbi_t3_neg_4_034_01    m/z 501.3387     0   2   4   6   8E+3   13   13.5  RT (min)      lcorbi_t3_neg_3_007_16    m/z 501.3714     0   1   2E+3   13   13.5  RT (min)      lcorbi_t3_neg_4_005_27    m/z 501.8241     0   1   2E+4   1   1.5  RT (min)      lcorbi_t3_neg_2_052_32    m/z 502.0728     0   1   2E+4   8   8.5  RT (min)      lcorbi_t3_neg_1_054_37    m/z 502.0908     0   2   4E+3   9.5   10  RT (min)      lcorbi_t3_neg_3_019_12    m/z 502.1288     0   2   4   6E+3   1   1.5  RT (min)      lcorbi_t3_neg_3_031_45    m/z 502.2265     0   1   2E+4   11   11.5  RT (min)      lcorbi_t3_neg_1_055_05    m/z 502.2341     0   1   2E+4   5   5.5  RT (min)      lcorbi_t3_neg_1_013_35    m/z 502.2703     0   1E+4   10.5   11  RT (min)      lcorbi_t3_neg_4_007_13    m/z 502.3159     0   1E+4   13.5   14  RT (min)      lcorbi_t3_neg_1_023_03    m/z 502.3242     0   2   4   6   8E+3   6.5   7  RT (min)      lcorbi_t3_neg_4_032_23    m/z 502.3283     0   2   4   6E+2   12.5   13  RT (min)      lcorbi_t3_neg_2_039_31    m/z 502.3351     0   1E+4   13   13.5  RT (min)      lcorbi_t3_neg_1_055_05    m/z 502.3589     0   1E+4   14   14.5  RT (min)      lcorbi_t3_neg_3_018_13    m/z 502.3621     0   1E+3   12   12.5  RT (min)      lcorbi_t3_neg_1_055_05    m/z 502.3790     0   1   2   3   4E+4   12   12.5  RT (min)      lcorbi_t3_neg_1_055_05    m/z 502.8494     0   1E+4   18   18.5  RT (min)      lcorbi_t3_neg_2_042_44    m/z 502.8590     0   1   2E+4   1   1.5  RT (min)      lcorbi_t3_neg_4_042_14    m/z 503.0460     0   1   2   3   4E+3   5   5.5  RT (min)      lcorbi_t3_neg_2_048_27    m/z 503.0938     0   1   2   3   4E+3   9.5   10  RT (min)      lcorbi_t3_neg_2_052_32    m/z 503.2040     0   1   2   3   4E+3   6   6.5  RT (min)      lcorbi_t3_neg_4_005_27    m/z 503.2211     0   2   4   6E+3   12.5   13  RT (min)      lcorbi_t3_neg_2_051_34    m/z 503.2388     0   2   4E+4   11.5   12  RT (min)      lcorbi_t3_neg_1_055_05    m/z 503.2480     0   2   4E+3   6   6.5  RT (min)      lcorbi_t3_neg_4_037_29    m/z 503.2709     0   1E+3   13   13.5  RT (min)      lcorbi_t3_neg_1_055_05    m/z 503.3000     0   2   4   6E+3   6.5   7  RT (min)      lcorbi_t3_neg_4_029_31    m/z 503.3224     0   1E+4   9   9.5  RT (min)      lcorbi_t3_neg_2_048_27    m/z 503.3314     0   1E+4   13   13.5  RT (min)      lcorbi_t3_neg_1_055_05    m/z 503.3547     0   1E+4   14   14.5  RT (min)      lcorbi_t3_neg_4_005_27    m/z 503.3587     0   1E+4   10   10.5  RT (min)      lcorbi_t3_neg_4_023_18    m/z 503.3713     0   1   2E+4   14   14.5  RT (min)      lcorbi_t3_neg_2_048_27    m/z 503.4364     0   2   4E+3   13   13.5  RT (min)      lcorbi_t3_neg_3_055_06    m/z 503.8812     0   1   2E+4   18   18.5  RT (min)      lcorbi_t3_neg_3_038_39    m/z 503.9808     0   2   4   6E+3   10   10.5  RT (min)      lcorbi_t3_neg_1_046_11    m/z 504.0666     0   1E+4   11   11.5  RT (min)      lcorbi_t3_neg_4_007_13    m/z 504.0698     0   1E+3   8   8.5  RT (min)      lcorbi_t3_neg_1_026_19    m/z 504.0838     0   1E+4   8   8.5  RT (min)      lcorbi_t3_neg_1_026_19    m/z 504.1047     0   1E+4   11   11.5  RT (min)      lcorbi_t3_neg_4_007_13    m/z 504.1574     0   2   4   6E+3   8   8.5  RT (min)      lcorbi_t3_neg_4_013_09    m/z 504.1807     0   1   2E+4   1   1.5  RT (min)      lcorbi_t3_neg_1_050_45    m/z 504.2705     0   1E+4   11.5   12  RT (min)      lcorbi_t3_neg_4_042_14    m/z 504.2837     0   1E+4   11.5   12  RT (min)      lcorbi_t3_neg_4_029_31    m/z 504.2999     0   1E+4   11.5   12  RT (min)      lcorbi_t3_neg_4_034_01    m/z 504.3549     0   1E+4   14   14.5  RT (min)      lcorbi_t3_neg_4_007_13    m/z 504.3804     0   1E+3   13   13.5  RT (min)      lcorbi_t3_neg_4_034_01    m/z 504.4138     0   2   4   6E+4   13.5   14  RT (min)      lcorbi_t3_neg_4_034_01    m/z 504.8642     0   2   4   6   8E+3   18   18.5  RT (min)      lcorbi_t3_neg_2_019_07    m/z 504.9363     0   1   2   3E+3   18   18.5  RT (min)      lcorbi_t3_neg_3_010_15    m/z 505.0633     0   2   4   6E+3   11   11.5  RT (min)      lcorbi_t3_neg_4_007_13    m/z 505.0920     0   2   4   6   8E+3   1.5   2  RT (min)      lcorbi_t3_neg_2_007_14    m/z 505.1018     0   2   4   6   8E+3   10.5   11  RT (min)      lcorbi_t3_neg_4_043_11    m/z 505.1139     0   1   2   3E+3   5   5.5  RT (min)      lcorbi_t3_neg_4_044_38    m/z 505.2075     0   2   4   6   8E+3   9   9.5  RT (min)      lcorbi_t3_neg_4_034_01    m/z 505.2436     0   1   9.5   10  RT (min)      lcorbi_t3_neg_4_032_23    m/z 505.2574     0   1   2   3E+3   11   11.5  RT (min)      lcorbi_t3_neg_3_028_35    m/z 505.2611     0   2   4   6E+3   6   6.5  RT (min)      lcorbi_t3_neg_2_048_27    m/z 505.2801     0   1E+4   11.5   12  RT (min)      lcorbi_t3_neg_4_029_31    m/z 505.2895     0   2   4   6E+2   12   12.5  RT (min)      lcorbi_t3_neg_1_044_23    m/z 505.3198     0   2   4   6E+2   12.5   13  RT (min)      lcorbi_t3_neg_4_007_13    m/z 505.3322     0   1   2   3E+4   14   14.5  RT (min)      lcorbi_t3_neg_1_055_05    m/z 505.3565     0   1   2E+3   13   13.5  RT (min)      lcorbi_t3_neg_2_052_32    m/z 505.4164     0   1E+4   13.5   14  RT (min)      lcorbi_t3_neg_4_034_01    m/z 505.7769     0   1   1   1.5  RT (min)      lcorbi_t3_neg_3_028_35    m/z 505.9777     0   2   4E+3   10   10.5  RT (min)      lcorbi_t3_neg_1_046_11    m/z 505.9963     0   2   4   6   8E+3   11   11.5  RT (min)      lcorbi_t3_neg_4_007_13    m/z 506.0235     0   2   4   6E+3   5.5   6  RT (min)      lcorbi_t3_neg_3_034_21    m/z 506.0806     0   1E+4   8   8.5  RT (min)      lcorbi_t3_neg_1_026_19    m/z 506.0851     0   2   4   6   8E+3   9.5   10  RT (min)      lcorbi_t3_neg_4_007_13    m/z 506.1243     0   1   2   3   4E+3   17.5   18  RT (min)      lcorbi_t3_neg_2_022_24    m/z 506.1519     0   2   4   6E+3   8   8.5  RT (min)      lcorbi_t3_neg_4_034_01    m/z 506.1664     0   1E+3   7   7.5  RT (min)      lcorbi_t3_neg_4_023_18    m/z 506.1792     0   1   2   3E+4   1   1.5  RT (min)      lcorbi_t3_neg_2_035_46    m/z 506.2326     0   1   2   3   4E+3   9   9.5  RT (min)      lcorbi_t3_neg_4_001_49    m/z 506.2343     0   2   4   6   8E+3   6   6.5  RT (min)      lcorbi_t3_neg_4_001_49    m/z 506.2394     0   2   4E+3   13   13.5  RT (min)      lcorbi_t3_neg_4_042_14    m/z 506.2589     0   2   4E+3   13   13.5  RT (min)      lcorbi_t3_neg_2_007_14    m/z 506.2651     0   1E+4   9   9.5  RT (min)      lcorbi_t3_neg_2_048_27    m/z 506.2707     0   2   4   6E+3   13   13.5  RT (min)      lcorbi_t3_neg_1_023_03    m/z 506.2708     0   2   4E+3   12   12.5  RT (min)      lcorbi_t3_neg_4_044_38    m/z 506.2859     0   1E+4   11.5   12  RT (min)      lcorbi_t3_neg_1_055_05    m/z 506.2927     0   1   2   3   4E+2   12   12.5  RT (min)      lcorbi_t3_neg_3_005_23    m/z 506.3386     0   1E+4   12.5   13  RT (min)      lcorbi_t3_neg_4_007_13    m/z 506.3538     0   1   2E+4   14   14.5  RT (min)      lcorbi_t3_neg_4_007_13    m/z 506.3963     0   1   2E+3   13.5   14  RT (min)      lcorbi_t3_neg_4_034_01    m/z 506.4293     0   1E+4   13.5   14  RT (min)      lcorbi_t3_neg_4_005_27    m/z 506.8611     0   1   2E+3   18   18.5  RT (min)      lcorbi_t3_neg_3_019_12    m/z 507.1191     0   1   2   3E+3   10   10.5  RT (min)      lcorbi_t3_neg_2_012_36    m/z 507.1209     0   1E+4   11.5   12  RT (min)      lcorbi_t3_neg_4_007_13    m/z 507.1601     0   1E+3   7   7.5  RT (min)      lcorbi_t3_neg_4_023_18    m/z 507.1687     0   2   4   6   8E+3   6   6.5  RT (min)      lcorbi_t3_neg_2_048_27    m/z 507.1947     0   1   2E+3   13   13.5  RT (min)      lcorbi_t3_neg_2_039_31    m/z 507.2127     0   1   2   3E+3   10.5   11  RT (min)      lcorbi_t3_neg_4_034_01    m/z 507.2357     0   2   4   6E+3   13   13.5  RT (min)      lcorbi_t3_neg_2_007_14    m/z 507.2552     0   2   4E+3   13   13.5  RT (min)      lcorbi_t3_neg_4_007_13    m/z 507.2595     0   1   2   3   4E+2   8   8.5  RT (min)      lcorbi_t3_neg_1_055_05    m/z 507.2724     0   2   4   6   8E+4   10.5   11  RT (min)      lcorbi_t3_neg_1_055_05    m/z 507.3184     0   1E+4   12.5   13  RT (min)      lcorbi_t3_neg_4_007_13    m/z 507.3201     0   1   2E+3   10.5   11  RT (min)      lcorbi_t3_neg_1_055_05    m/z 507.3687     0   1   2   3   4E+3   13   13.5  RT (min)      lcorbi_t3_neg_4_042_14    m/z 507.3869     0   1   2E+4   12   12.5  RT (min)      lcorbi_t3_neg_3_002_24    m/z 507.4417     0   2   4   6   8E+3   13.5   14  RT (min)      lcorbi_t3_neg_4_005_27    m/z 507.8756     0   1E+4   18   18.5  RT (min)      lcorbi_t3_neg_4_007_13    m/z 507.9548     0   1E+4   7.5   8  RT (min)      lcorbi_t3_neg_3_054_18    m/z 508.0550     0   1   2   3   4E+3   9   9.5  RT (min)      lcorbi_t3_neg_3_019_12    m/z 508.0856     0   2   4   6E+3   5   5.5  RT (min)      lcorbi_t3_neg_4_040_19    m/z 508.1080     0   2   4   6E+3   7.5   8  RT (min)      lcorbi_t3_neg_3_054_18    m/z 508.1212     0   1   2   3   4E+3   17.5   18  RT (min)      lcorbi_t3_neg_2_014_29    m/z 508.1635     0   2   4E+2   7   7.5  RT (min)      lcorbi_t3_neg_4_023_18    m/z 508.1692     0   2   4E+3   8   8.5  RT (min)      lcorbi_t3_neg_4_040_19    m/z 508.2011     0   1   2E+4   1   1.5  RT (min)      lcorbi_t3_neg_1_050_45    m/z 508.2138     0   2   4E+3   10   10.5  RT (min)      lcorbi_t3_neg_4_023_18    m/z 508.2551     0   2   4   6   8E+3   13   13.5  RT (min)      lcorbi_t3_neg_4_034_01    m/z 508.2620     0   2   4   6   8E+2   9   9.5  RT (min)      lcorbi_t3_neg_2_053_23    m/z 508.2676     0   2   4   6E+4   10.5   11  RT (min)      lcorbi_t3_neg_1_055_05    m/z 508.2945     0   1E+4   1   1.5  RT (min)      lcorbi_t3_neg_4_011_24    m/z 508.3000     0   1E+3   12   12.5  RT (min)      lcorbi_t3_neg_4_007_13    m/z 508.3107     0   2   4   6E+3   10   10.5  RT (min)      lcorbi_t3_neg_3_019_12    m/z 508.3276     0   1   9.5   10  RT (min)      lcorbi_t3_neg_4_021_25    m/z 508.3312     0   1E+4   11.5   12  RT (min)      lcorbi_t3_neg_4_007_13    m/z 508.3543     0   1E+4   13   13.5  RT (min)      lcorbi_t3_neg_4_007_13    m/z 508.7181     0   1   1   1.5  RT (min)      lcorbi_t3_neg_2_048_27    m/z 508.8758     0   2   4   6E+4   0.5   1  RT (min)      lcorbi_t3_neg_2_007_14    m/z 509.0181     0   1   9   9.5  RT (min)      lcorbi_t3_neg_2_053_23    m/z 509.0388     0   2   4   6   8E+3   4.5   5  RT (min)      lcorbi_t3_neg_3_024_26    m/z 509.0487     0   2   4   6   8E+3   9   9.5  RT (min)      lcorbi_t3_neg_3_019_12    m/z 509.2071     0   1   2   3E+3   7   7.5  RT (min)      lcorbi_t3_neg_2_004_28    m/z 509.2292     0   1   2   3E+3   10   10.5  RT (min)      lcorbi_t3_neg_4_034_01    m/z 509.2515     0   1E+4   13   13.5  RT (min)      lcorbi_t3_neg_4_034_01    m/z 509.2752     0   1   2   3E+2   8.5   9  RT (min)      lcorbi_t3_neg_4_021_25    m/z 509.2890     0   2   4   6   8E+3   12   12.5  RT (min)      lcorbi_t3_neg_4_034_01    m/z 509.2963     0   2   4   6   8E+3   10   10.5  RT (min)      lcorbi_t3_neg_4_029_31    m/z 509.3380     0   1   12   12.5  RT (min)      lcorbi_t3_neg_4_005_27    m/z 509.3630     0   2   4   6E+3   13   13.5  RT (min)      lcorbi_t3_neg_2_035_46    m/z 509.3845     0   1E+4   12   12.5  RT (min)      lcorbi_t3_neg_4_034_01    m/z 509.4320     0   1   12.5   13  RT (min)      lcorbi_t3_neg_4_034_01    m/z 509.8794     0   1   2   3   4E+3   18   18.5  RT (min)      lcorbi_t3_neg_2_019_07    m/z 509.9519     0   1E+4   7.5   8  RT (min)      lcorbi_t3_neg_3_054_18    m/z 510.0334     0   2   4E+3   9   9.5  RT (min)      lcorbi_t3_neg_4_007_13    m/z 510.0641     0   1   2   3   4E+3   6.5   7  RT (min)      lcorbi_t3_neg_4_001_49    m/z 510.1075     0   1E+4   6   6.5  RT (min)      lcorbi_t3_neg_2_030_47    m/z 510.1535     0   1E+4   13   13.5  RT (min)      lcorbi_t3_neg_4_040_19    m/z 510.1741     0   1   2   3E+4   13   13.5  RT (min)      lcorbi_t3_neg_4_044_38    m/z 510.1869     0   1   2E+4   7   7.5  RT (min)      lcorbi_t3_neg_3_024_26    m/z 510.2366     0   1E+4   11.5   12  RT (min)      lcorbi_t3_neg_1_055_05    m/z 510.2708     0   1E+4   13.5   14  RT (min)      lcorbi_t3_neg_4_040_19    m/z 510.2914     0   1E+4   12.5   13  RT (min)      lcorbi_t3_neg_3_024_26    m/z 510.2942     0   1   2E+4   5   5.5  RT (min)      lcorbi_t3_neg_2_048_27    m/z 510.2966     0   2   4E+3   10   10.5  RT (min)      lcorbi_t3_neg_3_019_12    m/z 510.3069     0   1E+3   13   13.5  RT (min)      lcorbi_t3_neg_4_034_01    m/z 510.3091     0   1   2   3E+4   10   10.5  RT (min)      lcorbi_t3_neg_3_019_12    m/z 510.3116     0   2   4   6E+3   5.5   6  RT (min)      lcorbi_t3_neg_4_044_38    m/z 510.3366     0   2   4   6E+2   11.5   12  RT (min)      lcorbi_t3_neg_2_048_27    m/z 510.4164     0   1   2E+4   14   14.5  RT (min)      lcorbi_t3_neg_4_032_23    m/z 510.8811     0   1E+4   12   12.5  RT (min)      lcorbi_t3_neg_2_018_40    m/z 510.8811     0   1E+4   10   10.5  RT (min)      lcorbi_t3_neg_2_003_42    m/z 510.9534     0   2   4E+3   18   18.5  RT (min)      lcorbi_t3_neg_4_042_14    m/z 510.9999     0   2   4   6E+3   8   8.5  RT (min)      lcorbi_t3_neg_4_023_18    m/z 511.0463     0   2   4   6   8E+3   7   7.5  RT (min)      lcorbi_t3_neg_2_002_26    m/z 511.0748     0   1   2E+4   8   8.5  RT (min)      lcorbi_t3_neg_4_040_19    m/z 511.0977     0   2   4   6   8E+3   10   10.5  RT (min)      lcorbi_t3_neg_4_023_18    m/z 511.1560     0   1   2E+3   13   13.5  RT (min)      lcorbi_t3_neg_4_044_38    m/z 511.1775     0   1E+4   13   13.5  RT (min)      lcorbi_t3_neg_4_044_38    m/z 511.1875     0   2   4E+3   8   8.5  RT (min)      lcorbi_t3_neg_2_048_27    m/z 511.2140     0   1E+3   13   13.5  RT (min)      lcorbi_t3_neg_4_047_30    m/z 511.2400     0   1   2   3E+3   11.5   12  RT (min)      lcorbi_t3_neg_1_055_05    m/z 511.2672     0   1E+4   13.5   14  RT (min)      lcorbi_t3_neg_4_040_19    m/z 511.2870     0   1   2   3   4E+4   11   11.5  RT (min)      lcorbi_t3_neg_4_005_27    m/z 511.3038     0   1   12   12.5  RT (min)      lcorbi_t3_neg_2_048_27    m/z 511.3096     0   1E+4   12.5   13  RT (min)      lcorbi_t3_neg_4_034_01    m/z 511.3258     0   1E+4   13   13.5  RT (min)      lcorbi_t3_neg_4_034_01    m/z 511.3362     0   2   4   6E+3   11   11.5  RT (min)      lcorbi_t3_neg_4_032_23    m/z 511.3375     0   2   4E+3   13   13.5  RT (min)      lcorbi_t3_neg_1_055_05    m/z 511.4000     0   1E+4   12.5   13  RT (min)      lcorbi_t3_neg_4_010_28    m/z 511.8155     0   1   2E+4   1   1.5  RT (min)      lcorbi_t3_neg_2_052_32    m/z 511.8846     0   2   4   6   8E+3   18   18.5  RT (min)      lcorbi_t3_neg_4_028_39    m/z 512.0669     0   1   2   3   4E+3   9   9.5  RT (min)      lcorbi_t3_neg_3_019_12    m/z 512.1018     0   1E+4   8   8.5  RT (min)      lcorbi_t3_neg_2_048_27    m/z 512.2189     0   2   4E+3   13   13.5  RT (min)      lcorbi_t3_neg_4_015_10    m/z 512.2337     0   2   4E+3   10   10.5  RT (min)      lcorbi_t3_neg_3_019_12    m/z 512.2704     0   1   13.5   14  RT (min)      lcorbi_t3_neg_3_005_23    m/z 512.2942     0   1E+4   8.5   9  RT (min)      lcorbi_t3_neg_2_048_27    m/z 512.2943     0   1E+4   11.5   12  RT (min)      lcorbi_t3_neg_4_005_27    m/z 512.3357     0   1E+4   11   11.5  RT (min)      lcorbi_t3_neg_4_042_14    m/z 512.3386     0   2   4   6   8E+3   14   14.5  RT (min)      lcorbi_t3_neg_2_048_27    m/z 512.3573     0   1   2E+4   12.5   13  RT (min)      lcorbi_t3_neg_4_010_28    m/z 512.3584     0   2   4E+4   10.5   11  RT (min)      lcorbi_t3_neg_1_055_05    m/z 512.3688     0   1   2E+4   12   12.5  RT (min)      lcorbi_t3_neg_4_007_13    m/z 512.3955     0   1   11.5   12  RT (min)      lcorbi_t3_neg_4_005_27    m/z 512.9622     0   1E+4   11   11.5  RT (min)      lcorbi_t3_neg_1_047_02    m/z 512.9809     0   1E+4   8   8.5  RT (min)      lcorbi_t3_neg_1_007_26    m/z 513.0433     0   2   4   6   8E+3   7   7.5  RT (min)      lcorbi_t3_neg_2_002_26    m/z 513.0883     0   1   2   3E+3   7.5   8  RT (min)      lcorbi_t3_neg_4_007_13    m/z 513.1117     0   2   4E+3   9.5   10  RT (min)      lcorbi_t3_neg_4_053_43    m/z 513.1912     0   1E+4   7   7.5  RT (min)      lcorbi_t3_neg_1_007_26    m/z 513.2055     0   2   4   6   8E+3   9   9.5  RT (min)      lcorbi_t3_neg_2_036_35    m/z 513.2153     0   1E+4   11.5   12  RT (min)      lcorbi_t3_neg_3_024_26    m/z 513.2321     0   1   9.5   10  RT (min)      lcorbi_t3_neg_4_105_61    m/z 513.3403     0   1E+3   12   12.5  RT (min)      lcorbi_t3_neg_4_023_18    m/z 513.3572     0   1E+4   13   13.5  RT (min)      lcorbi_t3_neg_4_042_14    m/z 513.3693     0   2   4   6   8E+3   9.5   10  RT (min)      lcorbi_t3_neg_3_024_26    m/z 513.3706     0   1   2E+4   11   11.5  RT (min)      lcorbi_t3_neg_1_055_05    m/z 513.8817     0   1   2   3E+3   18   18.5  RT (min)      lcorbi_t3_neg_2_027_13    m/z 514.0003     0   2   4   6   8E+3   9.5   10  RT (min)      lcorbi_t3_neg_2_048_27    m/z 514.0371     0   1E+4   7   7.5  RT (min)      lcorbi_t3_neg_2_002_26    m/z 514.1025     0   2   4   6   8E+3   5   5.5  RT (min)      lcorbi_t3_neg_4_040_19    m/z 514.1277     0   1   2   3E+3   9.5   10  RT (min)      lcorbi_t3_neg_4_050_02    m/z 514.1393     0   1   2   3   4E+3   10   10.5  RT (min)      lcorbi_t3_neg_4_045_35    m/z 514.1569     0   1   2   3   4E+3   9   9.5  RT (min)      lcorbi_t3_neg_2_046_20    m/z 514.2023     0   2   4   6E+3   9   9.5  RT (min)      lcorbi_t3_neg_2_051_34    m/z 514.2160     0   2   4   6E+3   11   11.5  RT (min)      lcorbi_t3_neg_4_034_01    m/z 514.2274     0   2   4   6E+3   12   12.5  RT (min)      lcorbi_t3_neg_4_005_27    m/z 514.2306     0   1E+3   10   10.5  RT (min)      lcorbi_t3_neg_1_055_05    m/z 514.2746     0   1   8.5   9  RT (min)      lcorbi_t3_neg_4_023_18    m/z 514.2755     0   1E+4   14   14.5  RT (min)      lcorbi_t3_neg_4_037_29    m/z 514.3069     0   1   2E+4   10.5   11  RT (min)      lcorbi_t3_neg_4_007_13    m/z 514.3216     0   1E+3   13   13.5  RT (min)      lcorbi_t3_neg_4_023_18    m/z 514.3789     0   1E+4   14   14.5  RT (min)      lcorbi_t3_neg_3_023_07    m/z 514.3840     0   1E+4   12   12.5  RT (min)      lcorbi_t3_neg_3_018_13    m/z 514.8928     0   1E+4   9.5   10  RT (min)      lcorbi_t3_neg_4_053_43    m/z 514.9593     0   2   4   6   8E+3   11   11.5  RT (min)      lcorbi_t3_neg_1_047_02    m/z 514.9779     0   2   4   6E+3   8   8.5  RT (min)      lcorbi_t3_neg_1_007_26    m/z 515.0117     0   1   2   3   4E+3   8   8.5  RT (min)      lcorbi_t3_neg_4_023_18    m/z 515.0695     0   1E+4   8   8.5  RT (min)      lcorbi_t3_neg_4_040_19    m/z 515.1186     0   1   2   3   4E+3   5   5.5  RT (min)      lcorbi_t3_neg_2_014_29    m/z 515.1215     0   1   9   9.5  RT (min)      lcorbi_t3_neg_4_019_08    m/z 515.1483     0   1E+4   10.5   11  RT (min)      lcorbi_t3_neg_4_011_24    m/z 515.2155     0   1   2E+4   10   10.5  RT (min)      lcorbi_t3_neg_3_019_12    m/z 515.2813     0   1E+3   13   13.5  RT (min)      lcorbi_t3_neg_2_007_14    m/z 515.2860     0   2   4   6E+5   8   8.5  RT (min)      lcorbi_t3_neg_4_023_18    m/z 515.3054     0   1E+4   12   12.5  RT (min)      lcorbi_t3_neg_4_023_18    m/z 515.3163     0   1   2   3E+3   11   11.5  RT (min)      lcorbi_t3_neg_1_055_05    m/z 515.3186     0   1E+3   13   13.5  RT (min)      lcorbi_t3_neg_4_034_01    m/z 515.3193     0   1   2   3E+3   14.5   15  RT (min)      lcorbi_t3_neg_1_005_07    m/z 515.3332     0   1   2   3E+3   13   13.5  RT (min)      lcorbi_t3_neg_4_007_13    m/z 515.3725     0   1   2E+4   13   13.5  RT (min)      lcorbi_t3_neg_4_034_01    m/z 515.4274     0   1E+4   12.5   13  RT (min)      lcorbi_t3_neg_4_039_32    m/z 515.9561     0   2   4   6   8E+3   11   11.5  RT (min)      lcorbi_t3_neg_1_047_02    m/z 516.0252     0   1   2E+4   11   11.5  RT (min)      lcorbi_t3_neg_4_007_13    m/z 516.0656     0   2   4   6E+3   1   1.5  RT (min)      lcorbi_t3_neg_3_053_14    m/z 516.0770     0   1E+3   9   9.5  RT (min)      lcorbi_t3_neg_3_019_12    m/z 516.1073     0   1E+4   12.5   13  RT (min)      lcorbi_t3_neg_4_047_30    m/z 516.1339     0   2   4   6E+3   10.5   11  RT (min)      lcorbi_t3_neg_4_043_11    m/z 516.1595     0   1E+5   1   1.5  RT (min)      lcorbi_t3_neg_2_035_46    m/z 516.2274     0   2   4   6E+3   12   12.5  RT (min)      lcorbi_t3_neg_4_007_13    m/z 516.2731     0   1E+4   11.5   12  RT (min)      lcorbi_t3_neg_3_007_16    m/z 516.2859     0   1   11   11.5  RT (min)      lcorbi_t3_neg_1_055_05    m/z 516.3301     0   1   2E+4   5.5   6  RT (min)      lcorbi_t3_neg_4_032_23    m/z 516.3348     0   1   2   3E+4   11   11.5  RT (min)      lcorbi_t3_neg_1_055_05    m/z 516.3369     0   1E+4   12   12.5  RT (min)      lcorbi_t3_neg_4_007_13    m/z 516.3520     0   1   12.5   13  RT (min)      lcorbi_t3_neg_3_007_16    m/z 516.3778     0   1   14   14.5  RT (min)      lcorbi_t3_neg_4_034_01    m/z 516.3898     0   1E+5   11.5   12  RT (min)      lcorbi_t3_neg_1_055_05    m/z 516.4137     0   1E+4   13.5   14  RT (min)      lcorbi_t3_neg_4_042_14    m/z 516.8976     0   2   4E+3   18   18.5  RT (min)      lcorbi_t3_neg_4_023_18    m/z 517.0071     0   2   4   6E+3   9   9.5  RT (min)      lcorbi_t3_neg_1_003_41    m/z 517.0387     0   1   2E+4   10   10.5  RT (min)      lcorbi_t3_neg_4_023_18    m/z 517.0879     0   1E+4   11   11.5  RT (min)      lcorbi_t3_neg_2_035_46    m/z 517.1014     0   1   2E+4   5   5.5  RT (min)      lcorbi_t3_neg_2_048_27    m/z 517.1502     0   2   4   6   8E+3   13   13.5  RT (min)      lcorbi_t3_neg_4_042_14    m/z 517.1804     0   2   4E+4   1   1.5  RT (min)      lcorbi_t3_neg_4_011_24    m/z 517.1872     0   2   4   6E+3   9   9.5  RT (min)      lcorbi_t3_neg_4_023_18    m/z 517.1898     0   1E+4   5   5.5  RT (min)      lcorbi_t3_neg_2_048_27    m/z 517.2062     0   1E+4   11   11.5  RT (min)      lcorbi_t3_neg_4_032_23    m/z 517.2762     0   1E+4   9.5   10  RT (min)      lcorbi_t3_neg_2_048_27    m/z 517.2974     0   1E+4   9   9.5  RT (min)      lcorbi_t3_neg_4_023_18    m/z 517.3075     0   1   2E+4   8   8.5  RT (min)      lcorbi_t3_neg_2_048_27    m/z 517.3212     0   2   4E+4   11.5   12  RT (min)      lcorbi_t3_neg_1_055_05    m/z 517.3320     0   1E+4   13.5   14  RT (min)      lcorbi_t3_neg_4_023_18    m/z 517.3514     0   1   2E+4   14   14.5  RT (min)      lcorbi_t3_neg_3_035_03    m/z 517.3719     0   1E+4   14   14.5  RT (min)      lcorbi_t3_neg_4_048_03    m/z 517.3894     0   1   2E+4   12.5   13  RT (min)      lcorbi_t3_neg_2_036_35    m/z 517.4260     0   1E+4   13.5   14  RT (min)      lcorbi_t3_neg_4_005_27    m/z 517.8704     0   1E+4   1   1.5  RT (min)      lcorbi_t3_neg_1_043_14    m/z 518.1042     0   1E+4   12.5   13  RT (min)      lcorbi_t3_neg_4_047_30    m/z 518.1291     0   2   4   6E+3   12.5   13  RT (min)      lcorbi_t3_neg_4_023_18    m/z 518.1359     0   1   2E+4   8   8.5  RT (min)      lcorbi_t3_neg_2_048_27    m/z 518.2800     0   1   2E+4   11.5   12  RT (min)      lcorbi_t3_neg_4_029_31    m/z 518.3121     0   2   4   6E+3   13   13.5  RT (min)      lcorbi_t3_neg_4_017_51    m/z 518.3196     0   1   2E+3   11.5   12  RT (min)      lcorbi_t3_neg_1_055_05    m/z 518.3343     0   2   4   6   8E+3   5.5   6  RT (min)      lcorbi_t3_neg_4_042_14    m/z 518.3353     0   2   4E+2   13.5   14  RT (min)      lcorbi_t3_neg_4_019_08    m/z 518.3525     0   1   2E+4   12.5   13  RT (min)      lcorbi_t3_neg_4_007_13    m/z 518.3926     0   1   2E+4   13   13.5  RT (min)      lcorbi_t3_neg_4_042_14    m/z 518.8316     0   1   2   3   4E+3   18   18.5  RT (min)      lcorbi_t3_neg_1_042_18    m/z 518.8859     0   1   2   3E+3   18   18.5  RT (min)      lcorbi_t3_neg_4_028_39    m/z 518.9633     0   2   4   6   8E+3   5   5.5  RT (min)      lcorbi_t3_neg_1_054_37    m/z 519.0043     0   1   2E+3   9   9.5  RT (min)      lcorbi_t3_neg_1_031_33    m/z 519.0464     0   1E+3   9.5   10  RT (min)      lcorbi_t3_neg_4_042_14    m/z 519.0780     0   1   11   11.5  RT (min)      lcorbi_t3_neg_4_043_11    m/z 519.0859     0   1   2   3E+3   9   9.5  RT (min)      lcorbi_t3_neg_3_019_12    m/z 519.1140     0   1E+4   7.5   8  RT (min)      lcorbi_t3_neg_4_037_29    m/z 519.1174     0   2   4   6   8E+3   11   11.5  RT (min)      lcorbi_t3_neg_4_043_11    m/z 519.1277     0   2   4   6E+3   6.5   7  RT (min)      lcorbi_t3_neg_4_034_01    m/z 519.1505     0   2   4   6   8E+3   13   13.5  RT (min)      lcorbi_t3_neg_4_007_13    m/z 519.1505     0   2   4   6   8E+3   10.5   11  RT (min)      lcorbi_t3_neg_4_042_14    m/z 519.2530     0   1   2E+4   11   11.5  RT (min)      lcorbi_t3_neg_1_055_05    m/z 519.2554     0   1   2   3   4E+4   10   10.5  RT (min)      lcorbi_t3_neg_1_055_05    m/z 519.2959     0   1E+4   11.5   12  RT (min)      lcorbi_t3_neg_4_032_23    m/z 519.3058     0   2   4   6E+3   13   13.5  RT (min)      lcorbi_t3_neg_4_017_51    m/z 519.3226     0   1E+4   12   12.5  RT (min)      lcorbi_t3_neg_1_055_05    m/z 519.3367     0   1   2E+4   12   12.5  RT (min)      lcorbi_t3_neg_1_055_05    m/z 519.3472     0   1   2   3E+4   11   11.5  RT (min)      lcorbi_t3_neg_1_055_05    m/z 519.3574     0   1   13   13.5  RT (min)      lcorbi_t3_neg_1_043_14    m/z 519.3949     0   2   4   6E+4   11.5   12  RT (min)      lcorbi_t3_neg_3_002_24    m/z 519.4051     0   1E+5   13   13.5  RT (min)      lcorbi_t3_neg_4_042_14    m/z 519.8206     0   1   2E+4   1   1.5  RT (min)      lcorbi_t3_neg_4_054_36    m/z 519.8674     0   1   2E+4   1   1.5  RT (min)      lcorbi_t3_neg_1_043_14    m/z 519.9954     0   1E+4   11   11.5  RT (min)      lcorbi_t3_neg_4_007_13    m/z 520.0495     0   1E+4   10.5   11  RT (min)      lcorbi_t3_neg_2_045_30    m/z 520.0977     0   2   4E+3   6.5   7  RT (min)      lcorbi_t3_neg_4_001_49    m/z 520.2266     0   2   4   6   8E+3   10   10.5  RT (min)      lcorbi_t3_neg_4_042_14    m/z 520.2379     0   2   4   6E+2   7.5   8  RT (min)      lcorbi_t3_neg_2_048_27    m/z 520.2576     0   2   4   6   8E+3   12   12.5  RT (min)      lcorbi_t3_neg_4_007_13    m/z 520.2849     0   1   2E+3   11   11.5  RT (min)      lcorbi_t3_neg_4_088_59    m/z 520.3457     0   1   2   3   4E+2   11   11.5  RT (min)      lcorbi_t3_neg_2_053_23    m/z 520.3507     0   2   4   6   8E+3   14   14.5  RT (min)      lcorbi_t3_neg_4_034_01    m/z 520.3854     0   1   2E+4   11.5   12  RT (min)      lcorbi_t3_neg_3_045_34    m/z 520.4085     0   1   2E+4   13   13.5  RT (min)      lcorbi_t3_neg_4_029_31    m/z 520.8321     0   1   2   3E+3   18   18.5  RT (min)      lcorbi_t3_neg_3_029_02    m/z 520.8492     0   2   4E+3   18   18.5  RT (min)      lcorbi_t3_neg_1_055_05    m/z 520.8870     0   1   2   3E+3   18   18.5  RT (min)      lcorbi_t3_neg_4_028_39    m/z 520.9098     0   1   2E+4   12   12.5  RT (min)      lcorbi_t3_neg_2_019_07    m/z 520.9098     0   1   2E+4   9.5   10  RT (min)      lcorbi_t3_neg_3_051_46    m/z 520.9330     0   1   2   3E+3   18   18.5  RT (min)      lcorbi_t3_neg_2_034_19    m/z 520.9603     0   2   4   6E+3   5   5.5  RT (min)      lcorbi_t3_neg_1_054_37    m/z 520.9943     0   1   2E+4   6   6.5  RT (min)      lcorbi_t3_neg_1_054_37    m/z 521.0752     0   2   4   6E+2   7   7.5  RT (min)      lcorbi_t3_neg_4_044_38    m/z 521.1083     0   1   2E+4   5   5.5  RT (min)      lcorbi_t3_neg_1_015_27    m/z 521.1142     0   2   4   6E+3   11   11.5  RT (min)      lcorbi_t3_neg_4_007_13    m/z 521.1476     0   2   4   6E+3   10.5   11  RT (min)      lcorbi_t3_neg_4_042_14    m/z 521.2343     0   2   4   6E+3   7.5   8  RT (min)      lcorbi_t3_neg_4_023_18    m/z 521.2393     0   1E+4   11   11.5  RT (min)      lcorbi_t3_neg_4_034_01    m/z 521.2538     0   1E+4   12   12.5  RT (min)      lcorbi_t3_neg_4_038_15    m/z 521.2815     0   1E+4   12.5   13  RT (min)      lcorbi_t3_neg_2_052_32    m/z 521.3270     0   2   4   6E+3   13   13.5  RT (min)      lcorbi_t3_neg_2_051_34    m/z 521.3321     0   2   4   6   8E+3   11   11.5  RT (min)      lcorbi_t3_neg_3_018_13    m/z 521.3442     0   1E+4   11.5   12  RT (min)      lcorbi_t3_neg_3_007_16    m/z 521.3925     0   2   4   6E+3   14.5   15  RT (min)      lcorbi_t3_neg_4_005_27    m/z 521.4110     0   1   2   3   4E+3   13   13.5  RT (min)      lcorbi_t3_neg_4_034_01    m/z 521.7732     0   1   2   3E+3   0   0.5  RT (min)      lcorbi_t3_neg_4_051_40    m/z 521.9542     0   1E+4   5   5.5  RT (min)      lcorbi_t3_neg_1_054_37    m/z 522.0335     0   2   4   6E+3   9   9.5  RT (min)      lcorbi_t3_neg_3_019_12    m/z 522.0560     0   1   2   3   4E+4   6   6.5  RT (min)      lcorbi_t3_neg_4_044_38    m/z 522.0688     0   1E+4   7   7.5  RT (min)      lcorbi_t3_neg_3_024_26    m/z 522.1302     0   1E+4   8   8.5  RT (min)      lcorbi_t3_neg_2_048_27    m/z 522.1436     0   1E+5   2.5   3  RT (min)       lcorbi_t3_neg_2_048_27    m/z 522.2065     0   1E+4   9   9.5  RT (min)      lcorbi_t3_neg_2_048_27    m/z 522.2736     0   1E+5   10.5   11  RT (min)      lcorbi_t3_neg_1_055_05    m/z 522.3006     0   1E+4   11.5   12  RT (min)      lcorbi_t3_neg_4_034_01    m/z 522.3200     0   2   4   6   8E+2   11.5   12  RT (min)      lcorbi_t3_neg_1_055_05    m/z 522.3653     0   1E+4   14   14.5  RT (min)      lcorbi_t3_neg_3_018_13    m/z 522.4023     0   1   2   3E+4   12   12.5  RT (min)      lcorbi_t3_neg_4_007_13    m/z 522.4243     0   1   2   3   4E+4   13   13.5  RT (min)      lcorbi_t3_neg_4_042_14    m/z 522.8368     0   1   2   3   4E+3   18   18.5  RT (min)      lcorbi_t3_neg_4_045_35    m/z 522.9914     0   1   2E+4   6   6.5  RT (min)      lcorbi_t3_neg_1_054_37    m/z 523.0254     0   1E+3   9   9.5  RT (min)      lcorbi_t3_neg_1_055_05    m/z 523.0366     0   1   2   3E+3   9   9.5  RT (min)      lcorbi_t3_neg_1_055_05    m/z 523.1735     0   1E+4   7   7.5  RT (min)      lcorbi_t3_neg_2_048_27    m/z 523.2126     0   2   4   6E+3   6   6.5  RT (min)      lcorbi_t3_neg_2_034_19    m/z 523.2545     0   2   4   6   8E+3   7.5   8  RT (min)      lcorbi_t3_neg_4_017_51    m/z 523.2659     0   1E+4   10   10.5  RT (min)      lcorbi_t3_neg_3_019_12    m/z 523.3018     0   1   2E+3   13   13.5  RT (min)      lcorbi_t3_neg_1_055_05    m/z 523.3213     0   1E+3   12.5   13  RT (min)      lcorbi_t3_neg_3_007_16    m/z 523.4027     0   1   2   3E+3   13   13.5  RT (min)      lcorbi_t3_neg_4_034_01    m/z 523.4243     0   1   2   3E+3   13.5   14  RT (min)      lcorbi_t3_neg_4_034_01    m/z 523.4366     0   1   2   3E+4   13.5   14  RT (min)      lcorbi_t3_neg_4_034_01    m/z 523.8622     0   1E+4   1   1.5  RT (min)      lcorbi_t3_neg_4_054_36    m/z 523.9852     0   1   2   3E+4   6   6.5  RT (min)      lcorbi_t3_neg_1_054_37    m/z 524.0428     0   2   4   6E+3   7.5   8  RT (min)      lcorbi_t3_neg_4_001_49    m/z 524.0435     0   2   4E+3   11   11.5  RT (min)      lcorbi_t3_neg_3_034_21    m/z 524.0581     0   1   2   3   4E+3   5   5.5  RT (min)      lcorbi_t3_neg_4_040_19    m/z 524.0760     0   1   2   3   4E+3   7.5   8  RT (min)      lcorbi_t3_neg_2_002_26    m/z 524.0824     0   2   4   6   8E+3   6   6.5  RT (min)      lcorbi_t3_neg_4_040_19    m/z 524.1266     0   1E+4   6   6.5  RT (min)      lcorbi_t3_neg_1_054_37    m/z 524.2550     0   1E+4   11   11.5  RT (min)      lcorbi_t3_neg_4_007_13    m/z 524.2576     0   1   2   3E+3   7.5   8  RT (min)      lcorbi_t3_neg_4_017_51    m/z 524.2595     0   1E+4   10   10.5  RT (min)      lcorbi_t3_neg_3_019_12    m/z 524.2894     0   1   11   11.5  RT (min)      lcorbi_t3_neg_4_034_01    m/z 524.3161     0   1E+3   12   12.5  RT (min)      lcorbi_t3_neg_1_055_05    m/z 524.3799     0   1E+4   13.5   14  RT (min)      lcorbi_t3_neg_4_034_01    m/z 524.4077     0   1E+4   14   14.5  RT (min)      lcorbi_t3_neg_4_023_18    m/z 524.4176     0   1E+4   12   12.5  RT (min)      lcorbi_t3_neg_3_018_13    m/z 524.8488     0   2   4   6E+3   18   18.5  RT (min)      lcorbi_t3_neg_4_053_43    m/z 524.9884     0   1   2   3E+4   11   11.5  RT (min)      lcorbi_t3_neg_4_007_13    m/z 525.2233     0   2   4   6E+3   11   11.5  RT (min)      lcorbi_t3_neg_4_042_14    m/z 525.2632     0   1   2   3E+3   10   10.5  RT (min)      lcorbi_t3_neg_3_019_12    m/z 525.2725     0   2   4   6   8E+3   13   13.5  RT (min)      lcorbi_t3_neg_2_007_14    m/z 525.2985     0   1E+4   13   13.5  RT (min)      lcorbi_t3_neg_3_005_23    m/z 525.3327     0   1E+4   12.5   13  RT (min)      lcorbi_t3_neg_2_053_23    m/z 525.3531     0   2   4   6   8E+3   13   13.5  RT (min)      lcorbi_t3_neg_1_055_05    m/z 525.3926     0   1E+4   13.5   14  RT (min)      lcorbi_t3_neg_4_034_01    m/z 525.4157     0   2   4   6   8E+3   13   13.5  RT (min)      lcorbi_t3_neg_4_007_13    m/z 525.9033     0   1E+4   1   1.5  RT (min)      lcorbi_t3_neg_2_007_14    m/z 525.9851     0   1E+4   11   11.5  RT (min)      lcorbi_t3_neg_4_007_13    m/z 526.0280     0   1E+4   9   9.5  RT (min)      lcorbi_t3_neg_4_007_13    m/z 526.0404     0   1   2   3E+3   11   11.5  RT (min)      lcorbi_t3_neg_3_034_21    m/z 526.0408     0   2   4E+3   7.5   8  RT (min)      lcorbi_t3_neg_4_001_49    m/z 526.2628     0   2   4   6E+3   12.5   13  RT (min)      lcorbi_t3_neg_4_007_13    m/z 526.2734     0   2   4   6E+3   7   7.5  RT (min)      lcorbi_t3_neg_4_034_01    m/z 526.2743     0   1E+4   10   10.5  RT (min)      lcorbi_t3_neg_3_019_12    m/z 526.3219     0   2   4   6E+3   13.5   14  RT (min)      lcorbi_t3_neg_4_040_19    m/z 526.3413     0   1   2E+3   11.5   12  RT (min)      lcorbi_t3_neg_4_007_13    m/z 526.3742     0   1   2   3   4E+4   10.5   11  RT (min)      lcorbi_t3_neg_1_055_05    m/z 526.3957     0   2   4E+3   13.5   14  RT (min)      lcorbi_t3_neg_4_005_27    m/z 526.4071     0   1E+4   14   14.5  RT (min)      lcorbi_t3_neg_4_040_19    m/z 526.8488     0   2   4   6E+3   18   18.5  RT (min)      lcorbi_t3_neg_4_024_07    m/z 526.8550     0   2   4   6E+3   18   18.5  RT (min)      lcorbi_t3_neg_2_031_25    m/z 527.0267     0   1   2   3   4E+3   1.5   2  RT (min)      lcorbi_t3_neg_2_007_14    m/z 527.0357     0   2   4   6E+3   9   9.5  RT (min)      lcorbi_t3_neg_3_019_12    m/z 527.0684     0   1E+4   10   10.5  RT (min)      lcorbi_t3_neg_4_029_31    m/z 527.0802     0   1E+4   7   7.5  RT (min)      lcorbi_t3_neg_4_005_27    m/z 527.0986     0   1   2   3   4E+3   10.5   11  RT (min)      lcorbi_t3_neg_1_002_16    m/z 527.1096     0   2   4   6   8E+3   11   11.5  RT (min)      lcorbi_t3_neg_4_007_13    m/z 527.1822     0   1   2   3   4E+3   13   13.5  RT (min)      lcorbi_t3_neg_4_020_44    m/z 527.2129     0   1E+4   7.5   8  RT (min)      lcorbi_t3_neg_2_048_27    m/z 527.2190     0   1   2   3   4E+4   9   9.5  RT (min)      lcorbi_t3_neg_2_048_27    m/z 527.2599     0   1E+4   10   10.5  RT (min)      lcorbi_t3_neg_2_048_27    m/z 527.2779     0   1E+3   10   10.5  RT (min)      lcorbi_t3_neg_4_005_27    m/z 527.2838     0   2   4   6   8E+3   12.5   13  RT (min)      lcorbi_t3_neg_4_007_13    m/z 527.2870     0   1E+4   11   11.5  RT (min)      lcorbi_t3_neg_3_007_16    m/z 527.3118     0   1E+3   13   13.5  RT (min)      lcorbi_t3_neg_4_017_51    m/z 527.3373     0   2   4   6   8E+3   13   13.5  RT (min)      lcorbi_t3_neg_4_017_51    m/z 527.3688     0   1E+4   13   13.5  RT (min)      lcorbi_t3_neg_2_022_24    m/z 527.3949     0   1   2   3   4E+3   11.5   12  RT (min)      lcorbi_t3_neg_4_052_26    m/z 527.3952     0   1E+3   13   13.5  RT (min)      lcorbi_t3_neg_4_039_32    m/z 527.4314     0   1   13   13.5  RT (min)      lcorbi_t3_neg_2_022_24    m/z 527.8713     0   2   4   6E+3   18   18.5  RT (min)      lcorbi_t3_neg_2_033_53    m/z 527.9028     0   2   4   6E+3   1   1.5  RT (min)      lcorbi_t3_neg_3_029_02    m/z 528.0943     0   2   4   6   8E+3   6   6.5  RT (min)      lcorbi_t3_neg_4_040_19    m/z 528.2164     0   1   2   3   4E+3   7.5   8  RT (min)      lcorbi_t3_neg_4_034_01    m/z 528.2404     0   2   4E+3   8   8.5  RT (min)      lcorbi_t3_neg_2_035_46    m/z 528.2657     0   1E+4   10   10.5  RT (min)      lcorbi_t3_neg_2_048_27    m/z 528.3030     0   1E+4   13   13.5  RT (min)      lcorbi_t3_neg_4_047_30    m/z 528.3207     0   1E+4   11   11.5  RT (min)      lcorbi_t3_neg_4_034_01    m/z 528.3215     0   2   4   6E+3   13.5   14  RT (min)      lcorbi_t3_neg_4_007_13    m/z 528.3360     0   1E+3   13   13.5  RT (min)      lcorbi_t3_neg_1_055_05    m/z 528.3551     0   1   2E+3   14   14.5  RT (min)      lcorbi_t3_neg_4_032_23    m/z 528.3901     0   2   4   6   8E+4   11   11.5  RT (min)      lcorbi_t3_neg_1_055_05    m/z 528.4134     0   1E+4   13.5   14  RT (min)      lcorbi_t3_neg_4_042_14    m/z 528.7031     0   1   1   1.5  RT (min)      lcorbi_t3_neg_2_048_27    m/z 528.8520     0   1   2   3E+3   18   18.5  RT (min)      lcorbi_t3_neg_4_024_07    m/z 528.8883     0   2   4E+3   8.5   9  RT (min)      lcorbi_t3_neg_4_053_43    m/z 529.0134     0   2   4E+3   5.5   6  RT (min)      lcorbi_t3_neg_4_023_18    m/z 529.0289     0   1   2   3E+3   7.5   8  RT (min)      lcorbi_t3_neg_4_034_01    m/z 529.0362     0   2   4E+3   10.5   11  RT (min)      lcorbi_t3_neg_2_011_04    m/z 529.0751     0   1E+4   9.5   10  RT (min)      lcorbi_t3_neg_4_043_11    m/z 529.1567     0   2   4E+3   8   8.5  RT (min)      lcorbi_t3_neg_4_040_19    m/z 529.1581     0   2   4   6E+4   1   1.5  RT (min)      lcorbi_t3_neg_2_048_27    m/z 529.2051     0   1E+4   10   10.5  RT (min)      lcorbi_t3_neg_4_034_01    m/z 529.2223     0   1E+4   8.5   9  RT (min)      lcorbi_t3_neg_4_017_51    m/z 529.2354     0   1E+4   13   13.5  RT (min)      lcorbi_t3_neg_4_007_13    m/z 529.2666     0   1E+4   8   8.5  RT (min)      lcorbi_t3_neg_4_034_01    m/z 529.2767     0   1E+4   11   11.5  RT (min)      lcorbi_t3_neg_4_042_14    m/z 529.2895     0   2   4   6E+3   10.5   11  RT (min)      lcorbi_t3_neg_4_007_13    m/z 529.2995     0   1   2E+4   13   13.5  RT (min)      lcorbi_t3_neg_4_047_30    m/z 529.3154     0   2   4   6   8E+2   14   14.5  RT (min)      lcorbi_t3_neg_2_048_27    m/z 529.3311     0   1E+3   12.5   13  RT (min)      lcorbi_t3_neg_4_007_13    m/z 529.3597     0   2   4E+3   11.5   12  RT (min)      lcorbi_t3_neg_4_034_01    m/z 529.3744     0   2   4   6   8E+2   13.5   14  RT (min)      lcorbi_t3_neg_4_005_27    m/z 530.0243     0   1   2   3E+4   11   11.5  RT (min)      lcorbi_t3_neg_4_007_13    m/z 530.0783     0   1   2E+3   9.5   10  RT (min)      lcorbi_t3_neg_4_053_43    m/z 530.1007     0   2   4E+3   7   7.5  RT (min)      lcorbi_t3_neg_3_045_34    m/z 530.1031     0   1   2E+4   11   11.5  RT (min)      lcorbi_t3_neg_4_007_13    m/z 530.1496     0   2   4   6E+3   11   11.5  RT (min)      lcorbi_t3_neg_4_043_11    m/z 530.2085     0   1   2   3E+3   10   10.5  RT (min)      lcorbi_t3_neg_4_034_01    m/z 530.2704     0   1   2E+4   14   14.5  RT (min)      lcorbi_t3_neg_4_007_13    m/z 530.2825     0   1   2   3E+3   14   14.5  RT (min)      lcorbi_t3_neg_4_007_13    m/z 530.2833     0   2   4E+3   10.5   11  RT (min)      lcorbi_t3_neg_4_016_05    m/z 530.2973     0   2   4   6   8E+3   12.5   13  RT (min)      lcorbi_t3_neg_4_047_30    m/z 530.3028     0   2   4E+3   13   13.5  RT (min)      lcorbi_t3_neg_4_047_30    m/z 530.3155     0   1E+4   11.5   12  RT (min)      lcorbi_t3_neg_4_007_13    m/z 530.3727     0   2   4   6   8E+3   14   14.5  RT (min)      lcorbi_t3_neg_4_034_01    m/z 530.4055     0   1   2   3   4E+4   11.5   12  RT (min)      lcorbi_t3_neg_1_055_05    m/z 530.4294     0   2   4E+4   13.5   14  RT (min)      lcorbi_t3_neg_4_034_01    m/z 530.8655     0   2   4   6E+3   9.5   10  RT (min)      lcorbi_t3_neg_1_034_04    m/z 530.8855     0   2   4E+2   8.5   9  RT (min)      lcorbi_t3_neg_4_022_41    m/z 531.0232     0   1E+4   6   6.5  RT (min)      lcorbi_t3_neg_1_054_37    m/z 531.0756     0   2   4E+3   9.5   10  RT (min)      lcorbi_t3_neg_4_029_31    m/z 531.1000     0   2   4   6   8E+3   11   11.5  RT (min)      lcorbi_t3_neg_4_007_13    m/z 531.1194     0   1   2E+4   11.5   12  RT (min)      lcorbi_t3_neg_4_007_13    m/z 531.1866     0   1   2   3   4E+4   7.5   8  RT (min)      lcorbi_t3_neg_3_019_12    m/z 531.1949     0   1   2   3   4E+3   13   13.5  RT (min)      lcorbi_t3_neg_4_034_01    m/z 531.2052     0   2   4   6   8E+3   8.5   9  RT (min)      lcorbi_t3_neg_4_017_51    m/z 531.2105     0   2   4   6E+3   10   10.5  RT (min)      lcorbi_t3_neg_3_019_12    m/z 531.2510     0   1   2E+4   13   13.5   14  RT (min)      lcorbi_t3_neg_4_007_13    m/z 531.2517     0   1   2   3E+3   9   9.5  RT (min)      lcorbi_t3_neg_4_001_49    m/z 531.2786     0   1E+4   12   12.5  RT (min)      lcorbi_t3_neg_4_035_33    m/z 531.2849     0   2   4   6E+3   10   10.5  RT (min)      lcorbi_t3_neg_3_019_12    m/z 531.2939     0   2   4   6E+3   12.5   13  RT (min)      lcorbi_t3_neg_4_047_30    m/z 531.3287     0   1   2   3   4E+3   11.5   12  RT (min)      lcorbi_t3_neg_4_044_38    m/z 531.3448     0   1E+5   9.5   10  RT (min)      lcorbi_t3_neg_3_002_24    m/z 531.3533     0   2   4   6E+4   10.5   11  RT (min)      lcorbi_t3_neg_1_055_05    m/z 531.3856     0   1E+4   13.5   14  RT (min)      lcorbi_t3_neg_4_029_31    m/z 531.8400     0   1   2   3E+3   18   18.5  RT (min)      lcorbi_t3_neg_4_003_21    m/z 532.0640     0   2   4   6   8E+3   8.5   9  RT (min)      lcorbi_t3_neg_1_049_55    m/z 532.0651     0   1E+4   10.5   11  RT (min)      lcorbi_t3_neg_1_051_15    m/z 532.0895     0   2   4E+3   6   6.5  RT (min)      lcorbi_t3_neg_4_006_47    m/z 532.1205     0   1   2E+4   11.5   12  RT (min)      lcorbi_t3_neg_4_007_13    m/z 532.1820     0   1   2E+4   7   7.5  RT (min)      lcorbi_t3_neg_4_040_19    m/z 532.1900     0   1E+4   7.5   8  RT (min)      lcorbi_t3_neg_3_019_12    m/z 532.2015     0   2   4   6E+2   7   7.5  RT (min)      lcorbi_t3_neg_4_040_19    m/z 532.2212     0   1E+4   10.5   11  RT (min)      lcorbi_t3_neg_2_020_05    m/z 532.2347     0   2   4   6E+3   7.5   8  RT (min)      lcorbi_t3_neg_4_034_01    m/z 532.2889     0   1   2E+5   8.5   9  RT (min)      lcorbi_t3_neg_4_023_18    m/z 532.3151     0   1   12.5   13  RT (min)      lcorbi_t3_neg_4_042_14    m/z 532.3405     0   1E+3   11.5   12  RT (min)      lcorbi_t3_neg_4_016_05    m/z 532.4121     0   2   4   6   8E+3   12.5   13  RT (min)      lcorbi_t3_neg_4_011_24    m/z 532.8425     0   1   2   3E+3   18   18.5  RT (min)      lcorbi_t3_neg_4_053_43    m/z 532.8658     0   1   2E+4   18   18.5  RT (min)      lcorbi_t3_neg_2_048_27    m/z 532.9203     0   1   1   1.5  RT (min)      lcorbi_t3_neg_3_022_27    m/z 532.9818     0   1   2   3   4E+3   8   8.5  RT (min)      lcorbi_t3_neg_4_023_18    m/z 533.0202     0   1E+5   11   11.5  RT (min)      lcorbi_t3_neg_4_007_13    m/z 533.0202     0   2   4   6E+3   6   6.5  RT (min)      lcorbi_t3_neg_1_054_37    m/z 533.0239     0   2   4E+3   8.5   9  RT (min)      lcorbi_t3_neg_4_034_01    m/z 533.0544     0   1E+4   9   9.5  RT (min)      lcorbi_t3_neg_2_048_27    m/z 533.1177     0   2   4E+4   11.5   12  RT (min)      lcorbi_t3_neg_4_007_13    m/z 533.1332     0   2   4   6   8E+3   11   11.5  RT (min)      lcorbi_t3_neg_4_043_11    m/z 533.1800     0   1E+4   8   8.5  RT (min)      lcorbi_t3_neg_4_034_01    m/z 533.1955     0   1   9.5   10  RT (min)      lcorbi_t3_neg_4_032_23    m/z 533.2365     0   1   2   3   4E+3   8.5   9  RT (min)      lcorbi_t3_neg_4_034_01    m/z 533.2924     0   2   4E+4   8.5   9  RT (min)      lcorbi_t3_neg_4_023_18    m/z 533.3016     0   2   4   6   8E+3   13.5   14  RT (min)      lcorbi_t3_neg_4_040_19    m/z 533.3204     0   1   2   3   4E+2   12.5   13  RT (min)      lcorbi_t3_neg_4_036_16    m/z 533.3606     0   1   2E+4   11   11.5  RT (min)      lcorbi_t3_neg_1_055_05    m/z 533.3690     0   1E+5   11   11.5  RT (min)      lcorbi_t3_neg_1_055_05    m/z 533.3841     0   2   4   6   8E+3   13   13.5  RT (min)      lcorbi_t3_neg_4_042_14    m/z 533.4242     0   1E+3   12   12.5  RT (min)      lcorbi_t3_neg_3_032_17    m/z 533.4575     0   1   2E+4   14   14.5  RT (min)      lcorbi_t3_neg_4_034_01    m/z 533.9710     0   2   4   6E+3   9   9.5  RT (min)      lcorbi_t3_neg_4_007_13    m/z 534.0049     0   1   2   3E+3   1.5   2  RT (min)      lcorbi_t3_neg_2_048_27    m/z 534.0171     0   2   4   6   8E+4   11   11.5  RT (min)      lcorbi_t3_neg_4_007_13    m/z 534.0610     0   2   4   6E+3   8.5   9  RT (min)      lcorbi_t3_neg_1_049_55    m/z 534.0707     0   1   2E+3   9   9.5  RT (min)      lcorbi_t3_neg_4_028_39    m/z 534.1179     0   1   2E+4   11.5   12  RT (min)      lcorbi_t3_neg_4_007_13    m/z 534.1536     0   1E+4   13   13.5  RT (min)      lcorbi_t3_neg_4_007_13    m/z 534.1555     0   1E+4   6   6.5  RT (min)      lcorbi_t3_neg_3_039_29    m/z 534.2037     0   2   4   6E+3   7   7.5  RT (min)      lcorbi_t3_neg_2_048_27    m/z 534.2043     0   1   2   3E+3   10   10.5  RT (min)      lcorbi_t3_neg_4_032_23    m/z 534.2503     0   1E+4   8.5   9  RT (min)      lcorbi_t3_neg_4_034_01    m/z 534.2539     0   1E+4   12   12.5  RT (min)      lcorbi_t3_neg_4_038_15    m/z 534.2810     0   2   4   6E+3   11.5   12  RT (min)      lcorbi_t3_neg_1_055_05    m/z 534.2976     0   2   4   6E+3   10   10.5  RT (min)      lcorbi_t3_neg_3_019_12    m/z 534.3164     0   1   2E+3   12.5   13  RT (min)      lcorbi_t3_neg_4_032_23    m/z 534.3561     0   1   2E+3   12   12.5  RT (min)      lcorbi_t3_neg_4_011_24    m/z 534.3729     0   1   2   3E+4   10.5   11  RT (min)      lcorbi_t3_neg_1_055_05    m/z 534.4610     0   1E+4   14   14.5  RT (min)      lcorbi_t3_neg_4_042_14    m/z 534.8037     0   1   2E+3   0   0.5  RT (min)      lcorbi_t3_neg_4_051_40    m/z 534.8612     0   2   4   6   8E+3   18   18.5  RT (min)      lcorbi_t3_neg_2_048_27    m/z 535.1141     0   1   2E+4   11.5   12  RT (min)      lcorbi_t3_neg_4_007_13    m/z 535.1146     0   1   2   3   4E+3   6   6.5  RT (min)      lcorbi_t3_neg_4_040_19    m/z 535.2302     0   1   2E+4   9   9.5  RT (min)      lcorbi_t3_neg_2_048_27    m/z 535.2402     0   2   4   6E+3   11   11.5  RT (min)      lcorbi_t3_neg_2_020_05    m/z 535.2504     0   1   2   3E+4   12   12.5  RT (min)      lcorbi_t3_neg_4_038_15    m/z 535.2872     0   1E+4   12.5   13  RT (min)      lcorbi_t3_neg_3_007_16    m/z 535.2901     0   1   2E+4   5.5   6  RT (min)      lcorbi_t3_neg_2_012_36    m/z 535.3168     0   2   4   6E+3   14   14.5  RT (min)      lcorbi_t3_neg_2_037_09    m/z 535.3426     0   2   4E+2   12.5   13  RT (min)      lcorbi_t3_neg_4_005_27    m/z 535.3792     0   2   4   6   8E+3   13   13.5  RT (min)      lcorbi_t3_neg_1_055_05    m/z 535.4002     0   1   2E+4   12.5   13  RT (min)      lcorbi_t3_neg_4_034_01    m/z 535.4365     0   1E+4   13   13.5  RT (min)      lcorbi_t3_neg_4_034_01    m/z 536.0396     0   2   4   6E+3   7   7.5  RT (min)      lcorbi_t3_neg_2_002_26    m/z 536.0659     0   1   2E+4   1   1.5  RT (min)      lcorbi_t3_neg_2_035_46    m/z 536.1693     0   2   4   6   8E+3   13   13.5  RT (min)      lcorbi_t3_neg_4_005_27    m/z 536.2338     0   2   4   6   8E+3   11   11.5  RT (min)      lcorbi_t3_neg_2_020_05    m/z 536.2942     0   1E+3   11.5   12  RT (min)      lcorbi_t3_neg_1_055_05    m/z 536.2990     0   1E+5   10.5   11  RT (min)      lcorbi_t3_neg_1_055_05    m/z 536.3573     0   2   4   6   8E+3   14   14.5  RT (min)      lcorbi_t3_neg_2_048_27    m/z 536.5048     0   1   2   3E+4   14   14.5  RT (min)      lcorbi_t3_neg_3_005_23    m/z 536.9936     0   2   4   6E+3   8   8.5  RT (min)      lcorbi_t3_neg_4_023_18    m/z 537.1718     0   1   2   3E+3   13   13.5  RT (min)      lcorbi_t3_neg_4_038_15    m/z 537.1821     0   2   4E+3   12   12.5  RT (min)      lcorbi_t3_neg_4_042_14    m/z 537.2505     0   2   4E+3   9   9.5  RT (min)      lcorbi_t3_neg_2_048_27    m/z 537.2556     0   2   4   6E+3   11.5   12  RT (min)      lcorbi_t3_neg_1_055_05    m/z 537.2716     0   1   2   3   4E+3   11   11.5  RT (min)      lcorbi_t3_neg_4_032_23    m/z 537.2963     0   1E+3   13   13.5  RT (min)      lcorbi_t3_neg_4_023_18    m/z 537.3026     0   1   2   3E+4   10.5   11  RT (min)      lcorbi_t3_neg_1_055_05    m/z 537.3278     0   1   2E+3   11   11.5  RT (min)      lcorbi_t3_neg_1_044_23    m/z 537.3350     0   1   2   3E+4   13   13.5  RT (min)      lcorbi_t3_neg_4_023_18    m/z 537.3522     0   1   11.5   12  RT (min)      lcorbi_t3_neg_2_022_24    m/z 537.3714     0   1   13.5   14  RT (min)      lcorbi_t3_neg_1_055_05    m/z 537.3974     0   1   2   3   4E+4   13   13.5  RT (min)      lcorbi_t3_neg_4_011_24    m/z 537.4521     0   1E+4   13.5   14  RT (min)      lcorbi_t3_neg_4_034_01    m/z 537.8873     0   1E+4   18   18.5  RT (min)      lcorbi_t3_neg_4_019_08    m/z 537.9119     0   1E+4   7.5   8  RT (min)      lcorbi_t3_neg_4_023_18    m/z 538.0365     0   2   4E+3   7   7.5  RT (min)      lcorbi_t3_neg_2_002_26    m/z 538.1005     0   2   4E+3   10   10.5  RT (min)      lcorbi_t3_neg_4_029_31    m/z 538.1480     0   1   2   3   4E+3   13   13.5  RT (min)      lcorbi_t3_neg_1_043_14    m/z 538.1764     0   1E+4   9   9.5  RT (min)      lcorbi_t3_neg_4_023_18    m/z 538.1849     0   1E+4   13.5   14  RT (min)      lcorbi_t3_neg_4_040_19    m/z 538.1928     0   1   2   3E+4   7.5   8  RT (min)      lcorbi_t3_neg_2_048_27    m/z 538.2346     0   1   2E+4   11   11.5  RT (min)      lcorbi_t3_neg_1_055_05    m/z 538.2485     0   1E+3   7.5   8  RT (min)      lcorbi_t3_neg_2_048_27    m/z 538.2493     0   2   4   6E+3   11.5   12  RT (min)      lcorbi_t3_neg_1_055_05    m/z 538.2904     0   1   2   3   4E+3   10.5   11  RT (min)      lcorbi_t3_neg_3_007_16    m/z 538.3217     0   2   4   6E+3   13   13.5  RT (min)      lcorbi_t3_neg_3_024_26    m/z 538.3392     0   1   2   3   4E+3   10.5   11  RT (min)      lcorbi_t3_neg_2_027_13    m/z 538.8590     0   1E+3   18   18.5  RT (min)      lcorbi_t3_neg_1_003_41    m/z 538.8822     0   2   4E+3   18   18.5  RT (min)      lcorbi_t3_neg_4_048_03    m/z 538.9891     0   1   2   3E+3   9   9.5  RT (min)      lcorbi_t3_neg_1_003_41    m/z 539.0527     0   1E+4   8   8.5  RT (min)      lcorbi_t3_neg_1_054_37    m/z 539.0990     0   1   2E+3   5   5.5  RT (min)      lcorbi_t3_neg_1_014_28    m/z 539.1863     0   1E+3   7.5   8  RT (min)      lcorbi_t3_neg_4_023_18    m/z 539.1874     0   1E+3   13.5   14  RT (min)      lcorbi_t3_neg_1_007_26    m/z 539.2713     0   1   2   3E+3   12   12.5  RT (min)      lcorbi_t3_neg_4_007_13    m/z 539.3328     0   2   4E+3   13   13.5  RT (min)      lcorbi_t3_neg_4_042_14    m/z 539.3549     0   1   2   3   4E+3   13.5   14  RT (min)      lcorbi_t3_neg_2_035_46    m/z 539.3610     0   1E+4   11.5   12  RT (min)      lcorbi_t3_neg_4_029_31    m/z 539.3871     0   1   2   3E+3   13.5   14  RT (min)      lcorbi_t3_neg_2_007_14    m/z 539.3947     0   1E+4   13   13.5  RT (min)      lcorbi_t3_neg_1_055_05    m/z 539.4216     0   1   2E+3   12.5   13  RT (min)      lcorbi_t3_neg_4_034_01    m/z 539.5137     0   1   2   3E+4   14   14.5  RT (min)      lcorbi_t3_neg_4_011_24    m/z 539.9095     0   1E+4   7.5   8  RT (min)      lcorbi_t3_neg_4_023_18    m/z 540.0487     0   1E+4   6   6.5  RT (min)      lcorbi_t3_neg_1_054_37    m/z 540.0685     0   1   2   3   4E+3   10.5   11  RT (min)      lcorbi_t3_neg_2_011_04    m/z 540.1051     0   2   4   6   8E+3   12   12.5  RT (min)      lcorbi_t3_neg_4_007_13    m/z 540.1294     0   2   4   6E+3   7   7.5  RT (min)      lcorbi_t3_neg_1_042_18    m/z 540.1315     0   2   4E+3   5   5.5  RT (min)      lcorbi_t3_neg_4_013_09    m/z 540.1637     0   2   4E+3   13   13.5  RT (min)      lcorbi_t3_neg_4_007_13    m/z 540.2339     0   1E+4   10   10.5  RT (min)      lcorbi_t3_neg_1_055_05    m/z 540.2527     0   1E+4   7   7.5  RT (min)      lcorbi_t3_neg_4_034_01    m/z 540.2633     0   1   2   3   4E+3   9   9.5  RT (min)      lcorbi_t3_neg_4_034_01    m/z 540.2650     0   1   2   3   4E+3   12   12.5  RT (min)      lcorbi_t3_neg_4_007_13    m/z 540.3355     0   1   2   3E+3   13   13.5  RT (min)      lcorbi_t3_neg_4_029_31    m/z 541.0621     0   2   4   6E+3   10.5   11  RT (min)      lcorbi_t3_neg_2_011_04    m/z 541.0996     0   1   2   3E+4   11.5   12  RT (min)      lcorbi_t3_neg_4_007_13    m/z 541.1560     0   2   4   6E+3   6   6.5  RT (min)      lcorbi_t3_neg_2_048_27    m/z 541.1675     0   2   4   6E+3   11   11.5  RT (min)      lcorbi_t3_neg_4_032_23    m/z 541.1977     0   1E+4   8   8.5  RT (min)      lcorbi_t3_neg_1_026_19    m/z 541.2391     0   1   2E+4   10   10.5  RT (min)      lcorbi_t3_neg_3_019_12    m/z 541.2693     0   1E+3   12   12.5  RT (min)      lcorbi_t3_neg_2_020_05    m/z 541.3052     0   2   4   6E+3   13.5   14  RT (min)      lcorbi_t3_neg_4_034_01    m/z 541.3233     0   2   4   6E+3   10.5   11  RT (min)      lcorbi_t3_neg_4_007_13    m/z 541.3339     0   1E+4   18   18.5  RT (min)      lcorbi_t3_neg_1_007_26    m/z 541.3353     0   2   4   6   8E+3   13   13.5  RT (min)      lcorbi_t3_neg_4_007_13    m/z 541.3872     0   1E+4   13   13.5  RT (min)      lcorbi_t3_neg_4_005_27    m/z 541.4376     0   2   4   6E+3   12.5   13  RT (min)      lcorbi_t3_neg_4_042_14    m/z 541.8756     0   2   4E+3   18   18.5  RT (min)      lcorbi_t3_neg_4_003_21    m/z 542.0436     0   1E+4   8   8.5  RT (min)      lcorbi_t3_neg_1_054_37    m/z 542.0533     0   1E+4   7   7.5  RT (min)      lcorbi_t3_neg_4_023_18    m/z 542.0601     0   1E+4   3   3.5  RT (min)      lcorbi_t3_neg_4_007_13    m/z 542.0927     0   2   4   6E+3   8.5   9  RT (min)      lcorbi_t3_neg_1_049_55    m/z 542.1001     0   1E+4   11.5   12  RT (min)      lcorbi_t3_neg_4_007_13    m/z 542.1230     0   1E+4   12.5   13  RT (min)      lcorbi_t3_neg_4_007_13    m/z 542.1793     0   1   2   3E+3   13.5   14  RT (min)      lcorbi_t3_neg_4_040_19    m/z 542.2094     0   2   4   6E+3   9   9.5  RT (min)      lcorbi_t3_neg_4_001_49    m/z 542.2684     0   1E+4   6.5   7  RT (min)      lcorbi_t3_neg_2_048_27    m/z 542.2838     0   2   4   6E+3   9   9.5  RT (min)      lcorbi_t3_neg_4_001_49    m/z 542.3418     0   1   2   3   4E+4   10.5   11  RT (min)      lcorbi_t3_neg_1_055_05    m/z 542.8405     0   1   2E+3   18   18.5  RT (min)      lcorbi_t3_neg_4_029_31    m/z 543.0770     0   2   4   6   8E+3   10.5   11  RT (min)      lcorbi_t3_neg_4_030_04    m/z 543.0969     0   1E+4   11.5   12  RT (min)      lcorbi_t3_neg_4_007_13    m/z 543.1283     0   1   2   3   4E+3   9.5   10  RT (min)      lcorbi_t3_neg_4_007_13    m/z 543.1619     0   1   2E+4   11   11.5  RT (min)      lcorbi_t3_neg_4_043_11    m/z 543.1772     0   1   2E+3   12   12.5  RT (min)      lcorbi_t3_neg_4_042_14    m/z 543.2529     0   2   4   6E+2   10.5   11  RT (min)      lcorbi_t3_neg_3_002_24    m/z 543.2549     0   1   2   3E+4   9.5   10  RT (min)      lcorbi_t3_neg_4_023_18    m/z 543.2759     0   2   4E+3   12.5   13  RT (min)      lcorbi_t3_neg_2_032_45    m/z 543.2807     0   2   4E+3   6.5   7  RT (min)      lcorbi_t3_neg_2_048_27    m/z 543.2920     0   1E+4   11   11.5  RT (min)      lcorbi_t3_neg_1_055_05    m/z 543.2941     0   2   4   6   8E+3   14   14.5  RT (min)      lcorbi_t3_neg_4_043_11    m/z 543.3128     0   2   4   6   8E+3   13.5   14  RT (min)      lcorbi_t3_neg_4_007_13    m/z 543.3281     0   1E+4   13   13.5  RT (min)      lcorbi_t3_neg_1_055_05    m/z 543.3444     0   1E+4   12   12.5  RT (min)      lcorbi_t3_neg_4_007_13    m/z 543.3622     0   1   2E+4   11   11.5  RT (min)      lcorbi_t3_neg_1_055_05    m/z 543.3866     0   1   2   3E+4   14   14.5  RT (min)      lcorbi_t3_neg_4_048_03    m/z 543.4035     0   2   4   6   8E+3   13   13.5  RT (min)      lcorbi_t3_neg_2_007_14    m/z 543.4415     0   1   2   3   4E+3   14   14.5  RT (min)      lcorbi_t3_neg_2_048_27    m/z 543.4427     0   1   2E+4   12.5   13  RT (min)      lcorbi_t3_neg_4_039_32    m/z 543.4499     0   1E+5   12   12.5  RT (min)      lcorbi_t3_neg_1_055_05    m/z 543.7276     0   1E+4   1   1.5  RT (min)      lcorbi_t3_neg_4_036_16    m/z 543.8738     0   1E+4   18   18.5  RT (min)      lcorbi_t3_neg_3_056_56    m/z 544.0140     0   1   2   3   4E+3   8   8.5  RT (min)      lcorbi_t3_neg_4_023_18    m/z 544.0385     0   1   2E+4   11.5   12  RT (min)      lcorbi_t3_neg_4_007_13    m/z 544.0513     0   1E+4   7   7.5  RT (min)      lcorbi_t3_neg_4_023_18    m/z 544.1008     0   1E+4   11   11.5  RT (min)      lcorbi_t3_neg_4_007_13    m/z 544.1185     0   1E+4   11   11.5  RT (min)      lcorbi_t3_neg_3_032_17    m/z 544.1201     0   1E+4   12.5   13  RT (min)      lcorbi_t3_neg_4_007_13    m/z 544.1387     0   2   4   6E+3   13   13.5  RT (min)      lcorbi_t3_neg_4_047_30    m/z 544.2411     0   1E+4   11.5   12  RT (min)      lcorbi_t3_neg_4_007_13    m/z 544.2472     0   1   10   10.5  RT (min)      lcorbi_t3_neg_4_019_08    m/z 544.2494     0   2   4   6   8E+3   13   13.5  RT (min)      lcorbi_t3_neg_4_042_14    m/z 544.2601     0   1E+4   12   12.5  RT (min)      lcorbi_t3_neg_4_007_13    m/z 544.3053     0   1   11.5   12  RT (min)      lcorbi_t3_neg_4_083_59    m/z 544.3351     0   1   12   12.5  RT (min)      lcorbi_t3_neg_4_019_08    m/z 544.3536     0   1   2   3E+3   12   13  RT (min)       lcorbi_t3_neg_4_090_60    m/z 544.3879     0   2   4E+3   14   14.5  RT (min)      lcorbi_t3_neg_3_029_02    m/z 544.9083     0   1E+4   18   18.5  RT (min)      lcorbi_t3_neg_1_003_41    m/z 545.0372     0   1   2E+4   11   11.5  RT (min)      lcorbi_t3_neg_4_007_13    m/z 545.0667     0   1   2   3   4E+3   9.5   10  RT (min)      lcorbi_t3_neg_2_006_10    m/z 545.0974     0   1   2   3   4E+4   11   11.5  RT (min)      lcorbi_t3_neg_4_007_13    m/z 545.1157     0   1E+4   11   11.5  RT (min)      lcorbi_t3_neg_3_007_16    m/z 545.1201     0   1E+4   8   8.5  RT (min)      lcorbi_t3_neg_1_026_19    m/z 545.2026     0   1E+4   8   8.5  RT (min)      lcorbi_t3_neg_4_001_49    m/z 545.2886     0   1E+3   11   11.5  RT (min)      lcorbi_t3_neg_4_016_05    m/z 545.3209     0   1   2   3   4E+2   11.5   12  RT (min)      lcorbi_t3_neg_1_055_05    m/z 545.3777     0   1   2E+4   11.5   12  RT (min)      lcorbi_t3_neg_1_055_05    m/z 545.3822     0   1   14   14.5  RT (min)      lcorbi_t3_neg_4_023_18    m/z 545.4208     0   1   2   3E+4   13   13.5  RT (min)      lcorbi_t3_neg_4_034_01    m/z 546.0337     0   1   2   3   4E+4   11   11.5  RT (min)      lcorbi_t3_neg_4_007_13    m/z 546.0980     0   1E+4   11   11.5  RT (min)      lcorbi_t3_neg_4_007_13    m/z 546.1147     0   2   4   6   8E+3   11   11.5  RT (min)      lcorbi_t3_neg_3_007_16    m/z 546.1234     0   1   2   3E+3   8   8.5  RT (min)      lcorbi_t3_neg_1_026_19    m/z 546.1357     0   2   4   6E+3   13   13.5  RT (min)      lcorbi_t3_neg_4_047_30    m/z 546.1672     0   2   4   6   8E+3   8   8.5  RT (min)      lcorbi_t3_neg_4_023_18    m/z 546.1897     0   1E+4   10.5   11  RT (min)      lcorbi_t3_neg_4_007_13    m/z 546.2713     0   2   4   6   8E+3   11   11.5  RT (min)      lcorbi_t3_neg_2_020_05    m/z 546.3111     0   1E+4   12.5   13  RT (min)      lcorbi_t3_neg_2_039_31    m/z 546.3197     0   2   4   6   8E+2   11.5   12  RT (min)      lcorbi_t3_neg_1_055_05    m/z 546.3525     0   2   4E+4   10.5   11  RT (min)      lcorbi_t3_neg_1_055_05    m/z 546.3927     0   2   4   6E+3   13.5   14  RT (min)      lcorbi_t3_neg_4_034_01    m/z 546.4243     0   1E+4   13   13.5  RT (min)      lcorbi_t3_neg_4_034_01    m/z 546.9094     0   2   4E+3   18   18.5  RT (min)      lcorbi_t3_neg_1_003_41    m/z 547.0030     0   1   2   3   4E+3   7.5   8  RT (min)      lcorbi_t3_neg_4_042_14    m/z 547.0338     0   1E+4   11   11.5  RT (min)      lcorbi_t3_neg_4_007_13    m/z 547.0636     0   1   2   3E+3   9.5   10  RT (min)      lcorbi_t3_neg_1_049_55    m/z 547.1116     0   2   4E+3   11   11.5  RT (min)      lcorbi_t3_neg_3_010_15    m/z 547.1208     0   1E+3   8   8.5  RT (min)      lcorbi_t3_neg_4_023_18    m/z 547.1249     0   1   2E+4   9   9.5  RT (min)      lcorbi_t3_neg_2_048_27    m/z 547.1448     0   1   2E+4   5   5.5  RT (min)      lcorbi_t3_neg_2_048_27    m/z 547.1687     0   2   4E+4   1   1.5  RT (min)      lcorbi_t3_neg_2_048_27    m/z 547.1833     0   1E+4   10.5   11  RT (min)      lcorbi_t3_neg_4_007_13    m/z 547.2426     0   2   4   6   8E+3   7.5   8  RT (min)      lcorbi_t3_neg_4_017_51    m/z 547.2943     0   2   4   6E+4   10   10.5  RT (min)      lcorbi_t3_neg_3_045_34    m/z 547.3076     0   1   2E+4   12.5   13  RT (min)      lcorbi_t3_neg_4_029_31    m/z 547.3253     0   2   4   6E+2   12.5   13  RT (min)      lcorbi_t3_neg_4_007_13    m/z 547.3610     0   1   2   3   4E+2   12   12.5  RT (min)      lcorbi_t3_neg_3_054_18    m/z 547.3767     0   1   2   3   4E+4   11.5   12  RT (min)      lcorbi_t3_neg_1_055_05    m/z 547.3811     0   1   14.5   15  RT (min)      lcorbi_t3_neg_4_023_18    m/z 547.4366     0   1   2   3E+4   13   13.5  RT (min)      lcorbi_t3_neg_4_042_14    m/z 547.7732     0   1   2E+4   5.5   6  RT (min)      lcorbi_t3_neg_4_042_14    m/z 548.0795     0   2   4   6E+3   6   6.5  RT (min)      lcorbi_t3_neg_1_054_37    m/z 548.0802     0   1E+4   3   3.5  RT (min)      lcorbi_t3_neg_4_007_13    m/z 548.2216     0   1   2E+4   7.5   8  RT (min)      lcorbi_t3_neg_4_037_29    m/z 548.2652     0   2   4   6E+3   10   10.5  RT (min)      lcorbi_t3_neg_3_019_12    m/z 548.2680     0   1E+4   12   12.5  RT (min)      lcorbi_t3_neg_4_007_13    m/z 548.2748     0   1E+4   5.5   6  RT (min)      lcorbi_t3_neg_2_048_27    m/z 548.2877     0   2   4   6E+2   11.5   12  RT (min)      lcorbi_t3_neg_2_022_24    m/z 548.2990     0   1E+5   10   10.5  RT (min)      lcorbi_t3_neg_1_055_05    m/z 548.3106     0   2   4E+3   11.5   12  RT (min)      lcorbi_t3_neg_1_055_05    m/z 548.3672     0   2   4   6E+4   13   13.5  RT (min)      lcorbi_t3_neg_4_023_18    m/z 548.3683     0   1   2E+4   11   11.5  RT (min)      lcorbi_t3_neg_1_055_05    m/z 548.3777     0   1E+4   13.5   14  RT (min)      lcorbi_t3_neg_4_007_13    m/z 548.4400     0   1E+4   13   13.5  RT (min)      lcorbi_t3_neg_4_042_14    m/z 548.6712     0   1   1   1.5  RT (min)      lcorbi_t3_neg_4_034_01    m/z 548.8042     0   2   4E+4   7.5   8  RT (min)      lcorbi_t3_neg_4_023_18    m/z 549.0622     0   1E+4   7   7.5  RT (min)      lcorbi_t3_neg_4_005_27    m/z 549.0752     0   2   4   6   8E+3   6.5   7  RT (min)      lcorbi_t3_neg_4_044_38    m/z 549.0796     0   1   2   3E+3   9.5   10  RT (min)      lcorbi_t3_neg_4_003_21    m/z 549.1206     0   2   4   6   8E+3   7.5   8  RT (min)      lcorbi_t3_neg_4_023_18    m/z 549.1218     0   1   2E+4   9   9.5  RT (min)      lcorbi_t3_neg_2_048_27    m/z 549.1748     0   2   4   6E+3   7   7.5  RT (min)      lcorbi_t3_neg_4_034_01    m/z 549.1856     0   1E+5   9.5   10  RT (min)      lcorbi_t3_neg_4_023_18    m/z 549.2655     0   2   4   6   8E+3   12.5   13  RT (min)      lcorbi_t3_neg_3_007_16    m/z 549.2816     0   1   2E+3   10   10.5  RT (min)      lcorbi_t3_neg_1_055_05    m/z 549.3195     0   1   2E+3   13   13.5  RT (min)      lcorbi_t3_neg_2_053_23    m/z 549.3694     0   2   4   6E+3   14.5   15  RT (min)      lcorbi_t3_neg_3_005_23    m/z 549.3806     0   1   2E+3   13.5   14  RT (min)      lcorbi_t3_neg_1_010_17    m/z 549.4234     0   1   14   14.5  RT (min)      lcorbi_t3_neg_4_007_13    m/z 549.4523     0   2   4   6E+4   13.5   14  RT (min)      lcorbi_t3_neg_4_042_14    m/z 549.5000     0   1   2   3   4E+3   14.5   15  RT (min)      lcorbi_t3_neg_4_050_02    m/z 550.0210     0   1   2E+3   9   9.5  RT (min)      lcorbi_t3_neg_3_019_12    m/z 550.0974     0   1E+4   10.5   11  RT (min)      lcorbi_t3_neg_4_030_04    m/z 550.1480     0   2   4   6   8E+3   7.5   8  RT (min)      lcorbi_t3_neg_3_045_34    m/z 550.2277     0   1   2   3   4E+3   13   13.5  RT (min)      lcorbi_t3_neg_4_042_14    m/z 550.2454     0   2   4E+3   8   8.5  RT (min)      lcorbi_t3_neg_4_034_01    m/z 550.2650     0   2   4   6E+3   12   12.5  RT (min)      lcorbi_t3_neg_1_055_05    m/z 550.2817     0   1E+4   12.5   13  RT (min)      lcorbi_t3_neg_4_007_13    m/z 550.3029     0   1   12   12.5  RT (min)      lcorbi_t3_neg_3_002_24    m/z 550.3148     0   1E+5   10.5   11  RT (min)      lcorbi_t3_neg_1_055_05    m/z 550.4558     0   1E+4   13.5   14  RT (min)      lcorbi_t3_neg_2_007_14    m/z 550.8643     0   1   2   3E+3   18   18.5  RT (min)      lcorbi_t3_neg_2_015_21    m/z 551.0185     0   1   2   3E+3   9   9.5  RT (min)      lcorbi_t3_neg_3_019_12    m/z 551.0264     0   1   2   3E+3   10   10.5  RT (min)      lcorbi_t3_neg_4_034_01    m/z 551.1259     0   1E+4   11.5   12  RT (min)      lcorbi_t3_neg_4_007_13    m/z 551.1835     0   1E+5   9.5   10  RT (min)      lcorbi_t3_neg_4_023_18    m/z 551.1951     0   1   2E+4   7   7.5  RT (min)      lcorbi_t3_neg_4_023_18    m/z 551.2084     0   1E+4   7   7.5  RT (min)      lcorbi_t3_neg_4_004_34    m/z 551.2210     0   1E+3   6   6.5  RT (min)      lcorbi_t3_neg_2_046_20    m/z 551.2233     0   1   2E+3   13   13.5  RT (min)      lcorbi_t3_neg_4_020_44    m/z 551.2604     0   1   2   3   4E+3   11   11.5  RT (min)      lcorbi_t3_neg_4_032_23    m/z 551.2849     0   1   2   3   4E+3   12.5   13  RT (min)      lcorbi_t3_neg_4_007_13    m/z 551.2923     0   2   4   6E+3   10.5   11  RT (min)      lcorbi_t3_neg_4_032_23    m/z 551.3103     0   1   12   12.5  RT (min)      lcorbi_t3_neg_1_055_05    m/z 551.3545     0   2   4   6E+3   11.5   12  RT (min)      lcorbi_t3_neg_4_005_27    m/z 551.3588     0   1   2E+3   13   13.5  RT (min)      lcorbi_t3_neg_4_009_50    m/z 551.3639     0   1   2E+3   14   14.5  RT (min)      lcorbi_t3_neg_3_023_07    m/z 551.4081     0   1E+4   13.5   14  RT (min)      lcorbi_t3_neg_4_042_14    m/z 551.4681     0   1   2E+4   13.5   14  RT (min)      lcorbi_t3_neg_4_038_15    m/z 551.9761     0   1   2   3   4E+3   9   9.5  RT (min)      lcorbi_t3_neg_1_044_23    m/z 552.0749     0   2   4   6E+3   12   12.5  RT (min)      lcorbi_t3_neg_3_014_19    m/z 552.1536     0   1E+4   9   9.5  RT (min)      lcorbi_t3_neg_4_023_18    m/z 552.1834     0   2   4   6   8E+3   13.5   14  RT (min)      lcorbi_t3_neg_4_041_54    m/z 552.2029     0   2   4E+3   11.5   12  RT (min)      lcorbi_t3_neg_1_055_05    m/z 552.2118     0   1   2   3E+3   7   7.5  RT (min)      lcorbi_t3_neg_3_054_18    m/z 552.2264     0   2   4   6   8E+3   14   14.5  RT (min)      lcorbi_t3_neg_4_040_19    m/z 552.2861     0   1   2   3   4E+4   10.5   11  RT (min)      lcorbi_t3_neg_1_055_05    m/z 552.2928     0   2   4   6   8E+3   12.5   13  RT (min)      lcorbi_t3_neg_1_023_03    m/z 552.3099     0   1E+3   11   11.5  RT (min)      lcorbi_t3_neg_1_055_05    m/z 552.3412     0   2   4   6E+3   12   12.5  RT (min)      lcorbi_t3_neg_4_010_28    m/z 552.3793     0   1   2E+3   11.5   12  RT (min)      lcorbi_t3_neg_1_055_05    m/z 552.4115     0   2   4E+3   13.5   14  RT (min)      lcorbi_t3_neg_4_042_14    m/z 552.8579     0   1   2   3   4E+3   18   18.5  RT (min)      lcorbi_t3_neg_4_053_43    m/z 552.9661     0   2   4   6E+3   8   8.5  RT (min)      lcorbi_t3_neg_4_023_18    m/z 553.0438     0   1   2E+3   9   9.5  RT (min)      lcorbi_t3_neg_4_051_40    m/z 553.1982     0   1   2   3E+3   13   13.5  RT (min)      lcorbi_t3_neg_4_007_13    m/z 553.2228     0   1E+4   6   6.5  RT (min)      lcorbi_t3_neg_1_012_20    m/z 553.2660     0   2   4   6   8E+3   10   10.5  RT (min)      lcorbi_t3_neg_4_029_31    m/z 553.2894     0   1E+4   10   10.5  RT (min)      lcorbi_t3_neg_4_029_31    m/z 553.3215     0   1E+4   6.5   7  RT (min)      lcorbi_t3_neg_4_032_23    m/z 553.3258     0   1   2E+3   10.5   11  RT (min)      lcorbi_t3_neg_1_055_05    m/z 553.3374     0   1E+4   14   14.5  RT (min)      lcorbi_t3_neg_2_048_27    m/z 553.3664     0   1   2   3   4E+4   12.5   13  RT (min)      lcorbi_t3_neg_4_034_01    m/z 553.3967     0   2   4   6E+3   7   7.5  RT (min)      lcorbi_t3_neg_4_032_23    m/z 553.4255     0   1E+4   13.5   14  RT (min)      lcorbi_t3_neg_4_034_01    m/z 553.4377     0   1E+4   13   13.5  RT (min)      lcorbi_t3_neg_4_034_01    m/z 553.4836     0   2   4   6   8E+3   13   13.5  RT (min)      lcorbi_t3_neg_4_042_14    m/z 554.0718     0   1   2   3   4E+3   12   12.5  RT (min)      lcorbi_t3_neg_3_014_19    m/z 554.1229     0   1E+4   1.5   2  RT (min)      lcorbi_t3_neg_4_032_23    m/z 554.1404     0   1E+4   7.5   8  RT (min)      lcorbi_t3_neg_2_048_27    m/z 554.2233     0   2   4   6   8E+3   14   14.5  RT (min)      lcorbi_t3_neg_4_040_19    m/z 554.2264     0   1E+4   11   11.5  RT (min)      lcorbi_t3_neg_1_055_05    m/z 554.2296     0   1E+4   13   13.5  RT (min)      lcorbi_t3_neg_4_005_27    m/z 554.2832     0   1E+4   10.5   11  RT (min)      lcorbi_t3_neg_1_055_05    m/z 554.3252     0   2   4   6E+3   11.5   12  RT (min)      lcorbi_t3_neg_4_034_01    m/z 554.3344     0   1   2E+4   14   14.5  RT (min)      lcorbi_t3_neg_4_032_23    m/z 554.3368     0   1   2   3E+3   11   11.5  RT (min)      lcorbi_t3_neg_1_013_35    m/z 554.4000     0   1E+4   14   14.5  RT (min)      lcorbi_t3_neg_3_023_07    m/z 554.4506     0   1   2   3E+4   13   13.5  RT (min)      lcorbi_t3_neg_4_034_01    m/z 554.8691     0   1E+4   18   18.5  RT (min)      lcorbi_t3_neg_1_006_40    m/z 555.0719     0   2   4   6E+3   6   6.5  RT (min)      lcorbi_t3_neg_1_054_37    m/z 555.1705     0   1E+4   13   13.5  RT (min)      lcorbi_t3_neg_4_007_13    m/z 555.1957     0   1   2E+3   13   13.5  RT (min)      lcorbi_t3_neg_4_007_13    m/z 555.2783     0   1E+4   12   12.5  RT (min)      lcorbi_t3_neg_4_023_18    m/z 555.2936     0   2   4   6   8E+3   12.5   13  RT (min)      lcorbi_t3_neg_4_017_51    m/z 555.3308     0   2   4E+4   14   14.5  RT (min)      lcorbi_t3_neg_4_032_23    m/z 555.3637     0   1   2   3   4E+3   12.5   13  RT (min)      lcorbi_t3_neg_4_034_01    m/z 555.3759     0   1   2   3E+3   12   12.5  RT (min)      lcorbi_t3_neg_1_021_31    m/z 555.3843     0   1   2   3   4E+3   14.5   15  RT (min)      lcorbi_t3_neg_4_046_37    m/z 556.0998     0   1   2   3   4E+3   9.5   10  RT (min)      lcorbi_t3_neg_2_040_43    m/z 556.1374     0   2   4   6   8E+3   7.5   8  RT (min)      lcorbi_t3_neg_2_048_27    m/z 556.1871     0   2   4E+3   9   9.5  RT (min)      lcorbi_t3_neg_4_023_18    m/z 556.2805     0   1   11.5   12  RT (min)      lcorbi_t3_neg_4_034_01    m/z 556.3341     0   1E+4   14   14.5  RT (min)      lcorbi_t3_neg_4_032_23    m/z 556.3625     0   1E+4   11.5   12  RT (min)      lcorbi_t3_neg_4_054_36    m/z 556.3886     0   1E+4   12.5   13  RT (min)      lcorbi_t3_neg_4_034_01    m/z 556.4664     0   1E+4   13.5   14  RT (min)      lcorbi_t3_neg_4_034_01    m/z 556.8811     0   2   4   6E+2   18   18.5  RT (min)      lcorbi_t3_neg_1_001_49    m/z 557.0935     0   2   4E+3   9.5   10  RT (min)      lcorbi_t3_neg_2_023_39    m/z 557.1311     0   1E+4   7.5   8  RT (min)      lcorbi_t3_neg_2_048_27    m/z 557.2216     0   1E+4   10.5   11  RT (min)      lcorbi_t3_neg_4_007_13    m/z 557.3092     0   1   2   3E+4   12   12.5  RT (min)      lcorbi_t3_neg_4_007_13    m/z 557.3289     0   1   2   3E+4   9   9.5  RT (min)      lcorbi_t3_neg_2_048_27    m/z 557.3291     0   2   4   6   8E+2   10.5   11  RT (min)      lcorbi_t3_neg_4_032_23    m/z 557.3322     0   1   2   3   4E+3   13   13.5  RT (min)      lcorbi_t3_neg_4_032_23    m/z 557.3549     0   1   2   3   4E+3   11.5   12  RT (min)      lcorbi_t3_neg_4_034_01    m/z 557.3641     0   1E+4   14   14.5  RT (min)      lcorbi_t3_neg_2_048_27    m/z 557.3660     0   2   4E+3   11.5   12  RT (min)      lcorbi_t3_neg_4_054_36    m/z 557.3977     0   1E+4   13   13.5  RT (min)      lcorbi_t3_neg_4_042_14    m/z 557.4459     0   2   4E+3   14   14.5  RT (min)      lcorbi_t3_neg_4_042_14    m/z 558.0036     0   2   4   6   8E+3   1.5   2  RT (min)      lcorbi_t3_neg_1_043_14    m/z 558.1081     0   2   4   6E+3   6   6.5  RT (min)      lcorbi_t3_neg_1_054_37    m/z 558.1537     0   1   2   3E+3   13   13.5  RT (min)      lcorbi_t3_neg_3_053_14    m/z 558.1617     0   1E+4   10   10.5  RT (min)      lcorbi_t3_neg_1_044_23    m/z 558.1868     0   2   4E+3   9   9.5  RT (min)      lcorbi_t3_neg_4_023_18    m/z 558.2412     0   2   4   6E+3   4.5   5  RT (min)      lcorbi_t3_neg_4_032_23    m/z 558.3052     0   1   2E+4   10.5   11  RT (min)      lcorbi_t3_neg_2_048_27    m/z 558.3239     0   2   4   6E+3   11.5   12  RT (min)      lcorbi_t3_neg_2_008_16    m/z 558.3285     0   1E+4   13   13.5  RT (min)      lcorbi_t3_neg_4_047_30    m/z 558.3400     0   2   4   6E+3   11   11.5  RT (min)      lcorbi_t3_neg_4_032_23    m/z 558.3468     0   2   4   6E+3   12.5   13  RT (min)      lcorbi_t3_neg_3_053_14    m/z 558.3641     0   1   2   3E+4   10   10.5  RT (min)      lcorbi_t3_neg_1_055_05    m/z 558.3648     0   1E+4   14   14.5  RT (min)      lcorbi_t3_neg_2_048_27    m/z 558.3797     0   2   4E+3   10.5   11  RT (min)      lcorbi_t3_neg_4_032_23    m/z 558.8110     0   2   4   6E+3   18   18.5  RT (min)      lcorbi_t3_neg_4_026_46    m/z 558.8297     0   2   4   6E+3   1   1.5  RT (min)      lcorbi_t3_neg_3_018_13    m/z 559.0466     0   1   2   3   4E+3   5.5   6  RT (min)      lcorbi_t3_neg_4_023_18    m/z 559.0495     0   2   4   6   8E+3   10.5   11  RT (min)      lcorbi_t3_neg_4_048_03    m/z 559.1748     0   2   4   6   8E+3   8   8.5  RT (min)      lcorbi_t3_neg_2_048_27    m/z 559.2403     0   2   4   6E+3   11   11.5  RT (min)      lcorbi_t3_neg_2_050_17    m/z 559.2911     0   1   2   3   4E+3   8   8.5  RT (min)      lcorbi_t3_neg_4_001_49    m/z 559.3252     0   1   2   3   4E+3   13   13.5  RT (min)      lcorbi_t3_neg_4_007_13    m/z 559.3462     0   1E+4   13.5   14  RT (min)      lcorbi_t3_neg_1_008_08    m/z 559.3588     0   2   4E+4   11.5   12  RT (min)      lcorbi_t3_neg_2_022_24    m/z 559.3607     0   1E+4   14   14.5  RT (min)      lcorbi_t3_neg_2_048_27    m/z 559.3762     0   1   2E+4   10.5   11  RT (min)      lcorbi_t3_neg_1_055_05    m/z 559.3810     0   2   4   6   8E+3   13.5   14  RT (min)      lcorbi_t3_neg_2_004_28    m/z 559.3982     0   2   4   6E+3   14.5   15  RT (min)      lcorbi_t3_neg_4_029_31    m/z 559.4639     0   1   2   3   4E+4   13.5   14  RT (min)      lcorbi_t3_neg_4_042_14    m/z 560.1118     0   1   2E+3   9.5   10  RT (min)      lcorbi_t3_neg_4_053_43    m/z 560.1585     0   1E+4   10   10.5  RT (min)      lcorbi_t3_neg_1_055_05    m/z 560.1853     0   2   4   6   8E+3   9.5   10  RT (min)      lcorbi_t3_neg_4_023_18    m/z 560.2342     0   2   4   6E+3   11   11.5  RT (min)      lcorbi_t3_neg_2_050_17    m/z 560.3217     0   2   4   6   8E+3   13.5   14  RT (min)      lcorbi_t3_neg_4_032_23    m/z 560.3462     0   1E+4   13   13.5  RT (min)      lcorbi_t3_neg_4_017_51    m/z 560.3817     0   1   2E+4   12   12.5  RT (min)      lcorbi_t3_neg_3_024_26    m/z 560.4012     0   1E+4   12.5   13  RT (min)      lcorbi_t3_neg_2_007_14    m/z 560.4768     0   1E+5   14   14.5  RT (min)      lcorbi_t3_neg_4_034_01    m/z 560.8082     0   2   4   6E+3   18   18.5  RT (min)      lcorbi_t3_neg_2_037_09    m/z 560.8309     0   1   2   3   4E+4   7.5   8  RT (min)      lcorbi_t3_neg_4_023_18    m/z 561.0550     0   2   4   6   8E+3   10   10.5  RT (min)      lcorbi_t3_neg_4_005_27    m/z 561.0917     0   1E+4   7.5   8  RT (min)      lcorbi_t3_neg_4_034_01    m/z 561.1127     0   1   2E+3   8   8.5  RT (min)      lcorbi_t3_neg_3_008_20    m/z 561.1312     0   1E+4   12   12.5  RT (min)      lcorbi_t3_neg_4_007_13    m/z 561.1520     0   1E+3   10   10.5  RT (min)      lcorbi_t3_neg_1_004_22    m/z 561.2331     0   2   4E+3   11   11.5  RT (min)      lcorbi_t3_neg_4_032_23    m/z 561.3226     0   1E+4   13   13.5  RT (min)      lcorbi_t3_neg_4_007_13    m/z 561.3273     0   1E+4   11.5   12  RT (min)      lcorbi_t3_neg_1_055_05    m/z 561.4798     0   1   2E+4   14   14.5  RT (min)      lcorbi_t3_neg_4_042_14    m/z 561.4891     0   2   4E+4   14   14.5  RT (min)      lcorbi_t3_neg_4_007_13    m/z 561.8399     0   2   4   6E+3   18   18.5  RT (min)      lcorbi_t3_neg_3_007_16    m/z 561.8880     0   2   4   6E+3   18   18.5  RT (min)      lcorbi_t3_neg_1_006_40    m/z 561.9305     0   2   4   6E+3   1   1.5  RT (min)      lcorbi_t3_neg_1_053_32    m/z 562.1115     0   2   4   6E+3   7   7.5  RT (min)      lcorbi_t3_neg_3_024_26    m/z 562.1551     0   1E+3   10   10.5  RT (min)      lcorbi_t3_neg_4_023_18    m/z 562.1601     0   1E+4   6   6.5  RT (min)      lcorbi_t3_neg_4_005_27    m/z 562.2243     0   1   2   3   4E+3   13.5   14  RT (min)      lcorbi_t3_neg_1_001_49    m/z 562.2315     0   1   2   3   4E+3   11.5   12  RT (min)      lcorbi_t3_neg_1_055_05    m/z 562.2511     0   1   2   3   4E+3   7.5   8  RT (min)      lcorbi_t3_neg_4_007_13    m/z 562.2995     0   2   4   6   8E+3   9   9.5  RT (min)      lcorbi_t3_neg_2_048_27    m/z 562.3261     0   2   4   6   8E+3   11.5   12  RT (min)      lcorbi_t3_neg_4_029_31    m/z 562.3618     0   1E+4   13   13.5  RT (min)      lcorbi_t3_neg_4_017_51    m/z 562.3752     0   1E+4   14   14.5  RT (min)      lcorbi_t3_neg_2_048_27    m/z 562.3828     0   1   2E+4   13   13.5  RT (min)      lcorbi_t3_neg_4_007_13    m/z 562.4171     0   2   4   6E+3   12.5   13  RT (min)      lcorbi_t3_neg_4_042_14    m/z 562.8836     0   1   2   3E+3   18   18.5  RT (min)      lcorbi_t3_neg_1_006_40    m/z 563.0729     0   2   4   6E+3   9   9.5  RT (min)      lcorbi_t3_neg_3_019_12    m/z 563.0886     0   1   2   3E+3   7.5   8  RT (min)      lcorbi_t3_neg_4_034_01    m/z 563.1074     0   1   2   3   4E+3   8   8.5  RT (min)      lcorbi_t3_neg_4_034_01    m/z 563.1472     0   2   4   6E+4   9   9.5  RT (min)      lcorbi_t3_neg_4_023_18    m/z 563.1660     0   1   2E+4   7.5   8  RT (min)      lcorbi_t3_neg_2_048_27    m/z 563.2210     0   2   4E+3   10   10.5  RT (min)      lcorbi_t3_neg_4_007_13    m/z 563.2966     0   1   2   3E+3   13   13.5  RT (min)      lcorbi_t3_neg_4_034_01    m/z 563.3330     0   2   4   6   8E+2   13   13.5  RT (min)      lcorbi_t3_neg_4_032_23    m/z 563.3718     0   1   2   3   4E+3   14   14.5  RT (min)      lcorbi_t3_neg_4_046_37    m/z 563.3930     0   1   14.5   15  RT (min)      lcorbi_t3_neg_4_011_24    m/z 563.3999     0   1   2   3E+4   10.5   11  RT (min)      lcorbi_t3_neg_1_055_05    m/z 563.9983     0   2   4   6E+3   6   6.5  RT (min)      lcorbi_t3_neg_1_054_37    m/z 564.0753     0   1   2E+3   9   9.5  RT (min)      lcorbi_t3_neg_3_019_12    m/z 564.0975     0   1E+4   6   6.5  RT (min)      lcorbi_t3_neg_1_054_37    m/z 564.1692     0   2   4   6E+3   7.5   8  RT (min)      lcorbi_t3_neg_2_048_27    m/z 564.2085     0   1   2E+4   7.5   8  RT (min)      lcorbi_t3_neg_4_037_29    m/z 564.2822     0   2   4   6   8E+3   11   11.5  RT (min)      lcorbi_t3_neg_3_007_16    m/z 564.3307     0   1E+4   18   18.5  RT (min)      lcorbi_t3_neg_2_008_16    m/z 564.3364     0   2   4E+3   13   13.5  RT (min)      lcorbi_t3_neg_4_034_01    m/z 564.3438     0   1E+4   14.5   15  RT (min)      lcorbi_t3_neg_4_046_37    m/z 564.3772     0   2   4   6E+3   13   13.5  RT (min)      lcorbi_t3_neg_4_044_38    m/z 564.3785     0   2   4   6   8E+3   11   11.5  RT (min)      lcorbi_t3_neg_4_034_01    m/z 564.4010     0   1E+4   14   14.5  RT (min)      lcorbi_t3_neg_3_023_07    m/z 564.9366     0   1   2   3E+3   18   18.5  RT (min)      lcorbi_t3_neg_4_026_46    m/z 565.0105     0   1   2   3   4E+3   9   9.5  RT (min)      lcorbi_t3_neg_2_015_21    m/z 565.0636     0   2   4   6E+3   5.5   6  RT (min)      lcorbi_t3_neg_4_023_18    m/z 565.0678     0   1   2   3   4E+3   10.5   11  RT (min)      lcorbi_t3_neg_4_030_04    m/z 565.1036     0   2   4   6E+3   7.5   8  RT (min)      lcorbi_t3_neg_4_034_01    m/z 565.1247     0   1   2   3E+3   8   8.5  RT (min)      lcorbi_t3_neg_2_034_19    m/z 565.1630     0   1E+4   7.5   8  RT (min)      lcorbi_t3_neg_2_048_27    m/z 565.2119     0   2   4   6E+3   7.5   8  RT (min)      lcorbi_t3_neg_4_037_29    m/z 565.2295     0   2   4   6E+3   12.5   13  RT (min)      lcorbi_t3_neg_3_007_16    m/z 565.2416     0   1E+3   7.5   8  RT (min)      lcorbi_t3_neg_4_037_29    m/z 565.3017     0   2   4   6   8E+3   8.5   9  RT (min)      lcorbi_t3_neg_2_048_27    m/z 565.3651     0   1E+4   13.5   14  RT (min)      lcorbi_t3_neg_4_039_32    m/z 565.3976     0   1   14   14.5  RT (min)      lcorbi_t3_neg_1_002_16    m/z 565.4473     0   1E+4   13   13.5  RT (min)      lcorbi_t3_neg_4_039_32    m/z 565.5397     0   1   2E+3   14.5   15  RT (min)      lcorbi_t3_neg_2_035_46    m/z 565.9953     0   2   4   6E+3   6   6.5  RT (min)      lcorbi_t3_neg_1_054_37    m/z 565.9977     0   2   4   6E+3   11   11.5  RT (min)      lcorbi_t3_neg_4_007_13    m/z 566.0486     0   2   4   6E+3   6   6.5  RT (min)      lcorbi_t3_neg_4_040_19    m/z 566.1657     0   2   4   6E+3   12.5   13  RT (min)      lcorbi_t3_neg_4_052_26    m/z 566.2886     0   1E+4   10.5   11  RT (min)      lcorbi_t3_neg_1_055_05    m/z 566.2948     0   1   2E+4   10   10.5  RT (min)      lcorbi_t3_neg_4_023_18    m/z 566.3800     0   1   2E+4   11.5   12  RT (min)      lcorbi_t3_neg_4_007_13    m/z 566.3946     0   2   4   6E+3   11.5   12  RT (min)      lcorbi_t3_neg_4_007_13    m/z 566.5232     0   1E+3   14.5   15  RT (min)      lcorbi_t3_neg_1_044_23    m/z 566.5517     0   1   2   3   4E+3   14.5   15  RT (min)      lcorbi_t3_neg_4_029_31    m/z 566.8357     0   2   4E+3   18   18.5  RT (min)      lcorbi_t3_neg_2_011_04    m/z 567.0076     0   1   2E+3   9   9.5  RT (min)      lcorbi_t3_neg_2_015_21    m/z 567.1309     0   1   2E+3   9.5   10  RT (min)      lcorbi_t3_neg_3_019_12    m/z 567.1773     0   2   4   6E+3   11   11.5  RT (min)      lcorbi_t3_neg_3_053_14    m/z 567.2532     0   2   4   6   8E+3   10.5   11  RT (min)      lcorbi_t3_neg_4_007_13    m/z 567.2920     0   1E+4   10.5   11  RT (min)      lcorbi_t3_neg_1_055_05    m/z 567.3111     0   2   4   6E+3   13.5   14  RT (min)      lcorbi_t3_neg_4_040_19    m/z 567.3829     0   1   11.5   12  RT (min)      lcorbi_t3_neg_4_077_58    m/z 567.4151     0   1E+4   14   14.5  RT (min)      lcorbi_t3_neg_3_023_07    m/z 567.4267     0   2   4   6E+3   13   13.5  RT (min)      lcorbi_t3_neg_3_002_24    m/z 567.5357     0   1   2E+3   13.5   14  RT (min)      lcorbi_t3_neg_4_010_28    m/z 568.1573     0   2   4   6E+3   13.5   14  RT (min)      lcorbi_t3_neg_4_040_19    m/z 568.2856     0   2   4   6E+3   11   11.5  RT (min)      lcorbi_t3_neg_3_007_16    m/z 568.3130     0   2   4   6   8E+3   12   12.5  RT (min)      lcorbi_t3_neg_4_012_17    m/z 568.3138     0   2   4   6   8E+3   13.5   14  RT (min)      lcorbi_t3_neg_4_040_19    m/z 568.3207     0   2   4   6E+3   9   9.5  RT (min)      lcorbi_t3_neg_2_048_27    m/z 568.3305     0   2   4E+3   13.5   14  RT (min)      lcorbi_t3_neg_4_040_19    m/z 568.4107     0   1   2E+4   12   12.5  RT (min)      lcorbi_t3_neg_4_038_15    m/z 568.4124     0   1E+4   14   14.5  RT (min)      lcorbi_t3_neg_3_023_07    m/z 568.4950     0   1E+4   14   14.5  RT (min)      lcorbi_t3_neg_3_005_23    m/z 568.7754     0   1   2E+3   1   1.5  RT (min)      lcorbi_t3_neg_3_029_02    m/z 568.9009     0   2   4   6   8E+3   18   18.5  RT (min)      lcorbi_t3_neg_1_004_22    m/z 569.0224     0   1   2   3E+3   9   9.5  RT (min)      lcorbi_t3_neg_3_019_12    m/z 569.1025     0   2   4   6E+3   10   10.5  RT (min)      lcorbi_t3_neg_4_043_11    m/z 569.2106     0   1   2   3E+4   11   11.5  RT (min)      lcorbi_t3_neg_2_020_05    m/z 569.2645     0   1E+4   5.5   6  RT (min)      lcorbi_t3_neg_3_037_36    m/z 569.2763     0   1   2E+4   10   10.5  RT (min)      lcorbi_t3_neg_4_004_34    m/z 569.2886     0   1E+4   8   8.5  RT (min)      lcorbi_t3_neg_2_048_27    m/z 569.3098     0   1E+4   13.5   14  RT (min)      lcorbi_t3_neg_4_040_19    m/z 569.3273     0   2   4   6   8E+3   13.5   14  RT (min)      lcorbi_t3_neg_4_040_19    m/z 569.3653     0   2   4   6   8E+3   13.5   14  RT (min)      lcorbi_t3_neg_4_040_19    m/z 569.3989     0   1E+4   12   12.5  RT (min)      lcorbi_t3_neg_4_005_27    m/z 569.4086     0   1E+4   14   14.5  RT (min)      lcorbi_t3_neg_3_023_07    m/z 569.4786     0   1E+4   13   13.5  RT (min)      lcorbi_t3_neg_4_007_13    m/z 570.0286     0   2   4   6E+3   5   5.5  RT (min)      lcorbi_t3_neg_3_014_19    m/z 570.0785     0   1   2   3E+3   9.5   10  RT (min)      lcorbi_t3_neg_4_053_43    m/z 570.1147     0   2   4   6E+3   11   11.5  RT (min)      lcorbi_t3_neg_4_026_46    m/z 570.1971     0   2   4   6   8E+3   13   13.5  RT (min)      lcorbi_t3_neg_4_042_14    m/z 570.2717     0   1   11   11.5  RT (min)      lcorbi_t3_neg_4_023_18    m/z 570.2810     0   1E+4   11.5   12  RT (min)      lcorbi_t3_neg_4_007_13    m/z 570.3281     0   1   13.5   14  RT (min)      lcorbi_t3_neg_3_002_24    m/z 570.3409     0   1E+3   12   12.5  RT (min)      lcorbi_t3_neg_3_002_24    m/z 570.3588     0   1   2E+4   11.5   12  RT (min)      lcorbi_t3_neg_1_055_05    m/z 570.3688     0   1   2E+3   14   14.5  RT (min)      lcorbi_t3_neg_1_001_49    m/z 570.4644     0   1   2E+4   14   14.5  RT (min)      lcorbi_t3_neg_3_005_23    m/z 571.0948     0   2   4   6E+3   10   10.5  RT (min)      lcorbi_t3_neg_4_054_36    m/z 571.1120     0   2   4   6E+3   11   11.5  RT (min)      lcorbi_t3_neg_4_026_46    m/z 571.1356     0   1E+4   6   6.5  RT (min)      lcorbi_t3_neg_1_054_37    m/z 571.1655     0   2   4   6   8E+3   9.5   10  RT (min)      lcorbi_t3_neg_4_053_43    m/z 571.2843     0   1   2E+4   4.5   5  RT (min)      lcorbi_t3_neg_4_038_15    m/z 571.3246     0   1E+4   13.5   14  RT (min)      lcorbi_t3_neg_4_040_19    m/z 571.3289     0   2   4   6E+3   12.5   13  RT (min)      lcorbi_t3_neg_3_007_16    m/z 571.4366     0   2   4E+4   13   13.5  RT (min)      lcorbi_t3_neg_4_034_01    m/z 571.4689     0   1   2E+4   11   11.5  RT (min)      lcorbi_t3_neg_2_039_31    m/z 571.5338     0   2   4   6E+3   15   15.5  RT (min)      lcorbi_t3_neg_4_020_44    m/z 571.8325     0   1   2E+3   18   18.5  RT (min)      lcorbi_t3_neg_2_049_55    m/z 571.8688     0   1E+4   18   18.5  RT (min)      lcorbi_t3_neg_3_050_40    m/z 571.9732     0   1E+4   11   11.5  RT (min)      lcorbi_t3_neg_4_007_13    m/z 572.1122     0   2   4E+3   11   11.5  RT (min)      lcorbi_t3_neg_4_026_46    m/z 572.1239     0   2   4   6E+3   7.5   8  RT (min)      lcorbi_t3_neg_4_034_01    m/z 572.1294     0   1E+4   6   6.5  RT (min)      lcorbi_t3_neg_1_054_37    m/z 572.1386     0   1E+4   9   9.5  RT (min)      lcorbi_t3_neg_4_023_18    m/z 572.1450     0   1   2   3   4E+3   8   8.5  RT (min)      lcorbi_t3_neg_2_046_20    m/z 572.1834     0   1   2   3E+4   1.5   2  RT (min)      lcorbi_t3_neg_2_022_24    m/z 572.2220     0   1   2E+4   9   9.5  RT (min)      lcorbi_t3_neg_2_048_27    m/z 572.2580     0   2   4   6   8E+3   11.5   12  RT (min)      lcorbi_t3_neg_4_042_14    m/z 572.2969     0   1   2E+4   11   11.5  RT (min)      lcorbi_t3_neg_4_016_05    m/z 572.3619     0   1E+4   18   18.5  RT (min)      lcorbi_t3_neg_1_014_28    m/z 572.4400     0   1E+4   13   13.5  RT (min)      lcorbi_t3_neg_4_042_14    m/z 572.5009     0   1   2E+3   14.5   15  RT (min)      lcorbi_t3_neg_3_007_16    m/z 572.8707     0   1   2   3E+3   18   18.5  RT (min)      lcorbi_t3_neg_2_018_40    m/z 572.9240     0   1   2   3E+3   18   18.5  RT (min)      lcorbi_t3_neg_3_002_24    m/z 573.0895     0   2   4   6E+3   10.5   11  RT (min)      lcorbi_t3_neg_4_043_11    m/z 573.1096     0   1   2   3   4E+3   11   11.5  RT (min)      lcorbi_t3_neg_4_026_46    m/z 573.1363     0   1E+4   8   8.5  RT (min)      lcorbi_t3_neg_4_034_01    m/z 573.1492     0   1   2E+4   11.5   12  RT (min)      lcorbi_t3_neg_4_007_13    m/z 573.2781     0   1   2E+4   10   10.5  RT (min)      lcorbi_t3_neg_4_023_18    m/z 573.2945     0   2   4   6   8E+3   10.5   11  RT (min)      lcorbi_t3_neg_1_055_05    m/z 573.3068     0   2   4   6E+3   8.5   9  RT (min)      lcorbi_t3_neg_4_001_49    m/z 573.3197     0   2   4   6E+3   14   14.5  RT (min)      lcorbi_t3_neg_4_023_18    m/z 573.3229     0   1   2   3   4E+3   12.5   13  RT (min)      lcorbi_t3_neg_4_007_13    m/z 573.3583     0   1   2E+4   18   18.5  RT (min)      lcorbi_t3_neg_1_010_17    m/z 573.3655     0   2   4   6   8E+3   7   7.5  RT (min)      lcorbi_t3_neg_4_032_23    m/z 573.3830     0   1E+5   10   10.5  RT (min)      lcorbi_t3_neg_1_055_05    m/z 573.4944     0   2   4   6   8E+4   14.5   15  RT (min)      lcorbi_t3_neg_4_036_16    m/z 574.0728     0   2   4   6E+3   9.5   10  RT (min)      lcorbi_t3_neg_4_007_13    m/z 574.1300     0   1E+4   11.5   12  RT (min)      lcorbi_t3_neg_4_007_13    m/z 574.1359     0   2   4E+3   9   9.5  RT (min)      lcorbi_t3_neg_4_023_18    m/z 574.1846     0   1   2E+4   9.5   10  RT (min)      lcorbi_t3_neg_4_007_13    m/z 574.2695     0   2   4   6   8E+3   10   10.5  RT (min)      lcorbi_t3_neg_4_023_18    m/z 574.2968     0   2   4   6   8E+3   14.5   15  RT (min)      lcorbi_t3_neg_4_056_56    m/z 574.3346     0   1E+4   10   10.5  RT (min)      lcorbi_t3_neg_4_032_23    m/z 574.8490     0   1   2   3E+3   18   18.5  RT (min)      lcorbi_t3_neg_3_050_40    m/z 575.1092     0   2   4   6E+3   11.5   12  RT (min)      lcorbi_t3_neg_4_007_13    m/z 575.1272     0   1   2E+4   11.5   12  RT (min)      lcorbi_t3_neg_4_007_13    m/z 575.1499     0   1E+4   13.5   14  RT (min)      lcorbi_t3_neg_4_040_19    m/z 575.1825     0   1   2   3E+3   13   13.5  RT (min)      lcorbi_t3_neg_3_053_14    m/z 575.2550     0   1E+4   12   12.5  RT (min)      lcorbi_t3_neg_4_023_18    m/z 575.2853     0   1E+4   10.5   11  RT (min)      lcorbi_t3_neg_4_007_13    m/z 575.3158     0   2   4   6E+3   11.5   12  RT (min)      lcorbi_t3_neg_4_034_01    m/z 575.3386     0   2   4E+3   13   13.5  RT (min)      lcorbi_t3_neg_4_007_13    m/z 575.3990     0   2   4   6   8E+4   10.5   11  RT (min)      lcorbi_t3_neg_1_055_05    m/z 575.8633     0   1E+4   18   18.5  RT (min)      lcorbi_t3_neg_2_048_27    m/z 576.0427     0   1   2   3E+3   9   9.5  RT (min)      lcorbi_t3_neg_2_015_21    m/z 576.1269     0   1   2E+4   11.5   12  RT (min)      lcorbi_t3_neg_4_007_13    m/z 576.1445     0   1   2   3   4E+3   12.5   13  RT (min)      lcorbi_t3_neg_4_040_19    m/z 576.1568     0   2   4E+3   8   8.5  RT (min)      lcorbi_t3_neg_4_014_20    m/z 576.2082     0   2   4   6   8E+3   11   11.5  RT (min)      lcorbi_t3_neg_2_053_23    m/z 576.2491     0   1E+4   12   12.5  RT (min)      lcorbi_t3_neg_4_023_18    m/z 576.2732     0   2   4   6E+3   14   14.5  RT (min)      lcorbi_t3_neg_1_023_03    m/z 576.2861     0   2   4   6E+3   10.5   11  RT (min)      lcorbi_t3_neg_4_007_13    m/z 576.2986     0   1   2E+3   10   10.5  RT (min)      lcorbi_t3_neg_4_011_24    m/z 576.3217     0   1E+4   11.5   12  RT (min)      lcorbi_t3_neg_2_039_31    m/z 576.4112     0   1   2   3   4E+4   11   11.5  RT (min)      lcorbi_t3_neg_1_055_05    m/z 576.4716     0   1   2   3   4E+4   13.5   14  RT (min)      lcorbi_t3_neg_4_046_37    m/z 577.0364     0   1   2   3   4E+3   9   9.5  RT (min)      lcorbi_t3_neg_3_019_12    m/z 577.0848     0   1E+4   11.5   12  RT (min)      lcorbi_t3_neg_4_007_13    m/z 577.1241     0   1   2   3   4E+4   11.5   12  RT (min)      lcorbi_t3_neg_4_007_13    m/z 577.1756     0   1E+4   9   9.5  RT (min)      lcorbi_t3_neg_4_023_18    m/z 577.2274     0   2   4   6E+3   10   10.5  RT (min)      lcorbi_t3_neg_4_032_23    m/z 577.2520     0   1   2   3E+3   12   12.5  RT (min)      lcorbi_t3_neg_3_013_28    m/z 577.2573     0   1E+4   9.5   10  RT (min)      lcorbi_t3_neg_4_001_49    m/z 577.2756     0   1E+4   12.5   13  RT (min)      lcorbi_t3_neg_4_052_26    m/z 577.2914     0   1E+4   13   13.5  RT (min)      lcorbi_t3_neg_4_032_23    m/z 577.3180     0   1E+3   13   13.5  RT (min)      lcorbi_t3_neg_4_032_23    m/z 577.3182     0   1   2   3   4E+4   11.5   12  RT (min)      lcorbi_t3_neg_4_029_31    m/z 577.3290     0   1   2   3E+4   8   8.5  RT (min)      lcorbi_t3_neg_2_048_27    m/z 577.3375     0   1   2E+4   11.5   12  RT (min)      lcorbi_t3_neg_3_020_31    m/z 577.3532     0   2   4   6E+3   12   12.5  RT (min)      lcorbi_t3_neg_3_013_28    m/z 577.3704     0   1E+4   10.5   11  RT (min)      lcorbi_t3_neg_1_055_05    m/z 577.4838     0   1E+5   13.5   14  RT (min)      lcorbi_t3_neg_4_005_27    m/z 577.8671     0   1   2   3   4E+3   18   18.5  RT (min)      lcorbi_t3_neg_2_023_39    m/z 578.1242     0   1E+4   11.5   12  RT (min)      lcorbi_t3_neg_4_007_13    m/z 578.1413     0   2   4   6E+3   13   13.5  RT (min)      lcorbi_t3_neg_4_020_44    m/z 578.1600     0   1   2E+4   1   1.5  RT (min)      lcorbi_t3_neg_2_035_46    m/z 578.2185     0   1E+4   11.5   12  RT (min)      lcorbi_t3_neg_1_055_05    m/z 578.2241     0   2   4   6E+3   11.5   12  RT (min)      lcorbi_t3_neg_2_020_05    m/z 578.2607     0   2   4   6E+2   9.5   10  RT (min)      lcorbi_t3_neg_1_055_05    m/z 578.2790     0   1   2   3   4E+3   12.5   13  RT (min)      lcorbi_t3_neg_4_052_26    m/z 578.2837     0   1E+5   10   10.5  RT (min)      lcorbi_t3_neg_4_078_58    m/z 578.3452     0   1E+4   11.5   12  RT (min)      lcorbi_t3_neg_4_029_31    m/z 578.3800     0   1   2E+4   12.5   13  RT (min)      lcorbi_t3_neg_4_007_13    m/z 578.4272     0   2   4   6   8E+3   13.5   14  RT (min)      lcorbi_t3_neg_2_007_14    m/z 578.4872     0   1   2   3   4E+4   13.5   14  RT (min)      lcorbi_t3_neg_4_034_01    m/z 578.8688     0   1E+4   9.5   10  RT (min)      lcorbi_t3_neg_4_020_44    m/z 578.9412     0   1   18   18.5  RT (min)      lcorbi_t3_neg_2_021_41    m/z 578.9877     0   1   2   3   4E+3   8   8.5  RT (min)      lcorbi_t3_neg_4_023_18    m/z 579.2250     0   2   4   6   8E+3   8.5   9  RT (min)      lcorbi_t3_neg_4_042_14    m/z 579.2543     0   1   2   3   4E+3   9.5   10  RT (min)      lcorbi_t3_neg_4_001_49    m/z 579.2548     0   1E+4   11.5   12  RT (min)      lcorbi_t3_neg_1_007_26    m/z 579.2550     0   1   2E+3   13.5   14  RT (min)      lcorbi_t3_neg_4_020_44    m/z 579.2785     0   2   4E+3   11.5   12  RT (min)      lcorbi_t3_neg_4_005_27    m/z 579.2787     0   1E+3   8.5   9  RT (min)      lcorbi_t3_neg_4_026_46    m/z 579.3300     0   1E+4   11.5   12  RT (min)      lcorbi_t3_neg_4_032_23    m/z 579.3414     0   2   4E+3   12.5   13  RT (min)      lcorbi_t3_neg_4_010_28    m/z 579.3516     0   2   4   6E+3   12   12.5  RT (min)      lcorbi_t3_neg_4_007_13    m/z 579.3690     0   2   4   6   8E+3   12.5   13  RT (min)      lcorbi_t3_neg_4_017_51    m/z 579.3860     0   2   4   6E+2   12   12.5  RT (min)      lcorbi_t3_neg_4_019_08    m/z 579.4299     0   1   2E+4   11.5   12  RT (min)      lcorbi_t3_neg_1_055_05    m/z 579.4994     0   1   2   3   4E+4   13.5   14  RT (min)      lcorbi_t3_neg_4_005_27    m/z 580.0546     0   1   2E+3   9   9.5  RT (min)      lcorbi_t3_neg_3_019_12    m/z 580.1612     0   1E+4   6   6.5  RT (min)      lcorbi_t3_neg_2_048_27    m/z 580.1814     0   1   13.5   14  RT (min)      lcorbi_t3_neg_4_041_54    m/z 580.2213     0   1   2   3   4E+3   10   10.5  RT (min)      lcorbi_t3_neg_3_019_12    m/z 580.2342     0   1   2   3E+3   12   12.5  RT (min)      lcorbi_t3_neg_4_042_14    m/z 580.2352     0   2   4   6   8E+3   10   10.5  RT (min)      lcorbi_t3_neg_4_007_13    m/z 580.3208     0   1E+4   8   8.5  RT (min)      lcorbi_t3_neg_2_048_27    m/z 580.3706     0   1   2E+4   10.5   11  RT (min)      lcorbi_t3_neg_1_055_05    m/z 580.3892     0   2   4   6E+4   11   11.5  RT (min)      lcorbi_t3_neg_1_055_05    m/z 580.4431     0   1E+4   14   14.5  RT (min)      lcorbi_t3_neg_3_053_14    m/z 581.0761     0   1   2   3E+3   7.5   8  RT (min)      lcorbi_t3_neg_4_042_14    m/z 581.0994     0   2   4E+3   9.5   10  RT (min)      lcorbi_t3_neg_4_023_18    m/z 581.1401     0   1E+4   12   12.5  RT (min)      lcorbi_t3_neg_4_007_13    m/z 581.2184     0   1E+4   13.5   14  RT (min)      lcorbi_t3_neg_2_041_54    m/z 581.2197     0   1   9.5   10  RT (min)      lcorbi_t3_neg_4_072_57    m/z 581.2526     0   1E+4   11.5   12  RT (min)      lcorbi_t3_neg_3_024_26    m/z 581.2707     0   1   2E+4   12   12.5  RT (min)      lcorbi_t3_neg_3_024_26    m/z 581.2901     0   1E+4   10.5   11  RT (min)      lcorbi_t3_neg_4_007_13    m/z 581.3025     0   2   4   6   8E+3   11   11.5  RT (min)      lcorbi_t3_neg_4_007_13    m/z 581.3095     0   1   2   3E+3   12.5   13  RT (min)      lcorbi_t3_neg_4_017_51    m/z 581.3250     0   1E+4   13.5   14  RT (min)      lcorbi_t3_neg_4_023_18    m/z 581.3290     0   2   4   6   8E+3   9.5   10  RT (min)      lcorbi_t3_neg_2_048_27    m/z 581.3652     0   2   4   6   8E+3   12   12.5  RT (min)      lcorbi_t3_neg_4_034_01    m/z 581.3945     0   1E+4   14   14.5  RT (min)      lcorbi_t3_neg_1_023_03    m/z 581.4017     0   2   4   6E+3   12.5   13  RT (min)      lcorbi_t3_neg_4_034_01    m/z 581.4557     0   1E+4   14   14.5  RT (min)      lcorbi_t3_neg_3_053_14    m/z 581.5152     0   1E+4   13.5   14  RT (min)      lcorbi_t3_neg_4_034_01    m/z 581.7069     0   1   2   3E+4   1   1.5  RT (min)      lcorbi_t3_neg_2_052_32    m/z 581.8695     0   1   2   3E+3   18   18.5  RT (min)      lcorbi_t3_neg_4_034_01    m/z 582.0024     0   1   2   3   4E+4   11   11.5  RT (min)      lcorbi_t3_neg_4_007_13    m/z 582.1270     0   1   2   3E+3   10   10.5  RT (min)      lcorbi_t3_neg_4_029_31    m/z 582.1785     0   2   4   6   8E+3   13.5   14  RT (min)      lcorbi_t3_neg_4_040_19    m/z 582.2183     0   1   2   3E+3   10   10.5  RT (min)      lcorbi_t3_neg_3_019_12    m/z 582.2370     0   1   11   11.5  RT (min)      lcorbi_t3_neg_4_023_18    m/z 582.2439     0   2   4   6E+4   9   9.5  RT (min)      lcorbi_t3_neg_4_007_13    m/z 582.2778     0   1E+4   11   11.5  RT (min)      lcorbi_t3_neg_1_055_05    m/z 582.3144     0   2   4   6E+2   11.5   12  RT (min)      lcorbi_t3_neg_2_020_05    m/z 582.3330     0   2   4   6   8E+2   11.5   12  RT (min)      lcorbi_t3_neg_1_055_05    m/z 582.3466     0   1E+5   14   14.5  RT (min)      lcorbi_t3_neg_4_032_23    m/z 582.3866     0   1   2E+4   11   11.5  RT (min)      lcorbi_t3_neg_1_055_05    m/z 582.4045     0   1   2   3   4E+4   11.5   12  RT (min)      lcorbi_t3_neg_1_055_05    m/z 582.4234     0   2   4   6   8E+3   5   5.5  RT (min)      lcorbi_t3_neg_4_054_36    m/z 582.4286     0   1E+4   14   14.5  RT (min)      lcorbi_t3_neg_4_016_05    m/z 582.5660     0   2   4   6   8E+3   15   15.5  RT (min)      lcorbi_t3_neg_4_029_31    m/z 582.8805     0   2   4   6   8E+3   9.5   10  RT (min)      lcorbi_t3_neg_3_050_40    m/z 583.1092     0   1   9   9.5  RT (min)      lcorbi_t3_neg_4_019_08    m/z 583.2232     0   1E+4   12   12.5  RT (min)      lcorbi_t3_neg_4_023_18    m/z 583.2502     0   1E+3   10   10.5  RT (min)      lcorbi_t3_neg_1_015_27    m/z 583.2687     0   1E+4   12   12.5  RT (min)      lcorbi_t3_neg_1_007_26    m/z 583.2887     0   1E+4   9.5   10  RT (min)      lcorbi_t3_neg_2_048_27    m/z 583.2963     0   2   4E+4   12   12.5  RT (min)      lcorbi_t3_neg_4_007_13    m/z 583.3124     0   2   4E+3   8   8.5  RT (min)      lcorbi_t3_neg_2_048_27    m/z 583.3214     0   1E+3   13   13.5  RT (min)      lcorbi_t3_neg_4_007_13    m/z 583.3361     0   2   4E+3   11.5   12  RT (min)      lcorbi_t3_neg_1_044_23    m/z 583.3415     0   1   2E+3   11   11.5  RT (min)      lcorbi_t3_neg_1_055_05    m/z 583.4167     0   1E+4   13   13.5  RT (min)      lcorbi_t3_neg_4_034_01    m/z 583.4249     0   1   2   3E+4   14   14.5  RT (min)      lcorbi_t3_neg_4_016_05    m/z 583.4752     0   2   4   6   8E+3   13   13.5  RT (min)      lcorbi_t3_neg_4_005_27    m/z 583.5331     0   1   2   3   4E+3   14.5   15  RT (min)      lcorbi_t3_neg_2_012_36    m/z 583.5502     0   1E+4   14.5   15  RT (min)      lcorbi_t3_neg_2_043_33    m/z 583.7041     0   1   1   1.5  RT (min)      lcorbi_t3_neg_2_048_27    m/z 584.1215     0   1   2   3   4E+3   10.5   11  RT (min)      lcorbi_t3_neg_4_043_11    m/z 584.1828     0   1E+4   7   7.5  RT (min)      lcorbi_t3_neg_4_040_19    m/z 584.2261     0   2   4   6E+3   12   12.5  RT (min)      lcorbi_t3_neg_4_023_18    m/z 584.2395     0   2   4E+3   11   11.5  RT (min)      lcorbi_t3_neg_2_054_15    m/z 584.2522     0   1   2E+3   11.5   12  RT (min)      lcorbi_t3_neg_1_001_49    m/z 584.2594     0   1   2E+4   7.5   8  RT (min)      lcorbi_t3_neg_2_048_27    m/z 584.2596     0   1   9.5   10  RT (min)      lcorbi_t3_neg_4_023_18    m/z 584.2732     0   1   11   11.5  RT (min)      lcorbi_t3_neg_1_055_05    m/z 584.2806     0   1   12   12.5  RT (min)      lcorbi_t3_neg_4_007_13    m/z 584.3100     0   1   2   3   4E+4   11.5   12  RT (min)      lcorbi_t3_neg_3_031_45    m/z 584.3402     0   2   4   6E+2   12   12.5  RT (min)      lcorbi_t3_neg_4_023_18    m/z 584.3457     0   2   4E+4   14   14.5  RT (min)      lcorbi_t3_neg_4_032_23    m/z 584.3568     0   1E+4   9.5   10  RT (min)      lcorbi_t3_neg_2_048_27    m/z 584.3815     0   1E+4   14.5   15  RT (min)      lcorbi_t3_neg_4_032_23    m/z 584.3822     0   2   4   6   8E+3   13   13.5  RT (min)      lcorbi_t3_neg_4_007_13    m/z 584.3985     0   1   14   14.5  RT (min)      lcorbi_t3_neg_4_032_23    m/z 584.8853     0   2   4   6E+3   18   18.5  RT (min)      lcorbi_t3_neg_3_011_04    m/z 585.0264     0   1E+4   10   10.5  RT (min)      lcorbi_t3_neg_4_023_18    m/z 585.1942     0   1   2   3   4E+3   11   11.5  RT (min)      lcorbi_t3_neg_4_032_23    m/z 585.2293     0   2   4   6   8E+3   10.5   11  RT (min)      lcorbi_t3_neg_4_034_01    m/z 585.2562     0   2   4   6   8E+2   11   11.5  RT (min)      lcorbi_t3_neg_1_055_05    m/z 585.2834     0   2   4   6E+2   12   12.5  RT (min)      lcorbi_t3_neg_4_007_13    m/z 585.2850     0   2   4   6   8E+3   9   9.5  RT (min)      lcorbi_t3_neg_4_023_18    m/z 585.2971     0   1   2E+4   11.5   12  RT (min)      lcorbi_t3_neg_4_007_13    m/z 585.3044     0   2   4   6   8E+3   11   11.5  RT (min)      lcorbi_t3_neg_1_055_05    m/z 585.3415     0   1E+4   11   11.5  RT (min)      lcorbi_t3_neg_1_043_14    m/z 585.3620     0   2   4E+4   14   14.5  RT (min)      lcorbi_t3_neg_4_032_23    m/z 585.3639     0   1E+4   11.5   12  RT (min)      lcorbi_t3_neg_4_007_13    m/z 585.3978     0   1   2E+3   13   13.5  RT (min)      lcorbi_t3_neg_4_034_01    m/z 585.4292     0   1   2E+4   13   13.5  RT (min)      lcorbi_t3_neg_4_034_01    m/z 586.0955     0   1   2   3   4E+4   2   2.5  RT (min)      lcorbi_t3_neg_4_029_31    m/z 586.1563     0   1E+4   1   1.5  RT (min)      lcorbi_t3_neg_2_051_34    m/z 586.2680     0   2   4E+3   11.5   12  RT (min)      lcorbi_t3_neg_4_007_13    m/z 586.3069     0   1   2E+3   11.5   12  RT (min)      lcorbi_t3_neg_2_020_05    m/z 586.3448     0   1E+4   14   14.5  RT (min)      lcorbi_t3_neg_4_029_31    m/z 586.3550     0   1E+4   12   12.5  RT (min)      lcorbi_t3_neg_4_034_01    m/z 586.3956     0   1E+5   10.5   11  RT (min)      lcorbi_t3_neg_1_055_05    m/z 586.6622     0   1   2   3E+4   1   1.5  RT (min)      lcorbi_t3_neg_2_048_27    m/z 586.8195     0   1   2   3   4E+3   18   18.5  RT (min)      lcorbi_t3_neg_1_042_18    m/z 586.8735     0   1   2   3E+3   18   18.5  RT (min)      lcorbi_t3_neg_2_038_22    m/z 586.9211     0   1   2   3   4E+3   18   18.5  RT (min)      lcorbi_t3_neg_2_002_26    m/z 586.9814     0   1   2E+4   11.5   12  RT (min)      lcorbi_t3_neg_4_007_13    m/z 587.0341     0   1E+3   9.5   10  RT (min)      lcorbi_t3_neg_4_042_14    m/z 587.0735     0   1   2E+3   9   9.5  RT (min)      lcorbi_t3_neg_4_051_40    m/z 587.1051     0   2   4   6E+3   11   11.5  RT (min)      lcorbi_t3_neg_4_043_11    m/z 587.1382     0   2   4E+3   10.5   11  RT (min)      lcorbi_t3_neg_4_042_14    m/z 587.2407     0   1E+4   11   11.5  RT (min)      lcorbi_t3_neg_4_021_25    m/z 587.3074     0   2   4   6E+3   9   9.5  RT (min)      lcorbi_t3_neg_4_044_38    m/z 587.3128     0   1E+4   12   12.5  RT (min)      lcorbi_t3_neg_4_007_13    m/z 587.3718     0   1E+3   11.5   12  RT (min)      lcorbi_t3_neg_3_002_24    m/z 587.3931     0   1E+3   13   13.5  RT (min)      lcorbi_t3_neg_4_034_01    m/z 587.4086     0   1   2   3   4E+4   13   13.5  RT (min)      lcorbi_t3_neg_4_034_01    m/z 587.4682     0   1E+4   13.5   14  RT (min)      lcorbi_t3_neg_4_012_17    m/z 587.5011     0   2   4   6   8E+3   14.5   15  RT (min)      lcorbi_t3_neg_2_053_23    m/z 588.1232     0   1E+4   5   5.5  RT (min)      lcorbi_t3_neg_1_004_22    m/z 588.1262     0   2   4   6   8E+3   11   11.5  RT (min)      lcorbi_t3_neg_3_032_17    m/z 588.2002     0   1E+4   10.5   11  RT (min)      lcorbi_t3_neg_4_007_13    m/z 588.2604     0   1   2   3E+3   11.5   12  RT (min)      lcorbi_t3_neg_4_007_13    m/z 588.2949     0   1E+4   11   11.5  RT (min)      lcorbi_t3_neg_4_007_13    m/z 588.3211     0   1E+4   10.5   11  RT (min)      lcorbi_t3_neg_4_007_13    m/z 588.3739     0   1   2   3   4E+4   11   11.5  RT (min)      lcorbi_t3_neg_1_055_05    m/z 588.3760     0   1   2E+4   14   14.5  RT (min)      lcorbi_t3_neg_3_035_03    m/z 588.4140     0   1   2E+4   13.5   14  RT (min)      lcorbi_t3_neg_3_024_26    m/z 588.5131     0   2   4   6   8E+3   14.5   15  RT (min)      lcorbi_t3_neg_2_053_23    m/z 588.8200     0   1   2   3   4E+3   18   18.5  RT (min)      lcorbi_t3_neg_3_035_03    m/z 588.8372     0   2   4E+3   18   18.5  RT (min)      lcorbi_t3_neg_1_055_05    m/z 588.8701     0   1   2   3E+3   18   18.5  RT (min)      lcorbi_t3_neg_2_034_19    m/z 588.8975     0   1E+4   12   12.5  RT (min)      lcorbi_t3_neg_2_019_07    m/z 588.8976     0   1   2E+4   10   10.5  RT (min)      lcorbi_t3_neg_3_035_03    m/z 588.9254     0   1   2   3E+3   18   18.5  RT (min)      lcorbi_t3_neg_2_023_39    m/z 588.9785     0   1   2E+4   11.5   12  RT (min)      lcorbi_t3_neg_4_007_13    m/z 589.0164     0   1   2   3E+3   8   8.5  RT (min)      lcorbi_t3_neg_2_015_21    m/z 589.1226     0   2   4   6E+3   11   11.5  RT (min)      lcorbi_t3_neg_3_007_16    m/z 589.1990     0   1   2E+4   5.5   6  RT (min)      lcorbi_t3_neg_2_053_23    m/z 589.2565     0   1E+3   11.5   12  RT (min)      lcorbi_t3_neg_4_069_57    m/z 589.2691     0   2   4E+3   12.5   13  RT (min)      lcorbi_t3_neg_4_039_32    m/z 589.3058     0   1E+4   11   11.5  RT (min)      lcorbi_t3_neg_4_007_13    m/z 589.3284     0   1E+4   12.5   13  RT (min)      lcorbi_t3_neg_4_007_13    m/z 589.3705     0   1E+4   12   12.5  RT (min)      lcorbi_t3_neg_3_024_26    m/z 589.3726     0   1   2   3E+3   13.5   14  RT (min)      lcorbi_t3_neg_4_039_32    m/z 589.3929     0   2   4   6   8E+3   6.5   7  RT (min)      lcorbi_t3_neg_4_032_23    m/z 589.4086     0   1   2   3E+3   13   13.5  RT (min)      lcorbi_t3_neg_4_029_31    m/z 589.4470     0   1E+4   12.5   13  RT (min)      lcorbi_t3_neg_4_046_37    m/z 589.4838     0   1   2E+4   13.5   14  RT (min)      lcorbi_t3_neg_4_007_13    m/z 589.9756     0   1E+4   11.5   12  RT (min)      lcorbi_t3_neg_4_007_13    m/z 590.0213     0   1   2   3E+3   9   9.5  RT (min)      lcorbi_t3_neg_3_019_12    m/z 590.0439     0   1E+4   11.5   12  RT (min)      lcorbi_t3_neg_4_007_13    m/z 590.0440     0   1E+4   6   6.5  RT (min)      lcorbi_t3_neg_4_044_38    m/z 590.1180     0   2   4   6   8E+3   8   8.5  RT (min)      lcorbi_t3_neg_2_048_27    m/z 590.1966     0   1   2   3E+4   14   14.5  RT (min)      lcorbi_t3_neg_2_048_27    m/z 590.2437     0   2   4   6   8E+3   6   6.5  RT (min)      lcorbi_t3_neg_2_034_19    m/z 590.2470     0   1E+4   12   12.5  RT (min)      lcorbi_t3_neg_3_054_18    m/z 590.2559     0   1E+4   13.5   14  RT (min)      lcorbi_t3_neg_4_017_51    m/z 590.2575     0   1   2   3E+3   11.5   12  RT (min)      lcorbi_t3_neg_4_007_13    m/z 590.3185     0   2   4   6   8E+3   11.5   12  RT (min)      lcorbi_t3_neg_4_007_13    m/z 590.3745     0   1   11.5   12  RT (min)      lcorbi_t3_neg_4_093_60    m/z 590.3867     0   2   4   6   8E+3   6.5   7  RT (min)      lcorbi_t3_neg_4_032_23    m/z 590.4063     0   1E+5   10.5   11  RT (min)      lcorbi_t3_neg_1_055_05    m/z 590.4266     0   1   2   3   4E+4   11.5   12  RT (min)      lcorbi_t3_neg_1_055_05    m/z 590.4329     0   2   4E+3   13.5   14  RT (min)      lcorbi_t3_neg_4_047_30    m/z 590.4958     0   2   4E+4   14   14.5  RT (min)      lcorbi_t3_neg_3_005_23    m/z 590.8245     0   2   4   6E+3   18   18.5  RT (min)      lcorbi_t3_neg_4_043_11    m/z 591.0246     0   1   2E+3   9   9.5  RT (min)      lcorbi_t3_neg_4_039_32    m/z 591.1539     0   1E+4   8   8.5  RT (min)      lcorbi_t3_neg_3_054_18    m/z 591.1927     0   1E+4   11   11.5  RT (min)      lcorbi_t3_neg_4_016_05    m/z 591.2187     0   2   4   6E+3   14.5   15  RT (min)      lcorbi_t3_neg_4_048_03    m/z 591.2266     0   2   4   6   8E+3   7.5   8  RT (min)      lcorbi_t3_neg_3_006_48    m/z 591.2486     0   2   4   6   8E+3   9.5   10  RT (min)      lcorbi_t3_neg_4_029_31    m/z 591.2537     0   2   4   6E+3   10   10.5  RT (min)      lcorbi_t3_neg_3_019_12    m/z 591.2888     0   1E+4   10   10.5  RT (min)      lcorbi_t3_neg_4_023_18    m/z 591.3295     0   1E+4   14   14.5  RT (min)      lcorbi_t3_neg_1_002_16    m/z 591.3645     0   1   2   3E+3   13.5   14  RT (min)      lcorbi_t3_neg_4_046_37    m/z 591.4421     0   2   4   6E+4   13.5   14  RT (min)      lcorbi_t3_neg_4_037_29    m/z 591.9055     0   1   2E+3   18   18.5  RT (min)      lcorbi_t3_neg_4_028_39    m/z 592.0459     0   1   2   3   4E+3   5   5.5  RT (min)      lcorbi_t3_neg_4_040_19    m/z 592.0701     0   2   4E+3   6   6.5  RT (min)      lcorbi_t3_neg_4_040_19    m/z 592.0854     0   1E+4   7.5   8  RT (min)      lcorbi_t3_neg_1_050_45    m/z 592.1565     0   1   2   3   4E+3   8   8.5  RT (min)      lcorbi_t3_neg_3_054_18    m/z 592.1935     0   1   2E+4   14   14.5  RT (min)      lcorbi_t3_neg_2_048_27    m/z 592.2292     0   1   2   3E+3   7.5   8  RT (min)      lcorbi_t3_neg_3_006_48    m/z 592.2436     0   1   2   3   4E+3   6   6.5  RT (min)      lcorbi_t3_neg_4_014_20    m/z 592.2472     0   2   4   6E+3   10   10.5  RT (min)      lcorbi_t3_neg_3_019_12    m/z 592.2538     0   2   4   6   8E+3   12.5   13  RT (min)      lcorbi_t3_neg_3_007_16    m/z 592.2927     0   1   2   3E+4   11.5   12  RT (min)      lcorbi_t3_neg_2_020_05    m/z 592.2929     0   1   2E+4   14   14.5  RT (min)      lcorbi_t3_neg_1_023_03    m/z 592.3037     0   1   12   12.5  RT (min)      lcorbi_t3_neg_4_023_18    m/z 592.3232     0   1E+4   11.5   12  RT (min)      lcorbi_t3_neg_4_005_27    m/z 592.3416     0   1   2   3   4E+3   11   11.5  RT (min)      lcorbi_t3_neg_4_038_15    m/z 592.4052     0   1   12   12.5  RT (min)      lcorbi_t3_neg_4_019_08    m/z 592.4455     0   1   2   3E+4   13.5   14  RT (min)      lcorbi_t3_neg_4_037_29    m/z 592.4663     0   2   4   6E+4   12.5   13  RT (min)      lcorbi_t3_neg_4_034_01    m/z 592.8369     0   2   4   6   8E+3   18   18.5  RT (min)      lcorbi_t3_neg_4_043_11    m/z 593.1758     0   1E+4   13   13.5  RT (min)      lcorbi_t3_neg_4_007_13    m/z 593.2106     0   2   4   6   8E+3   11   11.5  RT (min)      lcorbi_t3_neg_3_040_05    m/z 593.2157     0   2   4   6E+3   14.5   15  RT (min)      lcorbi_t3_neg_4_048_03    m/z 593.2472     0   2   4   6   8E+3   12.5   13  RT (min)      lcorbi_t3_neg_4_007_13    m/z 593.2603     0   1   12.5   13  RT (min)      lcorbi_t3_neg_4_032_23    m/z 593.3062     0   1   2   3   4E+3   7.5   8  RT (min)      lcorbi_t3_neg_2_048_27    m/z 593.3127     0   1E+3   10.5   11  RT (min)      lcorbi_t3_neg_3_053_14    m/z 593.3262     0   2   4   6E+3   11.5   12  RT (min)      lcorbi_t3_neg_4_042_14    m/z 593.3341     0   1E+4   12   12.5  RT (min)      lcorbi_t3_neg_4_023_18    m/z 593.3514     0   2   4   6E+3   13.5   14  RT (min)      lcorbi_t3_neg_4_005_27    m/z 593.3551     0   1E+4   11.5   12  RT (min)      lcorbi_t3_neg_4_007_13    m/z 593.3800     0   1   2   3E+3   13.5   14  RT (min)      lcorbi_t3_neg_4_007_13    m/z 593.3964     0   2   4E+3   13   13.5  RT (min)      lcorbi_t3_neg_2_027_13    m/z 593.4057     0   1E+3   10.5   11  RT (min)      lcorbi_t3_neg_4_011_24    m/z 593.4383     0   2   4   6E+4   11.5   12  RT (min)      lcorbi_t3_neg_1_055_05    m/z 593.4458     0   1   2   3   4E+4   12   12.5  RT (min)      lcorbi_t3_neg_1_055_05    m/z 593.5153     0   2   4   6   8E+3   13.5   14  RT (min)      lcorbi_t3_neg_3_024_26    m/z 593.5347     0   2   4   6   8E+3   14   14.5  RT (min)      lcorbi_t3_neg_4_032_23    m/z 594.0156     0   2   4E+3   9   9.5  RT (min)      lcorbi_t3_neg_4_007_13    m/z 594.0824     0   2   4   6   8E+3   7.5   8  RT (min)      lcorbi_t3_neg_1_050_45    m/z 594.2094     0   2   4   6E+3   14.5   15  RT (min)      lcorbi_t3_neg_4_048_03    m/z 594.2619     0   2   4   6   8E+3   10   10.5  RT (min)      lcorbi_t3_neg_3_019_12    m/z 594.2715     0   1   2E+3   11   11.5  RT (min)      lcorbi_t3_neg_1_050_45    m/z 594.2737     0   1E+4   8.5   9  RT (min)      lcorbi_t3_neg_4_023_18    m/z 594.3028     0   2   4   6   8E+2   7.5   8  RT (min)      lcorbi_t3_neg_1_055_05    m/z 594.3570     0   1   2E+4   10   10.5  RT (min)      lcorbi_t3_neg_1_055_05    m/z 594.3680     0   1E+3   11.5   12  RT (min)      lcorbi_t3_neg_1_055_05    m/z 594.4095     0   2   4   6E+4   13.5   14  RT (min)      lcorbi_t3_neg_4_007_13    m/z 594.4722     0   1   2   3E+4   13.5   14  RT (min)      lcorbi_t3_neg_3_024_26    m/z 594.5106     0   1   2E+4   14   14.5  RT (min)      lcorbi_t3_neg_3_005_23    m/z 594.5470     0   2   4   6E+3   14.5   15  RT (min)      lcorbi_t3_neg_4_029_31    m/z 594.8371     0   2   4   6E+3   18   18.5  RT (min)      lcorbi_t3_neg_4_036_16    m/z 595.0561     0   2   4   6E+3   10   10.5  RT (min)      lcorbi_t3_neg_4_029_31    m/z 595.0860     0   1   2   3   4E+3   10.5   11  RT (min)      lcorbi_t3_neg_4_030_04    m/z 595.2476     0   1E+4   10   10.5  RT (min)      lcorbi_t3_neg_4_038_15    m/z 595.2655     0   1E+3   10   10.5  RT (min)      lcorbi_t3_neg_1_032_13    m/z 595.2719     0   1   2E+4   7   7.5  RT (min)      lcorbi_t3_neg_4_023_18    m/z 595.3057     0   1E+3   11   11.5  RT (min)      lcorbi_t3_neg_1_055_05    m/z 595.3062     0   1   2   3E+2   7.5   8  RT (min)      lcorbi_t3_neg_1_055_05    m/z 595.3208     0   1   2   3   4E+4   5   5.5  RT (min)      lcorbi_t3_neg_2_048_27    m/z 595.4075     0   1   14   14.5  RT (min)      lcorbi_t3_neg_4_007_13    m/z 595.4346     0   1E+4   13   13.5  RT (min)      lcorbi_t3_neg_4_038_15    m/z 595.8591     0   2   4   6E+3   18   18.5  RT (min)      lcorbi_t3_neg_4_024_07    m/z 596.0811     0   2   4   6E+3   6   6.5  RT (min)      lcorbi_t3_neg_4_040_19    m/z 596.2335     0   2   4   6E+3   14   14.5  RT (min)      lcorbi_t3_neg_4_008_22    m/z 596.2528     0   1   2E+3   10   10.5  RT (min)      lcorbi_t3_neg_2_048_27    m/z 596.2736     0   1E+4   10.5   11  RT (min)      lcorbi_t3_neg_4_007_13    m/z 596.2753     0   1   7   7.5  RT (min)      lcorbi_t3_neg_4_032_23    m/z 596.3286     0   1   2   3   4E+3   13   13.5  RT (min)      lcorbi_t3_neg_4_037_29    m/z 596.3449     0   1   2E+3   11.5   12  RT (min)      lcorbi_t3_neg_2_027_13    m/z 596.3835     0   2   4   6E+3   12   12.5  RT (min)      lcorbi_t3_neg_4_034_01    m/z 596.4027     0   2   4   6   8E+3   5.5   6  RT (min)      lcorbi_t3_neg_4_032_23    m/z 596.4067     0   1   2   3E+4   11   11.5  RT (min)      lcorbi_t3_neg_4_029_31    m/z 596.4379     0   2   4   6E+3   13   13.5  RT (min)      lcorbi_t3_neg_4_034_01    m/z 596.8402     0   1   2   3E+3   18   18.5  RT (min)      lcorbi_t3_neg_4_015_10    m/z 596.8760     0   1   2   3   4E+3   8.5   9  RT (min)      lcorbi_t3_neg_4_053_43    m/z 597.0103     0   1   2   3   4E+4   11.5   12  RT (min)      lcorbi_t3_neg_4_007_13    m/z 597.0628     0   2   4   6   8E+3   9.5   10  RT (min)      lcorbi_t3_neg_4_053_43    m/z 597.1340     0   1E+4   11   11.5  RT (min)      lcorbi_t3_neg_4_043_11    m/z 597.1375     0   2   4   6E+3   9   9.5  RT (min)      lcorbi_t3_neg_4_023_18    m/z 597.1444     0   1   2   3E+3   8   8.5  RT (min)      lcorbi_t3_neg_4_040_19    m/z 597.1929     0   1E+4   10   10.5  RT (min)      lcorbi_t3_neg_4_023_18    m/z 597.2356     0   2   4E+3   14   14.5  RT (min)      lcorbi_t3_neg_1_042_18    m/z 597.2596     0   1   2E+4   5.5   6  RT (min)      lcorbi_t3_neg_3_037_36    m/z 597.2770     0   1   2E+3   10.5   11  RT (min)      lcorbi_t3_neg_1_039_38    m/z 597.2836     0   1   2E+4   8   8.5  RT (min)      lcorbi_t3_neg_2_048_27    m/z 597.3003     0   2   4   6E+3   11.5   12  RT (min)      lcorbi_t3_neg_1_055_05    m/z 597.3157     0   2   4   6E+3   12   12.5  RT (min)      lcorbi_t3_neg_4_010_28    m/z 597.3267     0   1   2E+3   10   10.5  RT (min)      lcorbi_t3_neg_3_043_25    m/z 597.3391     0   1E+4   14   14.5  RT (min)      lcorbi_t3_neg_2_053_23    m/z 597.3805     0   2   4   6   8E+3   11   11.5  RT (min)      lcorbi_t3_neg_2_039_31    m/z 597.4502     0   2   4   6E+3   13.5   14  RT (min)      lcorbi_t3_neg_1_020_36    m/z 597.5102     0   2   4E+3   13.5   14  RT (min)      lcorbi_t3_neg_4_042_14    m/z 597.8562     0   1   2   3E+3   18   18.5  RT (min)      lcorbi_t3_neg_4_014_20    m/z 597.9662     0   1E+4   9   9.5  RT (min)      lcorbi_t3_neg_4_007_13    m/z 598.0077     0   2   4   6E+3   6   6.5  RT (min)      lcorbi_t3_neg_1_054_37    m/z 598.1371     0   1   2   3   4E+3   11   11.5  RT (min)      lcorbi_t3_neg_4_043_11    m/z 598.1544     0   2   4   6E+3   9   9.5  RT (min)      lcorbi_t3_neg_4_023_18    m/z 598.2305     0   2   4   6E+3   14   14.5  RT (min)      lcorbi_t3_neg_4_008_22    m/z 598.2388     0   2   4   6   8E+3   8   8.5  RT (min)      lcorbi_t3_neg_4_042_14    m/z 598.2708     0   1E+3   10.5   11  RT (min)      lcorbi_t3_neg_2_022_24    m/z 598.2823     0   1   2   3E+3   11   11.5  RT (min)      lcorbi_t3_neg_4_007_13    m/z 598.2871     0   2   4   6E+3   8   8.5  RT (min)      lcorbi_t3_neg_2_048_27    m/z 598.2921     0   1E+3   13.5   14  RT (min)      lcorbi_t3_neg_1_026_19    m/z 598.3235     0   1   2E+3   10   10.5  RT (min)      lcorbi_t3_neg_3_007_16    m/z 598.3778     0   1E+4   14.5   15  RT (min)      lcorbi_t3_neg_4_029_31    m/z 598.3992     0   1   2   3E+3   12.5   13  RT (min)      lcorbi_t3_neg_4_034_01    m/z 598.4184     0   1E+4   6.5   7  RT (min)      lcorbi_t3_neg_4_032_23    m/z 598.4781     0   1   2   3   4E+4   14   14.5  RT (min)      lcorbi_t3_neg_2_053_23    m/z 598.4975     0   2   4   6   8E+3   14.5   15  RT (min)      lcorbi_t3_neg_3_045_34    m/z 598.8533     0   2   4   6E+3   9   9.5  RT (min)      lcorbi_t3_neg_3_048_08    m/z 599.0074     0   1   2E+4   11.5   12  RT (min)      lcorbi_t3_neg_4_007_13    m/z 599.0636     0   2   4E+3   9.5   10  RT (min)      lcorbi_t3_neg_4_029_31    m/z 599.1824     0   1   13   13.5  RT (min)      lcorbi_t3_neg_1_029_21    m/z 599.2726     0   1E+3   10   10.5  RT (min)      lcorbi_t3_neg_4_007_13    m/z 599.3108     0   1   2   3   4E+4   11   11.5  RT (min)      lcorbi_t3_neg_4_017_51    m/z 599.3202     0   2   4E+3   13.5   14  RT (min)      lcorbi_t3_neg_1_026_19    m/z 599.3802     0   1E+4   11   11.5  RT (min)      lcorbi_t3_neg_2_035_46    m/z 599.4657     0   1E+4   13   13.5  RT (min)      lcorbi_t3_neg_4_034_01    m/z 599.5652     0   1E+4   15   15.5  RT (min)      lcorbi_t3_neg_4_048_03    m/z 599.8279     0   1   2   3E+3   18   18.5  RT (min)      lcorbi_t3_neg_2_034_19    m/z 599.8578     0   1   2E+3   18   18.5  RT (min)      lcorbi_t3_neg_4_036_16    m/z 600.0067     0   2   4   6E+3   11   11.5  RT (min)      lcorbi_t3_neg_4_007_13    m/z 600.1274     0   1   2   3E+4   9   9.5  RT (min)      lcorbi_t3_neg_4_023_18    m/z 600.2033     0   1E+4   11   11.5  RT (min)      lcorbi_t3_neg_4_007_13    m/z 600.2766     0   1E+5   8.5   9  RT (min)      lcorbi_t3_neg_4_023_18    m/z 600.2847     0   2   4   6E+3   6.5   7  RT (min)      lcorbi_t3_neg_4_034_01    m/z 600.3205     0   2   4   6   8E+3   12.5   13  RT (min)      lcorbi_t3_neg_4_005_27    m/z 600.3612     0   1   2   3   4E+3   5.5   6  RT (min)      lcorbi_t3_neg_4_021_25    m/z 600.3773     0   2   4   6E+3   13.5   14  RT (min)      lcorbi_t3_neg_4_029_31    m/z 600.4185     0   2   4   6E+3   5.5   6   6.5  RT (min)      lcorbi_t3_neg_2_036_35    m/z 600.4938     0   1E+3   14.5   15  RT (min)      lcorbi_t3_neg_3_007_16    m/z 600.5322     0   2   4   6   8E+3   14.5   15  RT (min)      lcorbi_t3_neg_4_001_49    m/z 600.5485     0   1   14.5   15  RT (min)      lcorbi_t3_neg_4_007_13    m/z 600.5685     0   2   4E+3   15   15.5  RT (min)      lcorbi_t3_neg_4_048_03    m/z 600.8303     0   1E+3   18   18.5  RT (min)      lcorbi_t3_neg_4_019_08    m/z 600.9696     0   1   2   3   4E+3   8   8.5  RT (min)      lcorbi_t3_neg_4_023_18    m/z 601.0116     0   1   2   3   4E+3   8.5   9  RT (min)      lcorbi_t3_neg_4_034_01    m/z 601.0421     0   1   2   3   4E+3   9   9.5  RT (min)      lcorbi_t3_neg_3_019_12    m/z 601.1213     0   1   2E+4   9   9.5  RT (min)      lcorbi_t3_neg_4_023_18    m/z 601.1285     0   1   11.5   12  RT (min)      lcorbi_t3_neg_4_043_11    m/z 601.1458     0   1   2   3E+4   2   2.5  RT (min)      lcorbi_t3_neg_4_039_32    m/z 601.1677     0   2   4E+3   8   8.5  RT (min)      lcorbi_t3_neg_4_034_01    m/z 601.1831     0   1   9.5   10  RT (min)      lcorbi_t3_neg_4_032_23    m/z 601.1888     0   2   4   6E+3   10   10.5  RT (min)      lcorbi_t3_neg_4_029_31    m/z 601.2060     0   2   4   6E+3   11   11.5  RT (min)      lcorbi_t3_neg_4_007_13    m/z 601.2898     0   1E+4   11.5   12  RT (min)      lcorbi_t3_neg_2_052_32    m/z 601.3112     0   1E+4   10.5   11  RT (min)      lcorbi_t3_neg_4_017_51    m/z 601.3169     0   1E+4   13   13.5  RT (min)      lcorbi_t3_neg_4_005_27    m/z 601.3230     0   2   4   6   8E+3   9   9.5  RT (min)      lcorbi_t3_neg_4_044_38    m/z 601.3262     0   1E+4   11.5   12  RT (min)      lcorbi_t3_neg_3_007_16    m/z 601.3954     0   2   4   6E+3   13.5   14  RT (min)      lcorbi_t3_neg_3_024_26    m/z 601.4801     0   1   14.5   15  RT (min)      lcorbi_t3_neg_3_020_31    m/z 601.5260     0   1E+4   14.5   15  RT (min)      lcorbi_t3_neg_2_039_31    m/z 601.8569     0   1   2E+3   18   18.5  RT (min)      lcorbi_t3_neg_2_048_27    m/z 602.0989     0   1E+4   6   6.5  RT (min)      lcorbi_t3_neg_4_040_19    m/z 602.1142     0   2   4   6E+3   7.5   8  RT (min)      lcorbi_t3_neg_1_050_45    m/z 602.1412     0   1   2   3E+3   13   13.5  RT (min)      lcorbi_t3_neg_3_053_14    m/z 602.1934     0   1E+4   7   7.5  RT (min)      lcorbi_t3_neg_4_040_19    m/z 602.2242     0   1E+4   11   11.5  RT (min)      lcorbi_t3_neg_2_020_05    m/z 602.2854     0   1   2   3   4E+3   10   10.5  RT (min)      lcorbi_t3_neg_3_019_12    m/z 602.3016     0   1E+4   11   11.5  RT (min)      lcorbi_t3_neg_2_022_24    m/z 602.3135     0   2   4   6   8E+3   11   11.5  RT (min)      lcorbi_t3_neg_3_007_16    m/z 602.3200     0   2   4E+3   12.5   13  RT (min)      lcorbi_t3_neg_4_005_27    m/z 602.3264     0   1   2   3E+3   9   9.5  RT (min)      lcorbi_t3_neg_4_044_38    m/z 602.3399     0   2   4E+3   12   12.5  RT (min)      lcorbi_t3_neg_3_020_31    m/z 602.3657     0   2   4   6   8E+3   11   11.5  RT (min)      lcorbi_t3_neg_2_001_49    m/z 602.4736     0   1   2   3   4E+3   14.5   15  RT (min)      lcorbi_t3_neg_2_053_23    m/z 602.8491     0   2   4   6   8E+3   18   18.5  RT (min)      lcorbi_t3_neg_2_048_27    m/z 603.0792     0   1   2   3E+3   8   8.5  RT (min)      lcorbi_t3_neg_1_026_19    m/z 603.2833     0   1   2   3   4E+3   12   12.5  RT (min)      lcorbi_t3_neg_3_054_18    m/z 603.3050     0   1   2   3   4E+3   11   11.5  RT (min)      lcorbi_t3_neg_2_008_16    m/z 603.3176     0   1   2   3   4E+3   9   9.5  RT (min)      lcorbi_t3_neg_2_048_27    m/z 603.3542     0   2   4   6E+3   14   14.5  RT (min)      lcorbi_t3_neg_1_008_08    m/z 603.4249     0   2   4   6E+3   12   12.5  RT (min)      lcorbi_t3_neg_4_052_26    m/z 603.5397     0   1   2   3E+4   14.5   15  RT (min)      lcorbi_t3_neg_2_039_31    m/z 604.1222     0   1   2   3E+4   11.5   12  RT (min)      lcorbi_t3_neg_4_007_13    m/z 604.1569     0   1   2   3E+3   13   13.5  RT (min)      lcorbi_t3_neg_3_032_17    m/z 604.1738     0   1E+3   11.5   12  RT (min)      lcorbi_t3_neg_2_020_05    m/z 604.2400     0   1E+4   11.5   12  RT (min)      lcorbi_t3_neg_2_020_05    m/z 604.2640     0   1   2   3   4E+3   14   14.5  RT (min)      lcorbi_t3_neg_4_040_19    m/z 604.2924     0   2   4   6   8E+3   13   13.5  RT (min)      lcorbi_t3_neg_2_007_14    m/z 604.2991     0   2   4E+3   11   11.5  RT (min)      lcorbi_t3_neg_3_002_24    m/z 604.3108     0   2   4   6   8E+3   8.5   9  RT (min)      lcorbi_t3_neg_4_023_18    m/z 604.3256     0   1   2   3E+4   9.5   10  RT (min)      lcorbi_t3_neg_4_042_14    m/z 604.5257     0   1E+4   14.5   15  RT (min)      lcorbi_t3_neg_2_053_23    m/z 604.6331     0   2   4   6E+3   1   1.5  RT (min)      lcorbi_t3_neg_2_048_27    m/z 604.9814     0   2   4E+3   8   8.5  RT (min)      lcorbi_t3_neg_4_023_18    m/z 605.0849     0   1E+4   10   10.5  RT (min)      lcorbi_t3_neg_4_029_31    m/z 605.1194     0   1E+4   11.5   12  RT (min)      lcorbi_t3_neg_4_007_13    m/z 605.3052     0   1E+4   13   13.5  RT (min)      lcorbi_t3_neg_4_034_01    m/z 605.3207     0   2   4   6E+3   11   11.5  RT (min)      lcorbi_t3_neg_3_007_16    m/z 605.3290     0   1E+4   9.5   10  RT (min)      lcorbi_t3_neg_2_048_27    m/z 605.3389     0   1   2E+3   11.5   12  RT (min)      lcorbi_t3_neg_4_032_23    m/z 605.3852     0   1   2E+4   13   13.5  RT (min)      lcorbi_t3_neg_2_022_24    m/z 605.3947     0   1E+4   13.5   14  RT (min)      lcorbi_t3_neg_3_024_26    m/z 605.4786     0   1   2   3E+4   12.5   13  RT (min)      lcorbi_t3_neg_3_007_16    m/z 605.4788     0   1   2E+3   14.5   15  RT (min)      lcorbi_t3_neg_2_008_16    m/z 605.5293     0   2   4   6E+3   14.5   15  RT (min)      lcorbi_t3_neg_2_053_23    m/z 605.5926     0   1   2   3   4E+4   14   14.5  RT (min)      lcorbi_t3_neg_3_002_24    m/z 605.8198     0   2   4   6   8E+3   5.5   6  RT (min)      lcorbi_t3_neg_2_048_27    m/z 605.8445     0   2   4   6   8E+3   18   18.5  RT (min)      lcorbi_t3_neg_3_050_40    m/z 605.8749     0   1E+4   18   18.5  RT (min)      lcorbi_t3_neg_4_019_08    m/z 606.1359     0   1   2E+3   13   13.5  RT (min)      lcorbi_t3_neg_4_034_01    m/z 606.1640     0   1   2   3   4E+3   9   9.5  RT (min)      lcorbi_t3_neg_4_023_18    m/z 606.1727     0   2   4E+3   13.5   14  RT (min)      lcorbi_t3_neg_4_020_44    m/z 606.2555     0   2   4   6E+3   12   12.5  RT (min)      lcorbi_t3_neg_4_007_13    m/z 606.3284     0   2   4E+3   7.5   8  RT (min)      lcorbi_t3_neg_4_034_01    m/z 606.3330     0   2   4   6E+3   11.5   12  RT (min)      lcorbi_t3_neg_4_017_51    m/z 606.3412     0   2   4   6E+3   9.5   10  RT (min)      lcorbi_t3_neg_4_042_14    m/z 606.3524     0   1E+4   11.5   12  RT (min)      lcorbi_t3_neg_4_029_31    m/z 606.3684     0   1   2   3   4E+4   11   11.5  RT (min)      lcorbi_t3_neg_1_055_05    m/z 606.3763     0   1   2   3E+4   7   7.5  RT (min)      lcorbi_t3_neg_4_011_24    m/z 606.3775     0   2   4   6   8E+4   12   12.5  RT (min)      lcorbi_t3_neg_4_032_23    m/z 606.4819     0   1E+4   13   13.5  RT (min)      lcorbi_t3_neg_3_007_16    m/z 606.5278     0   1   2E+4   14.5   15  RT (min)      lcorbi_t3_neg_4_032_23    m/z 606.6300     0   1E+4   1   1.5  RT (min)      lcorbi_t3_neg_4_007_13    m/z 606.8467     0   2   4   6   8E+2   18   18.5  RT (min)      lcorbi_t3_neg_4_019_08    m/z 606.8705     0   1   2   3   4E+3   18   18.5  RT (min)      lcorbi_t3_neg_4_043_11    m/z 607.0393     0   1   2   3E+4   11.5   12  RT (min)      lcorbi_t3_neg_4_007_13    m/z 607.0528     0   1E+4   10.5   11  RT (min)      lcorbi_t3_neg_2_011_04    m/z 607.1369     0   1E+4   12   12.5  RT (min)      lcorbi_t3_neg_4_007_13    m/z 607.2444     0   1E+4   12.5   13  RT (min)      lcorbi_t3_neg_4_005_27    m/z 607.2845     0   2   4   6   8E+3   14   14.5  RT (min)      lcorbi_t3_neg_1_023_03    m/z 607.3411     0   1E+3   11   11.5  RT (min)      lcorbi_t3_neg_2_022_24    m/z 607.3487     0   2   4   6E+3   11   11.5  RT (min)      lcorbi_t3_neg_2_053_23    m/z 607.4140     0   1   2   3E+3   13.5   14  RT (min)      lcorbi_t3_neg_1_007_26    m/z 607.4733     0   1   2   3E+3   13   13.5  RT (min)      lcorbi_t3_neg_4_007_13    m/z 607.5310     0   1E+4   14.5   15  RT (min)      lcorbi_t3_neg_4_032_23    m/z 608.0363     0   1E+4   11.5   12  RT (min)      lcorbi_t3_neg_4_007_13    m/z 608.1514     0   1   2   3E+3   13   13.5  RT (min)      lcorbi_t3_neg_4_007_13    m/z 608.3158     0   1E+4   8   8.5  RT (min)      lcorbi_t3_neg_2_048_27    m/z 608.3441     0   2   4E+3   8.5   9  RT (min)      lcorbi_t3_neg_4_034_01    m/z 608.3565     0   1   2E+4   11   11.5  RT (min)      lcorbi_t3_neg_4_007_13    m/z 608.3584     0   2   4E+3   6.5   7  RT (min)      lcorbi_t3_neg_4_032_23    m/z 608.3841     0   2   4   6E+4   11.5   12  RT (min)      lcorbi_t3_neg_1_055_05    m/z 608.3884     0   1E+4   13   13.5  RT (min)      lcorbi_t3_neg_4_034_01    m/z 608.3962     0   1E+4   13   13.5  RT (min)      lcorbi_t3_neg_3_018_13    m/z 608.4979     0   1   2   3   4E+4   13.5   14  RT (min)      lcorbi_t3_neg_3_007_16    m/z 608.5626     0   2   4E+3   14.5   15  RT (min)      lcorbi_t3_neg_4_029_31    m/z 608.6116     0   1   2E+4   14   14.5  RT (min)      lcorbi_t3_neg_1_053_32    m/z 609.0498     0   1   2   3E+3   10.5   11  RT (min)      lcorbi_t3_neg_2_011_04    m/z 609.1549     0   1   2   3   4E+3   11   11.5  RT (min)      lcorbi_t3_neg_4_032_23    m/z 609.2525     0   1   2   3E+3   6.5   7  RT (min)      lcorbi_t3_neg_4_034_01    m/z 609.2972     0   1   11.5   12  RT (min)      lcorbi_t3_neg_4_069_57    m/z 609.3126     0   2   4   6E+3   8.5   9  RT (min)      lcorbi_t3_neg_4_007_13    m/z 609.3158     0   1E+5   12   12.5  RT (min)      lcorbi_t3_neg_4_007_13    m/z 609.3172     0   1E+4   5.5   6  RT (min)      lcorbi_t3_neg_3_037_36    m/z 609.3252     0   1   2   3E+3   13.5   14  RT (min)      lcorbi_t3_neg_2_034_19    m/z 609.3430     0   1E+4   13.5   14  RT (min)      lcorbi_t3_neg_4_040_19    m/z 609.3568     0   2   4E+3   11.5   12  RT (min)      lcorbi_t3_neg_4_034_01    m/z 609.3903     0   2   4E+3   13   13.5  RT (min)      lcorbi_t3_neg_4_007_13    m/z 609.8633     0   1   2   3   4E+3   18   18.5  RT (min)      lcorbi_t3_neg_3_014_19    m/z 610.1669     0   1   2   3   4E+3   13.5   14  RT (min)      lcorbi_t3_neg_2_023_39    m/z 610.3393     0   1   2E+4   13.5   14  RT (min)      lcorbi_t3_neg_4_040_19    m/z 610.3595     0   2   4E+3   8.5   9  RT (min)      lcorbi_t3_neg_2_048_27    m/z 610.3727     0   2   4E+2   11.5   12  RT (min)      lcorbi_t3_neg_4_019_08    m/z 610.3809     0   1E+4   11.5   12  RT (min)      lcorbi_t3_neg_1_055_05    m/z 610.4040     0   1E+4   11.5   12  RT (min)      lcorbi_t3_neg_4_011_24    m/z 610.8281     0   1   2   3E+3   18   18.5  RT (min)      lcorbi_t3_neg_1_055_05    m/z 610.9983     0   2   4E+3   8   8.5  RT (min)      lcorbi_t3_neg_2_015_21    m/z 611.0644     0   2   4   6E+3   10.5   11  RT (min)      lcorbi_t3_neg_4_030_04    m/z 611.1164     0   1   2E+3   9.5   10  RT (min)      lcorbi_t3_neg_4_053_43    m/z 611.1495     0   1E+4   11   11.5  RT (min)      lcorbi_t3_neg_4_043_11    m/z 611.2406     0   1E+4   10.5   11  RT (min)      lcorbi_t3_neg_4_021_25    m/z 611.2702     0   2   4E+3   8   8.5  RT (min)      lcorbi_t3_neg_4_005_27    m/z 611.3265     0   2   4E+2   11.5   12  RT (min)      lcorbi_t3_neg_4_032_23    m/z 611.3291     0   2   4   6E+3   8   8.5  RT (min)      lcorbi_t3_neg_2_034_19    m/z 611.3374     0   2   4E+3   13.5   14  RT (min)      lcorbi_t3_neg_3_042_22    m/z 611.3470     0   1   2E+4   9.5   10  RT (min)      lcorbi_t3_neg_3_019_12    m/z 611.3563     0   2   4   6E+3   11.5   12  RT (min)      lcorbi_t3_neg_4_007_13    m/z 611.3726     0   1E+4   12   12.5  RT (min)      lcorbi_t3_neg_4_034_01    m/z 611.3917     0   1E+4   11   11.5  RT (min)      lcorbi_t3_neg_1_055_05    m/z 611.4062     0   1E+3   11.5   12  RT (min)      lcorbi_t3_neg_4_011_24    m/z 611.4107     0   2   4   6   8E+3   13.5   14  RT (min)      lcorbi_t3_neg_4_047_30    m/z 611.4265     0   2   4   6E+3   13   13.5  RT (min)      lcorbi_t3_neg_4_044_38    m/z 611.4369     0   2   4   6   8E+4   14   14.5  RT (min)      lcorbi_t3_neg_4_016_05    m/z 611.4890     0   1   2   3   4E+3   12.5   13  RT (min)      lcorbi_t3_neg_4_042_14    m/z 611.5641     0   1   2   3   4E+3   14.5   15  RT (min)      lcorbi_t3_neg_4_035_33    m/z 611.6004     0   2   4E+3   15   15.5  RT (min)      lcorbi_t3_neg_4_046_37    m/z 612.2348     0   1   10   10.5  RT (min)      lcorbi_t3_neg_4_019_08    m/z 612.2605     0   1   2   3   4E+3   11   11.5  RT (min)      lcorbi_t3_neg_2_048_27    m/z 612.2730     0   1   2   3E+3   11   11.5  RT (min)      lcorbi_t3_neg_3_018_13    m/z 612.3049     0   2   4   6   8E+3   11.5   12  RT (min)      lcorbi_t3_neg_4_034_01    m/z 612.3412     0   2   4   6E+3   12.5   13  RT (min)      lcorbi_t3_neg_2_053_23    m/z 612.3522     0   1E+4   11.5   12  RT (min)      lcorbi_t3_neg_4_007_13    m/z 612.3739     0   1   2E+3   14.5   15  RT (min)      lcorbi_t3_neg_4_021_25    m/z 612.3879     0   1E+4   12   12.5  RT (min)      lcorbi_t3_neg_4_034_01    m/z 612.4146     0   1   14   14.5  RT (min)      lcorbi_t3_neg_3_024_26    m/z 612.4297     0   2   4   6E+3   13   13.5  RT (min)      lcorbi_t3_neg_4_042_14    m/z 612.4479     0   1   2   3E+4   13.5   14  RT (min)      lcorbi_t3_neg_4_034_01    m/z 612.4560     0   1   14   14.5  RT (min)      lcorbi_t3_neg_3_005_23    m/z 612.8632     0   1   2E+3   18   18.5  RT (min)      lcorbi_t3_neg_1_001_49    m/z 612.8958     0   1E+4   18   18.5  RT (min)      lcorbi_t3_neg_1_003_41    m/z 613.0552     0   1   2   3E+3   9.5   10  RT (min)      lcorbi_t3_neg_2_034_19    m/z 613.1077     0   1E+4   8   8.5  RT (min)      lcorbi_t3_neg_1_026_19    m/z 613.1519     0   2   4   6E+3   2   2.5  RT (min)      lcorbi_t3_neg_1_054_37    m/z 613.3081     0   1   2E+3   11.5   12  RT (min)      lcorbi_t3_neg_4_005_27    m/z 613.3162     0   1E+4   5.5   6  RT (min)      lcorbi_t3_neg_3_037_36    m/z 613.3551     0   1E+4   9.5   10  RT (min)      lcorbi_t3_neg_3_019_12    m/z 613.3702     0   2   4   6   8E+3   14.5   15  RT (min)      lcorbi_t3_neg_4_029_31    m/z 613.3955     0   1   2E+3   12.5   13  RT (min)      lcorbi_t3_neg_2_007_14    m/z 613.4268     0   2   4   6E+3   5.5   6  RT (min)      lcorbi_t3_neg_4_054_36    m/z 613.4447     0   1   2E+4   13   13.5  RT (min)      lcorbi_t3_neg_4_034_01    m/z 613.5046     0   2   4   6   8E+3   12.5   13  RT (min)      lcorbi_t3_neg_4_034_01    m/z 613.5518     0   1   2   3E+3   14.5   15  RT (min)      lcorbi_t3_neg_2_030_47    m/z 613.6056     0   1   2E+4   14   14.5  RT (min)      lcorbi_t3_neg_3_002_24    m/z 614.1437     0   1   2E+4   2   2.5  RT (min)      lcorbi_t3_neg_4_032_23    m/z 614.2581     0   2   4E+3   11   11.5  RT (min)      lcorbi_t3_neg_4_034_01    m/z 614.2763     0   2   4E+3   11.5   12  RT (min)      lcorbi_t3_neg_4_007_13    m/z 614.2824     0   1E+4   10   10.5  RT (min)      lcorbi_t3_neg_4_023_18    m/z 614.3020     0   2   4   6E+2   11.5   12  RT (min)      lcorbi_t3_neg_2_020_05    m/z 614.3164     0   2   4   6E+3   12.5   13  RT (min)      lcorbi_t3_neg_4_040_19    m/z 614.3342     0   2   4   6   8E+3   13   13.5  RT (min)      lcorbi_t3_neg_4_005_27    m/z 614.3627     0   1E+4   8   8.5  RT (min)      lcorbi_t3_neg_2_048_27    m/z 614.3736     0   1   2E+3   14.5   15  RT (min)      lcorbi_t3_neg_1_044_23    m/z 614.3925     0   1E+4   14.5   15  RT (min)      lcorbi_t3_neg_4_032_23    m/z 614.4481     0   2   4   6E+3   13   13.5  RT (min)      lcorbi_t3_neg_4_046_37    m/z 614.5081     0   1   2   3E+3   12.5   13  RT (min)      lcorbi_t3_neg_4_034_01    m/z 614.8968     0   2   4   6E+3   18   18.5  RT (min)      lcorbi_t3_neg_1_004_22    m/z 615.1084     0   1   2   3   4E+3   8   8.5  RT (min)      lcorbi_t3_neg_3_014_19    m/z 615.2459     0   2   4   6E+3   8.5   9  RT (min)      lcorbi_t3_neg_4_054_36    m/z 615.2721     0   1E+4   11.5   12  RT (min)      lcorbi_t3_neg_2_020_05    m/z 615.2788     0   1   2E+4   10   10.5  RT (min)      lcorbi_t3_neg_4_023_18    m/z 615.2823     0   1   11.5   12  RT (min)      lcorbi_t3_neg_4_032_23    m/z 615.2958     0   1   2   3   4E+4   12   12.5  RT (min)      lcorbi_t3_neg_4_032_23    m/z 615.3393     0   1   2   3   4E+3   11   11.5  RT (min)      lcorbi_t3_neg_2_054_15    m/z 615.3627     0   1E+3   11   11.5  RT (min)      lcorbi_t3_neg_4_034_01    m/z 615.4420     0   1   2   3E+4   13.5   14  RT (min)      lcorbi_t3_neg_4_037_29    m/z 615.4781     0   2   4   6   8E+3   14.5   15  RT (min)      lcorbi_t3_neg_4_011_24    m/z 615.5668     0   2   4E+3   14.5   15  RT (min)      lcorbi_t3_neg_4_029_31    m/z 616.2527     0   2   4   6E+3   10   10.5  RT (min)      lcorbi_t3_neg_3_019_12    m/z 616.2553     0   1   12   12.5  RT (min)      lcorbi_t3_neg_2_053_23    m/z 616.2732     0   2   4   6E+3   11.5   12  RT (min)      lcorbi_t3_neg_2_008_16    m/z 616.2812     0   2   4   6E+3   10   10.5  RT (min)      lcorbi_t3_neg_1_055_05    m/z 616.2856     0   1   2   3   4E+4   14   14.5  RT (min)      lcorbi_t3_neg_1_023_03    m/z 616.3054     0   2   4E+3   8   8.5  RT (min)      lcorbi_t3_neg_2_048_27    m/z 616.3162     0   2   4E+3   12.5   13  RT (min)      lcorbi_t3_neg_1_026_19    m/z 616.3254     0   1E+4   11   11.5  RT (min)      lcorbi_t3_neg_4_044_38    m/z 616.3731     0   1   2   3E+4   12   12.5  RT (min)      lcorbi_t3_neg_4_029_31    m/z 616.3886     0   1E+4   14.5   15  RT (min)      lcorbi_t3_neg_3_007_16    m/z 616.4085     0   1E+4   14   14.5  RT (min)      lcorbi_t3_neg_4_030_04    m/z 616.4893     0   1E+4   14   14.5  RT (min)      lcorbi_t3_neg_2_043_33    m/z 616.8397     0   1   2   3E+3   18   18.5  RT (min)      lcorbi_t3_neg_1_006_40    m/z 617.0055     0   1   2E+3   9   9.5  RT (min)      lcorbi_t3_neg_3_019_12    m/z 617.0498     0   2   4   6E+3   7   7.5  RT (min)      lcorbi_t3_neg_4_005_27    m/z 617.0670     0   1   2E+3   9.5   10  RT (min)      lcorbi_t3_neg_2_023_39    m/z 617.1407     0   2   4   6E+3   9   9.5  RT (min)      lcorbi_t3_neg_4_017_51    m/z 617.1977     0   1E+4   12   12.5  RT (min)      lcorbi_t3_neg_1_042_18    m/z 617.2117     0   1E+4   7.5   8  RT (min)      lcorbi_t3_neg_4_037_29    m/z 617.2530     0   1   12.5   13  RT (min)      lcorbi_t3_neg_4_032_23    m/z 617.2589     0   1E+4   9   9.5  RT (min)      lcorbi_t3_neg_4_023_18    m/z 617.2654     0   2   4   6   8E+3   13.5   14  RT (min)      lcorbi_t3_neg_4_023_18    m/z 617.2735     0   2   4   6   8E+3   10   10.5  RT (min)      lcorbi_t3_neg_4_023_18    m/z 617.2878     0   1E+4   12   12.5  RT (min)      lcorbi_t3_neg_4_034_01    m/z 617.2954     0   2   4   6E+3   12   12.5  RT (min)      lcorbi_t3_neg_2_008_16    m/z 617.3564     0   1E+3   12.5   13  RT (min)      lcorbi_t3_neg_4_007_13    m/z 617.3785     0   1   11.5   12  RT (min)      lcorbi_t3_neg_4_007_13    m/z 617.4037     0   1E+4   13.5   14  RT (min)      lcorbi_t3_neg_4_042_14    m/z 617.4654     0   1   2   3   4E+3   13   13.5  RT (min)      lcorbi_t3_neg_4_007_13    m/z 617.6371     0   1E+4   14.5   15  RT (min)      lcorbi_t3_neg_4_039_32    m/z 617.8771     0   1   2   3E+3   18   18.5  RT (min)      lcorbi_t3_neg_2_042_44    m/z 618.0848     0   2   4   6E+3   10.5   11  RT (min)      lcorbi_t3_neg_4_030_04    m/z 618.1194     0   1   2E+4   11.5   12  RT (min)      lcorbi_t3_neg_4_007_13    m/z 618.1373     0   2   4   6   8E+3   11.5   12  RT (min)      lcorbi_t3_neg_4_007_13    m/z 618.1545     0   1   2   3   4E+3   12.5   13  RT (min)      lcorbi_t3_neg_4_040_19    m/z 618.2384     0   1E+4   6.5   7  RT (min)      lcorbi_t3_neg_4_042_14    m/z 618.2523     0   2   4   6E+2   12   12.5  RT (min)      lcorbi_t3_neg_1_055_05    m/z 618.2600     0   2   4   6E+3   13.5   14  RT (min)      lcorbi_t3_neg_4_023_18    m/z 618.2870     0   1E+4   14   14.5  RT (min)      lcorbi_t3_neg_4_017_51    m/z 618.2888     0   2   4   6   8E+3   12   12.5  RT (min)      lcorbi_t3_neg_2_007_14    m/z 618.3079     0   2   4   6E+3   11.5   12  RT (min)      lcorbi_t3_neg_4_039_32    m/z 618.3410     0   2   4   6E+3   10.5   11  RT (min)      lcorbi_t3_neg_4_038_15    m/z 618.3607     0   2   4   6E+3   10   10.5  RT (min)      lcorbi_t3_neg_3_005_23    m/z 618.3774     0   1E+4   12   12.5  RT (min)      lcorbi_t3_neg_4_007_13    m/z 618.3879     0   2   4E+3   14.5   15  RT (min)      lcorbi_t3_neg_4_034_01    m/z 618.4067     0   1   2   3   4E+3   13.5   14  RT (min)      lcorbi_t3_neg_4_042_14    m/z 618.8515     0   1   2   3E+3   18   18.5  RT (min)      lcorbi_t3_neg_2_014_29    m/z 619.0060     0   1   2   3E+3   9   9.5  RT (min)      lcorbi_t3_neg_1_042_18    m/z 619.1947     0   1E+4   12   12.5  RT (min)      lcorbi_t3_neg_1_042_18    m/z 619.2953     0   1E+4   8.5   9  RT (min)      lcorbi_t3_neg_4_023_18    m/z 619.3705     0   1E+4   11.5   12  RT (min)      lcorbi_t3_neg_3_002_24    m/z 619.4819     0   2   4   6E+3   13   13.5  RT (min)      lcorbi_t3_neg_4_034_01    m/z 619.5275     0   1E+4   14.5   15  RT (min)      lcorbi_t3_neg_2_053_23    m/z 619.8828     0   1   2   3E+3   18   18.5  RT (min)      lcorbi_t3_neg_3_019_12    m/z 620.1147     0   2   4E+3   6   6.5  RT (min)      lcorbi_t3_neg_1_054_37    m/z 620.1885     0   2   4   6   8E+3   12   12.5  RT (min)      lcorbi_t3_neg_1_042_18    m/z 620.2735     0   1E+4   10.5   11  RT (min)      lcorbi_t3_neg_2_020_05    m/z 620.3180     0   1E+4   11   11.5  RT (min)      lcorbi_t3_neg_4_016_05    m/z 620.3481     0   1   2   3   4E+4   11.5   12  RT (min)      lcorbi_t3_neg_4_029_31    m/z 620.3636     0   2   4E+4   10   10.5  RT (min)      lcorbi_t3_neg_1_055_05    m/z 620.3931     0   1   2   3   4E+3   12.5   13  RT (min)      lcorbi_t3_neg_4_032_23    m/z 620.4223     0   1E+4   13   13.5  RT (min)      lcorbi_t3_neg_4_034_01    m/z 620.4464     0   1E+4   14   14.5  RT (min)      lcorbi_t3_neg_3_023_07    m/z 620.4557     0   1E+4   11.5   12  RT (min)      lcorbi_t3_neg_1_055_05    m/z 620.5206     0   1E+4   14.5   15  RT (min)      lcorbi_t3_neg_2_053_23    m/z 620.8454     0   1   2   3   4E+3   18   18.5  RT (min)      lcorbi_t3_neg_2_026_03    m/z 620.9537     0   1   2   3   4E+3   8   8.5  RT (min)      lcorbi_t3_neg_4_023_18    m/z 621.1136     0   1   2E+4   11.5   12  RT (min)      lcorbi_t3_neg_4_007_13    m/z 621.1917     0   1   2   3E+3   12   12.5  RT (min)      lcorbi_t3_neg_1_042_18    m/z 621.2106     0   2   4   6E+3   7.5   8  RT (min)      lcorbi_t3_neg_4_023_18    m/z 621.2119     0   1   2   3   4E+3   12.5   13  RT (min)      lcorbi_t3_neg_4_005_27    m/z 621.2535     0   1   2   3E+3   10   10.5  RT (min)      lcorbi_t3_neg_4_032_23    m/z 621.2745     0   1   2   3   4E+3   10   10.5  RT (min)      lcorbi_t3_neg_1_055_05    m/z 621.2950     0   1E+4   10.5   11  RT (min)      lcorbi_t3_neg_3_007_16    m/z 621.2976     0   2   4   6   8E+3   8   8.5  RT (min)      lcorbi_t3_neg_2_028_38    m/z 621.3245     0   2   4   6   8E+2   12   12.5  RT (min)      lcorbi_t3_neg_4_032_23    m/z 621.3279     0   1   2   3E+3   9.5   10  RT (min)      lcorbi_t3_neg_4_034_01    m/z 621.3760     0   1E+5   10.5   11  RT (min)      lcorbi_t3_neg_1_055_05    m/z 621.4374     0   2   4E+2   13.5   14  RT (min)      lcorbi_t3_neg_3_023_07    m/z 621.4907     0   1E+4   13   13.5  RT (min)      lcorbi_t3_neg_4_034_01    m/z 621.5003     0   1   2   3E+4   14   14.5  RT (min)      lcorbi_t3_neg_4_037_29    m/z 621.5099     0   2   4   6   8E+3   13   13.5  RT (min)      lcorbi_t3_neg_4_034_01    m/z 621.5577     0   1   2E+3   14.5   15  RT (min)      lcorbi_t3_neg_4_007_13    m/z 622.2171     0   1   2   3E+3   13   13.5  RT (min)      lcorbi_t3_neg_4_023_18    m/z 622.2819     0   1E+4   10   10.5  RT (min)      lcorbi_t3_neg_4_007_13    m/z 622.3078     0   1   2   3   4E+3   11   11.5  RT (min)      lcorbi_t3_neg_2_008_16    m/z 622.3484     0   1E+3   11.5   12  RT (min)      lcorbi_t3_neg_4_034_01    m/z 622.3603     0   1E+3   12   12.5  RT (min)      lcorbi_t3_neg_1_050_45    m/z 622.3710     0   1   2   3   4E+4   7   7.5  RT (min)      lcorbi_t3_neg_4_011_24    m/z 622.4063     0   1   14   14.5  RT (min)      lcorbi_t3_neg_4_019_08    m/z 622.4088     0   1   2E+3   12.5   13  RT (min)      lcorbi_t3_neg_1_055_05    m/z 622.4460     0   1E+4   14   14.5  RT (min)      lcorbi_t3_neg_3_023_07    m/z 622.4713     0   2   4   6E+4   12   12.5  RT (min)      lcorbi_t3_neg_1_055_05    m/z 622.4940     0   2   4   6   8E+3   13   13.5  RT (min)      lcorbi_t3_neg_4_034_01    m/z 622.5159     0   1   12.5   13  RT (min)      lcorbi_t3_neg_2_022_24    m/z 622.5780     0   2   4   6   8E+3   14.5   15  RT (min)      lcorbi_t3_neg_4_007_13    m/z 622.5908     0   1   2   3E+4   14   14.5  RT (min)      lcorbi_t3_neg_3_002_24    m/z 622.8566     0   2   4   6E+3   18   18.5  RT (min)      lcorbi_t3_neg_1_028_10    m/z 622.8919     0   2   4   6E+3   18   18.5  RT (min)      lcorbi_t3_neg_2_042_44    m/z 623.2025     0   1   2E+4   10.5   11  RT (min)      lcorbi_t3_neg_4_007_13    m/z 623.2363     0   1   2E+4   13.5   14  RT (min)      lcorbi_t3_neg_4_023_18    m/z 623.3737     0   2   4   6   8E+3   12.5   13  RT (min)      lcorbi_t3_neg_4_032_23    m/z 623.3916     0   1E+5   11   11.5  RT (min)      lcorbi_t3_neg_1_055_05    m/z 623.4028     0   1   14   14.5  RT (min)      lcorbi_t3_neg_4_019_08    m/z 623.4166     0   1E+4   13.5   14  RT (min)      lcorbi_t3_neg_4_007_13    m/z 623.4451     0   1E+4   13.5   14  RT (min)      lcorbi_t3_neg_3_007_16    m/z 623.4737     0   1E+4   13.5   14  RT (min)      lcorbi_t3_neg_2_030_47    m/z 623.4905     0   1E+4   13   13.5  RT (min)      lcorbi_t3_neg_3_018_13    m/z 623.5256     0   1E+4   13.5   14  RT (min)      lcorbi_t3_neg_4_042_14    m/z 623.5369     0   2   4E+3   13.5   14  RT (min)      lcorbi_t3_neg_2_030_47    m/z 623.9547     0   2   4   6E+3   11.5   12  RT (min)      lcorbi_t3_neg_4_007_13    m/z 624.0873     0   1   2E+3   9.5   10  RT (min)      lcorbi_t3_neg_4_053_43    m/z 624.2068     0   1E+4   10.5   11  RT (min)      lcorbi_t3_neg_4_007_13    m/z 624.2741     0   2   4   6E+2   10.5   11  RT (min)      lcorbi_t3_neg_2_022_24    m/z 624.3025     0   1   2   3   4E+2   8   8.5  RT (min)      lcorbi_t3_neg_4_034_01    m/z 624.3129     0   2   4   6   8E+3   10.5   11  RT (min)      lcorbi_t3_neg_4_007_13    m/z 624.3130     0   1E+4   13.5   14  RT (min)      lcorbi_t3_neg_4_023_18    m/z 624.3396     0   1   2E+3   11.5   12  RT (min)      lcorbi_t3_neg_2_008_16    m/z 624.3832     0   1   2E+4   10.5   11  RT (min)      lcorbi_t3_neg_1_055_05    m/z 624.4244     0   1E+4   13   13.5  RT (min)      lcorbi_t3_neg_4_034_01    m/z 624.5401     0   1   13.5   14  RT (min)      lcorbi_t3_neg_4_032_23    m/z 624.8620     0   1   2   3   4E+3   18   18.5  RT (min)      lcorbi_t3_neg_1_046_11    m/z 625.0810     0   1   2   3   4E+3   9.5   10  RT (min)      lcorbi_t3_neg_4_030_04    m/z 625.1498     0   2   4   6E+2   10   10.5  RT (min)      lcorbi_t3_neg_4_023_18    m/z 625.1602     0   1   2E+4   2.5   3  RT (min)      lcorbi_t3_neg_4_032_23    m/z 625.1605     0   2   4   6   8E+3   1   1.5  RT (min)      lcorbi_t3_neg_4_001_49    m/z 625.2348     0   1E+4   13.5   14  RT (min)      lcorbi_t3_neg_4_023_18    m/z 625.2900     0   2   4   6E+3   11   11.5  RT (min)      lcorbi_t3_neg_2_027_13    m/z 625.3066     0   1E+4   13.5   14  RT (min)      lcorbi_t3_neg_4_023_18    m/z 625.3308     0   1E+4   12.5   13  RT (min)      lcorbi_t3_neg_2_007_14    m/z 625.3412     0   1E+4   8   8.5  RT (min)      lcorbi_t3_neg_2_048_27    m/z 625.3512     0   1   2   3   4E+4   10.5   11  RT (min)      lcorbi_t3_neg_2_048_27    m/z 625.3593     0   1E+4   9   9.5  RT (min)      lcorbi_t3_neg_2_048_27    m/z 625.4072     0   1   2E+4   11.5   12  RT (min)      lcorbi_t3_neg_3_002_24    m/z 625.4280     0   1   13   13.5  RT (min)      lcorbi_t3_neg_3_040_05    m/z 625.4607     0   1E+3   14.5   15  RT (min)      lcorbi_t3_neg_2_050_17    m/z 625.5607     0   1E+4   14.5   15  RT (min)      lcorbi_t3_neg_2_039_31    m/z 625.8771     0   1   2E+3   18   18.5  RT (min)      lcorbi_t3_neg_3_003_09    m/z 626.1458     0   2   4   6   8E+3   12.5   13  RT (min)      lcorbi_t3_neg_4_007_13    m/z 626.2626     0   1   14   14.5  RT (min)      lcorbi_t3_neg_4_019_08    m/z 626.3343     0   1   2   3   4E+3   12.5   13  RT (min)      lcorbi_t3_neg_2_007_14    m/z 626.4442     0   2   4E+4   13   13.5  RT (min)      lcorbi_t3_neg_4_016_05    m/z 626.5095     0   1E+4   14.5   15  RT (min)      lcorbi_t3_neg_2_053_23    m/z 626.5477     0   1   2   3   4E+3   14.5   15  RT (min)      lcorbi_t3_neg_4_029_31    m/z 626.5731     0   1E+4   14.5   15  RT (min)      lcorbi_t3_neg_2_039_31    m/z 626.7986     0   2   4   6E+3   18   18.5  RT (min)      lcorbi_t3_neg_4_032_23    m/z 627.0370     0   2   4   6E+3   10.5   11  RT (min)      lcorbi_t3_neg_4_030_04    m/z 627.0958     0   2   4   6   8E+3   9.5   10  RT (min)      lcorbi_t3_neg_4_023_18    m/z 627.1624     0   1   2   3   4E+3   8   8.5  RT (min)      lcorbi_t3_neg_4_040_19    m/z 627.2197     0   2   4   6E+3   12.5   13  RT (min)      lcorbi_t3_neg_4_005_27    m/z 627.2275     0   1   11   11.5  RT (min)      lcorbi_t3_neg_4_019_08    m/z 627.2328     0   2   4   6E+3   13.5   14  RT (min)      lcorbi_t3_neg_1_012_20    m/z 627.2875     0   1E+4   12   12.5  RT (min)      lcorbi_t3_neg_3_007_16    m/z 627.3055     0   2   4   6E+2   11.5   12  RT (min)      lcorbi_t3_neg_3_040_05    m/z 627.3170     0   1E+4   11.5   12  RT (min)      lcorbi_t3_neg_2_020_05    m/z 627.3690     0   2   4   6   8E+3   13.5   14  RT (min)      lcorbi_t3_neg_4_040_19    m/z 627.3853     0   1E+4   12.5   13  RT (min)      lcorbi_t3_neg_4_042_14    m/z 627.4021     0   1E+4   10.5   11  RT (min)      lcorbi_t3_neg_4_029_31    m/z 627.4115     0   2   4   6E+3   11.5   12  RT (min)      lcorbi_t3_neg_2_035_46    m/z 627.4652     0   1   2   3E+3   14.5   15  RT (min)      lcorbi_t3_neg_4_045_35    m/z 627.4754     0   1E+4   13   13.5  RT (min)      lcorbi_t3_neg_2_052_32    m/z 627.5125     0   2   4   6E+3   14.5   15  RT (min)      lcorbi_t3_neg_2_053_23    m/z 627.5318     0   1   2   3   4E+4   14.5   15  RT (min)      lcorbi_t3_neg_4_019_08    m/z 627.5601     0   2   4   6E+3   14.5   15  RT (min)      lcorbi_t3_neg_1_030_47    m/z 628.0602     0   1   2   3   4E+3   5.5   6  RT (min)      lcorbi_t3_neg_4_001_49    m/z 628.0910     0   2   4   6   8E+3   6   6.5  RT (min)      lcorbi_t3_neg_3_034_21    m/z 628.1168     0   2   4   6E+4   8.5   9  RT (min)      lcorbi_t3_neg_2_048_27    m/z 628.1427     0   2   4   6E+3   12.5   13  RT (min)      lcorbi_t3_neg_4_007_13    m/z 628.1979     0   1E+4   10   10.5  RT (min)      lcorbi_t3_neg_3_045_34    m/z 628.2221     0   1   2   3E+3   11   11.5  RT (min)      lcorbi_t3_neg_2_010_02    m/z 628.2771     0   1E+4   13   13.5  RT (min)      lcorbi_t3_neg_1_023_03    m/z 628.2812     0   1E+4   12   12.5  RT (min)      lcorbi_t3_neg_3_007_16    m/z 628.3091     0   2   4   6   8E+3   13.5   14  RT (min)      lcorbi_t3_neg_1_023_03    m/z 628.3783     0   1E+4   8.5   9  RT (min)      lcorbi_t3_neg_2_048_27    m/z 628.3892     0   1   14.5   15  RT (min)      lcorbi_t3_neg_4_039_32    m/z 628.4796     0   1E+4   14.5   15  RT (min)      lcorbi_t3_neg_4_044_38    m/z 628.5249     0   1E+4   14.5   15  RT (min)      lcorbi_t3_neg_2_053_23    m/z 628.7957     0   2   4   6E+3   18   18.5  RT (min)      lcorbi_t3_neg_4_028_39    m/z 629.0426     0   2   4   6E+3   10   10.5  RT (min)      lcorbi_t3_neg_4_005_27    m/z 629.0791     0   2   4E+3   7.5   8  RT (min)      lcorbi_t3_neg_4_034_01    m/z 629.1002     0   1E+3   8   8.5  RT (min)      lcorbi_t3_neg_4_023_18    m/z 629.1693     0   1   2E+4   7   7.5  RT (min)      lcorbi_t3_neg_3_022_27    m/z 629.1756     0   1   2E+4   9   9.5  RT (min)      lcorbi_t3_neg_4_023_18    m/z 629.2052     0   1   2   3E+4   2   2.5  RT (min)      lcorbi_t3_neg_3_002_24    m/z 629.3668     0   1E+4   13   13.5  RT (min)      lcorbi_t3_neg_4_007_13    m/z 629.3861     0   1   2E+4   14   14.5  RT (min)      lcorbi_t3_neg_4_030_04    m/z 629.4009     0   2   4   6   8E+3   12.5   13  RT (min)      lcorbi_t3_neg_2_007_14    m/z 629.4144     0   1E+4   5.5   6   6.5  RT (min)      lcorbi_t3_neg_4_044_38    m/z 629.4732     0   2   4   6E+3   14.5   15  RT (min)      lcorbi_t3_neg_4_044_38    m/z 629.4806     0   1E+4   14.5   15  RT (min)      lcorbi_t3_neg_4_007_13    m/z 629.4918     0   2   4   6E+3   14.5   15  RT (min)      lcorbi_t3_neg_4_046_37    m/z 629.6553     0   1   2   3E+4   14   14.5  RT (min)      lcorbi_t3_neg_2_022_24    m/z 629.8275     0   2   4   6E+3   18   18.5  RT (min)      lcorbi_t3_neg_2_026_03    m/z 629.8756     0   2   4   6   8E+3   18   18.5  RT (min)      lcorbi_t3_neg_1_003_41    m/z 630.0583     0   2   4E+3   5.5   6  RT (min)      lcorbi_t3_neg_4_001_49    m/z 630.1719     0   2   4   6   8E+3   7   7.5  RT (min)      lcorbi_t3_neg_3_022_27    m/z 630.2360     0   1   2   3   4E+3   11   11.5  RT (min)      lcorbi_t3_neg_4_012_17    m/z 630.2966     0   1E+4   11.5   12  RT (min)      lcorbi_t3_neg_4_007_13    m/z 630.3215     0   1E+4   14   14.5  RT (min)      lcorbi_t3_neg_1_023_03    m/z 630.4043     0   2   4   6E+2   12.5   13  RT (min)      lcorbi_t3_neg_1_055_05    m/z 630.4842     0   1E+4   14.5   15  RT (min)      lcorbi_t3_neg_2_053_23    m/z 630.4953     0   1   2   3E+3   14.5   15  RT (min)      lcorbi_t3_neg_4_044_38    m/z 630.5234     0   1E+4   14.5   15  RT (min)      lcorbi_t3_neg_2_053_23    m/z 630.8714     0   2   4E+3   18   18.5  RT (min)      lcorbi_t3_neg_1_004_22    m/z 630.8847     0   1E+3   18   18.5  RT (min)      lcorbi_t3_neg_1_002_16    m/z 631.0602     0   1   2   3   4E+3   9   9.5  RT (min)      lcorbi_t3_neg_3_019_12    m/z 631.0949     0   1   2   3E+3   8   8.5  RT (min)      lcorbi_t3_neg_4_034_01    m/z 631.2423     0   1   2   3E+3   2   2.5  RT (min)      lcorbi_t3_neg_1_041_54    m/z 631.2912     0   2   4E+3   11   11.5  RT (min)      lcorbi_t3_neg_1_032_13    m/z 631.3203     0   2   4   6E+3   13   13.5  RT (min)      lcorbi_t3_neg_4_034_01    m/z 631.3413     0   2   4   6E+3   11   11.5  RT (min)      lcorbi_t3_neg_4_034_01    m/z 631.3824     0   1E+4   12   12.5  RT (min)      lcorbi_t3_neg_4_032_23    m/z 631.3953     0   1E+4   11.5   12  RT (min)      lcorbi_t3_neg_4_011_24    m/z 631.4235     0   2   4   6E+3   9   9.5  RT (min)      lcorbi_t3_neg_4_044_38    m/z 631.4345     0   1   2   3E+3   13.5   14  RT (min)      lcorbi_t3_neg_3_013_28    m/z 631.4586     0   1   2E+4   14   14.5  RT (min)      lcorbi_t3_neg_3_024_26    m/z 631.4732     0   1E+4   13   13.5  RT (min)      lcorbi_t3_neg_4_034_01    m/z 631.5085     0   2   4   6   8E+3   14   14.5  RT (min)      lcorbi_t3_neg_4_032_23    m/z 631.6437     0   1E+4   14.5   15  RT (min)      lcorbi_t3_neg_4_039_32    m/z 632.2050     0   1E+4   12   12.5  RT (min)      lcorbi_t3_neg_1_023_03    m/z 632.2429     0   2   4   6E+3   7.5   8  RT (min)      lcorbi_t3_neg_4_023_18    m/z 632.3105     0   1E+4   10.5   11  RT (min)      lcorbi_t3_neg_4_016_05    m/z 632.3365     0   1   2   3E+3   11   11.5  RT (min)      lcorbi_t3_neg_2_039_31    m/z 632.3843     0   2   4   6E+3   14.5   15  RT (min)      lcorbi_t3_neg_4_030_04    m/z 632.3930     0   1   2   3E+4   12   12.5  RT (min)      lcorbi_t3_neg_4_032_23    m/z 632.3990     0   2   4   6   8E+4   11.5   12  RT (min)      lcorbi_t3_neg_4_011_24    m/z 632.4747     0   2   4   6E+3   13   13.5  RT (min)      lcorbi_t3_neg_4_034_01    m/z 632.8170     0   1   2   3E+3   18   18.5  RT (min)      lcorbi_t3_neg_2_002_26    m/z 632.9241     0   1   2   3   4E+3   18   18.5  RT (min)      lcorbi_t3_neg_4_051_40    m/z 632.9980     0   1   2   3E+3   9   9.5  RT (min)      lcorbi_t3_neg_2_016_11    m/z 633.0511     0   1   2   3E+3   5.5   6  RT (min)      lcorbi_t3_neg_4_023_18    m/z 633.0553     0   1   2   3E+3   10.5   11  RT (min)      lcorbi_t3_neg_4_030_04    m/z 633.0910     0   2   4E+3   7.5   8  RT (min)      lcorbi_t3_neg_4_034_01    m/z 633.1120     0   1   2E+3   8   8.5  RT (min)      lcorbi_t3_neg_1_026_19    m/z 633.2895     0   1   2E+4   10   10.5  RT (min)      lcorbi_t3_neg_4_023_18    m/z 633.3407     0   1E+3   11   11.5  RT (min)      lcorbi_t3_neg_4_029_31    m/z 633.5102     0   1E+4   13.5   14  RT (min)      lcorbi_t3_neg_4_042_14    m/z 633.5298     0   1   2E+4   14   14.5  RT (min)      lcorbi_t3_neg_3_005_23    m/z 634.0360     0   1   2   3   4E+3   6   6.5  RT (min)      lcorbi_t3_neg_4_040_19    m/z 634.1162     0   2   4   6E+3   9.5   10  RT (min)      lcorbi_t3_neg_4_053_43    m/z 634.1293     0   1   2E+4   2   2.5  RT (min)      lcorbi_t3_neg_4_032_23    m/z 634.2959     0   2   4   6E+3   12.5   13  RT (min)      lcorbi_t3_neg_4_032_23    m/z 634.4271     0   2   4   6E+3   14   14.5  RT (min)      lcorbi_t3_neg_3_024_26    m/z 634.5228     0   1E+4   14.5   15  RT (min)      lcorbi_t3_neg_2_028_38    m/z 634.8232     0   2   4E+3   18   18.5  RT (min)      lcorbi_t3_neg_4_053_43    m/z 634.9951     0   1E+3   9   9.5  RT (min)      lcorbi_t3_neg_2_018_40    m/z 635.2799     0   1E+4   13.5   14  RT (min)      lcorbi_t3_neg_4_007_13    m/z 635.2906     0   1E+4   13   13.5  RT (min)      lcorbi_t3_neg_1_055_05    m/z 635.3795     0   1E+3   11.5   12  RT (min)      lcorbi_t3_neg_2_022_24    m/z 635.4144     0   1   13.5   14  RT (min)      lcorbi_t3_neg_4_040_19    m/z 635.4406     0   2   4   6E+2   13.5   14  RT (min)      lcorbi_t3_neg_1_029_21    m/z 635.4487     0   1E+4   13   13.5  RT (min)      lcorbi_t3_neg_4_034_01    m/z 635.5063     0   2   4   6   8E+3   13   13.5  RT (min)      lcorbi_t3_neg_1_052_24    m/z 635.5258     0   2   4   6E+3   14   14.5  RT (min)      lcorbi_t3_neg_2_053_23    m/z 635.6304     0   1E+4   14.5   15  RT (min)      lcorbi_t3_neg_2_052_32    m/z 636.2604     0   1   11   11.5  RT (min)      lcorbi_t3_neg_4_011_24    m/z 636.2925     0   1E+4   14   14.5  RT (min)      lcorbi_t3_neg_1_023_03    m/z 636.3119     0   2   4   6   8E+3   13.5   14  RT (min)      lcorbi_t3_neg_1_023_03    m/z 636.3333     0   1   2E+4   11   11.5  RT (min)      lcorbi_t3_neg_2_054_15    m/z 636.3881     0   2   4   6   8E+3   12   12.5  RT (min)      lcorbi_t3_neg_4_032_23    m/z 636.4485     0   1   2E+3   13   13.5  RT (min)      lcorbi_t3_neg_4_044_38    m/z 636.4781     0   2   4   6E+4   12   12.5  RT (min)      lcorbi_t3_neg_1_052_24    m/z 636.5009     0   1E+4   13   13.5  RT (min)      lcorbi_t3_neg_4_039_32    m/z 636.5573     0   2   4   6E+2   14.5   15  RT (min)      lcorbi_t3_neg_2_022_24    m/z 636.8887     0   2   4   6   8E+3   18   18.5  RT (min)      lcorbi_t3_neg_1_004_22    m/z 637.0098     0   1   2   3E+3   9   9.5  RT (min)      lcorbi_t3_neg_2_015_21    m/z 637.0899     0   1   2   3   4E+3   10   10.5  RT (min)      lcorbi_t3_neg_4_043_11    m/z 637.3202     0   1E+4   12   12.5  RT (min)      lcorbi_t3_neg_4_023_18    m/z 637.3585     0   1E+4   13.5   14  RT (min)      lcorbi_t3_neg_4_023_18    m/z 637.3749     0   1E+5   11.5   12  RT (min)      lcorbi_t3_neg_3_002_24    m/z 637.3915     0   1E+3   12   12.5  RT (min)      lcorbi_t3_neg_4_034_01    m/z 637.4084     0   1E+3   13   13.5  RT (min)      lcorbi_t3_neg_4_019_08    m/z 637.4321     0   1   2E+4   13.5   14  RT (min)      lcorbi_t3_neg_4_007_13    m/z 637.4855     0   1   2   3   4E+3   13   13.5  RT (min)      lcorbi_t3_neg_3_052_32    m/z 637.5050     0   2   4   6E+3   14.5   15  RT (min)      lcorbi_t3_neg_4_044_38    m/z 637.5415     0   1E+4   14.5   15  RT (min)      lcorbi_t3_neg_4_029_31    m/z 637.5622     0   1E+4   14.5   15  RT (min)      lcorbi_t3_neg_1_023_03    m/z 637.8306     0   1   2   3E+3   18   18.5  RT (min)      lcorbi_t3_neg_2_018_40    m/z 638.0161     0   1   2   3   4E+3   5   5.5  RT (min)      lcorbi_t3_neg_3_014_19    m/z 638.2372     0   1E+3   10.5   11  RT (min)      lcorbi_t3_neg_4_042_14    m/z 638.2589     0   1   2   3   4E+3   11   11.5  RT (min)      lcorbi_t3_neg_2_005_01    m/z 638.2689     0   1   11.5   12  RT (min)      lcorbi_t3_neg_4_011_24    m/z 638.2722     0   1E+4   14   14.5  RT (min)      lcorbi_t3_neg_4_007_13    m/z 638.2841     0   1   2E+4   14   14.5  RT (min)      lcorbi_t3_neg_4_007_13    m/z 638.3229     0   1   2   3   4E+3   12   12.5  RT (min)      lcorbi_t3_neg_4_023_18    m/z 638.3491     0   1E+4   11.5   12  RT (min)      lcorbi_t3_neg_4_011_24    m/z 638.3675     0   1   2   3E+4   11.5   12  RT (min)      lcorbi_t3_neg_4_007_13    m/z 638.4194     0   2   4   6   8E+3   14   14.5  RT (min)      lcorbi_t3_neg_3_023_07    m/z 638.4356     0   1E+4   13.5   14  RT (min)      lcorbi_t3_neg_4_007_13    m/z 638.4661     0   2   4E+3   13.5   14  RT (min)      lcorbi_t3_neg_4_034_01    m/z 638.5729     0   1E+4   14.5   15  RT (min)      lcorbi_t3_neg_4_007_13    m/z 638.5923     0   2   4   6   8E+3   14.5   15  RT (min)      lcorbi_t3_neg_4_039_32    m/z 638.6309     0   1E+4   14.5   15  RT (min)      lcorbi_t3_neg_2_052_32    m/z 638.6493     0   1   14.5   15  RT (min)      lcorbi_t3_neg_4_029_31    m/z 639.0492     0   2   4   6E+3   9   9.5  RT (min)      lcorbi_t3_neg_1_007_26    m/z 639.0823     0   1   2   3   4E+3   10   10.5  RT (min)      lcorbi_t3_neg_2_012_36    m/z 639.1291     0   1E+4   8   8.5  RT (min)      lcorbi_t3_neg_4_008_22    m/z 639.2761     0   2   4E+3   10.5   11  RT (min)      lcorbi_t3_neg_3_005_23    m/z 639.3107     0   1E+4   13.5   14  RT (min)      lcorbi_t3_neg_1_023_03    m/z 639.3689     0   2   4   6   8E+3   13.5   14  RT (min)      lcorbi_t3_neg_4_040_19    m/z 639.5012     0   1E+4   13   13.5  RT (min)      lcorbi_t3_neg_4_034_01    m/z 639.5571     0   2   4   6   8E+3   14.5   15  RT (min)      lcorbi_t3_neg_4_029_31    m/z 639.5684     0   2   4   6E+3   14.5   15  RT (min)      lcorbi_t3_neg_4_034_01    m/z 639.5764     0   1E+3   14.5   15  RT (min)      lcorbi_t3_neg_4_034_01    m/z 639.8201     0   1E+3   18   18.5  RT (min)      lcorbi_t3_neg_4_019_08    m/z 640.1112     0   1   2   3   4E+3   7.5   8  RT (min)      lcorbi_t3_neg_4_034_01    m/z 640.1262     0   2   4   6E+3   9   9.5  RT (min)      lcorbi_t3_neg_4_023_18    m/z 640.2523     0   1   2E+4   10.5   11  RT (min)      lcorbi_t3_neg_2_007_14    m/z 640.2701     0   1E+4   9.5   10  RT (min)      lcorbi_t3_neg_3_019_12    m/z 640.2922     0   1   12   12.5  RT (min)      lcorbi_t3_neg_2_020_05    m/z 640.3620     0   1E+4   11.5   12  RT (min)      lcorbi_t3_neg_4_007_13    m/z 640.5430     0   1E+4   14   14.5  RT (min)      lcorbi_t3_neg_1_055_05    m/z 640.5680     0   1E+4   10.5   11  RT (min)      lcorbi_t3_neg_4_046_37    m/z 640.6249     0   1   14   14.5  RT (min)      lcorbi_t3_neg_2_022_24    m/z 640.7109     0   1E+4   1   1.5  RT (min)      lcorbi_t3_neg_2_014_29    m/z 640.8218     0   2   4   6   8E+3   18   18.5  RT (min)      lcorbi_t3_neg_4_019_08    m/z 640.8582     0   1   2E+3   18   18.5  RT (min)      lcorbi_t3_neg_2_011_04    m/z 640.9115     0   1   2   3   4E+3   18   18.5  RT (min)      lcorbi_t3_neg_2_007_14    m/z 641.0768     0   1   2   3   4E+3   10.5   11  RT (min)      lcorbi_t3_neg_4_043_11    m/z 641.1070     0   1   2E+4   13   13.5  RT (min)      lcorbi_t3_neg_4_005_27    m/z 641.1238     0   2   4   6   8E+3   8   8.5  RT (min)      lcorbi_t3_neg_4_034_01    m/z 641.1686     0   2   4E+3   9.5   10  RT (min)      lcorbi_t3_neg_2_027_13    m/z 641.2147     0   1E+4   12.5   13  RT (min)      lcorbi_t3_neg_4_005_27    m/z 641.2903     0   1E+4   13.5   14  RT (min)      lcorbi_t3_neg_1_023_03    m/z 641.3186     0   1   2   3   4E+3   10   10.5  RT (min)      lcorbi_t3_neg_3_028_35    m/z 641.3541     0   1   2   3   4E+3   8.5   9  RT (min)      lcorbi_t3_neg_4_034_01    m/z 641.4420     0   1E+4   13.5   14  RT (min)      lcorbi_t3_neg_3_024_26    m/z 641.4909     0   1   2E+4   14   14.5  RT (min)      lcorbi_t3_neg_4_032_23    m/z 641.5838     0   1   2E+4   14.5   15  RT (min)      lcorbi_t3_neg_4_034_01    m/z 642.0600     0   1   2   3   4E+3   9.5   10  RT (min)      lcorbi_t3_neg_4_007_13    m/z 642.0735     0   1E+4   9   9.5  RT (min)      lcorbi_t3_neg_4_007_13    m/z 642.1075     0   1E+4   13   13.5  RT (min)      lcorbi_t3_neg_4_005_27    m/z 642.2181     0   1   2   3   4E+3   12.5   13  RT (min)      lcorbi_t3_neg_4_005_27    m/z 642.2866     0   2   4E+3   12   12.5  RT (min)      lcorbi_t3_neg_4_023_18    m/z 642.3774     0   2   4   6   8E+3   12   12.5  RT (min)      lcorbi_t3_neg_4_007_13    m/z 642.3846     0   1   2E+3   13.5   14  RT (min)      lcorbi_t3_neg_1_010_17    m/z 642.4223     0   1E+4   13.5   14  RT (min)      lcorbi_t3_neg_4_029_31    m/z 642.5679     0   2   4   6   8E+3   14.5   15  RT (min)      lcorbi_t3_neg_2_043_33    m/z 642.5872     0   1E+4   14.5   15  RT (min)      lcorbi_t3_neg_4_034_01    m/z 642.8365     0   1   2   3E+3   18   18.5  RT (min)      lcorbi_t3_neg_3_016_42    m/z 643.1171     0   1E+4   1   1.5  RT (min)      lcorbi_t3_neg_4_029_31    m/z 643.1345     0   1E+4   9   9.5  RT (min)      lcorbi_t3_neg_4_023_18    m/z 643.1409     0   1E+4   8   8.5  RT (min)      lcorbi_t3_neg_4_040_19    m/z 643.2119     0   1   2   3   4E+3   12.5   13  RT (min)      lcorbi_t3_neg_4_005_27    m/z 643.2884     0   1   2E+4   6   6.5  RT (min)      lcorbi_t3_neg_3_007_16    m/z 643.2916     0   1   2E+4   13.5   14  RT (min)      lcorbi_t3_neg_4_023_18    m/z 643.3809     0   1   2   3   4E+2   11.5   12  RT (min)      lcorbi_t3_neg_3_023_07    m/z 643.3981     0   2   4   6   8E+2   13   13.5  RT (min)      lcorbi_t3_neg_4_021_25    m/z 643.4240     0   1   2E+3   13.5   14  RT (min)      lcorbi_t3_neg_1_020_36    m/z 643.4711     0   2   4   6   8E+3   13.5   14  RT (min)      lcorbi_t3_neg_4_038_15    m/z 643.4906     0   1E+4   14.5   15  RT (min)      lcorbi_t3_neg_1_055_05    m/z 643.6438     0   1E+4   14.5   15  RT (min)      lcorbi_t3_neg_4_039_32    m/z 644.0113     0   1E+4   11.5   12  RT (min)      lcorbi_t3_neg_4_039_32    m/z 644.0301     0   1   2E+3   9   9.5  RT (min)      lcorbi_t3_neg_2_018_40    m/z 644.1156     0   2   4   6E+3   11.5   12  RT (min)      lcorbi_t3_neg_4_007_13    m/z 644.1198     0   1E+4   2.5   3  RT (min)      lcorbi_t3_neg_4_032_23    m/z 644.1287     0   1   2   3   4E+4   10.5   11  RT (min)      lcorbi_t3_neg_1_043_14    m/z 644.1380     0   2   4E+3   9   9.5  RT (min)      lcorbi_t3_neg_4_023_18    m/z 644.1442     0   1   2   3   4E+3   8   8.5  RT (min)      lcorbi_t3_neg_4_040_19    m/z 644.1957     0   2   4E+3   11   11.5  RT (min)      lcorbi_t3_neg_4_032_23    m/z 644.2735     0   2   4   6E+3   10.5   11  RT (min)      lcorbi_t3_neg_4_016_05    m/z 644.2917     0   2   4   6   8E+3   6   6.5  RT (min)      lcorbi_t3_neg_4_012_17    m/z 644.3930     0   1E+4   12   12.5  RT (min)      lcorbi_t3_neg_4_007_13    m/z 644.5202     0   1E+4   14.5   15  RT (min)      lcorbi_t3_neg_2_053_23    m/z 644.5393     0   2   4   6E+3   14.5   15  RT (min)      lcorbi_t3_neg_2_053_23    m/z 644.6188     0   1E+4   14.5   15  RT (min)      lcorbi_t3_neg_4_011_24    m/z 644.6559     0   1E+4   14.5   15  RT (min)      lcorbi_t3_neg_4_039_32    m/z 645.0239     0   1   2   3E+3   9   9.5  RT (min)      lcorbi_t3_neg_4_028_39    m/z 645.1126     0   1E+4   11.5   12  RT (min)      lcorbi_t3_neg_4_007_13    m/z 645.1318     0   2   4   6   8E+3   11.5   12  RT (min)      lcorbi_t3_neg_4_007_13    m/z 645.2149     0   1   2   3E+3   10   10.5  RT (min)      lcorbi_t3_neg_2_039_31    m/z 645.2272     0   1   2   3   4E+4   2.5   3  RT (min)      lcorbi_t3_neg_4_001_49    m/z 645.2445     0   2   4   6E+3   9.5   10  RT (min)      lcorbi_t3_neg_4_017_51    m/z 645.2507     0   1E+4   5   5.5  RT (min)      lcorbi_t3_neg_4_017_51    m/z 645.2630     0   1   2   3   4E+3   12.5   13  RT (min)      lcorbi_t3_neg_2_002_26    m/z 645.2942     0   2   4   6   8E+3   13.5   14  RT (min)      lcorbi_t3_neg_4_023_18    m/z 645.3044     0   2   4   6E+3   10.5   11  RT (min)      lcorbi_t3_neg_3_053_14    m/z 645.3617     0   1E+4   13   13.5  RT (min)      lcorbi_t3_neg_4_007_13    m/z 645.3887     0   1E+4   11   11.5  RT (min)      lcorbi_t3_neg_1_055_05    m/z 645.4008     0   2   4E+3   12.5   13  RT (min)      lcorbi_t3_neg_3_002_24    m/z 645.4108     0   1   13.5   14  RT (min)      lcorbi_t3_neg_4_019_08    m/z 645.4676     0   1   2E+5   14.5   15  RT (min)      lcorbi_t3_neg_4_052_26    m/z 645.4869     0   2   4   6E+2   13.5   14  RT (min)      lcorbi_t3_neg_1_022_34    m/z 645.5052     0   1E+4   14   14.5  RT (min)      lcorbi_t3_neg_1_055_05    m/z 645.8543     0   2   4   6E+3   18   18.5  RT (min)      lcorbi_t3_neg_2_003_42    m/z 646.0085     0   2   4   6E+3   11.5   12  RT (min)      lcorbi_t3_neg_4_039_32    m/z 646.1128     0   2   4   6E+3   11.5   12  RT (min)      lcorbi_t3_neg_4_007_13    m/z 646.1286     0   1   2E+3   13   13.5  RT (min)      lcorbi_t3_neg_4_007_13    m/z 646.2116     0   1   11.5   12  RT (min)      lcorbi_t3_neg_4_019_08    m/z 646.2206     0   1   2E+4   12   12.5  RT (min)      lcorbi_t3_neg_1_023_03    m/z 646.2573     0   2   4   6E+3   13.5   14  RT (min)      lcorbi_t3_neg_3_023_07    m/z 646.2707     0   2   4   6E+4   10   10.5  RT (min)      lcorbi_t3_neg_3_010_15    m/z 646.2976     0   1   2E+3   13.5   14  RT (min)      lcorbi_t3_neg_3_023_07    m/z 646.3606     0   1   2   3E+4   10.5   11  RT (min)      lcorbi_t3_neg_2_022_24    m/z 646.3652     0   1   2   3   4E+3   13   13.5  RT (min)      lcorbi_t3_neg_4_007_13    m/z 646.3920     0   1   2   3   4E+3   11   11.5  RT (min)      lcorbi_t3_neg_1_055_05    m/z 646.4383     0   2   4   6E+3   13.5   14  RT (min)      lcorbi_t3_neg_4_019_08    m/z 646.4903     0   1   13.5   14  RT (min)      lcorbi_t3_neg_1_022_34    m/z 646.5418     0   1   14   14.5  RT (min)      lcorbi_t3_neg_4_052_26    m/z 646.6175     0   2   4   6   8E+3   1   1.5  RT (min)      lcorbi_t3_neg_4_036_16    m/z 647.1289     0   2   4   6E+3   11.5   12  RT (min)      lcorbi_t3_neg_4_007_13    m/z 647.2241     0   2   4E+3   12   12.5  RT (min)      lcorbi_t3_neg_1_023_03    m/z 647.2700     0   2   4   6E+3   12.5   13  RT (min)      lcorbi_t3_neg_3_007_16    m/z 647.3047     0   2   4   6   8E+3   8   8.5  RT (min)      lcorbi_t3_neg_2_048_27    m/z 647.3198     0   2   4   6   8E+3   12.5   13  RT (min)      lcorbi_t3_neg_4_017_51    m/z 647.3831     0   1   14   14.5  RT (min)      lcorbi_t3_neg_2_024_37    m/z 647.4456     0   1E+4   14   14.5  RT (min)      lcorbi_t3_neg_2_028_38    m/z 647.4657     0   1   2   3E+3   14   14.5  RT (min)      lcorbi_t3_neg_2_055_18    m/z 647.4807     0   1   2   3E+4   14   14.5  RT (min)      lcorbi_t3_neg_3_024_26    m/z 647.6680     0   1   2E+4   14.5   15  RT (min)      lcorbi_t3_neg_2_052_32    m/z 647.6746     0   1   2E+4   14.5   15  RT (min)      lcorbi_t3_neg_4_039_32    m/z 648.0420     0   1   2E+3   9   9.5  RT (min)      lcorbi_t3_neg_4_055_06    m/z 648.1931     0   1   2E+4   10.5   11  RT (min)      lcorbi_t3_neg_1_043_14    m/z 648.2087     0   1   10   10.5  RT (min)      lcorbi_t3_neg_4_019_08    m/z 648.2229     0   1E+4   9   9.5  RT (min)      lcorbi_t3_neg_3_019_12    m/z 648.2483     0   1E+4   13.5   14  RT (min)      lcorbi_t3_neg_4_040_19    m/z 648.3047     0   2   4   6E+3   11.5   12  RT (min)      lcorbi_t3_neg_2_027_13    m/z 648.3490     0   1   2   3E+3   12   12.5  RT (min)      lcorbi_t3_neg_4_034_01    m/z 648.3578     0   2   4   6   8E+3   10.5   11  RT (min)      lcorbi_t3_neg_2_020_05    m/z 648.3764     0   1   2E+4   11   11.5  RT (min)      lcorbi_t3_neg_3_002_24    m/z 648.3868     0   1   2E+4   12   12.5  RT (min)      lcorbi_t3_neg_4_032_23    m/z 648.4322     0   2   4   6E+3   13   13.5  RT (min)      lcorbi_t3_neg_4_034_01    m/z 649.0635     0   1   2   3E+3   7.5   8  RT (min)      lcorbi_t3_neg_4_034_01    m/z 649.0780     0   1   2   3E+3   9   9.5  RT (min)      lcorbi_t3_neg_1_042_18    m/z 649.0866     0   1   2E+3   9.5   10  RT (min)      lcorbi_t3_neg_4_003_21    m/z 649.1111     0   1E+4   10   10.5  RT (min)      lcorbi_t3_neg_4_029_31    m/z 649.2072     0   1E+5   9.5   10  RT (min)      lcorbi_t3_neg_4_019_08    m/z 649.2089     0   1   2   3E+3   12.5   13  RT (min)      lcorbi_t3_neg_4_007_13    m/z 649.2864     0   1E+4   12   12.5  RT (min)      lcorbi_t3_neg_4_023_18    m/z 649.2942     0   1   2   3   4E+3   11   11.5  RT (min)      lcorbi_t3_neg_2_027_13    m/z 649.3088     0   1   2   3   4E+3   11   11.5  RT (min)      lcorbi_t3_neg_2_027_13    m/z 649.3122     0   2   4   6   8E+3   13.5   14  RT (min)      lcorbi_t3_neg_4_023_18    m/z 649.3488     0   1   13   13.5  RT (min)      lcorbi_t3_neg_1_026_19    m/z 649.4606     0   1E+3   14.5   15  RT (min)      lcorbi_t3_neg_4_034_01    m/z 649.8571     0   1   2   3   4E+3   18   18.5  RT (min)      lcorbi_t3_neg_2_045_30    m/z 650.1612     0   2   4   6E+3   8   8.5  RT (min)      lcorbi_t3_neg_4_040_19    m/z 650.2058     0   1   2E+3   10   10.5  RT (min)      lcorbi_t3_neg_2_011_04    m/z 650.2242     0   1   2   3E+3   11   11.5  RT (min)      lcorbi_t3_neg_2_010_02    m/z 650.2896     0   2   4   6E+3   12   12.5  RT (min)      lcorbi_t3_neg_4_007_13    m/z 650.3205     0   1   2   3E+3   11.5   12  RT (min)      lcorbi_t3_neg_4_034_01    m/z 650.3650     0   1   2E+3   12.5   13  RT (min)      lcorbi_t3_neg_2_053_23    m/z 650.3919     0   1   2E+4   11.5   12  RT (min)      lcorbi_t3_neg_1_055_05    m/z 650.4014     0   1E+3   13   13.5  RT (min)      lcorbi_t3_neg_2_053_23    m/z 650.4017     0   1E+4   10   10.5  RT (min)      lcorbi_t3_neg_4_039_32    m/z 650.4040     0   1E+4   11.5   12  RT (min)      lcorbi_t3_neg_4_032_23    m/z 650.4403     0   2   4   6   8E+3   13   13.5  RT (min)      lcorbi_t3_neg_4_007_13    m/z 650.4643     0   1E+4   14.5   15  RT (min)      lcorbi_t3_neg_1_055_05    m/z 650.5174     0   2   4   6E+4   14.5   15  RT (min)      lcorbi_t3_neg_3_005_23    m/z 650.6096     0   1   15   15.5  RT (min)      lcorbi_t3_neg_3_002_24    m/z 651.0965     0   1   9   9.5  RT (min)      lcorbi_t3_neg_4_019_08    m/z 651.1056     0   1E+4   10.5   11  RT (min)      lcorbi_t3_neg_4_043_11    m/z 651.1574     0   1   2E+4   9   9.5  RT (min)      lcorbi_t3_neg_4_023_18    m/z 651.2375     0   1   2   3E+3   10   10.5  RT (min)      lcorbi_t3_neg_3_019_12    m/z 651.2835     0   1   2   3   4E+3   12   12.5  RT (min)      lcorbi_t3_neg_4_023_18    m/z 651.3101     0   1   2   3   4E+3   11.5   12  RT (min)      lcorbi_t3_neg_4_044_38    m/z 651.3242     0   2   4E+3   11.5   12  RT (min)      lcorbi_t3_neg_4_007_13    m/z 651.3712     0   2   4   6E+4   11   11.5  RT (min)      lcorbi_t3_neg_4_032_23    m/z 651.4115     0   1E+4   13.5   14  RT (min)      lcorbi_t3_neg_3_007_16    m/z 651.4373     0   2   4   6E+3   13   13.5  RT (min)      lcorbi_t3_neg_2_007_14    m/z 651.4614     0   1   2E+4   13.5   14  RT (min)      lcorbi_t3_neg_4_040_19    m/z 651.5367     0   1   2E+4   14.5   15  RT (min)      lcorbi_t3_neg_2_053_23    m/z 651.5591     0   1   2E+4   14   14.5  RT (min)      lcorbi_t3_neg_1_044_23    m/z 651.6116     0   1   2   3E+4   14.5   15  RT (min)      lcorbi_t3_neg_3_002_24    m/z 652.1900     0   1E+4   2   2.5  RT (min)      lcorbi_t3_neg_2_022_24    m/z 652.2194     0   1   2   3E+4   10.5   11  RT (min)      lcorbi_t3_neg_4_042_14    m/z 652.2373     0   1   2   3E+3   12.5   13  RT (min)      lcorbi_t3_neg_3_007_16    m/z 652.2610     0   1   11   11.5  RT (min)      lcorbi_t3_neg_4_032_23    m/z 652.2682     0   1   12   12.5  RT (min)      lcorbi_t3_neg_4_012_17    m/z 652.2690     0   1E+4   9   9.5  RT (min)      lcorbi_t3_neg_4_023_18    m/z 652.2945     0   1E+4   5   5.5  RT (min)      lcorbi_t3_neg_4_044_38    m/z 652.2972     0   2   4   6E+3   11.5   12  RT (min)      lcorbi_t3_neg_2_007_14    m/z 652.3254     0   2   4   6   8E+3   11.5   12  RT (min)      lcorbi_t3_neg_4_004_34    m/z 652.3726     0   1   2   3E+3   18   18.5  RT (min)      lcorbi_t3_neg_4_030_04    m/z 652.4730     0   2   4   6E+4   11.5   12  RT (min)      lcorbi_t3_neg_3_002_24    m/z 652.5696     0   2   4   6   8E+3   14.5   15  RT (min)      lcorbi_t3_neg_2_053_23    m/z 652.5887     0   2   4E+3   14.5   15  RT (min)      lcorbi_t3_neg_4_007_13    m/z 652.8719     0   2   4E+3   18   18.5  RT (min)      lcorbi_t3_neg_4_022_41    m/z 652.9822     0   1   9   9.5  RT (min)      lcorbi_t3_neg_4_019_08    m/z 653.1744     0   1   9   9.5  RT (min)      lcorbi_t3_neg_1_023_03    m/z 653.2167     0   2   4   6   8E+3   10.5   11  RT (min)      lcorbi_t3_neg_4_034_01    m/z 653.2407     0   1   2   3   4E+4   9   9.5  RT (min)      lcorbi_t3_neg_3_019_12    m/z 653.2983     0   2   4   6   8E+3   12   12.5  RT (min)      lcorbi_t3_neg_4_023_18    m/z 653.3265     0   2   4   6   8E+3   12   12.5  RT (min)      lcorbi_t3_neg_4_034_01    m/z 653.3396     0   2   4   6   8E+3   12   12.5  RT (min)      lcorbi_t3_neg_4_038_15    m/z 653.3496     0   2   4   6E+3   13.5   14  RT (min)      lcorbi_t3_neg_4_040_19    m/z 653.3650     0   1   2   3E+3   13.5   14  RT (min)      lcorbi_t3_neg_4_023_18    m/z 653.4229     0   2   4   6E+3   12.5   13  RT (min)      lcorbi_t3_neg_4_005_27    m/z 653.4596     0   1E+4   13.5   14  RT (min)      lcorbi_t3_neg_4_007_13    m/z 653.4733     0   2   4   6E+3   14.5   15  RT (min)      lcorbi_t3_neg_1_055_05    m/z 653.5001     0   1E+4   14   14.5  RT (min)      lcorbi_t3_neg_4_032_23    m/z 653.5171     0   1   14   14.5  RT (min)      lcorbi_t3_neg_2_050_17    m/z 653.5553     0   2   4   6E+3   14.5   15  RT (min)      lcorbi_t3_neg_2_053_23    m/z 653.5843     0   1   2   3E+3   14.5   15  RT (min)      lcorbi_t3_neg_3_020_31    m/z 654.0589     0   2   4E+3   9   9.5  RT (min)      lcorbi_t3_neg_3_019_12    m/z 654.2697     0   2   4   6   8E+3   13.5   14  RT (min)      lcorbi_t3_neg_4_017_51    m/z 654.3349     0   2   4   6   8E+3   13.5   14  RT (min)      lcorbi_t3_neg_3_005_23    m/z 654.3463     0   2   4   6E+2   13.5   14  RT (min)      lcorbi_t3_neg_1_048_12    m/z 654.5488     0   2   4   6E+3   14.5   15  RT (min)      lcorbi_t3_neg_4_046_37    m/z 654.6449     0   1   14.5   15  RT (min)      lcorbi_t3_neg_4_044_38    m/z 654.8070     0   1   2   3   4E+3   18   18.5  RT (min)      lcorbi_t3_neg_4_028_39    m/z 654.8610     0   1   2   3E+3   18   18.5  RT (min)      lcorbi_t3_neg_4_028_39    m/z 654.9084     0   1   2   3   4E+3   18   18.5  RT (min)      lcorbi_t3_neg_4_050_02    m/z 655.0218     0   1   2   3   4E+3   9   9.5  RT (min)      lcorbi_t3_neg_4_042_14    m/z 655.0925     0   2   4   6E+3   11   11.5  RT (min)      lcorbi_t3_neg_4_043_11    m/z 655.1083     0   1   2E+4   2   2.5  RT (min)      lcorbi_t3_neg_4_032_23    m/z 655.1478     0   1   2   3   4E+3   9.5   10  RT (min)      lcorbi_t3_neg_4_046_37    m/z 655.1590     0   1   2   3   4E+4   12.5   13  RT (min)      lcorbi_t3_neg_4_045_35    m/z 655.1894     0   2   4   6   8E+3   2   2.5  RT (min)      lcorbi_t3_neg_2_022_24    m/z 655.2280     0   2   4   6E+3   11   11.5  RT (min)      lcorbi_t3_neg_4_032_23    m/z 655.2859     0   1   2E+5   14   14.5  RT (min)      lcorbi_t3_neg_4_019_08    m/z 655.3059     0   2   4   6   8E+3   10.5   11  RT (min)      lcorbi_t3_neg_4_007_13    m/z 655.3187     0   1   12.5   13  RT (min)      lcorbi_t3_neg_1_015_27    m/z 655.3432     0   2   4   6E+2   13.5   14  RT (min)      lcorbi_t3_neg_1_048_12    m/z 655.3861     0   1   2   3   4E+4   10.5   11  RT (min)      lcorbi_t3_neg_1_055_05    m/z 655.3866     0   2   4   6E+3   13.5   14  RT (min)      lcorbi_t3_neg_4_052_26    m/z 655.4890     0   1E+4   14.5   15  RT (min)      lcorbi_t3_neg_4_029_31    m/z 656.3632     0   2   4E+3   12   12.5  RT (min)      lcorbi_t3_neg_4_023_18    m/z 656.3767     0   2   4   6   8E+2   12   12.5  RT (min)      lcorbi_t3_neg_4_019_08    m/z 656.3960     0   2   4   6E+3   13   13.5  RT (min)      lcorbi_t3_neg_4_007_13    m/z 656.4382     0   1E+4   13.5   14  RT (min)      lcorbi_t3_neg_4_046_37    m/z 656.4925     0   2   4   6E+3   14.5   15  RT (min)      lcorbi_t3_neg_4_011_24    m/z 656.5755     0   1   2   3E+3   15   15.5  RT (min)      lcorbi_t3_neg_2_039_31    m/z 656.8074     0   1   2   3   4E+3   18   18.5  RT (min)      lcorbi_t3_neg_4_023_18    m/z 656.8246     0   1E+3   18   18.5  RT (min)      lcorbi_t3_neg_4_019_08    m/z 656.8849     0   2   4   6   8E+3   9.5   10  RT (min)      lcorbi_t3_neg_3_019_12    m/z 657.2215     0   1   2E+4   10.5   11  RT (min)      lcorbi_t3_neg_2_048_27    m/z 657.2440     0   2   4   6E+3   11.5   12  RT (min)      lcorbi_t3_neg_1_055_05    m/z 657.3018     0   1E+4   14   14.5  RT (min)      lcorbi_t3_neg_1_023_03    m/z 657.3115     0   1E+4   13.5   14  RT (min)      lcorbi_t3_neg_1_023_03    m/z 657.3312     0   1   2   3E+3   8.5   9  RT (min)      lcorbi_t3_neg_4_034_01    m/z 657.3600     0   1E+4   12   12.5  RT (min)      lcorbi_t3_neg_4_023_18    m/z 657.3852     0   2   4   6E+3   8   8.5  RT (min)      lcorbi_t3_neg_2_048_27    m/z 657.3868     0   2   4E+3   18   18.5  RT (min)      lcorbi_t3_neg_4_043_11    m/z 657.4018     0   1   2   3   4E+4   11   11.5  RT (min)      lcorbi_t3_neg_2_022_24    m/z 657.4679     0   1   14.5   15  RT (min)      lcorbi_t3_neg_4_039_32    m/z 657.4864     0   1E+4   14.5   15  RT (min)      lcorbi_t3_neg_4_011_24    m/z 657.4977     0   1   2   3   4E+3   13.5   14  RT (min)      lcorbi_t3_neg_1_022_34    m/z 657.5032     0   2   4   6   8E+3   14.5   15  RT (min)      lcorbi_t3_neg_1_055_05    m/z 658.1052     0   1E+3   8   8.5  RT (min)      lcorbi_t3_neg_3_054_18    m/z 658.2447     0   1   2   3E+3   11.5   12  RT (min)      lcorbi_t3_neg_2_026_03    m/z 658.2891     0   2   4E+4   14   14.5  RT (min)      lcorbi_t3_neg_4_030_04    m/z 658.3411     0   2   4   6E+3   13   13.5  RT (min)      lcorbi_t3_neg_4_005_27    m/z 658.3631     0   1   2   3E+3   12   12.5  RT (min)      lcorbi_t3_neg_4_023_18    m/z 658.3794     0   2   4   6E+3   12   12.5  RT (min)      lcorbi_t3_neg_4_052_26    m/z 658.3830     0   1E+4   13.5   14  RT (min)      lcorbi_t3_neg_3_024_26    m/z 658.4457     0   1E+4   13.5   14  RT (min)      lcorbi_t3_neg_2_039_31    m/z 658.4899     0   1E+3   13   13.5  RT (min)      lcorbi_t3_neg_2_022_24    m/z 658.5015     0   1   13.5   14  RT (min)      lcorbi_t3_neg_4_044_38    m/z 658.5154     0   2   4   6E+3   14.5   15  RT (min)      lcorbi_t3_neg_4_007_13    m/z 658.8120     0   2   4E+3   18   18.5  RT (min)      lcorbi_t3_neg_4_040_19    m/z 659.1065     0   1E+4   9   9.5  RT (min)      lcorbi_t3_neg_4_023_18    m/z 659.1190     0   2   4   6   8E+3   10   10.5  RT (min)      lcorbi_t3_neg_3_037_36    m/z 659.1284     0   1   2E+4   8   8.5  RT (min)      lcorbi_t3_neg_4_052_26    m/z 659.2083     0   1E+4   13.5   14  RT (min)      lcorbi_t3_neg_1_042_18    m/z 659.2408     0   1   2E+3   10   10.5  RT (min)      lcorbi_t3_neg_4_032_23    m/z 659.2805     0   1E+4   13.5   14  RT (min)      lcorbi_t3_neg_1_023_03    m/z 659.2840     0   1   2   3   4E+4   8   8.5  RT (min)      lcorbi_t3_neg_2_048_27    m/z 659.2925     0   1E+4   14   14.5  RT (min)      lcorbi_t3_neg_4_030_04    m/z 659.3267     0   1   11   11.5  RT (min)      lcorbi_t3_neg_1_055_05    m/z 659.3517     0   1   2   3   4E+3   13.5   14  RT (min)      lcorbi_t3_neg_4_042_14    m/z 659.4173     0   1   2E+4   11.5   12  RT (min)      lcorbi_t3_neg_1_055_05    m/z 659.4561     0   1   2   3   4E+4   13.5   14  RT (min)      lcorbi_t3_neg_4_019_08    m/z 659.4897     0   1E+3   14.5   15  RT (min)      lcorbi_t3_neg_4_039_32    m/z 659.8927     0   1   2E+3   18   18.5  RT (min)      lcorbi_t3_neg_2_011_04    m/z 660.1099     0   1   2   3E+3   9   9.5  RT (min)      lcorbi_t3_neg_4_023_18    m/z 660.2346     0   1   2   3E+3   10   10.5  RT (min)      lcorbi_t3_neg_3_019_12    m/z 660.2861     0   1E+4   14   14.5  RT (min)      lcorbi_t3_neg_4_030_04    m/z 660.3397     0   1   2   3   4E+3   11.5   12  RT (min)      lcorbi_t3_neg_3_018_13    m/z 660.3811     0   1E+4   13   13.5  RT (min)      lcorbi_t3_neg_4_034_01    m/z 660.4062     0   1E+4   13   13.5  RT (min)      lcorbi_t3_neg_4_042_14    m/z 660.4275     0   1E+4   13   13.5  RT (min)      lcorbi_t3_neg_4_007_13    m/z 660.4613     0   1E+4   13.5   14  RT (min)      lcorbi_t3_neg_4_019_08    m/z 660.5164     0   1   13.5   14  RT (min)      lcorbi_t3_neg_2_053_23    m/z 660.8246     0   2   4   6E+3   18   18.5  RT (min)      lcorbi_t3_neg_3_044_01    m/z 661.1956     0   1   2   3E+4   7.5   8  RT (min)      lcorbi_t3_neg_3_039_29    m/z 661.2054     0   1E+4   13.5   14  RT (min)      lcorbi_t3_neg_4_023_18    m/z 661.2377     0   2   4   6E+3   12.5   13  RT (min)      lcorbi_t3_neg_4_007_13    m/z 661.2396     0   2   4   6   8E+3   10.5   11  RT (min)      lcorbi_t3_neg_4_037_29    m/z 661.2476     0   1   2   3E+3   12.5   13  RT (min)      lcorbi_t3_neg_2_007_14    m/z 661.2565     0   2   4   6E+3   11   11.5  RT (min)      lcorbi_t3_neg_3_032_17    m/z 661.3214     0   2   4   6E+3   12   12.5  RT (min)      lcorbi_t3_neg_4_023_18    m/z 661.3527     0   2   4   6E+3   11.5   12  RT (min)      lcorbi_t3_neg_1_055_05    m/z 661.4083     0   2   4   6E+3   13   13.5  RT (min)      lcorbi_t3_neg_4_042_14    m/z 661.4307     0   1E+3   13   13.5  RT (min)      lcorbi_t3_neg_4_019_08    m/z 661.4724     0   1E+4   14   14.5  RT (min)      lcorbi_t3_neg_3_005_23    m/z 661.4834     0   2   4   6E+3   13   13.5  RT (min)      lcorbi_t3_neg_4_034_01    m/z 661.5051     0   2   4   6   8E+3   14.5   15  RT (min)      lcorbi_t3_neg_4_044_38    m/z 661.6471     0   1   2   3E+4   14   14.5  RT (min)      lcorbi_t3_neg_4_011_24    m/z 661.8636     0   1   2   3   4E+3   18   18.5  RT (min)      lcorbi_t3_neg_3_016_42    m/z 662.0030     0   1   2   3E+3   9   9.5  RT (min)      lcorbi_t3_neg_4_007_13    m/z 662.1803     0   1   2E+4   2.5   3  RT (min)      lcorbi_t3_neg_3_002_24    m/z 662.1983     0   1E+4   7.5   8  RT (min)      lcorbi_t3_neg_3_039_29    m/z 662.1992     0   1E+4   13.5   14  RT (min)      lcorbi_t3_neg_1_042_18    m/z 662.2492     0   2   4   6   8E+3   10   10.5  RT (min)      lcorbi_t3_neg_3_019_12    m/z 662.2687     0   1   2   3E+3   11.5   12  RT (min)      lcorbi_t3_neg_4_042_14    m/z 662.2897     0   1E+4   11   11.5  RT (min)      lcorbi_t3_neg_4_034_01    m/z 662.3311     0   1   2E+4   10.5   11  RT (min)      lcorbi_t3_neg_2_048_27    m/z 662.3869     0   2   4E+3   10   10.5  RT (min)      lcorbi_t3_neg_4_029_31    m/z 662.4363     0   1   2   3E+4   12   12.5  RT (min)      lcorbi_t3_neg_1_055_05    m/z 662.4767     0   1E+4   13.5   14  RT (min)      lcorbi_t3_neg_4_007_13    m/z 662.4848     0   2   4   6E+4   11.5   12  RT (min)      lcorbi_t3_neg_1_052_24    m/z 662.5731     0   1   2   3E+3   14.5   15  RT (min)      lcorbi_t3_neg_2_052_32    m/z 662.5851     0   2   4   6   8E+3   14.5   15  RT (min)      lcorbi_t3_neg_1_055_05    m/z 662.6093     0   1   2   3E+3   14.5   15  RT (min)      lcorbi_t3_neg_4_029_31    m/z 662.8293     0   2   4   6E+3   18   18.5  RT (min)      lcorbi_t3_neg_2_048_27    m/z 663.0111     0   1   2   3E+3   9   9.5  RT (min)      lcorbi_t3_neg_3_019_12    m/z 663.0435     0   1   2   3   4E+3   10   10.5  RT (min)      lcorbi_t3_neg_4_029_31    m/z 663.1840     0   1   2E+4   1   1.5  RT (min)      lcorbi_t3_neg_1_055_05    m/z 663.2529     0   1   2E+3   10   10.5  RT (min)      lcorbi_t3_neg_3_019_12    m/z 663.2595     0   1E+4   10.5   11  RT (min)      lcorbi_t3_neg_2_048_27    m/z 663.2723     0   1   11.5   12  RT (min)      lcorbi_t3_neg_1_055_05    m/z 663.2932     0   2   4E+3   11   11.5  RT (min)      lcorbi_t3_neg_4_034_01    m/z 663.3296     0   1E+4   11.5   12  RT (min)      lcorbi_t3_neg_4_042_14    m/z 663.3388     0   1E+4   8   8.5  RT (min)      lcorbi_t3_neg_2_048_27    m/z 663.3683     0   2   4   6E+3   12   12.5  RT (min)      lcorbi_t3_neg_4_034_01    m/z 663.3922     0   1   2E+3   13   13.5  RT (min)      lcorbi_t3_neg_4_054_36    m/z 663.4034     0   2   4   6E+2   13   13.5  RT (min)      lcorbi_t3_neg_4_034_01    m/z 663.4453     0   1   2   3E+3   13.5   14  RT (min)      lcorbi_t3_neg_4_005_27    m/z 663.4717     0   2   4   6E+3   12.5   13  RT (min)      lcorbi_t3_neg_4_007_13    m/z 663.4746     0   2   4   6E+3   14.5   15  RT (min)      lcorbi_t3_neg_4_001_49    m/z 663.5207     0   2   4   6E+3   14.5   15  RT (min)      lcorbi_t3_neg_4_044_38    m/z 663.5377     0   1   2E+4   12.5   13  RT (min)      lcorbi_t3_neg_1_055_05    m/z 663.5790     0   2   4   6   8E+3   14.5   15  RT (min)      lcorbi_t3_neg_1_023_03    m/z 663.5970     0   1E+4   14.5   15  RT (min)      lcorbi_t3_neg_3_002_24    m/z 663.8465     0   1E+4   18   18.5  RT (min)      lcorbi_t3_neg_4_019_08    m/z 663.9968     0   2   4   6E+3   11.5   12  RT (min)      lcorbi_t3_neg_4_007_13    m/z 664.0559     0   1   2E+4   2   2.5  RT (min)      lcorbi_t3_neg_1_044_23    m/z 664.0693     0   1   2   3   4E+3   6   6.5  RT (min)      lcorbi_t3_neg_4_040_19    m/z 664.1070     0   1E+4   1.5   2  RT (min)      lcorbi_t3_neg_2_012_36    m/z 664.1173     0   1   2   3E+4   2   2.5  RT (min)      lcorbi_t3_neg_4_044_38    m/z 664.2000     0   2   4   6E+3   12.5   13  RT (min)      lcorbi_t3_neg_4_005_27    m/z 664.2559     0   2   4   6E+3   11   11.5  RT (min)      lcorbi_t3_neg_3_037_36    m/z 664.2610     0   1E+4   10.5   11  RT (min)      lcorbi_t3_neg_4_007_13    m/z 664.2843     0   1E+4   11.5   12  RT (min)      lcorbi_t3_neg_4_034_01    m/z 664.3600     0   1   2   3E+4   10.5   11  RT (min)      lcorbi_t3_neg_4_011_24    m/z 664.3745     0   1   2   3E+4   11.5   12  RT (min)      lcorbi_t3_neg_2_039_31    m/z 664.3806     0   2   4   6E+2   12.5   13  RT (min)      lcorbi_t3_neg_3_005_23    m/z 664.3897     0   1E+4   4.5   5  RT (min)      lcorbi_t3_neg_4_042_14    m/z 664.4587     0   1E+4   13.5   14  RT (min)      lcorbi_t3_neg_4_040_19    m/z 664.4872     0   1E+3   13   13.5  RT (min)      lcorbi_t3_neg_2_031_25    m/z 664.6610     0   1E+4   14.5   15  RT (min)      lcorbi_t3_neg_1_053_32    m/z 664.8276     0   1   2   3E+3   18   18.5  RT (min)      lcorbi_t3_neg_2_006_10    m/z 664.8721     0   1   2E+3   18   18.5  RT (min)      lcorbi_t3_neg_1_020_36    m/z 665.0502     0   1   2   3   4E+3   9.5   10  RT (min)      lcorbi_t3_neg_4_040_19    m/z 665.0632     0   1   2E+4   2   2.5  RT (min)      lcorbi_t3_neg_1_044_23    m/z 665.1213     0   1E+4   11   11.5  RT (min)      lcorbi_t3_neg_4_043_11    m/z 665.1248     0   2   4E+3   9   9.5  RT (min)      lcorbi_t3_neg_4_023_18    m/z 665.1796     0   1E+4   2.5   3  RT (min)      lcorbi_t3_neg_3_002_24    m/z 665.1804     0   1   2   3   4E+3   10   10.5  RT (min)      lcorbi_t3_neg_4_034_01    m/z 665.1939     0   1   2   3   4E+3   12.5   13  RT (min)      lcorbi_t3_neg_4_005_27    m/z 665.2001     0   1E+4   9   9.5  RT (min)      lcorbi_t3_neg_3_019_12    m/z 665.2645     0   1   2   3E+3   10.5   11  RT (min)      lcorbi_t3_neg_4_007_13    m/z 665.2917     0   1   2   3E+3   11   11.5  RT (min)      lcorbi_t3_neg_4_029_31    m/z 665.3355     0   1E+4   8   8.5  RT (min)      lcorbi_t3_neg_2_048_27    m/z 665.3545     0   1   2E+4   8   8.5  RT (min)      lcorbi_t3_neg_2_048_27    m/z 665.3642     0   1E+4   18   18.5  RT (min)      lcorbi_t3_neg_2_028_38    m/z 665.3649     0   2   4E+4   13.5   14  RT (min)      lcorbi_t3_neg_2_036_35    m/z 665.3840     0   1   2   3E+3   12.5   13  RT (min)      lcorbi_t3_neg_4_042_14    m/z 665.4810     0   2   4   6E+3   14.5   15  RT (min)      lcorbi_t3_neg_2_020_05    m/z 665.5366     0   1   14.5   15  RT (min)      lcorbi_t3_neg_4_019_08    m/z 665.6036     0   1E+4   14.5   15  RT (min)      lcorbi_t3_neg_4_034_01    m/z 665.8438     0   2   4   6   8E+2   18   18.5  RT (min)      lcorbi_t3_neg_4_019_08    m/z 666.1475     0   2   4   6   8E+3   10   10.5  RT (min)      lcorbi_t3_neg_3_037_36    m/z 666.2582     0   1E+3   10.5   11  RT (min)      lcorbi_t3_neg_4_021_25    m/z 666.2958     0   1   11   11.5  RT (min)      lcorbi_t3_neg_2_022_24    m/z 666.3037     0   1E+4   14.5   15  RT (min)      lcorbi_t3_neg_4_017_51    m/z 666.3746     0   1   2E+4   11.5   12  RT (min)      lcorbi_t3_neg_4_029_31    m/z 666.3955     0   1   2   3   4E+3   12   12.5  RT (min)      lcorbi_t3_neg_1_021_31    m/z 666.4057     0   2   4   6E+3   6.5   7  RT (min)      lcorbi_t3_neg_4_029_31    m/z 666.4258     0   1E+3   11.5   12  RT (min)      lcorbi_t3_neg_2_039_31    m/z 666.4350     0   1E+4   13   13.5  RT (min)      lcorbi_t3_neg_3_005_23    m/z 666.5000     0   1E+4   14.5   15  RT (min)      lcorbi_t3_neg_4_044_38    m/z 666.5291     0   2   4E+3   14.5   15  RT (min)      lcorbi_t3_neg_4_029_31    m/z 666.5346     0   1   2E+4   13   13.5  RT (min)      lcorbi_t3_neg_3_018_13    m/z 666.5506     0   1   2   3   4E+3   13.5   14  RT (min)      lcorbi_t3_neg_4_032_23    m/z 666.5681     0   1   14.5   15  RT (min)      lcorbi_t3_neg_1_052_24    m/z 666.5974     0   1E+4   14.5   15  RT (min)      lcorbi_t3_neg_4_011_24    m/z 667.1508     0   1   2   3E+3   10   10.5  RT (min)      lcorbi_t3_neg_3_037_36    m/z 667.2802     0   1E+4   11.5   12  RT (min)      lcorbi_t3_neg_1_023_03    m/z 667.3079     0   2   4E+3   14.5   15  RT (min)      lcorbi_t3_neg_4_017_51    m/z 667.3657     0   1   2   3E+4   11   11.5  RT (min)      lcorbi_t3_neg_4_032_23    m/z 667.3984     0   1   2   3E+3   13   13.5  RT (min)      lcorbi_t3_neg_4_007_13    m/z 667.4251     0   1E+4   13   13.5  RT (min)      lcorbi_t3_neg_4_019_08    m/z 667.5098     0   1E+4   15   15.5  RT (min)      lcorbi_t3_neg_4_042_14    m/z 667.5321     0   1E+4   13   13.5  RT (min)      lcorbi_t3_neg_4_047_30    m/z 667.5521     0   1   14.5   15  RT (min)      lcorbi_t3_neg_4_011_24    m/z 667.6075     0   2   4   6   8E+3   15   15.5  RT (min)      lcorbi_t3_neg_4_007_13    m/z 667.8450     0   1   2E+3   18   18.5  RT (min)      lcorbi_t3_neg_4_037_29    m/z 668.1932     0   1   2   3   4E+3   10   10.5  RT (min)      lcorbi_t3_neg_4_032_23    m/z 668.2638     0   1   2   3   4E+4   8.5   9  RT (min)      lcorbi_t3_neg_4_023_18    m/z 668.2671     0   2   4   6   8E+4   5   5.5  RT (min)      lcorbi_t3_neg_2_048_27    m/z 668.2855     0   1E+4   12   12.5  RT (min)      lcorbi_t3_neg_4_042_14    m/z 668.3027     0   2   4   6   8E+3   9.5   10  RT (min)      lcorbi_t3_neg_3_019_12    m/z 668.4581     0   1   2E+4   13.5   14  RT (min)      lcorbi_t3_neg_4_040_19    m/z 668.4712     0   1   2   3   4E+3   14.5   15  RT (min)      lcorbi_t3_neg_4_007_13    m/z 668.4943     0   1   2E+4   13   13.5  RT (min)      lcorbi_t3_neg_3_018_13    m/z 668.4999     0   1E+4   14.5   15  RT (min)      lcorbi_t3_neg_4_044_38    m/z 668.5191     0   2   4   6   8E+3   14.5   15  RT (min)      lcorbi_t3_neg_2_039_31    m/z 668.5348     0   1E+4   13   13.5  RT (min)      lcorbi_t3_neg_3_018_13    m/z 668.5756     0   1E+4   14.5   15  RT (min)      lcorbi_t3_neg_1_055_05    m/z 668.5838     0   2   4E+3   14.5   15  RT (min)      lcorbi_t3_neg_4_054_36    m/z 668.6200     0   1   2   3   4E+3   14.5   15  RT (min)      lcorbi_t3_neg_4_011_24    m/z 669.0294     0   1   2   3E+3   9   9.5  RT (min)      lcorbi_t3_neg_4_027_12    m/z 669.1588     0   1E+4   2   2.5  RT (min)      lcorbi_t3_neg_2_022_24    m/z 669.1761     0   1   2   3   4E+3   10   10.5  RT (min)      lcorbi_t3_neg_4_029_31    m/z 669.2775     0   2   4E+3   11.5   12  RT (min)      lcorbi_t3_neg_1_023_03    m/z 669.4912     0   1E+3   13   13.5  RT (min)      lcorbi_t3_neg_3_005_23    m/z 669.5876     0   1E+4   14.5   15  RT (min)      lcorbi_t3_neg_4_034_01    m/z 669.6153     0   1   2E+4   15   15.5  RT (min)      lcorbi_t3_neg_4_034_01    m/z 669.6415     0   1E+4   14   14.5  RT (min)      lcorbi_t3_neg_3_002_24    m/z 669.8441     0   1   2E+3   18   18.5  RT (min)      lcorbi_t3_neg_3_024_26    m/z 670.0863     0   2   4   6   8E+3   6   6.5  RT (min)      lcorbi_t3_neg_4_040_19    m/z 670.2115     0   2   4   6E+3   11   11.5  RT (min)      lcorbi_t3_neg_2_053_23    m/z 670.2727     0   1   2E+3   10   10.5  RT (min)      lcorbi_t3_neg_3_019_12    m/z 670.3247     0   1   2   3E+4   14   14.5  RT (min)      lcorbi_t3_neg_1_023_03    m/z 670.4707     0   1E+4   14.5   15  RT (min)      lcorbi_t3_neg_2_039_31    m/z 670.4942     0   1E+3   13   13.5  RT (min)      lcorbi_t3_neg_4_032_23    m/z 670.5240     0   1   2E+4   14.5   15  RT (min)      lcorbi_t3_neg_3_020_31    m/z 670.6187     0   2   4   6   8E+3   15   15.5  RT (min)      lcorbi_t3_neg_4_034_01    m/z 670.6435     0   2   4   6   8E+3   14.5   15  RT (min)      lcorbi_t3_neg_4_011_24    m/z 670.6539     0   1E+4   14   14.5  RT (min)      lcorbi_t3_neg_4_011_24    m/z 670.6837     0   2   4   6   8E+3   1   1.5  RT (min)      lcorbi_t3_neg_3_032_17    m/z 670.8365     0   2   4   6   8E+3   18   18.5  RT (min)      lcorbi_t3_neg_2_048_27    m/z 671.1093     0   2   4   6   8E+3   11.5   12  RT (min)      lcorbi_t3_neg_4_007_13    m/z 671.1767     0   1E+4   9   9.5  RT (min)      lcorbi_t3_neg_4_023_18    m/z 671.2763     0   1E+4   13   13.5  RT (min)      lcorbi_t3_neg_2_007_14    m/z 671.2921     0   2   4   6   8E+3   11.5   12  RT (min)      lcorbi_t3_neg_1_043_14    m/z 671.2955     0   1E+4   8.5   9  RT (min)      lcorbi_t3_neg_4_023_18    m/z 671.3111     0   2   4E+2   9.5   10  RT (min)      lcorbi_t3_neg_4_038_15    m/z 671.3691     0   1   2   3E+4   18   18.5  RT (min)      lcorbi_t3_neg_2_028_38    m/z 671.3698     0   1E+5   14   14.5  RT (min)      lcorbi_t3_neg_2_036_35    m/z 671.4657     0   1E+4   18   18.5  RT (min)      lcorbi_t3_neg_4_046_37    m/z 671.5219     0   2   4   6   8E+3   14   14.5  RT (min)      lcorbi_t3_neg_4_046_37    m/z 671.8569     0   1   2E+4   18   18.5  RT (min)      lcorbi_t3_neg_1_022_34    m/z 671.8573     0   2   4   6E+4   13.5   14  RT (min)      lcorbi_t3_neg_2_012_36    m/z 671.8710     0   1E+4   18   18.5  RT (min)      lcorbi_t3_neg_2_036_35    m/z 671.8711     0   1   2   3   4E+4   14   14.5  RT (min)      lcorbi_t3_neg_2_012_36    m/z 672.2273     0   2   4   6   8E+3   11.5   12  RT (min)      lcorbi_t3_neg_2_020_05    m/z 672.2799     0   2   4E+3   13   13.5  RT (min)      lcorbi_t3_neg_2_007_14    m/z 672.2915     0   2   4   6   8E+3   11   11.5  RT (min)      lcorbi_t3_neg_4_034_01    m/z 672.3587     0   1E+4   18   18.5  RT (min)      lcorbi_t3_neg_2_028_38    m/z 672.3595     0   2   4E+4   13.5   14  RT (min)      lcorbi_t3_neg_2_028_38    m/z 672.4595     0   1E+5   13.5   14  RT (min)      lcorbi_t3_neg_4_019_08    m/z 672.4691     0   2   4   6   8E+3   18   18.5  RT (min)      lcorbi_t3_neg_4_046_37    m/z 672.4872     0   1E+4   13   13.5  RT (min)      lcorbi_t3_neg_4_034_01    m/z 672.5026     0   1E+4   14.5   15  RT (min)      lcorbi_t3_neg_2_051_34    m/z 672.5787     0   2   4   6   8E+3   14.5   15  RT (min)      lcorbi_t3_neg_2_053_23    m/z 672.6590     0   1E+4   14.5   15  RT (min)      lcorbi_t3_neg_3_002_24    m/z 672.9684     0   1   2   3E+3   8   8.5  RT (min)      lcorbi_t3_neg_4_023_18    m/z 673.0720     0   1E+4   10   10.5  RT (min)      lcorbi_t3_neg_4_029_31    m/z 673.1592     0   1E+4   9   9.5  RT (min)      lcorbi_t3_neg_4_023_18    m/z 673.2643     0   1E+4   12.5   13  RT (min)      lcorbi_t3_neg_1_055_05    m/z 673.2761     0   1   2   3E+4   14   14.5  RT (min)      lcorbi_t3_neg_3_011_04    m/z 673.3082     0   1   11   11.5  RT (min)      lcorbi_t3_neg_1_055_05    m/z 673.3867     0   1E+4   10   10.5  RT (min)      lcorbi_t3_neg_4_039_32    m/z 673.3931     0   1E+4   13.5   14  RT (min)      lcorbi_t3_neg_4_007_13    m/z 673.4188     0   1   2E+4   13   13.5  RT (min)      lcorbi_t3_neg_1_055_05    m/z 673.4811     0   2   4   6   8E+3   18   18.5  RT (min)      lcorbi_t3_neg_4_032_23    m/z 673.5050     0   1   2   3E+4   14.5   15  RT (min)      lcorbi_t3_neg_4_046_37    m/z 673.5278     0   1E+4   13.5   14  RT (min)      lcorbi_t3_neg_4_034_01    m/z 673.8316     0   2   4   6E+3   18   18.5  RT (min)      lcorbi_t3_neg_2_050_17    m/z 673.8621     0   1E+4   18   18.5  RT (min)      lcorbi_t3_neg_4_019_08    m/z 674.0180     0   1E+4   9   9.5  RT (min)      lcorbi_t3_neg_2_048_27    m/z 674.0256     0   2   4   6E+3   11.5   12  RT (min)      lcorbi_t3_neg_4_007_13    m/z 674.1598     0   1   2   3E+3   13.5   14  RT (min)      lcorbi_t3_neg_4_015_10    m/z 674.2427     0   2   4E+3   12   12.5  RT (min)      lcorbi_t3_neg_4_034_01    m/z 674.2518     0   2   4E+3   12.5   13  RT (min)      lcorbi_t3_neg_1_023_03    m/z 674.2637     0   1   2E+4   14   14.5  RT (min)      lcorbi_t3_neg_4_030_04    m/z 674.2678     0   1   2   3   4E+3   12.5   13  RT (min)      lcorbi_t3_neg_1_055_05    m/z 674.2719     0   1   2   3   4E+4   13.5   14  RT (min)      lcorbi_t3_neg_4_019_08    m/z 674.2836     0   1E+4   11.5   12  RT (min)      lcorbi_t3_neg_4_007_13    m/z 674.3202     0   1   2   3E+3   11.5   12  RT (min)      lcorbi_t3_neg_4_036_16    m/z 674.3495     0   1E+4   14.5   15  RT (min)      lcorbi_t3_neg_2_035_46    m/z 674.3963     0   2   4   6E+3   13.5   14  RT (min)      lcorbi_t3_neg_4_007_13    m/z 674.4014     0   1   12.5   13  RT (min)      lcorbi_t3_neg_1_055_05    m/z 674.4846     0   1   2   3   4E+3   18   18.5  RT (min)      lcorbi_t3_neg_4_046_37    m/z 674.5126     0   1   2   3E+5   14   14.5  RT (min)      lcorbi_t3_neg_1_055_05    m/z 674.5312     0   1   13.5   14  RT (min)      lcorbi_t3_neg_4_016_05    m/z 674.6290     0   2   4   6E+3   14.5   15  RT (min)      lcorbi_t3_neg_4_039_32    m/z 674.8337     0   1   2   3E+3   18   18.5  RT (min)      lcorbi_t3_neg_4_030_04    m/z 674.8587     0   1   2   3E+3   18   18.5  RT (min)      lcorbi_t3_neg_1_005_07    m/z 675.0399     0   2   4   6   8E+3   10.5   11  RT (min)      lcorbi_t3_neg_2_011_04    m/z 675.1477     0   1E+4   9   9.5  RT (min)       lcorbi_t3_neg_4_102_61    m/z 675.1753     0   1E+4   2   2.5  RT (min)      lcorbi_t3_neg_2_022_24    m/z 675.2314     0   1   2   3   4E+3   12.5   13  RT (min)      lcorbi_t3_neg_4_005_27    m/z 675.2889     0   2   4   6E+3   12   12.5  RT (min)      lcorbi_t3_neg_4_023_18    m/z 675.2995     0   1E+4   8   8.5  RT (min)      lcorbi_t3_neg_2_048_27    m/z 675.3527     0   1   2   3   4E+3   14.5   15  RT (min)      lcorbi_t3_neg_2_035_46    m/z 675.3915     0   2   4   6E+3   13.5   14  RT (min)      lcorbi_t3_neg_1_054_37    m/z 675.4346     0   2   4E+3   13   13.5  RT (min)      lcorbi_t3_neg_2_005_01    m/z 675.4607     0   2   4   6E+3   13.5   14  RT (min)      lcorbi_t3_neg_4_020_44    m/z 675.4760     0   1   2   3E+3   14.5   15  RT (min)      lcorbi_t3_neg_4_019_08    m/z 675.4961     0   1E+4   14.5   15  RT (min)      lcorbi_t3_neg_2_051_34    m/z 675.5217     0   1   13   13.5  RT (min)      lcorbi_t3_neg_1_055_05    m/z 675.5679     0   1E+4   14   14.5  RT (min)      lcorbi_t3_neg_1_034_04    m/z 676.0151     0   2   4   6   8E+3   9   9.5  RT (min)      lcorbi_t3_neg_2_048_27    m/z 676.1455     0   2   4E+4   9   9.5  RT (min)      lcorbi_t3_neg_4_023_18    m/z 676.2466     0   1E+4   9.5   10  RT (min)      lcorbi_t3_neg_3_023_07    m/z 676.2474     0   1E+4   14   14.5  RT (min)      lcorbi_t3_neg_3_011_04    m/z 676.3029     0   2   4   6E+3   8   8.5  RT (min)      lcorbi_t3_neg_2_048_27    m/z 676.3249     0   2   4E+3   11   11.5  RT (min)      lcorbi_t3_neg_3_028_35    m/z 676.4173     0   2   4   6E+2   13   13.5  RT (min)      lcorbi_t3_neg_4_032_23    m/z 676.4638     0   1   2   3E+3   13.5   14  RT (min)      lcorbi_t3_neg_2_050_17    m/z 676.4996     0   2   4   6   8E+3   14.5   15  RT (min)      lcorbi_t3_neg_4_004_34    m/z 676.5362     0   1   13.5   14  RT (min)      lcorbi_t3_neg_3_007_16    m/z 676.5713     0   1   2   3E+3   14   14.5  RT (min)      lcorbi_t3_neg_1_006_40    m/z 676.6092     0   1   2   3   4E+4   14   14.5  RT (min)      lcorbi_t3_neg_3_002_24    m/z 676.6249     0   1E+4   15   15.5  RT (min)      lcorbi_t3_neg_4_007_13    m/z 677.1401     0   1E+5   9   9.5  RT (min)      lcorbi_t3_neg_4_023_18    m/z 677.1406     0   1E+4   12   12.5  RT (min)      lcorbi_t3_neg_4_007_13    m/z 677.4918     0   1E+3   14.5   15  RT (min)      lcorbi_t3_neg_4_050_02    m/z 677.5812     0   2   4E+3   15   15.5  RT (min)      lcorbi_t3_neg_4_048_03    m/z 677.6282     0   2   4   6   8E+2   15   15.5  RT (min)      lcorbi_t3_neg_4_011_24    m/z 677.6762     0   1E+4   14.5   15  RT (min)      lcorbi_t3_neg_4_039_32    m/z 677.8508     0   1   2   3E+3   18   18.5  RT (min)      lcorbi_t3_neg_2_036_35    m/z 678.1656     0   2   4   6E+2   10.5   11  RT (min)      lcorbi_t3_neg_2_020_05    m/z 678.2216     0   2   4   6E+3   10   10.5  RT (min)      lcorbi_t3_neg_3_019_12    m/z 678.2246     0   1   2E+4   10.5   11  RT (min)      lcorbi_t3_neg_4_032_23    m/z 678.3407     0   2   4   6E+3   11.5   12  RT (min)      lcorbi_t3_neg_4_034_01    m/z 678.3772     0   2   4   6   8E+4   14   14.5  RT (min)      lcorbi_t3_neg_2_036_35    m/z 678.4216     0   1E+4   13   13.5  RT (min)      lcorbi_t3_neg_4_046_37    m/z 678.4326     0   2   4E+3   13.5   14  RT (min)      lcorbi_t3_neg_4_007_13    m/z 678.4714     0   2   4   6   8E+3   13.5   14  RT (min)      lcorbi_t3_neg_4_007_13    m/z 678.4842     0   1E+4   14.5   15  RT (min)      lcorbi_t3_neg_4_032_23    m/z 678.5183     0   1   2   3E+4   13   13.5  RT (min)      lcorbi_t3_neg_4_034_01    m/z 678.5490     0   1E+3   14   14.5  RT (min)      lcorbi_t3_neg_3_002_24    m/z 678.5687     0   2   4   6E+2   14.5   15  RT (min)      lcorbi_t3_neg_2_053_23    m/z 678.6406     0   1E+4   15   15.5  RT (min)      lcorbi_t3_neg_4_007_13    m/z 678.8155     0   1   2E+3   18   18.5  RT (min)      lcorbi_t3_neg_4_006_47    m/z 678.8783     0   1   2E+4   18   18.5  RT (min)      lcorbi_t3_neg_2_036_35    m/z 678.8788     0   1   2   3   4E+4   14   14.5  RT (min)      lcorbi_t3_neg_2_036_35    m/z 678.9855     0   2   4   6E+3   8   8.5  RT (min)      lcorbi_t3_neg_4_023_18    m/z 679.0516     0   1   2   3   4E+3   10.5   11  RT (min)      lcorbi_t3_neg_4_030_04    m/z 679.1368     0   1E+4   11   11.5  RT (min)      lcorbi_t3_neg_4_043_11    m/z 679.2253     0   1   2   3   4E+3   10   10.5  RT (min)      lcorbi_t3_neg_3_005_23    m/z 679.2799     0   1E+4   14   14.5  RT (min)      lcorbi_t3_neg_4_030_04    m/z 679.3123     0   2   4   6E+3   11   11.5  RT (min)      lcorbi_t3_neg_4_029_31    m/z 679.3290     0   2   4   6   8E+3   13.5   14  RT (min)      lcorbi_t3_neg_4_023_18    m/z 679.3342     0   2   4   6E+3   9.5   10  RT (min)      lcorbi_t3_neg_3_019_12    m/z 679.3599     0   1   2   3E+3   12   12.5  RT (min)      lcorbi_t3_neg_3_020_31    m/z 679.3698     0   1E+4   8   8.5  RT (min)      lcorbi_t3_neg_2_048_27    m/z 679.3863     0   1E+4   12   12.5  RT (min)      lcorbi_t3_neg_4_029_31    m/z 679.5156     0   1   2E+4   14   14.5  RT (min)      lcorbi_t3_neg_4_042_14    m/z 679.5533     0   1   2   3   4E+4   13.5   14  RT (min)      lcorbi_t3_neg_4_019_08    m/z 679.5884     0   2   4   6   8E+3   14.5   15  RT (min)      lcorbi_t3_neg_2_053_23    m/z 679.6076     0   1E+4   14.5   15  RT (min)      lcorbi_t3_neg_4_007_13    m/z 679.8814     0   1   2   3E+3   18   18.5  RT (min)      lcorbi_t3_neg_2_024_37    m/z 680.1024     0   2   4E+3   9   9.5  RT (min)      lcorbi_t3_neg_1_043_14    m/z 680.1609     0   1   2E+4   8.5   9  RT (min)      lcorbi_t3_neg_2_048_27    m/z 680.2223     0   1   2   3E+3   10   10.5  RT (min)      lcorbi_t3_neg_3_019_12    m/z 680.2920     0   2   4   6E+3   11.5   12  RT (min)      lcorbi_t3_neg_2_007_14    m/z 680.4116     0   1   2E+4   5.5   6   6.5  RT (min)      lcorbi_t3_neg_4_044_38    m/z 680.4142     0   1E+4   11.5   12  RT (min)      lcorbi_t3_neg_4_032_23    m/z 680.4269     0   1   14   14.5  RT (min)      lcorbi_t3_neg_4_019_08    m/z 680.5086     0   1E+5   14   14.5  RT (min)      lcorbi_t3_neg_4_029_31    m/z 680.5354     0   2   4   6E+3   14.5   15  RT (min)      lcorbi_t3_neg_2_028_38    m/z 680.5755     0   1E+4   14.5   15  RT (min)      lcorbi_t3_neg_4_007_13    m/z 680.5843     0   2   4   6E+3   14.5   15  RT (min)      lcorbi_t3_neg_4_029_31    m/z 680.8502     0   2   4   6   8E+3   18   18.5  RT (min)      lcorbi_t3_neg_2_029_12    m/z 680.8830     0   1E+4   18   18.5  RT (min)      lcorbi_t3_neg_1_001_49    m/z 681.0425     0   1   2E+3   9.5   10  RT (min)      lcorbi_t3_neg_2_011_04    m/z 681.0951     0   2   4E+3   8   8.5  RT (min)      lcorbi_t3_neg_1_026_19    m/z 681.1555     0   1   2E+4   12.5   13  RT (min)      lcorbi_t3_neg_4_045_35    m/z 681.2437     0   2   4   6E+3   11   11.5  RT (min)      lcorbi_t3_neg_4_032_23    m/z 681.2586     0   2   4   6E+3   11.5   12  RT (min)      lcorbi_t3_neg_4_050_02    m/z 681.2962     0   2   4   6   8E+3   14   14.5  RT (min)      lcorbi_t3_neg_3_039_29    m/z 681.3597     0   1   2   3   4E+3   12   12.5  RT (min)      lcorbi_t3_neg_3_030_33    m/z 681.4321     0   2   4E+3   13   13.5  RT (min)      lcorbi_t3_neg_4_045_35    m/z 681.4741     0   1   2   3   4E+3   13   13.5  RT (min)      lcorbi_t3_neg_4_007_13    m/z 681.5694     0   2   4   6   8E+3   13.5   14  RT (min)      lcorbi_t3_neg_1_032_13    m/z 681.5789     0   2   4   6E+3   14.5   15  RT (min)      lcorbi_t3_neg_4_007_13    m/z 681.6040     0   1   2   3   4E+4   14.5   15  RT (min)      lcorbi_t3_neg_1_055_05    m/z 681.6230     0   2   4   6   8E+3   15   15.5  RT (min)      lcorbi_t3_neg_4_007_13    m/z 681.8494     0   2   4E+3   18   18.5  RT (min)      lcorbi_t3_neg_4_018_42    m/z 682.1582     0   1E+4   8.5   9  RT (min)      lcorbi_t3_neg_2_048_27    m/z 682.1685     0   1   2E+4   1.5   2  RT (min)      lcorbi_t3_neg_1_053_32    m/z 682.3334     0   1E+4   11.5   12  RT (min)      lcorbi_t3_neg_4_007_13    m/z 682.3335     0   1E+4   13.5   14  RT (min)      lcorbi_t3_neg_1_023_03    m/z 682.3484     0   2   4   6E+3   5.5   6  RT (min)      lcorbi_t3_neg_4_046_37    m/z 682.5050     0   1   2E+3   14.5   15  RT (min)      lcorbi_t3_neg_1_055_05    m/z 682.5497     0   1   13.5   14  RT (min)      lcorbi_t3_neg_1_013_35    m/z 682.5996     0   1E+3   14.5   15  RT (min)      lcorbi_t3_neg_1_018_46    m/z 682.8557     0   1   2E+3   18   18.5  RT (min)      lcorbi_t3_neg_1_001_49    m/z 682.8842     0   2   4   6E+3   18   18.5  RT (min)      lcorbi_t3_neg_1_001_49    m/z 683.2596     0   2   4   6   8E+3   11.5   12  RT (min)      lcorbi_t3_neg_2_020_05    m/z 683.2876     0   1E+5   6   6.5  RT (min)      lcorbi_t3_neg_2_048_27    m/z 683.3022     0   2   4   6   8E+3   12.5   13  RT (min)      lcorbi_t3_neg_1_055_05    m/z 683.4232     0   1E+4   14   14.5  RT (min)      lcorbi_t3_neg_1_027_30    m/z 683.4836     0   1E+3   14.5   15  RT (min)      lcorbi_t3_neg_4_035_33    m/z 683.5945     0   1   15   15.5  RT (min)      lcorbi_t3_neg_1_044_23    m/z 684.1232     0   1   2E+4   2   2.5  RT (min)      lcorbi_t3_neg_1_052_24    m/z 684.1553     0   1   2   3E+4   10.5   11  RT (min)      lcorbi_t3_neg_1_043_14    m/z 684.2400     0   1   2   3   4E+3   10   10.5  RT (min)      lcorbi_t3_neg_3_019_12    m/z 684.2425     0   2   4   6E+3   12   12.5  RT (min)      lcorbi_t3_neg_4_007_13    m/z 684.2606     0   1   2   3E+3   11.5   12  RT (min)      lcorbi_t3_neg_4_034_01    m/z 684.2629     0   1E+3   13.5   14  RT (min)      lcorbi_t3_neg_1_023_03    m/z 684.3129     0   1   2   3E+4   13.5   14  RT (min)      lcorbi_t3_neg_1_023_03    m/z 684.3955     0   2   4   6   8E+3   13   13.5  RT (min)      lcorbi_t3_neg_4_042_14    m/z 684.4089     0   1E+4   11.5   12  RT (min)      lcorbi_t3_neg_4_007_13    m/z 684.4511     0   1   2E+4   13   13.5  RT (min)      lcorbi_t3_neg_4_034_01    m/z 684.4611     0   1   2   3   4E+4   13.5   14  RT (min)      lcorbi_t3_neg_4_029_31    m/z 684.4627     0   1   2   3E+4   11   11.5  RT (min)      lcorbi_t3_neg_1_052_24    m/z 684.5211     0   1E+4   14.5   15  RT (min)      lcorbi_t3_neg_4_044_38    m/z 684.5884     0   1   2E+3   15   15.5  RT (min)      lcorbi_t3_neg_4_045_35    m/z 684.6146     0   1   2E+3   14.5   15  RT (min)      lcorbi_t3_neg_2_028_38    m/z 684.8269     0   1   2   3E+3   18   18.5  RT (min)      lcorbi_t3_neg_1_006_40    m/z 684.8616     0   2   4   6   8E+3   18   18.5  RT (min)      lcorbi_t3_neg_4_023_18    m/z 684.9929     0   1   2E+3   9   9.5  RT (min)      lcorbi_t3_neg_3_048_08    m/z 685.1306     0   1   2E+4   2   2.5  RT (min)      lcorbi_t3_neg_1_052_24    m/z 685.1874     0   2   4E+4   2.5   3  RT (min)      lcorbi_t3_neg_1_053_32    m/z 685.2403     0   1   2   3   4E+3   12.5   13  RT (min)      lcorbi_t3_neg_3_007_16    m/z 685.2528     0   1   2E+4   11.5   12  RT (min)      lcorbi_t3_neg_4_007_13    m/z 685.2753     0   2   4E+3   12   12.5  RT (min)      lcorbi_t3_neg_4_034_01    m/z 685.2999     0   1E+4   13.5   14  RT (min)      lcorbi_t3_neg_3_024_26    m/z 685.3159     0   1E+4   13.5   14  RT (min)      lcorbi_t3_neg_1_023_03    m/z 685.3524     0   1E+4   11.5   12  RT (min)      lcorbi_t3_neg_4_007_13    m/z 685.4298     0   2   4E+4   14   14.5  RT (min)      lcorbi_t3_neg_4_052_26    m/z 685.4551     0   1   2   3E+4   11   11.5  RT (min)      lcorbi_t3_neg_4_029_31    m/z 685.4990     0   1   2E+3   14.5   15  RT (min)      lcorbi_t3_neg_4_007_13    m/z 685.5432     0   2   4   6   8E+3   15   15.5  RT (min)      lcorbi_t3_neg_4_029_31    m/z 685.6452     0   2   4   6   8E+2   14.5   15  RT (min)      lcorbi_t3_neg_3_002_24    m/z 685.8725     0   1E+4   18   18.5  RT (min)      lcorbi_t3_neg_2_036_35    m/z 685.8733     0   1   2E+4   14   14.5  RT (min)      lcorbi_t3_neg_2_036_35    m/z 686.0722     0   1   2   3   4E+3   10.5   11  RT (min)      lcorbi_t3_neg_4_030_04    m/z 686.2518     0   1   2   3E+4   13.5   14  RT (min)      lcorbi_t3_neg_4_048_03    m/z 686.2695     0   1E+4   14.5   15  RT (min)      lcorbi_t3_neg_4_020_44    m/z 686.2762     0   2   4E+3   12   12.5  RT (min)      lcorbi_t3_neg_1_010_17    m/z 686.3102     0   1   2   3E+4   10.5   11  RT (min)      lcorbi_t3_neg_4_042_14    m/z 686.3661     0   1   2   3   4E+3   18   18.5  RT (min)      lcorbi_t3_neg_4_015_10    m/z 686.4339     0   1E+4   12   12.5  RT (min)      lcorbi_t3_neg_3_002_24    m/z 686.4493     0   2   4E+3   7   7.5  RT (min)      lcorbi_t3_neg_4_032_23    m/z 686.4669     0   2   4   6E+2   13   13.5  RT (min)      lcorbi_t3_neg_4_019_08    m/z 686.5020     0   1   2   3   4E+3   14.5   15  RT (min)      lcorbi_t3_neg_2_051_34    m/z 686.5367     0   2   4E+3   14.5   15  RT (min)      lcorbi_t3_neg_4_044_38    m/z 686.5466     0   1   2   3E+3   15   15.5  RT (min)      lcorbi_t3_neg_2_039_31    m/z 686.5942     0   1E+4   14.5   15  RT (min)      lcorbi_t3_neg_2_020_05    m/z 686.6760     0   1   2   3E+4   14   14.5  RT (min)      lcorbi_t3_neg_1_052_24    m/z 686.8381     0   1   2   3E+3   18   18.5  RT (min)      lcorbi_t3_neg_2_034_19    m/z 686.8636     0   2   4   6E+3   18   18.5  RT (min)      lcorbi_t3_neg_4_053_43    m/z 687.2444     0   1E+4   13   13.5  RT (min)      lcorbi_t3_neg_4_039_32    m/z 687.2476     0   2   4E+3   8.5   9  RT (min)      lcorbi_t3_neg_2_048_27    m/z 687.2639     0   1E+4   11.5   12  RT (min)      lcorbi_t3_neg_3_037_36    m/z 687.2761     0   1E+4   10.5   11  RT (min)      lcorbi_t3_neg_4_007_13    m/z 687.3202     0   2   4   6   8E+3   6   6.5  RT (min)      lcorbi_t3_neg_4_042_14    m/z 687.4062     0   1E+4   13   13.5  RT (min)      lcorbi_t3_neg_4_034_01    m/z 687.4452     0   1E+4   13.5   14  RT (min)      lcorbi_t3_neg_4_042_14    m/z 687.5350     0   2   4   6   8E+3   14.5   15  RT (min)      lcorbi_t3_neg_4_038_15    m/z 687.5993     0   1E+4   14.5   15  RT (min)      lcorbi_t3_neg_4_039_32    m/z 688.2614     0   1   2E+4   11   11.5  RT (min)      lcorbi_t3_neg_1_043_14    m/z 688.2675     0   1   2E+4   13   13.5  RT (min)      lcorbi_t3_neg_4_046_37    m/z 688.3112     0   2   4   6E+3   12   12.5  RT (min)      lcorbi_t3_neg_3_052_32    m/z 688.3152     0   1   2   3E+4   14   14.5  RT (min)      lcorbi_t3_neg_4_030_04    m/z 688.3718     0   1   2   3   4E+3   18   18.5  RT (min)      lcorbi_t3_neg_2_003_42    m/z 688.4095     0   1   2   3   4E+3   13   13.5  RT (min)      lcorbi_t3_neg_3_032_17    m/z 688.4557     0   2   4   6   8E+3   13.5   14  RT (min)      lcorbi_t3_neg_4_038_15    m/z 688.5729     0   1E+4   14   14.5  RT (min)      lcorbi_t3_neg_1_053_32    m/z 688.5885     0   1E+4   14.5   15  RT (min)      lcorbi_t3_neg_4_007_13    m/z 688.8330     0   2   4E+3   18   18.5  RT (min)      lcorbi_t3_neg_2_019_07    m/z 689.2640     0   1   2   3E+3   10.5   11  RT (min)      lcorbi_t3_neg_2_020_05    m/z 689.2708     0   2   4   6E+3   13   13.5  RT (min)      lcorbi_t3_neg_1_023_03    m/z 689.3450     0   1E+4   12   12.5  RT (min)      lcorbi_t3_neg_4_034_01    m/z 689.4035     0   1E+4   8   8.5  RT (min)      lcorbi_t3_neg_4_017_51    m/z 689.4088     0   2   4   6E+3   14.5   15  RT (min)      lcorbi_t3_neg_1_055_05    m/z 689.4147     0   1E+4   11   11.5  RT (min)      lcorbi_t3_neg_1_055_05    m/z 689.4419     0   1E+4   14   14.5  RT (min)      lcorbi_t3_neg_4_023_18    m/z 689.4871     0   1   12.5   13  RT (min)      lcorbi_t3_neg_4_019_08    m/z 689.5503     0   1   2E+4   13.5   14  RT (min)      lcorbi_t3_neg_4_007_13    m/z 689.6154     0   1   2   3E+4   14.5   15  RT (min)      lcorbi_t3_neg_4_039_32    m/z 689.6765     0   1   2   3   4E+3   14.5   15  RT (min)      lcorbi_t3_neg_1_055_05    m/z 690.2581     0   1E+3   10.5   11  RT (min)      lcorbi_t3_neg_1_001_49    m/z 690.3665     0   1   2   3   4E+4   10.5   11  RT (min)      lcorbi_t3_neg_4_011_24    m/z 690.3955     0   2   4   6   8E+3   13   13.5  RT (min)      lcorbi_t3_neg_4_042_14    m/z 690.4069     0   1   2   3   4E+3   8   8.5  RT (min)      lcorbi_t3_neg_4_017_51    m/z 690.4641     0   2   4   6E+3   13.5   14  RT (min)      lcorbi_t3_neg_4_042_14    m/z 690.5081     0   1E+4   14   14.5  RT (min)      lcorbi_t3_neg_3_005_23    m/z 690.8438     0   2   4   6E+3   18   18.5  RT (min)      lcorbi_t3_neg_2_013_08    m/z 691.0712     0   2   4   6   8E+3   9.5   10  RT (min)      lcorbi_t3_neg_2_023_39    m/z 691.1210     0   1   2   3E+4   10.5   11  RT (min)      lcorbi_t3_neg_4_042_14    m/z 691.1933     0   1   2E+4   10.5   11  RT (min)      lcorbi_t3_neg_1_043_14    m/z 691.2585     0   2   4   6   8E+3   10.5   11  RT (min)      lcorbi_t3_neg_1_015_27    m/z 691.2823     0   1E+4   13.5   14  RT (min)      lcorbi_t3_neg_1_023_03    m/z 691.3242     0   1E+4   11.5   12  RT (min)      lcorbi_t3_neg_4_034_01    m/z 691.3607     0   2   4   6E+3   12.5   13  RT (min)      lcorbi_t3_neg_4_034_01    m/z 691.3804     0   1E+4   18   18.5  RT (min)      lcorbi_t3_neg_3_038_39    m/z 691.4524     0   1   2E+4   14   14.5  RT (min)      lcorbi_t3_neg_4_032_23    m/z 691.4767     0   1   2E+4   13.5   14  RT (min)      lcorbi_t3_neg_4_032_23    m/z 691.5629     0   1   2   3   4E+3   13.5   14  RT (min)      lcorbi_t3_neg_2_016_11    m/z 691.5901     0   1   2   3   4E+3   14   14.5  RT (min)      lcorbi_t3_neg_4_023_18    m/z 691.6193     0   1E+4   14.5   15  RT (min)      lcorbi_t3_neg_4_011_24    m/z 691.6286     0   1E+4   14   14.5  RT (min)      lcorbi_t3_neg_3_002_24    m/z 691.8813     0   2   4   6E+3   18   18.5  RT (min)      lcorbi_t3_neg_2_037_09    m/z 692.2759     0   1   2E+4   13.5   14  RT (min)      lcorbi_t3_neg_1_023_03    m/z 692.2759     0   1E+4   11   11.5  RT (min)      lcorbi_t3_neg_2_051_34    m/z 692.3002     0   1   2E+3   13.5   14  RT (min)      lcorbi_t3_neg_4_032_23    m/z 692.3821     0   1   2   3E+4   11   11.5  RT (min)      lcorbi_t3_neg_4_011_24    m/z 692.3958     0   2   4   6   8E+3   13   13.5  RT (min)      lcorbi_t3_neg_4_042_14    m/z 692.4505     0   1   2E+4   12   12.5  RT (min)      lcorbi_t3_neg_4_007_13    m/z 692.4710     0   1   2   3E+3   14.5   15  RT (min)      lcorbi_t3_neg_1_055_05    m/z 692.4748     0   2   4   6E+3   13   13.5  RT (min)      lcorbi_t3_neg_1_055_05    m/z 692.4998     0   1E+4   14.5   15  RT (min)      lcorbi_t3_neg_4_029_31    m/z 692.5155     0   2   4   6E+4   14.5   15  RT (min)      lcorbi_t3_neg_4_046_37    m/z 692.5661     0   1   2E+4   14   14.5  RT (min)      lcorbi_t3_neg_4_032_23    m/z 692.6101     0   1E+4   14.5   15  RT (min)      lcorbi_t3_neg_1_055_05    m/z 692.6953     0   1   2E+4   14.5   15  RT (min)      lcorbi_t3_neg_4_039_32    m/z 692.8492     0   2   4E+3   18   18.5  RT (min)      lcorbi_t3_neg_2_003_42    m/z 692.9564     0   1   2E+4   18   18.5  RT (min)      lcorbi_t3_neg_2_052_32    m/z 692.9573     0   1   2   3   4E+4   13.5   14  RT (min)      lcorbi_t3_neg_2_052_32    m/z 693.0684     0   1   2E+3   9.5   10  RT (min)      lcorbi_t3_neg_2_026_03    m/z 693.1346     0   1   2   3E+4   12   12.5  RT (min)      lcorbi_t3_neg_4_007_13    m/z 693.1371     0   1   2   3E+3   10   10.5  RT (min)      lcorbi_t3_neg_4_044_38    m/z 693.1951     0   2   4   6E+3   10.5   11  RT (min)      lcorbi_t3_neg_2_010_02    m/z 693.2580     0   1E+4   8   8.5  RT (min)      lcorbi_t3_neg_2_048_27    m/z 693.2659     0   2   4   6E+3   13.5   14  RT (min)      lcorbi_t3_neg_3_023_07    m/z 693.2771     0   2   4E+3   11   11.5  RT (min)      lcorbi_t3_neg_2_008_16    m/z 693.2941     0   2   4E+3   13.5   14  RT (min)      lcorbi_t3_neg_4_023_18    m/z 693.3181     0   2   4   6E+3   12.5   13  RT (min)      lcorbi_t3_neg_4_042_14    m/z 693.3622     0   2   4   6E+3   8   8.5  RT (min)      lcorbi_t3_neg_4_023_18    m/z 693.4022     0   2   4   6   8E+3   13   13.5  RT (min)      lcorbi_t3_neg_4_042_14    m/z 693.4404     0   1E+4   14.5   15  RT (min)      lcorbi_t3_neg_2_036_35    m/z 693.4539     0   1E+4   12   12.5  RT (min)      lcorbi_t3_neg_4_007_13    m/z 693.4714     0   1E+4   13   13.5  RT (min)      lcorbi_t3_neg_3_018_13    m/z 693.4923     0   2   4E+4   13.5   14  RT (min)      lcorbi_t3_neg_2_053_23    m/z 693.5094     0   1   2E+4   13   13.5  RT (min)      lcorbi_t3_neg_4_016_05    m/z 693.5185     0   2   4E+3   13   13.5  RT (min)      lcorbi_t3_neg_4_007_13    m/z 693.5842     0   1E+4   14.5   15  RT (min)      lcorbi_t3_neg_2_053_23    m/z 693.6353     0   1   14.5   15  RT (min)      lcorbi_t3_neg_4_044_38    m/z 693.8516     0   1E+3   18   18.5  RT (min)      lcorbi_t3_neg_4_019_08    m/z 694.1352     0   1E+4   12   12.5  RT (min)      lcorbi_t3_neg_4_007_13    m/z 694.2115     0   2   4   6E+3   11   11.5  RT (min)      lcorbi_t3_neg_2_010_02    m/z 694.2828     0   2   4   6E+3   13   13.5  RT (min)      lcorbi_t3_neg_4_005_27    m/z 694.3344     0   1E+4   14.5   15  RT (min)      lcorbi_t3_neg_4_007_13    m/z 694.4108     0   2   4   6E+3   13   13.5  RT (min)      lcorbi_t3_neg_4_042_14    m/z 694.4301     0   1   2E+3   13.5   14  RT (min)      lcorbi_t3_neg_4_019_08    m/z 694.4433     0   2   4   6E+3   14.5   15  RT (min)      lcorbi_t3_neg_2_036_35    m/z 694.4706     0   1   2   3E+4   14.5   15  RT (min)      lcorbi_t3_neg_1_055_05    m/z 694.4867     0   1   2   3E+3   14.5   15  RT (min)      lcorbi_t3_neg_3_040_05    m/z 694.5781     0   1E+4   13   13.5  RT (min)      lcorbi_t3_neg_4_034_01    m/z 694.6259     0   1   2E+4   15   15.5  RT (min)      lcorbi_t3_neg_1_055_05    m/z 694.7859     0   2   4   6E+3   18   18.5  RT (min)      lcorbi_t3_neg_2_034_19    m/z 694.8476     0   1   2   3E+3   18   18.5  RT (min)      lcorbi_t3_neg_4_051_40    m/z 695.0830     0   2   4E+3   9.5   10  RT (min)      lcorbi_t3_neg_4_043_11    m/z 695.1437     0   2   4   6E+3   9   9.5  RT (min)      lcorbi_t3_neg_4_023_18    m/z 695.1926     0   1   2   3   4E+3   10.5   11  RT (min)      lcorbi_t3_neg_1_023_03    m/z 695.2747     0   2   4   6E+2   12   12.5  RT (min)      lcorbi_t3_neg_4_034_01    m/z 695.2925     0   2   4   6E+3   11.5   12  RT (min)      lcorbi_t3_neg_2_027_13    m/z 695.3393     0   2   4E+3   14.5   15  RT (min)      lcorbi_t3_neg_4_007_13    m/z 695.3726     0   2   4   6E+3   12.5   13  RT (min)      lcorbi_t3_neg_4_042_14    m/z 695.4182     0   1E+4   13   13.5  RT (min)      lcorbi_t3_neg_2_007_14    m/z 695.4332     0   1   2   3E+4   11.5   12  RT (min)      lcorbi_t3_neg_4_032_23    m/z 695.5038     0   1   2E+3   14.5   15  RT (min)      lcorbi_t3_neg_4_029_31    m/z 695.5181     0   1E+4   14.5   15  RT (min)      lcorbi_t3_neg_2_053_23    m/z 695.5625     0   1   2E+4   13.5   14  RT (min)      lcorbi_t3_neg_4_040_19    m/z 695.5789     0   2   4   6E+2   13.5   14  RT (min)      lcorbi_t3_neg_3_005_23    m/z 695.6309     0   1E+4   15   15.5  RT (min)      lcorbi_t3_neg_4_034_01    m/z 695.8571     0   2   4E+3   18   18.5  RT (min)      lcorbi_t3_neg_2_011_04    m/z 696.2686     0   2   4   6E+2   12   12.5  RT (min)      lcorbi_t3_neg_4_034_01    m/z 696.2988     0   2   4   6   8E+2   14.5   15  RT (min)      lcorbi_t3_neg_1_029_21    m/z 696.3493     0   1E+4   13.5   14  RT (min)      lcorbi_t3_neg_1_023_03    m/z 696.4090     0   1   2   3E+4   11.5   12  RT (min)      lcorbi_t3_neg_4_032_23    m/z 696.4218     0   1E+3   13   13.5  RT (min)      lcorbi_t3_neg_4_032_23    m/z 696.4361     0   2   4   6   8E+3   11.5   12  RT (min)      lcorbi_t3_neg_4_032_23    m/z 696.5390     0   1   2E+4   14.5   15  RT (min)      lcorbi_t3_neg_4_032_23    m/z 696.5501     0   1E+4   14.5   15  RT (min)      lcorbi_t3_neg_4_029_31    m/z 696.6051     0   1   2   3E+4   14   14.5  RT (min)      lcorbi_t3_neg_4_039_32    m/z 696.6355     0   1   2E+4   14.5   15  RT (min)      lcorbi_t3_neg_3_002_24    m/z 696.7714     0   1E+4   6   6.5  RT (min)      lcorbi_t3_neg_2_048_27    m/z 696.7831     0   2   4   6E+3   18   18.5  RT (min)      lcorbi_t3_neg_2_013_08    m/z 697.0664     0   1   2   3   4E+3   7.5   8  RT (min)      lcorbi_t3_neg_4_034_01    m/z 697.1573     0   1   2   3E+3   2   2.5  RT (min)      lcorbi_t3_neg_2_022_24    m/z 697.2473     0   2   4   6E+3   11   11.5  RT (min)      lcorbi_t3_neg_4_008_22    m/z 697.2729     0   1E+4   6   6.5  RT (min)      lcorbi_t3_neg_2_048_27    m/z 697.3086     0   2   4   6E+3   12   12.5  RT (min)      lcorbi_t3_neg_2_050_17    m/z 697.3882     0   1   2   3   4E+3   12.5   13  RT (min)      lcorbi_t3_neg_4_005_27    m/z 697.4811     0   2   4E+3   18   18.5  RT (min)      lcorbi_t3_neg_4_046_37    m/z 697.5300     0   1E+5   11.5   12  RT (min)      lcorbi_t3_neg_3_002_24    m/z 697.6465     0   1E+4   15   15.5  RT (min)      lcorbi_t3_neg_4_034_01    m/z 697.6640     0   1E+4   14   14.5  RT (min)      lcorbi_t3_neg_1_055_05    m/z 697.8149     0   2   4E+3   18   18.5  RT (min)      lcorbi_t3_neg_3_013_28    m/z 697.8629     0   2   4   6E+3   18   18.5  RT (min)      lcorbi_t3_neg_1_006_40    m/z 698.2233     0   1   11   11.5  RT (min)      lcorbi_t3_neg_3_042_22    m/z 698.2285     0   1   2   3E+4   13.5   14  RT (min)      lcorbi_t3_neg_1_023_03    m/z 698.2412     0   1E+4   11   11.5  RT (min)      lcorbi_t3_neg_3_034_21    m/z 698.2764     0   1E+4   13   13.5  RT (min)      lcorbi_t3_neg_4_039_32    m/z 698.2862     0   1E+4   12.5   13  RT (min)      lcorbi_t3_neg_4_032_23    m/z 698.3700     0   1   2E+4   10.5   11  RT (min)      lcorbi_t3_neg_4_011_24    m/z 698.5470     0   2   4   6   8E+3   15   15.5  RT (min)      lcorbi_t3_neg_4_029_31    m/z 698.5865     0   1   14.5   15  RT (min)      lcorbi_t3_neg_3_002_24    m/z 698.6203     0   1E+4   14.5   15  RT (min)      lcorbi_t3_neg_4_039_32    m/z 698.6570     0   1   2   3   4E+3   15   15.5  RT (min)      lcorbi_t3_neg_4_034_01    m/z 698.8940     0   1   2E+3   18   18.5  RT (min)      lcorbi_t3_neg_2_016_11    m/z 699.0474     0   1   2   3E+3   9   9.5  RT (min)      lcorbi_t3_neg_4_028_39    m/z 699.2267     0   2   4   6E+3   7.5   8  RT (min)      lcorbi_t3_neg_4_023_18    m/z 699.2444     0   2   4E+3   11   11.5  RT (min)      lcorbi_t3_neg_4_008_22    m/z 699.2895     0   1   2   3   4E+3   12.5   13  RT (min)      lcorbi_t3_neg_4_032_23    m/z 699.2929     0   1E+4   10.5   11  RT (min)      lcorbi_t3_neg_2_020_05    m/z 699.3229     0   2   4   6   8E+3   12   12.5  RT (min)      lcorbi_t3_neg_1_043_14    m/z 699.3702     0   2   4E+3   13.5   14  RT (min)      lcorbi_t3_neg_4_023_18    m/z 699.3708     0   1   2   3   4E+3   18   18.5  RT (min)      lcorbi_t3_neg_4_040_19    m/z 699.4968     0   2   4   6   8E+3   18   18.5  RT (min)      lcorbi_t3_neg_4_046_37    m/z 699.5445     0   1E+3   14   14.5  RT (min)      lcorbi_t3_neg_4_046_37    m/z 699.5899     0   1   2   3   4E+3   14.5   15  RT (min)      lcorbi_t3_neg_2_053_23    m/z 699.6336     0   2   4   6   8E+3   14.5   15  RT (min)      lcorbi_t3_neg_2_043_33    m/z 700.2572     0   2   4   6   8E+3   13.5   14  RT (min)      lcorbi_t3_neg_4_017_51    m/z 700.2855     0   1   2E+4   5.5   6  RT (min)      lcorbi_t3_neg_4_032_23    m/z 700.3249     0   1   2E+3   11   11.5  RT (min)      lcorbi_t3_neg_4_029_31    m/z 700.3958     0   1E+4   12.5   13  RT (min)      lcorbi_t3_neg_3_005_23    m/z 700.5479     0   1   2E+3   14   14.5  RT (min)      lcorbi_t3_neg_2_053_23    m/z 700.6021     0   2   4   6   8E+3   15   15.5  RT (min)      lcorbi_t3_neg_2_020_05    m/z 700.6359     0   1E+3   14.5   15  RT (min)      lcorbi_t3_neg_4_044_38    m/z 700.6918     0   1   2E+4   14.5   15  RT (min)      lcorbi_t3_neg_4_011_24    m/z 700.8042     0   1   2   3E+3   18   18.5  RT (min)      lcorbi_t3_neg_2_016_11    m/z 700.9115     0   1   2   3E+3   18   18.5  RT (min)      lcorbi_t3_neg_2_011_04    m/z 700.9852     0   1   2   3E+3   9   9.5  RT (min)      lcorbi_t3_neg_2_016_11    m/z 701.0782     0   1   2   3   4E+3   7.5   8  RT (min)      lcorbi_t3_neg_4_034_01    m/z 701.3018     0   1E+4   10.5   11  RT (min)      lcorbi_t3_neg_4_011_24    m/z 701.3108     0   1E+3   13.5   14  RT (min)      lcorbi_t3_neg_3_042_22    m/z 701.3869     0   2   4E+3   13.5   14  RT (min)      lcorbi_t3_neg_4_023_18    m/z 701.4385     0   2   4   6   8E+3   8   8.5  RT (min)      lcorbi_t3_neg_4_017_51    m/z 701.4698     0   1   2   3E+4   18   18.5  RT (min)      lcorbi_t3_neg_2_043_33    m/z 701.4701     0   1E+3   14   14.5  RT (min)      lcorbi_t3_neg_3_005_23    m/z 701.4733     0   1E+4   6.5   7  RT (min)      lcorbi_t3_neg_4_032_23    m/z 701.5307     0   1   2   3E+3   15   15.5  RT (min)      lcorbi_t3_neg_4_039_32    m/z 701.6091     0   1   2E+5   14.5   15  RT (min)      lcorbi_t3_neg_2_020_05    m/z 701.8094     0   1   2E+3   18   18.5  RT (min)      lcorbi_t3_neg_1_047_02    m/z 701.9575     0   1E+4   18   18.5  RT (min)      lcorbi_t3_neg_2_043_33    m/z 701.9579     0   1   2   3   4E+4   13.5   14  RT (min)      lcorbi_t3_neg_2_043_33    m/z 702.1035     0   2   4E+3   9.5   10  RT (min)      lcorbi_t3_neg_4_003_21    m/z 702.2247     0   2   4   6   8E+3   10.5   11  RT (min)      lcorbi_t3_neg_4_034_01    m/z 702.3808     0   2   4E+3   14.5   15  RT (min)      lcorbi_t3_neg_2_035_46    m/z 702.3822     0   1   2E+4   10.5   11  RT (min)      lcorbi_t3_neg_4_007_13    m/z 702.4328     0   2   4   6E+3   13   13.5  RT (min)      lcorbi_t3_neg_4_044_38    m/z 702.5001     0   2   4   6E+4   14   14.5  RT (min)      lcorbi_t3_neg_4_029_31    m/z 702.5334     0   1   2   3   4E+3   15   15.5  RT (min)      lcorbi_t3_neg_4_029_31    m/z 702.6173     0   2   4   6   8E+3   15   15.5  RT (min)      lcorbi_t3_neg_4_039_32    m/z 702.6514     0   1E+4   14.5   15  RT (min)      lcorbi_t3_neg_4_039_32    m/z 702.6693     0   1E+4   14.5   15  RT (min)      lcorbi_t3_neg_4_011_24    m/z 702.8104     0   1   2   3   4E+3   18   18.5  RT (min)      lcorbi_t3_neg_2_002_26    m/z 703.2985     0   1   2   3E+4   14   14.5  RT (min)      lcorbi_t3_neg_4_030_04    m/z 703.3163     0   1   2   3E+4   11   11.5  RT (min)      lcorbi_t3_neg_4_011_24    m/z 703.3318     0   2   4   6E+3   13.5   14  RT (min)      lcorbi_t3_neg_4_023_18    m/z 703.4553     0   1E+5   13   13.5  RT (min)      lcorbi_t3_neg_4_032_23    m/z 703.5481     0   1   14.5   15  RT (min)      lcorbi_t3_neg_2_053_23    m/z 703.5555     0   1E+4   13.5   14  RT (min)      lcorbi_t3_neg_2_052_32    m/z 703.5990     0   1E+5   14   14.5  RT (min)      lcorbi_t3_neg_2_020_05    m/z 703.6170     0   1   2E+4   14.5   15  RT (min)      lcorbi_t3_neg_4_039_32    m/z 703.7104     0   1E+4   14.5   15  RT (min)      lcorbi_t3_neg_4_039_32    m/z 703.8799     0   1E+4   18   18.5  RT (min)      lcorbi_t3_neg_1_004_22    m/z 704.2274     0   1   2   3   4E+3   10.5   11  RT (min)      lcorbi_t3_neg_2_048_27    m/z 704.2477     0   1E+4   11   11.5  RT (min)      lcorbi_t3_neg_2_005_01    m/z 704.3737     0   2   4   6   8E+3   13   13.5  RT (min)      lcorbi_t3_neg_4_017_51    m/z 704.4503     0   1E+4   13.5   14  RT (min)      lcorbi_t3_neg_4_007_13    m/z 704.4506     0   2   4   6E+3   12   12.5  RT (min)      lcorbi_t3_neg_4_044_38    m/z 704.5687     0   1E+4   13.5   14  RT (min)      lcorbi_t3_neg_4_039_32    m/z 704.5758     0   2   4   6E+3   14.5   15  RT (min)      lcorbi_t3_neg_2_022_24    m/z 704.5971     0   1   2E+3   14.5   15  RT (min)      lcorbi_t3_neg_1_052_24    m/z 704.6673     0   1E+4   14.5   15  RT (min)      lcorbi_t3_neg_4_039_32    m/z 704.8767     0   2   4   6E+3   18   18.5  RT (min)      lcorbi_t3_neg_1_004_22    m/z 704.9972     0   1   2E+3   9   9.5  RT (min)      lcorbi_t3_neg_2_021_41    m/z 705.1132     0   1   2   3   4E+3   9   9.5  RT (min)      lcorbi_t3_neg_4_034_01    m/z 705.2214     0   2   4   6E+3   10.5   11  RT (min)      lcorbi_t3_neg_1_043_14    m/z 705.2438     0   1   2   3E+3   11   11.5  RT (min)      lcorbi_t3_neg_1_043_14    m/z 705.2657     0   1E+4   9   9.5  RT (min)      lcorbi_t3_neg_4_006_47    m/z 705.3066     0   1   2   3   4E+2   12   12.5  RT (min)      lcorbi_t3_neg_4_012_17    m/z 705.3324     0   1   2   3   4E+4   11.5   12  RT (min)      lcorbi_t3_neg_2_020_05    m/z 705.3970     0   2   4E+3   8   8.5  RT (min)      lcorbi_t3_neg_4_023_18    m/z 705.4008     0   2   4   6E+3   13   13.5  RT (min)      lcorbi_t3_neg_4_034_01    m/z 705.4097     0   2   4   6   8E+3   10   10.5  RT (min)      lcorbi_t3_neg_4_029_31    m/z 705.5908     0   1E+4   14.5   15  RT (min)      lcorbi_t3_neg_2_052_32    m/z 705.6003     0   1E+4   14.5   15  RT (min)      lcorbi_t3_neg_2_020_05    m/z 705.6327     0   1E+4   14.5   15  RT (min)      lcorbi_t3_neg_4_039_32    m/z 705.6444     0   1E+4   14   14.5  RT (min)      lcorbi_t3_neg_4_011_24    m/z 705.6985     0   2   4   6   8E+3   14.5   15  RT (min)      lcorbi_t3_neg_4_039_32    m/z 705.8174     0   1   2   3E+3   18   18.5  RT (min)      lcorbi_t3_neg_2_019_07    m/z 706.2456     0   1   2E+3   11   11.5  RT (min)      lcorbi_t3_neg_4_048_03    m/z 706.2567     0   1   11.5   12  RT (min)      lcorbi_t3_neg_3_002_24    m/z 706.3051     0   1E+4   14   14.5  RT (min)      lcorbi_t3_neg_4_048_03    m/z 706.4130     0   1   2   3E+3   10   10.5  RT (min)      lcorbi_t3_neg_4_029_31    m/z 706.4467     0   1E+4   13.5   14  RT (min)      lcorbi_t3_neg_4_029_31    m/z 706.6051     0   2   4   6   8E+3   14.5   15  RT (min)      lcorbi_t3_neg_3_002_24    m/z 706.6356     0   2   4   6   8E+3   15   15.5  RT (min)      lcorbi_t3_neg_4_007_13    m/z 706.7779     0   1   1   1.5  RT (min)      lcorbi_t3_neg_3_018_13    m/z 707.0953     0   2   4   6   8E+3   7.5   8  RT (min)      lcorbi_t3_neg_4_034_01    m/z 707.1164     0   2   4   6E+3   8   8.5  RT (min)      lcorbi_t3_neg_4_008_22    m/z 707.2634     0   1   2   3E+3   10.5   11  RT (min)      lcorbi_t3_neg_2_039_31    m/z 707.2739     0   1E+4   8   8.5  RT (min)      lcorbi_t3_neg_2_048_27    m/z 707.2744     0   1E+4   11.5   12  RT (min)      lcorbi_t3_neg_4_008_22    m/z 707.3450     0   2   4   6E+3   12   12.5  RT (min)      lcorbi_t3_neg_3_020_31    m/z 707.3933     0   1E+4   13   13.5  RT (min)      lcorbi_t3_neg_4_042_14    m/z 707.4426     0   1E+4   13.5   14  RT (min)      lcorbi_t3_neg_3_005_23    m/z 707.4497     0   2   4   6E+3   13.5   14  RT (min)      lcorbi_t3_neg_4_029_31    m/z 707.4900     0   1E+4   13   13.5  RT (min)      lcorbi_t3_neg_4_007_13    m/z 707.6361     0   2   4   6   8E+2   14.5   15  RT (min)      lcorbi_t3_neg_2_022_24    m/z 707.8071     0   1   2E+3   18   18.5  RT (min)      lcorbi_t3_neg_3_029_02    m/z 708.1881     0   2   4   6   8E+3   9   9.5  RT (min)      lcorbi_t3_neg_3_019_12    m/z 708.2428     0   1E+4   11   11.5  RT (min)      lcorbi_t3_neg_4_050_02    m/z 708.2764     0   2   4   6   8E+3   8   8.5  RT (min)      lcorbi_t3_neg_2_048_27    m/z 708.2770     0   2   4   6   8E+3   11.5   12  RT (min)      lcorbi_t3_neg_4_008_22    m/z 708.2984     0   1   2   3E+3   13.5   14  RT (min)      lcorbi_t3_neg_4_017_51    m/z 708.3377     0   2   4E+3   13   13.5  RT (min)      lcorbi_t3_neg_1_043_14    m/z 708.3611     0   1   2E+4   13   13.5  RT (min)      lcorbi_t3_neg_4_007_13    m/z 708.4560     0   1   2   3E+4   10.5   11  RT (min)      lcorbi_t3_neg_2_048_27    m/z 708.4772     0   1   2   3   4E+4   14   14.5  RT (min)      lcorbi_t3_neg_2_043_33    m/z 708.4775     0   1   2   3E+4   18   18.5  RT (min)      lcorbi_t3_neg_2_043_33    m/z 708.4932     0   1   2   3E+3   13   13.5  RT (min)      lcorbi_t3_neg_4_007_13    m/z 708.5213     0   1E+4   14.5   15  RT (min)      lcorbi_t3_neg_4_029_31    m/z 708.5395     0   2   4E+3   14.5   15  RT (min)      lcorbi_t3_neg_4_032_23    m/z 708.5701     0   1   2E+3   15   15.5  RT (min)      lcorbi_t3_neg_4_019_08    m/z 708.5808     0   1   13.5   14  RT (min)      lcorbi_t3_neg_4_022_41    m/z 708.5898     0   1E+4   18   18.5  RT (min)      lcorbi_t3_neg_3_002_24    m/z 708.6511     0   2   4   6   8E+3   15   15.5  RT (min)      lcorbi_t3_neg_4_034_01    m/z 708.8091     0   2   4   6   8E+3   18   18.5  RT (min)      lcorbi_t3_neg_4_019_08    m/z 708.8454     0   1E+3   18   18.5  RT (min)      lcorbi_t3_neg_4_039_32    m/z 708.8988     0   1   2   3E+3   18   18.5  RT (min)      lcorbi_t3_neg_2_011_04    m/z 708.9801     0   1E+5   14   14.5  RT (min)      lcorbi_t3_neg_2_043_33    m/z 709.1110     0   2   4E+3   8   8.5  RT (min)      lcorbi_t3_neg_4_034_01    m/z 709.1820     0   2   4   6   8E+3   9   9.5  RT (min)      lcorbi_t3_neg_3_019_12    m/z 709.2528     0   1E+4   13   13.5  RT (min)      lcorbi_t3_neg_4_019_08    m/z 709.2528     0   1E+4   11   11.5  RT (min)      lcorbi_t3_neg_4_007_13    m/z 709.2896     0   2   4   6E+3   12   12.5  RT (min)      lcorbi_t3_neg_4_038_15    m/z 709.4164     0   2   4E+4   11   11.5  RT (min)      lcorbi_t3_neg_4_029_31    m/z 709.4867     0   2   4   6E+3   5.5   6  RT (min)      lcorbi_t3_neg_4_044_38    m/z 709.5066     0   1E+3   14.5   15  RT (min)      lcorbi_t3_neg_2_052_32    m/z 709.5624     0   1   2E+4   14.5   15  RT (min)      lcorbi_t3_neg_4_029_31    m/z 710.2403     0   1E+4   10.5   11  RT (min)      lcorbi_t3_neg_2_007_14    m/z 710.2465     0   1E+4   13   13.5  RT (min)      lcorbi_t3_neg_4_019_08    m/z 710.2738     0   1   2   3   4E+3   12   12.5  RT (min)      lcorbi_t3_neg_2_055_18    m/z 710.2930     0   2   4   6E+3   12   12.5  RT (min)      lcorbi_t3_neg_4_039_32    m/z 710.4007     0   1   2   3E+3   11   11.5  RT (min)      lcorbi_t3_neg_3_005_23    m/z 710.4202     0   1   2   3   4E+4   11   11.5  RT (min)      lcorbi_t3_neg_4_032_23    m/z 710.4685     0   1E+3   13.5   14  RT (min)      lcorbi_t3_neg_4_007_13    m/z 710.4904     0   1E+4   13   13.5  RT (min)      lcorbi_t3_neg_2_007_14    m/z 710.5368     0   1   14.5   15  RT (min)      lcorbi_t3_neg_4_011_24    m/z 710.6054     0   1E+4   18   18.5  RT (min)      lcorbi_t3_neg_3_002_24    m/z 710.8246     0   1   2   3E+3   18   18.5  RT (min)      lcorbi_t3_neg_3_019_12    m/z 710.8666     0   2   4E+3   18   18.5  RT (min)      lcorbi_t3_neg_2_029_12    m/z 711.0141     0   2   4   6E+3   9   9.5  RT (min)      lcorbi_t3_neg_2_015_21    m/z 711.1218     0   2   4   6   8E+3   9   9.5  RT (min)      lcorbi_t3_neg_4_023_18    m/z 711.1282     0   2   4   6E+3   8   8.5  RT (min)      lcorbi_t3_neg_4_040_19    m/z 711.1691     0   1   2E+4   9   9.5  RT (min)      lcorbi_t3_neg_3_019_12    m/z 711.1966     0   2   4   6   8E+3   9   9.5  RT (min)      lcorbi_t3_neg_3_019_12    m/z 711.2683     0   1E+4   13   13.5  RT (min)      lcorbi_t3_neg_1_023_03    m/z 711.2685     0   1E+4   11.5   12  RT (min)      lcorbi_t3_neg_4_007_13    m/z 711.3210     0   1   2E+4   10.5   11  RT (min)      lcorbi_t3_neg_4_042_14    m/z 711.3514     0   2   4E+3   8.5   9  RT (min)      lcorbi_t3_neg_4_023_18    m/z 711.5022     0   2   4   6E+3   6.5   7  RT (min)      lcorbi_t3_neg_4_029_31    m/z 711.5221     0   1E+3   14.5   15  RT (min)      lcorbi_t3_neg_1_055_05    m/z 711.8379     0   1E+4   18   18.5  RT (min)      lcorbi_t3_neg_1_045_44    m/z 712.1831     0   1   2   3   4E+3   11   11.5  RT (min)      lcorbi_t3_neg_2_053_23    m/z 712.2616     0   1E+4   11   11.5  RT (min)      lcorbi_t3_neg_4_007_13    m/z 712.2621     0   1   2E+4   13   13.5  RT (min)      lcorbi_t3_neg_1_023_03    m/z 712.2759     0   1E+4   11.5   12  RT (min)      lcorbi_t3_neg_4_007_13    m/z 712.4403     0   1E+4   11.5   12  RT (min)      lcorbi_t3_neg_4_032_23    m/z 712.5066     0   1E+4   13.5   14  RT (min)      lcorbi_t3_neg_1_055_05    m/z 712.5374     0   2   4   6E+3   13.5   14  RT (min)      lcorbi_t3_neg_2_020_05    m/z 712.5529     0   1E+4   14.5   15  RT (min)      lcorbi_t3_neg_4_029_31    m/z 712.6202     0   2   4E+2   14.5   15  RT (min)      lcorbi_t3_neg_4_044_38    m/z 712.6732     0   1E+4   15   15.5  RT (min)      lcorbi_t3_neg_2_039_31    m/z 713.0114     0   1   2E+3   9   9.5  RT (min)      lcorbi_t3_neg_4_030_04    m/z 713.2518     0   1   2E+4   13.5   14  RT (min)      lcorbi_t3_neg_4_011_24    m/z 713.2817     0   1   13.5   14  RT (min)      lcorbi_t3_neg_1_052_24    m/z 713.2841     0   1   2E+4   12   12.5  RT (min)      lcorbi_t3_neg_4_007_13    m/z 713.3490     0   2   4   6   8E+3   13   13.5  RT (min)      lcorbi_t3_neg_4_007_13    m/z 713.4609     0   1   2   3E+4   14   14.5  RT (min)      lcorbi_t3_neg_3_024_26    m/z 713.4870     0   1E+5   13.5   14  RT (min)      lcorbi_t3_neg_4_019_08    m/z 713.5126     0   1   14   14.5  RT (min)      lcorbi_t3_neg_3_024_26    m/z 713.5543     0   1E+3   14.5   15  RT (min)      lcorbi_t3_neg_1_027_30    m/z 713.5746     0   1   2   3E+3   15   15.5  RT (min)      lcorbi_t3_neg_2_039_31    m/z 713.6050     0   1E+4   14.5   15  RT (min)      lcorbi_t3_neg_4_011_24    m/z 713.8414     0   2   4   6   8E+3   18   18.5  RT (min)      lcorbi_t3_neg_2_015_21    m/z 713.8751     0   2   4E+3   18   18.5  RT (min)      lcorbi_t3_neg_2_055_18    m/z 714.3009     0   1E+4   14.5   15  RT (min)      lcorbi_t3_neg_4_020_44    m/z 714.3479     0   1   2E+4   10.5   11  RT (min)      lcorbi_t3_neg_2_022_24    m/z 714.4074     0   1   13   13.5  RT (min)      lcorbi_t3_neg_1_055_05    m/z 714.4709     0   2   4   6E+4   14   14.5  RT (min)      lcorbi_t3_neg_2_028_38    m/z 714.5625     0   1   2   3E+4   14   14.5  RT (min)      lcorbi_t3_neg_1_044_23    m/z 714.6499     0   1   2E+4   14.5   15  RT (min)      lcorbi_t3_neg_4_039_32    m/z 714.7058     0   2   4   6E+3   14.5   15  RT (min)      lcorbi_t3_neg_4_039_32    m/z 715.0258     0   2   4E+3   9   9.5  RT (min)      lcorbi_t3_neg_3_019_12    m/z 715.1262     0   1E+4   13.5   14  RT (min)      lcorbi_t3_neg_4_045_35    m/z 715.2592     0   2   4   6   8E+3   10.5   11  RT (min)      lcorbi_t3_neg_3_029_02    m/z 715.2799     0   2   4   6   8E+3   11   11.5  RT (min)      lcorbi_t3_neg_2_005_01    m/z 715.2953     0   2   4   6   8E+3   12   12.5  RT (min)      lcorbi_t3_neg_4_042_14    m/z 715.4244     0   1E+4   14.5   15  RT (min)      lcorbi_t3_neg_1_054_37    m/z 715.4737     0   1   2   3   4E+4   14   14.5  RT (min)      lcorbi_t3_neg_3_005_23    m/z 715.4906     0   1E+5   14.5   15  RT (min)      lcorbi_t3_neg_4_019_08    m/z 715.5112     0   1E+4   18   18.5  RT (min)      lcorbi_t3_neg_3_005_23    m/z 715.5295     0   1   2   3   4E+3   14.5   15  RT (min)      lcorbi_t3_neg_3_045_34    m/z 715.5490     0   1   2   3   4E+4   14   14.5  RT (min)      lcorbi_t3_neg_1_044_23    m/z 716.1267     0   2   4   6   8E+3   13.5   14  RT (min)      lcorbi_t3_neg_2_036_35    m/z 716.1323     0   1E+4   9   9.5  RT (min)      lcorbi_t3_neg_3_019_12    m/z 716.1960     0   1   2   3E+3   10   10.5  RT (min)      lcorbi_t3_neg_2_015_21    m/z 716.2102     0   1   2   3E+3   9   9.5  RT (min)      lcorbi_t3_neg_3_019_12    m/z 716.2614     0   1E+4   10.5   11  RT (min)      lcorbi_t3_neg_2_027_13    m/z 716.3016     0   2   4   6E+3   11   11.5  RT (min)      lcorbi_t3_neg_4_023_18    m/z 716.3637     0   1   11   11.5  RT (min)      lcorbi_t3_neg_4_032_23    m/z 716.4142     0   1E+4   11   11.5  RT (min)      lcorbi_t3_neg_3_005_23    m/z 716.4274     0   2   4   6E+3   14.5   15  RT (min)      lcorbi_t3_neg_1_054_37    m/z 716.4512     0   1   13   13.5  RT (min)      lcorbi_t3_neg_2_022_24    m/z 716.4761     0   1E+4   14   14.5  RT (min)      lcorbi_t3_neg_4_044_38    m/z 716.4931     0   2   4E+4   14.5   15  RT (min)      lcorbi_t3_neg_4_054_36    m/z 716.5719     0   1E+4   14   14.5  RT (min)      lcorbi_t3_neg_3_047_37    m/z 716.6654     0   1E+4   14.5   15  RT (min)      lcorbi_t3_neg_4_039_32    m/z 717.0509     0   1   2E+3   7.5   8  RT (min)      lcorbi_t3_neg_3_032_17    m/z 717.0657     0   1   2E+3   9   9.5  RT (min)      lcorbi_t3_neg_4_023_18    m/z 717.0984     0   2   4   6E+3   10   10.5  RT (min)      lcorbi_t3_neg_4_029_31    m/z 717.1243     0   2   4   6   8E+3   13.5   14  RT (min)      lcorbi_t3_neg_2_036_35    m/z 717.1389     0   1E+4   9   9.5  RT (min)      lcorbi_t3_neg_1_023_03    m/z 717.1945     0   2   4   6E+4   9.5   10  RT (min)      lcorbi_t3_neg_4_019_08    m/z 717.2737     0   2   4   6   8E+3   12   12.5  RT (min)      lcorbi_t3_neg_4_023_18    m/z 717.2839     0   1E+4   8.5   9  RT (min)      lcorbi_t3_neg_2_048_27    m/z 717.2978     0   1E+4   13.5   14  RT (min)      lcorbi_t3_neg_1_023_03    m/z 717.3536     0   1   2E+4   6.5   7  RT (min)      lcorbi_t3_neg_2_048_27    m/z 717.3547     0   2   4   6E+3   9.5   10  RT (min)      lcorbi_t3_neg_4_029_31    m/z 717.4242     0   1   2E+3   14   14.5  RT (min)      lcorbi_t3_neg_2_040_43    m/z 717.4769     0   1E+4   6   6.5  RT (min)      lcorbi_t3_neg_4_044_38    m/z 717.4938     0   1   2   3   4E+4   14   14.5  RT (min)      lcorbi_t3_neg_3_005_23    m/z 717.5063     0   1E+5   14.5   15  RT (min)      lcorbi_t3_neg_4_019_08    m/z 717.6253     0   1E+4   14.5   15  RT (min)      lcorbi_t3_neg_4_039_32    m/z 717.8448     0   1   2   3E+3   18   18.5  RT (min)      lcorbi_t3_neg_2_037_09    m/z 717.8551     0   2   4   6E+3   6.5   7  RT (min)      lcorbi_t3_neg_4_042_14    m/z 718.1485     0   1   2   3   4E+3   8   8.5  RT (min)      lcorbi_t3_neg_4_013_09    m/z 718.2715     0   2   4   6   8E+2   11.5   12  RT (min)      lcorbi_t3_neg_4_044_38    m/z 718.2881     0   2   4   6E+2   11   11.5  RT (min)      lcorbi_t3_neg_4_019_08    m/z 718.2988     0   1   2E+4   13.5   14  RT (min)      lcorbi_t3_neg_3_005_23    m/z 718.3079     0   1   2E+3   11.5   12  RT (min)      lcorbi_t3_neg_2_005_01    m/z 718.3146     0   1E+4   14   14.5  RT (min)      lcorbi_t3_neg_4_030_04    m/z 718.3521     0   1   2   3   4E+3   12.5   13  RT (min)      lcorbi_t3_neg_4_034_01    m/z 718.4299     0   1   2   3   4E+4   11.5   12  RT (min)      lcorbi_t3_neg_4_032_23    m/z 718.4431     0   1E+4   14.5   15  RT (min)      lcorbi_t3_neg_2_039_31    m/z 718.5431     0   2   4   6E+3   14   14.5  RT (min)      lcorbi_t3_neg_2_016_11    m/z 718.5654     0   1E+3   13   13.5  RT (min)      lcorbi_t3_neg_1_055_05    m/z 718.5743     0   1   2E+4   13.5   14  RT (min)      lcorbi_t3_neg_4_039_32    m/z 718.6503     0   1E+3   14.5   15  RT (min)      lcorbi_t3_neg_2_012_36    m/z 719.0929     0   2   4   6E+3   10.5   11  RT (min)      lcorbi_t3_neg_4_043_11    m/z 719.1489     0   2   4   6   8E+4   13   13.5  RT (min)      lcorbi_t3_neg_4_045_35    m/z 719.2165     0   1   2   3E+3   13   13.5  RT (min)      lcorbi_t3_neg_4_021_25    m/z 719.2196     0   2   4   6E+3   9   9.5  RT (min)      lcorbi_t3_neg_3_019_12    m/z 719.2308     0   1   2   3   4E+3   12.5   13  RT (min)      lcorbi_t3_neg_4_005_27    m/z 719.2922     0   1   2   3E+3   11   11.5  RT (min)      lcorbi_t3_neg_4_037_29    m/z 719.3018     0   2   4   6E+3   13.5   14  RT (min)      lcorbi_t3_neg_3_023_07    m/z 719.3118     0   1   2   3   4E+3   11.5   12  RT (min)      lcorbi_t3_neg_4_034_01    m/z 719.3568     0   1   2   3E+3   18   18.5  RT (min)      lcorbi_t3_neg_2_003_42    m/z 719.3985     0   2   4   6E+3   13.5   14  RT (min)      lcorbi_t3_neg_4_054_36    m/z 719.4333     0   1E+4   11.5   12  RT (min)      lcorbi_t3_neg_4_032_23    m/z 719.4467     0   1   2   3   4E+4   7.5   8  RT (min)      lcorbi_t3_neg_4_023_18    m/z 719.4706     0   1E+3   13.5   14  RT (min)      lcorbi_t3_neg_4_007_13    m/z 719.4965     0   2   4   6E+3   13.5   14  RT (min)      lcorbi_t3_neg_4_005_27    m/z 719.5626     0   1   2E+4   13   13.5  RT (min)      lcorbi_t3_neg_4_034_01    m/z 719.5720     0   1E+4   13.5   14  RT (min)      lcorbi_t3_neg_1_052_24    m/z 719.5942     0   1E+4   14   14.5  RT (min)      lcorbi_t3_neg_1_055_05    m/z 720.1496     0   2   4E+4   13   13.5  RT (min)      lcorbi_t3_neg_4_045_35    m/z 720.2077     0   1   2   3E+3   10   10.5  RT (min)      lcorbi_t3_neg_4_023_18    m/z 720.2845     0   2   4E+3   11.5   12  RT (min)      lcorbi_t3_neg_2_007_14    m/z 720.3355     0   1E+3   14.5   15  RT (min)      lcorbi_t3_neg_4_042_14    m/z 720.3591     0   1   13   13.5  RT (min)      lcorbi_t3_neg_4_019_08    m/z 720.3599     0   2   4   6E+3   18   18.5  RT (min)      lcorbi_t3_neg_2_034_19    m/z 720.4455     0   2   4   6E+4   12   12.5  RT (min)      lcorbi_t3_neg_4_032_23    m/z 720.4819     0   1E+4   13   13.5  RT (min)      lcorbi_t3_neg_2_053_23    m/z 720.5167     0   1E+4   12.5   13  RT (min)      lcorbi_t3_neg_4_011_24    m/z 720.5265     0   1E+4   13   13.5  RT (min)      lcorbi_t3_neg_4_034_01    m/z 720.5907     0   1   2   3E+4   13.5   14  RT (min)      lcorbi_t3_neg_4_039_32    m/z 720.6401     0   1E+4   14.5   15  RT (min)      lcorbi_t3_neg_4_019_08    m/z 720.7175     0   1E+4   14.5   15  RT (min)      lcorbi_t3_neg_4_039_32    m/z 720.8582     0   1E+4   18   18.5  RT (min)      lcorbi_t3_neg_4_053_43    m/z 721.1467     0   2   4   6E+4   13   13.5  RT (min)      lcorbi_t3_neg_4_045_35    m/z 721.2039     0   1   2   3E+3   10.5   11  RT (min)      lcorbi_t3_neg_4_034_01    m/z 721.2853     0   2   4   6E+3   12   12.5  RT (min)      lcorbi_t3_neg_4_023_18    m/z 721.2889     0   1   2E+4   14   14.5  RT (min)      lcorbi_t3_neg_3_035_03    m/z 721.2894     0   2   4   6   8E+3   8   8.5  RT (min)      lcorbi_t3_neg_2_048_27    m/z 721.3131     0   2   4   6E+3   12   12.5  RT (min)      lcorbi_t3_neg_4_012_17    m/z 721.3272     0   2   4   6E+3   12   12.5  RT (min)      lcorbi_t3_neg_4_012_17    m/z 721.4904     0   1   2E+3   13.5   14  RT (min)      lcorbi_t3_neg_4_005_27    m/z 721.4990     0   1   2E+3   14.5   15  RT (min)      lcorbi_t3_neg_2_053_23    m/z 721.5064     0   2   4   6   8E+3   14.5   15  RT (min)      lcorbi_t3_neg_4_044_38    m/z 721.5242     0   1   2E+4   14   14.5  RT (min)      lcorbi_t3_neg_2_053_23    m/z 721.5853     0   1   13.5   14  RT (min)      lcorbi_t3_neg_4_007_13    m/z 721.6553     0   1E+4   14.5   15  RT (min)      lcorbi_t3_neg_4_039_32    m/z 721.9652     0   1E+4   14.5   15  RT (min)      lcorbi_t3_neg_4_046_37    m/z 722.0462     0   2   4E+3   9   9.5  RT (min)      lcorbi_t3_neg_4_051_40    m/z 722.2121     0   2   4   6   8E+3   11   11.5  RT (min)      lcorbi_t3_neg_4_032_23    m/z 722.2388     0   2   4   6   8E+3   14   14.5  RT (min)      lcorbi_t3_neg_4_053_43    m/z 722.2897     0   1E+4   10.5   11  RT (min)      lcorbi_t3_neg_4_007_13    m/z 722.3002     0   1   2E+4   14   14.5  RT (min)      lcorbi_t3_neg_4_048_03    m/z 722.3026     0   1E+4   12.5   13  RT (min)      lcorbi_t3_neg_3_007_16    m/z 722.3655     0   2   4   6E+3   18   18.5  RT (min)      lcorbi_t3_neg_3_016_42    m/z 722.3658     0   1   2   3   4E+3   14.5   15  RT (min)      lcorbi_t3_neg_2_043_33    m/z 722.3780     0   1E+4   14.5   15   15.5  RT (min)      lcorbi_t3_neg_4_035_33    m/z 722.3836     0   2   4   6   8E+3   8.5   9  RT (min)      lcorbi_t3_neg_4_023_18    m/z 722.4612     0   2   4   6   8E+2   12   12.5   13  RT (min)      lcorbi_t3_neg_4_019_08    m/z 722.4714     0   1   2   3E+3   14.5   15  RT (min)      lcorbi_t3_neg_4_009_50    m/z 722.5299     0   1   14   14.5  RT (min)      lcorbi_t3_neg_1_052_24    m/z 722.6058     0   2   4   6E+4   14   14.5  RT (min)      lcorbi_t3_neg_4_011_24    m/z 722.6161     0   1   2E+4   14.5   15  RT (min)      lcorbi_t3_neg_1_055_05    m/z 722.6303     0   2   4   6   8E+3   15   15.5  RT (min)      lcorbi_t3_neg_4_029_31    m/z 722.7945     0   1   2   3E+3   18   18.5  RT (min)      lcorbi_t3_neg_4_053_43    m/z 722.8958     0   1   2   3   4E+3   18   18.5  RT (min)      lcorbi_t3_neg_4_034_01    m/z 723.1303     0   1   2E+4   9   9.5  RT (min)      lcorbi_t3_neg_4_023_18    m/z 723.2085     0   1E+4   10.5   11  RT (min)      lcorbi_t3_neg_3_037_36    m/z 723.2930     0   2   4E+3   10.5   11  RT (min)      lcorbi_t3_neg_4_007_13    m/z 723.2995     0   2   4   6   8E+3   11.5   12  RT (min)      lcorbi_t3_neg_1_047_02    m/z 723.3735     0   1E+4   10.5   11  RT (min)      lcorbi_t3_neg_4_011_24    m/z 723.4645     0   1E+3   12   12.5  RT (min)      lcorbi_t3_neg_4_019_08    m/z 723.4775     0   1   2   3E+4   15   15.5  RT (min)      lcorbi_t3_neg_4_046_37    m/z 723.5043     0   1   2E+3   14.5   15  RT (min)      lcorbi_t3_neg_4_007_13    m/z 723.5220     0   2   4   6   8E+3   14.5   15  RT (min)      lcorbi_t3_neg_2_004_28    m/z 723.5459     0   1   2   3E+3   13.5   14  RT (min)      lcorbi_t3_neg_4_046_37    m/z 723.5579     0   1   2   3E+3   13.5   14  RT (min)      lcorbi_t3_neg_2_023_39    m/z 723.5779     0   1E+4   14.5   15  RT (min)      lcorbi_t3_neg_4_029_31    m/z 723.6334     0   2   4E+3   15   15.5  RT (min)      lcorbi_t3_neg_4_039_32    m/z 723.6513     0   1E+4   15   15.5  RT (min)      lcorbi_t3_neg_2_012_36    m/z 723.7728     0   1   2E+4   14.5   15  RT (min)      lcorbi_t3_neg_2_052_32    m/z 723.9802     0   2   4   6   8E+3   15   15.5  RT (min)      lcorbi_t3_neg_4_029_31    m/z 724.2117     0   2   4   6   8E+3   10.5   11  RT (min)      lcorbi_t3_neg_3_037_36    m/z 724.2284     0   2   4   6   8E+3   11.5   12  RT (min)      lcorbi_t3_neg_2_020_05    m/z 724.2349     0   2   4E+3   11.5   12  RT (min)      lcorbi_t3_neg_2_026_03    m/z 724.2442     0   2   4E+3   13   13.5  RT (min)      lcorbi_t3_neg_1_055_05    m/z 724.2752     0   2   4E+3   12   12.5  RT (min)      lcorbi_t3_neg_4_013_09    m/z 724.4406     0   1E+4   12   12.5  RT (min)      lcorbi_t3_neg_3_005_23    m/z 724.5047     0   1E+4   13.5   14  RT (min)      lcorbi_t3_neg_4_007_13    m/z 724.5198     0   1E+4   14.5   15  RT (min)      lcorbi_t3_neg_4_054_36    m/z 724.5575     0   1   13.5   14  RT (min)      lcorbi_t3_neg_1_052_24    m/z 724.5814     0   2   4   6   8E+3   14.5   15  RT (min)      lcorbi_t3_neg_4_029_31    m/z 724.5975     0   1   2E+4   13.5   14  RT (min)      lcorbi_t3_neg_4_019_08    m/z 724.6216     0   1E+4   14   14.5  RT (min)      lcorbi_t3_neg_3_002_24    m/z 724.6821     0   1   2   3   4E+3   15   15.5  RT (min)      lcorbi_t3_neg_2_012_36    m/z 724.7948     0   1   2   3E+3   18   18.5  RT (min)      lcorbi_t3_neg_4_008_22    m/z 724.8122     0   1   2   3   4E+3   18   18.5  RT (min)      lcorbi_t3_neg_1_055_05    m/z 724.9880     0   2   4   6E+3   15   15.5  RT (min)      lcorbi_t3_neg_2_039_31    m/z 725.1098     0   1   2E+4   8   8.5  RT (min)      lcorbi_t3_neg_4_023_18    m/z 725.2096     0   2   4   6E+3   10.5   11  RT (min)      lcorbi_t3_neg_1_047_02    m/z 725.2560     0   2   4   6E+3   9.5   10  RT (min)      lcorbi_t3_neg_2_048_27    m/z 725.3890     0   1E+4   11   11.5  RT (min)      lcorbi_t3_neg_4_011_24    m/z 725.4437     0   1E+4   12   12.5  RT (min)      lcorbi_t3_neg_4_032_23    m/z 725.4885     0   2   4E+3   14.5   15  RT (min)      lcorbi_t3_neg_2_024_37    m/z 725.4990     0   1E+3   14.5   15  RT (min)      lcorbi_t3_neg_3_005_23    m/z 725.5131     0   2   4E+3   14   14.5  RT (min)      lcorbi_t3_neg_4_005_27    m/z 725.5195     0   1E+4   14.5   15  RT (min)      lcorbi_t3_neg_4_044_38    m/z 725.5523     0   2   4   6E+4   11.5   12  RT (min)      lcorbi_t3_neg_4_011_24    m/z 725.5546     0   1   2E+4   13.5   14  RT (min)      lcorbi_t3_neg_4_040_19    m/z 725.6497     0   1   2   3E+3   15   15.5  RT (min)      lcorbi_t3_neg_1_055_05    m/z 726.2248     0   2   4   6   8E+3   10   10.5  RT (min)      lcorbi_t3_neg_3_019_12    m/z 726.3998     0   1E+4   5.5   6   6.5  RT (min)      lcorbi_t3_neg_3_037_36    m/z 726.4561     0   1   2   3E+4   11.5   12  RT (min)      lcorbi_t3_neg_4_007_13    m/z 726.5218     0   2   4   6   8E+3   14.5   15  RT (min)      lcorbi_t3_neg_4_048_03    m/z 726.5591     0   1E+4   15   15.5  RT (min)      lcorbi_t3_neg_4_032_23    m/z 726.6253     0   1   2E+4   14.5   15  RT (min)      lcorbi_t3_neg_2_056_56    m/z 726.6616     0   1E+4   15   15.5  RT (min)      lcorbi_t3_neg_2_039_31    m/z 726.7993     0   2   4   6E+3   18   18.5  RT (min)      lcorbi_t3_neg_2_048_27    m/z 726.8424     0   2   4   6E+3   18   18.5  RT (min)      lcorbi_t3_neg_2_011_04    m/z 727.0942     0   1E+4   9   9.5  RT (min)      lcorbi_t3_neg_4_023_18    m/z 727.2250     0   1E+4   12.5   13  RT (min)      lcorbi_t3_neg_3_007_16    m/z 727.2282     0   1E+3   10   10.5  RT (min)      lcorbi_t3_neg_4_029_31    m/z 727.2633     0   1   2E+4   13   13.5  RT (min)      lcorbi_t3_neg_1_023_03    m/z 727.2641     0   1E+4   11.5   12  RT (min)      lcorbi_t3_neg_3_032_17    m/z 727.3243     0   2   4   6E+3   11.5   12  RT (min)      lcorbi_t3_neg_2_027_13    m/z 727.3650     0   1E+4   13   13.5  RT (min)      lcorbi_t3_neg_4_007_13    m/z 727.4108     0   1   2   3   4E+3   13.5   14  RT (min)      lcorbi_t3_neg_2_040_43    m/z 727.4530     0   1   2   3   4E+3   14   14.5  RT (min)      lcorbi_t3_neg_4_046_37    m/z 727.4969     0   1   2   3   4E+3   14.5   15  RT (min)      lcorbi_t3_neg_4_054_36    m/z 727.5125     0   1   2   3E+4   11.5   12  RT (min)      lcorbi_t3_neg_4_011_24    m/z 727.5290     0   1   2E+3   14   14.5  RT (min)      lcorbi_t3_neg_2_052_32    m/z 727.5357     0   2   4   6   8E+3   14.5   15  RT (min)      lcorbi_t3_neg_4_029_31    m/z 727.5406     0   1E+5   11.5   12  RT (min)      lcorbi_t3_neg_4_011_24    m/z 727.6286     0   1   2E+4   14.5   15  RT (min)      lcorbi_t3_neg_4_039_32    m/z 727.6652     0   2   4   6E+3   15   15.5  RT (min)      lcorbi_t3_neg_2_012_36    m/z 727.8457     0   2   4   6E+3   18   18.5  RT (min)      lcorbi_t3_neg_2_029_12    m/z 728.0975     0   1   2   3   4E+3   9   9.5  RT (min)      lcorbi_t3_neg_4_023_18    m/z 728.1673     0   2   4   6   8E+3   9   9.5  RT (min)      lcorbi_t3_neg_4_007_13    m/z 728.2221     0   1   2E+3   10   10.5  RT (min)      lcorbi_t3_neg_3_019_12    m/z 728.2279     0   1   2   3E+3   12.5   13  RT (min)      lcorbi_t3_neg_3_007_16    m/z 728.2408     0   2   4   6   8E+3   11   11.5  RT (min)      lcorbi_t3_neg_3_029_02    m/z 728.2571     0   1   2E+4   13   13.5  RT (min)      lcorbi_t3_neg_1_023_03    m/z 728.3058     0   1E+4   12   12.5  RT (min)      lcorbi_t3_neg_4_023_18    m/z 728.3366     0   1E+4   11.5   12  RT (min)      lcorbi_t3_neg_4_034_01    m/z 728.3685     0   2   4E+3   13   13.5  RT (min)      lcorbi_t3_neg_4_007_13    m/z 728.4082     0   2   4   6E+4   11.5   12  RT (min)      lcorbi_t3_neg_4_016_05    m/z 728.5384     0   1   2   3   4E+3   14.5   15  RT (min)      lcorbi_t3_neg_4_007_13    m/z 728.6025     0   2   4   6E+3   14.5   15  RT (min)      lcorbi_t3_neg_2_043_33    m/z 728.6423     0   2   4E+4   1   1.5  RT (min)      lcorbi_t3_neg_4_047_30    m/z 728.6836     0   1   2   3E+4   14.5   15  RT (min)      lcorbi_t3_neg_4_011_24    m/z 728.7957     0   2   4   6E+3   18   18.5  RT (min)      lcorbi_t3_neg_4_019_08    m/z 728.8120     0   2   4   6   8E+3   18   18.5  RT (min)      lcorbi_t3_neg_4_040_19    m/z 729.0953     0   1   2   3E+3   9   9.5  RT (min)      lcorbi_t3_neg_4_023_18    m/z 729.1886     0   1   2   3E+4   1   1.5  RT (min)      lcorbi_t3_neg_1_055_05    m/z 729.2223     0   1   2   3   4E+3   12.5   13  RT (min)      lcorbi_t3_neg_2_008_16    m/z 729.2269     0   2   4   6   8E+3   10.5   11  RT (min)      lcorbi_t3_neg_4_050_02    m/z 729.2437     0   1   2E+3   11   11.5  RT (min)      lcorbi_t3_neg_3_029_02    m/z 729.2604     0   2   4   6   8E+3   13   13.5  RT (min)      lcorbi_t3_neg_1_023_03    m/z 729.3400     0   1   2   3   4E+3   11.5   12  RT (min)      lcorbi_t3_neg_4_034_01    m/z 729.4242     0   1E+4   13.5   14  RT (min)      lcorbi_t3_neg_2_040_43    m/z 729.4350     0   1   2   3E+4   10.5   11  RT (min)      lcorbi_t3_neg_2_048_27    m/z 729.4714     0   1   2E+4   13   13.5  RT (min)      lcorbi_t3_neg_3_005_23    m/z 729.4755     0   1   2   3E+4   7.5   8  RT (min)      lcorbi_t3_neg_4_023_18    m/z 729.4908     0   1   2E+4   13   13.5  RT (min)      lcorbi_t3_neg_4_032_23    m/z 729.5463     0   2   4   6E+3   14.5   15  RT (min)      lcorbi_t3_neg_4_029_31    m/z 729.8507     0   2   4E+3   18   18.5  RT (min)      lcorbi_t3_neg_4_040_19    m/z 730.0893     0   1E+4   1.5   2  RT (min)      lcorbi_t3_neg_3_020_31    m/z 730.2364     0   2   4E+3   10   10.5  RT (min)      lcorbi_t3_neg_3_019_12    m/z 730.2560     0   1E+3   11.5   12  RT (min)      lcorbi_t3_neg_4_042_14    m/z 730.2772     0   2   4   6E+3   11   11.5  RT (min)      lcorbi_t3_neg_4_034_01    m/z 730.3522     0   1E+4   12   12.5  RT (min)      lcorbi_t3_neg_4_034_01    m/z 730.3904     0   2   4E+3   13   13.5  RT (min)      lcorbi_t3_neg_4_042_14    m/z 730.4122     0   1   2E+3   15   15.5  RT (min)      lcorbi_t3_neg_2_035_46    m/z 730.4569     0   1   2   3E+4   13   13.5  RT (min)      lcorbi_t3_neg_4_019_08    m/z 730.5028     0   2   4   6E+3   13   13.5  RT (min)      lcorbi_t3_neg_1_055_05    m/z 730.5557     0   1   2E+3   13   13.5  RT (min)      lcorbi_t3_neg_3_040_05    m/z 730.5662     0   1   2   3   4E+3   14.5   15  RT (min)      lcorbi_t3_neg_4_039_32    m/z 730.7011     0   1   2E+4   14.5   15  RT (min)      lcorbi_t3_neg_4_011_24    m/z 730.8167     0   2   4   6   8E+3   18   18.5  RT (min)      lcorbi_t3_neg_4_003_21    m/z 730.8294     0   2   4   6   8E+3   18   18.5  RT (min)      lcorbi_t3_neg_1_055_05    m/z 730.8506     0   1   2E+3   18   18.5  RT (min)      lcorbi_t3_neg_2_003_42    m/z 730.9985     0   1E+3   9   9.5  RT (min)      lcorbi_t3_neg_2_007_14    m/z 731.0276     0   2   4E+3   9   9.5  RT (min)      lcorbi_t3_neg_4_042_14    m/z 731.1762     0   1E+5   13   13.5  RT (min)      lcorbi_t3_neg_4_045_35    m/z 731.1840     0   2   4   6   8E+3   12.5   13  RT (min)      lcorbi_t3_neg_4_005_27    m/z 731.3165     0   1E+4   5.5   6  RT (min)      lcorbi_t3_neg_4_042_14    m/z 731.3169     0   2   4   6E+3   11.5   12  RT (min)      lcorbi_t3_neg_4_042_14    m/z 731.3556     0   1   2   3E+3   12   12.5  RT (min)      lcorbi_t3_neg_4_034_01    m/z 731.5063     0   1   2E+4   13   13.5  RT (min)      lcorbi_t3_neg_4_032_23    m/z 731.5186     0   2   4   6E+3   14.5   15  RT (min)      lcorbi_t3_neg_4_034_01    m/z 731.5987     0   2   4   6E+4   14.5   15  RT (min)      lcorbi_t3_neg_1_055_05    m/z 731.8180     0   1E+4   5.5   6  RT (min)      lcorbi_t3_neg_4_042_14    m/z 731.8339     0   2   4   6E+3   18   18.5  RT (min)      lcorbi_t3_neg_2_029_12    m/z 732.1761     0   2   4   6E+4   13   13.5  RT (min)      lcorbi_t3_neg_4_045_35    m/z 732.1874     0   1   2   3E+3   12.5   13  RT (min)      lcorbi_t3_neg_3_022_27    m/z 732.2432     0   2   4   6   8E+3   11   11.5  RT (min)      lcorbi_t3_neg_4_034_01    m/z 732.2717     0   2   4   6E+3   11.5   12  RT (min)      lcorbi_t3_neg_4_034_01    m/z 732.3259     0   1   2   3   4E+3   8   8.5  RT (min)      lcorbi_t3_neg_4_040_19    m/z 732.3473     0   1E+4   10.5   11  RT (min)      lcorbi_t3_neg_4_011_24    m/z 732.3609     0   1   2E+4   13   13.5  RT (min)      lcorbi_t3_neg_4_007_13    m/z 732.3679     0   2   4   6E+3   12.5   13  RT (min)      lcorbi_t3_neg_4_034_01    m/z 732.4067     0   1E+4   11.5   12  RT (min)      lcorbi_t3_neg_4_032_23    m/z 732.5184     0   2   4E+3   13   13.5  RT (min)      lcorbi_t3_neg_4_044_38    m/z 732.5718     0   1E+4   13.5   14  RT (min)      lcorbi_t3_neg_4_017_51    m/z 732.6545     0   1E+4   14.5   15  RT (min)      lcorbi_t3_neg_4_039_32    m/z 732.8152     0   1E+3   18   18.5  RT (min)      lcorbi_t3_neg_4_019_08    m/z 733.0376     0   1   2   3E+3   9.5   10  RT (min)      lcorbi_t3_neg_4_043_11    m/z 733.1085     0   2   4   6E+3   11   11.5  RT (min)      lcorbi_t3_neg_4_043_11    m/z 733.1874     0   1   2   3   4E+3   9   9.5  RT (min)      lcorbi_t3_neg_3_019_12    m/z 733.3227     0   1E+4   8   8.5  RT (min)      lcorbi_t3_neg_4_040_19    m/z 733.3640     0   2   4E+3   18   18.5  RT (min)      lcorbi_t3_neg_2_042_44    m/z 733.4062     0   1   2   3   4E+3   13   13.5  RT (min)      lcorbi_t3_neg_4_029_31    m/z 733.4411     0   2   4E+3   11   11.5  RT (min)      lcorbi_t3_neg_1_055_05    m/z 733.4756     0   1E+4   13   13.5  RT (min)      lcorbi_t3_neg_4_044_38    m/z 733.5066     0   1E+3   14.5   15  RT (min)      lcorbi_t3_neg_2_044_06    m/z 733.5409     0   1E+4   14.5   15  RT (min)      lcorbi_t3_neg_4_044_38    m/z 733.5618     0   1E+4   14   14.5  RT (min)      lcorbi_t3_neg_4_005_27    m/z 733.6133     0   1   2E+4   14.5   15  RT (min)      lcorbi_t3_neg_2_052_32    m/z 733.8311     0   1   2   3E+3   18   18.5  RT (min)      lcorbi_t3_neg_2_048_27    m/z 733.8606     0   1   2   3E+3   18   18.5  RT (min)      lcorbi_t3_neg_3_008_20    m/z 734.2038     0   2   4E+3   9   9.5  RT (min)      lcorbi_t3_neg_3_019_12    m/z 734.2457     0   1   2E+3   10.5   11  RT (min)      lcorbi_t3_neg_4_007_13    m/z 734.2894     0   1E+4   14   14.5  RT (min)      lcorbi_t3_neg_3_011_04    m/z 734.3260     0   2   4   6E+3   8   8.5  RT (min)      lcorbi_t3_neg_4_040_19    m/z 734.5103     0   2   4E+3   14.5   15  RT (min)      lcorbi_t3_neg_4_044_38    m/z 734.5340     0   1   2   3E+3   14   14.5  RT (min)      lcorbi_t3_neg_4_034_01    m/z 734.5466     0   2   4   6   8E+3   14.5   15  RT (min)      lcorbi_t3_neg_4_007_13    m/z 734.6058     0   1   2   3E+4   14   14.5  RT (min)      lcorbi_t3_neg_4_011_24    m/z 735.2673     0   2   4   6E+3   11.5   12  RT (min)      lcorbi_t3_neg_1_023_03    m/z 735.2867     0   1E+4   9.5   10  RT (min)      lcorbi_t3_neg_3_019_12    m/z 735.2950     0   2   4E+3   11.5   12  RT (min)      lcorbi_t3_neg_4_052_26    m/z 735.3726     0   2   4   6E+3   7.5   8  RT (min)      lcorbi_t3_neg_4_023_18    m/z 735.4320     0   1E+4   14.5   15  RT (min)      lcorbi_t3_neg_1_054_37    m/z 735.4420     0   2   4   6E+3   13.5   14  RT (min)      lcorbi_t3_neg_1_020_36    m/z 735.5227     0   1E+4   14.5   15  RT (min)      lcorbi_t3_neg_4_046_37    m/z 735.5294     0   1   2E+5   13   13.5  RT (min)      lcorbi_t3_neg_4_007_13    m/z 735.5590     0   2   4E+3   15   15.5  RT (min)      lcorbi_t3_neg_2_012_36    m/z 735.6565     0   2   4E+4   14   14.5  RT (min)      lcorbi_t3_neg_3_002_24    m/z 735.6675     0   1   2   3   4E+3   15   15.5  RT (min)      lcorbi_t3_neg_2_043_33    m/z 735.8323     0   1   2   3E+3   18   18.5  RT (min)      lcorbi_t3_neg_4_027_12    m/z 736.1641     0   1   2E+4   8   8.5  RT (min)      lcorbi_t3_neg_2_048_27    m/z 736.2615     0   2   4   6   8E+3   11.5   12  RT (min)      lcorbi_t3_neg_4_048_03    m/z 736.2740     0   1E+3   12   12.5  RT (min)      lcorbi_t3_neg_4_019_08    m/z 736.2901     0   2   4   6E+2   9.5   10  RT (min)      lcorbi_t3_neg_1_051_15    m/z 736.3303     0   2   4   6E+3   13.5   14  RT (min)      lcorbi_t3_neg_3_005_23    m/z 736.4257     0   1E+4   13.5   14  RT (min)      lcorbi_t3_neg_2_053_23    m/z 736.4351     0   1   2E+3   14.5   15  RT (min)      lcorbi_t3_neg_4_046_37    m/z 736.5127     0   2   4   6   8E+3   13.5   14  RT (min)      lcorbi_t3_neg_2_053_23    m/z 736.6036     0   1E+5   13.5   14  RT (min)      lcorbi_t3_neg_4_019_08    m/z 736.6459     0   1E+4   15   15.5  RT (min)      lcorbi_t3_neg_4_029_31    m/z 737.0167     0   1   2E+3   9   9.5  RT (min)      lcorbi_t3_neg_4_028_39    m/z 737.1671     0   2   4   6E+3   8   8.5  RT (min)      lcorbi_t3_neg_2_048_27    m/z 737.2572     0   2   4   6E+3   10   10.5  RT (min)      lcorbi_t3_neg_3_019_12    m/z 737.2774     0   2   4E+2   12   12.5  RT (min)      lcorbi_t3_neg_4_019_08    m/z 737.2948     0   1   2E+4   10.5   11  RT (min)      lcorbi_t3_neg_3_053_14    m/z 737.3092     0   2   4   6E+3   14   14.5  RT (min)      lcorbi_t3_neg_3_005_23    m/z 737.4476     0   1E+3   14.5   15  RT (min)      lcorbi_t3_neg_1_044_23    m/z 737.5750     0   2   4   6   8E+2   15   15.5  RT (min)      lcorbi_t3_neg_1_055_05    m/z 737.6673     0   1   2E+3   15   15.5  RT (min)      lcorbi_t3_neg_1_053_32    m/z 737.8074     0   1   2E+3   18   18.5  RT (min)      lcorbi_t3_neg_3_029_02    m/z 737.8318     0   1   2   3E+3   18   18.5  RT (min)      lcorbi_t3_neg_4_014_20    m/z 738.0737     0   1   2   3   4E+3   6   6.5  RT (min)      lcorbi_t3_neg_4_040_19    m/z 738.1612     0   1   2E+4   8   8.5  RT (min)      lcorbi_t3_neg_2_048_27    m/z 738.1990     0   1   2   3   4E+3   11   11.5  RT (min)      lcorbi_t3_neg_3_005_23    m/z 738.2571     0   1E+4   12.5   13  RT (min)      lcorbi_t3_neg_3_007_16    m/z 738.2868     0   1E+4   10   10.5  RT (min)      lcorbi_t3_neg_2_048_27    m/z 738.3120     0   2   4   6   8E+3   8   8.5  RT (min)      lcorbi_t3_neg_2_048_27    m/z 738.3886     0   1   2E+4   7   7.5  RT (min)      lcorbi_t3_neg_4_023_18    m/z 738.4410     0   2   4   6E+3   13.5   14  RT (min)      lcorbi_t3_neg_4_030_04    m/z 738.4848     0   1E+6   13.5   14  RT (min)      lcorbi_t3_neg_4_019_08    m/z 738.5074     0   1E+4   18   18.5  RT (min)      lcorbi_t3_neg_3_005_23    m/z 738.5969     0   1   2E+3   14.5   15  RT (min)      lcorbi_t3_neg_4_019_08    m/z 738.6278     0   2   4   6   8E+3   18   18.5  RT (min)      lcorbi_t3_neg_1_052_24    m/z 738.8239     0   2   4   6E+3   18   18.5  RT (min)      lcorbi_t3_neg_2_048_27    m/z 739.2588     0   1   2   3   4E+3   12   12.5  RT (min)      lcorbi_t3_neg_1_042_18    m/z 739.2795     0   2   4E+3   11.5   12  RT (min)      lcorbi_t3_neg_1_023_03    m/z 739.2981     0   2   4   6E+3   9.5   10  RT (min)      lcorbi_t3_neg_3_019_12    m/z 739.3121     0   1E+4   8   8.5  RT (min)      lcorbi_t3_neg_1_015_27    m/z 739.4912     0   2   4   6E+4   14.5   15  RT (min)      lcorbi_t3_neg_4_019_08    m/z 739.5263     0   2   4   6E+3   13.5   14  RT (min)      lcorbi_t3_neg_1_039_38    m/z 739.5489     0   1   2   3E+4   14   14.5  RT (min)      lcorbi_t3_neg_4_032_23    m/z 739.6649     0   2   4   6E+3   15   15.5  RT (min)      lcorbi_t3_neg_2_056_56    m/z 739.7679     0   2   4   6   8E+3   14.5   15  RT (min)      lcorbi_t3_neg_1_053_32    m/z 739.7949     0   2   4   6   8E+3   15   15.5  RT (min)      lcorbi_t3_neg_2_052_32    m/z 740.2147     0   1   2   3   4E+3   11.5   12  RT (min)      lcorbi_t3_neg_2_020_05    m/z 740.2791     0   2   4   6   8E+3   11   11.5  RT (min)      lcorbi_t3_neg_4_038_15    m/z 740.3155     0   2   4   6E+3   8   8.5  RT (min)      lcorbi_t3_neg_1_026_19    m/z 740.3260     0   1E+4   14   14.5  RT (min)      lcorbi_t3_neg_4_017_51    m/z 740.4509     0   1   2   3E+3   13   13.5  RT (min)      lcorbi_t3_neg_4_032_23    m/z 740.5232     0   1E+4   18   18.5  RT (min)      lcorbi_t3_neg_3_005_23    m/z 740.5632     0   2   4   6E+3   14   14.5  RT (min)      lcorbi_t3_neg_3_002_24    m/z 740.5678     0   2   4   6E+4   14   14.5  RT (min)      lcorbi_t3_neg_3_005_23    m/z 740.5742     0   2   4   6E+2   14   14.5  RT (min)      lcorbi_t3_neg_2_022_24    m/z 740.5796     0   1E+5   13   13.5  RT (min)      lcorbi_t3_neg_3_002_24    m/z 740.6612     0   1E+4   15   15.5  RT (min)      lcorbi_t3_neg_4_011_24    m/z 740.7270     0   2   4   6E+3   14.5   15  RT (min)      lcorbi_t3_neg_2_022_24    m/z 740.7438     0   1E+4   14.5   15  RT (min)      lcorbi_t3_neg_4_039_32    m/z 741.0563     0   1E+4   9.5   10  RT (min)      lcorbi_t3_neg_4_042_14    m/z 741.2670     0   1E+4   13   13.5  RT (min)      lcorbi_t3_neg_4_030_04    m/z 741.3201     0   2   4E+3   14.5   15  RT (min)      lcorbi_t3_neg_4_020_44    m/z 741.3402     0   1   2E+4   5   5.5  RT (min)      lcorbi_t3_neg_2_048_27    m/z 741.3805     0   2   4   6E+3   13.5   14  RT (min)      lcorbi_t3_neg_4_007_13    m/z 741.4595     0   1   13.5   14  RT (min)      lcorbi_t3_neg_4_046_37    m/z 741.4711     0   1   13   13.5  RT (min)      lcorbi_t3_neg_2_040_43    m/z 741.4751     0   1   2E+4   12   12.5  RT (min)      lcorbi_t3_neg_4_007_13    m/z 741.4957     0   1   2E+4   6   6.5  RT (min)      lcorbi_t3_neg_4_044_38    m/z 741.5067     0   1E+5   14.5   15  RT (min)      lcorbi_t3_neg_4_019_08    m/z 741.5423     0   1   13.5   14  RT (min)      lcorbi_t3_neg_4_046_37    m/z 741.5710     0   1   2E+4   14.5   15  RT (min)      lcorbi_t3_neg_2_053_23    m/z 741.6451     0   1   2   3   4E+4   14.5   15  RT (min)      lcorbi_t3_neg_4_011_24    m/z 741.6807     0   1E+4   15   15.5  RT (min)      lcorbi_t3_neg_2_012_36    m/z 741.8191     0   1   18   18.5  RT (min)      lcorbi_t3_neg_4_019_08    m/z 741.8498     0   2   4   6   8E+3   18   18.5  RT (min)      lcorbi_t3_neg_4_019_08    m/z 742.0594     0   2   4   6E+3   9   9.5   10  RT (min)      lcorbi_t3_neg_4_042_14    m/z 742.2161     0   1E+4   12.5   13  RT (min)      lcorbi_t3_neg_4_005_27    m/z 742.2304     0   1   2   3E+3   12   12.5  RT (min)      lcorbi_t3_neg_2_008_16    m/z 742.2710     0   2   4E+3   11.5   12  RT (min)      lcorbi_t3_neg_3_007_16    m/z 742.2757     0   1   2   3E+4   14   14.5  RT (min)      lcorbi_t3_neg_4_030_04    m/z 742.3079     0   1   2E+3   11.5   12  RT (min)      lcorbi_t3_neg_4_036_16    m/z 742.3324     0   1E+4   14.5   15  RT (min)      lcorbi_t3_neg_4_020_44    m/z 742.3817     0   1E+4   5.5   6  RT (min)      lcorbi_t3_neg_4_042_14    m/z 742.5089     0   2   4E+4   14.5   15  RT (min)      lcorbi_t3_neg_4_029_31    m/z 742.5198     0   1   2E+4   14   14.5  RT (min)      lcorbi_t3_neg_4_044_38    m/z 742.5389     0   1E+4   18   18.5  RT (min)      lcorbi_t3_neg_4_032_23    m/z 742.5466     0   1   2E+5   13.5   14  RT (min)      lcorbi_t3_neg_4_007_13    m/z 742.5902     0   2   4   6E+4   14   14.5  RT (min)      lcorbi_t3_neg_1_055_05    m/z 742.8212     0   1   2   3E+3   18   18.5  RT (min)      lcorbi_t3_neg_4_028_39    m/z 742.8492     0   2   4   6E+3   18   18.5  RT (min)      lcorbi_t3_neg_1_006_40    m/z 743.0275     0   1   2   3   4E+3   10.5   11  RT (min)      lcorbi_t3_neg_2_011_04    m/z 743.2790     0   1E+4   14   14.5  RT (min)      lcorbi_t3_neg_4_030_04    m/z 743.2806     0   2   4   6E+3   10.5   11  RT (min)      lcorbi_t3_neg_3_018_13    m/z 743.3739     0   1E+4   7   7.5  RT (min)      lcorbi_t3_neg_4_023_18    m/z 743.5224     0   1   2E+5   14.5   15  RT (min)      lcorbi_t3_neg_4_019_08    m/z 743.5563     0   2   4   6   8E+3   15.5   16  RT (min)      lcorbi_t3_neg_2_007_14    m/z 743.5628     0   1   15   15.5  RT (min)      lcorbi_t3_neg_4_019_08    m/z 743.5777     0   1   2E+4   13.5   14  RT (min)      lcorbi_t3_neg_4_039_32    m/z 743.5831     0   2   4   6   8E+2   14.5   15  RT (min)      lcorbi_t3_neg_4_011_24    m/z 743.6121     0   1E+4   14.5   15  RT (min)      lcorbi_t3_neg_2_053_23    m/z 743.6322     0   1E+4   14.5   15  RT (min)      lcorbi_t3_neg_4_039_32    m/z 743.8229     0   1   2   3E+3   18   18.5  RT (min)      lcorbi_t3_neg_4_050_02    m/z 744.2853     0   1E+4   11   11.5  RT (min)      lcorbi_t3_neg_4_011_24    m/z 744.3126     0   2   4   6E+3   11   11.5  RT (min)      lcorbi_t3_neg_3_032_17    m/z 744.4731     0   1E+4   13   13.5  RT (min)      lcorbi_t3_neg_4_046_37    m/z 744.5060     0   1   2E+3   13   13.5  RT (min)      lcorbi_t3_neg_2_039_31    m/z 744.5246     0   1   2   3E+3   14.5   15  RT (min)      lcorbi_t3_neg_1_022_34    m/z 744.5375     0   2   4   6E+3   13.5   14  RT (min)      lcorbi_t3_neg_4_044_38    m/z 744.8217     0   1   2   3   4E+3   18   18.5  RT (min)      lcorbi_t3_neg_2_016_11    m/z 744.8602     0   2   4   6   8E+3   18   18.5  RT (min)      lcorbi_t3_neg_2_021_41    m/z 745.4686     0   1   2E+3   14.5   15  RT (min)      lcorbi_t3_neg_4_019_08    m/z 745.5765     0   1   2E+4   13.5   14  RT (min)      lcorbi_t3_neg_4_039_32    m/z 745.6476     0   1   2   3E+4   14.5   15  RT (min)      lcorbi_t3_neg_4_039_32    m/z 745.6673     0   1E+4   15   15.5  RT (min)      lcorbi_t3_neg_2_043_33    m/z 745.8386     0   1   2   3E+3   18   18.5  RT (min)      lcorbi_t3_neg_3_034_21    m/z 746.1928     0   1E+4   8   8.5  RT (min)      lcorbi_t3_neg_2_048_27    m/z 746.1977     0   1   12   12.5  RT (min)      lcorbi_t3_neg_4_019_08    m/z 746.2094     0   2   4   6E+3   10   10.5  RT (min)      lcorbi_t3_neg_2_053_23    m/z 746.2906     0   1E+4   13.5   14  RT (min)      lcorbi_t3_neg_1_023_03    m/z 746.3003     0   2   4   6E+3   12   12.5  RT (min)      lcorbi_t3_neg_3_012_47    m/z 746.3282     0   2   4   6   8E+3   11.5   12  RT (min)      lcorbi_t3_neg_4_034_01    m/z 746.4977     0   1   2   3   4E+3   18   18.5  RT (min)      lcorbi_t3_neg_4_032_23    m/z 746.5710     0   2   4   6E+3   14   14.5  RT (min)      lcorbi_t3_neg_4_007_13    m/z 746.6066     0   2   4   6   8E+2   14.5   15  RT (min)      lcorbi_t3_neg_1_052_24    m/z 746.6343     0   1E+4   14.5   15  RT (min)      lcorbi_t3_neg_2_012_36    m/z 747.2480     0   2   4   6   8E+3   13   13.5  RT (min)      lcorbi_t3_neg_4_017_51    m/z 747.3000     0   1   2   3   4E+3   11   11.5  RT (min)      lcorbi_t3_neg_3_022_27    m/z 747.3216     0   1E+3   9.5   10  RT (min)      lcorbi_t3_neg_4_044_38    m/z 747.4077     0   2   4E+3   7.5   8  RT (min)      lcorbi_t3_neg_4_023_18    m/z 747.4229     0   1E+4   7   7.5  RT (min)      lcorbi_t3_neg_4_023_18    m/z 747.5415     0   2   4E+4   14   14.5  RT (min)      lcorbi_t3_neg_4_007_13    m/z 747.5654     0   2   4   6   8E+3   18   18.5  RT (min)      lcorbi_t3_neg_4_034_01    m/z 747.6168     0   1E+4   13.5   14  RT (min)      lcorbi_t3_neg_2_014_29    m/z 747.6280     0   2   4   6   8E+3   13.5   14  RT (min)      lcorbi_t3_neg_2_004_28    m/z 747.6633     0   1   2   3E+4   14.5   15  RT (min)      lcorbi_t3_neg_4_039_32    m/z 747.8359     0   2   4   6   8E+3   18   18.5  RT (min)      lcorbi_t3_neg_3_034_21    m/z 747.8687     0   2   4   6E+3   18   18.5  RT (min)      lcorbi_t3_neg_4_048_03    m/z 748.1900     0   1E+4   8   8.5  RT (min)      lcorbi_t3_neg_2_048_27    m/z 748.2098     0   1   2E+3   10   10.5  RT (min)      lcorbi_t3_neg_3_019_12    m/z 748.2279     0   1E+4   11   11.5  RT (min)      lcorbi_t3_neg_4_032_23    m/z 748.2349     0   1E+4   11   11.5  RT (min)      lcorbi_t3_neg_3_040_05    m/z 748.2794     0   1   2   3   4E+3   11.5   12  RT (min)      lcorbi_t3_neg_3_053_14    m/z 748.3123     0   1E+4   11.5   12  RT (min)      lcorbi_t3_neg_4_007_13    m/z 748.3437     0   2   4   6E+3   12   12.5  RT (min)      lcorbi_t3_neg_3_022_27    m/z 748.4934     0   2   4   6   8E+3   13.5   14  RT (min)      lcorbi_t3_neg_4_046_37    m/z 748.5371     0   1   2E+4   13.5   14  RT (min)      lcorbi_t3_neg_4_007_13    m/z 748.6624     0   2   4E+4   14.5   15  RT (min)      lcorbi_t3_neg_4_019_08    m/z 748.6882     0   1   2   3E+3   15   15.5  RT (min)      lcorbi_t3_neg_1_030_47    m/z 748.8702     0   2   4   6E+3   18   18.5  RT (min)      lcorbi_t3_neg_1_003_41    m/z 749.0757     0   2   4   6E+3   9   9.5  RT (min)      lcorbi_t3_neg_4_023_18    m/z 749.0827     0   1   2   3   4E+3   8   8.5  RT (min)      lcorbi_t3_neg_4_014_20    m/z 749.2310     0   1   2   3E+3   11   11.5  RT (min)      lcorbi_t3_neg_2_053_23    m/z 749.2460     0   2   4   6E+3   11.5   12  RT (min)      lcorbi_t3_neg_3_029_02    m/z 749.3085     0   2   4   6E+3   11   11.5  RT (min)      lcorbi_t3_neg_4_036_16    m/z 749.3149     0   2   4   6E+3   11.5   12  RT (min)      lcorbi_t3_neg_4_007_13    m/z 749.3931     0   1   13.5   14  RT (min)      lcorbi_t3_neg_4_019_08    m/z 749.4357     0   2   4E+3   10   10.5  RT (min)      lcorbi_t3_neg_4_029_31    m/z 749.4880     0   2   4   6E+3   14.5   15  RT (min)      lcorbi_t3_neg_4_004_34    m/z 749.5446     0   2   4   6E+3   13.5   14  RT (min)      lcorbi_t3_neg_4_007_13    m/z 749.5618     0   1   13.5   14  RT (min)      lcorbi_t3_neg_1_052_24    m/z 749.7520     0   1E+4   14.5   15  RT (min)      lcorbi_t3_neg_4_039_32    m/z 749.7726     0   1E+3   14.5   15  RT (min)      lcorbi_t3_neg_3_002_24    m/z 749.8368     0   2   4   6E+3   18   18.5  RT (min)      lcorbi_t3_neg_2_040_43    m/z 750.1135     0   1   2   3   4E+3   9   9.5  RT (min)      lcorbi_t3_neg_3_019_12    m/z 750.2440     0   1E+4   11.5   12  RT (min)      lcorbi_t3_neg_4_016_05    m/z 750.2676     0   2   4   6E+3   12.5   13  RT (min)      lcorbi_t3_neg_1_023_03    m/z 750.2696     0   1E+4   14   14.5  RT (min)      lcorbi_t3_neg_3_011_04    m/z 750.2868     0   2   4   6   8E+3   12.5   13  RT (min)      lcorbi_t3_neg_2_053_23    m/z 750.3207     0   2   4   6E+3   11.5   12  RT (min)      lcorbi_t3_neg_2_008_16    m/z 750.4090     0   2   4   6E+3   15   15.5  RT (min)      lcorbi_t3_neg_4_035_33    m/z 750.4325     0   2   4   6E+3   13   13.5  RT (min)      lcorbi_t3_neg_4_029_31    m/z 750.4712     0   1E+4   13   13.5  RT (min)      lcorbi_t3_neg_4_032_23    m/z 750.6444     0   2   4   6E+3   15   15.5  RT (min)      lcorbi_t3_neg_2_039_31    m/z 750.7730     0   2   4   6   8E+2   14.5   15  RT (min)      lcorbi_t3_neg_1_055_05    m/z 750.8428     0   2   4E+3   18   18.5  RT (min)      lcorbi_t3_neg_4_022_41    m/z 750.8715     0   2   4E+3   18   18.5  RT (min)      lcorbi_t3_neg_1_004_22    m/z 751.2470     0   1   2   3   4E+3   11.5   12  RT (min)      lcorbi_t3_neg_2_020_05    m/z 751.2640     0   1   2   3E+3   11   11.5  RT (min)      lcorbi_t3_neg_2_045_30    m/z 751.3305     0   1E+4   12   12.5  RT (min)      lcorbi_t3_neg_3_010_15    m/z 751.4104     0   1E+4   14.5   15  RT (min)      lcorbi_t3_neg_1_002_16    m/z 751.4351     0   1   2E+4   13   13.5  RT (min)      lcorbi_t3_neg_4_034_01    m/z 751.5314     0   1   2   3   4E+4   11   11.5  RT (min)      lcorbi_t3_neg_1_052_24    m/z 751.5540     0   1   2   3   4E+5   14.5   15  RT (min)      lcorbi_t3_neg_1_039_38    m/z 751.5693     0   2   4   6E+3   15   15.5  RT (min)      lcorbi_t3_neg_4_042_14    m/z 751.6114     0   1E+4   14.5   15  RT (min)      lcorbi_t3_neg_4_019_08    m/z 751.6454     0   1   15   15.5  RT (min)      lcorbi_t3_neg_4_034_01    m/z 751.7678     0   1E+4   15   15.5  RT (min)      lcorbi_t3_neg_2_020_05    m/z 752.2273     0   1   2E+3   10   10.5  RT (min)      lcorbi_t3_neg_3_019_12    m/z 752.2298     0   1   2   3   4E+3   12   12.5  RT (min)      lcorbi_t3_neg_3_007_16    m/z 752.2481     0   1   2   3E+3   11.5   12  RT (min)      lcorbi_t3_neg_2_008_16    m/z 752.2597     0   1   12   12.5  RT (min)      lcorbi_t3_neg_3_014_19    m/z 752.2840     0   1E+3   12   12.5  RT (min)      lcorbi_t3_neg_3_014_19    m/z 752.2999     0   1   2E+4   13.5   14  RT (min)      lcorbi_t3_neg_1_023_03    m/z 752.3364     0   1E+4   11.5   12  RT (min)      lcorbi_t3_neg_3_007_16    m/z 752.3407     0   1E+4   14.5   15  RT (min)      lcorbi_t3_neg_4_032_23    m/z 752.4127     0   2   4   6E+3   14.5   15  RT (min)      lcorbi_t3_neg_1_007_26    m/z 752.4267     0   1E+4   11   11.5  RT (min)      lcorbi_t3_neg_2_039_31    m/z 752.4270     0   1   2E+4   6   6.5  RT (min)      lcorbi_t3_neg_4_044_38    m/z 752.4383     0   2   4   6E+3   13   13.5  RT (min)      lcorbi_t3_neg_2_007_14    m/z 752.4461     0   1E+4   11   11.5  RT (min)      lcorbi_t3_neg_3_020_31    m/z 752.4993     0   2   4   6   8E+3   14.5   15  RT (min)      lcorbi_t3_neg_4_029_31    m/z 752.5882     0   1E+4   15.5   16  RT (min)      lcorbi_t3_neg_4_042_14    m/z 752.5907     0   1   2E+4   14   14.5  RT (min)      lcorbi_t3_neg_2_052_32    m/z 752.6138     0   1E+5   14.5   15  RT (min)      lcorbi_t3_neg_4_019_08    m/z 752.6441     0   1   2   3E+5   14   14.5  RT (min)      lcorbi_t3_neg_3_002_24    m/z 752.6709     0   1   2   3E+4   13.5   14  RT (min)      lcorbi_t3_neg_4_039_32    m/z 752.6853     0   2   4E+3   14.5   15  RT (min)      lcorbi_t3_neg_2_053_23    m/z 752.8025     0   1   2E+3   18   18.5  RT (min)      lcorbi_t3_neg_4_013_09    m/z 752.8142     0   1   2E+3   18   18.5  RT (min)      lcorbi_t3_neg_2_006_10    m/z 753.0561     0   2   4   6   8E+3   10.5   11  RT (min)      lcorbi_t3_neg_4_030_04    m/z 753.2401     0   2   4   6   8E+3   11.5   12  RT (min)      lcorbi_t3_neg_1_047_02    m/z 753.2628     0   1   12   12.5  RT (min)      lcorbi_t3_neg_3_014_19    m/z 753.2783     0   2   4   6E+3   13   13.5  RT (min)      lcorbi_t3_neg_4_032_23    m/z 753.2797     0   1   2   3   4E+3   11.5   12  RT (min)      lcorbi_t3_neg_4_007_13    m/z 753.3032     0   2   4   6E+3   13.5   14  RT (min)      lcorbi_t3_neg_1_023_03    m/z 753.3097     0   1E+4   12   12.5  RT (min)      lcorbi_t3_neg_4_040_19    m/z 753.3398     0   2   4E+3   11.5   12  RT (min)      lcorbi_t3_neg_4_007_13    m/z 753.3439     0   2   4E+3   14.5   15  RT (min)      lcorbi_t3_neg_4_032_23    m/z 753.3505     0   1   2   3   4E+3   18   18.5  RT (min)      lcorbi_t3_neg_2_003_42    m/z 753.3935     0   1E+4   4.5   5  RT (min)      lcorbi_t3_neg_4_042_14    m/z 753.4234     0   1   2   3   4E+4   11   11.5  RT (min)      lcorbi_t3_neg_4_029_31    m/z 753.4684     0   1   14   14.5  RT (min)      lcorbi_t3_neg_4_029_31    m/z 753.4948     0   2   4   6   8E+3   14.5   15  RT (min)      lcorbi_t3_neg_4_029_31    m/z 753.5131     0   1E+3   14.5   15  RT (min)      lcorbi_t3_neg_4_019_08    m/z 753.5382     0   2   4   6E+3   13.5   14  RT (min)      lcorbi_t3_neg_4_054_36    m/z 753.5471     0   2   4   6   8E+4   11.5   12  RT (min)      lcorbi_t3_neg_1_052_24    m/z 753.5504     0   2   4   6E+3   14.5   15  RT (min)      lcorbi_t3_neg_4_039_32    m/z 753.6060     0   1E+3   15   15.5  RT (min)      lcorbi_t3_neg_3_002_24    m/z 753.6385     0   1E+5   18   18.5  RT (min)      lcorbi_t3_neg_3_002_24    m/z 753.6473     0   1   2E+6   14   14.5  RT (min)      lcorbi_t3_neg_3_002_24    m/z 753.8103     0   2   4   6E+3   15   15.5  RT (min)      lcorbi_t3_neg_2_052_32    m/z 754.2633     0   1   12   12.5  RT (min)      lcorbi_t3_neg_2_034_19    m/z 754.2723     0   1E+4   13   13.5  RT (min)      lcorbi_t3_neg_1_023_03    m/z 754.3379     0   2   4   6   8E+3   14.5   15  RT (min)      lcorbi_t3_neg_4_032_23    m/z 754.3534     0   2   4E+3   18   18.5  RT (min)      lcorbi_t3_neg_2_042_44    m/z 754.3565     0   2   4   6E+3   14.5   15  RT (min)      lcorbi_t3_neg_4_042_14    m/z 754.4268     0   1E+4   11   11.5  RT (min)      lcorbi_t3_neg_4_029_31    m/z 754.4460     0   1   2E+4   11   11.5  RT (min)      lcorbi_t3_neg_4_029_31    m/z 754.4543     0   1   2E+3   13   13.5  RT (min)      lcorbi_t3_neg_1_039_38    m/z 754.4621     0   2   4   6   8E+3   11   11.5  RT (min)      lcorbi_t3_neg_4_007_13    m/z 754.4872     0   1E+4   11.5   12  RT (min)      lcorbi_t3_neg_4_032_23    m/z 754.4975     0   1   2   3   4E+3   14.5   15  RT (min)      lcorbi_t3_neg_4_034_01    m/z 754.5155     0   1   2   3   4E+4   14.5   15  RT (min)      lcorbi_t3_neg_4_019_08    m/z 754.5538     0   1   2E+4   14.5   15  RT (min)      lcorbi_t3_neg_1_055_05    m/z 754.7867     0   1   2E+3   15   15.5  RT (min)      lcorbi_t3_neg_1_056_56    m/z 754.8243     0   1   2   3E+3   18   18.5  RT (min)      lcorbi_t3_neg_2_029_12    m/z 754.8510     0   2   4   6   8E+3   18   18.5  RT (min)      lcorbi_t3_neg_2_034_19    m/z 755.1092     0   1   2E+4   9   9.5  RT (min)      lcorbi_t3_neg_4_007_13    m/z 755.1744     0   2   4E+4   13   13.5  RT (min)      lcorbi_t3_neg_4_045_35    m/z 755.2631     0   2   4   6E+3   10.5   11  RT (min)      lcorbi_t3_neg_4_007_13    m/z 755.2952     0   1E+4   12   12.5  RT (min)      lcorbi_t3_neg_4_038_15    m/z 755.2991     0   1   2   3E+4   14   14.5  RT (min)      lcorbi_t3_neg_4_048_03    m/z 755.3067     0   1E+4   12   12.5  RT (min)      lcorbi_t3_neg_4_040_19    m/z 755.3937     0   2   4E+3   13   13.5  RT (min)      lcorbi_t3_neg_4_034_01    m/z 755.4877     0   2   4E+3   13.5   14  RT (min)      lcorbi_t3_neg_3_047_37    m/z 755.5291     0   1E+4   14.5   15  RT (min)      lcorbi_t3_neg_4_019_08    m/z 755.5627     0   1E+5   12   12.5  RT (min)      lcorbi_t3_neg_2_022_24    m/z 755.5696     0   1E+4   14.5   15  RT (min)      lcorbi_t3_neg_4_019_08    m/z 755.5913     0   2   4   6   8E+3   15.5   16  RT (min)      lcorbi_t3_neg_4_042_14    m/z 755.5968     0   1E+4   13   13.5  RT (min)      lcorbi_t3_neg_1_052_24    m/z 755.6541     0   2   4E+4   18   18.5  RT (min)      lcorbi_t3_neg_3_002_24    m/z 755.7627     0   2   4   6   8E+3   14.5   15  RT (min)      lcorbi_t3_neg_2_052_32    m/z 756.1038     0   2   4   6   8E+3   9   9.5  RT (min)      lcorbi_t3_neg_2_048_27    m/z 756.2480     0   2   4   6E+3   10.5   11  RT (min)      lcorbi_t3_neg_2_053_23    m/z 756.3537     0   2   4E+3   14.5   15  RT (min)      lcorbi_t3_neg_4_010_28    m/z 756.3590     0   2   4   6E+3   18   18.5  RT (min)      lcorbi_t3_neg_2_042_44    m/z 756.5343     0   2   4E+4   14   14.5  RT (min)      lcorbi_t3_neg_4_019_08    m/z 756.5459     0   1   2   3   4E+4   14   14.5  RT (min)      lcorbi_t3_neg_2_053_23    m/z 756.5893     0   1E+4   14.5   15  RT (min)      lcorbi_t3_neg_3_037_36    m/z 756.6108     0   1   2E+4   13.5   14  RT (min)      lcorbi_t3_neg_2_022_24    m/z 756.6415     0   1   14.5   15  RT (min)      lcorbi_t3_neg_4_029_31    m/z 756.8206     0   2   4E+3   18   18.5  RT (min)      lcorbi_t3_neg_3_026_44    m/z 757.3056     0   1E+4   7.5   8  RT (min)      lcorbi_t3_neg_2_048_27    m/z 757.3234     0   2   4   6E+3   14.5   15  RT (min)      lcorbi_t3_neg_4_012_17    m/z 757.3325     0   2   4E+3   12   12.5  RT (min)      lcorbi_t3_neg_4_042_14    m/z 757.4553     0   2   4   6   8E+3   14   14.5  RT (min)      lcorbi_t3_neg_2_040_43    m/z 757.4755     0   2   4   6E+3   13   13.5  RT (min)      lcorbi_t3_neg_4_044_38    m/z 757.4852     0   2   4   6   8E+3   18   18.5  RT (min)      lcorbi_t3_neg_4_046_37    m/z 757.5429     0   1   2E+4   14.5   15  RT (min)      lcorbi_t3_neg_4_019_08    m/z 757.5708     0   1   14   14.5  RT (min)      lcorbi_t3_neg_2_053_23    m/z 757.5833     0   1E+4   14.5   15  RT (min)      lcorbi_t3_neg_1_052_24    m/z 757.6007     0   1E+4   14.5   15  RT (min)      lcorbi_t3_neg_2_052_32    m/z 758.0462     0   2   4   6E+3   9   9.5  RT (min)      lcorbi_t3_neg_4_042_14    m/z 758.2704     0   1   2E+4   13.5   14  RT (min)      lcorbi_t3_neg_1_023_03    m/z 758.3446     0   2   4   6   8E+3   12.5   13  RT (min)      lcorbi_t3_neg_4_034_01    m/z 758.3538     0   1   2E+4   10.5   11  RT (min)      lcorbi_t3_neg_4_011_24    m/z 758.4455     0   1E+5   13.5   14  RT (min)      lcorbi_t3_neg_4_029_31    m/z 758.6244     0   1   2E+3   15   15.5  RT (min)      lcorbi_t3_neg_4_011_24    m/z 758.6510     0   1E+4   14.5   15  RT (min)      lcorbi_t3_neg_2_039_31    m/z 758.7459     0   1   14.5   15  RT (min)      lcorbi_t3_neg_4_044_38    m/z 758.8311     0   1E+3   18   18.5  RT (min)      lcorbi_t3_neg_4_019_08    m/z 759.0585     0   2   4E+3   9.5   10  RT (min)      lcorbi_t3_neg_2_029_12    m/z 759.2672     0   2   4   6   8E+3   11   11.5  RT (min)      lcorbi_t3_neg_2_008_16    m/z 759.2840     0   2   4   6E+3   13.5   14  RT (min)      lcorbi_t3_neg_4_023_18    m/z 759.3116     0   1   2   3   4E+3   11.5   12  RT (min)      lcorbi_t3_neg_3_053_14    m/z 759.3480     0   1   2   3E+3   12.5   13  RT (min)      lcorbi_t3_neg_4_034_01    m/z 759.3838     0   1E+4   7   7.5  RT (min)      lcorbi_t3_neg_4_042_14    m/z 759.4829     0   1   2   3   4E+3   14.5   15  RT (min)      lcorbi_t3_neg_3_037_36    m/z 759.5190     0   1E+4   13.5   14  RT (min)      lcorbi_t3_neg_4_032_23    m/z 759.6170     0   1   14.5   15  RT (min)      lcorbi_t3_neg_4_045_35    m/z 759.6268     0   1   2E+3   15   15.5  RT (min)      lcorbi_t3_neg_2_043_33    m/z 759.6467     0   1E+3   15   15.5  RT (min)      lcorbi_t3_neg_1_055_05    m/z 759.6544     0   1E+4   14.5   15  RT (min)      lcorbi_t3_neg_1_055_05    m/z 759.8687     0   2   4E+3   18   18.5  RT (min)      lcorbi_t3_neg_3_015_11    m/z 759.8854     0   2   4   6   8E+3   7   7.5  RT (min)      lcorbi_t3_neg_4_042_14    m/z 760.2132     0   1   2   3   4E+3   12   12.5  RT (min)      lcorbi_t3_neg_4_032_23    m/z 760.2466     0   2   4   6   8E+3   10.5   11  RT (min)      lcorbi_t3_neg_2_045_30    m/z 760.2498     0   1E+4   13.5   14  RT (min)      lcorbi_t3_neg_1_023_03    m/z 760.2634     0   1E+4   11   11.5  RT (min)      lcorbi_t3_neg_2_010_02    m/z 760.2942     0   1   2   3E+3   13.5   14  RT (min)      lcorbi_t3_neg_4_008_22    m/z 760.3695     0   1E+4   11   11.5  RT (min)      lcorbi_t3_neg_4_011_24    m/z 760.5308     0   2   4   6   8E+3   14   14.5  RT (min)      lcorbi_t3_neg_2_039_31    m/z 760.5468     0   2   4   6   8E+4   13.5   14  RT (min)      lcorbi_t3_neg_4_044_38    m/z 760.5856     0   1E+4   14.5   15  RT (min)      lcorbi_t3_neg_4_005_27    m/z 760.6045     0   1E+3   15   15.5  RT (min)      lcorbi_t3_neg_4_021_25    m/z 760.6267     0   2   4   6E+3   14.5   15  RT (min)      lcorbi_t3_neg_4_039_32    m/z 760.6464     0   2   4   6E+3   14.5   15  RT (min)      lcorbi_t3_neg_1_053_32    m/z 760.6591     0   2   4   6   8E+3   15   15.5  RT (min)      lcorbi_t3_neg_2_012_36    m/z 760.8364     0   2   4   6E+3   18   18.5  RT (min)      lcorbi_t3_neg_1_045_44    m/z 761.0922     0   1E+4   14.5   15  RT (min)      lcorbi_t3_neg_4_039_32    m/z 761.2829     0   2   4   6   8E+3   11.5   12  RT (min)      lcorbi_t3_neg_2_008_16    m/z 761.4282     0   2   4   6E+3   13.5   14  RT (min)      lcorbi_t3_neg_4_019_08    m/z 761.4473     0   1E+4   14.5   15  RT (min)      lcorbi_t3_neg_1_054_37    m/z 761.5493     0   1E+4   15   15.5  RT (min)      lcorbi_t3_neg_4_038_15    m/z 761.5720     0   2   4   6   8E+3   13.5   14  RT (min)      lcorbi_t3_neg_2_052_32    m/z 761.5973     0   2   4   6   8E+2   14.5   15  RT (min)      lcorbi_t3_neg_4_019_08    m/z 761.6327     0   1   2E+4   14.5   15  RT (min)      lcorbi_t3_neg_4_039_32    m/z 761.6624     0   2   4   6   8E+2   15   15.5  RT (min)      lcorbi_t3_neg_1_055_05    m/z 761.7468     0   2   4   6   8E+3   14.5   15  RT (min)      lcorbi_t3_neg_4_001_49    m/z 761.8388     0   2   4   6E+3   18   18.5  RT (min)      lcorbi_t3_neg_2_038_22    m/z 762.1989     0   1   2   3E+3   11   11.5  RT (min)      lcorbi_t3_neg_2_026_03    m/z 762.2586     0   1   12   12.5  RT (min)      lcorbi_t3_neg_4_040_19    m/z 762.2770     0   1E+4   11.5   12  RT (min)      lcorbi_t3_neg_2_008_16    m/z 762.2852     0   1E+4   14   14.5  RT (min)      lcorbi_t3_neg_3_011_04    m/z 762.3754     0   2   4   6   8E+3   14.5   15  RT (min)      lcorbi_t3_neg_4_029_31    m/z 762.4171     0   1E+4   11.5   12  RT (min)      lcorbi_t3_neg_4_032_23    m/z 762.4324     0   1   2   3E+4   9   9.5  RT (min)      lcorbi_t3_neg_2_048_27    m/z 762.4409     0   2   4   6E+3   13.5   14  RT (min)      lcorbi_t3_neg_4_030_04    m/z 762.4857     0   1E+4   13.5   14  RT (min)      lcorbi_t3_neg_2_053_23    m/z 762.5025     0   1E+3   14.5   15  RT (min)      lcorbi_t3_neg_4_019_08    m/z 762.5194     0   2   4   6E+3   18   18.5  RT (min)      lcorbi_t3_neg_4_029_31    m/z 762.5385     0   2   4   6   8E+3   13.5   14  RT (min)      lcorbi_t3_neg_3_017_51    m/z 762.5479     0   1   2E+4   14   14.5  RT (min)      lcorbi_t3_neg_4_007_13    m/z 762.6050     0   1   2E+4   15   15.5  RT (min)      lcorbi_t3_neg_4_039_32    m/z 762.6430     0   1E+4   14.5   15  RT (min)      lcorbi_t3_neg_4_011_24    m/z 762.7733     0   2   4   6E+3   18   18.5  RT (min)      lcorbi_t3_neg_2_034_19    m/z 762.8356     0   1   2   3E+3   18   18.5  RT (min)      lcorbi_t3_neg_2_026_03    m/z 763.0703     0   1   2   3E+3   9.5   10  RT (min)      lcorbi_t3_neg_4_023_18    m/z 763.1945     0   1   2   3   4E+3   12.5   13  RT (min)      lcorbi_t3_neg_4_005_27    m/z 763.2063     0   2   4E+3   9   9.5  RT (min)      lcorbi_t3_neg_3_019_12    m/z 763.2620     0   2   4   6E+2   12   12.5  RT (min)      lcorbi_t3_neg_4_040_19    m/z 763.2806     0   2   4   6E+3   11.5   12  RT (min)      lcorbi_t3_neg_3_024_26    m/z 763.3042     0   1E+3   13   13.5  RT (min)      lcorbi_t3_neg_2_005_01    m/z 763.3384     0   1E+4   12   12.5  RT (min)      lcorbi_t3_neg_4_040_19    m/z 763.3599     0   1   2   3E+3   12.5   13  RT (min)      lcorbi_t3_neg_4_005_27    m/z 763.4357     0   1E+4   9   9.5  RT (min)      lcorbi_t3_neg_2_048_27    m/z 763.4529     0   1   2E+4   13.5   14  RT (min)      lcorbi_t3_neg_4_032_23    m/z 763.4632     0   1E+3   14.5   15  RT (min)      lcorbi_t3_neg_1_021_31    m/z 763.5492     0   1   2   3E+4   14   14.5  RT (min)      lcorbi_t3_neg_1_043_14    m/z 763.6128     0   1   2   3E+2   15   15.5  RT (min)      lcorbi_t3_neg_4_019_08    m/z 763.6482     0   1E+4   14.5   15  RT (min)      lcorbi_t3_neg_4_039_32    m/z 763.6611     0   1   2   3E+3   15   15.5  RT (min)      lcorbi_t3_neg_2_039_31    m/z 763.6977     0   1   2   3   4E+3   15   15.5  RT (min)      lcorbi_t3_neg_2_039_31    m/z 763.8442     0   2   4   6   8E+3   18   18.5  RT (min)      lcorbi_t3_neg_4_015_10    m/z 764.1148     0   1E+4   15   15.5  RT (min)      lcorbi_t3_neg_4_039_32    m/z 764.2117     0   1   2E+4   10.5   11  RT (min)      lcorbi_t3_neg_3_053_14    m/z 764.2916     0   2   4   6   8E+3   11.5   12  RT (min)      lcorbi_t3_neg_4_029_31    m/z 764.3717     0   2   4   6   8E+3   14.5   15  RT (min)      lcorbi_t3_neg_1_021_31    m/z 764.4562     0   1E+4   13.5   14  RT (min)      lcorbi_t3_neg_4_044_38    m/z 764.4656     0   2   4E+3   14.5   15  RT (min)      lcorbi_t3_neg_1_054_37    m/z 764.4869     0   1   13.5   14  RT (min)      lcorbi_t3_neg_4_034_01    m/z 764.5081     0   2   4   6   8E+3   12   12.5  RT (min)      lcorbi_t3_neg_1_044_23    m/z 764.5635     0   1   2E+4   13.5   14  RT (min)      lcorbi_t3_neg_4_007_13    m/z 764.5711     0   1E+3   14   14.5  RT (min)      lcorbi_t3_neg_1_052_24    m/z 764.6205     0   2   4   6   8E+3   15   15.5  RT (min)      lcorbi_t3_neg_4_039_32    m/z 764.6363     0   1   2   3   4E+4   14   14.5  RT (min)      lcorbi_t3_neg_2_022_24    m/z 764.6583     0   1E+4   15   15.5  RT (min)      lcorbi_t3_neg_2_039_31    m/z 764.7705     0   2   4E+3   18   18.5  RT (min)      lcorbi_t3_neg_2_007_14    m/z 764.7886     0   2   4   6E+3   14.5   15  RT (min)      lcorbi_t3_neg_4_001_49    m/z 765.1225     0   1E+4   15   15.5  RT (min)      lcorbi_t3_neg_4_039_32    m/z 765.1433     0   1   2   3E+3   9   9.5  RT (min)      lcorbi_t3_neg_3_019_12    m/z 765.2798     0   2   4   6   8E+3   13.5   14  RT (min)      lcorbi_t3_neg_1_023_03    m/z 765.2958     0   2   4   6   8E+2   12   12.5  RT (min)      lcorbi_t3_neg_4_040_19    m/z 765.3041     0   1E+4   13   13.5  RT (min)      lcorbi_t3_neg_1_023_03    m/z 765.4516     0   2   4   6   8E+3   13   13.5  RT (min)      lcorbi_t3_neg_4_029_31    m/z 765.4693     0   1   14   14.5  RT (min)      lcorbi_t3_neg_4_007_13    m/z 765.4749     0   2   4   6E+3   14.5   15  RT (min)      lcorbi_t3_neg_4_054_36    m/z 765.4906     0   1   2   3   4E+3   13.5   14  RT (min)      lcorbi_t3_neg_4_054_36    m/z 765.5136     0   1   2   3E+2   14.5   15  RT (min)      lcorbi_t3_neg_4_019_08    m/z 765.5309     0   1   2   3E+3   15   15.5  RT (min)      lcorbi_t3_neg_1_044_23    m/z 765.5586     0   1E+4   13.5   14  RT (min)      lcorbi_t3_neg_3_053_14    m/z 765.5818     0   1E+4   12.5   13  RT (min)      lcorbi_t3_neg_4_007_13    m/z 765.6244     0   1E+4   15   15.5  RT (min)      lcorbi_t3_neg_4_029_31    m/z 765.6474     0   1E+5   14   14.5  RT (min)      lcorbi_t3_neg_1_052_24    m/z 765.6639     0   1E+4   14.5   15  RT (min)      lcorbi_t3_neg_4_011_24    m/z 765.6979     0   2   4   6   8E+3   15   15.5  RT (min)      lcorbi_t3_neg_2_012_36    m/z 765.8023     0   1   2   3E+3   18   18.5  RT (min)      lcorbi_t3_neg_2_020_05    m/z 765.8501     0   2   4   6E+3   18   18.5  RT (min)      lcorbi_t3_neg_1_003_41    m/z 766.3573     0   1E+4   10.5   11  RT (min)      lcorbi_t3_neg_4_011_24    m/z 766.4944     0   1   2E+4   14.5   15  RT (min)      lcorbi_t3_neg_4_007_13    m/z 766.5306     0   1E+5   13.5   14  RT (min)      lcorbi_t3_neg_4_044_38    m/z 766.5715     0   1   2E+4   14.5   15  RT (min)      lcorbi_t3_neg_4_042_14    m/z 766.5782     0   1   2   3   4E+4   13.5   14  RT (min)      lcorbi_t3_neg_4_039_32    m/z 766.5869     0   1   2   3E+4   13   13.5  RT (min)      lcorbi_t3_neg_4_039_32    m/z 766.6280     0   1   15   15.5  RT (min)      lcorbi_t3_neg_4_019_08    m/z 766.6511     0   2   4   6   8E+4   14   14.5  RT (min)      lcorbi_t3_neg_3_002_24    m/z 767.1406     0   1   2E+3   9   9.5  RT (min)      lcorbi_t3_neg_3_019_12    m/z 767.1535     0   1   2   3   4E+4   12.5   13  RT (min)      lcorbi_t3_neg_4_045_35    m/z 767.2803     0   2   4   6E+3   10.5   11  RT (min)      lcorbi_t3_neg_4_016_05    m/z 767.3108     0   2   4   6E+3   12   12.5  RT (min)      lcorbi_t3_neg_2_005_01    m/z 767.3575     0   2   4   6E+3   18   18.5  RT (min)      lcorbi_t3_neg_4_028_39    m/z 767.4196     0   1E+4   8   8.5  RT (min)      lcorbi_t3_neg_4_017_51    m/z 767.4851     0   2   4   6   8E+3   11   11.5  RT (min)      lcorbi_t3_neg_4_023_18    m/z 767.5013     0   1   2E+3   13.5   14  RT (min)      lcorbi_t3_neg_4_045_35    m/z 767.5199     0   1E+4   14   14.5  RT (min)      lcorbi_t3_neg_4_044_38    m/z 767.5313     0   2   4   6   8E+3   14   14.5  RT (min)      lcorbi_t3_neg_4_017_51    m/z 767.5562     0   1E+4   15.5   16  RT (min)      lcorbi_t3_neg_2_007_14    m/z 767.5820     0   2   4   6   8E+3   14.5   15  RT (min)      lcorbi_t3_neg_4_042_14    m/z 767.6073     0   1E+4   14.5   15  RT (min)      lcorbi_t3_neg_4_034_01    m/z 767.6178     0   2   4   6E+4   13   13.5  RT (min)      lcorbi_t3_neg_4_011_24    m/z 767.6902     0   1   2   3E+4   13.5   14  RT (min)      lcorbi_t3_neg_1_052_24    m/z 767.8195     0   1   2E+3   18   18.5  RT (min)      lcorbi_t3_neg_2_018_40    m/z 767.8376     0   1   2E+4   5.5   6  RT (min)      lcorbi_t3_neg_2_048_27    m/z 767.8543     0   1   2   3E+3   18   18.5  RT (min)      lcorbi_t3_neg_3_050_40    m/z 768.1540     0   1   2E+4   12.5   13  RT (min)      lcorbi_t3_neg_4_045_35    m/z 768.3133     0   1E+3   11   11.5  RT (min)      lcorbi_t3_neg_4_045_35    m/z 768.4230     0   1   2   3   4E+3   8   8.5  RT (min)      lcorbi_t3_neg_4_017_51    m/z 768.4939     0   2   4   6E+3   14.5   15  RT (min)      lcorbi_t3_neg_4_032_23    m/z 768.5029     0   2   4   6   8E+2   12.5   13  RT (min)      lcorbi_t3_neg_4_032_23    m/z 768.5345     0   1E+4   14   14.5  RT (min)      lcorbi_t3_neg_4_046_37    m/z 768.6031     0   2   4   6E+4   13.5   14  RT (min)      lcorbi_t3_neg_2_022_24    m/z 768.6103     0   1   2   3   4E+3   14.5   15  RT (min)      lcorbi_t3_neg_4_029_31    m/z 768.7661     0   1E+4   14.5   15  RT (min)      lcorbi_t3_neg_4_039_32    m/z 768.7915     0   1   2   3   4E+3   18   18.5  RT (min)      lcorbi_t3_neg_2_006_10    m/z 768.8988     0   1   2   3E+3   18   18.5  RT (min)      lcorbi_t3_neg_4_053_43    m/z 769.0874     0   2   4   6   8E+3   9.5   10  RT (min)      lcorbi_t3_neg_4_003_21    m/z 769.2783     0   1   2   3   4E+4   14   14.5  RT (min)      lcorbi_t3_neg_4_019_08    m/z 769.3514     0   2   4   6E+3   14.5   15  RT (min)      lcorbi_t3_neg_4_020_44    m/z 769.3602     0   2   4   6   8E+3   14   14.5  RT (min)      lcorbi_t3_neg_4_007_13    m/z 769.4676     0   1   14   14.5  RT (min)      lcorbi_t3_neg_4_034_01    m/z 769.5178     0   1   2   3   4E+3   14   14.5  RT (min)      lcorbi_t3_neg_4_044_38    m/z 769.5987     0   2   4   6E+4   13   13.5  RT (min)      lcorbi_t3_neg_4_039_32    m/z 769.6209     0   1E+4   14.5   15  RT (min)      lcorbi_t3_neg_3_007_16    m/z 769.6336     0   1   2   3   4E+4   13   13.5  RT (min)      lcorbi_t3_neg_4_011_24    m/z 770.0909     0   1   2E+3   9.5   10  RT (min)      lcorbi_t3_neg_4_015_10    m/z 770.2122     0   1   2   3   4E+3   10.5   11  RT (min)      lcorbi_t3_neg_3_007_16    m/z 770.2861     0   2   4E+4   2   2.5  RT (min)      lcorbi_t3_neg_2_022_24    m/z 770.2980     0   1E+4   11   11.5  RT (min)      lcorbi_t3_neg_4_011_24    m/z 770.3163     0   1   13.5   14  RT (min)      lcorbi_t3_neg_4_030_04    m/z 770.3635     0   2   4   6   8E+3   15   15.5  RT (min)      lcorbi_t3_neg_2_038_22    m/z 770.4873     0   2   4   6   8E+3   14.5   15  RT (min)      lcorbi_t3_neg_4_029_31    m/z 770.5527     0   1   2   3   4E+3   13.5   14  RT (min)      lcorbi_t3_neg_3_002_24    m/z 770.5981     0   2   4   6   8E+4   13   13.5  RT (min)      lcorbi_t3_neg_4_019_08    m/z 770.6233     0   2   4   6E+3   14.5   15  RT (min)      lcorbi_t3_neg_4_045_35    m/z 770.7977     0   1E+3   18   18.5  RT (min)      lcorbi_t3_neg_4_019_08    m/z 771.2115     0   2   4   6   8E+3   10.5   11  RT (min)      lcorbi_t3_neg_3_029_02    m/z 771.2941     0   1   2E+4   14   14.5  RT (min)      lcorbi_t3_neg_4_030_04    m/z 771.3002     0   2   4   6E+3   11   11.5  RT (min)      lcorbi_t3_neg_3_052_32    m/z 771.3782     0   2   4   6   8E+3   8   8.5  RT (min)      lcorbi_t3_neg_4_023_18    m/z 771.4647     0   1E+4   10.5   11  RT (min)      lcorbi_t3_neg_2_048_27    m/z 771.6044     0   1E+3   14.5   15  RT (min)      lcorbi_t3_neg_4_045_35    m/z 771.6280     0   1   2E+3   14.5   15  RT (min)      lcorbi_t3_neg_4_011_24    m/z 771.6797     0   1E+4   14   14.5  RT (min)      lcorbi_t3_neg_4_011_24    m/z 771.8673     0   2   4   6   8E+3   18   18.5  RT (min)      lcorbi_t3_neg_1_003_41    m/z 772.2150     0   1   2   3E+3   10.5   11  RT (min)      lcorbi_t3_neg_2_010_02    m/z 772.3143     0   2   4   6   8E+3   11.5   12  RT (min)      lcorbi_t3_neg_2_027_13    m/z 772.4354     0   1   2   3   4E+3   13.5   14  RT (min)      lcorbi_t3_neg_4_034_01    m/z 772.4767     0   2   4   6   8E+4   12.5   13  RT (min)      lcorbi_t3_neg_4_032_23    m/z 772.4874     0   1   2E+4   14.5   15  RT (min)      lcorbi_t3_neg_4_032_23    m/z 772.5933     0   2   4   6E+3   13.5   14  RT (min)      lcorbi_t3_neg_3_002_24    m/z 772.6035     0   1   13.5   14  RT (min)      lcorbi_t3_neg_4_007_13    m/z 772.6382     0   1   2E+4   14.5   15  RT (min)      lcorbi_t3_neg_2_012_36    m/z 772.6942     0   2   4   6E+3   15   15.5  RT (min)      lcorbi_t3_neg_2_012_36    m/z 772.7862     0   2   4   6   8E+3   14.5   15  RT (min)      lcorbi_t3_neg_2_052_32    m/z 772.8646     0   2   4E+3   18   18.5  RT (min)      lcorbi_t3_neg_1_004_22    m/z 773.2091     0   1   2   3E+3   10.5   11  RT (min)      lcorbi_t3_neg_1_023_03    m/z 773.2311     0   1   2   3E+3   11   11.5  RT (min)      lcorbi_t3_neg_2_010_02    m/z 773.2461     0   1   2   3   4E+3   11   11.5  RT (min)      lcorbi_t3_neg_3_028_35    m/z 773.2940     0   1   12   12.5  RT (min)      lcorbi_t3_neg_4_012_17    m/z 773.4802     0   1   2   3   4E+4   12.5   13  RT (min)      lcorbi_t3_neg_4_032_23    m/z 773.5062     0   1E+3   13   13.5  RT (min)      lcorbi_t3_neg_1_005_07    m/z 773.5165     0   2   4   6E+3   18   18.5  RT (min)      lcorbi_t3_neg_4_029_31    m/z 773.5328     0   2   4   6E+4   14   14.5  RT (min)      lcorbi_t3_neg_4_054_36    m/z 773.6170     0   1E+4   13.5   14  RT (min)      lcorbi_t3_neg_4_007_13    m/z 773.6424     0   1   2   3   4E+3   15   15.5  RT (min)      lcorbi_t3_neg_4_034_01    m/z 773.6624     0   1E+4   15   15.5  RT (min)      lcorbi_t3_neg_2_012_36    m/z 773.6705     0   1E+4   14.5   15  RT (min)      lcorbi_t3_neg_2_039_31    m/z 773.8048     0   1   2   3   4E+3   18   18.5  RT (min)      lcorbi_t3_neg_2_042_44    m/z 774.2290     0   1E+4   12.5   13  RT (min)      lcorbi_t3_neg_4_032_23    m/z 774.5282     0   2   4   6E+3   18   18.5  RT (min)      lcorbi_t3_neg_4_044_38    m/z 774.6380     0   1   15   15.5  RT (min)      lcorbi_t3_neg_4_034_01    m/z 774.6533     0   1E+4   14.5   15  RT (min)      lcorbi_t3_neg_4_039_32    m/z 774.6658     0   2   4   6E+3   15   15.5  RT (min)      lcorbi_t3_neg_2_012_36    m/z 774.6748     0   2   4   6   8E+3   14.5   15  RT (min)      lcorbi_t3_neg_4_039_32    m/z 774.7686     0   1   2   3   4E+3   15   15.5  RT (min)      lcorbi_t3_neg_4_001_49    m/z 774.8219     0   1   2E+4   5.5   6  RT (min)      lcorbi_t3_neg_2_048_27    m/z 774.8399     0   1   2E+3   18   18.5  RT (min)      lcorbi_t3_neg_2_037_09    m/z 775.0826     0   2   4E+3   7.5   8  RT (min)      lcorbi_t3_neg_4_034_01    m/z 775.1038     0   1   2   3   4E+3   8   8.5  RT (min)      lcorbi_t3_neg_4_013_09    m/z 775.2232     0   2   4   6   8E+3   10.5   11  RT (min)      lcorbi_t3_neg_1_047_02    m/z 775.2615     0   2   4   6E+3   11.5   12  RT (min)      lcorbi_t3_neg_4_005_27    m/z 775.2793     0   2   4   6E+3   13.5   14  RT (min)      lcorbi_t3_neg_4_034_01    m/z 775.3232     0   1E+4   5.5   6  RT (min)      lcorbi_t3_neg_2_048_27    m/z 775.4771     0   1   13   13.5  RT (min)      lcorbi_t3_neg_4_021_25    m/z 775.4958     0   1   2   3E+4   12.5   13  RT (min)      lcorbi_t3_neg_4_032_23    m/z 775.5234     0   1E+4   13.5   14  RT (min)      lcorbi_t3_neg_3_047_37    m/z 775.7234     0   1   2   3E+4   15   15.5  RT (min)      lcorbi_t3_neg_4_039_32    m/z 775.7719     0   1E+4   15   15.5  RT (min)      lcorbi_t3_neg_4_001_49    m/z 775.7943     0   1   2E+3   18   18.5  RT (min)      lcorbi_t3_neg_2_040_43    m/z 775.8310     0   2   4   6E+3   18   18.5  RT (min)      lcorbi_t3_neg_2_021_41    m/z 776.1753     0   1   2   3   4E+3   9   9.5  RT (min)      lcorbi_t3_neg_3_019_12    m/z 776.2300     0   2   4   6   8E+3   11.5   12  RT (min)      lcorbi_t3_neg_4_048_03    m/z 776.2644     0   1   2   3E+3   11.5   12  RT (min)      lcorbi_t3_neg_4_046_37    m/z 776.3331     0   1   2   3   4E+3   8   8.5  RT (min)      lcorbi_t3_neg_2_048_27    m/z 776.4861     0   1E+3   13   13.5  RT (min)      lcorbi_t3_neg_3_005_23    m/z 776.4984     0   2   4E+3   13   13.5  RT (min)      lcorbi_t3_neg_2_053_23    m/z 776.4992     0   1   2   3E+3   14.5   15  RT (min)      lcorbi_t3_neg_4_007_13    m/z 776.5243     0   2   4   6E+4   14   14.5  RT (min)      lcorbi_t3_neg_4_019_08    m/z 776.5527     0   1   2   3   4E+3   13   13.5  RT (min)      lcorbi_t3_neg_4_044_38    m/z 776.5882     0   1E+4   15.5   16  RT (min)      lcorbi_t3_neg_4_042_14    m/z 776.6690     0   1E+4   14.5   15  RT (min)      lcorbi_t3_neg_4_039_32    m/z 776.7478     0   1E+4   14.5   15  RT (min)      lcorbi_t3_neg_2_052_32    m/z 776.8863     0   1   2   3E+3   18   18.5  RT (min)      lcorbi_t3_neg_4_043_11    m/z 777.1697     0   1   2   3   4E+3   9   9.5  RT (min)      lcorbi_t3_neg_3_019_12    m/z 777.2400     0   2   4E+3   11   11.5  RT (min)      lcorbi_t3_neg_1_043_14    m/z 777.2480     0   1E+4   13.5   14  RT (min)      lcorbi_t3_neg_1_023_03    m/z 777.2772     0   2   4   6   8E+3   12   12.5  RT (min)      lcorbi_t3_neg_4_038_15    m/z 777.3298     0   2   4   6E+3   8   8.5  RT (min)      lcorbi_t3_neg_2_048_27    m/z 777.4955     0   1E+4   14.5   15  RT (min)      lcorbi_t3_neg_4_029_31    m/z 777.5114     0   1E+4   13   13.5  RT (min)      lcorbi_t3_neg_2_053_23    m/z 777.5172     0   1E+4   14.5   15  RT (min)      lcorbi_t3_neg_4_029_31    m/z 777.6132     0   1   2   3   4E+6   14   14.5  RT (min)      lcorbi_t3_neg_4_019_08    m/z 777.6388     0   1E+6   14   14.5  RT (min)      lcorbi_t3_neg_3_002_24    m/z 777.7134     0   1E+3   15   15.5  RT (min)      lcorbi_t3_neg_1_053_32    m/z 777.7688     0   2   4E+3   15   15.5  RT (min)      lcorbi_t3_neg_4_001_49    m/z 778.2477     0   1E+4   11   11.5  RT (min)      lcorbi_t3_neg_4_007_13    m/z 778.2805     0   2   4   6E+3   12   12.5  RT (min)      lcorbi_t3_neg_4_035_33    m/z 778.3566     0   2   4   6E+3   14.5   15  RT (min)      lcorbi_t3_neg_4_032_23    m/z 778.6065     0   1   2E+4   14   14.5  RT (min)      lcorbi_t3_neg_1_053_32    m/z 778.6512     0   1   14   14.5  RT (min)      lcorbi_t3_neg_4_034_01    m/z 778.7452     0   2   4E+3   14.5   15  RT (min)      lcorbi_t3_neg_2_052_32    m/z 778.7676     0   2   4   6E+3   14.5   15  RT (min)      lcorbi_t3_neg_4_039_32    m/z 778.8121     0   1   2   3E+3   18   18.5  RT (min)      lcorbi_t3_neg_2_037_09    m/z 778.8539     0   2   4   6E+3   18   18.5  RT (min)      lcorbi_t3_neg_2_003_42    m/z 779.0014     0   1   2   3E+3   9   9.5  RT (min)      lcorbi_t3_neg_2_015_21    m/z 779.1092     0   2   4   6   8E+3   9   9.5  RT (min)      lcorbi_t3_neg_4_023_18    m/z 779.1157     0   1   2   3   4E+3   8   8.5  RT (min)      lcorbi_t3_neg_4_013_09    m/z 779.1838     0   2   4   6   8E+3   9   9.5  RT (min)      lcorbi_t3_neg_3_019_12    m/z 779.2560     0   1E+4   11.5   12  RT (min)      lcorbi_t3_neg_4_007_13    m/z 779.3050     0   1E+4   14.5   15  RT (min)      lcorbi_t3_neg_1_023_03    m/z 779.4117     0   1   2E+4   8   8.5  RT (min)      lcorbi_t3_neg_2_048_27    m/z 779.4166     0   1   2E+4   4.5   5  RT (min)      lcorbi_t3_neg_2_048_27    m/z 779.4292     0   1E+4   8.5   9  RT (min)      lcorbi_t3_neg_4_017_51    m/z 779.4908     0   2   4E+2   13.5   14  RT (min)      lcorbi_t3_neg_3_055_06    m/z 779.5974     0   1E+4   12.5   13  RT (min)      lcorbi_t3_neg_4_007_13    m/z 779.6453     0   1   2E+4   18   18.5  RT (min)      lcorbi_t3_neg_4_011_24    m/z 779.6545     0   2   4   6E+5   14   14.5  RT (min)      lcorbi_t3_neg_4_011_24    m/z 779.7136     0   1   15   15.5  RT (min)      lcorbi_t3_neg_3_002_24    m/z 779.8251     0   2   4   6   8E+3   18   18.5  RT (min)      lcorbi_t3_neg_1_046_11    m/z 780.1873     0   1   2   3E+3   9   9.5  RT (min)      lcorbi_t3_neg_3_019_12    m/z 780.2627     0   2   4E+3   11.5   12  RT (min)      lcorbi_t3_neg_4_007_13    m/z 780.2957     0   1E+4   13   13.5  RT (min)      lcorbi_t3_neg_1_023_03    m/z 780.3537     0   2   4E+3   14.5   15  RT (min)      lcorbi_t3_neg_4_032_23    m/z 780.3722     0   2   4   6E+3   14.5   15  RT (min)      lcorbi_t3_neg_1_044_23    m/z 780.4696     0   1   2E+5   13.5   14  RT (min)      lcorbi_t3_neg_4_019_08    m/z 780.4936     0   2   4   6E+3   13.5   14  RT (min)      lcorbi_t3_neg_4_029_31    m/z 780.5180     0   1   2   3   4E+4   1   1.5  RT (min)      lcorbi_t3_neg_4_007_13    m/z 780.5274     0   1   2E+3   13   13.5  RT (min)      lcorbi_t3_neg_1_018_46    m/z 780.5832     0   1   2E+4   14.5   15  RT (min)      lcorbi_t3_neg_4_005_27    m/z 780.5941     0   1   2E+4   13   13.5  RT (min)      lcorbi_t3_neg_4_039_32    m/z 780.6363     0   2   4   6   8E+4   14.5   15  RT (min)      lcorbi_t3_neg_4_019_08    m/z 780.6666     0   1   2   3E+5   14   14.5  RT (min)      lcorbi_t3_neg_4_011_24    m/z 780.7166     0   2   4E+3   15   15.5  RT (min)      lcorbi_t3_neg_2_012_36    m/z 780.7933     0   1E+4   15   15.5  RT (min)      lcorbi_t3_neg_4_039_32    m/z 780.8111     0   1E+3   15   15.5  RT (min)      lcorbi_t3_neg_3_002_24    m/z 781.2717     0   1E+4   12   12.5  RT (min)      lcorbi_t3_neg_4_007_13    m/z 781.4271     0   2   4   6E+3   13.5   14  RT (min)      lcorbi_t3_neg_1_020_36    m/z 781.4831     0   2   4E+3   18   18.5  RT (min)      lcorbi_t3_neg_4_029_31    m/z 781.5100     0   2   4   6   8E+3   14.5   15  RT (min)      lcorbi_t3_neg_1_039_38    m/z 781.5221     0   1   2   3E+3   13.5   14  RT (min)      lcorbi_t3_neg_1_044_23    m/z 781.5622     0   2   4   6E+4   14.5   15  RT (min)      lcorbi_t3_neg_4_044_38    m/z 781.6610     0   2   4   6E+4   18   18.5  RT (min)      lcorbi_t3_neg_3_002_24    m/z 781.8287     0   1   18   18.5  RT (min)      lcorbi_t3_neg_4_019_08    m/z 781.8625     0   2   4   6E+3   18   18.5  RT (min)      lcorbi_t3_neg_2_026_03    m/z 782.2439     0   1E+4   10.5   11  RT (min)      lcorbi_t3_neg_4_050_02    m/z 782.2741     0   2   4   6   8E+3   13.5   14  RT (min)      lcorbi_t3_neg_1_023_03    m/z 782.2774     0   2   4   6E+3   12   12.5  RT (min)      lcorbi_t3_neg_4_046_37    m/z 782.2949     0   1   2   3E+3   14   14.5  RT (min)      lcorbi_t3_neg_2_037_09    m/z 782.3352     0   1E+4   10.5   11  RT (min)      lcorbi_t3_neg_1_052_24    m/z 782.3695     0   2   4   6E+3   14.5   15  RT (min)      lcorbi_t3_neg_1_044_23    m/z 782.4962     0   2   4   6   8E+3   18   18.5  RT (min)      lcorbi_t3_neg_4_029_31    m/z 782.5136     0   1E+3   14.5   15  RT (min)      lcorbi_t3_neg_1_044_23    m/z 782.5291     0   1E+4   14.5   15  RT (min)      lcorbi_t3_neg_3_045_34    m/z 782.5340     0   1E+4   13.5   14  RT (min)      lcorbi_t3_neg_2_053_23    m/z 782.5501     0   1E+3   14   14.5  RT (min)      lcorbi_t3_neg_4_019_08    m/z 782.5932     0   1   2   3E+3   14.5   15  RT (min)      lcorbi_t3_neg_4_042_14    m/z 782.6045     0   1   2   3E+3   14.5   15  RT (min)      lcorbi_t3_neg_3_053_14    m/z 782.6519     0   1   2E+5   14.5   15  RT (min)      lcorbi_t3_neg_4_019_08    m/z 782.8093     0   1   2E+4   15   15.5  RT (min)      lcorbi_t3_neg_3_002_24    m/z 783.0131     0   1   2   3   4E+3   9   9.5  RT (min)      lcorbi_t3_neg_4_028_39    m/z 783.2673     0   1   2   3   4E+3   11   11.5  RT (min)      lcorbi_t3_neg_2_010_02    m/z 783.4686     0   1E+3   13.5   14  RT (min)      lcorbi_t3_neg_4_046_37    m/z 783.4997     0   1   2   3   4E+3   18   18.5  RT (min)      lcorbi_t3_neg_4_029_31    m/z 783.5255     0   2   4   6E+3   14.5   15  RT (min)      lcorbi_t3_neg_4_007_13    m/z 783.5375     0   1E+4   13.5   14  RT (min)      lcorbi_t3_neg_4_044_38    m/z 783.6215     0   1   2E+4   14   14.5  RT (min)      lcorbi_t3_neg_4_035_33    m/z 783.6766     0   2   4E+4   18   18.5  RT (min)      lcorbi_t3_neg_4_011_24    m/z 783.7852     0   2   4   6   8E+3   14.5   15  RT (min)      lcorbi_t3_neg_2_052_32    m/z 783.7939     0   2   4E+3   15   15.5  RT (min)      lcorbi_t3_neg_2_052_32    m/z 784.1494     0   2   4   6   8E+3   4.5   5  RT (min)      lcorbi_t3_neg_4_046_37    m/z 784.2652     0   2   4   6E+3   11   11.5  RT (min)      lcorbi_t3_neg_2_027_13    m/z 784.2795     0   1   2E+3   11.5   12  RT (min)      lcorbi_t3_neg_1_043_14    m/z 784.4691     0   1E+4   13   13.5  RT (min)      lcorbi_t3_neg_2_036_35    m/z 784.4959     0   1   2   3   4E+4   13.5   14  RT (min)      lcorbi_t3_neg_1_039_38    m/z 784.5129     0   1E+4   18   18.5  RT (min)      lcorbi_t3_neg_4_046_37    m/z 784.5285     0   1E+3   14.5   15  RT (min)      lcorbi_t3_neg_2_053_23    m/z 784.6211     0   2   4   6   8E+3   14.5   15  RT (min)      lcorbi_t3_neg_4_039_32    m/z 784.6408     0   2   4   6   8E+3   14.5   15  RT (min)      lcorbi_t3_neg_2_022_24    m/z 784.7246     0   1E+4   14   14.5  RT (min)      lcorbi_t3_neg_4_016_05    m/z 785.0856     0   1   2   3   4E+3   10   10.5  RT (min)      lcorbi_t3_neg_4_029_31    m/z 785.1326     0   2   4   6   8E+3   8   8.5  RT (min)      lcorbi_t3_neg_4_008_22    m/z 785.2610     0   2   4   6E+3   12   12.5  RT (min)      lcorbi_t3_neg_2_015_21    m/z 785.3547     0   2   4   6   8E+3   14   14.5  RT (min)      lcorbi_t3_neg_1_023_03    m/z 785.3670     0   1   2   3E+4   6   6.5  RT (min)      lcorbi_t3_neg_2_048_27    m/z 785.4177     0   1E+4   9   9.5  RT (min)      lcorbi_t3_neg_2_048_27    m/z 785.4586     0   2   4E+3   13.5   14  RT (min)      lcorbi_t3_neg_1_055_05    m/z 785.5164     0   2   4E+3   18   18.5  RT (min)      lcorbi_t3_neg_4_046_37    m/z 785.5270     0   1   2   3E+3   14   14.5  RT (min)      lcorbi_t3_neg_1_044_23    m/z 785.5529     0   1E+4   14   14.5  RT (min)      lcorbi_t3_neg_4_039_32    m/z 785.5815     0   1   13.5   14  RT (min)      lcorbi_t3_neg_4_011_24    m/z 785.5945     0   2   4E+3   14.5   15  RT (min)      lcorbi_t3_neg_3_052_32    m/z 785.6180     0   1E+5   13.5   14  RT (min)      lcorbi_t3_neg_4_019_08    m/z 785.7070     0   2   4   6   8E+3   15   15.5  RT (min)      lcorbi_t3_neg_2_039_31    m/z 785.7355     0   1   2E+4   14.5   15  RT (min)      lcorbi_t3_neg_3_052_32    m/z 785.7488     0   1E+4   14.5   15  RT (min)      lcorbi_t3_neg_2_052_32    m/z 785.7734     0   1   2E+4   14.5   15  RT (min)      lcorbi_t3_neg_2_052_32    m/z 785.8327     0   1   2   3E+3   18   18.5  RT (min)      lcorbi_t3_neg_4_008_22    m/z 785.9966     0   1   2   3E+4   11   11.5  RT (min)      lcorbi_t3_neg_4_005_27    m/z 786.1361     0   1   2E+3   8   8.5  RT (min)      lcorbi_t3_neg_4_037_29    m/z 786.2586     0   1   11.5   12  RT (min)      lcorbi_t3_neg_4_044_38    m/z 786.2654     0   1   2   3   4E+4   14   14.5  RT (min)      lcorbi_t3_neg_4_048_03    m/z 786.2819     0   2   4   6   8E+3   11.5   12  RT (min)      lcorbi_t3_neg_4_034_01    m/z 786.5468     0   1   14   14.5  RT (min)      lcorbi_t3_neg_1_054_37    m/z 786.7492     0   1E+4   14.5   15  RT (min)      lcorbi_t3_neg_4_001_49    m/z 787.2069     0   2   4E+3   9   9.5  RT (min)      lcorbi_t3_neg_3_019_12    m/z 787.2886     0   1E+4   13.5   14  RT (min)      lcorbi_t3_neg_1_023_03    m/z 787.3439     0   1   2   3E+3   18   18.5  RT (min)      lcorbi_t3_neg_4_023_18    m/z 787.5117     0   1E+4   13   13.5  RT (min)      lcorbi_t3_neg_4_034_01    m/z 787.5578     0   1   2   3   4E+4   14   14.5  RT (min)      lcorbi_t3_neg_4_019_08    m/z 787.6258     0   1E+4   14.5   15  RT (min)      lcorbi_t3_neg_4_011_24    m/z 787.7720     0   2   4   6   8E+3   14.5   15  RT (min)      lcorbi_t3_neg_4_001_49    m/z 788.2718     0   1   2   3E+3   11.5   12  RT (min)      lcorbi_t3_neg_3_053_14    m/z 788.2981     0   1E+4   12   12.5  RT (min)      lcorbi_t3_neg_3_053_14    m/z 788.3256     0   1   2   3E+3   14   14.5  RT (min)      lcorbi_t3_neg_4_048_03    m/z 788.3473     0   2   4E+3   18   18.5  RT (min)      lcorbi_t3_neg_4_040_19    m/z 788.4249     0   2   4   6   8E+3   13   13.5  RT (min)      lcorbi_t3_neg_2_053_23    m/z 788.4483     0   2   4   6E+3   13   13.5  RT (min)      lcorbi_t3_neg_4_044_38    m/z 788.4637     0   2   4   6E+3   13   13.5  RT (min)      lcorbi_t3_neg_2_053_23    m/z 788.4715     0   1E+5   12.5   13  RT (min)      lcorbi_t3_neg_3_005_23    m/z 788.4823     0   1   13.5   14  RT (min)      lcorbi_t3_neg_4_007_13    m/z 788.5088     0   1   2E+4   13   13.5  RT (min)      lcorbi_t3_neg_1_035_25    m/z 788.5437     0   1E+4   18   18.5  RT (min)      lcorbi_t3_neg_1_044_23    m/z 788.5614     0   1E+4   13.5   14  RT (min)      lcorbi_t3_neg_4_019_08    m/z 788.5977     0   1   2E+3   14.5   15  RT (min)      lcorbi_t3_neg_4_011_24    m/z 788.6222     0   1E+4   14.5   15  RT (min)      lcorbi_t3_neg_4_011_24    m/z 788.6492     0   1E+4   14.5   15  RT (min)      lcorbi_t3_neg_4_039_32    m/z 788.6771     0   1E+4   14.5   15  RT (min)      lcorbi_t3_neg_1_053_32    m/z 788.7465     0   2   4E+3   14.5   15  RT (min)      lcorbi_t3_neg_4_001_49    m/z 788.7754     0   2   4   6E+3   14.5   15  RT (min)      lcorbi_t3_neg_2_052_32    m/z 788.8454     0   2   4   6   8E+3   18   18.5  RT (min)      lcorbi_t3_neg_4_053_43    m/z 789.0302     0   2   4   6E+3   9   9.5  RT (min)      lcorbi_t3_neg_3_019_12    m/z 789.2726     0   1   2   3   4E+3   12   12.5  RT (min)      lcorbi_t3_neg_2_015_21    m/z 789.4283     0   1   2   3E+3   13   13.5  RT (min)      lcorbi_t3_neg_1_044_23    m/z 789.4649     0   1E+4   14.5   15  RT (min)      lcorbi_t3_neg_4_011_24    m/z 789.4750     0   2   4E+4   12.5   13  RT (min)      lcorbi_t3_neg_4_032_23    m/z 789.4854     0   1E+3   13.5   14  RT (min)      lcorbi_t3_neg_4_007_13    m/z 789.5138     0   2   4   6E+3   14   14.5  RT (min)      lcorbi_t3_neg_3_005_23    m/z 789.5370     0   1   2E+4   13.5   14  RT (min)      lcorbi_t3_neg_1_021_31    m/z 789.5476     0   2   4E+3   18   18.5  RT (min)      lcorbi_t3_neg_1_044_23    m/z 789.5575     0   1   2E+4   13.5   14  RT (min)      lcorbi_t3_neg_4_007_13    m/z 789.5726     0   1   13   13.5  RT (min)      lcorbi_t3_neg_4_019_08    m/z 789.6314     0   1E+4   13.5   14  RT (min)      lcorbi_t3_neg_4_039_32    m/z 789.6668     0   1   2   3   4E+3   14.5   15  RT (min)      lcorbi_t3_neg_4_039_32    m/z 789.7877     0   1E+4   15   15.5  RT (min)      lcorbi_t3_neg_4_001_49    m/z 790.0335     0   1   2E+3   9   9.5  RT (min)      lcorbi_t3_neg_4_028_39    m/z 790.1997     0   2   4E+3   11   11.5  RT (min)      lcorbi_t3_neg_4_032_23    m/z 790.2768     0   2   4   6E+3   10.5   11  RT (min)      lcorbi_t3_neg_4_007_13    m/z 790.2853     0   2   4   6E+3   11.5   12  RT (min)      lcorbi_t3_neg_1_047_02    m/z 790.3527     0   2   4E+3   18   18.5  RT (min)      lcorbi_t3_neg_2_037_09    m/z 790.4440     0   1   13.5   14  RT (min)      lcorbi_t3_neg_4_007_13    m/z 790.4946     0   1E+3   14.5   15  RT (min)      lcorbi_t3_neg_4_019_08    m/z 790.5306     0   1E+4   13.5   14  RT (min)      lcorbi_t3_neg_4_029_31    m/z 790.6512     0   2   4   6E+3   15   15.5  RT (min)      lcorbi_t3_neg_2_012_36    m/z 790.6647     0   2   4   6   8E+3   14.5   15  RT (min)      lcorbi_t3_neg_3_052_32    m/z 790.6850     0   1E+4   14.5   15  RT (min)      lcorbi_t3_neg_4_039_32    m/z 790.8831     0   1   2   3   4E+3   18   18.5  RT (min)      lcorbi_t3_neg_4_022_41    m/z 791.1960     0   2   4   6E+3   10.5   11  RT (min)      lcorbi_t3_neg_4_050_02    m/z 791.3609     0   2   4   6   8E+3   10.5   11  RT (min)      lcorbi_t3_neg_4_011_24    m/z 791.4670     0   1E+4   13.5   14  RT (min)      lcorbi_t3_neg_4_044_38    m/z 791.5307     0   1E+3   14.5   15  RT (min)      lcorbi_t3_neg_4_011_24    m/z 791.6089     0   1   2   3   4E+4   13   13.5  RT (min)      lcorbi_t3_neg_3_002_24    m/z 791.6830     0   2   4E+3   14.5   15  RT (min)      lcorbi_t3_neg_3_002_24    m/z 791.8195     0   2   4E+3   14.5   15  RT (min)      lcorbi_t3_neg_4_001_49    m/z 792.2168     0   1   2   3E+3   11.5   12  RT (min)      lcorbi_t3_neg_2_026_03    m/z 792.4929     0   1   2E+4   6   6.5  RT (min)      lcorbi_t3_neg_4_044_38    m/z 792.4936     0   1E+5   13   13.5  RT (min)      lcorbi_t3_neg_2_053_23    m/z 792.4943     0   1E+3   14.5   15  RT (min)      lcorbi_t3_neg_4_019_08    m/z 792.5020     0   1E+5   13   13.5  RT (min)      lcorbi_t3_neg_4_032_23    m/z 792.6576     0   2   4   6   8E+4   14   14.5  RT (min)      lcorbi_t3_neg_3_002_24    m/z 792.6806     0   2   4   6   8E+3   14.5   15  RT (min)      lcorbi_t3_neg_4_011_24    m/z 792.7881     0   1E+4   15   15.5  RT (min)      lcorbi_t3_neg_4_001_49    m/z 792.7998     0   1   2E+3   18   18.5  RT (min)      lcorbi_t3_neg_4_019_08    m/z 792.8196     0   1E+4   14.5   15  RT (min)      lcorbi_t3_neg_4_001_49    m/z 793.0971     0   2   4   6   8E+3   8   8.5  RT (min)      lcorbi_t3_neg_4_023_18    m/z 793.3616     0   1E+4   18   18.5  RT (min)      lcorbi_t3_neg_2_046_20    m/z 793.4621     0   1   2   3E+3   10   10.5  RT (min)      lcorbi_t3_neg_4_029_31    m/z 793.4829     0   1   2E+4   13.5   14  RT (min)      lcorbi_t3_neg_2_053_23    m/z 793.5168     0   1E+4   13.5   14  RT (min)      lcorbi_t3_neg_1_013_35    m/z 793.5272     0   1   2   3   4E+3   14.5   15  RT (min)      lcorbi_t3_neg_4_038_15    m/z 793.5457     0   1   14.5   15  RT (min)      lcorbi_t3_neg_1_004_22    m/z 793.6159     0   1E+4   14.5   15  RT (min)      lcorbi_t3_neg_4_007_13    m/z 794.2121     0   2   4E+3   10   10.5  RT (min)      lcorbi_t3_neg_3_019_12    m/z 794.2985     0   2   4E+3   11   11.5  RT (min)      lcorbi_t3_neg_2_008_16    m/z 794.3089     0   1   2   3E+4   9   9.5  RT (min)      lcorbi_t3_neg_2_048_27    m/z 794.3800     0   1   2   3E+4   11   11.5  RT (min)      lcorbi_t3_neg_2_022_24    m/z 794.5097     0   1E+4   14.5   15  RT (min)      lcorbi_t3_neg_4_013_09    m/z 794.5258     0   1   2E+3   14.5   15  RT (min)      lcorbi_t3_neg_4_055_06    m/z 794.5834     0   1   14.5   15  RT (min)      lcorbi_t3_neg_4_019_08    m/z 794.7866     0   2   4   6   8E+2   18   18.5  RT (min)      lcorbi_t3_neg_4_019_08    m/z 794.8297     0   1   18   18.5  RT (min)      lcorbi_t3_neg_4_019_08    m/z 795.1565     0   2   4   6E+3   9   9.5  RT (min)      lcorbi_t3_neg_3_019_12    m/z 795.2107     0   1   2   3   4E+3   9.5   10  RT (min)      lcorbi_t3_neg_4_029_31    m/z 795.2123     0   2   4   6E+3   12.5   13  RT (min)      lcorbi_t3_neg_3_007_16    m/z 795.3118     0   1   2   3E+3   11.5   12  RT (min)      lcorbi_t3_neg_2_014_29    m/z 795.3523     0   2   4E+3   13   13.5  RT (min)      lcorbi_t3_neg_4_007_13    m/z 795.4404     0   2   4   6   8E+3   14   14.5  RT (min)      lcorbi_t3_neg_3_005_23    m/z 795.5125     0   1   2E+3   13   13.5  RT (min)      lcorbi_t3_neg_4_046_37    m/z 795.5742     0   1   2   3E+4   14.5   15  RT (min)      lcorbi_t3_neg_4_005_27    m/z 795.6475     0   1   2E+4   10   10.5  RT (min)      lcorbi_t3_neg_3_019_12    m/z 795.8327     0   2   4E+3   18   18.5  RT (min)      lcorbi_t3_neg_3_015_11    m/z 796.2285     0   1   2   3E+3   11   11.5  RT (min)      lcorbi_t3_neg_2_026_03    m/z 796.2760     0   1E+4   12.5   13  RT (min)      lcorbi_t3_neg_1_052_24    m/z 796.2931     0   2   4   6E+3   12   12.5  RT (min)      lcorbi_t3_neg_2_055_18    m/z 796.3089     0   1E+4   6.5   7  RT (min)      lcorbi_t3_neg_2_048_27    m/z 796.3161     0   1   12   12.5  RT (min)      lcorbi_t3_neg_4_023_18    m/z 796.3239     0   2   4   6E+3   11.5   12  RT (min)      lcorbi_t3_neg_4_034_01    m/z 796.3481     0   2   4   6E+3   14.5   15  RT (min)      lcorbi_t3_neg_4_032_23    m/z 796.3875     0   2   4   6   8E+3   14.5   15  RT (min)      lcorbi_t3_neg_4_044_38    m/z 796.3956     0   1   2E+4   11.5   12  RT (min)      lcorbi_t3_neg_4_016_05    m/z 796.4530     0   1E+4   11   11.5  RT (min)      lcorbi_t3_neg_1_021_31    m/z 796.4538     0   1   2E+4   13.5   14  RT (min)      lcorbi_t3_neg_4_044_38    m/z 796.5040     0   1   2   3E+4   13   13.5  RT (min)      lcorbi_t3_neg_4_029_31    m/z 796.5255     0   1E+4   14.5   15  RT (min)      lcorbi_t3_neg_4_007_13    m/z 796.5263     0   1E+4   13   13.5  RT (min)      lcorbi_t3_neg_4_045_35    m/z 796.5340     0   1E+4   13   13.5  RT (min)      lcorbi_t3_neg_4_054_36    m/z 796.5502     0   1   14   14.5  RT (min)      lcorbi_t3_neg_3_002_24    m/z 796.5658     0   1E+4   14   14.5  RT (min)      lcorbi_t3_neg_2_052_32    m/z 796.6183     0   1E+4   18   18.5  RT (min)      lcorbi_t3_neg_4_039_32    m/z 796.6516     0   1E+4   13.5   14  RT (min)      lcorbi_t3_neg_2_052_32    m/z 796.6710     0   1   2E+4   13.5   14  RT (min)      lcorbi_t3_neg_3_052_32    m/z 796.7830     0   2   4   6   8E+3   18   18.5  RT (min)      lcorbi_t3_neg_4_019_08    m/z 796.7994     0   2   4   6E+3   18   18.5  RT (min)      lcorbi_t3_neg_4_053_43    m/z 797.3196     0   1   2E+3   12   12.5  RT (min)      lcorbi_t3_neg_4_034_01    m/z 797.3826     0   1   2   3   4E+3   15   15.5  RT (min)      lcorbi_t3_neg_1_027_30    m/z 797.3910     0   1   2   3   4E+3   14.5   15  RT (min)      lcorbi_t3_neg_3_007_16    m/z 797.4495     0   1   2   3E+4   11   11.5  RT (min)      lcorbi_t3_neg_4_029_31    m/z 797.4630     0   1   2   3E+4   7.5   8  RT (min)      lcorbi_t3_neg_4_023_18    m/z 797.4693     0   1   2   3E+4   11   11.5  RT (min)      lcorbi_t3_neg_4_029_31    m/z 797.6211     0   1E+6   13.5   14  RT (min)      lcorbi_t3_neg_4_039_32    m/z 797.6427     0   2   4   6E+3   14.5   15  RT (min)      lcorbi_t3_neg_4_011_24    m/z 797.6645     0   2   4E+4   13.5   14  RT (min)      lcorbi_t3_neg_4_039_32    m/z 797.8009     0   1E+4   14.5   15  RT (min)      lcorbi_t3_neg_4_039_32    m/z 797.8380     0   2   4   6E+3   18   18.5  RT (min)      lcorbi_t3_neg_4_028_39    m/z 798.2237     0   1   2   3E+3   10   10.5  RT (min)      lcorbi_t3_neg_3_019_12    m/z 798.2432     0   1   2E+3   11.5   12  RT (min)      lcorbi_t3_neg_2_008_16    m/z 798.2647     0   1   2   3E+3   11   11.5  RT (min)      lcorbi_t3_neg_4_005_27    m/z 798.3008     0   2   4   6   8E+3   11.5   12  RT (min)      lcorbi_t3_neg_3_053_14    m/z 798.3396     0   1   2   3   4E+3   12   12.5  RT (min)      lcorbi_t3_neg_4_034_01    m/z 798.3454     0   2   4   6E+3   14.5   15  RT (min)      lcorbi_t3_neg_1_044_23    m/z 798.3641     0   1   2   3   4E+3   14.5   15  RT (min)      lcorbi_t3_neg_4_007_13    m/z 798.3951     0   2   4E+3   15   15.5  RT (min)      lcorbi_t3_neg_2_038_22    m/z 798.4538     0   1E+4   13   13.5  RT (min)      lcorbi_t3_neg_4_029_31    m/z 798.5032     0   1E+4   14.5   15  RT (min)      lcorbi_t3_neg_4_034_01    m/z 798.5527     0   1   2   3E+4   13.5   14  RT (min)      lcorbi_t3_neg_1_053_32    m/z 798.5672     0   2   4E+4   14   14.5  RT (min)      lcorbi_t3_neg_1_035_25    m/z 798.5991     0   2   4   6E+2   14   14.5  RT (min)      lcorbi_t3_neg_4_011_24    m/z 798.7266     0   1   2E+4   13.5   14  RT (min)      lcorbi_t3_neg_4_039_32    m/z 798.8041     0   2   4   6   8E+3   18   18.5  RT (min)      lcorbi_t3_neg_4_043_11    m/z 798.8129     0   1E+4   15   15.5  RT (min)      lcorbi_t3_neg_2_052_32    m/z 798.8166     0   1   2   3   4E+3   18   18.5  RT (min)      lcorbi_t3_neg_4_051_40    m/z 798.8385     0   1   2E+3   18   18.5  RT (min)      lcorbi_t3_neg_3_016_42    m/z 798.9860     0   1   2E+3   9   9.5  RT (min)      lcorbi_t3_neg_3_038_39    m/z 799.1713     0   1   2   3   4E+3   12.5   13  RT (min)      lcorbi_t3_neg_4_005_27    m/z 799.3042     0   1   2E+3   11.5   12  RT (min)      lcorbi_t3_neg_3_053_14    m/z 799.4571     0   2   4   6E+3   13   13.5  RT (min)      lcorbi_t3_neg_4_029_31    m/z 799.5151     0   1E+4   14.5   15  RT (min)      lcorbi_t3_neg_1_029_21    m/z 799.5574     0   2   4E+3   14   14.5  RT (min)      lcorbi_t3_neg_1_052_24    m/z 799.6266     0   2   4E+3   14.5   15  RT (min)      lcorbi_t3_neg_4_019_08    m/z 799.6661     0   1E+4   14.5   15  RT (min)      lcorbi_t3_neg_2_022_24    m/z 799.6715     0   1   2E+4   13   13.5  RT (min)      lcorbi_t3_neg_4_011_24    m/z 799.6882     0   1   2E+4   13.5   14  RT (min)      lcorbi_t3_neg_1_052_24    m/z 799.7076     0   1   2E+4   14.5   15  RT (min)      lcorbi_t3_neg_2_052_32    m/z 799.8213     0   1   2   3   4E+3   18   18.5  RT (min)      lcorbi_t3_neg_4_028_39    m/z 800.2354     0   2   4E+3   10.5   11  RT (min)      lcorbi_t3_neg_2_027_13    m/z 800.2591     0   1   2   3   4E+3   11.5   12  RT (min)      lcorbi_t3_neg_4_034_01    m/z 800.3346     0   2   4   6E+3   10.5   11  RT (min)      lcorbi_t3_neg_3_002_24    m/z 800.4126     0   1E+4   7.5   8  RT (min)      lcorbi_t3_neg_4_017_51    m/z 800.4823     0   2   4   6E+3   13.5   14  RT (min)      lcorbi_t3_neg_4_019_08    m/z 800.5003     0   2   4E+4   14   14.5  RT (min)      lcorbi_t3_neg_4_029_31    m/z 800.5183     0   1   2   3E+3   14.5   15  RT (min)      lcorbi_t3_neg_4_019_08    m/z 800.5255     0   1E+6   13.5   14  RT (min)      lcorbi_t3_neg_4_019_08    m/z 800.5355     0   1E+6   13.5   14  RT (min)      lcorbi_t3_neg_4_044_38    m/z 800.5442     0   1E+4   18   18.5  RT (min)      lcorbi_t3_neg_3_005_23    m/z 800.5611     0   2   4   6E+2   14.5   15  RT (min)      lcorbi_t3_neg_4_011_24    m/z 800.5782     0   1E+4   14.5   15  RT (min)      lcorbi_t3_neg_1_031_33    m/z 800.6154     0   1   15   15.5  RT (min)      lcorbi_t3_neg_4_019_08    m/z 800.6264     0   1   14.5   15  RT (min)      lcorbi_t3_neg_3_002_24    m/z 800.8024     0   1   2   3   4E+3   18   18.5  RT (min)      lcorbi_t3_neg_4_024_07    m/z 801.1748     0   1   2   3   4E+3   9   9.5  RT (min)      lcorbi_t3_neg_3_019_12    m/z 801.2881     0   2   4   6   8E+3   14   14.5  RT (min)      lcorbi_t3_neg_2_037_09    m/z 801.3513     0   2   4E+3   18   18.5  RT (min)      lcorbi_t3_neg_4_020_44    m/z 801.4515     0   2   4E+3   13   13.5  RT (min)      lcorbi_t3_neg_4_032_23    m/z 801.4682     0   1   2   3   4E+4   12   12.5  RT (min)      lcorbi_t3_neg_4_032_23    m/z 801.4732     0   2   4   6   8E+3   13.5   14  RT (min)      lcorbi_t3_neg_3_005_23    m/z 801.4921     0   2   4E+3   13   13.5  RT (min)      lcorbi_t3_neg_4_032_23    m/z 801.4965     0   1E+4   14.5   15  RT (min)      lcorbi_t3_neg_2_053_23    m/z 801.5175     0   1   14.5   15  RT (min)      lcorbi_t3_neg_4_019_08    m/z 801.5641     0   2   4   6   8E+3   14.5   15  RT (min)      lcorbi_t3_neg_2_052_32    m/z 801.6049     0   2   4   6   8E+3   14.5   15  RT (min)      lcorbi_t3_neg_4_047_30    m/z 801.6426     0   1E+4   14.5   15  RT (min)      lcorbi_t3_neg_4_011_24    m/z 801.7958     0   1E+4   14.5   15  RT (min)      lcorbi_t3_neg_2_052_32    m/z 801.8185     0   1   2   3E+3   18   18.5  RT (min)      lcorbi_t3_neg_4_040_19    m/z 801.8247     0   1E+4   15   15.5  RT (min)      lcorbi_t3_neg_2_052_32    m/z 802.2482     0   1E+4   11.5   12  RT (min)      lcorbi_t3_neg_3_035_03    m/z 802.3072     0   1E+4   13.5   14  RT (min)      lcorbi_t3_neg_2_045_30    m/z 802.4413     0   1E+4   13.5   14  RT (min)      lcorbi_t3_neg_4_019_08    m/z 802.4713     0   1   12   12.5  RT (min)      lcorbi_t3_neg_4_032_23    m/z 802.4955     0   2   4E+4   14   14.5  RT (min)      lcorbi_t3_neg_4_005_27    m/z 802.5249     0   1   2E+5   14   14.5  RT (min)      lcorbi_t3_neg_4_005_27    m/z 802.6717     0   1   2   3   4E+4   14   14.5  RT (min)      lcorbi_t3_neg_4_011_24    m/z 803.2549     0   2   4E+3   11.5   12  RT (min)      lcorbi_t3_neg_2_026_03    m/z 803.2640     0   2   4   6   8E+3   13.5   14  RT (min)      lcorbi_t3_neg_3_023_07    m/z 803.2740     0   2   4   6E+3   9.5   10  RT (min)      lcorbi_t3_neg_3_019_12    m/z 803.3296     0   2   4E+3   11   11.5  RT (min)      lcorbi_t3_neg_2_035_46    m/z 803.3706     0   2   4   6   8E+3   9   9.5  RT (min)      lcorbi_t3_neg_4_017_51    m/z 803.4430     0   2   4   6E+3   7.5   8  RT (min)      lcorbi_t3_neg_4_023_18    m/z 803.4541     0   1E+4   11   11.5  RT (min)      lcorbi_t3_neg_4_023_18    m/z 803.4954     0   1E+3   13.5   14  RT (min)      lcorbi_t3_neg_3_005_23    m/z 803.5791     0   2   4   6   8E+2   13   13.5  RT (min)      lcorbi_t3_neg_4_019_08    m/z 803.6206     0   2   4   6   8E+4   14   14.5  RT (min)      lcorbi_t3_neg_4_034_01    m/z 803.8198     0   1   2   3   4E+3   18   18.5  RT (min)      lcorbi_t3_neg_4_030_04    m/z 804.2409     0   2   4   6   8E+3   10   10.5  RT (min)      lcorbi_t3_neg_3_019_12    m/z 804.2491     0   2   4   6E+3   11.5   12  RT (min)      lcorbi_t3_neg_2_026_03    m/z 804.2614     0   1   12   12.5  RT (min)      lcorbi_t3_neg_4_019_08    m/z 804.3740     0   1   2   3   4E+3   9   9.5  RT (min)      lcorbi_t3_neg_4_017_51    m/z 804.4571     0   2   4   6   8E+3   11   11.5  RT (min)      lcorbi_t3_neg_4_023_18    m/z 804.4703     0   1E+4   13   13.5  RT (min)      lcorbi_t3_neg_4_029_31    m/z 804.5138     0   1E+4   14.5   15  RT (min)      lcorbi_t3_neg_2_053_23    m/z 804.5999     0   1   14   14.5  RT (min)      lcorbi_t3_neg_4_007_13    m/z 804.6488     0   2   4E+4   14   14.5  RT (min)      lcorbi_t3_neg_4_007_13    m/z 805.2445     0   1   2   3   4E+3   10   10.5  RT (min)      lcorbi_t3_neg_4_028_39    m/z 805.4273     0   1E+4   13.5   14  RT (min)      lcorbi_t3_neg_4_029_31    m/z 805.4686     0   2   4   6   8E+3   15   15.5  RT (min)      lcorbi_t3_neg_2_031_25    m/z 805.5096     0   1   2E+3   14.5   15  RT (min)      lcorbi_t3_neg_4_019_08    m/z 805.6123     0   1E+4   14   14.5  RT (min)      lcorbi_t3_neg_2_027_13    m/z 805.7947     0   1   2   3E+3   18   18.5  RT (min)      lcorbi_t3_neg_2_055_18    m/z 805.8202     0   2   4   6   8E+2   18   18.5  RT (min)      lcorbi_t3_neg_4_019_08    m/z 806.1949     0   1E+5   13.5   14  RT (min)      lcorbi_t3_neg_4_045_35    m/z 806.2445     0   2   4E+3   12.5   13  RT (min)      lcorbi_t3_neg_3_007_16    m/z 806.2632     0   2   4   6   8E+3   11.5   12  RT (min)      lcorbi_t3_neg_2_010_02    m/z 806.2826     0   1E+4   14   14.5  RT (min)      lcorbi_t3_neg_4_030_04    m/z 806.3181     0   1   2   3E+4   9   9.5  RT (min)      lcorbi_t3_neg_2_048_27    m/z 806.4305     0   2   4   6   8E+3   13.5   14  RT (min)      lcorbi_t3_neg_1_039_38    m/z 806.5105     0   1   2E+3   13.5   14  RT (min)      lcorbi_t3_neg_4_045_35    m/z 806.5455     0   1E+4   14.5   15  RT (min)      lcorbi_t3_neg_3_045_34    m/z 806.5910     0   1E+6   14.5   15  RT (min)      lcorbi_t3_neg_4_034_01    m/z 806.6068     0   1E+4   14.5   15  RT (min)      lcorbi_t3_neg_3_002_24    m/z 806.6155     0   1   2E+3   14   14.5  RT (min)      lcorbi_t3_neg_3_007_16    m/z 806.6468     0   1   2   3E+2   14.5   15  RT (min)      lcorbi_t3_neg_4_019_08    m/z 806.7861     0   2   4E+3   14.5   15  RT (min)      lcorbi_t3_neg_2_052_32    m/z 806.8113     0   2   4   6E+3   18   18.5  RT (min)      lcorbi_t3_neg_2_048_27    m/z 807.2325     0   1   2   3E+4   5   5.5  RT (min)      lcorbi_t3_neg_2_048_27    m/z 807.2857     0   2   4E+3   9.5   10  RT (min)      lcorbi_t3_neg_3_019_12    m/z 807.3217     0   1E+4   9   9.5  RT (min)      lcorbi_t3_neg_2_048_27    m/z 807.3362     0   1E+4   14.5   15  RT (min)      lcorbi_t3_neg_4_007_13    m/z 807.4403     0   1   2   3E+4   13.5   14  RT (min)      lcorbi_t3_neg_2_040_43    m/z 807.4823     0   1E+4   13   13.5  RT (min)      lcorbi_t3_neg_4_019_08    m/z 807.4890     0   1   2E+3   13   13.5  RT (min)      lcorbi_t3_neg_1_035_25    m/z 807.5251     0   1   2E+3   14.5   15  RT (min)      lcorbi_t3_neg_1_047_02    m/z 807.6157     0   1   2   3   4E+3   14.5   15  RT (min)      lcorbi_t3_neg_3_002_24    m/z 807.6286     0   2   4   6   8E+2   12.5   13  RT (min)      lcorbi_t3_neg_4_011_24    m/z 807.6405     0   1E+4   13.5   14  RT (min)      lcorbi_t3_neg_2_022_24    m/z 807.6764     0   1   2   3   4E+5   14   14.5  RT (min)      lcorbi_t3_neg_3_002_24    m/z 808.2661     0   1   2   3E+3   11   11.5  RT (min)      lcorbi_t3_neg_2_010_02    m/z 808.4438     0   1E+4   13.5   14  RT (min)      lcorbi_t3_neg_2_040_43    m/z 808.4979     0   1E+4   11.5   12  RT (min)      lcorbi_t3_neg_4_032_23    m/z 808.5027     0   1E+4   14   14.5  RT (min)      lcorbi_t3_neg_4_019_08    m/z 808.5270     0   1   2   3E+3   14.5   15  RT (min)      lcorbi_t3_neg_4_005_27    m/z 808.5449     0   1   2E+3   14.5   15  RT (min)      lcorbi_t3_neg_1_043_14    m/z 808.5622     0   1   2   3   4E+4   13.5   14  RT (min)      lcorbi_t3_neg_3_005_23    m/z 808.6226     0   2   4   6E+3   14.5   15  RT (min)      lcorbi_t3_neg_1_055_05    m/z 808.6305     0   1   2E+3   14   14.5  RT (min)      lcorbi_t3_neg_4_034_01    m/z 809.0436     0   2   4   6E+3   9   9.5  RT (min)      lcorbi_t3_neg_4_042_14    m/z 809.2002     0   1E+4   12.5   13  RT (min)      lcorbi_t3_neg_4_005_27    m/z 809.4560     0   1E+4   14   14.5  RT (min)      lcorbi_t3_neg_3_046_10    m/z 809.5015     0   2   4E+3   11.5   12  RT (min)      lcorbi_t3_neg_3_005_23    m/z 809.5415     0   2   4   6   8E+3   14.5   15  RT (min)      lcorbi_t3_neg_4_007_13    m/z 809.5533     0   1E+4   14   14.5  RT (min)      lcorbi_t3_neg_4_005_27    m/z 809.6146     0   1   2E+4   14.5   15  RT (min)      lcorbi_t3_neg_4_011_24    m/z 809.6731     0   1   2E+4   13.5   14  RT (min)      lcorbi_t3_neg_4_039_32    m/z 809.6920     0   1E+6   14   14.5  RT (min)      lcorbi_t3_neg_3_002_24    m/z 809.8063     0   1   2E+3   18   18.5  RT (min)      lcorbi_t3_neg_4_034_01    m/z 809.8372     0   2   4   6E+3   18   18.5  RT (min)      lcorbi_t3_neg_4_019_08    m/z 809.8503     0   2   4   6   8E+3   18   18.5  RT (min)      lcorbi_t3_neg_4_022_41    m/z 810.2034     0   2   4E+3   12.5   13  RT (min)      lcorbi_t3_neg_4_005_27    m/z 810.2757     0   2   4E+3   14.5   15  RT (min)      lcorbi_t3_neg_1_050_45    m/z 810.4069     0   2   4   6E+3   13   13.5  RT (min)      lcorbi_t3_neg_2_053_23    m/z 810.4545     0   1   2E+3   13   13.5  RT (min)      lcorbi_t3_neg_1_044_23    m/z 810.4725     0   1E+4   11   11.5  RT (min)      lcorbi_t3_neg_4_023_18    m/z 810.5079     0   1E+5   13   13.5  RT (min)      lcorbi_t3_neg_4_019_08    m/z 810.5417     0   1   2E+3   13   13.5  RT (min)      lcorbi_t3_neg_4_017_51    m/z 810.5445     0   1   2   3E+3   14.5   15  RT (min)      lcorbi_t3_neg_1_022_34    m/z 810.5495     0   1   2   3E+3   13.5   14  RT (min)      lcorbi_t3_neg_1_055_05    m/z 810.5761     0   1   13.5   14  RT (min)      lcorbi_t3_neg_4_011_24    m/z 810.8368     0   2   4   6E+3   18   18.5  RT (min)      lcorbi_t3_neg_1_003_41    m/z 811.2679     0   1   2   3   4E+3   10.5   11  RT (min)      lcorbi_t3_neg_3_018_13    m/z 811.5534     0   2   4   6   8E+2   13.5   14  RT (min)      lcorbi_t3_neg_1_053_32    m/z 811.6055     0   1   2   3E+3   14.5   15  RT (min)      lcorbi_t3_neg_4_039_32    m/z 811.6892     0   1   2   3E+4   13.5   14  RT (min)      lcorbi_t3_neg_4_039_32    m/z 812.2692     0   1E+4   7.5   8  RT (min)      lcorbi_t3_neg_2_048_27    m/z 812.2727     0   2   4   6E+3   11   11.5  RT (min)      lcorbi_t3_neg_4_011_24    m/z 812.3000     0   1   2   3   4E+3   11   11.5  RT (min)      lcorbi_t3_neg_4_036_16    m/z 812.3824     0   1E+4   14.5   15  RT (min)      lcorbi_t3_neg_4_034_01    m/z 812.4250     0   1   2   3   4E+3   13   13.5  RT (min)      lcorbi_t3_neg_2_012_36    m/z 812.4332     0   1E+4   10.5   11  RT (min)      lcorbi_t3_neg_4_005_27    m/z 812.4715     0   1   2E+4   13.5   14  RT (min)      lcorbi_t3_neg_3_005_23    m/z 812.5256     0   1   2   3E+5   13.5   14  RT (min)      lcorbi_t3_neg_4_019_08    m/z 812.5683     0   2   4   6E+3   13.5   14  RT (min)      lcorbi_t3_neg_4_039_32    m/z 812.6110     0   1E+5   12.5   13  RT (min)      lcorbi_t3_neg_3_002_24    m/z 812.6220     0   1E+4   14.5   15  RT (min)      lcorbi_t3_neg_4_039_32    m/z 812.6685     0   2   4E+3   14.5   15  RT (min)      lcorbi_t3_neg_4_039_32    m/z 812.8474     0   2   4E+3   18   18.5  RT (min)      lcorbi_t3_neg_2_021_41    m/z 813.2837     0   1E+4   11.5   12  RT (min)      lcorbi_t3_neg_4_048_03    m/z 813.3861     0   2   4   6E+3   14.5   15  RT (min)      lcorbi_t3_neg_4_034_01    m/z 813.3887     0   1E+4   7.5   8  RT (min)      lcorbi_t3_neg_4_023_18    m/z 813.4735     0   1E+4   13.5   14  RT (min)      lcorbi_t3_neg_3_005_23    m/z 813.4890     0   2   4E+3   13.5   14  RT (min)      lcorbi_t3_neg_4_007_13    m/z 813.5730     0   1   14   14.5  RT (min)      lcorbi_t3_neg_4_042_14    m/z 813.6223     0   1E+4   14.5   15  RT (min)      lcorbi_t3_neg_4_011_24    m/z 813.6336     0   1   2   3   4E+3   14.5   15  RT (min)      lcorbi_t3_neg_4_019_08    m/z 813.6507     0   2   4   6E+3   18   18.5  RT (min)      lcorbi_t3_neg_4_039_32    m/z 814.1968     0   2   4E+3   10   10.5  RT (min)      lcorbi_t3_neg_4_032_23    m/z 814.2848     0   1E+4   13.5   14  RT (min)      lcorbi_t3_neg_3_011_04    m/z 814.2859     0   2   4   6E+3   11.5   12  RT (min)      lcorbi_t3_neg_3_030_33    m/z 814.3061     0   2   4   6   8E+3   9.5   10  RT (min)      lcorbi_t3_neg_3_019_12    m/z 814.3152     0   1   2E+3   11.5   12  RT (min)      lcorbi_t3_neg_4_005_27    m/z 814.4851     0   1E+5   13   13.5  RT (min)      lcorbi_t3_neg_4_032_23    m/z 814.5574     0   2   4E+3   6.5   7  RT (min)      lcorbi_t3_neg_2_053_23    m/z 814.5762     0   2   4   6E+3   14   14.5  RT (min)      lcorbi_t3_neg_4_007_13    m/z 814.6197     0   2   4   6E+2   14.5   15  RT (min)      lcorbi_t3_neg_3_037_36    m/z 814.6310     0   1E+4   15   15.5  RT (min)      lcorbi_t3_neg_4_019_08    m/z 814.6529     0   1   2   3E+4   13.5   14  RT (min)      lcorbi_t3_neg_1_052_24    m/z 814.6847     0   1E+4   14.5   15  RT (min)      lcorbi_t3_neg_2_052_32    m/z 814.6991     0   1E+4   13.5   14  RT (min)      lcorbi_t3_neg_1_055_05    m/z 815.4716     0   2   4   6   8E+3   14.5   15  RT (min)      lcorbi_t3_neg_4_011_24    m/z 815.4916     0   1E+4   14.5   15  RT (min)      lcorbi_t3_neg_3_037_36    m/z 815.4963     0   1E+3   11   11.5  RT (min)      lcorbi_t3_neg_3_005_23    m/z 815.5374     0   1E+4   13.5   14  RT (min)      lcorbi_t3_neg_2_053_23    m/z 815.5439     0   1E+5   13.5   14  RT (min)      lcorbi_t3_neg_4_019_08    m/z 815.5639     0   1   2   3   4E+4   11.5   12  RT (min)      lcorbi_t3_neg_4_042_14    m/z 815.5792     0   2   4   6   8E+3   13   13.5  RT (min)      lcorbi_t3_neg_2_052_32    m/z 815.5910     0   1E+5   14   14.5  RT (min)      lcorbi_t3_neg_4_034_01    m/z 815.6456     0   2   4   6   8E+3   14.5   15  RT (min)      lcorbi_t3_neg_4_001_49    m/z 815.8116     0   2   4   6   8E+3   14.5   15  RT (min)      lcorbi_t3_neg_2_052_32    m/z 815.8234     0   2   4   6E+3   18   18.5  RT (min)      lcorbi_t3_neg_3_029_02    m/z 815.8389     0   2   4   6E+3   15   15.5  RT (min)      lcorbi_t3_neg_2_052_32    m/z 815.8564     0   2   4   6E+3   18   18.5  RT (min)      lcorbi_t3_neg_3_042_22    m/z 816.2155     0   2   4E+3   11   11.5  RT (min)      lcorbi_t3_neg_3_005_23    m/z 816.2668     0   1E+3   11.5   12  RT (min)      lcorbi_t3_neg_4_007_13    m/z 816.2804     0   1   2   3E+3   14   14.5  RT (min)      lcorbi_t3_neg_2_023_39    m/z 816.2926     0   2   4   6E+3   11   11.5  RT (min)      lcorbi_t3_neg_4_036_16    m/z 816.2999     0   2   4   6E+3   11.5   12  RT (min)      lcorbi_t3_neg_4_007_13    m/z 816.3314     0   2   4   6E+3   12   12.5  RT (min)      lcorbi_t3_neg_2_052_32    m/z 816.3568     0   1   2   3   4E+3   14   14.5  RT (min)      lcorbi_t3_neg_4_053_43    m/z 816.4387     0   1E+4   13.5   14  RT (min)      lcorbi_t3_neg_4_019_08    m/z 816.4565     0   1   13.5   14  RT (min)      lcorbi_t3_neg_4_007_13    m/z 816.5007     0   1   2   3   4E+3   13.5   14  RT (min)      lcorbi_t3_neg_1_035_25    m/z 816.5133     0   2   4   6E+4   14   14.5  RT (min)      lcorbi_t3_neg_4_019_08    m/z 816.5913     0   1E+4   13.5   14  RT (min)      lcorbi_t3_neg_2_022_24    m/z 816.6041     0   2   4   6   8E+3   14   14.5  RT (min)      lcorbi_t3_neg_4_032_23    m/z 816.6424     0   1   2   3   4E+4   13   13.5  RT (min)      lcorbi_t3_neg_1_052_24    m/z 816.6442     0   1E+3   14.5   15  RT (min)      lcorbi_t3_neg_4_034_01    m/z 816.6691     0   1   2   3E+4   13.5   14  RT (min)      lcorbi_t3_neg_4_011_24    m/z 816.6870     0   1E+4   13   13.5  RT (min)      lcorbi_t3_neg_4_039_32    m/z 816.7003     0   1E+3   14.5   15  RT (min)      lcorbi_t3_neg_3_023_07    m/z 816.7873     0   1   2   3   4E+3   15   15.5  RT (min)      lcorbi_t3_neg_4_001_49    m/z 816.8251     0   2   4   6   8E+3   18   18.5  RT (min)      lcorbi_t3_neg_2_029_12    m/z 816.8578     0   2   4   6E+3   18   18.5  RT (min)      lcorbi_t3_neg_1_004_22    m/z 817.1380     0   1   2   3E+3   9   9.5  RT (min)      lcorbi_t3_neg_3_019_12    m/z 817.2334     0   1   2   3   4E+3   11.5   12  RT (min)      lcorbi_t3_neg_3_035_03    m/z 817.3025     0   1   2   3   4E+3   11.5   12  RT (min)      lcorbi_t3_neg_3_022_27    m/z 817.4221     0   2   4   6E+3   7.5   8  RT (min)      lcorbi_t3_neg_4_023_18    m/z 817.4417     0   2   4   6E+3   13.5   14  RT (min)      lcorbi_t3_neg_4_019_08    m/z 817.4683     0   1   14.5   15  RT (min)      lcorbi_t3_neg_2_039_31    m/z 817.4871     0   1E+4   13   13.5  RT (min)      lcorbi_t3_neg_4_046_37    m/z 817.4873     0   1E+4   14.5   15  RT (min)      lcorbi_t3_neg_4_011_24    m/z 817.5438     0   2   4   6   8E+4   13.5   14  RT (min)      lcorbi_t3_neg_4_032_23    m/z 817.6079     0   1E+4   14   14.5  RT (min)      lcorbi_t3_neg_3_045_34    m/z 817.6258     0   2   4   6E+4   12.5   13  RT (min)      lcorbi_t3_neg_3_002_24    m/z 817.6294     0   1E+4   10   10.5  RT (min)      lcorbi_t3_neg_3_019_12    m/z 817.6360     0   1   2E+4   14.5   15  RT (min)      lcorbi_t3_neg_4_007_13    m/z 817.6547     0   1   2E+4   13   13.5  RT (min)      lcorbi_t3_neg_4_039_32    m/z 817.6767     0   1E+4   15   15.5  RT (min)      lcorbi_t3_neg_4_011_24    m/z 817.6847     0   1   2   3   4E+4   14   14.5  RT (min)      lcorbi_t3_neg_3_002_24    m/z 817.7155     0   2   4   6E+3   14.5   15  RT (min)      lcorbi_t3_neg_4_039_32    m/z 817.8247     0   2   4   6E+3   18   18.5  RT (min)      lcorbi_t3_neg_2_003_42    m/z 818.2325     0   2   4E+3   11.5   12  RT (min)      lcorbi_t3_neg_3_040_05    m/z 818.3154     0   1E+4   12   12.5  RT (min)      lcorbi_t3_neg_3_010_15    m/z 818.4382     0   2   4   6E+3   13   13.5  RT (min)      lcorbi_t3_neg_4_019_08    m/z 818.4914     0   1   2   3   4E+3   13   13.5  RT (min)      lcorbi_t3_neg_4_044_38    m/z 818.5728     0   2   4   6E+4   14   14.5  RT (min)      lcorbi_t3_neg_4_019_08    m/z 818.6623     0   2   4   6E+3   14.5   15  RT (min)      lcorbi_t3_neg_1_052_24    m/z 818.7155     0   1E+4   14.5   15  RT (min)      lcorbi_t3_neg_2_052_32    m/z 818.8304     0   1   2   3   4E+3   18   18.5  RT (min)      lcorbi_t3_neg_2_003_42    m/z 819.3182     0   2   4E+3   12   12.5  RT (min)      lcorbi_t3_neg_3_037_36    m/z 819.4226     0   2   4   6E+3   13   13.5  RT (min)      lcorbi_t3_neg_4_034_01    m/z 819.4544     0   1E+4   6   6.5  RT (min)      lcorbi_t3_neg_2_048_27    m/z 819.4845     0   1   14.5   15  RT (min)      lcorbi_t3_neg_4_007_13    m/z 819.5030     0   2   4   6E+3   14.5   15  RT (min)      lcorbi_t3_neg_1_055_05    m/z 819.5130     0   1E+3   14.5   15  RT (min)      lcorbi_t3_neg_4_019_08    m/z 819.5450     0   1   2E+4   10.5   11  RT (min)      lcorbi_t3_neg_4_039_32    m/z 819.6421     0   1   2   3E+4   13.5   14  RT (min)      lcorbi_t3_neg_4_039_32    m/z 819.6699     0   1E+4   13.5   14  RT (min)      lcorbi_t3_neg_2_052_32    m/z 819.6734     0   1   2   3   4E+3   14.5   15  RT (min)      lcorbi_t3_neg_3_002_24    m/z 820.2490     0   1   2   3   4E+3   12   12.5  RT (min)      lcorbi_t3_neg_2_054_15    m/z 820.3240     0   2   4   6E+3   11.5   12  RT (min)      lcorbi_t3_neg_3_007_16    m/z 820.4019     0   1   2   3E+4   10.5   11  RT (min)      lcorbi_t3_neg_4_029_31    m/z 820.5075     0   1   2E+4   13.5   14  RT (min)      lcorbi_t3_neg_4_046_37    m/z 820.5255     0   1E+3   14.5   15  RT (min)      lcorbi_t3_neg_4_019_08    m/z 820.5409     0   2   4   6   8E+3   14.5   15  RT (min)      lcorbi_t3_neg_4_007_13    m/z 820.6137     0   1E+4   18   18.5  RT (min)      lcorbi_t3_neg_4_039_32    m/z 820.6543     0   2   4   6E+3   14.5   15  RT (min)      lcorbi_t3_neg_4_034_01    m/z 820.6751     0   2   4   6   8E+3   14.5   15  RT (min)      lcorbi_t3_neg_4_039_32    m/z 820.7323     0   2   4   6   8E+3   15   15.5  RT (min)      lcorbi_t3_neg_4_039_32    m/z 820.7898     0   1   2E+3   18   18.5  RT (min)      lcorbi_t3_neg_3_046_10    m/z 820.8017     0   1   2E+3   18   18.5  RT (min)      lcorbi_t3_neg_2_015_21    m/z 820.8362     0   2   4   6   8E+3   18   18.5  RT (min)      lcorbi_t3_neg_2_040_43    m/z 821.0437     0   2   4E+3   10.5   11  RT (min)      lcorbi_t3_neg_4_030_04    m/z 821.2274     0   2   4   6   8E+4   5   5.5  RT (min)      lcorbi_t3_neg_2_048_27    m/z 821.4052     0   1E+4   10.5   11  RT (min)      lcorbi_t3_neg_4_029_31    m/z 821.5001     0   1E+3   14.5   15  RT (min)      lcorbi_t3_neg_4_029_31    m/z 821.5376     0   1   2E+4   13.5   14  RT (min)      lcorbi_t3_neg_4_035_33    m/z 821.5521     0   1   2   3E+4   10.5   11  RT (min)      lcorbi_t3_neg_4_039_32    m/z 821.5595     0   1E+5   14.5   15  RT (min)      lcorbi_t3_neg_4_042_14    m/z 822.2980     0   1   2   3   4E+4   7.5   8  RT (min)      lcorbi_t3_neg_2_048_27    m/z 822.3411     0   2   4E+3   18   18.5  RT (min)      lcorbi_t3_neg_2_021_41    m/z 822.3641     0   2   4E+3   14.5   15  RT (min)      lcorbi_t3_neg_4_032_23    m/z 822.4529     0   1E+4   13.5   14  RT (min)      lcorbi_t3_neg_1_054_37    m/z 822.5408     0   1E+4   14.5   15  RT (min)      lcorbi_t3_neg_4_007_13    m/z 822.5414     0   1   2   3E+5   13.5   14  RT (min)      lcorbi_t3_neg_2_043_33    m/z 822.6286     0   2   4   6E+3   18   18.5  RT (min)      lcorbi_t3_neg_4_011_24    m/z 822.8108     0   1   2   3E+3   18   18.5  RT (min)      lcorbi_t3_neg_2_013_08    m/z 822.8386     0   2   4   6E+3   18   18.5  RT (min)      lcorbi_t3_neg_3_011_04    m/z 822.8577     0   2   4   6E+3   15   15.5  RT (min)      lcorbi_t3_neg_4_001_49    m/z 823.3013     0   1   2E+4   7.5   8  RT (min)      lcorbi_t3_neg_2_048_27    m/z 823.4555     0   2   4   6   8E+3   13.5   14  RT (min)      lcorbi_t3_neg_3_021_38    m/z 823.5539     0   2   4   6   8E+3   15   15.5  RT (min)      lcorbi_t3_neg_4_054_36    m/z 823.5938     0   2   4   6   8E+3   14.5   15  RT (min)      lcorbi_t3_neg_2_033_53    m/z 823.6504     0   2   4   6   8E+2   14.5   15  RT (min)      lcorbi_t3_neg_4_011_24    m/z 824.2760     0   1E+4   7   7.5  RT (min)      lcorbi_t3_neg_3_045_34    m/z 824.3083     0   1   2   3   4E+3   14.5   15  RT (min)      lcorbi_t3_neg_1_023_03    m/z 824.3165     0   2   4   6   8E+3   12   12.5  RT (min)      lcorbi_t3_neg_4_034_01    m/z 824.3466     0   2   4E+3   18   18.5  RT (min)      lcorbi_t3_neg_2_029_12    m/z 824.3615     0   1   2   3   4E+3   14.5   15  RT (min)      lcorbi_t3_neg_4_007_13    m/z 824.3797     0   2   4E+3   14.5   15  RT (min)      lcorbi_t3_neg_4_007_13    m/z 824.4186     0   1E+3   14.5   15  RT (min)      lcorbi_t3_neg_4_032_23    m/z 824.4484     0   1   2E+4   12   12.5  RT (min)      lcorbi_t3_neg_4_032_23    m/z 824.5261     0   1   2   3E+6   13.5   14  RT (min)      lcorbi_t3_neg_4_019_08    m/z 824.5672     0   1   2E+3   14   14.5  RT (min)      lcorbi_t3_neg_3_005_23    m/z 824.5861     0   1   2   3   4E+3   6.5   7  RT (min)      lcorbi_t3_neg_4_029_31    m/z 824.6273     0   1   2   3E+5   13   13.5  RT (min)      lcorbi_t3_neg_4_019_08    m/z 824.6325     0   2   4   6   8E+3   14.5   15  RT (min)      lcorbi_t3_neg_2_007_14    m/z 824.6386     0   2   4   6   8E+3   18   18.5  RT (min)      lcorbi_t3_neg_4_039_32    m/z 824.8082     0   2   4E+3   18   18.5  RT (min)      lcorbi_t3_neg_2_016_11    m/z 825.3201     0   1   2   3E+3   12   12.5  RT (min)      lcorbi_t3_neg_4_034_01    m/z 825.4141     0   1   2   3   4E+3   15   15.5  RT (min)      lcorbi_t3_neg_2_038_22    m/z 825.4641     0   2   4   6E+3   11   11.5  RT (min)      lcorbi_t3_neg_4_023_18    m/z 825.4793     0   1   2   3   4E+4   6   6.5  RT (min)      lcorbi_t3_neg_2_048_27    m/z 825.4923     0   2   4   6   8E+3   13   13.5  RT (min)      lcorbi_t3_neg_4_029_31    m/z 825.5294     0   1E+6   13.5   14  RT (min)      lcorbi_t3_neg_4_019_08    m/z 825.5477     0   1   2E+4   18   18.5  RT (min)      lcorbi_t3_neg_1_020_36    m/z 825.5584     0   1   2E+4   14   14.5  RT (min)      lcorbi_t3_neg_4_007_13    m/z 825.5679     0   1   13.5   14  RT (min)      lcorbi_t3_neg_2_052_32    m/z 825.6070     0   1   2   3   4E+4   13.5   14  RT (min)      lcorbi_t3_neg_4_029_31    m/z 825.6507     0   2   4   6E+3   18   18.5  RT (min)      lcorbi_t3_neg_4_039_32    m/z 826.2957     0   2   4   6E+3   11.5   12  RT (min)      lcorbi_t3_neg_4_034_01    m/z 826.3321     0   2   4E+3   12.5   13  RT (min)      lcorbi_t3_neg_4_034_01    m/z 826.3412     0   1E+4   10.5   11  RT (min)      lcorbi_t3_neg_4_011_24    m/z 826.3771     0   1   2   3   4E+3   14.5   15  RT (min)      lcorbi_t3_neg_4_007_13    m/z 826.4407     0   1E+4   12.5   13  RT (min)      lcorbi_t3_neg_2_053_23    m/z 826.4767     0   1   2E+5   13.5   14  RT (min)      lcorbi_t3_neg_4_029_31    m/z 826.4828     0   1   2E+4   6   6.5  RT (min)      lcorbi_t3_neg_2_048_27    m/z 826.4957     0   2   4   6E+2   13   13.5  RT (min)      lcorbi_t3_neg_1_019_29    m/z 826.5414     0   1   2E+6   13.5   14  RT (min)      lcorbi_t3_neg_4_019_08    m/z 826.6195     0   2   4   6E+4   14   14.5  RT (min)      lcorbi_t3_neg_4_029_31    m/z 826.6325     0   1   2   3E+4   14   14.5  RT (min)      lcorbi_t3_neg_4_029_31    m/z 826.6545     0   1   2E+4   18   18.5  RT (min)      lcorbi_t3_neg_4_039_32    m/z 826.7007     0   1   2   3   4E+4   13.5   14  RT (min)      lcorbi_t3_neg_1_055_05    m/z 826.8185     0   1E+3   18   18.5  RT (min)      lcorbi_t3_neg_4_019_08    m/z 827.0460     0   1   2   3   4E+3   9.5   10  RT (min)      lcorbi_t3_neg_4_051_40    m/z 827.2547     0   2   4E+3   11   11.5  RT (min)      lcorbi_t3_neg_2_008_16    m/z 827.2716     0   1   2   3E+3   13.5   14  RT (min)      lcorbi_t3_neg_4_023_18    m/z 827.2991     0   1   2E+3   11.5   12  RT (min)      lcorbi_t3_neg_4_042_14    m/z 827.3554     0   2   4   6E+3   18   18.5  RT (min)      lcorbi_t3_neg_4_027_12    m/z 827.3689     0   1E+4   4.5   5  RT (min)      lcorbi_t3_neg_4_042_14    m/z 827.4779     0   1   2E+4   13   13.5  RT (min)      lcorbi_t3_neg_4_019_08    m/z 827.5439     0   1E+6   13.5   14  RT (min)      lcorbi_t3_neg_4_019_08    m/z 827.5891     0   2   4E+3   14.5   15  RT (min)      lcorbi_t3_neg_4_044_38    m/z 827.6458     0   2   4E+4   14.5   15  RT (min)      lcorbi_t3_neg_4_019_08    m/z 827.6663     0   1E+4   18   18.5  RT (min)      lcorbi_t3_neg_4_039_32    m/z 827.8562     0   1   2   3   4E+3   18   18.5  RT (min)      lcorbi_t3_neg_2_042_44    m/z 828.2495     0   1   2   3   4E+3   11   11.5  RT (min)      lcorbi_t3_neg_4_048_03    m/z 828.4126     0   2   4   6E+3   13.5   14  RT (min)      lcorbi_t3_neg_2_028_38    m/z 828.4828     0   1E+4   11.5   12  RT (min)      lcorbi_t3_neg_4_023_18    m/z 828.4907     0   1   2E+4   13.5   14  RT (min)      lcorbi_t3_neg_4_029_31    m/z 828.5921     0   1   14.5   15  RT (min)      lcorbi_t3_neg_4_007_13    m/z 828.6060     0   1E+5   12.5   13  RT (min)      lcorbi_t3_neg_2_022_24    m/z 828.6138     0   2   4   6   8E+4   14.5   15  RT (min)      lcorbi_t3_neg_4_034_01    m/z 828.6473     0   1E+4   14   14.5  RT (min)      lcorbi_t3_neg_3_007_16    m/z 828.7408     0   1   2   3E+3   14.5   15  RT (min)      lcorbi_t3_neg_3_030_33    m/z 828.8239     0   2   4   6E+3   18   18.5  RT (min)      lcorbi_t3_neg_2_029_12    m/z 829.1139     0   1E+5   11   11.5  RT (min)      lcorbi_t3_neg_4_007_13    m/z 829.1935     0   2   4E+4   13.5   14  RT (min)      lcorbi_t3_neg_4_045_35    m/z 829.2706     0   2   4E+3   11.5   12  RT (min)      lcorbi_t3_neg_2_005_01    m/z 829.4587     0   2   4   6E+3   7.5   8  RT (min)      lcorbi_t3_neg_4_023_18    m/z 829.4952     0   2   4   6   8E+3   10   10.5  RT (min)      lcorbi_t3_neg_4_023_18    m/z 829.5319     0   2   4   6   8E+4   14   14.5  RT (min)      lcorbi_t3_neg_4_007_13    m/z 829.5420     0   1   2E+4   13.5   14  RT (min)      lcorbi_t3_neg_2_039_31    m/z 829.5583     0   1E+4   6   6.5  RT (min)      lcorbi_t3_neg_4_044_38    m/z 829.5600     0   2   4   6E+5   13.5   14  RT (min)      lcorbi_t3_neg_4_019_08    m/z 829.6009     0   2   4E+3   13   13.5  RT (min)      lcorbi_t3_neg_1_052_24    m/z 829.6364     0   2   4E+5   14   14.5  RT (min)      lcorbi_t3_neg_4_034_01    m/z 829.6617     0   1   2E+4   14.5   15  RT (min)      lcorbi_t3_neg_4_019_08    m/z 829.6814     0   2   4   6   8E+3   18   18.5  RT (min)      lcorbi_t3_neg_3_002_24    m/z 829.6919     0   1E+4   14.5   15  RT (min)      lcorbi_t3_neg_4_011_24    m/z 829.6996     0   1E+4   14.5   15  RT (min)      lcorbi_t3_neg_4_011_24    m/z 829.8264     0   2   4   6E+3   18   18.5  RT (min)      lcorbi_t3_neg_3_011_04    m/z 830.1120     0   2   4   6   8E+4   11   11.5  RT (min)      lcorbi_t3_neg_4_007_13    m/z 830.2461     0   2   4   6E+3   12   12.5  RT (min)      lcorbi_t3_neg_3_007_16    m/z 830.2648     0   2   4E+3   11.5   12  RT (min)      lcorbi_t3_neg_2_050_17    m/z 830.3386     0   2   4   6   8E+3   14   14.5  RT (min)      lcorbi_t3_neg_2_023_39    m/z 830.4907     0   1E+4   13   13.5  RT (min)      lcorbi_t3_neg_4_044_38    m/z 830.6226     0   2   4   6E+5   13   13.5   14  RT (min)       lcorbi_t3_neg_4_021_25    m/z 830.7608     0   2   4E+3   18   18.5  RT (min)      lcorbi_t3_neg_2_015_21    m/z 830.8233     0   1   2   3E+3   18   18.5  RT (min)      lcorbi_t3_neg_3_042_22    m/z 830.8579     0   1E+4   15   15.5  RT (min)      lcorbi_t3_neg_2_052_32    m/z 831.1117     0   1   2   3   4E+4   11   11.5  RT (min)      lcorbi_t3_neg_4_007_13    m/z 831.1927     0   2   4   6E+3   9   9.5  RT (min)      lcorbi_t3_neg_3_019_12    m/z 831.1951     0   1E+4   4.5   5  RT (min)      lcorbi_t3_neg_2_048_27    m/z 831.2506     0   1   2E+4   9   9.5  RT (min)      lcorbi_t3_neg_4_023_18    m/z 831.2916     0   2   4   6   8E+3   13   13.5  RT (min)      lcorbi_t3_neg_1_023_03    m/z 831.4753     0   1   2   3   4E+4   7.5   8  RT (min)      lcorbi_t3_neg_2_048_27    m/z 831.5752     0   1   2E+5   14   14.5  RT (min)      lcorbi_t3_neg_4_019_08    m/z 831.5851     0   2   4   6E+5   14   14.5  RT (min)      lcorbi_t3_neg_4_007_13    m/z 831.7062     0   1E+4   13   13.5  RT (min)      lcorbi_t3_neg_4_039_32    m/z 831.7347     0   2   4   6E+3   14   14.5  RT (min)      lcorbi_t3_neg_3_040_05    m/z 831.8317     0   2   4   6E+3   18   18.5  RT (min)      lcorbi_t3_neg_4_028_39    m/z 832.2786     0   2   4   6E+3   11.5   12  RT (min)      lcorbi_t3_neg_4_034_01    m/z 832.4357     0   1E+4   13   13.5  RT (min)      lcorbi_t3_neg_4_029_31    m/z 832.4965     0   1   15.5   16  RT (min)      lcorbi_t3_neg_4_019_08    m/z 832.5265     0   2   4   6E+3   13   13.5  RT (min)      lcorbi_t3_neg_1_021_31    m/z 832.5344     0   1   2   3E+5   12.5   13  RT (min)      lcorbi_t3_neg_4_032_23    m/z 832.6824     0   1   15   15.5  RT (min)      lcorbi_t3_neg_4_019_08    m/z 832.7581     0   1   2   3   4E+3   18   18.5  RT (min)      lcorbi_t3_neg_2_029_12    m/z 833.1308     0   1   2E+3   9   9.5  RT (min)      lcorbi_t3_neg_2_023_39    m/z 833.2917     0   1E+4   13   13.5  RT (min)      lcorbi_t3_neg_1_023_03    m/z 833.4391     0   2   4   6E+3   13   13.5  RT (min)      lcorbi_t3_neg_3_020_31    m/z 833.4718     0   1   14.5   15  RT (min)      lcorbi_t3_neg_4_044_38    m/z 833.4910     0   2   4   6   8E+2   13.5   14  RT (min)      lcorbi_t3_neg_4_032_23    m/z 833.5093     0   2   4   6E+4   13.5   14  RT (min)      lcorbi_t3_neg_4_029_31    m/z 833.5301     0   1E+5   13   13.5  RT (min)      lcorbi_t3_neg_4_032_23    m/z 833.5910     0   1E+4   14   14.5  RT (min)      lcorbi_t3_neg_4_019_08    m/z 833.6575     0   1   2   3E+3   14.5   15  RT (min)      lcorbi_t3_neg_2_022_24    m/z 833.6775     0   2   4   6   8E+3   13.5   14  RT (min)      lcorbi_t3_neg_4_039_32    m/z 833.7900     0   1   2E+3   18   18.5  RT (min)      lcorbi_t3_neg_3_033_53    m/z 833.8375     0   1   2   3   4E+3   18   18.5  RT (min)      lcorbi_t3_neg_1_006_40    m/z 834.2645     0   1E+4   10.5   11  RT (min)      lcorbi_t3_neg_4_032_23    m/z 834.2708     0   1   2E+3   14   14.5  RT (min)      lcorbi_t3_neg_3_005_23    m/z 834.2943     0   2   4   6   8E+3   12   12.5  RT (min)      lcorbi_t3_neg_2_007_14    m/z 834.3842     0   2   4E+4   1   1.5  RT (min)      lcorbi_t3_neg_2_048_27    m/z 834.4435     0   2   4   6E+3   12   12.5  RT (min)      lcorbi_t3_neg_4_039_32    m/z 834.4857     0   2   4   6E+3   13.5   14  RT (min)      lcorbi_t3_neg_4_054_36    m/z 834.5026     0   1E+5   14   14.5  RT (min)      lcorbi_t3_neg_4_019_08    m/z 834.5121     0   1E+4   13.5   14  RT (min)      lcorbi_t3_neg_4_044_38    m/z 834.5325     0   1   2E+4   11   11.5  RT (min)      lcorbi_t3_neg_4_007_13    m/z 834.5684     0   1E+4   13.5   14  RT (min)      lcorbi_t3_neg_4_054_36    m/z 834.5769     0   1E+4   13.5   14  RT (min)      lcorbi_t3_neg_4_038_15    m/z 834.6919     0   2   4   6E+3   14.5   15  RT (min)      lcorbi_t3_neg_2_022_24    m/z 835.3448     0   2   4E+3   18   18.5  RT (min)      lcorbi_t3_neg_2_042_44    m/z 835.3644     0   1E+4   8   8.5  RT (min)      lcorbi_t3_neg_2_048_27    m/z 835.3673     0   2   4   6E+3   15   15.5  RT (min)      lcorbi_t3_neg_4_007_13    m/z 835.4545     0   1   2E+3   13.5   14  RT (min)      lcorbi_t3_neg_1_031_33    m/z 835.4714     0   1E+4   14   14.5  RT (min)      lcorbi_t3_neg_2_040_43    m/z 835.5071     0   1E+5   14   14.5  RT (min)      lcorbi_t3_neg_4_019_08    m/z 835.5531     0   1   2E+5   13   13.5  RT (min)      lcorbi_t3_neg_4_032_23    m/z 835.5740     0   1   2E+4   14   14.5  RT (min)      lcorbi_t3_neg_4_039_32    m/z 835.5881     0   1   2E+4   13.5   14  RT (min)      lcorbi_t3_neg_2_013_08    m/z 835.6286     0   1   14.5   15  RT (min)      lcorbi_t3_neg_2_022_24    m/z 835.6720     0   2   4E+3   14.5   15  RT (min)      lcorbi_t3_neg_2_022_24    m/z 835.7019     0   2   4E+4   14   14.5  RT (min)      lcorbi_t3_neg_3_002_24    m/z 835.8068     0   1   2   3E+3   18   18.5  RT (min)      lcorbi_t3_neg_3_048_08    m/z 836.4176     0   1   2   3E+4   13.5   14  RT (min)      lcorbi_t3_neg_1_054_37    m/z 836.4729     0   1E+4   13.5   14  RT (min)      lcorbi_t3_neg_4_032_23    m/z 836.4848     0   2   4E+3   11   11.5  RT (min)      lcorbi_t3_neg_2_053_23    m/z 836.5150     0   2   4   6E+3   11   11.5  RT (min)      lcorbi_t3_neg_1_023_03    m/z 836.5573     0   2   4   6E+2   13.5   14  RT (min)      lcorbi_t3_neg_3_002_24    m/z 836.6312     0   1E+4   14.5   15  RT (min)      lcorbi_t3_neg_4_019_08    m/z 836.6684     0   1E+4   13   13.5  RT (min)      lcorbi_t3_neg_2_052_32    m/z 836.6883     0   1   2   3E+4   14   14.5  RT (min)      lcorbi_t3_neg_1_052_24    m/z 836.6964     0   1E+4   13.5   14  RT (min)      lcorbi_t3_neg_2_052_32    m/z 836.7787     0   1   2   3   4E+3   18   18.5  RT (min)      lcorbi_t3_neg_2_006_10    m/z 836.8862     0   1   2   3E+3   18   18.5  RT (min)      lcorbi_t3_neg_4_018_42    m/z 837.0746     0   2   4E+3   9.5   10  RT (min)      lcorbi_t3_neg_4_003_21    m/z 837.3003     0   2   4   6   8E+3   13.5   14  RT (min)      lcorbi_t3_neg_4_023_18    m/z 837.4259     0   2   4   6   8E+3   13.5   14  RT (min)      lcorbi_t3_neg_2_053_23    m/z 837.4794     0   1   2E+4   6   6.5  RT (min)      lcorbi_t3_neg_3_037_36    m/z 837.5303     0   1   2E+4   13.5   14  RT (min)      lcorbi_t3_neg_4_044_38    m/z 837.5607     0   2   4   6   8E+3   13.5   14  RT (min)      lcorbi_t3_neg_1_044_23    m/z 837.6459     0   1   2   3E+3   14.5   15  RT (min)      lcorbi_t3_neg_3_002_24    m/z 838.2851     0   2   4   6   8E+3   11   11.5  RT (min)      lcorbi_t3_neg_4_011_24    m/z 838.3071     0   1E+4   14.5   15  RT (min)      lcorbi_t3_neg_2_032_45    m/z 838.4199     0   2   4   6   8E+3   13.5   14  RT (min)      lcorbi_t3_neg_4_019_08    m/z 838.4821     0   1E+4   6   6.5  RT (min)      lcorbi_t3_neg_3_037_36    m/z 838.4847     0   2   4   6E+4   13   13.5  RT (min)      lcorbi_t3_neg_3_005_23    m/z 838.4848     0   1E+4   11   11.5  RT (min)      lcorbi_t3_neg_4_023_18    m/z 838.5222     0   1E+4   13.5   14  RT (min)      lcorbi_t3_neg_4_032_23    m/z 838.5923     0   2   4   6   8E+3   6   6.5  RT (min)      lcorbi_t3_neg_4_032_23    m/z 838.6054     0   1   13.5   14  RT (min)      lcorbi_t3_neg_4_019_08    m/z 838.6429     0   1E+5   13   13.5  RT (min)      lcorbi_t3_neg_4_019_08    m/z 838.6476     0   1   2E+3   14.5   15  RT (min)      lcorbi_t3_neg_4_039_32    m/z 838.6750     0   1   2E+2   15   15.5  RT (min)      lcorbi_t3_neg_4_039_32    m/z 838.7850     0   2   4   6   8E+2   18   18.5  RT (min)      lcorbi_t3_neg_4_019_08    m/z 839.1989     0   2   4E+3   10.5   11  RT (min)      lcorbi_t3_neg_2_026_03    m/z 839.2919     0   2   4   6E+3   11   11.5  RT (min)      lcorbi_t3_neg_4_011_24    m/z 839.3098     0   1   2   3   4E+3   14.5   15  RT (min)      lcorbi_t3_neg_1_050_45    m/z 839.3115     0   2   4E+4   9   9.5  RT (min)      lcorbi_t3_neg_2_048_27    m/z 839.4437     0   2   4   6E+3   13   13.5  RT (min)      lcorbi_t3_neg_2_053_23    m/z 839.4654     0   1   2E+4   7.5   8  RT (min)      lcorbi_t3_neg_2_048_27    m/z 839.4882     0   1   2   3E+4   13   13.5  RT (min)      lcorbi_t3_neg_4_032_23    m/z 839.5192     0   1E+4   13.5   14  RT (min)      lcorbi_t3_neg_3_020_31    m/z 839.6270     0   2   4E+3   14.5   15  RT (min)      lcorbi_t3_neg_3_037_36    m/z 839.7219     0   1   2E+4   14   14.5  RT (min)      lcorbi_t3_neg_3_002_24    m/z 839.8546     0   2   4   6   8E+3   18   18.5  RT (min)      lcorbi_t3_neg_1_004_22    m/z 840.2153     0   2   4E+3   11   11.5  RT (min)      lcorbi_t3_neg_2_026_03    m/z 840.3016     0   1   2   3   4E+3   11.5   12  RT (min)      lcorbi_t3_neg_2_027_13    m/z 840.4371     0   1E+4   13.5   14  RT (min)      lcorbi_t3_neg_1_044_23    m/z 840.4564     0   1   2   3   4E+3   13.5   14  RT (min)      lcorbi_t3_neg_1_055_05    m/z 840.4926     0   1   2E+3   10.5   11  RT (min)      lcorbi_t3_neg_4_032_23    m/z 840.5391     0   1   2   3E+4   12.5   13  RT (min)      lcorbi_t3_neg_4_032_23    m/z 840.5670     0   2   4E+5   14   14.5  RT (min)      lcorbi_t3_neg_4_007_13    m/z 840.5757     0   1E+3   11.5   12  RT (min)      lcorbi_t3_neg_3_002_24    m/z 840.5904     0   1   2E+4   13.5   14  RT (min)      lcorbi_t3_neg_4_054_36    m/z 840.6251     0   2   4E+3   13.5   14  RT (min)      lcorbi_t3_neg_2_019_07    m/z 840.6584     0   1E+4   14.5   15  RT (min)      lcorbi_t3_neg_4_019_08    m/z 840.7339     0   1   2E+4   14   14.5  RT (min)      lcorbi_t3_neg_4_039_32    m/z 840.8523     0   1   2   3E+3   18   18.5  RT (min)      lcorbi_t3_neg_2_026_03    m/z 841.3076     0   1E+4   11.5   12  RT (min)      lcorbi_t3_neg_4_016_05    m/z 841.4595     0   2   4   6   8E+3   13.5   14  RT (min)      lcorbi_t3_neg_2_053_23    m/z 841.4756     0   1E+4   11   11.5  RT (min)      lcorbi_t3_neg_4_029_31    m/z 841.4930     0   2   4   6E+2   13   13.5  RT (min)      lcorbi_t3_neg_2_053_23    m/z 841.4953     0   2   4   6   8E+3   11   11.5  RT (min)      lcorbi_t3_neg_3_020_31    m/z 841.5053     0   1   2E+4   13.5   14  RT (min)      lcorbi_t3_neg_4_032_23    m/z 841.5594     0   2   4   6   8E+3   13.5   14  RT (min)      lcorbi_t3_neg_4_019_08    m/z 841.5791     0   2   4   6   8E+2   11.5   12  RT (min)      lcorbi_t3_neg_1_052_24    m/z 841.6067     0   2   4   6   8E+4   14   14.5  RT (min)      lcorbi_t3_neg_3_007_16    m/z 841.6612     0   1E+3   14.5   15  RT (min)      lcorbi_t3_neg_4_019_08    m/z 841.7922     0   1   2   3   4E+3   18   18.5  RT (min)      lcorbi_t3_neg_1_047_02    m/z 842.3174     0   1   2   3   4E+3   12   12.5  RT (min)      lcorbi_t3_neg_4_029_31    m/z 842.3836     0   1   2   3E+4   10.5   11  RT (min)      lcorbi_t3_neg_4_029_31    m/z 842.4646     0   1   2   3   4E+4   14   14.5  RT (min)      lcorbi_t3_neg_1_044_23    m/z 842.5981     0   1E+5   13.5   14  RT (min)      lcorbi_t3_neg_4_007_13    m/z 842.6051     0   1   2   3   4E+4   13.5   14  RT (min)      lcorbi_t3_neg_1_055_05    m/z 842.6852     0   1   2   3   4E+4   18   18.5  RT (min)      lcorbi_t3_neg_1_052_24    m/z 842.7957     0   2   4   6   8E+3   18   18.5  RT (min)      lcorbi_t3_neg_2_019_07    m/z 843.0699     0   1   2   3E+3   7.5   8  RT (min)      lcorbi_t3_neg_4_034_01    m/z 843.0910     0   1   2E+3   8   8.5  RT (min)      lcorbi_t3_neg_4_003_21    m/z 843.1594     0   2   4   6   8E+3   9   9.5  RT (min)      lcorbi_t3_neg_3_019_12    m/z 843.2106     0   2   4E+3   10.5   11  RT (min)      lcorbi_t3_neg_4_050_02    m/z 843.2487     0   1   2   3   4E+3   11.5   12  RT (min)      lcorbi_t3_neg_4_005_27    m/z 843.3871     0   1E+4   10.5   11  RT (min)      lcorbi_t3_neg_4_029_31    m/z 843.4666     0   1   2E+4   14   14.5  RT (min)      lcorbi_t3_neg_1_044_23    m/z 843.5105     0   1E+4   10   10.5  RT (min)      lcorbi_t3_neg_4_023_18    m/z 843.5313     0   1E+4   14   14.5  RT (min)      lcorbi_t3_neg_2_010_02    m/z 843.5751     0   1   14   14.5  RT (min)      lcorbi_t3_neg_2_053_23    m/z 843.5852     0   2   4   6E+4   14   14.5  RT (min)      lcorbi_t3_neg_4_005_27    m/z 843.6229     0   2   4   6E+3   14.5   15  RT (min)      lcorbi_t3_neg_4_011_24    m/z 843.6503     0   2   4   6E+4   14   14.5  RT (min)      lcorbi_t3_neg_2_022_24    m/z 843.7072     0   2   4E+3   14.5   15  RT (min)      lcorbi_t3_neg_4_007_13    m/z 843.7526     0   1E+4   14.5   15  RT (min)      lcorbi_t3_neg_2_052_32    m/z 843.8184     0   2   4   6E+3   18   18.5  RT (min)      lcorbi_t3_neg_3_036_41    m/z 844.1626     0   1   2   3E+3   9   9.5  RT (min)      lcorbi_t3_neg_3_019_12    m/z 844.3879     0   2   4E+3   14.5   15  RT (min)      lcorbi_t3_neg_2_023_39    m/z 844.4075     0   2   4   6   8E+3   13.5   14  RT (min)      lcorbi_t3_neg_4_054_36    m/z 844.4803     0   1E+4   14.5   15  RT (min)      lcorbi_t3_neg_1_044_23    m/z 844.4822     0   2   4E+4   10.5   11  RT (min)      lcorbi_t3_neg_4_039_32    m/z 844.5066     0   2   4   6   8E+3   13   13.5  RT (min)      lcorbi_t3_neg_4_046_37    m/z 844.5162     0   1   2E+4   11   11.5  RT (min)      lcorbi_t3_neg_4_007_13    m/z 844.5224     0   1   2E+4   13.5   14  RT (min)      lcorbi_t3_neg_4_007_13    m/z 844.5451     0   1   2E+4   14   14.5  RT (min)      lcorbi_t3_neg_1_002_16    m/z 844.6177     0   1E+3   13   13.5  RT (min)      lcorbi_t3_neg_4_042_14    m/z 844.8012     0   2   4   6   8E+3   18   18.5  RT (min)      lcorbi_t3_neg_2_011_04    m/z 844.8736     0   1   2   3E+3   18   18.5  RT (min)      lcorbi_t3_neg_4_024_07    m/z 845.1574     0   1   2   3E+3   9   9.5  RT (min)      lcorbi_t3_neg_4_040_19    m/z 845.2649     0   2   4   6E+3   12   12.5  RT (min)      lcorbi_t3_neg_3_053_14    m/z 845.4389     0   2   4   6   8E+3   13   13.5  RT (min)      lcorbi_t3_neg_3_028_35    m/z 845.5494     0   1E+4   14.5   15  RT (min)      lcorbi_t3_neg_2_004_28    m/z 845.5721     0   1E+4   15.5   16  RT (min)      lcorbi_t3_neg_4_042_14    m/z 845.7217     0   1   2   3E+4   13.5   14  RT (min)      lcorbi_t3_neg_4_039_32    m/z 846.2351     0   2   4   6E+3   11   11.5  RT (min)      lcorbi_t3_neg_4_036_16    m/z 846.2673     0   2   4E+3   12   12.5  RT (min)      lcorbi_t3_neg_4_007_13    m/z 846.2895     0   2   4E+3   8.5   9  RT (min)      lcorbi_t3_neg_4_001_49    m/z 846.3964     0   1E+4   8   8.5  RT (min)      lcorbi_t3_neg_4_040_19    m/z 846.4837     0   1   2   3   4E+3   13.5   14  RT (min)      lcorbi_t3_neg_4_029_31    m/z 846.5107     0   2   4   6   8E+3   13   13.5  RT (min)      lcorbi_t3_neg_2_043_33    m/z 846.6509     0   2   4   6E+3   14.5   15  RT (min)      lcorbi_t3_neg_3_010_15    m/z 846.6588     0   1E+3   14.5   15  RT (min)      lcorbi_t3_neg_3_023_07    m/z 846.6702     0   1E+5   14.5   15  RT (min)      lcorbi_t3_neg_2_052_32    m/z 846.6803     0   1   13.5   14  RT (min)      lcorbi_t3_neg_4_034_01    m/z 846.6978     0   1   2   3   4E+2   15   15.5  RT (min)      lcorbi_t3_neg_4_019_08    m/z 846.7553     0   1   2   3   4E+3   15   15.5  RT (min)      lcorbi_t3_neg_4_029_31    m/z 846.7996     0   1   2   3   4E+3   18   18.5  RT (min)      lcorbi_t3_neg_2_015_21    m/z 846.8413     0   1   2   3   4E+3   18   18.5  RT (min)      lcorbi_t3_neg_3_003_09    m/z 846.9887     0   1   2E+3   9   9.5  RT (min)      lcorbi_t3_neg_2_018_40    m/z 847.0967     0   1   2   3   4E+3   9   9.5  RT (min)      lcorbi_t3_neg_4_023_18    m/z 847.1710     0   2   4   6E+3   9   9.5  RT (min)      lcorbi_t3_neg_3_019_12    m/z 847.4784     0   1   13.5   14  RT (min)      lcorbi_t3_neg_4_032_23    m/z 847.5251     0   1E+5   13.5   14  RT (min)      lcorbi_t3_neg_4_032_23    m/z 847.5533     0   1   14   14.5  RT (min)      lcorbi_t3_neg_1_052_24    m/z 847.6547     0   1E+4   13.5   14  RT (min)      lcorbi_t3_neg_3_043_25    m/z 847.6631     0   2   4   6E+3   14.5   15  RT (min)      lcorbi_t3_neg_4_007_13    m/z 847.6650     0   1   2   3   4E+3   14.5   15  RT (min)      lcorbi_t3_neg_2_052_32    m/z 847.8121     0   2   4   6   8E+3   18   18.5  RT (min)      lcorbi_t3_neg_1_046_11    m/z 848.1745     0   1   2   3E+3   9   9.5  RT (min)      lcorbi_t3_neg_3_019_12    m/z 848.2506     0   2   4E+3   11.5   12  RT (min)      lcorbi_t3_neg_4_007_13    m/z 848.2832     0   1E+4   14   14.5  RT (min)      lcorbi_t3_neg_1_023_03    m/z 848.4575     0   1   2E+4   13.5   14  RT (min)      lcorbi_t3_neg_2_053_23    m/z 848.4647     0   1   2E+4   11   11.5  RT (min)      lcorbi_t3_neg_4_023_18    m/z 848.4666     0   2   4   6   8E+3   18   18.5  RT (min)      lcorbi_t3_neg_3_005_23    m/z 848.4812     0   2   4   6E+2   13.5   14  RT (min)      lcorbi_t3_neg_3_045_34    m/z 848.5268     0   1   2E+4   13   13.5  RT (min)      lcorbi_t3_neg_2_053_23    m/z 848.6021     0   2   4E+2   14   14.5  RT (min)      lcorbi_t3_neg_4_011_24    m/z 848.6196     0   1   2   3   4E+4   13   13.5  RT (min)      lcorbi_t3_neg_4_019_08    m/z 848.6318     0   1   2   3E+4   11.5   12  RT (min)      lcorbi_t3_neg_1_052_24    m/z 848.6575     0   1   2   3   4E+3   13.5   14  RT (min)      lcorbi_t3_neg_2_052_32    m/z 848.6681     0   1   2E+4   13   13.5  RT (min)      lcorbi_t3_neg_2_022_24    m/z 848.6710     0   1   2   3E+4   13.5   14  RT (min)      lcorbi_t3_neg_4_039_32    m/z 848.6858     0   1   2E+3   14.5   15  RT (min)      lcorbi_t3_neg_1_030_47    m/z 848.7326     0   1E+4   14.5   15  RT (min)      lcorbi_t3_neg_2_022_24    m/z 849.2278     0   1E+4   10.5   11  RT (min)      lcorbi_t3_neg_3_029_02    m/z 849.2588     0   2   4   6E+3   12   12.5  RT (min)      lcorbi_t3_neg_1_043_14    m/z 849.3757     0   2   4E+3   8   8.5  RT (min)      lcorbi_t3_neg_4_017_51    m/z 849.4597     0   1E+4   11   11.5  RT (min)      lcorbi_t3_neg_4_023_18    m/z 849.4700     0   1   2E+4   13.5   14  RT (min)      lcorbi_t3_neg_3_005_23    m/z 849.4701     0   1   2   3   4E+3   18   18.5  RT (min)      lcorbi_t3_neg_4_046_37    m/z 849.5018     0   2   4E+4   7.5   8  RT (min)      lcorbi_t3_neg_2_048_27    m/z 849.5127     0   2   4E+3   12.5   13  RT (min)      lcorbi_t3_neg_4_032_23    m/z 849.5397     0   1   2   3   4E+4   13.5   14  RT (min)      lcorbi_t3_neg_3_047_37    m/z 849.6349     0   1   2   3   4E+3   14.5   15  RT (min)      lcorbi_t3_neg_2_027_13    m/z 849.8158     0   2   4   6E+3   18   18.5  RT (min)      lcorbi_t3_neg_2_018_40    m/z 849.8497     0   1   2   3E+3   18   18.5  RT (min)      lcorbi_t3_neg_3_014_19    m/z 850.2313     0   2   4E+3   10.5   11  RT (min)      lcorbi_t3_neg_3_029_02    m/z 850.2513     0   2   4   6E+3   11   11.5  RT (min)      lcorbi_t3_neg_2_026_03    m/z 850.2658     0   1E+4   13   13.5  RT (min)      lcorbi_t3_neg_3_005_23    m/z 850.3225     0   2   4   6E+3   10.5   11  RT (min)      lcorbi_t3_neg_2_022_24    m/z 850.5370     0   1E+4   13   13.5  RT (min)      lcorbi_t3_neg_2_053_23    m/z 850.5505     0   1   2E+5   13.5   14  RT (min)      lcorbi_t3_neg_4_044_38    m/z 850.5599     0   1E+4   18   18.5  RT (min)      lcorbi_t3_neg_1_014_28    m/z 850.5931     0   1   2E+4   14.5   15  RT (min)      lcorbi_t3_neg_4_007_13    m/z 850.6006     0   1   2   3   4E+3   14.5   15  RT (min)      lcorbi_t3_neg_4_011_24    m/z 850.6350     0   1   2   3E+4   13.5   14  RT (min)      lcorbi_t3_neg_1_055_05    m/z 851.0003     0   1   2E+3   9   9.5  RT (min)      lcorbi_t3_neg_4_028_39    m/z 851.2546     0   1   2E+3   11   11.5  RT (min)      lcorbi_t3_neg_2_026_03    m/z 851.4695     0   2   4   6E+3   13.5   14  RT (min)      lcorbi_t3_neg_4_019_08    m/z 851.4889     0   2   4   6   8E+3   13.5   14  RT (min)      lcorbi_t3_neg_3_048_08    m/z 851.5483     0   2   4   6   8E+3   13.5   14  RT (min)      lcorbi_t3_neg_2_053_23    m/z 851.5546     0   1   2   3E+6   13.5   14  RT (min)      lcorbi_t3_neg_4_007_13    m/z 851.5952     0   1   2   3E+3   14.5   15  RT (min)      lcorbi_t3_neg_4_038_15    m/z 851.6060     0   1   13.5   14  RT (min)      lcorbi_t3_neg_4_019_08    m/z 851.6511     0   1   2   3   4E+3   14.5   15  RT (min)      lcorbi_t3_neg_3_052_32    m/z 852.2527     0   2   4E+3   11   11.5  RT (min)      lcorbi_t3_neg_3_032_17    m/z 852.4016     0   2   4   6E+3   9   9.5  RT (min)      lcorbi_t3_neg_2_048_27    m/z 852.4560     0   2   4E+3   13   13.5  RT (min)      lcorbi_t3_neg_2_012_36    m/z 852.5263     0   2   4   6E+4   14   14.5  RT (min)      lcorbi_t3_neg_4_007_13    m/z 852.5752     0   1E+4   18   18.5  RT (min)      lcorbi_t3_neg_1_014_28    m/z 852.6126     0   1E+4   14.5   15  RT (min)      lcorbi_t3_neg_3_013_28    m/z 853.1195     0   1   2   3   4E+3   8   8.5  RT (min)      lcorbi_t3_neg_4_014_20    m/z 853.2380     0   1   2E+4   5   5.5  RT (min)      lcorbi_t3_neg_2_048_27    m/z 853.2483     0   1   2   3E+3   12   12.5  RT (min)      lcorbi_t3_neg_3_054_18    m/z 853.5041     0   1   2E+3   13.5   14  RT (min)      lcorbi_t3_neg_4_024_07    m/z 853.5142     0   1E+4   14   14.5  RT (min)      lcorbi_t3_neg_2_053_23    m/z 853.5427     0   1E+4   13.5   14  RT (min)      lcorbi_t3_neg_4_029_31    m/z 853.5763     0   1E+4   6   6.5  RT (min)      lcorbi_t3_neg_4_044_38    m/z 853.6170     0   1E+4   14.5   15  RT (min)      lcorbi_t3_neg_3_032_17    m/z 853.6614     0   2   4   6E+3   14.5   15  RT (min)      lcorbi_t3_neg_4_024_07    m/z 853.7881     0   1E+4   15   15.5  RT (min)      lcorbi_t3_neg_4_001_49    m/z 853.8201     0   1   2   3   4E+3   18   18.5  RT (min)      lcorbi_t3_neg_2_034_19    m/z 854.1916     0   1E+4   9   9.5  RT (min)      lcorbi_t3_neg_3_019_12    m/z 854.2628     0   2   4E+3   11   11.5  RT (min)      lcorbi_t3_neg_2_008_16    m/z 854.2694     0   2   4E+3   11.5   12  RT (min)      lcorbi_t3_neg_4_005_27    m/z 854.4718     0   1E+4   13.5   14  RT (min)      lcorbi_t3_neg_3_005_23    m/z 854.4965     0   2   4   6   8E+3   13   13.5  RT (min)      lcorbi_t3_neg_4_007_13    m/z 854.5185     0   1   2   3   4E+5   12.5   13  RT (min)      lcorbi_t3_neg_4_032_23    m/z 854.5551     0   1   2E+3   13.5   14  RT (min)      lcorbi_t3_neg_4_042_14    m/z 854.5795     0   2   4   6E+3   6   6.5  RT (min)      lcorbi_t3_neg_4_044_38    m/z 854.7640     0   1E+4   14.5   15  RT (min)      lcorbi_t3_neg_2_052_32    m/z 855.0333     0   1E+4   9   9.5  RT (min)      lcorbi_t3_neg_4_042_14    m/z 855.1942     0   1   2E+3   9   9.5  RT (min)      lcorbi_t3_neg_4_048_03    m/z 855.3329     0   1   2E+4   9   9.5  RT (min)      lcorbi_t3_neg_2_048_27    m/z 855.4754     0   1   13.5   14  RT (min)      lcorbi_t3_neg_4_007_13    m/z 856.0359     0   2   4   6E+3   9   9.5  RT (min)      lcorbi_t3_neg_4_042_14    m/z 856.2854     0   2   4E+3   12   12.5  RT (min)      lcorbi_t3_neg_3_052_32    m/z 856.3183     0   1E+4   8.5   9  RT (min)      lcorbi_t3_neg_2_048_27    m/z 856.3346     0   2   4E+3   18   18.5  RT (min)      lcorbi_t3_neg_4_028_39    m/z 856.4874     0   2   4   6E+3   13.5   14  RT (min)      lcorbi_t3_neg_2_013_08    m/z 856.5260     0   1E+6   13   13.5  RT (min)      lcorbi_t3_neg_4_032_23    m/z 856.5421     0   1   2   3   4E+3   14   14.5  RT (min)      lcorbi_t3_neg_1_044_23    m/z 856.5576     0   1   2   3E+4   14.5   15  RT (min)      lcorbi_t3_neg_3_018_13    m/z 856.5874     0   1   2E+4   14   14.5  RT (min)      lcorbi_t3_neg_1_005_07    m/z 856.6054     0   2   4   6   8E+3   18   18.5  RT (min)      lcorbi_t3_neg_4_034_01    m/z 856.8325     0   2   4   6E+3   18   18.5  RT (min)      lcorbi_t3_neg_2_021_41    m/z 857.0176     0   1   2   3   4E+3   9   9.5  RT (min)      lcorbi_t3_neg_4_028_39    m/z 857.3216     0   2   4   6   8E+3   8.5   9  RT (min)      lcorbi_t3_neg_2_048_27    m/z 857.5598     0   2   4   6E+3   13   13.5  RT (min)      lcorbi_t3_neg_4_007_13    m/z 857.6372     0   1E+3   15   15.5  RT (min)      lcorbi_t3_neg_4_007_13    m/z 857.6423     0   1E+4   13.5   14  RT (min)      lcorbi_t3_neg_2_022_24    m/z 857.7375     0   2   4   6E+3   14.5   15  RT (min)      lcorbi_t3_neg_3_028_35    m/z 857.7503     0   1E+4   14   14.5  RT (min)      lcorbi_t3_neg_4_011_24    m/z 858.2643     0   1   2   3   4E+3   10.5   11  RT (min)      lcorbi_t3_neg_2_027_13    m/z 858.3403     0   1   2   3   4E+3   18   18.5  RT (min)      lcorbi_t3_neg_2_034_19    m/z 858.4090     0   1   2E+4   13.5   14  RT (min)      lcorbi_t3_neg_4_019_08    m/z 858.4510     0   1E+4   13.5   14  RT (min)      lcorbi_t3_neg_2_024_37    m/z 858.4887     0   2   4   6E+3   13.5   14  RT (min)      lcorbi_t3_neg_4_019_08    m/z 858.5320     0   1E+4   11   11.5  RT (min)      lcorbi_t3_neg_4_007_13    m/z 858.5380     0   2   4   6E+3   14   14.5  RT (min)      lcorbi_t3_neg_1_035_25    m/z 858.5474     0   1   2E+4   14.5   15  RT (min)      lcorbi_t3_neg_2_053_23    m/z 858.6528     0   2   4   6E+2   14.5   15  RT (min)      lcorbi_t3_neg_4_034_01    m/z 858.8703     0   1   2   3   4E+3   18   18.5  RT (min)      lcorbi_t3_neg_4_036_16    m/z 859.1835     0   1   2   3   4E+3   10.5   11  RT (min)      lcorbi_t3_neg_4_050_02    m/z 859.4266     0   2   4   6E+4   13.5   14  RT (min)      lcorbi_t3_neg_1_054_37    m/z 859.4545     0   1E+3   13.5   14  RT (min)      lcorbi_t3_neg_2_043_33    m/z 859.4951     0   1   2E+3   13.5   14  RT (min)      lcorbi_t3_neg_4_029_31    m/z 859.5071     0   1   2   3E+3   13.5   14  RT (min)      lcorbi_t3_neg_4_034_01    m/z 859.5513     0   1   14.5   15  RT (min)      lcorbi_t3_neg_4_007_13    m/z 859.5602     0   2   4E+3   13.5   14  RT (min)      lcorbi_t3_neg_4_038_15    m/z 859.6369     0   1   2E+4   13.5   14  RT (min)      lcorbi_t3_neg_4_039_32    m/z 859.6721     0   2   4E+4   14.5   15  RT (min)      lcorbi_t3_neg_4_007_13    m/z 860.3052     0   2   4   6E+3   9   9.5  RT (min)      lcorbi_t3_neg_4_001_49    m/z 860.3515     0   1E+4   10.5   11  RT (min)      lcorbi_t3_neg_4_011_24    m/z 860.4225     0   1   2   3   4E+4   13   13.5  RT (min)      lcorbi_t3_neg_1_044_23    m/z 860.5020     0   1   2   3   4E+3   14   14.5  RT (min)      lcorbi_t3_neg_3_032_17    m/z 860.5273     0   1E+4   13.5   14  RT (min)      lcorbi_t3_neg_1_039_38    m/z 860.5568     0   2   4E+5   13.5   14  RT (min)      lcorbi_t3_neg_4_032_23    m/z 860.5795     0   1E+3   13.5   14  RT (min)      lcorbi_t3_neg_4_038_15    m/z 860.5922     0   1   2E+3   13.5   14  RT (min)      lcorbi_t3_neg_4_019_08    m/z 860.6678     0   1   2   3E+5   13.5   14  RT (min)      lcorbi_t3_neg_3_002_24    m/z 860.7247     0   1   2   3E+4   14   14.5  RT (min)      lcorbi_t3_neg_3_002_24    m/z 860.7319     0   1   2   3E+4   14   14.5  RT (min)      lcorbi_t3_neg_3_002_24    m/z 860.7419     0   1   2   3E+4   13.5   14  RT (min)      lcorbi_t3_neg_2_022_24    m/z 860.7858     0   1   2   3E+3   18   18.5  RT (min)      lcorbi_t3_neg_4_053_43    m/z 861.2835     0   2   4   6   8E+3   11   11.5  RT (min)      lcorbi_t3_neg_3_029_02    m/z 861.3489     0   2   4   6   8E+3   18   18.5  RT (min)      lcorbi_t3_neg_3_046_10    m/z 861.4257     0   1E+4   13   13.5  RT (min)      lcorbi_t3_neg_2_053_23    m/z 861.4938     0   2   4   6E+3   14.5   15  RT (min)      lcorbi_t3_neg_4_011_24    m/z 861.6323     0   2   4   6E+4   14.5   15  RT (min)      lcorbi_t3_neg_4_007_13    m/z 861.6622     0   1   2E+3   14.5   15  RT (min)      lcorbi_t3_neg_4_044_38    m/z 861.6880     0   1   2   3E+4   14.5   15  RT (min)      lcorbi_t3_neg_4_034_01    m/z 862.1994     0   1   2   3E+3   10   10.5  RT (min)      lcorbi_t3_neg_4_030_04    m/z 862.2861     0   1   2E+3   11   11.5  RT (min)      lcorbi_t3_neg_3_029_02    m/z 862.2957     0   2   4   6E+3   11.5   12  RT (min)      lcorbi_t3_neg_3_018_13    m/z 862.3672     0   1E+4   11   11.5  RT (min)      lcorbi_t3_neg_4_011_24    m/z 862.3954     0   2   4   6   8E+3   15   15.5  RT (min)      lcorbi_t3_neg_4_008_22    m/z 862.4240     0   2   4   6   8E+3   13.5   14  RT (min)      lcorbi_t3_neg_2_053_23    m/z 862.4486     0   1E+4   10.5   11  RT (min)      lcorbi_t3_neg_4_029_31    m/z 862.5061     0   2   4E+3   13   13.5  RT (min)      lcorbi_t3_neg_1_044_23    m/z 862.5624     0   1E+3   13   13.5  RT (min)      lcorbi_t3_neg_4_021_25    m/z 862.5810     0   2   4   6   8E+4   13.5   14  RT (min)      lcorbi_t3_neg_3_005_23    m/z 862.6074     0   1   2   3E+3   14   14.5  RT (min)      lcorbi_t3_neg_4_005_27    m/z 862.6353     0   2   4   6E+3   14.5   15  RT (min)      lcorbi_t3_neg_4_039_32    m/z 862.6647     0   1   2E+3   14.5   15  RT (min)      lcorbi_t3_neg_2_052_32    m/z 862.6841     0   2   4   6E+4   13.5   14  RT (min)      lcorbi_t3_neg_2_052_32    m/z 862.6884     0   1   2   3E+3   14.5   15  RT (min)      lcorbi_t3_neg_4_038_15    m/z 862.7228     0   1   14.5   15  RT (min)      lcorbi_t3_neg_4_034_01    m/z 862.7738     0   1E+3   18   18.5  RT (min)      lcorbi_t3_neg_4_019_08    m/z 862.8163     0   1E+3   18   18.5  RT (min)      lcorbi_t3_neg_4_019_08    m/z 863.1996     0   1   2   3   4E+3   12.5   13  RT (min)      lcorbi_t3_neg_2_008_16    m/z 863.2771     0   2   4   6   8E+3   12   12.5  RT (min)      lcorbi_t3_neg_4_023_18    m/z 863.3988     0   1   2   3E+3   15   15.5  RT (min)      lcorbi_t3_neg_2_038_22    m/z 863.4838     0   2   4E+4   7.5   8  RT (min)      lcorbi_t3_neg_2_048_27    m/z 863.4910     0   2   4   6E+3   14.5   15  RT (min)      lcorbi_t3_neg_4_011_24    m/z 863.5097     0   1   2   3   4E+3   14.5   15  RT (min)      lcorbi_t3_neg_1_055_05    m/z 863.5097     0   1   13   13.5  RT (min)      lcorbi_t3_neg_2_022_24    m/z 863.5844     0   1   2   3   4E+4   13.5   14  RT (min)      lcorbi_t3_neg_3_005_23    m/z 863.7059     0   1   2E+4   14   14.5  RT (min)      lcorbi_t3_neg_1_052_24    m/z 863.8200     0   1   2   3   4E+3   18   18.5  RT (min)      lcorbi_t3_neg_4_015_10    m/z 864.2806     0   1   2   3E+3   12   12.5  RT (min)      lcorbi_t3_neg_3_054_18    m/z 864.3112     0   1   2   3   4E+3   11.5   12  RT (min)      lcorbi_t3_neg_4_034_01    m/z 864.3745     0   1   2   3   4E+3   14.5   15  RT (min)      lcorbi_t3_neg_2_050_17    m/z 864.4916     0   2   4E+3   13   13.5  RT (min)      lcorbi_t3_neg_2_019_07    m/z 864.5356     0   1   2E+4   14   14.5  RT (min)      lcorbi_t3_neg_3_011_04    m/z 864.5391     0   1   2E+4   13.5   14  RT (min)      lcorbi_t3_neg_4_044_38    m/z 864.5825     0   1   2E+4   12   12.5  RT (min)      lcorbi_t3_neg_4_011_24    m/z 864.6604     0   1   2E+4   13.5   14  RT (min)      lcorbi_t3_neg_1_053_32    m/z 864.7705     0   2   4E+3   18   18.5  RT (min)      lcorbi_t3_neg_4_019_08    m/z 864.7867     0   2   4   6   8E+3   18   18.5  RT (min)      lcorbi_t3_neg_4_053_43    m/z 865.3489     0   1   2   3E+4   10.5   11  RT (min)      lcorbi_t3_neg_2_048_27    m/z 865.4628     0   1   2E+3   13.5   14  RT (min)      lcorbi_t3_neg_4_029_31    m/z 865.4717     0   2   4   6E+3   10   10.5  RT (min)      lcorbi_t3_neg_4_023_18    m/z 865.5621     0   1   13.5   14  RT (min)      lcorbi_t3_neg_4_011_24    m/z 865.6974     0   2   4   6   8E+3   13.5   14  RT (min)      lcorbi_t3_neg_4_055_06    m/z 865.7235     0   1E+4   13.5   14  RT (min)      lcorbi_t3_neg_4_035_33    m/z 865.7432     0   1E+4   15   15.5  RT (min)      lcorbi_t3_neg_4_039_32    m/z 865.8255     0   2   4   6   8E+3   18   18.5  RT (min)      lcorbi_t3_neg_4_048_03    m/z 866.2111     0   1   2E+3   10   10.5  RT (min)      lcorbi_t3_neg_4_023_18    m/z 866.2882     0   2   4E+3   11.5   12  RT (min)      lcorbi_t3_neg_2_007_14    m/z 866.4718     0   1E+4   13.5   14  RT (min)      lcorbi_t3_neg_2_012_36    m/z 866.5071     0   1   2E+4   13.5   14  RT (min)      lcorbi_t3_neg_3_020_31    m/z 866.5550     0   1E+5   13   13.5  RT (min)      lcorbi_t3_neg_4_042_14    m/z 866.6183     0   1   2   3   4E+5   14   14.5  RT (min)      lcorbi_t3_neg_4_005_27    m/z 866.6946     0   1   2E+6   14   14.5  RT (min)      lcorbi_t3_neg_3_002_24    m/z 866.7062     0   1E+4   13.5   14  RT (min)      lcorbi_t3_neg_4_019_08    m/z 866.7866     0   2   4   6E+3   18   18.5  RT (min)      lcorbi_t3_neg_4_050_02    m/z 866.7923     0   2   4   6   8E+3   18   18.5  RT (min)      lcorbi_t3_neg_4_028_39    m/z 866.8028     0   2   4   6   8E+3   18   18.5  RT (min)      lcorbi_t3_neg_1_055_05    m/z 867.1864     0   1   2   3E+4   13.5   14  RT (min)      lcorbi_t3_neg_4_045_35    m/z 867.3409     0   2   4   6   8E+3   14.5   15  RT (min)      lcorbi_t3_neg_1_050_45    m/z 867.3586     0   1E+4   12   12.5  RT (min)      lcorbi_t3_neg_2_007_14    m/z 867.4652     0   1   2   3   4E+3   13.5   14  RT (min)      lcorbi_t3_neg_4_007_13    m/z 867.4754     0   1   2   3   4E+3   13.5   14  RT (min)      lcorbi_t3_neg_1_044_23    m/z 867.5106     0   2   4   6E+4   13.5   14  RT (min)      lcorbi_t3_neg_2_053_23    m/z 867.5299     0   2   4   6E+3   14   14.5  RT (min)      lcorbi_t3_neg_3_016_42    m/z 867.5391     0   1   13.5   14  RT (min)      lcorbi_t3_neg_3_045_34    m/z 867.5584     0   2   4   6E+4   13   13.5  RT (min)      lcorbi_t3_neg_4_042_14    m/z 867.6585     0   2   4   6E+2   14.5   15  RT (min)      lcorbi_t3_neg_4_019_08    m/z 867.8038     0   2   4   6E+3   15   15.5  RT (min)      lcorbi_t3_neg_4_001_49    m/z 867.8086     0   1   2   3E+3   18   18.5  RT (min)      lcorbi_t3_neg_4_053_43    m/z 868.2228     0   1   2   3E+3   10.5   11  RT (min)      lcorbi_t3_neg_2_027_13    m/z 868.3456     0   2   4   6E+3   9   9.5  RT (min)      lcorbi_t3_neg_4_017_51    m/z 868.4355     0   1E+4   13   13.5  RT (min)      lcorbi_t3_neg_2_039_31    m/z 868.5716     0   2   4   6   8E+4   13.5   14  RT (min)      lcorbi_t3_neg_2_007_14    m/z 868.7698     0   1   2   3   4E+4   13.5   14  RT (min)      lcorbi_t3_neg_1_053_32    m/z 868.7899     0   1E+3   18   18.5  RT (min)      lcorbi_t3_neg_4_019_08    m/z 868.8341     0   2   4   6E+3   18   18.5  RT (min)      lcorbi_t3_neg_2_042_44    m/z 869.1621     0   1   2E+3   9   9.5  RT (min)      lcorbi_t3_neg_3_019_12    m/z 869.1848     0   1   2E+4   13.5   14  RT (min)      lcorbi_t3_neg_4_045_35    m/z 869.3386     0   1   2   3E+3   18   18.5  RT (min)      lcorbi_t3_neg_2_018_40    m/z 869.4388     0   2   4   6E+3   13   13.5  RT (min)      lcorbi_t3_neg_4_029_31    m/z 869.4434     0   1   2E+4   11   11.5  RT (min)      lcorbi_t3_neg_4_023_18    m/z 869.5248     0   1   2E+4   13.5   14  RT (min)      lcorbi_t3_neg_4_007_13    m/z 869.5550     0   1   2E+4   13.5   14  RT (min)      lcorbi_t3_neg_4_007_13    m/z 869.5675     0   1E+4   14   14.5  RT (min)      lcorbi_t3_neg_4_005_27    m/z 869.5740     0   2   4E+4   13   13.5  RT (min)      lcorbi_t3_neg_4_042_14    m/z 869.5996     0   1   2E+3   14   14.5  RT (min)      lcorbi_t3_neg_4_005_27    m/z 869.6545     0   2   4E+4   14   14.5  RT (min)      lcorbi_t3_neg_4_039_32    m/z 870.2386     0   2   4   6   8E+3   11.5   12  RT (min)      lcorbi_t3_neg_2_026_03    m/z 870.3951     0   1   2E+4   11   11.5  RT (min)      lcorbi_t3_neg_4_036_16    m/z 870.4468     0   1E+4   11   11.5  RT (min)      lcorbi_t3_neg_4_023_18    m/z 870.5385     0   1E+5   13.5   14  RT (min)      lcorbi_t3_neg_3_018_13    m/z 870.5585     0   2   4   6   8E+3   13.5   14  RT (min)      lcorbi_t3_neg_1_043_14    m/z 870.6155     0   1E+4   13   13.5  RT (min)      lcorbi_t3_neg_4_039_32    m/z 870.6430     0   1E+4   13   13.5  RT (min)      lcorbi_t3_neg_1_052_24    m/z 870.7853     0   1   2E+4   14   14.5  RT (min)      lcorbi_t3_neg_3_030_33    m/z 871.1838     0   2   4   6   8E+4   13.5   14  RT (min)      lcorbi_t3_neg_4_045_35    m/z 871.2612     0   2   4E+3   9.5   10  RT (min)      lcorbi_t3_neg_3_019_12    m/z 871.4420     0   2   4   6E+3   11   11.5  RT (min)      lcorbi_t3_neg_4_023_18    m/z 871.4830     0   1E+3   13.5   14  RT (min)      lcorbi_t3_neg_4_046_37    m/z 871.4869     0   1   2   3E+4   7.5   8  RT (min)      lcorbi_t3_neg_2_048_27    m/z 871.5816     0   1E+4   14   14.5  RT (min)      lcorbi_t3_neg_3_037_36    m/z 871.6160     0   1   2E+4   14.5   15  RT (min)      lcorbi_t3_neg_4_005_27    m/z 871.6250     0   1   2E+4   14.5   15  RT (min)      lcorbi_t3_neg_4_005_27    m/z 871.6304     0   1   2E+4   15   15.5  RT (min)      lcorbi_t3_neg_4_034_01    m/z 871.6526     0   1E+4   15   15.5  RT (min)      lcorbi_t3_neg_1_044_23    m/z 871.7001     0   2   4   6E+5   13.5   14  RT (min)      lcorbi_t3_neg_4_019_08    m/z 871.8073     0   1   2   3   4E+3   18   18.5  RT (min)      lcorbi_t3_neg_4_005_27    m/z 872.1845     0   2   4   6E+4   13.5   14  RT (min)      lcorbi_t3_neg_4_045_35    m/z 872.2283     0   2   4   6E+3   10   10.5  RT (min)      lcorbi_t3_neg_4_030_04    m/z 872.3404     0   1   2E+4   8.5   9  RT (min)      lcorbi_t3_neg_4_023_18    m/z 872.4573     0   2   4   6E+3   13   13.5  RT (min)      lcorbi_t3_neg_1_020_36    m/z 872.4904     0   1   2E+4   7.5   8  RT (min)      lcorbi_t3_neg_2_048_27    m/z 872.4995     0   1   2E+3   13.5   14  RT (min)      lcorbi_t3_neg_2_053_23    m/z 872.5286     0   1E+3   13.5   14  RT (min)      lcorbi_t3_neg_3_023_07    m/z 872.5380     0   1   2E+4   13.5   14  RT (min)      lcorbi_t3_neg_4_007_13    m/z 872.5711     0   1E+3   13.5   14  RT (min)      lcorbi_t3_neg_4_039_32    m/z 872.6196     0   2   4   6E+3   14.5   15  RT (min)      lcorbi_t3_neg_4_005_27    m/z 872.6378     0   1E+4   14.5   15  RT (min)      lcorbi_t3_neg_3_002_24    m/z 872.6503     0   1E+4   14.5   15  RT (min)      lcorbi_t3_neg_4_016_05    m/z 872.6676     0   1E+4   13.5   14  RT (min)      lcorbi_t3_neg_3_002_24    m/z 872.6832     0   1   2   3   4E+4   14.5   15  RT (min)      lcorbi_t3_neg_4_011_24    m/z 872.7232     0   2   4E+4   18   18.5  RT (min)      lcorbi_t3_neg_1_052_24    m/z 872.7693     0   2   4   6E+4   14   14.5  RT (min)      lcorbi_t3_neg_4_011_24    m/z 873.1825     0   2   4E+4   13.5   14  RT (min)      lcorbi_t3_neg_4_045_35    m/z 873.2286     0   2   4   6E+3   12.5   13  RT (min)      lcorbi_t3_neg_3_007_16    m/z 873.2318     0   1   2E+3   10   10.5  RT (min)      lcorbi_t3_neg_4_027_12    m/z 873.3701     0   1   2E+4   8.5   9  RT (min)      lcorbi_t3_neg_4_023_18    m/z 873.4017     0   1   2E+4   10.5   11  RT (min)      lcorbi_t3_neg_4_042_14    m/z 873.4744     0   1E+4   13   13.5  RT (min)      lcorbi_t3_neg_4_054_36    m/z 873.4946     0   2   4   6   8E+3   13.5   14  RT (min)      lcorbi_t3_neg_2_027_13    m/z 873.5115     0   1   2E+4   13.5   14  RT (min)      lcorbi_t3_neg_2_010_02    m/z 873.5325     0   1   2   3E+3   13.5   14  RT (min)      lcorbi_t3_neg_1_035_25    m/z 873.5471     0   1   2E+4   13.5   14  RT (min)      lcorbi_t3_neg_2_010_02    m/z 873.5557     0   1   2   3E+4   14   14.5  RT (min)      lcorbi_t3_neg_2_048_27    m/z 873.5809     0   1   10   10.5  RT (min)      lcorbi_t3_neg_2_022_24    m/z 873.5817     0   1   2   3   4E+3   13.5   14  RT (min)      lcorbi_t3_neg_4_035_33    m/z 873.5971     0   1   2E+4   14.5   15  RT (min)      lcorbi_t3_neg_2_005_01    m/z 873.6418     0   2   4   6E+3   14.5   15  RT (min)      lcorbi_t3_neg_2_036_35    m/z 873.6782     0   1E+4   14.5   15  RT (min)      lcorbi_t3_neg_4_007_13    m/z 873.6971     0   1   2   3E+4   14.5   15  RT (min)      lcorbi_t3_neg_4_034_01    m/z 873.7062     0   2   4   6E+2   14.5   15  RT (min)      lcorbi_t3_neg_4_019_08    m/z 873.8076     0   1   2E+3   18   18.5  RT (min)      lcorbi_t3_neg_4_003_21    m/z 874.2503     0   2   4E+3   11.5   12  RT (min)      lcorbi_t3_neg_4_048_03    m/z 874.2553     0   1   2E+4   6   6.5  RT (min)      lcorbi_t3_neg_1_054_37    m/z 874.3735     0   1E+4   8.5   9  RT (min)      lcorbi_t3_neg_4_023_18    m/z 874.4660     0   2   4   6E+3   13   13.5  RT (min)      lcorbi_t3_neg_2_053_23    m/z 874.4822     0   1   2E+4   14   14.5  RT (min)      lcorbi_t3_neg_2_013_08    m/z 874.4974     0   2   4   6E+3   13.5   14  RT (min)      lcorbi_t3_neg_1_044_23    m/z 874.5152     0   1E+4   13.5   14  RT (min)      lcorbi_t3_neg_3_029_02    m/z 874.5444     0   1E+4   13.5   14  RT (min)      lcorbi_t3_neg_4_032_23    m/z 874.5812     0   1   2   3E+4   13.5   14  RT (min)      lcorbi_t3_neg_4_044_38    m/z 874.5937     0   2   4   6   8E+3   14.5   15  RT (min)      lcorbi_t3_neg_2_008_16    m/z 874.6028     0   2   4E+3   14   14.5  RT (min)      lcorbi_t3_neg_3_044_01    m/z 874.6098     0   1E+4   14   14.5  RT (min)      lcorbi_t3_neg_4_042_14    m/z 874.6816     0   2   4E+3   14.5   15  RT (min)      lcorbi_t3_neg_4_038_15    m/z 874.6900     0   2   4E+2   15   15.5  RT (min)      lcorbi_t3_neg_3_002_24    m/z 874.7003     0   1E+4   14.5   15  RT (min)      lcorbi_t3_neg_3_002_24    m/z 874.7866     0   1   2   3E+2   15.5   16  RT (min)      lcorbi_t3_neg_2_022_24    m/z 874.7991     0   1   2   3   4E+3   18   18.5  RT (min)      lcorbi_t3_neg_3_038_39    m/z 875.2200     0   1E+4   5   5.5  RT (min)      lcorbi_t3_neg_2_048_27    m/z 875.2733     0   1   2   3   4E+3   9.5   10  RT (min)      lcorbi_t3_neg_3_054_18    m/z 875.2986     0   2   4E+3   13   13.5  RT (min)      lcorbi_t3_neg_1_023_03    m/z 875.4895     0   1E+4   13.5   14  RT (min)      lcorbi_t3_neg_4_007_13    m/z 875.5103     0   1   2   3   4E+3   13.5   14  RT (min)      lcorbi_t3_neg_4_007_13    m/z 875.5534     0   2   4   6   8E+3   13.5   14  RT (min)      lcorbi_t3_neg_4_032_23    m/z 875.5854     0   2   4   6E+3   13.5   14  RT (min)      lcorbi_t3_neg_4_011_24    m/z 875.6090     0   1   13.5   14  RT (min)      lcorbi_t3_neg_4_019_08    m/z 875.7128     0   1   2E+3   14.5   15  RT (min)      lcorbi_t3_neg_2_052_32    m/z 876.2532     0   1E+4   6   6.5  RT (min)      lcorbi_t3_neg_1_054_37    m/z 876.2955     0   1E+4   13   13.5  RT (min)      lcorbi_t3_neg_1_023_03    m/z 876.3138     0   1E+4   14   14.5  RT (min)      lcorbi_t3_neg_1_023_03    m/z 876.5146     0   2   4   6E+3   13   13.5  RT (min)      lcorbi_t3_neg_2_007_14    m/z 876.5213     0   1   2   3   4E+3   13   13.5  RT (min)      lcorbi_t3_neg_2_031_25    m/z 876.5982     0   2   4E+4   13   13.5  RT (min)      lcorbi_t3_neg_3_002_24    m/z 876.6625     0   1E+4   18   18.5  RT (min)      lcorbi_t3_neg_3_052_32    m/z 876.6700     0   1E+4   14.5   15  RT (min)      lcorbi_t3_neg_2_052_32    m/z 876.6805     0   1E+4   14.5   15  RT (min)      lcorbi_t3_neg_4_034_01    m/z 876.6891     0   2   4   6E+2   14.5   15  RT (min)      lcorbi_t3_neg_3_002_24    m/z 877.1876     0   1   2   3   4E+3   12.5   13  RT (min)      lcorbi_t3_neg_4_005_27    m/z 877.4565     0   2   4   6E+3   11   11.5  RT (min)      lcorbi_t3_neg_4_023_18    m/z 877.5444     0   1E+4   13   13.5  RT (min)      lcorbi_t3_neg_4_042_14    m/z 877.5659     0   1   2E+4   14   14.5  RT (min)      lcorbi_t3_neg_4_029_31    m/z 877.6147     0   1E+4   13   13.5  RT (min)      lcorbi_t3_neg_1_055_05    m/z 877.6898     0   1   2E+3   14.5   15  RT (min)      lcorbi_t3_neg_2_052_32    m/z 877.7937     0   1   2E+3   18   18.5  RT (min)      lcorbi_t3_neg_2_021_41    m/z 877.8248     0   2   4   6E+3   18   18.5  RT (min)      lcorbi_t3_neg_2_042_44    m/z 877.8374     0   2   4   6E+3   18   18.5  RT (min)      lcorbi_t3_neg_4_018_42    m/z 878.2518     0   2   4   6E+3   10.5   11  RT (min)      lcorbi_t3_neg_3_018_13    m/z 878.3205     0   1   2   3   4E+3   9   9.5  RT (min)      lcorbi_t3_neg_4_017_51    m/z 878.5298     0   1E+4   13.5   14  RT (min)      lcorbi_t3_neg_3_005_23    m/z 878.5479     0   1   2   3E+3   13   13.5  RT (min)      lcorbi_t3_neg_2_053_23    m/z 878.5540     0   2   4   6   8E+2   13.5   14  RT (min)      lcorbi_t3_neg_4_007_13    m/z 878.5758     0   2   4   6E+3   13.5   14  RT (min)      lcorbi_t3_neg_4_011_24    m/z 878.6190     0   1   2   3   4E+4   13   13.5  RT (min)      lcorbi_t3_neg_4_011_24    m/z 878.6315     0   1   15   15.5  RT (min)      lcorbi_t3_neg_4_011_24    m/z 878.7592     0   1   2E+4   14.5   15  RT (min)      lcorbi_t3_neg_2_022_24    m/z 878.8238     0   2   4   6E+3   18   18.5  RT (min)      lcorbi_t3_neg_1_006_40    m/z 879.2551     0   1   2E+3   10.5   11  RT (min)      lcorbi_t3_neg_4_034_01    m/z 879.5170     0   1   2E+4   14   14.5  RT (min)      lcorbi_t3_neg_4_032_23    m/z 879.5412     0   1E+4   13.5   14  RT (min)      lcorbi_t3_neg_1_044_23    m/z 879.5798     0   1E+4   13.5   14  RT (min)      lcorbi_t3_neg_4_035_33    m/z 879.5883     0   2   4   6   8E+3   13.5   14  RT (min)      lcorbi_t3_neg_4_007_13    m/z 879.6288     0   2   4   6E+3   13.5   14  RT (min)      lcorbi_t3_neg_2_052_32    m/z 879.6716     0   2   4   6E+3   13.5   14  RT (min)      lcorbi_t3_neg_1_055_05    m/z 880.3692     0   2   4E+3   14.5   15  RT (min)      lcorbi_t3_neg_2_050_17    m/z 880.4067     0   1E+4   13   13.5  RT (min)      lcorbi_t3_neg_4_044_38    m/z 880.5811     0   1E+4   13   13.5  RT (min)      lcorbi_t3_neg_2_052_32    m/z 880.6435     0   1E+3   14.5   15  RT (min)      lcorbi_t3_neg_1_030_47    m/z 880.8344     0   1   2   3E+3   18   18.5  RT (min)      lcorbi_t3_neg_3_008_20    m/z 881.2124     0   1   2E+5   13.5   14  RT (min)      lcorbi_t3_neg_4_045_35    m/z 881.2711     0   2   4   6   8E+3   11.5   12  RT (min)      lcorbi_t3_neg_2_026_03    m/z 881.2900     0   1E+4   9.5   10  RT (min)      lcorbi_t3_neg_3_019_12    m/z 881.4243     0   1E+4   10.5   11  RT (min)      lcorbi_t3_neg_3_010_15    m/z 881.5743     0   1E+4   13.5   14  RT (min)      lcorbi_t3_neg_4_038_15    m/z 881.5936     0   2   4E+3   13.5   14  RT (min)      lcorbi_t3_neg_2_022_24    m/z 881.6467     0   1   2   3E+3   14.5   15  RT (min)      lcorbi_t3_neg_4_038_15    m/z 881.6592     0   2   4E+4   13   13.5  RT (min)      lcorbi_t3_neg_4_039_32    m/z 881.6925     0   1   2E+4   13.5   14  RT (min)      lcorbi_t3_neg_4_039_32    m/z 881.7022     0   1E+5   14   14.5  RT (min)      lcorbi_t3_neg_4_016_05    m/z 882.2719     0   1   2   3   4E+3   11   11.5  RT (min)      lcorbi_t3_neg_3_013_28    m/z 882.2935     0   2   4   6E+3   9.5   10  RT (min)      lcorbi_t3_neg_3_019_12    m/z 882.4221     0   1E+4   13.5   14  RT (min)      lcorbi_t3_neg_4_054_36    m/z 882.4813     0   1E+3   11   11.5  RT (min)      lcorbi_t3_neg_1_044_23    m/z 882.5323     0   2   4   6E+3   11   11.5  RT (min)      lcorbi_t3_neg_1_044_23    m/z 882.5776     0   1   2   3E+3   13   13.5  RT (min)      lcorbi_t3_neg_4_032_23    m/z 882.5860     0   1   2E+4   13.5   14  RT (min)      lcorbi_t3_neg_4_038_15    m/z 882.5973     0   1E+4   13.5   14  RT (min)      lcorbi_t3_neg_1_052_24    m/z 882.6592     0   2   4   6   8E+3   14.5   15  RT (min)      lcorbi_t3_neg_4_007_13    m/z 882.7014     0   1   2E+4   13.5   14  RT (min)      lcorbi_t3_neg_2_052_32    m/z 883.5063     0   1   2   3E+3   14   14.5  RT (min)      lcorbi_t3_neg_3_032_17    m/z 883.5892     0   1E+4   13.5   14  RT (min)      lcorbi_t3_neg_4_038_15    m/z 883.6018     0   1   14   14.5  RT (min)      lcorbi_t3_neg_4_005_27    m/z 883.6541     0   1   2   3E+4   13.5   14  RT (min)      lcorbi_t3_neg_4_011_24    m/z 883.6652     0   1E+4   13.5   14  RT (min)      lcorbi_t3_neg_4_039_32    m/z 883.6791     0   1E+3   14.5   15  RT (min)      lcorbi_t3_neg_2_031_25    m/z 883.8107     0   2   4   6E+3   18   18.5  RT (min)      lcorbi_t3_neg_3_042_22    m/z 883.8436     0   1   2   3E+3   18   18.5  RT (min)      lcorbi_t3_neg_3_046_10    m/z 884.1065     0   1E+4   4.5   5  RT (min)      lcorbi_t3_neg_2_048_27    m/z 884.2868     0   1   2   3   4E+3   11.5   12  RT (min)      lcorbi_t3_neg_3_004_30    m/z 884.4872     0   1   2   3E+3   13.5   14  RT (min)      lcorbi_t3_neg_4_005_27    m/z 884.5031     0   1   2   3E+3   13.5   14  RT (min)      lcorbi_t3_neg_3_053_14    m/z 884.5698     0   1E+4   13   13.5  RT (min)      lcorbi_t3_neg_4_039_32    m/z 884.5791     0   1E+4   13.5   14  RT (min)      lcorbi_t3_neg_3_002_24    m/z 884.6133     0   1   2E+4   10   10.5  RT (min)      lcorbi_t3_neg_3_019_12    m/z 884.6662     0   1   2E+4   14   14.5  RT (min)      lcorbi_t3_neg_4_011_24    m/z 884.7225     0   1   2   3E+4   14   14.5  RT (min)      lcorbi_t3_neg_3_002_24    m/z 884.7591     0   1E+4   14.5   15  RT (min)      lcorbi_t3_neg_1_053_32    m/z 884.8125     0   2   4   6E+3   18   18.5  RT (min)      lcorbi_t3_neg_3_014_19    m/z 884.8454     0   2   4   6E+3   18   18.5  RT (min)      lcorbi_t3_neg_1_003_41    m/z 885.5495     0   2   4   6E+3   18   18.5  RT (min)      lcorbi_t3_neg_4_034_01    m/z 885.5664     0   2   4   6   8E+3   14   14.5  RT (min)      lcorbi_t3_neg_4_044_38    m/z 885.6028     0   2   4   6E+3   14.5   15  RT (min)      lcorbi_t3_neg_4_007_13    m/z 885.6345     0   1E+4   13   13.5  RT (min)      lcorbi_t3_neg_4_039_32    m/z 885.6970     0   1E+4   14.5   15  RT (min)      lcorbi_t3_neg_4_034_01    m/z 885.8118     0   2   4E+3   18   18.5  RT (min)      lcorbi_t3_neg_2_029_12    m/z 886.2194     0   1   2E+3   11.5   12  RT (min)      lcorbi_t3_neg_4_016_05    m/z 886.2725     0   2   4   6   8E+3   10   10.5  RT (min)      lcorbi_t3_neg_3_037_36    m/z 886.3026     0   2   4E+3   12   12.5  RT (min)      lcorbi_t3_neg_1_054_37    m/z 886.6048     0   1E+4   13.5   14  RT (min)      lcorbi_t3_neg_4_007_13    m/z 886.6146     0   2   4   6   8E+3   14.5   15  RT (min)      lcorbi_t3_neg_3_010_15    m/z 886.6292     0   2   4E+2   14.5   15  RT (min)      lcorbi_t3_neg_2_019_07    m/z 886.6354     0   1E+4   14.5   15  RT (min)      lcorbi_t3_neg_3_032_17    m/z 886.6903     0   1   2E+3   14.5   15  RT (min)      lcorbi_t3_neg_4_007_13    m/z 886.7094     0   1E+4   14.5   15  RT (min)      lcorbi_t3_neg_4_007_13    m/z 886.7389     0   1   2E+4   14   14.5  RT (min)      lcorbi_t3_neg_3_002_24    m/z 886.8175     0   1   2E+3   18   18.5  RT (min)      lcorbi_t3_neg_1_003_41    m/z 887.2758     0   2   4E+3   10   10.5  RT (min)      lcorbi_t3_neg_3_034_21    m/z 887.4900     0   2   4E+3   13.5   14  RT (min)      lcorbi_t3_neg_2_039_31    m/z 887.5224     0   1E+4   14.5   15  RT (min)      lcorbi_t3_neg_1_034_04    m/z 887.5891     0   2   4   6E+3   13.5   14  RT (min)      lcorbi_t3_neg_4_032_23    m/z 887.6022     0   1   2   3   4E+3   14.5   15  RT (min)      lcorbi_t3_neg_2_019_07    m/z 887.6187     0   2   4   6E+3   14.5   15  RT (min)      lcorbi_t3_neg_4_034_01    m/z 887.6310     0   1   2   3E+3   14.5   15  RT (min)      lcorbi_t3_neg_3_007_16    m/z 887.7206     0   1   14.5   15  RT (min)      lcorbi_t3_neg_4_019_08    m/z 888.3112     0   1   2   3E+3   11.5   12  RT (min)      lcorbi_t3_neg_3_018_13    m/z 888.4088     0   2   4   6E+3   15   15.5  RT (min)      lcorbi_t3_neg_1_039_38    m/z 888.5064     0   2   4   6E+3   13.5   14  RT (min)      lcorbi_t3_neg_4_007_13    m/z 888.5231     0   2   4   6E+3   13.5   14  RT (min)      lcorbi_t3_neg_4_046_37    m/z 888.5360     0   1   2E+4   10.5   11  RT (min)      lcorbi_t3_neg_4_039_32    m/z 888.5376     0   1E+4   14   14.5  RT (min)      lcorbi_t3_neg_3_011_04    m/z 888.5493     0   1   13.5   14  RT (min)      lcorbi_t3_neg_4_019_08    m/z 888.6011     0   2   4   6   8E+3   18   18.5  RT (min)      lcorbi_t3_neg_4_039_32    m/z 888.6127     0   2   4   6E+3   14   14.5  RT (min)      lcorbi_t3_neg_2_048_27    m/z 888.6625     0   2   4   6E+2   14.5   15  RT (min)      lcorbi_t3_neg_3_002_24    m/z 888.6690     0   2   4   6   8E+3   14.5   15  RT (min)      lcorbi_t3_neg_4_044_38    m/z 888.7118     0   2   4E+3   14   14.5  RT (min)      lcorbi_t3_neg_3_002_24    m/z 888.7195     0   2   4   6E+2   14.5   15  RT (min)      lcorbi_t3_neg_4_029_31    m/z 888.7548     0   1E+4   14.5   15  RT (min)      lcorbi_t3_neg_2_022_24    m/z 888.7902     0   1E+4   14.5   15  RT (min)      lcorbi_t3_neg_2_022_24    m/z 888.8235     0   2   4   6E+3   18   18.5  RT (min)      lcorbi_t3_neg_3_026_44    m/z 889.4264     0   2   4   6E+3   13   13.5  RT (min)      lcorbi_t3_neg_3_020_31    m/z 889.4612     0   2   4   6   8E+3   13.5   14  RT (min)      lcorbi_t3_neg_2_053_23    m/z 889.5168     0   1   2E+4   7.5   8  RT (min)      lcorbi_t3_neg_2_048_27    m/z 889.5624     0   2   4E+4   13.5   14  RT (min)      lcorbi_t3_neg_4_034_01    m/z 889.6002     0   1   2   3   4E+3   13.5   14  RT (min)      lcorbi_t3_neg_4_019_08    m/z 889.6178     0   1   2   3E+4   14.5   15  RT (min)      lcorbi_t3_neg_3_048_08    m/z 889.6724     0   1   2   3   4E+3   14.5   15  RT (min)      lcorbi_t3_neg_4_007_13    m/z 889.7081     0   1   2   3E+4   14   14.5  RT (min)      lcorbi_t3_neg_4_037_29    m/z 890.2851     0   1   2   3E+4   7.5   8  RT (min)      lcorbi_t3_neg_2_048_27    m/z 890.3284     0   1   2   3E+3   18   18.5  RT (min)      lcorbi_t3_neg_2_046_20    m/z 890.4064     0   2   4E+3   15   15.5  RT (min)      lcorbi_t3_neg_1_039_38    m/z 890.4270     0   2   4E+3   15   15.5  RT (min)      lcorbi_t3_neg_2_038_22    m/z 890.4701     0   1   2E+3   13.5   14  RT (min)      lcorbi_t3_neg_3_047_37    m/z 890.5742     0   1   14   14.5  RT (min)      lcorbi_t3_neg_2_022_24    m/z 890.6006     0   1E+4   13.5   14  RT (min)      lcorbi_t3_neg_2_022_24    m/z 890.6103     0   1   2   3E+4   13   13.5  RT (min)      lcorbi_t3_neg_2_019_07    m/z 890.6420     0   1   2E+4   12.5   13  RT (min)      lcorbi_t3_neg_4_011_24    m/z 890.6975     0   1   2E+4   14   14.5  RT (min)      lcorbi_t3_neg_2_022_24    m/z 890.7100     0   1E+4   14   14.5  RT (min)      lcorbi_t3_neg_1_052_24    m/z 890.7968     0   1   2   3E+3   18   18.5  RT (min)      lcorbi_t3_neg_2_018_40    m/z 890.8263     0   1   2   3   4E+3   18   18.5  RT (min)      lcorbi_t3_neg_2_011_04    m/z 891.2886     0   1E+4   7.5   8  RT (min)      lcorbi_t3_neg_2_048_27    m/z 891.3641     0   2   4   6E+4   10.5   11  RT (min)      lcorbi_t3_neg_1_043_14    m/z 891.5601     0   1   13.5   14  RT (min)      lcorbi_t3_neg_1_055_05    m/z 891.5692     0   2   4E+3   14.5   15  RT (min)      lcorbi_t3_neg_4_034_01    m/z 891.7735     0   2   4   6   8E+3   14.5   15  RT (min)      lcorbi_t3_neg_4_039_32    m/z 891.8321     0   2   4   6   8E+3   18   18.5  RT (min)      lcorbi_t3_neg_4_027_12    m/z 892.3039     0   2   4E+3   12   12.5  RT (min)      lcorbi_t3_neg_3_053_14    m/z 892.4568     0   1E+4   13.5   14  RT (min)      lcorbi_t3_neg_2_053_23    m/z 892.4810     0   1   2   3E+3   13.5   14  RT (min)      lcorbi_t3_neg_3_037_36    m/z 892.4890     0   2   4E+3   13.5   14  RT (min)      lcorbi_t3_neg_2_028_38    m/z 892.5457     0   1   2   3   4E+3   13.5   14  RT (min)      lcorbi_t3_neg_4_021_25    m/z 892.5628     0   1   2E+3   14.5   15  RT (min)      lcorbi_t3_neg_4_007_13    m/z 892.5646     0   1   2E+3   14.5   15  RT (min)      lcorbi_t3_neg_4_007_13    m/z 892.6073     0   1E+4   14.5   15  RT (min)      lcorbi_t3_neg_3_022_27    m/z 892.6851     0   2   4E+4   13.5   14  RT (min)      lcorbi_t3_neg_3_002_24    m/z 892.7957     0   1   2   3   4E+3   18   18.5  RT (min)      lcorbi_t3_neg_3_016_42    m/z 893.3072     0   1   2   3E+4   9   9.5  RT (min)      lcorbi_t3_neg_2_048_27    m/z 893.4603     0   2   4   6   8E+3   13.5   14  RT (min)      lcorbi_t3_neg_3_005_23    m/z 893.4793     0   1   2   3E+3   13.5   14  RT (min)      lcorbi_t3_neg_2_012_36    m/z 893.6108     0   2   4   6E+3   14.5   15  RT (min)      lcorbi_t3_neg_4_044_38    m/z 893.6381     0   1E+4   13.5   14  RT (min)      lcorbi_t3_neg_1_052_24    m/z 893.6854     0   1E+4   13.5   14  RT (min)      lcorbi_t3_neg_2_022_24    m/z 894.2388     0   2   4   6E+3   11   11.5  RT (min)      lcorbi_t3_neg_2_010_02    m/z 894.2831     0   1   2   3   4E+3   11.5   12  RT (min)      lcorbi_t3_neg_4_042_14    m/z 894.4986     0   2   4   6   8E+2   13.5   14  RT (min)      lcorbi_t3_neg_2_053_23    m/z 894.5864     0   2   4   6E+4   13.5   14  RT (min)      lcorbi_t3_neg_2_007_14    m/z 894.5965     0   1E+3   13   13.5  RT (min)      lcorbi_t3_neg_4_042_14    m/z 894.6224     0   1   2E+4   14.5   15  RT (min)      lcorbi_t3_neg_4_005_27    m/z 894.6862     0   1   2   3   4E+4   13   13.5  RT (min)      lcorbi_t3_neg_2_022_24    m/z 894.8058     0   1   2   3   4E+3   18   18.5  RT (min)      lcorbi_t3_neg_3_024_26    m/z 895.3429     0   2   4E+3   18   18.5  RT (min)      lcorbi_t3_neg_4_014_20    m/z 895.3725     0   2   4   6   8E+3   15   15.5  RT (min)      lcorbi_t3_neg_1_050_45    m/z 895.4027     0   2   4   6E+3   8   8.5  RT (min)      lcorbi_t3_neg_4_023_18    m/z 895.4223     0   1E+4   10.5   11  RT (min)      lcorbi_t3_neg_4_042_14    m/z 895.5405     0   1   2   3E+4   13.5   14  RT (min)      lcorbi_t3_neg_2_012_36    m/z 895.5897     0   1   2   3E+4   13.5   14  RT (min)      lcorbi_t3_neg_4_042_14    m/z 895.6268     0   2   4   6E+3   14.5   15  RT (min)      lcorbi_t3_neg_2_007_14    m/z 895.6535     0   1   2E+4   14   14.5  RT (min)      lcorbi_t3_neg_1_052_24    m/z 896.2545     0   2   4   6   8E+3   11.5   12  RT (min)      lcorbi_t3_neg_2_027_13    m/z 896.3319     0   1   2E+4   9   9.5  RT (min)      lcorbi_t3_neg_2_048_27    m/z 896.4992     0   2   4   6   8E+3   13.5   14  RT (min)      lcorbi_t3_neg_1_043_14    m/z 896.6011     0   1   2   3   4E+4   13.5   14  RT (min)      lcorbi_t3_neg_4_042_14    m/z 896.6122     0   1   13.5   14  RT (min)      lcorbi_t3_neg_2_007_14    m/z 896.6311     0   2   4   6E+3   13   13.5  RT (min)      lcorbi_t3_neg_4_039_32    m/z 896.6383     0   2   4   6   8E+3   14.5   15  RT (min)      lcorbi_t3_neg_3_022_27    m/z 896.6675     0   1E+5   14.5   15  RT (min)      lcorbi_t3_neg_4_007_13    m/z 896.6748     0   1E+5   14.5   15  RT (min)      lcorbi_t3_neg_4_007_13    m/z 896.6983     0   1   2   3E+4   13.5   14  RT (min)      lcorbi_t3_neg_4_011_24    m/z 896.8111     0   2   4   6E+3   18   18.5  RT (min)      lcorbi_t3_neg_2_046_20    m/z 897.2580     0   1   2E+3   11.5   12  RT (min)      lcorbi_t3_neg_2_027_13    m/z 897.4989     0   1   13.5   14  RT (min)      lcorbi_t3_neg_4_019_08    m/z 897.5116     0   1   2   3E+3   13.5   14  RT (min)      lcorbi_t3_neg_4_005_27    m/z 897.5562     0   1E+4   14   14.5  RT (min)      lcorbi_t3_neg_1_055_05    m/z 897.6050     0   1   2E+4   13.5   14  RT (min)      lcorbi_t3_neg_4_042_14    m/z 897.6122     0   1   2   3E+4   11.5   12  RT (min)      lcorbi_t3_neg_4_007_13    m/z 897.6506     0   1   2E+4   13   13.5  RT (min)      lcorbi_t3_neg_4_039_32    m/z 897.7100     0   1E+4   13.5   14  RT (min)      lcorbi_t3_neg_4_019_08    m/z 897.8140     0   1   2   3   4E+3   18   18.5  RT (min)      lcorbi_t3_neg_3_011_04    m/z 898.2505     0   2   4E+3   11   11.5  RT (min)      lcorbi_t3_neg_2_027_13    m/z 898.2702     0   2   4   6   8E+3   12   12.5  RT (min)      lcorbi_t3_neg_2_005_01    m/z 898.2765     0   1   2E+4   9   9.5  RT (min)      lcorbi_t3_neg_4_040_19    m/z 898.3962     0   1   2   3E+4   9   9.5  RT (min)      lcorbi_t3_neg_2_048_27    m/z 898.4131     0   1E+4   13.5   14  RT (min)      lcorbi_t3_neg_1_044_23    m/z 898.5698     0   1   2   3   4E+5   14   14.5  RT (min)      lcorbi_t3_neg_2_048_27    m/z 898.6100     0   1E+4   13.5   14  RT (min)      lcorbi_t3_neg_2_007_14    m/z 898.6276     0   2   4   6E+3   13.5   14  RT (min)      lcorbi_t3_neg_1_052_24    m/z 898.7026     0   1E+4   13   13.5  RT (min)      lcorbi_t3_neg_3_002_24    m/z 898.7481     0   1   2   3   4E+3   18   18.5  RT (min)      lcorbi_t3_neg_2_038_22    m/z 898.8105     0   1   2   3   4E+3   18   18.5  RT (min)      lcorbi_t3_neg_2_019_07    m/z 898.9710     0   1E+4   11   11.5  RT (min)      lcorbi_t3_neg_2_041_54    m/z 899.1807     0   1   2E+3   9   9.5  RT (min)      lcorbi_t3_neg_3_019_12    m/z 899.2738     0   1   2E+3   12   12.5  RT (min)      lcorbi_t3_neg_2_014_29    m/z 899.4715     0   1E+4   7.5   8  RT (min)      lcorbi_t3_neg_4_017_51    m/z 899.5059     0   1   2   3E+3   13   13.5  RT (min)      lcorbi_t3_neg_1_044_23    m/z 899.5504     0   1   2   3E+4   13.5   14  RT (min)      lcorbi_t3_neg_4_044_38    m/z 899.6666     0   1   2   3E+4   13.5   14  RT (min)      lcorbi_t3_neg_4_039_32    m/z 899.6853     0   2   4E+4   14.5   15  RT (min)      lcorbi_t3_neg_2_022_24    m/z 899.8188     0   1   2   3   4E+3   18   18.5  RT (min)      lcorbi_t3_neg_2_029_12    m/z 900.2661     0   1   2   3   4E+3   11.5   12  RT (min)      lcorbi_t3_neg_2_027_13    m/z 900.4276     0   1   2E+4   13.5   14  RT (min)      lcorbi_t3_neg_1_044_23    m/z 900.4748     0   2   4   6E+3   7.5   8  RT (min)      lcorbi_t3_neg_4_017_51    m/z 900.4750     0   1E+4   13.5   14  RT (min)      lcorbi_t3_neg_2_027_13    m/z 900.5124     0   1   2   3   4E+3   13   13.5  RT (min)      lcorbi_t3_neg_2_051_34    m/z 900.5582     0   1E+4   13.5   14  RT (min)      lcorbi_t3_neg_4_038_15    m/z 900.5745     0   1   2   3E+4   14   14.5  RT (min)      lcorbi_t3_neg_4_034_01    m/z 900.5852     0   2   4   6   8E+4   14   14.5  RT (min)      lcorbi_t3_neg_2_048_27    m/z 900.6870     0   2   4E+3   14.5   15  RT (min)      lcorbi_t3_neg_2_054_15    m/z 900.6976     0   1E+4   15   15.5  RT (min)      lcorbi_t3_neg_4_007_13    m/z 900.7059     0   1E+4   15   15.5  RT (min)      lcorbi_t3_neg_4_034_01    m/z 900.7178     0   2   4   6E+3   13.5   14  RT (min)      lcorbi_t3_neg_4_039_32    m/z 900.7456     0   1   2   3E+3   18   18.5  RT (min)      lcorbi_t3_neg_2_033_53    m/z 901.4045     0   1E+4   13.5   14  RT (min)      lcorbi_t3_neg_1_044_23    m/z 901.4785     0   2   4E+3   13.5   14  RT (min)      lcorbi_t3_neg_2_027_13    m/z 901.4962     0   2   4   6   8E+3   13.5   14  RT (min)      lcorbi_t3_neg_4_042_14    m/z 901.5215     0   1E+4   13.5   14  RT (min)      lcorbi_t3_neg_4_032_23    m/z 901.5447     0   2   4   6E+4   13   13.5  RT (min)      lcorbi_t3_neg_2_007_14    m/z 901.5612     0   2   4   6   8E+3   13.5   14  RT (min)      lcorbi_t3_neg_4_038_15    m/z 901.5870     0   1   2E+4   14   14.5  RT (min)      lcorbi_t3_neg_2_048_27    m/z 901.6189     0   1   2   3E+3   14.5   15  RT (min)      lcorbi_t3_neg_4_034_01    m/z 901.6378     0   1E+3   14.5   15  RT (min)      lcorbi_t3_neg_4_048_03    m/z 901.7098     0   2   4   6E+3   15   15.5  RT (min)      lcorbi_t3_neg_4_034_01    m/z 901.7774     0   1   2E+3   18   18.5  RT (min)      lcorbi_t3_neg_3_055_06    m/z 901.8249     0   2   4   6E+3   18   18.5  RT (min)      lcorbi_t3_neg_1_005_07    m/z 902.2519     0   2   4   6E+3   10.5   11  RT (min)      lcorbi_t3_neg_4_032_23    m/z 902.2815     0   2   4E+3   12   12.5  RT (min)      lcorbi_t3_neg_1_024_01    m/z 902.3275     0   1   2E+4   8   8.5  RT (min)      lcorbi_t3_neg_4_023_18    m/z 902.4387     0   2   4   6E+3   13.5   14  RT (min)      lcorbi_t3_neg_3_005_23    m/z 902.4527     0   1E+4   6   6.5  RT (min)      lcorbi_t3_neg_2_048_27    m/z 902.5241     0   1   2E+3   13.5   14  RT (min)      lcorbi_t3_neg_4_042_14    m/z 902.5481     0   1E+3   13   13.5  RT (min)      lcorbi_t3_neg_4_032_23    m/z 902.5624     0   1E+3   13.5   14  RT (min)      lcorbi_t3_neg_4_019_08    m/z 902.6128     0   1   2   3E+4   14   14.5  RT (min)      lcorbi_t3_neg_4_054_36    m/z 902.6217     0   1   2E+3   14.5   15  RT (min)      lcorbi_t3_neg_4_007_13    m/z 902.6403     0   2   4   6   8E+4   13   13.5  RT (min)      lcorbi_t3_neg_4_019_08    m/z 902.6581     0   1   2E+4   13   13.5  RT (min)      lcorbi_t3_neg_4_039_32    m/z 902.6858     0   1E+4   14.5   15  RT (min)      lcorbi_t3_neg_4_034_01    m/z 902.7498     0   1E+4   14   14.5  RT (min)      lcorbi_t3_neg_3_002_24    m/z 902.8290     0   2   4   6E+3   18   18.5  RT (min)      lcorbi_t3_neg_2_016_11    m/z 903.2124     0   2   4E+4   13.5   14  RT (min)      lcorbi_t3_neg_2_036_35    m/z 903.3520     0   1   2E+3   8   8.5  RT (min)      lcorbi_t3_neg_1_036_09    m/z 903.4198     0   2   4E+4   13.5   14  RT (min)      lcorbi_t3_neg_1_044_23    m/z 903.4556     0   2   4   6   8E+3   5.5   6  RT (min)      lcorbi_t3_neg_2_048_27    m/z 903.5407     0   1E+4   13.5   14  RT (min)      lcorbi_t3_neg_4_032_23    m/z 903.5543     0   1E+4   13   13.5  RT (min)      lcorbi_t3_neg_4_042_14    m/z 903.6039     0   1   2   3   4E+2   14.5   15  RT (min)      lcorbi_t3_neg_1_029_21    m/z 903.6221     0   1   2E+4   13   13.5  RT (min)      lcorbi_t3_neg_4_039_32    m/z 903.6340     0   2   4   6E+3   14.5   15  RT (min)      lcorbi_t3_neg_3_010_15    m/z 903.6913     0   1   2   3   4E+4   13.5   14  RT (min)      lcorbi_t3_neg_3_002_24    m/z 903.7730     0   1   2E+4   14.5   15  RT (min)      lcorbi_t3_neg_3_002_24    m/z 904.2128     0   1   2   3E+4   13.5   14  RT (min)      lcorbi_t3_neg_3_028_35    m/z 904.4098     0   2   4   6   8E+3   13.5   14  RT (min)      lcorbi_t3_neg_3_005_23    m/z 904.5638     0   2   4E+3   13   13.5  RT (min)      lcorbi_t3_neg_2_022_24    m/z 904.5739     0   1   2E+4   6   6.5  RT (min)      lcorbi_t3_neg_4_044_38    m/z 904.5789     0   2   4E+3   13.5   14  RT (min)      lcorbi_t3_neg_2_013_08    m/z 904.7028     0   1E+3   13   13.5  RT (min)      lcorbi_t3_neg_2_043_33    m/z 905.2712     0   2   4   6   8E+3   11   11.5  RT (min)      lcorbi_t3_neg_2_027_13    m/z 905.4132     0   1   2   3E+3   13.5   14  RT (min)      lcorbi_t3_neg_2_053_23    m/z 905.4478     0   1E+4   7.5   8  RT (min)      lcorbi_t3_neg_2_048_27    m/z 905.4692     0   1E+4   11   11.5  RT (min)      lcorbi_t3_neg_4_023_18    m/z 905.5281     0   2   4E+3   13.5   14  RT (min)      lcorbi_t3_neg_4_007_13    m/z 905.5575     0   1   13.5   14  RT (min)      lcorbi_t3_neg_4_039_32    m/z 905.6048     0   1   2   3E+4   14.5   15  RT (min)      lcorbi_t3_neg_4_007_13    m/z 905.6097     0   1   2   3E+4   14   14.5  RT (min)      lcorbi_t3_neg_4_039_32    m/z 905.6255     0   2   4   6E+3   13   13.5  RT (min)      lcorbi_t3_neg_3_002_24    m/z 905.7604     0   2   4E+3   15   15.5  RT (min)      lcorbi_t3_neg_4_038_15    m/z 906.2721     0   1   2   3   4E+3   11   11.5  RT (min)      lcorbi_t3_neg_2_022_24    m/z 906.3369     0   1E+4   6   6.5  RT (min)      lcorbi_t3_neg_3_014_19    m/z 906.4723     0   1   2E+4   13   13.5  RT (min)      lcorbi_t3_neg_4_032_23    m/z 906.4898     0   1E+4   13.5   14  RT (min)      lcorbi_t3_neg_4_045_35    m/z 906.5476     0   1   11   11.5  RT (min)      lcorbi_t3_neg_1_044_23    m/z 906.6899     0   1E+5   13.5   14  RT (min)      lcorbi_t3_neg_4_039_32    m/z 906.7568     0   2   4E+3   14.5   15  RT (min)      lcorbi_t3_neg_4_001_49    m/z 906.7728     0   1   2   3E+3   18   18.5  RT (min)      lcorbi_t3_neg_2_026_03    m/z 906.7929     0   1E+4   14.5   15  RT (min)      lcorbi_t3_neg_4_039_32    m/z 907.2867     0   2   4   6E+3   11.5   12  RT (min)      lcorbi_t3_neg_3_044_01    m/z 907.5376     0   1E+4   13.5   14  RT (min)      lcorbi_t3_neg_4_007_13    m/z 907.6029     0   2   4   6E+3   14.5   15  RT (min)      lcorbi_t3_neg_2_013_08    m/z 907.8423     0   2   4   6E+3   18   18.5  RT (min)      lcorbi_t3_neg_1_001_49    m/z 908.5070     0   1   2E+4   13.5   14  RT (min)      lcorbi_t3_neg_1_020_36    m/z 908.5184     0   1E+4   14   14.5  RT (min)      lcorbi_t3_neg_1_034_04    m/z 908.5403     0   1E+4   13.5   14  RT (min)      lcorbi_t3_neg_4_007_13    m/z 908.5633     0   1E+4   11.5   12  RT (min)      lcorbi_t3_neg_4_042_14    m/z 908.5635     0   1E+4   14   14.5  RT (min)      lcorbi_t3_neg_2_026_03    m/z 908.6524     0   1   2   3   4E+5   12.5   13  RT (min)      lcorbi_t3_neg_3_002_24    m/z 908.6750     0   2   4   6   8E+3   14.5   15  RT (min)      lcorbi_t3_neg_4_007_13    m/z 908.7995     0   1E+4   14.5   15  RT (min)      lcorbi_t3_neg_2_022_24    m/z 908.8401     0   1   2   3E+3   18   18.5  RT (min)      lcorbi_t3_neg_2_003_42    m/z 909.3025     0   1E+4   12   12.5  RT (min)      lcorbi_t3_neg_3_044_01    m/z 909.3124     0   1E+4   8.5   9  RT (min)      lcorbi_t3_neg_4_040_19    m/z 909.5224     0   2   4   6E+3   14   14.5  RT (min)      lcorbi_t3_neg_2_048_27    m/z 909.5608     0   2   4   6   8E+3   13.5   14  RT (min)      lcorbi_t3_neg_4_046_37    m/z 909.6164     0   1E+3   13.5   14  RT (min)      lcorbi_t3_neg_1_005_07    m/z 909.6305     0   1   2   3E+3   14   14.5  RT (min)      lcorbi_t3_neg_4_044_38    m/z 910.3289     0   1E+4   8.5   9  RT (min)      lcorbi_t3_neg_2_048_27    m/z 910.5209     0   1E+4   14   14.5  RT (min)      lcorbi_t3_neg_2_011_04    m/z 910.5426     0   2   4   6E+4   13   13.5  RT (min)      lcorbi_t3_neg_2_007_14    m/z 910.5771     0   2   4   6E+3   13   13.5  RT (min)      lcorbi_t3_neg_3_052_32    m/z 910.6171     0   1   2   3   4E+3   14   14.5  RT (min)      lcorbi_t3_neg_2_024_37    m/z 910.7746     0   2   4   6   8E+3   18   18.5  RT (min)      lcorbi_t3_neg_2_038_22    m/z 910.7833     0   2   4   6   8E+3   18   18.5  RT (min)      lcorbi_t3_neg_2_013_08    m/z 911.1471     0   1   2   3   4E+3   9   9.5  RT (min)      lcorbi_t3_neg_3_019_12    m/z 911.3062     0   1E+4   9   9.5  RT (min)      lcorbi_t3_neg_2_048_27    m/z 911.3325     0   2   4   6E+3   8.5   9  RT (min)      lcorbi_t3_neg_2_048_27    m/z 911.4365     0   1   2E+4   10.5   11  RT (min)      lcorbi_t3_neg_4_017_51    m/z 911.4692     0   2   4   6   8E+2   13   13.5  RT (min)      lcorbi_t3_neg_4_042_14    m/z 911.5461     0   1   2   3E+4   13   13.5  RT (min)      lcorbi_t3_neg_4_042_14    m/z 911.5887     0   1E+4   13   13.5  RT (min)      lcorbi_t3_neg_3_052_32    m/z 911.6182     0   2   4E+3   14.5   15  RT (min)      lcorbi_t3_neg_2_025_52    m/z 911.7133     0   1   2E+4   12   12.5  RT (min)      lcorbi_t3_neg_4_023_18    m/z 911.8061     0   1   2   3   4E+3   18   18.5  RT (min)      lcorbi_t3_neg_3_036_41    m/z 912.1503     0   1   2E+3   9   9.5  RT (min)      lcorbi_t3_neg_4_051_40    m/z 912.3988     0   1   2   3   4E+4   9   9.5  RT (min)      lcorbi_t3_neg_2_048_27    m/z 912.4232     0   2   4   6   8E+3   13   13.5  RT (min)      lcorbi_t3_neg_3_021_38    m/z 912.4727     0   2   4   6   8E+3   13   13.5  RT (min)      lcorbi_t3_neg_2_053_23    m/z 912.5094     0   2   4   6E+3   13.5   14  RT (min)      lcorbi_t3_neg_2_027_13    m/z 912.5945     0   2   4E+3   14.5   15  RT (min)      lcorbi_t3_neg_2_052_32    m/z 912.6300     0   2   4   6   8E+3   14.5   15  RT (min)      lcorbi_t3_neg_4_034_01    m/z 912.6731     0   1   2   3   4E+4   13   13.5  RT (min)      lcorbi_t3_neg_4_039_32    m/z 912.7151     0   1   2   3   4E+2   14.5   15  RT (min)      lcorbi_t3_neg_4_007_13    m/z 912.7890     0   2   4   6   8E+3   18   18.5  RT (min)      lcorbi_t3_neg_2_003_42    m/z 913.1454     0   1   2E+3   9   9.5  RT (min)      lcorbi_t3_neg_2_023_39    m/z 913.3805     0   1   2E+4   8   8.5  RT (min)      lcorbi_t3_neg_4_040_19    m/z 913.4269     0   1   2   3   4E+3   13   13.5  RT (min)      lcorbi_t3_neg_3_047_37    m/z 913.4677     0   1E+3   13.5   14  RT (min)      lcorbi_t3_neg_4_046_37    m/z 913.4847     0   1E+4   18   18.5  RT (min)      lcorbi_t3_neg_1_022_34    m/z 913.5613     0   1E+4   13   13.5  RT (min)      lcorbi_t3_neg_4_042_14    m/z 913.5721     0   1   14   14.5  RT (min)      lcorbi_t3_neg_2_013_08    m/z 913.5804     0   1   2   3E+4   14   14.5  RT (min)      lcorbi_t3_neg_4_019_08    m/z 913.6048     0   1   2   3   4E+4   13.5   14  RT (min)      lcorbi_t3_neg_2_013_08    m/z 913.6362     0   1   2E+4   14   14.5  RT (min)      lcorbi_t3_neg_1_052_24    m/z 913.6849     0   1E+4   13.5   14  RT (min)      lcorbi_t3_neg_4_011_24    m/z 913.7011     0   1   2E+4   14   14.5  RT (min)      lcorbi_t3_neg_3_002_24    m/z 913.7181     0   1   14.5   15  RT (min)      lcorbi_t3_neg_4_007_13    m/z 913.7278     0   1   2E+4   14   14.5  RT (min)      lcorbi_t3_neg_2_048_27    m/z 914.3677     0   1   2E+4   6   6.5  RT (min)      lcorbi_t3_neg_3_014_19    m/z 914.3840     0   1E+4   8   8.5  RT (min)      lcorbi_t3_neg_4_040_19    m/z 914.4711     0   1   13.5   14  RT (min)      lcorbi_t3_neg_4_045_35    m/z 914.4882     0   2   4   6E+3   18   18.5  RT (min)      lcorbi_t3_neg_1_022_34    m/z 914.4884     0   1   2E+4   13   13.5  RT (min)      lcorbi_t3_neg_4_046_37    m/z 914.5238     0   2   4   6E+3   14   14.5  RT (min)      lcorbi_t3_neg_1_047_02    m/z 914.5656     0   1   2E+4   14.5   15  RT (min)      lcorbi_t3_neg_2_026_03    m/z 914.6211     0   2   4   6   8E+3   14.5   15  RT (min)      lcorbi_t3_neg_1_047_02    m/z 914.7330     0   1E+4   14   14.5  RT (min)      lcorbi_t3_neg_2_052_32    m/z 914.7877     0   1   2   3E+3   18   18.5  RT (min)      lcorbi_t3_neg_2_015_21    m/z 915.1587     0   1   2   3   4E+3   9   9.5  RT (min)      lcorbi_t3_neg_3_019_12    m/z 915.1913     0   1   2E+4   13.5   14  RT (min)      lcorbi_t3_neg_4_045_35    m/z 915.4764     0   1   13.5   14  RT (min)      lcorbi_t3_neg_4_046_37    m/z 915.4914     0   1   2   3   4E+4   13   13.5  RT (min)      lcorbi_t3_neg_4_046_37    m/z 915.4997     0   2   4   6   8E+3   18   18.5  RT (min)      lcorbi_t3_neg_1_022_34    m/z 915.5603     0   1E+4   13.5   14  RT (min)      lcorbi_t3_neg_4_038_15    m/z 915.5697     0   1   2E+3   14.5   15  RT (min)      lcorbi_t3_neg_4_042_14    m/z 915.6164     0   1   2   3E+4   14.5   15  RT (min)      lcorbi_t3_neg_2_013_08    m/z 915.6530     0   2   4   6E+3   13.5   14  RT (min)      lcorbi_t3_neg_1_055_05    m/z 915.6631     0   1   2   3E+3   14.5   15  RT (min)      lcorbi_t3_neg_4_044_38    m/z 915.6695     0   2   4   6   8E+3   14.5   15  RT (min)      lcorbi_t3_neg_4_034_01    m/z 915.6886     0   1E+4   15   15.5  RT (min)      lcorbi_t3_neg_4_029_31    m/z 915.7997     0   2   4   6E+3   18   18.5  RT (min)      lcorbi_t3_neg_1_045_44    m/z 916.4453     0   2   4   6E+3   13.5   14  RT (min)      lcorbi_t3_neg_4_032_23    m/z 916.4543     0   2   4   6E+3   18   18.5  RT (min)      lcorbi_t3_neg_4_029_31    m/z 916.4557     0   1E+4   13   13.5  RT (min)      lcorbi_t3_neg_2_013_08    m/z 916.4799     0   1   13.5   14  RT (min)      lcorbi_t3_neg_3_048_08    m/z 916.4869     0   1   2   3E+4   7.5   8  RT (min)      lcorbi_t3_neg_2_048_27    m/z 916.5036     0   2   4   6E+3   18   18.5  RT (min)      lcorbi_t3_neg_1_022_34    m/z 916.5038     0   1   2E+4   13.5   14  RT (min)      lcorbi_t3_neg_4_046_37    m/z 916.5670     0   2   4E+3   14.5   15  RT (min)      lcorbi_t3_neg_4_007_13    m/z 916.5811     0   1   2E+3   14.5   15  RT (min)      lcorbi_t3_neg_2_043_33    m/z 916.6001     0   1E+4   14   14.5  RT (min)      lcorbi_t3_neg_4_019_08    m/z 916.7007     0   2   4   6E+3   15   15.5  RT (min)      lcorbi_t3_neg_4_029_31    m/z 916.8218     0   2   4   6E+3   14.5   15  RT (min)      lcorbi_t3_neg_4_039_32    m/z 917.2155     0   2   4   6E+3   10.5   11  RT (min)      lcorbi_t3_neg_3_035_03    m/z 917.4582     0   2   4   6E+3   13.5   14  RT (min)      lcorbi_t3_neg_2_013_08    m/z 917.4773     0   1   13.5   14  RT (min)      lcorbi_t3_neg_3_040_05    m/z 917.4926     0   1   13.5   14  RT (min)      lcorbi_t3_neg_2_013_08    m/z 917.5067     0   2   4   6   8E+3   13.5   14  RT (min)      lcorbi_t3_neg_2_053_23    m/z 917.5354     0   1   2E+4   13.5   14  RT (min)      lcorbi_t3_neg_1_047_02    m/z 917.5846     0   1   2   3E+3   14.5   15  RT (min)      lcorbi_t3_neg_4_007_13    m/z 917.6036     0   1   2E+3   14   14.5  RT (min)      lcorbi_t3_neg_2_013_08    m/z 917.6557     0   1E+4   13   13.5  RT (min)      lcorbi_t3_neg_4_039_32    m/z 917.6805     0   1E+4   13.5   14  RT (min)      lcorbi_t3_neg_3_002_24    m/z 917.7100     0   2   4   6E+3   14   14.5  RT (min)      lcorbi_t3_neg_3_002_24    m/z 917.8036     0   2   4E+3   18   18.5  RT (min)      lcorbi_t3_neg_2_006_10    m/z 918.2389     0   1   2   3E+3   11   11.5  RT (min)      lcorbi_t3_neg_4_048_03    m/z 918.3291     0   1   2   3E+4   9   9.5  RT (min)      lcorbi_t3_neg_2_048_27    m/z 918.5343     0   1   2   3   4E+3   13   13.5  RT (min)      lcorbi_t3_neg_3_010_15    m/z 918.6351     0   1   2E+3   14.5   15  RT (min)      lcorbi_t3_neg_3_052_32    m/z 918.6539     0   1   13   13.5  RT (min)      lcorbi_t3_neg_1_014_28    m/z 918.9889     0   2   4E+3   12.5   13  RT (min)      lcorbi_t3_neg_4_017_51    m/z 919.3488     0   1   2   3   4E+3   8.5   9  RT (min)      lcorbi_t3_neg_4_040_19    m/z 919.5080     0   1   2   3   4E+3   13.5   14  RT (min)      lcorbi_t3_neg_4_050_02    m/z 919.6863     0   1E+4   13   13.5  RT (min)      lcorbi_t3_neg_4_017_51    m/z 919.8087     0   2   4   6   8E+3   18   18.5  RT (min)      lcorbi_t3_neg_2_038_22    m/z 920.3895     0   1   2   3E+3   9   9.5  RT (min)      lcorbi_t3_neg_2_048_27    m/z 920.4266     0   1E+4   10.5   11  RT (min)      lcorbi_t3_neg_4_005_27    m/z 920.6497     0   1   13.5   14  RT (min)      lcorbi_t3_neg_1_021_31    m/z 920.6637     0   2   4   6   8E+3   12.5   13  RT (min)      lcorbi_t3_neg_3_002_24    m/z 920.6642     0   1   2E+4   14.5   15  RT (min)      lcorbi_t3_neg_2_022_24    m/z 920.6896     0   1   2   3E+4   13   13.5  RT (min)      lcorbi_t3_neg_4_017_51    m/z 920.7086     0   1   2E+3   14   14.5  RT (min)      lcorbi_t3_neg_1_005_07    m/z 920.7358     0   1   2E+4   14   14.5  RT (min)      lcorbi_t3_neg_3_002_24    m/z 921.4099     0   1   2   3E+3   13   13.5  RT (min)      lcorbi_t3_neg_3_028_35    m/z 921.4817     0   2   4   6   8E+3   13   13.5  RT (min)      lcorbi_t3_neg_4_007_13    m/z 921.4897     0   2   4   6   8E+3   12.5   13  RT (min)      lcorbi_t3_neg_4_046_37    m/z 921.5551     0   1   2E+4   13.5   14  RT (min)      lcorbi_t3_neg_3_020_31    m/z 921.6614     0   1E+4   13.5   14  RT (min)      lcorbi_t3_neg_1_052_24    m/z 921.7020     0   2   4E+3   13.5   14  RT (min)      lcorbi_t3_neg_4_017_51    m/z 921.8075     0   1   2E+3   18   18.5  RT (min)      lcorbi_t3_neg_2_042_44    m/z 922.1793     0   2   4E+3   9   9.5  RT (min)      lcorbi_t3_neg_3_019_12    m/z 922.2568     0   1   2   3E+3   11.5   12  RT (min)      lcorbi_t3_neg_2_027_13    m/z 922.4014     0   1E+4   15   15.5  RT (min)      lcorbi_t3_neg_2_038_22    m/z 922.4910     0   1E+3   12.5   13  RT (min)      lcorbi_t3_neg_4_032_23    m/z 922.5136     0   2   4   6E+3   10.5   11  RT (min)      lcorbi_t3_neg_3_005_23    m/z 922.5473     0   1E+4   13   13.5  RT (min)      lcorbi_t3_neg_4_039_32    m/z 922.6580     0   2   4   6   8E+3   14.5   15  RT (min)      lcorbi_t3_neg_4_005_27    m/z 923.3280     0   1   2   3E+4   9   9.5  RT (min)      lcorbi_t3_neg_4_040_19    m/z 923.3343     0   1   2   3   4E+3   13   13.5  RT (min)      lcorbi_t3_neg_4_038_15    m/z 923.4041     0   2   4   6E+3   15   15.5  RT (min)      lcorbi_t3_neg_2_038_22    m/z 923.4052     0   1E+4   10.5   11  RT (min)      lcorbi_t3_neg_2_007_14    m/z 923.4257     0   1E+4   13.5   14  RT (min)      lcorbi_t3_neg_1_044_23    m/z 923.5056     0   2   4   6   8E+3   13   13.5  RT (min)      lcorbi_t3_neg_4_032_23    m/z 923.6610     0   1   2   3   4E+4   13   13.5  RT (min)      lcorbi_t3_neg_4_039_32    m/z 923.8206     0   2   4   6   8E+3   18   18.5  RT (min)      lcorbi_t3_neg_4_015_10    m/z 924.2728     0   1   2E+3   12   12.5  RT (min)      lcorbi_t3_neg_3_029_02    m/z 924.3300     0   1E+4   9   9.5  RT (min)      lcorbi_t3_neg_4_040_19    m/z 924.3445     0   2   4   6   8E+3   9.5   10  RT (min)      lcorbi_t3_neg_2_048_27    m/z 924.5101     0   1E+3   13   13.5  RT (min)      lcorbi_t3_neg_3_005_23    m/z 924.5628     0   2   4   6E+3   13.5   14  RT (min)      lcorbi_t3_neg_3_032_17    m/z 924.6545     0   2   4E+3   18   18.5  RT (min)      lcorbi_t3_neg_1_013_35    m/z 924.6698     0   1E+4   14.5   15  RT (min)      lcorbi_t3_neg_4_005_27    m/z 924.8202     0   2   4E+3   18   18.5  RT (min)      lcorbi_t3_neg_2_018_40    m/z 925.3377     0   1   2   3E+4   10.5   11  RT (min)      lcorbi_t3_neg_1_043_14    m/z 925.5056     0   1   2   3E+4   9.5   10  RT (min)      lcorbi_t3_neg_3_019_12    m/z 925.5231     0   1   2   3   4E+4   13   13.5  RT (min)      lcorbi_t3_neg_4_032_23    m/z 925.5616     0   1E+3   13.5   14  RT (min)      lcorbi_t3_neg_4_044_38    m/z 925.6766     0   1   2E+4   13.5   14  RT (min)      lcorbi_t3_neg_4_039_32    m/z 925.7064     0   1E+4   15   15.5  RT (min)      lcorbi_t3_neg_4_007_13    m/z 925.7096     0   1E+4   15   15.5  RT (min)      lcorbi_t3_neg_4_007_13    m/z 925.8255     0   2   4   6E+3   18   18.5  RT (min)      lcorbi_t3_neg_3_026_44    m/z 926.4390     0   1   2   3E+3   13.5   14  RT (min)      lcorbi_t3_neg_4_032_23    m/z 926.5091     0   1E+4   9.5   10  RT (min)      lcorbi_t3_neg_3_019_12    m/z 926.5505     0   1E+4   13   13.5  RT (min)      lcorbi_t3_neg_3_052_32    m/z 926.7146     0   2   4   6E+3   15   15.5  RT (min)      lcorbi_t3_neg_4_007_13    m/z 926.7363     0   1E+3   13.5   14  RT (min)      lcorbi_t3_neg_2_013_08    m/z 926.8582     0   1   2   3E+3   18   18.5  RT (min)      lcorbi_t3_neg_2_042_44    m/z 927.5002     0   1E+4   13.5   14  RT (min)      lcorbi_t3_neg_2_028_38    m/z 927.5090     0   1E+4   13.5   14  RT (min)      lcorbi_t3_neg_4_005_27    m/z 927.5794     0   1   2E+4   13.5   14  RT (min)      lcorbi_t3_neg_4_005_27    m/z 927.6920     0   1   2   3   4E+4   13   13.5  RT (min)      lcorbi_t3_neg_4_039_32    m/z 928.2678     0   2   4   6   8E+3   11   11.5  RT (min)      lcorbi_t3_neg_4_048_03    m/z 928.3392     0   2   4   6   8E+3   10.5   11  RT (min)      lcorbi_t3_neg_4_011_24    m/z 928.4100     0   2   4E+3   13   13.5  RT (min)      lcorbi_t3_neg_2_053_23    m/z 928.5621     0   2   4   6E+3   13.5   14  RT (min)      lcorbi_t3_neg_4_005_27    m/z 928.6175     0   2   4   6   8E+3   14.5   15  RT (min)      lcorbi_t3_neg_3_010_15    m/z 928.6254     0   1E+4   14.5   15  RT (min)      lcorbi_t3_neg_3_007_16    m/z 928.7736     0   1   2E+3   18   18.5  RT (min)      lcorbi_t3_neg_3_055_06    m/z 929.2713     0   1   2   3   4E+3   11   11.5  RT (min)      lcorbi_t3_neg_2_010_02    m/z 929.3367     0   1   2   3E+3   18   18.5  RT (min)      lcorbi_t3_neg_3_011_04    m/z 929.4506     0   1E+4   7.5   8  RT (min)      lcorbi_t3_neg_2_048_27    m/z 929.5261     0   2   4E+3   13.5   14  RT (min)      lcorbi_t3_neg_4_007_13    m/z 929.5566     0   1   2   3   4E+4   13.5   14  RT (min)      lcorbi_t3_neg_2_053_23    m/z 929.5955     0   2   4   6E+4   13.5   14  RT (min)      lcorbi_t3_neg_4_005_27    m/z 929.6291     0   2   4   6E+3   14.5   15  RT (min)      lcorbi_t3_neg_3_007_16    m/z 929.6309     0   1E+4   13.5   14  RT (min)      lcorbi_t3_neg_3_002_24    m/z 929.6850     0   1E+4   14.5   15  RT (min)      lcorbi_t3_neg_4_034_01    m/z 929.7014     0   2   4E+3   15   15.5  RT (min)      lcorbi_t3_neg_2_007_14    m/z 930.2832     0   1   2   3E+3   11.5   12  RT (min)      lcorbi_t3_neg_2_027_13    m/z 930.5612     0   1E+4   13.5   14  RT (min)      lcorbi_t3_neg_3_022_27    m/z 930.6628     0   1   2   3E+3   13.5   14  RT (min)      lcorbi_t3_neg_4_019_08    m/z 930.6877     0   1   14.5   15  RT (min)      lcorbi_t3_neg_2_013_08    m/z 930.7052     0   1   2   3   4E+3   15   15.5  RT (min)      lcorbi_t3_neg_2_007_14    m/z 930.7617     0   1   2E+3   18   18.5  RT (min)      lcorbi_t3_neg_2_039_31    m/z 930.7840     0   1   2E+4   14.5   15  RT (min)      lcorbi_t3_neg_4_039_32    m/z 930.8396     0   2   4   6   8E+3   18   18.5  RT (min)      lcorbi_t3_neg_3_015_11    m/z 931.1319     0   1   2E+3   9   9.5  RT (min)      lcorbi_t3_neg_3_019_12    m/z 931.2649     0   1   2   3   4E+3   12   12.5  RT (min)      lcorbi_t3_neg_3_054_18    m/z 931.4760     0   2   4   6   8E+3   13   13.5  RT (min)      lcorbi_t3_neg_4_044_38    m/z 931.5449     0   1E+4   13   13.5  RT (min)      lcorbi_t3_neg_4_046_37    m/z 931.6028     0   1   13   13.5  RT (min)      lcorbi_t3_neg_4_032_23    m/z 931.7157     0   1E+4   13.5   14  RT (min)      lcorbi_t3_neg_4_011_24    m/z 931.7168     0   2   4E+3   15   15.5  RT (min)      lcorbi_t3_neg_4_042_14    m/z 931.7245     0   1   14.5   15  RT (min)      lcorbi_t3_neg_1_052_24    m/z 931.7551     0   1E+4   16   16.5  RT (min)      lcorbi_t3_neg_1_007_26    m/z 931.7963     0   2   4   6   8E+3   14.5   15  RT (min)      lcorbi_t3_neg_2_052_32    m/z 931.8076     0   1   2   3E+3   18   18.5  RT (min)      lcorbi_t3_neg_4_015_10    m/z 932.3081     0   1   2   3E+4   9   9.5  RT (min)      lcorbi_t3_neg_2_048_27    m/z 932.3447     0   1E+4   9   9.5  RT (min)      lcorbi_t3_neg_2_048_27    m/z 932.3654     0   1   2   3E+4   10.5   11  RT (min)      lcorbi_t3_neg_1_043_14    m/z 932.5488     0   2   4   6E+3   13   13.5  RT (min)      lcorbi_t3_neg_4_044_38    m/z 932.5906     0   1E+4   13.5   14  RT (min)      lcorbi_t3_neg_4_011_24    m/z 932.6000     0   2   4E+3   14.5   15  RT (min)      lcorbi_t3_neg_1_053_32    m/z 932.6145     0   1   2E+4   14   14.5  RT (min)      lcorbi_t3_neg_4_005_27    m/z 932.7235     0   1   2   3   4E+3   15   15.5  RT (min)      lcorbi_t3_neg_4_042_14    m/z 932.7744     0   1   2   3   4E+3   18   18.5  RT (min)      lcorbi_t3_neg_4_051_40    m/z 932.7996     0   1   2E+4   14.5   15  RT (min)      lcorbi_t3_neg_4_039_32    m/z 932.8444     0   1E+4   14.5   15  RT (min)      lcorbi_t3_neg_4_039_32    m/z 933.3300     0   2   4   6E+3   8.5   9  RT (min)      lcorbi_t3_neg_4_023_18    m/z 933.3715     0   1   2   3E+3   13   13.5  RT (min)      lcorbi_t3_neg_4_034_01    m/z 933.4440     0   1E+4   13.5   14  RT (min)      lcorbi_t3_neg_4_054_36    m/z 933.6035     0   2   4   6E+3   14.5   15  RT (min)      lcorbi_t3_neg_4_011_24    m/z 933.6183     0   1   13.5   14  RT (min)      lcorbi_t3_neg_4_019_08    m/z 933.7203     0   1   2   3   4E+3   15   15.5  RT (min)      lcorbi_t3_neg_4_042_14    m/z 933.7321     0   2   4E+3   15   15.5  RT (min)      lcorbi_t3_neg_4_044_38    m/z 934.4597     0   1E+3   13.5   14  RT (min)      lcorbi_t3_neg_4_042_14    m/z 934.5427     0   1   2E+4   13   13.5  RT (min)      lcorbi_t3_neg_4_042_14    m/z 934.6037     0   1   2E+4   13.5   14  RT (min)      lcorbi_t3_neg_4_019_08    m/z 934.6538     0   1   2E+3   14.5   15  RT (min)      lcorbi_t3_neg_4_019_08    m/z 934.6882     0   1E+4   13.5   14  RT (min)      lcorbi_t3_neg_4_011_24    m/z 934.7360     0   1   2   3   4E+3   15   15.5  RT (min)      lcorbi_t3_neg_4_029_31    m/z 934.7810     0   2   4   6E+3   18   18.5  RT (min)      lcorbi_t3_neg_2_003_42    m/z 934.7897     0   2   4   6E+3   18   18.5  RT (min)      lcorbi_t3_neg_2_010_02    m/z 934.8154     0   1   2E+4   14.5   15  RT (min)      lcorbi_t3_neg_4_039_32    m/z 935.4669     0   1   13   13.5  RT (min)      lcorbi_t3_neg_2_007_14    m/z 935.5043     0   1E+4   13.5   14  RT (min)      lcorbi_t3_neg_3_018_13    m/z 935.5461     0   1E+4   13   13.5  RT (min)      lcorbi_t3_neg_2_007_14    m/z 935.6580     0   2   4   6E+2   14.5   15  RT (min)      lcorbi_t3_neg_4_019_08    m/z 935.7017     0   1E+4   12   12.5  RT (min)      lcorbi_t3_neg_2_022_24    m/z 936.2852     0   1E+5   9   9.5  RT (min)      lcorbi_t3_neg_2_048_27    m/z 936.4234     0   2   4E+3   13   13.5  RT (min)      lcorbi_t3_neg_4_054_36    m/z 936.4705     0   1   2E+3   13   13.5  RT (min)      lcorbi_t3_neg_3_005_23    m/z 936.5070     0   1   2   3E+4   13.5   14  RT (min)      lcorbi_t3_neg_1_054_37    m/z 936.5575     0   1E+4   13   13.5  RT (min)      lcorbi_t3_neg_2_007_14    m/z 936.6186     0   1E+4   13.5   14  RT (min)      lcorbi_t3_neg_1_052_24    m/z 936.6692     0   1E+4   14.5   15  RT (min)      lcorbi_t3_neg_4_007_13    m/z 936.7776     0   1   2E+3   18   18.5  RT (min)      lcorbi_t3_neg_3_016_42    m/z 936.8218     0   2   4E+3   18   18.5  RT (min)      lcorbi_t3_neg_3_015_11    m/z 937.2886     0   2   4   6   8E+4   9   9.5  RT (min)      lcorbi_t3_neg_2_048_27    m/z 937.4310     0   1E+4   11   11.5  RT (min)      lcorbi_t3_neg_4_023_18    m/z 937.4824     0   1E+3   13.5   14  RT (min)      lcorbi_t3_neg_3_052_32    m/z 937.5124     0   1E+4   13.5   14  RT (min)      lcorbi_t3_neg_1_054_37    m/z 937.5228     0   1E+4   13.5   14  RT (min)      lcorbi_t3_neg_3_029_02    m/z 937.5443     0   1E+4   13.5   14  RT (min)      lcorbi_t3_neg_3_002_24    m/z 937.5613     0   2   4   6   8E+3   13   13.5  RT (min)      lcorbi_t3_neg_4_042_14    m/z 937.5731     0   2   4E+3   15   15.5  RT (min)      lcorbi_t3_neg_4_017_51    m/z 937.6631     0   1   2E+3   14.5   15  RT (min)      lcorbi_t3_neg_4_019_08    m/z 937.6711     0   1E+3   14.5   15  RT (min)      lcorbi_t3_neg_4_019_08    m/z 938.2264     0   1   2   3E+3   11.5   12  RT (min)      lcorbi_t3_neg_4_030_04    m/z 938.3216     0   2   4   6E+3   8.5   9  RT (min)      lcorbi_t3_neg_4_023_18    m/z 938.3727     0   2   4   6E+3   10.5   11  RT (min)      lcorbi_t3_neg_1_024_01    m/z 938.5159     0   1   2E+4   13.5   14  RT (min)      lcorbi_t3_neg_4_019_08    m/z 938.5237     0   1   2   3E+4   13.5   14  RT (min)      lcorbi_t3_neg_1_054_37    m/z 938.5740     0   1   2   3   4E+4   13.5   14  RT (min)      lcorbi_t3_neg_2_007_14    m/z 938.6341     0   1E+3   13.5   14  RT (min)      lcorbi_t3_neg_4_042_14    m/z 938.6449     0   1E+4   14.5   15  RT (min)      lcorbi_t3_neg_4_034_01    m/z 938.6511     0   1E+4   14.5   15  RT (min)      lcorbi_t3_neg_4_019_08    m/z 938.6837     0   1E+4   15   15.5  RT (min)      lcorbi_t3_neg_4_034_01    m/z 938.7003     0   1E+4   13   13.5  RT (min)      lcorbi_t3_neg_1_052_24    m/z 939.2488     0   1   2   3E+3   9.5   10  RT (min)      lcorbi_t3_neg_3_019_12    m/z 939.4422     0   2   4   6   8E+3   13   13.5  RT (min)      lcorbi_t3_neg_2_053_23    m/z 939.4751     0   1E+4   7.5   8  RT (min)      lcorbi_t3_neg_4_040_19    m/z 939.4835     0   1   13.5   14  RT (min)      lcorbi_t3_neg_4_019_08    m/z 939.5004     0   2   4E+3   13   13.5  RT (min)      lcorbi_t3_neg_4_046_37    m/z 939.5216     0   1   2   3E+4   13.5   14  RT (min)      lcorbi_t3_neg_1_054_37    m/z 939.5602     0   1   2   3   4E+4   13.5   14  RT (min)      lcorbi_t3_neg_4_011_24    m/z 939.5702     0   2   4E+3   15   15.5  RT (min)      lcorbi_t3_neg_4_017_51    m/z 939.5774     0   1   2E+4   13.5   14  RT (min)      lcorbi_t3_neg_2_007_14    m/z 939.6413     0   1E+4   6.5   7  RT (min)      lcorbi_t3_neg_4_029_31    m/z 939.6426     0   1   2   3   4E+3   13   13.5  RT (min)      lcorbi_t3_neg_2_022_24    m/z 939.6492     0   1   2   3E+3   14.5   15  RT (min)      lcorbi_t3_neg_3_053_14    m/z 939.7950     0   1   2E+3   18   18.5  RT (min)      lcorbi_t3_neg_4_018_42    m/z 940.2157     0   1   2   3E+3   10   10.5  RT (min)      lcorbi_t3_neg_4_023_18    m/z 940.3398     0   1   2   3E+4   9   9.5  RT (min)      lcorbi_t3_neg_2_048_27    m/z 940.3857     0   2   4   6   8E+3   10.5   11  RT (min)      lcorbi_t3_neg_1_027_30    m/z 940.4530     0   1   2E+4   6   6.5  RT (min)      lcorbi_t3_neg_2_048_27    m/z 940.4548     0   2   4   6E+3   13.5   14  RT (min)      lcorbi_t3_neg_1_008_08    m/z 940.4866     0   2   4E+3   13.5   14  RT (min)      lcorbi_t3_neg_2_027_13    m/z 940.5252     0   1   2   3E+3   13.5   14  RT (min)      lcorbi_t3_neg_2_050_17    m/z 940.5381     0   1   2   3E+4   14   14.5  RT (min)      lcorbi_t3_neg_2_011_04    m/z 940.5684     0   1E+4   14   14.5  RT (min)      lcorbi_t3_neg_1_052_24    m/z 940.5814     0   1   2E+3   14.5   15  RT (min)      lcorbi_t3_neg_4_042_14    m/z 940.6185     0   1E+4   13   13.5  RT (min)      lcorbi_t3_neg_4_039_32    m/z 940.6278     0   1E+4   13.5   14  RT (min)      lcorbi_t3_neg_4_039_32    m/z 940.6445     0   2   4   6   8E+3   6.5   7  RT (min)      lcorbi_t3_neg_4_044_38    m/z 940.6812     0   1   2   3E+4   14.5   15  RT (min)      lcorbi_t3_neg_3_005_23    m/z 940.6997     0   1E+4   14   14.5  RT (min)      lcorbi_t3_neg_4_011_24    m/z 940.7908     0   2   4   6   8E+3   18   18.5  RT (min)      lcorbi_t3_neg_4_003_21    m/z 941.2161     0   1   2   3   4E+3   12.5   13  RT (min)      lcorbi_t3_neg_3_007_16    m/z 941.3406     0   2   4E+4   9   9.5  RT (min)      lcorbi_t3_neg_2_048_27    m/z 941.4560     0   1E+4   6   6.5  RT (min)      lcorbi_t3_neg_2_048_27    m/z 941.4816     0   1E+3   13.5   14  RT (min)      lcorbi_t3_neg_4_046_37    m/z 941.4911     0   2   4   6E+3   13.5   14  RT (min)      lcorbi_t3_neg_2_027_13    m/z 941.5160     0   1E+4   13.5   14  RT (min)      lcorbi_t3_neg_4_046_37    m/z 941.5407     0   1E+4   14   14.5  RT (min)      lcorbi_t3_neg_3_011_04    m/z 941.5682     0   2   4   6E+3   10   10.5  RT (min)      lcorbi_t3_neg_3_019_12    m/z 941.6357     0   1   14.5   15  RT (min)      lcorbi_t3_neg_2_022_24    m/z 941.6406     0   2   4E+3   6.5   7  RT (min)      lcorbi_t3_neg_4_044_38    m/z 941.6935     0   1   2   3E+3   14.5   15  RT (min)      lcorbi_t3_neg_4_019_08    m/z 942.5881     0   1   2E+4   14   14.5  RT (min)      lcorbi_t3_neg_2_022_24    m/z 942.6157     0   2   4   6   8E+3   14   14.5  RT (min)      lcorbi_t3_neg_4_019_08    m/z 942.6546     0   1   2   3E+3   14.5   15  RT (min)      lcorbi_t3_neg_4_039_32    m/z 942.7167     0   1   2   3   4E+3   15   15.5  RT (min)      lcorbi_t3_neg_4_029_31    m/z 942.7867     0   1   2   3   4E+3   18   18.5  RT (min)      lcorbi_t3_neg_2_048_27    m/z 943.4453     0   1   2E+4   10.5   11  RT (min)      lcorbi_t3_neg_4_042_14    m/z 943.4771     0   2   4E+3   13.5   14  RT (min)      lcorbi_t3_neg_4_007_13    m/z 943.5510     0   1   2E+4   14   14.5  RT (min)      lcorbi_t3_neg_1_052_24    m/z 943.5665     0   1   13.5   14  RT (min)      lcorbi_t3_neg_1_052_24    m/z 943.6208     0   2   4   6E+3   14.5   15  RT (min)      lcorbi_t3_neg_1_047_02    m/z 943.6574     0   1   2   3   4E+3   14.5   15  RT (min)      lcorbi_t3_neg_1_053_32    m/z 943.7247     0   1E+4   13.5   14  RT (min)      lcorbi_t3_neg_3_002_24    m/z 943.7386     0   1   2   3E+4   14.5   15  RT (min)      lcorbi_t3_neg_2_022_24    m/z 943.9971     0   1E+4   6.5   7  RT (min)      lcorbi_t3_neg_2_048_27    m/z 944.3261     0   1E+4   9.5   10  RT (min)      lcorbi_t3_neg_2_048_27    m/z 944.4986     0   1E+4   6.5   7  RT (min)      lcorbi_t3_neg_2_048_27    m/z 944.5006     0   1   13   13.5  RT (min)      lcorbi_t3_neg_4_040_19    m/z 944.5112     0   1E+4   13.5   14  RT (min)      lcorbi_t3_neg_2_010_02    m/z 944.5443     0   1   2E+3   13.5   14  RT (min)      lcorbi_t3_neg_4_029_31    m/z 944.5617     0   1E+4   14   14.5  RT (min)      lcorbi_t3_neg_2_048_27    m/z 944.5700     0   1   2   3E+4   14   14.5  RT (min)      lcorbi_t3_neg_1_047_02    m/z 944.6609     0   2   4   6E+4   13   13.5  RT (min)      lcorbi_t3_neg_4_039_32    m/z 944.6995     0   1   2E+4   13.5   14  RT (min)      lcorbi_t3_neg_1_052_24    m/z 944.7107     0   1   14   14.5  RT (min)      lcorbi_t3_neg_4_011_24    m/z 945.0002     0   2   4   6   8E+3   6.5   7  RT (min)      lcorbi_t3_neg_2_048_27    m/z 945.2026     0   1   2   3   4E+4   13.5   14  RT (min)      lcorbi_t3_neg_2_036_35    m/z 945.3267     0   1E+4   10.5   11  RT (min)      lcorbi_t3_neg_1_043_14    m/z 945.4138     0   1   2E+4   7.5   8  RT (min)      lcorbi_t3_neg_2_048_27    m/z 945.5083     0   2   4   6   8E+3   13.5   14  RT (min)      lcorbi_t3_neg_1_002_16    m/z 945.5607     0   1   13   13.5  RT (min)      lcorbi_t3_neg_4_039_32    m/z 945.6364     0   1   2   3   4E+2   14.5   15  RT (min)      lcorbi_t3_neg_4_019_08    m/z 945.6427     0   1E+4   13   13.5  RT (min)      lcorbi_t3_neg_4_039_32    m/z 945.8127     0   2   4E+3   18   18.5  RT (min)      lcorbi_t3_neg_3_046_10    m/z 945.8243     0   1   2   3   4E+3   18   18.5  RT (min)      lcorbi_t3_neg_4_027_12    m/z 946.2033     0   1   2E+4   13.5   14  RT (min)      lcorbi_t3_neg_4_045_35    m/z 946.2392     0   1   2   3E+3   10.5   11  RT (min)      lcorbi_t3_neg_2_027_13    m/z 946.2590     0   1E+4   9   9.5  RT (min)      lcorbi_t3_neg_2_048_27    m/z 946.4607     0   2   4   6E+3   11   11.5  RT (min)      lcorbi_t3_neg_4_008_22    m/z 946.4927     0   1   2E+4   13.5   14  RT (min)      lcorbi_t3_neg_4_040_19    m/z 946.6066     0   1E+4   13   13.5  RT (min)      lcorbi_t3_neg_4_011_24    m/z 946.6545     0   1   2E+4   13.5   14  RT (min)      lcorbi_t3_neg_1_052_24    m/z 946.6764     0   1   2E+4   13.5   14  RT (min)      lcorbi_t3_neg_2_052_32    m/z 946.8112     0   1   2   3E+3   18   18.5  RT (min)      lcorbi_t3_neg_1_003_41    m/z 947.2017     0   1   2E+4   13.5   14  RT (min)      lcorbi_t3_neg_2_036_35    m/z 947.3456     0   1   2E+4   9   9.5  RT (min)      lcorbi_t3_neg_4_040_19    m/z 947.3523     0   1E+4   13   13.5  RT (min)      lcorbi_t3_neg_4_005_27    m/z 947.4061     0   1E+4   10.5   11  RT (min)      lcorbi_t3_neg_3_024_26    m/z 947.6106     0   1E+4   11.5   12  RT (min)      lcorbi_t3_neg_4_029_31    m/z 947.7424     0   1E+5   14   14.5  RT (min)      lcorbi_t3_neg_4_030_04    m/z 947.8633     0   1E+4   14.5   15  RT (min)      lcorbi_t3_neg_2_052_32    m/z 948.2551     0   2   4   6   8E+3   11.5   12  RT (min)      lcorbi_t3_neg_3_035_03    m/z 948.2569     0   1E+4   9   9.5  RT (min)      lcorbi_t3_neg_2_048_27    m/z 948.3556     0   1   2   3   4E+3   13   13.5  RT (min)      lcorbi_t3_neg_4_005_27    m/z 948.5170     0   1E+4   14.5   15  RT (min)      lcorbi_t3_neg_3_005_23    m/z 948.5581     0   1E+3   13.5   14  RT (min)      lcorbi_t3_neg_2_011_04    m/z 948.6139     0   1E+3   11.5   12  RT (min)      lcorbi_t3_neg_2_022_24    m/z 949.2774     0   2   4   6E+3   9.5   10  RT (min)      lcorbi_t3_neg_4_053_43    m/z 949.4657     0   2   4   6   8E+3   11   11.5  RT (min)      lcorbi_t3_neg_4_023_18    m/z 950.2808     0   1   2E+3   9.5   10  RT (min)      lcorbi_t3_neg_3_019_12    m/z 950.3004     0   1E+5   9   9.5  RT (min)      lcorbi_t3_neg_2_048_27    m/z 950.4323     0   2   4   6   8E+3   15   15.5  RT (min)      lcorbi_t3_neg_1_049_55    m/z 950.4913     0   1   13   13.5  RT (min)      lcorbi_t3_neg_1_034_04    m/z 950.5066     0   1   2E+4   14   14.5  RT (min)      lcorbi_t3_neg_4_048_03    m/z 950.6372     0   1E+4   13.5   14  RT (min)      lcorbi_t3_neg_1_052_24    m/z 950.6734     0   2   4   6   8E+3   6.5   7  RT (min)      lcorbi_t3_neg_4_029_31    m/z 950.6849     0   2   4   6E+3   14.5   15  RT (min)      lcorbi_t3_neg_4_005_27    m/z 951.4351     0   2   4E+3   15   15.5  RT (min)      lcorbi_t3_neg_1_049_55    m/z 951.4947     0   2   4   6   8E+2   13   13.5  RT (min)      lcorbi_t3_neg_1_034_04    m/z 951.5971     0   1   2E+4   10   10.5  RT (min)      lcorbi_t3_neg_3_019_12    m/z 951.6796     0   2   4   6   8E+3   14.5   15  RT (min)      lcorbi_t3_neg_3_007_16    m/z 951.6896     0   1E+4   13.5   14  RT (min)      lcorbi_t3_neg_4_013_09    m/z 951.7251     0   1E+4   15   15.5  RT (min)      lcorbi_t3_neg_4_034_01    m/z 951.7983     0   1   2   3   4E+3   18   18.5  RT (min)      lcorbi_t3_neg_3_046_10    m/z 952.5071     0   1E+4   14   14.5  RT (min)      lcorbi_t3_neg_4_048_03    m/z 952.5571     0   2   4   6   8E+3   13   13.5  RT (min)      lcorbi_t3_neg_3_052_32    m/z 952.5915     0   1   2E+4   13.5   14  RT (min)      lcorbi_t3_neg_4_005_27    m/z 952.6007     0   1E+4   10   10.5  RT (min)      lcorbi_t3_neg_3_019_12    m/z 952.6183     0   2   4   6   8E+3   14.5   15  RT (min)      lcorbi_t3_neg_3_032_17    m/z 952.6313     0   2   4   6   8E+3   13.5   14  RT (min)      lcorbi_t3_neg_4_019_08    m/z 952.6538     0   1E+4   13.5   14  RT (min)      lcorbi_t3_neg_3_002_24    m/z 952.7293     0   1   2   3E+3   15   15.5  RT (min)      lcorbi_t3_neg_4_034_01    m/z 952.7801     0   2   4E+4   14   14.5  RT (min)      lcorbi_t3_neg_3_023_07    m/z 952.8000     0   2   4E+3   18   18.5  RT (min)      lcorbi_t3_neg_2_003_42    m/z 953.5264     0   2   4   6   8E+3   13.5   14  RT (min)      lcorbi_t3_neg_4_007_13    m/z 953.5640     0   1   2   3   4E+4   11   11.5  RT (min)      lcorbi_t3_neg_1_055_05    m/z 953.5696     0   2   4   6   8E+3   13.5   14  RT (min)      lcorbi_t3_neg_2_022_24    m/z 953.5973     0   1   2E+4   12.5   13  RT (min)      lcorbi_t3_neg_4_007_13    m/z 953.6211     0   1   2E+4   14.5   15  RT (min)      lcorbi_t3_neg_2_023_39    m/z 954.3549     0   1   2E+4   9.5   10  RT (min)      lcorbi_t3_neg_2_048_27    m/z 954.6009     0   1E+4   12.5   13  RT (min)      lcorbi_t3_neg_4_007_13    m/z 954.6077     0   1   2E+4   14   14.5  RT (min)      lcorbi_t3_neg_4_005_27    m/z 954.6396     0   1   2E+3   14.5   15  RT (min)      lcorbi_t3_neg_2_004_28    m/z 954.6689     0   1   2E+4   14   14.5  RT (min)      lcorbi_t3_neg_3_005_23    m/z 954.7816     0   1   2   3   4E+4   14   14.5  RT (min)      lcorbi_t3_neg_3_023_07    m/z 955.3553     0   1E+4   10.5   11  RT (min)      lcorbi_t3_neg_4_038_15    m/z 955.5334     0   1E+4   13   13.5  RT (min)      lcorbi_t3_neg_4_042_14    m/z 955.5906     0   2   4   6E+2   14.5   15  RT (min)      lcorbi_t3_neg_1_055_05    m/z 955.6254     0   2   4   6E+3   18   18.5  RT (min)      lcorbi_t3_neg_1_053_32    m/z 955.6258     0   1   2E+4   13   13.5  RT (min)      lcorbi_t3_neg_4_039_32    m/z 955.7061     0   1   2   3E+4   13   13.5  RT (min)      lcorbi_t3_neg_4_039_32    m/z 955.7165     0   2   4   6   8E+3   15   15.5  RT (min)      lcorbi_t3_neg_4_032_23    m/z 956.2307     0   2   4   6E+4   13.5   14  RT (min)      lcorbi_t3_neg_4_045_35    m/z 956.3524     0   2   4E+4   9   9.5  RT (min)      lcorbi_t3_neg_2_048_27    m/z 956.3711     0   1E+4   9   9.5  RT (min)      lcorbi_t3_neg_2_048_27    m/z 956.4104     0   2   4   6E+3   13   13.5  RT (min)      lcorbi_t3_neg_3_020_31    m/z 956.4894     0   2   4E+2   11   11.5  RT (min)      lcorbi_t3_neg_4_032_23    m/z 956.5461     0   2   4   6   8E+3   13.5   14  RT (min)      lcorbi_t3_neg_3_004_30    m/z 956.5553     0   2   4   6   8E+3   14   14.5  RT (min)      lcorbi_t3_neg_3_004_30    m/z 956.5885     0   1E+4   13.5   14  RT (min)      lcorbi_t3_neg_2_019_07    m/z 956.6290     0   2   4   6   8E+4   13   13.5  RT (min)      lcorbi_t3_neg_4_039_32    m/z 956.6291     0   1   2E+4   18   18.5  RT (min)      lcorbi_t3_neg_1_053_32    m/z 956.8110     0   2   4   6E+3   18   18.5  RT (min)      lcorbi_t3_neg_4_013_09    m/z 956.8204     0   1E+4   14   14.5  RT (min)      lcorbi_t3_neg_2_022_24    m/z 957.3554     0   1   2E+4   9   9.5  RT (min)      lcorbi_t3_neg_2_048_27    m/z 957.4927     0   2   4   6   8E+3   11   11.5  RT (min)      lcorbi_t3_neg_4_008_22    m/z 957.5821     0   1E+4   13.5   14  RT (min)      lcorbi_t3_neg_4_046_37    m/z 957.6046     0   2   4   6E+3   14.5   15  RT (min)      lcorbi_t3_neg_2_013_08    m/z 957.6366     0   1   2   3   4E+3   14   14.5  RT (min)      lcorbi_t3_neg_3_052_32    m/z 957.6519     0   1E+3   13.5   14  RT (min)      lcorbi_t3_neg_4_029_31    m/z 957.7214     0   1   2E+3   15   15.5  RT (min)      lcorbi_t3_neg_2_053_23    m/z 957.7507     0   1E+4   14.5   15  RT (min)      lcorbi_t3_neg_4_034_01    m/z 957.8141     0   2   4   6E+3   18   18.5  RT (min)      lcorbi_t3_neg_2_011_04    m/z 958.4964     0   1   2   3   4E+4   13   13.5  RT (min)      lcorbi_t3_neg_4_046_37    m/z 958.5533     0   2   4   6   8E+3   14.5   15  RT (min)      lcorbi_t3_neg_2_026_03    m/z 958.5868     0   1   2E+3   13.5   14  RT (min)      lcorbi_t3_neg_1_053_32    m/z 958.6113     0   1   13   13.5  RT (min)      lcorbi_t3_neg_1_054_37    m/z 958.6389     0   2   4   6E+3   14   14.5  RT (min)      lcorbi_t3_neg_3_003_09    m/z 958.6443     0   1E+4   18   18.5  RT (min)      lcorbi_t3_neg_1_053_32    m/z 958.6445     0   2   4E+4   13.5   14  RT (min)      lcorbi_t3_neg_1_053_32    m/z 958.7395     0   1   2   3   4E+3   15   15.5  RT (min)      lcorbi_t3_neg_4_042_14    m/z 958.7543     0   2   4   6E+3   14.5   15  RT (min)      lcorbi_t3_neg_4_034_01    m/z 958.8137     0   1   2   3E+3   18   18.5  RT (min)      lcorbi_t3_neg_4_048_03    m/z 959.2896     0   1   2   3E+4   9   9.5  RT (min)      lcorbi_t3_neg_2_048_27    m/z 959.4651     0   1E+3   13.5   14  RT (min)      lcorbi_t3_neg_2_013_08    m/z 959.6247     0   1   13.5   14  RT (min)      lcorbi_t3_neg_1_029_21    m/z 959.7371     0   1   2E+4   15   15.5  RT (min)      lcorbi_t3_neg_4_042_14    m/z 959.7662     0   2   4   6   8E+3   15   15.5  RT (min)      lcorbi_t3_neg_4_034_01    m/z 959.8195     0   2   4E+3   18   18.5  RT (min)      lcorbi_t3_neg_2_023_39    m/z 960.2746     0   1E+4   9   9.5  RT (min)      lcorbi_t3_neg_2_048_27    m/z 960.4706     0   2   4   6   8E+4   7.5   8  RT (min)      lcorbi_t3_neg_2_048_27    m/z 960.6659     0   2   4E+4   12   12.5  RT (min)      lcorbi_t3_neg_3_024_26    m/z 960.7407     0   1E+4   15   15.5  RT (min)      lcorbi_t3_neg_4_042_14    m/z 960.7700     0   2   4E+3   15   15.5  RT (min)      lcorbi_t3_neg_4_034_01    m/z 960.7831     0   1   2   3   4E+3   18   18.5  RT (min)      lcorbi_t3_neg_4_015_10    m/z 961.2783     0   2   4   6   8E+3   9   9.5  RT (min)      lcorbi_t3_neg_2_048_27    m/z 961.3734     0   1   2   3   4E+3   10.5   11  RT (min)      lcorbi_t3_neg_4_050_02    m/z 961.6345     0   1E+4   14.5   15  RT (min)      lcorbi_t3_neg_4_007_13    m/z 961.6690     0   1   2E+4   12   12.5  RT (min)      lcorbi_t3_neg_4_010_28    m/z 961.7524     0   1E+4   15   15.5  RT (min)      lcorbi_t3_neg_4_029_31    m/z 962.2262     0   1   2E+3   11   11.5  RT (min)      lcorbi_t3_neg_2_026_03    m/z 962.2731     0   2   4   6   8E+3   9   9.5  RT (min)      lcorbi_t3_neg_2_048_27    m/z 962.3893     0   2   4   6   8E+3   10.5   11  RT (min)      lcorbi_t3_neg_3_010_15    m/z 962.4470     0   1E+4   6.5   7  RT (min)      lcorbi_t3_neg_4_042_14    m/z 962.5183     0   1   2E+3   13   13.5  RT (min)      lcorbi_t3_neg_4_019_08    m/z 962.5279     0   2   4E+4   13.5   14  RT (min)      lcorbi_t3_neg_4_044_38    m/z 962.6372     0   2   4E+4   13.5   14  RT (min)      lcorbi_t3_neg_4_054_36    m/z 962.6718     0   1E+4   13   13.5  RT (min)      lcorbi_t3_neg_1_052_24    m/z 962.7564     0   2   4   6E+3   15   15.5  RT (min)      lcorbi_t3_neg_4_029_31    m/z 962.7930     0   1   2   3   4E+3   18   18.5  RT (min)      lcorbi_t3_neg_2_044_06    m/z 962.8286     0   2   4   6   8E+3   18   18.5  RT (min)      lcorbi_t3_neg_2_019_07    m/z 962.8378     0   1E+4   14.5   15  RT (min)      lcorbi_t3_neg_4_039_32    m/z 963.3304     0   1   2   3E+3   18   18.5  RT (min)      lcorbi_t3_neg_2_027_13    m/z 963.4502     0   2   4   6E+3   6.5   7  RT (min)      lcorbi_t3_neg_4_042_14    m/z 963.5308     0   1   2   3E+4   13.5   14  RT (min)      lcorbi_t3_neg_4_030_04    m/z 963.6376     0   2   4E+4   13.5   14  RT (min)      lcorbi_t3_neg_4_054_36    m/z 963.7638     0   1   2   3E+3   15   15.5  RT (min)      lcorbi_t3_neg_4_029_31    m/z 963.8499     0   2   4E+3   15   15.5  RT (min)      lcorbi_t3_neg_4_039_32    m/z 963.8857     0   2   4   6E+3   14.5   15  RT (min)      lcorbi_t3_neg_4_039_32    m/z 964.2418     0   1   2   3E+3   11.5   12  RT (min)      lcorbi_t3_neg_2_014_29    m/z 964.5068     0   2   4   6   8E+3   13   13.5  RT (min)      lcorbi_t3_neg_4_046_37    m/z 964.5398     0   1E+4   14   14.5  RT (min)      lcorbi_t3_neg_4_030_04    m/z 964.5512     0   2   4   6E+4   14   14.5  RT (min)      lcorbi_t3_neg_2_011_04    m/z 964.6002     0   1E+3   13.5   14  RT (min)      lcorbi_t3_neg_2_007_14    m/z 964.6480     0   1E+4   13.5   14  RT (min)      lcorbi_t3_neg_1_020_36    m/z 964.6618     0   1E+4   14.5   15  RT (min)      lcorbi_t3_neg_4_007_13    m/z 964.7986     0   1   2   3   4E+3   18   18.5  RT (min)      lcorbi_t3_neg_1_045_44    m/z 964.8533     0   1E+4   15   15.5  RT (min)      lcorbi_t3_neg_4_039_32    m/z 965.3978     0   2   4   6E+3   13.5   14  RT (min)      lcorbi_t3_neg_1_044_23    m/z 965.5100     0   2   4   6E+3   13   13.5  RT (min)      lcorbi_t3_neg_4_046_37    m/z 965.5671     0   1E+4   13.5   14  RT (min)      lcorbi_t3_neg_1_052_24    m/z 965.6193     0   1E+4   14.5   15  RT (min)      lcorbi_t3_neg_3_035_03    m/z 965.6655     0   1E+4   14.5   15  RT (min)      lcorbi_t3_neg_4_007_13    m/z 965.7014     0   1   2E+4   13.5   14  RT (min)      lcorbi_t3_neg_4_011_24    m/z 965.8013     0   1   2   3   4E+3   18   18.5  RT (min)      lcorbi_t3_neg_2_021_41    m/z 966.2577     0   1   2   3   4E+3   12   12.5  RT (min)      lcorbi_t3_neg_2_010_02    m/z 966.2578     0   1E+4   9   9.5  RT (min)      lcorbi_t3_neg_4_040_19    m/z 966.3750     0   1   2   3E+4   9   9.5  RT (min)      lcorbi_t3_neg_2_048_27    m/z 966.5666     0   2   4   6   8E+4   14   14.5  RT (min)      lcorbi_t3_neg_3_011_04    m/z 966.6228     0   2   4   6E+3   14.5   15  RT (min)      lcorbi_t3_neg_4_048_03    m/z 966.6399     0   2   4   6   8E+3   12.5   13  RT (min)      lcorbi_t3_neg_1_055_05    m/z 966.6769     0   2   4   6   8E+3   15   15.5  RT (min)      lcorbi_t3_neg_4_034_01    m/z 966.7356     0   1   2   3E+3   18   18.5  RT (min)      lcorbi_t3_neg_2_007_14    m/z 966.8688     0   2   4   6E+3   15   15.5  RT (min)      lcorbi_t3_neg_4_039_32    m/z 967.5697     0   1   2   3   4E+4   14   14.5  RT (min)      lcorbi_t3_neg_3_011_04    m/z 967.6809     0   2   4E+3   15   15.5  RT (min)      lcorbi_t3_neg_4_007_13    m/z 967.7049     0   1   14.5   15  RT (min)      lcorbi_t3_neg_1_052_24    m/z 967.7370     0   1   15   15.5  RT (min)      lcorbi_t3_neg_4_007_13    m/z 968.4620     0   1   2   3E+4   13.5   14  RT (min)      lcorbi_t3_neg_4_054_36    m/z 968.4809     0   2   4   6   8E+3   13.5   14  RT (min)      lcorbi_t3_neg_3_005_23    m/z 968.5456     0   2   4   6   8E+2   13.5   14  RT (min)      lcorbi_t3_neg_4_054_36    m/z 968.5705     0   1   2E+4   14   14.5  RT (min)      lcorbi_t3_neg_3_011_04    m/z 968.6377     0   1E+4   14.5   15  RT (min)      lcorbi_t3_neg_3_010_15    m/z 968.6546     0   2   4E+4   13.5   14  RT (min)      lcorbi_t3_neg_2_052_32    m/z 968.6811     0   2   4   6E+4   14   14.5  RT (min)      lcorbi_t3_neg_2_053_23    m/z 968.7330     0   1   2E+3   18   18.5  RT (min)      lcorbi_t3_neg_2_037_09    m/z 968.8100     0   1E+4   18   18.5  RT (min)      lcorbi_t3_neg_1_004_22    m/z 969.6371     0   1E+4   14.5   15  RT (min)      lcorbi_t3_neg_1_047_02    m/z 969.6703     0   1   2E+3   14.5   15  RT (min)      lcorbi_t3_neg_2_052_32    m/z 969.6843     0   1   14   14.5  RT (min)      lcorbi_t3_neg_3_002_24    m/z 969.7526     0   2   4   6E+2   15   15.5  RT (min)      lcorbi_t3_neg_2_022_24    m/z 970.2594     0   1   2   3E+4   9   9.5  RT (min)      lcorbi_t3_neg_2_048_27    m/z 970.4447     0   2   4   6E+4   10.5   11  RT (min)      lcorbi_t3_neg_4_042_14    m/z 970.5503     0   1   13   13.5  RT (min)      lcorbi_t3_neg_4_019_08    m/z 970.5859     0   1E+4   14.5   15  RT (min)      lcorbi_t3_neg_1_047_02    m/z 970.6101     0   1   2   3E+4   14   14.5  RT (min)      lcorbi_t3_neg_3_002_24    m/z 970.6399     0   2   4   6   8E+3   14.5   15  RT (min)      lcorbi_t3_neg_1_047_02    m/z 970.7490     0   1   2E+4   14   14.5  RT (min)      lcorbi_t3_neg_2_022_24    m/z 970.8166     0   1   2   3   4E+3   18   18.5  RT (min)      lcorbi_t3_neg_2_030_47    m/z 971.2618     0   1   2E+4   9   9.5  RT (min)      lcorbi_t3_neg_2_048_27    m/z 971.4364     0   1   2E+4   7.5   8  RT (min)      lcorbi_t3_neg_4_040_19    m/z 971.4993     0   1   2   3E+3   13.5   14  RT (min)      lcorbi_t3_neg_4_044_38    m/z 971.5088     0   1   2   3E+3   13.5   14  RT (min)      lcorbi_t3_neg_4_046_37    m/z 971.5880     0   2   4   6   8E+3   14.5   15  RT (min)      lcorbi_t3_neg_1_047_02    m/z 971.6100     0   2   4   6E+3   13   13.5  RT (min)      lcorbi_t3_neg_4_039_32    m/z 972.2551     0   2   4   6E+3   11   11.5  RT (min)      lcorbi_t3_neg_3_029_02    m/z 972.4398     0   1E+4   7.5   8  RT (min)      lcorbi_t3_neg_4_040_19    m/z 972.5124     0   2   4   6   8E+2   13.5   14  RT (min)      lcorbi_t3_neg_1_013_35    m/z 972.5877     0   2   4   6E+3   13.5   14  RT (min)      lcorbi_t3_neg_4_007_13    m/z 972.6011     0   1   2E+3   14.5   15  RT (min)      lcorbi_t3_neg_2_022_24    m/z 972.6256     0   1E+4   14.5   15  RT (min)      lcorbi_t3_neg_2_022_24    m/z 972.6722     0   1   2   3E+4   13.5   14  RT (min)      lcorbi_t3_neg_1_052_24    m/z 972.7646     0   1E+4   14.5   15  RT (min)      lcorbi_t3_neg_4_011_24    m/z 972.8153     0   2   4   6   8E+3   18   18.5  RT (min)      lcorbi_t3_neg_2_052_32    m/z 973.2587     0   1   2   3E+3   11   11.5  RT (min)      lcorbi_t3_neg_4_050_02    m/z 973.3051     0   1   2E+4   9   9.5  RT (min)      lcorbi_t3_neg_2_048_27    m/z 973.6259     0   1E+4   13.5   14  RT (min)      lcorbi_t3_neg_2_052_32    m/z 973.6264     0   1E+4   11.5   12  RT (min)      lcorbi_t3_neg_4_007_13    m/z 973.6576     0   2   4   6   8E+3   14.5   15  RT (min)      lcorbi_t3_neg_1_051_15    m/z 973.6680     0   2   4   6E+3   14.5   15  RT (min)      lcorbi_t3_neg_1_051_15    m/z 973.8204     0   1   2E+4   14   14.5  RT (min)      lcorbi_t3_neg_3_002_24    m/z 974.2707     0   2   4   6   8E+3   11.5   12  RT (min)      lcorbi_t3_neg_4_037_29    m/z 974.2983     0   1   2   3   4E+3   8.5   9  RT (min)      lcorbi_t3_neg_4_040_19    m/z 974.5715     0   1E+4   13   13.5   14  RT (min)      lcorbi_t3_neg_4_039_32    m/z 974.6032     0   1E+4   13.5   14  RT (min)      lcorbi_t3_neg_4_019_08    m/z 974.6296     0   1E+3   11.5   12  RT (min)      lcorbi_t3_neg_2_020_05    m/z 974.6712     0   2   4   6E+2   14.5   15  RT (min)      lcorbi_t3_neg_4_019_08    m/z 974.8201     0   1E+4   14   14.5  RT (min)      lcorbi_t3_neg_3_002_24    m/z 975.2743     0   1   2   3E+3   11.5   12  RT (min)      lcorbi_t3_neg_2_014_29    m/z 975.4481     0   1   2E+4   7.5   8  RT (min)      lcorbi_t3_neg_2_048_27    m/z 975.4945     0   1   2E+4   13   13.5  RT (min)      lcorbi_t3_neg_4_046_37    m/z 975.5275     0   2   4E+3   13.5   14  RT (min)      lcorbi_t3_neg_3_018_13    m/z 975.5842     0   1   2   3E+4   13.5   14  RT (min)      lcorbi_t3_neg_1_052_24    m/z 975.6068     0   2   4   6E+3   13.5   14  RT (min)      lcorbi_t3_neg_2_050_17    m/z 975.8297     0   1   2   3   4E+3   18   18.5  RT (min)      lcorbi_t3_neg_1_003_41    m/z 976.2865     0   1E+4   12   12.5  RT (min)      lcorbi_t3_neg_3_029_02    m/z 976.5065     0   2   4   6   8E+3   18   18.5  RT (min)      lcorbi_t3_neg_1_020_36    m/z 976.5508     0   2   4   6   8E+3   11.5   12  RT (min)      lcorbi_t3_neg_1_055_05    m/z 976.5872     0   1   14.5   15  RT (min)      lcorbi_t3_neg_1_052_24    m/z 976.6255     0   1   14.5   15  RT (min)      lcorbi_t3_neg_4_019_08    m/z 976.6409     0   1E+5   11   11.5  RT (min)      lcorbi_t3_neg_1_055_05    m/z 976.6449     0   2   4   6   8E+3   13.5   14  RT (min)      lcorbi_t3_neg_3_002_24    m/z 976.7368     0   2   4E+3   15   15.5  RT (min)      lcorbi_t3_neg_2_007_14    m/z 976.7644     0   2   4   6   8E+3   18   18.5  RT (min)      lcorbi_t3_neg_2_029_12    m/z 977.2316     0   1   2E+4   13.5   14  RT (min)      lcorbi_t3_neg_2_036_35    m/z 977.2898     0   2   4E+3   12   12.5  RT (min)      lcorbi_t3_neg_3_029_02    m/z 977.5099     0   1   2   3   4E+3   18   18.5  RT (min)      lcorbi_t3_neg_1_013_35    m/z 977.5373     0   2   4   6E+3   13.5   14  RT (min)      lcorbi_t3_neg_3_029_02    m/z 977.5638     0   1   2E+4   11   11.5  RT (min)      lcorbi_t3_neg_4_007_13    m/z 977.5901     0   1   14.5   15  RT (min)      lcorbi_t3_neg_1_052_24    m/z 977.6048     0   2   4   6E+3   13.5   14  RT (min)      lcorbi_t3_neg_4_039_32    m/z 977.6291     0   1   2E+3   14.5   15  RT (min)      lcorbi_t3_neg_4_038_15    m/z 978.2317     0   1   2E+4   13.5   14  RT (min)      lcorbi_t3_neg_3_028_35    m/z 978.5218     0   1E+4   18   18.5  RT (min)      lcorbi_t3_neg_1_022_34    m/z 978.5672     0   1E+4   11   11.5  RT (min)      lcorbi_t3_neg_4_007_13    m/z 978.6108     0   1   2E+4   13   13.5  RT (min)      lcorbi_t3_neg_4_039_32    m/z 978.6355     0   1   14.5   15  RT (min)      lcorbi_t3_neg_4_019_08    m/z 978.7530     0   1   2E+4   15   15.5  RT (min)      lcorbi_t3_neg_2_007_14    m/z 978.7620     0   2   4   6E+3   18   18.5  RT (min)      lcorbi_t3_neg_2_029_12    m/z 978.7705     0   1   2   3   4E+3   18   18.5  RT (min)      lcorbi_t3_neg_4_051_40    m/z 979.1345     0   1   2E+3   9   9.5  RT (min)      lcorbi_t3_neg_3_019_12    m/z 979.4948     0   2   4   6E+2   13.5   14  RT (min)      lcorbi_t3_neg_2_053_23    m/z 979.5254     0   2   4E+3   18   18.5  RT (min)      lcorbi_t3_neg_1_021_31    m/z 979.5662     0   1E+4   13   13.5  RT (min)      lcorbi_t3_neg_4_044_38    m/z 979.5760     0   2   4   6E+3   13   13.5  RT (min)      lcorbi_t3_neg_2_052_32    m/z 979.7568     0   1E+4   15   15.5  RT (min)      lcorbi_t3_neg_2_007_14    m/z 979.7936     0   1   2   3E+3   18   18.5  RT (min)      lcorbi_t3_neg_4_006_47    m/z 980.5289     0   1   2   3E+4   13.5   14  RT (min)      lcorbi_t3_neg_4_046_37    m/z 980.5383     0   1   2   3E+3   14.5   15  RT (min)      lcorbi_t3_neg_2_024_37    m/z 980.6259     0   1   2   3E+3   13.5   14  RT (min)      lcorbi_t3_neg_4_011_24    m/z 980.6378     0   2   4   6E+3   14.5   15  RT (min)      lcorbi_t3_neg_3_053_14    m/z 980.6495     0   2   4   6   8E+3   14.5   15  RT (min)      lcorbi_t3_neg_4_007_13    m/z 980.6544     0   1   2   3E+4   13.5   14  RT (min)      lcorbi_t3_neg_4_019_08    m/z 980.7672     0   2   4   6   8E+3   15   15.5  RT (min)      lcorbi_t3_neg_2_007_14    m/z 980.7765     0   1   2   3   4E+3   18   18.5  RT (min)      lcorbi_t3_neg_2_015_21    m/z 981.3679     0   1E+4   8   8.5  RT (min)      lcorbi_t3_neg_4_040_19    m/z 981.4548     0   1   13.5   14  RT (min)      lcorbi_t3_neg_3_021_38    m/z 981.6062     0   2   4   6E+3   14.5   15  RT (min)      lcorbi_t3_neg_2_048_27    m/z 981.6502     0   2   4   6   8E+3   14.5   15  RT (min)      lcorbi_t3_neg_3_010_15    m/z 981.7174     0   1   2   3   4E+3   14.5   15  RT (min)      lcorbi_t3_neg_2_053_23    m/z 981.7720     0   1   2   3   4E+3   15   15.5  RT (min)      lcorbi_t3_neg_2_007_14    m/z 982.2883     0   1   2   3E+4   9   9.5  RT (min)      lcorbi_t3_neg_2_048_27    m/z 982.3714     0   2   4E+3   8   8.5  RT (min)      lcorbi_t3_neg_4_040_19    m/z 982.4261     0   1E+4   13.5   14  RT (min)      lcorbi_t3_neg_2_053_23    m/z 982.4938     0   2   4   6   8E+3   13   13.5  RT (min)      lcorbi_t3_neg_4_044_38    m/z 982.5608     0   1E+4   13.5   14  RT (min)      lcorbi_t3_neg_4_042_14    m/z 982.5865     0   1   2E+3   13   13.5  RT (min)      lcorbi_t3_neg_3_053_14    m/z 982.5956     0   1E+4   13.5   14  RT (min)      lcorbi_t3_neg_4_039_32    m/z 982.6532     0   1   2   3E+2   14.5   15  RT (min)      lcorbi_t3_neg_4_019_08    m/z 982.6704     0   1   2   3E+4   13.5   14  RT (min)      lcorbi_t3_neg_1_053_32    m/z 983.1459     0   1   2E+3   9   9.5  RT (min)      lcorbi_t3_neg_3_019_12    m/z 983.4295     0   2   4   6   8E+3   13.5   14  RT (min)      lcorbi_t3_neg_3_005_23    m/z 983.4439     0   1   2E+4   10.5   11  RT (min)      lcorbi_t3_neg_2_048_27    m/z 983.4714     0   1E+4   7.5   8  RT (min)      lcorbi_t3_neg_4_040_19    m/z 983.7661     0   1E+3   15   15.5  RT (min)      lcorbi_t3_neg_2_007_14    m/z 983.7872     0   2   4   6E+3   18   18.5  RT (min)      lcorbi_t3_neg_1_046_11    m/z 983.8129     0   2   4E+3   14   14.5  RT (min)      lcorbi_t3_neg_4_030_04    m/z 984.3439     0   1   2   3E+4   9   9.5  RT (min)      lcorbi_t3_neg_2_048_27    m/z 984.6511     0   1   2E+4   13   13.5  RT (min)      lcorbi_t3_neg_4_039_32    m/z 984.6863     0   1   2   3E+4   14   14.5  RT (min)      lcorbi_t3_neg_1_052_24    m/z 985.2780     0   2   4E+4   9   9.5  RT (min)      lcorbi_t3_neg_2_048_27    m/z 985.4746     0   1E+4   13.5   14  RT (min)      lcorbi_t3_neg_4_042_14    m/z 985.6211     0   2   4   6   8E+3   14.5   15  RT (min)      lcorbi_t3_neg_2_013_08    m/z 985.6366     0   1   2   3   4E+3   14.5   15  RT (min)      lcorbi_t3_neg_2_007_14    m/z 985.6671     0   1   2   3E+3   15   15.5  RT (min)      lcorbi_t3_neg_3_032_17    m/z 985.7907     0   1   2   3E+3   18   18.5  RT (min)      lcorbi_t3_neg_4_053_43    m/z 986.3333     0   1E+4   8.5   9  RT (min)      lcorbi_t3_neg_4_023_18    m/z 986.6175     0   1E+4   13.5   14  RT (min)      lcorbi_t3_neg_3_005_23    m/z 986.6283     0   2   4   6   8E+3   13.5   14  RT (min)      lcorbi_t3_neg_1_020_36    m/z 986.6462     0   2   4   6   8E+3   14.5   15  RT (min)      lcorbi_t3_neg_4_034_01    m/z 986.6670     0   2   4   6   8E+3   18   18.5  RT (min)      lcorbi_t3_neg_1_053_32    m/z 986.6670     0   1   2   3E+4   13.5   14  RT (min)      lcorbi_t3_neg_1_053_32    m/z 986.8160     0   1E+4   14.5   15  RT (min)      lcorbi_t3_neg_4_011_24    m/z 986.8316     0   1   2E+4   14   14.5  RT (min)      lcorbi_t3_neg_3_002_24    m/z 987.3401     0   1E+4   9   9.5  RT (min)      lcorbi_t3_neg_3_019_12    m/z 987.5388     0   1   2E+4   14   14.5  RT (min)      lcorbi_t3_neg_2_011_04    m/z 987.6503     0   1E+3   14.5   15  RT (min)      lcorbi_t3_neg_1_002_16    m/z 987.7676     0   2   4   6   8E+3   15   15.5  RT (min)      lcorbi_t3_neg_4_029_31    m/z 987.7960     0   2   4   6E+3   18   18.5  RT (min)      lcorbi_t3_neg_4_003_21    m/z 988.2984     0   1E+4   9   9.5  RT (min)      lcorbi_t3_neg_2_048_27    m/z 988.6504     0   1E+4   13.5   14  RT (min)      lcorbi_t3_neg_2_012_36    m/z 988.6769     0   1   2   3E+3   13   13.5  RT (min)      lcorbi_t3_neg_3_003_09    m/z 988.7720     0   2   4   6E+3   15   15.5  RT (min)      lcorbi_t3_neg_4_039_32    m/z 989.1632     0   2   4   6E+3   9   9.5  RT (min)      lcorbi_t3_neg_3_019_12    m/z 989.6545     0   1E+4   14.5   15  RT (min)      lcorbi_t3_neg_4_011_24    m/z 989.6754     0   1E+4   13.5   14  RT (min)      lcorbi_t3_neg_1_052_24    m/z 989.7776     0   1   2   3   4E+3   14.5   15  RT (min)      lcorbi_t3_neg_2_039_31    m/z 990.1665     0   1   2E+3   9   9.5  RT (min)      lcorbi_t3_neg_3_019_12    m/z 990.5226     0   1E+4   13.5   14  RT (min)      lcorbi_t3_neg_4_046_37    m/z 990.5662     0   1   2   3   4E+4   14   14.5  RT (min)      lcorbi_t3_neg_3_011_04    m/z 990.7878     0   1   2   3   4E+3   15   15.5  RT (min)      lcorbi_t3_neg_4_029_31    m/z 991.4777     0   2   4E+3   13   13.5  RT (min)      lcorbi_t3_neg_4_029_31    m/z 991.5103     0   1E+4   14   14.5  RT (min)      lcorbi_t3_neg_1_023_03    m/z 991.5695     0   1   2E+4   14   14.5  RT (min)      lcorbi_t3_neg_3_011_04    m/z 991.6535     0   1E+4   12   12.5  RT (min)      lcorbi_t3_neg_4_023_18    m/z 991.8078     0   2   4   6   8E+3   18   18.5  RT (min)      lcorbi_t3_neg_2_006_10    m/z 992.4874     0   2   4   6E+3   13.5   14  RT (min)      lcorbi_t3_neg_3_018_13    m/z 992.6570     0   2   4   6E+3   12   12.5  RT (min)      lcorbi_t3_neg_4_023_18    m/z 992.6863     0   1E+4   15   15.5  RT (min)      lcorbi_t3_neg_4_007_13    m/z 992.8073     0   1   2   3E+3   18   18.5  RT (min)      lcorbi_t3_neg_4_034_01    m/z 993.4928     0   1E+4   9.5   10  RT (min)      lcorbi_t3_neg_3_019_12    m/z 993.5854     0   1E+4   14.5   15  RT (min)      lcorbi_t3_neg_2_026_03    m/z 993.8127     0   1   2   3   4E+3   18   18.5  RT (min)      lcorbi_t3_neg_2_034_19    m/z 994.4967     0   1E+4   13.5   14  RT (min)      lcorbi_t3_neg_4_007_13    m/z 994.5378     0   1   2   3   4E+3   13   13.5  RT (min)      lcorbi_t3_neg_3_052_32    m/z 994.5880     0   1E+4   14.5   15  RT (min)      lcorbi_t3_neg_2_053_23    m/z 994.5968     0   2   4   6   8E+3   14.5   15  RT (min)      lcorbi_t3_neg_3_018_13    m/z 994.6381     0   1E+4   13   13.5  RT (min)      lcorbi_t3_neg_4_039_32    m/z 994.6599     0   1   2E+4   13.5   14  RT (min)      lcorbi_t3_neg_4_005_27    m/z 995.5070     0   1E+4   13.5   14  RT (min)      lcorbi_t3_neg_2_027_13    m/z 995.6012     0   1   2   3   4E+3   14.5   15  RT (min)      lcorbi_t3_neg_3_029_02    m/z 995.6568     0   1E+4   14.5   15  RT (min)      lcorbi_t3_neg_2_054_15    m/z 995.6654     0   1E+4   14.5   15  RT (min)      lcorbi_t3_neg_4_034_01    m/z 995.7685     0   1   2   3   4E+3   15   15.5  RT (min)      lcorbi_t3_neg_2_032_45    m/z 996.2552     0   1   2   3   4E+3   11   11.5  RT (min)      lcorbi_t3_neg_4_048_03    m/z 996.3034     0   1E+4   9   9.5  RT (min)      lcorbi_t3_neg_2_048_27    m/z 996.4733     0   1   2   3   4E+3   13   13.5  RT (min)      lcorbi_t3_neg_2_053_23    m/z 996.5109     0   1   2   3   4E+3   13.5   14  RT (min)      lcorbi_t3_neg_3_018_13    m/z 996.5793     0   1E+4   13.5   14  RT (min)      lcorbi_t3_neg_4_005_27    m/z 996.6025     0   1   2E+4   14   14.5  RT (min)      lcorbi_t3_neg_2_053_23    m/z 996.6535     0   2   4   6   8E+3   13.5   14  RT (min)      lcorbi_t3_neg_2_013_08    m/z 996.6691     0   2   4   6   8E+3   14.5   15  RT (min)      lcorbi_t3_neg_4_034_01    m/z 996.7595     0   1   2E+3   18   18.5  RT (min)      lcorbi_t3_neg_4_053_43    m/z 996.8149     0   1E+4   14.5   15  RT (min)      lcorbi_t3_neg_2_022_24    m/z 997.3461     0   2   4E+3   13   13.5  RT (min)      lcorbi_t3_neg_4_012_17    m/z 997.3643     0   2   4   6   8E+3   10.5   11  RT (min)      lcorbi_t3_neg_4_039_32    m/z 997.6185     0   1   2   3   4E+3   13.5   14  RT (min)      lcorbi_t3_neg_2_022_24    m/z 997.8270     0   2   4   6   8E+3   14.5   15  RT (min)      lcorbi_t3_neg_2_052_32    m/z 998.2908     0   2   4E+4   9   9.5  RT (min)      lcorbi_t3_neg_2_048_27    m/z 998.3497     0   1   2   3   4E+3   13   13.5  RT (min)      lcorbi_t3_neg_4_005_27    m/z 998.4597     0   1   2   3   4E+3   13   13.5  RT (min)      lcorbi_t3_neg_4_046_37    m/z 998.5272     0   2   4   6   8E+3   13.5   14  RT (min)      lcorbi_t3_neg_1_047_02    m/z 998.6205     0   1   2   3   4E+4   11   11.5  RT (min)      lcorbi_t3_neg_1_055_05    m/z 998.6297     0   2   4   6   8E+2   11.5   12  RT (min)      lcorbi_t3_neg_2_022_24    m/z 998.8267     0   2   4E+3   18   18.5  RT (min)      lcorbi_t3_neg_4_027_12    m/z 998.8304     0   1   2E+4   14.5   15  RT (min)      lcorbi_t3_neg_2_052_32    m/z 999.2572     0   1   2   3E+4   9   9.5  RT (min)      lcorbi_t3_neg_2_048_27    m/z 999.2935     0   1   2   3E+4   9   9.5  RT (min)      lcorbi_t3_neg_2_048_27    m/z 999.4923     0   1   2E+4   13   13.5  RT (min)      lcorbi_t3_neg_4_046_37    m/z 999.5457     0   1E+4   11   11.5  RT (min)      lcorbi_t3_neg_4_007_13    m/z 999.6420     0   1   2E+4   11.5   12  RT (min)      lcorbi_t3_neg_4_034_01    m/z 1000.2576     0   1   2E+4   9   9.5  RT (min)      lcorbi_t3_neg_2_048_27    m/z 1000.3492     0   1   2   3   4E+4   9   9.5  RT (min)      lcorbi_t3_neg_4_040_19    m/z 1000.4753     0   1   2   3E+3   13.5   14  RT (min)      lcorbi_t3_neg_4_046_37    m/z 1000.5047     0   1E+4   13.5   14  RT (min)      lcorbi_t3_neg_4_054_36    m/z 1000.5492     0   2   4   6E+3   11   11.5  RT (min)      lcorbi_t3_neg_4_007_13    m/z 1000.5933     0   2   4   6   8E+3   13.5   14  RT (min)      lcorbi_t3_neg_3_052_32    m/z 1000.6453     0   1E+4   11.5   12  RT (min)      lcorbi_t3_neg_4_034_01    m/z 1000.6650     0   1   2E+4   14.5   15  RT (min)      lcorbi_t3_neg_1_043_14    m/z 1000.7615     0   1   2   3E+3   18   18.5  RT (min)      lcorbi_t3_neg_4_020_44    m/z 1000.8469     0   1   2E+4   14.5   15  RT (min)      lcorbi_t3_neg_2_052_32    m/z 1001.5896     0   2   4   6E+3   14.5   15  RT (min)      lcorbi_t3_neg_4_042_14    m/z 1002.4901     0   1   2   3E+3   13.5   14  RT (min)      lcorbi_t3_neg_2_010_02    m/z 1002.5227     0   1E+4   13.5   14  RT (min)      lcorbi_t3_neg_4_046_37    m/z 1002.5301     0   2   4   6   8E+3   13   13.5  RT (min)      lcorbi_t3_neg_4_042_14    m/z 1002.5999     0   2   4   6   8E+3   14.5   15  RT (min)      lcorbi_t3_neg_1_043_14    m/z 1002.7531     0   1   2E+4   15   15.5  RT (min)      lcorbi_t3_neg_4_042_14    m/z 1002.7691     0   2   4E+3   18   18.5  RT (min)      lcorbi_t3_neg_2_014_29    m/z 1003.3582     0   2   4   6   8E+3   10.5   11  RT (min)      lcorbi_t3_neg_2_010_02    m/z 1003.5262     0   1E+4   13.5   14  RT (min)      lcorbi_t3_neg_4_046_37    m/z 1003.5706     0   2   4   6   8E+3   11   11.5  RT (min)      lcorbi_t3_neg_4_034_01    m/z 1003.6041     0   1E+4   14.5   15  RT (min)      lcorbi_t3_neg_4_037_29    m/z 1003.6645     0   1   2E+4   13.5   14  RT (min)      lcorbi_t3_neg_2_013_08    m/z 1003.6733     0   1E+4   12   12.5  RT (min)      lcorbi_t3_neg_4_034_01    m/z 1003.6891     0   2   4   6E+3   12   12.5  RT (min)      lcorbi_t3_neg_4_023_18    m/z 1003.7564     0   1E+4   15   15.5  RT (min)      lcorbi_t3_neg_4_042_14    m/z 1003.8298     0   1   2E+4   14   14.5  RT (min)      lcorbi_t3_neg_3_002_24    m/z 1003.8584     0   1E+4   14.5   15  RT (min)      lcorbi_t3_neg_2_052_32    m/z 1003.9744     0   1E+4   6.5   7  RT (min)      lcorbi_t3_neg_2_048_27    m/z 1004.3615     0   2   4   6E+3   10.5   11  RT (min)      lcorbi_t3_neg_2_010_02    m/z 1004.4759     0   1   2E+4   6.5   7  RT (min)      lcorbi_t3_neg_2_048_27    m/z 1004.5382     0   1   2   3   4E+4   13.5   14  RT (min)      lcorbi_t3_neg_4_046_37    m/z 1004.6162     0   1E+4   14.5   15  RT (min)      lcorbi_t3_neg_4_038_15    m/z 1004.6196     0   1   2   3   4E+3   13   13.5  RT (min)      lcorbi_t3_neg_2_043_33    m/z 1004.6446     0   2   4   6   8E+4   13.5   14  RT (min)      lcorbi_t3_neg_4_019_08    m/z 1004.6766     0   1E+3   12   12.5  RT (min)      lcorbi_t3_neg_2_022_24    m/z 1004.9773     0   1E+4   6.5   7  RT (min)      lcorbi_t3_neg_2_048_27    m/z 1005.4996     0   1   13.5   14  RT (min)      lcorbi_t3_neg_4_054_36    m/z 1005.6069     0   2   4   6   8E+3   14.5   15  RT (min)      lcorbi_t3_neg_1_043_14    m/z 1005.6629     0   1   2E+4   13.5   14  RT (min)      lcorbi_t3_neg_4_039_32    m/z 1005.7165     0   1E+4   14.5   15  RT (min)      lcorbi_t3_neg_4_029_31    m/z 1006.4260     0   2   4   6E+3   13   13.5  RT (min)      lcorbi_t3_neg_2_012_36    m/z 1006.6096     0   2   4   6   8E+3   14.5   15  RT (min)      lcorbi_t3_neg_1_043_14    m/z 1006.7198     0   1E+4   14.5   15  RT (min)      lcorbi_t3_neg_4_029_31    m/z 1006.7787     0   2   4   6   8E+3   18   18.5  RT (min)      lcorbi_t3_neg_4_014_20    m/z 1007.3696     0   2   4   6E+3   10.5   11  RT (min)      lcorbi_t3_neg_2_002_26    m/z 1007.4296     0   1   2   3E+3   13   13.5  RT (min)      lcorbi_t3_neg_2_053_23    m/z 1007.4705     0   1   2   3E+3   13.5   14  RT (min)      lcorbi_t3_neg_2_027_13    m/z 1007.7879     0   1   2E+4   15   15.5  RT (min)      lcorbi_t3_neg_2_007_14    m/z 1007.8715     0   2   4E+3   15   15.5  RT (min)      lcorbi_t3_neg_2_035_46    m/z 1008.6061     0   2   4   6E+3   13   13.5  RT (min)      lcorbi_t3_neg_4_039_32    m/z 1008.6825     0   2   4   6E+4   13.5   14  RT (min)      lcorbi_t3_neg_4_039_32    m/z 1008.7782     0   2   4E+3   18   18.5  RT (min)      lcorbi_t3_neg_3_036_41    m/z 1008.7971     0   2   4   6   8E+3   15   15.5  RT (min)      lcorbi_t3_neg_2_007_14    m/z 1009.2076     0   1   2   3   4E+3   13   13.5  RT (min)      lcorbi_t3_neg_4_017_51    m/z 1009.3273     0   1   2   3E+4   9   9.5  RT (min)      lcorbi_t3_neg_2_048_27    m/z 1009.5557     0   1   2   3   4E+3   10   10.5  RT (min)      lcorbi_t3_neg_3_019_12    m/z 1009.6184     0   1E+4   13   13.5  RT (min)      lcorbi_t3_neg_3_052_32    m/z 1009.8872     0   1   2   3   4E+3   15   15.5  RT (min)      lcorbi_t3_neg_2_032_45    m/z 1010.7206     0   1E+4   14.5   15  RT (min)      lcorbi_t3_neg_3_048_08    m/z 1011.7629     0   1E+4   14.5   15  RT (min)      lcorbi_t3_neg_4_046_37    m/z 1012.2698     0   1   2   3E+4   9   9.5  RT (min)      lcorbi_t3_neg_2_048_27    m/z 1012.4752     0   2   4   6E+3   13.5   14  RT (min)      lcorbi_t3_neg_4_004_34    m/z 1012.6618     0   2   4E+3   14.5   15  RT (min)      lcorbi_t3_neg_4_007_13    m/z 1012.7700     0   2   4   6   8E+3   14.5   15  RT (min)      lcorbi_t3_neg_4_054_36    m/z 1013.1652     0   1   2E+4   9   9.5  RT (min)      lcorbi_t3_neg_2_048_27    m/z 1013.2730     0   1   2E+4   9   9.5  RT (min)      lcorbi_t3_neg_2_048_27    m/z 1013.4942     0   2   4   6   8E+3   14   14.5  RT (min)      lcorbi_t3_neg_3_011_04    m/z 1013.7751     0   2   4E+3   14.5   15  RT (min)      lcorbi_t3_neg_4_054_36    m/z 1013.8009     0   1   2   3E+3   18   18.5  RT (min)      lcorbi_t3_neg_3_008_20    m/z 1013.8110     0   1   2   3   4E+3   18   18.5  RT (min)      lcorbi_t3_neg_4_050_02    m/z 1014.1685     0   1E+4   9   9.5  RT (min)      lcorbi_t3_neg_2_048_27    m/z 1014.6288     0   1   2E+4   13.5   14  RT (min)      lcorbi_t3_neg_4_005_27    m/z 1014.6782     0   2   4   6E+3   14.5   15  RT (min)      lcorbi_t3_neg_4_007_13    m/z 1014.7102     0   2   4   6   8E+3   14   14.5  RT (min)      lcorbi_t3_neg_4_034_01    m/z 1014.7888     0   2   4   6E+3   14.5   15  RT (min)      lcorbi_t3_neg_4_039_32    m/z 1015.1631     0   1   2E+4   9   9.5  RT (min)      lcorbi_t3_neg_2_048_27    m/z 1015.3936     0   1   2   3   4E+3   10.5   11  RT (min)      lcorbi_t3_neg_4_050_02    m/z 1015.4470     0   1E+4   11   11.5  RT (min)      lcorbi_t3_neg_4_023_18    m/z 1015.5984     0   2   4   6E+3   11.5   12  RT (min)      lcorbi_t3_neg_4_029_31    m/z 1015.6816     0   1   2   3   4E+3   14.5   15  RT (min)      lcorbi_t3_neg_4_007_13    m/z 1015.7929     0   1   2   3   4E+3   14.5   15   15.5  RT (min)      lcorbi_t3_neg_4_039_32    m/z 1016.1658     0   1E+4   9   9.5  RT (min)      lcorbi_t3_neg_2_048_27    m/z 1016.2429     0   2   4E+3   11.5   12  RT (min)      lcorbi_t3_neg_3_011_04    m/z 1016.4504     0   2   4   6   8E+3   11   11.5  RT (min)      lcorbi_t3_neg_4_023_18    m/z 1016.5806     0   2   4   6E+3   14.5   15  RT (min)      lcorbi_t3_neg_3_011_04    m/z 1016.6301     0   1   2E+4   13.5   14  RT (min)      lcorbi_t3_neg_4_039_32    m/z 1016.6935     0   2   4   6   8E+3   15   15.5  RT (min)      lcorbi_t3_neg_4_007_13    m/z 1016.7255     0   1E+4   14.5   15  RT (min)      lcorbi_t3_neg_4_007_13    m/z 1017.2649     0   1   2   3   4E+3   9.5   10  RT (min)      lcorbi_t3_neg_4_053_43    m/z 1017.3164     0   1   2E+4   9   9.5  RT (min)      lcorbi_t3_neg_2_048_27    m/z 1017.5262     0   1E+4   14   14.5  RT (min)      lcorbi_t3_neg_2_010_02    m/z 1017.6479     0   1   2   3E+4   13.5   14  RT (min)      lcorbi_t3_neg_4_005_27    m/z 1017.6972     0   2   4E+3   15   15.5  RT (min)      lcorbi_t3_neg_4_007_13    m/z 1017.7289     0   1E+4   14.5   15  RT (min)      lcorbi_t3_neg_4_007_13    m/z 1017.8455     0   1   2E+4   14   14.5  RT (min)      lcorbi_t3_neg_4_055_06    m/z 1017.8727     0   1E+4   14.5   15  RT (min)      lcorbi_t3_neg_2_052_32    m/z 1018.5027     0   1E+4   13.5   14  RT (min)      lcorbi_t3_neg_2_027_13    m/z 1018.5969     0   2   4E+3   14.5   15  RT (min)      lcorbi_t3_neg_3_010_15    m/z 1018.6603     0   1   2   3   4E+4   13.5   14  RT (min)      lcorbi_t3_neg_4_005_27    m/z 1018.6880     0   1   2   3   4E+4   11   11.5  RT (min)      lcorbi_t3_neg_4_039_32    m/z 1018.7092     0   1   2   3   4E+3   15   15.5  RT (min)      lcorbi_t3_neg_4_007_13    m/z 1019.2217     0   1   2E+4   13.5   14  RT (min)      lcorbi_t3_neg_2_036_35    m/z 1019.5087     0   1E+4   13.5   14  RT (min)      lcorbi_t3_neg_2_027_13    m/z 1019.5410     0   2   4   6E+3   14.5   15  RT (min)      lcorbi_t3_neg_2_008_16    m/z 1019.5845     0   1E+4   10   10.5  RT (min)      lcorbi_t3_neg_3_019_12    m/z 1019.6001     0   1   2   3   4E+3   14.5   15  RT (min)      lcorbi_t3_neg_2_014_29    m/z 1019.7228     0   1   2E+4   13.5   14  RT (min)      lcorbi_t3_neg_4_023_18    m/z 1020.2222     0   1E+4   13.5   14  RT (min)      lcorbi_t3_neg_4_045_35    m/z 1020.5122     0   2   4   6E+3   13.5   14  RT (min)      lcorbi_t3_neg_3_018_13    m/z 1020.5437     0   2   4   6E+3   14.5   15  RT (min)      lcorbi_t3_neg_2_008_16    m/z 1020.5882     0   2   4   6E+3   10   10.5  RT (min)      lcorbi_t3_neg_3_019_12    m/z 1020.6456     0   1   2   3E+4   13   13.5  RT (min)      lcorbi_t3_neg_4_039_32    m/z 1020.6500     0   1   2   3E+4   13   13.5  RT (min)      lcorbi_t3_neg_2_052_32    m/z 1021.2207     0   1E+4   13.5   14  RT (min)      lcorbi_t3_neg_4_045_35    m/z 1021.5852     0   2   4   6   8E+3   12.5   13  RT (min)      lcorbi_t3_neg_3_007_16    m/z 1022.5175     0   2   4   6   8E+3   13   13.5  RT (min)      lcorbi_t3_neg_2_007_14    m/z 1022.5605     0   2   4   6   8E+3   13.5   14  RT (min)      lcorbi_t3_neg_2_022_24    m/z 1022.6207     0   1   2   3E+4   11   11.5  RT (min)      lcorbi_t3_neg_4_007_13    m/z 1022.6299     0   1   2   3   4E+4   11.5   12  RT (min)      lcorbi_t3_neg_4_034_01    m/z 1023.1939     0   1E+4   9   9.5  RT (min)      lcorbi_t3_neg_2_048_27    m/z 1023.2233     0   1   2   3   4E+3   13   13.5  RT (min)      lcorbi_t3_neg_4_017_51    m/z 1023.3432     0   2   4   6E+3   10.5   11  RT (min)      lcorbi_t3_neg_3_037_36    m/z 1023.5209     0   1   2   3   4E+3   13   13.5  RT (min)      lcorbi_t3_neg_2_007_14    m/z 1023.5316     0   2   4   6E+3   13.5   14  RT (min)      lcorbi_t3_neg_3_029_02    m/z 1023.6422     0   1   2E+4   11.5   12  RT (min)      lcorbi_t3_neg_4_007_13    m/z 1023.6738     0   2   4E+4   13.5   14  RT (min)      lcorbi_t3_neg_2_052_32    m/z 1024.5493     0   2   4   6   8E+3   11   11.5  RT (min)      lcorbi_t3_neg_4_034_01    m/z 1025.3404     0   2   4E+4   9   9.5  RT (min)      lcorbi_t3_neg_2_048_27    m/z 1025.8017     0   2   4E+3   18   18.5  RT (min)      lcorbi_t3_neg_4_051_40    m/z 1026.2749     0   1   2E+4   9   9.5  RT (min)      lcorbi_t3_neg_2_048_27    m/z 1026.4776     0   1E+4   13.5   14  RT (min)      lcorbi_t3_neg_4_019_08    m/z 1026.5671     0   1E+4   11   11.5  RT (min)      lcorbi_t3_neg_4_034_01    m/z 1026.6800     0   1   2   3E+4   14.5   15  RT (min)      lcorbi_t3_neg_1_043_14    m/z 1026.7533     0   2   4   6   8E+3   15   15.5  RT (min)      lcorbi_t3_neg_2_007_14    m/z 1026.8528     0   1E+4   14.5   15  RT (min)      lcorbi_t3_neg_2_052_32    m/z 1027.1811     0   1E+4   9   9.5  RT (min)      lcorbi_t3_neg_2_048_27    m/z 1027.2772     0   2   4   6E+3   9   9.5  RT (min)      lcorbi_t3_neg_2_048_27    m/z 1027.7566     0   2   4E+3   15   15.5  RT (min)      lcorbi_t3_neg_4_042_14    m/z 1028.2863     0   1   2E+4   9   9.5  RT (min)      lcorbi_t3_neg_2_048_27    m/z 1028.6185     0   1   2   3E+3   14.5   15  RT (min)      lcorbi_t3_neg_3_004_30    m/z 1028.6364     0   1   11   11.5  RT (min)      lcorbi_t3_neg_2_022_24    m/z 1028.6481     0   1E+4   13   13.5  RT (min)      lcorbi_t3_neg_4_039_32    m/z 1028.6960     0   1E+4   14.5   15  RT (min)      lcorbi_t3_neg_1_043_14    m/z 1028.7706     0   1   2E+3   18   18.5  RT (min)      lcorbi_t3_neg_4_028_39    m/z 1028.8685     0   1E+4   14.5   15  RT (min)      lcorbi_t3_neg_2_052_32    m/z 1029.1790     0   1E+4   9   9.5  RT (min)      lcorbi_t3_neg_2_048_27    m/z 1029.3292     0   2   4   6E+4   9   9.5  RT (min)      lcorbi_t3_neg_2_048_27    m/z 1029.7723     0   1E+4   15   15.5  RT (min)      lcorbi_t3_neg_4_042_14    m/z 1030.3169     0   2   4   6E+3   8.5   9  RT (min)      lcorbi_t3_neg_4_010_28    m/z 1030.3293     0   1   2   3   4E+4   9   9.5  RT (min)      lcorbi_t3_neg_2_048_27    m/z 1030.3697     0   1E+4   9   9.5  RT (min)      lcorbi_t3_neg_4_023_18    m/z 1030.5226     0   1E+4   14   14.5  RT (min)      lcorbi_t3_neg_3_011_04    m/z 1030.5545     0   1E+4   12   12.5  RT (min)      lcorbi_t3_neg_4_040_19    m/z 1030.6341     0   2   4   6E+3   14.5   15  RT (min)      lcorbi_t3_neg_2_027_13    m/z 1030.7805     0   1   2   3E+3   18   18.5  RT (min)      lcorbi_t3_neg_1_004_22    m/z 1030.8162     0   2   4   6E+3   18   18.5  RT (min)      lcorbi_t3_neg_2_037_09    m/z 1031.3321     0   1E+4   9   9.5  RT (min)      lcorbi_t3_neg_2_048_27    m/z 1031.6373     0   1   2   3   4E+3   14.5   15  RT (min)      lcorbi_t3_neg_4_007_13    m/z 1032.5467     0   1E+4   13.5   14  RT (min)      lcorbi_t3_neg_4_040_19    m/z 1032.6498     0   1   2   3E+3   14.5   15  RT (min)      lcorbi_t3_neg_4_007_13    m/z 1032.7640     0   1E+4   13.5   14  RT (min)      lcorbi_t3_neg_2_052_32    m/z 1032.8917     0   1E+4   14.5   15  RT (min)      lcorbi_t3_neg_2_052_32    m/z 1033.8037     0   1   2E+4   15   15.5  RT (min)      lcorbi_t3_neg_2_007_14    m/z 1034.6130     0   1E+4   13   13.5  RT (min)      lcorbi_t3_neg_4_005_27    m/z 1034.7792     0   1E+4   13.5   14  RT (min)      lcorbi_t3_neg_3_002_24    m/z 1034.8108     0   2   4   6E+3   15   15.5  RT (min)      lcorbi_t3_neg_1_043_14    m/z 1035.4655     0   2   4E+3   13   13.5  RT (min)      lcorbi_t3_neg_2_053_23    m/z 1035.6245     0   1E+4   13   13.5  RT (min)      lcorbi_t3_neg_4_005_27    m/z 1035.9029     0   1   2   3   4E+3   15   15.5  RT (min)      lcorbi_t3_neg_2_035_46    m/z 1036.5692     0   2   4   6E+3   14.5   15  RT (min)      lcorbi_t3_neg_4_030_04    m/z 1036.5977     0   2   4   6E+3   13.5   14  RT (min)      lcorbi_t3_neg_4_044_38    m/z 1036.6685     0   1   2   3E+3   14.5   15  RT (min)      lcorbi_t3_neg_4_007_13    m/z 1036.6754     0   2   4   6E+3   12   12.5  RT (min)      lcorbi_t3_neg_2_002_26    m/z 1036.7975     0   2   4E+3   18   18.5  RT (min)      lcorbi_t3_neg_4_051_40    m/z 1037.3126     0   1E+4   9   9.5  RT (min)      lcorbi_t3_neg_2_048_27    m/z 1037.4802     0   2   4   6E+3   13.5   14  RT (min)      lcorbi_t3_neg_3_007_16    m/z 1037.6412     0   2   4   6   8E+3   13.5   14  RT (min)      lcorbi_t3_neg_4_011_24    m/z 1038.4835     0   1   2   3E+3   13.5   14  RT (min)      lcorbi_t3_neg_3_018_13    m/z 1038.7152     0   2   4E+3   13.5   14  RT (min)      lcorbi_t3_neg_2_035_46    m/z 1038.9169     0   1E+4   14.5   15  RT (min)      lcorbi_t3_neg_4_039_32    m/z 1039.2240     0   2   4   6E+3   13   13.5  RT (min)      lcorbi_t3_neg_1_023_03    m/z 1039.2941     0   1   2E+4   9   9.5  RT (min)      lcorbi_t3_neg_2_048_27    m/z 1039.6400     0   2   4   6E+3   14.5   15  RT (min)      lcorbi_t3_neg_1_051_15    m/z 1039.9204     0   1   2E+4   14.5   15  RT (min)      lcorbi_t3_neg_4_039_32    m/z 1040.2428     0   1   2   3E+3   11   11.5  RT (min)      lcorbi_t3_neg_3_029_02    m/z 1040.2909     0   1E+4   9   9.5  RT (min)      lcorbi_t3_neg_2_048_27    m/z 1040.6010     0   2   4   6   8E+3   13   13.5  RT (min)      lcorbi_t3_neg_4_039_32    m/z 1040.6433     0   1   2   3E+3   14.5   15  RT (min)      lcorbi_t3_neg_4_005_27    m/z 1040.9324     0   1E+4   14.5   15  RT (min)      lcorbi_t3_neg_1_053_32    m/z 1041.2924     0   1E+4   9   9.5  RT (min)      lcorbi_t3_neg_2_048_27    m/z 1041.6139     0   1E+4   11.5   12  RT (min)      lcorbi_t3_neg_1_055_05    m/z 1042.2582     0   1   2   3E+3   11.5   12  RT (min)      lcorbi_t3_neg_3_022_27    m/z 1042.3014     0   1E+4   9   9.5  RT (min)      lcorbi_t3_neg_2_048_27    m/z 1042.6169     0   2   4E+3   13.5   14  RT (min)      lcorbi_t3_neg_3_048_08    m/z 1042.8063     0   2   4   6E+3   15   15.5  RT (min)      lcorbi_t3_neg_4_042_14    m/z 1042.8147     0   2   4E+3   18   18.5  RT (min)      lcorbi_t3_neg_3_015_11    m/z 1043.2927     0   1E+4   9   9.5  RT (min)      lcorbi_t3_neg_3_019_12    m/z 1043.5404     0   1   2   3   4E+3   14.5   15  RT (min)      lcorbi_t3_neg_1_047_02    m/z 1043.6398     0   1E+4   13   13.5  RT (min)      lcorbi_t3_neg_4_039_32    m/z 1043.8097     0   1   2   3   4E+3   15   15.5  RT (min)      lcorbi_t3_neg_2_007_14    m/z 1044.2739     0   2   4E+3   12   12.5  RT (min)      lcorbi_t3_neg_3_029_02    m/z 1044.2957     0   2   4   6E+3   9   9.5  RT (min)      lcorbi_t3_neg_3_019_12    m/z 1044.3328     0   2   4   6E+3   9   9.5  RT (min)      lcorbi_t3_neg_2_048_27    m/z 1044.4811     0   1   2E+4   8   8.5  RT (min)      lcorbi_t3_neg_4_023_18    m/z 1044.6087     0   1E+4   14.5   15  RT (min)      lcorbi_t3_neg_3_032_17    m/z 1044.6431     0   1   2   3   4E+4   13   13.5  RT (min)      lcorbi_t3_neg_4_039_32    m/z 1044.7519     0   2   4E+3   18   18.5  RT (min)      lcorbi_t3_neg_2_055_18    m/z 1045.3415     0   2   4   6E+3   13   13.5  RT (min)      lcorbi_t3_neg_1_054_37    m/z 1045.6117     0   2   4   6   8E+3   14.5   15  RT (min)      lcorbi_t3_neg_4_007_13    m/z 1045.6175     0   1E+4   11   11.5  RT (min)      lcorbi_t3_neg_4_034_01    m/z 1045.6476     0   1E+4   11   11.5  RT (min)      lcorbi_t3_neg_4_007_13    m/z 1046.4789     0   2   4E+3   13.5   14  RT (min)      lcorbi_t3_neg_3_018_13    m/z 1046.5174     0   2   4   6E+3   13   13.5  RT (min)      lcorbi_t3_neg_3_053_14    m/z 1046.6221     0   1   2   3   4E+3   14.5   15  RT (min)      lcorbi_t3_neg_2_005_01    m/z 1046.7497     0   1   2   3   4E+3   18   18.5  RT (min)      lcorbi_t3_neg_2_002_26    m/z 1047.1220     0   1   2E+3   9   9.5  RT (min)      lcorbi_t3_neg_4_048_03    m/z 1047.5459     0   2   4   6E+3   11   11.5  RT (min)      lcorbi_t3_neg_4_034_01    m/z 1047.6218     0   1   2E+3   14.5   15  RT (min)      lcorbi_t3_neg_3_004_30    m/z 1047.6421     0   1   2   3E+4   11.5   12  RT (min)      lcorbi_t3_neg_4_007_13    m/z 1047.7074     0   1   2   3E+4   12   12.5  RT (min)      lcorbi_t3_neg_3_024_26    m/z 1048.5493     0   2   4   6E+3   11   11.5  RT (min)      lcorbi_t3_neg_4_034_01    m/z 1048.5670     0   2   4   6E+3   13   13.5  RT (min)      lcorbi_t3_neg_3_052_32    m/z 1048.6366     0   1   2   3E+3   15   15.5  RT (min)      lcorbi_t3_neg_4_007_13    m/z 1048.6454     0   1E+4   11.5   12  RT (min)      lcorbi_t3_neg_4_007_13    m/z 1048.6677     0   1E+4   13   13.5  RT (min)      lcorbi_t3_neg_4_039_32    m/z 1048.7029     0   2   4   6E+3   11.5   12  RT (min)      lcorbi_t3_neg_4_034_01    m/z 1048.7106     0   1E+4   12   12.5  RT (min)      lcorbi_t3_neg_3_024_26    m/z 1048.7637     0   1   2   3E+3   18   18.5  RT (min)      lcorbi_t3_neg_3_034_21    m/z 1049.3555     0   1   2   3   4E+3   8   8.5  RT (min)      lcorbi_t3_neg_4_040_19    m/z 1049.6349     0   2   4E+3   11   11.5  RT (min)      lcorbi_t3_neg_4_005_27    m/z 1049.6805     0   1E+4   13.5   14  RT (min)      lcorbi_t3_neg_4_039_32    m/z 1049.7710     0   1E+4   13   13.5  RT (min)      lcorbi_t3_neg_2_052_32    m/z 1050.2743     0   1E+4   9   9.5  RT (min)      lcorbi_t3_neg_2_048_27    m/z 1050.3258     0   1   2E+4   9   9.5  RT (min)      lcorbi_t3_neg_2_048_27    m/z 1050.4137     0   1   2   3E+3   13.5   14  RT (min)      lcorbi_t3_neg_3_048_08    m/z 1050.4559     0   1E+4   7.5   8  RT (min)      lcorbi_t3_neg_4_040_19    m/z 1050.5447     0   1   2E+4   4.5   5  RT (min)      lcorbi_t3_neg_2_048_27    m/z 1050.6836     0   1   2   3   4E+4   13.5   14  RT (min)      lcorbi_t3_neg_4_039_32    m/z 1050.7745     0   1   2   3E+4   13   13.5  RT (min)      lcorbi_t3_neg_2_052_32    m/z 1051.1334     0   1E+3   9   9.5  RT (min)      lcorbi_t3_neg_4_030_04    m/z 1051.3051     0   2   4   6E+3   14.5   15  RT (min)      lcorbi_t3_neg_4_007_13    m/z 1051.6087     0   2   4E+3   14.5   15  RT (min)      lcorbi_t3_neg_4_050_02    m/z 1051.6506     0   2   4E+3   11.5   12  RT (min)      lcorbi_t3_neg_3_007_16    m/z 1051.7744     0   1   2   3   4E+3   18   18.5  RT (min)      lcorbi_t3_neg_1_048_12    m/z 1052.3145     0   1E+4   8.5   9  RT (min)      lcorbi_t3_neg_4_023_18    m/z 1052.6536     0   1E+4   11.5   12  RT (min)      lcorbi_t3_neg_3_010_15    m/z 1052.6992     0   1   2   3E+4   13.5   14  RT (min)      lcorbi_t3_neg_2_052_32    m/z 1052.7749     0   2   4   6E+3   18   18.5  RT (min)      lcorbi_t3_neg_4_051_40    m/z 1053.5698     0   2   4   6E+3   11   11.5  RT (min)      lcorbi_t3_neg_2_027_13    m/z 1054.3247     0   1E+4   9   9.5  RT (min)      lcorbi_t3_neg_3_019_12    m/z 1054.5645     0   1E+4   4.5   5  RT (min)      lcorbi_t3_neg_4_042_14    m/z 1054.6811     0   2   4   6E+3   14.5   15  RT (min)      lcorbi_t3_neg_2_053_23    m/z 1054.7849     0   1E+4   15   15.5  RT (min)      lcorbi_t3_neg_2_007_14    m/z 1055.5430     0   1   2E+4   6.5   7  RT (min)      lcorbi_t3_neg_2_048_27    m/z 1055.6204     0   1E+4   13.5   14  RT (min)      lcorbi_t3_neg_1_035_25    m/z 1055.7835     0   1   2   3   4E+3   18   18.5  RT (min)      lcorbi_t3_neg_2_023_39    m/z 1055.7885     0   2   4   6E+3   15   15.5  RT (min)      lcorbi_t3_neg_2_007_14    m/z 1056.6499     0   2   4   6E+3   14.5   15  RT (min)      lcorbi_t3_neg_4_007_13    m/z 1056.7794     0   2   4E+3   18   18.5  RT (min)      lcorbi_t3_neg_2_003_42    m/z 1056.8000     0   2   4   6   8E+3   15   15.5  RT (min)      lcorbi_t3_neg_2_007_14    m/z 1057.1508     0   1   2   3E+3   9   9.5  RT (min)      lcorbi_t3_neg_4_028_39    m/z 1057.6415     0   1E+4   14   14.5  RT (min)      lcorbi_t3_neg_4_011_24    m/z 1057.6476     0   1E+4   11   11.5  RT (min)      lcorbi_t3_neg_4_007_13    m/z 1057.6532     0   1   2   3   4E+3   14.5   15  RT (min)      lcorbi_t3_neg_4_034_01    m/z 1057.8043     0   2   4E+3   15   15.5  RT (min)      lcorbi_t3_neg_2_007_14    m/z 1058.5539     0   1E+4   5   5.5  RT (min)      lcorbi_t3_neg_2_048_27    m/z 1058.6164     0   1   2   3E+4   13.5   14  RT (min)      lcorbi_t3_neg_4_005_27    m/z 1058.6656     0   1E+4   14.5   15   15.5  RT (min)      lcorbi_t3_neg_4_007_13    m/z 1058.6769     0   1E+4   13.5   14  RT (min)      lcorbi_t3_neg_4_032_23    m/z 1058.8149     0   1   2   3   4E+3   15   15.5  RT (min)      lcorbi_t3_neg_2_007_14    m/z 1059.3777     0   2   4   6   8E+3   10.5   11  RT (min)      lcorbi_t3_neg_2_002_26    m/z 1059.4715     0   2   4E+3   13.5   14  RT (min)      lcorbi_t3_neg_3_018_13    m/z 1059.5566     0   2   4   6   8E+3   5   5.5  RT (min)      lcorbi_t3_neg_2_048_27    m/z 1059.6218     0   1E+3   14.5   15  RT (min)      lcorbi_t3_neg_3_022_27    m/z 1059.6412     0   2   4E+3   12   12.5  RT (min)      lcorbi_t3_neg_4_023_18    m/z 1059.6690     0   2   4   6E+3   15   15.5  RT (min)      lcorbi_t3_neg_4_007_13    m/z 1059.7440     0   1   11   11.5  RT (min)      lcorbi_t3_neg_4_007_13    m/z 1059.7953     0   1   2   3   4E+3   18   18.5  RT (min)      lcorbi_t3_neg_2_011_04    m/z 1061.4802     0   2   4   6   8E+3   9.5   10  RT (min)      lcorbi_t3_neg_3_019_12    m/z 1061.4806     0   2   4   6E+3   13.5   14  RT (min)      lcorbi_t3_neg_2_027_13    m/z 1062.6408     0   1E+4   14.5   15  RT (min)      lcorbi_t3_neg_4_037_29    m/z 1063.3372     0   1E+4   9   9.5  RT (min)      lcorbi_t3_neg_4_023_18    m/z 1063.6531     0   1   2   3E+3   14.5   15  RT (min)      lcorbi_t3_neg_4_042_14    m/z 1063.9116     0   1E+4   14.5   15  RT (min)      lcorbi_t3_neg_4_039_32    m/z 1064.2883     0   1E+4   9   9.5  RT (min)      lcorbi_t3_neg_2_048_27    m/z 1064.3408     0   2   4   6   8E+3   9   9.5  RT (min)      lcorbi_t3_neg_4_023_18    m/z 1064.7174     0   1E+4   14.5   15  RT (min)      lcorbi_t3_neg_1_043_14    m/z 1064.9239     0   1E+4   14.5   15  RT (min)      lcorbi_t3_neg_3_002_24    m/z 1065.6050     0   2   4   6   8E+3   11   11.5  RT (min)      lcorbi_t3_neg_3_007_16    m/z 1065.8133     0   2   4   6   8E+3   18   18.5  RT (min)      lcorbi_t3_neg_4_020_44    m/z 1066.2419     0   1   2E+4   9   9.5  RT (min)      lcorbi_t3_neg_2_048_27    m/z 1066.4767     0   1E+4   13   13.5  RT (min)      lcorbi_t3_neg_4_046_37    m/z 1066.6532     0   1   2   3   4E+3   13.5   14  RT (min)      lcorbi_t3_neg_4_007_13    m/z 1066.8065     0   1E+4   15   15.5  RT (min)      lcorbi_t3_neg_4_042_14    m/z 1066.8146     0   1   2   3E+3   18   18.5  RT (min)      lcorbi_t3_neg_3_034_21    m/z 1066.9394     0   1E+4   14.5   15  RT (min)      lcorbi_t3_neg_4_011_24    m/z 1067.2454     0   1E+4   9   9.5  RT (min)      lcorbi_t3_neg_2_048_27    m/z 1067.4801     0   2   4   6   8E+3   13   13.5  RT (min)      lcorbi_t3_neg_2_051_34    m/z 1067.6300     0   1E+4   11.5   12  RT (min)      lcorbi_t3_neg_4_034_01    m/z 1067.8100     0   1E+4   15   15.5  RT (min)      lcorbi_t3_neg_2_007_14    m/z 1067.9431     0   1   2E+4   14.5   15  RT (min)      lcorbi_t3_neg_4_011_24    m/z 1068.6148     0   1E+4   11   11.5  RT (min)      lcorbi_t3_neg_4_007_13    m/z 1068.6692     0   1E+4   13.5   14  RT (min)      lcorbi_t3_neg_4_007_13    m/z 1068.8206     0   1E+4   15   15.5  RT (min)      lcorbi_t3_neg_4_042_14    m/z 1068.9551     0   2   4   6E+3   14.5   15  RT (min)      lcorbi_t3_neg_2_022_24    m/z 1069.6194     0   1E+4   11   11.5  RT (min)      lcorbi_t3_neg_4_007_13    m/z 1069.6701     0   1   2E+4   12   12.5  RT (min)      lcorbi_t3_neg_4_023_18    m/z 1069.6727     0   1E+4   13.5   14  RT (min)      lcorbi_t3_neg_4_007_13    m/z 1069.7291     0   1E+4   14.5   15  RT (min)      lcorbi_t3_neg_2_053_23    m/z 1069.8251     0   2   4   6E+3   15   15.5  RT (min)      lcorbi_t3_neg_2_007_14    m/z 1069.9588     0   1E+4   14.5   15  RT (min)      lcorbi_t3_neg_3_002_24    m/z 1070.3018     0   2   4   6E+3   9   9.5  RT (min)      lcorbi_t3_neg_4_040_19    m/z 1070.5957     0   1E+4   10.5   11  RT (min)      lcorbi_t3_neg_4_034_01    m/z 1070.6736     0   1E+4   12   12.5  RT (min)      lcorbi_t3_neg_4_023_18    m/z 1070.7328     0   2   4E+3   14.5   15  RT (min)      lcorbi_t3_neg_2_053_23    m/z 1070.7575     0   1   2   3   4E+3   18   18.5  RT (min)      lcorbi_t3_neg_2_040_43    m/z 1070.8371     0   1   15   15.5  RT (min)      lcorbi_t3_neg_4_055_06    m/z 1071.3459     0   1   2   3   4E+3   10.5   11  RT (min)      lcorbi_t3_neg_1_023_03    m/z 1071.6767     0   2   4E+3   12   12.5  RT (min)      lcorbi_t3_neg_4_023_18    m/z 1071.7164     0   1   2E+4   14   14.5  RT (min)      lcorbi_t3_neg_1_044_23    m/z 1073.6959     0   1   2E+4   14   14.5  RT (min)      lcorbi_t3_neg_3_005_23    m/z 1074.6383     0   1E+4   11   11.5  RT (min)      lcorbi_t3_neg_4_007_13    m/z 1074.7666     0   2   4E+3   18   18.5  RT (min)      lcorbi_t3_neg_4_048_03    m/z 1075.5195     0   1E+4   6.5   7  RT (min)      lcorbi_t3_neg_4_042_14    m/z 1075.6352     0   1E+4   11   11.5  RT (min)      lcorbi_t3_neg_4_005_27    m/z 1075.6949     0   1E+4   11.5   12  RT (min)      lcorbi_t3_neg_4_034_01    m/z 1076.5225     0   2   4   6E+3   6.5   7  RT (min)      lcorbi_t3_neg_4_042_14    m/z 1076.6987     0   2   4   6E+3   11.5   12  RT (min)      lcorbi_t3_neg_4_034_01    m/z 1076.7660     0   1   2   3   4E+3   18   18.5  RT (min)      lcorbi_t3_neg_4_022_41    m/z 1077.5702     0   2   4   6E+3   11   11.5  RT (min)      lcorbi_t3_neg_2_050_17    m/z 1077.6060     0   1   2   3   4E+3   13   13.5  RT (min)      lcorbi_t3_neg_2_052_32    m/z 1078.6096     0   1E+4   13   13.5  RT (min)      lcorbi_t3_neg_2_052_32    m/z 1078.6696     0   1E+4   11.5   12  RT (min)      lcorbi_t3_neg_4_007_13    m/z 1079.6713     0   1E+4   11   11.5  RT (min)      lcorbi_t3_neg_2_054_15    m/z 1080.2576     0   1E+4   9   9.5  RT (min)      lcorbi_t3_neg_2_048_27    m/z 1080.6743     0   1E+4   11   11.5   12  RT (min)      lcorbi_t3_neg_4_042_14    m/z 1081.3748     0   1E+4   10.5   11  RT (min)      lcorbi_t3_neg_3_029_02    m/z 1081.7007     0   1E+4   14.5   15  RT (min)      lcorbi_t3_neg_2_039_31    m/z 1082.0253     0   1E+4   6   6.5  RT (min)      lcorbi_t3_neg_2_048_27    m/z 1082.3785     0   2   4E+3   10.5   11  RT (min)      lcorbi_t3_neg_3_029_02    m/z 1082.5267     0   1   2E+4   6   6.5  RT (min)      lcorbi_t3_neg_2_048_27    m/z 1083.0281     0   1E+4   6   6.5  RT (min)      lcorbi_t3_neg_2_048_27    m/z 1083.4348     0   2   4   6E+3   11   11.5  RT (min)      lcorbi_t3_neg_4_023_18    m/z 1083.5890     0   2   4E+3   11.5   12  RT (min)      lcorbi_t3_neg_1_047_02    m/z 1083.7167     0   1E+4   14.5   15  RT (min)      lcorbi_t3_neg_2_053_23    m/z 1084.6994     0   1E+4   12   12.5  RT (min)      lcorbi_t3_neg_2_007_14    m/z 1084.8692     0   1   2E+4   15.5   16  RT (min)      lcorbi_t3_neg_4_005_27    m/z 1085.4872     0   1   2   3   4E+3   13.5   14  RT (min)      lcorbi_t3_neg_2_027_13    m/z 1085.7041     0   1   2   3E+4   13.5   14  RT (min)      lcorbi_t3_neg_4_023_18    m/z 1085.7240     0   1E+4   14.5   15  RT (min)      lcorbi_t3_neg_2_053_23    m/z 1085.8726     0   1E+4   15.5   16  RT (min)      lcorbi_t3_neg_4_005_27    m/z 1086.8797     0   2   4   6E+3   15.5   16  RT (min)      lcorbi_t3_neg_4_005_27    m/z 1087.4968     0   1   2   3E+3   13.5   14  RT (min)      lcorbi_t3_neg_2_027_13    m/z 1087.5723     0   2   4E+3   10   10.5  RT (min)      lcorbi_t3_neg_3_019_12    m/z 1087.6785     0   2   4   6E+3   11   11.5  RT (min)      lcorbi_t3_neg_1_055_05    m/z 1089.6049     0   2   4   6E+3   11   11.5  RT (min)      lcorbi_t3_neg_4_034_01    m/z 1089.6145     0   2   4   6   8E+3   11.5   12  RT (min)      lcorbi_t3_neg_3_039_29    m/z 1090.5485     0   1   2   3E+3   13.5   14  RT (min)      lcorbi_t3_neg_3_048_08    m/z 1090.7044     0   1E+4   11.5   12  RT (min)      lcorbi_t3_neg_4_007_13    m/z 1090.7206     0   1   2   3E+4   14   14.5  RT (min)      lcorbi_t3_neg_2_053_23    m/z 1090.8066     0   1E+4   15   15.5  RT (min)      lcorbi_t3_neg_4_042_14    m/z 1091.5339     0   1   2   3   4E+3   11   11.5  RT (min)      lcorbi_t3_neg_2_050_17    m/z 1091.8100     0   1E+4   15   15.5  RT (min)      lcorbi_t3_neg_4_042_14    m/z 1092.2627     0   1E+4   9   9.5  RT (min)      lcorbi_t3_neg_2_048_27    m/z 1093.2408     0   1   2   3   4E+4   14   14.5  RT (min)      lcorbi_t3_neg_4_045_35    m/z 1093.2665     0   2   4   6E+3   9   9.5  RT (min)      lcorbi_t3_neg_4_040_19    m/z 1093.8256     0   1E+4   15   15.5  RT (min)      lcorbi_t3_neg_2_007_14    m/z 1094.2417     0   2   4E+4   14   14.5  RT (min)      lcorbi_t3_neg_4_045_35    m/z 1094.2921     0   1E+4   14.5   15  RT (min)      lcorbi_t3_neg_4_048_03    m/z 1094.5957     0   1   2   3E+4   10.5   11  RT (min)      lcorbi_t3_neg_4_029_31    m/z 1094.7132     0   2   4   6E+3   14.5   15  RT (min)      lcorbi_t3_neg_2_053_23    m/z 1094.8355     0   2   4   6   8E+3   15   15.5  RT (min)      lcorbi_t3_neg_2_007_14    m/z 1095.2404     0   1   2   3E+4   14   14.5  RT (min)      lcorbi_t3_neg_4_045_35    m/z 1095.3483     0   1   2E+4   9   9.5  RT (min)      lcorbi_t3_neg_4_023_18    m/z 1095.5992     0   1   2E+4   10.5   11  RT (min)      lcorbi_t3_neg_4_029_31    m/z 1095.7166     0   1   2   3   4E+3   14.5   15  RT (min)      lcorbi_t3_neg_3_005_23    m/z 1095.7386     0   1E+4   12   12.5  RT (min)      lcorbi_t3_neg_4_034_01    m/z 1095.8408     0   2   4   6E+3   15   15.5  RT (min)      lcorbi_t3_neg_4_042_14    m/z 1096.3514     0   1E+4   9   9.5  RT (min)      lcorbi_t3_neg_4_023_18    m/z 1096.5546     0   2   4   6   8E+3   5.5   6  RT (min)      lcorbi_t3_neg_2_048_27    m/z 1096.6242     0   2   4   6   8E+3   11.5   12  RT (min)      lcorbi_t3_neg_1_015_27    m/z 1096.7286     0   2   4E+3   15   15.5  RT (min)      lcorbi_t3_neg_2_053_23    m/z 1097.6351     0   2   4   6   8E+3   11   11.5  RT (min)      lcorbi_t3_neg_4_007_13    m/z 1098.6383     0   2   4E+3   11   11.5  RT (min)      lcorbi_t3_neg_3_007_16    m/z 1098.7448     0   1   2   3E+4   14   14.5  RT (min)      lcorbi_t3_neg_2_053_23    m/z 1098.8040     0   2   4E+3   18   18.5  RT (min)      lcorbi_t3_neg_3_003_09    m/z 1099.7480     0   1E+4   14   14.5  RT (min)      lcorbi_t3_neg_2_053_23    m/z 1101.7266     0   1E+4   18   18.5  RT (min)      lcorbi_t3_neg_1_013_35    m/z 1102.7020     0   2   4   6   8E+3   18   18.5  RT (min)      lcorbi_t3_neg_1_039_38    m/z 1103.2947     0   1   2E+4   9   9.5  RT (min)      lcorbi_t3_neg_2_048_27    m/z 1103.7055     0   2   4E+3   18   18.5  RT (min)      lcorbi_t3_neg_1_020_36    m/z 1104.2695     0   2   4   6   8E+4   14   14.5  RT (min)      lcorbi_t3_neg_4_045_35    m/z 1104.2976     0   1E+4   9   9.5  RT (min)      lcorbi_t3_neg_2_048_27    m/z 1104.6163     0   2   4   6   8E+3   13   13.5  RT (min)      lcorbi_t3_neg_3_052_32    m/z 1104.6975     0   2   4   6E+3   14.5   15  RT (min)      lcorbi_t3_neg_2_039_31    m/z 1105.7007     0   1   2   3   4E+3   14.5   15  RT (min)      lcorbi_t3_neg_2_039_31    m/z 1106.2782     0   2   4   6   8E+3   9   9.5  RT (min)      lcorbi_t3_neg_2_048_27    m/z 1106.6320     0   2   4   6E+3   13.5   14  RT (min)      lcorbi_t3_neg_2_022_24    m/z 1106.7129     0   1E+4   14.5   15  RT (min)      lcorbi_t3_neg_2_039_31    m/z 1106.8537     0   2   4   6E+3   15.5   16  RT (min)      lcorbi_t3_neg_4_005_27    m/z 1107.5867     0   2   4   6E+3   11   11.5  RT (min)      lcorbi_t3_neg_2_039_31    m/z 1107.7164     0   2   4   6E+3   14.5   15  RT (min)      lcorbi_t3_neg_2_039_31    m/z 1107.8595     0   1   14.5   15  RT (min)      lcorbi_t3_neg_4_005_27    m/z 1108.7160     0   2   4   6E+3   12   12.5  RT (min)      lcorbi_t3_neg_4_038_15    m/z 1109.3028     0   1   2E+4   14   14.5  RT (min)      lcorbi_t3_neg_3_035_03    m/z 1109.6704     0   1   2   3E+4   11   11.5  RT (min)      lcorbi_t3_neg_4_007_13    m/z 1109.6797     0   1   2   3E+4   11.5   12  RT (min)      lcorbi_t3_neg_4_007_13    m/z 1110.3059     0   1E+4   14   14.5  RT (min)      lcorbi_t3_neg_2_011_04    m/z 1110.6734     0   1   2E+4   11   11.5  RT (min)      lcorbi_t3_neg_4_007_13    m/z 1110.6827     0   1   2E+4   11.5   12  RT (min)      lcorbi_t3_neg_4_007_13    m/z 1110.7359     0   2   4   6E+3   14.5   15  RT (min)      lcorbi_t3_neg_2_053_23    m/z 1110.7546     0   1   2   3E+4   11   11.5  RT (min)      lcorbi_t3_neg_1_055_05    m/z 1110.8025     0   2   4E+3   18   18.5  RT (min)      lcorbi_t3_neg_2_042_44    m/z 1110.8517     0   1E+4   13.5   14  RT (min)      lcorbi_t3_neg_1_052_24    m/z 1111.2805     0   1   2   3   4E+3   9   9.5  RT (min)      lcorbi_t3_neg_3_019_12    m/z 1111.6941     0   1E+4   11.5   12  RT (min)      lcorbi_t3_neg_4_007_13    m/z 1111.8859     0   1   2E+4   15.5   16  RT (min)      lcorbi_t3_neg_4_005_27    m/z 1112.6023     0   1E+4   11   11.5  RT (min)      lcorbi_t3_neg_4_007_13    m/z 1112.6311     0   1E+4   13   13.5  RT (min)      lcorbi_t3_neg_2_052_32    m/z 1112.7396     0   1   2   3   4E+3   18   18.5  RT (min)      lcorbi_t3_neg_2_018_40    m/z 1112.8919     0   1   2E+4   15.5   16  RT (min)      lcorbi_t3_neg_4_005_27    m/z 1112.9004     0   1   2E+4   16   16.5  RT (min)      lcorbi_t3_neg_1_015_27    m/z 1113.6045     0   2   4E+3   11   11.5  RT (min)      lcorbi_t3_neg_3_007_16    m/z 1113.7822     0   1E+4   11.5   12  RT (min)      lcorbi_t3_neg_1_055_05    m/z 1113.9039     0   1   2E+4   16   16.5  RT (min)      lcorbi_t3_neg_4_005_27    m/z 1114.6922     0   1   2E+4   12   12.5  RT (min)      lcorbi_t3_neg_3_024_26    m/z 1114.9083     0   2   4   6   8E+3   16   16.5  RT (min)      lcorbi_t3_neg_4_005_27    m/z 1115.6299     0   2   4E+3   11.5   12  RT (min)      lcorbi_t3_neg_4_032_23    m/z 1115.6872     0   2   4   6E+3   11.5   12  RT (min)      lcorbi_t3_neg_4_034_01    m/z 1115.6957     0   1   2E+4   12   12.5  RT (min)      lcorbi_t3_neg_3_024_26    m/z 1116.3066     0   1   2E+4   9   9.5  RT (min)      lcorbi_t3_neg_2_048_27    m/z 1116.6990     0   2   4   6E+3   12   12.5  RT (min)      lcorbi_t3_neg_4_004_34    m/z 1116.7465     0   1E+4   18   18.5  RT (min)      lcorbi_t3_neg_1_013_35    m/z 1116.8223     0   1   2   3   4E+3   15   15.5  RT (min)      lcorbi_t3_neg_4_042_14    m/z 1117.3097     0   1E+4   9   9.5  RT (min)      lcorbi_t3_neg_2_048_27    m/z 1117.3303     0   1E+4   9   9.5  RT (min)      lcorbi_t3_neg_4_023_18    m/z 1117.4402     0   1E+4   7.5   8  RT (min)      lcorbi_t3_neg_4_040_19    m/z 1117.6227     0   1   2   3E+3   11   11.5  RT (min)      lcorbi_t3_neg_2_051_34    m/z 1117.7225     0   1   2   3E+4   14   14.5  RT (min)      lcorbi_t3_neg_1_044_23    m/z 1117.8258     0   1   2   3E+3   15   15.5  RT (min)      lcorbi_t3_neg_2_007_14    m/z 1118.2599     0   1E+4   9   9.5  RT (min)      lcorbi_t3_neg_2_048_27    m/z 1118.5959     0   1   2E+4   10.5   11  RT (min)      lcorbi_t3_neg_4_034_01    m/z 1118.8378     0   2   4   6E+3   15   15.5  RT (min)      lcorbi_t3_neg_2_007_14    m/z 1118.8536     0   1E+4   15.5   16  RT (min)      lcorbi_t3_neg_4_029_31    m/z 1119.6257     0   1E+4   11   11.5  RT (min)      lcorbi_t3_neg_4_042_14    m/z 1119.8415     0   2   4E+3   15   15.5  RT (min)      lcorbi_t3_neg_2_007_14    m/z 1119.8570     0   2   4   6E+3   15.5   16  RT (min)      lcorbi_t3_neg_4_029_31    m/z 1120.6353     0   2   4   6   8E+3   11.5   12  RT (min)      lcorbi_t3_neg_1_043_14    m/z 1120.6830     0   2   4E+4   10.5   11  RT (min)      lcorbi_t3_neg_4_039_32    m/z 1120.7628     0   1   2   3   4E+3   18   18.5  RT (min)      lcorbi_t3_neg_2_015_21    m/z 1120.8521     0   1   2   3   4E+3   15   15.5  RT (min)      lcorbi_t3_neg_4_042_14    m/z 1120.8652     0   1   2   3   4E+3   15.5   16  RT (min)      lcorbi_t3_neg_4_029_31    m/z 1121.3093     0   1E+4   9   9.5  RT (min)      lcorbi_t3_neg_3_019_12    m/z 1121.6351     0   2   4   6E+3   11   11.5  RT (min)      lcorbi_t3_neg_4_007_13    m/z 1121.8568     0   1   2E+3   15   15.5  RT (min)      lcorbi_t3_neg_4_042_14    m/z 1122.3126     0   2   4   6   8E+3   9   9.5  RT (min)      lcorbi_t3_neg_3_019_12    m/z 1122.7444     0   1   15   15.5  RT (min)      lcorbi_t3_neg_1_029_21    m/z 1122.8847     0   1E+4   15.5   16  RT (min)      lcorbi_t3_neg_4_005_27    m/z 1123.7799     0   2   4   6E+3   14.5   15  RT (min)      lcorbi_t3_neg_4_005_27    m/z 1123.8883     0   2   4   6   8E+3   15.5   16  RT (min)      lcorbi_t3_neg_4_005_27    m/z 1123.9103     0   2   4   6E+3   14.5   15  RT (min)      lcorbi_t3_neg_2_052_32    m/z 1124.7674     0   1   2   3   4E+3   18   18.5  RT (min)      lcorbi_t3_neg_2_016_11    m/z 1124.7836     0   2   4E+3   14.5   15  RT (min)      lcorbi_t3_neg_4_005_27    m/z 1124.8946     0   1   2E+4   14   14.5  RT (min)      lcorbi_t3_neg_3_002_24    m/z 1125.1386     0   1   2E+3   9   9.5  RT (min)      lcorbi_t3_neg_4_051_40    m/z 1125.8704     0   1E+4   14   14.5  RT (min)      lcorbi_t3_neg_2_022_24    m/z 1126.7799     0   1E+4   18   18.5  RT (min)      lcorbi_t3_neg_2_034_19    m/z 1128.4799     0   1   2   3E+4   7   7.5  RT (min)      lcorbi_t3_neg_2_048_27    m/z 1129.4828     0   1E+4   7   7.5  RT (min)      lcorbi_t3_neg_2_048_27    m/z 1129.6252     0   1   2   3E+3   14.5   15  RT (min)      lcorbi_t3_neg_3_029_02    m/z 1129.6577     0   1   2E+4   11   11.5  RT (min)      lcorbi_t3_neg_4_046_37    m/z 1130.7334     0   1E+4   18   18.5  RT (min)      lcorbi_t3_neg_1_039_38    m/z 1130.7622     0   1E+4   14.5   15  RT (min)      lcorbi_t3_neg_2_053_23    m/z 1132.2740     0   2   4   6   8E+3   9   9.5  RT (min)      lcorbi_t3_neg_4_040_19    m/z 1132.7409     0   1   2   3   4E+4   14   14.5  RT (min)      lcorbi_t3_neg_2_053_23    m/z 1133.6288     0   1   2   3   4E+4   13.5   14  RT (min)      lcorbi_t3_neg_4_036_16    m/z 1133.6702     0   1   2E+4   11   11.5  RT (min)      lcorbi_t3_neg_4_034_01    m/z 1133.7511     0   1   11   11.5  RT (min)      lcorbi_t3_neg_4_007_13    m/z 1133.8009     0   1   2   3   4E+3   18   18.5  RT (min)      lcorbi_t3_neg_2_040_43    m/z 1134.2295     0   1E+4   9   9.5  RT (min)      lcorbi_t3_neg_2_048_27    m/z 1134.4644     0   1   2   3   4E+3   13   13.5  RT (min)      lcorbi_t3_neg_3_021_38    m/z 1134.8785     0   1E+4   14.5   15  RT (min)      lcorbi_t3_neg_4_011_24    m/z 1135.5990     0   1E+4   11   11.5  RT (min)      lcorbi_t3_neg_4_007_13    m/z 1135.6201     0   2   4   6E+3   11.5   12  RT (min)      lcorbi_t3_neg_4_034_01    m/z 1136.6021     0   1   2   3   4E+3   11   11.5  RT (min)      lcorbi_t3_neg_3_007_16    m/z 1136.7375     0   2   4   6   8E+3   12   12.5  RT (min)      lcorbi_t3_neg_4_034_01    m/z 1136.7602     0   2   4E+3   15   15.5  RT (min)      lcorbi_t3_neg_2_053_23    m/z 1136.8945     0   1   2   3   4E+2   14.5   15  RT (min)      lcorbi_t3_neg_4_046_37    m/z 1137.6577     0   1E+4   12   12.5  RT (min)      lcorbi_t3_neg_4_023_18    m/z 1137.7103     0   1E+4   11.5   12  RT (min)      lcorbi_t3_neg_4_034_01    m/z 1137.8792     0   1   2E+4   14   14.5  RT (min)      lcorbi_t3_neg_3_002_24    m/z 1137.8981     0   1   14.5   15  RT (min)      lcorbi_t3_neg_4_042_14    m/z 1138.6611     0   2   4   6   8E+3   12   12.5  RT (min)      lcorbi_t3_neg_4_023_18    m/z 1138.7137     0   2   4   6   8E+3   11.5   12  RT (min)      lcorbi_t3_neg_4_034_01    m/z 1139.6547     0   2   4   6E+3   12   12.5  RT (min)      lcorbi_t3_neg_2_050_17    m/z 1139.8854     0   1   2E+4   18   18.5  RT (min)      lcorbi_t3_neg_1_053_32    m/z 1139.9057     0   2   4   6   8E+3   14.5   15  RT (min)      lcorbi_t3_neg_2_052_32    m/z 1139.9129     0   1E+4   16   16.5  RT (min)      lcorbi_t3_neg_4_005_27    m/z 1142.5182     0   1E+4   10.5   11  RT (min)      lcorbi_t3_neg_4_029_31    m/z 1142.6260     0   1E+4   11   11.5  RT (min)      lcorbi_t3_neg_2_027_13    m/z 1142.8536     0   1   2   3   4E+3   15.5   16  RT (min)      lcorbi_t3_neg_4_029_31    m/z 1143.6233     0   2   4   6E+3   11   11.5  RT (min)      lcorbi_t3_neg_3_018_13    m/z 1143.6822     0   2   4   6E+3   11.5   12  RT (min)      lcorbi_t3_neg_4_034_01    m/z 1143.7742     0   1   2E+4   14.5   15  RT (min)      lcorbi_t3_neg_2_053_23    m/z 1143.8570     0   1   2   3E+3   15.5   16  RT (min)      lcorbi_t3_neg_4_029_31    m/z 1144.8688     0   1E+4   15.5   16  RT (min)      lcorbi_t3_neg_4_029_31    m/z 1145.8726     0   2   4   6   8E+3   15.5   16  RT (min)      lcorbi_t3_neg_4_029_31    m/z 1146.5972     0   1   2   3   4E+3   13   13.5  RT (min)      lcorbi_t3_neg_3_052_32    m/z 1146.6570     0   1E+4   11.5   12  RT (min)      lcorbi_t3_neg_4_007_13    m/z 1146.8836     0   1E+4   15.5   16  RT (min)      lcorbi_t3_neg_2_039_31    m/z 1147.6544     0   2   4   6   8E+3   11.5   12  RT (min)      lcorbi_t3_neg_4_007_13    m/z 1147.7800     0   2   4   6   8E+3   14.5   15  RT (min)      lcorbi_t3_neg_4_005_27    m/z 1147.8881     0   2   4   6   8E+3   15.5   16  RT (min)      lcorbi_t3_neg_2_039_31    m/z 1148.2453     0   2   4   6   8E+3   9   9.5  RT (min)      lcorbi_t3_neg_2_048_27    m/z 1148.7713     0   2   4   6E+3   18   18.5  RT (min)      lcorbi_t3_neg_2_042_44    m/z 1148.7834     0   2   4E+3   14.5   15  RT (min)      lcorbi_t3_neg_4_035_33    m/z 1148.8949     0   1   2   3   4E+3   15.5   16  RT (min)      lcorbi_t3_neg_4_029_31    m/z 1149.3626     0   2   4E+3   10.5   11  RT (min)      lcorbi_t3_neg_3_029_02    m/z 1149.6422     0   2   4   6   8E+3   11   11.5  RT (min)      lcorbi_t3_neg_3_010_15    m/z 1149.6906     0   1E+4   13   13.5  RT (min)      lcorbi_t3_neg_4_007_13    m/z 1149.7946     0   1   2   3E+3   14.5   15  RT (min)      lcorbi_t3_neg_4_005_27    m/z 1150.6450     0   2   4   6   8E+3   11   11.5  RT (min)      lcorbi_t3_neg_2_005_01    m/z 1150.6941     0   1E+4   13   13.5  RT (min)      lcorbi_t3_neg_4_007_13    m/z 1152.6871     0   1E+4   12   12.5  RT (min)      lcorbi_t3_neg_2_007_14    m/z 1153.6669     0   1   2   3E+4   11.5   12  RT (min)      lcorbi_t3_neg_4_007_13    m/z 1153.6914     0   1E+4   13.5   14  RT (min)      lcorbi_t3_neg_4_023_18    m/z 1153.7130     0   2   4   6E+3   13   13.5  RT (min)      lcorbi_t3_neg_4_034_01    m/z 1154.6613     0   1E+4   11   11.5  RT (min)      lcorbi_t3_neg_4_039_32    m/z 1154.6704     0   1E+4   11.5   12  RT (min)      lcorbi_t3_neg_2_027_13    m/z 1154.6948     0   1E+4   18   18.5  RT (min)      lcorbi_t3_neg_1_020_36    m/z 1154.6953     0   1   2E+4   13.5   14  RT (min)      lcorbi_t3_neg_1_044_23    m/z 1154.8774     0   1   2E+4   14   14.5  RT (min)      lcorbi_t3_neg_4_019_08    m/z 1155.6642     0   1E+4   11   11.5  RT (min)      lcorbi_t3_neg_4_007_13    m/z 1155.9081     0   1E+4   18   18.5  RT (min)      lcorbi_t3_neg_1_053_32    m/z 1156.5902     0   1E+4   11   11.5  RT (min)      lcorbi_t3_neg_4_034_01    m/z 1156.6254     0   1   2   3E+4   13.5   14  RT (min)      lcorbi_t3_neg_4_036_16    m/z 1156.6674     0   1E+4   11   11.5  RT (min)      lcorbi_t3_neg_4_007_13    m/z 1156.8836     0   2   4   6   8E+3   18   18.5  RT (min)      lcorbi_t3_neg_1_053_32    m/z 1156.9134     0   1E+4   14   14.5  RT (min)      lcorbi_t3_neg_3_002_24    m/z 1157.5558     0   1   2E+4   10.5   11  RT (min)      lcorbi_t3_neg_2_007_14    m/z 1157.5942     0   1   2   3   4E+3   11   11.5  RT (min)      lcorbi_t3_neg_4_034_01    m/z 1158.6919     0   2   4   6E+3   11.5   12  RT (min)      lcorbi_t3_neg_4_039_32    m/z 1158.8851     0   1   2E+4   15.5   16  RT (min)      lcorbi_t3_neg_4_005_27    m/z 1159.8884     0   1   2E+4   15.5   16  RT (min)      lcorbi_t3_neg_4_005_27    m/z 1161.8885     0   2   4   6   8E+3   14.5   15  RT (min)      lcorbi_t3_neg_1_053_32    m/z 1161.9039     0   1   2E+4   15.5   16  RT (min)      lcorbi_t3_neg_4_005_27    m/z 1162.6132     0   1E+4   11.5   12  RT (min)      lcorbi_t3_neg_1_053_32    m/z 1162.9009     0   1   14.5   15  RT (min)      lcorbi_t3_neg_4_042_14    m/z 1162.9130     0   1   2E+4   16   16.5  RT (min)      lcorbi_t3_neg_4_005_27    m/z 1163.6102     0   1E+4   11.5   12  RT (min)      lcorbi_t3_neg_2_010_02    m/z 1163.7510     0   2   4   6E+3   14.5   15  RT (min)      lcorbi_t3_neg_3_005_23    m/z 1163.9057     0   1E+4   14.5   15  RT (min)      lcorbi_t3_neg_4_011_24    m/z 1163.9187     0   1E+4   16   16.5  RT (min)      lcorbi_t3_neg_4_005_27    m/z 1164.6119     0   1   11.5   12  RT (min)      lcorbi_t3_neg_2_022_24    m/z 1164.9171     0   1   2E+4   14.5   15  RT (min)      lcorbi_t3_neg_3_002_24    m/z 1164.9280     0   2   4E+3   16   16.5  RT (min)      lcorbi_t3_neg_4_005_27    m/z 1165.6208     0   1E+4   11   11.5  RT (min)      lcorbi_t3_neg_3_044_01    m/z 1165.7665     0   1   2   3   4E+3   14.5   15  RT (min)      lcorbi_t3_neg_4_032_23    m/z 1166.5187     0   1   2   3   4E+4   10.5   11  RT (min)      lcorbi_t3_neg_1_043_14    m/z 1166.5988     0   1   2E+4   10.5   11  RT (min)      lcorbi_t3_neg_4_039_32    m/z 1167.2597     0   1   2E+4   14   14.5  RT (min)      lcorbi_t3_neg_4_045_35    m/z 1167.5200     0   2   4   6E+3   12.5   13  RT (min)      lcorbi_t3_neg_4_005_27    m/z 1167.6245     0   2   4   6E+3   11   11.5  RT (min)      lcorbi_t3_neg_4_007_13    m/z 1168.2606     0   1   2E+4   14   14.5  RT (min)      lcorbi_t3_neg_4_045_35    m/z 1169.2594     0   1   2   3E+4   14   14.5  RT (min)      lcorbi_t3_neg_2_036_35    m/z 1169.9235     0   1E+4   18   18.5  RT (min)      lcorbi_t3_neg_1_053_32    m/z 1170.2790     0   2   4   6   8E+3   9   9.5  RT (min)      lcorbi_t3_neg_2_048_27    m/z 1171.2823     0   2   4   6E+3   9   9.5  RT (min)      lcorbi_t3_neg_4_040_19    m/z 1171.7801     0   2   4E+3   14.5   15  RT (min)      lcorbi_t3_neg_4_005_27    m/z 1172.4046     0   1   2   3E+4   9   9.5  RT (min)      lcorbi_t3_neg_2_048_27    m/z 1172.7834     0   1   2   3E+3   14.5   15  RT (min)      lcorbi_t3_neg_2_043_33    m/z 1172.9320     0   1   2   3E+4   14   14.5  RT (min)      lcorbi_t3_neg_4_011_24    m/z 1173.5356     0   1   2E+4   10.5   11  RT (min)      lcorbi_t3_neg_4_042_14    m/z 1174.4854     0   1   2E+4   7   7.5  RT (min)      lcorbi_t3_neg_2_048_27    m/z 1174.6479     0   2   4   6   8E+3   11.5   12  RT (min)      lcorbi_t3_neg_3_013_28    m/z 1174.9316     0   1E+4   16   16.5  RT (min)      lcorbi_t3_neg_4_005_27    m/z 1175.4882     0   2   4   6   8E+3   7   7.5  RT (min)      lcorbi_t3_neg_2_048_27    m/z 1175.6875     0   1E+4   12   12.5  RT (min)      lcorbi_t3_neg_4_034_01    m/z 1175.7772     0   1E+4   14.5   15  RT (min)      lcorbi_t3_neg_4_053_43    m/z 1175.9384     0   2   4   6E+3   16   16.5  RT (min)      lcorbi_t3_neg_4_005_27    m/z 1176.6970     0   2   4E+3   12   12.5  RT (min)      lcorbi_t3_neg_3_053_14    m/z 1176.8569     0   1E+4   15.5   16  RT (min)      lcorbi_t3_neg_4_042_14    m/z 1177.6579     0   1   11   11.5  RT (min)      lcorbi_t3_neg_1_055_05    m/z 1177.6668     0   2   4   6   8E+3   11.5   12  RT (min)      lcorbi_t3_neg_4_007_13    m/z 1177.7220     0   2   4   6   8E+3   13   13.5  RT (min)      lcorbi_t3_neg_4_007_13    m/z 1177.7386     0   1   2   3E+4   11   11.5  RT (min)      lcorbi_t3_neg_1_055_05    m/z 1177.8603     0   1E+4   15.5   16  RT (min)      lcorbi_t3_neg_4_042_14    m/z 1178.2887     0   2   4   6E+4   14   14.5  RT (min)      lcorbi_t3_neg_4_045_35    m/z 1178.7253     0   2   4E+3   13   13.5  RT (min)      lcorbi_t3_neg_4_007_13    m/z 1179.2682     0   1   2   3E+3   9   9.5  RT (min)      lcorbi_t3_neg_3_019_12    m/z 1179.5868     0   2   4   6   8E+3   11   11.5  RT (min)      lcorbi_t3_neg_3_032_17    m/z 1179.7076     0   1E+5   8   8.5  RT (min)      lcorbi_t3_neg_2_048_27    m/z 1179.7379     0   2   4   6   8E+3   13.5   14  RT (min)      lcorbi_t3_neg_4_034_01    m/z 1180.4658     0   2   4   6E+4   9   9.5  RT (min)      lcorbi_t3_neg_2_048_27    m/z 1180.5901     0   2   4E+3   11   11.5  RT (min)      lcorbi_t3_neg_2_050_17    m/z 1180.7411     0   2   4   6E+3   13.5   14  RT (min)      lcorbi_t3_neg_4_034_01    m/z 1180.9394     0   2   4E+3   15   15.5  RT (min)      lcorbi_t3_neg_3_002_24    m/z 1181.4691     0   1   2   3   4E+4   9   9.5  RT (min)      lcorbi_t3_neg_2_048_27    m/z 1182.6798     0   1E+4   12   12.5  RT (min)      lcorbi_t3_neg_4_052_26    m/z 1182.7267     0   2   4   6E+3   18   18.5  RT (min)      lcorbi_t3_neg_1_054_37    m/z 1182.9061     0   1   2   3   4E+3   16   16.5  RT (min)      lcorbi_t3_neg_4_029_31    m/z 1183.6829     0   1E+4   12   12.5  RT (min)      lcorbi_t3_neg_4_004_34    m/z 1183.7202     0   1   2E+4   14   14.5  RT (min)      lcorbi_t3_neg_2_053_23    m/z 1183.9097     0   1   2   3E+3   16   16.5  RT (min)      lcorbi_t3_neg_4_029_31    m/z 1184.7233     0   1E+4   14   14.5  RT (min)      lcorbi_t3_neg_3_005_23    m/z 1184.9430     0   1E+4   14.5   15  RT (min)      lcorbi_t3_neg_2_052_32    m/z 1185.4172     0   1   2   3E+4   9   9.5  RT (min)      lcorbi_t3_neg_4_023_18    m/z 1185.7354     0   1   2   3   4E+4   14   14.5  RT (min)      lcorbi_t3_neg_4_032_23    m/z 1185.9467     0   1   2E+4   14.5   15  RT (min)      lcorbi_t3_neg_2_052_32    m/z 1186.4203     0   1E+4   9   9.5  RT (min)      lcorbi_t3_neg_4_023_18    m/z 1186.7391     0   1   2E+4   14   14.5  RT (min)      lcorbi_t3_neg_4_032_23    m/z 1186.8935     0   1   16   16.5  RT (min)      lcorbi_t3_neg_4_019_08    m/z 1186.9216     0   1E+4   18   18.5  RT (min)      lcorbi_t3_neg_1_053_32    m/z 1187.6133     0   2   4   6   8E+3   11.5   12  RT (min)      lcorbi_t3_neg_3_029_02    m/z 1187.7508     0   1   2E+4   14   14.5  RT (min)      lcorbi_t3_neg_4_032_23    m/z 1187.8968     0   2   4E+3   16   16.5  RT (min)      lcorbi_t3_neg_2_007_14    m/z 1188.5056     0   1   2   3   4E+4   10.5   11  RT (min)      lcorbi_t3_neg_1_043_14    m/z 1188.7549     0   1E+4   14   14.5  RT (min)      lcorbi_t3_neg_4_032_23    m/z 1189.2968     0   2   4   6   8E+3   9   9.5  RT (min)      lcorbi_t3_neg_3_019_12    m/z 1189.5050     0   1   2   3   4E+4   10.5   11  RT (min)      lcorbi_t3_neg_1_043_14    m/z 1189.6227     0   1   2   3   4E+3   11   11.5  RT (min)      lcorbi_t3_neg_2_050_17    m/z 1189.7654     0   2   4   6   8E+3   14.5   15  RT (min)      lcorbi_t3_neg_4_032_23    m/z 1190.3002     0   1   2   3   4E+3   9   9.5  RT (min)      lcorbi_t3_neg_3_019_12    m/z 1190.5065     0   1   2   3E+4   10.5   11  RT (min)      lcorbi_t3_neg_1_043_14    m/z 1190.7561     0   2   4   6E+3   18   18.5  RT (min)      lcorbi_t3_neg_4_028_39    m/z 1191.6346     0   1   2E+4   8   8.5  RT (min)      lcorbi_t3_neg_2_048_27    m/z 1194.5238     0   1   2E+4   10.5   11  RT (min)      lcorbi_t3_neg_4_042_14    m/z 1194.7676     0   1   2   3   4E+3   18   18.5  RT (min)      lcorbi_t3_neg_2_006_10    m/z 1194.8592     0   2   4   6E+3   16   16.5  RT (min)      lcorbi_t3_neg_2_007_14    m/z 1195.9584     0   2   4   6   8E+3   15   15.5  RT (min)      lcorbi_t3_neg_3_002_24    m/z 1196.6427     0   2   4   6E+3   11   11.5  RT (min)      lcorbi_t3_neg_4_045_35    m/z 1197.6447     0   2   4   6   8E+3   11   11.5  RT (min)      lcorbi_t3_neg_3_022_27    m/z 1197.6544     0   1E+4   11.5   12  RT (min)      lcorbi_t3_neg_4_036_16    m/z 1198.6578     0   2   4   6   8E+3   11.5   12  RT (min)      lcorbi_t3_neg_2_027_13    m/z 1199.5741     0   2   4   6E+3   11   11.5  RT (min)      lcorbi_t3_neg_4_006_47    m/z 1199.6651     0   2   4   6E+4   8   8.5  RT (min)      lcorbi_t3_neg_2_048_27    m/z 1199.9384     0   1E+4   16   16.5  RT (min)      lcorbi_t3_neg_4_005_27    m/z 1199.9621     0   1E+4   14.5   15  RT (min)      lcorbi_t3_neg_2_052_32   
