## Supplementary material for "The Everything Bagel Feature Finder: Ultra-fast automated feature finding for untargeted metabolomics": SI Data File: tool_unique_xic_st_XCMS_ST004501.html

XCMS-unique features (not detected by the other two tools) - ST004501

### XCMS-unique features (not detected by the other two tools) - ST004501

3893 features · orbitrap · XIC window m/z ± 10 ppm (≥ 0.002 Da) · ± 0.5 min

Every panel is a feature **XCMS** detected that neither of the other two tools detected — three-way unique across EB (standard mode (min\_detection\_rate=0.1)), Masscube and XCMS, under the standard matching rule (m/z ± 10 ppm; apex within the reference RT window ± 0.2 min). It shows the MS1 extracted-ion chromatogram at the feature's m/z, drawn from the raw mzML named in `alignment_reference_file`, zoomed to ± 0.5 min around the feature's reported apex RT. **Gray solid line** = the feature's reported apex RT; **red dashed line** = XCMS's assigned MS/MS — `RTINSECONDS` in its `aligned_features.mgf`; the panel widens to keep an off-apex MS/MS on-screen. Features with no assigned MS/MS show only the gray line. y is per-cell autoscaled. Rows keep the table order (m/z ascending).

m/z 50.3714

0123E+433.5RT (min)

lcorbi\_hilic\_pos\_4\_088\_59

m/z 53.2748

024E+355.5RT (min)

lcorbi\_hilic\_pos\_2\_053\_27

m/z 53.6004

01E+50.51RT (min)

lcorbi\_hilic\_pos\_4\_018\_31

m/z 54.0488

01E+322.5RT (min)

lcorbi\_hilic\_pos\_4\_013\_23

m/z 54.8035

024E+433.5RT (min)

lcorbi\_hilic\_pos\_1\_031\_36

m/z 56.0495

0123E+444.5RT (min)

lcorbi\_hilic\_pos\_4\_109\_62

m/z 56.0495

01E+411.5RT (min)

lcorbi\_hilic\_pos\_1\_052\_35

m/z 56.7377

02468E+35.56RT (min)

lcorbi\_hilic\_pos\_4\_031\_17

m/z 56.9648

01E+499.5RT (min)

lcorbi\_hilic\_pos\_4\_086\_59

m/z 57.0335

0123E+40.51RT (min)

lcorbi\_hilic\_pos\_3\_046\_41

m/z 57.5618

01E+42.53RT (min)

lcorbi\_hilic\_pos\_1\_054\_23

m/z 57.9267

0246E+344.5RT (min)

lcorbi\_hilic\_pos\_4\_071\_57

m/z 58.0650

01E+545RT (min)

lcorbi\_hilic\_pos\_4\_070\_57

m/z 58.2873

0123E+311.5RT (min)

lcorbi\_hilic\_pos\_1\_044\_17

m/z 58.3749

02468E+344.5RT (min)

lcorbi\_hilic\_pos\_4\_078\_58

m/z 58.4925

0246E+40.51RT (min)

lcorbi\_hilic\_pos\_4\_087\_59

m/z 58.9327

01234E+388.5RT (min)

lcorbi\_hilic\_pos\_4\_105\_61

m/z 59.0604

01E+444.5RT (min)

lcorbi\_hilic\_pos\_4\_077\_58

m/z 59.0604

01E+41.52RT (min)

lcorbi\_hilic\_pos\_1\_054\_23

m/z 59.7755

02468E+311.5RT (min)

lcorbi\_hilic\_pos\_4\_020\_37

m/z 60.0444

02468E+41313.5RT (min)

lcorbi\_hilic\_pos\_1\_044\_17

m/z 60.0444

0246E+41111.5RT (min)

lcorbi\_hilic\_pos\_1\_044\_17

m/z 60.0444

0123E+499.5RT (min)

lcorbi\_hilic\_pos\_1\_025\_52

m/z 60.0444

01234E+46.57RT (min)

lcorbi\_hilic\_pos\_1\_023\_28

m/z 60.0525

01E+40.51RT (min)

lcorbi\_hilic\_pos\_1\_038\_21

m/z 60.0556

024E+37.58RT (min)

lcorbi\_hilic\_pos\_4\_083\_59

m/z 60.0556

01E+44.55RT (min)

lcorbi\_hilic\_pos\_4\_087\_59

m/z 60.0808

01E+43.54RT (min)

lcorbi\_hilic\_pos\_4\_073\_57

m/z 60.2818

012E+333.5RT (min)

lcorbi\_hilic\_pos\_2\_038\_48

m/z 60.4824

01E+444.5RT (min)

lcorbi\_hilic\_pos\_4\_066\_57

m/z 61.0284

01E+433.5RT (min)

lcorbi\_hilic\_pos\_4\_104\_61

m/z 61.0416

01E+622.5RT (min)

lcorbi\_hilic\_pos\_1\_007\_16

m/z 61.0478

01234E+43.54RT (min)

lcorbi\_hilic\_pos\_4\_086\_59

m/z 61.0590

012E+47.58RT (min)

lcorbi\_hilic\_pos\_4\_093\_60

m/z 61.2685

01234E+355.5RT (min)

lcorbi\_hilic\_pos\_4\_020\_37

m/z 62.0151

01E+43.54RT (min)

lcorbi\_hilic\_pos\_4\_105\_61

m/z 62.0511

0123E+46.57RT (min)

lcorbi\_hilic\_pos\_4\_094\_60

m/z 62.7615

012E+40.51RT (min)

lcorbi\_hilic\_pos\_2\_040\_14

m/z 62.9366

012E+43.54RT (min)

lcorbi\_hilic\_pos\_4\_083\_59

m/z 63.0439

01E+422.5RT (min)

lcorbi\_hilic\_pos\_4\_076\_58

m/z 63.5221

0123E+40.51RT (min)

lcorbi\_hilic\_pos\_2\_047\_22

m/z 64.5274

02468E+36.57RT (min)

lcorbi\_hilic\_pos\_4\_020\_37

m/z 65.0260

0123E+434RT (min)

lcorbi\_hilic\_pos\_4\_082\_59

m/z 65.0260

0123E+40.51RT (min)

lcorbi\_hilic\_pos\_4\_076\_58

m/z 65.5592

01E+433.5RT (min)

lcorbi\_hilic\_pos\_1\_034\_37

m/z 66.0851

01E+455.5RT (min)

lcorbi\_hilic\_pos\_4\_087\_59

m/z 68.0470

0123E+34.55RT (min)

lcorbi\_hilic\_pos\_4\_075\_58

m/z 68.5683

0246E+42.53RT (min)

lcorbi\_hilic\_pos\_3\_037\_20

m/z 69.0335

012E+444.5RT (min)

lcorbi\_hilic\_pos\_4\_089\_59

m/z 69.0698

0123E+20.51RT (min)

lcorbi\_hilic\_pos\_2\_044\_26

m/z 69.0699

0246E+355.5RT (min)

lcorbi\_hilic\_pos\_4\_077\_58

m/z 70.0651

01234E+47.58RT (min)

lcorbi\_hilic\_pos\_4\_088\_59

m/z 71.0685

0246E+35.56RT (min)

lcorbi\_hilic\_pos\_4\_076\_58

m/z 72.0808

01E+51212.5RT (min)

lcorbi\_hilic\_pos\_3\_039\_29

m/z 72.0888

012E+40.51RT (min)

lcorbi\_hilic\_pos\_4\_075\_58

m/z 72.5671

02468E+32.53RT (min)

lcorbi\_hilic\_pos\_3\_019\_43

m/z 73.0648

012E+40.51RT (min)

lcorbi\_hilic\_pos\_1\_046\_30

m/z 73.7823

0123E+344.5RT (min)

lcorbi\_hilic\_pos\_1\_037\_18

m/z 74.0785

01E+31.52RT (min)

lcorbi\_hilic\_pos\_4\_110\_62

m/z 74.0953

01E+61314RT (min)

lcorbi\_hilic\_pos\_4\_100\_61

m/z 74.0964

01E+65.56RT (min)

lcorbi\_hilic\_pos\_4\_106\_62

m/z 74.0976

0246E+600.5RT (min)

lcorbi\_hilic\_pos\_4\_066\_57

m/z 74.5241

01234E+322.5RT (min)

lcorbi\_hilic\_pos\_2\_013\_36

m/z 75.0626

02468E+344.5RT (min)

lcorbi\_hilic\_pos\_4\_092\_60

m/z 75.0997

0123E+41212.5RT (min)

lcorbi\_hilic\_pos\_3\_044\_10

m/z 75.0998

0123E+488.5RT (min)

lcorbi\_hilic\_pos\_4\_097\_60

m/z 75.0998

0246E+45.56RT (min)

lcorbi\_hilic\_pos\_4\_102\_61

m/z 75.0998

01E+43.54RT (min)

lcorbi\_hilic\_pos\_4\_087\_59

m/z 76.0303

01E+41.52RT (min)

lcorbi\_hilic\_pos\_4\_096\_60

m/z 76.0767

0246E+544.5RT (min)

lcorbi\_hilic\_pos\_4\_072\_57

m/z 76.0942

01E+55.56RT (min)

lcorbi\_hilic\_pos\_1\_003\_24

m/z 77.0382

012E+40.51RT (min)

lcorbi\_hilic\_pos\_4\_074\_58

m/z 77.0701

012E+41.52RT (min)

lcorbi\_hilic\_pos\_4\_094\_60

m/z 77.0790

012E+444.5RT (min)

lcorbi\_hilic\_pos\_4\_066\_57

m/z 77.0901

01E+344.5RT (min)

lcorbi\_hilic\_pos\_4\_020\_37

m/z 77.2836

012E+322.5RT (min)

lcorbi\_hilic\_pos\_2\_003\_33

m/z 78.2894

0123E+444.5RT (min)

lcorbi\_hilic\_pos\_2\_012\_37

m/z 79.2753

024E+40.51RT (min)

lcorbi\_hilic\_pos\_4\_103\_61

m/z 80.9478

0246E+400.5RT (min)

lcorbi\_hilic\_pos\_3\_013\_30

m/z 80.9478

024E+45.56RT (min)

lcorbi\_hilic\_pos\_1\_015\_44

m/z 81.0447

012E+433.54RT (min)

lcorbi\_hilic\_pos\_2\_040\_14

m/z 81.0447

012E+40.51RT (min)

lcorbi\_hilic\_pos\_4\_074\_58

m/z 81.0447

024E+46.57RT (min)

lcorbi\_hilic\_pos\_2\_020\_31

m/z 81.0699

0123E+40.51RT (min)

lcorbi\_hilic\_pos\_3\_019\_43

m/z 82.5371

01234E+48.59RT (min)

lcorbi\_hilic\_pos\_4\_078\_58

m/z 82.9449

012E+455.5RT (min)

lcorbi\_hilic\_pos\_4\_098\_61

m/z 83.0604

01234E+30.51RT (min)

lcorbi\_hilic\_pos\_2\_028\_13

m/z 83.0855

02468E+344.5RT (min)

lcorbi\_hilic\_pos\_1\_001\_49

m/z 84.0612

024E+33.54RT (min)

lcorbi\_hilic\_pos\_4\_082\_59

m/z 84.0613

0246E+311.5RT (min)

lcorbi\_hilic\_pos\_4\_096\_60

m/z 84.0808

01E+455.5RT (min)

lcorbi\_hilic\_pos\_4\_072\_57

m/z 84.0808

01E+41.52RT (min)

lcorbi\_hilic\_pos\_4\_071\_57

m/z 84.0900

01E+444.5RT (min)

lcorbi\_hilic\_pos\_3\_036\_33

m/z 84.2860

01E+40.51RT (min)

lcorbi\_hilic\_pos\_4\_073\_57

m/z 84.3967

01E+43.54RT (min)

lcorbi\_hilic\_pos\_4\_068\_57

m/z 84.3975

02468E+30.51RT (min)

lcorbi\_hilic\_pos\_4\_069\_57

m/z 84.5091

024E+33.54RT (min)

lcorbi\_hilic\_pos\_4\_068\_57

m/z 84.5100

0246E+30.51RT (min)

lcorbi\_hilic\_pos\_4\_069\_57

m/z 84.9597

012E+499.5RT (min)

lcorbi\_hilic\_pos\_4\_071\_57

m/z 85.0284

0123E+456RT (min)

lcorbi\_hilic\_pos\_1\_019\_01

m/z 85.0396

012E+40.51RT (min)

lcorbi\_hilic\_pos\_2\_027\_38

m/z 85.0396

012E+42.53RT (min)

lcorbi\_hilic\_pos\_2\_048\_04

m/z 85.0522

012E+344.5RT (min)

lcorbi\_hilic\_pos\_4\_092\_60

m/z 85.0651

0123E+511.5RT (min)

lcorbi\_hilic\_pos\_1\_036\_32

m/z 86.0316

01E+44.55RT (min)

lcorbi\_hilic\_pos\_4\_070\_57

m/z 86.0503

01E+444.5RT (min)

lcorbi\_hilic\_pos\_4\_097\_60

m/z 86.0600

0123E+344.5RT (min)

lcorbi\_hilic\_pos\_4\_092\_60

m/z 86.0950

01234E+555.5RT (min)

lcorbi\_hilic\_pos\_2\_013\_36

m/z 86.5039

0246E+33.54RT (min)

lcorbi\_hilic\_pos\_4\_089\_59

m/z 86.7279

01E+43.54RT (min)

lcorbi\_hilic\_pos\_4\_082\_59

m/z 86.9519

01E+433.5RT (min)

lcorbi\_hilic\_pos\_4\_066\_57

m/z 86.9527

010.51RT (min)

lcorbi\_hilic\_pos\_4\_104\_61

m/z 87.0440

012E+44.555.5RT (min)

lcorbi\_hilic\_pos\_4\_089\_59

m/z 87.0633

024E+333.5RT (min)

lcorbi\_hilic\_pos\_4\_066\_57

m/z 87.0738

01E+46.57RT (min)

lcorbi\_hilic\_pos\_4\_090\_60

m/z 87.0804

010.51RT (min)

lcorbi\_hilic\_pos\_4\_046\_21

m/z 87.1758

01234E+33.54RT (min)

lcorbi\_hilic\_pos\_4\_066\_57

m/z 87.1759

0123E+20.51RT (min)

lcorbi\_hilic\_pos\_1\_037\_18

m/z 87.2547

012E+37.58RT (min)

lcorbi\_hilic\_pos\_1\_029\_14

m/z 87.3998

01E+43.54RT (min)

lcorbi\_hilic\_pos\_4\_071\_57

m/z 87.3998

010.51RT (min)

lcorbi\_hilic\_pos\_1\_038\_21

m/z 87.5034

0123E+37.58RT (min)

lcorbi\_hilic\_pos\_1\_029\_14

m/z 87.5113

01234E+33.54RT (min)

lcorbi\_hilic\_pos\_4\_066\_57

m/z 88.0216

0246E+31.52RT (min)

lcorbi\_hilic\_pos\_4\_084\_59

m/z 88.0216

01E+388.5RT (min)

lcorbi\_hilic\_pos\_4\_018\_31

m/z 88.0577

0246E+4234RT (min)

lcorbi\_hilic\_pos\_4\_090\_60

m/z 88.0757

0246E+455.5RT (min)

lcorbi\_hilic\_pos\_3\_002\_04

m/z 89.0386

012E+43.54RT (min)

lcorbi\_hilic\_pos\_4\_103\_61

m/z 89.0418

01E+46.57RT (min)

lcorbi\_hilic\_pos\_4\_091\_60

m/z 89.0614

01234E+511.5RT (min)

lcorbi\_hilic\_pos\_1\_052\_35

m/z 89.0790

01234E+355.5RT (min)

lcorbi\_hilic\_pos\_4\_068\_57

m/z 89.1073

01234E+400.5RT (min)

lcorbi\_hilic\_pos\_3\_013\_30

m/z 89.1154

024E+333.5RT (min)

lcorbi\_hilic\_pos\_4\_107\_62

m/z 90.0549

012E+344.5RT (min)

lcorbi\_hilic\_pos\_4\_020\_37

m/z 90.0631

01E+411.5RT (min)

lcorbi\_hilic\_pos\_4\_069\_57

m/z 90.0653

012E+46.57RT (min)

lcorbi\_hilic\_pos\_4\_094\_60

m/z 90.0665

0133.5RT (min)

lcorbi\_hilic\_pos\_4\_093\_60

m/z 90.0735

0123E+45.56RT (min)

lcorbi\_hilic\_pos\_4\_095\_60

m/z 90.0913

012E+544.5RT (min)

lcorbi\_hilic\_pos\_4\_007\_05

m/z 90.1779

01234E+333.5RT (min)

lcorbi\_hilic\_pos\_4\_071\_57

m/z 90.5070

02468E+388.5RT (min)

lcorbi\_hilic\_pos\_4\_070\_57

m/z 90.5259

01234E+499.5RT (min)

lcorbi\_hilic\_pos\_4\_079\_58

m/z 90.9733

01E+45.56RT (min)

lcorbi\_hilic\_pos\_3\_031\_36

m/z 91.0291

024E+44.55RT (min)

lcorbi\_hilic\_pos\_3\_018\_01

m/z 91.0316

01E+388.5RT (min)

lcorbi\_hilic\_pos\_4\_105\_61

m/z 91.0493

01E+33.54RT (min)

lcorbi\_hilic\_pos\_4\_077\_58

m/z 91.0542

01234E+50.51RT (min)

lcorbi\_hilic\_pos\_4\_020\_37

m/z 91.0558

02468E+63.54RT (min)

lcorbi\_hilic\_pos\_4\_098\_61

m/z 91.0573

01234E+45.56RT (min)

lcorbi\_hilic\_pos\_4\_096\_60

m/z 91.0616

01E+43.54RT (min)

lcorbi\_hilic\_pos\_4\_104\_61

m/z 91.1132

01E+42.53RT (min)

lcorbi\_hilic\_pos\_4\_094\_60

m/z 91.1229

01E+500.5RT (min)

lcorbi\_hilic\_pos\_4\_066\_57

m/z 91.1230

02468E+34.55RT (min)

lcorbi\_hilic\_pos\_4\_074\_58

m/z 91.5047

0123E+388.5RT (min)

lcorbi\_hilic\_pos\_1\_013\_42

m/z 91.6332

02468E+33.54RT (min)

lcorbi\_hilic\_pos\_4\_095\_60

m/z 91.8571

0246E+33.54RT (min)

lcorbi\_hilic\_pos\_4\_092\_60

m/z 92.0369

012E+41212.5RT (min)

lcorbi\_hilic\_pos\_1\_047\_47

m/z 92.0369

0246E+499.5RT (min)

lcorbi\_hilic\_pos\_1\_013\_42

m/z 92.0369

0123E+50.51RT (min)

lcorbi\_hilic\_pos\_4\_067\_57

m/z 92.0369

012E+53.54RT (min)

lcorbi\_hilic\_pos\_2\_007\_28

m/z 92.0369

01E+56.57RT (min)

lcorbi\_hilic\_pos\_2\_040\_14

m/z 92.0891

01E+444.5RT (min)

lcorbi\_hilic\_pos\_1\_003\_24

m/z 92.9808

015.56RT (min)

lcorbi\_hilic\_pos\_4\_112\_62

m/z 93.0402

01E+422.5RT (min)

lcorbi\_hilic\_pos\_4\_097\_60

m/z 93.0609

012E+43.54RT (min)

lcorbi\_hilic\_pos\_4\_102\_61

m/z 93.1207

01E+477.5RT (min)

lcorbi\_hilic\_pos\_4\_067\_57

m/z 94.0651

0123E+41.52RT (min)

lcorbi\_hilic\_pos\_4\_077\_58

m/z 94.1047

0123E+544.5RT (min)

lcorbi\_hilic\_pos\_4\_007\_05

m/z 94.8051

01E+422.5RT (min)

lcorbi\_hilic\_pos\_4\_014\_27

m/z 95.0561

01E+322.5RT (min)

lcorbi\_hilic\_pos\_3\_035\_22

m/z 95.0603

0123E+344.5RT (min)

lcorbi\_hilic\_pos\_4\_020\_37

m/z 95.0855

01E+40.51RT (min)

lcorbi\_hilic\_pos\_4\_075\_58

m/z 96.0355

0111.5RT (min)

lcorbi\_hilic\_pos\_4\_073\_57

m/z 96.0443

02468E+34.55RT (min)

lcorbi\_hilic\_pos\_4\_113\_62

m/z 96.0710

012E+35.56RT (min)

lcorbi\_hilic\_pos\_4\_079\_58

m/z 97.0284

0166.5RT (min)

lcorbi\_hilic\_pos\_4\_089\_59

m/z 97.0396

01234E+322.5RT (min)

lcorbi\_hilic\_pos\_4\_077\_58

m/z 97.0477

012E+311.5RT (min)

lcorbi\_hilic\_pos\_4\_020\_37

m/z 97.0648

0123E+40.51RT (min)

lcorbi\_hilic\_pos\_2\_002\_11

m/z 97.0743

0246E+35.56RT (min)

lcorbi\_hilic\_pos\_4\_102\_61

m/z 97.2803

01E+40.51RT (min)

lcorbi\_hilic\_pos\_3\_053\_21

m/z 97.9913

012E+499.5RT (min)

lcorbi\_hilic\_pos\_1\_001\_49

m/z 98.0600

01E+34.55RT (min)

lcorbi\_hilic\_pos\_4\_104\_61

m/z 98.0600

02468E+311.5RT (min)

lcorbi\_hilic\_pos\_3\_004\_27

m/z 98.0777

0123E+355.5RT (min)

lcorbi\_hilic\_pos\_4\_090\_60

m/z 98.0793

01E+43.54RT (min)

lcorbi\_hilic\_pos\_4\_086\_59

m/z 98.9841

02468E+46.57RT (min)

lcorbi\_hilic\_pos\_1\_003\_24

m/z 99.0440

0123E+444.5RT (min)

lcorbi\_hilic\_pos\_4\_086\_59

m/z 99.0804

01E+40.51RT (min)

lcorbi\_hilic\_pos\_3\_023\_18

m/z 99.0917

01E+411.5RT (min)

lcorbi\_hilic\_pos\_4\_091\_60

m/z 99.9691

024E+388.5RT (min)

lcorbi\_hilic\_pos\_4\_070\_57

m/z 100.0243

0246E+38.59RT (min)

lcorbi\_hilic\_pos\_4\_102\_61

m/z 100.0497

0123E+422.5RT (min)

lcorbi\_hilic\_pos\_4\_091\_60

m/z 100.0505

01E+344.5RT (min)

lcorbi\_hilic\_pos\_4\_109\_62

m/z 100.0756

012E+433.5RT (min)

lcorbi\_hilic\_pos\_4\_067\_57

m/z 100.0769

024E+511.5RT (min)

lcorbi\_hilic\_pos\_1\_044\_17

m/z 100.0861

01E+43.54RT (min)

lcorbi\_hilic\_pos\_4\_091\_60

m/z 100.0869

02468E+277.5RT (min)

lcorbi\_hilic\_pos\_4\_071\_57

m/z 100.1121

01234E+45.56RT (min)

lcorbi\_hilic\_pos\_4\_045\_33

m/z 100.5100

02468E+388.5RT (min)

lcorbi\_hilic\_pos\_4\_072\_57

m/z 100.9567

01E+488.5RT (min)

lcorbi\_hilic\_pos\_4\_082\_59

m/z 101.0709

0123E+46.57RT (min)

lcorbi\_hilic\_pos\_1\_017\_51

m/z 101.0790

0123E+211.5RT (min)

lcorbi\_hilic\_pos\_1\_013\_42

m/z 101.0961

01234E+30.51RT (min)

lcorbi\_hilic\_pos\_2\_018\_12

m/z 101.1154

0246E+244.5RT (min)

lcorbi\_hilic\_pos\_1\_039\_41

m/z 102.9702

01E+499.5RT (min)

lcorbi\_hilic\_pos\_4\_085\_59

m/z 103.0542

01E+44.55RT (min)

lcorbi\_hilic\_pos\_4\_101\_61

m/z 103.0542

01E+311.5RT (min)

lcorbi\_hilic\_pos\_1\_036\_32

m/z 103.0687

01234E+333.5RT (min)

lcorbi\_hilic\_pos\_4\_018\_31

m/z 103.0768

01E+433.5RT (min)

lcorbi\_hilic\_pos\_4\_094\_60

m/z 104.0237

012E+400.5RT (min)

lcorbi\_hilic\_pos\_4\_106\_62

m/z 104.0237

012E+34.55RT (min)

lcorbi\_hilic\_pos\_4\_110\_62

m/z 104.0369

0123E+444.5RT (min)

lcorbi\_hilic\_pos\_3\_038\_15

m/z 104.0369

024E+40.51RT (min)

lcorbi\_hilic\_pos\_4\_081\_58

m/z 104.0528

0246E+355.5RT (min)

lcorbi\_hilic\_pos\_4\_069\_57

m/z 104.1035

012E+43.54RT (min)

lcorbi\_hilic\_pos\_2\_027\_38

m/z 104.1098

012E+43.54RT (min)

lcorbi\_hilic\_pos\_4\_068\_57

m/z 105.0335

01E+30.51RT (min)

lcorbi\_hilic\_pos\_4\_102\_61

m/z 105.0447

0123E+444.5RT (min)

lcorbi\_hilic\_pos\_3\_043\_42

m/z 105.0447

024E+40.51RT (min)

lcorbi\_hilic\_pos\_4\_076\_58

m/z 105.0650

01E+46.57RT (min)

lcorbi\_hilic\_pos\_2\_030\_05

m/z 105.0698

01E+411.5RT (min)

lcorbi\_hilic\_pos\_4\_076\_58

m/z 105.1103

01E+36.57RT (min)

lcorbi\_hilic\_pos\_4\_097\_60

m/z 105.9537

024E+40.51RT (min)

lcorbi\_hilic\_pos\_1\_006\_11

m/z 106.0525

012E+40.51RT (min)

lcorbi\_hilic\_pos\_4\_067\_57

m/z 106.0525

012E+42.53RT (min)

lcorbi\_hilic\_pos\_4\_070\_57

m/z 106.0652

01234E+311.52RT (min)

lcorbi\_hilic\_pos\_4\_092\_60

m/z 106.0683

01E+344.5RT (min)

lcorbi\_hilic\_pos\_4\_104\_61

m/z 106.0773

013.54RT (min)

lcorbi\_hilic\_pos\_4\_069\_57

m/z 106.0784

0166.5RT (min)

lcorbi\_hilic\_pos\_4\_107\_62

m/z 106.1112

012E+33.54RT (min)

lcorbi\_hilic\_pos\_4\_084\_59

m/z 107.0491

0246E+311.5RT (min)

lcorbi\_hilic\_pos\_4\_111\_62

m/z 107.0603

01E+46.57RT (min)

lcorbi\_hilic\_pos\_4\_088\_59

m/z 107.0855

01234E+30.51RT (min)

lcorbi\_hilic\_pos\_4\_086\_59

m/z 107.0894

012E+444.55RT (min)

lcorbi\_hilic\_pos\_4\_034\_36

m/z 107.9668

02468E+311.5RT (min)

lcorbi\_hilic\_pos\_3\_019\_43

m/z 108.0444

02468E+31.52RT (min)

lcorbi\_hilic\_pos\_4\_020\_37

m/z 108.0665

01234E+355.5RT (min)

lcorbi\_hilic\_pos\_1\_012\_19

m/z 108.0777

012E+33.54RT (min)

lcorbi\_hilic\_pos\_2\_030\_05

m/z 109.0284

01E+477.5RT (min)

lcorbi\_hilic\_pos\_4\_020\_37

m/z 109.0648

01E+42.53RT (min)

lcorbi\_hilic\_pos\_4\_111\_62

m/z 109.0760

0123E+311.5RT (min)

lcorbi\_hilic\_pos\_4\_087\_59

m/z 109.1011

0246E+30.51RT (min)

lcorbi\_hilic\_pos\_2\_016\_18

m/z 110.0349

01234E+444.5RT (min)

lcorbi\_hilic\_pos\_4\_020\_37

m/z 110.0600

012E+42.53RT (min)

lcorbi\_hilic\_pos\_4\_069\_57

m/z 110.0600

01234E+411.5RT (min)

lcorbi\_hilic\_pos\_4\_079\_58

m/z 110.0600

01E+444.5RT (min)

lcorbi\_hilic\_pos\_4\_066\_57

m/z 110.0713

0123E+34.55RT (min)

lcorbi\_hilic\_pos\_4\_013\_23

m/z 111.0440

02468E+322.5RT (min)

lcorbi\_hilic\_pos\_4\_089\_59

m/z 111.0648

01E+411.5RT (min)

lcorbi\_hilic\_pos\_4\_097\_60

m/z 111.0746

01E+37.58RT (min)

lcorbi\_hilic\_pos\_4\_071\_57

m/z 111.0809

0246E+311.5RT (min)

lcorbi\_hilic\_pos\_2\_002\_11

m/z 111.1168

01E+30.51RT (min)

lcorbi\_hilic\_pos\_3\_023\_18

m/z 111.5219

012E+388.5RT (min)

lcorbi\_hilic\_pos\_1\_026\_39

m/z 112.0179

01E+488.5RT (min)

lcorbi\_hilic\_pos\_4\_090\_60

m/z 112.0392

01234E+333.5RT (min)

lcorbi\_hilic\_pos\_4\_068\_57

m/z 112.0564

012E+311.5RT (min)

lcorbi\_hilic\_pos\_2\_020\_31

m/z 112.0757

012E+45.56RT (min)

lcorbi\_hilic\_pos\_4\_083\_59

m/z 112.0869

01E+34.55RT (min)

lcorbi\_hilic\_pos\_4\_076\_58

m/z 112.0869

02468E+37.58RT (min)

lcorbi\_hilic\_pos\_4\_082\_59

m/z 112.0950

02468E+300.5RT (min)

lcorbi\_hilic\_pos\_4\_066\_57

m/z 112.1120

01E+43.54RT (min)

lcorbi\_hilic\_pos\_4\_082\_59

m/z 113.0643

0123E+433.5RT (min)

lcorbi\_hilic\_pos\_4\_090\_60

m/z 113.0805

01E+36.57RT (min)

lcorbi\_hilic\_pos\_4\_104\_61

m/z 113.1073

02468E+45.56RT (min)

lcorbi\_hilic\_pos\_4\_082\_59

m/z 114.0483

012E+444.5RT (min)

lcorbi\_hilic\_pos\_1\_016\_33

m/z 114.0549

0123E+31.52RT (min)

lcorbi\_hilic\_pos\_4\_089\_59

m/z 114.0581

012E+43.54RT (min)

lcorbi\_hilic\_pos\_4\_079\_58

m/z 114.0662

01234E+40.51RT (min)

lcorbi\_hilic\_pos\_1\_019\_01

m/z 114.0731

012E+43.54RT (min)

lcorbi\_hilic\_pos\_4\_068\_57

m/z 114.1026

017.58RT (min)

lcorbi\_hilic\_pos\_1\_013\_42

m/z 115.0502

01E+32.53RT (min)

lcorbi\_hilic\_pos\_4\_112\_62

m/z 115.0542

01E+355.5RT (min)

lcorbi\_hilic\_pos\_4\_106\_62

m/z 115.0542

02468E+20.51RT (min)

lcorbi\_hilic\_pos\_4\_043\_42

m/z 115.0866

012E+355.5RT (min)

lcorbi\_hilic\_pos\_4\_072\_57

m/z 115.0866

01234E+47.58RT (min)

lcorbi\_hilic\_pos\_4\_090\_60

m/z 115.0946

0246E+300.5RT (min)

lcorbi\_hilic\_pos\_4\_070\_57

m/z 115.0946

0123E+31.52RT (min)

lcorbi\_hilic\_pos\_4\_108\_62

m/z 116.0019

0123E+499.5RT (min)

lcorbi\_hilic\_pos\_4\_079\_58

m/z 116.0107

0177.5RT (min)

lcorbi\_hilic\_pos\_4\_073\_57

m/z 116.0342

01234E+46.57RT (min)

lcorbi\_hilic\_pos\_1\_031\_36

m/z 116.0369

012E+433.54RT (min)

lcorbi\_hilic\_pos\_2\_037\_24

m/z 116.0369

01E+40.51RT (min)

lcorbi\_hilic\_pos\_4\_043\_42

m/z 116.0470

0155.5RT (min)

lcorbi\_hilic\_pos\_4\_103\_61

m/z 116.0801

012E+41.52RT (min)

lcorbi\_hilic\_pos\_4\_092\_60

m/z 116.0818

01E+34.55RT (min)

lcorbi\_hilic\_pos\_4\_066\_57

m/z 116.8908

0246E+355.5RT (min)

lcorbi\_hilic\_pos\_4\_076\_58

m/z 116.9719

02468E+400.5RT (min)

lcorbi\_hilic\_pos\_1\_015\_44

m/z 116.9922

0155.5RT (min)

lcorbi\_hilic\_pos\_4\_087\_59

m/z 117.0442

01234E+366.5RT (min)

lcorbi\_hilic\_pos\_4\_066\_57

m/z 117.0465

01E+43.54RT (min)

lcorbi\_hilic\_pos\_1\_016\_33

m/z 117.0572

012E+344.5RT (min)

lcorbi\_hilic\_pos\_4\_088\_59

m/z 117.0739

01E+311.5RT (min)

lcorbi\_hilic\_pos\_4\_020\_37

m/z 117.0739

017.58RT (min)

lcorbi\_hilic\_pos\_4\_104\_61

m/z 117.0743

01E+43.54RT (min)

lcorbi\_hilic\_pos\_4\_104\_61

m/z 117.0815

0246E+43.54RT (min)

lcorbi\_hilic\_pos\_4\_106\_62

m/z 117.0939

0123E+43.54RT (min)

lcorbi\_hilic\_pos\_4\_098\_61

m/z 117.1037

012E+47.58RT (min)

lcorbi\_hilic\_pos\_4\_096\_60

m/z 117.1236

01234E+33.54RT (min)

lcorbi\_hilic\_pos\_4\_100\_61

m/z 117.1385

01234E+400.5RT (min)

lcorbi\_hilic\_pos\_4\_020\_37

m/z 118.0819

01E+555.5RT (min)

lcorbi\_hilic\_pos\_4\_070\_57

m/z 118.0885

02468E+455.5RT (min)

lcorbi\_hilic\_pos\_3\_042\_17

m/z 118.0974

01E+44.555.5RT (min)

lcorbi\_hilic\_pos\_4\_069\_57

m/z 118.1226

0246E+433.5RT (min)

lcorbi\_hilic\_pos\_4\_018\_31

m/z 118.9776

01234E+388.5RT (min)

lcorbi\_hilic\_pos\_4\_074\_58

m/z 119.0161

0246E+35.56RT (min)

lcorbi\_hilic\_pos\_4\_072\_57

m/z 119.0352

01E+533.5RT (min)

lcorbi\_hilic\_pos\_3\_029\_23

m/z 119.0491

024E+35.56RT (min)

lcorbi\_hilic\_pos\_4\_076\_58

m/z 119.0685

024E+311.5RT (min)

lcorbi\_hilic\_pos\_4\_052\_32

m/z 119.0717

01E+355.5RT (min)

lcorbi\_hilic\_pos\_1\_048\_02

m/z 119.0855

024E+30.51RT (min)

lcorbi\_hilic\_pos\_1\_046\_30

m/z 119.0915

0123E+43.54RT (min)

lcorbi\_hilic\_pos\_4\_101\_61

m/z 119.1086

01234E+33.54RT (min)

lcorbi\_hilic\_pos\_4\_102\_61

m/z 119.1193

012E+47.58RT (min)

lcorbi\_hilic\_pos\_4\_097\_60

m/z 119.9752

0246E+388.5RT (min)

lcorbi\_hilic\_pos\_4\_094\_60

m/z 120.0476

01E+577.5RT (min)

lcorbi\_hilic\_pos\_1\_003\_24

m/z 120.0556

01234E+31.52RT (min)

lcorbi\_hilic\_pos\_4\_082\_59

m/z 120.0655

01E+422.5RT (min)

lcorbi\_hilic\_pos\_1\_020\_13

m/z 120.0840

0123E+47.58RT (min)

lcorbi\_hilic\_pos\_4\_097\_60

m/z 120.0927

02468E+355.5RT (min)

lcorbi\_hilic\_pos\_4\_074\_58

m/z 120.1034

02468E+31.52RT (min)

lcorbi\_hilic\_pos\_3\_055\_24

m/z 120.9808

0246E+399.5RT (min)

lcorbi\_hilic\_pos\_2\_047\_22

m/z 121.0284

012E+37.58RT (min)

lcorbi\_hilic\_pos\_4\_084\_59

m/z 121.0551

012E+466.5RT (min)

lcorbi\_hilic\_pos\_3\_010\_32

m/z 121.0648

02468E+344.5RT (min)

lcorbi\_hilic\_pos\_1\_037\_18

m/z 121.0648

012E+433.5RT (min)

lcorbi\_hilic\_pos\_3\_023\_18

m/z 121.0648

01E+411.5RT (min)

lcorbi\_hilic\_pos\_4\_066\_57

m/z 121.0689

0166.5RT (min)

lcorbi\_hilic\_pos\_4\_094\_60

m/z 121.0874

0123E+422.5RT (min)

lcorbi\_hilic\_pos\_1\_003\_24

m/z 121.1011

0246E+30.51RT (min)

lcorbi\_hilic\_pos\_4\_078\_58

m/z 121.1052

024E+34.55RT (min)

lcorbi\_hilic\_pos\_4\_093\_60

m/z 121.9661

0123E+412.513RT (min)

lcorbi\_hilic\_pos\_3\_039\_29

m/z 122.0099

0246E+266.5RT (min)

lcorbi\_hilic\_pos\_1\_001\_49

m/z 122.0521

0166.5RT (min)

lcorbi\_hilic\_pos\_1\_003\_24

m/z 122.0600

012E+311.5RT (min)

lcorbi\_hilic\_pos\_4\_088\_59

m/z 122.0714

0246E+266.5RT (min)

lcorbi\_hilic\_pos\_1\_002\_29

m/z 122.0786

02468E+35RT (min)

lcorbi\_hilic\_pos\_4\_091\_60

m/z 122.9271

01234E+355.5RT (min)

lcorbi\_hilic\_pos\_4\_077\_58

m/z 123.0440

0246E+35.56RT (min)

lcorbi\_hilic\_pos\_4\_080\_58

m/z 123.0819

0246E+355.5RT (min)

lcorbi\_hilic\_pos\_4\_091\_60

m/z 123.0917

01234E+433.5RT (min)

lcorbi\_hilic\_pos\_1\_054\_23

m/z 123.1030

0246E+45.56RT (min)

lcorbi\_hilic\_pos\_4\_092\_60

m/z 123.1168

02468E+30.51RT (min)

lcorbi\_hilic\_pos\_4\_079\_58

m/z 123.9278

02468E+355.5RT (min)

lcorbi\_hilic\_pos\_4\_075\_58

m/z 124.0393

01E+311.5RT (min)

lcorbi\_hilic\_pos\_1\_001\_49

m/z 124.0586

015.56RT (min)

lcorbi\_hilic\_pos\_4\_105\_61

m/z 124.1120

012.53RT (min)

lcorbi\_hilic\_pos\_4\_112\_62

m/z 124.3353

01234E+344.5RT (min)

lcorbi\_hilic\_pos\_4\_074\_58

m/z 124.8973

0123E+32.53RT (min)

lcorbi\_hilic\_pos\_4\_079\_58

m/z 124.9998

0246E+36.57RT (min)

lcorbi\_hilic\_pos\_4\_074\_58

m/z 125.0594

012E+411.5RT (min)

lcorbi\_hilic\_pos\_4\_020\_37

m/z 125.0709

01234E+42RT (min)

lcorbi\_hilic\_pos\_4\_083\_59

m/z 125.0841

012E+355.5RT (min)

lcorbi\_hilic\_pos\_1\_017\_51

m/z 125.0902

01E+40.51RT (min)

lcorbi\_hilic\_pos\_4\_067\_57

m/z 125.0961

01234E+30.51RT (min)

lcorbi\_hilic\_pos\_2\_038\_48

m/z 125.0975

0246E+355.5RT (min)

lcorbi\_hilic\_pos\_4\_097\_60

m/z 125.1073

01E+333.5RT (min)

lcorbi\_hilic\_pos\_4\_102\_61

m/z 125.1187

01E+544.5RT (min)

lcorbi\_hilic\_pos\_4\_092\_60

m/z 125.9722

02468E+388.5RT (min)

lcorbi\_hilic\_pos\_4\_105\_61

m/z 126.0549

01E+44.55RT (min)

lcorbi\_hilic\_pos\_1\_034\_37

m/z 126.0661

01234E+44.55RT (min)

lcorbi\_hilic\_pos\_4\_020\_37

m/z 126.0743

024E+36.57RT (min)

lcorbi\_hilic\_pos\_3\_003\_37

m/z 126.0757

01E+41.52RT (min)

lcorbi\_hilic\_pos\_4\_097\_60

m/z 126.1026

0246E+377.5RT (min)

lcorbi\_hilic\_pos\_4\_020\_37

m/z 126.9207

024E+355.5RT (min)

lcorbi\_hilic\_pos\_4\_077\_58

m/z 127.0389

01E+47.58RT (min)

lcorbi\_hilic\_pos\_4\_086\_59

m/z 127.0688

0111.5RT (min)

lcorbi\_hilic\_pos\_4\_020\_37

m/z 127.0726

01E+344.5RT (min)

lcorbi\_hilic\_pos\_4\_112\_62

m/z 127.0866

012E+34.55RT (min)

lcorbi\_hilic\_pos\_4\_080\_58

m/z 127.0947

0123E+31RT (min)

lcorbi\_hilic\_pos\_4\_066\_57

m/z 127.0996

02468E+24.55RT (min)

lcorbi\_hilic\_pos\_1\_045\_12

m/z 127.1117

024E+30.51RT (min)

lcorbi\_hilic\_pos\_2\_007\_28

m/z 127.1229

0123E+444.5RT (min)

lcorbi\_hilic\_pos\_3\_037\_20

m/z 127.9677

01234E+388.5RT (min)

lcorbi\_hilic\_pos\_4\_069\_57

m/z 128.0177

024E+45.56RT (min)

lcorbi\_hilic\_pos\_4\_020\_37

m/z 128.0192

024E+48.59RT (min)

lcorbi\_hilic\_pos\_4\_091\_60

m/z 128.0343

014.55RT (min)

lcorbi\_hilic\_pos\_4\_096\_60

m/z 128.0454

012E+444.5RT (min)

lcorbi\_hilic\_pos\_4\_086\_59

m/z 128.0620

01E+40.51RT (min)

lcorbi\_hilic\_pos\_4\_111\_62

m/z 128.0995

0111.5RT (min)

lcorbi\_hilic\_pos\_4\_053\_01

m/z 128.1069

0123E+43.54RT (min)

lcorbi\_hilic\_pos\_4\_085\_59

m/z 128.1070

012E+411.5RT (min)

lcorbi\_hilic\_pos\_4\_042\_02

m/z 128.1075

012E+455.5RT (min)

lcorbi\_hilic\_pos\_4\_092\_60

m/z 129.0200

012E+48.59RT (min)

lcorbi\_hilic\_pos\_4\_081\_58

m/z 129.0658

012E+322.5RT (min)

lcorbi\_hilic\_pos\_4\_097\_60

m/z 129.0738

0123E+611.5RT (min)

lcorbi\_hilic\_pos\_2\_003\_33

m/z 129.0739

01E+35RT (min)

lcorbi\_hilic\_pos\_1\_044\_17

m/z 129.0786

01E+411.5RT (min)

lcorbi\_hilic\_pos\_4\_090\_60

m/z 130.0168

02468E+38.59RT (min)

lcorbi\_hilic\_pos\_4\_108\_62

m/z 130.0611

024E+30.51RT (min)

lcorbi\_hilic\_pos\_1\_031\_36

m/z 130.0651

01E+44.555.5RT (min)

lcorbi\_hilic\_pos\_4\_101\_61

m/z 130.0692

0246E+366.5RT (min)

lcorbi\_hilic\_pos\_4\_074\_58

m/z 130.0837

0123E+344.5RT (min)

lcorbi\_hilic\_pos\_1\_028\_34

m/z 130.1151

01234E+433.5RT (min)

lcorbi\_hilic\_pos\_4\_052\_32

m/z 130.1618

012E+32.53RT (min)

lcorbi\_hilic\_pos\_4\_097\_60

m/z 131.0491

01E+455.5RT (min)

lcorbi\_hilic\_pos\_4\_020\_37

m/z 131.0702

01E+411.5RT (min)

lcorbi\_hilic\_pos\_4\_020\_37

m/z 131.0796

01E+411.5RT (min)

lcorbi\_hilic\_pos\_4\_091\_60

m/z 131.0895

024E+411.5RT (min)

lcorbi\_hilic\_pos\_1\_050\_05

m/z 131.0896

02468E+478RT (min)

lcorbi\_hilic\_pos\_4\_092\_60

m/z 131.0910

0246E+42.53RT (min)

lcorbi\_hilic\_pos\_4\_024\_24

m/z 131.0921

01E+344.5RT (min)

lcorbi\_hilic\_pos\_2\_018\_12

m/z 131.1008

017.58RT (min)

lcorbi\_hilic\_pos\_1\_048\_02

m/z 131.1066

0246E+30.51RT (min)

lcorbi\_hilic\_pos\_4\_019\_30

m/z 132.0540

02468E+36.57RT (min)

lcorbi\_hilic\_pos\_4\_101\_61

m/z 132.0588

01E+444.5RT (min)

lcorbi\_hilic\_pos\_4\_097\_60

m/z 132.0655

01234E+322.5RT (min)

lcorbi\_hilic\_pos\_4\_020\_37

m/z 132.0737

024E+366.5RT (min)

lcorbi\_hilic\_pos\_1\_036\_32

m/z 132.0795

0246E+366.5RT (min)

lcorbi\_hilic\_pos\_4\_078\_58

m/z 132.0904

012E+344.5RT (min)

lcorbi\_hilic\_pos\_4\_097\_60

m/z 132.0993

01E+355.5RT (min)

lcorbi\_hilic\_pos\_4\_020\_37

m/z 132.1019

012E+42.53RT (min)

lcorbi\_hilic\_pos\_4\_083\_59

m/z 132.9569

0123E+388.5RT (min)

lcorbi\_hilic\_pos\_1\_045\_12

m/z 133.0522

02468E+255.5RT (min)

lcorbi\_hilic\_pos\_4\_074\_58

m/z 133.0522

01E+311.5RT (min)

lcorbi\_hilic\_pos\_1\_023\_28

m/z 133.0634

01E+42.53RT (min)

lcorbi\_hilic\_pos\_3\_050\_44

m/z 133.0737

01234E+466.5RT (min)

lcorbi\_hilic\_pos\_4\_020\_37

m/z 133.1011

012E+30.511.5RT (min)

lcorbi\_hilic\_pos\_4\_082\_59

m/z 134.0324

012E+366.5RT (min)

lcorbi\_hilic\_pos\_1\_029\_14

m/z 134.0632

012E+433.5RT (min)

lcorbi\_hilic\_pos\_1\_036\_32

m/z 134.0713

01E+344.5RT (min)

lcorbi\_hilic\_pos\_4\_082\_59

m/z 134.0793

01E+311.5RT (min)

lcorbi\_hilic\_pos\_4\_020\_37

m/z 134.0811

024E+444.5RT (min)

lcorbi\_hilic\_pos\_4\_082\_59

m/z 134.0825

012E+466.5RT (min)

lcorbi\_hilic\_pos\_2\_024\_32

m/z 134.0892

02468E+311.5RT (min)

lcorbi\_hilic\_pos\_2\_012\_37

m/z 134.0996

02468E+33.54RT (min)

lcorbi\_hilic\_pos\_4\_090\_60

m/z 134.1005

02468E+37.58RT (min)

lcorbi\_hilic\_pos\_4\_084\_59

m/z 134.1108

02468E+344.5RT (min)

lcorbi\_hilic\_pos\_4\_097\_60

m/z 134.1175

01E+444.5RT (min)

lcorbi\_hilic\_pos\_4\_087\_59

m/z 134.1464

0123E+444.5RT (min)

lcorbi\_hilic\_pos\_4\_052\_32

m/z 134.9725

0246E+388.5RT (min)

lcorbi\_hilic\_pos\_4\_101\_61

m/z 135.0028

02468E+355.5RT (min)

lcorbi\_hilic\_pos\_4\_081\_58

m/z 135.0481

01E+46.57RT (min)

lcorbi\_hilic\_pos\_1\_031\_36

m/z 135.0633

0123E+411.5RT (min)

lcorbi\_hilic\_pos\_4\_076\_58

m/z 135.0755

011.52RT (min)

lcorbi\_hilic\_pos\_1\_019\_01

m/z 135.0916

0111.5RT (min)

lcorbi\_hilic\_pos\_4\_077\_58

m/z 135.0955

0123E+366.5RT (min)

lcorbi\_hilic\_pos\_4\_090\_60

m/z 135.1030

01E+366.5RT (min)

lcorbi\_hilic\_pos\_1\_016\_33

m/z 135.1168

01E+333.5RT (min)

lcorbi\_hilic\_pos\_4\_052\_32

m/z 135.1168

02468E+30.51RT (min)

lcorbi\_hilic\_pos\_4\_079\_58

m/z 135.1223

01E+37.58RT (min)

lcorbi\_hilic\_pos\_2\_050\_06

m/z 136.0393

01E+41.52RT (min)

lcorbi\_hilic\_pos\_4\_074\_58

m/z 136.0757

01E+322.5RT (min)

lcorbi\_hilic\_pos\_4\_097\_60

m/z 136.0968

01E+44.55RT (min)

lcorbi\_hilic\_pos\_1\_044\_17

m/z 136.1063

012E+45.56RT (min)

lcorbi\_hilic\_pos\_4\_091\_60

m/z 136.1151

01E+43.54RT (min)

lcorbi\_hilic\_pos\_3\_023\_18

m/z 137.0458

01E+577.5RT (min)

lcorbi\_hilic\_pos\_4\_020\_37

m/z 137.0548

024E+36.57RT (min)

lcorbi\_hilic\_pos\_4\_097\_60

m/z 137.0596

01234E+433.5RT (min)

lcorbi\_hilic\_pos\_1\_054\_23

m/z 137.0693

01E+555.5RT (min)

lcorbi\_hilic\_pos\_4\_073\_57

m/z 137.0709

012E+411.5RT (min)

lcorbi\_hilic\_pos\_4\_081\_58

m/z 137.0912

024E+444.5RT (min)

lcorbi\_hilic\_pos\_4\_090\_60

m/z 137.1073

02468E+30.51RT (min)

lcorbi\_hilic\_pos\_1\_039\_41

m/z 137.1090

0246E+33.54RT (min)

lcorbi\_hilic\_pos\_2\_002\_11

m/z 137.1106

012E+47.58RT (min)

lcorbi\_hilic\_pos\_4\_094\_60

m/z 137.1324

0246E+30.51RT (min)

lcorbi\_hilic\_pos\_4\_110\_62

m/z 137.9831

01E+37.58RT (min)

lcorbi\_hilic\_pos\_4\_032\_35

m/z 138.0404

01E+488.5RT (min)

lcorbi\_hilic\_pos\_4\_090\_60

m/z 138.0583

01E+344.5RT (min)

lcorbi\_hilic\_pos\_1\_045\_12

m/z 138.0742

014.55RT (min)

lcorbi\_hilic\_pos\_4\_106\_62

m/z 138.0913

01234E+344.5RT (min)

lcorbi\_hilic\_pos\_4\_090\_60

m/z 138.9064

01234E+34.55RT (min)

lcorbi\_hilic\_pos\_2\_055\_42

m/z 139.0179

01234E+499.5RT (min)

lcorbi\_hilic\_pos\_4\_079\_58

m/z 139.0390

01234E+37.58RT (min)

lcorbi\_hilic\_pos\_4\_083\_59

m/z 139.0582

01234E+31.52RT (min)

lcorbi\_hilic\_pos\_4\_088\_59

m/z 139.0865

01E+41.52RT (min)

lcorbi\_hilic\_pos\_4\_083\_59

m/z 139.0866

01E+34.55RT (min)

lcorbi\_hilic\_pos\_4\_066\_57

m/z 139.0978

0166.5RT (min)

lcorbi\_hilic\_pos\_4\_070\_57

m/z 139.1117

01E+30.51RT (min)

lcorbi\_hilic\_pos\_4\_093\_60

m/z 139.1229

02468E+32.53RT (min)

lcorbi\_hilic\_pos\_4\_071\_57

m/z 139.1310

02468E+32.53RT (min)

lcorbi\_hilic\_pos\_4\_079\_58

m/z 139.9879

02468E+400.5RT (min)

lcorbi\_hilic\_pos\_1\_013\_42

m/z 139.9915

01234E+388.5RT (min)

lcorbi\_hilic\_pos\_4\_071\_57

m/z 140.0430

0246E+266.5RT (min)

lcorbi\_hilic\_pos\_1\_001\_49

m/z 140.0535

0246E+422.5RT (min)

lcorbi\_hilic\_pos\_3\_035\_22

m/z 140.0706

012E+41.52RT (min)

lcorbi\_hilic\_pos\_4\_018\_31

m/z 140.0819

024E+366.5RT (min)

lcorbi\_hilic\_pos\_4\_096\_60

m/z 140.1070

02468E+311.5RT (min)

lcorbi\_hilic\_pos\_2\_021\_17

m/z 140.1263

012E+400.5RT (min)

lcorbi\_hilic\_pos\_4\_106\_62

m/z 140.9035

012E+34.55RT (min)

lcorbi\_hilic\_pos\_3\_023\_18

m/z 141.0401

0246E+322.5RT (min)

lcorbi\_hilic\_pos\_3\_010\_32

m/z 141.0509

01E+48.59RT (min)

lcorbi\_hilic\_pos\_4\_079\_58

m/z 141.0785

01E+477.5RT (min)

lcorbi\_hilic\_pos\_4\_096\_60

m/z 141.0910

01E+411.5RT (min)

lcorbi\_hilic\_pos\_2\_002\_11

m/z 141.1109

012E+355.5RT (min)

lcorbi\_hilic\_pos\_1\_020\_13

m/z 141.9586

01234E+41213RT (min)

lcorbi\_hilic\_pos\_4\_083\_59

m/z 141.9935

0246E+388.5RT (min)

lcorbi\_hilic\_pos\_3\_008\_14

m/z 141.9991

01E+44.55RT (min)

lcorbi\_hilic\_pos\_2\_013\_36

m/z 142.0474

01E+466.5RT (min)

lcorbi\_hilic\_pos\_2\_012\_37

m/z 142.0499

01E+31.52RT (min)

lcorbi\_hilic\_pos\_4\_081\_58

m/z 142.0650

01E+34RT (min)

lcorbi\_hilic\_pos\_1\_036\_32

m/z 142.0755

0133.5RT (min)

lcorbi\_hilic\_pos\_4\_074\_58

m/z 142.0756

02468E+322.5RT (min)

lcorbi\_hilic\_pos\_2\_024\_32

m/z 142.1227

01234E+31.52RT (min)

lcorbi\_hilic\_pos\_4\_020\_37

m/z 142.9836

0246E+388.5RT (min)

lcorbi\_hilic\_pos\_4\_029\_41

m/z 143.0729

0155.5RT (min)

lcorbi\_hilic\_pos\_4\_088\_59

m/z 143.0815

01E+45.56RT (min)

lcorbi\_hilic\_pos\_4\_020\_37

m/z 143.0815

01234E+30.51RT (min)

lcorbi\_hilic\_pos\_1\_047\_47

m/z 143.0874

01E+40.51RT (min)

lcorbi\_hilic\_pos\_2\_015\_39

m/z 143.1067

02468E+30.51RT (min)

lcorbi\_hilic\_pos\_2\_047\_22

m/z 143.1067

01234E+444.5RT (min)

lcorbi\_hilic\_pos\_4\_112\_62

m/z 143.1179

02468E+344.5RT (min)

lcorbi\_hilic\_pos\_4\_072\_57

m/z 143.1179

01E+411.5RT (min)

lcorbi\_hilic\_pos\_4\_073\_57

m/z 143.1182

012E+433.5RT (min)

lcorbi\_hilic\_pos\_4\_024\_24

m/z 143.1543

0123E+45.56RT (min)

lcorbi\_hilic\_pos\_4\_105\_61

m/z 143.6099

01234E+377.5RT (min)

lcorbi\_hilic\_pos\_4\_086\_59

m/z 144.0058

01234E+44.55RT (min)

lcorbi\_hilic\_pos\_1\_036\_32

m/z 144.0420

0177.5RT (min)

lcorbi\_hilic\_pos\_1\_002\_29

m/z 144.0808

024E+22.53RT (min)

lcorbi\_hilic\_pos\_2\_051\_15

m/z 144.0986

0246E+455.5RT (min)

lcorbi\_hilic\_pos\_4\_073\_57

m/z 144.1747

0246E+32.53RT (min)

lcorbi\_hilic\_pos\_4\_054\_45

m/z 144.1772

01E+444.5RT (min)

lcorbi\_hilic\_pos\_4\_077\_58

m/z 144.9886

0123E+388.5RT (min)

lcorbi\_hilic\_pos\_1\_003\_24

m/z 145.0495

011.52RT (min)

lcorbi\_hilic\_pos\_1\_003\_24

m/z 145.0608

012E+311.5RT (min)

lcorbi\_hilic\_pos\_4\_020\_37

m/z 145.0609

01E+44.55RT (min)

lcorbi\_hilic\_pos\_4\_034\_36

m/z 145.0688

014.55RT (min)

lcorbi\_hilic\_pos\_4\_097\_60

m/z 145.0689

024E+366.5RT (min)

lcorbi\_hilic\_pos\_4\_084\_59

m/z 145.0785

012E+41.52RT (min)

lcorbi\_hilic\_pos\_3\_055\_24

m/z 145.0840

012E+34.55RT (min)

lcorbi\_hilic\_pos\_4\_093\_60

m/z 145.0971

01E+422.5RT (min)

lcorbi\_hilic\_pos\_4\_089\_59

m/z 145.0989

0246E+355.5RT (min)

lcorbi\_hilic\_pos\_3\_020\_31

m/z 145.1011

02468E+30.51RT (min)

lcorbi\_hilic\_pos\_1\_054\_23

m/z 145.1179

02468E+366.5RT (min)

lcorbi\_hilic\_pos\_1\_036\_32

m/z 145.1335

01234E+34.55RT (min)

lcorbi\_hilic\_pos\_2\_013\_36

m/z 145.1416

024E+300.5RT (min)

lcorbi\_hilic\_pos\_4\_091\_60

m/z 145.1416

01234E+32.53RT (min)

lcorbi\_hilic\_pos\_4\_080\_58

m/z 145.1529

024E+322.5RT (min)

lcorbi\_hilic\_pos\_4\_090\_60

m/z 145.9805

01E+488.5RT (min)

lcorbi\_hilic\_pos\_4\_070\_57

m/z 146.0298

01E+48.59RT (min)

lcorbi\_hilic\_pos\_1\_045\_12

m/z 146.0351

02468E+355.5RT (min)

lcorbi\_hilic\_pos\_4\_099\_61

m/z 146.0812

01E+46.57RT (min)

lcorbi\_hilic\_pos\_4\_109\_62

m/z 146.1061

02468E+245RT (min)

lcorbi\_hilic\_pos\_4\_097\_60

m/z 146.1101

011.52RT (min)

lcorbi\_hilic\_pos\_4\_020\_37

m/z 146.1175

02468E+42.53RT (min)

lcorbi\_hilic\_pos\_4\_020\_37

m/z 147.0275

0246E+255.5RT (min)

lcorbi\_hilic\_pos\_1\_001\_49

m/z 147.0441

01E+35.56RT (min)

lcorbi\_hilic\_pos\_1\_001\_49

m/z 147.0553

0246E+21.52RT (min)

lcorbi\_hilic\_pos\_1\_056\_56

m/z 147.0634

0123E+355.5RT (min)

lcorbi\_hilic\_pos\_4\_074\_58

m/z 147.0731

01E+46.57RT (min)

lcorbi\_hilic\_pos\_3\_048\_38

m/z 147.0766

0246E+311.5RT (min)

lcorbi\_hilic\_pos\_4\_073\_57

m/z 147.0798

02468E+36.57RT (min)

lcorbi\_hilic\_pos\_4\_089\_59

m/z 147.0845

01E+444.5RT (min)

lcorbi\_hilic\_pos\_4\_091\_60

m/z 147.0917

01234E+311.5RT (min)

lcorbi\_hilic\_pos\_4\_089\_59

m/z 147.0956

0123E+444.5RT (min)

lcorbi\_hilic\_pos\_2\_022\_35

m/z 147.1168

01E+40.51RT (min)

lcorbi\_hilic\_pos\_3\_019\_43

m/z 147.9519

0123E+388.5RT (min)

lcorbi\_hilic\_pos\_4\_073\_57

m/z 148.0039

01E+55.56RT (min)

lcorbi\_hilic\_pos\_4\_020\_37

m/z 148.0345

01E+45.56RT (min)

lcorbi\_hilic\_pos\_4\_087\_59

m/z 148.0427

024E+35.56RT (min)

lcorbi\_hilic\_pos\_4\_013\_23

m/z 148.0604

0246E+333.5RT (min)

lcorbi\_hilic\_pos\_4\_086\_59

m/z 148.0756

01234E+311.5RT (min)

lcorbi\_hilic\_pos\_4\_077\_58

m/z 148.0757

016.57RT (min)

lcorbi\_hilic\_pos\_4\_090\_60

m/z 148.0937

01E+444.5RT (min)

lcorbi\_hilic\_pos\_1\_034\_37

m/z 148.1049

012E+311.5RT (min)

lcorbi\_hilic\_pos\_4\_014\_27

m/z 148.1139

0246E+23.54RT (min)

lcorbi\_hilic\_pos\_2\_037\_24

m/z 148.1145

01234E+46.57RT (min)

lcorbi\_hilic\_pos\_1\_016\_33

m/z 148.1332

012E+344.5RT (min)

lcorbi\_hilic\_pos\_2\_020\_31

m/z 149.0266

01E+46.57RT (min)

lcorbi\_hilic\_pos\_1\_031\_36

m/z 149.0271

0246E+40.51RT (min)

lcorbi\_hilic\_pos\_3\_034\_26

m/z 149.0575

01E+46.57RT (min)

lcorbi\_hilic\_pos\_1\_016\_33

m/z 149.0597

0155.5RT (min)

lcorbi\_hilic\_pos\_4\_096\_60

m/z 149.0597

02468E+233.5RT (min)

lcorbi\_hilic\_pos\_2\_037\_24

m/z 149.0597

0123E+40.51RT (min)

lcorbi\_hilic\_pos\_4\_086\_59

m/z 149.0806

01E+36.57RT (min)

lcorbi\_hilic\_pos\_4\_092\_60

m/z 149.0823

024E+333.5RT (min)

lcorbi\_hilic\_pos\_1\_043\_25

m/z 149.0823

012E+411.5RT (min)

lcorbi\_hilic\_pos\_3\_001\_49

m/z 149.0921

01E+422.5RT (min)

lcorbi\_hilic\_pos\_4\_031\_17

m/z 149.0961

01E+30.51RT (min)

lcorbi\_hilic\_pos\_4\_086\_59

m/z 149.1016

012E+455.5RT (min)

lcorbi\_hilic\_pos\_4\_096\_60

m/z 149.1325

0246E+30.51RT (min)

lcorbi\_hilic\_pos\_4\_086\_59

m/z 149.9928

01234E+366.5RT (min)

lcorbi\_hilic\_pos\_4\_078\_58

m/z 150.0549

01E+433.5RT (min)

lcorbi\_hilic\_pos\_4\_099\_61

m/z 150.0662

02468E+300.5RT (min)

lcorbi\_hilic\_pos\_4\_106\_62

m/z 150.0774

01E+35.56RT (min)

lcorbi\_hilic\_pos\_4\_091\_60

m/z 150.1010

01234E+344.5RT (min)

lcorbi\_hilic\_pos\_2\_039\_09

m/z 150.1251

0123E+455.5RT (min)

lcorbi\_hilic\_pos\_4\_097\_60

m/z 150.1277

01E+433.5RT (min)

lcorbi\_hilic\_pos\_1\_049\_55

m/z 150.6334

01E+477.5RT (min)

lcorbi\_hilic\_pos\_4\_093\_60

m/z 151.0389

0246E+32.53RT (min)

lcorbi\_hilic\_pos\_4\_107\_62

m/z 151.0617

024E+35.56RT (min)

lcorbi\_hilic\_pos\_2\_003\_33

m/z 151.0753

0246E+32.53RT (min)

lcorbi\_hilic\_pos\_4\_020\_37

m/z 151.0753

012E+311.5RT (min)

lcorbi\_hilic\_pos\_1\_024\_03

m/z 151.0898

0144.5RT (min)

lcorbi\_hilic\_pos\_4\_087\_59

m/z 151.0931

024E+311.5RT (min)

lcorbi\_hilic\_pos\_1\_052\_35

m/z 151.1092

01E+44.55RT (min)

lcorbi\_hilic\_pos\_2\_037\_24

m/z 151.1117

0246E+30.51RT (min)

lcorbi\_hilic\_pos\_4\_082\_59

m/z 152.0220

01E+43.54RT (min)

lcorbi\_hilic\_pos\_4\_074\_58

m/z 152.0479

01E+45.56RT (min)

lcorbi\_hilic\_pos\_4\_092\_60

m/z 152.0707

012E+356RT (min)

lcorbi\_hilic\_pos\_4\_094\_60

m/z 152.0897

01E+36.57RT (min)

lcorbi\_hilic\_pos\_4\_095\_60

m/z 152.1296

01E+466.5RT (min)

lcorbi\_hilic\_pos\_4\_097\_60

m/z 152.1296

01E+47.58RT (min)

lcorbi\_hilic\_pos\_4\_094\_60

m/z 152.1377

012.53RT (min)

lcorbi\_hilic\_pos\_2\_044\_26

m/z 152.9471

01E+499.5RT (min)

lcorbi\_hilic\_pos\_4\_082\_59

m/z 153.0328

01E+48.59RT (min)

lcorbi\_hilic\_pos\_4\_088\_59

m/z 153.0407

024E+44.55RT (min)

lcorbi\_hilic\_pos\_4\_020\_37

m/z 153.0537

01E+44.55RT (min)

lcorbi\_hilic\_pos\_4\_082\_59

m/z 153.0634

0123E+35.56RT (min)

lcorbi\_hilic\_pos\_4\_087\_59

m/z 153.0738

012E+311.5RT (min)

lcorbi\_hilic\_pos\_4\_074\_58

m/z 153.0901

012E+36.57RT (min)

lcorbi\_hilic\_pos\_3\_033\_53

m/z 153.1007

0246E+311.5RT (min)

lcorbi\_hilic\_pos\_4\_066\_57

m/z 153.1066

0246E+255.5RT (min)

lcorbi\_hilic\_pos\_1\_042\_40

m/z 153.1136

01E+444.5RT (min)

lcorbi\_hilic\_pos\_4\_096\_60

m/z 153.1410

0246E+34.55RT (min)

lcorbi\_hilic\_pos\_4\_024\_24

m/z 153.9999

01234E+366.5RT (min)

lcorbi\_hilic\_pos\_4\_112\_62

m/z 154.0434

01234E+46.57RT (min)

lcorbi\_hilic\_pos\_1\_043\_25

m/z 154.0692

02468E+31.52RT (min)

lcorbi\_hilic\_pos\_4\_069\_57

m/z 154.0742

0123E+46.57RT (min)

lcorbi\_hilic\_pos\_4\_020\_37

m/z 154.0973

02468E+34.55RT (min)

lcorbi\_hilic\_pos\_4\_066\_57

m/z 155.0427

01234E+46.57RT (min)

lcorbi\_hilic\_pos\_2\_027\_38

m/z 155.0532

0123E+31.52RT (min)

lcorbi\_hilic\_pos\_4\_071\_57

m/z 155.0620

0246E+366.5RT (min)

lcorbi\_hilic\_pos\_4\_020\_37

m/z 155.0703

0123E+311.5RT (min)

lcorbi\_hilic\_pos\_4\_067\_57

m/z 155.0751

012E+455.5RT (min)

lcorbi\_hilic\_pos\_1\_003\_24

m/z 155.0791

012E+47.58RT (min)

lcorbi\_hilic\_pos\_4\_085\_59

m/z 155.1066

024E+311.5RT (min)

lcorbi\_hilic\_pos\_4\_067\_57

m/z 156.0267

0246E+477.5RT (min)

lcorbi\_hilic\_pos\_1\_054\_23

m/z 156.0509

0246E+344.5RT (min)

lcorbi\_hilic\_pos\_2\_020\_31

m/z 156.0573

012E+444.5RT (min)

lcorbi\_hilic\_pos\_4\_006\_14

m/z 156.0655

02468E+31.52RT (min)

lcorbi\_hilic\_pos\_1\_034\_37

m/z 156.0701

024E+44.55RT (min)

lcorbi\_hilic\_pos\_4\_097\_60

m/z 156.1576

01E+42.53RT (min)

lcorbi\_hilic\_pos\_4\_091\_60

m/z 156.9542

02468E+311.5RT (min)

lcorbi\_hilic\_pos\_4\_069\_57

m/z 156.9637

012E+466.5RT (min)

lcorbi\_hilic\_pos\_4\_097\_60

m/z 157.0647

01E+344.5RT (min)

lcorbi\_hilic\_pos\_3\_039\_29

m/z 157.0801

01E+47.58RT (min)

lcorbi\_hilic\_pos\_4\_020\_37

m/z 157.0891

02468E+344.5RT (min)

lcorbi\_hilic\_pos\_4\_097\_60

m/z 157.0977

01E+411.5RT (min)

lcorbi\_hilic\_pos\_3\_010\_32

m/z 157.1084

024E+27.58RT (min)

lcorbi\_hilic\_pos\_2\_021\_17

m/z 157.1335

01E+400.5RT (min)

lcorbi\_hilic\_pos\_4\_113\_62

m/z 157.1416

012E+433.5RT (min)

lcorbi\_hilic\_pos\_4\_080\_58

m/z 158.0575

01E+54.55RT (min)

lcorbi\_hilic\_pos\_4\_090\_60

m/z 158.0704

012E+44.55RT (min)

lcorbi\_hilic\_pos\_4\_097\_60

m/z 158.0818

01E+511.5RT (min)

lcorbi\_hilic\_pos\_4\_020\_37

m/z 158.0958

012E+344.5RT (min)

lcorbi\_hilic\_pos\_3\_055\_24

m/z 158.1175

0246E+311.5RT (min)

lcorbi\_hilic\_pos\_1\_036\_32

m/z 158.1733

02468E+36.57RT (min)

lcorbi\_hilic\_pos\_4\_069\_57

m/z 158.9220

01234E+377.5RT (min)

lcorbi\_hilic\_pos\_1\_010\_07

m/z 158.9683

012E+35.56RT (min)

lcorbi\_hilic\_pos\_3\_002\_04

m/z 159.0561

016.57RT (min)

lcorbi\_hilic\_pos\_2\_023\_34

m/z 159.0652

024E+344.5RT (min)

lcorbi\_hilic\_pos\_4\_066\_57

m/z 159.0764

01E+44.55RT (min)

lcorbi\_hilic\_pos\_4\_109\_62

m/z 159.0764

01234E+322.5RT (min)

lcorbi\_hilic\_pos\_4\_068\_57

m/z 159.0958

02468E+37.58RT (min)

lcorbi\_hilic\_pos\_4\_087\_59

m/z 159.1208

02468E+355.5RT (min)

lcorbi\_hilic\_pos\_4\_069\_57

m/z 159.1490

01E+44.55RT (min)

lcorbi\_hilic\_pos\_4\_066\_57

m/z 159.1492

0246E+26.57RT (min)

lcorbi\_hilic\_pos\_1\_031\_36

m/z 160.0401

02468E+477.5RT (min)

lcorbi\_hilic\_pos\_1\_036\_32

m/z 160.0556

0166.5RT (min)

lcorbi\_hilic\_pos\_4\_090\_60

m/z 160.0604

01E+41.52RT (min)

lcorbi\_hilic\_pos\_3\_001\_49

m/z 160.0669

01E+466.5RT (min)

lcorbi\_hilic\_pos\_4\_020\_37

m/z 160.0798

01E+46.57RT (min)

lcorbi\_hilic\_pos\_4\_085\_59

m/z 160.0968

01E+41.52RT (min)

lcorbi\_hilic\_pos\_1\_024\_03

m/z 160.0968

01E+54.555.5RT (min)

lcorbi\_hilic\_pos\_2\_046\_46

m/z 160.1176

01E+422.5RT (min)

lcorbi\_hilic\_pos\_4\_097\_60

m/z 160.1176

01E+466.5RT (min)

lcorbi\_hilic\_pos\_4\_091\_60

m/z 160.1374

0123E+344.5RT (min)

lcorbi\_hilic\_pos\_2\_004\_01

m/z 160.1808

0123E+400.5RT (min)

lcorbi\_hilic\_pos\_4\_020\_37

m/z 160.1808

0123E+31313.5RT (min)

lcorbi\_hilic\_pos\_4\_022\_08

m/z 160.9648

0123E+322.5RT (min)

lcorbi\_hilic\_pos\_4\_077\_58

m/z 160.9683

01E+45.56RT (min)

lcorbi\_hilic\_pos\_4\_072\_57

m/z 160.9855

0246E+388.5RT (min)

lcorbi\_hilic\_pos\_4\_087\_59

m/z 161.0378

01E+45.56RT (min)

lcorbi\_hilic\_pos\_3\_011\_34

m/z 161.0597

024E+32.53RT (min)

lcorbi\_hilic\_pos\_4\_111\_62

m/z 161.0934

02468E+377.5RT (min)

lcorbi\_hilic\_pos\_4\_096\_60

m/z 161.1002

024E+344.5RT (min)

lcorbi\_hilic\_pos\_2\_009\_50

m/z 161.1172

01E+411.5RT (min)

lcorbi\_hilic\_pos\_3\_028\_07

m/z 161.1324

01E+40.51RT (min)

lcorbi\_hilic\_pos\_4\_074\_58

m/z 161.1729

0123E+333.5RT (min)

lcorbi\_hilic\_pos\_3\_015\_11

m/z 162.0074

012E+499.5RT (min)

lcorbi\_hilic\_pos\_4\_073\_57

m/z 162.0249

02468E+366.5RT (min)

lcorbi\_hilic\_pos\_4\_069\_57

m/z 162.0743

024E+32.53RT (min)

lcorbi\_hilic\_pos\_2\_023\_34

m/z 162.0954

024E+255.5RT (min)

lcorbi\_hilic\_pos\_3\_002\_04

m/z 162.1103

0123E+555.5RT (min)

lcorbi\_hilic\_pos\_1\_051\_26

m/z 162.1277

02468E+35.56RT (min)

lcorbi\_hilic\_pos\_1\_019\_01

m/z 162.1277

01E+411.5RT (min)

lcorbi\_hilic\_pos\_4\_079\_58

m/z 162.1488

01E+43.54RT (min)

lcorbi\_hilic\_pos\_4\_101\_61

m/z 162.9069

02468E+34.55RT (min)

lcorbi\_hilic\_pos\_2\_035\_44

m/z 162.9069

01E+46.57RT (min)

lcorbi\_hilic\_pos\_4\_084\_59

m/z 162.9809

0123E+388.5RT (min)

lcorbi\_hilic\_pos\_1\_043\_25

m/z 163.0390

012E+411.5RT (min)

lcorbi\_hilic\_pos\_2\_054\_29

m/z 163.0534

0123E+35.56RT (min)

lcorbi\_hilic\_pos\_4\_072\_57

m/z 163.0617

01E+41.522.5RT (min)

lcorbi\_hilic\_pos\_2\_007\_28

m/z 163.0794

0123E+25.56RT (min)

lcorbi\_hilic\_pos\_2\_021\_17

m/z 163.0866

02468E+31.52RT (min)

lcorbi\_hilic\_pos\_2\_003\_33

m/z 163.0965

01E+411.5RT (min)

lcorbi\_hilic\_pos\_3\_019\_43

m/z 163.1092

01E+47.58RT (min)

lcorbi\_hilic\_pos\_4\_094\_60

m/z 163.1229

012E+43.54RT (min)

lcorbi\_hilic\_pos\_4\_073\_57

m/z 163.1285

012E+47.58RT (min)

lcorbi\_hilic\_pos\_4\_097\_60

m/z 163.1481

0123E+30.51RT (min)

lcorbi\_hilic\_pos\_4\_088\_59

m/z 163.9707

01234E+388.5RT (min)

lcorbi\_hilic\_pos\_4\_082\_59

m/z 164.0376

02468E+31.52RT (min)

lcorbi\_hilic\_pos\_1\_044\_17

m/z 164.0541

012E+322.5RT (min)

lcorbi\_hilic\_pos\_4\_066\_57

m/z 164.0566

01E+43.54RT (min)

lcorbi\_hilic\_pos\_4\_082\_59

m/z 164.0705

01E+311.5RT (min)

lcorbi\_hilic\_pos\_2\_001\_49

m/z 164.0739

01234E+411.52RT (min)

lcorbi\_hilic\_pos\_4\_068\_57

m/z 164.0794

01E+36.57RT (min)

lcorbi\_hilic\_pos\_4\_106\_62

m/z 164.0917

02468E+34.55RT (min)

lcorbi\_hilic\_pos\_2\_023\_34

m/z 164.1069

01E+31.52RT (min)

lcorbi\_hilic\_pos\_3\_029\_23

m/z 164.1190

0123E+455.5RT (min)

lcorbi\_hilic\_pos\_1\_051\_26

m/z 164.1206

01E+41.52RT (min)

lcorbi\_hilic\_pos\_4\_096\_60

m/z 164.1281

01234E+54.55RT (min)

lcorbi\_hilic\_pos\_1\_040\_06

m/z 164.1370

010.51RT (min)

lcorbi\_hilic\_pos\_1\_037\_18

m/z 165.0134

02468E+355.5RT (min)

lcorbi\_hilic\_pos\_4\_066\_57

m/z 165.0327

0246E+366.5RT (min)

lcorbi\_hilic\_pos\_4\_067\_57

m/z 165.0740

0246E+333.5RT (min)

lcorbi\_hilic\_pos\_4\_105\_61

m/z 165.0772

012E+423RT (min)

lcorbi\_hilic\_pos\_4\_094\_60

m/z 165.0910

0123E+40.51RT (min)

lcorbi\_hilic\_pos\_4\_070\_57

m/z 165.1274

01E+40.51RT (min)

lcorbi\_hilic\_pos\_4\_101\_61

m/z 165.9625

024E+388.5RT (min)

lcorbi\_hilic\_pos\_4\_067\_57

m/z 165.9878

0246E+36.57RT (min)

lcorbi\_hilic\_pos\_4\_071\_57

m/z 166.0144

02468E+37.58RT (min)

lcorbi\_hilic\_pos\_4\_084\_59

m/z 166.0532

01E+44.55RT (min)

lcorbi\_hilic\_pos\_4\_020\_37

m/z 166.0580

0246E+35.56RT (min)

lcorbi\_hilic\_pos\_4\_081\_58

m/z 166.0839

024E+455.5RT (min)

lcorbi\_hilic\_pos\_4\_021\_13

m/z 166.0862

0123E+42.53RT (min)

lcorbi\_hilic\_pos\_4\_083\_59

m/z 166.0862

0246E+411.5RT (min)

lcorbi\_hilic\_pos\_4\_020\_37

m/z 166.0975

01E+411.5RT (min)

lcorbi\_hilic\_pos\_3\_034\_26

m/z 166.1226

012E+33.54RT (min)

lcorbi\_hilic\_pos\_2\_038\_48

m/z 167.0508

02468E+333.5RT (min)

lcorbi\_hilic\_pos\_4\_083\_59

m/z 167.0563

01E+444.5RT (min)

lcorbi\_hilic\_pos\_4\_020\_37

m/z 167.0563

0246E+322.5RT (min)

lcorbi\_hilic\_pos\_4\_067\_57

m/z 167.0566

02468E+36.57RT (min)

lcorbi\_hilic\_pos\_1\_034\_37

m/z 167.0669

012E+45.56RT (min)

lcorbi\_hilic\_pos\_4\_096\_60

m/z 167.0757

01E+444.5RT (min)

lcorbi\_hilic\_pos\_4\_020\_37

m/z 167.0815

01E+355.5RT (min)

lcorbi\_hilic\_pos\_2\_001\_49

m/z 167.0927

01234E+455.5RT (min)

lcorbi\_hilic\_pos\_4\_095\_60

m/z 167.1066

012E+30.51RT (min)

lcorbi\_hilic\_pos\_2\_044\_26

m/z 167.1122

02468E+36.57RT (min)

lcorbi\_hilic\_pos\_4\_093\_60

m/z 168.0113

01234E+411.5RT (min)

lcorbi\_hilic\_pos\_2\_023\_34

m/z 168.0574

01E+42.53RT (min)

lcorbi\_hilic\_pos\_4\_072\_57

m/z 168.0767

02468E+30.51RT (min)

lcorbi\_hilic\_pos\_2\_027\_38

m/z 168.1131

0123E+45.56RT (min)

lcorbi\_hilic\_pos\_4\_020\_37

m/z 168.1383

012E+322.5RT (min)

lcorbi\_hilic\_pos\_4\_072\_57

m/z 169.0090

01E+433.5RT (min)

lcorbi\_hilic\_pos\_2\_005\_23

m/z 169.0332

024E+300.5RT (min)

lcorbi\_hilic\_pos\_4\_032\_35

m/z 169.0458

0123E+48.59RT (min)

lcorbi\_hilic\_pos\_1\_047\_47

m/z 169.0624

01E+322.5RT (min)

lcorbi\_hilic\_pos\_4\_070\_57

m/z 169.0735

02468E+23.54RT (min)

lcorbi\_hilic\_pos\_4\_020\_37

m/z 169.0759

02468E+21.52RT (min)

lcorbi\_hilic\_pos\_3\_042\_17

m/z 169.0859

01E+311.5RT (min)

lcorbi\_hilic\_pos\_1\_007\_16

m/z 169.1011

0111.5RT (min)

lcorbi\_hilic\_pos\_4\_072\_57

m/z 169.1223

01E+40.51RT (min)

lcorbi\_hilic\_pos\_4\_110\_62

m/z 169.1252

01E+455.5RT (min)

lcorbi\_hilic\_pos\_4\_099\_61

m/z 169.1335

01E+32.53RT (min)

lcorbi\_hilic\_pos\_4\_067\_57

m/z 170.0038

01E+433.5RT (min)

lcorbi\_hilic\_pos\_2\_005\_23

m/z 170.0326

01E+466.5RT (min)

lcorbi\_hilic\_pos\_4\_020\_37

m/z 170.0787

0246E+43.54RT (min)

lcorbi\_hilic\_pos\_4\_093\_60

m/z 170.0924

01E+31.52RT (min)

lcorbi\_hilic\_pos\_4\_020\_37

m/z 170.1539

01234E+32.53RT (min)

lcorbi\_hilic\_pos\_4\_078\_58

m/z 170.1539

01E+40.51RT (min)

lcorbi\_hilic\_pos\_3\_019\_43

m/z 170.1732

012E+400.5RT (min)

lcorbi\_hilic\_pos\_4\_089\_59

m/z 170.1903

0123E+411.5RT (min)

lcorbi\_hilic\_pos\_4\_032\_35

m/z 170.9793

01234E+366.5RT (min)

lcorbi\_hilic\_pos\_4\_091\_60

m/z 171.0166

024E+36.57RT (min)

lcorbi\_hilic\_pos\_2\_027\_38

m/z 171.0457

0246E+36.57RT (min)

lcorbi\_hilic\_pos\_4\_087\_59

m/z 171.0764

01E+43.54RT (min)

lcorbi\_hilic\_pos\_4\_020\_37

m/z 171.0764

01E+31.52RT (min)

lcorbi\_hilic\_pos\_4\_090\_60

m/z 171.0916

02468E+40.51RT (min)

lcorbi\_hilic\_pos\_4\_104\_61

m/z 171.0957

01E+477.5RT (min)

lcorbi\_hilic\_pos\_4\_020\_37

m/z 171.1030

02468E+355.5RT (min)

lcorbi\_hilic\_pos\_4\_103\_61

m/z 171.1128

024E+44.55RT (min)

lcorbi\_hilic\_pos\_4\_020\_37

m/z 171.1492

01234E+30.51RT (min)

lcorbi\_hilic\_pos\_3\_011\_34

m/z 171.1855

0246E+400.5RT (min)

lcorbi\_hilic\_pos\_4\_105\_61

m/z 171.1855

01E+41313.5RT (min)

lcorbi\_hilic\_pos\_4\_101\_61

m/z 172.0007

01E+46.57RT (min)

lcorbi\_hilic\_pos\_1\_054\_23

m/z 172.0191

01E+433.5RT (min)

lcorbi\_hilic\_pos\_4\_095\_60

m/z 172.0604

01E+344.5RT (min)

lcorbi\_hilic\_pos\_2\_024\_32

m/z 172.0675

01234E+433.5RT (min)

lcorbi\_hilic\_pos\_4\_095\_60

m/z 172.0757

0144.5RT (min)

lcorbi\_hilic\_pos\_4\_071\_57

m/z 172.1080

02468E+366.5RT (min)

lcorbi\_hilic\_pos\_3\_001\_49

m/z 172.1167

01E+411.5RT (min)

lcorbi\_hilic\_pos\_3\_010\_32

m/z 172.1193

01E+54.55RT (min)

lcorbi\_hilic\_pos\_1\_032\_31

m/z 172.1332

012E+422.5RT (min)

lcorbi\_hilic\_pos\_4\_111\_62

m/z 172.1332

01E+40.51RT (min)

lcorbi\_hilic\_pos\_1\_054\_23

m/z 172.1444

024E+400.5RT (min)

lcorbi\_hilic\_pos\_1\_005\_45

m/z 173.0209

02468E+366.5RT (min)

lcorbi\_hilic\_pos\_4\_020\_37

m/z 173.0557

01E+41.52RT (min)

lcorbi\_hilic\_pos\_4\_066\_57

m/z 173.0658

012E+46.57RT (min)

lcorbi\_hilic\_pos\_1\_016\_33

m/z 173.0718

01E+46.57RT (min)

lcorbi\_hilic\_pos\_4\_097\_60

m/z 173.0808

01234E+344.5RT (min)

lcorbi\_hilic\_pos\_4\_070\_57

m/z 173.0921

012E+44.55RT (min)

lcorbi\_hilic\_pos\_1\_009\_50

m/z 173.0979

012E+311.5RT (min)

lcorbi\_hilic\_pos\_1\_037\_18

m/z 173.1097

02468E+32.53RT (min)

lcorbi\_hilic\_pos\_4\_024\_24

m/z 173.1097

024E+355.5RT (min)

lcorbi\_hilic\_pos\_4\_104\_61

m/z 173.1284

0246E+233.5RT (min)

lcorbi\_hilic\_pos\_4\_020\_37

m/z 173.1365

024E+344.5RT (min)

lcorbi\_hilic\_pos\_4\_068\_57

m/z 173.1536

0123E+30.51RT (min)

lcorbi\_hilic\_pos\_2\_016\_18

m/z 173.1729

024E+333.5RT (min)

lcorbi\_hilic\_pos\_4\_105\_61

m/z 173.2093

02468E+322.5RT (min)

lcorbi\_hilic\_pos\_4\_054\_45

m/z 174.0137

02468E+333.5RT (min)

lcorbi\_hilic\_pos\_4\_075\_58

m/z 174.0331

024E+366.5RT (min)

lcorbi\_hilic\_pos\_4\_076\_58

m/z 174.0501

0246E+35.56RT (min)

lcorbi\_hilic\_pos\_4\_068\_57

m/z 174.0761

0111.5RT (min)

lcorbi\_hilic\_pos\_1\_014\_27

m/z 174.0932

02468E+24.55RT (min)

lcorbi\_hilic\_pos\_4\_052\_32

m/z 174.1125

0246E+344.5RT (min)

lcorbi\_hilic\_pos\_1\_043\_25

m/z 174.1277

01E+30.51RT (min)

lcorbi\_hilic\_pos\_3\_019\_43

m/z 174.1324

01234E+444.5RT (min)

lcorbi\_hilic\_pos\_4\_020\_37

m/z 174.1488

0246E+311.5RT (min)

lcorbi\_hilic\_pos\_1\_028\_34

m/z 174.1687

024E+333.5RT (min)

lcorbi\_hilic\_pos\_2\_007\_28

m/z 174.1852

0123E+411.5RT (min)

lcorbi\_hilic\_pos\_1\_045\_12

m/z 174.2028

02468E+333.5RT (min)

lcorbi\_hilic\_pos\_4\_050\_16

m/z 174.9978

02468E+36.57RT (min)

lcorbi\_hilic\_pos\_4\_082\_59

m/z 175.0206

012E+433.5RT (min)

lcorbi\_hilic\_pos\_4\_095\_60

m/z 175.0866

01E+444.5RT (min)

lcorbi\_hilic\_pos\_4\_105\_61

m/z 175.0965

0246E+30.51RT (min)

lcorbi\_hilic\_pos\_2\_036\_08

m/z 175.1078

01234E+45.566.5RT (min)

lcorbi\_hilic\_pos\_4\_096\_60

m/z 175.1157

012E+47.58RT (min)

lcorbi\_hilic\_pos\_1\_054\_23

m/z 175.1164

0123E+455.5RT (min)

lcorbi\_hilic\_pos\_4\_092\_60

m/z 175.1252

01E+47.58RT (min)

lcorbi\_hilic\_pos\_4\_087\_59

m/z 175.1481

024E+40.51RT (min)

lcorbi\_hilic\_pos\_3\_019\_43

m/z 175.9546

01E+488.5RT (min)

lcorbi\_hilic\_pos\_4\_071\_57

m/z 176.0108

0246E+355.5RT (min)

lcorbi\_hilic\_pos\_4\_068\_57

m/z 176.0141

02468E+36.57RT (min)

lcorbi\_hilic\_pos\_3\_010\_32

m/z 176.0819

024E+31.52RT (min)

lcorbi\_hilic\_pos\_1\_031\_36

m/z 176.0917

01E+411.52RT (min)

lcorbi\_hilic\_pos\_4\_001\_49

m/z 176.1028

01E+44.55RT (min)

lcorbi\_hilic\_pos\_1\_049\_55

m/z 176.1125

01E+54.55RT (min)

lcorbi\_hilic\_pos\_4\_020\_37

m/z 176.1281

0246E+355.5RT (min)

lcorbi\_hilic\_pos\_4\_066\_57

m/z 176.1362

01E+40.51RT (min)

lcorbi\_hilic\_pos\_4\_101\_61

m/z 176.1893

01E+333.5RT (min)

lcorbi\_hilic\_pos\_4\_105\_61

m/z 177.0392

014.55RT (min)

lcorbi\_hilic\_pos\_4\_078\_58

m/z 177.0588

012E+433.5RT (min)

lcorbi\_hilic\_pos\_4\_003\_26

m/z 177.0610

024E+47.58RT (min)

lcorbi\_hilic\_pos\_4\_068\_57

m/z 177.0659

01E+41.52RT (min)

lcorbi\_hilic\_pos\_4\_020\_37

m/z 177.0980

02468E+311.5RT (min)

lcorbi\_hilic\_pos\_2\_013\_36

m/z 177.1063

01E+46.57RT (min)

lcorbi\_hilic\_pos\_4\_020\_37

m/z 177.1191

0123E+37.58RT (min)

lcorbi\_hilic\_pos\_4\_020\_37

m/z 177.1240

01E+34.55RT (min)

lcorbi\_hilic\_pos\_3\_009\_50

m/z 177.1638

01E+40.51RT (min)

lcorbi\_hilic\_pos\_3\_007\_19

m/z 178.0280

0123E+36.57RT (min)

lcorbi\_hilic\_pos\_4\_084\_59

m/z 178.0406

01E+44.55RT (min)

lcorbi\_hilic\_pos\_4\_070\_57

m/z 178.0498

012E+34.55RT (min)

lcorbi\_hilic\_pos\_4\_079\_58

m/z 178.0862

01E+455.5RT (min)

lcorbi\_hilic\_pos\_4\_071\_57

m/z 178.0896

012E+411.5RT (min)

lcorbi\_hilic\_pos\_1\_048\_02

m/z 178.1074

02468E+356RT (min)

lcorbi\_hilic\_pos\_1\_048\_02

m/z 178.1206

0123E+377.5RT (min)

lcorbi\_hilic\_pos\_4\_087\_59

m/z 179.0120

0123E+377.5RT (min)

lcorbi\_hilic\_pos\_2\_022\_35

m/z 179.0302

012E+466.5RT (min)

lcorbi\_hilic\_pos\_4\_072\_57

m/z 179.0453

014.55RT (min)

lcorbi\_hilic\_pos\_4\_080\_58

m/z 179.0711

01E+444.5RT (min)

lcorbi\_hilic\_pos\_4\_097\_60

m/z 179.0817

0246E+311.5RT (min)

lcorbi\_hilic\_pos\_1\_048\_02

m/z 179.0839

012E+444.5RT (min)

lcorbi\_hilic\_pos\_4\_097\_60

m/z 179.0903

017.58RT (min)

lcorbi\_hilic\_pos\_4\_094\_60

m/z 179.0928

01234E+411.5RT (min)

lcorbi\_hilic\_pos\_4\_090\_60

m/z 179.1090

01E+311.5RT (min)

lcorbi\_hilic\_pos\_2\_002\_11

m/z 179.1179

01E+34.55RT (min)

lcorbi\_hilic\_pos\_2\_030\_05

m/z 179.1234

01E+466.5RT (min)

lcorbi\_hilic\_pos\_4\_091\_60

m/z 179.1285

01E+577.5RT (min)

lcorbi\_hilic\_pos\_4\_020\_37

m/z 179.1404

013.54RT (min)

lcorbi\_hilic\_pos\_1\_052\_35

m/z 179.1430

012E+30.51RT (min)

lcorbi\_hilic\_pos\_4\_029\_41

m/z 180.0033

01E+466.5RT (min)

lcorbi\_hilic\_pos\_4\_069\_57

m/z 180.0066

0246E+35.56RT (min)

lcorbi\_hilic\_pos\_4\_067\_57

m/z 180.0376

01E+44.55RT (min)

lcorbi\_hilic\_pos\_4\_067\_57

m/z 180.0468

0123E+34.55RT (min)

lcorbi\_hilic\_pos\_4\_020\_37

m/z 180.0631

015.56RT (min)

lcorbi\_hilic\_pos\_4\_092\_60

m/z 180.0655

0123E+311.5RT (min)

lcorbi\_hilic\_pos\_1\_036\_32

m/z 180.0744

024E+27.58RT (min)

lcorbi\_hilic\_pos\_1\_036\_32

m/z 180.0866

0246E+35.56RT (min)

lcorbi\_hilic\_pos\_4\_066\_57

m/z 180.0879

0123E+433.5RT (min)

lcorbi\_hilic\_pos\_4\_020\_37

m/z 180.1312

012E+311.5RT (min)

lcorbi\_hilic\_pos\_1\_001\_49

m/z 180.1495

01E+400.5RT (min)

lcorbi\_hilic\_pos\_1\_005\_45

m/z 180.1747

01E+411.5RT (min)

lcorbi\_hilic\_pos\_3\_019\_43

m/z 180.8731

0123E+34.55RT (min)

lcorbi\_hilic\_pos\_4\_077\_58

m/z 180.8944

02468E+37.58RT (min)

lcorbi\_hilic\_pos\_4\_102\_61

m/z 180.9039

024E+377.5RT (min)

lcorbi\_hilic\_pos\_2\_036\_08

m/z 180.9780

01E+488.5RT (min)

lcorbi\_hilic\_pos\_4\_066\_57

m/z 181.0145

01234E+388.5RT (min)

lcorbi\_hilic\_pos\_4\_073\_57

m/z 181.0260

02468E+311.5RT (min)

lcorbi\_hilic\_pos\_1\_034\_37

m/z 181.0496

0246E+311.5RT (min)

lcorbi\_hilic\_pos\_4\_113\_62

m/z 181.0859

012E+311.5RT (min)

lcorbi\_hilic\_pos\_1\_032\_31

m/z 181.0970

01E+455.5RT (min)

lcorbi\_hilic\_pos\_4\_082\_59

m/z 181.1055

0123E+311.5RT (min)

lcorbi\_hilic\_pos\_2\_049\_55

m/z 181.1085

01234E+422.5RT (min)

lcorbi\_hilic\_pos\_3\_001\_49

m/z 181.1197

01E+46.57RT (min)

lcorbi\_hilic\_pos\_4\_090\_60

m/z 181.1223

01E+30.51RT (min)

lcorbi\_hilic\_pos\_3\_037\_20

m/z 181.1327

01234E+37.58RT (min)

lcorbi\_hilic\_pos\_2\_036\_08

m/z 181.1335

024E+31.52RT (min)

lcorbi\_hilic\_pos\_1\_028\_34

m/z 181.1416

012E+32.53RT (min)

lcorbi\_hilic\_pos\_4\_031\_17

m/z 181.1487

012E+47.58RT (min)

lcorbi\_hilic\_pos\_4\_094\_60

m/z 181.1583

0246E+37.58RT (min)

lcorbi\_hilic\_pos\_4\_090\_60

m/z 181.1587

01234E+40.51RT (min)

lcorbi\_hilic\_pos\_3\_019\_43

m/z 181.9988

01234E+34.55RT (min)

lcorbi\_hilic\_pos\_3\_042\_17

m/z 182.0346

0123E+34.55RT (min)

lcorbi\_hilic\_pos\_4\_071\_57

m/z 182.0599

01E+355.5RT (min)

lcorbi\_hilic\_pos\_4\_080\_58

m/z 182.0754

01E+411.5RT (min)

lcorbi\_hilic\_pos\_4\_020\_37

m/z 182.0754

01234E+33.54RT (min)

lcorbi\_hilic\_pos\_1\_019\_01

m/z 182.0788

01E+45.56RT (min)

lcorbi\_hilic\_pos\_4\_020\_37

m/z 182.0811

01E+433.5RT (min)

lcorbi\_hilic\_pos\_4\_104\_61

m/z 182.0812

024E+31.52RT (min)

lcorbi\_hilic\_pos\_1\_019\_01

m/z 182.1005

01E+311.5RT (min)

lcorbi\_hilic\_pos\_3\_001\_49

m/z 182.1312

0123E+466.5RT (min)

lcorbi\_hilic\_pos\_4\_096\_60

m/z 182.1387

01E+411.5RT (min)

lcorbi\_hilic\_pos\_1\_024\_03

m/z 182.1452

012E+47.58RT (min)

lcorbi\_hilic\_pos\_4\_097\_60

m/z 182.9852

0123E+41213RT (min)

lcorbi\_hilic\_pos\_4\_091\_60

m/z 183.0247

0246E+333.5RT (min)

lcorbi\_hilic\_pos\_4\_083\_59

m/z 183.0611

024E+355.5RT (min)

lcorbi\_hilic\_pos\_4\_079\_58

m/z 183.0740

02468E+34.55RT (min)

lcorbi\_hilic\_pos\_4\_067\_57

m/z 183.0764

024E+31.52RT (min)

lcorbi\_hilic\_pos\_1\_032\_31

m/z 183.0845

015.56RT (min)

lcorbi\_hilic\_pos\_4\_094\_60

m/z 183.1016

01E+411.5RT (min)

lcorbi\_hilic\_pos\_4\_110\_62

m/z 183.1064

01E+422.5RT (min)

lcorbi\_hilic\_pos\_2\_024\_32

m/z 183.1169

010.51RT (min)

lcorbi\_hilic\_pos\_3\_054\_16

m/z 183.1380

02468E+20.51RT (min)

lcorbi\_hilic\_pos\_2\_032\_30

m/z 183.1710

0246E+377.5RT (min)

lcorbi\_hilic\_pos\_2\_037\_24

m/z 183.1937

0246E+32.53RT (min)

lcorbi\_hilic\_pos\_4\_096\_60

m/z 183.9959

024E+34.55RT (min)

lcorbi\_hilic\_pos\_3\_018\_01

m/z 184.0168

01E+466.5RT (min)

lcorbi\_hilic\_pos\_4\_091\_60

m/z 184.0195

01E+433.5RT (min)

lcorbi\_hilic\_pos\_4\_086\_59

m/z 184.0290

01234E+366.5RT (min)

lcorbi\_hilic\_pos\_4\_074\_58

m/z 184.0559

024E+355.5RT (min)

lcorbi\_hilic\_pos\_4\_081\_58

m/z 184.0604

01E+31.52RT (min)

lcorbi\_hilic\_pos\_3\_054\_16

m/z 184.0652

01E+46.57RT (min)

lcorbi\_hilic\_pos\_2\_024\_32

m/z 184.0734

010.51RT (min)

lcorbi\_hilic\_pos\_4\_046\_21

m/z 184.0968

012E+41.52RT (min)

lcorbi\_hilic\_pos\_4\_020\_37

m/z 184.1081

01E+444.5RT (min)

lcorbi\_hilic\_pos\_4\_084\_59

m/z 184.1162

02468E+344.5RT (min)

lcorbi\_hilic\_pos\_4\_089\_59

m/z 184.1332

0123E+40.51RT (min)

lcorbi\_hilic\_pos\_4\_020\_37

m/z 184.1696

0246E+32.53RT (min)

lcorbi\_hilic\_pos\_4\_086\_59

m/z 184.8889

012E+45.56RT (min)

lcorbi\_hilic\_pos\_4\_100\_61

m/z 184.9906

012E+388.5RT (min)

lcorbi\_hilic\_pos\_4\_071\_57

m/z 185.0234

012E+499.5RT (min)

lcorbi\_hilic\_pos\_4\_107\_62

m/z 185.0921

024E+311.5RT (min)

lcorbi\_hilic\_pos\_3\_029\_23

m/z 185.1033

012E+311.5RT (min)

lcorbi\_hilic\_pos\_4\_075\_58

m/z 185.1104

024E+27.58RT (min)

lcorbi\_hilic\_pos\_4\_020\_37

m/z 185.1172

012E+311.5RT (min)

lcorbi\_hilic\_pos\_1\_002\_29

m/z 185.1202

012E+44.55RT (min)

lcorbi\_hilic\_pos\_4\_091\_60

m/z 185.1284

0246E+377.5RT (min)

lcorbi\_hilic\_pos\_4\_018\_31

m/z 185.1648

01E+444.5RT (min)

lcorbi\_hilic\_pos\_4\_078\_58

m/z 185.1729

0246E+30.51RT (min)

lcorbi\_hilic\_pos\_2\_047\_22

m/z 185.1729

012E+444.5RT (min)

lcorbi\_hilic\_pos\_4\_085\_59

m/z 185.2093

01E+433.5RT (min)

lcorbi\_hilic\_pos\_3\_027\_05

m/z 186.0526

0144.5RT (min)

lcorbi\_hilic\_pos\_4\_071\_57

m/z 186.0697

0111.5RT (min)

lcorbi\_hilic\_pos\_4\_020\_37

m/z 186.0944

0246E+37.58RT (min)

lcorbi\_hilic\_pos\_4\_091\_60

m/z 186.1125

024E+322.5RT (min)

lcorbi\_hilic\_pos\_4\_020\_37

m/z 186.1237

024E+36.57RT (min)

lcorbi\_hilic\_pos\_4\_088\_59

m/z 186.1489

01E+42.53RT (min)

lcorbi\_hilic\_pos\_4\_077\_58

m/z 186.1489

0123E+40.51RT (min)

lcorbi\_hilic\_pos\_4\_099\_61

m/z 186.2217

012E+30.51RT (min)

lcorbi\_hilic\_pos\_1\_044\_17

m/z 186.6491

0246E+377.5RT (min)

lcorbi\_hilic\_pos\_4\_092\_60

m/z 186.8959

0123E+36.57RT (min)

lcorbi\_hilic\_pos\_4\_103\_61

m/z 186.9119

01234E+377.5RT (min)

lcorbi\_hilic\_pos\_2\_004\_01

m/z 186.9543

01E+34.55RT (min)

lcorbi\_hilic\_pos\_2\_019\_16

m/z 187.0304

01234E+36.57RT (min)

lcorbi\_hilic\_pos\_4\_085\_59

m/z 187.0577

01E+41.52RT (min)

lcorbi\_hilic\_pos\_1\_024\_03

m/z 187.0753

0133.5RT (min)

lcorbi\_hilic\_pos\_1\_036\_32

m/z 187.0785

0246E+45RT (min)

lcorbi\_hilic\_pos\_4\_090\_60

m/z 187.0979

01E+35.56RT (min)

lcorbi\_hilic\_pos\_4\_052\_32

m/z 187.1157

01E+355.5RT (min)

lcorbi\_hilic\_pos\_4\_085\_59

m/z 187.1230

0246E+244.5RT (min)

lcorbi\_hilic\_pos\_3\_012\_13

m/z 187.1329

01E+40.51RT (min)

lcorbi\_hilic\_pos\_2\_011\_03

m/z 187.1441

01234E+36.57RT (min)

lcorbi\_hilic\_pos\_4\_072\_57

m/z 187.1441

012E+411.5RT (min)

lcorbi\_hilic\_pos\_4\_020\_37

m/z 187.1886

012E+433.5RT (min)

lcorbi\_hilic\_pos\_1\_044\_17

m/z 187.2250

01E+433.5RT (min)

lcorbi\_hilic\_pos\_4\_091\_60

m/z 187.9843

024E+388.5RT (min)

lcorbi\_hilic\_pos\_1\_026\_39

m/z 188.0103

01E+50.51RT (min)

lcorbi\_hilic\_pos\_1\_037\_18

m/z 188.0568

012E+44.55RT (min)

lcorbi\_hilic\_pos\_4\_083\_59

m/z 188.0706

01E+41.52RT (min)

lcorbi\_hilic\_pos\_2\_019\_16

m/z 188.0837

01234E+311.5RT (min)

lcorbi\_hilic\_pos\_4\_020\_37

m/z 188.0909

024E+34.55RT (min)

lcorbi\_hilic\_pos\_2\_001\_49

m/z 188.0974

0123E+311.5RT (min)

lcorbi\_hilic\_pos\_4\_052\_32

m/z 188.1111

0246E+25.56RT (min)

lcorbi\_hilic\_pos\_1\_036\_32

m/z 188.1206

0246E+411.5RT (min)

lcorbi\_hilic\_pos\_2\_024\_32

m/z 188.1282

02468E+411.5RT (min)

lcorbi\_hilic\_pos\_4\_020\_37

m/z 188.1645

01E+43.54RT (min)

lcorbi\_hilic\_pos\_4\_105\_61

m/z 188.1645

024E+411.5RT (min)

lcorbi\_hilic\_pos\_1\_027\_10

m/z 188.1757

01E+46.57RT (min)

lcorbi\_hilic\_pos\_4\_068\_57

m/z 189.0037

024E+366.5RT (min)

lcorbi\_hilic\_pos\_4\_076\_58

m/z 189.0134

01E+466.5RT (min)

lcorbi\_hilic\_pos\_4\_083\_59

m/z 189.0734

01234E+32.53RT (min)

lcorbi\_hilic\_pos\_4\_086\_59

m/z 189.0870

01234E+444.5RT (min)

lcorbi\_hilic\_pos\_4\_020\_37

m/z 189.0901

01E+47.58RT (min)

lcorbi\_hilic\_pos\_4\_097\_60

m/z 189.0910

012E+411.5RT (min)

lcorbi\_hilic\_pos\_4\_108\_62

m/z 189.1022

012E+433.5RT (min)

lcorbi\_hilic\_pos\_4\_113\_62

m/z 189.1022

02468E+31.52RT (min)

lcorbi\_hilic\_pos\_4\_075\_58

m/z 189.1022

01E+344.5RT (min)

lcorbi\_hilic\_pos\_4\_092\_60

m/z 189.1047

0123E+45.56RT (min)

lcorbi\_hilic\_pos\_4\_020\_37

m/z 189.1122

0123E+40.51RT (min)

lcorbi\_hilic\_pos\_4\_102\_61

m/z 189.1234

0246E+366.5RT (min)

lcorbi\_hilic\_pos\_4\_086\_59

m/z 189.1240

0111.5RT (min)

lcorbi\_hilic\_pos\_4\_113\_62

m/z 189.1321

01E+411.5RT (min)

lcorbi\_hilic\_pos\_4\_097\_60

m/z 189.2042

01E+42.53RT (min)

lcorbi\_hilic\_pos\_4\_054\_45

m/z 189.9876

0246E+333.5RT (min)

lcorbi\_hilic\_pos\_4\_080\_58

m/z 190.0491

0123E+46.57RT (min)

lcorbi\_hilic\_pos\_4\_095\_60

m/z 190.0499

024E+40.51RT (min)

lcorbi\_hilic\_pos\_4\_079\_58

m/z 190.0532

02468E+34.55RT (min)

lcorbi\_hilic\_pos\_4\_113\_62

m/z 190.0863

01E+344.5RT (min)

lcorbi\_hilic\_pos\_4\_091\_60

m/z 190.1186

01E+34.55RT (min)

lcorbi\_hilic\_pos\_3\_029\_23

m/z 190.1281

014.55RT (min)

lcorbi\_hilic\_pos\_1\_054\_23

m/z 190.1363

01E+444.5RT (min)

lcorbi\_hilic\_pos\_4\_024\_24

m/z 190.1380

02468E+366.5RT (min)

lcorbi\_hilic\_pos\_2\_001\_49

m/z 190.1437

0246E+20.51RT (min)

lcorbi\_hilic\_pos\_3\_023\_18

m/z 190.1631

02468E+377.5RT (min)

lcorbi\_hilic\_pos\_4\_020\_37

m/z 190.1753

012E+333.5RT (min)

lcorbi\_hilic\_pos\_4\_098\_61

m/z 190.1801

02468E+311.5RT (min)

lcorbi\_hilic\_pos\_4\_028\_10

m/z 190.1874

0246E+333.5RT (min)

lcorbi\_hilic\_pos\_1\_011\_09

m/z 190.9119

01E+455.5RT (min)

lcorbi\_hilic\_pos\_4\_091\_60

m/z 190.9781

0246E+388.5RT (min)

lcorbi\_hilic\_pos\_4\_071\_57

m/z 190.9927

024E+34.55RT (min)

lcorbi\_hilic\_pos\_1\_029\_14

m/z 191.0104

0246E+37.58RT (min)

lcorbi\_hilic\_pos\_4\_084\_59

m/z 191.0316

012E+444.5RT (min)

lcorbi\_hilic\_pos\_4\_089\_59

m/z 191.0485

012E+46.57RT (min)

lcorbi\_hilic\_pos\_4\_089\_59

m/z 191.0703

012E+411.5RT (min)

lcorbi\_hilic\_pos\_4\_020\_37

m/z 191.0849

01E+411.5RT (min)

lcorbi\_hilic\_pos\_4\_086\_59

m/z 191.0896

012E+42.53RT (min)

lcorbi\_hilic\_pos\_3\_048\_38

m/z 191.1066

01E+311.5RT (min)

lcorbi\_hilic\_pos\_3\_019\_43

m/z 191.1107

014.55RT (min)

lcorbi\_hilic\_pos\_4\_110\_62

m/z 191.1133

0246E+355.5RT (min)

lcorbi\_hilic\_pos\_4\_113\_62

m/z 191.1238

01E+455.5RT (min)

lcorbi\_hilic\_pos\_4\_028\_10

m/z 191.1430

01E+50.51RT (min)

lcorbi\_hilic\_pos\_3\_019\_43

m/z 191.1772

0123E+333.5RT (min)

lcorbi\_hilic\_pos\_4\_099\_61

m/z 191.1795

01E+40.51RT (min)

lcorbi\_hilic\_pos\_4\_066\_57

m/z 192.0146

0246E+366.5RT (min)

lcorbi\_hilic\_pos\_4\_081\_58

m/z 192.0397

02468E+355.5RT (min)

lcorbi\_hilic\_pos\_4\_074\_58

m/z 192.0437

01E+46.57RT (min)

lcorbi\_hilic\_pos\_4\_071\_57

m/z 192.0519

011.52RT (min)

lcorbi\_hilic\_pos\_4\_110\_62

m/z 192.0744

01E+377.5RT (min)

lcorbi\_hilic\_pos\_4\_020\_37

m/z 192.0866

01E+466.5RT (min)

lcorbi\_hilic\_pos\_1\_050\_05

m/z 192.0964

01E+411.5RT (min)

lcorbi\_hilic\_pos\_2\_046\_46

m/z 192.1019

01E+355.5RT (min)

lcorbi\_hilic\_pos\_4\_020\_37

m/z 192.1230

01234E+43.54RT (min)

lcorbi\_hilic\_pos\_1\_042\_40

m/z 192.1382

0246E+40.51RT (min)

lcorbi\_hilic\_pos\_4\_112\_62

m/z 192.1430

012E+444.5RT (min)

lcorbi\_hilic\_pos\_4\_024\_24

m/z 192.1438

012E+46.57RT (min)

lcorbi\_hilic\_pos\_4\_090\_60

m/z 192.1519

01E+444.5RT (min)

lcorbi\_hilic\_pos\_4\_024\_24

m/z 192.1868

0246E+333.5RT (min)

lcorbi\_hilic\_pos\_4\_036\_09

m/z 192.9613

01E+466.5RT (min)

lcorbi\_hilic\_pos\_4\_094\_60

m/z 192.9986

0246E+36.57RT (min)

lcorbi\_hilic\_pos\_4\_083\_59

m/z 193.0277

02468E+36.57RT (min)

lcorbi\_hilic\_pos\_4\_082\_59

m/z 193.0496

012E+411.5RT (min)

lcorbi\_hilic\_pos\_2\_027\_38

m/z 193.0723

01E+411.5RT (min)

lcorbi\_hilic\_pos\_4\_008\_19

m/z 193.0737

02468E+44.55RT (min)

lcorbi\_hilic\_pos\_1\_036\_32

m/z 193.0819

0246E+377.5RT (min)

lcorbi\_hilic\_pos\_4\_078\_58

m/z 193.0972

01E+42.53RT (min)

lcorbi\_hilic\_pos\_4\_073\_57

m/z 193.1189

01E+44.55RT (min)

lcorbi\_hilic\_pos\_4\_097\_60

m/z 193.1270

01E+41.52RT (min)

lcorbi\_hilic\_pos\_4\_093\_60

m/z 193.1386

0123E+40.51RT (min)

lcorbi\_hilic\_pos\_1\_042\_40

m/z 193.1586

01234E+40.51RT (min)

lcorbi\_hilic\_pos\_3\_019\_43

m/z 193.1628

02468E+33.54RT (min)

lcorbi\_hilic\_pos\_4\_097\_60

m/z 193.9826

024E+377.5RT (min)

lcorbi\_hilic\_pos\_2\_023\_34

m/z 194.0659

01E+41.52RT (min)

lcorbi\_hilic\_pos\_4\_104\_61

m/z 194.0812

01E+32.53RT (min)

lcorbi\_hilic\_pos\_1\_054\_23

m/z 194.0925

01E+42.53RT (min)

lcorbi\_hilic\_pos\_1\_021\_46

m/z 194.1029

01E+34.55RT (min)

lcorbi\_hilic\_pos\_4\_066\_57

m/z 194.1029

01E+433.5RT (min)

lcorbi\_hilic\_pos\_2\_001\_49

m/z 194.1231

0144.5RT (min)

lcorbi\_hilic\_pos\_4\_020\_37

m/z 194.1515

01E+466.5RT (min)

lcorbi\_hilic\_pos\_4\_092\_60

m/z 194.9196

01234E+366.5RT (min)

lcorbi\_hilic\_pos\_1\_043\_25

m/z 195.0029

016.57RT (min)

lcorbi\_hilic\_pos\_1\_037\_18

m/z 195.0061

01E+44.55RT (min)

lcorbi\_hilic\_pos\_1\_053\_08

m/z 195.0293

0246E+44.55RT (min)

lcorbi\_hilic\_pos\_1\_003\_24

m/z 195.0479

016.57RT (min)

lcorbi\_hilic\_pos\_4\_113\_62

m/z 195.0764

012E+41.52RT (min)

lcorbi\_hilic\_pos\_4\_083\_59

m/z 195.0877

0123E+311.5RT (min)

lcorbi\_hilic\_pos\_2\_016\_18

m/z 195.1379

0246E+30.51RT (min)

lcorbi\_hilic\_pos\_4\_086\_59

m/z 195.1477

01E+40.51RT (min)

lcorbi\_hilic\_pos\_4\_107\_62

m/z 196.0377

02468E+35.56RT (min)

lcorbi\_hilic\_pos\_4\_091\_60

m/z 196.0484

01E+444.5RT (min)

lcorbi\_hilic\_pos\_3\_014\_08

m/z 196.0605

024E+31.52RT (min)

lcorbi\_hilic\_pos\_4\_020\_37

m/z 196.0718

0133.5RT (min)

lcorbi\_hilic\_pos\_4\_098\_61

m/z 196.0735

0123E+255.5RT (min)

lcorbi\_hilic\_pos\_1\_042\_40

m/z 196.0829

0144.5RT (min)

lcorbi\_hilic\_pos\_1\_036\_32

m/z 196.0912

01E+311.5RT (min)

lcorbi\_hilic\_pos\_1\_025\_52

m/z 196.1525

02468E+433.5RT (min)

lcorbi\_hilic\_pos\_2\_044\_26

m/z 196.1580

01E+366.5RT (min)

lcorbi\_hilic\_pos\_4\_024\_24

m/z 196.1672

0123E+333.5RT (min)

lcorbi\_hilic\_pos\_4\_080\_58

m/z 196.8571

0155.5RT (min)

lcorbi\_hilic\_pos\_2\_035\_44

m/z 197.0120

0246E+36.57RT (min)

lcorbi\_hilic\_pos\_4\_090\_60

m/z 197.0687

0246E+31.52RT (min)

lcorbi\_hilic\_pos\_4\_090\_60

m/z 197.0897

01E+466.5RT (min)

lcorbi\_hilic\_pos\_3\_031\_36

m/z 197.0944

012E+433.54RT (min)

lcorbi\_hilic\_pos\_4\_089\_59

m/z 197.1034

0166.5RT (min)

lcorbi\_hilic\_pos\_4\_020\_37

m/z 197.1272

01E+466.5RT (min)

lcorbi\_hilic\_pos\_3\_024\_35

m/z 197.1365

0246E+333.5RT (min)

lcorbi\_hilic\_pos\_4\_101\_61

m/z 197.1502

01E+466.5RT (min)

lcorbi\_hilic\_pos\_4\_094\_60

m/z 197.1536

0246E+20.51RT (min)

lcorbi\_hilic\_pos\_4\_036\_09

m/z 197.1774

02468E+31.52RT (min)

lcorbi\_hilic\_pos\_1\_044\_17

m/z 197.1900

010.51RT (min)

lcorbi\_hilic\_pos\_4\_036\_09

m/z 197.2012

012E+400.5RT (min)

lcorbi\_hilic\_pos\_4\_105\_61

m/z 197.9728

012E+34.55RT (min)

lcorbi\_hilic\_pos\_1\_049\_55

m/z 197.9960

01234E+377.5RT (min)

lcorbi\_hilic\_pos\_2\_003\_33

m/z 198.0130

01E+46.57RT (min)

lcorbi\_hilic\_pos\_1\_050\_05

m/z 198.0526

012E+36.57RT (min)

lcorbi\_hilic\_pos\_4\_045\_33

m/z 198.0641

01E+44.55RT (min)

lcorbi\_hilic\_pos\_4\_078\_58

m/z 198.0737

012E+45.56RT (min)

lcorbi\_hilic\_pos\_4\_088\_59

m/z 198.0954

011.52RT (min)

lcorbi\_hilic\_pos\_2\_002\_11

m/z 198.0972

01E+34.55RT (min)

lcorbi\_hilic\_pos\_4\_020\_37

m/z 198.1042

01E+455.5RT (min)

lcorbi\_hilic\_pos\_4\_101\_61

m/z 198.1043

02468E+27.58RT (min)

lcorbi\_hilic\_pos\_4\_003\_26

m/z 198.1084

02468E+311.5RT (min)

lcorbi\_hilic\_pos\_2\_024\_32

m/z 198.1237

024E+36.57RT (min)

lcorbi\_hilic\_pos\_1\_008\_38

m/z 198.1488

02468E+311.5RT (min)

lcorbi\_hilic\_pos\_4\_110\_62

m/z 198.1906

01234E+322.5RT (min)

lcorbi\_hilic\_pos\_2\_007\_28

m/z 198.8621

0246E+355.5RT (min)

lcorbi\_hilic\_pos\_4\_098\_61

m/z 199.0213

01E+41.52RT (min)

lcorbi\_hilic\_pos\_4\_101\_61

m/z 199.0390

01E+455.5RT (min)

lcorbi\_hilic\_pos\_2\_037\_24

m/z 199.0672

0246E+377.5RT (min)

lcorbi\_hilic\_pos\_1\_028\_34

m/z 199.0689

01E+411.5RT (min)

lcorbi\_hilic\_pos\_4\_050\_16

m/z 199.0827

012.53RT (min)

lcorbi\_hilic\_pos\_4\_094\_60

m/z 199.0883

012E+46.57RT (min)

lcorbi\_hilic\_pos\_4\_087\_59

m/z 199.1013

012E+43.54RT (min)

lcorbi\_hilic\_pos\_4\_095\_60

m/z 199.1077

01234E+31.52RT (min)

lcorbi\_hilic\_pos\_4\_069\_57

m/z 199.1441

02468E+36.57RT (min)

lcorbi\_hilic\_pos\_1\_032\_31

m/z 199.1465

0123E+44.55RT (min)

lcorbi\_hilic\_pos\_4\_090\_60

m/z 199.1805

0123E+30.51RT (min)

lcorbi\_hilic\_pos\_2\_039\_09

m/z 199.9699

012E+34.55RT (min)

lcorbi\_hilic\_pos\_1\_001\_49

m/z 200.1281

01E+40.51RT (min)

lcorbi\_hilic\_pos\_3\_054\_16

m/z 200.1453

02468E+311.5RT (min)

lcorbi\_hilic\_pos\_4\_099\_61

m/z 200.1645

0246E+333.5RT (min)

lcorbi\_hilic\_pos\_3\_047\_28

m/z 200.1645

024E+40.51RT (min)

lcorbi\_hilic\_pos\_3\_019\_43

m/z 200.1920

012E+322.5RT (min)

lcorbi\_hilic\_pos\_4\_066\_57

m/z 201.0440

02468E+377.5RT (min)

lcorbi\_hilic\_pos\_1\_020\_13

m/z 201.0546

012E+411.5RT (min)

lcorbi\_hilic\_pos\_4\_073\_57

m/z 201.0693

012E+44.55RT (min)

lcorbi\_hilic\_pos\_1\_032\_31

m/z 201.0870

02468E+344.5RT (min)

lcorbi\_hilic\_pos\_4\_085\_59

m/z 201.0871

0123E+31.52RT (min)

lcorbi\_hilic\_pos\_4\_069\_57

m/z 201.1234

024E+344.5RT (min)

lcorbi\_hilic\_pos\_4\_083\_59

m/z 201.1240

0246E+211.5RT (min)

lcorbi\_hilic\_pos\_1\_024\_03

m/z 201.1316

024E+34.55RT (min)

lcorbi\_hilic\_pos\_3\_055\_24

m/z 201.1485

0111.5RT (min)

lcorbi\_hilic\_pos\_4\_108\_62

m/z 201.1598

02468E+311.5RT (min)

lcorbi\_hilic\_pos\_4\_020\_37

m/z 201.2325

012E+400.5RT (min)

lcorbi\_hilic\_pos\_4\_107\_62

m/z 201.6681

01E+46.57RT (min)

lcorbi\_hilic\_pos\_4\_090\_60

m/z 202.0451

01E+45.56RT (min)

lcorbi\_hilic\_pos\_4\_083\_59

m/z 202.0532

0111.5RT (min)

lcorbi\_hilic\_pos\_3\_048\_38

m/z 202.0777

012E+500.5RT (min)

lcorbi\_hilic\_pos\_4\_066\_57

m/z 202.0862

02468E+311.5RT (min)

lcorbi\_hilic\_pos\_3\_054\_16

m/z 202.0863

02468E+333.5RT (min)

lcorbi\_hilic\_pos\_2\_001\_49

m/z 202.1074

01234E+366.5RT (min)

lcorbi\_hilic\_pos\_4\_084\_59

m/z 202.1074

01234E+41.52RT (min)

lcorbi\_hilic\_pos\_4\_020\_37

m/z 202.1177

01234E+37.58RT (min)

lcorbi\_hilic\_pos\_4\_092\_60

m/z 202.1313

012E+44.55RT (min)

lcorbi\_hilic\_pos\_4\_090\_60

m/z 202.1438

0123E+33.54RT (min)

lcorbi\_hilic\_pos\_4\_004\_38

m/z 202.1438

012E+311.5RT (min)

lcorbi\_hilic\_pos\_3\_054\_16

m/z 202.2440

0246E+222.5RT (min)

lcorbi\_hilic\_pos\_1\_019\_01

m/z 202.9961

02468E+388.5RT (min)

lcorbi\_hilic\_pos\_4\_066\_57

m/z 203.0436

01E+43.54RT (min)

lcorbi\_hilic\_pos\_4\_072\_57

m/z 203.0680

01234E+311.5RT (min)

lcorbi\_hilic\_pos\_4\_090\_60

m/z 203.0694

01E+46.57RT (min)

lcorbi\_hilic\_pos\_4\_092\_60

m/z 203.0815

012E+311.5RT (min)

lcorbi\_hilic\_pos\_3\_053\_21

m/z 203.1026

012E+46.57RT (min)

lcorbi\_hilic\_pos\_4\_093\_60

m/z 203.1026

02468E+333.5RT (min)

lcorbi\_hilic\_pos\_4\_083\_59

m/z 203.1029

012E+311.5RT (min)

lcorbi\_hilic\_pos\_4\_096\_60

m/z 203.1278

0123E+40.51RT (min)

lcorbi\_hilic\_pos\_1\_012\_19

m/z 203.1396

0123E+411.5RT (min)

lcorbi\_hilic\_pos\_4\_090\_60

m/z 203.1794

01E+40.51RT (min)

lcorbi\_hilic\_pos\_4\_107\_62

m/z 204.0372

012E+50.51RT (min)

lcorbi\_hilic\_pos\_1\_037\_18

m/z 204.0866

01E+322.5RT (min)

lcorbi\_hilic\_pos\_4\_096\_60

m/z 204.1138

01E+44.55RT (min)

lcorbi\_hilic\_pos\_4\_019\_30

m/z 204.1242

01234E+311.5RT (min)

lcorbi\_hilic\_pos\_4\_052\_32

m/z 204.1388

0246E+34.55RT (min)

lcorbi\_hilic\_pos\_2\_004\_01

m/z 204.1536

02468E+36.57RT (min)

lcorbi\_hilic\_pos\_1\_052\_35

m/z 204.2206

0123E+333.5RT (min)

lcorbi\_hilic\_pos\_1\_008\_38

m/z 204.9917

0123E+388.5RT (min)

lcorbi\_hilic\_pos\_4\_073\_57

m/z 205.0748

012E+34.55RT (min)

lcorbi\_hilic\_pos\_1\_020\_13

m/z 205.0819

02468E+377.5RT (min)

lcorbi\_hilic\_pos\_2\_013\_36

m/z 205.0899

0145RT (min)

lcorbi\_hilic\_pos\_1\_019\_01

m/z 205.0905

012E+411.5RT (min)

lcorbi\_hilic\_pos\_3\_055\_24

m/z 205.0923

012E+477.5RT (min)

lcorbi\_hilic\_pos\_4\_072\_57

m/z 205.1189

01E+31.52RT (min)

lcorbi\_hilic\_pos\_4\_020\_37

m/z 205.1464

014.55RT (min)

lcorbi\_hilic\_pos\_4\_070\_57

m/z 205.1587

01E+50.51RT (min)

lcorbi\_hilic\_pos\_4\_030\_18

m/z 206.0608

0123E+47.58RT (min)

lcorbi\_hilic\_pos\_1\_003\_24

m/z 206.0812

0123E+41.52RT (min)

lcorbi\_hilic\_pos\_4\_020\_37

m/z 206.0907

0246E+333.5RT (min)

lcorbi\_hilic\_pos\_2\_049\_55

m/z 206.1175

01E+344.5RT (min)

lcorbi\_hilic\_pos\_4\_106\_62

m/z 206.1222

011.52RT (min)

lcorbi\_hilic\_pos\_3\_035\_22

m/z 206.1387

024E+411.5RT (min)

lcorbi\_hilic\_pos\_3\_035\_22

m/z 206.1468

012E+40.51RT (min)

lcorbi\_hilic\_pos\_1\_043\_25

m/z 206.2026

0246E+333.5RT (min)

lcorbi\_hilic\_pos\_1\_031\_36

m/z 206.6536

01E+46.57RT (min)

lcorbi\_hilic\_pos\_4\_020\_37

m/z 207.0053

0246E+37.58RT (min)

lcorbi\_hilic\_pos\_4\_084\_59

m/z 207.0143

01E+46.57RT (min)

lcorbi\_hilic\_pos\_4\_070\_57

m/z 207.0256

02468E+355.5RT (min)

lcorbi\_hilic\_pos\_4\_098\_61

m/z 207.0433

0166.5RT (min)

lcorbi\_hilic\_pos\_4\_067\_57

m/z 207.0718

01234E+34.55RT (min)

lcorbi\_hilic\_pos\_4\_066\_57

m/z 207.0863

012.53RT (min)

lcorbi\_hilic\_pos\_4\_066\_57

m/z 207.0893

0123E+44.55RT (min)

lcorbi\_hilic\_pos\_1\_036\_32

m/z 207.1016

010.51RT (min)

lcorbi\_hilic\_pos\_4\_091\_60

m/z 207.1128

01234E+544.5RT (min)

lcorbi\_hilic\_pos\_4\_020\_37

m/z 207.1128

0246E+411.5RT (min)

lcorbi\_hilic\_pos\_4\_091\_60

m/z 207.1240

0246E+30.51RT (min)

lcorbi\_hilic\_pos\_4\_091\_60

m/z 207.1321

01E+44.55RT (min)

lcorbi\_hilic\_pos\_1\_050\_05

m/z 207.1380

012E+30.51RT (min)

lcorbi\_hilic\_pos\_4\_036\_09

m/z 207.1784

02468E+333.5RT (min)

lcorbi\_hilic\_pos\_4\_050\_16

m/z 207.8893

02468E+355.5RT (min)

lcorbi\_hilic\_pos\_4\_075\_58

m/z 207.9673

02468E+36.57RT (min)

lcorbi\_hilic\_pos\_4\_089\_59

m/z 207.9983

02468E+36.57RT (min)

lcorbi\_hilic\_pos\_4\_089\_59

m/z 208.0137

01E+355.5RT (min)

lcorbi\_hilic\_pos\_4\_103\_61

m/z 208.0638

01E+411.5RT (min)

lcorbi\_hilic\_pos\_4\_110\_62

m/z 208.0944

02468E+355.5RT (min)

lcorbi\_hilic\_pos\_4\_072\_57

m/z 208.1332

0123E+344.5RT (min)

lcorbi\_hilic\_pos\_4\_020\_37

m/z 208.1543

01E+30.51RT (min)

lcorbi\_hilic\_pos\_1\_034\_37

m/z 208.1626

010.51RT (min)

lcorbi\_hilic\_pos\_2\_031\_21

m/z 208.1696

0123E+322.5RT (min)

lcorbi\_hilic\_pos\_4\_102\_61

m/z 208.8778

0246E+36.57RT (min)

lcorbi\_hilic\_pos\_4\_103\_61

m/z 209.0259

01E+46.57RT (min)

lcorbi\_hilic\_pos\_4\_096\_60

m/z 209.0671

01E+44.55RT (min)

lcorbi\_hilic\_pos\_4\_113\_62

m/z 209.0920

01E+35.56RT (min)

lcorbi\_hilic\_pos\_4\_083\_59

m/z 209.1138

02468E+344.5RT (min)

lcorbi\_hilic\_pos\_4\_052\_32

m/z 209.1285

0123E+34.55RT (min)

lcorbi\_hilic\_pos\_2\_012\_37

m/z 209.1309

024E+311.5RT (min)

lcorbi\_hilic\_pos\_1\_037\_18

m/z 209.1776

012E+46.57RT (min)

lcorbi\_hilic\_pos\_4\_096\_60

m/z 210.0171

0246E+36.57RT (min)

lcorbi\_hilic\_pos\_4\_089\_59

m/z 210.0595

02468E+34.55RT (min)

lcorbi\_hilic\_pos\_4\_113\_62

m/z 210.0762

01E+31.52RT (min)

lcorbi\_hilic\_pos\_4\_091\_60

m/z 210.0873

01E+34.55RT (min)

lcorbi\_hilic\_pos\_4\_020\_37

m/z 210.0873

0246E+300.5RT (min)

lcorbi\_hilic\_pos\_4\_022\_08

m/z 210.1102

024E+411.5RT (min)

lcorbi\_hilic\_pos\_4\_020\_37

m/z 210.1125

024E+333.5RT (min)

lcorbi\_hilic\_pos\_2\_012\_37

m/z 210.1488

0246E+33.54RT (min)

lcorbi\_hilic\_pos\_4\_077\_58

m/z 210.1618

01E+41.52RT (min)

lcorbi\_hilic\_pos\_4\_086\_59

m/z 210.1852

0123E+311.5RT (min)

lcorbi\_hilic\_pos\_4\_013\_23

m/z 210.1852

02468E+333.5RT (min)

lcorbi\_hilic\_pos\_3\_010\_32

m/z 210.2046

01E+422.5RT (min)

lcorbi\_hilic\_pos\_4\_092\_60

m/z 210.8821

0246E+355.5RT (min)

lcorbi\_hilic\_pos\_4\_077\_58

m/z 210.9777

012E+466.5RT (min)

lcorbi\_hilic\_pos\_4\_067\_57

m/z 211.0577

0123E+31.52RT (min)

lcorbi\_hilic\_pos\_4\_075\_58

m/z 211.0965

01E+42.53RT (min)

lcorbi\_hilic\_pos\_4\_110\_62

m/z 211.1053

012E+466.5RT (min)

lcorbi\_hilic\_pos\_4\_089\_59

m/z 211.1294

02468E+333.5RT (min)

lcorbi\_hilic\_pos\_4\_093\_60

m/z 211.1376

0166.5RT (min)

lcorbi\_hilic\_pos\_4\_089\_59

m/z 211.1426

012E+31.52RT (min)

lcorbi\_hilic\_pos\_2\_020\_31

m/z 211.1693

0123E+40.51RT (min)

lcorbi\_hilic\_pos\_3\_019\_43

m/z 211.1771

01234E+46.57RT (min)

lcorbi\_hilic\_pos\_1\_003\_24

m/z 212.0601

0123E+466.5RT (min)

lcorbi\_hilic\_pos\_4\_095\_60

m/z 212.0917

02468E+233.5RT (min)

lcorbi\_hilic\_pos\_4\_082\_59

m/z 212.1031

01E+42.53RT (min)

lcorbi\_hilic\_pos\_3\_021\_46

m/z 212.1053

012E+377.5RT (min)

lcorbi\_hilic\_pos\_4\_020\_37

m/z 212.1134

024E+44.55RT (min)

lcorbi\_hilic\_pos\_4\_096\_60

m/z 212.1135

01E+488.5RT (min)

lcorbi\_hilic\_pos\_2\_024\_32

m/z 212.1135

0123E+466.5RT (min)

lcorbi\_hilic\_pos\_4\_092\_60

m/z 212.1136

0246E+32.53RT (min)

lcorbi\_hilic\_pos\_4\_023\_03

m/z 212.1281

024E+33.54RT (min)

lcorbi\_hilic\_pos\_1\_044\_17

m/z 212.1493

01E+411.5RT (min)

lcorbi\_hilic\_pos\_4\_067\_57

m/z 212.1497

012E+44.55RT (min)

lcorbi\_hilic\_pos\_2\_024\_32

m/z 212.1645

012E+411.5RT (min)

lcorbi\_hilic\_pos\_3\_019\_43

m/z 212.2009

024E+322.5RT (min)

lcorbi\_hilic\_pos\_4\_087\_59

m/z 212.2373

012E+42.53RT (min)

lcorbi\_hilic\_pos\_3\_019\_43

m/z 212.9746

01E+34.55RT (min)

lcorbi\_hilic\_pos\_4\_075\_58

m/z 213.0789

01E+444.5RT (min)

lcorbi\_hilic\_pos\_4\_066\_57

m/z 213.0847

0155.5RT (min)

lcorbi\_hilic\_pos\_1\_014\_27

m/z 213.0888

0246E+40.51RT (min)

lcorbi\_hilic\_pos\_4\_008\_19

m/z 213.1005

0111.5RT (min)

lcorbi\_hilic\_pos\_4\_071\_57

m/z 213.1217

012E+433.5RT (min)

lcorbi\_hilic\_pos\_4\_023\_03

m/z 213.1234

01E+411.5RT (min)

lcorbi\_hilic\_pos\_4\_020\_37

m/z 213.1234

02468E+366.5RT (min)

lcorbi\_hilic\_pos\_4\_085\_59

m/z 213.1598

012E+311.5RT (min)

lcorbi\_hilic\_pos\_1\_051\_26

m/z 213.1961

01234E+411.5RT (min)

lcorbi\_hilic\_pos\_2\_022\_35

m/z 213.2042

012E+43.54RT (min)

lcorbi\_hilic\_pos\_2\_023\_34

m/z 213.9876

0123E+377.5RT (min)

lcorbi\_hilic\_pos\_4\_086\_59

m/z 214.0530

012E+411.5RT (min)

lcorbi\_hilic\_pos\_2\_029\_19

m/z 214.0711

01234E+322.5RT (min)

lcorbi\_hilic\_pos\_4\_088\_59

m/z 214.1438

012E+43.54RT (min)

lcorbi\_hilic\_pos\_4\_073\_57

m/z 214.1579

0123E+411.5RT (min)

lcorbi\_hilic\_pos\_2\_024\_32

m/z 215.0096

01E+36.57RT (min)

lcorbi\_hilic\_pos\_1\_050\_05

m/z 215.0339

01E+46.57RT (min)

lcorbi\_hilic\_pos\_4\_094\_60

m/z 215.0669

01234E+355.5RT (min)

lcorbi\_hilic\_pos\_4\_096\_60

m/z 215.0698

01E+41.52RT (min)

lcorbi\_hilic\_pos\_4\_104\_61

m/z 215.0849

0246E+344.5RT (min)

lcorbi\_hilic\_pos\_4\_084\_59

m/z 215.0927

014.55RT (min)

lcorbi\_hilic\_pos\_4\_090\_60

m/z 215.1026

0246E+36.57RT (min)

lcorbi\_hilic\_pos\_4\_087\_59

m/z 215.1032

024E+31.52RT (min)

lcorbi\_hilic\_pos\_4\_094\_60

m/z 215.1065

01E+455.5RT (min)

lcorbi\_hilic\_pos\_4\_014\_27

m/z 215.1307

024E+355.5RT (min)

lcorbi\_hilic\_pos\_4\_014\_27

m/z 215.1390

0123E+44.55RT (min)

lcorbi\_hilic\_pos\_4\_007\_05

m/z 215.2199

0123E+30.51RT (min)

lcorbi\_hilic\_pos\_4\_012\_29

m/z 215.9645

0123E+377.5RT (min)

lcorbi\_hilic\_pos\_1\_034\_37

m/z 216.0146

02468E+377.5RT (min)

lcorbi\_hilic\_pos\_1\_020\_13

m/z 216.0689

0246E+31.52RT (min)

lcorbi\_hilic\_pos\_1\_007\_16

m/z 216.0866

01E+322.5RT (min)

lcorbi\_hilic\_pos\_4\_082\_59

m/z 216.1067

0123E+41.52RT (min)

lcorbi\_hilic\_pos\_4\_092\_60

m/z 216.1110

0166.5RT (min)

lcorbi\_hilic\_pos\_4\_020\_37

m/z 216.1594

012E+411.5RT (min)

lcorbi\_hilic\_pos\_1\_032\_31

m/z 216.1594

01E+43.54RT (min)

lcorbi\_hilic\_pos\_4\_073\_57

m/z 216.1788

0123E+411.5RT (min)

lcorbi\_hilic\_pos\_4\_089\_59

m/z 216.1855

012E+46.57RT (min)

lcorbi\_hilic\_pos\_4\_091\_60

m/z 216.1901

01234E+322.5RT (min)

lcorbi\_hilic\_pos\_3\_054\_16

m/z 216.2021

01234E+322.5RT (min)

lcorbi\_hilic\_pos\_3\_037\_20

m/z 216.8368

0246E+355.5RT (min)

lcorbi\_hilic\_pos\_4\_092\_60

m/z 216.9227

01234E+45.56RT (min)

lcorbi\_hilic\_pos\_2\_036\_08

m/z 216.9880

0155.5RT (min)

lcorbi\_hilic\_pos\_4\_103\_61

m/z 217.0103

02468E+33.54RT (min)

lcorbi\_hilic\_pos\_4\_106\_62

m/z 217.0736

012E+355.5RT (min)

lcorbi\_hilic\_pos\_4\_091\_60

m/z 217.0819

0123E+366.5RT (min)

lcorbi\_hilic\_pos\_4\_078\_58

m/z 217.0854

0123E+311.5RT (min)

lcorbi\_hilic\_pos\_4\_020\_37

m/z 217.0972

0246E+344.5RT (min)

lcorbi\_hilic\_pos\_4\_102\_61

m/z 217.1295

0246E+366.5RT (min)

lcorbi\_hilic\_pos\_4\_020\_37

m/z 217.1295

01E+40.51RT (min)

lcorbi\_hilic\_pos\_4\_035\_43

m/z 217.1547

0246E+35.56RT (min)

lcorbi\_hilic\_pos\_4\_072\_57

m/z 217.1547

0123E+311.5RT (min)

lcorbi\_hilic\_pos\_1\_019\_01

m/z 217.1910

0123E+411.5RT (min)

lcorbi\_hilic\_pos\_2\_007\_28

m/z 217.1959

0246E+333.5RT (min)

lcorbi\_hilic\_pos\_3\_044\_10

m/z 218.0812

01E+433.5RT (min)

lcorbi\_hilic\_pos\_2\_049\_55

m/z 218.1014

02468E+455.5RT (min)

lcorbi\_hilic\_pos\_4\_096\_60

m/z 218.1080

01E+311.5RT (min)

lcorbi\_hilic\_pos\_2\_003\_33

m/z 218.1135

0123E+344.5RT (min)

lcorbi\_hilic\_pos\_4\_077\_58

m/z 218.1175

01234E+311.5RT (min)

lcorbi\_hilic\_pos\_4\_089\_59

m/z 218.1497

01E+344.5RT (min)

lcorbi\_hilic\_pos\_4\_020\_37

m/z 218.2213

01E+533.5RT (min)

lcorbi\_hilic\_pos\_1\_002\_29

m/z 218.2286

0246E+433.5RT (min)

lcorbi\_hilic\_pos\_1\_054\_23

m/z 218.9197

01E+45.56RT (min)

lcorbi\_hilic\_pos\_1\_022\_43

m/z 218.9876

0246E+31.52RT (min)

lcorbi\_hilic\_pos\_4\_074\_58

m/z 219.0716

0166.5RT (min)

lcorbi\_hilic\_pos\_4\_092\_60

m/z 219.0763

0133.5RT (min)

lcorbi\_hilic\_pos\_1\_051\_26

m/z 219.0975

02468E+36.57RT (min)

lcorbi\_hilic\_pos\_4\_085\_59

m/z 219.1128

024E+34.55RT (min)

lcorbi\_hilic\_pos\_4\_089\_59

m/z 219.1322

0123E+466.5RT (min)

lcorbi\_hilic\_pos\_4\_094\_60

m/z 219.1339

017.58RT (min)

lcorbi\_hilic\_pos\_4\_066\_57

m/z 219.1346

01E+444.5RT (min)

lcorbi\_hilic\_pos\_4\_090\_60

m/z 219.1346

011.52RT (min)

lcorbi\_hilic\_pos\_4\_106\_62

m/z 219.1452

01234E+46.57RT (min)

lcorbi\_hilic\_pos\_4\_007\_05

m/z 219.1743

01E+333.5RT (min)

lcorbi\_hilic\_pos\_1\_034\_37

m/z 219.1743

01E+50.51RT (min)

lcorbi\_hilic\_pos\_4\_112\_62

m/z 219.1955

0246E+333.5RT (min)

lcorbi\_hilic\_pos\_4\_012\_29

m/z 219.2107

010.51RT (min)

lcorbi\_hilic\_pos\_4\_106\_62

m/z 219.9780

0177.5RT (min)

lcorbi\_hilic\_pos\_1\_028\_34

m/z 220.0087

02468E+27.58RT (min)

lcorbi\_hilic\_pos\_4\_099\_61

m/z 220.0299

0246E+355.5RT (min)

lcorbi\_hilic\_pos\_4\_077\_58

m/z 220.0458

012E+477.5RT (min)

lcorbi\_hilic\_pos\_1\_051\_26

m/z 220.0643

0246E+433.5RT (min)

lcorbi\_hilic\_pos\_4\_003\_26

m/z 220.0686

01E+311.5RT (min)

lcorbi\_hilic\_pos\_1\_030\_48

m/z 220.0789

016.57RT (min)

lcorbi\_hilic\_pos\_4\_083\_59

m/z 220.0831

0123E+411.52RT (min)

lcorbi\_hilic\_pos\_4\_087\_59

m/z 220.0862

01E+47.58RT (min)

lcorbi\_hilic\_pos\_4\_070\_57

m/z 220.1057

01E+36.57RT (min)

lcorbi\_hilic\_pos\_2\_048\_04

m/z 220.1096

0246E+322.5RT (min)

lcorbi\_hilic\_pos\_1\_043\_25

m/z 220.1191

0246E+34.55RT (min)

lcorbi\_hilic\_pos\_4\_074\_58

m/z 220.1331

012E+433.5RT (min)

lcorbi\_hilic\_pos\_4\_079\_58

m/z 220.1463

01E+40.51RT (min)

lcorbi\_hilic\_pos\_2\_030\_05

m/z 220.1625

01E+50.51RT (min)

lcorbi\_hilic\_pos\_3\_013\_30

m/z 220.9346

01E+499.5RT (min)

lcorbi\_hilic\_pos\_4\_081\_58

m/z 221.0247

02468E+355.5RT (min)

lcorbi\_hilic\_pos\_4\_078\_58

m/z 221.0590

02468E+35.56RT (min)

lcorbi\_hilic\_pos\_4\_088\_59

m/z 221.0702

01E+36.57RT (min)

lcorbi\_hilic\_pos\_4\_097\_60

m/z 221.1213

0123E+41.52RT (min)

lcorbi\_hilic\_pos\_4\_020\_37

m/z 221.1384

01234E+311.5RT (min)

lcorbi\_hilic\_pos\_1\_012\_19

m/z 221.1413

01E+444.5RT (min)

lcorbi\_hilic\_pos\_2\_024\_32

m/z 221.1536

01E+30.51RT (min)

lcorbi\_hilic\_pos\_4\_043\_42

m/z 221.1940

01E+433.5RT (min)

lcorbi\_hilic\_pos\_1\_044\_17

m/z 221.2186

024E+433.5RT (min)

lcorbi\_hilic\_pos\_1\_054\_23

m/z 221.9880

0246E+366.5RT (min)

lcorbi\_hilic\_pos\_4\_073\_57

m/z 221.9962

0123E+455.5RT (min)

lcorbi\_hilic\_pos\_2\_016\_18

m/z 222.0139

0166.5RT (min)

lcorbi\_hilic\_pos\_4\_073\_57

m/z 222.0724

01E+466.5RT (min)

lcorbi\_hilic\_pos\_4\_066\_57

m/z 222.0985

01234E+433.5RT (min)

lcorbi\_hilic\_pos\_4\_005\_34

m/z 222.1125

012E+333.5RT (min)

lcorbi\_hilic\_pos\_1\_031\_36

m/z 222.1125

02468E+311.5RT (min)

lcorbi\_hilic\_pos\_4\_020\_37

m/z 222.1700

010.51RT (min)

lcorbi\_hilic\_pos\_4\_032\_35

m/z 222.1852

0123E+322.5RT (min)

lcorbi\_hilic\_pos\_1\_044\_17

m/z 222.8886

0123E+377.5RT (min)

lcorbi\_hilic\_pos\_1\_018\_04

m/z 223.0748

01E+42.53RT (min)

lcorbi\_hilic\_pos\_4\_042\_02

m/z 223.0965

0246E+41.52RT (min)

lcorbi\_hilic\_pos\_4\_020\_37

m/z 223.1570

01E+45.56RT (min)

lcorbi\_hilic\_pos\_4\_097\_60

m/z 223.1804

0122.5RT (min)

lcorbi\_hilic\_pos\_4\_094\_60

m/z 223.9890

012E+48.59RT (min)

lcorbi\_hilic\_pos\_4\_079\_58

m/z 224.0706

012E+40.51RT (min)

lcorbi\_hilic\_pos\_4\_072\_57

m/z 224.0781

02468E+388.5RT (min)

lcorbi\_hilic\_pos\_4\_084\_59

m/z 224.0959

01E+45.56RT (min)

lcorbi\_hilic\_pos\_4\_072\_57

m/z 224.1281

012E+411.5RT (min)

lcorbi\_hilic\_pos\_3\_037\_20

m/z 224.1282

01E+33.54RT (min)

lcorbi\_hilic\_pos\_4\_093\_60

m/z 224.1856

0246E+30.51RT (min)

lcorbi\_hilic\_pos\_1\_011\_09

m/z 225.0159

01E+47.58RT (min)

lcorbi\_hilic\_pos\_4\_086\_59

m/z 225.0705

0246E+388.5RT (min)

lcorbi\_hilic\_pos\_4\_083\_59

m/z 225.0757

012E+411.5RT (min)

lcorbi\_hilic\_pos\_1\_014\_27

m/z 225.0836

0246E+36.57RT (min)

lcorbi\_hilic\_pos\_4\_094\_60

m/z 225.0845

02468E+333.5RT (min)

lcorbi\_hilic\_pos\_2\_005\_23

m/z 225.1234

01E+444.5RT (min)

lcorbi\_hilic\_pos\_4\_082\_59

m/z 225.1485

0246E+30.51RT (min)

lcorbi\_hilic\_pos\_1\_038\_21

m/z 225.1598

0122.5RT (min)

lcorbi\_hilic\_pos\_2\_051\_15

m/z 225.9866

02468E+38.59RT (min)

lcorbi\_hilic\_pos\_4\_081\_58

m/z 226.0710

01E+433.5RT (min)

lcorbi\_hilic\_pos\_4\_088\_59

m/z 226.0911

0166.5RT (min)

lcorbi\_hilic\_pos\_4\_022\_08

m/z 226.1046

01E+44.55RT (min)

lcorbi\_hilic\_pos\_1\_051\_26

m/z 226.1285

01E+44.55RT (min)

lcorbi\_hilic\_pos\_4\_076\_58

m/z 226.1404

02468E+344.5RT (min)

lcorbi\_hilic\_pos\_4\_052\_32

m/z 226.1550

0246E+355.5RT (min)

lcorbi\_hilic\_pos\_4\_084\_59

m/z 226.1914

012E+444.5RT (min)

lcorbi\_hilic\_pos\_4\_084\_59

m/z 226.2530

0111.5RT (min)

lcorbi\_hilic\_pos\_4\_074\_58

m/z 226.9430

012E+45.56RT (min)

lcorbi\_hilic\_pos\_3\_054\_16

m/z 226.9676

01E+45.56RT (min)

lcorbi\_hilic\_pos\_4\_066\_57

m/z 227.0429

0246E+344.5RT (min)

lcorbi\_hilic\_pos\_1\_008\_38

m/z 227.0484

015.56RT (min)

lcorbi\_hilic\_pos\_4\_101\_61

m/z 227.0639

024E+377.5RT (min)

lcorbi\_hilic\_pos\_2\_020\_31

m/z 227.0791

012E+455.5RT (min)

lcorbi\_hilic\_pos\_4\_096\_60

m/z 227.1027

0246E+344.5RT (min)

lcorbi\_hilic\_pos\_4\_086\_59

m/z 227.1027

01E+41.52RT (min)

lcorbi\_hilic\_pos\_4\_020\_37

m/z 227.1139

0123E+477.5RT (min)

lcorbi\_hilic\_pos\_4\_020\_37

m/z 227.1140

024E+344.5RT (min)

lcorbi\_hilic\_pos\_4\_051\_48

m/z 227.1244

0246E+36.57RT (min)

lcorbi\_hilic\_pos\_4\_090\_60

m/z 227.1390

0123E+422.53RT (min)

lcorbi\_hilic\_pos\_3\_010\_32

m/z 227.1405

02468E+344.5RT (min)

lcorbi\_hilic\_pos\_1\_056\_56

m/z 227.1663

024E+31.52RT (min)

lcorbi\_hilic\_pos\_1\_049\_55

m/z 227.2118

0246E+333.5RT (min)

lcorbi\_hilic\_pos\_3\_010\_32

m/z 227.2118

01E+411.5RT (min)

lcorbi\_hilic\_pos\_4\_071\_57

m/z 228.0268

01E+433.5RT (min)

lcorbi\_hilic\_pos\_1\_054\_23

m/z 228.0882

01E+422.5RT (min)

lcorbi\_hilic\_pos\_4\_090\_60

m/z 228.1231

0246E+30.51RT (min)

lcorbi\_hilic\_pos\_1\_029\_14

m/z 228.1285

01E+43.54RT (min)

lcorbi\_hilic\_pos\_4\_075\_58

m/z 228.1448

02468E+31.52RT (min)

lcorbi\_hilic\_pos\_4\_091\_60

m/z 228.1594

01E+30.51RT (min)

lcorbi\_hilic\_pos\_4\_105\_61

m/z 228.1594

01E+34.55RT (min)

lcorbi\_hilic\_pos\_4\_100\_61

m/z 228.1958

01E+42.53RT (min)

lcorbi\_hilic\_pos\_3\_009\_50

m/z 228.2686

0246E+422.5RT (min)

lcorbi\_hilic\_pos\_4\_016\_28

m/z 229.0253

02468E+366.5RT (min)

lcorbi\_hilic\_pos\_4\_084\_59

m/z 229.0496

01E+50.51RT (min)

lcorbi\_hilic\_pos\_4\_104\_61

m/z 229.0617

011.52RT (min)

lcorbi\_hilic\_pos\_4\_090\_60

m/z 229.0807

01E+422.5RT (min)

lcorbi\_hilic\_pos\_4\_020\_37

m/z 229.0949

02468E+388.5RT (min)

lcorbi\_hilic\_pos\_4\_091\_60

m/z 229.1183

02468E+34.55RT (min)

lcorbi\_hilic\_pos\_4\_089\_59

m/z 229.1183

01E+466.5RT (min)

lcorbi\_hilic\_pos\_4\_020\_37

m/z 229.1278

01E+422.5RT (min)

lcorbi\_hilic\_pos\_4\_092\_60

m/z 229.1296

02468E+366.5RT (min)

lcorbi\_hilic\_pos\_1\_050\_05

m/z 229.1991

01234E+40.51RT (min)

lcorbi\_hilic\_pos\_3\_019\_43

m/z 229.2275

01234E+42.53RT (min)

lcorbi\_hilic\_pos\_4\_083\_59

m/z 229.2356

0123E+30.51RT (min)

lcorbi\_hilic\_pos\_2\_012\_37

m/z 229.9802

01234E+36.57RT (min)

lcorbi\_hilic\_pos\_4\_079\_58

m/z 230.0211

012E+50.51RT (min)

lcorbi\_hilic\_pos\_1\_051\_26

m/z 230.0375

024E+35.56RT (min)

lcorbi\_hilic\_pos\_4\_101\_61

m/z 230.0458

012E+411.5RT (min)

lcorbi\_hilic\_pos\_2\_029\_19

m/z 230.0772

01E+455.5RT (min)

lcorbi\_hilic\_pos\_3\_004\_27

m/z 230.1104

012E+42.53RT (min)

lcorbi\_hilic\_pos\_3\_007\_19

m/z 230.1240

01E+44.55RT (min)

lcorbi\_hilic\_pos\_4\_096\_60

m/z 230.1400

01E+40.51RT (min)

lcorbi\_hilic\_pos\_4\_073\_57

m/z 230.1751

01E+411.5RT (min)

lcorbi\_hilic\_pos\_4\_098\_61

m/z 230.1751

01234E+33.54RT (min)

lcorbi\_hilic\_pos\_4\_077\_58

m/z 230.1863

0246E+34.55RT (min)

lcorbi\_hilic\_pos\_4\_071\_57

m/z 230.2115

012E+322.5RT (min)

lcorbi\_hilic\_pos\_1\_023\_28

m/z 231.0360

0246E+37.58RT (min)

lcorbi\_hilic\_pos\_4\_090\_60

m/z 231.0619

02468E+35.56RT (min)

lcorbi\_hilic\_pos\_4\_105\_61

m/z 231.0993

0123E+311.5RT (min)

lcorbi\_hilic\_pos\_4\_020\_37

m/z 231.1340

01E+433.5RT (min)

lcorbi\_hilic\_pos\_4\_102\_61

m/z 231.1591

012E+40.51RT (min)

lcorbi\_hilic\_pos\_4\_018\_31

m/z 231.1703

0246E+45.56RT (min)

lcorbi\_hilic\_pos\_4\_020\_37

m/z 231.2148

024E+30.51RT (min)

lcorbi\_hilic\_pos\_4\_018\_31

m/z 231.2453

012E+42.53RT (min)

lcorbi\_hilic\_pos\_3\_019\_43

m/z 232.0169

0123E+366.5RT (min)

lcorbi\_hilic\_pos\_4\_068\_57

m/z 232.0816

0123E+322.5RT (min)

lcorbi\_hilic\_pos\_3\_048\_38

m/z 232.0940

02468E+388.5RT (min)

lcorbi\_hilic\_pos\_4\_096\_60

m/z 232.1030

010.51RT (min)

lcorbi\_hilic\_pos\_2\_031\_21

m/z 232.1193

02468E+30.51RT (min)

lcorbi\_hilic\_pos\_4\_105\_61

m/z 232.1397

0246E+466.5RT (min)

lcorbi\_hilic\_pos\_4\_093\_60

m/z 232.1404

024E+37.58RT (min)

lcorbi\_hilic\_pos\_4\_020\_37

m/z 232.1543

02468E+222.5RT (min)

lcorbi\_hilic\_pos\_4\_012\_29

m/z 232.8108

01E+455.5RT (min)

lcorbi\_hilic\_pos\_4\_088\_59

m/z 232.8966

01E+455.5RT (min)

lcorbi\_hilic\_pos\_4\_090\_60

m/z 232.9174

01E+377.5RT (min)

lcorbi\_hilic\_pos\_4\_101\_61

m/z 232.9282

0246E+499.5RT (min)

lcorbi\_hilic\_pos\_4\_081\_58

m/z 233.0387

01E+45.56RT (min)

lcorbi\_hilic\_pos\_4\_095\_60

m/z 233.0421

02468E+377.5RT (min)

lcorbi\_hilic\_pos\_4\_083\_59

m/z 233.0591

01E+46.57RT (min)

lcorbi\_hilic\_pos\_2\_020\_31

m/z 233.0632

0246E+32.53RT (min)

lcorbi\_hilic\_pos\_4\_080\_58

m/z 233.0685

0123E+35.56RT (min)

lcorbi\_hilic\_pos\_4\_112\_62

m/z 233.0768

0246E+444.5RT (min)

lcorbi\_hilic\_pos\_4\_005\_34

m/z 233.0787

01234E+311.5RT (min)

lcorbi\_hilic\_pos\_1\_051\_26

m/z 233.0921

0246E+211.5RT (min)

lcorbi\_hilic\_pos\_1\_037\_18

m/z 233.0985

0246E+377.5RT (min)

lcorbi\_hilic\_pos\_4\_066\_57

m/z 233.1034

01E+44.55RT (min)

lcorbi\_hilic\_pos\_4\_007\_05

m/z 233.1132

01234E+366.5RT (min)

lcorbi\_hilic\_pos\_4\_086\_59

m/z 233.1138

011.52RT (min)

lcorbi\_hilic\_pos\_4\_113\_62

m/z 233.1284

012E+41.52RT (min)

lcorbi\_hilic\_pos\_4\_110\_62

m/z 233.1432

02468E+311.5RT (min)

lcorbi\_hilic\_pos\_1\_051\_26

m/z 233.1565

02468E+311.5RT (min)

lcorbi\_hilic\_pos\_1\_044\_17

m/z 233.1748

01E+30.51RT (min)

lcorbi\_hilic\_pos\_4\_016\_28

m/z 233.1941

012E+411.5RT (min)

lcorbi\_hilic\_pos\_3\_002\_04

m/z 233.9993

024E+366.5RT (min)

lcorbi\_hilic\_pos\_4\_081\_58

m/z 234.0664

0144.5RT (min)

lcorbi\_hilic\_pos\_4\_020\_37

m/z 234.0873

012E+477.5RT (min)

lcorbi\_hilic\_pos\_4\_088\_59

m/z 234.1336

012E+411.5RT (min)

lcorbi\_hilic\_pos\_2\_046\_46

m/z 234.1467

01E+411.5RT (min)

lcorbi\_hilic\_pos\_1\_044\_17

m/z 234.1598

012E+33.54RT (min)

lcorbi\_hilic\_pos\_4\_012\_29

m/z 234.1873

024E+333.5RT (min)

lcorbi\_hilic\_pos\_4\_031\_17

m/z 234.1998

01E+533.5RT (min)

lcorbi\_hilic\_pos\_2\_019\_16

m/z 234.2064

0246E+31.52RT (min)

lcorbi\_hilic\_pos\_4\_028\_10

m/z 234.2152

0246E+433.5RT (min)

lcorbi\_hilic\_pos\_2\_020\_31

m/z 234.8089

01234E+355.5RT (min)

lcorbi\_hilic\_pos\_4\_097\_60

m/z 235.0485

012E+50.51RT (min)

lcorbi\_hilic\_pos\_4\_046\_21

m/z 235.0568

01E+47.58RT (min)

lcorbi\_hilic\_pos\_4\_070\_57

m/z 235.0753

01E+45.56RT (min)

lcorbi\_hilic\_pos\_4\_097\_60

m/z 235.0802

012E+42.53RT (min)

lcorbi\_hilic\_pos\_4\_073\_57

m/z 235.0925

0123E+377.5RT (min)

lcorbi\_hilic\_pos\_4\_004\_38

m/z 235.1441

01234E+411.5RT (min)

lcorbi\_hilic\_pos\_4\_086\_59

m/z 235.1652

01E+455.5RT (min)

lcorbi\_hilic\_pos\_4\_071\_57

m/z 235.1693

01E+30.51RT (min)

lcorbi\_hilic\_pos\_4\_015\_20

m/z 235.2037

01234E+433.5RT (min)

lcorbi\_hilic\_pos\_2\_022\_35

m/z 235.2644

01234E+40.51RT (min)

lcorbi\_hilic\_pos\_4\_090\_60

m/z 235.6341

0246E+377.5RT (min)

lcorbi\_hilic\_pos\_4\_082\_59

m/z 235.9848

01234E+35.56RT (min)

lcorbi\_hilic\_pos\_4\_080\_58

m/z 236.0379

01E+44.55RT (min)

lcorbi\_hilic\_pos\_4\_068\_57

m/z 236.0408

01E+46.57RT (min)

lcorbi\_hilic\_pos\_4\_084\_59

m/z 236.0505

012E+466.5RT (min)

lcorbi\_hilic\_pos\_2\_030\_05

m/z 236.0877

01E+533.5RT (min)

lcorbi\_hilic\_pos\_3\_034\_26

m/z 236.1029

0246E+36.57RT (min)

lcorbi\_hilic\_pos\_1\_036\_32

m/z 236.1045

01E+433.5RT (min)

lcorbi\_hilic\_pos\_1\_036\_32

m/z 236.1111

01E+344.5RT (min)

lcorbi\_hilic\_pos\_4\_004\_38

m/z 236.1222

014.55RT (min)

lcorbi\_hilic\_pos\_4\_104\_61

m/z 236.1281

01E+411.5RT (min)

lcorbi\_hilic\_pos\_4\_020\_37

m/z 236.1493

01E+444.5RT (min)

lcorbi\_hilic\_pos\_3\_028\_07

m/z 236.1574

01234E+40.51RT (min)

lcorbi\_hilic\_pos\_3\_007\_19

m/z 236.1603

012E+44.55RT (min)

lcorbi\_hilic\_pos\_4\_097\_60

m/z 236.9043

0123E+366.5RT (min)

lcorbi\_hilic\_pos\_1\_043\_25

m/z 236.9475

02468E+37.58RT (min)

lcorbi\_hilic\_pos\_4\_113\_62

m/z 237.0279

0123E+355.5RT (min)

lcorbi\_hilic\_pos\_1\_018\_04

m/z 237.0458

0246E+377.5RT (min)

lcorbi\_hilic\_pos\_4\_079\_58

m/z 237.0794

024E+40.51RT (min)

lcorbi\_hilic\_pos\_4\_073\_57

m/z 237.0870

01E+433.5RT (min)

lcorbi\_hilic\_pos\_4\_088\_59

m/z 237.1362

012E+422.5RT (min)

lcorbi\_hilic\_pos\_4\_091\_60

m/z 237.1451

0123E+44.55RT (min)

lcorbi\_hilic\_pos\_2\_037\_24

m/z 237.1849

012E+40.51RT (min)

lcorbi\_hilic\_pos\_3\_019\_43

m/z 237.9732

0246E+25.56RT (min)

lcorbi\_hilic\_pos\_1\_037\_18

m/z 238.0589

01234E+477.5RT (min)

lcorbi\_hilic\_pos\_4\_093\_60

m/z 238.1074

01E+311.5RT (min)

lcorbi\_hilic\_pos\_4\_076\_58

m/z 238.1186

0246E+444.5RT (min)

lcorbi\_hilic\_pos\_4\_097\_60

m/z 238.1438

01E+433.5RT (min)

lcorbi\_hilic\_pos\_4\_077\_58

m/z 238.1730

010.51RT (min)

lcorbi\_hilic\_pos\_4\_106\_62

m/z 238.2529

01E+40.51RT (min)

lcorbi\_hilic\_pos\_1\_007\_16

m/z 239.0915

0246E+32.53RT (min)

lcorbi\_hilic\_pos\_4\_020\_37

m/z 239.1026

01E+433.5RT (min)

lcorbi\_hilic\_pos\_4\_020\_37

m/z 239.1122

0246E+333.5RT (min)

lcorbi\_hilic\_pos\_4\_049\_55

m/z 239.1519

02468E+322.5RT (min)

lcorbi\_hilic\_pos\_1\_003\_24

m/z 239.1681

01234E+311.5RT (min)

lcorbi\_hilic\_pos\_4\_093\_60

m/z 239.2046

0123E+40.51RT (min)

lcorbi\_hilic\_pos\_2\_034\_41

m/z 239.2118

0246E+322.5RT (min)

lcorbi\_hilic\_pos\_4\_070\_57

m/z 239.9666

012E+499.5RT (min)

lcorbi\_hilic\_pos\_4\_077\_58

m/z 240.0963

012E+433.5RT (min)

lcorbi\_hilic\_pos\_4\_073\_57

m/z 240.0997

024E+311.52RT (min)

lcorbi\_hilic\_pos\_4\_085\_59

m/z 240.1230

0246E+311.5RT (min)

lcorbi\_hilic\_pos\_2\_027\_38

m/z 240.1357

01E+42.53RT (min)

lcorbi\_hilic\_pos\_4\_096\_60

m/z 240.1789

02468E+33.54RT (min)

lcorbi\_hilic\_pos\_4\_078\_58

m/z 240.1933

02468E+433.5RT (min)

lcorbi\_hilic\_pos\_2\_052\_10

m/z 240.2322

01E+40.51RT (min)

lcorbi\_hilic\_pos\_3\_019\_43

m/z 240.2322

012E+422.5RT (min)

lcorbi\_hilic\_pos\_4\_082\_59

m/z 240.2686

012E+444.5RT (min)

lcorbi\_hilic\_pos\_1\_020\_13

m/z 240.9250

012E+477.5RT (min)

lcorbi\_hilic\_pos\_4\_083\_59

m/z 241.0311

0246E+388.5RT (min)

lcorbi\_hilic\_pos\_4\_066\_57

m/z 241.0584

01234E+333.5RT (min)

lcorbi\_hilic\_pos\_1\_003\_24

m/z 241.0674

024E+466.5RT (min)

lcorbi\_hilic\_pos\_4\_093\_60

m/z 241.0796

0246E+36.57RT (min)

lcorbi\_hilic\_pos\_4\_088\_59

m/z 241.0908

0177.5RT (min)

lcorbi\_hilic\_pos\_4\_096\_60

m/z 241.1053

0144.5RT (min)

lcorbi\_hilic\_pos\_4\_092\_60

m/z 241.1071

01E+411.5RT (min)

lcorbi\_hilic\_pos\_4\_083\_59

m/z 241.1183

01E+44.55RT (min)

lcorbi\_hilic\_pos\_4\_082\_59

m/z 241.1537

0246E+311.5RT (min)

lcorbi\_hilic\_pos\_4\_089\_59

m/z 241.1675

0246E+31.52RT (min)

lcorbi\_hilic\_pos\_4\_096\_60

m/z 241.2036

01E+54.55RT (min)

lcorbi\_hilic\_pos\_4\_020\_37

m/z 241.2720

01234E+344.5RT (min)

lcorbi\_hilic\_pos\_2\_028\_13

m/z 241.9980

01E+455.5RT (min)

lcorbi\_hilic\_pos\_1\_016\_33

m/z 241.9996

0123E+48.59RT (min)

lcorbi\_hilic\_pos\_4\_075\_58

m/z 242.0277

01E+477.5RT (min)

lcorbi\_hilic\_pos\_1\_037\_18

m/z 242.0464

012E+433.5RT (min)

lcorbi\_hilic\_pos\_4\_003\_26

m/z 242.0609

01E+46.57RT (min)

lcorbi\_hilic\_pos\_4\_092\_60

m/z 242.0772

01E+444.5RT (min)

lcorbi\_hilic\_pos\_4\_088\_59

m/z 242.1000

012E+41.52RT (min)

lcorbi\_hilic\_pos\_4\_020\_37

m/z 242.1023

01E+433.5RT (min)

lcorbi\_hilic\_pos\_4\_020\_37

m/z 242.1135

0123E+344.5RT (min)

lcorbi\_hilic\_pos\_4\_020\_37

m/z 242.1136

0111.5RT (min)

lcorbi\_hilic\_pos\_4\_020\_37

m/z 242.1249

02468E+333.5RT (min)

lcorbi\_hilic\_pos\_3\_056\_56

m/z 242.1387

0123E+31.52RT (min)

lcorbi\_hilic\_pos\_2\_027\_38

m/z 242.1580

01E+455.5RT (min)

lcorbi\_hilic\_pos\_4\_020\_37

m/z 242.1751

01E+411.5RT (min)

lcorbi\_hilic\_pos\_3\_019\_43

m/z 242.1751

012.53RT (min)

lcorbi\_hilic\_pos\_3\_019\_43

m/z 242.2479

01E+42.53RT (min)

lcorbi\_hilic\_pos\_1\_034\_37

m/z 242.9255

01E+455.5RT (min)

lcorbi\_hilic\_pos\_4\_101\_61

m/z 243.0005

02468E+38.59RT (min)

lcorbi\_hilic\_pos\_4\_077\_58

m/z 243.0265

01E+47.58RT (min)

lcorbi\_hilic\_pos\_4\_087\_59

m/z 243.0410

0246E+35.56RT (min)

lcorbi\_hilic\_pos\_4\_089\_59

m/z 243.0531

02468E+355.5RT (min)

lcorbi\_hilic\_pos\_4\_075\_58

m/z 243.0798

01E+422.5RT (min)

lcorbi\_hilic\_pos\_4\_001\_49

m/z 243.0839

02468E+40.51RT (min)

lcorbi\_hilic\_pos\_4\_089\_59

m/z 243.1321

012E+333.5RT (min)

lcorbi\_hilic\_pos\_4\_081\_58

m/z 243.1637

0111.5RT (min)

lcorbi\_hilic\_pos\_4\_040\_39

m/z 243.1704

0246E+33.54RT (min)

lcorbi\_hilic\_pos\_4\_077\_58

m/z 243.1829

01E+44.55RT (min)

lcorbi\_hilic\_pos\_2\_005\_23

m/z 243.1955

0123E+40.51RT (min)

lcorbi\_hilic\_pos\_4\_006\_14

m/z 243.2067

02468E+333.5RT (min)

lcorbi\_hilic\_pos\_4\_112\_62

m/z 243.2107

024E+20.51RT (min)

lcorbi\_hilic\_pos\_2\_016\_18

m/z 243.2512

01E+40.51RT (min)

lcorbi\_hilic\_pos\_3\_019\_43

m/z 243.2876

0123E+32.53RT (min)

lcorbi\_hilic\_pos\_4\_086\_59

m/z 243.9959

0246E+366.5RT (min)

lcorbi\_hilic\_pos\_4\_086\_59

m/z 243.9972

01E+48.59RT (min)

lcorbi\_hilic\_pos\_4\_075\_58

m/z 244.0169

012E+45.56RT (min)

lcorbi\_hilic\_pos\_4\_083\_59

m/z 244.0281

02468E+366.5RT (min)

lcorbi\_hilic\_pos\_4\_077\_58

m/z 244.0394

012E+47.58RT (min)

lcorbi\_hilic\_pos\_2\_024\_32

m/z 244.1115

0123E+32.53RT (min)

lcorbi\_hilic\_pos\_2\_024\_32

m/z 244.1292

01E+46.57RT (min)

lcorbi\_hilic\_pos\_4\_020\_37

m/z 244.1379

01E+411.5RT (min)

lcorbi\_hilic\_pos\_4\_097\_60

m/z 244.1402

0123E+44.55RT (min)

lcorbi\_hilic\_pos\_4\_091\_60

m/z 244.1544

01E+311.5RT (min)

lcorbi\_hilic\_pos\_4\_105\_61

m/z 244.1544

01E+444.5RT (min)

lcorbi\_hilic\_pos\_4\_066\_57

m/z 244.1656

02468E+377.5RT (min)

lcorbi\_hilic\_pos\_4\_087\_59

m/z 244.2060

01E+411.5RT (min)

lcorbi\_hilic\_pos\_4\_082\_59

m/z 244.2078

01E+433.5RT (min)

lcorbi\_hilic\_pos\_4\_012\_29

m/z 244.8638

024E+37.58RT (min)

lcorbi\_hilic\_pos\_4\_103\_61

m/z 244.8964

01E+477.5RT (min)

lcorbi\_hilic\_pos\_1\_053\_08

m/z 245.0676

01E+455.5RT (min)

lcorbi\_hilic\_pos\_4\_080\_58

m/z 245.0825

01E+455.5RT (min)

lcorbi\_hilic\_pos\_4\_109\_62

m/z 245.0921

02468E+344.5RT (min)

lcorbi\_hilic\_pos\_4\_105\_61

m/z 245.1019

024E+31.52RT (min)

lcorbi\_hilic\_pos\_2\_046\_46

m/z 245.1035

0144.5RT (min)

lcorbi\_hilic\_pos\_4\_092\_60

m/z 245.1227

01E+433.5RT (min)

lcorbi\_hilic\_pos\_4\_052\_32

m/z 245.1360

0246E+411.5RT (min)

lcorbi\_hilic\_pos\_2\_052\_10

m/z 245.1496

011.522.5RT (min)

lcorbi\_hilic\_pos\_4\_092\_60

m/z 245.1496

014.55RT (min)

lcorbi\_hilic\_pos\_4\_107\_62

m/z 245.1608

02468E+355.5RT (min)

lcorbi\_hilic\_pos\_4\_020\_37

m/z 245.1649

01E+43.54RT (min)

lcorbi\_hilic\_pos\_4\_111\_62

m/z 245.1860

01E+46.57RT (min)

lcorbi\_hilic\_pos\_4\_068\_57

m/z 245.1860

02468E+444.55RT (min)

lcorbi\_hilic\_pos\_4\_083\_59

m/z 245.2054

0246E+355.5RT (min)

lcorbi\_hilic\_pos\_4\_083\_59

m/z 245.2224

01E+411.5RT (min)

lcorbi\_hilic\_pos\_2\_007\_28

m/z 245.9493

024E+366.5RT (min)

lcorbi\_hilic\_pos\_4\_073\_57

m/z 246.0196

01E+44.55RT (min)

lcorbi\_hilic\_pos\_1\_008\_38

m/z 246.0601

01234E+388.5RT (min)

lcorbi\_hilic\_pos\_4\_085\_59

m/z 246.0715

0246E+35.56RT (min)

lcorbi\_hilic\_pos\_4\_073\_57

m/z 246.0948

012E+466.5RT (min)

lcorbi\_hilic\_pos\_4\_072\_57

m/z 246.0972

0123E+444.5RT (min)

lcorbi\_hilic\_pos\_4\_088\_59

m/z 246.0988

01E+422.5RT (min)

lcorbi\_hilic\_pos\_4\_091\_60

m/z 246.1447

0246E+311.5RT (min)

lcorbi\_hilic\_pos\_4\_094\_60

m/z 246.1558

01234E+44.55RT (min)

lcorbi\_hilic\_pos\_4\_097\_60

m/z 246.1561

024E+37.58RT (min)

lcorbi\_hilic\_pos\_4\_082\_59

m/z 246.1693

01E+477.5RT (min)

lcorbi\_hilic\_pos\_4\_091\_60

m/z 246.1893

01E+44.55RT (min)

lcorbi\_hilic\_pos\_4\_085\_59

m/z 246.2064

012E+40.51RT (min)

lcorbi\_hilic\_pos\_1\_024\_03

m/z 246.2701

012E+42.53RT (min)

lcorbi\_hilic\_pos\_4\_104\_61

m/z 247.0688

02468E+31.52RT (min)

lcorbi\_hilic\_pos\_4\_082\_59

m/z 247.0900

01E+455.5RT (min)

lcorbi\_hilic\_pos\_3\_003\_37

m/z 247.0913

0122.5RT (min)

lcorbi\_hilic\_pos\_3\_023\_18

m/z 247.0925

02468E+377.5RT (min)

lcorbi\_hilic\_pos\_2\_003\_33

m/z 247.1053

0246E+21.52RT (min)

lcorbi\_hilic\_pos\_3\_029\_23

m/z 247.1078

01E+422.5RT (min)

lcorbi\_hilic\_pos\_4\_066\_57

m/z 247.1402

01E+42.53RT (min)

lcorbi\_hilic\_pos\_3\_021\_46

m/z 247.1666

0123E+40.51RT (min)

lcorbi\_hilic\_pos\_4\_076\_58

m/z 247.2268

0246E+30.51RT (min)

lcorbi\_hilic\_pos\_4\_106\_62

m/z 248.0182

01E+455.5RT (min)

lcorbi\_hilic\_pos\_1\_003\_24

m/z 248.0590

024E+30.51RT (min)

lcorbi\_hilic\_pos\_3\_023\_18

m/z 248.0821

01E+44.55RT (min)

lcorbi\_hilic\_pos\_2\_009\_50

m/z 248.0918

01E+422.5RT (min)

lcorbi\_hilic\_pos\_4\_070\_57

m/z 248.1031

0246E+311.5RT (min)

lcorbi\_hilic\_pos\_2\_009\_50

m/z 248.1223

01E+444.5RT (min)

lcorbi\_hilic\_pos\_1\_036\_32

m/z 248.1328

0123E+41.52RT (min)

lcorbi\_hilic\_pos\_4\_093\_60

m/z 248.1347

0166.5RT (min)

lcorbi\_hilic\_pos\_4\_092\_60

m/z 248.1643

01234E+333.5RT (min)

lcorbi\_hilic\_pos\_4\_084\_59

m/z 248.1856

0123E+333.5RT (min)

lcorbi\_hilic\_pos\_4\_012\_29

m/z 248.1856

01E+411.5RT (min)

lcorbi\_hilic\_pos\_1\_044\_17

m/z 248.2027

0133.5RT (min)

lcorbi\_hilic\_pos\_4\_012\_29

m/z 248.2373

012E+31.52RT (min)

lcorbi\_hilic\_pos\_4\_113\_62

m/z 248.8706

01E+455.5RT (min)

lcorbi\_hilic\_pos\_4\_082\_59

m/z 249.0336

012E+45.56RT (min)

lcorbi\_hilic\_pos\_4\_095\_60

m/z 249.0458

01E+377.5RT (min)

lcorbi\_hilic\_pos\_4\_094\_60

m/z 249.0546

010.51RT (min)

lcorbi\_hilic\_pos\_4\_074\_58

m/z 249.0703

01E+44.55RT (min)

lcorbi\_hilic\_pos\_4\_068\_57

m/z 249.0870

0246E+344.5RT (min)

lcorbi\_hilic\_pos\_1\_031\_36

m/z 249.1064

012E+46.57RT (min)

lcorbi\_hilic\_pos\_1\_050\_05

m/z 249.1409

01234E+466.5RT (min)

lcorbi\_hilic\_pos\_4\_069\_57

m/z 249.1445

0123E+44.55RT (min)

lcorbi\_hilic\_pos\_1\_044\_17

m/z 249.1849

01E+411.5RT (min)

lcorbi\_hilic\_pos\_4\_081\_58

m/z 249.2253

01234E+333.5RT (min)

lcorbi\_hilic\_pos\_1\_044\_17

m/z 250.0087

024E+333.5RT (min)

lcorbi\_hilic\_pos\_1\_054\_23

m/z 250.0298

0123E+34.55RT (min)

lcorbi\_hilic\_pos\_2\_020\_31

m/z 250.0644

01E+455.5RT (min)

lcorbi\_hilic\_pos\_4\_077\_58

m/z 250.0658

02468E+40.51RT (min)

lcorbi\_hilic\_pos\_4\_076\_58

m/z 250.1187

01E+41.52RT (min)

lcorbi\_hilic\_pos\_2\_046\_46

m/z 250.1199

0123E+466.5RT (min)

lcorbi\_hilic\_pos\_4\_069\_57

m/z 250.1285

0246E+35.56RT (min)

lcorbi\_hilic\_pos\_1\_021\_46

m/z 250.1404

0144.5RT (min)

lcorbi\_hilic\_pos\_4\_020\_37

m/z 250.1649

01E+44.55RT (min)

lcorbi\_hilic\_pos\_2\_001\_49

m/z 250.1730

0123E+40.51RT (min)

lcorbi\_hilic\_pos\_2\_052\_10

m/z 250.2094

01E+40.51RT (min)

lcorbi\_hilic\_pos\_3\_019\_43

m/z 250.2166

01234E+322.5RT (min)

lcorbi\_hilic\_pos\_3\_027\_05

m/z 250.8680

0246E+355.5RT (min)

lcorbi\_hilic\_pos\_4\_082\_59

m/z 251.0088

012E+477.5RT (min)

lcorbi\_hilic\_pos\_4\_081\_58

m/z 251.0395

010.51RT (min)

lcorbi\_hilic\_pos\_4\_074\_58

m/z 251.0544

024E+40.51RT (min)

lcorbi\_hilic\_pos\_3\_008\_14

m/z 251.0674

01E+34.55RT (min)

lcorbi\_hilic\_pos\_4\_020\_37

m/z 251.0945

0155.5RT (min)

lcorbi\_hilic\_pos\_4\_080\_58

m/z 251.1027

01E+42.53RT (min)

lcorbi\_hilic\_pos\_4\_086\_59

m/z 251.1067

0111.5RT (min)

lcorbi\_hilic\_pos\_4\_015\_20

m/z 251.1116

0246E+466.5RT (min)

lcorbi\_hilic\_pos\_1\_050\_05

m/z 251.1301

0246E+31.52RT (min)

lcorbi\_hilic\_pos\_4\_096\_60

m/z 251.1366

01E+455.5RT (min)

lcorbi\_hilic\_pos\_4\_080\_58

m/z 251.1490

01E+40.51RT (min)

lcorbi\_hilic\_pos\_2\_029\_19

m/z 251.1601

01E+455.5RT (min)

lcorbi\_hilic\_pos\_1\_044\_17

m/z 251.2006

012E+40.51RT (min)

lcorbi\_hilic\_pos\_4\_081\_58

m/z 252.0068

01E+411.5RT (min)

lcorbi\_hilic\_pos\_4\_106\_62

m/z 252.0277

01E+44.55RT (min)

lcorbi\_hilic\_pos\_4\_106\_62

m/z 252.1060

0123E+44.55RT (min)

lcorbi\_hilic\_pos\_2\_003\_33

m/z 252.1231

02468E+311.5RT (min)

lcorbi\_hilic\_pos\_4\_020\_37

m/z 252.1259

02468E+33.54RT (min)

lcorbi\_hilic\_pos\_1\_024\_03

m/z 252.1383

02468E+40.51RT (min)

lcorbi\_hilic\_pos\_4\_083\_59

m/z 252.1806

0246E+30.51RT (min)

lcorbi\_hilic\_pos\_1\_051\_26

m/z 252.1887

012E+40.51RT (min)

lcorbi\_hilic\_pos\_4\_100\_61

m/z 252.8764

01E+45.56RT (min)

lcorbi\_hilic\_pos\_4\_103\_61

m/z 253.0070

01E+477.5RT (min)

lcorbi\_hilic\_pos\_4\_081\_58

m/z 253.0520

02468E+40.51RT (min)

lcorbi\_hilic\_pos\_4\_075\_58

m/z 253.0779

0123E+466.5RT (min)

lcorbi\_hilic\_pos\_4\_092\_60

m/z 253.1028

01234E+40.51RT (min)

lcorbi\_hilic\_pos\_2\_016\_18

m/z 253.1183

0123E+444.5RT (min)

lcorbi\_hilic\_pos\_4\_088\_59

m/z 253.1281

012E+31.52RT (min)

lcorbi\_hilic\_pos\_4\_106\_62

m/z 253.1479

01E+466.5RT (min)

lcorbi\_hilic\_pos\_4\_068\_57

m/z 253.1798

01E+30.51RT (min)

lcorbi\_hilic\_pos\_4\_034\_36

m/z 253.1910

01E+433.5RT (min)

lcorbi\_hilic\_pos\_4\_109\_62

m/z 253.1950

012E+455.5RT (min)

lcorbi\_hilic\_pos\_4\_090\_60

m/z 253.2162

02468E+20.51RT (min)

lcorbi\_hilic\_pos\_4\_034\_36

m/z 254.0200

02468E+366.5RT (min)

lcorbi\_hilic\_pos\_4\_067\_57

m/z 254.1154

0246E+311.5RT (min)

lcorbi\_hilic\_pos\_4\_082\_59

m/z 254.1234

0123E+566.5RT (min)

lcorbi\_hilic\_pos\_1\_003\_24

m/z 254.1388

0123E+311.5RT (min)

lcorbi\_hilic\_pos\_4\_106\_62

m/z 254.1598

0246E+31.52RT (min)

lcorbi\_hilic\_pos\_1\_054\_23

m/z 254.1679

01E+411.5RT (min)

lcorbi\_hilic\_pos\_4\_102\_61

m/z 254.2114

0246E+30.51RT (min)

lcorbi\_hilic\_pos\_4\_099\_61

m/z 254.2308

01E+422.5RT (min)

lcorbi\_hilic\_pos\_4\_092\_60

m/z 254.8157

0155.5RT (min)

lcorbi\_hilic\_pos\_4\_098\_61

m/z 254.8237

0246E+355.5RT (min)

lcorbi\_hilic\_pos\_4\_098\_61

m/z 254.9662

01E+43.54RT (min)

lcorbi\_hilic\_pos\_4\_108\_62

m/z 255.0240

01E+477.5RT (min)

lcorbi\_hilic\_pos\_4\_082\_59

m/z 255.0264

01E+47.58RT (min)

lcorbi\_hilic\_pos\_4\_084\_59

m/z 255.0976

012E+433.5RT (min)

lcorbi\_hilic\_pos\_4\_088\_59

m/z 255.1187

012E+466.5RT (min)

lcorbi\_hilic\_pos\_4\_096\_60

m/z 255.1236

0111.5RT (min)

lcorbi\_hilic\_pos\_4\_031\_17

m/z 255.1339

0144.5RT (min)

lcorbi\_hilic\_pos\_4\_097\_60

m/z 255.1453

0123E+45.56RT (min)

lcorbi\_hilic\_pos\_2\_030\_05

m/z 255.1955

01E+30.51RT (min)

lcorbi\_hilic\_pos\_4\_084\_59

m/z 255.2025

02468E+377.5RT (min)

lcorbi\_hilic\_pos\_4\_097\_60

m/z 255.2068

02468E+311.5RT (min)

lcorbi\_hilic\_pos\_1\_044\_17

m/z 255.2319

01E+40.51RT (min)

lcorbi\_hilic\_pos\_4\_077\_58

m/z 255.2512

012E+422.5RT (min)

lcorbi\_hilic\_pos\_4\_104\_61

m/z 255.9781

01E+466.5RT (min)

lcorbi\_hilic\_pos\_4\_073\_57

m/z 256.0693

02468E+377.5RT (min)

lcorbi\_hilic\_pos\_4\_020\_37

m/z 256.0904

02468E+366.5RT (min)

lcorbi\_hilic\_pos\_2\_013\_36

m/z 256.1099

012E+433.5RT (min)

lcorbi\_hilic\_pos\_4\_068\_57

m/z 256.1180

012E+44.55RT (min)

lcorbi\_hilic\_pos\_4\_052\_32

m/z 256.1292

0246E+366.5RT (min)

lcorbi\_hilic\_pos\_4\_091\_60

m/z 256.1481

01E+43.54RT (min)

lcorbi\_hilic\_pos\_4\_103\_61

m/z 256.1597

0246E+355.5RT (min)

lcorbi\_hilic\_pos\_4\_092\_60

m/z 256.1907

0246E+32.53RT (min)

lcorbi\_hilic\_pos\_4\_092\_60

m/z 256.2271

0123E+40.51RT (min)

lcorbi\_hilic\_pos\_3\_019\_43

m/z 256.2271

024E+32.53RT (min)

lcorbi\_hilic\_pos\_4\_093\_60

m/z 256.2635

012E+40.51RT (min)

lcorbi\_hilic\_pos\_1\_054\_23

m/z 256.2999

02468E+41.52RT (min)

lcorbi\_hilic\_pos\_3\_020\_31

m/z 256.8131

0155.5RT (min)

lcorbi\_hilic\_pos\_4\_103\_61

m/z 256.9475

0246E+399.5RT (min)

lcorbi\_hilic\_pos\_4\_079\_58

m/z 257.0533

0246E+34.55RT (min)

lcorbi\_hilic\_pos\_2\_024\_32

m/z 257.0744

0123E+444.5RT (min)

lcorbi\_hilic\_pos\_4\_082\_59

m/z 257.0806

01E+50.51RT (min)

lcorbi\_hilic\_pos\_4\_100\_61

m/z 257.1023

01234E+411.52RT (min)

lcorbi\_hilic\_pos\_4\_020\_37

m/z 257.1132

012E+433.5RT (min)

lcorbi\_hilic\_pos\_3\_011\_34

m/z 257.1496

02468E+311.5RT (min)

lcorbi\_hilic\_pos\_4\_082\_59

m/z 257.1609

01E+45.56RT (min)

lcorbi\_hilic\_pos\_3\_021\_46

m/z 257.1683

0123E+466.5RT (min)

lcorbi\_hilic\_pos\_4\_050\_16

m/z 257.1789

01E+411.5RT (min)

lcorbi\_hilic\_pos\_1\_025\_52

m/z 257.2111

010.51RT (min)

lcorbi\_hilic\_pos\_4\_082\_59

m/z 257.2264

0246E+344.5RT (min)

lcorbi\_hilic\_pos\_4\_074\_58

m/z 257.2588

024E+42.53RT (min)

lcorbi\_hilic\_pos\_4\_087\_59

m/z 257.2669

024E+32.53RT (min)

lcorbi\_hilic\_pos\_1\_046\_30

m/z 257.9393

0246E+38.59RT (min)

lcorbi\_hilic\_pos\_4\_098\_61

m/z 257.9493

02468E+377.5RT (min)

lcorbi\_hilic\_pos\_4\_018\_31

m/z 257.9917

012E+36.57RT (min)

lcorbi\_hilic\_pos\_4\_074\_58

m/z 258.0298

01E+466.5RT (min)

lcorbi\_hilic\_pos\_4\_076\_58

m/z 258.1023

01E+36.57RT (min)

lcorbi\_hilic\_pos\_4\_081\_58

m/z 258.1084

0246E+33.54RT (min)

lcorbi\_hilic\_pos\_4\_068\_57

m/z 258.1238

02468E+33.54RT (min)

lcorbi\_hilic\_pos\_4\_070\_57

m/z 258.1337

01E+511.5RT (min)

lcorbi\_hilic\_pos\_4\_020\_37

m/z 258.1464

02468E+32.53RT (min)

lcorbi\_hilic\_pos\_4\_095\_60

m/z 258.1700

0246E+311.5RT (min)

lcorbi\_hilic\_pos\_4\_102\_61

m/z 258.1812

0246E+433.5RT (min)

lcorbi\_hilic\_pos\_3\_029\_23

m/z 258.2063

01E+40.51RT (min)

lcorbi\_hilic\_pos\_1\_020\_13

m/z 258.2064

01E+33.54RT (min)

lcorbi\_hilic\_pos\_2\_048\_04

m/z 258.2428

024E+33.54RT (min)

lcorbi\_hilic\_pos\_1\_019\_01

m/z 258.8994

01E+455.5RT (min)

lcorbi\_hilic\_pos\_4\_090\_60

m/z 258.9294

01E+47.58RT (min)

lcorbi\_hilic\_pos\_4\_104\_61

m/z 258.9373

0246E+38.59RT (min)

lcorbi\_hilic\_pos\_4\_098\_61

m/z 259.0788

0123E+311.5RT (min)

lcorbi\_hilic\_pos\_3\_001\_49

m/z 259.0901

0123E+444.5RT (min)

lcorbi\_hilic\_pos\_4\_073\_57

m/z 259.0946

02468E+355.5RT (min)

lcorbi\_hilic\_pos\_4\_102\_61

m/z 259.1071

0246E+36.57RT (min)

lcorbi\_hilic\_pos\_4\_077\_58

m/z 259.1288

01234E+277.5RT (min)

lcorbi\_hilic\_pos\_2\_037\_24

m/z 259.1289

02468E+34.55RT (min)

lcorbi\_hilic\_pos\_4\_085\_59

m/z 259.1323

0123E+40.51RT (min)

lcorbi\_hilic\_pos\_4\_043\_42

m/z 259.1516

012E+40.51RT (min)

lcorbi\_hilic\_pos\_4\_042\_02

m/z 259.1700

012E+466.5RT (min)

lcorbi\_hilic\_pos\_4\_096\_60

m/z 259.1804

024E+322.5RT (min)

lcorbi\_hilic\_pos\_4\_107\_62

m/z 259.2063

01234E+433.5RT (min)

lcorbi\_hilic\_pos\_3\_047\_28

m/z 259.2461

02468E+40.51RT (min)

lcorbi\_hilic\_pos\_1\_020\_13

m/z 259.2825

0246E+322.5RT (min)

lcorbi\_hilic\_pos\_1\_054\_23

m/z 259.9369

0246E+399.5RT (min)

lcorbi\_hilic\_pos\_4\_078\_58

m/z 259.9463

0123E+377.5RT (min)

lcorbi\_hilic\_pos\_4\_018\_31

m/z 259.9649

0246E+366.5RT (min)

lcorbi\_hilic\_pos\_4\_075\_58

m/z 260.0408

01234E+477.5RT (min)

lcorbi\_hilic\_pos\_4\_096\_60

m/z 260.0531

012E+44.55RT (min)

lcorbi\_hilic\_pos\_4\_110\_62

m/z 260.0747

01E+34.55RT (min)

lcorbi\_hilic\_pos\_4\_102\_61

m/z 260.1144

01E+311.5RT (min)

lcorbi\_hilic\_pos\_4\_020\_37

m/z 260.1259

01E+411.5RT (min)

lcorbi\_hilic\_pos\_2\_042\_02

m/z 260.1322

0246E+36.57RT (min)

lcorbi\_hilic\_pos\_4\_087\_59

m/z 260.1492

012E+41.52RT (min)

lcorbi\_hilic\_pos\_4\_082\_59

m/z 260.1493

014.55RT (min)

lcorbi\_hilic\_pos\_4\_099\_61

m/z 260.1621

012E+55.56RT (min)

lcorbi\_hilic\_pos\_4\_097\_60

m/z 260.1686

0122.5RT (min)

lcorbi\_hilic\_pos\_4\_092\_60

m/z 260.1857

01E+411.5RT (min)

lcorbi\_hilic\_pos\_4\_100\_61

m/z 260.9249

024E+388.5RT (min)

lcorbi\_hilic\_pos\_4\_102\_61

m/z 261.0280

01234E+36.57RT (min)

lcorbi\_hilic\_pos\_4\_080\_58

m/z 261.0288

024E+311.5RT (min)

lcorbi\_hilic\_pos\_1\_046\_30

m/z 261.0370

0246E+38.59RT (min)

lcorbi\_hilic\_pos\_4\_086\_59

m/z 261.0458

0123E+466.5RT (min)

lcorbi\_hilic\_pos\_4\_080\_58

m/z 261.0573

01234E+31.52RT (min)

lcorbi\_hilic\_pos\_4\_105\_61

m/z 261.0878

01E+41.52RT (min)

lcorbi\_hilic\_pos\_1\_020\_13

m/z 261.1116

0246E+211.5RT (min)

lcorbi\_hilic\_pos\_4\_020\_37

m/z 261.1445

0123E+35.56RT (min)

lcorbi\_hilic\_pos\_4\_020\_37

m/z 261.1891

01E+43.54RT (min)

lcorbi\_hilic\_pos\_4\_068\_57

m/z 261.2061

012E+433.5RT (min)

lcorbi\_hilic\_pos\_3\_047\_28

m/z 261.2173

024E+333.5RT (min)

lcorbi\_hilic\_pos\_4\_020\_37

m/z 261.2251

0246E+233.5RT (min)

lcorbi\_hilic\_pos\_3\_031\_36

m/z 261.2618

0123E+42.53RT (min)

lcorbi\_hilic\_pos\_1\_054\_23

m/z 261.9626

02468E+377.5RT (min)

lcorbi\_hilic\_pos\_1\_010\_07

m/z 262.0358

0246E+37.58RT (min)

lcorbi\_hilic\_pos\_4\_081\_58

m/z 262.0380

024E+366.5RT (min)

lcorbi\_hilic\_pos\_4\_074\_58

m/z 262.0380

0246E+311.5RT (min)

lcorbi\_hilic\_pos\_2\_053\_27

m/z 262.0517

01E+455.5RT (min)

lcorbi\_hilic\_pos\_4\_077\_58

m/z 262.0898

0246E+377.5RT (min)

lcorbi\_hilic\_pos\_4\_083\_59

m/z 262.0937

01E+344.5RT (min)

lcorbi\_hilic\_pos\_4\_020\_37

m/z 262.1115

0246E+366.5RT (min)

lcorbi\_hilic\_pos\_4\_085\_59

m/z 262.1397

0123E+388.5RT (min)

lcorbi\_hilic\_pos\_4\_020\_37

m/z 262.1510

01234E+388.5RT (min)

lcorbi\_hilic\_pos\_4\_020\_37

m/z 262.1649

0123E+44.55RT (min)

lcorbi\_hilic\_pos\_4\_020\_37

m/z 262.2013

011.52RT (min)

lcorbi\_hilic\_pos\_4\_110\_62

m/z 262.2299

01E+42.53RT (min)

lcorbi\_hilic\_pos\_2\_020\_31

m/z 262.2457

012E+42.53RT (min)

lcorbi\_hilic\_pos\_2\_019\_16

m/z 262.8879

01E+47.58RT (min)

lcorbi\_hilic\_pos\_4\_103\_61

m/z 263.0466

01E+355.5RT (min)

lcorbi\_hilic\_pos\_1\_009\_50

m/z 263.0861

01234E+244.5RT (min)

lcorbi\_hilic\_pos\_3\_004\_27

m/z 263.0874

0123E+377.5RT (min)

lcorbi\_hilic\_pos\_4\_089\_59

m/z 263.1027

01E+444.5RT (min)

lcorbi\_hilic\_pos\_4\_104\_61

m/z 263.1126

0246E+31.52RT (min)

lcorbi\_hilic\_pos\_4\_048\_46

m/z 263.1518

012E+444.5RT (min)

lcorbi\_hilic\_pos\_1\_003\_24

m/z 263.2006

01E+30.51RT (min)

lcorbi\_hilic\_pos\_4\_030\_18

m/z 263.2348

01E+42.53RT (min)

lcorbi\_hilic\_pos\_1\_046\_30

m/z 263.8948

012E+388.5RT (min)

lcorbi\_hilic\_pos\_1\_039\_41

m/z 263.9598

0123E+377.5RT (min)

lcorbi\_hilic\_pos\_2\_010\_07

m/z 263.9720

0246E+366.5RT (min)

lcorbi\_hilic\_pos\_4\_110\_62

m/z 263.9809

0246E+355.5RT (min)

lcorbi\_hilic\_pos\_4\_103\_61

m/z 264.0009

0177.5RT (min)

lcorbi\_hilic\_pos\_2\_044\_26

m/z 264.0097

01E+477.5RT (min)

lcorbi\_hilic\_pos\_2\_016\_18

m/z 264.0956

012E+411.5RT (min)

lcorbi\_hilic\_pos\_2\_046\_46

m/z 264.0957

01234E+32.53RT (min)

lcorbi\_hilic\_pos\_3\_041\_54

m/z 264.1013

0246E+31.52RT (min)

lcorbi\_hilic\_pos\_4\_077\_58

m/z 264.1108

02468E+366.5RT (min)

lcorbi\_hilic\_pos\_4\_092\_60

m/z 264.1173

024E+466.5RT (min)

lcorbi\_hilic\_pos\_4\_020\_37

m/z 264.1442

01E+41.52RT (min)

lcorbi\_hilic\_pos\_1\_046\_30

m/z 264.1887

01E+50.51RT (min)

lcorbi\_hilic\_pos\_2\_032\_30

m/z 264.1958

01E+42.53RT (min)

lcorbi\_hilic\_pos\_4\_073\_57

m/z 264.2170

01234E+32.53RT (min)

lcorbi\_hilic\_pos\_1\_034\_37

m/z 264.2322

0123E+322.5RT (min)

lcorbi\_hilic\_pos\_1\_054\_23

m/z 264.2443

01234E+433.5RT (min)

lcorbi\_hilic\_pos\_1\_034\_37

m/z 264.2686

012E+40.51RT (min)

lcorbi\_hilic\_pos\_4\_076\_58

m/z 264.8553

0123E+377.5RT (min)

lcorbi\_hilic\_pos\_1\_037\_18

m/z 264.8927

0123E+388.5RT (min)

lcorbi\_hilic\_pos\_4\_066\_57

m/z 265.0156

0123E+48.59RT (min)

lcorbi\_hilic\_pos\_4\_077\_58

m/z 265.1013

0144.5RT (min)

lcorbi\_hilic\_pos\_3\_023\_18

m/z 265.1112

012E+444.5RT (min)

lcorbi\_hilic\_pos\_3\_029\_23

m/z 265.1312

01E+41.52RT (min)

lcorbi\_hilic\_pos\_4\_082\_59

m/z 265.1585

02468E+344.5RT (min)

lcorbi\_hilic\_pos\_1\_003\_24

m/z 265.1758

0123E+355.5RT (min)

lcorbi\_hilic\_pos\_1\_016\_33

m/z 265.1981

0123E+455.5RT (min)

lcorbi\_hilic\_pos\_4\_096\_60

m/z 265.2162

012E+30.51RT (min)

lcorbi\_hilic\_pos\_4\_015\_20

m/z 266.0165

01E+48.59RT (min)

lcorbi\_hilic\_pos\_4\_075\_58

m/z 266.0323

024E+377.5RT (min)

lcorbi\_hilic\_pos\_4\_081\_58

m/z 266.0764

012E+466.5RT (min)

lcorbi\_hilic\_pos\_3\_031\_36

m/z 266.1345

0246E+35.56RT (min)

lcorbi\_hilic\_pos\_4\_096\_60

m/z 266.1387

0123E+444.5RT (min)

lcorbi\_hilic\_pos\_4\_020\_37

m/z 266.1501

01E+311.5RT (min)

lcorbi\_hilic\_pos\_3\_019\_43

m/z 266.1727

012E+411.5RT (min)

lcorbi\_hilic\_pos\_3\_019\_43

m/z 266.1839

0123E+455.5RT (min)

lcorbi\_hilic\_pos\_4\_082\_59

m/z 266.2842

01E+444.5RT (min)

lcorbi\_hilic\_pos\_4\_070\_57

m/z 266.8420

01E+455.5RT (min)

lcorbi\_hilic\_pos\_4\_082\_59

m/z 267.0132

01E+48.59RT (min)

lcorbi\_hilic\_pos\_4\_076\_58

m/z 267.0386

01E+488.5RT (min)

lcorbi\_hilic\_pos\_4\_088\_59

m/z 267.0451

012E+34.55RT (min)

lcorbi\_hilic\_pos\_3\_012\_13

m/z 267.0572

01E+48.59RT (min)

lcorbi\_hilic\_pos\_4\_093\_60

m/z 267.0588

01E+44.55RT (min)

lcorbi\_hilic\_pos\_4\_089\_59

m/z 267.0775

01E+422.5RT (min)

lcorbi\_hilic\_pos\_4\_090\_60

m/z 267.1057

0111.5RT (min)

lcorbi\_hilic\_pos\_4\_106\_62

m/z 267.1227

012E+40.51RT (min)

lcorbi\_hilic\_pos\_4\_052\_32

m/z 267.1339

0111.5RT (min)

lcorbi\_hilic\_pos\_4\_108\_62

m/z 267.1551

012E+44.55RT (min)

lcorbi\_hilic\_pos\_4\_113\_62

m/z 267.1632

012E+466.5RT (min)

lcorbi\_hilic\_pos\_4\_073\_57

m/z 267.1703

01234E+44.55RT (min)

lcorbi\_hilic\_pos\_4\_069\_57

m/z 267.2108

010.51RT (min)

lcorbi\_hilic\_pos\_3\_019\_43

m/z 267.2319

01E+30.51RT (min)

lcorbi\_hilic\_pos\_2\_014\_20

m/z 267.8897

0123E+388.5RT (min)

lcorbi\_hilic\_pos\_4\_066\_57

m/z 267.9781

0246E+377.5RT (min)

lcorbi\_hilic\_pos\_4\_018\_31

m/z 268.0016

0246E+34.55RT (min)

lcorbi\_hilic\_pos\_4\_106\_62

m/z 268.0696

012.53RT (min)

lcorbi\_hilic\_pos\_4\_020\_37

m/z 268.0767

02468E+211.5RT (min)

lcorbi\_hilic\_pos\_4\_106\_62

m/z 268.0968

01234E+411.5RT (min)

lcorbi\_hilic\_pos\_4\_108\_62

m/z 268.1214

012E+344.5RT (min)

lcorbi\_hilic\_pos\_4\_020\_37

m/z 268.1247

01E+466.5RT (min)

lcorbi\_hilic\_pos\_3\_010\_32

m/z 268.1405

01E+466.5RT (min)

lcorbi\_hilic\_pos\_1\_021\_46

m/z 268.1500

0123E+433.5RT (min)

lcorbi\_hilic\_pos\_3\_010\_32

m/z 268.1502

01E+355.5RT (min)

lcorbi\_hilic\_pos\_4\_066\_57

m/z 268.1737

02468E+34.55RT (min)

lcorbi\_hilic\_pos\_4\_072\_57

m/z 268.2119

0123E+40.51RT (min)

lcorbi\_hilic\_pos\_4\_104\_61

m/z 268.2271

01E+30.51RT (min)

lcorbi\_hilic\_pos\_4\_104\_61

m/z 268.2635

02468E+311.5RT (min)

lcorbi\_hilic\_pos\_1\_054\_23

m/z 268.2999

012E+444.5RT (min)

lcorbi\_hilic\_pos\_4\_068\_57

m/z 268.8737

0123E+388.5RT (min)

lcorbi\_hilic\_pos\_4\_069\_57

m/z 269.0611

0111.5RT (min)

lcorbi\_hilic\_pos\_3\_048\_38

m/z 269.1132

024E+34.55RT (min)

lcorbi\_hilic\_pos\_4\_084\_59

m/z 269.1243

012E+466.5RT (min)

lcorbi\_hilic\_pos\_1\_043\_25

m/z 269.1496

012.53RT (min)

lcorbi\_hilic\_pos\_4\_112\_62

m/z 269.1608

012E+366.5RT (min)

lcorbi\_hilic\_pos\_4\_020\_37

m/z 269.1722

01E+45.56RT (min)

lcorbi\_hilic\_pos\_4\_007\_05

m/z 269.1940

010.51RT (min)

lcorbi\_hilic\_pos\_1\_051\_26

m/z 269.2588

0111.5RT (min)

lcorbi\_hilic\_pos\_4\_079\_58

m/z 269.2669

012E+43.54RT (min)

lcorbi\_hilic\_pos\_1\_054\_23

m/z 269.9937

01E+466.5RT (min)

lcorbi\_hilic\_pos\_4\_079\_58

m/z 270.0407

01E+41.52RT (min)

lcorbi\_hilic\_pos\_1\_044\_17

m/z 270.0842

0246E+40.51RT (min)

lcorbi\_hilic\_pos\_1\_012\_19

m/z 270.1166

01E+42.53RT (min)

lcorbi\_hilic\_pos\_2\_012\_37

m/z 270.1169

01E+477.5RT (min)

lcorbi\_hilic\_pos\_4\_097\_60

m/z 270.1279

0123E+444.5RT (min)

lcorbi\_hilic\_pos\_1\_050\_05

m/z 270.1383

0246E+366.5RT (min)

lcorbi\_hilic\_pos\_4\_094\_60

m/z 270.1576

0246E+34.55RT (min)

lcorbi\_hilic\_pos\_4\_024\_24

m/z 270.1657

01E+46.57RT (min)

lcorbi\_hilic\_pos\_4\_092\_60

m/z 270.1700

0123E+30.51RT (min)

lcorbi\_hilic\_pos\_4\_069\_57

m/z 270.1911

01E+411.5RT (min)

lcorbi\_hilic\_pos\_3\_019\_43

m/z 270.2038

0246E+37.58RT (min)

lcorbi\_hilic\_pos\_1\_050\_05

m/z 270.2216

0111.5RT (min)

lcorbi\_hilic\_pos\_4\_068\_57

m/z 270.2428

01E+40.51RT (min)

lcorbi\_hilic\_pos\_1\_020\_13

m/z 270.2792

012E+40.51RT (min)

lcorbi\_hilic\_pos\_4\_085\_59

m/z 270.7896

02468E+355.5RT (min)

lcorbi\_hilic\_pos\_4\_067\_57

m/z 270.8692

01234E+388.5RT (min)

lcorbi\_hilic\_pos\_4\_066\_57

m/z 271.0278

0246E+377.5RT (min)

lcorbi\_hilic\_pos\_4\_083\_59

m/z 271.0400

0123E+355.5RT (min)

lcorbi\_hilic\_pos\_4\_074\_58

m/z 271.0586

0155.5RT (min)

lcorbi\_hilic\_pos\_4\_052\_32

m/z 271.0835

016.57RT (min)

lcorbi\_hilic\_pos\_4\_074\_58

m/z 271.0901

024E+34.55RT (min)

lcorbi\_hilic\_pos\_4\_088\_59

m/z 271.0925

013.54RT (min)

lcorbi\_hilic\_pos\_2\_024\_32

m/z 271.1037

02468E+37.58RT (min)

lcorbi\_hilic\_pos\_4\_088\_59

m/z 271.1152

01E+30.51RT (min)

lcorbi\_hilic\_pos\_1\_051\_26

m/z 271.1187

02468E+33.54RT (min)

lcorbi\_hilic\_pos\_1\_004\_22

m/z 271.1654

01E+41.52RT (min)

lcorbi\_hilic\_pos\_4\_079\_58

m/z 271.2420

0123E+322.5RT (min)

lcorbi\_hilic\_pos\_4\_074\_58

m/z 271.2631

01234E+40.51RT (min)

lcorbi\_hilic\_pos\_4\_084\_59

m/z 271.2744

01E+42.53RT (min)

lcorbi\_hilic\_pos\_4\_086\_59

m/z 271.9915

024E+377.5RT (min)

lcorbi\_hilic\_pos\_2\_010\_07

m/z 272.0530

0123E+388.5RT (min)

lcorbi\_hilic\_pos\_4\_084\_59

m/z 272.1105

0123E+35.56RT (min)

lcorbi\_hilic\_pos\_1\_021\_46

m/z 272.1257

0246E+34.55RT (min)

lcorbi\_hilic\_pos\_1\_054\_23

m/z 272.1281

010.51RT (min)

lcorbi\_hilic\_pos\_4\_030\_18

m/z 272.1506

0123E+40.51RT (min)

lcorbi\_hilic\_pos\_4\_034\_36

m/z 272.1718

02468E+37.58RT (min)

lcorbi\_hilic\_pos\_4\_088\_59

m/z 272.1857

0246E+33.54RT (min)

lcorbi\_hilic\_pos\_4\_020\_37

m/z 272.2391

01E+433.5RT (min)

lcorbi\_hilic\_pos\_4\_086\_59

m/z 272.3222

01E+32.53RT (min)

lcorbi\_hilic\_pos\_1\_023\_28

m/z 272.7870

0246E+355.5RT (min)

lcorbi\_hilic\_pos\_4\_067\_57

m/z 272.9869

01E+51.52RT (min)

lcorbi\_hilic\_pos\_3\_034\_26

m/z 273.0038

01E+466.5RT (min)

lcorbi\_hilic\_pos\_4\_094\_60

m/z 273.0484

012E+444.5RT (min)

lcorbi\_hilic\_pos\_4\_082\_59

m/z 273.0758

01E+50.51RT (min)

lcorbi\_hilic\_pos\_2\_016\_18

m/z 273.0904

01E+41.52RT (min)

lcorbi\_hilic\_pos\_1\_032\_31

m/z 273.0904

0123E+46.57RT (min)

lcorbi\_hilic\_pos\_4\_091\_60

m/z 273.0999

0166.5RT (min)

lcorbi\_hilic\_pos\_4\_050\_16

m/z 273.1080

012E+433.5RT (min)

lcorbi\_hilic\_pos\_4\_088\_59

m/z 273.1194

01E+47.58RT (min)

lcorbi\_hilic\_pos\_1\_050\_05

m/z 273.1212

01E+455.5RT (min)

lcorbi\_hilic\_pos\_4\_080\_58

m/z 273.2003

0122.5RT (min)

lcorbi\_hilic\_pos\_4\_075\_58

m/z 273.2061

01E+40.51RT (min)

lcorbi\_hilic\_pos\_4\_031\_17

m/z 273.2173

0246E+31.52RT (min)

lcorbi\_hilic\_pos\_1\_032\_31

m/z 273.2537

01E+411.5RT (min)

lcorbi\_hilic\_pos\_3\_019\_43

m/z 273.2982

02468E+32.53RT (min)

lcorbi\_hilic\_pos\_1\_034\_37

m/z 273.9620

02468E+36.57RT (min)

lcorbi\_hilic\_pos\_4\_066\_57

m/z 273.9836

012E+411.5RT (min)

lcorbi\_hilic\_pos\_4\_113\_62

m/z 274.0289

024E+355.5RT (min)

lcorbi\_hilic\_pos\_1\_055\_20

m/z 274.0621

01E+488.5RT (min)

lcorbi\_hilic\_pos\_4\_093\_60

m/z 274.1413

01E+56.577.5RT (min)

lcorbi\_hilic\_pos\_4\_096\_60

m/z 274.1439

0246E+311.5RT (min)

lcorbi\_hilic\_pos\_2\_019\_16

m/z 274.1454

0246E+244.5RT (min)

lcorbi\_hilic\_pos\_4\_024\_24

m/z 274.1842

024E+31.52RT (min)

lcorbi\_hilic\_pos\_4\_089\_59

m/z 274.2013

012E+333.5RT (min)

lcorbi\_hilic\_pos\_2\_036\_08

m/z 274.2377

010.51RT (min)

lcorbi\_hilic\_pos\_4\_106\_62

m/z 274.2548

0123E+42.53RT (min)

lcorbi\_hilic\_pos\_3\_037\_20

m/z 274.2917

024E+42.53RT (min)

lcorbi\_hilic\_pos\_3\_037\_20

m/z 275.0598

0123E+466.5RT (min)

lcorbi\_hilic\_pos\_4\_092\_60

m/z 275.0738

012E+444.5RT (min)

lcorbi\_hilic\_pos\_4\_066\_57

m/z 275.0850

024E+466.5RT (min)

lcorbi\_hilic\_pos\_2\_054\_29

m/z 275.1002

012E+411.5RT (min)

lcorbi\_hilic\_pos\_1\_031\_36

m/z 275.1101

012E+31.52RT (min)

lcorbi\_hilic\_pos\_4\_030\_18

m/z 275.1853

0111.5RT (min)

lcorbi\_hilic\_pos\_4\_109\_62

m/z 275.1966

0246E+344.5RT (min)

lcorbi\_hilic\_pos\_3\_019\_43

m/z 275.2005

02468E+30.51RT (min)

lcorbi\_hilic\_pos\_4\_077\_58

m/z 275.2206

024E+47.58RT (min)

lcorbi\_hilic\_pos\_1\_003\_24

m/z 275.2330

02468E+322.5RT (min)

lcorbi\_hilic\_pos\_3\_004\_27

m/z 275.2411

0123E+433.5RT (min)

lcorbi\_hilic\_pos\_2\_036\_08

m/z 275.2711

01E+32.53RT (min)

lcorbi\_hilic\_pos\_4\_105\_61

m/z 276.0011

0246E+36.57RT (min)

lcorbi\_hilic\_pos\_4\_082\_59

m/z 276.0718

01E+55RT (min)

lcorbi\_hilic\_pos\_4\_020\_37

m/z 276.1079

0123E+322.5RT (min)

lcorbi\_hilic\_pos\_4\_085\_59

m/z 276.1191

01E+477.5RT (min)

lcorbi\_hilic\_pos\_4\_082\_59

m/z 276.1442

012E+45RT (min)

lcorbi\_hilic\_pos\_4\_095\_60

m/z 276.1463

02468E+311.5RT (min)

lcorbi\_hilic\_pos\_2\_024\_32

m/z 276.1554

01234E+37.58RT (min)

lcorbi\_hilic\_pos\_4\_086\_59

m/z 276.1557

012E+46.57RT (min)

lcorbi\_hilic\_pos\_4\_093\_60

m/z 276.1601

0123E+311.5RT (min)

lcorbi\_hilic\_pos\_4\_050\_16

m/z 276.1806

01E+44.55RT (min)

lcorbi\_hilic\_pos\_1\_021\_46

m/z 276.1806

024E+311.5RT (min)

lcorbi\_hilic\_pos\_4\_105\_61

m/z 276.2170

01E+411.5RT (min)

lcorbi\_hilic\_pos\_1\_045\_12

m/z 276.8715

0246E+355.5RT (min)

lcorbi\_hilic\_pos\_4\_090\_60

m/z 276.9478

01234E+38.59RT (min)

lcorbi\_hilic\_pos\_4\_111\_62

m/z 277.0383

01E+46.57RT (min)

lcorbi\_hilic\_pos\_4\_070\_57

m/z 277.0723

01234E+344.5RT (min)

lcorbi\_hilic\_pos\_4\_086\_59

m/z 277.0860

0123E+40.51RT (min)

lcorbi\_hilic\_pos\_4\_084\_59

m/z 277.0894

01234E+44.555.5RT (min)

lcorbi\_hilic\_pos\_4\_073\_57

m/z 277.1007

0123E+355.5RT (min)

lcorbi\_hilic\_pos\_4\_071\_57

m/z 277.1048

01234E+311.5RT (min)

lcorbi\_hilic\_pos\_4\_070\_57

m/z 277.1273

01E+45.56RT (min)

lcorbi\_hilic\_pos\_1\_050\_05

m/z 277.1282

02468E+40.51RT (min)

lcorbi\_hilic\_pos\_1\_040\_06

m/z 277.1566

01E+411.5RT (min)

lcorbi\_hilic\_pos\_2\_020\_31

m/z 277.1758

012E+411.5RT (min)

lcorbi\_hilic\_pos\_2\_007\_28

m/z 277.2565

012E+32.53RT (min)

lcorbi\_hilic\_pos\_3\_037\_20

m/z 277.9340

0246E+377.5RT (min)

lcorbi\_hilic\_pos\_3\_029\_23

m/z 278.0143

0246E+35.56RT (min)

lcorbi\_hilic\_pos\_4\_083\_59

m/z 278.0255

0123E+36.57RT (min)

lcorbi\_hilic\_pos\_4\_071\_57

m/z 278.1024

012E+344.5RT (min)

lcorbi\_hilic\_pos\_3\_024\_35

m/z 278.1218

012E+444.5RT (min)

lcorbi\_hilic\_pos\_2\_030\_05

m/z 278.1264

01234E+31.52RT (min)

lcorbi\_hilic\_pos\_3\_001\_49

m/z 278.1377

012E+43.54RT (min)

lcorbi\_hilic\_pos\_4\_093\_60

m/z 278.1388

0111.5RT (min)

lcorbi\_hilic\_pos\_1\_040\_06

m/z 278.1428

01E+47.58RT (min)

lcorbi\_hilic\_pos\_4\_086\_59

m/z 278.1708

01E+411.5RT (min)

lcorbi\_hilic\_pos\_1\_036\_32

m/z 278.2043

010.51RT (min)

lcorbi\_hilic\_pos\_3\_034\_26

m/z 278.2133

02468E+333.5RT (min)

lcorbi\_hilic\_pos\_1\_034\_37

m/z 278.2196

01E+40.51RT (min)

lcorbi\_hilic\_pos\_4\_080\_58

m/z 278.2479

012E+40.51RT (min)

lcorbi\_hilic\_pos\_3\_019\_43

m/z 278.8758

02468E+377.5RT (min)

lcorbi\_hilic\_pos\_1\_043\_25

m/z 278.9944

02468E+388.5RT (min)

lcorbi\_hilic\_pos\_4\_082\_59

m/z 279.1340

012E+411.5RT (min)

lcorbi\_hilic\_pos\_4\_072\_57

m/z 279.1468

0246E+32.53RT (min)

lcorbi\_hilic\_pos\_4\_024\_24

m/z 279.1704

01E+444.5RT (min)

lcorbi\_hilic\_pos\_4\_080\_58

m/z 279.1743

0246E+311.5RT (min)

lcorbi\_hilic\_pos\_4\_076\_58

m/z 279.2167

0246E+30.51RT (min)

lcorbi\_hilic\_pos\_4\_077\_58

m/z 279.2360

01E+433.5RT (min)

lcorbi\_hilic\_pos\_4\_012\_29

m/z 280.0485

016.57RT (min)

lcorbi\_hilic\_pos\_4\_091\_60

m/z 280.0605

01E+47.58RT (min)

lcorbi\_hilic\_pos\_4\_095\_60

m/z 280.0876

01234E+366.5RT (min)

lcorbi\_hilic\_pos\_4\_066\_57

m/z 280.1203

0246E+34.55RT (min)

lcorbi\_hilic\_pos\_4\_095\_60

m/z 280.1391

01E+45.56RT (min)

lcorbi\_hilic\_pos\_4\_070\_57

m/z 280.2636

01E+43.54RT (min)

lcorbi\_hilic\_pos\_4\_080\_58

m/z 281.0512

01E+40.51RT (min)

lcorbi\_hilic\_pos\_4\_105\_61

m/z 281.0858

0144.5RT (min)

lcorbi\_hilic\_pos\_1\_003\_24

m/z 281.1108

012E+46.57RT (min)

lcorbi\_hilic\_pos\_4\_083\_59

m/z 281.1214

024E+311.5RT (min)

lcorbi\_hilic\_pos\_4\_071\_57

m/z 281.1345

01E+46.57RT (min)

lcorbi\_hilic\_pos\_4\_084\_59

m/z 281.1474

02468E+322.5RT (min)

lcorbi\_hilic\_pos\_1\_032\_31

m/z 281.1579

013.54RT (min)

lcorbi\_hilic\_pos\_4\_093\_60

m/z 281.2013

012E+422.5RT (min)

lcorbi\_hilic\_pos\_4\_104\_61

m/z 281.9474

01234E+377.5RT (min)

lcorbi\_hilic\_pos\_3\_010\_32

m/z 281.9938

01E+46.57RT (min)

lcorbi\_hilic\_pos\_4\_069\_57

m/z 282.0133

0246E+455.5RT (min)

lcorbi\_hilic\_pos\_4\_096\_60

m/z 282.0228

01E+477.5RT (min)

lcorbi\_hilic\_pos\_1\_003\_24

m/z 282.0777

01E+433.5RT (min)

lcorbi\_hilic\_pos\_4\_089\_59

m/z 282.0963

0246E+36.57RT (min)

lcorbi\_hilic\_pos\_4\_078\_58

m/z 282.1184

0246E+477.5RT (min)

lcorbi\_hilic\_pos\_4\_088\_59

m/z 282.1198

0246E+322.5RT (min)

lcorbi\_hilic\_pos\_2\_005\_23

m/z 282.1198

01E+45.56RT (min)

lcorbi\_hilic\_pos\_1\_032\_31

m/z 282.1295

02468E+33.54RT (min)

lcorbi\_hilic\_pos\_4\_096\_60

m/z 282.1313

0246E+31.52RT (min)

lcorbi\_hilic\_pos\_4\_083\_59

m/z 282.1441

01E+55.56RT (min)

lcorbi\_hilic\_pos\_4\_097\_60

m/z 282.1912

012E+41.52RT (min)

lcorbi\_hilic\_pos\_3\_019\_43

m/z 282.2008

024E+34.55RT (min)

lcorbi\_hilic\_pos\_1\_021\_46

m/z 282.2047

02468E+322.5RT (min)

lcorbi\_hilic\_pos\_4\_103\_61

m/z 282.2429

0123E+30.51RT (min)

lcorbi\_hilic\_pos\_4\_081\_58

m/z 282.2622

01E+422.5RT (min)

lcorbi\_hilic\_pos\_4\_091\_60

m/z 282.8161

01E+455.5RT (min)

lcorbi\_hilic\_pos\_4\_077\_58

m/z 282.9051

01E+499.5RT (min)

lcorbi\_hilic\_pos\_4\_103\_61

m/z 283.1078

024E+30.51RT (min)

lcorbi\_hilic\_pos\_4\_019\_30

m/z 283.1176

01E+411.5RT (min)

lcorbi\_hilic\_pos\_4\_004\_38

m/z 283.1219

0246E+377.5RT (min)

lcorbi\_hilic\_pos\_4\_087\_59

m/z 283.1403

02468E+444.5RT (min)

lcorbi\_hilic\_pos\_4\_007\_05

m/z 283.1530

01E+47.58RT (min)

lcorbi\_hilic\_pos\_4\_092\_60

m/z 283.2131

02468E+344.5RT (min)

lcorbi\_hilic\_pos\_3\_056\_56

m/z 283.2745

012E+444.5RT (min)

lcorbi\_hilic\_pos\_4\_021\_13

m/z 283.2826

0123E+411.5RT (min)

lcorbi\_hilic\_pos\_1\_054\_23

m/z 284.0094

02468E+35.56RT (min)

lcorbi\_hilic\_pos\_1\_051\_26

m/z 284.0472

024E+34.55RT (min)

lcorbi\_hilic\_pos\_4\_072\_57

m/z 284.0531

02468E+377.5RT (min)

lcorbi\_hilic\_pos\_4\_087\_59

m/z 284.0985

0246E+366.5RT (min)

lcorbi\_hilic\_pos\_4\_052\_32

m/z 284.1030

01E+411.5RT (min)

lcorbi\_hilic\_pos\_4\_068\_57

m/z 284.1065

0246E+36.57RT (min)

lcorbi\_hilic\_pos\_4\_077\_58

m/z 284.1281

011.52RT (min)

lcorbi\_hilic\_pos\_4\_093\_60

m/z 284.1449

0133.5RT (min)

lcorbi\_hilic\_pos\_3\_037\_20

m/z 284.1493

0246E+31.52RT (min)

lcorbi\_hilic\_pos\_3\_048\_38

m/z 284.1857

013.54RT (min)

lcorbi\_hilic\_pos\_4\_085\_59

m/z 284.2192

01E+43.54RT (min)

lcorbi\_hilic\_pos\_4\_082\_59

m/z 284.2585

024E+30.51RT (min)

lcorbi\_hilic\_pos\_1\_054\_23

m/z 284.2948

012E+40.51RT (min)

lcorbi\_hilic\_pos\_1\_054\_23

m/z 284.3313

01E+51.52RT (min)

lcorbi\_hilic\_pos\_1\_040\_06

m/z 284.3313

01234E+300.5RT (min)

lcorbi\_hilic\_pos\_1\_001\_49

m/z 284.8139

024E+355.5RT (min)

lcorbi\_hilic\_pos\_4\_101\_61

m/z 284.8699

01E+47.58RT (min)

lcorbi\_hilic\_pos\_4\_103\_61

m/z 284.8889

01E+477.5RT (min)

lcorbi\_hilic\_pos\_1\_053\_08

m/z 285.0047

02468E+366.5RT (min)

lcorbi\_hilic\_pos\_4\_070\_57

m/z 285.0946

01234E+31.52RT (min)

lcorbi\_hilic\_pos\_1\_021\_46

m/z 285.0956

01E+366.5RT (min)

lcorbi\_hilic\_pos\_4\_020\_37

m/z 285.1234

01E+433.5RT (min)

lcorbi\_hilic\_pos\_4\_079\_58

m/z 285.1338

01E+411.5RT (min)

lcorbi\_hilic\_pos\_4\_097\_60

m/z 285.1849

0246E+311.5RT (min)

lcorbi\_hilic\_pos\_4\_100\_61

m/z 285.2102

0246E+311.5RT (min)

lcorbi\_hilic\_pos\_4\_096\_60

m/z 285.2213

0123E+411.5RT (min)

lcorbi\_hilic\_pos\_4\_098\_61

m/z 285.2791

0123E+40.51RT (min)

lcorbi\_hilic\_pos\_4\_088\_59

m/z 285.2901

01E+40.51RT (min)

lcorbi\_hilic\_pos\_4\_082\_59

m/z 285.9638

0123E+37.58RT (min)

lcorbi\_hilic\_pos\_4\_100\_61

m/z 285.9806

01E+466.5RT (min)

lcorbi\_hilic\_pos\_4\_089\_59

m/z 285.9887

024E+36.57RT (min)

lcorbi\_hilic\_pos\_3\_035\_22

m/z 286.0179

02468E+388.5RT (min)

lcorbi\_hilic\_pos\_4\_092\_60

m/z 286.0442

0246E+34.55RT (min)

lcorbi\_hilic\_pos\_4\_070\_57

m/z 286.1286

02468E+34.55RT (min)

lcorbi\_hilic\_pos\_4\_068\_57

m/z 286.1513

0123E+46.57RT (min)

lcorbi\_hilic\_pos\_1\_050\_05

m/z 286.1762

01234E+31.52RT (min)

lcorbi\_hilic\_pos\_4\_082\_59

m/z 286.2126

02468E+322.5RT (min)

lcorbi\_hilic\_pos\_4\_087\_59

m/z 286.2126

01E+477.5RT (min)

lcorbi\_hilic\_pos\_2\_020\_31

m/z 286.2377

01234E+20.51RT (min)

lcorbi\_hilic\_pos\_2\_044\_26

m/z 286.2741

01E+411.52RT (min)

lcorbi\_hilic\_pos\_1\_042\_40

m/z 286.3016

012.53RT (min)

lcorbi\_hilic\_pos\_4\_019\_30

m/z 286.3603

02468E+333.5RT (min)

lcorbi\_hilic\_pos\_1\_036\_32

m/z 286.8373

0123E+377.5RT (min)

lcorbi\_hilic\_pos\_1\_055\_20

m/z 287.1108

0123E+411.5RT (min)

lcorbi\_hilic\_pos\_4\_020\_37

m/z 287.1156

01E+46.57RT (min)

lcorbi\_hilic\_pos\_1\_050\_05

m/z 287.1237

01234E+433.5RT (min)

lcorbi\_hilic\_pos\_1\_054\_23

m/z 287.1426

0144.5RT (min)

lcorbi\_hilic\_pos\_1\_029\_14

m/z 287.1447

0122.5RT (min)

lcorbi\_hilic\_pos\_4\_066\_57

m/z 287.1578

012E+455.5RT (min)

lcorbi\_hilic\_pos\_1\_048\_02

m/z 287.2330

01E+41.52RT (min)

lcorbi\_hilic\_pos\_1\_014\_27

m/z 287.2775

0123E+311.5RT (min)

lcorbi\_hilic\_pos\_1\_028\_34

m/z 287.2967

01E+422.5RT (min)

lcorbi\_hilic\_pos\_4\_088\_59

m/z 287.3139

01E+51.52RT (min)

lcorbi\_hilic\_pos\_4\_016\_28

m/z 287.9389

01E+36.57RT (min)

lcorbi\_hilic\_pos\_4\_089\_59

m/z 287.9776

01234E+366.5RT (min)

lcorbi\_hilic\_pos\_4\_088\_59

m/z 288.0054

0246E+34.55RT (min)

lcorbi\_hilic\_pos\_4\_067\_57

m/z 288.0228

02468E+35.56RT (min)

lcorbi\_hilic\_pos\_4\_096\_60

m/z 288.0584

01E+466.5RT (min)

lcorbi\_hilic\_pos\_4\_084\_59

m/z 288.1037

01E+422.5RT (min)

lcorbi\_hilic\_pos\_4\_066\_57

m/z 288.1055

024E+444.5RT (min)

lcorbi\_hilic\_pos\_1\_054\_23

m/z 288.1271

0246E+333.5RT (min)

lcorbi\_hilic\_pos\_4\_084\_59

m/z 288.1503

010.51RT (min)

lcorbi\_hilic\_pos\_3\_034\_26

m/z 288.1594

01E+30.51RT (min)

lcorbi\_hilic\_pos\_3\_034\_26

m/z 288.2031

01E+477.5RT (min)

lcorbi\_hilic\_pos\_1\_052\_35

m/z 288.2080

02468E+33.54RT (min)

lcorbi\_hilic\_pos\_4\_076\_58

m/z 288.2170

0123E+43.54RT (min)

lcorbi\_hilic\_pos\_4\_074\_58

m/z 288.9221

02468E+31010.5RT (min)

lcorbi\_hilic\_pos\_2\_041\_54

m/z 289.0206

0123E+388.5RT (min)

lcorbi\_hilic\_pos\_4\_087\_59

m/z 289.0392

01E+47.58RT (min)

lcorbi\_hilic\_pos\_4\_093\_60

m/z 289.1125

0246E+344.5RT (min)

lcorbi\_hilic\_pos\_4\_092\_60

m/z 289.1507

012E+377.5RT (min)

lcorbi\_hilic\_pos\_3\_029\_23

m/z 289.1620

0246E+388.5RT (min)

lcorbi\_hilic\_pos\_3\_031\_36

m/z 289.1759

0246E+211.5RT (min)

lcorbi\_hilic\_pos\_3\_051\_25

m/z 289.2486

02468E+311.5RT (min)

lcorbi\_hilic\_pos\_3\_002\_04

m/z 289.9755

0246E+366.5RT (min)

lcorbi\_hilic\_pos\_4\_067\_57

m/z 290.0025

01E+44.55RT (min)

lcorbi\_hilic\_pos\_4\_078\_58

m/z 290.0844

01E+477.5RT (min)

lcorbi\_hilic\_pos\_2\_005\_23

m/z 290.1044

0133.5RT (min)

lcorbi\_hilic\_pos\_4\_066\_57

m/z 290.1114

01E+41.52RT (min)

lcorbi\_hilic\_pos\_1\_056\_56

m/z 290.1249

01E+31.52RT (min)

lcorbi\_hilic\_pos\_3\_012\_13

m/z 290.1460

01E+488.5RT (min)

lcorbi\_hilic\_pos\_4\_083\_59

m/z 290.1963

014.55RT (min)

lcorbi\_hilic\_pos\_1\_020\_13

m/z 290.2599

01E+42.53RT (min)

lcorbi\_hilic\_pos\_2\_019\_16

m/z 290.2792

01E+42.53RT (min)

lcorbi\_hilic\_pos\_1\_040\_06

m/z 290.2964

01E+322.5RT (min)

lcorbi\_hilic\_pos\_1\_033\_53

m/z 291.0382

01E+44.55RT (min)

lcorbi\_hilic\_pos\_4\_070\_57

m/z 291.0478

01234E+344.5RT (min)

lcorbi\_hilic\_pos\_4\_073\_57

m/z 291.0647

016.57RT (min)

lcorbi\_hilic\_pos\_4\_091\_60

m/z 291.0840

02468E+311.5RT (min)

lcorbi\_hilic\_pos\_2\_004\_01

m/z 291.1188

01E+466.5RT (min)

lcorbi\_hilic\_pos\_4\_066\_57

m/z 291.1381

01E+46.57RT (min)

lcorbi\_hilic\_pos\_4\_083\_59

m/z 291.1551

01E+377.5RT (min)

lcorbi\_hilic\_pos\_4\_096\_60

m/z 291.1633

0123E+34.55RT (min)

lcorbi\_hilic\_pos\_4\_066\_57

m/z 291.1680

01E+56.57RT (min)

lcorbi\_hilic\_pos\_4\_091\_60

m/z 291.1823

01E+466.5RT (min)

lcorbi\_hilic\_pos\_3\_003\_37

m/z 291.1956

01E+411.5RT (min)

lcorbi\_hilic\_pos\_4\_079\_58

m/z 291.2319

01234E+20.51RT (min)

lcorbi\_hilic\_pos\_3\_037\_20

m/z 291.2531

01E+32.53RT (min)

lcorbi\_hilic\_pos\_4\_019\_30

m/z 291.8507

01E+455.5RT (min)

lcorbi\_hilic\_pos\_4\_075\_58

m/z 291.9497

016.57RT (min)

lcorbi\_hilic\_pos\_4\_094\_60

m/z 291.9995

0246E+34.55RT (min)

lcorbi\_hilic\_pos\_4\_079\_58

m/z 292.0659

012E+43.54RT (min)

lcorbi\_hilic\_pos\_1\_002\_29

m/z 292.1099

01E+444.5RT (min)

lcorbi\_hilic\_pos\_3\_027\_05

m/z 292.1334

01234E+37.58RT (min)

lcorbi\_hilic\_pos\_4\_087\_59

m/z 292.1414

02468E+32.53RT (min)

lcorbi\_hilic\_pos\_4\_072\_57

m/z 292.1832

0246E+466.5RT (min)

lcorbi\_hilic\_pos\_3\_031\_36

m/z 292.2111

024E+333.5RT (min)

lcorbi\_hilic\_pos\_2\_025\_52

m/z 293.0550

01E+41.52RT (min)

lcorbi\_hilic\_pos\_4\_071\_57

m/z 293.1474

024E+21.52RT (min)

lcorbi\_hilic\_pos\_1\_036\_32

m/z 293.1567

01E+422.5RT (min)

lcorbi\_hilic\_pos\_4\_092\_60

m/z 293.1958

0246E+30.51RT (min)

lcorbi\_hilic\_pos\_3\_051\_25

m/z 293.2112

0246E+30.51RT (min)

lcorbi\_hilic\_pos\_3\_054\_16

m/z 293.2952

01E+41.52RT (min)

lcorbi\_hilic\_pos\_4\_105\_61

m/z 293.8488

0246E+355.5RT (min)

lcorbi\_hilic\_pos\_4\_080\_58

m/z 294.0326

024E+35.56RT (min)

lcorbi\_hilic\_pos\_4\_078\_58

m/z 294.1490

024E+36.57RT (min)

lcorbi\_hilic\_pos\_2\_013\_36

m/z 294.1813

024E+36.57RT (min)

lcorbi\_hilic\_pos\_4\_077\_58

m/z 294.1913

0246E+33.54RT (min)

lcorbi\_hilic\_pos\_1\_048\_02

m/z 294.1966

01E+422.5RT (min)

lcorbi\_hilic\_pos\_4\_094\_60

m/z 294.2640

01234E+30.51RT (min)

lcorbi\_hilic\_pos\_1\_004\_22

m/z 294.8435

01E+455.5RT (min)

lcorbi\_hilic\_pos\_4\_075\_58

m/z 294.8832

01E+488.5RT (min)

lcorbi\_hilic\_pos\_4\_071\_57

m/z 294.9381

01234E+377.5RT (min)

lcorbi\_hilic\_pos\_1\_050\_05

m/z 295.0779

01E+47.58RT (min)

lcorbi\_hilic\_pos\_4\_084\_59

m/z 295.0850

0246E+333.54RT (min)

lcorbi\_hilic\_pos\_4\_089\_59

m/z 295.1022

01234E+477.5RT (min)

lcorbi\_hilic\_pos\_2\_030\_05

m/z 295.1197

01234E+444.5RT (min)

lcorbi\_hilic\_pos\_4\_090\_60

m/z 295.1372

0111.5RT (min)

lcorbi\_hilic\_pos\_4\_096\_60

m/z 295.1390

0246E+344.5RT (min)

lcorbi\_hilic\_pos\_1\_046\_30

m/z 295.1806

02468E+33.54RT (min)

lcorbi\_hilic\_pos\_4\_106\_62

m/z 295.1946

01E+444.5RT (min)

lcorbi\_hilic\_pos\_2\_036\_08

m/z 295.2106

010.51RT (min)

lcorbi\_hilic\_pos\_4\_077\_58

m/z 295.2131

01E+44.55RT (min)

lcorbi\_hilic\_pos\_3\_021\_46

m/z 295.2633

01E+30.51RT (min)

lcorbi\_hilic\_pos\_2\_037\_24

m/z 295.2766

02468E+40.51RT (min)

lcorbi\_hilic\_pos\_4\_090\_60

m/z 295.8815

0246E+388.5RT (min)

lcorbi\_hilic\_pos\_4\_069\_57

m/z 295.9319

0123E+35.56RT (min)

lcorbi\_hilic\_pos\_4\_074\_58

m/z 295.9425

01E+45.56RT (min)

lcorbi\_hilic\_pos\_4\_071\_57

m/z 296.0441

0123E+388.5RT (min)

lcorbi\_hilic\_pos\_4\_095\_60

m/z 296.0813

0144.5RT (min)

lcorbi\_hilic\_pos\_2\_044\_26

m/z 296.1174

016.57RT (min)

lcorbi\_hilic\_pos\_4\_093\_60

m/z 296.1219

012E+41.52RT (min)

lcorbi\_hilic\_pos\_1\_030\_48

m/z 296.1494

014.55RT (min)

lcorbi\_hilic\_pos\_4\_102\_61

m/z 296.1606

02468E+355.5RT (min)

lcorbi\_hilic\_pos\_4\_082\_59

m/z 296.1705

012E+45.56RT (min)

lcorbi\_hilic\_pos\_1\_021\_46

m/z 296.1903

012E+466.5RT (min)

lcorbi\_hilic\_pos\_4\_069\_57

m/z 296.2145

012E+455.5RT (min)

lcorbi\_hilic\_pos\_4\_079\_58

m/z 296.2302

01E+411.5RT (min)

lcorbi\_hilic\_pos\_4\_077\_58

m/z 296.7162

01E+455.5RT (min)

lcorbi\_hilic\_pos\_4\_080\_58

m/z 296.9665

01E+46.57RT (min)

lcorbi\_hilic\_pos\_4\_093\_60

m/z 297.0129

0123E+35.56RT (min)

lcorbi\_hilic\_pos\_4\_083\_59

m/z 297.0419

0123E+466.5RT (min)

lcorbi\_hilic\_pos\_4\_092\_60

m/z 297.0558

0246E+45.56RT (min)

lcorbi\_hilic\_pos\_4\_067\_57

m/z 297.0625

01E+366.5RT (min)

lcorbi\_hilic\_pos\_4\_080\_58

m/z 297.0799

015.56RT (min)

lcorbi\_hilic\_pos\_1\_003\_24

m/z 297.0826

012E+40.51RT (min)

lcorbi\_hilic\_pos\_4\_105\_61

m/z 297.0829

01E+43.54RT (min)

lcorbi\_hilic\_pos\_1\_003\_24

m/z 297.0912

02468E+36.57RT (min)

lcorbi\_hilic\_pos\_2\_024\_32

m/z 297.1082

02468E+322.5RT (min)

lcorbi\_hilic\_pos\_4\_085\_59

m/z 297.1251

01E+43.54RT (min)

lcorbi\_hilic\_pos\_4\_069\_57

m/z 297.1446

0123E+311.5RT (min)

lcorbi\_hilic\_pos\_4\_020\_37

m/z 297.1543

01E+31.52RT (min)

lcorbi\_hilic\_pos\_4\_007\_05

m/z 297.1810

0246E+344.5RT (min)

lcorbi\_hilic\_pos\_4\_068\_57

m/z 297.1851

0111.5RT (min)

lcorbi\_hilic\_pos\_3\_019\_43

m/z 297.2983

01E+33.54RT (min)

lcorbi\_hilic\_pos\_4\_017\_51

m/z 297.9676

012E+477.5RT (min)

lcorbi\_hilic\_pos\_4\_018\_31

m/z 298.0518

01E+433.5RT (min)

lcorbi\_hilic\_pos\_4\_088\_59

m/z 298.0642

01234E+36.57RT (min)

lcorbi\_hilic\_pos\_4\_076\_58

m/z 298.0879

02468E+322.5RT (min)

lcorbi\_hilic\_pos\_2\_027\_38

m/z 298.1011

01E+477.5RT (min)

lcorbi\_hilic\_pos\_4\_088\_59

m/z 298.1071

01E+422.5RT (min)

lcorbi\_hilic\_pos\_1\_032\_31

m/z 298.1148

01E+444.5RT (min)

lcorbi\_hilic\_pos\_4\_084\_59

m/z 298.1390

01E+42.53RT (min)

lcorbi\_hilic\_pos\_4\_096\_60

m/z 298.1391

01E+566.5RT (min)

lcorbi\_hilic\_pos\_4\_097\_60

m/z 298.1625

01E+46.57RT (min)

lcorbi\_hilic\_pos\_2\_003\_33

m/z 298.1862

01E+41.52RT (min)

lcorbi\_hilic\_pos\_4\_034\_36

m/z 298.2378

0246E+40.51RT (min)

lcorbi\_hilic\_pos\_4\_086\_59

m/z 298.2742

01234E+40.51RT (min)

lcorbi\_hilic\_pos\_4\_020\_37

m/z 298.3106

01E+40.51RT (min)

lcorbi\_hilic\_pos\_1\_054\_23

m/z 298.3106

0246E+32.53RT (min)

lcorbi\_hilic\_pos\_4\_082\_59

m/z 298.3240

02468E+33.54RT (min)

lcorbi\_hilic\_pos\_4\_096\_60

m/z 298.3470

012E+41.52RT (min)

lcorbi\_hilic\_pos\_4\_045\_33

m/z 298.3470

01E+400.5RT (min)

lcorbi\_hilic\_pos\_1\_027\_10

m/z 299.0285

01234E+35.56RT (min)

lcorbi\_hilic\_pos\_4\_102\_61

m/z 299.0464

0144.5RT (min)

lcorbi\_hilic\_pos\_4\_020\_37

m/z 299.0564

0246E+366.5RT (min)

lcorbi\_hilic\_pos\_4\_079\_58

m/z 299.0619

01E+40.51RT (min)

lcorbi\_hilic\_pos\_4\_105\_61

m/z 299.0851

024E+36.57RT (min)

lcorbi\_hilic\_pos\_4\_089\_59

m/z 299.1150

012E+577.5RT (min)

lcorbi\_hilic\_pos\_2\_037\_24

m/z 299.2007

0123E+40.51RT (min)

lcorbi\_hilic\_pos\_1\_053\_08

m/z 299.2007

012E+422.5RT (min)

lcorbi\_hilic\_pos\_4\_102\_61

m/z 299.9160

024E+377.5RT (min)

lcorbi\_hilic\_pos\_2\_022\_35

m/z 299.9257

0166.5RT (min)

lcorbi\_hilic\_pos\_2\_027\_38

m/z 299.9639

0246E+38.59RT (min)

lcorbi\_hilic\_pos\_4\_099\_61

m/z 300.0043

01E+466.5RT (min)

lcorbi\_hilic\_pos\_4\_067\_57

m/z 300.0244

0166.5RT (min)

lcorbi\_hilic\_pos\_4\_072\_57

m/z 300.0528

0246E+35.56RT (min)

lcorbi\_hilic\_pos\_4\_106\_62

m/z 300.0724

012E+366.5RT (min)

lcorbi\_hilic\_pos\_4\_010\_47

m/z 300.1066

01E+433.5RT (min)

lcorbi\_hilic\_pos\_4\_007\_05

m/z 300.1231

012E+411.5RT (min)

lcorbi\_hilic\_pos\_1\_033\_53

m/z 300.1365

01234E+344.5RT (min)

lcorbi\_hilic\_pos\_1\_040\_06

m/z 300.1425

010.51RT (min)

lcorbi\_hilic\_pos\_3\_053\_21

m/z 300.1555

0246E+34.55RT (min)

lcorbi\_hilic\_pos\_4\_067\_57

m/z 300.1684

01E+46.57RT (min)

lcorbi\_hilic\_pos\_4\_096\_60

m/z 300.1784

012E+43.54RT (min)

lcorbi\_hilic\_pos\_4\_020\_37

m/z 300.1796

01E+488.5RT (min)

lcorbi\_hilic\_pos\_4\_093\_60

m/z 300.2283

024E+31.52RT (min)

lcorbi\_hilic\_pos\_4\_087\_59

m/z 300.2434

01E+477.5RT (min)

lcorbi\_hilic\_pos\_2\_037\_24

m/z 300.2616

0123E+411.5RT (min)

lcorbi\_hilic\_pos\_4\_081\_58

m/z 300.3262

0246E+31.52RT (min)

lcorbi\_hilic\_pos\_3\_019\_43

m/z 300.8578

02468E+377.5RT (min)

lcorbi\_hilic\_pos\_4\_088\_59

m/z 301.0962

0122.5RT (min)

lcorbi\_hilic\_pos\_4\_098\_61

m/z 301.0981

01E+47.58RT (min)

lcorbi\_hilic\_pos\_4\_095\_60

m/z 301.1120

01E+455.5RT (min)

lcorbi\_hilic\_pos\_4\_067\_57

m/z 301.1184

01E+411.5RT (min)

lcorbi\_hilic\_pos\_4\_072\_57

m/z 301.1411

01E+50.51RT (min)

lcorbi\_hilic\_pos\_4\_079\_58

m/z 301.1636

012E+47.58RT (min)

lcorbi\_hilic\_pos\_4\_094\_60

m/z 301.1717

0246E+44.55RT (min)

lcorbi\_hilic\_pos\_4\_093\_60

m/z 301.1800

01E+30.51RT (min)

lcorbi\_hilic\_pos\_4\_006\_14

m/z 301.2052

01234E+411.5RT (min)

lcorbi\_hilic\_pos\_3\_019\_43

m/z 301.2849

0123E+433.5RT (min)

lcorbi\_hilic\_pos\_4\_036\_09

m/z 301.2932

012E+411.5RT (min)

lcorbi\_hilic\_pos\_4\_089\_59

m/z 301.9659

015.56RT (min)

lcorbi\_hilic\_pos\_4\_088\_59

m/z 301.9811

0177.5RT (min)

lcorbi\_hilic\_pos\_4\_074\_58

m/z 301.9933

01E+45.56RT (min)

lcorbi\_hilic\_pos\_4\_084\_59

m/z 302.0379

01E+44.55RT (min)

lcorbi\_hilic\_pos\_4\_111\_62

m/z 302.1000

01E+311.5RT (min)

lcorbi\_hilic\_pos\_4\_020\_37

m/z 302.1114

024E+311.5RT (min)

lcorbi\_hilic\_pos\_3\_021\_46

m/z 302.1445

0123E+40.51RT (min)

lcorbi\_hilic\_pos\_4\_080\_58

m/z 302.1907

0246E+35.56RT (min)

lcorbi\_hilic\_pos\_1\_021\_46

m/z 302.2197

010.51RT (min)

lcorbi\_hilic\_pos\_4\_046\_21

m/z 302.2753

02468E+36.57RT (min)

lcorbi\_hilic\_pos\_4\_096\_60

m/z 302.3055

0246E+30.51RT (min)

lcorbi\_hilic\_pos\_1\_015\_44

m/z 302.9463

01E+477.5RT (min)

lcorbi\_hilic\_pos\_1\_003\_24

m/z 303.0231

01E+466.5RT (min)

lcorbi\_hilic\_pos\_4\_074\_58

m/z 303.0331

0246E+35.56RT (min)

lcorbi\_hilic\_pos\_4\_072\_57

m/z 303.0419

015.56RT (min)

lcorbi\_hilic\_pos\_4\_067\_57

m/z 303.0477

024E+37.58RT (min)

lcorbi\_hilic\_pos\_1\_052\_35

m/z 303.0743

012E+422.5RT (min)

lcorbi\_hilic\_pos\_4\_066\_57

m/z 303.0928

02468E+36.57RT (min)

lcorbi\_hilic\_pos\_4\_084\_59

m/z 303.1075

01E+422.5RT (min)

lcorbi\_hilic\_pos\_4\_048\_46

m/z 303.1164

01E+46.57RT (min)

lcorbi\_hilic\_pos\_4\_082\_59

m/z 303.1188

01E+444.5RT (min)

lcorbi\_hilic\_pos\_4\_084\_59

m/z 303.1375

0246E+311.5RT (min)

lcorbi\_hilic\_pos\_4\_082\_59

m/z 303.1793

0123E+46.57RT (min)

lcorbi\_hilic\_pos\_4\_097\_60

m/z 303.1916

01234E+211.5RT (min)

lcorbi\_hilic\_pos\_3\_014\_08

m/z 303.1997

012E+444.5RT (min)

lcorbi\_hilic\_pos\_4\_076\_58

m/z 303.2167

0111.5RT (min)

lcorbi\_hilic\_pos\_4\_081\_58

m/z 303.2360

01234E+333.5RT (min)

lcorbi\_hilic\_pos\_4\_080\_58

m/z 303.2551

012E+30.51RT (min)

lcorbi\_hilic\_pos\_3\_019\_43

m/z 303.2724

0246E+333.5RT (min)

lcorbi\_hilic\_pos\_2\_036\_08

m/z 303.9294

01234E+377.5RT (min)

lcorbi\_hilic\_pos\_1\_040\_06

m/z 303.9473

01234E+366.5RT (min)

lcorbi\_hilic\_pos\_4\_066\_57

m/z 304.0178

0246E+366.5RT (min)

lcorbi\_hilic\_pos\_4\_094\_60

m/z 304.0350

0246E+34.55RT (min)

lcorbi\_hilic\_pos\_4\_106\_62

m/z 304.1004

02468E+477.5RT (min)

lcorbi\_hilic\_pos\_2\_005\_23

m/z 304.1018

0246E+35.56RT (min)

lcorbi\_hilic\_pos\_4\_068\_57

m/z 304.1545

01234E+422.5RT (min)

lcorbi\_hilic\_pos\_4\_067\_57

m/z 304.1617

01E+466.5RT (min)

lcorbi\_hilic\_pos\_4\_020\_37

m/z 304.1756

01E+455.5RT (min)

lcorbi\_hilic\_pos\_4\_070\_57

m/z 304.1756

0246E+312RT (min)

lcorbi\_hilic\_pos\_4\_092\_60

m/z 304.2353

01E+411.5RT (min)

lcorbi\_hilic\_pos\_4\_075\_58

m/z 304.2484

0246E+311.5RT (min)

lcorbi\_hilic\_pos\_1\_034\_37

m/z 304.3120

013.54RT (min)

lcorbi\_hilic\_pos\_3\_037\_20

m/z 305.0010

01234E+27.58RT (min)

lcorbi\_hilic\_pos\_1\_029\_14

m/z 305.0180

012E+466.5RT (min)

lcorbi\_hilic\_pos\_4\_080\_58

m/z 305.0180

01E+47.58RT (min)

lcorbi\_hilic\_pos\_3\_008\_14

m/z 305.0280

01E+47.58RT (min)

lcorbi\_hilic\_pos\_4\_081\_58

m/z 305.0544

0246E+344.5RT (min)

lcorbi\_hilic\_pos\_4\_082\_59

m/z 305.1472

0123E+46.57RT (min)

lcorbi\_hilic\_pos\_4\_093\_60

m/z 305.1532

01E+444.5RT (min)

lcorbi\_hilic\_pos\_3\_010\_32

m/z 305.1578

02468E+322.5RT (min)

lcorbi\_hilic\_pos\_4\_069\_57

m/z 305.2154

01E+444.5RT (min)

lcorbi\_hilic\_pos\_4\_077\_58

m/z 305.2477

02468E+30.51RT (min)

lcorbi\_hilic\_pos\_4\_078\_58

m/z 305.3034

012E+41.52RT (min)

lcorbi\_hilic\_pos\_4\_020\_37

m/z 305.3034

0246E+300.5RT (min)

lcorbi\_hilic\_pos\_4\_098\_61

m/z 305.9193

01234E+388.5RT (min)

lcorbi\_hilic\_pos\_4\_071\_57

m/z 306.0731

0123E+366.5RT (min)

lcorbi\_hilic\_pos\_4\_067\_57

m/z 306.1103

024E+36.57RT (min)

lcorbi\_hilic\_pos\_1\_028\_34

m/z 306.1270

0123E+433.5RT (min)

lcorbi\_hilic\_pos\_4\_097\_60

m/z 306.1297

024E+377.5RT (min)

lcorbi\_hilic\_pos\_4\_086\_59

m/z 306.1512

02468E+32.53RT (min)

lcorbi\_hilic\_pos\_3\_034\_26

m/z 306.2447

01E+433.5RT (min)

lcorbi\_hilic\_pos\_1\_044\_17

m/z 306.2639

0123E+433.5RT (min)

lcorbi\_hilic\_pos\_3\_019\_43

m/z 306.8519

02468E+37.58RT (min)

lcorbi\_hilic\_pos\_4\_111\_62

m/z 306.8709

0123E+377.5RT (min)

lcorbi\_hilic\_pos\_1\_053\_08

m/z 306.8949

02468E+37.58RT (min)

lcorbi\_hilic\_pos\_4\_102\_61

m/z 306.9925

02468E+35.56RT (min)

lcorbi\_hilic\_pos\_4\_091\_60

m/z 307.0198

02468E+355.5RT (min)

lcorbi\_hilic\_pos\_4\_069\_57

m/z 307.0234

024E+37.58RT (min)

lcorbi\_hilic\_pos\_4\_080\_58

m/z 307.0292

0123E+37.58RT (min)

lcorbi\_hilic\_pos\_1\_029\_14

m/z 307.0691

0246E+366.5RT (min)

lcorbi\_hilic\_pos\_4\_083\_59

m/z 307.0790

01E+477.5RT (min)

lcorbi\_hilic\_pos\_4\_082\_59

m/z 307.0835

0246E+388.5RT (min)

lcorbi\_hilic\_pos\_4\_083\_59

m/z 307.1001

01E+455.5RT (min)

lcorbi\_hilic\_pos\_4\_072\_57

m/z 307.1182

012E+444.5RT (min)

lcorbi\_hilic\_pos\_4\_094\_60

m/z 307.1272

024E+322.5RT (min)

lcorbi\_hilic\_pos\_2\_024\_32

m/z 307.1561

012E+466.5RT (min)

lcorbi\_hilic\_pos\_3\_031\_36

m/z 307.2115

01E+40.51RT (min)

lcorbi\_hilic\_pos\_1\_023\_28

m/z 307.2481

01234E+40.51RT (min)

lcorbi\_hilic\_pos\_4\_018\_31

m/z 308.1220

024E+32.53RT (min)

lcorbi\_hilic\_pos\_3\_001\_49

m/z 308.1344

01234E+46.57RT (min)

lcorbi\_hilic\_pos\_1\_050\_05

m/z 308.1857

01234E+33.54RT (min)

lcorbi\_hilic\_pos\_4\_071\_57

m/z 308.2433

01E+40.51RT (min)

lcorbi\_hilic\_pos\_2\_016\_18

m/z 308.3143

0246E+322.5RT (min)

lcorbi\_hilic\_pos\_4\_084\_59

m/z 309.0047

0246E+366.5RT (min)

lcorbi\_hilic\_pos\_4\_079\_58

m/z 309.0832

01E+466.5RT (min)

lcorbi\_hilic\_pos\_1\_036\_32

m/z 309.1240

01E+422.5RT (min)

lcorbi\_hilic\_pos\_3\_036\_33

m/z 309.1447

02468E+31.52RT (min)

lcorbi\_hilic\_pos\_1\_001\_49

m/z 309.1599

012E+444.5RT (min)

lcorbi\_hilic\_pos\_4\_111\_62

m/z 309.1604

0246E+366.5RT (min)

lcorbi\_hilic\_pos\_3\_003\_37

m/z 309.2426

0246E+20.51RT (min)

lcorbi\_hilic\_pos\_3\_053\_21

m/z 309.2902

01E+444.5RT (min)

lcorbi\_hilic\_pos\_4\_021\_13

m/z 309.9004

024E+388.5RT (min)

lcorbi\_hilic\_pos\_4\_071\_57

m/z 310.0222

01E+477.58RT (min)

lcorbi\_hilic\_pos\_4\_078\_58

m/z 310.0957

012E+36.57RT (min)

lcorbi\_hilic\_pos\_1\_054\_23

m/z 310.1091

012E+433.5RT (min)

lcorbi\_hilic\_pos\_4\_084\_59

m/z 310.1135

017RT (min)

lcorbi\_hilic\_pos\_4\_090\_60

m/z 310.1486

0166.5RT (min)

lcorbi\_hilic\_pos\_4\_093\_60

m/z 310.1584

01E+47.58RT (min)

lcorbi\_hilic\_pos\_1\_003\_24

m/z 310.1633

0246E+344.5RT (min)

lcorbi\_hilic\_pos\_4\_112\_62

m/z 310.1763

01E+477.5RT (min)

lcorbi\_hilic\_pos\_4\_074\_58

m/z 310.2226

01E+40.51RT (min)

lcorbi\_hilic\_pos\_4\_019\_30

m/z 310.2741

0246E+30.51RT (min)

lcorbi\_hilic\_pos\_3\_019\_43

m/z 310.3106

01E+50.51RT (min)

lcorbi\_hilic\_pos\_3\_019\_43

m/z 310.8983

0246E+388.5RT (min)

lcorbi\_hilic\_pos\_4\_073\_57

m/z 311.0285

012E+35.56RT (min)

lcorbi\_hilic\_pos\_1\_028\_34

m/z 311.0640

012E+433.5RT (min)

lcorbi\_hilic\_pos\_4\_087\_59

m/z 311.0739

024E+42.53RT (min)

lcorbi\_hilic\_pos\_4\_112\_62

m/z 311.0908

0123E+377.5RT (min)

lcorbi\_hilic\_pos\_4\_089\_59

m/z 311.1382

0123E+322.5RT (min)

lcorbi\_hilic\_pos\_4\_093\_60

m/z 311.1669

0123E+466.5RT (min)

lcorbi\_hilic\_pos\_3\_036\_33

m/z 311.2048

012E+43.54RT (min)

lcorbi\_hilic\_pos\_4\_074\_58

m/z 311.2065

01E+511.5RT (min)

lcorbi\_hilic\_pos\_4\_018\_31

m/z 311.2582

01E+40.51RT (min)

lcorbi\_hilic\_pos\_2\_054\_29

m/z 311.2717

024E+20.51RT (min)

lcorbi\_hilic\_pos\_2\_021\_17

m/z 311.3059

02468E+344.5RT (min)

lcorbi\_hilic\_pos\_4\_021\_13

m/z 311.3274

02468E+333.5RT (min)

lcorbi\_hilic\_pos\_2\_037\_24

m/z 311.5487

02468E+34.55RT (min)

lcorbi\_hilic\_pos\_4\_081\_58

m/z 311.8961

01E+488.5RT (min)

lcorbi\_hilic\_pos\_4\_067\_57

m/z 311.9646

0123E+377.5RT (min)

lcorbi\_hilic\_pos\_1\_043\_25

m/z 312.0044

012E+45.56RT (min)

lcorbi\_hilic\_pos\_4\_082\_59

m/z 312.0156

01E+466.5RT (min)

lcorbi\_hilic\_pos\_4\_075\_58

m/z 312.0204

0246E+377.5RT (min)

lcorbi\_hilic\_pos\_4\_080\_58

m/z 312.0408

0246E+355.5RT (min)

lcorbi\_hilic\_pos\_4\_066\_57

m/z 312.0868

01234E+311.5RT (min)

lcorbi\_hilic\_pos\_4\_017\_51

m/z 312.0974

01E+544.5RT (min)

lcorbi\_hilic\_pos\_1\_003\_24

m/z 312.1340

01234E+444.5RT (min)

lcorbi\_hilic\_pos\_4\_014\_27

m/z 312.1418

01234E+544.5RT (min)

lcorbi\_hilic\_pos\_3\_048\_38

m/z 312.1641

012E+466.5RT (min)

lcorbi\_hilic\_pos\_4\_068\_57

m/z 312.2171

0246E+33.54RT (min)

lcorbi\_hilic\_pos\_4\_074\_58

m/z 312.2899

0123E+30.51RT (min)

lcorbi\_hilic\_pos\_1\_050\_05

m/z 312.3262

0246E+32.53RT (min)

lcorbi\_hilic\_pos\_4\_087\_59

m/z 312.3626

01E+41.52RT (min)

lcorbi\_hilic\_pos\_3\_050\_44

m/z 312.5472

01E+44.55RT (min)

lcorbi\_hilic\_pos\_4\_078\_58

m/z 312.9996

01234E+36.57RT (min)

lcorbi\_hilic\_pos\_4\_067\_57

m/z 313.0361

012E+466.5RT (min)

lcorbi\_hilic\_pos\_4\_081\_58

m/z 313.0709

0123E+411.5RT (min)

lcorbi\_hilic\_pos\_3\_023\_18

m/z 313.0784

0246E+34.55RT (min)

lcorbi\_hilic\_pos\_4\_089\_59

m/z 313.0854

012E+333.5RT (min)

lcorbi\_hilic\_pos\_2\_042\_02

m/z 313.0966

01E+466.5RT (min)

lcorbi\_hilic\_pos\_4\_091\_60

m/z 313.1035

02468E+33.54RT (min)

lcorbi\_hilic\_pos\_4\_098\_61

m/z 313.1120

02468E+37.58RT (min)

lcorbi\_hilic\_pos\_4\_082\_59

m/z 313.1175

012E+411.5RT (min)

lcorbi\_hilic\_pos\_4\_097\_60

m/z 313.1201

0144.5RT (min)

lcorbi\_hilic\_pos\_4\_013\_23

m/z 313.1202

02468E+36.57RT (min)

lcorbi\_hilic\_pos\_4\_086\_59

m/z 313.1500

012E+56.57RT (min)

lcorbi\_hilic\_pos\_4\_091\_60

m/z 313.1630

010.51RT (min)

lcorbi\_hilic\_pos\_1\_004\_22

m/z 313.2375

01E+411.5RT (min)

lcorbi\_hilic\_pos\_4\_080\_58

m/z 313.2487

012E+30.51RT (min)

lcorbi\_hilic\_pos\_1\_004\_22

m/z 313.2739

01234E+444.5RT (min)

lcorbi\_hilic\_pos\_2\_030\_05

m/z 313.3295

024E+40.51RT (min)

lcorbi\_hilic\_pos\_1\_054\_23

m/z 313.3579

0246E+355.5RT (min)

lcorbi\_hilic\_pos\_1\_010\_07

m/z 313.9317

016.57RT (min)

lcorbi\_hilic\_pos\_4\_090\_60

m/z 314.0330

024E+37.58RT (min)

lcorbi\_hilic\_pos\_4\_084\_59

m/z 314.0701

01E+466.5RT (min)

lcorbi\_hilic\_pos\_1\_044\_17

m/z 314.1534

0246E+34.55RT (min)

lcorbi\_hilic\_pos\_4\_109\_62

m/z 314.1600

012E+411.5RT (min)

lcorbi\_hilic\_pos\_1\_029\_14

m/z 314.1772

0123E+466.5RT (min)

lcorbi\_hilic\_pos\_3\_010\_32

m/z 314.1811

01E+31.52RT (min)

lcorbi\_hilic\_pos\_4\_096\_60

m/z 314.2175

01234E+40.51RT (min)

lcorbi\_hilic\_pos\_3\_019\_43

m/z 314.2439

02468E+31.52RT (min)

lcorbi\_hilic\_pos\_4\_086\_59

m/z 314.3055

01234E+322.5RT (min)

lcorbi\_hilic\_pos\_1\_054\_23

m/z 314.3055

01E+344.5RT (min)

lcorbi\_hilic\_pos\_3\_027\_05

m/z 314.3055

0246E+30.51RT (min)

lcorbi\_hilic\_pos\_1\_054\_23

m/z 314.3249

0123E+422.5RT (min)

lcorbi\_hilic\_pos\_4\_083\_59

m/z 314.3419

02468E+31.52RT (min)

lcorbi\_hilic\_pos\_4\_100\_61

m/z 315.0590

0177.5RT (min)

lcorbi\_hilic\_pos\_2\_037\_24

m/z 315.0860

01E+422.5RT (min)

lcorbi\_hilic\_pos\_3\_054\_16

m/z 315.1058

0123E+466.5RT (min)

lcorbi\_hilic\_pos\_4\_093\_60

m/z 315.1259

012E+433.5RT (min)

lcorbi\_hilic\_pos\_4\_007\_05

m/z 315.1487

0246E+344.5RT (min)

lcorbi\_hilic\_pos\_4\_071\_57

m/z 315.1741

0166.5RT (min)

lcorbi\_hilic\_pos\_1\_050\_05

m/z 315.1780

01E+411.5RT (min)

lcorbi\_hilic\_pos\_4\_088\_59

m/z 315.1956

012E+411.5RT (min)

lcorbi\_hilic\_pos\_4\_022\_08

m/z 315.1986

01E+47.58RT (min)

lcorbi\_hilic\_pos\_2\_030\_05

m/z 315.1986

012E+466.5RT (min)

lcorbi\_hilic\_pos\_4\_052\_32

m/z 315.2280

01E+411.5RT (min)

lcorbi\_hilic\_pos\_4\_085\_59

m/z 315.2280

02468E+344.5RT (min)

lcorbi\_hilic\_pos\_4\_020\_37

m/z 315.2531

0123E+411.5RT (min)

lcorbi\_hilic\_pos\_4\_081\_58

m/z 316.0808

02468E+233.5RT (min)

lcorbi\_hilic\_pos\_4\_042\_02

m/z 316.0834

0246E+377.5RT (min)

lcorbi\_hilic\_pos\_1\_050\_05

m/z 316.0866

0123E+433.5RT (min)

lcorbi\_hilic\_pos\_1\_048\_02

m/z 316.0877

012E+444.5RT (min)

lcorbi\_hilic\_pos\_4\_083\_59

m/z 316.1142

014.55RT (min)

lcorbi\_hilic\_pos\_4\_088\_59

m/z 316.1157

02468E+311.5RT (min)

lcorbi\_hilic\_pos\_1\_029\_14

m/z 316.1311

02468E+36.57RT (min)

lcorbi\_hilic\_pos\_4\_069\_57

m/z 316.1367

01E+411.5RT (min)

lcorbi\_hilic\_pos\_1\_042\_40

m/z 316.1480

01E+411.5RT (min)

lcorbi\_hilic\_pos\_4\_091\_60

m/z 316.1755

0246E+34.55RT (min)

lcorbi\_hilic\_pos\_2\_036\_08

m/z 316.1811

012.53RT (min)

lcorbi\_hilic\_pos\_4\_045\_33

m/z 316.2393

012E+333.5RT (min)

lcorbi\_hilic\_pos\_4\_066\_57

m/z 316.2655

01234E+42.53RT (min)

lcorbi\_hilic\_pos\_1\_034\_37

m/z 316.2848

0123E+31.52RT (min)

lcorbi\_hilic\_pos\_4\_026\_11

m/z 316.9201

0123E+377.5RT (min)

lcorbi\_hilic\_pos\_4\_082\_59

m/z 317.0213

015.56RT (min)

lcorbi\_hilic\_pos\_4\_020\_37

m/z 317.0483

01E+45.56RT (min)

lcorbi\_hilic\_pos\_4\_091\_60

m/z 317.0562

02468E+45.56RT (min)

lcorbi\_hilic\_pos\_1\_008\_38

m/z 317.0958

0144.55RT (min)

lcorbi\_hilic\_pos\_3\_023\_18

m/z 317.1028

0166.5RT (min)

lcorbi\_hilic\_pos\_1\_002\_29

m/z 317.1150

010.51RT (min)

lcorbi\_hilic\_pos\_4\_099\_61

m/z 317.1171

0188.5RT (min)

lcorbi\_hilic\_pos\_1\_029\_14

m/z 317.1215

016.57RT (min)

lcorbi\_hilic\_pos\_1\_029\_14

m/z 317.1507

01E+444.5RT (min)

lcorbi\_hilic\_pos\_3\_036\_33

m/z 317.1667

02468E+377.5RT (min)

lcorbi\_hilic\_pos\_4\_096\_60

m/z 317.2154

012E+422.5RT (min)

lcorbi\_hilic\_pos\_4\_090\_60

m/z 317.2186

0246E+36.57RT (min)

lcorbi\_hilic\_pos\_4\_089\_59

m/z 317.2437

02468E+344.5RT (min)

lcorbi\_hilic\_pos\_4\_100\_61

m/z 317.2475

02468E+30.51RT (min)

lcorbi\_hilic\_pos\_4\_080\_58

m/z 317.2689

01234E+32.53RT (min)

lcorbi\_hilic\_pos\_1\_054\_23

m/z 317.2800

024E+30.51RT (min)

lcorbi\_hilic\_pos\_3\_008\_14

m/z 317.3245

012E+422.5RT (min)

lcorbi\_hilic\_pos\_1\_054\_23

m/z 317.3469

0133.5RT (min)

lcorbi\_hilic\_pos\_4\_066\_57

m/z 318.0339

024E+34.55RT (min)

lcorbi\_hilic\_pos\_4\_080\_58

m/z 318.0568

01234E+355.5RT (min)

lcorbi\_hilic\_pos\_2\_012\_37

m/z 318.0650

024E+26.57RT (min)

lcorbi\_hilic\_pos\_2\_024\_32

m/z 318.1337

0246E+311.5RT (min)

lcorbi\_hilic\_pos\_4\_077\_58

m/z 318.1913

01E+444.5RT (min)

lcorbi\_hilic\_pos\_4\_070\_57

m/z 318.2722

01E+411.5RT (min)

lcorbi\_hilic\_pos\_1\_035\_15

m/z 318.3503

01E+411.5RT (min)

lcorbi\_hilic\_pos\_4\_096\_60

m/z 318.8508

024E+388.5RT (min)

lcorbi\_hilic\_pos\_4\_067\_57

m/z 318.9375

024E+38.59RT (min)

lcorbi\_hilic\_pos\_4\_109\_62

m/z 319.1114

0144.5RT (min)

lcorbi\_hilic\_pos\_4\_105\_61

m/z 319.1291

01E+411.5RT (min)

lcorbi\_hilic\_pos\_4\_020\_37

m/z 319.1654

01E+344.5RT (min)

lcorbi\_hilic\_pos\_3\_023\_18

m/z 319.2269

012E+40.51RT (min)

lcorbi\_hilic\_pos\_2\_040\_14

m/z 319.2844

0246E+42.53RT (min)

lcorbi\_hilic\_pos\_3\_037\_20

m/z 319.3038

02468E+42.53RT (min)

lcorbi\_hilic\_pos\_3\_037\_20

m/z 319.3627

012E+43.54RT (min)

lcorbi\_hilic\_pos\_4\_096\_60

m/z 319.9497

01E+477.5RT (min)

lcorbi\_hilic\_pos\_4\_018\_31

m/z 319.9996

0246E+377.5RT (min)

lcorbi\_hilic\_pos\_1\_035\_15

m/z 320.0532

0123E+47.58RT (min)

lcorbi\_hilic\_pos\_4\_095\_60

m/z 320.0831

0246E+377.5RT (min)

lcorbi\_hilic\_pos\_4\_069\_57

m/z 320.1206

01E+46.57RT (min)

lcorbi\_hilic\_pos\_1\_036\_32

m/z 320.1319

012E+455.5RT (min)

lcorbi\_hilic\_pos\_4\_070\_57

m/z 320.1690

0144.5RT (min)

lcorbi\_hilic\_pos\_4\_095\_60

m/z 320.2140

012E+57.58RT (min)

lcorbi\_hilic\_pos\_4\_093\_60

m/z 320.2222

012E+31.52RT (min)

lcorbi\_hilic\_pos\_4\_093\_60

m/z 320.2628

01E+43.54RT (min)

lcorbi\_hilic\_pos\_1\_044\_17

m/z 320.2949

024E+33.54RT (min)

lcorbi\_hilic\_pos\_4\_084\_59

m/z 321.0420

012E+433.5RT (min)

lcorbi\_hilic\_pos\_2\_042\_02

m/z 321.0502

024E+366.5RT (min)

lcorbi\_hilic\_pos\_4\_113\_62

m/z 321.0808

01E+46.57RT (min)

lcorbi\_hilic\_pos\_1\_028\_34

m/z 321.0969

0246E+477.5RT (min)

lcorbi\_hilic\_pos\_4\_091\_60

m/z 321.1035

01E+47.58RT (min)

lcorbi\_hilic\_pos\_1\_054\_23

m/z 321.1370

02468E+366.5RT (min)

lcorbi\_hilic\_pos\_3\_021\_46

m/z 321.1599

012E+43.54RT (min)

lcorbi\_hilic\_pos\_4\_109\_62

m/z 321.2426

012E+41.52RT (min)

lcorbi\_hilic\_pos\_4\_100\_61

m/z 321.3154

01E+30.51RT (min)

lcorbi\_hilic\_pos\_4\_083\_59

m/z 321.3266

01E+41.52RT (min)

lcorbi\_hilic\_pos\_4\_095\_60

m/z 322.0984

01E+42.53RT (min)

lcorbi\_hilic\_pos\_4\_101\_61

m/z 322.1876

02468E+36.57RT (min)

lcorbi\_hilic\_pos\_1\_032\_31

m/z 322.2378

01E+30.51RT (min)

lcorbi\_hilic\_pos\_3\_037\_20

m/z 322.2952

0123E+40.51RT (min)

lcorbi\_hilic\_pos\_4\_082\_59

m/z 322.3300

01234E+31.52RT (min)

lcorbi\_hilic\_pos\_4\_094\_60

m/z 323.0801

0246E+37.58RT (min)

lcorbi\_hilic\_pos\_2\_010\_07

m/z 323.0918

024E+411.5RT (min)

lcorbi\_hilic\_pos\_4\_107\_62

m/z 323.1299

0166.5RT (min)

lcorbi\_hilic\_pos\_1\_050\_05

m/z 323.1642

01234E+411.5RT (min)

lcorbi\_hilic\_pos\_4\_097\_60

m/z 323.2525

01E+422.5RT (min)

lcorbi\_hilic\_pos\_4\_086\_59

m/z 323.3139

0246E+33.54RT (min)

lcorbi\_hilic\_pos\_4\_086\_59

m/z 323.9631

0177.5RT (min)

lcorbi\_hilic\_pos\_4\_074\_58

m/z 324.0457

0177.5RT (min)

lcorbi\_hilic\_pos\_4\_074\_58

m/z 324.1255

02468E+444.5RT (min)

lcorbi\_hilic\_pos\_4\_091\_60

m/z 324.1408

0123E+50.51RT (min)

lcorbi\_hilic\_pos\_4\_032\_35

m/z 324.1655

0246E+466.5RT (min)

lcorbi\_hilic\_pos\_4\_069\_57

m/z 324.1675

02468E+411.5RT (min)

lcorbi\_hilic\_pos\_4\_098\_61

m/z 324.2171

02468E+40.51RT (min)

lcorbi\_hilic\_pos\_4\_081\_58

m/z 324.2233

01E+44.55RT (min)

lcorbi\_hilic\_pos\_4\_094\_60

m/z 324.2382

02468E+30.51RT (min)

lcorbi\_hilic\_pos\_4\_081\_58

m/z 324.2534

02468E+30.51RT (min)

lcorbi\_hilic\_pos\_1\_026\_39

m/z 324.2899

012E+40.51RT (min)

lcorbi\_hilic\_pos\_1\_044\_17

m/z 324.3263

02468E+33.54RT (min)

lcorbi\_hilic\_pos\_4\_089\_59

m/z 324.9283

0246E+377.5RT (min)

lcorbi\_hilic\_pos\_1\_010\_07

m/z 325.0050

01234E+366.5RT (min)

lcorbi\_hilic\_pos\_4\_076\_58

m/z 325.0434

01E+477.5RT (min)

lcorbi\_hilic\_pos\_1\_054\_23

m/z 325.0797

012E+433.5RT (min)

lcorbi\_hilic\_pos\_4\_085\_59

m/z 325.1131

0123E+455.5RT (min)

lcorbi\_hilic\_pos\_4\_110\_62

m/z 325.1218

0246E+50.51RT (min)

lcorbi\_hilic\_pos\_1\_051\_26

m/z 325.1342

01234E+366.5RT (min)

lcorbi\_hilic\_pos\_3\_003\_37

m/z 325.1613

01234E+46.57RT (min)

lcorbi\_hilic\_pos\_4\_090\_60

m/z 325.2276

01E+422.5RT (min)

lcorbi\_hilic\_pos\_4\_096\_60

m/z 325.9293

01234E+366.5RT (min)

lcorbi\_hilic\_pos\_4\_066\_57

m/z 326.0824

0123E+477.5RT (min)

lcorbi\_hilic\_pos\_4\_087\_59

m/z 326.0953

02468E+344.5RT (min)

lcorbi\_hilic\_pos\_1\_053\_08

m/z 326.1263

012E+444.5RT (min)

lcorbi\_hilic\_pos\_4\_093\_60

m/z 326.1538

012E+466.5RT (min)

lcorbi\_hilic\_pos\_2\_046\_46

m/z 326.1947

01E+43.54RT (min)

lcorbi\_hilic\_pos\_4\_112\_62

m/z 326.2076

01E+411.5RT (min)

lcorbi\_hilic\_pos\_4\_083\_59

m/z 326.2175

0123E+41.52RT (min)

lcorbi\_hilic\_pos\_3\_019\_43

m/z 326.2439

011.52RT (min)

lcorbi\_hilic\_pos\_3\_019\_43

m/z 326.2561

0123E+40.51RT (min)

lcorbi\_hilic\_pos\_4\_008\_19

m/z 326.2692

012E+322.5RT (min)

lcorbi\_hilic\_pos\_4\_087\_59

m/z 326.3055

01E+40.51RT (min)

lcorbi\_hilic\_pos\_3\_019\_43

m/z 326.3250

0246E+322.5RT (min)

lcorbi\_hilic\_pos\_4\_084\_59

m/z 326.3420

01E+400.5RT (min)

lcorbi\_hilic\_pos\_1\_027\_10

m/z 326.3783

012E+51.52RT (min)

lcorbi\_hilic\_pos\_1\_040\_06

m/z 327.0000

02468E+366.5RT (min)

lcorbi\_hilic\_pos\_4\_078\_58

m/z 327.0454

0177.5RT (min)

lcorbi\_hilic\_pos\_4\_074\_58

m/z 327.0541

01E+466.5RT (min)

lcorbi\_hilic\_pos\_4\_089\_59

m/z 327.1247

01234E+444.5RT (min)

lcorbi\_hilic\_pos\_4\_090\_60

m/z 327.1293

012E+46.57RT (min)

lcorbi\_hilic\_pos\_4\_097\_60

m/z 327.1382

0144.5RT (min)

lcorbi\_hilic\_pos\_4\_082\_59

m/z 327.1408

01E+466.5RT (min)

lcorbi\_hilic\_pos\_3\_036\_33

m/z 327.1592

01E+411.5RT (min)

lcorbi\_hilic\_pos\_1\_053\_08

m/z 327.2320

012E+411.5RT (min)

lcorbi\_hilic\_pos\_4\_078\_58

m/z 327.2531

01E+40.51RT (min)

lcorbi\_hilic\_pos\_1\_044\_17

m/z 327.2645

01E+40.51RT (min)

lcorbi\_hilic\_pos\_4\_096\_60

m/z 327.3372

0246E+30.51RT (min)

lcorbi\_hilic\_pos\_1\_030\_48

m/z 327.3782

01E+422.5RT (min)

lcorbi\_hilic\_pos\_4\_095\_60

m/z 327.9473

01E+466.5RT (min)

lcorbi\_hilic\_pos\_4\_083\_59

m/z 328.0856

0123E+355.5RT (min)

lcorbi\_hilic\_pos\_4\_101\_61

m/z 328.1117

01234E+377.5RT (min)

lcorbi\_hilic\_pos\_4\_082\_59

m/z 328.1379

01E+466.5RT (min)

lcorbi\_hilic\_pos\_3\_038\_15

m/z 328.2049

012E+411.5RT (min)

lcorbi\_hilic\_pos\_3\_019\_43

m/z 328.2332

0123E+411.5RT (min)

lcorbi\_hilic\_pos\_3\_019\_43

m/z 328.2848

02468E+20.51RT (min)

lcorbi\_hilic\_pos\_4\_018\_31

m/z 328.3211

01E+50.51RT (min)

lcorbi\_hilic\_pos\_4\_018\_31

m/z 328.3212

0133.5RT (min)

lcorbi\_hilic\_pos\_3\_037\_20

m/z 329.0610

01E+477.5RT (min)

lcorbi\_hilic\_pos\_4\_082\_59

m/z 329.0923

01E+466.5RT (min)

lcorbi\_hilic\_pos\_4\_020\_37

m/z 329.1491

01234E+411.5RT (min)

lcorbi\_hilic\_pos\_4\_042\_02

m/z 329.2517

024E+333.5RT (min)

lcorbi\_hilic\_pos\_4\_075\_58

m/z 329.9973

01E+455.5RT (min)

lcorbi\_hilic\_pos\_3\_042\_17

m/z 330.1314

012E+45.56RT (min)

lcorbi\_hilic\_pos\_1\_048\_02

m/z 330.1509

012E+466.5RT (min)

lcorbi\_hilic\_pos\_3\_010\_32

m/z 330.1702

0123E+44.55RT (min)

lcorbi\_hilic\_pos\_4\_070\_57

m/z 330.1913

012E+34.55RT (min)

lcorbi\_hilic\_pos\_4\_072\_57

m/z 330.2277

024E+31.52RT (min)

lcorbi\_hilic\_pos\_4\_006\_14

m/z 330.2277

0144.5RT (min)

lcorbi\_hilic\_pos\_4\_013\_23

m/z 330.2464

0246E+33.54RT (min)

lcorbi\_hilic\_pos\_4\_075\_58

m/z 330.2470

02468E+311.5RT (min)

lcorbi\_hilic\_pos\_4\_082\_59

m/z 330.2640

012E+433.5RT (min)

lcorbi\_hilic\_pos\_4\_081\_58

m/z 330.3369

01234E+52.53RT (min)

lcorbi\_hilic\_pos\_3\_037\_20

m/z 330.6719

01E+44.55RT (min)

lcorbi\_hilic\_pos\_4\_073\_57

m/z 331.0603

012E+46.57RT (min)

lcorbi\_hilic\_pos\_1\_054\_23

m/z 331.0766

01E+46.57RT (min)

lcorbi\_hilic\_pos\_4\_083\_59

m/z 331.0992

012E+477.5RT (min)

lcorbi\_hilic\_pos\_4\_007\_05

m/z 331.1405

01E+47.58RT (min)

lcorbi\_hilic\_pos\_4\_090\_60

m/z 331.1479

024E+366.5RT (min)

lcorbi\_hilic\_pos\_4\_010\_47

m/z 331.2229

012E+44.55RT (min)

lcorbi\_hilic\_pos\_4\_096\_60

m/z 331.2268

0123E+30.51RT (min)

lcorbi\_hilic\_pos\_1\_023\_28

m/z 331.2481

0123E+411.5RT (min)

lcorbi\_hilic\_pos\_4\_077\_58

m/z 331.2593

0111.5RT (min)

lcorbi\_hilic\_pos\_2\_012\_37

m/z 331.2593

012E+444.5RT (min)

lcorbi\_hilic\_pos\_4\_012\_29

m/z 331.3402

02468E+42.53RT (min)

lcorbi\_hilic\_pos\_3\_037\_20

m/z 331.9685

0246E+355.5RT (min)

lcorbi\_hilic\_pos\_4\_098\_61

m/z 331.9865

0246E+355.5RT (min)

lcorbi\_hilic\_pos\_1\_054\_23

m/z 332.0690

016.57RT (min)

lcorbi\_hilic\_pos\_4\_092\_60

m/z 332.0954

0177.5RT (min)

lcorbi\_hilic\_pos\_4\_092\_60

m/z 332.1431

02468E+344.5RT (min)

lcorbi\_hilic\_pos\_2\_036\_08

m/z 332.2070

024E+311.5RT (min)

lcorbi\_hilic\_pos\_1\_048\_02

m/z 332.2969

01E+322.5RT (min)

lcorbi\_hilic\_pos\_4\_066\_57

m/z 332.3069

0111.5RT (min)

lcorbi\_hilic\_pos\_4\_031\_17

m/z 332.3161

0122.5RT (min)

lcorbi\_hilic\_pos\_4\_074\_58

m/z 332.3313

01234E+433.5RT (min)

lcorbi\_hilic\_pos\_4\_086\_59

m/z 332.3934

01E+42.53RT (min)

lcorbi\_hilic\_pos\_4\_091\_60

m/z 332.9165

02468E+388.5RT (min)

lcorbi\_hilic\_pos\_4\_107\_62

m/z 333.0283

01234E+477.5RT (min)

lcorbi\_hilic\_pos\_2\_020\_31

m/z 333.1447

012E+311.5RT (min)

lcorbi\_hilic\_pos\_4\_020\_37

m/z 333.1632

01E+411.5RT (min)

lcorbi\_hilic\_pos\_1\_042\_40

m/z 333.2062

02468E+311.5RT (min)

lcorbi\_hilic\_pos\_4\_098\_61

m/z 333.2468

02468E+33.54RT (min)

lcorbi\_hilic\_pos\_4\_066\_57

m/z 333.2750

01E+43.54RT (min)

lcorbi\_hilic\_pos\_1\_048\_02

m/z 333.2790

01234E+411.5RT (min)

lcorbi\_hilic\_pos\_3\_034\_26

m/z 333.3348

01E+50.51RT (min)

lcorbi\_hilic\_pos\_4\_106\_62

m/z 333.7061

02468E+36.57RT (min)

lcorbi\_hilic\_pos\_4\_074\_58

m/z 333.9143

01E+488.5RT (min)

lcorbi\_hilic\_pos\_4\_068\_57

m/z 334.0735

012E+477.5RT (min)

lcorbi\_hilic\_pos\_4\_097\_60

m/z 334.0883

0123E+444.5RT (min)

lcorbi\_hilic\_pos\_1\_037\_18

m/z 334.0987

01E+46.57RT (min)

lcorbi\_hilic\_pos\_4\_087\_59

m/z 334.1597

012E+44.55RT (min)

lcorbi\_hilic\_pos\_4\_109\_62

m/z 334.1699

0246E+34.55RT (min)

lcorbi\_hilic\_pos\_4\_109\_62

m/z 334.2460

0123E+40.51RT (min)

lcorbi\_hilic\_pos\_3\_029\_23

m/z 334.2760

01E+42.53RT (min)

lcorbi\_hilic\_pos\_1\_034\_37

m/z 334.2844

01E+411.5RT (min)

lcorbi\_hilic\_pos\_4\_024\_24

m/z 334.2953

01E+42.53RT (min)

lcorbi\_hilic\_pos\_1\_034\_37

m/z 334.3226

012E+444.5RT (min)

lcorbi\_hilic\_pos\_4\_005\_34

m/z 334.6902

02468E+366.5RT (min)

lcorbi\_hilic\_pos\_4\_080\_58

m/z 335.0116

01E+47.58RT (min)

lcorbi\_hilic\_pos\_4\_084\_59

m/z 335.0641

024E+37.58RT (min)

lcorbi\_hilic\_pos\_4\_087\_59

m/z 335.0738

01E+46.57RT (min)

lcorbi\_hilic\_pos\_1\_043\_25

m/z 335.0831

01E+444.5RT (min)

lcorbi\_hilic\_pos\_1\_037\_18

m/z 335.1603

01E+411.5RT (min)

lcorbi\_hilic\_pos\_4\_033\_53

m/z 335.1631

0123E+34.55RT (min)

lcorbi\_hilic\_pos\_4\_107\_62

m/z 335.1735

01E+444.5RT (min)

lcorbi\_hilic\_pos\_3\_045\_06

m/z 335.2219

012E+40.51RT (min)

lcorbi\_hilic\_pos\_2\_052\_10

m/z 335.9097

024E+388.5RT (min)

lcorbi\_hilic\_pos\_4\_106\_62

m/z 336.0869

0123E+411.5RT (min)

lcorbi\_hilic\_pos\_2\_044\_26

m/z 336.1155

0123E+444.5RT (min)

lcorbi\_hilic\_pos\_4\_013\_23

m/z 336.1160

024E+477.5RT (min)

lcorbi\_hilic\_pos\_4\_093\_60

m/z 336.1232

0246E+322.5RT (min)

lcorbi\_hilic\_pos\_4\_100\_61

m/z 336.2536

024E+31.52RT (min)

lcorbi\_hilic\_pos\_4\_090\_60

m/z 336.2828

012E+40.51RT (min)

lcorbi\_hilic\_pos\_4\_086\_59

m/z 336.8944

02468E+377.5RT (min)

lcorbi\_hilic\_pos\_1\_003\_24

m/z 337.0148

01E+466.5RT (min)

lcorbi\_hilic\_pos\_4\_077\_58

m/z 337.0417

0123E+477.5RT (min)

lcorbi\_hilic\_pos\_1\_040\_06

m/z 337.0514

01E+44.55RT (min)

lcorbi\_hilic\_pos\_4\_078\_58

m/z 337.0878

012E+466.5RT (min)

lcorbi\_hilic\_pos\_4\_091\_60

m/z 337.1427

01234E+34.55RT (min)

lcorbi\_hilic\_pos\_4\_078\_58

m/z 337.1599

01E+444.5RT (min)

lcorbi\_hilic\_pos\_4\_022\_08

m/z 337.1721

01E+466.5RT (min)

lcorbi\_hilic\_pos\_1\_021\_46

m/z 337.1913

01E+422.5RT (min)

lcorbi\_hilic\_pos\_4\_102\_61

m/z 337.2739

0246E+444.5RT (min)

lcorbi\_hilic\_pos\_3\_027\_05

m/z 337.2739

0246E+30.51RT (min)

lcorbi\_hilic\_pos\_4\_083\_59

m/z 337.2835

010.51RT (min)

lcorbi\_hilic\_pos\_4\_088\_59

m/z 337.8524

024E+37.58RT (min)

lcorbi\_hilic\_pos\_4\_099\_61

m/z 338.0509

0123E+36.57RT (min)

lcorbi\_hilic\_pos\_4\_077\_58

m/z 338.0962

01E+44.55RT (min)

lcorbi\_hilic\_pos\_1\_021\_46

m/z 338.1141

01E+466.5RT (min)

lcorbi\_hilic\_pos\_1\_021\_46

m/z 338.1560

02468E+37.58RT (min)

lcorbi\_hilic\_pos\_4\_068\_57

m/z 338.1789

0246E+31.52RT (min)

lcorbi\_hilic\_pos\_4\_100\_61

m/z 338.2198

01E+411.5RT (min)

lcorbi\_hilic\_pos\_4\_113\_62

m/z 338.2328

01E+433.5RT (min)

lcorbi\_hilic\_pos\_4\_078\_58

m/z 338.3055

0246E+30.51RT (min)

lcorbi\_hilic\_pos\_1\_054\_23

m/z 338.3419

024E+444.5RT (min)

lcorbi\_hilic\_pos\_4\_083\_59

m/z 338.3650

0123E+444.5RT (min)

lcorbi\_hilic\_pos\_3\_036\_33

m/z 338.8729

0123E+388.5RT (min)

lcorbi\_hilic\_pos\_4\_098\_61

m/z 338.9021

02468E+377.5RT (min)

lcorbi\_hilic\_pos\_4\_083\_59

m/z 339.0484

01E+44.55RT (min)

lcorbi\_hilic\_pos\_4\_078\_58

m/z 339.0871

0123E+46.57RT (min)

lcorbi\_hilic\_pos\_1\_003\_24

m/z 339.1034

024E+467RT (min)

lcorbi\_hilic\_pos\_4\_096\_60

m/z 339.1867

0111.5RT (min)

lcorbi\_hilic\_pos\_1\_053\_08

m/z 339.1940

0144.5RT (min)

lcorbi\_hilic\_pos\_4\_004\_38

m/z 339.2645

01E+411.5RT (min)

lcorbi\_hilic\_pos\_1\_044\_17

m/z 339.2757

02468E+344.5RT (min)

lcorbi\_hilic\_pos\_4\_088\_59

m/z 339.3453

024E+344.5RT (min)

lcorbi\_hilic\_pos\_4\_044\_25

m/z 339.9086

024E+377.5RT (min)

lcorbi\_hilic\_pos\_4\_082\_59

m/z 340.0048

024E+377.5RT (min)

lcorbi\_hilic\_pos\_1\_051\_26

m/z 340.0876

01E+433.5RT (min)

lcorbi\_hilic\_pos\_1\_037\_18

m/z 340.1019

0144.5RT (min)

lcorbi\_hilic\_pos\_4\_098\_61

m/z 340.1182

01E+422.5RT (min)

lcorbi\_hilic\_pos\_4\_105\_61

m/z 340.1505

01E+45.56RT (min)

lcorbi\_hilic\_pos\_4\_086\_59

m/z 340.1739

0123E+444.5RT (min)

lcorbi\_hilic\_pos\_1\_012\_19

m/z 340.2481

0123E+422.5RT (min)

lcorbi\_hilic\_pos\_2\_005\_23

m/z 340.2718

01E+322.5RT (min)

lcorbi\_hilic\_pos\_4\_084\_59

m/z 340.2789

01E+422.5RT (min)

lcorbi\_hilic\_pos\_2\_005\_23

m/z 340.3213

01E+40.51RT (min)

lcorbi\_hilic\_pos\_3\_019\_43

m/z 340.3405

010.51RT (min)

lcorbi\_hilic\_pos\_3\_019\_43

m/z 340.3486

01E+50.51RT (min)

lcorbi\_hilic\_pos\_3\_019\_43

m/z 340.3576

0246E+40.51RT (min)

lcorbi\_hilic\_pos\_3\_019\_43

m/z 340.7613

01E+444.5RT (min)

lcorbi\_hilic\_pos\_2\_023\_34

m/z 341.0451

02468E+34.55RT (min)

lcorbi\_hilic\_pos\_4\_068\_57

m/z 341.1232

02468E+32.53RT (min)

lcorbi\_hilic\_pos\_4\_110\_62

m/z 341.1242

01E+466.5RT (min)

lcorbi\_hilic\_pos\_1\_021\_46

m/z 341.1323

01234E+344.5RT (min)

lcorbi\_hilic\_pos\_3\_045\_06

m/z 341.1862

01E+444.5RT (min)

lcorbi\_hilic\_pos\_3\_021\_46

m/z 341.3529

01234E+40.51RT (min)

lcorbi\_hilic\_pos\_1\_054\_23

m/z 341.3529

01234E+422.5RT (min)

lcorbi\_hilic\_pos\_4\_084\_59

m/z 341.9973

01E+455.5RT (min)

lcorbi\_hilic\_pos\_4\_098\_61

m/z 342.0154

012E+455.5RT (min)

lcorbi\_hilic\_pos\_1\_054\_23

m/z 342.0352

024E+37.58RT (min)

lcorbi\_hilic\_pos\_1\_010\_07

m/z 342.0650

0123E+36.57RT (min)

lcorbi\_hilic\_pos\_4\_084\_59

m/z 342.1397

01E+366.5RT (min)

lcorbi\_hilic\_pos\_2\_049\_55

m/z 342.1538

012E+377.5RT (min)

lcorbi\_hilic\_pos\_3\_009\_50

m/z 342.1563

02468E+366.5RT (min)

lcorbi\_hilic\_pos\_4\_020\_37

m/z 342.1610

02468E+311.5RT (min)

lcorbi\_hilic\_pos\_4\_103\_61

m/z 342.1913

0246E+40.51RT (min)

lcorbi\_hilic\_pos\_2\_001\_49

m/z 342.3369

0123E+422.5RT (min)

lcorbi\_hilic\_pos\_2\_030\_05

m/z 342.3562

01E+422.5RT (min)

lcorbi\_hilic\_pos\_4\_087\_59

m/z 343.0058

024E+34.55RT (min)

lcorbi\_hilic\_pos\_4\_066\_57

m/z 343.0403

0123E+37.58RT (min)

lcorbi\_hilic\_pos\_4\_084\_59

m/z 343.0766

02468E+36.57RT (min)

lcorbi\_hilic\_pos\_2\_005\_23

m/z 343.1309

012E+466.5RT (min)

lcorbi\_hilic\_pos\_3\_021\_46

m/z 343.1979

01E+455.5RT (min)

lcorbi\_hilic\_pos\_1\_021\_46

m/z 343.2094

01E+411.5RT (min)

lcorbi\_hilic\_pos\_4\_082\_59

m/z 343.2269

012E+30.51RT (min)

lcorbi\_hilic\_pos\_4\_084\_59

m/z 343.2674

0123E+433.5RT (min)

lcorbi\_hilic\_pos\_4\_068\_57

m/z 343.2859

01E+444.5RT (min)

lcorbi\_hilic\_pos\_4\_068\_57

m/z 343.7879

012E+43.544.5RT (min)

lcorbi\_hilic\_pos\_4\_003\_26

m/z 343.9221

0123E+377.5RT (min)

lcorbi\_hilic\_pos\_2\_010\_07

m/z 344.0416

01E+44.55RT (min)

lcorbi\_hilic\_pos\_4\_068\_57

m/z 344.1042

01E+47.58RT (min)

lcorbi\_hilic\_pos\_1\_050\_05

m/z 344.1320

0166.5RT (min)

lcorbi\_hilic\_pos\_4\_097\_60

m/z 344.1455

0144.5RT (min)

lcorbi\_hilic\_pos\_4\_020\_37

m/z 344.1576

0246E+30.51RT (min)

lcorbi\_hilic\_pos\_1\_047\_47

m/z 344.2546

012E+42.53RT (min)

lcorbi\_hilic\_pos\_4\_088\_59

m/z 344.2620

024E+344.5RT (min)

lcorbi\_hilic\_pos\_1\_020\_13

m/z 344.3102

01E+411.5RT (min)

lcorbi\_hilic\_pos\_3\_029\_23

m/z 344.3161

01E+444.5RT (min)

lcorbi\_hilic\_pos\_1\_054\_23

m/z 344.3355

01E+433.5RT (min)

lcorbi\_hilic\_pos\_4\_087\_59

m/z 344.8757

01E+499.5RT (min)

lcorbi\_hilic\_pos\_4\_076\_58

m/z 345.0029

012E+34.55RT (min)

lcorbi\_hilic\_pos\_4\_083\_59

m/z 345.0760

01234E+36.57RT (min)

lcorbi\_hilic\_pos\_4\_086\_59

m/z 345.0873

01E+31.52RT (min)

lcorbi\_hilic\_pos\_4\_113\_62

m/z 345.1357

01E+46.57RT (min)

lcorbi\_hilic\_pos\_4\_090\_60

m/z 345.1714

01E+50.51RT (min)

lcorbi\_hilic\_pos\_3\_011\_34

m/z 345.2314

0123E+411.5RT (min)

lcorbi\_hilic\_pos\_3\_019\_43

m/z 345.2386

01E+311.5RT (min)

lcorbi\_hilic\_pos\_1\_053\_08

m/z 345.2640

01E+40.51RT (min)

lcorbi\_hilic\_pos\_3\_054\_16

m/z 345.2830

01234E+433.5RT (min)

lcorbi\_hilic\_pos\_4\_072\_57

m/z 346.0097

012E+56.57RT (min)

lcorbi\_hilic\_pos\_1\_056\_56

m/z 346.0256

01E+355.5RT (min)

lcorbi\_hilic\_pos\_4\_087\_59

m/z 346.0472

016.57RT (min)

lcorbi\_hilic\_pos\_4\_074\_58

m/z 346.1010

012E+47.58RT (min)

lcorbi\_hilic\_pos\_1\_003\_24

m/z 346.1797

01E+411.5RT (min)

lcorbi\_hilic\_pos\_2\_001\_49

m/z 347.0254

0177.5RT (min)

lcorbi\_hilic\_pos\_4\_074\_58

m/z 347.0318

0246E+35.56RT (min)

lcorbi\_hilic\_pos\_4\_093\_60

m/z 347.0803

01E+466.5RT (min)

lcorbi\_hilic\_pos\_4\_088\_59

m/z 347.1067

024E+34.55RT (min)

lcorbi\_hilic\_pos\_4\_071\_57

m/z 347.1434

01E+46.57RT (min)

lcorbi\_hilic\_pos\_4\_096\_60

m/z 347.2907

01E+43.54RT (min)

lcorbi\_hilic\_pos\_3\_008\_14

m/z 347.2988

0123E+32.53RT (min)

lcorbi\_hilic\_pos\_3\_040\_45

m/z 348.0141

0246E+366.5RT (min)

lcorbi\_hilic\_pos\_4\_079\_58

m/z 348.0352

01E+45.56RT (min)

lcorbi\_hilic\_pos\_4\_093\_60

m/z 348.1477

02468E+466.5RT (min)

lcorbi\_hilic\_pos\_4\_092\_60

m/z 348.1897

011.52RT (min)

lcorbi\_hilic\_pos\_4\_082\_59

m/z 348.2019

0246E+30.51RT (min)

lcorbi\_hilic\_pos\_1\_012\_19

m/z 348.3609

012E+411.5RT (min)

lcorbi\_hilic\_pos\_4\_090\_60

m/z 349.0053

012E+47.58RT (min)

lcorbi\_hilic\_pos\_4\_076\_58

m/z 349.0079

012E+466.5RT (min)

lcorbi\_hilic\_pos\_4\_080\_58

m/z 349.0298

017.58RT (min)

lcorbi\_hilic\_pos\_4\_079\_58

m/z 349.0342

01E+37.58RT (min)

lcorbi\_hilic\_pos\_4\_081\_58

m/z 349.0740

01E+477.5RT (min)

lcorbi\_hilic\_pos\_4\_020\_37

m/z 349.1109

01E+466.5RT (min)

lcorbi\_hilic\_pos\_4\_092\_60

m/z 349.2318

0111.5RT (min)

lcorbi\_hilic\_pos\_4\_100\_61

m/z 349.2950

012E+40.51RT (min)

lcorbi\_hilic\_pos\_4\_088\_59

m/z 349.3508

02468E+40.51RT (min)

lcorbi\_hilic\_pos\_1\_032\_31

m/z 349.3580

012E+41.52RT (min)

lcorbi\_hilic\_pos\_4\_092\_60

m/z 350.0039

012E+47.58RT (min)

lcorbi\_hilic\_pos\_4\_078\_58

m/z 350.0274

017.58RT (min)

lcorbi\_hilic\_pos\_4\_081\_58

m/z 350.0321

017.58RT (min)

lcorbi\_hilic\_pos\_4\_081\_58

m/z 350.0849

02468E+355.5RT (min)

lcorbi\_hilic\_pos\_4\_071\_57

m/z 350.0897

0123E+444.5RT (min)

lcorbi\_hilic\_pos\_1\_054\_23

m/z 350.2175

01E+411.5RT (min)

lcorbi\_hilic\_pos\_1\_048\_02

m/z 350.2343

0246E+37.58RT (min)

lcorbi\_hilic\_pos\_1\_034\_37

m/z 350.3613

024E+31.52RT (min)

lcorbi\_hilic\_pos\_4\_095\_60

m/z 351.0313

01234E+366.5RT (min)

lcorbi\_hilic\_pos\_4\_088\_59

m/z 351.0430

01E+477.5RT (min)

lcorbi\_hilic\_pos\_4\_083\_59

m/z 351.9913

0246E+35.56RT (min)

lcorbi\_hilic\_pos\_4\_081\_58

m/z 352.0454

012E+455.5RT (min)

lcorbi\_hilic\_pos\_4\_106\_62

m/z 352.1069

01E+46.57RT (min)

lcorbi\_hilic\_pos\_2\_024\_32

m/z 352.1658

01E+444.5RT (min)

lcorbi\_hilic\_pos\_4\_066\_57

m/z 352.1871

0111.5RT (min)

lcorbi\_hilic\_pos\_1\_037\_18

m/z 352.2848

010.51RT (min)

lcorbi\_hilic\_pos\_3\_035\_22

m/z 352.2849

024E+322.5RT (min)

lcorbi\_hilic\_pos\_4\_038\_04

m/z 352.8978

01E+45.56RT (min)

lcorbi\_hilic\_pos\_4\_109\_62

m/z 353.0178

01E+37.58RT (min)

lcorbi\_hilic\_pos\_4\_076\_58

m/z 353.0458

01E+45.56RT (min)

lcorbi\_hilic\_pos\_1\_032\_31

m/z 353.0586

02468E+36.57RT (min)

lcorbi\_hilic\_pos\_4\_083\_59

m/z 353.1941

01E+34.55RT (min)

lcorbi\_hilic\_pos\_4\_020\_37

m/z 353.2366

01E+43.54RT (min)

lcorbi\_hilic\_pos\_2\_036\_08

m/z 353.2688

010.51RT (min)

lcorbi\_hilic\_pos\_4\_094\_60

m/z 353.2801

0123E+411.5RT (min)

lcorbi\_hilic\_pos\_3\_042\_17

m/z 353.3097

0246E+40.51RT (min)

lcorbi\_hilic\_pos\_4\_090\_60

m/z 353.3247

024E+30.51RT (min)

lcorbi\_hilic\_pos\_4\_094\_60

m/z 353.3528

01E+422.5RT (min)

lcorbi\_hilic\_pos\_4\_088\_59

m/z 353.9243

024E+36.57RT (min)

lcorbi\_hilic\_pos\_4\_069\_57

m/z 354.0123

01E+47.58RT (min)

lcorbi\_hilic\_pos\_4\_081\_58

m/z 354.0702

02468E+34.55RT (min)

lcorbi\_hilic\_pos\_1\_021\_46

m/z 354.0926

01E+411.5RT (min)

lcorbi\_hilic\_pos\_1\_042\_40

m/z 354.1662

012E+45.56RT (min)

lcorbi\_hilic\_pos\_4\_088\_59

m/z 354.2389

02468E+31.52RT (min)

lcorbi\_hilic\_pos\_2\_005\_23

m/z 354.3369

0246E+30.51RT (min)

lcorbi\_hilic\_pos\_3\_029\_23

m/z 355.0871

0123E+511.5RT (min)

lcorbi\_hilic\_pos\_2\_044\_26

m/z 355.1066

01E+444.5RT (min)

lcorbi\_hilic\_pos\_3\_045\_06

m/z 355.1308

012E+43.54RT (min)

lcorbi\_hilic\_pos\_4\_068\_57

m/z 355.1493

012E+44.55RT (min)

lcorbi\_hilic\_pos\_4\_112\_62

m/z 355.2170

01E+311.5RT (min)

lcorbi\_hilic\_pos\_4\_008\_19

m/z 355.2342

0246E+377.5RT (min)

lcorbi\_hilic\_pos\_4\_085\_59

m/z 355.2822

0246E+30.51RT (min)

lcorbi\_hilic\_pos\_1\_054\_23

m/z 355.3493

01E+433.5RT (min)

lcorbi\_hilic\_pos\_4\_013\_23

m/z 356.0105

012E+377.5RT (min)

lcorbi\_hilic\_pos\_1\_054\_23

m/z 356.0419

0246E+36.57RT (min)

lcorbi\_hilic\_pos\_4\_073\_57

m/z 356.0556

01E+477.5RT (min)

lcorbi\_hilic\_pos\_4\_097\_60

m/z 356.0807

0246E+36.57RT (min)

lcorbi\_hilic\_pos\_4\_089\_59

m/z 356.1417

024E+34.55RT (min)

lcorbi\_hilic\_pos\_4\_112\_62

m/z 356.2546

01E+411.5RT (min)

lcorbi\_hilic\_pos\_1\_032\_31

m/z 356.2951

010.51RT (min)

lcorbi\_hilic\_pos\_4\_082\_59

m/z 356.3526

01E+52.53RT (min)

lcorbi\_hilic\_pos\_3\_037\_20

m/z 357.0086

012E+47.58RT (min)

lcorbi\_hilic\_pos\_1\_001\_49

m/z 357.1008

012E+477.5RT (min)

lcorbi\_hilic\_pos\_4\_091\_60

m/z 357.2499

01E+577.5RT (min)

lcorbi\_hilic\_pos\_4\_020\_37

m/z 357.2831

01E+41.52RT (min)

lcorbi\_hilic\_pos\_4\_101\_61

m/z 358.0115

0246E+35.56RT (min)

lcorbi\_hilic\_pos\_4\_107\_62

m/z 358.0214

024E+366.5RT (min)

lcorbi\_hilic\_pos\_4\_081\_58

m/z 358.0472

01E+46.57RT (min)

lcorbi\_hilic\_pos\_1\_054\_23

m/z 358.2074

01E+41.52RT (min)

lcorbi\_hilic\_pos\_1\_034\_37

m/z 358.2108

024E+34.55RT (min)

lcorbi\_hilic\_pos\_1\_049\_55

m/z 358.2338

0123E+433.5RT (min)

lcorbi\_hilic\_pos\_3\_029\_23

m/z 358.2591

02468E+33.54RT (min)

lcorbi\_hilic\_pos\_4\_072\_57

m/z 358.2744

012E+30.51RT (min)

lcorbi\_hilic\_pos\_4\_026\_11

m/z 358.2777

013.54RT (min)

lcorbi\_hilic\_pos\_4\_107\_62

m/z 358.2897

01E+455.5RT (min)

lcorbi\_hilic\_pos\_4\_088\_59

m/z 358.2954

01E+433.5RT (min)

lcorbi\_hilic\_pos\_4\_067\_57

m/z 358.3683

0123E+422.5RT (min)

lcorbi\_hilic\_pos\_3\_037\_20

m/z 358.8349

0246E+355.5RT (min)

lcorbi\_hilic\_pos\_4\_082\_59

m/z 358.8764

01234E+377.5RT (min)

lcorbi\_hilic\_pos\_1\_003\_24

m/z 358.9968

01E+466.5RT (min)

lcorbi\_hilic\_pos\_4\_079\_58

m/z 359.0074

01E+47.58RT (min)

lcorbi\_hilic\_pos\_1\_001\_49

m/z 359.0237

01E+477.5RT (min)

lcorbi\_hilic\_pos\_1\_030\_48

m/z 359.0699

01E+46.577.5RT (min)

lcorbi\_hilic\_pos\_4\_091\_60

m/z 359.0973

012E+311.5RT (min)

lcorbi\_hilic\_pos\_4\_108\_62

m/z 359.1248

01234E+366.5RT (min)

lcorbi\_hilic\_pos\_1\_003\_24

m/z 359.1524

01E+43.54RT (min)

lcorbi\_hilic\_pos\_4\_109\_62

m/z 359.2292

0246E+355.5RT (min)

lcorbi\_hilic\_pos\_4\_087\_59

m/z 359.2542

01E+411.5RT (min)

lcorbi\_hilic\_pos\_1\_036\_32

m/z 359.2737

012E+44.55RT (min)

lcorbi\_hilic\_pos\_4\_083\_59

m/z 359.3352

01E+411.5RT (min)

lcorbi\_hilic\_pos\_4\_088\_59

m/z 359.3522

024E+40.51RT (min)

lcorbi\_hilic\_pos\_1\_032\_31

m/z 359.8774

01234E+388.5RT (min)

lcorbi\_hilic\_pos\_4\_073\_57

m/z 360.0757

01E+46.57RT (min)

lcorbi\_hilic\_pos\_4\_084\_59

m/z 360.0951

02468E+36.57RT (min)

lcorbi\_hilic\_pos\_4\_066\_57

m/z 360.1080

01E+411.5RT (min)

lcorbi\_hilic\_pos\_2\_007\_28

m/z 360.1164

0246E+344.5RT (min)

lcorbi\_hilic\_pos\_1\_016\_33

m/z 360.1504

0123E+46.57RT (min)

lcorbi\_hilic\_pos\_4\_068\_57

m/z 360.1571

01E+311.5RT (min)

lcorbi\_hilic\_pos\_4\_020\_37

m/z 360.1808

012E+40.51RT (min)

lcorbi\_hilic\_pos\_4\_086\_59

m/z 360.2244

0246E+377.5RT (min)

lcorbi\_hilic\_pos\_4\_083\_59

m/z 360.2678

02468E+37.58RT (min)

lcorbi\_hilic\_pos\_1\_003\_24

m/z 360.2748

01E+43.54RT (min)

lcorbi\_hilic\_pos\_4\_068\_57

m/z 360.3628

0123E+400.51RT (min)

lcorbi\_hilic\_pos\_4\_079\_58

m/z 360.8841

01234E+377.5RT (min)

lcorbi\_hilic\_pos\_2\_010\_07

m/z 360.9277

01234E+35.56RT (min)

lcorbi\_hilic\_pos\_1\_020\_13

m/z 361.1391

02468E+32.53RT (min)

lcorbi\_hilic\_pos\_4\_111\_62

m/z 361.2352

012E+41.52RT (min)

lcorbi\_hilic\_pos\_4\_101\_61

m/z 361.2482

02468E+311.5RT (min)

lcorbi\_hilic\_pos\_4\_076\_58

m/z 361.2740

012E+444.5RT (min)

lcorbi\_hilic\_pos\_4\_075\_58

m/z 361.3043

012E+311.5RT (min)

lcorbi\_hilic\_pos\_1\_023\_28

m/z 361.3733

012E+411.5RT (min)

lcorbi\_hilic\_pos\_4\_092\_60

m/z 361.8731

0246E+388.5RT (min)

lcorbi\_hilic\_pos\_4\_067\_57

m/z 362.0631

0123E+36.57RT (min)

lcorbi\_hilic\_pos\_4\_090\_60

m/z 362.2207

01E+311.5RT (min)

lcorbi\_hilic\_pos\_2\_002\_11

m/z 362.2773

0246E+344.5RT (min)

lcorbi\_hilic\_pos\_4\_075\_58

m/z 362.2903

0123E+433.5RT (min)

lcorbi\_hilic\_pos\_1\_048\_02

m/z 362.8863

01E+499.5RT (min)

lcorbi\_hilic\_pos\_4\_080\_58

m/z 363.0235

0123E+35.56RT (min)

lcorbi\_hilic\_pos\_4\_077\_58

m/z 363.0286

01E+366.5RT (min)

lcorbi\_hilic\_pos\_4\_074\_58

m/z 363.0372

012E+477.5RT (min)

lcorbi\_hilic\_pos\_2\_024\_32

m/z 363.0890

01E+433.5RT (min)

lcorbi\_hilic\_pos\_2\_042\_02

m/z 363.1179

0246E+344.5RT (min)

lcorbi\_hilic\_pos\_4\_055\_06

m/z 363.2203

02468E+311.5RT (min)

lcorbi\_hilic\_pos\_3\_040\_45

m/z 363.2451

01E+422.5RT (min)

lcorbi\_hilic\_pos\_1\_032\_31

m/z 363.2644

02468E+311.5RT (min)

lcorbi\_hilic\_pos\_4\_080\_58

m/z 363.3802

01E+411.5RT (min)

lcorbi\_hilic\_pos\_4\_024\_24

m/z 364.0553

0123E+46.57RT (min)

lcorbi\_hilic\_pos\_4\_083\_59

m/z 364.0827

01E+45.56RT (min)

lcorbi\_hilic\_pos\_1\_036\_32

m/z 364.1085

012E+46.57RT (min)

lcorbi\_hilic\_pos\_1\_010\_07

m/z 364.9311

01E+45.56RT (min)

lcorbi\_hilic\_pos\_4\_071\_57

m/z 364.9786

01234E+355.5RT (min)

lcorbi\_hilic\_pos\_4\_071\_57

m/z 364.9846

012E+40.51RT (min)

lcorbi\_hilic\_pos\_4\_068\_57

m/z 365.0290

01E+46.57RT (min)

lcorbi\_hilic\_pos\_1\_044\_17

m/z 365.0926

01E+36.57RT (min)

lcorbi\_hilic\_pos\_4\_112\_62

m/z 365.1710

0123E+411.5RT (min)

lcorbi\_hilic\_pos\_1\_029\_14

m/z 365.2074

012E+44.55RT (min)

lcorbi\_hilic\_pos\_4\_107\_62

m/z 365.3165

0246E+333.5RT (min)

lcorbi\_hilic\_pos\_2\_012\_37

m/z 365.3956

01E+444.5RT (min)

lcorbi\_hilic\_pos\_1\_036\_32

m/z 365.9964

01234E+366.5RT (min)

lcorbi\_hilic\_pos\_4\_079\_58

m/z 366.2641

01E+433.5RT (min)

lcorbi\_hilic\_pos\_4\_068\_57

m/z 366.2812

02468E+33.54RT (min)

lcorbi\_hilic\_pos\_4\_018\_31

m/z 366.2925

0111.5RT (min)

lcorbi\_hilic\_pos\_4\_013\_23

m/z 366.3217

012E+40.51RT (min)

lcorbi\_hilic\_pos\_4\_089\_59

m/z 366.3370

01234E+41.52RT (min)

lcorbi\_hilic\_pos\_2\_030\_05

m/z 366.3489

0246E+40.51RT (min)

lcorbi\_hilic\_pos\_4\_085\_59

m/z 367.0419

0123E+366.5RT (min)

lcorbi\_hilic\_pos\_1\_036\_32

m/z 367.1103

0123E+366.5RT (min)

lcorbi\_hilic\_pos\_4\_110\_62

m/z 367.1177

01234E+36.57RT (min)

lcorbi\_hilic\_pos\_4\_097\_60

m/z 367.2706

012E+311.5RT (min)

lcorbi\_hilic\_pos\_4\_018\_31

m/z 367.3686

0123E+422.5RT (min)

lcorbi\_hilic\_pos\_4\_088\_59

m/z 368.0194

0123E+355.5RT (min)

lcorbi\_hilic\_pos\_4\_095\_60

m/z 368.0293

016.57RT (min)

lcorbi\_hilic\_pos\_4\_074\_58

m/z 368.0768

0111.5RT (min)

lcorbi\_hilic\_pos\_4\_020\_37

m/z 368.2219

02468E+344.5RT (min)

lcorbi\_hilic\_pos\_4\_022\_08

m/z 368.2434

01E+411.5RT (min)

lcorbi\_hilic\_pos\_4\_098\_61

m/z 368.3162

010.51RT (min)

lcorbi\_hilic\_pos\_4\_018\_31

m/z 368.3526

01E+41.52RT (min)

lcorbi\_hilic\_pos\_2\_030\_05

m/z 368.4254

01E+51.52RT (min)

lcorbi\_hilic\_pos\_3\_035\_22

m/z 368.4255

01E+400.5RT (min)

lcorbi\_hilic\_pos\_1\_001\_49

m/z 368.9949

0246E+366.5RT (min)

lcorbi\_hilic\_pos\_4\_079\_58

m/z 369.0074

02468E+377.5RT (min)

lcorbi\_hilic\_pos\_4\_084\_59

m/z 369.0624

01E+466.5RT (min)

lcorbi\_hilic\_pos\_4\_086\_59

m/z 369.1253

012E+46.57RT (min)

lcorbi\_hilic\_pos\_4\_095\_60

m/z 369.1253

01234E+40.51RT (min)

lcorbi\_hilic\_pos\_1\_032\_31

m/z 369.1449

011.52RT (min)

lcorbi\_hilic\_pos\_4\_103\_61

m/z 369.2500

01E+477.5RT (min)

lcorbi\_hilic\_pos\_4\_020\_37

m/z 369.2830

012E+433.5RT (min)

lcorbi\_hilic\_pos\_4\_073\_57

m/z 369.3843

0123E+422.5RT (min)

lcorbi\_hilic\_pos\_1\_040\_06

m/z 369.4288

012E+41.52RT (min)

lcorbi\_hilic\_pos\_3\_035\_22

m/z 370.0527

0123E+477.5RT (min)

lcorbi\_hilic\_pos\_4\_083\_59

m/z 370.2200

012E+45.56RT (min)

lcorbi\_hilic\_pos\_4\_093\_60

m/z 370.2211

024E+32.53RT (min)

lcorbi\_hilic\_pos\_4\_033\_53

m/z 370.2341

024E+31.52RT (min)

lcorbi\_hilic\_pos\_4\_086\_59

m/z 370.2756

01234E+46.57RT (min)

lcorbi\_hilic\_pos\_4\_092\_60

m/z 370.2777

01E+411.5RT (min)

lcorbi\_hilic\_pos\_4\_020\_37

m/z 370.3684

02468E+322.5RT (min)

lcorbi\_hilic\_pos\_4\_109\_62

m/z 370.9844

0123E+36.57RT (min)

lcorbi\_hilic\_pos\_2\_022\_35

m/z 371.0933

024E+46.57RT (min)

lcorbi\_hilic\_pos\_4\_095\_60

m/z 371.1234

024E+34.55RT (min)

lcorbi\_hilic\_pos\_4\_108\_62

m/z 371.1397

012E+411.5RT (min)

lcorbi\_hilic\_pos\_4\_104\_61

m/z 371.2129

012E+47.58RT (min)

lcorbi\_hilic\_pos\_1\_008\_38

m/z 371.2230

02468E+33.54RT (min)

lcorbi\_hilic\_pos\_4\_111\_62

m/z 371.2279

01E+411.5RT (min)

lcorbi\_hilic\_pos\_3\_019\_43

m/z 371.2989

01234E+30.51RT (min)

lcorbi\_hilic\_pos\_2\_007\_28

m/z 371.3159

0123E+40.51RT (min)

lcorbi\_hilic\_pos\_3\_003\_37

m/z 371.3635

01E+433.5RT (min)

lcorbi\_hilic\_pos\_4\_088\_59

m/z 372.0669

02468E+355.5RT (min)

lcorbi\_hilic\_pos\_4\_066\_57

m/z 372.1136

024E+40.51RT (min)

lcorbi\_hilic\_pos\_2\_044\_26

m/z 372.1382

02468E+366.5RT (min)

lcorbi\_hilic\_pos\_1\_056\_56

m/z 372.1907

01E+477.5RT (min)

lcorbi\_hilic\_pos\_4\_093\_60

m/z 372.2913

01E+477.5RT (min)

lcorbi\_hilic\_pos\_4\_092\_60

m/z 372.2933

01E+411.5RT (min)

lcorbi\_hilic\_pos\_4\_020\_37

m/z 372.3111

012E+411.5RT (min)

lcorbi\_hilic\_pos\_3\_031\_36

m/z 372.3305

024E+40.51RT (min)

lcorbi\_hilic\_pos\_4\_028\_10

m/z 372.3476

012E+41.52RT (min)

lcorbi\_hilic\_pos\_4\_086\_59

m/z 373.0012

01E+47.58RT (min)

lcorbi\_hilic\_pos\_4\_081\_58

m/z 373.0573

01E+466.5RT (min)

lcorbi\_hilic\_pos\_4\_067\_57

m/z 373.0986

01E+30.51RT (min)

lcorbi\_hilic\_pos\_1\_051\_26

m/z 373.1623

01E+422.5RT (min)

lcorbi\_hilic\_pos\_4\_073\_57

m/z 373.1649

01E+50.51RT (min)

lcorbi\_hilic\_pos\_2\_044\_26

m/z 373.1809

012E+46.57RT (min)

lcorbi\_hilic\_pos\_1\_031\_36

m/z 373.2530

012E+433.5RT (min)

lcorbi\_hilic\_pos\_4\_020\_37

m/z 373.3145

0133.5RT (min)

lcorbi\_hilic\_pos\_4\_087\_59

m/z 373.3337

01E+43.54RT (min)

lcorbi\_hilic\_pos\_4\_088\_59

m/z 373.3552

02468E+40.51RT (min)

lcorbi\_hilic\_pos\_3\_055\_24

m/z 373.9484

01234E+35.56RT (min)

lcorbi\_hilic\_pos\_4\_071\_57

m/z 374.0997

0246E+40.51RT (min)

lcorbi\_hilic\_pos\_2\_032\_30

m/z 374.1025

01E+46.57RT (min)

lcorbi\_hilic\_pos\_4\_083\_59

m/z 374.1560

012E+422.5RT (min)

lcorbi\_hilic\_pos\_4\_073\_57

m/z 374.1673

0177.5RT (min)

lcorbi\_hilic\_pos\_1\_048\_02

m/z 374.3268

01E+422.5RT (min)

lcorbi\_hilic\_pos\_1\_040\_06

m/z 374.8089

01E+455.5RT (min)

lcorbi\_hilic\_pos\_4\_075\_58

m/z 374.9929

0123E+37.58RT (min)

lcorbi\_hilic\_pos\_4\_074\_58

m/z 375.0075

0123E+355.5RT (min)

lcorbi\_hilic\_pos\_4\_069\_57

m/z 375.2241

01234E+36.57RT (min)

lcorbi\_hilic\_pos\_4\_082\_59

m/z 375.3222

01E+400.5RT (min)

lcorbi\_hilic\_pos\_4\_068\_57

m/z 375.3895

010.51RT (min)

lcorbi\_hilic\_pos\_4\_090\_60

m/z 375.7636

012E+44.55RT (min)

lcorbi\_hilic\_pos\_4\_075\_58

m/z 375.9942

01E+477.5RT (min)

lcorbi\_hilic\_pos\_4\_081\_58

m/z 376.1280

012E+44.55RT (min)

lcorbi\_hilic\_pos\_4\_106\_62

m/z 376.3003

01E+422.5RT (min)

lcorbi\_hilic\_pos\_4\_048\_46

m/z 376.3446

011.52RT (min)

lcorbi\_hilic\_pos\_3\_020\_31

m/z 376.3789

024E+40.51RT (min)

lcorbi\_hilic\_pos\_2\_020\_31

m/z 376.8070

024E+355.5RT (min)

lcorbi\_hilic\_pos\_4\_080\_58

m/z 376.9924

024E+377.5RT (min)

lcorbi\_hilic\_pos\_2\_027\_38

m/z 377.1461

024E+477.5RT (min)

lcorbi\_hilic\_pos\_4\_091\_60

m/z 377.1830

01E+45.56RT (min)

lcorbi\_hilic\_pos\_4\_091\_60

m/z 377.2332

01E+47.58RT (min)

lcorbi\_hilic\_pos\_1\_009\_50

m/z 377.3202

01E+40.51RT (min)

lcorbi\_hilic\_pos\_4\_086\_59

m/z 377.3894

01E+41.52RT (min)

lcorbi\_hilic\_pos\_4\_090\_60

m/z 377.9924

0246E+377.5RT (min)

lcorbi\_hilic\_pos\_4\_081\_58

m/z 378.0127

0246E+35.56RT (min)

lcorbi\_hilic\_pos\_4\_083\_59

m/z 378.0376

01E+477.5RT (min)

lcorbi\_hilic\_pos\_4\_094\_60

m/z 378.1375

012E+566.5RT (min)

lcorbi\_hilic\_pos\_4\_089\_59

m/z 378.1658

01E+433.5RT (min)

lcorbi\_hilic\_pos\_4\_102\_61

m/z 378.2083

0246E+36.57RT (min)

lcorbi\_hilic\_pos\_4\_074\_58

m/z 378.2157

02468E+322.5RT (min)

lcorbi\_hilic\_pos\_1\_032\_31

m/z 378.2661

012E+30.51RT (min)

lcorbi\_hilic\_pos\_4\_082\_59

m/z 378.3003

01E+40.51RT (min)

lcorbi\_hilic\_pos\_4\_082\_59

m/z 379.0002

012E+40.51RT (min)

lcorbi\_hilic\_pos\_4\_067\_57

m/z 379.1214

01E+455.5RT (min)

lcorbi\_hilic\_pos\_4\_067\_57

m/z 379.2010

01E+46.57RT (min)

lcorbi\_hilic\_pos\_1\_016\_33

m/z 379.2414

0122.5RT (min)

lcorbi\_hilic\_pos\_4\_111\_62

m/z 379.2496

02468E+322.5RT (min)

lcorbi\_hilic\_pos\_1\_014\_27

m/z 379.2683

01E+455.5RT (min)

lcorbi\_hilic\_pos\_4\_082\_59

m/z 379.2833

010.51RT (min)

lcorbi\_hilic\_pos\_4\_082\_59

m/z 379.3055

01E+40.51RT (min)

lcorbi\_hilic\_pos\_4\_088\_59

m/z 380.0177

02468E+36.57RT (min)

lcorbi\_hilic\_pos\_4\_081\_58

m/z 380.1239

0177.5RT (min)

lcorbi\_hilic\_pos\_4\_095\_60

m/z 380.1449

012E+40.51RT (min)

lcorbi\_hilic\_pos\_4\_010\_47

m/z 380.1989

0246E+40.51RT (min)

lcorbi\_hilic\_pos\_4\_076\_58

m/z 380.2564

0246E+34.55RT (min)

lcorbi\_hilic\_pos\_4\_078\_58

m/z 380.8584

0123E+377.5RT (min)

lcorbi\_hilic\_pos\_1\_010\_07

m/z 381.0058

012E+377.5RT (min)

lcorbi\_hilic\_pos\_1\_040\_06

m/z 381.0798

01E+466.5RT (min)

lcorbi\_hilic\_pos\_4\_066\_57

m/z 381.2042

012E+411.5RT (min)

lcorbi\_hilic\_pos\_4\_076\_58

m/z 381.2518

012E+43.54RT (min)

lcorbi\_hilic\_pos\_4\_018\_31

m/z 381.2630

01E+411.5RT (min)

lcorbi\_hilic\_pos\_1\_008\_38

m/z 381.9865

0246E+366.5RT (min)

lcorbi\_hilic\_pos\_4\_077\_58

m/z 382.0577

01E+46.57RT (min)

lcorbi\_hilic\_pos\_4\_084\_59

m/z 382.1193

024E+32.53RT (min)

lcorbi\_hilic\_pos\_4\_113\_62

m/z 382.1766

01E+511.5RT (min)

lcorbi\_hilic\_pos\_4\_105\_61

m/z 382.2314

012E+422.5RT (min)

lcorbi\_hilic\_pos\_4\_109\_62

m/z 382.2323

012E+455.5RT (min)

lcorbi\_hilic\_pos\_4\_113\_62

m/z 382.3167

01E+50.51RT (min)

lcorbi\_hilic\_pos\_4\_018\_31

m/z 382.8374

01234E+37.58RT (min)

lcorbi\_hilic\_pos\_4\_068\_57

m/z 382.8661

012E+477.5RT (min)

lcorbi\_hilic\_pos\_1\_053\_08

m/z 383.1022

0123E+344.5RT (min)

lcorbi\_hilic\_pos\_1\_053\_08

m/z 383.1411

012E+40.51RT (min)

lcorbi\_hilic\_pos\_1\_032\_31

m/z 383.1544

02468E+466.5RT (min)

lcorbi\_hilic\_pos\_4\_044\_25

m/z 383.2334

02468E+322.5RT (min)

lcorbi\_hilic\_pos\_3\_001\_49

m/z 383.3271

0123E+433.5RT (min)

lcorbi\_hilic\_pos\_4\_087\_59

m/z 383.3314

012E+40.51RT (min)

lcorbi\_hilic\_pos\_4\_084\_59

m/z 383.9658

01234E+366.5RT (min)

lcorbi\_hilic\_pos\_4\_074\_58

m/z 384.0709

01E+46.57RT (min)

lcorbi\_hilic\_pos\_4\_052\_32

m/z 384.1198

01E+455.5RT (min)

lcorbi\_hilic\_pos\_4\_068\_57

m/z 384.1300

01E+311.5RT (min)

lcorbi\_hilic\_pos\_4\_110\_62

m/z 384.1573

0246E+36.57RT (min)

lcorbi\_hilic\_pos\_2\_024\_32

m/z 384.1744

0123E+42.53RT (min)

lcorbi\_hilic\_pos\_4\_073\_57

m/z 384.2749

0246E+33.54RT (min)

lcorbi\_hilic\_pos\_4\_075\_58

m/z 384.3112

01E+41.52RT (min)

lcorbi\_hilic\_pos\_4\_101\_61

m/z 384.3839

01E+40.51RT (min)

lcorbi\_hilic\_pos\_1\_031\_36

m/z 384.7725

0246E+34.55RT (min)

lcorbi\_hilic\_pos\_4\_088\_59

m/z 385.0159

0123E+36.57RT (min)

lcorbi\_hilic\_pos\_2\_010\_07

m/z 385.0364

01234E+355.5RT (min)

lcorbi\_hilic\_pos\_4\_066\_57

m/z 385.1292

0246E+36.57RT (min)

lcorbi\_hilic\_pos\_4\_084\_59

m/z 385.1304

02468E+32.53RT (min)

lcorbi\_hilic\_pos\_2\_014\_20

m/z 385.1779

02468E+32.53RT (min)

lcorbi\_hilic\_pos\_4\_073\_57

m/z 385.2212

024E+46.57RT (min)

lcorbi\_hilic\_pos\_4\_094\_60

m/z 385.2928

012E+40.51RT (min)

lcorbi\_hilic\_pos\_4\_018\_31

m/z 385.3793

0246E+30.51RT (min)

lcorbi\_hilic\_pos\_4\_101\_61

m/z 385.9024

01E+499.5RT (min)

lcorbi\_hilic\_pos\_4\_077\_58

m/z 386.0135

01E+46.57RT (min)

lcorbi\_hilic\_pos\_1\_044\_17

m/z 386.0374

01E+46.57RT (min)

lcorbi\_hilic\_pos\_4\_083\_59

m/z 386.0444

0246E+355.5RT (min)

lcorbi\_hilic\_pos\_4\_101\_61

m/z 386.1644

012E+466.5RT (min)

lcorbi\_hilic\_pos\_4\_044\_25

m/z 386.2542

0246E+300.5RT (min)

lcorbi\_hilic\_pos\_1\_022\_43

m/z 386.2654

0246E+322.5RT (min)

lcorbi\_hilic\_pos\_4\_089\_59

m/z 386.2906

01E+43.54RT (min)

lcorbi\_hilic\_pos\_4\_067\_57

m/z 386.3211

012E+422.5RT (min)

lcorbi\_hilic\_pos\_4\_089\_59

m/z 386.9739

0123E+37.58RT (min)

lcorbi\_hilic\_pos\_4\_077\_58

m/z 387.0843

0123E+50.51RT (min)

lcorbi\_hilic\_pos\_4\_030\_18

m/z 387.0878

0246E+366.5RT (min)

lcorbi\_hilic\_pos\_4\_069\_57

m/z 387.1528

024E+411.5RT (min)

lcorbi\_hilic\_pos\_1\_052\_35

m/z 387.1935

01E+40.51RT (min)

lcorbi\_hilic\_pos\_3\_003\_37

m/z 387.3104

01E+477.5RT (min)

lcorbi\_hilic\_pos\_4\_096\_60

m/z 387.3586

01E+422.5RT (min)

lcorbi\_hilic\_pos\_2\_007\_28

m/z 387.9718

024E+37.58RT (min)

lcorbi\_hilic\_pos\_4\_075\_58

m/z 387.9784

0123E+366.5RT (min)

lcorbi\_hilic\_pos\_4\_077\_58

m/z 388.0586

01E+46.57RT (min)

lcorbi\_hilic\_pos\_3\_045\_06

m/z 388.1540

01E+43.54RT (min)

lcorbi\_hilic\_pos\_4\_069\_57

m/z 388.1818

02468E+36.57RT (min)

lcorbi\_hilic\_pos\_4\_069\_57

m/z 388.1839

01234E+40.51RT (min)

lcorbi\_hilic\_pos\_3\_003\_37

m/z 388.2122

01E+50.51RT (min)

lcorbi\_hilic\_pos\_4\_084\_59

m/z 388.3062

01E+43.54RT (min)

lcorbi\_hilic\_pos\_4\_073\_57

m/z 388.3621

0123E+322.5RT (min)

lcorbi\_hilic\_pos\_2\_007\_28

m/z 389.0567

024E+34.55RT (min)

lcorbi\_hilic\_pos\_4\_078\_58

m/z 389.0910

012E+466.5RT (min)

lcorbi\_hilic\_pos\_4\_086\_59

m/z 389.1631

02468E+32.53RT (min)

lcorbi\_hilic\_pos\_4\_066\_57

m/z 389.2580

01E+411.5RT (min)

lcorbi\_hilic\_pos\_3\_019\_43

m/z 389.2691

02468E+20.51RT (min)

lcorbi\_hilic\_pos\_3\_027\_05

m/z 390.1574

024E+377.5RT (min)

lcorbi\_hilic\_pos\_1\_005\_45

m/z 390.3217

012E+433.5RT (min)

lcorbi\_hilic\_pos\_1\_048\_02

m/z 390.3372

01E+31.52RT (min)

lcorbi\_hilic\_pos\_4\_094\_60

m/z 390.9770

024E+366.5RT (min)

lcorbi\_hilic\_pos\_4\_075\_58

m/z 391.0538

01E+44.55RT (min)

lcorbi\_hilic\_pos\_4\_080\_58

m/z 391.0677

01E+466.5RT (min)

lcorbi\_hilic\_pos\_4\_077\_58

m/z 391.0738

0246E+30.51RT (min)

lcorbi\_hilic\_pos\_4\_103\_61

m/z 391.1605

01E+42.53RT (min)

lcorbi\_hilic\_pos\_2\_029\_19

m/z 391.1812

0123E+566.5RT (min)

lcorbi\_hilic\_pos\_4\_097\_60

m/z 391.2696

01E+50.51RT (min)

lcorbi\_hilic\_pos\_2\_040\_14

m/z 391.3574

01E+40.51RT (min)

lcorbi\_hilic\_pos\_4\_075\_58

m/z 392.0347

01E+477.5RT (min)

lcorbi\_hilic\_pos\_4\_087\_59

m/z 392.1021

01E+44.55RT (min)

lcorbi\_hilic\_pos\_4\_110\_62

m/z 392.1314

024E+40.51RT (min)

lcorbi\_hilic\_pos\_4\_100\_61

m/z 392.9153

01234E+377.5RT (min)

lcorbi\_hilic\_pos\_1\_050\_05

m/z 392.9720

024E+366.5RT (min)

lcorbi\_hilic\_pos\_4\_080\_58

m/z 392.9820

01E+47.58RT (min)

lcorbi\_hilic\_pos\_4\_076\_58

m/z 393.0161

01E+40.51RT (min)

lcorbi\_hilic\_pos\_4\_069\_57

m/z 393.0510

01E+44.55RT (min)

lcorbi\_hilic\_pos\_4\_068\_57

m/z 393.0754

012E+46.57RT (min)

lcorbi\_hilic\_pos\_4\_093\_60

m/z 393.1949

02468E+37.58RT (min)

lcorbi\_hilic\_pos\_4\_068\_57

m/z 393.2387

01E+50.51RT (min)

lcorbi\_hilic\_pos\_4\_106\_62

m/z 393.2406

012E+444.5RT (min)

lcorbi\_hilic\_pos\_2\_030\_05

m/z 393.2987

012E+42.53RT (min)

lcorbi\_hilic\_pos\_4\_109\_62

m/z 393.3094

02468E+33.54RT (min)

lcorbi\_hilic\_pos\_4\_018\_31

m/z 394.3108

01E+72.53RT (min)

lcorbi\_hilic\_pos\_2\_016\_18

m/z 394.9774

0123E+37.58RT (min)

lcorbi\_hilic\_pos\_2\_004\_01

m/z 395.0479

0246E+34.55RT (min)

lcorbi\_hilic\_pos\_4\_068\_57

m/z 395.1629

01E+46.57RT (min)

lcorbi\_hilic\_pos\_4\_084\_59

m/z 395.1924

01E+433.5RT (min)

lcorbi\_hilic\_pos\_4\_104\_61

m/z 395.2929

0246E+30.51RT (min)

lcorbi\_hilic\_pos\_3\_029\_23

m/z 395.4000

0246E+32.53RT (min)

lcorbi\_hilic\_pos\_4\_074\_58

m/z 396.0732

01E+455.5RT (min)

lcorbi\_hilic\_pos\_4\_105\_61

m/z 396.1477

012E+444.5RT (min)

lcorbi\_hilic\_pos\_4\_094\_60

m/z 397.0518

0246E+344.5RT (min)

lcorbi\_hilic\_pos\_4\_098\_61

m/z 397.0900

01E+42.53RT (min)

lcorbi\_hilic\_pos\_4\_106\_62

m/z 397.2952

0246E+40.51RT (min)

lcorbi\_hilic\_pos\_2\_022\_35

m/z 397.3035

012E+444.5RT (min)

lcorbi\_hilic\_pos\_4\_087\_59

m/z 397.3299

02468E+32.53RT (min)

lcorbi\_hilic\_pos\_4\_106\_62

m/z 397.8050

01E+444.5RT (min)

lcorbi\_hilic\_pos\_4\_084\_59

m/z 397.9762

01E+477.5RT (min)

lcorbi\_hilic\_pos\_4\_078\_58

m/z 398.2672

01234E+355.5RT (min)

lcorbi\_hilic\_pos\_4\_083\_59

m/z 398.3268

01E+411.5RT (min)

lcorbi\_hilic\_pos\_4\_069\_57

m/z 398.3827

0123E+40.51RT (min)

lcorbi\_hilic\_pos\_2\_007\_28

m/z 398.3997

02468E+322.5RT (min)

lcorbi\_hilic\_pos\_4\_074\_58

m/z 399.1569

01E+433.5RT (min)

lcorbi\_hilic\_pos\_4\_068\_57

m/z 399.2354

01E+45.56RT (min)

lcorbi\_hilic\_pos\_4\_077\_58

m/z 399.3083

010.51RT (min)

lcorbi\_hilic\_pos\_4\_102\_61

m/z 399.3302

02468E+32.53RT (min)

lcorbi\_hilic\_pos\_4\_070\_57

m/z 399.3585

024E+40.51RT (min)

lcorbi\_hilic\_pos\_4\_086\_59

m/z 400.0863

024E+366.5RT (min)

lcorbi\_hilic\_pos\_1\_009\_50

m/z 400.1692

01E+433.5RT (min)

lcorbi\_hilic\_pos\_4\_072\_57

m/z 400.2446

024E+31.52RT (min)

lcorbi\_hilic\_pos\_4\_083\_59

m/z 400.2545

012E+40.51RT (min)

lcorbi\_hilic\_pos\_4\_067\_57

m/z 400.3789

01E+422.5RT (min)

lcorbi\_hilic\_pos\_4\_091\_60

m/z 400.3790

0123E+300.5RT (min)

lcorbi\_hilic\_pos\_1\_054\_23

m/z 400.3938

01E+50.51RT (min)

lcorbi\_hilic\_pos\_4\_052\_32

m/z 401.1830

0123E+36.57RT (min)

lcorbi\_hilic\_pos\_2\_003\_33

m/z 401.2900

0123E+40.51RT (min)

lcorbi\_hilic\_pos\_1\_054\_23

m/z 401.3650

01E+43.54RT (min)

lcorbi\_hilic\_pos\_4\_083\_59

m/z 401.3779

0123E+40.51RT (min)

lcorbi\_hilic\_pos\_4\_024\_24

m/z 402.0683

012E+477.5RT (min)

lcorbi\_hilic\_pos\_4\_097\_60

m/z 402.2084

01E+422.5RT (min)

lcorbi\_hilic\_pos\_4\_112\_62

m/z 402.2337

01E+31.52RT (min)

lcorbi\_hilic\_pos\_4\_093\_60

m/z 402.3219

01E+41.52RT (min)

lcorbi\_hilic\_pos\_1\_052\_35

m/z 402.3489

0123E+32.53RT (min)

lcorbi\_hilic\_pos\_4\_068\_57

m/z 402.3582

01E+522.5RT (min)

lcorbi\_hilic\_pos\_1\_040\_06

m/z 403.1651

02468E+344.5RT (min)

lcorbi\_hilic\_pos\_4\_087\_59

m/z 403.1862

02468E+311.5RT (min)

lcorbi\_hilic\_pos\_4\_079\_58

m/z 403.2306

0246E+35.56RT (min)

lcorbi\_hilic\_pos\_4\_082\_59

m/z 404.1919

012E+42.53RT (min)

lcorbi\_hilic\_pos\_4\_111\_62

m/z 404.2947

0246E+344.5RT (min)

lcorbi\_hilic\_pos\_1\_020\_13

m/z 405.0237

0246E+36.57RT (min)

lcorbi\_hilic\_pos\_4\_071\_57

m/z 405.1106

02468E+311.5RT (min)

lcorbi\_hilic\_pos\_3\_023\_18

m/z 405.2173

01E+455.5RT (min)

lcorbi\_hilic\_pos\_4\_106\_62

m/z 405.3153

01E+455.5RT (min)

lcorbi\_hilic\_pos\_4\_083\_59

m/z 405.3578

01E+40.51RT (min)

lcorbi\_hilic\_pos\_4\_092\_60

m/z 405.3732

010.51RT (min)

lcorbi\_hilic\_pos\_4\_092\_60

m/z 405.4208

0246E+31.52RT (min)

lcorbi\_hilic\_pos\_4\_091\_60

m/z 405.9477

01234E+366.5RT (min)

lcorbi\_hilic\_pos\_4\_076\_58

m/z 406.2746

02468E+33.54RT (min)

lcorbi\_hilic\_pos\_4\_113\_62

m/z 407.1170

0246E+36.57RT (min)

lcorbi\_hilic\_pos\_4\_080\_58

m/z 407.2032

01E+46.57RT (min)

lcorbi\_hilic\_pos\_4\_097\_60

m/z 407.3887

01234E+30.51RT (min)

lcorbi\_hilic\_pos\_1\_036\_32

m/z 407.4302

02468E+40.51RT (min)

lcorbi\_hilic\_pos\_2\_024\_32

m/z 408.2902

012E+433.54RT (min)

lcorbi\_hilic\_pos\_4\_110\_62

m/z 408.3265

02468E+32.53RT (min)

lcorbi\_hilic\_pos\_4\_109\_62

m/z 409.1902

012E+42.53RT (min)

lcorbi\_hilic\_pos\_2\_029\_19

m/z 409.1931

0123E+30.51RT (min)

lcorbi\_hilic\_pos\_4\_075\_58

m/z 409.2336

01E+50.51RT (min)

lcorbi\_hilic\_pos\_4\_075\_58

m/z 409.2783

01234E+411.5RT (min)

lcorbi\_hilic\_pos\_4\_018\_31

m/z 409.2833

01E+43.54RT (min)

lcorbi\_hilic\_pos\_1\_034\_37

m/z 409.2936

02468E+33.54RT (min)

lcorbi\_hilic\_pos\_4\_110\_62

m/z 410.1171

01E+433.5RT (min)

lcorbi\_hilic\_pos\_2\_005\_23

m/z 410.3480

02468E+20.51RT (min)

lcorbi\_hilic\_pos\_1\_037\_18

m/z 411.0265

01E+40.51RT (min)

lcorbi\_hilic\_pos\_4\_067\_57

m/z 411.0730

01E+466.5RT (min)

lcorbi\_hilic\_pos\_4\_083\_59

m/z 411.0843

01E+50.51RT (min)

lcorbi\_hilic\_pos\_1\_051\_26

m/z 411.1054

01E+50.51RT (min)

lcorbi\_hilic\_pos\_3\_035\_22

m/z 411.1628

0246E+311.5RT (min)

lcorbi\_hilic\_pos\_4\_029\_41

m/z 412.0471

02468E+366.5RT (min)

lcorbi\_hilic\_pos\_4\_076\_58

m/z 412.2099

01E+44.55RT (min)

lcorbi\_hilic\_pos\_3\_051\_25

m/z 412.3061

012E+43.54RT (min)

lcorbi\_hilic\_pos\_2\_030\_05

m/z 412.3282

01E+40.51RT (min)

lcorbi\_hilic\_pos\_1\_054\_23

m/z 412.8633

01E+499.5RT (min)

lcorbi\_hilic\_pos\_4\_077\_58

m/z 413.0497

02468E+366.5RT (min)

lcorbi\_hilic\_pos\_4\_077\_58

m/z 414.1486

013.54RT (min)

lcorbi\_hilic\_pos\_4\_113\_62

m/z 414.2337

012E+311.5RT (min)

lcorbi\_hilic\_pos\_4\_020\_37

m/z 414.2856

02468E+300.5RT (min)

lcorbi\_hilic\_pos\_1\_001\_49

m/z 414.3008

013.54RT (min)

lcorbi\_hilic\_pos\_4\_108\_62

m/z 414.3218

0123E+33.54RT (min)

lcorbi\_hilic\_pos\_4\_074\_58

m/z 414.3371

01E+433.5RT (min)

lcorbi\_hilic\_pos\_4\_107\_62

m/z 414.3582

0123E+32.53RT (min)

lcorbi\_hilic\_pos\_4\_068\_57

m/z 414.8973

0246E+377.5RT (min)

lcorbi\_hilic\_pos\_1\_043\_25

m/z 414.9639

012E+37.58RT (min)

lcorbi\_hilic\_pos\_1\_048\_02

m/z 415.1199

01E+433.5RT (min)

lcorbi\_hilic\_pos\_4\_103\_61

m/z 415.3251

0246E+40.51RT (min)

lcorbi\_hilic\_pos\_2\_032\_30

m/z 415.3706

01234E+433.5RT (min)

lcorbi\_hilic\_pos\_3\_047\_28

m/z 415.3900

012E+322.5RT (min)

lcorbi\_hilic\_pos\_1\_040\_06

m/z 415.9618

01E+47.58RT (min)

lcorbi\_hilic\_pos\_1\_024\_03

m/z 416.2284

01E+411.5RT (min)

lcorbi\_hilic\_pos\_2\_020\_31

m/z 416.2680

02468E+20.51RT (min)

lcorbi\_hilic\_pos\_1\_012\_19

m/z 416.3164

01E+41.52RT (min)

lcorbi\_hilic\_pos\_4\_073\_57

m/z 416.3375

0123E+33.54RT (min)

lcorbi\_hilic\_pos\_4\_066\_57

m/z 416.3740

0123E+400.5RT (min)

lcorbi\_hilic\_pos\_1\_001\_49

m/z 416.7986

01234E+355.5RT (min)

lcorbi\_hilic\_pos\_4\_082\_59

m/z 416.8610

01E+477.5RT (min)

lcorbi\_hilic\_pos\_1\_003\_24

m/z 417.1224

012E+45.56RT (min)

lcorbi\_hilic\_pos\_4\_088\_59

m/z 417.1449

01E+46.57RT (min)

lcorbi\_hilic\_pos\_4\_087\_59

m/z 417.2406

0123E+444.5RT (min)

lcorbi\_hilic\_pos\_1\_050\_05

m/z 417.3775

012E+40.51RT (min)

lcorbi\_hilic\_pos\_4\_007\_05

m/z 418.1654

0246E+322.5RT (min)

lcorbi\_hilic\_pos\_4\_102\_61

m/z 418.1826

02468E+322.5RT (min)

lcorbi\_hilic\_pos\_4\_106\_62

m/z 418.2957

01E+411.5RT (min)

lcorbi\_hilic\_pos\_4\_106\_62

m/z 419.3735

0123E+40.51RT (min)

lcorbi\_hilic\_pos\_4\_083\_59

m/z 419.3938

0123E+40.51RT (min)

lcorbi\_hilic\_pos\_4\_024\_24

m/z 420.3193

012E+30.51RT (min)

lcorbi\_hilic\_pos\_4\_083\_59

m/z 420.3325

02468E+322.5RT (min)

lcorbi\_hilic\_pos\_4\_097\_60

m/z 420.4200

01E+40.51RT (min)

lcorbi\_hilic\_pos\_4\_097\_60

m/z 420.4289

0246E+40.51RT (min)

lcorbi\_hilic\_pos\_4\_097\_60

m/z 420.8854

0246E+35.56RT (min)

lcorbi\_hilic\_pos\_1\_010\_07

m/z 421.0420

012E+366.5RT (min)

lcorbi\_hilic\_pos\_4\_006\_14

m/z 421.2234

012E+40.51RT (min)

lcorbi\_hilic\_pos\_2\_022\_35

m/z 421.3377

01E+433.5RT (min)

lcorbi\_hilic\_pos\_4\_091\_60

m/z 421.3429

0111.5RT (min)

lcorbi\_hilic\_pos\_4\_085\_59

m/z 421.3472

01E+444.5RT (min)

lcorbi\_hilic\_pos\_4\_112\_62

m/z 422.1005

0123E+411.5RT (min)

lcorbi\_hilic\_pos\_4\_015\_20

m/z 422.2219

012E+433.5RT (min)

lcorbi\_hilic\_pos\_4\_100\_61

m/z 422.3059

02468E+322.5RT (min)

lcorbi\_hilic\_pos\_4\_106\_62

m/z 422.3268

0246E+32.53RT (min)

lcorbi\_hilic\_pos\_4\_078\_58

m/z 423.0646

01E+466.5RT (min)

lcorbi\_hilic\_pos\_4\_102\_61

m/z 423.1650

01234E+36.57RT (min)

lcorbi\_hilic\_pos\_3\_033\_53

m/z 423.2729

0123E+43.54RT (min)

lcorbi\_hilic\_pos\_4\_012\_29

m/z 423.3587

0246E+40.51RT (min)

lcorbi\_hilic\_pos\_4\_087\_59

m/z 424.1025

0123E+377.5RT (min)

lcorbi\_hilic\_pos\_4\_084\_59

m/z 424.2851

012E+43.54RT (min)

lcorbi\_hilic\_pos\_4\_004\_38

m/z 424.3626

02468E+33.54RT (min)

lcorbi\_hilic\_pos\_4\_083\_59

m/z 425.0422

012E+40.51RT (min)

lcorbi\_hilic\_pos\_2\_019\_16

m/z 426.1618

0246E+36.57RT (min)

lcorbi\_hilic\_pos\_4\_074\_58

m/z 426.2766

0144.5RT (min)

lcorbi\_hilic\_pos\_4\_020\_37

m/z 426.3089

01234E+50.51RT (min)

lcorbi\_hilic\_pos\_4\_018\_31

m/z 426.3219

0123E+41.52RT (min)

lcorbi\_hilic\_pos\_2\_016\_18

m/z 426.3430

010.51RT (min)

lcorbi\_hilic\_pos\_4\_018\_31

m/z 426.4141

012E+40.51RT (min)

lcorbi\_hilic\_pos\_2\_007\_28

m/z 426.4310

02468E+322.5RT (min)

lcorbi\_hilic\_pos\_4\_074\_58

m/z 427.3209

0246E+411.52RT (min)

lcorbi\_hilic\_pos\_3\_029\_23

m/z 427.3325

012E+444.5RT (min)

lcorbi\_hilic\_pos\_4\_071\_57

m/z 427.3535

0123E+40.51RT (min)

lcorbi\_hilic\_pos\_3\_019\_43

m/z 427.3806

013.54RT (min)

lcorbi\_hilic\_pos\_4\_082\_59

m/z 428.3363

01E+444.5RT (min)

lcorbi\_hilic\_pos\_4\_073\_57

m/z 428.3647

012E+32.53RT (min)

lcorbi\_hilic\_pos\_4\_070\_57

m/z 428.3819

0246E+40.51RT (min)

lcorbi\_hilic\_pos\_3\_020\_31

m/z 428.3933

01E+43.54RT (min)

lcorbi\_hilic\_pos\_4\_082\_59

m/z 429.1162

01234E+42.53RT (min)

lcorbi\_hilic\_pos\_1\_012\_19

m/z 429.1851

01E+46.57RT (min)

lcorbi\_hilic\_pos\_4\_090\_60

m/z 429.2043

01E+411.5RT (min)

lcorbi\_hilic\_pos\_4\_111\_62

m/z 429.2425

02468E+36.57RT (min)

lcorbi\_hilic\_pos\_4\_083\_59

m/z 429.3439

014.55RT (min)

lcorbi\_hilic\_pos\_4\_097\_60

m/z 429.3772

012E+42.53RT (min)

lcorbi\_hilic\_pos\_1\_040\_06

m/z 430.0078

0123E+35.56RT (min)

lcorbi\_hilic\_pos\_4\_079\_58

m/z 430.2874

01E+333.5RT (min)

lcorbi\_hilic\_pos\_3\_023\_18

m/z 430.3104

02468E+344.5RT (min)

lcorbi\_hilic\_pos\_4\_007\_05

m/z 430.3895

01E+522.5RT (min)

lcorbi\_hilic\_pos\_1\_040\_06

m/z 431.2908

0246E+333.5RT (min)

lcorbi\_hilic\_pos\_3\_021\_46

m/z 431.2949

01E+50.51RT (min)

lcorbi\_hilic\_pos\_4\_008\_19

m/z 431.3371

0111.5RT (min)

lcorbi\_hilic\_pos\_4\_089\_59

m/z 431.3525

01234E+20.51RT (min)

lcorbi\_hilic\_pos\_3\_019\_43

m/z 431.3845

01E+50.51RT (min)

lcorbi\_hilic\_pos\_3\_019\_43

m/z 431.9848

0246E+35.56RT (min)

lcorbi\_hilic\_pos\_4\_084\_59

m/z 432.2385

0123E+40.51RT (min)

lcorbi\_hilic\_pos\_4\_084\_59

m/z 432.2844

02468E+333.5RT (min)

lcorbi\_hilic\_pos\_4\_048\_46

m/z 432.3113

01E+43.54RT (min)

lcorbi\_hilic\_pos\_4\_074\_58

m/z 432.9188

01E+45.56RT (min)

lcorbi\_hilic\_pos\_4\_066\_57

m/z 433.1516

012E+455.5RT (min)

lcorbi\_hilic\_pos\_4\_107\_62

m/z 433.2588

01E+50.51RT (min)

lcorbi\_hilic\_pos\_4\_112\_62

m/z 433.3316

0246E+40.51RT (min)

lcorbi\_hilic\_pos\_1\_032\_31

m/z 433.3680

02468E+30.51RT (min)

lcorbi\_hilic\_pos\_4\_080\_58

m/z 434.0003

024E+35.56RT (min)

lcorbi\_hilic\_pos\_4\_099\_61

m/z 434.0291

02468E+366.5RT (min)

lcorbi\_hilic\_pos\_4\_080\_58

m/z 435.1231

016.57RT (min)

lcorbi\_hilic\_pos\_4\_091\_60

m/z 435.3884

010.51RT (min)

lcorbi\_hilic\_pos\_4\_084\_59

m/z 436.0279

0123E+35.56RT (min)

lcorbi\_hilic\_pos\_4\_100\_61

m/z 436.2182

012E+42.53RT (min)

lcorbi\_hilic\_pos\_4\_107\_62

m/z 436.2350

0123E+43.54RT (min)

lcorbi\_hilic\_pos\_4\_014\_27

m/z 436.8793

01234E+377.5RT (min)

lcorbi\_hilic\_pos\_4\_088\_59

m/z 437.4039

0246E+40.51RT (min)

lcorbi\_hilic\_pos\_4\_096\_60

m/z 438.1495

02468E+377.5RT (min)

lcorbi\_hilic\_pos\_4\_092\_60

m/z 438.2984

0144.5RT (min)

lcorbi\_hilic\_pos\_2\_030\_05

m/z 438.3694

01E+444.5RT (min)

lcorbi\_hilic\_pos\_4\_072\_57

m/z 438.8429

01E+477.5RT (min)

lcorbi\_hilic\_pos\_1\_003\_24

m/z 438.9442

0246E+35.56RT (min)

lcorbi\_hilic\_pos\_1\_020\_13

m/z 439.1044

01E+45.56RT (min)

lcorbi\_hilic\_pos\_4\_083\_59

m/z 439.3211

0246E+30.51RT (min)

lcorbi\_hilic\_pos\_3\_046\_41

m/z 439.3576

01234E+444.5RT (min)

lcorbi\_hilic\_pos\_4\_107\_62

m/z 439.4263

010.51RT (min)

lcorbi\_hilic\_pos\_3\_003\_37

m/z 440.3143

01E+444.5RT (min)

lcorbi\_hilic\_pos\_4\_077\_58

m/z 440.3611

01E+444.5RT (min)

lcorbi\_hilic\_pos\_4\_109\_62

m/z 440.4103

012E+41.52RT (min)

lcorbi\_hilic\_pos\_4\_092\_60

m/z 441.1113

024E+388.5RT (min)

lcorbi\_hilic\_pos\_4\_088\_59

m/z 441.9359

024E+35.56RT (min)

lcorbi\_hilic\_pos\_4\_075\_58

m/z 442.3036

01234E+40.51RT (min)

lcorbi\_hilic\_pos\_4\_018\_31

m/z 442.3279

01E+41.52RT (min)

lcorbi\_hilic\_pos\_4\_082\_59

m/z 442.3474

01E+41.52RT (min)

lcorbi\_hilic\_pos\_4\_086\_59

m/z 442.7965

0246E+355.5RT (min)

lcorbi\_hilic\_pos\_4\_097\_60

m/z 443.9868

0123E+377.5RT (min)

lcorbi\_hilic\_pos\_1\_016\_33

m/z 444.0571

0246E+34.55RT (min)

lcorbi\_hilic\_pos\_4\_068\_57

m/z 444.3323

024E+40.51RT (min)

lcorbi\_hilic\_pos\_4\_087\_59

m/z 444.4051

0246E+40.51RT (min)

lcorbi\_hilic\_pos\_1\_032\_31

m/z 445.1205

01E+40.51RT (min)

lcorbi\_hilic\_pos\_4\_105\_61

m/z 445.3277

012E+411.5RT (min)

lcorbi\_hilic\_pos\_4\_069\_57

m/z 445.3681

01234E+40.51RT (min)

lcorbi\_hilic\_pos\_2\_005\_23

m/z 446.0002

024E+35.56RT (min)

lcorbi\_hilic\_pos\_4\_084\_59

m/z 446.0542

01E+44.55RT (min)

lcorbi\_hilic\_pos\_4\_068\_57

m/z 446.1101

02468E+366.5RT (min)

lcorbi\_hilic\_pos\_4\_069\_57

m/z 446.2600

02468E+31.52RT (min)

lcorbi\_hilic\_pos\_4\_079\_58

m/z 447.3440

012E+444.5RT (min)

lcorbi\_hilic\_pos\_2\_013\_36

m/z 447.9325

0123E+366.5RT (min)

lcorbi\_hilic\_pos\_3\_002\_04

m/z 448.2911

02468E+34.55RT (min)

lcorbi\_hilic\_pos\_4\_082\_59

m/z 448.2912

01E+400.5RT (min)

lcorbi\_hilic\_pos\_4\_076\_58

m/z 448.3063

02468E+31.52RT (min)

lcorbi\_hilic\_pos\_4\_066\_57

m/z 448.3426

02468E+32.53RT (min)

lcorbi\_hilic\_pos\_4\_077\_58

m/z 448.3638

01234E+322.5RT (min)

lcorbi\_hilic\_pos\_4\_022\_08

m/z 448.4513

02468E+40.51RT (min)

lcorbi\_hilic\_pos\_4\_092\_60

m/z 448.7416

01234E+355.5RT (min)

lcorbi\_hilic\_pos\_4\_076\_58

m/z 448.8396

01234E+40.51RT (min)

lcorbi\_hilic\_pos\_4\_089\_59

m/z 449.1255

024E+355.5RT (min)

lcorbi\_hilic\_pos\_1\_014\_27

m/z 449.3056

012E+40.51RT (min)

lcorbi\_hilic\_pos\_4\_008\_19

m/z 449.3638

01234E+30.51RT (min)

lcorbi\_hilic\_pos\_4\_096\_60

m/z 450.3130

02468E+36.57RT (min)

lcorbi\_hilic\_pos\_4\_093\_60

m/z 450.3583

024E+32.53RT (min)

lcorbi\_hilic\_pos\_4\_078\_58

m/z 450.7392

0123E+355.5RT (min)

lcorbi\_hilic\_pos\_4\_081\_58

m/z 451.9896

024E+35.56RT (min)

lcorbi\_hilic\_pos\_4\_072\_57

m/z 452.1644

0246E+433.5RT (min)

lcorbi\_hilic\_pos\_4\_082\_59

m/z 452.2778

01E+444.5RT (min)

lcorbi\_hilic\_pos\_4\_082\_59

m/z 452.3740

01234E+32.53RT (min)

lcorbi\_hilic\_pos\_4\_020\_37

m/z 453.1605

024E+36.57RT (min)

lcorbi\_hilic\_pos\_2\_024\_32

m/z 453.3439

0246E+52.53RT (min)

lcorbi\_hilic\_pos\_4\_007\_05

m/z 454.2570

02468E+311.5RT (min)

lcorbi\_hilic\_pos\_4\_113\_62

m/z 454.3298

0144.5RT (min)

lcorbi\_hilic\_pos\_4\_082\_59

m/z 454.3744

02468E+30.51RT (min)

lcorbi\_hilic\_pos\_4\_088\_59

m/z 454.3896

01234E+32.53RT (min)

lcorbi\_hilic\_pos\_4\_020\_37

m/z 454.4454

01E+40.51RT (min)

lcorbi\_hilic\_pos\_4\_026\_11

m/z 454.4624

024E+31.52RT (min)

lcorbi\_hilic\_pos\_4\_079\_58

m/z 455.2241

0246E+455.5RT (min)

lcorbi\_hilic\_pos\_4\_110\_62

m/z 455.3502

02468E+433.5RT (min)

lcorbi\_hilic\_pos\_4\_087\_59

m/z 455.3636

01E+444.5RT (min)

lcorbi\_hilic\_pos\_4\_067\_57

m/z 455.4151

01E+50.51RT (min)

lcorbi\_hilic\_pos\_4\_090\_60

m/z 456.0112

0246E+366.5RT (min)

lcorbi\_hilic\_pos\_4\_080\_58

m/z 456.3326

02468E+33.54RT (min)

lcorbi\_hilic\_pos\_4\_069\_57

m/z 457.3390

0246E+33.54RT (min)

lcorbi\_hilic\_pos\_4\_082\_59

m/z 457.3684

0123E+444.5RT (min)

lcorbi\_hilic\_pos\_4\_112\_62

m/z 457.4308

02468E+40.51RT (min)

lcorbi\_hilic\_pos\_4\_091\_60

m/z 457.9613

01234E+366.5RT (min)

lcorbi\_hilic\_pos\_2\_048\_04

m/z 458.0100

01234E+35.56RT (min)

lcorbi\_hilic\_pos\_4\_105\_61

m/z 458.2727

012E+477.5RT (min)

lcorbi\_hilic\_pos\_4\_079\_58

m/z 458.3481

01E+50.51RT (min)

lcorbi\_hilic\_pos\_2\_007\_28

m/z 458.3482

02468E+33.54RT (min)

lcorbi\_hilic\_pos\_4\_081\_58

m/z 458.3718

01E+444.5RT (min)

lcorbi\_hilic\_pos\_4\_107\_62

m/z 460.2701

01E+400.5RT (min)

lcorbi\_hilic\_pos\_4\_071\_57

m/z 460.7686

012E+455.5RT (min)

lcorbi\_hilic\_pos\_4\_080\_58

m/z 460.8249

01234E+377.5RT (min)

lcorbi\_hilic\_pos\_1\_043\_25

m/z 461.1085

0123E+43.54RT (min)

lcorbi\_hilic\_pos\_1\_012\_19

m/z 461.3128

0246E+31.52RT (min)

lcorbi\_hilic\_pos\_4\_084\_59

m/z 461.3631

012E+40.51RT (min)

lcorbi\_hilic\_pos\_4\_020\_37

m/z 461.8574

01E+455.5RT (min)

lcorbi\_hilic\_pos\_4\_089\_59

m/z 462.1119

02468E+33.54RT (min)

lcorbi\_hilic\_pos\_1\_012\_19

m/z 462.3494

01E+455.5RT (min)

lcorbi\_hilic\_pos\_4\_085\_59

m/z 463.3000

02468E+31.52RT (min)

lcorbi\_hilic\_pos\_4\_085\_59

m/z 463.3431

024E+40.51RT (min)

lcorbi\_hilic\_pos\_2\_027\_38

m/z 463.3787

010.51RT (min)

lcorbi\_hilic\_pos\_2\_039\_09

m/z 464.3140

012E+444.5RT (min)

lcorbi\_hilic\_pos\_2\_013\_36

m/z 464.3565

012E+40.51RT (min)

lcorbi\_hilic\_pos\_1\_032\_31

m/z 465.0682

01E+466.5RT (min)

lcorbi\_hilic\_pos\_4\_079\_58

m/z 466.3168

01E+53.54RT (min)

lcorbi\_hilic\_pos\_3\_023\_18

m/z 466.3297

01E+444.5RT (min)

lcorbi\_hilic\_pos\_4\_076\_58

m/z 466.3321

0123E+32.53RT (min)

lcorbi\_hilic\_pos\_3\_047\_28

m/z 466.3742

01234E+40.51RT (min)

lcorbi\_hilic\_pos\_4\_087\_59

m/z 466.3826

024E+40.51RT (min)

lcorbi\_hilic\_pos\_4\_088\_59

m/z 467.1599

012E+45.56RT (min)

lcorbi\_hilic\_pos\_4\_070\_57

m/z 467.3484

02468E+31.52RT (min)

lcorbi\_hilic\_pos\_4\_079\_58

m/z 467.3738

02468E+30.51RT (min)

lcorbi\_hilic\_pos\_4\_084\_59

m/z 467.4140

01E+50.51RT (min)

lcorbi\_hilic\_pos\_4\_084\_59

m/z 467.4576

01E+40.51RT (min)

lcorbi\_hilic\_pos\_2\_007\_28

m/z 468.3454

01234E+344.5RT (min)

lcorbi\_hilic\_pos\_1\_016\_33

m/z 468.4053

0123E+40.51RT (min)

lcorbi\_hilic\_pos\_4\_088\_59

m/z 468.4415

02468E+31.52RT (min)

lcorbi\_hilic\_pos\_4\_095\_60

m/z 468.9675

0246E+37.58RT (min)

lcorbi\_hilic\_pos\_4\_080\_58

m/z 469.3350

02468E+311.5RT (min)

lcorbi\_hilic\_pos\_4\_099\_61

m/z 469.3389

01E+44.55RT (min)

lcorbi\_hilic\_pos\_4\_082\_59

m/z 469.8405

01E+44.55RT (min)

lcorbi\_hilic\_pos\_4\_086\_59

m/z 470.3151

024E+344.5RT (min)

lcorbi\_hilic\_pos\_4\_067\_57

m/z 470.3692

01E+40.51RT (min)

lcorbi\_hilic\_pos\_4\_084\_59

m/z 470.3703

012E+433.5RT (min)

lcorbi\_hilic\_pos\_3\_029\_23

m/z 470.3706

012E+455.5RT (min)

lcorbi\_hilic\_pos\_4\_087\_59

m/z 470.3958

01234E+40.51RT (min)

lcorbi\_hilic\_pos\_4\_083\_59

m/z 470.8527

024E+40.51RT (min)

lcorbi\_hilic\_pos\_4\_089\_59

m/z 471.3184

01E+43.54RT (min)

lcorbi\_hilic\_pos\_4\_087\_59

m/z 471.3546

012E+455.5RT (min)

lcorbi\_hilic\_pos\_4\_089\_59

m/z 471.4099

0123E+40.51RT (min)

lcorbi\_hilic\_pos\_4\_091\_60

m/z 472.3039

01E+41.52RT (min)

lcorbi\_hilic\_pos\_4\_081\_58

m/z 472.3580

024E+355.5RT (min)

lcorbi\_hilic\_pos\_4\_085\_59

m/z 472.4001

01E+411.5RT (min)

lcorbi\_hilic\_pos\_4\_082\_59

m/z 474.2676

01E+433.5RT (min)

lcorbi\_hilic\_pos\_3\_039\_29

m/z 475.4061

012E+40.51RT (min)

lcorbi\_hilic\_pos\_3\_034\_26

m/z 476.3070

01E+411.5RT (min)

lcorbi\_hilic\_pos\_4\_089\_59

m/z 476.3823

01234E+444.5RT (min)

lcorbi\_hilic\_pos\_4\_112\_62

m/z 477.2811

012E+444.5RT (min)

lcorbi\_hilic\_pos\_4\_082\_59

m/z 478.0552

02468E+366.5RT (min)

lcorbi\_hilic\_pos\_4\_076\_58

m/z 478.3300

0144.5RT (min)

lcorbi\_hilic\_pos\_4\_086\_59

m/z 480.1077

0123E+466.5RT (min)

lcorbi\_hilic\_pos\_1\_008\_38

m/z 480.1927

01E+411.5RT (min)

lcorbi\_hilic\_pos\_4\_106\_62

m/z 480.4262

024E+40.51RT (min)

lcorbi\_hilic\_pos\_4\_083\_59

m/z 481.1080

01234E+37.58RT (min)

lcorbi\_hilic\_pos\_3\_048\_38

m/z 481.3125

012E+444.5RT (min)

lcorbi\_hilic\_pos\_4\_087\_59

m/z 481.3489

01E+444.5RT (min)

lcorbi\_hilic\_pos\_4\_074\_58

m/z 481.4733

0246E+40.51RT (min)

lcorbi\_hilic\_pos\_4\_026\_11

m/z 484.3499

0144.5RT (min)

lcorbi\_hilic\_pos\_3\_023\_18

m/z 484.3675

02468E+344.5RT (min)

lcorbi\_hilic\_pos\_4\_074\_58

m/z 484.3850

02468E+30.51RT (min)

lcorbi\_hilic\_pos\_4\_088\_59

m/z 486.1249

0246E+37.58RT (min)

lcorbi\_hilic\_pos\_3\_010\_32

m/z 486.3373

01E+43.54RT (min)

lcorbi\_hilic\_pos\_4\_083\_59

m/z 486.4014

01E+50.51RT (min)

lcorbi\_hilic\_pos\_4\_090\_60

m/z 486.4206

01E+40.51RT (min)

lcorbi\_hilic\_pos\_4\_096\_60

m/z 486.9879

0123E+366.5RT (min)

lcorbi\_hilic\_pos\_4\_076\_58

m/z 487.0501

02468E+366.5RT (min)

lcorbi\_hilic\_pos\_4\_077\_58

m/z 487.3610

0246E+30.51RT (min)

lcorbi\_hilic\_pos\_4\_093\_60

m/z 488.3198

01E+466.5RT (min)

lcorbi\_hilic\_pos\_4\_081\_58

m/z 488.3255

01234E+40.51RT (min)

lcorbi\_hilic\_pos\_4\_083\_59

m/z 490.1069

01E+455.5RT (min)

lcorbi\_hilic\_pos\_4\_113\_62

m/z 490.2915

02468E+344.5RT (min)

lcorbi\_hilic\_pos\_2\_030\_05

m/z 491.2466

01E+477.5RT (min)

lcorbi\_hilic\_pos\_4\_083\_59

m/z 492.3092

0123E+444.5RT (min)

lcorbi\_hilic\_pos\_1\_050\_05

m/z 492.4781

01234E+40.51RT (min)

lcorbi\_hilic\_pos\_4\_068\_57

m/z 492.8658

0123E+40.51RT (min)

lcorbi\_hilic\_pos\_4\_085\_59

m/z 493.0632

012E+46.57RT (min)

lcorbi\_hilic\_pos\_1\_056\_56

m/z 495.3799

01E+41.52RT (min)

lcorbi\_hilic\_pos\_4\_075\_58

m/z 495.4890

0123E+40.51RT (min)

lcorbi\_hilic\_pos\_4\_014\_27

m/z 496.1216

01E+47.58RT (min)

lcorbi\_hilic\_pos\_4\_077\_58

m/z 496.3064

02468E+344.5RT (min)

lcorbi\_hilic\_pos\_4\_006\_14

m/z 496.3737

0123E+444.5RT (min)

lcorbi\_hilic\_pos\_4\_076\_58

m/z 496.3801

01234E+444.5RT (min)

lcorbi\_hilic\_pos\_4\_076\_58

m/z 498.2834

0123E+455.5RT (min)

lcorbi\_hilic\_pos\_4\_087\_59

m/z 498.3657

01234E+33.54RT (min)

lcorbi\_hilic\_pos\_4\_082\_59

m/z 498.4017

024E+433.5RT (min)

lcorbi\_hilic\_pos\_4\_085\_59

m/z 499.2867

02468E+355.5RT (min)

lcorbi\_hilic\_pos\_4\_087\_59

m/z 499.9054

01E+45.56RT (min)

lcorbi\_hilic\_pos\_4\_068\_57

m/z 500.2773

01E+444.5RT (min)

lcorbi\_hilic\_pos\_4\_074\_58

m/z 500.3674

01234E+444.5RT (min)

lcorbi\_hilic\_pos\_4\_092\_60

m/z 500.9064

0246E+35.56RT (min)

lcorbi\_hilic\_pos\_4\_066\_57

m/z 501.9878

01234E+35.56RT (min)

lcorbi\_hilic\_pos\_4\_101\_61

m/z 502.3745

0246E+30.51RT (min)

lcorbi\_hilic\_pos\_4\_018\_31

m/z 503.8862

01E+444.5RT (min)

lcorbi\_hilic\_pos\_4\_082\_59

m/z 504.2994

02468E+344.5RT (min)

lcorbi\_hilic\_pos\_4\_078\_58

m/z 504.3093

01E+444.5RT (min)

lcorbi\_hilic\_pos\_2\_030\_05

m/z 504.3205

0246E+40.51RT (min)

lcorbi\_hilic\_pos\_1\_031\_36

m/z 505.5097

012E+40.51RT (min)

lcorbi\_hilic\_pos\_2\_021\_17

m/z 506.0502

01E+46.57RT (min)

lcorbi\_hilic\_pos\_4\_074\_58

m/z 506.3248

0123E+43.54RT (min)

lcorbi\_hilic\_pos\_4\_007\_05

m/z 506.3611

01234E+444.5RT (min)

lcorbi\_hilic\_pos\_3\_031\_36

m/z 506.9316

01234E+35.56RT (min)

lcorbi\_hilic\_pos\_1\_014\_27

m/z 508.5458

01E+40.51RT (min)

lcorbi\_hilic\_pos\_1\_007\_16

m/z 509.2176

0123E+411.5RT (min)

lcorbi\_hilic\_pos\_4\_109\_62

m/z 509.3439

02468E+344.5RT (min)

lcorbi\_hilic\_pos\_4\_084\_59

m/z 510.3040

02468E+355.5RT (min)

lcorbi\_hilic\_pos\_4\_078\_58

m/z 511.2249

01E+40.51RT (min)

lcorbi\_hilic\_pos\_1\_020\_13

m/z 511.3956

0123E+344.5RT (min)

lcorbi\_hilic\_pos\_4\_071\_57

m/z 512.3353

012E+44.55RT (min)

lcorbi\_hilic\_pos\_4\_070\_57

m/z 512.3622

024E+344.5RT (min)

lcorbi\_hilic\_pos\_4\_079\_58

m/z 512.4680

024E+32.53RT (min)

lcorbi\_hilic\_pos\_4\_067\_57

m/z 512.5044

0246E+30.51RT (min)

lcorbi\_hilic\_pos\_2\_023\_34

m/z 513.3387

0246E+34.55RT (min)

lcorbi\_hilic\_pos\_4\_072\_57

m/z 514.3772

0123E+40.51RT (min)

lcorbi\_hilic\_pos\_4\_089\_59

m/z 514.4521

01E+50.51RT (min)

lcorbi\_hilic\_pos\_4\_096\_60

m/z 515.0451

0246E+36.57RT (min)

lcorbi\_hilic\_pos\_4\_074\_58

m/z 515.2135

01E+46.57RT (min)

lcorbi\_hilic\_pos\_4\_020\_37

m/z 515.3099

012E+43.54RT (min)

lcorbi\_hilic\_pos\_4\_069\_57

m/z 516.3035

012E+43.54RT (min)

lcorbi\_hilic\_pos\_4\_068\_57

m/z 516.3897

012E+444.5RT (min)

lcorbi\_hilic\_pos\_2\_030\_05

m/z 517.4019

01234E+444.5RT (min)

lcorbi\_hilic\_pos\_4\_096\_60

m/z 517.4258

0246E+30.51RT (min)

lcorbi\_hilic\_pos\_3\_054\_16

m/z 518.1037

0246E+37.58RT (min)

lcorbi\_hilic\_pos\_4\_078\_58

m/z 519.4415

010.51RT (min)

lcorbi\_hilic\_pos\_4\_018\_31

m/z 519.4608

02468E+40.51RT (min)

lcorbi\_hilic\_pos\_4\_098\_61

m/z 519.4891

024E+322.5RT (min)

lcorbi\_hilic\_pos\_4\_072\_57

m/z 519.4972

01E+30.51RT (min)

lcorbi\_hilic\_pos\_4\_102\_61

m/z 520.0463

012E+40.51RT (min)

lcorbi\_hilic\_pos\_4\_072\_57

m/z 520.3332

0246E+311.5RT (min)

lcorbi\_hilic\_pos\_4\_078\_58

m/z 520.3572

01234E+455.5RT (min)

lcorbi\_hilic\_pos\_4\_091\_60

m/z 522.2833

01234E+455.5RT (min)

lcorbi\_hilic\_pos\_1\_050\_05

m/z 522.4240

01234E+40.51RT (min)

lcorbi\_hilic\_pos\_4\_006\_14

m/z 523.4362

012E+40.51RT (min)

lcorbi\_hilic\_pos\_4\_075\_58

m/z 524.1280

01E+47.58RT (min)

lcorbi\_hilic\_pos\_4\_020\_37

m/z 524.2990

01E+455.5RT (min)

lcorbi\_hilic\_pos\_4\_084\_59

m/z 524.3479

01E+444.5RT (min)

lcorbi\_hilic\_pos\_1\_029\_14

m/z 524.3949

0144.5RT (min)

lcorbi\_hilic\_pos\_4\_078\_58

m/z 526.2653

02468E+33.54RT (min)

lcorbi\_hilic\_pos\_4\_079\_58

m/z 526.7579

01234E+355.5RT (min)

lcorbi\_hilic\_pos\_4\_091\_60

m/z 527.1431

01E+46.57RT (min)

lcorbi\_hilic\_pos\_4\_069\_57

m/z 527.2967

01E+444.5RT (min)

lcorbi\_hilic\_pos\_4\_072\_57

m/z 527.3804

01E+444.5RT (min)

lcorbi\_hilic\_pos\_4\_076\_58

m/z 527.4790

012E+411.5RT (min)

lcorbi\_hilic\_pos\_4\_088\_59

m/z 528.3091

01E+444.5RT (min)

lcorbi\_hilic\_pos\_4\_067\_57

m/z 529.3123

024E+344.5RT (min)

lcorbi\_hilic\_pos\_4\_069\_57

m/z 530.1480

01E+47.58RT (min)

lcorbi\_hilic\_pos\_4\_052\_32

m/z 530.3247

01E+444.5RT (min)

lcorbi\_hilic\_pos\_4\_071\_57

m/z 530.7887

01E+477.5RT (min)

lcorbi\_hilic\_pos\_4\_074\_58

m/z 531.0141

01234E+366.5RT (min)

lcorbi\_hilic\_pos\_4\_078\_58

m/z 531.4086

012E+50.51RT (min)

lcorbi\_hilic\_pos\_4\_020\_37

m/z 531.4087

0123E+43.54RT (min)

lcorbi\_hilic\_pos\_4\_007\_05

m/z 532.4627

012E+50.51RT (min)

lcorbi\_hilic\_pos\_4\_090\_60

m/z 532.7031

0123E+355.5RT (min)

lcorbi\_hilic\_pos\_4\_075\_58

m/z 533.4207

01234E+40.51RT (min)

lcorbi\_hilic\_pos\_3\_047\_28

m/z 533.4243

0123E+43.54RT (min)

lcorbi\_hilic\_pos\_4\_091\_60

m/z 533.5410

0123E+40.51RT (min)

lcorbi\_hilic\_pos\_4\_039\_15

m/z 534.7007

0123E+355.5RT (min)

lcorbi\_hilic\_pos\_4\_080\_58

m/z 535.2998

024E+344.5RT (min)

lcorbi\_hilic\_pos\_4\_076\_58

m/z 536.3355

0246E+344.5RT (min)

lcorbi\_hilic\_pos\_4\_045\_33

m/z 536.5043

01234E+40.51RT (min)

lcorbi\_hilic\_pos\_2\_022\_35

m/z 536.5773

024E+30.51RT (min)

lcorbi\_hilic\_pos\_1\_053\_08

m/z 537.1955

0123E+46.57RT (min)

lcorbi\_hilic\_pos\_1\_029\_14

m/z 537.4522

02468E+30.51RT (min)

lcorbi\_hilic\_pos\_4\_066\_57

m/z 537.5362

0246E+30.51RT (min)

lcorbi\_hilic\_pos\_3\_044\_10

m/z 537.5725

01E+40.51RT (min)

lcorbi\_hilic\_pos\_1\_044\_17

m/z 538.3874

0144.5RT (min)

lcorbi\_hilic\_pos\_4\_109\_62

m/z 538.4472

01234E+40.51RT (min)

lcorbi\_hilic\_pos\_1\_029\_14

m/z 539.1639

0246E+40.51RT (min)

lcorbi\_hilic\_pos\_4\_105\_61

m/z 539.5232

01234E+40.51RT (min)

lcorbi\_hilic\_pos\_1\_031\_36

m/z 540.3065

02468E+344.5RT (min)

lcorbi\_hilic\_pos\_4\_082\_59

m/z 540.3668

0246E+344.5RT (min)

lcorbi\_hilic\_pos\_4\_045\_33

m/z 540.4628

010.51RT (min)

lcorbi\_hilic\_pos\_4\_089\_59

m/z 540.4993

024E+322.5RT (min)

lcorbi\_hilic\_pos\_4\_071\_57

m/z 540.5357

01E+40.51RT (min)

lcorbi\_hilic\_pos\_2\_023\_34

m/z 541.4509

0246E+40.51RT (min)

lcorbi\_hilic\_pos\_1\_049\_55

m/z 542.3380

01E+43.54RT (min)

lcorbi\_hilic\_pos\_4\_068\_57

m/z 542.7319

0123E+455.5RT (min)

lcorbi\_hilic\_pos\_4\_075\_58

m/z 543.3275

02468E+344.5RT (min)

lcorbi\_hilic\_pos\_4\_067\_57

m/z 544.7301

01E+455.5RT (min)

lcorbi\_hilic\_pos\_4\_074\_58

m/z 546.3453

012E+444.5RT (min)

lcorbi\_hilic\_pos\_4\_018\_31

m/z 546.3639

0123E+455.5RT (min)

lcorbi\_hilic\_pos\_4\_091\_60

m/z 546.5252

024E+40.51RT (min)

lcorbi\_hilic\_pos\_4\_068\_57

m/z 547.4399

0246E+444.5RT (min)

lcorbi\_hilic\_pos\_4\_091\_60

m/z 548.3719

0144.55RT (min)

lcorbi\_hilic\_pos\_4\_091\_60

m/z 549.3750

024E+344.5RT (min)

lcorbi\_hilic\_pos\_4\_076\_58

m/z 550.3875

012E+444.5RT (min)

lcorbi\_hilic\_pos\_4\_078\_58

m/z 550.4591

01234E+444.5RT (min)

lcorbi\_hilic\_pos\_4\_092\_60

m/z 550.5200

0246E+40.51RT (min)

lcorbi\_hilic\_pos\_1\_052\_35

m/z 550.6293

01E+300.5RT (min)

lcorbi\_hilic\_pos\_4\_036\_09

m/z 551.2283

0246E+311.5RT (min)

lcorbi\_hilic\_pos\_4\_113\_62

m/z 551.3906

0246E+344.5RT (min)

lcorbi\_hilic\_pos\_4\_079\_58

m/z 552.0608

01E+44.55RT (min)

lcorbi\_hilic\_pos\_4\_072\_57

m/z 552.4031

01E+444.5RT (min)

lcorbi\_hilic\_pos\_3\_036\_33

m/z 553.0594

01E+44.55RT (min)

lcorbi\_hilic\_pos\_4\_071\_57

m/z 554.0577

02468E+34.55RT (min)

lcorbi\_hilic\_pos\_4\_078\_58

m/z 554.1766

012E+40.51RT (min)

lcorbi\_hilic\_pos\_4\_105\_61

m/z 554.4283

01E+42.53RT (min)

lcorbi\_hilic\_pos\_4\_082\_59

m/z 554.4502

010.51RT (min)

lcorbi\_hilic\_pos\_4\_056\_56

m/z 554.5148

012E+40.51RT (min)

lcorbi\_hilic\_pos\_4\_088\_59

m/z 557.4460

02468E+40.51RT (min)

lcorbi\_hilic\_pos\_4\_018\_31

m/z 558.2962

0246E+344.5RT (min)

lcorbi\_hilic\_pos\_4\_082\_59

m/z 560.3355

01E+444.5RT (min)

lcorbi\_hilic\_pos\_4\_081\_58

m/z 561.3389

0123E+344.5RT (min)

lcorbi\_hilic\_pos\_3\_020\_31

m/z 561.4267

02468E+40.51RT (min)

lcorbi\_hilic\_pos\_3\_042\_17

m/z 562.3275

0246E+344.5RT (min)

lcorbi\_hilic\_pos\_4\_074\_58

m/z 565.5392

01E+50.51RT (min)

lcorbi\_hilic\_pos\_3\_011\_34

m/z 565.5676

01E+40.51RT (min)

lcorbi\_hilic\_pos\_3\_054\_16

m/z 566.3226

0144.5RT (min)

lcorbi\_hilic\_pos\_4\_089\_59

m/z 566.6073

01E+40.51RT (min)

lcorbi\_hilic\_pos\_1\_053\_08

m/z 567.3260

0144.5RT (min)

lcorbi\_hilic\_pos\_4\_082\_59

m/z 568.4350

012E+43.54RT (min)

lcorbi\_hilic\_pos\_4\_084\_59

m/z 568.5307

024E+40.51RT (min)

lcorbi\_hilic\_pos\_4\_032\_35

m/z 568.5670

024E+40.51RT (min)

lcorbi\_hilic\_pos\_4\_083\_59

m/z 569.2389

01234E+411.5RT (min)

lcorbi\_hilic\_pos\_4\_106\_62

m/z 569.3439

012E+444.5RT (min)

lcorbi\_hilic\_pos\_4\_066\_57

m/z 570.4737

010.51RT (min)

lcorbi\_hilic\_pos\_3\_024\_35

m/z 571.3596

0123E+444.5RT (min)

lcorbi\_hilic\_pos\_4\_034\_36

m/z 574.3380

02468E+40.51RT (min)

lcorbi\_hilic\_pos\_1\_051\_26

m/z 574.3744

012E+40.51RT (min)

lcorbi\_hilic\_pos\_4\_010\_47

m/z 575.4677

012E+40.51RT (min)

lcorbi\_hilic\_pos\_4\_076\_58

m/z 575.5042

01E+444.5RT (min)

lcorbi\_hilic\_pos\_4\_083\_59

m/z 576.4666

024E+300.5RT (min)

lcorbi\_hilic\_pos\_4\_111\_62

m/z 577.5198

02468E+322.5RT (min)

lcorbi\_hilic\_pos\_4\_082\_59

m/z 577.5198

01E+444.5RT (min)

lcorbi\_hilic\_pos\_4\_084\_59

m/z 578.4176

012E+544.5RT (min)

lcorbi\_hilic\_pos\_4\_090\_60

m/z 580.5308

02468E+40.51RT (min)

lcorbi\_hilic\_pos\_1\_052\_35

m/z 582.2082

012E+40.51RT (min)

lcorbi\_hilic\_pos\_4\_073\_57

m/z 582.2968

0144.5RT (min)

lcorbi\_hilic\_pos\_4\_087\_59

m/z 582.5942

0246E+300.5RT (min)

lcorbi\_hilic\_pos\_4\_072\_57

m/z 583.2558

012E+41.52RT (min)

lcorbi\_hilic\_pos\_4\_066\_57

m/z 583.4552

01E+43.54RT (min)

lcorbi\_hilic\_pos\_4\_084\_59

m/z 583.5497

01E+50.51RT (min)

lcorbi\_hilic\_pos\_3\_011\_34

m/z 584.2637

0123E+411.52RT (min)

lcorbi\_hilic\_pos\_2\_019\_16

m/z 584.4392

012E+455.5RT (min)

lcorbi\_hilic\_pos\_4\_020\_37

m/z 584.4740

02468E+40.51RT (min)

lcorbi\_hilic\_pos\_1\_049\_55

m/z 585.2715

01E+433.5RT (min)

lcorbi\_hilic\_pos\_4\_074\_58

m/z 588.4106

01234E+43.54RT (min)

lcorbi\_hilic\_pos\_3\_048\_38

m/z 589.2514

01E+444.5RT (min)

lcorbi\_hilic\_pos\_2\_016\_18

m/z 589.4581

01234E+40.51RT (min)

lcorbi\_hilic\_pos\_4\_039\_15

m/z 592.4710

01E+444.5RT (min)

lcorbi\_hilic\_pos\_4\_018\_31

m/z 595.4940

01234E+40.51RT (min)

lcorbi\_hilic\_pos\_4\_076\_58

m/z 600.4690

02468E+40.51RT (min)

lcorbi\_hilic\_pos\_3\_011\_34

m/z 600.6568

0123E+40.51RT (min)

lcorbi\_hilic\_pos\_3\_010\_32

m/z 602.5881

012E+40.51RT (min)

lcorbi\_hilic\_pos\_4\_067\_57

m/z 604.3855

01E+400.5RT (min)

lcorbi\_hilic\_pos\_1\_015\_44

m/z 608.0897

01E+488.5RT (min)

lcorbi\_hilic\_pos\_1\_052\_35

m/z 608.3932

0246E+444.5RT (min)

lcorbi\_hilic\_pos\_4\_090\_60

m/z 608.5259

024E+40.51RT (min)

lcorbi\_hilic\_pos\_4\_004\_38

m/z 611.4865

01E+43.54RT (min)

lcorbi\_hilic\_pos\_4\_085\_59

m/z 612.5204

01E+411.5RT (min)

lcorbi\_hilic\_pos\_4\_078\_58

m/z 612.6283

012E+444.5RT (min)

lcorbi\_hilic\_pos\_4\_090\_60

m/z 613.1604

01E+488.5RT (min)

lcorbi\_hilic\_pos\_4\_083\_59

m/z 613.1832

0123E+40.51RT (min)

lcorbi\_hilic\_pos\_1\_014\_27

m/z 614.6439

012E+444.5RT (min)

lcorbi\_hilic\_pos\_4\_095\_60

m/z 615.4374

0123E+50.51RT (min)

lcorbi\_hilic\_pos\_4\_036\_09

m/z 616.4908

0246E+40.51RT (min)

lcorbi\_hilic\_pos\_1\_049\_55

m/z 616.6516

012E+40.51RT (min)

lcorbi\_hilic\_pos\_4\_052\_32

m/z 618.6671

0246E+40.51RT (min)

lcorbi\_hilic\_pos\_3\_001\_49

m/z 628.1956

012E+40.51RT (min)

lcorbi\_hilic\_pos\_1\_028\_34

m/z 628.6034

0123E+40.51RT (min)

lcorbi\_hilic\_pos\_4\_068\_57

m/z 630.6189

012E+40.51RT (min)

lcorbi\_hilic\_pos\_4\_068\_57

m/z 632.4168

0246E+300.5RT (min)

lcorbi\_hilic\_pos\_1\_008\_38

m/z 633.2275

0123E+488.5RT (min)

lcorbi\_hilic\_pos\_4\_094\_60

m/z 634.8769

02468E+35.56RT (min)

lcorbi\_hilic\_pos\_4\_071\_57

m/z 635.5447

010.51RT (min)

lcorbi\_hilic\_pos\_4\_018\_31

m/z 637.3062

01234E+40.51RT (min)

lcorbi\_hilic\_pos\_2\_027\_38

m/z 637.5603

01234E+40.51RT (min)

lcorbi\_hilic\_pos\_4\_004\_38

m/z 638.5726

0246E+40.51RT (min)

lcorbi\_hilic\_pos\_1\_032\_31

m/z 646.2571

01E+40.51RT (min)

lcorbi\_hilic\_pos\_1\_044\_17

m/z 647.5771

0123E+40.51RT (min)

lcorbi\_hilic\_pos\_1\_029\_14

m/z 647.6654

0246E+40.51RT (min)

lcorbi\_hilic\_pos\_1\_036\_32

m/z 655.3362

0246E+40.51RT (min)

lcorbi\_hilic\_pos\_4\_004\_38

m/z 664.1172

01E+47.58RT (min)

lcorbi\_hilic\_pos\_2\_012\_37

m/z 669.6409

0246E+40.51RT (min)

lcorbi\_hilic\_pos\_2\_024\_32

m/z 671.5771

0246E+40.51RT (min)

lcorbi\_hilic\_pos\_4\_006\_14

m/z 671.6566

01E+50.51RT (min)

lcorbi\_hilic\_pos\_3\_010\_32

m/z 672.5802

0246E+40.51RT (min)

lcorbi\_hilic\_pos\_1\_029\_14

m/z 673.5287

01E+444.5RT (min)

lcorbi\_hilic\_pos\_4\_068\_57

m/z 673.6720

01E+50.51RT (min)

lcorbi\_hilic\_pos\_3\_010\_32

m/z 675.2915

012E+40.51RT (min)

lcorbi\_hilic\_pos\_1\_054\_23

m/z 675.6877

01E+50.51RT (min)

lcorbi\_hilic\_pos\_3\_010\_32

m/z 677.7033

01E+50.51RT (min)

lcorbi\_hilic\_pos\_3\_010\_32

m/z 681.3209

012E+53.54RT (min)

lcorbi\_hilic\_pos\_3\_004\_27

m/z 684.6508

0123E+40.51RT (min)

lcorbi\_hilic\_pos\_4\_089\_59

m/z 686.2020

0123E+40.51RT (min)

lcorbi\_hilic\_pos\_1\_014\_27

m/z 690.5076

01E+40.51RT (min)

lcorbi\_hilic\_pos\_4\_089\_59

m/z 690.5635

01234E+444.5RT (min)

lcorbi\_hilic\_pos\_2\_054\_29

m/z 691.3147

01E+40.51RT (min)

lcorbi\_hilic\_pos\_1\_054\_23

m/z 702.5370

01E+50.51RT (min)

lcorbi\_hilic\_pos\_4\_049\_55

m/z 704.5234

01E+43.54RT (min)

lcorbi\_hilic\_pos\_4\_087\_59

m/z 706.5390

0123E+43.54RT (min)

lcorbi\_hilic\_pos\_4\_066\_57

m/z 706.6139

01234E+40.51RT (min)

lcorbi\_hilic\_pos\_2\_040\_14

m/z 707.2604

02468E+50.51RT (min)

lcorbi\_hilic\_pos\_1\_052\_35

m/z 707.5424

01E+43.54RT (min)

lcorbi\_hilic\_pos\_4\_070\_57

m/z 708.2639

012E+50.51RT (min)

lcorbi\_hilic\_pos\_4\_003\_26

m/z 708.5124

012E+40.51RT (min)

lcorbi\_hilic\_pos\_4\_089\_59

m/z 713.0673

01E+44.55RT (min)

lcorbi\_hilic\_pos\_4\_078\_58

m/z 714.5077

01234E+433.5RT (min)

lcorbi\_hilic\_pos\_4\_083\_59

m/z 714.5078

01E+40.51RT (min)

lcorbi\_hilic\_pos\_4\_005\_34

m/z 715.5112

012E+433.5RT (min)

lcorbi\_hilic\_pos\_4\_084\_59

m/z 717.5267

02468E+40.51RT (min)

lcorbi\_hilic\_pos\_4\_032\_35

m/z 718.5389

01E+40.51RT (min)

lcorbi\_hilic\_pos\_4\_084\_59

m/z 719.5424

024E+433.5RT (min)

lcorbi\_hilic\_pos\_4\_083\_59

m/z 719.5424

0246E+40.51RT (min)

lcorbi\_hilic\_pos\_2\_023\_34

m/z 720.5453

01E+433.5RT (min)

lcorbi\_hilic\_pos\_4\_086\_59

m/z 720.5911

01234E+33.54RT (min)

lcorbi\_hilic\_pos\_4\_066\_57

m/z 726.5078

0246E+33.54RT (min)

lcorbi\_hilic\_pos\_4\_082\_59

m/z 728.5235

012E+43.54RT (min)

lcorbi\_hilic\_pos\_4\_083\_59

m/z 728.5235

02468E+30.51RT (min)

lcorbi\_hilic\_pos\_4\_088\_59

m/z 729.5270

01E+43.54RT (min)

lcorbi\_hilic\_pos\_4\_089\_59

m/z 730.5391

012E+40.51RT (min)

lcorbi\_hilic\_pos\_4\_087\_59

m/z 730.5950

01234E+444.5RT (min)

lcorbi\_hilic\_pos\_1\_046\_30

m/z 731.5426

012E+43.54RT (min)

lcorbi\_hilic\_pos\_4\_089\_59

m/z 731.5426

01E+40.51RT (min)

lcorbi\_hilic\_pos\_4\_089\_59

m/z 731.6066

0246E+43.54RT (min)

lcorbi\_hilic\_pos\_2\_054\_29

m/z 733.5582

01E+40.51RT (min)

lcorbi\_hilic\_pos\_4\_073\_57

m/z 734.5701

012E+40.51RT (min)

lcorbi\_hilic\_pos\_4\_020\_37

m/z 735.5738

01E+43.54RT (min)

lcorbi\_hilic\_pos\_4\_066\_57

m/z 736.4923

01E+433.5RT (min)

lcorbi\_hilic\_pos\_4\_083\_59

m/z 736.5771

024E+33.54RT (min)

lcorbi\_hilic\_pos\_4\_070\_57

m/z 737.3496

012E+422.5RT (min)

lcorbi\_hilic\_pos\_4\_109\_62

m/z 739.5112

024E+40.51RT (min)

lcorbi\_hilic\_pos\_2\_012\_37

m/z 740.5232

02468E+40.51RT (min)

lcorbi\_hilic\_pos\_2\_013\_36

m/z 741.5269

01234E+40.51RT (min)

lcorbi\_hilic\_pos\_2\_012\_37

m/z 742.5390

0123E+40.51RT (min)

lcorbi\_hilic\_pos\_4\_083\_59

m/z 743.5426

01E+40.51RT (min)

lcorbi\_hilic\_pos\_4\_089\_59

m/z 744.5913

013.54RT (min)

lcorbi\_hilic\_pos\_4\_082\_59

m/z 744.5916

01E+50.51RT (min)

lcorbi\_hilic\_pos\_4\_084\_59

m/z 745.5581

0123E+40.51RT (min)

lcorbi\_hilic\_pos\_4\_084\_59

m/z 745.5947

013.54RT (min)

lcorbi\_hilic\_pos\_4\_089\_59

m/z 745.5961

0246E+40.51RT (min)

lcorbi\_hilic\_pos\_4\_084\_59

m/z 746.5696

0123E+40.51RT (min)

lcorbi\_hilic\_pos\_4\_089\_59

m/z 746.5701

02468E+33.54RT (min)

lcorbi\_hilic\_pos\_4\_082\_59

m/z 746.6066

013.54RT (min)

lcorbi\_hilic\_pos\_4\_082\_59

m/z 746.6068

01E+40.51RT (min)

lcorbi\_hilic\_pos\_4\_083\_59

m/z 747.5733

01E+40.51RT (min)

lcorbi\_hilic\_pos\_4\_088\_59

m/z 747.5738

024E+33.54RT (min)

lcorbi\_hilic\_pos\_4\_089\_59

m/z 750.5077

01E+43.54RT (min)

lcorbi\_hilic\_pos\_4\_087\_59

m/z 752.5233

01E+40.51RT (min)

lcorbi\_hilic\_pos\_4\_087\_59

m/z 753.5266

0246E+30.51RT (min)

lcorbi\_hilic\_pos\_4\_086\_59

m/z 753.5267

012E+43.54RT (min)

lcorbi\_hilic\_pos\_4\_089\_59

m/z 753.6390

0123E+433.5RT (min)

lcorbi\_hilic\_pos\_4\_090\_60

m/z 754.6506

01E+40.51RT (min)

lcorbi\_hilic\_pos\_4\_097\_60

m/z 754.6507

024E+433.5RT (min)

lcorbi\_hilic\_pos\_4\_095\_60

m/z 756.5987

0246E+41.52RT (min)

lcorbi\_hilic\_pos\_4\_087\_59

m/z 756.6658

012E+433.5RT (min)

lcorbi\_hilic\_pos\_4\_090\_60

m/z 757.6697

012E+40.51RT (min)

lcorbi\_hilic\_pos\_4\_097\_60

m/z 758.2224

01234E+40.51RT (min)

lcorbi\_hilic\_pos\_1\_014\_27

m/z 758.4974

012E+444.5RT (min)

lcorbi\_hilic\_pos\_4\_088\_59

m/z 758.5294

01E+33.54RT (min)

lcorbi\_hilic\_pos\_4\_069\_57

m/z 758.6107

01E+33.54RT (min)

lcorbi\_hilic\_pos\_4\_069\_57

m/z 759.2230

0123E+40.51RT (min)

lcorbi\_hilic\_pos\_1\_032\_31

m/z 760.2209

0123E+40.51RT (min)

lcorbi\_hilic\_pos\_1\_014\_27

m/z 761.5164

0246E+444.5RT (min)

lcorbi\_hilic\_pos\_4\_082\_59

m/z 762.5276

024E+444.5RT (min)

lcorbi\_hilic\_pos\_4\_082\_59

m/z 763.5952

02468E+33.54RT (min)

lcorbi\_hilic\_pos\_4\_068\_57

m/z 763.5954

0246E+30.51RT (min)

lcorbi\_hilic\_pos\_4\_073\_57

m/z 764.5232

012E+43.54RT (min)

lcorbi\_hilic\_pos\_4\_089\_59

m/z 764.5233

02468E+30.51RT (min)

lcorbi\_hilic\_pos\_3\_029\_23

m/z 765.5266

01E+43.54RT (min)

lcorbi\_hilic\_pos\_4\_087\_59

m/z 766.5386

0123E+43.54RT (min)

lcorbi\_hilic\_pos\_4\_082\_59

m/z 766.5389

012E+40.51RT (min)

lcorbi\_hilic\_pos\_4\_085\_59

m/z 766.5753

0123E+433.5RT (min)

lcorbi\_hilic\_pos\_1\_036\_32

m/z 767.5422

01E+43.54RT (min)

lcorbi\_hilic\_pos\_4\_082\_59

m/z 767.5422

01E+40.51RT (min)

lcorbi\_hilic\_pos\_4\_087\_59

m/z 767.5787

01E+433.5RT (min)

lcorbi\_hilic\_pos\_1\_036\_32

m/z 767.6541

0246E+33.54RT (min)

lcorbi\_hilic\_pos\_4\_096\_60

m/z 768.6576

012E+43.54RT (min)

lcorbi\_hilic\_pos\_4\_096\_60

m/z 769.5578

0123E+40.51RT (min)

lcorbi\_hilic\_pos\_4\_089\_59

m/z 769.5579

012E+43.54RT (min)

lcorbi\_hilic\_pos\_4\_082\_59

m/z 769.5943

010.51RT (min)

lcorbi\_hilic\_pos\_4\_087\_59

m/z 769.6696

01E+43.54RT (min)

lcorbi\_hilic\_pos\_4\_096\_60

m/z 770.5698

01234E+43.54RT (min)

lcorbi\_hilic\_pos\_4\_082\_59

m/z 770.6056

0133.5RT (min)

lcorbi\_hilic\_pos\_4\_082\_59

m/z 770.6057

010.51RT (min)

lcorbi\_hilic\_pos\_4\_082\_59

m/z 770.6731

024E+43.54RT (min)

lcorbi\_hilic\_pos\_4\_092\_60

m/z 770.6732

012E+40.51RT (min)

lcorbi\_hilic\_pos\_4\_090\_60

m/z 771.5734

012E+43.54RT (min)

lcorbi\_hilic\_pos\_4\_086\_59

m/z 771.5735

012E+40.51RT (min)

lcorbi\_hilic\_pos\_4\_085\_59

m/z 771.6096

0133.5RT (min)

lcorbi\_hilic\_pos\_4\_086\_59

m/z 772.5855

012E+43.54RT (min)

lcorbi\_hilic\_pos\_4\_082\_59

m/z 772.6217

0133.5RT (min)

lcorbi\_hilic\_pos\_4\_086\_59

m/z 772.6887

01E+40.51RT (min)

lcorbi\_hilic\_pos\_4\_090\_60

m/z 772.6887

012E+43.54RT (min)

lcorbi\_hilic\_pos\_4\_093\_60

m/z 773.5891

01E+43.54RT (min)

lcorbi\_hilic\_pos\_4\_086\_59

m/z 773.5892

01E+40.51RT (min)

lcorbi\_hilic\_pos\_4\_083\_59

m/z 774.6006

024E+33.54RT (min)

lcorbi\_hilic\_pos\_4\_069\_57

m/z 774.6007

012E+40.51RT (min)

lcorbi\_hilic\_pos\_3\_004\_27

m/z 778.6419

01E+40.51RT (min)

lcorbi\_hilic\_pos\_4\_097\_60

m/z 778.6419

01234E+433.5RT (min)

lcorbi\_hilic\_pos\_2\_037\_24

m/z 780.6572

012E+433.5RT (min)

lcorbi\_hilic\_pos\_1\_003\_24

m/z 781.6609

012E+40.51RT (min)

lcorbi\_hilic\_pos\_4\_090\_60

m/z 782.6730

01E+40.51RT (min)

lcorbi\_hilic\_pos\_4\_097\_60

m/z 782.6731

01E+433.5RT (min)

lcorbi\_hilic\_pos\_4\_052\_32

m/z 783.6767

0123E+40.51RT (min)

lcorbi\_hilic\_pos\_4\_090\_60

m/z 784.6885

012E+40.51RT (min)

lcorbi\_hilic\_pos\_4\_096\_60

m/z 784.6887

012E+433.5RT (min)

lcorbi\_hilic\_pos\_3\_010\_32

m/z 785.6921

0246E+40.51RT (min)

lcorbi\_hilic\_pos\_4\_097\_60

m/z 787.7076

01E+40.51RT (min)

lcorbi\_hilic\_pos\_4\_090\_60

m/z 787.7078

01E+433.5RT (min)

lcorbi\_hilic\_pos\_3\_010\_32

m/z 789.6195

012E+433.5RT (min)

lcorbi\_hilic\_pos\_4\_066\_57

m/z 789.6196

012E+40.51RT (min)

lcorbi\_hilic\_pos\_4\_069\_57

m/z 790.6230

0246E+333.5RT (min)

lcorbi\_hilic\_pos\_4\_072\_57

m/z 792.5910

0246E+333.5RT (min)

lcorbi\_hilic\_pos\_4\_020\_37

m/z 792.6493

0246E+33.54RT (min)

lcorbi\_hilic\_pos\_4\_091\_60

m/z 793.5943

01234E+333.5RT (min)

lcorbi\_hilic\_pos\_4\_020\_37

m/z 793.6610

01E+43.54RT (min)

lcorbi\_hilic\_pos\_4\_092\_60

m/z 793.7576

02468E+40.51RT (min)

lcorbi\_hilic\_pos\_2\_037\_24

m/z 794.6651

01E+40.51RT (min)

lcorbi\_hilic\_pos\_1\_003\_24

m/z 795.6100

01234E+40.51RT (min)

lcorbi\_hilic\_pos\_4\_014\_27

m/z 795.6766

012E+40.51RT (min)

lcorbi\_hilic\_pos\_1\_003\_24

m/z 796.5856

01E+433.5RT (min)

lcorbi\_hilic\_pos\_4\_050\_16

m/z 796.5857

012E+40.51RT (min)

lcorbi\_hilic\_pos\_1\_014\_27

m/z 796.6217

012E+40.51RT (min)

lcorbi\_hilic\_pos\_1\_014\_27

m/z 797.5892

012E+333.5RT (min)

lcorbi\_hilic\_pos\_2\_001\_49

m/z 797.6255

02468E+30.51RT (min)

lcorbi\_hilic\_pos\_4\_069\_57

m/z 798.5651

012E+43.54RT (min)

lcorbi\_hilic\_pos\_3\_008\_14

m/z 798.5884

0123E+50.51RT (min)

lcorbi\_hilic\_pos\_2\_040\_14

m/z 798.6004

012E+333.5RT (min)

lcorbi\_hilic\_pos\_4\_071\_57

m/z 798.6301

01E+333.5RT (min)

lcorbi\_hilic\_pos\_4\_090\_60

m/z 799.5685

01E+43.54RT (min)

lcorbi\_hilic\_pos\_3\_008\_14

m/z 799.5917

01E+50.51RT (min)

lcorbi\_hilic\_pos\_2\_040\_14

m/z 799.7073

01234E+40.51RT (min)

lcorbi\_hilic\_pos\_1\_003\_24

m/z 800.6473

0246E+444.5RT (min)

lcorbi\_hilic\_pos\_4\_090\_60

m/z 801.7206

01E+43.54RT (min)

lcorbi\_hilic\_pos\_4\_095\_60

m/z 802.7265

012E+40.51RT (min)

lcorbi\_hilic\_pos\_4\_091\_60

m/z 802.7265

012E+43.54RT (min)

lcorbi\_hilic\_pos\_4\_090\_60

m/z 804.5539

01234E+333.5RT (min)

lcorbi\_hilic\_pos\_4\_078\_58

m/z 807.6677

01E+43.54RT (min)

lcorbi\_hilic\_pos\_4\_090\_60

m/z 809.6834

01E+43.54RT (min)

lcorbi\_hilic\_pos\_4\_092\_60

m/z 809.6835

012E+40.51RT (min)

lcorbi\_hilic\_pos\_1\_003\_24

m/z 810.6953

02468E+33.54RT (min)

lcorbi\_hilic\_pos\_4\_091\_60

m/z 810.6954

01E+40.51RT (min)

lcorbi\_hilic\_pos\_1\_003\_24

m/z 811.6989

012E+43.54RT (min)

lcorbi\_hilic\_pos\_4\_092\_60

m/z 811.6990

01234E+40.51RT (min)

lcorbi\_hilic\_pos\_1\_003\_24

m/z 812.7109

01E+40.51RT (min)

lcorbi\_hilic\_pos\_1\_003\_24

m/z 813.6200

012E+40.51RT (min)

lcorbi\_hilic\_pos\_4\_070\_57

m/z 813.7145

01234E+40.51RT (min)

lcorbi\_hilic\_pos\_3\_055\_24

m/z 813.7145

012E+43.54RT (min)

lcorbi\_hilic\_pos\_4\_096\_60

m/z 814.6249

01E+433.5RT (min)

lcorbi\_hilic\_pos\_4\_071\_57

m/z 814.6253

01E+50.51RT (min)

lcorbi\_hilic\_pos\_1\_038\_21

m/z 815.7300

012E+40.51RT (min)

lcorbi\_hilic\_pos\_4\_090\_60

m/z 815.7301

01E+43.54RT (min)

lcorbi\_hilic\_pos\_4\_093\_60

m/z 818.6066

0246E+333.5RT (min)

lcorbi\_hilic\_pos\_4\_072\_57

m/z 820.6220

02468E+333.5RT (min)

lcorbi\_hilic\_pos\_4\_069\_57

m/z 820.6797

01E+43.54RT (min)

lcorbi\_hilic\_pos\_4\_090\_60

m/z 821.6256

024E+333.5RT (min)

lcorbi\_hilic\_pos\_4\_070\_57

m/z 821.6834

01234E+40.51RT (min)

lcorbi\_hilic\_pos\_1\_003\_24

m/z 822.5651

0246E+33.54RT (min)

lcorbi\_hilic\_pos\_2\_040\_14

m/z 822.6376

01E+433.5RT (min)

lcorbi\_hilic\_pos\_4\_066\_57

m/z 823.6413

0246E+333.5RT (min)

lcorbi\_hilic\_pos\_4\_069\_57

m/z 824.5804

0246E+33.54RT (min)

lcorbi\_hilic\_pos\_3\_008\_14

m/z 824.6528

0133.5RT (min)

lcorbi\_hilic\_pos\_4\_093\_60

m/z 826.5963

01E+43.54RT (min)

lcorbi\_hilic\_pos\_3\_008\_14

m/z 827.5997

02468E+33.54RT (min)

lcorbi\_hilic\_pos\_3\_008\_14

m/z 827.7294

01234E+40.51RT (min)

lcorbi\_hilic\_pos\_1\_003\_24

m/z 829.7443

0123E+433.5RT (min)

lcorbi\_hilic\_pos\_4\_094\_60

m/z 830.5701

012E+433.5RT (min)

lcorbi\_hilic\_pos\_4\_020\_37

m/z 830.7490

01234E+40.51RT (min)

lcorbi\_hilic\_pos\_1\_003\_24

m/z 832.2414

0246E+40.51RT (min)

lcorbi\_hilic\_pos\_2\_020\_31

m/z 832.5855

0246E+30.51RT (min)

lcorbi\_hilic\_pos\_4\_070\_57

m/z 832.5856

01E+433.5RT (min)

lcorbi\_hilic\_pos\_4\_020\_37

m/z 833.2422

024E+40.51RT (min)

lcorbi\_hilic\_pos\_2\_020\_31

m/z 833.5891

01E+433.5RT (min)

lcorbi\_hilic\_pos\_4\_020\_37

m/z 834.2401

01234E+40.51RT (min)

lcorbi\_hilic\_pos\_1\_032\_31

m/z 834.6015

02468E+40.51RT (min)

lcorbi\_hilic\_pos\_3\_004\_27

m/z 835.6050

01E+40.51RT (min)

lcorbi\_hilic\_pos\_4\_070\_57

m/z 836.6161

01E+40.51RT (min)

lcorbi\_hilic\_pos\_4\_070\_57

m/z 837.6199

01E+433.5RT (min)

lcorbi\_hilic\_pos\_4\_071\_57

m/z 837.6200

0246E+30.51RT (min)

lcorbi\_hilic\_pos\_4\_067\_57

m/z 838.6314

01E+433.5RT (min)

lcorbi\_hilic\_pos\_4\_070\_57

m/z 839.6355

02468E+333.5RT (min)

lcorbi\_hilic\_pos\_4\_066\_57

m/z 870.7547

012E+40.51RT (min)

lcorbi\_hilic\_pos\_2\_020\_31

m/z 872.7700

012E+40.51RT (min)

lcorbi\_hilic\_pos\_1\_032\_31

m/z 874.7852

01E+40.51RT (min)

lcorbi\_hilic\_pos\_4\_004\_38

m/z 894.7547

012E+40.51RT (min)

lcorbi\_hilic\_pos\_4\_089\_59

m/z 896.7701

012E+40.51RT (min)

lcorbi\_hilic\_pos\_4\_089\_59

m/z 897.7740

01E+40.51RT (min)

lcorbi\_hilic\_pos\_4\_089\_59

m/z 898.7855

0123E+40.51RT (min)

lcorbi\_hilic\_pos\_4\_089\_59

m/z 899.7896

012E+40.51RT (min)

lcorbi\_hilic\_pos\_4\_004\_38

m/z 900.8006

012E+40.51RT (min)

lcorbi\_hilic\_pos\_4\_089\_59

m/z 906.2600

012E+40.51RT (min)

lcorbi\_hilic\_pos\_1\_032\_31
