## Supplementary material for "The Everything Bagel Feature Finder: Ultra-fast automated feature finding for untargeted metabolomics": SI Data File: tool_unique_xic_st_XCMS_ST004502.html

 
 
 XCMS-unique features (not detected by the other two tools) - ST004502 
 
 XCMS-unique features (not detected by the other two tools) - ST004502 
 8974 features · orbitrap · XIC window m/z ± 10 ppm (≥ 0.002 Da) · ± 0.5 min 
 Every panel is a feature  XCMS  detected that neither of the other two tools detected — three-way unique across EB (standard mode (min_detection_rate=0.1)), Masscube and XCMS, under the standard matching rule (m/z ± 10 ppm; apex within the reference RT window ± 0.2 min). It shows the MS1 extracted-ion chromatogram at the feature's m/z, drawn from the raw mzML named in  alignment_reference_file , zoomed to ± 0.5 min around the feature's reported apex RT.  Gray solid line  = the feature's reported apex RT;  red dashed line  = XCMS's assigned MS/MS —  RTINSECONDS  in its  aligned_features.mgf ; the panel widens to keep an off-apex MS/MS on-screen. Features with no assigned MS/MS show only the gray line. y is per-cell autoscaled. Rows keep the table order (m/z ascending). 
   m/z 50.0228     0   1   2   3E+3   5   5.5  RT (min)      lcorbi_hilic_neg_2_020_27    m/z 50.2029     0   1   2   3E+3   4.5   5  RT (min)      lcorbi_hilic_neg_4_055_38    m/z 50.2523     0   1E+4   5   5.5  RT (min)      lcorbi_hilic_neg_2_020_27    m/z 50.2590     0   1   2   3E+3   3.5   4  RT (min)      lcorbi_hilic_neg_4_044_01    m/z 50.2644     0   1   2E+3   2   2.5  RT (min)      lcorbi_hilic_neg_4_055_38    m/z 50.4623     0   1E+4   4.5   5  RT (min)      lcorbi_hilic_neg_4_107_62    m/z 50.5099     0   2   4   6E+3   2   2.5  RT (min)      lcorbi_hilic_neg_4_075_58    m/z 50.7589     0   2   4   6   8E+3   2   2.5  RT (min)      lcorbi_hilic_neg_4_068_57    m/z 50.9473     0   1E+4   1   1.5  RT (min)      lcorbi_hilic_neg_4_011_19    m/z 50.9902     0   2   4   6E+3   3.5   4  RT (min)      lcorbi_hilic_neg_2_020_27    m/z 51.2482     0   1E+3   3.5   4  RT (min)      lcorbi_hilic_neg_2_050_17    m/z 51.6455     0   2   4   6E+3   4.5   5  RT (min)      lcorbi_hilic_neg_4_078_58    m/z 51.6949     0   2   4   6E+3   6   6.5  RT (min)      lcorbi_hilic_neg_4_089_59    m/z 51.8124     0   1   2E+4   0.5   1  RT (min)      lcorbi_hilic_neg_2_050_17    m/z 51.9057     0   1   2E+4   0.5   1  RT (min)      lcorbi_hilic_neg_2_019_21    m/z 52.2669     0   2   4   6   8E+3   3.5   4  RT (min)      lcorbi_hilic_neg_2_020_27    m/z 52.3501     0   2   4   6E+4   1   1.5  RT (min)      lcorbi_hilic_neg_1_027_18    m/z 52.5920     0   2   4   6   8E+3   5   5.5  RT (min)      lcorbi_hilic_neg_2_020_27    m/z 52.7298     0   1   2   3   4E+3   5   5.5  RT (min)      lcorbi_hilic_neg_2_020_27    m/z 52.7506     0   2   4   6E+3   1.5   2  RT (min)      lcorbi_hilic_neg_1_027_18    m/z 53.0008     0   2   4   6E+3   1   1.5  RT (min)      lcorbi_hilic_neg_1_048_35    m/z 53.1442     0   1E+4   4   4.5  RT (min)      lcorbi_hilic_neg_1_047_05    m/z 53.2499     0   1   2E+3   1.5   2  RT (min)      lcorbi_hilic_neg_1_018_21    m/z 53.2683     0   1   2   3   4E+3   1   1.5  RT (min)      lcorbi_hilic_neg_4_014_18    m/z 53.3227     0   2   4E+3   6.5   7  RT (min)      lcorbi_hilic_neg_4_023_34    m/z 53.3680     0   1E+4   4   4.5  RT (min)      lcorbi_hilic_neg_2_012_14    m/z 53.3685     0   1E+4   4   4.5  RT (min)      lcorbi_hilic_neg_2_012_14    m/z 53.5416     0   1E+4   6   6.5  RT (min)      lcorbi_hilic_neg_4_023_34    m/z 54.0072     0   1E+4   5   5.5  RT (min)      lcorbi_hilic_neg_2_020_27    m/z 54.0093     0   1   2   3   4E+3   5   5.5  RT (min)      lcorbi_hilic_neg_1_015_30    m/z 54.2577     0   1E+4   5   5.5  RT (min)      lcorbi_hilic_neg_2_054_13    m/z 54.2634     0   1   2E+3   2   2.5  RT (min)      lcorbi_hilic_neg_2_020_27    m/z 54.5262     0   1E+3   1.5   2  RT (min)      lcorbi_hilic_neg_2_040_35    m/z 54.6382     0   1   2   3   4E+3   5   5.5  RT (min)      lcorbi_hilic_neg_2_020_27    m/z 54.6441     0   1E+4   6   6.5  RT (min)      lcorbi_hilic_neg_1_012_32    m/z 54.8920     0   2   4E+3   1   1.5  RT (min)      lcorbi_hilic_neg_1_046_19    m/z 54.9906     0   2   4   6   8E+3   5   5.5  RT (min)      lcorbi_hilic_neg_2_034_01    m/z 55.0177     0   2   4E+3   1   1.5  RT (min)      lcorbi_hilic_neg_2_020_27    m/z 55.0189     0   1   2E+4   6   6.5  RT (min)      lcorbi_hilic_neg_1_005_36    m/z 55.0898     0   1E+4   1   1.5  RT (min)      lcorbi_hilic_neg_1_027_18    m/z 55.2472     0   2   4   6E+3   0.5   1  RT (min)      lcorbi_hilic_neg_4_008_35    m/z 55.2480     0   1   2   3E+3   5   5.5  RT (min)      lcorbi_hilic_neg_2_020_27    m/z 55.2602     0   1E+4   4   4.5  RT (min)      lcorbi_hilic_neg_1_047_05    m/z 55.2889     0   1   2E+4   0.5   1  RT (min)      lcorbi_hilic_neg_3_007_07    m/z 55.4843     0   1   2   3   4E+4   4   4.5  RT (min)      lcorbi_hilic_neg_1_047_05    m/z 55.6916     0   1   2   3   4E+3   3   3.5  RT (min)      lcorbi_hilic_neg_3_034_34    m/z 55.6987     0   1   4   4.5  RT (min)      lcorbi_hilic_neg_1_047_05    m/z 55.8863     0   1E+4   1   1.5  RT (min)      lcorbi_hilic_neg_1_036_09    m/z 56.0159     0   1   2E+3   3   3.5  RT (min)      lcorbi_hilic_neg_3_053_13    m/z 56.0814     0   2   4   6E+3   1   1.5  RT (min)      lcorbi_hilic_neg_3_010_18    m/z 56.1910     0   1   2   3E+3   3   3.5  RT (min)      lcorbi_hilic_neg_2_031_34    m/z 56.2657     0   1E+4   5   5.5  RT (min)      lcorbi_hilic_neg_3_024_27    m/z 57.1317     0   1E+4   1   1.5  RT (min)      lcorbi_hilic_neg_2_013_18    m/z 57.2384     0   1E+4   1   1.5  RT (min)      lcorbi_hilic_neg_2_019_21    m/z 57.2624     0   1   2   3E+3   2   2.5  RT (min)      lcorbi_hilic_neg_4_004_13    m/z 57.9910     0   1   2E+3   5   5.5  RT (min)      lcorbi_hilic_neg_2_020_27    m/z 58.0289     0   1   2   3E+4   6   6.5  RT (min)      lcorbi_hilic_neg_4_005_32    m/z 58.2535     0   1E+4   4   4.5  RT (min)      lcorbi_hilic_neg_4_068_57    m/z 58.2713     0   1E+4   2   2.5  RT (min)      lcorbi_hilic_neg_4_008_35    m/z 58.7147     0   1   2   3E+3   5   5.5  RT (min)      lcorbi_hilic_neg_2_020_27    m/z 58.9121     0   1E+4   1   1.5  RT (min)      lcorbi_hilic_neg_2_004_09    m/z 58.9229     0   1E+4   4   4.5  RT (min)      lcorbi_hilic_neg_4_079_58    m/z 59.0138     0   1E+4   1.5   2  RT (min)      lcorbi_hilic_neg_4_099_61    m/z 59.0266     0   2   4   6   8E+3   0.5   1  RT (min)      lcorbi_hilic_neg_4_100_61    m/z 59.1268     0   1   2E+3   7   7.5  RT (min)      lcorbi_hilic_neg_3_011_24    m/z 59.7670     0   2   4   6E+3   0.5   1  RT (min)      lcorbi_hilic_neg_2_053_03    m/z 59.7697     0   1   2E+3   2   2.5  RT (min)      lcorbi_hilic_neg_4_051_16    m/z 59.8132     0   2   4   6E+3   4   4.5  RT (min)      lcorbi_hilic_neg_4_070_57    m/z 60.0172     0   1E+4   5   5.5  RT (min)      lcorbi_hilic_neg_4_076_58    m/z 60.2600     0   2   4   6   8E+3   4   4.5  RT (min)      lcorbi_hilic_neg_4_077_58    m/z 60.5203     0   1   2   3E+3   6   6.5  RT (min)      lcorbi_hilic_neg_4_079_58    m/z 61.0205     0   1E+4   1.5   2  RT (min)      lcorbi_hilic_neg_4_096_60    m/z 61.2612     0   1   2   3E+3   6   6.5  RT (min)      lcorbi_hilic_neg_1_029_31    m/z 61.8020     0   2   4   6   8E+3   1   1.5  RT (min)      lcorbi_hilic_neg_2_004_09    m/z 61.9404     0   2   4E+4   0.5   1  RT (min)      lcorbi_hilic_neg_1_003_14    m/z 62.2402     0   1E+4   5   5.5  RT (min)      lcorbi_hilic_neg_3_019_01    m/z 62.8154     0   1E+4   4   4.5  RT (min)      lcorbi_hilic_neg_4_070_57    m/z 63.0394     0   1E+4   4   4.5  RT (min)      lcorbi_hilic_neg_4_070_57    m/z 63.6902     0   1   2E+3   5   5.5  RT (min)      lcorbi_hilic_neg_2_020_27    m/z 64.0151     0   2   4   6   8E+3   4.5   5  RT (min)      lcorbi_hilic_neg_2_020_27    m/z 64.2638     0   1   2   3   4E+3   5.5   6  RT (min)      lcorbi_hilic_neg_4_073_57    m/z 64.5144     0   1   2E+3   4.5   5  RT (min)      lcorbi_hilic_neg_1_039_38    m/z 65.3281     0   1   2   3E+3   1   1.5  RT (min)      lcorbi_hilic_neg_4_019_09    m/z 65.4821     0   1E+4   4   4.5  RT (min)      lcorbi_hilic_neg_4_068_57    m/z 66.0194     0   2   4E+3   2.5   3  RT (min)      lcorbi_hilic_neg_1_037_46    m/z 66.0296     0   2   4   6   8E+3   4   4.5  RT (min)      lcorbi_hilic_neg_4_005_32    m/z 66.5381     0   2   4E+3   0.5   1  RT (min)      lcorbi_hilic_neg_4_037_31    m/z 66.7106     0   1E+4   3.5   4  RT (min)      lcorbi_hilic_neg_4_087_59    m/z 66.7710     0   2   4   6   8E+3   5.5   6  RT (min)      lcorbi_hilic_neg_4_071_57    m/z 66.9440     0   1E+4   0.5   1  RT (min)      lcorbi_hilic_neg_4_105_61    m/z 67.9385     0   1E+3   5   5.5  RT (min)      lcorbi_hilic_neg_3_018_38    m/z 68.5263     0   2   4E+3   2.5   3  RT (min)      lcorbi_hilic_neg_1_037_46    m/z 68.9957     0   1   2   3E+5   4   5   6  RT (min)       lcorbi_hilic_neg_1_021_28    m/z 68.9957     0   1   2E+4   9   9.5  RT (min)      lcorbi_hilic_neg_1_002_26    m/z 68.9957     0   1E+4   12.5   13  RT (min)      lcorbi_hilic_neg_1_032_02    m/z 68.9957     0   1   2   3E+4   11   11.5  RT (min)      lcorbi_hilic_neg_1_002_26    m/z 69.4117     0   2   4   6E+4   0.5   1  RT (min)      lcorbi_hilic_neg_4_078_58    m/z 69.6593     0   2   4E+3   0.5   1  RT (min)      lcorbi_hilic_neg_4_087_59    m/z 69.9714     0   2   4   6   8E+3   5   5.5  RT (min)      lcorbi_hilic_neg_2_020_27    m/z 70.5361     0   1   2E+4   1   1.5  RT (min)      lcorbi_hilic_neg_1_002_26    m/z 70.9460     0   1   2E+4   0.5   1  RT (min)      lcorbi_hilic_neg_1_008_39    m/z 70.9470     0   2   4   6   8E+5   4.5   5  RT (min)       lcorbi_hilic_neg_3_023_36    m/z 70.9485     0   1   2E+4   4.5   5  RT (min)      lcorbi_hilic_neg_4_031_14    m/z 71.0138     0   2   4E+4   1   1.5  RT (min)       lcorbi_hilic_neg_4_084_59    m/z 71.0502     0   1E+4   5   5.5  RT (min)      lcorbi_hilic_neg_2_020_27    m/z 71.4740     0   2   4   6   8E+3   0.5   1  RT (min)      lcorbi_hilic_neg_4_089_59    m/z 72.0305     0   1   2   3   4E+3   6.5   7  RT (min)      lcorbi_hilic_neg_3_028_23    m/z 72.2590     0   1   2E+3   7.5   8  RT (min)      lcorbi_hilic_neg_2_014_31    m/z 72.7728     0   2   4   6E+3   2   2.5  RT (min)      lcorbi_hilic_neg_4_051_16    m/z 72.9431     0   1   2E+4   0.5   1  RT (min)      lcorbi_hilic_neg_1_008_39    m/z 72.9442     0   1E+4   4.5   5  RT (min)       lcorbi_hilic_neg_1_028_45    m/z 72.9882     0   1E+3   5   5.5  RT (min)      lcorbi_hilic_neg_1_001_49    m/z 73.0117     0   1   2E+4   5   5.5  RT (min)       lcorbi_hilic_neg_4_105_61    m/z 73.0205     0   2   4E+3   2   2.5  RT (min)      lcorbi_hilic_neg_4_072_57    m/z 73.0480     0   1   2E+4   2   2.5  RT (min)       lcorbi_hilic_neg_4_095_60    m/z 73.2310     0   1   2   3   4E+3   7   7.5  RT (min)      lcorbi_hilic_neg_1_047_05    m/z 73.2787     0   2   4E+3   4   4.5  RT (min)      lcorbi_hilic_neg_4_099_61    m/z 73.2947     0   1E+4   0.5   1  RT (min)      lcorbi_hilic_neg_1_030_04    m/z 73.5452     0   1   2   3   4E+3   0.5   1  RT (min)      lcorbi_hilic_neg_3_007_07    m/z 74.0328     0   1   2E+3   1.5   2  RT (min)      lcorbi_hilic_neg_4_093_60    m/z 74.2207     0   1   2   3   4E+3   4.5   5  RT (min)      lcorbi_hilic_neg_4_075_58    m/z 74.5738     0   2   4   6   8E+3   1   1.5  RT (min)      lcorbi_hilic_neg_4_014_18    m/z 74.7359     0   2   4E+3   0.5   1  RT (min)      lcorbi_hilic_neg_4_108_62    m/z 75.3805     0   2   4E+3   4   4.5  RT (min)      lcorbi_hilic_neg_3_044_32    m/z 76.1145     0   2   4   6E+3   0.5   1  RT (min)      lcorbi_hilic_neg_4_087_59    m/z 76.4817     0   1   2   3   4E+3   5   5.5  RT (min)      lcorbi_hilic_neg_2_020_27    m/z 76.5193     0   2   4   6E+3   6.5   7  RT (min)      lcorbi_hilic_neg_1_029_31    m/z 77.0392     0   1E+5   0.5   1  RT (min)      lcorbi_hilic_neg_4_079_58    m/z 77.2937     0   1E+4   0.5   1  RT (min)      lcorbi_hilic_neg_4_089_59    m/z 77.5354     0   1E+4   3.5   4  RT (min)      lcorbi_hilic_neg_4_107_62    m/z 77.8111     0   2   4   6E+3   4   4.5  RT (min)      lcorbi_hilic_neg_3_044_32    m/z 78.0349     0   2   4E+3   4.5   5  RT (min)      lcorbi_hilic_neg_2_020_27    m/z 78.7708     0   1E+3   6   6.5  RT (min)      lcorbi_hilic_neg_3_028_23    m/z 78.9174     0   1   2   3E+6   3   3.5  RT (min)      lcorbi_hilic_neg_4_073_57    m/z 78.9188     0   1   2   3   4E+4   0.5   1  RT (min)       lcorbi_hilic_neg_4_102_61    m/z 78.9200     0   1   2E+6   3   3.5  RT (min)       lcorbi_hilic_neg_4_080_58    m/z 78.9518     0   2   4E+3   7   7.5  RT (min)      lcorbi_hilic_neg_4_097_60    m/z 79.2371     0   1E+4   5   5.5  RT (min)      lcorbi_hilic_neg_4_004_13    m/z 79.2671     0   1   2   3E+4   1   1.5  RT (min)      lcorbi_hilic_neg_1_027_18    m/z 79.3904     0   1   2E+4   3   3.5  RT (min)      lcorbi_hilic_neg_4_089_59    m/z 79.3904     0   1E+4   0.5   1  RT (min)      lcorbi_hilic_neg_1_023_23    m/z 79.5018     0   1E+4   3   3.5  RT (min)      lcorbi_hilic_neg_4_084_59    m/z 79.5179     0   1   2   3E+3   1   1.5  RT (min)      lcorbi_hilic_neg_4_014_18    m/z 79.6142     0   2   4   6   8E+3   3   3.5  RT (min)      lcorbi_hilic_neg_3_028_23    m/z 79.9573     0   1E+4   4   4.5  RT (min)      lcorbi_hilic_neg_4_090_60    m/z 79.9573     0   1   2E+4   5.5   6  RT (min)       lcorbi_hilic_neg_4_081_58    m/z 79.9632     0   1   2   3E+3   7   7.5  RT (min)      lcorbi_hilic_neg_1_055_24    m/z 80.7657     0   1   2E+3   6   6.5  RT (min)      lcorbi_hilic_neg_4_084_59    m/z 80.9168     0   1   2   3E+4   0.5   1  RT (min)       lcorbi_hilic_neg_4_098_61    m/z 80.9198     0   1   2E+4   3   3.5  RT (min)       lcorbi_hilic_neg_3_051_46    m/z 80.9483     0   1   2   3   4E+3   6   6.5  RT (min)      lcorbi_hilic_neg_4_070_57    m/z 80.9651     0   1E+3   5   5.5  RT (min)      lcorbi_hilic_neg_2_006_45    m/z 80.9652     0   1E+4   1   1.5  RT (min)      lcorbi_hilic_neg_4_097_60    m/z 81.0345     0   1E+4   1   1.5  RT (min)      lcorbi_hilic_neg_4_104_61    m/z 81.8331     0   1   2E+4   3   3.5  RT (min)      lcorbi_hilic_neg_4_089_59    m/z 81.8338     0   1   2E+4   3   3.5  RT (min)      lcorbi_hilic_neg_4_089_59    m/z 81.9446     0   2   4   6E+3   3   3.5  RT (min)      lcorbi_hilic_neg_4_087_59    m/z 82.0298     0   1   2   3   4E+3   6.5   7  RT (min)      lcorbi_hilic_neg_4_086_59    m/z 82.0299     0   2   4   6   8E+3   3   3.5  RT (min)      lcorbi_hilic_neg_4_105_61    m/z 82.0570     0   1E+4   3   3.5  RT (min)      lcorbi_hilic_neg_4_083_59    m/z 82.0986     0   1   2   3   4E+3   5   5.5  RT (min)      lcorbi_hilic_neg_2_020_27    m/z 82.2810     0   1E+4   3   3.5  RT (min)      lcorbi_hilic_neg_4_053_23    m/z 82.5049     0   1E+4   3   3.5  RT (min)      lcorbi_hilic_neg_4_053_23    m/z 82.5375     0   1E+4   4.5   5  RT (min)      lcorbi_hilic_neg_2_020_27    m/z 82.9542     0   1   2   3E+4   7.5   8  RT (min)       lcorbi_hilic_neg_2_040_35    m/z 83.0250     0   1E+3   5.5   6  RT (min)       lcorbi_hilic_neg_4_055_38    m/z 83.0332     0   1   2E+3   1   1.5  RT (min)      lcorbi_hilic_neg_4_034_45    m/z 83.0502     0   1E+4   1   1.5  RT (min)      lcorbi_hilic_neg_4_082_59    m/z 83.7382     0   1   2E+4   3   3.5  RT (min)      lcorbi_hilic_neg_1_055_24    m/z 83.7391     0   1   2E+4   3   3.5  RT (min)      lcorbi_hilic_neg_4_015_24    m/z 83.9308     0   1   2E+4   0.5   1  RT (min)       lcorbi_hilic_neg_4_098_61    m/z 84.0091     0   2   4   6   8E+3   5.5   6  RT (min)      lcorbi_hilic_neg_4_072_57    m/z 84.0455     0   1E+4   1   1.5  RT (min)      lcorbi_hilic_neg_4_110_62    m/z 84.0455     0   1   2   3   4E+4   6.5   7  RT (min)      lcorbi_hilic_neg_1_016_37    m/z 84.2780     0   2   4   6E+4   4   4.5  RT (min)      lcorbi_hilic_neg_4_087_59    m/z 84.2790     0   2   4   6E+4   4   4.5  RT (min)      lcorbi_hilic_neg_4_087_59    m/z 84.3016     0   2   4E+3   0.5   1  RT (min)      lcorbi_hilic_neg_4_089_59    m/z 84.3897     0   1   2E+4   4   4.5  RT (min)      lcorbi_hilic_neg_4_084_59    m/z 84.8084     0   2   4   6   8E+4   0.5   1  RT (min)      lcorbi_hilic_neg_4_004_13    m/z 84.9387     0   1E+4   6.5   7  RT (min)      lcorbi_hilic_neg_4_067_57    m/z 84.9888     0   1   2   3   4E+4   5.5   6  RT (min)       lcorbi_hilic_neg_1_022_48    m/z 84.9906     0   2   4   6   8E+4   0.5   1  RT (min)      lcorbi_hilic_neg_1_001_49    m/z 85.9821     0   2   4   6   8E+3   6   6.5  RT (min)      lcorbi_hilic_neg_4_067_57    m/z 85.9997     0   2   4   6E+3   6   6.5  RT (min)      lcorbi_hilic_neg_1_003_14    m/z 86.2802     0   1   2   3   4E+3   3   3.5  RT (min)      lcorbi_hilic_neg_4_104_61    m/z 86.4044     0   1   2E+4   3   3.5  RT (min)      lcorbi_hilic_neg_4_015_24    m/z 86.6279     0   1E+4   3   3.5  RT (min)      lcorbi_hilic_neg_4_015_24    m/z 86.7510     0   2   4   6E+3   7.5   8  RT (min)      lcorbi_hilic_neg_4_080_58    m/z 86.8519     0   1E+4   3   3.5  RT (min)      lcorbi_hilic_neg_1_055_24    m/z 86.9498     0   1E+4   8   8.5  RT (min)      lcorbi_hilic_neg_4_066_57    m/z 87.0004     0   1   7.5   8  RT (min)      lcorbi_hilic_neg_1_006_08    m/z 87.0271     0   1   4   4.5  RT (min)       lcorbi_hilic_neg_1_005_36    m/z 87.0273     0   1E+3   2   2.5  RT (min)      lcorbi_hilic_neg_1_002_26    m/z 87.0318     0   1   5   5.5  RT (min)      lcorbi_hilic_neg_1_007_10    m/z 87.0759     0   1E+4   3   3.5  RT (min)      lcorbi_hilic_neg_3_011_24    m/z 87.1176     0   1   2   3E+3   5   5.5  RT (min)      lcorbi_hilic_neg_2_020_27    m/z 87.2499     0   1   2   3E+3   7.5   8  RT (min)      lcorbi_hilic_neg_4_079_58    m/z 87.9742     0   1E+4   4.5   5  RT (min)      lcorbi_hilic_neg_1_028_45    m/z 88.0121     0   2   4   6   8E+3   5   5.5  RT (min)      lcorbi_hilic_neg_4_067_57    m/z 88.0122     0   1E+4   2   2.5  RT (min)      lcorbi_hilic_neg_4_080_58    m/z 88.0589     0   1E+4   6.5   7  RT (min)      lcorbi_hilic_neg_4_094_60    m/z 88.3006     0   1E+4   0.5   1  RT (min)      lcorbi_hilic_neg_2_038_36    m/z 88.5270     0   2   4   6E+3   6   6.5  RT (min)      lcorbi_hilic_neg_3_051_46    m/z 88.7262     0   1   2   3   4E+3   3   3.5  RT (min)      lcorbi_hilic_neg_4_089_59    m/z 88.9501     0   1E+3   3   3.5  RT (min)      lcorbi_hilic_neg_4_074_58    m/z 88.9697     0   1E+3   4.5   5  RT (min)      lcorbi_hilic_neg_1_001_49    m/z 88.9879     0   2   4   6   8E+3   4   4.5  RT (min)      lcorbi_hilic_neg_1_001_49    m/z 89.0052     0   1   2E+3   2   2.5  RT (min)      lcorbi_hilic_neg_3_034_34    m/z 89.0257     0   1   2   3   4E+6   5   5.5  RT (min)       lcorbi_hilic_neg_1_003_14    m/z 89.0259     0   2   4   6   8E+6   2   2.5  RT (min)       lcorbi_hilic_neg_3_048_33    m/z 89.0259     0   1E+6   11.5   12  RT (min)      lcorbi_hilic_neg_1_001_49    m/z 89.0295     0   1E+4   5   5.5  RT (min)      lcorbi_hilic_neg_1_003_14    m/z 89.0308     0   1E+4   1   1.5  RT (min)      lcorbi_hilic_neg_2_022_37    m/z 89.0437     0   2   4E+2   5.5   6  RT (min)      lcorbi_hilic_neg_1_007_10    m/z 89.0616     0   1   2   3E+3   3   3.5  RT (min)      lcorbi_hilic_neg_4_088_59    m/z 89.1733     0   2   4   6E+4   3   3.5  RT (min)      lcorbi_hilic_neg_4_070_57    m/z 89.1741     0   1   2   3   4E+4   0.5   1  RT (min)      lcorbi_hilic_neg_4_067_57    m/z 89.3987     0   1   2E+4   3   3.5  RT (min)      lcorbi_hilic_neg_4_004_13    m/z 89.5095     0   2   4   6   8E+3   3   3.5  RT (min)      lcorbi_hilic_neg_1_020_13    m/z 89.9257     0   1   2   3   4E+3   7.5   8  RT (min)      lcorbi_hilic_neg_4_082_59    m/z 89.9410     0   2   4   6E+3   1   1.5  RT (min)      lcorbi_hilic_neg_4_082_59    m/z 89.9709     0   2   4   6E+5   4.5   5  RT (min)       lcorbi_hilic_neg_4_077_58    m/z 90.0189     0   1   2E+3   7   7.5  RT (min)       lcorbi_hilic_neg_1_002_26    m/z 90.9236     0   1   2E+3   8   8.5  RT (min)      lcorbi_hilic_neg_2_052_41    m/z 91.0036     0   1E+4   0.5   1  RT (min)      lcorbi_hilic_neg_4_113_62    m/z 91.0220     0   1E+4   0.5   1  RT (min)      lcorbi_hilic_neg_4_113_62    m/z 91.0286     0   1E+3   5   5.5  RT (min)      lcorbi_hilic_neg_4_091_60    m/z 91.0286     0   1   2   3   4E+4   2   2.5  RT (min)       lcorbi_hilic_neg_4_081_58    m/z 91.0517     0   1   2   3   4E+6   1   1.5  RT (min)       lcorbi_hilic_neg_4_067_57    m/z 91.0545     0   1E+5   1   1.5  RT (min)      lcorbi_hilic_neg_4_070_57    m/z 91.3928     0   2   4   6E+3   3   3.5  RT (min)      lcorbi_hilic_neg_4_089_59    m/z 91.3937     0   2   4   6E+3   3   3.5  RT (min)      lcorbi_hilic_neg_4_086_59    m/z 91.6168     0   1E+4   3   3.5  RT (min)      lcorbi_hilic_neg_1_023_23    m/z 91.7283     0   2   4E+3   3   3.5  RT (min)      lcorbi_hilic_neg_4_089_59    m/z 91.7292     0   1   2   3   4E+3   3   3.5  RT (min)      lcorbi_hilic_neg_4_088_59    m/z 91.7899     0   1   2   3   4E+3   1   1.5  RT (min)      lcorbi_hilic_neg_2_040_35    m/z 91.9392     0   1   2   3E+3   1   1.5  RT (min)      lcorbi_hilic_neg_4_092_60    m/z 91.9523     0   1E+4   3   3.5  RT (min)      lcorbi_hilic_neg_4_069_57    m/z 92.0254     0   1   2   3E+3   3   3.5  RT (min)      lcorbi_hilic_neg_4_097_60    m/z 92.0474     0   1E+4   1   1.5  RT (min)      lcorbi_hilic_neg_4_067_57    m/z 92.0474     0   1E+4   6   6.5  RT (min)      lcorbi_hilic_neg_4_096_60    m/z 92.0646     0   1E+4   3   3.5  RT (min)      lcorbi_hilic_neg_1_020_13    m/z 92.1762     0   2   4   6E+3   3   3.5  RT (min)      lcorbi_hilic_neg_4_071_57    m/z 92.2895     0   1   2E+4   3   3.5  RT (min)      lcorbi_hilic_neg_4_004_13    m/z 92.4001     0   1E+4   3   3.5  RT (min)      lcorbi_hilic_neg_2_018_16    m/z 92.8421     0   1   2   3   4E+3   1   1.5  RT (min)      lcorbi_hilic_neg_3_010_18    m/z 92.9290     0   1E+5   8   8.5  RT (min)      lcorbi_hilic_neg_1_007_10    m/z 92.9297     0   1E+6   4.5   5  RT (min)      lcorbi_hilic_neg_4_097_60    m/z 93.0458     0   1   2   3E+5   1   1.5  RT (min)       lcorbi_hilic_neg_2_038_36    m/z 93.0458     0   1   2   3E+3   7   7.5  RT (min)      lcorbi_hilic_neg_4_070_57    m/z 93.0546     0   1E+4   1   1.5  RT (min)      lcorbi_hilic_neg_4_089_59    m/z 93.5556     0   1   2E+3   1.5   2  RT (min)      lcorbi_hilic_neg_4_089_59    m/z 93.6325     0   1E+4   3   3.5  RT (min)      lcorbi_hilic_neg_4_015_24    m/z 93.9470     0   1   2E+4   8   8.5  RT (min)      lcorbi_hilic_neg_4_066_57    m/z 93.9711     0   1   2   3E+3   5   5.5  RT (min)      lcorbi_hilic_neg_2_020_27    m/z 94.0379     0   1   2   3E+3   1   1.5  RT (min)      lcorbi_hilic_neg_4_005_32    m/z 94.2716     0   2   4E+3   6   6.5  RT (min)      lcorbi_hilic_neg_2_020_27    m/z 94.3023     0   2   4   6E+3   2   2.5  RT (min)      lcorbi_hilic_neg_4_112_62    m/z 94.5074     0   2   4E+3   3   3.5  RT (min)      lcorbi_hilic_neg_4_066_57    m/z 94.5531     0   1   2E+3   2   2.5  RT (min)      lcorbi_hilic_neg_1_046_19    m/z 94.9554     0   1   3   3.5  RT (min)      lcorbi_hilic_neg_4_103_61    m/z 94.9807     0   1E+5   1  RT (min)       lcorbi_hilic_neg_4_067_57    m/z 94.9950     0   1   2E+4   1   1.5  RT (min)       lcorbi_hilic_neg_2_036_42    m/z 94.9950     0   1   2E+4   3   3.5   4  RT (min)       lcorbi_hilic_neg_4_003_08    m/z 95.0250     0   1   2   3   4E+3   6.5   7  RT (min)      lcorbi_hilic_neg_4_067_57    m/z 95.0669     0   2   4   6   8E+3   3   3.5  RT (min)      lcorbi_hilic_neg_4_066_57    m/z 95.9761     0   2   4   6E+3   5   5.5  RT (min)      lcorbi_hilic_neg_4_086_59    m/z 96.0172     0   1E+4   1.5   2  RT (min)      lcorbi_hilic_neg_4_101_61    m/z 96.2340     0   2   4   6E+3   5   5.5  RT (min)      lcorbi_hilic_neg_3_024_27    m/z 96.2982     0   1E+4   3   3.5  RT (min)      lcorbi_hilic_neg_2_055_32    m/z 96.5221     0   1E+4   3   3.5  RT (min)      lcorbi_hilic_neg_4_015_24    m/z 96.7461     0   1E+4   3   3.5  RT (min)      lcorbi_hilic_neg_2_055_32    m/z 96.9221     0   1E+4   6.5   7  RT (min)      lcorbi_hilic_neg_2_007_08    m/z 96.9221     0   1   2E+4   0   0.5  RT (min)       lcorbi_hilic_neg_4_076_58    m/z 96.9237     0   1E+5   4.5   5  RT (min)      lcorbi_hilic_neg_4_066_57    m/z 96.9415     0   1   7   7.5  RT (min)      lcorbi_hilic_neg_4_108_62    m/z 96.9513     0   1   2   3   4E+3   7   7.5  RT (min)      lcorbi_hilic_neg_4_096_60    m/z 96.9616     0   1   2   3   4E+6   7.5   8  RT (min)       lcorbi_hilic_neg_4_074_58    m/z 96.9617     0   1   2E+5   4.5   5  RT (min)      lcorbi_hilic_neg_2_035_22    m/z 96.9622     0   2   4   6E+5   12   12.5  RT (min)      lcorbi_hilic_neg_2_024_11    m/z 96.9694     0   1E+5   11.5   12  RT (min)      lcorbi_hilic_neg_4_096_60    m/z 96.9865     0   1   2   3   4E+2   7   7.5  RT (min)      lcorbi_hilic_neg_3_005_08    m/z 96.9972     0   2   4E+3   7   7.5  RT (min)      lcorbi_hilic_neg_4_096_60    m/z 97.0043     0   1   2   3   4E+2   4   4.5  RT (min)      lcorbi_hilic_neg_1_047_05    m/z 97.0144     0   1E+4   6   6.5  RT (min)      lcorbi_hilic_neg_2_027_02    m/z 97.9480     0   2   4   6   8E+2   1   1.5  RT (min)      lcorbi_hilic_neg_4_082_59    m/z 97.9650     0   1E+4   1   1.5  RT (min)      lcorbi_hilic_neg_1_052_41    m/z 97.9736     0   1E+4   7   7.5  RT (min)      lcorbi_hilic_neg_4_092_60    m/z 98.0247     0   1   2E+4   1.5   2  RT (min)       lcorbi_hilic_neg_4_086_59    m/z 98.0359     0   2   4   6E+3   4   4.5  RT (min)      lcorbi_hilic_neg_4_072_57    m/z 98.0611     0   1   2   3E+4   1   1.5  RT (min)       lcorbi_hilic_neg_4_110_62    m/z 98.0611     0   1   6   6.5  RT (min)      lcorbi_hilic_neg_1_012_32    m/z 98.2742     0   2   4E+3   2   2.5  RT (min)      lcorbi_hilic_neg_4_110_62    m/z 98.7736     0   1   2E+3   2   2.5  RT (min)      lcorbi_hilic_neg_2_020_27    m/z 98.9047     0   2   4   6   8E+2   12   12.5  RT (min)      lcorbi_hilic_neg_3_051_46    m/z 98.9047     0   2   4   6   8E+2   4.5   5  RT (min)      lcorbi_hilic_neg_4_098_61    m/z 98.9068     0   2   4   6   8E+4   0.5   1  RT (min)      lcorbi_hilic_neg_3_050_03    m/z 98.9391     0   2   4   6   8E+4   7.5   8  RT (min)      lcorbi_hilic_neg_4_074_58    m/z 98.9643     0   1E+3   7.5   8  RT (min)       lcorbi_hilic_neg_2_032_25    m/z 99.0160     0   1   2   3E+2   6   6.5  RT (min)       lcorbi_hilic_neg_1_003_14    m/z 99.0563     0   1   2   3E+3   6.5   7  RT (min)      lcorbi_hilic_neg_4_048_37    m/z 99.3069     0   2   4   6E+3   0.5   1  RT (min)      lcorbi_hilic_neg_1_039_38    m/z 99.5164     0   1   2   3   4E+3   2   2.5  RT (min)      lcorbi_hilic_neg_4_011_19    m/z 99.9258     0   1   2   3E+5   7   7.5  RT (min)      lcorbi_hilic_neg_4_023_34    m/z 99.9496     0   1   6.5   7  RT (min)      lcorbi_hilic_neg_4_106_62    m/z 99.9620     0   1   2   3   4E+3   1   1.5  RT (min)      lcorbi_hilic_neg_1_034_43    m/z 100.0041     0   1E+3   2   2.5  RT (min)      lcorbi_hilic_neg_1_029_31    m/z 100.0312     0   1   2   3E+3   5   5.5  RT (min)      lcorbi_hilic_neg_4_004_13    m/z 100.0404     0   1E+4   4.5   5  RT (min)      lcorbi_hilic_neg_2_020_27    m/z 100.0405     0   2   4   6   8E+3   6   6.5  RT (min)      lcorbi_hilic_neg_4_088_59    m/z 100.7815     0   1E+4   2   2.5  RT (min)      lcorbi_hilic_neg_4_109_62    m/z 100.9037     0   1   2E+4   12.5   13  RT (min)      lcorbi_hilic_neg_3_046_39    m/z 100.9039     0   2   4   6E+3   4   4.5  RT (min)      lcorbi_hilic_neg_4_098_61    m/z 100.9263     0   1E+5   1   1.5  RT (min)       lcorbi_hilic_neg_4_109_62    m/z 100.9314     0   2   4   6E+5   4.5   5  RT (min)      lcorbi_hilic_neg_1_055_24    m/z 100.9314     0   1E+4   7   7.5  RT (min)      lcorbi_hilic_neg_4_074_58    m/z 100.9336     0   1   2E+6   6   6.5  RT (min)       lcorbi_hilic_neg_4_072_57    m/z 100.9458     0   2   4   6   8E+3   0   0.5  RT (min)      lcorbi_hilic_neg_4_082_59    m/z 100.9608     0   1   2   3E+3   1   1.5  RT (min)      lcorbi_hilic_neg_4_083_59    m/z 101.0052     0   1E+3   2   2.5  RT (min)      lcorbi_hilic_neg_3_028_23    m/z 101.0221     0   1E+4   5   5.5  RT (min)      lcorbi_hilic_neg_3_031_14    m/z 101.0245     0   1E+5   2   2.5  RT (min)       lcorbi_hilic_neg_2_013_18    m/z 101.9241     0   1   2   3E+4   0.5   1  RT (min)      lcorbi_hilic_neg_4_106_62    m/z 101.9337     0   1   2   3E+5   6   6.5  RT (min)      lcorbi_hilic_neg_4_072_57    m/z 102.0188     0   2   4   6   8E+4   1.5   2  RT (min)       lcorbi_hilic_neg_3_007_07    m/z 102.0381     0   1E+4   1.5   2  RT (min)      lcorbi_hilic_neg_4_091_60    m/z 102.0461     0   1   2E+4   4   4.5  RT (min)       lcorbi_hilic_neg_4_091_60    m/z 102.0560     0   1E+4   3   3.5  RT (min)      lcorbi_hilic_neg_4_088_59    m/z 102.0592     0   1   2E+4   6   6.5  RT (min)      lcorbi_hilic_neg_1_047_05    m/z 102.5309     0   1   2   3   4E+3   3   3.5  RT (min)      lcorbi_hilic_neg_4_105_61    m/z 102.9018     0   1   2   3E+3   4   4.5  RT (min)      lcorbi_hilic_neg_4_098_61    m/z 102.9020     0   1E+4   0.5   1  RT (min)      lcorbi_hilic_neg_4_106_62    m/z 102.9319     0   2   4E+4   5.5   6  RT (min)      lcorbi_hilic_neg_4_106_62    m/z 102.9681     0   1   6   6.5  RT (min)      lcorbi_hilic_neg_4_109_62    m/z 103.0036     0   1E+4   4   4.5  RT (min)       lcorbi_hilic_neg_4_112_62    m/z 103.0036     0   1   2   3   4E+3   1.5   2  RT (min)       lcorbi_hilic_neg_4_055_38    m/z 103.0220     0   1E+3   6   6.5  RT (min)       lcorbi_hilic_neg_4_107_62    m/z 103.0243     0   2   4   6E+2   1.5   2  RT (min)       lcorbi_hilic_neg_4_048_37    m/z 103.0377     0   1   5   5.5  RT (min)      lcorbi_hilic_neg_1_015_30    m/z 103.0401     0   1   2E+5   12.5   13  RT (min)      lcorbi_hilic_neg_4_054_30    m/z 103.0531     0   1   2E+3   6   6.5  RT (min)      lcorbi_hilic_neg_4_053_23    m/z 103.9201     0   2   4   6E+3   1   1.5  RT (min)      lcorbi_hilic_neg_4_097_60    m/z 103.9894     0   2   4   6E+3   8   8.5  RT (min)      lcorbi_hilic_neg_4_103_61    m/z 104.0344     0   1   2   3   4E+3   4   4.5  RT (min)       lcorbi_hilic_neg_1_006_08    m/z 104.0367     0   1E+6   6.5   7  RT (min)       lcorbi_hilic_neg_4_082_59    m/z 104.0393     0   1E+4   6.5   7  RT (min)      lcorbi_hilic_neg_3_013_37    m/z 104.0434     0   2   4E+4   1   1.5  RT (min)      lcorbi_hilic_neg_1_012_32    m/z 104.0537     0   1E+3   5   5.5  RT (min)      lcorbi_hilic_neg_4_096_60    m/z 104.0537     0   2   4   6E+3   3.5   4  RT (min)      lcorbi_hilic_neg_4_067_57    m/z 104.0603     0   1   6   6.5  RT (min)      lcorbi_hilic_neg_4_091_60    m/z 104.0618     0   1   2E+4   1.5   2  RT (min)       lcorbi_hilic_neg_3_044_32    m/z 104.2153     0   1   2E+3   5   5.5  RT (min)      lcorbi_hilic_neg_2_020_27    m/z 104.9538     0   1E+4   0   0.5  RT (min)      lcorbi_hilic_neg_4_036_36    m/z 104.9862     0   2   4   6   8E+3   8   8.5  RT (min)      lcorbi_hilic_neg_4_076_58    m/z 105.0323     0   1   2E+3   6.5   7  RT (min)      lcorbi_hilic_neg_4_079_58    m/z 105.0661     0   1   2E+4   6   6.5  RT (min)       lcorbi_hilic_neg_4_094_60    m/z 105.0854     0   1   2E+4   6.5   7  RT (min)       lcorbi_hilic_neg_4_091_60    m/z 105.9470     0   1E+4   8   8.5  RT (min)      lcorbi_hilic_neg_4_066_57    m/z 105.9702     0   1E+4   6   6.5  RT (min)      lcorbi_hilic_neg_4_073_57    m/z 106.0396     0   1   2E+3   6.5   7  RT (min)       lcorbi_hilic_neg_4_091_60    m/z 106.0423     0   2   4   6   8E+3   1   1.5  RT (min)      lcorbi_hilic_neg_4_110_62    m/z 106.0694     0   1E+4   3   3.5  RT (min)      lcorbi_hilic_neg_4_097_60    m/z 106.0711     0   2   4   6   8E+5   6   6.5  RT (min)       lcorbi_hilic_neg_1_055_24    m/z 106.2808     0   1   2   3E+3   2.5   3  RT (min)      lcorbi_hilic_neg_4_110_62    m/z 106.9119     0   1E+4   6.5   7  RT (min)      lcorbi_hilic_neg_4_105_61    m/z 106.9247     0   1   2E+5   4.5   5  RT (min)      lcorbi_hilic_neg_4_077_58    m/z 106.9265     0   2   4   6E+3   8   8.5  RT (min)      lcorbi_hilic_neg_4_066_57    m/z 107.0213     0   2   4   6   8E+3   2   2.5  RT (min)      lcorbi_hilic_neg_4_083_59    m/z 107.0362     0   1   2   3E+4   3   3.5  RT (min)       lcorbi_hilic_neg_4_085_59    m/z 107.0376     0   1   2   3E+4   1   1.5  RT (min)      lcorbi_hilic_neg_1_026_34    m/z 107.0473     0   1E+5   6.5   7  RT (min)       lcorbi_hilic_neg_4_003_08    m/z 107.0614     0   1   1.5   2  RT (min)      lcorbi_hilic_neg_4_106_62    m/z 108.0216     0   1E+4   1   1.5  RT (min)      lcorbi_hilic_neg_1_010_22    m/z 108.0293     0   1E+5   6   6.5  RT (min)       lcorbi_hilic_neg_2_055_32    m/z 108.7879     0   2   4   6   8E+3   2.5   3  RT (min)      lcorbi_hilic_neg_4_111_62    m/z 108.7891     0   2   4   6   8E+3   2.5   3  RT (min)      lcorbi_hilic_neg_4_111_62    m/z 108.9046     0   1E+3   4.5   5  RT (min)      lcorbi_hilic_neg_1_028_45    m/z 108.9356     0   1   2   3   4E+4   0.5   1  RT (min)      lcorbi_hilic_neg_2_036_42    m/z 109.0059     0   2   4   6E+3   4.5   5  RT (min)      lcorbi_hilic_neg_4_083_59    m/z 109.0294     0   2   4   6E+3   0.5   1  RT (min)      lcorbi_hilic_neg_4_108_62    m/z 109.8974     0   1   2   3   4E+3   8   8.5  RT (min)      lcorbi_hilic_neg_4_082_59    m/z 109.9339     0   2   4   6   8E+3   6   6.5  RT (min)      lcorbi_hilic_neg_4_078_58    m/z 109.9419     0   2   4   6   8E+3   8   8.5  RT (min)      lcorbi_hilic_neg_4_082_59    m/z 110.0247     0   1E+4   4   4.5  RT (min)      lcorbi_hilic_neg_4_073_57    m/z 110.0359     0   2   4   6E+3   6   6.5  RT (min)      lcorbi_hilic_neg_4_103_61    m/z 110.0440     0   2   4   6E+3   5.5   6  RT (min)      lcorbi_hilic_neg_4_083_59    m/z 110.0724     0   1   2   3E+3   7   7.5  RT (min)      lcorbi_hilic_neg_2_020_27    m/z 110.9041     0   1E+4   4.5   5  RT (min)      lcorbi_hilic_neg_2_028_33    m/z 110.9338     0   2   4   6E+3   0.5   1  RT (min)      lcorbi_hilic_neg_3_004_41    m/z 110.9580     0   1   2E+3   1   1.5  RT (min)      lcorbi_hilic_neg_4_051_16    m/z 110.9805     0   2   4   6E+3   1   1.5  RT (min)      lcorbi_hilic_neg_2_046_47    m/z 111.0176     0   1   2   3E+4   4   5   6  RT (min)       lcorbi_hilic_neg_3_001_49    m/z 111.0200     0   1   2   3E+4   0   0.5  RT (min)      lcorbi_hilic_neg_4_089_59    m/z 111.0451     0   2   4   6E+3   1   1.5  RT (min)      lcorbi_hilic_neg_4_071_57    m/z 111.9789     0   2   4E+4   1   1.5  RT (min)       lcorbi_hilic_neg_4_068_57    m/z 112.0121     0   1E+3   2   3  RT (min)       lcorbi_hilic_neg_4_073_57    m/z 112.0404     0   1   2E+3   1   1.5  RT (min)      lcorbi_hilic_neg_2_050_17    m/z 112.0404     0   2   4E+4   3   3.5  RT (min)      lcorbi_hilic_neg_1_002_26    m/z 112.0516     0   2   4   6E+3   5.5   6  RT (min)      lcorbi_hilic_neg_4_075_58    m/z 112.6282     0   2   4   6   8E+3   5   5.5  RT (min)      lcorbi_hilic_neg_4_066_57    m/z 112.8961     0   2   4   6E+3   0.5   1  RT (min)      lcorbi_hilic_neg_4_082_59    m/z 112.8988     0   1E+3   4.5   5  RT (min)      lcorbi_hilic_neg_1_051_33    m/z 112.9138     0   2   4E+3   5   5.5  RT (min)      lcorbi_hilic_neg_4_081_58    m/z 112.9244     0   1   2E+3   5   5.5  RT (min)      lcorbi_hilic_neg_4_066_57    m/z 112.9317     0   1   2   3   4E+3   5   5.5  RT (min)      lcorbi_hilic_neg_4_066_57    m/z 112.9498     0   2   4   6E+3   5   5.5  RT (min)      lcorbi_hilic_neg_4_074_58    m/z 112.9587     0   1   2   3   4E+3   5   5.5  RT (min)      lcorbi_hilic_neg_4_066_57    m/z 112.9691     0   2   4   6   8E+3   5   5.5  RT (min)      lcorbi_hilic_neg_4_075_58    m/z 112.9831     0   2   4E+4   5   5.5  RT (min)      lcorbi_hilic_neg_4_075_58    m/z 112.9874     0   1E+6   12   12.5  RT (min)      lcorbi_hilic_neg_1_021_28    m/z 112.9874     0   1E+6   9   9.5  RT (min)      lcorbi_hilic_neg_1_035_29    m/z 112.9874     0   1   2   3   4E+6   1   1.5  RT (min)       lcorbi_hilic_neg_1_023_23    m/z 112.9874     0   1   2E+7   5   5.5  RT (min)       lcorbi_hilic_neg_4_071_57    m/z 113.0015     0   2   4   6   8E+3   1   1.5  RT (min)       lcorbi_hilic_neg_1_003_14    m/z 113.0033     0   2   4   6   8E+3   5   5.5  RT (min)      lcorbi_hilic_neg_4_076_58    m/z 113.0128     0   1   2   3   4E+3   5   5.5  RT (min)      lcorbi_hilic_neg_4_074_58    m/z 113.0260     0   1   2E+3   5   5.5  RT (min)       lcorbi_hilic_neg_1_010_22    m/z 113.0353     0   2   4   6   8E+2   1   1.5  RT (min)      lcorbi_hilic_neg_1_008_39    m/z 113.0356     0   1   2   3   4E+3   5   5.5  RT (min)      lcorbi_hilic_neg_4_072_57    m/z 113.0506     0   1   2   3E+3   5   5.5  RT (min)      lcorbi_hilic_neg_4_066_57    m/z 113.0580     0   1   5   5.5  RT (min)      lcorbi_hilic_neg_4_097_60    m/z 113.0607     0   1E+4   1   1.5  RT (min)      lcorbi_hilic_neg_4_085_59    m/z 113.0701     0   2   4   6   8E+3   5.5   6  RT (min)      lcorbi_hilic_neg_4_105_61    m/z 113.2310     0   2   4   6   8E+3   5   5.5  RT (min)      lcorbi_hilic_neg_4_044_01    m/z 113.3445     0   2   4   6   8E+3   5   5.5  RT (min)      lcorbi_hilic_neg_4_066_57    m/z 113.9261     0   1   2   3   4E+3   0.5   1  RT (min)      lcorbi_hilic_neg_4_041_54    m/z 113.9864     0   2   4   6E+2   9   9.5  RT (min)      lcorbi_hilic_neg_1_002_26    m/z 113.9864     0   1   12.5   13  RT (min)      lcorbi_hilic_neg_1_001_49    m/z 114.0299     0   1   4   4.5  RT (min)      lcorbi_hilic_neg_1_007_10    m/z 114.0299     0   1   1   1.5  RT (min)      lcorbi_hilic_neg_1_006_08    m/z 114.0389     0   1E+3   6.5   7  RT (min)      lcorbi_hilic_neg_1_002_26    m/z 114.0558     0   1   2E+5   5.5   6  RT (min)       lcorbi_hilic_neg_4_097_60    m/z 114.8942     0   1   4.5   5  RT (min)      lcorbi_hilic_neg_4_102_61    m/z 114.8955     0   1   3   3.5  RT (min)      lcorbi_hilic_neg_4_113_62    m/z 114.9340     0   2   4E+4   7   7.5  RT (min)       lcorbi_hilic_neg_4_077_58    m/z 114.9491     0   1   2E+4   5   5.5  RT (min)       lcorbi_hilic_neg_4_074_58    m/z 114.9886     0   1E+4   8   8.5  RT (min)      lcorbi_hilic_neg_4_074_58    m/z 115.0285     0   2   4   6   8E+3   1.5   2  RT (min)       lcorbi_hilic_neg_4_071_57    m/z 115.0308     0   1   2E+3   5   5.5   6  RT (min)       lcorbi_hilic_neg_1_012_32    m/z 115.0400     0   2   4   6   8E+3   2   2.5  RT (min)       lcorbi_hilic_neg_2_011_29    m/z 115.0593     0   2   4E+3   2   2.5  RT (min)      lcorbi_hilic_neg_4_004_13    m/z 115.9028     0   1   5.5   6  RT (min)      lcorbi_hilic_neg_4_045_25    m/z 115.9028     0   1   1   1.5  RT (min)      lcorbi_hilic_neg_4_082_59    m/z 115.9207     0   1   2E+5   5.5   6  RT (min)       lcorbi_hilic_neg_4_055_38    m/z 115.9444     0   1   2   2.5  RT (min)      lcorbi_hilic_neg_4_090_60    m/z 115.9930     0   2   4   6   8E+2   5   5.5  RT (min)      lcorbi_hilic_neg_1_010_22    m/z 116.0038     0   1E+4   1   2  RT (min)       lcorbi_hilic_neg_1_009_50    m/z 116.0039     0   1   2   3E+3   4.5   5  RT (min)      lcorbi_hilic_neg_4_079_58    m/z 116.0504     0   1E+4   1   1.5  RT (min)      lcorbi_hilic_neg_4_076_58    m/z 116.3264     0   2   4   6   8E+3   0.5   1  RT (min)      lcorbi_hilic_neg_4_072_57    m/z 116.8934     0   1E+3   3   3.5  RT (min)      lcorbi_hilic_neg_4_108_62    m/z 116.9286     0   2   4   6E+5   7   7.5  RT (min)      lcorbi_hilic_neg_3_031_14    m/z 116.9305     0   2   4   6E+5   4.5   5  RT (min)      lcorbi_hilic_neg_1_008_39    m/z 116.9306     0   1   2   3E+4   1   1.5  RT (min)      lcorbi_hilic_neg_4_110_62    m/z 116.9957     0   2   4   6E+4   1   1.5  RT (min)       lcorbi_hilic_neg_2_032_25    m/z 116.9957     0   1   2   3E+4   3   4  RT (min)       lcorbi_hilic_neg_2_030_43    m/z 117.0087     0   2   4   6E+3   1.5   2  RT (min)      lcorbi_hilic_neg_4_085_59    m/z 117.0167     0   2   4   6   8E+2   4   5   6  RT (min)       lcorbi_hilic_neg_1_003_14    m/z 117.0212     0   1E+6   1.5   2  RT (min)       lcorbi_hilic_neg_4_075_58    m/z 117.0378     0   1   2E+3   1.5   2  RT (min)       lcorbi_hilic_neg_1_008_39    m/z 117.0538     0   1   2   3E+3   5   5.5  RT (min)      lcorbi_hilic_neg_1_016_37    m/z 118.0008     0   2   4   6   8E+3   0   0.5  RT (min)      lcorbi_hilic_neg_4_083_59    m/z 118.0010     0   1   2   3   4E+2   4.5   5  RT (min)      lcorbi_hilic_neg_1_028_45    m/z 118.0159     0   2   4   6   8E+2   1.5   2  RT (min)       lcorbi_hilic_neg_4_089_59    m/z 118.0298     0   1   2E+3   1   1.5  RT (min)      lcorbi_hilic_neg_1_001_49    m/z 118.0411     0   2   4E+2   6   6.5  RT (min)      lcorbi_hilic_neg_4_035_48    m/z 118.0435     0   1   2   3E+3   2   2.5  RT (min)      lcorbi_hilic_neg_4_051_16    m/z 118.0525     0   1E+6   6   6.5  RT (min)       lcorbi_hilic_neg_3_023_36    m/z 118.0545     0   1E+4   6   6.5  RT (min)      lcorbi_hilic_neg_3_030_30    m/z 118.0589     0   1   2   3   4E+4   1   1.5  RT (min)      lcorbi_hilic_neg_1_012_32    m/z 118.9267     0   1   2   3E+4   6.5   7   7.5  RT (min)       lcorbi_hilic_neg_4_083_59    m/z 118.9346     0   1E+3   4.5   5  RT (min)      lcorbi_hilic_neg_1_051_33    m/z 118.9662     0   1   2   3   4E+4   8.5   9  RT (min)       lcorbi_hilic_neg_4_074_58    m/z 118.9899     0   1   2E+3   5.5   6  RT (min)      lcorbi_hilic_neg_1_002_26    m/z 118.9925     0   1   2E+5   0.5   1  RT (min)      lcorbi_hilic_neg_1_002_26    m/z 119.0452     0   1   1.5   2  RT (min)      lcorbi_hilic_neg_1_002_26    m/z 119.0502     0   1   2   3   4E+4   2   3  RT (min)       lcorbi_hilic_neg_4_100_61    m/z 119.0726     0   1   2   3   4E+4   5.5   6  RT (min)       lcorbi_hilic_neg_4_091_60    m/z 119.9074     0   2   4   6   8E+3   6   6.5  RT (min)       lcorbi_hilic_neg_4_098_61    m/z 119.9242     0   2   4   6E+3   0   0.5  RT (min)      lcorbi_hilic_neg_4_072_57    m/z 119.9262     0   1   2   3   4E+3   8   8.5  RT (min)      lcorbi_hilic_neg_4_069_57    m/z 119.9341     0   1   2E+3   4.5   5  RT (min)      lcorbi_hilic_neg_4_103_61    m/z 119.9372     0   1   2E+3   8   8.5  RT (min)      lcorbi_hilic_neg_3_002_15    m/z 119.9492     0   1   2   3E+3   1.5   2  RT (min)      lcorbi_hilic_neg_4_103_61    m/z 120.0123     0   2   4   6   8E+3   5.5   6  RT (min)      lcorbi_hilic_neg_4_037_31    m/z 120.0211     0   1   2E+3   2   2.5  RT (min)      lcorbi_hilic_neg_2_039_07    m/z 120.0382     0   1   2   3E+4   5   5.5  RT (min)       lcorbi_hilic_neg_4_075_58    m/z 120.0382     0   2   4E+3   2   2.5  RT (min)      lcorbi_hilic_neg_4_093_60    m/z 120.0550     0   2   4E+3   6   6.5  RT (min)      lcorbi_hilic_neg_4_078_58    m/z 120.9307     0   1E+3   4.5   5  RT (min)      lcorbi_hilic_neg_2_041_54    m/z 120.9547     0   1E+4   8   8.5  RT (min)      lcorbi_hilic_neg_4_074_58    m/z 121.0325     0   2   4   6E+2   6   6.5  RT (min)      lcorbi_hilic_neg_4_005_32    m/z 121.0366     0   1E+5   1.5   2  RT (min)      lcorbi_hilic_neg_4_092_60    m/z 121.0391     0   2   4   6E+3   5   5.5  RT (min)      lcorbi_hilic_neg_4_075_58    m/z 121.0609     0   1E+4   6   6.5  RT (min)      lcorbi_hilic_neg_4_095_60    m/z 121.0657     0   2   4   6   8E+3   0.5   1  RT (min)      lcorbi_hilic_neg_4_077_58    m/z 121.0882     0   1E+4   5.5   6  RT (min)      lcorbi_hilic_neg_4_097_60    m/z 121.9045     0   1   2E+3   6   6.5  RT (min)       lcorbi_hilic_neg_4_102_61    m/z 121.9046     0   1   2   3E+4   8   8.5  RT (min)       lcorbi_hilic_neg_4_075_58    m/z 121.9311     0   1E+3   4.5   5  RT (min)      lcorbi_hilic_neg_4_102_61    m/z 121.9439     0   1   2E+4   6.5   7  RT (min)      lcorbi_hilic_neg_4_101_61    m/z 121.9440     0   2   4E+4   1  RT (min)       lcorbi_hilic_neg_2_008_39    m/z 121.9440     0   1   2   3E+4   4   4.5  RT (min)       lcorbi_hilic_neg_4_109_62    m/z 122.0610     0   1   2E+4   3.5   4  RT (min)      lcorbi_hilic_neg_2_026_23    m/z 122.0610     0   1E+4   1   1.5  RT (min)      lcorbi_hilic_neg_4_089_59    m/z 122.9365     0   2   4E+3   5.5   6   6.5  RT (min)      lcorbi_hilic_neg_4_072_57    m/z 123.0482     0   2   4   6E+4   4   4.5  RT (min)       lcorbi_hilic_neg_4_095_60    m/z 123.9019     0   1E+4   5   5.5  RT (min)       lcorbi_hilic_neg_4_075_58    m/z 123.9112     0   1   2   3   4E+4   0.5   1  RT (min)      lcorbi_hilic_neg_2_012_14    m/z 123.9709     0   1E+4   1.5   2  RT (min)      lcorbi_hilic_neg_1_005_36    m/z 124.0102     0   1   5.5   6  RT (min)       lcorbi_hilic_neg_4_082_59    m/z 124.0161     0   1   2E+4   0.5   1  RT (min)      lcorbi_hilic_neg_4_105_61    m/z 124.0403     0   1   2E+3   1   1.5  RT (min)      lcorbi_hilic_neg_1_002_26    m/z 124.0530     0   2   4E+4   3.5   4  RT (min)       lcorbi_hilic_neg_4_095_60    m/z 124.9645     0   2   4   6E+3   7   7.5  RT (min)      lcorbi_hilic_neg_4_096_60    m/z 124.9879     0   1E+3   6   6.5  RT (min)      lcorbi_hilic_neg_4_092_60    m/z 124.9940     0   2   4   6   8E+3   1   1.5  RT (min)      lcorbi_hilic_neg_4_068_57    m/z 125.0106     0   1   5.5   6  RT (min)      lcorbi_hilic_neg_4_106_62    m/z 125.0470     0   2   4   6   8E+3   3   3.5  RT (min)      lcorbi_hilic_neg_4_082_59    m/z 125.0702     0   1E+4   5.5   6  RT (min)      lcorbi_hilic_neg_3_037_12    m/z 125.0713     0   1   2E+3   4   4.5  RT (min)      lcorbi_hilic_neg_4_069_57    m/z 125.0971     0   1   2   3   4E+4   0.5   1  RT (min)      lcorbi_hilic_neg_4_076_58    m/z 125.8731     0   1   2E+4   9   9.5  RT (min)      lcorbi_hilic_neg_4_066_57    m/z 125.8731     0   1E+4   1.5   2  RT (min)      lcorbi_hilic_neg_4_097_60    m/z 125.8999     0   1E+4   1   1.5  RT (min)      lcorbi_hilic_neg_4_089_59    m/z 125.9287     0   1E+3   1   1.5  RT (min)      lcorbi_hilic_neg_4_076_58    m/z 125.9489     0   1   2   3   4E+3   4.5   5  RT (min)      lcorbi_hilic_neg_4_079_58    m/z 125.9946     0   1E+4   1   1.5  RT (min)      lcorbi_hilic_neg_4_068_57    m/z 125.9947     0   1E+3   4   4.5  RT (min)      lcorbi_hilic_neg_1_001_49    m/z 126.0195     0   2   4   6   8E+3   2.5   3  RT (min)      lcorbi_hilic_neg_4_072_57    m/z 126.0277     0   1E+4   5   5.5  RT (min)      lcorbi_hilic_neg_4_075_58    m/z 126.0308     0   1   2   3E+4   1   1.5  RT (min)       lcorbi_hilic_neg_4_081_58    m/z 126.0389     0   1   2E+4   1.5   2  RT (min)      lcorbi_hilic_neg_4_040_17    m/z 126.0434     0   2   4   6E+3   4   4.5  RT (min)      lcorbi_hilic_neg_4_101_61    m/z 126.8809     0   1   2   3E+4   8.5   9   9.5  RT (min)       lcorbi_hilic_neg_4_076_58    m/z 126.8810     0   1   2   3   4E+4   11  RT (min)       lcorbi_hilic_neg_4_078_58    m/z 126.9527     0   1E+4   1   1.5  RT (min)      lcorbi_hilic_neg_4_077_58    m/z 126.9871     0   1   2E+4   1   1.5  RT (min)       lcorbi_hilic_neg_4_068_57    m/z 126.9871     0   1E+4   4   4.5  RT (min)      lcorbi_hilic_neg_2_038_36    m/z 127.0012     0   1E+4   2   2.5  RT (min)      lcorbi_hilic_neg_4_073_57    m/z 127.0024     0   1   2E+3   6   6.5  RT (min)      lcorbi_hilic_neg_1_001_49    m/z 127.0262     0   2   4E+4   4.5   5  RT (min)       lcorbi_hilic_neg_4_101_61    m/z 127.0512     0   1E+4   2   2.5  RT (min)      lcorbi_hilic_neg_4_072_57    m/z 127.0539     0   2   4   6E+3   6.5   7  RT (min)      lcorbi_hilic_neg_2_026_23    m/z 127.8701     0   2   4   6   8E+3   1.5   2  RT (min)      lcorbi_hilic_neg_4_097_60    m/z 127.8703     0   1E+4   8   8.5  RT (min)       lcorbi_hilic_neg_4_075_58    m/z 127.8972     0   2   4   6E+3   1   1.5  RT (min)      lcorbi_hilic_neg_4_082_59    m/z 127.9941     0   2   4E+3   3   3.5  RT (min)      lcorbi_hilic_neg_4_088_59    m/z 128.0063     0   1   2   3E+3   3   3.5  RT (min)      lcorbi_hilic_neg_3_018_38    m/z 128.0128     0   1   2   3E+3   3   3.5  RT (min)       lcorbi_hilic_neg_1_029_31    m/z 128.0248     0   1E+4   3   3.5  RT (min)      lcorbi_hilic_neg_4_088_59    m/z 128.0326     0   1   2   3   4E+2   12.5   13  RT (min)      lcorbi_hilic_neg_4_017_51    m/z 128.0396     0   1   2E+3   6   6.5  RT (min)      lcorbi_hilic_neg_3_029_25    m/z 128.0426     0   1   2   3   4E+2   2   3  RT (min)       lcorbi_hilic_neg_2_019_21    m/z 128.0526     0   1E+4   3   3.5  RT (min)      lcorbi_hilic_neg_4_088_59    m/z 128.0764     0   2   4E+3   3   3.5  RT (min)      lcorbi_hilic_neg_4_088_59    m/z 128.2946     0   2   4E+3   1   1.5  RT (min)      lcorbi_hilic_neg_4_101_61    m/z 128.8813     0   2   4E+4   8   8.5  RT (min)       lcorbi_hilic_neg_4_080_58    m/z 128.8922     0   1E+4   4.5   5  RT (min)       lcorbi_hilic_neg_4_069_57    m/z 128.9048     0   2   4   6E+3   4.5   5  RT (min)      lcorbi_hilic_neg_4_076_58    m/z 128.9182     0   1   4.5   5  RT (min)      lcorbi_hilic_neg_4_113_62    m/z 128.9285     0   1   2E+4   6  RT (min)       lcorbi_hilic_neg_4_098_61    m/z 128.9285     0   1   2   3E+4   0.5   1  RT (min)       lcorbi_hilic_neg_4_111_62    m/z 128.9400     0   1   4.5   5  RT (min)      lcorbi_hilic_neg_4_113_62    m/z 128.9490     0   1   4.5   5  RT (min)      lcorbi_hilic_neg_4_113_62    m/z 128.9765     0   2   4   6E+3   4.5   5  RT (min)      lcorbi_hilic_neg_4_079_58    m/z 128.9850     0   1   2   3   4E+3   4.5   5  RT (min)      lcorbi_hilic_neg_4_080_58    m/z 128.9957     0   1E+4   1   1.5  RT (min)      lcorbi_hilic_neg_4_089_59    m/z 129.0010     0   1   2   3   4E+3   4.5   5  RT (min)      lcorbi_hilic_neg_4_080_58    m/z 129.0071     0   1   2.5   3  RT (min)       lcorbi_hilic_neg_3_039_31    m/z 129.0323     0   2   4E+4   2.5   3  RT (min)       lcorbi_hilic_neg_4_089_59    m/z 129.0325     0   1   2E+3   6   6.5  RT (min)      lcorbi_hilic_neg_3_018_38    m/z 129.0409     0   1E+4   3   3.5  RT (min)      lcorbi_hilic_neg_3_034_34    m/z 129.0510     0   1E+6   1.5   2  RT (min)       lcorbi_hilic_neg_4_069_57    m/z 129.0749     0   1E+4   1.5   2  RT (min)      lcorbi_hilic_neg_4_067_57    m/z 129.0845     0   1   2E+4   1   1.5  RT (min)       lcorbi_hilic_neg_4_092_60    m/z 129.0845     0   1E+4   3.5   4  RT (min)      lcorbi_hilic_neg_4_097_60    m/z 129.9326     0   2   4E+3   4   4.5  RT (min)      lcorbi_hilic_neg_3_001_49    m/z 129.9362     0   1   2E+4   1   2  RT (min)       lcorbi_hilic_neg_4_113_62    m/z 129.9363     0   2   4   6E+4   8   8.5  RT (min)       lcorbi_hilic_neg_4_081_58    m/z 129.9376     0   2   4   6   8E+3   5   6  RT (min)       lcorbi_hilic_neg_4_053_23    m/z 129.9601     0   1   6   6.5  RT (min)      lcorbi_hilic_neg_4_111_62    m/z 129.9944     0   1E+5   4.5   5  RT (min)       lcorbi_hilic_neg_4_089_59    m/z 130.0046     0   1   6   6.5  RT (min)      lcorbi_hilic_neg_4_113_62    m/z 130.0338     0   1E+3   1.5   2  RT (min)      lcorbi_hilic_neg_4_106_62    m/z 130.0397     0   1E+3   6   6.5  RT (min)      lcorbi_hilic_neg_1_051_33    m/z 130.0410     0   1E+5   3   3.5  RT (min)      lcorbi_hilic_neg_4_088_59    m/z 130.0660     0   1E+3   1   1.5  RT (min)      lcorbi_hilic_neg_2_054_13    m/z 130.0660     0   1   5   5.5  RT (min)      lcorbi_hilic_neg_4_005_32    m/z 130.2278     0   2   4   6E+3   5   5.5  RT (min)      lcorbi_hilic_neg_2_020_27    m/z 130.2294     0   2   4   6E+3   5   5.5  RT (min)      lcorbi_hilic_neg_2_020_27    m/z 130.8784     0   1   2E+4   8   8.5  RT (min)       lcorbi_hilic_neg_4_074_58    m/z 130.8894     0   1E+4   4.5   5  RT (min)       lcorbi_hilic_neg_4_071_57    m/z 130.9019     0   2   4   6E+3   4.5   5  RT (min)      lcorbi_hilic_neg_4_078_58    m/z 130.9290     0   2   4   6E+3   4.5   5  RT (min)      lcorbi_hilic_neg_4_075_58    m/z 130.9442     0   1E+3   4.5   5  RT (min)      lcorbi_hilic_neg_4_105_61    m/z 130.9465     0   1E+4   1   1.5  RT (min)       lcorbi_hilic_neg_4_110_62    m/z 130.9493     0   2   4   6   8E+3   1   1.5  RT (min)      lcorbi_hilic_neg_2_004_09    m/z 130.9573     0   1E+4   7.5   8  RT (min)       lcorbi_hilic_neg_2_027_02    m/z 130.9601     0   1E+4   4.5   5  RT (min)       lcorbi_hilic_neg_4_074_58    m/z 130.9924     0   2   4   6   8E+3   5   5.5  RT (min)       lcorbi_hilic_neg_4_031_14    m/z 130.9925     0   1   2   3E+4   3   3.5  RT (min)       lcorbi_hilic_neg_4_048_37    m/z 131.0235     0   1   1.5   2  RT (min)      lcorbi_hilic_neg_4_095_60    m/z 131.0459     0   1E+3   2   3  RT (min)       lcorbi_hilic_neg_4_088_59    m/z 131.0494     0   1E+3   6.5   7  RT (min)      lcorbi_hilic_neg_1_051_33    m/z 131.0542     0   1E+4   1.5   2  RT (min)      lcorbi_hilic_neg_4_074_58    m/z 131.0653     0   2   4   6   8E+3   5.5   6  RT (min)      lcorbi_hilic_neg_4_095_60    m/z 131.0855     0   2   4   6E+3   6.5   7  RT (min)      lcorbi_hilic_neg_4_089_59    m/z 131.9374     0   2   4   6   8E+3   8   8.5  RT (min)      lcorbi_hilic_neg_4_092_60    m/z 131.9608     0   1   2E+4   4.5   5  RT (min)       lcorbi_hilic_neg_4_079_58    m/z 132.0202     0   1E+4   6   6.5  RT (min)      lcorbi_hilic_neg_4_091_60    m/z 132.0203     0   1E+3   1.5   2  RT (min)       lcorbi_hilic_neg_4_083_59    m/z 132.0362     0   1E+4   6.5   7  RT (min)      lcorbi_hilic_neg_2_040_35    m/z 132.0406     0   2   4   6E+2   1.5   2  RT (min)      lcorbi_hilic_neg_2_005_24    m/z 132.0931     0   1   2   3E+4   1   1.5  RT (min)       lcorbi_hilic_neg_4_096_60    m/z 132.8651     0   1   2E+3   7   7.5  RT (min)      lcorbi_hilic_neg_1_008_39    m/z 132.8702     0   1E+4   12   12.5  RT (min)      lcorbi_hilic_neg_4_005_32    m/z 132.8703     0   2   4   6E+4   0.5   1  RT (min)      lcorbi_hilic_neg_4_099_61    m/z 132.8739     0   2   4   6E+4   13   13.5  RT (min)      lcorbi_hilic_neg_4_099_61    m/z 132.8866     0   1   2   3E+3   4.5   5  RT (min)      lcorbi_hilic_neg_1_042_25    m/z 132.9234     0   2   4E+2   8   8.5  RT (min)       lcorbi_hilic_neg_1_007_10    m/z 132.9351     0   2   4   6E+3   8   8.5  RT (min)      lcorbi_hilic_neg_4_052_43    m/z 132.9369     0   2   4   6E+3   4.5   5  RT (min)      lcorbi_hilic_neg_1_055_24    m/z 132.9463     0   1E+3   7   7.5  RT (min)       lcorbi_hilic_neg_1_007_10    m/z 132.9483     0   1   1   1.5  RT (min)      lcorbi_hilic_neg_3_004_41    m/z 132.9588     0   1   4.5   5  RT (min)      lcorbi_hilic_neg_1_036_09    m/z 132.9877     0   2   4E+3   8   8.5  RT (min)      lcorbi_hilic_neg_4_078_58    m/z 132.9950     0   1   2   3   4E+2   4.5   5  RT (min)       lcorbi_hilic_neg_1_005_36    m/z 133.0244     0   2   4E+3   7   7.5  RT (min)      lcorbi_hilic_neg_2_040_35    m/z 133.0391     0   1   1   1.5  RT (min)      lcorbi_hilic_neg_4_107_62    m/z 133.0436     0   1   2   3E+4   3   3.5  RT (min)      lcorbi_hilic_neg_4_095_60    m/z 133.0549     0   1   2   3E+3   6   6.5  RT (min)      lcorbi_hilic_neg_4_025_52    m/z 133.0620     0   1   2   3E+3   3   3.5  RT (min)      lcorbi_hilic_neg_4_056_56    m/z 133.0717     0   1E+4   3   3.5  RT (min)      lcorbi_hilic_neg_4_092_60    m/z 133.8733     0   1E+4   8   8.5  RT (min)      lcorbi_hilic_neg_4_082_59    m/z 133.9236     0   1E+4   4.5   5  RT (min)      lcorbi_hilic_neg_4_071_57    m/z 133.9913     0   1   2   3   4E+3   4   4.5  RT (min)      lcorbi_hilic_neg_4_112_62    m/z 134.0068     0   1E+3   1   1.5  RT (min)      lcorbi_hilic_neg_2_020_27    m/z 134.0142     0   1E+4   2   2.5  RT (min)      lcorbi_hilic_neg_4_106_62    m/z 134.0176     0   2   4   6   8E+4   7   7.5  RT (min)       lcorbi_hilic_neg_3_012_35    m/z 134.0281     0   1   7.5   8  RT (min)      lcorbi_hilic_neg_4_112_62    m/z 134.0344     0   1E+3   6.5   7   7.5  RT (min)       lcorbi_hilic_neg_4_100_61    m/z 134.0562     0   1   2   3   4E+4   6.5   7  RT (min)       lcorbi_hilic_neg_4_097_60    m/z 134.0742     0   2   4   6E+3   4   4.5  RT (min)      lcorbi_hilic_neg_4_073_57    m/z 134.8692     0   1E+4   4.5   5  RT (min)      lcorbi_hilic_neg_4_091_60    m/z 134.8697     0   1   2   3   4E+4   0.5   1  RT (min)      lcorbi_hilic_neg_4_019_09    m/z 134.8704     0   1E+5   13   13.5  RT (min)      lcorbi_hilic_neg_4_091_60    m/z 134.8848     0   1   2E+4   4.5   5  RT (min)      lcorbi_hilic_neg_4_092_60    m/z 134.8915     0   2   4   6E+3   0.5   1  RT (min)      lcorbi_hilic_neg_3_037_12    m/z 134.8990     0   2   4E+4   7.5   8  RT (min)      lcorbi_hilic_neg_4_106_62    m/z 134.8998     0   2   4   6E+3   4.5   5  RT (min)      lcorbi_hilic_neg_3_022_06    m/z 134.9232     0   2   4   6E+3   4.5   5  RT (min)      lcorbi_hilic_neg_4_092_60    m/z 134.9332     0   1   2E+3   4.5   5  RT (min)      lcorbi_hilic_neg_4_090_60    m/z 134.9428     0   2   4E+3   4.5   5  RT (min)      lcorbi_hilic_neg_4_109_62    m/z 134.9432     0   1   2   3   4E+3   7   7.5  RT (min)      lcorbi_hilic_neg_4_081_58    m/z 135.0183     0   1E+3   7   7.5  RT (min)       lcorbi_hilic_neg_2_012_14    m/z 135.0339     0   2   4   6E+3   3   3.5  RT (min)      lcorbi_hilic_neg_4_070_57    m/z 135.0512     0   1   2   3E+4   1   1.5  RT (min)      lcorbi_hilic_neg_4_113_62    m/z 135.0673     0   1E+5   5.5   6  RT (min)       lcorbi_hilic_neg_4_015_24    m/z 135.0675     0   1   3.5   4  RT (min)      lcorbi_hilic_neg_4_085_59    m/z 135.0813     0   1E+3   0.5   1  RT (min)      lcorbi_hilic_neg_4_097_60    m/z 135.0829     0   1E+4   4   4.5  RT (min)      lcorbi_hilic_neg_4_075_58    m/z 135.0846     0   1E+4   4   4.5  RT (min)      lcorbi_hilic_neg_4_075_58    m/z 135.0926     0   1E+3   1.5   2  RT (min)      lcorbi_hilic_neg_4_097_60    m/z 135.0958     0   1   2E+4   7   7.5  RT (min)       lcorbi_hilic_neg_4_097_60    m/z 135.2090     0   2   4   6E+3   4   4.5  RT (min)      lcorbi_hilic_neg_4_079_58    m/z 135.3337     0   2   4E+3   4   4.5  RT (min)      lcorbi_hilic_neg_4_079_58    m/z 135.8917     0   2   4   6E+5   4.5   5  RT (min)      lcorbi_hilic_neg_3_033_53    m/z 135.9207     0   1   2E+4   0.5   1   1.5  RT (min)       lcorbi_hilic_neg_4_093_60    m/z 135.9708     0   1   2   3   4E+4   0.5   1  RT (min)       lcorbi_hilic_neg_4_075_58    m/z 136.0249     0   1   2E+3   7   7.5  RT (min)       lcorbi_hilic_neg_1_028_45    m/z 136.0281     0   1   3   3.5  RT (min)      lcorbi_hilic_neg_4_107_62    m/z 136.0303     0   1   2   2.5  RT (min)      lcorbi_hilic_neg_4_106_62    m/z 136.0344     0   1   3   3.5  RT (min)       lcorbi_hilic_neg_4_107_62    m/z 136.0407     0   1   2   3E+4   7   7.5  RT (min)      lcorbi_hilic_neg_4_097_60    m/z 136.0461     0   2   4   6   8E+4   7   7.5  RT (min)       lcorbi_hilic_neg_4_093_60    m/z 136.0564     0   1   2   3   4E+3   6.5   7  RT (min)      lcorbi_hilic_neg_2_005_24    m/z 136.0963     0   1E+3   7   7.5  RT (min)      lcorbi_hilic_neg_4_090_60    m/z 136.8650     0   2   4   6E+3   12.5   13  RT (min)      lcorbi_hilic_neg_4_006_06    m/z 136.8651     0   2   4E+3   0.5   1  RT (min)      lcorbi_hilic_neg_3_033_53    m/z 136.8947     0   1E+4   1   1.5  RT (min)      lcorbi_hilic_neg_4_019_09    m/z 136.8998     0   1   2   3E+4   4.5   5  RT (min)      lcorbi_hilic_neg_4_090_60    m/z 136.9398     0   1   2E+4   12.5   13   13.5  RT (min)       lcorbi_hilic_neg_4_070_57    m/z 136.9399     0   1E+5   10  RT (min)       lcorbi_hilic_neg_4_109_62    m/z 136.9912     0   1   2E+3   1.5   2  RT (min)      lcorbi_hilic_neg_4_072_57    m/z 137.0228     0   2   4   6E+3   3   3.5  RT (min)      lcorbi_hilic_neg_1_012_32    m/z 137.0304     0   2   4E+3   6.5   7  RT (min)      lcorbi_hilic_neg_3_022_06    m/z 137.0400     0   1   2E+4   7   7.5  RT (min)       lcorbi_hilic_neg_4_095_60    m/z 137.0428     0   2   4   6   8E+2   3.5   4  RT (min)      lcorbi_hilic_neg_1_001_49    m/z 137.0495     0   1E+4   1   1.5  RT (min)      lcorbi_hilic_neg_4_108_62    m/z 137.0517     0   1   2.5   3  RT (min)      lcorbi_hilic_neg_4_097_60    m/z 137.0606     0   2   4   6   8E+3   0.5   1  RT (min)      lcorbi_hilic_neg_2_020_27    m/z 137.0636     0   2   4   6   8E+3   6.5   7  RT (min)      lcorbi_hilic_neg_4_096_60    m/z 137.8953     0   2   4   6   8E+3   4.5   5  RT (min)      lcorbi_hilic_neg_4_082_59    m/z 137.9178     0   1   2E+4   1   1.5  RT (min)       lcorbi_hilic_neg_4_100_61    m/z 137.9407     0   1E+4   8   8.5  RT (min)      lcorbi_hilic_neg_4_081_58    m/z 137.9911     0   1   2   3   4E+3   2   2.5  RT (min)      lcorbi_hilic_neg_1_001_49    m/z 138.0348     0   1   4   4.5  RT (min)       lcorbi_hilic_neg_1_007_10    m/z 138.0388     0   1E+3   1   1.5  RT (min)       lcorbi_hilic_neg_1_001_49    m/z 138.0476     0   1   7   7.5  RT (min)       lcorbi_hilic_neg_1_002_26    m/z 138.8600     0   1E+5   11.5   12  RT (min)      lcorbi_hilic_neg_4_048_37    m/z 138.8691     0   2   4E+3   8.5   9  RT (min)       lcorbi_hilic_neg_1_007_10    m/z 138.8899     0   1E+4   7   7.5  RT (min)       lcorbi_hilic_neg_4_109_62    m/z 138.8900     0   2   4   6   8E+3   0.5   1  RT (min)      lcorbi_hilic_neg_4_032_22    m/z 138.9209     0   1   2   3E+3   8   8.5  RT (min)       lcorbi_hilic_neg_1_007_10    m/z 138.9209     0   1   2   3E+3   4.5   5  RT (min)      lcorbi_hilic_neg_4_096_60    m/z 138.9375     0   2   4E+3   8   8.5  RT (min)      lcorbi_hilic_neg_1_007_10    m/z 138.9523     0   2   4   6   8E+2   1   1.5  RT (min)      lcorbi_hilic_neg_1_001_49    m/z 138.9706     0   1   6   6.5  RT (min)      lcorbi_hilic_neg_2_005_24    m/z 138.9847     0   1   2E+4   7   7.5  RT (min)       lcorbi_hilic_neg_4_097_60    m/z 138.9966     0   1E+3   6   6.5  RT (min)      lcorbi_hilic_neg_1_001_49    m/z 139.0036     0   1E+3   1   1.5  RT (min)      lcorbi_hilic_neg_4_069_57    m/z 139.0259     0   1E+3   4.5   5  RT (min)      lcorbi_hilic_neg_1_001_49    m/z 139.0285     0   2   4   6E+2   1   1.5  RT (min)      lcorbi_hilic_neg_1_001_49    m/z 139.0317     0   1   2   3E+4   6.5   7  RT (min)       lcorbi_hilic_neg_3_011_24    m/z 139.0512     0   1E+3   4.5   5  RT (min)      lcorbi_hilic_neg_4_093_60    m/z 139.0592     0   1   2   3   4E+3   1.5   2  RT (min)      lcorbi_hilic_neg_3_033_53    m/z 139.0608     0   1   2E+3   4   4.5  RT (min)      lcorbi_hilic_neg_4_096_60    m/z 139.0962     0   1   2   3E+3   5   5.5  RT (min)      lcorbi_hilic_neg_4_093_60    m/z 139.8706     0   1   2   3E+4   4.5   5  RT (min)      lcorbi_hilic_neg_4_066_57    m/z 139.8894     0   1   2   3   4E+3   8   8.5  RT (min)      lcorbi_hilic_neg_4_082_59    m/z 139.8968     0   2   4   6   8E+3   4.5   5  RT (min)      lcorbi_hilic_neg_4_091_60    m/z 139.9292     0   1   2   3E+3   1   1.5  RT (min)      lcorbi_hilic_neg_4_089_59    m/z 139.9811     0   2   4   6   8E+3   6   6.5  RT (min)      lcorbi_hilic_neg_4_087_59    m/z 139.9974     0   1   2   3E+4   2   2.5  RT (min)       lcorbi_hilic_neg_4_085_59    m/z 140.0100     0   1   2.5   3  RT (min)      lcorbi_hilic_neg_4_083_59    m/z 140.0100     0   1E+4   4.5   5  RT (min)      lcorbi_hilic_neg_4_073_57    m/z 140.0352     0   1E+3   2   2.5  RT (min)      lcorbi_hilic_neg_4_098_61    m/z 140.0478     0   1   2E+4   3   3.5  RT (min)       lcorbi_hilic_neg_4_093_60    m/z 140.0539     0   1E+4   6   6.5  RT (min)      lcorbi_hilic_neg_4_070_57    m/z 140.0608     0   1E+5   3   3.5  RT (min)       lcorbi_hilic_neg_4_095_60    m/z 140.8679     0   1E+4   8.5   9  RT (min)       lcorbi_hilic_neg_4_105_61    m/z 140.8724     0   1E+4   2.5   3  RT (min)       lcorbi_hilic_neg_4_073_57    m/z 140.9128     0   1E+3   4.5   5  RT (min)      lcorbi_hilic_neg_4_072_57    m/z 140.9180     0   1E+4   8   8.5  RT (min)      lcorbi_hilic_neg_4_077_58    m/z 140.9789     0   1   2E+4   4.5   5  RT (min)      lcorbi_hilic_neg_4_070_57    m/z 140.9957     0   1   2   3   4E+4   1   2  RT (min)       lcorbi_hilic_neg_4_091_60    m/z 141.0127     0   1E+3   5.5   6  RT (min)      lcorbi_hilic_neg_4_093_60    m/z 141.0193     0   1E+4   1   1.5  RT (min)      lcorbi_hilic_neg_4_084_59    m/z 141.0304     0   1   2E+3   2.5   3  RT (min)       lcorbi_hilic_neg_4_097_60    m/z 141.0379     0   1   6   6.5  RT (min)      lcorbi_hilic_neg_4_081_58    m/z 141.0555     0   2   4   6   8E+3   1   1.5  RT (min)      lcorbi_hilic_neg_4_088_59    m/z 141.0668     0   2   4   6   8E+3   5   5.5  RT (min)      lcorbi_hilic_neg_4_082_59    m/z 141.0668     0   2   4   6E+3   1   1.5  RT (min)      lcorbi_hilic_neg_4_079_58    m/z 141.0668     0   2   4   6   8E+3   3   3.5  RT (min)      lcorbi_hilic_neg_4_080_58    m/z 141.0829     0   2   4E+3   4   4.5  RT (min)      lcorbi_hilic_neg_4_068_57    m/z 141.0847     0   1   2   3E+3   4   4.5  RT (min)      lcorbi_hilic_neg_4_068_57    m/z 141.5867     0   2   4E+3   4   4.5  RT (min)      lcorbi_hilic_neg_4_070_57    m/z 141.5886     0   1   2   3E+3   4   4.5  RT (min)      lcorbi_hilic_neg_4_072_57    m/z 141.8680     0   1E+3   1   1.5  RT (min)      lcorbi_hilic_neg_4_067_57    m/z 141.9050     0   1E+4   8   8.5  RT (min)      lcorbi_hilic_neg_4_069_57    m/z 141.9144     0   1E+3   2.5   3  RT (min)      lcorbi_hilic_neg_4_101_61    m/z 141.9727     0   2   4   6   8E+3   4.5   5  RT (min)      lcorbi_hilic_neg_4_076_58    m/z 141.9802     0   2   4   6E+2   4   4.5  RT (min)      lcorbi_hilic_neg_1_012_32    m/z 141.9944     0   1E+4   2   2.5  RT (min)       lcorbi_hilic_neg_4_084_59    m/z 142.0153     0   1   6.5   7  RT (min)       lcorbi_hilic_neg_4_096_60    m/z 142.0661     0   1   2   3   4E+3   1   1.5  RT (min)      lcorbi_hilic_neg_2_020_27    m/z 142.0872     0   2   4   6   8E+3   1   1.5  RT (min)      lcorbi_hilic_neg_2_028_33    m/z 142.0907     0   1   4   4.5  RT (min)      lcorbi_hilic_neg_4_087_59    m/z 142.8561     0   1E+3   3   3.5  RT (min)      lcorbi_hilic_neg_4_098_61    m/z 142.9097     0   2   4   6   8E+3   0   0.5  RT (min)       lcorbi_hilic_neg_4_090_60    m/z 142.9249     0   1   2   3E+4   1   2  RT (min)       lcorbi_hilic_neg_4_091_60    m/z 142.9288     0   1   2E+4   7.5   8  RT (min)       lcorbi_hilic_neg_4_103_61    m/z 142.9604     0   1E+3   6.5   7  RT (min)      lcorbi_hilic_neg_4_073_57    m/z 142.9750     0   1   2   3E+4   7   7.5  RT (min)       lcorbi_hilic_neg_4_096_60    m/z 142.9760     0   1   2E+4   4.5   5  RT (min)      lcorbi_hilic_neg_4_077_58    m/z 142.9820     0   1   2   3E+4   2   2.5  RT (min)       lcorbi_hilic_neg_4_096_60    m/z 142.9924     0   1   2E+4   1.5   2  RT (min)       lcorbi_hilic_neg_4_083_59    m/z 143.0220     0   1   6.5   7  RT (min)      lcorbi_hilic_neg_4_067_57    m/z 143.0227     0   1E+3   1   1.5  RT (min)      lcorbi_hilic_neg_4_077_58    m/z 143.0349     0   2   4E+4   1   1.5  RT (min)       lcorbi_hilic_neg_4_088_59    m/z 143.0460     0   2   4E+3   5  RT (min)       lcorbi_hilic_neg_2_055_32    m/z 143.0461     0   2   4E+2   2.5   3  RT (min)       lcorbi_hilic_neg_2_013_18    m/z 143.0520     0   1   2E+4   1   1.5  RT (min)       lcorbi_hilic_neg_4_080_58    m/z 143.8651     0   2   4E+3   1   1.5  RT (min)      lcorbi_hilic_neg_4_097_60    m/z 143.9326     0   1   2E+4   2   2.5  RT (min)       lcorbi_hilic_neg_4_082_59    m/z 143.9668     0   1   2E+4   6.5   7  RT (min)       lcorbi_hilic_neg_4_097_60    m/z 143.9698     0   2   4   6E+3   4.5   5  RT (min)      lcorbi_hilic_neg_4_079_58    m/z 143.9866     0   1   2E+4   4   4.5  RT (min)      lcorbi_hilic_neg_1_012_32    m/z 143.9879     0   1   7   7.5  RT (min)      lcorbi_hilic_neg_1_048_35    m/z 144.0301     0   1   2E+4   3   4  RT (min)       lcorbi_hilic_neg_4_089_59    m/z 144.0301     0   1   2   3E+4   1   1.5  RT (min)       lcorbi_hilic_neg_4_075_58    m/z 144.0454     0   1E+3   4.5   5  RT (min)      lcorbi_hilic_neg_4_005_32    m/z 144.0551     0   1   2E+4   3   3.5  RT (min)      lcorbi_hilic_neg_4_089_59    m/z 144.1029     0   1E+4   4   4.5  RT (min)      lcorbi_hilic_neg_4_073_57    m/z 144.8542     0   1E+3   3   3.5  RT (min)      lcorbi_hilic_neg_4_098_61    m/z 144.8663     0   1   2   3   4E+3   8   8.5  RT (min)      lcorbi_hilic_neg_4_079_58    m/z 144.8696     0   1   2   3E+3   4.5   5  RT (min)      lcorbi_hilic_neg_4_090_60    m/z 144.8787     0   1   2E+3   4.5   5  RT (min)      lcorbi_hilic_neg_4_108_62    m/z 144.9204     0   2   4E+3   5   5.5  RT (min)       lcorbi_hilic_neg_4_091_60    m/z 144.9204     0   1E+3   7   7.5  RT (min)       lcorbi_hilic_neg_4_105_61    m/z 144.9234     0   2   4   6E+5   5   6  RT (min)       lcorbi_hilic_neg_4_098_61    m/z 144.9735     0   1   2   3E+3   4.5   5  RT (min)      lcorbi_hilic_neg_1_016_37    m/z 144.9759     0   1E+3   7   7.5  RT (min)      lcorbi_hilic_neg_1_002_26    m/z 144.9886     0   1   5   6   7  RT (min)       lcorbi_hilic_neg_4_095_60    m/z 144.9917     0   1   2   3   4E+4   3   3.5  RT (min)      lcorbi_hilic_neg_4_056_56    m/z 145.0147     0   2   4E+4   6   6.5  RT (min)       lcorbi_hilic_neg_4_048_37    m/z 145.0312     0   2   4   6   8E+3   1.5   2  RT (min)       lcorbi_hilic_neg_4_077_58    m/z 145.0391     0   1   5   5.5  RT (min)      lcorbi_hilic_neg_4_094_60    m/z 145.0532     0   1   2   3E+4   6.5   7  RT (min)      lcorbi_hilic_neg_4_088_59    m/z 145.0646     0   1E+5   6.5   7  RT (min)       lcorbi_hilic_neg_4_023_34    m/z 145.0714     0   1   2   3E+3   6.5   7  RT (min)      lcorbi_hilic_neg_2_032_25    m/z 145.0808     0   1E+4   6.5   7  RT (min)      lcorbi_hilic_neg_1_016_37    m/z 145.0868     0   1   2   3E+3   1   1.5  RT (min)      lcorbi_hilic_neg_4_037_31    m/z 145.0981     0   1   2E+4   7   7.5  RT (min)       lcorbi_hilic_neg_4_073_57    m/z 145.5207     0   2   4   6E+3   6   6.5  RT (min)      lcorbi_hilic_neg_4_082_59    m/z 145.9235     0   2   4   6E+4   4.5   5  RT (min)       lcorbi_hilic_neg_4_100_61    m/z 145.9389     0   1   6   6.5  RT (min)      lcorbi_hilic_neg_4_074_58    m/z 145.9495     0   1   4.5   5  RT (min)      lcorbi_hilic_neg_4_113_62    m/z 145.9496     0   2   4   6   8E+3   1   1.5  RT (min)      lcorbi_hilic_neg_4_106_62    m/z 145.9606     0   1   2   3E+3   6   6.5  RT (min)      lcorbi_hilic_neg_4_087_59    m/z 145.9950     0   2   4   6E+3   1   1.5  RT (min)      lcorbi_hilic_neg_4_088_59    m/z 145.9988     0   1   2E+4   6   6.5  RT (min)       lcorbi_hilic_neg_4_078_58    m/z 146.0127     0   1E+3   4.5   5  RT (min)      lcorbi_hilic_neg_1_003_14    m/z 146.0148     0   2   4   6E+3   6   6.5  RT (min)      lcorbi_hilic_neg_4_087_59    m/z 146.0174     0   1   2   3   4E+4   2   2.5  RT (min)      lcorbi_hilic_neg_4_037_31    m/z 146.0243     0   2   4   6   8E+3   6   6.5  RT (min)      lcorbi_hilic_neg_4_089_59    m/z 146.0339     0   1E+4   6   6.5  RT (min)      lcorbi_hilic_neg_4_087_59    m/z 146.0539     0   1E+4   1   1.5  RT (min)       lcorbi_hilic_neg_4_084_59    m/z 146.0609     0   1E+4   4   4.5  RT (min)      lcorbi_hilic_neg_4_105_61    m/z 146.0642     0   1   2E+3   1   1.5  RT (min)      lcorbi_hilic_neg_4_072_57    m/z 146.0724     0   1   6   6.5  RT (min)      lcorbi_hilic_neg_3_053_13    m/z 146.0821     0   1E+4   1.5   2  RT (min)      lcorbi_hilic_neg_4_089_59    m/z 146.0830     0   2   4   6E+2   6   6.5  RT (min)      lcorbi_hilic_neg_3_053_13    m/z 146.0997     0   1   2   3E+3   6   6.5  RT (min)       lcorbi_hilic_neg_4_088_59    m/z 146.1526     0   2   4   6E+3   6   6.5  RT (min)      lcorbi_hilic_neg_2_038_36    m/z 146.5737     0   2   4   6E+3   6   6.5  RT (min)      lcorbi_hilic_neg_4_082_59    m/z 146.8668     0   1E+3   4.5   5  RT (min)      lcorbi_hilic_neg_2_020_27    m/z 146.8758     0   2   4   6   8E+4   4.5   5  RT (min)       lcorbi_hilic_neg_3_016_16    m/z 146.9062     0   1   2E+4   2.5   3  RT (min)      lcorbi_hilic_neg_4_073_57    m/z 146.9082     0   1   1   1.5  RT (min)      lcorbi_hilic_neg_4_100_61    m/z 146.9386     0   1   2   3E+4   11   11.5  RT (min)      lcorbi_hilic_neg_2_040_35    m/z 146.9644     0   2   4   6E+3   8.5   9  RT (min)      lcorbi_hilic_neg_4_112_62    m/z 146.9658     0   1   2E+3   6.5   7  RT (min)      lcorbi_hilic_neg_1_026_34    m/z 146.9691     0   1   2E+4   1   1.5  RT (min)      lcorbi_hilic_neg_4_105_61    m/z 146.9825     0   2   4   6   8E+2   7   7.5  RT (min)      lcorbi_hilic_neg_1_026_34    m/z 146.9828     0   1   2   3E+3   4.5   5  RT (min)      lcorbi_hilic_neg_4_108_62    m/z 147.0086     0   1   2   3   4E+2   0.5   1  RT (min)      lcorbi_hilic_neg_4_002_41    m/z 147.0183     0   1   2   2.5  RT (min)      lcorbi_hilic_neg_4_002_41    m/z 147.0297     0   2   4E+3   1.5   2  RT (min)       lcorbi_hilic_neg_1_027_18    m/z 147.0298     0   2   4   6E+5   6.5   7  RT (min)       lcorbi_hilic_neg_1_026_34    m/z 147.0491     0   1   3   3.5  RT (min)      lcorbi_hilic_neg_3_003_44    m/z 147.0676     0   2   4   6   8E+4   3   3.5  RT (min)       lcorbi_hilic_neg_4_097_60    m/z 147.0813     0   1E+4   0.5   1  RT (min)      lcorbi_hilic_neg_4_080_58    m/z 147.0854     0   1E+4   5   5.5  RT (min)      lcorbi_hilic_neg_4_090_60    m/z 147.2249     0   1   2   3   4E+3   5   5.5  RT (min)      lcorbi_hilic_neg_2_020_27    m/z 147.9245     0   1   2   3   4E+3   8   8.5  RT (min)      lcorbi_hilic_neg_4_083_59    m/z 147.9461     0   1   2   3   4E+3   4   4.5  RT (min)      lcorbi_hilic_neg_4_088_59    m/z 147.9469     0   1   2E+4   8.5   9  RT (min)       lcorbi_hilic_neg_4_074_58    m/z 147.9628     0   1E+4   1   1.5  RT (min)      lcorbi_hilic_neg_4_074_58    m/z 147.9701     0   1E+3   1   1.5  RT (min)      lcorbi_hilic_neg_4_106_62    m/z 147.9959     0   1   6   6.5  RT (min)      lcorbi_hilic_neg_4_005_32    m/z 148.0151     0   2   4   6E+2   6   6.5  RT (min)       lcorbi_hilic_neg_4_005_32    m/z 148.0153     0   1E+5   3   3.5  RT (min)      lcorbi_hilic_neg_2_005_24    m/z 148.0244     0   1   2   3   4E+4   6   6.5  RT (min)       lcorbi_hilic_neg_4_072_57    m/z 148.0402     0   1E+3   1   1.5  RT (min)      lcorbi_hilic_neg_4_106_62    m/z 148.0404     0   1   5   5.5  RT (min)      lcorbi_hilic_neg_1_026_34    m/z 148.0435     0   1   3.5   4  RT (min)      lcorbi_hilic_neg_3_011_24    m/z 148.0483     0   1E+4   1   1.5  RT (min)       lcorbi_hilic_neg_2_020_27    m/z 148.0623     0   1E+4   4   4.5  RT (min)       lcorbi_hilic_neg_4_096_60    m/z 148.0709     0   1   2E+3   6.5   7  RT (min)      lcorbi_hilic_neg_4_037_31    m/z 148.0744     0   2   4   6E+3   3   3.5  RT (min)      lcorbi_hilic_neg_3_001_49    m/z 148.8728     0   1   2   3E+4   4.5   5  RT (min)      lcorbi_hilic_neg_2_020_27    m/z 148.9041     0   1   2   3E+4   2.5   3  RT (min)       lcorbi_hilic_neg_4_070_57    m/z 148.9244     0   2   4E+3   8   8.5  RT (min)      lcorbi_hilic_neg_4_090_60    m/z 148.9389     0   2   4   6E+3   8   8.5  RT (min)       lcorbi_hilic_neg_2_015_19    m/z 148.9700     0   1   2E+4   1   1.5  RT (min)       lcorbi_hilic_neg_4_074_58    m/z 148.9800     0   1   2E+4   1.5   2  RT (min)       lcorbi_hilic_neg_4_081_58    m/z 149.0030     0   1   2   3E+4   1   1.5  RT (min)       lcorbi_hilic_neg_4_095_60    m/z 149.0275     0   1   2   3E+3   6   6.5  RT (min)       lcorbi_hilic_neg_1_003_14    m/z 149.0309     0   2   4E+2   4.5   5  RT (min)      lcorbi_hilic_neg_1_022_48    m/z 149.0469     0   1   2E+3   1   1.5  RT (min)      lcorbi_hilic_neg_4_104_61    m/z 149.0549     0   2   4E+4   1.5   2  RT (min)       lcorbi_hilic_neg_4_092_60    m/z 149.0559     0   1   6   6.5  RT (min)       lcorbi_hilic_neg_2_005_24    m/z 149.0622     0   1   2   3E+4   1   1.5  RT (min)       lcorbi_hilic_neg_4_091_60    m/z 149.0719     0   1   2E+4   1   1.5  RT (min)      lcorbi_hilic_neg_2_055_32    m/z 149.0970     0   1   2   3E+4   0.5   1  RT (min)       lcorbi_hilic_neg_4_077_58    m/z 149.1082     0   2   4   6   8E+3   1   1.5  RT (min)      lcorbi_hilic_neg_4_082_59    m/z 149.9191     0   1   2   3   4E+3   8   8.5  RT (min)      lcorbi_hilic_neg_4_088_59    m/z 149.9399     0   1E+4   8   8.5  RT (min)      lcorbi_hilic_neg_4_066_57    m/z 150.0060     0   2   4E+2   6   6.5  RT (min)      lcorbi_hilic_neg_1_026_34    m/z 150.0090     0   1   2E+3   0.5   1  RT (min)      lcorbi_hilic_neg_4_106_62    m/z 150.0229     0   1   6   6.5  RT (min)      lcorbi_hilic_neg_4_097_60    m/z 150.0252     0   1   3.5   4  RT (min)      lcorbi_hilic_neg_1_019_06    m/z 150.0343     0   2   4E+3   2   2.5  RT (min)      lcorbi_hilic_neg_3_033_53    m/z 150.0488     0   1   5   5.5  RT (min)      lcorbi_hilic_neg_1_051_33    m/z 150.0489     0   1   3   3.5  RT (min)      lcorbi_hilic_neg_4_113_62    m/z 150.0714     0   2   4   6E+4   6.5   7  RT (min)      lcorbi_hilic_neg_4_093_60    m/z 150.0752     0   1E+4   2.5   3  RT (min)      lcorbi_hilic_neg_4_095_60    m/z 150.0858     0   1   2E+3   6.5   7  RT (min)      lcorbi_hilic_neg_3_022_06    m/z 150.0866     0   1   1   1.5  RT (min)      lcorbi_hilic_neg_1_020_13    m/z 150.0866     0   1   2   3E+4   3   3.5  RT (min)       lcorbi_hilic_neg_4_093_60    m/z 150.8865     0   1   2E+4   0   0.5  RT (min)       lcorbi_hilic_neg_4_076_58    m/z 150.9163     0   1   2E+4   8   8.5  RT (min)      lcorbi_hilic_neg_4_090_60    m/z 150.9401     0   1E+4   8   8.5  RT (min)      lcorbi_hilic_neg_4_066_57    m/z 150.9481     0   1E+3   0.5   1  RT (min)       lcorbi_hilic_neg_4_071_57    m/z 151.0069     0   1E+3   3.5   4  RT (min)      lcorbi_hilic_neg_4_103_61    m/z 151.0069     0   1   2E+3   1   1.5  RT (min)       lcorbi_hilic_neg_1_027_18    m/z 151.0199     0   1   4   4.5  RT (min)      lcorbi_hilic_neg_4_005_32    m/z 151.0393     0   2   4   6E+3   4   4.5  RT (min)      lcorbi_hilic_neg_4_005_32    m/z 151.0402     0   1E+4   6   6.5  RT (min)      lcorbi_hilic_neg_4_092_60    m/z 151.0471     0   1E+5   4   4.5  RT (min)       lcorbi_hilic_neg_3_037_12    m/z 151.0493     0   1   2E+4   6   6.5  RT (min)      lcorbi_hilic_neg_4_097_60    m/z 151.0512     0   1E+3   1.5   2  RT (min)      lcorbi_hilic_neg_4_089_59    m/z 151.0610     0   1E+4   1   1.5  RT (min)       lcorbi_hilic_neg_3_028_23    m/z 151.0821     0   1E+4   6   6.5  RT (min)      lcorbi_hilic_neg_4_090_60    m/z 151.0827     0   1   2   3E+4   4   4.5  RT (min)      lcorbi_hilic_neg_4_083_59    m/z 151.0935     0   1E+4   4   4.5  RT (min)      lcorbi_hilic_neg_4_089_59    m/z 151.1127     0   1   1   1.5  RT (min)      lcorbi_hilic_neg_4_097_60    m/z 151.1137     0   1E+4   4   4.5  RT (min)      lcorbi_hilic_neg_4_089_59    m/z 151.1148     0   1   2E+3   6   6.5  RT (min)      lcorbi_hilic_neg_3_048_33    m/z 151.8740     0   1E+4   0.5   1  RT (min)      lcorbi_hilic_neg_4_109_62    m/z 151.8741     0   1   2   3   4E+4   7.5   8   8.5  RT (min)       lcorbi_hilic_neg_4_066_57    m/z 151.8741     0   1   2   3   4E+4   4.5   5  RT (min)       lcorbi_hilic_neg_4_088_59    m/z 151.9352     0   1   2.5   3  RT (min)      lcorbi_hilic_neg_4_082_59    m/z 151.9410     0   1   2   3E+3   8   8.5  RT (min)      lcorbi_hilic_neg_4_067_57    m/z 152.0231     0   2   4   6E+3   4   4.5  RT (min)      lcorbi_hilic_neg_1_012_32    m/z 152.0293     0   1   2   3   4E+3   4   4.5  RT (min)      lcorbi_hilic_neg_1_019_06    m/z 152.0465     0   1   2E+4   1.5   2  RT (min)       lcorbi_hilic_neg_4_094_60    m/z 152.0465     0   2   4E+5   6.5   7   7.5  RT (min)       lcorbi_hilic_neg_1_012_32    m/z 152.0482     0   1   2E+5   4   4.5  RT (min)      lcorbi_hilic_neg_1_012_32    m/z 152.0577     0   2   4   6E+2   3.5   4  RT (min)      lcorbi_hilic_neg_2_041_54    m/z 152.0666     0   2   4   6   8E+3   6   6.5  RT (min)      lcorbi_hilic_neg_4_005_32    m/z 152.0668     0   2   4   6E+4   4   4.5  RT (min)      lcorbi_hilic_neg_3_028_23    m/z 152.0716     0   1   2E+4   1   1.5  RT (min)       lcorbi_hilic_neg_4_094_60    m/z 152.0996     0   2   4   6   8E+3   5   5.5  RT (min)      lcorbi_hilic_neg_4_109_62    m/z 152.1022     0   1   2E+4   1.5   2  RT (min)       lcorbi_hilic_neg_4_094_60    m/z 152.8513     0   1   2E+4   2.5   3  RT (min)       lcorbi_hilic_neg_4_067_57    m/z 152.8836     0   1E+4   0   0.5  RT (min)       lcorbi_hilic_neg_4_099_61    m/z 152.9176     0   2   4   6   8E+3   8   8.5  RT (min)       lcorbi_hilic_neg_1_054_27    m/z 152.9176     0   2   4   6   8E+4   12   13  RT (min)       lcorbi_hilic_neg_4_080_58    m/z 152.9618     0   1E+4   1.5   2  RT (min)      lcorbi_hilic_neg_2_014_31    m/z 152.9862     0   1   2E+4   1   1.5  RT (min)      lcorbi_hilic_neg_4_076_58    m/z 152.9863     0   1   2   3   4E+4   4.5   5  RT (min)       lcorbi_hilic_neg_4_096_60    m/z 152.9966     0   1   2E+4   1   1.5  RT (min)      lcorbi_hilic_neg_4_073_57    m/z 153.0071     0   1   3.5   4  RT (min)      lcorbi_hilic_neg_4_096_60    m/z 153.0919     0   1E+3   1   1.5  RT (min)       lcorbi_hilic_neg_4_090_60    m/z 153.8744     0   1E+4   4.5   5  RT (min)      lcorbi_hilic_neg_4_066_57    m/z 153.9331     0   1   3   3.5  RT (min)      lcorbi_hilic_neg_4_082_59    m/z 154.0120     0   2   4   6   8E+2   7   7.5  RT (min)      lcorbi_hilic_neg_3_055_26    m/z 154.0177     0   1E+3   2   2.5  RT (min)      lcorbi_hilic_neg_1_003_14    m/z 154.0257     0   2   4   6E+3   2   2.5  RT (min)      lcorbi_hilic_neg_1_004_17    m/z 154.0274     0   1   2E+3   6   6.5  RT (min)      lcorbi_hilic_neg_3_055_26    m/z 154.0620     0   1   2E+4   5.5   6  RT (min)       lcorbi_hilic_neg_4_094_60    m/z 154.0622     0   1   2   3   4E+4   2   2.5  RT (min)       lcorbi_hilic_neg_4_090_60    m/z 154.0656     0   2   4   6   8E+3   7.5   8  RT (min)      lcorbi_hilic_neg_4_081_58    m/z 154.0701     0   1   2E+4   1   1.5  RT (min)      lcorbi_hilic_neg_1_029_31    m/z 154.8499     0   1E+3   4.5   5  RT (min)      lcorbi_hilic_neg_1_017_51    m/z 154.8654     0   1   4   4.5  RT (min)      lcorbi_hilic_neg_4_103_61    m/z 154.8687     0   1   2   3   4E+4   0.5   1  RT (min)       lcorbi_hilic_neg_4_112_62    m/z 154.8806     0   1   2   3E+3   0   0.5  RT (min)      lcorbi_hilic_neg_1_005_36    m/z 154.8807     0   2   4   6E+3   4.5   5  RT (min)       lcorbi_hilic_neg_1_009_50    m/z 154.8954     0   1   2   3   4E+3   8   8.5  RT (min)      lcorbi_hilic_neg_4_075_58    m/z 154.9062     0   1   2E+4   8.5   9  RT (min)       lcorbi_hilic_neg_4_101_61    m/z 154.9248     0   2   4E+2   7.5   8  RT (min)       lcorbi_hilic_neg_4_003_08    m/z 154.9507     0   1   2E+3   6   6.5  RT (min)      lcorbi_hilic_neg_4_073_57    m/z 154.9702     0   1E+4   6   6.5  RT (min)      lcorbi_hilic_neg_4_072_57    m/z 154.9924     0   1E+4   0.5   1  RT (min)      lcorbi_hilic_neg_4_069_57    m/z 154.9962     0   1E+4   1.5   2   2.5  RT (min)       lcorbi_hilic_neg_1_016_37    m/z 155.0116     0   2   4E+3   1.5   2  RT (min)      lcorbi_hilic_neg_4_094_60    m/z 155.0209     0   1E+3   2.5   3  RT (min)       lcorbi_hilic_neg_4_091_60    m/z 155.0267     0   1   0.5   1  RT (min)       lcorbi_hilic_neg_4_084_59    m/z 155.0461     0   1   1.5   2  RT (min)      lcorbi_hilic_neg_4_094_60    m/z 155.0781     0   2   4   6E+4   4   4.5  RT (min)       lcorbi_hilic_neg_4_091_60    m/z 155.0824     0   1   1   1.5  RT (min)      lcorbi_hilic_neg_4_096_60    m/z 155.0862     0   1E+4   4   4.5  RT (min)      lcorbi_hilic_neg_4_005_32    m/z 155.1439     0   1E+3   0.5   1  RT (min)      lcorbi_hilic_neg_4_097_60    m/z 155.8693     0   1   2   3E+3   7.5   8  RT (min)      lcorbi_hilic_neg_4_067_57    m/z 155.8695     0   1E+4   4.5   5  RT (min)      lcorbi_hilic_neg_4_066_57    m/z 155.9064     0   1   2E+3   1   1.5  RT (min)      lcorbi_hilic_neg_1_054_27    m/z 155.9520     0   1E+4   2   2.5  RT (min)      lcorbi_hilic_neg_3_039_31    m/z 155.9600     0   1   2E+4   4   4.5  RT (min)      lcorbi_hilic_neg_4_093_60    m/z 156.0057     0   1   2E+4   6   6.5  RT (min)      lcorbi_hilic_neg_4_095_60    m/z 156.0208     0   1   2E+4   4   4.5  RT (min)      lcorbi_hilic_neg_4_093_60    m/z 156.0276     0   1   2   3E+4   6   6.5  RT (min)       lcorbi_hilic_neg_4_080_58    m/z 156.0302     0   2   4   6E+4   4   4.5  RT (min)       lcorbi_hilic_neg_4_082_59    m/z 156.0542     0   1   4   4.5  RT (min)      lcorbi_hilic_neg_4_103_61    m/z 156.0682     0   2   4E+5   4   4.5  RT (min)      lcorbi_hilic_neg_4_095_60    m/z 156.0780     0   1   2E+5   1.5   2  RT (min)      lcorbi_hilic_neg_2_005_24    m/z 156.0977     0   1   4   4.5  RT (min)      lcorbi_hilic_neg_4_103_61    m/z 156.1072     0   1   2   3E+4   4   4.5  RT (min)      lcorbi_hilic_neg_4_093_60    m/z 156.1177     0   1   2E+4   4   4.5  RT (min)      lcorbi_hilic_neg_3_044_32    m/z 156.1328     0   1   2   3E+4   4   4.5  RT (min)      lcorbi_hilic_neg_4_095_60    m/z 156.1955     0   1   2E+4   4   4.5  RT (min)      lcorbi_hilic_neg_3_011_24    m/z 156.6606     0   2   4   6   8E+4   4   4.5  RT (min)      lcorbi_hilic_neg_4_095_60    m/z 156.8465     0   2   4   6   8E+3   2.5   3  RT (min)      lcorbi_hilic_neg_4_069_57    m/z 156.8473     0   1E+4   4.5   5  RT (min)      lcorbi_hilic_neg_4_076_58    m/z 156.8657     0   1   2E+4   0.5   1  RT (min)       lcorbi_hilic_neg_4_106_62    m/z 156.8908     0   2   4   6   8E+3   8   8.5  RT (min)      lcorbi_hilic_neg_4_107_62    m/z 156.9033     0   1E+4   8.5   9  RT (min)      lcorbi_hilic_neg_4_110_62    m/z 156.9156     0   1E+4   0   0.5  RT (min)      lcorbi_hilic_neg_2_003_20    m/z 156.9157     0   1E+4   4.5   5  RT (min)      lcorbi_hilic_neg_4_082_59    m/z 156.9267     0   1E+3   4.5   5  RT (min)      lcorbi_hilic_neg_4_105_61    m/z 156.9267     0   1   2   3E+4   7.5   8  RT (min)       lcorbi_hilic_neg_4_087_59    m/z 156.9363     0   1   7.5   8  RT (min)      lcorbi_hilic_neg_4_112_62    m/z 156.9513     0   1   2   3   4E+3   6   6.5  RT (min)      lcorbi_hilic_neg_4_079_58    m/z 156.9673     0   1   6   6.5  RT (min)      lcorbi_hilic_neg_4_078_58    m/z 156.9758     0   2   4E+2   6   6.5  RT (min)      lcorbi_hilic_neg_1_008_39    m/z 157.0118     0   1   2E+4   5.5   6  RT (min)      lcorbi_hilic_neg_4_076_58    m/z 157.0770     0   1   2E+4   4   4.5  RT (min)      lcorbi_hilic_neg_2_005_24    m/z 157.0987     0   1   2   2.5  RT (min)       lcorbi_hilic_neg_4_004_13    m/z 157.8627     0   1   2   3E+3   0.5   1  RT (min)       lcorbi_hilic_neg_4_055_38    m/z 157.8719     0   2   4   6   8E+5   0.5   1  RT (min)       lcorbi_hilic_neg_1_003_14    m/z 157.9311     0   1   2E+3   1   1.5  RT (min)      lcorbi_hilic_neg_1_012_32    m/z 157.9675     0   1   2   3   4E+3   1   1.5  RT (min)       lcorbi_hilic_neg_4_091_60    m/z 158.0205     0   1E+4   3.5   4  RT (min)      lcorbi_hilic_neg_4_073_57    m/z 158.0254     0   1E+4   7   7.5  RT (min)      lcorbi_hilic_neg_4_096_60    m/z 158.0314     0   2   4   6   8E+4   2   2.5  RT (min)       lcorbi_hilic_neg_4_075_58    m/z 158.0422     0   1   4   4.5  RT (min)      lcorbi_hilic_neg_1_005_36    m/z 158.0457     0   1   2   3   4E+3   1   1.5  RT (min)      lcorbi_hilic_neg_4_067_57    m/z 158.0610     0   1   2   3E+3   1   1.5  RT (min)      lcorbi_hilic_neg_4_078_58    m/z 158.0650     0   1   2E+4   5.5   6  RT (min)      lcorbi_hilic_neg_4_089_59    m/z 158.0707     0   1   2E+4   1.5   2  RT (min)      lcorbi_hilic_neg_4_082_59    m/z 158.8462     0   2   4   6E+4   4.5   5  RT (min)      lcorbi_hilic_neg_4_066_57    m/z 158.8467     0   2   4   6   8E+4   0   0.5  RT (min)      lcorbi_hilic_neg_4_083_59    m/z 158.8467     0   2   4E+4   13.5   14  RT (min)      lcorbi_hilic_neg_4_094_60    m/z 158.8627     0   2   4   6   8E+3   4.5   5  RT (min)      lcorbi_hilic_neg_4_080_58    m/z 158.9009     0   1   2   3E+4   1   1.5  RT (min)      lcorbi_hilic_neg_4_005_32    m/z 158.9391     0   1   2   3   4E+4   8   8.5  RT (min)       lcorbi_hilic_neg_4_090_60    m/z 158.9503     0   1   2   2.5  RT (min)       lcorbi_hilic_neg_4_084_59    m/z 158.9521     0   1E+4   4.5   5  RT (min)      lcorbi_hilic_neg_4_077_58    m/z 158.9699     0   1   2E+4   8   8.5  RT (min)      lcorbi_hilic_neg_4_084_59    m/z 158.9787     0   1   2E+6   9   10  RT (min)       lcorbi_hilic_neg_3_039_31    m/z 158.9852     0   1   2   3E+4   8   8.5  RT (min)      lcorbi_hilic_neg_4_079_58    m/z 158.9856     0   1   2   3E+3   6   6.5  RT (min)      lcorbi_hilic_neg_4_111_62    m/z 159.0073     0   1   2E+4   3.5   4  RT (min)       lcorbi_hilic_neg_4_069_57    m/z 159.0074     0   1E+5   1   1.5   2  RT (min)       lcorbi_hilic_neg_4_083_59    m/z 159.0280     0   1E+5   2   2.5  RT (min)       lcorbi_hilic_neg_4_076_58    m/z 159.0298     0   2   4   6   8E+4   4   4.5  RT (min)       lcorbi_hilic_neg_1_016_37    m/z 159.0376     0   1   6   6.5  RT (min)      lcorbi_hilic_neg_4_101_61    m/z 159.0377     0   1E+3   1   1.5  RT (min)      lcorbi_hilic_neg_1_046_19    m/z 159.0410     0   1E+3   3.5   4  RT (min)       lcorbi_hilic_neg_4_091_60    m/z 159.0772     0   1   6   6.5  RT (min)      lcorbi_hilic_neg_2_005_24    m/z 159.0788     0   1E+4   7.5   8  RT (min)       lcorbi_hilic_neg_4_092_60    m/z 159.0856     0   1   2   3E+3   1   1.5  RT (min)      lcorbi_hilic_neg_4_085_59    m/z 159.1025     0   1   2   3   4E+3   1   1.5  RT (min)      lcorbi_hilic_neg_4_088_59    m/z 159.8451     0   1E+4   4.5   5  RT (min)      lcorbi_hilic_neg_4_100_61    m/z 159.8452     0   2   4   6   8E+4   8.5   9  RT (min)       lcorbi_hilic_neg_4_080_58    m/z 159.8693     0   1E+5   0.5   1  RT (min)      lcorbi_hilic_neg_1_003_14    m/z 159.8866     0   1   2E+4   1   1.5  RT (min)      lcorbi_hilic_neg_4_004_13    m/z 159.8966     0   1   0.5   1  RT (min)      lcorbi_hilic_neg_4_090_60    m/z 159.9191     0   1E+4   1   1.5  RT (min)      lcorbi_hilic_neg_4_004_13    m/z 159.9391     0   1   2   3E+4   2   2.5  RT (min)       lcorbi_hilic_neg_4_113_62    m/z 159.9391     0   1   2E+3   5   5.5  RT (min)      lcorbi_hilic_neg_4_109_62    m/z 159.9837     0   1   2E+3   8   8.5  RT (min)      lcorbi_hilic_neg_4_066_57    m/z 160.0072     0   1   6.5   7  RT (min)      lcorbi_hilic_neg_4_076_58    m/z 160.0348     0   2   4   6E+3   6   6.5  RT (min)      lcorbi_hilic_neg_4_101_61    m/z 160.0657     0   2   4   6E+2   6   6.5  RT (min)      lcorbi_hilic_neg_3_027_45    m/z 160.0695     0   2   4   6   8E+3   1   1.5  RT (min)      lcorbi_hilic_neg_4_084_59    m/z 160.0820     0   1E+4   5.5   6  RT (min)      lcorbi_hilic_neg_4_096_60    m/z 160.0903     0   1   2E+4   4.5   5  RT (min)      lcorbi_hilic_neg_4_093_60    m/z 160.8422     0   1   2E+6   8   10  RT (min)       lcorbi_hilic_neg_4_040_17    m/z 160.8458     0   2   4E+4   10   10.5  RT (min)      lcorbi_hilic_neg_4_079_58    m/z 160.9153     0   1E+4   5   5.5  RT (min)      lcorbi_hilic_neg_4_073_57    m/z 160.9553     0   1   5.5   6  RT (min)      lcorbi_hilic_neg_4_099_61    m/z 160.9565     0   1   0.5   1  RT (min)      lcorbi_hilic_neg_4_085_59    m/z 160.9708     0   1E+3   6.5   7  RT (min)      lcorbi_hilic_neg_1_048_35    m/z 160.9913     0   1   1.5   2  RT (min)       lcorbi_hilic_neg_4_106_62    m/z 160.9985     0   1   2E+4   6.5   7  RT (min)      lcorbi_hilic_neg_4_104_61    m/z 161.0069     0   1   2   3   4E+3   2   2.5  RT (min)      lcorbi_hilic_neg_4_080_58    m/z 161.0090     0   1   6   6.5  RT (min)      lcorbi_hilic_neg_4_110_62    m/z 161.0420     0   2   4E+5   5.5   6  RT (min)      lcorbi_hilic_neg_2_025_52    m/z 161.0567     0   1E+3   5.5   6  RT (min)      lcorbi_hilic_neg_3_029_25    m/z 161.0648     0   2   4E+3   2   2.5  RT (min)      lcorbi_hilic_neg_4_073_57    m/z 161.0818     0   1   2E+4   1   1.5  RT (min)      lcorbi_hilic_neg_4_091_60    m/z 161.8423     0   2   4   6   8E+4   9   10  RT (min)       lcorbi_hilic_neg_4_081_58    m/z 161.8423     0   1   2   3E+4   4.5   5  RT (min)       lcorbi_hilic_neg_4_110_62    m/z 161.8848     0   1E+3   1   1.5  RT (min)      lcorbi_hilic_neg_4_026_33    m/z 161.9356     0   1   2E+4   2   2.5  RT (min)       lcorbi_hilic_neg_4_106_62    m/z 161.9747     0   2   4E+4   1   1.5  RT (min)      lcorbi_hilic_neg_4_100_61    m/z 161.9773     0   1   6.5   7  RT (min)       lcorbi_hilic_neg_4_068_57    m/z 161.9795     0   1E+4   8   8.5  RT (min)      lcorbi_hilic_neg_4_077_58    m/z 162.0014     0   1   2E+4   1   1.5  RT (min)      lcorbi_hilic_neg_4_101_61    m/z 162.0400     0   1   2   3   4E+2   2   2.5  RT (min)      lcorbi_hilic_neg_2_019_21    m/z 162.0418     0   1   5   5.5  RT (min)      lcorbi_hilic_neg_4_079_58    m/z 162.0488     0   2   4E+4   5   5.5  RT (min)      lcorbi_hilic_neg_4_074_58    m/z 162.0503     0   2   4   6   8E+3   2.5   3   3.5  RT (min)       lcorbi_hilic_neg_4_090_60    m/z 162.0656     0   2   4   6   8E+3   6   6.5  RT (min)      lcorbi_hilic_neg_4_083_59    m/z 162.0866     0   2   4E+4   1.5   2  RT (min)       lcorbi_hilic_neg_4_090_60    m/z 162.8396     0   1E+6   8   8.5  RT (min)       lcorbi_hilic_neg_1_003_14    m/z 162.8441     0   1E+4   8.5   9  RT (min)      lcorbi_hilic_neg_1_003_14    m/z 162.8801     0   1   2.5   3  RT (min)      lcorbi_hilic_neg_4_108_62    m/z 162.9027     0   1   2   3E+3   1   1.5  RT (min)      lcorbi_hilic_neg_4_066_57    m/z 162.9124     0   1E+4   5   5.5  RT (min)      lcorbi_hilic_neg_4_070_57    m/z 162.9137     0   1   2   3E+3   1   1.5  RT (min)       lcorbi_hilic_neg_4_105_61    m/z 162.9569     0   1   2   3   4E+3   1.5   2  RT (min)       lcorbi_hilic_neg_1_002_26    m/z 162.9821     0   1   2   3   4E+5   6.5   7  RT (min)      lcorbi_hilic_neg_1_002_26    m/z 162.9822     0   1E+5   12   12.5  RT (min)      lcorbi_hilic_neg_1_011_01    m/z 162.9823     0   1E+3   9   9.5  RT (min)      lcorbi_hilic_neg_2_005_24    m/z 163.0055     0   2   4   6   8E+2   2   2.5  RT (min)      lcorbi_hilic_neg_3_016_16    m/z 163.0248     0   1   2   3   4E+3   6   6.5  RT (min)      lcorbi_hilic_neg_1_001_49    m/z 163.0340     0   1   2E+4   4   4.5  RT (min)      lcorbi_hilic_neg_1_001_49    m/z 163.0362     0   2   4   6E+2   6   6.5  RT (min)      lcorbi_hilic_neg_1_001_49    m/z 163.0875     0   2   4   6   8E+3   3.5   4  RT (min)      lcorbi_hilic_neg_4_095_60    m/z 163.0945     0   1   2E+3   1.5   2  RT (min)       lcorbi_hilic_neg_1_022_48    m/z 163.1127     0   1E+5   0.5   1  RT (min)       lcorbi_hilic_neg_4_066_57    m/z 163.1239     0   1   2E+4   1   1.5  RT (min)       lcorbi_hilic_neg_4_089_59    m/z 163.8395     0   1   2E+4   4.5   5  RT (min)       lcorbi_hilic_neg_4_110_62    m/z 163.8448     0   1   2   3E+4   8   8.5  RT (min)      lcorbi_hilic_neg_4_112_62    m/z 163.8877     0   1   2   3E+3   7.5   8  RT (min)      lcorbi_hilic_neg_4_074_58    m/z 163.9933     0   1   6   6.5  RT (min)       lcorbi_hilic_neg_4_081_58    m/z 164.0576     0   2   4   6   8E+3   3.5   4  RT (min)      lcorbi_hilic_neg_4_077_58    m/z 164.0740     0   1E+5   0.5   1  RT (min)       lcorbi_hilic_neg_2_055_32    m/z 164.0841     0   1E+4   0.5   1  RT (min)      lcorbi_hilic_neg_4_084_59    m/z 164.8780     0   1E+4   2.5   3  RT (min)      lcorbi_hilic_neg_4_071_57    m/z 164.9378     0   1   2E+4   1   1.5  RT (min)       lcorbi_hilic_neg_4_097_60    m/z 164.9441     0   1   1   1.5  RT (min)      lcorbi_hilic_neg_2_003_20    m/z 164.9473     0   1E+4   4   4.5  RT (min)       lcorbi_hilic_neg_4_088_59    m/z 164.9475     0   2   4E+4   8   8.5  RT (min)       lcorbi_hilic_neg_4_076_58    m/z 165.0193     0   1   2   3E+4   3.5   4  RT (min)       lcorbi_hilic_neg_4_076_58    m/z 165.0250     0   1E+4   1   1.5  RT (min)      lcorbi_hilic_neg_3_009_50    m/z 165.0403     0   2   4   6E+3   5   5.5  RT (min)      lcorbi_hilic_neg_1_005_36    m/z 165.0495     0   1   2E+5   4   4.5  RT (min)      lcorbi_hilic_neg_1_012_32    m/z 165.0707     0   1   2E+3   1   1.5  RT (min)      lcorbi_hilic_neg_1_003_14    m/z 165.0749     0   2   4   6E+3   5   5.5  RT (min)      lcorbi_hilic_neg_4_076_58    m/z 165.0767     0   1E+4   3.5   4  RT (min)      lcorbi_hilic_neg_4_077_58    m/z 165.0919     0   2   4   6E+3   1   1.5  RT (min)      lcorbi_hilic_neg_1_002_26    m/z 165.1055     0   1   1   1.5  RT (min)      lcorbi_hilic_neg_1_004_17    m/z 165.8418     0   1   2   3E+4   8   8.5  RT (min)      lcorbi_hilic_neg_4_101_61    m/z 165.8943     0   1   2E+4   5.5   6   6.5  RT (min)       lcorbi_hilic_neg_4_076_58    m/z 165.8943     0   1   2E+4   8   8.5  RT (min)       lcorbi_hilic_neg_4_031_14    m/z 165.9028     0   1E+4   1   1.5  RT (min)      lcorbi_hilic_neg_3_044_32    m/z 165.9356     0   1   2   3E+3   8   8.5  RT (min)      lcorbi_hilic_neg_3_002_15    m/z 165.9601     0   1   6   6.5  RT (min)       lcorbi_hilic_neg_1_008_39    m/z 165.9789     0   1   2E+3   0.5   1  RT (min)      lcorbi_hilic_neg_1_001_49    m/z 165.9967     0   1   1   1.5  RT (min)      lcorbi_hilic_neg_3_002_15    m/z 166.0091     0   1   2   3   4E+2   3   3.5  RT (min)       lcorbi_hilic_neg_1_008_39    m/z 166.0141     0   2   4E+4   4   4.5  RT (min)      lcorbi_hilic_neg_1_022_48    m/z 166.0274     0   1E+3   6   6.5  RT (min)       lcorbi_hilic_neg_1_006_08    m/z 166.0403     0   1E+3   0.5   1  RT (min)      lcorbi_hilic_neg_1_003_14    m/z 166.0622     0   2   4   6E+3   1.5   2  RT (min)      lcorbi_hilic_neg_4_015_24    m/z 166.8332     0   1   2   3   4E+2   10.5   11  RT (min)       lcorbi_hilic_neg_4_005_32    m/z 166.8345     0   1   2   3E+4   4.5   5  RT (min)      lcorbi_hilic_neg_4_066_57    m/z 166.8604     0   1E+4   0   0.5  RT (min)      lcorbi_hilic_neg_4_082_59    m/z 166.8605     0   1E+4   4.5   5  RT (min)      lcorbi_hilic_neg_4_099_61    m/z 166.9155     0   2   4E+4   0   0.5  RT (min)      lcorbi_hilic_neg_4_079_58    m/z 166.9243     0   1   2   3E+4   8.5   9  RT (min)       lcorbi_hilic_neg_4_080_58    m/z 166.9323     0   1E+4   8   8.5  RT (min)      lcorbi_hilic_neg_4_074_58    m/z 166.9751     0   1E+4   6   6.5  RT (min)      lcorbi_hilic_neg_4_089_59    m/z 166.9924     0   1E+4   3   3.5  RT (min)      lcorbi_hilic_neg_4_105_61    m/z 166.9924     0   2   4   6   8E+3   1   1.5  RT (min)       lcorbi_hilic_neg_4_014_18    m/z 167.0008     0   1   2   3E+2   6.5   7  RT (min)      lcorbi_hilic_neg_1_023_23    m/z 167.0115     0   1   2   3E+3   1   1.5  RT (min)       lcorbi_hilic_neg_2_013_18    m/z 167.0127     0   1   2E+4   5.5   6  RT (min)      lcorbi_hilic_neg_4_079_58    m/z 167.0296     0   1   2   3   4E+3   5.5   6  RT (min)      lcorbi_hilic_neg_2_016_30    m/z 167.0407     0   2   4   6E+3   1   1.5  RT (min)       lcorbi_hilic_neg_2_049_55    m/z 167.0461     0   1E+3   4.5   5  RT (min)      lcorbi_hilic_neg_4_098_61    m/z 167.0461     0   1E+3   3   3.5  RT (min)       lcorbi_hilic_neg_4_103_61    m/z 167.1440     0   1E+4   0.5   1  RT (min)      lcorbi_hilic_neg_4_106_62    m/z 167.8367     0   1   2E+5   12.5   13  RT (min)      lcorbi_hilic_neg_4_097_60    m/z 167.8396     0   1   2E+4   8.5   9  RT (min)      lcorbi_hilic_neg_4_102_61    m/z 167.8407     0   1E+4   4.5   5  RT (min)      lcorbi_hilic_neg_1_008_39    m/z 167.9970     0   2   4E+3   6.5   7  RT (min)      lcorbi_hilic_neg_4_087_59    m/z 168.0068     0   1E+4   7   7.5  RT (min)      lcorbi_hilic_neg_4_085_59    m/z 168.0526     0   1   2   2.5  RT (min)      lcorbi_hilic_neg_4_071_57    m/z 168.0777     0   1   2E+4   2.5   3  RT (min)       lcorbi_hilic_neg_4_071_57    m/z 168.8362     0   1E+4   4.5   5  RT (min)      lcorbi_hilic_neg_4_096_60    m/z 168.8575     0   1E+4   0   0.5  RT (min)      lcorbi_hilic_neg_4_066_57    m/z 168.8577     0   1E+4   4.5   5  RT (min)      lcorbi_hilic_neg_4_104_61    m/z 168.9437     0   1   2E+4   6   6.5  RT (min)       lcorbi_hilic_neg_4_097_60    m/z 168.9893     0   1E+4   3   3.5  RT (min)      lcorbi_hilic_neg_4_095_60    m/z 168.9993     0   1E+3   1.5   2  RT (min)       lcorbi_hilic_neg_4_071_57    m/z 169.0175     0   2   4   6E+4   1   1.5  RT (min)       lcorbi_hilic_neg_4_014_18    m/z 169.0250     0   1E+3   6  RT (min)       lcorbi_hilic_neg_1_007_10    m/z 169.0253     0   1E+3   1.5   2  RT (min)       lcorbi_hilic_neg_4_028_03    m/z 169.0311     0   1   6   6.5  RT (min)      lcorbi_hilic_neg_1_008_39    m/z 169.0557     0   1   2   3E+5   5.5   6   6.5  RT (min)       lcorbi_hilic_neg_4_094_60    m/z 169.0617     0   1E+4   1   1.5  RT (min)       lcorbi_hilic_neg_4_074_58    m/z 169.0698     0   1   2E+3   1.5   2  RT (min)      lcorbi_hilic_neg_3_011_24    m/z 169.0811     0   1   2   3E+3   7   7.5  RT (min)      lcorbi_hilic_neg_4_074_58    m/z 169.0869     0   1   2   3E+5   1   1.5  RT (min)       lcorbi_hilic_neg_3_028_23    m/z 169.0980     0   1   1   1.5  RT (min)      lcorbi_hilic_neg_2_026_23    m/z 169.8342     0   1   2E+5   12.5   13  RT (min)      lcorbi_hilic_neg_4_030_05    m/z 169.8388     0   1E+4   4.5   5  RT (min)      lcorbi_hilic_neg_1_008_39    m/z 169.8494     0   1   2   3   4E+2   3   3.5  RT (min)      lcorbi_hilic_neg_1_008_39    m/z 169.8739     0   2   4E+4   8.5   9  RT (min)       lcorbi_hilic_neg_4_077_58    m/z 169.9085     0   1   2   3   4E+3   8   8.5  RT (min)      lcorbi_hilic_neg_4_066_57    m/z 169.9286     0   1E+4   0   0.5  RT (min)       lcorbi_hilic_neg_4_081_58    m/z 169.9735     0   1   2   3E+4   6   6.5  RT (min)       lcorbi_hilic_neg_4_094_60    m/z 169.9846     0   1   2E+4   1.5   2  RT (min)       lcorbi_hilic_neg_4_069_57    m/z 169.9851     0   2   4E+4   6   6.5  RT (min)       lcorbi_hilic_neg_4_096_60    m/z 170.0256     0   1   2E+4   3.5   4   4.5  RT (min)       lcorbi_hilic_neg_4_083_59    m/z 170.0433     0   1   5.5   6  RT (min)      lcorbi_hilic_neg_4_095_60    m/z 170.0457     0   1E+4   1   1.5  RT (min)       lcorbi_hilic_neg_4_084_59    m/z 170.0822     0   1   2E+4   1   1.5  RT (min)       lcorbi_hilic_neg_4_088_59    m/z 170.8331     0   1   2E+4   4.5   5  RT (min)      lcorbi_hilic_neg_4_100_61    m/z 170.8480     0   2   4   6   8E+3   4.5   5  RT (min)      lcorbi_hilic_neg_4_104_61    m/z 170.8708     0   2   4   6   8E+4   10   10.5   11  RT (min)       lcorbi_hilic_neg_4_106_62    m/z 170.8925     0   2   4E+3   0   0.5  RT (min)      lcorbi_hilic_neg_1_006_08    m/z 170.8926     0   1   2   3   4E+3   4.5   5  RT (min)      lcorbi_hilic_neg_4_098_61    m/z 170.9313     0   1   2   3E+4   1   1.5  RT (min)       lcorbi_hilic_neg_4_066_57    m/z 170.9441     0   1   2E+4   0   0.5  RT (min)      lcorbi_hilic_neg_1_004_17    m/z 170.9628     0   1   6   6.5  RT (min)      lcorbi_hilic_neg_4_099_61    m/z 170.9769     0   1   2   3E+4   1.5   2  RT (min)       lcorbi_hilic_neg_4_069_57    m/z 170.9874     0   1E+4   1   1.5  RT (min)       lcorbi_hilic_neg_4_076_58    m/z 170.9910     0   2   4E+3   6   6.5  RT (min)      lcorbi_hilic_neg_1_047_05    m/z 170.9911     0   2   4   6   8E+3   7.5   8  RT (min)      lcorbi_hilic_neg_4_087_59    m/z 171.0064     0   1   2E+5   6   6.5  RT (min)      lcorbi_hilic_neg_1_047_05    m/z 171.0077     0   1   2   3E+4   3   3.5  RT (min)       lcorbi_hilic_neg_4_081_58    m/z 171.0258     0   1   2   3E+4   3   3.5  RT (min)       lcorbi_hilic_neg_4_095_60    m/z 171.0274     0   1   2   3E+3   1   1.5  RT (min)       lcorbi_hilic_neg_4_088_59    m/z 171.0511     0   1E+4   6   6.5  RT (min)      lcorbi_hilic_neg_4_091_60    m/z 171.0773     0   1   2   3   4E+3   5.5   6  RT (min)      lcorbi_hilic_neg_4_083_59    m/z 171.0814     0   1E+4   0.5   1  RT (min)      lcorbi_hilic_neg_4_081_58    m/z 171.1508     0   1   2   3   4E+4   1   1.5  RT (min)       lcorbi_hilic_neg_4_090_60    m/z 171.8315     0   1E+5   13   13.5  RT (min)      lcorbi_hilic_neg_4_108_62    m/z 171.8355     0   1   2   3   4E+3   2   2.5  RT (min)      lcorbi_hilic_neg_4_089_59    m/z 171.8355     0   1   2E+4   8.5   9  RT (min)       lcorbi_hilic_neg_1_043_40    m/z 171.8458     0   2   4   6E+2   4.5   5  RT (min)      lcorbi_hilic_neg_1_008_39    m/z 171.8710     0   1   2   3E+4   8.5   9  RT (min)       lcorbi_hilic_neg_4_077_58    m/z 171.9469     0   2   4   6E+4   2.5   3   3.5  RT (min)       lcorbi_hilic_neg_4_081_58    m/z 171.9809     0   1   2E+3   6.5   7  RT (min)       lcorbi_hilic_neg_4_073_57    m/z 172.0181     0   1E+4   1.5   2  RT (min)      lcorbi_hilic_neg_2_055_32    m/z 172.0208     0   1E+4   3   3.5  RT (min)      lcorbi_hilic_neg_4_084_59    m/z 172.0299     0   2   4   6E+4   1.5   2   2.5  RT (min)       lcorbi_hilic_neg_4_090_60    m/z 172.0423     0   1   2   3   4E+4   3   3.5  RT (min)       lcorbi_hilic_neg_4_084_59    m/z 172.0468     0   2   4   6E+3   5.5   6  RT (min)      lcorbi_hilic_neg_4_069_57    m/z 172.0556     0   1E+4   3.5   4  RT (min)      lcorbi_hilic_neg_4_073_57    m/z 172.0766     0   1   2   3   4E+3   4   4.5  RT (min)      lcorbi_hilic_neg_4_104_61    m/z 172.0808     0   1E+4   1.5   2  RT (min)       lcorbi_hilic_neg_4_071_57    m/z 172.0978     0   1   2   3   4E+3   1   1.5  RT (min)       lcorbi_hilic_neg_4_091_60    m/z 172.8424     0   1   2   3E+5   4   4.5  RT (min)       lcorbi_hilic_neg_4_066_57    m/z 172.8678     0   2   4E+4   9.5   10  RT (min)       lcorbi_hilic_neg_4_066_57    m/z 172.8857     0   1E+4   8   8.5  RT (min)      lcorbi_hilic_neg_4_069_57    m/z 172.9410     0   2   4E+3   0.5   1  RT (min)      lcorbi_hilic_neg_1_003_14    m/z 172.9411     0   1E+4   5.5   6  RT (min)      lcorbi_hilic_neg_1_007_10    m/z 172.9577     0   2   4   6E+3   0.5   1  RT (min)      lcorbi_hilic_neg_4_081_58    m/z 172.9618     0   1   2   3E+3   6   6.5  RT (min)      lcorbi_hilic_neg_1_005_36    m/z 172.9622     0   2   4   6   8E+2   2   2.5  RT (min)      lcorbi_hilic_neg_1_003_14    m/z 172.9856     0   2   4E+2   4.5   5  RT (min)      lcorbi_hilic_neg_1_007_10    m/z 172.9872     0   1E+4   1   1.5  RT (min)       lcorbi_hilic_neg_4_038_27    m/z 172.9999     0   1   2E+3   1.5   2  RT (min)      lcorbi_hilic_neg_4_023_34    m/z 173.0178     0   1   2   3   4E+2   6   6.5  RT (min)      lcorbi_hilic_neg_1_012_32    m/z 173.0294     0   1   6.5   7  RT (min)       lcorbi_hilic_neg_1_006_08    m/z 173.0388     0   1   1   1.5  RT (min)       lcorbi_hilic_neg_4_084_59    m/z 173.0566     0   2   4   6   8E+3   4   4.5  RT (min)      lcorbi_hilic_neg_4_078_58    m/z 173.0628     0   2   4E+3   2   2.5  RT (min)      lcorbi_hilic_neg_4_087_59    m/z 173.1082     0   1E+4   4   4.5  RT (min)      lcorbi_hilic_neg_4_101_61    m/z 173.1107     0   1   2E+3   7   7.5  RT (min)      lcorbi_hilic_neg_3_029_25    m/z 173.1181     0   1   2   3   4E+3   1   1.5  RT (min)      lcorbi_hilic_neg_4_037_31    m/z 173.1575     0   1E+4   1   1.5  RT (min)      lcorbi_hilic_neg_4_090_60    m/z 173.4167     0   1   2E+4   5   5.5  RT (min)      lcorbi_hilic_neg_4_091_60    m/z 173.8426     0   1   2E+3   4.5   5  RT (min)       lcorbi_hilic_neg_1_034_43    m/z 173.9254     0   1E+4   4   4.5  RT (min)      lcorbi_hilic_neg_4_096_60    m/z 173.9260     0   1   2   3   4E+4   1   2  RT (min)       lcorbi_hilic_neg_4_087_59    m/z 173.9261     0   1   2   3E+4   6   6.5  RT (min)       lcorbi_hilic_neg_4_076_58    m/z 173.9261     0   1E+5   8   8.5   9  RT (min)       lcorbi_hilic_neg_4_066_57    m/z 173.9490     0   2   4   6E+3   2   2.5  RT (min)      lcorbi_hilic_neg_4_089_59    m/z 173.9500     0   1E+3   6   6.5  RT (min)       lcorbi_hilic_neg_4_025_52    m/z 173.9625     0   1   2E+4   2   2.5  RT (min)       lcorbi_hilic_neg_4_055_38    m/z 173.9710     0   1   5.5   6  RT (min)      lcorbi_hilic_neg_4_085_59    m/z 173.9865     0   1   2   3   4E+4   1   1.5  RT (min)      lcorbi_hilic_neg_2_020_27    m/z 173.9940     0   1   1.5   2  RT (min)      lcorbi_hilic_neg_4_108_62    m/z 174.0069     0   2   4   6E+2   3.5   4  RT (min)      lcorbi_hilic_neg_4_014_18    m/z 174.0164     0   1   2E+5   6.5   7  RT (min)       lcorbi_hilic_neg_1_047_05    m/z 174.0263     0   1E+3   1   2  RT (min)       lcorbi_hilic_neg_4_082_59    m/z 174.0302     0   1   5   5.5  RT (min)      lcorbi_hilic_neg_1_028_45    m/z 174.0421     0   2   4   6E+2   1.5   2  RT (min)       lcorbi_hilic_neg_4_031_14    m/z 174.0558     0   1E+4   1   1.5  RT (min)       lcorbi_hilic_neg_2_012_14    m/z 174.0621     0   1   2   3E+4   3.5   4  RT (min)       lcorbi_hilic_neg_4_071_57    m/z 174.0795     0   1E+4   5.5   6  RT (min)       lcorbi_hilic_neg_4_089_59    m/z 174.0851     0   1E+4   1   1.5  RT (min)      lcorbi_hilic_neg_4_091_60    m/z 174.8357     0   1   2E+3   4.5   5  RT (min)      lcorbi_hilic_neg_4_110_62    m/z 174.8394     0   1   2E+4   0.5   1  RT (min)      lcorbi_hilic_neg_4_066_57    m/z 174.8395     0   1E+5   4   4.5  RT (min)       lcorbi_hilic_neg_4_098_61    m/z 174.8678     0   1   2E+4   8   8.5  RT (min)       lcorbi_hilic_neg_4_079_58    m/z 174.8869     0   1E+4   0   0.5  RT (min)      lcorbi_hilic_neg_4_098_61    m/z 174.8870     0   1   2E+4   6.5   7  RT (min)      lcorbi_hilic_neg_4_106_62    m/z 174.8870     0   1   2E+4   4.5   5  RT (min)       lcorbi_hilic_neg_4_112_62    m/z 174.9374     0   1E+4   4.5   5  RT (min)      lcorbi_hilic_neg_4_075_58    m/z 174.9468     0   2   4E+2   7   7.5  RT (min)      lcorbi_hilic_neg_1_022_48    m/z 174.9469     0   1   2   3   4E+3   4.5   5  RT (min)      lcorbi_hilic_neg_4_100_61    m/z 174.9539     0   2   4   6E+3   4.5   5  RT (min)      lcorbi_hilic_neg_1_008_39    m/z 174.9560     0   1E+5   7.5   8  RT (min)      lcorbi_hilic_neg_4_077_58    m/z 174.9562     0   1E+6   11.5   12  RT (min)      lcorbi_hilic_neg_4_108_62    m/z 174.9609     0   2   4   6   8E+4   8.5   9  RT (min)      lcorbi_hilic_neg_4_080_58    m/z 174.9864     0   1E+3   6   6.5  RT (min)      lcorbi_hilic_neg_4_101_61    m/z 174.9871     0   1   0.5   1  RT (min)      lcorbi_hilic_neg_2_015_19    m/z 175.0011     0   1E+4   1   1.5  RT (min)      lcorbi_hilic_neg_4_098_61    m/z 175.0132     0   1   2   3   4E+3   1.5   2  RT (min)      lcorbi_hilic_neg_4_101_61    m/z 175.0430     0   2   4E+3   6   6.5  RT (min)      lcorbi_hilic_neg_4_109_62    m/z 175.0430     0   1E+4   0.5   1  RT (min)       lcorbi_hilic_neg_1_046_19    m/z 175.0611     0   1   2E+5   6.5   7  RT (min)       lcorbi_hilic_neg_3_034_34    m/z 175.0615     0   1   2   3E+5   2   2.5  RT (min)      lcorbi_hilic_neg_1_005_36    m/z 175.0723     0   1E+3   6.5   7  RT (min)       lcorbi_hilic_neg_1_023_23    m/z 175.0803     0   2   4   6E+2   1.5   2  RT (min)      lcorbi_hilic_neg_1_001_49    m/z 175.0879     0   1   6.5   7  RT (min)      lcorbi_hilic_neg_4_087_59    m/z 175.1085     0   1E+4   7   7.5  RT (min)      lcorbi_hilic_neg_4_085_59    m/z 175.8259     0   1E+4   4.5   5  RT (min)      lcorbi_hilic_neg_4_111_62    m/z 175.8259     0   2   4   6E+3   0.5   1  RT (min)      lcorbi_hilic_neg_4_083_59    m/z 175.8281     0   1   2   3E+4   8.5   9  RT (min)       lcorbi_hilic_neg_4_091_60    m/z 175.8397     0   1   2E+4   4.5   5  RT (min)       lcorbi_hilic_neg_4_099_61    m/z 175.8716     0   1E+4   3   3.5  RT (min)       lcorbi_hilic_neg_4_069_57    m/z 175.9085     0   1   2E+4   6.5   7   7.5  RT (min)       lcorbi_hilic_neg_3_002_15    m/z 175.9086     0   1   2E+4   4.5   5  RT (min)       lcorbi_hilic_neg_4_104_61    m/z 175.9369     0   1   2E+3   7.5   8  RT (min)      lcorbi_hilic_neg_4_012_20    m/z 175.9684     0   2   4E+3   8   8.5  RT (min)      lcorbi_hilic_neg_4_089_59    m/z 175.9811     0   1   2   3E+3   1.5   2  RT (min)      lcorbi_hilic_neg_4_073_57    m/z 175.9909     0   1   1.5   2  RT (min)      lcorbi_hilic_neg_4_108_62    m/z 176.0137     0   1   2   3   4E+3   6   6.5  RT (min)      lcorbi_hilic_neg_4_081_58    m/z 176.0224     0   2   4   6E+4   1.5   2  RT (min)       lcorbi_hilic_neg_4_100_61    m/z 176.0387     0   1E+3   1   1.5  RT (min)       lcorbi_hilic_neg_4_107_62    m/z 176.0427     0   1   2E+4   3.5   4  RT (min)       lcorbi_hilic_neg_4_067_57    m/z 176.0595     0   1   2E+3   2.5   3  RT (min)      lcorbi_hilic_neg_4_090_60    m/z 176.0812     0   1E+4   5.5   6  RT (min)      lcorbi_hilic_neg_4_089_59    m/z 176.0841     0   2   4   6E+3   0.5   1  RT (min)      lcorbi_hilic_neg_4_095_60    m/z 176.0925     0   2   4   6   8E+3   6.5   7  RT (min)      lcorbi_hilic_neg_4_089_59    m/z 176.8172     0   2   4   6E+3   0.5   1  RT (min)      lcorbi_hilic_neg_4_107_62    m/z 176.8216     0   2   4   6E+3   2.5   3  RT (min)      lcorbi_hilic_neg_4_068_57    m/z 176.8365     0   1E+4   8   8.5  RT (min)      lcorbi_hilic_neg_4_066_57    m/z 176.8496     0   1E+3   3   3.5  RT (min)      lcorbi_hilic_neg_3_050_03    m/z 176.8748     0   1E+5   0.5   1  RT (min)       lcorbi_hilic_neg_4_055_38    m/z 176.8842     0   2   4E+3   0.5   1  RT (min)      lcorbi_hilic_neg_4_082_59    m/z 176.8843     0   2   4   6E+3   4   4.5  RT (min)      lcorbi_hilic_neg_4_098_61    m/z 176.8953     0   1   2E+4   0.5   1  RT (min)      lcorbi_hilic_neg_4_074_58    m/z 176.9169     0   1E+4   7.5   8  RT (min)      lcorbi_hilic_neg_4_090_60    m/z 176.9273     0   1   2   3E+3   7   7.5  RT (min)      lcorbi_hilic_neg_4_015_24    m/z 176.9293     0   1   2   3E+3   5   5.5  RT (min)      lcorbi_hilic_neg_4_098_61    m/z 176.9337     0   2   4   6   8E+4   7   7.5  RT (min)       lcorbi_hilic_neg_1_047_05    m/z 176.9401     0   2   4E+3   7   7.5  RT (min)      lcorbi_hilic_neg_2_005_24    m/z 176.9513     0   1   0.5   1  RT (min)      lcorbi_hilic_neg_4_107_62    m/z 176.9519     0   1   2   3   4E+4   8.5   9  RT (min)       lcorbi_hilic_neg_4_077_58    m/z 176.9611     0   1E+4   2   2.5  RT (min)      lcorbi_hilic_neg_4_083_59    m/z 176.9979     0   2   4E+4   3   3.5  RT (min)       lcorbi_hilic_neg_4_111_62    m/z 177.0127     0   1E+3   6   6.5  RT (min)      lcorbi_hilic_neg_4_095_60    m/z 177.0250     0   1E+3   5.5   6  RT (min)      lcorbi_hilic_neg_4_078_58    m/z 177.0288     0   1E+3   1.5   2  RT (min)      lcorbi_hilic_neg_4_084_59    m/z 177.0338     0   1   6.5   7  RT (min)      lcorbi_hilic_neg_4_090_60    m/z 177.0403     0   2   4   6E+4   2   2.5   3  RT (min)       lcorbi_hilic_neg_4_075_58    m/z 177.0599     0   2   4   6E+3   1   1.5  RT (min)      lcorbi_hilic_neg_4_001_49    m/z 177.0628     0   2   4   6E+3   6   6.5  RT (min)      lcorbi_hilic_neg_4_089_59    m/z 177.0692     0   2   4E+4   3   3.5  RT (min)       lcorbi_hilic_neg_4_093_60    m/z 177.0692     0   1   1   1.5  RT (min)       lcorbi_hilic_neg_4_011_19    m/z 177.1179     0   1E+4   7   7.5  RT (min)      lcorbi_hilic_neg_4_090_60    m/z 177.8368     0   2   4   6   8E+3   4.5   5  RT (min)      lcorbi_hilic_neg_4_110_62    m/z 177.8631     0   1   2   3E+3   8   8.5  RT (min)       lcorbi_hilic_neg_2_002_28    m/z 177.8882     0   1E+3   7.5   8  RT (min)      lcorbi_hilic_neg_1_015_30    m/z 177.9129     0   1   2E+3   0   0.5  RT (min)      lcorbi_hilic_neg_1_011_01    m/z 177.9400     0   1   7   7.5  RT (min)      lcorbi_hilic_neg_3_002_15    m/z 177.9709     0   2   4E+3   5.5   6  RT (min)      lcorbi_hilic_neg_3_048_33    m/z 177.9879     0   1   2   3   4E+4   1.5   2  RT (min)      lcorbi_hilic_neg_4_047_07    m/z 177.9968     0   1   2   3E+3   2   2.5  RT (min)      lcorbi_hilic_neg_4_107_62    m/z 178.0182     0   1E+4   2   2.5  RT (min)      lcorbi_hilic_neg_4_084_59    m/z 178.0258     0   1E+3   2   3  RT (min)       lcorbi_hilic_neg_4_067_57    m/z 178.0376     0   1   2E+4   2   2.5  RT (min)       lcorbi_hilic_neg_4_075_58    m/z 178.0397     0   1E+3   3.5   4  RT (min)      lcorbi_hilic_neg_4_089_59    m/z 178.0660     0   2   4   6E+3   3.5   4  RT (min)      lcorbi_hilic_neg_4_099_61    m/z 178.0953     0   2   4   6   8E+3   0.5   1  RT (min)      lcorbi_hilic_neg_4_095_60    m/z 178.6273     0   1   2   3E+3   3   3.5  RT (min)      lcorbi_hilic_neg_4_089_59    m/z 178.8151     0   1E+4   0.5   1  RT (min)      lcorbi_hilic_neg_4_104_61    m/z 178.8151     0   1   2E+4   13.5   14  RT (min)      lcorbi_hilic_neg_4_085_59    m/z 178.8196     0   1E+4   3   3.5  RT (min)      lcorbi_hilic_neg_4_069_57    m/z 178.8338     0   2   4   6E+3   4.5   5  RT (min)      lcorbi_hilic_neg_4_101_61    m/z 178.8485     0   1   2   3E+3   3   3.5  RT (min)      lcorbi_hilic_neg_1_001_49    m/z 178.8722     0   1   2   3   4E+3   0.5   1  RT (min)      lcorbi_hilic_neg_2_048_38    m/z 178.8847     0   2   4E+3   0.5   1  RT (min)      lcorbi_hilic_neg_4_037_31    m/z 178.9030     0   1   2   3E+4   7.5   8  RT (min)      lcorbi_hilic_neg_4_029_02    m/z 178.9484     0   1   0.5   1  RT (min)      lcorbi_hilic_neg_4_098_61    m/z 178.9836     0   1E+4   5.5   6  RT (min)       lcorbi_hilic_neg_4_083_59    m/z 178.9926     0   1   2   3E+4   3   3.5  RT (min)       lcorbi_hilic_neg_4_086_59    m/z 178.9927     0   1   2   3   4E+4   1   1.5  RT (min)       lcorbi_hilic_neg_4_107_62    m/z 179.0078     0   1   2E+3   3.5   4  RT (min)      lcorbi_hilic_neg_4_091_60    m/z 179.0146     0   1E+3   2   2.5  RT (min)       lcorbi_hilic_neg_3_011_24    m/z 179.0210     0   2   4   6E+2   4   4.5  RT (min)      lcorbi_hilic_neg_1_047_05    m/z 179.0517     0   1   2   3E+4   5   5.5  RT (min)      lcorbi_hilic_neg_2_020_27    m/z 179.0655     0   2   4   6   8E+3   5   5.5  RT (min)      lcorbi_hilic_neg_3_002_15    m/z 179.0824     0   1   6   6.5  RT (min)      lcorbi_hilic_neg_4_108_62    m/z 179.0826     0   1   2E+3   1.5   2  RT (min)       lcorbi_hilic_neg_1_012_32    m/z 179.1152     0   2   4E+3   7   7.5  RT (min)      lcorbi_hilic_neg_3_042_05    m/z 179.1188     0   1   2.5   3  RT (min)      lcorbi_hilic_neg_4_014_18    m/z 179.1319     0   2   4   6E+4   7   7.5  RT (min)      lcorbi_hilic_neg_4_097_60    m/z 179.1441     0   1   2   3E+4   1   1.5  RT (min)       lcorbi_hilic_neg_4_071_57    m/z 179.8598     0   1   2E+4   4.5   5  RT (min)       lcorbi_hilic_neg_4_070_57    m/z 179.8675     0   1   2E+4   2.5   3  RT (min)       lcorbi_hilic_neg_4_069_57    m/z 179.8773     0   1   0.5   1  RT (min)      lcorbi_hilic_neg_4_102_61    m/z 179.8825     0   1   2E+4   0.5   1  RT (min)      lcorbi_hilic_neg_4_055_38    m/z 179.8919     0   1   7.5   8  RT (min)       lcorbi_hilic_neg_3_022_06    m/z 179.9005     0   1   2E+4   6   6.5  RT (min)      lcorbi_hilic_neg_4_075_58    m/z 179.9006     0   1E+4   4.5   5  RT (min)      lcorbi_hilic_neg_4_069_57    m/z 179.9024     0   2   4   6   8E+3   0   0.5  RT (min)      lcorbi_hilic_neg_4_075_58    m/z 179.9093     0   1E+4   2   2.5  RT (min)      lcorbi_hilic_neg_4_084_59    m/z 179.9850     0   1   1.5   2  RT (min)      lcorbi_hilic_neg_1_045_42    m/z 180.0229     0   1   6   6.5  RT (min)       lcorbi_hilic_neg_1_052_41    m/z 180.0289     0   2   4   6   8E+2   3   3.5  RT (min)       lcorbi_hilic_neg_1_043_40    m/z 180.0376     0   1E+5   5.5   6  RT (min)      lcorbi_hilic_neg_2_005_24    m/z 180.0414     0   2   4   6E+4   1   1.5  RT (min)      lcorbi_hilic_neg_2_009_50    m/z 180.0525     0   1   2   3   4E+2   4   4.5  RT (min)      lcorbi_hilic_neg_3_035_48    m/z 180.0549     0   1   1   1.5  RT (min)      lcorbi_hilic_neg_1_047_05    m/z 180.0664     0   1   2   3   4E+4   1   1.5  RT (min)       lcorbi_hilic_neg_4_089_59    m/z 180.0697     0   2   4   6E+4   5.5   6  RT (min)       lcorbi_hilic_neg_4_095_60    m/z 180.0745     0   1E+3   0.5   1  RT (min)       lcorbi_hilic_neg_4_095_60    m/z 180.1001     0   1   2E+3   6.5   7  RT (min)      lcorbi_hilic_neg_1_042_25    m/z 180.1247     0   1   2E+4   1   1.5  RT (min)       lcorbi_hilic_neg_4_091_60    m/z 180.1304     0   1   7   7.5  RT (min)      lcorbi_hilic_neg_4_005_32    m/z 180.8174     0   1   2E+3   2.5   3  RT (min)      lcorbi_hilic_neg_4_004_13    m/z 180.8468     0   1E+4   3   3.5  RT (min)      lcorbi_hilic_neg_4_070_57    m/z 180.8995     0   1   2   3   4E+4   0   0.5  RT (min)       lcorbi_hilic_neg_4_099_61    m/z 180.8999     0   1E+4   7   7.5  RT (min)      lcorbi_hilic_neg_4_087_59    m/z 180.9124     0   2   4E+4   8.5   9  RT (min)       lcorbi_hilic_neg_4_080_58    m/z 180.9213     0   2   4   6   8E+2   4.5   5  RT (min)      lcorbi_hilic_neg_1_031_20    m/z 180.9310     0   1   2E+4   7   7.5  RT (min)       lcorbi_hilic_neg_4_088_59    m/z 180.9388     0   2   4   6E+3   2   2.5  RT (min)      lcorbi_hilic_neg_4_096_60    m/z 180.9653     0   2   4   6   8E+3   5   5.5  RT (min)      lcorbi_hilic_neg_3_002_15    m/z 180.9893     0   1E+4   3   3.5  RT (min)      lcorbi_hilic_neg_1_003_14    m/z 180.9920     0   1E+3   6   6.5   7  RT (min)      lcorbi_hilic_neg_2_055_32    m/z 181.0003     0   1E+3   2   2.5  RT (min)      lcorbi_hilic_neg_1_028_45    m/z 181.0093     0   2   4   6   8E+3   1   1.5  RT (min)       lcorbi_hilic_neg_4_070_57    m/z 181.0332     0   1E+4   1.5   2  RT (min)      lcorbi_hilic_neg_4_093_60    m/z 181.0448     0   1   2   3E+3   6.5   7  RT (min)       lcorbi_hilic_neg_4_005_32    m/z 181.0617     0   1E+3   1.5   2  RT (min)       lcorbi_hilic_neg_4_096_60    m/z 181.0749     0   1   2   3E+2   7   7.5  RT (min)      lcorbi_hilic_neg_3_011_24    m/z 181.0967     0   1E+3   5.5   6  RT (min)      lcorbi_hilic_neg_4_090_60    m/z 181.0982     0   1E+4   4   4.5  RT (min)      lcorbi_hilic_neg_4_085_59    m/z 181.1005     0   1   1.5   2  RT (min)      lcorbi_hilic_neg_4_079_58    m/z 181.1014     0   1   6.5   7  RT (min)       lcorbi_hilic_neg_4_097_60    m/z 181.1285     0   1   2   3E+4   7   7.5  RT (min)       lcorbi_hilic_neg_4_097_60    m/z 181.8572     0   2   4   6   8E+3   4.5   5  RT (min)      lcorbi_hilic_neg_4_069_57    m/z 181.8682     0   1E+4   2.5   3  RT (min)      lcorbi_hilic_neg_4_080_58    m/z 181.8784     0   1E+4   7.5   8  RT (min)      lcorbi_hilic_neg_4_066_57    m/z 181.8816     0   1   0.5   1  RT (min)      lcorbi_hilic_neg_3_004_41    m/z 181.8868     0   1   2   3   4E+3   7.5   8  RT (min)      lcorbi_hilic_neg_4_066_57    m/z 181.8983     0   2   4E+3   7.5   8  RT (min)      lcorbi_hilic_neg_4_067_57    m/z 181.9658     0   1   2E+3   5.5   6  RT (min)      lcorbi_hilic_neg_4_083_59    m/z 181.9707     0   1E+4   1   1.5  RT (min)      lcorbi_hilic_neg_2_048_38    m/z 181.9797     0   2   4   6E+4   1   1.5  RT (min)      lcorbi_hilic_neg_4_108_62    m/z 181.9892     0   1   6   6.5  RT (min)      lcorbi_hilic_neg_4_078_58    m/z 181.9982     0   1   1   1.5  RT (min)      lcorbi_hilic_neg_4_098_61    m/z 182.0081     0   1   2   3   4E+4   1   1.5  RT (min)      lcorbi_hilic_neg_4_108_62    m/z 182.0175     0   2   4   6   8E+3   1   1.5  RT (min)      lcorbi_hilic_neg_3_018_38    m/z 182.0200     0   1   2E+3   6   6.5  RT (min)       lcorbi_hilic_neg_4_089_59    m/z 182.0281     0   1   3   4  RT (min)       lcorbi_hilic_neg_4_111_62    m/z 182.0318     0   1   5.5   6  RT (min)      lcorbi_hilic_neg_4_089_59    m/z 182.0456     0   2   4   6E+3   9   9.5  RT (min)      lcorbi_hilic_neg_2_017_51    m/z 182.0457     0   1E+4   11   12  RT (min)       lcorbi_hilic_neg_4_087_59    m/z 182.0457     0   1   2   3   4E+4   6   6.5  RT (min)       lcorbi_hilic_neg_1_036_09    m/z 182.0542     0   1E+3   1.5   2  RT (min)      lcorbi_hilic_neg_1_009_50    m/z 182.0649     0   2   4E+2   1.5   2  RT (min)      lcorbi_hilic_neg_2_055_32    m/z 182.0710     0   1E+3   5   5.5  RT (min)      lcorbi_hilic_neg_1_051_33    m/z 182.0820     0   1E+3   1   1.5  RT (min)      lcorbi_hilic_neg_4_094_60    m/z 182.0846     0   1   2E+5   6.5   7  RT (min)       lcorbi_hilic_neg_4_097_60    m/z 182.1124     0   1E+4   6.5   7  RT (min)      lcorbi_hilic_neg_4_097_60    m/z 182.8297     0   2   4   6E+3   3   3.5  RT (min)      lcorbi_hilic_neg_4_070_57    m/z 182.8395     0   2   4   6E+3   4.5   5  RT (min)      lcorbi_hilic_neg_1_029_31    m/z 182.8402     0   1   2E+3   3   3.5  RT (min)      lcorbi_hilic_neg_4_029_02    m/z 182.8573     0   1   2E+4   4.5   5  RT (min)       lcorbi_hilic_neg_4_068_57    m/z 182.8713     0   2   4   6   8E+3   4.5   5  RT (min)      lcorbi_hilic_neg_4_072_57    m/z 182.8792     0   2   4E+3   0   0.5  RT (min)       lcorbi_hilic_neg_2_012_14    m/z 182.8999     0   1E+4   8   8.5  RT (min)      lcorbi_hilic_neg_4_069_57    m/z 182.9369     0   1   2   3E+3   2   2.5  RT (min)      lcorbi_hilic_neg_4_092_60    m/z 182.9435     0   1   4.5   5  RT (min)      lcorbi_hilic_neg_4_087_59    m/z 182.9498     0   1E+3   1   1.5  RT (min)       lcorbi_hilic_neg_4_106_62    m/z 182.9499     0   1   4   4.5  RT (min)      lcorbi_hilic_neg_4_107_62    m/z 182.9789     0   1   2   3E+3   1   1.5  RT (min)      lcorbi_hilic_neg_2_020_27    m/z 182.9911     0   2   4   6E+4   1   2  RT (min)       lcorbi_hilic_neg_4_106_62    m/z 182.9916     0   1   2   3   4E+3   6.5   7  RT (min)      lcorbi_hilic_neg_4_088_59    m/z 183.0071     0   2   4   6   8E+3   1   1.5  RT (min)       lcorbi_hilic_neg_2_020_27    m/z 183.0345     0   1E+3   2.5   3  RT (min)      lcorbi_hilic_neg_1_001_49    m/z 183.0465     0   1   6   6.5  RT (min)      lcorbi_hilic_neg_4_106_62    m/z 183.0522     0   2   4   6   8E+3   3.5   4  RT (min)      lcorbi_hilic_neg_4_073_57    m/z 183.0604     0   1   2   3E+4   2.5   3  RT (min)       lcorbi_hilic_neg_4_096_60    m/z 183.0605     0   1E+5   1   1.5  RT (min)       lcorbi_hilic_neg_4_093_60    m/z 183.0774     0   1   1   1.5  RT (min)      lcorbi_hilic_neg_4_097_60    m/z 183.1025     0   2   4   6E+3   1   1.5  RT (min)      lcorbi_hilic_neg_4_081_58    m/z 183.1054     0   1   2E+4   4   4.5  RT (min)       lcorbi_hilic_neg_4_093_60    m/z 183.1388     0   1E+4   1   1.5  RT (min)      lcorbi_hilic_neg_4_068_57    m/z 183.9628     0   1   5.5   6  RT (min)      lcorbi_hilic_neg_4_095_60    m/z 183.9867     0   1   2E+4   6.5   7  RT (min)      lcorbi_hilic_neg_4_086_59    m/z 183.9900     0   1   2   3E+3   6.5   7  RT (min)      lcorbi_hilic_neg_4_083_59    m/z 183.9980     0   1E+4   1   1.5  RT (min)      lcorbi_hilic_neg_4_108_62    m/z 184.0167     0   1   1   1.5  RT (min)      lcorbi_hilic_neg_1_007_10    m/z 184.0227     0   1   2E+4   1.5   2  RT (min)       lcorbi_hilic_neg_4_073_57    m/z 184.0281     0   1   2   3   4E+3   6   6.5  RT (min)      lcorbi_hilic_neg_4_074_58    m/z 184.0339     0   2   4   6   8E+3   6   6.5  RT (min)      lcorbi_hilic_neg_4_077_58    m/z 184.0477     0   1   3   3.5  RT (min)      lcorbi_hilic_neg_4_113_62    m/z 184.0590     0   1   2   3E+4   5   5.5  RT (min)       lcorbi_hilic_neg_4_100_61    m/z 184.0614     0   1   2E+4   1.5   2  RT (min)       lcorbi_hilic_neg_4_089_59    m/z 184.0765     0   1   2E+4   6   6.5  RT (min)       lcorbi_hilic_neg_4_093_60    m/z 184.0977     0   2   4   6E+3   1   1.5  RT (min)      lcorbi_hilic_neg_4_072_57    m/z 184.8280     0   1   2E+3   3   3.5  RT (min)      lcorbi_hilic_neg_4_074_58    m/z 184.8367     0   1   2   3   4E+3   4.5   5  RT (min)      lcorbi_hilic_neg_2_034_01    m/z 184.8576     0   1   2E+4   8.5   9  RT (min)       lcorbi_hilic_neg_4_105_61    m/z 184.8586     0   2   4E+3   4.5   5  RT (min)      lcorbi_hilic_neg_4_111_62    m/z 184.8765     0   1   2   3   4E+4   8.5   9  RT (min)       lcorbi_hilic_neg_1_003_14    m/z 184.8765     0   1   2   3E+4   0   0.5  RT (min)       lcorbi_hilic_neg_4_106_62    m/z 184.8765     0   1E+5   4.5   5  RT (min)       lcorbi_hilic_neg_4_066_57    m/z 184.9082     0   1   2   3E+4   1   1.5  RT (min)       lcorbi_hilic_neg_4_066_57    m/z 184.9469     0   1E+4   1   1.5  RT (min)      lcorbi_hilic_neg_4_066_57    m/z 184.9582     0   1E+3   1.5   2  RT (min)      lcorbi_hilic_neg_4_083_59    m/z 184.9744     0   2   4E+3   6   6.5  RT (min)      lcorbi_hilic_neg_4_087_59    m/z 184.9852     0   1   2E+4   6.5   7  RT (min)       lcorbi_hilic_neg_4_083_59    m/z 185.0089     0   1   2   3   4E+3   1.5   2  RT (min)       lcorbi_hilic_neg_4_097_60    m/z 185.0092     0   1   6   6.5  RT (min)       lcorbi_hilic_neg_4_097_60    m/z 185.0221     0   1E+4   3.5   4  RT (min)      lcorbi_hilic_neg_4_081_58    m/z 185.0222     0   1   2E+3   2   2.5  RT (min)      lcorbi_hilic_neg_4_097_60    m/z 185.0429     0   1   1   1.5  RT (min)      lcorbi_hilic_neg_4_106_62    m/z 185.0566     0   1   6.5   7  RT (min)      lcorbi_hilic_neg_4_094_60    m/z 185.0567     0   2   4   6   8E+3   4.5   5  RT (min)      lcorbi_hilic_neg_4_073_57    m/z 185.0693     0   1   3   3.5  RT (min)       lcorbi_hilic_neg_4_074_58    m/z 185.0760     0   1E+4   7   7.5  RT (min)      lcorbi_hilic_neg_4_091_60    m/z 185.0761     0   2   4   6   8E+4   5   5.5  RT (min)       lcorbi_hilic_neg_4_094_60    m/z 185.0817     0   2   4   6   8E+3   1   1.5  RT (min)       lcorbi_hilic_neg_4_096_60    m/z 185.1012     0   1   5   5.5  RT (min)      lcorbi_hilic_neg_4_092_60    m/z 185.8948     0   1E+4   8   8.5  RT (min)      lcorbi_hilic_neg_4_067_57    m/z 185.9111     0   1   2E+3   1   1.5  RT (min)      lcorbi_hilic_neg_4_074_58    m/z 185.9624     0   2   4   6   8E+3   1   1.5  RT (min)       lcorbi_hilic_neg_4_106_62    m/z 185.9691     0   1   6   6.5  RT (min)      lcorbi_hilic_neg_4_093_60    m/z 185.9748     0   1E+3   6   6.5  RT (min)      lcorbi_hilic_neg_4_076_58    m/z 185.9897     0   1   2   3E+4   6   6.5  RT (min)       lcorbi_hilic_neg_4_097_60    m/z 186.0018     0   1   2E+4   1   1.5  RT (min)       lcorbi_hilic_neg_4_090_60    m/z 186.0075     0   1E+3   2   2.5  RT (min)      lcorbi_hilic_neg_4_077_58    m/z 186.0285     0   1   2E+3   2   2.5  RT (min)      lcorbi_hilic_neg_3_018_38    m/z 186.0408     0   1E+3   3.5   4  RT (min)       lcorbi_hilic_neg_4_108_62    m/z 186.0560     0   1E+4   3   3.5  RT (min)      lcorbi_hilic_neg_4_083_59    m/z 186.0715     0   1   2E+4   3.5   4  RT (min)      lcorbi_hilic_neg_3_042_05    m/z 186.0964     0   1   5   5.5  RT (min)      lcorbi_hilic_neg_4_112_62    m/z 186.1060     0   1   2E+4   1.5   2  RT (min)       lcorbi_hilic_neg_4_093_60    m/z 186.1216     0   1   1   1.5  RT (min)      lcorbi_hilic_neg_4_087_59    m/z 186.8560     0   1E+4   0   0.5  RT (min)      lcorbi_hilic_neg_4_109_62    m/z 186.8735     0   1E+4   4.5   5  RT (min)      lcorbi_hilic_neg_4_104_61    m/z 186.8974     0   1E+3   1   1.5  RT (min)      lcorbi_hilic_neg_4_066_57    m/z 186.9055     0   1E+4   1   1.5  RT (min)      lcorbi_hilic_neg_4_066_57    m/z 186.9180     0   1E+4   5   5.5  RT (min)      lcorbi_hilic_neg_4_080_58    m/z 186.9181     0   1   2E+3   0.5   1  RT (min)      lcorbi_hilic_neg_2_022_37    m/z 186.9293     0   1   2E+4   7.5   8  RT (min)       lcorbi_hilic_neg_4_102_61    m/z 186.9340     0   1E+4   2   2.5  RT (min)      lcorbi_hilic_neg_4_079_58    m/z 186.9649     0   2   4   6   8E+3   7   7.5  RT (min)      lcorbi_hilic_neg_4_085_59    m/z 186.9815     0   2   4E+3   6.5   7  RT (min)       lcorbi_hilic_neg_4_075_58    m/z 186.9821     0   1E+4   1.5   2  RT (min)      lcorbi_hilic_neg_4_082_59    m/z 186.9961     0   2   4E+3   0.5   1  RT (min)      lcorbi_hilic_neg_4_008_35    m/z 187.0024     0   2   4   6   8E+3   4   4.5  RT (min)      lcorbi_hilic_neg_3_028_23    m/z 187.0049     0   1   2   3   4E+4   1   1.5  RT (min)       lcorbi_hilic_neg_3_024_27    m/z 187.0119     0   1E+4   0.5   1  RT (min)      lcorbi_hilic_neg_2_054_13    m/z 187.0244     0   1E+4   4   4.5  RT (min)      lcorbi_hilic_neg_1_039_38    m/z 187.0416     0   2   4   6E+3   6   6.5  RT (min)      lcorbi_hilic_neg_4_084_59    m/z 187.0523     0   2   4   6   8E+4   4   4.5  RT (min)      lcorbi_hilic_neg_4_088_59    m/z 187.0593     0   2   4   6E+4   1.5   2   2.5  RT (min)       lcorbi_hilic_neg_4_083_59    m/z 187.0722     0   1   2   3E+3   2   2.5  RT (min)      lcorbi_hilic_neg_4_078_58    m/z 187.0804     0   1   1   1.5  RT (min)      lcorbi_hilic_neg_4_076_58    m/z 187.1199     0   1E+4   7   7.5  RT (min)      lcorbi_hilic_neg_4_072_57    m/z 187.8747     0   2   4   6   8E+3   8   8.5  RT (min)      lcorbi_hilic_neg_4_066_57    m/z 187.9409     0   1   2E+4   4   4.5  RT (min)      lcorbi_hilic_neg_4_099_61    m/z 187.9418     0   2   4   6   8E+3   6   6.5  RT (min)       lcorbi_hilic_neg_1_019_06    m/z 187.9578     0   1   6.5   7  RT (min)      lcorbi_hilic_neg_4_094_60    m/z 187.9654     0   1E+3   6   6.5  RT (min)       lcorbi_hilic_neg_4_109_62    m/z 187.9657     0   1   2E+4   2.5   3  RT (min)       lcorbi_hilic_neg_4_105_61    m/z 187.9997     0   2   4   6E+3   6.5   7  RT (min)       lcorbi_hilic_neg_4_026_33    m/z 188.0044     0   1E+4   0.5   1  RT (min)      lcorbi_hilic_neg_4_079_58    m/z 188.0094     0   1   6   6.5  RT (min)      lcorbi_hilic_neg_4_112_62    m/z 188.0206     0   1   2   3   4E+2   5.5   6  RT (min)       lcorbi_hilic_neg_1_023_23    m/z 188.0352     0   1E+4   4.5   5  RT (min)      lcorbi_hilic_neg_3_022_06    m/z 188.0522     0   2   4E+4   6.5   7  RT (min)       lcorbi_hilic_neg_4_092_60    m/z 188.0608     0   2   4   6E+4   3   3.5  RT (min)       lcorbi_hilic_neg_4_088_59    m/z 188.0636     0   2   4E+4   6.5   7  RT (min)       lcorbi_hilic_neg_4_090_60    m/z 188.0645     0   2   4   6E+3   1   1.5  RT (min)      lcorbi_hilic_neg_4_086_59    m/z 188.0666     0   1   2E+3   3   3.5  RT (min)      lcorbi_hilic_neg_4_030_05    m/z 188.0926     0   2   4   6   8E+3   5   5.5  RT (min)      lcorbi_hilic_neg_4_073_57    m/z 188.0928     0   2   4   6E+3   1   1.5  RT (min)      lcorbi_hilic_neg_4_067_57    m/z 188.1008     0   1   2   3E+4   1   1.5  RT (min)      lcorbi_hilic_neg_4_088_59    m/z 188.1041     0   1E+3   6.5   7  RT (min)      lcorbi_hilic_neg_4_096_60    m/z 188.1127     0   1   2   3E+4   1.5   2  RT (min)       lcorbi_hilic_neg_4_090_60    m/z 188.1371     0   2   4E+3   1   1.5  RT (min)      lcorbi_hilic_neg_4_081_58    m/z 188.8266     0   1   2   3   4E+4   4.5   5  RT (min)      lcorbi_hilic_neg_2_020_27    m/z 188.8440     0   1   2   3   4E+3   2.5   3  RT (min)      lcorbi_hilic_neg_4_068_57    m/z 188.9026     0   1   2E+4   0.5   1   1.5  RT (min)       lcorbi_hilic_neg_4_106_62    m/z 188.9152     0   2   4E+3   5   5.5  RT (min)      lcorbi_hilic_neg_4_077_58    m/z 188.9361     0   1   2   2.5  RT (min)      lcorbi_hilic_neg_4_074_58    m/z 188.9563     0   1E+4   1   1.5  RT (min)      lcorbi_hilic_neg_4_072_57    m/z 188.9817     0   1   2E+3   7.5   8  RT (min)      lcorbi_hilic_neg_1_016_37    m/z 188.9829     0   2   4   6E+4   6   6.5  RT (min)      lcorbi_hilic_neg_4_093_60    m/z 188.9977     0   1   2E+4   3   3.5  RT (min)      lcorbi_hilic_neg_4_071_57    m/z 188.9978     0   1E+4   0.5   1  RT (min)       lcorbi_hilic_neg_3_024_27    m/z 189.0200     0   2   4E+4   1.5   2  RT (min)       lcorbi_hilic_neg_4_090_60    m/z 189.0384     0   1   2   3E+4   2   2.5  RT (min)       lcorbi_hilic_neg_4_004_13    m/z 189.0515     0   1   2E+4   6.5   7  RT (min)       lcorbi_hilic_neg_4_089_59    m/z 189.0516     0   1E+4   4.5   5  RT (min)      lcorbi_hilic_neg_4_069_57    m/z 189.0548     0   2   4   6E+3   6   6.5  RT (min)      lcorbi_hilic_neg_1_012_32    m/z 189.0645     0   1   2E+3   3   3.5  RT (min)      lcorbi_hilic_neg_4_089_59    m/z 189.0758     0   1E+3   5.5   6  RT (min)      lcorbi_hilic_neg_3_013_37    m/z 189.0879     0   1E+3   6   6.5  RT (min)      lcorbi_hilic_neg_4_089_59    m/z 189.0885     0   2   4E+2   1.5   2  RT (min)      lcorbi_hilic_neg_4_044_01    m/z 189.1027     0   1E+4   4   4.5  RT (min)      lcorbi_hilic_neg_4_099_61    m/z 189.1030     0   1E+3   1   1.5  RT (min)      lcorbi_hilic_neg_1_005_36    m/z 189.8613     0   1E+4   8   8.5  RT (min)      lcorbi_hilic_neg_4_066_57    m/z 189.9497     0   1   1.5   2  RT (min)      lcorbi_hilic_neg_4_090_60    m/z 189.9677     0   2   4   6E+2   6.5   7  RT (min)      lcorbi_hilic_neg_4_005_32    m/z 189.9677     0   1   1.5   2  RT (min)      lcorbi_hilic_neg_4_090_60    m/z 189.9759     0   1E+3   1.5   2  RT (min)      lcorbi_hilic_neg_4_106_62    m/z 189.9824     0   1E+4   6   6.5  RT (min)      lcorbi_hilic_neg_4_090_60    m/z 189.9895     0   1E+4   0.5   1  RT (min)      lcorbi_hilic_neg_4_079_58    m/z 190.0060     0   1   2E+4   0.5   1  RT (min)      lcorbi_hilic_neg_2_020_27    m/z 190.0065     0   1   2   3   4E+3   6   6.5  RT (min)      lcorbi_hilic_neg_4_076_58    m/z 190.0175     0   1E+4   5.5   6  RT (min)      lcorbi_hilic_neg_4_078_58    m/z 190.0322     0   1   2E+4   4.5   5  RT (min)       lcorbi_hilic_neg_4_097_60    m/z 190.0368     0   2   4   6   8E+3   1   1.5  RT (min)      lcorbi_hilic_neg_2_020_27    m/z 190.0437     0   1   2   3   4E+2   1   1.5   2  RT (min)       lcorbi_hilic_neg_2_005_24    m/z 190.0478     0   2   4   6   8E+3   4   4.5  RT (min)      lcorbi_hilic_neg_2_026_23    m/z 190.0517     0   1E+5   6   6.5  RT (min)       lcorbi_hilic_neg_4_096_60    m/z 190.0606     0   1   2   3E+4   3   3.5  RT (min)      lcorbi_hilic_neg_4_089_59    m/z 190.0719     0   1   6   6.5  RT (min)      lcorbi_hilic_neg_4_090_60    m/z 190.0721     0   1   2   3E+3   1   1.5  RT (min)       lcorbi_hilic_neg_1_005_36    m/z 190.0814     0   2   4   6E+3   1   1.5  RT (min)      lcorbi_hilic_neg_1_054_27    m/z 190.0872     0   1   2.5   3  RT (min)      lcorbi_hilic_neg_4_113_62    m/z 190.0998     0   1   2E+4   0.5   1  RT (min)      lcorbi_hilic_neg_4_107_62    m/z 190.8428     0   1   2   3   4E+4   0.5   1  RT (min)      lcorbi_hilic_neg_3_044_32    m/z 190.8527     0   1   2   3E+4   8   8.5  RT (min)       lcorbi_hilic_neg_4_098_61    m/z 190.8611     0   2   4   6E+3   1   1.5  RT (min)       lcorbi_hilic_neg_3_032_42    m/z 190.8947     0   1E+4   0   0.5  RT (min)       lcorbi_hilic_neg_4_097_60    m/z 190.9240     0   1E+3   4.5   5  RT (min)      lcorbi_hilic_neg_3_050_03    m/z 190.9325     0   1   2.5   3  RT (min)      lcorbi_hilic_neg_4_111_62    m/z 190.9754     0   1   6   6.5  RT (min)       lcorbi_hilic_neg_4_073_57    m/z 190.9820     0   1   0.5   1  RT (min)      lcorbi_hilic_neg_4_092_60    m/z 190.9932     0   1   2   3E+4   1   1.5  RT (min)       lcorbi_hilic_neg_4_081_58    m/z 190.9932     0   1   2E+4   3   3.5  RT (min)      lcorbi_hilic_neg_4_093_60    m/z 191.0017     0   1E+3   6.5   7  RT (min)      lcorbi_hilic_neg_1_003_14    m/z 191.0136     0   1   2E+4   0.5   1  RT (min)       lcorbi_hilic_neg_4_080_58    m/z 191.0196     0   1   2   3E+4   3.5   4  RT (min)       lcorbi_hilic_neg_4_038_27    m/z 191.0347     0   1E+3   1   1.5  RT (min)      lcorbi_hilic_neg_4_104_61    m/z 191.0542     0   2   4E+3   1   1.5  RT (min)      lcorbi_hilic_neg_4_107_62    m/z 191.0623     0   1E+4   1   1.5  RT (min)      lcorbi_hilic_neg_4_107_62    m/z 191.0673     0   2   4E+4   7   7.5  RT (min)       lcorbi_hilic_neg_4_081_58    m/z 191.0765     0   2   4   6E+3   1   1.5  RT (min)       lcorbi_hilic_neg_4_107_62    m/z 191.1076     0   1E+4   3   3.5  RT (min)       lcorbi_hilic_neg_4_097_60    m/z 191.1189     0   1   1   1.5  RT (min)      lcorbi_hilic_neg_1_010_22    m/z 191.8164     0   1E+4   4.5   5  RT (min)      lcorbi_hilic_neg_4_067_57    m/z 191.8550     0   1   2   3   4E+3   1   1.5  RT (min)      lcorbi_hilic_neg_4_081_58    m/z 191.8579     0   2   4   6   8E+3   8   8.5  RT (min)      lcorbi_hilic_neg_4_099_61    m/z 191.9187     0   2   4   6E+3   0.5   1  RT (min)      lcorbi_hilic_neg_4_106_62    m/z 191.9475     0   1E+3   3   3.5  RT (min)      lcorbi_hilic_neg_4_074_58    m/z 191.9518     0   1   2E+4   1   1.5  RT (min)       lcorbi_hilic_neg_4_072_57    m/z 191.9653     0   1   1.5   2  RT (min)      lcorbi_hilic_neg_4_090_60    m/z 191.9736     0   2   4   6   8E+3   2   2.5  RT (min)      lcorbi_hilic_neg_4_089_59    m/z 191.9859     0   1   2E+4   7   7.5  RT (min)       lcorbi_hilic_neg_4_081_58    m/z 191.9995     0   1   2   3E+3   5.5   6  RT (min)      lcorbi_hilic_neg_4_077_58    m/z 192.0017     0   1E+4   2   2.5  RT (min)      lcorbi_hilic_neg_4_081_58    m/z 192.0228     0   1   2   3E+4   6.5   7  RT (min)      lcorbi_hilic_neg_1_003_14    m/z 192.0342     0   2   4   6   8E+3   4   4.5  RT (min)      lcorbi_hilic_neg_4_005_32    m/z 192.0357     0   1   2   3   4E+3   7.5   8  RT (min)      lcorbi_hilic_neg_4_083_59    m/z 192.0410     0   2   4   6   8E+3   1.5   2  RT (min)       lcorbi_hilic_neg_4_094_60    m/z 192.0445     0   2   4   6E+4   4   4.5  RT (min)      lcorbi_hilic_neg_1_055_24    m/z 192.0446     0   1E+4   6.5   7  RT (min)      lcorbi_hilic_neg_3_002_15    m/z 192.0546     0   1   2   3E+3   1   1.5  RT (min)      lcorbi_hilic_neg_4_110_62    m/z 192.0655     0   2   4   6   8E+2   1   1.5  RT (min)      lcorbi_hilic_neg_3_040_19    m/z 192.0690     0   1   2   3   4E+3   5.5   6  RT (min)      lcorbi_hilic_neg_4_095_60    m/z 192.0707     0   2   4   6E+4   4   4.5  RT (min)      lcorbi_hilic_neg_4_097_60    m/z 192.0809     0   2   4   6E+3   3.5   4  RT (min)      lcorbi_hilic_neg_3_056_56    m/z 192.0881     0   1E+4   1.5   2  RT (min)      lcorbi_hilic_neg_4_094_60    m/z 192.1202     0   2   4   6   8E+3   0.5   1  RT (min)      lcorbi_hilic_neg_2_029_10    m/z 192.8403     0   1   2E+4   0.5   1  RT (min)      lcorbi_hilic_neg_3_044_32    m/z 192.8562     0   1   8   8.5  RT (min)       lcorbi_hilic_neg_2_032_25    m/z 192.8632     0   1   4.5   5  RT (min)      lcorbi_hilic_neg_4_104_61    m/z 192.9112     0   2   4E+3   0.5   1  RT (min)      lcorbi_hilic_neg_4_106_62    m/z 192.9114     0   1   2E+4   4.5   5  RT (min)       lcorbi_hilic_neg_3_003_44    m/z 192.9302     0   1E+4   2.5   3  RT (min)      lcorbi_hilic_neg_4_076_58    m/z 192.9335     0   1E+4   8   8.5  RT (min)      lcorbi_hilic_neg_4_082_59    m/z 192.9490     0   1   2   3E+4   1.5   2  RT (min)       lcorbi_hilic_neg_4_095_60    m/z 192.9599     0   1   3.5   4  RT (min)      lcorbi_hilic_neg_4_095_60    m/z 192.9789     0   1   2   3E+3   1   1.5  RT (min)      lcorbi_hilic_neg_4_091_60    m/z 192.9892     0   1   2E+4   0.5   1  RT (min)       lcorbi_hilic_neg_4_075_58    m/z 192.9892     0   1E+4   3   3.5  RT (min)      lcorbi_hilic_neg_4_093_60    m/z 192.9980     0   1   5.5   6  RT (min)      lcorbi_hilic_neg_4_084_59    m/z 193.0072     0   1E+4   5.5   6  RT (min)      lcorbi_hilic_neg_4_072_57    m/z 193.0091     0   1E+4   1   1.5  RT (min)      lcorbi_hilic_neg_4_089_59    m/z 193.0132     0   1   2   3   4E+2   3.5   4   4.5  RT (min)       lcorbi_hilic_neg_2_055_32    m/z 193.0261     0   2   4E+2   5   5.5  RT (min)      lcorbi_hilic_neg_1_023_23    m/z 193.0351     0   1E+3   5   5.5   6  RT (min)       lcorbi_hilic_neg_2_026_23    m/z 193.0504     0   1E+3   1   1.5  RT (min)       lcorbi_hilic_neg_4_015_24    m/z 193.0600     0   1E+5   4   4.5  RT (min)       lcorbi_hilic_neg_4_090_60    m/z 193.0616     0   1E+3   1   1.5   2  RT (min)       lcorbi_hilic_neg_4_090_60    m/z 193.0868     0   1E+4   3   3.5  RT (min)      lcorbi_hilic_neg_4_113_62    m/z 193.0982     0   1   1   1.5  RT (min)       lcorbi_hilic_neg_4_093_60    m/z 193.1197     0   1   2   3E+4   5   5.5  RT (min)       lcorbi_hilic_neg_2_055_32    m/z 193.1596     0   1E+4   0.5   1  RT (min)      lcorbi_hilic_neg_4_089_59    m/z 193.8155     0   1   2   3E+4   8   8.5  RT (min)      lcorbi_hilic_neg_4_066_57    m/z 193.8156     0   2   4E+4   13   13.5  RT (min)       lcorbi_hilic_neg_4_082_59    m/z 193.8795     0   2   4   6E+3   0.5   1  RT (min)      lcorbi_hilic_neg_4_068_57    m/z 193.9080     0   2   4   6   8E+3   4.5   5  RT (min)      lcorbi_hilic_neg_4_101_61    m/z 193.9312     0   1E+4   8   8.5  RT (min)      lcorbi_hilic_neg_4_087_59    m/z 193.9864     0   2   4   6E+2   5.5   6  RT (min)      lcorbi_hilic_neg_2_005_24    m/z 194.0012     0   1   2E+4   5.5   6  RT (min)       lcorbi_hilic_neg_4_097_60    m/z 194.0304     0   1E+3   2   2.5  RT (min)      lcorbi_hilic_neg_2_005_24    m/z 194.0458     0   1E+3   1.5   2  RT (min)       lcorbi_hilic_neg_4_095_60    m/z 194.0464     0   2   4   6E+6   4   4.5  RT (min)       lcorbi_hilic_neg_2_041_54    m/z 194.0530     0   2   4   6E+3   1.5   2  RT (min)       lcorbi_hilic_neg_1_037_46    m/z 194.0626     0   1E+5   4   4.5  RT (min)      lcorbi_hilic_neg_4_091_60    m/z 194.0764     0   1E+5   2.5   3  RT (min)       lcorbi_hilic_neg_3_011_24    m/z 194.0875     0   1   2E+4   6.5   7  RT (min)       lcorbi_hilic_neg_4_097_60    m/z 194.1184     0   1E+3   1   1.5  RT (min)      lcorbi_hilic_neg_4_095_60    m/z 194.8359     0   2   4E+3   2.5   3  RT (min)      lcorbi_hilic_neg_4_074_58    m/z 194.8555     0   2   4   6   8E+3   1   1.5  RT (min)       lcorbi_hilic_neg_4_084_59    m/z 194.8603     0   1   4.5   5  RT (min)      lcorbi_hilic_neg_4_107_62    m/z 194.9013     0   1   2E+3   4.5   5  RT (min)      lcorbi_hilic_neg_4_098_61    m/z 194.9205     0   1E+4   7.5   8  RT (min)      lcorbi_hilic_neg_3_048_33    m/z 194.9273     0   2   4E+4   0   0.5  RT (min)      lcorbi_hilic_neg_2_005_24    m/z 194.9370     0   1   1   1.5  RT (min)      lcorbi_hilic_neg_1_008_39    m/z 194.9468     0   1   2   3   4E+2   2   2.5  RT (min)      lcorbi_hilic_neg_1_045_42    m/z 194.9595     0   1E+4   7   7.5  RT (min)      lcorbi_hilic_neg_4_094_60    m/z 194.9652     0   1   1.5   2  RT (min)      lcorbi_hilic_neg_4_084_59    m/z 194.9674     0   1   2   3E+3   7   7.5  RT (min)      lcorbi_hilic_neg_3_042_05    m/z 194.9760     0   1   4.5   5  RT (min)      lcorbi_hilic_neg_4_092_60    m/z 194.9842     0   1   6   6.5  RT (min)      lcorbi_hilic_neg_4_093_60    m/z 194.9967     0   1   2E+3   3   3.5  RT (min)      lcorbi_hilic_neg_4_090_60    m/z 194.9968     0   1   2E+3   0.5   1  RT (min)      lcorbi_hilic_neg_4_093_60    m/z 195.0136     0   1   2E+3   1.5   2  RT (min)      lcorbi_hilic_neg_4_083_59    m/z 195.0150     0   1   6.5   7  RT (min)       lcorbi_hilic_neg_4_070_57    m/z 195.0177     0   1   3.5   4  RT (min)      lcorbi_hilic_neg_4_095_60    m/z 195.0265     0   2   4   6   8E+3   1.5   2  RT (min)      lcorbi_hilic_neg_4_094_60    m/z 195.0355     0   1E+3   2   2.5  RT (min)      lcorbi_hilic_neg_4_097_60    m/z 195.0384     0   1   2   3E+2   6   6.5  RT (min)       lcorbi_hilic_neg_1_023_23    m/z 195.0492     0   1   2E+3   2   3  RT (min)       lcorbi_hilic_neg_4_113_62    m/z 195.0660     0   1   2E+3   2   2.5  RT (min)      lcorbi_hilic_neg_2_028_33    m/z 195.0716     0   2   4   6   8E+3   4.5   5  RT (min)      lcorbi_hilic_neg_4_097_60    m/z 195.0717     0   1   2   3E+4   6.5   7  RT (min)       lcorbi_hilic_neg_4_095_60    m/z 195.0774     0   2   4   6E+3   3   3.5  RT (min)       lcorbi_hilic_neg_1_055_24    m/z 195.0848     0   1   2   3   4E+3   3   3.5  RT (min)      lcorbi_hilic_neg_3_005_08    m/z 195.0855     0   1E+4   0.5   1  RT (min)      lcorbi_hilic_neg_4_110_62    m/z 195.1066     0   1   3.5   4  RT (min)      lcorbi_hilic_neg_4_090_60    m/z 195.8112     0   2   4   6   8E+5   12.5   13  RT (min)      lcorbi_hilic_neg_4_108_62    m/z 195.8160     0   1   2   3E+4   4.5   5  RT (min)      lcorbi_hilic_neg_4_107_62    m/z 195.8766     0   2   4E+3   0.5   1  RT (min)      lcorbi_hilic_neg_4_082_59    m/z 195.9057     0   1   2E+4   0.5   1  RT (min)      lcorbi_hilic_neg_4_082_59    m/z 195.9364     0   1   7   7.5  RT (min)      lcorbi_hilic_neg_2_016_30    m/z 195.9506     0   1   7   7.5  RT (min)      lcorbi_hilic_neg_4_050_26    m/z 195.9809     0   1   4.5   5  RT (min)      lcorbi_hilic_neg_4_077_58    m/z 195.9921     0   2   4   6E+2   7   7.5  RT (min)      lcorbi_hilic_neg_4_050_26    m/z 195.9991     0   1   7   7.5  RT (min)      lcorbi_hilic_neg_2_045_26    m/z 196.0283     0   2   4   6E+3   4   4.5  RT (min)      lcorbi_hilic_neg_2_055_32    m/z 196.0402     0   1   1   1.5  RT (min)      lcorbi_hilic_neg_4_071_57    m/z 196.0523     0   1   2   3E+4   1.5   2  RT (min)       lcorbi_hilic_neg_4_096_60    m/z 196.0555     0   1E+5   3   3.5   4  RT (min)       lcorbi_hilic_neg_4_097_60    m/z 196.0588     0   1   7.5   8  RT (min)      lcorbi_hilic_neg_2_005_24    m/z 196.0613     0   1E+3   1   1.5  RT (min)      lcorbi_hilic_neg_4_097_60    m/z 196.0744     0   1E+4   5   5.5  RT (min)      lcorbi_hilic_neg_1_021_28    m/z 196.0768     0   1   3   3.5  RT (min)      lcorbi_hilic_neg_4_067_57    m/z 196.0799     0   1   1.5   2  RT (min)      lcorbi_hilic_neg_4_091_60    m/z 196.0919     0   1   1   1.5  RT (min)      lcorbi_hilic_neg_4_089_59    m/z 196.0977     0   2   4   6E+3   6.5   7  RT (min)      lcorbi_hilic_neg_4_089_59    m/z 196.1058     0   1   1   1.5  RT (min)      lcorbi_hilic_neg_4_097_60    m/z 196.1090     0   2   4E+3   6   6.5  RT (min)      lcorbi_hilic_neg_4_098_61    m/z 196.1422     0   2   4   6E+4   0.5   1  RT (min)      lcorbi_hilic_neg_2_019_21    m/z 196.8112     0   1   2   3E+4   0.5   1  RT (min)       lcorbi_hilic_neg_4_109_62    m/z 196.8113     0   1   2   3E+4   4   4.5  RT (min)       lcorbi_hilic_neg_4_099_61    m/z 196.8113     0   2   4   6   8E+3   8   8.5  RT (min)      lcorbi_hilic_neg_4_067_57    m/z 196.8336     0   1   2   3   4E+3   2.5   3  RT (min)      lcorbi_hilic_neg_4_068_57    m/z 196.8531     0   1E+4   1   1.5  RT (min)       lcorbi_hilic_neg_4_085_59    m/z 196.8769     0   1   2E+3   0   0.5  RT (min)      lcorbi_hilic_neg_2_005_24    m/z 196.8771     0   1   2   3E+4   4.5   5  RT (min)       lcorbi_hilic_neg_4_107_62    m/z 196.8852     0   1E+4   0.5   1  RT (min)      lcorbi_hilic_neg_4_106_62    m/z 196.8995     0   1   4.5   5  RT (min)       lcorbi_hilic_neg_4_082_59    m/z 196.9015     0   1E+4   0.5   1  RT (min)      lcorbi_hilic_neg_4_066_57    m/z 196.9071     0   1   2   3E+4   7   7.5  RT (min)      lcorbi_hilic_neg_4_083_59    m/z 196.9097     0   1E+4   4.5   5  RT (min)      lcorbi_hilic_neg_4_110_62    m/z 196.9598     0   1E+3   1.5   2  RT (min)      lcorbi_hilic_neg_3_053_13    m/z 196.9730     0   1E+3   4.5   5  RT (min)      lcorbi_hilic_neg_4_088_59    m/z 196.9840     0   1E+4   1   1.5  RT (min)      lcorbi_hilic_neg_4_071_57    m/z 197.0030     0   1E+4   1   1.5  RT (min)      lcorbi_hilic_neg_4_075_58    m/z 197.0222     0   2   4   6E+4   3.5   4  RT (min)       lcorbi_hilic_neg_4_069_57    m/z 197.0332     0   1E+3   3.5   4  RT (min)      lcorbi_hilic_neg_4_071_57    m/z 197.0398     0   2   4   6E+4   7   7.5  RT (min)       lcorbi_hilic_neg_1_012_32    m/z 197.0428     0   2   4   6   8E+2   1   1.5  RT (min)      lcorbi_hilic_neg_1_003_14    m/z 197.0445     0   1   4   4.5  RT (min)       lcorbi_hilic_neg_4_097_60    m/z 197.0486     0   1   1.5   2  RT (min)      lcorbi_hilic_neg_4_077_58    m/z 197.0524     0   1   6.5   7  RT (min)      lcorbi_hilic_neg_4_088_59    m/z 197.0608     0   1   2   3   4E+4   4   4.5  RT (min)      lcorbi_hilic_neg_3_048_33    m/z 197.0839     0   1   2E+3   3   3.5  RT (min)       lcorbi_hilic_neg_4_089_59    m/z 197.0929     0   1E+3   6   6.5  RT (min)      lcorbi_hilic_neg_4_092_60    m/z 197.1294     0   1E+3   2   2.5  RT (min)      lcorbi_hilic_neg_4_089_59    m/z 197.1545     0   1E+4   0.5   1  RT (min)      lcorbi_hilic_neg_4_113_62    m/z 197.8027     0   1   2E+4   4.5   5  RT (min)      lcorbi_hilic_neg_4_110_62    m/z 197.8029     0   1   2E+3   0   0.5  RT (min)      lcorbi_hilic_neg_4_074_58    m/z 197.8081     0   2   4   6E+5   12   12.5  RT (min)      lcorbi_hilic_neg_2_050_17    m/z 197.8138     0   1   2E+4   13.5   14  RT (min)      lcorbi_hilic_neg_4_104_61    m/z 197.8155     0   1   2E+4   4.5   5  RT (min)      lcorbi_hilic_neg_4_107_62    m/z 197.8179     0   2   4E+3   3   3.5  RT (min)      lcorbi_hilic_neg_4_108_62    m/z 197.9114     0   1E+4   1   1.5  RT (min)      lcorbi_hilic_neg_4_108_62    m/z 197.9430     0   1   4   4.5  RT (min)      lcorbi_hilic_neg_4_112_62    m/z 197.9490     0   1   1   1.5  RT (min)      lcorbi_hilic_neg_4_108_62    m/z 197.9630     0   1E+4   0.5   1  RT (min)      lcorbi_hilic_neg_1_034_43    m/z 197.9630     0   1E+4   6   6.5  RT (min)      lcorbi_hilic_neg_4_002_41    m/z 197.9862     0   1   6   6.5  RT (min)      lcorbi_hilic_neg_1_034_43    m/z 197.9865     0   1   2   2.5  RT (min)      lcorbi_hilic_neg_1_004_17    m/z 198.0170     0   1E+4   7.5   8  RT (min)      lcorbi_hilic_neg_4_085_59    m/z 198.0171     0   1   2   3E+2   6   6.5  RT (min)      lcorbi_hilic_neg_1_018_21    m/z 198.0185     0   2   4   6   8E+3   3.5   4  RT (min)      lcorbi_hilic_neg_1_012_32    m/z 198.0256     0   1   2E+4   2   2.5  RT (min)      lcorbi_hilic_neg_4_004_13    m/z 198.0381     0   1   2   3E+2   5.5   6   6.5  RT (min)       lcorbi_hilic_neg_1_018_21    m/z 198.0516     0   1   2   3E+4   6   6.5  RT (min)       lcorbi_hilic_neg_2_020_27    m/z 198.0537     0   1   2E+5   2   2.5  RT (min)      lcorbi_hilic_neg_1_012_32    m/z 198.0746     0   1   5   6  RT (min)       lcorbi_hilic_neg_4_097_60    m/z 198.0883     0   2   4   6   8E+2   1.5   2  RT (min)      lcorbi_hilic_neg_3_011_24    m/z 198.1133     0   2   4   6   8E+3   1   1.5  RT (min)      lcorbi_hilic_neg_4_073_57    m/z 198.8083     0   1   2   3E+4   0.5   1  RT (min)       lcorbi_hilic_neg_4_108_62    m/z 198.8083     0   1   2   3E+4   4.5   5  RT (min)       lcorbi_hilic_neg_4_103_61    m/z 198.8083     0   1E+4   8   8.5  RT (min)      lcorbi_hilic_neg_4_067_57    m/z 198.8506     0   1E+4   1   1.5  RT (min)      lcorbi_hilic_neg_4_066_57    m/z 198.8643     0   1   2   3E+3   0.5   1  RT (min)      lcorbi_hilic_neg_4_069_57    m/z 198.8681     0   2   4   6E+3   3   3.5  RT (min)      lcorbi_hilic_neg_4_073_57    m/z 198.8741     0   1   2E+3   7.5   8  RT (min)      lcorbi_hilic_neg_4_082_59    m/z 198.8741     0   1   2   3   4E+4   4.5   5  RT (min)      lcorbi_hilic_neg_4_066_57    m/z 198.8812     0   1E+4   8   8.5  RT (min)       lcorbi_hilic_neg_4_070_57    m/z 198.9179     0   1   2E+4   7.5   8  RT (min)       lcorbi_hilic_neg_4_036_36    m/z 198.9275     0   2   4   6E+2   7.5   8  RT (min)      lcorbi_hilic_neg_1_036_09    m/z 198.9315     0   2   4E+3   1   1.5  RT (min)      lcorbi_hilic_neg_4_106_62    m/z 198.9343     0   1   2E+4   4.5   5  RT (min)       lcorbi_hilic_neg_4_074_58    m/z 198.9448     0   1E+4   2   2.5  RT (min)      lcorbi_hilic_neg_4_090_60    m/z 198.9449     0   1   2   3E+3   4.5   5  RT (min)      lcorbi_hilic_neg_4_070_57    m/z 198.9543     0   1E+4   4.5   5  RT (min)      lcorbi_hilic_neg_4_068_57    m/z 198.9649     0   2   4   6   8E+2   2   2.5  RT (min)      lcorbi_hilic_neg_4_097_60    m/z 198.9704     0   1   2   3   4E+3   5.5   6  RT (min)      lcorbi_hilic_neg_2_012_14    m/z 198.9823     0   1E+4   0.5   1  RT (min)      lcorbi_hilic_neg_2_020_27    m/z 199.0025     0   1   2   3E+3   6.5   7  RT (min)      lcorbi_hilic_neg_1_020_13    m/z 199.0068     0   1   2E+5   1   1.5  RT (min)       lcorbi_hilic_neg_3_018_38    m/z 199.0107     0   1   4   4.5  RT (min)      lcorbi_hilic_neg_4_087_59    m/z 199.0192     0   2   4   6E+3   3.5   4  RT (min)       lcorbi_hilic_neg_4_079_58    m/z 199.0245     0   1   6   6.5  RT (min)      lcorbi_hilic_neg_4_071_57    m/z 199.0280     0   1   1   1.5  RT (min)      lcorbi_hilic_neg_4_095_60    m/z 199.0433     0   2   4   6E+4   1   1.5  RT (min)      lcorbi_hilic_neg_2_005_24    m/z 199.0452     0   1   2E+4   3.5   4  RT (min)      lcorbi_hilic_neg_4_066_57    m/z 199.0527     0   1   2   3E+3   6.5   7  RT (min)      lcorbi_hilic_neg_4_111_62    m/z 199.0582     0   1E+3   3.5   4  RT (min)      lcorbi_hilic_neg_4_031_14    m/z 199.0722     0   1E+3   1.5   2  RT (min)      lcorbi_hilic_neg_4_090_60    m/z 199.0875     0   1E+4   4   4.5  RT (min)      lcorbi_hilic_neg_4_102_61    m/z 199.0973     0   1E+4   1   1.5  RT (min)       lcorbi_hilic_neg_4_083_59    m/z 199.1338     0   1E+4   0.5   1  RT (min)      lcorbi_hilic_neg_4_089_59    m/z 199.1814     0   1E+4   4   4.5  RT (min)      lcorbi_hilic_neg_4_026_33    m/z 199.7996     0   1   2E+3   0   0.5  RT (min)      lcorbi_hilic_neg_4_074_58    m/z 199.8006     0   1E+4   4.5   5  RT (min)      lcorbi_hilic_neg_4_110_62    m/z 199.8157     0   1   2   3   4E+3   3   3.5  RT (min)      lcorbi_hilic_neg_4_111_62    m/z 199.8509     0   1E+4   4.5   5  RT (min)       lcorbi_hilic_neg_4_109_62    m/z 199.8509     0   1   2   3   4E+4   7.5   8  RT (min)       lcorbi_hilic_neg_4_108_62    m/z 199.8811     0   1E+5   0.5   1  RT (min)      lcorbi_hilic_neg_4_055_38    m/z 199.8923     0   2   4E+3   8   8.5  RT (min)      lcorbi_hilic_neg_4_100_61    m/z 199.8990     0   1   7   7.5  RT (min)      lcorbi_hilic_neg_4_074_58    m/z 199.9492     0   1   4   4.5  RT (min)      lcorbi_hilic_neg_4_110_62    m/z 199.9642     0   1   2   3   4E+3   2   2.5  RT (min)      lcorbi_hilic_neg_4_075_58    m/z 199.9690     0   1E+4   6   6.5  RT (min)      lcorbi_hilic_neg_4_070_57    m/z 199.9868     0   1E+4   6   6.5  RT (min)      lcorbi_hilic_neg_4_087_59    m/z 200.0021     0   1   2E+5   2   2.5  RT (min)       lcorbi_hilic_neg_4_004_13    m/z 200.0102     0   1   2E+4   0.5   1  RT (min)      lcorbi_hilic_neg_2_020_27    m/z 200.0225     0   1E+3   2   2.5  RT (min)      lcorbi_hilic_neg_4_075_58    m/z 200.0381     0   2   4   6E+4   6   6.5  RT (min)       lcorbi_hilic_neg_4_071_57    m/z 200.0456     0   2   4E+4   2   2.5  RT (min)       lcorbi_hilic_neg_4_079_58    m/z 200.0571     0   2   4E+3   6.5   7  RT (min)      lcorbi_hilic_neg_4_107_62    m/z 200.0673     0   1   3.5   4   4.5  RT (min)       lcorbi_hilic_neg_1_002_26    m/z 200.0715     0   1   2   2.5  RT (min)       lcorbi_hilic_neg_4_075_58    m/z 200.0742     0   2   4   6E+2   4   4.5  RT (min)       lcorbi_hilic_neg_2_019_21    m/z 200.1290     0   1E+4   1   1.5  RT (min)      lcorbi_hilic_neg_4_068_57    m/z 200.6404     0   2   4   6E+3   3   3.5  RT (min)      lcorbi_hilic_neg_4_070_57    m/z 200.6435     0   1   2   3E+3   3   3.5  RT (min)      lcorbi_hilic_neg_4_070_57    m/z 200.8053     0   1   2E+4   0.5   1  RT (min)      lcorbi_hilic_neg_4_109_62    m/z 200.8054     0   1   2E+4   4.5   5  RT (min)       lcorbi_hilic_neg_4_106_62    m/z 200.8660     0   1   3   3.5  RT (min)      lcorbi_hilic_neg_4_082_59    m/z 200.8806     0   1   2E+4   8   8.5  RT (min)      lcorbi_hilic_neg_4_106_62    m/z 200.8897     0   2   4   6   8E+3   4.5   5  RT (min)      lcorbi_hilic_neg_4_091_60    m/z 200.8912     0   1E+3   3   3.5  RT (min)      lcorbi_hilic_neg_4_106_62    m/z 200.8941     0   1   3   3.5  RT (min)      lcorbi_hilic_neg_4_106_62    m/z 200.9116     0   2   4E+3   8   8.5  RT (min)      lcorbi_hilic_neg_4_066_57    m/z 200.9314     0   1   2   3E+4   4.5   5  RT (min)       lcorbi_hilic_neg_4_074_58    m/z 200.9343     0   2   4E+3   5   5.5  RT (min)      lcorbi_hilic_neg_4_075_58    m/z 200.9527     0   1E+4   4.5   5  RT (min)      lcorbi_hilic_neg_4_070_57    m/z 200.9605     0   2   4   6E+3   2   2.5  RT (min)      lcorbi_hilic_neg_4_075_58    m/z 200.9708     0   1E+3   1   1.5  RT (min)      lcorbi_hilic_neg_2_020_27    m/z 200.9979     0   1   2   3E+4   0.5   1  RT (min)       lcorbi_hilic_neg_4_076_58    m/z 200.9979     0   1   2E+4   3   3.5  RT (min)      lcorbi_hilic_neg_4_093_60    m/z 201.0099     0   1   2   2.5  RT (min)      lcorbi_hilic_neg_4_112_62    m/z 201.0264     0   1E+4   3.5   4  RT (min)      lcorbi_hilic_neg_4_081_58    m/z 201.0382     0   1   2   3   4E+4   2   2.5  RT (min)       lcorbi_hilic_neg_4_075_58    m/z 201.0516     0   1   3.5   4  RT (min)      lcorbi_hilic_neg_2_005_24    m/z 201.0542     0   2   4   6E+4   6  RT (min)       lcorbi_hilic_neg_4_091_60    m/z 201.0647     0   1   3.5   4  RT (min)      lcorbi_hilic_neg_4_105_61    m/z 201.0756     0   1   2E+3   4   4.5  RT (min)       lcorbi_hilic_neg_4_094_60    m/z 201.0768     0   1   2E+4   1   1.5  RT (min)       lcorbi_hilic_neg_4_088_59    m/z 201.0880     0   1   3   3.5  RT (min)      lcorbi_hilic_neg_4_077_58    m/z 201.0880     0   2   4   6   8E+2   2   2.5  RT (min)       lcorbi_hilic_neg_4_046_15    m/z 201.1242     0   1   5   5.5  RT (min)       lcorbi_hilic_neg_4_094_60    m/z 201.1355     0   1E+4   6.5   7  RT (min)      lcorbi_hilic_neg_4_071_57    m/z 201.1436     0   1E+3   3   3.5  RT (min)      lcorbi_hilic_neg_4_075_58    m/z 201.1494     0   1E+4   1   1.5  RT (min)      lcorbi_hilic_neg_4_089_59    m/z 201.8020     0   2   4   6   8E+4   13   13.5  RT (min)      lcorbi_hilic_neg_3_038_21    m/z 201.8130     0   1E+3   3   3.5  RT (min)      lcorbi_hilic_neg_4_113_62    m/z 201.8589     0   1E+4   0.5   1  RT (min)      lcorbi_hilic_neg_4_102_61    m/z 201.9059     0   1   2   3E+4   1  RT (min)       lcorbi_hilic_neg_4_106_62    m/z 201.9393     0   1E+4   6   6.5  RT (min)      lcorbi_hilic_neg_4_078_58    m/z 201.9426     0   1   2E+4   2.5   3  RT (min)       lcorbi_hilic_neg_4_105_61    m/z 201.9671     0   1   4.5   5  RT (min)      lcorbi_hilic_neg_4_090_60    m/z 202.0065     0   1   2.5   3  RT (min)       lcorbi_hilic_neg_4_076_58    m/z 202.0121     0   1E+4   6   6.5  RT (min)      lcorbi_hilic_neg_4_096_60    m/z 202.0154     0   2   4   6E+3   5.5   6  RT (min)      lcorbi_hilic_neg_4_078_58    m/z 202.0246     0   1   2E+4   1.5   2  RT (min)       lcorbi_hilic_neg_4_085_59    m/z 202.0306     0   1   2E+4   7.5   8  RT (min)       lcorbi_hilic_neg_4_093_60    m/z 202.0424     0   1   2E+4   6   6.5  RT (min)      lcorbi_hilic_neg_4_072_57    m/z 202.0453     0   1   2   3   4E+3   3.5   4  RT (min)      lcorbi_hilic_neg_4_078_58    m/z 202.0477     0   1   2E+3   2   2.5  RT (min)      lcorbi_hilic_neg_3_024_27    m/z 202.0544     0   2   4   6E+3   1   1.5  RT (min)       lcorbi_hilic_neg_4_048_37    m/z 202.0581     0   1   6   6.5  RT (min)      lcorbi_hilic_neg_4_105_61    m/z 202.0627     0   1   2E+4   4   4.5  RT (min)      lcorbi_hilic_neg_4_005_32    m/z 202.0722     0   1   2.5   3  RT (min)      lcorbi_hilic_neg_4_110_62    m/z 202.0800     0   1E+4   4.5   5  RT (min)      lcorbi_hilic_neg_1_042_25    m/z 202.0899     0   1   2   3   4E+4   0   0.5  RT (min)      lcorbi_hilic_neg_4_077_58    m/z 202.0917     0   1E+3   2   2.5  RT (min)      lcorbi_hilic_neg_4_077_58    m/z 202.1057     0   1E+4   4   4.5  RT (min)      lcorbi_hilic_neg_3_044_32    m/z 202.1082     0   1E+3   1   1.5  RT (min)      lcorbi_hilic_neg_4_090_60    m/z 202.1557     0   1   2E+4   1   1.5  RT (min)       lcorbi_hilic_neg_4_096_60    m/z 202.8819     0   2   4   6E+3   0   0.5  RT (min)      lcorbi_hilic_neg_4_105_61    m/z 202.9033     0   1E+4   0   0.5  RT (min)      lcorbi_hilic_neg_4_071_57    m/z 202.9288     0   2   4   6   8E+3   0   0.5  RT (min)       lcorbi_hilic_neg_4_113_62    m/z 202.9288     0   2   4E+4   8   8.5  RT (min)       lcorbi_hilic_neg_4_082_59    m/z 202.9347     0   1   2E+3   5.5   6  RT (min)      lcorbi_hilic_neg_4_070_57    m/z 202.9652     0   2   4E+4   6.5   7  RT (min)       lcorbi_hilic_neg_4_070_57    m/z 202.9814     0   1   6   6.5  RT (min)      lcorbi_hilic_neg_4_069_57    m/z 202.9933     0   1E+4   1   1.5  RT (min)      lcorbi_hilic_neg_4_077_58    m/z 202.9971     0   1   2   3E+3   6   6.5  RT (min)      lcorbi_hilic_neg_4_069_57    m/z 203.0294     0   2   4   6   8E+3   2   2.5  RT (min)      lcorbi_hilic_neg_4_077_58    m/z 203.0389     0   2   4E+4   2   2.5  RT (min)       lcorbi_hilic_neg_4_081_58    m/z 203.0425     0   1   6.5   7  RT (min)      lcorbi_hilic_neg_4_090_60    m/z 203.0503     0   1   2E+3   3.5   4  RT (min)      lcorbi_hilic_neg_4_074_58    m/z 203.0514     0   1E+3   6  RT (min)       lcorbi_hilic_neg_4_097_60    m/z 203.0616     0   1E+4   2   2.5  RT (min)      lcorbi_hilic_neg_4_080_58    m/z 203.0732     0   1   2   3E+4   1.5   2  RT (min)      lcorbi_hilic_neg_4_075_58    m/z 203.0780     0   2   4   6E+2   5   5.5  RT (min)      lcorbi_hilic_neg_2_020_27    m/z 203.1286     0   1E+4   1   1.5  RT (min)      lcorbi_hilic_neg_4_088_59    m/z 203.7989     0   1E+4   0.5   1  RT (min)      lcorbi_hilic_neg_4_107_62    m/z 203.7990     0   1E+4   4.5   5  RT (min)      lcorbi_hilic_neg_4_109_62    m/z 203.8898     0   2   4   6   8E+3   4.5   5  RT (min)      lcorbi_hilic_neg_4_080_58    m/z 203.9289     0   1E+4   5   5.5  RT (min)      lcorbi_hilic_neg_4_077_58    m/z 203.9367     0   1   2E+4   0   0.5  RT (min)      lcorbi_hilic_neg_4_066_57    m/z 203.9460     0   1   2E+4   2   2.5  RT (min)       lcorbi_hilic_neg_4_098_61    m/z 203.9602     0   1   2E+4   6.5   7  RT (min)       lcorbi_hilic_neg_4_081_58    m/z 203.9650     0   1E+4   3   3.5  RT (min)       lcorbi_hilic_neg_4_086_59    m/z 203.9652     0   1   2E+4   1   1.5  RT (min)      lcorbi_hilic_neg_4_066_57    m/z 203.9870     0   1   7   7.5  RT (min)      lcorbi_hilic_neg_2_055_32    m/z 203.9945     0   2   4   6E+3   2   2.5  RT (min)      lcorbi_hilic_neg_4_107_62    m/z 203.9967     0   1   2   3E+4   5   5.5  RT (min)       lcorbi_hilic_neg_4_089_59    m/z 204.0051     0   1   2E+4   0.5   1  RT (min)       lcorbi_hilic_neg_4_078_58    m/z 204.0301     0   2   4   6E+3   4.5   5  RT (min)      lcorbi_hilic_neg_4_077_58    m/z 204.0387     0   1   2E+4   3.5   4  RT (min)       lcorbi_hilic_neg_4_093_60    m/z 204.0539     0   1   2   3E+3   1.5   2  RT (min)      lcorbi_hilic_neg_4_077_58    m/z 204.0543     0   2   4E+3   7   7.5  RT (min)      lcorbi_hilic_neg_4_077_58    m/z 204.0858     0   1E+6   4   5  RT (min)       lcorbi_hilic_neg_4_094_60    m/z 204.0875     0   2   4E+3   2   2.5  RT (min)      lcorbi_hilic_neg_4_089_59    m/z 204.1069     0   1E+4   4   4.5  RT (min)      lcorbi_hilic_neg_4_092_60    m/z 204.1082     0   1   2   2.5  RT (min)      lcorbi_hilic_neg_4_089_59    m/z 204.1239     0   1   3.5   4  RT (min)      lcorbi_hilic_neg_1_055_24    m/z 204.1239     0   1E+5   1   1.5  RT (min)      lcorbi_hilic_neg_2_056_56    m/z 204.1474     0   1   2   3E+4   0.5   1  RT (min)      lcorbi_hilic_neg_4_076_58    m/z 204.8766     0   1E+4   1   1.5  RT (min)      lcorbi_hilic_neg_4_106_62    m/z 204.9245     0   1E+4   1.5   2  RT (min)      lcorbi_hilic_neg_4_106_62    m/z 204.9269     0   1   2   3E+3   5   5.5  RT (min)      lcorbi_hilic_neg_4_074_58    m/z 204.9424     0   1   2   3E+4   1.5   2  RT (min)       lcorbi_hilic_neg_4_082_59    m/z 204.9893     0   1   2   3   4E+4   3   3.5   4  RT (min)       lcorbi_hilic_neg_4_089_59    m/z 204.9962     0   1   5.5   6  RT (min)      lcorbi_hilic_neg_2_007_08    m/z 205.0067     0   2   4   6   8E+3   1.5   2  RT (min)      lcorbi_hilic_neg_4_070_57    m/z 205.0091     0   1   2E+4   1.5   2  RT (min)       lcorbi_hilic_neg_4_083_59    m/z 205.0389     0   1   2E+3   2   2.5  RT (min)       lcorbi_hilic_neg_4_087_59    m/z 205.0537     0   2   4   6   8E+3   7   7.5  RT (min)      lcorbi_hilic_neg_4_072_57    m/z 205.0561     0   1E+4   3.5   4  RT (min)      lcorbi_hilic_neg_4_071_57    m/z 205.0650     0   2   4   6   8E+3   1.5   2  RT (min)       lcorbi_hilic_neg_4_089_59    m/z 205.0827     0   2   4E+3   6.5   7  RT (min)      lcorbi_hilic_neg_4_074_58    m/z 205.0870     0   1   2E+3   1   1.5  RT (min)       lcorbi_hilic_neg_4_092_60    m/z 205.8396     0   2   4   6E+3   4.5   5  RT (min)      lcorbi_hilic_neg_4_086_59    m/z 205.8396     0   2   4   6E+4   13   13.5  RT (min)       lcorbi_hilic_neg_4_075_58    m/z 205.8396     0   1E+5   9.5   10  RT (min)       lcorbi_hilic_neg_4_075_58    m/z 205.8869     0   1E+4   4.5   5  RT (min)      lcorbi_hilic_neg_4_082_59    m/z 205.8981     0   2   4   6E+3   7.5   8  RT (min)      lcorbi_hilic_neg_2_045_26    m/z 205.9830     0   1   2   3   4E+3   6   6.5  RT (min)      lcorbi_hilic_neg_4_070_57    m/z 205.9843     0   2   4   6   8E+3   0.5   1  RT (min)      lcorbi_hilic_neg_4_081_58    m/z 205.9996     0   1E+3   0.5   1  RT (min)      lcorbi_hilic_neg_4_076_58    m/z 206.0013     0   1   6   6.5  RT (min)       lcorbi_hilic_neg_4_090_60    m/z 206.0109     0   1   2E+4   6.5   7  RT (min)      lcorbi_hilic_neg_4_091_60    m/z 206.0219     0   1E+4   2   2.5  RT (min)      lcorbi_hilic_neg_4_083_59    m/z 206.0253     0   2   4   6E+3   6   6.5  RT (min)      lcorbi_hilic_neg_4_091_60    m/z 206.0296     0   2   4E+4   2.5   3   3.5  RT (min)       lcorbi_hilic_neg_4_091_60    m/z 206.0489     0   1   2   3   4E+3   1   1.5  RT (min)      lcorbi_hilic_neg_4_093_60    m/z 206.0682     0   1E+3   2   2.5  RT (min)      lcorbi_hilic_neg_4_088_59    m/z 206.0821     0   1   2E+4   1   1.5  RT (min)       lcorbi_hilic_neg_4_089_59    m/z 206.1231     0   1E+4   6   6.5  RT (min)      lcorbi_hilic_neg_4_097_60    m/z 206.1630     0   2   4E+4   0.5   1  RT (min)      lcorbi_hilic_neg_4_075_58    m/z 206.6404     0   1   2   3E+3   3   3.5  RT (min)      lcorbi_hilic_neg_4_068_57    m/z 206.8872     0   2   4E+3   4.5   5  RT (min)      lcorbi_hilic_neg_4_076_58    m/z 206.9214     0   2   4E+3   1.5   2  RT (min)      lcorbi_hilic_neg_4_106_62    m/z 206.9433     0   2   4E+4   0.5   1  RT (min)      lcorbi_hilic_neg_2_020_27    m/z 206.9581     0   1   6   6.5  RT (min)      lcorbi_hilic_neg_4_090_60    m/z 206.9703     0   1   2E+4   6.5   7  RT (min)       lcorbi_hilic_neg_4_091_60    m/z 206.9757     0   1   2   3E+6   0   0.5  RT (min)      lcorbi_hilic_neg_1_005_36    m/z 206.9840     0   1   2E+4   6   6.5  RT (min)       lcorbi_hilic_neg_4_070_57    m/z 206.9862     0   1   2   3   4E+4   3   3.5  RT (min)       lcorbi_hilic_neg_4_093_60    m/z 206.9881     0   1   2   3E+4   0.5   1  RT (min)       lcorbi_hilic_neg_4_066_57    m/z 206.9948     0   1   2   3E+4   6   6.5  RT (min)      lcorbi_hilic_neg_4_091_60    m/z 206.9968     0   2   4   6   8E+2   1   1.5  RT (min)      lcorbi_hilic_neg_4_099_61    m/z 207.0144     0   2   4   6E+3   7   7.5  RT (min)      lcorbi_hilic_neg_4_074_58    m/z 207.0181     0   1E+4   2   2.5  RT (min)       lcorbi_hilic_neg_4_073_57    m/z 207.0299     0   2   4   6   8E+3   1.5   2  RT (min)      lcorbi_hilic_neg_4_084_59    m/z 207.0417     0   1   2E+4   6   6.5  RT (min)       lcorbi_hilic_neg_4_090_60    m/z 207.0522     0   1   2   3E+4   4.5   5  RT (min)      lcorbi_hilic_neg_4_107_62    m/z 207.0653     0   1   2E+3   5.5   6  RT (min)      lcorbi_hilic_neg_4_077_58    m/z 207.0707     0   1   4   4.5  RT (min)      lcorbi_hilic_neg_4_082_59    m/z 207.0871     0   1   2E+3   1.5   2  RT (min)      lcorbi_hilic_neg_4_073_57    m/z 207.1067     0   1   5   5.5  RT (min)      lcorbi_hilic_neg_4_084_59    m/z 207.1137     0   1E+4   1   1.5  RT (min)       lcorbi_hilic_neg_4_093_60    m/z 207.6480     0   1   2E+3   3   3.5  RT (min)      lcorbi_hilic_neg_4_073_57    m/z 207.8366     0   1   2   3E+4   0.5   1  RT (min)       lcorbi_hilic_neg_4_097_60    m/z 207.8367     0   1E+5   9.5   10  RT (min)       lcorbi_hilic_neg_4_077_58    m/z 207.8367     0   2   4   6E+4   13   13.5  RT (min)       lcorbi_hilic_neg_4_100_61    m/z 207.8840     0   2   4   6E+3   4.5   5  RT (min)      lcorbi_hilic_neg_4_082_59    m/z 207.9184     0   1   2E+3   1   1.5  RT (min)      lcorbi_hilic_neg_4_106_62    m/z 207.9944     0   1   2E+4   2   2.5  RT (min)       lcorbi_hilic_neg_4_101_61    m/z 208.0050     0   1E+5   1   1.5  RT (min)       lcorbi_hilic_neg_4_097_60    m/z 208.0282     0   1   2E+4   3   3.5   4  RT (min)       lcorbi_hilic_neg_4_097_60    m/z 208.0450     0   1E+4   4.5   5  RT (min)      lcorbi_hilic_neg_4_113_62    m/z 208.0574     0   1   6.5   7  RT (min)      lcorbi_hilic_neg_4_112_62    m/z 208.0614     0   1   2E+3   1   1.5  RT (min)       lcorbi_hilic_neg_4_090_60    m/z 208.0694     0   1   2   3E+4   1   1.5  RT (min)      lcorbi_hilic_neg_4_076_58    m/z 208.0831     0   1   2E+4   2.5   3  RT (min)       lcorbi_hilic_neg_4_097_60    m/z 208.1114     0   1   2   2.5  RT (min)      lcorbi_hilic_neg_4_068_57    m/z 208.7863     0   1   2E+4   2.5   3  RT (min)       lcorbi_hilic_neg_4_081_58    m/z 208.8450     0   1E+4   0   0.5  RT (min)      lcorbi_hilic_neg_4_075_58    m/z 208.9346     0   2   4   6E+3   1   1.5  RT (min)      lcorbi_hilic_neg_2_014_31    m/z 208.9404     0   2   4   6E+3   4.5   5  RT (min)      lcorbi_hilic_neg_1_002_26    m/z 208.9648     0   1   3   3.5  RT (min)      lcorbi_hilic_neg_4_090_60    m/z 208.9734     0   1   2E+6   0   0.5  RT (min)      lcorbi_hilic_neg_1_008_39    m/z 208.9809     0   1   6   6.5  RT (min)      lcorbi_hilic_neg_4_093_60    m/z 208.9845     0   1   2   3   4E+3   5.5   6  RT (min)      lcorbi_hilic_neg_4_092_60    m/z 208.9907     0   2   4   6   8E+2   2   2.5  RT (min)       lcorbi_hilic_neg_2_055_32    m/z 209.0211     0   2   4   6   8E+4   6   6.5  RT (min)       lcorbi_hilic_neg_4_092_60    m/z 209.0259     0   1   3.5   4  RT (min)       lcorbi_hilic_neg_4_094_60    m/z 209.0446     0   1E+3   3.5   4  RT (min)      lcorbi_hilic_neg_4_095_60    m/z 209.0550     0   1   2   3E+3   3.5   4  RT (min)      lcorbi_hilic_neg_2_020_27    m/z 209.0676     0   1   2   3E+4   1.5   2  RT (min)       lcorbi_hilic_neg_4_067_57    m/z 209.0720     0   1E+3   3.5   4  RT (min)       lcorbi_hilic_neg_4_078_58    m/z 209.0722     0   1E+3   6   6.5  RT (min)      lcorbi_hilic_neg_4_097_60    m/z 209.0760     0   2   4   6E+3   1.5   2  RT (min)       lcorbi_hilic_neg_4_073_57    m/z 209.0852     0   1   2   3E+5   0.5   1  RT (min)       lcorbi_hilic_neg_2_004_09    m/z 209.0859     0   2   4   6   8E+3   3.5   4  RT (min)      lcorbi_hilic_neg_4_081_58    m/z 209.1147     0   1E+4   3   3.5  RT (min)       lcorbi_hilic_neg_4_091_60    m/z 209.8337     0   2   4   6   8E+3   0.5   1  RT (min)      lcorbi_hilic_neg_4_113_62    m/z 209.8338     0   1   2   3E+4   10   10.5   11  RT (min)       lcorbi_hilic_neg_4_074_58    m/z 209.9488     0   1E+4   1   1.5  RT (min)      lcorbi_hilic_neg_4_067_57    m/z 209.9500     0   1   2E+4   3.5   4  RT (min)       lcorbi_hilic_neg_4_085_59    m/z 209.9927     0   1E+4   2   2.5  RT (min)      lcorbi_hilic_neg_4_101_61    m/z 209.9974     0   1E+4   7.5   8  RT (min)      lcorbi_hilic_neg_4_081_58    m/z 210.0020     0   2   4   6E+4   1   1.5  RT (min)       lcorbi_hilic_neg_4_096_60    m/z 210.0252     0   1   1.5   2  RT (min)       lcorbi_hilic_neg_4_093_60    m/z 210.0341     0   1   2   3   4E+4   6.5   7  RT (min)       lcorbi_hilic_neg_4_090_60    m/z 210.0522     0   1   2   3E+3   2   2.5  RT (min)      lcorbi_hilic_neg_4_097_60    m/z 210.0606     0   1   6.5   7  RT (min)      lcorbi_hilic_neg_4_089_59    m/z 210.0760     0   1E+3   3.5   4  RT (min)      lcorbi_hilic_neg_4_076_58    m/z 210.0770     0   1E+4   1   1.5  RT (min)      lcorbi_hilic_neg_1_039_38    m/z 210.0882     0   1   6.5   7  RT (min)       lcorbi_hilic_neg_3_036_17    m/z 210.0901     0   2   4E+4   4   4.5  RT (min)       lcorbi_hilic_neg_4_090_60    m/z 210.0988     0   1   2E+4   2.5   3  RT (min)       lcorbi_hilic_neg_4_095_60    m/z 210.1579     0   1E+4   0.5   1  RT (min)      lcorbi_hilic_neg_4_108_62    m/z 210.8421     0   1E+4   0   0.5  RT (min)      lcorbi_hilic_neg_4_068_57    m/z 210.9009     0   2   4   6E+4   1   1.5  RT (min)       lcorbi_hilic_neg_4_040_17    m/z 210.9313     0   1   2   3   4E+3   0.5   1  RT (min)      lcorbi_hilic_neg_4_073_57    m/z 210.9381     0   2   4E+3   4.5   5  RT (min)      lcorbi_hilic_neg_1_002_26    m/z 210.9469     0   1E+4   0.5   1  RT (min)      lcorbi_hilic_neg_4_073_57    m/z 210.9499     0   1E+3   8   8.5  RT (min)       lcorbi_hilic_neg_1_025_52    m/z 210.9579     0   1   2   3E+4   3   3.5  RT (min)       lcorbi_hilic_neg_4_085_59    m/z 210.9683     0   1   2   3E+3   4   4.5  RT (min)      lcorbi_hilic_neg_4_086_59    m/z 210.9708     0   1   2   3   4E+5   0   0.5  RT (min)      lcorbi_hilic_neg_1_004_17    m/z 210.9893     0   1   2E+3   1.5   2  RT (min)      lcorbi_hilic_neg_3_010_18    m/z 210.9920     0   1E+3   6   6.5  RT (min)      lcorbi_hilic_neg_3_029_25    m/z 210.9995     0   2   4   6E+5   1   1.5  RT (min)       lcorbi_hilic_neg_4_099_61    m/z 211.0011     0   2   4   6E+2   7   7.5  RT (min)       lcorbi_hilic_neg_1_022_48    m/z 211.0080     0   1   2   3   4E+3   1.5   2  RT (min)      lcorbi_hilic_neg_4_016_29    m/z 211.0182     0   1   2E+4   6   6.5  RT (min)       lcorbi_hilic_neg_4_094_60    m/z 211.0246     0   1   1   1.5  RT (min)      lcorbi_hilic_neg_4_094_60    m/z 211.0361     0   1   2   3E+3   0.5   1  RT (min)      lcorbi_hilic_neg_4_101_61    m/z 211.0464     0   1   4   4.5  RT (min)      lcorbi_hilic_neg_4_105_61    m/z 211.0488     0   1   2   2.5  RT (min)      lcorbi_hilic_neg_4_098_61    m/z 211.0567     0   1E+4   4   4.5  RT (min)      lcorbi_hilic_neg_3_011_24    m/z 211.0683     0   1   2E+4   6.5   7  RT (min)       lcorbi_hilic_neg_4_088_59    m/z 211.0778     0   1E+3   5   5.5   6  RT (min)       lcorbi_hilic_neg_4_026_33    m/z 211.0780     0   1E+4   7   7.5  RT (min)      lcorbi_hilic_neg_4_085_59    m/z 211.0823     0   1E+4   2.5   3  RT (min)      lcorbi_hilic_neg_4_073_57    m/z 211.0836     0   2   4   6   8E+2   7   7.5  RT (min)      lcorbi_hilic_neg_1_012_32    m/z 211.0984     0   1E+4   1   1.5  RT (min)      lcorbi_hilic_neg_4_089_59    m/z 211.8740     0   1E+4   4.5   5  RT (min)      lcorbi_hilic_neg_4_066_57    m/z 211.8851     0   1E+4   6   6.5  RT (min)      lcorbi_hilic_neg_4_075_58    m/z 211.9310     0   2   4   6   8E+3   4.5   5  RT (min)      lcorbi_hilic_neg_1_034_43    m/z 211.9413     0   2   4   6   8E+3   8   8.5  RT (min)      lcorbi_hilic_neg_4_080_58    m/z 211.9478     0   1   2   3E+3   3.5   4  RT (min)      lcorbi_hilic_neg_4_080_58    m/z 211.9705     0   1   2   3E+4   6.5   7  RT (min)       lcorbi_hilic_neg_4_082_59    m/z 211.9766     0   1   3   3.5   4  RT (min)       lcorbi_hilic_neg_4_105_61    m/z 212.0199     0   1   2   3   4E+4   1.5   2  RT (min)       lcorbi_hilic_neg_4_093_60    m/z 212.0329     0   1E+4   6   6.5  RT (min)      lcorbi_hilic_neg_4_076_58    m/z 212.0331     0   2   4   6   8E+3   2.5   3  RT (min)      lcorbi_hilic_neg_4_048_37    m/z 212.0385     0   1   1.5   2  RT (min)      lcorbi_hilic_neg_1_023_23    m/z 212.0741     0   1   3.5   4  RT (min)      lcorbi_hilic_neg_4_088_59    m/z 212.0927     0   1   2   3   4E+4   0.5   1  RT (min)      lcorbi_hilic_neg_4_075_58    m/z 212.1430     0   1   1   1.5  RT (min)      lcorbi_hilic_neg_4_089_59    m/z 212.8393     0   2   4   6E+3   0   0.5  RT (min)      lcorbi_hilic_neg_4_068_57    m/z 212.8394     0   1   2   3E+4   4.5   5  RT (min)       lcorbi_hilic_neg_4_104_61    m/z 212.8772     0   2   4   6E+3   0   0.5  RT (min)      lcorbi_hilic_neg_1_007_10    m/z 212.8773     0   1E+4   4.5   5  RT (min)      lcorbi_hilic_neg_4_103_61    m/z 212.8867     0   2   4   6E+3   7.5   8  RT (min)      lcorbi_hilic_neg_4_103_61    m/z 212.9106     0   1E+4   0   0.5   1  RT (min)      lcorbi_hilic_neg_4_098_61    m/z 212.9162     0   1   4.5   5  RT (min)      lcorbi_hilic_neg_4_074_58    m/z 212.9242     0   1   2   3   4E+3   4.5   5  RT (min)       lcorbi_hilic_neg_4_075_58    m/z 212.9377     0   1E+4   8   8.5  RT (min)      lcorbi_hilic_neg_4_087_59    m/z 212.9440     0   1   0.5   1  RT (min)      lcorbi_hilic_neg_4_090_60    m/z 212.9689     0   1E+3   3.5   4  RT (min)      lcorbi_hilic_neg_1_015_30    m/z 212.9791     0   1E+5   3   3.5  RT (min)       lcorbi_hilic_neg_1_012_32    m/z 213.0049     0   1E+4   1.5   2  RT (min)       lcorbi_hilic_neg_2_013_18    m/z 213.0129     0   1   2E+3   3   3.5  RT (min)       lcorbi_hilic_neg_1_012_32    m/z 213.0133     0   1E+3   6.5   7  RT (min)      lcorbi_hilic_neg_1_008_39    m/z 213.0168     0   1E+4   2.5   3  RT (min)       lcorbi_hilic_neg_4_083_59    m/z 213.0264     0   1E+3   2   2.5  RT (min)      lcorbi_hilic_neg_1_001_49    m/z 213.0372     0   2   4E+2   2   2.5  RT (min)       lcorbi_hilic_neg_1_001_49    m/z 213.0653     0   2   4   6   8E+3   6.5   7  RT (min)      lcorbi_hilic_neg_4_088_59    m/z 213.0709     0   1   2E+4   1.5   2  RT (min)       lcorbi_hilic_neg_4_097_60    m/z 213.0855     0   1E+4   7   7.5  RT (min)      lcorbi_hilic_neg_4_071_57    m/z 213.0879     0   2   4   6E+3   2   2.5  RT (min)      lcorbi_hilic_neg_4_081_58    m/z 213.1130     0   1E+4   1   1.5  RT (min)      lcorbi_hilic_neg_4_084_59    m/z 213.1160     0   2   4   6E+3   5   5.5  RT (min)      lcorbi_hilic_neg_4_105_61    m/z 213.1242     0   1E+4   6.5   7  RT (min)      lcorbi_hilic_neg_4_089_59    m/z 213.1495     0   1E+4   1   1.5  RT (min)      lcorbi_hilic_neg_4_077_58    m/z 213.8711     0   2   4   6E+3   4.5   5  RT (min)      lcorbi_hilic_neg_4_067_57    m/z 213.8797     0   1   2E+4   0   0.5  RT (min)       lcorbi_hilic_neg_4_075_58    m/z 213.9241     0   1   4.5   5  RT (min)       lcorbi_hilic_neg_4_107_62    m/z 213.9678     0   1E+4   6.5   7  RT (min)      lcorbi_hilic_neg_4_083_59    m/z 213.9824     0   1   2   3E+3   0.5   1  RT (min)      lcorbi_hilic_neg_4_076_58    m/z 214.0062     0   2   4   6E+3   1   1.5  RT (min)      lcorbi_hilic_neg_1_015_30    m/z 214.0177     0   2   4   6   8E+3   7.5   8  RT (min)      lcorbi_hilic_neg_4_067_57    m/z 214.0179     0   1E+4   3   3.5  RT (min)      lcorbi_hilic_neg_4_078_58    m/z 214.0217     0   1E+4   4   4.5  RT (min)      lcorbi_hilic_neg_4_080_58    m/z 214.0335     0   2   4   6E+4   2   2.5  RT (min)       lcorbi_hilic_neg_4_080_58    m/z 214.0490     0   2   4E+3   3.5   4  RT (min)      lcorbi_hilic_neg_4_005_32    m/z 214.0508     0   1E+4   0.5   1  RT (min)       lcorbi_hilic_neg_3_024_27    m/z 214.0544     0   1E+4   6.5   7  RT (min)      lcorbi_hilic_neg_3_012_35    m/z 214.0767     0   1E+4   2   2.5  RT (min)      lcorbi_hilic_neg_4_074_58    m/z 214.0830     0   1   4   4.5  RT (min)      lcorbi_hilic_neg_4_085_59    m/z 214.1447     0   2   4   6E+4   1   1.5  RT (min)       lcorbi_hilic_neg_4_034_45    m/z 214.8100     0   1E+5   8   8.5  RT (min)       lcorbi_hilic_neg_4_111_62    m/z 214.8421     0   1   2E+3   3   3.5  RT (min)      lcorbi_hilic_neg_4_004_13    m/z 214.8516     0   1E+3   7.5   8  RT (min)      lcorbi_hilic_neg_4_083_59    m/z 214.8593     0   1E+4   7.5   8  RT (min)      lcorbi_hilic_neg_4_091_60    m/z 214.8747     0   2   4   6E+3   4.5   5  RT (min)       lcorbi_hilic_neg_4_098_61    m/z 214.8796     0   1   2E+4   7   7.5  RT (min)       lcorbi_hilic_neg_4_077_58    m/z 214.9085     0   1E+5   4.5   5  RT (min)       lcorbi_hilic_neg_2_020_27    m/z 214.9249     0   1E+3   4.5   5  RT (min)      lcorbi_hilic_neg_4_005_32    m/z 214.9288     0   1   2   3   4E+4   1   2  RT (min)       lcorbi_hilic_neg_4_082_59    m/z 214.9460     0   1   2E+4   2   2.5  RT (min)       lcorbi_hilic_neg_4_075_58    m/z 214.9526     0   1   7   7.5  RT (min)      lcorbi_hilic_neg_4_078_58    m/z 214.9666     0   1   2E+3   3   3.5   4  RT (min)       lcorbi_hilic_neg_4_075_58    m/z 215.0046     0   1   2E+4   1   1.5  RT (min)       lcorbi_hilic_neg_4_113_62    m/z 215.0101     0   1E+4   5   5.5  RT (min)      lcorbi_hilic_neg_2_020_27    m/z 215.0182     0   1E+3   1   1.5  RT (min)       lcorbi_hilic_neg_4_082_59    m/z 215.0196     0   1   2E+4   5   5.5  RT (min)      lcorbi_hilic_neg_1_004_17    m/z 215.0325     0   2   4   6   8E+4   1.5   2  RT (min)       lcorbi_hilic_neg_3_028_23    m/z 215.0463     0   1E+4   5   5.5  RT (min)      lcorbi_hilic_neg_1_012_32    m/z 215.0512     0   1E+4   0   1   2  RT (min)       lcorbi_hilic_neg_4_081_58    m/z 215.0670     0   1   2E+4   6   6.5  RT (min)       lcorbi_hilic_neg_4_081_58    m/z 215.0707     0   2   4E+3   5   5.5  RT (min)      lcorbi_hilic_neg_4_070_57    m/z 215.0815     0   1   6.5   7  RT (min)      lcorbi_hilic_neg_4_084_59    m/z 215.0826     0   1   4   4.5  RT (min)      lcorbi_hilic_neg_4_086_59    m/z 215.1035     0   1   2E+3   3.5   4  RT (min)      lcorbi_hilic_neg_4_108_62    m/z 215.1147     0   1   5.5   6  RT (min)      lcorbi_hilic_neg_4_113_62    m/z 215.1222     0   2   4   6E+3   4.5   5  RT (min)      lcorbi_hilic_neg_4_110_62    m/z 215.1651     0   1   2   3   4E+4   1   1.5  RT (min)      lcorbi_hilic_neg_1_011_01    m/z 215.8112     0   1E+5   8   8.5  RT (min)       lcorbi_hilic_neg_4_111_62    m/z 215.8637     0   1   8.5   9  RT (min)      lcorbi_hilic_neg_4_071_57    m/z 215.8766     0   2   4   6   8E+3   0   0.5  RT (min)      lcorbi_hilic_neg_4_098_61    m/z 215.8767     0   1   2E+4   7.5   8  RT (min)       lcorbi_hilic_neg_1_033_53    m/z 215.8768     0   1   2   3   4E+4   4.5   5  RT (min)       lcorbi_hilic_neg_4_107_62    m/z 215.9221     0   1   4.5   5  RT (min)      lcorbi_hilic_neg_4_107_62    m/z 215.9365     0   1E+3   6.5   7  RT (min)      lcorbi_hilic_neg_3_002_15    m/z 215.9607     0   1   1.5   2  RT (min)      lcorbi_hilic_neg_4_090_60    m/z 215.9817     0   2   4   6E+3   1   1.5  RT (min)      lcorbi_hilic_neg_4_093_60    m/z 215.9826     0   1   2E+4   7   7.5  RT (min)       lcorbi_hilic_neg_4_090_60    m/z 215.9832     0   1   3   3.5  RT (min)       lcorbi_hilic_neg_4_095_60    m/z 215.9967     0   2   4E+2   6   6.5  RT (min)      lcorbi_hilic_neg_1_027_18    m/z 216.0061     0   1   3.5   4  RT (min)      lcorbi_hilic_neg_4_074_58    m/z 216.0125     0   1   1   1.5  RT (min)      lcorbi_hilic_neg_4_071_57    m/z 216.0131     0   1E+5   6   7  RT (min)       lcorbi_hilic_neg_4_005_32    m/z 216.0236     0   1E+4   6.5   7  RT (min)      lcorbi_hilic_neg_4_089_59    m/z 216.0245     0   1   2   3   4E+3   1   1.5  RT (min)       lcorbi_hilic_neg_4_082_59    m/z 216.0335     0   1E+3   2   2.5  RT (min)       lcorbi_hilic_neg_4_083_59    m/z 216.0415     0   1   2E+3   1   2  RT (min)       lcorbi_hilic_neg_4_067_57    m/z 216.0419     0   1E+4   5   5.5  RT (min)      lcorbi_hilic_neg_2_020_27    m/z 216.0525     0   1   6.5   7  RT (min)      lcorbi_hilic_neg_4_090_60    m/z 216.0698     0   1E+4   3.5   4  RT (min)      lcorbi_hilic_neg_4_096_60    m/z 216.0876     0   1E+3   2   2.5  RT (min)      lcorbi_hilic_neg_4_068_57    m/z 216.0897     0   1   2   3   4E+4   5   5.5  RT (min)       lcorbi_hilic_neg_4_097_60    m/z 216.0986     0   1   2   3   4E+3   4   4.5  RT (min)      lcorbi_hilic_neg_4_075_58    m/z 216.0988     0   1   2   3E+3   6.5   7  RT (min)      lcorbi_hilic_neg_4_097_60    m/z 216.1351     0   2   4   6   8E+3   7.5   8  RT (min)      lcorbi_hilic_neg_4_077_58    m/z 216.1439     0   1   2   3   4E+4   1   1.5  RT (min)       lcorbi_hilic_neg_4_090_60    m/z 216.8094     0   1   2   3E+5   8   8.5  RT (min)       lcorbi_hilic_neg_4_111_62    m/z 216.8400     0   1   2   3E+3   3   3.5  RT (min)      lcorbi_hilic_neg_4_071_57    m/z 216.8766     0   1E+4   6   6.5  RT (min)      lcorbi_hilic_neg_4_075_58    m/z 216.8769     0   1E+4   1   1.5  RT (min)      lcorbi_hilic_neg_4_097_60    m/z 216.8891     0   2   4   6E+3   8.5   9  RT (min)      lcorbi_hilic_neg_4_078_58    m/z 216.9055     0   1   4.5   5  RT (min)       lcorbi_hilic_neg_4_098_61    m/z 216.9092     0   1   2E+4   0   0.5  RT (min)      lcorbi_hilic_neg_3_010_18    m/z 216.9154     0   2   4E+3   2   2.5  RT (min)      lcorbi_hilic_neg_4_092_60    m/z 216.9236     0   1E+5   7.5   8  RT (min)       lcorbi_hilic_neg_2_005_24    m/z 216.9283     0   1   2   3E+4   0.5   1  RT (min)      lcorbi_hilic_neg_4_099_61    m/z 216.9810     0   1   5.5   6  RT (min)      lcorbi_hilic_neg_4_070_57    m/z 217.0133     0   1   2E+3   3   3.5  RT (min)       lcorbi_hilic_neg_2_045_26    m/z 217.0197     0   1E+4   5   5.5  RT (min)      lcorbi_hilic_neg_2_020_27    m/z 217.0204     0   2   4   6E+3   8   8.5  RT (min)      lcorbi_hilic_neg_4_080_58    m/z 217.0462     0   1   1.5   2  RT (min)      lcorbi_hilic_neg_4_084_59    m/z 217.0507     0   1   2E+5   1.5   2  RT (min)       lcorbi_hilic_neg_4_108_62    m/z 217.0577     0   1   7   7.5  RT (min)      lcorbi_hilic_neg_4_091_60    m/z 217.0616     0   1   3   3.5  RT (min)      lcorbi_hilic_neg_4_106_62    m/z 217.0941     0   1   7   7.5  RT (min)       lcorbi_hilic_neg_4_091_60    m/z 217.1081     0   1   2   3   4E+3   1   1.5  RT (min)      lcorbi_hilic_neg_4_107_62    m/z 217.1190     0   1   1   1.5  RT (min)       lcorbi_hilic_neg_1_046_19    m/z 217.1191     0   2   4   6   8E+3   7.5   8  RT (min)      lcorbi_hilic_neg_4_089_59    m/z 217.1444     0   1   2E+3   1   1.5  RT (min)      lcorbi_hilic_neg_1_031_20    m/z 217.7790     0   1   2   3   4E+4   3   3.5  RT (min)       lcorbi_hilic_neg_4_067_57    m/z 217.8099     0   2   4   6   8E+4   8   8.5  RT (min)       lcorbi_hilic_neg_4_113_62    m/z 217.8500     0   2   4E+4   8.5   9  RT (min)       lcorbi_hilic_neg_4_066_57    m/z 217.8500     0   1   2E+4   4.5   5  RT (min)       lcorbi_hilic_neg_4_103_61    m/z 217.8537     0   1E+4   1   1.5  RT (min)      lcorbi_hilic_neg_4_088_59    m/z 217.8654     0   1   2   3E+4   8.5   9  RT (min)       lcorbi_hilic_neg_4_074_58    m/z 217.8799     0   1E+4   1   1.5  RT (min)      lcorbi_hilic_neg_4_106_62    m/z 217.9242     0   2   4   6   8E+2   8   8.5  RT (min)      lcorbi_hilic_neg_1_003_14    m/z 217.9312     0   2   4   6   8E+3   3   3.5  RT (min)      lcorbi_hilic_neg_4_085_59    m/z 217.9442     0   1   1.5   2  RT (min)      lcorbi_hilic_neg_4_090_60    m/z 217.9522     0   2   4   6   8E+3   7   7.5  RT (min)      lcorbi_hilic_neg_4_078_58    m/z 217.9619     0   1   2E+3   1.5   2  RT (min)      lcorbi_hilic_neg_4_090_60    m/z 217.9707     0   1   2   3E+4   1.5   2  RT (min)       lcorbi_hilic_neg_4_082_59    m/z 217.9779     0   1E+4   1   1.5  RT (min)      lcorbi_hilic_neg_4_068_57    m/z 217.9805     0   1E+3   2.5   3  RT (min)      lcorbi_hilic_neg_4_092_60    m/z 218.0021     0   2   4   6   8E+2   5   5.5  RT (min)       lcorbi_hilic_neg_2_020_27    m/z 218.0128     0   1   1   1.5  RT (min)      lcorbi_hilic_neg_4_093_60    m/z 218.0138     0   1E+4   2   2.5  RT (min)      lcorbi_hilic_neg_4_066_57    m/z 218.0163     0   1   6   6.5  RT (min)       lcorbi_hilic_neg_4_090_60    m/z 218.0207     0   1   0.5   1  RT (min)      lcorbi_hilic_neg_4_092_60    m/z 218.0396     0   2   4   6   8E+3   5   6  RT (min)       lcorbi_hilic_neg_1_048_35    m/z 218.0513     0   1E+4   5   5.5  RT (min)      lcorbi_hilic_neg_4_070_57    m/z 218.0619     0   1   2   3E+4   6.5   7  RT (min)       lcorbi_hilic_neg_4_091_60    m/z 218.0781     0   1   6.5   7  RT (min)       lcorbi_hilic_neg_4_097_60    m/z 218.0821     0   1E+3   1   1.5  RT (min)      lcorbi_hilic_neg_4_108_62    m/z 218.1013     0   1   4   4.5  RT (min)      lcorbi_hilic_neg_2_005_24    m/z 218.1055     0   2   4E+4   5.5   6  RT (min)       lcorbi_hilic_neg_4_097_60    m/z 218.1506     0   1E+4   1   1.5  RT (min)      lcorbi_hilic_neg_4_090_60    m/z 218.7896     0   1E+4   2.5   3  RT (min)      lcorbi_hilic_neg_4_066_57    m/z 218.8523     0   2   4   6E+3   7.5   8  RT (min)      lcorbi_hilic_neg_4_072_57    m/z 218.8671     0   2   4E+3   0.5   1  RT (min)      lcorbi_hilic_neg_4_112_62    m/z 218.8738     0   1   2E+4   5   5.5  RT (min)      lcorbi_hilic_neg_4_110_62    m/z 218.9026     0   2   4E+4   4.5   5  RT (min)      lcorbi_hilic_neg_2_020_27    m/z 218.9051     0   1E+4   7.5   8  RT (min)       lcorbi_hilic_neg_4_109_62    m/z 218.9135     0   1   2E+3   2.5   3  RT (min)      lcorbi_hilic_neg_4_106_62    m/z 218.9632     0   1   2   3   4E+3   0.5   1  RT (min)      lcorbi_hilic_neg_4_077_58    m/z 218.9883     0   1E+4   0.5   1  RT (min)       lcorbi_hilic_neg_4_002_41    m/z 218.9883     0   1   2   3E+4   3   3.5  RT (min)      lcorbi_hilic_neg_4_086_59    m/z 218.9942     0   1   6.5   7  RT (min)      lcorbi_hilic_neg_4_005_32    m/z 219.0048     0   2   4   6   8E+3   1   1.5  RT (min)       lcorbi_hilic_neg_4_083_59    m/z 219.0099     0   1   2E+4   2   2.5  RT (min)       lcorbi_hilic_neg_4_090_60    m/z 219.0175     0   1   6.5   7  RT (min)      lcorbi_hilic_neg_4_072_57    m/z 219.0371     0   1   2   3E+3   2  RT (min)       lcorbi_hilic_neg_4_083_59    m/z 219.0540     0   1   2   3   4E+4   1   1.5  RT (min)      lcorbi_hilic_neg_4_107_62    m/z 219.0662     0   1   2E+5   4   5   6  RT (min)       lcorbi_hilic_neg_4_113_62    m/z 219.0861     0   1E+4   1   1.5  RT (min)      lcorbi_hilic_neg_4_107_62    m/z 219.0980     0   1   2   3E+3   0.5   1  RT (min)      lcorbi_hilic_neg_3_040_19    m/z 219.1104     0   1   7   7.5  RT (min)       lcorbi_hilic_neg_4_106_62    m/z 219.8084     0   1E+3   8   8.5  RT (min)       lcorbi_hilic_neg_1_004_17    m/z 219.8407     0   2   4E+3   4.5   5  RT (min)      lcorbi_hilic_neg_4_066_57    m/z 219.8454     0   2   4   6   8E+4   13   13.5  RT (min)      lcorbi_hilic_neg_4_090_60    m/z 219.8515     0   1   2   3   4E+3   8.5   9  RT (min)      lcorbi_hilic_neg_1_010_22    m/z 219.9425     0   1E+3   1.5   2   2.5  RT (min)       lcorbi_hilic_neg_1_018_21    m/z 219.9500     0   1   2E+4   6   6.5  RT (min)      lcorbi_hilic_neg_3_044_32    m/z 219.9545     0   2   4E+4   1.5   2  RT (min)       lcorbi_hilic_neg_3_007_07    m/z 219.9627     0   2   4E+3   6   6.5  RT (min)      lcorbi_hilic_neg_4_095_60    m/z 219.9685     0   1   2   3E+3   2   2.5  RT (min)      lcorbi_hilic_neg_4_106_62    m/z 219.9815     0   1   2E+4   4   4.5  RT (min)      lcorbi_hilic_neg_4_091_60    m/z 219.9916     0   1E+4   5.5   6  RT (min)       lcorbi_hilic_neg_4_103_61    m/z 219.9959     0   2   4   6   8E+3   7   7.5  RT (min)      lcorbi_hilic_neg_4_091_60    m/z 219.9992     0   1   5   5.5  RT (min)      lcorbi_hilic_neg_4_005_32    m/z 220.0131     0   1E+5   3   4  RT (min)       lcorbi_hilic_neg_4_005_32    m/z 220.0170     0   1   6   6.5  RT (min)      lcorbi_hilic_neg_4_093_60    m/z 220.0276     0   1   7.5   8  RT (min)      lcorbi_hilic_neg_4_096_60    m/z 220.0368     0   1   5   5.5  RT (min)      lcorbi_hilic_neg_4_092_60    m/z 220.0369     0   1E+4   6.5   7  RT (min)      lcorbi_hilic_neg_4_097_60    m/z 220.0487     0   2   4   6   8E+3   5   5.5  RT (min)      lcorbi_hilic_neg_1_055_24    m/z 220.0757     0   1E+4   1   1.5  RT (min)      lcorbi_hilic_neg_1_039_38    m/z 220.0839     0   1E+3   3   3.5  RT (min)      lcorbi_hilic_neg_4_107_62    m/z 220.7730     0   1   2   3   4E+3   3   3.5  RT (min)      lcorbi_hilic_neg_2_040_35    m/z 220.8073     0   1   2E+5   8   8.5  RT (min)       lcorbi_hilic_neg_4_111_62    m/z 220.8451     0   1E+4   0   0.5  RT (min)      lcorbi_hilic_neg_4_098_61    m/z 220.8452     0   2   4E+4   8.5   9  RT (min)       lcorbi_hilic_neg_4_066_57    m/z 220.8452     0   2   4E+4   4.5   5  RT (min)       lcorbi_hilic_neg_4_106_62    m/z 220.8709     0   1   2E+4   5   5.5  RT (min)       lcorbi_hilic_neg_4_106_62    m/z 220.8984     0   2   4   6   8E+3   1   1.5  RT (min)      lcorbi_hilic_neg_4_106_62    m/z 220.9600     0   1E+3   7   7.5  RT (min)      lcorbi_hilic_neg_4_067_57    m/z 220.9707     0   1   2E+4   6.5   7  RT (min)      lcorbi_hilic_neg_4_075_58    m/z 220.9709     0   1   4.5   5  RT (min)      lcorbi_hilic_neg_1_051_33    m/z 220.9879     0   1E+3   6.5   7  RT (min)      lcorbi_hilic_neg_4_107_62    m/z 220.9927     0   2   4E+3   2   2.5  RT (min)      lcorbi_hilic_neg_4_107_62    m/z 221.0038     0   1   2E+4   1   1.5  RT (min)       lcorbi_hilic_neg_4_090_60    m/z 221.0209     0   1E+4   7   7.5  RT (min)      lcorbi_hilic_neg_4_090_60    m/z 221.0305     0   2   4   6   8E+3   5.5   6  RT (min)      lcorbi_hilic_neg_4_088_59    m/z 221.0341     0   1   2   3   4E+3   1.5   2   2.5  RT (min)       lcorbi_hilic_neg_4_098_61    m/z 221.0529     0   1E+5   1   1.5  RT (min)       lcorbi_hilic_neg_3_011_24    m/z 221.0818     0   1   2   3E+3   1   1.5  RT (min)       lcorbi_hilic_neg_1_010_22    m/z 221.1060     0   1   4   4.5  RT (min)      lcorbi_hilic_neg_4_092_60    m/z 221.1462     0   2   4E+4   0.5   1  RT (min)      lcorbi_hilic_neg_1_046_19    m/z 221.1607     0   2   4   6E+4   0.5   1  RT (min)       lcorbi_hilic_neg_4_111_62    m/z 221.8003     0   2   4   6E+4   1   1.5  RT (min)      lcorbi_hilic_neg_3_038_21    m/z 221.8424     0   2   4E+4   13   13.5  RT (min)      lcorbi_hilic_neg_4_077_58    m/z 221.8485     0   1E+4   8.5   9  RT (min)      lcorbi_hilic_neg_4_066_57    m/z 221.8488     0   1E+4   4.5   5  RT (min)      lcorbi_hilic_neg_4_066_57    m/z 221.8744     0   2   4E+4   1   1.5  RT (min)       lcorbi_hilic_neg_4_097_60    m/z 221.9026     0   1   2E+4   7.5   8  RT (min)      lcorbi_hilic_neg_1_003_14    m/z 221.9027     0   1   2   3E+4   4   4.5  RT (min)       lcorbi_hilic_neg_4_101_61    m/z 221.9027     0   2   4   6   8E+4   0   0.5  RT (min)       lcorbi_hilic_neg_4_066_57    m/z 221.9444     0   1   0.5   1  RT (min)      lcorbi_hilic_neg_4_090_60    m/z 221.9621     0   1   1   1.5  RT (min)      lcorbi_hilic_neg_4_090_60    m/z 221.9770     0   2   4E+4   3   3.5  RT (min)      lcorbi_hilic_neg_2_040_35    m/z 221.9841     0   1   2   3   4E+4   1.5   2  RT (min)       lcorbi_hilic_neg_4_096_60    m/z 221.9893     0   1   2E+4   6   6.5  RT (min)       lcorbi_hilic_neg_4_096_60    m/z 222.0096     0   2   4   6   8E+3   2.5   3   3.5  RT (min)      lcorbi_hilic_neg_3_044_32    m/z 222.0252     0   2   4   6   8E+3   2   2.5  RT (min)      lcorbi_hilic_neg_4_082_59    m/z 222.0301     0   1   5   5.5  RT (min)      lcorbi_hilic_neg_4_095_60    m/z 222.0463     0   1   2E+4   1.5   2  RT (min)       lcorbi_hilic_neg_4_096_60    m/z 222.0631     0   2   4   6   8E+3   3.5   4  RT (min)      lcorbi_hilic_neg_4_094_60    m/z 222.0634     0   1   2E+4   7.5   8  RT (min)      lcorbi_hilic_neg_4_083_59    m/z 222.0769     0   2   4   6   8E+3   1.5   2  RT (min)      lcorbi_hilic_neg_4_073_57    m/z 222.0811     0   1E+4   5.5   6  RT (min)      lcorbi_hilic_neg_4_066_57    m/z 222.0981     0   1E+4   6   6.5  RT (min)      lcorbi_hilic_neg_4_087_59    m/z 222.1134     0   1E+4   1   1.5  RT (min)      lcorbi_hilic_neg_4_005_32    m/z 222.7568     0   1E+4   3   3.5  RT (min)      lcorbi_hilic_neg_3_012_35    m/z 222.7721     0   1E+4   3   3.5  RT (min)      lcorbi_hilic_neg_4_073_57    m/z 222.8045     0   2   4   6E+4   8   8.5  RT (min)       lcorbi_hilic_neg_4_112_62    m/z 222.8296     0   1   2   3   4E+4   1   1.5  RT (min)       lcorbi_hilic_neg_4_097_60    m/z 222.8410     0   2   4E+3   0.5   1  RT (min)      lcorbi_hilic_neg_4_076_58    m/z 222.8422     0   1   2   3E+4   8.5   9   9.5  RT (min)       lcorbi_hilic_neg_4_066_57    m/z 222.8423     0   1   2   3E+4   4.5   5  RT (min)       lcorbi_hilic_neg_4_107_62    m/z 222.8716     0   1E+3   0   0.5  RT (min)      lcorbi_hilic_neg_4_066_57    m/z 222.8986     0   1   2   3   4E+3   0.5   1  RT (min)      lcorbi_hilic_neg_4_072_57    m/z 222.8989     0   1E+4   8.5   9  RT (min)      lcorbi_hilic_neg_4_103_61    m/z 222.9007     0   1E+4   4.5   5  RT (min)      lcorbi_hilic_neg_4_073_57    m/z 222.9154     0   2   4   6E+3   7.5   8  RT (min)      lcorbi_hilic_neg_4_070_57    m/z 222.9577     0   1   2E+4   6   6.5  RT (min)       lcorbi_hilic_neg_4_073_57    m/z 222.9680     0   1   4.5   5  RT (min)      lcorbi_hilic_neg_4_005_32    m/z 222.9702     0   1   1   1.5  RT (min)      lcorbi_hilic_neg_4_076_58    m/z 222.9800     0   1   1   1.5  RT (min)       lcorbi_hilic_neg_4_070_57    m/z 222.9994     0   1   2   3E+4   0.5   1  RT (min)       lcorbi_hilic_neg_4_097_60    m/z 222.9994     0   1   2E+4   3   3.5  RT (min)      lcorbi_hilic_neg_4_080_58    m/z 223.0086     0   1   2E+3   2   2.5  RT (min)      lcorbi_hilic_neg_3_023_36    m/z 223.0197     0   1   2E+4   3   3.5  RT (min)       lcorbi_hilic_neg_4_072_57    m/z 223.0378     0   2   4E+3   1.5   2  RT (min)      lcorbi_hilic_neg_4_075_58    m/z 223.0459     0   1E+3   3   3.5  RT (min)      lcorbi_hilic_neg_4_091_60    m/z 223.0497     0   1E+5   1.5   2  RT (min)       lcorbi_hilic_neg_4_093_60    m/z 223.0558     0   1E+4   7.5   8  RT (min)      lcorbi_hilic_neg_4_089_59    m/z 223.0684     0   1   2E+3   5   5.5  RT (min)       lcorbi_hilic_neg_4_005_32    m/z 223.0974     0   2   4   6   8E+3   1   1.5  RT (min)       lcorbi_hilic_neg_4_106_62    m/z 223.1086     0   2   4   6E+2   2.5   3  RT (min)      lcorbi_hilic_neg_4_013_46    m/z 223.1110     0   1   1   1.5  RT (min)      lcorbi_hilic_neg_2_020_27    m/z 223.1702     0   2   4E+3   0.5   1  RT (min)      lcorbi_hilic_neg_4_108_62    m/z 223.1703     0   1   2   3E+3   2.5   3  RT (min)      lcorbi_hilic_neg_4_078_58    m/z 223.7982     0   1   2   3E+4   12   13  RT (min)       lcorbi_hilic_neg_4_096_60    m/z 223.8393     0   1   2   3E+4   0   0.5  RT (min)       lcorbi_hilic_neg_4_098_61    m/z 223.8394     0   1E+5   4.5   5  RT (min)       lcorbi_hilic_neg_4_107_62    m/z 223.8394     0   1E+5   9   10  RT (min)       lcorbi_hilic_neg_4_066_57    m/z 223.8723     0   1   2   3E+4   1   1.5  RT (min)       lcorbi_hilic_neg_4_097_60    m/z 223.8999     0   1   2E+4   0   0.5  RT (min)      lcorbi_hilic_neg_4_066_57    m/z 223.9416     0   1   0.5   1  RT (min)      lcorbi_hilic_neg_4_090_60    m/z 223.9542     0   2   4   6   8E+3   2   2.5  RT (min)      lcorbi_hilic_neg_4_100_61    m/z 223.9656     0   1   2   3   4E+3   3   3.5  RT (min)      lcorbi_hilic_neg_3_023_36    m/z 223.9749     0   1   2   3   4E+4   3   3.5  RT (min)      lcorbi_hilic_neg_2_040_35    m/z 223.9941     0   1   6   6.5  RT (min)      lcorbi_hilic_neg_4_090_60    m/z 223.9952     0   1   2   2.5  RT (min)      lcorbi_hilic_neg_4_107_62    m/z 224.0111     0   2   4   6   8E+3   6   6.5  RT (min)      lcorbi_hilic_neg_4_093_60    m/z 224.0278     0   2   4   6E+4   0.5   1  RT (min)      lcorbi_hilic_neg_4_113_62    m/z 224.0303     0   1   2   3E+4   4.5   5  RT (min)       lcorbi_hilic_neg_4_094_60    m/z 224.0401     0   1E+4   6   6.5  RT (min)      lcorbi_hilic_neg_4_096_60    m/z 224.0411     0   1E+3   1.5   2  RT (min)      lcorbi_hilic_neg_4_091_60    m/z 224.0561     0   2   4   6E+4   1.5   2  RT (min)       lcorbi_hilic_neg_4_082_59    m/z 224.0644     0   1E+4   1   1.5   2  RT (min)       lcorbi_hilic_neg_4_095_60    m/z 224.0682     0   1E+3   3   3.5  RT (min)      lcorbi_hilic_neg_4_070_57    m/z 224.0764     0   1   2   3E+4   1.5   2  RT (min)       lcorbi_hilic_neg_4_083_59    m/z 224.0855     0   1E+4   2.5   3  RT (min)      lcorbi_hilic_neg_1_037_46    m/z 224.0926     0   1   2   3   4E+3   1.5   2  RT (min)      lcorbi_hilic_neg_4_089_59    m/z 224.1249     0   1   2E+4   4   4.5  RT (min)       lcorbi_hilic_neg_4_095_60    m/z 224.1336     0   1   3.5   4  RT (min)      lcorbi_hilic_neg_4_066_57    m/z 224.1370     0   1E+3   1   1.5  RT (min)      lcorbi_hilic_neg_4_094_60    m/z 224.7568     0   2   4   6   8E+3   3   3.5  RT (min)      lcorbi_hilic_neg_3_012_35    m/z 224.7692     0   2   4   6   8E+3   3   3.5  RT (min)      lcorbi_hilic_neg_4_023_34    m/z 224.7891     0   1E+3   3   3.5  RT (min)      lcorbi_hilic_neg_3_039_31    m/z 224.8191     0   1E+4   4.5   5  RT (min)      lcorbi_hilic_neg_4_099_61    m/z 224.8192     0   2   4   6   8E+3   0.5   1   1.5  RT (min)       lcorbi_hilic_neg_4_068_57    m/z 224.9086     0   1   2   3   4E+3   1   1.5  RT (min)      lcorbi_hilic_neg_4_097_60    m/z 224.9242     0   1E+4   8   8.5  RT (min)      lcorbi_hilic_neg_4_089_59    m/z 224.9353     0   1E+4   0.5   1  RT (min)      lcorbi_hilic_neg_4_078_58    m/z 224.9506     0   1E+4   2   2.5  RT (min)       lcorbi_hilic_neg_4_083_59    m/z 224.9547     0   1   6   6.5  RT (min)      lcorbi_hilic_neg_1_003_14    m/z 224.9748     0   1   2   3   4E+3   1   1.5  RT (min)      lcorbi_hilic_neg_4_113_62    m/z 224.9877     0   1E+3   2   2.5  RT (min)      lcorbi_hilic_neg_4_107_62    m/z 225.0240     0   1   4   4.5  RT (min)       lcorbi_hilic_neg_4_089_59    m/z 225.0242     0   1   2E+3   2.5   3  RT (min)      lcorbi_hilic_neg_4_097_60    m/z 225.0249     0   2   4E+4   0.5   1  RT (min)      lcorbi_hilic_neg_4_113_62    m/z 225.0361     0   1E+4   1   1.5  RT (min)      lcorbi_hilic_neg_4_094_60    m/z 225.0491     0   1   6.5   7  RT (min)      lcorbi_hilic_neg_3_049_55    m/z 225.0532     0   1   2E+3   1.5   2  RT (min)      lcorbi_hilic_neg_2_054_13    m/z 225.0553     0   1   2   3E+4   5   5.5  RT (min)      lcorbi_hilic_neg_2_020_27    m/z 225.0721     0   1E+4   5   5.5  RT (min)      lcorbi_hilic_neg_2_012_14    m/z 225.0766     0   1E+3   1   1.5  RT (min)      lcorbi_hilic_neg_4_097_60    m/z 225.0879     0   1   2   3   4E+2   1.5   2  RT (min)      lcorbi_hilic_neg_1_020_13    m/z 225.0978     0   1   2   2.5  RT (min)      lcorbi_hilic_neg_2_013_18    m/z 225.0992     0   1   2   3E+4   7   7.5  RT (min)       lcorbi_hilic_neg_4_078_58    m/z 225.1097     0   1   3.5   4  RT (min)       lcorbi_hilic_neg_4_068_57    m/z 225.1097     0   1   2   3E+4   6.5   7  RT (min)       lcorbi_hilic_neg_4_090_60    m/z 225.1298     0   1   2   3   4E+4   7   7.5  RT (min)       lcorbi_hilic_neg_4_096_60    m/z 225.1494     0   1E+4   0.5   1  RT (min)      lcorbi_hilic_neg_4_089_59    m/z 225.1606     0   1   2   3E+4   1.5   2  RT (min)       lcorbi_hilic_neg_4_084_59    m/z 225.1859     0   1   2   3   4E+3   2.5   3  RT (min)      lcorbi_hilic_neg_4_069_57    m/z 225.2221     0   1E+4   1   1.5  RT (min)      lcorbi_hilic_neg_4_077_58    m/z 225.5613     0   1E+4   3   3.5  RT (min)      lcorbi_hilic_neg_4_071_57    m/z 225.7725     0   1E+4   3   3.5  RT (min)      lcorbi_hilic_neg_4_066_57    m/z 225.7958     0   2   4E+4   1   1.5  RT (min)      lcorbi_hilic_neg_3_026_09    m/z 225.8308     0   2   4   6E+3   8   8.5  RT (min)      lcorbi_hilic_neg_4_074_58    m/z 225.8972     0   1E+4   8.5   9  RT (min)      lcorbi_hilic_neg_4_075_58    m/z 225.9084     0   1E+4   3   3.5  RT (min)      lcorbi_hilic_neg_4_105_61    m/z 225.9264     0   1E+4   1   1.5  RT (min)      lcorbi_hilic_neg_4_070_57    m/z 225.9338     0   1   3   3.5  RT (min)      lcorbi_hilic_neg_4_104_61    m/z 225.9661     0   1   2   3E+3   3   3.5  RT (min)      lcorbi_hilic_neg_4_082_59    m/z 226.0070     0   1   2   3   4E+4   0.5   1  RT (min)       lcorbi_hilic_neg_4_113_62    m/z 226.0121     0   1   2   3   4E+4   2.5   3  RT (min)       lcorbi_hilic_neg_4_083_59    m/z 226.0128     0   1   2   3E+4   7   7.5  RT (min)       lcorbi_hilic_neg_4_090_60    m/z 226.0363     0   2   4   6E+3   7   7.5  RT (min)      lcorbi_hilic_neg_4_071_57    m/z 226.0486     0   1   1   1.5  RT (min)      lcorbi_hilic_neg_4_107_62    m/z 226.0567     0   1   2   3   4E+4   3.5   4  RT (min)      lcorbi_hilic_neg_4_068_57    m/z 226.0660     0   2   4   6   8E+3   1.5   2  RT (min)      lcorbi_hilic_neg_4_045_25    m/z 226.0674     0   1E+4   3.5   4  RT (min)      lcorbi_hilic_neg_4_068_57    m/z 226.0721     0   2   4   6   8E+2   3.5   4  RT (min)      lcorbi_hilic_neg_1_037_46    m/z 226.0856     0   1E+4   6.5   7  RT (min)      lcorbi_hilic_neg_4_091_60    m/z 226.1055     0   1   6   6.5  RT (min)      lcorbi_hilic_neg_4_070_57    m/z 226.1196     0   1   2E+4   6.5   7  RT (min)       lcorbi_hilic_neg_4_086_59    m/z 226.1448     0   1E+4   4.5   5  RT (min)      lcorbi_hilic_neg_4_096_60    m/z 226.1493     0   1E+4   7.5   8  RT (min)      lcorbi_hilic_neg_4_094_60    m/z 226.8161     0   1E+4   0.5   1  RT (min)      lcorbi_hilic_neg_4_068_57    m/z 226.8162     0   1   2E+4   4.5   5  RT (min)       lcorbi_hilic_neg_4_104_61    m/z 226.8240     0   1E+4   1   1.5  RT (min)      lcorbi_hilic_neg_4_097_60    m/z 226.9023     0   2   4   6E+3   6   6.5  RT (min)      lcorbi_hilic_neg_4_091_60    m/z 226.9272     0   1E+4   8.5   9  RT (min)       lcorbi_hilic_neg_4_081_58    m/z 226.9444     0   1   2   3   4E+3   2   2.5  RT (min)      lcorbi_hilic_neg_4_093_60    m/z 226.9765     0   1E+3   5.5   6  RT (min)      lcorbi_hilic_neg_4_093_60    m/z 226.9782     0   2   4   6   8E+3   1   1.5  RT (min)      lcorbi_hilic_neg_4_091_60    m/z 226.9844     0   1E+3   1.5   2  RT (min)      lcorbi_hilic_neg_4_085_59    m/z 227.0116     0   1E+4   0.5   1  RT (min)       lcorbi_hilic_neg_4_097_60    m/z 227.0211     0   1E+3   1   1.5  RT (min)      lcorbi_hilic_neg_4_085_59    m/z 227.0261     0   1E+3   4  RT (min)       lcorbi_hilic_neg_4_082_59    m/z 227.0325     0   1   2E+3   3   3.5   4  RT (min)       lcorbi_hilic_neg_4_092_60    m/z 227.0343     0   1   2E+3   1   1.5  RT (min)       lcorbi_hilic_neg_4_084_59    m/z 227.0438     0   1E+3   4.5   5  RT (min)       lcorbi_hilic_neg_4_098_61    m/z 227.0593     0   1   2   3E+3   1.5   2  RT (min)      lcorbi_hilic_neg_2_055_32    m/z 227.0672     0   1E+4   2   2.5  RT (min)      lcorbi_hilic_neg_4_089_59    m/z 227.0769     0   1   2E+4   4.5   5  RT (min)       lcorbi_hilic_neg_4_079_58    m/z 227.0801     0   1   2E+4   7.5   8  RT (min)       lcorbi_hilic_neg_4_097_60    m/z 227.0922     0   2   4   6E+3   1   1.5  RT (min)      lcorbi_hilic_neg_4_073_57    m/z 227.1035     0   1   2E+4   6   6.5  RT (min)       lcorbi_hilic_neg_4_068_57    m/z 227.1036     0   1   2.5   3  RT (min)      lcorbi_hilic_neg_4_108_62    m/z 227.1147     0   2   4   6E+2   6.5   7  RT (min)      lcorbi_hilic_neg_4_030_05    m/z 227.1399     0   1   4.5   5  RT (min)      lcorbi_hilic_neg_4_080_58    m/z 227.7705     0   1   2   3E+4   3   3.5  RT (min)       lcorbi_hilic_neg_4_066_57    m/z 227.8279     0   1   2   3E+3   8   8.5  RT (min)      lcorbi_hilic_neg_4_082_59    m/z 227.9062     0   1E+4   3   3.5  RT (min)      lcorbi_hilic_neg_4_103_61    m/z 228.0123     0   1E+3   1   1.5  RT (min)      lcorbi_hilic_neg_4_106_62    m/z 228.0275     0   1   2E+4   7.5   8  RT (min)       lcorbi_hilic_neg_4_089_59    m/z 228.0414     0   1E+3   0.5   1  RT (min)      lcorbi_hilic_neg_4_069_57    m/z 228.0487     0   2   4   6E+3   1.5   2  RT (min)      lcorbi_hilic_neg_4_081_58    m/z 228.0489     0   1E+4   5   5.5  RT (min)       lcorbi_hilic_neg_3_037_12    m/z 228.0641     0   1   6.5   7   7.5  RT (min)       lcorbi_hilic_neg_4_091_60    m/z 228.0900     0   1E+4   1.5   2  RT (min)      lcorbi_hilic_neg_4_091_60    m/z 228.0988     0   1E+5   6   6.5   7  RT (min)       lcorbi_hilic_neg_3_028_23    m/z 228.1319     0   1   2   3   4E+3   1   1.5  RT (min)      lcorbi_hilic_neg_4_089_59    m/z 228.1603     0   1   2   3E+4   1   1.5  RT (min)       lcorbi_hilic_neg_4_087_59    m/z 228.7985     0   1E+4   1   1.5  RT (min)      lcorbi_hilic_neg_1_001_49    m/z 228.8130     0   1E+4   4.5   5  RT (min)      lcorbi_hilic_neg_4_070_57    m/z 228.8134     0   2   4   6E+3   0.5   1  RT (min)      lcorbi_hilic_neg_4_074_58    m/z 228.9026     0   1   2   3   4E+4   5.5   6  RT (min)       lcorbi_hilic_neg_4_106_62    m/z 228.9027     0   2   4   6E+3   0   0.5  RT (min)      lcorbi_hilic_neg_4_068_57    m/z 228.9376     0   1E+4   8   8.5  RT (min)       lcorbi_hilic_neg_4_093_60    m/z 228.9593     0   1   2   3E+4   0.5   1  RT (min)      lcorbi_hilic_neg_4_104_61    m/z 228.9607     0   1E+3   3   3.5  RT (min)      lcorbi_hilic_neg_2_027_02    m/z 228.9609     0   1E+3   6   6.5  RT (min)      lcorbi_hilic_neg_4_076_58    m/z 228.9899     0   1   2   3   4E+4   0.5   1  RT (min)       lcorbi_hilic_neg_4_076_58    m/z 229.0002     0   1   1   1.5  RT (min)      lcorbi_hilic_neg_4_084_59    m/z 229.0095     0   2   4   6E+3   0.5   1  RT (min)      lcorbi_hilic_neg_4_007_21    m/z 229.0116     0   1   2E+4   6.5   7  RT (min)      lcorbi_hilic_neg_4_089_59    m/z 229.0230     0   1E+4   6.5   7  RT (min)      lcorbi_hilic_neg_4_073_57    m/z 229.0230     0   1E+5   4.5   5  RT (min)       lcorbi_hilic_neg_3_011_24    m/z 229.0295     0   1   2E+3   3.5   4  RT (min)       lcorbi_hilic_neg_4_093_60    m/z 229.0313     0   1   2E+3   1   1.5  RT (min)       lcorbi_hilic_neg_3_017_51    m/z 229.0659     0   1   2E+5   1.5   2  RT (min)       lcorbi_hilic_neg_4_094_60    m/z 229.0827     0   1E+4   5.5   6  RT (min)      lcorbi_hilic_neg_4_088_59    m/z 229.0889     0   1   1.5   2  RT (min)      lcorbi_hilic_neg_4_091_60    m/z 229.0958     0   1   2E+4   3   4  RT (min)       lcorbi_hilic_neg_4_090_60    m/z 229.1375     0   1E+3   1.5   2  RT (min)      lcorbi_hilic_neg_4_041_54    m/z 229.1595     0   1E+4   0.5   1  RT (min)      lcorbi_hilic_neg_4_011_19    m/z 229.1807     0   1   2E+4   1   1.5  RT (min)       lcorbi_hilic_neg_4_089_59    m/z 229.7684     0   1E+4   3   3.5  RT (min)      lcorbi_hilic_neg_4_066_57    m/z 229.8740     0   1   2   3E+3   0.5   1  RT (min)      lcorbi_hilic_neg_4_106_62    m/z 229.9010     0   1   2E+4   1   1.5  RT (min)      lcorbi_hilic_neg_4_074_58    m/z 229.9180     0   1E+3   2.5   3  RT (min)      lcorbi_hilic_neg_4_076_58    m/z 229.9371     0   2   4   6   8E+3   0.5   1  RT (min)      lcorbi_hilic_neg_4_066_57    m/z 229.9556     0   2   4   6   8E+3   4.5   5  RT (min)      lcorbi_hilic_neg_4_079_58    m/z 229.9591     0   2   4   6   8E+3   3   3.5  RT (min)      lcorbi_hilic_neg_4_089_59    m/z 229.9812     0   1E+3   6.5   7  RT (min)      lcorbi_hilic_neg_4_053_23    m/z 229.9982     0   1   2E+4   6   7  RT (min)       lcorbi_hilic_neg_4_083_59    m/z 230.0236     0   1E+4   6.5   7  RT (min)      lcorbi_hilic_neg_4_089_59    m/z 230.0237     0   2   4   6E+3   2   2.5  RT (min)      lcorbi_hilic_neg_4_100_61    m/z 230.0337     0   1   1   1.5  RT (min)      lcorbi_hilic_neg_3_034_34    m/z 230.0392     0   1   2   3   4E+4   6.5   7  RT (min)       lcorbi_hilic_neg_4_069_57    m/z 230.0473     0   1   2   3   4E+3   5   5.5  RT (min)      lcorbi_hilic_neg_3_037_12    m/z 230.0571     0   1   2   3   4E+4   0.5   1  RT (min)      lcorbi_hilic_neg_4_081_58    m/z 230.0645     0   1E+3   6   6.5  RT (min)      lcorbi_hilic_neg_4_102_61    m/z 230.0670     0   2   4   6   8E+2   4   4.5  RT (min)      lcorbi_hilic_neg_4_097_60    m/z 230.0782     0   2   4   6E+4   1.5   2  RT (min)       lcorbi_hilic_neg_4_095_60    m/z 230.0783     0   1   2E+4   7.5   8  RT (min)       lcorbi_hilic_neg_4_096_60    m/z 230.1031     0   1   2.5   3  RT (min)      lcorbi_hilic_neg_4_085_59    m/z 230.1248     0   1E+5   6   6.5  RT (min)       lcorbi_hilic_neg_4_095_60    m/z 230.1257     0   1   7.5   8  RT (min)      lcorbi_hilic_neg_4_097_60    m/z 230.1589     0   2   4   6E+3   4   4.5  RT (min)      lcorbi_hilic_neg_4_076_58    m/z 230.1759     0   2   4   6   8E+3   3.5   4  RT (min)      lcorbi_hilic_neg_4_089_59    m/z 230.7662     0   1   2E+5   0.5   1  RT (min)      lcorbi_hilic_neg_2_052_41    m/z 230.7664     0   1   2   3   4E+5   12   12.5  RT (min)      lcorbi_hilic_neg_4_087_59    m/z 230.8013     0   2   4   6E+3   4.5   5  RT (min)      lcorbi_hilic_neg_4_097_60    m/z 230.8693     0   1E+3   1   1.5  RT (min)       lcorbi_hilic_neg_2_031_34    m/z 230.8997     0   1   2E+4   5   5.5  RT (min)       lcorbi_hilic_neg_4_098_61    m/z 230.9249     0   1E+3   1   1.5  RT (min)       lcorbi_hilic_neg_2_025_52    m/z 230.9423     0   2   4   6   8E+2   1   1.5  RT (min)      lcorbi_hilic_neg_2_031_34    m/z 230.9675     0   1E+3   1.5   2  RT (min)      lcorbi_hilic_neg_4_070_57    m/z 230.9698     0   1E+3   6.5   7  RT (min)      lcorbi_hilic_neg_3_010_18    m/z 230.9815     0   1E+4   7.5   8  RT (min)      lcorbi_hilic_neg_4_085_59    m/z 230.9829     0   1E+3   3   3.5  RT (min)       lcorbi_hilic_neg_3_043_02    m/z 230.9859     0   1   2E+4   0.5   1  RT (min)       lcorbi_hilic_neg_4_113_62    m/z 230.9911     0   2   4   6E+4   7   7.5  RT (min)       lcorbi_hilic_neg_4_087_59    m/z 231.0054     0   2   4   6   8E+3   3   3.5  RT (min)       lcorbi_hilic_neg_2_032_25    m/z 231.0055     0   1   2   3E+4   1   1.5  RT (min)       lcorbi_hilic_neg_4_076_58    m/z 231.0162     0   1   2   2.5  RT (min)      lcorbi_hilic_neg_4_077_58    m/z 231.0272     0   1E+5   6.5   7  RT (min)       lcorbi_hilic_neg_4_088_59    m/z 231.0306     0   1   2   3E+3   2   2.5  RT (min)       lcorbi_hilic_neg_1_020_13    m/z 231.0330     0   1E+3   0.5   1  RT (min)       lcorbi_hilic_neg_4_093_60    m/z 231.0516     0   1   2E+3   6   6.5  RT (min)      lcorbi_hilic_neg_4_095_60    m/z 231.0620     0   2   4   6   8E+3   4.5   5  RT (min)      lcorbi_hilic_neg_4_081_58    m/z 231.0679     0   1E+4   1.5   2  RT (min)       lcorbi_hilic_neg_4_110_62    m/z 231.0815     0   2   4   6E+4   5   5.5  RT (min)       lcorbi_hilic_neg_4_097_60    m/z 231.0984     0   1E+4   6   6.5  RT (min)      lcorbi_hilic_neg_4_089_59    m/z 231.1235     0   1   2   3E+3   1   1.5  RT (min)      lcorbi_hilic_neg_4_112_62    m/z 231.1347     0   1   5.5   6  RT (min)      lcorbi_hilic_neg_4_092_60    m/z 231.1348     0   2   4   6E+3   7.5   8  RT (min)      lcorbi_hilic_neg_4_088_59    m/z 231.1753     0   1E+4   0.5   1  RT (min)      lcorbi_hilic_neg_4_109_62    m/z 231.8028     0   2   4   6E+3   4.5   5  RT (min)      lcorbi_hilic_neg_4_096_60    m/z 231.8696     0   1   2   3E+3   0.5   1  RT (min)      lcorbi_hilic_neg_4_113_62    m/z 231.8955     0   1   2E+3   1   1.5  RT (min)      lcorbi_hilic_neg_4_090_60    m/z 231.9160     0   1E+3   2.5   3  RT (min)      lcorbi_hilic_neg_4_100_61    m/z 231.9316     0   2   4E+4   3.5   4   4.5  RT (min)       lcorbi_hilic_neg_1_006_08    m/z 231.9316     0   1E+5   7   7.5  RT (min)       lcorbi_hilic_neg_1_003_14    m/z 231.9526     0   1E+4   4.5   5  RT (min)      lcorbi_hilic_neg_4_077_58    m/z 231.9740     0   1E+4   3   3.5  RT (min)      lcorbi_hilic_neg_4_093_60    m/z 231.9856     0   2   4   6E+3   6   6.5  RT (min)      lcorbi_hilic_neg_4_073_57    m/z 231.9919     0   1E+4   1   1.5  RT (min)      lcorbi_hilic_neg_4_075_58    m/z 232.0084     0   1   2E+3   1   1.5  RT (min)      lcorbi_hilic_neg_4_082_59    m/z 232.0207     0   1   4   4.5  RT (min)      lcorbi_hilic_neg_4_096_60    m/z 232.0283     0   1   3   3.5   4  RT (min)       lcorbi_hilic_neg_4_091_60    m/z 232.0283     0   1E+3   1.5   2  RT (min)      lcorbi_hilic_neg_4_109_62    m/z 232.0307     0   1E+3   7   7.5  RT (min)      lcorbi_hilic_neg_4_055_38    m/z 232.0490     0   2   4   6E+3   2   2.5  RT (min)       lcorbi_hilic_neg_4_055_38    m/z 232.0588     0   1   2E+4   5   5.5  RT (min)       lcorbi_hilic_neg_4_078_58    m/z 232.0591     0   1   2E+4   3.5   4  RT (min)      lcorbi_hilic_neg_1_039_38    m/z 232.0686     0   1E+4   1   1.5  RT (min)      lcorbi_hilic_neg_4_090_60    m/z 232.0724     0   2   4   6E+3   6.5   7  RT (min)      lcorbi_hilic_neg_4_092_60    m/z 232.0937     0   1E+3   6.5   7  RT (min)      lcorbi_hilic_neg_4_042_28    m/z 232.1024     0   1   2   3E+4   1.5   2  RT (min)       lcorbi_hilic_neg_4_093_60    m/z 232.1189     0   1   1.5   2  RT (min)      lcorbi_hilic_neg_4_092_60    m/z 232.1424     0   1   2   3   4E+4   0.5   1  RT (min)      lcorbi_hilic_neg_4_076_58    m/z 232.7639     0   2   4   6   8E+5   12   12.5  RT (min)      lcorbi_hilic_neg_4_106_62    m/z 232.8009     0   2   4   6E+3   4.5   5  RT (min)      lcorbi_hilic_neg_4_086_59    m/z 232.8536     0   2   4   6   8E+3   8.5   9  RT (min)      lcorbi_hilic_neg_4_104_61    m/z 232.8537     0   1   2E+3   4.5   5  RT (min)      lcorbi_hilic_neg_1_009_50    m/z 232.8743     0   1E+4   8.5   9  RT (min)      lcorbi_hilic_neg_4_090_60    m/z 232.8834     0   1E+4   7.5   8  RT (min)      lcorbi_hilic_neg_4_085_59    m/z 232.9023     0   1   7   7.5  RT (min)      lcorbi_hilic_neg_1_020_13    m/z 232.9222     0   1   2E+4   8.5   9  RT (min)       lcorbi_hilic_neg_4_103_61    m/z 232.9350     0   2   4   6E+3   0.5   1  RT (min)      lcorbi_hilic_neg_4_112_62    m/z 232.9442     0   1   2   3   4E+3   7   7.5  RT (min)       lcorbi_hilic_neg_1_020_13    m/z 232.9525     0   1   0.5   1  RT (min)      lcorbi_hilic_neg_4_096_60    m/z 232.9554     0   1   7   7.5  RT (min)      lcorbi_hilic_neg_4_068_57    m/z 232.9759     0   1   1   1.5  RT (min)      lcorbi_hilic_neg_4_091_60    m/z 232.9773     0   1E+5   6   6.5  RT (min)       lcorbi_hilic_neg_4_093_60    m/z 232.9842     0   1   2   3E+4   3   3.5  RT (min)       lcorbi_hilic_neg_4_093_60    m/z 233.0006     0   1E+4   6.5   7  RT (min)      lcorbi_hilic_neg_4_096_60    m/z 233.0040     0   1   2E+4   0.5   1  RT (min)       lcorbi_hilic_neg_4_075_58    m/z 233.0131     0   1   4   4.5  RT (min)      lcorbi_hilic_neg_4_092_60    m/z 233.0253     0   2   4   6   8E+3   4   4.5  RT (min)      lcorbi_hilic_neg_4_085_59    m/z 233.0294     0   1   2   3E+2   2   2.5  RT (min)      lcorbi_hilic_neg_1_006_08    m/z 233.0445     0   1   3.5   4  RT (min)      lcorbi_hilic_neg_4_083_59    m/z 233.0445     0   1E+4   7.5   8  RT (min)      lcorbi_hilic_neg_4_094_60    m/z 233.0463     0   2   4E+4   2   2.5  RT (min)       lcorbi_hilic_neg_4_080_58    m/z 233.0599     0   1E+4   7   7.5  RT (min)      lcorbi_hilic_neg_4_067_57    m/z 233.0665     0   1E+3   1.5   2  RT (min)       lcorbi_hilic_neg_4_092_60    m/z 233.0669     0   1E+4   4   4.5  RT (min)      lcorbi_hilic_neg_1_039_38    m/z 233.0778     0   2   4   6E+3   7   7.5  RT (min)      lcorbi_hilic_neg_4_083_59    m/z 233.0819     0   2   4E+6   1.5   2  RT (min)       lcorbi_hilic_neg_4_055_38    m/z 233.0929     0   1   2E+3   2   2.5  RT (min)      lcorbi_hilic_neg_4_040_17    m/z 233.0932     0   1   5   5.5  RT (min)       lcorbi_hilic_neg_4_090_60    m/z 233.7999     0   2   4E+3   4.5   5  RT (min)      lcorbi_hilic_neg_4_093_60    m/z 233.8610     0   1E+4   0.5   1  RT (min)      lcorbi_hilic_neg_4_106_62    m/z 233.9181     0   1   2   3E+3   0   0.5  RT (min)      lcorbi_hilic_neg_1_005_36    m/z 233.9241     0   1   2E+4   5   5.5  RT (min)      lcorbi_hilic_neg_2_008_39    m/z 233.9301     0   1   2   3   4E+3   0.5   1  RT (min)      lcorbi_hilic_neg_4_066_57    m/z 233.9319     0   1E+4   3   3.5  RT (min)      lcorbi_hilic_neg_3_053_13    m/z 233.9433     0   1E+4   3   3.5  RT (min)      lcorbi_hilic_neg_4_089_59    m/z 233.9496     0   1   2E+3   4.5   5  RT (min)      lcorbi_hilic_neg_1_047_05    m/z 233.9679     0   2   4   6E+3   6   6.5  RT (min)      lcorbi_hilic_neg_4_071_57    m/z 233.9741     0   2   4   6   8E+3   6   6.5  RT (min)      lcorbi_hilic_neg_4_113_62    m/z 233.9775     0   1   2   3   4E+4   3   3.5  RT (min)      lcorbi_hilic_neg_4_094_60    m/z 233.9932     0   1   6.5   7  RT (min)      lcorbi_hilic_neg_4_095_60    m/z 234.0036     0   1E+4   3   3.5  RT (min)      lcorbi_hilic_neg_4_093_60    m/z 234.0076     0   2   4   6E+3   1   1.5  RT (min)      lcorbi_hilic_neg_2_020_27    m/z 234.0108     0   1   2   3E+4   7.5   8  RT (min)       lcorbi_hilic_neg_4_091_60    m/z 234.0260     0   1E+4   2   2.5  RT (min)      lcorbi_hilic_neg_4_099_61    m/z 234.0310     0   1E+4   5.5   6  RT (min)      lcorbi_hilic_neg_4_089_59    m/z 234.0438     0   1E+4   6.5   7  RT (min)       lcorbi_hilic_neg_4_089_59    m/z 234.0478     0   1E+4   3.5   4  RT (min)      lcorbi_hilic_neg_4_068_57    m/z 234.0488     0   1E+3   2   2.5  RT (min)      lcorbi_hilic_neg_4_100_61    m/z 234.0696     0   1   2E+4   5.5   6   6.5  RT (min)       lcorbi_hilic_neg_1_033_53    m/z 234.0730     0   1   2   3E+4   1.5   2  RT (min)       lcorbi_hilic_neg_4_073_57    m/z 234.0811     0   2   4   6   8E+3   4   4.5  RT (min)      lcorbi_hilic_neg_4_088_59    m/z 234.0891     0   1   6   6.5  RT (min)       lcorbi_hilic_neg_4_096_60    m/z 234.0982     0   1   2E+4   2.5   3  RT (min)       lcorbi_hilic_neg_4_082_59    m/z 234.1133     0   1   1   1.5  RT (min)       lcorbi_hilic_neg_4_111_62    m/z 234.7613     0   1E+5   0.5   1  RT (min)      lcorbi_hilic_neg_4_109_62    m/z 234.7615     0   2   4   6E+5   12   12.5  RT (min)      lcorbi_hilic_neg_4_106_62    m/z 234.8025     0   2   4   6E+3   4.5   5  RT (min)      lcorbi_hilic_neg_4_068_57    m/z 234.8197     0   1E+4   0.5   1  RT (min)      lcorbi_hilic_neg_4_112_62    m/z 234.8198     0   1   2E+4   7.5   8  RT (min)       lcorbi_hilic_neg_4_106_62    m/z 234.8478     0   2   4   6E+3   5   5.5  RT (min)      lcorbi_hilic_neg_4_086_59    m/z 234.8590     0   2   4E+3   0   0.5  RT (min)      lcorbi_hilic_neg_1_001_49    m/z 234.8592     0   1E+4   4.5   5  RT (min)      lcorbi_hilic_neg_4_100_61    m/z 234.9342     0   2   4   6E+3   2   2.5  RT (min)      lcorbi_hilic_neg_4_098_61    m/z 234.9601     0   1   2   3E+4   3   3.5  RT (min)      lcorbi_hilic_neg_1_012_32    m/z 234.9779     0   1   1   1.5  RT (min)      lcorbi_hilic_neg_4_092_60    m/z 234.9832     0   1   2   3   4E+4   0.5   1  RT (min)       lcorbi_hilic_neg_4_078_58    m/z 234.9843     0   1   2E+4   6.5   7  RT (min)       lcorbi_hilic_neg_4_090_60    m/z 234.9950     0   1   2   3E+4   7.5   8  RT (min)       lcorbi_hilic_neg_4_091_60    m/z 234.9998     0   1E+4   0.5   1  RT (min)      lcorbi_hilic_neg_4_090_60    m/z 235.0044     0   1E+5   7   7.5  RT (min)       lcorbi_hilic_neg_4_092_60    m/z 235.0150     0   1E+4   6   6.5  RT (min)      lcorbi_hilic_neg_4_096_60    m/z 235.0307     0   1E+5   4.5   5  RT (min)       lcorbi_hilic_neg_4_096_60    m/z 235.0403     0   1   2E+4   6.5   7  RT (min)       lcorbi_hilic_neg_4_095_60    m/z 235.0415     0   1E+3   3.5   4  RT (min)      lcorbi_hilic_neg_4_074_58    m/z 235.0489     0   1E+3   3.5   4  RT (min)      lcorbi_hilic_neg_4_088_59    m/z 235.0568     0   1E+4   6.5   7  RT (min)      lcorbi_hilic_neg_4_091_60    m/z 235.0683     0   1E+4   1.5   2  RT (min)      lcorbi_hilic_neg_4_093_60    m/z 235.0756     0   1E+4   6   6.5  RT (min)      lcorbi_hilic_neg_4_080_58    m/z 235.0854     0   1   2   2.5  RT (min)       lcorbi_hilic_neg_4_088_59    m/z 235.0934     0   1   2   3E+4   3   3.5  RT (min)       lcorbi_hilic_neg_4_073_57    m/z 235.0974     0   2   4   6E+4   1   1.5  RT (min)      lcorbi_hilic_neg_4_066_57    m/z 235.0978     0   2   4E+3   5   5.5  RT (min)      lcorbi_hilic_neg_4_100_61    m/z 235.1214     0   1   2   3   4E+4   3.5   4  RT (min)       lcorbi_hilic_neg_4_097_60    m/z 235.1214     0   1   2E+4   2   2.5  RT (min)       lcorbi_hilic_neg_4_096_60    m/z 235.1349     0   1   2E+4   1   1.5  RT (min)      lcorbi_hilic_neg_4_105_61    m/z 235.1611     0   1   2   3E+4   0.5   1  RT (min)      lcorbi_hilic_neg_4_079_58    m/z 235.8581     0   2   4   6E+3   0.5   1  RT (min)      lcorbi_hilic_neg_4_104_61    m/z 235.9185     0   1E+4   1.5   2  RT (min)      lcorbi_hilic_neg_1_027_18    m/z 235.9300     0   1   3   3.5  RT (min)      lcorbi_hilic_neg_4_108_62    m/z 235.9534     0   1   4.5   5  RT (min)      lcorbi_hilic_neg_2_005_24    m/z 235.9652     0   1   6.5   7  RT (min)       lcorbi_hilic_neg_2_055_32    m/z 236.0054     0   1E+3   5.5   6  RT (min)      lcorbi_hilic_neg_3_011_24    m/z 236.0253     0   1   2   3E+3   4   4.5  RT (min)      lcorbi_hilic_neg_1_017_51    m/z 236.0499     0   1   2   3E+3   2   2.5  RT (min)      lcorbi_hilic_neg_4_075_58    m/z 236.0530     0   1E+4   3.5   4  RT (min)      lcorbi_hilic_neg_4_096_60    m/z 236.0587     0   2   4   6   8E+3   2.5   3  RT (min)      lcorbi_hilic_neg_4_103_61    m/z 236.0650     0   2   4   6   8E+3   7   7.5  RT (min)      lcorbi_hilic_neg_4_068_57    m/z 236.0774     0   2   4E+3   7   7.5  RT (min)      lcorbi_hilic_neg_4_066_57    m/z 236.0775     0   1   1   1.5  RT (min)      lcorbi_hilic_neg_4_073_57    m/z 236.0788     0   1E+4   4.5   5  RT (min)      lcorbi_hilic_neg_4_087_59    m/z 236.0943     0   2   4   6   8E+3   1   2  RT (min)       lcorbi_hilic_neg_4_094_60    m/z 236.1145     0   1   2   3   4E+4   0.5   1  RT (min)      lcorbi_hilic_neg_2_029_10    m/z 236.1289     0   2   4E+2   1   1.5  RT (min)      lcorbi_hilic_neg_4_004_13    m/z 236.1448     0   1   2E+4   6.5   7  RT (min)       lcorbi_hilic_neg_4_096_60    m/z 236.7587     0   2   4   6E+4   0.5   1  RT (min)      lcorbi_hilic_neg_3_037_12    m/z 236.7589     0   1   2E+5   12   12.5  RT (min)      lcorbi_hilic_neg_4_106_62    m/z 236.7992     0   2   4   6E+3   4.5   5  RT (min)      lcorbi_hilic_neg_4_086_59    m/z 236.8171     0   1E+4   7.5   8  RT (min)      lcorbi_hilic_neg_4_106_62    m/z 236.8448     0   2   4   6E+3   5   5.5  RT (min)      lcorbi_hilic_neg_4_073_57    m/z 236.8561     0   1   2   3   4E+3   0   0.5  RT (min)      lcorbi_hilic_neg_1_010_22    m/z 236.8562     0   2   4   6   8E+3   4.5   5  RT (min)      lcorbi_hilic_neg_4_066_57    m/z 236.8835     0   1   2   3   4E+2   7.5   8  RT (min)      lcorbi_hilic_neg_4_005_32    m/z 236.9310     0   1   2   3E+2   6   6.5  RT (min)      lcorbi_hilic_neg_2_009_50    m/z 236.9495     0   1E+3   1   1.5  RT (min)      lcorbi_hilic_neg_4_075_58    m/z 236.9666     0   1   2   3E+3   2   2.5   3  RT (min)       lcorbi_hilic_neg_4_077_58    m/z 236.9684     0   1   2E+3   0.5   1  RT (min)      lcorbi_hilic_neg_4_081_58    m/z 236.9689     0   1   7   7.5  RT (min)      lcorbi_hilic_neg_4_098_61    m/z 236.9790     0   1   2E+4   0.5   1  RT (min)       lcorbi_hilic_neg_4_077_58    m/z 236.9791     0   1E+4   3   3.5  RT (min)      lcorbi_hilic_neg_4_090_60    m/z 236.9846     0   1   7.5   8  RT (min)      lcorbi_hilic_neg_4_099_61    m/z 236.9954     0   1   2E+4   0.5   1  RT (min)       lcorbi_hilic_neg_4_078_58    m/z 236.9955     0   1E+4   3   3.5  RT (min)       lcorbi_hilic_neg_4_092_60    m/z 237.0272     0   1   4   4.5  RT (min)      lcorbi_hilic_neg_4_098_61    m/z 237.0338     0   1E+3   4   5  RT (min)       lcorbi_hilic_neg_4_098_61    m/z 237.0457     0   1E+3   3.5   4  RT (min)      lcorbi_hilic_neg_4_085_59    m/z 237.0491     0   1   6   6.5  RT (min)      lcorbi_hilic_neg_4_098_61    m/z 237.0614     0   1   2   3   4E+4   3.5   4  RT (min)       lcorbi_hilic_neg_4_089_59    m/z 237.0615     0   1E+3   7.5   8  RT (min)       lcorbi_hilic_neg_1_005_36    m/z 237.0616     0   1   2   3   4E+4   1.5   2  RT (min)      lcorbi_hilic_neg_2_014_31    m/z 237.0727     0   1   2E+4   4.5   5  RT (min)      lcorbi_hilic_neg_4_066_57    m/z 237.0766     0   1E+3   1   1.5  RT (min)       lcorbi_hilic_neg_4_092_60    m/z 237.0822     0   1   2   2.5  RT (min)       lcorbi_hilic_neg_4_082_59    m/z 237.0830     0   1E+3   6.5   7  RT (min)       lcorbi_hilic_neg_4_091_60    m/z 237.0912     0   2   4   6E+2   1.5   2  RT (min)      lcorbi_hilic_neg_1_029_31    m/z 237.1041     0   1   2E+5   2   2.5  RT (min)      lcorbi_hilic_neg_4_005_32    m/z 237.1289     0   1E+5   6   6.5  RT (min)       lcorbi_hilic_neg_4_097_60    m/z 237.1371     0   2   4   6E+4   2   2.5  RT (min)       lcorbi_hilic_neg_3_011_24    m/z 237.1493     0   1   2E+3   1   1.5  RT (min)       lcorbi_hilic_neg_2_029_10    m/z 237.8351     0   1   2E+5   4.5   5  RT (min)      lcorbi_hilic_neg_2_020_27    m/z 237.8880     0   1E+4   7.5   8  RT (min)      lcorbi_hilic_neg_4_085_59    m/z 237.8976     0   2   4E+3   7   7.5  RT (min)      lcorbi_hilic_neg_1_005_36    m/z 237.9289     0   1   2   3E+3   4.5   5  RT (min)      lcorbi_hilic_neg_4_074_58    m/z 237.9525     0   1   2E+4   5   5.5  RT (min)      lcorbi_hilic_neg_4_099_61    m/z 237.9733     0   1   2E+3   7   7.5  RT (min)      lcorbi_hilic_neg_4_090_60    m/z 237.9813     0   1E+3   6   6.5  RT (min)      lcorbi_hilic_neg_4_072_57    m/z 238.0065     0   1E+3   1   1.5  RT (min)      lcorbi_hilic_neg_2_005_24    m/z 238.0144     0   1   7   7.5  RT (min)      lcorbi_hilic_neg_4_092_60    m/z 238.0381     0   2   4   6E+4   4   5  RT (min)       lcorbi_hilic_neg_4_097_60    m/z 238.0381     0   1   2E+4   2   2.5  RT (min)       lcorbi_hilic_neg_4_097_60    m/z 238.0486     0   1E+3   2.5   3  RT (min)      lcorbi_hilic_neg_4_088_59    m/z 238.1096     0   1   2.5   3  RT (min)      lcorbi_hilic_neg_4_074_58    m/z 238.1129     0   1E+4   5.5   6  RT (min)      lcorbi_hilic_neg_4_094_60    m/z 238.1198     0   1E+4   0.5   1  RT (min)      lcorbi_hilic_neg_2_038_36    m/z 238.1294     0   2   4   6E+3   3.5   4  RT (min)      lcorbi_hilic_neg_4_076_58    m/z 238.7564     0   1   2   3   4E+4   12   12.5  RT (min)       lcorbi_hilic_neg_4_069_57    m/z 238.8787     0   1   2E+4   8.5   9  RT (min)       lcorbi_hilic_neg_4_088_59    m/z 238.9129     0   2   4   6E+3   2   2.5  RT (min)      lcorbi_hilic_neg_4_067_57    m/z 238.9516     0   1   1   1.5  RT (min)      lcorbi_hilic_neg_4_084_59    m/z 238.9601     0   1   2   3   4E+4   3   3.5  RT (min)       lcorbi_hilic_neg_4_094_60    m/z 238.9687     0   1E+3   1   1.5  RT (min)      lcorbi_hilic_neg_4_071_57    m/z 238.9943     0   1   2E+4   2.5   3   3.5  RT (min)       lcorbi_hilic_neg_4_078_58    m/z 239.0035     0   1   2E+3   2   2.5  RT (min)       lcorbi_hilic_neg_4_092_60    m/z 239.0074     0   1   2   3   4E+4   4   4.5  RT (min)      lcorbi_hilic_neg_1_020_13    m/z 239.0163     0   1E+4   8   8.5  RT (min)      lcorbi_hilic_neg_4_070_57    m/z 239.0232     0   1   2   3E+3   0.5   1  RT (min)       lcorbi_hilic_neg_4_083_59    m/z 239.0284     0   1   2   3E+3   5.5   6  RT (min)      lcorbi_hilic_neg_4_067_57    m/z 239.0341     0   1   2   3E+4   1   1.5  RT (min)      lcorbi_hilic_neg_4_013_46    m/z 239.0395     0   1   7.5   8  RT (min)      lcorbi_hilic_neg_4_107_62    m/z 239.0441     0   2   4   6   8E+4   4.5   5  RT (min)       lcorbi_hilic_neg_4_073_57    m/z 239.0530     0   1   2E+3   1   1.5  RT (min)      lcorbi_hilic_neg_1_012_32    m/z 239.0550     0   2   4   6   8E+3   4.5   5  RT (min)      lcorbi_hilic_neg_1_005_36    m/z 239.0647     0   1   2E+4   6.5   7  RT (min)       lcorbi_hilic_neg_4_085_59    m/z 239.0671     0   1E+4   3.5   4  RT (min)      lcorbi_hilic_neg_4_084_59    m/z 239.0962     0   1E+4   4   4.5  RT (min)      lcorbi_hilic_neg_4_090_60    m/z 239.1117     0   2   4   6E+2   1   1.5  RT (min)      lcorbi_hilic_neg_4_004_13    m/z 239.1528     0   1   2E+4   1.5   2  RT (min)       lcorbi_hilic_neg_3_011_24    m/z 239.1650     0   1   1   1.5  RT (min)      lcorbi_hilic_neg_4_091_60    m/z 239.5868     0   1   2E+4   2.5   3  RT (min)      lcorbi_hilic_neg_4_077_58    m/z 239.8990     0   1   2   3   4E+3   6.5   7  RT (min)      lcorbi_hilic_neg_4_070_57    m/z 239.9133     0   1E+4   2   2.5  RT (min)      lcorbi_hilic_neg_4_090_60    m/z 239.9318     0   1   2E+4   1.5   2  RT (min)      lcorbi_hilic_neg_4_090_60    m/z 240.0037     0   1   1.5   2  RT (min)      lcorbi_hilic_neg_4_087_59    m/z 240.0221     0   1E+4   5.5   6  RT (min)      lcorbi_hilic_neg_4_081_58    m/z 240.0281     0   1   2E+4   1.5   2  RT (min)       lcorbi_hilic_neg_4_082_59    m/z 240.0302     0   2   4   6E+3   7   7.5  RT (min)      lcorbi_hilic_neg_4_085_59    m/z 240.0361     0   1E+3   1   1.5  RT (min)      lcorbi_hilic_neg_4_106_62    m/z 240.0489     0   1   2E+4   3   3.5  RT (min)       lcorbi_hilic_neg_4_076_58    m/z 240.0591     0   2   4   6E+3   0.5   1  RT (min)       lcorbi_hilic_neg_4_083_59    m/z 240.0639     0   1   2E+4   6.5   7  RT (min)       lcorbi_hilic_neg_4_097_60    m/z 240.0876     0   1E+3   6.5   7  RT (min)      lcorbi_hilic_neg_4_091_60    m/z 240.0958     0   1E+3   1   1.5  RT (min)      lcorbi_hilic_neg_3_029_25    m/z 240.1164     0   1   2.5   3  RT (min)      lcorbi_hilic_neg_4_066_57    m/z 240.1205     0   1   2   3E+3   6.5   7  RT (min)      lcorbi_hilic_neg_3_011_24    m/z 240.1285     0   1E+4   6   6.5  RT (min)      lcorbi_hilic_neg_4_093_60    m/z 240.1286     0   2   4   6E+3   3   3.5  RT (min)      lcorbi_hilic_neg_4_094_60    m/z 240.7834     0   2   4   6   8E+3   2.5   3  RT (min)      lcorbi_hilic_neg_4_066_57    m/z 240.7929     0   1E+4   4.5   5  RT (min)      lcorbi_hilic_neg_4_073_57    m/z 240.9019     0   1E+4   1   1.5  RT (min)      lcorbi_hilic_neg_4_097_60    m/z 240.9286     0   1   2   3   4E+4   5.5   6  RT (min)       lcorbi_hilic_neg_4_104_61    m/z 240.9899     0   1   2E+4   0.5   1  RT (min)       lcorbi_hilic_neg_4_108_62    m/z 240.9899     0   1   2E+4   3   3.5  RT (min)      lcorbi_hilic_neg_4_078_58    m/z 241.0296     0   1   2   3E+3   1.5   2  RT (min)       lcorbi_hilic_neg_4_097_60    m/z 241.0380     0   1   5   6  RT (min)       lcorbi_hilic_neg_4_091_60    m/z 241.0434     0   2   4E+3   1.5   2  RT (min)       lcorbi_hilic_neg_4_088_59    m/z 241.0462     0   1   2E+3   3.5   4   4.5  RT (min)       lcorbi_hilic_neg_4_091_60    m/z 241.0561     0   1   2E+4   0.5   1  RT (min)      lcorbi_hilic_neg_4_113_62    m/z 241.0594     0   1   2E+4   6.5   7  RT (min)       lcorbi_hilic_neg_4_085_59    m/z 241.0595     0   1E+3   4   4.5  RT (min)      lcorbi_hilic_neg_4_036_36    m/z 241.0652     0   1   2   3E+2   3.5   4  RT (min)       lcorbi_hilic_neg_4_004_13    m/z 241.0713     0   1   2E+4   0.5   1  RT (min)      lcorbi_hilic_neg_4_107_62    m/z 241.0824     0   1   2E+3   2   2.5  RT (min)      lcorbi_hilic_neg_4_074_58    m/z 241.0826     0   1E+4   6   6.5  RT (min)      lcorbi_hilic_neg_4_088_59    m/z 241.0940     0   1   2   3   4E+4   7.5   8  RT (min)      lcorbi_hilic_neg_4_079_58    m/z 241.1080     0   2   4   6E+4   1   1.5  RT (min)       lcorbi_hilic_neg_2_032_25    m/z 241.1444     0   1   2   3E+4   0.5   1  RT (min)      lcorbi_hilic_neg_4_113_62    m/z 241.1807     0   1E+4   1   1.5  RT (min)      lcorbi_hilic_neg_4_075_58    m/z 241.7673     0   1   2   3E+3   3   3.5  RT (min)      lcorbi_hilic_neg_3_054_29    m/z 241.8563     0   2   4   6E+3   8   8.5  RT (min)      lcorbi_hilic_neg_4_066_57    m/z 241.8842     0   1E+4   1.5   2  RT (min)      lcorbi_hilic_neg_4_082_59    m/z 241.8988     0   2   4E+4   1   1.5  RT (min)       lcorbi_hilic_neg_4_097_60    m/z 241.9166     0   1   2   3E+3   0.5   1  RT (min)      lcorbi_hilic_neg_4_004_13    m/z 241.9471     0   2   4E+4   1   1.5  RT (min)       lcorbi_hilic_neg_4_106_62    m/z 241.9603     0   2   4   6E+3   6   6.5  RT (min)      lcorbi_hilic_neg_4_071_57    m/z 241.9758     0   2   4E+4   2   3  RT (min)       lcorbi_hilic_neg_4_083_59    m/z 241.9761     0   2   4   6   8E+3   6   6.5  RT (min)      lcorbi_hilic_neg_4_023_34    m/z 241.9868     0   2   4   6E+3   6.5   7  RT (min)      lcorbi_hilic_neg_4_090_60    m/z 241.9960     0   1   2E+3   1   1.5  RT (min)      lcorbi_hilic_neg_3_028_23    m/z 241.9989     0   1   2E+4   6   6.5  RT (min)       lcorbi_hilic_neg_4_085_59    m/z 242.0259     0   1   2E+4   4.5   5  RT (min)       lcorbi_hilic_neg_4_113_62    m/z 242.0282     0   2   4   6   8E+3   3   3.5  RT (min)      lcorbi_hilic_neg_4_085_59    m/z 242.0426     0   1E+4   1   1.5  RT (min)      lcorbi_hilic_neg_4_053_23    m/z 242.0515     0   1E+4   3.5   4  RT (min)      lcorbi_hilic_neg_4_078_58    m/z 242.0646     0   1E+3   6   6.5  RT (min)       lcorbi_hilic_neg_4_046_15    m/z 242.0668     0   1E+4   1   1.5  RT (min)      lcorbi_hilic_neg_4_089_59    m/z 242.0782     0   1   2   3E+3   1.5   2  RT (min)      lcorbi_hilic_neg_4_090_60    m/z 242.0866     0   1   6   6.5  RT (min)      lcorbi_hilic_neg_4_074_58    m/z 242.0951     0   1E+4   7   7.5  RT (min)      lcorbi_hilic_neg_4_086_59    m/z 242.0966     0   1E+3   6   6.5  RT (min)       lcorbi_hilic_neg_4_089_59    m/z 242.0975     0   1E+5   1   1.5  RT (min)       lcorbi_hilic_neg_4_096_60    m/z 242.1144     0   1   2   3   4E+2   6   6.5  RT (min)      lcorbi_hilic_neg_3_031_14    m/z 242.1145     0   1   2   3   4E+3   2.5   3  RT (min)      lcorbi_hilic_neg_4_013_46    m/z 242.1249     0   1   2   3   4E+4   2   2.5  RT (min)       lcorbi_hilic_neg_4_096_60    m/z 242.1524     0   2   4   6   8E+3   1.5   2  RT (min)      lcorbi_hilic_neg_4_093_60    m/z 242.7901     0   1   2E+4   4.5   5  RT (min)      lcorbi_hilic_neg_4_072_57    m/z 242.9434     0   1E+5   8.5   9  RT (min)      lcorbi_hilic_neg_4_075_58    m/z 242.9860     0   1   2   3   4E+4   0.5   1  RT (min)       lcorbi_hilic_neg_4_100_61    m/z 242.9861     0   1   2   3E+4   3   4  RT (min)       lcorbi_hilic_neg_4_086_59    m/z 242.9925     0   1   6.5   7  RT (min)      lcorbi_hilic_neg_4_108_62    m/z 242.9965     0   1E+3   0.5   1  RT (min)      lcorbi_hilic_neg_4_092_60    m/z 242.9994     0   2   4   6E+3   4.5   5  RT (min)      lcorbi_hilic_neg_4_083_59    m/z 243.0053     0   1E+4   0.5   1  RT (min)      lcorbi_hilic_neg_4_097_60    m/z 243.0175     0   1   2   3E+4   4.5   5   5.5  RT (min)       lcorbi_hilic_neg_4_069_57    m/z 243.0177     0   1E+5   2   2.5  RT (min)       lcorbi_hilic_neg_4_004_13    m/z 243.0221     0   1   6.5   7  RT (min)      lcorbi_hilic_neg_4_107_62    m/z 243.0330     0   1   0.5   1  RT (min)       lcorbi_hilic_neg_4_092_60    m/z 243.0386     0   1   6.5   7  RT (min)      lcorbi_hilic_neg_4_108_62    m/z 243.0442     0   1   1.5   2  RT (min)      lcorbi_hilic_neg_4_096_60    m/z 243.0448     0   1E+3   3.5   4  RT (min)       lcorbi_hilic_neg_4_085_59    m/z 243.0540     0   1E+3   1   1.5  RT (min)      lcorbi_hilic_neg_4_076_58    m/z 243.0622     0   1   2   3E+5   2.5   3  RT (min)       lcorbi_hilic_neg_4_079_58    m/z 243.0720     0   1   4   4.5  RT (min)       lcorbi_hilic_neg_4_105_61    m/z 243.0932     0   1   2   3E+4   3.5   4  RT (min)       lcorbi_hilic_neg_4_095_60    m/z 243.0983     0   1E+3   6   6.5  RT (min)      lcorbi_hilic_neg_4_080_58    m/z 243.1036     0   1   2E+4   6.5   7  RT (min)       lcorbi_hilic_neg_4_096_60    m/z 243.1349     0   1   2   2.5  RT (min)       lcorbi_hilic_neg_3_011_24    m/z 243.1458     0   2   4   6   8E+2   5.5   6  RT (min)      lcorbi_hilic_neg_4_076_58    m/z 243.1792     0   1E+4   1   1.5  RT (min)      lcorbi_hilic_neg_4_086_59    m/z 243.7653     0   1E+3   3   3.5  RT (min)       lcorbi_hilic_neg_3_050_03    m/z 243.8534     0   1E+3   8   8.5  RT (min)      lcorbi_hilic_neg_1_003_14    m/z 243.8634     0   1E+4   7   7.5  RT (min)      lcorbi_hilic_neg_1_001_49    m/z 243.8816     0   1   2   3E+4   0   0.5  RT (min)       lcorbi_hilic_neg_4_111_62    m/z 243.8817     0   2   4   6E+4   4   4.5  RT (min)       lcorbi_hilic_neg_4_066_57    m/z 243.8817     0   1   2E+4   9  RT (min)       lcorbi_hilic_neg_4_066_57    m/z 243.8991     0   1   2   3   4E+4   1   1.5  RT (min)       lcorbi_hilic_neg_4_097_60    m/z 243.9272     0   1   2   3E+2   3   3.5  RT (min)      lcorbi_hilic_neg_3_032_42    m/z 243.9524     0   1   2   3   4E+3   3   3.5  RT (min)      lcorbi_hilic_neg_4_074_58    m/z 243.9748     0   2   4   6E+3   2   2.5  RT (min)       lcorbi_hilic_neg_4_044_01    m/z 243.9766     0   1   2E+4   6   6.5  RT (min)       lcorbi_hilic_neg_4_088_59    m/z 243.9967     0   1E+4   6   6.5  RT (min)      lcorbi_hilic_neg_4_072_57    m/z 244.0228     0   1E+4   4.5   5  RT (min)      lcorbi_hilic_neg_4_113_62    m/z 244.0397     0   1   2   3E+3   1   1.5  RT (min)      lcorbi_hilic_neg_3_051_46    m/z 244.0454     0   2   4   6E+3   7.5   8  RT (min)      lcorbi_hilic_neg_4_086_59    m/z 244.0494     0   1E+5   1   2  RT (min)       lcorbi_hilic_neg_1_037_46    m/z 244.0597     0   1   2   3   4E+4   2   2.5  RT (min)       lcorbi_hilic_neg_4_080_58    m/z 244.0655     0   1E+4   2   3  RT (min)       lcorbi_hilic_neg_4_106_62    m/z 244.0655     0   1E+4   4.5   5  RT (min)      lcorbi_hilic_neg_4_069_57    m/z 244.0758     0   2   4E+3   5.5   6  RT (min)      lcorbi_hilic_neg_4_066_57    m/z 244.0837     0   1E+3   6   7   8  RT (min)       lcorbi_hilic_neg_4_092_60    m/z 244.0839     0   1   2E+3   2   2.5  RT (min)      lcorbi_hilic_neg_4_075_58    m/z 244.0847     0   1   2   3E+4   3.5   4  RT (min)       lcorbi_hilic_neg_4_096_60    m/z 244.0938     0   1   1.5   2  RT (min)      lcorbi_hilic_neg_4_108_62    m/z 244.1124     0   2   4   6E+4   2.5   3  RT (min)       lcorbi_hilic_neg_3_011_24    m/z 244.1188     0   1E+4   1.5   2  RT (min)       lcorbi_hilic_neg_4_088_59    m/z 244.1301     0   1   2E+3   6.5   7  RT (min)      lcorbi_hilic_neg_4_076_58    m/z 244.1745     0   1E+4   4   4.5  RT (min)      lcorbi_hilic_neg_4_076_58    m/z 244.7873     0   1E+4   4.5   5  RT (min)      lcorbi_hilic_neg_4_072_57    m/z 244.8765     0   2   4E+3   5   5.5  RT (min)      lcorbi_hilic_neg_4_113_62    m/z 244.8973     0   2   4   6   8E+3   7   7.5  RT (min)      lcorbi_hilic_neg_4_067_57    m/z 244.9012     0   1   2E+3   0.5   1  RT (min)      lcorbi_hilic_neg_4_079_58    m/z 244.9147     0   1E+3   0.5   1  RT (min)      lcorbi_hilic_neg_4_076_58    m/z 244.9629     0   2   4E+3   6.5   7  RT (min)      lcorbi_hilic_neg_4_068_57    m/z 244.9638     0   2   4   6E+3   8   8.5  RT (min)      lcorbi_hilic_neg_4_079_58    m/z 244.9680     0   1E+4   2   2.5  RT (min)      lcorbi_hilic_neg_4_067_57    m/z 244.9867     0   2   4   6   8E+4   6.5   7  RT (min)       lcorbi_hilic_neg_4_071_57    m/z 244.9996     0   2   4E+3   6.5   7  RT (min)      lcorbi_hilic_neg_4_073_57    m/z 245.0022     0   1E+4   0.5   1  RT (min)      lcorbi_hilic_neg_4_076_58    m/z 245.0122     0   1   2E+4   1   1.5  RT (min)      lcorbi_hilic_neg_4_081_58    m/z 245.0211     0   1   2E+3   5   5.5  RT (min)      lcorbi_hilic_neg_4_110_62    m/z 245.0233     0   1   2E+3   7   7.5  RT (min)      lcorbi_hilic_neg_4_107_62    m/z 245.0245     0   1E+4   2   2.5  RT (min)       lcorbi_hilic_neg_4_073_57    m/z 245.0500     0   1E+4   6   6.5  RT (min)      lcorbi_hilic_neg_4_087_59    m/z 245.0779     0   2   4   6E+2   6   6.5  RT (min)      lcorbi_hilic_neg_3_029_25    m/z 245.0882     0   2   4   6   8E+2   5.5   6  RT (min)      lcorbi_hilic_neg_1_013_47    m/z 245.0914     0   1   2   2.5  RT (min)      lcorbi_hilic_neg_4_107_62    m/z 245.0971     0   1   1.5   2  RT (min)       lcorbi_hilic_neg_4_108_62    m/z 245.1140     0   2   4   6E+3   6   6.5  RT (min)      lcorbi_hilic_neg_4_085_59    m/z 245.1142     0   1   2   2.5  RT (min)      lcorbi_hilic_neg_4_106_62    m/z 245.1293     0   1E+4   4   4.5  RT (min)      lcorbi_hilic_neg_4_069_57    m/z 245.1393     0   1   2   3E+3   1   1.5  RT (min)      lcorbi_hilic_neg_4_086_59    m/z 245.1506     0   1E+4   1   1.5  RT (min)      lcorbi_hilic_neg_4_078_58    m/z 245.7522     0   1E+4   3   3.5  RT (min)       lcorbi_hilic_neg_4_080_58    m/z 245.7633     0   1   2E+4   3   3.5  RT (min)      lcorbi_hilic_neg_4_066_57    m/z 245.8786     0   1E+4   0   0.5  RT (min)      lcorbi_hilic_neg_4_113_62    m/z 245.8787     0   1   2E+4   4   4.5  RT (min)       lcorbi_hilic_neg_4_100_61    m/z 245.8787     0   1E+4   8.5   9  RT (min)      lcorbi_hilic_neg_4_066_57    m/z 245.8946     0   1E+4   1   1.5  RT (min)       lcorbi_hilic_neg_4_097_60    m/z 245.9173     0   1E+4   3   3.5  RT (min)      lcorbi_hilic_neg_2_031_34    m/z 245.9242     0   1   3   3.5  RT (min)      lcorbi_hilic_neg_4_081_58    m/z 245.9476     0   1E+3   1   1.5  RT (min)      lcorbi_hilic_neg_4_095_60    m/z 245.9568     0   1   2   3E+4   3   3.5  RT (min)      lcorbi_hilic_neg_3_034_34    m/z 245.9664     0   2   4   6E+3   2   2.5  RT (min)      lcorbi_hilic_neg_4_075_58    m/z 245.9835     0   1E+4   3.5   4  RT (min)      lcorbi_hilic_neg_4_079_58    m/z 245.9835     0   1   2E+4   1.5   2  RT (min)      lcorbi_hilic_neg_4_075_58    m/z 245.9869     0   1E+3   6.5   7  RT (min)      lcorbi_hilic_neg_4_045_25    m/z 245.9942     0   1   2   3   4E+3   6   6.5  RT (min)      lcorbi_hilic_neg_2_055_32    m/z 246.0075     0   2   4E+4   1   1.5  RT (min)       lcorbi_hilic_neg_1_002_26    m/z 246.0149     0   1   2E+4   3   3.5  RT (min)      lcorbi_hilic_neg_1_023_23    m/z 246.0219     0   1   2E+4   6   6.5  RT (min)      lcorbi_hilic_neg_4_090_60    m/z 246.0231     0   1   1   1.5   2  RT (min)       lcorbi_hilic_neg_4_082_59    m/z 246.0280     0   1   6   6.5  RT (min)       lcorbi_hilic_neg_4_089_59    m/z 246.0423     0   2   4   6E+3   5.5   6  RT (min)      lcorbi_hilic_neg_4_110_62    m/z 246.0464     0   1   2   3   4E+5   1   1.5   2  RT (min)       lcorbi_hilic_neg_4_069_57    m/z 246.0574     0   1E+5   1.5   2  RT (min)       lcorbi_hilic_neg_4_097_60    m/z 246.0593     0   1E+4   6   6.5  RT (min)      lcorbi_hilic_neg_4_081_58    m/z 246.0619     0   1   2E+4   4.5   5  RT (min)       lcorbi_hilic_neg_4_082_59    m/z 246.0662     0   1   2   3   4E+4   1.5   2  RT (min)      lcorbi_hilic_neg_4_013_46    m/z 246.0729     0   1E+4   7   7.5  RT (min)      lcorbi_hilic_neg_4_089_59    m/z 246.0812     0   1   2   3   4E+3   3   3.5  RT (min)      lcorbi_hilic_neg_4_077_58    m/z 246.0882     0   2   4   6   8E+3   5   5.5  RT (min)      lcorbi_hilic_neg_2_020_27    m/z 246.0883     0   1   2E+3   1.5   2  RT (min)      lcorbi_hilic_neg_4_096_60    m/z 246.1074     0   1   2E+3   1.5   2  RT (min)      lcorbi_hilic_neg_4_096_60    m/z 246.1086     0   2   4   6E+3   7   7.5  RT (min)      lcorbi_hilic_neg_4_097_60    m/z 246.1090     0   2   4   6   8E+4   4   4.5  RT (min)      lcorbi_hilic_neg_3_044_32    m/z 246.1181     0   1E+3   1   1.5  RT (min)      lcorbi_hilic_neg_2_044_46    m/z 246.1204     0   1E+4   6.5   7  RT (min)      lcorbi_hilic_neg_4_095_60    m/z 246.5764     0   1   2E+4   3.5   4  RT (min)      lcorbi_hilic_neg_4_079_58    m/z 246.8737     0   1E+4   5   5.5  RT (min)      lcorbi_hilic_neg_4_078_58    m/z 246.8987     0   2   4   6   8E+3   1   1.5  RT (min)      lcorbi_hilic_neg_4_069_57    m/z 246.9099     0   1E+4   0.5   1  RT (min)       lcorbi_hilic_neg_4_077_58    m/z 246.9345     0   1   2   3E+4   3   3.5  RT (min)       lcorbi_hilic_neg_4_085_59    m/z 246.9364     0   1   2   3E+4   1   1.5  RT (min)      lcorbi_hilic_neg_4_040_17    m/z 246.9505     0   1   1   1.5  RT (min)      lcorbi_hilic_neg_4_084_59    m/z 246.9598     0   2   4   6E+3   8   8.5  RT (min)      lcorbi_hilic_neg_4_076_58    m/z 246.9641     0   1E+3   1   1.5  RT (min)       lcorbi_hilic_neg_4_084_59    m/z 246.9710     0   2   4E+4   6   6.5  RT (min)      lcorbi_hilic_neg_4_087_59    m/z 246.9810     0   1   2E+4   1   1.5  RT (min)       lcorbi_hilic_neg_4_084_59    m/z 246.9916     0   1E+5   1   1.5  RT (min)       lcorbi_hilic_neg_4_031_14    m/z 247.0003     0   2   4E+4   1   1.5  RT (min)       lcorbi_hilic_neg_4_081_58    m/z 247.0219     0   1E+4   2   2.5  RT (min)       lcorbi_hilic_neg_4_067_57    m/z 247.0317     0   1   2   3   4E+4   6   6.5  RT (min)       lcorbi_hilic_neg_4_096_60    m/z 247.0473     0   1E+3   6   6.5  RT (min)      lcorbi_hilic_neg_4_103_61    m/z 247.0492     0   1E+3   4   4.5  RT (min)      lcorbi_hilic_neg_4_076_58    m/z 247.0547     0   1   7   7.5  RT (min)      lcorbi_hilic_neg_4_093_60    m/z 247.0586     0   1E+3   5.5   6  RT (min)      lcorbi_hilic_neg_2_055_32    m/z 247.0713     0   2   4   6   8E+3   6   6.5  RT (min)      lcorbi_hilic_neg_4_037_31    m/z 247.0725     0   2   4   6E+3   3.5   4  RT (min)      lcorbi_hilic_neg_4_096_60    m/z 247.0807     0   1   5.5   6  RT (min)      lcorbi_hilic_neg_2_025_52    m/z 247.0883     0   1   2   2.5  RT (min)      lcorbi_hilic_neg_4_055_38    m/z 247.0932     0   1E+4   6.5   7  RT (min)       lcorbi_hilic_neg_4_092_60    m/z 247.1037     0   1E+3   5.5   6  RT (min)       lcorbi_hilic_neg_4_005_32    m/z 247.1040     0   2   4   6   8E+3   1.5   2  RT (min)      lcorbi_hilic_neg_4_094_60    m/z 247.8096     0   1   2   3   4E+4   1   1.5  RT (min)      lcorbi_hilic_neg_2_055_32    m/z 247.9152     0   1E+4   3   3.5  RT (min)      lcorbi_hilic_neg_2_031_34    m/z 247.9418     0   1   1   1.5  RT (min)      lcorbi_hilic_neg_4_092_60    m/z 247.9693     0   1E+4   6   6.5  RT (min)      lcorbi_hilic_neg_4_087_59    m/z 247.9846     0   1E+3   3   3.5  RT (min)      lcorbi_hilic_neg_4_095_60    m/z 247.9866     0   1E+3   6   6.5  RT (min)      lcorbi_hilic_neg_4_076_58    m/z 247.9990     0   1   1.5   2  RT (min)      lcorbi_hilic_neg_4_090_60    m/z 247.9991     0   1   3.5   4  RT (min)      lcorbi_hilic_neg_4_095_60    m/z 248.0088     0   2   4   6E+3   6   6.5  RT (min)      lcorbi_hilic_neg_4_090_60    m/z 248.0207     0   2   4   6E+3   6   6.5  RT (min)      lcorbi_hilic_neg_4_090_60    m/z 248.0307     0   2   4   6   8E+3   5.5   6  RT (min)      lcorbi_hilic_neg_1_047_05    m/z 248.0400     0   1E+5   3   3.5  RT (min)       lcorbi_hilic_neg_1_023_23    m/z 248.0437     0   1   6   6.5  RT (min)      lcorbi_hilic_neg_4_079_58    m/z 248.0609     0   1   2E+3   5.5   6  RT (min)      lcorbi_hilic_neg_4_078_58    m/z 248.0620     0   1E+3   2   2.5  RT (min)      lcorbi_hilic_neg_1_037_46    m/z 248.0829     0   2   4   6   8E+3   6.5   7  RT (min)       lcorbi_hilic_neg_4_015_24    m/z 248.0968     0   1E+4   4.5   5  RT (min)      lcorbi_hilic_neg_4_089_59    m/z 248.1003     0   2   4   6   8E+3   6   6.5  RT (min)      lcorbi_hilic_neg_4_091_60    m/z 248.1007     0   1   2E+4   1   1.5  RT (min)      lcorbi_hilic_neg_4_077_58    m/z 248.1139     0   1   4.5   5  RT (min)      lcorbi_hilic_neg_4_107_62    m/z 248.1248     0   2   4   6E+4   1   1.5  RT (min)      lcorbi_hilic_neg_4_090_60    m/z 248.5815     0   2   4   6   8E+3   1   1.5  RT (min)      lcorbi_hilic_neg_4_095_60    m/z 248.7408     0   1E+4   3   3.5  RT (min)      lcorbi_hilic_neg_3_055_26    m/z 248.8840     0   2   4   6E+3   6   6.5  RT (min)      lcorbi_hilic_neg_4_092_60    m/z 248.8936     0   2   4   6   8E+4   1   1.5  RT (min)       lcorbi_hilic_neg_4_004_13    m/z 248.8998     0   1   2E+4   8.5   9  RT (min)       lcorbi_hilic_neg_4_083_59    m/z 248.9095     0   2   4E+3   0.5   1  RT (min)      lcorbi_hilic_neg_4_080_58    m/z 248.9216     0   1   2   3E+4   0   0.5  RT (min)       lcorbi_hilic_neg_4_066_57    m/z 248.9299     0   2   4   6   8E+3   5   5.5  RT (min)      lcorbi_hilic_neg_4_068_57    m/z 248.9402     0   2   4   6E+3   5   5.5  RT (min)      lcorbi_hilic_neg_4_009_50    m/z 248.9790     0   1E+4   1   1.5  RT (min)       lcorbi_hilic_neg_4_091_60    m/z 248.9791     0   1E+4   3   3.5  RT (min)      lcorbi_hilic_neg_4_071_57    m/z 248.9793     0   2   4   6E+3   5   5.5  RT (min)       lcorbi_hilic_neg_3_050_03    m/z 248.9935     0   2   4E+3   5   6  RT (min)       lcorbi_hilic_neg_4_044_01    m/z 248.9956     0   1   2   3E+4   3   3.5  RT (min)       lcorbi_hilic_neg_4_069_57    m/z 249.0072     0   1   2E+3   1   1.5  RT (min)       lcorbi_hilic_neg_4_084_59    m/z 249.0078     0   1   3.5   4  RT (min)      lcorbi_hilic_neg_4_092_60    m/z 249.0205     0   1E+4   1   1.5   2  RT (min)       lcorbi_hilic_neg_4_082_59    m/z 249.0320     0   1   2   3E+3   1   1.5  RT (min)       lcorbi_hilic_neg_4_084_59    m/z 249.0377     0   2   4   6   8E+3   4.5   5  RT (min)      lcorbi_hilic_neg_4_077_58    m/z 249.0377     0   1E+3   6   6.5  RT (min)      lcorbi_hilic_neg_4_113_62    m/z 249.0460     0   1E+3   2   2.5  RT (min)      lcorbi_hilic_neg_4_082_59    m/z 249.0570     0   1E+4   5.5   6  RT (min)      lcorbi_hilic_neg_4_079_58    m/z 249.0765     0   2   4E+4   1.5   2  RT (min)       lcorbi_hilic_neg_4_091_60    m/z 249.0777     0   1E+5   4   4.5  RT (min)      lcorbi_hilic_neg_4_096_60    m/z 249.0876     0   1   2E+4   5.5   6  RT (min)       lcorbi_hilic_neg_4_095_60    m/z 249.0979     0   1E+4   2   2.5  RT (min)       lcorbi_hilic_neg_4_089_59    m/z 249.1007     0   1E+4   3   3.5  RT (min)      lcorbi_hilic_neg_4_095_60    m/z 249.1089     0   1   2   3E+3   1.5   2  RT (min)      lcorbi_hilic_neg_4_090_60    m/z 249.1193     0   1   2E+4   6   6.5  RT (min)       lcorbi_hilic_neg_4_095_60    m/z 249.1242     0   1   2E+4   4   4.5  RT (min)       lcorbi_hilic_neg_4_103_61    m/z 249.1858     0   1   2E+4   0.5   1  RT (min)      lcorbi_hilic_neg_4_029_02    m/z 249.9053     0   1E+4   0.5   1  RT (min)      lcorbi_hilic_neg_4_077_58    m/z 249.9217     0   2   4   6   8E+3   1   1.5  RT (min)      lcorbi_hilic_neg_4_066_57    m/z 249.9532     0   2   4   6   8E+2   5.5   6  RT (min)      lcorbi_hilic_neg_4_005_32    m/z 249.9711     0   1   5   5.5  RT (min)      lcorbi_hilic_neg_1_051_33    m/z 250.0037     0   1E+5   6.5   7  RT (min)       lcorbi_hilic_neg_4_090_60    m/z 250.0413     0   2   4E+3   2   2.5  RT (min)      lcorbi_hilic_neg_4_099_61    m/z 250.0598     0   1   3.5   4  RT (min)      lcorbi_hilic_neg_2_014_31    m/z 250.0600     0   1   2   3E+4   5.5   6  RT (min)       lcorbi_hilic_neg_4_097_60    m/z 250.0717     0   1   2   3E+3   1   1.5  RT (min)      lcorbi_hilic_neg_4_073_57    m/z 250.0765     0   1   2E+4   3.5   4  RT (min)      lcorbi_hilic_neg_3_011_24    m/z 250.0800     0   2   4   6   8E+3   6.5   7  RT (min)      lcorbi_hilic_neg_4_095_60    m/z 250.1044     0   1E+4   6.5   7  RT (min)      lcorbi_hilic_neg_4_085_59    m/z 250.1082     0   2   4E+3   1   1.5  RT (min)      lcorbi_hilic_neg_3_039_31    m/z 250.1318     0   2   4   6E+3   5.5   6  RT (min)      lcorbi_hilic_neg_4_093_60    m/z 250.8216     0   2   4   6   8E+3   4.5   5  RT (min)      lcorbi_hilic_neg_4_078_58    m/z 250.8330     0   2   4E+3   0   0.5  RT (min)      lcorbi_hilic_neg_1_002_26    m/z 250.8332     0   2   4   6E+3   4.5   5  RT (min)      lcorbi_hilic_neg_4_104_61    m/z 250.8471     0   2   4   6   8E+3   8   8.5  RT (min)      lcorbi_hilic_neg_4_111_62    m/z 250.8565     0   2   4   6E+3   7.5   8  RT (min)      lcorbi_hilic_neg_4_083_59    m/z 250.8929     0   1   2   3E+4   1   1.5  RT (min)      lcorbi_hilic_neg_4_004_13    m/z 250.8991     0   1   0.5   1  RT (min)      lcorbi_hilic_neg_4_098_61    m/z 250.9203     0   2   4   6   8E+3   8   8.5  RT (min)      lcorbi_hilic_neg_4_074_58    m/z 250.9341     0   2   4   6   8E+3   3   3.5  RT (min)      lcorbi_hilic_neg_1_012_32    m/z 250.9512     0   1   4   4.5  RT (min)      lcorbi_hilic_neg_4_086_59    m/z 250.9683     0   1   1   1.5  RT (min)      lcorbi_hilic_neg_4_085_59    m/z 250.9814     0   1   2   3E+3   1   1.5  RT (min)      lcorbi_hilic_neg_4_085_59    m/z 250.9945     0   1   2   3   4E+4   0.5   1  RT (min)       lcorbi_hilic_neg_4_097_60    m/z 250.9946     0   1   2   3E+4   3   3.5  RT (min)      lcorbi_hilic_neg_4_068_57    m/z 251.0049     0   1   2   3   4E+3   2   2.5  RT (min)      lcorbi_hilic_neg_4_081_58    m/z 251.0200     0   2   4   6   8E+3   5   5.5  RT (min)      lcorbi_hilic_neg_1_008_39    m/z 251.0381     0   1   0.5   1  RT (min)       lcorbi_hilic_neg_4_093_60    m/z 251.0634     0   1   2.5   3  RT (min)      lcorbi_hilic_neg_4_071_57    m/z 251.0835     0   1   2   3   4E+4   2   2.5  RT (min)      lcorbi_hilic_neg_3_044_32    m/z 251.0882     0   1   5.5   6  RT (min)      lcorbi_hilic_neg_4_078_58    m/z 251.1109     0   1   2E+3   1   1.5  RT (min)      lcorbi_hilic_neg_4_082_59    m/z 251.1162     0   1   2E+4   6   6.5  RT (min)       lcorbi_hilic_neg_4_096_60    m/z 251.1481     0   1   2   3E+5   0.5   1  RT (min)      lcorbi_hilic_neg_4_113_62    m/z 251.1651     0   1E+3   0.5   1  RT (min)      lcorbi_hilic_neg_4_095_60    m/z 251.1964     0   2   4   6E+4   1   1.5  RT (min)       lcorbi_hilic_neg_3_011_24    m/z 251.8381     0   2   4E+3   0.5   1  RT (min)      lcorbi_hilic_neg_4_068_57    m/z 251.8449     0   2   4   6   8E+3   8   8.5  RT (min)      lcorbi_hilic_neg_4_098_61    m/z 251.8543     0   1   2E+3   8   8.5  RT (min)      lcorbi_hilic_neg_4_074_58    m/z 251.8851     0   1E+4   8   8.5  RT (min)      lcorbi_hilic_neg_4_066_57    m/z 251.9037     0   1   2E+4   7.5   8  RT (min)       lcorbi_hilic_neg_4_066_57    m/z 251.9177     0   1   2   3E+3   6   6.5  RT (min)      lcorbi_hilic_neg_4_078_58    m/z 251.9513     0   1   2   3E+4   3   3.5  RT (min)       lcorbi_hilic_neg_4_093_60    m/z 251.9606     0   1   2E+3   7   7.5  RT (min)      lcorbi_hilic_neg_4_088_59    m/z 251.9677     0   2   4E+3   5   5.5  RT (min)      lcorbi_hilic_neg_4_071_57    m/z 251.9795     0   2   4   6E+3   7   7.5  RT (min)       lcorbi_hilic_neg_1_023_23    m/z 251.9833     0   1E+4   1   1.5  RT (min)      lcorbi_hilic_neg_4_077_58    m/z 251.9865     0   1   2   3E+4   4   4.5  RT (min)      lcorbi_hilic_neg_4_086_59    m/z 252.0016     0   1   2   3   4E+4   1.5   2  RT (min)       lcorbi_hilic_neg_4_099_61    m/z 252.0251     0   1E+3   6   6.5  RT (min)      lcorbi_hilic_neg_4_096_60    m/z 252.0389     0   2   4   6E+4   7   7.5  RT (min)      lcorbi_hilic_neg_1_016_37    m/z 252.0489     0   1   2   2.5  RT (min)      lcorbi_hilic_neg_4_098_61    m/z 252.0517     0   2   4E+4   5.5   6  RT (min)       lcorbi_hilic_neg_4_071_57    m/z 252.0567     0   2   4   6   8E+2   4   4.5  RT (min)      lcorbi_hilic_neg_4_090_60    m/z 252.0664     0   1E+3   4.5   5  RT (min)       lcorbi_hilic_neg_4_005_32    m/z 252.0752     0   1   2E+3   5   5.5  RT (min)      lcorbi_hilic_neg_4_072_57    m/z 252.0875     0   1E+3   1.5   2  RT (min)      lcorbi_hilic_neg_4_095_60    m/z 252.0993     0   2   4   6E+4   3.5   4  RT (min)       lcorbi_hilic_neg_4_090_60    m/z 252.1239     0   1E+3   1   1.5  RT (min)      lcorbi_hilic_neg_3_028_23    m/z 252.1477     0   1E+4   6   6.5  RT (min)      lcorbi_hilic_neg_4_094_60    m/z 252.1735     0   2   4   6   8E+3   2.5   3  RT (min)      lcorbi_hilic_neg_4_089_59    m/z 252.7364     0   1   2   3E+4   3   3.5  RT (min)      lcorbi_hilic_neg_3_055_26    m/z 252.8188     0   2   4   6   8E+3   4.5   5  RT (min)      lcorbi_hilic_neg_4_076_58    m/z 252.8383     0   2   4E+3   4.5   5  RT (min)      lcorbi_hilic_neg_4_068_57    m/z 252.8425     0   1   2   3E+3   7.5   8  RT (min)      lcorbi_hilic_neg_4_067_57    m/z 252.9146     0   1E+4   7.5   8  RT (min)      lcorbi_hilic_neg_4_085_59    m/z 252.9147     0   2   4E+3   4.5   5  RT (min)      lcorbi_hilic_neg_4_068_57    m/z 252.9321     0   2   4   6   8E+3   3   3.5  RT (min)      lcorbi_hilic_neg_1_012_32    m/z 252.9735     0   1E+3   4.5   5  RT (min)       lcorbi_hilic_neg_1_010_22    m/z 252.9785     0   1E+4   4.5   5  RT (min)      lcorbi_hilic_neg_4_100_61    m/z 252.9902     0   1E+4   0.5   1  RT (min)      lcorbi_hilic_neg_4_079_58    m/z 252.9973     0   1   2   3E+3   6   6.5  RT (min)      lcorbi_hilic_neg_4_079_58    m/z 252.9978     0   1   2   3E+4   2   2.5  RT (min)       lcorbi_hilic_neg_4_099_61    m/z 253.0103     0   1E+4   0.5   1  RT (min)      lcorbi_hilic_neg_4_070_57    m/z 253.0115     0   1E+4   6.5   7  RT (min)      lcorbi_hilic_neg_4_089_59    m/z 253.0270     0   1   2E+3   1   1.5  RT (min)      lcorbi_hilic_neg_4_091_60    m/z 253.0384     0   2   4   6   8E+2   3   3.5  RT (min)      lcorbi_hilic_neg_4_050_26    m/z 253.0417     0   2   4   6E+2   1   1.5  RT (min)      lcorbi_hilic_neg_4_055_38    m/z 253.0498     0   2   4   6E+4   1   1.5  RT (min)       lcorbi_hilic_neg_4_113_62    m/z 253.0526     0   1E+5   6   6.5  RT (min)       lcorbi_hilic_neg_4_071_57    m/z 253.0596     0   1   3.5   4  RT (min)      lcorbi_hilic_neg_4_079_58    m/z 253.0705     0   2   4   6E+3   5.5   6  RT (min)      lcorbi_hilic_neg_4_105_61    m/z 253.0748     0   1E+4   1   1.5  RT (min)      lcorbi_hilic_neg_1_037_46    m/z 253.0805     0   2   4   6   8E+3   6   6.5  RT (min)      lcorbi_hilic_neg_4_088_59    m/z 253.0829     0   1   2   3   4E+4   3   3.5  RT (min)       lcorbi_hilic_neg_4_088_59    m/z 253.0927     0   1   2   3E+4   1.5   2  RT (min)       lcorbi_hilic_neg_4_088_59    m/z 253.1081     0   1E+3   1   1.5  RT (min)      lcorbi_hilic_neg_4_095_60    m/z 253.1192     0   1   2   2.5  RT (min)      lcorbi_hilic_neg_4_105_61    m/z 253.1304     0   2   4   6   8E+3   5.5   6  RT (min)      lcorbi_hilic_neg_4_092_60    m/z 253.1343     0   1   1.5   2  RT (min)       lcorbi_hilic_neg_4_091_60    m/z 253.2172     0   1   0.5   1  RT (min)      lcorbi_hilic_neg_4_091_60    m/z 253.8266     0   1   2E+4   8.5   9  RT (min)      lcorbi_hilic_neg_4_066_57    m/z 253.8267     0   1   2E+4   4.5   5  RT (min)       lcorbi_hilic_neg_4_107_62    m/z 253.8351     0   2   4E+3   0.5   1  RT (min)      lcorbi_hilic_neg_4_066_57    m/z 253.9021     0   2   4   6   8E+3   8   8.5  RT (min)      lcorbi_hilic_neg_4_075_58    m/z 253.9146     0   2   4E+3   6   6.5  RT (min)      lcorbi_hilic_neg_4_075_58    m/z 253.9560     0   1   3   3.5  RT (min)      lcorbi_hilic_neg_4_094_60    m/z 253.9808     0   2   4   6E+3   1   1.5  RT (min)      lcorbi_hilic_neg_1_026_34    m/z 253.9983     0   2   4   6E+3   1.5   2  RT (min)      lcorbi_hilic_neg_4_099_61    m/z 254.0039     0   2   4   6E+3   1.5   2  RT (min)      lcorbi_hilic_neg_4_093_60    m/z 254.0076     0   1E+4   6.5   7  RT (min)      lcorbi_hilic_neg_2_045_26    m/z 254.0265     0   2   4   6E+4   3   3.5  RT (min)      lcorbi_hilic_neg_1_002_26    m/z 254.0305     0   1E+4   1.5   2  RT (min)      lcorbi_hilic_neg_4_084_59    m/z 254.0602     0   1   2E+3   5.5   6  RT (min)      lcorbi_hilic_neg_4_091_60    m/z 254.0667     0   2   4   6E+3   1   1.5  RT (min)      lcorbi_hilic_neg_4_087_59    m/z 254.0845     0   1E+3   1.5   2  RT (min)       lcorbi_hilic_neg_4_089_59    m/z 254.0960     0   1E+3   3   3.5   4  RT (min)      lcorbi_hilic_neg_2_044_46    m/z 254.0999     0   1E+4   7   7.5  RT (min)      lcorbi_hilic_neg_4_096_60    m/z 254.1079     0   1   2E+4   3   3.5  RT (min)       lcorbi_hilic_neg_4_091_60    m/z 254.1146     0   1E+4   2   2.5  RT (min)       lcorbi_hilic_neg_4_097_60    m/z 254.1478     0   2   4   6E+3   1   1.5  RT (min)      lcorbi_hilic_neg_2_044_46    m/z 254.2487     0   1E+4   0.5   1  RT (min)      lcorbi_hilic_neg_3_008_04    m/z 254.7339     0   2   4   6E+3   2.5   3  RT (min)      lcorbi_hilic_neg_3_055_26    m/z 254.8983     0   1   2   3E+4   8   8.5  RT (min)       lcorbi_hilic_neg_4_075_58    m/z 254.9054     0   1   2   3E+4   5   5.5  RT (min)       lcorbi_hilic_neg_4_089_59    m/z 254.9133     0   1   7   7.5  RT (min)      lcorbi_hilic_neg_4_100_61    m/z 254.9167     0   1E+4   5   5.5  RT (min)      lcorbi_hilic_neg_4_101_61    m/z 254.9261     0   2   4E+3   7   7.5  RT (min)      lcorbi_hilic_neg_1_001_49    m/z 254.9600     0   1   5.5   6  RT (min)      lcorbi_hilic_neg_4_090_60    m/z 254.9814     0   1   1   1.5  RT (min)      lcorbi_hilic_neg_4_096_60    m/z 254.9861     0   1   2   3E+4   0.5   1  RT (min)       lcorbi_hilic_neg_2_017_51    m/z 254.9943     0   1   2E+3   0.5   1  RT (min)      lcorbi_hilic_neg_4_113_62    m/z 255.0059     0   1E+4   1   1.5  RT (min)       lcorbi_hilic_neg_4_091_60    m/z 255.0208     0   1   5.5   6  RT (min)      lcorbi_hilic_neg_4_094_60    m/z 255.0260     0   1   2E+4   0.5   1  RT (min)       lcorbi_hilic_neg_4_089_59    m/z 255.0289     0   1   2E+3   6   6.5  RT (min)      lcorbi_hilic_neg_4_095_60    m/z 255.0428     0   1E+3   1   1.5  RT (min)       lcorbi_hilic_neg_4_084_59    m/z 255.0718     0   1E+4   2.5   3  RT (min)      lcorbi_hilic_neg_4_080_58    m/z 255.0816     0   1   6.5   7  RT (min)      lcorbi_hilic_neg_4_091_60    m/z 255.0905     0   1E+5   3.5   4  RT (min)       lcorbi_hilic_neg_4_096_60    m/z 255.1096     0   2   4   6E+3   7.5   8  RT (min)      lcorbi_hilic_neg_4_079_58    m/z 255.1137     0   1   2   3E+4   1   1.5  RT (min)      lcorbi_hilic_neg_4_066_57    m/z 255.1349     0   1   1.5   2  RT (min)      lcorbi_hilic_neg_4_091_60    m/z 255.1600     0   1E+3   1   1.5  RT (min)       lcorbi_hilic_neg_4_092_60    m/z 255.2328     0   1   2   3E+4   4   4.5  RT (min)       lcorbi_hilic_neg_4_076_58    m/z 255.8221     0   2   4E+4   13.5   14  RT (min)       lcorbi_hilic_neg_4_103_61    m/z 255.8222     0   1   2E+5   10   12  RT (min)       lcorbi_hilic_neg_2_037_15    m/z 255.8290     0   1   8.5   9  RT (min)      lcorbi_hilic_neg_4_066_57    m/z 255.8373     0   2   4E+3   4.5   5  RT (min)      lcorbi_hilic_neg_4_067_57    m/z 255.8802     0   1   2   3E+4   6.5   7  RT (min)       lcorbi_hilic_neg_1_011_01    m/z 255.8803     0   1E+4   4.5   5  RT (min)       lcorbi_hilic_neg_4_098_61    m/z 255.8992     0   2   4   6   8E+3   8   8.5  RT (min)      lcorbi_hilic_neg_4_081_58    m/z 255.9241     0   1   2   3   4E+4   1   1.5  RT (min)       lcorbi_hilic_neg_4_106_62    m/z 255.9265     0   2   4   6E+3   6   6.5  RT (min)      lcorbi_hilic_neg_4_066_57    m/z 255.9757     0   1   6   6.5  RT (min)      lcorbi_hilic_neg_4_096_60    m/z 255.9916     0   1E+5   1.5   2  RT (min)       lcorbi_hilic_neg_4_082_59    m/z 255.9926     0   1   2   3E+4   7   7.5  RT (min)       lcorbi_hilic_neg_4_095_60    m/z 256.0024     0   2   4   6   8E+4   7   7.5  RT (min)       lcorbi_hilic_neg_4_091_60    m/z 256.0147     0   1   2   3E+4   6   6.5  RT (min)       lcorbi_hilic_neg_4_075_58    m/z 256.0162     0   1   3.5   4  RT (min)      lcorbi_hilic_neg_2_021_05    m/z 256.0233     0   2   4   6   8E+3   3   3.5  RT (min)      lcorbi_hilic_neg_4_075_58    m/z 256.0436     0   1E+4   3   3.5  RT (min)      lcorbi_hilic_neg_4_083_59    m/z 256.0436     0   1   2E+4   1   1.5  RT (min)       lcorbi_hilic_neg_4_106_62    m/z 256.0462     0   1E+4   4.5   5  RT (min)      lcorbi_hilic_neg_4_107_62    m/z 256.0507     0   1E+4   6   6.5  RT (min)      lcorbi_hilic_neg_4_091_60    m/z 256.0667     0   1   2   3E+4   4.5   5  RT (min)      lcorbi_hilic_neg_4_108_62    m/z 256.0769     0   1   2   3E+4   1.5   2  RT (min)       lcorbi_hilic_neg_4_088_59    m/z 256.0844     0   2   4E+4   1   1.5  RT (min)       lcorbi_hilic_neg_4_091_60    m/z 256.0855     0   1E+4   6   6.5  RT (min)      lcorbi_hilic_neg_4_081_58    m/z 256.0937     0   2   4   6   8E+3   5.5   6  RT (min)      lcorbi_hilic_neg_4_071_57    m/z 256.0938     0   2   4E+3   7   7.5  RT (min)      lcorbi_hilic_neg_4_089_59    m/z 256.0938     0   1   2E+4   1   1.5  RT (min)      lcorbi_hilic_neg_2_044_46    m/z 256.0938     0   1   2   3   4E+4   3.5   4  RT (min)      lcorbi_hilic_neg_1_047_05    m/z 256.1237     0   2   4E+2   4.5   5  RT (min)       lcorbi_hilic_neg_1_046_19    m/z 256.2361     0   2   4   6E+3   4   4.5  RT (min)      lcorbi_hilic_neg_4_081_58    m/z 256.2362     0   1E+5   0.5   1  RT (min)       lcorbi_hilic_neg_4_113_62    m/z 256.7573     0   1   2.5   3  RT (min)      lcorbi_hilic_neg_4_108_62    m/z 256.8219     0   1   2   3E+4   8.5   9  RT (min)       lcorbi_hilic_neg_4_066_57    m/z 256.8220     0   1   2E+4   4.5   5  RT (min)       lcorbi_hilic_neg_4_113_62    m/z 256.8359     0   1   2E+3   4.5   5  RT (min)      lcorbi_hilic_neg_2_020_27    m/z 256.8616     0   1E+4   7.5   8  RT (min)      lcorbi_hilic_neg_4_085_59    m/z 256.8901     0   2   4E+3   2   2.5  RT (min)      lcorbi_hilic_neg_4_090_60    m/z 256.8957     0   1E+4   8   8.5  RT (min)      lcorbi_hilic_neg_4_077_58    m/z 256.9026     0   1E+4   5   5.5  RT (min)      lcorbi_hilic_neg_4_076_58    m/z 256.9388     0   1   2E+4   1   1.5  RT (min)      lcorbi_hilic_neg_4_072_57    m/z 256.9608     0   1   2   3   4E+4   5   5.5  RT (min)      lcorbi_hilic_neg_1_010_22    m/z 256.9665     0   1   2.5   3  RT (min)      lcorbi_hilic_neg_4_093_60    m/z 256.9849     0   1   2E+4   0.5   1  RT (min)       lcorbi_hilic_neg_4_075_58    m/z 256.9849     0   1   2E+4   3   3.5  RT (min)       lcorbi_hilic_neg_4_073_57    m/z 256.9939     0   2   4E+3   0.5   1  RT (min)      lcorbi_hilic_neg_4_084_59    m/z 257.0029     0   1   2   3   4E+4   4   4.5  RT (min)      lcorbi_hilic_neg_4_091_60    m/z 257.0100     0   1E+3   2   2.5  RT (min)      lcorbi_hilic_neg_4_108_62    m/z 257.0122     0   1E+3   0.5   1  RT (min)      lcorbi_hilic_neg_4_102_61    m/z 257.0431     0   1   2   3   4E+4   4   4.5  RT (min)       lcorbi_hilic_neg_4_083_59    m/z 257.0443     0   2   4E+3   5.5   6  RT (min)      lcorbi_hilic_neg_3_027_45    m/z 257.0657     0   1E+4   7   7.5  RT (min)      lcorbi_hilic_neg_4_085_59    m/z 257.0673     0   1E+4   5.5   6  RT (min)      lcorbi_hilic_neg_4_081_58    m/z 257.0776     0   1   2   3E+3   1.5   2  RT (min)      lcorbi_hilic_neg_4_094_60    m/z 257.0839     0   1E+4   5.5   6  RT (min)      lcorbi_hilic_neg_4_090_60    m/z 257.1141     0   1   2   3   4E+5   7   7.5  RT (min)       lcorbi_hilic_neg_4_088_59    m/z 257.1392     0   1   2   3   4E+3   0.5   1  RT (min)      lcorbi_hilic_neg_4_095_60    m/z 257.1756     0   1   2   3E+4   1   1.5  RT (min)      lcorbi_hilic_neg_4_075_58    m/z 257.8191     0   2   4E+4   13   13.5  RT (min)       lcorbi_hilic_neg_4_086_59    m/z 257.8259     0   1E+4   8.5   9  RT (min)      lcorbi_hilic_neg_4_066_57    m/z 257.8448     0   2   4E+2   7.5   8  RT (min)      lcorbi_hilic_neg_1_003_14    m/z 257.9213     0   1E+4   1   1.5  RT (min)      lcorbi_hilic_neg_4_106_62    m/z 257.9684     0   1   6   6.5  RT (min)      lcorbi_hilic_neg_4_090_60    m/z 257.9836     0   1E+4   1.5   2  RT (min)      lcorbi_hilic_neg_4_066_57    m/z 257.9871     0   1   2   3   4E+2   5.5   6  RT (min)      lcorbi_hilic_neg_3_010_18    m/z 257.9936     0   1   2E+4   5.5   6  RT (min)      lcorbi_hilic_neg_4_066_57    m/z 258.0053     0   1E+3   6   6.5  RT (min)      lcorbi_hilic_neg_4_090_60    m/z 258.0077     0   1   2E+3   2   2.5  RT (min)      lcorbi_hilic_neg_4_069_57    m/z 258.0381     0   1E+4   8   8.5  RT (min)      lcorbi_hilic_neg_1_047_05    m/z 258.0389     0   1   2E+4   6   6.5  RT (min)       lcorbi_hilic_neg_4_097_60    m/z 258.0491     0   2   4E+5   4.5   5  RT (min)      lcorbi_hilic_neg_1_028_45    m/z 258.0666     0   1   2   2.5  RT (min)      lcorbi_hilic_neg_4_090_60    m/z 258.0813     0   1E+3   2   2.5  RT (min)      lcorbi_hilic_neg_4_089_59    m/z 258.0910     0   1   4.5   5  RT (min)       lcorbi_hilic_neg_2_020_27    m/z 258.1093     0   1   4.5   5  RT (min)      lcorbi_hilic_neg_4_109_62    m/z 258.1344     0   1   2   3E+4   1.5   2  RT (min)       lcorbi_hilic_neg_4_089_59    m/z 258.1456     0   1   2E+3   5.5   6  RT (min)       lcorbi_hilic_neg_4_073_57    m/z 258.1539     0   1E+4   2   2.5  RT (min)      lcorbi_hilic_neg_4_088_59    m/z 258.8188     0   1   2E+4   8.5   9  RT (min)       lcorbi_hilic_neg_4_066_57    m/z 258.8189     0   1E+4   4.5   5  RT (min)       lcorbi_hilic_neg_2_020_27    m/z 258.8367     0   1   2   3E+3   4.5   5  RT (min)      lcorbi_hilic_neg_2_020_27    m/z 258.9129     0   1E+4   6   6.5  RT (min)      lcorbi_hilic_neg_4_096_60    m/z 258.9173     0   2   4   6E+3   9   9.5  RT (min)      lcorbi_hilic_neg_4_098_61    m/z 258.9210     0   1   2E+4   4.5   5  RT (min)       lcorbi_hilic_neg_4_069_57    m/z 258.9357     0   1E+4   1   1.5  RT (min)      lcorbi_hilic_neg_4_073_57    m/z 258.9644     0   1E+3   3   3.5  RT (min)      lcorbi_hilic_neg_4_083_59    m/z 258.9789     0   2   4   6E+3   6   6.5  RT (min)      lcorbi_hilic_neg_4_073_57    m/z 258.9810     0   1E+4   1   1.5  RT (min)      lcorbi_hilic_neg_4_091_60    m/z 258.9915     0   1   1   1.5  RT (min)      lcorbi_hilic_neg_4_091_60    m/z 258.9973     0   1E+4   6   6.5  RT (min)      lcorbi_hilic_neg_4_070_57    m/z 259.0007     0   2   4   6   8E+4   0.5   1  RT (min)       lcorbi_hilic_neg_4_091_60    m/z 259.0028     0   1   2   3   4E+3   7.5   8  RT (min)      lcorbi_hilic_neg_3_028_23    m/z 259.0116     0   1   2   3   4E+3   2   2.5  RT (min)      lcorbi_hilic_neg_4_108_62    m/z 259.0155     0   1   2E+3   7.5   8  RT (min)      lcorbi_hilic_neg_4_048_37    m/z 259.0279     0   1   0.5   1  RT (min)      lcorbi_hilic_neg_4_092_60    m/z 259.0302     0   1   7.5   8  RT (min)      lcorbi_hilic_neg_4_082_59    m/z 259.0356     0   1E+4   4.5   5  RT (min)      lcorbi_hilic_neg_4_070_57    m/z 259.0358     0   1E+4   0.5   1  RT (min)      lcorbi_hilic_neg_4_113_62    m/z 259.0439     0   1   2   3E+5   5.5   6  RT (min)      lcorbi_hilic_neg_2_009_50    m/z 259.0448     0   1E+3   1   1.5  RT (min)      lcorbi_hilic_neg_4_091_60    m/z 259.0489     0   2   4E+3   2   2.5  RT (min)      lcorbi_hilic_neg_4_097_60    m/z 259.0781     0   2   4   6E+2   6   6.5  RT (min)      lcorbi_hilic_neg_1_004_17    m/z 259.0976     0   1   2E+3   6   6.5  RT (min)      lcorbi_hilic_neg_2_020_27    m/z 259.1016     0   1E+5   1   1.5  RT (min)       lcorbi_hilic_neg_4_094_60    m/z 259.1039     0   2   4   6E+2   3.5   4  RT (min)       lcorbi_hilic_neg_3_001_49    m/z 259.1047     0   1   6.5   7  RT (min)      lcorbi_hilic_neg_4_095_60    m/z 259.1296     0   1E+3   5.5   6  RT (min)      lcorbi_hilic_neg_1_030_04    m/z 259.1299     0   2   4   6E+3   2   2.5  RT (min)      lcorbi_hilic_neg_3_013_37    m/z 259.1409     0   2   4   6   8E+3   7.5   8  RT (min)      lcorbi_hilic_neg_4_083_59    m/z 259.1661     0   1E+4   0.5   1  RT (min)      lcorbi_hilic_neg_4_084_59    m/z 259.7602     0   1   3   3.5  RT (min)      lcorbi_hilic_neg_4_108_62    m/z 259.8160     0   1   2   3   4E+4   0   0.5  RT (min)       lcorbi_hilic_neg_4_078_58    m/z 259.8161     0   1E+5   4.5   5  RT (min)       lcorbi_hilic_neg_4_068_57    m/z 259.8464     0   2   4   6   8E+3   7.5   8  RT (min)      lcorbi_hilic_neg_4_106_62    m/z 259.8849     0   1   2E+5   1   1.5  RT (min)       lcorbi_hilic_neg_2_019_21    m/z 259.9012     0   2   4E+3   3   3.5  RT (min)      lcorbi_hilic_neg_4_098_61    m/z 259.9327     0   1E+4   3   3.5  RT (min)      lcorbi_hilic_neg_3_034_34    m/z 259.9555     0   1   2E+3   1.5   2  RT (min)      lcorbi_hilic_neg_4_069_57    m/z 260.0068     0   2   4   6   8E+3   5.5   6  RT (min)      lcorbi_hilic_neg_4_075_58    m/z 260.0072     0   1   2   3E+4   7.5   8  RT (min)       lcorbi_hilic_neg_4_090_60    m/z 260.0255     0   1   2   3E+4   7.5   8  RT (min)      lcorbi_hilic_neg_4_089_59    m/z 260.0378     0   1E+4   1   1.5  RT (min)       lcorbi_hilic_neg_4_082_59    m/z 260.0390     0   2   4   6   8E+3   5.5   6  RT (min)      lcorbi_hilic_neg_4_078_58    m/z 260.0623     0   2   4E+4   2.5   3  RT (min)      lcorbi_hilic_neg_1_037_46    m/z 260.0718     0   1   2E+3   4.5   5  RT (min)      lcorbi_hilic_neg_1_009_50    m/z 260.0731     0   1   2E+4   1   1.5  RT (min)       lcorbi_hilic_neg_4_082_59    m/z 260.0942     0   1   2E+4   6   6.5  RT (min)      lcorbi_hilic_neg_4_089_59    m/z 260.0965     0   1   3.5   4  RT (min)      lcorbi_hilic_neg_4_095_60    m/z 260.1251     0   1E+4   7.5   8  RT (min)      lcorbi_hilic_neg_4_082_59    m/z 260.1337     0   1   2   3E+4   1   1.5  RT (min)       lcorbi_hilic_neg_4_095_60    m/z 260.1362     0   1   2E+3   7.5   8  RT (min)      lcorbi_hilic_neg_4_088_59    m/z 260.8505     0   1   2E+4   4.5   5  RT (min)       lcorbi_hilic_neg_4_078_58    m/z 260.8730     0   1E+4   0   0.5  RT (min)      lcorbi_hilic_neg_2_003_20    m/z 260.8826     0   2   4   6E+3   7.5   8  RT (min)      lcorbi_hilic_neg_4_011_19    m/z 260.8920     0   1   2E+3   7   7.5  RT (min)      lcorbi_hilic_neg_4_112_62    m/z 260.9161     0   1   2   3   4E+3   8   8.5  RT (min)       lcorbi_hilic_neg_4_080_58    m/z 260.9390     0   1   7   7.5  RT (min)      lcorbi_hilic_neg_4_112_62    m/z 260.9461     0   2   4   6E+2   0.5   1  RT (min)      lcorbi_hilic_neg_2_003_20    m/z 260.9583     0   1   2E+4   7   7.5  RT (min)      lcorbi_hilic_neg_1_001_49    m/z 260.9621     0   1E+3   3   3.5  RT (min)      lcorbi_hilic_neg_4_082_59    m/z 260.9669     0   2   4E+3   0.5   1  RT (min)       lcorbi_hilic_neg_4_086_59    m/z 260.9790     0   1E+4   0.5   1  RT (min)      lcorbi_hilic_neg_4_091_60    m/z 260.9790     0   1   2E+4   6.5   7  RT (min)       lcorbi_hilic_neg_4_069_57    m/z 260.9957     0   1   2E+4   0.5   1  RT (min)       lcorbi_hilic_neg_4_069_57    m/z 261.0052     0   1E+3   6.5   7  RT (min)      lcorbi_hilic_neg_4_073_57    m/z 261.0263     0   2   4   6   8E+3   7.5   8  RT (min)      lcorbi_hilic_neg_4_086_59    m/z 261.0282     0   2   4   6   8E+3   5.5   6  RT (min)      lcorbi_hilic_neg_4_076_58    m/z 261.0556     0   2   4   6   8E+3   5.5   6  RT (min)      lcorbi_hilic_neg_4_081_58    m/z 261.0713     0   1   2E+4   3.5   4  RT (min)      lcorbi_hilic_neg_4_096_60    m/z 261.0724     0   1E+4   5   5.5  RT (min)      lcorbi_hilic_neg_4_089_59    m/z 261.0725     0   1E+4   6.5   7  RT (min)      lcorbi_hilic_neg_4_084_59    m/z 261.0744     0   2   4E+3   1   1.5  RT (min)       lcorbi_hilic_neg_4_089_59    m/z 261.0823     0   1   3.5   4  RT (min)       lcorbi_hilic_neg_4_101_61    m/z 261.0847     0   1E+5   6   6.5  RT (min)      lcorbi_hilic_neg_4_030_05    m/z 261.0976     0   1E+3   1.5   2  RT (min)      lcorbi_hilic_neg_4_094_60    m/z 261.1192     0   1   4   4.5  RT (min)      lcorbi_hilic_neg_1_002_26    m/z 261.1195     0   1   1.5   2  RT (min)       lcorbi_hilic_neg_4_083_59    m/z 261.8130     0   1   2E+4   4.5   5  RT (min)      lcorbi_hilic_neg_4_107_62    m/z 261.8130     0   1   2E+4   8.5   9  RT (min)      lcorbi_hilic_neg_4_066_57    m/z 261.8353     0   1   2E+4   7.5   8  RT (min)      lcorbi_hilic_neg_4_106_62    m/z 261.8438     0   1   2   3E+3   7.5   8  RT (min)      lcorbi_hilic_neg_4_106_62    m/z 261.8825     0   1E+5   0.5   1  RT (min)       lcorbi_hilic_neg_3_010_18    m/z 261.8922     0   1   2E+4   0   0.5  RT (min)       lcorbi_hilic_neg_4_083_59    m/z 261.8922     0   1   2E+4   8.5   9  RT (min)       lcorbi_hilic_neg_4_066_57    m/z 261.8922     0   2   4E+4   4   4.5  RT (min)       lcorbi_hilic_neg_4_066_57    m/z 261.9139     0   1   2   3E+4   8   8.5  RT (min)       lcorbi_hilic_neg_4_066_57    m/z 261.9307     0   1   3   3.5  RT (min)      lcorbi_hilic_neg_4_112_62    m/z 261.9783     0   1   2   3E+4   5   5.5   6  RT (min)       lcorbi_hilic_neg_4_066_57    m/z 261.9888     0   2   4E+3   3   3.5  RT (min)      lcorbi_hilic_neg_2_026_23    m/z 262.0120     0   1E+4   1   1.5  RT (min)      lcorbi_hilic_neg_4_079_58    m/z 262.0444     0   1   5   5.5  RT (min)      lcorbi_hilic_neg_2_009_50    m/z 262.0506     0   1   2E+3   1.5   2  RT (min)      lcorbi_hilic_neg_3_015_47    m/z 262.0594     0   1   2   3E+3   4.5   5  RT (min)      lcorbi_hilic_neg_4_098_61    m/z 262.0598     0   1E+3   6   6.5  RT (min)      lcorbi_hilic_neg_4_087_59    m/z 262.0786     0   1E+4   2   2.5  RT (min)      lcorbi_hilic_neg_4_073_57    m/z 262.0825     0   1E+4   6   6.5  RT (min)      lcorbi_hilic_neg_3_042_05    m/z 262.0947     0   1E+3   4   4.5  RT (min)       lcorbi_hilic_neg_4_103_61    m/z 262.1012     0   2   4   6E+3   6   6.5  RT (min)      lcorbi_hilic_neg_4_072_57    m/z 262.1067     0   2   4   6   8E+2   1.5   2  RT (min)      lcorbi_hilic_neg_4_017_51    m/z 262.1321     0   1E+4   3.5   4  RT (min)      lcorbi_hilic_neg_4_094_60    m/z 262.1405     0   1   2   3E+4   1   1.5  RT (min)       lcorbi_hilic_neg_4_082_59    m/z 262.7590     0   1E+5   4.5   5  RT (min)       lcorbi_hilic_neg_2_020_27    m/z 262.8476     0   1E+4   4.5   5  RT (min)      lcorbi_hilic_neg_4_079_58    m/z 262.8831     0   2   4E+3   1.5   2  RT (min)      lcorbi_hilic_neg_4_104_61    m/z 262.9023     0   2   4   6   8E+3   8   8.5  RT (min)      lcorbi_hilic_neg_4_092_60    m/z 262.9076     0   1E+3   1   1.5  RT (min)      lcorbi_hilic_neg_4_098_61    m/z 262.9594     0   1   2   3E+3   2   2.5  RT (min)      lcorbi_hilic_neg_3_037_12    m/z 262.9761     0   1E+4   5.5   6  RT (min)      lcorbi_hilic_neg_4_090_60    m/z 262.9863     0   2   4   6   8E+4   3.5   4  RT (min)       lcorbi_hilic_neg_4_074_58    m/z 262.9943     0   1   2E+4   0.5   1  RT (min)       lcorbi_hilic_neg_4_075_58    m/z 262.9944     0   1   2E+4   3   3.5  RT (min)       lcorbi_hilic_neg_4_075_58    m/z 263.0066     0   1   2   3E+4   5.5   6  RT (min)       lcorbi_hilic_neg_4_097_60    m/z 263.0121     0   1E+4   1   1.5  RT (min)      lcorbi_hilic_neg_4_083_59    m/z 263.0172     0   1   2E+4   5   5.5  RT (min)      lcorbi_hilic_neg_3_053_13    m/z 263.0230     0   1E+3   1.5   2  RT (min)       lcorbi_hilic_neg_4_084_59    m/z 263.0259     0   1   6   6.5  RT (min)      lcorbi_hilic_neg_4_098_61    m/z 263.0379     0   1E+4   5   5.5  RT (min)      lcorbi_hilic_neg_4_079_58    m/z 263.0480     0   1   2   3E+3   2.5   3  RT (min)       lcorbi_hilic_neg_3_006_28    m/z 263.0670     0   1E+3   3   3.5  RT (min)      lcorbi_hilic_neg_1_002_26    m/z 263.0742     0   1E+3   2   2.5  RT (min)       lcorbi_hilic_neg_2_009_50    m/z 263.0747     0   1   2   3E+4   5.5   6  RT (min)       lcorbi_hilic_neg_4_095_60    m/z 263.0827     0   1   2   3   4E+4   4.5   5  RT (min)       lcorbi_hilic_neg_4_093_60    m/z 263.0857     0   1E+4   7   7.5  RT (min)      lcorbi_hilic_neg_4_093_60    m/z 263.0938     0   1   2E+3   2   2.5  RT (min)       lcorbi_hilic_neg_4_073_57    m/z 263.1163     0   1   2   3   4E+2   3   3.5  RT (min)      lcorbi_hilic_neg_1_002_26    m/z 263.1321     0   2   4   6E+4   1   1.5  RT (min)       lcorbi_hilic_neg_3_028_23    m/z 263.1651     0   1E+4   0.5   1  RT (min)      lcorbi_hilic_neg_4_112_62    m/z 263.1992     0   1   2E+4   5   5.5  RT (min)       lcorbi_hilic_neg_4_095_60    m/z 263.7266     0   2   4   6E+3   3   3.5  RT (min)      lcorbi_hilic_neg_4_067_57    m/z 263.7948     0   1   2   3   4E+3   2.5   3  RT (min)      lcorbi_hilic_neg_3_055_26    m/z 263.8892     0   1E+4   8   8.5  RT (min)      lcorbi_hilic_neg_4_075_58    m/z 263.8893     0   1   2   3E+4   4   4.5  RT (min)       lcorbi_hilic_neg_4_109_62    m/z 263.8906     0   2   4   6   8E+3   1   1.5  RT (min)      lcorbi_hilic_neg_4_113_62    m/z 263.9173     0   1   2   3E+3   8   8.5  RT (min)      lcorbi_hilic_neg_4_077_58    m/z 263.9309     0   1   2   3E+4   8   8.5  RT (min)       lcorbi_hilic_neg_4_079_58    m/z 263.9609     0   1E+4   6.5   7  RT (min)      lcorbi_hilic_neg_4_076_58    m/z 263.9626     0   2   4   6   8E+4   3   3.5  RT (min)      lcorbi_hilic_neg_1_012_32    m/z 263.9849     0   1   2   3E+2   3   3.5  RT (min)      lcorbi_hilic_neg_1_012_32    m/z 264.0077     0   2   4   6E+3   4.5   5  RT (min)      lcorbi_hilic_neg_4_107_62    m/z 264.0101     0   2   4   6E+2   3   3.5  RT (min)      lcorbi_hilic_neg_1_012_32    m/z 264.0173     0   1E+4   7.5   8  RT (min)      lcorbi_hilic_neg_4_081_58    m/z 264.0236     0   1E+4   2   2.5  RT (min)      lcorbi_hilic_neg_1_051_33    m/z 264.0277     0   1   5.5   6  RT (min)      lcorbi_hilic_neg_3_042_05    m/z 264.0344     0   1E+4   2   2.5  RT (min)      lcorbi_hilic_neg_4_098_61    m/z 264.0365     0   1   6.5   7  RT (min)      lcorbi_hilic_neg_2_032_25    m/z 264.0554     0   2   4   6E+4   3   3.5  RT (min)      lcorbi_hilic_neg_3_055_26    m/z 264.0584     0   1E+5   7.5   8  RT (min)       lcorbi_hilic_neg_1_047_05    m/z 264.0619     0   2   4   6E+3   5   5.5  RT (min)      lcorbi_hilic_neg_4_070_57    m/z 264.0640     0   1E+4   1.5   2  RT (min)      lcorbi_hilic_neg_4_091_60    m/z 264.0710     0   1E+3   6.5   7  RT (min)      lcorbi_hilic_neg_4_089_59    m/z 264.0871     0   1E+3   1.5   2  RT (min)      lcorbi_hilic_neg_1_037_46    m/z 264.0947     0   1   6   6.5  RT (min)      lcorbi_hilic_neg_4_085_59    m/z 264.0956     0   1E+3   1.5   2  RT (min)      lcorbi_hilic_neg_4_108_62    m/z 264.1076     0   2   4   6   8E+4   4   4.5  RT (min)      lcorbi_hilic_neg_3_028_23    m/z 264.1114     0   1   2E+4   6.5   7  RT (min)      lcorbi_hilic_neg_1_012_32    m/z 264.1377     0   1E+4   1.5   2  RT (min)      lcorbi_hilic_neg_4_082_59    m/z 264.8185     0   2   4E+3   8   8.5  RT (min)      lcorbi_hilic_neg_4_066_57    m/z 264.8369     0   1E+4   0.5   1  RT (min)      lcorbi_hilic_neg_4_070_57    m/z 264.9242     0   1   4.5   5  RT (min)      lcorbi_hilic_neg_4_005_32    m/z 264.9443     0   2   4   6   8E+4   1   1.5  RT (min)       lcorbi_hilic_neg_4_106_62    m/z 264.9448     0   1E+4   6   6.5  RT (min)      lcorbi_hilic_neg_4_076_58    m/z 264.9680     0   1E+3   1   1.5  RT (min)      lcorbi_hilic_neg_4_085_59    m/z 264.9909     0   1   2E+4   0.5   1  RT (min)      lcorbi_hilic_neg_4_002_41    m/z 265.0101     0   1E+4   0.5   1  RT (min)      lcorbi_hilic_neg_4_113_62    m/z 265.0109     0   2   4E+3   6.5   7  RT (min)      lcorbi_hilic_neg_4_108_62    m/z 265.0188     0   1   4   4.5  RT (min)      lcorbi_hilic_neg_4_107_62    m/z 265.0192     0   1E+3   2   2.5  RT (min)      lcorbi_hilic_neg_4_095_60    m/z 265.0205     0   1   6.5   7  RT (min)      lcorbi_hilic_neg_4_110_62    m/z 265.0328     0   1E+3   5.5   6  RT (min)      lcorbi_hilic_neg_4_111_62    m/z 265.0403     0   2   4E+3   2   2.5  RT (min)      lcorbi_hilic_neg_4_111_62    m/z 265.0443     0   1   5.5   6  RT (min)      lcorbi_hilic_neg_4_110_62    m/z 265.0675     0   1   2E+4   7   7.5  RT (min)       lcorbi_hilic_neg_4_087_59    m/z 265.0699     0   2   4   6E+4   4   4.5  RT (min)       lcorbi_hilic_neg_4_096_60    m/z 265.0830     0   1   2   3E+2   6   6.5  RT (min)      lcorbi_hilic_neg_4_030_05    m/z 265.0927     0   1   2   3   4E+4   3   4  RT (min)       lcorbi_hilic_neg_4_073_57    m/z 265.0931     0   1E+3   6   6.5  RT (min)       lcorbi_hilic_neg_4_097_60    m/z 265.1239     0   1   3   3.5  RT (min)      lcorbi_hilic_neg_4_070_57    m/z 265.1479     0   2   4   6E+3   4   4.5  RT (min)       lcorbi_hilic_neg_1_016_37    m/z 265.1571     0   1E+4   3.5   4  RT (min)      lcorbi_hilic_neg_4_073_57    m/z 265.1800     0   1   0.5   1  RT (min)       lcorbi_hilic_neg_4_092_60    m/z 265.7925     0   1   2.5   3  RT (min)      lcorbi_hilic_neg_4_108_62    m/z 265.8521     0   1   2E+4   3   3.5  RT (min)      lcorbi_hilic_neg_4_066_57    m/z 265.9223     0   1E+3   8   8.5  RT (min)       lcorbi_hilic_neg_4_076_58    m/z 265.9529     0   1E+4   1   1.5  RT (min)      lcorbi_hilic_neg_4_112_62    m/z 265.9605     0   2   4   6   8E+4   3   3.5  RT (min)      lcorbi_hilic_neg_1_012_32    m/z 265.9838     0   1   2E+4   1   1.5  RT (min)       lcorbi_hilic_neg_4_106_62    m/z 265.9951     0   1E+4   6   6.5  RT (min)      lcorbi_hilic_neg_4_089_59    m/z 266.0047     0   2   4   6E+3   6   6.5  RT (min)      lcorbi_hilic_neg_4_083_59    m/z 266.0269     0   2   4   6   8E+3   5.5   6  RT (min)      lcorbi_hilic_neg_4_105_61    m/z 266.0368     0   1   2   3E+4   5.5   6  RT (min)       lcorbi_hilic_neg_4_076_58    m/z 266.0392     0   2   4   6   8E+3   7.5   8  RT (min)      lcorbi_hilic_neg_4_092_60    m/z 266.0423     0   2   4   6E+3   3   3.5  RT (min)      lcorbi_hilic_neg_4_066_57    m/z 266.0444     0   1   1.5   2  RT (min)      lcorbi_hilic_neg_4_106_62    m/z 266.0570     0   1   7.5   8  RT (min)      lcorbi_hilic_neg_4_067_57    m/z 266.0618     0   1   2   3E+3   1.5   2  RT (min)      lcorbi_hilic_neg_4_094_60    m/z 266.0782     0   2   4   6   8E+4   1.5   2  RT (min)      lcorbi_hilic_neg_4_013_46    m/z 266.0790     0   1E+4   2.5   3  RT (min)      lcorbi_hilic_neg_2_044_46    m/z 266.0880     0   2   4   6E+4   6.5   7  RT (min)       lcorbi_hilic_neg_4_089_59    m/z 266.1082     0   2   4E+4   3   3.5  RT (min)       lcorbi_hilic_neg_4_095_60    m/z 266.1115     0   1   2E+5   1   1.5  RT (min)      lcorbi_hilic_neg_1_035_29    m/z 266.1145     0   1   3   3.5   4  RT (min)       lcorbi_hilic_neg_4_093_60    m/z 266.1244     0   2   4E+3   6.5   7  RT (min)      lcorbi_hilic_neg_4_095_60    m/z 266.1245     0   2   4   6E+2   2.5   3  RT (min)      lcorbi_hilic_neg_4_009_50    m/z 266.1398     0   1   2   3   4E+3   0.5   1  RT (min)      lcorbi_hilic_neg_4_105_61    m/z 266.8039     0   1   2   3   4E+4   4.5   5  RT (min)       lcorbi_hilic_neg_4_105_61    m/z 266.8726     0   1E+4   8   8.5  RT (min)      lcorbi_hilic_neg_4_113_62    m/z 266.9151     0   2   4   6   8E+3   1   1.5  RT (min)      lcorbi_hilic_neg_4_066_57    m/z 266.9246     0   1E+4   8   8.5  RT (min)      lcorbi_hilic_neg_4_081_58    m/z 266.9460     0   2   4   6E+3   5.5   6  RT (min)      lcorbi_hilic_neg_4_075_58    m/z 266.9599     0   1   1   1.5  RT (min)      lcorbi_hilic_neg_4_091_60    m/z 266.9860     0   1   2   3E+4   3   3.5  RT (min)      lcorbi_hilic_neg_4_088_59    m/z 267.0095     0   1E+4   0.5   1  RT (min)      lcorbi_hilic_neg_4_079_58    m/z 267.0207     0   1E+4   6.5   7  RT (min)      lcorbi_hilic_neg_4_089_59    m/z 267.0222     0   1   2E+3   1.5   2  RT (min)      lcorbi_hilic_neg_4_085_59    m/z 267.0328     0   1   0.5   1  RT (min)      lcorbi_hilic_neg_4_092_60    m/z 267.0565     0   1   2.5   3  RT (min)      lcorbi_hilic_neg_4_082_59    m/z 267.0664     0   1   2   3E+4   5.5   6  RT (min)      lcorbi_hilic_neg_4_070_57    m/z 267.0670     0   2   4   6   8E+4   1.5   2  RT (min)       lcorbi_hilic_neg_4_093_60    m/z 267.0671     0   1   4   4.5  RT (min)       lcorbi_hilic_neg_4_083_59    m/z 267.0694     0   2   4E+2   0.5   1  RT (min)      lcorbi_hilic_neg_1_023_23    m/z 267.0723     0   1   2E+3   5.5   6  RT (min)      lcorbi_hilic_neg_4_035_48    m/z 267.0731     0   1E+4   4   4.5  RT (min)      lcorbi_hilic_neg_4_083_59    m/z 267.0871     0   1E+4   1   1.5  RT (min)      lcorbi_hilic_neg_4_068_57    m/z 267.0915     0   1   2E+3   5.5   6  RT (min)       lcorbi_hilic_neg_4_071_57    m/z 267.1112     0   1E+4   1.5   2  RT (min)      lcorbi_hilic_neg_4_096_60    m/z 267.1195     0   1   2E+4   6.5   7  RT (min)      lcorbi_hilic_neg_2_020_27    m/z 267.1459     0   1   6   6.5  RT (min)      lcorbi_hilic_neg_4_073_57    m/z 267.1598     0   2   4   6E+3   0.5   1  RT (min)      lcorbi_hilic_neg_4_055_38    m/z 267.1964     0   2   4E+4   1   1.5  RT (min)      lcorbi_hilic_neg_4_045_25    m/z 267.8501     0   1   2E+4   3   3.5  RT (min)      lcorbi_hilic_neg_4_066_57    m/z 267.8703     0   1E+4   8   8.5  RT (min)      lcorbi_hilic_neg_4_066_57    m/z 267.9088     0   2   4E+3   1   1.5  RT (min)      lcorbi_hilic_neg_4_081_58    m/z 267.9806     0   2   4   6   8E+3   1.5   2  RT (min)      lcorbi_hilic_neg_4_092_60    m/z 267.9825     0   1E+4   3   3.5  RT (min)      lcorbi_hilic_neg_1_029_31    m/z 267.9968     0   1   2E+3   3   3.5  RT (min)      lcorbi_hilic_neg_1_029_31    m/z 268.0165     0   2   4   6E+4   1   1.5  RT (min)      lcorbi_hilic_neg_2_050_17    m/z 268.0253     0   1   2E+4   5.5   6   6.5  RT (min)       lcorbi_hilic_neg_4_071_57    m/z 268.0309     0   1E+4   1.5   2  RT (min)      lcorbi_hilic_neg_4_073_57    m/z 268.0374     0   1E+4   3.5   4  RT (min)      lcorbi_hilic_neg_4_110_62    m/z 268.0480     0   1E+4   1.5   2  RT (min)      lcorbi_hilic_neg_4_066_57    m/z 268.0548     0   1   2E+4   7.5   8  RT (min)      lcorbi_hilic_neg_1_011_01    m/z 268.0607     0   1   2E+3   1.5   2  RT (min)      lcorbi_hilic_neg_4_089_59    m/z 268.0702     0   1   6   6.5  RT (min)       lcorbi_hilic_neg_4_095_60    m/z 268.0800     0   1   2E+3   5.5   6  RT (min)      lcorbi_hilic_neg_4_073_57    m/z 268.0952     0   1   1   1.5  RT (min)      lcorbi_hilic_neg_4_106_62    m/z 268.1034     0   1   6   6.5  RT (min)       lcorbi_hilic_neg_4_095_60    m/z 268.1038     0   1   3.5   4  RT (min)      lcorbi_hilic_neg_4_089_59    m/z 268.1176     0   1E+3   6   6.5  RT (min)      lcorbi_hilic_neg_4_090_60    m/z 268.1234     0   2   4   6E+3   3.5   4  RT (min)      lcorbi_hilic_neg_3_011_24    m/z 268.1234     0   1   2   3E+4   6   6.5  RT (min)       lcorbi_hilic_neg_4_096_60    m/z 268.1429     0   2   4   6   8E+4   4.5   5  RT (min)       lcorbi_hilic_neg_3_011_24    m/z 268.1509     0   1E+5   7   7.5  RT (min)       lcorbi_hilic_neg_4_096_60    m/z 268.1549     0   1E+3   0.5   1  RT (min)      lcorbi_hilic_neg_4_095_60    m/z 268.1683     0   1E+4   3   3.5  RT (min)      lcorbi_hilic_neg_4_108_62    m/z 268.8007     0   1E+4   0   0.5  RT (min)      lcorbi_hilic_neg_4_075_58    m/z 268.8856     0   1E+4   0.5   1  RT (min)      lcorbi_hilic_neg_4_077_58    m/z 268.8863     0   1E+4   8.5   9  RT (min)      lcorbi_hilic_neg_4_112_62    m/z 268.9012     0   1E+4   4.5   5  RT (min)      lcorbi_hilic_neg_4_102_61    m/z 268.9417     0   1E+4   6   6.5  RT (min)      lcorbi_hilic_neg_4_096_60    m/z 268.9633     0   1   3   3.5  RT (min)       lcorbi_hilic_neg_4_106_62    m/z 268.9851     0   1   2E+4   3   3.5  RT (min)      lcorbi_hilic_neg_4_083_59    m/z 269.0018     0   1E+4   0.5   1  RT (min)      lcorbi_hilic_neg_4_087_59    m/z 269.0123     0   2   4E+3   0.5   1  RT (min)      lcorbi_hilic_neg_4_089_59    m/z 269.0432     0   1   2   3   4E+3   1   1.5  RT (min)       lcorbi_hilic_neg_4_084_59    m/z 269.0446     0   1E+4   6.5   7  RT (min)      lcorbi_hilic_neg_4_089_59    m/z 269.0489     0   1   0.5   1  RT (min)      lcorbi_hilic_neg_4_089_59    m/z 269.0600     0   2   4E+3   2   2.5  RT (min)      lcorbi_hilic_neg_4_083_59    m/z 269.0778     0   1   2   3   4E+2   3.5   4  RT (min)       lcorbi_hilic_neg_3_011_24    m/z 269.0830     0   1   7.5   8  RT (min)      lcorbi_hilic_neg_1_016_37    m/z 269.0876     0   1E+5   5   5.5  RT (min)       lcorbi_hilic_neg_4_074_58    m/z 269.0956     0   2   4   6E+4   6   6.5  RT (min)       lcorbi_hilic_neg_4_092_60    m/z 269.1030     0   1   2   3   4E+4   1   1.5  RT (min)       lcorbi_hilic_neg_4_090_60    m/z 269.1310     0   1   2   3   4E+3   0.5   1   1.5  RT (min)       lcorbi_hilic_neg_4_089_59    m/z 269.1393     0   2   4   6E+3   0.5   1  RT (min)      lcorbi_hilic_neg_4_084_59    m/z 269.1505     0   1   2E+4   1.5   2  RT (min)       lcorbi_hilic_neg_4_073_57    m/z 269.1744     0   1E+4   7.5   8  RT (min)      lcorbi_hilic_neg_4_096_60    m/z 269.1756     0   1E+4   1   1.5  RT (min)      lcorbi_hilic_neg_4_075_58    m/z 269.2484     0   1E+4   1   1.5  RT (min)      lcorbi_hilic_neg_3_048_33    m/z 269.8378     0   2   4   6E+3   0.5   1  RT (min)      lcorbi_hilic_neg_3_052_40    m/z 269.9293     0   1E+4   7   7.5  RT (min)      lcorbi_hilic_neg_4_082_59    m/z 269.9301     0   2   4E+3   3   3.5  RT (min)      lcorbi_hilic_neg_4_103_61    m/z 269.9606     0   1E+4   1   1.5  RT (min)      lcorbi_hilic_neg_4_074_58    m/z 269.9802     0   2   4   6E+2   3   3.5  RT (min)      lcorbi_hilic_neg_2_011_29    m/z 269.9843     0   1E+3   1   1.5  RT (min)      lcorbi_hilic_neg_4_069_57    m/z 269.9914     0   1E+3   5.5   6  RT (min)      lcorbi_hilic_neg_4_076_58    m/z 270.0357     0   1   2E+4   5.5   6  RT (min)       lcorbi_hilic_neg_4_077_58    m/z 270.0479     0   1   2E+3   5.5   6  RT (min)      lcorbi_hilic_neg_4_079_58    m/z 270.0576     0   1E+3   4   4.5  RT (min)      lcorbi_hilic_neg_4_093_60    m/z 270.0608     0   2   4   6E+3   1   1.5  RT (min)      lcorbi_hilic_neg_4_092_60    m/z 270.0647     0   1   4   4.5  RT (min)      lcorbi_hilic_neg_4_093_60    m/z 270.0674     0   2   4   6   8E+3   6   6.5  RT (min)      lcorbi_hilic_neg_4_005_32    m/z 270.0746     0   1E+4   3   3.5   4  RT (min)       lcorbi_hilic_neg_1_012_32    m/z 270.0811     0   1E+4   0.5   1  RT (min)      lcorbi_hilic_neg_4_083_59    m/z 270.0910     0   2   4E+4   2.5   3  RT (min)      lcorbi_hilic_neg_1_037_46    m/z 270.0910     0   1   2   3   4E+3   5   5.5  RT (min)      lcorbi_hilic_neg_4_097_60    m/z 270.0926     0   2   4   6E+3   7.5   8  RT (min)      lcorbi_hilic_neg_2_032_25    m/z 270.1063     0   2   4   6   8E+3   1   1.5  RT (min)      lcorbi_hilic_neg_4_091_60    m/z 270.1204     0   1E+3   6.5   7  RT (min)      lcorbi_hilic_neg_4_096_60    m/z 270.1280     0   1   4.5   5  RT (min)      lcorbi_hilic_neg_4_090_60    m/z 270.1572     0   1E+4   7   7.5  RT (min)      lcorbi_hilic_neg_4_091_60    m/z 270.1660     0   2   4   6   8E+3   2   2.5  RT (min)      lcorbi_hilic_neg_4_097_60    m/z 270.7978     0   1   2   3   4E+4   4.5   5  RT (min)       lcorbi_hilic_neg_4_100_61    m/z 270.8836     0   2   4   6E+3   8.5   9  RT (min)      lcorbi_hilic_neg_4_106_62    m/z 270.8923     0   2   4E+3   4.5   5  RT (min)      lcorbi_hilic_neg_4_082_59    m/z 270.9080     0   2   4   6E+3   1   1.5  RT (min)      lcorbi_hilic_neg_4_081_58    m/z 270.9253     0   1E+4   7.5   8  RT (min)      lcorbi_hilic_neg_4_085_59    m/z 270.9458     0   1   2E+4   3   3.5  RT (min)       lcorbi_hilic_neg_4_073_57    m/z 270.9669     0   1E+3   1.5   2  RT (min)      lcorbi_hilic_neg_3_034_34    m/z 270.9809     0   1   2E+4   1   1.5  RT (min)       lcorbi_hilic_neg_4_067_57    m/z 271.0005     0   1E+4   3   3.5  RT (min)      lcorbi_hilic_neg_4_073_57    m/z 271.0006     0   1   2E+4   0.5   1  RT (min)       lcorbi_hilic_neg_4_068_57    m/z 271.0077     0   1   2E+4   6.5   7  RT (min)       lcorbi_hilic_neg_4_069_57    m/z 271.0223     0   1E+3   7.5   8  RT (min)      lcorbi_hilic_neg_4_094_60    m/z 271.0280     0   1   0.5   1  RT (min)      lcorbi_hilic_neg_4_085_59    m/z 271.0344     0   1E+4   6.5   7  RT (min)      lcorbi_hilic_neg_4_090_60    m/z 271.0402     0   1E+4   1   1.5  RT (min)       lcorbi_hilic_neg_4_085_59    m/z 271.0546     0   1   2   3E+4   5.5   6  RT (min)      lcorbi_hilic_neg_1_023_23    m/z 271.0592     0   1E+4   1.5   2  RT (min)       lcorbi_hilic_neg_4_004_13    m/z 271.0683     0   1E+4   1   1.5  RT (min)      lcorbi_hilic_neg_4_013_46    m/z 271.0747     0   1   5   5.5  RT (min)      lcorbi_hilic_neg_4_076_58    m/z 271.0909     0   1   2   3   4E+2   6   6.5  RT (min)      lcorbi_hilic_neg_1_047_05    m/z 271.0951     0   1E+5   2   3  RT (min)       lcorbi_hilic_neg_4_089_59    m/z 271.1136     0   1   2E+4   6   6.5  RT (min)      lcorbi_hilic_neg_4_030_05    m/z 271.1139     0   1   4   4.5  RT (min)      lcorbi_hilic_neg_4_087_59    m/z 271.1295     0   1   1.5   2  RT (min)      lcorbi_hilic_neg_4_093_60    m/z 271.1297     0   1E+4   6.5   7  RT (min)      lcorbi_hilic_neg_4_088_59    m/z 271.1579     0   1   2   3E+3   1   1.5  RT (min)      lcorbi_hilic_neg_4_084_59    m/z 271.1913     0   2   4E+3   1   1.5  RT (min)      lcorbi_hilic_neg_1_051_33    m/z 271.2197     0   2   4   6E+3   1   1.5  RT (min)      lcorbi_hilic_neg_1_026_34    m/z 271.8348     0   1   2   3   4E+3   0.5   1  RT (min)      lcorbi_hilic_neg_3_052_40    m/z 271.9069     0   1   2   3   4E+3   1   1.5  RT (min)      lcorbi_hilic_neg_3_016_16    m/z 271.9845     0   1E+4   0.5   1  RT (min)      lcorbi_hilic_neg_3_024_27    m/z 272.0021     0   1E+4   1.5   2  RT (min)      lcorbi_hilic_neg_4_089_59    m/z 272.0069     0   1   2   3E+4   6.5   7  RT (min)       lcorbi_hilic_neg_4_086_59    m/z 272.0175     0   1E+4   1   1.5  RT (min)      lcorbi_hilic_neg_4_106_62    m/z 272.0176     0   1   2   3E+3   7   7.5  RT (min)      lcorbi_hilic_neg_1_012_32    m/z 272.0413     0   1E+4   3   3.5  RT (min)      lcorbi_hilic_neg_4_089_59    m/z 272.0542     0   1E+3   6   6.5  RT (min)      lcorbi_hilic_neg_4_090_60    m/z 272.0548     0   1   4   4.5  RT (min)      lcorbi_hilic_neg_4_096_60    m/z 272.0622     0   1   2   2.5  RT (min)      lcorbi_hilic_neg_4_090_60    m/z 272.0927     0   1   1.5   2  RT (min)      lcorbi_hilic_neg_4_089_59    m/z 272.1076     0   1   6.5   7  RT (min)      lcorbi_hilic_neg_1_029_31    m/z 272.1261     0   2   4   6   8E+4   6   6.5  RT (min)       lcorbi_hilic_neg_4_096_60    m/z 272.1265     0   1   2   3E+5   3.5   4  RT (min)       lcorbi_hilic_neg_4_093_60    m/z 272.1290     0   2   4   6   8E+3   1   1.5  RT (min)      lcorbi_hilic_neg_4_070_57    m/z 272.1345     0   1   2   3   4E+4   4   4.5  RT (min)       lcorbi_hilic_neg_4_096_60    m/z 272.7879     0   1   4.5   5  RT (min)      lcorbi_hilic_neg_4_098_61    m/z 272.7950     0   1E+4   4.5   5  RT (min)      lcorbi_hilic_neg_4_100_61    m/z 272.8830     0   1   2   3E+3   0.5   1  RT (min)      lcorbi_hilic_neg_1_001_49    m/z 272.9125     0   1E+4   1   1.5  RT (min)      lcorbi_hilic_neg_4_073_57    m/z 272.9287     0   1E+3   5.5   6  RT (min)      lcorbi_hilic_neg_4_101_61    m/z 272.9342     0   1E+4   1.5   2  RT (min)      lcorbi_hilic_neg_4_074_58    m/z 272.9438     0   1   2E+4   3   3.5  RT (min)       lcorbi_hilic_neg_4_073_57    m/z 272.9588     0   1E+4   12   12.5  RT (min)      lcorbi_hilic_neg_4_104_61    m/z 272.9702     0   1E+3   1   1.5  RT (min)      lcorbi_hilic_neg_4_091_60    m/z 272.9769     0   2   4   6   8E+3   1   1.5  RT (min)       lcorbi_hilic_neg_4_069_57    m/z 272.9879     0   1   1   1.5  RT (min)      lcorbi_hilic_neg_4_091_60    m/z 272.9963     0   1   2E+4   1   1.5  RT (min)       lcorbi_hilic_neg_4_093_60    m/z 272.9971     0   1E+5   3   3.5  RT (min)       lcorbi_hilic_neg_4_069_57    m/z 273.0048     0   1   6.5   7  RT (min)      lcorbi_hilic_neg_4_113_62    m/z 273.0165     0   1   2   3E+4   3   3.5  RT (min)       lcorbi_hilic_neg_4_067_57    m/z 273.0285     0   1   2E+4   2   2.5  RT (min)       lcorbi_hilic_neg_4_073_57    m/z 273.0375     0   2   4E+3   2   2.5  RT (min)      lcorbi_hilic_neg_4_072_57    m/z 273.0378     0   1E+4   6.5   7  RT (min)      lcorbi_hilic_neg_3_028_23    m/z 273.0480     0   1   1.5   2  RT (min)      lcorbi_hilic_neg_4_084_59    m/z 273.0587     0   1   2   3   4E+3   1.5   2   2.5  RT (min)       lcorbi_hilic_neg_4_077_58    m/z 273.0675     0   1E+3   2   2.5  RT (min)       lcorbi_hilic_neg_4_080_58    m/z 273.0703     0   1   2E+4   6.5   7  RT (min)       lcorbi_hilic_neg_4_089_59    m/z 273.0765     0   1E+3   1   1.5  RT (min)      lcorbi_hilic_neg_4_089_59    m/z 273.0920     0   2   4E+4   6.5   7  RT (min)       lcorbi_hilic_neg_4_096_60    m/z 273.1011     0   2   4   6E+4   2.5   3  RT (min)      lcorbi_hilic_neg_1_037_46    m/z 273.1089     0   1E+3   6   6.5  RT (min)       lcorbi_hilic_neg_4_030_05    m/z 273.1092     0   2   4   6E+3   3.5   4  RT (min)      lcorbi_hilic_neg_4_088_59    m/z 273.1202     0   2   4   6   8E+4   6.5   7  RT (min)       lcorbi_hilic_neg_4_030_05    m/z 273.1493     0   1E+4   0.5   1  RT (min)      lcorbi_hilic_neg_2_004_09    m/z 273.1720     0   1E+4   1   1.5  RT (min)       lcorbi_hilic_neg_4_084_59    m/z 273.2341     0   1E+4   1   1.5  RT (min)      lcorbi_hilic_neg_4_076_58    m/z 273.8363     0   1   2   3E+3   8   8.5  RT (min)      lcorbi_hilic_neg_4_070_57    m/z 273.8829     0   1E+4   6   6.5  RT (min)      lcorbi_hilic_neg_4_100_61    m/z 273.8907     0   1   2   3E+3   6.5   7  RT (min)      lcorbi_hilic_neg_4_098_61    m/z 273.9075     0   2   4   6E+3   1   1.5  RT (min)      lcorbi_hilic_neg_4_081_58    m/z 273.9425     0   1   7   7.5  RT (min)      lcorbi_hilic_neg_4_074_58    m/z 273.9802     0   1   2   3E+4   3   3.5  RT (min)       lcorbi_hilic_neg_4_093_60    m/z 274.0025     0   2   4   6   8E+3   2   2.5  RT (min)      lcorbi_hilic_neg_4_080_58    m/z 274.0042     0   1   2E+3   6.5   7  RT (min)      lcorbi_hilic_neg_4_078_58    m/z 274.0208     0   1   2E+3   7.5   8  RT (min)      lcorbi_hilic_neg_4_053_23    m/z 274.0332     0   2   4   6E+3   2   2.5  RT (min)      lcorbi_hilic_neg_4_100_61    m/z 274.0388     0   1E+3   0.5   1  RT (min)      lcorbi_hilic_neg_4_077_58    m/z 274.0698     0   2   4   6E+3   7   7.5  RT (min)      lcorbi_hilic_neg_4_083_59    m/z 274.0712     0   1E+3   2   2.5  RT (min)      lcorbi_hilic_neg_4_077_58    m/z 274.0749     0   1E+4   5   5.5  RT (min)      lcorbi_hilic_neg_4_110_62    m/z 274.0785     0   1   2   3E+4   1   1.5  RT (min)       lcorbi_hilic_neg_4_096_60    m/z 274.0794     0   1E+3   2.5   3  RT (min)      lcorbi_hilic_neg_3_051_46    m/z 274.0890     0   2   4E+4   1   1.5  RT (min)      lcorbi_hilic_neg_4_089_59    m/z 274.0906     0   1E+4   5   5.5  RT (min)      lcorbi_hilic_neg_4_066_57    m/z 274.0932     0   1E+4   2   2.5  RT (min)      lcorbi_hilic_neg_1_009_50    m/z 274.1043     0   2   4   6E+4   7   7.5  RT (min)       lcorbi_hilic_neg_4_088_59    m/z 274.1045     0   2   4   6   8E+3   4.5   5  RT (min)      lcorbi_hilic_neg_4_089_59    m/z 274.1133     0   1E+4   2.5   3  RT (min)      lcorbi_hilic_neg_4_013_46    m/z 274.1145     0   2   4E+4   4.5   5  RT (min)       lcorbi_hilic_neg_4_090_60    m/z 274.1406     0   1E+4   7   8  RT (min)       lcorbi_hilic_neg_4_085_59    m/z 274.4824     0   1E+4   4.5   5  RT (min)      lcorbi_hilic_neg_4_079_58    m/z 274.8245     0   1   2   3E+4   8   8.5  RT (min)       lcorbi_hilic_neg_4_106_62    m/z 274.8720     0   1   2E+4   7.5   8  RT (min)       lcorbi_hilic_neg_4_095_60    m/z 274.8803     0   2   4   6   8E+3   4.5   5  RT (min)      lcorbi_hilic_neg_4_086_59    m/z 274.8869     0   1E+4   7   7.5  RT (min)      lcorbi_hilic_neg_4_082_59    m/z 274.9498     0   1   2E+4   1.5   2  RT (min)       lcorbi_hilic_neg_4_104_61    m/z 274.9658     0   1   2E+4   1.5   2  RT (min)       lcorbi_hilic_neg_4_082_59    m/z 274.9713     0   1E+4   6   6.5  RT (min)       lcorbi_hilic_neg_4_072_57    m/z 274.9737     0   1   2E+4   6   6.5  RT (min)      lcorbi_hilic_neg_4_071_57    m/z 274.9757     0   1E+4   1   1.5  RT (min)      lcorbi_hilic_neg_4_099_61    m/z 274.9859     0   1   2E+4   1   1.5  RT (min)      lcorbi_hilic_neg_4_050_26    m/z 274.9950     0   1   2E+4   0.5   1  RT (min)       lcorbi_hilic_neg_4_070_57    m/z 274.9956     0   1E+5   3   3.5  RT (min)       lcorbi_hilic_neg_4_072_57    m/z 275.0195     0   1   2   3   4E+3   6.5   7  RT (min)      lcorbi_hilic_neg_4_084_59    m/z 275.0204     0   1   3   3.5  RT (min)      lcorbi_hilic_neg_4_069_57    m/z 275.0227     0   1   1   1.5  RT (min)       lcorbi_hilic_neg_4_091_60    m/z 275.0437     0   1E+5   1   1.5  RT (min)       lcorbi_hilic_neg_4_082_59    m/z 275.0537     0   1   2   3   4E+4   5   5.5  RT (min)       lcorbi_hilic_neg_4_087_59    m/z 275.0557     0   1E+4   7   7.5  RT (min)      lcorbi_hilic_neg_4_070_57    m/z 275.0648     0   1E+3   1.5   2  RT (min)      lcorbi_hilic_neg_4_074_58    m/z 275.0882     0   1E+4   5   5.5  RT (min)      lcorbi_hilic_neg_4_082_59    m/z 275.0927     0   1   4   4.5  RT (min)       lcorbi_hilic_neg_1_002_26    m/z 275.0962     0   1E+3   6.5   7  RT (min)      lcorbi_hilic_neg_4_091_60    m/z 275.1034     0   1E+3   2.5   3  RT (min)       lcorbi_hilic_neg_1_037_46    m/z 275.1146     0   2   4E+4   5.5   6  RT (min)      lcorbi_hilic_neg_3_016_16    m/z 275.1357     0   1   6   6.5  RT (min)      lcorbi_hilic_neg_4_005_32    m/z 275.4809     0   1E+4   4.5   5  RT (min)      lcorbi_hilic_neg_4_077_58    m/z 275.9094     0   1   2   3   4E+3   1   1.5  RT (min)      lcorbi_hilic_neg_1_028_45    m/z 275.9258     0   2   4E+4   4   4.5  RT (min)       lcorbi_hilic_neg_4_083_59    m/z 275.9352     0   1E+4   4.5   5  RT (min)      lcorbi_hilic_neg_4_071_57    m/z 275.9577     0   1E+4   8   8.5  RT (min)      lcorbi_hilic_neg_4_080_58    m/z 275.9611     0   2   4   6   8E+3   7   7.5  RT (min)      lcorbi_hilic_neg_4_090_60    m/z 276.0003     0   1E+4   1   1.5  RT (min)      lcorbi_hilic_neg_1_002_26    m/z 276.0378     0   1E+4   2   2.5  RT (min)      lcorbi_hilic_neg_4_100_61    m/z 276.0502     0   2   4   6E+2   1.5   2  RT (min)      lcorbi_hilic_neg_4_037_31    m/z 276.0617     0   1   2   3E+4   1.5   2  RT (min)       lcorbi_hilic_neg_4_086_59    m/z 276.0756     0   1E+4   2   2.5  RT (min)      lcorbi_hilic_neg_2_041_54    m/z 276.0769     0   1   2   3   4E+2   4   4.5  RT (min)      lcorbi_hilic_neg_1_027_18    m/z 276.0969     0   1E+3   6   6.5  RT (min)      lcorbi_hilic_neg_4_005_32    m/z 276.1057     0   1   2   3   4E+3   4   4.5   5  RT (min)       lcorbi_hilic_neg_3_011_24    m/z 276.1085     0   1   2   3E+4   2.5   3  RT (min)      lcorbi_hilic_neg_1_037_46    m/z 276.1244     0   1E+3   6   6.5  RT (min)      lcorbi_hilic_neg_4_091_60    m/z 276.1479     0   1E+4   3   3.5  RT (min)      lcorbi_hilic_neg_4_095_60    m/z 276.1719     0   1   2   3E+5   1   1.5  RT (min)      lcorbi_hilic_neg_4_113_62    m/z 276.4794     0   1E+4   4.5   5  RT (min)      lcorbi_hilic_neg_4_077_58    m/z 276.8210     0   1   2E+3   7.5   8  RT (min)      lcorbi_hilic_neg_3_026_09    m/z 276.8517     0   1   2   3   4E+4   7   7.5  RT (min)       lcorbi_hilic_neg_4_083_59    m/z 276.8613     0   2   4   6E+3   7   7.5  RT (min)      lcorbi_hilic_neg_4_093_60    m/z 276.9316     0   1   2   3   4E+4   4.5   5  RT (min)       lcorbi_hilic_neg_4_069_57    m/z 276.9357     0   1   2   3   4E+3   8   8.5  RT (min)      lcorbi_hilic_neg_1_002_26    m/z 276.9539     0   2   4   6   8E+3   8   8.5  RT (min)      lcorbi_hilic_neg_4_080_58    m/z 276.9558     0   1   1   1.5  RT (min)      lcorbi_hilic_neg_4_091_60    m/z 276.9683     0   1E+3   6   6.5  RT (min)      lcorbi_hilic_neg_1_012_32    m/z 276.9779     0   1   2   3   4E+3   4.5   5  RT (min)      lcorbi_hilic_neg_4_106_62    m/z 276.9974     0   1   1   2  RT (min)       lcorbi_hilic_neg_4_080_58    m/z 277.0151     0   1   1.5   2  RT (min)      lcorbi_hilic_neg_4_092_60    m/z 277.0236     0   1   2   3   4E+4   6.5   7  RT (min)       lcorbi_hilic_neg_4_096_60    m/z 277.0249     0   1   2E+3   1   1.5  RT (min)      lcorbi_hilic_neg_4_091_60    m/z 277.0345     0   2   4   6   8E+3   2   2.5  RT (min)      lcorbi_hilic_neg_4_113_62    m/z 277.0507     0   1   2E+3   5.5   6  RT (min)      lcorbi_hilic_neg_4_081_58    m/z 277.0594     0   1   2E+4   6   6.5  RT (min)       lcorbi_hilic_neg_4_093_60    m/z 277.0595     0   1   2E+3   7.5   8  RT (min)       lcorbi_hilic_neg_4_015_24    m/z 277.0595     0   2   4   6E+3   1.5   2  RT (min)      lcorbi_hilic_neg_4_077_58    m/z 277.0735     0   1E+3   2   2.5  RT (min)      lcorbi_hilic_neg_4_082_59    m/z 277.0736     0   1   5.5   6  RT (min)      lcorbi_hilic_neg_4_066_57    m/z 277.0979     0   1   2   3   4E+4   0.5   1  RT (min)      lcorbi_hilic_neg_4_031_14    m/z 277.1055     0   1   4   4.5  RT (min)       lcorbi_hilic_neg_4_078_58    m/z 277.1070     0   1E+3   6   6.5  RT (min)      lcorbi_hilic_neg_4_095_60    m/z 277.1098     0   1   2   3   4E+4   0.5   1  RT (min)      lcorbi_hilic_neg_3_031_14    m/z 277.1200     0   1E+5   0.5   1  RT (min)      lcorbi_hilic_neg_4_046_15    m/z 277.1226     0   2   4   6   8E+3   6   6.5  RT (min)       lcorbi_hilic_neg_4_095_60    m/z 277.1227     0   1   2E+3   2.5   3  RT (min)      lcorbi_hilic_neg_4_080_58    m/z 277.1302     0   2   4   6E+5   0.5   1  RT (min)      lcorbi_hilic_neg_4_078_58    m/z 277.1319     0   1   2.5   3  RT (min)      lcorbi_hilic_neg_4_080_58    m/z 277.1443     0   2   4E+4   4.5   5  RT (min)       lcorbi_hilic_neg_4_081_58    m/z 277.1443     0   1   2   3   4E+4   3   3.5  RT (min)      lcorbi_hilic_neg_4_078_58    m/z 277.1554     0   1   2E+4   4   4.5  RT (min)       lcorbi_hilic_neg_4_077_58    m/z 277.1841     0   2   4E+4   0.5   1  RT (min)      lcorbi_hilic_neg_3_031_14    m/z 277.2167     0   1   2   3E+4   0.5   1  RT (min)      lcorbi_hilic_neg_4_031_14    m/z 277.4779     0   1E+4   4.5   5  RT (min)      lcorbi_hilic_neg_4_080_58    m/z 277.8422     0   2   4   6E+3   7.5   8  RT (min)      lcorbi_hilic_neg_4_106_62    m/z 277.8608     0   1E+4   1.5   2  RT (min)      lcorbi_hilic_neg_4_082_59    m/z 277.9230     0   1   2   3   4E+4   4   4.5  RT (min)       lcorbi_hilic_neg_4_082_59    m/z 277.9747     0   2   4E+3   7   7.5  RT (min)      lcorbi_hilic_neg_4_093_60    m/z 277.9761     0   1   2   3E+4   1   1.5  RT (min)       lcorbi_hilic_neg_4_066_57    m/z 277.9841     0   1   2   3E+4   7   7.5  RT (min)      lcorbi_hilic_neg_4_094_60    m/z 277.9977     0   1   2E+4   2   2.5  RT (min)       lcorbi_hilic_neg_4_098_61    m/z 278.0360     0   1   2E+4   2   2.5  RT (min)       lcorbi_hilic_neg_4_091_60    m/z 278.0587     0   2   4E+3   2   2.5  RT (min)      lcorbi_hilic_neg_4_078_58    m/z 278.0701     0   1   4   4.5  RT (min)      lcorbi_hilic_neg_4_074_58    m/z 278.0726     0   1   2E+4   2   2.5  RT (min)       lcorbi_hilic_neg_4_092_60    m/z 278.0880     0   1   2   3   4E+4   4.5   5  RT (min)       lcorbi_hilic_neg_4_087_59    m/z 278.0912     0   1E+4   2   2.5  RT (min)      lcorbi_hilic_neg_1_037_46    m/z 278.1018     0   2   4E+4   3   3.5  RT (min)       lcorbi_hilic_neg_4_090_60    m/z 278.1032     0   1E+3   1   1.5  RT (min)       lcorbi_hilic_neg_4_096_60    m/z 278.1146     0   2   4E+3   5.5   6  RT (min)      lcorbi_hilic_neg_4_093_60    m/z 278.1592     0   1   4   4.5  RT (min)      lcorbi_hilic_neg_4_088_59    m/z 278.1759     0   1E+3   0.5   1  RT (min)      lcorbi_hilic_neg_3_031_14    m/z 278.1842     0   1   2E+4   0.5   1  RT (min)      lcorbi_hilic_neg_4_079_58    m/z 278.8095     0   1   2   3   4E+4   7   7.5  RT (min)       lcorbi_hilic_neg_4_086_59    m/z 278.8296     0   1   2E+4   5   5.5  RT (min)       lcorbi_hilic_neg_4_104_61    m/z 278.8380     0   1   2E+3   7.5   8  RT (min)      lcorbi_hilic_neg_4_090_60    m/z 278.8569     0   1   2   3   4E+3   1.5   2  RT (min)      lcorbi_hilic_neg_4_066_57    m/z 278.8582     0   1   2   3   4E+3   7.5   8  RT (min)      lcorbi_hilic_neg_1_019_06    m/z 278.8798     0   2   4   6E+3   8.5   9  RT (min)      lcorbi_hilic_neg_4_079_58    m/z 278.9075     0   1   2   3E+4   8.5   9  RT (min)       lcorbi_hilic_neg_4_075_58    m/z 278.9157     0   2   4   6   8E+3   0.5   1  RT (min)      lcorbi_hilic_neg_4_078_58    m/z 278.9246     0   2   4E+3   8   8.5  RT (min)      lcorbi_hilic_neg_4_081_58    m/z 278.9496     0   1   7   7.5  RT (min)      lcorbi_hilic_neg_4_076_58    m/z 278.9811     0   2   4E+4   5   5.5  RT (min)       lcorbi_hilic_neg_4_071_57    m/z 278.9863     0   1   2E+4   0.5   1  RT (min)       lcorbi_hilic_neg_4_071_57    m/z 279.0061     0   1   2E+4   1   1.5  RT (min)       lcorbi_hilic_neg_4_071_57    m/z 279.0091     0   1   2   3   4E+4   3   3.5  RT (min)       lcorbi_hilic_neg_4_104_61    m/z 279.0212     0   1   7   7.5  RT (min)      lcorbi_hilic_neg_4_075_58    m/z 279.0325     0   1   2E+4   2   2.5  RT (min)       lcorbi_hilic_neg_4_091_60    m/z 279.0387     0   2   4   6   8E+4   6   6.5   7  RT (min)       lcorbi_hilic_neg_4_074_58    m/z 279.0522     0   1E+4   2   2.5  RT (min)       lcorbi_hilic_neg_4_076_58    m/z 279.0688     0   1E+4   1.5   2  RT (min)       lcorbi_hilic_neg_4_080_58    m/z 279.0781     0   1   2   3E+4   4   4.5  RT (min)       lcorbi_hilic_neg_4_096_60    m/z 279.0863     0   1   2   3   4E+4   6.5   7  RT (min)      lcorbi_hilic_neg_4_030_05    m/z 279.0985     0   1   3.5   4  RT (min)      lcorbi_hilic_neg_4_090_60    m/z 279.0986     0   1   1   1.5  RT (min)      lcorbi_hilic_neg_4_104_61    m/z 279.1112     0   2   4   6E+2   1   1.5  RT (min)       lcorbi_hilic_neg_1_002_26    m/z 279.1224     0   1E+5   7.5   8  RT (min)       lcorbi_hilic_neg_1_055_24    m/z 279.1237     0   1E+5   1   1.5  RT (min)       lcorbi_hilic_neg_4_100_61    m/z 279.1478     0   1   4   4.5  RT (min)      lcorbi_hilic_neg_4_095_60    m/z 279.2327     0   1E+4   4   4.5  RT (min)      lcorbi_hilic_neg_4_069_57    m/z 279.2328     0   2   4   6   8E+3   2.5   3  RT (min)      lcorbi_hilic_neg_4_079_58    m/z 279.8585     0   1   2   3   4E+4   2   2.5  RT (min)       lcorbi_hilic_neg_4_090_60    m/z 279.9161     0   1E+4   8   8.5  RT (min)      lcorbi_hilic_neg_4_081_58    m/z 279.9503     0   2   4   6E+3   1   1.5  RT (min)      lcorbi_hilic_neg_4_111_62    m/z 279.9580     0   1E+4   7   7.5  RT (min)      lcorbi_hilic_neg_4_083_59    m/z 279.9590     0   2   4   6E+3   3   3.5  RT (min)      lcorbi_hilic_neg_4_098_61    m/z 279.9894     0   1E+4   1   1.5  RT (min)      lcorbi_hilic_neg_4_079_58    m/z 280.0106     0   1E+3   6   6.5  RT (min)      lcorbi_hilic_neg_4_080_58    m/z 280.0157     0   1   2E+4   2   2.5  RT (min)       lcorbi_hilic_neg_4_081_58    m/z 280.0203     0   1   2   3E+2   5.5   6  RT (min)      lcorbi_hilic_neg_4_005_32    m/z 280.0284     0   2   4   6   8E+3   3.5   4  RT (min)      lcorbi_hilic_neg_4_076_58    m/z 280.0336     0   2   4   6E+4   6.5   7  RT (min)       lcorbi_hilic_neg_4_097_60    m/z 280.0452     0   1   2E+3   1   1.5  RT (min)       lcorbi_hilic_neg_4_068_57    m/z 280.0574     0   1   3   3.5  RT (min)      lcorbi_hilic_neg_4_085_59    m/z 280.0705     0   1   1.5   2  RT (min)      lcorbi_hilic_neg_4_004_13    m/z 280.0751     0   1E+4   3.5   4  RT (min)      lcorbi_hilic_neg_4_089_59    m/z 280.0752     0   1E+4   6   6.5  RT (min)      lcorbi_hilic_neg_4_083_59    m/z 280.0991     0   1   2   3   4E+4   6.5   7  RT (min)      lcorbi_hilic_neg_3_029_25    m/z 280.1244     0   1E+4   1   1.5  RT (min)      lcorbi_hilic_neg_4_077_58    m/z 280.1542     0   1E+4   7   7.5  RT (min)       lcorbi_hilic_neg_4_091_60    m/z 280.1666     0   1   2   3   4E+5   0.5   1  RT (min)      lcorbi_hilic_neg_4_073_57    m/z 280.2361     0   2   4   6E+3   0.5   1  RT (min)      lcorbi_hilic_neg_4_073_57    m/z 280.6119     0   1   2E+4   4   4.5  RT (min)      lcorbi_hilic_neg_4_037_31    m/z 280.9126     0   1   2E+4   8   8.5  RT (min)       lcorbi_hilic_neg_4_076_58    m/z 280.9162     0   1   6   6.5  RT (min)      lcorbi_hilic_neg_1_003_14    m/z 280.9308     0   1   2E+4   1   1.5  RT (min)       lcorbi_hilic_neg_4_066_57    m/z 280.9423     0   1   2   3E+4   4   4.5  RT (min)      lcorbi_hilic_neg_4_047_07    m/z 280.9619     0   1   4   4.5  RT (min)       lcorbi_hilic_neg_4_088_59    m/z 280.9836     0   1   2   3E+4   3   3.5  RT (min)      lcorbi_hilic_neg_4_069_57    m/z 280.9839     0   1   2   3E+4   0.5   1  RT (min)       lcorbi_hilic_neg_4_111_62    m/z 280.9929     0   1   2   3   4E+3   4.5   5  RT (min)      lcorbi_hilic_neg_4_078_58    m/z 281.0018     0   1   2E+4   0.5   1  RT (min)       lcorbi_hilic_neg_4_111_62    m/z 281.0018     0   1   2E+4   3   3.5  RT (min)       lcorbi_hilic_neg_4_071_57    m/z 281.0104     0   1   6.5   7  RT (min)      lcorbi_hilic_neg_4_073_57    m/z 281.0122     0   1   2E+4   2   2.5  RT (min)       lcorbi_hilic_neg_4_076_58    m/z 281.0358     0   2   4   6   8E+3   3   3.5  RT (min)       lcorbi_hilic_neg_4_088_59    m/z 281.0362     0   1   2   3E+3   2   2.5  RT (min)       lcorbi_hilic_neg_4_081_58    m/z 281.0363     0   2   4E+4   5   6   7  RT (min)       lcorbi_hilic_neg_4_067_57    m/z 281.0518     0   1   2   3   4E+3   5.5   6  RT (min)      lcorbi_hilic_neg_4_098_61    m/z 281.0545     0   1   2E+3   3   3.5   4  RT (min)       lcorbi_hilic_neg_4_104_61    m/z 281.0876     0   1   2   3E+3   6   6.5  RT (min)      lcorbi_hilic_neg_4_098_61    m/z 281.0987     0   1E+3   3.5   4  RT (min)       lcorbi_hilic_neg_4_014_18    m/z 281.0997     0   1   2   3   4E+3   1.5   2  RT (min)       lcorbi_hilic_neg_4_105_61    m/z 281.1028     0   2   4E+3   5   5.5  RT (min)      lcorbi_hilic_neg_4_105_61    m/z 281.1073     0   1   2E+3   3   3.5  RT (min)      lcorbi_hilic_neg_4_105_61    m/z 281.1098     0   1   6.5   7  RT (min)      lcorbi_hilic_neg_4_014_18    m/z 281.1199     0   1   2E+4   1   1.5  RT (min)      lcorbi_hilic_neg_1_048_35    m/z 281.1204     0   1   6   6.5  RT (min)       lcorbi_hilic_neg_4_098_61    m/z 281.1236     0   2   4   6E+3   4   4.5  RT (min)      lcorbi_hilic_neg_4_098_61    m/z 281.1381     0   2   4E+4   7.5   8  RT (min)       lcorbi_hilic_neg_4_098_61    m/z 281.1563     0   2   4   6   8E+4   1   1.5  RT (min)      lcorbi_hilic_neg_4_099_61    m/z 281.1755     0   1E+4   0.5   1  RT (min)      lcorbi_hilic_neg_4_100_61    m/z 281.2484     0   2   4   6   8E+3   4   4.5  RT (min)      lcorbi_hilic_neg_4_072_57    m/z 281.2484     0   1   2.5   3  RT (min)      lcorbi_hilic_neg_4_105_61    m/z 281.8556     0   1   2   3   4E+4   2   2.5  RT (min)       lcorbi_hilic_neg_4_090_60    m/z 281.8743     0   1   2E+4   1.5   2  RT (min)       lcorbi_hilic_neg_4_097_60    m/z 281.9115     0   2   4   6   8E+3   4.5   5  RT (min)      lcorbi_hilic_neg_4_090_60    m/z 281.9167     0   1   2   3   4E+4   1   1.5  RT (min)      lcorbi_hilic_neg_4_106_62    m/z 281.9335     0   1E+3   2   2.5  RT (min)      lcorbi_hilic_neg_3_052_40    m/z 282.0051     0   1E+4   5.5   6  RT (min)      lcorbi_hilic_neg_4_076_58    m/z 282.0358     0   1   2   3   4E+4   6.5   7  RT (min)       lcorbi_hilic_neg_4_083_59    m/z 282.0417     0   1E+5   1   1.5  RT (min)      lcorbi_hilic_neg_1_037_46    m/z 282.0487     0   2   4   6E+4   1   1.5  RT (min)       lcorbi_hilic_neg_4_106_62    m/z 282.0609     0   1E+3   3.5   4  RT (min)      lcorbi_hilic_neg_4_098_61    m/z 282.0842     0   1   2E+4   5   5.5  RT (min)       lcorbi_hilic_neg_4_111_62    m/z 282.0965     0   1   2E+5   7   7.5  RT (min)       lcorbi_hilic_neg_4_094_60    m/z 282.1029     0   1   2   2.5  RT (min)       lcorbi_hilic_neg_4_105_61    m/z 282.1306     0   1   3   3.5  RT (min)       lcorbi_hilic_neg_4_104_61    m/z 282.1341     0   1E+4   1   1.5  RT (min)      lcorbi_hilic_neg_2_045_26    m/z 282.2518     0   1   0.5   1  RT (min)      lcorbi_hilic_neg_3_037_12    m/z 282.7778     0   2   4   6E+3   0.5   1  RT (min)      lcorbi_hilic_neg_4_068_57    m/z 282.7778     0   1E+4   4.5   5  RT (min)      lcorbi_hilic_neg_4_100_61    m/z 282.8890     0   1   2E+3   1.5   2  RT (min)      lcorbi_hilic_neg_4_113_62    m/z 282.9062     0   1   2E+4   4.5   5  RT (min)      lcorbi_hilic_neg_4_090_60    m/z 282.9134     0   2   4   6E+3   6   6.5  RT (min)      lcorbi_hilic_neg_4_071_57    m/z 282.9394     0   1   2E+4   4   4.5  RT (min)      lcorbi_hilic_neg_4_047_07    m/z 282.9481     0   2   4   6E+2   0.5   1  RT (min)      lcorbi_hilic_neg_3_055_26    m/z 282.9574     0   1E+3   5.5   6  RT (min)      lcorbi_hilic_neg_4_106_62    m/z 282.9810     0   1   2   3   4E+4   0.5   1  RT (min)       lcorbi_hilic_neg_4_101_61    m/z 282.9810     0   1   2E+4   3   3.5  RT (min)      lcorbi_hilic_neg_4_108_62    m/z 282.9910     0   1   4.5   5  RT (min)      lcorbi_hilic_neg_4_094_60    m/z 283.0006     0   1   2E+4   0.5   1  RT (min)       lcorbi_hilic_neg_4_076_58    m/z 283.0007     0   1   2E+4   3   3.5  RT (min)      lcorbi_hilic_neg_4_076_58    m/z 283.0200     0   1E+4   5   5.5  RT (min)       lcorbi_hilic_neg_4_071_57    m/z 283.0285     0   1   0.5   1  RT (min)      lcorbi_hilic_neg_4_091_60    m/z 283.0514     0   1E+3   3   3.5  RT (min)      lcorbi_hilic_neg_4_102_61    m/z 283.0610     0   1E+3   3.5   4  RT (min)       lcorbi_hilic_neg_4_085_59    m/z 283.0649     0   1   1   1.5  RT (min)       lcorbi_hilic_neg_4_096_60    m/z 283.1032     0   2   4   6E+4   5   5.5  RT (min)       lcorbi_hilic_neg_4_071_57    m/z 283.1046     0   2   4   6E+4   6   7  RT (min)       lcorbi_hilic_neg_4_074_58    m/z 283.1175     0   1   2E+4   6   6.5  RT (min)       lcorbi_hilic_neg_4_091_60    m/z 283.1372     0   1   1   1.5  RT (min)      lcorbi_hilic_neg_1_002_26    m/z 283.1702     0   2   4   6   8E+3   0.5   1  RT (min)      lcorbi_hilic_neg_4_072_57    m/z 283.1912     0   1   2E+4   1   1.5  RT (min)      lcorbi_hilic_neg_4_080_58    m/z 283.2276     0   2   4   6E+3   1   1.5  RT (min)      lcorbi_hilic_neg_4_113_62    m/z 283.2640     0   1E+4   4   4.5  RT (min)      lcorbi_hilic_neg_4_073_57    m/z 283.2754     0   2   4   6E+3   3.5   4  RT (min)      lcorbi_hilic_neg_4_088_59    m/z 283.8526     0   1E+4   2   2.5  RT (min)      lcorbi_hilic_neg_4_090_60    m/z 283.8713     0   1   2E+4   1.5   2  RT (min)       lcorbi_hilic_neg_4_097_60    m/z 283.8851     0   1   2E+4   1   1.5  RT (min)      lcorbi_hilic_neg_4_110_62    m/z 283.9449     0   1E+4   6   6.5  RT (min)      lcorbi_hilic_neg_4_072_57    m/z 283.9713     0   1E+4   7   7.5  RT (min)      lcorbi_hilic_neg_4_092_60    m/z 283.9992     0   2   4   6E+3   6   6.5  RT (min)      lcorbi_hilic_neg_4_072_57    m/z 284.0347     0   2   4   6E+4   1.5   2  RT (min)      lcorbi_hilic_neg_4_046_15    m/z 284.0673     0   1   5.5   6  RT (min)      lcorbi_hilic_neg_4_068_57    m/z 284.0714     0   2   4   6   8E+2   1   1.5  RT (min)      lcorbi_hilic_neg_1_002_26    m/z 284.0902     0   2   4E+3   5.5   6  RT (min)       lcorbi_hilic_neg_4_068_57    m/z 284.0924     0   2   4   6E+3   0.5   1  RT (min)      lcorbi_hilic_neg_2_009_50    m/z 284.0985     0   1   6.5   7  RT (min)       lcorbi_hilic_neg_4_097_60    m/z 284.1069     0   2   4   6E+2   2   2.5  RT (min)      lcorbi_hilic_neg_1_001_49    m/z 284.1249     0   2   4   6   8E+3   3.5   4  RT (min)      lcorbi_hilic_neg_4_079_58    m/z 284.1253     0   1   2E+4   2   2.5  RT (min)      lcorbi_hilic_neg_4_001_49    m/z 284.1364     0   2   4   6E+4   6.5   7  RT (min)       lcorbi_hilic_neg_4_096_60    m/z 284.1378     0   1   4.5   5  RT (min)      lcorbi_hilic_neg_4_106_62    m/z 284.1470     0   2   4   6E+4   1   1.5  RT (min)      lcorbi_hilic_neg_1_002_26    m/z 284.2229     0   1   2E+4   1   1.5  RT (min)       lcorbi_hilic_neg_4_085_59    m/z 284.2595     0   1   2   3   4E+3   0.5   1  RT (min)       lcorbi_hilic_neg_4_073_57    m/z 284.7331     0   2   4   6   8E+3   3   3.5  RT (min)      lcorbi_hilic_neg_4_066_57    m/z 284.7748     0   1E+4   0.5   1  RT (min)      lcorbi_hilic_neg_4_068_57    m/z 284.7749     0   1   2E+4   4.5   5  RT (min)       lcorbi_hilic_neg_4_100_61    m/z 284.8718     0   2   4E+3   8   8.5  RT (min)      lcorbi_hilic_neg_4_089_59    m/z 284.9157     0   2   4   6   8E+3   7   7.5  RT (min)      lcorbi_hilic_neg_3_005_08    m/z 284.9338     0   1E+4   1.5   2  RT (min)      lcorbi_hilic_neg_4_098_61    m/z 284.9531     0   1E+3   0.5   1  RT (min)      lcorbi_hilic_neg_4_067_57    m/z 284.9744     0   2   4   6E+4   0.5   1  RT (min)       lcorbi_hilic_neg_4_113_62    m/z 284.9964     0   1   2E+4   3   3.5  RT (min)      lcorbi_hilic_neg_4_072_57    m/z 284.9964     0   1   2E+4   0.5   1  RT (min)      lcorbi_hilic_neg_4_096_60    m/z 285.0057     0   2   4   6E+2   1.5   2  RT (min)      lcorbi_hilic_neg_1_027_18    m/z 285.0166     0   2   4   6E+3   5   5.5  RT (min)      lcorbi_hilic_neg_4_073_57    m/z 285.0453     0   1   2   2.5  RT (min)      lcorbi_hilic_neg_1_027_18    m/z 285.0628     0   1E+5   0.5   1  RT (min)      lcorbi_hilic_neg_1_006_08    m/z 285.0766     0   1   1   1.5  RT (min)      lcorbi_hilic_neg_4_078_58    m/z 285.0811     0   1   5.5   6   6.5  RT (min)       lcorbi_hilic_neg_4_096_60    m/z 285.0919     0   1   3   3.5  RT (min)      lcorbi_hilic_neg_4_095_60    m/z 285.0920     0   1   2   3E+4   1.5   2  RT (min)       lcorbi_hilic_neg_4_092_60    m/z 285.0959     0   1E+4   6   6.5  RT (min)      lcorbi_hilic_neg_4_095_60    m/z 285.1091     0   2   4   6E+4   3   3.5  RT (min)       lcorbi_hilic_neg_4_089_59    m/z 285.1164     0   1   2E+4   0.5   1  RT (min)       lcorbi_hilic_neg_4_107_62    m/z 285.1202     0   1   2   3E+4   4   4.5  RT (min)       lcorbi_hilic_neg_1_047_05    m/z 285.1454     0   1   1   1.5  RT (min)      lcorbi_hilic_neg_4_079_58    m/z 285.1581     0   1E+4   7.5   8  RT (min)      lcorbi_hilic_neg_4_095_60    m/z 285.1606     0   2   4E+3   2   2.5  RT (min)      lcorbi_hilic_neg_4_109_62    m/z 285.1775     0   1E+4   7.5   8  RT (min)      lcorbi_hilic_neg_4_093_60    m/z 285.1858     0   1E+4   0.5   1  RT (min)      lcorbi_hilic_neg_4_105_61    m/z 285.2070     0   1   2   3   4E+4   1   1.5  RT (min)       lcorbi_hilic_neg_4_112_62    m/z 285.9420     0   2   4   6   8E+3   6   6.5  RT (min)      lcorbi_hilic_neg_4_073_57    m/z 285.9726     0   1E+4   7   7.5  RT (min)      lcorbi_hilic_neg_4_091_60    m/z 285.9899     0   1   2E+4   6.5   7  RT (min)      lcorbi_hilic_neg_1_016_37    m/z 286.0416     0   1   2E+4   2   2.5  RT (min)       lcorbi_hilic_neg_4_099_61    m/z 286.0469     0   2   4   6E+3   6.5   7  RT (min)      lcorbi_hilic_neg_2_005_24    m/z 286.0527     0   1   2E+4   3.5   4  RT (min)      lcorbi_hilic_neg_2_045_26    m/z 286.0533     0   1   6.5   7  RT (min)      lcorbi_hilic_neg_4_068_57    m/z 286.0778     0   1   2E+3   3.5   4  RT (min)      lcorbi_hilic_neg_1_037_46    m/z 286.0874     0   1E+4   5.5   6  RT (min)      lcorbi_hilic_neg_4_089_59    m/z 286.0909     0   1E+4   1.5   2  RT (min)      lcorbi_hilic_neg_4_089_59    m/z 286.1043     0   2   4   6   8E+3   6.5   7  RT (min)      lcorbi_hilic_neg_4_089_59    m/z 286.1044     0   1E+4   1   1.5  RT (min)      lcorbi_hilic_neg_3_001_49    m/z 286.1238     0   1E+4   7   7.5  RT (min)      lcorbi_hilic_neg_4_030_05    m/z 286.1293     0   1   1   1.5  RT (min)      lcorbi_hilic_neg_4_090_60    m/z 286.1409     0   1   1.5   2  RT (min)      lcorbi_hilic_neg_4_077_58    m/z 286.1516     0   2   4   6E+4   5.5   6  RT (min)      lcorbi_hilic_neg_3_052_40    m/z 286.1593     0   2   4   6E+3   4.5   5  RT (min)      lcorbi_hilic_neg_4_110_62    m/z 286.1882     0   1E+4   6.5   7  RT (min)      lcorbi_hilic_neg_4_085_59    m/z 286.2468     0   2   4   6E+3   1   1.5  RT (min)      lcorbi_hilic_neg_1_029_31    m/z 286.7310     0   2   4   6   8E+3   2.5   3  RT (min)      lcorbi_hilic_neg_4_066_57    m/z 286.7720     0   2   4   6E+3   0.5   1  RT (min)      lcorbi_hilic_neg_4_068_57    m/z 286.7720     0   1E+4   4.5   5  RT (min)      lcorbi_hilic_neg_4_102_61    m/z 286.8614     0   1   2E+4   5   5.5   6  RT (min)       lcorbi_hilic_neg_4_099_61    m/z 286.8837     0   2   4E+3   1   1.5  RT (min)      lcorbi_hilic_neg_4_074_58    m/z 286.9118     0   2   4   6   8E+4   6   6.5  RT (min)       lcorbi_hilic_neg_1_004_17    m/z 286.9311     0   1E+4   1.5   2  RT (min)      lcorbi_hilic_neg_4_098_61    m/z 286.9500     0   1   1.5   2  RT (min)      lcorbi_hilic_neg_4_094_60    m/z 286.9506     0   1   2   3   4E+4   6.5   7  RT (min)      lcorbi_hilic_neg_1_015_30    m/z 286.9507     0   1   2   3E+4   4.5   5  RT (min)      lcorbi_hilic_neg_1_008_39    m/z 286.9741     0   1E+3   6   6.5  RT (min)      lcorbi_hilic_neg_1_008_39    m/z 286.9759     0   1E+4   0.5   1  RT (min)      lcorbi_hilic_neg_4_112_62    m/z 286.9923     0   2   4   6E+4   1   1.5  RT (min)       lcorbi_hilic_neg_4_097_60    m/z 287.0119     0   1E+4   0.5   1  RT (min)      lcorbi_hilic_neg_4_071_57    m/z 287.0229     0   1   2E+3   0.5   1  RT (min)      lcorbi_hilic_neg_4_111_62    m/z 287.0319     0   2   4   6   8E+3   3   3.5  RT (min)      lcorbi_hilic_neg_4_081_58    m/z 287.0357     0   2   4   6E+3   5   5.5  RT (min)      lcorbi_hilic_neg_4_096_60    m/z 287.0363     0   1   2   3   4E+4   1.5   2  RT (min)       lcorbi_hilic_neg_4_099_61    m/z 287.0465     0   2   4   6   8E+4   3.5   4  RT (min)      lcorbi_hilic_neg_2_045_26    m/z 287.0537     0   2   4   6   8E+3   1   1.5  RT (min)      lcorbi_hilic_neg_4_100_61    m/z 287.0578     0   1E+4   2   2.5  RT (min)      lcorbi_hilic_neg_4_076_58    m/z 287.0802     0   1   2   3   4E+4   1   1.5  RT (min)       lcorbi_hilic_neg_4_088_59    m/z 287.0884     0   1E+3   6   6.5  RT (min)       lcorbi_hilic_neg_1_023_23    m/z 287.0891     0   1E+4   1.5   2  RT (min)      lcorbi_hilic_neg_4_096_60    m/z 287.0989     0   1   3   3.5  RT (min)       lcorbi_hilic_neg_4_069_57    m/z 287.1093     0   1E+3   4   4.5  RT (min)      lcorbi_hilic_neg_4_070_57    m/z 287.1215     0   2   4   6   8E+4   0.5   1  RT (min)      lcorbi_hilic_neg_3_024_27    m/z 287.1246     0   2   4   6E+3   2.5   3  RT (min)      lcorbi_hilic_neg_4_099_61    m/z 287.1359     0   1   2   3   4E+4   6   6.5  RT (min)       lcorbi_hilic_neg_4_087_59    m/z 287.1472     0   1E+4   7.5   8  RT (min)      lcorbi_hilic_neg_4_081_58    m/z 287.1499     0   1E+4   0.5   1  RT (min)      lcorbi_hilic_neg_4_104_61    m/z 287.2226     0   1   2   3   4E+3   1   1.5  RT (min)      lcorbi_hilic_neg_4_098_61    m/z 287.2813     0   1   2   3   4E+4   1   1.5  RT (min)       lcorbi_hilic_neg_4_091_60    m/z 287.6115     0   1   2   3E+3   3   3.5  RT (min)      lcorbi_hilic_neg_1_003_14    m/z 287.8568     0   1E+4   4.5   5  RT (min)      lcorbi_hilic_neg_4_078_58    m/z 287.9114     0   1E+4   4.5   5  RT (min)      lcorbi_hilic_neg_1_010_22    m/z 287.9261     0   1   2   3   4E+4   3.5   4  RT (min)       lcorbi_hilic_neg_4_081_58    m/z 287.9398     0   2   4   6E+2   6.5   7  RT (min)      lcorbi_hilic_neg_1_006_08    m/z 287.9527     0   2   4   6E+3   1   1.5  RT (min)      lcorbi_hilic_neg_4_107_62    m/z 287.9771     0   1E+4   6   6.5  RT (min)      lcorbi_hilic_neg_4_088_59    m/z 287.9816     0   2   4E+4   3   3.5  RT (min)       lcorbi_hilic_neg_4_088_59    m/z 287.9850     0   1   2E+4   1   1.5  RT (min)      lcorbi_hilic_neg_4_106_62    m/z 288.0044     0   1   1   1.5  RT (min)      lcorbi_hilic_neg_4_106_62    m/z 288.0490     0   1   2   3E+4   8   8.5  RT (min)       lcorbi_hilic_neg_4_087_59    m/z 288.0551     0   1   2   3E+4   5.5   6  RT (min)      lcorbi_hilic_neg_4_088_59    m/z 288.0660     0   1E+3   6   6.5  RT (min)       lcorbi_hilic_neg_4_095_60    m/z 288.0755     0   1E+3   2   2.5  RT (min)       lcorbi_hilic_neg_4_095_60    m/z 288.0811     0   1   2E+4   6.5   7  RT (min)      lcorbi_hilic_neg_1_023_23    m/z 288.0906     0   1   2   3   4E+4   3   3.5  RT (min)       lcorbi_hilic_neg_4_094_60    m/z 288.1020     0   1   5.5   6  RT (min)      lcorbi_hilic_neg_4_089_59    m/z 288.1202     0   1   2   3E+4   5   5.5  RT (min)       lcorbi_hilic_neg_4_097_60    m/z 288.1295     0   2   4   6E+3   6.5   7  RT (min)      lcorbi_hilic_neg_4_055_38    m/z 288.1816     0   2   4   6   8E+3   1   1.5  RT (min)      lcorbi_hilic_neg_4_067_57    m/z 288.2049     0   1   2E+4   1   1.5  RT (min)      lcorbi_hilic_neg_3_055_26    m/z 288.6196     0   2   4   6   8E+3   4.5   5  RT (min)      lcorbi_hilic_neg_4_079_58    m/z 288.8031     0   2   4   6E+3   8   8.5  RT (min)      lcorbi_hilic_neg_4_108_62    m/z 288.8491     0   2   4   6E+3   8   8.5  RT (min)      lcorbi_hilic_neg_4_074_58    m/z 288.8584     0   1E+4   5.5   6  RT (min)      lcorbi_hilic_neg_3_004_41    m/z 288.8771     0   1   2   3   4E+4   3   3.5  RT (min)       lcorbi_hilic_neg_4_085_59    m/z 288.8971     0   1E+4   1.5   2  RT (min)      lcorbi_hilic_neg_3_003_44    m/z 288.9088     0   1   2   3E+4   6   6.5  RT (min)       lcorbi_hilic_neg_1_003_14    m/z 288.9364     0   1   2E+5   9   10   11  RT (min)       lcorbi_hilic_neg_4_070_57    m/z 288.9652     0   1   5   5.5  RT (min)      lcorbi_hilic_neg_4_090_60    m/z 288.9754     0   1   2E+3   3   3.5  RT (min)      lcorbi_hilic_neg_4_082_59    m/z 288.9909     0   1   2E+4   0.5   1  RT (min)       lcorbi_hilic_neg_4_073_57    m/z 288.9975     0   1   2E+3   7.5   8  RT (min)      lcorbi_hilic_neg_1_029_31    m/z 289.0007     0   2   4   6E+4   1   1.5  RT (min)       lcorbi_hilic_neg_1_044_03    m/z 289.0211     0   1   7   7.5  RT (min)      lcorbi_hilic_neg_4_087_59    m/z 289.0309     0   1   2   3   4E+4   2   2.5  RT (min)       lcorbi_hilic_neg_4_082_59    m/z 289.0419     0   2   4   6E+3   7.5   8  RT (min)      lcorbi_hilic_neg_4_082_59    m/z 289.0453     0   2   4   6E+3   7.5   8  RT (min)      lcorbi_hilic_neg_4_008_35    m/z 289.0649     0   1E+4   5.5   6  RT (min)       lcorbi_hilic_neg_4_095_60    m/z 289.0789     0   1   1.5   2  RT (min)      lcorbi_hilic_neg_4_074_58    m/z 289.0805     0   1E+4   2.5   3  RT (min)      lcorbi_hilic_neg_4_089_59    m/z 289.0952     0   2   4   6E+2   6   6.5  RT (min)      lcorbi_hilic_neg_4_005_32    m/z 289.1056     0   1   2   3E+4   1   1.5  RT (min)      lcorbi_hilic_neg_4_066_57    m/z 289.1146     0   1   1   1.5  RT (min)       lcorbi_hilic_neg_4_089_59    m/z 289.1154     0   1   2   3   4E+4   7.5   8  RT (min)       lcorbi_hilic_neg_4_086_59    m/z 289.1400     0   1   7.5   8  RT (min)       lcorbi_hilic_neg_4_093_60    m/z 289.1597     0   2   4E+4   1.5   2  RT (min)      lcorbi_hilic_neg_4_089_59    m/z 289.7027     0   1   2E+4   3   3.5  RT (min)       lcorbi_hilic_neg_4_080_58    m/z 289.8544     0   2   4   6   8E+3   4.5   5  RT (min)      lcorbi_hilic_neg_4_076_58    m/z 289.8841     0   1   1   1.5  RT (min)       lcorbi_hilic_neg_4_082_59    m/z 289.9021     0   2   4   6E+3   1.5   2  RT (min)      lcorbi_hilic_neg_4_074_58    m/z 289.9083     0   1   2   3   4E+3   4.5   5  RT (min)      lcorbi_hilic_neg_1_010_22    m/z 289.9231     0   1   2E+4   3.5   4  RT (min)       lcorbi_hilic_neg_4_081_58    m/z 289.9379     0   1   2   3E+4   8.5   9  RT (min)      lcorbi_hilic_neg_4_077_58    m/z 289.9491     0   2   4   6   8E+3   8   8.5  RT (min)      lcorbi_hilic_neg_4_078_58    m/z 289.9504     0   1   2   3   4E+3   1   1.5  RT (min)      lcorbi_hilic_neg_3_024_27    m/z 289.9870     0   1   2E+4   7   7.5  RT (min)       lcorbi_hilic_neg_4_082_59    m/z 290.0055     0   2   4   6   8E+3   6.5   7  RT (min)      lcorbi_hilic_neg_4_083_59    m/z 290.0326     0   1E+4   2   2.5  RT (min)      lcorbi_hilic_neg_4_098_61    m/z 290.0524     0   2   4   6   8E+2   2   2.5  RT (min)       lcorbi_hilic_neg_4_051_16    m/z 290.0542     0   1   2   3E+4   3.5   4  RT (min)       lcorbi_hilic_neg_4_070_57    m/z 290.0629     0   1   2   3E+4   1   1.5  RT (min)      lcorbi_hilic_neg_4_088_59    m/z 290.0654     0   1E+3   6   6.5  RT (min)      lcorbi_hilic_neg_2_040_35    m/z 290.0710     0   1   2E+3   3   3.5  RT (min)      lcorbi_hilic_neg_4_112_62    m/z 290.0846     0   1E+3   3.5   4  RT (min)       lcorbi_hilic_neg_4_082_59    m/z 290.0880     0   1   2   3E+4   5.5   6  RT (min)       lcorbi_hilic_neg_3_023_36    m/z 290.1043     0   1E+4   6   6.5  RT (min)      lcorbi_hilic_neg_4_092_60    m/z 290.1064     0   2   4   6   8E+3   3   3.5  RT (min)      lcorbi_hilic_neg_4_091_60    m/z 290.1309     0   1   2   3E+4   6.5   7  RT (min)       lcorbi_hilic_neg_4_097_60    m/z 290.1371     0   1   2E+5   4   4.5  RT (min)       lcorbi_hilic_neg_4_090_60    m/z 290.1396     0   1E+3   0.5   1  RT (min)      lcorbi_hilic_neg_2_005_24    m/z 290.1760     0   1   2E+4   0.5   1  RT (min)       lcorbi_hilic_neg_4_105_61    m/z 290.8738     0   1   2E+4   1   1.5  RT (min)      lcorbi_hilic_neg_3_032_42    m/z 290.8742     0   1   2   3   4E+4   3   3.5  RT (min)       lcorbi_hilic_neg_4_085_59    m/z 290.8937     0   2   4E+3   4.5   5  RT (min)      lcorbi_hilic_neg_4_106_62    m/z 290.9050     0   1   2E+3   4.5   5  RT (min)      lcorbi_hilic_neg_4_098_61    m/z 290.9277     0   1   8.5   9  RT (min)      lcorbi_hilic_neg_4_080_58    m/z 290.9340     0   1   2   3E+4   8.5   9  RT (min)      lcorbi_hilic_neg_4_080_58    m/z 290.9452     0   1   2E+4   6   6.5  RT (min)       lcorbi_hilic_neg_4_072_57    m/z 290.9867     0   1E+4   1   1.5  RT (min)      lcorbi_hilic_neg_4_067_57    m/z 290.9943     0   1   3   3.5  RT (min)      lcorbi_hilic_neg_4_072_57    m/z 291.0071     0   1   2   3   4E+4   3   3.5  RT (min)       lcorbi_hilic_neg_4_088_59    m/z 291.0209     0   1E+3   3.5   4  RT (min)      lcorbi_hilic_neg_4_096_60    m/z 291.0366     0   1   0.5   1  RT (min)       lcorbi_hilic_neg_4_107_62    m/z 291.0373     0   1   2E+4   7.5   8  RT (min)      lcorbi_hilic_neg_4_089_59    m/z 291.0489     0   1E+5   1.5   2  RT (min)       lcorbi_hilic_neg_4_068_57    m/z 291.0566     0   1E+3   4.5   5  RT (min)      lcorbi_hilic_neg_4_090_60    m/z 291.0662     0   1   2   3E+4   1.5   2  RT (min)       lcorbi_hilic_neg_4_111_62    m/z 291.0663     0   1   2E+4   4  RT (min)       lcorbi_hilic_neg_4_004_13    m/z 291.0979     0   1   2   3   4E+2   5.5   6  RT (min)      lcorbi_hilic_neg_1_047_05    m/z 291.1025     0   1   2   3E+3   2   2.5  RT (min)      lcorbi_hilic_neg_2_050_17    m/z 291.1041     0   1E+5   5.5   6  RT (min)      lcorbi_hilic_neg_2_005_24    m/z 291.1048     0   2   4   6E+4   4.5   5  RT (min)       lcorbi_hilic_neg_4_090_60    m/z 291.1237     0   2   4   6E+4   1   1.5  RT (min)       lcorbi_hilic_neg_4_075_58    m/z 291.1601     0   1   2   3   4E+3   0.5   1  RT (min)       lcorbi_hilic_neg_1_042_25    m/z 291.1996     0   1   2   3   4E+3   0.5   1  RT (min)      lcorbi_hilic_neg_4_045_25    m/z 291.8438     0   2   4E+3   8   8.5  RT (min)      lcorbi_hilic_neg_4_098_61    m/z 291.8811     0   1E+4   1   1.5  RT (min)       lcorbi_hilic_neg_4_081_58    m/z 291.9267     0   1   2   3   4E+3   6   6.5  RT (min)      lcorbi_hilic_neg_4_072_57    m/z 291.9761     0   1   2E+4   1   1.5  RT (min)      lcorbi_hilic_neg_4_106_62    m/z 291.9891     0   1   2   3   4E+4   5.5   6  RT (min)       lcorbi_hilic_neg_4_108_62    m/z 292.0026     0   2   4E+3   6.5   7  RT (min)      lcorbi_hilic_neg_4_088_59    m/z 292.0356     0   1   1.5   2  RT (min)      lcorbi_hilic_neg_4_098_61    m/z 292.0704     0   1   2   3   4E+4   1   1.5  RT (min)      lcorbi_hilic_neg_4_013_46    m/z 292.0706     0   1E+3   3.5   4  RT (min)      lcorbi_hilic_neg_4_074_58    m/z 292.0722     0   2   4E+4   6   6.5  RT (min)       lcorbi_hilic_neg_4_079_58    m/z 292.0866     0   1   3   3.5  RT (min)      lcorbi_hilic_neg_4_105_61    m/z 292.0868     0   1E+4   7   7.5  RT (min)      lcorbi_hilic_neg_1_005_36    m/z 292.1037     0   1   2E+3   1.5   2  RT (min)      lcorbi_hilic_neg_4_097_60    m/z 292.1038     0   1E+5   3.5   4  RT (min)       lcorbi_hilic_neg_4_074_58    m/z 292.1046     0   1   2   3E+3   5   5.5  RT (min)       lcorbi_hilic_neg_4_091_60    m/z 292.1150     0   1E+5   6.5   7  RT (min)       lcorbi_hilic_neg_1_016_37    m/z 292.1421     0   1   2E+4   1   1.5  RT (min)       lcorbi_hilic_neg_4_090_60    m/z 292.1665     0   1E+4   6.5   7  RT (min)      lcorbi_hilic_neg_4_078_58    m/z 292.6908     0   1   2E+4   3   3.5  RT (min)      lcorbi_hilic_neg_3_034_34    m/z 292.8066     0   1   2   3   4E+3   5   5.5  RT (min)      lcorbi_hilic_neg_4_107_62    m/z 292.8710     0   1E+4   3   3.5  RT (min)      lcorbi_hilic_neg_4_085_59    m/z 292.8777     0   1   2E+3   7   7.5  RT (min)      lcorbi_hilic_neg_1_047_05    m/z 292.9421     0   1   6   6.5  RT (min)      lcorbi_hilic_neg_1_003_14    m/z 292.9653     0   1   7   7.5  RT (min)      lcorbi_hilic_neg_4_106_62    m/z 292.9670     0   1E+3   5.5   6  RT (min)      lcorbi_hilic_neg_1_003_14    m/z 292.9972     0   1   3   3.5  RT (min)      lcorbi_hilic_neg_4_090_60    m/z 293.0154     0   1   2   3   4E+2   6   6.5  RT (min)      lcorbi_hilic_neg_1_027_18    m/z 293.0335     0   1   1   1.5  RT (min)      lcorbi_hilic_neg_4_084_59    m/z 293.0490     0   1   3.5   4  RT (min)      lcorbi_hilic_neg_4_088_59    m/z 293.0545     0   1   2E+4   2.5   3  RT (min)      lcorbi_hilic_neg_4_074_58    m/z 293.0867     0   1   2E+3   7.5   8  RT (min)      lcorbi_hilic_neg_3_023_36    m/z 293.0940     0   1   2   3E+4   2   2.5  RT (min)      lcorbi_hilic_neg_4_051_16    m/z 293.1021     0   1E+3   6.5   7  RT (min)       lcorbi_hilic_neg_4_096_60    m/z 293.1022     0   1E+3   4   4.5  RT (min)      lcorbi_hilic_neg_4_104_61    m/z 293.1141     0   2   4   6   8E+2   5.5   6  RT (min)      lcorbi_hilic_neg_3_011_24    m/z 293.1144     0   1E+4   2.5   3  RT (min)      lcorbi_hilic_neg_4_089_59    m/z 293.1225     0   2   4   6   8E+3   1   1.5  RT (min)      lcorbi_hilic_neg_4_076_58    m/z 293.1238     0   1   4   4.5  RT (min)      lcorbi_hilic_neg_4_103_61    m/z 293.1338     0   2   4   6E+4   5   5.5  RT (min)       lcorbi_hilic_neg_4_097_60    m/z 293.1576     0   2   4   6   8E+3   7.5   8  RT (min)      lcorbi_hilic_neg_4_093_60    m/z 293.8911     0   1E+4   4.5   5  RT (min)      lcorbi_hilic_neg_1_010_22    m/z 293.8941     0   1   2   3   4E+4   3   3.5  RT (min)       lcorbi_hilic_neg_4_093_60    m/z 293.9276     0   2   4   6   8E+3   7   7.5  RT (min)      lcorbi_hilic_neg_4_097_60    m/z 293.9733     0   1   2E+4   1.5   2  RT (min)       lcorbi_hilic_neg_4_095_60    m/z 293.9737     0   1E+4   6   6.5  RT (min)      lcorbi_hilic_neg_4_071_57    m/z 293.9863     0   1E+4   0.5   1  RT (min)      lcorbi_hilic_neg_1_044_03    m/z 294.0002     0   1E+4   7   7.5  RT (min)      lcorbi_hilic_neg_4_090_60    m/z 294.0126     0   1   2E+3   6   6.5  RT (min)      lcorbi_hilic_neg_4_089_59    m/z 294.0359     0   2   4   6   8E+3   6   6.5  RT (min)      lcorbi_hilic_neg_4_087_59    m/z 294.0675     0   1E+4   1   1.5  RT (min)      lcorbi_hilic_neg_4_013_46    m/z 294.0680     0   1   4.5   5  RT (min)       lcorbi_hilic_neg_4_101_61    m/z 294.0691     0   1   6.5   7  RT (min)      lcorbi_hilic_neg_4_090_60    m/z 294.0790     0   1   2E+4   3   3.5  RT (min)      lcorbi_hilic_neg_4_085_59    m/z 294.1010     0   1E+4   6.5   7  RT (min)      lcorbi_hilic_neg_4_095_60    m/z 294.1199     0   2   4E+3   5.5   6  RT (min)       lcorbi_hilic_neg_4_095_60    m/z 294.1416     0   1E+4   6   6.5  RT (min)      lcorbi_hilic_neg_4_097_60    m/z 294.1425     0   2   4   6   8E+3   1   1.5  RT (min)      lcorbi_hilic_neg_4_074_58    m/z 294.1708     0   1   2E+3   1   1.5  RT (min)      lcorbi_hilic_neg_4_004_13    m/z 294.6193     0   2   4   6   8E+3   3   3.5  RT (min)      lcorbi_hilic_neg_4_078_58    m/z 294.6886     0   2   4   6E+4   3   3.5  RT (min)      lcorbi_hilic_neg_3_034_34    m/z 294.8036     0   2   4   6   8E+3   5   5.5  RT (min)      lcorbi_hilic_neg_4_066_57    m/z 294.8149     0   1E+4   4.5   5  RT (min)      lcorbi_hilic_neg_4_109_62    m/z 294.8623     0   1   2   3   4E+3   7   7.5  RT (min)      lcorbi_hilic_neg_1_042_25    m/z 294.8851     0   2   4   6   8E+3   12.5   13  RT (min)      lcorbi_hilic_neg_4_101_61    m/z 294.8852     0   1   2   3   4E+4   9   10  RT (min)       lcorbi_hilic_neg_4_076_58    m/z 294.8884     0   2   4   6E+3   4.5   5  RT (min)      lcorbi_hilic_neg_1_043_40    m/z 294.9390     0   2   4E+3   1.5   2  RT (min)      lcorbi_hilic_neg_4_108_62    m/z 294.9626     0   1   1.5   2  RT (min)      lcorbi_hilic_neg_4_084_59    m/z 294.9818     0   1   2E+4   3   3.5  RT (min)       lcorbi_hilic_neg_4_085_59    m/z 294.9818     0   1   2   3E+4   0.5   1  RT (min)       lcorbi_hilic_neg_4_075_58    m/z 295.0007     0   1E+4   1   1.5  RT (min)       lcorbi_hilic_neg_4_068_57    m/z 295.0054     0   1   3   3.5  RT (min)       lcorbi_hilic_neg_4_087_59    m/z 295.0279     0   2   4E+4   1   1.5  RT (min)      lcorbi_hilic_neg_1_005_36    m/z 295.0390     0   1   7.5   8  RT (min)      lcorbi_hilic_neg_4_067_57    m/z 295.0525     0   1   2   3E+4   6.5   7   7.5  RT (min)       lcorbi_hilic_neg_2_055_32    m/z 295.0645     0   1E+3   5   6  RT (min)       lcorbi_hilic_neg_1_055_24    m/z 295.0676     0   2   4   6E+3   2   2.5  RT (min)      lcorbi_hilic_neg_4_089_59    m/z 295.0726     0   1   2E+3   7.5   8  RT (min)       lcorbi_hilic_neg_1_012_32    m/z 295.0782     0   1   3.5   4  RT (min)      lcorbi_hilic_neg_4_110_62    m/z 295.0822     0   1E+4   1   1.5  RT (min)      lcorbi_hilic_neg_1_001_49    m/z 295.1033     0   1   2   3   4E+2   4   4.5  RT (min)      lcorbi_hilic_neg_3_040_19    m/z 295.1122     0   1   3.5   4  RT (min)       lcorbi_hilic_neg_4_107_62    m/z 295.1183     0   1   2E+4   1   1.5  RT (min)      lcorbi_hilic_neg_4_107_62    m/z 295.1189     0   1E+4   1   1.5  RT (min)       lcorbi_hilic_neg_4_094_60    m/z 295.1199     0   1E+5   4   4.5  RT (min)      lcorbi_hilic_neg_4_102_61    m/z 295.1375     0   1   2   3   4E+4   1   1.5  RT (min)       lcorbi_hilic_neg_4_112_62    m/z 295.8313     0   2   4   6   8E+3   7.5   8  RT (min)      lcorbi_hilic_neg_4_107_62    m/z 295.8719     0   1E+4   7   7.5  RT (min)      lcorbi_hilic_neg_4_088_59    m/z 295.8911     0   1   2   3E+4   3   3.5  RT (min)       lcorbi_hilic_neg_4_093_60    m/z 295.8937     0   2   4E+4   7.5   8  RT (min)       lcorbi_hilic_neg_4_089_59    m/z 295.9434     0   1   2E+4   7   7.5  RT (min)       lcorbi_hilic_neg_4_082_59    m/z 295.9841     0   1   2   3   4E+3   5.5   6  RT (min)      lcorbi_hilic_neg_4_078_58    m/z 296.0083     0   1E+4   2   2.5  RT (min)      lcorbi_hilic_neg_4_099_61    m/z 296.0231     0   1   6.5   7  RT (min)      lcorbi_hilic_neg_4_103_61    m/z 296.0294     0   1   2E+4   6   6.5  RT (min)       lcorbi_hilic_neg_4_092_60    m/z 296.0376     0   1E+4   3   3.5  RT (min)      lcorbi_hilic_neg_4_089_59    m/z 296.0437     0   2   4   6E+3   4.5   5  RT (min)      lcorbi_hilic_neg_4_071_57    m/z 296.0492     0   1   2   2.5  RT (min)      lcorbi_hilic_neg_4_083_59    m/z 296.0557     0   1   3.5   4  RT (min)      lcorbi_hilic_neg_4_103_61    m/z 296.0605     0   1   2E+4   7   7.5  RT (min)       lcorbi_hilic_neg_4_091_60    m/z 296.0714     0   1E+3   4   4.5  RT (min)       lcorbi_hilic_neg_4_097_60    m/z 296.0757     0   1E+3   6   6.5  RT (min)      lcorbi_hilic_neg_4_094_60    m/z 296.0821     0   1   2   3E+4   1.5   2  RT (min)       lcorbi_hilic_neg_4_089_59    m/z 296.0839     0   2   4E+4   3.5   4  RT (min)       lcorbi_hilic_neg_4_096_60    m/z 296.0858     0   1   2E+4   6   6.5   7  RT (min)       lcorbi_hilic_neg_4_089_59    m/z 296.0995     0   2   4   6E+3   1.5   2  RT (min)      lcorbi_hilic_neg_4_096_60    m/z 296.1206     0   1   2E+5   4   5  RT (min)       lcorbi_hilic_neg_4_097_60    m/z 296.1218     0   1E+3   1   1.5  RT (min)      lcorbi_hilic_neg_4_096_60    m/z 296.1476     0   1E+5   6.5   7  RT (min)      lcorbi_hilic_neg_1_012_32    m/z 296.1616     0   1E+4   1.5   2  RT (min)      lcorbi_hilic_neg_4_092_60    m/z 296.1845     0   1   2   3   4E+2   0.5   1  RT (min)      lcorbi_hilic_neg_4_031_14    m/z 296.1997     0   2   4   6E+3   2.5   3  RT (min)      lcorbi_hilic_neg_4_110_62    m/z 296.2310     0   1E+5   1   1.5  RT (min)      lcorbi_hilic_neg_3_031_14    m/z 296.2593     0   1   2   3E+3   0.5   1  RT (min)      lcorbi_hilic_neg_4_091_60    m/z 296.6867     0   2   4   6E+4   3   3.5  RT (min)      lcorbi_hilic_neg_3_034_34    m/z 296.8525     0   2   4   6E+3   7.5   8  RT (min)      lcorbi_hilic_neg_4_098_61    m/z 296.8818     0   1E+4   8.5   9  RT (min)      lcorbi_hilic_neg_4_082_59    m/z 296.8901     0   1   2   3E+4   5.5   6  RT (min)       lcorbi_hilic_neg_4_101_61    m/z 296.9047     0   1E+4   1   1.5  RT (min)      lcorbi_hilic_neg_4_070_57    m/z 296.9913     0   1   4   4.5  RT (min)      lcorbi_hilic_neg_4_085_59    m/z 296.9969     0   1   2   3E+4   3   3.5  RT (min)      lcorbi_hilic_neg_4_078_58    m/z 296.9970     0   1   2   3E+4   0.5   1  RT (min)       lcorbi_hilic_neg_4_076_58    m/z 297.0051     0   1   2   3   4E+3   2   2.5  RT (min)      lcorbi_hilic_neg_4_089_59    m/z 297.0316     0   1   3.5   4  RT (min)       lcorbi_hilic_neg_4_068_57    m/z 297.0492     0   1   6   6.5  RT (min)      lcorbi_hilic_neg_4_103_61    m/z 297.0550     0   1E+4   1.5   2  RT (min)      lcorbi_hilic_neg_4_075_58    m/z 297.0703     0   1E+4   6.5   7  RT (min)      lcorbi_hilic_neg_4_088_59    m/z 297.0744     0   1E+3   1.5   2  RT (min)      lcorbi_hilic_neg_4_077_58    m/z 297.0937     0   1E+5   6   6.5  RT (min)      lcorbi_hilic_neg_3_054_29    m/z 297.0962     0   1E+3   4   4.5  RT (min)      lcorbi_hilic_neg_4_070_57    m/z 297.1200     0   1   2E+4   3.5   4  RT (min)      lcorbi_hilic_neg_4_014_18    m/z 297.1219     0   1   2E+4   1.5   2  RT (min)       lcorbi_hilic_neg_4_096_60    m/z 297.1517     0   1   2   3E+4   4   4.5  RT (min)       lcorbi_hilic_neg_4_093_60    m/z 297.2068     0   1   1   1.5  RT (min)      lcorbi_hilic_neg_4_095_60    m/z 297.2930     0   1   2E+5   0.5   1  RT (min)      lcorbi_hilic_neg_1_047_05    m/z 297.8204     0   1   2E+4   7   7.5  RT (min)      lcorbi_hilic_neg_4_092_60    m/z 297.8229     0   2   4   6E+3   4.5   5  RT (min)      lcorbi_hilic_neg_1_052_41    m/z 297.8504     0   1   2   3E+3   8   8.5  RT (min)      lcorbi_hilic_neg_4_074_58    m/z 297.8717     0   1   2   3E+3   8   8.5  RT (min)      lcorbi_hilic_neg_4_098_61    m/z 297.9091     0   1   2   3E+4   1   1.5  RT (min)      lcorbi_hilic_neg_4_040_17    m/z 297.9093     0   1E+4   7.5   8  RT (min)      lcorbi_hilic_neg_4_074_58    m/z 297.9605     0   1E+4   6   6.5  RT (min)      lcorbi_hilic_neg_4_083_59    m/z 297.9686     0   1E+4   6.5   7  RT (min)      lcorbi_hilic_neg_4_073_57    m/z 298.0096     0   1E+4   6.5   7  RT (min)      lcorbi_hilic_neg_4_075_58    m/z 298.0466     0   2   4   6   8E+4   0.5   1  RT (min)      lcorbi_hilic_neg_4_113_62    m/z 298.0528     0   1   2   3E+3   3.5   4  RT (min)      lcorbi_hilic_neg_4_082_59    m/z 298.0604     0   1E+4   7.5   8  RT (min)      lcorbi_hilic_neg_4_092_60    m/z 298.0632     0   1   5.5   6  RT (min)      lcorbi_hilic_neg_4_089_59    m/z 298.0696     0   1   2   2.5  RT (min)      lcorbi_hilic_neg_4_005_32    m/z 298.0890     0   2   4   6E+3   3   3.5  RT (min)      lcorbi_hilic_neg_2_021_05    m/z 298.0992     0   1   2E+4   6.5   7  RT (min)      lcorbi_hilic_neg_1_012_32    m/z 298.1100     0   2   4E+3   7.5   8  RT (min)      lcorbi_hilic_neg_2_005_24    m/z 298.1126     0   1E+4   2   2.5  RT (min)      lcorbi_hilic_neg_3_044_32    m/z 298.1142     0   1   6   6.5  RT (min)       lcorbi_hilic_neg_4_096_60    m/z 298.1409     0   1E+3   3   3.5  RT (min)      lcorbi_hilic_neg_4_090_60    m/z 298.1444     0   1E+4   1   1.5  RT (min)      lcorbi_hilic_neg_4_097_60    m/z 298.1470     0   2   4E+4   6.5   7  RT (min)      lcorbi_hilic_neg_4_005_32    m/z 298.1538     0   2   4   6E+4   6.5   7  RT (min)       lcorbi_hilic_neg_4_091_60    m/z 298.1648     0   1E+4   7.5   8  RT (min)      lcorbi_hilic_neg_4_097_60    m/z 298.1727     0   1   2E+4   6   6.5  RT (min)       lcorbi_hilic_neg_4_097_60    m/z 298.2466     0   1   2E+4   1   1.5  RT (min)      lcorbi_hilic_neg_4_113_62    m/z 298.6849     0   1   2E+4   3   3.5  RT (min)      lcorbi_hilic_neg_3_034_34    m/z 298.7517     0   1E+4   4.5   5  RT (min)      lcorbi_hilic_neg_4_073_57    m/z 298.7796     0   1E+4   8   8.5  RT (min)      lcorbi_hilic_neg_4_066_57    m/z 298.8652     0   2   4   6E+3   7.5   8  RT (min)      lcorbi_hilic_neg_4_066_57    m/z 298.8785     0   1   2E+4   8   8.5  RT (min)       lcorbi_hilic_neg_4_082_59    m/z 298.8870     0   1E+4   5   5.5  RT (min)       lcorbi_hilic_neg_4_075_58    m/z 298.9334     0   1   0.5   1  RT (min)      lcorbi_hilic_neg_4_077_58    m/z 298.9525     0   1E+4   0.5   1  RT (min)      lcorbi_hilic_neg_4_101_61    m/z 298.9759     0   1E+4   0.5   1  RT (min)      lcorbi_hilic_neg_4_112_62    m/z 298.9927     0   1   2E+4   1   1.5  RT (min)      lcorbi_hilic_neg_4_084_59    m/z 298.9936     0   1   2E+4   1   1.5  RT (min)       lcorbi_hilic_neg_4_071_57    m/z 298.9938     0   1E+4   5   5.5  RT (min)      lcorbi_hilic_neg_4_076_58    m/z 299.0158     0   1E+4   0.5   1  RT (min)      lcorbi_hilic_neg_4_113_62    m/z 299.0438     0   2   4   6E+3   0.5   1  RT (min)      lcorbi_hilic_neg_4_095_60    m/z 299.0502     0   1E+3   7   7.5  RT (min)      lcorbi_hilic_neg_4_090_60    m/z 299.0538     0   1E+3   1.5   2  RT (min)       lcorbi_hilic_neg_4_075_58    m/z 299.0688     0   1   4   4.5  RT (min)       lcorbi_hilic_neg_4_096_60    m/z 299.0803     0   1   2E+4   2   2.5  RT (min)       lcorbi_hilic_neg_4_066_57    m/z 299.0811     0   1E+4   0.5   1  RT (min)      lcorbi_hilic_neg_4_109_62    m/z 299.0882     0   1E+4   5.5   6  RT (min)       lcorbi_hilic_neg_4_089_59    m/z 299.0962     0   1E+3   3   3.5  RT (min)      lcorbi_hilic_neg_4_105_61    m/z 299.0974     0   1   2E+4   5.5   6  RT (min)       lcorbi_hilic_neg_4_067_57    m/z 299.1131     0   1E+4   1   1.5  RT (min)      lcorbi_hilic_neg_4_001_49    m/z 299.1146     0   1   2E+4   6   6.5  RT (min)       lcorbi_hilic_neg_4_095_60    m/z 299.1247     0   1   4   4.5  RT (min)      lcorbi_hilic_neg_4_082_59    m/z 299.1360     0   1   2E+4   6.5   7  RT (min)      lcorbi_hilic_neg_4_030_05    m/z 299.1487     0   1E+5   1   1.5  RT (min)      lcorbi_hilic_neg_4_113_62    m/z 299.1489     0   1   2   3E+4   7.5   8  RT (min)       lcorbi_hilic_neg_4_094_60    m/z 299.1651     0   2   4   6E+4   0.5   1  RT (min)      lcorbi_hilic_neg_4_014_18    m/z 299.1862     0   2   4E+3   1   1.5  RT (min)      lcorbi_hilic_neg_4_097_60    m/z 299.2225     0   1   2   3E+3   1   1.5  RT (min)      lcorbi_hilic_neg_4_096_60    m/z 299.7718     0   2   4   6   8E+3   2.5   3  RT (min)      lcorbi_hilic_neg_4_080_58    m/z 299.8198     0   1   2E+4   4.5   5  RT (min)      lcorbi_hilic_neg_4_066_57    m/z 299.8660     0   1   2E+4   4   4.5  RT (min)       lcorbi_hilic_neg_4_109_62    m/z 299.8755     0   2   4   6E+4   8   8.5  RT (min)       lcorbi_hilic_neg_4_066_57    m/z 299.8837     0   1E+4   4.5   5  RT (min)       lcorbi_hilic_neg_4_078_58    m/z 299.9071     0   2   4   6   8E+4   7.5   8  RT (min)       lcorbi_hilic_neg_4_091_60    m/z 299.9569     0   1   6.5   7  RT (min)      lcorbi_hilic_neg_4_074_58    m/z 299.9662     0   1   7.5   8  RT (min)       lcorbi_hilic_neg_1_003_14    m/z 299.9731     0   2   4   6E+3   1   1.5  RT (min)      lcorbi_hilic_neg_4_107_62    m/z 300.0182     0   2   4E+3   2   2.5  RT (min)      lcorbi_hilic_neg_4_077_58    m/z 300.0257     0   2   4   6E+4   0.5   1  RT (min)       lcorbi_hilic_neg_4_113_62    m/z 300.0344     0   1E+3   6.5   7  RT (min)      lcorbi_hilic_neg_1_003_14    m/z 300.0393     0   1E+4   4   5  RT (min)       lcorbi_hilic_neg_4_067_57    m/z 300.0492     0   1   2   3   4E+2   7   7.5  RT (min)       lcorbi_hilic_neg_1_003_14    m/z 300.0672     0   1   2E+3   6.5   7  RT (min)      lcorbi_hilic_neg_4_084_59    m/z 300.0778     0   1   2.5   3  RT (min)      lcorbi_hilic_neg_4_108_62    m/z 300.0849     0   1   2   3E+4   6   6.5  RT (min)       lcorbi_hilic_neg_4_086_59    m/z 300.0852     0   1E+5   4   4.5  RT (min)       lcorbi_hilic_neg_4_082_59    m/z 300.0856     0   1   2.5   3  RT (min)      lcorbi_hilic_neg_4_111_62    m/z 300.1151     0   2   4E+4   3   3.5   4  RT (min)       lcorbi_hilic_neg_4_105_61    m/z 300.1217     0   1   1.5   2  RT (min)       lcorbi_hilic_neg_4_075_58    m/z 300.1313     0   1   4.5   5  RT (min)      lcorbi_hilic_neg_4_089_59    m/z 300.1565     0   1E+4   1   1.5  RT (min)      lcorbi_hilic_neg_3_001_49    m/z 300.7488     0   2   4   6E+3   4.5   5  RT (min)       lcorbi_hilic_neg_4_101_61    m/z 300.7766     0   2   4   6E+3   8   8.5  RT (min)      lcorbi_hilic_neg_4_066_57    m/z 300.8662     0   1   2E+4   7   7.5  RT (min)       lcorbi_hilic_neg_4_084_59    m/z 300.8784     0   1   2E+4   4.5   5  RT (min)      lcorbi_hilic_neg_4_075_58    m/z 300.9097     0   2   4   6   8E+3   8.5   9  RT (min)      lcorbi_hilic_neg_4_111_62    m/z 300.9472     0   1   2E+3   0.5   1  RT (min)      lcorbi_hilic_neg_1_033_53    m/z 300.9681     0   1E+3   4.5   5  RT (min)      lcorbi_hilic_neg_4_098_61    m/z 300.9740     0   1   2   3E+4   6   6.5  RT (min)       lcorbi_hilic_neg_4_073_57    m/z 300.9794     0   1   2E+3   1.5   2  RT (min)      lcorbi_hilic_neg_4_088_59    m/z 300.9841     0   2   4   6E+3   4.5   5  RT (min)      lcorbi_hilic_neg_4_006_06    m/z 300.9913     0   1   2   3E+4   3   3.5  RT (min)       lcorbi_hilic_neg_4_077_58    m/z 300.9914     0   1   2E+4   0.5   1  RT (min)       lcorbi_hilic_neg_4_078_58    m/z 301.0083     0   1   2E+4   6.5   7  RT (min)      lcorbi_hilic_neg_4_075_58    m/z 301.0231     0   2   4E+4   0.5   1  RT (min)       lcorbi_hilic_neg_4_113_62    m/z 301.0332     0   1   2E+4   7   7.5  RT (min)       lcorbi_hilic_neg_4_089_59    m/z 301.0395     0   1   2E+3   2   2.5  RT (min)      lcorbi_hilic_neg_4_095_60    m/z 301.0472     0   1E+3   7   7.5  RT (min)      lcorbi_hilic_neg_4_099_61    m/z 301.0516     0   1   2   3   4E+4   1.5   2  RT (min)       lcorbi_hilic_neg_4_075_58    m/z 301.0661     0   1   2E+3   1.5   2  RT (min)      lcorbi_hilic_neg_4_068_57    m/z 301.0698     0   2   4   6E+4   2   3  RT (min)       lcorbi_hilic_neg_2_027_02    m/z 301.0806     0   1   2E+4   6.5   7  RT (min)       lcorbi_hilic_neg_4_087_59    m/z 301.0972     0   1   4   4.5  RT (min)      lcorbi_hilic_neg_1_048_35    m/z 301.0997     0   1   2   3   4E+4   1   1.5  RT (min)       lcorbi_hilic_neg_4_096_60    m/z 301.1017     0   1E+4   6   6.5  RT (min)      lcorbi_hilic_neg_4_089_59    m/z 301.1027     0   1   3.5   4   4.5  RT (min)       lcorbi_hilic_neg_4_072_57    m/z 301.1113     0   1E+3   0.5   1  RT (min)      lcorbi_hilic_neg_4_092_60    m/z 301.1189     0   2   4   6E+3   3   3.5  RT (min)      lcorbi_hilic_neg_4_105_61    m/z 301.1247     0   1   2E+4   5   5.5  RT (min)       lcorbi_hilic_neg_4_096_60    m/z 301.1516     0   1   5.5   6  RT (min)      lcorbi_hilic_neg_2_005_24    m/z 301.1644     0   1   2   3E+5   6.5   7  RT (min)       lcorbi_hilic_neg_4_015_24    m/z 301.1655     0   1   2E+4   1   1.5  RT (min)       lcorbi_hilic_neg_4_083_59    m/z 301.1805     0   1   2E+4   7.5   8  RT (min)       lcorbi_hilic_neg_4_095_60    m/z 301.1807     0   2   4   6E+2   0.5   1  RT (min)      lcorbi_hilic_neg_2_012_14    m/z 301.1920     0   1E+4   2   2.5  RT (min)      lcorbi_hilic_neg_4_107_62    m/z 301.2018     0   1   2   3E+3   1   1.5  RT (min)      lcorbi_hilic_neg_4_083_59    m/z 301.2651     0   1   2   3   4E+4   1   1.5  RT (min)      lcorbi_hilic_neg_4_113_62    m/z 301.6220     0   2   4   6   8E+3   2   2.5  RT (min)      lcorbi_hilic_neg_4_001_49    m/z 301.7693     0   1E+4   2.5   3  RT (min)      lcorbi_hilic_neg_4_080_58    m/z 301.8727     0   1   2   3   4E+3   8   8.5  RT (min)      lcorbi_hilic_neg_4_098_61    m/z 301.9418     0   2   4E+3   0.5   1  RT (min)      lcorbi_hilic_neg_4_068_57    m/z 301.9554     0   1E+4   6   6.5  RT (min)      lcorbi_hilic_neg_4_067_57    m/z 301.9886     0   1   2E+4   1   1.5  RT (min)      lcorbi_hilic_neg_4_106_62    m/z 301.9929     0   1E+3   6   6.5  RT (min)      lcorbi_hilic_neg_4_106_62    m/z 302.0035     0   1E+4   5.5   6  RT (min)      lcorbi_hilic_neg_4_088_59    m/z 302.0315     0   1   2   3   4E+3   6.5   7  RT (min)      lcorbi_hilic_neg_4_088_59    m/z 302.0423     0   2   4   6E+3   1   1.5  RT (min)      lcorbi_hilic_neg_4_042_28    m/z 302.0492     0   1   2   3   4E+3   3   3.5  RT (min)      lcorbi_hilic_neg_1_023_23    m/z 302.0551     0   1E+3   1.5   2  RT (min)      lcorbi_hilic_neg_4_097_60    m/z 302.0660     0   1   2   3E+4   6   6.5  RT (min)       lcorbi_hilic_neg_4_104_61    m/z 302.0705     0   1   1.5   2  RT (min)      lcorbi_hilic_neg_4_097_60    m/z 302.0768     0   2   4   6   8E+3   5.5   6  RT (min)      lcorbi_hilic_neg_4_069_57    m/z 302.0818     0   1   4   4.5  RT (min)       lcorbi_hilic_neg_4_110_62    m/z 302.0911     0   1   6   6.5  RT (min)      lcorbi_hilic_neg_4_078_58    m/z 302.0912     0   1E+3   1.5   2  RT (min)      lcorbi_hilic_neg_4_069_57    m/z 302.0956     0   1   4   4.5  RT (min)       lcorbi_hilic_neg_4_113_62    m/z 302.1091     0   2   4   6   8E+4   2.5   3  RT (min)       lcorbi_hilic_neg_4_093_60    m/z 302.1107     0   1   6.5   7  RT (min)      lcorbi_hilic_neg_4_076_58    m/z 302.1298     0   1   2E+5   0.5   1  RT (min)       lcorbi_hilic_neg_4_099_61    m/z 302.1356     0   1   5.5   6  RT (min)      lcorbi_hilic_neg_4_099_61    m/z 302.1468     0   1   7.5   8  RT (min)      lcorbi_hilic_neg_4_106_62    m/z 302.1469     0   1   6   6.5  RT (min)       lcorbi_hilic_neg_3_031_14    m/z 302.1484     0   1E+4   1.5   2  RT (min)      lcorbi_hilic_neg_4_096_60    m/z 302.2206     0   1   2   3   4E+4   0.5   1  RT (min)      lcorbi_hilic_neg_2_012_14    m/z 302.7401     0   1E+4   8   8.5  RT (min)      lcorbi_hilic_neg_4_066_57    m/z 302.7460     0   1E+4   4.5   5  RT (min)      lcorbi_hilic_neg_4_069_57    m/z 302.8383     0   2   4   6   8E+3   4.5   5  RT (min)      lcorbi_hilic_neg_4_066_57    m/z 302.9811     0   1   2E+4   4.5   5  RT (min)       lcorbi_hilic_neg_4_077_58    m/z 302.9872     0   1   2E+4   3   3.5  RT (min)       lcorbi_hilic_neg_4_069_57    m/z 302.9872     0   1   2E+4   0.5   1  RT (min)       lcorbi_hilic_neg_4_076_58    m/z 302.9964     0   2   4   6   8E+3   6   6.5  RT (min)       lcorbi_hilic_neg_4_066_57    m/z 303.0130     0   1   2   3E+4   7   8  RT (min)       lcorbi_hilic_neg_4_074_58    m/z 303.0238     0   2   4   6E+2   7   7.5  RT (min)      lcorbi_hilic_neg_3_001_49    m/z 303.0558     0   1E+4   2   2.5  RT (min)      lcorbi_hilic_neg_4_083_59    m/z 303.0621     0   1E+4   7   7.5  RT (min)      lcorbi_hilic_neg_4_088_59    m/z 303.0671     0   1   2   3   4E+4   1.5   2  RT (min)       lcorbi_hilic_neg_4_073_57    m/z 303.0790     0   1   6.5   7  RT (min)      lcorbi_hilic_neg_3_023_36    m/z 303.0834     0   2   4   6   8E+2   2.5   3  RT (min)       lcorbi_hilic_neg_2_042_06    m/z 303.1382     0   1   2   3E+4   1   1.5  RT (min)       lcorbi_hilic_neg_4_089_59    m/z 303.1672     0   2   4E+3   7.5   8  RT (min)      lcorbi_hilic_neg_4_089_59    m/z 303.2328     0   1   2E+4   4   4.5  RT (min)       lcorbi_hilic_neg_4_073_57    m/z 303.2763     0   1   2E+4   1   1.5  RT (min)      lcorbi_hilic_neg_4_092_60    m/z 303.7388     0   1E+4   8   8.5  RT (min)      lcorbi_hilic_neg_4_072_57    m/z 303.7665     0   2   4   6E+3   2.5   3  RT (min)      lcorbi_hilic_neg_4_080_58    m/z 303.9525     0   1   6   6.5  RT (min)      lcorbi_hilic_neg_4_101_61    m/z 304.0022     0   1   2E+4   6   6.5  RT (min)       lcorbi_hilic_neg_4_083_59    m/z 304.0024     0   2   4E+4   0.5   1  RT (min)      lcorbi_hilic_neg_4_073_57    m/z 304.0109     0   1   2E+4   7.5   8  RT (min)       lcorbi_hilic_neg_4_075_58    m/z 304.0270     0   1   6  RT (min)       lcorbi_hilic_neg_4_106_62    m/z 304.0518     0   1   2E+3   7   7.5  RT (min)      lcorbi_hilic_neg_4_069_57    m/z 304.0649     0   1   2   3E+4   6   6.5  RT (min)       lcorbi_hilic_neg_4_072_57    m/z 304.0692     0   1   2E+4   1.5   2  RT (min)      lcorbi_hilic_neg_4_030_05    m/z 304.0707     0   1   5.5   6  RT (min)      lcorbi_hilic_neg_3_001_49    m/z 304.0786     0   1   2   3   4E+2   1   1.5  RT (min)      lcorbi_hilic_neg_1_001_49    m/z 304.0887     0   1   2   3   4E+2   5   5.5  RT (min)      lcorbi_hilic_neg_2_032_25    m/z 304.1057     0   1E+4   6   6.5  RT (min)      lcorbi_hilic_neg_4_079_58    m/z 304.1085     0   1E+3   2.5   3  RT (min)       lcorbi_hilic_neg_4_093_60    m/z 304.1151     0   1E+4   7   7.5  RT (min)      lcorbi_hilic_neg_4_083_59    m/z 304.1302     0   2   4   6E+4   1   1.5  RT (min)      lcorbi_hilic_neg_3_001_49    m/z 304.1345     0   1   6.5   7  RT (min)      lcorbi_hilic_neg_4_069_57    m/z 304.1526     0   1   2   3   4E+5   5.5   6  RT (min)       lcorbi_hilic_neg_4_097_60    m/z 304.2361     0   1E+5   0.5   1  RT (min)      lcorbi_hilic_neg_4_077_58    m/z 304.2362     0   2   4E+3   4   4.5  RT (min)      lcorbi_hilic_neg_4_073_57    m/z 304.7371     0   1   2E+4   8   8.5  RT (min)       lcorbi_hilic_neg_4_068_57    m/z 304.8316     0   2   4   6E+3   8   8.5  RT (min)      lcorbi_hilic_neg_4_066_57    m/z 304.8324     0   2   4   6   8E+3   5   5.5  RT (min)      lcorbi_hilic_neg_4_071_57    m/z 304.8711     0   2   4   6E+3   1.5   2  RT (min)      lcorbi_hilic_neg_3_003_44    m/z 304.9046     0   1E+4   8.5   9  RT (min)      lcorbi_hilic_neg_4_082_59    m/z 304.9140     0   2   4   6   8E+5   10   12  RT (min)       lcorbi_hilic_neg_4_111_62    m/z 304.9780     0   1E+4   4.5   5  RT (min)      lcorbi_hilic_neg_4_079_58    m/z 304.9832     0   1   2E+4   0.5   1  RT (min)       lcorbi_hilic_neg_4_100_61    m/z 304.9916     0   1   1.5   2  RT (min)      lcorbi_hilic_neg_4_086_59    m/z 305.0026     0   1   2E+4   0.5   1  RT (min)       lcorbi_hilic_neg_4_078_58    m/z 305.0027     0   1   2E+4   3   3.5  RT (min)       lcorbi_hilic_neg_4_070_57    m/z 305.0085     0   1E+4   7.5   8  RT (min)      lcorbi_hilic_neg_4_074_58    m/z 305.0183     0   2   4   6   8E+3   5.5   6  RT (min)      lcorbi_hilic_neg_4_081_58    m/z 305.0226     0   1   2E+4   0.5   1  RT (min)       lcorbi_hilic_neg_4_075_58    m/z 305.0226     0   1E+4   3   3.5  RT (min)      lcorbi_hilic_neg_4_078_58    m/z 305.0390     0   1E+4   5.5   6  RT (min)      lcorbi_hilic_neg_4_067_57    m/z 305.0400     0   1   2E+3   2   2.5  RT (min)      lcorbi_hilic_neg_4_083_59    m/z 305.0544     0   1   3.5   4  RT (min)      lcorbi_hilic_neg_3_042_05    m/z 305.0645     0   1   2E+5   6.5   7  RT (min)       lcorbi_hilic_neg_4_089_59    m/z 305.0680     0   1   2E+5   3   3.5  RT (min)       lcorbi_hilic_neg_4_095_60    m/z 305.0815     0   2   4   6E+3   3   3.5  RT (min)      lcorbi_hilic_neg_1_047_05    m/z 305.1004     0   1   6   6.5  RT (min)       lcorbi_hilic_neg_4_083_59    m/z 305.1005     0   2   4   6E+3   3.5   4  RT (min)      lcorbi_hilic_neg_4_093_60    m/z 305.1050     0   1   2   3   4E+4   2   2.5  RT (min)       lcorbi_hilic_neg_4_051_16    m/z 305.1757     0   1   2   3E+3   1   1.5  RT (min)      lcorbi_hilic_neg_4_092_60    m/z 305.2154     0   2   4   6E+4   0.5   1  RT (min)      lcorbi_hilic_neg_2_038_36    m/z 305.2484     0   1   2   3E+3   0.5   1  RT (min)      lcorbi_hilic_neg_2_032_25    m/z 305.8759     0   2   4   6E+3   1.5   2  RT (min)      lcorbi_hilic_neg_4_074_58    m/z 305.9173     0   2   4E+4   8.5   9  RT (min)      lcorbi_hilic_neg_4_075_58    m/z 305.9717     0   1E+4   1   1.5  RT (min)      lcorbi_hilic_neg_4_106_62    m/z 305.9767     0   1   2   3E+4   4.5   5  RT (min)       lcorbi_hilic_neg_4_079_58    m/z 306.0099     0   1   2E+3   2.5   3  RT (min)      lcorbi_hilic_neg_4_075_58    m/z 306.0124     0   1E+4   5.5   6  RT (min)      lcorbi_hilic_neg_4_075_58    m/z 306.0479     0   1   6.5   7  RT (min)      lcorbi_hilic_neg_4_073_57    m/z 306.0549     0   1   6.5   7  RT (min)      lcorbi_hilic_neg_2_049_55    m/z 306.0578     0   2   4   6   8E+2   1   1.5  RT (min)       lcorbi_hilic_neg_3_011_24    m/z 306.0666     0   1   2   3E+3   5   5.5  RT (min)      lcorbi_hilic_neg_4_098_61    m/z 306.0787     0   1   3.5   4  RT (min)      lcorbi_hilic_neg_4_092_60    m/z 306.0852     0   1E+4   0   0.5  RT (min)      lcorbi_hilic_neg_4_067_57    m/z 306.0861     0   1   2   3   4E+4   2   2.5  RT (min)       lcorbi_hilic_neg_4_067_57    m/z 306.0862     0   1   6.5   7  RT (min)      lcorbi_hilic_neg_4_089_59    m/z 306.0929     0   2   4   6E+3   6.5   7  RT (min)      lcorbi_hilic_neg_3_012_35    m/z 306.1082     0   1   2   3E+3   2   2.5  RT (min)      lcorbi_hilic_neg_4_071_57    m/z 306.1197     0   1   6.5   7  RT (min)       lcorbi_hilic_neg_4_082_59    m/z 306.1400     0   2   4E+4   3.5   4  RT (min)       lcorbi_hilic_neg_4_095_60    m/z 306.1418     0   1E+3   6.5   7  RT (min)      lcorbi_hilic_neg_1_016_37    m/z 306.1458     0   1E+4   1   2  RT (min)       lcorbi_hilic_neg_4_103_61    m/z 306.7341     0   1E+4   8   8.5  RT (min)      lcorbi_hilic_neg_4_066_57    m/z 306.8609     0   2   4   6   8E+4   0.5   1  RT (min)       lcorbi_hilic_neg_2_019_21    m/z 306.8663     0   1E+4   8.5   9  RT (min)      lcorbi_hilic_neg_4_066_57    m/z 306.9095     0   1   2   3E+4   8   8.5  RT (min)       lcorbi_hilic_neg_4_082_59    m/z 306.9433     0   2   4   6   8E+3   3.5   4  RT (min)      lcorbi_hilic_neg_4_112_62    m/z 306.9821     0   1E+4   0.5   1  RT (min)      lcorbi_hilic_neg_4_073_57    m/z 307.0006     0   1   2E+4   2   3  RT (min)       lcorbi_hilic_neg_4_075_58    m/z 307.0363     0   1E+3   5.5   6  RT (min)      lcorbi_hilic_neg_4_112_62    m/z 307.0514     0   1   3.5   4  RT (min)      lcorbi_hilic_neg_4_090_60    m/z 307.0561     0   1   6   6.5  RT (min)      lcorbi_hilic_neg_4_096_60    m/z 307.0717     0   2   4   6   8E+4   1.5   2  RT (min)       lcorbi_hilic_neg_4_093_60    m/z 307.0800     0   2   4   6   8E+3   2   2.5  RT (min)       lcorbi_hilic_neg_4_100_61    m/z 307.0997     0   1   2   3   4E+4   6   6.5  RT (min)       lcorbi_hilic_neg_4_095_60    m/z 307.0998     0   1E+5   3.5   4  RT (min)       lcorbi_hilic_neg_4_092_60    m/z 307.1115     0   2   4   6   8E+3   1.5   2  RT (min)      lcorbi_hilic_neg_4_092_60    m/z 307.1190     0   1   1.5   2  RT (min)       lcorbi_hilic_neg_4_105_61    m/z 307.1357     0   2   4   6E+4   2   2.5  RT (min)       lcorbi_hilic_neg_4_098_61    m/z 307.1913     0   1E+3   1   1.5  RT (min)      lcorbi_hilic_neg_1_043_40    m/z 307.2043     0   1   2   3E+3   1   1.5  RT (min)      lcorbi_hilic_neg_4_087_59    m/z 307.2640     0   1   0.5   1  RT (min)      lcorbi_hilic_neg_4_095_60    m/z 307.9072     0   2   4   6E+4   8   8.5  RT (min)       lcorbi_hilic_neg_4_090_60    m/z 307.9685     0   1E+4   1   1.5  RT (min)      lcorbi_hilic_neg_4_106_62    m/z 307.9737     0   2   4   6   8E+3   4.5   5  RT (min)      lcorbi_hilic_neg_4_077_58    m/z 307.9826     0   1E+3   6   6.5  RT (min)      lcorbi_hilic_neg_4_087_59    m/z 307.9971     0   1E+4   6.5   7  RT (min)      lcorbi_hilic_neg_4_071_57    m/z 308.0095     0   2   4E+3   5.5   6  RT (min)      lcorbi_hilic_neg_4_076_58    m/z 308.0556     0   2   4   6E+4   4   4.5  RT (min)      lcorbi_hilic_neg_1_037_46    m/z 308.0655     0   1   2E+4   1.5   2  RT (min)       lcorbi_hilic_neg_4_092_60    m/z 308.0840     0   1   2E+3   2.5   3  RT (min)      lcorbi_hilic_neg_4_093_60    m/z 308.0990     0   1   2E+3   4   4.5  RT (min)       lcorbi_hilic_neg_4_094_60    m/z 308.1077     0   1   2   3   4E+3   1.5   2  RT (min)      lcorbi_hilic_neg_4_095_60    m/z 308.1111     0   2   4   6E+4   4   4.5  RT (min)       lcorbi_hilic_neg_4_090_60    m/z 308.1350     0   2   4   6E+3   6   6.5  RT (min)      lcorbi_hilic_neg_4_083_59    m/z 308.1371     0   2   4   6E+3   1   1.5  RT (min)       lcorbi_hilic_neg_4_098_61    m/z 308.1436     0   1   2   3E+4   7.5   8  RT (min)       lcorbi_hilic_neg_4_095_60    m/z 308.1615     0   1   2   3E+4   7   7.5  RT (min)       lcorbi_hilic_neg_4_077_58    m/z 308.1616     0   1E+4   0.5   1  RT (min)      lcorbi_hilic_neg_4_097_60    m/z 308.1925     0   2   4   6   8E+4   5   5.5  RT (min)       lcorbi_hilic_neg_4_091_60    m/z 308.1981     0   1E+4   0.5   1  RT (min)      lcorbi_hilic_neg_4_071_57    m/z 308.7804     0   2   4   6E+3   4.5   5  RT (min)      lcorbi_hilic_neg_4_075_58    m/z 308.8402     0   1   2   3E+4   0.5   1  RT (min)       lcorbi_hilic_neg_4_077_58    m/z 308.9068     0   1   2   3   4E+4   8.5   9  RT (min)       lcorbi_hilic_neg_4_080_58    m/z 308.9160     0   1   2   3E+4   5.5   6  RT (min)       lcorbi_hilic_neg_4_106_62    m/z 308.9535     0   1   1   1.5  RT (min)      lcorbi_hilic_neg_4_084_59    m/z 308.9976     0   1   2   3E+4   3   3.5  RT (min)       lcorbi_hilic_neg_4_106_62    m/z 308.9976     0   2   4E+4   0.5   1  RT (min)       lcorbi_hilic_neg_4_073_57    m/z 309.0207     0   1   2   3   4E+4   6.5   7  RT (min)       lcorbi_hilic_neg_4_076_58    m/z 309.0270     0   1   2.5   3  RT (min)      lcorbi_hilic_neg_4_093_60    m/z 309.0318     0   2   4   6   8E+4   7   7.5  RT (min)       lcorbi_hilic_neg_1_039_38    m/z 309.0359     0   1E+3   5   5.5  RT (min)       lcorbi_hilic_neg_4_085_59    m/z 309.0496     0   2   4   6E+4   3   3.5  RT (min)       lcorbi_hilic_neg_4_085_59    m/z 309.0597     0   1E+4   7   7.5  RT (min)      lcorbi_hilic_neg_4_095_60    m/z 309.0606     0   2   4E+4   1   1.5  RT (min)       lcorbi_hilic_neg_4_013_46    m/z 309.0681     0   1E+4   5.5   6  RT (min)      lcorbi_hilic_neg_4_072_57    m/z 309.0690     0   1   2E+3   1.5   2  RT (min)       lcorbi_hilic_neg_2_044_46    m/z 309.0791     0   1   2E+4   7   7.5  RT (min)       lcorbi_hilic_neg_4_073_57    m/z 309.0820     0   2   4   6E+3   4   4.5  RT (min)      lcorbi_hilic_neg_1_037_46    m/z 309.0831     0   1   2E+3   1.5   2  RT (min)       lcorbi_hilic_neg_4_091_60    m/z 309.0890     0   1E+3   3.5   4  RT (min)       lcorbi_hilic_neg_4_093_60    m/z 309.1002     0   1   2   3E+3   1.5   2  RT (min)       lcorbi_hilic_neg_4_093_60    m/z 309.1091     0   1E+4   3.5   4  RT (min)      lcorbi_hilic_neg_4_087_59    m/z 309.1092     0   1E+4   2   2.5  RT (min)      lcorbi_hilic_neg_4_089_59    m/z 309.1094     0   1   2E+4   6   6.5  RT (min)       lcorbi_hilic_neg_4_093_60    m/z 309.1315     0   1   6   6.5  RT (min)      lcorbi_hilic_neg_4_087_59    m/z 309.1530     0   2   4E+3   1   1.5  RT (min)      lcorbi_hilic_neg_1_002_26    m/z 309.1686     0   1E+4   6   6.5  RT (min)      lcorbi_hilic_neg_4_096_60    m/z 309.1874     0   1   0.5   1  RT (min)      lcorbi_hilic_neg_4_102_61    m/z 309.1950     0   2   4   6E+3   0.5   1  RT (min)      lcorbi_hilic_neg_2_014_31    m/z 309.2070     0   2   4E+3   1   1.5  RT (min)       lcorbi_hilic_neg_4_103_61    m/z 309.2795     0   1   0.5   1  RT (min)      lcorbi_hilic_neg_4_015_24    m/z 309.9049     0   2   4   6   8E+3   8   8.5  RT (min)      lcorbi_hilic_neg_4_098_61    m/z 309.9589     0   1E+4   6   6.5  RT (min)      lcorbi_hilic_neg_4_083_59    m/z 310.0034     0   1   2E+4   1   1.5  RT (min)       lcorbi_hilic_neg_4_108_62    m/z 310.0351     0   1   2   3   4E+3   7   7.5  RT (min)      lcorbi_hilic_neg_1_039_38    m/z 310.0564     0   1   6.5   7  RT (min)      lcorbi_hilic_neg_4_093_60    m/z 310.0632     0   1   2E+3   6.5   7  RT (min)       lcorbi_hilic_neg_4_015_24    m/z 310.1048     0   1   1.5   2  RT (min)      lcorbi_hilic_neg_4_066_57    m/z 310.1108     0   1   2   3E+5   6.5   7  RT (min)       lcorbi_hilic_neg_4_097_60    m/z 310.1187     0   1   2   3E+3   3.5   4  RT (min)       lcorbi_hilic_neg_1_018_21    m/z 310.1214     0   1   2   3   4E+5   6   6.5  RT (min)       lcorbi_hilic_neg_4_097_60    m/z 310.1271     0   1   2E+3   3.5   4  RT (min)      lcorbi_hilic_neg_1_014_12    m/z 310.1287     0   1   6   6.5  RT (min)      lcorbi_hilic_neg_2_026_23    m/z 310.1619     0   2   4E+4   3.5   4  RT (min)      lcorbi_hilic_neg_4_108_62    m/z 310.2104     0   2   4   6   8E+3   1   1.5  RT (min)      lcorbi_hilic_neg_4_085_59    m/z 310.7775     0   2   4   6E+3   4.5   5  RT (min)      lcorbi_hilic_neg_4_071_57    m/z 310.8569     0   2   4   6   8E+4   0.5   1  RT (min)       lcorbi_hilic_neg_2_019_21    m/z 310.8704     0   2   4   6   8E+3   9   10   11  RT (min)       lcorbi_hilic_neg_2_025_52    m/z 310.8704     0   1E+4   13   13.5  RT (min)      lcorbi_hilic_neg_4_084_59    m/z 310.8799     0   1E+4   8.5   9  RT (min)       lcorbi_hilic_neg_4_073_57    m/z 310.9309     0   1   2   3E+3   9   9.5  RT (min)      lcorbi_hilic_neg_3_012_35    m/z 310.9505     0   1   1   1.5  RT (min)      lcorbi_hilic_neg_4_084_59    m/z 311.0111     0   1E+4   6   6.5  RT (min)      lcorbi_hilic_neg_4_067_57    m/z 311.0505     0   1E+4   7   7.5  RT (min)      lcorbi_hilic_neg_1_012_32    m/z 311.0579     0   1   1   1.5  RT (min)      lcorbi_hilic_neg_4_071_57    m/z 311.0803     0   1   4   4.5  RT (min)      lcorbi_hilic_neg_4_085_59    m/z 311.0807     0   1E+3   1.5   2  RT (min)       lcorbi_hilic_neg_4_071_57    m/z 311.0972     0   2   4   6E+4   7   8  RT (min)       lcorbi_hilic_neg_4_093_60    m/z 311.0983     0   1E+5   1.5   2  RT (min)       lcorbi_hilic_neg_4_072_57    m/z 311.0983     0   1   2   3E+4   3   3.5  RT (min)       lcorbi_hilic_neg_4_070_57    m/z 311.1350     0   1E+4   4.5   5  RT (min)      lcorbi_hilic_neg_1_047_05    m/z 311.1541     0   1   5   5.5  RT (min)       lcorbi_hilic_neg_1_041_54    m/z 311.1685     0   2   4E+4   3   4  RT (min)       lcorbi_hilic_neg_4_084_59    m/z 311.7710     0   1   2E+4   8   8.5  RT (min)       lcorbi_hilic_neg_4_066_57    m/z 311.8888     0   1   2   3E+4   7.5   8  RT (min)      lcorbi_hilic_neg_2_045_26    m/z 311.9683     0   1E+4   2   2.5  RT (min)      lcorbi_hilic_neg_4_103_61    m/z 311.9761     0   1E+4   5.5   6  RT (min)      lcorbi_hilic_neg_4_082_59    m/z 311.9843     0   1   2E+4   6   6.5  RT (min)       lcorbi_hilic_neg_4_068_57    m/z 312.0326     0   1E+4   5.5   6  RT (min)      lcorbi_hilic_neg_4_105_61    m/z 312.0759     0   2   4   6   8E+2   4   4.5  RT (min)      lcorbi_hilic_neg_2_014_31    m/z 312.0872     0   1   2   3E+2   6.5   7  RT (min)      lcorbi_hilic_neg_4_004_13    m/z 312.0874     0   1   1   1.5  RT (min)      lcorbi_hilic_neg_4_090_60    m/z 312.0949     0   1   2   3E+4   5.5   6  RT (min)       lcorbi_hilic_neg_4_104_61    m/z 312.0991     0   1   3.5   4  RT (min)       lcorbi_hilic_neg_4_086_59    m/z 312.1094     0   2   4E+4   6   6.5  RT (min)       lcorbi_hilic_neg_4_097_60    m/z 312.1112     0   1   2E+4   1.5   2  RT (min)       lcorbi_hilic_neg_4_097_60    m/z 312.1160     0   1E+5   6   6.5   7  RT (min)       lcorbi_hilic_neg_4_093_60    m/z 312.1181     0   1   3   3.5  RT (min)      lcorbi_hilic_neg_4_083_59    m/z 312.1288     0   1E+4   4.5   5  RT (min)      lcorbi_hilic_neg_4_110_62    m/z 312.1307     0   1   2   3E+4   0.5   1  RT (min)       lcorbi_hilic_neg_4_110_62    m/z 312.1494     0   1   2E+3   4.5   5  RT (min)      lcorbi_hilic_neg_4_110_62    m/z 312.1686     0   1   2E+4   4.5   5  RT (min)       lcorbi_hilic_neg_4_093_60    m/z 312.1815     0   2   4   6   8E+3   0.5   1  RT (min)      lcorbi_hilic_neg_4_111_62    m/z 312.2259     0   1   2   3   4E+3   0.5   1  RT (min)      lcorbi_hilic_neg_4_112_62    m/z 312.2542     0   1   1   1.5  RT (min)      lcorbi_hilic_neg_4_091_60    m/z 312.2847     0   1E+5   0.5   1  RT (min)       lcorbi_hilic_neg_4_097_60    m/z 312.7749     0   1   2   3   4E+3   4.5   5  RT (min)      lcorbi_hilic_neg_4_076_58    m/z 312.8361     0   1   2   3E+4   0.5   1  RT (min)       lcorbi_hilic_neg_4_077_58    m/z 312.8599     0   2   4   6   8E+3   7   7.5  RT (min)      lcorbi_hilic_neg_4_082_59    m/z 312.8642     0   1E+4   5   5.5  RT (min)      lcorbi_hilic_neg_4_087_59    m/z 312.9084     0   1E+4   4.5   5  RT (min)      lcorbi_hilic_neg_4_068_57    m/z 312.9648     0   1E+4   2   2.5  RT (min)      lcorbi_hilic_neg_4_104_61    m/z 312.9913     0   1   2E+4   0.5   1  RT (min)       lcorbi_hilic_neg_4_089_59    m/z 312.9914     0   1E+4   3   3.5  RT (min)       lcorbi_hilic_neg_4_067_57    m/z 313.0306     0   1E+3   6   6.5  RT (min)       lcorbi_hilic_neg_1_028_45    m/z 313.0418     0   1E+3   0.5   1  RT (min)      lcorbi_hilic_neg_1_030_04    m/z 313.0805     0   2   4   6E+3   7.5   8  RT (min)      lcorbi_hilic_neg_4_090_60    m/z 313.0959     0   1   2   3E+4   3.5   4  RT (min)       lcorbi_hilic_neg_4_089_59    m/z 313.1114     0   1E+3   2.5   3  RT (min)      lcorbi_hilic_neg_4_093_60    m/z 313.1155     0   1E+5   4   4.5  RT (min)      lcorbi_hilic_neg_1_047_05    m/z 313.1186     0   1   2E+3   1.5   2  RT (min)       lcorbi_hilic_neg_4_013_46    m/z 313.1410     0   1E+3   1.5   2  RT (min)      lcorbi_hilic_neg_4_072_57    m/z 313.1481     0   1   3.5   4  RT (min)      lcorbi_hilic_neg_4_094_60    m/z 313.1748     0   1E+5   1   1.5  RT (min)      lcorbi_hilic_neg_4_113_62    m/z 313.1769     0   1   2E+4   6   6.5  RT (min)      lcorbi_hilic_neg_2_005_24    m/z 313.1837     0   1   2   3   4E+4   6   6.5  RT (min)       lcorbi_hilic_neg_4_097_60    m/z 313.1837     0   1E+4   7.5   8  RT (min)      lcorbi_hilic_neg_4_096_60    m/z 313.2170     0   1E+3   0.5   1  RT (min)      lcorbi_hilic_neg_4_095_60    m/z 313.2940     0   2   4   6   8E+3   0.5   1  RT (min)      lcorbi_hilic_neg_3_008_04    m/z 313.7676     0   1   2E+4   8   8.5  RT (min)       lcorbi_hilic_neg_4_066_57    m/z 313.8557     0   2   4   6   8E+3   8.5   9  RT (min)      lcorbi_hilic_neg_4_090_60    m/z 313.9309     0   1E+3   4.5   5  RT (min)      lcorbi_hilic_neg_4_092_60    m/z 313.9729     0   1E+4   5.5   6  RT (min)      lcorbi_hilic_neg_4_082_59    m/z 313.9742     0   1   2E+4   6   6.5  RT (min)       lcorbi_hilic_neg_4_082_59    m/z 313.9761     0   1E+4   0.5   1  RT (min)      lcorbi_hilic_neg_4_089_59    m/z 314.0173     0   1   5.5   6  RT (min)      lcorbi_hilic_neg_4_091_60    m/z 314.0296     0   2   4E+3   5.5   6  RT (min)      lcorbi_hilic_neg_4_102_61    m/z 314.0726     0   2   4   6   8E+4   3   3.5  RT (min)       lcorbi_hilic_neg_4_068_57    m/z 314.0779     0   1   2   3E+4   0.5   1  RT (min)      lcorbi_hilic_neg_4_113_62    m/z 314.0847     0   2   4   6   8E+2   6   6.5  RT (min)      lcorbi_hilic_neg_2_005_24    m/z 314.0881     0   1   2   3E+4   1.5   2  RT (min)      lcorbi_hilic_neg_1_012_32    m/z 314.0990     0   1   2   3   4E+4   4   4.5  RT (min)      lcorbi_hilic_neg_1_037_46    m/z 314.1001     0   1   0.5   1  RT (min)      lcorbi_hilic_neg_1_047_05    m/z 314.1159     0   1E+4   3   3.5  RT (min)      lcorbi_hilic_neg_4_066_57    m/z 314.1228     0   1E+4   1.5   2  RT (min)      lcorbi_hilic_neg_4_067_57    m/z 314.1446     0   1E+4   4.5   5  RT (min)      lcorbi_hilic_neg_4_111_62    m/z 314.1677     0   2   4   6E+4   1   1.5  RT (min)      lcorbi_hilic_neg_4_112_62    m/z 314.1678     0   1E+4   6.5   7  RT (min)      lcorbi_hilic_neg_4_096_60    m/z 314.2416     0   2   4   6   8E+3   1   1.5  RT (min)      lcorbi_hilic_neg_4_067_57    m/z 314.7257     0   1E+4   4.5   5  RT (min)      lcorbi_hilic_neg_4_072_57    m/z 314.8291     0   1E+4   7.5   8  RT (min)      lcorbi_hilic_neg_4_074_58    m/z 314.8606     0   2   4   6   8E+3   7.5   8  RT (min)      lcorbi_hilic_neg_4_106_62    m/z 314.8613     0   1   2   3   4E+3   5   5.5  RT (min)      lcorbi_hilic_neg_4_098_61    m/z 314.8769     0   1   2E+4   7.5   8  RT (min)       lcorbi_hilic_neg_4_102_61    m/z 314.9489     0   1   2E+4   5.5   6  RT (min)       lcorbi_hilic_neg_4_072_57    m/z 314.9874     0   1   2E+4   0.5   1  RT (min)       lcorbi_hilic_neg_4_078_58    m/z 314.9874     0   1   2E+4   3   3.5  RT (min)       lcorbi_hilic_neg_4_081_58    m/z 315.0070     0   1E+4   0.5   1  RT (min)      lcorbi_hilic_neg_4_068_57    m/z 315.0130     0   2   4   6   8E+3   5.5   6  RT (min)      lcorbi_hilic_neg_4_081_58    m/z 315.0508     0   1   2   3   4E+3   3.5   4  RT (min)      lcorbi_hilic_neg_1_047_05    m/z 315.0524     0   1   2E+4   1   2  RT (min)       lcorbi_hilic_neg_4_067_57    m/z 315.0653     0   1   2E+3   6   6.5  RT (min)      lcorbi_hilic_neg_4_091_60    m/z 315.0710     0   1   2E+3   6   6.5  RT (min)      lcorbi_hilic_neg_4_089_59    m/z 315.0751     0   1   2E+4   0.5   1  RT (min)      lcorbi_hilic_neg_4_113_62    m/z 315.0831     0   1   2E+3   3.5   4  RT (min)      lcorbi_hilic_neg_2_055_32    m/z 315.0852     0   1E+3   2   2.5  RT (min)      lcorbi_hilic_neg_4_075_58    m/z 315.0927     0   2   4   6   8E+3   0.5   1  RT (min)      lcorbi_hilic_neg_4_110_62    m/z 315.1176     0   1   2   3   4E+3   6   6.5  RT (min)      lcorbi_hilic_neg_4_083_59    m/z 315.1269     0   1E+4   0.5   1  RT (min)      lcorbi_hilic_neg_4_113_62    m/z 315.1269     0   1   2E+3   3.5   4  RT (min)      lcorbi_hilic_neg_4_069_57    m/z 315.1520     0   1E+4   7   7.5  RT (min)       lcorbi_hilic_neg_4_094_60    m/z 315.1572     0   2   4   6E+3   1.5   2  RT (min)      lcorbi_hilic_neg_4_018_12    m/z 315.1671     0   1E+4   5.5   6  RT (min)      lcorbi_hilic_neg_4_089_59    m/z 315.1811     0   1   2   3   4E+4   1   1.5  RT (min)       lcorbi_hilic_neg_4_084_59    m/z 315.1929     0   1E+5   1.5   2  RT (min)      lcorbi_hilic_neg_1_029_31    m/z 315.2036     0   1E+4   6.5   7  RT (min)      lcorbi_hilic_neg_4_089_59    m/z 315.3036     0   1E+5   0.5   1  RT (min)       lcorbi_hilic_neg_4_097_60    m/z 315.7642     0   1E+4   8   8.5  RT (min)      lcorbi_hilic_neg_4_102_61    m/z 315.8333     0   1   2   3E+4   8   8.5  RT (min)       lcorbi_hilic_neg_4_066_57    m/z 315.8334     0   2   4   6E+4   4.5   5  RT (min)       lcorbi_hilic_neg_4_067_57    m/z 315.9269     0   1   2   3E+3   1   1.5  RT (min)      lcorbi_hilic_neg_4_104_61    m/z 315.9298     0   2   4   6E+3   5   5.5  RT (min)      lcorbi_hilic_neg_4_099_61    m/z 315.9975     0   1   2   3E+3   6   6.5  RT (min)      lcorbi_hilic_neg_4_074_58    m/z 316.0177     0   1   6   6.5  RT (min)      lcorbi_hilic_neg_4_081_58    m/z 316.0284     0   1E+4   1   1.5  RT (min)      lcorbi_hilic_neg_4_107_62    m/z 316.0374     0   1E+4   5.5   6  RT (min)      lcorbi_hilic_neg_3_033_53    m/z 316.0880     0   1   2E+4   2   2.5  RT (min)      lcorbi_hilic_neg_4_055_38    m/z 316.0971     0   2   4   6E+4   1.5   2  RT (min)       lcorbi_hilic_neg_4_090_60    m/z 316.1217     0   1   2   3E+4   2.5   3  RT (min)       lcorbi_hilic_neg_4_105_61    m/z 316.1666     0   1E+3   4   4.5  RT (min)      lcorbi_hilic_neg_4_005_32    m/z 316.1752     0   1   2   3   4E+4   6.5   7  RT (min)       lcorbi_hilic_neg_4_096_60    m/z 316.1820     0   1E+4   1   1.5  RT (min)      lcorbi_hilic_neg_2_055_32    m/z 316.2572     0   1E+3   1   1.5  RT (min)      lcorbi_hilic_neg_4_092_60    m/z 316.3159     0   1E+5   1   1.5  RT (min)      lcorbi_hilic_neg_4_095_60    m/z 316.7228     0   1   2E+4   4.5   5  RT (min)       lcorbi_hilic_neg_4_073_57    m/z 316.7390     0   2   4   6E+4   1   1.5  RT (min)      lcorbi_hilic_neg_1_026_34    m/z 316.7628     0   1E+4   8   8.5  RT (min)      lcorbi_hilic_neg_4_069_57    m/z 316.8269     0   1E+3   8   8.5  RT (min)      lcorbi_hilic_neg_3_044_32    m/z 316.8692     0   1   7   7.5  RT (min)      lcorbi_hilic_neg_4_106_62    m/z 316.9083     0   1   2   3   4E+4   1.5   2  RT (min)       lcorbi_hilic_neg_4_088_59    m/z 316.9337     0   2   4E+3   5   5.5  RT (min)      lcorbi_hilic_neg_3_008_04    m/z 316.9477     0   2   4   6E+6   5.5   6  RT (min)       lcorbi_hilic_neg_2_020_27    m/z 316.9647     0   1   2   3E+3   1   1.5  RT (min)      lcorbi_hilic_neg_4_072_57    m/z 316.9680     0   1   2   3   4E+4   5   5.5  RT (min)      lcorbi_hilic_neg_4_067_57    m/z 316.9802     0   1   2E+3   5   5.5  RT (min)      lcorbi_hilic_neg_3_024_27    m/z 316.9811     0   1   2   3E+3   7.5   8  RT (min)      lcorbi_hilic_neg_4_081_58    m/z 317.0008     0   1E+4   4   4.5  RT (min)      lcorbi_hilic_neg_4_089_59    m/z 317.0036     0   1E+4   0.5   1  RT (min)      lcorbi_hilic_neg_4_109_62    m/z 317.0100     0   1   5.5   6  RT (min)      lcorbi_hilic_neg_4_092_60    m/z 317.0342     0   1   2E+4   3.5   4  RT (min)       lcorbi_hilic_neg_4_087_59    m/z 317.0452     0   1   2   3E+4   1   1.5   2  RT (min)       lcorbi_hilic_neg_4_067_57    m/z 317.0515     0   1E+5   7   7.5  RT (min)       lcorbi_hilic_neg_2_005_24    m/z 317.0586     0   1E+5   7   7.5  RT (min)      lcorbi_hilic_neg_1_012_32    m/z 317.0666     0   1E+5   3   3.5   4  RT (min)       lcorbi_hilic_neg_4_103_61    m/z 317.0872     0   2   4   6E+3   1   1.5  RT (min)      lcorbi_hilic_neg_4_014_18    m/z 317.1000     0   1   5.5   6  RT (min)      lcorbi_hilic_neg_4_005_32    m/z 317.1009     0   1   3.5   4  RT (min)      lcorbi_hilic_neg_2_019_21    m/z 317.1060     0   1   2E+3   0.5   1   1.5  RT (min)       lcorbi_hilic_neg_1_024_11    m/z 317.1253     0   1E+5   6   6.5  RT (min)      lcorbi_hilic_neg_1_037_46    m/z 317.1273     0   1   2E+3   3   4  RT (min)       lcorbi_hilic_neg_4_098_61    m/z 317.1674     0   1   2E+4   1   1.5  RT (min)       lcorbi_hilic_neg_4_096_60    m/z 317.1869     0   1   2   3   4E+3   2   2.5  RT (min)      lcorbi_hilic_neg_4_109_62    m/z 317.8303     0   1   2E+4   8   8.5  RT (min)       lcorbi_hilic_neg_4_066_57    m/z 317.9406     0   2   4   6   8E+3   5.5   6  RT (min)      lcorbi_hilic_neg_4_080_58    m/z 317.9837     0   2   4   6   8E+3   1   1.5  RT (min)      lcorbi_hilic_neg_4_106_62    m/z 318.0181     0   1   2E+4   6   6.5  RT (min)       lcorbi_hilic_neg_4_082_59    m/z 318.0285     0   1   2E+4   1.5   2  RT (min)       lcorbi_hilic_neg_4_089_59    m/z 318.0503     0   1E+3   5.5   6  RT (min)      lcorbi_hilic_neg_4_013_46    m/z 318.0514     0   1   2   3   4E+3   1   1.5  RT (min)      lcorbi_hilic_neg_4_098_61    m/z 318.0611     0   1E+4   4.5   5  RT (min)      lcorbi_hilic_neg_4_087_59    m/z 318.0660     0   1E+3   6.5   7  RT (min)      lcorbi_hilic_neg_4_089_59    m/z 318.0873     0   1   2   2.5  RT (min)      lcorbi_hilic_neg_4_098_61    m/z 318.0926     0   1   2   3E+2   6.5   7  RT (min)      lcorbi_hilic_neg_4_035_48    m/z 318.1037     0   2   4   6   8E+4   6.5   7  RT (min)      lcorbi_hilic_neg_1_012_32    m/z 318.1191     0   1E+4   1   1.5  RT (min)       lcorbi_hilic_neg_2_045_26    m/z 318.1198     0   1   2E+5   6   6.5  RT (min)      lcorbi_hilic_neg_2_044_46    m/z 318.1412     0   1   5   5.5  RT (min)      lcorbi_hilic_neg_4_097_60    m/z 318.1424     0   2   4E+3   0.5   1  RT (min)      lcorbi_hilic_neg_4_094_60    m/z 318.1596     0   1   2E+5   4.5   5  RT (min)       lcorbi_hilic_neg_4_093_60    m/z 318.1790     0   1   0.5   1  RT (min)      lcorbi_hilic_neg_3_031_14    m/z 318.7200     0   1E+4   4.5   5  RT (min)      lcorbi_hilic_neg_4_070_57    m/z 318.8091     0   2   4   6   8E+3   4.5   5  RT (min)      lcorbi_hilic_neg_4_080_58    m/z 318.8270     0   2   4   6   8E+3   7.5   8  RT (min)      lcorbi_hilic_neg_4_104_61    m/z 318.9053     0   1   2   3E+4   1.5   2  RT (min)       lcorbi_hilic_neg_4_089_59    m/z 318.9493     0   1   3   3.5  RT (min)      lcorbi_hilic_neg_4_090_60    m/z 318.9624     0   1   1   1.5  RT (min)      lcorbi_hilic_neg_4_067_57    m/z 318.9812     0   1   2E+4   3   3.5  RT (min)       lcorbi_hilic_neg_4_070_57    m/z 318.9814     0   1   2   3   4E+4   0.5   1  RT (min)       lcorbi_hilic_neg_4_079_58    m/z 318.9904     0   1   2   3E+4   6.5   7  RT (min)       lcorbi_hilic_neg_4_095_60    m/z 318.9914     0   1   1   1.5  RT (min)       lcorbi_hilic_neg_4_076_58    m/z 319.0005     0   1   2E+4   1   1.5  RT (min)       lcorbi_hilic_neg_4_085_59    m/z 319.0270     0   1   2   3   4E+4   6.5   7  RT (min)       lcorbi_hilic_neg_4_071_57    m/z 319.0311     0   1   3.5   4  RT (min)      lcorbi_hilic_neg_4_069_57    m/z 319.0369     0   1   2E+4   7.5   8  RT (min)      lcorbi_hilic_neg_1_051_33    m/z 319.0519     0   2   4   6E+2   6   6.5  RT (min)       lcorbi_hilic_neg_1_037_46    m/z 319.0649     0   1   2   3   4E+4   5   5.5  RT (min)       lcorbi_hilic_neg_4_067_57    m/z 319.0886     0   1E+4   7.5   8  RT (min)      lcorbi_hilic_neg_4_085_59    m/z 319.0955     0   1E+3   6   6.5  RT (min)       lcorbi_hilic_neg_3_049_55    m/z 319.0961     0   2   4   6   8E+4   1   1.5  RT (min)       lcorbi_hilic_neg_4_107_62    m/z 319.0971     0   1   3   3.5  RT (min)      lcorbi_hilic_neg_4_104_61    m/z 319.1094     0   1E+5   6.5   7  RT (min)      lcorbi_hilic_neg_3_029_25    m/z 319.1162     0   1E+4   3.5   4  RT (min)      lcorbi_hilic_neg_4_105_61    m/z 319.1251     0   1   2E+3   1.5   2  RT (min)       lcorbi_hilic_neg_4_105_61    m/z 319.1265     0   1   2E+5   6.5   7  RT (min)      lcorbi_hilic_neg_2_044_46    m/z 319.1339     0   1E+5   6   6.5  RT (min)      lcorbi_hilic_neg_1_009_50    m/z 319.1340     0   1   2.5   3  RT (min)       lcorbi_hilic_neg_4_105_61    m/z 319.1549     0   2   4   6   8E+2   0.5   1  RT (min)      lcorbi_hilic_neg_4_031_14    m/z 319.9375     0   1   5.5   6  RT (min)      lcorbi_hilic_neg_4_093_60    m/z 319.9555     0   1   5   5.5  RT (min)      lcorbi_hilic_neg_4_102_61    m/z 320.0272     0   2   4E+3   6.5   7  RT (min)      lcorbi_hilic_neg_4_072_57    m/z 320.0388     0   1   2E+4   5.5   6  RT (min)       lcorbi_hilic_neg_4_071_57    m/z 320.0468     0   2   4   6E+3   7.5   8  RT (min)      lcorbi_hilic_neg_4_088_59    m/z 320.0492     0   1   1   1.5  RT (min)      lcorbi_hilic_neg_4_099_61    m/z 320.0514     0   2   4   6   8E+3   5.5   6  RT (min)      lcorbi_hilic_neg_4_083_59    m/z 320.0581     0   1   2   3   4E+3   4.5   5  RT (min)      lcorbi_hilic_neg_4_086_59    m/z 320.0668     0   1E+4   3   3.5  RT (min)      lcorbi_hilic_neg_1_012_32    m/z 320.0721     0   1   2   3E+4   6.5   7  RT (min)      lcorbi_hilic_neg_4_089_59    m/z 320.0741     0   1E+4   1   1.5  RT (min)       lcorbi_hilic_neg_4_101_61    m/z 320.0850     0   2   4E+2   6   6.5  RT (min)      lcorbi_hilic_neg_1_051_33    m/z 320.0864     0   1   2   3E+4   3   3.5  RT (min)       lcorbi_hilic_neg_4_095_60    m/z 320.0921     0   1   2   3   4E+5   6   6.5  RT (min)      lcorbi_hilic_neg_3_029_25    m/z 320.1020     0   1E+5   2   2.5  RT (min)      lcorbi_hilic_neg_2_001_49    m/z 320.1164     0   1E+4   1   1.5  RT (min)      lcorbi_hilic_neg_4_105_61    m/z 320.1175     0   1   6   6.5  RT (min)      lcorbi_hilic_neg_2_047_48    m/z 320.1280     0   1   6   6.5  RT (min)      lcorbi_hilic_neg_2_007_08    m/z 320.1354     0   1E+3   6.5   7  RT (min)       lcorbi_hilic_neg_4_003_08    m/z 320.1727     0   1E+4   6.5   7  RT (min)      lcorbi_hilic_neg_4_076_58    m/z 320.8062     0   1E+4   4.5   5  RT (min)      lcorbi_hilic_neg_4_080_58    m/z 320.8277     0   1E+4   8   8.5  RT (min)       lcorbi_hilic_neg_4_077_58    m/z 320.9765     0   2   4E+3   5.5   6  RT (min)      lcorbi_hilic_neg_4_068_57    m/z 320.9778     0   1E+4   1   1.5  RT (min)       lcorbi_hilic_neg_4_068_57    m/z 320.9974     0   1   2E+4   3   3.5  RT (min)       lcorbi_hilic_neg_4_078_58    m/z 321.0029     0   1E+4   5.5   6  RT (min)      lcorbi_hilic_neg_4_073_57    m/z 321.0085     0   1   2   3   4E+3   2   2.5  RT (min)      lcorbi_hilic_neg_4_096_60    m/z 321.0400     0   2   4   6E+4   6   6.5  RT (min)       lcorbi_hilic_neg_4_071_57    m/z 321.0623     0   1   2   3E+4   2   2.5  RT (min)       lcorbi_hilic_neg_4_066_57    m/z 321.0713     0   1E+4   7   7.5  RT (min)      lcorbi_hilic_neg_4_097_60    m/z 321.0817     0   1E+4   2   2.5  RT (min)      lcorbi_hilic_neg_4_073_57    m/z 321.0972     0   1   6   6.5  RT (min)      lcorbi_hilic_neg_4_090_60    m/z 321.1164     0   1   6.5   7  RT (min)      lcorbi_hilic_neg_4_094_60    m/z 321.1717     0   1   2E+3   1   1.5  RT (min)      lcorbi_hilic_neg_3_055_26    m/z 321.2197     0   1   0.5   1  RT (min)      lcorbi_hilic_neg_4_111_62    m/z 321.2343     0   1   2E+4   0.5   1  RT (min)      lcorbi_hilic_neg_4_080_58    m/z 321.7997     0   1   2E+4   8   8.5  RT (min)      lcorbi_hilic_neg_4_066_57    m/z 321.9037     0   1E+4   4.5   5  RT (min)      lcorbi_hilic_neg_4_107_62    m/z 321.9453     0   2   4   6   8E+3   1   1.5  RT (min)      lcorbi_hilic_neg_4_106_62    m/z 321.9639     0   1   2E+3   2   2.5  RT (min)      lcorbi_hilic_neg_4_106_62    m/z 321.9981     0   2   4   6E+3   5.5   6  RT (min)      lcorbi_hilic_neg_4_082_59    m/z 322.0010     0   2   4E+3   0.5   1  RT (min)      lcorbi_hilic_neg_4_082_59    m/z 322.0048     0   1   2E+4   2   2.5  RT (min)       lcorbi_hilic_neg_4_099_61    m/z 322.0129     0   1   2E+4   6   6.5  RT (min)       lcorbi_hilic_neg_4_068_57    m/z 322.0430     0   1   2   3E+3   2   2.5  RT (min)      lcorbi_hilic_neg_4_106_62    m/z 322.0445     0   1   2   3   4E+4   7.5   8  RT (min)       lcorbi_hilic_neg_1_047_05    m/z 322.0598     0   1E+4   2.5   3  RT (min)      lcorbi_hilic_neg_4_073_57    m/z 322.0656     0   1E+4   1   1.5  RT (min)      lcorbi_hilic_neg_1_037_46    m/z 322.0712     0   2   4   6E+3   3.5   4  RT (min)      lcorbi_hilic_neg_4_086_59    m/z 322.0909     0   1   4   4.5  RT (min)      lcorbi_hilic_neg_4_082_59    m/z 322.0990     0   1   2E+4   2   2.5  RT (min)      lcorbi_hilic_neg_2_001_49    m/z 322.1028     0   1E+4   6   6.5  RT (min)      lcorbi_hilic_neg_3_035_48    m/z 322.1142     0   1   2   3   4E+4   4   4.5  RT (min)       lcorbi_hilic_neg_4_082_59    m/z 322.1284     0   1   2E+4   3.5   4  RT (min)       lcorbi_hilic_neg_4_097_60    m/z 322.2468     0   2   4E+4   0.5   1  RT (min)      lcorbi_hilic_neg_4_031_14    m/z 322.8298     0   1   8   8.5  RT (min)      lcorbi_hilic_neg_3_035_48    m/z 322.8440     0   1E+4   9   9.5  RT (min)       lcorbi_hilic_neg_4_072_57    m/z 322.8867     0   1E+4   8.5   9  RT (min)      lcorbi_hilic_neg_4_090_60    m/z 322.8929     0   1   2E+4   5   5.5  RT (min)      lcorbi_hilic_neg_4_082_59    m/z 322.9413     0   1E+4   8   8.5  RT (min)      lcorbi_hilic_neg_4_080_58    m/z 322.9745     0   1   2E+3   5.5   6  RT (min)      lcorbi_hilic_neg_4_095_60    m/z 322.9749     0   2   4   6   8E+4   1.5   2  RT (min)      lcorbi_hilic_neg_3_030_30    m/z 322.9933     0   1   2E+4   3   3.5  RT (min)      lcorbi_hilic_neg_4_068_57    m/z 322.9934     0   1   2E+4   0.5   1  RT (min)       lcorbi_hilic_neg_4_071_57    m/z 323.0131     0   1   2   3E+4   0.5   1  RT (min)       lcorbi_hilic_neg_4_067_57    m/z 323.0132     0   1   2   3E+4   3   3.5  RT (min)       lcorbi_hilic_neg_4_079_58    m/z 323.0397     0   1   2   3E+3   2   2.5  RT (min)      lcorbi_hilic_neg_4_090_60    m/z 323.0480     0   1   6.5   7  RT (min)      lcorbi_hilic_neg_4_078_58    m/z 323.0592     0   1   2   2.5  RT (min)       lcorbi_hilic_neg_4_110_62    m/z 323.0650     0   2   4E+4   2.5   3  RT (min)       lcorbi_hilic_neg_4_085_59    m/z 323.0764     0   1   2   3E+3   6   6.5  RT (min)      lcorbi_hilic_neg_4_084_59    m/z 323.0835     0   1   2E+4   6   6.5  RT (min)       lcorbi_hilic_neg_4_089_59    m/z 323.0911     0   2   4   6   8E+4   1   1.5  RT (min)      lcorbi_hilic_neg_4_014_18    m/z 323.0981     0   1E+3   6   6.5  RT (min)      lcorbi_hilic_neg_4_084_59    m/z 323.0982     0   1   2E+4   4   4.5  RT (min)      lcorbi_hilic_neg_3_043_02    m/z 323.1146     0   1E+4   1.5   2  RT (min)      lcorbi_hilic_neg_4_076_58    m/z 323.1177     0   1   4   4.5  RT (min)       lcorbi_hilic_neg_4_090_60    m/z 323.1562     0   2   4E+4   3.5   4  RT (min)       lcorbi_hilic_neg_4_095_60    m/z 323.1562     0   2   4   6E+4   2   2.5  RT (min)       lcorbi_hilic_neg_4_096_60    m/z 323.1686     0   1   2E+5   0.5   1  RT (min)       lcorbi_hilic_neg_4_113_62    m/z 323.2061     0   2   4   6   8E+3   0.5   1  RT (min)      lcorbi_hilic_neg_4_068_57    m/z 323.2225     0   1   2E+5   1   1.5  RT (min)       lcorbi_hilic_neg_4_083_59    m/z 323.2590     0   1   2   3E+4   1   1.5  RT (min)       lcorbi_hilic_neg_4_066_57    m/z 323.7962     0   1   2E+4   8   8.5  RT (min)      lcorbi_hilic_neg_4_066_57    m/z 323.8759     0   1E+4   2   2.5  RT (min)       lcorbi_hilic_neg_4_096_60    m/z 323.8846     0   1E+4   8.5   9  RT (min)      lcorbi_hilic_neg_4_082_59    m/z 323.9029     0   1E+4   3.5   4  RT (min)      lcorbi_hilic_neg_4_081_58    m/z 323.9424     0   2   4   6E+3   1   1.5  RT (min)      lcorbi_hilic_neg_4_106_62    m/z 323.9674     0   1   6   6.5  RT (min)      lcorbi_hilic_neg_4_097_60    m/z 323.9746     0   1E+4   6   6.5  RT (min)      lcorbi_hilic_neg_4_087_59    m/z 323.9966     0   1E+4   5.5   6  RT (min)      lcorbi_hilic_neg_4_075_58    m/z 324.0190     0   2   4   6   8E+4   1   1.5  RT (min)      lcorbi_hilic_neg_4_042_28    m/z 324.0317     0   1   2E+4   6.5   7  RT (min)      lcorbi_hilic_neg_4_089_59    m/z 324.0442     0   2   4   6   8E+3   6.5   7  RT (min)      lcorbi_hilic_neg_4_091_60    m/z 324.0572     0   1   1.5   2  RT (min)      lcorbi_hilic_neg_4_109_62    m/z 324.0789     0   1E+4   3   3.5  RT (min)      lcorbi_hilic_neg_4_092_60    m/z 324.0965     0   1   2   3E+4   5   5.5  RT (min)      lcorbi_hilic_neg_1_047_05    m/z 324.1010     0   1   2   2.5  RT (min)      lcorbi_hilic_neg_4_091_60    m/z 324.1086     0   2   4   6E+3   3   3.5  RT (min)      lcorbi_hilic_neg_4_101_61    m/z 324.1115     0   2   4   6E+4   6   6.5  RT (min)       lcorbi_hilic_neg_4_094_60    m/z 324.1196     0   1   2   3E+3   6.5   7  RT (min)      lcorbi_hilic_neg_4_005_32    m/z 324.1330     0   2   4   6E+4   3.5   4  RT (min)       lcorbi_hilic_neg_4_095_60    m/z 324.1386     0   1   6   6.5  RT (min)      lcorbi_hilic_neg_2_038_36    m/z 324.2309     0   2   4   6   8E+3   2   3  RT (min)       lcorbi_hilic_neg_4_106_62    m/z 324.2543     0   1   2   3E+3   0.5   1  RT (min)      lcorbi_hilic_neg_4_066_57    m/z 324.7545     0   1   4.5   5  RT (min)      lcorbi_hilic_neg_4_106_62    m/z 324.7946     0   1   2   3   4E+4   8   8.5  RT (min)       lcorbi_hilic_neg_4_066_57    m/z 324.9806     0   1E+5   1   1.5  RT (min)      lcorbi_hilic_neg_1_027_18    m/z 324.9910     0   1   2E+4   1   1.5  RT (min)       lcorbi_hilic_neg_4_075_58    m/z 324.9967     0   2   4E+3   7   7.5  RT (min)      lcorbi_hilic_neg_1_029_31    m/z 325.0092     0   1E+4   0.5   1  RT (min)       lcorbi_hilic_neg_4_081_58    m/z 325.0552     0   2   4   6   8E+3   5   5.5  RT (min)      lcorbi_hilic_neg_4_066_57    m/z 325.0665     0   1   6   6.5  RT (min)      lcorbi_hilic_neg_4_073_57    m/z 325.0757     0   1E+4   0.5   1  RT (min)      lcorbi_hilic_neg_4_091_60    m/z 325.0822     0   1E+4   3.5   4  RT (min)       lcorbi_hilic_neg_4_091_60    m/z 325.1032     0   1   2E+4   0.5   1  RT (min)       lcorbi_hilic_neg_4_099_61    m/z 325.1141     0   1E+6   6   6.5  RT (min)       lcorbi_hilic_neg_4_093_60    m/z 325.1143     0   1   2E+3   2   2.5  RT (min)      lcorbi_hilic_neg_2_044_46    m/z 325.1152     0   1   3.5   4  RT (min)      lcorbi_hilic_neg_4_092_60    m/z 325.1252     0   1   2   3E+3   6.5   7  RT (min)       lcorbi_hilic_neg_4_097_60    m/z 325.1479     0   1   2   3   4E+4   1   1.5  RT (min)      lcorbi_hilic_neg_4_111_62    m/z 325.1609     0   1   2E+4   5.5   6  RT (min)       lcorbi_hilic_neg_4_096_60    m/z 325.1841     0   1   2   3   4E+4   3   3.5   4  RT (min)       lcorbi_hilic_neg_4_079_58    m/z 325.2016     0   1   2E+4   1   1.5  RT (min)      lcorbi_hilic_neg_4_097_60    m/z 325.2170     0   1E+4   0.5   1  RT (min)       lcorbi_hilic_neg_4_111_62    m/z 325.2381     0   1   2E+3   1   1.5  RT (min)      lcorbi_hilic_neg_4_095_60    m/z 325.2746     0   1   2E+5   0.5   1  RT (min)       lcorbi_hilic_neg_4_113_62    m/z 325.7927     0   2   4   6   8E+3   8   8.5  RT (min)      lcorbi_hilic_neg_4_066_57    m/z 325.8608     0   1   2   3   4E+4   1   1.5  RT (min)       lcorbi_hilic_neg_4_107_62    m/z 325.8729     0   1E+4   2   2.5  RT (min)      lcorbi_hilic_neg_4_097_60    m/z 325.9001     0   1E+4   3.5   4  RT (min)      lcorbi_hilic_neg_4_081_58    m/z 325.9775     0   2   4E+3   1   1.5  RT (min)      lcorbi_hilic_neg_4_108_62    m/z 325.9810     0   1E+4   5.5   6  RT (min)      lcorbi_hilic_neg_4_072_57    m/z 325.9913     0   1   2   3   4E+4   6.5   7  RT (min)      lcorbi_hilic_neg_3_039_31    m/z 326.0157     0   2   4   6   8E+3   6   6.5  RT (min)      lcorbi_hilic_neg_4_098_61    m/z 326.0265     0   1E+4   6   6.5  RT (min)      lcorbi_hilic_neg_4_090_60    m/z 326.0290     0   2   4   6E+3   1   1.5  RT (min)      lcorbi_hilic_neg_4_106_62    m/z 326.0453     0   2   4   6E+3   5.5   6  RT (min)      lcorbi_hilic_neg_4_069_57    m/z 326.0511     0   2   4E+4   7.5   8  RT (min)       lcorbi_hilic_neg_4_095_60    m/z 326.0607     0   1   2E+4   7.5   8  RT (min)      lcorbi_hilic_neg_1_012_32    m/z 326.0653     0   1E+4   4.5   5  RT (min)      lcorbi_hilic_neg_4_073_57    m/z 326.0667     0   1   2E+3   2.5   3  RT (min)      lcorbi_hilic_neg_4_049_55    m/z 326.0759     0   1   2E+3   6.5   7  RT (min)       lcorbi_hilic_neg_4_090_60    m/z 326.0993     0   1E+4   2   2.5  RT (min)      lcorbi_hilic_neg_4_066_57    m/z 326.1139     0   1   2E+5   6.5   7  RT (min)       lcorbi_hilic_neg_4_097_60    m/z 326.1165     0   1E+5   2.5   3  RT (min)      lcorbi_hilic_neg_3_051_46    m/z 326.1284     0   1   2   3   4E+3   5.5   6   6.5  RT (min)       lcorbi_hilic_neg_4_095_60    m/z 326.1481     0   1   2   3   4E+4   6   6.5  RT (min)       lcorbi_hilic_neg_4_093_60    m/z 326.1722     0   2   4   6   8E+3   0.5   1  RT (min)      lcorbi_hilic_neg_4_111_62    m/z 326.4872     0   1E+4   4.5   5  RT (min)      lcorbi_hilic_neg_4_080_58    m/z 326.7515     0   1   4.5   5  RT (min)      lcorbi_hilic_neg_4_098_61    m/z 326.7599     0   1E+4   4.5   5  RT (min)      lcorbi_hilic_neg_4_100_61    m/z 326.7746     0   2   4E+3   8   8.5  RT (min)      lcorbi_hilic_neg_4_067_57    m/z 326.7917     0   2   4   6   8E+3   8   8.5  RT (min)      lcorbi_hilic_neg_4_066_57    m/z 326.9005     0   2   4   6   8E+3   6   6.5  RT (min)      lcorbi_hilic_neg_4_090_60    m/z 326.9855     0   1E+4   6   6.5  RT (min)      lcorbi_hilic_neg_4_072_57    m/z 326.9882     0   1   2E+4   3   3.5  RT (min)       lcorbi_hilic_neg_4_078_58    m/z 326.9882     0   1   2E+4   0.5   1  RT (min)       lcorbi_hilic_neg_4_076_58    m/z 327.0097     0   1E+4   7.5   8  RT (min)      lcorbi_hilic_neg_4_087_59    m/z 327.0387     0   1E+4   5.5   6  RT (min)       lcorbi_hilic_neg_4_085_59    m/z 327.0587     0   1   2   3E+4   1.5   2  RT (min)       lcorbi_hilic_neg_4_097_60    m/z 327.0630     0   1   2E+4   3.5   4  RT (min)       lcorbi_hilic_neg_4_072_57    m/z 327.0633     0   1   7.5   8  RT (min)      lcorbi_hilic_neg_2_055_32    m/z 327.0703     0   2   4   6E+4   7.5   8  RT (min)       lcorbi_hilic_neg_4_096_60    m/z 327.0759     0   2   4E+3   2   2.5  RT (min)      lcorbi_hilic_neg_4_097_60    m/z 327.0784     0   2   4   6   8E+3   6   6.5  RT (min)      lcorbi_hilic_neg_4_071_57    m/z 327.0835     0   1E+4   1.5   2  RT (min)      lcorbi_hilic_neg_4_089_59    m/z 327.0912     0   2   4   6E+3   5   5.5  RT (min)      lcorbi_hilic_neg_4_068_57    m/z 327.0988     0   1   2   3   4E+3   2   2.5  RT (min)      lcorbi_hilic_neg_4_079_58    m/z 327.1044     0   1E+4   5.5   6  RT (min)      lcorbi_hilic_neg_2_002_28    m/z 327.1157     0   1E+4   6.5   7  RT (min)      lcorbi_hilic_neg_4_097_60    m/z 327.1266     0   1   2E+4   0.5   1  RT (min)      lcorbi_hilic_neg_4_066_57    m/z 327.1304     0   1   4   4.5  RT (min)      lcorbi_hilic_neg_4_070_57    m/z 327.1634     0   1E+5   1   1.5  RT (min)      lcorbi_hilic_neg_4_113_62    m/z 327.1873     0   2   4   6   8E+3   3.5   4  RT (min)      lcorbi_hilic_neg_4_090_60    m/z 327.2175     0   2   4   6E+3   1   1.5  RT (min)      lcorbi_hilic_neg_4_111_62    m/z 327.2328     0   1   2   3   4E+3   4   4.5  RT (min)      lcorbi_hilic_neg_4_069_57    m/z 327.4859     0   1   2   3   4E+4   4.5   5  RT (min)       lcorbi_hilic_neg_4_077_58    m/z 327.6598     0   1E+4   3   3.5  RT (min)      lcorbi_hilic_neg_4_081_58    m/z 327.8577     0   1   2E+4   1   1.5  RT (min)       lcorbi_hilic_neg_4_106_62    m/z 327.9583     0   1   6   6.5  RT (min)      lcorbi_hilic_neg_4_104_61    m/z 327.9943     0   1   2E+3   5.5   6  RT (min)      lcorbi_hilic_neg_4_084_59    m/z 328.0243     0   1E+4   2.5   3  RT (min)      lcorbi_hilic_neg_4_013_46    m/z 328.0451     0   1   2   3E+4   5.5   6  RT (min)       lcorbi_hilic_neg_4_098_61    m/z 328.0706     0   1E+3   1.5   2  RT (min)      lcorbi_hilic_neg_4_066_57    m/z 328.0805     0   1   3.5   4  RT (min)      lcorbi_hilic_neg_2_005_24    m/z 328.0884     0   1   2E+4   1.5   2  RT (min)      lcorbi_hilic_neg_4_066_57    m/z 328.1073     0   2   4E+4   5   5.5  RT (min)      lcorbi_hilic_neg_1_005_36    m/z 328.1107     0   2   4   6   8E+4   6.5   7  RT (min)       lcorbi_hilic_neg_4_095_60    m/z 328.1238     0   1E+3   6.5   7  RT (min)       lcorbi_hilic_neg_1_016_37    m/z 328.1376     0   1   2E+4   3   3.5  RT (min)       lcorbi_hilic_neg_4_092_60    m/z 328.1513     0   1   2   3   4E+3   4   4.5  RT (min)      lcorbi_hilic_neg_2_005_24    m/z 328.1626     0   1   2   3   4E+4   4  RT (min)       lcorbi_hilic_neg_4_096_60    m/z 328.1832     0   2   4   6   8E+4   0.5   1  RT (min)       lcorbi_hilic_neg_4_113_62    m/z 328.2361     0   2   4   6   8E+4   0.5   1  RT (min)      lcorbi_hilic_neg_4_077_58    m/z 328.4845     0   1E+5   4.5   5  RT (min)       lcorbi_hilic_neg_4_077_58    m/z 328.7567     0   1E+4   4.5   5  RT (min)      lcorbi_hilic_neg_4_098_61    m/z 328.7717     0   1   2   3E+3   8   8.5  RT (min)      lcorbi_hilic_neg_4_068_57    m/z 328.8100     0   1   2E+4   1   1.5  RT (min)      lcorbi_hilic_neg_4_082_59    m/z 328.8379     0   1E+4   4.5   5  RT (min)      lcorbi_hilic_neg_4_075_58    m/z 328.9588     0   1E+4   5   5.5  RT (min)      lcorbi_hilic_neg_4_104_61    m/z 328.9664     0   2   4   6E+3   6.5   7  RT (min)      lcorbi_hilic_neg_4_071_57    m/z 328.9857     0   1E+4   1   1.5  RT (min)      lcorbi_hilic_neg_4_072_57    m/z 328.9974     0   1E+4   5   5.5  RT (min)      lcorbi_hilic_neg_1_008_39    m/z 329.0032     0   1E+4   0.5   1  RT (min)      lcorbi_hilic_neg_4_076_58    m/z 329.0049     0   1   2E+4   7   7.5  RT (min)       lcorbi_hilic_neg_4_085_59    m/z 329.0285     0   1E+4   5   5.5  RT (min)      lcorbi_hilic_neg_4_070_57    m/z 329.0399     0   2   4   6   8E+3   7   7.5  RT (min)      lcorbi_hilic_neg_4_082_59    m/z 329.0483     0   1   1   1.5  RT (min)      lcorbi_hilic_neg_4_098_61    m/z 329.0740     0   2   4   6E+4   1.5   2  RT (min)      lcorbi_hilic_neg_1_047_05    m/z 329.0757     0   1   2E+5   4.5   5  RT (min)       lcorbi_hilic_neg_4_073_57    m/z 329.0850     0   2   4   6   8E+2   1   1.5  RT (min)      lcorbi_hilic_neg_1_047_05    m/z 329.1089     0   1   5   5.5  RT (min)       lcorbi_hilic_neg_4_096_60    m/z 329.1130     0   1   6.5   7  RT (min)      lcorbi_hilic_neg_4_082_59    m/z 329.1132     0   1   2   3   4E+4   1   1.5  RT (min)       lcorbi_hilic_neg_4_092_60    m/z 329.1243     0   1E+4   1.5   2  RT (min)      lcorbi_hilic_neg_4_091_60    m/z 329.1310     0   1   2E+3   6.5   7  RT (min)      lcorbi_hilic_neg_4_085_59    m/z 329.1388     0   1   5   5.5  RT (min)      lcorbi_hilic_neg_2_005_24    m/z 329.1395     0   2   4   6   8E+4   1   1.5  RT (min)      lcorbi_hilic_neg_2_020_27    m/z 329.1757     0   1E+4   1   1.5  RT (min)      lcorbi_hilic_neg_4_075_58    m/z 329.1864     0   2   4   6   8E+3   1   1.5  RT (min)      lcorbi_hilic_neg_4_081_58    m/z 329.2484     0   1   0.5   1  RT (min)      lcorbi_hilic_neg_1_024_11    m/z 329.9015     0   2   4   6E+3   8   8.5  RT (min)      lcorbi_hilic_neg_4_067_57    m/z 329.9720     0   1E+4   1   1.5  RT (min)      lcorbi_hilic_neg_4_106_62    m/z 329.9761     0   1   5.5   6  RT (min)      lcorbi_hilic_neg_4_081_58    m/z 330.0366     0   1E+3   5   5.5  RT (min)      lcorbi_hilic_neg_1_047_05    m/z 330.0486     0   1E+4   2   2.5  RT (min)      lcorbi_hilic_neg_4_098_61    m/z 330.0541     0   1   2E+3   6.5   7  RT (min)       lcorbi_hilic_neg_4_091_60    m/z 330.0620     0   2   4   6   8E+3   6.5   7  RT (min)      lcorbi_hilic_neg_3_028_23    m/z 330.0711     0   2   4   6   8E+2   6.5   7  RT (min)      lcorbi_hilic_neg_2_005_24    m/z 330.0822     0   2   4   6E+4   5   6  RT (min)       lcorbi_hilic_neg_4_097_60    m/z 330.1053     0   1E+4   3.5   4  RT (min)      lcorbi_hilic_neg_4_030_05    m/z 330.1056     0   1   2   3E+3   7   7.5  RT (min)      lcorbi_hilic_neg_2_014_31    m/z 330.1056     0   1   2   3   4E+4   2   2.5  RT (min)      lcorbi_hilic_neg_4_055_38    m/z 330.1172     0   2   4E+3   1.5   2  RT (min)      lcorbi_hilic_neg_3_013_37    m/z 330.1261     0   2   4E+3   4.5   5  RT (min)      lcorbi_hilic_neg_2_020_27    m/z 330.1344     0   1   4.5   5  RT (min)      lcorbi_hilic_neg_4_112_62    m/z 330.1417     0   1E+4   6.5   7  RT (min)      lcorbi_hilic_neg_3_042_05    m/z 330.1612     0   1E+3   4.5   5  RT (min)      lcorbi_hilic_neg_4_013_46    m/z 330.1628     0   2   4   6   8E+3   7   7.5  RT (min)      lcorbi_hilic_neg_4_096_60    m/z 330.1670     0   2   4   6E+3   2.5   3  RT (min)      lcorbi_hilic_neg_4_089_59    m/z 330.1743     0   1   2E+4   5   5.5  RT (min)      lcorbi_hilic_neg_4_096_60    m/z 330.2365     0   2   4E+4   1   1.5  RT (min)      lcorbi_hilic_neg_4_075_58    m/z 330.2517     0   1   0.5   1  RT (min)      lcorbi_hilic_neg_4_015_24    m/z 330.4816     0   2   4   6   8E+4   4.5   5  RT (min)       lcorbi_hilic_neg_4_079_58    m/z 330.7003     0   1   2E+5   4.5   5  RT (min)       lcorbi_hilic_neg_2_020_27    m/z 330.8308     0   1   2   3   4E+3   8   8.5  RT (min)      lcorbi_hilic_neg_4_106_62    m/z 330.8351     0   1E+4   4.5   5  RT (min)      lcorbi_hilic_neg_4_078_58    m/z 330.8420     0   1E+4   7.5   8  RT (min)      lcorbi_hilic_neg_4_105_61    m/z 330.8721     0   1   2   3E+3   7   7.5  RT (min)      lcorbi_hilic_neg_4_082_59    m/z 330.8795     0   1E+4   7   7.5  RT (min)      lcorbi_hilic_neg_4_083_59    m/z 330.9192     0   1E+4   4.5   5  RT (min)      lcorbi_hilic_neg_4_069_57    m/z 330.9529     0   1   2   3E+4   7   7.5  RT (min)       lcorbi_hilic_neg_4_082_59    m/z 330.9559     0   1E+3   4.5   5  RT (min)      lcorbi_hilic_neg_4_084_59    m/z 330.9812     0   1   2   3E+4   0.5   1  RT (min)       lcorbi_hilic_neg_4_097_60    m/z 330.9878     0   1   2E+4   7.5   8  RT (min)       lcorbi_hilic_neg_4_074_58    m/z 331.0002     0   1E+4   1   1.5  RT (min)      lcorbi_hilic_neg_4_113_62    m/z 331.0163     0   2   4   6E+3   0.5   1  RT (min)      lcorbi_hilic_neg_4_076_58    m/z 331.0394     0   2   4E+4   3   3.5  RT (min)      lcorbi_hilic_neg_1_002_26    m/z 331.0451     0   1   2   3E+3   1   1.5   2  RT (min)       lcorbi_hilic_neg_4_098_61    m/z 331.0467     0   1E+4   6   6.5  RT (min)      lcorbi_hilic_neg_4_088_59    m/z 331.0606     0   1E+4   1.5   2  RT (min)      lcorbi_hilic_neg_4_073_57    m/z 331.0675     0   1   7.5   8  RT (min)      lcorbi_hilic_neg_4_090_60    m/z 331.0726     0   2   4   6E+4   4.5   5  RT (min)       lcorbi_hilic_neg_4_073_57    m/z 331.0747     0   1E+5   7.5   8  RT (min)       lcorbi_hilic_neg_4_097_60    m/z 331.0836     0   1E+3   6   6.5  RT (min)      lcorbi_hilic_neg_4_089_59    m/z 331.0911     0   1   2   2.5  RT (min)      lcorbi_hilic_neg_4_106_62    m/z 331.1018     0   1E+5   2   2.5  RT (min)      lcorbi_hilic_neg_4_055_38    m/z 331.1104     0   1   2E+4   5   5.5  RT (min)       lcorbi_hilic_neg_4_093_60    m/z 331.1135     0   1   7   7.5  RT (min)      lcorbi_hilic_neg_4_097_60    m/z 331.1164     0   1   2   2.5  RT (min)      lcorbi_hilic_neg_4_106_62    m/z 331.1375     0   2   4   6   8E+3   3   3.5  RT (min)      lcorbi_hilic_neg_1_012_32    m/z 331.1621     0   1E+4   6   6.5  RT (min)      lcorbi_hilic_neg_4_089_59    m/z 331.2042     0   1   0.5   1  RT (min)      lcorbi_hilic_neg_4_007_21    m/z 331.2276     0   2   4E+4   0.5   1  RT (min)       lcorbi_hilic_neg_4_089_59    m/z 331.2488     0   1   2   3   4E+3   0.5   1  RT (min)      lcorbi_hilic_neg_4_023_34    m/z 331.2640     0   1   0.5   1  RT (min)      lcorbi_hilic_neg_4_015_24    m/z 331.4800     0   2   4E+4   4.5   5  RT (min)       lcorbi_hilic_neg_4_077_58    m/z 331.7856     0   2   4E+3   8   8.5  RT (min)      lcorbi_hilic_neg_4_082_59    m/z 331.8286     0   1   2   3E+4   8   8.5  RT (min)       lcorbi_hilic_neg_4_066_57    m/z 331.9840     0   1   2E+4   1.5   2  RT (min)       lcorbi_hilic_neg_4_104_61    m/z 331.9955     0   1   6.5   7  RT (min)      lcorbi_hilic_neg_4_096_60    m/z 332.0153     0   1   5.5   6  RT (min)      lcorbi_hilic_neg_4_082_59    m/z 332.0171     0   1E+4   7   7.5  RT (min)      lcorbi_hilic_neg_4_083_59    m/z 332.0234     0   1   2   3   4E+4   3   3.5  RT (min)       lcorbi_hilic_neg_4_066_57    m/z 332.0239     0   1   6.5   7  RT (min)      lcorbi_hilic_neg_4_083_59    m/z 332.0444     0   1   3.5   4  RT (min)      lcorbi_hilic_neg_4_084_59    m/z 332.0729     0   1E+4   2.5   3  RT (min)      lcorbi_hilic_neg_1_037_46    m/z 332.0842     0   1   2E+3   1.5   2  RT (min)       lcorbi_hilic_neg_4_107_62    m/z 332.0899     0   2   4E+3   6.5   7  RT (min)      lcorbi_hilic_neg_4_086_59    m/z 332.0918     0   1E+4   4   4.5  RT (min)      lcorbi_hilic_neg_4_103_61    m/z 332.0963     0   1   2E+4   1.5   2  RT (min)      lcorbi_hilic_neg_4_084_59    m/z 332.1250     0   1E+3   5.5   6  RT (min)      lcorbi_hilic_neg_4_030_05    m/z 332.1368     0   1   2E+3   6.5   7  RT (min)      lcorbi_hilic_neg_4_091_60    m/z 332.1527     0   1E+4   1   1.5  RT (min)       lcorbi_hilic_neg_4_097_60    m/z 332.1737     0   1   2E+3   0.5   1  RT (min)      lcorbi_hilic_neg_4_105_61    m/z 332.1799     0   1   2   2.5  RT (min)      lcorbi_hilic_neg_4_097_60    m/z 332.4785     0   1   2E+4   4.5   5  RT (min)       lcorbi_hilic_neg_4_077_58    m/z 332.8265     0   1   2   3   4E+4   8   8.5  RT (min)      lcorbi_hilic_neg_4_066_57    m/z 332.9216     0   1E+4   5   5.5  RT (min)       lcorbi_hilic_neg_4_108_62    m/z 332.9977     0   1   2E+4   3   3.5  RT (min)       lcorbi_hilic_neg_4_076_58    m/z 332.9977     0   1   2E+4   0.5   1  RT (min)       lcorbi_hilic_neg_4_076_58    m/z 333.0082     0   1   3.5   4  RT (min)      lcorbi_hilic_neg_4_070_57    m/z 333.0237     0   2   4E+3   7   7.5  RT (min)      lcorbi_hilic_neg_1_012_32    m/z 333.0373     0   1E+3   5.5   6  RT (min)      lcorbi_hilic_neg_4_099_61    m/z 333.0380     0   1   2E+4   2   2.5  RT (min)       lcorbi_hilic_neg_4_098_61    m/z 333.0520     0   2   4   6E+2   7   7.5  RT (min)      lcorbi_hilic_neg_1_023_23    m/z 333.0734     0   2   4   6E+3   5.5   6  RT (min)      lcorbi_hilic_neg_3_016_16    m/z 333.0765     0   1E+5   2   2.5  RT (min)       lcorbi_hilic_neg_4_107_62    m/z 333.0877     0   1E+4   4   4.5  RT (min)      lcorbi_hilic_neg_1_017_51    m/z 333.0882     0   1   6   6.5  RT (min)      lcorbi_hilic_neg_2_055_32    m/z 333.0938     0   1E+4   3   3.5  RT (min)      lcorbi_hilic_neg_4_078_58    m/z 333.0952     0   2   4   6E+4   6   6.5  RT (min)       lcorbi_hilic_neg_4_097_60    m/z 333.1000     0   1E+3   2   2.5  RT (min)      lcorbi_hilic_neg_4_107_62    m/z 333.1486     0   1E+4   4.5   5  RT (min)      lcorbi_hilic_neg_4_107_62    m/z 333.2069     0   2   4E+4   0.5   1  RT (min)      lcorbi_hilic_neg_4_079_58    m/z 333.2433     0   1   2   3   4E+4   0.5   1  RT (min)      lcorbi_hilic_neg_4_081_58    m/z 333.3143     0   1   2E+4   0.5   1  RT (min)      lcorbi_hilic_neg_4_095_60    m/z 333.6530     0   1E+4   3   3.5  RT (min)      lcorbi_hilic_neg_4_081_58    m/z 333.8248     0   1   2E+4   8   8.5  RT (min)      lcorbi_hilic_neg_4_066_57    m/z 333.9978     0   1E+4   6.5   7  RT (min)      lcorbi_hilic_neg_4_085_59    m/z 333.9999     0   2   4   6E+3   5   5.5  RT (min)      lcorbi_hilic_neg_4_074_58    m/z 334.0143     0   1   2   3   4E+2   5.5   6  RT (min)      lcorbi_hilic_neg_2_005_24    m/z 334.0213     0   1E+5   7   7.5  RT (min)       lcorbi_hilic_neg_4_097_60    m/z 334.0243     0   1   2E+4   5.5   6  RT (min)       lcorbi_hilic_neg_4_076_58    m/z 334.0627     0   1   7.5   8  RT (min)      lcorbi_hilic_neg_4_093_60    m/z 334.0630     0   2   4   6E+2   6   6.5   7  RT (min)       lcorbi_hilic_neg_4_005_32    m/z 334.0756     0   1E+5   4.5   5  RT (min)       lcorbi_hilic_neg_4_106_62    m/z 334.0812     0   1   1.5   2  RT (min)      lcorbi_hilic_neg_4_097_60    m/z 334.0906     0   1   2   3   4E+5   1.5   2  RT (min)       lcorbi_hilic_neg_4_097_60    m/z 334.1002     0   1   2   2.5  RT (min)      lcorbi_hilic_neg_4_107_62    m/z 334.1007     0   1E+4   4   4.5  RT (min)      lcorbi_hilic_neg_2_017_51    m/z 334.1042     0   1   6.5   7  RT (min)      lcorbi_hilic_neg_4_090_60    m/z 334.1192     0   1   2   3E+4   3.5   4  RT (min)       lcorbi_hilic_neg_4_075_58    m/z 334.1365     0   1   6.5   7  RT (min)      lcorbi_hilic_neg_4_094_60    m/z 334.1438     0   1   6.5   7  RT (min)      lcorbi_hilic_neg_4_096_60    m/z 334.1537     0   2   4   6E+4   4   4.5  RT (min)       lcorbi_hilic_neg_4_090_60    m/z 334.2104     0   1E+4   0.5   1  RT (min)      lcorbi_hilic_neg_4_077_58    m/z 334.2235     0   2   4   6E+4   0.5   1  RT (min)      lcorbi_hilic_neg_4_031_14    m/z 334.7831     0   2   4   6   8E+3   4.5   5  RT (min)      lcorbi_hilic_neg_4_076_58    m/z 334.8234     0   1E+4   8   8.5  RT (min)      lcorbi_hilic_neg_4_066_57    m/z 334.8242     0   2   4   6E+3   3   3.5  RT (min)      lcorbi_hilic_neg_4_079_58    m/z 334.8799     0   1   2E+4   7   7.5  RT (min)       lcorbi_hilic_neg_1_023_23    m/z 334.9383     0   1   2E+4   1   1.5  RT (min)      lcorbi_hilic_neg_2_012_14    m/z 334.9938     0   1   2E+4   0.5   1  RT (min)       lcorbi_hilic_neg_4_101_61    m/z 335.0133     0   1E+4   0.5   1  RT (min)      lcorbi_hilic_neg_4_079_58    m/z 335.0198     0   1E+3   6.5   7  RT (min)      lcorbi_hilic_neg_1_019_06    m/z 335.0245     0   1   3   3.5  RT (min)      lcorbi_hilic_neg_4_083_59    m/z 335.0599     0   1   3   3.5  RT (min)      lcorbi_hilic_neg_4_106_62    m/z 335.0599     0   2   4E+3   7   7.5  RT (min)      lcorbi_hilic_neg_4_090_60    m/z 335.0811     0   1   6   6.5  RT (min)      lcorbi_hilic_neg_4_098_61    m/z 335.1040     0   1   4   4.5  RT (min)      lcorbi_hilic_neg_1_027_18    m/z 335.1158     0   1E+5   3.5   4  RT (min)       lcorbi_hilic_neg_4_075_58    m/z 335.1335     0   1E+5   6.5   7  RT (min)       lcorbi_hilic_neg_4_093_60    m/z 335.1468     0   1E+4   4   4.5  RT (min)      lcorbi_hilic_neg_4_091_60    m/z 335.1586     0   1E+5   6.5   7  RT (min)       lcorbi_hilic_neg_4_097_60    m/z 335.1896     0   1E+3   0.5   1  RT (min)       lcorbi_hilic_neg_2_012_14    m/z 335.2172     0   1E+5   0.5   1  RT (min)      lcorbi_hilic_neg_2_012_14    m/z 335.2260     0   1E+4   0.5   1  RT (min)      lcorbi_hilic_neg_4_083_59    m/z 335.2355     0   2   4   6E+4   1   1.5  RT (min)       lcorbi_hilic_neg_4_110_62    m/z 335.8057     0   2   4   6E+3   8   8.5  RT (min)      lcorbi_hilic_neg_4_066_57    m/z 335.8580     0   2   4   6E+3   8   8.5  RT (min)      lcorbi_hilic_neg_4_067_57    m/z 335.9697     0   1   2E+5   7   7.5  RT (min)       lcorbi_hilic_neg_4_091_60    m/z 336.0130     0   1E+3   6   6.5  RT (min)      lcorbi_hilic_neg_1_026_34    m/z 336.0193     0   1   2   3   4E+4   3   3.5  RT (min)       lcorbi_hilic_neg_4_066_57    m/z 336.0528     0   1   5.5   6  RT (min)      lcorbi_hilic_neg_4_097_60    m/z 336.0605     0   1E+4   4   4.5  RT (min)      lcorbi_hilic_neg_1_042_25    m/z 336.0642     0   1E+3   6   6.5  RT (min)      lcorbi_hilic_neg_2_026_23    m/z 336.0910     0   1   6   6.5  RT (min)      lcorbi_hilic_neg_4_005_32    m/z 336.1028     0   1   2E+4   6   6.5  RT (min)       lcorbi_hilic_neg_4_097_60    m/z 336.1063     0   1E+3   1   1.5  RT (min)       lcorbi_hilic_neg_4_096_60    m/z 336.1302     0   1E+3   4   4.5  RT (min)      lcorbi_hilic_neg_1_017_51    m/z 336.1312     0   1   2   3   4E+2   6   6.5  RT (min)      lcorbi_hilic_neg_3_011_24    m/z 336.1333     0   1   2E+4   2   2.5  RT (min)      lcorbi_hilic_neg_2_001_49    m/z 336.1412     0   1   2   3E+4   7.5   8  RT (min)       lcorbi_hilic_neg_4_072_57    m/z 336.1567     0   2   4   6E+3   1.5   2  RT (min)      lcorbi_hilic_neg_4_017_51    m/z 336.2260     0   1E+4   1   1.5  RT (min)      lcorbi_hilic_neg_3_031_14    m/z 336.6922     0   1E+5   4.5   5  RT (min)       lcorbi_hilic_neg_2_020_27    m/z 336.7804     0   1E+4   4.5   5  RT (min)      lcorbi_hilic_neg_4_080_58    m/z 336.8035     0   1E+4   8   8.5  RT (min)      lcorbi_hilic_neg_4_066_57    m/z 336.8686     0   2   4E+3   7.5   8  RT (min)      lcorbi_hilic_neg_4_102_61    m/z 336.8986     0   2   4   6   8E+3   4.5   5  RT (min)      lcorbi_hilic_neg_4_068_57    m/z 336.9293     0   1E+4   6   6.5  RT (min)      lcorbi_hilic_neg_4_090_60    m/z 336.9912     0   1   2E+4   0.5   1  RT (min)       lcorbi_hilic_neg_4_075_58    m/z 337.0041     0   1   2.5   3  RT (min)      lcorbi_hilic_neg_4_088_59    m/z 337.0139     0   1   2E+4   6   6.5  RT (min)       lcorbi_hilic_neg_4_075_58    m/z 337.0189     0   1   3   3.5  RT (min)      lcorbi_hilic_neg_4_069_57    m/z 337.0290     0   1E+4   0.5   1  RT (min)      lcorbi_hilic_neg_4_092_60    m/z 337.0362     0   1   2   2.5  RT (min)      lcorbi_hilic_neg_4_097_60    m/z 337.0554     0   1E+5   6.5   7  RT (min)       lcorbi_hilic_neg_4_006_06    m/z 337.0722     0   1E+4   5.5   6  RT (min)       lcorbi_hilic_neg_4_091_60    m/z 337.0753     0   1E+3   2.5   3  RT (min)      lcorbi_hilic_neg_4_109_62    m/z 337.0886     0   1   6.5   7  RT (min)      lcorbi_hilic_neg_4_082_59    m/z 337.0955     0   1   2   3E+3   3   3.5  RT (min)      lcorbi_hilic_neg_4_105_61    m/z 337.1129     0   1   2E+3   6.5   7  RT (min)       lcorbi_hilic_neg_4_083_59    m/z 337.1226     0   1   2   3E+4   1   1.5  RT (min)       lcorbi_hilic_neg_4_067_57    m/z 337.1337     0   1E+3   2   2.5  RT (min)      lcorbi_hilic_neg_4_097_60    m/z 337.1425     0   2   4   6E+3   4.5   5  RT (min)      lcorbi_hilic_neg_4_097_60    m/z 337.1718     0   1   1   1.5  RT (min)      lcorbi_hilic_neg_4_054_30    m/z 337.1843     0   1E+5   0.5   1  RT (min)       lcorbi_hilic_neg_4_113_62    m/z 337.1944     0   1   0.5   1  RT (min)      lcorbi_hilic_neg_4_085_59    m/z 337.2383     0   2   4E+3   0.5   1  RT (min)      lcorbi_hilic_neg_4_113_62    m/z 337.8021     0   1   2E+3   8   8.5  RT (min)      lcorbi_hilic_neg_4_072_57    m/z 337.9795     0   2   4   6E+2   5.5   6  RT (min)      lcorbi_hilic_neg_3_044_32    m/z 337.9898     0   1E+4   6   6.5  RT (min)      lcorbi_hilic_neg_4_089_59    m/z 338.0086     0   2   4   6E+3   2.5   3  RT (min)      lcorbi_hilic_neg_1_037_46    m/z 338.0192     0   1   5.5   6  RT (min)       lcorbi_hilic_neg_4_110_62    m/z 338.0352     0   1   2E+3   5.5   6  RT (min)      lcorbi_hilic_neg_4_079_58    m/z 338.0664     0   2   4E+4   6   6.5   7  RT (min)       lcorbi_hilic_neg_4_088_59    m/z 338.0788     0   1   2E+4   1   1.5  RT (min)      lcorbi_hilic_neg_1_052_41    m/z 338.1030     0   2   4   6   8E+3   1   1.5  RT (min)      lcorbi_hilic_neg_4_013_46    m/z 338.1217     0   1   5.5   6  RT (min)      lcorbi_hilic_neg_4_089_59    m/z 338.1303     0   1   2   2.5  RT (min)      lcorbi_hilic_neg_4_090_60    m/z 338.1572     0   2   4   6E+4   6.5   7  RT (min)       lcorbi_hilic_neg_4_097_60    m/z 338.7776     0   1   4.5   5  RT (min)      lcorbi_hilic_neg_4_098_61    m/z 338.7828     0   1   2   3   4E+3   8   8.5  RT (min)      lcorbi_hilic_neg_4_106_62    m/z 338.8004     0   1   2E+3   8   8.5  RT (min)      lcorbi_hilic_neg_4_107_62    m/z 338.8168     0   2   4   6   8E+3   7.5   8  RT (min)      lcorbi_hilic_neg_4_111_62    m/z 338.8547     0   2   4   6E+3   7.5   8  RT (min)      lcorbi_hilic_neg_4_099_61    m/z 338.8666     0   1   2E+4   4.5   5  RT (min)       lcorbi_hilic_neg_4_082_59    m/z 338.9882     0   1   2E+4   3   3.5  RT (min)      lcorbi_hilic_neg_4_076_58    m/z 338.9882     0   1   2E+4   0.5   1  RT (min)       lcorbi_hilic_neg_4_073_57    m/z 338.9952     0   1   6.5   7  RT (min)      lcorbi_hilic_neg_4_107_62    m/z 339.0023     0   1   2.5   3  RT (min)      lcorbi_hilic_neg_4_066_57    m/z 339.0073     0   1   2E+4   0.5   1  RT (min)      lcorbi_hilic_neg_4_066_57    m/z 339.0134     0   1   2E+4   5.5   6  RT (min)       lcorbi_hilic_neg_4_076_58    m/z 339.0227     0   1   2   3   4E+3   5.5   6  RT (min)      lcorbi_hilic_neg_4_066_57    m/z 339.0329     0   1E+5   7   7.5  RT (min)      lcorbi_hilic_neg_4_005_32    m/z 339.0573     0   1E+4   5   5.5  RT (min)      lcorbi_hilic_neg_4_071_57    m/z 339.0706     0   2   4   6E+3   5   5.5  RT (min)      lcorbi_hilic_neg_4_069_57    m/z 339.0800     0   1E+3   0.5   1  RT (min)      lcorbi_hilic_neg_2_052_41    m/z 339.0812     0   2   4   6E+4   3   3.5  RT (min)       lcorbi_hilic_neg_4_085_59    m/z 339.0976     0   2   4   6   8E+3   6   6.5  RT (min)      lcorbi_hilic_neg_4_096_60    m/z 339.0981     0   1   2   3E+4   1.5   2  RT (min)       lcorbi_hilic_neg_4_092_60    m/z 339.1044     0   1E+5   4.5   5  RT (min)       lcorbi_hilic_neg_4_066_57    m/z 339.1080     0   1   2   3   4E+2   6.5   7  RT (min)      lcorbi_hilic_neg_2_005_24    m/z 339.1202     0   1   2E+4   6   6.5  RT (min)       lcorbi_hilic_neg_4_091_60    m/z 339.1390     0   1   2   3E+4   5.5   6  RT (min)       lcorbi_hilic_neg_4_092_60    m/z 339.1536     0   1   2   3   4E+4   1.5   2  RT (min)       lcorbi_hilic_neg_4_084_59    m/z 339.1635     0   1E+4   1   1.5  RT (min)      lcorbi_hilic_neg_4_096_60    m/z 339.1839     0   1E+4   0.5   1  RT (min)      lcorbi_hilic_neg_3_053_13    m/z 339.2000     0   1   2   3E+4   4   4.5  RT (min)       lcorbi_hilic_neg_4_094_60    m/z 339.2139     0   1E+4   0.5   1  RT (min)      lcorbi_hilic_neg_4_009_50    m/z 339.2481     0   1   0.5   1  RT (min)      lcorbi_hilic_neg_4_066_57    m/z 339.2539     0   1E+5   1   1.5  RT (min)      lcorbi_hilic_neg_4_069_57    m/z 339.3013     0   2   4   6E+3   0.5   1  RT (min)      lcorbi_hilic_neg_4_111_62    m/z 339.9126     0   2   4   6E+4   0.5   1  RT (min)      lcorbi_hilic_neg_4_038_27    m/z 339.9929     0   1E+4   1   1.5  RT (min)      lcorbi_hilic_neg_4_042_28    m/z 340.0002     0   1E+4   5.5   6  RT (min)      lcorbi_hilic_neg_4_081_58    m/z 340.0510     0   1E+3   6.5   7  RT (min)      lcorbi_hilic_neg_4_087_59    m/z 340.0520     0   2   4   6   8E+3   1   1.5  RT (min)      lcorbi_hilic_neg_4_098_61    m/z 340.0653     0   1E+3   5   5.5  RT (min)      lcorbi_hilic_neg_4_050_26    m/z 340.0755     0   1E+3   7   7.5  RT (min)       lcorbi_hilic_neg_4_097_60    m/z 340.0764     0   1   1.5   2  RT (min)      lcorbi_hilic_neg_4_096_60    m/z 340.0880     0   1   2E+4   5.5   6  RT (min)      lcorbi_hilic_neg_1_047_05    m/z 340.1068     0   1E+4   1   1.5  RT (min)      lcorbi_hilic_neg_1_001_49    m/z 340.1102     0   1E+4   3.5   4  RT (min)      lcorbi_hilic_neg_4_091_60    m/z 340.1124     0   2   4   6E+4   5.5   6  RT (min)       lcorbi_hilic_neg_4_096_60    m/z 340.1265     0   1E+3   1.5   2  RT (min)      lcorbi_hilic_neg_1_026_34    m/z 340.1339     0   1   5.5   6   6.5  RT (min)       lcorbi_hilic_neg_4_094_60    m/z 340.1400     0   1E+4   0.5   1  RT (min)      lcorbi_hilic_neg_1_003_14    m/z 340.1412     0   1E+3   6   6.5  RT (min)      lcorbi_hilic_neg_1_009_50    m/z 340.1620     0   1   0.5   1  RT (min)       lcorbi_hilic_neg_4_066_57    m/z 340.7287     0   1   2E+4   4.5   5  RT (min)       lcorbi_hilic_neg_4_070_57    m/z 340.8595     0   1   2E+4   8   8.5  RT (min)      lcorbi_hilic_neg_4_066_57    m/z 340.8640     0   1E+4   4.5   5  RT (min)      lcorbi_hilic_neg_4_074_58    m/z 340.9466     0   1E+4   8   8.5  RT (min)      lcorbi_hilic_neg_4_079_58    m/z 340.9767     0   1   1   1.5  RT (min)      lcorbi_hilic_neg_4_090_60    m/z 340.9843     0   1E+4   1   1.5  RT (min)      lcorbi_hilic_neg_4_084_59    m/z 341.0037     0   1   2   3E+4   3   3.5  RT (min)      lcorbi_hilic_neg_4_087_59    m/z 341.0037     0   1   2   3   4E+4   0.5   1  RT (min)       lcorbi_hilic_neg_4_069_57    m/z 341.0098     0   1   2   3E+4   7.5   8  RT (min)       lcorbi_hilic_neg_4_086_59    m/z 341.0295     0   1E+4   7.5   8  RT (min)      lcorbi_hilic_neg_4_093_60    m/z 341.0527     0   1E+4   2   2.5  RT (min)      lcorbi_hilic_neg_4_073_57    m/z 341.0616     0   1   6.5   7  RT (min)      lcorbi_hilic_neg_4_069_57    m/z 341.0697     0   1   6   6.5  RT (min)      lcorbi_hilic_neg_4_014_18    m/z 341.0771     0   1   2   3E+4   4   4.5  RT (min)       lcorbi_hilic_neg_4_092_60    m/z 341.0781     0   1   2E+3   6   6.5  RT (min)      lcorbi_hilic_neg_4_073_57    m/z 341.0817     0   1E+3   0.5   1  RT (min)      lcorbi_hilic_neg_4_097_60    m/z 341.0990     0   1   1   1.5  RT (min)      lcorbi_hilic_neg_4_092_60    m/z 341.1091     0   1E+4   3.5   4  RT (min)      lcorbi_hilic_neg_4_073_57    m/z 341.1200     0   2   4   6   8E+3   5.5   6  RT (min)      lcorbi_hilic_neg_4_066_57    m/z 341.1264     0   1   2   3   4E+4   1.5   2  RT (min)       lcorbi_hilic_neg_1_012_32    m/z 341.1453     0   1   2   3   4E+3   2   2.5  RT (min)      lcorbi_hilic_neg_4_035_48    m/z 341.1628     0   1E+3   0.5   1  RT (min)       lcorbi_hilic_neg_3_053_13    m/z 341.1790     0   1   2   3   4E+3   0.5   1  RT (min)       lcorbi_hilic_neg_4_066_57    m/z 341.2694     0   1E+4   0.5   1  RT (min)      lcorbi_hilic_neg_4_081_58    m/z 341.5423     0   1   2E+3   7   7.5  RT (min)      lcorbi_hilic_neg_4_015_24    m/z 341.8143     0   2   4E+3   8   8.5  RT (min)      lcorbi_hilic_neg_4_101_61    m/z 341.9097     0   2   4   6   8E+4   0.5   1  RT (min)      lcorbi_hilic_neg_3_024_27    m/z 341.9851     0   1   2   3   4E+4   0.5   1  RT (min)      lcorbi_hilic_neg_4_107_62    m/z 341.9996     0   2   4   6   8E+4   7.5   8  RT (min)       lcorbi_hilic_neg_4_075_58    m/z 342.0623     0   1   2   3E+3   5   5.5  RT (min)      lcorbi_hilic_neg_4_110_62    m/z 342.0973     0   1E+4   5.5   6  RT (min)      lcorbi_hilic_neg_4_079_58    m/z 342.1047     0   1   7   7.5  RT (min)      lcorbi_hilic_neg_4_082_59    m/z 342.1123     0   2   4   6   8E+4   6.5   7  RT (min)      lcorbi_hilic_neg_4_014_18    m/z 342.1193     0   1   7   7.5  RT (min)      lcorbi_hilic_neg_4_092_60    m/z 342.1395     0   2   4   6E+3   6.5   7  RT (min)      lcorbi_hilic_neg_4_088_59    m/z 342.1531     0   1   0.5   1  RT (min)      lcorbi_hilic_neg_3_010_18    m/z 342.1612     0   1E+4   7.5   8  RT (min)      lcorbi_hilic_neg_1_037_46    m/z 342.1668     0   2   4   6   8E+3   3.5   4  RT (min)      lcorbi_hilic_neg_4_075_58    m/z 342.1988     0   1   2   3   4E+3   0.5   1  RT (min)      lcorbi_hilic_neg_2_019_21    m/z 342.2422     0   1   0.5   1  RT (min)       lcorbi_hilic_neg_1_007_10    m/z 342.2646     0   1   1   1.5  RT (min)      lcorbi_hilic_neg_4_091_60    m/z 342.2863     0   1   2   3E+5   1   1.5  RT (min)       lcorbi_hilic_neg_4_091_60    m/z 342.7258     0   1   2   3E+4   4.5   5  RT (min)       lcorbi_hilic_neg_4_073_57    m/z 342.7336     0   1E+4   4.5   5  RT (min)      lcorbi_hilic_neg_4_101_61    m/z 342.9438     0   1   2   3   4E+3   0.5   1  RT (min)      lcorbi_hilic_neg_4_108_62    m/z 342.9588     0   1E+4   6   6.5  RT (min)      lcorbi_hilic_neg_4_071_57    m/z 342.9999     0   1   2E+4   1   1.5  RT (min)       lcorbi_hilic_neg_4_071_57    m/z 343.0030     0   1   2   3E+4   6.5   7  RT (min)       lcorbi_hilic_neg_4_074_58    m/z 343.0381     0   1   5   5.5  RT (min)      lcorbi_hilic_neg_1_002_26    m/z 343.0457     0   2   4E+4   5   5.5  RT (min)      lcorbi_hilic_neg_1_004_17    m/z 343.0557     0   1   2E+4   7.5   8  RT (min)       lcorbi_hilic_neg_4_096_60    m/z 343.0742     0   1E+4   4.5   5  RT (min)      lcorbi_hilic_neg_4_091_60    m/z 343.0789     0   1   7   7.5  RT (min)      lcorbi_hilic_neg_4_096_60    m/z 343.0847     0   1   6.5   7  RT (min)      lcorbi_hilic_neg_4_072_57    m/z 343.0851     0   1E+4   1   1.5  RT (min)      lcorbi_hilic_neg_4_091_60    m/z 343.1031     0   1E+3   6   6.5   7  RT (min)       lcorbi_hilic_neg_3_016_16    m/z 343.1135     0   1   2E+4   6   6.5  RT (min)       lcorbi_hilic_neg_4_089_59    m/z 343.1241     0   1E+4   6.5   7  RT (min)      lcorbi_hilic_neg_4_088_59    m/z 343.1375     0   2   4   6E+3   0.5   1  RT (min)      lcorbi_hilic_neg_2_020_27    m/z 343.1563     0   1E+3   0.5   1  RT (min)      lcorbi_hilic_neg_3_031_14    m/z 343.1873     0   1E+4   0.5   1  RT (min)       lcorbi_hilic_neg_2_020_27    m/z 343.1985     0   2   4   6   8E+3   5   5.5  RT (min)      lcorbi_hilic_neg_4_089_59    m/z 343.2487     0   1E+4   0.5   1  RT (min)      lcorbi_hilic_neg_4_077_58    m/z 343.2851     0   1   1   1.5  RT (min)      lcorbi_hilic_neg_4_095_60    m/z 343.2965     0   1   2E+4   3.5   4  RT (min)       lcorbi_hilic_neg_4_089_59    m/z 343.8528     0   1E+4   8   8.5  RT (min)      lcorbi_hilic_neg_4_066_57    m/z 343.9244     0   1E+4   7.5   8  RT (min)      lcorbi_hilic_neg_4_107_62    m/z 343.9309     0   2   4   6E+4   1   1.5  RT (min)       lcorbi_hilic_neg_4_107_62    m/z 343.9788     0   1   2E+4   6   6.5  RT (min)      lcorbi_hilic_neg_4_074_58    m/z 343.9874     0   1E+4   7.5   8  RT (min)      lcorbi_hilic_neg_4_074_58    m/z 343.9926     0   2   4   6E+4   0.5   1  RT (min)      lcorbi_hilic_neg_4_050_26    m/z 344.0060     0   1E+3   6   6.5  RT (min)      lcorbi_hilic_neg_4_074_58    m/z 344.0160     0   1   2   3E+4   6   6.5  RT (min)      lcorbi_hilic_neg_4_074_58    m/z 344.0317     0   1   2   3E+3   6.5   7  RT (min)       lcorbi_hilic_neg_4_074_58    m/z 344.0834     0   1   1.5   2  RT (min)      lcorbi_hilic_neg_4_098_61    m/z 344.0981     0   1   3   3.5  RT (min)      lcorbi_hilic_neg_4_105_61    m/z 344.1065     0   1   2E+3   6   6.5  RT (min)       lcorbi_hilic_neg_2_005_24    m/z 344.1178     0   1   2E+3   6.5   7  RT (min)       lcorbi_hilic_neg_1_055_24    m/z 344.1275     0   1   5   5.5  RT (min)      lcorbi_hilic_neg_4_080_58    m/z 344.1809     0   1   0.5   1  RT (min)       lcorbi_hilic_neg_2_012_14    m/z 344.2158     0   1E+3   1   1.5  RT (min)      lcorbi_hilic_neg_4_097_60    m/z 344.2804     0   1E+4   0.5   1  RT (min)      lcorbi_hilic_neg_3_008_04    m/z 344.7232     0   1   2E+4   4.5   5  RT (min)       lcorbi_hilic_neg_4_073_57    m/z 344.7852     0   2   4   6E+3   8   8.5  RT (min)      lcorbi_hilic_neg_4_066_57    m/z 344.8118     0   1E+4   4.5   5  RT (min)      lcorbi_hilic_neg_4_077_58    m/z 344.8514     0   1   2   3   4E+3   8   8.5  RT (min)      lcorbi_hilic_neg_4_066_57    m/z 344.9516     0   1   6.5   7  RT (min)      lcorbi_hilic_neg_1_003_14    m/z 344.9789     0   1   2E+4   0.5   1  RT (min)      lcorbi_hilic_neg_4_097_60    m/z 344.9789     0   1E+4   3   3.5  RT (min)      lcorbi_hilic_neg_4_070_57    m/z 344.9976     0   1E+4   0.5   1  RT (min)      lcorbi_hilic_neg_4_073_57    m/z 344.9980     0   1E+5   7.5   8  RT (min)      lcorbi_hilic_neg_4_075_58    m/z 345.0097     0   2   4   6   8E+2   6.5   7  RT (min)      lcorbi_hilic_neg_1_003_14    m/z 345.0292     0   1   6.5   7  RT (min)      lcorbi_hilic_neg_1_003_14    m/z 345.0318     0   1   2E+4   0.5   1  RT (min)      lcorbi_hilic_neg_4_113_62    m/z 345.0433     0   1   6   6.5  RT (min)      lcorbi_hilic_neg_4_005_32    m/z 345.0761     0   1   7   7.5  RT (min)      lcorbi_hilic_neg_2_005_24    m/z 345.0799     0   1   2E+3   1   1.5  RT (min)      lcorbi_hilic_neg_4_050_26    m/z 345.0929     0   1   2   3   4E+4   1.5   2  RT (min)       lcorbi_hilic_neg_1_012_32    m/z 345.1068     0   1   3   3.5  RT (min)      lcorbi_hilic_neg_4_099_61    m/z 345.1080     0   1E+5   5.5   6  RT (min)       lcorbi_hilic_neg_4_095_60    m/z 345.1082     0   2   4   6   8E+4   1   1.5  RT (min)       lcorbi_hilic_neg_4_090_60    m/z 345.1146     0   1E+4   6   6.5  RT (min)      lcorbi_hilic_neg_4_097_60    m/z 345.1293     0   1   2.5   3  RT (min)      lcorbi_hilic_neg_4_100_61    m/z 345.1553     0   2   4   6E+3   4   4.5  RT (min)      lcorbi_hilic_neg_4_072_57    m/z 345.1819     0   1   4.5   5  RT (min)      lcorbi_hilic_neg_4_053_23    m/z 345.1842     0   1E+3   0.5   1  RT (min)      lcorbi_hilic_neg_4_031_14    m/z 345.2069     0   2   4   6E+2   0.5   1  RT (min)      lcorbi_hilic_neg_4_031_14    m/z 345.7193     0   1   2   3   4E+3   2.5   3  RT (min)      lcorbi_hilic_neg_3_053_13    m/z 345.8344     0   1   2   3   4E+3   8   8.5  RT (min)      lcorbi_hilic_neg_4_066_57    m/z 345.9281     0   2   4E+4   1   1.5  RT (min)       lcorbi_hilic_neg_4_107_62    m/z 345.9905     0   2   4   6E+4   0.5   1  RT (min)      lcorbi_hilic_neg_4_050_26    m/z 346.0448     0   1   2E+5   6.5   7  RT (min)      lcorbi_hilic_neg_3_029_25    m/z 346.0608     0   2   4E+3   3.5   4  RT (min)      lcorbi_hilic_neg_4_090_60    m/z 346.0721     0   2   4   6E+3   0.5   1  RT (min)      lcorbi_hilic_neg_1_034_43    m/z 346.0808     0   1E+3   2   2.5  RT (min)       lcorbi_hilic_neg_1_037_46    m/z 346.0856     0   1E+3   7   7.5  RT (min)      lcorbi_hilic_neg_4_087_59    m/z 346.0933     0   1   2   3E+4   6   6.5  RT (min)       lcorbi_hilic_neg_4_094_60    m/z 346.1048     0   1   3   3.5  RT (min)      lcorbi_hilic_neg_4_014_18    m/z 346.1124     0   1   2E+5   6.5   7  RT (min)       lcorbi_hilic_neg_4_091_60    m/z 346.1350     0   1   0.5   1  RT (min)      lcorbi_hilic_neg_4_087_59    m/z 346.1362     0   1   4   4.5  RT (min)      lcorbi_hilic_neg_4_107_62    m/z 346.1440     0   2   4   6E+3   5.5   6  RT (min)      lcorbi_hilic_neg_4_109_62    m/z 346.1621     0   1E+4   2   2.5  RT (min)      lcorbi_hilic_neg_3_035_48    m/z 346.1853     0   1E+4   2   2.5  RT (min)      lcorbi_hilic_neg_4_109_62    m/z 346.2467     0   1E+5   0.5   1  RT (min)      lcorbi_hilic_neg_4_031_14    m/z 346.2790     0   1   2   3   4E+4   0.5   1  RT (min)      lcorbi_hilic_neg_4_067_57    m/z 346.8092     0   1E+4   4.5   5  RT (min)      lcorbi_hilic_neg_4_078_58    m/z 346.8160     0   2   4   6E+3   8   8.5  RT (min)      lcorbi_hilic_neg_4_098_61    m/z 346.8323     0   2   4   6E+3   8   8.5  RT (min)      lcorbi_hilic_neg_4_066_57    m/z 346.8785     0   1E+4   5   5.5  RT (min)      lcorbi_hilic_neg_1_010_22    m/z 346.9026     0   1E+4   8   8.5  RT (min)      lcorbi_hilic_neg_4_105_61    m/z 346.9241     0   2   4E+3   1   1.5  RT (min)      lcorbi_hilic_neg_4_081_58    m/z 346.9625     0   1   2   3   4E+5   1   1.5  RT (min)       lcorbi_hilic_neg_1_027_18    m/z 346.9840     0   1E+4   7   7.5   8  RT (min)       lcorbi_hilic_neg_4_081_58    m/z 346.9938     0   1E+4   0.5   1  RT (min)      lcorbi_hilic_neg_4_071_57    m/z 347.0191     0   1   2   3E+3   7.5   8  RT (min)      lcorbi_hilic_neg_1_028_45    m/z 347.0282     0   1   2   3E+3   7.5   8  RT (min)      lcorbi_hilic_neg_3_002_15    m/z 347.0400     0   2   4   6E+2   7   7.5  RT (min)      lcorbi_hilic_neg_4_031_14    m/z 347.0530     0   1   4   4.5  RT (min)      lcorbi_hilic_neg_4_093_60    m/z 347.0558     0   1   0.5   1  RT (min)       lcorbi_hilic_neg_4_087_59    m/z 347.0593     0   2   4   6E+4   6.5   7  RT (min)      lcorbi_hilic_neg_4_089_59    m/z 347.0649     0   1   2   3   4E+2   0.5   1  RT (min)      lcorbi_hilic_neg_4_014_18    m/z 347.0684     0   1   4   4.5  RT (min)      lcorbi_hilic_neg_4_090_60    m/z 347.0750     0   1E+4   6   6.5  RT (min)      lcorbi_hilic_neg_4_072_57    m/z 347.0778     0   1   2   3E+4   1.5   2  RT (min)       lcorbi_hilic_neg_4_071_57    m/z 347.0934     0   1   3   3.5  RT (min)      lcorbi_hilic_neg_4_101_61    m/z 347.1135     0   1   2   3   4E+4   1   2  RT (min)       lcorbi_hilic_neg_4_104_61    m/z 347.1207     0   1   2E+4   3.5   4  RT (min)       lcorbi_hilic_neg_4_110_62    m/z 347.1346     0   1E+4   1   1.5  RT (min)      lcorbi_hilic_neg_4_088_59    m/z 347.1481     0   2   4E+4   0.5   1  RT (min)      lcorbi_hilic_neg_4_087_59    m/z 347.1626     0   2   4   6   8E+3   3.5   4  RT (min)      lcorbi_hilic_neg_4_076_58    m/z 347.1862     0   1   2   3E+3   0.5   1  RT (min)      lcorbi_hilic_neg_4_076_58    m/z 347.2438     0   2   4   6E+4   1   1.5  RT (min)       lcorbi_hilic_neg_4_077_58    m/z 347.8138     0   1E+4   8   8.5  RT (min)      lcorbi_hilic_neg_4_066_57    m/z 347.8607     0   2   4   6   8E+3   8   8.5  RT (min)      lcorbi_hilic_neg_4_071_57    m/z 347.9232     0   2   4E+3   1   1.5  RT (min)      lcorbi_hilic_neg_4_081_58    m/z 347.9664     0   1E+4   7.5   8  RT (min)      lcorbi_hilic_neg_4_080_58    m/z 347.9808     0   1   2E+3   7.5   8  RT (min)      lcorbi_hilic_neg_4_031_14    m/z 347.9883     0   1   7.5   8  RT (min)      lcorbi_hilic_neg_4_074_58    m/z 348.0077     0   2   4   6   8E+3   5.5   6  RT (min)      lcorbi_hilic_neg_4_087_59    m/z 348.0158     0   1E+4   3   3.5  RT (min)      lcorbi_hilic_neg_4_072_57    m/z 348.0276     0   1   2E+3   7.5   8  RT (min)       lcorbi_hilic_neg_4_046_15    m/z 348.0372     0   2   4E+3   2.5   3  RT (min)      lcorbi_hilic_neg_1_037_46    m/z 348.0538     0   1E+4   4   4.5  RT (min)      lcorbi_hilic_neg_4_082_59    m/z 348.0603     0   1E+4   6.5   7  RT (min)      lcorbi_hilic_neg_4_086_59    m/z 348.0701     0   1E+4   5   5.5  RT (min)      lcorbi_hilic_neg_4_067_57    m/z 348.0813     0   1E+4   2   2.5  RT (min)      lcorbi_hilic_neg_4_066_57    m/z 348.0958     0   2   4E+4   7   7.5  RT (min)       lcorbi_hilic_neg_4_090_60    m/z 348.1047     0   1   2E+3   6.5   7  RT (min)      lcorbi_hilic_neg_4_096_60    m/z 348.1267     0   2   4E+3   3   3.5  RT (min)      lcorbi_hilic_neg_1_037_46    m/z 348.1410     0   1E+4   6   6.5  RT (min)      lcorbi_hilic_neg_4_001_49    m/z 348.2026     0   1   2E+4   1   1.5  RT (min)      lcorbi_hilic_neg_4_079_58    m/z 348.2202     0   1E+4   7   7.5  RT (min)      lcorbi_hilic_neg_4_089_59    m/z 348.2473     0   1E+4   1   1.5  RT (min)      lcorbi_hilic_neg_4_081_58    m/z 348.2905     0   1E+5   1   1.5  RT (min)      lcorbi_hilic_neg_1_002_26    m/z 348.8138     0   1E+4   8   8.5  RT (min)      lcorbi_hilic_neg_4_098_61    m/z 348.8689     0   1   2E+4   1   1.5  RT (min)      lcorbi_hilic_neg_4_066_57    m/z 348.8955     0   1   2   3   4E+4   4.5   5  RT (min)       lcorbi_hilic_neg_4_090_60    m/z 348.9540     0   1   2   3E+4   0.5   1  RT (min)       lcorbi_hilic_neg_4_076_58    m/z 348.9793     0   2   4E+3   7.5   8  RT (min)      lcorbi_hilic_neg_4_074_58    m/z 348.9868     0   1   7.5   8  RT (min)      lcorbi_hilic_neg_4_074_58    m/z 348.9912     0   1E+4   0.5   1  RT (min)      lcorbi_hilic_neg_4_112_62    m/z 349.0241     0   1E+4   6.5   7  RT (min)      lcorbi_hilic_neg_4_069_57    m/z 349.0560     0   2   4   6   8E+3   2   2.5  RT (min)      lcorbi_hilic_neg_4_098_61    m/z 349.0632     0   1   2   3E+4   4   4.5  RT (min)       lcorbi_hilic_neg_4_029_02    m/z 349.0632     0   1   2E+3   6   6.5  RT (min)       lcorbi_hilic_neg_4_035_48    m/z 349.0727     0   1   2   3   4E+4   3.5   4  RT (min)       lcorbi_hilic_neg_4_089_59    m/z 349.0830     0   1E+6   6   6.5  RT (min)      lcorbi_hilic_neg_2_044_46    m/z 349.1010     0   1   2E+3   6   6.5  RT (min)      lcorbi_hilic_neg_2_044_46    m/z 349.1091     0   1   2   3   4E+4   0.5   1   1.5  RT (min)       lcorbi_hilic_neg_4_096_60    m/z 349.1179     0   2   4   6   8E+3   3.5   4  RT (min)      lcorbi_hilic_neg_4_107_62    m/z 349.1395     0   1E+5   6.5   7  RT (min)      lcorbi_hilic_neg_1_047_05    m/z 349.1515     0   1   2   3   4E+3   5   5.5  RT (min)      lcorbi_hilic_neg_4_077_58    m/z 349.2018     0   2   4   6E+4   1   1.5  RT (min)      lcorbi_hilic_neg_4_003_08    m/z 349.2513     0   2   4E+4   3.5   4  RT (min)       lcorbi_hilic_neg_4_086_59    m/z 349.2745     0   1   1   1.5  RT (min)      lcorbi_hilic_neg_1_002_26    m/z 349.8119     0   1E+4   8   8.5  RT (min)      lcorbi_hilic_neg_4_066_57    m/z 349.9237     0   2   4   6E+3   1   1.5  RT (min)      lcorbi_hilic_neg_4_081_58    m/z 350.0308     0   1   6.5   7  RT (min)      lcorbi_hilic_neg_4_083_59    m/z 350.0403     0   1   2   3E+4   6.5   7  RT (min)       lcorbi_hilic_neg_4_083_59    m/z 350.0519     0   1E+4   2   2.5  RT (min)      lcorbi_hilic_neg_4_098_61    m/z 350.0682     0   1E+3   4   4.5  RT (min)      lcorbi_hilic_neg_1_037_46    m/z 350.0868     0   1E+5   6.5   7  RT (min)       lcorbi_hilic_neg_2_044_46    m/z 350.0941     0   1E+5   5   6  RT (min)       lcorbi_hilic_neg_4_013_46    m/z 350.0986     0   1   2E+4   1   1.5  RT (min)      lcorbi_hilic_neg_4_067_57    m/z 350.1059     0   2   4   6   8E+3   7   7.5  RT (min)      lcorbi_hilic_neg_4_085_59    m/z 350.1141     0   2   4E+3   3   3.5  RT (min)      lcorbi_hilic_neg_4_076_58    m/z 350.1252     0   1   2E+3   0.5   1  RT (min)      lcorbi_hilic_neg_4_076_58    m/z 350.1359     0   1   2   3E+4   1   1.5  RT (min)      lcorbi_hilic_neg_2_001_49    m/z 350.1390     0   1   2E+4   3.5   4  RT (min)       lcorbi_hilic_neg_4_091_60    m/z 350.1428     0   2   4   6E+4   6.5   7  RT (min)      lcorbi_hilic_neg_1_047_05    m/z 350.1608     0   2   4   6E+4   0.5   1  RT (min)      lcorbi_hilic_neg_4_031_14    m/z 350.1877     0   2   4   6E+3   6.5   7  RT (min)      lcorbi_hilic_neg_4_089_59    m/z 350.2450     0   1   2   3E+4   0.5   1  RT (min)      lcorbi_hilic_neg_2_056_56    m/z 350.8100     0   2   4   6E+3   8   8.5  RT (min)      lcorbi_hilic_neg_4_112_62    m/z 350.8604     0   1E+4   6   6.5  RT (min)      lcorbi_hilic_neg_4_087_59    m/z 350.8604     0   1   2E+4   4.5   5  RT (min)       lcorbi_hilic_neg_4_104_61    m/z 350.8935     0   1   4.5   5  RT (min)      lcorbi_hilic_neg_4_104_61    m/z 350.8960     0   1   2E+4   1   1.5  RT (min)       lcorbi_hilic_neg_4_107_62    m/z 350.9448     0   1   5.5   6  RT (min)      lcorbi_hilic_neg_4_100_61    m/z 350.9883     0   1E+4   3   3.5  RT (min)      lcorbi_hilic_neg_4_066_57    m/z 350.9884     0   1   2E+4   0.5   1  RT (min)       lcorbi_hilic_neg_4_111_62    m/z 351.0025     0   1   7.5   8  RT (min)      lcorbi_hilic_neg_4_074_58    m/z 351.0075     0   1E+4   0.5   1  RT (min)      lcorbi_hilic_neg_4_071_57    m/z 351.0152     0   1E+5   6.5   7  RT (min)       lcorbi_hilic_neg_4_076_58    m/z 351.0446     0   2   4   6   8E+3   5.5   6  RT (min)      lcorbi_hilic_neg_4_067_57    m/z 351.0577     0   1E+4   1.5   2  RT (min)      lcorbi_hilic_neg_4_100_61    m/z 351.0672     0   1E+4   6.5   7  RT (min)      lcorbi_hilic_neg_1_048_35    m/z 351.0795     0   1   4   4.5  RT (min)      lcorbi_hilic_neg_4_101_61    m/z 351.0955     0   1E+3   6   6.5  RT (min)      lcorbi_hilic_neg_2_044_46    m/z 351.1216     0   2   4E+4   3   3.5  RT (min)       lcorbi_hilic_neg_4_093_60    m/z 351.1585     0   1E+4   5   5.5  RT (min)      lcorbi_hilic_neg_4_097_60    m/z 351.1634     0   1E+4   0.5   1  RT (min)      lcorbi_hilic_neg_4_080_58    m/z 351.2182     0   1   2   3   4E+4   0.5   1  RT (min)       lcorbi_hilic_neg_4_072_57    m/z 351.2536     0   1E+3   0.5   1  RT (min)      lcorbi_hilic_neg_4_092_60    m/z 351.2618     0   1E+5   1   1.5  RT (min)       lcorbi_hilic_neg_4_093_60    m/z 351.2803     0   2   4   6   8E+4   0.5   1  RT (min)       lcorbi_hilic_neg_4_097_60    m/z 351.2903     0   1E+4   1   1.5  RT (min)      lcorbi_hilic_neg_4_071_57    m/z 351.8602     0   2   4   6E+3   4.5   5  RT (min)      lcorbi_hilic_neg_4_099_61    m/z 351.8823     0   1   2   3   4E+3   6   6.5  RT (min)      lcorbi_hilic_neg_4_072_57    m/z 351.9588     0   1   7   7.5  RT (min)      lcorbi_hilic_neg_4_076_58    m/z 352.0144     0   2   4   6   8E+4   6.5   7  RT (min)       lcorbi_hilic_neg_4_076_58    m/z 352.0640     0   2   4   6   8E+3   6   6.5  RT (min)      lcorbi_hilic_neg_4_085_59    m/z 352.0857     0   1   2   3   4E+4   0.5   1  RT (min)       lcorbi_hilic_neg_4_080_58    m/z 352.0910     0   1E+3   6   6.5  RT (min)      lcorbi_hilic_neg_2_044_46    m/z 352.0956     0   2   4   6E+3   1   1.5  RT (min)      lcorbi_hilic_neg_4_067_57    m/z 352.1044     0   2   4E+3   2   2.5  RT (min)      lcorbi_hilic_neg_4_080_58    m/z 352.1218     0   2   4   6   8E+3   1.5   2  RT (min)      lcorbi_hilic_neg_4_097_60    m/z 352.1313     0   1E+4   4.5   5  RT (min)      lcorbi_hilic_neg_4_097_60    m/z 352.1376     0   1   2   3E+3   1   1.5  RT (min)      lcorbi_hilic_neg_4_079_58    m/z 352.1383     0   1   7   7.5  RT (min)      lcorbi_hilic_neg_4_090_60    m/z 352.1460     0   1   2E+4   7   7.5  RT (min)       lcorbi_hilic_neg_4_097_60    m/z 352.1667     0   1   2   3E+4   3   3.5  RT (min)      lcorbi_hilic_neg_4_072_57    m/z 352.1823     0   1   2E+4   5.5   6  RT (min)       lcorbi_hilic_neg_4_095_60    m/z 352.2210     0   2   4   6   8E+3   1.5   2  RT (min)      lcorbi_hilic_neg_4_105_61    m/z 352.2855     0   2   4   6   8E+3   0.5   1  RT (min)      lcorbi_hilic_neg_4_107_62    m/z 352.2981     0   2   4   6E+4   0.5   1  RT (min)      lcorbi_hilic_neg_4_095_60    m/z 352.7736     0   2   4   6   8E+3   4.5   5  RT (min)      lcorbi_hilic_neg_4_099_61    m/z 352.8329     0   2   4   6   8E+3   7.5   8  RT (min)      lcorbi_hilic_neg_4_110_62    m/z 352.8576     0   1   2   3   4E+3   4.5   5  RT (min)      lcorbi_hilic_neg_4_102_61    m/z 352.8595     0   1E+4   8.5   9  RT (min)      lcorbi_hilic_neg_4_071_57    m/z 352.8928     0   1   2E+4   1   1.5  RT (min)       lcorbi_hilic_neg_4_107_62    m/z 352.9488     0   1E+4   0.5   1  RT (min)      lcorbi_hilic_neg_4_080_58    m/z 352.9846     0   1E+4   1   1.5  RT (min)      lcorbi_hilic_neg_4_067_57    m/z 352.9874     0   1   2E+4   6   6.5  RT (min)       lcorbi_hilic_neg_4_071_57    m/z 352.9891     0   1E+4   4.5   5  RT (min)      lcorbi_hilic_neg_4_079_58    m/z 352.9929     0   1   3   3.5  RT (min)      lcorbi_hilic_neg_4_074_58    m/z 353.0027     0   1   2E+4   7.5   8  RT (min)      lcorbi_hilic_neg_4_076_58    m/z 353.0039     0   1   2E+4   0.5   1  RT (min)       lcorbi_hilic_neg_4_076_58    m/z 353.0105     0   2   4   6   8E+5   7  RT (min)       lcorbi_hilic_neg_3_031_14    m/z 353.0209     0   1   2   3E+3   6   6.5  RT (min)      lcorbi_hilic_neg_4_074_58    m/z 353.0279     0   1E+5   1   1.5  RT (min)      lcorbi_hilic_neg_4_077_58    m/z 353.0337     0   1   0.5   1  RT (min)      lcorbi_hilic_neg_4_066_57    m/z 353.0370     0   1E+4   6   6.5  RT (min)       lcorbi_hilic_neg_4_074_58    m/z 353.0741     0   1   2   3E+4   1   1.5  RT (min)       lcorbi_hilic_neg_4_079_58    m/z 353.0892     0   1E+3   0.5   1  RT (min)      lcorbi_hilic_neg_4_068_57    m/z 353.0963     0   1E+5   6   6.5  RT (min)      lcorbi_hilic_neg_2_044_46    m/z 353.1074     0   2   4   6E+2   2   2.5  RT (min)      lcorbi_hilic_neg_3_011_24    m/z 353.1312     0   1   2E+3   1   1.5  RT (min)       lcorbi_hilic_neg_1_012_32    m/z 353.1425     0   1   2   3E+3   0.5   1  RT (min)       lcorbi_hilic_neg_4_103_61    m/z 353.1591     0   2   4   6E+3   3   3.5  RT (min)      lcorbi_hilic_neg_4_071_57    m/z 353.1643     0   1   2   3   4E+3   0.5   1  RT (min)       lcorbi_hilic_neg_2_022_37    m/z 353.1765     0   2   4   6E+3   1   1.5  RT (min)      lcorbi_hilic_neg_4_079_58    m/z 353.1793     0   2   4   6   8E+3   3   3.5  RT (min)      lcorbi_hilic_neg_4_070_57    m/z 353.1871     0   1   0.5   1  RT (min)      lcorbi_hilic_neg_4_089_59    m/z 353.2331     0   1E+4   0.5   1  RT (min)      lcorbi_hilic_neg_4_113_62    m/z 353.2657     0   1   2   3E+4   3.5   4  RT (min)       lcorbi_hilic_neg_4_082_59    m/z 353.2693     0   1E+4   0.5   1  RT (min)      lcorbi_hilic_neg_4_113_62    m/z 353.3056     0   1E+5   0.5   1  RT (min)       lcorbi_hilic_neg_4_113_62    m/z 353.7703     0   2   4E+3   8   8.5  RT (min)      lcorbi_hilic_neg_4_098_61    m/z 353.9771     0   1E+4   5   5.5  RT (min)      lcorbi_hilic_neg_4_100_61    m/z 353.9875     0   1   2E+4   4.5   5  RT (min)       lcorbi_hilic_neg_4_079_58    m/z 354.0127     0   1E+5   6.5   7  RT (min)      lcorbi_hilic_neg_4_078_58    m/z 354.0408     0   1E+3   6.5   7  RT (min)       lcorbi_hilic_neg_4_076_58    m/z 354.0656     0   1E+3   6.5   7  RT (min)      lcorbi_hilic_neg_4_088_59    m/z 354.0736     0   1E+4   4   4.5  RT (min)      lcorbi_hilic_neg_4_101_61    m/z 354.0824     0   2   4   6E+2   6.5   7  RT (min)      lcorbi_hilic_neg_4_005_32    m/z 354.0897     0   1E+4   3.5   4  RT (min)      lcorbi_hilic_neg_2_021_05    m/z 354.0903     0   1   2   3   4E+3   6   6.5  RT (min)      lcorbi_hilic_neg_3_056_56    m/z 354.1164     0   1E+4   6   6.5  RT (min)      lcorbi_hilic_neg_3_051_46    m/z 354.1311     0   1   6   6.5  RT (min)      lcorbi_hilic_neg_4_083_59    m/z 354.1776     0   1   0.5   1  RT (min)       lcorbi_hilic_neg_2_014_31    m/z 354.2395     0   1   0.5   1  RT (min)      lcorbi_hilic_neg_4_113_62    m/z 354.2594     0   2   4E+4   3.5   4  RT (min)       lcorbi_hilic_neg_4_082_59    m/z 354.7521     0   1E+3   4.5   5  RT (min)       lcorbi_hilic_neg_4_084_59    m/z 354.7681     0   2   4E+3   8   8.5  RT (min)      lcorbi_hilic_neg_4_066_57    m/z 354.8387     0   1E+4   7   7.5  RT (min)      lcorbi_hilic_neg_4_082_59    m/z 354.8407     0   1   2   3E+4   4.5   5  RT (min)       lcorbi_hilic_neg_4_074_58    m/z 354.8898     0   2   4   6E+3   1   1.5  RT (min)      lcorbi_hilic_neg_4_111_62    m/z 354.9861     0   2   4   6E+4   4.5   5  RT (min)       lcorbi_hilic_neg_4_077_58    m/z 355.0001     0   1E+4   0.5   1  RT (min)       lcorbi_hilic_neg_4_079_58    m/z 355.0070     0   2   4   6E+4   6.5   7  RT (min)      lcorbi_hilic_neg_4_076_58    m/z 355.0148     0   2   4   6   8E+3   6.5   7  RT (min)       lcorbi_hilic_neg_4_076_58    m/z 355.0195     0   1   2E+4   3   3.5  RT (min)       lcorbi_hilic_neg_4_077_58    m/z 355.0195     0   1   2E+4   0.5   1  RT (min)       lcorbi_hilic_neg_4_072_57    m/z 355.0316     0   1   5   5.5  RT (min)      lcorbi_hilic_neg_4_090_60    m/z 355.0485     0   1   2E+3   1.5   2  RT (min)      lcorbi_hilic_neg_4_088_59    m/z 355.0553     0   1   2E+4   3   3.5  RT (min)       lcorbi_hilic_neg_4_085_59    m/z 355.0931     0   2   4E+3   3.5   4  RT (min)      lcorbi_hilic_neg_1_047_05    m/z 355.1100     0   1E+4   1   1.5  RT (min)      lcorbi_hilic_neg_4_089_59    m/z 355.1105     0   2   4   6   8E+2   5.5   6  RT (min)      lcorbi_hilic_neg_2_027_02    m/z 355.1202     0   1   2E+4   3.5   4   4.5  RT (min)       lcorbi_hilic_neg_1_012_32    m/z 355.1208     0   1   2   3E+3   1.5   2  RT (min)      lcorbi_hilic_neg_1_032_02    m/z 355.1244     0   2   4   6E+4   5   5.5  RT (min)       lcorbi_hilic_neg_2_027_02    m/z 355.1363     0   1   6   6.5  RT (min)      lcorbi_hilic_neg_4_089_59    m/z 355.1440     0   1E+3   6   6.5  RT (min)      lcorbi_hilic_neg_2_005_24    m/z 355.1581     0   2   4   6E+3   5   5.5  RT (min)      lcorbi_hilic_neg_4_074_58    m/z 355.1964     0   1   6.5   7  RT (min)      lcorbi_hilic_neg_4_090_60    m/z 355.2041     0   1   6.5   7  RT (min)      lcorbi_hilic_neg_4_090_60    m/z 355.2270     0   1   2E+3   0.5   1  RT (min)      lcorbi_hilic_neg_4_111_62    m/z 355.2487     0   2   4   6E+3   1   1.5  RT (min)      lcorbi_hilic_neg_4_097_60    m/z 355.9196     0   2   4   6   8E+3   6   6.5  RT (min)      lcorbi_hilic_neg_4_083_59    m/z 355.9769     0   1   4.5   5  RT (min)      lcorbi_hilic_neg_4_077_58    m/z 355.9848     0   2   4   6E+4   4.5   5  RT (min)       lcorbi_hilic_neg_4_077_58    m/z 355.9957     0   1   2E+4   7.5   8  RT (min)       lcorbi_hilic_neg_4_081_58    m/z 356.0063     0   1   2   3   4E+4   6   7  RT (min)       lcorbi_hilic_neg_4_091_60    m/z 356.0321     0   1E+4   6.5   7  RT (min)       lcorbi_hilic_neg_4_089_59    m/z 356.0450     0   1E+4   2   2.5  RT (min)      lcorbi_hilic_neg_4_099_61    m/z 356.0868     0   2   4   6   8E+3   3.5   4  RT (min)      lcorbi_hilic_neg_4_095_60    m/z 356.1100     0   1E+3   4   4.5  RT (min)       lcorbi_hilic_neg_4_095_60    m/z 356.1142     0   1   2   3   4E+3   6   6.5  RT (min)      lcorbi_hilic_neg_4_013_46    m/z 356.1278     0   1   5.5   6  RT (min)      lcorbi_hilic_neg_3_001_49    m/z 356.2081     0   1   2   3   4E+3   0.5   1  RT (min)      lcorbi_hilic_neg_4_111_62    m/z 356.2203     0   1   2E+4   3   3.5  RT (min)      lcorbi_hilic_neg_4_082_59    m/z 356.2264     0   1   0.5   1  RT (min)      lcorbi_hilic_neg_4_113_62    m/z 356.2563     0   1E+4   4.5   5  RT (min)      lcorbi_hilic_neg_4_089_59    m/z 356.2885     0   1E+4   0.5   1  RT (min)      lcorbi_hilic_neg_4_076_58    m/z 356.3248     0   1E+4   0.5   1  RT (min)      lcorbi_hilic_neg_4_113_62    m/z 356.8382     0   1E+4   4.5   5  RT (min)      lcorbi_hilic_neg_4_074_58    m/z 356.9238     0   1E+4   8.5   9  RT (min)      lcorbi_hilic_neg_4_092_60    m/z 356.9790     0   1E+4   0.5   1  RT (min)      lcorbi_hilic_neg_4_090_60    m/z 356.9832     0   2   4   6   8E+4   4.5   5  RT (min)       lcorbi_hilic_neg_4_077_58    m/z 356.9978     0   1   2E+4   1   1.5  RT (min)       lcorbi_hilic_neg_4_097_60    m/z 357.0203     0   2   4   6   8E+2   7.5   8  RT (min)       lcorbi_hilic_neg_2_055_32    m/z 357.0396     0   1   2E+3   6.5   7  RT (min)      lcorbi_hilic_neg_4_067_57    m/z 357.0413     0   1   2E+4   2   2.5  RT (min)       lcorbi_hilic_neg_4_098_61    m/z 357.0548     0   1E+3   3   3.5  RT (min)      lcorbi_hilic_neg_4_090_60    m/z 357.0613     0   1E+4   5   5.5  RT (min)      lcorbi_hilic_neg_4_071_57    m/z 357.0616     0   1E+4   1   1.5  RT (min)       lcorbi_hilic_neg_4_075_58    m/z 357.0701     0   1   6.5   7  RT (min)      lcorbi_hilic_neg_4_068_57    m/z 357.0861     0   1   2   3E+4   6.5   7  RT (min)       lcorbi_hilic_neg_4_094_60    m/z 357.1148     0   1E+4   5   6  RT (min)       lcorbi_hilic_neg_4_092_60    m/z 357.1425     0   2   4E+4   3   3.5   4  RT (min)       lcorbi_hilic_neg_4_013_46    m/z 357.1566     0   1   2E+5   6.5   7  RT (min)       lcorbi_hilic_neg_4_097_60    m/z 357.1747     0   1E+4   1   1.5  RT (min)      lcorbi_hilic_neg_4_089_59    m/z 357.2070     0   1   2E+4   1   1.5  RT (min)      lcorbi_hilic_neg_2_031_34    m/z 357.2182     0   1   2E+4   0.5   1  RT (min)      lcorbi_hilic_neg_3_017_51    m/z 357.2418     0   1   2E+4   0.5   1  RT (min)      lcorbi_hilic_neg_4_093_60    m/z 357.2797     0   1E+4   0.5   1  RT (min)      lcorbi_hilic_neg_2_011_29    m/z 357.7432     0   1   2   3   4E+3   5   5.5  RT (min)      lcorbi_hilic_neg_2_020_27    m/z 357.9162     0   2   4   6E+3   6   6.5  RT (min)      lcorbi_hilic_neg_4_083_59    m/z 357.9817     0   1   2   3E+4   4.5   5  RT (min)      lcorbi_hilic_neg_4_079_58    m/z 357.9957     0   2   4   6E+3   7.5   8  RT (min)      lcorbi_hilic_neg_4_081_58    m/z 358.0346     0   1   2E+4   5.5   6  RT (min)      lcorbi_hilic_neg_4_069_57    m/z 358.0378     0   1   2E+4   2   2.5  RT (min)      lcorbi_hilic_neg_4_099_61    m/z 358.0460     0   1E+4   6   6.5  RT (min)      lcorbi_hilic_neg_3_022_06    m/z 358.0464     0   2   4   6E+4   1.5   2  RT (min)       lcorbi_hilic_neg_4_100_61    m/z 358.0547     0   1E+4   6   6.5  RT (min)      lcorbi_hilic_neg_2_039_07    m/z 358.0660     0   1E+3   2.5   3  RT (min)      lcorbi_hilic_neg_1_001_49    m/z 358.0824     0   1   2   3E+4   4   4.5  RT (min)       lcorbi_hilic_neg_4_089_59    m/z 358.0909     0   1   2   3E+4   3.5   4  RT (min)       lcorbi_hilic_neg_4_070_57    m/z 358.0934     0   1   2   3E+4   6.5   7  RT (min)       lcorbi_hilic_neg_4_096_60    m/z 358.1089     0   1E+4   6   6.5  RT (min)      lcorbi_hilic_neg_1_037_46    m/z 358.1225     0   1   2E+4   2   2.5  RT (min)      lcorbi_hilic_neg_4_107_62    m/z 358.1246     0   1   2   3E+4   7   7.5  RT (min)       lcorbi_hilic_neg_4_092_60    m/z 358.1411     0   1E+4   3   3.5  RT (min)      lcorbi_hilic_neg_1_001_49    m/z 358.1536     0   1E+4   6   6.5  RT (min)      lcorbi_hilic_neg_4_045_25    m/z 358.2100     0   1E+4   7   7.5  RT (min)      lcorbi_hilic_neg_4_082_59    m/z 358.8284     0   1E+3   8   8.5  RT (min)      lcorbi_hilic_neg_4_006_06    m/z 358.8409     0   1   2   3   4E+4   7.5   8  RT (min)       lcorbi_hilic_neg_4_112_62    m/z 358.8506     0   2   4E+3   7.5   8  RT (min)      lcorbi_hilic_neg_4_102_61    m/z 358.9326     0   1E+4   6   6.5  RT (min)      lcorbi_hilic_neg_4_069_57    m/z 358.9804     0   1   2E+4   4.5   5  RT (min)       lcorbi_hilic_neg_4_077_58    m/z 358.9943     0   2   4E+4   0.5   1  RT (min)       lcorbi_hilic_neg_4_112_62    m/z 358.9944     0   1   2E+4   3   3.5  RT (min)       lcorbi_hilic_neg_4_067_57    m/z 359.0376     0   1E+4   6   6.5  RT (min)      lcorbi_hilic_neg_4_087_59    m/z 359.0750     0   1E+4   5   5.5  RT (min)      lcorbi_hilic_neg_4_070_57    m/z 359.0804     0   2   4   6E+3   0.5   1  RT (min)      lcorbi_hilic_neg_4_070_57    m/z 359.0866     0   1   2   3E+4   4   4.5  RT (min)       lcorbi_hilic_neg_4_073_57    m/z 359.0946     0   1   3.5   4  RT (min)      lcorbi_hilic_neg_4_089_59    m/z 359.0962     0   1   6   6.5  RT (min)      lcorbi_hilic_neg_4_083_59    m/z 359.1116     0   2   4   6E+2   6   6.5  RT (min)      lcorbi_hilic_neg_1_037_46    m/z 359.1249     0   1E+3   2.5   3  RT (min)      lcorbi_hilic_neg_4_069_57    m/z 359.1499     0   1   2E+3   1   1.5  RT (min)      lcorbi_hilic_neg_1_001_49    m/z 359.1608     0   1   2   3   4E+3   6   6.5  RT (min)      lcorbi_hilic_neg_4_017_51    m/z 359.1977     0   1   2E+4   2   2.5  RT (min)       lcorbi_hilic_neg_4_107_62    m/z 359.2227     0   1   2E+4   1   1.5  RT (min)      lcorbi_hilic_neg_4_031_14    m/z 359.2339     0   1E+4   1   1.5  RT (min)      lcorbi_hilic_neg_4_075_58    m/z 359.2549     0   1   2E+4   3.5   4  RT (min)       lcorbi_hilic_neg_4_084_59    m/z 359.2572     0   1E+5   1   1.5  RT (min)       lcorbi_hilic_neg_4_097_60    m/z 359.2914     0   2   4E+3   0.5   1  RT (min)       lcorbi_hilic_neg_4_094_60    m/z 359.9141     0   2   4   6E+3   6   6.5  RT (min)      lcorbi_hilic_neg_4_071_57    m/z 360.0463     0   1   2   3   4E+4   1.5   2  RT (min)      lcorbi_hilic_neg_4_100_61    m/z 360.0487     0   1   6   6.5  RT (min)      lcorbi_hilic_neg_4_095_60    m/z 360.0559     0   1E+4   2   2.5  RT (min)      lcorbi_hilic_neg_4_066_57    m/z 360.0630     0   1   6   6.5  RT (min)      lcorbi_hilic_neg_4_095_60    m/z 360.0733     0   1   6   6.5  RT (min)      lcorbi_hilic_neg_2_005_24    m/z 360.0879     0   1   3.5   4  RT (min)      lcorbi_hilic_neg_4_094_60    m/z 360.1097     0   2   4   6E+4   3.5   4  RT (min)       lcorbi_hilic_neg_4_095_60    m/z 360.1173     0   1E+6   5.5   6  RT (min)      lcorbi_hilic_neg_2_005_24    m/z 360.1290     0   2   4   6E+3   2.5   3  RT (min)      lcorbi_hilic_neg_4_106_62    m/z 360.1394     0   1E+3   6   6.5  RT (min)      lcorbi_hilic_neg_2_005_24    m/z 360.1527     0   1   6.5   7  RT (min)      lcorbi_hilic_neg_3_029_25    m/z 360.1604     0   2   4E+4   6   6.5  RT (min)      lcorbi_hilic_neg_4_045_25    m/z 360.2007     0   2   4E+3   2   2.5  RT (min)      lcorbi_hilic_neg_4_109_62    m/z 360.8365     0   2   4   6E+3   7.5   8  RT (min)      lcorbi_hilic_neg_4_107_62    m/z 360.9767     0   2   4   6E+3   4.5   5  RT (min)      lcorbi_hilic_neg_4_079_58    m/z 360.9912     0   1   2E+4   1   1.5  RT (min)      lcorbi_hilic_neg_4_097_60    m/z 361.0096     0   1   2E+4   0.5   1  RT (min)      lcorbi_hilic_neg_4_077_58    m/z 361.0362     0   1E+4   5   5.5  RT (min)      lcorbi_hilic_neg_4_070_57    m/z 361.0506     0   2   4   6   8E+3   7   7.5  RT (min)      lcorbi_hilic_neg_4_083_59    m/z 361.0520     0   1E+4   1.5   2  RT (min)      lcorbi_hilic_neg_4_067_57    m/z 361.0656     0   2   4   6E+4   3   3.5  RT (min)       lcorbi_hilic_neg_4_023_34    m/z 361.0906     0   1E+4   4   4.5  RT (min)      lcorbi_hilic_neg_4_070_57    m/z 361.0939     0   1   2   3   4E+4   1   1.5  RT (min)      lcorbi_hilic_neg_2_005_24    m/z 361.1209     0   1   2   3E+4   6   6.5  RT (min)       lcorbi_hilic_neg_4_097_60    m/z 361.1361     0   1E+4   5.5   6  RT (min)      lcorbi_hilic_neg_4_097_60    m/z 361.1622     0   1   2   3   4E+2   6   6.5  RT (min)      lcorbi_hilic_neg_3_029_25    m/z 361.2234     0   2   4E+4   1.5   2  RT (min)       lcorbi_hilic_neg_4_079_58    m/z 361.2242     0   1E+5   6.5   7  RT (min)       lcorbi_hilic_neg_4_097_60    m/z 361.2382     0   2   4E+3   1   1.5  RT (min)      lcorbi_hilic_neg_4_107_62    m/z 361.2495     0   1   2   3   4E+4   1   1.5  RT (min)       lcorbi_hilic_neg_4_077_58    m/z 361.2624     0   2   4   6E+4   0.5   1  RT (min)      lcorbi_hilic_neg_4_067_57    m/z 361.9112     0   2   4E+3   6   6.5  RT (min)      lcorbi_hilic_neg_4_072_57    m/z 361.9611     0   1E+4   6   6.5  RT (min)      lcorbi_hilic_neg_4_070_57    m/z 361.9924     0   1E+4   6   6.5  RT (min)      lcorbi_hilic_neg_4_098_61    m/z 362.0129     0   1   2   3E+4   1   1.5  RT (min)      lcorbi_hilic_neg_1_002_26    m/z 362.0508     0   1   2E+4   7.5   8  RT (min)       lcorbi_hilic_neg_4_089_59    m/z 362.0524     0   1   2.5   3  RT (min)      lcorbi_hilic_neg_4_100_61    m/z 362.0865     0   1   2   3   4E+4   6.5   7  RT (min)       lcorbi_hilic_neg_4_091_60    m/z 362.0940     0   1   2   3E+3   6   6.5  RT (min)      lcorbi_hilic_neg_4_094_60    m/z 362.1071     0   1E+5   3   3.5   4  RT (min)       lcorbi_hilic_neg_4_105_61    m/z 362.1091     0   1E+4   1   1.5  RT (min)      lcorbi_hilic_neg_4_078_58    m/z 362.1220     0   2   4E+4   1   1.5  RT (min)      lcorbi_hilic_neg_4_079_58    m/z 362.1596     0   1   2   3E+4   3   3.5  RT (min)      lcorbi_hilic_neg_4_017_51    m/z 362.1666     0   1   2   3E+4   0.5   1  RT (min)      lcorbi_hilic_neg_1_034_43    m/z 362.2215     0   1   0.5   1  RT (min)      lcorbi_hilic_neg_4_079_58    m/z 362.2372     0   1E+4   1   1.5  RT (min)      lcorbi_hilic_neg_4_095_60    m/z 362.9366     0   2   4   6   8E+4   1   1.5  RT (min)      lcorbi_hilic_neg_4_098_61    m/z 362.9883     0   1E+4   0.5   1  RT (min)      lcorbi_hilic_neg_4_076_58    m/z 362.9915     0   1E+4   7.5   8  RT (min)       lcorbi_hilic_neg_4_083_59    m/z 363.0039     0   1   2   3E+3   7   7.5  RT (min)      lcorbi_hilic_neg_4_090_60    m/z 363.0228     0   2   4E+2   6.5   7  RT (min)      lcorbi_hilic_neg_1_006_08    m/z 363.0626     0   1   2   3   4E+3   6   6.5  RT (min)      lcorbi_hilic_neg_4_091_60    m/z 363.0639     0   2   4   6E+4   3   3.5  RT (min)      lcorbi_hilic_neg_4_023_34    m/z 363.0707     0   2   4   6E+4   6   6.5  RT (min)       lcorbi_hilic_neg_4_094_60    m/z 363.0754     0   1   2   3   4E+3   1   1.5  RT (min)      lcorbi_hilic_neg_4_074_58    m/z 363.0875     0   1   2E+4   6   6.5  RT (min)       lcorbi_hilic_neg_4_069_57    m/z 363.1008     0   1E+4   6   6.5  RT (min)      lcorbi_hilic_neg_4_095_60    m/z 363.1125     0   1   2E+4   2.5   3  RT (min)      lcorbi_hilic_neg_4_104_61    m/z 363.1255     0   1E+3   1   1.5  RT (min)      lcorbi_hilic_neg_4_055_38    m/z 363.1311     0   1   2E+4   6   6.5  RT (min)      lcorbi_hilic_neg_3_051_46    m/z 363.1419     0   1   3   3.5  RT (min)      lcorbi_hilic_neg_4_101_61    m/z 363.1927     0   1   2E+4   4.5   5  RT (min)       lcorbi_hilic_neg_4_107_62    m/z 363.2209     0   1   0.5   1  RT (min)       lcorbi_hilic_neg_4_097_60    m/z 363.2539     0   1   0.5   1  RT (min)      lcorbi_hilic_neg_4_113_62    m/z 363.2782     0   2   4   6   8E+4   0.5   1  RT (min)       lcorbi_hilic_neg_4_051_16    m/z 363.2946     0   1E+4   3.5   4  RT (min)      lcorbi_hilic_neg_4_082_59    m/z 363.9691     0   2   4   6E+4   0.5   1  RT (min)       lcorbi_hilic_neg_4_075_58    m/z 363.9716     0   1E+3   5.5   6  RT (min)      lcorbi_hilic_neg_1_006_08    m/z 363.9896     0   2   4E+3   6   6.5  RT (min)      lcorbi_hilic_neg_4_104_61    m/z 364.0060     0   1E+4   5   5.5  RT (min)      lcorbi_hilic_neg_4_105_61    m/z 364.0168     0   2   4   6   8E+3   6.5   7  RT (min)      lcorbi_hilic_neg_4_081_58    m/z 364.0250     0   2   4   6E+3   5   5.5  RT (min)      lcorbi_hilic_neg_4_087_59    m/z 364.0439     0   1E+4   7.5   8  RT (min)      lcorbi_hilic_neg_4_096_60    m/z 364.0735     0   1E+3   6.5   7  RT (min)      lcorbi_hilic_neg_2_026_23    m/z 364.1613     0   1   2E+4   5   5.5  RT (min)       lcorbi_hilic_neg_4_078_58    m/z 364.2623     0   1   2   3E+4   1   1.5  RT (min)       lcorbi_hilic_neg_4_082_59    m/z 364.7810     0   1   2   3E+4   4.5   5  RT (min)       lcorbi_hilic_neg_4_078_58    m/z 364.8774     0   1E+4   5   5.5  RT (min)       lcorbi_hilic_neg_4_090_60    m/z 364.8894     0   1E+4   4.5   5  RT (min)      lcorbi_hilic_neg_1_007_10    m/z 364.9280     0   1   5   5.5  RT (min)      lcorbi_hilic_neg_4_090_60    m/z 364.9663     0   1   2   3   4E+4   0.5   1  RT (min)       lcorbi_hilic_neg_4_076_58    m/z 364.9676     0   2   4E+4   7.5   8  RT (min)       lcorbi_hilic_neg_4_095_60    m/z 365.0040     0   1E+4   0.5   1  RT (min)      lcorbi_hilic_neg_4_094_60    m/z 365.0115     0   2   4   6   8E+3   6.5   7  RT (min)      lcorbi_hilic_neg_4_081_58    m/z 365.0661     0   1   1   1.5  RT (min)       lcorbi_hilic_neg_4_082_59    m/z 365.0677     0   1   2E+3   5.5   6  RT (min)      lcorbi_hilic_neg_4_030_05    m/z 365.0700     0   2   4E+4   3.5   4  RT (min)       lcorbi_hilic_neg_1_020_13    m/z 365.0900     0   1   2E+4   1   1.5  RT (min)       lcorbi_hilic_neg_4_096_60    m/z 365.1046     0   2   4   6   8E+3   2   2.5  RT (min)      lcorbi_hilic_neg_3_011_24    m/z 365.1055     0   1   2E+4   6.5   7  RT (min)       lcorbi_hilic_neg_4_089_59    m/z 365.1139     0   2   4   6   8E+3   1.5   2  RT (min)      lcorbi_hilic_neg_4_096_60    m/z 365.1359     0   2   4   6E+4   6   6.5  RT (min)       lcorbi_hilic_neg_1_037_46    m/z 365.1427     0   1E+4   1   1.5  RT (min)      lcorbi_hilic_neg_4_111_62    m/z 365.2100     0   1E+4   1   1.5  RT (min)      lcorbi_hilic_neg_4_094_60    m/z 365.2695     0   1   0.5   1  RT (min)       lcorbi_hilic_neg_4_097_60    m/z 365.8727     0   2   4   6   8E+3   4.5   5  RT (min)      lcorbi_hilic_neg_4_090_60    m/z 365.8952     0   2   4   6   8E+3   1   1.5  RT (min)      lcorbi_hilic_neg_4_092_60    m/z 365.9561     0   2   4   6   8E+3   6.5   7  RT (min)      lcorbi_hilic_neg_4_067_57    m/z 366.0117     0   1E+4   2   2.5  RT (min)      lcorbi_hilic_neg_4_099_61    m/z 366.0128     0   1E+3   6   6.5  RT (min)      lcorbi_hilic_neg_4_101_61    m/z 366.0682     0   1   2E+3   5   5.5  RT (min)      lcorbi_hilic_neg_4_075_58    m/z 366.0872     0   2   4E+3   6   6.5  RT (min)      lcorbi_hilic_neg_4_089_59    m/z 366.0979     0   2   4   6E+3   6   6.5  RT (min)      lcorbi_hilic_neg_4_086_59    m/z 366.1001     0   1   2   3   4E+3   4   4.5  RT (min)      lcorbi_hilic_neg_3_049_55    m/z 366.1091     0   1   5   5.5  RT (min)      lcorbi_hilic_neg_1_028_45    m/z 366.1400     0   1E+4   6   6.5  RT (min)      lcorbi_hilic_neg_3_051_46    m/z 366.1677     0   2   4   6   8E+3   2   2.5  RT (min)      lcorbi_hilic_neg_4_013_46    m/z 366.1805     0   1E+5   6   6.5  RT (min)       lcorbi_hilic_neg_4_097_60    m/z 366.2399     0   1   2   3E+4   0.5   1  RT (min)       lcorbi_hilic_neg_4_089_59    m/z 366.2780     0   2   4   6   8E+3   3.5   4  RT (min)      lcorbi_hilic_neg_4_088_59    m/z 366.8677     0   1   2E+4   4.5   5  RT (min)       lcorbi_hilic_neg_4_092_60    m/z 366.8861     0   1   2   3   4E+3   4.5   5  RT (min)      lcorbi_hilic_neg_1_007_10    m/z 366.9006     0   2   4   6E+3   0.5   1  RT (min)      lcorbi_hilic_neg_2_020_27    m/z 366.9675     0   2   4E+4   1   1.5  RT (min)      lcorbi_hilic_neg_3_026_09    m/z 366.9818     0   1   2   3E+3   5   5.5  RT (min)      lcorbi_hilic_neg_4_093_60    m/z 367.0376     0   1   2   3E+3   1.5   2  RT (min)      lcorbi_hilic_neg_4_089_59    m/z 367.0378     0   1E+4   7   7.5  RT (min)      lcorbi_hilic_neg_4_071_57    m/z 367.0470     0   1   2   3   4E+4   4   4.5  RT (min)       lcorbi_hilic_neg_4_103_61    m/z 367.0709     0   1   2   3   4E+3   7   7.5  RT (min)      lcorbi_hilic_neg_4_090_60    m/z 367.0775     0   1E+3   6   6.5  RT (min)       lcorbi_hilic_neg_4_087_59    m/z 367.1055     0   1   2   3   4E+4   5   5.5  RT (min)       lcorbi_hilic_neg_4_075_58    m/z 367.1219     0   1E+4   5.5   6  RT (min)      lcorbi_hilic_neg_1_047_05    m/z 367.1325     0   1   2   3E+4   0.5   1  RT (min)       lcorbi_hilic_neg_4_098_61    m/z 367.1448     0   1E+4   1   1.5  RT (min)      lcorbi_hilic_neg_3_002_15    m/z 367.1692     0   1   2   3   4E+3   1   1.5  RT (min)      lcorbi_hilic_neg_4_040_17    m/z 367.1760     0   1   6   6.5  RT (min)      lcorbi_hilic_neg_4_095_60    m/z 367.1794     0   2   4   6   8E+3   0.5   1  RT (min)      lcorbi_hilic_neg_4_066_57    m/z 367.2011     0   1E+4   5.5   6  RT (min)      lcorbi_hilic_neg_4_094_60    m/z 367.2311     0   1   2   3E+3   0.5   1  RT (min)       lcorbi_hilic_neg_4_068_57    m/z 367.2482     0   1E+3   1   1.5  RT (min)      lcorbi_hilic_neg_2_054_13    m/z 367.8668     0   1   2   3E+3   1   1.5  RT (min)      lcorbi_hilic_neg_4_066_57    m/z 368.0218     0   1E+4   6.5   7  RT (min)      lcorbi_hilic_neg_4_070_57    m/z 368.0506     0   2   4   6E+3   4   4.5  RT (min)      lcorbi_hilic_neg_4_102_61    m/z 368.0608     0   1   2E+4   6.5   7  RT (min)       lcorbi_hilic_neg_4_088_59    m/z 368.0844     0   1E+5   7   7.5   8  RT (min)       lcorbi_hilic_neg_1_047_05    m/z 368.1117     0   1E+3   1   1.5  RT (min)      lcorbi_hilic_neg_1_033_53    m/z 368.1141     0   1   4.5   5  RT (min)      lcorbi_hilic_neg_2_020_27    m/z 368.1220     0   2   4   6E+3   1   1.5  RT (min)      lcorbi_hilic_neg_2_033_53    m/z 368.1599     0   1   4   4.5  RT (min)      lcorbi_hilic_neg_4_091_60    m/z 368.1970     0   2   4   6   8E+4   5.5   6  RT (min)      lcorbi_hilic_neg_4_005_32    m/z 368.2355     0   1   2E+4   0.5   1  RT (min)      lcorbi_hilic_neg_4_069_57    m/z 368.2607     0   1   2E+4   6.5   7  RT (min)       lcorbi_hilic_neg_4_090_60    m/z 368.8976     0   2   4   6   8E+3   0.5   1  RT (min)      lcorbi_hilic_neg_2_020_27    m/z 368.9614     0   1   2E+3   6   6.5  RT (min)      lcorbi_hilic_neg_4_081_58    m/z 368.9979     0   1   2   3E+4   0.5   1  RT (min)      lcorbi_hilic_neg_4_078_58    m/z 369.0051     0   2   4   6E+3   7.5   8  RT (min)       lcorbi_hilic_neg_1_004_17    m/z 369.0054     0   1   2   3   4E+4   6   6.5  RT (min)       lcorbi_hilic_neg_4_074_58    m/z 369.0284     0   1   1   1.5  RT (min)      lcorbi_hilic_neg_4_084_59    m/z 369.0442     0   1E+4   4   4.5  RT (min)      lcorbi_hilic_neg_4_088_59    m/z 369.0588     0   1   2   3E+4   7   7.5  RT (min)       lcorbi_hilic_neg_4_086_59    m/z 369.0793     0   1   2E+5   6   6.5  RT (min)       lcorbi_hilic_neg_4_091_60    m/z 369.0986     0   1   6.5   7  RT (min)      lcorbi_hilic_neg_4_068_57    m/z 369.1026     0   2   4   6   8E+3   4   4.5  RT (min)      lcorbi_hilic_neg_4_109_62    m/z 369.1058     0   2   4E+3   1   1.5  RT (min)      lcorbi_hilic_neg_4_097_60    m/z 369.1175     0   2   4   6   8E+4   4   4.5  RT (min)      lcorbi_hilic_neg_4_095_60    m/z 369.1265     0   1   4.5   5  RT (min)       lcorbi_hilic_neg_4_095_60    m/z 369.1275     0   2   4E+3   1   1.5  RT (min)      lcorbi_hilic_neg_4_069_57    m/z 369.1301     0   1   2E+4   6.5   7  RT (min)       lcorbi_hilic_neg_4_073_57    m/z 369.1642     0   1   0.5   1  RT (min)      lcorbi_hilic_neg_4_068_57    m/z 369.1875     0   1   2   3E+4   6   6.5  RT (min)       lcorbi_hilic_neg_4_095_60    m/z 369.1959     0   2   4E+4   0.5   1  RT (min)      lcorbi_hilic_neg_2_022_37    m/z 369.2281     0   2   4   6   8E+2   1   1.5  RT (min)       lcorbi_hilic_neg_1_020_13    m/z 369.3008     0   1E+4   0.5   1  RT (min)      lcorbi_hilic_neg_4_113_62    m/z 369.3371     0   1   2E+4   0.5   1  RT (min)      lcorbi_hilic_neg_4_113_62    m/z 369.9428     0   1E+4   6   6.5  RT (min)      lcorbi_hilic_neg_4_068_57    m/z 369.9888     0   2   4E+3   7.5   8  RT (min)      lcorbi_hilic_neg_4_079_58    m/z 369.9931     0   2   4   6E+3   6   6.5  RT (min)      lcorbi_hilic_neg_1_005_36    m/z 370.0004     0   1   2E+4   1   1.5  RT (min)      lcorbi_hilic_neg_4_038_27    m/z 370.0156     0   1   7.5   8  RT (min)       lcorbi_hilic_neg_4_080_58    m/z 370.0571     0   1   6.5   7  RT (min)      lcorbi_hilic_neg_4_076_58    m/z 370.0644     0   1   6   6.5  RT (min)      lcorbi_hilic_neg_4_094_60    m/z 370.0774     0   1   2E+4   5.5   6  RT (min)       lcorbi_hilic_neg_4_092_60    m/z 370.0883     0   1   2   3   4E+4   6   6.5  RT (min)       lcorbi_hilic_neg_4_079_58    m/z 370.0942     0   1   6   6.5  RT (min)      lcorbi_hilic_neg_4_078_58    m/z 370.0964     0   1E+4   3.5   4  RT (min)      lcorbi_hilic_neg_4_108_62    m/z 370.1048     0   1   2   3E+5   1   1.5  RT (min)      lcorbi_hilic_neg_1_052_41    m/z 370.1190     0   2   4E+3   4.5   5  RT (min)      lcorbi_hilic_neg_4_107_62    m/z 370.1442     0   1   2E+3   5.5   6  RT (min)       lcorbi_hilic_neg_4_095_60    m/z 370.1560     0   1E+5   6   6.5  RT (min)       lcorbi_hilic_neg_4_093_60    m/z 370.1575     0   1E+4   1   1.5  RT (min)      lcorbi_hilic_neg_4_071_57    m/z 370.1843     0   1   0.5   1  RT (min)      lcorbi_hilic_neg_4_091_60    m/z 370.2025     0   1E+4   7   7.5  RT (min)      lcorbi_hilic_neg_4_084_59    m/z 370.2089     0   1E+4   0.5   1  RT (min)      lcorbi_hilic_neg_4_090_60    m/z 370.2316     0   1   1   1.5  RT (min)      lcorbi_hilic_neg_4_004_13    m/z 370.2961     0   1   2   3E+4   1   1.5  RT (min)      lcorbi_hilic_neg_4_037_31    m/z 370.3225     0   1   2   3E+4   3.5   4  RT (min)      lcorbi_hilic_neg_4_090_60    m/z 370.7965     0   1E+4   8   8.5  RT (min)      lcorbi_hilic_neg_4_073_57    m/z 370.8695     0   2   4E+3   4.5   5  RT (min)      lcorbi_hilic_neg_3_004_41    m/z 370.9864     0   1   2   3E+4   7.5   8  RT (min)      lcorbi_hilic_neg_4_076_58    m/z 370.9946     0   1E+4   0.5   1  RT (min)      lcorbi_hilic_neg_4_071_57    m/z 371.0268     0   1E+3   1   1.5  RT (min)      lcorbi_hilic_neg_4_084_59    m/z 371.1047     0   1E+4   4.5   5  RT (min)      lcorbi_hilic_neg_1_019_06    m/z 371.1055     0   2   4E+3   6   6.5  RT (min)      lcorbi_hilic_neg_4_073_57    m/z 371.1192     0   2   4E+4   6   6.5  RT (min)       lcorbi_hilic_neg_4_069_57    m/z 371.1236     0   1   2   3   4E+4   4   4.5  RT (min)       lcorbi_hilic_neg_4_092_60    m/z 371.1256     0   1   6.5   7  RT (min)      lcorbi_hilic_neg_4_076_58    m/z 371.1327     0   1   2   3E+4   1   1.5  RT (min)       lcorbi_hilic_neg_4_095_60    m/z 371.1434     0   2   4   6   8E+4   5.5   6  RT (min)       lcorbi_hilic_neg_4_089_59    m/z 371.1575     0   1   2.5   3  RT (min)      lcorbi_hilic_neg_4_083_59    m/z 371.1799     0   1   2E+4   1   1.5  RT (min)      lcorbi_hilic_neg_4_073_57    m/z 371.2116     0   1E+5   0.5   1  RT (min)       lcorbi_hilic_neg_4_090_60    m/z 371.2204     0   1   2E+3   1   1.5  RT (min)      lcorbi_hilic_neg_4_069_57    m/z 371.2381     0   1E+4   2.5   3  RT (min)      lcorbi_hilic_neg_4_089_59    m/z 371.3166     0   1E+4   1   1.5  RT (min)      lcorbi_hilic_neg_4_036_36    m/z 371.6097     0   1E+4   3   3.5  RT (min)      lcorbi_hilic_neg_3_034_34    m/z 371.7824     0   2   4E+3   8   8.5  RT (min)      lcorbi_hilic_neg_4_102_61    m/z 371.7946     0   2   4   6   8E+3   8   8.5  RT (min)      lcorbi_hilic_neg_4_070_57    m/z 371.8184     0   1E+4   4.5   5  RT (min)      lcorbi_hilic_neg_4_075_58    m/z 371.9318     0   2   4   6E+3   5.5   6  RT (min)      lcorbi_hilic_neg_4_082_59    m/z 371.9400     0   1E+3   6   6.5  RT (min)      lcorbi_hilic_neg_3_028_23    m/z 371.9831     0   1E+4   7.5   8  RT (min)      lcorbi_hilic_neg_4_074_58    m/z 371.9870     0   1E+4   7.5   8  RT (min)      lcorbi_hilic_neg_4_074_58    m/z 371.9942     0   1E+4   1   1.5  RT (min)      lcorbi_hilic_neg_4_100_61    m/z 372.0212     0   1E+4   6   6.5  RT (min)      lcorbi_hilic_neg_4_103_61    m/z 372.0552     0   2   4   6E+3   6.5   7  RT (min)      lcorbi_hilic_neg_4_097_60    m/z 372.0660     0   1   2   3   4E+4   0.5   1  RT (min)       lcorbi_hilic_neg_4_113_62    m/z 372.0755     0   1   6.5   7  RT (min)      lcorbi_hilic_neg_2_005_24    m/z 372.0845     0   2   4E+4   7.5   8  RT (min)       lcorbi_hilic_neg_4_097_60    m/z 372.0994     0   1   2E+4   6   7  RT (min)       lcorbi_hilic_neg_4_095_60    m/z 372.1524     0   1E+4   7   7.5  RT (min)      lcorbi_hilic_neg_4_082_59    m/z 372.1695     0   1E+4   6.5   7  RT (min)      lcorbi_hilic_neg_1_012_32    m/z 372.1730     0   2   4   6   8E+3   1   1.5  RT (min)      lcorbi_hilic_neg_4_071_57    m/z 372.1810     0   2   4   6   8E+4   3.5   4  RT (min)       lcorbi_hilic_neg_4_091_60    m/z 372.1931     0   1E+4   1   1.5  RT (min)      lcorbi_hilic_neg_4_074_58    m/z 372.2503     0   1E+4   0.5   1  RT (min)      lcorbi_hilic_neg_4_084_59    m/z 372.2948     0   1   2   3E+4   0.5   1  RT (min)      lcorbi_hilic_neg_4_067_57    m/z 372.7803     0   1   2E+4   8   8.5  RT (min)       lcorbi_hilic_neg_4_066_57    m/z 372.7923     0   2   4   6   8E+3   7.5   8  RT (min)      lcorbi_hilic_neg_4_100_61    m/z 372.9013     0   2   4   6E+3   12.5   13  RT (min)      lcorbi_hilic_neg_4_070_57    m/z 372.9013     0   1E+4   10.5   11  RT (min)      lcorbi_hilic_neg_4_103_61    m/z 372.9014     0   1   2   3   4E+4   8.5   9  RT (min)       lcorbi_hilic_neg_4_080_58    m/z 372.9778     0   1E+4   7.5   8  RT (min)      lcorbi_hilic_neg_4_074_58    m/z 372.9910     0   1E+4   0.5   1  RT (min)      lcorbi_hilic_neg_4_113_62    m/z 373.0100     0   1   2E+4   0.5   1  RT (min)       lcorbi_hilic_neg_4_112_62    m/z 373.0101     0   1   2E+4   3   3.5  RT (min)       lcorbi_hilic_neg_4_079_58    m/z 373.0451     0   1   2E+5   0.5   1  RT (min)       lcorbi_hilic_neg_4_113_62    m/z 373.0538     0   1   2   3E+4   6   6.5  RT (min)       lcorbi_hilic_neg_4_072_57    m/z 373.0586     0   1   2   3   4E+4   4   4.5  RT (min)      lcorbi_hilic_neg_4_030_05    m/z 373.0687     0   1   2E+4   6.5   7  RT (min)       lcorbi_hilic_neg_4_072_57    m/z 373.0918     0   2   4   6   8E+3   7.5   8  RT (min)      lcorbi_hilic_neg_4_090_60    m/z 373.0961     0   2   4   6E+3   0.5   1  RT (min)      lcorbi_hilic_neg_4_068_57    m/z 373.1000     0   1E+3   5.5   6  RT (min)      lcorbi_hilic_neg_1_047_05    m/z 373.1208     0   1E+4   4   4.5  RT (min)      lcorbi_hilic_neg_4_093_60    m/z 373.1324     0   1   2E+4   1   1.5  RT (min)       lcorbi_hilic_neg_4_077_58    m/z 373.1364     0   2   4   6E+2   4   4.5  RT (min)      lcorbi_hilic_neg_4_030_05    m/z 373.2092     0   1E+4   6.5   7  RT (min)      lcorbi_hilic_neg_4_089_59    m/z 373.2673     0   1E+4   4.5   5  RT (min)      lcorbi_hilic_neg_4_097_60    m/z 373.7789     0   2   4   6E+3   8   8.5  RT (min)      lcorbi_hilic_neg_4_066_57    m/z 373.8160     0   2   4   6   8E+3   4.5   5  RT (min)      lcorbi_hilic_neg_4_076_58    m/z 373.8748     0   1   2   3   4E+3   1   1.5  RT (min)      lcorbi_hilic_neg_3_003_44    m/z 373.9791     0   1   2   3   4E+4   7.5   8  RT (min)       lcorbi_hilic_neg_4_080_58    m/z 373.9921     0   1   5.5   6  RT (min)      lcorbi_hilic_neg_4_080_58    m/z 373.9999     0   1E+4   5.5   6  RT (min)      lcorbi_hilic_neg_4_080_58    m/z 374.0416     0   1   6.5   7  RT (min)      lcorbi_hilic_neg_4_077_58    m/z 374.0453     0   2   4   6E+4   0.5   1  RT (min)      lcorbi_hilic_neg_4_113_62    m/z 374.0539     0   2   4   6E+3   5   5.5  RT (min)      lcorbi_hilic_neg_4_113_62    m/z 374.0777     0   2   4E+3   1   1.5  RT (min)      lcorbi_hilic_neg_4_096_60    m/z 374.1220     0   2   4   6E+3   5.5   6  RT (min)      lcorbi_hilic_neg_3_043_02    m/z 374.1323     0   2   4   6   8E+4   0.5   1  RT (min)       lcorbi_hilic_neg_4_113_62    m/z 374.1332     0   1E+4   5.5   6  RT (min)      lcorbi_hilic_neg_4_089_59    m/z 374.1568     0   1E+4   4   4.5  RT (min)      lcorbi_hilic_neg_4_085_59    m/z 374.2084     0   1   1.5   2  RT (min)      lcorbi_hilic_neg_4_089_59    m/z 374.2911     0   1   0.5   1  RT (min)      lcorbi_hilic_neg_4_004_13    m/z 374.3102     0   1   2   3E+4   0.5   1  RT (min)      lcorbi_hilic_neg_4_067_57    m/z 374.7773     0   1E+4   8   8.5  RT (min)      lcorbi_hilic_neg_4_066_57    m/z 374.8059     0   2   4E+3   7.5   8  RT (min)      lcorbi_hilic_neg_4_102_61    m/z 374.8103     0   1   2   3E+4   4.5   5  RT (min)       lcorbi_hilic_neg_4_076_58    m/z 374.8150     0   1E+4   7.5   8  RT (min)      lcorbi_hilic_neg_4_102_61    m/z 374.8386     0   2   4   6E+3   7   7.5  RT (min)      lcorbi_hilic_neg_4_088_59    m/z 374.9063     0   2   4   6E+4   5.5   6  RT (min)       lcorbi_hilic_neg_4_106_62    m/z 374.9787     0   1E+4   7.5   8  RT (min)      lcorbi_hilic_neg_4_081_58    m/z 374.9885     0   1E+4   0.5   1  RT (min)      lcorbi_hilic_neg_4_068_57    m/z 374.9923     0   1E+5   6.5   7  RT (min)       lcorbi_hilic_neg_4_076_58    m/z 375.0065     0   1   2   3   4E+3   0.5   1  RT (min)      lcorbi_hilic_neg_3_031_14    m/z 375.0192     0   2   4   6   8E+4   6   6.5  RT (min)       lcorbi_hilic_neg_4_087_59    m/z 375.0498     0   1   2   3E+2   7.5   8  RT (min)      lcorbi_hilic_neg_4_015_24    m/z 375.0518     0   2   4   6E+3   5   5.5  RT (min)      lcorbi_hilic_neg_4_071_57    m/z 375.0625     0   1   2E+4   4   4.5  RT (min)       lcorbi_hilic_neg_4_066_57    m/z 375.0707     0   1   2E+3   7   7.5  RT (min)       lcorbi_hilic_neg_4_097_60    m/z 375.0752     0   1E+5   0.5   1  RT (min)      lcorbi_hilic_neg_4_074_58    m/z 375.0794     0   1   2   3   4E+4   7   7.5  RT (min)       lcorbi_hilic_neg_4_090_60    m/z 375.1010     0   2   4   6   8E+3   5.5   6  RT (min)      lcorbi_hilic_neg_4_086_59    m/z 375.1179     0   1   2   2.5  RT (min)      lcorbi_hilic_neg_4_106_62    m/z 375.1284     0   2   4   6E+2   3.5   4  RT (min)      lcorbi_hilic_neg_4_051_16    m/z 375.1660     0   1   2E+3   1   1.5  RT (min)      lcorbi_hilic_neg_4_079_58    m/z 375.1676     0   1   2   3   4E+4   5.5   6  RT (min)       lcorbi_hilic_neg_4_096_60    m/z 375.2130     0   1   1.5   2  RT (min)       lcorbi_hilic_neg_4_084_59    m/z 375.2187     0   1E+4   7   7.5  RT (min)      lcorbi_hilic_neg_4_017_51    m/z 375.2387     0   1E+3   1   1.5  RT (min)       lcorbi_hilic_neg_4_089_59    m/z 375.2539     0   1E+4   0.5   1  RT (min)      lcorbi_hilic_neg_3_055_26    m/z 375.2783     0   1E+5   0.5   1  RT (min)       lcorbi_hilic_neg_4_066_57    m/z 375.6055     0   1E+5   3   3.5  RT (min)       lcorbi_hilic_neg_3_034_34    m/z 375.9773     0   1   7.5   8  RT (min)      lcorbi_hilic_neg_4_093_60    m/z 375.9848     0   1E+3   6.5   7  RT (min)      lcorbi_hilic_neg_4_077_58    m/z 375.9967     0   1   2E+3   5.5   6  RT (min)      lcorbi_hilic_neg_4_048_37    m/z 376.0286     0   1   2E+4   1   1.5  RT (min)       lcorbi_hilic_neg_4_098_61    m/z 376.0817     0   1   1.5   2  RT (min)       lcorbi_hilic_neg_4_107_62    m/z 376.0891     0   1E+4   6.5   7  RT (min)      lcorbi_hilic_neg_4_096_60    m/z 376.1094     0   1   2   3E+3   5.5   6   6.5  RT (min)       lcorbi_hilic_neg_4_093_60    m/z 376.1222     0   1   2E+5   6   6.5  RT (min)       lcorbi_hilic_neg_4_097_60    m/z 376.1299     0   1E+3   2   2.5  RT (min)      lcorbi_hilic_neg_4_101_61    m/z 376.1513     0   1   2E+4   3.5   4  RT (min)       lcorbi_hilic_neg_4_077_58    m/z 376.1722     0   1   2E+4   6.5   7  RT (min)      lcorbi_hilic_neg_1_044_03    m/z 376.1856     0   1   2E+4   3   3.5  RT (min)       lcorbi_hilic_neg_4_096_60    m/z 376.2263     0   1E+4   4.5   5  RT (min)      lcorbi_hilic_neg_4_079_58    m/z 376.2815     0   1   2E+4   0.5   1  RT (min)      lcorbi_hilic_neg_4_066_57    m/z 376.9035     0   1   2E+4   5.5   6  RT (min)       lcorbi_hilic_neg_4_106_62    m/z 376.9108     0   2   4   6E+3   8   8.5  RT (min)      lcorbi_hilic_neg_4_075_58    m/z 376.9755     0   2   4   6   8E+3   7.5   8  RT (min)      lcorbi_hilic_neg_4_081_58    m/z 377.0039     0   2   4E+3   0.5   1  RT (min)      lcorbi_hilic_neg_1_044_03    m/z 377.0107     0   1   5   5.5  RT (min)      lcorbi_hilic_neg_4_100_61    m/z 377.0221     0   1   2   3   4E+4   0.5   1  RT (min)       lcorbi_hilic_neg_4_098_61    m/z 377.0376     0   1   2E+3   3   3.5  RT (min)      lcorbi_hilic_neg_4_066_57    m/z 377.0410     0   1   6   6.5  RT (min)      lcorbi_hilic_neg_4_089_59    m/z 377.0539     0   2   4   6E+3   3   3.5  RT (min)      lcorbi_hilic_neg_2_040_35    m/z 377.0665     0   1E+3   7   7.5  RT (min)      lcorbi_hilic_neg_1_005_36    m/z 377.0743     0   1E+4   6   6.5  RT (min)      lcorbi_hilic_neg_2_020_27    m/z 377.0758     0   1E+4   4   4.5  RT (min)      lcorbi_hilic_neg_4_101_61    m/z 377.0999     0   1E+4   6   6.5  RT (min)      lcorbi_hilic_neg_2_048_38    m/z 377.1135     0   1E+4   7.5   8  RT (min)      lcorbi_hilic_neg_4_088_59    m/z 377.1141     0   2   4   6E+3   6.5   7  RT (min)       lcorbi_hilic_neg_4_073_57    m/z 377.1267     0   1E+3   2   2.5  RT (min)      lcorbi_hilic_neg_4_107_62    m/z 377.1374     0   1   2   3E+3   0.5   1  RT (min)      lcorbi_hilic_neg_4_080_58    m/z 377.1568     0   2   4   6   8E+3   7   7.5  RT (min)      lcorbi_hilic_neg_4_088_59    m/z 377.1857     0   1   2   3   4E+3   2   2.5  RT (min)      lcorbi_hilic_neg_4_032_22    m/z 377.1861     0   1   2E+4   6.5   7  RT (min)      lcorbi_hilic_neg_1_047_05    m/z 377.1941     0   1   2   2.5  RT (min)      lcorbi_hilic_neg_4_107_62    m/z 377.2222     0   1   1   2  RT (min)       lcorbi_hilic_neg_4_108_62    m/z 377.2697     0   1   2E+3   1   1.5  RT (min)      lcorbi_hilic_neg_4_051_16    m/z 377.3954     0   2   4   6   8E+4   0.5   1  RT (min)       lcorbi_hilic_neg_4_097_60    m/z 377.9883     0   1E+3   0.5   1  RT (min)      lcorbi_hilic_neg_3_031_14    m/z 377.9885     0   2   4   6E+3   6   6.5  RT (min)      lcorbi_hilic_neg_4_080_58    m/z 378.0256     0   2   4   6   8E+3   1   1.5  RT (min)      lcorbi_hilic_neg_4_098_61    m/z 378.0509     0   2   4   6E+4   6   6.5  RT (min)       lcorbi_hilic_neg_4_082_59    m/z 378.0923     0   1E+4   4   4.5  RT (min)      lcorbi_hilic_neg_4_101_61    m/z 378.0960     0   1   2   3E+4   1   1.5  RT (min)       lcorbi_hilic_neg_4_079_58    m/z 378.1016     0   1   2E+4   0.5   1  RT (min)       lcorbi_hilic_neg_4_081_58    m/z 378.1074     0   1E+4   2   2.5  RT (min)      lcorbi_hilic_neg_4_073_57    m/z 378.1098     0   1   7   7.5  RT (min)      lcorbi_hilic_neg_2_005_24    m/z 378.1194     0   1   6   6.5  RT (min)       lcorbi_hilic_neg_4_109_62    m/z 378.1345     0   1   6   6.5  RT (min)      lcorbi_hilic_neg_4_073_57    m/z 378.1534     0   1E+4   1   1.5  RT (min)      lcorbi_hilic_neg_4_108_62    m/z 378.1613     0   1   2E+3   2   2.5  RT (min)      lcorbi_hilic_neg_4_108_62    m/z 378.2116     0   2   4   6   8E+5   1.5   2  RT (min)      lcorbi_hilic_neg_4_106_62    m/z 378.2420     0   1   2E+4   4.5   5  RT (min)       lcorbi_hilic_neg_4_078_58    m/z 378.2610     0   1E+4   3.5   4  RT (min)      lcorbi_hilic_neg_4_084_59    m/z 378.2779     0   1E+4   0.5   1  RT (min)      lcorbi_hilic_neg_4_084_59    m/z 378.8578     0   1   2E+4   10   11  RT (min)       lcorbi_hilic_neg_4_109_62    m/z 378.8578     0   1E+4   12.5   13  RT (min)      lcorbi_hilic_neg_4_109_62    m/z 378.9082     0   2   4   6E+3   8   8.5  RT (min)      lcorbi_hilic_neg_4_081_58    m/z 378.9721     0   1   2   3E+4   7.5   8  RT (min)       lcorbi_hilic_neg_4_089_59    m/z 378.9822     0   2   4   6   8E+3   0.5   1  RT (min)       lcorbi_hilic_neg_4_097_60    m/z 379.0003     0   1E+4   0.5   1  RT (min)       lcorbi_hilic_neg_4_112_62    m/z 379.0131     0   1   2   3   4E+4   5.5   6  RT (min)       lcorbi_hilic_neg_4_067_57    m/z 379.0385     0   2   4   6   8E+3   0.5   1  RT (min)      lcorbi_hilic_neg_4_104_61    m/z 379.0509     0   2   4   6E+3   2.5   3  RT (min)      lcorbi_hilic_neg_4_004_13    m/z 379.0820     0   1   0.5   1  RT (min)      lcorbi_hilic_neg_4_074_58    m/z 379.0900     0   2   4   6   8E+3   6   6.5  RT (min)      lcorbi_hilic_neg_2_020_27    m/z 379.0911     0   1E+4   1   1.5  RT (min)      lcorbi_hilic_neg_4_068_57    m/z 379.0973     0   1   2E+4   3   3.5  RT (min)      lcorbi_hilic_neg_4_098_61    m/z 379.1010     0   1   6   6.5  RT (min)       lcorbi_hilic_neg_4_104_61    m/z 379.1039     0   1E+4   2   2.5  RT (min)       lcorbi_hilic_neg_4_068_57    m/z 379.1317     0   1E+4   3.5   4  RT (min)      lcorbi_hilic_neg_4_112_62    m/z 379.1790     0   2   4   6   8E+3   0.5   1  RT (min)      lcorbi_hilic_neg_4_077_58    m/z 379.1893     0   1   2E+4   1   1.5  RT (min)       lcorbi_hilic_neg_4_075_58    m/z 379.1952     0   1E+3   2.5   3  RT (min)       lcorbi_hilic_neg_4_106_62    m/z 379.2451     0   2   4E+3   4.5   5  RT (min)      lcorbi_hilic_neg_4_077_58    m/z 379.2547     0   1   2E+4   3.5   4  RT (min)       lcorbi_hilic_neg_4_088_59    m/z 379.2909     0   1   2   3E+4   0.5   1  RT (min)      lcorbi_hilic_neg_4_066_57    m/z 379.6013     0   1E+4   3   3.5  RT (min)      lcorbi_hilic_neg_3_034_34    m/z 379.9082     0   1   2E+4   1   1.5  RT (min)       lcorbi_hilic_neg_4_109_62    m/z 379.9558     0   1   5.5   6  RT (min)      lcorbi_hilic_neg_4_082_59    m/z 379.9716     0   1E+4   6   6.5  RT (min)      lcorbi_hilic_neg_4_068_57    m/z 380.0120     0   1   5.5   6  RT (min)      lcorbi_hilic_neg_4_079_58    m/z 380.0201     0   1E+4   5.5   6  RT (min)      lcorbi_hilic_neg_4_104_61    m/z 380.0361     0   1   6   6.5  RT (min)       lcorbi_hilic_neg_4_074_58    m/z 380.0496     0   1   2E+4   6   6.5  RT (min)      lcorbi_hilic_neg_4_084_59    m/z 380.0773     0   1E+3   6.5   7  RT (min)      lcorbi_hilic_neg_4_073_57    m/z 380.1605     0   2   4   6E+4   0.5   1  RT (min)      lcorbi_hilic_neg_4_074_58    m/z 380.1982     0   2   4   6E+3   2.5   3  RT (min)      lcorbi_hilic_neg_4_075_58    m/z 380.2164     0   1E+4   2   2.5  RT (min)      lcorbi_hilic_neg_4_106_62    m/z 380.8228     0   1   2E+4   7.5   8  RT (min)       lcorbi_hilic_neg_4_102_61    m/z 380.8303     0   1   4.5   5  RT (min)      lcorbi_hilic_neg_4_080_58    m/z 380.9971     0   1E+4   0.5   1  RT (min)      lcorbi_hilic_neg_4_072_57    m/z 381.0055     0   2   4   6   8E+4   6   6.5  RT (min)       lcorbi_hilic_neg_4_074_58    m/z 381.0423     0   1   6   6.5  RT (min)      lcorbi_hilic_neg_4_078_58    m/z 381.0511     0   1E+3   6   6.5  RT (min)      lcorbi_hilic_neg_4_068_57    m/z 381.0648     0   1   1   1.5  RT (min)      lcorbi_hilic_neg_2_020_27    m/z 381.0680     0   1   2E+3   3   3.5  RT (min)      lcorbi_hilic_neg_4_098_61    m/z 381.0684     0   1E+4   6   6.5  RT (min)      lcorbi_hilic_neg_4_083_59    m/z 381.0784     0   1   2   3   4E+2   3.5   4  RT (min)      lcorbi_hilic_neg_2_032_25    m/z 381.0876     0   1E+3   6   6.5  RT (min)      lcorbi_hilic_neg_4_013_46    m/z 381.0921     0   1E+4   4   4.5  RT (min)      lcorbi_hilic_neg_4_053_23    m/z 381.1263     0   2   4   6E+4   6   6.5  RT (min)      lcorbi_hilic_neg_4_013_46    m/z 381.1472     0   1E+4   2   2.5  RT (min)      lcorbi_hilic_neg_4_001_49    m/z 381.1525     0   1   2   3E+4   4   4.5  RT (min)       lcorbi_hilic_neg_4_092_60    m/z 381.1862     0   2   4   6E+3   1   1.5  RT (min)      lcorbi_hilic_neg_4_077_58    m/z 381.1948     0   2   4   6   8E+3   0.5   1  RT (min)       lcorbi_hilic_neg_4_087_59    m/z 381.2467     0   2   4   6E+3   0.5   1  RT (min)       lcorbi_hilic_neg_4_089_59    m/z 381.3373     0   1   2E+3   1   1.5  RT (min)      lcorbi_hilic_neg_4_095_60    m/z 381.4879     0   1E+4   4.5   5  RT (min)      lcorbi_hilic_neg_4_080_58    m/z 381.8112     0   2   4   6   8E+3   8   8.5  RT (min)      lcorbi_hilic_neg_4_066_57    m/z 381.9051     0   1   2E+4   1   1.5  RT (min)      lcorbi_hilic_neg_4_109_62    m/z 381.9606     0   1E+4   5.5   6  RT (min)      lcorbi_hilic_neg_4_083_59    m/z 381.9687     0   2   4   6E+3   6   6.5  RT (min)      lcorbi_hilic_neg_4_068_57    m/z 382.0129     0   1   2E+4   1.5   2  RT (min)      lcorbi_hilic_neg_3_030_30    m/z 382.0171     0   2   4E+3   5.5   6  RT (min)      lcorbi_hilic_neg_4_104_61    m/z 382.0415     0   2   4   6E+4   3   3.5  RT (min)      lcorbi_hilic_neg_4_023_34    m/z 382.0997     0   1   5.5   6  RT (min)      lcorbi_hilic_neg_4_072_57    m/z 382.1340     0   1   6.5   7  RT (min)      lcorbi_hilic_neg_4_092_60    m/z 382.1425     0   1   2E+4   6.5   7  RT (min)       lcorbi_hilic_neg_4_096_60    m/z 382.1763     0   1   2   3   4E+3   0.5   1  RT (min)      lcorbi_hilic_neg_4_087_59    m/z 382.4864     0   1   2E+4   4.5   5  RT (min)       lcorbi_hilic_neg_4_077_58    m/z 382.8092     0   1E+4   8   8.5  RT (min)      lcorbi_hilic_neg_4_066_57    m/z 382.9364     0   2   4   6   8E+3   5.5   6  RT (min)      lcorbi_hilic_neg_4_070_57    m/z 382.9554     0   1   0.5   1  RT (min)      lcorbi_hilic_neg_4_084_59    m/z 382.9948     0   1E+4   0.5   1  RT (min)      lcorbi_hilic_neg_4_070_57    m/z 383.0005     0   2   4   6E+3   5.5   6  RT (min)      lcorbi_hilic_neg_4_076_58    m/z 383.0215     0   1   2.5   3  RT (min)      lcorbi_hilic_neg_4_088_59    m/z 383.0323     0   1   2   3   4E+3   7   7.5  RT (min)      lcorbi_hilic_neg_4_070_57    m/z 383.0431     0   1   2.5   3  RT (min)      lcorbi_hilic_neg_4_088_59    m/z 383.0563     0   1E+4   1   1.5  RT (min)      lcorbi_hilic_neg_1_005_36    m/z 383.0751     0   1E+4   3   3.5  RT (min)      lcorbi_hilic_neg_4_104_61    m/z 383.0840     0   1   2   3E+3   6.5   7  RT (min)       lcorbi_hilic_neg_2_022_37    m/z 383.0977     0   1   2E+4   3.5   4  RT (min)       lcorbi_hilic_neg_4_111_62    m/z 383.1061     0   1   6.5   7  RT (min)      lcorbi_hilic_neg_4_089_59    m/z 383.1214     0   1   2E+3   1   1.5  RT (min)      lcorbi_hilic_neg_4_107_62    m/z 383.1460     0   2   4E+3   2   2.5  RT (min)      lcorbi_hilic_neg_4_001_49    m/z 383.1895     0   2   4   6E+3   5   5.5  RT (min)      lcorbi_hilic_neg_4_082_59    m/z 383.1938     0   1   7   7.5  RT (min)       lcorbi_hilic_neg_2_005_24    m/z 383.2062     0   1E+5   6.5   7  RT (min)       lcorbi_hilic_neg_4_097_60    m/z 383.2314     0   1E+4   4.5   5  RT (min)      lcorbi_hilic_neg_4_078_58    m/z 383.2372     0   2   4   6   8E+3   0.5   1  RT (min)      lcorbi_hilic_neg_4_101_61    m/z 383.2916     0   2   4   6E+3   1   1.5  RT (min)      lcorbi_hilic_neg_4_074_58    m/z 383.3165     0   2   4   6   8E+3   0.5   1  RT (min)      lcorbi_hilic_neg_4_113_62    m/z 383.4851     0   1   2E+4   4.5   5  RT (min)      lcorbi_hilic_neg_4_077_58    m/z 383.8075     0   2   4   6E+3   8   8.5  RT (min)      lcorbi_hilic_neg_4_066_57    m/z 383.8453     0   1E+4   4.5   5  RT (min)      lcorbi_hilic_neg_4_075_58    m/z 383.9744     0   2   4   6   8E+3   6   6.5  RT (min)      lcorbi_hilic_neg_4_098_61    m/z 383.9872     0   1E+4   0.5   1  RT (min)      lcorbi_hilic_neg_4_004_13    m/z 384.0108     0   1   2E+4   1.5   2  RT (min)      lcorbi_hilic_neg_4_054_30    m/z 384.0289     0   1   2E+4   5.5   6  RT (min)       lcorbi_hilic_neg_4_081_58    m/z 384.0708     0   2   4   6   8E+3   6.5   7  RT (min)      lcorbi_hilic_neg_4_093_60    m/z 384.0736     0   1E+4   3.5   4  RT (min)      lcorbi_hilic_neg_4_108_62    m/z 384.0907     0   2   4   6E+3   1.5   2  RT (min)      lcorbi_hilic_neg_4_112_62    m/z 384.1254     0   1   5.5   6  RT (min)      lcorbi_hilic_neg_4_097_60    m/z 384.1498     0   1E+4   5   5.5  RT (min)      lcorbi_hilic_neg_2_020_27    m/z 384.1795     0   2   4   6E+4   1   1.5  RT (min)      lcorbi_hilic_neg_4_076_58    m/z 384.2304     0   1   2   3   4E+3   0.5   1  RT (min)      lcorbi_hilic_neg_4_105_61    m/z 384.3565     0   1   2   3E+4   1   1.5  RT (min)      lcorbi_hilic_neg_1_047_05    m/z 384.4834     0   1E+4   4.5   5  RT (min)      lcorbi_hilic_neg_4_077_58    m/z 384.8062     0   2   4   6E+3   8   8.5  RT (min)      lcorbi_hilic_neg_4_066_57    m/z 384.8401     0   1   2E+4   4.5   5  RT (min)       lcorbi_hilic_neg_4_075_58    m/z 384.9059     0   1   2   3E+3   5   5.5  RT (min)      lcorbi_hilic_neg_2_020_27    m/z 384.9163     0   2   4   6   8E+3   5   5.5  RT (min)      lcorbi_hilic_neg_4_042_28    m/z 384.9523     0   1   0.5   1  RT (min)      lcorbi_hilic_neg_4_111_62    m/z 384.9590     0   1   2E+4   7.5   8  RT (min)       lcorbi_hilic_neg_4_075_58    m/z 384.9646     0   1   2E+4   5   5.5  RT (min)      lcorbi_hilic_neg_4_067_57    m/z 384.9962     0   2   4   6E+3   5.5   6  RT (min)      lcorbi_hilic_neg_4_072_57    m/z 385.0104     0   1E+4   0.5   1  RT (min)      lcorbi_hilic_neg_4_071_57    m/z 385.0371     0   1E+4   6   6.5  RT (min)      lcorbi_hilic_neg_3_029_25    m/z 385.0463     0   1E+3   6   6.5  RT (min)       lcorbi_hilic_neg_4_094_60    m/z 385.0675     0   1   2E+4   5.5   6  RT (min)      lcorbi_hilic_neg_1_011_01    m/z 385.0718     0   1   2   3E+4   0.5   1  RT (min)      lcorbi_hilic_neg_3_031_14    m/z 385.0930     0   1E+3   1   1.5  RT (min)      lcorbi_hilic_neg_4_102_61    m/z 385.0947     0   1E+4   3.5   4   4.5  RT (min)      lcorbi_hilic_neg_4_109_62    m/z 385.1212     0   1   6   6.5  RT (min)      lcorbi_hilic_neg_4_072_57    m/z 385.1476     0   2   4   6   8E+3   6   6.5  RT (min)      lcorbi_hilic_neg_4_068_57    m/z 385.1492     0   1E+3   1   1.5  RT (min)      lcorbi_hilic_neg_4_098_61    m/z 385.1592     0   1   1   1.5  RT (min)      lcorbi_hilic_neg_4_089_59    m/z 385.1906     0   2   4   6E+3   7   7.5  RT (min)      lcorbi_hilic_neg_4_083_59    m/z 385.2179     0   1   1   1.5  RT (min)       lcorbi_hilic_neg_4_071_57    m/z 385.9199     0   1   2E+3   5.5   6  RT (min)      lcorbi_hilic_neg_4_071_57    m/z 385.9282     0   1E+5   5.5   6  RT (min)      lcorbi_hilic_neg_4_005_32    m/z 385.9530     0   2   4E+3   5.5   6  RT (min)      lcorbi_hilic_neg_4_071_57    m/z 385.9568     0   1   2   3   4E+4   7.5   8  RT (min)       lcorbi_hilic_neg_4_074_58    m/z 386.0058     0   1E+4   6   6.5  RT (min)      lcorbi_hilic_neg_4_082_59    m/z 386.0576     0   1E+4   1   1.5  RT (min)      lcorbi_hilic_neg_4_102_61    m/z 386.0577     0   1   2   3   4E+2   6   6.5  RT (min)      lcorbi_hilic_neg_4_001_49    m/z 386.0710     0   1   4   4.5  RT (min)      lcorbi_hilic_neg_4_090_60    m/z 386.0883     0   1   3.5   4  RT (min)      lcorbi_hilic_neg_4_090_60    m/z 386.1145     0   1   1.5   2  RT (min)       lcorbi_hilic_neg_4_071_57    m/z 386.1174     0   1   2   3   4E+2   6   6.5  RT (min)      lcorbi_hilic_neg_2_044_46    m/z 386.1335     0   1E+4   5.5   6  RT (min)       lcorbi_hilic_neg_4_097_60    m/z 386.1351     0   1E+4   1   1.5  RT (min)      lcorbi_hilic_neg_4_066_57    m/z 386.1724     0   1E+4   1.5   2  RT (min)      lcorbi_hilic_neg_4_075_58    m/z 386.1820     0   1   2   3   4E+4   1   1.5  RT (min)       lcorbi_hilic_neg_4_101_61    m/z 386.2371     0   2   4E+3   1   1.5  RT (min)      lcorbi_hilic_neg_4_067_57    m/z 386.2564     0   1E+4   1.5   2  RT (min)      lcorbi_hilic_neg_4_089_59    m/z 386.7160     0   2   4   6E+3   4.5   5  RT (min)      lcorbi_hilic_neg_4_101_61    m/z 386.7995     0   1   2   3E+3   4.5   5  RT (min)      lcorbi_hilic_neg_4_077_58    m/z 386.8623     0   2   4E+3   7.5   8  RT (min)      lcorbi_hilic_neg_4_099_61    m/z 386.9312     0   1   5   5.5  RT (min)      lcorbi_hilic_neg_4_077_58    m/z 386.9546     0   1E+4   7.5   8  RT (min)      lcorbi_hilic_neg_4_078_58    m/z 386.9637     0   1   2E+3   7.5   8  RT (min)      lcorbi_hilic_neg_1_016_37    m/z 386.9889     0   1E+4   0.5   1  RT (min)      lcorbi_hilic_neg_4_095_60    m/z 387.0006     0   2   4E+3   7.5   8  RT (min)      lcorbi_hilic_neg_4_083_59    m/z 387.0259     0   1E+4   0.5   1  RT (min)      lcorbi_hilic_neg_4_079_58    m/z 387.0464     0   1E+4   6   6.5  RT (min)      lcorbi_hilic_neg_4_072_57    m/z 387.0609     0   1E+4   0.5   1  RT (min)      lcorbi_hilic_neg_4_097_60    m/z 387.0725     0   1   2E+4   1   1.5  RT (min)       lcorbi_hilic_neg_4_092_60    m/z 387.0831     0   2   4   6   8E+3   7.5   8  RT (min)      lcorbi_hilic_neg_4_086_59    m/z 387.1847     0   1E+4   1.5   2  RT (min)      lcorbi_hilic_neg_4_067_57    m/z 387.1943     0   1   1   1.5  RT (min)      lcorbi_hilic_neg_4_098_61    m/z 387.2542     0   1E+4   1   1.5  RT (min)      lcorbi_hilic_neg_1_002_26    m/z 387.9251     0   1   2   3   4E+3   5.5   6  RT (min)      lcorbi_hilic_neg_4_069_57    m/z 387.9431     0   2   4E+3   5   5.5  RT (min)      lcorbi_hilic_neg_4_068_57    m/z 387.9763     0   1   2   3   4E+4   1   1.5  RT (min)      lcorbi_hilic_neg_4_106_62    m/z 388.0058     0   1   6.5   7  RT (min)      lcorbi_hilic_neg_4_088_59    m/z 388.0143     0   1   2E+4   6.5   7  RT (min)       lcorbi_hilic_neg_4_082_59    m/z 388.0270     0   2   4   6E+3   5.5   6  RT (min)      lcorbi_hilic_neg_4_072_57    m/z 388.0455     0   2   4   6E+3   6.5   7  RT (min)      lcorbi_hilic_neg_4_090_60    m/z 388.0548     0   1E+4   1   1.5  RT (min)      lcorbi_hilic_neg_4_088_59    m/z 388.1112     0   1E+4   2   2.5  RT (min)      lcorbi_hilic_neg_4_051_16    m/z 388.1180     0   1   2E+4   6.5   7  RT (min)       lcorbi_hilic_neg_4_071_57    m/z 388.1271     0   1   2E+4   6   6.5  RT (min)       lcorbi_hilic_neg_4_091_60    m/z 388.1352     0   1   2E+4   7  RT (min)       lcorbi_hilic_neg_4_095_60    m/z 388.1728     0   1E+4   4   4.5  RT (min)      lcorbi_hilic_neg_4_083_59    m/z 388.1796     0   1E+4   1   1.5  RT (min)      lcorbi_hilic_neg_4_102_61    m/z 388.1839     0   2   4   6E+3   8   8.5  RT (min)      lcorbi_hilic_neg_4_093_60    m/z 388.2527     0   1   2E+3   1   1.5  RT (min)      lcorbi_hilic_neg_1_039_38    m/z 388.2621     0   1   2E+4   0.5   1  RT (min)       lcorbi_hilic_neg_4_089_59    m/z 388.3020     0   1   2E+4   1   1.5  RT (min)      lcorbi_hilic_neg_1_048_35    m/z 388.7940     0   1   2   3   4E+3   5   5.5  RT (min)      lcorbi_hilic_neg_4_075_58    m/z 388.9723     0   1   2E+4   6   6.5   7  RT (min)       lcorbi_hilic_neg_4_083_59    m/z 388.9855     0   1E+4   0.5   1  RT (min)      lcorbi_hilic_neg_4_099_61    m/z 389.0044     0   1E+4   0.5   1  RT (min)      lcorbi_hilic_neg_4_073_57    m/z 389.0278     0   1   2E+4   6   6.5  RT (min)       lcorbi_hilic_neg_4_071_57    m/z 389.0485     0   1   2E+4   1   1.5  RT (min)       lcorbi_hilic_neg_4_069_57    m/z 389.0519     0   1E+4   7   7.5  RT (min)      lcorbi_hilic_neg_1_051_33    m/z 389.0609     0   1E+4   7   7.5  RT (min)      lcorbi_hilic_neg_1_005_36    m/z 389.0795     0   1E+4   6   6.5  RT (min)       lcorbi_hilic_neg_4_090_60    m/z 389.0913     0   1E+4   1   1.5  RT (min)      lcorbi_hilic_neg_4_074_58    m/z 389.1076     0   1   2   3   4E+4   2   2.5  RT (min)      lcorbi_hilic_neg_4_051_16    m/z 389.1117     0   1E+4   4.5   5  RT (min)      lcorbi_hilic_neg_4_111_62    m/z 389.1447     0   1   6.5   7  RT (min)      lcorbi_hilic_neg_4_092_60    m/z 389.1458     0   1   2   3   4E+4   2.5   3  RT (min)      lcorbi_hilic_neg_4_023_34    m/z 389.2060     0   2   4   6E+3   4   4.5  RT (min)      lcorbi_hilic_neg_4_096_60    m/z 389.2137     0   1   2   3   4E+3   7   7.5  RT (min)      lcorbi_hilic_neg_4_001_49    m/z 389.2406     0   1   2   3   4E+4   1   1.5  RT (min)       lcorbi_hilic_neg_4_089_59    m/z 389.2628     0   1E+4   6.5   7  RT (min)      lcorbi_hilic_neg_4_096_60    m/z 389.2698     0   1E+4   1   1.5  RT (min)      lcorbi_hilic_neg_4_078_58    m/z 389.2926     0   1E+4   3.5   4  RT (min)      lcorbi_hilic_neg_4_088_59    m/z 389.8489     0   2   4E+3   1   1.5   2  RT (min)      lcorbi_hilic_neg_4_066_57    m/z 389.9387     0   2   4   6E+3   8   8.5  RT (min)      lcorbi_hilic_neg_4_077_58    m/z 389.9596     0   1E+4   7.5   8  RT (min)      lcorbi_hilic_neg_3_018_38    m/z 389.9925     0   1   2   3E+4   5.5   6  RT (min)       lcorbi_hilic_neg_4_088_59    m/z 390.0007     0   1E+4   6   6.5  RT (min)      lcorbi_hilic_neg_4_068_57    m/z 390.0174     0   1E+4   6.5   7  RT (min)      lcorbi_hilic_neg_4_097_60    m/z 390.0407     0   1   5.5   6  RT (min)      lcorbi_hilic_neg_4_082_59    m/z 390.0591     0   1   2E+4   3.5   4  RT (min)      lcorbi_hilic_neg_4_090_60    m/z 390.0830     0   1E+4   1   1.5  RT (min)      lcorbi_hilic_neg_4_098_61    m/z 390.0976     0   1   2E+4   6   6.5  RT (min)       lcorbi_hilic_neg_4_089_59    m/z 390.1464     0   1   2   3   4E+4   5.5   6  RT (min)      lcorbi_hilic_neg_2_044_46    m/z 390.1866     0   1E+4   5.5   6  RT (min)       lcorbi_hilic_neg_4_097_60    m/z 390.2399     0   1   2E+3   0.5   1  RT (min)      lcorbi_hilic_neg_4_088_59    m/z 390.8538     0   2   4E+3   1   1.5  RT (min)      lcorbi_hilic_neg_4_074_58    m/z 391.0009     0   1   2E+4   3   3.5  RT (min)       lcorbi_hilic_neg_4_081_58    m/z 391.0009     0   1   2   3   4E+4   0.5   1  RT (min)       lcorbi_hilic_neg_4_103_61    m/z 391.0093     0   1   7   7.5  RT (min)      lcorbi_hilic_neg_4_087_59    m/z 391.0125     0   1E+3   2   2.5  RT (min)      lcorbi_hilic_neg_4_097_60    m/z 391.0409     0   1   2   3E+4   7   7.5  RT (min)       lcorbi_hilic_neg_4_084_59    m/z 391.0494     0   1   2E+4   5.5   6  RT (min)       lcorbi_hilic_neg_4_089_59    m/z 391.0631     0   1E+4   6.5   7  RT (min)      lcorbi_hilic_neg_4_090_60    m/z 391.0716     0   1E+4   6   6.5  RT (min)      lcorbi_hilic_neg_4_089_59    m/z 391.0797     0   1   2E+4   1   1.5  RT (min)       lcorbi_hilic_neg_4_107_62    m/z 391.0949     0   1E+4   1   1.5  RT (min)      lcorbi_hilic_neg_3_024_27    m/z 391.1013     0   1   2   3E+4   5  RT (min)       lcorbi_hilic_neg_4_053_23    m/z 391.1084     0   1   4.5   5  RT (min)      lcorbi_hilic_neg_4_073_57    m/z 391.1411     0   1   6   6.5  RT (min)       lcorbi_hilic_neg_4_083_59    m/z 391.2257     0   1E+4   2   2.5  RT (min)      lcorbi_hilic_neg_4_073_57    m/z 391.2484     0   1   2   3   4E+4   5   5.5  RT (min)       lcorbi_hilic_neg_4_089_59    m/z 391.2620     0   1   2E+4   0.5   1  RT (min)      lcorbi_hilic_neg_4_069_57    m/z 391.2967     0   1   2   3   4E+5   1   1.5  RT (min)      lcorbi_hilic_neg_1_002_26    m/z 391.8404     0   1   2   3   4E+3   8   8.5  RT (min)      lcorbi_hilic_neg_4_066_57    m/z 391.9733     0   2   4E+3   5.5   6  RT (min)      lcorbi_hilic_neg_4_104_61    m/z 391.9896     0   1E+4   5.5   6  RT (min)      lcorbi_hilic_neg_4_088_59    m/z 392.0082     0   1E+4   2   2.5  RT (min)      lcorbi_hilic_neg_4_099_61    m/z 392.0438     0   1E+4   3   3.5  RT (min)      lcorbi_hilic_neg_4_066_57    m/z 392.0573     0   2   4E+2   6   6.5  RT (min)      lcorbi_hilic_neg_4_053_23    m/z 392.1049     0   1   2E+3   5   5.5  RT (min)      lcorbi_hilic_neg_3_029_25    m/z 392.2295     0   1   2E+3   2   2.5  RT (min)      lcorbi_hilic_neg_4_027_47    m/z 392.2898     0   1   2   3   4E+4   1   1.5  RT (min)       lcorbi_hilic_neg_4_068_57    m/z 392.3003     0   2   4   6E+4   1   1.5  RT (min)      lcorbi_hilic_neg_1_002_26    m/z 392.7787     0   1E+4   8   8.5  RT (min)      lcorbi_hilic_neg_4_106_62    m/z 392.8384     0   2   4E+3   8   8.5  RT (min)      lcorbi_hilic_neg_4_106_62    m/z 392.8493     0   1E+4   7   7.5  RT (min)      lcorbi_hilic_neg_4_088_59    m/z 392.8967     0   1E+4   8   8.5  RT (min)      lcorbi_hilic_neg_4_079_58    m/z 392.9052     0   2   4   6   8E+3   7   7.5  RT (min)      lcorbi_hilic_neg_4_096_60    m/z 392.9076     0   1E+3   5   5.5  RT (min)      lcorbi_hilic_neg_4_082_59    m/z 393.0295     0   1   2E+4   5.5   6  RT (min)       lcorbi_hilic_neg_4_071_57    m/z 393.0493     0   1   2E+4   3   3.5  RT (min)       lcorbi_hilic_neg_4_105_61    m/z 393.0764     0   2   4   6E+3   2   2.5  RT (min)      lcorbi_hilic_neg_3_021_20    m/z 393.0813     0   1   2   2.5  RT (min)      lcorbi_hilic_neg_4_107_62    m/z 393.0982     0   1   5.5   6  RT (min)      lcorbi_hilic_neg_1_023_23    m/z 393.1014     0   1   2E+4   1   1.5  RT (min)      lcorbi_hilic_neg_4_082_59    m/z 393.1090     0   2   4   6E+3   8   8.5  RT (min)      lcorbi_hilic_neg_4_083_59    m/z 393.1111     0   1E+4   2   2.5  RT (min)      lcorbi_hilic_neg_4_011_19    m/z 393.1173     0   1E+5   6   6.5  RT (min)       lcorbi_hilic_neg_4_089_59    m/z 393.1200     0   2   4   6   8E+3   2   2.5  RT (min)      lcorbi_hilic_neg_4_109_62    m/z 393.1256     0   1E+5   5.5   6  RT (min)       lcorbi_hilic_neg_4_089_59    m/z 393.1354     0   1   2   3E+2   0.5   1  RT (min)      lcorbi_hilic_neg_1_024_11    m/z 393.1454     0   1E+4   4.5   5  RT (min)      lcorbi_hilic_neg_4_106_62    m/z 393.1718     0   1   2   3   4E+4   1   1.5  RT (min)       lcorbi_hilic_neg_4_113_62    m/z 393.2227     0   1   2   3   4E+4   7   7.5  RT (min)       lcorbi_hilic_neg_4_085_59    m/z 393.2316     0   1   2   3   4E+3   0.5   1  RT (min)      lcorbi_hilic_neg_4_078_58    m/z 393.3011     0   1E+4   1   1.5  RT (min)      lcorbi_hilic_neg_4_078_58    m/z 393.7767     0   2   4   6E+3   8   8.5  RT (min)      lcorbi_hilic_neg_4_108_62    m/z 393.8969     0   2   4   6E+3   8   8.5  RT (min)      lcorbi_hilic_neg_4_079_58    m/z 393.9699     0   1   2   3   4E+3   5.5   6  RT (min)      lcorbi_hilic_neg_4_068_57    m/z 393.9804     0   1   7   7.5  RT (min)      lcorbi_hilic_neg_4_097_60    m/z 394.0032     0   1E+4   6   6.5  RT (min)      lcorbi_hilic_neg_4_100_61    m/z 394.0296     0   1   2   3E+4   7.5   8  RT (min)      lcorbi_hilic_neg_4_097_60    m/z 394.0418     0   1E+4   4   4.5  RT (min)      lcorbi_hilic_neg_4_103_61    m/z 394.0427     0   1   2E+4   3   3.5  RT (min)      lcorbi_hilic_neg_4_066_57    m/z 394.1023     0   1E+4   3.5   4  RT (min)      lcorbi_hilic_neg_4_108_62    m/z 394.1105     0   1   4   4.5  RT (min)      lcorbi_hilic_neg_4_095_60    m/z 394.1191     0   1   2   3   4E+4   6   6.5  RT (min)      lcorbi_hilic_neg_3_029_25    m/z 394.1275     0   2   4   6   8E+3   1   1.5  RT (min)      lcorbi_hilic_neg_4_106_62    m/z 394.1327     0   1E+5   4.5   5  RT (min)       lcorbi_hilic_neg_4_095_60    m/z 394.1527     0   2   4E+4   2   2.5  RT (min)       lcorbi_hilic_neg_4_101_61    m/z 394.1833     0   1E+4   5   5.5  RT (min)      lcorbi_hilic_neg_4_078_58    m/z 394.2531     0   1E+4   1   1.5  RT (min)      lcorbi_hilic_neg_4_084_59    m/z 394.2680     0   1   2E+4   0.5   1  RT (min)      lcorbi_hilic_neg_4_105_61    m/z 394.8462     0   1E+4   7   7.5  RT (min)      lcorbi_hilic_neg_4_003_08    m/z 394.8937     0   2   4   6E+3   8   8.5  RT (min)      lcorbi_hilic_neg_4_074_58    m/z 395.0268     0   1E+4   5.5   6  RT (min)      lcorbi_hilic_neg_4_066_57    m/z 395.0356     0   1   2   3   4E+4   1   1.5  RT (min)       lcorbi_hilic_neg_4_107_62    m/z 395.0464     0   1   2E+3   3   3.5  RT (min)      lcorbi_hilic_neg_4_090_60    m/z 395.0539     0   1   2E+3   3.5   4  RT (min)      lcorbi_hilic_neg_4_101_61    m/z 395.0639     0   1   2   2.5  RT (min)      lcorbi_hilic_neg_4_099_61    m/z 395.0662     0   1E+3   6   6.5  RT (min)      lcorbi_hilic_neg_2_005_24    m/z 395.0741     0   2   4   6E+4   7   7.5   8  RT (min)       lcorbi_hilic_neg_4_091_60    m/z 395.0847     0   2   4E+3   7.5   8  RT (min)      lcorbi_hilic_neg_4_093_60    m/z 395.0962     0   1   2E+4   5   5.5  RT (min)      lcorbi_hilic_neg_1_015_30    m/z 395.1041     0   1   2   2.5  RT (min)      lcorbi_hilic_neg_4_103_61    m/z 395.1042     0   1E+4   6.5   7  RT (min)      lcorbi_hilic_neg_2_005_24    m/z 395.1253     0   1E+6   6   6.5  RT (min)      lcorbi_hilic_neg_3_029_25    m/z 395.1626     0   1   2   3   4E+4   5.5   6  RT (min)      lcorbi_hilic_neg_3_037_12    m/z 395.1641     0   1E+3   0.5   1  RT (min)      lcorbi_hilic_neg_1_024_11    m/z 395.1899     0   2   4   6E+3   3.5   4  RT (min)      lcorbi_hilic_neg_4_079_58    m/z 395.2028     0   1   2   3   4E+3   0.5   1  RT (min)       lcorbi_hilic_neg_1_007_10    m/z 395.2205     0   1   0.5   1  RT (min)      lcorbi_hilic_neg_1_007_10    m/z 395.2339     0   1   2   3E+4   7   7.5  RT (min)       lcorbi_hilic_neg_4_090_60    m/z 395.2470     0   1E+3   0.5   1  RT (min)       lcorbi_hilic_neg_1_024_11    m/z 395.3107     0   1   2E+4   1   1.5  RT (min)      lcorbi_hilic_neg_4_087_59    m/z 395.3530     0   1   2E+4   0.5   1  RT (min)      lcorbi_hilic_neg_4_077_58    m/z 395.8107     0   2   4   6E+3   5   5.5  RT (min)      lcorbi_hilic_neg_2_020_27    m/z 395.9454     0   1   2   3   4E+3   6   6.5  RT (min)      lcorbi_hilic_neg_4_067_57    m/z 395.9817     0   2   4   6   8E+4   7.5   8  RT (min)       lcorbi_hilic_neg_2_055_32    m/z 395.9819     0   1E+4   5.5   6  RT (min)      lcorbi_hilic_neg_4_071_57    m/z 396.0326     0   2   4   6E+3   5.5   6  RT (min)      lcorbi_hilic_neg_4_072_57    m/z 396.0348     0   1E+4   6.5   7  RT (min)      lcorbi_hilic_neg_4_082_59    m/z 396.0402     0   1E+4   3   3.5  RT (min)      lcorbi_hilic_neg_4_066_57    m/z 396.0437     0   1E+5   0.5   1  RT (min)       lcorbi_hilic_neg_4_074_58    m/z 396.0484     0   1   2E+4   6.5   7  RT (min)       lcorbi_hilic_neg_4_082_59    m/z 396.1154     0   1   5.5   6  RT (min)      lcorbi_hilic_neg_4_096_60    m/z 396.1474     0   1   2E+3   1.5   2  RT (min)      lcorbi_hilic_neg_4_101_61    m/z 396.1934     0   1E+4   1   1.5  RT (min)      lcorbi_hilic_neg_1_026_34    m/z 396.2518     0   1   2   3E+4   4.5   5  RT (min)       lcorbi_hilic_neg_4_085_59    m/z 396.3121     0   1   2E+3   1   1.5  RT (min)      lcorbi_hilic_neg_4_074_58    m/z 396.7973     0   2   4   6E+3   7.5   8  RT (min)      lcorbi_hilic_neg_4_112_62    m/z 396.8256     0   2   4   6E+3   5   5.5  RT (min)      lcorbi_hilic_neg_4_076_58    m/z 396.8912     0   2   4E+3   7.5   8  RT (min)      lcorbi_hilic_neg_4_098_61    m/z 396.9912     0   1   2E+4   0.5   1  RT (min)      lcorbi_hilic_neg_4_066_57    m/z 396.9993     0   2   4   6   8E+3   7   7.5  RT (min)      lcorbi_hilic_neg_4_090_60    m/z 397.0164     0   2   4   6   8E+3   5   5.5  RT (min)      lcorbi_hilic_neg_4_070_57    m/z 397.0610     0   1   2E+4   7   7.5  RT (min)       lcorbi_hilic_neg_4_090_60    m/z 397.0714     0   1E+3   1   1.5  RT (min)      lcorbi_hilic_neg_4_087_59    m/z 397.0946     0   1   6   6.5  RT (min)      lcorbi_hilic_neg_3_029_25    m/z 397.0985     0   1E+4   2   2.5  RT (min)      lcorbi_hilic_neg_4_108_62    m/z 397.1084     0   1E+4   0   0.5  RT (min)      lcorbi_hilic_neg_4_069_57    m/z 397.1196     0   1   4   4.5  RT (min)       lcorbi_hilic_neg_4_110_62    m/z 397.1242     0   1   2E+4   1   1.5  RT (min)      lcorbi_hilic_neg_4_107_62    m/z 397.1415     0   1   2E+3   5.5   6  RT (min)      lcorbi_hilic_neg_4_095_60    m/z 397.1499     0   1   2   3E+4   5.5   6  RT (min)       lcorbi_hilic_neg_4_095_60    m/z 397.1553     0   1   2   3   4E+3   1   1.5  RT (min)      lcorbi_hilic_neg_4_044_01    m/z 397.1614     0   1   2   3E+4   6.5   7  RT (min)       lcorbi_hilic_neg_4_097_60    m/z 397.1692     0   1E+4   1   1.5  RT (min)      lcorbi_hilic_neg_4_077_58    m/z 397.1789     0   1   2E+4   1   1.5  RT (min)       lcorbi_hilic_neg_4_079_58    m/z 397.1955     0   2   4   6   8E+3   6.5   7  RT (min)      lcorbi_hilic_neg_4_095_60    m/z 397.2119     0   1   2E+4   6.5   7  RT (min)       lcorbi_hilic_neg_4_096_60    m/z 397.2204     0   2   4   6E+4   5.5   6  RT (min)       lcorbi_hilic_neg_4_081_58    m/z 397.2555     0   2   4   6E+3   4.5   5  RT (min)      lcorbi_hilic_neg_4_085_59    m/z 397.2748     0   1   2E+4   0.5   1  RT (min)       lcorbi_hilic_neg_4_090_60    m/z 397.3321     0   1   2   3   4E+3   0.5   1  RT (min)      lcorbi_hilic_neg_4_093_60    m/z 397.9790     0   2   4E+3   5.5   6  RT (min)      lcorbi_hilic_neg_4_072_57    m/z 398.0171     0   1   2   3   4E+4   6.5   7  RT (min)       lcorbi_hilic_neg_4_089_59    m/z 398.0242     0   1   2   3   4E+3   5   5.5  RT (min)      lcorbi_hilic_neg_4_103_61    m/z 398.0322     0   1   2   3E+3   6.5   7  RT (min)      lcorbi_hilic_neg_4_085_59    m/z 398.0408     0   2   4E+4   0.5   1  RT (min)      lcorbi_hilic_neg_4_074_58    m/z 398.0477     0   2   4   6E+3   6.5   7  RT (min)      lcorbi_hilic_neg_4_091_60    m/z 398.0838     0   1E+4   1   1.5  RT (min)      lcorbi_hilic_neg_4_089_59    m/z 398.0965     0   1E+4   0.5   1  RT (min)      lcorbi_hilic_neg_3_037_12    m/z 398.0971     0   1E+4   7.5   8  RT (min)      lcorbi_hilic_neg_3_042_05    m/z 398.1048     0   1E+3   6   6.5  RT (min)      lcorbi_hilic_neg_3_042_05    m/z 398.1142     0   1   2E+4   4   4.5  RT (min)      lcorbi_hilic_neg_4_013_46    m/z 398.1149     0   1   2   3   4E+3   6   6.5  RT (min)      lcorbi_hilic_neg_3_042_05    m/z 398.1238     0   2   4E+3   1   1.5  RT (min)      lcorbi_hilic_neg_4_108_62    m/z 398.1347     0   1   2   3   4E+5   6   6.5  RT (min)      lcorbi_hilic_neg_3_029_25    m/z 398.1513     0   1   2E+3   6.5   7  RT (min)       lcorbi_hilic_neg_4_056_56    m/z 398.2239     0   1E+4   5.5   6  RT (min)      lcorbi_hilic_neg_4_075_58    m/z 398.2782     0   1   2   3   4E+3   0.5   1  RT (min)      lcorbi_hilic_neg_4_089_59    m/z 398.2910     0   1   2E+4   0.5   1  RT (min)       lcorbi_hilic_neg_4_089_59    m/z 398.3235     0   1   2E+4   1   1.5  RT (min)      lcorbi_hilic_neg_4_083_59    m/z 398.3720     0   1   2   3E+4   0.5   1  RT (min)      lcorbi_hilic_neg_4_080_58    m/z 398.8227     0   2   4   6E+3   5   5.5  RT (min)      lcorbi_hilic_neg_4_074_58    m/z 398.8523     0   2   4   6E+3   8   8.5  RT (min)      lcorbi_hilic_neg_4_074_58    m/z 399.0016     0   2   4   6   8E+3   7   7.5  RT (min)      lcorbi_hilic_neg_4_083_59    m/z 399.0073     0   1E+4   0.5   1  RT (min)      lcorbi_hilic_neg_4_066_57    m/z 399.0392     0   1   2E+4   3   3.5  RT (min)       lcorbi_hilic_neg_4_104_61    m/z 399.0466     0   1E+3   6.5   7  RT (min)      lcorbi_hilic_neg_4_095_60    m/z 399.0674     0   1   2E+4   6   6.5  RT (min)       lcorbi_hilic_neg_4_110_62    m/z 399.0831     0   2   4   6   8E+3   5.5   6  RT (min)      lcorbi_hilic_neg_4_071_57    m/z 399.1028     0   1   2   3E+4   3   3.5  RT (min)       lcorbi_hilic_neg_3_028_23    m/z 399.1151     0   2   4   6   8E+3   5.5   6  RT (min)      lcorbi_hilic_neg_4_097_60    m/z 399.1269     0   2   4E+4   4   4.5  RT (min)      lcorbi_hilic_neg_1_042_25    m/z 399.1454     0   1   2E+3   6   6.5  RT (min)      lcorbi_hilic_neg_2_005_24    m/z 399.1526     0   1E+4   1   1.5  RT (min)      lcorbi_hilic_neg_4_078_58    m/z 399.1848     0   1E+4   1   1.5  RT (min)      lcorbi_hilic_neg_4_074_58    m/z 399.2014     0   2   4   6   8E+3   1   1.5  RT (min)      lcorbi_hilic_neg_4_077_58    m/z 399.2425     0   1   7   7.5  RT (min)      lcorbi_hilic_neg_4_096_60    m/z 399.3228     0   1E+5   4   4.5  RT (min)       lcorbi_hilic_neg_4_072_57    m/z 399.3270     0   1E+4   1   1.5  RT (min)      lcorbi_hilic_neg_4_074_58    m/z 399.9400     0   1E+4   0.5   1  RT (min)      lcorbi_hilic_neg_4_068_57    m/z 400.0137     0   1   6.5   7  RT (min)      lcorbi_hilic_neg_2_032_25    m/z 400.0522     0   1E+4   2   2.5  RT (min)      lcorbi_hilic_neg_4_099_61    m/z 400.1308     0   1E+3   4   4.5  RT (min)      lcorbi_hilic_neg_4_107_62    m/z 400.1327     0   1   6   6.5  RT (min)      lcorbi_hilic_neg_4_089_59    m/z 400.1412     0   1   6   6.5  RT (min)       lcorbi_hilic_neg_4_089_59    m/z 400.2246     0   1   2E+4   1   1.5  RT (min)       lcorbi_hilic_neg_4_074_58    m/z 400.2344     0   1E+4   0.5   1  RT (min)      lcorbi_hilic_neg_4_106_62    m/z 400.3070     0   1E+4   1   1.5  RT (min)      lcorbi_hilic_neg_4_088_59    m/z 400.3262     0   1   2E+4   4   4.5  RT (min)       lcorbi_hilic_neg_4_072_57    m/z 400.9092     0   1E+4   5   5.5  RT (min)      lcorbi_hilic_neg_4_083_59    m/z 400.9326     0   2   4   6   8E+3   0.5   1  RT (min)      lcorbi_hilic_neg_4_068_57    m/z 400.9953     0   2   4   6E+4   5.5   6  RT (min)       lcorbi_hilic_neg_4_073_57    m/z 401.0040     0   1E+4   0.5   1  RT (min)      lcorbi_hilic_neg_4_095_60    m/z 401.0263     0   1   2   3E+3   7   7.5  RT (min)      lcorbi_hilic_neg_4_070_57    m/z 401.0280     0   1   2E+3   1   1.5  RT (min)      lcorbi_hilic_neg_4_085_59    m/z 401.0477     0   1E+4   2   2.5  RT (min)      lcorbi_hilic_neg_4_099_61    m/z 401.0642     0   1E+3   6   6.5  RT (min)      lcorbi_hilic_neg_4_069_57    m/z 401.0762     0   1E+4   5   5.5  RT (min)      lcorbi_hilic_neg_2_020_27    m/z 401.0969     0   1   2E+4   6   6.5  RT (min)       lcorbi_hilic_neg_4_069_57    m/z 401.1091     0   1E+4   2   2.5  RT (min)      lcorbi_hilic_neg_4_108_62    m/z 401.1300     0   2   4   6   8E+2   5   5.5  RT (min)       lcorbi_hilic_neg_2_009_50    m/z 401.1502     0   1E+3   4   4.5  RT (min)      lcorbi_hilic_neg_1_037_46    m/z 401.1509     0   1E+4   6   6.5  RT (min)      lcorbi_hilic_neg_4_095_60    m/z 401.1972     0   1   2   3   4E+4   1   1.5  RT (min)       lcorbi_hilic_neg_4_105_61    m/z 401.2155     0   1E+4   5.5   6  RT (min)       lcorbi_hilic_neg_4_086_59    m/z 401.8988     0   1   2E+4   1   1.5  RT (min)      lcorbi_hilic_neg_4_074_58    m/z 401.9536     0   1   2   3E+4   6   6.5  RT (min)       lcorbi_hilic_neg_4_074_58    m/z 401.9914     0   1   6.5   7  RT (min)      lcorbi_hilic_neg_4_074_58    m/z 402.0020     0   1E+4   5.5   6  RT (min)      lcorbi_hilic_neg_4_105_61    m/z 402.0066     0   2   4   6   8E+3   0.5   1  RT (min)      lcorbi_hilic_neg_4_093_60    m/z 402.0118     0   1E+4   5.5   6  RT (min)      lcorbi_hilic_neg_4_075_58    m/z 402.0657     0   1   1   1.5  RT (min)      lcorbi_hilic_neg_1_052_41    m/z 402.0862     0   1   2   3E+2   6   6.5  RT (min)       lcorbi_hilic_neg_2_005_24    m/z 402.1334     0   1E+4   5   5.5  RT (min)      lcorbi_hilic_neg_4_094_60    m/z 402.1448     0   1E+4   2   2.5  RT (min)      lcorbi_hilic_neg_4_013_46    m/z 402.1636     0   1   2   3E+4   0.5   1  RT (min)       lcorbi_hilic_neg_4_112_62    m/z 402.1993     0   1   2   3E+4   5.5   6  RT (min)       lcorbi_hilic_neg_4_089_59    m/z 402.2803     0   1   2E+4   4   4.5  RT (min)       lcorbi_hilic_neg_4_071_57    m/z 402.8049     0   1E+4   7.5   8  RT (min)      lcorbi_hilic_neg_4_101_61    m/z 402.9467     0   1E+5   1   1.5  RT (min)       lcorbi_hilic_neg_4_106_62    m/z 403.0011     0   1E+4   0.5   1  RT (min)      lcorbi_hilic_neg_4_109_62    m/z 403.0182     0   2   4   6   8E+3   0.5   1  RT (min)      lcorbi_hilic_neg_4_067_57    m/z 403.0476     0   1   2E+4   1   1.5  RT (min)      lcorbi_hilic_neg_4_098_61    m/z 403.0845     0   1   2E+3   2   2.5  RT (min)      lcorbi_hilic_neg_4_038_27    m/z 403.1040     0   1   2   3   4E+3   2   2.5  RT (min)      lcorbi_hilic_neg_3_056_56    m/z 403.1232     0   1E+3   4   4.5  RT (min)      lcorbi_hilic_neg_1_037_46    m/z 403.1309     0   1   4   4.5  RT (min)      lcorbi_hilic_neg_4_097_60    m/z 403.1353     0   2   4   6E+2   1   1.5  RT (min)      lcorbi_hilic_neg_1_052_41    m/z 403.1404     0   1   6   6.5  RT (min)      lcorbi_hilic_neg_3_023_36    m/z 403.1405     0   2   4   6E+4   4   4.5  RT (min)       lcorbi_hilic_neg_4_095_60    m/z 403.1494     0   1   2   3E+3   2   2.5  RT (min)      lcorbi_hilic_neg_4_048_37    m/z 403.1581     0   1   2   3   4E+4   0.5   1  RT (min)      lcorbi_hilic_neg_4_074_58    m/z 403.1897     0   2   4   6E+3   1   1.5  RT (min)      lcorbi_hilic_neg_3_010_18    m/z 403.2046     0   1   2E+4   2   2.5  RT (min)       lcorbi_hilic_neg_4_106_62    m/z 403.2488     0   1   2E+3   0.5   1  RT (min)      lcorbi_hilic_neg_4_097_60    m/z 403.2851     0   1E+4   1   1.5  RT (min)      lcorbi_hilic_neg_1_029_31    m/z 403.9507     0   1E+4   6   6.5  RT (min)      lcorbi_hilic_neg_4_074_58    m/z 403.9994     0   2   4E+3   5.5   6  RT (min)      lcorbi_hilic_neg_4_105_61    m/z 404.0025     0   1   2E+4   1   1.5  RT (min)      lcorbi_hilic_neg_1_025_52    m/z 404.0146     0   1   2   3E+4   6.5   7  RT (min)       lcorbi_hilic_neg_4_089_59    m/z 404.0616     0   1   5.5   6  RT (min)      lcorbi_hilic_neg_2_005_24    m/z 404.0682     0   1   2   3E+4   6   6.5  RT (min)       lcorbi_hilic_neg_4_091_60    m/z 404.0825     0   1   4.5   5  RT (min)      lcorbi_hilic_neg_4_097_60    m/z 404.1123     0   1   2   2.5  RT (min)      lcorbi_hilic_neg_4_107_62    m/z 404.1177     0   2   4E+4   7.5   8  RT (min)       lcorbi_hilic_neg_4_094_60    m/z 404.1415     0   1   2E+4   2   2.5  RT (min)      lcorbi_hilic_neg_4_005_32    m/z 404.1447     0   1   6   6.5  RT (min)      lcorbi_hilic_neg_4_073_57    m/z 404.1541     0   1E+5   6   6.5  RT (min)       lcorbi_hilic_neg_4_095_60    m/z 404.2264     0   1E+4   3.5   4  RT (min)      lcorbi_hilic_neg_3_011_24    m/z 404.7679     0   2   4E+3   4.5   5  RT (min)      lcorbi_hilic_neg_4_079_58    m/z 404.9392     0   1   1   1.5  RT (min)      lcorbi_hilic_neg_4_106_62    m/z 404.9517     0   1E+4   6   6.5  RT (min)      lcorbi_hilic_neg_4_074_58    m/z 404.9983     0   1E+4   0.5   1  RT (min)      lcorbi_hilic_neg_4_111_62    m/z 405.0015     0   2   4E+3   6   6.5  RT (min)      lcorbi_hilic_neg_4_095_60    m/z 405.0166     0   1   2E+4   3   3.5  RT (min)       lcorbi_hilic_neg_4_071_57    m/z 405.0166     0   1E+4   0.5   1  RT (min)       lcorbi_hilic_neg_4_071_57    m/z 405.0663     0   1   2E+4   6.5   7  RT (min)       lcorbi_hilic_neg_4_089_59    m/z 405.1044     0   1   1   1.5  RT (min)      lcorbi_hilic_neg_4_107_62    m/z 405.1045     0   1   5.5   6  RT (min)      lcorbi_hilic_neg_2_005_24    m/z 405.1352     0   1E+4   5.5   6  RT (min)      lcorbi_hilic_neg_4_089_59    m/z 405.1928     0   1E+4   0.5   1  RT (min)      lcorbi_hilic_neg_4_113_62    m/z 405.2278     0   1E+4   3.5   4  RT (min)      lcorbi_hilic_neg_4_082_59    m/z 405.2624     0   1E+5   7   7.5  RT (min)       lcorbi_hilic_neg_4_096_60    m/z 405.2892     0   1   2   3   4E+4   3.5   4  RT (min)       lcorbi_hilic_neg_4_089_59    m/z 405.3008     0   1E+4   1   1.5  RT (min)      lcorbi_hilic_neg_1_029_31    m/z 405.8228     0   2   4E+3   1   1.5  RT (min)      lcorbi_hilic_neg_4_066_57    m/z 405.9598     0   2   4   6E+3   7.5   8  RT (min)      lcorbi_hilic_neg_4_080_58    m/z 405.9925     0   2   4   6   8E+3   4.5   5  RT (min)      lcorbi_hilic_neg_4_080_58    m/z 406.0023     0   1   2   3   4E+4   6.5   7  RT (min)      lcorbi_hilic_neg_3_039_31    m/z 406.0109     0   1   2   3E+4   5.5   6  RT (min)       lcorbi_hilic_neg_4_072_57    m/z 406.0226     0   1E+3   5.5   6  RT (min)      lcorbi_hilic_neg_4_076_58    m/z 406.0710     0   1E+4   6.5   7  RT (min)      lcorbi_hilic_neg_4_091_60    m/z 406.0795     0   1   4.5   5  RT (min)      lcorbi_hilic_neg_4_096_60    m/z 406.0799     0   1E+4   6.5   7  RT (min)      lcorbi_hilic_neg_1_012_32    m/z 406.0974     0   2   4E+3   1   1.5  RT (min)      lcorbi_hilic_neg_4_110_62    m/z 406.1112     0   2   4   6   8E+3   3   3.5  RT (min)      lcorbi_hilic_neg_2_031_34    m/z 406.1327     0   1   2E+4   2   2.5  RT (min)       lcorbi_hilic_neg_4_111_62    m/z 406.1410     0   1   2   3E+5   4   4.5  RT (min)       lcorbi_hilic_neg_4_090_60    m/z 406.1544     0   1E+4   1   1.5  RT (min)      lcorbi_hilic_neg_4_071_57    m/z 406.1826     0   1E+5   1   1.5  RT (min)      lcorbi_hilic_neg_1_003_14    m/z 406.2776     0   2   4E+3   0.5   1  RT (min)      lcorbi_hilic_neg_4_108_62    m/z 406.2925     0   2   4   6   8E+3   3.5   4  RT (min)      lcorbi_hilic_neg_4_088_59    m/z 406.6311     0   2   4   6   8E+4   4.5   5  RT (min)      lcorbi_hilic_neg_2_020_27    m/z 406.8544     0   1E+4   5   5.5  RT (min)      lcorbi_hilic_neg_4_094_60    m/z 406.9627     0   1   1   1.5  RT (min)      lcorbi_hilic_neg_4_083_59    m/z 406.9950     0   1   2   3E+4   0.5   1  RT (min)      lcorbi_hilic_neg_4_097_60    m/z 407.0454     0   1E+4   5   5.5  RT (min)      lcorbi_hilic_neg_4_066_57    m/z 407.0689     0   2   4E+3   1   1.5  RT (min)      lcorbi_hilic_neg_4_038_27    m/z 407.0920     0   1   2   3E+4   6   6.5  RT (min)       lcorbi_hilic_neg_4_090_60    m/z 407.1478     0   1   2E+4   1   1.5  RT (min)      lcorbi_hilic_neg_4_068_57    m/z 407.1573     0   2   4   6E+3   5.5   6  RT (min)       lcorbi_hilic_neg_4_092_60    m/z 407.1823     0   1E+5   6   6.5  RT (min)      lcorbi_hilic_neg_1_012_32    m/z 407.1861     0   1E+3   0.5   1  RT (min)       lcorbi_hilic_neg_1_030_04    m/z 407.2105     0   2   4E+2   0.5   1  RT (min)      lcorbi_hilic_neg_1_006_08    m/z 407.2206     0   2   4   6   8E+4   3.5   4  RT (min)      lcorbi_hilic_neg_1_047_05    m/z 407.2808     0   1E+5   1.5   2  RT (min)       lcorbi_hilic_neg_4_014_18    m/z 407.2861     0   1E+4   3.5   4  RT (min)      lcorbi_hilic_neg_4_089_59    m/z 407.3530     0   1   2   3E+4   0.5   1  RT (min)      lcorbi_hilic_neg_4_076_58    m/z 407.4337     0   1   2E+5   1   1.5  RT (min)       lcorbi_hilic_neg_1_047_05    m/z 407.9900     0   1   2E+4   4.5   5  RT (min)       lcorbi_hilic_neg_4_077_58    m/z 407.9982     0   1E+4   1   1.5  RT (min)      lcorbi_hilic_neg_4_105_61    m/z 407.9991     0   2   4   6   8E+3   6.5   7  RT (min)      lcorbi_hilic_neg_4_083_59    m/z 408.0460     0   1   2   3   4E+4   6.5   7  RT (min)       lcorbi_hilic_neg_4_088_59    m/z 408.0530     0   1E+4   5   5.5  RT (min)      lcorbi_hilic_neg_4_108_62    m/z 408.0612     0   1E+4   6.5   7  RT (min)      lcorbi_hilic_neg_4_071_57    m/z 408.1092     0   1   2   3E+3   7   7.5  RT (min)      lcorbi_hilic_neg_4_089_59    m/z 408.1426     0   2   4   6E+4   4.5   5  RT (min)      lcorbi_hilic_neg_3_027_45    m/z 408.1673     0   2   4   6E+4   6   6.5  RT (min)      lcorbi_hilic_neg_2_055_32    m/z 408.1723     0   1   2E+4   0.5   1  RT (min)      lcorbi_hilic_neg_1_030_04    m/z 408.1798     0   1   2   3E+4   1   1.5  RT (min)      lcorbi_hilic_neg_4_075_58    m/z 408.2029     0   2   4   6E+3   1   1.5  RT (min)      lcorbi_hilic_neg_4_107_62    m/z 408.2139     0   1   2E+4   0.5   1  RT (min)      lcorbi_hilic_neg_4_076_58    m/z 408.2840     0   1   2E+4   1.5   2  RT (min)      lcorbi_hilic_neg_3_010_18    m/z 408.6284     0   2   4   6   8E+4   4.5   5  RT (min)       lcorbi_hilic_neg_2_020_27    m/z 408.8518     0   2   4E+3   5   5.5  RT (min)      lcorbi_hilic_neg_4_090_60    m/z 408.9341     0   2   4   6E+3   8   8.5  RT (min)      lcorbi_hilic_neg_4_070_57    m/z 408.9823     0   1   7.5   8  RT (min)      lcorbi_hilic_neg_4_082_59    m/z 408.9887     0   2   4   6   8E+3   4.5   5  RT (min)      lcorbi_hilic_neg_4_096_60    m/z 408.9918     0   1   2E+4   0.5   1  RT (min)      lcorbi_hilic_neg_4_076_58    m/z 409.0112     0   2   4   6   8E+4   0.5   1  RT (min)       lcorbi_hilic_neg_4_079_58    m/z 409.0235     0   1   2   3E+4   3   3.5  RT (min)       lcorbi_hilic_neg_4_105_61    m/z 409.0322     0   1   2   3E+4   4   4.5  RT (min)       lcorbi_hilic_neg_4_102_61    m/z 409.0964     0   1   2   3   4E+4   6   6.5  RT (min)       lcorbi_hilic_neg_4_112_62    m/z 409.0991     0   1   2E+4   1   2  RT (min)       lcorbi_hilic_neg_4_085_59    m/z 409.1133     0   1   2E+4   3.5   4  RT (min)       lcorbi_hilic_neg_4_107_62    m/z 409.1967     0   1E+4   1   1.5  RT (min)      lcorbi_hilic_neg_4_107_62    m/z 409.2097     0   1E+4   1   1.5  RT (min)      lcorbi_hilic_neg_4_088_59    m/z 409.2264     0   1   2E+4   0.5   1  RT (min)      lcorbi_hilic_neg_3_016_16    m/z 409.2729     0   2   4E+3   1   1.5  RT (min)      lcorbi_hilic_neg_2_006_45    m/z 409.3328     0   2   4   6E+4   0.5   1  RT (min)       lcorbi_hilic_neg_4_091_60    m/z 409.9870     0   1   2   3   4E+4   4.5   5  RT (min)       lcorbi_hilic_neg_4_080_58    m/z 409.9957     0   2   4   6   8E+3   7.5   8  RT (min)      lcorbi_hilic_neg_4_089_59    m/z 410.0118     0   1E+4   6.5   7  RT (min)      lcorbi_hilic_neg_4_085_59    m/z 410.0324     0   1E+4   6   6.5  RT (min)      lcorbi_hilic_neg_4_075_58    m/z 410.0501     0   1   2   3E+3   5   5.5  RT (min)      lcorbi_hilic_neg_4_107_62    m/z 410.0666     0   2   4   6   8E+3   5   5.5  RT (min)      lcorbi_hilic_neg_4_066_57    m/z 410.0996     0   1   2   3E+4   6   6.5   7  RT (min)       lcorbi_hilic_neg_4_096_60    m/z 410.1062     0   1   3   3.5  RT (min)      lcorbi_hilic_neg_4_101_61    m/z 410.1283     0   2   4   6   8E+3   1   1.5  RT (min)      lcorbi_hilic_neg_4_107_62    m/z 410.1383     0   1   3   3.5  RT (min)      lcorbi_hilic_neg_4_105_61    m/z 410.1564     0   1E+4   4.5   5  RT (min)      lcorbi_hilic_neg_4_107_62    m/z 410.1642     0   1E+4   0.5   1  RT (min)      lcorbi_hilic_neg_4_081_58    m/z 410.2216     0   1   2   3E+4   0.5   1  RT (min)      lcorbi_hilic_neg_4_106_62    m/z 410.2395     0   1   2   3   4E+4   4   4.5  RT (min)      lcorbi_hilic_neg_1_047_05    m/z 410.2664     0   1E+4   0.5   1  RT (min)      lcorbi_hilic_neg_4_089_59    m/z 410.3149     0   1   2E+3   0.5   1  RT (min)      lcorbi_hilic_neg_4_089_59    m/z 410.3721     0   1   2E+4   0.5   1  RT (min)      lcorbi_hilic_neg_4_076_58    m/z 410.7329     0   2   4   6   8E+3   4.5   5  RT (min)      lcorbi_hilic_neg_4_101_61    m/z 410.9466     0   2   4   6E+3   6   6.5  RT (min)      lcorbi_hilic_neg_4_070_57    m/z 410.9854     0   1   2   3   4E+3   4.5   5  RT (min)      lcorbi_hilic_neg_4_090_60    m/z 411.0071     0   1E+4   0.5   1  RT (min)       lcorbi_hilic_neg_4_107_62    m/z 411.0207     0   1E+4   3   3.5  RT (min)       lcorbi_hilic_neg_4_105_61    m/z 411.0291     0   1   2E+4   4   4.5  RT (min)       lcorbi_hilic_neg_4_102_61    m/z 411.1047     0   2   4   6   8E+3   6   6.5  RT (min)      lcorbi_hilic_neg_4_087_59    m/z 411.1103     0   2   4   6E+3   4   4.5  RT (min)      lcorbi_hilic_neg_4_113_62    m/z 411.1247     0   1   2E+4   6  RT (min)       lcorbi_hilic_neg_4_088_59    m/z 411.1590     0   1E+3   0.5   1  RT (min)      lcorbi_hilic_neg_4_068_57    m/z 411.1621     0   1   4   4.5  RT (min)      lcorbi_hilic_neg_4_082_59    m/z 411.1774     0   1   2   3E+4   6   6.5  RT (min)      lcorbi_hilic_neg_1_047_05    m/z 411.1848     0   1E+4   1   1.5  RT (min)      lcorbi_hilic_neg_4_076_58    m/z 411.2034     0   1   2   3   4E+3   2   2.5  RT (min)      lcorbi_hilic_neg_4_074_58    m/z 411.2257     0   2   4E+3   0.5   1  RT (min)      lcorbi_hilic_neg_4_066_57    m/z 411.2420     0   1   2E+5   0.5   1  RT (min)       lcorbi_hilic_neg_4_073_57    m/z 411.2522     0   2   4E+4   1   1.5  RT (min)       lcorbi_hilic_neg_4_088_59    m/z 411.2623     0   2   4   6   8E+3   4   4.5  RT (min)      lcorbi_hilic_neg_4_075_58    m/z 411.2979     0   1E+4   3.5   4  RT (min)      lcorbi_hilic_neg_4_073_57    m/z 411.3298     0   1   2E+4   0.5   1  RT (min)      lcorbi_hilic_neg_4_091_60    m/z 411.3479     0   1   2E+4   0.5   1  RT (min)      lcorbi_hilic_neg_4_080_58    m/z 411.8592     0   2   4   6E+3   8   8.5  RT (min)      lcorbi_hilic_neg_4_098_61    m/z 411.9360     0   1E+4   5   5.5  RT (min)      lcorbi_hilic_neg_4_104_61    m/z 411.9824     0   1   2E+4   6   6.5  RT (min)       lcorbi_hilic_neg_4_074_58    m/z 411.9840     0   1   2E+4   4.5   5  RT (min)       lcorbi_hilic_neg_4_077_58    m/z 411.9894     0   2   4   6   8E+3   1   1.5  RT (min)      lcorbi_hilic_neg_4_100_61    m/z 412.0223     0   1   5.5   6  RT (min)      lcorbi_hilic_neg_4_113_62    m/z 412.1127     0   2   4E+3   6   6.5  RT (min)      lcorbi_hilic_neg_1_047_05    m/z 412.1380     0   1   2E+3   5   5.5  RT (min)       lcorbi_hilic_neg_4_097_60    m/z 412.1688     0   1   2   3E+4   6   7  RT (min)       lcorbi_hilic_neg_4_090_60    m/z 412.1738     0   1E+4   2   2.5  RT (min)      lcorbi_hilic_neg_4_013_46    m/z 412.1981     0   1   2   3   4E+4   1   1.5  RT (min)       lcorbi_hilic_neg_4_101_61    m/z 412.2185     0   2   4E+3   0.5   1  RT (min)      lcorbi_hilic_neg_4_073_57    m/z 412.3877     0   1   2   3   4E+4   0.5   1  RT (min)      lcorbi_hilic_neg_4_080_58    m/z 412.7994     0   2   4   6   8E+3   4.5   5  RT (min)      lcorbi_hilic_neg_4_080_58    m/z 412.8592     0   2   4E+3   8   8.5  RT (min)      lcorbi_hilic_neg_4_066_57    m/z 412.8696     0   1   2   3E+3   7   7.5  RT (min)      lcorbi_hilic_neg_4_089_59    m/z 412.9669     0   1E+3   0.5   1  RT (min)      lcorbi_hilic_neg_4_086_59    m/z 413.0225     0   1E+4   7   7.5  RT (min)      lcorbi_hilic_neg_4_084_59    m/z 413.0644     0   1E+4   4   4.5  RT (min)      lcorbi_hilic_neg_4_099_61    m/z 413.1070     0   1E+4   3   3.5  RT (min)       lcorbi_hilic_neg_4_105_61    m/z 413.1643     0   1   2   3   4E+3   1   1.5  RT (min)      lcorbi_hilic_neg_4_077_58    m/z 413.1894     0   1E+5   6   6.5  RT (min)       lcorbi_hilic_neg_4_095_60    m/z 413.2184     0   1   2   3E+4   1   1.5  RT (min)       lcorbi_hilic_neg_4_101_61    m/z 413.2491     0   1   1   1.5  RT (min)      lcorbi_hilic_neg_4_074_58    m/z 413.3138     0   1   2   3E+4   1   1.5  RT (min)      lcorbi_hilic_neg_1_048_35    m/z 413.3637     0   1E+4   0.5   1  RT (min)      lcorbi_hilic_neg_4_076_58    m/z 413.9805     0   2   4   6E+3   4.5   5  RT (min)      lcorbi_hilic_neg_4_080_58    m/z 414.0312     0   2   4   6E+3   1   1.5  RT (min)      lcorbi_hilic_neg_3_033_53    m/z 414.0441     0   2   4E+4   6.5   7  RT (min)       lcorbi_hilic_neg_4_090_60    m/z 414.0798     0   2   4   6E+4   6   6.5  RT (min)       lcorbi_hilic_neg_4_089_59    m/z 414.1250     0   1E+4   3   3.5  RT (min)      lcorbi_hilic_neg_4_101_61    m/z 414.2779     0   1   2E+4   0.5   1  RT (min)      lcorbi_hilic_neg_4_087_59    m/z 414.2898     0   1E+4   1   1.5  RT (min)      lcorbi_hilic_neg_2_006_45    m/z 414.3176     0   2   4   6   8E+3   1   1.5  RT (min)      lcorbi_hilic_neg_1_039_38    m/z 414.7607     0   1E+4   7.5   8  RT (min)      lcorbi_hilic_neg_4_105_61    m/z 414.7967     0   1   4.5   5  RT (min)      lcorbi_hilic_neg_4_098_61    m/z 414.8041     0   1   2   3E+3   7.5   8  RT (min)      lcorbi_hilic_neg_4_105_61    m/z 414.8904     0   1E+4   8.5   9  RT (min)      lcorbi_hilic_neg_4_069_57    m/z 414.9472     0   2   4E+3   5.5   6  RT (min)      lcorbi_hilic_neg_4_070_57    m/z 414.9654     0   1E+3   1   1.5  RT (min)      lcorbi_hilic_neg_4_084_59    m/z 414.9783     0   1E+4   7.5   8  RT (min)       lcorbi_hilic_neg_4_078_58    m/z 414.9808     0   1E+4   6   6.5  RT (min)      lcorbi_hilic_neg_4_075_58    m/z 415.0013     0   1E+4   0.5   1  RT (min)      lcorbi_hilic_neg_4_110_62    m/z 415.0403     0   1E+3   6   6.5  RT (min)      lcorbi_hilic_neg_4_078_58    m/z 415.1056     0   1E+5   5.5   6  RT (min)       lcorbi_hilic_neg_4_089_59    m/z 415.1794     0   1   2E+4   0.5   1  RT (min)      lcorbi_hilic_neg_4_080_58    m/z 415.1964     0   2   4   6   8E+3   1   1.5  RT (min)      lcorbi_hilic_neg_4_074_58    m/z 415.2063     0   2   4   6   8E+3   1   1.5  RT (min)      lcorbi_hilic_neg_4_078_58    m/z 415.2165     0   1E+4   1.5   2  RT (min)       lcorbi_hilic_neg_4_074_58    m/z 415.2258     0   1   1.5   2  RT (min)      lcorbi_hilic_neg_4_108_62    m/z 415.3295     0   2   4   6E+4   1   1.5  RT (min)      lcorbi_hilic_neg_1_029_31    m/z 415.8018     0   2   4   6E+3   8   8.5  RT (min)      lcorbi_hilic_neg_4_066_57    m/z 415.8463     0   2   4E+3   7   7.5  RT (min)      lcorbi_hilic_neg_1_023_23    m/z 415.9761     0   2   4   6   8E+3   7.5   8  RT (min)      lcorbi_hilic_neg_4_080_58    m/z 415.9954     0   2   4   6   8E+3   5.5   6  RT (min)      lcorbi_hilic_neg_4_089_59    m/z 416.0158     0   2   4E+4   7.5   8  RT (min)       lcorbi_hilic_neg_4_097_60    m/z 416.0292     0   1   2   3   4E+3   1   1.5  RT (min)      lcorbi_hilic_neg_4_025_52    m/z 416.0362     0   1E+3   6.5   7  RT (min)      lcorbi_hilic_neg_4_092_60    m/z 416.0451     0   1E+4   6.5   7  RT (min)      lcorbi_hilic_neg_4_097_60    m/z 416.0746     0   2   4   6   8E+4   0.5   1  RT (min)       lcorbi_hilic_neg_4_113_62    m/z 416.1112     0   1E+4   5.5   6  RT (min)      lcorbi_hilic_neg_4_089_59    m/z 416.1448     0   1E+4   6.5   7  RT (min)      lcorbi_hilic_neg_4_095_60    m/z 416.1836     0   1   6   6.5  RT (min)      lcorbi_hilic_neg_4_091_60    m/z 416.1858     0   1E+4   2   2.5  RT (min)      lcorbi_hilic_neg_4_112_62    m/z 416.1940     0   2   4   6   8E+4   6   6.5  RT (min)       lcorbi_hilic_neg_4_094_60    m/z 416.2196     0   2   4   6E+3   1   1.5  RT (min)      lcorbi_hilic_neg_4_079_58    m/z 416.2869     0   2   4E+3   1   1.5  RT (min)      lcorbi_hilic_neg_2_006_45    m/z 416.3214     0   1   2   3   4E+4   3.5   4  RT (min)       lcorbi_hilic_neg_4_088_59    m/z 416.7996     0   2   4   6E+3   8   8.5  RT (min)      lcorbi_hilic_neg_4_066_57    m/z 416.8832     0   1E+4   5   5.5  RT (min)      lcorbi_hilic_neg_4_090_60    m/z 416.9862     0   2   4E+4   1.5   2  RT (min)      lcorbi_hilic_neg_4_054_30    m/z 417.0038     0   1   2E+4   6   6.5  RT (min)       lcorbi_hilic_neg_4_071_57    m/z 417.0116     0   1   2E+3   7.5   8  RT (min)      lcorbi_hilic_neg_4_074_58    m/z 417.0167     0   1E+4   0.5   1  RT (min)      lcorbi_hilic_neg_4_101_61    m/z 417.0659     0   1E+4   7.5   8  RT (min)      lcorbi_hilic_neg_4_089_59    m/z 417.0739     0   2   4   6   8E+3   5.5   6  RT (min)      lcorbi_hilic_neg_4_071_57    m/z 417.1207     0   1   6.5   7  RT (min)      lcorbi_hilic_neg_4_095_60    m/z 417.1360     0   1E+5   6   6.5  RT (min)       lcorbi_hilic_neg_4_089_59    m/z 417.1447     0   1E+4   5.5   6  RT (min)      lcorbi_hilic_neg_3_037_12    m/z 417.1742     0   1   2E+3   6.5   7  RT (min)      lcorbi_hilic_neg_4_097_60    m/z 417.1964     0   1   4   4.5  RT (min)      lcorbi_hilic_neg_2_045_26    m/z 417.2109     0   1   2   3E+4   1   1.5  RT (min)       lcorbi_hilic_neg_4_095_60    m/z 417.2132     0   2   4   6   8E+3   3   3.5  RT (min)      lcorbi_hilic_neg_4_072_57    m/z 417.2414     0   1   1.5   2  RT (min)      lcorbi_hilic_neg_1_018_21    m/z 417.3012     0   1   2   3E+4   0.5   1  RT (min)      lcorbi_hilic_neg_4_081_58    m/z 417.3124     0   1E+5   1   1.5  RT (min)      lcorbi_hilic_neg_1_002_26    m/z 417.8730     0   1   2   3E+4   1   1.5  RT (min)       lcorbi_hilic_neg_4_074_58    m/z 417.9944     0   1   2   3   4E+4   1   1.5  RT (min)      lcorbi_hilic_neg_4_098_61    m/z 418.0021     0   1E+4   6.5   7  RT (min)      lcorbi_hilic_neg_3_039_31    m/z 418.0055     0   1E+4   6.5   7  RT (min)      lcorbi_hilic_neg_4_084_59    m/z 418.0284     0   1E+4   6.5   7  RT (min)      lcorbi_hilic_neg_4_086_59    m/z 418.0716     0   2   4   6   8E+4   0.5   1  RT (min)       lcorbi_hilic_neg_4_113_62    m/z 418.0819     0   1E+4   5   5.5  RT (min)       lcorbi_hilic_neg_4_108_62    m/z 418.0959     0   1E+4   2   2.5  RT (min)      lcorbi_hilic_neg_4_106_62    m/z 418.1124     0   1   6   6.5  RT (min)      lcorbi_hilic_neg_4_089_59    m/z 418.2086     0   1   2E+4   0.5   1  RT (min)      lcorbi_hilic_neg_4_098_61    m/z 418.2320     0   1E+4   1   1.5  RT (min)      lcorbi_hilic_neg_4_108_62    m/z 418.7446     0   2   4   6E+3   4.5   5  RT (min)      lcorbi_hilic_neg_4_066_57    m/z 418.8947     0   2   4   6E+3   12   12.5  RT (min)      lcorbi_hilic_neg_4_089_59    m/z 418.8950     0   2   4   6E+4   8.5   9  RT (min)       lcorbi_hilic_neg_4_077_58    m/z 418.9770     0   1   2E+3   6.5   7  RT (min)      lcorbi_hilic_neg_4_066_57    m/z 418.9841     0   2   4   6   8E+4   1.5   2  RT (min)      lcorbi_hilic_neg_4_054_30    m/z 418.9949     0   1E+4   0.5   1  RT (min)      lcorbi_hilic_neg_4_105_61    m/z 419.0425     0   2   4   6   8E+3   7   7.5  RT (min)      lcorbi_hilic_neg_4_090_60    m/z 419.0446     0   2   4   6   8E+3   6   6.5  RT (min)      lcorbi_hilic_neg_1_023_23    m/z 419.0507     0   2   4   6   8E+4   0.5   1  RT (min)       lcorbi_hilic_neg_4_113_62    m/z 419.0520     0   1E+4   3   3.5  RT (min)      lcorbi_hilic_neg_4_103_61    m/z 419.0612     0   1E+4   4   4.5  RT (min)      lcorbi_hilic_neg_4_098_61    m/z 419.0926     0   1   2   3   4E+4   2   2.5  RT (min)       lcorbi_hilic_neg_4_106_62    m/z 419.1074     0   1E+5   7   7.5  RT (min)       lcorbi_hilic_neg_2_054_13    m/z 419.1328     0   1   5   5.5  RT (min)      lcorbi_hilic_neg_4_095_60    m/z 419.1532     0   1   2   3E+4   1   1.5  RT (min)      lcorbi_hilic_neg_4_076_58    m/z 419.2571     0   2   4   6E+3   1.5   2  RT (min)      lcorbi_hilic_neg_4_071_57    m/z 419.3017     0   1   1   1.5  RT (min)      lcorbi_hilic_neg_4_074_58    m/z 419.3168     0   2   4E+4   0.5   1  RT (min)       lcorbi_hilic_neg_4_075_58    m/z 419.3282     0   1E+5   1   1.5  RT (min)      lcorbi_hilic_neg_1_002_26    m/z 419.3532     0   1   0.5   1  RT (min)      lcorbi_hilic_neg_4_094_60    m/z 419.3618     0   2   4E+4   0.5   1  RT (min)       lcorbi_hilic_neg_4_091_60    m/z 419.8964     0   1E+4   8.5   9  RT (min)      lcorbi_hilic_neg_4_080_58    m/z 419.9513     0   1   2   3   4E+3   6   6.5  RT (min)      lcorbi_hilic_neg_4_103_61    m/z 420.0756     0   1   2E+4   1   1.5  RT (min)      lcorbi_hilic_neg_4_106_62    m/z 420.1110     0   1   2   3   4E+4   7   7.5  RT (min)      lcorbi_hilic_neg_4_071_57    m/z 420.1344     0   1E+4   3   3.5  RT (min)      lcorbi_hilic_neg_4_007_21    m/z 420.1397     0   2   4   6   8E+4   6   6.5  RT (min)       lcorbi_hilic_neg_4_095_60    m/z 420.1679     0   1   3   3.5  RT (min)      lcorbi_hilic_neg_4_103_61    m/z 420.1773     0   1   2E+4   3   3.5  RT (min)      lcorbi_hilic_neg_4_105_61    m/z 420.7420     0   2   4   6E+3   4.5   5  RT (min)      lcorbi_hilic_neg_4_073_57    m/z 420.7604     0   1   2   3   4E+3   8   8.5  RT (min)      lcorbi_hilic_neg_4_108_62    m/z 420.8469     0   2   4   6E+3   8.5   9  RT (min)      lcorbi_hilic_neg_4_100_61    m/z 420.8899     0   2   4   6   8E+3   8   8.5  RT (min)      lcorbi_hilic_neg_4_082_59    m/z 420.9753     0   2   4   6   8E+3   6   6.5  RT (min)      lcorbi_hilic_neg_4_072_57    m/z 421.0466     0   2   4E+3   1   1.5  RT (min)      lcorbi_hilic_neg_4_069_57    m/z 421.0691     0   1   2E+4   1   1.5  RT (min)      lcorbi_hilic_neg_4_107_62    m/z 421.0765     0   1   2   3E+4   8.5   9  RT (min)       lcorbi_hilic_neg_4_084_59    m/z 421.0890     0   1E+4   2   2.5  RT (min)      lcorbi_hilic_neg_4_109_62    m/z 421.1047     0   2   4   6E+4   7   7.5  RT (min)      lcorbi_hilic_neg_4_067_57    m/z 421.1187     0   1E+4   2   2.5  RT (min)      lcorbi_hilic_neg_4_111_62    m/z 421.1502     0   1   2   3E+4   1   1.5  RT (min)       lcorbi_hilic_neg_4_077_58    m/z 421.1877     0   1   0.5   1  RT (min)       lcorbi_hilic_neg_3_054_29    m/z 421.2540     0   1   2E+4   0.5   1  RT (min)      lcorbi_hilic_neg_4_105_61    m/z 421.2811     0   2   4E+4   1   1.5  RT (min)       lcorbi_hilic_neg_4_083_59    m/z 421.7583     0   2   4   6E+3   8   8.5  RT (min)      lcorbi_hilic_neg_4_106_62    m/z 421.8885     0   1E+4   8   8.5  RT (min)      lcorbi_hilic_neg_4_082_59    m/z 421.9649     0   1E+4   5   5.5  RT (min)       lcorbi_hilic_neg_4_105_61    m/z 421.9848     0   2   4E+3   5.5   6  RT (min)      lcorbi_hilic_neg_4_077_58    m/z 422.0793     0   1   2   3   4E+3   8   8.5  RT (min)      lcorbi_hilic_neg_4_089_59    m/z 422.0951     0   1E+4   6.5   7  RT (min)      lcorbi_hilic_neg_1_037_46    m/z 422.1442     0   1   6   6.5  RT (min)      lcorbi_hilic_neg_4_093_60    m/z 422.1538     0   1   2E+4   6   6.5  RT (min)       lcorbi_hilic_neg_4_093_60    m/z 422.1635     0   1   2   3   4E+4   6.5   7  RT (min)       lcorbi_hilic_neg_4_096_60    m/z 422.1784     0   1E+4   6.5   7  RT (min)      lcorbi_hilic_neg_4_089_59    m/z 422.2301     0   1E+5   0.5   1  RT (min)      lcorbi_hilic_neg_3_054_29    m/z 422.2846     0   1E+4   1   1.5  RT (min)      lcorbi_hilic_neg_4_082_59    m/z 422.3360     0   1E+4   1   1.5  RT (min)      lcorbi_hilic_neg_4_044_01    m/z 422.3722     0   1   2E+4   0.5   1  RT (min)      lcorbi_hilic_neg_4_074_58    m/z 422.7561     0   2   4   6E+3   8   8.5  RT (min)      lcorbi_hilic_neg_4_098_61    m/z 422.8283     0   1   4.5   5  RT (min)      lcorbi_hilic_neg_4_106_62    m/z 422.8874     0   2   4   6   8E+3   8   8.5  RT (min)      lcorbi_hilic_neg_4_066_57    m/z 422.9900     0   1E+4   0.5   1  RT (min)      lcorbi_hilic_neg_4_095_60    m/z 423.0072     0   1   2E+4   3   3.5  RT (min)       lcorbi_hilic_neg_4_080_58    m/z 423.0074     0   1   2E+4   0.5   1  RT (min)       lcorbi_hilic_neg_4_081_58    m/z 423.0197     0   1E+3   5   5.5  RT (min)      lcorbi_hilic_neg_4_102_61    m/z 423.0747     0   1   2   3E+4   0.5   1  RT (min)      lcorbi_hilic_neg_3_043_02    m/z 423.0823     0   1   7.5   8  RT (min)      lcorbi_hilic_neg_4_082_59    m/z 423.0916     0   1E+5   7.5   8  RT (min)       lcorbi_hilic_neg_4_088_59    m/z 423.1073     0   1E+4   2   2.5  RT (min)      lcorbi_hilic_neg_4_113_62    m/z 423.1433     0   1E+4   1   1.5  RT (min)      lcorbi_hilic_neg_4_066_57    m/z 423.1592     0   1E+4   3   3.5  RT (min)      lcorbi_hilic_neg_4_101_61    m/z 423.2516     0   1   0.5   1  RT (min)      lcorbi_hilic_neg_4_074_58    m/z 424.0322     0   2   4E+4   5   5.5  RT (min)       lcorbi_hilic_neg_4_066_57    m/z 424.0680     0   1   6.5   7  RT (min)      lcorbi_hilic_neg_4_091_60    m/z 424.0771     0   2   4   6   8E+4   6.5   7  RT (min)       lcorbi_hilic_neg_4_094_60    m/z 424.3069     0   1   2   3E+4   0.5   1  RT (min)       lcorbi_hilic_neg_4_089_59    m/z 424.3878     0   1E+4   0.5   1  RT (min)      lcorbi_hilic_neg_4_074_58    m/z 424.8257     0   1   4.5   5  RT (min)      lcorbi_hilic_neg_4_106_62    m/z 424.8517     0   1E+4   8.5   9  RT (min)      lcorbi_hilic_neg_4_113_62    m/z 425.0064     0   2   4   6E+3   4   4.5  RT (min)      lcorbi_hilic_neg_4_070_57    m/z 425.0192     0   1E+4   5.5   6  RT (min)      lcorbi_hilic_neg_4_079_58    m/z 425.0701     0   1   2   3E+4   6   6.5  RT (min)       lcorbi_hilic_neg_4_072_57    m/z 425.0815     0   1E+4   8   8.5  RT (min)      lcorbi_hilic_neg_4_089_59    m/z 425.1048     0   2   4   6E+3   2.5   3  RT (min)      lcorbi_hilic_neg_2_011_29    m/z 425.1172     0   1   1   1.5  RT (min)      lcorbi_hilic_neg_4_113_62    m/z 425.1238     0   1   2   3   4E+3   1   1.5  RT (min)      lcorbi_hilic_neg_4_066_57    m/z 425.1317     0   1   2E+4   6   6.5  RT (min)       lcorbi_hilic_neg_4_079_58    m/z 425.1440     0   1   2.5   3   3.5  RT (min)       lcorbi_hilic_neg_4_107_62    m/z 425.1676     0   2   4   6E+3   6   6.5  RT (min)      lcorbi_hilic_neg_4_045_25    m/z 425.2006     0   1E+4   1   1.5  RT (min)      lcorbi_hilic_neg_4_070_57    m/z 425.2160     0   1E+4   6.5   7  RT (min)      lcorbi_hilic_neg_4_077_58    m/z 425.3220     0   1   2   3   4E+3   1   1.5  RT (min)      lcorbi_hilic_neg_2_046_47    m/z 425.3386     0   1   2   3   4E+4   4   4.5  RT (min)       lcorbi_hilic_neg_4_072_57    m/z 425.3999     0   1E+4   0.5   1  RT (min)      lcorbi_hilic_neg_4_079_58    m/z 425.9652     0   2   4   6   8E+4   0.5   1  RT (min)       lcorbi_hilic_neg_4_099_61    m/z 426.0263     0   2   4   6E+3   6.5   7  RT (min)      lcorbi_hilic_neg_4_082_59    m/z 426.1290     0   1   2   3   4E+4   6   6.5  RT (min)      lcorbi_hilic_neg_1_037_46    m/z 426.1493     0   2   4   6E+3   4.5   5  RT (min)      lcorbi_hilic_neg_4_073_57    m/z 426.3672     0   2   4   6E+4   0.5   1  RT (min)      lcorbi_hilic_neg_4_080_58    m/z 426.6880     0   1E+4   4.5   5  RT (min)      lcorbi_hilic_neg_4_073_57    m/z 426.9204     0   2   4   6   8E+3   6   6.5  RT (min)      lcorbi_hilic_neg_4_071_57    m/z 426.9439     0   1E+4   7.5   8  RT (min)      lcorbi_hilic_neg_4_079_58    m/z 427.0339     0   1E+4   5.5   6  RT (min)      lcorbi_hilic_neg_3_056_56    m/z 427.0440     0   2   4   6E+3   2   2.5  RT (min)      lcorbi_hilic_neg_4_099_61    m/z 427.1071     0   2   4   6   8E+4   6   6.5  RT (min)       lcorbi_hilic_neg_1_037_46    m/z 427.1125     0   1   1   1.5  RT (min)      lcorbi_hilic_neg_1_002_26    m/z 427.1329     0   1   2E+4   1   1.5  RT (min)       lcorbi_hilic_neg_4_092_60    m/z 427.1409     0   1   4.5   5  RT (min)      lcorbi_hilic_neg_4_096_60    m/z 427.1458     0   1   2   2.5  RT (min)      lcorbi_hilic_neg_1_027_18    m/z 427.1584     0   2   4   6E+3   7.5   8  RT (min)      lcorbi_hilic_neg_4_089_59    m/z 427.1948     0   1   6.5   7  RT (min)       lcorbi_hilic_neg_4_090_60    m/z 427.2162     0   2   4   6   8E+3   1   1.5  RT (min)      lcorbi_hilic_neg_4_113_62    m/z 427.2533     0   1   2   3E+4   0.5   1  RT (min)      lcorbi_hilic_neg_4_089_59    m/z 427.2636     0   1E+4   0.5   1  RT (min)      lcorbi_hilic_neg_4_088_59    m/z 427.2886     0   1   2   2.5  RT (min)      lcorbi_hilic_neg_4_053_23    m/z 427.3129     0   1   2   3   4E+3   0.5   1  RT (min)      lcorbi_hilic_neg_4_084_59    m/z 427.3543     0   2   4E+4   4   4.5  RT (min)       lcorbi_hilic_neg_4_069_57    m/z 427.3794     0   1E+4   0.5   1  RT (min)      lcorbi_hilic_neg_4_076_58    m/z 427.9419     0   1   2   3   4E+4   7.5   8  RT (min)       lcorbi_hilic_neg_4_076_58    m/z 427.9620     0   2   4   6E+4   0.5   1  RT (min)      lcorbi_hilic_neg_4_098_61    m/z 428.0049     0   1   2   3   4E+3   5.5   6  RT (min)      lcorbi_hilic_neg_4_098_61    m/z 428.0392     0   1   2   3E+4   6.5   7  RT (min)       lcorbi_hilic_neg_4_095_60    m/z 428.0995     0   1   7.5   8  RT (min)      lcorbi_hilic_neg_1_047_05    m/z 428.1087     0   1   2   3   4E+2   7.5   8  RT (min)      lcorbi_hilic_neg_2_032_25    m/z 428.1111     0   1E+4   6   6.5  RT (min)      lcorbi_hilic_neg_4_094_60    m/z 428.1319     0   2   4E+3   3   3.5  RT (min)       lcorbi_hilic_neg_4_107_62    m/z 428.2083     0   1E+4   0.5   1  RT (min)      lcorbi_hilic_neg_1_031_20    m/z 428.2569     0   1   2E+3   0.5   1  RT (min)      lcorbi_hilic_neg_4_095_60    m/z 428.3339     0   1   2   3E+4   4.5   5  RT (min)      lcorbi_hilic_neg_4_073_57    m/z 428.7735     0   1E+4   4.5   5  RT (min)      lcorbi_hilic_neg_4_074_58    m/z 428.8477     0   2   4   6E+3   7   7.5  RT (min)      lcorbi_hilic_neg_2_032_25    m/z 428.8573     0   2   4   6   8E+3   7   7.5  RT (min)      lcorbi_hilic_neg_4_082_59    m/z 428.9401     0   1E+4   7.5   8  RT (min)      lcorbi_hilic_neg_4_081_58    m/z 429.0063     0   1E+4   6.5   7  RT (min)      lcorbi_hilic_neg_4_081_58    m/z 429.0311     0   1   2   3E+4   6.5   7  RT (min)       lcorbi_hilic_neg_4_072_57    m/z 429.0813     0   1E+4   3   3.5  RT (min)      lcorbi_hilic_neg_4_103_61    m/z 429.0901     0   1E+4   4   4.5  RT (min)      lcorbi_hilic_neg_4_098_61    m/z 429.1222     0   2   4E+4   2.5   3  RT (min)       lcorbi_hilic_neg_4_107_62    m/z 429.1251     0   1   2   3E+3   6.5   7  RT (min)       lcorbi_hilic_neg_4_071_57    m/z 429.1363     0   1E+5   7   7.5  RT (min)      lcorbi_hilic_neg_4_072_57    m/z 429.1601     0   1   2   3   4E+4   3   3.5  RT (min)      lcorbi_hilic_neg_2_031_34    m/z 429.1609     0   1E+4   5   5.5  RT (min)      lcorbi_hilic_neg_4_109_62    m/z 429.1957     0   1E+4   1   1.5   2  RT (min)      lcorbi_hilic_neg_4_077_58    m/z 429.1991     0   1   2E+4   5.5   6  RT (min)       lcorbi_hilic_neg_4_089_59    m/z 429.2813     0   1E+4   1.5   2  RT (min)      lcorbi_hilic_neg_2_014_31    m/z 429.3011     0   1E+4   1   1.5  RT (min)      lcorbi_hilic_neg_4_082_59    m/z 429.9389     0   1E+4   7.5   8  RT (min)      lcorbi_hilic_neg_4_081_58    m/z 430.0097     0   1   2E+4   3   3.5  RT (min)      lcorbi_hilic_neg_4_098_61    m/z 430.0863     0   1   2   3E+4   3   3.5  RT (min)       lcorbi_hilic_neg_4_083_59    m/z 430.1148     0   1E+4   7.5   8  RT (min)      lcorbi_hilic_neg_1_047_05    m/z 430.1799     0   1E+5   6   6.5  RT (min)       lcorbi_hilic_neg_4_095_60    m/z 430.2967     0   1E+4   1   1.5  RT (min)      lcorbi_hilic_neg_4_072_57    m/z 430.3118     0   1E+4   4   4.5  RT (min)      lcorbi_hilic_neg_4_071_57    m/z 430.7709     0   2   4   6   8E+3   4.5   5  RT (min)      lcorbi_hilic_neg_4_079_58    m/z 430.9558     0   1E+4   7.5   8  RT (min)      lcorbi_hilic_neg_4_076_58    m/z 430.9881     0   2   4E+4   7.5   8  RT (min)       lcorbi_hilic_neg_4_075_58    m/z 431.0284     0   1   2E+4   6.5   7  RT (min)       lcorbi_hilic_neg_4_072_57    m/z 431.0772     0   1E+4   4.5   5  RT (min)      lcorbi_hilic_neg_4_106_62    m/z 431.1221     0   2   4   6   8E+3   6.5   7  RT (min)      lcorbi_hilic_neg_1_037_46    m/z 431.1411     0   1E+4   5.5   6  RT (min)      lcorbi_hilic_neg_4_013_46    m/z 431.1712     0   2   4E+4   0.5   1  RT (min)      lcorbi_hilic_neg_4_105_61    m/z 431.1842     0   1   2   3E+4   0.5   1  RT (min)       lcorbi_hilic_neg_4_091_60    m/z 431.2288     0   1   2   3   4E+3   1   1.5  RT (min)      lcorbi_hilic_neg_4_091_60    m/z 431.3168     0   2   4E+4   0.5   1  RT (min)      lcorbi_hilic_neg_4_076_58    m/z 431.3532     0   1E+4   0.5   1  RT (min)      lcorbi_hilic_neg_4_077_58    m/z 431.9381     0   1E+4   7.5   8  RT (min)       lcorbi_hilic_neg_4_080_58    m/z 431.9537     0   1E+4   7.5   8  RT (min)      lcorbi_hilic_neg_4_081_58    m/z 431.9589     0   2   4   6E+3   5.5   6  RT (min)      lcorbi_hilic_neg_4_077_58    m/z 431.9841     0   1   2E+4   7   7.5  RT (min)       lcorbi_hilic_neg_4_097_60    m/z 431.9938     0   1E+4   5   5.5  RT (min)      lcorbi_hilic_neg_4_098_61    m/z 432.0129     0   2   4   6E+3   5   5.5  RT (min)      lcorbi_hilic_neg_4_088_59    m/z 432.0183     0   1E+5   1   1.5  RT (min)       lcorbi_hilic_neg_4_099_61    m/z 432.1726     0   1   2   3E+4   6   6.5  RT (min)      lcorbi_hilic_neg_1_029_31    m/z 432.3125     0   1   2E+4   1   1.5  RT (min)       lcorbi_hilic_neg_4_069_57    m/z 432.3473     0   2   4   6   8E+3   4.5   5  RT (min)      lcorbi_hilic_neg_1_037_46    m/z 432.8570     0   1   2E+4   4.5   5  RT (min)       lcorbi_hilic_neg_4_090_60    m/z 432.8651     0   1E+4   5.5   6  RT (min)       lcorbi_hilic_neg_3_037_12    m/z 432.9514     0   1   2   3E+4   6.5   7  RT (min)       lcorbi_hilic_neg_4_074_58    m/z 432.9920     0   1E+4   0.5   1  RT (min)      lcorbi_hilic_neg_4_113_62    m/z 433.0477     0   2   4   6E+3   3   3.5  RT (min)      lcorbi_hilic_neg_4_068_57    m/z 433.0686     0   1   2   3E+3   1   1.5  RT (min)      lcorbi_hilic_neg_4_078_58    m/z 433.1073     0   1   2   3E+4   8   8.5   9  RT (min)      lcorbi_hilic_neg_2_005_24    m/z 433.1168     0   2   4E+4   8.5   9  RT (min)       lcorbi_hilic_neg_4_091_60    m/z 433.1272     0   1   2E+4   6   6.5  RT (min)      lcorbi_hilic_neg_1_037_46    m/z 433.1354     0   1   2   3   4E+4   1.5   2  RT (min)       lcorbi_hilic_neg_4_107_62    m/z 433.1685     0   1   0.5   1  RT (min)      lcorbi_hilic_neg_4_068_57    m/z 433.1743     0   1   2   3   4E+4   3   3.5  RT (min)      lcorbi_hilic_neg_4_034_45    m/z 433.1760     0   2   4E+3   6   6.5  RT (min)      lcorbi_hilic_neg_4_097_60    m/z 433.2266     0   1   2   3E+4   0.5   1  RT (min)      lcorbi_hilic_neg_3_016_16    m/z 433.2808     0   1   2   3   4E+3   2   2.5  RT (min)      lcorbi_hilic_neg_4_071_57    m/z 433.2973     0   1E+4   1   1.5  RT (min)      lcorbi_hilic_neg_4_068_57    m/z 433.3207     0   1   2E+4   3.5   4  RT (min)       lcorbi_hilic_neg_4_113_62    m/z 433.3688     0   1   2E+4   0.5   1  RT (min)      lcorbi_hilic_neg_4_074_58    m/z 433.8468     0   1   2E+4   1   1.5  RT (min)       lcorbi_hilic_neg_4_074_58    m/z 433.9357     0   1E+4   7.5   8  RT (min)      lcorbi_hilic_neg_4_081_58    m/z 434.0097     0   2   4   6   8E+3   6   6.5  RT (min)      lcorbi_hilic_neg_4_110_62    m/z 434.0557     0   2   4E+3   5   5.5  RT (min)      lcorbi_hilic_neg_4_090_60    m/z 434.1392     0   1E+4   2   2.5  RT (min)      lcorbi_hilic_neg_4_107_62    m/z 434.2395     0   1   2E+5   4   4.5  RT (min)      lcorbi_hilic_neg_3_042_05    m/z 434.2909     0   1E+4   1   1.5  RT (min)      lcorbi_hilic_neg_4_082_59    m/z 434.3721     0   2   4   6E+3   0.5   1  RT (min)      lcorbi_hilic_neg_4_074_58    m/z 434.4913     0   2   4   6   8E+3   4.5   5  RT (min)      lcorbi_hilic_neg_4_079_58    m/z 434.7194     0   1   2E+4   4.5   5  RT (min)       lcorbi_hilic_neg_4_078_58    m/z 434.8270     0   2   4   6E+3   7.5   8  RT (min)      lcorbi_hilic_neg_4_102_61    m/z 434.8622     0   1E+4   5   5.5  RT (min)      lcorbi_hilic_neg_4_111_62    m/z 435.0074     0   1E+4   0.5   1  RT (min)      lcorbi_hilic_neg_4_073_57    m/z 435.0257     0   1E+4   7   7.5  RT (min)      lcorbi_hilic_neg_4_072_57    m/z 435.0443     0   1E+4   6   6.5  RT (min)      lcorbi_hilic_neg_4_111_62    m/z 435.1034     0   2   4   6   8E+3   3   3.5  RT (min)      lcorbi_hilic_neg_2_021_05    m/z 435.1226     0   1E+3   8   8.5  RT (min)      lcorbi_hilic_neg_4_015_24    m/z 435.1279     0   2   4   6E+3   2.5   3  RT (min)      lcorbi_hilic_neg_1_046_19    m/z 435.1294     0   1   2   3E+4   1   1.5  RT (min)       lcorbi_hilic_neg_4_075_58    m/z 435.1322     0   1E+5   7.5   8  RT (min)       lcorbi_hilic_neg_4_096_60    m/z 435.1332     0   1E+4   2.5   3  RT (min)      lcorbi_hilic_neg_1_046_19    m/z 435.1452     0   2   4   6E+2   6.5   7  RT (min)      lcorbi_hilic_neg_2_005_24    m/z 435.1461     0   1   2E+4   1   1.5  RT (min)      lcorbi_hilic_neg_4_077_58    m/z 435.1734     0   1   2   3   4E+3   2.5   3  RT (min)      lcorbi_hilic_neg_2_022_37    m/z 435.2953     0   1   2   3   4E+4   1   1.5  RT (min)      lcorbi_hilic_neg_3_042_05    m/z 435.3122     0   1E+4   0.5   1  RT (min)      lcorbi_hilic_neg_4_076_58    m/z 435.3844     0   1E+4   0.5   1  RT (min)      lcorbi_hilic_neg_4_076_58    m/z 435.4901     0   1E+4   4.5   5  RT (min)      lcorbi_hilic_neg_4_080_58    m/z 435.8759     0   1E+4   8.5   9  RT (min)      lcorbi_hilic_neg_4_079_58    m/z 436.0096     0   2   4   6   8E+3   6.5   7  RT (min)      lcorbi_hilic_neg_4_070_57    m/z 436.0644     0   1   2   3E+4   6   6.5  RT (min)       lcorbi_hilic_neg_4_083_59    m/z 436.0870     0   2   4E+3   3.5   4  RT (min)      lcorbi_hilic_neg_4_106_62    m/z 436.1260     0   1   7   7.5  RT (min)      lcorbi_hilic_neg_4_091_60    m/z 436.1355     0   1   2   3E+4   7   7.5  RT (min)       lcorbi_hilic_neg_4_095_60    m/z 436.1709     0   1   2E+4   2   2.5  RT (min)      lcorbi_hilic_neg_4_106_62    m/z 436.2553     0   1E+5   4   4.5  RT (min)      lcorbi_hilic_neg_4_030_05    m/z 436.2835     0   1   4   4.5  RT (min)       lcorbi_hilic_neg_1_047_05    m/z 436.3517     0   1   2E+4   0.5   1  RT (min)      lcorbi_hilic_neg_4_091_60    m/z 436.4887     0   1E+4   4.5   5  RT (min)      lcorbi_hilic_neg_4_079_58    m/z 436.7168     0   1   2   3E+4   4.5   5  RT (min)       lcorbi_hilic_neg_4_075_58    m/z 436.8287     0   2   4   6E+3   7   7.5  RT (min)      lcorbi_hilic_neg_4_091_60    m/z 436.9492     0   1   2E+4   6   6.5  RT (min)      lcorbi_hilic_neg_4_069_57    m/z 436.9609     0   2   4E+3   0.5   1  RT (min)      lcorbi_hilic_neg_4_076_58    m/z 436.9723     0   1   5.5   6  RT (min)      lcorbi_hilic_neg_4_091_60    m/z 436.9727     0   1   2   3E+4   7.5   8  RT (min)       lcorbi_hilic_neg_4_078_58    m/z 436.9877     0   2   4   6E+3   0.5   1  RT (min)       lcorbi_hilic_neg_4_090_60    m/z 437.0229     0   1E+4   0.5   1  RT (min)      lcorbi_hilic_neg_4_108_62    m/z 437.0414     0   1E+4   6   6.5  RT (min)      lcorbi_hilic_neg_4_107_62    m/z 437.0553     0   1   2E+4   6   6.5  RT (min)      lcorbi_hilic_neg_1_023_23    m/z 437.0868     0   1E+4   5.5   6  RT (min)       lcorbi_hilic_neg_4_086_59    m/z 437.1029     0   1   2E+4   2   2.5  RT (min)       lcorbi_hilic_neg_4_106_62    m/z 437.1323     0   2   4   6   8E+4   0.5   1  RT (min)       lcorbi_hilic_neg_4_097_60    m/z 437.1418     0   2   4   6   8E+4   3   3.5  RT (min)       lcorbi_hilic_neg_4_103_61    m/z 437.1642     0   1   2E+4   2   2.5  RT (min)       lcorbi_hilic_neg_4_096_60    m/z 437.1855     0   2   4   6E+2   7   7.5  RT (min)      lcorbi_hilic_neg_2_009_50    m/z 437.1894     0   1   2E+4   6   7  RT (min)       lcorbi_hilic_neg_4_095_60    m/z 437.2153     0   1   1.5   2  RT (min)      lcorbi_hilic_neg_4_093_60    m/z 437.2579     0   1   2E+3   0.5   1  RT (min)      lcorbi_hilic_neg_4_075_58    m/z 437.2676     0   1   2E+4   4   4.5  RT (min)      lcorbi_hilic_neg_3_042_05    m/z 437.2680     0   2   4E+4   1   1.5  RT (min)       lcorbi_hilic_neg_4_070_57    m/z 437.2867     0   1   4   4.5  RT (min)      lcorbi_hilic_neg_1_047_05    m/z 437.2912     0   2   4E+4   1   1.5  RT (min)       lcorbi_hilic_neg_4_070_57    m/z 437.3048     0   1E+3   1   1.5  RT (min)      lcorbi_hilic_neg_4_094_60    m/z 437.3551     0   2   4   6   8E+4   0.5   1  RT (min)       lcorbi_hilic_neg_4_091_60    m/z 437.4002     0   1   0.5   1  RT (min)      lcorbi_hilic_neg_4_093_60    m/z 437.4872     0   1E+4   4.5   5  RT (min)      lcorbi_hilic_neg_4_079_58    m/z 437.9306     0   2   4   6   8E+3   6   6.5  RT (min)      lcorbi_hilic_neg_4_072_57    m/z 437.9536     0   2   4   6E+3   0.5   1  RT (min)      lcorbi_hilic_neg_4_104_61    m/z 437.9789     0   2   4   6E+3   5.5   6  RT (min)      lcorbi_hilic_neg_4_113_62    m/z 437.9862     0   1   2   3   4E+5   0.5   1  RT (min)       lcorbi_hilic_neg_3_031_14    m/z 438.0761     0   2   4   6   8E+3   6   6.5  RT (min)      lcorbi_hilic_neg_4_075_58    m/z 438.1439     0   2   4E+4   6   6.5  RT (min)       lcorbi_hilic_neg_4_095_60    m/z 438.1453     0   1E+4   3   3.5  RT (min)      lcorbi_hilic_neg_4_102_61    m/z 438.1596     0   2   4   6E+3   2.5   3  RT (min)       lcorbi_hilic_neg_4_101_61    m/z 438.2318     0   1E+5   6.5   7  RT (min)       lcorbi_hilic_neg_4_097_60    m/z 438.2579     0   1E+4   4   4.5  RT (min)      lcorbi_hilic_neg_4_073_57    m/z 438.2963     0   1   2E+4   1   1.5  RT (min)      lcorbi_hilic_neg_1_047_05    m/z 438.2996     0   1   2   3   4E+4   4   4.5  RT (min)       lcorbi_hilic_neg_4_081_58    m/z 438.4854     0   2   4   6   8E+3   4.5   5  RT (min)      lcorbi_hilic_neg_4_077_58    m/z 438.7145     0   1E+4   4.5   5  RT (min)      lcorbi_hilic_neg_4_075_58    m/z 438.8024     0   1   2E+4   4.5   5  RT (min)       lcorbi_hilic_neg_4_074_58    m/z 438.8651     0   1E+4   8.5   9  RT (min)      lcorbi_hilic_neg_4_074_58    m/z 438.9683     0   1E+4   7.5   8  RT (min)      lcorbi_hilic_neg_4_075_58    m/z 438.9708     0   1E+4   5.5   6  RT (min)      lcorbi_hilic_neg_4_070_57    m/z 438.9892     0   2   4   6E+4   0.5   1  RT (min)      lcorbi_hilic_neg_3_031_14    m/z 439.0599     0   1   2E+4   6.5   7  RT (min)       lcorbi_hilic_neg_4_072_57    m/z 439.0639     0   1E+4   3   3.5  RT (min)      lcorbi_hilic_neg_4_103_61    m/z 439.0770     0   2   4   6E+3   7   7.5  RT (min)      lcorbi_hilic_neg_1_023_23    m/z 439.0974     0   1   2E+4   8   8.5  RT (min)       lcorbi_hilic_neg_4_088_59    m/z 439.1071     0   1E+4   6   6.5  RT (min)      lcorbi_hilic_neg_4_068_57    m/z 439.1410     0   1E+4   1   1.5  RT (min)      lcorbi_hilic_neg_4_111_62    m/z 439.1540     0   1E+4   6.5   7  RT (min)      lcorbi_hilic_neg_4_088_59    m/z 439.1612     0   2   4   6E+3   3.5   4  RT (min)      lcorbi_hilic_neg_4_079_58    m/z 439.2260     0   1   1.5   2  RT (min)      lcorbi_hilic_neg_4_101_61    m/z 439.2736     0   1   2E+4   0.5   1  RT (min)      lcorbi_hilic_neg_4_004_13    m/z 439.3027     0   1E+4   4   4.5  RT (min)      lcorbi_hilic_neg_4_081_58    m/z 439.3523     0   1   0.5   1  RT (min)       lcorbi_hilic_neg_4_076_58    m/z 439.3706     0   1   2E+4   0.5   1  RT (min)       lcorbi_hilic_neg_4_091_60    m/z 439.9276     0   2   4   6E+3   6   6.5  RT (min)      lcorbi_hilic_neg_4_072_57    m/z 439.9760     0   1   2   3E+3   5.5   6  RT (min)      lcorbi_hilic_neg_4_084_59    m/z 439.9832     0   1   2E+5   0.5   1  RT (min)      lcorbi_hilic_neg_4_046_15    m/z 440.0434     0   1   2E+4   6.5   7  RT (min)       lcorbi_hilic_neg_4_075_58    m/z 440.0530     0   2   4   6E+3   3.5   4  RT (min)      lcorbi_hilic_neg_4_112_62    m/z 440.0747     0   1E+4   0.5   1  RT (min)      lcorbi_hilic_neg_4_077_58    m/z 440.0904     0   1E+4   7   7.5  RT (min)      lcorbi_hilic_neg_4_085_59    m/z 440.1203     0   2   4   6   8E+3   3   3.5  RT (min)      lcorbi_hilic_neg_2_021_05    m/z 440.1376     0   1E+4   6.5   7  RT (min)      lcorbi_hilic_neg_4_089_59    m/z 440.1433     0   1E+4   1.5   2  RT (min)      lcorbi_hilic_neg_4_089_59    m/z 440.3057     0   2   4   6   8E+3   1   1.5  RT (min)      lcorbi_hilic_neg_2_006_45    m/z 440.3133     0   1   2E+5   1   1.5  RT (min)      lcorbi_hilic_neg_1_047_05    m/z 440.3828     0   1   2E+4   0.5   1  RT (min)      lcorbi_hilic_neg_4_077_58    m/z 440.8000     0   1E+4   4.5   5  RT (min)      lcorbi_hilic_neg_4_074_58    m/z 440.8291     0   1   2   3E+4   8.5   9  RT (min)       lcorbi_hilic_neg_4_081_58    m/z 440.8888     0   2   4   6E+3   11.5   12  RT (min)      lcorbi_hilic_neg_4_066_57    m/z 440.9980     0   1E+4   3.5   4  RT (min)      lcorbi_hilic_neg_4_080_58    m/z 440.9982     0   1   2   3E+4   0.5   1  RT (min)       lcorbi_hilic_neg_4_072_57    m/z 441.0413     0   1   2E+4   6   6.5  RT (min)       lcorbi_hilic_neg_4_072_57    m/z 441.0569     0   2   4   6   8E+3   6.5   7  RT (min)      lcorbi_hilic_neg_4_070_57    m/z 441.0812     0   1   2   3   4E+3   5.5   6  RT (min)      lcorbi_hilic_neg_4_077_58    m/z 441.1124     0   1E+4   1   1.5  RT (min)      lcorbi_hilic_neg_4_074_58    m/z 441.1654     0   1   2   3   4E+4   6.5   7  RT (min)       lcorbi_hilic_neg_4_096_60    m/z 441.3166     0   1   2E+4   1   1.5  RT (min)       lcorbi_hilic_neg_1_047_05    m/z 441.9669     0   1E+4   7.5   8  RT (min)      lcorbi_hilic_neg_4_080_58    m/z 441.9805     0   1E+4   0.5   1  RT (min)      lcorbi_hilic_neg_4_105_61    m/z 441.9877     0   1E+4   5.5   6  RT (min)      lcorbi_hilic_neg_4_077_58    m/z 442.0416     0   2   4   6   8E+3   5   5.5  RT (min)      lcorbi_hilic_neg_4_106_62    m/z 442.0502     0   2   4   6E+3   3.5   4  RT (min)      lcorbi_hilic_neg_4_107_62    m/z 442.0701     0   1   2E+4   7   7.5  RT (min)      lcorbi_hilic_neg_1_023_23    m/z 442.0716     0   1E+4   0.5   1  RT (min)      lcorbi_hilic_neg_4_076_58    m/z 442.0902     0   1E+4   0.5   1  RT (min)      lcorbi_hilic_neg_4_076_58    m/z 442.1374     0   1E+4   1.5   2  RT (min)      lcorbi_hilic_neg_4_077_58    m/z 442.1439     0   1E+4   4.5   5  RT (min)      lcorbi_hilic_neg_4_097_60    m/z 442.1730     0   1   2E+5   3   3.5  RT (min)      lcorbi_hilic_neg_1_026_34    m/z 442.3049     0   1   2E+3   1   1.5  RT (min)      lcorbi_hilic_neg_4_083_59    m/z 442.3985     0   1   2   3E+4   0.5   1  RT (min)      lcorbi_hilic_neg_4_076_58    m/z 442.6779     0   1E+4   4.5   5  RT (min)      lcorbi_hilic_neg_4_098_61    m/z 442.8940     0   2   4   6E+4   5.5   6  RT (min)       lcorbi_hilic_neg_4_106_62    m/z 442.9397     0   1E+4   1   1.5  RT (min)      lcorbi_hilic_neg_4_100_61    m/z 442.9802     0   1   2   3   4E+4   6.5   7  RT (min)       lcorbi_hilic_neg_4_077_58    m/z 442.9961     0   2   4   6   8E+3   6   6.5  RT (min)      lcorbi_hilic_neg_4_078_58    m/z 443.1562     0   1   2   3E+4   3.5   4  RT (min)       lcorbi_hilic_neg_4_068_57    m/z 443.1758     0   2   4E+4   3   3.5  RT (min)      lcorbi_hilic_neg_1_026_34    m/z 443.2098     0   2   4E+3   0.5   1  RT (min)      lcorbi_hilic_neg_4_072_57    m/z 443.2584     0   1E+3   0.5   1  RT (min)      lcorbi_hilic_neg_2_013_18    m/z 443.3169     0   1   2   3   4E+4   1   1.5  RT (min)      lcorbi_hilic_neg_4_077_58    m/z 443.9160     0   1E+4   7.5   8  RT (min)      lcorbi_hilic_neg_4_075_58    m/z 443.9652     0   1E+3   6.5   7  RT (min)      lcorbi_hilic_neg_4_081_58    m/z 443.9765     0   1E+4   6.5   7  RT (min)      lcorbi_hilic_neg_4_082_59    m/z 443.9785     0   1E+4   0.5   1  RT (min)      lcorbi_hilic_neg_4_099_61    m/z 443.9847     0   1   2   3   4E+3   5.5   6  RT (min)      lcorbi_hilic_neg_4_067_57    m/z 444.0531     0   1   2   3E+4   7.5   8  RT (min)       lcorbi_hilic_neg_4_089_59    m/z 444.0874     0   1   0.5   1  RT (min)      lcorbi_hilic_neg_4_111_62    m/z 444.0895     0   1E+4   4   4.5  RT (min)      lcorbi_hilic_neg_4_084_59    m/z 444.1020     0   1   2E+4   3   3.5  RT (min)       lcorbi_hilic_neg_4_083_59    m/z 444.1060     0   1   2E+4   0.5   1  RT (min)       lcorbi_hilic_neg_4_113_62    m/z 444.1339     0   1   2   3E+5   1   1.5  RT (min)      lcorbi_hilic_neg_1_027_18    m/z 444.1414     0   1   2   3   4E+3   4.5   5  RT (min)      lcorbi_hilic_neg_4_090_60    m/z 444.1545     0   1   2E+4   7   7.5  RT (min)      lcorbi_hilic_neg_2_021_05    m/z 444.3201     0   1E+4   1   1.5  RT (min)      lcorbi_hilic_neg_1_047_05    m/z 444.3532     0   1   2E+4   3.5   4  RT (min)       lcorbi_hilic_neg_4_089_59    m/z 444.6749     0   1   2E+4   4.5   5  RT (min)      lcorbi_hilic_neg_4_098_61    m/z 444.8911     0   1   2E+4   5.5   6  RT (min)       lcorbi_hilic_neg_4_106_62    m/z 444.9366     0   1E+4   1   1.5  RT (min)      lcorbi_hilic_neg_4_100_61    m/z 444.9805     0   1   2   3E+3   7.5   8  RT (min)      lcorbi_hilic_neg_4_081_58    m/z 445.0321     0   2   4   6E+3   0.5   1  RT (min)      lcorbi_hilic_neg_4_093_60    m/z 445.0356     0   1   2E+4   6   6.5  RT (min)       lcorbi_hilic_neg_4_083_59    m/z 445.0542     0   1E+4   7   7.5  RT (min)      lcorbi_hilic_neg_4_068_57    m/z 445.0683     0   1E+3   3.5   4  RT (min)      lcorbi_hilic_neg_4_096_60    m/z 445.0731     0   1   2E+4   6   6.5  RT (min)       lcorbi_hilic_neg_4_071_57    m/z 445.1741     0   1   6.5   7  RT (min)      lcorbi_hilic_neg_4_087_59    m/z 445.2067     0   2   4   6E+3   0.5   1  RT (min)      lcorbi_hilic_neg_4_067_57    m/z 445.9691     0   1   2   3E+4   7.5   8  RT (min)      lcorbi_hilic_neg_4_080_58    m/z 445.9785     0   1E+4   7.5   8  RT (min)      lcorbi_hilic_neg_2_055_32    m/z 446.0279     0   1   2E+4   0.5   1  RT (min)       lcorbi_hilic_neg_4_099_61    m/z 446.0383     0   1E+4   6.5   7  RT (min)      lcorbi_hilic_neg_4_085_59    m/z 446.0606     0   1   2E+4   3   3.5  RT (min)       lcorbi_hilic_neg_4_085_59    m/z 446.0679     0   1   2E+4   4.5   5  RT (min)       lcorbi_hilic_neg_4_107_62    m/z 446.1030     0   1   2E+4   0.5   1  RT (min)      lcorbi_hilic_neg_4_113_62    m/z 446.1157     0   2   4   6E+3   4   4.5  RT (min)      lcorbi_hilic_neg_4_108_62    m/z 446.1399     0   1   2   2.5  RT (min)      lcorbi_hilic_neg_4_096_60    m/z 446.1524     0   2   4E+4   6.5   7  RT (min)       lcorbi_hilic_neg_4_069_57    m/z 446.1723     0   1E+4   1   1.5  RT (min)      lcorbi_hilic_neg_4_075_58    m/z 446.1983     0   2   4   6   8E+3   7   7.5  RT (min)      lcorbi_hilic_neg_4_089_59    m/z 446.1998     0   1   2E+4   2   2.5  RT (min)      lcorbi_hilic_neg_4_106_62    m/z 446.2187     0   2   4   6E+3   0.5   1  RT (min)      lcorbi_hilic_neg_2_053_03    m/z 446.2924     0   1   2E+4   1.5   2  RT (min)       lcorbi_hilic_neg_4_066_57    m/z 446.6719     0   1E+4   4.5   5  RT (min)      lcorbi_hilic_neg_4_098_61    m/z 446.8452     0   2   4   6E+3   11.5   12  RT (min)      lcorbi_hilic_neg_4_111_62    m/z 446.9286     0   2   4   6   8E+3   7.5   8  RT (min)      lcorbi_hilic_neg_4_074_58    m/z 446.9667     0   1   2E+4   7.5   8  RT (min)       lcorbi_hilic_neg_4_079_58    m/z 446.9759     0   1   2E+3   7.5   8  RT (min)      lcorbi_hilic_neg_4_075_58    m/z 446.9871     0   2   4   6   8E+3   5.5   6  RT (min)      lcorbi_hilic_neg_4_072_57    m/z 446.9940     0   1E+4   1   1.5  RT (min)      lcorbi_hilic_neg_4_079_58    m/z 447.0014     0   2   4   6   8E+3   5.5   6  RT (min)      lcorbi_hilic_neg_4_069_57    m/z 447.0643     0   1E+5   0.5   1  RT (min)       lcorbi_hilic_neg_4_113_62    m/z 447.0652     0   1E+4   3.5   4  RT (min)      lcorbi_hilic_neg_4_108_62    m/z 447.1013     0   1   5.5   6   6.5  RT (min)       lcorbi_hilic_neg_2_013_18    m/z 447.1207     0   1E+4   7   7.5  RT (min)      lcorbi_hilic_neg_4_089_59    m/z 447.1286     0   1   2   3   4E+4   4.5   5  RT (min)      lcorbi_hilic_neg_4_107_62    m/z 447.1345     0   1E+4   0.5   1  RT (min)      lcorbi_hilic_neg_4_084_59    m/z 447.1615     0   1   2E+5   0.5   1  RT (min)       lcorbi_hilic_neg_4_097_60    m/z 447.2758     0   2   4   6E+4   1   1.5  RT (min)       lcorbi_hilic_neg_4_074_58    m/z 447.2964     0   2   4   6   8E+4   1   1.5  RT (min)       lcorbi_hilic_neg_4_070_57    m/z 447.3118     0   1   2   3E+4   0.5   1  RT (min)      lcorbi_hilic_neg_4_080_58    m/z 447.3613     0   1E+3   0.5   1  RT (min)      lcorbi_hilic_neg_4_093_60    m/z 447.3841     0   1E+5   0.5   1  RT (min)       lcorbi_hilic_neg_4_091_60    m/z 447.9267     0   1E+4   7.5   8  RT (min)      lcorbi_hilic_neg_4_074_58    m/z 447.9514     0   1E+4   5.5   6  RT (min)      lcorbi_hilic_neg_4_089_59    m/z 447.9595     0   1E+4   6   6.5  RT (min)      lcorbi_hilic_neg_4_067_57    m/z 448.0079     0   1E+4   5.5   6  RT (min)      lcorbi_hilic_neg_4_102_61    m/z 448.0254     0   1   2E+4   0.5   1  RT (min)      lcorbi_hilic_neg_4_099_61    m/z 448.0647     0   1E+4   4.5   5  RT (min)      lcorbi_hilic_neg_4_111_62    m/z 448.0647     0   1   2   3   4E+4   0.5   1  RT (min)       lcorbi_hilic_neg_4_113_62    m/z 448.0918     0   1   2E+4   6   6.5  RT (min)      lcorbi_hilic_neg_1_055_24    m/z 448.1380     0   2   4   6E+3   3   3.5  RT (min)      lcorbi_hilic_neg_4_082_59    m/z 448.1932     0   2   4   6E+3   3   3.5  RT (min)      lcorbi_hilic_neg_1_025_52    m/z 448.2999     0   1   2E+4   1   1.5  RT (min)      lcorbi_hilic_neg_4_070_57    m/z 448.7433     0   1E+4   4.5   5  RT (min)       lcorbi_hilic_neg_4_075_58    m/z 448.9282     0   1E+4   7.5   8  RT (min)      lcorbi_hilic_neg_4_076_58    m/z 448.9341     0   1E+4   0.5   1  RT (min)      lcorbi_hilic_neg_4_068_57    m/z 449.0888     0   1   2E+4   6.5   7  RT (min)       lcorbi_hilic_neg_4_073_57    m/z 449.0944     0   1   2E+4   1   1.5  RT (min)       lcorbi_hilic_neg_4_070_57    m/z 449.1071     0   2   4   6   8E+4   6   6.5  RT (min)       lcorbi_hilic_neg_4_070_57    m/z 449.1312     0   1   2   3E+4   3   3.5  RT (min)       lcorbi_hilic_neg_4_066_57    m/z 449.1501     0   1   2   3   4E+4   0.5   1  RT (min)       lcorbi_hilic_neg_4_096_60    m/z 449.1525     0   1   2   3   4E+4   3   3.5  RT (min)       lcorbi_hilic_neg_4_094_60    m/z 449.1623     0   2   4E+4   6.5   7  RT (min)      lcorbi_hilic_neg_2_005_24    m/z 449.1715     0   1E+5   7   7.5  RT (min)       lcorbi_hilic_neg_4_090_60    m/z 449.2190     0   2   4   6   8E+3   0.5   1  RT (min)      lcorbi_hilic_neg_4_113_62    m/z 449.3998     0   1   2   3   4E+4   0.5   1  RT (min)       lcorbi_hilic_neg_4_091_60    m/z 449.8206     0   1E+4   1   1.5  RT (min)      lcorbi_hilic_neg_4_074_58    m/z 449.9265     0   1E+4   7.5   8  RT (min)      lcorbi_hilic_neg_4_074_58    m/z 449.9489     0   1   5.5   6  RT (min)      lcorbi_hilic_neg_4_070_57    m/z 449.9753     0   1   2E+3   7.5   8  RT (min)      lcorbi_hilic_neg_4_056_56    m/z 450.0050     0   2   4E+3   5.5   6  RT (min)      lcorbi_hilic_neg_4_102_61    m/z 450.0865     0   1E+4   1   1.5  RT (min)      lcorbi_hilic_neg_4_040_17    m/z 450.0973     0   2   4   6   8E+3   6.5   7  RT (min)      lcorbi_hilic_neg_1_019_06    m/z 450.0996     0   1   4.5   5  RT (min)      lcorbi_hilic_neg_1_047_05    m/z 450.1344     0   1E+4   3   3.5  RT (min)      lcorbi_hilic_neg_4_066_57    m/z 450.2290     0   1   2E+4   1   1.5  RT (min)      lcorbi_hilic_neg_3_001_49    m/z 450.2591     0   1   2   3E+4   2   2.5  RT (min)       lcorbi_hilic_neg_4_079_58    m/z 450.3144     0   1   2E+4   1.5   2  RT (min)      lcorbi_hilic_neg_4_077_58    m/z 450.3341     0   1E+4   0.5   1  RT (min)      lcorbi_hilic_neg_3_015_47    m/z 450.3671     0   2   4   6   8E+4   0.5   1  RT (min)      lcorbi_hilic_neg_4_080_58    m/z 450.7925     0   2   4   6E+3   7.5   8  RT (min)      lcorbi_hilic_neg_4_099_61    m/z 450.8295     0   1E+4   4.5   5  RT (min)      lcorbi_hilic_neg_4_090_60    m/z 450.8393     0   2   4   6   8E+3   7   7.5  RT (min)      lcorbi_hilic_neg_4_082_59    m/z 450.9242     0   2   4   6E+3   7.5   8  RT (min)      lcorbi_hilic_neg_4_080_58    m/z 450.9242     0   1E+4   5.5   6  RT (min)      lcorbi_hilic_neg_4_073_57    m/z 450.9578     0   1E+4   5   5.5  RT (min)      lcorbi_hilic_neg_2_020_27    m/z 451.0550     0   2   4   6   8E+3   2   2.5  RT (min)      lcorbi_hilic_neg_4_108_62    m/z 451.0776     0   1   3   3.5  RT (min)      lcorbi_hilic_neg_4_106_62    m/z 451.0872     0   1   2   3E+4   6.5   7  RT (min)       lcorbi_hilic_neg_4_088_59    m/z 451.1245     0   1   1.5   2  RT (min)      lcorbi_hilic_neg_4_107_62    m/z 451.1855     0   1   2   3   4E+4   0.5   1  RT (min)      lcorbi_hilic_neg_3_055_26    m/z 451.2144     0   1   2E+4   7   7.5  RT (min)      lcorbi_hilic_neg_1_047_05    m/z 451.2614     0   2   4E+3   2   2.5  RT (min)      lcorbi_hilic_neg_4_081_58    m/z 451.2663     0   1   2E+4   4   4.5  RT (min)      lcorbi_hilic_neg_1_047_05    m/z 451.3429     0   1   2E+4   0.5   1  RT (min)      lcorbi_hilic_neg_4_077_58    m/z 451.9964     0   2   4   6   8E+3   6.5   7  RT (min)      lcorbi_hilic_neg_4_084_59    m/z 452.0167     0   1   2E+4   5.5   6  RT (min)       lcorbi_hilic_neg_4_081_58    m/z 452.0584     0   2   4E+3   6.5   7  RT (min)      lcorbi_hilic_neg_4_092_60    m/z 452.1361     0   1E+4   7   7.5  RT (min)      lcorbi_hilic_neg_4_089_59    m/z 452.1448     0   1E+4   2   2.5  RT (min)      lcorbi_hilic_neg_4_106_62    m/z 452.2391     0   2   4   6E+4   1.5   2  RT (min)      lcorbi_hilic_neg_1_005_36    m/z 452.3494     0   1E+4   0.5   1  RT (min)      lcorbi_hilic_neg_3_015_47    m/z 452.8888     0   2   4E+3   5   5.5  RT (min)      lcorbi_hilic_neg_2_020_27    m/z 452.8987     0   2   4   6   8E+3   5   5.5  RT (min)      lcorbi_hilic_neg_1_011_01    m/z 452.9364     0   1   2   3   4E+4   7.5   8  RT (min)       lcorbi_hilic_neg_4_081_58    m/z 452.9469     0   1   7.5   8  RT (min)      lcorbi_hilic_neg_4_075_58    m/z 452.9470     0   2   4   6E+3   5   5.5  RT (min)      lcorbi_hilic_neg_1_002_26    m/z 452.9600     0   1   2E+4   5   5.5  RT (min)      lcorbi_hilic_neg_4_072_57    m/z 452.9980     0   1E+4   0.5   1  RT (min)      lcorbi_hilic_neg_4_068_57    m/z 453.0250     0   1E+4   6   6.5  RT (min)      lcorbi_hilic_neg_4_071_57    m/z 453.0421     0   2   4   6   8E+3   1   1.5  RT (min)      lcorbi_hilic_neg_4_098_61    m/z 453.0520     0   2   4   6E+3   2   2.5  RT (min)      lcorbi_hilic_neg_4_111_62    m/z 453.1214     0   1E+3   5.5   6  RT (min)      lcorbi_hilic_neg_4_015_24    m/z 453.1387     0   1E+4   2   2.5  RT (min)      lcorbi_hilic_neg_4_106_62    m/z 453.1445     0   1   2E+4   8   8.5  RT (min)       lcorbi_hilic_neg_4_093_60    m/z 453.1448     0   1E+4   2.5   3  RT (min)      lcorbi_hilic_neg_4_111_62    m/z 453.1836     0   1   2   3   4E+4   0.5   1  RT (min)      lcorbi_hilic_neg_3_055_26    m/z 453.2330     0   1   2E+4   1   1.5  RT (min)      lcorbi_hilic_neg_4_107_62    m/z 453.2423     0   2   4   6E+3   1.5   2  RT (min)      lcorbi_hilic_neg_4_084_59    m/z 453.2633     0   1   2E+4   1   1.5  RT (min)       lcorbi_hilic_neg_4_066_57    m/z 453.2892     0   1   2E+4   0.5   1  RT (min)       lcorbi_hilic_neg_4_087_59    m/z 453.3224     0   1   2E+4   0.5   1  RT (min)      lcorbi_hilic_neg_4_076_58    m/z 453.3467     0   2   4   6E+4   4.5   5  RT (min)       lcorbi_hilic_neg_4_073_57    m/z 453.3587     0   1   2E+4   0.5   1  RT (min)      lcorbi_hilic_neg_4_081_58    m/z 453.3953     0   1   2E+4   0.5   1  RT (min)      lcorbi_hilic_neg_4_081_58    m/z 453.9158     0   1   2E+5   5.5   6  RT (min)      lcorbi_hilic_neg_1_051_33    m/z 453.9448     0   1E+4   7.5   8  RT (min)      lcorbi_hilic_neg_4_078_58    m/z 453.9935     0   1   2E+3   5   5.5  RT (min)      lcorbi_hilic_neg_4_083_59    m/z 454.0476     0   1E+4   1   1.5  RT (min)      lcorbi_hilic_neg_4_107_62    m/z 454.0594     0   1   2E+3   6   6.5  RT (min)      lcorbi_hilic_neg_4_096_60    m/z 454.0729     0   1E+4   3   3.5  RT (min)      lcorbi_hilic_neg_4_098_61    m/z 454.1030     0   1   2E+3   6   6.5  RT (min)      lcorbi_hilic_neg_2_005_24    m/z 454.1110     0   1   2E+4   7   7.5  RT (min)       lcorbi_hilic_neg_1_012_32    m/z 454.1411     0   2   4E+3   2   2.5  RT (min)      lcorbi_hilic_neg_3_024_27    m/z 454.1439     0   1   5.5   6  RT (min)      lcorbi_hilic_neg_3_035_48    m/z 454.2846     0   1E+4   4   4.5  RT (min)      lcorbi_hilic_neg_4_083_59    m/z 454.2923     0   1   2   3E+3   0.5   1  RT (min)      lcorbi_hilic_neg_1_047_05    m/z 454.9283     0   2   4   6   8E+3   8   8.5  RT (min)      lcorbi_hilic_neg_4_080_58    m/z 454.9756     0   1E+3   5   5.5  RT (min)      lcorbi_hilic_neg_4_094_60    m/z 455.0134     0   1E+4   3   3.5  RT (min)      lcorbi_hilic_neg_4_075_58    m/z 455.0138     0   1E+4   0.5   1  RT (min)      lcorbi_hilic_neg_4_073_57    m/z 455.0872     0   1E+4   2   2.5  RT (min)      lcorbi_hilic_neg_4_108_62    m/z 455.0901     0   1   2E+4   7   7.5  RT (min)       lcorbi_hilic_neg_4_096_60    m/z 455.1084     0   1   5.5   6  RT (min)      lcorbi_hilic_neg_1_005_36    m/z 455.1376     0   1   2E+4   2   2.5  RT (min)      lcorbi_hilic_neg_4_055_38    m/z 455.1589     0   1   2.5   3  RT (min)      lcorbi_hilic_neg_4_097_60    m/z 455.2293     0   1   3   3.5  RT (min)      lcorbi_hilic_neg_4_093_60    m/z 455.2479     0   1   2E+4   1   1.5  RT (min)       lcorbi_hilic_neg_4_066_57    m/z 455.2685     0   2   4   6E+4   0.5   1  RT (min)       lcorbi_hilic_neg_4_069_57    m/z 455.2796     0   2   4   6E+3   1   1.5  RT (min)      lcorbi_hilic_neg_4_091_60    m/z 455.3170     0   2   4   6   8E+3   0.5   1  RT (min)      lcorbi_hilic_neg_4_068_57    m/z 455.3626     0   2   4   6E+4   4.5   5  RT (min)       lcorbi_hilic_neg_4_071_57    m/z 455.7807     0   2   4   6   8E+3   4.5   5  RT (min)      lcorbi_hilic_neg_4_078_58    m/z 455.9128     0   2   4   6E+3   5.5   6  RT (min)      lcorbi_hilic_neg_4_072_57    m/z 455.9307     0   2   4   6E+3   5   5.5  RT (min)      lcorbi_hilic_neg_4_072_57    m/z 456.0147     0   1   2E+4   6   7  RT (min)       lcorbi_hilic_neg_4_095_60    m/z 456.0697     0   1E+4   6   6.5  RT (min)      lcorbi_hilic_neg_1_023_23    m/z 456.0708     0   1E+4   3   3.5  RT (min)      lcorbi_hilic_neg_4_098_61    m/z 456.0944     0   2   4   6E+3   3   3.5  RT (min)      lcorbi_hilic_neg_2_021_05    m/z 456.0962     0   1E+4   4.5   5  RT (min)      lcorbi_hilic_neg_4_106_62    m/z 456.1053     0   1E+4   6   6.5  RT (min)      lcorbi_hilic_neg_4_071_57    m/z 456.1619     0   1   2   3E+4   4.5   5  RT (min)      lcorbi_hilic_neg_4_106_62    m/z 456.2744     0   1   2E+4   0.5   1  RT (min)      lcorbi_hilic_neg_4_070_57    m/z 456.3571     0   1   2E+4   1   1.5  RT (min)       lcorbi_hilic_neg_4_104_61    m/z 456.3661     0   1E+4   4.5   5  RT (min)      lcorbi_hilic_neg_4_071_57    m/z 456.3673     0   1   2E+5   1   1.5  RT (min)       lcorbi_hilic_neg_1_047_05    m/z 456.7815     0   1   2   3   4E+3   5   5.5  RT (min)      lcorbi_hilic_neg_4_084_59    m/z 456.8471     0   1   2E+4   7   7.5  RT (min)       lcorbi_hilic_neg_4_047_07    m/z 456.9598     0   1   6   6.5  RT (min)      lcorbi_hilic_neg_4_083_59    m/z 457.0154     0   1E+4   5.5   6   6.5  RT (min)      lcorbi_hilic_neg_4_072_57    m/z 457.0449     0   1E+4   2   2.5  RT (min)      lcorbi_hilic_neg_4_108_62    m/z 457.1058     0   1E+4   6   6.5   7  RT (min)       lcorbi_hilic_neg_4_083_59    m/z 457.1202     0   1   2E+4   6   6.5  RT (min)       lcorbi_hilic_neg_4_085_59    m/z 457.1554     0   1E+4   6.5   7  RT (min)      lcorbi_hilic_neg_4_030_05    m/z 457.2029     0   2   4E+4   0.5   1  RT (min)       lcorbi_hilic_neg_4_113_62    m/z 457.2347     0   2   4   6   8E+4   7   7.5  RT (min)       lcorbi_hilic_neg_4_096_60    m/z 457.2615     0   1E+4   7   7.5  RT (min)      lcorbi_hilic_neg_4_081_58    m/z 457.2788     0   1   2   3   4E+3   0.5   1  RT (min)      lcorbi_hilic_neg_4_077_58    m/z 457.9708     0   1   5.5   6  RT (min)      lcorbi_hilic_neg_1_003_14    m/z 457.9805     0   1   2E+4   5.5   6  RT (min)       lcorbi_hilic_neg_4_088_59    m/z 457.9886     0   1E+4   6   6.5  RT (min)      lcorbi_hilic_neg_4_070_57    m/z 458.1104     0   1   7   7.5  RT (min)      lcorbi_hilic_neg_4_097_60    m/z 458.2398     0   1   2E+4   4   4.5  RT (min)      lcorbi_hilic_neg_4_081_58    m/z 458.6517     0   1E+4   4.5   5  RT (min)      lcorbi_hilic_neg_4_100_61    m/z 458.7724     0   1   2E+4   4.5   5  RT (min)       lcorbi_hilic_neg_4_076_58    m/z 458.8053     0   2   4   6   8E+3   7.5   8  RT (min)      lcorbi_hilic_neg_4_099_61    m/z 458.8801     0   2   4E+3   5   5.5  RT (min)      lcorbi_hilic_neg_4_109_62    m/z 458.9957     0   1   4.5   5  RT (min)      lcorbi_hilic_neg_4_090_60    m/z 459.0453     0   1E+4   6   6.5  RT (min)      lcorbi_hilic_neg_4_067_57    m/z 459.0598     0   2   4   6   8E+3   6   6.5  RT (min)      lcorbi_hilic_neg_4_083_59    m/z 459.1385     0   1E+4   6   6.5  RT (min)      lcorbi_hilic_neg_4_072_57    m/z 459.1456     0   1E+5   1   1.5  RT (min)      lcorbi_hilic_neg_1_002_26    m/z 459.1720     0   2   4   6E+3   5.5   6  RT (min)      lcorbi_hilic_neg_4_110_62    m/z 459.1796     0   1E+4   2.5   3  RT (min)      lcorbi_hilic_neg_1_037_46    m/z 459.2184     0   2   4   6E+4   0.5   1  RT (min)       lcorbi_hilic_neg_4_105_61    m/z 459.2512     0   1   1   1.5  RT (min)      lcorbi_hilic_neg_4_004_13    m/z 459.2519     0   2   4   6E+3   4   4.5  RT (min)      lcorbi_hilic_neg_4_081_58    m/z 459.2908     0   1E+5   0.5   1  RT (min)       lcorbi_hilic_neg_4_066_57    m/z 459.3119     0   1E+3   0.5   1  RT (min)      lcorbi_hilic_neg_4_004_13    m/z 459.3274     0   1E+3   0.5   1  RT (min)      lcorbi_hilic_neg_4_066_57    m/z 459.3368     0   1E+4   3.5   4  RT (min)      lcorbi_hilic_neg_4_088_59    m/z 459.3483     0   2   4   6E+3   0.5   1  RT (min)      lcorbi_hilic_neg_3_053_13    m/z 459.9130     0   2   4   6   8E+3   6   6.5  RT (min)      lcorbi_hilic_neg_4_074_58    m/z 459.9607     0   2   4   6   8E+3   5.5   6  RT (min)      lcorbi_hilic_neg_4_088_59    m/z 459.9773     0   1   5.5   6  RT (min)      lcorbi_hilic_neg_4_074_58    m/z 460.0764     0   1E+4   3   3.5  RT (min)      lcorbi_hilic_neg_4_085_59    m/z 460.1316     0   2   4   6E+3   6   6.5  RT (min)      lcorbi_hilic_neg_4_089_59    m/z 460.1510     0   1E+4   0.5   1  RT (min)      lcorbi_hilic_neg_4_091_60    m/z 460.1757     0   2   4   6E+4   2.5   3  RT (min)      lcorbi_hilic_neg_1_037_46    m/z 460.2219     0   1   0.5   1  RT (min)      lcorbi_hilic_neg_4_091_60    m/z 460.2402     0   2   4   6E+3   4   4.5  RT (min)      lcorbi_hilic_neg_1_047_05    m/z 460.2824     0   1E+4   1   1.5  RT (min)       lcorbi_hilic_neg_4_082_59    m/z 460.6487     0   1E+4   4.5   5  RT (min)      lcorbi_hilic_neg_4_111_62    m/z 460.8847     0   1E+4   8   8.5  RT (min)      lcorbi_hilic_neg_4_081_58    m/z 460.9341     0   1E+4   0.5   1  RT (min)      lcorbi_hilic_neg_4_070_57    m/z 460.9598     0   1E+4   7.5   8  RT (min)      lcorbi_hilic_neg_4_083_59    m/z 460.9937     0   1   2   3E+4   4.5   5  RT (min)       lcorbi_hilic_neg_4_077_58    m/z 461.0172     0   2   4   6   8E+3   5.5   6  RT (min)      lcorbi_hilic_neg_4_068_57    m/z 461.0577     0   1   2E+4   1   1.5  RT (min)      lcorbi_hilic_neg_4_106_62    m/z 461.0839     0   1E+4   2   2.5  RT (min)      lcorbi_hilic_neg_4_108_62    m/z 461.0998     0   1E+4   3.5   4  RT (min)      lcorbi_hilic_neg_4_111_62    m/z 461.1082     0   2   4   6   8E+3   6   6.5  RT (min)      lcorbi_hilic_neg_4_079_58    m/z 461.1706     0   2   4E+4   2   2.5  RT (min)       lcorbi_hilic_neg_4_106_62    m/z 461.1708     0   1E+4   6   6.5  RT (min)      lcorbi_hilic_neg_4_073_57    m/z 461.1794     0   2   4   6E+3   2.5   3  RT (min)      lcorbi_hilic_neg_1_037_46    m/z 461.3274     0   1   2E+4   0.5   1  RT (min)      lcorbi_hilic_neg_4_078_58    m/z 461.3640     0   2   4E+4   0.5   1  RT (min)      lcorbi_hilic_neg_4_080_58    m/z 461.8847     0   2   4E+3   8   8.5  RT (min)      lcorbi_hilic_neg_4_075_58    m/z 461.9564     0   2   4E+3   5.5   6  RT (min)      lcorbi_hilic_neg_4_076_58    m/z 461.9907     0   2   4   6E+3   6   6.5  RT (min)      lcorbi_hilic_neg_4_104_61    m/z 461.9921     0   1E+4   4.5   5  RT (min)       lcorbi_hilic_neg_4_080_58    m/z 462.0256     0   1E+4   6.5   7  RT (min)      lcorbi_hilic_neg_4_078_58    m/z 462.0415     0   1E+4   4.5   5  RT (min)      lcorbi_hilic_neg_4_107_62    m/z 462.1259     0   1   2   3E+3   6   6.5  RT (min)      lcorbi_hilic_neg_3_029_25    m/z 462.1385     0   1   2   3E+3   6   6.5  RT (min)      lcorbi_hilic_neg_1_013_47    m/z 462.1480     0   1E+4   0.5   1  RT (min)      lcorbi_hilic_neg_4_091_60    m/z 462.1733     0   2   4   6   8E+3   2   2.5  RT (min)      lcorbi_hilic_neg_4_106_62    m/z 462.1775     0   1E+4   3.5   4  RT (min)      lcorbi_hilic_neg_4_075_58    m/z 462.1867     0   1E+4   2   2.5  RT (min)      lcorbi_hilic_neg_4_011_19    m/z 462.2996     0   1   2E+4   4   4.5  RT (min)       lcorbi_hilic_neg_4_078_58    m/z 462.6457     0   2   4   6   8E+3   4.5   5  RT (min)      lcorbi_hilic_neg_4_104_61    m/z 462.8814     0   2   4   6E+3   8   8.5  RT (min)      lcorbi_hilic_neg_4_080_58    m/z 462.9103     0   2   4E+3   6   6.5  RT (min)      lcorbi_hilic_neg_4_076_58    m/z 462.9607     0   2   4   6E+3   5.5   6   6.5  RT (min)      lcorbi_hilic_neg_4_069_57    m/z 462.9907     0   2   4   6E+4   4.5   5  RT (min)       lcorbi_hilic_neg_4_077_58    m/z 463.0225     0   2   4   6   8E+3   0.5   1  RT (min)      lcorbi_hilic_neg_4_091_60    m/z 463.0540     0   1E+4   6   6.5  RT (min)      lcorbi_hilic_neg_4_070_57    m/z 463.0544     0   2   4   6   8E+3   1   1.5  RT (min)      lcorbi_hilic_neg_4_106_62    m/z 463.1958     0   1   2   3E+4   1   1.5  RT (min)       lcorbi_hilic_neg_1_002_26    m/z 463.2313     0   1E+4   7   7.5  RT (min)      lcorbi_hilic_neg_4_080_58    m/z 463.2911     0   2   4   6E+3   0.5   1  RT (min)       lcorbi_hilic_neg_4_074_58    m/z 463.3070     0   1   2E+4   4.5   5  RT (min)       lcorbi_hilic_neg_4_073_57    m/z 463.3079     0   2   4E+3   1.5   2  RT (min)      lcorbi_hilic_neg_4_077_58    m/z 463.3793     0   1   2E+4   0.5   1  RT (min)      lcorbi_hilic_neg_4_079_58    m/z 463.9336     0   2   4   6   8E+2   6   6.5  RT (min)      lcorbi_hilic_neg_4_005_32    m/z 463.9697     0   1E+4   5.5   6  RT (min)      lcorbi_hilic_neg_4_073_57    m/z 463.9891     0   1   2E+4   4.5   5  RT (min)       lcorbi_hilic_neg_4_079_58    m/z 464.0387     0   2   4   6   8E+3   4.5   5  RT (min)      lcorbi_hilic_neg_4_111_62    m/z 464.0525     0   2   4   6   8E+3   7   7.5  RT (min)      lcorbi_hilic_neg_1_039_38    m/z 464.0781     0   1E+5   0.5   1  RT (min)      lcorbi_hilic_neg_4_074_58    m/z 464.1267     0   1E+4   6   6.5  RT (min)      lcorbi_hilic_neg_4_089_59    m/z 464.1778     0   1   2E+4   3   3.5  RT (min)      lcorbi_hilic_neg_4_023_34    m/z 464.1964     0   1   2E+4   2.5   3  RT (min)      lcorbi_hilic_neg_1_037_46    m/z 464.2153     0   2   4   6   8E+3   7   7.5  RT (min)      lcorbi_hilic_neg_4_080_58    m/z 464.2783     0   2   4E+4   4   4.5  RT (min)       lcorbi_hilic_neg_1_047_05    m/z 464.3465     0   2   4   6E+2   0.5   1  RT (min)      lcorbi_hilic_neg_2_021_05    m/z 464.8132     0   2   4E+3   5   5.5  RT (min)      lcorbi_hilic_neg_4_093_60    m/z 464.9734     0   1   2   3   4E+3   7.5   8  RT (min)      lcorbi_hilic_neg_4_081_58    m/z 464.9774     0   1   2E+3   4.5   5  RT (min)      lcorbi_hilic_neg_4_077_58    m/z 464.9877     0   1   2   3   4E+4   4.5   5  RT (min)       lcorbi_hilic_neg_4_077_58    m/z 465.0044     0   1E+3   5   5.5  RT (min)      lcorbi_hilic_neg_4_090_60    m/z 465.0480     0   1E+4   1   1.5  RT (min)      lcorbi_hilic_neg_4_050_26    m/z 465.0686     0   1E+4   1   1.5  RT (min)      lcorbi_hilic_neg_4_087_59    m/z 465.0802     0   1   2   3   4E+4   0.5   1  RT (min)      lcorbi_hilic_neg_3_031_14    m/z 465.1270     0   1   2   3   4E+4   1   1.5  RT (min)       lcorbi_hilic_neg_4_091_60    m/z 465.1371     0   1E+4   5.5   6  RT (min)      lcorbi_hilic_neg_4_096_60    m/z 465.2143     0   1E+4   0.5   1  RT (min)      lcorbi_hilic_neg_4_085_59    m/z 465.2497     0   1E+4   3   3.5  RT (min)      lcorbi_hilic_neg_4_077_58    m/z 465.2869     0   1   2E+3   1.5   2  RT (min)      lcorbi_hilic_neg_2_018_16    m/z 465.3066     0   1   2   3   4E+5   3.5   4  RT (min)      lcorbi_hilic_neg_4_014_18    m/z 465.3186     0   1E+4   4   4.5  RT (min)      lcorbi_hilic_neg_4_081_58    m/z 465.3237     0   1   2   3E+4   0.5   1  RT (min)      lcorbi_hilic_neg_3_016_16    m/z 465.8092     0   1   2E+4   4.5   5  RT (min)       lcorbi_hilic_neg_4_076_58    m/z 465.9714     0   2   4E+3   7.5   8  RT (min)      lcorbi_hilic_neg_4_078_58    m/z 466.0118     0   2   4   6E+3   5   5.5  RT (min)      lcorbi_hilic_neg_4_110_62    m/z 466.2638     0   2   4   6   8E+3   0.5   1  RT (min)      lcorbi_hilic_neg_4_069_57    m/z 466.2944     0   1E+4   4   4.5  RT (min)       lcorbi_hilic_neg_4_089_59    m/z 466.3307     0   1   2E+4   4   4.5  RT (min)       lcorbi_hilic_neg_4_075_58    m/z 466.3622     0   2   4E+4   0.5   1  RT (min)      lcorbi_hilic_neg_4_075_58    m/z 466.8103     0   1E+3   5   5.5  RT (min)      lcorbi_hilic_neg_1_047_05    m/z 466.8401     0   2   4   6   8E+3   8   8.5  RT (min)      lcorbi_hilic_neg_4_105_61    m/z 466.9529     0   1   2   3E+4   7.5   8  RT (min)       lcorbi_hilic_neg_4_080_58    m/z 466.9628     0   2   4   6E+3   7.5   8  RT (min)      lcorbi_hilic_neg_4_075_58    m/z 466.9850     0   1   4.5   5  RT (min)       lcorbi_hilic_neg_4_105_61    m/z 467.0296     0   2   4   6   8E+3   2   2.5  RT (min)      lcorbi_hilic_neg_4_110_62    m/z 467.0366     0   1   2E+4   7   7.5  RT (min)       lcorbi_hilic_neg_4_086_59    m/z 467.0548     0   1E+4   6   6.5  RT (min)      lcorbi_hilic_neg_4_109_62    m/z 467.0989     0   2   4   6   8E+3   1   1.5  RT (min)      lcorbi_hilic_neg_4_100_61    m/z 467.1426     0   1   2E+4   6.5   7  RT (min)       lcorbi_hilic_neg_4_094_60    m/z 467.1767     0   1E+4   7   7.5  RT (min)      lcorbi_hilic_neg_4_087_59    m/z 467.2216     0   1   2   3   4E+4   1   1.5  RT (min)       lcorbi_hilic_neg_4_107_62    m/z 467.2410     0   1E+4   2   2.5  RT (min)      lcorbi_hilic_neg_3_018_38    m/z 467.3341     0   2   4   6   8E+3   4   4.5  RT (min)      lcorbi_hilic_neg_4_080_58    m/z 467.3503     0   1E+4   3.5   4  RT (min)      lcorbi_hilic_neg_4_086_59    m/z 467.8072     0   1E+4   4.5   5  RT (min)      lcorbi_hilic_neg_4_076_58    m/z 467.9506     0   1   2E+4   7.5   8  RT (min)      lcorbi_hilic_neg_4_078_58    m/z 467.9992     0   1   5.5   6  RT (min)      lcorbi_hilic_neg_2_048_38    m/z 468.0090     0   1   2   3E+4   5.5   6  RT (min)       lcorbi_hilic_neg_4_089_59    m/z 468.0817     0   1E+4   3   3.5  RT (min)      lcorbi_hilic_neg_4_103_61    m/z 468.1239     0   1E+3   6   6.5  RT (min)      lcorbi_hilic_neg_1_023_23    m/z 468.1339     0   1   2E+5   6   6.5  RT (min)       lcorbi_hilic_neg_1_023_23    m/z 468.2249     0   1E+4   1   1.5  RT (min)      lcorbi_hilic_neg_4_107_62    m/z 468.2474     0   2   4   6   8E+3   7   7.5  RT (min)      lcorbi_hilic_neg_4_086_59    m/z 468.3144     0   1   0.5   1  RT (min)      lcorbi_hilic_neg_4_071_57    m/z 468.6918     0   2   4E+3   4.5   5  RT (min)      lcorbi_hilic_neg_4_100_61    m/z 468.7649     0   2   4   6   8E+3   7.5   8  RT (min)      lcorbi_hilic_neg_4_106_62    m/z 468.8021     0   1   2E+4   4.5   5  RT (min)       lcorbi_hilic_neg_4_078_58    m/z 468.8967     0   1   2E+4   5   5.5  RT (min)       lcorbi_hilic_neg_4_113_62    m/z 468.9481     0   1E+4   7.5   8  RT (min)      lcorbi_hilic_neg_4_081_58    m/z 468.9578     0   1   2   3E+3   7.5   8  RT (min)      lcorbi_hilic_neg_4_078_58    m/z 468.9638     0   1   2E+3   7.5   8  RT (min)      lcorbi_hilic_neg_3_034_34    m/z 468.9831     0   1   5.5   6  RT (min)      lcorbi_hilic_neg_4_098_61    m/z 468.9891     0   1E+4   4   4.5  RT (min)      lcorbi_hilic_neg_4_102_61    m/z 469.0294     0   1E+4   0.5   1  RT (min)      lcorbi_hilic_neg_4_076_58    m/z 469.0859     0   1E+4   6.5   7  RT (min)      lcorbi_hilic_neg_4_084_59    m/z 469.1176     0   1   2   3E+4   4.5   5   5.5  RT (min)       lcorbi_hilic_neg_4_093_60    m/z 469.2180     0   1E+4   1   1.5  RT (min)      lcorbi_hilic_neg_4_106_62    m/z 469.2264     0   2   4   6   8E+3   1   1.5  RT (min)      lcorbi_hilic_neg_4_075_58    m/z 469.2307     0   1   2   3   4E+4   4   4.5  RT (min)       lcorbi_hilic_neg_4_087_59    m/z 469.2579     0   1   2E+3   4   4.5  RT (min)      lcorbi_hilic_neg_1_047_05    m/z 469.3039     0   1E+4   3.5   4  RT (min)      lcorbi_hilic_neg_4_083_59    m/z 469.3330     0   1   2   3   4E+4   1   1.5  RT (min)      lcorbi_hilic_neg_1_002_26    m/z 469.3397     0   1E+4   4.5   5  RT (min)      lcorbi_hilic_neg_4_089_59    m/z 469.3649     0   2   4   6E+3   0.5   1  RT (min)      lcorbi_hilic_neg_4_085_59    m/z 469.3904     0   1   2   3   4E+4   0.5   1  RT (min)      lcorbi_hilic_neg_4_097_60    m/z 469.9418     0   1E+4   6   6.5  RT (min)      lcorbi_hilic_neg_4_075_58    m/z 469.9786     0   2   4   6   8E+4   0.5   1  RT (min)       lcorbi_hilic_neg_4_076_58    m/z 469.9898     0   1E+4   5.5   6  RT (min)      lcorbi_hilic_neg_4_105_61    m/z 470.0763     0   1E+4   2.5   3  RT (min)      lcorbi_hilic_neg_4_038_27    m/z 470.1907     0   1   2E+4   0.5   1  RT (min)      lcorbi_hilic_neg_4_113_62    m/z 470.2343     0   1E+4   4   4.5  RT (min)      lcorbi_hilic_neg_4_083_59    m/z 470.3355     0   1   2E+3   1   1.5  RT (min)      lcorbi_hilic_neg_1_047_05    m/z 470.3692     0   1   2E+4   3.5   4  RT (min)       lcorbi_hilic_neg_4_089_59    m/z 470.3855     0   1E+4   1   1.5  RT (min)      lcorbi_hilic_neg_4_084_59    m/z 470.7587     0   1   4.5   5  RT (min)      lcorbi_hilic_neg_4_098_61    m/z 471.0239     0   2   4   6   8E+3   4   4.5  RT (min)      lcorbi_hilic_neg_4_103_61    m/z 471.0459     0   1E+4   5.5   6  RT (min)      lcorbi_hilic_neg_4_066_57    m/z 471.0755     0   2   4   6E+3   7   7.5  RT (min)      lcorbi_hilic_neg_4_091_60    m/z 471.0853     0   1E+4   6.5   7  RT (min)       lcorbi_hilic_neg_4_095_60    m/z 471.0864     0   2   4   6   8E+3   1   1.5  RT (min)      lcorbi_hilic_neg_4_106_62    m/z 471.1474     0   1   2   3   4E+4   5   6   7  RT (min)       lcorbi_hilic_neg_4_111_62    m/z 471.2435     0   1E+4   1.5   2  RT (min)      lcorbi_hilic_neg_4_072_57    m/z 471.2795     0   1   0.5   1  RT (min)      lcorbi_hilic_neg_4_073_57    m/z 471.2904     0   1   2E+3   0.5   1  RT (min)      lcorbi_hilic_neg_4_073_57    m/z 471.2942     0   1E+4   6.5   7  RT (min)      lcorbi_hilic_neg_4_078_58    m/z 471.3318     0   1E+5   4   4.5  RT (min)       lcorbi_hilic_neg_4_095_60    m/z 471.3488     0   1   2   3   4E+4   1   1.5  RT (min)      lcorbi_hilic_neg_4_075_58    m/z 471.3815     0   1   2   3E+4   3.5   4  RT (min)       lcorbi_hilic_neg_4_088_59    m/z 471.9571     0   2   4   6   8E+3   7.5   8  RT (min)      lcorbi_hilic_neg_4_079_58    m/z 471.9870     0   1   2   3   4E+3   5.5   6  RT (min)      lcorbi_hilic_neg_4_100_61    m/z 472.0034     0   2   4   6   8E+3   6.5   7  RT (min)      lcorbi_hilic_neg_4_088_59    m/z 472.0741     0   1E+4   2.5   3  RT (min)      lcorbi_hilic_neg_4_038_27    m/z 472.0788     0   1   2   3   4E+2   6.5   7  RT (min)       lcorbi_hilic_neg_1_012_32    m/z 472.1162     0   1E+4   2   2.5  RT (min)      lcorbi_hilic_neg_4_109_62    m/z 472.1551     0   1E+4   6.5   7  RT (min)      lcorbi_hilic_neg_4_097_60    m/z 472.1671     0   2   4   6E+3   6   6.5  RT (min)      lcorbi_hilic_neg_4_095_60    m/z 472.2473     0   2   4   6E+3   1   1.5  RT (min)      lcorbi_hilic_neg_4_067_57    m/z 472.2709     0   2   4   6   8E+3   0.5   1  RT (min)      lcorbi_hilic_neg_4_068_57    m/z 472.3449     0   2   4   6E+4   4   4.5  RT (min)       lcorbi_hilic_neg_4_092_60    m/z 472.3455     0   1E+5   0.5   1  RT (min)      lcorbi_hilic_neg_1_047_05    m/z 472.7557     0   1   4.5   5  RT (min)      lcorbi_hilic_neg_4_102_61    m/z 472.7837     0   1E+4   7.5   8  RT (min)      lcorbi_hilic_neg_4_102_61    m/z 472.9394     0   2   4   6   8E+3   6   6.5  RT (min)      lcorbi_hilic_neg_4_075_58    m/z 472.9892     0   2   4   6   8E+3   6   6.5  RT (min)      lcorbi_hilic_neg_4_066_57    m/z 473.0042     0   1E+4   0.5   1  RT (min)      lcorbi_hilic_neg_4_073_57    m/z 473.0567     0   1   2E+4   7   7.5  RT (min)      lcorbi_hilic_neg_4_097_60    m/z 473.0666     0   1   2   3   4E+4   5.5   6  RT (min)       lcorbi_hilic_neg_4_089_59    m/z 473.1483     0   1E+4   2   2.5  RT (min)      lcorbi_hilic_neg_4_055_38    m/z 473.1507     0   1   6   6.5  RT (min)      lcorbi_hilic_neg_2_032_25    m/z 473.1631     0   2   4   6E+3   6.5   7  RT (min)      lcorbi_hilic_neg_4_005_32    m/z 473.1738     0   1   2   3   4E+4   6   6.5  RT (min)       lcorbi_hilic_neg_4_095_60    m/z 473.1799     0   1   2   3   4E+3   4   4.5  RT (min)      lcorbi_hilic_neg_2_016_30    m/z 473.2055     0   1E+4   1   1.5  RT (min)      lcorbi_hilic_neg_2_019_21    m/z 473.3122     0   1   2   3   4E+3   1   1.5  RT (min)      lcorbi_hilic_neg_1_018_21    m/z 473.3495     0   2   4E+4   1   1.5  RT (min)       lcorbi_hilic_neg_4_069_57    m/z 473.9547     0   1   5.5   6  RT (min)      lcorbi_hilic_neg_4_068_57    m/z 473.9554     0   2   4E+3   7.5   8  RT (min)      lcorbi_hilic_neg_4_080_58    m/z 473.9611     0   1   2E+4   1   1.5  RT (min)      lcorbi_hilic_neg_4_109_62    m/z 473.9986     0   1   2E+4   5.5   6  RT (min)       lcorbi_hilic_neg_4_080_58    m/z 474.0102     0   1E+3   5.5   6  RT (min)      lcorbi_hilic_neg_4_078_58    m/z 474.3525     0   1E+4   1   1.5  RT (min)      lcorbi_hilic_neg_4_066_57    m/z 474.3671     0   1E+5   0.5   1  RT (min)      lcorbi_hilic_neg_4_076_58    m/z 474.8421     0   2   4   6   8E+3   5   5.5  RT (min)      lcorbi_hilic_neg_4_091_60    m/z 474.9667     0   1   2   3   4E+3   5.5   6  RT (min)      lcorbi_hilic_neg_4_066_57    m/z 475.0334     0   1E+4   5   5.5  RT (min)      lcorbi_hilic_neg_4_071_57    m/z 475.0558     0   2   4   6   8E+3   2   2.5  RT (min)      lcorbi_hilic_neg_4_110_62    m/z 475.1474     0   1   2   3   4E+3   2   2.5  RT (min)      lcorbi_hilic_neg_4_042_28    m/z 475.1500     0   1E+4   7   7.5  RT (min)      lcorbi_hilic_neg_4_085_59    m/z 475.2660     0   1E+5   4   4.5  RT (min)       lcorbi_hilic_neg_1_047_05    m/z 475.3063     0   1E+4   0.5   1  RT (min)      lcorbi_hilic_neg_4_082_59    m/z 475.3280     0   2   4E+4   0.5   1  RT (min)      lcorbi_hilic_neg_4_066_57    m/z 475.3425     0   2   4   6   8E+2   0.5   1  RT (min)      lcorbi_hilic_neg_1_013_47    m/z 475.9582     0   1   2E+4   1   1.5  RT (min)      lcorbi_hilic_neg_4_107_62    m/z 476.0405     0   1E+4   5   5.5  RT (min)      lcorbi_hilic_neg_4_110_62    m/z 476.0913     0   2   4   6E+3   6   6.5  RT (min)      lcorbi_hilic_neg_1_012_32    m/z 476.1508     0   1E+4   6   6.5  RT (min)      lcorbi_hilic_neg_2_032_25    m/z 476.1608     0   1E+4   6   6.5  RT (min)      lcorbi_hilic_neg_1_047_05    m/z 476.1686     0   2   4   6   8E+3   2   2.5  RT (min)      lcorbi_hilic_neg_4_108_62    m/z 476.1745     0   1E+4   6.5   7  RT (min)      lcorbi_hilic_neg_4_088_59    m/z 476.2923     0   1E+4   0.5   1  RT (min)      lcorbi_hilic_neg_1_028_45    m/z 476.3313     0   1E+4   1   1.5  RT (min)      lcorbi_hilic_neg_4_068_57    m/z 476.3830     0   2   4   6   8E+4   0.5   1  RT (min)      lcorbi_hilic_neg_4_031_14    m/z 477.0319     0   1E+4   1   1.5  RT (min)      lcorbi_hilic_neg_4_106_62    m/z 477.0577     0   2   4E+3   2   2.5  RT (min)      lcorbi_hilic_neg_4_098_61    m/z 477.0838     0   1E+4   6   6.5  RT (min)       lcorbi_hilic_neg_4_112_62    m/z 477.1342     0   2   4   6E+3   2   2.5  RT (min)      lcorbi_hilic_neg_4_110_62    m/z 477.1633     0   2   4   6E+3   2.5   3  RT (min)      lcorbi_hilic_neg_4_080_58    m/z 477.1770     0   1E+5   0.5   1  RT (min)      lcorbi_hilic_neg_4_105_61    m/z 477.2498     0   1   2E+4   0.5   1  RT (min)      lcorbi_hilic_neg_4_107_62    m/z 477.2635     0   1E+4   1   1.5  RT (min)      lcorbi_hilic_neg_4_066_57    m/z 477.3042     0   1   2   3E+3   0.5   1  RT (min)      lcorbi_hilic_neg_4_086_59    m/z 477.3226     0   2   4   6E+3   1   1.5  RT (min)      lcorbi_hilic_neg_4_074_58    m/z 477.3582     0   2   4   6   8E+3   0.5   1  RT (min)      lcorbi_hilic_neg_4_080_58    m/z 477.9092     0   1E+4   1   1.5  RT (min)      lcorbi_hilic_neg_4_098_61    m/z 478.0377     0   2   4E+3   5   5.5  RT (min)      lcorbi_hilic_neg_4_109_62    m/z 478.1475     0   2   4E+4   2   2.5  RT (min)      lcorbi_hilic_neg_4_106_62    m/z 478.1605     0   2   4   6E+3   2   2.5  RT (min)      lcorbi_hilic_neg_4_011_19    m/z 478.1608     0   1   6.5   7   7.5  RT (min)       lcorbi_hilic_neg_4_085_59    m/z 478.1806     0   1   2E+4   0.5   1  RT (min)      lcorbi_hilic_neg_4_105_61    m/z 478.2835     0   1   4   4.5  RT (min)      lcorbi_hilic_neg_4_070_57    m/z 478.3985     0   2   4   6E+4   0.5   1  RT (min)      lcorbi_hilic_neg_4_031_14    m/z 478.7008     0   2   4E+3   4.5   5  RT (min)      lcorbi_hilic_neg_4_080_58    m/z 478.7912     0   1E+4   7.5   8  RT (min)      lcorbi_hilic_neg_4_102_61    m/z 478.9343     0   2   4E+3   5.5   6   6.5  RT (min)      lcorbi_hilic_neg_2_053_03    m/z 479.0290     0   2   4   6E+3   1   1.5  RT (min)      lcorbi_hilic_neg_4_111_62    m/z 479.0746     0   1   2   3E+3   6.5   7  RT (min)      lcorbi_hilic_neg_4_088_59    m/z 479.0890     0   1   2E+4   0.5   1  RT (min)      lcorbi_hilic_neg_1_012_32    m/z 479.0961     0   1   5.5   6  RT (min)      lcorbi_hilic_neg_4_072_57    m/z 479.1399     0   2   4   6E+4   5.5   6   6.5  RT (min)       lcorbi_hilic_neg_4_072_57    m/z 479.1782     0   2   4   6   8E+4   2.5   3  RT (min)      lcorbi_hilic_neg_4_055_38    m/z 479.2169     0   1E+4   7   7.5  RT (min)      lcorbi_hilic_neg_4_096_60    m/z 479.2903     0   2   4   6   8E+4   1.5   2  RT (min)       lcorbi_hilic_neg_4_072_57    m/z 479.3381     0   1E+4   0.5   1  RT (min)      lcorbi_hilic_neg_4_080_58    m/z 479.3741     0   1   2E+4   0.5   1  RT (min)      lcorbi_hilic_neg_4_031_14    m/z 479.4107     0   1   2   3   4E+4   0.5   1  RT (min)      lcorbi_hilic_neg_4_078_58    m/z 479.9062     0   1E+4   1   1.5  RT (min)      lcorbi_hilic_neg_4_098_61    m/z 479.9234     0   2   4   6   8E+3   5   5.5  RT (min)      lcorbi_hilic_neg_4_101_61    m/z 479.9707     0   1E+4   6   6.5  RT (min)      lcorbi_hilic_neg_4_074_58    m/z 480.0603     0   2   4   6E+3   6   6.5  RT (min)      lcorbi_hilic_neg_4_048_37    m/z 480.0740     0   1   2E+4   6.5   7  RT (min)      lcorbi_hilic_neg_2_031_34    m/z 480.0884     0   1   5.5   6  RT (min)      lcorbi_hilic_neg_4_072_57    m/z 480.1089     0   1   5.5   6  RT (min)      lcorbi_hilic_neg_2_005_24    m/z 480.1430     0   1E+4   5.5   6  RT (min)      lcorbi_hilic_neg_4_068_57    m/z 480.1816     0   1E+4   2.5   3  RT (min)      lcorbi_hilic_neg_4_055_38    m/z 480.1840     0   1E+4   6.5   7  RT (min)      lcorbi_hilic_neg_4_013_46    m/z 480.2874     0   1   2   3   4E+3   0.5   1  RT (min)      lcorbi_hilic_neg_4_066_57    m/z 480.3235     0   1E+4   0.5   1  RT (min)      lcorbi_hilic_neg_2_012_14    m/z 480.5646     0   2   4   6   8E+4   4.5   5  RT (min)      lcorbi_hilic_neg_2_020_27    m/z 480.7872     0   1   4.5   5  RT (min)      lcorbi_hilic_neg_4_098_61    m/z 480.9785     0   1E+3   5   5.5  RT (min)      lcorbi_hilic_neg_4_092_60    m/z 481.0136     0   1E+4   0.5   1  RT (min)      lcorbi_hilic_neg_4_112_62    m/z 481.0317     0   1   2E+4   1   1.5  RT (min)      lcorbi_hilic_neg_4_082_59    m/z 481.0598     0   1E+3   6   6.5  RT (min)      lcorbi_hilic_neg_3_031_14    m/z 481.1373     0   1   2E+4   5.5   6  RT (min)       lcorbi_hilic_neg_4_073_57    m/z 481.1551     0   1   2   3   4E+4   0.5   1  RT (min)       lcorbi_hilic_neg_4_104_61    m/z 481.1772     0   1E+5   6   6.5  RT (min)       lcorbi_hilic_neg_4_095_60    m/z 481.2366     0   1   2E+4   4   4.5  RT (min)       lcorbi_hilic_neg_4_076_58    m/z 481.2452     0   2   4E+3   3.5   4  RT (min)      lcorbi_hilic_neg_4_076_58    m/z 481.2574     0   1   2E+3   4   4.5  RT (min)      lcorbi_hilic_neg_1_032_02    m/z 481.2830     0   1   2E+4   2   2.5  RT (min)      lcorbi_hilic_neg_4_071_57    m/z 481.2999     0   2   4   6   8E+3   0.5   1  RT (min)      lcorbi_hilic_neg_4_080_58    m/z 481.3538     0   1   2E+4   0.5   1  RT (min)      lcorbi_hilic_neg_4_080_58    m/z 481.3902     0   1E+4   0.5   1  RT (min)      lcorbi_hilic_neg_4_073_57    m/z 482.0343     0   1E+4   6.5   7  RT (min)      lcorbi_hilic_neg_4_089_59    m/z 482.0726     0   1E+4   6.5   7  RT (min)      lcorbi_hilic_neg_4_076_58    m/z 482.0953     0   1E+4   6.5   7  RT (min)      lcorbi_hilic_neg_4_084_59    m/z 482.1487     0   1   6.5   7  RT (min)      lcorbi_hilic_neg_4_076_58    m/z 482.1679     0   1E+4   6.5   7  RT (min)      lcorbi_hilic_neg_4_072_57    m/z 482.1798     0   1   6.5   7  RT (min)      lcorbi_hilic_neg_4_089_59    m/z 482.1904     0   1   2   3E+4   6.5   7  RT (min)       lcorbi_hilic_neg_4_083_59    m/z 482.3158     0   1   2   3E+4   4   4.5  RT (min)      lcorbi_hilic_neg_4_079_58    m/z 482.3360     0   1   2E+3   0.5   1  RT (min)      lcorbi_hilic_neg_2_021_05    m/z 482.5618     0   1E+5   4.5   5  RT (min)       lcorbi_hilic_neg_2_020_27    m/z 482.7846     0   1   2   3E+3   4.5   5  RT (min)      lcorbi_hilic_neg_4_066_57    m/z 482.9657     0   1E+4   4   4.5  RT (min)      lcorbi_hilic_neg_4_102_61    m/z 482.9684     0   2   4   6   8E+3   6   6.5  RT (min)      lcorbi_hilic_neg_4_079_58    m/z 483.0448     0   2   4   6E+3   3   3.5  RT (min)      lcorbi_hilic_neg_4_077_58    m/z 483.0581     0   1E+4   6   6.5  RT (min)      lcorbi_hilic_neg_3_023_36    m/z 483.0669     0   1   2   3   4E+3   6.5   7  RT (min)      lcorbi_hilic_neg_4_076_58    m/z 483.0910     0   1E+4   6   6.5  RT (min)       lcorbi_hilic_neg_4_079_58    m/z 483.3197     0   1   2   3   4E+4   3.5   4  RT (min)       lcorbi_hilic_neg_4_086_59    m/z 483.3239     0   1   2   3   4E+4   1   1.5  RT (min)      lcorbi_hilic_neg_1_021_28    m/z 483.3696     0   1   2E+4   0.5   1  RT (min)      lcorbi_hilic_neg_4_077_58    m/z 483.9832     0   2   4   6E+3   5.5   6  RT (min)      lcorbi_hilic_neg_4_086_59    m/z 484.0527     0   1   6   6.5  RT (min)      lcorbi_hilic_neg_4_089_59    m/z 484.0559     0   1   2E+4   3   3.5  RT (min)       lcorbi_hilic_neg_4_101_61    m/z 484.2772     0   1E+4   3.5   4  RT (min)      lcorbi_hilic_neg_4_086_59    m/z 484.2851     0   1E+4   1.5   2  RT (min)      lcorbi_hilic_neg_4_076_58    m/z 484.3236     0   1E+4   3.5   4  RT (min)      lcorbi_hilic_neg_4_089_59    m/z 484.5593     0   2   4   6E+4   4.5   5  RT (min)       lcorbi_hilic_neg_2_020_27    m/z 484.8708     0   1E+4   5   5.5  RT (min)      lcorbi_hilic_neg_4_090_60    m/z 484.9629     0   1E+4   4   4.5  RT (min)      lcorbi_hilic_neg_4_105_61    m/z 484.9801     0   1   2E+4   1   1.5  RT (min)      lcorbi_hilic_neg_4_098_61    m/z 485.1674     0   2   4   6   8E+3   3.5   4  RT (min)      lcorbi_hilic_neg_4_075_58    m/z 485.2731     0   1   0.5   1  RT (min)       lcorbi_hilic_neg_2_032_25    m/z 485.3347     0   1   0.5   1  RT (min)      lcorbi_hilic_neg_4_113_62    m/z 485.3899     0   1   2   3E+4   0.5   1  RT (min)      lcorbi_hilic_neg_4_097_60    m/z 485.9010     0   2   4E+3   7.5   8  RT (min)      lcorbi_hilic_neg_4_077_58    m/z 486.0346     0   2   4   6   8E+3   7   7.5  RT (min)      lcorbi_hilic_neg_1_016_37    m/z 486.0529     0   1E+4   3   3.5  RT (min)      lcorbi_hilic_neg_4_101_61    m/z 486.0697     0   1   5   5.5  RT (min)       lcorbi_hilic_neg_4_005_32    m/z 486.0877     0   1   6.5   7  RT (min)      lcorbi_hilic_neg_4_005_32    m/z 486.1095     0   1E+4   6   6.5  RT (min)      lcorbi_hilic_neg_4_083_59    m/z 486.1716     0   1E+4   6.5   7  RT (min)      lcorbi_hilic_neg_4_091_60    m/z 486.1823     0   1   6.5   7  RT (min)      lcorbi_hilic_neg_4_090_60    m/z 486.1880     0   1E+4   7   7.5  RT (min)      lcorbi_hilic_neg_4_030_05    m/z 486.3052     0   2   4E+4   6   6.5  RT (min)       lcorbi_hilic_neg_4_080_58    m/z 486.3126     0   1E+3   0.5   1  RT (min)      lcorbi_hilic_neg_4_068_57    m/z 486.7323     0   2   4   6E+3   4.5   5  RT (min)      lcorbi_hilic_neg_4_078_58    m/z 487.0199     0   1E+4   3   3.5  RT (min)      lcorbi_hilic_neg_4_090_60    m/z 487.0202     0   1E+4   0.5   1  RT (min)      lcorbi_hilic_neg_4_069_57    m/z 487.0726     0   2   4   6   8E+4   6   6.5  RT (min)      lcorbi_hilic_neg_3_044_32    m/z 487.0870     0   1   2E+4   2   2.5  RT (min)       lcorbi_hilic_neg_4_108_62    m/z 487.1256     0   1E+4   1.5   2  RT (min)      lcorbi_hilic_neg_4_104_61    m/z 487.2498     0   1   2E+4   0.5   1  RT (min)      lcorbi_hilic_neg_4_111_62    m/z 487.2746     0   1   2   3E+4   0.5   1  RT (min)      lcorbi_hilic_neg_3_039_31    m/z 487.2914     0   1E+4   0.5   1  RT (min)       lcorbi_hilic_neg_4_073_57    m/z 487.3069     0   1E+5   2   2.5  RT (min)       lcorbi_hilic_neg_4_089_59    m/z 487.3086     0   1E+4   6   6.5  RT (min)      lcorbi_hilic_neg_4_078_58    m/z 487.3290     0   1   2   2.5  RT (min)      lcorbi_hilic_neg_4_089_59    m/z 488.0760     0   2   4E+3   1   1.5  RT (min)      lcorbi_hilic_neg_4_107_62    m/z 488.1264     0   1E+4   5.5   6  RT (min)      lcorbi_hilic_neg_3_015_47    m/z 488.1393     0   1   2   3E+4   3   3.5  RT (min)       lcorbi_hilic_neg_4_100_61    m/z 488.1635     0   1   2E+4   6   6.5  RT (min)      lcorbi_hilic_neg_4_023_34    m/z 488.1828     0   1   2   3E+4   6.5   7  RT (min)      lcorbi_hilic_neg_4_071_57    m/z 488.4936     0   1E+4   4.5   5  RT (min)      lcorbi_hilic_neg_4_079_58    m/z 488.7297     0   2   4   6   8E+3   4.5   5  RT (min)      lcorbi_hilic_neg_4_080_58    m/z 488.9350     0   1E+4   7.5   8  RT (min)       lcorbi_hilic_neg_4_077_58    m/z 488.9449     0   2   4   6E+3   7.5   8  RT (min)      lcorbi_hilic_neg_4_075_58    m/z 488.9631     0   2   4   6E+3   6   6.5  RT (min)      lcorbi_hilic_neg_4_066_57    m/z 489.0187     0   1   2E+4   7   7.5  RT (min)       lcorbi_hilic_neg_4_086_59    m/z 489.1589     0   1E+4   7   7.5  RT (min)      lcorbi_hilic_neg_4_082_59    m/z 489.1686     0   1   2E+4   5.5   6  RT (min)       lcorbi_hilic_neg_4_072_57    m/z 489.2178     0   1   2   3   4E+4   6.5   7  RT (min)       lcorbi_hilic_neg_4_095_60    m/z 489.2233     0   1E+4   2   2.5  RT (min)      lcorbi_hilic_neg_4_038_27    m/z 489.3043     0   1   2E+5   2   2.5   3  RT (min)       lcorbi_hilic_neg_4_082_59    m/z 489.3190     0   1E+5   1   1.5  RT (min)       lcorbi_hilic_neg_4_066_57    m/z 489.3588     0   2   4E+4   0.5   1  RT (min)       lcorbi_hilic_neg_4_074_58    m/z 489.4925     0   1E+4   4.5   5  RT (min)      lcorbi_hilic_neg_4_079_58    m/z 489.9323     0   1E+4   7.5   8  RT (min)      lcorbi_hilic_neg_4_081_58    m/z 489.9328     0   2   4   6E+3   5.5   6  RT (min)      lcorbi_hilic_neg_4_073_57    m/z 489.9427     0   1E+4   7.5   8  RT (min)      lcorbi_hilic_neg_4_080_58    m/z 489.9526     0   1E+4   5   5.5  RT (min)      lcorbi_hilic_neg_4_103_61    m/z 489.9718     0   1   2   3   4E+3   5   5.5  RT (min)      lcorbi_hilic_neg_4_113_62    m/z 490.0828     0   2   4E+3   5.5   6  RT (min)      lcorbi_hilic_neg_1_037_46    m/z 490.2013     0   1E+5   7   7.5  RT (min)       lcorbi_hilic_neg_4_097_60    m/z 490.2356     0   1E+4   6.5   7  RT (min)      lcorbi_hilic_neg_4_087_59    m/z 490.2958     0   1   2   3E+4   3.5   4  RT (min)       lcorbi_hilic_neg_4_089_59    m/z 490.3073     0   2   4   6E+4   2.5   3  RT (min)       lcorbi_hilic_neg_4_089_59    m/z 490.3623     0   1   2E+4   0.5   1  RT (min)      lcorbi_hilic_neg_4_076_58    m/z 490.4909     0   1E+4   4.5   5  RT (min)      lcorbi_hilic_neg_4_079_58    m/z 490.7871     0   2   4   6E+3   8   8.5  RT (min)      lcorbi_hilic_neg_4_066_57    m/z 490.8160     0   1   4.5   5  RT (min)      lcorbi_hilic_neg_4_106_62    m/z 490.9308     0   2   4   6   8E+3   7.5   8  RT (min)      lcorbi_hilic_neg_4_078_58    m/z 491.0075     0   1E+3   5   5.5  RT (min)      lcorbi_hilic_neg_4_105_61    m/z 491.1882     0   1E+3   6.5   7  RT (min)      lcorbi_hilic_neg_3_044_32    m/z 491.1994     0   1   2E+4   6.5   7  RT (min)      lcorbi_hilic_neg_3_044_32    m/z 491.3205     0   1   2E+4   0.5   1  RT (min)      lcorbi_hilic_neg_4_069_57    m/z 491.4892     0   1E+4   4.5   5  RT (min)      lcorbi_hilic_neg_4_081_58    m/z 491.7851     0   2   4   6E+3   8   8.5  RT (min)      lcorbi_hilic_neg_4_066_57    m/z 492.0147     0   2   4E+3   5   5.5  RT (min)      lcorbi_hilic_neg_4_093_60    m/z 492.0546     0   1E+4   6   6.5  RT (min)      lcorbi_hilic_neg_4_072_57    m/z 492.0647     0   1E+4   6.5   7  RT (min)      lcorbi_hilic_neg_4_091_60    m/z 492.1966     0   2   4   6   8E+3   6.5   7  RT (min)      lcorbi_hilic_neg_1_037_46    m/z 492.2737     0   1   2   3E+3   4   4.5  RT (min)      lcorbi_hilic_neg_4_088_59    m/z 492.3101     0   1E+4   3.5   4  RT (min)       lcorbi_hilic_neg_4_084_59    m/z 492.3461     0   2   4   6E+3   4   4.5  RT (min)      lcorbi_hilic_neg_4_079_58    m/z 492.3781     0   1   2   3E+4   0.5   1  RT (min)      lcorbi_hilic_neg_4_074_58    m/z 492.8136     0   1   4.5   5  RT (min)      lcorbi_hilic_neg_4_108_62    m/z 493.0041     0   1   2   3   4E+3   6   6.5  RT (min)      lcorbi_hilic_neg_4_109_62    m/z 493.0556     0   1   2   3E+4   6   6.5  RT (min)       lcorbi_hilic_neg_4_072_57    m/z 493.0974     0   1E+4   6   6.5  RT (min)      lcorbi_hilic_neg_4_089_59    m/z 493.1194     0   1E+4   6   6.5  RT (min)      lcorbi_hilic_neg_4_079_58    m/z 493.1264     0   2   4   6E+3   2   2.5  RT (min)      lcorbi_hilic_neg_4_108_62    m/z 493.1815     0   2   4E+3   6   6.5  RT (min)      lcorbi_hilic_neg_4_017_51    m/z 493.9392     0   2   4   6   8E+3   7.5   8  RT (min)      lcorbi_hilic_neg_4_081_58    m/z 494.0848     0   1E+4   3   3.5  RT (min)      lcorbi_hilic_neg_4_101_61    m/z 494.2455     0   1E+4   3.5   4  RT (min)       lcorbi_hilic_neg_4_068_57    m/z 494.3394     0   1   2E+4   0.5   1  RT (min)      lcorbi_hilic_neg_4_073_57    m/z 494.6749     0   2   4   6   8E+3   4.5   5  RT (min)      lcorbi_hilic_neg_4_071_57    m/z 494.9082     0   2   4   6   8E+3   6   6.5  RT (min)      lcorbi_hilic_neg_4_072_57    m/z 494.9318     0   1   2   3   4E+3   7.5   8  RT (min)      lcorbi_hilic_neg_4_078_58    m/z 494.9474     0   1E+4   5.5   6  RT (min)      lcorbi_hilic_neg_4_073_57    m/z 495.0714     0   1E+4   3.5   4  RT (min)      lcorbi_hilic_neg_4_107_62    m/z 495.1834     0   1   2   3   4E+3   6   6.5  RT (min)      lcorbi_hilic_neg_4_106_62    m/z 495.2260     0   1E+4   4.5   5  RT (min)      lcorbi_hilic_neg_4_071_57    m/z 495.2731     0   2   4   6E+4   0.5   1  RT (min)      lcorbi_hilic_neg_3_036_17    m/z 495.2982     0   1E+4   3.5   4  RT (min)      lcorbi_hilic_neg_4_074_58    m/z 495.3692     0   1E+3   0.5   1  RT (min)      lcorbi_hilic_neg_2_012_14    m/z 495.9299     0   2   4   6   8E+3   7.5   8  RT (min)      lcorbi_hilic_neg_4_078_58    m/z 495.9924     0   1   2   3E+3   5.5   6  RT (min)      lcorbi_hilic_neg_4_106_62    m/z 496.3193     0   1   2E+4   0.5   1  RT (min)      lcorbi_hilic_neg_4_078_58    m/z 496.3882     0   2   4E+4   4   4.5  RT (min)       lcorbi_hilic_neg_4_070_57    m/z 496.7611     0   2   4   6   8E+3   4.5   5  RT (min)      lcorbi_hilic_neg_4_078_58    m/z 497.0045     0   2   4   6E+3   3   3.5  RT (min)      lcorbi_hilic_neg_4_068_57    m/z 497.0236     0   2   4   6E+3   3   3.5  RT (min)      lcorbi_hilic_neg_4_068_57    m/z 497.0607     0   1   2   3E+3   3   3.5  RT (min)      lcorbi_hilic_neg_4_108_62    m/z 497.0950     0   1   2E+4   6.5   7  RT (min)       lcorbi_hilic_neg_4_083_59    m/z 497.1389     0   1E+4   6.5   7  RT (min)      lcorbi_hilic_neg_4_072_57    m/z 497.1676     0   2   4   6   8E+3   1   1.5  RT (min)      lcorbi_hilic_neg_4_101_61    m/z 497.1676     0   1E+5   4   4.5  RT (min)       lcorbi_hilic_neg_4_097_60    m/z 497.2342     0   1   2   3   4E+4   2   2.5  RT (min)       lcorbi_hilic_neg_4_107_62    m/z 497.2723     0   2   4   6   8E+4   5   5.5  RT (min)      lcorbi_hilic_neg_4_083_59    m/z 497.2758     0   1   3   3.5  RT (min)      lcorbi_hilic_neg_4_088_59    m/z 497.3156     0   1E+4   0.5   1  RT (min)      lcorbi_hilic_neg_4_087_59    m/z 497.3853     0   2   4E+3   0.5   1  RT (min)      lcorbi_hilic_neg_4_014_18    m/z 498.0117     0   2   4   6E+3   3.5   4  RT (min)      lcorbi_hilic_neg_4_107_62    m/z 498.0747     0   2   4   6   8E+3   7   7.5  RT (min)      lcorbi_hilic_neg_1_012_32    m/z 498.0863     0   2   4   6   8E+3   6.5   7  RT (min)      lcorbi_hilic_neg_2_020_27    m/z 498.0959     0   2   4   6   8E+4   1   1.5  RT (min)       lcorbi_hilic_neg_4_098_61    m/z 498.1712     0   1   2E+4   4   4.5  RT (min)      lcorbi_hilic_neg_4_096_60    m/z 498.1786     0   1   2E+4   1   1.5  RT (min)       lcorbi_hilic_neg_4_074_58    m/z 498.1886     0   1   2E+4   3.5   4  RT (min)      lcorbi_hilic_neg_4_005_32    m/z 498.2371     0   1E+4   2   2.5  RT (min)      lcorbi_hilic_neg_4_107_62    m/z 498.2744     0   1E+4   5   5.5  RT (min)      lcorbi_hilic_neg_4_084_59    m/z 498.7586     0   2   4   6   8E+3   4.5   5  RT (min)      lcorbi_hilic_neg_4_078_58    m/z 499.0011     0   2   4   6   8E+3   5.5   6  RT (min)      lcorbi_hilic_neg_4_072_57    m/z 499.0199     0   2   4   6   8E+3   3   3.5  RT (min)      lcorbi_hilic_neg_4_076_58    m/z 499.0558     0   1E+4   2.5   3  RT (min)      lcorbi_hilic_neg_4_110_62    m/z 499.0996     0   1E+4   1   1.5  RT (min)      lcorbi_hilic_neg_4_098_61    m/z 499.2933     0   2   4   6E+4   3.5   4  RT (min)      lcorbi_hilic_neg_4_014_18    m/z 499.3413     0   1E+4   2.5   3  RT (min)      lcorbi_hilic_neg_4_088_59    m/z 499.3656     0   1E+3   0.5   1  RT (min)       lcorbi_hilic_neg_4_031_14    m/z 499.3799     0   1E+5   0.5   1  RT (min)      lcorbi_hilic_neg_4_031_14    m/z 499.9345     0   1   2   3E+4   0.5   1  RT (min)      lcorbi_hilic_neg_4_068_57    m/z 499.9413     0   2   4   6E+3   7.5   8  RT (min)      lcorbi_hilic_neg_4_081_58    m/z 499.9465     0   1   2   3   4E+3   5.5   6  RT (min)      lcorbi_hilic_neg_4_077_58    m/z 499.9813     0   1E+4   5   5.5  RT (min)      lcorbi_hilic_neg_4_098_61    m/z 500.0007     0   1E+4   5   5.5  RT (min)      lcorbi_hilic_neg_4_086_59    m/z 500.0094     0   2   4   6E+3   3.5   4   4.5  RT (min)      lcorbi_hilic_neg_4_112_62    m/z 500.0749     0   1   2E+4   5.5   6  RT (min)      lcorbi_hilic_neg_3_018_38    m/z 500.1117     0   1   5.5   6  RT (min)      lcorbi_hilic_neg_3_018_38    m/z 500.2564     0   1E+3   4   4.5  RT (min)      lcorbi_hilic_neg_4_073_57    m/z 500.2673     0   1   4   4.5  RT (min)      lcorbi_hilic_neg_4_070_57    m/z 500.3001     0   1   4   4.5  RT (min)      lcorbi_hilic_neg_4_069_57    m/z 500.3678     0   1E+5   4   4.5  RT (min)       lcorbi_hilic_neg_4_092_60    m/z 500.6369     0   1E+4   4.5   5  RT (min)      lcorbi_hilic_neg_4_098_61    m/z 500.8527     0   1   2   3E+4   5.5   6  RT (min)       lcorbi_hilic_neg_4_098_61    m/z 500.9217     0   1E+4   8   8.5  RT (min)      lcorbi_hilic_neg_4_066_57    m/z 500.9271     0   1E+4   0.5   1  RT (min)      lcorbi_hilic_neg_4_072_57    m/z 500.9985     0   2   4   6E+3   3   3.5  RT (min)      lcorbi_hilic_neg_4_072_57    m/z 501.0355     0   2   4   6   8E+3   3   3.5  RT (min)      lcorbi_hilic_neg_4_069_57    m/z 501.0771     0   1E+4   6.5   7  RT (min)      lcorbi_hilic_neg_4_083_59    m/z 501.1202     0   1   2E+4   6.5   7  RT (min)       lcorbi_hilic_neg_4_089_59    m/z 501.2141     0   1E+4   1   1.5  RT (min)      lcorbi_hilic_neg_4_098_61    m/z 501.9228     0   2   4   6E+3   8   8.5  RT (min)      lcorbi_hilic_neg_4_079_58    m/z 502.0433     0   2   4E+3   5   5.5  RT (min)      lcorbi_hilic_neg_4_106_62    m/z 502.0869     0   1   2E+4   5   5.5  RT (min)       lcorbi_hilic_neg_4_107_62    m/z 502.2714     0   1   2   3E+3   4   4.5  RT (min)      lcorbi_hilic_neg_4_068_57    m/z 502.3799     0   1   0.5   1  RT (min)      lcorbi_hilic_neg_3_031_14    m/z 502.6340     0   1E+4   4.5   5  RT (min)      lcorbi_hilic_neg_4_098_61    m/z 502.7063     0   2   4   6   8E+3   4.5   5  RT (min)      lcorbi_hilic_neg_4_071_57    m/z 502.8498     0   1   2E+4   5.5   6  RT (min)       lcorbi_hilic_neg_4_102_61    m/z 502.8506     0   1   2E+4   1   1.5  RT (min)      lcorbi_hilic_neg_4_074_58    m/z 502.8607     0   1   2   3   4E+4   8.5   9  RT (min)       lcorbi_hilic_neg_4_080_58    m/z 502.9192     0   2   4   6   8E+3   8   8.5  RT (min)      lcorbi_hilic_neg_4_066_57    m/z 502.9948     0   2   4   6E+3   3   3.5  RT (min)      lcorbi_hilic_neg_4_081_58    m/z 503.0213     0   1E+3   7.5   8  RT (min)      lcorbi_hilic_neg_4_005_32    m/z 503.0320     0   2   4   6   8E+3   6   6.5  RT (min)      lcorbi_hilic_neg_1_012_32    m/z 503.0471     0   1E+4   1   1.5  RT (min)      lcorbi_hilic_neg_4_103_61    m/z 503.1396     0   1   2E+4   2.5   3  RT (min)       lcorbi_hilic_neg_4_111_62    m/z 503.1620     0   1E+4   7   7.5  RT (min)       lcorbi_hilic_neg_4_071_57    m/z 503.1897     0   2   4   6   8E+4   3   3.5  RT (min)      lcorbi_hilic_neg_1_026_34    m/z 503.2866     0   1E+3   4   4.5  RT (min)      lcorbi_hilic_neg_4_069_57    m/z 503.3875     0   2   4   6E+4   4   4.5  RT (min)       lcorbi_hilic_neg_4_095_60    m/z 503.9196     0   1   2   3E+3   8   8.5  RT (min)      lcorbi_hilic_neg_4_066_57    m/z 503.9972     0   2   4   6E+3   6.5   7  RT (min)      lcorbi_hilic_neg_4_070_57    m/z 504.0161     0   2   4   6E+3   6.5   7  RT (min)      lcorbi_hilic_neg_4_088_59    m/z 504.0265     0   1E+4   4.5   5  RT (min)      lcorbi_hilic_neg_4_112_62    m/z 504.0841     0   1E+4   5   5.5  RT (min)      lcorbi_hilic_neg_4_107_62    m/z 504.1133     0   1   2E+4   3   3.5  RT (min)       lcorbi_hilic_neg_4_101_61    m/z 504.1933     0   1E+4   3   3.5  RT (min)      lcorbi_hilic_neg_1_026_34    m/z 504.2688     0   1   4   4.5  RT (min)      lcorbi_hilic_neg_4_097_60    m/z 504.3331     0   1   2   3   4E+4   5   5.5  RT (min)       lcorbi_hilic_neg_4_089_59    m/z 504.6307     0   1E+4   4.5   5  RT (min)      lcorbi_hilic_neg_4_098_61    m/z 504.7038     0   1E+4   4.5   5  RT (min)      lcorbi_hilic_neg_4_066_57    m/z 504.9607     0   2   4   6E+3   7.5   8  RT (min)      lcorbi_hilic_neg_4_079_58    m/z 505.0105     0   1E+4   3.5   4  RT (min)      lcorbi_hilic_neg_4_096_60    m/z 505.0108     0   1E+4   0.5   1  RT (min)      lcorbi_hilic_neg_4_101_61    m/z 505.0765     0   2   4   6E+4   1   1.5  RT (min)      lcorbi_hilic_neg_2_045_26    m/z 505.0990     0   1   2   3   4E+3   6   6.5  RT (min)      lcorbi_hilic_neg_4_072_57    m/z 505.1340     0   2   4   6   8E+3   6.5   7  RT (min)      lcorbi_hilic_neg_4_086_59    m/z 505.2091     0   1E+4   3.5   4  RT (min)      lcorbi_hilic_neg_4_080_58    m/z 505.3477     0   2   4   6   8E+4   3.5   4  RT (min)       lcorbi_hilic_neg_4_096_60    m/z 505.3540     0   1E+4   1   1.5  RT (min)      lcorbi_hilic_neg_4_076_58    m/z 505.3904     0   2   4   6E+3   0.5   1  RT (min)      lcorbi_hilic_neg_2_020_27    m/z 505.9181     0   2   4   6E+3   6   6.5  RT (min)      lcorbi_hilic_neg_4_073_57    m/z 505.9587     0   1E+4   7.5   8  RT (min)      lcorbi_hilic_neg_4_077_58    m/z 505.9665     0   2   4   6E+3   5.5   6  RT (min)      lcorbi_hilic_neg_4_105_61    m/z 506.0236     0   1   4.5   5  RT (min)      lcorbi_hilic_neg_4_091_60    m/z 506.0796     0   1E+4   1   1.5  RT (min)      lcorbi_hilic_neg_3_055_26    m/z 506.1004     0   1   2E+4   6   6.5  RT (min)       lcorbi_hilic_neg_4_084_59    m/z 506.1675     0   1E+5   0.5   1  RT (min)      lcorbi_hilic_neg_4_106_62    m/z 506.2352     0   1E+4   2   2.5  RT (min)      lcorbi_hilic_neg_4_001_49    m/z 506.2674     0   1   2E+4   1.5   2  RT (min)       lcorbi_hilic_neg_4_073_57    m/z 506.3394     0   1   2E+4   0.5   1  RT (min)      lcorbi_hilic_neg_4_073_57    m/z 506.4299     0   2   4E+4   0.5   1  RT (min)      lcorbi_hilic_neg_4_031_14    m/z 506.7012     0   2   4   6E+3   4.5   5  RT (min)      lcorbi_hilic_neg_4_066_57    m/z 506.7899     0   1E+4   4.5   5  RT (min)      lcorbi_hilic_neg_4_086_59    m/z 506.8767     0   2   4   6E+3   8   8.5  RT (min)      lcorbi_hilic_neg_4_104_61    m/z 506.9583     0   2   4   6   8E+3   5.5   6  RT (min)      lcorbi_hilic_neg_4_079_58    m/z 506.9701     0   1E+4   7.5   8  RT (min)      lcorbi_hilic_neg_4_089_59    m/z 507.0992     0   1E+4   2   2.5  RT (min)      lcorbi_hilic_neg_4_112_62    m/z 507.2259     0   2   4   6E+3   1.5   2  RT (min)      lcorbi_hilic_neg_4_080_58    m/z 507.2704     0   2   4   6   8E+3   1.5   2  RT (min)      lcorbi_hilic_neg_4_073_57    m/z 507.2738     0   1   2   3E+4   3.5   4  RT (min)       lcorbi_hilic_neg_4_088_59    m/z 507.3288     0   1   2E+4   4   4.5  RT (min)       lcorbi_hilic_neg_4_068_57    m/z 507.3517     0   2   4E+4   0.5   1  RT (min)      lcorbi_hilic_neg_4_073_57    m/z 507.3694     0   1E+4   0.5   1  RT (min)      lcorbi_hilic_neg_4_075_58    m/z 507.4059     0   2   4   6E+3   0.5   1  RT (min)      lcorbi_hilic_neg_4_072_57    m/z 507.4420     0   1   2   3E+4   0.5   1  RT (min)      lcorbi_hilic_neg_2_012_14    m/z 507.8770     0   2   4E+3   8   8.5  RT (min)      lcorbi_hilic_neg_4_106_62    m/z 507.9635     0   1   2   3   4E+3   5.5   6  RT (min)      lcorbi_hilic_neg_4_108_62    m/z 508.0290     0   1E+3   6   6.5  RT (min)      lcorbi_hilic_neg_1_039_38    m/z 508.1650     0   1   2   3   4E+4   0.5   1  RT (min)      lcorbi_hilic_neg_4_106_62    m/z 508.1801     0   1   2   3E+3   5   5.5  RT (min)      lcorbi_hilic_neg_4_096_60    m/z 508.2067     0   2   4   6   8E+3   3   3.5  RT (min)      lcorbi_hilic_neg_4_001_49    m/z 508.2283     0   1   2E+4   3.5   4  RT (min)      lcorbi_hilic_neg_4_081_58    m/z 508.2643     0   2   4   6E+3   1.5   2  RT (min)      lcorbi_hilic_neg_4_080_58    m/z 508.2958     0   2   4   6E+4   6.5   7  RT (min)       lcorbi_hilic_neg_4_095_60    m/z 508.3554     0   1E+4   0.5   1  RT (min)      lcorbi_hilic_neg_4_073_57    m/z 508.6526     0   1E+4   4.5   5  RT (min)      lcorbi_hilic_neg_4_069_57    m/z 508.7874     0   1E+4   4.5   5  RT (min)      lcorbi_hilic_neg_4_082_59    m/z 508.8162     0   1E+4   8.5   9  RT (min)      lcorbi_hilic_neg_4_078_58    m/z 509.0294     0   1   2E+4   5.5   6   6.5  RT (min)       lcorbi_hilic_neg_4_072_57    m/z 509.1004     0   2   4E+3   2   2.5  RT (min)      lcorbi_hilic_neg_4_107_62    m/z 509.1489     0   1   2   3E+4   6.5   7  RT (min)       lcorbi_hilic_neg_4_095_60    m/z 509.2718     0   2   4   6   8E+3   4   4.5  RT (min)      lcorbi_hilic_neg_4_053_23    m/z 509.2896     0   1   2   3E+4   3.5   4  RT (min)       lcorbi_hilic_neg_4_088_59    m/z 509.2941     0   1E+4   5   5.5  RT (min)      lcorbi_hilic_neg_4_080_58    m/z 509.3039     0   1   2E+4   4   4.5  RT (min)       lcorbi_hilic_neg_4_089_59    m/z 509.3853     0   1   2   3E+4   0.5   1  RT (min)       lcorbi_hilic_neg_4_075_58    m/z 509.4577     0   1E+5   0.5   1  RT (min)       lcorbi_hilic_neg_4_113_62    m/z 509.9755     0   1E+4   5.5   6  RT (min)      lcorbi_hilic_neg_4_077_58    m/z 510.0293     0   2   4   6E+3   5   5.5  RT (min)      lcorbi_hilic_neg_4_106_62    m/z 510.0969     0   1E+4   6.5   7  RT (min)      lcorbi_hilic_neg_4_076_58    m/z 510.1084     0   1   2   3   4E+3   5.5   6  RT (min)      lcorbi_hilic_neg_2_006_45    m/z 510.1881     0   1   2   3E+3   2   2.5  RT (min)      lcorbi_hilic_neg_1_046_19    m/z 510.2382     0   2   4   6   8E+3   4   4.5  RT (min)      lcorbi_hilic_neg_4_073_57    m/z 510.2848     0   1   2E+4   4.5   5  RT (min)       lcorbi_hilic_neg_4_083_59    m/z 510.3351     0   1   2   3E+4   0.5   1  RT (min)       lcorbi_hilic_neg_4_077_58    m/z 510.3471     0   2   4   6   8E+3   4   4.5  RT (min)      lcorbi_hilic_neg_4_078_58    m/z 510.3887     0   1E+4   0.5   1  RT (min)      lcorbi_hilic_neg_4_080_58    m/z 510.6498     0   1   2E+4   4.5   5  RT (min)       lcorbi_hilic_neg_4_073_57    m/z 510.9173     0   2   4   6E+3   7.5   8  RT (min)      lcorbi_hilic_neg_4_079_58    m/z 510.9839     0   2   4   6E+3   6   6.5  RT (min)      lcorbi_hilic_neg_4_070_57    m/z 511.0200     0   2   4   6E+3   3   3.5  RT (min)      lcorbi_hilic_neg_4_081_58    m/z 511.1171     0   1   2   3   4E+3   6   6.5  RT (min)      lcorbi_hilic_neg_1_047_05    m/z 511.1475     0   2   4   6   8E+3   6.5   7  RT (min)      lcorbi_hilic_neg_4_092_60    m/z 511.2582     0   1   2   3E+4   1.5   2  RT (min)      lcorbi_hilic_neg_4_075_58    m/z 511.2915     0   1   0.5   1  RT (min)      lcorbi_hilic_neg_4_083_59    m/z 511.2917     0   1   2E+4   4   4.5  RT (min)       lcorbi_hilic_neg_4_079_58    m/z 511.3048     0   2   4   6   8E+3   1.5   2  RT (min)      lcorbi_hilic_neg_4_070_57    m/z 511.3165     0   1E+4   1   1.5  RT (min)      lcorbi_hilic_neg_4_066_57    m/z 511.3669     0   1   2   3   4E+4   4   4.5  RT (min)       lcorbi_hilic_neg_4_070_57    m/z 511.4008     0   1   2E+4   0.5   1  RT (min)      lcorbi_hilic_neg_4_076_58    m/z 512.1157     0   1   2   3   4E+4   5   5.5  RT (min)       lcorbi_hilic_neg_4_106_62    m/z 512.1972     0   2   4E+5   1   1.5  RT (min)       lcorbi_hilic_neg_1_027_18    m/z 512.3694     0   1E+4   4   4.5  RT (min)      lcorbi_hilic_neg_4_071_57    m/z 512.6475     0   1E+4   4.5   5  RT (min)      lcorbi_hilic_neg_4_072_57    m/z 512.7352     0   1E+4   4.5   5  RT (min)      lcorbi_hilic_neg_4_076_58    m/z 512.8788     0   1   2   3E+4   5.5   6  RT (min)       lcorbi_hilic_neg_4_110_62    m/z 512.9805     0   1   2   3   4E+3   6   6.5  RT (min)      lcorbi_hilic_neg_4_070_57    m/z 513.0316     0   1   2E+4   6   6.5  RT (min)      lcorbi_hilic_neg_4_023_34    m/z 513.0356     0   2   4E+3   3   3.5  RT (min)      lcorbi_hilic_neg_4_081_58    m/z 513.0607     0   2   4   6   8E+3   6   6.5  RT (min)      lcorbi_hilic_neg_4_066_57    m/z 513.2005     0   2   4   6E+4   1   1.5  RT (min)      lcorbi_hilic_neg_4_102_61    m/z 513.2742     0   1   2E+4   2   2.5  RT (min)      lcorbi_hilic_neg_4_066_57    m/z 513.3825     0   1   2   3E+4   4   4.5  RT (min)       lcorbi_hilic_neg_4_073_57    m/z 514.0467     0   2   4   6E+3   6.5   7  RT (min)      lcorbi_hilic_neg_4_090_60    m/z 514.0702     0   1   2   3E+4   1   1.5  RT (min)      lcorbi_hilic_neg_4_098_61    m/z 514.1129     0   1E+4   5   5.5  RT (min)      lcorbi_hilic_neg_4_107_62    m/z 514.1654     0   2   4   6E+3   6.5   7  RT (min)      lcorbi_hilic_neg_3_042_05    m/z 514.3065     0   1E+4   4   4.5  RT (min)      lcorbi_hilic_neg_4_073_57    m/z 514.3260     0   1   2   3E+5   2.5   3  RT (min)       lcorbi_hilic_neg_4_089_59    m/z 514.3755     0   2   4E+4   3.5   4  RT (min)       lcorbi_hilic_neg_4_096_60    m/z 514.3853     0   1E+4   4   4.5  RT (min)      lcorbi_hilic_neg_4_071_57    m/z 514.7327     0   1E+4   4.5   5  RT (min)      lcorbi_hilic_neg_4_074_58    m/z 514.9751     0   2   4   6   8E+3   5.5   6  RT (min)      lcorbi_hilic_neg_1_051_33    m/z 514.9891     0   1   2E+4   5.5   6  RT (min)       lcorbi_hilic_neg_4_073_57    m/z 515.0140     0   2   4   6E+3   3   3.5  RT (min)      lcorbi_hilic_neg_4_081_58    m/z 515.0511     0   1   2   3   4E+3   3   3.5  RT (min)      lcorbi_hilic_neg_4_079_58    m/z 515.0648     0   1E+4   6   6.5  RT (min)      lcorbi_hilic_neg_4_075_58    m/z 515.1298     0   2   4E+3   7   7.5  RT (min)      lcorbi_hilic_neg_3_011_24    m/z 515.2224     0   1   2   3E+4   7   7.5  RT (min)       lcorbi_hilic_neg_4_089_59    m/z 515.3021     0   2   4E+4   0.5   1  RT (min)       lcorbi_hilic_neg_4_108_62    m/z 515.3294     0   2   4   6   8E+4   2   3  RT (min)       lcorbi_hilic_neg_4_089_59    m/z 515.3747     0   1   2E+4   0.5   1  RT (min)      lcorbi_hilic_neg_2_037_15    m/z 515.4109     0   1E+5   0.5   1  RT (min)      lcorbi_hilic_neg_4_031_14    m/z 515.9468     0   1E+4   6   6.5  RT (min)      lcorbi_hilic_neg_4_072_57    m/z 515.9827     0   2   4   6E+3   4   4.5  RT (min)      lcorbi_hilic_neg_4_112_62    m/z 515.9847     0   1   4.5   5  RT (min)      lcorbi_hilic_neg_4_077_58    m/z 515.9954     0   1E+4   5.5   6  RT (min)      lcorbi_hilic_neg_4_104_61    m/z 516.1865     0   1   2E+4   1   1.5  RT (min)       lcorbi_hilic_neg_4_100_61    m/z 516.2058     0   1   2E+4   6.5   7  RT (min)       lcorbi_hilic_neg_4_087_59    m/z 516.2737     0   2   4   6E+4   4   4.5  RT (min)       lcorbi_hilic_neg_4_078_58    m/z 516.2869     0   2   4   6   8E+3   4   4.5  RT (min)      lcorbi_hilic_neg_4_075_58    m/z 516.2962     0   1E+4   1.5   2  RT (min)      lcorbi_hilic_neg_4_073_57    m/z 516.3236     0   1   2   3   4E+4   4   4.5   5  RT (min)       lcorbi_hilic_neg_4_081_58    m/z 516.3908     0   1   2   3   4E+4   3.5   4  RT (min)       lcorbi_hilic_neg_4_096_60    m/z 516.7303     0   2   4   6E+3   4.5   5  RT (min)      lcorbi_hilic_neg_4_077_58    m/z 516.8186     0   1   2   3E+4   4.5   5  RT (min)       lcorbi_hilic_neg_4_082_59    m/z 517.0106     0   2   4   6   8E+3   3   3.5  RT (min)      lcorbi_hilic_neg_4_071_57    m/z 517.0533     0   1E+4   3.5   4  RT (min)      lcorbi_hilic_neg_4_109_62    m/z 517.1990     0   1   2   3   4E+4   0.5   1  RT (min)      lcorbi_hilic_neg_4_113_62    m/z 517.2772     0   1   2E+4   4   4.5  RT (min)      lcorbi_hilic_neg_4_078_58    m/z 517.3267     0   1E+4   4   4.5  RT (min)      lcorbi_hilic_neg_4_095_60    m/z 517.3392     0   1E+5   5   5.5  RT (min)       lcorbi_hilic_neg_4_097_60    m/z 517.9923     0   1   2E+3   5.5   6  RT (min)      lcorbi_hilic_neg_4_079_58    m/z 518.0176     0   1   2E+4   1   1.5  RT (min)       lcorbi_hilic_neg_4_101_61    m/z 518.0494     0   1   6   6.5  RT (min)      lcorbi_hilic_neg_4_005_32    m/z 518.0609     0   1   2E+3   4.5   5  RT (min)      lcorbi_hilic_neg_4_108_62    m/z 518.1010     0   1   2E+4   1   1.5  RT (min)      lcorbi_hilic_neg_4_011_19    m/z 518.1893     0   2   4   6E+3   2   2.5  RT (min)      lcorbi_hilic_neg_2_020_27    m/z 518.2457     0   2   4   6E+4   3.5   4  RT (min)       lcorbi_hilic_neg_4_070_57    m/z 518.2527     0   1   5   5.5  RT (min)       lcorbi_hilic_neg_4_097_60    m/z 518.6074     0   2   4   6   8E+3   4.5   5  RT (min)      lcorbi_hilic_neg_4_100_61    m/z 518.6814     0   1   2   3E+4   4.5   5  RT (min)       lcorbi_hilic_neg_4_076_58    m/z 518.8169     0   1E+4   4.5   5  RT (min)      lcorbi_hilic_neg_4_082_59    m/z 518.8244     0   1   2E+4   1   1.5  RT (min)      lcorbi_hilic_neg_4_074_58    m/z 518.9120     0   1E+4   5.5   6  RT (min)      lcorbi_hilic_neg_4_073_57    m/z 519.0262     0   1E+4   3   3.5  RT (min)      lcorbi_hilic_neg_4_076_58    m/z 519.0771     0   1   2E+4   6.5   7  RT (min)       lcorbi_hilic_neg_4_083_59    m/z 519.0980     0   2   4   6E+3   5.5   6  RT (min)      lcorbi_hilic_neg_4_081_58    m/z 519.1135     0   1E+4   2.5   3  RT (min)      lcorbi_hilic_neg_4_111_62    m/z 519.1594     0   2   4   6   8E+3   3.5   4  RT (min)      lcorbi_hilic_neg_4_096_60    m/z 519.2162     0   1   2E+4   2   2.5  RT (min)       lcorbi_hilic_neg_4_107_62    m/z 519.2249     0   1   2E+4   3   3.5  RT (min)      lcorbi_hilic_neg_4_074_58    m/z 519.2496     0   1   3.5   4  RT (min)      lcorbi_hilic_neg_4_084_59    m/z 520.0044     0   1E+4   5.5   6  RT (min)      lcorbi_hilic_neg_4_079_58    m/z 520.1046     0   1E+4   6.5   7  RT (min)      lcorbi_hilic_neg_4_088_59    m/z 520.2429     0   1   2E+4   3.5   4  RT (min)      lcorbi_hilic_neg_4_071_57    m/z 520.2855     0   2   4   6   8E+4   0.5   1  RT (min)      lcorbi_hilic_neg_3_036_17    m/z 520.6787     0   1   2   3   4E+4   4.5   5  RT (min)       lcorbi_hilic_neg_4_076_58    m/z 520.9335     0   1E+4   5   5.5  RT (min)      lcorbi_hilic_neg_2_031_34    m/z 520.9402     0   2   4   6E+3   5   5.5  RT (min)      lcorbi_hilic_neg_3_024_27    m/z 521.0127     0   2   4   6   8E+3   6   6.5  RT (min)      lcorbi_hilic_neg_4_068_57    m/z 521.0627     0   1   2E+4   7  RT (min)       lcorbi_hilic_neg_4_087_59    m/z 521.1863     0   2   4   6E+4   1   1.5  RT (min)       lcorbi_hilic_neg_4_107_62    m/z 521.2404     0   1   2E+4   1   1.5  RT (min)       lcorbi_hilic_neg_4_079_58    m/z 521.2720     0   1E+5   4.5   5  RT (min)      lcorbi_hilic_neg_4_089_59    m/z 521.3283     0   1   2   3E+4   4   4.5  RT (min)       lcorbi_hilic_neg_4_070_57    m/z 521.3302     0   1   2E+4   7   7.5  RT (min)       lcorbi_hilic_neg_4_086_59    m/z 521.3451     0   1E+4   1   1.5  RT (min)      lcorbi_hilic_neg_4_068_57    m/z 521.9034     0   1E+5   5.5   6  RT (min)      lcorbi_hilic_neg_1_051_33    m/z 521.9813     0   1E+3   5   5.5  RT (min)      lcorbi_hilic_neg_4_083_59    m/z 521.9974     0   2   4   6E+3   4.5   5  RT (min)      lcorbi_hilic_neg_4_107_62    m/z 522.0470     0   1E+4   6   6.5  RT (min)      lcorbi_hilic_neg_4_074_58    m/z 522.0489     0   2   4   6E+4   1   1.5  RT (min)      lcorbi_hilic_neg_2_003_20    m/z 522.0570     0   1E+4   5.5   6  RT (min)      lcorbi_hilic_neg_3_018_38    m/z 522.1655     0   2   4   6   8E+3   2   2.5  RT (min)      lcorbi_hilic_neg_4_108_62    m/z 522.2414     0   1E+4   1.5   2  RT (min)      lcorbi_hilic_neg_4_071_57    m/z 522.2633     0   1E+3   4   4.5  RT (min)      lcorbi_hilic_neg_3_010_18    m/z 522.2734     0   1   4.5   5  RT (min)      lcorbi_hilic_neg_4_088_59    m/z 522.2843     0   1   2   3E+5   4.5   5  RT (min)       lcorbi_hilic_neg_1_047_05    m/z 522.3010     0   2   4   6   8E+4   0.5   1  RT (min)      lcorbi_hilic_neg_1_004_17    m/z 522.4038     0   1E+4   4   4.5  RT (min)       lcorbi_hilic_neg_4_071_57    m/z 522.6763     0   1   2E+4   4.5   5  RT (min)       lcorbi_hilic_neg_4_077_58    m/z 522.7642     0   1   2E+4   4.5   5  RT (min)       lcorbi_hilic_neg_4_074_58    m/z 523.0011     0   2   4   6E+3   3   3.5  RT (min)      lcorbi_hilic_neg_4_090_60    m/z 523.0896     0   1E+4   6   6.5  RT (min)      lcorbi_hilic_neg_4_105_61    m/z 523.1480     0   1   2E+4   4.5   5   5.5  RT (min)      lcorbi_hilic_neg_4_106_62    m/z 523.2875     0   1   2   3E+4   4.5   5  RT (min)       lcorbi_hilic_neg_4_083_59    m/z 523.4014     0   1E+3   0.5   1  RT (min)      lcorbi_hilic_neg_3_031_14    m/z 523.9008     0   2   4   6E+3   5.5   6  RT (min)      lcorbi_hilic_neg_4_070_57    m/z 523.9185     0   2   4   6   8E+3   5   5.5  RT (min)      lcorbi_hilic_neg_4_072_57    m/z 524.2101     0   2   4E+3   7   7.5  RT (min)      lcorbi_hilic_neg_4_106_62    m/z 524.2998     0   1   2E+4   4.5   5  RT (min)       lcorbi_hilic_neg_4_085_59    m/z 524.4195     0   1E+4   4   4.5  RT (min)      lcorbi_hilic_neg_4_072_57    m/z 524.4405     0   1   2   3E+4   0.5   1  RT (min)      lcorbi_hilic_neg_4_078_58    m/z 524.7619     0   1   2E+4   4.5   5  RT (min)       lcorbi_hilic_neg_4_082_59    m/z 525.0032     0   1E+4   5.5   6   6.5  RT (min)      lcorbi_hilic_neg_4_072_57    m/z 525.0144     0   2   4   6   8E+3   6   6.5  RT (min)      lcorbi_hilic_neg_4_107_62    m/z 525.0534     0   2   4   6E+3   7.5   8  RT (min)      lcorbi_hilic_neg_4_077_58    m/z 525.1180     0   2   4   6E+3   5.5   6  RT (min)      lcorbi_hilic_neg_4_073_57    m/z 525.1376     0   2   4   6E+3   6.5   7  RT (min)      lcorbi_hilic_neg_4_087_59    m/z 525.2147     0   2   4E+4   0.5   1  RT (min)      lcorbi_hilic_neg_4_111_62    m/z 525.3401     0   1   2E+3   4   4.5  RT (min)       lcorbi_hilic_neg_2_018_16    m/z 525.4167     0   1E+4   0.5   1  RT (min)      lcorbi_hilic_neg_4_073_57    m/z 525.9681     0   1E+4   5.5   6  RT (min)      lcorbi_hilic_neg_4_088_59    m/z 525.9761     0   2   4   6   8E+3   6   6.5  RT (min)      lcorbi_hilic_neg_4_068_57    m/z 526.0243     0   1E+4   5.5   6  RT (min)      lcorbi_hilic_neg_4_100_61    m/z 526.0407     0   1E+4   3   3.5  RT (min)      lcorbi_hilic_neg_4_098_61    m/z 526.0871     0   1E+4   5   5.5  RT (min)      lcorbi_hilic_neg_4_109_62    m/z 526.1902     0   2   4   6E+3   7   7.5  RT (min)      lcorbi_hilic_neg_1_009_50    m/z 526.7595     0   2   4E+3   4.5   5  RT (min)      lcorbi_hilic_neg_4_083_59    m/z 526.8557     0   1E+4   5.5   6  RT (min)       lcorbi_hilic_neg_4_099_61    m/z 527.0967     0   1E+4   6.5   7  RT (min)      lcorbi_hilic_neg_4_088_59    m/z 527.2136     0   2   4   6E+3   3   3.5  RT (min)      lcorbi_hilic_neg_4_077_58    m/z 527.3192     0   1   2   3E+4   4   4.5  RT (min)       lcorbi_hilic_neg_4_080_58    m/z 527.3502     0   1   2   3   4E+3   7   7.5  RT (min)      lcorbi_hilic_neg_1_047_05    m/z 527.3788     0   1   2   3E+4   3.5   4  RT (min)       lcorbi_hilic_neg_4_093_60    m/z 527.4109     0   2   4   6E+4   0.5   1  RT (min)      lcorbi_hilic_neg_4_031_14    m/z 528.0378     0   1E+4   3   3.5  RT (min)      lcorbi_hilic_neg_4_098_61    m/z 528.0815     0   2   4E+4   6   6.5  RT (min)       lcorbi_hilic_neg_4_087_59    m/z 528.0898     0   1E+4   4.5   5  RT (min)      lcorbi_hilic_neg_4_038_27    m/z 528.2742     0   1   2E+3   3.5   4  RT (min)      lcorbi_hilic_neg_4_074_58    m/z 528.2871     0   1   2   3E+4   4   4.5  RT (min)       lcorbi_hilic_neg_4_072_57    m/z 528.3585     0   1E+4   4   4.5  RT (min)      lcorbi_hilic_neg_4_077_58    m/z 528.9454     0   2   4   6E+4   0.5   1  RT (min)       lcorbi_hilic_neg_4_071_57    m/z 528.9523     0   1E+4   7.5   8  RT (min)      lcorbi_hilic_neg_4_089_59    m/z 529.0107     0   2   4   6E+3   3   3.5  RT (min)      lcorbi_hilic_neg_4_075_58    m/z 529.0300     0   1   2   3   4E+3   3   3.5  RT (min)      lcorbi_hilic_neg_4_075_58    m/z 529.2066     0   2   4   6   8E+3   3.5   4  RT (min)      lcorbi_hilic_neg_4_075_58    m/z 529.2687     0   1   2   3E+4   3.5   4  RT (min)      lcorbi_hilic_neg_4_081_58    m/z 529.2903     0   2   4   6   8E+3   4   4.5  RT (min)      lcorbi_hilic_neg_4_070_57    m/z 529.3951     0   1E+5   3.5   4  RT (min)       lcorbi_hilic_neg_4_093_60    m/z 529.9438     0   2   4   6   8E+3   5.5   6  RT (min)      lcorbi_hilic_neg_4_072_57    m/z 529.9535     0   1E+4   6   6.5  RT (min)      lcorbi_hilic_neg_4_074_58    m/z 530.1118     0   1   2E+4   1   1.5  RT (min)       lcorbi_hilic_neg_4_100_61    m/z 530.1296     0   1E+4   6   6.5  RT (min)      lcorbi_hilic_neg_4_098_61    m/z 530.3091     0   1E+5   0.5   1  RT (min)      lcorbi_hilic_neg_3_039_31    m/z 530.3261     0   1   2   3   4E+3   4   4.5  RT (min)      lcorbi_hilic_neg_4_077_58    m/z 530.9553     0   2   4   6   8E+3   7   7.5  RT (min)      lcorbi_hilic_neg_1_016_37    m/z 531.0258     0   1E+4   5.5   6  RT (min)      lcorbi_hilic_neg_4_082_59    m/z 531.0262     0   2   4   6E+3   3   3.5  RT (min)      lcorbi_hilic_neg_4_076_58    m/z 531.0419     0   2   4   6E+3   6   6.5  RT (min)      lcorbi_hilic_neg_4_070_57    m/z 531.1152     0   2   4   6E+3   1   1.5  RT (min)      lcorbi_hilic_neg_4_098_61    m/z 531.1379     0   2   4   6E+3   5.5   6  RT (min)      lcorbi_hilic_neg_3_037_12    m/z 531.1887     0   1   2E+4   1.5   2  RT (min)       lcorbi_hilic_neg_4_103_61    m/z 531.3037     0   1   2E+4   0.5   1  RT (min)      lcorbi_hilic_neg_4_089_59    m/z 531.3546     0   1   2E+4   3.5   4  RT (min)       lcorbi_hilic_neg_4_092_60    m/z 531.4066     0   1E+3   0.5   1  RT (min)      lcorbi_hilic_neg_3_031_14    m/z 531.4108     0   2   4   6   8E+4   3.5   4  RT (min)       lcorbi_hilic_neg_4_097_60    m/z 531.9211     0   1   5.5   6  RT (min)      lcorbi_hilic_neg_4_079_58    m/z 531.9575     0   2   4   6E+3   5.5   6  RT (min)      lcorbi_hilic_neg_4_071_57    m/z 532.1844     0   1   2   3   4E+4   0.5   1  RT (min)      lcorbi_hilic_neg_4_106_62    m/z 532.2026     0   1   2   3E+4   1   1.5  RT (min)      lcorbi_hilic_neg_4_036_36    m/z 532.2369     0   1E+4   1.5   2  RT (min)      lcorbi_hilic_neg_4_078_58    m/z 532.3656     0   2   4   6   8E+3   4   4.5  RT (min)      lcorbi_hilic_neg_4_094_60    m/z 532.4457     0   2   4   6   8E+4   0.5   1  RT (min)      lcorbi_hilic_neg_4_031_14    m/z 532.7052     0   1   2   3E+4   4.5   5  RT (min)       lcorbi_hilic_neg_4_076_58    m/z 532.8006     0   2   4   6E+3   5   5.5  RT (min)      lcorbi_hilic_neg_4_090_60    m/z 532.8429     0   1   2   3E+4   7   7.5  RT (min)       lcorbi_hilic_neg_4_086_59    m/z 533.0047     0   2   4   6E+3   3   3.5  RT (min)      lcorbi_hilic_neg_4_069_57    m/z 533.0418     0   2   4   6E+3   3   3.5  RT (min)      lcorbi_hilic_neg_4_069_57    m/z 533.0997     0   2   4   6E+3   6   6.5  RT (min)      lcorbi_hilic_neg_4_003_08    m/z 533.2925     0   1   2   3   4E+4   0.5   1  RT (min)       lcorbi_hilic_neg_4_071_57    m/z 533.3293     0   1   1   1.5  RT (min)      lcorbi_hilic_neg_4_086_59    m/z 533.3703     0   2   4   6E+4   3.5   4  RT (min)       lcorbi_hilic_neg_4_094_60    m/z 533.3704     0   2   4   6E+3   7   7.5  RT (min)      lcorbi_hilic_neg_2_028_33    m/z 533.3853     0   1   0.5   1  RT (min)      lcorbi_hilic_neg_4_091_60    m/z 533.4218     0   1E+4   0.5   1  RT (min)      lcorbi_hilic_neg_4_071_57    m/z 533.7960     0   2   4   6E+3   4.5   5  RT (min)      lcorbi_hilic_neg_4_090_60    m/z 533.9996     0   2   4E+3   5   5.5  RT (min)      lcorbi_hilic_neg_4_106_62    m/z 534.2382     0   1   2   3E+4   4   4.5  RT (min)       lcorbi_hilic_neg_4_069_57    m/z 534.2528     0   2   4   6E+3   1.5   2  RT (min)      lcorbi_hilic_neg_4_066_57    m/z 534.2660     0   2   4   6   8E+4   6.5   7  RT (min)       lcorbi_hilic_neg_4_094_60    m/z 534.2960     0   1E+4   0.5   1  RT (min)      lcorbi_hilic_neg_4_069_57    m/z 534.3057     0   2   4   6   8E+3   4   4.5  RT (min)      lcorbi_hilic_neg_4_070_57    m/z 534.4612     0   1   2   3   4E+4   0.5   1  RT (min)      lcorbi_hilic_neg_2_037_15    m/z 534.7915     0   1   2E+4   4.5   5  RT (min)       lcorbi_hilic_neg_4_090_60    m/z 534.8063     0   2   4E+3   7   7.5  RT (min)      lcorbi_hilic_neg_1_023_23    m/z 535.0013     0   2   4   6E+3   3   3.5  RT (min)      lcorbi_hilic_neg_4_078_58    m/z 535.0134     0   2   4   6   8E+3   6   6.5  RT (min)      lcorbi_hilic_neg_4_074_58    m/z 535.0424     0   1E+4   6   6.5  RT (min)      lcorbi_hilic_neg_4_112_62    m/z 535.1900     0   1E+4   2   2.5  RT (min)      lcorbi_hilic_neg_4_107_62    m/z 535.2605     0   1   2E+4   3   3.5  RT (min)      lcorbi_hilic_neg_2_031_34    m/z 535.3067     0   2   4E+3   0.5   1  RT (min)      lcorbi_hilic_neg_4_099_61    m/z 535.4375     0   1   2E+4   0.5   1  RT (min)       lcorbi_hilic_neg_4_080_58    m/z 535.4739     0   2   4E+4   0.5   1  RT (min)      lcorbi_hilic_neg_1_003_14    m/z 535.9968     0   1   2E+4   5.5   6  RT (min)       lcorbi_hilic_neg_4_083_59    m/z 536.1078     0   2   4E+3   6   6.5  RT (min)      lcorbi_hilic_neg_4_089_59    m/z 536.2535     0   1   2E+3   4   4.5  RT (min)      lcorbi_hilic_neg_1_047_05    m/z 536.3004     0   1   2E+4   4   4.5  RT (min)       lcorbi_hilic_neg_4_089_59    m/z 536.4043     0   1E+4   0.5   1  RT (min)      lcorbi_hilic_neg_4_081_58    m/z 536.5054     0   2   4   6E+3   0.5   1  RT (min)      lcorbi_hilic_neg_4_083_59    m/z 536.8843     0   1   2   3E+4   5.5   6  RT (min)       lcorbi_hilic_neg_4_109_62    m/z 536.9711     0   1E+4   5.5   6  RT (min)      lcorbi_hilic_neg_4_068_57    m/z 537.0169     0   2   4   6   8E+3   3   3.5  RT (min)      lcorbi_hilic_neg_4_076_58    m/z 537.1415     0   1   2E+4   1   1.5  RT (min)       lcorbi_hilic_neg_4_102_61    m/z 537.3359     0   2   4E+4   0.5   1  RT (min)       lcorbi_hilic_neg_4_101_61    m/z 537.4533     0   1   2E+3   0.5   1  RT (min)       lcorbi_hilic_neg_4_105_61    m/z 537.7574     0   2   4E+3   7   7.5  RT (min)      lcorbi_hilic_neg_1_047_05    m/z 537.9774     0   1E+4   5.5   6  RT (min)      lcorbi_hilic_neg_4_082_59    m/z 538.0231     0   1   2E+4   1   1.5  RT (min)      lcorbi_hilic_neg_3_026_09    m/z 538.0898     0   1E+4   5   5.5  RT (min)      lcorbi_hilic_neg_2_020_27    m/z 538.1185     0   2   4E+4   4.5   5  RT (min)      lcorbi_hilic_neg_3_018_38    m/z 538.1449     0   2   4   6E+3   1   1.5  RT (min)      lcorbi_hilic_neg_4_102_61    m/z 538.3362     0   1E+4   1   1.5  RT (min)       lcorbi_hilic_neg_4_067_57    m/z 538.4202     0   1   2   3   4E+4   0.5   1  RT (min)      lcorbi_hilic_neg_3_031_14    m/z 538.4926     0   2   4   6E+4   0.5   1  RT (min)      lcorbi_hilic_neg_4_113_62    m/z 539.0333     0   2   4   6E+3   5.5   6  RT (min)      lcorbi_hilic_neg_4_073_57    m/z 539.1221     0   2   4   6   8E+3   4.5   5  RT (min)      lcorbi_hilic_neg_3_018_38    m/z 539.1239     0   1   2   3   4E+4   6.5   7  RT (min)       lcorbi_hilic_neg_4_072_57    m/z 539.1412     0   1E+4   6.5   7   7.5  RT (min)       lcorbi_hilic_neg_4_004_13    m/z 539.2521     0   1E+4   4.5   5  RT (min)      lcorbi_hilic_neg_4_073_57    m/z 539.3904     0   2   4   6   8E+4   7   7.5  RT (min)       lcorbi_hilic_neg_4_096_60    m/z 539.7424     0   1E+4   4.5   5  RT (min)      lcorbi_hilic_neg_4_076_58    m/z 540.0555     0   2   4E+4   7.5   8  RT (min)      lcorbi_hilic_neg_1_016_37    m/z 540.2649     0   1E+4   1   1.5  RT (min)      lcorbi_hilic_neg_4_077_58    m/z 540.2679     0   1   2E+4   4   4.5  RT (min)       lcorbi_hilic_neg_4_077_58    m/z 540.2729     0   1   2E+4   4   4.5  RT (min)       lcorbi_hilic_neg_4_070_57    m/z 540.2913     0   1   2   3   4E+4   4   4.5  RT (min)      lcorbi_hilic_neg_4_078_58    m/z 540.2992     0   1E+3   4   4.5  RT (min)      lcorbi_hilic_neg_4_077_58    m/z 540.3598     0   1   4   4.5  RT (min)      lcorbi_hilic_neg_4_077_58    m/z 540.3689     0   2   4E+4   4   4.5  RT (min)      lcorbi_hilic_neg_4_077_58    m/z 540.3778     0   2   4   6E+4   4   4.5  RT (min)      lcorbi_hilic_neg_4_077_58    m/z 540.3897     0   1   2E+4   4   4.5  RT (min)      lcorbi_hilic_neg_4_069_57    m/z 540.4357     0   2   4   6E+4   0.5   1  RT (min)      lcorbi_hilic_neg_4_076_58    m/z 540.9771     0   2   4E+3   6   6.5  RT (min)      lcorbi_hilic_neg_4_067_57    m/z 541.0541     0   1E+4   5.5   6  RT (min)      lcorbi_hilic_neg_4_089_59    m/z 541.2516     0   1E+4   2   2.5  RT (min)      lcorbi_hilic_neg_4_109_62    m/z 541.2684     0   1E+4   4.5   5  RT (min)      lcorbi_hilic_neg_4_080_58    m/z 541.3021     0   1   2E+3   4   4.5  RT (min)      lcorbi_hilic_neg_4_077_58    m/z 541.3091     0   1   4   4.5  RT (min)      lcorbi_hilic_neg_4_074_58    m/z 541.3582     0   1   4   4.5  RT (min)      lcorbi_hilic_neg_4_077_58    m/z 541.3652     0   1   4   4.5  RT (min)      lcorbi_hilic_neg_4_077_58    m/z 541.7401     0   2   4   6   8E+3   4.5   5  RT (min)      lcorbi_hilic_neg_4_077_58    m/z 541.9864     0   1E+4   5.5   6  RT (min)      lcorbi_hilic_neg_4_070_57    m/z 541.9980     0   1E+3   5.5   6  RT (min)      lcorbi_hilic_neg_4_071_57    m/z 542.0142     0   2   4   6   8E+3   3   3.5  RT (min)      lcorbi_hilic_neg_4_101_61    m/z 542.2485     0   1   2   3E+3   4   4.5  RT (min)      lcorbi_hilic_neg_4_073_57    m/z 542.2893     0   1E+4   0.5   1  RT (min)      lcorbi_hilic_neg_4_079_58    m/z 542.2908     0   1   2E+4   3   3.5   4  RT (min)       lcorbi_hilic_neg_4_079_58    m/z 542.7343     0   1   2   3E+4   4.5   5  RT (min)       lcorbi_hilic_neg_4_076_58    m/z 542.8296     0   2   4   6   8E+3   5   5.5  RT (min)      lcorbi_hilic_neg_4_092_60    m/z 543.0265     0   1   2   3E+3   3   3.5  RT (min)      lcorbi_hilic_neg_4_081_58    m/z 543.0457     0   1E+4   6.5   7  RT (min)       lcorbi_hilic_neg_4_075_58    m/z 543.1163     0   1   2E+4   6.5   7  RT (min)       lcorbi_hilic_neg_4_088_59    m/z 543.1724     0   1   2   3   4E+4   7   7.5  RT (min)       lcorbi_hilic_neg_4_086_59    m/z 543.2944     0   1E+4   4   4.5  RT (min)      lcorbi_hilic_neg_4_089_59    m/z 543.3348     0   1   4.5   5  RT (min)      lcorbi_hilic_neg_4_097_60    m/z 544.0118     0   2   4   6   8E+3   3   3.5  RT (min)      lcorbi_hilic_neg_4_105_61    m/z 544.0281     0   1   5   5.5  RT (min)      lcorbi_hilic_neg_4_074_58    m/z 544.0291     0   1E+4   6   6.5  RT (min)      lcorbi_hilic_neg_4_048_37    m/z 544.2662     0   1   2E+4   4   4.5  RT (min)       lcorbi_hilic_neg_4_073_57    m/z 544.3175     0   1   4   4.5  RT (min)      lcorbi_hilic_neg_4_082_59    m/z 545.0420     0   1   2E+3   3   3.5  RT (min)      lcorbi_hilic_neg_4_080_58    m/z 545.0715     0   1   2E+4   6   6.5  RT (min)       lcorbi_hilic_neg_4_112_62    m/z 545.1292     0   2   4E+3   5.5   6  RT (min)      lcorbi_hilic_neg_4_082_59    m/z 545.3610     0   2   4   6E+4   4.5   5  RT (min)       lcorbi_hilic_neg_4_097_60    m/z 546.0255     0   2   4E+3   5   5.5  RT (min)      lcorbi_hilic_neg_4_101_61    m/z 546.0861     0   2   4   6   8E+3   1   1.5  RT (min)      lcorbi_hilic_neg_4_100_61    m/z 546.2816     0   2   4   6   8E+3   4   4.5  RT (min)      lcorbi_hilic_neg_4_083_59    m/z 546.2977     0   2   4   6   8E+3   4   4.5  RT (min)      lcorbi_hilic_neg_4_069_57    m/z 547.0014     0   2   4E+3   3   3.5  RT (min)      lcorbi_hilic_neg_4_077_58    m/z 547.0206     0   1   2   3   4E+3   3   3.5  RT (min)      lcorbi_hilic_neg_4_070_57    m/z 547.0575     0   1   2   3E+3   3   3.5  RT (min)      lcorbi_hilic_neg_4_072_57    m/z 547.2192     0   2   4   6E+4   0.5   1  RT (min)       lcorbi_hilic_neg_4_112_62    m/z 547.3427     0   1E+4   4   4.5  RT (min)      lcorbi_hilic_neg_4_070_57    m/z 547.3649     0   1E+5   0.5   1  RT (min)       lcorbi_hilic_neg_4_104_61    m/z 547.9066     0   2   4   6E+3   5.5   6  RT (min)      lcorbi_hilic_neg_4_105_61    m/z 548.0062     0   1E+4   5.5   6  RT (min)      lcorbi_hilic_neg_4_105_61    m/z 548.2698     0   1   2E+4   3.5   4  RT (min)       lcorbi_hilic_neg_4_070_57    m/z 548.2996     0   1E+4   4   4.5  RT (min)      lcorbi_hilic_neg_4_089_59    m/z 548.3801     0   2   4   6E+4   4.5   5  RT (min)       lcorbi_hilic_neg_4_097_60    m/z 548.4405     0   2   4E+4   0.5   1  RT (min)      lcorbi_hilic_neg_4_031_14    m/z 548.7748     0   1   2   3   4E+2   5   5.5  RT (min)      lcorbi_hilic_neg_1_047_05    m/z 548.8163     0   1E+4   7   7.5  RT (min)      lcorbi_hilic_neg_4_082_59    m/z 548.8539     0   1E+4   8.5   9  RT (min)      lcorbi_hilic_neg_4_077_58    m/z 549.0169     0   2   4   6E+3   3   3.5  RT (min)      lcorbi_hilic_neg_4_076_58    m/z 549.0640     0   2   4   6E+3   7.5   8  RT (min)      lcorbi_hilic_neg_4_089_59    m/z 549.0841     0   1   2   3E+4   0.5   1  RT (min)       lcorbi_hilic_neg_4_107_62    m/z 549.1526     0   1   2   3E+4   6.5   7  RT (min)       lcorbi_hilic_neg_4_073_57    m/z 549.1800     0   1   7   7.5  RT (min)      lcorbi_hilic_neg_4_073_57    m/z 549.1924     0   1   7   7.5  RT (min)      lcorbi_hilic_neg_4_072_57    m/z 549.2670     0   1   2E+4   0.5   1  RT (min)      lcorbi_hilic_neg_4_069_57    m/z 549.3584     0   1E+4   4   4.5  RT (min)      lcorbi_hilic_neg_4_073_57    m/z 549.4169     0   1   2   3   4E+4   0.5   1  RT (min)       lcorbi_hilic_neg_4_074_58    m/z 549.7712     0   1   2E+4   4.5   5  RT (min)       lcorbi_hilic_neg_4_093_60    m/z 550.0874     0   1E+4   0.5   1  RT (min)      lcorbi_hilic_neg_4_110_62    m/z 550.1585     0   1E+4   1   1.5  RT (min)      lcorbi_hilic_neg_4_098_61    m/z 550.2114     0   2   4   6   8E+3   4   4.5  RT (min)      lcorbi_hilic_neg_4_073_57    m/z 550.2472     0   1E+4   3.5   4  RT (min)      lcorbi_hilic_neg_4_076_58    m/z 550.2669     0   1E+4   3.5   4  RT (min)      lcorbi_hilic_neg_4_067_57    m/z 550.3524     0   1E+4   4   4.5  RT (min)      lcorbi_hilic_neg_4_080_58    m/z 550.4203     0   1E+4   0.5   1  RT (min)      lcorbi_hilic_neg_4_076_58    m/z 551.0326     0   2   4   6E+3   3   3.5  RT (min)      lcorbi_hilic_neg_4_077_58    m/z 551.0835     0   1E+4   5.5   6  RT (min)      lcorbi_hilic_neg_4_083_59    m/z 551.1264     0   2   4E+3   2   2.5  RT (min)      lcorbi_hilic_neg_3_024_27    m/z 551.1522     0   1E+4   1   1.5  RT (min)      lcorbi_hilic_neg_4_098_61    m/z 551.1973     0   1   2   3   4E+4   1.5   2  RT (min)       lcorbi_hilic_neg_4_098_61    m/z 551.2410     0   1E+4   4.5   5  RT (min)      lcorbi_hilic_neg_4_106_62    m/z 551.7689     0   1E+4   4.5   5  RT (min)      lcorbi_hilic_neg_4_091_60    m/z 552.2289     0   1   2E+4   0.5   1  RT (min)      lcorbi_hilic_neg_1_010_22    m/z 552.2868     0   2   4   6E+4   4   4.5  RT (min)       lcorbi_hilic_neg_4_085_59    m/z 552.5003     0   1   0.5   1  RT (min)      lcorbi_hilic_neg_2_012_14    m/z 552.7641     0   1   2E+4   4.5   5  RT (min)       lcorbi_hilic_neg_4_093_60    m/z 553.2419     0   1E+4   1.5   2  RT (min)      lcorbi_hilic_neg_1_002_26    m/z 553.3341     0   2   4   6E+3   4   4.5  RT (min)      lcorbi_hilic_neg_4_069_57    m/z 553.3707     0   1   2   3E+4   4   4.5  RT (min)      lcorbi_hilic_neg_4_076_58    m/z 554.0571     0   1E+4   5   5.5  RT (min)      lcorbi_hilic_neg_4_106_62    m/z 554.1410     0   2   4   6   8E+3   6   6.5  RT (min)      lcorbi_hilic_neg_4_098_61    m/z 554.4517     0   2   4E+4   0.5   1  RT (min)       lcorbi_hilic_neg_4_074_58    m/z 554.8250     0   1E+4   7   7.5  RT (min)      lcorbi_hilic_neg_4_085_59    m/z 555.0075     0   2   4   6   8E+3   3   3.5  RT (min)      lcorbi_hilic_neg_4_071_57    m/z 555.1056     0   1   2   3E+4   7.5   8  RT (min)      lcorbi_hilic_neg_1_012_32    m/z 555.1364     0   1E+4   7   7.5  RT (min)      lcorbi_hilic_neg_4_098_61    m/z 555.1715     0   1   2   3E+4   0.5   1  RT (min)      lcorbi_hilic_neg_4_004_13    m/z 555.2124     0   1   2   3   4E+4   7   7.5  RT (min)       lcorbi_hilic_neg_4_090_60    m/z 555.2157     0   1   2   3E+4   0.5   1  RT (min)      lcorbi_hilic_neg_4_069_57    m/z 555.3831     0   1E+4   0.5   1  RT (min)      lcorbi_hilic_neg_4_081_58    m/z 555.3862     0   2   4   6E+3   4   4.5  RT (min)      lcorbi_hilic_neg_4_069_57    m/z 555.4635     0   1E+5   0.5   1  RT (min)      lcorbi_hilic_neg_2_012_14    m/z 556.2191     0   1E+4   0.5   1  RT (min)      lcorbi_hilic_neg_4_067_57    m/z 556.3002     0   2   4E+5   0.5   1  RT (min)      lcorbi_hilic_neg_4_031_14    m/z 556.3527     0   2   4   6   8E+3   4   4.5  RT (min)      lcorbi_hilic_neg_4_079_58    m/z 556.4664     0   1   2   3E+4   0.5   1  RT (min)      lcorbi_hilic_neg_3_031_14    m/z 556.7175     0   2   4   6E+3   4.5   5  RT (min)      lcorbi_hilic_neg_4_078_58    m/z 557.0237     0   1   2   3E+3   3   3.5  RT (min)      lcorbi_hilic_neg_4_072_57    m/z 557.3024     0   2   4   6   8E+3   4   4.5  RT (min)      lcorbi_hilic_neg_4_075_58    m/z 557.3030     0   1E+5   0.5   1  RT (min)      lcorbi_hilic_neg_4_031_14    m/z 557.3070     0   1   2   3E+4   7   7.5  RT (min)       lcorbi_hilic_neg_4_085_59    m/z 557.3296     0   1E+4   4   4.5  RT (min)      lcorbi_hilic_neg_4_070_57    m/z 557.3987     0   1   2   3E+4   0.5   1  RT (min)      lcorbi_hilic_neg_4_076_58    m/z 558.2246     0   1   2   3E+4   4.5   5  RT (min)       lcorbi_hilic_neg_4_082_59    m/z 558.2378     0   1   2   3E+4   4   4.5  RT (min)       lcorbi_hilic_neg_4_066_57    m/z 558.2973     0   1E+4   0.5   1  RT (min)      lcorbi_hilic_neg_4_072_57    m/z 558.3154     0   1   2E+4   4   4.5  RT (min)      lcorbi_hilic_neg_4_079_58    m/z 558.3402     0   1   2E+3   0.5   1  RT (min)       lcorbi_hilic_neg_4_073_57    m/z 558.8037     0   1   4.5   5  RT (min)      lcorbi_hilic_neg_4_108_62    m/z 559.3020     0   1   2   3   4E+4   3.5   4  RT (min)      lcorbi_hilic_neg_4_070_57    m/z 559.3186     0   2   4E+3   4   4.5  RT (min)      lcorbi_hilic_neg_4_079_58    m/z 559.3356     0   2   4   6   8E+3   0.5   1  RT (min)      lcorbi_hilic_neg_4_073_57    m/z 560.0022     0   1   2   3   4E+3   5   5.5  RT (min)      lcorbi_hilic_neg_4_109_62    m/z 560.2396     0   1   2   3E+3   4   4.5  RT (min)      lcorbi_hilic_neg_4_082_59    m/z 560.2945     0   2   4   6E+3   0.5   1  RT (min)      lcorbi_hilic_neg_4_072_57    m/z 560.2958     0   1   2   3   4E+4   3.5   4  RT (min)       lcorbi_hilic_neg_4_070_57    m/z 560.9950     0   2   4   6   8E+3   5.5   6  RT (min)      lcorbi_hilic_neg_4_079_58    m/z 561.0171     0   1   2   3   4E+3   3   3.5  RT (min)      lcorbi_hilic_neg_4_081_58    m/z 561.0437     0   1E+4   6   6.5  RT (min)      lcorbi_hilic_neg_4_071_57    m/z 561.3079     0   2   4E+4   0.5   1  RT (min)       lcorbi_hilic_neg_4_108_62    m/z 561.3346     0   1   2E+4   4   4.5  RT (min)       lcorbi_hilic_neg_4_081_58    m/z 561.3820     0   1E+4   2   3  RT (min)       lcorbi_hilic_neg_4_073_57    m/z 562.0893     0   1E+4   6.5   7  RT (min)      lcorbi_hilic_neg_4_072_57    m/z 562.1995     0   1   2   3E+3   8   8.5  RT (min)      lcorbi_hilic_neg_4_082_59    m/z 562.2528     0   2   4   6   8E+2   4   4.5  RT (min)      lcorbi_hilic_neg_1_047_05    m/z 562.3374     0   1   4   4.5  RT (min)      lcorbi_hilic_neg_4_082_59    m/z 562.3961     0   1E+5   4   4.5  RT (min)       lcorbi_hilic_neg_4_091_60    m/z 562.9352     0   1E+4   5.5   6  RT (min)      lcorbi_hilic_neg_4_070_57    m/z 563.0326     0   1   2   3E+3   3   3.5  RT (min)      lcorbi_hilic_neg_4_077_58    m/z 563.2334     0   1   2   3   4E+3   2   2.5  RT (min)      lcorbi_hilic_neg_3_054_29    m/z 563.2874     0   1   2   3E+5   1   1.5  RT (min)      lcorbi_hilic_neg_1_002_26    m/z 563.4324     0   1   2   3E+3   0.5   1  RT (min)      lcorbi_hilic_neg_4_097_60    m/z 563.5054     0   1   2   3   4E+4   0.5   1  RT (min)      lcorbi_hilic_neg_4_044_01    m/z 564.3046     0   1   4   4.5  RT (min)      lcorbi_hilic_neg_4_071_57    m/z 564.3572     0   1   4   4.5  RT (min)      lcorbi_hilic_neg_3_042_05    m/z 564.4111     0   2   4   6   8E+4   4   4.5  RT (min)       lcorbi_hilic_neg_4_095_60    m/z 564.4149     0   1   2E+4   0.5   1  RT (min)      lcorbi_hilic_neg_4_054_30    m/z 564.7487     0   2   4   6E+3   4.5   5  RT (min)      lcorbi_hilic_neg_4_079_58    m/z 564.8317     0   1   2   3E+4   8.5   9  RT (min)       lcorbi_hilic_neg_4_080_58    m/z 565.0499     0   1E+4   8   8.5  RT (min)      lcorbi_hilic_neg_4_089_59    m/z 565.0984     0   1   2E+4   6.5   7  RT (min)       lcorbi_hilic_neg_4_088_59    m/z 565.1545     0   1   2E+4   7   7.5  RT (min)       lcorbi_hilic_neg_4_085_59    m/z 565.3229     0   1   2E+4   2.5   3  RT (min)       lcorbi_hilic_neg_4_082_59    m/z 565.3524     0   2   4   6   8E+4   0.5   1  RT (min)      lcorbi_hilic_neg_3_010_18    m/z 565.5204     0   1   2   3E+4   0.5   1  RT (min)       lcorbi_hilic_neg_4_105_61    m/z 566.0108     0   2   4   6   8E+3   6.5   7  RT (min)      lcorbi_hilic_neg_4_079_58    m/z 566.2206     0   1   2   3   4E+3   5.5   6  RT (min)      lcorbi_hilic_neg_4_082_59    m/z 566.3198     0   1   4   4.5  RT (min)      lcorbi_hilic_neg_4_069_57    m/z 566.3341     0   2   4   6   8E+4   4.5   5  RT (min)       lcorbi_hilic_neg_2_041_54    m/z 566.4512     0   1   0.5   1  RT (min)      lcorbi_hilic_neg_4_087_59    m/z 566.5158     0   1   2   3E+4   0.5   1  RT (min)      lcorbi_hilic_neg_4_082_59    m/z 566.7462     0   2   4   6E+3   4.5   5  RT (min)      lcorbi_hilic_neg_4_083_59    m/z 567.0075     0   1   2   3   4E+3   3   3.5  RT (min)      lcorbi_hilic_neg_4_080_58    m/z 567.2292     0   1   2E+4   0.5   1  RT (min)      lcorbi_hilic_neg_4_056_56    m/z 567.3195     0   1E+4   4   4.5  RT (min)      lcorbi_hilic_neg_4_068_57    m/z 567.3361     0   1E+4   7   7.5  RT (min)      lcorbi_hilic_neg_4_086_59    m/z 567.3427     0   1   2   3E+3   4   4.5  RT (min)      lcorbi_hilic_neg_4_070_57    m/z 567.4540     0   2   4   6   8E+4   0.5   1  RT (min)      lcorbi_hilic_neg_4_042_28    m/z 567.9882     0   2   4   6E+3   5   5.5  RT (min)      lcorbi_hilic_neg_4_083_59    m/z 568.2669     0   2   4   6   8E+2   4   4.5  RT (min)       lcorbi_hilic_neg_2_041_54    m/z 568.3219     0   1   2E+3   4   4.5  RT (min)      lcorbi_hilic_neg_2_041_54    m/z 568.3502     0   2   4E+5   4   4.5  RT (min)      lcorbi_hilic_neg_2_041_54    m/z 568.3628     0   1E+5   1   1.5  RT (min)       lcorbi_hilic_neg_4_073_57    m/z 568.3882     0   1   4   4.5  RT (min)      lcorbi_hilic_neg_4_080_58    m/z 568.4577     0   1   2E+4   0.5   1  RT (min)      lcorbi_hilic_neg_3_055_26    m/z 568.4671     0   1   2   3E+4   0.5   1  RT (min)      lcorbi_hilic_neg_4_031_14    m/z 568.8322     0   2   4   6E+3   5   5.5  RT (min)      lcorbi_hilic_neg_3_022_06    m/z 568.8403     0   1   2E+4   5.5   6  RT (min)       lcorbi_hilic_neg_4_112_62    m/z 569.0232     0   2   4   6E+3   3   3.5  RT (min)      lcorbi_hilic_neg_4_070_57    m/z 569.2260     0   1   2E+4   0.5   1  RT (min)      lcorbi_hilic_neg_1_049_55    m/z 569.3473     0   2   4   6E+3   7   7.5  RT (min)      lcorbi_hilic_neg_1_001_49    m/z 569.3661     0   1   2   3E+4   1   1.5  RT (min)      lcorbi_hilic_neg_4_073_57    m/z 569.3662     0   1E+6   3.5   4  RT (min)       lcorbi_hilic_neg_4_067_57    m/z 569.4793     0   2   4E+4   0.5   1  RT (min)      lcorbi_hilic_neg_2_012_14    m/z 570.0306     0   2   4E+3   5   5.5  RT (min)      lcorbi_hilic_neg_4_108_62    m/z 570.1360     0   1E+4   6.5   7  RT (min)      lcorbi_hilic_neg_4_106_62    m/z 570.4667     0   1   2E+4   0.5   1  RT (min)       lcorbi_hilic_neg_4_087_59    m/z 570.7873     0   1E+4   8.5   9  RT (min)      lcorbi_hilic_neg_4_113_62    m/z 570.8375     0   1E+4   5.5   6  RT (min)       lcorbi_hilic_neg_4_112_62    m/z 571.1372     0   1E+4   1   1.5  RT (min)      lcorbi_hilic_neg_4_098_61    m/z 571.1745     0   1   7   7.5  RT (min)      lcorbi_hilic_neg_3_022_06    m/z 571.2856     0   1   2   3   4E+4   6.5   7  RT (min)      lcorbi_hilic_neg_1_016_37    m/z 571.2903     0   2   4E+4   5   5.5  RT (min)       lcorbi_hilic_neg_4_087_59    m/z 571.3285     0   1   2E+5   0.5   1  RT (min)       lcorbi_hilic_neg_2_012_14    m/z 571.3711     0   1   2   3E+4   4   4.5  RT (min)       lcorbi_hilic_neg_4_081_58    m/z 571.4728     0   1   2E+4   0.5   1  RT (min)      lcorbi_hilic_neg_3_016_16    m/z 572.0714     0   2   4   6E+3   6.5   7  RT (min)      lcorbi_hilic_neg_4_078_58    m/z 572.1161     0   2   4E+3   6   6.5  RT (min)      lcorbi_hilic_neg_4_066_57    m/z 572.3319     0   1E+5   0.5   1  RT (min)      lcorbi_hilic_neg_2_012_14    m/z 572.4405     0   1   2   3E+4   0.5   1  RT (min)      lcorbi_hilic_neg_4_031_14    m/z 572.9984     0   1   2   3   4E+3   3   3.5  RT (min)      lcorbi_hilic_neg_4_068_57    m/z 573.1819     0   1E+4   0.5   1  RT (min)      lcorbi_hilic_neg_3_053_13    m/z 573.1824     0   1E+4   3   3.5  RT (min)      lcorbi_hilic_neg_4_072_57    m/z 573.2376     0   2   4E+4   1   1.5  RT (min)      lcorbi_hilic_neg_4_108_62    m/z 573.3323     0   1E+5   0.5   1  RT (min)      lcorbi_hilic_neg_3_039_31    m/z 573.9541     0   2   4E+3   5.5   6  RT (min)      lcorbi_hilic_neg_4_104_61    m/z 574.2831     0   1   2   3E+4   4   4.5  RT (min)       lcorbi_hilic_neg_4_071_57    m/z 574.4796     0   1   2   3   4E+4   0.5   1  RT (min)      lcorbi_hilic_neg_4_087_59    m/z 574.4981     0   1   2E+4   0.5   1  RT (min)      lcorbi_hilic_neg_4_086_59    m/z 574.7775     0   1E+4   4.5   5  RT (min)      lcorbi_hilic_neg_4_083_59    m/z 575.3238     0   2   4   6E+4   0.5   1  RT (min)       lcorbi_hilic_neg_2_019_21    m/z 575.3674     0   1   2   3E+4   7   7.5  RT (min)       lcorbi_hilic_neg_4_091_60    m/z 575.4829     0   1   0.5   1  RT (min)      lcorbi_hilic_neg_4_075_58    m/z 575.5017     0   1   2E+3   0.5   1  RT (min)      lcorbi_hilic_neg_4_075_58    m/z 577.1775     0   1   2E+4   1   1.5  RT (min)      lcorbi_hilic_neg_4_102_61    m/z 577.1944     0   1   2E+4   7   7.5  RT (min)      lcorbi_hilic_neg_4_090_60    m/z 577.3769     0   1   2E+4   1.5   2  RT (min)       lcorbi_hilic_neg_4_075_58    m/z 577.4229     0   2   4E+5   0.5   1  RT (min)       lcorbi_hilic_neg_4_046_15    m/z 577.9632     0   2   4   6   8E+3   5.5   6  RT (min)      lcorbi_hilic_neg_4_074_58    m/z 578.2822     0   1   2   3E+5   0.5   1  RT (min)      lcorbi_hilic_neg_2_012_14    m/z 578.3743     0   1   2   3   4E+4   4   4.5  RT (min)       lcorbi_hilic_neg_2_021_05    m/z 578.3799     0   2   4   6   8E+3   1.5   2  RT (min)      lcorbi_hilic_neg_4_077_58    m/z 578.4262     0   1E+5   0.5   1  RT (min)      lcorbi_hilic_neg_2_037_15    m/z 578.5161     0   1   0.5   1  RT (min)      lcorbi_hilic_neg_4_031_14    m/z 578.8694     0   2   4   6E+4   5.5   6  RT (min)       lcorbi_hilic_neg_4_069_57    m/z 579.0079     0   1   2   3E+3   3   3.5  RT (min)      lcorbi_hilic_neg_4_076_58    m/z 579.0273     0   1   2   3E+3   3   3.5  RT (min)      lcorbi_hilic_neg_4_077_58    m/z 579.0748     0   1E+4   7.5   8  RT (min)      lcorbi_hilic_neg_4_089_59    m/z 579.1476     0   2   4   6   8E+3   6.5   7  RT (min)      lcorbi_hilic_neg_4_098_61    m/z 579.2853     0   2   4   6   8E+4   0.5   1  RT (min)      lcorbi_hilic_neg_2_012_14    m/z 579.4402     0   2   4   6E+4   0.5   1  RT (min)      lcorbi_hilic_neg_1_003_14    m/z 579.4642     0   1   2E+3   0.5   1  RT (min)      lcorbi_hilic_neg_3_002_15    m/z 580.1034     0   1E+4   5   5.5  RT (min)      lcorbi_hilic_neg_4_109_62    m/z 580.2157     0   1E+4   0.5   1  RT (min)      lcorbi_hilic_neg_4_101_61    m/z 580.3201     0   1E+4   4   4.5  RT (min)      lcorbi_hilic_neg_4_068_57    m/z 580.3627     0   1   2E+3   4   4.5  RT (min)      lcorbi_hilic_neg_4_091_60    m/z 580.3729     0   1   2E+4   3   3.5  RT (min)       lcorbi_hilic_neg_4_088_59    m/z 580.3910     0   1   2E+4   4   4.5  RT (min)       lcorbi_hilic_neg_4_015_24    m/z 580.5037     0   1   0.5   1  RT (min)      lcorbi_hilic_neg_4_082_59    m/z 580.7227     0   2   4   6E+3   4.5   5  RT (min)      lcorbi_hilic_neg_4_074_58    m/z 580.8666     0   1   2E+4   5   5.5  RT (min)       lcorbi_hilic_neg_4_105_61    m/z 581.0232     0   1   2   3   4E+3   3   3.5  RT (min)      lcorbi_hilic_neg_4_076_58    m/z 581.1661     0   1   2E+4   1   1.5  RT (min)       lcorbi_hilic_neg_4_098_61    m/z 581.2416     0   1E+5   1.5   2  RT (min)       lcorbi_hilic_neg_4_072_57    m/z 581.3033     0   1E+4   4   4.5  RT (min)       lcorbi_hilic_neg_4_068_57    m/z 581.3217     0   2   4   6   8E+3   4   4.5  RT (min)      lcorbi_hilic_neg_4_072_57    m/z 581.3297     0   1   2   3   4E+3   4   4.5  RT (min)      lcorbi_hilic_neg_4_080_58    m/z 581.4798     0   1   0.5   1  RT (min)      lcorbi_hilic_neg_4_084_59    m/z 581.4991     0   1E+4   0.5   1  RT (min)      lcorbi_hilic_neg_4_083_59    m/z 581.5351     0   2   4   6E+4   0.5   1  RT (min)      lcorbi_hilic_neg_4_084_59    m/z 582.1694     0   2   4   6E+3   1   1.5  RT (min)      lcorbi_hilic_neg_4_098_61    m/z 582.2974     0   1E+4   0.5   1  RT (min)      lcorbi_hilic_neg_4_072_57    m/z 582.2974     0   2   4   6E+4   3.5   4  RT (min)       lcorbi_hilic_neg_4_071_57    m/z 582.2992     0   2   4   6E+4   4   5  RT (min)       lcorbi_hilic_neg_4_090_60    m/z 582.3210     0   1   3   3.5  RT (min)      lcorbi_hilic_neg_4_068_57    m/z 582.3786     0   1E+4   4   4.5  RT (min)      lcorbi_hilic_neg_4_079_58    m/z 582.4245     0   1E+4   4.5   5  RT (min)      lcorbi_hilic_neg_4_088_59    m/z 582.7202     0   2   4   6   8E+3   4.5   5  RT (min)      lcorbi_hilic_neg_4_078_58    m/z 582.9768     0   1E+4   5.5   6  RT (min)      lcorbi_hilic_neg_4_070_57    m/z 583.0389     0   2   4E+3   3   3.5  RT (min)      lcorbi_hilic_neg_4_075_58    m/z 583.1297     0   1E+4   9   9.5  RT (min)      lcorbi_hilic_neg_4_084_59    m/z 583.3020     0   1   2E+4   3.5   4  RT (min)      lcorbi_hilic_neg_4_071_57    m/z 583.3344     0   1E+4   3.5   4  RT (min)      lcorbi_hilic_neg_4_084_59    m/z 583.9830     0   1E+4   5.5   6  RT (min)      lcorbi_hilic_neg_4_105_61    m/z 584.2958     0   1E+4   3.5   4  RT (min)      lcorbi_hilic_neg_4_070_57    m/z 584.3147     0   2   4E+4   4   5  RT (min)       lcorbi_hilic_neg_4_092_60    m/z 584.3575     0   1E+4   4   4.5  RT (min)      lcorbi_hilic_neg_4_069_57    m/z 584.3599     0   1   2E+5   0.5   1  RT (min)      lcorbi_hilic_neg_3_055_26    m/z 584.8062     0   1E+4   4.5   5  RT (min)      lcorbi_hilic_neg_4_082_59    m/z 585.2635     0   1E+4   0.5   1  RT (min)      lcorbi_hilic_neg_2_020_27    m/z 585.3527     0   2   4   6   8E+3   1   1.5  RT (min)      lcorbi_hilic_neg_4_073_57    m/z 585.3529     0   1E+4   4   4.5  RT (min)      lcorbi_hilic_neg_4_077_58    m/z 585.3632     0   1E+5   0.5   1  RT (min)      lcorbi_hilic_neg_3_030_30    m/z 585.3951     0   1E+3   0.5   1  RT (min)      lcorbi_hilic_neg_2_020_27    m/z 585.5303     0   1   2E+4   0.5   1  RT (min)       lcorbi_hilic_neg_4_089_59    m/z 586.1583     0   2   4   6   8E+3   8   8.5  RT (min)      lcorbi_hilic_neg_4_084_59    m/z 586.3160     0   2   4   6E+4   4   4.5  RT (min)       lcorbi_hilic_neg_4_072_57    m/z 586.3303     0   1   2   3E+4   3.5   4  RT (min)       lcorbi_hilic_neg_4_070_57    m/z 586.4981     0   1   2E+4   0.5   1  RT (min)       lcorbi_hilic_neg_4_087_59    m/z 586.8560     0   1E+4   7   7.5  RT (min)      lcorbi_hilic_neg_4_088_59    m/z 586.8995     0   1E+4   5.5   6  RT (min)      lcorbi_hilic_neg_4_068_57    m/z 587.0139     0   2   4E+3   3   3.5  RT (min)      lcorbi_hilic_neg_4_070_57    m/z 587.0632     0   1   2   3E+3   7   7.5  RT (min)      lcorbi_hilic_neg_4_095_60    m/z 587.0857     0   2   4E+3   5.5   6  RT (min)      lcorbi_hilic_neg_4_078_58    m/z 587.3190     0   1E+4   4   4.5  RT (min)       lcorbi_hilic_neg_4_070_57    m/z 587.3600     0   1   2E+5   0.5   1  RT (min)       lcorbi_hilic_neg_3_007_07    m/z 587.3998     0   2   4   6   8E+4   4   4.5  RT (min)       lcorbi_hilic_neg_1_047_05    m/z 587.4690     0   1   2E+4   0.5   1  RT (min)       lcorbi_hilic_neg_4_051_16    m/z 587.9921     0   1E+4   5.5   6  RT (min)      lcorbi_hilic_neg_4_079_58    m/z 588.2611     0   2   4   6   8E+3   4   4.5  RT (min)      lcorbi_hilic_neg_4_072_57    m/z 588.3462     0   2   4   6E+3   5   5.5  RT (min)      lcorbi_hilic_neg_1_029_31    m/z 588.3636     0   2   4   6   8E+4   0.5   1  RT (min)      lcorbi_hilic_neg_1_034_43    m/z 588.4140     0   1   2   3E+4   0.5   1  RT (min)      lcorbi_hilic_neg_4_074_58    m/z 588.4769     0   2   4   6E+4   0.5   1  RT (min)      lcorbi_hilic_neg_4_082_59    m/z 588.6654     0   2   4   6   8E+3   4.5   5  RT (min)      lcorbi_hilic_neg_4_066_57    m/z 589.4152     0   2   4E+4   4   4.5  RT (min)       lcorbi_hilic_neg_4_094_60    m/z 589.4260     0   2   4   6E+4   0.5   1  RT (min)      lcorbi_hilic_neg_1_003_14    m/z 589.4834     0   1   2   3E+4   0.5   1  RT (min)       lcorbi_hilic_neg_4_051_16    m/z 589.8911     0   1E+5   5.5   6  RT (min)      lcorbi_hilic_neg_1_051_33    m/z 589.9689     0   2   4E+3   5   5.5  RT (min)      lcorbi_hilic_neg_4_089_59    m/z 590.1322     0   1E+4   4.5   5   5.5  RT (min)      lcorbi_hilic_neg_4_106_62    m/z 590.2876     0   1E+4   0.5   1  RT (min)      lcorbi_hilic_neg_4_090_60    m/z 590.4512     0   1   2E+4   0.5   1  RT (min)      lcorbi_hilic_neg_4_066_57    m/z 590.7515     0   1E+4   4.5   5  RT (min)      lcorbi_hilic_neg_4_082_59    m/z 591.2737     0   2   4   6E+5   0.5   1  RT (min)       lcorbi_hilic_neg_1_027_18    m/z 591.5003     0   2   4   6E+2   0.5   1  RT (min)      lcorbi_hilic_neg_2_012_14    m/z 591.9069     0   2   4E+3   5   5.5  RT (min)      lcorbi_hilic_neg_4_072_57    m/z 592.2772     0   1   2E+5   0.5   1  RT (min)      lcorbi_hilic_neg_1_027_18    m/z 592.2935     0   1E+4   4   4.5  RT (min)      lcorbi_hilic_neg_4_075_58    m/z 592.3631     0   1   2   3E+3   4   4.5  RT (min)       lcorbi_hilic_neg_4_090_60    m/z 592.3961     0   1   2   3E+4   1.5   2  RT (min)      lcorbi_hilic_neg_4_075_58    m/z 592.4348     0   1E+5   4   4.5  RT (min)       lcorbi_hilic_neg_4_095_60    m/z 592.4420     0   1   2   3E+5   0.5   1  RT (min)      lcorbi_hilic_neg_2_037_15    m/z 592.7492     0   1E+4   4.5   5  RT (min)      lcorbi_hilic_neg_4_082_59    m/z 593.1055     0   2   4   6E+3   5.5   6  RT (min)      lcorbi_hilic_neg_4_067_57    m/z 593.2529     0   1   2   3E+5   0.5   1  RT (min)      lcorbi_hilic_neg_2_012_14    m/z 593.2708     0   1   0.5   1  RT (min)      lcorbi_hilic_neg_4_076_58    m/z 593.3662     0   1E+4   4   4.5  RT (min)      lcorbi_hilic_neg_4_081_58    m/z 593.4699     0   1   2E+4   0.5   1  RT (min)      lcorbi_hilic_neg_4_042_28    m/z 594.0118     0   1E+4   5.5   6  RT (min)      lcorbi_hilic_neg_4_105_61    m/z 594.2563     0   1E+5   0.5   1  RT (min)      lcorbi_hilic_neg_2_037_15    m/z 594.3792     0   1   2   3   4E+4   4   4.5  RT (min)       lcorbi_hilic_neg_4_079_58    m/z 595.3819     0   1E+4   4   4.5  RT (min)      lcorbi_hilic_neg_4_079_58    m/z 595.4857     0   1   2   3   4E+4   0.5   1  RT (min)      lcorbi_hilic_neg_4_016_29    m/z 596.2930     0   1E+4   5   5.5  RT (min)      lcorbi_hilic_neg_4_085_59    m/z 596.3739     0   1   2E+5   5   5.5  RT (min)       lcorbi_hilic_neg_4_097_60    m/z 596.3946     0   1E+4   4   4.5  RT (min)      lcorbi_hilic_neg_4_079_58    m/z 596.4039     0   1E+4   3.5   4  RT (min)      lcorbi_hilic_neg_4_089_59    m/z 596.4895     0   1   2   3   4E+4   0.5   1  RT (min)      lcorbi_hilic_neg_4_089_59    m/z 596.6968     0   1E+4   4.5   5  RT (min)      lcorbi_hilic_neg_4_078_58    m/z 596.9287     0   1   2   3E+4   5.5   6  RT (min)       lcorbi_hilic_neg_4_073_57    m/z 597.2365     0   1   2E+4   3   3.5  RT (min)       lcorbi_hilic_neg_4_076_58    m/z 597.2612     0   1   2   3   4E+4   2.5   3  RT (min)       lcorbi_hilic_neg_4_082_59    m/z 597.2898     0   2   4E+3   7   7.5  RT (min)      lcorbi_hilic_neg_3_029_25    m/z 597.3027     0   2   4   6E+3   7   7.5  RT (min)      lcorbi_hilic_neg_3_011_24    m/z 597.3051     0   1   2   3E+4   4.5   5  RT (min)       lcorbi_hilic_neg_4_085_59    m/z 597.3803     0   2   4   6E+5   0.5   1  RT (min)      lcorbi_hilic_neg_4_031_14    m/z 597.5014     0   2   4   6   8E+4   0.5   1  RT (min)      lcorbi_hilic_neg_4_080_58    m/z 597.5305     0   1   2E+4   0.5   1  RT (min)      lcorbi_hilic_neg_4_087_59    m/z 598.2900     0   1   2E+4   4   4.5  RT (min)       lcorbi_hilic_neg_4_069_57    m/z 598.3083     0   1E+4   4.5   5  RT (min)      lcorbi_hilic_neg_4_085_59    m/z 598.3839     0   1   2E+5   0.5   1  RT (min)      lcorbi_hilic_neg_4_031_14    m/z 598.4982     0   1   2   3   4E+3   0.5   1  RT (min)      lcorbi_hilic_neg_2_032_25    m/z 598.5428     0   1   2   3   4E+3   0.5   1  RT (min)      lcorbi_hilic_neg_2_032_25    m/z 598.6944     0   1E+4   4.5   5  RT (min)      lcorbi_hilic_neg_4_078_58    m/z 599.0140     0   1   2   3E+3   3   3.5  RT (min)      lcorbi_hilic_neg_4_077_58    m/z 599.3213     0   2   4   6E+4   4.5   5  RT (min)       lcorbi_hilic_neg_4_079_58    m/z 599.5095     0   2   4   6E+4   0.5   1  RT (min)      lcorbi_hilic_neg_4_084_59    m/z 599.5383     0   1E+4   0.5   1  RT (min)      lcorbi_hilic_neg_4_074_58    m/z 599.5668     0   1E+4   0.5   1  RT (min)      lcorbi_hilic_neg_4_098_61    m/z 599.9202     0   1   2E+4   5.5   6  RT (min)       lcorbi_hilic_neg_4_072_57    m/z 600.3244     0   1   2E+4   4.5   5  RT (min)      lcorbi_hilic_neg_4_078_58    m/z 600.7803     0   1   2E+4   4.5   5  RT (min)       lcorbi_hilic_neg_4_082_59    m/z 601.0295     0   1   2   3   4E+3   3   3.5  RT (min)      lcorbi_hilic_neg_4_069_57    m/z 601.5173     0   1   2   3   4E+4   0.5   1  RT (min)      lcorbi_hilic_neg_4_087_59    m/z 601.5336     0   1   2   3E+4   0.5   1  RT (min)       lcorbi_hilic_neg_4_083_59    m/z 602.3031     0   1   2E+4   4   4.5  RT (min)       lcorbi_hilic_neg_4_067_57    m/z 602.4928     0   1   2   3   4E+4   0.5   1  RT (min)      lcorbi_hilic_neg_4_083_59    m/z 602.5110     0   1E+4   0.5   1  RT (min)      lcorbi_hilic_neg_4_084_59    m/z 602.5296     0   1E+4   0.5   1  RT (min)      lcorbi_hilic_neg_4_083_59    m/z 602.6429     0   1E+4   4.5   5  RT (min)      lcorbi_hilic_neg_4_079_58    m/z 603.0301     0   2   4   6   8E+3   6   6.5  RT (min)      lcorbi_hilic_neg_4_112_62    m/z 603.3922     0   2   4   6E+4   3.5   4  RT (min)       lcorbi_hilic_neg_4_088_59    m/z 603.4310     0   1   2E+4   0.5   1  RT (min)      lcorbi_hilic_neg_4_042_28    m/z 603.4386     0   1   2   3   4E+5   0.5   1  RT (min)       lcorbi_hilic_neg_3_036_17    m/z 603.4966     0   1   2   3E+4   0.5   1  RT (min)      lcorbi_hilic_neg_3_039_31    m/z 603.5142     0   2   4   6   8E+3   0.5   1  RT (min)      lcorbi_hilic_neg_4_083_59    m/z 603.5325     0   1   0.5   1  RT (min)      lcorbi_hilic_neg_4_074_58    m/z 603.8490     0   1   2   3E+4   3   3.5  RT (min)      lcorbi_hilic_neg_4_070_57    m/z 604.3238     0   1   2   3E+4   4   4.5  RT (min)       lcorbi_hilic_neg_4_079_58    m/z 604.3520     0   1E+4   3   3.5  RT (min)      lcorbi_hilic_neg_4_069_57    m/z 604.3949     0   1E+4   3.5   4  RT (min)      lcorbi_hilic_neg_4_085_59    m/z 604.4419     0   1E+5   0.5   1  RT (min)      lcorbi_hilic_neg_3_036_17    m/z 604.5813     0   1   2   3E+4   0.5   1  RT (min)      lcorbi_hilic_neg_4_091_60    m/z 604.6405     0   1   2E+4   4.5   5  RT (min)       lcorbi_hilic_neg_4_076_58    m/z 604.8720     0   1   2E+4   5   5.5  RT (min)       lcorbi_hilic_neg_4_105_61    m/z 604.9585     0   2   4   6   8E+3   5.5   6  RT (min)      lcorbi_hilic_neg_4_073_57    m/z 605.2395     0   1E+4   0.5   1  RT (min)      lcorbi_hilic_neg_4_038_27    m/z 605.3223     0   1E+4   4   4.5  RT (min)      lcorbi_hilic_neg_4_067_57    m/z 605.4078     0   1   2   3E+4   1.5   2  RT (min)       lcorbi_hilic_neg_4_066_57    m/z 605.4542     0   2   4   6E+5   0.5   1  RT (min)       lcorbi_hilic_neg_2_037_15    m/z 605.4797     0   1E+4   0.5   1  RT (min)       lcorbi_hilic_neg_3_036_17    m/z 606.2252     0   1   2   3   4E+4   8   8.5  RT (min)       lcorbi_hilic_neg_4_096_60    m/z 606.2990     0   1   2   3   4E+4   3.5   4  RT (min)       lcorbi_hilic_neg_4_067_57    m/z 606.3345     0   1E+4   4   4.5  RT (min)      lcorbi_hilic_neg_4_072_57    m/z 606.3784     0   2   4   6   8E+4   3.5   4  RT (min)       lcorbi_hilic_neg_4_088_59    m/z 606.4109     0   1E+4   1.5   2  RT (min)      lcorbi_hilic_neg_4_066_57    m/z 606.4576     0   1   2E+5   0.5   1  RT (min)      lcorbi_hilic_neg_3_036_17    m/z 606.4832     0   1   2   3E+3   0.5   1  RT (min)      lcorbi_hilic_neg_4_044_01    m/z 606.5968     0   2   4   6   8E+4   0.5   1  RT (min)       lcorbi_hilic_neg_4_091_60    m/z 606.7257     0   1   2E+4   4.5   5  RT (min)       lcorbi_hilic_neg_4_082_59    m/z 607.0798     0   1   2E+4   8   8.5  RT (min)      lcorbi_hilic_neg_4_089_59    m/z 607.3025     0   1E+5   0.5   1  RT (min)      lcorbi_hilic_neg_3_037_12    m/z 607.4148     0   2   4   6E+4   0.5   1  RT (min)      lcorbi_hilic_neg_2_045_26    m/z 607.4602     0   2   4E+4   0.5   1  RT (min)      lcorbi_hilic_neg_2_037_15    m/z 608.0816     0   2   4   6E+3   8   8.5  RT (min)      lcorbi_hilic_neg_4_088_59    m/z 608.3066     0   2   4E+4   0.5   1  RT (min)      lcorbi_hilic_neg_3_037_12    m/z 608.3849     0   1E+4   3.5   4  RT (min)      lcorbi_hilic_neg_4_094_60    m/z 608.4182     0   1   2E+4   0.5   1  RT (min)      lcorbi_hilic_neg_2_045_26    m/z 608.4299     0   2   4   6E+3   4   4.5  RT (min)      lcorbi_hilic_neg_4_095_60    m/z 608.4990     0   1   2   3   4E+4   0.5   1  RT (min)       lcorbi_hilic_neg_4_073_57    m/z 608.5271     0   1   0.5   1  RT (min)       lcorbi_hilic_neg_4_073_57    m/z 608.5938     0   1   2E+4   0.5   1  RT (min)      lcorbi_hilic_neg_4_091_60    m/z 608.7232     0   1E+4   4.5   5  RT (min)      lcorbi_hilic_neg_4_082_59    m/z 608.8381     0   1E+4   7   7.5  RT (min)      lcorbi_hilic_neg_4_086_59    m/z 609.3224     0   1   2   3E+4   4   4.5  RT (min)      lcorbi_hilic_neg_4_074_58    m/z 609.4298     0   2   4   6E+4   0.5   1  RT (min)      lcorbi_hilic_neg_4_076_58    m/z 609.4744     0   1   2   3E+3   0.5   1  RT (min)      lcorbi_hilic_neg_4_004_13    m/z 609.5956     0   1E+4   0.5   1  RT (min)      lcorbi_hilic_neg_4_098_61    m/z 610.4804     0   1   2   3E+3   7.5   8  RT (min)      lcorbi_hilic_neg_1_003_14    m/z 610.5062     0   1   2E+4   0.5   1  RT (min)      lcorbi_hilic_neg_4_087_59    m/z 610.5142     0   1   2E+3   0.5   1  RT (min)       lcorbi_hilic_neg_4_089_59    m/z 610.8017     0   1E+4   7   7.5  RT (min)      lcorbi_hilic_neg_1_023_23    m/z 611.3726     0   1   2   3   4E+5   0.5   1  RT (min)      lcorbi_hilic_neg_4_031_14    m/z 611.4237     0   1   2   3   4E+4   3.5   4  RT (min)      lcorbi_hilic_neg_4_085_59    m/z 611.4454     0   1   2   3   4E+4   0.5   1  RT (min)      lcorbi_hilic_neg_4_031_14    m/z 611.5265     0   1   0.5   1  RT (min)      lcorbi_hilic_neg_4_087_59    m/z 612.0158     0   2   4   6E+3   5   5.5  RT (min)      lcorbi_hilic_neg_4_110_62    m/z 612.1499     0   2   4   6   8E+4   8   8.5  RT (min)      lcorbi_hilic_neg_4_084_59    m/z 612.3758     0   1E+5   0.5   1  RT (min)      lcorbi_hilic_neg_4_031_14    m/z 612.5216     0   1E+5   0.5   1  RT (min)       lcorbi_hilic_neg_3_039_31    m/z 612.6716     0   1   2   3E+4   4.5   5  RT (min)       lcorbi_hilic_neg_4_079_58    m/z 613.0587     0   2   4   6   8E+3   6   6.5  RT (min)      lcorbi_hilic_neg_4_112_62    m/z 613.1430     0   1   2   3E+4   8   8.5  RT (min)      lcorbi_hilic_neg_4_085_59    m/z 613.5251     0   2   4E+4   0.5   1  RT (min)      lcorbi_hilic_neg_4_083_59    m/z 613.5614     0   1   2E+4   0.5   1  RT (min)       lcorbi_hilic_neg_4_087_59    m/z 614.4929     0   2   4E+4   0.5   1  RT (min)      lcorbi_hilic_neg_4_083_59    m/z 614.6100     0   1   2   3   4E+4   0.5   1  RT (min)       lcorbi_hilic_neg_4_092_60    m/z 614.6370     0   1E+5   0.5   1  RT (min)      lcorbi_hilic_neg_4_097_60    m/z 614.6691     0   1   2   3E+4   4.5   5  RT (min)       lcorbi_hilic_neg_4_076_58    m/z 614.8431     0   2   4E+3   3   3.5  RT (min)      lcorbi_hilic_neg_4_069_57    m/z 615.2906     0   2   4   6E+3   5.5   6  RT (min)      lcorbi_hilic_neg_4_103_61    m/z 615.3500     0   1   2   3   4E+4   4   4.5  RT (min)       lcorbi_hilic_neg_4_072_57    m/z 615.4962     0   1   2E+4   0.5   1  RT (min)      lcorbi_hilic_neg_4_083_59    m/z 615.6222     0   1   2   3E+4   0.5   1  RT (min)       lcorbi_hilic_neg_4_091_60    m/z 615.8503     0   2   4   6E+3   3   3.5  RT (min)      lcorbi_hilic_neg_4_069_57    m/z 616.1436     0   2   4   6E+3   1   1.5  RT (min)      lcorbi_hilic_neg_4_106_62    m/z 616.3525     0   1   2   3   4E+3   3   3.5  RT (min)      lcorbi_hilic_neg_4_067_57    m/z 616.6255     0   1E+5   0.5   1  RT (min)       lcorbi_hilic_neg_4_091_60    m/z 616.6523     0   1E+5   0.5   1  RT (min)      lcorbi_hilic_neg_4_097_60    m/z 616.6663     0   1   4.5   5  RT (min)      lcorbi_hilic_neg_4_006_06    m/z 616.7545     0   1   2   3E+4   4.5   5  RT (min)       lcorbi_hilic_neg_4_090_60    m/z 616.8584     0   1E+4   3   3.5  RT (min)      lcorbi_hilic_neg_4_068_57    m/z 617.2185     0   1E+5   0.5   1  RT (min)      lcorbi_hilic_neg_3_031_14    m/z 617.3580     0   1E+4   0.5   1  RT (min)      lcorbi_hilic_neg_3_012_35    m/z 617.3656     0   1   2   3E+4   3.5   4  RT (min)      lcorbi_hilic_neg_4_073_57    m/z 617.4434     0   1E+5   0.5   1  RT (min)      lcorbi_hilic_neg_3_055_26    m/z 617.5119     0   1   0.5   1  RT (min)      lcorbi_hilic_neg_3_036_17    m/z 618.4467     0   2   4   6E+4   0.5   1  RT (min)      lcorbi_hilic_neg_3_055_26    m/z 618.4574     0   1   2E+5   0.5   1  RT (min)      lcorbi_hilic_neg_3_036_17    m/z 618.5055     0   1   0.5   1  RT (min)      lcorbi_hilic_neg_3_036_17    m/z 618.6410     0   1   2E+4   0.5   1  RT (min)      lcorbi_hilic_neg_4_091_60    m/z 618.7527     0   1E+4   4.5   5  RT (min)      lcorbi_hilic_neg_4_090_60    m/z 619.0200     0   1   2   3   4E+3   3   3.5  RT (min)      lcorbi_hilic_neg_4_078_58    m/z 619.2900     0   1   2   3   4E+4   4.5   5  RT (min)       lcorbi_hilic_neg_4_068_57    m/z 619.3078     0   1   2   3   4E+4   7   7.5  RT (min)       lcorbi_hilic_neg_4_085_59    m/z 619.5091     0   1   0.5   1  RT (min)      lcorbi_hilic_neg_2_037_15    m/z 620.2930     0   1E+4   4.5   5  RT (min)      lcorbi_hilic_neg_4_073_57    m/z 620.4731     0   1   2   3E+5   0.5   1  RT (min)      lcorbi_hilic_neg_2_037_15    m/z 620.5757     0   2   4   6   8E+4   0.5   1  RT (min)      lcorbi_hilic_neg_4_095_60    m/z 621.4759     0   2   4   6E+4   0.5   1  RT (min)      lcorbi_hilic_neg_3_036_17    m/z 622.0447     0   2   4   6   8E+3   5   5.5  RT (min)      lcorbi_hilic_neg_4_106_62    m/z 622.3677     0   1   2E+3   4.5   5  RT (min)       lcorbi_hilic_neg_4_015_24    m/z 622.3798     0   1   2E+5   4.5   5  RT (min)       lcorbi_hilic_neg_4_097_60    m/z 622.5914     0   2   4   6E+4   0.5   1  RT (min)      lcorbi_hilic_neg_4_095_60    m/z 622.8927     0   1E+4   5   5.5  RT (min)      lcorbi_hilic_neg_4_069_57    m/z 623.1333     0   2   4   6   8E+4   8   8.5  RT (min)       lcorbi_hilic_neg_4_095_60    m/z 623.3352     0   1   2E+4   0.5   1  RT (min)      lcorbi_hilic_neg_4_111_62    m/z 623.3923     0   1   2   3   4E+4   5   5.5  RT (min)       lcorbi_hilic_neg_4_097_60    m/z 624.2922     0   1E+4   4   4.5  RT (min)      lcorbi_hilic_neg_4_070_57    m/z 624.3957     0   1   2E+5   4.5   5  RT (min)       lcorbi_hilic_neg_4_091_60    m/z 624.6979     0   1   2   3E+4   4.5   5  RT (min)       lcorbi_hilic_neg_4_075_58    m/z 625.5252     0   1   2   3   4E+4   0.5   1  RT (min)      lcorbi_hilic_neg_4_083_59    m/z 626.4106     0   1   2   3   4E+4   4.5   5  RT (min)       lcorbi_hilic_neg_4_091_60    m/z 626.6956     0   1E+4   4.5   5  RT (min)      lcorbi_hilic_neg_4_077_58    m/z 627.2465     0   1   2E+5   1   1.5  RT (min)       lcorbi_hilic_neg_4_066_57    m/z 627.2757     0   1   2   3E+4   0.5   1  RT (min)      lcorbi_hilic_neg_4_004_13    m/z 627.4109     0   2   4   6E+4   3.5   4  RT (min)       lcorbi_hilic_neg_4_089_59    m/z 627.5698     0   1E+4   0.5   1  RT (min)      lcorbi_hilic_neg_4_074_58    m/z 628.0584     0   2   4   6E+3   8   8.5  RT (min)      lcorbi_hilic_neg_4_089_59    m/z 628.2501     0   2   4   6E+4   1   1.5  RT (min)      lcorbi_hilic_neg_4_066_57    m/z 628.4139     0   1   2E+4   3.5   4  RT (min)      lcorbi_hilic_neg_4_085_59    m/z 628.5090     0   1E+4   0.5   1  RT (min)      lcorbi_hilic_neg_4_083_59    m/z 628.5435     0   2   4   6   8E+4   0.5   1  RT (min)      lcorbi_hilic_neg_4_083_59    m/z 628.5539     0   2   4E+4   0.5   1  RT (min)      lcorbi_hilic_neg_4_026_33    m/z 628.9821     0   1E+4   5.5   6  RT (min)      lcorbi_hilic_neg_4_073_57    m/z 630.2105     0   1E+5   8   8.5  RT (min)       lcorbi_hilic_neg_4_094_60    m/z 630.6045     0   1   2   3E+5   0.5   1  RT (min)       lcorbi_hilic_neg_4_097_60    m/z 630.8199     0   1E+4   7   7.5  RT (min)      lcorbi_hilic_neg_4_082_59    m/z 631.4616     0   1E+4   0.5   1  RT (min)      lcorbi_hilic_neg_4_016_29    m/z 632.3190     0   1   2E+4   4   4.5  RT (min)       lcorbi_hilic_neg_4_066_57    m/z 632.5270     0   1E+4   0.5   1  RT (min)      lcorbi_hilic_neg_4_085_59    m/z 632.6202     0   1   2E+5   0.5   1  RT (min)       lcorbi_hilic_neg_4_097_60    m/z 632.8179     0   1E+4   8.5   9  RT (min)      lcorbi_hilic_neg_4_081_58    m/z 633.1162     0   2   4   6E+3   8   8.5  RT (min)      lcorbi_hilic_neg_4_083_59    m/z 633.2097     0   2   4   6   8E+4   8   8.5  RT (min)       lcorbi_hilic_neg_4_094_60    m/z 633.7318     0   1E+4   4.5   5  RT (min)      lcorbi_hilic_neg_4_093_60    m/z 636.3509     0   1   2   3E+4   4   4.5  RT (min)       lcorbi_hilic_neg_4_074_58    m/z 636.6349     0   2   4E+4   0.5   1  RT (min)       lcorbi_hilic_neg_4_091_60    m/z 636.7252     0   1E+4   4.5   5  RT (min)      lcorbi_hilic_neg_4_092_60    m/z 636.8278     0   1E+4   5.5   6  RT (min)      lcorbi_hilic_neg_4_111_62    m/z 637.2814     0   1E+4   0.5   1  RT (min)      lcorbi_hilic_neg_4_099_61    m/z 638.3087     0   1E+4   0.5   1  RT (min)      lcorbi_hilic_neg_4_093_60    m/z 638.5376     0   1   2   3E+4   0.5   1  RT (min)      lcorbi_hilic_neg_4_084_59    m/z 641.5857     0   1   2E+4   0.5   1  RT (min)       lcorbi_hilic_neg_4_074_58    m/z 643.4592     0   1   2   3   4E+4   0.5   1  RT (min)      lcorbi_hilic_neg_3_055_26    m/z 643.4715     0   1   2E+4   0.5   1  RT (min)       lcorbi_hilic_neg_3_036_17    m/z 645.4509     0   2   4E+4   3.5   4  RT (min)       lcorbi_hilic_neg_4_106_62    m/z 645.4749     0   1   0.5   1  RT (min)      lcorbi_hilic_neg_4_068_57    m/z 645.4863     0   2   4   6E+4   0.5   1  RT (min)      lcorbi_hilic_neg_2_037_15    m/z 646.4542     0   1   2E+4   3.5   4  RT (min)      lcorbi_hilic_neg_4_107_62    m/z 646.5411     0   1   5   5.5  RT (min)      lcorbi_hilic_neg_4_108_62    m/z 646.6637     0   2   4   6   8E+4   0.5   1  RT (min)       lcorbi_hilic_neg_4_091_60    m/z 646.8568     0   1   2   3   4E+4   5   5.5  RT (min)       lcorbi_hilic_neg_4_108_62    m/z 648.6795     0   1   2   3E+4   0.5   1  RT (min)       lcorbi_hilic_neg_4_091_60    m/z 648.8543     0   1E+4   5   5.5  RT (min)      lcorbi_hilic_neg_4_112_62    m/z 649.2289     0   2   4   6E+4   1   1.5  RT (min)      lcorbi_hilic_neg_4_071_57    m/z 649.5533     0   2   4   6E+4   3.5   4  RT (min)       lcorbi_hilic_neg_4_113_62    m/z 649.5534     0   1   2   3   4E+4   0.5   1  RT (min)      lcorbi_hilic_neg_4_067_57    m/z 650.5567     0   1   2E+4   3.5   4  RT (min)      lcorbi_hilic_neg_4_109_62    m/z 650.5567     0   1E+4   0.5   1  RT (min)      lcorbi_hilic_neg_4_069_57    m/z 650.6138     0   1   2E+5   0.5   1  RT (min)       lcorbi_hilic_neg_4_097_60    m/z 650.9639     0   1E+4   5.5   6  RT (min)      lcorbi_hilic_neg_4_067_57    m/z 651.6259     0   2   4   6E+4   0.5   1  RT (min)      lcorbi_hilic_neg_4_005_32    m/z 651.8711     0   2   4   6   8E+3   5   5.5  RT (min)      lcorbi_hilic_neg_4_113_62    m/z 651.9704     0   2   4   6E+3   5.5   6  RT (min)      lcorbi_hilic_neg_4_105_61    m/z 652.6294     0   1   2E+5   0.5   1  RT (min)       lcorbi_hilic_neg_1_012_32    m/z 654.6263     0   2   4   6E+4   0.5   1  RT (min)      lcorbi_hilic_neg_1_012_32    m/z 656.3103     0   1E+4   0.5   1  RT (min)      lcorbi_hilic_neg_4_090_60    m/z 656.3189     0   1   2E+4   4   4.5  RT (min)       lcorbi_hilic_neg_4_068_57    m/z 657.3884     0   2   4   6E+3   5   5.5  RT (min)      lcorbi_hilic_neg_4_069_57    m/z 657.8772     0   2   4   6   8E+3   5   5.5  RT (min)      lcorbi_hilic_neg_4_070_57    m/z 658.1278     0   1   2   3E+4   1   1.5  RT (min)      lcorbi_hilic_neg_4_105_61    m/z 658.2910     0   1E+4   0.5   1  RT (min)      lcorbi_hilic_neg_4_105_61    m/z 658.8908     0   2   4   6   8E+3   5   5.5  RT (min)      lcorbi_hilic_neg_4_067_57    m/z 659.4665     0   1   2E+4   3   3.5  RT (min)       lcorbi_hilic_neg_4_106_62    m/z 659.6395     0   1E+5   0.5   1  RT (min)      lcorbi_hilic_neg_4_097_60    m/z 660.4698     0   1E+4   3   3.5  RT (min)      lcorbi_hilic_neg_4_112_62    m/z 660.4698     0   1   2E+4   0.5   1  RT (min)      lcorbi_hilic_neg_4_111_62    m/z 661.6548     0   1   2E+5   0.5   1  RT (min)       lcorbi_hilic_neg_4_097_60    m/z 661.9991     0   2   4   6E+3   5.5   6  RT (min)      lcorbi_hilic_neg_4_104_61    m/z 662.1034     0   1E+4   7.5   8  RT (min)      lcorbi_hilic_neg_4_083_59    m/z 663.3302     0   1E+5   1   1.5  RT (min)      lcorbi_hilic_neg_1_002_26    m/z 664.4201     0   1   2   3E+4   3.5   4  RT (min)       lcorbi_hilic_neg_4_089_59    m/z 664.9158     0   1E+4   5.5   6  RT (min)      lcorbi_hilic_neg_4_066_57    m/z 665.3127     0   1E+4   0.5   1  RT (min)      lcorbi_hilic_neg_4_105_61    m/z 666.3594     0   1   2   3   4E+4   6   6.5  RT (min)       lcorbi_hilic_neg_4_078_58    m/z 667.4351     0   2   4   6   8E+3   0.5   1  RT (min)      lcorbi_hilic_neg_4_084_59    m/z 667.9078     0   1E+4   5.5   6  RT (min)      lcorbi_hilic_neg_4_072_57    m/z 669.4513     0   1E+5   2.5   3  RT (min)       lcorbi_hilic_neg_4_082_59    m/z 670.4541     0   1   2E+4   0.5   1  RT (min)      lcorbi_hilic_neg_4_086_59    m/z 670.4544     0   2   4   6E+4   3   3.5  RT (min)       lcorbi_hilic_neg_4_082_59    m/z 671.2691     0   1   2E+4   0.5   1  RT (min)      lcorbi_hilic_neg_4_082_59    m/z 672.4700     0   1   2   3   4E+4   3   3.5  RT (min)      lcorbi_hilic_neg_4_109_62    m/z 673.0981     0   1E+4   1   1.5  RT (min)      lcorbi_hilic_neg_4_103_61    m/z 673.4817     0   1   2E+5   3   3.5  RT (min)      lcorbi_hilic_neg_4_087_59    m/z 674.4852     0   1   2   3   4E+4   0.5   1  RT (min)       lcorbi_hilic_neg_4_084_59    m/z 674.4855     0   2   4   6E+4   2.5   3  RT (min)       lcorbi_hilic_neg_4_083_59    m/z 675.4612     0   2   4E+3   0.5   1  RT (min)      lcorbi_hilic_neg_4_068_57    m/z 675.6613     0   1   2   3E+5   0.5   1  RT (min)       lcorbi_hilic_neg_4_097_60    m/z 681.2978     0   1   2   3   4E+4   0.5   1  RT (min)       lcorbi_hilic_neg_4_082_59    m/z 684.4616     0   2   4   6   8E+3   3   3.5  RT (min)      lcorbi_hilic_neg_4_089_59    m/z 685.4819     0   1   2E+4   0.5   1  RT (min)      lcorbi_hilic_neg_4_082_59    m/z 685.4820     0   1   2   3E+4   3   3.5  RT (min)       lcorbi_hilic_neg_4_082_59    m/z 685.5298     0   2   4   6   8E+3   3.5   4  RT (min)      lcorbi_hilic_neg_4_105_61    m/z 685.8649     0   2   4   6E+3   5   5.5  RT (min)      lcorbi_hilic_neg_4_113_62    m/z 686.4777     0   1E+4   0.5   1  RT (min)      lcorbi_hilic_neg_4_084_59    m/z 687.4806     0   1   2E+4   3   3.5  RT (min)       lcorbi_hilic_neg_4_087_59    m/z 688.4930     0   1   2   3E+4   0.5   1  RT (min)       lcorbi_hilic_neg_4_085_59    m/z 689.4962     0   1E+4   0.5   1  RT (min)      lcorbi_hilic_neg_4_082_59    m/z 690.4992     0   2   4   6   8E+3   3   3.5  RT (min)      lcorbi_hilic_neg_4_084_59    m/z 690.6868     0   1E+4   4.5   5  RT (min)      lcorbi_hilic_neg_4_089_59    m/z 690.8798     0   1E+4   5   5.5  RT (min)      lcorbi_hilic_neg_4_004_13    m/z 691.4349     0   1E+4   0.5   1  RT (min)      lcorbi_hilic_neg_4_084_59    m/z 691.4353     0   1   2   3   4E+4   3   3.5  RT (min)       lcorbi_hilic_neg_4_082_59    m/z 692.6214     0   2   4E+4   0.5   1  RT (min)      lcorbi_hilic_neg_1_020_13    m/z 693.4506     0   1   2   3E+4   0.5   1  RT (min)       lcorbi_hilic_neg_4_084_59    m/z 693.4511     0   2   4   6E+4   2.5   3  RT (min)       lcorbi_hilic_neg_4_088_59    m/z 694.4307     0   1   2   3E+4   3.5   4  RT (min)       lcorbi_hilic_neg_4_088_59    m/z 694.4540     0   1E+4   0.5   1  RT (min)      lcorbi_hilic_neg_4_084_59    m/z 695.4661     0   1   2   3   4E+4   0.5   1  RT (min)       lcorbi_hilic_neg_4_084_59    m/z 695.4665     0   2   4   6E+4   2.5   3  RT (min)       lcorbi_hilic_neg_4_088_59    m/z 695.8772     0   1E+4   5   5.5  RT (min)      lcorbi_hilic_neg_2_020_27    m/z 696.4696     0   1E+4   0.5   1  RT (min)      lcorbi_hilic_neg_4_084_59    m/z 697.4823     0   2   4   6   8E+4   2.5   3  RT (min)       lcorbi_hilic_neg_4_087_59    m/z 698.4774     0   1E+4   3   3.5  RT (min)      lcorbi_hilic_neg_4_088_59    m/z 698.4850     0   1E+4   0.5   1  RT (min)      lcorbi_hilic_neg_4_084_59    m/z 698.5135     0   1E+4   3   3.5  RT (min)      lcorbi_hilic_neg_1_012_32    m/z 699.4976     0   1   2   3E+4   3   3.5  RT (min)       lcorbi_hilic_neg_4_111_62    m/z 699.5170     0   1   3   3.5  RT (min)      lcorbi_hilic_neg_4_109_62    m/z 700.1892     0   1   2   3   4E+3   5   5.5  RT (min)      lcorbi_hilic_neg_2_020_27    m/z 700.4999     0   1   2   3   4E+4   0.5   1  RT (min)      lcorbi_hilic_neg_4_085_59    m/z 700.5007     0   1E+4   3   3.5  RT (min)       lcorbi_hilic_neg_4_108_62    m/z 700.7156     0   1E+4   4.5   5  RT (min)      lcorbi_hilic_neg_4_090_60    m/z 701.4964     0   1   2E+4   3   3.5  RT (min)       lcorbi_hilic_neg_4_084_59    m/z 702.5087     0   1   2E+4   0.5   1  RT (min)      lcorbi_hilic_neg_4_084_59    m/z 703.5118     0   1   2E+4   3   3.5  RT (min)      lcorbi_hilic_neg_4_087_59    m/z 703.5121     0   2   4   6   8E+3   0.5   1  RT (min)      lcorbi_hilic_neg_4_082_59    m/z 704.5148     0   2   4E+3   3   3.5  RT (min)      lcorbi_hilic_neg_4_084_59    m/z 706.5751     0   1E+4   0.5   1  RT (min)      lcorbi_hilic_neg_4_090_60    m/z 706.6608     0   1E+4   4.5   5  RT (min)      lcorbi_hilic_neg_4_078_58    m/z 707.5871     0   1   2   3E+4   0.5   1  RT (min)      lcorbi_hilic_neg_4_091_60    m/z 708.4616     0   1E+4   3   3.5  RT (min)      lcorbi_hilic_neg_4_089_59    m/z 708.5906     0   1   2E+5   3   3.5  RT (min)       lcorbi_hilic_neg_4_090_60    m/z 708.6585     0   1E+4   4.5   5  RT (min)      lcorbi_hilic_neg_4_080_58    m/z 709.4649     0   2   4   6   8E+3   3   3.5  RT (min)      lcorbi_hilic_neg_4_089_59    m/z 709.6022     0   1   2E+4   0.5   1  RT (min)      lcorbi_hilic_neg_4_091_60    m/z 710.4773     0   1E+4   0.5   1  RT (min)      lcorbi_hilic_neg_4_082_59    m/z 710.6061     0   2   4   6E+4   0.5   1  RT (min)       lcorbi_hilic_neg_4_091_60    m/z 710.6061     0   1E+5   3   3.5  RT (min)       lcorbi_hilic_neg_4_091_60    m/z 711.4805     0   2   4   6E+3   0.5   1  RT (min)      lcorbi_hilic_neg_4_085_59    m/z 711.4805     0   1   2   3E+4   3   3.5  RT (min)       lcorbi_hilic_neg_4_089_59    m/z 714.4708     0   1   2   3   4E+3   3   3.5  RT (min)      lcorbi_hilic_neg_2_014_31    m/z 714.5451     0   1E+3   3   3.5  RT (min)      lcorbi_hilic_neg_4_084_59    m/z 714.8441     0   1   2E+4   5   5.5  RT (min)       lcorbi_hilic_neg_4_108_62    m/z 716.4482     0   1   2E+3   7   7.5  RT (min)      lcorbi_hilic_neg_3_011_24    m/z 716.6896     0   1   2   3E+4   4.5   5  RT (min)       lcorbi_hilic_neg_4_093_60    m/z 718.5036     0   1   2   3   4E+3   0.5   1  RT (min)      lcorbi_hilic_neg_4_082_59    m/z 718.5302     0   1   2   3   4E+4   3   3.5  RT (min)       lcorbi_hilic_neg_4_089_59    m/z 718.5305     0   1E+4   0.5   1  RT (min)      lcorbi_hilic_neg_4_082_59    m/z 718.6879     0   1E+4   4.5   5  RT (min)      lcorbi_hilic_neg_4_093_60    m/z 718.9449     0   1   2   3E+3   3.5   4  RT (min)      lcorbi_hilic_neg_1_010_22    m/z 719.5345     0   1E+5   3   3.5   4  RT (min)       lcorbi_hilic_neg_4_067_57    m/z 719.5351     0   1   2   3   4E+4   0.5   1  RT (min)       lcorbi_hilic_neg_4_070_57    m/z 720.4612     0   2   4   6E+3   3   3.5  RT (min)      lcorbi_hilic_neg_4_089_59    m/z 720.5385     0   1E+4   0.5   1  RT (min)      lcorbi_hilic_neg_4_067_57    m/z 721.2215     0   2   4   6   8E+3   1   1.5  RT (min)      lcorbi_hilic_neg_2_015_19    m/z 722.4772     0   1E+4   0.5   1  RT (min)      lcorbi_hilic_neg_4_085_59    m/z 722.5139     0   1   3   3.5  RT (min)      lcorbi_hilic_neg_4_082_59    m/z 722.5981     0   2   4   6E+3   3   3.5  RT (min)      lcorbi_hilic_neg_4_095_60    m/z 723.4803     0   1   2E+4   3   3.5  RT (min)      lcorbi_hilic_neg_4_084_59    m/z 723.4805     0   1   2   3   4E+3   0.5   1  RT (min)      lcorbi_hilic_neg_2_014_31    m/z 723.5184     0   1   3   3.5  RT (min)      lcorbi_hilic_neg_4_082_59    m/z 723.6015     0   1   2E+4   3   3.5  RT (min)       lcorbi_hilic_neg_4_093_60    m/z 723.6023     0   2   4   6   8E+3   0.5   1  RT (min)      lcorbi_hilic_neg_4_091_60    m/z 724.4924     0   1   2   3E+4   3   3.5  RT (min)       lcorbi_hilic_neg_4_089_59    m/z 724.4928     0   1E+4   0.5   1  RT (min)      lcorbi_hilic_neg_4_082_59    m/z 724.5288     0   1   0.5   1  RT (min)      lcorbi_hilic_neg_2_026_23    m/z 724.5293     0   1   3   3.5  RT (min)      lcorbi_hilic_neg_4_082_59    m/z 724.6134     0   1E+4   3   3.5  RT (min)      lcorbi_hilic_neg_4_093_60    m/z 725.4960     0   1E+4   3   3.5  RT (min)      lcorbi_hilic_neg_4_085_59    m/z 725.4962     0   1   0.5   1  RT (min)      lcorbi_hilic_neg_4_092_60    m/z 725.5327     0   1   2   3   4E+4   3   3.5  RT (min)       lcorbi_hilic_neg_4_096_60    m/z 725.6171     0   1E+4   0.5   1  RT (min)      lcorbi_hilic_neg_4_091_60    m/z 725.8761     0   2   4   6E+3   5   5.5  RT (min)      lcorbi_hilic_neg_4_112_62    m/z 726.5084     0   1   2E+4   0.5   1  RT (min)      lcorbi_hilic_neg_4_084_59    m/z 726.5445     0   1   3   3.5  RT (min)      lcorbi_hilic_neg_4_082_59    m/z 727.5117     0   1   2E+4   3   3.5  RT (min)       lcorbi_hilic_neg_4_089_59    m/z 727.5119     0   1E+4   0.5   1  RT (min)      lcorbi_hilic_neg_4_083_59    m/z 728.6417     0   1   2   3E+4   0.5   1  RT (min)      lcorbi_hilic_neg_4_084_59    m/z 729.4943     0   2   4   6E+3   4   4.5  RT (min)      lcorbi_hilic_neg_3_044_32    m/z 729.5276     0   1   2E+4   0.5   1  RT (min)      lcorbi_hilic_neg_4_084_59    m/z 729.6454     0   1   2   3E+4   0.5   1  RT (min)      lcorbi_hilic_neg_4_083_59    m/z 730.4671     0   1   2   3   4E+4   4   4.5  RT (min)       lcorbi_hilic_neg_4_084_59    m/z 730.5391     0   1   2E+4   3   3.5  RT (min)      lcorbi_hilic_neg_4_089_59    m/z 730.5393     0   1E+4   0.5   1  RT (min)      lcorbi_hilic_neg_4_084_59    m/z 731.4710     0   1E+4   4   4.5  RT (min)      lcorbi_hilic_neg_4_084_59    m/z 731.5428     0   1E+4   3   3.5  RT (min)      lcorbi_hilic_neg_4_088_59    m/z 732.4821     0   2   4   6E+4   4   4.5  RT (min)       lcorbi_hilic_neg_4_084_59    m/z 733.4862     0   1   2   3E+4   4   4.5  RT (min)      lcorbi_hilic_neg_4_084_59    m/z 733.5509     0   1E+4   0.5   1  RT (min)      lcorbi_hilic_neg_4_073_57    m/z 735.4807     0   1E+4   0.5   1  RT (min)      lcorbi_hilic_neg_4_085_59    m/z 735.5221     0   1   0.5   1  RT (min)      lcorbi_hilic_neg_4_082_59    m/z 736.5312     0   1   3   3.5  RT (min)      lcorbi_hilic_neg_4_085_59    m/z 736.6128     0   1   2E+4   0.5   1  RT (min)      lcorbi_hilic_neg_4_091_60    m/z 737.5376     0   1   0.5   1  RT (min)       lcorbi_hilic_neg_4_082_59    m/z 737.6175     0   1E+4   3   3.5  RT (min)      lcorbi_hilic_neg_4_093_60    m/z 737.6244     0   1E+4   0.5   1  RT (min)      lcorbi_hilic_neg_4_091_60    m/z 738.5411     0   1   0.5   1  RT (min)      lcorbi_hilic_neg_4_082_59    m/z 738.6209     0   2   4   6E+4   3   3.5  RT (min)      lcorbi_hilic_neg_4_095_60    m/z 738.6281     0   2   4E+4   0.5   1  RT (min)      lcorbi_hilic_neg_4_091_60    m/z 740.6361     0   1   2   3E+4   3   3.5  RT (min)      lcorbi_hilic_neg_4_095_60    m/z 740.6423     0   1E+4   0.5   1  RT (min)      lcorbi_hilic_neg_4_091_60    m/z 743.4875     0   2   4   6E+3   0.5   1  RT (min)      lcorbi_hilic_neg_3_042_05    m/z 744.4823     0   2   4   6E+4   4   4.5  RT (min)       lcorbi_hilic_neg_4_087_59    m/z 744.5483     0   1   2   3E+4   0.5   1  RT (min)      lcorbi_hilic_neg_4_087_59    m/z 745.4862     0   1   2E+4   4   4.5  RT (min)       lcorbi_hilic_neg_4_084_59    m/z 745.5560     0   1E+4   3   3.5  RT (min)       lcorbi_hilic_neg_4_067_57    m/z 746.4978     0   2   4   6   8E+4   4  RT (min)       lcorbi_hilic_neg_4_087_59    m/z 746.5064     0   1   2   3E+4   1   2  RT (min)       lcorbi_hilic_neg_4_084_59    m/z 746.5545     0   1   2   3E+4   0.5   1  RT (min)       lcorbi_hilic_neg_4_072_57    m/z 746.5602     0   1E+5   3   3.5   4  RT (min)       lcorbi_hilic_neg_4_067_57    m/z 747.5016     0   1   2E+4   4   4.5  RT (min)      lcorbi_hilic_neg_4_087_59    m/z 747.5660     0   1E+6   3   3.5   4  RT (min)       lcorbi_hilic_neg_4_067_57    m/z 747.5929     0   1   2   3E+2   3   3.5  RT (min)      lcorbi_hilic_neg_2_020_27    m/z 748.5140     0   1E+4   3   3.5  RT (min)      lcorbi_hilic_neg_4_084_59    m/z 748.5222     0   2   4   6E+4   0.5   1  RT (min)      lcorbi_hilic_neg_4_085_59    m/z 748.5293     0   1E+4   3   3.5  RT (min)      lcorbi_hilic_neg_4_072_57    m/z 748.5695     0   1   2   3   4E+4   3   3.5  RT (min)      lcorbi_hilic_neg_4_067_57    m/z 748.6047     0   1E+4   3   3.5  RT (min)      lcorbi_hilic_neg_3_011_24    m/z 749.5251     0   2   4   6   8E+3   0.5   1  RT (min)      lcorbi_hilic_neg_4_089_59    m/z 749.5327     0   2   4   6E+3   3   3.5  RT (min)      lcorbi_hilic_neg_4_067_57    m/z 749.5735     0   1   2E+4   0.5   1  RT (min)      lcorbi_hilic_neg_4_067_57    m/z 749.6084     0   1   0.5   1  RT (min)      lcorbi_hilic_neg_4_072_57    m/z 750.5081     0   1   0.5   1  RT (min)      lcorbi_hilic_neg_1_012_32    m/z 750.5297     0   1   2   3   4E+4   3   3.5  RT (min)       lcorbi_hilic_neg_4_070_57    m/z 750.5446     0   1   2E+4   3   3.5  RT (min)      lcorbi_hilic_neg_4_074_58    m/z 750.5447     0   2   4   6   8E+3   0.5   1  RT (min)      lcorbi_hilic_neg_1_012_32    m/z 750.6206     0   1E+4   0.5   1  RT (min)      lcorbi_hilic_neg_2_005_24    m/z 751.5330     0   1E+4   3   3.5  RT (min)      lcorbi_hilic_neg_4_096_60    m/z 751.5483     0   2   4   6E+3   3   3.5  RT (min)      lcorbi_hilic_neg_4_069_57    m/z 752.5236     0   2   4   6E+3   0.5   1  RT (min)      lcorbi_hilic_neg_4_084_59    m/z 752.5598     0   1   3   3.5  RT (min)      lcorbi_hilic_neg_4_090_60    m/z 754.4671     0   1E+4   4   4.5  RT (min)      lcorbi_hilic_neg_4_084_59    m/z 754.5395     0   1E+4   0.5   1  RT (min)      lcorbi_hilic_neg_4_087_59    m/z 754.5397     0   1   2E+4   2.5   3   3.5  RT (min)       lcorbi_hilic_neg_4_089_59    m/z 754.6508     0   1   2   3   4E+4   0.5   1  RT (min)       lcorbi_hilic_neg_4_092_60    m/z 756.5549     0   1E+4   0.5   1  RT (min)      lcorbi_hilic_neg_4_088_59    m/z 756.6599     0   1E+4   3   3.5  RT (min)      lcorbi_hilic_neg_4_095_60    m/z 757.6590     0   2   4E+3   3   3.5  RT (min)      lcorbi_hilic_neg_4_091_60    m/z 761.6078     0   2   4   6   8E+3   3   3.5  RT (min)      lcorbi_hilic_neg_2_055_32    m/z 762.5083     0   1   2   3E+3   0.5   1  RT (min)      lcorbi_hilic_neg_4_088_59    m/z 762.5191     0   2   4   6E+4   4   4.5  RT (min)       lcorbi_hilic_neg_4_087_59    m/z 762.5658     0   1E+3   3   3.5  RT (min)      lcorbi_hilic_neg_3_044_32    m/z 762.6112     0   1   2   3E+4   3   3.5  RT (min)      lcorbi_hilic_neg_2_055_32    m/z 762.6113     0   2   4   6   8E+3   0.5   1  RT (min)      lcorbi_hilic_neg_2_005_24    m/z 763.5117     0   1E+4   3   3.5  RT (min)      lcorbi_hilic_neg_4_067_57    m/z 763.5118     0   1   2   3E+3   0.5   1  RT (min)      lcorbi_hilic_neg_4_085_59    m/z 763.5231     0   1E+4   4   4.5  RT (min)      lcorbi_hilic_neg_4_087_59    m/z 763.6229     0   1   2   3E+3   3   3.5  RT (min)      lcorbi_hilic_neg_1_055_24    m/z 764.5007     0   2   4E+3   3   3.5  RT (min)      lcorbi_hilic_neg_4_070_57    m/z 764.5239     0   2   4E+4   0.5   1   1.5  RT (min)       lcorbi_hilic_neg_4_087_59    m/z 764.5816     0   1   3   3.5  RT (min)      lcorbi_hilic_neg_1_055_24    m/z 764.6269     0   1E+4   3   3.5  RT (min)      lcorbi_hilic_neg_1_055_24    m/z 764.6271     0   2   4   6   8E+3   0.5   1  RT (min)      lcorbi_hilic_neg_1_055_24    m/z 765.5272     0   2   4   6E+3   0.5   1  RT (min)      lcorbi_hilic_neg_4_089_59    m/z 765.5273     0   1E+4   3   3.5  RT (min)      lcorbi_hilic_neg_4_068_57    m/z 765.6390     0   1   2   3   4E+3   3   3.5  RT (min)      lcorbi_hilic_neg_1_055_24    m/z 766.5164     0   1E+4   3   3.5  RT (min)      lcorbi_hilic_neg_4_070_57    m/z 766.6426     0   1E+4   3   3.5  RT (min)      lcorbi_hilic_neg_1_055_24    m/z 766.6429     0   1E+4   0.5   1  RT (min)      lcorbi_hilic_neg_1_055_24    m/z 767.5200     0   2   4   6E+3   3   3.5  RT (min)      lcorbi_hilic_neg_4_070_57    m/z 767.5429     0   2   4   6   8E+3   0.5   1  RT (min)      lcorbi_hilic_neg_4_089_59    m/z 767.6547     0   2   4   6   8E+3   3   3.5  RT (min)      lcorbi_hilic_neg_4_092_60    m/z 767.6549     0   1E+4   0.5   1  RT (min)      lcorbi_hilic_neg_2_005_24    m/z 768.4827     0   1   2   3   4E+4   3.5   4   4.5  RT (min)       lcorbi_hilic_neg_4_088_59    m/z 768.5473     0   1E+4   3   3.5  RT (min)      lcorbi_hilic_neg_4_069_57    m/z 768.5546     0   1E+4   3   3.5  RT (min)      lcorbi_hilic_neg_4_070_57    m/z 768.5551     0   2   4   6E+4   0.5   1   1.5  RT (min)       lcorbi_hilic_neg_4_089_59    m/z 768.6582     0   1   2   3E+4   2.5   3   3.5  RT (min)       lcorbi_hilic_neg_4_015_24    m/z 768.6584     0   1   2E+4   0.5   1  RT (min)      lcorbi_hilic_neg_1_055_24    m/z 769.5573     0   1   2   3   4E+3   3   3.5  RT (min)      lcorbi_hilic_neg_4_069_57    m/z 769.5586     0   1E+4   0.5   1  RT (min)      lcorbi_hilic_neg_4_088_59    m/z 770.4980     0   1   2   3   4E+4   3.5   4   4.5  RT (min)       lcorbi_hilic_neg_4_087_59    m/z 770.5702     0   1   2   3   4E+3   3   3.5  RT (min)      lcorbi_hilic_neg_4_068_57    m/z 770.5703     0   1E+4   0.5   1  RT (min)      lcorbi_hilic_neg_4_089_59    m/z 770.6082     0   1   2   3E+4   4   4.5  RT (min)       lcorbi_hilic_neg_4_093_60    m/z 770.6738     0   2   4   6   8E+3   3   3.5  RT (min)      lcorbi_hilic_neg_4_015_24    m/z 771.5016     0   1E+4   4   4.5  RT (min)      lcorbi_hilic_neg_4_086_59    m/z 772.5139     0   1   2E+4   0.5   1  RT (min)      lcorbi_hilic_neg_3_028_23    m/z 772.5584     0   1   2   3   4E+4   1  RT (min)       lcorbi_hilic_neg_4_089_59    m/z 773.5172     0   1   2E+4   3   3.5  RT (min)       lcorbi_hilic_neg_4_089_59    m/z 773.5821     0   1   2E+4   0.5   1  RT (min)       lcorbi_hilic_neg_4_067_57    m/z 774.5296     0   1   2   3E+4   0.5   1  RT (min)      lcorbi_hilic_neg_3_028_23    m/z 774.5855     0   1E+4   0.5   1  RT (min)      lcorbi_hilic_neg_4_073_57    m/z 774.6121     0   1   2E+4   3   3.5  RT (min)      lcorbi_hilic_neg_2_055_32    m/z 774.6122     0   1   0.5   1  RT (min)      lcorbi_hilic_neg_4_068_57    m/z 775.5330     0   1E+4   0.5   1  RT (min)      lcorbi_hilic_neg_2_026_23    m/z 775.5970     0   1   0.5   1  RT (min)      lcorbi_hilic_neg_2_005_24    m/z 775.6168     0   1   2   3E+4   0.5   1  RT (min)      lcorbi_hilic_neg_2_005_24    m/z 777.5487     0   1   2   3E+4   0.5   1  RT (min)      lcorbi_hilic_neg_4_067_57    m/z 777.5878     0   1   2   3   4E+3   3   3.5  RT (min)      lcorbi_hilic_neg_2_005_24    m/z 778.5599     0   1   2   3E+2   1  RT (min)       lcorbi_hilic_neg_2_005_24    m/z 779.5639     0   1   3   3.5  RT (min)       lcorbi_hilic_neg_1_055_24    m/z 780.4810     0   2   4   6E+4   4   4.5  RT (min)       lcorbi_hilic_neg_4_082_59    m/z 780.5670     0   1   3   3.5  RT (min)      lcorbi_hilic_neg_4_086_59    m/z 781.4851     0   1   2   3E+4   4   4.5  RT (min)       lcorbi_hilic_neg_4_082_59    m/z 782.8310     0   1E+4   5   5.5  RT (min)      lcorbi_hilic_neg_4_108_62    m/z 783.5005     0   2   4E+4   4   4.5  RT (min)       lcorbi_hilic_neg_4_088_59    m/z 783.6122     0   1   2E+4   3.5   4   4.5  RT (min)       lcorbi_hilic_neg_4_096_60    m/z 784.5136     0   1E+5   3   3.5   4  RT (min)       lcorbi_hilic_neg_4_089_59    m/z 784.6336     0   2   4   6   8E+3   0.5   1  RT (min)      lcorbi_hilic_neg_4_096_60    m/z 784.6867     0   1   2   3E+4   3   3.5  RT (min)       lcorbi_hilic_neg_4_091_60    m/z 784.6870     0   1E+4   0.5   1  RT (min)      lcorbi_hilic_neg_1_055_24    m/z 785.5172     0   2   4E+4   4   4.5  RT (min)       lcorbi_hilic_neg_4_087_59    m/z 785.6374     0   2   4   6E+4   1   2  RT (min)       lcorbi_hilic_neg_4_090_60    m/z 785.6928     0   1   2   3   4E+4   0.5   1  RT (min)      lcorbi_hilic_neg_2_005_24    m/z 786.4856     0   2   4   6E+3   3   3.5  RT (min)      lcorbi_hilic_neg_4_086_59    m/z 786.5296     0   1E+4   0.5   1  RT (min)      lcorbi_hilic_neg_3_028_23    m/z 786.5658     0   1E+4   3   3.5  RT (min)      lcorbi_hilic_neg_3_044_32    m/z 786.5740     0   2   4   6   8E+4   0.5   1   1.5  RT (min)       lcorbi_hilic_neg_4_083_59    m/z 786.6492     0   1   2E+4   0.5   1  RT (min)      lcorbi_hilic_neg_4_090_60    m/z 787.5329     0   1E+4   3   3.5  RT (min)      lcorbi_hilic_neg_4_089_59    m/z 787.5693     0   2   4   6E+3   3   3.5  RT (min)      lcorbi_hilic_neg_2_055_32    m/z 787.5774     0   1   2   3E+4   0.5   1   1.5  RT (min)       lcorbi_hilic_neg_4_086_59    m/z 787.6529     0   2   4   6E+4   1   2  RT (min)       lcorbi_hilic_neg_4_015_24    m/z 788.5009     0   1   3   3.5  RT (min)      lcorbi_hilic_neg_4_070_57    m/z 788.5242     0   1   3   3.5  RT (min)      lcorbi_hilic_neg_4_082_59    m/z 788.5453     0   1E+4   0.5   1  RT (min)      lcorbi_hilic_neg_1_023_23    m/z 788.5824     0   2   4E+3   0.5   1  RT (min)      lcorbi_hilic_neg_4_088_59    m/z 789.5043     0   2   4   6   8E+3   3   3.5  RT (min)      lcorbi_hilic_neg_4_089_59    m/z 789.5486     0   2   4   6   8E+3   0.5   1  RT (min)      lcorbi_hilic_neg_1_023_23    m/z 789.5487     0   1E+4   3   3.5  RT (min)      lcorbi_hilic_neg_4_089_59    m/z 789.5850     0   1E+4   3   3.5  RT (min)      lcorbi_hilic_neg_4_069_57    m/z 789.5861     0   1   2E+4   0.5   1   1.5  RT (min)       lcorbi_hilic_neg_4_083_59    m/z 790.5165     0   1E+4   3   3.5  RT (min)       lcorbi_hilic_neg_4_089_59    m/z 790.5167     0   2   4   6E+3   0.5   1  RT (min)      lcorbi_hilic_neg_4_085_59    m/z 790.5397     0   1   2E+4   3   3.5  RT (min)      lcorbi_hilic_neg_4_072_57    m/z 790.5605     0   1   2E+4   3   3.5  RT (min)      lcorbi_hilic_neg_4_068_57    m/z 790.5607     0   2   4   6   8E+4   0.5   1   1.5  RT (min)       lcorbi_hilic_neg_4_086_59    m/z 790.5968     0   1   2E+4   3   3.5  RT (min)      lcorbi_hilic_neg_4_073_57    m/z 790.5968     0   2   4   6   8E+2   0.5   1  RT (min)      lcorbi_hilic_neg_2_026_23    m/z 791.5199     0   2   4   6   8E+3   3   3.5  RT (min)      lcorbi_hilic_neg_4_089_59    m/z 791.5431     0   1E+4   3   3.5  RT (min)      lcorbi_hilic_neg_4_068_57    m/z 791.5643     0   2   4   6E+3   3   3.5  RT (min)      lcorbi_hilic_neg_4_070_57    m/z 791.5644     0   1   2   3   4E+3   0.5   1  RT (min)      lcorbi_hilic_neg_4_086_59    m/z 791.6006     0   2   4   6   8E+3   3   3.5  RT (min)      lcorbi_hilic_neg_4_069_57    m/z 792.5323     0   1   2E+4   0.5   1  RT (min)       lcorbi_hilic_neg_4_068_57    m/z 792.5547     0   1   3   3.5  RT (min)      lcorbi_hilic_neg_4_086_59    m/z 792.6494     0   1E+4   0.5   1  RT (min)      lcorbi_hilic_neg_1_055_24    m/z 793.5357     0   1E+4   0.5   1  RT (min)      lcorbi_hilic_neg_4_066_57    m/z 794.4984     0   1   3   3.5  RT (min)      lcorbi_hilic_neg_4_070_57    m/z 794.5295     0   1E+4   0.5   1  RT (min)      lcorbi_hilic_neg_4_069_57    m/z 794.5477     0   1E+4   0.5   1  RT (min)      lcorbi_hilic_neg_4_071_57    m/z 794.5705     0   1   3   3.5  RT (min)      lcorbi_hilic_neg_4_086_59    m/z 795.5016     0   1   2E+3   3   3.5  RT (min)      lcorbi_hilic_neg_4_070_57    m/z 795.5329     0   1E+4   3   3.5  RT (min)       lcorbi_hilic_neg_4_070_57    m/z 795.5515     0   1E+4   3   3.5  RT (min)      lcorbi_hilic_neg_4_070_57    m/z 795.6121     0   1   2   3E+4   4   4.5  RT (min)       lcorbi_hilic_neg_4_090_60    m/z 796.5141     0   1   2E+4   0.5   1  RT (min)      lcorbi_hilic_neg_3_028_23    m/z 796.5452     0   1   3   3.5  RT (min)      lcorbi_hilic_neg_4_089_59    m/z 796.6159     0   1E+5   4   4.5  RT (min)       lcorbi_hilic_neg_4_090_60    m/z 796.6808     0   1E+4   0.5   1  RT (min)      lcorbi_hilic_neg_4_094_60    m/z 797.5175     0   1E+4   0.5   1  RT (min)      lcorbi_hilic_neg_3_028_23    m/z 797.5489     0   2   4   6E+2   3   3.5  RT (min)      lcorbi_hilic_neg_4_036_36    m/z 799.5332     0   2   4   6E+4   0.5   1  RT (min)       lcorbi_hilic_neg_3_028_23    m/z 799.6413     0   1E+5   4   4.5  RT (min)       lcorbi_hilic_neg_4_090_60    m/z 800.5900     0   1   2E+5   0.5   1   1.5  RT (min)       lcorbi_hilic_neg_4_086_59    m/z 801.5933     0   1E+4   0.5   1  RT (min)      lcorbi_hilic_neg_4_086_59    m/z 801.6136     0   1   0.5   1  RT (min)      lcorbi_hilic_neg_3_028_23    m/z 802.5162     0   1   3   3.5  RT (min)      lcorbi_hilic_neg_4_067_57    m/z 802.5249     0   1   3   3.5  RT (min)      lcorbi_hilic_neg_4_067_57    m/z 802.6049     0   1   3   3.5  RT (min)      lcorbi_hilic_neg_4_066_57    m/z 802.6174     0   2   4E+3   3   3.5  RT (min)      lcorbi_hilic_neg_1_004_17    m/z 802.6298     0   1E+4   3   3.5  RT (min)      lcorbi_hilic_neg_4_070_57    m/z 803.5194     0   2   4   6E+3   3   3.5  RT (min)      lcorbi_hilic_neg_1_004_17    m/z 803.6083     0   1   3   3.5  RT (min)      lcorbi_hilic_neg_4_070_57    m/z 803.6273     0   1   2E+4   3.5   4  RT (min)      lcorbi_hilic_neg_4_073_57    m/z 804.5287     0   2   4   6E+3   3   3.5  RT (min)      lcorbi_hilic_neg_3_053_13    m/z 805.5243     0   1E+4   3   3.5  RT (min)      lcorbi_hilic_neg_4_089_59    m/z 807.5857     0   1E+4   0.5   1  RT (min)      lcorbi_hilic_neg_4_067_57    m/z 807.6686     0   1   2E+4   0.5   1   1.5  RT (min)       lcorbi_hilic_neg_4_096_60    m/z 808.5139     0   1E+4   0.5   1  RT (min)      lcorbi_hilic_neg_3_028_23    m/z 808.5881     0   1   2   3   4E+3   3   3.5  RT (min)      lcorbi_hilic_neg_4_070_57    m/z 808.6805     0   2   4   6   8E+3   0.5   1  RT (min)      lcorbi_hilic_neg_4_090_60    m/z 809.5172     0   1E+4   3   3.5  RT (min)      lcorbi_hilic_neg_4_088_59    m/z 809.6843     0   1   2E+4   0.5   1   1.5  RT (min)       lcorbi_hilic_neg_4_094_60    m/z 810.5295     0   1   2E+4   0.5   1  RT (min)      lcorbi_hilic_neg_3_028_23    m/z 810.5661     0   1   2E+4   3   3.5  RT (min)      lcorbi_hilic_neg_1_012_32    m/z 810.6961     0   1E+4   0.5   1  RT (min)      lcorbi_hilic_neg_4_090_60    m/z 811.5327     0   1E+4   3   3.5  RT (min)      lcorbi_hilic_neg_4_089_59    m/z 811.5329     0   1E+4   0.5   1  RT (min)      lcorbi_hilic_neg_3_028_23    m/z 811.5697     0   1E+4   3   3.5  RT (min)      lcorbi_hilic_neg_1_012_32    m/z 811.6997     0   1   2   3E+4   0.5   1   1.5  RT (min)       lcorbi_hilic_neg_4_092_60    m/z 812.5007     0   1E+4   3   3.5  RT (min)      lcorbi_hilic_neg_4_088_59    m/z 812.5011     0   1   2   3E+3   0.5   1  RT (min)      lcorbi_hilic_neg_4_082_59    m/z 812.5451     0   2   4   6E+4   0.5   1  RT (min)      lcorbi_hilic_neg_1_023_23    m/z 812.5817     0   1   2   3E+4   0.5   1  RT (min)       lcorbi_hilic_neg_4_067_57    m/z 812.6484     0   1   0.5   1  RT (min)      lcorbi_hilic_neg_4_069_57    m/z 812.6484     0   1   2E+4   3   3.5  RT (min)       lcorbi_hilic_neg_4_092_60    m/z 813.5040     0   2   4   6   8E+3   3   3.5  RT (min)      lcorbi_hilic_neg_4_089_59    m/z 813.5487     0   1   2   3E+4   0.5   1  RT (min)      lcorbi_hilic_neg_1_023_23    m/z 813.5851     0   1   2E+4   0.5   1  RT (min)      lcorbi_hilic_neg_4_067_57    m/z 813.6502     0   1   2   3   4E+4   3.5   4   4.5  RT (min)       lcorbi_hilic_neg_4_090_60    m/z 813.6601     0   1   2   3   4E+4   3   3.5  RT (min)      lcorbi_hilic_neg_4_090_60    m/z 813.6960     0   1   2   3E+4   1  RT (min)       lcorbi_hilic_neg_4_096_60    m/z 813.7154     0   1   2E+4   0.5   1  RT (min)      lcorbi_hilic_neg_4_090_60    m/z 814.5164     0   1E+4   3   3.5  RT (min)      lcorbi_hilic_neg_4_089_59    m/z 814.5168     0   1E+4   0.5   1  RT (min)      lcorbi_hilic_neg_4_085_59    m/z 814.5604     0   1   2   3   4E+4   0.5   1  RT (min)      lcorbi_hilic_neg_3_028_23    m/z 814.5959     0   2   4   6E+2   0.5   1  RT (min)      lcorbi_hilic_neg_2_026_23    m/z 814.6639     0   1   2   3E+4   0.5   1  RT (min)      lcorbi_hilic_neg_2_005_24    m/z 815.5197     0   1E+4   3   3.5  RT (min)      lcorbi_hilic_neg_4_089_59    m/z 815.5642     0   1   2E+4   0.5   1  RT (min)      lcorbi_hilic_neg_1_023_23    m/z 815.6003     0   1   3   3.5  RT (min)      lcorbi_hilic_neg_4_089_59    m/z 815.6748     0   1   2   3   4E+4   3   3.5  RT (min)      lcorbi_hilic_neg_4_092_60    m/z 816.5129     0   1   3   3.5  RT (min)      lcorbi_hilic_neg_1_003_14    m/z 816.5325     0   1E+4   0.5   1  RT (min)      lcorbi_hilic_neg_4_067_57    m/z 816.5761     0   1   2E+4   0.5   1  RT (min)       lcorbi_hilic_neg_4_066_57    m/z 816.6123     0   1   3   3.5  RT (min)      lcorbi_hilic_neg_4_089_59    m/z 816.6793     0   1   2   3E+4   3   3.5  RT (min)      lcorbi_hilic_neg_3_011_24    m/z 817.5357     0   1   2   3   4E+3   0.5   1  RT (min)      lcorbi_hilic_neg_4_084_59    m/z 817.5798     0   2   4E+3   0.5   1  RT (min)      lcorbi_hilic_neg_4_086_59    m/z 817.6162     0   2   4   6   8E+3   3   3.5  RT (min)      lcorbi_hilic_neg_4_094_60    m/z 818.5297     0   1   2E+4   3   3.5  RT (min)      lcorbi_hilic_neg_4_078_58    m/z 818.5490     0   2   4   6   8E+3   0.5   1  RT (min)      lcorbi_hilic_neg_4_066_57    m/z 818.5908     0   1E+4   3   3.5  RT (min)      lcorbi_hilic_neg_4_069_57    m/z 819.5329     0   1E+4   3   3.5  RT (min)      lcorbi_hilic_neg_4_075_58    m/z 819.5511     0   1E+4   3   3.5  RT (min)      lcorbi_hilic_neg_4_073_57    m/z 819.5518     0   1   2   3E+3   0.5   1  RT (min)      lcorbi_hilic_neg_4_084_59    m/z 819.5953     0   2   4E+3   3   3.5  RT (min)      lcorbi_hilic_neg_4_077_58    m/z 820.5442     0   1   2   3E+3   3   3.5  RT (min)      lcorbi_hilic_neg_4_084_59    m/z 820.5635     0   1E+4   0.5   1  RT (min)      lcorbi_hilic_neg_4_066_57    m/z 821.5671     0   2   4   6   8E+3   0.5   1  RT (min)      lcorbi_hilic_neg_4_068_57    m/z 822.5616     0   1E+4   3   3.5  RT (min)      lcorbi_hilic_neg_4_070_57    m/z 823.5332     0   1E+5   0.5   1  RT (min)       lcorbi_hilic_neg_3_028_23    m/z 823.5647     0   2   4   6E+3   3   3.5  RT (min)      lcorbi_hilic_neg_4_073_57    m/z 824.6377     0   2   4   6E+4   4   4.5  RT (min)       lcorbi_hilic_neg_4_090_60    m/z 825.6516     0   1   2   3E+4   3   3.5   4  RT (min)       lcorbi_hilic_neg_4_090_60    m/z 826.5142     0   2   4   6E+3   3   3.5  RT (min)      lcorbi_hilic_neg_3_053_13    m/z 826.6069     0   1   2E+3   3   3.5  RT (min)      lcorbi_hilic_neg_3_053_13    m/z 826.6534     0   1E+5   4   4.5  RT (min)       lcorbi_hilic_neg_1_012_32    m/z 827.6672     0   1E+4   3   3.5   4  RT (min)       lcorbi_hilic_neg_2_055_32    m/z 828.6686     0   1   2E+5   4   4.5  RT (min)       lcorbi_hilic_neg_1_012_32    m/z 828.7153     0   1   2   3   4E+4   0.5   1   1.5  RT (min)       lcorbi_hilic_neg_4_096_60    m/z 829.6399     0   2   4   6   8E+2   3   3.5  RT (min)       lcorbi_hilic_neg_1_002_26    m/z 829.6441     0   2   4E+3   0.5   1  RT (min)      lcorbi_hilic_neg_4_037_31    m/z 829.6826     0   2   4   6E+3   3   3.5  RT (min)      lcorbi_hilic_neg_4_093_60    m/z 830.5440     0   2   4   6E+3   3   3.5  RT (min)      lcorbi_hilic_neg_4_004_13    m/z 830.6408     0   2   4E+3   3   3.5  RT (min)      lcorbi_hilic_neg_2_054_13    m/z 830.6441     0   1   2   3   4E+3   0.5   1  RT (min)      lcorbi_hilic_neg_3_016_16    m/z 831.6540     0   1E+4   3   3.5  RT (min)      lcorbi_hilic_neg_4_072_57    m/z 831.6983     0   2   4E+3   3   3.5  RT (min)      lcorbi_hilic_neg_4_015_24    m/z 832.6016     0   1E+5   0.5   1  RT (min)      lcorbi_hilic_neg_4_067_57    m/z 832.6574     0   1E+4   3   3.5  RT (min)      lcorbi_hilic_neg_1_055_24    m/z 833.5189     0   2   4   6E+4   4   4.5  RT (min)       lcorbi_hilic_neg_4_086_59    m/z 833.6095     0   1   2   3   4E+4   0.5   1  RT (min)       lcorbi_hilic_neg_4_067_57    m/z 834.6132     0   1E+4   3   3.5  RT (min)      lcorbi_hilic_neg_4_068_57    m/z 834.6729     0   2   4   6   8E+3   3   3.5  RT (min)      lcorbi_hilic_neg_1_055_24    m/z 835.5329     0   2   4E+4   4   4.5  RT (min)       lcorbi_hilic_neg_4_086_59    m/z 835.6157     0   1   2   3E+3   3   3.5  RT (min)      lcorbi_hilic_neg_4_070_57    m/z 836.5815     0   1E+4   3   3.5  RT (min)      lcorbi_hilic_neg_4_066_57    m/z 837.5850     0   2   4   6E+3   3   3.5  RT (min)      lcorbi_hilic_neg_4_066_57    m/z 837.6520     0   1E+4   3   3.5  RT (min)      lcorbi_hilic_neg_2_055_32    m/z 838.5595     0   1E+3   3   3.5  RT (min)      lcorbi_hilic_neg_1_005_36    m/z 838.5969     0   1   2E+4   0.5   1  RT (min)       lcorbi_hilic_neg_3_053_13    m/z 838.6559     0   1   2E+4   0.5   1  RT (min)      lcorbi_hilic_neg_1_055_24    m/z 839.6003     0   1E+4   0.5   1  RT (min)      lcorbi_hilic_neg_3_053_13    m/z 839.6670     0   1   2   3   4E+4   0.5   1  RT (min)      lcorbi_hilic_neg_1_055_24    m/z 840.5319     0   1E+4   3   3.5  RT (min)      lcorbi_hilic_neg_4_067_57    m/z 840.6117     0   1E+4   0.5   1  RT (min)      lcorbi_hilic_neg_4_066_57    m/z 840.6707     0   1E+5   0.5   1  RT (min)       lcorbi_hilic_neg_1_055_24    m/z 841.5352     0   1E+3   3   3.5  RT (min)      lcorbi_hilic_neg_4_084_59    m/z 841.5795     0   1E+4   3   3.5  RT (min)      lcorbi_hilic_neg_4_068_57    m/z 841.6159     0   1   2   3E+4   3   3.5  RT (min)       lcorbi_hilic_neg_4_094_60    m/z 841.6824     0   2   4   6   8E+4   0.5   1  RT (min)      lcorbi_hilic_neg_1_055_24    m/z 842.5471     0   1E+4   3   3.5  RT (min)      lcorbi_hilic_neg_4_068_57    m/z 842.5908     0   2   4   6E+3   3   3.5  RT (min)      lcorbi_hilic_neg_2_020_27    m/z 842.6228     0   2   4   6E+3   3   3.5  RT (min)      lcorbi_hilic_neg_4_094_60    m/z 842.7307     0   1   2   3E+3   0.5   1   1.5  RT (min)       lcorbi_hilic_neg_4_030_05    m/z 843.5510     0   2   4   6   8E+3   3   3.5  RT (min)      lcorbi_hilic_neg_4_068_57    m/z 843.5947     0   1   2   3E+3   3   3.5  RT (min)      lcorbi_hilic_neg_2_020_27    m/z 843.6975     0   2   4   6E+4   0.5   1  RT (min)      lcorbi_hilic_neg_1_055_24    m/z 843.7342     0   1   2E+4   0.5   1   1.5  RT (min)       lcorbi_hilic_neg_4_090_60    m/z 844.6067     0   1   2E+3   3   3.5  RT (min)      lcorbi_hilic_neg_1_001_49    m/z 844.7016     0   1   2E+5   0.5   1  RT (min)       lcorbi_hilic_neg_1_055_24    m/z 845.5666     0   1   2E+4   3   3.5  RT (min)      lcorbi_hilic_neg_4_066_57    m/z 845.6107     0   1E+3   3   3.5  RT (min)      lcorbi_hilic_neg_1_011_01    m/z 845.7093     0   1E+4   3   3.5  RT (min)      lcorbi_hilic_neg_4_015_24    m/z 846.5610     0   1E+4   3   3.5  RT (min)      lcorbi_hilic_neg_4_068_57    m/z 846.5784     0   2   4   6E+3   3   3.5  RT (min)      lcorbi_hilic_neg_4_077_58    m/z 846.5864     0   2   4   6   8E+3   3   3.5  RT (min)      lcorbi_hilic_neg_4_078_58    m/z 846.7167     0   1   2   3E+4   3   3.5  RT (min)       lcorbi_hilic_neg_4_015_24    m/z 846.7169     0   1   2   3E+4   0.5   1  RT (min)      lcorbi_hilic_neg_2_005_24    m/z 847.5640     0   2   4   6E+3   3   3.5  RT (min)      lcorbi_hilic_neg_4_070_57    m/z 847.5825     0   2   4   6E+3   3   3.5  RT (min)      lcorbi_hilic_neg_4_078_58    m/z 848.5448     0   1   2   3   4E+3   3   3.5  RT (min)      lcorbi_hilic_neg_2_018_16    m/z 849.5482     0   1   2E+3   3   3.5  RT (min)      lcorbi_hilic_neg_1_004_17    m/z 850.5608     0   1E+5   0.5   1  RT (min)       lcorbi_hilic_neg_3_053_13    m/z 851.5642     0   2   4   6E+4   0.5   1  RT (min)       lcorbi_hilic_neg_3_053_13    m/z 851.6584     0   1   2E+4   3   3.5  RT (min)      lcorbi_hilic_neg_2_055_32    m/z 852.5758     0   1E+5   0.5   1  RT (min)       lcorbi_hilic_neg_2_054_13    m/z 852.6700     0   2   4   6E+3   3   3.5  RT (min)      lcorbi_hilic_neg_1_012_32    m/z 853.5795     0   2   4   6E+4   0.5   1  RT (min)       lcorbi_hilic_neg_3_053_13    m/z 853.6739     0   1   2E+4   3   3.5  RT (min)      lcorbi_hilic_neg_2_055_32    m/z 854.6858     0   1E+4   3   3.5  RT (min)      lcorbi_hilic_neg_3_011_24    m/z 855.5954     0   1E+5   0.5   1  RT (min)      lcorbi_hilic_neg_3_053_13    m/z 855.6448     0   1   3   3.5  RT (min)      lcorbi_hilic_neg_1_003_14    m/z 855.6599     0   1   0.5   1  RT (min)      lcorbi_hilic_neg_1_055_24    m/z 855.6894     0   1   2   3   4E+4   3   3.5  RT (min)      lcorbi_hilic_neg_3_011_24    m/z 855.6895     0   1   2   3E+4   0.5   1  RT (min)      lcorbi_hilic_neg_1_055_24    m/z 856.6051     0   2   4   6   8E+4   0.5   1  RT (min)      lcorbi_hilic_neg_4_066_57    m/z 856.6486     0   1   3   3.5  RT (min)      lcorbi_hilic_neg_1_003_14    m/z 856.6632     0   2   4   6E+3   0.5   1  RT (min)      lcorbi_hilic_neg_4_069_57    m/z 856.7011     0   1E+4   3   3.5  RT (min)      lcorbi_hilic_neg_3_011_24    m/z 857.6102     0   1   2   3   4E+4   0.5   1  RT (min)       lcorbi_hilic_neg_2_054_13    m/z 857.6606     0   1   2E+3   3   3.5  RT (min)      lcorbi_hilic_neg_4_095_60    m/z 857.6750     0   1   0.5   1  RT (min)      lcorbi_hilic_neg_1_055_24    m/z 857.7049     0   1   2   3E+4   3   3.5  RT (min)      lcorbi_hilic_neg_4_015_24    m/z 857.7050     0   1   2E+4   0.5   1  RT (min)      lcorbi_hilic_neg_1_055_24    m/z 858.6641     0   1E+4   3   3.5  RT (min)      lcorbi_hilic_neg_1_055_24    m/z 859.6228     0   2   4   6   8E+3   3   3.5  RT (min)      lcorbi_hilic_neg_4_073_57    m/z 859.6759     0   2   4   6E+3   3   3.5  RT (min)      lcorbi_hilic_neg_1_055_24    m/z 859.7204     0   1E+4   3   3.5  RT (min)      lcorbi_hilic_neg_4_015_24    m/z 860.6605     0   1   3   3.5  RT (min)      lcorbi_hilic_neg_4_094_60    m/z 860.6797     0   1   2E+4   3   3.5  RT (min)      lcorbi_hilic_neg_1_055_24    m/z 860.8471     0   2   4   6   8E+3   5   5.5  RT (min)      lcorbi_hilic_neg_4_110_62    m/z 862.6954     0   1E+4   3   3.5  RT (min)      lcorbi_hilic_neg_1_055_24    m/z 864.5474     0   2   4   6   8E+3   1   1.5  RT (min)      lcorbi_hilic_neg_1_003_14    m/z 864.5761     0   1   2E+3   3   3.5  RT (min)      lcorbi_hilic_neg_3_016_16    m/z 864.6128     0   2   4   6E+3   3   3.5  RT (min)      lcorbi_hilic_neg_4_068_57    m/z 864.6704     0   1   2E+4   3   3.5  RT (min)      lcorbi_hilic_neg_2_055_32    m/z 865.6741     0   2   4   6   8E+4   0.5   1  RT (min)      lcorbi_hilic_neg_1_055_24    m/z 866.6279     0   2   4   6   8E+3   3   3.5  RT (min)      lcorbi_hilic_neg_4_068_57    m/z 866.6777     0   1   2   3E+5   0.5   1  RT (min)       lcorbi_hilic_neg_1_055_24    m/z 867.6893     0   1E+5   0.5   1  RT (min)      lcorbi_hilic_neg_1_055_24    m/z 868.5629     0   2   4   6E+3   3   3.5  RT (min)      lcorbi_hilic_neg_4_071_57    m/z 869.7043     0   1E+5   0.5   1  RT (min)      lcorbi_hilic_neg_1_055_24    m/z 870.5481     0   1E+4   3   3.5  RT (min)      lcorbi_hilic_neg_4_066_57    m/z 870.5865     0   1E+4   3   3.5  RT (min)      lcorbi_hilic_neg_4_075_58    m/z 870.7084     0   2   4E+5   0.5   1  RT (min)       lcorbi_hilic_neg_1_055_24    m/z 871.5515     0   2   4   6E+3   3   3.5  RT (min)      lcorbi_hilic_neg_4_066_57    m/z 871.5842     0   2   4E+3   3   3.5  RT (min)      lcorbi_hilic_neg_4_067_57    m/z 871.7196     0   2   4   6   8E+4   0.5   1  RT (min)      lcorbi_hilic_neg_1_055_24    m/z 872.5625     0   2   4E+3   3   3.5  RT (min)      lcorbi_hilic_neg_4_070_57    m/z 872.7240     0   1   2E+5   0.5   1  RT (min)       lcorbi_hilic_neg_1_055_24    m/z 873.7337     0   1   2   3   4E+4   3   3.5  RT (min)      lcorbi_hilic_neg_4_015_24    m/z 874.7395     0   2   4   6E+4   0.5   1  RT (min)      lcorbi_hilic_neg_1_055_24    m/z 876.5764     0   1   2E+4   3   3.5  RT (min)       lcorbi_hilic_neg_4_068_57    m/z 876.7547     0   2   4   6   8E+3   3   3.5  RT (min)      lcorbi_hilic_neg_4_015_24    m/z 877.5799     0   1E+4   3   3.5  RT (min)      lcorbi_hilic_neg_4_066_57    m/z 878.5919     0   1   2   3E+4   0.5   1  RT (min)      lcorbi_hilic_neg_3_053_13    m/z 881.6108     0   1   2E+4   3   3.5  RT (min)       lcorbi_hilic_neg_4_066_57    m/z 882.6218     0   1   2E+4   3   3.5  RT (min)      lcorbi_hilic_neg_4_066_57    m/z 883.6263     0   1E+4   3   3.5  RT (min)      lcorbi_hilic_neg_4_066_57    m/z 886.5222     0   1   2   3   4E+3   3   3.5  RT (min)      lcorbi_hilic_neg_4_070_57    m/z 894.5480     0   2   4E+3   3   3.5  RT (min)      lcorbi_hilic_neg_1_003_14    m/z 895.5516     0   1   2   3E+3   3   3.5  RT (min)      lcorbi_hilic_neg_1_004_17    m/z 896.5627     0   1   2   3E+3   3   3.5  RT (min)      lcorbi_hilic_neg_1_003_14    m/z 898.5795     0   2   4E+3   3   3.5  RT (min)      lcorbi_hilic_neg_1_003_14    m/z 898.7312     0   2   4E+3   3   3.5  RT (min)      lcorbi_hilic_neg_3_044_32    m/z 900.7465     0   1E+4   3   3.5  RT (min)      lcorbi_hilic_neg_3_044_32    m/z 902.7622     0   2   4   6E+3   3   3.5  RT (min)      lcorbi_hilic_neg_1_012_32    m/z 925.7469     0   1   2E+4   2   2.5  RT (min)      lcorbi_hilic_neg_2_055_32    m/z 936.8596     0   2   4E+3   2   2.5  RT (min)      lcorbi_hilic_neg_2_055_32    m/z 978.1975     0   1   2   3E+3   1   1.5  RT (min)      lcorbi_hilic_neg_1_046_19    m/z 1017.9630     0   1   2E+3   5   5.5  RT (min)      lcorbi_hilic_neg_4_050_26    m/z 1023.7532     0   1   2E+3   5   5.5  RT (min)      lcorbi_hilic_neg_3_019_01    m/z 1036.2656     0   1   2E+3   4.5   5  RT (min)      lcorbi_hilic_neg_4_055_38    m/z 1090.2994     0   2   4   6E+3   7   7.5  RT (min)      lcorbi_hilic_neg_4_095_60    m/z 1109.4914     0   2   4E+3   4.5   5  RT (min)      lcorbi_hilic_neg_3_044_32    m/z 1114.1552     0   2   4E+3   1.5   2  RT (min)      lcorbi_hilic_neg_1_038_15    m/z 1116.2397     0   2   4   6   8E+3   7.5   8  RT (min)      lcorbi_hilic_neg_3_023_36    m/z 1116.3389     0   1E+4   7   7.5  RT (min)      lcorbi_hilic_neg_3_011_24    m/z 1177.7533     0   2   4E+3   4   4.5  RT (min)      lcorbi_hilic_neg_3_044_32    m/z 1179.9842     0   2   4   6E+3   6.5   7  RT (min)      lcorbi_hilic_neg_3_023_36   
