## Supplementary material for "The Everything Bagel Feature Finder: Ultra-fast automated feature finding for untargeted metabolomics": SI Data File: tool_unique_xic_st_XCMS_ST004503.html

XCMS-unique features (not detected by the other two tools) - ST004503

### XCMS-unique features (not detected by the other two tools) - ST004503

429 features · qtof · XIC window m/z ± 25 ppm (≥ 0.002 Da) · ± 0.5 min

Every panel is a feature **XCMS** detected that neither of the other two tools detected — three-way unique across EB (standard mode (min\_detection\_rate=0.1)), Masscube and XCMS, under the standard matching rule (m/z ± 25 ppm; apex within the reference RT window ± 0.2 min). It shows the MS1 extracted-ion chromatogram at the feature's m/z, drawn from the raw mzML named in `alignment_reference_file`, zoomed to ± 0.5 min around the feature's reported apex RT. **Gray solid line** = the feature's reported apex RT; **red dashed line** = XCMS's assigned MS/MS — `RTINSECONDS` in its `aligned_features.mgf`; the panel widens to keep an off-apex MS/MS on-screen. Features with no assigned MS/MS show only the gray line. y is per-cell autoscaled. Rows keep the table order (m/z ascending).

m/z 80.0506

01E+417.518RT (min)

lcqtof\_t3\_pos\_3\_069\_57

m/z 81.0706

01E+41414.5RT (min)

lcqtof\_t3\_pos\_2\_024\_45

m/z 81.0708

01E+47.58RT (min)

lcqtof\_t3\_pos\_4\_021\_01

m/z 82.0667

0123E+317.518RT (min)

lcqtof\_t3\_pos\_3\_069\_57

m/z 82.0675

0246E+31616.5RT (min)

lcqtof\_t3\_pos\_2\_015\_13

m/z 83.0486

01234E+388.5RT (min)

lcqtof\_t3\_pos\_4\_101\_61

m/z 83.0611

012E+41212.5RT (min)

lcqtof\_t3\_pos\_1\_003\_33

m/z 83.0613

0246E+417.518RT (min)

lcqtof\_t3\_pos\_3\_069\_57

m/z 84.0642

012E+316.517RT (min)

lcqtof\_t3\_pos\_3\_056\_56

m/z 84.0823

024E+316.517RT (min)

lcqtof\_t3\_pos\_2\_008\_31

m/z 85.0772

0123E+316.517RT (min)

lcqtof\_t3\_pos\_2\_006\_04

m/z 88.0768

024E+31212.5RT (min)

lcqtof\_t3\_pos\_1\_011\_43

m/z 94.0662

012E+317.518RT (min)

lcqtof\_t3\_pos\_1\_009\_50

m/z 95.0609

0123E+31616.5RT (min)

lcqtof\_t3\_pos\_2\_038\_21

m/z 95.0864

01E+41010.5RT (min)

lcqtof\_t3\_pos\_4\_111\_62

m/z 96.0812

01234E+41212.5RT (min)

lcqtof\_t3\_pos\_1\_001\_49

m/z 97.0768

0123E+544.5RT (min)

lcqtof\_t3\_pos\_1\_011\_43

m/z 98.0798

012E+41414.5RT (min)

lcqtof\_t3\_pos\_2\_008\_31

m/z 98.0798

01234E+417.518RT (min)

lcqtof\_t3\_pos\_3\_069\_57

m/z 98.0799

01E+41212.5RT (min)

lcqtof\_t3\_pos\_2\_002\_37

m/z 98.9853

01E+41313.5RT (min)

lcqtof\_t3\_pos\_2\_028\_34

m/z 99.0833

0246E+316.517RT (min)

lcqtof\_t3\_pos\_1\_005\_46

m/z 102.0923

0123E+317.518RT (min)

lcqtof\_t3\_pos\_1\_053\_18

m/z 109.0769

01E+417.518RT (min)

lcqtof\_t3\_pos\_3\_066\_57

m/z 112.0773

01E+41717.5RT (min)

lcqtof\_t3\_pos\_2\_004\_42

m/z 113.1079

01E+31212.5RT (min)

lcqtof\_t3\_pos\_4\_076\_58

m/z 113.1083

0123E+414.515RT (min)

lcqtof\_t3\_pos\_2\_004\_42

m/z 114.1365

01234E+31616.5RT (min)

lcqtof\_t3\_pos\_2\_011\_19

m/z 116.1080

012E+417.518RT (min)

lcqtof\_t3\_pos\_2\_003\_16

m/z 117.0703

0246E+313.514RT (min)

lcqtof\_t3\_pos\_2\_020\_27

m/z 117.1115

01234E+31212.513RT (min)

lcqtof\_t3\_pos\_1\_015\_04

m/z 119.0896

0123E+44.55RT (min)

lcqtof\_t3\_pos\_4\_101\_61

m/z 124.1133

012E+217.518RT (min)

lcqtof\_t3\_pos\_4\_017\_51

m/z 125.1070

01234E+315.516RT (min)

lcqtof\_t3\_pos\_2\_012\_35

m/z 125.1083

024E+317.518RT (min)

lcqtof\_t3\_pos\_2\_007\_07

m/z 125.9871

02468E+318.519RT (min)

lcqtof\_t3\_pos\_2\_006\_04

m/z 126.0923

01234E+315.516RT (min)

lcqtof\_t3\_pos\_2\_015\_13

m/z 127.0868

011616.5RT (min)

lcqtof\_t3\_pos\_4\_110\_62

m/z 128.1077

0123E+41212.5RT (min)

lcqtof\_t3\_pos\_1\_020\_35

m/z 130.0588

0123E+312.513RT (min)

lcqtof\_t3\_pos\_4\_099\_61

m/z 133.0652

02468E+355.5RT (min)

lcqtof\_t3\_pos\_4\_029\_37

m/z 133.1016

01234E+313.514RT (min)

lcqtof\_t3\_pos\_4\_112\_62

m/z 138.0281

01E+41717.5RT (min)

lcqtof\_t3\_pos\_4\_101\_61

m/z 138.0992

01E+413.514RT (min)

lcqtof\_t3\_pos\_4\_014\_24

m/z 138.1284

01E+317.518RT (min)

lcqtof\_t3\_pos\_4\_098\_61

m/z 139.0736

01E+444.5RT (min)

lcqtof\_t3\_pos\_4\_085\_59

m/z 139.0873

01E+217.518RT (min)

lcqtof\_t3\_pos\_4\_109\_62

m/z 140.0693

0123E+318.519RT (min)

lcqtof\_t3\_pos\_4\_106\_62

m/z 140.0821

012E+211.5RT (min)

lcqtof\_t3\_pos\_2\_015\_13

m/z 141.1030

0123E+317.518RT (min)

lcqtof\_t3\_pos\_4\_080\_58

m/z 141.9596

01234E+42.53RT (min)

lcqtof\_t3\_pos\_4\_099\_61

m/z 142.1233

0246E+49.510RT (min)

lcqtof\_t3\_pos\_1\_014\_14

m/z 142.1233

0246E+412.513RT (min)

lcqtof\_t3\_pos\_1\_002\_36

m/z 142.1235

01234E+444.5RT (min)

lcqtof\_t3\_pos\_1\_001\_49

m/z 145.0179

02468E+312.513RT (min)

lcqtof\_t3\_pos\_1\_032\_13

m/z 145.0180

02468E+31010.5RT (min)

lcqtof\_t3\_pos\_3\_070\_57

m/z 145.0180

01E+456RT (min)

lcqtof\_t3\_pos\_2\_032\_47

m/z 145.0185

01E+46.57RT (min)

lcqtof\_t3\_pos\_1\_010\_27

m/z 151.0730

0246E+414.515RT (min)

lcqtof\_t3\_pos\_4\_096\_60

m/z 151.0730

024E+417RT (min)

lcqtof\_t3\_pos\_4\_078\_58

m/z 152.0571

012E+311.5RT (min)

lcqtof\_t3\_pos\_4\_092\_60

m/z 152.0740

0246E+31515.5RT (min)

lcqtof\_t3\_pos\_4\_086\_59

m/z 153.0992

024E+313.514RT (min)

lcqtof\_t3\_pos\_2\_041\_54

m/z 155.0538

01E+41010.5RT (min)

lcqtof\_t3\_pos\_4\_085\_59

m/z 155.0541

01E+41818.5RT (min)

lcqtof\_t3\_pos\_4\_096\_60

m/z 155.1183

0246E+21616.5RT (min)

lcqtof\_t3\_pos\_4\_106\_62

m/z 156.1390

01E+41212.5RT (min)

lcqtof\_t3\_pos\_1\_008\_31

m/z 156.1391

012E+417.518RT (min)

lcqtof\_t3\_pos\_1\_007\_07

m/z 163.0389

01234E+31717.5RT (min)

lcqtof\_t3\_pos\_2\_016\_39

m/z 164.0488

0246E+36.57RT (min)

lcqtof\_t3\_pos\_4\_029\_37

m/z 164.0777

0246E+244.5RT (min)

lcqtof\_t3\_pos\_4\_040\_28

m/z 166.0483

0123E+444.5RT (min)

lcqtof\_t3\_pos\_4\_029\_37

m/z 167.1183

024E+317.518RT (min)

lcqtof\_t3\_pos\_3\_068\_57

m/z 169.1347

01234E+317.518RT (min)

lcqtof\_t3\_pos\_3\_069\_57

m/z 171.1162

01234E+113.514RT (min)

lcqtof\_t3\_pos\_4\_106\_62

m/z 174.0865

012E+211.5RT (min)

lcqtof\_t3\_pos\_4\_021\_01

m/z 181.0970

0123E+315.516RT (min)

lcqtof\_t3\_pos\_2\_003\_16

m/z 183.0734

0123E+41617RT (min)

lcqtof\_t3\_pos\_4\_108\_62

m/z 183.1162

0246E+113.514RT (min)

lcqtof\_t3\_pos\_4\_098\_61

m/z 186.0446

0246E+34.55RT (min)

lcqtof\_t3\_pos\_2\_005\_17

m/z 187.0401

0246E+377.5RT (min)

lcqtof\_t3\_pos\_4\_042\_18

m/z 189.0979

02468E+317.518RT (min)

lcqtof\_t3\_pos\_2\_045\_10

m/z 189.1641

01234E+213.514RT (min)

lcqtof\_t3\_pos\_4\_047\_13

m/z 191.0714

01234E+31717.5RT (min)

lcqtof\_t3\_pos\_2\_009\_50

m/z 191.0717

0246E+314.515RT (min)

lcqtof\_t3\_pos\_2\_009\_50

m/z 191.0878

012E+41414.5RT (min)

lcqtof\_t3\_pos\_4\_014\_24

m/z 192.1109

024E+312.513RT (min)

lcqtof\_t3\_pos\_4\_024\_20

m/z 195.1748

024E+312.513RT (min)

lcqtof\_t3\_pos\_4\_042\_18

m/z 197.1325

02468E+313.514RT (min)

lcqtof\_t3\_pos\_2\_011\_19

m/z 201.0548

0123E+37.58RT (min)

lcqtof\_t3\_pos\_4\_072\_57

m/z 205.0754

02468E+34.55RT (min)

lcqtof\_t3\_pos\_2\_034\_24

m/z 205.0758

01E+41212.5RT (min)

lcqtof\_t3\_pos\_2\_005\_17

m/z 209.0820

0246E+31717.5RT (min)

lcqtof\_t3\_pos\_2\_004\_42

m/z 213.1106

01E+466.5RT (min)

lcqtof\_t3\_pos\_2\_008\_31

m/z 213.1107

02468E+31212.5RT (min)

lcqtof\_t3\_pos\_3\_017\_51

m/z 213.1107

01E+41717.5RT (min)

lcqtof\_t3\_pos\_2\_023\_05

m/z 213.1107

01E+414.515RT (min)

lcqtof\_t3\_pos\_2\_014\_03

m/z 213.1615

01E+47.58RT (min)

lcqtof\_t3\_pos\_4\_014\_24

m/z 215.1426

0246E+313.514RT (min)

lcqtof\_t3\_pos\_2\_011\_19

m/z 216.9238

02468E+31717.5RT (min)

lcqtof\_t3\_pos\_2\_011\_19

m/z 220.9353

0246E+36.57RT (min)

lcqtof\_t3\_pos\_2\_011\_19

m/z 220.9354

0246E+399.5RT (min)

lcqtof\_t3\_pos\_2\_030\_33

m/z 222.1132

012E+31818.5RT (min)

lcqtof\_t3\_pos\_4\_087\_59

m/z 223.1295

0246E+315.516RT (min)

lcqtof\_t3\_pos\_4\_073\_57

m/z 226.1225

0123E+34.55RT (min)

lcqtof\_t3\_pos\_4\_006\_41

m/z 227.1434

0113.514RT (min)

lcqtof\_t3\_pos\_4\_113\_62

m/z 228.1612

01E+455.5RT (min)

lcqtof\_t3\_pos\_4\_029\_37

m/z 231.0943

0123E+31616.5RT (min)

lcqtof\_t3\_pos\_2\_001\_49

m/z 232.0653

0246E+311.52RT (min)

lcqtof\_t3\_pos\_4\_104\_61

m/z 240.1121

02468E+316.517RT (min)

lcqtof\_t3\_pos\_4\_112\_62

m/z 242.2761

0246E+311.512RT (min)

lcqtof\_t3\_pos\_4\_021\_01

m/z 244.0667

02468E+36.57RT (min)

lcqtof\_t3\_pos\_4\_029\_37

m/z 251.1095

01234E+37.58RT (min)

lcqtof\_t3\_pos\_4\_049\_55

m/z 251.1291

024E+314.515RT (min)

lcqtof\_t3\_pos\_2\_028\_34

m/z 251.1291

01234E+31717.5RT (min)

lcqtof\_t3\_pos\_2\_011\_19

m/z 252.0704

0246E+31717.5RT (min)

lcqtof\_t3\_pos\_4\_093\_60

m/z 252.0711

01E+444.5RT (min)

lcqtof\_t3\_pos\_4\_074\_58

m/z 252.1648

02468E+377.5RT (min)

lcqtof\_t3\_pos\_4\_021\_01

m/z 254.1099

02468E+316.517RT (min)

lcqtof\_t3\_pos\_4\_109\_62

m/z 256.1087

0246E+316.517RT (min)

lcqtof\_t3\_pos\_4\_095\_60

m/z 259.1913

0123E+31717.5RT (min)

lcqtof\_t3\_pos\_1\_010\_27

m/z 261.2415

0123E+210.511RT (min)

lcqtof\_t3\_pos\_4\_003\_05

m/z 264.2694

01E+41717.5RT (min)

lcqtof\_t3\_pos\_1\_006\_29

m/z 266.1238

012E+422.5RT (min)

lcqtof\_t3\_pos\_4\_001\_49

m/z 266.2854

01E+31717.5RT (min)

lcqtof\_t3\_pos\_4\_007\_14

m/z 268.2164

01234E+313.514RT (min)

lcqtof\_t3\_pos\_4\_020\_19

m/z 268.3000

0246E+31010.5RT (min)

lcqtof\_t3\_pos\_4\_029\_37

m/z 269.1221

012E+41717.5RT (min)

lcqtof\_t3\_pos\_4\_081\_58

m/z 271.2413

01234E+214.515RT (min)

lcqtof\_t3\_pos\_2\_004\_42

m/z 272.1407

024E+31717.5RT (min)

lcqtof\_t3\_pos\_4\_104\_61

m/z 272.2326

0246E+388.5RT (min)

lcqtof\_t3\_pos\_4\_021\_01

m/z 272.2960

0246E+399.5RT (min)

lcqtof\_t3\_pos\_4\_108\_62

m/z 278.2485

0246E+31717.5RT (min)

lcqtof\_t3\_pos\_1\_001\_49

m/z 280.2635

01E+41717.5RT (min)

lcqtof\_t3\_pos\_1\_004\_05

m/z 281.2301

0246E+313.514RT (min)

lcqtof\_t3\_pos\_4\_106\_62

m/z 282.1459

012E+34.55RT (min)

lcqtof\_t3\_pos\_4\_001\_49

m/z 282.2801

0246E+31717.5RT (min)

lcqtof\_t3\_pos\_2\_004\_42

m/z 283.2667

02468E+31717.5RT (min)

lcqtof\_t3\_pos\_1\_001\_49

m/z 284.1356

02468E+316.517RT (min)

lcqtof\_t3\_pos\_4\_105\_61

m/z 284.2936

01E+416.517RT (min)

lcqtof\_t3\_pos\_4\_077\_58

m/z 285.0117

012E+513.514RT (min)

lcqtof\_t3\_pos\_4\_024\_20

m/z 285.1547

0123E+300.5RT (min)

lcqtof\_t3\_pos\_4\_106\_62

m/z 285.2562

01E+314.515RT (min)

lcqtof\_t3\_pos\_4\_021\_01

m/z 285.3337

02468E+31717.5RT (min)

lcqtof\_t3\_pos\_4\_113\_62

m/z 289.1574

01E+31010.5RT (min)

lcqtof\_t3\_pos\_4\_042\_18

m/z 294.3148

0246E+31212.5RT (min)

lcqtof\_t3\_pos\_4\_021\_01

m/z 296.2956

01E+41717.5RT (min)

lcqtof\_t3\_pos\_4\_088\_59

m/z 297.2929

02468E+316.517RT (min)

lcqtof\_t3\_pos\_4\_111\_62

m/z 298.1830

024E+388.5RT (min)

lcqtof\_t3\_pos\_2\_006\_04

m/z 300.1713

0246E+31717.5RT (min)

lcqtof\_t3\_pos\_4\_093\_60

m/z 310.3113

0246E+31717.5RT (min)

lcqtof\_t3\_pos\_4\_072\_57

m/z 321.3164

0246E+31717.5RT (min)

lcqtof\_t3\_pos\_4\_042\_18

m/z 324.0472

024E+35.56RT (min)

lcqtof\_t3\_pos\_4\_005\_30

m/z 325.3215

01E+41717.5RT (min)

lcqtof\_t3\_pos\_4\_107\_62

m/z 338.3904

0112.513RT (min)

lcqtof\_t3\_pos\_4\_049\_55

m/z 339.1391

0246E+316.517RT (min)

lcqtof\_t3\_pos\_4\_096\_60

m/z 340.1586

01E+41717.5RT (min)

lcqtof\_t3\_pos\_4\_094\_60

m/z 340.3526

02468E+31717.5RT (min)

lcqtof\_t3\_pos\_4\_079\_58

m/z 341.1751

01E+41717.5RT (min)

lcqtof\_t3\_pos\_4\_112\_62

m/z 345.0970

0246E+37.58RT (min)

lcqtof\_t3\_pos\_4\_042\_18

m/z 345.2178

012E+31212.5RT (min)

lcqtof\_t3\_pos\_2\_002\_37

m/z 348.2862

024E+31212.5RT (min)

lcqtof\_t3\_pos\_4\_006\_41

m/z 352.2130

024E+39.510RT (min)

lcqtof\_t3\_pos\_1\_014\_14

m/z 353.2667

012E+41717.5RT (min)

lcqtof\_t3\_pos\_2\_004\_42

m/z 353.2668

012E+41515.5RT (min)

lcqtof\_t3\_pos\_2\_006\_04

m/z 354.2778

0246E+31616.5RT (min)

lcqtof\_t3\_pos\_2\_003\_16

m/z 361.3285

01234E+31717.5RT (min)

lcqtof\_t3\_pos\_4\_089\_59

m/z 367.8833

012E+37.58RT (min)

lcqtof\_t3\_pos\_2\_037\_46

m/z 369.2107

024E+31212.5RT (min)

lcqtof\_t3\_pos\_2\_013\_29

m/z 369.2782

02468E+211.512RT (min)

lcqtof\_t3\_pos\_4\_021\_01

m/z 371.1634

0246E+314.515RT (min)

lcqtof\_t3\_pos\_2\_049\_55

m/z 371.1639

0246E+31717.5RT (min)

lcqtof\_t3\_pos\_4\_092\_60

m/z 372.1998

01E+41010.5RT (min)

lcqtof\_t3\_pos\_1\_010\_27

m/z 372.3202

01E+41717.5RT (min)

lcqtof\_t3\_pos\_2\_004\_42

m/z 373.0580

01E+51313.5RT (min)

lcqtof\_t3\_pos\_4\_024\_20

m/z 378.3106

01E+411.512RT (min)

lcqtof\_t3\_pos\_4\_034\_23

m/z 379.1175

01234E+399.5RT (min)

lcqtof\_t3\_pos\_4\_029\_37

m/z 380.3259

01234E+49.510RT (min)

lcqtof\_t3\_pos\_4\_029\_37

m/z 385.3312

0246E+31717.5RT (min)

lcqtof\_t3\_pos\_2\_027\_38

m/z 386.1830

0246E+31010.5RT (min)

lcqtof\_t3\_pos\_4\_012\_12

m/z 386.3419

0246E+31717.5RT (min)

lcqtof\_t3\_pos\_2\_020\_27

m/z 391.4845

024E+414.515RT (min)

lcqtof\_t3\_pos\_2\_012\_35

m/z 391.4849

024E+41616.5RT (min)

lcqtof\_t3\_pos\_2\_009\_50

m/z 391.6707

0246E+317.518RT (min)

lcqtof\_t3\_pos\_4\_110\_62

m/z 392.0812

01234E+36.57RT (min)

lcqtof\_t3\_pos\_2\_010\_26

m/z 392.3628

0246E+31717.5RT (min)

lcqtof\_t3\_pos\_4\_072\_57

m/z 398.2896

0246E+29.510RT (min)

lcqtof\_t3\_pos\_4\_021\_01

m/z 403.2668

0123E+21111.5RT (min)

lcqtof\_t3\_pos\_4\_025\_52

m/z 407.6817

024E+317.518RT (min)

lcqtof\_t3\_pos\_4\_104\_61

m/z 410.2763

0112.513RT (min)

lcqtof\_t3\_pos\_4\_020\_19

m/z 411.1938

024E+31717.5RT (min)

lcqtof\_t3\_pos\_4\_095\_60

m/z 411.3095

0123E+388.5RT (min)

lcqtof\_t3\_pos\_4\_029\_37

m/z 415.2730

012E+417.518RT (min)

lcqtof\_t3\_pos\_2\_003\_16

m/z 416.2764

0123E+31717.5RT (min)

lcqtof\_t3\_pos\_2\_032\_47

m/z 429.3400

0246E+310.511RT (min)

lcqtof\_t3\_pos\_4\_029\_37

m/z 429.3771

0246E+310.511RT (min)

lcqtof\_t3\_pos\_4\_021\_01

m/z 435.3222

02468E+414.515RT (min)

lcqtof\_t3\_pos\_2\_004\_42

m/z 435.3473

01E+41212.5RT (min)

lcqtof\_t3\_pos\_4\_021\_01

m/z 436.3308

0123E+415.516RT (min)

lcqtof\_t3\_pos\_2\_006\_04

m/z 436.3311

0123E+41414.5RT (min)

lcqtof\_t3\_pos\_2\_004\_42

m/z 436.3313

01E+417.518RT (min)

lcqtof\_t3\_pos\_2\_010\_26

m/z 439.2990

01E+313.514RT (min)

lcqtof\_t3\_pos\_4\_042\_18

m/z 441.3075

01234E+355.5RT (min)

lcqtof\_t3\_pos\_4\_029\_37

m/z 442.3524

0246E+31111.5RT (min)

lcqtof\_t3\_pos\_4\_003\_05

m/z 442.4278

0246E+31212.5RT (min)

lcqtof\_t3\_pos\_4\_029\_37

m/z 443.3062

0123E+312.513RT (min)

lcqtof\_t3\_pos\_4\_030\_08

m/z 449.2705

01234E+37.58RT (min)

lcqtof\_t3\_pos\_4\_029\_37

m/z 451.3615

012E+41414.5RT (min)

lcqtof\_t3\_pos\_3\_067\_57

m/z 451.3618

01234E+417.518RT (min)

lcqtof\_t3\_pos\_2\_003\_16

m/z 453.1675

0246E+31010.5RT (min)

lcqtof\_t3\_pos\_4\_012\_12

m/z 457.3478

0246E+310.511RT (min)

lcqtof\_t3\_pos\_4\_003\_05

m/z 462.1409

012E+29.510RT (min)

lcqtof\_t3\_pos\_4\_012\_12

m/z 463.3554

01234E+212.513RT (min)

lcqtof\_t3\_pos\_4\_021\_01

m/z 464.3229

0246E+277.5RT (min)

lcqtof\_t3\_pos\_4\_042\_18

m/z 475.0162

0123E+31111.5RT (min)

lcqtof\_t3\_pos\_4\_001\_49

m/z 475.2101

0246E+310.511RT (min)

lcqtof\_t3\_pos\_2\_031\_01

m/z 476.3571

012E+41717.5RT (min)

lcqtof\_t3\_pos\_2\_004\_42

m/z 478.3542

02468E+31717.5RT (min)

lcqtof\_t3\_pos\_1\_012\_42

m/z 482.8625

01E+41111.5RT (min)

lcqtof\_t3\_pos\_2\_036\_20

m/z 485.3857

0123E+39.510RT (min)

lcqtof\_t3\_pos\_4\_093\_60

m/z 489.3046

01E+39.510RT (min)

lcqtof\_t3\_pos\_3\_005\_18

m/z 492.3684

01E+417.518RT (min)

lcqtof\_t3\_pos\_2\_004\_42

m/z 495.0136

01E+38.59RT (min)

lcqtof\_t3\_pos\_3\_019\_06

m/z 495.4069

02468E+311.512RT (min)

lcqtof\_t3\_pos\_2\_022\_32

m/z 500.3361

01234E+388.5RT (min)

lcqtof\_t3\_pos\_4\_072\_57

m/z 503.3855

012E+41717.5RT (min)

lcqtof\_t3\_pos\_2\_004\_42

m/z 506.3849

0123E+415.516RT (min)

lcqtof\_t3\_pos\_2\_009\_50

m/z 506.3850

0123E+417.518RT (min)

lcqtof\_t3\_pos\_2\_003\_16

m/z 507.1394

01234E+31010.5RT (min)

lcqtof\_t3\_pos\_4\_012\_12

m/z 510.4391

012E+314.515RT (min)

lcqtof\_t3\_pos\_4\_029\_37

m/z 515.2615

024E+31010.5RT (min)

lcqtof\_t3\_pos\_4\_034\_23

m/z 517.3892

01E+31212.5RT (min)

lcqtof\_t3\_pos\_4\_021\_01

m/z 518.2601

012E+31717.5RT (min)

lcqtof\_t3\_pos\_2\_025\_52

m/z 518.7303

0246E+410.511RT (min)

lcqtof\_t3\_pos\_4\_003\_05

m/z 533.2937

024E+399.5RT (min)

lcqtof\_t3\_pos\_2\_027\_38

m/z 538.4858

0123E+312.513RT (min)

lcqtof\_t3\_pos\_4\_071\_57

m/z 545.2683

02468E+29.510RT (min)

lcqtof\_t3\_pos\_4\_005\_30

m/z 549.2762

0123E+31010.5RT (min)

lcqtof\_t3\_pos\_4\_072\_57

m/z 552.2451

012E+31717.5RT (min)

lcqtof\_t3\_pos\_2\_011\_19

m/z 552.3070

01234E+310.511RT (min)

lcqtof\_t3\_pos\_4\_021\_01

m/z 556.3695

0123E+41717.5RT (min)

lcqtof\_t3\_pos\_4\_092\_60

m/z 558.3132

012E+388.5RT (min)

lcqtof\_t3\_pos\_4\_103\_61

m/z 576.4695

01E+313.514RT (min)

lcqtof\_t3\_pos\_4\_003\_05

m/z 581.3797

024E+31212.5RT (min)

lcqtof\_t3\_pos\_2\_039\_23

m/z 582.4563

024E+312.513RT (min)

lcqtof\_t3\_pos\_4\_074\_58

m/z 584.9300

012E+31111.5RT (min)

lcqtof\_t3\_pos\_4\_008\_15

m/z 587.2426

0246E+37.58RT (min)

lcqtof\_t3\_pos\_4\_100\_61

m/z 589.3268

012E+312.513RT (min)

lcqtof\_t3\_pos\_4\_103\_61

m/z 589.3371

0246E+35.56RT (min)

lcqtof\_t3\_pos\_4\_106\_62

m/z 592.3724

012E+39.510RT (min)

lcqtof\_t3\_pos\_4\_049\_55

m/z 592.5613

02468E+313.514RT (min)

lcqtof\_t3\_pos\_2\_039\_23

m/z 594.5515

01E+312.513RT (min)

lcqtof\_t3\_pos\_4\_021\_01

m/z 599.1173

024E+38.59RT (min)

lcqtof\_t3\_pos\_4\_029\_37

m/z 606.1651

01234E+388.5RT (min)

lcqtof\_t3\_pos\_4\_042\_18

m/z 606.6020

0246E+312.513RT (min)

lcqtof\_t3\_pos\_4\_014\_24

m/z 609.5096

02468E+312.513RT (min)

lcqtof\_t3\_pos\_4\_021\_01

m/z 611.3298

024E+312.513RT (min)

lcqtof\_t3\_pos\_2\_055\_18

m/z 617.5422

012E+314.515RT (min)

lcqtof\_t3\_pos\_4\_029\_37

m/z 621.3119

024E+211.512RT (min)

lcqtof\_t3\_pos\_3\_007\_32

m/z 624.6266

01E+31313.5RT (min)

lcqtof\_t3\_pos\_1\_007\_07

m/z 631.9744

024E+35.56RT (min)

lcqtof\_t3\_pos\_4\_029\_37

m/z 633.2162

01234E+39.510RT (min)

lcqtof\_t3\_pos\_2\_050\_14

m/z 633.2271

02468E+51.52RT (min)

lcqtof\_t3\_pos\_4\_014\_24

m/z 634.4221

01234E+399.5RT (min)

lcqtof\_t3\_pos\_4\_015\_16

m/z 641.4045

01234E+31212.5RT (min)

lcqtof\_t3\_pos\_2\_002\_37

m/z 643.2782

01E+41717.5RT (min)

lcqtof\_t3\_pos\_4\_107\_62

m/z 646.6332

02468E+31717.5RT (min)

lcqtof\_t3\_pos\_2\_030\_33

m/z 648.4237

012E+41010.5RT (min)

lcqtof\_t3\_pos\_4\_021\_01

m/z 649.2828

0123E+310.511RT (min)

lcqtof\_t3\_pos\_2\_020\_27

m/z 650.6454

011313.5RT (min)

lcqtof\_t3\_pos\_4\_093\_60

m/z 652.4983

012E+31111.5RT (min)

lcqtof\_t3\_pos\_4\_003\_05

m/z 654.2064

01E+41.52RT (min)

lcqtof\_t3\_pos\_4\_014\_24

m/z 658.5967

01E+41717.5RT (min)

lcqtof\_t3\_pos\_2\_011\_19

m/z 663.6404

01E+41717.5RT (min)

lcqtof\_t3\_pos\_2\_050\_14

m/z 665.4105

024E+31212.5RT (min)

lcqtof\_t3\_pos\_2\_027\_38

m/z 665.5106

0246E+311.512RT (min)

lcqtof\_t3\_pos\_4\_003\_05

m/z 669.6257

01E+41717.5RT (min)

lcqtof\_t3\_pos\_2\_006\_04

m/z 671.1773

02468E+322.5RT (min)

lcqtof\_t3\_pos\_4\_014\_24

m/z 674.6972

012E+315.516RT (min)

lcqtof\_t3\_pos\_4\_111\_62

m/z 676.6361

012E+315.516RT (min)

lcqtof\_t3\_pos\_4\_086\_59

m/z 677.3148

0123E+31111.5RT (min)

lcqtof\_t3\_pos\_2\_020\_27

m/z 680.4257

024E+39.510RT (min)

lcqtof\_t3\_pos\_4\_029\_37

m/z 681.4949

0246E+317.518RT (min)

lcqtof\_t3\_pos\_4\_109\_62

m/z 681.6867

0112.513RT (min)

lcqtof\_t3\_pos\_3\_052\_08

m/z 683.1027

01E+377.5RT (min)

lcqtof\_t3\_pos\_2\_055\_18

m/z 684.2720

0123E+310.511RT (min)

lcqtof\_t3\_pos\_2\_031\_01

m/z 684.3970

01234E+31212.5RT (min)

lcqtof\_t3\_pos\_4\_052\_06

m/z 686.6276

02468E+41717.5RT (min)

lcqtof\_t3\_pos\_2\_003\_16

m/z 691.4185

012E+39.510RT (min)

lcqtof\_t3\_pos\_3\_011\_15

m/z 691.4307

024E+312.513RT (min)

lcqtof\_t3\_pos\_2\_039\_23

m/z 696.4379

01234E+317.518RT (min)

lcqtof\_t3\_pos\_4\_106\_62

m/z 699.1726

0246E+322.5RT (min)

lcqtof\_t3\_pos\_4\_014\_24

m/z 700.2807

01234E+310.511RT (min)

lcqtof\_t3\_pos\_2\_003\_16

m/z 701.4655

01234E+312.513RT (min)

lcqtof\_t3\_pos\_2\_002\_37

m/z 701.4860

02468E+31414.5RT (min)

lcqtof\_t3\_pos\_2\_008\_31

m/z 702.6217

012E+515.516RT (min)

lcqtof\_t3\_pos\_2\_009\_50

m/z 703.2189

012E+39.510RT (min)

lcqtof\_t3\_pos\_3\_011\_15

m/z 703.4777

0123E+312.513RT (min)

lcqtof\_t3\_pos\_4\_012\_12

m/z 703.6265

0246E+41616.5RT (min)

lcqtof\_t3\_pos\_2\_005\_17

m/z 706.7741

0246E+31313.5RT (min)

lcqtof\_t3\_pos\_2\_022\_32

m/z 708.4327

012E+37.58RT (min)

lcqtof\_t3\_pos\_1\_010\_27

m/z 711.1802

0123E+38.59RT (min)

lcqtof\_t3\_pos\_4\_021\_01

m/z 713.6588

0246E+31717.5RT (min)

lcqtof\_t3\_pos\_2\_004\_42

m/z 719.2837

02468E+317.518RT (min)

lcqtof\_t3\_pos\_4\_017\_51

m/z 720.6276

012E+31111.5RT (min)

lcqtof\_t3\_pos\_2\_004\_42

m/z 720.7649

01E+31414.5RT (min)

lcqtof\_t3\_pos\_4\_049\_55

m/z 724.6044

012E+41515.5RT (min)

lcqtof\_t3\_pos\_2\_002\_37

m/z 727.4062

024E+399.5RT (min)

lcqtof\_t3\_pos\_4\_014\_24

m/z 730.6624

02468E+31717.5RT (min)

lcqtof\_t3\_pos\_2\_006\_04

m/z 732.4561

012E+31313.5RT (min)

lcqtof\_t3\_pos\_3\_068\_57

m/z 732.6093

01E+31414.5RT (min)

lcqtof\_t3\_pos\_2\_050\_14

m/z 735.3000

0123E+388.5RT (min)

lcqtof\_t3\_pos\_4\_029\_37

m/z 736.3089

0246E+317.518RT (min)

lcqtof\_t3\_pos\_4\_106\_62

m/z 737.1937

0123E+39.510RT (min)

lcqtof\_t3\_pos\_3\_011\_15

m/z 739.6361

0246E+312.513RT (min)

lcqtof\_t3\_pos\_4\_093\_60

m/z 743.1961

012E+39.510RT (min)

lcqtof\_t3\_pos\_3\_011\_15

m/z 744.6475

024E+111.512RT (min)

lcqtof\_t3\_pos\_4\_082\_59

m/z 745.2997

01234E+31111.5RT (min)

lcqtof\_t3\_pos\_2\_050\_14

m/z 747.4100

0123E+37.58RT (min)

lcqtof\_t3\_pos\_4\_110\_62

m/z 747.6972

0246E+313.514RT (min)

lcqtof\_t3\_pos\_2\_034\_24

m/z 753.1647

012E+38.59RT (min)

lcqtof\_t3\_pos\_4\_006\_41

m/z 754.4335

012E+312.513RT (min)

lcqtof\_t3\_pos\_4\_089\_59

m/z 756.6024

024E+31414.5RT (min)

lcqtof\_t3\_pos\_4\_086\_59

m/z 762.7826

0246E+31414.5RT (min)

lcqtof\_t3\_pos\_2\_022\_32

m/z 769.4758

0123E+312.513RT (min)

lcqtof\_t3\_pos\_4\_087\_59

m/z 773.6874

0123E+31212.5RT (min)

lcqtof\_t3\_pos\_1\_022\_28

m/z 776.4453

0246E+21212.5RT (min)

lcqtof\_t3\_pos\_4\_012\_12

m/z 776.5579

0246E+31515.5RT (min)

lcqtof\_t3\_pos\_2\_020\_27

m/z 781.6308

01E+214.515RT (min)

lcqtof\_t3\_pos\_2\_050\_14

m/z 790.9917

0246E+31010.5RT (min)

lcqtof\_t3\_pos\_4\_047\_13

m/z 791.3145

0246E+31212.5RT (min)

lcqtof\_t3\_pos\_4\_084\_59

m/z 792.5317

02468E+312.513RT (min)

lcqtof\_t3\_pos\_4\_048\_11

m/z 793.6829

01E+31414.5RT (min)

lcqtof\_t3\_pos\_4\_049\_55

m/z 795.3935

02468E+313.514RT (min)

lcqtof\_t3\_pos\_2\_005\_17

m/z 795.5360

01E+411.512RT (min)

lcqtof\_t3\_pos\_4\_029\_37

m/z 797.2874

0123E+313.514RT (min)

lcqtof\_t3\_pos\_2\_005\_17

m/z 800.4871

0246E+310.511RT (min)

lcqtof\_t3\_pos\_4\_037\_32

m/z 803.4457

0246E+37.58RT (min)

lcqtof\_t3\_pos\_4\_110\_62

m/z 804.3369

012E+31212.5RT (min)

lcqtof\_t3\_pos\_2\_034\_24

m/z 808.1064

024E+313.514RT (min)

lcqtof\_t3\_pos\_4\_012\_12

m/z 811.5327

01E+312.513RT (min)

lcqtof\_t3\_pos\_4\_086\_59

m/z 812.2854

012E+31111.5RT (min)

lcqtof\_t3\_pos\_2\_008\_31

m/z 813.4930

012E+313.514RT (min)

lcqtof\_t3\_pos\_4\_097\_60

m/z 814.3536

01234E+313.514RT (min)

lcqtof\_t3\_pos\_4\_021\_01

m/z 837.5014

01234E+212.513RT (min)

lcqtof\_t3\_pos\_4\_003\_05

m/z 837.5162

012E+38.59RT (min)

lcqtof\_t3\_pos\_4\_104\_61

m/z 839.3384

01234E+388.5RT (min)

lcqtof\_t3\_pos\_4\_029\_37

m/z 841.7787

012E+311.512RT (min)

lcqtof\_t3\_pos\_1\_001\_49

m/z 844.6785

01E+31414.5RT (min)

lcqtof\_t3\_pos\_1\_016\_47

m/z 846.5970

0246E+312.513RT (min)

lcqtof\_t3\_pos\_4\_004\_33

m/z 848.4470

0246E+31212.5RT (min)

lcqtof\_t3\_pos\_2\_039\_23

m/z 853.5524

0123E+31313.5RT (min)

lcqtof\_t3\_pos\_3\_011\_15

m/z 854.5845

024E+39.510RT (min)

lcqtof\_t3\_pos\_4\_023\_39

m/z 856.5039

024E+31212.5RT (min)

lcqtof\_t3\_pos\_4\_034\_23

m/z 856.8245

024E+313.514RT (min)

lcqtof\_t3\_pos\_4\_097\_60

m/z 862.7436

01E+311.512RT (min)

lcqtof\_t3\_pos\_4\_003\_05

m/z 865.6880

02468E+31414.5RT (min)

lcqtof\_t3\_pos\_2\_050\_14

m/z 866.5950

01E+41212.5RT (min)

lcqtof\_t3\_pos\_2\_019\_08

m/z 872.7199

01E+213.514RT (min)

lcqtof\_t3\_pos\_3\_069\_57

m/z 874.4538

0246E+21010.5RT (min)

lcqtof\_t3\_pos\_4\_040\_28

m/z 877.4813

024E+31212.5RT (min)

lcqtof\_t3\_pos\_2\_039\_23

m/z 879.5435

012E+38.59RT (min)

lcqtof\_t3\_pos\_4\_108\_62

m/z 880.3306

0123E+388.5RT (min)

lcqtof\_t3\_pos\_4\_029\_37

m/z 880.6968

0123E+312.513RT (min)

lcqtof\_t3\_pos\_4\_095\_60

m/z 881.5140

0123E+31010.5RT (min)

lcqtof\_t3\_pos\_4\_074\_58

m/z 882.6880

012E+312.513RT (min)

lcqtof\_t3\_pos\_4\_111\_62

m/z 884.5358

01234E+312.513RT (min)

lcqtof\_t3\_pos\_2\_015\_13

m/z 884.6308

0246E+312.513RT (min)

lcqtof\_t3\_pos\_3\_013\_24

m/z 886.6161

01234E+313.514RT (min)

lcqtof\_t3\_pos\_2\_003\_16

m/z 890.6553

0123E+313.514RT (min)

lcqtof\_t3\_pos\_4\_021\_01

m/z 892.5964

0246E+313.514RT (min)

lcqtof\_t3\_pos\_3\_003\_14

m/z 893.6767

01E+31414.5RT (min)

lcqtof\_t3\_pos\_2\_044\_25

m/z 896.6375

01E+113.514RT (min)

lcqtof\_t3\_pos\_4\_092\_60

m/z 904.5014

0123E+312.513RT (min)

lcqtof\_t3\_pos\_2\_015\_13

m/z 906.6201

0123E+313.514RT (min)

lcqtof\_t3\_pos\_2\_003\_16

m/z 907.8518

0246E+31212.5RT (min)

lcqtof\_t3\_pos\_4\_021\_01

m/z 914.6194

024E+312.513RT (min)

lcqtof\_t3\_pos\_2\_034\_24

m/z 916.7428

01E+413.514RT (min)

lcqtof\_t3\_pos\_4\_086\_59

m/z 930.5146

0123E+312.513RT (min)

lcqtof\_t3\_pos\_2\_015\_13

m/z 935.5497

0123E+31313.5RT (min)

lcqtof\_t3\_pos\_2\_031\_01

m/z 943.6692

024E+31313.5RT (min)

lcqtof\_t3\_pos\_2\_039\_23

m/z 947.7183

02468E+312.513RT (min)

lcqtof\_t3\_pos\_4\_014\_24

m/z 956.7067

0246E+313.514RT (min)

lcqtof\_t3\_pos\_2\_027\_38

m/z 957.4132

024E+39.510RT (min)

lcqtof\_t3\_pos\_4\_008\_15

m/z 957.7496

0246E+314.515RT (min)

lcqtof\_t3\_pos\_4\_021\_01

m/z 960.7713

01E+313.514RT (min)

lcqtof\_t3\_pos\_4\_055\_47

m/z 962.5254

024E+31212.5RT (min)

lcqtof\_t3\_pos\_4\_029\_37

m/z 962.8714

0246E+31515.5RT (min)

lcqtof\_t3\_pos\_4\_109\_62

m/z 965.7161

02468E+41515.5RT (min)

lcqtof\_t3\_pos\_4\_045\_04

m/z 966.6934

024E+39.510RT (min)

lcqtof\_t3\_pos\_4\_003\_05

m/z 966.9104

01E+415.516RT (min)

lcqtof\_t3\_pos\_4\_101\_61

m/z 968.6190

01E+412.513RT (min)

lcqtof\_t3\_pos\_2\_020\_27

m/z 970.5777

012E+313.514RT (min)

lcqtof\_t3\_pos\_2\_014\_03

m/z 981.4052

01234E+388.5RT (min)

lcqtof\_t3\_pos\_4\_029\_37

m/z 987.8632

012E+31515.5RT (min)

lcqtof\_t3\_pos\_4\_113\_62

m/z 989.8862

0246E+315.516RT (min)

lcqtof\_t3\_pos\_4\_105\_61

m/z 991.8710

024E+315.516RT (min)

lcqtof\_t3\_pos\_4\_107\_62

m/z 1001.5057

0123E+31212.5RT (min)

lcqtof\_t3\_pos\_1\_003\_33

m/z 1002.5183

024E+312.513RT (min)

lcqtof\_t3\_pos\_2\_027\_38

m/z 1003.9012

01234E+315.516RT (min)

lcqtof\_t3\_pos\_4\_100\_61

m/z 1006.8594

024E+316.517RT (min)

lcqtof\_t3\_pos\_2\_025\_52

m/z 1007.2818

0123E+388.5RT (min)

lcqtof\_t3\_pos\_2\_020\_27

m/z 1009.3968

0246E+388.5RT (min)

lcqtof\_t3\_pos\_4\_029\_37

m/z 1009.7428

012E+31111.5RT (min)

lcqtof\_t3\_pos\_4\_011\_10

m/z 1010.9081

0123E+315.516RT (min)

lcqtof\_t3\_pos\_4\_086\_59

m/z 1019.8579

02468E+31313.5RT (min)

lcqtof\_t3\_pos\_2\_022\_32

m/z 1019.9022

012E+315.516RT (min)

lcqtof\_t3\_pos\_4\_072\_57

m/z 1022.4844

024E+31212.5RT (min)

lcqtof\_t3\_pos\_2\_039\_23

m/z 1023.6729

02468E+31212.5RT (min)

lcqtof\_t3\_pos\_4\_014\_24

m/z 1025.8460

024E+313.514RT (min)

lcqtof\_t3\_pos\_2\_034\_24

m/z 1028.1572

0246E+310.511RT (min)

lcqtof\_t3\_pos\_4\_047\_13

m/z 1037.3977

0246E+388.5RT (min)

lcqtof\_t3\_pos\_4\_029\_37

m/z 1038.7768

012E+311.512RT (min)

lcqtof\_t3\_pos\_4\_008\_15

m/z 1042.5931

0123E+31010.5RT (min)

lcqtof\_t3\_pos\_4\_074\_58

m/z 1047.9253

0123E+315.516RT (min)

lcqtof\_t3\_pos\_4\_105\_61

m/z 1048.8298

024E+313.514RT (min)

lcqtof\_t3\_pos\_2\_034\_24

m/z 1052.7665

024E+310.511RT (min)

lcqtof\_t3\_pos\_4\_003\_05

m/z 1057.9929

0123E+315.516RT (min)

lcqtof\_t3\_pos\_4\_090\_60

m/z 1071.9300

01234E+31414.5RT (min)

lcqtof\_t3\_pos\_2\_034\_24

m/z 1114.7215

0246E+31313.5RT (min)

lcqtof\_t3\_pos\_4\_089\_59

m/z 1127.6327

0246E+310.511RT (min)

lcqtof\_t3\_pos\_1\_014\_14

m/z 1144.8700

0246E+31010.5RT (min)

lcqtof\_t3\_pos\_2\_039\_23

m/z 1147.4675

02468E+31010.5RT (min)

lcqtof\_t3\_pos\_4\_029\_37

m/z 1147.6681

0246E+31010.5RT (min)

lcqtof\_t3\_pos\_4\_029\_37

m/z 1168.8477

024E+31111.5RT (min)

lcqtof\_t3\_pos\_4\_003\_05

m/z 1169.8517

0123E+31111.5RT (min)

lcqtof\_t3\_pos\_4\_003\_05

m/z 1170.9332

0246E+313.514RT (min)

lcqtof\_t3\_pos\_2\_022\_32

m/z 1192.8233

01234E+31313.5RT (min)

lcqtof\_t3\_pos\_2\_040\_28

m/z 1193.3707

024E+31313.5RT (min)

lcqtof\_t3\_pos\_2\_003\_16
