## Supplementary material for "The Everything Bagel Feature Finder: Ultra-fast automated feature finding for untargeted metabolomics": SI Data File: tool_unique_xic_st_XCMS_ST004504.html

 
 
 XCMS-unique features (not detected by the other two tools) - ST004504 
 
 XCMS-unique features (not detected by the other two tools) - ST004504 
 9423 features · orbitrap · XIC window m/z ± 10 ppm (≥ 0.002 Da) · ± 0.5 min 
 Every panel is a feature  XCMS  detected that neither of the other two tools detected — three-way unique across EB (standard mode (min_detection_rate=0.1)), Masscube and XCMS, under the standard matching rule (m/z ± 10 ppm; apex within the reference RT window ± 0.2 min). It shows the MS1 extracted-ion chromatogram at the feature's m/z, drawn from the raw mzML named in  alignment_reference_file , zoomed to ± 0.5 min around the feature's reported apex RT.  Gray solid line  = the feature's reported apex RT;  red dashed line  = XCMS's assigned MS/MS —  RTINSECONDS  in its  aligned_features.mgf ; the panel widens to keep an off-apex MS/MS on-screen. Features with no assigned MS/MS show only the gray line. y is per-cell autoscaled. Rows keep the table order (m/z ascending). 
   m/z 50.4829     0   1   2E+4   5.5   6  RT (min)      lcorbi_t3_pos_4_018_01    m/z 50.5540     0   1E+4   8   8.5  RT (min)      lcorbi_t3_pos_2_044_21    m/z 50.5821     0   1   7.5   8  RT (min)      lcorbi_t3_pos_4_099_61    m/z 50.7630     0   1   2   3   4E+4   1   1.5  RT (min)      lcorbi_t3_pos_1_046_22    m/z 51.0306     0   1E+4   1   1.5  RT (min)      lcorbi_t3_pos_3_012_31    m/z 51.8065     0   1   2   3E+4   10   10.5  RT (min)      lcorbi_t3_pos_4_080_58    m/z 52.0342     0   1   2   3E+4   10   10.5  RT (min)      lcorbi_t3_pos_4_072_57    m/z 53.1462     0   1   2   3E+4   10   10.5  RT (min)      lcorbi_t3_pos_4_077_58    m/z 53.6005     0   1   2E+5   11   11.5  RT (min)      lcorbi_t3_pos_2_020_22    m/z 53.6967     0   1E+4   10   10.5  RT (min)      lcorbi_t3_pos_4_079_58    m/z 53.8077     0   1   2E+4   9   9.5  RT (min)      lcorbi_t3_pos_4_072_57    m/z 54.2305     0   2   4   6   8E+3   1   1.5  RT (min)      lcorbi_t3_pos_1_020_31    m/z 54.2475     0   2   4   6   8E+3   7   7.5  RT (min)      lcorbi_t3_pos_4_113_62    m/z 54.6778     0   1   2   3E+4   6.5   7  RT (min)      lcorbi_t3_pos_2_044_21    m/z 54.7115     0   1E+5   13   13.5  RT (min)      lcorbi_t3_pos_4_076_58    m/z 54.7706     0   2   4   6   8E+3   1   1.5  RT (min)      lcorbi_t3_pos_1_034_34    m/z 54.7752     0   1   2   3   4E+3   6   6.5  RT (min)      lcorbi_t3_pos_3_048_30    m/z 54.7934     0   1   2E+4   10   10.5  RT (min)      lcorbi_t3_pos_1_011_44    m/z 54.8035     0   2   4   6E+4   6.5   7  RT (min)      lcorbi_t3_pos_4_038_38    m/z 54.9248     0   2   4   6E+4   10.5   11  RT (min)      lcorbi_t3_pos_4_069_57    m/z 55.2616     0   1   2   3E+4   0   0.5  RT (min)      lcorbi_t3_pos_1_007_07    m/z 55.3716     0   2   4   6   8E+4   11   11.5  RT (min)      lcorbi_t3_pos_4_078_58    m/z 55.8103     0   1   2   3E+4   11   11.5  RT (min)      lcorbi_t3_pos_4_073_57    m/z 56.2756     0   2   4   6   8E+3   1   1.5  RT (min)      lcorbi_t3_pos_1_036_08    m/z 56.7061     0   1   2   3E+4   11.5   12  RT (min)      lcorbi_t3_pos_4_079_58    m/z 56.7377     0   1   2   3E+4   1   1.5  RT (min)      lcorbi_t3_pos_1_044_25    m/z 56.7937     0   1E+4   4.5   5  RT (min)      lcorbi_t3_pos_2_012_01    m/z 56.9648     0   2   4E+4   18   18.5  RT (min)       lcorbi_t3_pos_4_084_59    m/z 56.9850     0   2   4   6   8E+4   13   13.5  RT (min)       lcorbi_t3_pos_1_029_29    m/z 57.0335     0   1   2   3   4E+4   1   1.5  RT (min)       lcorbi_t3_pos_4_086_59    m/z 57.2083     0   1   2   3   4E+4   1   1.5  RT (min)      lcorbi_t3_pos_4_077_58    m/z 57.5618     0   1   2   3E+4   9   9.5  RT (min)      lcorbi_t3_pos_2_044_21    m/z 57.5913     0   1   2   3   4E+4   11   11.5  RT (min)      lcorbi_t3_pos_4_007_46    m/z 57.7570     0   1E+4   6.5   7  RT (min)      lcorbi_t3_pos_2_023_26    m/z 58.0650     0   2   4E+4   1   1.5  RT (min)      lcorbi_t3_pos_4_071_57    m/z 58.0732     0   2   4   6   8E+4   13   13.5  RT (min)      lcorbi_t3_pos_4_039_35    m/z 58.2609     0   1E+4   6.5   7  RT (min)      lcorbi_t3_pos_3_011_42    m/z 58.4921     0   1   2E+5   11   11.5  RT (min)      lcorbi_t3_pos_4_089_59    m/z 58.5514     0   1   2   3   4E+4   7   7.5  RT (min)      lcorbi_t3_pos_1_019_16    m/z 58.8022     0   2   4   6E+3   7   7.5  RT (min)      lcorbi_t3_pos_4_053_19    m/z 59.0491     0   1E+6   15.5   16  RT (min)      lcorbi_t3_pos_1_050_13    m/z 59.0491     0   2   4E+5   11   11.5  RT (min)      lcorbi_t3_pos_1_013_10    m/z 59.1567     0   2   4   6E+4   13.5   14  RT (min)      lcorbi_t3_pos_4_077_58    m/z 59.8376     0   2   4E+4   14   14.5  RT (min)      lcorbi_t3_pos_4_098_61    m/z 60.0436     0   1E+5   1   1.5  RT (min)       lcorbi_t3_pos_4_097_60    m/z 60.5936     0   2   4   6   8E+4   11   11.5  RT (min)      lcorbi_t3_pos_4_072_57    m/z 60.7061     0   2   4E+4   11   11.5  RT (min)      lcorbi_t3_pos_4_068_57    m/z 60.7312     0   2   4   6   8E+3   1   1.5  RT (min)      lcorbi_t3_pos_4_052_06    m/z 61.0396     0   1   2E+4   1   1.5  RT (min)      lcorbi_t3_pos_1_029_29    m/z 61.0648     0   1E+6   10   10.5  RT (min)      lcorbi_t3_pos_1_032_26    m/z 61.5605     0   1   2   3   4E+4   8   8.5  RT (min)      lcorbi_t3_pos_1_041_54    m/z 61.8113     0   1E+4   8   8.5  RT (min)      lcorbi_t3_pos_2_044_21    m/z 62.0157     0   1   2   3E+4   7   7.5  RT (min)      lcorbi_t3_pos_2_016_23    m/z 62.0173     0   2   4E+3   4.5   5  RT (min)      lcorbi_t3_pos_1_029_29    m/z 62.0681     0   1E+5   15   15.5  RT (min)      lcorbi_t3_pos_2_016_23    m/z 62.2781     0   1E+5   14.5   15  RT (min)      lcorbi_t3_pos_4_099_61    m/z 62.7615     0   1   2   3   4E+4   11.5   12  RT (min)      lcorbi_t3_pos_1_029_29    m/z 62.8000     0   1E+4   10.5   11  RT (min)      lcorbi_t3_pos_2_053_28    m/z 62.9363     0   1E+5   5.5   6  RT (min)      lcorbi_t3_pos_3_024_07    m/z 62.9521     0   2   4E+4   14.5   15  RT (min)      lcorbi_t3_pos_4_082_59    m/z 62.9876     0   1E+5   1   1.5  RT (min)      lcorbi_t3_pos_3_012_31    m/z 63.0479     0   1E+4   5.5   6  RT (min)      lcorbi_t3_pos_4_034_42    m/z 63.1488     0   1   2   3E+4   11   11.5  RT (min)      lcorbi_t3_pos_4_072_57    m/z 63.3841     0   1E+5   11   11.5  RT (min)      lcorbi_t3_pos_4_088_59    m/z 63.5221     0   2   4E+4   8.5   9  RT (min)      lcorbi_t3_pos_4_030_05    m/z 63.8152     0   1E+4   8.5   9  RT (min)      lcorbi_t3_pos_4_099_61    m/z 64.2600     0   1   2E+4   8   8.5  RT (min)      lcorbi_t3_pos_4_108_62    m/z 64.5273     0   1   2   3E+4   1   1.5  RT (min)      lcorbi_t3_pos_4_013_36    m/z 64.5277     0   1   2   3   4E+4   1   1.5  RT (min)      lcorbi_t3_pos_4_005_33    m/z 64.6059     0   2   4E+4   14   14.5  RT (min)      lcorbi_t3_pos_4_098_61    m/z 64.7625     0   1   2   3E+4   1   1.5  RT (min)      lcorbi_t3_pos_4_075_58    m/z 65.5592     0   1E+4   8   8.5  RT (min)      lcorbi_t3_pos_2_004_42    m/z 66.0403     0   1   2   3E+4   8   8.5  RT (min)      lcorbi_t3_pos_4_109_62    m/z 66.1519     0   1E+4   8   8.5  RT (min)      lcorbi_t3_pos_4_107_62    m/z 66.5909     0   1   2   3E+4   8   8.5  RT (min)      lcorbi_t3_pos_4_109_62    m/z 66.7025     0   1E+4   8   8.5  RT (min)      lcorbi_t3_pos_4_107_62    m/z 67.0542     0   1E+4   1   1.5  RT (min)       lcorbi_t3_pos_1_020_31    m/z 67.0542     0   2   4   6E+4   11   11.5  RT (min)      lcorbi_t3_pos_1_040_27    m/z 67.6138     0   2   4E+4   14   14.5  RT (min)      lcorbi_t3_pos_4_088_59    m/z 67.9492     0   1   14.5   15  RT (min)      lcorbi_t3_pos_4_113_62    m/z 68.0494     0   1   2E+4   1   1.5  RT (min)      lcorbi_t3_pos_1_028_32    m/z 68.0617     0   2   4   6   8E+4   14.5   15  RT (min)      lcorbi_t3_pos_3_020_35    m/z 68.1288     0   2   4   6   8E+2   2   2.5  RT (min)      lcorbi_t3_pos_4_082_59    m/z 68.3659     0   2   4   6   8E+3   8   8.5  RT (min)      lcorbi_t3_pos_4_107_62    m/z 68.5683     0   1   2   3E+4   8.5   9  RT (min)      lcorbi_t3_pos_4_014_48    m/z 68.9971     0   1   11   11.5  RT (min)      lcorbi_t3_pos_1_001_49    m/z 69.0447     0   1   2   3   4E+6   1   1.5  RT (min)      lcorbi_t3_pos_1_001_49    m/z 69.0447     0   2   4   6E+5   3.5   4  RT (min)      lcorbi_t3_pos_1_052_38    m/z 69.0447     0   1   2   3   4E+6   6   6.5  RT (min)      lcorbi_t3_pos_1_001_49    m/z 69.0447     0   2   4   6   8E+6   12   12.5  RT (min)      lcorbi_t3_pos_1_004_24    m/z 69.0698     0   1E+6   15.5   16  RT (min)       lcorbi_t3_pos_1_033_53    m/z 69.0698     0   1   2E+5   11.5   12  RT (min)      lcorbi_t3_pos_1_056_56    m/z 69.0698     0   2   4   6   8E+4   6.5   7  RT (min)      lcorbi_t3_pos_1_009_50    m/z 69.7896     0   1   2   3E+4   10   10.5  RT (min)      lcorbi_t3_pos_1_029_29    m/z 70.0368     0   2   4E+3   6.5   7  RT (min)      lcorbi_t3_pos_1_007_07    m/z 70.0399     0   1E+5   0   0.5  RT (min)      lcorbi_t3_pos_1_001_49    m/z 70.2805     0   2   4   6E+4   14   14.5  RT (min)      lcorbi_t3_pos_4_085_59    m/z 70.3585     0   1   2   2.5  RT (min)      lcorbi_t3_pos_4_090_60    m/z 70.5044     0   1   2E+5   14.5   15  RT (min)      lcorbi_t3_pos_3_020_35    m/z 70.5418     0   1E+5   1   1.5  RT (min)      lcorbi_t3_pos_4_068_57    m/z 70.6159     0   2   4E+4   14.5   15  RT (min)      lcorbi_t3_pos_4_082_59    m/z 70.8399     0   2   4   6E+4   14.5   15  RT (min)      lcorbi_t3_pos_4_083_59    m/z 70.9804     0   2   4E+3   1   1.5  RT (min)      lcorbi_t3_pos_1_028_32    m/z 71.0127     0   1   2E+6   15  RT (min)       lcorbi_t3_pos_1_046_22    m/z 71.0491     0   1   2E+5   15   15.5  RT (min)      lcorbi_t3_pos_1_047_41    m/z 71.0491     0   1   2E+5   12   12.5  RT (min)      lcorbi_t3_pos_1_047_41    m/z 71.0491     0   2   4   6   8E+4   6.5   7  RT (min)      lcorbi_t3_pos_1_046_22    m/z 71.0491     0   1E+5   1   1.5  RT (min)      lcorbi_t3_pos_4_083_59    m/z 71.0638     0   1   2E+4   14.5   15  RT (min)      lcorbi_t3_pos_4_074_58    m/z 71.0684     0   2   4   6   8E+4   1   1.5  RT (min)      lcorbi_t3_pos_4_079_58    m/z 71.2558     0   2   4   6   8E+4   14   14.5  RT (min)      lcorbi_t3_pos_4_104_61    m/z 71.9627     0   1   2   3E+5   14.5   15  RT (min)      lcorbi_t3_pos_1_036_08    m/z 72.0435     0   2   4   6E+4   1   1.5  RT (min)       lcorbi_t3_pos_1_006_15    m/z 72.0807     0   1E+5   1.5   2  RT (min)       lcorbi_t3_pos_2_030_14    m/z 72.0807     0   1   2E+5   12.5   13  RT (min)      lcorbi_t3_pos_1_009_50    m/z 72.0888     0   1   10   10.5  RT (min)      lcorbi_t3_pos_1_026_17    m/z 72.3167     0   1   2E+4   1   1.5  RT (min)      lcorbi_t3_pos_4_081_58    m/z 72.5670     0   1   2   3E+4   9   9.5  RT (min)      lcorbi_t3_pos_2_044_21    m/z 72.7035     0   2   4E+4   12   12.5  RT (min)      lcorbi_t3_pos_4_086_59    m/z 72.9371     0   2   4E+5   0.5   1  RT (min)      lcorbi_t3_pos_1_007_07    m/z 73.0106     0   1   2E+4   2.5   3   3.5  RT (min)       lcorbi_t3_pos_4_098_61    m/z 73.0284     0   1   2E+5   11.5   12  RT (min)      lcorbi_t3_pos_4_068_57    m/z 73.0648     0   1   2   3E+5   12   12.5  RT (min)      lcorbi_t3_pos_1_008_36    m/z 73.0752     0   1E+5   1.5   2  RT (min)       lcorbi_t3_pos_4_093_60    m/z 73.0841     0   2   4   6E+4   1   1.5  RT (min)      lcorbi_t3_pos_4_075_58    m/z 73.0841     0   1E+5   18   18.5  RT (min)       lcorbi_t3_pos_4_098_61    m/z 73.7177     0   1E+5   14.5   15  RT (min)      lcorbi_t3_pos_4_083_59    m/z 73.7820     0   1   2E+3   5   5.5  RT (min)      lcorbi_t3_pos_3_034_26    m/z 73.8284     0   2   4   6   8E+4   14.5   15  RT (min)      lcorbi_t3_pos_4_082_59    m/z 74.0600     0   1E+5   1   1.5  RT (min)       lcorbi_t3_pos_4_093_60    m/z 74.0600     0   1   2E+5   6   6.5  RT (min)      lcorbi_t3_pos_1_001_49    m/z 74.0600     0   1   2E+5   3.5   4  RT (min)      lcorbi_t3_pos_1_001_49    m/z 74.0600     0   1   2E+5   12.5   13  RT (min)      lcorbi_t3_pos_1_002_01    m/z 74.0600     0   1   2E+5   9   9.5  RT (min)      lcorbi_t3_pos_1_001_49    m/z 74.0866     0   1E+3   2.5   3  RT (min)       lcorbi_t3_pos_1_002_01    m/z 74.0964     0   1   2E+5   1   1.5  RT (min)      lcorbi_t3_pos_4_091_60    m/z 74.0964     0   2   4   6   8E+5   6   6.5  RT (min)      lcorbi_t3_pos_1_001_49    m/z 74.0964     0   1E+6   10   12  RT (min)       lcorbi_t3_pos_1_014_03    m/z 74.4818     0   1   2   3E+4   14   14.5  RT (min)      lcorbi_t3_pos_4_107_62    m/z 74.5240     0   2   4   6E+3   4.5   5  RT (min)      lcorbi_t3_pos_2_013_33    m/z 74.6284     0   1E+5   14   14.5  RT (min)      lcorbi_t3_pos_4_052_06    m/z 74.8523     0   1   2   3E+5   14.5   15  RT (min)      lcorbi_t3_pos_1_036_08    m/z 74.9209     0   2   4E+4   14   14.5  RT (min)      lcorbi_t3_pos_4_101_61    m/z 75.0064     0   2   4   6   8E+2   2.5   3  RT (min)      lcorbi_t3_pos_1_044_25    m/z 75.0440     0   1E+5   15   15.5  RT (min)      lcorbi_t3_pos_2_021_09    m/z 75.0440     0   1   2   3E+4   6.5   7  RT (min)      lcorbi_t3_pos_1_017_51    m/z 75.0440     0   1E+5   11.5   12  RT (min)      lcorbi_t3_pos_4_094_60    m/z 75.0606     0   1   13   13.5  RT (min)      lcorbi_t3_pos_1_027_43    m/z 75.0908     0   1   2   3E+3   1   1.5  RT (min)      lcorbi_t3_pos_3_027_27    m/z 75.0998     0   1E+5   16   18  RT (min)       lcorbi_t3_pos_1_014_03    m/z 75.2850     0   1E+4   10   10.5  RT (min)      lcorbi_t3_pos_4_006_12    m/z 75.3896     0   2   4   6E+4   13.5   14  RT (min)      lcorbi_t3_pos_4_098_61    m/z 75.5013     0   2   4   6   8E+3   6   6.5  RT (min)      lcorbi_t3_pos_4_084_59    m/z 75.6088     0   1E+5   14.5   15  RT (min)      lcorbi_t3_pos_4_082_59    m/z 76.0358     0   1E+4   9   9.5  RT (min)      lcorbi_t3_pos_1_004_24    m/z 76.0393     0   1   2   3E+5   1   1.5  RT (min)       lcorbi_t3_pos_4_096_60    m/z 76.0838     0   1   2E+5   15   15.5  RT (min)      lcorbi_t3_pos_4_099_61    m/z 76.0838     0   1E+5   13   13.5  RT (min)      lcorbi_t3_pos_1_025_52    m/z 76.9733     0   1E+4   7   7.5  RT (min)      lcorbi_t3_pos_3_023_05    m/z 77.0553     0   1   2E+4   9.5   10  RT (min)      lcorbi_t3_pos_1_028_32    m/z 77.0701     0   1   2E+5   1   1.5  RT (min)       lcorbi_t3_pos_4_095_60    m/z 77.2831     0   2   4   6E+3   4.5   5  RT (min)      lcorbi_t3_pos_3_010_44    m/z 77.5964     0   2   4   6E+4   14.5   15  RT (min)      lcorbi_t3_pos_4_107_62    m/z 78.0355     0   2   4E+4   14.5   15  RT (min)      lcorbi_t3_pos_4_100_61    m/z 78.0625     0   2   4   6   8E+4   13.5   14  RT (min)      lcorbi_t3_pos_4_067_57    m/z 78.0960     0   2   4E+4   1   1.5  RT (min)      lcorbi_t3_pos_4_068_57    m/z 78.2894     0   1E+4   5   5.5  RT (min)      lcorbi_t3_pos_2_029_43    m/z 79.0542     0   1E+5   6.5   7  RT (min)      lcorbi_t3_pos_4_090_60    m/z 79.0542     0   1   2   3E+5   12.5   13  RT (min)      lcorbi_t3_pos_1_001_49    m/z 79.0753     0   2   4   6E+5   11.5   12  RT (min)      lcorbi_t3_pos_1_002_01    m/z 79.2751     0   1   2E+4   6.5   7  RT (min)      lcorbi_t3_pos_4_005_33    m/z 79.5748     0   1E+4   10   10.5  RT (min)      lcorbi_t3_pos_1_054_48    m/z 79.7251     0   1E+5   14   14.5  RT (min)      lcorbi_t3_pos_4_089_59    m/z 79.8375     0   1   2E+5   14   14.5  RT (min)      lcorbi_t3_pos_4_088_59    m/z 80.0494     0   1   2E+5   15   15.5  RT (min)      lcorbi_t3_pos_1_020_31    m/z 80.0494     0   1   2   3E+5   12   12.5  RT (min)      lcorbi_t3_pos_1_056_56    m/z 80.7110     0   2   4   6E+4   14.5   15  RT (min)      lcorbi_t3_pos_4_110_62    m/z 80.8407     0   1E+5   15   15.5  RT (min)      lcorbi_t3_pos_4_088_59    m/z 80.9189     0   1E+4   4.5   5  RT (min)      lcorbi_t3_pos_4_107_62    m/z 80.9462     0   2   4E+6   1   1.5  RT (min)      lcorbi_t3_pos_3_039_03    m/z 80.9478     0   1E+4   3.5   4  RT (min)      lcorbi_t3_pos_2_019_13    m/z 80.9478     0   1   2E+4   5   5.5  RT (min)      lcorbi_t3_pos_3_016_21    m/z 80.9478     0   2   4   6   8E+4   11   11.5  RT (min)      lcorbi_t3_pos_3_021_39    m/z 81.0334     0   2   4   6   8E+4   1   1.5  RT (min)       lcorbi_t3_pos_4_083_59    m/z 81.0443     0   2   4E+4   4.5   5  RT (min)      lcorbi_t3_pos_3_004_24    m/z 81.0447     0   2   4   6E+4   0   1  RT (min)       lcorbi_t3_pos_4_084_59    m/z 81.0699     0   2   4   6E+5   6.5   7  RT (min)      lcorbi_t3_pos_2_012_01    m/z 81.1557     0   1E+4   5.5   6  RT (min)      lcorbi_t3_pos_3_053_48    m/z 81.1709     0   1E+5   13.5   14  RT (min)      lcorbi_t3_pos_4_082_59    m/z 81.2812     0   1   2   3E+4   5.5   6  RT (min)      lcorbi_t3_pos_4_104_61    m/z 81.5063     0   1   2E+5   13.5   14  RT (min)      lcorbi_t3_pos_4_071_57    m/z 81.6187     0   1   2   3   4E+4   14   14.5  RT (min)      lcorbi_t3_pos_4_078_58    m/z 82.0263     0   2   4   6   8E+3   1   1.5  RT (min)      lcorbi_t3_pos_1_032_26    m/z 82.0732     0   1E+5   15   15.5  RT (min)      lcorbi_t3_pos_1_027_43    m/z 82.0732     0   1E+5   11   12  RT (min)       lcorbi_t3_pos_1_054_48    m/z 82.1678     0   1E+5   13.5   14  RT (min)      lcorbi_t3_pos_4_083_59    m/z 82.2803     0   1   2E+5   13.5   14  RT (min)      lcorbi_t3_pos_4_089_59    m/z 82.3918     0   2   4   6   8E+4   13.5   14  RT (min)      lcorbi_t3_pos_4_083_59    m/z 82.5043     0   1E+5   14   14.5  RT (min)      lcorbi_t3_pos_4_087_59    m/z 82.5507     0   2   4   6   8E+3   9.5   10  RT (min)      lcorbi_t3_pos_3_023_05    m/z 82.7323     0   1   14   14.5  RT (min)      lcorbi_t3_pos_4_088_59    m/z 82.9449     0   2   4   6   8E+4   15   15.5  RT (min)      lcorbi_t3_pos_4_094_60    m/z 82.9564     0   1   14.5   15  RT (min)      lcorbi_t3_pos_1_011_44    m/z 83.0467     0   2   4E+5   18   18.5  RT (min)      lcorbi_t3_pos_4_091_60    m/z 83.0467     0   1   2E+4   1   1.5  RT (min)      lcorbi_t3_pos_1_046_22    m/z 83.0467     0   2   4   6E+4   6.5   7  RT (min)      lcorbi_t3_pos_1_028_32    m/z 83.0491     0   1   2E+5   13   13.5  RT (min)      lcorbi_t3_pos_1_044_25    m/z 83.0679     0   1   14.5   15  RT (min)      lcorbi_t3_pos_1_020_31    m/z 83.6137     0   1E+5   13   13.5  RT (min)      lcorbi_t3_pos_4_086_59    m/z 83.8256     0   2   4   6E+4   14.5   15  RT (min)      lcorbi_t3_pos_4_086_59    m/z 83.9615     0   1   14   14.5  RT (min)      lcorbi_t3_pos_1_045_12    m/z 84.0444     0   2   4   6   8E+5   6   6.5  RT (min)      lcorbi_t3_pos_1_023_02    m/z 84.0444     0   1   2   3   4E+5   11.5   12  RT (min)      lcorbi_t3_pos_1_003_45    m/z 84.0807     0   1   2   3E+5   16   18  RT (min)       lcorbi_t3_pos_1_011_44    m/z 84.0807     0   1   2   3E+5   12   12.5  RT (min)      lcorbi_t3_pos_1_017_51    m/z 84.0888     0   2   4   6E+5   10.5   11  RT (min)      lcorbi_t3_pos_1_021_18    m/z 84.1855     0   1   2E+5   14   14.5  RT (min)      lcorbi_t3_pos_2_037_27    m/z 84.5851     0   2   4E+4   13   13.5  RT (min)      lcorbi_t3_pos_1_018_35    m/z 84.7253     0   1E+5   13.5   14  RT (min)      lcorbi_t3_pos_4_083_59    m/z 84.7324     0   1E+5   14   14.5  RT (min)      lcorbi_t3_pos_4_075_58    m/z 84.8359     0   1   2   3   4E+4   13   13.5  RT (min)      lcorbi_t3_pos_4_085_59    m/z 84.9472     0   2   4   6   8E+4   13   13.5  RT (min)      lcorbi_t3_pos_4_084_59    m/z 84.9597     0   1   2   3E+5   4.5   5  RT (min)      lcorbi_t3_pos_1_008_36    m/z 84.9597     0   1E+5   6.5   7  RT (min)      lcorbi_t3_pos_2_038_03    m/z 85.0477     0   1   2   3E+4   1   1.5  RT (min)      lcorbi_t3_pos_4_074_58    m/z 85.0581     0   1   2   3E+3   1   1.5  RT (min)       lcorbi_t3_pos_1_002_01    m/z 85.1712     0   1E+5   13.5   14  RT (min)      lcorbi_t3_pos_4_089_59    m/z 85.3986     0   1   2E+5   14   14.5  RT (min)      lcorbi_t3_pos_4_074_58    m/z 85.5102     0   1   14   14.5  RT (min)      lcorbi_t3_pos_4_082_59    m/z 85.6190     0   1   2E+5   13.5   14  RT (min)      lcorbi_t3_pos_4_082_59    m/z 85.6303     0   1E+5   13.5   14  RT (min)      lcorbi_t3_pos_4_091_60    m/z 85.7303     0   1E+5   13.5   14  RT (min)      lcorbi_t3_pos_4_083_59    m/z 85.8428     0   2   4   6   8E+4   14   14.5  RT (min)      lcorbi_t3_pos_4_088_59    m/z 85.9542     0   1   2   3   4E+4   14   14.5  RT (min)      lcorbi_t3_pos_4_089_59    m/z 86.0317     0   1   2   3E+4   1   1.5  RT (min)      lcorbi_t3_pos_1_029_29    m/z 86.0396     0   2   4   6E+4   18   18.5  RT (min)       lcorbi_t3_pos_4_083_59    m/z 86.0503     0   1   2E+4   1   1.5  RT (min)       lcorbi_t3_pos_1_008_36    m/z 86.0600     0   1   2   3E+5   1   2   3  RT (min)       lcorbi_t3_pos_1_054_48    m/z 86.0713     0   1E+4   1   1.5  RT (min)       lcorbi_t3_pos_1_001_49    m/z 86.0978     0   2   4   6E+5   2.5   3  RT (min)      lcorbi_t3_pos_1_016_05    m/z 86.1045     0   1   2   3E+5   10.5   11  RT (min)      lcorbi_t3_pos_1_014_03    m/z 86.3381     0   1E+4   5   5.5  RT (min)      lcorbi_t3_pos_4_014_48    m/z 86.8512     0   1E+5   13.5   14  RT (min)      lcorbi_t3_pos_4_092_60    m/z 87.0553     0   1E+4   1   1.5  RT (min)       lcorbi_t3_pos_1_001_49    m/z 87.0738     0   1   2   3   4E+4   1   1.5  RT (min)      lcorbi_t3_pos_4_093_60    m/z 87.0804     0   1   2E+5   11.5   12  RT (min)      lcorbi_t3_pos_3_017_51    m/z 87.0934     0   1E+4   2.5   3  RT (min)      lcorbi_t3_pos_1_028_32    m/z 87.0998     0   2   4   6E+4   2.5   3  RT (min)       lcorbi_t3_pos_1_034_34    m/z 87.1777     0   1   2E+6   14   14.5  RT (min)      lcorbi_t3_pos_1_020_31    m/z 87.8013     0   2   4   6E+4   7   7.5  RT (min)      lcorbi_t3_pos_4_098_61    m/z 88.0215     0   1   2   3E+3   1   1.5  RT (min)      lcorbi_t3_pos_1_001_49    m/z 88.0216     0   1   2   3   4E+3   8   8.5  RT (min)      lcorbi_t3_pos_1_002_01    m/z 88.0344     0   1   2   3   4E+3   6.5   7  RT (min)      lcorbi_t3_pos_1_001_49    m/z 88.0474     0   1E+4   6.5   7  RT (min)       lcorbi_t3_pos_4_106_62    m/z 88.0529     0   2   4E+3   7   7.5  RT (min)      lcorbi_t3_pos_1_025_52    m/z 88.0578     0   2   4   6   8E+4   1   1.5   2  RT (min)       lcorbi_t3_pos_1_022_20    m/z 88.1120     0   1   2   3   4E+7   15   15.5  RT (min)      lcorbi_t3_pos_2_023_26    m/z 88.1135     0   1   2   3   4E+7   12   13  RT (min)       lcorbi_t3_pos_1_055_33    m/z 88.1136     0   2   4   6E+7   16  RT (min)       lcorbi_t3_pos_4_081_58    m/z 88.2694     0   1E+4   8   8.5  RT (min)      lcorbi_t3_pos_3_034_26    m/z 88.2897     0   1E+5   14   14.5  RT (min)      lcorbi_t3_pos_4_075_58    m/z 88.4010     0   2   4   6E+4   14   14.5  RT (min)      lcorbi_t3_pos_4_075_58    m/z 88.6253     0   1   14   14.5  RT (min)      lcorbi_t3_pos_4_091_60    m/z 88.9468     0   1E+5   13   13.5  RT (min)      lcorbi_t3_pos_4_087_59    m/z 88.9525     0   1   2E+4   0.5   1  RT (min)       lcorbi_t3_pos_1_006_15    m/z 88.9909     0   1E+4   1   1.5  RT (min)      lcorbi_t3_pos_1_009_50    m/z 89.0233     0   2   4E+5   15.5   16  RT (min)      lcorbi_t3_pos_1_046_22    m/z 89.0233     0   1   2   3   4E+5   11.5   12  RT (min)      lcorbi_t3_pos_1_047_41    m/z 89.0418     0   2   4E+4   1   1.5  RT (min)      lcorbi_t3_pos_4_095_60    m/z 89.0596     0   1   2   3E+5   15   15.5  RT (min)      lcorbi_t3_pos_1_006_15    m/z 89.0597     0   1   2E+5   12   12.5  RT (min)      lcorbi_t3_pos_1_026_17    m/z 89.0597     0   1   2   3   4E+4   1   1.5  RT (min)      lcorbi_t3_pos_1_036_08    m/z 89.0901     0   1E+4   1   1.5  RT (min)       lcorbi_t3_pos_1_008_36    m/z 89.0975     0   1   1   1.5  RT (min)       lcorbi_t3_pos_1_002_01    m/z 89.1072     0   1   2   3E+5   18.5   19  RT (min)      lcorbi_t3_pos_4_099_61    m/z 89.1090     0   1   2   3E+5   15.5   16  RT (min)      lcorbi_t3_pos_1_028_32    m/z 89.1091     0   1E+5   12   12.5  RT (min)      lcorbi_t3_pos_1_046_22    m/z 89.1154     0   1   2   3E+5   6.5   7  RT (min)      lcorbi_t3_pos_2_053_28    m/z 89.1154     0   1E+5   4   4.5  RT (min)      lcorbi_t3_pos_4_071_57    m/z 89.1154     0   2   4   6E+4   1   1.5  RT (min)      lcorbi_t3_pos_2_024_15    m/z 89.1183     0   1   2E+5   14.5   15  RT (min)       lcorbi_t3_pos_1_023_02    m/z 89.1853     0   1   2E+5   13.5   14  RT (min)      lcorbi_t3_pos_4_092_60    m/z 89.5500     0   1E+4   9.5   10  RT (min)      lcorbi_t3_pos_4_054_18    m/z 89.7304     0   1E+5   13.5   14  RT (min)      lcorbi_t3_pos_4_076_58    m/z 89.9433     0   2   4E+4   15   15.5  RT (min)      lcorbi_t3_pos_4_068_57    m/z 89.9544     0   1E+5   14   14.5  RT (min)      lcorbi_t3_pos_4_074_58    m/z 89.9636     0   1   2   3   4E+5   0.5   1  RT (min)      lcorbi_t3_pos_1_007_07    m/z 90.0630     0   2   4E+3   2.5   3  RT (min)      lcorbi_t3_pos_1_001_49    m/z 90.0661     0   2   4E+2   1   1.5  RT (min)       lcorbi_t3_pos_3_029_13    m/z 90.1188     0   1E+5   16   18  RT (min)       lcorbi_t3_pos_4_074_58    m/z 90.2972     0   1   2   3E+4   14   14.5  RT (min)      lcorbi_t3_pos_1_034_34    m/z 90.4095     0   1   14.5   15  RT (min)      lcorbi_t3_pos_4_091_60    m/z 90.5150     0   2   4E+4   14   14.5  RT (min)      lcorbi_t3_pos_4_072_57    m/z 90.7312     0   2   4   6   8E+3   1   1.5  RT (min)      lcorbi_t3_pos_3_012_31    m/z 90.9476     0   1   2E+4   18.5   19  RT (min)       lcorbi_t3_pos_4_090_60    m/z 90.9476     0   1   2E+6   0   1  RT (min)       lcorbi_t3_pos_1_006_15    m/z 90.9721     0   1E+4   1   1.5  RT (min)      lcorbi_t3_pos_1_034_34    m/z 90.9766     0   1   2E+4   5.5   6  RT (min)      lcorbi_t3_pos_2_007_02    m/z 90.9766     0   2   4E+4   9   9.5  RT (min)      lcorbi_t3_pos_1_037_19    m/z 91.0240     0   1   2E+3   1   1.5  RT (min)      lcorbi_t3_pos_1_008_36    m/z 91.0513     0   1   2   3E+4   4.5   5  RT (min)      lcorbi_t3_pos_4_099_61    m/z 91.0542     0   2   4   6   8E+4   12   12.5  RT (min)       lcorbi_t3_pos_1_035_28    m/z 91.0542     0   2   4   6   8E+4   15   15.5  RT (min)      lcorbi_t3_pos_4_099_61    m/z 91.0556     0   2   4   6E+6   4.5   5  RT (min)       lcorbi_t3_pos_4_099_61    m/z 91.1132     0   1E+5   2.5   3  RT (min)       lcorbi_t3_pos_4_095_60    m/z 91.1869     0   1E+5   13   13.5   14  RT (min)      lcorbi_t3_pos_1_036_08    m/z 91.9799     0   2   4   6E+4   1   1.5  RT (min)       lcorbi_t3_pos_1_023_02    m/z 92.0826     0   1E+6   14   14.5  RT (min)      lcorbi_t3_pos_2_037_27    m/z 92.0891     0   2   4   6E+4   1   1.5  RT (min)       lcorbi_t3_pos_4_093_60    m/z 92.4711     0   1E+5   14.5   15  RT (min)      lcorbi_t3_pos_4_105_61    m/z 92.7336     0   2   4   6   8E+4   14   14.5  RT (min)      lcorbi_t3_pos_4_074_58    m/z 92.9577     0   2   4   6E+4   14   14.5  RT (min)      lcorbi_t3_pos_4_075_58    m/z 92.9809     0   1   1   1.5  RT (min)      lcorbi_t3_pos_1_045_12    m/z 93.0214     0   2   4   6   8E+4   0.5   1  RT (min)      lcorbi_t3_pos_1_048_47    m/z 93.0609     0   1   2   3   4E+3   4.5   5  RT (min)      lcorbi_t3_pos_1_002_01    m/z 93.1817     0   2   4   6E+4   14   14.5  RT (min)      lcorbi_t3_pos_4_067_57    m/z 93.2908     0   2   4   6E+3   8   8.5  RT (min)      lcorbi_t3_pos_3_023_05    m/z 94.0527     0   2   4E+3   8   8.5  RT (min)      lcorbi_t3_pos_1_049_55    m/z 94.8053     0   1E+4   7.5   8  RT (min)      lcorbi_t3_pos_4_110_62    m/z 94.9267     0   1   2E+4   8.5   9  RT (min)      lcorbi_t3_pos_4_111_62    m/z 95.0491     0   1E+5   13   13.5  RT (min)      lcorbi_t3_pos_4_070_57    m/z 95.0491     0   2   4   6E+4   6.5   7  RT (min)      lcorbi_t3_pos_4_093_60    m/z 95.0603     0   1   2   3   4E+4   0   1  RT (min)       lcorbi_t3_pos_1_001_49    m/z 95.0855     0   2   4   6   8E+4   14   14.5  RT (min)      lcorbi_t3_pos_4_067_57    m/z 96.0356     0   1   5   5.5  RT (min)      lcorbi_t3_pos_1_003_45    m/z 96.2865     0   1E+4   5   5.5  RT (min)      lcorbi_t3_pos_4_014_48    m/z 96.5267     0   1E+5   15.5   16  RT (min)      lcorbi_t3_pos_4_086_59    m/z 96.7506     0   1E+5   15.5   16  RT (min)      lcorbi_t3_pos_4_089_59    m/z 96.8621     0   2   4   6   8E+4   15.5   16  RT (min)      lcorbi_t3_pos_4_084_59    m/z 96.9236     0   1E+6   1   1.5  RT (min)      lcorbi_t3_pos_1_002_01    m/z 96.9746     0   1   2E+4   15.5   16  RT (min)      lcorbi_t3_pos_1_047_41    m/z 96.9951     0   1E+4   0   0.5  RT (min)      lcorbi_t3_pos_1_001_49    m/z 97.0076     0   1   2E+4   18.5   19  RT (min)       lcorbi_t3_pos_4_107_62    m/z 97.0076     0   1   2   3E+4   0.5   1  RT (min)      lcorbi_t3_pos_1_046_22    m/z 97.0648     0   1   2   3   4E+4   1   1.5  RT (min)      lcorbi_t3_pos_1_052_38    m/z 97.0648     0   1   2E+5   13   13.5  RT (min)      lcorbi_t3_pos_1_013_10    m/z 97.0648     0   1E+5   6.5   7  RT (min)      lcorbi_t3_pos_2_044_21    m/z 97.0760     0   1   2E+4   1   2  RT (min)       lcorbi_t3_pos_1_001_49    m/z 97.0861     0   1   15.5   16  RT (min)      lcorbi_t3_pos_1_013_10    m/z 97.2805     0   2   4E+3   8   8.5  RT (min)      lcorbi_t3_pos_2_023_26    m/z 97.3101     0   2   4E+4   15.5   16  RT (min)      lcorbi_t3_pos_4_089_59    m/z 97.8116     0   1E+4   6   6.5  RT (min)      lcorbi_t3_pos_4_106_62    m/z 97.8125     0   1E+4   6   6.5  RT (min)      lcorbi_t3_pos_4_113_62    m/z 97.9661     0   2   4   6   8E+3   0.5   1  RT (min)      lcorbi_t3_pos_4_066_57    m/z 97.9744     0   1   2   3E+4   1   1.5  RT (min)      lcorbi_t3_pos_4_073_57    m/z 98.0039     0   2   4   6E+4   18   18.5  RT (min)       lcorbi_t3_pos_4_076_58    m/z 98.0964     0   1E+4   1.5   2  RT (min)      lcorbi_t3_pos_1_001_49    m/z 98.8280     0   1   2   3E+4   7   7.5  RT (min)      lcorbi_t3_pos_4_111_62    m/z 98.9841     0   1E+5   18   18.5  RT (min)      lcorbi_t3_pos_4_097_60    m/z 99.0032     0   1E+4   18.5   19  RT (min)       lcorbi_t3_pos_4_106_62    m/z 99.0032     0   2   4E+5   0   0.5  RT (min)      lcorbi_t3_pos_1_001_49    m/z 99.0263     0   1E+3   2.5   3  RT (min)      lcorbi_t3_pos_1_001_49    m/z 99.0804     0   1   2E+5   12.5   13  RT (min)      lcorbi_t3_pos_2_003_19    m/z 99.0804     0   1E+5   6.5   7  RT (min)      lcorbi_t3_pos_2_012_01    m/z 99.1933     0   1E+5   15.5   16  RT (min)      lcorbi_t3_pos_4_086_59    m/z 99.4173     0   1   2E+5   15.5   16  RT (min)      lcorbi_t3_pos_4_089_59    m/z 99.5288     0   1E+5   15.5   16  RT (min)      lcorbi_t3_pos_4_089_59    m/z 99.6413     0   1   2E+5   15.5   16  RT (min)      lcorbi_t3_pos_4_083_59    m/z 99.7528     0   1E+5   15.5   16  RT (min)      lcorbi_t3_pos_4_083_59    m/z 99.8652     0   1   2E+5   15.5   16  RT (min)      lcorbi_t3_pos_4_084_59    m/z 99.9634     0   2   4E+3   0.5   1  RT (min)      lcorbi_t3_pos_1_001_49    m/z 99.9767     0   1E+5   15.5   16  RT (min)      lcorbi_t3_pos_4_087_59    m/z 100.0474     0   1E+5   1   1.5  RT (min)       lcorbi_t3_pos_4_084_59    m/z 100.0757     0   1   2   3E+5   12.5   13  RT (min)      lcorbi_t3_pos_1_036_08    m/z 100.0757     0   1E+5   8.5   9  RT (min)      lcorbi_t3_pos_2_032_39    m/z 100.0892     0   1   15.5   16  RT (min)      lcorbi_t3_pos_1_001_49    m/z 100.1121     0   1E+5   14.5   15  RT (min)      lcorbi_t3_pos_4_101_61    m/z 100.2007     0   1E+5   15.5   16  RT (min)      lcorbi_t3_pos_4_087_59    m/z 100.6955     0   1E+5   15   15.5  RT (min)      lcorbi_t3_pos_4_087_59    m/z 100.8066     0   2   4   6   8E+4   15   15.5  RT (min)      lcorbi_t3_pos_4_087_59    m/z 100.8079     0   1E+4   11   11.5  RT (min)      lcorbi_t3_pos_4_007_46    m/z 100.9169     0   1   2   3E+4   1   1.5  RT (min)      lcorbi_t3_pos_1_029_29    m/z 100.9820     0   1   2   3E+4   1   1.5  RT (min)      lcorbi_t3_pos_4_094_60    m/z 100.9989     0   2   4   6E+5   0   0.5  RT (min)      lcorbi_t3_pos_1_001_49    m/z 101.0611     0   1E+5   2.5   3  RT (min)       lcorbi_t3_pos_1_001_49    m/z 101.9762     0   1   2   3   4E+4   0   0.5  RT (min)      lcorbi_t3_pos_1_009_50    m/z 102.0464     0   2   4   6   8E+3   2.5   3  RT (min)      lcorbi_t3_pos_1_008_36    m/z 102.0550     0   1   2E+5   1  RT (min)       lcorbi_t3_pos_1_011_44    m/z 102.0927     0   1   2E+5   1   1.5  RT (min)       lcorbi_t3_pos_1_002_01    m/z 102.0995     0   1   2   3E+4   6.5   7  RT (min)      lcorbi_t3_pos_4_090_60    m/z 102.0995     0   1E+5   12.5   13  RT (min)      lcorbi_t3_pos_1_041_54    m/z 102.0995     0   1   2E+5   15   15.5  RT (min)      lcorbi_t3_pos_1_008_36    m/z 102.1221     0   1E+4   4.5   5  RT (min)      lcorbi_t3_pos_1_010_46    m/z 102.1292     0   1E+8   15   16   17  RT (min)       lcorbi_t3_pos_1_015_11    m/z 102.1292     0   1   2   3   4E+7   10   12  RT (min)       lcorbi_t3_pos_1_044_25    m/z 102.1295     0   2   4   6   8E+6   4.5   5  RT (min)      lcorbi_t3_pos_1_013_10    m/z 102.1339     0   1   2E+3   2.5   3  RT (min)      lcorbi_t3_pos_1_012_14    m/z 102.9702     0   1   2E+5   0.5   1  RT (min)       lcorbi_t3_pos_4_106_62    m/z 102.9702     0   1   2E+5   4   4.5  RT (min)      lcorbi_t3_pos_1_006_15    m/z 103.0542     0   2   4E+3   5   5.5  RT (min)       lcorbi_t3_pos_1_013_10    m/z 103.0542     0   2   4   6E+3   2.5   3  RT (min)       lcorbi_t3_pos_1_012_14    m/z 103.0575     0   1   2E+4   1   1.5  RT (min)       lcorbi_t3_pos_1_002_01    m/z 103.0753     0   1   2E+4   4.5   5  RT (min)      lcorbi_t3_pos_4_082_59    m/z 103.0753     0   1   2E+5   13   13.5  RT (min)      lcorbi_t3_pos_1_015_11    m/z 103.1133     0   1   2   3E+7   15   15.5  RT (min)      lcorbi_t3_pos_1_003_45    m/z 103.1247     0   1   2E+5   12.5   13  RT (min)      lcorbi_t3_pos_1_012_14    m/z 103.1247     0   1   2   3E+4   4.5   5  RT (min)      lcorbi_t3_pos_1_019_16    m/z 103.1338     0   1   2E+5   15   16   17  RT (min)       lcorbi_t3_pos_1_008_36    m/z 103.1338     0   1   2E+4   4.5   5  RT (min)      lcorbi_t3_pos_1_029_29    m/z 103.1338     0   1E+5   12   12.5   13  RT (min)       lcorbi_t3_pos_1_017_51    m/z 103.1339     0   1   2E+4   6.5   7  RT (min)      lcorbi_t3_pos_1_041_54    m/z 103.3071     0   2   4   6E+3   6   6.5  RT (min)      lcorbi_t3_pos_2_053_28    m/z 103.3162     0   2   4   6E+4   13   13.5  RT (min)      lcorbi_t3_pos_2_021_09    m/z 104.0004     0   2   4E+4   0   0.5  RT (min)      lcorbi_t3_pos_1_009_50    m/z 104.0494     0   1   2E+4   4.5   5  RT (min)       lcorbi_t3_pos_4_098_61    m/z 104.0494     0   1E+3   1   1.5  RT (min)      lcorbi_t3_pos_4_082_59    m/z 104.0528     0   1   2   3   4E+3   4.5   5  RT (min)       lcorbi_t3_pos_1_003_45    m/z 104.0576     0   1   2E+3   3   3.5  RT (min)      lcorbi_t3_pos_1_001_49    m/z 104.1150     0   2   4E+5   10.5   11  RT (min)      lcorbi_t3_pos_4_080_58    m/z 104.1343     0   1   2E+5   15   15.5  RT (min)      lcorbi_t3_pos_1_004_24    m/z 104.1344     0   1E+5   12   12.5  RT (min)      lcorbi_t3_pos_1_004_24    m/z 104.1344     0   1E+4   4.5   5  RT (min)      lcorbi_t3_pos_1_001_49    m/z 104.3284     0   2   4   6E+4   14   14.5  RT (min)      lcorbi_t3_pos_1_008_36    m/z 104.9078     0   2   4   6E+3   0.5   1  RT (min)      lcorbi_t3_pos_4_080_58    m/z 104.9633     0   2   4   6E+3   1   1.5  RT (min)      lcorbi_t3_pos_4_088_59    m/z 104.9923     0   1   1   1.5  RT (min)      lcorbi_t3_pos_4_082_59    m/z 105.0032     0   1E+5   4   4.5  RT (min)       lcorbi_t3_pos_1_003_45    m/z 105.0182     0   1   11   11.5  RT (min)      lcorbi_t3_pos_1_001_49    m/z 105.0334     0   1E+5   9   9.5  RT (min)       lcorbi_t3_pos_1_001_49    m/z 105.0334     0   2   4E+4   5   5.5  RT (min)       lcorbi_t3_pos_1_002_01    m/z 105.0650     0   1E+3   1.5   2  RT (min)       lcorbi_t3_pos_1_052_38    m/z 105.0698     0   2   4   6   8E+4   8.5   9  RT (min)      lcorbi_t3_pos_1_036_08    m/z 105.0731     0   2   4   6E+3   1   1.5  RT (min)      lcorbi_t3_pos_1_003_45    m/z 105.0924     0   2   4   6   8E+2   3   3.5  RT (min)      lcorbi_t3_pos_4_078_58    m/z 105.1040     0   2   4   6   8E+4   1   1.5  RT (min)      lcorbi_t3_pos_4_100_61    m/z 105.1103     0   1   11   11.5  RT (min)       lcorbi_t3_pos_1_004_24    m/z 106.0040     0   1E+4   1   1.5  RT (min)      lcorbi_t3_pos_1_054_48    m/z 106.0263     0   2   4   6E+2   1   1.5  RT (min)      lcorbi_t3_pos_3_029_13    m/z 106.0602     0   1   1.5   2  RT (min)      lcorbi_t3_pos_1_052_38    m/z 106.0862     0   2   4   6E+4   6.5   7  RT (min)       lcorbi_t3_pos_1_041_54    m/z 106.0883     0   2   4   6   8E+5   1   1.5  RT (min)      lcorbi_t3_pos_1_044_25    m/z 106.3034     0   1E+4   11   11.5  RT (min)      lcorbi_t3_pos_3_006_38    m/z 107.0016     0   1   2   3E+4   2   3  RT (min)       lcorbi_t3_pos_4_106_62    m/z 107.0491     0   1   2   3E+5   12.5   13   13.5  RT (min)       lcorbi_t3_pos_4_113_62    m/z 107.0603     0   2   4   6   8E+3   1   1.5  RT (min)       lcorbi_t3_pos_1_001_49    m/z 107.0806     0   1E+4   9.5   10  RT (min)       lcorbi_t3_pos_4_080_58    m/z 107.0832     0   1   1   1.5  RT (min)       lcorbi_t3_pos_1_002_01    m/z 107.0896     0   2   4   6   8E+4   1   1.5  RT (min)      lcorbi_t3_pos_1_047_41    m/z 107.1006     0   2   4   6E+3   1   1.5  RT (min)      lcorbi_t3_pos_1_001_49    m/z 107.7281     0   1E+4   1   1.5  RT (min)      lcorbi_t3_pos_4_073_57    m/z 107.9715     0   1   2E+3   1   1.5  RT (min)      lcorbi_t3_pos_1_044_25    m/z 108.0665     0   1E+3   3   3.5  RT (min)      lcorbi_t3_pos_1_013_10    m/z 108.0777     0   1   2   3   4E+3   1.5   2  RT (min)      lcorbi_t3_pos_2_037_27    m/z 108.0904     0   2   4   6E+3   1   1.5  RT (min)       lcorbi_t3_pos_4_090_60    m/z 108.1204     0   1   2   3E+5   1   1.5  RT (min)      lcorbi_t3_pos_1_050_13    m/z 108.9138     0   1   2   3   4E+4   1   1.5  RT (min)      lcorbi_t3_pos_1_007_07    m/z 108.9487     0   1   2E+4   1   1.5  RT (min)      lcorbi_t3_pos_1_036_08    m/z 108.9582     0   1E+4   18.5   19  RT (min)      lcorbi_t3_pos_4_081_58    m/z 108.9582     0   1E+5   0.5   1  RT (min)       lcorbi_t3_pos_1_013_10    m/z 109.0284     0   2   4E+3   1   1.5  RT (min)       lcorbi_t3_pos_1_006_15    m/z 109.0508     0   2   4   6E+2   1.5   2  RT (min)      lcorbi_t3_pos_2_037_27    m/z 109.0522     0   1   2E+3   4.5   5  RT (min)      lcorbi_t3_pos_1_002_01    m/z 109.0648     0   1E+4   1   1.5   2  RT (min)       lcorbi_t3_pos_4_093_60    m/z 109.0648     0   1   2   3   4E+4   5.5   6  RT (min)       lcorbi_t3_pos_2_029_43    m/z 109.0648     0   2   4   6E+4   7.5   8  RT (min)       lcorbi_t3_pos_1_001_49    m/z 109.0648     0   1E+5   15   16  RT (min)       lcorbi_t3_pos_4_080_58    m/z 109.0743     0   1   2   3E+4   2   3  RT (min)       lcorbi_t3_pos_1_013_10    m/z 109.1012     0   1E+5   10.5   11  RT (min)      lcorbi_t3_pos_4_111_62    m/z 110.0349     0   1E+4   1   1.5  RT (min)      lcorbi_t3_pos_4_067_57    m/z 110.0362     0   2   4E+4   5.5   6  RT (min)       lcorbi_t3_pos_4_099_61    m/z 110.0362     0   2   4   6   8E+4   10   10.5  RT (min)       lcorbi_t3_pos_4_074_58    m/z 110.0964     0   1   2   3E+4   1   1.5  RT (min)      lcorbi_t3_pos_1_031_40    m/z 111.0108     0   2   4   6   8E+3   0   0.5  RT (min)      lcorbi_t3_pos_1_030_21    m/z 111.0205     0   1   2E+3   2.5   3  RT (min)      lcorbi_t3_pos_1_048_47    m/z 111.0232     0   2   4   6E+4   1   1.5  RT (min)      lcorbi_t3_pos_1_001_49    m/z 111.0440     0   2   4   6E+4   1   1.5  RT (min)      lcorbi_t3_pos_2_003_19    m/z 111.0440     0   1   2   3E+5   8   9   10  RT (min)       lcorbi_t3_pos_1_047_41    m/z 111.0648     0   2   4   6E+3   2.5   3  RT (min)      lcorbi_t3_pos_1_001_49    m/z 111.1168     0   1   2E+5   5.5   6  RT (min)       lcorbi_t3_pos_4_113_62    m/z 111.7216     0   2   4E+3   1   1.5  RT (min)      lcorbi_t3_pos_1_029_29    m/z 112.0369     0   1   2   3E+3   1   1.5  RT (min)      lcorbi_t3_pos_1_045_12    m/z 112.0505     0   1   2   3E+5   4.5   5  RT (min)      lcorbi_t3_pos_1_028_32    m/z 112.0505     0   1   2   3   4E+5   10.5   11  RT (min)      lcorbi_t3_pos_1_008_36    m/z 112.0869     0   1   2   3   4E+3   1   1.5  RT (min)       lcorbi_t3_pos_4_075_58    m/z 112.1120     0   1E+3   1   1.5  RT (min)       lcorbi_t3_pos_4_095_60    m/z 112.1202     0   1   2E+4   9.5   10  RT (min)       lcorbi_t3_pos_4_091_60    m/z 112.9998     0   1E+5   1   1.5  RT (min)       lcorbi_t3_pos_4_097_60    m/z 113.0189     0   1E+4   1   1.5  RT (min)      lcorbi_t3_pos_1_003_45    m/z 113.0345     0   2   4   6   8E+3   1   1.5  RT (min)      lcorbi_t3_pos_1_003_45    m/z 113.0597     0   1   2E+5   1.5   2  RT (min)      lcorbi_t3_pos_1_002_01    m/z 113.0597     0   1E+6   11   12  RT (min)       lcorbi_t3_pos_1_046_22    m/z 113.0804     0   2   4   6E+3   1   1.5  RT (min)       lcorbi_t3_pos_1_002_01    m/z 113.0961     0   2   4   6E+4   11.5   12  RT (min)      lcorbi_t3_pos_1_046_22    m/z 113.1073     0   1   2E+4   1   1.5  RT (min)      lcorbi_t3_pos_1_003_45    m/z 113.1324     0   1E+5   17.5   18  RT (min)      lcorbi_t3_pos_4_074_58    m/z 113.3356     0   2   4   6   8E+3   5.5   6  RT (min)      lcorbi_t3_pos_4_088_59    m/z 113.9791     0   1   2E+3   1   1.5  RT (min)      lcorbi_t3_pos_1_003_45    m/z 113.9966     0   2   4   6   8E+4   0   0.5  RT (min)      lcorbi_t3_pos_1_009_50    m/z 114.0684     0   1E+5   1   1.5  RT (min)       lcorbi_t3_pos_1_048_47    m/z 114.0913     0   1   2E+5   1.5   2  RT (min)      lcorbi_t3_pos_1_023_02    m/z 114.0913     0   2   4   6   8E+5   8   8.5  RT (min)      lcorbi_t3_pos_1_001_49    m/z 114.0913     0   1E+6   12   12.5  RT (min)      lcorbi_t3_pos_1_001_49    m/z 114.1025     0   1   2   3   4E+3   5   5.5  RT (min)      lcorbi_t3_pos_1_001_49    m/z 114.1026     0   2   4   6E+2   1   1.5  RT (min)       lcorbi_t3_pos_2_040_25    m/z 114.9708     0   1E+4   18.5   19  RT (min)      lcorbi_t3_pos_4_075_58    m/z 114.9708     0   1   2   3E+4   0.5   1  RT (min)      lcorbi_t3_pos_1_037_19    m/z 114.9950     0   2   4   6E+3   0   0.5  RT (min)      lcorbi_t3_pos_1_040_27    m/z 115.0057     0   1E+4   0.5   1  RT (min)      lcorbi_t3_pos_1_022_20    m/z 115.0366     0   1   2   3   4E+3   1   1.5  RT (min)       lcorbi_t3_pos_1_054_48    m/z 115.0470     0   1   1   1.5  RT (min)      lcorbi_t3_pos_1_008_36    m/z 115.0542     0   2   4   6E+4   7   7.5  RT (min)      lcorbi_t3_pos_1_002_01    m/z 115.0542     0   1E+5   16   18  RT (min)       lcorbi_t3_pos_1_004_24    m/z 115.0542     0   2   4   6E+4   10   10.5  RT (min)      lcorbi_t3_pos_1_051_09    m/z 115.0753     0   1   2E+4   1   1.5  RT (min)       lcorbi_t3_pos_4_077_58    m/z 115.0753     0   1E+5   16   16.5  RT (min)      lcorbi_t3_pos_4_067_57    m/z 115.0754     0   2   4   6   8E+4   12   12.5  RT (min)      lcorbi_t3_pos_4_069_57    m/z 115.0880     0   2   4E+4   1   1.5  RT (min)       lcorbi_t3_pos_1_002_01    m/z 115.9700     0   1E+4   18.5   19  RT (min)      lcorbi_t3_pos_4_086_59    m/z 115.9700     0   1   2E+4   0.5   1  RT (min)      lcorbi_t3_pos_1_008_36    m/z 115.9919     0   1   2   3   4E+4   18.5   19  RT (min)       lcorbi_t3_pos_4_077_58    m/z 115.9919     0   1E+6   0.5   1  RT (min)      lcorbi_t3_pos_1_007_07    m/z 115.9919     0   2   4E+4   4.5   5  RT (min)       lcorbi_t3_pos_1_007_07    m/z 116.0441     0   1E+3   1   1.5  RT (min)      lcorbi_t3_pos_1_054_48    m/z 116.0528     0   1E+3   2.5   3  RT (min)      lcorbi_t3_pos_4_083_59    m/z 116.0681     0   1E+4   1   1.5  RT (min)       lcorbi_t3_pos_4_078_58    m/z 116.0832     0   1E+5   13   13.5  RT (min)      lcorbi_t3_pos_1_015_11    m/z 116.0832     0   2   4   6E+4   8   8.5  RT (min)      lcorbi_t3_pos_4_090_60    m/z 116.1070     0   1   2E+4   1   1.5   2  RT (min)       lcorbi_t3_pos_2_005_11    m/z 116.1454     0   1   2E+5   14.5   15  RT (min)      lcorbi_t3_pos_3_013_19    m/z 116.8909     0   1   2E+4   1   1.5  RT (min)      lcorbi_t3_pos_1_029_29    m/z 116.9473     0   1   2   3E+5   0   0.5  RT (min)      lcorbi_t3_pos_1_005_42    m/z 116.9661     0   1   2   3E+5   1   1.5  RT (min)      lcorbi_t3_pos_1_013_10    m/z 116.9661     0   1   2E+5   4.5   5  RT (min)      lcorbi_t3_pos_1_050_13    m/z 116.9859     0   1E+4   4.5   5  RT (min)      lcorbi_t3_pos_1_038_30    m/z 116.9859     0   1E+4   1   1.5  RT (min)      lcorbi_t3_pos_1_047_41    m/z 117.0077     0   1   1   1.5  RT (min)      lcorbi_t3_pos_1_022_20    m/z 117.0223     0   1   2   3E+5   5   5.5  RT (min)      lcorbi_t3_pos_4_107_62    m/z 117.0223     0   1   2E+5   6.5   7  RT (min)      lcorbi_t3_pos_3_044_46    m/z 117.0573     0   2   4   6E+3   4.5   5  RT (min)       lcorbi_t3_pos_4_082_59    m/z 117.0677     0   1E+4   1   1.5  RT (min)       lcorbi_t3_pos_1_003_45    m/z 117.0779     0   1   2   3E+4   1   1.5  RT (min)       lcorbi_t3_pos_4_106_62    m/z 117.0873     0   2   4E+5   1   1.5  RT (min)       lcorbi_t3_pos_4_087_59    m/z 117.0949     0   1E+4   1   1.5  RT (min)      lcorbi_t3_pos_1_056_56    m/z 117.1103     0   1   2E+3   1   1.5  RT (min)       lcorbi_t3_pos_3_029_13    m/z 117.1404     0   2   4   6E+4   15.5   16  RT (min)      lcorbi_t3_pos_3_038_18    m/z 117.1467     0   2   4E+5   11   11.5  RT (min)      lcorbi_t3_pos_1_052_38    m/z 117.1467     0   1E+5   6.5   7  RT (min)      lcorbi_t3_pos_2_054_29    m/z 117.1622     0   1E+4   1   1.5  RT (min)       lcorbi_t3_pos_1_039_23    m/z 117.9586     0   1E+5   0.5   1  RT (min)      lcorbi_t3_pos_1_006_15    m/z 117.9660     0   2   4   6   8E+3   18.5   19  RT (min)      lcorbi_t3_pos_4_075_58    m/z 117.9739     0   1E+4   18.5   19  RT (min)      lcorbi_t3_pos_4_098_61    m/z 117.9740     0   1   2E+4   0.5   1  RT (min)      lcorbi_t3_pos_1_038_30    m/z 117.9923     0   1   2   3   4E+4   0   0.5  RT (min)      lcorbi_t3_pos_1_004_24    m/z 118.0301     0   2   4E+4   18   18.5  RT (min)       lcorbi_t3_pos_4_081_58    m/z 118.0498     0   2   4E+4   1   1.5  RT (min)       lcorbi_t3_pos_1_032_26    m/z 118.0611     0   2   4   6   8E+2   1   1.5  RT (min)       lcorbi_t3_pos_2_016_23    m/z 118.0793     0   1E+3   1   1.5  RT (min)       lcorbi_t3_pos_4_092_60    m/z 118.0881     0   1   2E+7   1   1.5  RT (min)       lcorbi_t3_pos_4_068_57    m/z 118.0943     0   2   4   6E+3   5   5.5  RT (min)       lcorbi_t3_pos_1_002_01    m/z 118.0957     0   1E+4   1   1.5  RT (min)      lcorbi_t3_pos_1_014_03    m/z 118.0958     0   2   4E+3   3   3.5  RT (min)      lcorbi_t3_pos_1_002_01    m/z 118.3495     0   2   4   6E+4   14.5   15  RT (min)      lcorbi_t3_pos_4_103_61    m/z 118.9426     0   1   2   3   4E+6   0   0.5  RT (min)      lcorbi_t3_pos_1_006_15    m/z 118.9630     0   2   4   6   8E+3   18.5   19  RT (min)      lcorbi_t3_pos_4_093_60    m/z 118.9630     0   1   2E+4   0.5   1  RT (min)      lcorbi_t3_pos_1_026_17    m/z 118.9926     0   1   1   1.5  RT (min)      lcorbi_t3_pos_1_054_48    m/z 119.0103     0   1E+3   4.5   5  RT (min)      lcorbi_t3_pos_1_024_04    m/z 119.0161     0   2   4   6   8E+3   1   1.5  RT (min)       lcorbi_t3_pos_4_090_60    m/z 119.0189     0   2   4   6E+4   1   1.5  RT (min)       lcorbi_t3_pos_4_082_59    m/z 119.0189     0   1   2   3   4E+4   4   4.5  RT (min)       lcorbi_t3_pos_4_074_58    m/z 119.0352     0   1   2E+4   4.5   5  RT (min)       lcorbi_t3_pos_4_074_58    m/z 119.0553     0   2   4   6   8E+3   1   1.5  RT (min)      lcorbi_t3_pos_4_073_57    m/z 119.0684     0   1E+3   4.5   5  RT (min)       lcorbi_t3_pos_4_096_60    m/z 119.0717     0   2   4E+3   1   1.5  RT (min)       lcorbi_t3_pos_1_020_31    m/z 119.0729     0   1   2   3E+4   2.5   3  RT (min)       lcorbi_t3_pos_4_091_60    m/z 119.0798     0   1E+5   2.5   3  RT (min)       lcorbi_t3_pos_4_097_60    m/z 119.0832     0   2   4   6   8E+3   1   1.5  RT (min)       lcorbi_t3_pos_4_095_60    m/z 119.0855     0   2   4   6E+4   10.5   11  RT (min)       lcorbi_t3_pos_1_053_39    m/z 119.0880     0   2   4   6E+3   4.5   5  RT (min)      lcorbi_t3_pos_4_113_62    m/z 119.1193     0   1   1   1.5  RT (min)       lcorbi_t3_pos_2_040_25    m/z 119.9877     0   1   2   3E+4   0   0.5  RT (min)      lcorbi_t3_pos_1_005_42    m/z 120.0032     0   1   1   1.5  RT (min)      lcorbi_t3_pos_2_008_16    m/z 120.0197     0   2   4E+3   4   4.5  RT (min)      lcorbi_t3_pos_1_004_24    m/z 120.0386     0   2   4   6   8E+2   1.5   2  RT (min)      lcorbi_t3_pos_2_037_27    m/z 120.0904     0   1E+3   1   1.5  RT (min)       lcorbi_t3_pos_4_091_60    m/z 120.1019     0   1E+6   1   1.5  RT (min)       lcorbi_t3_pos_4_092_60    m/z 120.1100     0   1E+5   15   15.5  RT (min)       lcorbi_t3_pos_1_050_13    m/z 120.9808     0   1   2E+5   4   4.5  RT (min)      lcorbi_t3_pos_4_101_61    m/z 120.9872     0   1   2E+3   1   1.5  RT (min)      lcorbi_t3_pos_4_082_59    m/z 121.0165     0   2   4   6E+3   4   4.5  RT (min)      lcorbi_t3_pos_4_106_62    m/z 121.0396     0   1E+4   1.5   2  RT (min)      lcorbi_t3_pos_4_082_59    m/z 121.0396     0   1E+5   13   13.5  RT (min)       lcorbi_t3_pos_1_021_18    m/z 121.0690     0   1   2E+4   1   1.5  RT (min)       lcorbi_t3_pos_1_007_07    m/z 121.0841     0   2   4   6   8E+3   2.5   3  RT (min)       lcorbi_t3_pos_1_001_49    m/z 121.0848     0   2   4   6   8E+4   1.5   2  RT (min)       lcorbi_t3_pos_4_092_60    m/z 121.0886     0   2   4   6   8E+3   4.5   5  RT (min)      lcorbi_t3_pos_1_008_36    m/z 121.1012     0   1   2   3E+4   1   1.5  RT (min)      lcorbi_t3_pos_4_097_60    m/z 121.1012     0   2   4   6E+4   10.5   11  RT (min)      lcorbi_t3_pos_1_001_49    m/z 121.1223     0   2   4E+7   15   15.5   16  RT (min)       lcorbi_t3_pos_1_010_46    m/z 121.1223     0   1E+6   5.5   6  RT (min)       lcorbi_t3_pos_1_028_32    m/z 121.1223     0   1   2   3   4E+7   12.5   13  RT (min)       lcorbi_t3_pos_1_001_49    m/z 121.1223     0   1E+7   9   10  RT (min)       lcorbi_t3_pos_2_013_33    m/z 121.1246     0   2   4   6E+5   14   14.5  RT (min)      lcorbi_t3_pos_4_106_62    m/z 122.0270     0   1   2   3E+4   1.5   2  RT (min)      lcorbi_t3_pos_1_032_26    m/z 122.0576     0   1E+3   3.5   4  RT (min)      lcorbi_t3_pos_1_001_49    m/z 122.0576     0   1   1   1.5  RT (min)      lcorbi_t3_pos_3_027_27    m/z 122.0811     0   2   4   6E+3   7   7.5  RT (min)      lcorbi_t3_pos_1_002_01    m/z 122.0844     0   1E+4   1.5   2  RT (min)       lcorbi_t3_pos_1_017_51    m/z 122.0964     0   2   4E+4   1.5   2  RT (min)       lcorbi_t3_pos_1_002_01    m/z 122.1284     0   1   2E+5   14.5   15  RT (min)      lcorbi_t3_pos_1_011_44    m/z 122.8569     0   1   2   3   4E+4   14.5   15  RT (min)      lcorbi_t3_pos_4_103_61    m/z 122.9739     0   2   4   6E+3   1   1.5  RT (min)      lcorbi_t3_pos_4_081_58    m/z 123.0295     0   2   4   6   8E+2   1.5   2  RT (min)       lcorbi_t3_pos_4_082_59    m/z 123.0396     0   2   4E+4   9.5   10  RT (min)       lcorbi_t3_pos_4_098_61    m/z 123.0578     0   2   4   6E+3   2.5   3  RT (min)      lcorbi_t3_pos_1_005_42    m/z 123.0916     0   1   2E+5   14.5   15  RT (min)      lcorbi_t3_pos_1_005_42    m/z 123.0997     0   1   2E+4   4.5   5  RT (min)      lcorbi_t3_pos_1_003_45    m/z 123.1168     0   1   2   3E+4   5.5   6  RT (min)      lcorbi_t3_pos_1_002_01    m/z 123.1265     0   1   2E+5   15   15.5  RT (min)      lcorbi_t3_pos_1_035_28    m/z 123.1265     0   1   2E+5   13   13.5  RT (min)      lcorbi_t3_pos_1_011_44    m/z 123.1290     0   1E+5   15   15.5  RT (min)      lcorbi_t3_pos_1_001_49    m/z 124.0146     0   2   4   6   8E+3   0.5   1  RT (min)       lcorbi_t3_pos_1_001_49    m/z 124.0146     0   1E+4   4   4.5  RT (min)       lcorbi_t3_pos_1_001_49    m/z 124.0369     0   1   2   3   4E+3   1   1.5  RT (min)      lcorbi_t3_pos_4_086_59    m/z 124.0506     0   1   2   3E+3   4.5   5  RT (min)       lcorbi_t3_pos_1_004_24    m/z 124.0520     0   2   4   6   8E+5   1.5   2  RT (min)       lcorbi_t3_pos_2_037_27    m/z 124.0586     0   1E+5   2.5   3  RT (min)       lcorbi_t3_pos_4_087_59    m/z 124.0844     0   2   4   6E+3   4.5   5  RT (min)      lcorbi_t3_pos_1_001_49    m/z 124.0853     0   2   4   6E+3   3   3.5  RT (min)      lcorbi_t3_pos_1_013_10    m/z 124.0949     0   2   4   6   8E+4   4.5   5  RT (min)      lcorbi_t3_pos_1_001_49    m/z 124.0950     0   2   4   6E+4   1.5   2  RT (min)      lcorbi_t3_pos_1_003_45    m/z 124.1121     0   1   2E+4   2.5   3  RT (min)      lcorbi_t3_pos_1_001_49    m/z 124.1121     0   1   2E+4   4.5   5  RT (min)      lcorbi_t3_pos_1_014_03    m/z 124.2099     0   1   2E+4   11   11.5  RT (min)      lcorbi_t3_pos_4_006_12    m/z 124.3354     0   2   4   6E+4   11   11.5  RT (min)      lcorbi_t3_pos_1_029_29    m/z 124.7250     0   1E+4   1   1.5  RT (min)      lcorbi_t3_pos_3_012_31    m/z 124.9226     0   1E+5   1   1.5  RT (min)       lcorbi_t3_pos_4_090_60    m/z 124.9375     0   1E+3   1   1.5  RT (min)      lcorbi_t3_pos_2_037_27    m/z 124.9998     0   1E+3   1   1.5  RT (min)       lcorbi_t3_pos_1_004_24    m/z 125.0114     0   1E+4   0.5   1  RT (min)       lcorbi_t3_pos_4_105_61    m/z 125.0114     0   1   2E+4   2.5   3  RT (min)      lcorbi_t3_pos_4_100_61    m/z 125.0114     0   1E+4   4   4.5  RT (min)       lcorbi_t3_pos_1_001_49    m/z 125.0264     0   1   0   0.5  RT (min)      lcorbi_t3_pos_1_001_49    m/z 125.0389     0   1   2   3   4E+3   1   1.5  RT (min)      lcorbi_t3_pos_4_111_62    m/z 125.0709     0   1E+5   0   1  RT (min)       lcorbi_t3_pos_1_032_26    m/z 125.0841     0   1   2E+3   4.5   5  RT (min)       lcorbi_t3_pos_1_013_10    m/z 125.0902     0   1   2   3E+3   1   1.5  RT (min)      lcorbi_t3_pos_1_002_01    m/z 125.0975     0   1   2E+4   3   3.5  RT (min)      lcorbi_t3_pos_1_013_10    m/z 125.1073     0   1   2E+4   4.5   5  RT (min)       lcorbi_t3_pos_1_002_01    m/z 125.1170     0   1E+5   1   1.5  RT (min)       lcorbi_t3_pos_4_093_60    m/z 125.1325     0   1   2E+4   0   0.5  RT (min)      lcorbi_t3_pos_1_004_24    m/z 125.9528     0   1E+4   0.5   1  RT (min)      lcorbi_t3_pos_1_046_22    m/z 126.0219     0   1   2E+5   1   1.5  RT (min)       lcorbi_t3_pos_3_052_47    m/z 126.0526     0   1E+4   1   1.5  RT (min)      lcorbi_t3_pos_4_068_57    m/z 126.0662     0   1   2E+4   1.5   2  RT (min)       lcorbi_t3_pos_4_083_59    m/z 126.0662     0   1   2E+4   4.5   5  RT (min)       lcorbi_t3_pos_1_037_19    m/z 126.0743     0   1E+3   1   1.5  RT (min)      lcorbi_t3_pos_4_078_58    m/z 126.1277     0   1E+4   0.5   1  RT (min)      lcorbi_t3_pos_4_079_58    m/z 126.9244     0   1   2E+4   0.5   1  RT (min)      lcorbi_t3_pos_1_007_07    m/z 126.9981     0   1   2E+4   18   18.5  RT (min)       lcorbi_t3_pos_4_095_60    m/z 126.9982     0   2   4   6E+4   0.5   1  RT (min)       lcorbi_t3_pos_1_001_49    m/z 127.0155     0   1   1.5   2  RT (min)      lcorbi_t3_pos_1_020_31    m/z 127.0687     0   1E+3   2.5   3  RT (min)      lcorbi_t3_pos_1_001_49    m/z 127.0802     0   1E+3   1   1.5  RT (min)       lcorbi_t3_pos_1_001_49    m/z 127.0841     0   2   4E+4   18   18.5  RT (min)       lcorbi_t3_pos_4_098_61    m/z 127.0842     0   1   2E+3   1   1.5  RT (min)       lcorbi_t3_pos_1_003_45    m/z 127.1042     0   1   2E+3   3   3.5  RT (min)      lcorbi_t3_pos_1_001_49    m/z 127.1481     0   1E+5   13.5   14  RT (min)       lcorbi_t3_pos_4_085_59    m/z 128.0108     0   1E+4   1   1.5  RT (min)      lcorbi_t3_pos_1_032_26    m/z 128.0318     0   1E+5   1   1.5  RT (min)      lcorbi_t3_pos_1_031_40    m/z 128.0620     0   1   2   3E+4   6   7  RT (min)       lcorbi_t3_pos_1_001_49    m/z 128.0682     0   1E+5   1   2  RT (min)       lcorbi_t3_pos_1_028_32    m/z 128.0995     0   1   2E+3   5   5.5  RT (min)       lcorbi_t3_pos_1_002_01    m/z 128.9865     0   1E+3   1   1.5  RT (min)      lcorbi_t3_pos_4_080_58    m/z 129.0182     0   1   2E+4   1.5   2  RT (min)      lcorbi_t3_pos_1_029_29    m/z 129.0395     0   2   4   6E+3   0.5   1  RT (min)      lcorbi_t3_pos_3_012_31    m/z 129.0395     0   2   4   6E+4   18.5   19  RT (min)       lcorbi_t3_pos_4_091_60    m/z 129.0443     0   2   4   6E+3   2.5   3  RT (min)      lcorbi_t3_pos_1_020_31    m/z 129.0699     0   1E+4   6   7  RT (min)       lcorbi_t3_pos_1_001_49    m/z 129.0779     0   1   2   3   4E+4   2.5   3  RT (min)       lcorbi_t3_pos_4_092_60    m/z 129.1022     0   1   2   3E+4   1   1.5  RT (min)       lcorbi_t3_pos_1_029_29    m/z 129.9712     0   1E+3   0   0.5  RT (min)      lcorbi_t3_pos_4_082_59    m/z 129.9856     0   2   4   6E+3   1   1.5  RT (min)      lcorbi_t3_pos_2_005_11    m/z 130.0080     0   1   2E+3   0   0.5  RT (min)      lcorbi_t3_pos_4_082_59    m/z 130.0216     0   1   11   11.5  RT (min)      lcorbi_t3_pos_1_004_24    m/z 130.0527     0   1   2   3   4E+4   1   2  RT (min)       lcorbi_t3_pos_1_043_06    m/z 130.0651     0   2   4   6E+4   7  RT (min)       lcorbi_t3_pos_1_019_16    m/z 130.0651     0   1   2E+4   4.5   5  RT (min)       lcorbi_t3_pos_1_001_49    m/z 130.0692     0   1E+3   1   2  RT (min)       lcorbi_t3_pos_1_038_30    m/z 130.0846     0   2   4   6E+4   11   11.5  RT (min)      lcorbi_t3_pos_1_020_31    m/z 130.0890     0   2   4E+3   1   1.5  RT (min)      lcorbi_t3_pos_1_001_49    m/z 130.1151     0   1   1.5   2  RT (min)      lcorbi_t3_pos_1_052_38    m/z 130.1308     0   2   4   6E+4   9.5   10  RT (min)       lcorbi_t3_pos_1_039_23    m/z 130.1559     0   2   4   6   8E+3   4.5   5  RT (min)      lcorbi_t3_pos_4_082_59    m/z 130.1590     0   1   2   3   4E+5   6.5   7  RT (min)      lcorbi_t3_pos_2_053_28    m/z 130.1617     0   2   4E+3   4.5   5  RT (min)      lcorbi_t3_pos_2_022_38    m/z 130.3356     0   1   2E+4   11   11.5  RT (min)      lcorbi_t3_pos_4_067_57    m/z 130.5886     0   2   4E+4   11   11.5  RT (min)      lcorbi_t3_pos_1_023_02    m/z 130.8395     0   1E+4   11   11.5  RT (min)      lcorbi_t3_pos_4_006_12    m/z 130.9818     0   1   2E+4   18.5   19  RT (min)       lcorbi_t3_pos_4_113_62    m/z 130.9818     0   2   4   6   8E+4   0.5   1  RT (min)       lcorbi_t3_pos_1_039_23    m/z 131.0028     0   1   2E+4   1   1.5  RT (min)       lcorbi_t3_pos_1_034_34    m/z 131.0028     0   1   2   3E+4   4   4.5  RT (min)       lcorbi_t3_pos_1_015_11    m/z 131.0339     0   1E+3   1   1.5  RT (min)       lcorbi_t3_pos_4_071_57    m/z 131.0419     0   1E+3   1   1.5  RT (min)       lcorbi_t3_pos_1_027_43    m/z 131.0522     0   1E+4   4.5   5  RT (min)      lcorbi_t3_pos_1_001_49    m/z 131.0685     0   1   2   3   4E+4   7  RT (min)       lcorbi_t3_pos_1_001_49    m/z 131.0686     0   2   4   6E+4   4.5   5  RT (min)      lcorbi_t3_pos_1_001_49    m/z 131.0796     0   1E+4   2   2.5  RT (min)      lcorbi_t3_pos_1_001_49    m/z 131.0896     0   1   2E+4   1   1.5  RT (min)       lcorbi_t3_pos_2_012_01    m/z 131.1040     0   1   1   1.5  RT (min)      lcorbi_t3_pos_4_073_57    m/z 131.1179     0   2   4   6   8E+4   1  RT (min)       lcorbi_t3_pos_1_054_48    m/z 131.1624     0   1E+5   13.5   14  RT (min)      lcorbi_t3_pos_4_067_57    m/z 131.1652     0   1E+4   4.5   5  RT (min)      lcorbi_t3_pos_4_106_62    m/z 131.3434     0   1   2   3E+4   11.5   12  RT (min)      lcorbi_t3_pos_1_029_29    m/z 131.9532     0   2   4   6   8E+4   0.5   1  RT (min)      lcorbi_t3_pos_1_045_12    m/z 131.9743     0   1E+3   0   0.5  RT (min)      lcorbi_t3_pos_1_047_41    m/z 131.9818     0   2   4E+3   1   1.5  RT (min)      lcorbi_t3_pos_1_046_22    m/z 131.9897     0   1E+3   1   1.5  RT (min)      lcorbi_t3_pos_1_045_12    m/z 132.0142     0   1E+4   0.5   1  RT (min)      lcorbi_t3_pos_1_047_41    m/z 132.0444     0   1   2E+3   4.5   5  RT (min)      lcorbi_t3_pos_1_045_12    m/z 132.0541     0   1E+4   1   1.5  RT (min)      lcorbi_t3_pos_1_029_29    m/z 132.0751     0   1E+7   1   1.5  RT (min)       lcorbi_t3_pos_4_079_58    m/z 132.0797     0   1   2E+4   1   1.5  RT (min)      lcorbi_t3_pos_1_030_21    m/z 132.0808     0   1   2E+4   4.5   5  RT (min)       lcorbi_t3_pos_1_001_49    m/z 132.1050     0   1E+4   2.5   3  RT (min)      lcorbi_t3_pos_1_001_49    m/z 132.1657     0   2   4   6E+3   4.5   5  RT (min)      lcorbi_t3_pos_4_090_60    m/z 132.8518     0   1   13.5   14  RT (min)      lcorbi_t3_pos_1_002_01    m/z 132.8895     0   1   1   1.5  RT (min)      lcorbi_t3_pos_1_003_45    m/z 132.9027     0   1   2E+4   0   0.5  RT (min)      lcorbi_t3_pos_1_002_01    m/z 132.9658     0   1   2E+3   0.5   1  RT (min)      lcorbi_t3_pos_1_012_14    m/z 132.9787     0   1   2   3   4E+3   1   1.5  RT (min)      lcorbi_t3_pos_1_003_45    m/z 132.9946     0   2   4   6E+4   2  RT (min)       lcorbi_t3_pos_1_012_14    m/z 133.0172     0   1E+5   18   18.5  RT (min)      lcorbi_t3_pos_4_068_57    m/z 133.0318     0   1E+3   4.5   5  RT (min)       lcorbi_t3_pos_4_077_58    m/z 133.0345     0   1   2   3   4E+3   1.5   2  RT (min)      lcorbi_t3_pos_1_003_45    m/z 133.0522     0   1   2E+3   1.5   2  RT (min)       lcorbi_t3_pos_3_038_18    m/z 133.0648     0   1   2   3   4E+4   5   5.5  RT (min)       lcorbi_t3_pos_1_008_36    m/z 133.0760     0   1   2   3E+4   4.5   5  RT (min)       lcorbi_t3_pos_1_016_05    m/z 133.0859     0   1   2   3E+4   2.5   3  RT (min)       lcorbi_t3_pos_1_028_32    m/z 133.0859     0   1   2E+4   4.5   5  RT (min)       lcorbi_t3_pos_1_003_45    m/z 133.0955     0   1   2E+3   4.5   5  RT (min)      lcorbi_t3_pos_1_004_24    m/z 133.0989     0   1   2   3   4E+3   2.5   3  RT (min)      lcorbi_t3_pos_1_001_49    m/z 133.1012     0   2   4E+4   7   7.5  RT (min)       lcorbi_t3_pos_4_098_61    m/z 133.9908     0   1   2   3E+4   0   0.5  RT (min)      lcorbi_t3_pos_1_034_34    m/z 134.0026     0   1   2   3   4E+5   0   0.5  RT (min)      lcorbi_t3_pos_1_006_15    m/z 134.0189     0   2   4   6   8E+2   1   1.5  RT (min)       lcorbi_t3_pos_3_014_29    m/z 134.0325     0   1E+4   1   1.5  RT (min)       lcorbi_t3_pos_1_054_48    m/z 134.0600     0   2   4   6   8E+4   9   9.5  RT (min)       lcorbi_t3_pos_1_037_19    m/z 134.0633     0   1   2E+5   1   1.5  RT (min)       lcorbi_t3_pos_2_037_27    m/z 134.0634     0   1   2E+4   4.5   5  RT (min)       lcorbi_t3_pos_1_013_10    m/z 134.0907     0   1E+3   1.5   2  RT (min)       lcorbi_t3_pos_2_042_45    m/z 134.0964     0   1   2   3   4E+4   4.5   5  RT (min)       lcorbi_t3_pos_1_002_01    m/z 134.1005     0   1   1   1.5  RT (min)      lcorbi_t3_pos_1_037_19    m/z 134.1062     0   1   2   3E+3   2.5   3  RT (min)      lcorbi_t3_pos_1_010_46    m/z 134.1175     0   1   2E+4   1   1.5  RT (min)      lcorbi_t3_pos_4_090_60    m/z 134.1175     0   1   2   3E+4   4   4.5  RT (min)      lcorbi_t3_pos_1_001_49    m/z 134.1464     0   1   1   1.5  RT (min)       lcorbi_t3_pos_1_054_48    m/z 134.3837     0   1E+4   14   14.5  RT (min)      lcorbi_t3_pos_4_037_26    m/z 134.8865     0   1   2   3E+4   1   1.5  RT (min)      lcorbi_t3_pos_4_071_57    m/z 134.9578     0   1   2E+3   0   0.5  RT (min)      lcorbi_t3_pos_4_101_61    m/z 134.9965     0   2   4E+4   4   4.5  RT (min)       lcorbi_t3_pos_4_090_60    m/z 134.9965     0   2   4   6   8E+4   1   1.5  RT (min)       lcorbi_t3_pos_4_066_57    m/z 135.0028     0   2   4   6E+2   1.5   2  RT (min)      lcorbi_t3_pos_2_016_23    m/z 135.0328     0   1   2E+5   0   0.5   1  RT (min)       lcorbi_t3_pos_4_110_62    m/z 135.0474     0   1   2   3E+3   5  RT (min)       lcorbi_t3_pos_1_043_06    m/z 135.0699     0   1   2   3E+4   1   1.5  RT (min)      lcorbi_t3_pos_3_051_12    m/z 135.0804     0   1   2   3   4E+5   9.5   10  RT (min)       lcorbi_t3_pos_4_098_61    m/z 135.0845     0   1   2   3E+3   1   1.5  RT (min)      lcorbi_t3_pos_3_051_12    m/z 135.0956     0   1   2E+4   1   1.5  RT (min)       lcorbi_t3_pos_1_008_36    m/z 135.1016     0   1E+5   4.5   5  RT (min)       lcorbi_t3_pos_1_002_01    m/z 135.1030     0   1   2   3E+4   1.5   2  RT (min)       lcorbi_t3_pos_2_005_11    m/z 135.1224     0   2   4   6E+3   1   1.5  RT (min)      lcorbi_t3_pos_1_016_05    m/z 135.1379     0   1E+5   13   13.5  RT (min)      lcorbi_t3_pos_4_090_60    m/z 135.9563     0   1   2E+4   0   0.5  RT (min)      lcorbi_t3_pos_1_039_23    m/z 135.9691     0   1E+5   0   0.5   1  RT (min)       lcorbi_t3_pos_1_030_21    m/z 135.9692     0   2   4E+4   4   4.5  RT (min)       lcorbi_t3_pos_1_002_01    m/z 136.0029     0   1E+4   0   0.5  RT (min)      lcorbi_t3_pos_1_046_22    m/z 136.0362     0   2   4   6E+3   0.5   1  RT (min)      lcorbi_t3_pos_1_003_45    m/z 136.0427     0   1E+3   1.5   2  RT (min)      lcorbi_t3_pos_1_004_24    m/z 136.0685     0   1   2E+5   1   1.5  RT (min)      lcorbi_t3_pos_4_084_59    m/z 136.0733     0   2   4E+3   5   5.5  RT (min)      lcorbi_t3_pos_1_051_09    m/z 136.0790     0   1E+3   1   1.5  RT (min)       lcorbi_t3_pos_4_085_59    m/z 136.0838     0   2   4   6   8E+4   15   15.5  RT (min)      lcorbi_t3_pos_4_088_59    m/z 136.0861     0   2   4   6   8E+3   11   11.5  RT (min)      lcorbi_t3_pos_4_106_62    m/z 136.0870     0   1E+3   2.5   3  RT (min)       lcorbi_t3_pos_4_082_59    m/z 136.0968     0   2   4   6   8E+2   1   1.5  RT (min)      lcorbi_t3_pos_4_084_59    m/z 136.1064     0   1   2E+3   1   1.5  RT (min)       lcorbi_t3_pos_1_054_48    m/z 136.1153     0   1   2.5   3  RT (min)      lcorbi_t3_pos_4_082_59    m/z 136.1163     0   2   4   6   8E+3   4.5   5  RT (min)      lcorbi_t3_pos_1_012_14    m/z 136.1459     0   1   1   1.5  RT (min)      lcorbi_t3_pos_1_054_48    m/z 136.9401     0   1   2   3E+4   1   1.5  RT (min)      lcorbi_t3_pos_4_090_60    m/z 136.9531     0   2   4   6E+6   0   0.5  RT (min)       lcorbi_t3_pos_1_006_15    m/z 136.9895     0   1   2E+3   1   1.5  RT (min)      lcorbi_t3_pos_1_044_25    m/z 137.0249     0   1   2   3E+4   7.5   8  RT (min)      lcorbi_t3_pos_1_004_24    m/z 137.0294     0   2   4E+4   1  RT (min)       lcorbi_t3_pos_4_066_57    m/z 137.0295     0   1   2   3E+4   4   4.5  RT (min)       lcorbi_t3_pos_4_090_60    m/z 137.0548     0   1   2E+5   1   1.5  RT (min)       lcorbi_t3_pos_4_090_60    m/z 137.0626     0   1E+5   9.5   10  RT (min)       lcorbi_t3_pos_4_098_61    m/z 137.0652     0   1E+4   4.5   5  RT (min)      lcorbi_t3_pos_1_001_49    m/z 137.0685     0   1   2E+4   1   1.5  RT (min)       lcorbi_t3_pos_4_013_36    m/z 137.0912     0   2   4   6E+2   1.5   2  RT (min)      lcorbi_t3_pos_3_029_13    m/z 137.0961     0   1E+5   14.5   15  RT (min)      lcorbi_t3_pos_1_009_50    m/z 137.0961     0   2   4   6E+4   10   10.5  RT (min)      lcorbi_t3_pos_4_113_62    m/z 137.1091     0   1   2E+4   4.5   5  RT (min)       lcorbi_t3_pos_1_008_36    m/z 137.1106     0   1   1   1.5  RT (min)       lcorbi_t3_pos_1_054_48    m/z 137.1187     0   1   2E+3   2   3  RT (min)       lcorbi_t3_pos_1_010_46    m/z 137.1380     0   1   2E+3   1.5   2  RT (min)      lcorbi_t3_pos_2_050_18    m/z 137.9536     0   1E+5   0   0.5  RT (min)      lcorbi_t3_pos_1_003_45    m/z 137.9695     0   1   2E+4   0   0.5  RT (min)       lcorbi_t3_pos_4_110_62    m/z 137.9983     0   1   2E+4   0   0.5  RT (min)      lcorbi_t3_pos_1_012_14    m/z 138.0259     0   1   1   1.5  RT (min)      lcorbi_t3_pos_4_082_59    m/z 138.0410     0   1   2   3   4E+3   4.5   5  RT (min)      lcorbi_t3_pos_4_090_60    m/z 138.0491     0   1E+4   1.5   2  RT (min)      lcorbi_t3_pos_4_094_60    m/z 138.0549     0   2   4   6E+4   4.5   5  RT (min)       lcorbi_t3_pos_1_001_49    m/z 138.0946     0   1   1.5   2  RT (min)       lcorbi_t3_pos_1_052_38    m/z 138.1187     0   1   4.5   5  RT (min)      lcorbi_t3_pos_1_003_45    m/z 138.1220     0   2   4E+5   1   2   3  RT (min)       lcorbi_t3_pos_1_036_08    m/z 138.1277     0   1E+5   15   15.5  RT (min)      lcorbi_t3_pos_1_056_56    m/z 138.9674     0   1   2   3E+3   0.5   1  RT (min)       lcorbi_t3_pos_4_074_58    m/z 138.9914     0   1   2   3E+4   0.5   1  RT (min)      lcorbi_t3_pos_4_108_62    m/z 138.9914     0   2   4   6E+4   4   4.5  RT (min)      lcorbi_t3_pos_4_083_59    m/z 139.0257     0   2   4   6E+3   0.5   1  RT (min)      lcorbi_t3_pos_4_105_61    m/z 139.0271     0   1E+4   4   4.5  RT (min)      lcorbi_t3_pos_4_066_57    m/z 139.0365     0   1   2.5   3  RT (min)      lcorbi_t3_pos_4_074_58    m/z 139.0421     0   1   0   0.5  RT (min)      lcorbi_t3_pos_4_082_59    m/z 139.0545     0   1   2E+4   18.5   19  RT (min)       lcorbi_t3_pos_4_077_58    m/z 139.0545     0   1   2E+4   1   1.5  RT (min)       lcorbi_t3_pos_1_001_49    m/z 139.0640     0   1E+4   9.5   10  RT (min)      lcorbi_t3_pos_1_001_49    m/z 139.0947     0   1   2   3   4E+3   2.5   3  RT (min)      lcorbi_t3_pos_1_001_49    m/z 139.0953     0   2   4   6   8E+3   4.5   5  RT (min)      lcorbi_t3_pos_1_001_49    m/z 139.1230     0   1E+7   0   0.5  RT (min)      lcorbi_t3_pos_1_001_49    m/z 139.1311     0   1   2   3E+4   4.5   5  RT (min)      lcorbi_t3_pos_4_066_57    m/z 139.1311     0   2   4   6E+3   2.5   3  RT (min)      lcorbi_t3_pos_1_001_49    m/z 140.0131     0   1E+4   18.5   19  RT (min)       lcorbi_t3_pos_4_100_61    m/z 140.0318     0   1E+4   1   1.5  RT (min)      lcorbi_t3_pos_1_028_32    m/z 140.0423     0   1E+5   10   10.5  RT (min)      lcorbi_t3_pos_4_075_58    m/z 140.0625     0   2   4   6   8E+3   1   1.5  RT (min)       lcorbi_t3_pos_4_045_08    m/z 140.0752     0   2   4   6E+3   4.5   5  RT (min)      lcorbi_t3_pos_1_001_49    m/z 140.1182     0   1   2   3E+3   0.5   1  RT (min)      lcorbi_t3_pos_1_001_49    m/z 140.1434     0   1E+4   4.5   5  RT (min)      lcorbi_t3_pos_1_045_12    m/z 140.9971     0   1   2E+4   18.5   19  RT (min)       lcorbi_t3_pos_4_099_61    m/z 141.0138     0   2   4   6E+3   1   1.5  RT (min)      lcorbi_t3_pos_1_003_45    m/z 141.0311     0   1E+4   4.5   5  RT (min)      lcorbi_t3_pos_1_001_49    m/z 141.0401     0   1   2E+5   1.5   2  RT (min)       lcorbi_t3_pos_1_036_08    m/z 141.0502     0   1E+4   18   18.5  RT (min)       lcorbi_t3_pos_4_105_61    m/z 141.0699     0   2   4   6   8E+3   6   6.5  RT (min)      lcorbi_t3_pos_1_001_49    m/z 141.0699     0   1E+5   14   14.5  RT (min)       lcorbi_t3_pos_1_014_03    m/z 141.0886     0   1   2   3E+4   3.5   4  RT (min)       lcorbi_t3_pos_1_020_31    m/z 141.1109     0   1E+3   4.5   5  RT (min)      lcorbi_t3_pos_1_013_10    m/z 141.1135     0   1   2   3E+3   0   0.5  RT (min)       lcorbi_t3_pos_1_001_49    m/z 141.6066     0   1E+4   5.5   6  RT (min)      lcorbi_t3_pos_4_032_40    m/z 141.7219     0   1E+4   1   1.5  RT (min)      lcorbi_t3_pos_3_012_31    m/z 141.9586     0   1   2E+6   0   0.5  RT (min)      lcorbi_t3_pos_1_002_01    m/z 142.0625     0   1   2E+5   1.5   2  RT (min)       lcorbi_t3_pos_2_037_27    m/z 142.0651     0   1   4.5   5  RT (min)       lcorbi_t3_pos_1_004_24    m/z 142.0756     0   1   2E+3   4.5   5  RT (min)      lcorbi_t3_pos_1_002_01    m/z 142.0958     0   1E+4   1   2  RT (min)       lcorbi_t3_pos_1_007_07    m/z 142.1010     0   1E+4   0   1  RT (min)       lcorbi_t3_pos_1_017_51    m/z 142.1092     0   1   2   3   4E+3   1   1.5  RT (min)      lcorbi_t3_pos_4_005_33    m/z 142.1226     0   1E+4   5.5   6  RT (min)       lcorbi_t3_pos_1_002_01    m/z 142.1585     0   1   2E+4   6.5   7  RT (min)      lcorbi_t3_pos_4_091_60    m/z 142.9183     0   1E+4   1   1.5  RT (min)      lcorbi_t3_pos_1_036_08    m/z 142.9385     0   1   2   3E+5   18   18.5  RT (min)      lcorbi_t3_pos_4_089_59    m/z 142.9385     0   1   2E+4   5.5   6  RT (min)      lcorbi_t3_pos_4_019_47    m/z 142.9385     0   1E+5   10.5   11  RT (min)      lcorbi_t3_pos_1_031_40    m/z 142.9386     0   1E+4   3   3.5  RT (min)      lcorbi_t3_pos_4_006_12    m/z 142.9903     0   1   1   1.5  RT (min)      lcorbi_t3_pos_4_006_12    m/z 143.0016     0   1E+5   0.5   1  RT (min)      lcorbi_t3_pos_1_046_22    m/z 143.0315     0   1E+3   1.5   2  RT (min)      lcorbi_t3_pos_1_045_12    m/z 143.0459     0   1E+3   0   0.5  RT (min)      lcorbi_t3_pos_1_053_39    m/z 143.0581     0   1   2   3   4E+3   1   1.5  RT (min)      lcorbi_t3_pos_1_034_34    m/z 143.0679     0   1   2E+3   2.5   3  RT (min)      lcorbi_t3_pos_1_001_49    m/z 143.0694     0   1   2   3E+4   1   1.5  RT (min)       lcorbi_t3_pos_2_020_22    m/z 143.0729     0   1   4.5   5  RT (min)       lcorbi_t3_pos_4_106_62    m/z 143.0945     0   1   0   1  RT (min)       lcorbi_t3_pos_1_002_01    m/z 143.1066     0   1E+4   1   1.5  RT (min)      lcorbi_t3_pos_4_084_59    m/z 143.1259     0   1   2   3E+3   1.5   2  RT (min)      lcorbi_t3_pos_1_036_08    m/z 143.1430     0   2   4   6   8E+3   0   0.5  RT (min)      lcorbi_t3_pos_1_001_49    m/z 143.9634     0   2   4   6   8E+3   0.5   1  RT (min)      lcorbi_t3_pos_1_041_54    m/z 143.9968     0   2   4   6E+3   0.5   1  RT (min)      lcorbi_t3_pos_1_007_07    m/z 144.0444     0   1   4.5   5  RT (min)      lcorbi_t3_pos_1_003_45    m/z 144.0570     0   1   2   3E+3   6   6.5  RT (min)      lcorbi_t3_pos_1_003_45    m/z 144.0688     0   1   2.5   3  RT (min)      lcorbi_t3_pos_4_082_59    m/z 144.0764     0   1   2   3E+3   4.5   5  RT (min)      lcorbi_t3_pos_4_066_57    m/z 144.0848     0   1   2   3E+3   1.5   2  RT (min)      lcorbi_t3_pos_1_052_38    m/z 144.1383     0   1   2E+4   2   3  RT (min)       lcorbi_t3_pos_2_001_49    m/z 144.1383     0   1   2   3E+4   8.5   9  RT (min)       lcorbi_t3_pos_1_004_24    m/z 144.1383     0   2   4E+4   4.5   5  RT (min)       lcorbi_t3_pos_1_002_01    m/z 144.1495     0   1   4.5   5  RT (min)       lcorbi_t3_pos_4_066_57    m/z 144.1772     0   2   4   6E+3   1   1.5  RT (min)      lcorbi_t3_pos_4_110_62    m/z 144.3340     0   1E+4   8   8.5  RT (min)      lcorbi_t3_pos_1_014_03    m/z 144.9474     0   1   2E+4   0   0.5  RT (min)      lcorbi_t3_pos_1_001_49    m/z 144.9921     0   1   2E+3   1   1.5  RT (min)      lcorbi_t3_pos_1_006_15    m/z 144.9975     0   1   2   3E+3   1   1.5  RT (min)      lcorbi_t3_pos_3_038_18    m/z 145.0172     0   1   2E+5   0.5   1  RT (min)      lcorbi_t3_pos_4_092_60    m/z 145.0284     0   1   6.5   7   7.5  RT (min)       lcorbi_t3_pos_1_015_11    m/z 145.0345     0   1E+5   18.5   19  RT (min)      lcorbi_t3_pos_4_112_62    m/z 145.0471     0   1E+4   1.5   2  RT (min)      lcorbi_t3_pos_4_006_12    m/z 145.0625     0   1   2E+3   2.5   3  RT (min)      lcorbi_t3_pos_4_082_59    m/z 145.0679     0   2   4E+4   4.5   5  RT (min)       lcorbi_t3_pos_4_082_59    m/z 145.0688     0   2   4   6E+3   2.5   3  RT (min)      lcorbi_t3_pos_1_003_45    m/z 145.0784     0   1E+3   1.5   2  RT (min)      lcorbi_t3_pos_2_028_24    m/z 145.0841     0   1   2   3   4E+4   4.5   5  RT (min)      lcorbi_t3_pos_1_003_45    m/z 145.0859     0   2   4   6E+4   8   8.5  RT (min)       lcorbi_t3_pos_1_002_01    m/z 145.0989     0   1E+4   1   1.5  RT (min)      lcorbi_t3_pos_3_003_33    m/z 145.1012     0   1E+4   7   7.5  RT (min)       lcorbi_t3_pos_1_001_49    m/z 145.1157     0   1   1   1.5  RT (min)      lcorbi_t3_pos_1_044_25    m/z 145.1223     0   1E+4   10   10.5  RT (min)      lcorbi_t3_pos_1_001_49    m/z 145.1416     0   1   2E+3   2.5   3  RT (min)      lcorbi_t3_pos_2_001_49    m/z 145.1416     0   1   2   3   4E+3   4.5   5  RT (min)      lcorbi_t3_pos_1_001_49    m/z 145.9603     0   1E+4   0.5   1  RT (min)      lcorbi_t3_pos_4_040_21    m/z 146.0205     0   2   4E+4   18.5   19  RT (min)      lcorbi_t3_pos_4_071_57    m/z 146.0205     0   1   2   3E+4   0.5   1  RT (min)       lcorbi_t3_pos_4_075_58    m/z 146.0228     0   1   2E+4   6.5   7  RT (min)      lcorbi_t3_pos_4_074_58    m/z 146.0352     0   1   2E+4   2.5   3  RT (min)       lcorbi_t3_pos_2_022_38    m/z 146.0353     0   2   4   6E+3   0.5   1  RT (min)       lcorbi_t3_pos_3_012_31    m/z 146.0448     0   1   2   3E+3   1   1.5  RT (min)      lcorbi_t3_pos_1_006_15    m/z 146.0529     0   2   4   6E+3   1   1.5  RT (min)       lcorbi_t3_pos_1_054_48    m/z 146.0600     0   1   2   3E+4   6.5   7  RT (min)       lcorbi_t3_pos_4_091_60    m/z 146.0662     0   2   4   6E+3   1   1.5  RT (min)      lcorbi_t3_pos_4_071_57    m/z 146.0697     0   1   2E+3   2.5   3  RT (min)      lcorbi_t3_pos_1_003_45    m/z 146.1061     0   2   4   6E+3   1   1.5  RT (min)      lcorbi_t3_pos_1_001_49    m/z 146.1101     0   2   4E+3   1   1.5  RT (min)      lcorbi_t3_pos_4_012_37    m/z 146.1175     0   1E+4   4.5   5  RT (min)      lcorbi_t3_pos_2_053_28    m/z 146.1347     0   1   2   3   4E+3   0   0.5  RT (min)      lcorbi_t3_pos_1_047_41    m/z 146.1539     0   2   4   6   8E+4   4.5   5  RT (min)       lcorbi_t3_pos_4_066_57    m/z 146.1539     0   1   2   3   4E+3   2.5   3  RT (min)      lcorbi_t3_pos_4_085_59    m/z 146.9443     0   1   2   3   4E+3   0   0.5  RT (min)      lcorbi_t3_pos_1_008_36    m/z 147.0288     0   2   4   6E+3   1   2  RT (min)       lcorbi_t3_pos_1_001_49    m/z 147.0324     0   1   2   3   4E+4   18.5   19  RT (min)       lcorbi_t3_pos_4_084_59    m/z 147.0634     0   1E+4   4.5   5  RT (min)      lcorbi_t3_pos_4_106_62    m/z 147.0764     0   1   2   3E+7   1   1.5  RT (min)       lcorbi_t3_pos_4_083_59    m/z 147.0811     0   2   4   6E+4   1   1.5  RT (min)       lcorbi_t3_pos_1_023_02    m/z 147.0957     0   2   4   6E+3   1   1.5  RT (min)      lcorbi_t3_pos_3_029_13    m/z 147.1016     0   1E+4   4.5   5  RT (min)       lcorbi_t3_pos_1_012_14    m/z 147.1573     0   2   4   6   8E+3   4.5   5  RT (min)      lcorbi_t3_pos_4_066_57    m/z 147.9917     0   1   2E+4   0.5   1  RT (min)       lcorbi_t3_pos_1_045_12    m/z 148.0039     0   1   2   3E+4   1   1.5  RT (min)      lcorbi_t3_pos_3_054_15    m/z 148.0198     0   1   2   3   4E+3   6.5   7  RT (min)      lcorbi_t3_pos_1_004_24    m/z 148.0660     0   1E+4   1   1.5  RT (min)      lcorbi_t3_pos_1_008_36    m/z 148.0950     0   1E+4   4.5   5  RT (min)      lcorbi_t3_pos_1_034_34    m/z 148.1000     0   1   2E+3   1   1.5  RT (min)      lcorbi_t3_pos_1_036_08    m/z 148.1121     0   1E+4   4.5   5  RT (min)      lcorbi_t3_pos_1_023_02    m/z 148.1332     0   1   2   3E+3   1   1.5  RT (min)      lcorbi_t3_pos_1_034_34    m/z 148.9757     0   1E+5   2.5   3  RT (min)      lcorbi_t3_pos_4_082_59    m/z 148.9896     0   1   2   3   4E+3   0   0.5  RT (min)      lcorbi_t3_pos_4_098_61    m/z 149.0101     0   2   4   6   8E+3   0.5   1  RT (min)      lcorbi_t3_pos_1_001_49    m/z 149.0186     0   1   2   3   4E+3   1   1.5  RT (min)       lcorbi_t3_pos_3_010_44    m/z 149.0195     0   1   2   3   4E+5   13   13.5  RT (min)      lcorbi_t3_pos_4_080_58    m/z 149.0427     0   1E+4   4.5   5  RT (min)      lcorbi_t3_pos_4_077_58    m/z 149.0427     0   1E+3   2.5   3  RT (min)      lcorbi_t3_pos_4_106_62    m/z 149.0544     0   2   4   6E+3   1   1.5  RT (min)       lcorbi_t3_pos_4_104_61    m/z 149.0743     0   2   4E+3   4.5   5  RT (min)       lcorbi_t3_pos_1_003_45    m/z 149.0806     0   1E+4   1.5   2  RT (min)       lcorbi_t3_pos_1_003_45    m/z 149.0920     0   2   4E+3   1.5   2  RT (min)      lcorbi_t3_pos_1_002_01    m/z 149.1073     0   1   2E+4   4.5   5  RT (min)       lcorbi_t3_pos_1_001_49    m/z 149.1187     0   2   4   6E+3   2.5   3  RT (min)       lcorbi_t3_pos_1_002_01    m/z 149.1238     0   1   1   1.5  RT (min)      lcorbi_t3_pos_4_103_61    m/z 150.0063     0   1E+4   0.5   1  RT (min)       lcorbi_t3_pos_2_013_33    m/z 150.0200     0   2   4   6E+4   18   18.5  RT (min)       lcorbi_t3_pos_4_075_58    m/z 150.0247     0   1E+4   0.5   1  RT (min)      lcorbi_t3_pos_4_076_58    m/z 150.0385     0   1E+3   1.5   2  RT (min)      lcorbi_t3_pos_1_045_12    m/z 150.0583     0   1   2E+4   1.5   2  RT (min)       lcorbi_t3_pos_4_085_59    m/z 150.0631     0   1   2E+4   6.5   7  RT (min)      lcorbi_t3_pos_1_001_49    m/z 150.0774     0   2   4E+4   1.5   2  RT (min)       lcorbi_t3_pos_4_067_57    m/z 150.0774     0   1   2   3   4E+4   4.5   5  RT (min)       lcorbi_t3_pos_4_106_62    m/z 150.0937     0   1   2E+3   1   2   3  RT (min)       lcorbi_t3_pos_1_002_01    m/z 150.1125     0   1   2E+4   1   1.5  RT (min)       lcorbi_t3_pos_1_002_01    m/z 150.1218     0   1   1   1.5  RT (min)       lcorbi_t3_pos_4_077_58    m/z 150.1277     0   1   2   3   4E+4   4.5   5  RT (min)       lcorbi_t3_pos_1_001_49    m/z 150.6334     0   1   2   3E+4   1   1.5  RT (min)       lcorbi_t3_pos_4_094_60    m/z 150.9001     0   1E+4   1   1.5  RT (min)      lcorbi_t3_pos_1_002_01    m/z 150.9323     0   2   4   6E+4   1   1.5  RT (min)      lcorbi_t3_pos_1_002_01    m/z 150.9688     0   2   4   6E+3   4   4.5  RT (min)      lcorbi_t3_pos_4_074_58    m/z 150.9688     0   2   4   6   8E+3   1   1.5  RT (min)      lcorbi_t3_pos_1_034_34    m/z 150.9900     0   1E+4   0   0.5  RT (min)      lcorbi_t3_pos_1_006_15    m/z 151.0052     0   1E+5   0.5   1  RT (min)      lcorbi_t3_pos_1_006_15    m/z 151.0087     0   2   4E+4   1.5   2   2.5  RT (min)       lcorbi_t3_pos_4_098_61    m/z 151.0087     0   1   2   3E+4   4   4.5  RT (min)       lcorbi_t3_pos_4_075_58    m/z 151.0276     0   1E+5   10   10.5  RT (min)       lcorbi_t3_pos_4_076_58    m/z 151.0300     0   1   2E+5   13   13.5  RT (min)      lcorbi_t3_pos_4_079_58    m/z 151.0324     0   2   4   6E+3   5   5.5  RT (min)      lcorbi_t3_pos_1_006_15    m/z 151.0390     0   2   4   6E+3   5   5.5  RT (min)       lcorbi_t3_pos_1_006_15    m/z 151.0615     0   1E+4   2.5   3  RT (min)       lcorbi_t3_pos_1_001_49    m/z 151.0616     0   1   2   3   4E+3   4.5   5  RT (min)      lcorbi_t3_pos_1_001_49    m/z 151.0705     0   1E+4   9.5   10  RT (min)       lcorbi_t3_pos_1_002_01    m/z 151.0753     0   1   2E+4   5   5.5  RT (min)       lcorbi_t3_pos_1_001_49    m/z 151.0787     0   1   2E+3   2.5   3  RT (min)      lcorbi_t3_pos_1_001_49    m/z 151.0866     0   1E+4   1   1.5  RT (min)       lcorbi_t3_pos_1_002_01    m/z 151.0965     0   2   4E+4   13   13.5  RT (min)      lcorbi_t3_pos_4_007_46    m/z 151.0965     0   2   4E+5   10   11  RT (min)       lcorbi_t3_pos_1_032_26    m/z 151.0999     0   2   4   6E+3   2.5   3  RT (min)      lcorbi_t3_pos_1_001_49    m/z 151.1091     0   1   1.5   2  RT (min)       lcorbi_t3_pos_1_002_01    m/z 151.1230     0   1   2E+4   4.5   5  RT (min)      lcorbi_t3_pos_1_002_01    m/z 152.0014     0   1   2   3E+4   0   0.5  RT (min)      lcorbi_t3_pos_1_001_49    m/z 152.0221     0   2   4   6E+2   1   1.5  RT (min)       lcorbi_t3_pos_3_029_13    m/z 152.0456     0   2   4   6   8E+2   1.5   2  RT (min)      lcorbi_t3_pos_2_020_22    m/z 152.0620     0   1E+5   11.5   12  RT (min)       lcorbi_t3_pos_4_078_58    m/z 152.0986     0   2   4   6E+3   4.5   5  RT (min)      lcorbi_t3_pos_4_066_57    m/z 152.0998     0   1   2   3   4E+4   2.5   3  RT (min)      lcorbi_t3_pos_1_001_49    m/z 152.1102     0   1   1.5   2  RT (min)       lcorbi_t3_pos_2_005_11    m/z 152.1281     0   1   2E+4   4.5   5  RT (min)       lcorbi_t3_pos_1_007_07    m/z 152.1296     0   1E+4   1   1.5  RT (min)       lcorbi_t3_pos_1_054_48    m/z 152.1434     0   2   4   6E+3   4.5   5  RT (min)      lcorbi_t3_pos_4_070_57    m/z 152.1434     0   1E+4   7   7.5  RT (min)      lcorbi_t3_pos_4_083_59    m/z 152.8971     0   1E+4   1   1.5  RT (min)      lcorbi_t3_pos_2_040_25    m/z 152.9974     0   1   2   3   4E+4   0.5   1  RT (min)      lcorbi_t3_pos_1_001_49    m/z 152.9974     0   1   2E+4   18   18.5  RT (min)       lcorbi_t3_pos_4_082_59    m/z 153.0068     0   2   4   6E+3   4   4.5  RT (min)      lcorbi_t3_pos_1_001_49    m/z 153.0069     0   1E+4   1   1.5  RT (min)      lcorbi_t3_pos_1_001_49    m/z 153.0095     0   1   2E+3   1   1.5  RT (min)      lcorbi_t3_pos_1_004_24    m/z 153.0238     0   2   4   6   8E+4   1   1.5  RT (min)      lcorbi_t3_pos_2_024_15    m/z 153.0352     0   1E+3   1   1.5  RT (min)      lcorbi_t3_pos_1_036_08    m/z 153.0407     0   2   4   6E+3   1   1.5   2  RT (min)       lcorbi_t3_pos_1_002_01    m/z 153.0434     0   2   4   6E+4   4.5   5  RT (min)      lcorbi_t3_pos_1_051_09    m/z 153.0522     0   1E+4   1   1.5  RT (min)      lcorbi_t3_pos_1_054_48    m/z 153.0522     0   1   2E+3   2.5   3  RT (min)       lcorbi_t3_pos_4_069_57    m/z 153.0600     0   1E+4   1   1.5   2  RT (min)       lcorbi_t3_pos_4_068_57    m/z 153.0646     0   1   2E+4   4.5   5  RT (min)      lcorbi_t3_pos_1_002_01    m/z 153.0699     0   1E+5   11.5   12  RT (min)       lcorbi_t3_pos_4_075_58    m/z 153.0739     0   2   4   6   8E+3   4.5   5  RT (min)      lcorbi_t3_pos_1_002_01    m/z 153.0807     0   1   2E+4   1   1.5  RT (min)      lcorbi_t3_pos_2_006_47    m/z 153.0903     0   1   2   3E+4   1   1.5  RT (min)       lcorbi_t3_pos_1_054_48    m/z 153.1007     0   2   4   6E+3   1   2   3  RT (min)       lcorbi_t3_pos_4_071_57    m/z 153.1065     0   1   4.5   5  RT (min)      lcorbi_t3_pos_1_015_11    m/z 153.1135     0   1   1.5   2  RT (min)      lcorbi_t3_pos_2_005_11    m/z 153.1410     0   1   2   3E+3   4.5   5  RT (min)      lcorbi_t3_pos_1_001_49    m/z 153.9559     0   1   2   3E+4   0.5   1  RT (min)      lcorbi_t3_pos_1_007_07    m/z 154.0265     0   1   1.5   2  RT (min)       lcorbi_t3_pos_2_037_27    m/z 154.0474     0   1E+3   1   1.5  RT (min)       lcorbi_t3_pos_4_090_60    m/z 154.0587     0   1E+3   1   1.5  RT (min)       lcorbi_t3_pos_3_029_13    m/z 154.0692     0   1   2E+3   1   2   3  RT (min)       lcorbi_t3_pos_1_006_15    m/z 154.0838     0   1E+5   2.5   3  RT (min)       lcorbi_t3_pos_4_006_12    m/z 154.0974     0   1E+5   1   1.5  RT (min)       lcorbi_t3_pos_4_090_60    m/z 154.1143     0   2   4   6E+4   4.5   5  RT (min)       lcorbi_t3_pos_4_106_62    m/z 154.1169     0   1   2.5   3  RT (min)       lcorbi_t3_pos_4_032_40    m/z 154.8804     0   2   4   6E+4   1   1.5  RT (min)       lcorbi_t3_pos_4_078_58    m/z 154.9417     0   1E+5   1   1.5  RT (min)       lcorbi_t3_pos_4_082_59    m/z 154.9637     0   1   2   3E+5   0   0.5  RT (min)      lcorbi_t3_pos_1_008_36    m/z 155.0338     0   2   4   6E+3   1   1.5  RT (min)       lcorbi_t3_pos_2_027_05    m/z 155.0409     0   1E+5   1   1.5  RT (min)       lcorbi_t3_pos_1_034_34    m/z 155.0791     0   1   1   1.5  RT (min)       lcorbi_t3_pos_1_004_24    m/z 155.0872     0   2   4E+3   2.5   3  RT (min)      lcorbi_t3_pos_1_001_49    m/z 155.0902     0   2   4   6E+3   6.5   7  RT (min)       lcorbi_t3_pos_1_002_01    m/z 155.0902     0   1   2   3E+3   4.5   5  RT (min)       lcorbi_t3_pos_1_013_10    m/z 155.0928     0   1   2   3   4E+3   1   1.5  RT (min)       lcorbi_t3_pos_1_029_29    m/z 155.1260     0   1   2   3E+3   2.5   3  RT (min)      lcorbi_t3_pos_1_001_49    m/z 155.1260     0   1E+4   4.5   5  RT (min)      lcorbi_t3_pos_1_002_01    m/z 155.1260     0   2   4   6   8E+3   7   7.5  RT (min)      lcorbi_t3_pos_1_002_01    m/z 155.1543     0   1E+5   4.5   5  RT (min)       lcorbi_t3_pos_4_090_60    m/z 155.9809     0   2   4   6E+3   1   1.5  RT (min)      lcorbi_t3_pos_2_023_26    m/z 156.0421     0   2   4E+3   1   1.5  RT (min)       lcorbi_t3_pos_3_029_13    m/z 156.0478     0   1   2E+4   4.5   5  RT (min)      lcorbi_t3_pos_4_090_60    m/z 156.0573     0   2   4   6E+3   1.5   2  RT (min)      lcorbi_t3_pos_1_020_31    m/z 156.0631     0   1E+5   1   1.5  RT (min)       lcorbi_t3_pos_2_051_35    m/z 156.0736     0   1E+4   6.5   7  RT (min)      lcorbi_t3_pos_1_001_49    m/z 156.0824     0   1E+4   1   1.5  RT (min)      lcorbi_t3_pos_1_034_34    m/z 156.0849     0   1   2E+3   2.5   3  RT (min)      lcorbi_t3_pos_4_076_58    m/z 156.1383     0   1   2   3   4E+4   3  RT (min)       lcorbi_t3_pos_1_016_05    m/z 156.1576     0   2   4   6   8E+3   4.5   5  RT (min)      lcorbi_t3_pos_4_096_60    m/z 156.1747     0   1   2E+4   8.5   9  RT (min)      lcorbi_t3_pos_2_005_11    m/z 156.5926     0   1E+4   1   1.5  RT (min)      lcorbi_t3_pos_4_075_58    m/z 156.8775     0   1   2   3   4E+4   1   1.5  RT (min)      lcorbi_t3_pos_4_078_58    m/z 156.9747     0   2   4   6   8E+4   1   1.5  RT (min)       lcorbi_t3_pos_4_082_59    m/z 156.9975     0   1E+4   0.5   1  RT (min)      lcorbi_t3_pos_2_056_56    m/z 157.0108     0   1   1.5   2  RT (min)      lcorbi_t3_pos_1_036_08    m/z 157.0301     0   1E+3   1   1.5  RT (min)      lcorbi_t3_pos_3_010_44    m/z 157.0486     0   1   2E+5   1.5   2  RT (min)       lcorbi_t3_pos_2_024_15    m/z 157.0536     0   2   4   6   8E+4   18.5   19  RT (min)       lcorbi_t3_pos_4_087_59    m/z 157.0608     0   1   2   3E+3   1   1.5  RT (min)       lcorbi_t3_pos_4_092_60    m/z 157.0648     0   2   4E+3   5.5   6  RT (min)      lcorbi_t3_pos_1_001_49    m/z 157.0760     0   2   4E+3   4.5   5  RT (min)      lcorbi_t3_pos_1_002_01    m/z 157.0835     0   1E+4   4.5   5  RT (min)       lcorbi_t3_pos_1_001_49    m/z 157.1012     0   1   2   3E+4   14   14.5  RT (min)       lcorbi_t3_pos_1_002_01    m/z 157.1084     0   1   2E+4   1   1.5  RT (min)       lcorbi_t3_pos_4_092_60    m/z 157.1416     0   1   2   3E+3   2.5   3  RT (min)      lcorbi_t3_pos_1_001_49    m/z 157.1699     0   1   2E+4   7.5   8  RT (min)       lcorbi_t3_pos_4_066_57    m/z 157.1699     0   2   4E+4   1   1.5  RT (min)       lcorbi_t3_pos_4_086_59    m/z 157.1780     0   1   2E+4   7   7.5  RT (min)      lcorbi_t3_pos_1_021_18    m/z 158.0124     0   1E+3   0.5   1  RT (min)      lcorbi_t3_pos_1_002_01    m/z 158.0165     0   2   4   6   8E+3   11   11.5  RT (min)      lcorbi_t3_pos_4_050_31    m/z 158.0270     0   1   2   3   4E+3   8   8.5  RT (min)      lcorbi_t3_pos_1_001_49    m/z 158.0388     0   2   4   6E+4   18.5   19  RT (min)       lcorbi_t3_pos_4_079_58    m/z 158.0389     0   1   2   3E+5   0   0.5  RT (min)      lcorbi_t3_pos_1_008_36    m/z 158.0575     0   1   1   1.5   2  RT (min)       lcorbi_t3_pos_2_005_11    m/z 158.0600     0   1E+3   4.5   5  RT (min)      lcorbi_t3_pos_4_082_59    m/z 158.0705     0   1   2E+3   1.5   2  RT (min)      lcorbi_t3_pos_2_024_15    m/z 158.0726     0   1   6   6.5  RT (min)      lcorbi_t3_pos_4_096_60    m/z 158.0838     0   2   4   6E+3   4.5   5  RT (min)       lcorbi_t3_pos_4_082_59    m/z 158.1101     0   1   5.5   6  RT (min)      lcorbi_t3_pos_4_097_60    m/z 158.1175     0   1   2E+4   5   5.5  RT (min)       lcorbi_t3_pos_4_096_60    m/z 158.1539     0   1E+5   6.5   7  RT (min)      lcorbi_t3_pos_1_001_49    m/z 158.1540     0   1E+5   1   1.5  RT (min)      lcorbi_t3_pos_1_001_49    m/z 158.1576     0   1E+4   8.5   9  RT (min)      lcorbi_t3_pos_4_082_59    m/z 158.1733     0   1   2   3E+3   1   1.5  RT (min)      lcorbi_t3_pos_1_003_45    m/z 158.1903     0   1   2E+4   8   8.5  RT (min)       lcorbi_t3_pos_1_005_42    m/z 158.9125     0   1E+4   1   1.5  RT (min)       lcorbi_t3_pos_1_016_05    m/z 158.9278     0   1   2E+3   0.5   1  RT (min)      lcorbi_t3_pos_4_091_60    m/z 158.9599     0   1   2E+4   1   1.5  RT (min)      lcorbi_t3_pos_1_034_34    m/z 158.9676     0   2   4E+4   1   1.5  RT (min)       lcorbi_t3_pos_4_069_57    m/z 158.9739     0   2   4E+5   0.5   1  RT (min)      lcorbi_t3_pos_1_014_03    m/z 158.9942     0   1   2   3E+5   0   0.5  RT (min)      lcorbi_t3_pos_1_006_15    m/z 159.0131     0   1   2E+4   18.5   19  RT (min)       lcorbi_t3_pos_4_098_61    m/z 159.0131     0   1E+4   0.5   1  RT (min)       lcorbi_t3_pos_1_007_07    m/z 159.0277     0   1   2E+4   1.5   2  RT (min)       lcorbi_t3_pos_4_073_57    m/z 159.0421     0   2   4E+3   0   0.5  RT (min)      lcorbi_t3_pos_1_020_31    m/z 159.0505     0   1   2   3   4E+3   1   1.5  RT (min)       lcorbi_t3_pos_3_029_13    m/z 159.0692     0   1E+5   18.5   19  RT (min)      lcorbi_t3_pos_4_091_60    m/z 159.0724     0   1   2.5   3  RT (min)       lcorbi_t3_pos_1_002_01    m/z 159.0823     0   1   2   3   4E+3   2.5   3  RT (min)      lcorbi_t3_pos_4_087_59    m/z 159.0837     0   2   4E+3   5   5.5  RT (min)       lcorbi_t3_pos_1_003_45    m/z 159.0961     0   1   2   3E+3   1   1.5  RT (min)       lcorbi_t3_pos_1_002_01    m/z 159.1168     0   1E+5   10.5   11  RT (min)       lcorbi_t3_pos_4_106_62    m/z 159.1209     0   1   2   3   4E+3   1.5   2  RT (min)      lcorbi_t3_pos_1_052_38    m/z 159.1492     0   1   2E+3   2.5   3  RT (min)      lcorbi_t3_pos_1_001_49    m/z 159.1493     0   1E+3   0.5   1  RT (min)      lcorbi_t3_pos_4_085_59    m/z 159.9927     0   1   2   3E+4   0   0.5  RT (min)      lcorbi_t3_pos_1_023_02    m/z 160.0155     0   1   2   3E+4   6.5   7   7.5  RT (min)       lcorbi_t3_pos_1_028_32    m/z 160.0210     0   1   2   3   4E+3   0.5   1  RT (min)      lcorbi_t3_pos_4_092_60    m/z 160.0345     0   1   2E+4   1   1.5  RT (min)       lcorbi_t3_pos_1_036_08    m/z 160.0427     0   1   2E+4   1   1.5  RT (min)      lcorbi_t3_pos_1_036_08    m/z 160.0519     0   1   2E+3   6   6.5  RT (min)      lcorbi_t3_pos_1_001_49    m/z 160.0660     0   1   1   1.5  RT (min)       lcorbi_t3_pos_1_036_08    m/z 160.0757     0   1   2E+4   4.5   5  RT (min)       lcorbi_t3_pos_1_002_01    m/z 160.0771     0   2   4   6E+4   18   18.5  RT (min)      lcorbi_t3_pos_4_090_60    m/z 160.0798     0   2   4   6   8E+3   1   1.5  RT (min)      lcorbi_t3_pos_1_036_08    m/z 160.0854     0   1   2E+3   3   3.5  RT (min)      lcorbi_t3_pos_1_001_49    m/z 160.0950     0   2   4E+4   4.5   5  RT (min)      lcorbi_t3_pos_1_001_49    m/z 160.1040     0   2   4   6E+3   2.5   3  RT (min)      lcorbi_t3_pos_1_008_36    m/z 160.1121     0   1E+4   4.5   5  RT (min)      lcorbi_t3_pos_1_001_49    m/z 160.1177     0   1E+3   1   1.5  RT (min)      lcorbi_t3_pos_3_054_15    m/z 160.1332     0   1E+4   5   5.5  RT (min)       lcorbi_t3_pos_1_001_49    m/z 160.1696     0   1   2E+4   6   6.5  RT (min)       lcorbi_t3_pos_3_026_04    m/z 160.1808     0   1   2   3E+4   1   1.5  RT (min)      lcorbi_t3_pos_4_012_37    m/z 160.9202     0   1   1   1.5  RT (min)      lcorbi_t3_pos_2_040_25    m/z 160.9289     0   2   4   6E+4   1   1.5  RT (min)       lcorbi_t3_pos_4_066_57    m/z 160.9608     0   1   2   3E+3   1   1.5  RT (min)      lcorbi_t3_pos_1_047_41    m/z 160.9683     0   2   4   6   8E+2   1   1.5  RT (min)      lcorbi_t3_pos_1_055_33    m/z 160.9896     0   1   2   3   4E+6   0   0.5  RT (min)      lcorbi_t3_pos_1_007_07    m/z 161.0121     0   1   2E+5   6.5   7  RT (min)      lcorbi_t3_pos_3_009_50    m/z 161.0233     0   1   2   3   4E+3   7   7.5  RT (min)       lcorbi_t3_pos_4_107_62    m/z 161.0242     0   1   1   1.5  RT (min)       lcorbi_t3_pos_1_045_12    m/z 161.0659     0   1   2   3E+4   4.5   5  RT (min)       lcorbi_t3_pos_1_001_49    m/z 161.0659     0   2   4E+4   1   1.5  RT (min)       lcorbi_t3_pos_1_006_15    m/z 161.0709     0   1   2E+4   5.5   6  RT (min)       lcorbi_t3_pos_1_001_49    m/z 161.0709     0   1E+4   9   9.5  RT (min)      lcorbi_t3_pos_1_001_49    m/z 161.0904     0   2   4   6E+3   5   5.5  RT (min)      lcorbi_t3_pos_1_002_01    m/z 161.0961     0   2   4   6   8E+4   15   15.5  RT (min)      lcorbi_t3_pos_4_080_58    m/z 161.0961     0   2   4   6E+4   9   9.5  RT (min)      lcorbi_t3_pos_4_097_60    m/z 161.1285     0   1   2E+3   1   1.5  RT (min)       lcorbi_t3_pos_1_003_45    m/z 161.1841     0   1   2   3E+4   18   18.5  RT (min)       lcorbi_t3_pos_4_111_62    m/z 161.9848     0   1E+4   0.5   1  RT (min)      lcorbi_t3_pos_1_006_15    m/z 161.9929     0   1   2E+5   0   0.5  RT (min)      lcorbi_t3_pos_1_005_42    m/z 162.0074     0   2   4   6   8E+3   0.5   1  RT (min)      lcorbi_t3_pos_1_028_32    m/z 162.0155     0   2   4E+4   18.5   19  RT (min)       lcorbi_t3_pos_4_066_57    m/z 162.0155     0   2   4   6   8E+3   0.5   1  RT (min)      lcorbi_t3_pos_2_041_54    m/z 162.0220     0   1   1.5   2  RT (min)      lcorbi_t3_pos_1_048_47    m/z 162.0501     0   1   2   3E+3   1   1.5  RT (min)      lcorbi_t3_pos_1_030_21    m/z 162.0798     0   1   1   1.5  RT (min)      lcorbi_t3_pos_3_039_03    m/z 162.0925     0   1E+4   1   1.5  RT (min)       lcorbi_t3_pos_3_034_26    m/z 162.0947     0   1   2E+4   5.5   6  RT (min)       lcorbi_t3_pos_4_098_61    m/z 162.1050     0   2   4E+4   1   1.5  RT (min)       lcorbi_t3_pos_1_031_40    m/z 162.1203     0   1   2   3E+4   1   1.5  RT (min)      lcorbi_t3_pos_1_051_09    m/z 162.1277     0   2   4   6E+4   4.5   5  RT (min)       lcorbi_t3_pos_1_014_03    m/z 162.1489     0   1   2   3E+3   2.5   3  RT (min)      lcorbi_t3_pos_1_046_22    m/z 162.1489     0   1E+3   4.5   5  RT (min)      lcorbi_t3_pos_1_004_24    m/z 162.9259     0   1   2E+4   1   1.5  RT (min)      lcorbi_t3_pos_4_066_57    m/z 162.9939     0   1E+3   0.5   1  RT (min)      lcorbi_t3_pos_4_112_62    m/z 163.0026     0   2   4E+3   1   1.5  RT (min)       lcorbi_t3_pos_1_011_44    m/z 163.0221     0   1E+4   1   1.5  RT (min)       lcorbi_t3_pos_1_026_17    m/z 163.0278     0   1   2   3   4E+5   11.5   12  RT (min)      lcorbi_t3_pos_2_054_29    m/z 163.0554     0   1   2.5   3  RT (min)      lcorbi_t3_pos_1_003_45    m/z 163.0616     0   1   2   3E+3   2.5   3  RT (min)       lcorbi_t3_pos_1_003_45    m/z 163.0794     0   1E+4   1.5   2  RT (min)      lcorbi_t3_pos_3_038_18    m/z 163.0899     0   1   2   3E+3   5   5.5  RT (min)      lcorbi_t3_pos_1_001_49    m/z 163.0964     0   2   4   6   8E+3   2.5   3  RT (min)      lcorbi_t3_pos_1_002_01    m/z 163.0978     0   1   2E+3   1   1.5  RT (min)      lcorbi_t3_pos_2_040_25    m/z 163.1117     0   1   2   3E+4   10   10.5  RT (min)       lcorbi_t3_pos_1_006_15    m/z 163.1188     0   1   2   3   4E+4   1   1.5  RT (min)      lcorbi_t3_pos_1_034_34    m/z 163.1230     0   1   2E+4   4.5   5  RT (min)      lcorbi_t3_pos_1_001_49    m/z 163.1328     0   1E+5   1   1.5  RT (min)       lcorbi_t3_pos_4_077_58    m/z 163.1328     0   1   2E+5   8.5   9  RT (min)      lcorbi_t3_pos_1_036_08    m/z 163.1329     0   1E+5   12   12.5  RT (min)      lcorbi_t3_pos_4_068_57    m/z 163.9640     0   1   2E+3   0.5   1  RT (min)      lcorbi_t3_pos_4_081_58    m/z 164.0423     0   1   2E+3   6.5   7  RT (min)      lcorbi_t3_pos_4_075_58    m/z 164.0424     0   1   10   10.5  RT (min)       lcorbi_t3_pos_4_090_60    m/z 164.0541     0   2   4   6E+4   2.5   3  RT (min)      lcorbi_t3_pos_1_046_22    m/z 164.0626     0   1   2   3   4E+3   2.5   3  RT (min)      lcorbi_t3_pos_1_028_32    m/z 164.0672     0   1E+3   1   1.5  RT (min)      lcorbi_t3_pos_1_012_14    m/z 164.0706     0   2   4E+3   4.5   5  RT (min)       lcorbi_t3_pos_4_082_59    m/z 164.0739     0   1E+3   2.5   3  RT (min)      lcorbi_t3_pos_4_095_60    m/z 164.0787     0   1   2   3   4E+4   10   10.5  RT (min)       lcorbi_t3_pos_1_032_26    m/z 164.0794     0   1   2E+3   1   1.5  RT (min)       lcorbi_t3_pos_4_012_37    m/z 164.0832     0   1   2   3E+3   6.5   7  RT (min)      lcorbi_t3_pos_1_001_49    m/z 164.0931     0   1E+5   4.5   5  RT (min)       lcorbi_t3_pos_4_090_60    m/z 164.0931     0   2   4   6E+3   2.5   3  RT (min)      lcorbi_t3_pos_4_086_59    m/z 164.1013     0   2   4   6   8E+2   4.5   5  RT (min)      lcorbi_t3_pos_4_066_57    m/z 164.1167     0   1   2E+3   1   1.5  RT (min)       lcorbi_t3_pos_3_029_13    m/z 164.1281     0   1E+4   1   1.5  RT (min)      lcorbi_t3_pos_1_054_48    m/z 164.9092     0   2   4   6E+3   1   1.5  RT (min)      lcorbi_t3_pos_4_045_08    m/z 164.9205     0   1   2   3E+4   5.5   6  RT (min)      lcorbi_t3_pos_1_051_09    m/z 164.9205     0   1E+4   3   3.5  RT (min)      lcorbi_t3_pos_4_003_39    m/z 164.9205     0   1   2   3   4E+4   6.5   7  RT (min)      lcorbi_t3_pos_4_004_14    m/z 164.9205     0   1   2E+5   11   11.5  RT (min)      lcorbi_t3_pos_1_041_54    m/z 164.9338     0   1   2E+4   1   1.5  RT (min)      lcorbi_t3_pos_1_034_34    m/z 165.0050     0   1E+4   0.5   1  RT (min)      lcorbi_t3_pos_4_076_58    m/z 165.0134     0   1   2   3   4E+4   1   1.5  RT (min)      lcorbi_t3_pos_2_053_28    m/z 165.0319     0   1   2E+4   0.5   1  RT (min)      lcorbi_t3_pos_4_077_58    m/z 165.0426     0   1E+4   0.5   1  RT (min)      lcorbi_t3_pos_4_080_58    m/z 165.0546     0   1E+4   9.5   10  RT (min)      lcorbi_t3_pos_4_083_59    m/z 165.0546     0   1   2   3E+3   2   2.5  RT (min)       lcorbi_t3_pos_4_098_61    m/z 165.0757     0   1   2   3E+3   1   1.5  RT (min)      lcorbi_t3_pos_3_010_44    m/z 165.0772     0   1E+3   2.5   3  RT (min)      lcorbi_t3_pos_4_106_62    m/z 165.0965     0   1   2   3   4E+3   4.5   5  RT (min)      lcorbi_t3_pos_1_024_04    m/z 165.1386     0   1   2   3   4E+6   0   0.5  RT (min)      lcorbi_t3_pos_1_001_49    m/z 165.9876     0   1   1   1.5  RT (min)      lcorbi_t3_pos_4_090_60    m/z 166.0532     0   1E+6   1   2  RT (min)       lcorbi_t3_pos_4_084_59    m/z 166.0783     0   1E+3   1.5   2  RT (min)      lcorbi_t3_pos_2_005_11    m/z 166.0821     0   1   2   3E+3   2.5   3  RT (min)      lcorbi_t3_pos_1_031_40    m/z 166.0884     0   2   4   6   8E+3   3   4  RT (min)       lcorbi_t3_pos_4_080_58    m/z 166.0944     0   2   4   6   8E+4   14.5   15  RT (min)       lcorbi_t3_pos_4_104_61    m/z 166.0959     0   2   4   6E+3   1   1.5  RT (min)      lcorbi_t3_pos_4_077_58    m/z 166.1155     0   2   4E+3   4.5   5  RT (min)       lcorbi_t3_pos_1_028_32    m/z 166.1533     0   2   4   6E+4   5   5.5  RT (min)       lcorbi_t3_pos_4_090_60    m/z 166.1590     0   1   2   3   4E+4   7.5   8  RT (min)       lcorbi_t3_pos_4_098_61    m/z 166.9163     0   1E+4   1   1.5  RT (min)      lcorbi_t3_pos_4_042_09    m/z 166.9342     0   1   2E+4   1   1.5  RT (min)      lcorbi_t3_pos_1_028_32    m/z 166.9487     0   2   4   6E+4   0   0.5  RT (min)      lcorbi_t3_pos_1_007_07    m/z 166.9863     0   2   4   6E+4   4   4.5  RT (min)       lcorbi_t3_pos_4_107_62    m/z 167.0206     0   2   4   6   8E+4   18.5   19  RT (min)      lcorbi_t3_pos_4_098_61    m/z 167.0206     0   1   2   3E+4   1   1.5  RT (min)      lcorbi_t3_pos_4_082_59    m/z 167.0298     0   1   2E+4   10   10.5  RT (min)      lcorbi_t3_pos_4_077_58    m/z 167.0339     0   1   2   3E+5   17.5   18  RT (min)       lcorbi_t3_pos_4_090_60    m/z 167.0372     0   1   1   1.5  RT (min)       lcorbi_t3_pos_4_098_61    m/z 167.0494     0   1   2   3   4E+3   1   1.5  RT (min)      lcorbi_t3_pos_3_001_49    m/z 167.0532     0   1   2E+3   2.5   3  RT (min)      lcorbi_t3_pos_4_106_62    m/z 167.0532     0   2   4   6E+4   4.5   5  RT (min)       lcorbi_t3_pos_4_090_60    m/z 167.0586     0   2   4   6E+4   18   18.5  RT (min)       lcorbi_t3_pos_4_090_60    m/z 167.0756     0   2   4   6E+3   1   1.5  RT (min)      lcorbi_t3_pos_3_027_27    m/z 167.0757     0   2   4E+3   4.5   5  RT (min)       lcorbi_t3_pos_1_002_01    m/z 167.0815     0   2   4   6E+3   1.5   2  RT (min)      lcorbi_t3_pos_4_054_18    m/z 167.0833     0   1   2E+4   2.5   3  RT (min)      lcorbi_t3_pos_1_031_40    m/z 167.0855     0   2   4   6   8E+4   9   9.5  RT (min)       lcorbi_t3_pos_4_066_57    m/z 167.0896     0   1   2   3E+3   3   3.5  RT (min)      lcorbi_t3_pos_1_003_45    m/z 167.1095     0   1   2   3E+3   4.5   5  RT (min)      lcorbi_t3_pos_1_001_49    m/z 167.1179     0   2   4   6E+3   2.5   3  RT (min)       lcorbi_t3_pos_1_004_24    m/z 167.1486     0   1   1   1.5  RT (min)      lcorbi_t3_pos_1_002_01    m/z 168.0429     0   2   4   6E+4   1   1.5  RT (min)      lcorbi_t3_pos_1_036_08    m/z 168.0477     0   1E+3   6   6.5  RT (min)      lcorbi_t3_pos_1_047_41    m/z 168.0540     0   1   2E+3   2.5   3  RT (min)      lcorbi_t3_pos_1_024_04    m/z 168.0541     0   1   2   3E+3   4.5   5  RT (min)      lcorbi_t3_pos_1_001_49    m/z 168.0574     0   1   2E+3   6   6.5  RT (min)      lcorbi_t3_pos_1_001_49    m/z 168.0632     0   1   2   3   4E+3   2.5   3  RT (min)      lcorbi_t3_pos_1_005_42    m/z 168.0655     0   1   2E+4   1   1.5  RT (min)       lcorbi_t3_pos_4_093_60    m/z 168.0840     0   2   4E+3   1   1.5  RT (min)      lcorbi_t3_pos_1_001_49    m/z 168.0848     0   1   2   3E+3   5   5.5  RT (min)      lcorbi_t3_pos_1_001_49    m/z 168.0889     0   1   9   9.5  RT (min)      lcorbi_t3_pos_1_003_45    m/z 168.0905     0   1E+3   2.5   3  RT (min)      lcorbi_t3_pos_1_002_01    m/z 168.0936     0   1E+3   3   4  RT (min)       lcorbi_t3_pos_1_003_45    m/z 168.1051     0   1E+4   1   1.5  RT (min)      lcorbi_t3_pos_1_003_45    m/z 168.1131     0   1E+3   1   1.5  RT (min)       lcorbi_t3_pos_4_093_60    m/z 168.1174     0   1   4.5   5  RT (min)      lcorbi_t3_pos_1_002_01    m/z 168.9195     0   1   2   3   4E+4   0.5   1  RT (min)       lcorbi_t3_pos_1_005_42    m/z 168.9210     0   2   4E+4   18   18.5  RT (min)      lcorbi_t3_pos_4_079_58    m/z 168.9712     0   1   2   3   4E+3   1   1.5  RT (min)      lcorbi_t3_pos_4_079_58    m/z 168.9837     0   2   4   6E+3   0   0.5  RT (min)      lcorbi_t3_pos_1_005_42    m/z 169.0356     0   1   2E+5   1.5   2  RT (min)       lcorbi_t3_pos_1_020_31    m/z 169.0381     0   1E+5   10   10.5  RT (min)       lcorbi_t3_pos_4_074_58    m/z 169.0382     0   1E+5   13   13.5  RT (min)      lcorbi_t3_pos_4_077_58    m/z 169.0451     0   1   2E+4   18   18.5  RT (min)       lcorbi_t3_pos_4_078_58    m/z 169.0471     0   1E+4   2   2.5  RT (min)      lcorbi_t3_pos_4_074_58    m/z 169.0648     0   1   2E+3   5.5   6  RT (min)      lcorbi_t3_pos_1_001_49    m/z 169.0683     0   1E+4   1   1.5  RT (min)       lcorbi_t3_pos_2_023_26    m/z 169.0760     0   2   4   6   8E+4   5   5.5  RT (min)       lcorbi_t3_pos_4_098_61    m/z 169.0860     0   2   4   6E+4   4  RT (min)       lcorbi_t3_pos_1_001_49    m/z 169.1012     0   2   4E+3   7   7.5  RT (min)       lcorbi_t3_pos_1_001_49    m/z 169.1052     0   2   4   6E+3   1   1.5  RT (min)      lcorbi_t3_pos_1_031_40    m/z 169.1053     0   2   4   6E+3   3.5   4  RT (min)      lcorbi_t3_pos_1_001_49    m/z 169.1252     0   2   4   6   8E+3   4.5   5  RT (min)       lcorbi_t3_pos_1_001_49    m/z 169.1264     0   1E+4   2.5   3  RT (min)      lcorbi_t3_pos_1_001_49    m/z 169.1335     0   1E+4   4.5   5  RT (min)      lcorbi_t3_pos_1_001_49    m/z 169.1360     0   1   1   1.5  RT (min)       lcorbi_t3_pos_1_034_34    m/z 169.9858     0   1   1   1.5  RT (min)      lcorbi_t3_pos_1_054_48    m/z 170.0212     0   1E+4   5   5.5  RT (min)      lcorbi_t3_pos_4_074_58    m/z 170.0255     0   2   4   6   8E+3   1   1.5  RT (min)      lcorbi_t3_pos_2_023_26    m/z 170.0327     0   1E+4   1.5   2  RT (min)      lcorbi_t3_pos_1_028_32    m/z 170.0448     0   1E+4   4.5   5  RT (min)      lcorbi_t3_pos_4_066_57    m/z 170.0560     0   2   4   6E+4   9   9.5  RT (min)       lcorbi_t3_pos_4_074_58    m/z 170.0600     0   2   4   6E+4   4.5   5  RT (min)       lcorbi_t3_pos_4_074_58    m/z 170.0617     0   1   2E+3   1   1.5  RT (min)       lcorbi_t3_pos_1_010_46    m/z 170.0768     0   1   2.5   3  RT (min)       lcorbi_t3_pos_4_090_60    m/z 170.0788     0   2   4E+4   1   1.5  RT (min)       lcorbi_t3_pos_1_028_32    m/z 170.0793     0   2   4   6E+4   5   5.5  RT (min)       lcorbi_t3_pos_4_066_57    m/z 170.0996     0   1   2E+4   1   1.5  RT (min)       lcorbi_t3_pos_4_084_59    m/z 170.1036     0   2   4   6E+3   2.5   3  RT (min)      lcorbi_t3_pos_4_094_60    m/z 170.1092     0   2   4E+3   4.5   5  RT (min)      lcorbi_t3_pos_1_002_01    m/z 170.1539     0   1   2E+4   8.5   9  RT (min)       lcorbi_t3_pos_4_018_01    m/z 170.8544     0   1   2   3E+4   1   1.5  RT (min)      lcorbi_t3_pos_1_029_29    m/z 170.9903     0   1E+3   1.5   2  RT (min)      lcorbi_t3_pos_4_104_61    m/z 171.0095     0   1E+4   1   1.5  RT (min)       lcorbi_t3_pos_2_003_19    m/z 171.0279     0   1   2E+5   1.5   2  RT (min)      lcorbi_t3_pos_1_029_29    m/z 171.0447     0   1   2   3   4E+3   4.5   5  RT (min)      lcorbi_t3_pos_4_067_57    m/z 171.0457     0   2   4   6E+2   1   1.5  RT (min)      lcorbi_t3_pos_2_048_36    m/z 171.0530     0   1   0   1  RT (min)       lcorbi_t3_pos_4_090_60    m/z 171.0628     0   1   2E+3   2.5   3  RT (min)      lcorbi_t3_pos_1_003_45    m/z 171.0632     0   1E+4   4.5   5  RT (min)      lcorbi_t3_pos_1_001_49    m/z 171.0747     0   2   4   6E+3   1   2  RT (min)       lcorbi_t3_pos_4_090_60    m/z 171.0839     0   1   2E+3   2.5   3  RT (min)      lcorbi_t3_pos_1_001_49    m/z 171.0851     0   1   2   3E+3   4.5   5  RT (min)      lcorbi_t3_pos_1_002_01    m/z 171.0958     0   1E+3   1   1.5  RT (min)      lcorbi_t3_pos_2_008_16    m/z 171.1030     0   2   4   6E+3   2.5   3  RT (min)       lcorbi_t3_pos_4_111_62    m/z 171.1057     0   1E+3   1   1.5  RT (min)       lcorbi_t3_pos_2_037_27    m/z 171.1166     0   1   2   3   4E+3   2.5   3  RT (min)      lcorbi_t3_pos_1_001_49    m/z 171.1168     0   2   4E+4   14   14.5  RT (min)       lcorbi_t3_pos_1_002_01    m/z 171.1209     0   2   4   6   8E+3   5.5   6  RT (min)       lcorbi_t3_pos_1_005_42    m/z 171.1379     0   1E+5   12   12.5  RT (min)       lcorbi_t3_pos_1_001_49    m/z 171.9610     0   1   2E+3   1   1.5  RT (min)      lcorbi_t3_pos_1_054_48    m/z 172.0007     0   1E+5   1   1.5  RT (min)       lcorbi_t3_pos_2_005_11    m/z 172.0281     0   1   2.5   3  RT (min)      lcorbi_t3_pos_4_082_59    m/z 172.0385     0   1E+4   1   1.5  RT (min)      lcorbi_t3_pos_1_043_06    m/z 172.0541     0   2   4   6E+4   1   1.5  RT (min)       lcorbi_t3_pos_1_036_08    m/z 172.0676     0   1E+3   2.5   3  RT (min)       lcorbi_t3_pos_1_002_01    m/z 172.0757     0   2   4   6E+3   5   5.5  RT (min)       lcorbi_t3_pos_1_001_49    m/z 172.0797     0   1E+3   2.5   3  RT (min)      lcorbi_t3_pos_1_002_01    m/z 172.0925     0   1   1.5   2  RT (min)      lcorbi_t3_pos_2_020_22    m/z 172.0950     0   1   2   3E+4   9   9.5  RT (min)      lcorbi_t3_pos_1_001_49    m/z 172.1081     0   1E+4   1   1.5  RT (min)      lcorbi_t3_pos_4_071_57    m/z 172.1193     0   1   1.5   2  RT (min)      lcorbi_t3_pos_1_036_08    m/z 172.1332     0   1   2   3   4E+4   15   15.5  RT (min)      lcorbi_t3_pos_4_039_35    m/z 172.1696     0   1   2   3E+4   5   5.5  RT (min)       lcorbi_t3_pos_1_001_49    m/z 172.2059     0   2   4   6   8E+4   8   8.5  RT (min)       lcorbi_t3_pos_3_003_33    m/z 173.0057     0   1   2   3E+3   4.5   5  RT (min)      lcorbi_t3_pos_4_090_60    m/z 173.0057     0   1E+3   2.5   3  RT (min)      lcorbi_t3_pos_4_090_60    m/z 173.0288     0   2   4E+3   1   1.5  RT (min)      lcorbi_t3_pos_2_005_11    m/z 173.0422     0   1   2E+3   1   1.5  RT (min)      lcorbi_t3_pos_3_029_13    m/z 173.0485     0   2   4   6   8E+4   18   18.5  RT (min)       lcorbi_t3_pos_4_066_57    m/z 173.0668     0   1   2E+5   1   1.5  RT (min)       lcorbi_t3_pos_4_090_60    m/z 173.0709     0   1   2   3   4E+3   4.5   5  RT (min)      lcorbi_t3_pos_1_001_49    m/z 173.0784     0   1E+4   1   1.5  RT (min)      lcorbi_t3_pos_1_003_45    m/z 173.0921     0   1E+4   1   1.5  RT (min)       lcorbi_t3_pos_1_002_01    m/z 173.1097     0   1   3   3.5  RT (min)      lcorbi_t3_pos_1_013_10    m/z 173.1325     0   1E+4   10.5   11  RT (min)       lcorbi_t3_pos_2_002_06    m/z 173.1365     0   2   4   6   8E+3   7.5   8  RT (min)      lcorbi_t3_pos_1_001_49    m/z 173.1365     0   1   2   3   4E+4   5.5   6  RT (min)       lcorbi_t3_pos_4_098_61    m/z 173.1648     0   1E+4   8.5   9  RT (min)      lcorbi_t3_pos_4_027_07    m/z 173.1729     0   1   2   3   4E+4   9.5   10  RT (min)       lcorbi_t3_pos_1_023_02    m/z 173.2093     0   1E+4   8   8.5  RT (min)      lcorbi_t3_pos_4_027_07    m/z 173.4124     0   1   2E+5   1   1.5  RT (min)      lcorbi_t3_pos_1_028_32    m/z 174.0138     0   1   2E+4   1   1.5  RT (min)      lcorbi_t3_pos_2_023_26    m/z 174.0315     0   1E+4   10.5   11  RT (min)      lcorbi_t3_pos_4_108_62    m/z 174.0338     0   1   2E+4   1   1.5  RT (min)       lcorbi_t3_pos_1_008_36    m/z 174.0542     0   2   4   6E+3   4.5   5  RT (min)       lcorbi_t3_pos_4_111_62    m/z 174.0611     0   2   4   6E+3   0.5   1  RT (min)      lcorbi_t3_pos_4_091_60    m/z 174.0675     0   2   4E+4   6   6.5  RT (min)       lcorbi_t3_pos_1_014_03    m/z 174.0761     0   1   2E+4   2.5   3  RT (min)       lcorbi_t3_pos_1_001_49    m/z 174.0818     0   2   4E+3   1   2   3  RT (min)       lcorbi_t3_pos_1_001_49    m/z 174.0842     0   1   2E+4   6.5   7  RT (min)      lcorbi_t3_pos_1_001_49    m/z 174.0932     0   1   2E+3   1   1.5  RT (min)       lcorbi_t3_pos_2_003_19    m/z 174.1050     0   1   2E+3   5   5.5  RT (min)      lcorbi_t3_pos_1_001_49    m/z 174.1140     0   1   2E+5   1   1.5  RT (min)       lcorbi_t3_pos_4_093_60    m/z 174.1277     0   1E+4   1   1.5  RT (min)       lcorbi_t3_pos_4_035_17    m/z 174.1488     0   1E+4   1.5   2  RT (min)       lcorbi_t3_pos_2_003_19    m/z 174.1792     0   1   2   3   4E+4   7   7.5  RT (min)      lcorbi_t3_pos_1_003_45    m/z 174.1852     0   2   4   6   8E+4   4.5   5  RT (min)      lcorbi_t3_pos_4_075_58    m/z 174.1894     0   2   4E+4   7   7.5  RT (min)       lcorbi_t3_pos_1_043_06    m/z 174.1941     0   1E+4   7   7.5  RT (min)      lcorbi_t3_pos_1_023_02    m/z 174.8705     0   2   4E+3   1   1.5  RT (min)      lcorbi_t3_pos_4_054_18    m/z 174.8865     0   1   1   1.5  RT (min)       lcorbi_t3_pos_1_029_29    m/z 174.8960     0   1   1   1.5  RT (min)       lcorbi_t3_pos_1_029_29    m/z 174.9505     0   2   4   6   8E+3   1   1.5  RT (min)      lcorbi_t3_pos_1_054_48    m/z 174.9725     0   1   2E+4   0   0.5  RT (min)      lcorbi_t3_pos_1_006_15    m/z 174.9853     0   1E+5   1   1.5  RT (min)       lcorbi_t3_pos_4_083_59    m/z 175.0017     0   1   2E+3   1.5   2  RT (min)      lcorbi_t3_pos_1_001_49    m/z 175.0155     0   1   2E+4   9   9.5  RT (min)      lcorbi_t3_pos_4_074_58    m/z 175.0237     0   1E+3   1.5   2  RT (min)      lcorbi_t3_pos_2_005_11    m/z 175.0263     0   1   2E+5   0   0.5   1  RT (min)       lcorbi_t3_pos_1_007_07    m/z 175.0397     0   1E+3   2.5   3  RT (min)      lcorbi_t3_pos_4_069_57    m/z 175.0451     0   1E+4   0.5   1  RT (min)       lcorbi_t3_pos_4_111_62    m/z 175.0478     0   1   2.5   3  RT (min)       lcorbi_t3_pos_3_019_17    m/z 175.0530     0   1   2   3   4E+4   1   1.5  RT (min)      lcorbi_t3_pos_2_005_11    m/z 175.0642     0   1E+5   13.5   14  RT (min)      lcorbi_t3_pos_2_051_35    m/z 175.0642     0   1   2E+5   7.5   8  RT (min)      lcorbi_t3_pos_4_046_44    m/z 175.0642     0   1   2E+5   11   11.5  RT (min)      lcorbi_t3_pos_1_011_44    m/z 175.0753     0   2   4   6   8E+3   6   6.5  RT (min)       lcorbi_t3_pos_4_105_61    m/z 175.0794     0   1   2E+3   2.5   3  RT (min)      lcorbi_t3_pos_1_003_45    m/z 175.0866     0   1   2E+3   5   5.5  RT (min)       lcorbi_t3_pos_4_068_57    m/z 175.1083     0   1   2E+5   5   5.5  RT (min)       lcorbi_t3_pos_1_036_08    m/z 175.1113     0   2   4E+3   1   1.5  RT (min)       lcorbi_t3_pos_1_053_39    m/z 175.1164     0   1   2.5   3   3.5  RT (min)       lcorbi_t3_pos_4_090_60    m/z 175.1329     0   2   4   6E+3   5   5.5  RT (min)       lcorbi_t3_pos_2_013_33    m/z 175.1440     0   1   2   3   4E+3   1   1.5  RT (min)      lcorbi_t3_pos_2_030_14    m/z 175.1481     0   1   2   3E+3   7   7.5  RT (min)       lcorbi_t3_pos_4_097_60    m/z 175.1823     0   1   2E+4   7   7.5  RT (min)       lcorbi_t3_pos_3_026_04    m/z 175.1886     0   1   2   3E+4   8.5   9  RT (min)      lcorbi_t3_pos_3_044_46    m/z 175.1886     0   1E+4   4.5   5  RT (min)      lcorbi_t3_pos_4_077_58    m/z 175.8936     0   2   4   6E+4   13.5   14  RT (min)      lcorbi_t3_pos_4_072_57    m/z 175.9414     0   1E+4   1   1.5  RT (min)      lcorbi_t3_pos_4_075_58    m/z 175.9876     0   1   2   3E+4   0   0.5  RT (min)      lcorbi_t3_pos_1_009_50    m/z 176.0141     0   1   2E+5   1   1.5  RT (min)       lcorbi_t3_pos_4_097_60    m/z 176.0294     0   1E+4   1   1.5  RT (min)      lcorbi_t3_pos_1_003_45    m/z 176.0423     0   2   4   6E+3   6.5   7   7.5  RT (min)       lcorbi_t3_pos_1_001_49    m/z 176.0494     0   2   4E+4   0   0.5  RT (min)      lcorbi_t3_pos_1_007_07    m/z 176.0541     0   1   2   3E+3   4.5   5  RT (min)      lcorbi_t3_pos_4_056_56    m/z 176.0658     0   1   2E+3   2.5   3  RT (min)      lcorbi_t3_pos_1_001_49    m/z 176.0698     0   2   4   6E+3   1   1.5  RT (min)       lcorbi_t3_pos_4_020_41    m/z 176.0740     0   2   4   6E+3   4.5   5  RT (min)      lcorbi_t3_pos_1_001_49    m/z 176.0740     0   1   2   3   4E+3   2.5   3  RT (min)      lcorbi_t3_pos_1_001_49    m/z 176.0961     0   1E+4   1.5   2  RT (min)      lcorbi_t3_pos_1_030_21    m/z 176.1362     0   1   5   5.5  RT (min)      lcorbi_t3_pos_1_001_49    m/z 176.1645     0   1E+3   4   4.5  RT (min)      lcorbi_t3_pos_1_041_54    m/z 176.1919     0   1   2   3   4E+4   7   7.5  RT (min)       lcorbi_t3_pos_4_013_36    m/z 176.8677     0   1E+4   1   1.5  RT (min)      lcorbi_t3_pos_4_012_37    m/z 176.9383     0   2   4E+3   0.5   1  RT (min)       lcorbi_t3_pos_4_088_59    m/z 176.9739     0   1E+4   1   1.5  RT (min)      lcorbi_t3_pos_1_031_40    m/z 176.9845     0   1E+6   0.5   1  RT (min)      lcorbi_t3_pos_1_014_03    m/z 177.0083     0   2   4   6E+4   0   1  RT (min)       lcorbi_t3_pos_1_005_42    m/z 177.0083     0   1   2   3E+4   4   4.5  RT (min)      lcorbi_t3_pos_4_102_61    m/z 177.0244     0   1E+5   1   1.5  RT (min)      lcorbi_t3_pos_1_028_32    m/z 177.0338     0   1   2E+4   0.5   1  RT (min)      lcorbi_t3_pos_1_008_36    m/z 177.0338     0   1   2   3E+4   18.5   19  RT (min)       lcorbi_t3_pos_4_082_59    m/z 177.0434     0   1   2E+4   0.5   1  RT (min)      lcorbi_t3_pos_1_011_44    m/z 177.0434     0   2   4   6   8E+4   18   18.5  RT (min)       lcorbi_t3_pos_4_074_58    m/z 177.0700     0   1E+4   9   9.5  RT (min)      lcorbi_t3_pos_4_083_59    m/z 177.0758     0   1   2E+3   1   1.5  RT (min)      lcorbi_t3_pos_3_027_27    m/z 177.0798     0   1   2   3E+5   13   13.5  RT (min)      lcorbi_t3_pos_2_004_42    m/z 177.0888     0   2   4   6   8E+3   1.5   2  RT (min)       lcorbi_t3_pos_1_017_51    m/z 177.0909     0   1E+4   6.5   7  RT (min)       lcorbi_t3_pos_1_042_37    m/z 177.1063     0   1   2E+5   1   1.5  RT (min)       lcorbi_t3_pos_4_082_59    m/z 177.1121     0   1   2   3   4E+3   4.5   5  RT (min)      lcorbi_t3_pos_4_004_14    m/z 177.1193     0   1E+4   1   1.5  RT (min)      lcorbi_t3_pos_1_032_26    m/z 177.1239     0   1   2   3E+3   1   1.5  RT (min)       lcorbi_t3_pos_4_006_12    m/z 177.1274     0   1   2   3E+4   12.5   13  RT (min)       lcorbi_t3_pos_1_013_10    m/z 177.1360     0   1   1.5   2  RT (min)      lcorbi_t3_pos_2_005_11    m/z 177.1638     0   2   4   6E+4   14   14.5  RT (min)       lcorbi_t3_pos_4_108_62    m/z 177.9564     0   1   2E+4   0   0.5  RT (min)      lcorbi_t3_pos_1_006_15    m/z 177.9923     0   1E+4   0.5   1  RT (min)      lcorbi_t3_pos_1_027_43    m/z 178.0009     0   1   2   3   4E+4   0   0.5  RT (min)      lcorbi_t3_pos_1_003_45    m/z 178.0032     0   1   2   3E+4   0.5   1  RT (min)       lcorbi_t3_pos_1_007_07    m/z 178.0161     0   1E+5   0.5   1  RT (min)      lcorbi_t3_pos_1_007_07    m/z 178.0474     0   1E+3   2.5   3  RT (min)      lcorbi_t3_pos_1_006_15    m/z 178.0498     0   2   4   6E+4   18   18.5  RT (min)       lcorbi_t3_pos_4_106_62    m/z 178.0580     0   1   2E+4   8.5   9  RT (min)       lcorbi_t3_pos_1_024_04    m/z 178.0587     0   1   2   3E+5   1   1.5  RT (min)       lcorbi_t3_pos_1_031_40    m/z 178.0681     0   1   1   1.5  RT (min)       lcorbi_t3_pos_3_012_31    m/z 178.0724     0   1   2   3E+3   4.5   5  RT (min)      lcorbi_t3_pos_1_034_34    m/z 178.0862     0   1E+4   1   1.5   2  RT (min)       lcorbi_t3_pos_2_005_11    m/z 178.0896     0   1E+4   4.5   5  RT (min)       lcorbi_t3_pos_1_002_01    m/z 178.0943     0   2   4E+3   6.5   7  RT (min)      lcorbi_t3_pos_4_111_62    m/z 178.0983     0   1   2   3E+3   1.5   2  RT (min)      lcorbi_t3_pos_4_084_59    m/z 178.0993     0   2   4   6E+3   7.5   8  RT (min)      lcorbi_t3_pos_4_075_58    m/z 178.1056     0   1E+5   2.5   3  RT (min)      lcorbi_t3_pos_4_073_57    m/z 178.1120     0   1   2E+3   5   5.5  RT (min)      lcorbi_t3_pos_4_066_57    m/z 178.1283     0   1   2E+3   1   1.5  RT (min)      lcorbi_t3_pos_1_034_34    m/z 178.1339     0   2   4E+3   4.5   5  RT (min)      lcorbi_t3_pos_1_002_01    m/z 178.9394     0   1E+4   1   1.5  RT (min)      lcorbi_t3_pos_4_042_09    m/z 178.9849     0   2   4   6E+4   0   0.5  RT (min)      lcorbi_t3_pos_1_005_42    m/z 179.0001     0   1E+5   18.5   19  RT (min)      lcorbi_t3_pos_4_082_59    m/z 179.0001     0   1   2E+5   4.5   5  RT (min)      lcorbi_t3_pos_1_002_01    m/z 179.0082     0   1   2E+4   0   0.5  RT (min)      lcorbi_t3_pos_1_009_50    m/z 179.0193     0   1   2   3   4E+3   1   1.5  RT (min)      lcorbi_t3_pos_1_008_36    m/z 179.0226     0   1E+5   18.5   19  RT (min)      lcorbi_t3_pos_4_066_57    m/z 179.0485     0   1   1.5   2  RT (min)      lcorbi_t3_pos_1_006_15    m/z 179.0554     0   2   4   6E+4   1   1.5  RT (min)       lcorbi_t3_pos_4_094_60    m/z 179.0570     0   2   4   6   8E+4   18   18.5  RT (min)      lcorbi_t3_pos_4_078_58    m/z 179.0637     0   1   2   3E+4   7   7.5  RT (min)      lcorbi_t3_pos_4_090_60    m/z 179.0759     0   1   2   3   4E+4   18   18.5  RT (min)      lcorbi_t3_pos_4_082_59    m/z 179.0764     0   1E+4   1   1.5  RT (min)       lcorbi_t3_pos_1_036_08    m/z 179.0811     0   1   2   3   4E+4   9   9.5  RT (min)       lcorbi_t3_pos_4_077_58    m/z 179.0903     0   1E+3   1   1.5  RT (min)       lcorbi_t3_pos_4_090_60    m/z 179.0915     0   1E+4   4   5  RT (min)       lcorbi_t3_pos_1_001_49    m/z 179.1089     0   1E+3   2.5   3  RT (min)      lcorbi_t3_pos_4_091_60    m/z 179.1107     0   2   4   6E+3   1   1.5  RT (min)      lcorbi_t3_pos_1_016_05    m/z 179.1202     0   2   4   6E+2   5   5.5  RT (min)      lcorbi_t3_pos_4_038_38    m/z 179.1234     0   1E+3   1   1.5  RT (min)      lcorbi_t3_pos_1_016_05    m/z 179.1324     0   1   1   1.5  RT (min)       lcorbi_t3_pos_1_031_40    m/z 179.9936     0   1E+4   6.5   7  RT (min)      lcorbi_t3_pos_4_082_59    m/z 179.9963     0   1   2E+4   0   0.5  RT (min)      lcorbi_t3_pos_1_006_15    m/z 180.0034     0   2   4   6   8E+5   0   0.5  RT (min)      lcorbi_t3_pos_1_006_15    m/z 180.0165     0   2   4   6E+4   0   0.5  RT (min)      lcorbi_t3_pos_1_003_45    m/z 180.0524     0   1E+4   1   1.5  RT (min)      lcorbi_t3_pos_1_016_05    m/z 180.0631     0   1   2E+3   3   3.5  RT (min)       lcorbi_t3_pos_4_081_58    m/z 180.0637     0   1   2E+4   1   1.5  RT (min)      lcorbi_t3_pos_3_026_04    m/z 180.0655     0   1   2E+4   5   5.5  RT (min)       lcorbi_t3_pos_2_006_47    m/z 180.0736     0   1   5   5.5  RT (min)       lcorbi_t3_pos_4_073_57    m/z 180.0821     0   2   4   6   8E+3   1   1.5  RT (min)       lcorbi_t3_pos_4_080_58    m/z 180.1019     0   1   2   3E+4   4.5   5  RT (min)       lcorbi_t3_pos_1_028_32    m/z 180.1050     0   1E+3   1   1.5  RT (min)       lcorbi_t3_pos_1_010_46    m/z 180.1099     0   1E+4   8   8.5  RT (min)      lcorbi_t3_pos_1_002_01    m/z 180.1155     0   1   2   3   4E+3   1.5   2  RT (min)       lcorbi_t3_pos_2_003_19    m/z 180.1243     0   2   4   6E+3   1   1.5   2  RT (min)       lcorbi_t3_pos_2_041_54    m/z 180.1311     0   2   4   6E+3   4.5   5  RT (min)      lcorbi_t3_pos_1_031_40    m/z 180.1383     0   1   2   3E+4   6.5   7  RT (min)      lcorbi_t3_pos_1_023_02    m/z 180.1551     0   2   4E+3   1   1.5  RT (min)      lcorbi_t3_pos_1_001_49    m/z 180.8944     0   1   2   3E+3   1   1.5  RT (min)      lcorbi_t3_pos_1_004_24    m/z 180.8944     0   2   4   6E+4   12   12.5  RT (min)      lcorbi_t3_pos_1_043_06    m/z 180.9803     0   1   2E+4   0   0.5  RT (min)      lcorbi_t3_pos_1_006_15    m/z 180.9958     0   1   1.5   2  RT (min)      lcorbi_t3_pos_4_110_62    m/z 181.0005     0   1   2   3   4E+3   0   0.5  RT (min)      lcorbi_t3_pos_4_069_57    m/z 181.0043     0   2   4   6E+3   0   0.5  RT (min)      lcorbi_t3_pos_1_034_34    m/z 181.0144     0   2   4   6E+3   1   1.5  RT (min)       lcorbi_t3_pos_4_078_58    m/z 181.0285     0   1E+4   18   18.5  RT (min)       lcorbi_t3_pos_4_074_58    m/z 181.0383     0   1   2E+5   5   5.5  RT (min)       lcorbi_t3_pos_4_098_61    m/z 181.0383     0   1   2E+5   6.5   7  RT (min)      lcorbi_t3_pos_4_005_33    m/z 181.0495     0   1E+4   10   10.5  RT (min)       lcorbi_t3_pos_4_100_61    m/z 181.0529     0   1   2   3E+3   1   1.5  RT (min)       lcorbi_t3_pos_1_003_45    m/z 181.0806     0   1E+4   1   1.5  RT (min)      lcorbi_t3_pos_2_008_16    m/z 181.0922     0   1E+4   9   10  RT (min)       lcorbi_t3_pos_1_029_29    m/z 181.0970     0   1   2E+4   4.5   5  RT (min)       lcorbi_t3_pos_1_014_03    m/z 181.1014     0   2   4E+4   0.5   1  RT (min)      lcorbi_t3_pos_1_003_45    m/z 181.1014     0   1   2   3   4E+4   18.5   19  RT (min)       lcorbi_t3_pos_4_090_60    m/z 181.1052     0   1   2   3E+3   4.5   5  RT (min)      lcorbi_t3_pos_4_073_57    m/z 181.1197     0   1   1   1.5  RT (min)       lcorbi_t3_pos_2_040_25    m/z 181.1295     0   2   4   6   8E+3   1   1.5  RT (min)      lcorbi_t3_pos_1_030_21    m/z 181.1346     0   1E+4   1   1.5  RT (min)       lcorbi_t3_pos_4_091_60    m/z 181.1455     0   1E+5   1   1.5  RT (min)      lcorbi_t3_pos_2_037_27    m/z 181.1780     0   1   2   3E+4   8   8.5  RT (min)       lcorbi_t3_pos_4_090_60    m/z 181.9984     0   1   2E+4   0.5   1  RT (min)       lcorbi_t3_pos_3_052_47    m/z 182.0093     0   1   2   3   4E+4   9.5   10  RT (min)       lcorbi_t3_pos_4_098_61    m/z 182.0119     0   1   2   3E+4   0   0.5  RT (min)      lcorbi_t3_pos_1_011_44    m/z 182.0214     0   1E+3   2.5   3  RT (min)      lcorbi_t3_pos_4_012_37    m/z 182.0417     0   1   2   3   4E+4   18   18.5  RT (min)       lcorbi_t3_pos_4_098_61    m/z 182.0495     0   1   2   3   4E+3   4.5   5  RT (min)      lcorbi_t3_pos_4_112_62    m/z 182.0529     0   1E+5   13   13.5  RT (min)      lcorbi_t3_pos_4_075_58    m/z 182.0600     0   1   2E+4   5   5.5  RT (min)       lcorbi_t3_pos_4_105_61    m/z 182.0690     0   1E+3   4.5   5  RT (min)      lcorbi_t3_pos_1_024_04    m/z 182.0754     0   1   2   3E+3   4.5   5  RT (min)       lcorbi_t3_pos_1_024_04    m/z 182.0789     0   2   4   6E+5   1   1.5   2  RT (min)       lcorbi_t3_pos_1_036_08    m/z 182.0893     0   1E+5   13   13.5  RT (min)       lcorbi_t3_pos_4_081_58    m/z 182.0909     0   1E+4   1   1.5  RT (min)       lcorbi_t3_pos_1_036_08    m/z 182.1005     0   1   5.5   6  RT (min)      lcorbi_t3_pos_4_098_61    m/z 182.1118     0   1   1.5   2  RT (min)       lcorbi_t3_pos_2_051_35    m/z 182.1230     0   1   2   3   4E+6   1   1.5  RT (min)       lcorbi_t3_pos_4_094_60    m/z 182.1256     0   2   4E+3   7.5   8  RT (min)      lcorbi_t3_pos_3_054_15    m/z 182.1453     0   1   1   1.5  RT (min)       lcorbi_t3_pos_4_012_37    m/z 182.1651     0   1   2E+6   0   0.5  RT (min)      lcorbi_t3_pos_1_001_49    m/z 182.1903     0   2   4   6E+4   5.5   6  RT (min)       lcorbi_t3_pos_4_090_60    m/z 182.9022     0   1   2   3E+4   1   1.5  RT (min)      lcorbi_t3_pos_1_031_40    m/z 182.9023     0   1   2E+4   18   18.5  RT (min)       lcorbi_t3_pos_4_095_60    m/z 182.9263     0   1   2   3E+5   0   0.5  RT (min)      lcorbi_t3_pos_1_007_07    m/z 182.9807     0   1   2   3   4E+3   6.5   7  RT (min)      lcorbi_t3_pos_4_108_62    m/z 182.9959     0   1E+4   0   0.5  RT (min)      lcorbi_t3_pos_1_003_45    m/z 183.0652     0   1E+4   9.5   10  RT (min)      lcorbi_t3_pos_1_002_01    m/z 183.0685     0   1   2E+3   1   1.5  RT (min)      lcorbi_t3_pos_4_054_18    m/z 183.0781     0   2   4   6   8E+3   6.5   7  RT (min)      lcorbi_t3_pos_1_001_49    m/z 183.0845     0   1E+3   2   2.5  RT (min)      lcorbi_t3_pos_1_003_45    m/z 183.0925     0   1E+4   1   1.5  RT (min)      lcorbi_t3_pos_4_093_60    m/z 183.0971     0   1   2E+4   18   18.5  RT (min)       lcorbi_t3_pos_4_082_59    m/z 183.0992     0   1E+4   4.5   5  RT (min)      lcorbi_t3_pos_1_002_01    m/z 183.1056     0   1   2   3   4E+3   1   1.5  RT (min)      lcorbi_t3_pos_3_027_27    m/z 183.1128     0   1   2E+3   1   1.5  RT (min)       lcorbi_t3_pos_4_093_60    m/z 183.1168     0   1   2   3   4E+3   7   7.5  RT (min)      lcorbi_t3_pos_1_001_49    m/z 183.1168     0   1   2E+4   14   14.5  RT (min)       lcorbi_t3_pos_1_003_45    m/z 183.1355     0   2   4E+4   8   8.5  RT (min)       lcorbi_t3_pos_4_074_58    m/z 183.1379     0   2   4   6E+4   9   9.5  RT (min)       lcorbi_t3_pos_1_044_25    m/z 183.1383     0   2   4   6E+4   13.5   14  RT (min)      lcorbi_t3_pos_1_002_01    m/z 183.1420     0   1   2E+3   4.5   5  RT (min)      lcorbi_t3_pos_1_002_01    m/z 183.1433     0   1E+3   1   1.5  RT (min)       lcorbi_t3_pos_1_004_24    m/z 183.1856     0   1   2   3   4E+4   5   5.5  RT (min)       lcorbi_t3_pos_4_106_62    m/z 183.1937     0   2   4   6   8E+3   5.5   6  RT (min)      lcorbi_t3_pos_4_090_60    m/z 183.9959     0   1   2   3E+4   1   1.5  RT (min)      lcorbi_t3_pos_4_070_57    m/z 184.0290     0   1E+5   18.5   19  RT (min)       lcorbi_t3_pos_4_091_60    m/z 184.0403     0   2   4   6   8E+4   2.5   3  RT (min)       lcorbi_t3_pos_3_034_26    m/z 184.0580     0   1E+4   1   1.5  RT (min)       lcorbi_t3_pos_1_043_06    m/z 184.0733     0   2   4   6E+4   11   11.5  RT (min)       lcorbi_t3_pos_4_067_57    m/z 184.0733     0   1E+5   13.5   14  RT (min)       lcorbi_t3_pos_1_051_09    m/z 184.0838     0   1   9   9.5  RT (min)       lcorbi_t3_pos_4_080_58    m/z 184.0992     0   1   2   3E+4   2  RT (min)       lcorbi_t3_pos_2_024_15    m/z 184.0993     0   1   2E+3   5   5.5  RT (min)      lcorbi_t3_pos_4_066_57    m/z 184.1180     0   2   4   6   8E+3   1   1.5  RT (min)      lcorbi_t3_pos_3_029_13    m/z 184.1332     0   2   4   6E+4   6.5   7  RT (min)       lcorbi_t3_pos_4_106_62    m/z 184.1444     0   1E+4   4.5   5  RT (min)      lcorbi_t3_pos_4_106_62    m/z 184.1889     0   2   4   6E+3   5   5.5  RT (min)      lcorbi_t3_pos_4_074_58    m/z 184.2059     0   2   4   6   8E+4   8   8.5  RT (min)       lcorbi_t3_pos_2_044_21    m/z 184.6266     0   1   2   3E+4   13.5   14  RT (min)      lcorbi_t3_pos_4_083_59    m/z 184.9563     0   1   2   3   4E+3   2.5   3  RT (min)      lcorbi_t3_pos_4_068_57    m/z 185.0420     0   1   2   3E+3   1.5   2  RT (min)       lcorbi_t3_pos_2_005_11    m/z 185.0422     0   1   2   3   4E+4   18   18.5  RT (min)       lcorbi_t3_pos_4_109_62    m/z 185.0604     0   2   4E+3   4.5   5  RT (min)      lcorbi_t3_pos_1_001_49    m/z 185.0638     0   1   2   3E+3   2.5   3  RT (min)      lcorbi_t3_pos_4_082_59    m/z 185.0638     0   1   2   3   4E+4   4.5   5  RT (min)      lcorbi_t3_pos_4_098_61    m/z 185.0658     0   2   4   6E+4   18   18.5  RT (min)       lcorbi_t3_pos_4_100_61    m/z 185.0659     0   1   2E+3   0.5   1  RT (min)       lcorbi_t3_pos_1_001_49    m/z 185.0687     0   1   2   3E+4   1   1.5  RT (min)       lcorbi_t3_pos_4_087_59    m/z 185.0785     0   1   2   3E+3   4.5   5  RT (min)      lcorbi_t3_pos_1_002_01    m/z 185.0857     0   2   4E+3   1   1.5  RT (min)       lcorbi_t3_pos_1_003_45    m/z 185.0961     0   1E+4   8.5   9  RT (min)       lcorbi_t3_pos_1_001_49    m/z 185.0977     0   1E+5   1   1.5  RT (min)       lcorbi_t3_pos_4_093_60    m/z 185.1147     0   1   2   3E+3   1   1.5  RT (min)       lcorbi_t3_pos_1_004_24    m/z 185.1201     0   1   4.5   5  RT (min)       lcorbi_t3_pos_1_002_01    m/z 185.1284     0   1E+4   4.5   5  RT (min)       lcorbi_t3_pos_1_003_45    m/z 185.1365     0   1   2   3E+3   6.5   7  RT (min)      lcorbi_t3_pos_1_002_01    m/z 185.1648     0   2   4   6   8E+3   1.5   2  RT (min)      lcorbi_t3_pos_1_001_49    m/z 185.1649     0   1   2   3   4E+3   6.5   7  RT (min)      lcorbi_t3_pos_1_046_22    m/z 185.1729     0   1   7.5   8  RT (min)      lcorbi_t3_pos_1_046_22    m/z 185.1729     0   1   9.5   10  RT (min)      lcorbi_t3_pos_1_044_25    m/z 185.2093     0   1   2   3   4E+3   8   8.5  RT (min)      lcorbi_t3_pos_1_045_12    m/z 185.9948     0   1E+4   18.5   19  RT (min)       lcorbi_t3_pos_4_073_57    m/z 185.9951     0   1   2   3   4E+3   5   5.5  RT (min)      lcorbi_t3_pos_4_066_57    m/z 186.0083     0   1E+4   1   1.5  RT (min)       lcorbi_t3_pos_2_005_11    m/z 186.0084     0   1   2E+4   4.5   5  RT (min)      lcorbi_t3_pos_4_074_58    m/z 186.0261     0   2   4   6E+4   18   18.5  RT (min)       lcorbi_t3_pos_4_106_62    m/z 186.0357     0   1E+4   1   1.5  RT (min)      lcorbi_t3_pos_3_029_13    m/z 186.0526     0   1   2   3E+3   5.5   6  RT (min)      lcorbi_t3_pos_1_001_49    m/z 186.0528     0   1   2   3E+3   1   1.5  RT (min)      lcorbi_t3_pos_1_055_33    m/z 186.0647     0   1   2E+3   4.5   5  RT (min)      lcorbi_t3_pos_1_001_49    m/z 186.0696     0   1   2   3   4E+3   2.5   3  RT (min)      lcorbi_t3_pos_1_001_49    m/z 186.0842     0   1E+4   11   11.5  RT (min)       lcorbi_t3_pos_1_001_49    m/z 186.0945     0   1E+3   1   1.5  RT (min)       lcorbi_t3_pos_2_029_43    m/z 186.1060     0   1   2E+3   0   0.5   1  RT (min)       lcorbi_t3_pos_4_073_57    m/z 186.1220     0   1   2E+3   4.5   5  RT (min)      lcorbi_t3_pos_4_075_58    m/z 186.1237     0   1   1   1.5  RT (min)      lcorbi_t3_pos_2_054_29    m/z 186.1318     0   1   2   3E+3   4.5   5  RT (min)      lcorbi_t3_pos_1_001_49    m/z 186.1481     0   2   4   6E+4   15   15.5  RT (min)      lcorbi_t3_pos_4_088_59    m/z 186.1489     0   2   4   6E+4   6.5   7  RT (min)       lcorbi_t3_pos_4_098_61    m/z 186.1489     0   1E+4   2.5   3  RT (min)      lcorbi_t3_pos_1_001_49    m/z 186.1852     0   2   4   6   8E+4   10   10.5  RT (min)      lcorbi_t3_pos_1_002_01    m/z 186.2216     0   2   4   6   8E+4   9   9.5  RT (min)       lcorbi_t3_pos_2_044_21    m/z 186.9300     0   1   2   3E+3   1   1.5  RT (min)       lcorbi_t3_pos_3_011_42    m/z 186.9356     0   1   2E+3   2.5   3  RT (min)      lcorbi_t3_pos_1_011_44    m/z 186.9543     0   2   4E+4   1   1.5  RT (min)      lcorbi_t3_pos_4_021_25    m/z 186.9679     0   1   2E+3   1   1.5  RT (min)      lcorbi_t3_pos_4_021_25    m/z 186.9943     0   2   4E+4   18   18.5  RT (min)       lcorbi_t3_pos_4_106_62    m/z 186.9983     0   1E+4   1   1.5  RT (min)      lcorbi_t3_pos_4_012_37    m/z 187.0304     0   1   2E+4   1   1.5  RT (min)      lcorbi_t3_pos_2_013_33    m/z 187.0444     0   1E+3   1.5   2  RT (min)      lcorbi_t3_pos_2_037_27    m/z 187.0536     0   1E+3   2.5   3  RT (min)      lcorbi_t3_pos_4_012_37    m/z 187.0713     0   1   2E+4   1   1.5  RT (min)       lcorbi_t3_pos_1_044_25    m/z 187.0713     0   1E+3   6   6.5  RT (min)      lcorbi_t3_pos_1_048_47    m/z 187.0815     0   1E+5   6.5   7  RT (min)      lcorbi_t3_pos_3_025_52    m/z 187.0815     0   1E+5   8.5   9  RT (min)      lcorbi_t3_pos_2_021_09    m/z 187.0815     0   1E+5   5   5.5  RT (min)       lcorbi_t3_pos_3_003_33    m/z 187.0815     0   1E+5   10.5   11  RT (min)      lcorbi_t3_pos_2_003_19    m/z 187.0815     0   1E+5   13   13.5  RT (min)      lcorbi_t3_pos_4_066_57    m/z 187.0815     0   2   4   6   8E+4   3   3.5  RT (min)      lcorbi_t3_pos_3_016_21    m/z 187.0941     0   2   4   6E+3   4.5   5  RT (min)       lcorbi_t3_pos_1_020_31    m/z 187.1115     0   1   2   3E+3   1   1.5   2  RT (min)       lcorbi_t3_pos_4_084_59    m/z 187.1230     0   2   4   6   8E+3   4.5   5  RT (min)      lcorbi_t3_pos_1_055_33    m/z 187.1441     0   2   4   6   8E+3   1   1.5  RT (min)       lcorbi_t3_pos_1_051_09    m/z 187.1522     0   2   4   6E+3   6.5   7  RT (min)       lcorbi_t3_pos_1_028_32    m/z 187.1886     0   2   4E+4   7   7.5  RT (min)      lcorbi_t3_pos_4_053_19    m/z 187.2250     0   1E+4   9   9.5  RT (min)      lcorbi_t3_pos_2_044_21    m/z 187.2250     0   2   4   6E+3   5.5   6  RT (min)      lcorbi_t3_pos_4_074_58    m/z 187.9916     0   1   2E+4   1   1.5  RT (min)       lcorbi_t3_pos_1_034_34    m/z 188.0054     0   1   2E+4   1   1.5  RT (min)       lcorbi_t3_pos_4_075_58    m/z 188.0104     0   1   2   3   4E+4   6.5   7  RT (min)       lcorbi_t3_pos_1_028_32    m/z 188.0318     0   1E+4   4.5   5  RT (min)       lcorbi_t3_pos_4_075_58    m/z 188.0318     0   1   2.5   3  RT (min)       lcorbi_t3_pos_4_082_59    m/z 188.0351     0   2   4   6   8E+4   1   1.5  RT (min)      lcorbi_t3_pos_1_052_38    m/z 188.0610     0   2   4E+3   1   1.5  RT (min)      lcorbi_t3_pos_1_001_49    m/z 188.0682     0   1E+3   2.5   3  RT (min)       lcorbi_t3_pos_4_081_58    m/z 188.0823     0   1E+5   18.5   19  RT (min)       lcorbi_t3_pos_4_074_58    m/z 188.0849     0   1   2   3   4E+3   0.5   1  RT (min)       lcorbi_t3_pos_4_071_57    m/z 188.0918     0   1   2   3E+4   18   18.5  RT (min)       lcorbi_t3_pos_4_098_61    m/z 188.0974     0   1E+3   4.5   5  RT (min)      lcorbi_t3_pos_1_052_38    m/z 188.1111     0   1   2E+3   1   1.5  RT (min)       lcorbi_t3_pos_1_002_01    m/z 188.1206     0   1   2E+3   5.5   6  RT (min)       lcorbi_t3_pos_4_082_59    m/z 188.1281     0   2   4   6E+4   1   1.5  RT (min)       lcorbi_t3_pos_1_023_02    m/z 188.1362     0   1   2E+4   8   8.5  RT (min)      lcorbi_t3_pos_1_003_45    m/z 188.1757     0   1E+4   1   1.5  RT (min)      lcorbi_t3_pos_4_067_57    m/z 188.2009     0   1E+5   7   7.5   8  RT (min)       lcorbi_t3_pos_3_003_33    m/z 188.6345     0   1E+5   13.5   14  RT (min)      lcorbi_t3_pos_1_032_26    m/z 188.8852     0   2   4   6E+4   13.5   14  RT (min)      lcorbi_t3_pos_1_028_32    m/z 188.9246     0   2   4   6   8E+3   1   1.5  RT (min)       lcorbi_t3_pos_1_004_24    m/z 188.9513     0   1   2   3E+4   1   1.5  RT (min)      lcorbi_t3_pos_1_019_16    m/z 188.9683     0   1E+4   1   1.5  RT (min)      lcorbi_t3_pos_1_044_25    m/z 188.9912     0   1   2   3E+4   18   18.5  RT (min)       lcorbi_t3_pos_4_088_59    m/z 189.0070     0   1E+4   1   1.5  RT (min)       lcorbi_t3_pos_1_044_25    m/z 189.0159     0   1   2   3   4E+3   5   5.5  RT (min)      lcorbi_t3_pos_4_081_58    m/z 189.0192     0   2   4   6   8E+2   1.5   2  RT (min)      lcorbi_t3_pos_4_021_25    m/z 189.0394     0   1   2   2.5  RT (min)      lcorbi_t3_pos_4_107_62    m/z 189.0522     0   1E+4   1   1.5  RT (min)       lcorbi_t3_pos_1_025_52    m/z 189.0524     0   1E+4   2.5   3  RT (min)       lcorbi_t3_pos_1_005_42    m/z 189.0717     0   1   2E+3   2.5   3  RT (min)      lcorbi_t3_pos_1_003_45    m/z 189.0732     0   2   4   6E+3   6.5   7  RT (min)      lcorbi_t3_pos_1_001_49    m/z 189.0733     0   1   2   3E+3   1   1.5  RT (min)       lcorbi_t3_pos_1_006_15    m/z 189.0740     0   1E+4   4.5   5  RT (min)       lcorbi_t3_pos_1_012_14    m/z 189.0791     0   1E+5   18.5   19  RT (min)       lcorbi_t3_pos_4_095_60    m/z 189.0871     0   1   2E+3   1   1.5  RT (min)       lcorbi_t3_pos_1_015_11    m/z 189.0971     0   1   2   3   4E+5   1   1.5  RT (min)      lcorbi_t3_pos_3_012_31    m/z 189.0971     0   2   4   6   8E+5   6   6.5   7  RT (min)       lcorbi_t3_pos_2_051_35    m/z 189.0971     0   1   2   3   4E+5   2.5   3  RT (min)      lcorbi_t3_pos_2_024_15    m/z 189.0972     0   2   4E+5   10.5   11  RT (min)      lcorbi_t3_pos_3_021_39    m/z 189.1120     0   1   2   3E+4   2   2.5  RT (min)      lcorbi_t3_pos_1_005_42    m/z 189.1233     0   1   2   3E+3   1.5   2  RT (min)       lcorbi_t3_pos_1_015_11    m/z 189.1314     0   2   4   6   8E+3   5.5   6  RT (min)       lcorbi_t3_pos_1_012_14    m/z 189.1346     0   1   0   1  RT (min)       lcorbi_t3_pos_1_011_44    m/z 189.2042     0   1   2   3   4E+3   7.5   8  RT (min)      lcorbi_t3_pos_4_027_07    m/z 189.3892     0   2   4   6   8E+4   13.5   14  RT (min)      lcorbi_t3_pos_4_083_59    m/z 189.7675     0   1   2   3   4E+4   13.5   14  RT (min)       lcorbi_t3_pos_1_020_31    m/z 189.9887     0   1   2   3E+3   1   1.5  RT (min)      lcorbi_t3_pos_2_037_27    m/z 190.0126     0   1   2E+3   1   1.5  RT (min)      lcorbi_t3_pos_1_003_45    m/z 190.0215     0   1   2   3E+4   7   7.5  RT (min)      lcorbi_t3_pos_4_098_61    m/z 190.0247     0   1E+4   1   1.5  RT (min)      lcorbi_t3_pos_4_054_18    m/z 190.0386     0   1   2   3   4E+3   0.5   1  RT (min)      lcorbi_t3_pos_4_092_60    m/z 190.0491     0   1   2   3E+5   1   1.5  RT (min)       lcorbi_t3_pos_2_037_27    m/z 190.0499     0   1   2   3E+3   6   6.5  RT (min)       lcorbi_t3_pos_1_026_17    m/z 190.0668     0   1   2   3   4E+4   6.5   7  RT (min)       lcorbi_t3_pos_4_106_62    m/z 190.0710     0   2   4   6E+3   1.5   2  RT (min)       lcorbi_t3_pos_1_001_49    m/z 190.0773     0   1   4.5   5  RT (min)      lcorbi_t3_pos_1_001_49    m/z 190.0783     0   1   1.5   2  RT (min)      lcorbi_t3_pos_1_002_01    m/z 190.0862     0   1E+4   5   5.5  RT (min)       lcorbi_t3_pos_1_025_52    m/z 190.0862     0   2   4   6E+3   7.5   8  RT (min)       lcorbi_t3_pos_1_039_23    m/z 190.0863     0   2   4   6E+3   2.5   3  RT (min)      lcorbi_t3_pos_1_001_49    m/z 190.0882     0   1   2E+3   1   1.5  RT (min)      lcorbi_t3_pos_2_040_25    m/z 190.0979     0   2   4   6   8E+4   6   6.5  RT (min)      lcorbi_t3_pos_4_042_09    m/z 190.1187     0   1   2   3   4E+3   1.5   2  RT (min)      lcorbi_t3_pos_1_002_01    m/z 190.1273     0   1   2E+3   5.5   6  RT (min)       lcorbi_t3_pos_1_037_19    m/z 190.1280     0   1   13.5   14  RT (min)      lcorbi_t3_pos_1_022_20    m/z 190.1380     0   2   4   6E+3   1   1.5  RT (min)      lcorbi_t3_pos_1_034_34    m/z 190.1631     0   1E+4   1   1.5  RT (min)      lcorbi_t3_pos_1_052_38    m/z 190.1671     0   1   2E+4   10.5   11  RT (min)      lcorbi_t3_pos_1_004_24    m/z 190.1801     0   1   2E+3   7   7.5  RT (min)      lcorbi_t3_pos_1_006_15    m/z 190.2714     0   2   4   6   8E+3   14   14.5  RT (min)      lcorbi_t3_pos_4_072_57    m/z 190.3969     0   1E+5   14   14.5  RT (min)      lcorbi_t3_pos_1_020_31    m/z 190.6477     0   2   4   6   8E+3   14   14.5  RT (min)      lcorbi_t3_pos_4_068_57    m/z 190.8481     0   2   4   6E+4   1   1.5  RT (min)      lcorbi_t3_pos_4_070_57    m/z 190.9201     0   1   1   1.5  RT (min)      lcorbi_t3_pos_1_029_29    m/z 190.9628     0   2   4   6   8E+3   1   1.5  RT (min)       lcorbi_t3_pos_4_070_57    m/z 190.9901     0   1   2   3E+3   0   0.5  RT (min)      lcorbi_t3_pos_4_094_60    m/z 190.9901     0   1   2   3E+4   18   18.5  RT (min)       lcorbi_t3_pos_4_093_60    m/z 191.0176     0   1E+3   1.5   2  RT (min)       lcorbi_t3_pos_2_005_11    m/z 191.0550     0   1   2   3E+3   1.5   2  RT (min)      lcorbi_t3_pos_2_029_43    m/z 191.0767     0   1   2E+3   1   1.5  RT (min)      lcorbi_t3_pos_4_008_30    m/z 191.0848     0   1   2   3E+3   5   5.5  RT (min)      lcorbi_t3_pos_1_002_01    m/z 191.0915     0   1   2   3E+3   2.5   3  RT (min)      lcorbi_t3_pos_1_002_01    m/z 191.0947     0   2   4E+4   3   3.5  RT (min)      lcorbi_t3_pos_3_009_50    m/z 191.0947     0   1E+5   5   5.5  RT (min)      lcorbi_t3_pos_3_026_04    m/z 191.0947     0   1E+5   7   7.5  RT (min)      lcorbi_t3_pos_2_012_01    m/z 191.0947     0   1E+5   13   13.5  RT (min)      lcorbi_t3_pos_1_029_29    m/z 191.0947     0   2   4   6   8E+4   9.5   10  RT (min)      lcorbi_t3_pos_4_025_52    m/z 191.0948     0   2   4   6   8E+4   11.5   12  RT (min)      lcorbi_t3_pos_4_091_60    m/z 191.1013     0   1   2E+4   0.5   1  RT (min)      lcorbi_t3_pos_4_111_62    m/z 191.1066     0   2   4E+5   9.5   10  RT (min)      lcorbi_t3_pos_4_075_58    m/z 191.1178     0   1E+7   1.5   2  RT (min)       lcorbi_t3_pos_4_112_62    m/z 191.1244     0   2   4E+4   1.5   2  RT (min)      lcorbi_t3_pos_1_031_40    m/z 191.1277     0   1   2E+4   5   5.5  RT (min)      lcorbi_t3_pos_1_019_16    m/z 191.1542     0   1   2   3E+4   4.5   5  RT (min)       lcorbi_t3_pos_4_090_60    m/z 191.1771     0   1   2E+3   5   5.5  RT (min)      lcorbi_t3_pos_1_046_22    m/z 191.9097     0   2   4   6E+4   1   1.5  RT (min)       lcorbi_t3_pos_4_070_57    m/z 191.9775     0   1   2   3   4E+3   1   1.5  RT (min)      lcorbi_t3_pos_2_006_47    m/z 192.0079     0   1   2   3E+3   4   4.5  RT (min)      lcorbi_t3_pos_1_010_46    m/z 192.0325     0   1   2   3E+3   1   1.5  RT (min)      lcorbi_t3_pos_1_017_51    m/z 192.0437     0   2   4   6E+4   1   1.5  RT (min)      lcorbi_t3_pos_1_052_38    m/z 192.0631     0   1   2.5   3  RT (min)      lcorbi_t3_pos_1_001_49    m/z 192.0669     0   1E+5   18   18.5  RT (min)       lcorbi_t3_pos_4_070_57    m/z 192.0736     0   1E+4   10   10.5  RT (min)      lcorbi_t3_pos_1_047_41    m/z 192.0743     0   1   1   1.5  RT (min)      lcorbi_t3_pos_4_015_45    m/z 192.0801     0   2   4   6E+2   1   1.5  RT (min)       lcorbi_t3_pos_3_037_32    m/z 192.0889     0   1E+4   10.5   11  RT (min)      lcorbi_t3_pos_1_044_25    m/z 192.0981     0   2   4E+3   0.5   1  RT (min)      lcorbi_t3_pos_4_111_62    m/z 192.1100     0   2   4E+4   9.5   10  RT (min)      lcorbi_t3_pos_1_047_41    m/z 192.1149     0   1E+5   1.5   2  RT (min)       lcorbi_t3_pos_4_110_62    m/z 192.1311     0   1   2   3E+3   5   5.5  RT (min)      lcorbi_t3_pos_1_046_22    m/z 192.1383     0   1   2E+4   4.5   5  RT (min)      lcorbi_t3_pos_1_001_49    m/z 192.1430     0   1   2.5   3  RT (min)      lcorbi_t3_pos_1_001_49    m/z 192.1438     0   1   1   1.5  RT (min)       lcorbi_t3_pos_4_008_30    m/z 192.1593     0   1E+5   1   1.5  RT (min)       lcorbi_t3_pos_2_044_21    m/z 192.1675     0   1E+4   8   8.5  RT (min)      lcorbi_t3_pos_4_098_61    m/z 192.1868     0   1   5   5.5  RT (min)      lcorbi_t3_pos_4_098_61    m/z 192.6071     0   1   2   3   4E+4   13.5   14  RT (min)       lcorbi_t3_pos_1_023_02    m/z 192.7296     0   1   2E+4   13.5   14  RT (min)      lcorbi_t3_pos_4_077_58    m/z 192.8452     0   2   4   6E+4   1   1.5  RT (min)      lcorbi_t3_pos_4_072_57    m/z 192.8529     0   1   2E+4   14   14.5  RT (min)      lcorbi_t3_pos_1_031_40    m/z 192.8811     0   1E+4   1   1.5  RT (min)      lcorbi_t3_pos_4_012_37    m/z 192.9101     0   2   4   6E+3   1   1.5  RT (min)       lcorbi_t3_pos_1_006_15    m/z 192.9470     0   1   2E+4   1   1.5  RT (min)       lcorbi_t3_pos_4_068_57    m/z 192.9642     0   1   2   3E+3   4.5   5  RT (min)      lcorbi_t3_pos_4_082_59    m/z 192.9642     0   1   2   3E+3   2.5   3  RT (min)      lcorbi_t3_pos_1_001_49    m/z 193.0019     0   1E+4   1.5   2  RT (min)      lcorbi_t3_pos_4_074_58    m/z 193.0158     0   1E+5   0   0.5  RT (min)      lcorbi_t3_pos_1_005_42    m/z 193.0342     0   2   4E+4   1   1.5  RT (min)       lcorbi_t3_pos_4_075_58    m/z 193.0351     0   1E+4   4.5   5  RT (min)      lcorbi_t3_pos_4_066_57    m/z 193.0521     0   1   2E+3   1   1.5  RT (min)       lcorbi_t3_pos_4_103_61    m/z 193.0557     0   1   2E+4   4   4.5   5  RT (min)       lcorbi_t3_pos_1_001_49    m/z 193.0706     0   1   2E+3   1   1.5  RT (min)       lcorbi_t3_pos_1_003_45    m/z 193.0721     0   1   2   3   4E+3   4.5   5  RT (min)      lcorbi_t3_pos_1_003_45    m/z 193.0737     0   1E+3   2.5   3  RT (min)      lcorbi_t3_pos_1_002_01    m/z 193.0835     0   1   2   3E+3   6.5   7  RT (min)       lcorbi_t3_pos_1_001_49    m/z 193.0859     0   2   4   6   8E+3   5   5.5  RT (min)       lcorbi_t3_pos_1_025_52    m/z 193.0971     0   1E+4   2  RT (min)       lcorbi_t3_pos_1_001_49    m/z 193.1070     0   2   4   6E+3   4.5   5  RT (min)      lcorbi_t3_pos_1_002_01    m/z 193.1083     0   2   4E+3   2.5   3  RT (min)      lcorbi_t3_pos_1_002_01    m/z 193.1199     0   2   4   6   8E+3   8   8.5  RT (min)      lcorbi_t3_pos_1_001_49    m/z 193.1244     0   1E+4   1.5   2  RT (min)      lcorbi_t3_pos_1_007_07    m/z 193.1547     0   1E+5   1   1.5  RT (min)       lcorbi_t3_pos_4_068_57    m/z 193.1627     0   2   4E+3   1   1.5  RT (min)      lcorbi_t3_pos_1_001_49    m/z 193.1699     0   1E+7   0   0.5  RT (min)      lcorbi_t3_pos_1_001_49    m/z 193.3570     0   2   4   6   8E+3   14   14.5  RT (min)      lcorbi_t3_pos_4_072_57    m/z 193.9067     0   2   4   6   8E+4   1   1.5  RT (min)       lcorbi_t3_pos_4_068_57    m/z 193.9826     0   2   4   6   8E+4   1   1.5  RT (min)       lcorbi_t3_pos_1_028_32    m/z 194.0326     0   2   4   6E+4   0   1  RT (min)       lcorbi_t3_pos_4_089_59    m/z 194.0647     0   2   4   6   8E+4   4.5   5  RT (min)      lcorbi_t3_pos_1_012_14    m/z 194.0659     0   1E+5   1   1.5  RT (min)       lcorbi_t3_pos_4_103_61    m/z 194.0811     0   2   4   6E+3   5.5   6  RT (min)       lcorbi_t3_pos_4_098_61    m/z 194.0883     0   1E+4   3   3.5  RT (min)      lcorbi_t3_pos_4_113_62    m/z 194.0925     0   2   4   6   8E+4   1.5   2  RT (min)      lcorbi_t3_pos_3_021_39    m/z 194.1023     0   1   2   3   4E+3   5   5.5  RT (min)      lcorbi_t3_pos_1_025_52    m/z 194.1151     0   2   4   6E+4   5.5   6  RT (min)       lcorbi_t3_pos_4_098_61    m/z 194.1175     0   1   2   3E+3   1.5   2  RT (min)      lcorbi_t3_pos_4_030_05    m/z 194.1288     0   2   4   6   8E+3   4.5   5  RT (min)      lcorbi_t3_pos_1_001_49    m/z 194.1288     0   1   2   3   4E+3   2.5   3  RT (min)      lcorbi_t3_pos_1_001_49    m/z 194.1514     0   1E+4   1   1.5  RT (min)      lcorbi_t3_pos_4_095_60    m/z 194.1515     0   1   2E+4   9.5   10  RT (min)      lcorbi_t3_pos_4_027_07    m/z 194.8783     0   1E+4   1   1.5  RT (min)      lcorbi_t3_pos_2_054_29    m/z 194.8851     0   1E+5   13.5   14  RT (min)      lcorbi_t3_pos_1_031_40    m/z 194.8879     0   2   4E+4   13.5   14  RT (min)      lcorbi_t3_pos_1_034_34    m/z 194.9282     0   1E+4   1   1.5  RT (min)      lcorbi_t3_pos_2_019_13    m/z 194.9441     0   2   4E+3   1   1.5  RT (min)      lcorbi_t3_pos_1_006_15    m/z 194.9801     0   1   2E+3   1   1.5  RT (min)       lcorbi_t3_pos_4_113_62    m/z 195.0188     0   1   2E+4   0.5   1  RT (min)       lcorbi_t3_pos_1_018_35    m/z 195.0293     0   1   2   3   4E+4   1.5   2  RT (min)       lcorbi_t3_pos_4_092_60    m/z 195.0444     0   1E+4   0.5   1  RT (min)       lcorbi_t3_pos_1_008_36    m/z 195.0446     0   1   2   3   4E+4   18   18.5  RT (min)       lcorbi_t3_pos_4_068_57    m/z 195.0626     0   1   2   3E+3   2.5   3  RT (min)      lcorbi_t3_pos_1_003_45    m/z 195.0626     0   2   4   6   8E+3   4.5   5  RT (min)      lcorbi_t3_pos_1_047_41    m/z 195.0764     0   1   2   3E+3   1.5   2  RT (min)      lcorbi_t3_pos_1_001_49    m/z 195.0764     0   2   4   6E+3   4.5   5  RT (min)       lcorbi_t3_pos_1_002_01    m/z 195.0904     0   1   2E+5   10   10.5  RT (min)      lcorbi_t3_pos_4_105_61    m/z 195.0991     0   2   4   6E+3   1   1.5  RT (min)       lcorbi_t3_pos_1_003_45    m/z 195.1128     0   2   4   6   8E+3   4.5   5  RT (min)       lcorbi_t3_pos_1_004_24    m/z 195.1145     0   2   4   6   8E+3   8.5   9  RT (min)      lcorbi_t3_pos_1_001_49    m/z 195.1181     0   1   2   3E+3   5.5   6   6.5  RT (min)       lcorbi_t3_pos_1_001_49    m/z 195.1346     0   1   1.5   2  RT (min)       lcorbi_t3_pos_1_002_01    m/z 195.1409     0   1E+5   13.5   14  RT (min)      lcorbi_t3_pos_4_082_59    m/z 195.1491     0   1E+4   5   5.5  RT (min)      lcorbi_t3_pos_1_001_49    m/z 195.1856     0   1   2E+4   6.5   7  RT (min)       lcorbi_t3_pos_4_106_62    m/z 195.3892     0   1E+5   13.5   14  RT (min)      lcorbi_t3_pos_4_082_59    m/z 195.8931     0   1E+5   13.5   14  RT (min)      lcorbi_t3_pos_1_028_32    m/z 195.9038     0   1   2   3E+4   1   1.5  RT (min)      lcorbi_t3_pos_4_070_57    m/z 196.0028     0   1   2   3E+4   0.5   1  RT (min)      lcorbi_t3_pos_1_012_14    m/z 196.0250     0   1   1   1.5  RT (min)      lcorbi_t3_pos_4_090_60    m/z 196.0396     0   1E+4   1   1.5  RT (min)       lcorbi_t3_pos_2_003_19    m/z 196.0523     0   2   4   6   8E+3   8   8.5  RT (min)      lcorbi_t3_pos_1_018_35    m/z 196.0579     0   2   4   6   8E+5   1   1.5  RT (min)      lcorbi_t3_pos_1_036_08    m/z 196.0661     0   2   4E+3   6.5   7  RT (min)      lcorbi_t3_pos_1_005_42    m/z 196.0718     0   1   2E+3   4.5   5  RT (min)      lcorbi_t3_pos_1_002_01    m/z 196.0847     0   1E+4   4.5   5  RT (min)      lcorbi_t3_pos_1_007_07    m/z 196.0910     0   1E+5   4.5   5  RT (min)       lcorbi_t3_pos_1_020_31    m/z 196.0937     0   2   4   6   8E+4   18   18.5  RT (min)       lcorbi_t3_pos_4_082_59    m/z 196.0987     0   1   2E+5   1  RT (min)       lcorbi_t3_pos_4_084_59    m/z 196.1104     0   1E+3   1   1.5  RT (min)       lcorbi_t3_pos_1_002_01    m/z 196.1160     0   1E+4   4.5   5  RT (min)      lcorbi_t3_pos_4_082_59    m/z 196.1194     0   2   4   6   8E+3   1   1.5  RT (min)       lcorbi_t3_pos_1_036_08    m/z 196.1260     0   1E+4   4.5   5  RT (min)      lcorbi_t3_pos_1_003_45    m/z 196.1332     0   1   2E+4   2.5   3  RT (min)       lcorbi_t3_pos_1_002_01    m/z 196.1543     0   1   2   3E+4   4.5   5  RT (min)       lcorbi_t3_pos_1_014_03    m/z 196.1543     0   1E+4   2.5   3  RT (min)      lcorbi_t3_pos_1_001_49    m/z 196.1581     0   1   2   3   4E+4   1   1.5  RT (min)       lcorbi_t3_pos_4_091_60    m/z 196.3967     0   2   4   6E+4   14   14.5  RT (min)      lcorbi_t3_pos_4_084_59    m/z 196.3995     0   1   2   3   4E+4   14   14.5  RT (min)      lcorbi_t3_pos_1_020_31    m/z 196.6528     0   1E+5   14   14.5  RT (min)      lcorbi_t3_pos_1_007_07    m/z 196.7754     0   1   2E+4   14   14.5  RT (min)      lcorbi_t3_pos_1_034_34    m/z 196.8779     0   1   1   1.5  RT (min)       lcorbi_t3_pos_3_039_03    m/z 196.9325     0   1   2   3E+3   1   1.5  RT (min)      lcorbi_t3_pos_1_044_25    m/z 196.9613     0   2   4   6E+3   1   1.5  RT (min)      lcorbi_t3_pos_1_029_29    m/z 196.9734     0   1   2   3   4E+3   1.5   2  RT (min)      lcorbi_t3_pos_2_003_19    m/z 197.0119     0   1   2E+3   1   1.5  RT (min)      lcorbi_t3_pos_1_006_15    m/z 197.0271     0   2   4E+4   1   1.5  RT (min)      lcorbi_t3_pos_2_009_50    m/z 197.0411     0   2   4   6   8E+5   1   1.5  RT (min)      lcorbi_t3_pos_1_036_08    m/z 197.0789     0   1E+4   1   1.5  RT (min)       lcorbi_t3_pos_1_008_36    m/z 197.0921     0   1E+4   4.5   5  RT (min)       lcorbi_t3_pos_1_002_01    m/z 197.0945     0   1   2   3   4E+3   1   1.5   2  RT (min)       lcorbi_t3_pos_1_030_21    m/z 197.1009     0   1E+6   1   1.5  RT (min)       lcorbi_t3_pos_1_031_40    m/z 197.1115     0   1   5   6  RT (min)       lcorbi_t3_pos_4_111_62    m/z 197.1269     0   1   2   3   4E+3   3   3.5  RT (min)      lcorbi_t3_pos_1_003_45    m/z 197.1284     0   1   2   3   4E+3   5   5.5  RT (min)       lcorbi_t3_pos_3_007_28    m/z 197.1325     0   2   4   6E+4   14   14.5  RT (min)       lcorbi_t3_pos_2_027_05    m/z 197.1365     0   1   2E+3   2.5   3  RT (min)      lcorbi_t3_pos_1_001_49    m/z 197.1502     0   1E+5   1.5   2  RT (min)       lcorbi_t3_pos_4_093_60    m/z 197.1518     0   2   4   6E+4   14   14.5  RT (min)      lcorbi_t3_pos_1_019_16    m/z 197.1648     0   2   4   6E+3   4.5   5  RT (min)       lcorbi_t3_pos_1_001_49    m/z 197.9960     0   1E+5   1   1.5  RT (min)       lcorbi_t3_pos_1_036_08    m/z 198.0042     0   1   2   3   4E+4   6.5   7  RT (min)       lcorbi_t3_pos_4_082_59    m/z 198.0278     0   2   4E+4   18   18.5  RT (min)      lcorbi_t3_pos_4_076_58    m/z 198.0305     0   1   2E+3   1   1.5  RT (min)      lcorbi_t3_pos_4_054_18    m/z 198.0465     0   1   2E+5   1   1.5  RT (min)       lcorbi_t3_pos_1_036_08    m/z 198.0518     0   2   4   6E+2   1   1.5  RT (min)       lcorbi_t3_pos_4_012_37    m/z 198.0549     0   1   2E+3   6.5   7  RT (min)      lcorbi_t3_pos_4_100_61    m/z 198.0842     0   2   4   6   8E+4   15   15.5  RT (min)      lcorbi_t3_pos_4_084_59    m/z 198.1041     0   1   4.5   5  RT (min)       lcorbi_t3_pos_1_022_20    m/z 198.1043     0   2   4   6E+4   1   1.5  RT (min)      lcorbi_t3_pos_1_002_01    m/z 198.1237     0   1E+5   18.5   19  RT (min)      lcorbi_t3_pos_4_095_60    m/z 198.1349     0   2   4   6E+3   5   5.5  RT (min)      lcorbi_t3_pos_4_110_62    m/z 198.1601     0   2   4   6E+3   5.5   6  RT (min)      lcorbi_t3_pos_1_003_45    m/z 198.1899     0   1E+5   1   1.5  RT (min)       lcorbi_t3_pos_4_091_60    m/z 198.8760     0   1   1   1.5  RT (min)      lcorbi_t3_pos_2_040_25    m/z 198.9398     0   1E+3   1   1.5  RT (min)      lcorbi_t3_pos_4_073_57    m/z 198.9570     0   1   2E+3   1   1.5  RT (min)      lcorbi_t3_pos_4_099_61    m/z 198.9743     0   2   4   6E+4   1   1.5  RT (min)      lcorbi_t3_pos_4_077_58    m/z 199.0075     0   1   2E+3   6.5   7  RT (min)      lcorbi_t3_pos_1_002_01    m/z 199.0216     0   1E+5   1   1.5  RT (min)       lcorbi_t3_pos_4_098_61    m/z 199.0247     0   2   4   6   8E+4   18   18.5  RT (min)      lcorbi_t3_pos_4_090_60    m/z 199.0390     0   1E+4   7   7.5  RT (min)      lcorbi_t3_pos_1_001_49    m/z 199.0502     0   1   2   3E+3   4.5   5  RT (min)       lcorbi_t3_pos_1_001_49    m/z 199.0544     0   1   1.5   2  RT (min)      lcorbi_t3_pos_1_002_01    m/z 199.0577     0   1   2E+3   5   5.5  RT (min)      lcorbi_t3_pos_1_001_49    m/z 199.0671     0   1   1.5   2  RT (min)      lcorbi_t3_pos_3_004_24    m/z 199.0761     0   1E+4   2  RT (min)       lcorbi_t3_pos_1_023_02    m/z 199.0942     0   1   2E+4   1   1.5  RT (min)      lcorbi_t3_pos_2_023_26    m/z 199.0965     0   1   2E+4   5   5.5  RT (min)       lcorbi_t3_pos_1_001_49    m/z 199.1006     0   2   4   6   8E+5   1   1.5  RT (min)       lcorbi_t3_pos_4_078_58    m/z 199.1077     0   1E+4   1.5   2  RT (min)       lcorbi_t3_pos_1_014_03    m/z 199.1077     0   1E+4   2.5   3  RT (min)       lcorbi_t3_pos_1_001_49    m/z 199.1304     0   1   2   3   4E+3   5.5   6  RT (min)      lcorbi_t3_pos_1_003_45    m/z 199.1304     0   2   4E+3   1   1.5  RT (min)      lcorbi_t3_pos_4_075_58    m/z 199.1465     0   2   4   6E+2   1   1.5  RT (min)       lcorbi_t3_pos_4_012_37    m/z 199.1522     0   1   2   3E+3   7   7.5  RT (min)      lcorbi_t3_pos_1_001_49    m/z 199.1724     0   1   2   3   4E+4   13.5   14  RT (min)      lcorbi_t3_pos_1_003_45    m/z 199.6162     0   1   2E+4   14   14.5  RT (min)      lcorbi_t3_pos_1_044_25    m/z 199.6735     0   1   2E+5   13.5   14  RT (min)      lcorbi_t3_pos_1_036_08    m/z 200.0000     0   2   4E+3   6.5   7  RT (min)      lcorbi_t3_pos_4_074_58    m/z 200.0230     0   1E+4   1   1.5  RT (min)       lcorbi_t3_pos_1_034_34    m/z 200.0319     0   1E+4   1   1.5  RT (min)       lcorbi_t3_pos_2_012_01    m/z 200.0528     0   1   2E+3   1   1.5  RT (min)       lcorbi_t3_pos_4_012_37    m/z 200.0684     0   1   2E+4   1   1.5  RT (min)      lcorbi_t3_pos_3_034_26    m/z 200.0706     0   1   2   3   4E+4   5   5.5  RT (min)       lcorbi_t3_pos_4_104_61    m/z 200.0740     0   1   2   3E+3   7   7.5  RT (min)      lcorbi_t3_pos_1_001_49    m/z 200.0766     0   1   2   3E+3   2.5   3  RT (min)      lcorbi_t3_pos_1_001_49    m/z 200.0893     0   1   2E+3   1   1.5  RT (min)      lcorbi_t3_pos_4_012_37    m/z 200.0917     0   1E+4   2.5   3  RT (min)      lcorbi_t3_pos_1_002_01    m/z 200.0930     0   1   2E+4   8   8.5  RT (min)      lcorbi_t3_pos_4_066_57    m/z 200.0998     0   2   4E+3   5   5.5  RT (min)      lcorbi_t3_pos_1_002_01    m/z 200.1191     0   1E+3   1.5   2  RT (min)       lcorbi_t3_pos_1_020_31    m/z 200.1452     0   1   2E+3   2.5   3  RT (min)      lcorbi_t3_pos_1_001_49    m/z 200.1726     0   1   2   3   4E+4   9.5   10  RT (min)      lcorbi_t3_pos_1_001_49    m/z 200.1726     0   1   2   3E+4   11   11.5  RT (min)      lcorbi_t3_pos_1_001_49    m/z 200.1775     0   1E+5   13.5   14  RT (min)      lcorbi_t3_pos_4_096_60    m/z 200.1839     0   2   4   6   8E+3   8.5   9  RT (min)      lcorbi_t3_pos_1_003_45    m/z 200.1921     0   1   2E+4   6.5   7  RT (min)      lcorbi_t3_pos_2_044_21    m/z 200.8769     0   1   2   3   4E+4   1   1.5  RT (min)      lcorbi_t3_pos_4_068_57    m/z 200.9067     0   1   1   1.5  RT (min)      lcorbi_t3_pos_2_028_24    m/z 200.9473     0   1E+4   1   1.5  RT (min)      lcorbi_t3_pos_1_051_09    m/z 200.9940     0   1   2   3E+4   1   1.5  RT (min)      lcorbi_t3_pos_1_052_38    m/z 201.0434     0   1E+5   18.5   19  RT (min)       lcorbi_t3_pos_4_066_57    m/z 201.0439     0   1E+4   1   1.5  RT (min)      lcorbi_t3_pos_1_046_22    m/z 201.0464     0   1   2   3   4E+3   8   8.5  RT (min)      lcorbi_t3_pos_3_025_52    m/z 201.0601     0   2   4   6   8E+3   0.5   1  RT (min)       lcorbi_t3_pos_4_092_60    m/z 201.0601     0   1E+4   18.5   19  RT (min)       lcorbi_t3_pos_4_098_61    m/z 201.0733     0   1   2   3E+3   2.5   3  RT (min)       lcorbi_t3_pos_1_001_49    m/z 201.0739     0   1   2   3   4E+3   5   5.5  RT (min)      lcorbi_t3_pos_1_001_49    m/z 201.0870     0   1   2   3E+3   1   1.5  RT (min)      lcorbi_t3_pos_1_004_24    m/z 201.0910     0   1E+5   6   6.5  RT (min)       lcorbi_t3_pos_2_053_28    m/z 201.1022     0   2   4   6   8E+2   1.5   2  RT (min)      lcorbi_t3_pos_3_055_10    m/z 201.1022     0   2   4   6E+4   4.5   5  RT (min)      lcorbi_t3_pos_1_001_49    m/z 201.1097     0   1E+4   4.5   5  RT (min)       lcorbi_t3_pos_1_001_49    m/z 201.1258     0   2   4   6E+3   5   5.5  RT (min)      lcorbi_t3_pos_1_001_49    m/z 201.1314     0   1E+3   1   1.5  RT (min)       lcorbi_t3_pos_3_027_27    m/z 201.1597     0   2   4   6E+4   18   18.5  RT (min)      lcorbi_t3_pos_4_083_59    m/z 201.1678     0   1   2E+4   7   7.5  RT (min)      lcorbi_t3_pos_4_098_61    m/z 201.2043     0   1   2E+4   10.5   11  RT (min)      lcorbi_t3_pos_4_107_62    m/z 201.2406     0   1   2   3   4E+4   9   9.5  RT (min)      lcorbi_t3_pos_1_023_02    m/z 202.0161     0   1   2   3E+6   0   0.5  RT (min)       lcorbi_t3_pos_1_001_49    m/z 202.0261     0   1   7.5   8  RT (min)       lcorbi_t3_pos_4_090_60    m/z 202.0473     0   2   4E+3   5   5.5  RT (min)      lcorbi_t3_pos_4_103_61    m/z 202.0476     0   1   2   3E+5   1   1.5  RT (min)       lcorbi_t3_pos_3_052_47    m/z 202.0685     0   1E+5   1   1.5  RT (min)      lcorbi_t3_pos_3_055_10    m/z 202.0777     0   1   2E+4   15.5   16  RT (min)      lcorbi_t3_pos_1_002_01    m/z 202.0777     0   1E+4   7   7.5  RT (min)      lcorbi_t3_pos_4_076_58    m/z 202.0777     0   2   4   6   8E+5   0   0.5  RT (min)      lcorbi_t3_pos_1_048_47    m/z 202.0863     0   2   4   6   8E+3   4.5   5  RT (min)      lcorbi_t3_pos_1_001_49    m/z 202.0944     0   1   2E+3   5.5   6  RT (min)      lcorbi_t3_pos_1_001_49    m/z 202.1009     0   1   2E+3   1   1.5  RT (min)       lcorbi_t3_pos_1_004_24    m/z 202.1186     0   1   2E+3   1   1.5  RT (min)       lcorbi_t3_pos_1_004_24    m/z 202.1186     0   2   4   6   8E+4   4.5   5  RT (min)       lcorbi_t3_pos_4_066_57    m/z 202.1186     0   1E+3   2.5   3  RT (min)       lcorbi_t3_pos_4_082_59    m/z 202.1227     0   1E+4   5.5   6  RT (min)       lcorbi_t3_pos_1_001_49    m/z 202.1282     0   1   2E+3   1   1.5  RT (min)      lcorbi_t3_pos_1_003_45    m/z 202.1438     0   1   2   3   4E+4   6.5   7  RT (min)       lcorbi_t3_pos_1_001_49    m/z 202.1461     0   1   2   3   4E+4   14   14.5  RT (min)      lcorbi_t3_pos_4_078_58    m/z 202.1671     0   1   2   3   4E+4   10   10.5  RT (min)      lcorbi_t3_pos_4_113_62    m/z 202.1801     0   1   2   3E+4   4.5   5  RT (min)       lcorbi_t3_pos_1_025_52    m/z 202.2109     0   1   2   3E+4   8   8.5  RT (min)      lcorbi_t3_pos_1_018_35    m/z 202.2165     0   2   4   6E+3   6   6.5  RT (min)      lcorbi_t3_pos_1_002_01    m/z 202.2276     0   2   4   6   8E+3   8   8.5  RT (min)      lcorbi_t3_pos_2_044_21    m/z 202.8562     0   1   2   3E+4   1   1.5  RT (min)      lcorbi_t3_pos_1_051_09    m/z 202.8740     0   1   2   3E+4   1   1.5  RT (min)      lcorbi_t3_pos_1_002_01    m/z 202.8859     0   2   4   6E+4   1   1.5  RT (min)      lcorbi_t3_pos_1_036_08    m/z 202.9009     0   2   4   6E+4   14   14.5  RT (min)      lcorbi_t3_pos_1_023_02    m/z 202.9758     0   2   4E+4   18   18.5  RT (min)       lcorbi_t3_pos_4_102_61    m/z 202.9759     0   1   2   3E+4   1   1.5  RT (min)       lcorbi_t3_pos_4_073_57    m/z 203.0007     0   2   4   6E+2   1   1.5  RT (min)      lcorbi_t3_pos_4_012_37    m/z 203.0317     0   2   4   6E+4   1   1.5  RT (min)       lcorbi_t3_pos_4_074_58    m/z 203.0591     0   1E+5   1   1.5  RT (min)      lcorbi_t3_pos_1_044_25    m/z 203.0591     0   1   2E+5   5   5.5  RT (min)      lcorbi_t3_pos_2_019_13    m/z 203.0591     0   2   4   6E+4   3   3.5  RT (min)      lcorbi_t3_pos_3_009_50    m/z 203.0591     0   1   2E+5   6.5   7  RT (min)      lcorbi_t3_pos_2_012_01    m/z 203.0591     0   1E+5   9.5   10  RT (min)      lcorbi_t3_pos_4_034_42    m/z 203.0591     0   2   4   6   8E+4   13   13.5  RT (min)      lcorbi_t3_pos_3_009_50    m/z 203.0678     0   2   4   6E+3   7   7.5  RT (min)      lcorbi_t3_pos_1_051_09    m/z 203.0680     0   1   2E+3   4.5   5  RT (min)      lcorbi_t3_pos_1_052_38    m/z 203.0811     0   2   4   6E+4   0   0.5  RT (min)      lcorbi_t3_pos_4_068_57    m/z 203.0855     0   1   2E+4   18.5   19  RT (min)       lcorbi_t3_pos_4_066_57    m/z 203.0914     0   2   4   6   8E+3   11   11.5  RT (min)      lcorbi_t3_pos_1_053_39    m/z 203.1026     0   1   2E+3   2.5   3  RT (min)      lcorbi_t3_pos_4_110_62    m/z 203.1210     0   1   2E+3   1   1.5  RT (min)       lcorbi_t3_pos_2_040_25    m/z 203.1220     0   2   4E+3   4.5   5  RT (min)      lcorbi_t3_pos_4_113_62    m/z 203.1278     0   1   2   3   4E+4   7.5   8  RT (min)       lcorbi_t3_pos_1_054_48    m/z 203.1390     0   1   2   3   4E+3   1.5   2  RT (min)       lcorbi_t3_pos_2_048_36    m/z 203.1471     0   1   2   3E+3   6.5   7  RT (min)       lcorbi_t3_pos_1_051_09    m/z 203.1477     0   1   2E+3   5   5.5  RT (min)       lcorbi_t3_pos_1_055_33    m/z 203.1541     0   2   4   6E+4   14   14.5  RT (min)      lcorbi_t3_pos_1_007_07    m/z 203.1754     0   2   4E+4   18   18.5  RT (min)      lcorbi_t3_pos_4_103_61    m/z 203.1754     0   1   2   3   4E+4   9   9.5  RT (min)       lcorbi_t3_pos_4_106_62    m/z 203.1754     0   2   4   6   8E+3   1   1.5  RT (min)      lcorbi_t3_pos_4_018_01    m/z 203.1835     0   1   2   3E+3   1.5   2  RT (min)       lcorbi_t3_pos_4_021_25    m/z 203.1835     0   1   2E+3   4.5   5  RT (min)      lcorbi_t3_pos_4_082_59    m/z 203.2138     0   1E+4   8   8.5  RT (min)       lcorbi_t3_pos_4_069_57    m/z 203.2199     0   1   2   3   4E+4   9.5   10  RT (min)      lcorbi_t3_pos_4_104_61    m/z 204.0179     0   2   4   6   8E+4   1   1.5  RT (min)       lcorbi_t3_pos_4_082_59    m/z 204.0367     0   1   2E+4   1   1.5  RT (min)       lcorbi_t3_pos_3_010_44    m/z 204.0421     0   1   2.5   3  RT (min)      lcorbi_t3_pos_4_098_61    m/z 204.0457     0   1   1   1.5  RT (min)      lcorbi_t3_pos_1_016_05    m/z 204.0625     0   1E+5   18.5   19  RT (min)      lcorbi_t3_pos_4_091_60    m/z 204.0717     0   1   2   3   4E+3   1   1.5  RT (min)      lcorbi_t3_pos_1_021_18    m/z 204.0867     0   1E+4   1.5   2  RT (min)       lcorbi_t3_pos_4_066_57    m/z 204.0979     0   1   6   6.5  RT (min)      lcorbi_t3_pos_4_083_59    m/z 204.1066     0   1   2   3   4E+3   3.5   4   4.5  RT (min)       lcorbi_t3_pos_1_001_49    m/z 204.1174     0   1   2E+4   1   1.5  RT (min)       lcorbi_t3_pos_2_021_09    m/z 204.1294     0   1   7   7.5  RT (min)      lcorbi_t3_pos_4_105_61    m/z 204.1340     0   1E+4   1   1.5  RT (min)      lcorbi_t3_pos_1_051_09    m/z 204.1536     0   1   2   3   4E+4   1   1.5  RT (min)      lcorbi_t3_pos_3_024_07    m/z 204.1594     0   2   4   6E+4   8   8.5  RT (min)       lcorbi_t3_pos_1_001_49    m/z 204.1898     0   1   1   1.5  RT (min)      lcorbi_t3_pos_1_001_49    m/z 204.2207     0   2   4   6   8E+3   8   8.5  RT (min)      lcorbi_t3_pos_1_002_01    m/z 204.2232     0   1E+4   8   8.5  RT (min)      lcorbi_t3_pos_1_012_14    m/z 204.9333     0   1   2E+4   0.5   1  RT (min)       lcorbi_t3_pos_4_076_58    m/z 204.9570     0   1   2E+5   1   1.5  RT (min)       lcorbi_t3_pos_4_079_58    m/z 204.9570     0   2   4   6E+4   11.5   12  RT (min)      lcorbi_t3_pos_4_106_62    m/z 204.9783     0   1E+3   1   1.5  RT (min)      lcorbi_t3_pos_1_014_03    m/z 205.0158     0   2   4   6E+3   0.5   1  RT (min)      lcorbi_t3_pos_4_076_58    m/z 205.0384     0   2   4   6   8E+3   0.5   1  RT (min)       lcorbi_t3_pos_4_080_58    m/z 205.0593     0   1   2E+4   1   1.5  RT (min)      lcorbi_t3_pos_1_052_38    m/z 205.0642     0   1   2E+3   2.5   3  RT (min)      lcorbi_t3_pos_1_002_01    m/z 205.0747     0   1E+6   1   1.5  RT (min)      lcorbi_t3_pos_4_075_58    m/z 205.0747     0   1   2   3E+5   3   3.5  RT (min)      lcorbi_t3_pos_4_017_51    m/z 205.0747     0   1   2   3   4E+5   13   13.5  RT (min)      lcorbi_t3_pos_2_054_29    m/z 205.0819     0   1   2   3E+3   1   1.5  RT (min)       lcorbi_t3_pos_1_015_11    m/z 205.0899     0   2   4E+3   1   1.5  RT (min)      lcorbi_t3_pos_1_006_15    m/z 205.0920     0   1   2E+4   8   8.5  RT (min)      lcorbi_t3_pos_1_019_16    m/z 205.1336     0   1   2   3E+3   1   1.5  RT (min)      lcorbi_t3_pos_1_011_44    m/z 205.1434     0   2   4   6E+3   5.5   6   6.5  RT (min)       lcorbi_t3_pos_1_004_24    m/z 205.1434     0   1E+4   7.5   8  RT (min)       lcorbi_t3_pos_1_011_44    m/z 205.1463     0   1   1.5   2  RT (min)      lcorbi_t3_pos_1_012_14    m/z 205.1547     0   1   2   3E+3   1   1.5  RT (min)      lcorbi_t3_pos_1_003_45    m/z 205.1587     0   1   2E+4   7   7.5  RT (min)       lcorbi_t3_pos_1_037_19    m/z 205.1951     0   1   2   3E+4   10.5   11  RT (min)       lcorbi_t3_pos_4_112_62    m/z 205.1991     0   2   4   6   8E+4   6   6.5  RT (min)       lcorbi_t3_pos_2_044_21    m/z 205.4207     0   1E+5   13.5   14  RT (min)      lcorbi_t3_pos_1_036_08    m/z 205.9245     0   2   4   6E+4   13.5   14  RT (min)      lcorbi_t3_pos_4_091_60    m/z 205.9578     0   1   2E+3   1   1.5  RT (min)      lcorbi_t3_pos_4_090_60    m/z 206.0022     0   1   2   3E+3   1   1.5  RT (min)      lcorbi_t3_pos_3_055_10    m/z 206.0110     0   1   2   3E+4   0   0.5  RT (min)      lcorbi_t3_pos_1_002_01    m/z 206.0222     0   1   2.5   3  RT (min)      lcorbi_t3_pos_4_082_59    m/z 206.0336     0   2   4   6E+4   1   1.5  RT (min)       lcorbi_t3_pos_4_094_60    m/z 206.0424     0   1   2.5   3  RT (min)      lcorbi_t3_pos_4_082_59    m/z 206.0424     0   1   2E+4   4.5   5  RT (min)       lcorbi_t3_pos_4_074_58    m/z 206.0481     0   1E+4   5.5   6  RT (min)      lcorbi_t3_pos_1_014_03    m/z 206.0482     0   1   2.5   3  RT (min)      lcorbi_t3_pos_4_082_59    m/z 206.0554     0   1   2E+3   1   1.5  RT (min)      lcorbi_t3_pos_1_036_08    m/z 206.0672     0   2   4   6E+3   4.5   5  RT (min)      lcorbi_t3_pos_1_008_36    m/z 206.0708     0   1E+5   1.5   2  RT (min)      lcorbi_t3_pos_4_047_28    m/z 206.0812     0   1   2.5   3  RT (min)       lcorbi_t3_pos_4_077_58    m/z 206.0845     0   2   4   6   8E+2   5   5.5  RT (min)      lcorbi_t3_pos_4_038_38    m/z 206.0888     0   1   1   1.5  RT (min)      lcorbi_t3_pos_2_051_35    m/z 206.0893     0   1E+4   10.5   11  RT (min)      lcorbi_t3_pos_1_006_15    m/z 206.0943     0   2   4E+3   4.5   5  RT (min)      lcorbi_t3_pos_4_080_58    m/z 206.1005     0   1E+5   4.5   5  RT (min)       lcorbi_t3_pos_4_074_58    m/z 206.1022     0   1   2   3E+4   18   18.5  RT (min)       lcorbi_t3_pos_4_093_60    m/z 206.1141     0   2   4   6   8E+2   1.5   2  RT (min)       lcorbi_t3_pos_3_014_29    m/z 206.1273     0   1   1   1.5   2  RT (min)       lcorbi_t3_pos_4_047_28    m/z 206.1467     0   1   2   3E+3   5.5   6  RT (min)      lcorbi_t3_pos_1_002_01    m/z 206.1519     0   1E+5   1   1.5  RT (min)       lcorbi_t3_pos_3_029_13    m/z 206.1750     0   1   2   3E+3   3   3.5  RT (min)      lcorbi_t3_pos_1_001_49    m/z 206.4285     0   2   4E+4   13.5   14  RT (min)      lcorbi_t3_pos_4_090_60    m/z 206.6536     0   1   2   3   4E+4   1   1.5  RT (min)       lcorbi_t3_pos_4_087_59    m/z 206.8859     0   1   2E+3   1   1.5  RT (min)       lcorbi_t3_pos_1_019_16    m/z 206.9546     0   1   2E+3   1   1.5  RT (min)       lcorbi_t3_pos_1_054_48    m/z 206.9546     0   2   4E+4   18   18.5  RT (min)       lcorbi_t3_pos_4_089_59    m/z 206.9627     0   1   0   0.5  RT (min)      lcorbi_t3_pos_4_074_58    m/z 207.0142     0   2   4   6   8E+4   1   1.5  RT (min)       lcorbi_t3_pos_1_028_32    m/z 207.0499     0   1   2E+3   2.5   3  RT (min)      lcorbi_t3_pos_1_002_01    m/z 207.0507     0   2   4   6E+3   1   1.5  RT (min)      lcorbi_t3_pos_4_070_57    m/z 207.0520     0   1   2   3   4E+4   18   18.5  RT (min)      lcorbi_t3_pos_4_066_57    m/z 207.0707     0   2   4E+4   18.5   19  RT (min)      lcorbi_t3_pos_4_072_57    m/z 207.0709     0   1   2   3   4E+3   0.5   1  RT (min)      lcorbi_t3_pos_4_085_59    m/z 207.0764     0   1   4.5   5  RT (min)      lcorbi_t3_pos_4_112_62    m/z 207.0792     0   1   2   3   4E+4   18.5   19  RT (min)      lcorbi_t3_pos_4_067_57    m/z 207.0837     0   2   4   6E+3   7   7.5  RT (min)      lcorbi_t3_pos_1_033_53    m/z 207.0863     0   1   1.5   2  RT (min)      lcorbi_t3_pos_2_028_24    m/z 207.0878     0   1   2E+3   5   5.5  RT (min)      lcorbi_t3_pos_1_013_10    m/z 207.0992     0   1   2   3   4E+3   4.5   5  RT (min)      lcorbi_t3_pos_4_014_48    m/z 207.1077     0   1   2   3E+5   6   7  RT (min)       lcorbi_t3_pos_2_004_42    m/z 207.1077     0   1   2   3   4E+4   2.5   3   3.5  RT (min)      lcorbi_t3_pos_4_021_25    m/z 207.1077     0   1   2E+5   9.5   10  RT (min)      lcorbi_t3_pos_4_080_58    m/z 207.1078     0   1E+5   13   13.5  RT (min)      lcorbi_t3_pos_2_003_19    m/z 207.1227     0   1   2   3E+3   4.5   5  RT (min)      lcorbi_t3_pos_4_046_44    m/z 207.1339     0   1   2   3E+4   1   1.5  RT (min)      lcorbi_t3_pos_4_077_58    m/z 207.1458     0   1   1   1.5  RT (min)      lcorbi_t3_pos_4_070_57    m/z 207.1592     0   1   2   3E+4   6.5   7  RT (min)      lcorbi_t3_pos_1_001_49    m/z 207.1784     0   1   2E+4   5.5   6  RT (min)      lcorbi_t3_pos_1_019_16    m/z 207.1833     0   1E+5   14   14.5  RT (min)      lcorbi_t3_pos_3_055_10    m/z 207.9724     0   2   4   6E+3   1   1.5  RT (min)      lcorbi_t3_pos_2_025_52    m/z 207.9984     0   1   2E+5   1   1.5  RT (min)       lcorbi_t3_pos_1_034_34    m/z 208.0482     0   1E+4   1   1.5  RT (min)      lcorbi_t3_pos_4_079_58    m/z 208.0638     0   1   2   3   4E+5   1.5   2  RT (min)       lcorbi_t3_pos_3_048_30    m/z 208.0638     0   1   2   3E+4   4.5   5  RT (min)       lcorbi_t3_pos_4_012_37    m/z 208.0716     0   1E+4   9   9.5  RT (min)      lcorbi_t3_pos_1_027_43    m/z 208.0798     0   1   2E+3   4.5   5  RT (min)      lcorbi_t3_pos_4_048_24    m/z 208.0815     0   1   2   3   4E+3   1   1.5  RT (min)      lcorbi_t3_pos_3_003_33    m/z 208.0987     0   1   2E+3   1   1.5  RT (min)      lcorbi_t3_pos_3_003_33    m/z 208.1049     0   2   4E+4   14.5   15  RT (min)       lcorbi_t3_pos_2_016_23    m/z 208.1085     0   1   2   3   4E+4   9.5   10  RT (min)      lcorbi_t3_pos_1_035_28    m/z 208.1085     0   1   2E+4   4.5   5  RT (min)      lcorbi_t3_pos_4_048_24    m/z 208.1086     0   1   2   3   4E+4   6   7  RT (min)       lcorbi_t3_pos_3_002_16    m/z 208.1087     0   2   4   6   8E+4   18   18.5  RT (min)       lcorbi_t3_pos_4_067_57    m/z 208.1298     0   1   1.5   2  RT (min)      lcorbi_t3_pos_3_048_30    m/z 208.1308     0   2   4E+3   6.5   7  RT (min)      lcorbi_t3_pos_4_034_42    m/z 208.1379     0   1   1   1.5  RT (min)       lcorbi_t3_pos_3_003_33    m/z 208.1379     0   1   2   3E+4   2.5   3  RT (min)      lcorbi_t3_pos_1_036_08    m/z 208.1543     0   1   2   3E+4   5   5.5  RT (min)       lcorbi_t3_pos_1_006_15    m/z 208.1696     0   1   2   3E+4   8.5   9  RT (min)      lcorbi_t3_pos_4_106_62    m/z 208.1907     0   2   4   6   8E+3   8   8.5  RT (min)      lcorbi_t3_pos_1_001_49    m/z 208.2060     0   2   4   6   8E+3   9   9.5  RT (min)      lcorbi_t3_pos_1_001_49    m/z 208.8587     0   1   2E+4   1   1.5  RT (min)      lcorbi_t3_pos_1_034_34    m/z 208.9549     0   2   4E+3   0.5   1  RT (min)      lcorbi_t3_pos_4_083_59    m/z 208.9801     0   1   2.5   3  RT (min)      lcorbi_t3_pos_1_002_01    m/z 209.0089     0   1   0.5   1  RT (min)      lcorbi_t3_pos_4_078_58    m/z 209.0332     0   1E+5   0   1  RT (min)       lcorbi_t3_pos_2_005_11    m/z 209.0333     0   2   4   6E+4   4.5   5  RT (min)       lcorbi_t3_pos_4_106_62    m/z 209.0556     0   2   4   6E+3   1.5   2  RT (min)      lcorbi_t3_pos_2_027_05    m/z 209.0727     0   1E+5   1   1.5  RT (min)       lcorbi_t3_pos_1_036_08    m/z 209.0808     0   1E+4   7.5   8  RT (min)      lcorbi_t3_pos_4_090_60    m/z 209.0809     0   1   2   3E+5   17.5   18  RT (min)      lcorbi_t3_pos_4_074_58    m/z 209.0921     0   1   2E+3   2.5   3  RT (min)      lcorbi_t3_pos_1_003_45    m/z 209.0964     0   1   2   3E+4   0.5   1  RT (min)      lcorbi_t3_pos_1_001_49    m/z 209.0964     0   1   2   3E+4   18   18.5  RT (min)       lcorbi_t3_pos_4_076_58    m/z 209.1020     0   2   4   6E+2   2.5   3  RT (min)      lcorbi_t3_pos_3_023_05    m/z 209.1038     0   1E+4   4.5   5  RT (min)      lcorbi_t3_pos_1_050_13    m/z 209.1054     0   2   4   6E+4   6.5   7   7.5  RT (min)       lcorbi_t3_pos_2_041_54    m/z 209.1172     0   2   4E+5   8   9  RT (min)       lcorbi_t3_pos_2_053_28    m/z 209.1172     0   2   4E+5   12.5   13  RT (min)      lcorbi_t3_pos_4_107_62    m/z 209.1201     0   1   5   5.5  RT (min)      lcorbi_t3_pos_1_001_49    m/z 209.1291     0   1E+4   1   1.5  RT (min)      lcorbi_t3_pos_1_001_49    m/z 209.1325     0   1   14   14.5  RT (min)       lcorbi_t3_pos_4_091_60    m/z 209.1703     0   1   1   1.5  RT (min)       lcorbi_t3_pos_1_002_01    m/z 209.2012     0   2   4   6   8E+4   6   6.5  RT (min)       lcorbi_t3_pos_4_090_60    m/z 209.9405     0   2   4   6E+3   1   1.5  RT (min)      lcorbi_t3_pos_4_098_61    m/z 209.9769     0   1   2   3   4E+3   1   1.5  RT (min)      lcorbi_t3_pos_3_011_42    m/z 210.0042     0   1E+4   7   7.5  RT (min)      lcorbi_t3_pos_4_082_59    m/z 210.0501     0   1   2.5   3  RT (min)      lcorbi_t3_pos_4_044_32    m/z 210.0761     0   1   2E+3   4.5   5  RT (min)       lcorbi_t3_pos_1_019_16    m/z 210.0832     0   2   4E+3   2   2.5  RT (min)      lcorbi_t3_pos_1_028_32    m/z 210.0842     0   2   4   6E+3   7.5   8  RT (min)      lcorbi_t3_pos_1_044_25    m/z 210.0873     0   1   2   3   4E+4   10   10.5  RT (min)       lcorbi_t3_pos_4_076_58    m/z 210.0891     0   1   2E+4   5.5   6  RT (min)       lcorbi_t3_pos_4_106_62    m/z 210.0972     0   1   2   3E+3   1   1.5  RT (min)       lcorbi_t3_pos_4_081_58    m/z 210.1100     0   1   2   3   4E+4   5.5   6  RT (min)       lcorbi_t3_pos_4_098_61    m/z 210.1206     0   1E+4   0   0.5  RT (min)      lcorbi_t3_pos_1_005_42    m/z 210.1206     0   1E+5   13.5   14  RT (min)      lcorbi_t3_pos_4_074_58    m/z 210.1206     0   1E+5   15   15.5  RT (min)      lcorbi_t3_pos_4_055_02    m/z 210.1373     0   1   1   1.5   2  RT (min)       lcorbi_t3_pos_4_079_58    m/z 210.1417     0   1   2   3E+4   4.5   5  RT (min)      lcorbi_t3_pos_2_054_29    m/z 210.1488     0   1   2E+4   4.5   5  RT (min)       lcorbi_t3_pos_1_031_40    m/z 210.1852     0   2   4   6E+3   6   6.5  RT (min)      lcorbi_t3_pos_1_001_49    m/z 210.1934     0   1E+4   10   10.5  RT (min)      lcorbi_t3_pos_1_003_45    m/z 210.1965     0   2   4   6E+6   0   0.5  RT (min)      lcorbi_t3_pos_1_001_49    m/z 210.2046     0   2   4E+3   6   6.5  RT (min)      lcorbi_t3_pos_1_001_49    m/z 210.8558     0   2   4E+4   1   1.5  RT (min)      lcorbi_t3_pos_4_067_57    m/z 210.9057     0   2   4   6E+4   18   18.5  RT (min)       lcorbi_t3_pos_4_113_62    m/z 210.9058     0   1   2   3E+4   1   1.5  RT (min)      lcorbi_t3_pos_4_070_57    m/z 210.9058     0   1   2   3E+4   11   11.5  RT (min)      lcorbi_t3_pos_2_002_06    m/z 210.9213     0   1   2E+5   0   0.5  RT (min)      lcorbi_t3_pos_1_007_07    m/z 210.9576     0   1   2   3   4E+4   1   1.5  RT (min)      lcorbi_t3_pos_4_073_57    m/z 210.9777     0   1E+5   1   1.5  RT (min)       lcorbi_t3_pos_4_067_57    m/z 210.9889     0   1   2   3   4E+3   1   1.5  RT (min)      lcorbi_t3_pos_4_012_37    m/z 211.0217     0   1   1   1.5  RT (min)      lcorbi_t3_pos_1_006_15    m/z 211.0367     0   1E+4   6.5   7  RT (min)      lcorbi_t3_pos_4_066_57    m/z 211.0577     0   1   2E+3   4.5   5  RT (min)      lcorbi_t3_pos_1_001_49    m/z 211.0642     0   1   1.5   2  RT (min)      lcorbi_t3_pos_1_004_24    m/z 211.0812     0   2   4E+3   1   1.5  RT (min)      lcorbi_t3_pos_1_001_49    m/z 211.0851     0   1   2E+4   13   13.5  RT (min)      lcorbi_t3_pos_1_004_24    m/z 211.0920     0   1   2E+4   18   18.5  RT (min)       lcorbi_t3_pos_4_075_58    m/z 211.0997     0   1   2E+3   1   1.5  RT (min)      lcorbi_t3_pos_1_001_49    m/z 211.1077     0   1E+4   4.5   5  RT (min)      lcorbi_t3_pos_1_001_49    m/z 211.1077     0   2   4   6   8E+3   2.5   3  RT (min)       lcorbi_t3_pos_1_001_49    m/z 211.1189     0   1   2E+3   1.5   2  RT (min)       lcorbi_t3_pos_1_002_01    m/z 211.1295     0   1   2E+3   2.5   3  RT (min)      lcorbi_t3_pos_1_001_49    m/z 211.1376     0   1   2E+3   2.5   3  RT (min)      lcorbi_t3_pos_1_001_49    m/z 211.1441     0   2   4   6   8E+3   5.5   6  RT (min)       lcorbi_t3_pos_1_001_49    m/z 211.1481     0   1   2   3   4E+3   7   7.5  RT (min)      lcorbi_t3_pos_1_001_49    m/z 211.1733     0   1   2   3E+4   5   5.5  RT (min)      lcorbi_t3_pos_4_099_61    m/z 211.1771     0   1E+5   1   1.5  RT (min)       lcorbi_t3_pos_4_091_60    m/z 211.9376     0   1E+4   1   1.5  RT (min)      lcorbi_t3_pos_4_106_62    m/z 212.0310     0   2   4   6E+4   1   1.5  RT (min)       lcorbi_t3_pos_1_036_08    m/z 212.0529     0   1   1.5   2  RT (min)      lcorbi_t3_pos_3_016_21    m/z 212.0701     0   2   4   6E+2   1   1.5  RT (min)      lcorbi_t3_pos_3_001_49    m/z 212.0893     0   1   2E+3   4.5   5  RT (min)      lcorbi_t3_pos_1_047_41    m/z 212.0917     0   1   2   3E+3   2.5   3  RT (min)      lcorbi_t3_pos_4_094_60    m/z 212.0999     0   2   4   6E+3   6.5   7  RT (min)      lcorbi_t3_pos_4_110_62    m/z 212.1182     0   2   4E+4   5   5.5  RT (min)       lcorbi_t3_pos_4_082_59    m/z 212.1281     0   1   2E+4   8   8.5  RT (min)      lcorbi_t3_pos_1_023_02    m/z 212.1492     0   1   2   3E+4   3   3.5  RT (min)      lcorbi_t3_pos_1_002_01    m/z 212.1757     0   1   2   3   4E+3   6   6.5  RT (min)      lcorbi_t3_pos_1_001_49    m/z 212.2009     0   1E+5   8.5   9  RT (min)       lcorbi_t3_pos_4_085_59    m/z 212.2009     0   1   2   3   4E+4   13   13.5  RT (min)      lcorbi_t3_pos_2_044_21    m/z 212.2373     0   2   4   6   8E+4   9   9.5  RT (min)       lcorbi_t3_pos_2_044_21    m/z 212.8525     0   1E+5   1   1.5  RT (min)       lcorbi_t3_pos_4_093_60    m/z 212.9028     0   2   4   6   8E+3   1   1.5  RT (min)      lcorbi_t3_pos_4_012_37    m/z 212.9547     0   1   2   3E+4   1   1.5  RT (min)      lcorbi_t3_pos_4_077_58    m/z 213.0010     0   1   2E+3   1.5   2  RT (min)      lcorbi_t3_pos_1_025_52    m/z 213.0151     0   1E+5   1   1.5  RT (min)      lcorbi_t3_pos_1_036_08    m/z 213.0564     0   2   4E+4   18   18.5  RT (min)      lcorbi_t3_pos_4_070_57    m/z 213.0565     0   2   4   6E+3   1   1.5  RT (min)      lcorbi_t3_pos_3_029_13    m/z 213.0635     0   2   4E+3   6   6.5  RT (min)      lcorbi_t3_pos_1_002_01    m/z 213.0788     0   1   3   3.5  RT (min)      lcorbi_t3_pos_1_002_01    m/z 213.0870     0   2   4   6   8E+2   1.5   2  RT (min)       lcorbi_t3_pos_4_014_48    m/z 213.0888     0   2   4   6   8E+3   5   5.5  RT (min)      lcorbi_t3_pos_1_002_01    m/z 213.0916     0   2   4   6   8E+3   6.5   7  RT (min)      lcorbi_t3_pos_1_001_49    m/z 213.0998     0   1E+5   2  RT (min)       lcorbi_t3_pos_4_106_62    m/z 213.1022     0   1   2E+4   5   5.5  RT (min)       lcorbi_t3_pos_4_082_59    m/z 213.1097     0   1   2   3   4E+3   5   5.5  RT (min)       lcorbi_t3_pos_1_001_49    m/z 213.1162     0   1   1   1.5  RT (min)      lcorbi_t3_pos_3_029_13    m/z 213.1216     0   2   4E+3   5   5.5  RT (min)      lcorbi_t3_pos_1_002_01    m/z 213.1233     0   1   2   3   4E+3   1.5   2  RT (min)       lcorbi_t3_pos_2_053_28    m/z 213.1526     0   2   4E+3   4.5   5  RT (min)      lcorbi_t3_pos_1_001_49    m/z 213.1532     0   1E+5   1   1.5   2  RT (min)       lcorbi_t3_pos_4_091_60    m/z 213.1679     0   1   2E+4   7.5   8  RT (min)      lcorbi_t3_pos_4_093_60    m/z 213.1961     0   1   2   3E+4   8   8.5  RT (min)       lcorbi_t3_pos_4_066_57    m/z 213.2042     0   1   2   3   4E+3   8.5   9  RT (min)      lcorbi_t3_pos_1_001_49    m/z 213.9991     0   2   4   6   8E+3   6.5   7  RT (min)      lcorbi_t3_pos_4_106_62    m/z 214.0063     0   1E+6   1   1.5  RT (min)       lcorbi_t3_pos_1_031_40    m/z 214.0079     0   2   4   6E+4   18   18.5  RT (min)      lcorbi_t3_pos_4_094_60    m/z 214.0508     0   1E+5   4.5   5  RT (min)       lcorbi_t3_pos_4_014_48    m/z 214.0562     0   1   1   1.5  RT (min)      lcorbi_t3_pos_4_096_60    m/z 214.0589     0   2   4   6   8E+4   1   1.5  RT (min)      lcorbi_t3_pos_1_031_40    m/z 214.0685     0   1   2   3E+3   1   1.5  RT (min)       lcorbi_t3_pos_2_040_25    m/z 214.0782     0   1   1   1.5  RT (min)      lcorbi_t3_pos_4_090_60    m/z 214.0791     0   1   11   11.5  RT (min)      lcorbi_t3_pos_1_006_15    m/z 214.1017     0   1   2   3E+3   2.5   3  RT (min)      lcorbi_t3_pos_1_009_50    m/z 214.1032     0   1E+3   1.5   2  RT (min)      lcorbi_t3_pos_1_028_32    m/z 214.1169     0   1   2   3E+3   6   6.5  RT (min)      lcorbi_t3_pos_1_002_01    m/z 214.1269     0   1   2   3   4E+4   1   1.5  RT (min)       lcorbi_t3_pos_4_094_60    m/z 214.1437     0   1E+4   1   1.5  RT (min)      lcorbi_t3_pos_1_034_34    m/z 214.1581     0   1E+4   6.5   7  RT (min)      lcorbi_t3_pos_4_052_06    m/z 214.1849     0   1E+4   1   1.5  RT (min)      lcorbi_t3_pos_4_091_60    m/z 214.2165     0   2   4   6   8E+4   8   8.5  RT (min)       lcorbi_t3_pos_4_113_62    m/z 214.9176     0   2   4   6   8E+4   0.5   1  RT (min)       lcorbi_t3_pos_4_075_58    m/z 214.9176     0   1E+5   18.5   19  RT (min)       lcorbi_t3_pos_4_081_58    m/z 214.9896     0   2   4E+4   18   18.5  RT (min)      lcorbi_t3_pos_4_087_59    m/z 214.9897     0   1E+4   0.5   1  RT (min)      lcorbi_t3_pos_4_087_59    m/z 215.0097     0   1   1   1.5  RT (min)      lcorbi_t3_pos_3_029_13    m/z 215.0162     0   1E+5   1.5   2  RT (min)       lcorbi_t3_pos_1_029_29    m/z 215.0339     0   1E+5   8   8.5  RT (min)       lcorbi_t3_pos_1_051_09    m/z 215.0339     0   2   4E+4   6.5   7   7.5  RT (min)       lcorbi_t3_pos_1_028_32    m/z 215.0513     0   2   4   6   8E+3   1   1.5  RT (min)       lcorbi_t3_pos_1_008_36    m/z 215.0541     0   1   2   3E+3   4.5   5  RT (min)      lcorbi_t3_pos_1_004_24    m/z 215.0696     0   1E+5   2.5   3  RT (min)       lcorbi_t3_pos_4_091_60    m/z 215.0731     0   1   1   1.5  RT (min)      lcorbi_t3_pos_1_002_01    m/z 215.0849     0   1   2   3E+3   2.5   3  RT (min)      lcorbi_t3_pos_1_003_45    m/z 215.0877     0   1E+4   1   1.5  RT (min)      lcorbi_t3_pos_2_040_25    m/z 215.0890     0   2   4   6   8E+4   4.5   5  RT (min)       lcorbi_t3_pos_2_021_09    m/z 215.0928     0   2   4   6   8E+3   8   8.5  RT (min)      lcorbi_t3_pos_1_002_01    m/z 215.0955     0   2   4E+4   18   18.5  RT (min)      lcorbi_t3_pos_4_068_57    m/z 215.1026     0   1   2E+4   1.5   2  RT (min)       lcorbi_t3_pos_3_048_30    m/z 215.1067     0   1   2   3E+4   7   7.5  RT (min)       lcorbi_t3_pos_4_005_33    m/z 215.1108     0   1   2   3E+4   4.5   5  RT (min)      lcorbi_t3_pos_4_098_61    m/z 215.1179     0   1   2   3   4E+3   7.5   8  RT (min)      lcorbi_t3_pos_1_001_49    m/z 215.1254     0   2   4   6   8E+4   1   1.5  RT (min)      lcorbi_t3_pos_1_002_01    m/z 215.1307     0   1   2E+4   4.5   5  RT (min)       lcorbi_t3_pos_4_106_62    m/z 215.1390     0   1E+4   1.5   2  RT (min)      lcorbi_t3_pos_1_008_36    m/z 215.1754     0   2   4   6   8E+3   8.5   9  RT (min)      lcorbi_t3_pos_1_002_01    m/z 215.1835     0   1E+4   7.5   8  RT (min)      lcorbi_t3_pos_1_002_01    m/z 215.2199     0   1E+4   8   8.5  RT (min)      lcorbi_t3_pos_1_001_49    m/z 215.2199     0   1E+4   10.5   11  RT (min)      lcorbi_t3_pos_4_112_62    m/z 215.2563     0   2   4   6E+4   9   9.5  RT (min)      lcorbi_t3_pos_4_086_59    m/z 215.9646     0   2   4   6E+4   1   1.5  RT (min)      lcorbi_t3_pos_2_020_22    m/z 215.9784     0   2   4   6E+3   5   5.5  RT (min)      lcorbi_t3_pos_4_082_59    m/z 216.0146     0   1   2   3   4E+4   1   1.5  RT (min)       lcorbi_t3_pos_1_032_26    m/z 216.0196     0   2   4   6   8E+2   1.5   2  RT (min)      lcorbi_t3_pos_2_005_11    m/z 216.0353     0   1E+5   1   1.5  RT (min)       lcorbi_t3_pos_4_085_59    m/z 216.0416     0   2   4   6   8E+3   6.5   7  RT (min)      lcorbi_t3_pos_4_098_61    m/z 216.0448     0   1   2E+3   0   0.5  RT (min)      lcorbi_t3_pos_4_096_60    m/z 216.0543     0   2   4E+4   2.5   3  RT (min)       lcorbi_t3_pos_4_006_12    m/z 216.0632     0   1   2E+5   1   1.5  RT (min)       lcorbi_t3_pos_4_089_59    m/z 216.0703     0   1   2   3E+3   3   3.5  RT (min)      lcorbi_t3_pos_1_001_49    m/z 216.0810     0   2   4E+4   1.5   2  RT (min)       lcorbi_t3_pos_4_052_06    m/z 216.0879     0   1   2   3   4E+3   5   5.5  RT (min)      lcorbi_t3_pos_1_017_51    m/z 216.1066     0   1   2E+3   4.5   5  RT (min)      lcorbi_t3_pos_1_001_49    m/z 216.1230     0   1E+4   1   1.5  RT (min)      lcorbi_t3_pos_3_029_13    m/z 216.1288     0   2   4   6   8E+4   5   5.5  RT (min)       lcorbi_t3_pos_4_103_61    m/z 216.1788     0   1   2   3E+3   5.5   6  RT (min)      lcorbi_t3_pos_1_001_49    m/z 216.1828     0   2   4   6   8E+4   10.5   11  RT (min)      lcorbi_t3_pos_4_106_62    m/z 216.1958     0   1E+6   7   7.5  RT (min)      lcorbi_t3_pos_2_018_04    m/z 216.6872     0   1   2E+5   1   1.5  RT (min)       lcorbi_t3_pos_4_095_60    m/z 216.8332     0   1   2   3   4E+4   1   1.5  RT (min)      lcorbi_t3_pos_4_075_58    m/z 216.9227     0   2   4E+5   17.5   18  RT (min)      lcorbi_t3_pos_4_074_58    m/z 216.9227     0   1   2   3   4E+4   12.5   13  RT (min)      lcorbi_t3_pos_3_029_13    m/z 216.9506     0   1   2   3E+4   0.5   1  RT (min)       lcorbi_t3_pos_4_098_61    m/z 217.0322     0   1   2E+3   1   1.5  RT (min)      lcorbi_t3_pos_3_008_20    m/z 217.0577     0   1   2E+3   2.5   3  RT (min)      lcorbi_t3_pos_1_001_49    m/z 217.0683     0   1E+6   1.5   2  RT (min)       lcorbi_t3_pos_3_016_21    m/z 217.0696     0   1   2   3E+3   4.5   5  RT (min)      lcorbi_t3_pos_1_001_49    m/z 217.0836     0   2   4E+3   4.5   5  RT (min)      lcorbi_t3_pos_1_001_49    m/z 217.0855     0   1   2   3E+3   2.5   3  RT (min)      lcorbi_t3_pos_1_001_49    m/z 217.0972     0   1   2   3   4E+3   5   5.5  RT (min)      lcorbi_t3_pos_1_001_49    m/z 217.1044     0   1   8   8.5  RT (min)      lcorbi_t3_pos_1_001_49    m/z 217.1046     0   2   4   6E+3   1   1.5  RT (min)      lcorbi_t3_pos_3_054_15    m/z 217.1182     0   2   4E+3   4.5   5  RT (min)      lcorbi_t3_pos_4_112_62    m/z 217.1183     0   1   2   3E+3   1.5   2  RT (min)       lcorbi_t3_pos_2_042_45    m/z 217.1198     0   2   4   6   8E+3   7.5   8  RT (min)      lcorbi_t3_pos_1_001_49    m/z 217.1295     0   1   2   3   4E+3   1.5   2  RT (min)       lcorbi_t3_pos_1_003_45    m/z 217.1296     0   1   5   5.5  RT (min)      lcorbi_t3_pos_1_001_49    m/z 217.1546     0   1   2E+4   1   1.5  RT (min)       lcorbi_t3_pos_4_075_58    m/z 217.1628     0   1E+4   7   7.5  RT (min)      lcorbi_t3_pos_4_082_59    m/z 217.1911     0   1   2E+4   8.5   9  RT (min)      lcorbi_t3_pos_3_044_46    m/z 217.1958     0   1E+4   7   7.5  RT (min)       lcorbi_t3_pos_1_027_43    m/z 217.2355     0   1E+4   8   8.5  RT (min)      lcorbi_t3_pos_4_027_07    m/z 217.2556     0   1   2   3   4E+4   6.5   7  RT (min)      lcorbi_t3_pos_4_014_48    m/z 217.9261     0   2   4   6E+4   1   1.5  RT (min)      lcorbi_t3_pos_1_020_31    m/z 217.9972     0   2   4   6   8E+4   1   1.5  RT (min)       lcorbi_t3_pos_4_082_59    m/z 218.0209     0   1E+5   7   7.5  RT (min)       lcorbi_t3_pos_4_102_61    m/z 218.0236     0   1E+4   0.5   1  RT (min)      lcorbi_t3_pos_4_088_59    m/z 218.0345     0   1   2   3E+4   1   1.5  RT (min)      lcorbi_t3_pos_4_086_59    m/z 218.0377     0   1E+4   11   11.5  RT (min)      lcorbi_t3_pos_3_013_19    m/z 218.0439     0   1E+4   0.5   1  RT (min)      lcorbi_t3_pos_4_083_59    m/z 218.0529     0   1   2   3   4E+4   7.5   8  RT (min)      lcorbi_t3_pos_4_098_61    m/z 218.0537     0   1   4   4.5  RT (min)       lcorbi_t3_pos_4_074_58    m/z 218.0716     0   2   4   6   8E+4   1   1.5  RT (min)      lcorbi_t3_pos_3_016_21    m/z 218.0813     0   1   2   3   4E+3   4.5   5  RT (min)      lcorbi_t3_pos_1_001_49    m/z 218.0874     0   1E+3   2.5   3  RT (min)      lcorbi_t3_pos_1_003_45    m/z 218.0918     0   2   4   6   8E+4   1   1.5  RT (min)       lcorbi_t3_pos_4_094_60    m/z 218.0981     0   1   2E+4   8   8.5  RT (min)       lcorbi_t3_pos_4_074_58    m/z 218.1023     0   2   4   6E+3   1.5   2  RT (min)       lcorbi_t3_pos_3_012_31    m/z 218.1080     0   1E+3   3   3.5  RT (min)      lcorbi_t3_pos_1_001_49    m/z 218.1080     0   2   4E+3   4.5   5  RT (min)      lcorbi_t3_pos_1_001_49    m/z 218.1104     0   1   2E+4   5.5   6  RT (min)      lcorbi_t3_pos_1_001_49    m/z 218.1257     0   1E+4   9   9.5  RT (min)      lcorbi_t3_pos_1_001_49    m/z 218.1330     0   1   2   3E+3   1.5   2  RT (min)      lcorbi_t3_pos_1_020_31    m/z 218.1387     0   1E+4   5.5   6  RT (min)       lcorbi_t3_pos_4_092_60    m/z 218.1387     0   2   4   6   8E+4   3  RT (min)       lcorbi_t3_pos_1_016_05    m/z 218.1497     0   1   1.5   2  RT (min)      lcorbi_t3_pos_3_012_31    m/z 218.1586     0   2   4   6   8E+3   6.5   7  RT (min)      lcorbi_t3_pos_1_001_49    m/z 218.1832     0   1E+4   9.5   10  RT (min)       lcorbi_t3_pos_4_014_48    m/z 218.1846     0   2   4   6E+4   6.5   7  RT (min)      lcorbi_t3_pos_1_046_22    m/z 218.2264     0   1E+4   6.5   7  RT (min)      lcorbi_t3_pos_4_011_22    m/z 218.2354     0   1E+5   6.5   7  RT (min)      lcorbi_t3_pos_1_019_16    m/z 218.2454     0   2   4E+4   6.5   7  RT (min)      lcorbi_t3_pos_2_044_21    m/z 218.4079     0   2   4   6E+4   6.5   7  RT (min)      lcorbi_t3_pos_1_041_54    m/z 218.9482     0   2   4   6E+3   0.5   1  RT (min)      lcorbi_t3_pos_4_069_57    m/z 218.9577     0   1E+3   1   1.5  RT (min)      lcorbi_t3_pos_4_012_37    m/z 218.9837     0   1E+4   0.5   1  RT (min)      lcorbi_t3_pos_4_069_57    m/z 219.0053     0   1   1   1.5  RT (min)       lcorbi_t3_pos_2_040_25    m/z 219.0244     0   1   6.5   7   7.5  RT (min)       lcorbi_t3_pos_4_047_28    m/z 219.0474     0   1   2   3   4E+4   7.5   8  RT (min)      lcorbi_t3_pos_1_001_49    m/z 219.0518     0   1E+6   0   0.5  RT (min)      lcorbi_t3_pos_1_006_15    m/z 219.0612     0   1   2   3E+5   1.5   2  RT (min)      lcorbi_t3_pos_2_044_21    m/z 219.0652     0   2   4   6   8E+3   6   6.5  RT (min)       lcorbi_t3_pos_1_014_03    m/z 219.0798     0   1   2E+4   2   2.5  RT (min)       lcorbi_t3_pos_4_077_58    m/z 219.0829     0   1   2   3   4E+6   1   1.5  RT (min)       lcorbi_t3_pos_1_034_34    m/z 219.0854     0   1   4.5   5  RT (min)      lcorbi_t3_pos_4_106_62    m/z 219.0951     0   1E+4   9   10  RT (min)       lcorbi_t3_pos_1_042_37    m/z 219.0975     0   1   1   1.5  RT (min)       lcorbi_t3_pos_4_091_60    m/z 219.1079     0   2   4   6E+3   1   1.5  RT (min)      lcorbi_t3_pos_1_032_26    m/z 219.1128     0   2   4E+4   4.5   5  RT (min)      lcorbi_t3_pos_4_113_62    m/z 219.1257     0   1   2.5   3  RT (min)      lcorbi_t3_pos_4_090_60    m/z 219.1268     0   1E+5   8  RT (min)       lcorbi_t3_pos_3_026_04    m/z 219.1268     0   1E+5   10.5   11  RT (min)      lcorbi_t3_pos_4_070_57    m/z 219.1268     0   1E+5   13   13.5  RT (min)      lcorbi_t3_pos_3_027_27    m/z 219.1420     0   1   2   3E+3   2.5   3  RT (min)      lcorbi_t3_pos_1_003_45    m/z 219.1426     0   1   2E+3   4.5   5  RT (min)      lcorbi_t3_pos_4_106_62    m/z 219.1440     0   1E+5   18   18.5  RT (min)       lcorbi_t3_pos_4_083_59    m/z 219.1452     0   1   2E+3   1   1.5  RT (min)       lcorbi_t3_pos_1_034_34    m/z 219.1591     0   1   2   3E+4   6.5   7  RT (min)      lcorbi_t3_pos_4_027_07    m/z 219.1591     0   1   2   3E+4   9   9.5  RT (min)       lcorbi_t3_pos_1_030_21    m/z 219.1743     0   1   2E+5   7.5   8  RT (min)       lcorbi_t3_pos_1_012_14    m/z 219.1744     0   1E+5   12   12.5  RT (min)      lcorbi_t3_pos_4_109_62    m/z 219.1872     0   1E+4   9.5   10  RT (min)      lcorbi_t3_pos_2_014_37    m/z 219.1956     0   1   2E+4   7   7.5  RT (min)      lcorbi_t3_pos_1_019_16    m/z 219.9231     0   1   2   3E+4   1   1.5  RT (min)      lcorbi_t3_pos_1_023_02    m/z 219.9665     0   1   1.5   2  RT (min)      lcorbi_t3_pos_1_011_44    m/z 219.9780     0   1   1   1.5  RT (min)      lcorbi_t3_pos_1_011_44    m/z 220.0034     0   1   2E+4   0   0.5  RT (min)       lcorbi_t3_pos_1_012_14    m/z 220.0058     0   1   2E+3   4.5   5  RT (min)      lcorbi_t3_pos_1_012_14    m/z 220.0502     0   1   2   3   4E+5   0   0.5   1  RT (min)       lcorbi_t3_pos_1_002_01    m/z 220.0669     0   1E+5   1   1.5  RT (min)      lcorbi_t3_pos_1_034_34    m/z 220.0686     0   1E+3   6   6.5  RT (min)      lcorbi_t3_pos_2_053_28    m/z 220.0803     0   1E+5   1   1.5  RT (min)       lcorbi_t3_pos_1_036_08    m/z 220.0829     0   1   2   3E+3   5   5.5  RT (min)      lcorbi_t3_pos_1_001_49    m/z 220.0968     0   1   2   3   4E+3   6   6.5  RT (min)      lcorbi_t3_pos_1_034_34    m/z 220.1056     0   2   4E+3   1   1.5  RT (min)       lcorbi_t3_pos_4_093_60    m/z 220.1180     0   1E+5   4.5   5  RT (min)       lcorbi_t3_pos_4_082_59    m/z 220.1180     0   2   4   6E+3   1   1.5   2  RT (min)       lcorbi_t3_pos_1_036_08    m/z 220.1413     0   1   2   3   4E+4   6.5   7  RT (min)      lcorbi_t3_pos_4_066_57    m/z 220.1458     0   1E+4   8.5   9  RT (min)      lcorbi_t3_pos_1_002_01    m/z 220.1463     0   1E+3   2.5   3  RT (min)      lcorbi_t3_pos_1_002_01    m/z 220.1543     0   1E+4   1   1.5  RT (min)       lcorbi_t3_pos_2_024_15    m/z 220.1543     0   1   2E+4   4   4.5   5  RT (min)       lcorbi_t3_pos_1_001_49    m/z 220.1625     0   1   6.5   7  RT (min)      lcorbi_t3_pos_1_001_49    m/z 220.1777     0   1   2   3   4E+4   7.5   8  RT (min)      lcorbi_t3_pos_1_012_14    m/z 220.1907     0   2   4   6   8E+4   6.5   7  RT (min)       lcorbi_t3_pos_4_053_19    m/z 220.2117     0   1   6.5   7  RT (min)      lcorbi_t3_pos_4_081_58    m/z 220.7016     0   2   4   6E+3   4.5   5  RT (min)      lcorbi_t3_pos_1_050_13    m/z 220.8964     0   1   1   1.5  RT (min)      lcorbi_t3_pos_1_046_22    m/z 220.9056     0   1E+5   0   0.5  RT (min)      lcorbi_t3_pos_1_009_50    m/z 220.9346     0   2   4   6E+5   1   1.5  RT (min)       lcorbi_t3_pos_4_078_58    m/z 220.9864     0   1E+3   1   1.5  RT (min)       lcorbi_t3_pos_1_006_15    m/z 221.0318     0   2   4   6E+4   0   0.5  RT (min)      lcorbi_t3_pos_1_006_15    m/z 221.0470     0   1E+7   0.5   1  RT (min)       lcorbi_t3_pos_1_005_42    m/z 221.0471     0   1E+5   18  RT (min)       lcorbi_t3_pos_4_077_58    m/z 221.0471     0   2   4   6   8E+5   14.5   15  RT (min)       lcorbi_t3_pos_1_006_15    m/z 221.0527     0   1E+5   0   0.5  RT (min)      lcorbi_t3_pos_1_006_15    m/z 221.0631     0   2   4   6   8E+3   8.5   9  RT (min)       lcorbi_t3_pos_1_037_19    m/z 221.0784     0   1   2   3E+4   5   5.5  RT (min)       lcorbi_t3_pos_4_079_58    m/z 221.0800     0   1E+5   1   1.5  RT (min)       lcorbi_t3_pos_4_093_60    m/z 221.0931     0   1E+5   1   1.5  RT (min)       lcorbi_t3_pos_4_091_60    m/z 221.1020     0   2   4   6E+3   4.5   5  RT (min)      lcorbi_t3_pos_1_002_01    m/z 221.1056     0   1E+4   6   6.5  RT (min)      lcorbi_t3_pos_4_113_62    m/z 221.1078     0   1   1   1.5  RT (min)       lcorbi_t3_pos_4_089_59    m/z 221.1084     0   1   2   3E+3   6   6.5  RT (min)      lcorbi_t3_pos_1_004_24    m/z 221.1148     0   1E+4   8.5   9  RT (min)      lcorbi_t3_pos_4_085_59    m/z 221.1384     0   1E+4   4.5   5  RT (min)      lcorbi_t3_pos_1_001_49    m/z 221.1598     0   1E+5   18   18.5  RT (min)      lcorbi_t3_pos_4_075_58    m/z 221.1699     0   1   2   3E+3   4.5   5  RT (min)      lcorbi_t3_pos_1_001_49    m/z 221.1747     0   1E+4   7   7.5  RT (min)      lcorbi_t3_pos_2_056_56    m/z 221.1810     0   1E+5   10   10.5  RT (min)      lcorbi_t3_pos_4_108_62    m/z 221.1941     0   1E+4   6.5   7  RT (min)      lcorbi_t3_pos_1_019_16    m/z 221.2188     0   1E+5   6.5   7  RT (min)       lcorbi_t3_pos_4_038_38    m/z 221.9380     0   1E+4   1   1.5  RT (min)      lcorbi_t3_pos_1_035_28    m/z 221.9880     0   1   1   1.5  RT (min)      lcorbi_t3_pos_2_036_12    m/z 222.0028     0   2   4E+3   4.5   5  RT (min)      lcorbi_t3_pos_4_074_58    m/z 222.0106     0   2   4   6E+4   1   1.5  RT (min)       lcorbi_t3_pos_4_090_60    m/z 222.0163     0   1   2   3   4E+3   4.5   5  RT (min)      lcorbi_t3_pos_4_111_62    m/z 222.0164     0   1   2E+3   2.5   3  RT (min)      lcorbi_t3_pos_3_038_18    m/z 222.0347     0   1   1   1.5  RT (min)       lcorbi_t3_pos_4_082_59    m/z 222.0437     0   2   4E+3   3   3.5  RT (min)      lcorbi_t3_pos_4_080_58    m/z 222.0504     0   1   2   3E+4   0.5   1  RT (min)      lcorbi_t3_pos_1_008_36    m/z 222.0587     0   1   2.5   3  RT (min)      lcorbi_t3_pos_4_013_36    m/z 222.0664     0   1   1   1.5  RT (min)      lcorbi_t3_pos_1_002_01    m/z 222.0794     0   1E+4   2.5   3  RT (min)      lcorbi_t3_pos_3_001_49    m/z 222.0817     0   1   5   5.5  RT (min)      lcorbi_t3_pos_1_006_15    m/z 222.0838     0   1   2   3   4E+4   7   7.5  RT (min)       lcorbi_t3_pos_4_106_62    m/z 222.0850     0   2   4E+2   1.5   2  RT (min)      lcorbi_t3_pos_3_004_24    m/z 222.0973     0   1   2   3E+3   4.5   5  RT (min)      lcorbi_t3_pos_1_006_15    m/z 222.1292     0   1   1.5   2  RT (min)      lcorbi_t3_pos_2_053_28    m/z 222.1359     0   1   2E+4   9   9.5  RT (min)      lcorbi_t3_pos_4_107_62    m/z 222.1464     0   1   2   3   4E+4   7   7.5  RT (min)       lcorbi_t3_pos_4_098_61    m/z 222.1640     0   1   2E+4   1   1.5  RT (min)      lcorbi_t3_pos_4_047_28    m/z 222.1700     0   1   2   3E+4   5.5   6  RT (min)       lcorbi_t3_pos_4_027_07    m/z 222.1828     0   1E+4   10.5   11  RT (min)      lcorbi_t3_pos_3_020_35    m/z 222.8791     0   1E+5   1   1.5  RT (min)      lcorbi_t3_pos_4_073_57    m/z 222.9196     0   1E+4   0.5   1  RT (min)      lcorbi_t3_pos_4_081_58    m/z 222.9291     0   1   2E+4   1   1.5  RT (min)      lcorbi_t3_pos_2_012_01    m/z 222.9390     0   1   2   3E+3   1   1.5  RT (min)      lcorbi_t3_pos_1_032_26    m/z 223.0272     0   1E+4   0   0.5  RT (min)      lcorbi_t3_pos_1_018_35    m/z 223.0475     0   2   4E+3   0   0.5  RT (min)      lcorbi_t3_pos_4_084_59    m/z 223.0536     0   1   2   3E+3   5   5.5  RT (min)      lcorbi_t3_pos_4_014_48    m/z 223.0754     0   2   4   6E+4   9   9.5  RT (min)       lcorbi_t3_pos_1_039_23    m/z 223.0884     0   1   2   3   4E+3   1.5   2  RT (min)      lcorbi_t3_pos_2_028_24    m/z 223.0965     0   2   4   6E+4   13   13.5  RT (min)       lcorbi_t3_pos_1_011_44    m/z 223.1176     0   2   4   6   8E+3   4.5   5  RT (min)       lcorbi_t3_pos_1_001_49    m/z 223.1197     0   2   4E+4   18   18.5  RT (min)       lcorbi_t3_pos_4_106_62    m/z 223.1329     0   1   2   3E+4   8.5   9  RT (min)       lcorbi_t3_pos_1_016_05    m/z 223.1484     0   1E+4   18   18.5  RT (min)       lcorbi_t3_pos_4_092_60    m/z 223.1569     0   1E+5   1   1.5  RT (min)       lcorbi_t3_pos_4_091_60    m/z 223.1733     0   1   5.5   6  RT (min)      lcorbi_t3_pos_1_001_49    m/z 223.2169     0   2   4   6E+3   7.5   8  RT (min)      lcorbi_t3_pos_1_001_49    m/z 224.0453     0   2   4   6E+3   0.5   1  RT (min)      lcorbi_t3_pos_1_001_49    m/z 224.0632     0   2   4   6E+4   10   10.5  RT (min)      lcorbi_t3_pos_4_113_62    m/z 224.0765     0   1   2   3   4E+3   2   2.5  RT (min)      lcorbi_t3_pos_4_087_59    m/z 224.0787     0   2   4   6   8E+3   9   9.5  RT (min)      lcorbi_t3_pos_1_039_23    m/z 224.0887     0   1   2E+4   6.5   7  RT (min)      lcorbi_t3_pos_4_052_06    m/z 224.0887     0   1   2E+4   4.5   5  RT (min)      lcorbi_t3_pos_2_053_28    m/z 224.0887     0   1E+5   18   18.5  RT (min)       lcorbi_t3_pos_4_075_58    m/z 224.0917     0   2   4   6E+4   1   1.5   2  RT (min)       lcorbi_t3_pos_4_052_06    m/z 224.0999     0   1   2E+4   8.5   9  RT (min)      lcorbi_t3_pos_1_029_29    m/z 224.1013     0   1   2E+3   4.5   5  RT (min)      lcorbi_t3_pos_1_021_18    m/z 224.1129     0   1E+4   4.5   5  RT (min)      lcorbi_t3_pos_2_053_28    m/z 224.1257     0   1E+4   6.5   7  RT (min)      lcorbi_t3_pos_4_106_62    m/z 224.1328     0   1   1.5   2  RT (min)      lcorbi_t3_pos_4_021_25    m/z 224.1362     0   2   4   6E+3   8.5   9  RT (min)      lcorbi_t3_pos_1_039_23    m/z 224.1410     0   2   4   6E+2   1   1.5  RT (min)      lcorbi_t3_pos_4_021_25    m/z 224.1493     0   1   2   3   4E+3   4.5   5  RT (min)      lcorbi_t3_pos_1_021_18    m/z 224.1530     0   2   4   6E+4   1   1.5   2  RT (min)       lcorbi_t3_pos_4_091_60    m/z 224.1574     0   1E+4   5   5.5  RT (min)      lcorbi_t3_pos_4_014_48    m/z 224.1726     0   1   2   3E+4   10   10.5  RT (min)       lcorbi_t3_pos_4_027_07    m/z 224.8761     0   1E+4   1   1.5  RT (min)      lcorbi_t3_pos_1_034_34    m/z 224.8857     0   1E+5   1   1.5  RT (min)      lcorbi_t3_pos_1_031_40    m/z 224.9275     0   1E+4   1   1.5  RT (min)      lcorbi_t3_pos_4_077_58    m/z 224.9563     0   1   2   3   4E+3   1   1.5  RT (min)      lcorbi_t3_pos_1_029_29    m/z 225.0429     0   1E+5   0   0.5  RT (min)      lcorbi_t3_pos_1_002_01    m/z 225.0429     0   1   2   3E+5   10   10.5   11  RT (min)       lcorbi_t3_pos_4_113_62    m/z 225.0641     0   2   4   6E+3   1   1.5  RT (min)      lcorbi_t3_pos_3_010_44    m/z 225.0757     0   2   4E+4   4.5   5  RT (min)       lcorbi_t3_pos_4_112_62    m/z 225.0845     0   1   1.5   2  RT (min)      lcorbi_t3_pos_2_024_15    m/z 225.0870     0   2   4   6E+4   4.5   5  RT (min)       lcorbi_t3_pos_4_082_59    m/z 225.0910     0   1   2   3E+4   9.5   10  RT (min)      lcorbi_t3_pos_1_004_24    m/z 225.0968     0   1E+4   1   1.5  RT (min)      lcorbi_t3_pos_2_003_19    m/z 225.1094     0   1   2E+4   1   1.5  RT (min)       lcorbi_t3_pos_1_036_08    m/z 225.1097     0   2   4   6   8E+3   4.5   5  RT (min)       lcorbi_t3_pos_1_034_34    m/z 225.1234     0   1   2   3E+3   2.5   3  RT (min)      lcorbi_t3_pos_1_001_49    m/z 225.1234     0   2   4   6   8E+3   4.5   5  RT (min)      lcorbi_t3_pos_1_002_01    m/z 225.1653     0   1E+6   1   1.5  RT (min)       lcorbi_t3_pos_4_097_60    m/z 225.1654     0   1   8.5   9  RT (min)      lcorbi_t3_pos_4_076_58    m/z 225.1962     0   1E+5   15   15.5   16  RT (min)       lcorbi_t3_pos_4_020_41    m/z 225.1962     0   1E+5   13.5   14  RT (min)      lcorbi_t3_pos_4_071_57    m/z 225.1962     0   1E+5   17   17.5   18  RT (min)       lcorbi_t3_pos_4_073_57    m/z 225.2073     0   1   2E+6   0   0.5  RT (min)      lcorbi_t3_pos_1_001_49    m/z 225.2092     0   1E+5   12.5   13  RT (min)      lcorbi_t3_pos_2_006_47    m/z 225.9148     0   1E+4   1   1.5  RT (min)      lcorbi_t3_pos_1_009_50    m/z 226.0288     0   1E+5   1   1.5  RT (min)       lcorbi_t3_pos_4_090_60    m/z 226.0451     0   1   2   3E+3   2   2.5  RT (min)      lcorbi_t3_pos_4_074_58    m/z 226.0839     0   1   2E+3   5.5   6  RT (min)      lcorbi_t3_pos_1_001_49    m/z 226.0904     0   1   2E+3   4.5   5  RT (min)      lcorbi_t3_pos_1_001_49    m/z 226.0921     0   1   1   1.5  RT (min)       lcorbi_t3_pos_1_003_45    m/z 226.0975     0   2   4   6   8E+4   11   11.5  RT (min)       lcorbi_t3_pos_1_002_01    m/z 226.1048     0   2   4   6   8E+3   1   1.5  RT (min)      lcorbi_t3_pos_2_023_26    m/z 226.1187     0   1   2   3E+3   2.5   3  RT (min)      lcorbi_t3_pos_1_001_49    m/z 226.1226     0   1   2   3   4E+4   9   9.5  RT (min)       lcorbi_t3_pos_4_082_59    m/z 226.1286     0   1E+4   2.5   3  RT (min)       lcorbi_t3_pos_1_002_01    m/z 226.1486     0   1E+3   1.5   2  RT (min)       lcorbi_t3_pos_3_028_11    m/z 226.1519     0   1   2E+4   8.5   9  RT (min)       lcorbi_t3_pos_1_006_15    m/z 226.1550     0   2   4E+3   1   2   3  RT (min)       lcorbi_t3_pos_1_001_49    m/z 226.1649     0   1   2   3   4E+4   4.5   5  RT (min)       lcorbi_t3_pos_1_023_02    m/z 226.1914     0   1   2   3   4E+6   0   0.5  RT (min)      lcorbi_t3_pos_1_001_49    m/z 226.1914     0   1   2   3E+4   7   7.5  RT (min)       lcorbi_t3_pos_1_001_49    m/z 226.1914     0   1   2E+4   4.5   5  RT (min)      lcorbi_t3_pos_1_001_49    m/z 226.1932     0   1E+4   9   9.5  RT (min)      lcorbi_t3_pos_1_012_14    m/z 226.1994     0   1   2   3E+4   7   7.5  RT (min)      lcorbi_t3_pos_4_052_06    m/z 226.8622     0   2   4   6E+3   1   1.5  RT (min)      lcorbi_t3_pos_3_012_31    m/z 226.9004     0   1E+3   1   1.5  RT (min)      lcorbi_t3_pos_1_007_07    m/z 226.9229     0   1E+5   0   0.5  RT (min)      lcorbi_t3_pos_1_001_49    m/z 226.9344     0   2   4E+3   1   1.5  RT (min)      lcorbi_t3_pos_1_044_25    m/z 226.9429     0   2   4   6E+4   1   1.5  RT (min)      lcorbi_t3_pos_3_004_24    m/z 226.9515     0   1   2   3   4E+4   12.5   13  RT (min)      lcorbi_t3_pos_2_017_51    m/z 226.9619     0   1   2   3E+4   1   1.5  RT (min)      lcorbi_t3_pos_1_002_01    m/z 227.0166     0   1   2E+3   1   1.5  RT (min)      lcorbi_t3_pos_1_020_31    m/z 227.0222     0   1E+5   10   10.5  RT (min)       lcorbi_t3_pos_4_113_62    m/z 227.0526     0   1   4.5   5  RT (min)      lcorbi_t3_pos_4_074_58    m/z 227.0682     0   2   4   6   8E+2   1   1.5  RT (min)      lcorbi_t3_pos_3_038_18    m/z 227.0791     0   2   4   6   8E+4   4.5   5  RT (min)       lcorbi_t3_pos_4_074_58    m/z 227.0833     0   1E+5   0   0.5  RT (min)      lcorbi_t3_pos_4_010_43    m/z 227.1026     0   1E+3   1.5   2  RT (min)       lcorbi_t3_pos_2_037_27    m/z 227.1027     0   1   2   3E+3   4.5   5  RT (min)      lcorbi_t3_pos_4_077_58    m/z 227.1067     0   2   4   6E+4   7   7.5  RT (min)       lcorbi_t3_pos_4_005_33    m/z 227.1139     0   1E+4   1   1.5  RT (min)      lcorbi_t3_pos_4_012_37    m/z 227.1140     0   1   4.5   5  RT (min)      lcorbi_t3_pos_4_074_58    m/z 227.1253     0   1E+4   5.5   6  RT (min)      lcorbi_t3_pos_4_027_07    m/z 227.1325     0   1   1.5   2  RT (min)      lcorbi_t3_pos_1_052_38    m/z 227.1647     0   1E+4   4.5   5  RT (min)      lcorbi_t3_pos_3_048_30    m/z 227.1814     0   1   2   3E+3   4.5   5  RT (min)      lcorbi_t3_pos_4_084_59    m/z 227.2030     0   1   2   3E+4   9   9.5  RT (min)      lcorbi_t3_pos_1_001_49    m/z 227.2118     0   2   4   6E+3   6   6.5  RT (min)      lcorbi_t3_pos_1_001_49    m/z 227.2199     0   2   4   6E+4   8   8.5  RT (min)       lcorbi_t3_pos_3_044_46    m/z 228.0219     0   1   2E+3   1   1.5  RT (min)      lcorbi_t3_pos_2_009_50    m/z 228.0417     0   1E+4   18.5   19  RT (min)      lcorbi_t3_pos_4_101_61    m/z 228.0645     0   1E+3   3   3.5  RT (min)       lcorbi_t3_pos_4_066_57    m/z 228.0710     0   1E+4   1   1.5  RT (min)      lcorbi_t3_pos_4_098_61    m/z 228.0824     0   1   4.5   5  RT (min)      lcorbi_t3_pos_1_001_49    m/z 228.0882     0   1   5   5.5  RT (min)      lcorbi_t3_pos_1_002_01    m/z 228.0979     0   1   2E+3   1   1.5  RT (min)       lcorbi_t3_pos_4_030_05    m/z 228.1078     0   2   4   6   8E+4   1   1.5  RT (min)       lcorbi_t3_pos_4_110_62    m/z 228.1724     0   1   4.5   5  RT (min)       lcorbi_t3_pos_1_002_01    m/z 228.1959     0   2   4   6   8E+4   15   15.5  RT (min)      lcorbi_t3_pos_1_001_49    m/z 228.2322     0   2   4   6E+4   8   8.5  RT (min)      lcorbi_t3_pos_1_034_34    m/z 228.2686     0   1   2   3E+4   9.5   10  RT (min)      lcorbi_t3_pos_1_032_26    m/z 228.8130     0   1E+4   1   1.5  RT (min)      lcorbi_t3_pos_4_006_12    m/z 228.9240     0   1   2   3   4E+4   12   12.5  RT (min)      lcorbi_t3_pos_2_047_31    m/z 228.9241     0   2   4   6   8E+4   18   18.5  RT (min)       lcorbi_t3_pos_4_106_62    m/z 228.9770     0   2   4   6E+4   0   0.5  RT (min)      lcorbi_t3_pos_1_002_01    m/z 229.0412     0   2   4E+4   18   18.5  RT (min)       lcorbi_t3_pos_4_101_61    m/z 229.0413     0   2   4   6E+4   0   0.5   1  RT (min)       lcorbi_t3_pos_1_001_49    m/z 229.0496     0   1   2   3E+4   6.5   7  RT (min)       lcorbi_t3_pos_4_005_33    m/z 229.0550     0   2   4E+3   0.5   1  RT (min)       lcorbi_t3_pos_4_075_58    m/z 229.0616     0   1   2.5   3  RT (min)      lcorbi_t3_pos_4_106_62    m/z 229.0682     0   1   2   3E+4   8   8.5  RT (min)       lcorbi_t3_pos_4_086_59    m/z 229.0682     0   1E+3   2.5   3  RT (min)      lcorbi_t3_pos_4_084_59    m/z 229.0738     0   2   4   6E+2   1.5   2  RT (min)       lcorbi_t3_pos_4_021_25    m/z 229.0861     0   1   2   3   4E+3   4.5   5  RT (min)      lcorbi_t3_pos_4_090_60    m/z 229.0914     0   1   2   3   4E+3   1   1.5  RT (min)      lcorbi_t3_pos_4_050_31    m/z 229.1183     0   1   2   3   4E+3   4.5   5  RT (min)      lcorbi_t3_pos_4_090_60    m/z 229.1223     0   1   2E+4   7   7.5  RT (min)      lcorbi_t3_pos_3_011_42    m/z 229.1295     0   1   2   3E+3   1   1.5  RT (min)       lcorbi_t3_pos_4_090_60    m/z 229.1482     0   1E+5   4.5   5  RT (min)       lcorbi_t3_pos_4_090_60    m/z 229.1547     0   2   4   6E+5   1.5   2  RT (min)       lcorbi_t3_pos_4_039_35    m/z 229.1821     0   2   4E+4   4.5   5  RT (min)      lcorbi_t3_pos_4_086_59    m/z 229.1910     0   1   2   3   4E+3   6.5   7  RT (min)      lcorbi_t3_pos_1_003_45    m/z 229.2275     0   1   2   3E+4   7   7.5  RT (min)       lcorbi_t3_pos_4_085_59    m/z 229.2356     0   2   4   6   8E+4   11.5   12  RT (min)      lcorbi_t3_pos_1_032_26    m/z 229.9578     0   2   4   6E+3   4.5   5  RT (min)      lcorbi_t3_pos_4_090_60    m/z 229.9580     0   1   2   3E+3   1   1.5  RT (min)      lcorbi_t3_pos_1_034_34    m/z 229.9802     0   1E+5   1   1.5  RT (min)      lcorbi_t3_pos_1_034_34    m/z 230.0210     0   1E+5   8   8.5  RT (min)       lcorbi_t3_pos_1_051_09    m/z 230.0211     0   1   2E+4   6.5   7  RT (min)       lcorbi_t3_pos_2_023_26    m/z 230.0248     0   1   2E+4   4.5   5  RT (min)       lcorbi_t3_pos_4_111_62    m/z 230.0333     0   1E+4   2.5   3  RT (min)      lcorbi_t3_pos_3_034_26    m/z 230.0457     0   1E+5   1   1.5  RT (min)       lcorbi_t3_pos_3_048_30    m/z 230.0771     0   2   4   6E+3   1   1.5  RT (min)      lcorbi_t3_pos_4_055_02    m/z 230.0958     0   1   2   3E+3   0.5   1  RT (min)       lcorbi_t3_pos_4_074_58    m/z 230.1023     0   2   4   6   8E+3   2.5   3  RT (min)      lcorbi_t3_pos_1_002_01    m/z 230.1023     0   1E+4   4.5   5  RT (min)      lcorbi_t3_pos_1_001_49    m/z 230.1104     0   2   4   6E+4   5   5.5  RT (min)      lcorbi_t3_pos_4_008_30    m/z 230.1169     0   2   4   6   8E+3   8.5   9  RT (min)      lcorbi_t3_pos_1_001_49    m/z 230.1241     0   1   2   3E+4   4.5   5  RT (min)       lcorbi_t3_pos_4_090_60    m/z 230.1241     0   1   2E+3   1   1.5  RT (min)       lcorbi_t3_pos_2_040_25    m/z 230.1580     0   1E+5   1.5   2  RT (min)      lcorbi_t3_pos_4_077_58    m/z 230.2412     0   1   2   3E+4   9   9.5  RT (min)      lcorbi_t3_pos_1_032_26    m/z 230.2545     0   1   2   3   4E+4   9   9.5  RT (min)      lcorbi_t3_pos_4_038_38    m/z 230.8899     0   1   2   3   4E+5   0   0.5  RT (min)      lcorbi_t3_pos_1_006_15    m/z 230.9652     0   1   2   3E+4   9   9.5  RT (min)      lcorbi_t3_pos_2_039_30    m/z 230.9652     0   2   4   6   8E+3   5   5.5   6  RT (min)      lcorbi_t3_pos_2_025_52    m/z 230.9662     0   1   2E+3   1   1.5  RT (min)      lcorbi_t3_pos_3_029_13    m/z 230.9716     0   2   4   6E+4   0   0.5  RT (min)      lcorbi_t3_pos_1_006_15    m/z 230.9902     0   2   4   6E+3   1.5   2  RT (min)      lcorbi_t3_pos_1_052_38    m/z 231.0221     0   1   2E+4   6.5   7  RT (min)      lcorbi_t3_pos_4_104_61    m/z 231.0352     0   1   2   3   4E+3   6.5   7  RT (min)      lcorbi_t3_pos_4_104_61    m/z 231.0381     0   1   2E+4   18   18.5  RT (min)       lcorbi_t3_pos_4_106_62    m/z 231.0476     0   1E+5   1   1.5  RT (min)       lcorbi_t3_pos_1_028_32    m/z 231.0628     0   1E+4   7.5   8  RT (min)      lcorbi_t3_pos_4_075_58    m/z 231.0652     0   2   4E+3   1   1.5  RT (min)      lcorbi_t3_pos_4_012_37    m/z 231.0839     0   1   2   3   4E+4   2.5   3  RT (min)       lcorbi_t3_pos_4_034_42    m/z 231.0840     0   1E+3   4.5   5  RT (min)      lcorbi_t3_pos_4_037_26    m/z 231.0976     0   1   2E+4   1.5   2  RT (min)      lcorbi_t3_pos_4_084_59    m/z 231.0993     0   1   2   3   4E+4   5   5.5  RT (min)       lcorbi_t3_pos_4_103_61    m/z 231.1016     0   2   4E+4   8.5   9  RT (min)       lcorbi_t3_pos_4_072_57    m/z 231.1036     0   1E+3   1   1.5  RT (min)      lcorbi_t3_pos_3_016_21    m/z 231.1128     0   1   2   3E+3   4.5   5  RT (min)      lcorbi_t3_pos_2_044_21    m/z 231.1339     0   2   4   6   8E+2   4.5   5  RT (min)      lcorbi_t3_pos_4_024_27    m/z 231.1340     0   1   2   3E+4   1   1.5  RT (min)      lcorbi_t3_pos_3_011_42    m/z 231.1380     0   1E+5   11   11.5  RT (min)       lcorbi_t3_pos_1_028_32    m/z 231.1473     0   1E+5   1   1.5  RT (min)      lcorbi_t3_pos_4_074_58    m/z 231.1703     0   2   4   6   8E+2   1   1.5  RT (min)       lcorbi_t3_pos_2_037_27    m/z 231.1703     0   1   2E+4   4.5   5  RT (min)      lcorbi_t3_pos_3_020_35    m/z 231.1784     0   1   2   3E+4   7.5   8  RT (min)       lcorbi_t3_pos_4_106_62    m/z 231.2452     0   1   2   3   4E+4   9   9.5  RT (min)      lcorbi_t3_pos_2_044_21    m/z 231.2512     0   1   2   3   4E+4   10.5   11  RT (min)      lcorbi_t3_pos_4_110_62    m/z 231.9556     0   1   2E+3   1.5   2  RT (min)      lcorbi_t3_pos_2_005_11    m/z 231.9556     0   1   2   3   4E+3   4.5   5  RT (min)      lcorbi_t3_pos_1_056_56    m/z 231.9661     0   2   4   6   8E+3   4.5   5  RT (min)      lcorbi_t3_pos_1_037_19    m/z 232.0129     0   2   4   6   8E+4   1   1.5  RT (min)       lcorbi_t3_pos_4_089_59    m/z 232.0271     0   1E+4   2.5   3  RT (min)      lcorbi_t3_pos_1_008_36    m/z 232.0653     0   1   2   2.5  RT (min)      lcorbi_t3_pos_1_001_49    m/z 232.0751     0   2   4E+3   4.5   5  RT (min)      lcorbi_t3_pos_1_021_18    m/z 232.0757     0   1   2   3E+4   12   12.5  RT (min)      lcorbi_t3_pos_4_083_59    m/z 232.0968     0   1   2   3E+3   4.5   5  RT (min)      lcorbi_t3_pos_1_001_49    m/z 232.1050     0   1   2   3   4E+4   11.5   12  RT (min)      lcorbi_t3_pos_4_080_58    m/z 232.1179     0   1E+4   1.5   2  RT (min)      lcorbi_t3_pos_2_037_27    m/z 232.1237     0   1E+4   4.5   5  RT (min)      lcorbi_t3_pos_4_038_38    m/z 232.1293     0   1   2E+3   1.5   2  RT (min)      lcorbi_t3_pos_1_028_32    m/z 232.1332     0   1   2E+4   7.5   8  RT (min)       lcorbi_t3_pos_4_074_58    m/z 232.1404     0   1   2E+5   1   1.5  RT (min)       lcorbi_t3_pos_4_090_60    m/z 232.1484     0   2   4   6   8E+4   1   1.5  RT (min)       lcorbi_t3_pos_4_082_59    m/z 232.1544     0   1E+5   3   4  RT (min)       lcorbi_t3_pos_4_066_57    m/z 232.2546     0   1E+5   9   9.5  RT (min)      lcorbi_t3_pos_2_044_21    m/z 232.9079     0   1   2E+3   1   1.5  RT (min)       lcorbi_t3_pos_4_082_59    m/z 232.9282     0   1E+5   18.5   19  RT (min)      lcorbi_t3_pos_4_107_62    m/z 232.9282     0   2   4E+4   0   0.5  RT (min)       lcorbi_t3_pos_4_080_58    m/z 232.9631     0   1E+4   5.5   6  RT (min)      lcorbi_t3_pos_1_025_52    m/z 232.9631     0   1   2   3E+4   9   9.5  RT (min)      lcorbi_t3_pos_1_011_44    m/z 232.9855     0   1   5.5   6  RT (min)      lcorbi_t3_pos_2_025_52    m/z 233.0002     0   2   4   6   8E+4   18   18.5  RT (min)       lcorbi_t3_pos_4_084_59    m/z 233.0002     0   1E+4   0.5   1  RT (min)      lcorbi_t3_pos_4_083_59    m/z 233.0099     0   1   2E+4   1   1.5  RT (min)       lcorbi_t3_pos_1_031_40    m/z 233.0362     0   2   4   6E+4   11.5   12  RT (min)       lcorbi_t3_pos_4_081_58    m/z 233.0365     0   2   4E+3   6.5   7   7.5  RT (min)       lcorbi_t3_pos_4_107_62    m/z 233.0422     0   1E+5   1   1.5  RT (min)       lcorbi_t3_pos_3_016_21    m/z 233.0506     0   2   4   6E+3   1.5   2  RT (min)       lcorbi_t3_pos_1_034_34    m/z 233.0768     0   1E+3   1   1.5  RT (min)      lcorbi_t3_pos_4_007_46    m/z 233.0785     0   2   4   6E+4   6.5   7  RT (min)       lcorbi_t3_pos_1_009_50    m/z 233.0786     0   1   2   3E+3   3   3.5  RT (min)      lcorbi_t3_pos_2_040_25    m/z 233.0786     0   1   2E+4   4.5   5  RT (min)       lcorbi_t3_pos_4_082_59    m/z 233.0921     0   1   6.5   7  RT (min)      lcorbi_t3_pos_4_079_58    m/z 233.1033     0   1   2E+3   2.5   3  RT (min)      lcorbi_t3_pos_1_002_01    m/z 233.1061     0   2   4   6   8E+4   18   18.5  RT (min)       lcorbi_t3_pos_4_095_60    m/z 233.1107     0   1E+4   9.5   10  RT (min)      lcorbi_t3_pos_4_082_59    m/z 233.1138     0   2   4   6   8E+3   5   5.5  RT (min)      lcorbi_t3_pos_4_090_60    m/z 233.1285     0   1E+3   4.5   5  RT (min)      lcorbi_t3_pos_4_066_57    m/z 233.1577     0   1   2E+3   3   3.5  RT (min)      lcorbi_t3_pos_1_001_49    m/z 233.1747     0   2   4   6E+4   9   9.5  RT (min)       lcorbi_t3_pos_1_002_01    m/z 233.1941     0   2   4E+4   7.5   8   8.5  RT (min)       lcorbi_t3_pos_2_005_11    m/z 233.2112     0   1   2E+4   10   10.5  RT (min)      lcorbi_t3_pos_1_001_49    m/z 233.2264     0   1E+4   10   10.5  RT (min)      lcorbi_t3_pos_1_001_49    m/z 233.2305     0   2   4   6E+4   7.5   8  RT (min)      lcorbi_t3_pos_2_044_21    m/z 233.9000     0   1   2E+4   1   1.5  RT (min)      lcorbi_t3_pos_1_029_29    m/z 233.9315     0   1   2E+4   0.5   1  RT (min)      lcorbi_t3_pos_4_078_58    m/z 234.0549     0   2   4   6   8E+3   1   1.5  RT (min)      lcorbi_t3_pos_1_008_36    m/z 234.0612     0   1   2   3   4E+3   3   3.5  RT (min)      lcorbi_t3_pos_1_001_49    m/z 234.0649     0   1   2.5   3  RT (min)       lcorbi_t3_pos_1_003_45    m/z 234.0817     0   2   4   6E+3   6.5   7  RT (min)      lcorbi_t3_pos_1_002_01    m/z 234.0851     0   1   2.5   3  RT (min)      lcorbi_t3_pos_1_002_01    m/z 234.1237     0   1   2   3   4E+3   4.5   5  RT (min)      lcorbi_t3_pos_1_001_49    m/z 234.1336     0   1E+5   5.5   6  RT (min)       lcorbi_t3_pos_1_001_49    m/z 234.1530     0   1   2   3E+4   3   3.5  RT (min)      lcorbi_t3_pos_4_024_27    m/z 234.1598     0   1   2E+3   1   1.5  RT (min)      lcorbi_t3_pos_3_042_14    m/z 234.1619     0   2   4   6E+4   10   10.5  RT (min)       lcorbi_t3_pos_4_071_57    m/z 234.1661     0   2   4   6   8E+4   1   1.5  RT (min)       lcorbi_t3_pos_4_047_28    m/z 234.1700     0   2   4   6E+3   4.5   5  RT (min)      lcorbi_t3_pos_1_001_49    m/z 234.1700     0   2   4   6E+3   2.5   3  RT (min)      lcorbi_t3_pos_1_002_01    m/z 234.1871     0   1E+4   5   5.5  RT (min)      lcorbi_t3_pos_4_038_38    m/z 234.1904     0   1E+4   7   7.5  RT (min)      lcorbi_t3_pos_1_019_16    m/z 234.1974     0   1   2   3E+4   7   7.5  RT (min)      lcorbi_t3_pos_4_102_61    m/z 234.2064     0   1   2E+4   5   5.5  RT (min)      lcorbi_t3_pos_2_044_21    m/z 234.2064     0   1E+4   9.5   10  RT (min)      lcorbi_t3_pos_1_013_10    m/z 234.2153     0   1   2   3E+4   7   7.5  RT (min)      lcorbi_t3_pos_1_030_21    m/z 234.2242     0   2   4   6   8E+3   7   7.5  RT (min)      lcorbi_t3_pos_1_019_16    m/z 234.8263     0   1   2E+4   1   1.5  RT (min)      lcorbi_t3_pos_4_019_47    m/z 234.9241     0   1E+3   0.5   1  RT (min)      lcorbi_t3_pos_4_071_57    m/z 234.9612     0   1   2   3E+4   4.5   5  RT (min)       lcorbi_t3_pos_4_106_62    m/z 234.9612     0   1   2   3E+4   0.5   1  RT (min)       lcorbi_t3_pos_4_098_61    m/z 234.9612     0   1   2   3   4E+4   18   18.5  RT (min)      lcorbi_t3_pos_4_094_60    m/z 234.9815     0   1   1   1.5  RT (min)      lcorbi_t3_pos_4_082_59    m/z 234.9970     0   1E+3   1   1.5  RT (min)      lcorbi_t3_pos_4_082_59    m/z 235.0134     0   2   4   6E+4   1   1.5  RT (min)      lcorbi_t3_pos_1_028_32    m/z 235.0492     0   2   4E+4   6.5   7   7.5  RT (min)       lcorbi_t3_pos_4_099_61    m/z 235.0795     0   2   4   6E+4   4.5   5  RT (min)       lcorbi_t3_pos_4_090_60    m/z 235.0803     0   2   4   6   8E+4   2.5   3  RT (min)       lcorbi_t3_pos_4_067_57    m/z 235.0925     0   1E+4   1   1.5  RT (min)       lcorbi_t3_pos_1_031_40    m/z 235.0941     0   1   2E+3   5   5.5  RT (min)      lcorbi_t3_pos_2_016_23    m/z 235.0965     0   1E+5   8  RT (min)       lcorbi_t3_pos_4_106_62    m/z 235.1190     0   2   4   6   8E+3   1.5   2  RT (min)       lcorbi_t3_pos_4_044_32    m/z 235.1205     0   1E+5   4.5   5  RT (min)       lcorbi_t3_pos_1_036_08    m/z 235.1369     0   1   5.5   6  RT (min)      lcorbi_t3_pos_4_097_60    m/z 235.1441     0   2   4   6E+2   5.5   6  RT (min)       lcorbi_t3_pos_4_030_05    m/z 235.1540     0   1   2E+4   5.5   6  RT (min)       lcorbi_t3_pos_1_046_22    m/z 235.1904     0   2   4   6E+3   8.5   9  RT (min)      lcorbi_t3_pos_1_001_49    m/z 235.2038     0   1   2E+4   7   7.5  RT (min)      lcorbi_t3_pos_4_108_62    m/z 235.6341     0   1   2E+4   1.5   2  RT (min)       lcorbi_t3_pos_4_087_59    m/z 235.9673     0   1E+5   1   1.5  RT (min)       lcorbi_t3_pos_1_034_34    m/z 236.0329     0   1   2E+4   4.5   5  RT (min)       lcorbi_t3_pos_4_111_62    m/z 236.0457     0   1   1   1.5  RT (min)      lcorbi_t3_pos_1_034_34    m/z 236.0601     0   1   2   3   4E+3   1   1.5  RT (min)      lcorbi_t3_pos_1_051_09    m/z 236.0716     0   1   2   3E+4   8   8.5  RT (min)      lcorbi_t3_pos_1_001_49    m/z 236.0796     0   1   2E+4   1   1.5  RT (min)       lcorbi_t3_pos_4_091_60    m/z 236.0877     0   1E+5   1.5   2  RT (min)      lcorbi_t3_pos_3_034_26    m/z 236.0999     0   1   2E+4   7   7.5  RT (min)      lcorbi_t3_pos_4_109_62    m/z 236.1062     0   2   4   6E+3   1.5   2  RT (min)      lcorbi_t3_pos_1_023_02    m/z 236.1070     0   2   4   6E+4   4.5   5  RT (min)       lcorbi_t3_pos_3_048_30    m/z 236.1129     0   1   2E+4   5.5   6  RT (min)       lcorbi_t3_pos_1_002_01    m/z 236.1129     0   2   4   6   8E+3   1.5   2  RT (min)       lcorbi_t3_pos_1_051_09    m/z 236.1282     0   1   2   3   4E+3   6.5   7  RT (min)      lcorbi_t3_pos_1_001_49    m/z 236.1504     0   2   4   6E+3   1   1.5  RT (min)      lcorbi_t3_pos_1_051_09    m/z 236.1574     0   2   4E+3   5.5   6  RT (min)      lcorbi_t3_pos_1_001_49    m/z 236.1602     0   1   2   3E+3   1.5   2  RT (min)      lcorbi_t3_pos_1_001_49    m/z 236.1621     0   2   4   6E+4   7.5   8  RT (min)       lcorbi_t3_pos_4_106_62    m/z 236.1758     0   2   4E+3   4.5   5  RT (min)      lcorbi_t3_pos_1_001_49    m/z 236.1856     0   1E+4   6.5   7  RT (min)      lcorbi_t3_pos_4_027_07    m/z 236.2009     0   1   2E+4   9   9.5  RT (min)      lcorbi_t3_pos_4_074_58    m/z 236.2121     0   1E+5   7   7.5  RT (min)      lcorbi_t3_pos_4_053_19    m/z 236.8918     0   2   4   6   8E+3   1   1.5  RT (min)      lcorbi_t3_pos_1_009_50    m/z 236.9082     0   1   2   3   4E+4   1   1.5  RT (min)      lcorbi_t3_pos_1_001_49    m/z 236.9982     0   1   2   3E+3   1.5   2  RT (min)       lcorbi_t3_pos_1_001_49    m/z 237.0159     0   1   1   1.5  RT (min)       lcorbi_t3_pos_1_001_49    m/z 237.0524     0   1   5   5.5  RT (min)      lcorbi_t3_pos_4_036_11    m/z 237.0545     0   1   8.5   9  RT (min)       lcorbi_t3_pos_1_039_23    m/z 237.0549     0   1   1   1.5  RT (min)      lcorbi_t3_pos_1_003_45    m/z 237.0658     0   1E+4   0.5   1  RT (min)      lcorbi_t3_pos_1_013_10    m/z 237.0728     0   1   2E+3   4.5   5  RT (min)      lcorbi_t3_pos_1_001_49    m/z 237.0854     0   1   2E+4   1   1.5  RT (min)       lcorbi_t3_pos_4_094_60    m/z 237.0870     0   1E+3   4.5   5  RT (min)      lcorbi_t3_pos_4_008_30    m/z 237.0913     0   1E+4   18   18.5  RT (min)       lcorbi_t3_pos_4_074_58    m/z 237.0963     0   1   2   3E+4   1   1.5  RT (min)       lcorbi_t3_pos_4_095_60    m/z 237.1122     0   2   4   6   8E+3   5.5   6  RT (min)       lcorbi_t3_pos_4_030_05    m/z 237.1122     0   2   4E+4   15   15.5  RT (min)       lcorbi_t3_pos_1_007_07    m/z 237.1452     0   1   2E+4   2.5   3  RT (min)      lcorbi_t3_pos_4_024_27    m/z 237.1573     0   1E+4   8.5   9  RT (min)      lcorbi_t3_pos_1_001_49    m/z 237.1573     0   1   2   3E+3   5.5   6  RT (min)      lcorbi_t3_pos_1_001_49    m/z 237.1651     0   2   4   6E+4   1   1.5   2  RT (min)       lcorbi_t3_pos_4_090_60    m/z 237.1654     0   2   4   6   8E+3   7.5   8  RT (min)      lcorbi_t3_pos_4_106_62    m/z 237.1697     0   1   2E+4   5.5   6  RT (min)       lcorbi_t3_pos_4_074_58    m/z 237.1849     0   1   2E+5   7.5   8  RT (min)       lcorbi_t3_pos_1_012_14    m/z 237.1919     0   1   2.5   3  RT (min)      lcorbi_t3_pos_1_002_01    m/z 237.1961     0   2   4   6E+3   6   6.5  RT (min)      lcorbi_t3_pos_1_001_49    m/z 237.2138     0   1   2E+3   7   7.5  RT (min)      lcorbi_t3_pos_1_002_01    m/z 237.9644     0   1   1   1.5  RT (min)      lcorbi_t3_pos_3_010_44    m/z 238.0206     0   1   1   1.5  RT (min)      lcorbi_t3_pos_4_112_62    m/z 238.0331     0   1   2   3   4E+5   1   1.5  RT (min)      lcorbi_t3_pos_1_036_08    m/z 238.0363     0   1   2.5   3  RT (min)      lcorbi_t3_pos_1_012_14    m/z 238.0498     0   2   4E+4   1   1.5  RT (min)       lcorbi_t3_pos_1_005_42    m/z 238.0498     0   2   4   6E+4   4.5   5  RT (min)       lcorbi_t3_pos_4_113_62    m/z 238.0589     0   1E+5   1   1.5  RT (min)       lcorbi_t3_pos_4_091_60    m/z 238.0922     0   2   4E+2   1.5   2  RT (min)      lcorbi_t3_pos_4_042_09    m/z 238.0993     0   1   2   3E+4   6   6.5  RT (min)       lcorbi_t3_pos_4_073_57    m/z 238.1092     0   1   2   3E+4   1.5   2  RT (min)      lcorbi_t3_pos_4_042_09    m/z 238.1160     0   1   2   3E+4   4.5   5  RT (min)       lcorbi_t3_pos_4_106_62    m/z 238.1189     0   1   2   3E+4   1   1.5  RT (min)      lcorbi_t3_pos_4_095_60    m/z 238.1204     0   2   4E+3   7   7.5  RT (min)      lcorbi_t3_pos_2_016_23    m/z 238.1438     0   1   2   3   4E+4   5   5.5  RT (min)       lcorbi_t3_pos_4_073_57    m/z 238.1563     0   1E+4   8.5   9  RT (min)      lcorbi_t3_pos_4_066_57    m/z 238.1649     0   1   2E+4   4.5   5  RT (min)       lcorbi_t3_pos_4_086_59    m/z 238.1778     0   1   2E+4   7.5   8  RT (min)       lcorbi_t3_pos_4_106_62    m/z 238.1883     0   1   2   3E+4   7.5   8  RT (min)       lcorbi_t3_pos_4_008_30    m/z 238.2013     0   1E+4   5   5.5  RT (min)      lcorbi_t3_pos_4_098_61    m/z 238.2359     0   1E+4   8   8.5  RT (min)      lcorbi_t3_pos_4_082_59    m/z 238.8418     0   2   4E+4   1   1.5  RT (min)      lcorbi_t3_pos_1_029_29    m/z 238.8531     0   1E+4   1   1.5  RT (min)      lcorbi_t3_pos_3_010_44    m/z 238.8626     0   2   4   6   8E+4   1   1.5  RT (min)      lcorbi_t3_pos_1_036_08    m/z 238.9029     0   1   1   1.5  RT (min)      lcorbi_t3_pos_3_011_42    m/z 239.0168     0   1   2E+4   1   1.5  RT (min)      lcorbi_t3_pos_4_074_58    m/z 239.0316     0   1   2   3   4E+3   1   1.5  RT (min)      lcorbi_t3_pos_4_080_58    m/z 239.0434     0   2   4   6E+4   18   18.5  RT (min)       lcorbi_t3_pos_4_100_61    m/z 239.0505     0   1   1   1.5  RT (min)       lcorbi_t3_pos_1_001_49    m/z 239.0658     0   1   8.5   9  RT (min)       lcorbi_t3_pos_1_003_45    m/z 239.0890     0   1E+4   5.5   6  RT (min)      lcorbi_t3_pos_1_001_49    m/z 239.0914     0   1   7.5   8  RT (min)      lcorbi_t3_pos_1_001_49    m/z 239.1027     0   1   2   3E+3   1.5   2  RT (min)      lcorbi_t3_pos_1_001_49    m/z 239.1067     0   1   2E+4   9   9.5  RT (min)       lcorbi_t3_pos_1_001_49    m/z 239.1430     0   1   2   3   4E+3   7   7.5  RT (min)      lcorbi_t3_pos_1_001_49    m/z 239.1472     0   2   4   6   8E+3   5   5.5  RT (min)      lcorbi_t3_pos_1_001_49    m/z 239.1754     0   1   2.5   3  RT (min)      lcorbi_t3_pos_4_081_58    m/z 239.2249     0   1   2E+4   4.5   5  RT (min)      lcorbi_t3_pos_4_040_21    m/z 239.2249     0   1E+3   2.5   3  RT (min)      lcorbi_t3_pos_1_002_01    m/z 239.2482     0   1E+4   5   5.5  RT (min)      lcorbi_t3_pos_4_106_62    m/z 239.9807     0   1E+5   1   1.5  RT (min)      lcorbi_t3_pos_1_036_08    m/z 240.0178     0   1   2E+4   1   1.5  RT (min)      lcorbi_t3_pos_1_036_08    m/z 240.0301     0   1E+5   1   1.5  RT (min)      lcorbi_t3_pos_1_036_08    m/z 240.0736     0   1   1   1.5  RT (min)       lcorbi_t3_pos_3_052_47    m/z 240.0813     0   1   2   3   4E+3   3   3.5  RT (min)      lcorbi_t3_pos_1_001_49    m/z 240.0963     0   1E+4   6   6.5  RT (min)      lcorbi_t3_pos_4_071_57    m/z 240.1019     0   2   4   6E+4   8.5   9  RT (min)       lcorbi_t3_pos_4_107_62    m/z 240.1019     0   1   2.5   3  RT (min)       lcorbi_t3_pos_4_074_58    m/z 240.1060     0   1   2E+4   5   5.5  RT (min)      lcorbi_t3_pos_4_087_59    m/z 240.1078     0   1   2E+4   4.5   5  RT (min)      lcorbi_t3_pos_4_082_59    m/z 240.1078     0   1   2.5   3  RT (min)      lcorbi_t3_pos_4_076_58    m/z 240.1190     0   2   4E+4   1   1.5  RT (min)      lcorbi_t3_pos_4_069_57    m/z 240.1243     0   1   2   3E+3   4.5   5  RT (min)      lcorbi_t3_pos_4_066_57    m/z 240.1278     0   1   2E+4   1.5   2  RT (min)      lcorbi_t3_pos_4_097_60    m/z 240.1311     0   1   2E+4   8   8.5  RT (min)      lcorbi_t3_pos_1_001_49    m/z 240.1361     0   1E+4   8.5   9  RT (min)      lcorbi_t3_pos_4_106_62    m/z 240.1442     0   1   2E+4   4.5   5  RT (min)      lcorbi_t3_pos_4_082_59    m/z 240.1495     0   1   2   3E+4   5.5   6  RT (min)       lcorbi_t3_pos_4_074_58    m/z 240.1523     0   1   2   3E+4   4.5   5  RT (min)      lcorbi_t3_pos_4_026_20    m/z 240.1594     0   2   4   6   8E+4   7.5   8  RT (min)       lcorbi_t3_pos_4_066_57    m/z 240.1672     0   1   2   3E+3   0.5   1  RT (min)      lcorbi_t3_pos_1_001_49    m/z 240.1754     0   1   2E+4   1.5   2  RT (min)      lcorbi_t3_pos_4_095_60    m/z 240.1806     0   1E+4   5   5.5  RT (min)       lcorbi_t3_pos_4_073_57    m/z 240.1934     0   2   4E+4   6.5   7  RT (min)      lcorbi_t3_pos_4_038_38    m/z 240.2071     0   1   2E+4   7.5   8  RT (min)       lcorbi_t3_pos_4_074_58    m/z 240.2322     0   1   2E+4   5   5.5  RT (min)       lcorbi_t3_pos_4_106_62    m/z 240.2322     0   2   4   6E+4   13   13.5  RT (min)       lcorbi_t3_pos_2_044_21    m/z 240.8390     0   1   2   3E+4   1   1.5  RT (min)      lcorbi_t3_pos_1_029_29    m/z 240.8598     0   1   2   3   4E+4   1   1.5  RT (min)      lcorbi_t3_pos_1_036_08    m/z 240.9249     0   1   2   3   4E+4   1   1.5  RT (min)      lcorbi_t3_pos_1_036_08    m/z 240.9673     0   2   4E+3   1   1.5  RT (min)      lcorbi_t3_pos_2_006_47    m/z 241.0417     0   1E+5   1   1.5  RT (min)      lcorbi_t3_pos_3_034_26    m/z 241.0432     0   1E+5   1   1.5  RT (min)       lcorbi_t3_pos_1_051_09    m/z 241.0591     0   1   2E+5   18.5   19  RT (min)       lcorbi_t3_pos_4_079_58    m/z 241.0591     0   1E+4   0.5   1  RT (min)      lcorbi_t3_pos_4_094_60    m/z 241.0770     0   2   4   6   8E+5   1   1.5  RT (min)       lcorbi_t3_pos_4_091_60    m/z 241.0914     0   2   4   6E+3   0.5   1  RT (min)      lcorbi_t3_pos_4_094_60    m/z 241.1046     0   1   2   3E+3   4.5   5  RT (min)      lcorbi_t3_pos_2_041_54    m/z 241.1053     0   2   4   6E+2   2.5   3  RT (min)      lcorbi_t3_pos_4_024_27    m/z 241.1053     0   2   4E+3   8.5   9  RT (min)      lcorbi_t3_pos_1_042_37    m/z 241.1183     0   1E+3   2.5   3  RT (min)      lcorbi_t3_pos_4_012_37    m/z 241.1224     0   1   2   3   4E+4   7   7.5  RT (min)       lcorbi_t3_pos_4_098_61    m/z 241.1303     0   2   4   6E+4   18   18.5  RT (min)       lcorbi_t3_pos_4_074_58    m/z 241.1408     0   1   2   3   4E+3   6.5   7  RT (min)      lcorbi_t3_pos_4_078_58    m/z 241.1531     0   1E+3   5.5   6  RT (min)      lcorbi_t3_pos_1_047_41    m/z 241.1547     0   1E+3   1   1.5  RT (min)       lcorbi_t3_pos_1_045_12    m/z 241.1563     0   1   10   10.5  RT (min)       lcorbi_t3_pos_4_092_60    m/z 241.1628     0   1   2   3E+3   7.5   8  RT (min)      lcorbi_t3_pos_1_047_41    m/z 241.1675     0   1E+5   1   1.5   2  RT (min)       lcorbi_t3_pos_4_090_60    m/z 241.1733     0   1E+5   15   15.5  RT (min)       lcorbi_t3_pos_4_068_57    m/z 241.1733     0   1E+5   17.5   18  RT (min)       lcorbi_t3_pos_4_067_57    m/z 241.2037     0   1   2E+4   6   6.5  RT (min)      lcorbi_t3_pos_1_001_49    m/z 241.2219     0   1   2E+3   4.5   5  RT (min)      lcorbi_t3_pos_1_047_41    m/z 241.2355     0   1   2E+3   5   5.5  RT (min)      lcorbi_t3_pos_1_046_22    m/z 241.2638     0   1E+5   10   10.5  RT (min)      lcorbi_t3_pos_1_005_42    m/z 241.7053     0   2   4   6E+3   6   6.5  RT (min)      lcorbi_t3_pos_1_001_49    m/z 241.9778     0   1   1   1.5  RT (min)      lcorbi_t3_pos_3_045_43    m/z 242.0310     0   1   1   1.5  RT (min)      lcorbi_t3_pos_1_003_45    m/z 242.0893     0   1   2E+4   9   9.5  RT (min)      lcorbi_t3_pos_1_001_49    m/z 242.1003     0   2   4   6E+4   4.5   5  RT (min)       lcorbi_t3_pos_4_082_59    m/z 242.1042     0   1   2E+3   1   1.5  RT (min)      lcorbi_t3_pos_2_005_11    m/z 242.1136     0   1   2   3E+3   1   1.5  RT (min)      lcorbi_t3_pos_1_021_18    m/z 242.1138     0   1   2E+3   2.5   3  RT (min)      lcorbi_t3_pos_1_042_37    m/z 242.1231     0   1E+5   0.5   1  RT (min)       lcorbi_t3_pos_4_086_59    m/z 242.1306     0   1E+4   6   6.5  RT (min)      lcorbi_t3_pos_4_068_57    m/z 242.1312     0   1   4.5   5  RT (min)      lcorbi_t3_pos_4_082_59    m/z 242.1434     0   1   1   1.5   2  RT (min)       lcorbi_t3_pos_2_005_11    m/z 242.1580     0   1   1.5   2  RT (min)      lcorbi_t3_pos_3_055_10    m/z 242.1766     0   1   2   3   4E+4   9.5   10  RT (min)      lcorbi_t3_pos_4_074_58    m/z 242.2227     0   1E+4   6   6.5  RT (min)      lcorbi_t3_pos_4_106_62    m/z 242.2479     0   2   4   6   8E+4   9   9.5  RT (min)       lcorbi_t3_pos_4_113_62    m/z 242.2842     0   1   2E+4   6   6.5  RT (min)       lcorbi_t3_pos_4_091_60    m/z 242.2843     0   1   2E+5   10   11  RT (min)       lcorbi_t3_pos_4_086_59    m/z 242.9186     0   1E+4   1   1.5  RT (min)      lcorbi_t3_pos_1_034_34    m/z 242.9328     0   1E+4   1   1.5  RT (min)      lcorbi_t3_pos_1_029_29    m/z 242.9942     0   1   2   3   4E+4   1   1.5  RT (min)       lcorbi_t3_pos_4_086_59    m/z 243.0037     0   1   1.5   2  RT (min)      lcorbi_t3_pos_3_055_10    m/z 243.0265     0   2   4   6E+4   1   1.5  RT (min)       lcorbi_t3_pos_4_088_59    m/z 243.0326     0   2   4E+3   1   1.5  RT (min)      lcorbi_t3_pos_2_008_16    m/z 243.0476     0   1   2E+3   1   1.5  RT (min)      lcorbi_t3_pos_2_005_11    m/z 243.0531     0   1   4.5   5  RT (min)      lcorbi_t3_pos_1_027_43    m/z 243.0660     0   1   2E+3   1   1.5  RT (min)      lcorbi_t3_pos_1_015_11    m/z 243.0750     0   2   4   6E+4   1   1.5  RT (min)       lcorbi_t3_pos_4_091_60    m/z 243.0799     0   1E+5   2.5   3  RT (min)       lcorbi_t3_pos_1_036_08    m/z 243.0839     0   2   4   6E+4   9   9.5  RT (min)       lcorbi_t3_pos_4_082_59    m/z 243.0938     0   1   1   1.5  RT (min)       lcorbi_t3_pos_1_034_34    m/z 243.0976     0   1   2E+3   4.5   5  RT (min)      lcorbi_t3_pos_3_048_30    m/z 243.1070     0   1   2E+4   18.5   19  RT (min)       lcorbi_t3_pos_4_081_58    m/z 243.1071     0   1   2   3E+4   0   1  RT (min)       lcorbi_t3_pos_1_036_08    m/z 243.1128     0   1   2   3E+3   6.5   7  RT (min)      lcorbi_t3_pos_3_048_30    m/z 243.1203     0   1   2   3E+3   4.5   5  RT (min)       lcorbi_t3_pos_1_027_43    m/z 243.1269     0   1   2E+3   1   1.5  RT (min)      lcorbi_t3_pos_1_022_20    m/z 243.1639     0   1   4.5   5  RT (min)      lcorbi_t3_pos_4_082_59    m/z 243.1691     0   1E+4   9.5   10  RT (min)      lcorbi_t3_pos_1_003_45    m/z 243.1704     0   1   2E+3   6   6.5  RT (min)      lcorbi_t3_pos_1_001_49    m/z 243.1829     0   2   4E+3   4.5   5  RT (min)      lcorbi_t3_pos_1_001_49    m/z 243.2067     0   2   4   6   8E+3   7   7.5  RT (min)      lcorbi_t3_pos_1_001_49    m/z 243.2067     0   1   2   3E+3   5   5.5  RT (min)      lcorbi_t3_pos_1_003_45    m/z 243.2108     0   1E+4   10   10.5  RT (min)      lcorbi_t3_pos_1_001_49    m/z 243.2108     0   2   4E+4   15   15.5  RT (min)      lcorbi_t3_pos_2_040_25    m/z 243.2513     0   1   2E+4   11   11.5  RT (min)      lcorbi_t3_pos_3_020_35    m/z 243.2876     0   1   2   3E+4   9.5   10  RT (min)      lcorbi_t3_pos_4_088_59    m/z 243.9289     0   1E+5   1   1.5  RT (min)       lcorbi_t3_pos_4_075_58    m/z 243.9927     0   2   4E+3   1   1.5  RT (min)      lcorbi_t3_pos_1_003_45    m/z 244.0483     0   1   2E+4   1   1.5  RT (min)       lcorbi_t3_pos_4_093_60    m/z 244.0662     0   1   2E+4   6.5   7  RT (min)      lcorbi_t3_pos_1_012_14    m/z 244.0805     0   1   2   3E+3   4.5   5  RT (min)      lcorbi_t3_pos_4_090_60    m/z 244.0910     0   1   2E+4   6   6.5  RT (min)      lcorbi_t3_pos_4_082_59    m/z 244.0945     0   1E+6   1  RT (min)       lcorbi_t3_pos_4_083_59    m/z 244.1292     0   1   1   1.5  RT (min)       lcorbi_t3_pos_2_054_29    m/z 244.1398     0   2   4   6E+2   1   1.5  RT (min)       lcorbi_t3_pos_3_029_13    m/z 244.1402     0   1   2E+4   2.5   3  RT (min)      lcorbi_t3_pos_4_095_60    m/z 244.1544     0   1   2   3   4E+4   5.5   6  RT (min)       lcorbi_t3_pos_4_098_61    m/z 244.1626     0   1E+4   8.5   9  RT (min)      lcorbi_t3_pos_1_001_49    m/z 244.2020     0   2   4E+4   2.5   3  RT (min)       lcorbi_t3_pos_4_089_59    m/z 244.2272     0   1   2   3   4E+4   10   10.5  RT (min)       lcorbi_t3_pos_4_111_62    m/z 244.8706     0   2   4   6   8E+3   1   1.5  RT (min)      lcorbi_t3_pos_4_039_35    m/z 244.8964     0   1E+3   1   1.5  RT (min)      lcorbi_t3_pos_3_004_24    m/z 244.9149     0   2   4   6   8E+3   1   1.5  RT (min)      lcorbi_t3_pos_1_001_49    m/z 244.9236     0   1   2E+5   1   1.5  RT (min)       lcorbi_t3_pos_4_077_58    m/z 244.9297     0   1   2   3E+4   1   1.5  RT (min)      lcorbi_t3_pos_3_010_44    m/z 244.9639     0   1   1.5   2  RT (min)      lcorbi_t3_pos_2_037_27    m/z 245.0271     0   2   4   6E+4   1   1.5  RT (min)       lcorbi_t3_pos_4_084_59    m/z 245.0445     0   1   2   3E+4   6.5   7  RT (min)       lcorbi_t3_pos_1_028_32    m/z 245.0486     0   1   2   3E+4   1   1.5  RT (min)       lcorbi_t3_pos_1_032_26    m/z 245.0632     0   1   2E+4   1   1.5  RT (min)      lcorbi_t3_pos_3_012_31    m/z 245.0632     0   2   4   6   8E+4   4.5   5  RT (min)       lcorbi_t3_pos_4_090_60    m/z 245.0632     0   1   2E+3   2.5   3  RT (min)      lcorbi_t3_pos_4_092_60    m/z 245.0745     0   2   4   6E+6   1   1.5  RT (min)       lcorbi_t3_pos_4_070_57    m/z 245.0769     0   2   4E+3   4.5   5  RT (min)      lcorbi_t3_pos_4_082_59    m/z 245.0785     0   1E+4   8.5   9  RT (min)      lcorbi_t3_pos_4_074_58    m/z 245.0825     0   1   2   3E+4   1.5   2  RT (min)      lcorbi_t3_pos_1_020_31    m/z 245.0921     0   1   2   3   4E+4   4.5   5  RT (min)      lcorbi_t3_pos_3_047_40    m/z 245.0979     0   1   1   1.5  RT (min)      lcorbi_t3_pos_3_012_31    m/z 245.0995     0   1   2   3E+3   4.5   5  RT (min)       lcorbi_t3_pos_1_050_13    m/z 245.0996     0   1   2.5   3  RT (min)      lcorbi_t3_pos_4_092_60    m/z 245.1132     0   1   2   3   4E+3   1   1.5   2  RT (min)       lcorbi_t3_pos_4_021_25    m/z 245.1242     0   1   2.5   3  RT (min)      lcorbi_t3_pos_4_082_59    m/z 245.1245     0   1   1.5   2  RT (min)      lcorbi_t3_pos_4_021_25    m/z 245.1360     0   2   4   6E+4   5   5.5  RT (min)       lcorbi_t3_pos_4_044_32    m/z 245.1384     0   1   2E+4   11   11.5  RT (min)      lcorbi_t3_pos_3_028_11    m/z 245.1413     0   1   2   3E+4   5.5   6  RT (min)       lcorbi_t3_pos_4_094_60    m/z 245.1495     0   2   4E+3   1   1.5  RT (min)       lcorbi_t3_pos_1_023_02    m/z 245.1496     0   1   2   3   4E+3   4.5   5  RT (min)      lcorbi_t3_pos_4_090_60    m/z 245.1649     0   1   1   1.5  RT (min)      lcorbi_t3_pos_4_021_25    m/z 245.1649     0   1   2.5   3  RT (min)      lcorbi_t3_pos_4_106_62    m/z 245.1748     0   1   2   3E+4   9   9.5  RT (min)       lcorbi_t3_pos_4_090_60    m/z 245.1860     0   1   2   3   4E+4   1.5   2  RT (min)       lcorbi_t3_pos_4_067_57    m/z 245.1860     0   2   4   6E+4   4.5   5  RT (min)       lcorbi_t3_pos_4_082_59    m/z 245.2054     0   1   2.5   3  RT (min)      lcorbi_t3_pos_4_111_62    m/z 245.2224     0   1   2E+3   6   6.5  RT (min)      lcorbi_t3_pos_1_002_01    m/z 245.2264     0   2   4   6   8E+4   12   12.5  RT (min)       lcorbi_t3_pos_1_020_31    m/z 245.2270     0   1E+4   8   8.5  RT (min)      lcorbi_t3_pos_4_082_59    m/z 245.2305     0   2   4   6   8E+4   8   8.5  RT (min)       lcorbi_t3_pos_3_044_46    m/z 245.9271     0   2   4   6   8E+3   1   1.5  RT (min)      lcorbi_t3_pos_4_077_58    m/z 245.9836     0   1   2E+4   1   1.5  RT (min)      lcorbi_t3_pos_1_031_40    m/z 245.9891     0   1   2E+4   1   1.5  RT (min)      lcorbi_t3_pos_4_074_58    m/z 246.0108     0   1   2E+4   2.5   3  RT (min)      lcorbi_t3_pos_4_019_47    m/z 246.0285     0   1   2.5   3  RT (min)      lcorbi_t3_pos_4_104_61    m/z 246.0479     0   1   2   3E+4   8   8.5  RT (min)      lcorbi_t3_pos_3_034_26    m/z 246.0715     0   1   2   3E+4   1   1.5  RT (min)      lcorbi_t3_pos_1_002_01    m/z 246.0808     0   2   4   6E+4   9.5   10  RT (min)       lcorbi_t3_pos_4_082_59    m/z 246.0907     0   2   4   6   8E+4   1   1.5  RT (min)       lcorbi_t3_pos_4_088_59    m/z 246.0955     0   1E+3   4.5   5  RT (min)      lcorbi_t3_pos_4_028_04    m/z 246.1085     0   1E+4   1   1.5  RT (min)      lcorbi_t3_pos_2_023_26    m/z 246.1393     0   1   2   3   4E+3   5   6  RT (min)       lcorbi_t3_pos_1_012_14    m/z 246.1489     0   1   2E+3   5.5   6  RT (min)      lcorbi_t3_pos_1_005_42    m/z 246.1559     0   1   1   1.5   2  RT (min)       lcorbi_t3_pos_3_053_48    m/z 246.1695     0   1E+4   1.5   2  RT (min)      lcorbi_t3_pos_1_036_08    m/z 246.1812     0   1E+5   1   1.5  RT (min)      lcorbi_t3_pos_3_052_47    m/z 246.1893     0   1   2E+3   4.5   5  RT (min)      lcorbi_t3_pos_1_001_49    m/z 246.1976     0   1   2   3E+3   8   8.5  RT (min)      lcorbi_t3_pos_1_019_16    m/z 246.2064     0   2   4   6   8E+3   9.5   10  RT (min)      lcorbi_t3_pos_2_044_21    m/z 246.2235     0   2   4   6   8E+4   8   8.5  RT (min)       lcorbi_t3_pos_4_079_58    m/z 246.2331     0   1   2   3   4E+4   8   8.5  RT (min)      lcorbi_t3_pos_1_017_51    m/z 246.2524     0   1E+5   8   8.5  RT (min)      lcorbi_t3_pos_2_025_52    m/z 246.2578     0   1   2E+4   8   8.5  RT (min)      lcorbi_t3_pos_1_019_16    m/z 246.8677     0   1   2   3   4E+4   1   1.5  RT (min)      lcorbi_t3_pos_1_034_34    m/z 246.9052     0   1E+5   0   0.5  RT (min)      lcorbi_t3_pos_1_009_50    m/z 247.0215     0   1   2   3   4E+4   1   1.5  RT (min)      lcorbi_t3_pos_1_028_32    m/z 247.0327     0   2   4   6E+4   7   7.5  RT (min)      lcorbi_t3_pos_4_101_61    m/z 247.0719     0   1E+4   7   7.5  RT (min)      lcorbi_t3_pos_1_006_15    m/z 247.0926     0   2   4   6   8E+3   1   1.5  RT (min)      lcorbi_t3_pos_1_056_56    m/z 247.0940     0   2   4   6E+3   8   8.5  RT (min)      lcorbi_t3_pos_4_105_61    m/z 247.0942     0   1   2   3E+3   4.5   5  RT (min)      lcorbi_t3_pos_1_020_31    m/z 247.1288     0   1   3   3.5  RT (min)      lcorbi_t3_pos_4_105_61    m/z 247.1289     0   1   2   3E+3   1.5   2  RT (min)      lcorbi_t3_pos_4_049_55    m/z 247.1370     0   2   4   6   8E+4   5.5   6  RT (min)      lcorbi_t3_pos_4_111_62    m/z 247.1402     0   2   4   6E+4   1   1.5  RT (min)      lcorbi_t3_pos_1_049_55    m/z 247.1553     0   1   2   3E+3   4.5   5  RT (min)      lcorbi_t3_pos_1_001_49    m/z 247.1652     0   2   4   6   8E+2   1   1.5  RT (min)       lcorbi_t3_pos_4_090_60    m/z 247.1733     0   1   2   3   4E+3   7.5   8  RT (min)      lcorbi_t3_pos_1_001_49    m/z 247.1734     0   1   2E+3   4.5   5  RT (min)      lcorbi_t3_pos_1_003_45    m/z 247.1781     0   1   2   3E+4   9   9.5  RT (min)      lcorbi_t3_pos_4_108_62    m/z 247.2098     0   2   4E+4   7   7.5  RT (min)      lcorbi_t3_pos_4_090_60    m/z 247.2268     0   1E+4   8   8.5  RT (min)      lcorbi_t3_pos_1_019_16    m/z 247.2400     0   1   2   3E+4   8   8.5  RT (min)      lcorbi_t3_pos_4_074_58    m/z 247.9869     0   2   4E+4   1   1.5  RT (min)       lcorbi_t3_pos_1_003_45    m/z 248.0434     0   1E+4   4.5   5  RT (min)      lcorbi_t3_pos_4_048_24    m/z 248.0494     0   2   4   6   8E+4   1   1.5  RT (min)      lcorbi_t3_pos_1_036_08    m/z 248.0561     0   1   2E+4   7   7.5  RT (min)      lcorbi_t3_pos_4_101_61    m/z 248.0619     0   2   4   6   8E+4   1   1.5  RT (min)      lcorbi_t3_pos_1_036_08    m/z 248.0707     0   1   7   7.5  RT (min)      lcorbi_t3_pos_4_101_61    m/z 248.0822     0   1   2   3   4E+4   1.5   2  RT (min)      lcorbi_t3_pos_2_009_50    m/z 248.0936     0   1   1   1.5  RT (min)       lcorbi_t3_pos_4_070_57    m/z 248.1030     0   2   4E+3   1.5   2  RT (min)      lcorbi_t3_pos_1_041_54    m/z 248.1129     0   2   4   6   8E+3   1   1.5  RT (min)      lcorbi_t3_pos_1_056_56    m/z 248.1129     0   1   2E+3   4.5   5  RT (min)      lcorbi_t3_pos_4_052_06    m/z 248.1239     0   1   2   3E+3   1.5   2  RT (min)       lcorbi_t3_pos_1_009_50    m/z 248.1281     0   2   4   6   8E+3   6   6.5  RT (min)      lcorbi_t3_pos_4_074_58    m/z 248.1353     0   1   2E+4   1   1.5  RT (min)      lcorbi_t3_pos_2_037_27    m/z 248.1475     0   2   4E+3   4.5   5  RT (min)      lcorbi_t3_pos_1_005_42    m/z 248.1492     0   1   2   3   4E+4   1   1.5  RT (min)       lcorbi_t3_pos_4_070_57    m/z 248.1856     0   1   2   3   4E+3   4.5   5  RT (min)      lcorbi_t3_pos_4_096_60    m/z 248.1857     0   2   4E+3   7.5   8  RT (min)      lcorbi_t3_pos_3_016_21    m/z 248.2028     0   2   4   6   8E+3   6   6.5  RT (min)      lcorbi_t3_pos_4_038_38    m/z 248.2220     0   2   4   6E+3   6   6.5  RT (min)      lcorbi_t3_pos_2_044_21    m/z 248.2221     0   2   4   6E+4   7.5   8  RT (min)       lcorbi_t3_pos_4_053_19    m/z 248.2302     0   1E+4   10.5   11  RT (min)      lcorbi_t3_pos_4_018_01    m/z 248.8068     0   1   2   3   4E+4   1   1.5  RT (min)      lcorbi_t3_pos_4_072_57    m/z 248.8706     0   1   2E+5   1   1.5  RT (min)       lcorbi_t3_pos_1_029_29    m/z 248.8819     0   1   1   1.5  RT (min)      lcorbi_t3_pos_1_006_15    m/z 248.8914     0   1   2E+3   1   1.5  RT (min)      lcorbi_t3_pos_2_051_35    m/z 248.9006     0   2   4   6E+4   0.5   1  RT (min)      lcorbi_t3_pos_1_006_15    m/z 248.9769     0   2   4E+3   1   1.5  RT (min)      lcorbi_t3_pos_2_040_25    m/z 248.9891     0   1   2   3   4E+4   12   12.5  RT (min)      lcorbi_t3_pos_4_039_35    m/z 248.9989     0   1E+5   1   1.5  RT (min)      lcorbi_t3_pos_1_028_32    m/z 249.0282     0   1E+4   1   1.5  RT (min)       lcorbi_t3_pos_3_004_24    m/z 249.0394     0   1E+5   6.5   7   7.5  RT (min)       lcorbi_t3_pos_4_005_33    m/z 249.0456     0   2   4   6E+4   1   1.5  RT (min)       lcorbi_t3_pos_1_044_25    m/z 249.0547     0   1E+5   10   10.5  RT (min)       lcorbi_t3_pos_4_089_59    m/z 249.0611     0   1   2E+4   4.5   5  RT (min)       lcorbi_t3_pos_4_079_58    m/z 249.0670     0   1E+4   4.5   5  RT (min)      lcorbi_t3_pos_4_077_58    m/z 249.0793     0   1   2   3E+4   1   1.5  RT (min)      lcorbi_t3_pos_1_020_31    m/z 249.0911     0   1   9   9.5  RT (min)      lcorbi_t3_pos_1_029_29    m/z 249.1122     0   2   4   6E+4   9   9.5  RT (min)       lcorbi_t3_pos_1_029_29    m/z 249.1139     0   1   2   3E+4   1   1.5  RT (min)       lcorbi_t3_pos_4_094_60    m/z 249.1234     0   2   4E+5   4.5   5  RT (min)       lcorbi_t3_pos_4_113_62    m/z 249.1362     0   1   2E+3   5   5.5  RT (min)       lcorbi_t3_pos_4_113_62    m/z 249.1446     0   1   2   3E+4   2.5   3  RT (min)      lcorbi_t3_pos_4_024_27    m/z 249.1557     0   1   2   3   4E+3   1   1.5  RT (min)      lcorbi_t3_pos_1_001_49    m/z 249.1697     0   1E+4   6   6.5  RT (min)       lcorbi_t3_pos_1_001_49    m/z 249.1961     0   2   4   6   8E+3   7.5   8  RT (min)      lcorbi_t3_pos_1_002_01    m/z 249.2061     0   2   4E+4   9   9.5  RT (min)       lcorbi_t3_pos_1_029_29    m/z 249.2254     0   1   7.5   8  RT (min)      lcorbi_t3_pos_1_007_07    m/z 249.9830     0   1E+5   1   1.5  RT (min)      lcorbi_t3_pos_1_028_32    m/z 249.9971     0   1   1   1.5  RT (min)      lcorbi_t3_pos_1_034_34    m/z 250.0096     0   1   1   1.5  RT (min)      lcorbi_t3_pos_1_034_34    m/z 250.0387     0   1   2.5   3  RT (min)       lcorbi_t3_pos_4_081_58    m/z 250.0428     0   1E+4   6.5   7  RT (min)      lcorbi_t3_pos_3_023_05    m/z 250.0561     0   1   2   2.5  RT (min)      lcorbi_t3_pos_1_001_49    m/z 250.0628     0   2   4   6   8E+3   7.5   8  RT (min)       lcorbi_t3_pos_4_105_61    m/z 250.0653     0   1E+5   1   1.5  RT (min)      lcorbi_t3_pos_1_036_08    m/z 250.0658     0   1E+5   4.5   5  RT (min)      lcorbi_t3_pos_4_077_58    m/z 250.0742     0   2   4   6E+3   4.5   5  RT (min)      lcorbi_t3_pos_1_034_34    m/z 250.0800     0   1   1.5   2  RT (min)      lcorbi_t3_pos_2_005_11    m/z 250.0863     0   1   2   3   4E+4   12   12.5  RT (min)       lcorbi_t3_pos_4_086_59    m/z 250.0935     0   1   2   3E+3   2.5   3  RT (min)      lcorbi_t3_pos_3_021_39    m/z 250.1121     0   1   2E+4   8.5   9  RT (min)      lcorbi_t3_pos_4_082_59    m/z 250.1155     0   1E+5   11.5   12  RT (min)      lcorbi_t3_pos_4_080_58    m/z 250.1188     0   1   4.5   5  RT (min)      lcorbi_t3_pos_4_113_62    m/z 250.1199     0   1E+3   1   1.5  RT (min)      lcorbi_t3_pos_1_036_08    m/z 250.1286     0   2   4   6   8E+4   1   1.5  RT (min)       lcorbi_t3_pos_4_083_59    m/z 250.1396     0   1   5   5.5  RT (min)      lcorbi_t3_pos_4_113_62    m/z 250.1397     0   1   2   3E+3   1.5   2  RT (min)      lcorbi_t3_pos_4_048_24    m/z 250.1503     0   1   2   3E+5   1   1.5  RT (min)       lcorbi_t3_pos_4_094_60    m/z 250.1607     0   1E+4   4.5   5  RT (min)      lcorbi_t3_pos_4_086_59    m/z 250.1650     0   1E+4   2.5   3  RT (min)      lcorbi_t3_pos_1_056_56    m/z 250.1802     0   2   4   6E+3   4.5   5  RT (min)      lcorbi_t3_pos_1_002_01    m/z 250.1953     0   1   2E+4   4.5   5  RT (min)       lcorbi_t3_pos_4_090_60    m/z 250.2384     0   1   4.5   5  RT (min)      lcorbi_t3_pos_4_090_60    m/z 250.4753     0   1   2E+4   4.5   5  RT (min)      lcorbi_t3_pos_1_036_08    m/z 250.7261     0   1E+4   4.5   5  RT (min)      lcorbi_t3_pos_1_056_56    m/z 250.8039     0   1   2   3   4E+4   1   1.5  RT (min)      lcorbi_t3_pos_4_068_57    m/z 250.8679     0   1   2   3E+4   1   1.5  RT (min)      lcorbi_t3_pos_2_007_02    m/z 250.9388     0   1   2E+5   18.5   19  RT (min)       lcorbi_t3_pos_4_075_58    m/z 250.9388     0   1   2E+5   0.5   1  RT (min)       lcorbi_t3_pos_4_074_58    m/z 250.9388     0   1   2   3   4E+4   4.5   5  RT (min)      lcorbi_t3_pos_4_106_62    m/z 250.9960     0   1   1   1.5  RT (min)      lcorbi_t3_pos_4_044_32    m/z 251.0308     0   1E+5   1   1.5  RT (min)       lcorbi_t3_pos_4_089_59    m/z 251.0394     0   1   2   3E+4   11.5   12  RT (min)      lcorbi_t3_pos_4_074_58    m/z 251.0479     0   2   4   6E+4   18   18.5  RT (min)       lcorbi_t3_pos_4_069_57    m/z 251.0534     0   1   2   3E+4   4.5   5  RT (min)       lcorbi_t3_pos_4_090_60    m/z 251.0541     0   1E+5   11.5   12  RT (min)      lcorbi_t3_pos_4_006_12    m/z 251.0922     0   2   4   6   8E+2   1.5   2  RT (min)      lcorbi_t3_pos_2_037_27    m/z 251.1067     0   1   2E+4   8.5   9  RT (min)      lcorbi_t3_pos_4_074_58    m/z 251.1139     0   1   2   3   4E+3   1   1.5  RT (min)      lcorbi_t3_pos_3_010_44    m/z 251.1254     0   1   2E+4   7.5   8  RT (min)       lcorbi_t3_pos_4_090_60    m/z 251.1278     0   2   4   6   8E+4   9.5   10  RT (min)       lcorbi_t3_pos_4_074_58    m/z 251.1301     0   2   4   6   8E+4   4.5   5  RT (min)       lcorbi_t3_pos_4_090_60    m/z 251.1617     0   1   2E+4   8.5   9  RT (min)      lcorbi_t3_pos_1_001_49    m/z 251.1642     0   1   2E+5   13.5   14  RT (min)      lcorbi_t3_pos_2_055_46    m/z 251.1643     0   1   2E+5   17.5   18  RT (min)      lcorbi_t3_pos_4_089_59    m/z 251.1730     0   2   4   6   8E+3   6.5   7  RT (min)      lcorbi_t3_pos_1_001_49    m/z 251.1812     0   1E+4   8   8.5  RT (min)      lcorbi_t3_pos_1_004_24    m/z 251.1853     0   1   2   3   4E+4   6   6.5  RT (min)       lcorbi_t3_pos_1_002_01    m/z 251.2370     0   1E+4   9.5   10  RT (min)      lcorbi_t3_pos_1_001_49    m/z 251.2482     0   1E+4   8.5   9  RT (min)      lcorbi_t3_pos_1_002_01    m/z 251.9414     0   1   2   3E+4   1   1.5  RT (min)      lcorbi_t3_pos_1_034_34    m/z 252.0152     0   1   2   3   4E+3   2.5   3  RT (min)      lcorbi_t3_pos_1_051_09    m/z 252.0718     0   1   3   3.5  RT (min)      lcorbi_t3_pos_4_082_59    m/z 252.0778     0   2   4   6E+4   8.5   9  RT (min)       lcorbi_t3_pos_4_100_61    m/z 252.0802     0   2   4   6E+4   1   1.5  RT (min)       lcorbi_t3_pos_1_036_08    m/z 252.0924     0   2   4   6E+4   5.5   6  RT (min)      lcorbi_t3_pos_4_106_62    m/z 252.0956     0   2   4   6E+5   1.5   2  RT (min)      lcorbi_t3_pos_2_046_07    m/z 252.1060     0   1   1   1.5  RT (min)      lcorbi_t3_pos_1_012_14    m/z 252.1091     0   2   4E+4   1   1.5  RT (min)       lcorbi_t3_pos_4_071_57    m/z 252.1091     0   2   4   6   8E+4   4.5   5  RT (min)       lcorbi_t3_pos_4_066_57    m/z 252.1092     0   1   2E+3   2.5   3  RT (min)       lcorbi_t3_pos_4_082_59    m/z 252.1231     0   1   2E+3   4.5   5  RT (min)      lcorbi_t3_pos_4_110_62    m/z 252.1231     0   1   2   3E+4   9   9.5  RT (min)      lcorbi_t3_pos_1_003_45    m/z 252.1361     0   2   4   6E+4   7.5   8  RT (min)       lcorbi_t3_pos_4_083_59    m/z 252.1594     0   1E+4   7.5   8  RT (min)      lcorbi_t3_pos_1_001_49    m/z 252.1806     0   1E+4   5.5   6  RT (min)      lcorbi_t3_pos_1_019_16    m/z 252.1887     0   2   4   6E+3   6   6.5  RT (min)      lcorbi_t3_pos_1_001_49    m/z 252.8009     0   1   2E+4   1   1.5  RT (min)      lcorbi_t3_pos_4_070_57    m/z 252.8825     0   2   4E+4   1   1.5  RT (min)       lcorbi_t3_pos_4_090_60    m/z 252.9186     0   2   4E+4   1   1.5  RT (min)      lcorbi_t3_pos_4_082_59    m/z 252.9880     0   1   1.5   2  RT (min)      lcorbi_t3_pos_1_004_24    m/z 253.0108     0   1   2   3E+4   1   1.5  RT (min)      lcorbi_t3_pos_2_023_26    m/z 253.0472     0   1E+4   1   1.5  RT (min)      lcorbi_t3_pos_4_084_59    m/z 253.0509     0   1   2E+5   11.5   12  RT (min)      lcorbi_t3_pos_4_079_58    m/z 253.0538     0   1   2E+5   11.5   12  RT (min)      lcorbi_t3_pos_4_079_58    m/z 253.0654     0   1   2E+5   8.5   9  RT (min)      lcorbi_t3_pos_4_100_61    m/z 253.0901     0   1   1   1.5   2  RT (min)       lcorbi_t3_pos_4_113_62    m/z 253.0990     0   2   4E+4   1   1.5  RT (min)       lcorbi_t3_pos_4_091_60    m/z 253.1008     0   1   2E+4   8.5   9  RT (min)      lcorbi_t3_pos_1_034_34    m/z 253.1125     0   1   2   3E+3   4.5   5  RT (min)      lcorbi_t3_pos_4_082_59    m/z 253.1182     0   1   2.5   3  RT (min)      lcorbi_t3_pos_4_106_62    m/z 253.1184     0   2   4   6E+3   9.5   10  RT (min)      lcorbi_t3_pos_4_103_61    m/z 253.1282     0   1   2   3E+3   4.5   5  RT (min)      lcorbi_t3_pos_4_104_61    m/z 253.1310     0   1   1.5   2  RT (min)      lcorbi_t3_pos_4_105_61    m/z 253.1411     0   2   4   6E+3   7   7.5  RT (min)      lcorbi_t3_pos_4_104_61    m/z 253.1603     0   2   4   6E+3   7.5   8  RT (min)      lcorbi_t3_pos_1_001_49    m/z 253.1646     0   1   2   3E+4   4.5   5  RT (min)       lcorbi_t3_pos_4_014_48    m/z 253.1798     0   1   2   3   4E+4   6.5   7  RT (min)       lcorbi_t3_pos_1_008_36    m/z 253.1911     0   2   4E+3   4.5   5  RT (min)      lcorbi_t3_pos_1_001_49    m/z 253.1950     0   1   2   3   4E+3   1.5   2  RT (min)      lcorbi_t3_pos_1_001_49    m/z 253.1968     0   1   9.5   10  RT (min)      lcorbi_t3_pos_1_001_49    m/z 253.2073     0   2   4   6   8E+4   10.5   11  RT (min)      lcorbi_t3_pos_4_106_62    m/z 253.2275     0   2   4   6E+4   8   8.5  RT (min)       lcorbi_t3_pos_1_023_02    m/z 253.2406     0   1   2E+3   4.5   5  RT (min)      lcorbi_t3_pos_2_044_21    m/z 253.9097     0   1E+4   1   1.5  RT (min)      lcorbi_t3_pos_1_007_07    m/z 253.9981     0   2   4   6E+4   1   1.5  RT (min)      lcorbi_t3_pos_1_036_08    m/z 254.0159     0   2   4E+4   1   1.5  RT (min)      lcorbi_t3_pos_1_036_08    m/z 254.0817     0   1E+5   8.5   9  RT (min)      lcorbi_t3_pos_3_023_05    m/z 254.0968     0   1   8.5   9  RT (min)      lcorbi_t3_pos_4_104_61    m/z 254.1057     0   2   4   6   8E+3   1.5   2  RT (min)      lcorbi_t3_pos_3_055_10    m/z 254.1121     0   1   2E+4   9.5   10  RT (min)      lcorbi_t3_pos_4_102_61    m/z 254.1151     0   1E+4   1.5   2  RT (min)      lcorbi_t3_pos_2_037_27    m/z 254.1175     0   1E+4   9   9.5  RT (min)       lcorbi_t3_pos_4_098_61    m/z 254.1217     0   1   2   3   4E+4   5.5   6  RT (min)      lcorbi_t3_pos_4_087_59    m/z 254.1234     0   1   2   3E+4   1   1.5  RT (min)      lcorbi_t3_pos_3_055_10    m/z 254.1387     0   2   4E+4   7.5   8  RT (min)       lcorbi_t3_pos_4_067_57    m/z 254.1599     0   1E+4   4.5   5  RT (min)      lcorbi_t3_pos_4_103_61    m/z 254.1612     0   2   4   6   8E+4   2   3  RT (min)       lcorbi_t3_pos_4_097_60    m/z 254.1726     0   2   4   6E+3   5.5   6  RT (min)      lcorbi_t3_pos_4_106_62    m/z 254.1832     0   1E+4   6.5   7  RT (min)      lcorbi_t3_pos_4_108_62    m/z 254.1962     0   1   2   3E+3   5.5   6  RT (min)      lcorbi_t3_pos_3_049_55    m/z 254.2308     0   1   2   3   4E+3   8   8.5  RT (min)      lcorbi_t3_pos_2_005_11    m/z 254.2479     0   1E+5   11.5   12  RT (min)       lcorbi_t3_pos_2_044_21    m/z 254.2479     0   2   4   6E+4   13   13.5  RT (min)       lcorbi_t3_pos_2_044_21    m/z 254.8885     0   2   4   6   8E+5   0   0.5  RT (min)      lcorbi_t3_pos_1_007_07    m/z 254.9316     0   1   2   3E+3   1   1.5  RT (min)      lcorbi_t3_pos_3_011_42    m/z 254.9517     0   2   4E+4   1   1.5  RT (min)      lcorbi_t3_pos_4_082_59    m/z 254.9998     0   1E+5   1   1.5  RT (min)      lcorbi_t3_pos_1_036_08    m/z 255.0210     0   1E+5   18.5   19  RT (min)       lcorbi_t3_pos_4_092_60    m/z 255.0210     0   2   4E+3   0.5   1  RT (min)      lcorbi_t3_pos_4_083_59    m/z 255.0839     0   1   2   3   4E+4   11   11.5  RT (min)       lcorbi_t3_pos_2_037_27    m/z 255.1128     0   2   4   6E+3   7   7.5  RT (min)      lcorbi_t3_pos_4_082_59    m/z 255.1274     0   1   2   3E+3   1   1.5  RT (min)      lcorbi_t3_pos_1_050_13    m/z 255.1279     0   1   2   3   4E+4   6   6.5  RT (min)       lcorbi_t3_pos_4_090_60    m/z 255.1316     0   2   4E+3   2.5   3  RT (min)      lcorbi_t3_pos_4_037_26    m/z 255.1339     0   1E+4   4.5   5  RT (min)      lcorbi_t3_pos_4_082_59    m/z 255.1380     0   2   4   6E+4   7   7.5  RT (min)       lcorbi_t3_pos_1_028_32    m/z 255.1454     0   1   2   3   4E+5   1   1.5  RT (min)       lcorbi_t3_pos_2_030_14    m/z 255.1570     0   1   2   3   4E+4   9   9.5  RT (min)       lcorbi_t3_pos_1_009_50    m/z 255.1760     0   1E+4   8   8.5  RT (min)      lcorbi_t3_pos_1_001_49    m/z 255.1817     0   1   2.5   3  RT (min)      lcorbi_t3_pos_1_002_01    m/z 255.2067     0   2   4E+3   7   7.5  RT (min)      lcorbi_t3_pos_1_002_01    m/z 255.2148     0   1   9   9.5  RT (min)      lcorbi_t3_pos_1_003_45    m/z 255.2193     0   2   4E+3   5.5   6  RT (min)      lcorbi_t3_pos_1_001_49    m/z 255.2431     0   1   2   3E+3   5   5.5  RT (min)      lcorbi_t3_pos_1_001_49    m/z 255.2512     0   2   4E+4   9   9.5  RT (min)       lcorbi_t3_pos_2_044_21    m/z 255.2795     0   1   2   3   4E+4   10.5   11  RT (min)      lcorbi_t3_pos_1_038_30    m/z 255.9548     0   2   4   6E+4   1   1.5  RT (min)      lcorbi_t3_pos_1_036_08    m/z 255.9781     0   1   2   3E+4   1   1.5  RT (min)      lcorbi_t3_pos_4_072_57    m/z 255.9952     0   1   2E+4   1   1.5  RT (min)      lcorbi_t3_pos_1_036_08    m/z 256.0468     0   1   2   3   4E+4   1   2   3  RT (min)       lcorbi_t3_pos_4_093_60    m/z 256.0589     0   2   4   6   8E+3   3   3.5  RT (min)      lcorbi_t3_pos_4_103_61    m/z 256.0816     0   2   4   6E+3   1.5   2  RT (min)       lcorbi_t3_pos_4_090_60    m/z 256.0933     0   1E+5   8.5   9  RT (min)       lcorbi_t3_pos_4_102_61    m/z 256.1014     0   1   1   1.5  RT (min)      lcorbi_t3_pos_4_095_60    m/z 256.1099     0   1E+4   6   6.5  RT (min)      lcorbi_t3_pos_4_069_57    m/z 256.1180     0   2   4   6   8E+2   1.5   2  RT (min)      lcorbi_t3_pos_2_037_27    m/z 256.1294     0   1E+5   1   1.5  RT (min)       lcorbi_t3_pos_4_095_60    m/z 256.1391     0   1E+4   9.5   10  RT (min)      lcorbi_t3_pos_4_027_07    m/z 256.1413     0   1E+4   7   7.5  RT (min)      lcorbi_t3_pos_4_030_05    m/z 256.1544     0   1E+5   10   10.5  RT (min)       lcorbi_t3_pos_4_106_62    m/z 256.1625     0   1E+4   8.5   9  RT (min)      lcorbi_t3_pos_1_001_49    m/z 256.2271     0   2   4   6E+3   5   5.5  RT (min)      lcorbi_t3_pos_1_001_49    m/z 256.2271     0   1   2E+4   9   9.5  RT (min)      lcorbi_t3_pos_1_001_49    m/z 256.2272     0   1   2   3E+4   11   11.5  RT (min)       lcorbi_t3_pos_4_110_62    m/z 256.8951     0   2   4   6   8E+3   0.5   1  RT (min)      lcorbi_t3_pos_2_051_35    m/z 257.0171     0   1   2   3   4E+4   1   1.5  RT (min)      lcorbi_t3_pos_3_034_26    m/z 257.0367     0   2   4E+5   18.5   19  RT (min)      lcorbi_t3_pos_4_099_61    m/z 257.0367     0   1E+4   0.5   1  RT (min)      lcorbi_t3_pos_2_006_47    m/z 257.0460     0   2   4   6   8E+4   18   18.5  RT (min)       lcorbi_t3_pos_4_097_60    m/z 257.0508     0   2   4   6   8E+5   1   1.5  RT (min)       lcorbi_t3_pos_4_091_60    m/z 257.0537     0   2   4   6E+4   18   18.5  RT (min)      lcorbi_t3_pos_4_084_59    m/z 257.0632     0   1   4.5   5  RT (min)      lcorbi_t3_pos_1_015_11    m/z 257.0745     0   2   4   6   8E+4   1   1.5   2  RT (min)       lcorbi_t3_pos_2_022_38    m/z 257.0808     0   2   4   6E+4   6.5   7  RT (min)       lcorbi_t3_pos_3_011_42    m/z 257.0851     0   1   2   3E+3   1   1.5  RT (min)      lcorbi_t3_pos_4_007_46    m/z 257.0995     0   2   4   6E+4   9.5   10  RT (min)       lcorbi_t3_pos_4_082_59    m/z 257.1024     0   2   4E+3   4.5   5  RT (min)      lcorbi_t3_pos_1_014_03    m/z 257.1133     0   1E+4   2.5   3  RT (min)      lcorbi_t3_pos_1_052_38    m/z 257.1227     0   1   2   3E+4   1.5   2  RT (min)       lcorbi_t3_pos_1_042_37    m/z 257.1227     0   1E+4   18.5   19  RT (min)       lcorbi_t3_pos_4_068_57    m/z 257.1360     0   1E+4   5.5   6  RT (min)      lcorbi_t3_pos_4_110_62    m/z 257.1789     0   1E+4   4.5   5  RT (min)      lcorbi_t3_pos_4_026_20    m/z 257.2182     0   1E+4   1   1.5  RT (min)      lcorbi_t3_pos_4_094_60    m/z 257.2224     0   2   4   6E+3   7.5   8  RT (min)      lcorbi_t3_pos_1_003_45    m/z 257.2476     0   2   4E+4   11.5   12  RT (min)       lcorbi_t3_pos_3_020_35    m/z 257.2587     0   2   4   6E+4   8   8.5  RT (min)       lcorbi_t3_pos_4_086_59    m/z 257.9493     0   1   2   3E+5   1   1.5  RT (min)      lcorbi_t3_pos_1_034_34    m/z 258.0046     0   2   4   6E+4   1   1.5  RT (min)       lcorbi_t3_pos_4_090_60    m/z 258.0174     0   1E+5   1   1.5  RT (min)      lcorbi_t3_pos_1_003_45    m/z 258.0298     0   1   2   3   4E+4   1   1.5  RT (min)       lcorbi_t3_pos_4_074_58    m/z 258.0842     0   1E+4   6.5   7  RT (min)      lcorbi_t3_pos_4_104_61    m/z 258.0949     0   1E+4   1   1.5  RT (min)       lcorbi_t3_pos_1_034_34    m/z 258.1101     0   2   4   6   8E+4   18.5   19  RT (min)      lcorbi_t3_pos_4_075_58    m/z 258.1186     0   1E+5   1   1.5  RT (min)      lcorbi_t3_pos_1_034_34    m/z 258.1515     0   2   4   6E+4   10   10.5  RT (min)      lcorbi_t3_pos_4_106_62    m/z 258.1570     0   1E+5   7   7.5  RT (min)      lcorbi_t3_pos_4_005_33    m/z 258.1700     0   2   4   6   8E+3   4.5   5  RT (min)      lcorbi_t3_pos_1_001_49    m/z 258.1812     0   1   2E+4   4.5   5  RT (min)      lcorbi_t3_pos_4_082_59    m/z 258.2064     0   1   2   3   4E+4   8.5   9  RT (min)       lcorbi_t3_pos_4_082_59    m/z 258.2621     0   1E+4   5   5.5  RT (min)      lcorbi_t3_pos_4_106_62    m/z 258.3066     0   1E+5   9.5   10  RT (min)      lcorbi_t3_pos_1_001_49    m/z 258.8653     0   2   4   6   8E+3   1   1.5  RT (min)      lcorbi_t3_pos_4_006_12    m/z 258.9976     0   2   4E+5   18.5   19  RT (min)       lcorbi_t3_pos_4_089_59    m/z 258.9976     0   1E+5   0.5   1  RT (min)       lcorbi_t3_pos_4_075_58    m/z 259.0102     0   1   2   3   4E+4   1   1.5  RT (min)       lcorbi_t3_pos_2_020_22    m/z 259.0278     0   1   2E+4   1   1.5  RT (min)      lcorbi_t3_pos_1_031_40    m/z 259.0368     0   1   2E+4   6.5   7  RT (min)       lcorbi_t3_pos_1_008_36    m/z 259.0642     0   1   1   1.5  RT (min)       lcorbi_t3_pos_3_027_27    m/z 259.0697     0   1   2   3   4E+4   11.5   12  RT (min)      lcorbi_t3_pos_3_019_17    m/z 259.0789     0   1E+4   2.5   3  RT (min)      lcorbi_t3_pos_3_012_31    m/z 259.0826     0   1   2   3E+3   5   5.5  RT (min)      lcorbi_t3_pos_4_069_57    m/z 259.0925     0   2   4   6E+3   1   1.5  RT (min)       lcorbi_t3_pos_3_029_13    m/z 259.1071     0   1   2E+4   1   1.5  RT (min)      lcorbi_t3_pos_1_030_21    m/z 259.1112     0   2   4   6E+3   6   6.5  RT (min)      lcorbi_t3_pos_4_102_61    m/z 259.1289     0   1   2   3E+3   1.5   2  RT (min)      lcorbi_t3_pos_4_094_60    m/z 259.1324     0   1   2E+4   5   5.5  RT (min)       lcorbi_t3_pos_4_103_61    m/z 259.1385     0   1   2E+3   4.5   5  RT (min)      lcorbi_t3_pos_4_090_60    m/z 259.1403     0   1   2.5   3  RT (min)      lcorbi_t3_pos_4_069_57    m/z 259.1516     0   2   4   6E+3   5.5   6  RT (min)      lcorbi_t3_pos_1_005_42    m/z 259.1653     0   2   4E+4   4.5   5  RT (min)       lcorbi_t3_pos_4_034_42    m/z 259.1669     0   2   4   6   8E+4   7.5   8  RT (min)       lcorbi_t3_pos_4_008_30    m/z 259.1699     0   1   2   3   4E+4   1   1.5  RT (min)       lcorbi_t3_pos_4_097_60    m/z 259.2016     0   1   5.5   6  RT (min)      lcorbi_t3_pos_1_001_49    m/z 259.2064     0   2   4   6E+4   7   7.5  RT (min)       lcorbi_t3_pos_4_053_19    m/z 259.2825     0   2   4   6   8E+4   9.5   10  RT (min)      lcorbi_t3_pos_2_044_21    m/z 259.9028     0   1   2E+4   1   1.5  RT (min)      lcorbi_t3_pos_4_081_58    m/z 259.9463     0   2   4   6   8E+4   1   1.5  RT (min)      lcorbi_t3_pos_1_028_32    m/z 259.9991     0   1   2E+4   1   1.5  RT (min)      lcorbi_t3_pos_4_020_41    m/z 260.0270     0   1   2   3   4E+4   1   1.5  RT (min)      lcorbi_t3_pos_2_020_22    m/z 260.0408     0   1   2   3   4E+4   1   1.5  RT (min)       lcorbi_t3_pos_4_096_60    m/z 260.0438     0   1E+4   4.5   5  RT (min)      lcorbi_t3_pos_3_048_30    m/z 260.0531     0   2   4   6E+4   1   1.5  RT (min)      lcorbi_t3_pos_2_027_05    m/z 260.0740     0   2   4   6   8E+3   4.5   5  RT (min)      lcorbi_t3_pos_4_087_59    m/z 260.0918     0   1   2   3   4E+4   5   5.5  RT (min)       lcorbi_t3_pos_4_104_61    m/z 260.0980     0   2   4   6E+3   4.5   5  RT (min)      lcorbi_t3_pos_4_098_61    m/z 260.1064     0   1   2   3E+3   1.5   2  RT (min)      lcorbi_t3_pos_3_048_30    m/z 260.1149     0   1   2   3E+3   4.5   5  RT (min)      lcorbi_t3_pos_1_001_49    m/z 260.1242     0   2   4   6   8E+2   1.5   2  RT (min)      lcorbi_t3_pos_3_004_24    m/z 260.1352     0   1E+3   1   1.5  RT (min)      lcorbi_t3_pos_3_029_13    m/z 260.1574     0   2   4E+4   11   11.5  RT (min)      lcorbi_t3_pos_4_090_60    m/z 260.1621     0   1   2   3E+6   1   1.5  RT (min)       lcorbi_t3_pos_4_097_60    m/z 260.1646     0   1   6.5   7  RT (min)      lcorbi_t3_pos_4_008_30    m/z 260.1686     0   2   4   6E+3   4.5   5  RT (min)      lcorbi_t3_pos_4_086_59    m/z 260.1857     0   1   2E+4   5.5   6  RT (min)       lcorbi_t3_pos_1_007_07    m/z 260.1969     0   1   1   1.5  RT (min)      lcorbi_t3_pos_4_090_60    m/z 260.2221     0   1   2   3E+4   7   7.5  RT (min)       lcorbi_t3_pos_2_044_21    m/z 260.2495     0   1   2   3E+4   10.5   11  RT (min)      lcorbi_t3_pos_4_099_61    m/z 260.2584     0   1   2   3E+5   8.5   9  RT (min)       lcorbi_t3_pos_2_044_21    m/z 260.7203     0   2   4E+3   4.5   5  RT (min)      lcorbi_t3_pos_1_054_48    m/z 260.8149     0   1E+5   1   1.5  RT (min)       lcorbi_t3_pos_1_020_31    m/z 260.8445     0   1   2E+3   1   1.5  RT (min)      lcorbi_t3_pos_3_039_03    m/z 260.8623     0   2   4E+4   1   1.5  RT (min)      lcorbi_t3_pos_1_028_32    m/z 260.8975     0   2   4   6E+4   1   1.5  RT (min)      lcorbi_t3_pos_3_008_20    m/z 260.9769     0   1   2E+3   1.5   2  RT (min)      lcorbi_t3_pos_4_039_35    m/z 260.9952     0   2   4E+3   0.5   1  RT (min)      lcorbi_t3_pos_4_083_59    m/z 261.0093     0   2   4E+5   1   1.5  RT (min)       lcorbi_t3_pos_1_023_02    m/z 261.0143     0   1   2   3E+3   1.5   2  RT (min)      lcorbi_t3_pos_2_020_22    m/z 261.0306     0   2   4   6E+3   0.5   1  RT (min)       lcorbi_t3_pos_4_071_57    m/z 261.0547     0   1   2E+3   4.5   5  RT (min)      lcorbi_t3_pos_4_106_62    m/z 261.0790     0   1   2.5   3  RT (min)      lcorbi_t3_pos_1_036_08    m/z 261.0796     0   1   1   1.5  RT (min)      lcorbi_t3_pos_3_011_42    m/z 261.0846     0   1E+4   5   5.5  RT (min)      lcorbi_t3_pos_4_087_59    m/z 261.0904     0   2   4   6   8E+4   2.5   3  RT (min)       lcorbi_t3_pos_1_036_08    m/z 261.0952     0   1   5   5.5  RT (min)      lcorbi_t3_pos_4_090_60    m/z 261.1082     0   1E+5   1   1.5  RT (min)       lcorbi_t3_pos_3_029_13    m/z 261.1098     0   1   2   3   4E+4   8   8.5  RT (min)       lcorbi_t3_pos_1_013_10    m/z 261.1116     0   1   2E+3   4.5   5  RT (min)       lcorbi_t3_pos_2_027_05    m/z 261.1268     0   1E+4   4.5   5  RT (min)      lcorbi_t3_pos_4_098_61    m/z 261.1445     0   1   5   5.5  RT (min)      lcorbi_t3_pos_1_003_45    m/z 261.1588     0   1   2E+4   1   1.5  RT (min)      lcorbi_t3_pos_4_090_60    m/z 261.1809     0   1E+4   4.5   5  RT (min)      lcorbi_t3_pos_4_066_57    m/z 261.1809     0   2   4E+3   2.5   3  RT (min)      lcorbi_t3_pos_4_095_60    m/z 261.1890     0   1   5.5   6  RT (min)      lcorbi_t3_pos_4_094_60    m/z 261.2062     0   1   2   3   4E+4   7   7.5  RT (min)      lcorbi_t3_pos_4_053_19    m/z 261.2173     0   1E+4   6.5   7  RT (min)      lcorbi_t3_pos_2_044_21    m/z 261.2214     0   2   4E+4   9.5   10  RT (min)       lcorbi_t3_pos_2_010_08    m/z 261.2425     0   1E+5   11   11.5  RT (min)       lcorbi_t3_pos_1_032_26    m/z 261.2521     0   1   2   3   4E+4   12.5   13  RT (min)       lcorbi_t3_pos_4_102_61    m/z 261.2618     0   1   2   3   4E+4   8.5   9  RT (min)      lcorbi_t3_pos_2_044_21    m/z 261.9626     0   1   2   3   4E+5   1   1.5  RT (min)      lcorbi_t3_pos_1_036_08    m/z 262.0146     0   2   4E+3   1   1.5  RT (min)      lcorbi_t3_pos_4_054_18    m/z 262.0314     0   1   2E+3   1   1.5  RT (min)      lcorbi_t3_pos_4_072_57    m/z 262.0503     0   1E+5   0   0.5  RT (min)       lcorbi_t3_pos_1_002_01    m/z 262.0898     0   1   2   3   4E+4   1   1.5  RT (min)       lcorbi_t3_pos_4_087_59    m/z 262.0936     0   1E+4   2.5   3  RT (min)      lcorbi_t3_pos_1_036_08    m/z 262.1115     0   1   1   1.5  RT (min)      lcorbi_t3_pos_3_054_15    m/z 262.1131     0   1E+4   8   8.5  RT (min)      lcorbi_t3_pos_1_022_20    m/z 262.1162     0   1   4.5   5  RT (min)      lcorbi_t3_pos_4_111_62    m/z 262.1285     0   1   2   3E+3   1.5   2  RT (min)       lcorbi_t3_pos_3_029_13    m/z 262.1343     0   1   2E+4   4.5   5  RT (min)      lcorbi_t3_pos_4_109_62    m/z 262.1485     0   1   2E+4   5.5   6  RT (min)       lcorbi_t3_pos_4_093_60    m/z 262.1650     0   2   4E+4   2   3  RT (min)       lcorbi_t3_pos_4_095_60    m/z 262.2013     0   2   4E+4   10   10.5  RT (min)      lcorbi_t3_pos_1_015_11    m/z 262.2247     0   1E+4   10   10.5  RT (min)      lcorbi_t3_pos_1_039_23    m/z 262.2251     0   1   2   3   4E+4   6.5   7  RT (min)      lcorbi_t3_pos_1_046_22    m/z 262.2458     0   1   2   3E+4   8   8.5  RT (min)      lcorbi_t3_pos_2_044_21    m/z 262.8880     0   2   4   6E+4   12   12.5  RT (min)      lcorbi_t3_pos_4_009_50    m/z 262.8975     0   1E+4   1   1.5  RT (min)      lcorbi_t3_pos_1_036_08    m/z 262.9069     0   1   2E+5   1   1.5  RT (min)      lcorbi_t3_pos_1_036_08    m/z 262.9157     0   2   4   6E+4   18   18.5  RT (min)       lcorbi_t3_pos_4_099_61    m/z 262.9157     0   1   2E+4   1   1.5  RT (min)      lcorbi_t3_pos_2_023_26    m/z 262.9526     0   2   4E+4   0   0.5  RT (min)      lcorbi_t3_pos_1_006_15    m/z 263.0282     0   1   2   3   4E+3   1   1.5  RT (min)      lcorbi_t3_pos_1_051_09    m/z 263.0466     0   1E+4   1   1.5  RT (min)      lcorbi_t3_pos_2_054_29    m/z 263.0539     0   1   2E+4   1   1.5  RT (min)      lcorbi_t3_pos_2_009_50    m/z 263.0562     0   1   2E+5   9   9.5  RT (min)       lcorbi_t3_pos_4_113_62    m/z 263.0653     0   2   4   6E+4   18   18.5  RT (min)       lcorbi_t3_pos_4_087_59    m/z 263.0679     0   2   4   6E+3   9   9.5  RT (min)      lcorbi_t3_pos_4_113_62    m/z 263.0791     0   1E+4   6.5   7  RT (min)      lcorbi_t3_pos_4_082_59    m/z 263.0815     0   2   4   6E+4   8.5   9  RT (min)      lcorbi_t3_pos_1_041_54    m/z 263.0815     0   1   0   0.5  RT (min)      lcorbi_t3_pos_4_032_40    m/z 263.1027     0   1E+4   5   5.5  RT (min)      lcorbi_t3_pos_4_101_61    m/z 263.1238     0   1   2   3   4E+4   1   1.5  RT (min)       lcorbi_t3_pos_3_029_13    m/z 263.1519     0   1   2E+5   5.5   6  RT (min)       lcorbi_t3_pos_4_094_60    m/z 263.1542     0   1   2E+5   1   1.5  RT (min)       lcorbi_t3_pos_3_029_13    m/z 263.1853     0   2   4   6E+3   6.5   7  RT (min)      lcorbi_t3_pos_1_002_01    m/z 263.2346     0   1E+4   8   8.5  RT (min)      lcorbi_t3_pos_4_112_62    m/z 263.9598     0   1E+5   1   1.5  RT (min)      lcorbi_t3_pos_1_036_08    m/z 264.0270     0   1E+5   1   1.5  RT (min)       lcorbi_t3_pos_1_036_08    m/z 264.0716     0   2   4E+3   6.5   7  RT (min)      lcorbi_t3_pos_4_066_57    m/z 264.0956     0   1   2E+4   4.5   5  RT (min)      lcorbi_t3_pos_4_041_54    m/z 264.0957     0   1E+4   2.5   3  RT (min)      lcorbi_t3_pos_1_041_54    m/z 264.1079     0   2   4   6E+4   1   1.5   2  RT (min)       lcorbi_t3_pos_4_087_59    m/z 264.1189     0   1   1   1.5  RT (min)      lcorbi_t3_pos_4_013_36    m/z 264.1320     0   1E+4   4.5   5  RT (min)      lcorbi_t3_pos_3_001_49    m/z 264.1553     0   1   5.5   6  RT (min)      lcorbi_t3_pos_4_105_61    m/z 264.1555     0   2   4   6   8E+4   1   1.5   2  RT (min)       lcorbi_t3_pos_4_084_59    m/z 264.1959     0   1E+4   10.5   11  RT (min)      lcorbi_t3_pos_4_074_58    m/z 264.2170     0   1   2E+3   5   5.5  RT (min)      lcorbi_t3_pos_1_001_49    m/z 264.2404     0   1   2   3E+4   12   12.5  RT (min)       lcorbi_t3_pos_1_052_38    m/z 264.2444     0   2   4   6   8E+4   8   8.5  RT (min)      lcorbi_t3_pos_4_111_62    m/z 264.2534     0   2   4   6E+4   10.5   11  RT (min)       lcorbi_t3_pos_1_032_26    m/z 264.8446     0   1   2E+4   1   1.5  RT (min)      lcorbi_t3_pos_4_090_60    m/z 264.8729     0   2   4   6E+6   0   0.5  RT (min)      lcorbi_t3_pos_1_007_07    m/z 264.9128     0   2   4E+3   1   1.5  RT (min)      lcorbi_t3_pos_4_012_37    m/z 264.9609     0   2   4   6   8E+2   1.5   2  RT (min)      lcorbi_t3_pos_2_035_34    m/z 264.9666     0   1   2E+4   10.5   11  RT (min)      lcorbi_t3_pos_4_043_13    m/z 265.0439     0   1   1   1.5  RT (min)      lcorbi_t3_pos_2_040_25    m/z 265.0526     0   1   2   3E+3   1   1.5  RT (min)      lcorbi_t3_pos_1_050_13    m/z 265.0696     0   1   2   3E+4   6   6.5  RT (min)       lcorbi_t3_pos_4_082_59    m/z 265.0796     0   1   2   3E+4   4.5   5  RT (min)      lcorbi_t3_pos_4_082_59    m/z 265.0796     0   1E+4   2.5   3  RT (min)      lcorbi_t3_pos_4_032_40    m/z 265.0885     0   1E+3   1   1.5  RT (min)       lcorbi_t3_pos_4_091_60    m/z 265.1013     0   2   4   6   8E+3   1.5   2  RT (min)      lcorbi_t3_pos_1_050_13    m/z 265.1071     0   1E+4   4.5   5  RT (min)      lcorbi_t3_pos_4_098_61    m/z 265.1183     0   1   2   3   4E+4   5   5.5  RT (min)       lcorbi_t3_pos_4_066_57    m/z 265.1282     0   1   2E+3   4.5   5  RT (min)      lcorbi_t3_pos_4_089_59    m/z 265.1336     0   2   4   6   8E+5   1   1.5  RT (min)       lcorbi_t3_pos_4_090_60    m/z 265.1410     0   1   2   3E+3   5   5.5  RT (min)      lcorbi_t3_pos_1_001_49    m/z 265.1565     0   2   4   6E+4   10   10.5  RT (min)       lcorbi_t3_pos_4_099_61    m/z 265.1587     0   1   2E+4   1.5   2  RT (min)       lcorbi_t3_pos_4_052_06    m/z 265.1646     0   1   2   3E+3   6   6.5  RT (min)       lcorbi_t3_pos_1_036_08    m/z 265.1759     0   2   4   6   8E+3   1   1.5  RT (min)      lcorbi_t3_pos_1_002_01    m/z 265.1887     0   2   4E+3   7   7.5  RT (min)      lcorbi_t3_pos_1_001_49    m/z 265.1981     0   1   2.5   3  RT (min)       lcorbi_t3_pos_1_002_01    m/z 265.2639     0   2   4   6E+3   9   9.5  RT (min)      lcorbi_t3_pos_1_001_49    m/z 266.0336     0   2   4   6E+3   2   2.5  RT (min)      lcorbi_t3_pos_4_074_58    m/z 266.0730     0   1   2   3   4E+3   6   6.5  RT (min)      lcorbi_t3_pos_4_085_59    m/z 266.0763     0   2   4   6E+3   1   1.5  RT (min)      lcorbi_t3_pos_1_003_45    m/z 266.0884     0   1   2   3   4E+3   4.5   5  RT (min)      lcorbi_t3_pos_4_082_59    m/z 266.0943     0   1E+4   7.5   8  RT (min)      lcorbi_t3_pos_4_066_57    m/z 266.1071     0   1   2E+3   6.5   7  RT (min)      lcorbi_t3_pos_4_096_60    m/z 266.1114     0   1   2   3E+3   2.5   3  RT (min)      lcorbi_t3_pos_4_091_60    m/z 266.1218     0   1   1   1.5  RT (min)       lcorbi_t3_pos_1_036_08    m/z 266.1247     0   1E+4   4.5   5  RT (min)      lcorbi_t3_pos_4_082_59    m/z 266.1302     0   1   2   3   4E+4   9.5   10  RT (min)      lcorbi_t3_pos_4_098_61    m/z 266.1362     0   1E+4   7   7.5  RT (min)      lcorbi_t3_pos_4_066_57    m/z 266.1370     0   1E+7   1   1.5  RT (min)       lcorbi_t3_pos_3_027_27    m/z 266.1389     0   1E+3   4.5   5  RT (min)      lcorbi_t3_pos_3_010_44    m/z 266.1517     0   1   2E+4   8   8.5  RT (min)       lcorbi_t3_pos_4_108_62    m/z 266.1599     0   1   2E+5   1   1.5  RT (min)       lcorbi_t3_pos_4_070_57    m/z 266.1962     0   1E+4   6   6.5  RT (min)      lcorbi_t3_pos_1_023_02    m/z 266.2091     0   1   2E+4   10.5   11  RT (min)      lcorbi_t3_pos_4_082_59    m/z 266.2228     0   2   4   6E+3   6.5   7  RT (min)      lcorbi_t3_pos_4_066_57    m/z 266.2560     0   1   2E+4   12   12.5  RT (min)      lcorbi_t3_pos_4_080_58    m/z 266.8421     0   2   4E+3   1   1.5  RT (min)      lcorbi_t3_pos_1_035_28    m/z 266.8783     0   1   1   1.5  RT (min)      lcorbi_t3_pos_1_007_07    m/z 266.9112     0   1E+6   0   0.5  RT (min)      lcorbi_t3_pos_1_007_07    m/z 266.9275     0   1   2   3   4E+4   0.5   1  RT (min)       lcorbi_t3_pos_4_075_58    m/z 266.9743     0   2   4   6E+4   0   0.5  RT (min)      lcorbi_t3_pos_1_006_15    m/z 267.0228     0   1   1   1.5  RT (min)       lcorbi_t3_pos_1_031_40    m/z 267.0387     0   1   2   3   4E+4   1   1.5  RT (min)      lcorbi_t3_pos_2_027_05    m/z 267.0587     0   2   4E+4   1   1.5  RT (min)       lcorbi_t3_pos_1_050_13    m/z 267.0630     0   2   4   6E+4   5.5   6  RT (min)       lcorbi_t3_pos_4_110_62    m/z 267.0993     0   1E+4   7.5   8  RT (min)      lcorbi_t3_pos_4_066_57    m/z 267.1016     0   2   4   6E+3   8.5   9  RT (min)       lcorbi_t3_pos_1_028_32    m/z 267.1075     0   2   4   6E+4   4.5   5  RT (min)       lcorbi_t3_pos_4_090_60    m/z 267.1186     0   1   1.5   2  RT (min)       lcorbi_t3_pos_2_040_25    m/z 267.1228     0   1   2E+4   8.5   9  RT (min)      lcorbi_t3_pos_1_028_32    m/z 267.1316     0   2   4   6E+3   4.5   5  RT (min)      lcorbi_t3_pos_4_094_60    m/z 267.1340     0   1E+4   6.5   7  RT (min)      lcorbi_t3_pos_4_041_54    m/z 267.1380     0   1E+4   8   8.5  RT (min)      lcorbi_t3_pos_4_104_61    m/z 267.1429     0   2   4E+4   1   1.5  RT (min)       lcorbi_t3_pos_4_090_60    m/z 267.1453     0   2   4   6E+2   2.5   3  RT (min)      lcorbi_t3_pos_4_021_25    m/z 267.1533     0   1E+4   10   10.5  RT (min)      lcorbi_t3_pos_4_082_59    m/z 267.1633     0   1   2   3E+4   1   1.5  RT (min)      lcorbi_t3_pos_3_016_21    m/z 267.1704     0   1   2   3E+4   4.5   5  RT (min)       lcorbi_t3_pos_4_066_57    m/z 267.1704     0   1E+4   2.5   3  RT (min)      lcorbi_t3_pos_4_021_25    m/z 267.1720     0   2   4   6E+4   10.5   11  RT (min)       lcorbi_t3_pos_4_066_57    m/z 267.1760     0   1   2   3E+4   8   8.5  RT (min)      lcorbi_t3_pos_4_082_59    m/z 267.1803     0   1   2E+4   5   5.5  RT (min)       lcorbi_t3_pos_2_044_21    m/z 267.2108     0   1   2   3E+3   7.5   8  RT (min)      lcorbi_t3_pos_1_008_36    m/z 267.2108     0   1   2E+5   11.5   12  RT (min)       lcorbi_t3_pos_4_077_58    m/z 267.2108     0   2   4   6E+4   14   14.5  RT (min)       lcorbi_t3_pos_3_048_30    m/z 267.2562     0   1   4.5   5  RT (min)      lcorbi_t3_pos_1_003_45    m/z 267.2877     0   1E+5   12.5   13  RT (min)       lcorbi_t3_pos_2_021_09    m/z 267.6471     0   2   4E+4   11   11.5  RT (min)       lcorbi_t3_pos_4_081_58    m/z 267.9781     0   1   2   3   4E+4   1   1.5  RT (min)      lcorbi_t3_pos_1_032_26    m/z 267.9928     0   1   2.5   3  RT (min)      lcorbi_t3_pos_4_098_61    m/z 268.0281     0   1   2E+4   1   1.5  RT (min)      lcorbi_t3_pos_1_031_40    m/z 268.0663     0   1E+4   5.5   6  RT (min)      lcorbi_t3_pos_4_110_62    m/z 268.0696     0   1   1   1.5  RT (min)      lcorbi_t3_pos_4_007_46    m/z 268.0969     0   2   4   6E+4   8.5   9  RT (min)       lcorbi_t3_pos_4_100_61    m/z 268.1041     0   1E+5   1   1.5  RT (min)       lcorbi_t3_pos_4_071_57    m/z 268.1041     0   1E+5   2.5   3  RT (min)       lcorbi_t3_pos_3_052_47    m/z 268.1294     0   1   2E+3   4.5   5  RT (min)      lcorbi_t3_pos_4_082_59    m/z 268.1339     0   1   1.5   2  RT (min)      lcorbi_t3_pos_4_042_09    m/z 268.1391     0   1E+4   4.5   5  RT (min)      lcorbi_t3_pos_1_003_45    m/z 268.1408     0   1E+5   1   1.5  RT (min)       lcorbi_t3_pos_1_050_13    m/z 268.1521     0   2   4E+3   6.5   7  RT (min)      lcorbi_t3_pos_4_090_60    m/z 268.1544     0   1   2   3E+4   8.5   9  RT (min)      lcorbi_t3_pos_1_010_46    m/z 268.1710     0   1E+5   1   1.5  RT (min)       lcorbi_t3_pos_3_055_10    m/z 268.1755     0   1   2E+4   5   5.5  RT (min)       lcorbi_t3_pos_4_088_59    m/z 268.1897     0   1   2E+4   8   8.5  RT (min)       lcorbi_t3_pos_4_066_57    m/z 268.2142     0   2   4   6E+4   11.5   12  RT (min)      lcorbi_t3_pos_4_077_58    m/z 268.2636     0   2   4   6   8E+4   13   13.5  RT (min)       lcorbi_t3_pos_2_044_21    m/z 268.2999     0   1   2E+4   10.5   11  RT (min)      lcorbi_t3_pos_3_029_13    m/z 268.8467     0   2   4E+4   1   1.5  RT (min)      lcorbi_t3_pos_2_028_24    m/z 268.8644     0   1   2E+4   1   1.5  RT (min)      lcorbi_t3_pos_1_023_02    m/z 268.8942     0   1   2E+4   1   1.5  RT (min)      lcorbi_t3_pos_1_036_08    m/z 268.9406     0   1   2E+5   0   0.5  RT (min)      lcorbi_t3_pos_1_005_42    m/z 268.9605     0   2   4   6E+4   18   18.5  RT (min)      lcorbi_t3_pos_4_096_60    m/z 269.0611     0   2   4E+3   5.5   6  RT (min)      lcorbi_t3_pos_4_108_62    m/z 269.0744     0   1E+3   1.5   2  RT (min)      lcorbi_t3_pos_4_007_46    m/z 269.0995     0   1   2E+4   11.5   12  RT (min)       lcorbi_t3_pos_4_111_62    m/z 269.1002     0   1E+4   8.5   9  RT (min)      lcorbi_t3_pos_4_103_61    m/z 269.1020     0   1   2   3E+4   5   5.5  RT (min)       lcorbi_t3_pos_4_074_58    m/z 269.1072     0   1   2   3   4E+3   1   1.5  RT (min)      lcorbi_t3_pos_3_027_27    m/z 269.1072     0   2   4   6   8E+3   2.5   3  RT (min)      lcorbi_t3_pos_2_030_14    m/z 269.1133     0   2   4   6E+2   5   5.5  RT (min)      lcorbi_t3_pos_4_006_12    m/z 269.1133     0   1   2E+3   2.5   3  RT (min)      lcorbi_t3_pos_1_054_48    m/z 269.1172     0   1   2E+4   8   8.5  RT (min)      lcorbi_t3_pos_1_033_53    m/z 269.1245     0   1E+5   1.5   2  RT (min)       lcorbi_t3_pos_3_027_27    m/z 269.1259     0   1   2   3E+3   4.5   5  RT (min)      lcorbi_t3_pos_4_090_60    m/z 269.1497     0   1E+3   2.5   3  RT (min)      lcorbi_t3_pos_1_054_48    m/z 269.1537     0   2   4   6E+3   7   7.5  RT (min)       lcorbi_t3_pos_1_001_49    m/z 269.1609     0   1E+3   2.5   3  RT (min)      lcorbi_t3_pos_1_001_49    m/z 269.1722     0   2   4   6E+4   1.5   2  RT (min)      lcorbi_t3_pos_3_051_12    m/z 269.1723     0   1E+4   8   8.5  RT (min)      lcorbi_t3_pos_1_003_45    m/z 269.2669     0   2   4   6E+4   10   10.5  RT (min)      lcorbi_t3_pos_4_087_59    m/z 269.2952     0   1   2   3E+4   10.5   11  RT (min)      lcorbi_t3_pos_2_045_40    m/z 269.9687     0   1E+5   1   1.5  RT (min)       lcorbi_t3_pos_4_088_59    m/z 270.0455     0   1   2   3   4E+4   7   7.5  RT (min)      lcorbi_t3_pos_4_101_61    m/z 270.0493     0   1   2   3E+3   2.5   3  RT (min)      lcorbi_t3_pos_3_008_20    m/z 270.0520     0   1E+5   4.5   5  RT (min)       lcorbi_t3_pos_4_077_58    m/z 270.0755     0   1   1   1.5  RT (min)      lcorbi_t3_pos_3_054_15    m/z 270.1055     0   1   5   5.5  RT (min)      lcorbi_t3_pos_4_104_61    m/z 270.1279     0   2   4   6E+3   1.5   2  RT (min)      lcorbi_t3_pos_3_012_31    m/z 270.1383     0   1   1.5   2  RT (min)      lcorbi_t3_pos_3_012_31    m/z 270.1418     0   1E+4   11   11.5  RT (min)      lcorbi_t3_pos_4_111_62    m/z 270.1466     0   1E+4   8   8.5  RT (min)      lcorbi_t3_pos_2_005_11    m/z 270.1548     0   1   2   3   4E+3   4.5   5  RT (min)      lcorbi_t3_pos_1_001_49    m/z 270.1548     0   1   2   3E+3   1.5   2  RT (min)      lcorbi_t3_pos_3_055_10    m/z 270.1658     0   1   2   3E+4   1   1.5   2  RT (min)       lcorbi_t3_pos_4_091_60    m/z 270.1912     0   1E+4   4.5   5  RT (min)      lcorbi_t3_pos_4_110_62    m/z 270.2064     0   1   2E+4   8.5   9  RT (min)      lcorbi_t3_pos_4_082_59    m/z 270.2339     0   2   4E+4   10.5   11  RT (min)      lcorbi_t3_pos_4_106_62    m/z 270.2428     0   1E+4   9.5   10  RT (min)      lcorbi_t3_pos_2_002_06    m/z 270.3156     0   1E+4   9   9.5  RT (min)      lcorbi_t3_pos_1_001_49    m/z 270.8615     0   2   4   6E+3   1   1.5  RT (min)      lcorbi_t3_pos_1_036_08    m/z 270.8734     0   2   4   6E+4   1   1.5  RT (min)      lcorbi_t3_pos_2_024_15    m/z 270.9292     0   2   4   6E+4   1   1.5  RT (min)      lcorbi_t3_pos_4_085_59    m/z 270.9605     0   1E+4   0.5   1  RT (min)      lcorbi_t3_pos_1_056_56    m/z 270.9710     0   1   2E+4   12   12.5  RT (min)      lcorbi_t3_pos_4_021_25    m/z 270.9711     0   2   4   6E+4   18   18.5  RT (min)       lcorbi_t3_pos_4_102_61    m/z 270.9986     0   1   2E+4   1.5   2  RT (min)       lcorbi_t3_pos_4_073_57    m/z 271.0554     0   2   4   6E+3   4.5   5  RT (min)      lcorbi_t3_pos_1_014_03    m/z 271.0611     0   1E+3   2.5   3  RT (min)      lcorbi_t3_pos_1_022_20    m/z 271.0633     0   1   2E+4   8.5   9  RT (min)       lcorbi_t3_pos_4_073_57    m/z 271.0876     0   1   2   3E+4   9.5   10  RT (min)      lcorbi_t3_pos_4_102_61    m/z 271.0926     0   1   2E+3   4.5   5  RT (min)      lcorbi_t3_pos_1_012_14    m/z 271.0926     0   1E+3   2.5   3  RT (min)      lcorbi_t3_pos_1_026_17    m/z 271.0965     0   2   4   6E+4   6.5   7   7.5  RT (min)       lcorbi_t3_pos_4_005_33    m/z 271.1020     0   1   2   3E+4   0.5   1  RT (min)      lcorbi_t3_pos_1_027_43    m/z 271.1152     0   2   4E+4   10   10.5  RT (min)       lcorbi_t3_pos_4_082_59    m/z 271.1217     0   1   1   1.5  RT (min)      lcorbi_t3_pos_3_027_27    m/z 271.1289     0   1   2E+3   4.5   5  RT (min)      lcorbi_t3_pos_1_026_17    m/z 271.1289     0   1E+3   2.5   3  RT (min)      lcorbi_t3_pos_1_024_04    m/z 271.1330     0   1   2E+4   9   9.5  RT (min)      lcorbi_t3_pos_4_074_58    m/z 271.1515     0   2   4E+3   1   1.5  RT (min)      lcorbi_t3_pos_1_002_01    m/z 271.1517     0   1   2   3E+3   6   6.5  RT (min)      lcorbi_t3_pos_1_001_49    m/z 271.2856     0   1   2   3   4E+4   11.5   12  RT (min)       lcorbi_t3_pos_4_092_60    m/z 271.9915     0   1   1   1.5  RT (min)      lcorbi_t3_pos_1_034_34    m/z 272.0207     0   2   4   6   8E+3   3   3.5  RT (min)      lcorbi_t3_pos_4_080_58    m/z 272.0538     0   1   2   3E+3   2   2.5  RT (min)      lcorbi_t3_pos_4_085_59    m/z 272.0916     0   1   2   3   4E+3   4.5   5  RT (min)      lcorbi_t3_pos_4_066_57    m/z 272.0999     0   2   4   6E+3   6.5   7  RT (min)      lcorbi_t3_pos_4_030_05    m/z 272.1242     0   1   2E+4   4.5   5  RT (min)      lcorbi_t3_pos_4_066_57    m/z 272.1242     0   1   2.5   3  RT (min)      lcorbi_t3_pos_4_082_59    m/z 272.1242     0   1   2E+4   1   1.5  RT (min)      lcorbi_t3_pos_2_037_27    m/z 272.1340     0   1E+4   1   1.5  RT (min)      lcorbi_t3_pos_4_070_57    m/z 272.1718     0   2   4   6E+4   1   1.5  RT (min)       lcorbi_t3_pos_2_020_22    m/z 272.1727     0   1   2   3E+4   7   7.5  RT (min)      lcorbi_t3_pos_4_005_33    m/z 272.1969     0   1   2   3E+4   4.5   5  RT (min)       lcorbi_t3_pos_3_048_30    m/z 272.2333     0   1   2   3   4E+3   4.5   5  RT (min)      lcorbi_t3_pos_1_002_01    m/z 272.2948     0   1E+5   10   10.5  RT (min)       lcorbi_t3_pos_2_044_21    m/z 272.9595     0   1   2E+3   1   1.5  RT (min)      lcorbi_t3_pos_4_087_59    m/z 273.0090     0   1   2E+5   0   0.5  RT (min)      lcorbi_t3_pos_1_006_15    m/z 273.0221     0   1E+5   1   1.5  RT (min)       lcorbi_t3_pos_4_087_59    m/z 273.0370     0   1E+5   1   1.5  RT (min)       lcorbi_t3_pos_4_084_59    m/z 273.0484     0   1   2   3E+4   1   1.5  RT (min)      lcorbi_t3_pos_2_020_22    m/z 273.0581     0   1   2E+3   1.5   2  RT (min)      lcorbi_t3_pos_1_028_32    m/z 273.0693     0   1   2   3   4E+5   1.5   2  RT (min)      lcorbi_t3_pos_1_028_32    m/z 273.0721     0   1E+4   4.5   5  RT (min)      lcorbi_t3_pos_4_081_58    m/z 273.0758     0   2   4   6   8E+4   6.5   7   7.5  RT (min)       lcorbi_t3_pos_4_005_33    m/z 273.0945     0   1E+4   9   9.5  RT (min)      lcorbi_t3_pos_4_066_57    m/z 273.0945     0   1E+4   4.5   5  RT (min)      lcorbi_t3_pos_4_066_57    m/z 273.1195     0   2   4   6E+4   1   1.5  RT (min)       lcorbi_t3_pos_1_050_13    m/z 273.1308     0   1E+4   5   5.5  RT (min)      lcorbi_t3_pos_4_088_59    m/z 273.1421     0   1   1   1.5  RT (min)       lcorbi_t3_pos_4_093_60    m/z 273.1673     0   1E+4   6   6.5  RT (min)       lcorbi_t3_pos_1_001_49    m/z 273.1810     0   1   2   3E+3   5   5.5  RT (min)      lcorbi_t3_pos_1_001_49    m/z 273.1890     0   1E+4   8.5   9  RT (min)      lcorbi_t3_pos_1_001_49    m/z 273.2213     0   1E+4   7.5   8  RT (min)      lcorbi_t3_pos_1_001_49    m/z 273.2255     0   1E+4   9   9.5  RT (min)      lcorbi_t3_pos_1_001_49    m/z 273.2618     0   2   4   6   8E+4   11   11.5  RT (min)       lcorbi_t3_pos_4_111_62    m/z 273.2982     0   1   2   3   4E+4   10   10.5  RT (min)      lcorbi_t3_pos_2_044_21    m/z 273.9233     0   2   4   6   8E+4   1   1.5  RT (min)      lcorbi_t3_pos_1_028_32    m/z 273.9453     0   1   2E+4   1   1.5  RT (min)      lcorbi_t3_pos_4_075_58    m/z 273.9620     0   1E+5   1   1.5  RT (min)      lcorbi_t3_pos_1_028_32    m/z 273.9821     0   1   1   1.5  RT (min)       lcorbi_t3_pos_1_031_40    m/z 273.9973     0   1   1   1.5  RT (min)      lcorbi_t3_pos_1_052_38    m/z 274.0483     0   1   2E+4   1   1.5  RT (min)      lcorbi_t3_pos_3_010_44    m/z 274.0621     0   1E+4   1   1.5  RT (min)      lcorbi_t3_pos_4_090_60    m/z 274.0727     0   1   1.5   2  RT (min)      lcorbi_t3_pos_2_037_27    m/z 274.0791     0   1E+4   6.5   7  RT (min)      lcorbi_t3_pos_4_005_33    m/z 274.0913     0   2   4E+3   4.5   5  RT (min)      lcorbi_t3_pos_4_106_62    m/z 274.1003     0   1   2   3E+4   11   11.5  RT (min)      lcorbi_t3_pos_4_038_38    m/z 274.1209     0   1   2   3E+4   4.5   5  RT (min)       lcorbi_t3_pos_4_096_60    m/z 274.1286     0   1   2E+4   4.5   5  RT (min)      lcorbi_t3_pos_4_082_59    m/z 274.1286     0   1E+3   2.5   3  RT (min)      lcorbi_t3_pos_4_090_60    m/z 274.1414     0   2   4   6   8E+5   1.5   2  RT (min)       lcorbi_t3_pos_4_093_60    m/z 274.1704     0   1   2   3E+3   6   6.5  RT (min)      lcorbi_t3_pos_1_002_01    m/z 274.1890     0   1   2E+5   1.5   2  RT (min)       lcorbi_t3_pos_2_037_27    m/z 274.2185     0   1E+4   7   7.5  RT (min)      lcorbi_t3_pos_4_053_19    m/z 274.2377     0   1E+5   8   8.5  RT (min)       lcorbi_t3_pos_4_091_60    m/z 274.2549     0   1   2E+4   9   9.5  RT (min)       lcorbi_t3_pos_1_013_10    m/z 274.2571     0   1E+4   5   5.5  RT (min)      lcorbi_t3_pos_4_106_62    m/z 274.2655     0   2   4E+4   9   9.5  RT (min)      lcorbi_t3_pos_4_003_39    m/z 274.2828     0   2   4   6E+4   8.5   9  RT (min)      lcorbi_t3_pos_1_019_16    m/z 274.7919     0   2   4   6   8E+4   1   1.5  RT (min)       lcorbi_t3_pos_1_029_29    m/z 274.8392     0   1   1   1.5  RT (min)      lcorbi_t3_pos_2_051_35    m/z 274.8735     0   1   2E+3   1   1.5  RT (min)       lcorbi_t3_pos_1_020_31    m/z 274.9057     0   2   4E+3   0.5   1  RT (min)      lcorbi_t3_pos_4_094_60    m/z 274.9420     0   1   2E+4   1   1.5  RT (min)      lcorbi_t3_pos_4_080_58    m/z 274.9752     0   1E+5   0.5   1  RT (min)      lcorbi_t3_pos_4_080_58    m/z 274.9926     0   1   2E+3   1   1.5  RT (min)      lcorbi_t3_pos_3_039_03    m/z 274.9927     0   2   4   6E+4   18   18.5  RT (min)       lcorbi_t3_pos_4_078_58    m/z 275.0472     0   1   2   3   4E+5   18.5   19  RT (min)       lcorbi_t3_pos_4_101_61    m/z 275.0472     0   1   2   3   4E+4   6.5   7  RT (min)      lcorbi_t3_pos_1_041_54    m/z 275.0473     0   1   2   3E+4   10.5   11  RT (min)      lcorbi_t3_pos_3_022_02    m/z 275.0738     0   2   4   6E+5   1   1.5   2  RT (min)       lcorbi_t3_pos_2_028_24    m/z 275.0850     0   1E+5   1   1.5  RT (min)      lcorbi_t3_pos_2_054_29    m/z 275.0890     0   1E+4   8.5   9  RT (min)      lcorbi_t3_pos_4_106_62    m/z 275.1002     0   1E+4   5.5   6  RT (min)      lcorbi_t3_pos_4_084_59    m/z 275.1101     0   1   2   3   4E+4   4.5   5  RT (min)      lcorbi_t3_pos_4_066_57    m/z 275.1102     0   2   4E+4   2.5   3  RT (min)      lcorbi_t3_pos_4_030_05    m/z 275.1238     0   1   2   3E+3   1   1.5   2  RT (min)       lcorbi_t3_pos_2_028_24    m/z 275.1254     0   1E+4   6.5   7  RT (min)      lcorbi_t3_pos_4_066_57    m/z 275.1273     0   1   2E+3   4.5   5  RT (min)      lcorbi_t3_pos_1_001_49    m/z 275.1350     0   1   1.5   2  RT (min)      lcorbi_t3_pos_3_012_31    m/z 275.1465     0   1   2E+4   4.5   5  RT (min)       lcorbi_t3_pos_4_106_62    m/z 275.1618     0   1   2E+4   6.5   7  RT (min)       lcorbi_t3_pos_2_023_26    m/z 275.1643     0   1E+5   13.5   14  RT (min)       lcorbi_t3_pos_4_085_59    m/z 275.1854     0   1   8.5   9  RT (min)       lcorbi_t3_pos_4_090_60    m/z 275.2330     0   1   2   3E+4   7   8  RT (min)       lcorbi_t3_pos_2_052_48    m/z 275.2582     0   1E+4   9   9.5  RT (min)      lcorbi_t3_pos_1_019_16    m/z 275.2693     0   2   4   6E+3   5   5.5  RT (min)      lcorbi_t3_pos_4_106_62    m/z 275.2710     0   1   2   3E+4   8.5   9   9.5  RT (min)      lcorbi_t3_pos_1_019_16    m/z 275.8917     0   1E+4   1   1.5  RT (min)      lcorbi_t3_pos_1_009_50    m/z 275.9205     0   1   2   3E+4   1   1.5  RT (min)      lcorbi_t3_pos_1_034_34    m/z 275.9785     0   1   2E+4   1   1.5  RT (min)      lcorbi_t3_pos_1_032_26    m/z 275.9945     0   1   1   1.5  RT (min)      lcorbi_t3_pos_1_028_32    m/z 276.0689     0   1   4.5   5  RT (min)      lcorbi_t3_pos_4_090_60    m/z 276.0770     0   2   4   6E+4   1   1.5  RT (min)      lcorbi_t3_pos_3_012_31    m/z 276.1044     0   1   1   1.5  RT (min)      lcorbi_t3_pos_4_067_57    m/z 276.1078     0   1E+5   4.5   5  RT (min)       lcorbi_t3_pos_4_090_60    m/z 276.1079     0   1   2   3   4E+4   2.5   3  RT (min)       lcorbi_t3_pos_4_032_40    m/z 276.1191     0   1   2   3E+4   1   1.5  RT (min)      lcorbi_t3_pos_4_084_59    m/z 276.1442     0   2   4E+2   1   1.5  RT (min)       lcorbi_t3_pos_3_029_13    m/z 276.1443     0   2   4   6   8E+3   2.5   3  RT (min)      lcorbi_t3_pos_1_034_34    m/z 276.1806     0   1E+4   4.5   5  RT (min)      lcorbi_t3_pos_4_074_58    m/z 276.1806     0   2   4   6E+2   1   1.5  RT (min)      lcorbi_t3_pos_4_007_46    m/z 276.1959     0   1   2   3E+4   9   9.5  RT (min)      lcorbi_t3_pos_4_090_60    m/z 276.2534     0   1E+5   8.5   9  RT (min)       lcorbi_t3_pos_4_113_62    m/z 276.2807     0   1   2   3E+5   8.5   9  RT (min)      lcorbi_t3_pos_2_044_21    m/z 276.7891     0   1   2E+4   1   1.5  RT (min)      lcorbi_t3_pos_1_029_29    m/z 276.8364     0   1   2   3E+4   1   1.5  RT (min)      lcorbi_t3_pos_1_036_08    m/z 276.8715     0   1   2   3E+4   1   1.5  RT (min)       lcorbi_t3_pos_4_090_60    m/z 276.9545     0   1   1.5   2  RT (min)      lcorbi_t3_pos_4_110_62    m/z 276.9867     0   1E+4   1.5   2  RT (min)      lcorbi_t3_pos_1_036_08    m/z 277.0082     0   1E+6   18.5   19  RT (min)       lcorbi_t3_pos_4_086_59    m/z 277.0082     0   1   2E+5   0.5   1  RT (min)       lcorbi_t3_pos_4_077_58    m/z 277.0707     0   1   7   7.5  RT (min)      lcorbi_t3_pos_4_110_62    m/z 277.0860     0   2   4   6E+4   10   10.5  RT (min)       lcorbi_t3_pos_4_088_59    m/z 277.0894     0   1   2   3   4E+3   1   1.5  RT (min)       lcorbi_t3_pos_3_029_13    m/z 277.1049     0   1   2   3   4E+4   4.5   5  RT (min)       lcorbi_t3_pos_4_112_62    m/z 277.1099     0   1   2E+3   6   6.5  RT (min)      lcorbi_t3_pos_4_098_61    m/z 277.1112     0   1   2E+3   2.5   3  RT (min)      lcorbi_t3_pos_4_070_57    m/z 277.1183     0   2   4E+4   5   5.5  RT (min)       lcorbi_t3_pos_4_104_61    m/z 277.1282     0   1E+4   7   7.5  RT (min)      lcorbi_t3_pos_4_103_61    m/z 277.1330     0   1   2E+4   10   10.5  RT (min)      lcorbi_t3_pos_4_108_62    m/z 277.1410     0   1E+4   8.5   9  RT (min)      lcorbi_t3_pos_4_090_60    m/z 277.1566     0   1   3   3.5  RT (min)      lcorbi_t3_pos_1_001_49    m/z 277.1675     0   1   2E+4   6.5   7  RT (min)       lcorbi_t3_pos_3_001_49    m/z 277.1799     0   2   4E+4   9.5   10  RT (min)      lcorbi_t3_pos_4_074_58    m/z 277.1887     0   1E+5   1   1.5  RT (min)       lcorbi_t3_pos_1_003_45    m/z 277.2010     0   1E+5   8.5   9  RT (min)       lcorbi_t3_pos_4_086_59    m/z 277.2139     0   2   4   6E+3   1   1.5  RT (min)      lcorbi_t3_pos_1_003_45    m/z 277.2141     0   1   2   3E+4   11.5   12  RT (min)       lcorbi_t3_pos_4_080_58    m/z 277.2374     0   1   10   10.5  RT (min)       lcorbi_t3_pos_4_074_58    m/z 277.2568     0   1   2E+4   8.5   9  RT (min)      lcorbi_t3_pos_4_112_62    m/z 277.2649     0   2   4   6   8E+4   11.5   12  RT (min)      lcorbi_t3_pos_4_087_59    m/z 277.9367     0   1E+5   1   1.5  RT (min)      lcorbi_t3_pos_1_036_08    m/z 278.0294     0   2   4   6   8E+3   1   1.5  RT (min)      lcorbi_t3_pos_1_003_45    m/z 278.0389     0   1   2E+5   1   1.5  RT (min)       lcorbi_t3_pos_4_093_60    m/z 278.0408     0   2   4E+3   3   3.5  RT (min)      lcorbi_t3_pos_4_102_61    m/z 278.0637     0   1   2   3   4E+4   1   1.5  RT (min)       lcorbi_t3_pos_4_084_59    m/z 278.0707     0   1E+4   5   5.5  RT (min)      lcorbi_t3_pos_4_066_57    m/z 278.0924     0   1   1   1.5  RT (min)      lcorbi_t3_pos_3_029_13    m/z 278.1114     0   2   4   6E+3   4.5   5  RT (min)      lcorbi_t3_pos_3_001_49    m/z 278.1217     0   2   4   6E+3   5   5.5  RT (min)      lcorbi_t3_pos_4_099_61    m/z 278.1235     0   1   2   3E+4   1   1.5  RT (min)       lcorbi_t3_pos_1_031_40    m/z 278.1377     0   2   4   6E+4   2.5   3  RT (min)       lcorbi_t3_pos_1_036_08    m/z 278.1444     0   1   2E+4   6.5   7  RT (min)       lcorbi_t3_pos_4_066_57    m/z 278.1477     0   1E+4   4.5   5  RT (min)      lcorbi_t3_pos_3_001_49    m/z 278.1751     0   1   4.5   5  RT (min)      lcorbi_t3_pos_4_104_61    m/z 278.2133     0   1   2E+4   7   7.5  RT (min)      lcorbi_t3_pos_4_053_19    m/z 278.2326     0   2   4   6E+4   7   7.5  RT (min)       lcorbi_t3_pos_4_053_19    m/z 278.2455     0   1E+5   12   12.5  RT (min)       lcorbi_t3_pos_4_082_59    m/z 278.2691     0   2   4   6E+4   11   11.5  RT (min)       lcorbi_t3_pos_1_032_26    m/z 278.8809     0   2   4E+4   1   1.5  RT (min)      lcorbi_t3_pos_1_036_08    m/z 278.8932     0   2   4E+4   18   18.5  RT (min)       lcorbi_t3_pos_4_079_58    m/z 278.8932     0   2   4   6E+4   1   1.5  RT (min)       lcorbi_t3_pos_1_020_31    m/z 278.8933     0   2   4E+4   12   12.5  RT (min)      lcorbi_t3_pos_2_002_06    m/z 278.9231     0   1   2E+3   1   1.5  RT (min)      lcorbi_t3_pos_2_054_29    m/z 278.9653     0   1E+3   1   1.5  RT (min)      lcorbi_t3_pos_1_003_45    m/z 278.9681     0   2   4   6E+4   18   18.5  RT (min)       lcorbi_t3_pos_4_085_59    m/z 278.9788     0   2   4   6   8E+4   1   1.5  RT (min)      lcorbi_t3_pos_1_036_08    m/z 278.9875     0   1E+3   1.5   2  RT (min)      lcorbi_t3_pos_1_001_49    m/z 279.0319     0   2   4   6E+3   1   1.5  RT (min)      lcorbi_t3_pos_1_001_49    m/z 279.0388     0   1E+4   1   1.5  RT (min)      lcorbi_t3_pos_1_036_08    m/z 279.0934     0   1   2   3E+4   8.5   9  RT (min)       lcorbi_t3_pos_4_087_59    m/z 279.1187     0   2   4   6   8E+4   1   1.5  RT (min)      lcorbi_t3_pos_4_089_59    m/z 279.1452     0   2   4   6E+3   8.5   9  RT (min)      lcorbi_t3_pos_4_087_59    m/z 279.1503     0   1E+5   10   10.5  RT (min)      lcorbi_t3_pos_4_077_58    m/z 279.1592     0   1   2   3E+5   17.5   18  RT (min)       lcorbi_t3_pos_4_075_58    m/z 279.1705     0   1   2   3E+3   5.5   6  RT (min)      lcorbi_t3_pos_1_002_01    m/z 279.1707     0   2   4   6E+4   10   10.5  RT (min)      lcorbi_t3_pos_1_029_29    m/z 279.1843     0   2   4   6   8E+4   18   18.5  RT (min)       lcorbi_t3_pos_4_106_62    m/z 279.1996     0   1   4.5   5  RT (min)      lcorbi_t3_pos_1_001_49    m/z 279.2043     0   2   4E+3   7.5   8  RT (min)      lcorbi_t3_pos_1_001_49    m/z 279.2360     0   1   2   3E+3   7   7.5  RT (min)      lcorbi_t3_pos_1_001_49    m/z 279.2795     0   1   2E+4   9.5   10  RT (min)      lcorbi_t3_pos_2_030_14    m/z 279.9339     0   2   4E+4   1   1.5  RT (min)      lcorbi_t3_pos_1_036_08    m/z 280.0605     0   1   1   1.5  RT (min)       lcorbi_t3_pos_4_006_12    m/z 280.0908     0   1E+4   4.5   5  RT (min)      lcorbi_t3_pos_4_074_58    m/z 280.0968     0   2   4   6   8E+3   0.5   1  RT (min)       lcorbi_t3_pos_4_075_58    m/z 280.1076     0   1E+3   1   1.5   2  RT (min)       lcorbi_t3_pos_1_052_38    m/z 280.1180     0   1   4.5   5  RT (min)      lcorbi_t3_pos_4_106_62    m/z 280.1374     0   1   2   3E+4   4.5   5  RT (min)      lcorbi_t3_pos_4_066_57    m/z 280.1391     0   2   4   6   8E+3   1   1.5  RT (min)      lcorbi_t3_pos_1_052_38    m/z 280.1544     0   1   2   3   4E+3   6   6.5  RT (min)      lcorbi_t3_pos_1_001_49    m/z 280.1544     0   1   2   3E+4   4.5   5  RT (min)       lcorbi_t3_pos_4_067_57    m/z 280.2119     0   2   4   6E+3   6.5   7  RT (min)      lcorbi_t3_pos_4_077_58    m/z 280.2272     0   1E+4   8.5   9  RT (min)      lcorbi_t3_pos_3_047_40    m/z 280.2384     0   2   4   6E+3   6   6.5  RT (min)      lcorbi_t3_pos_4_082_59    m/z 280.2636     0   2   4E+4   10   10.5  RT (min)       lcorbi_t3_pos_2_051_35    m/z 280.8265     0   1   2E+4   1   1.5  RT (min)      lcorbi_t3_pos_4_045_08    m/z 280.8380     0   1E+4   1   1.5  RT (min)      lcorbi_t3_pos_1_034_34    m/z 280.8902     0   1   2E+3   1   1.5  RT (min)      lcorbi_t3_pos_1_004_24    m/z 281.0535     0   1   1   1.5  RT (min)       lcorbi_t3_pos_2_040_25    m/z 281.0745     0   1   2   3   4E+3   4.5   5  RT (min)      lcorbi_t3_pos_1_021_18    m/z 281.0746     0   1   2.5   3  RT (min)      lcorbi_t3_pos_4_090_60    m/z 281.0955     0   1   2   3   4E+5   1   1.5  RT (min)      lcorbi_t3_pos_4_075_58    m/z 281.1020     0   1E+4   8.5   9  RT (min)      lcorbi_t3_pos_4_052_06    m/z 281.1065     0   2   4E+3   4.5   5  RT (min)      lcorbi_t3_pos_4_072_57    m/z 281.1133     0   1E+4   5.5   6  RT (min)      lcorbi_t3_pos_4_076_58    m/z 281.1344     0   1E+5   1   1.5  RT (min)       lcorbi_t3_pos_1_054_48    m/z 281.1496     0   2   4   6   8E+4   4.5   5  RT (min)       lcorbi_t3_pos_4_082_59    m/z 281.1577     0   1   2E+3   4.5   5  RT (min)      lcorbi_t3_pos_1_001_49    m/z 281.1658     0   1   2E+5   10   10.5  RT (min)      lcorbi_t3_pos_4_080_58    m/z 281.1884     0   1E+4   7   7.5  RT (min)      lcorbi_t3_pos_4_053_19    m/z 281.1959     0   1   2   3E+3   5.5   6  RT (min)      lcorbi_t3_pos_1_002_01    m/z 281.1974     0   1   2   3   4E+4   1.5   2  RT (min)      lcorbi_t3_pos_4_049_55    m/z 281.2281     0   2   4   6E+4   10.5   11  RT (min)       lcorbi_t3_pos_4_113_62    m/z 281.2588     0   2   4   6E+4   9   9.5  RT (min)       lcorbi_t3_pos_2_044_21    m/z 281.2952     0   1   2E+5   10   11   12  RT (min)       lcorbi_t3_pos_1_050_13    m/z 281.2952     0   1   2E+4   14   14.5  RT (min)      lcorbi_t3_pos_2_040_25    m/z 281.9045     0   1E+3   1   1.5  RT (min)      lcorbi_t3_pos_1_006_15    m/z 282.0277     0   1E+4   7.5   8  RT (min)      lcorbi_t3_pos_4_074_58    m/z 282.0963     0   2   4   6E+4   1   1.5  RT (min)      lcorbi_t3_pos_4_078_58    m/z 282.1126     0   1E+4   9   9.5  RT (min)      lcorbi_t3_pos_2_053_28    m/z 282.1196     0   1E+4   4.5   5  RT (min)      lcorbi_t3_pos_4_082_59    m/z 282.1196     0   1   2E+5   1  RT (min)       lcorbi_t3_pos_3_029_13    m/z 282.1466     0   1   2E+4   8   8.5  RT (min)      lcorbi_t3_pos_4_082_59    m/z 282.1530     0   2   4   6   8E+3   4.5   5  RT (min)      lcorbi_t3_pos_1_001_49    m/z 282.1701     0   1   2E+4   9   9.5  RT (min)      lcorbi_t3_pos_1_010_46    m/z 282.1848     0   2   4   6E+4   7   7.5  RT (min)       lcorbi_t3_pos_4_053_19    m/z 282.1912     0   1   2E+3   4.5   5  RT (min)      lcorbi_t3_pos_1_003_45    m/z 282.1912     0   1   2E+3   2.5   3  RT (min)      lcorbi_t3_pos_1_003_45    m/z 282.2428     0   1   10.5   11  RT (min)      lcorbi_t3_pos_1_005_42    m/z 282.2621     0   1E+4   9   9.5  RT (min)      lcorbi_t3_pos_4_038_38    m/z 282.8236     0   1   2E+4   1   1.5  RT (min)      lcorbi_t3_pos_4_045_08    m/z 282.8348     0   1   2   3   4E+3   1   1.5  RT (min)      lcorbi_t3_pos_2_023_26    m/z 282.8443     0   2   4   6   8E+4   1   1.5  RT (min)      lcorbi_t3_pos_1_052_38    m/z 282.8835     0   2   4E+6   0   0.5  RT (min)      lcorbi_t3_pos_1_007_07    m/z 282.9051     0   1E+5   0.5   1  RT (min)      lcorbi_t3_pos_4_075_58    m/z 282.9569     0   2   4   6E+4   18   18.5  RT (min)      lcorbi_t3_pos_4_072_57    m/z 282.9912     0   1   2   3   4E+3   1.5   2  RT (min)      lcorbi_t3_pos_3_008_20    m/z 283.0004     0   1   2   3E+3   1   1.5  RT (min)       lcorbi_t3_pos_1_034_34    m/z 283.0214     0   1   2   3E+4   7.5   8  RT (min)       lcorbi_t3_pos_4_098_61    m/z 283.0617     0   1   2   3E+4   1   1.5  RT (min)      lcorbi_t3_pos_1_036_08    m/z 283.0724     0   1   2.5   3  RT (min)      lcorbi_t3_pos_1_045_12    m/z 283.0901     0   1   1.5   2  RT (min)      lcorbi_t3_pos_4_052_06    m/z 283.1021     0   1   1   1.5  RT (min)      lcorbi_t3_pos_1_045_12    m/z 283.1153     0   1   2   3   4E+3   4.5   5  RT (min)      lcorbi_t3_pos_1_050_13    m/z 283.1155     0   1   2E+3   6   6.5  RT (min)      lcorbi_t3_pos_1_054_48    m/z 283.1230     0   1E+3   1   1.5  RT (min)      lcorbi_t3_pos_1_045_12    m/z 283.1384     0   1   2   3E+4   0.5   1  RT (min)      lcorbi_t3_pos_1_045_12    m/z 283.1587     0   1   2E+4   18.5   19  RT (min)       lcorbi_t3_pos_4_100_61    m/z 283.1587     0   1E+4   0.5   1  RT (min)       lcorbi_t3_pos_3_011_42    m/z 283.1689     0   2   4E+4   9.5   10  RT (min)       lcorbi_t3_pos_1_039_23    m/z 283.1752     0   1E+5   4.5   5  RT (min)       lcorbi_t3_pos_2_044_21    m/z 283.2074     0   1E+4   9   9.5  RT (min)      lcorbi_t3_pos_4_103_61    m/z 283.2087     0   1   2   3E+4   1   1.5  RT (min)      lcorbi_t3_pos_1_036_08    m/z 283.2251     0   2   4   6E+4   13   13.5  RT (min)       lcorbi_t3_pos_4_101_61    m/z 283.2826     0   1E+4   10   10.5  RT (min)      lcorbi_t3_pos_1_001_49    m/z 283.3109     0   1   2E+4   11   12  RT (min)       lcorbi_t3_pos_1_001_49    m/z 283.7175     0   1   2   3E+4   5.5   6  RT (min)       lcorbi_t3_pos_4_034_42    m/z 283.9843     0   1E+5   1   1.5  RT (min)       lcorbi_t3_pos_4_085_59    m/z 284.0482     0   1   2E+3   1.5   2  RT (min)      lcorbi_t3_pos_2_005_11    m/z 284.0990     0   2   4E+3   4.5   5  RT (min)      lcorbi_t3_pos_4_069_57    m/z 284.1065     0   1E+5   1   1.5  RT (min)       lcorbi_t3_pos_4_074_58    m/z 284.1090     0   1   2   3   4E+3   4.5   5  RT (min)      lcorbi_t3_pos_4_106_62    m/z 284.1340     0   1E+5   4.5   5  RT (min)       lcorbi_t3_pos_4_066_57    m/z 284.1341     0   2   4   6E+4   2.5   3  RT (min)       lcorbi_t3_pos_1_048_47    m/z 284.1578     0   1   2E+4   0.5   1  RT (min)      lcorbi_t3_pos_1_032_26    m/z 284.1705     0   2   4   6   8E+3   4.5   5  RT (min)       lcorbi_t3_pos_1_055_33    m/z 284.1785     0   1   2E+3   4.5   5  RT (min)      lcorbi_t3_pos_1_015_11    m/z 284.2068     0   1   2   3   4E+3   5   5.5  RT (min)      lcorbi_t3_pos_1_012_14    m/z 284.2221     0   1   2E+4   9   9.5  RT (min)      lcorbi_t3_pos_4_074_58    m/z 284.2949     0   2   4   6E+4   15   15.5  RT (min)      lcorbi_t3_pos_3_042_14    m/z 284.8497     0   2   4E+4   11.5   12  RT (min)      lcorbi_t3_pos_3_026_04    m/z 284.8592     0   1   1   1.5  RT (min)      lcorbi_t3_pos_1_020_31    m/z 284.8794     0   1   2   3E+4   1   1.5  RT (min)      lcorbi_t3_pos_1_028_32    m/z 284.8889     0   1E+3   1   1.5  RT (min)      lcorbi_t3_pos_1_020_31    m/z 284.9102     0   1   2   3E+5   1   1.5  RT (min)       lcorbi_t3_pos_1_002_01    m/z 284.9381     0   1   2   3E+5   18   18.5  RT (min)       lcorbi_t3_pos_4_078_58    m/z 284.9381     0   1   2E+4   8   8.5  RT (min)      lcorbi_t3_pos_2_020_22    m/z 284.9381     0   2   4   6   8E+3   0.5   1  RT (min)       lcorbi_t3_pos_4_067_57    m/z 284.9581     0   1   2   3   4E+4   1   1.5  RT (min)      lcorbi_t3_pos_3_020_35    m/z 284.9612     0   1E+4   7.5   8  RT (min)      lcorbi_t3_pos_1_032_26    m/z 285.0160     0   2   4E+4   7   7.5  RT (min)       lcorbi_t3_pos_4_030_05    m/z 285.0945     0   1   2E+4   4.5   5  RT (min)      lcorbi_t3_pos_4_048_24    m/z 285.1122     0   1   2   3E+4   10   10.5  RT (min)      lcorbi_t3_pos_1_001_49    m/z 285.1193     0   1   2E+4   1   1.5  RT (min)      lcorbi_t3_pos_4_089_59    m/z 285.1338     0   1E+4   5.5   6  RT (min)      lcorbi_t3_pos_4_092_60    m/z 285.1373     0   1   2   3   4E+3   4.5   5  RT (min)      lcorbi_t3_pos_4_048_24    m/z 285.1446     0   1E+4   9.5   10  RT (min)      lcorbi_t3_pos_4_090_60    m/z 285.1446     0   1E+4   4.5   5  RT (min)      lcorbi_t3_pos_4_090_60    m/z 285.1446     0   1   2.5   3  RT (min)      lcorbi_t3_pos_4_082_59    m/z 285.1540     0   1E+5   6.5   7  RT (min)      lcorbi_t3_pos_1_036_08    m/z 285.1540     0   1E+5   0.5   1  RT (min)       lcorbi_t3_pos_4_089_59    m/z 285.1540     0   1   2E+5   4.5   5  RT (min)      lcorbi_t3_pos_1_033_53    m/z 285.1672     0   1E+3   6.5   7  RT (min)      lcorbi_t3_pos_1_028_32    m/z 285.1687     0   1E+3   1   1.5  RT (min)       lcorbi_t3_pos_1_038_30    m/z 285.1809     0   1   4.5   5  RT (min)      lcorbi_t3_pos_1_043_06    m/z 285.2213     0   1   2   3E+3   7.5   8  RT (min)       lcorbi_t3_pos_1_022_20    m/z 285.2286     0   1   4.5   5  RT (min)      lcorbi_t3_pos_1_012_14    m/z 285.2425     0   1E+5   14.5   15  RT (min)       lcorbi_t3_pos_1_031_40    m/z 285.2982     0   2   4   6E+4   10.5   11  RT (min)       lcorbi_t3_pos_1_008_36    m/z 285.3346     0   1   2   3E+4   10   10.5  RT (min)      lcorbi_t3_pos_2_009_50    m/z 285.9394     0   2   4   6E+4   18   18.5  RT (min)      lcorbi_t3_pos_4_091_60    m/z 286.0259     0   1E+5   4.5   5  RT (min)       lcorbi_t3_pos_4_081_58    m/z 286.0269     0   1   2.5   3  RT (min)      lcorbi_t3_pos_4_083_59    m/z 286.0899     0   1   2E+4   1   1.5  RT (min)       lcorbi_t3_pos_4_084_59    m/z 286.1399     0   2   4   6   8E+2   2.5   3  RT (min)      lcorbi_t3_pos_2_020_22    m/z 286.1538     0   1E+5   0.5   1  RT (min)       lcorbi_t3_pos_1_050_13    m/z 286.1574     0   1E+4   0.5   1  RT (min)      lcorbi_t3_pos_3_024_07    m/z 286.2126     0   1   2   3E+4   1   1.5  RT (min)       lcorbi_t3_pos_4_082_59    m/z 286.2247     0   1E+4   11.5   12  RT (min)       lcorbi_t3_pos_4_052_06    m/z 286.2247     0   2   4   6   8E+3   7.5   8  RT (min)      lcorbi_t3_pos_1_038_30    m/z 286.2489     0   2   4   6   8E+2   4.5   5  RT (min)      lcorbi_t3_pos_1_042_37    m/z 286.2741     0   2   4   6E+4   9   9.5  RT (min)       lcorbi_t3_pos_1_019_16    m/z 286.2741     0   1E+5   11.5   12  RT (min)       lcorbi_t3_pos_2_044_21    m/z 286.2935     0   1E+5   9   9.5  RT (min)       lcorbi_t3_pos_4_083_59    m/z 286.3105     0   1   2E+4   9   9.5  RT (min)       lcorbi_t3_pos_1_032_26    m/z 286.7717     0   1   2   3E+4   1   1.5  RT (min)      lcorbi_t3_pos_2_040_25    m/z 286.9072     0   2   4   6   8E+4   1   1.5  RT (min)      lcorbi_t3_pos_1_002_01    m/z 286.9355     0   1   2   3   4E+4   18   18.5  RT (min)       lcorbi_t3_pos_4_099_61    m/z 286.9486     0   2   4   6E+4   18   18.5  RT (min)       lcorbi_t3_pos_4_071_57    m/z 286.9551     0   1   2E+4   1.5   2  RT (min)      lcorbi_t3_pos_4_004_14    m/z 286.9582     0   1E+4   7.5   8  RT (min)      lcorbi_t3_pos_1_032_26    m/z 287.0051     0   2   4   6   8E+3   1   1.5  RT (min)      lcorbi_t3_pos_3_048_30    m/z 287.0294     0   2   4   6   8E+3   4.5   5  RT (min)      lcorbi_t3_pos_4_078_58    m/z 287.0704     0   1E+5   1   1.5  RT (min)       lcorbi_t3_pos_2_022_38    m/z 287.0808     0   2   4   6   8E+3   4.5   5  RT (min)      lcorbi_t3_pos_4_074_58    m/z 287.0890     0   2   4   6   8E+3   6   6.5  RT (min)      lcorbi_t3_pos_2_056_56    m/z 287.1003     0   2   4   6   8E+3   5   5.5  RT (min)      lcorbi_t3_pos_4_073_57    m/z 287.1108     0   1E+4   8.5   9  RT (min)      lcorbi_t3_pos_4_109_62    m/z 287.1155     0   1E+4   1   1.5  RT (min)      lcorbi_t3_pos_2_008_16    m/z 287.1238     0   1   2   3E+3   1.5   2  RT (min)      lcorbi_t3_pos_1_050_13    m/z 287.1352     0   2   4E+5   1   1.5  RT (min)      lcorbi_t3_pos_1_050_13    m/z 287.1464     0   1   2E+3   6   6.5  RT (min)      lcorbi_t3_pos_3_001_49    m/z 287.1472     0   2   4   6   8E+4   9   9.5  RT (min)      lcorbi_t3_pos_4_078_58    m/z 287.1509     0   2   4   6E+3   0.5   1  RT (min)      lcorbi_t3_pos_1_001_49    m/z 287.1618     0   1E+4   9.5   10  RT (min)      lcorbi_t3_pos_1_001_49    m/z 287.1918     0   1E+4   7   7.5  RT (min)      lcorbi_t3_pos_4_098_61    m/z 287.2047     0   1   6.5   7  RT (min)       lcorbi_t3_pos_3_023_05    m/z 287.2330     0   2   4   6E+4   7.5   8  RT (min)       lcorbi_t3_pos_4_082_59    m/z 287.2775     0   1E+4   9   9.5  RT (min)      lcorbi_t3_pos_3_015_06    m/z 287.9389     0   1E+5   1   1.5  RT (min)      lcorbi_t3_pos_2_020_22    m/z 288.0156     0   2   4   6E+4   1   2  RT (min)       lcorbi_t3_pos_4_090_60    m/z 288.1055     0   1E+5   1   2  RT (min)       lcorbi_t3_pos_3_029_13    m/z 288.1189     0   1E+6   1   1.5  RT (min)       lcorbi_t3_pos_1_036_08    m/z 288.1764     0   1E+5   1   1.5  RT (min)       lcorbi_t3_pos_4_090_60    m/z 288.2031     0   2   4E+4   1   1.5   2  RT (min)       lcorbi_t3_pos_2_020_22    m/z 288.2039     0   1   2E+4   7   7.5  RT (min)       lcorbi_t3_pos_4_005_33    m/z 288.2040     0   2   4   6   8E+4   13.5   14  RT (min)      lcorbi_t3_pos_1_051_09    m/z 288.2251     0   1   2   3E+4   11   11.5  RT (min)      lcorbi_t3_pos_1_004_24    m/z 288.2341     0   2   4   6E+4   8   8.5  RT (min)       lcorbi_t3_pos_4_053_19    m/z 288.2534     0   1E+4   6   6.5  RT (min)      lcorbi_t3_pos_1_001_49    m/z 288.2705     0   1E+4   9   9.5  RT (min)      lcorbi_t3_pos_2_044_21    m/z 288.7688     0   1   2   3   4E+4   1   1.5  RT (min)      lcorbi_t3_pos_2_040_25    m/z 288.9220     0   2   4   6E+5   1   1.5  RT (min)       lcorbi_t3_pos_1_029_29    m/z 288.9522     0   2   4   6E+4   0   0.5  RT (min)      lcorbi_t3_pos_1_007_07    m/z 288.9903     0   1   2   3E+3   1.5   2  RT (min)      lcorbi_t3_pos_1_028_32    m/z 288.9998     0   1   2E+5   1   1.5  RT (min)       lcorbi_t3_pos_4_084_59    m/z 289.0206     0   1   2   3E+4   1   1.5  RT (min)      lcorbi_t3_pos_2_012_01    m/z 289.0392     0   1   1   1.5  RT (min)      lcorbi_t3_pos_3_029_13    m/z 289.0496     0   1   2E+4   4.5   5  RT (min)       lcorbi_t3_pos_4_079_58    m/z 289.0894     0   1   2   3   4E+5   4.5   5  RT (min)       lcorbi_t3_pos_4_066_57    m/z 289.0895     0   1   2E+5   2.5   3  RT (min)       lcorbi_t3_pos_4_021_25    m/z 289.1273     0   1   2.5   3  RT (min)      lcorbi_t3_pos_4_021_25    m/z 289.1296     0   2   4   6E+3   4.5   5  RT (min)      lcorbi_t3_pos_4_082_59    m/z 289.1429     0   1   2E+3   5   5.5  RT (min)      lcorbi_t3_pos_4_051_34    m/z 289.1510     0   1E+5   1   1.5  RT (min)       lcorbi_t3_pos_4_086_59    m/z 289.1565     0   1E+5   10.5   11  RT (min)       lcorbi_t3_pos_4_108_62    m/z 289.1605     0   1E+4   1   2  RT (min)       lcorbi_t3_pos_2_023_26    m/z 289.1622     0   1   2E+4   5   5.5  RT (min)      lcorbi_t3_pos_4_014_48    m/z 289.2073     0   1E+5   11   11.5  RT (min)      lcorbi_t3_pos_4_098_61    m/z 289.2122     0   1E+4   4.5   5  RT (min)      lcorbi_t3_pos_4_066_57    m/z 289.2163     0   1E+4   8.5   9  RT (min)      lcorbi_t3_pos_1_007_07    m/z 289.2203     0   1E+4   7   7.5  RT (min)       lcorbi_t3_pos_4_075_58    m/z 289.2374     0   1E+4   8   8.5  RT (min)      lcorbi_t3_pos_4_053_19    m/z 289.2486     0   1   5   5.5  RT (min)      lcorbi_t3_pos_2_021_09    m/z 289.2534     0   1E+5   8   8.5  RT (min)       lcorbi_t3_pos_4_014_48    m/z 289.2568     0   1E+5   10   10.5  RT (min)       lcorbi_t3_pos_2_044_21    m/z 289.9255     0   1   2   3E+4   1   1.5  RT (min)      lcorbi_t3_pos_1_035_28    m/z 289.9749     0   1   1   1.5  RT (min)      lcorbi_t3_pos_1_031_40    m/z 290.0013     0   2   4   6   8E+4   1   1.5  RT (min)      lcorbi_t3_pos_1_031_40    m/z 290.0361     0   1   2   3E+4   1   1.5  RT (min)      lcorbi_t3_pos_4_094_60    m/z 290.0638     0   1   1   1.5  RT (min)      lcorbi_t3_pos_4_093_60    m/z 290.0860     0   1E+4   4.5   5  RT (min)      lcorbi_t3_pos_4_074_58    m/z 290.0861     0   1   2.5   3  RT (min)      lcorbi_t3_pos_4_082_59    m/z 290.0928     0   1   2   3E+3   4.5   5  RT (min)      lcorbi_t3_pos_1_021_18    m/z 290.0928     0   1E+4   2.5   3  RT (min)      lcorbi_t3_pos_3_012_31    m/z 290.1159     0   1   2   3   4E+3   4.5   5  RT (min)       lcorbi_t3_pos_4_048_24    m/z 290.1235     0   1   2E+3   4.5   5  RT (min)      lcorbi_t3_pos_4_048_24    m/z 290.1347     0   2   4   6   8E+4   1   1.5   2  RT (min)       lcorbi_t3_pos_4_088_59    m/z 290.1459     0   1E+4   1   1.5  RT (min)       lcorbi_t3_pos_4_018_01    m/z 290.1599     0   1   2   3E+4   4.5   5  RT (min)      lcorbi_t3_pos_4_066_57    m/z 290.1599     0   1   1   2   3  RT (min)       lcorbi_t3_pos_4_090_60    m/z 290.2567     0   1E+4   9   9.5  RT (min)      lcorbi_t3_pos_4_111_62    m/z 290.2772     0   1   2E+5   11.5   12  RT (min)       lcorbi_t3_pos_1_032_26    m/z 290.2894     0   2   4E+4   9   9.5  RT (min)      lcorbi_t3_pos_4_107_62    m/z 290.7660     0   1   2E+4   1   1.5  RT (min)      lcorbi_t3_pos_1_029_29    m/z 290.7879     0   2   4   6   8E+3   1   1.5  RT (min)      lcorbi_t3_pos_1_029_29    m/z 290.8553     0   2   4   6E+4   1   1.5  RT (min)      lcorbi_t3_pos_1_029_29    m/z 290.8666     0   1E+4   1   1.5  RT (min)      lcorbi_t3_pos_1_016_05    m/z 290.8762     0   1   2   3   4E+4   1   1.5  RT (min)      lcorbi_t3_pos_1_036_08    m/z 290.8945     0   1   2   3   4E+3   0.5   1  RT (min)      lcorbi_t3_pos_1_005_42    m/z 290.9169     0   2   4   6E+3   1   1.5  RT (min)      lcorbi_t3_pos_1_034_34    m/z 290.9263     0   2   4   6   8E+3   1   1.5  RT (min)      lcorbi_t3_pos_1_008_36    m/z 290.9475     0   1   2E+5   0.5   1  RT (min)      lcorbi_t3_pos_1_006_15    m/z 291.0477     0   2   4E+4   1   1.5  RT (min)       lcorbi_t3_pos_3_012_31    m/z 291.0602     0   1   2   3   4E+4   18   18.5  RT (min)      lcorbi_t3_pos_4_066_57    m/z 291.0628     0   1   2   3   4E+4   9.5   10  RT (min)       lcorbi_t3_pos_2_039_30    m/z 291.0839     0   1   5   5.5  RT (min)       lcorbi_t3_pos_4_034_42    m/z 291.1065     0   1   2   3   4E+4   1.5   2  RT (min)      lcorbi_t3_pos_4_007_46    m/z 291.1187     0   2   4E+4   1   1.5  RT (min)       lcorbi_t3_pos_4_067_57    m/z 291.1204     0   2   4   6   8E+3   4.5   5  RT (min)      lcorbi_t3_pos_4_106_62    m/z 291.1300     0   1   2E+5   1   1.5  RT (min)       lcorbi_t3_pos_4_082_59    m/z 291.1381     0   1E+4   1.5   2  RT (min)      lcorbi_t3_pos_4_082_59    m/z 291.1421     0   1E+5   5   5.5  RT (min)      lcorbi_t3_pos_4_073_57    m/z 291.1550     0   2   4   6E+3   1   1.5  RT (min)       lcorbi_t3_pos_2_008_16    m/z 291.1737     0   1   2   3E+4   5   5.5  RT (min)       lcorbi_t3_pos_4_106_62    m/z 291.1955     0   2   4   6   8E+4   10   10.5  RT (min)       lcorbi_t3_pos_3_054_15    m/z 291.2108     0   2   4   6E+3   6   6.5  RT (min)      lcorbi_t3_pos_2_053_28    m/z 291.2319     0   1   7.5   8  RT (min)      lcorbi_t3_pos_4_081_58    m/z 291.2531     0   1E+5   10.5   11  RT (min)       lcorbi_t3_pos_4_084_59    m/z 291.2531     0   2   4   6   8E+4   8   8.5  RT (min)      lcorbi_t3_pos_2_044_21    m/z 291.2642     0   2   4   6E+3   5   5.5  RT (min)      lcorbi_t3_pos_4_106_62    m/z 291.9513     0   1E+4   1.5   2  RT (min)      lcorbi_t3_pos_1_029_29    m/z 291.9719     0   1   2   3E+4   1   1.5  RT (min)      lcorbi_t3_pos_1_036_08    m/z 292.0452     0   1   2   3E+4   8   8.5  RT (min)      lcorbi_t3_pos_4_109_62    m/z 292.0874     0   1   2E+3   5   5.5  RT (min)      lcorbi_t3_pos_4_070_57    m/z 292.0874     0   1E+3   1   1.5  RT (min)      lcorbi_t3_pos_3_027_27    m/z 292.1026     0   1   2   3   4E+4   1   1.5  RT (min)       lcorbi_t3_pos_4_087_59    m/z 292.1028     0   1E+4   4.5   5  RT (min)      lcorbi_t3_pos_4_082_59    m/z 292.1180     0   2   4   6E+4   7   7.5  RT (min)       lcorbi_t3_pos_1_004_24    m/z 292.1270     0   1   2.5   3  RT (min)      lcorbi_t3_pos_4_007_46    m/z 292.1391     0   2   4   6   8E+3   1.5   2  RT (min)      lcorbi_t3_pos_4_007_46    m/z 292.1453     0   1E+4   5   5.5  RT (min)      lcorbi_t3_pos_4_066_57    m/z 292.1519     0   1E+6   1   1.5  RT (min)       lcorbi_t3_pos_4_090_60    m/z 292.1695     0   1   1   1.5  RT (min)       lcorbi_t3_pos_2_037_27    m/z 292.1755     0   1E+4   7.5   8  RT (min)      lcorbi_t3_pos_4_027_07    m/z 292.2308     0   1E+4   4.5   5  RT (min)       lcorbi_t3_pos_4_089_59    m/z 292.2756     0   1E+5   9   9.5  RT (min)      lcorbi_t3_pos_4_113_62    m/z 292.7324     0   2   4   6E+3   4.5   5  RT (min)      lcorbi_t3_pos_4_089_59    m/z 292.7850     0   1   2   3E+4   1   1.5  RT (min)      lcorbi_t3_pos_4_073_57    m/z 292.8525     0   2   4E+4   1   1.5  RT (min)      lcorbi_t3_pos_1_029_29    m/z 292.9150     0   1E+4   1   1.5  RT (min)      lcorbi_t3_pos_4_006_12    m/z 292.9558     0   2   4   6   8E+4   1   1.5  RT (min)      lcorbi_t3_pos_1_036_08    m/z 292.9857     0   2   4   6E+4   0.5   1  RT (min)      lcorbi_t3_pos_1_006_15    m/z 293.0745     0   1   2   3E+3   4.5   5  RT (min)      lcorbi_t3_pos_1_055_33    m/z 293.0745     0   1   2.5   3  RT (min)      lcorbi_t3_pos_4_074_58    m/z 293.0808     0   1   2E+4   7.5   8  RT (min)      lcorbi_t3_pos_4_041_54    m/z 293.0840     0   1E+4   1   1.5  RT (min)      lcorbi_t3_pos_4_077_58    m/z 293.0980     0   2   4   6   8E+5   1   1.5   2  RT (min)       lcorbi_t3_pos_4_075_58    m/z 293.1150     0   1E+4   6.5   7  RT (min)      lcorbi_t3_pos_4_082_59    m/z 293.1237     0   1   1.5   2  RT (min)      lcorbi_t3_pos_4_006_12    m/z 293.1335     0   1   1   1.5  RT (min)      lcorbi_t3_pos_4_006_12    m/z 293.1384     0   2   4   6E+3   8   8.5  RT (min)      lcorbi_t3_pos_3_056_56    m/z 293.1566     0   2   4   6   8E+2   2.5   3  RT (min)      lcorbi_t3_pos_4_090_60    m/z 293.1810     0   1   2E+4   4.5   5  RT (min)      lcorbi_t3_pos_4_090_60    m/z 293.1901     0   2   4   6   8E+4   14   14.5  RT (min)       lcorbi_t3_pos_2_027_05    m/z 293.1910     0   1   2   3E+3   4.5   5  RT (min)      lcorbi_t3_pos_1_002_01    m/z 293.2476     0   2   4   6   8E+4   12.5   13  RT (min)       lcorbi_t3_pos_3_042_14    m/z 293.2952     0   1   2   3E+4   9   9.5  RT (min)      lcorbi_t3_pos_4_027_07    m/z 293.9891     0   2   4   6   8E+4   18   18.5  RT (min)      lcorbi_t3_pos_4_084_59    m/z 294.0070     0   1   2E+4   1.5   2  RT (min)      lcorbi_t3_pos_4_096_60    m/z 294.0165     0   1   2   3E+5   1   1.5  RT (min)       lcorbi_t3_pos_4_097_60    m/z 294.1061     0   1E+4   4.5   5  RT (min)      lcorbi_t3_pos_4_090_60    m/z 294.1062     0   1   2E+3   2.5   3  RT (min)      lcorbi_t3_pos_4_076_58    m/z 294.1184     0   1E+4   4.5   5  RT (min)      lcorbi_t3_pos_4_090_60    m/z 294.1296     0   2   4   6E+4   1   1.5  RT (min)       lcorbi_t3_pos_4_067_57    m/z 294.1336     0   1   2   3E+4   5   5.5  RT (min)       lcorbi_t3_pos_4_080_58    m/z 294.1489     0   1E+5   0   0.5  RT (min)      lcorbi_t3_pos_1_005_42    m/z 294.1489     0   1   2E+4   7   7.5  RT (min)      lcorbi_t3_pos_4_100_61    m/z 294.1548     0   1E+3   2.5   3  RT (min)      lcorbi_t3_pos_4_092_60    m/z 294.1569     0   1E+4   4.5   5  RT (min)      lcorbi_t3_pos_4_052_06    m/z 294.1700     0   1   4.5   5  RT (min)      lcorbi_t3_pos_1_006_15    m/z 294.1707     0   1   2   3   4E+4   1   1.5  RT (min)      lcorbi_t3_pos_4_090_60    m/z 294.1782     0   1   2   3   4E+4   9.5   10  RT (min)      lcorbi_t3_pos_4_066_57    m/z 294.1912     0   1   2.5   3  RT (min)       lcorbi_t3_pos_4_090_60    m/z 294.2041     0   1E+4   9   9.5  RT (min)      lcorbi_t3_pos_1_032_26    m/z 294.2152     0   2   4   6E+3   6   6.5  RT (min)      lcorbi_t3_pos_4_074_58    m/z 294.2276     0   1E+5   8.5   9  RT (min)       lcorbi_t3_pos_1_032_26    m/z 294.2405     0   2   4   6E+4   11   11.5  RT (min)       lcorbi_t3_pos_4_110_62    m/z 294.2541     0   1   2   3   4E+3   7.5   8  RT (min)      lcorbi_t3_pos_4_006_12    m/z 294.2640     0   1   2   3   4E+4   10   10.5  RT (min)      lcorbi_t3_pos_4_085_59    m/z 294.7821     0   1   2E+4   1   1.5  RT (min)      lcorbi_t3_pos_4_067_57    m/z 294.8671     0   1E+4   1   1.5  RT (min)      lcorbi_t3_pos_3_010_44    m/z 294.8785     0   2   4E+4   11   11.5  RT (min)      lcorbi_t3_pos_1_030_21    m/z 294.9656     0   2   4E+3   1   1.5  RT (min)      lcorbi_t3_pos_2_013_33    m/z 294.9869     0   1   2E+4   1   1.5  RT (min)      lcorbi_t3_pos_2_027_05    m/z 295.0169     0   1E+5   1   1.5  RT (min)      lcorbi_t3_pos_3_048_30    m/z 295.0540     0   1   4.5   5  RT (min)      lcorbi_t3_pos_4_013_36    m/z 295.0846     0   2   4   6   8E+4   1   1.5  RT (min)      lcorbi_t3_pos_4_055_02    m/z 295.0860     0   1   2E+3   4.5   5  RT (min)      lcorbi_t3_pos_4_090_60    m/z 295.1136     0   1E+5   1   1.5  RT (min)       lcorbi_t3_pos_4_087_59    m/z 295.1329     0   1   2   3   4E+4   8   8.5  RT (min)       lcorbi_t3_pos_4_098_61    m/z 295.1514     0   2   4   6E+3   1   1.5  RT (min)       lcorbi_t3_pos_4_097_60    m/z 295.1672     0   1   2   3E+4   1   1.5  RT (min)      lcorbi_t3_pos_1_019_16    m/z 295.1728     0   2   4   6E+4   13   13.5  RT (min)       lcorbi_t3_pos_1_036_08    m/z 295.1728     0   1   2E+4   9.5   10  RT (min)      lcorbi_t3_pos_4_074_58    m/z 295.1947     0   1   2.5   3  RT (min)      lcorbi_t3_pos_4_048_24    m/z 295.2131     0   1E+4   2.5   3  RT (min)      lcorbi_t3_pos_4_048_24    m/z 295.2268     0   2   4   6E+4   9   9.5  RT (min)       lcorbi_t3_pos_4_074_58    m/z 295.2269     0   1E+5   15.5   16  RT (min)       lcorbi_t3_pos_4_081_58    m/z 295.9424     0   2   4   6E+4   1   1.5  RT (min)      lcorbi_t3_pos_1_020_31    m/z 295.9587     0   1   2   3   4E+3   5   5.5  RT (min)      lcorbi_t3_pos_4_066_57    m/z 295.9640     0   1E+4   1   1.5  RT (min)      lcorbi_t3_pos_1_036_08    m/z 295.9792     0   1   2E+4   1   1.5  RT (min)      lcorbi_t3_pos_2_024_15    m/z 296.0467     0   2   4E+4   1   1.5  RT (min)       lcorbi_t3_pos_4_075_58    m/z 296.1217     0   2   4   6   8E+4   7   7.5  RT (min)       lcorbi_t3_pos_4_072_57    m/z 296.1233     0   1   2E+5   1.5   2  RT (min)       lcorbi_t3_pos_4_091_60    m/z 296.1324     0   1   2   3E+3   1.5   2  RT (min)       lcorbi_t3_pos_1_052_38    m/z 296.1361     0   2   4   6   8E+3   8   8.5  RT (min)      lcorbi_t3_pos_4_103_61    m/z 296.1493     0   1   2   3   4E+4   5.5   6  RT (min)       lcorbi_t3_pos_4_079_58    m/z 296.1523     0   1E+5   1   1.5  RT (min)       lcorbi_t3_pos_1_036_08    m/z 296.1606     0   1   2   3   4E+4   4.5   5  RT (min)       lcorbi_t3_pos_4_082_59    m/z 296.1705     0   1   2E+4   1   1.5  RT (min)      lcorbi_t3_pos_2_046_07    m/z 296.1717     0   1   2E+3   4.5   5  RT (min)      lcorbi_t3_pos_4_074_58    m/z 296.1857     0   2   4E+4   6.5   7  RT (min)       lcorbi_t3_pos_4_099_61    m/z 296.1938     0   1E+4   9.5   10  RT (min)      lcorbi_t3_pos_4_077_58    m/z 296.1988     0   1   2   3   4E+4   10.5   11  RT (min)       lcorbi_t3_pos_4_113_62    m/z 296.2068     0   1   2E+4   5.5   6  RT (min)      lcorbi_t3_pos_4_010_43    m/z 296.2144     0   1E+4   4.5   5  RT (min)      lcorbi_t3_pos_4_039_35    m/z 296.2221     0   1   2   3   4E+4   10.5   11  RT (min)       lcorbi_t3_pos_4_090_60    m/z 296.2949     0   2   4   6   8E+4   11   11.5  RT (min)       lcorbi_t3_pos_4_088_59    m/z 296.2950     0   2   4E+4   13   13.5  RT (min)       lcorbi_t3_pos_2_002_06    m/z 296.7162     0   2   4E+3   4.5   5  RT (min)      lcorbi_t3_pos_4_039_35    m/z 296.8121     0   1E+4   1   1.5  RT (min)      lcorbi_t3_pos_2_023_26    m/z 296.8213     0   2   4E+4   1   1.5  RT (min)      lcorbi_t3_pos_1_036_08    m/z 296.9432     0   1   2E+4   1   1.5  RT (min)      lcorbi_t3_pos_1_007_07    m/z 296.9754     0   1   2   3   4E+4   1   1.5  RT (min)      lcorbi_t3_pos_4_079_58    m/z 296.9786     0   2   4   6E+4   18   18.5  RT (min)       lcorbi_t3_pos_4_085_59    m/z 297.0299     0   1   2   3   4E+3   0.5   1  RT (min)       lcorbi_t3_pos_3_013_19    m/z 297.0694     0   1   2E+5   1   1.5  RT (min)      lcorbi_t3_pos_4_006_12    m/z 297.0826     0   1   2E+5   10.5   11  RT (min)       lcorbi_t3_pos_4_113_62    m/z 297.0881     0   1E+3   4.5   5  RT (min)      lcorbi_t3_pos_3_029_13    m/z 297.1163     0   1   2   3E+4   2.5   3  RT (min)       lcorbi_t3_pos_4_091_60    m/z 297.1250     0   1E+4   7   7.5  RT (min)      lcorbi_t3_pos_4_073_57    m/z 297.1320     0   2   4E+4   5   5.5  RT (min)       lcorbi_t3_pos_4_106_62    m/z 297.1388     0   1E+4   4.5   5  RT (min)      lcorbi_t3_pos_4_066_57    m/z 297.1486     0   1   2   3E+4   8   8.5  RT (min)       lcorbi_t3_pos_4_099_61    m/z 297.1527     0   1   2E+3   5   5.5  RT (min)      lcorbi_t3_pos_1_002_01    m/z 297.1850     0   2   4E+4   9   9.5  RT (min)       lcorbi_t3_pos_4_102_61    m/z 297.1908     0   1   2E+3   4.5   5  RT (min)      lcorbi_t3_pos_1_001_49    m/z 297.2650     0   1   4.5   5  RT (min)      lcorbi_t3_pos_1_001_49    m/z 297.2901     0   2   4   6   8E+4   12.5   13  RT (min)       lcorbi_t3_pos_4_072_57    m/z 297.2923     0   1E+5   10.5   11  RT (min)       lcorbi_t3_pos_4_092_60    m/z 297.9677     0   1E+5   1   1.5  RT (min)      lcorbi_t3_pos_1_034_34    m/z 298.0031     0   1   2E+4   4.5   5  RT (min)      lcorbi_t3_pos_4_014_48    m/z 298.0436     0   2   4   6E+3   1   1.5  RT (min)      lcorbi_t3_pos_1_028_32    m/z 298.0642     0   1E+5   1   1.5  RT (min)      lcorbi_t3_pos_4_006_12    m/z 298.0898     0   2   4   6E+4   4.5   5  RT (min)       lcorbi_t3_pos_4_090_60    m/z 298.0898     0   1   2   3E+4   2.5   3  RT (min)      lcorbi_t3_pos_4_032_40    m/z 298.1147     0   1E+4   2.5   3  RT (min)      lcorbi_t3_pos_4_032_40    m/z 298.1205     0   1E+4   7.5   8  RT (min)      lcorbi_t3_pos_4_090_60    m/z 298.1285     0   2   4E+3   5.5   6  RT (min)      lcorbi_t3_pos_4_111_62    m/z 298.1398     0   1   2   3E+3   4.5   5  RT (min)      lcorbi_t3_pos_1_032_26    m/z 298.1496     0   1   2E+3   1.5   2  RT (min)      lcorbi_t3_pos_4_052_06    m/z 298.1587     0   1E+4   7   7.5  RT (min)      lcorbi_t3_pos_4_053_19    m/z 298.1860     0   1   2   3   4E+3   5   5.5  RT (min)      lcorbi_t3_pos_4_026_20    m/z 298.2225     0   1   2   3E+3   5.5   6  RT (min)      lcorbi_t3_pos_4_038_38    m/z 298.2435     0   1E+5   11.5   12  RT (min)      lcorbi_t3_pos_4_084_59    m/z 298.3469     0   2   4   6E+4   9.5   10  RT (min)      lcorbi_t3_pos_2_055_46    m/z 298.8184     0   1   2   3   4E+4   1   1.5  RT (min)      lcorbi_t3_pos_2_024_15    m/z 298.8775     0   1E+4   0.5   1  RT (min)      lcorbi_t3_pos_1_006_15    m/z 298.9688     0   2   4E+3   1.5   2  RT (min)      lcorbi_t3_pos_3_008_20    m/z 299.0618     0   1   2   3E+5   10.5   11  RT (min)       lcorbi_t3_pos_4_113_62    m/z 299.1214     0   2   4   6E+3   1   1.5  RT (min)       lcorbi_t3_pos_4_095_60    m/z 299.1391     0   1   2   3   4E+4   12.5   13  RT (min)       lcorbi_t3_pos_2_053_28    m/z 299.1619     0   1   2   3   4E+4   9.5   10  RT (min)      lcorbi_t3_pos_1_038_30    m/z 299.1829     0   1   2   3   4E+4   8.5   9  RT (min)       lcorbi_t3_pos_4_089_59    m/z 299.1916     0   1   2   3E+4   7   7.5  RT (min)      lcorbi_t3_pos_4_101_61    m/z 299.2006     0   2   4   6E+3   8   8.5  RT (min)       lcorbi_t3_pos_1_053_39    m/z 299.2205     0   1E+4   5   5.5  RT (min)      lcorbi_t3_pos_4_087_59    m/z 299.2404     0   2   4   6   8E+4   12   12.5  RT (min)       lcorbi_t3_pos_4_082_59    m/z 299.3058     0   1   10.5   11  RT (min)      lcorbi_t3_pos_1_002_01    m/z 299.3139     0   1E+5   13   13.5  RT (min)      lcorbi_t3_pos_2_044_21    m/z 299.7124     0   1E+4   5.5   6  RT (min)      lcorbi_t3_pos_4_018_01    m/z 300.0037     0   1   2   3   4E+4   1.5   2  RT (min)      lcorbi_t3_pos_1_036_08    m/z 300.0209     0   1   2E+4   1   1.5  RT (min)      lcorbi_t3_pos_1_041_54    m/z 300.0334     0   1   2   3E+5   1   1.5  RT (min)      lcorbi_t3_pos_1_036_08    m/z 300.0723     0   2   4   6E+3   8   8.5  RT (min)      lcorbi_t3_pos_3_045_43    m/z 300.0844     0   2   4   6E+3   2.5   3  RT (min)      lcorbi_t3_pos_4_089_59    m/z 300.0865     0   1E+4   4.5   5  RT (min)      lcorbi_t3_pos_4_098_61    m/z 300.0927     0   1E+5   4.5   5  RT (min)      lcorbi_t3_pos_3_048_30    m/z 300.0990     0   2   4   6   8E+4   1.5   2  RT (min)       lcorbi_t3_pos_4_093_60    m/z 300.1021     0   1E+4   4.5   5  RT (min)      lcorbi_t3_pos_3_048_30    m/z 300.1231     0   2   4   6   8E+4   9   9.5  RT (min)       lcorbi_t3_pos_4_107_62    m/z 300.1555     0   1   1.5   2  RT (min)      lcorbi_t3_pos_3_055_10    m/z 300.1683     0   1E+4   1.5   2  RT (min)      lcorbi_t3_pos_3_055_10    m/z 300.1796     0   1   2E+4   1   1.5  RT (min)      lcorbi_t3_pos_4_094_60    m/z 300.1877     0   1E+5   1   1.5  RT (min)       lcorbi_t3_pos_4_095_60    m/z 300.2017     0   1E+5   4.5   5  RT (min)       lcorbi_t3_pos_2_044_21    m/z 300.2282     0   1E+4   7   7.5  RT (min)      lcorbi_t3_pos_4_090_60    m/z 300.2434     0   1   2E+4   1.5   2  RT (min)      lcorbi_t3_pos_4_091_60    m/z 300.2596     0   2   4   6E+4   1   1.5  RT (min)       lcorbi_t3_pos_4_093_60    m/z 300.2645     0   1   2   3   4E+3   4.5   5  RT (min)      lcorbi_t3_pos_1_001_49    m/z 300.3396     0   1   10   10.5  RT (min)       lcorbi_t3_pos_1_008_36    m/z 300.7005     0   2   4   6E+3   4.5   5  RT (min)      lcorbi_t3_pos_4_052_06    m/z 300.8628     0   1   1   1.5  RT (min)      lcorbi_t3_pos_1_014_03    m/z 300.8841     0   1E+5   1   1.5  RT (min)       lcorbi_t3_pos_1_029_29    m/z 300.8954     0   2   4   6E+4   1   1.5  RT (min)       lcorbi_t3_pos_4_067_57    m/z 300.9049     0   1E+5   1   1.5  RT (min)      lcorbi_t3_pos_2_024_15    m/z 300.9158     0   1   2   3   4E+3   5   5.5  RT (min)      lcorbi_t3_pos_3_019_17    m/z 300.9923     0   1   2   3E+4   11   11.5  RT (min)      lcorbi_t3_pos_4_098_61    m/z 301.0932     0   1E+4   5.5   6  RT (min)      lcorbi_t3_pos_4_104_61    m/z 301.1166     0   2   4E+4   18   18.5  RT (min)       lcorbi_t3_pos_4_085_59    m/z 301.1193     0   1   2E+4   4.5   5  RT (min)      lcorbi_t3_pos_4_074_58    m/z 301.1265     0   1   9   9.5  RT (min)      lcorbi_t3_pos_2_040_25    m/z 301.1283     0   1E+4   4.5   5  RT (min)      lcorbi_t3_pos_4_098_61    m/z 301.1635     0   2   4   6   8E+4   1   1.5  RT (min)       lcorbi_t3_pos_4_090_60    m/z 301.1717     0   1E+5   1   1.5   2  RT (min)       lcorbi_t3_pos_4_095_60    m/z 301.1872     0   2   4   6   8E+4   1   1.5  RT (min)      lcorbi_t3_pos_2_046_07    m/z 301.2051     0   1   2E+4   4.5   5  RT (min)      lcorbi_t3_pos_2_044_21    m/z 301.2163     0   1E+5   15   15.5  RT (min)       lcorbi_t3_pos_3_008_20    m/z 301.2534     0   1   2E+4   7   7.5  RT (min)      lcorbi_t3_pos_4_053_19    m/z 301.2850     0   1   2E+4   9   9.5  RT (min)       lcorbi_t3_pos_4_089_59    m/z 301.2932     0   1   2   3   4E+4   12   12.5  RT (min)      lcorbi_t3_pos_4_018_01    m/z 301.3519     0   2   4   6E+4   10.5   11  RT (min)       lcorbi_t3_pos_4_093_60    m/z 301.9810     0   1   2E+5   1   1.5  RT (min)      lcorbi_t3_pos_1_036_08    m/z 302.0007     0   1E+3   2.5   3  RT (min)      lcorbi_t3_pos_1_005_42    m/z 302.0196     0   1E+5   1   1.5  RT (min)       lcorbi_t3_pos_4_006_12    m/z 302.0399     0   1E+3   2.5   3  RT (min)      lcorbi_t3_pos_4_074_58    m/z 302.0999     0   1E+3   4.5   5  RT (min)      lcorbi_t3_pos_4_048_24    m/z 302.1113     0   1E+4   4.5   5  RT (min)      lcorbi_t3_pos_4_048_24    m/z 302.1289     0   2   4   6   8E+4   9   9.5  RT (min)       lcorbi_t3_pos_4_098_61    m/z 302.1556     0   2   4   6   8E+4   1   1.5  RT (min)      lcorbi_t3_pos_2_037_27    m/z 302.1599     0   1   2   3   4E+3   4.5   5  RT (min)      lcorbi_t3_pos_4_074_58    m/z 302.1600     0   1   2E+4   2.5   3  RT (min)      lcorbi_t3_pos_4_001_49    m/z 302.1963     0   2   4   6   8E+4   4.5   5  RT (min)       lcorbi_t3_pos_4_074_58    m/z 302.1963     0   1E+4   2.5   3  RT (min)      lcorbi_t3_pos_4_039_35    m/z 302.2326     0   1E+4   7   7.5  RT (min)      lcorbi_t3_pos_4_076_58    m/z 302.2498     0   1E+5   7   7.5  RT (min)       lcorbi_t3_pos_4_053_19    m/z 302.2520     0   1E+4   9.5   10  RT (min)      lcorbi_t3_pos_1_046_22    m/z 302.2802     0   2   4   6E+3   5.5   6  RT (min)      lcorbi_t3_pos_4_088_59    m/z 302.6910     0   1   2E+4   5.5   6  RT (min)      lcorbi_t3_pos_4_034_42    m/z 302.8293     0   2   4   6E+4   1   1.5  RT (min)      lcorbi_t3_pos_2_020_22    m/z 302.8813     0   1   2   3   4E+4   1   1.5  RT (min)      lcorbi_t3_pos_1_029_29    m/z 302.9361     0   2   4E+3   1   1.5  RT (min)      lcorbi_t3_pos_1_035_28    m/z 303.0063     0   1E+4   7.5   8  RT (min)      lcorbi_t3_pos_4_090_60    m/z 303.0273     0   1   2   3E+3   1.5   2  RT (min)      lcorbi_t3_pos_4_075_58    m/z 303.0444     0   1   2   3E+4   1   1.5  RT (min)       lcorbi_t3_pos_4_091_60    m/z 303.0776     0   1   2   3E+5   18.5   19  RT (min)       lcorbi_t3_pos_4_089_59    m/z 303.0776     0   2   4E+3   1   1.5  RT (min)       lcorbi_t3_pos_4_097_60    m/z 303.0862     0   2   4   6   8E+2   6   6.5  RT (min)      lcorbi_t3_pos_4_039_35    m/z 303.1017     0   1   2E+4   4.5   5  RT (min)       lcorbi_t3_pos_4_066_57    m/z 303.1052     0   1   2   3E+4   2.5   3  RT (min)      lcorbi_t3_pos_4_051_34    m/z 303.1322     0   2   4E+4   5.5   6  RT (min)      lcorbi_t3_pos_1_051_09    m/z 303.1322     0   1   9   9.5  RT (min)      lcorbi_t3_pos_1_014_03    m/z 303.1374     0   1   2   3   4E+3   5   5.5  RT (min)      lcorbi_t3_pos_1_030_21    m/z 303.1380     0   1E+4   7   7.5  RT (min)      lcorbi_t3_pos_1_012_14    m/z 303.1453     0   1E+3   2.5   3  RT (min)      lcorbi_t3_pos_4_051_34    m/z 303.1477     0   1E+5   10   10.5  RT (min)      lcorbi_t3_pos_4_077_58    m/z 303.1551     0   2   4   6E+3   4.5   5  RT (min)      lcorbi_t3_pos_4_075_58    m/z 303.1552     0   1E+4   2.5   3  RT (min)      lcorbi_t3_pos_4_004_14    m/z 303.1779     0   2   4E+3   5.5   6  RT (min)      lcorbi_t3_pos_4_106_62    m/z 303.1955     0   2   4   6   8E+4   7   7.5  RT (min)       lcorbi_t3_pos_4_005_33    m/z 303.1996     0   1E+4   4.5   5  RT (min)      lcorbi_t3_pos_4_074_58    m/z 303.2167     0   1   2E+4   9.5   10  RT (min)      lcorbi_t3_pos_4_087_59    m/z 303.2319     0   1   2   3   4E+4   7.5   8  RT (min)       lcorbi_t3_pos_4_100_61    m/z 303.2531     0   1   2E+4   7   7.5  RT (min)      lcorbi_t3_pos_4_053_19    m/z 303.2690     0   2   4E+4   6.5   7  RT (min)      lcorbi_t3_pos_2_044_21    m/z 303.2724     0   1   2   3   4E+4   10.5   11  RT (min)      lcorbi_t3_pos_4_112_62    m/z 304.0790     0   1   2E+3   1   1.5  RT (min)      lcorbi_t3_pos_4_097_60    m/z 304.0929     0   2   4   6   8E+5   1   1.5  RT (min)       lcorbi_t3_pos_4_007_46    m/z 304.1504     0   1   2.5   3  RT (min)      lcorbi_t3_pos_1_002_01    m/z 304.1504     0   1   1   1.5  RT (min)      lcorbi_t3_pos_1_002_01    m/z 304.1544     0   1   2   3E+3   5.5   6  RT (min)       lcorbi_t3_pos_1_001_49    m/z 304.1616     0   2   4   6   8E+4   1   1.5   2  RT (min)       lcorbi_t3_pos_4_089_59    m/z 304.1755     0   2   4   6   8E+3   8   8.5  RT (min)      lcorbi_t3_pos_1_002_01    m/z 304.1762     0   1   2   3   4E+3   1.5   2  RT (min)      lcorbi_t3_pos_1_001_49    m/z 304.1765     0   2   4E+3   4.5   5  RT (min)      lcorbi_t3_pos_1_001_49    m/z 304.1989     0   1   7   7.5  RT (min)      lcorbi_t3_pos_1_001_49    m/z 304.2119     0   1   2E+3   6   6.5  RT (min)       lcorbi_t3_pos_1_003_45    m/z 304.2353     0   1   2   3E+3   7.5   8  RT (min)      lcorbi_t3_pos_1_001_49    m/z 304.2353     0   1   2   3E+4   15   15.5  RT (min)      lcorbi_t3_pos_1_001_49    m/z 304.2613     0   1   2   3   4E+4   12   12.5  RT (min)       lcorbi_t3_pos_4_088_59    m/z 304.2654     0   1E+5   6.5   7  RT (min)       lcorbi_t3_pos_2_044_21    m/z 304.3082     0   2   4E+4   13   13.5  RT (min)      lcorbi_t3_pos_2_002_06    m/z 304.3709     0   1E+5   10.5   11  RT (min)       lcorbi_t3_pos_1_036_08    m/z 304.8263     0   2   4E+4   1   1.5  RT (min)      lcorbi_t3_pos_1_032_26    m/z 304.8960     0   2   4   6E+4   18   18.5  RT (min)      lcorbi_t3_pos_4_103_61    m/z 305.0009     0   2   4   6E+4   1   1.5  RT (min)      lcorbi_t3_pos_1_028_32    m/z 305.0634     0   2   4E+4   4.5   5  RT (min)       lcorbi_t3_pos_4_066_57    m/z 305.0634     0   1   2E+4   2.5   3  RT (min)      lcorbi_t3_pos_4_042_09    m/z 305.0751     0   1   2E+3   1   1.5  RT (min)      lcorbi_t3_pos_4_095_60    m/z 305.0857     0   1E+3   2.5   3  RT (min)      lcorbi_t3_pos_3_016_21    m/z 305.0996     0   1E+3   6   6.5  RT (min)      lcorbi_t3_pos_3_034_26    m/z 305.1068     0   2   4   6E+4   1   1.5  RT (min)      lcorbi_t3_pos_2_020_22    m/z 305.1379     0   2   4E+3   4.5   5  RT (min)      lcorbi_t3_pos_4_066_57    m/z 305.1419     0   2   4   6E+3   9   9.5  RT (min)      lcorbi_t3_pos_4_099_61    m/z 305.1479     0   2   4   6E+4   6   6.5  RT (min)      lcorbi_t3_pos_3_034_26    m/z 305.1571     0   1   2   3   4E+4   4.5   5  RT (min)       lcorbi_t3_pos_4_113_62    m/z 305.1595     0   1E+4   8.5   9  RT (min)      lcorbi_t3_pos_4_090_60    m/z 305.1894     0   2   4E+3   5   5.5  RT (min)      lcorbi_t3_pos_4_066_57    m/z 305.2111     0   2   4   6E+4   10.5   11  RT (min)       lcorbi_t3_pos_1_037_19    m/z 305.2153     0   2   4E+3   6   6.5  RT (min)      lcorbi_t3_pos_3_042_14    m/z 305.2291     0   1   2   3E+4   6.5   7  RT (min)      lcorbi_t3_pos_4_012_37    m/z 305.2452     0   1E+3   1   1.5  RT (min)      lcorbi_t3_pos_4_093_60    m/z 305.2517     0   1E+4   6.5   7  RT (min)      lcorbi_t3_pos_1_001_49    m/z 305.2517     0   1   2E+4   11   12  RT (min)       lcorbi_t3_pos_1_029_29    m/z 305.2687     0   2   4   6   8E+3   11   11.5  RT (min)       lcorbi_t3_pos_2_044_21    m/z 305.2880     0   1E+5   9.5   10  RT (min)       lcorbi_t3_pos_3_018_34    m/z 305.8993     0   2   4   6E+3   1   1.5  RT (min)      lcorbi_t3_pos_4_006_12    m/z 305.9662     0   2   4   6E+4   1   1.5  RT (min)      lcorbi_t3_pos_4_084_59    m/z 306.0239     0   2   4E+3   2.5   3  RT (min)      lcorbi_t3_pos_4_068_57    m/z 306.0496     0   1E+3   1   1.5  RT (min)      lcorbi_t3_pos_3_003_33    m/z 306.0600     0   1   2   3   4E+3   4.5   5  RT (min)      lcorbi_t3_pos_1_021_18    m/z 306.0600     0   1   2.5   3  RT (min)      lcorbi_t3_pos_1_012_14    m/z 306.0809     0   1   2   3E+3   4.5   5  RT (min)      lcorbi_t3_pos_4_048_24    m/z 306.1336     0   2   4   6   8E+3   4.5   5  RT (min)      lcorbi_t3_pos_4_072_57    m/z 306.1511     0   1   2   3E+3   5.5   6  RT (min)      lcorbi_t3_pos_2_001_49    m/z 306.1511     0   2   4   6   8E+4   9   9.5  RT (min)      lcorbi_t3_pos_4_099_61    m/z 306.2254     0   2   4   6   8E+4   6.5   7  RT (min)       lcorbi_t3_pos_2_044_21    m/z 306.2639     0   1   2   3E+4   8   8.5  RT (min)       lcorbi_t3_pos_4_112_62    m/z 306.2768     0   1   2E+4   10.5   11  RT (min)      lcorbi_t3_pos_3_020_35    m/z 306.2769     0   2   4   6E+4   12.5   13  RT (min)       lcorbi_t3_pos_4_087_59    m/z 306.2904     0   2   4   6E+3   8   8.5  RT (min)      lcorbi_t3_pos_4_113_62    m/z 306.3067     0   1   2   3E+4   9   9.5  RT (min)      lcorbi_t3_pos_1_017_51    m/z 306.7655     0   1   2   3   4E+4   1   1.5  RT (min)      lcorbi_t3_pos_4_068_57    m/z 306.8518     0   2   4   6   8E+3   1   1.5  RT (min)      lcorbi_t3_pos_1_001_49    m/z 306.8614     0   1E+4   1   1.5  RT (min)      lcorbi_t3_pos_2_023_26    m/z 306.8721     0   2   4   6   8E+3   0.5   1  RT (min)      lcorbi_t3_pos_2_010_08    m/z 306.8943     0   1E+4   1   1.5  RT (min)      lcorbi_t3_pos_1_001_49    m/z 306.9199     0   1   2E+6   0   0.5  RT (min)      lcorbi_t3_pos_1_006_15    m/z 306.9400     0   1   2   3E+4   1.5   2  RT (min)      lcorbi_t3_pos_1_052_38    m/z 306.9577     0   1   2E+4   1   1.5  RT (min)      lcorbi_t3_pos_2_020_22    m/z 307.0489     0   1   2   3E+3   6   6.5  RT (min)      lcorbi_t3_pos_1_001_49    m/z 307.0579     0   1E+4   5   5.5  RT (min)      lcorbi_t3_pos_4_079_58    m/z 307.0691     0   1   2.5   3  RT (min)       lcorbi_t3_pos_4_082_59    m/z 307.0834     0   1   2   2.5  RT (min)       lcorbi_t3_pos_4_082_59    m/z 307.0941     0   1E+4   10   10.5  RT (min)      lcorbi_t3_pos_1_002_01    m/z 307.1001     0   1   1   1.5  RT (min)      lcorbi_t3_pos_4_077_58    m/z 307.1137     0   1E+5   1.5   2  RT (min)       lcorbi_t3_pos_4_080_58    m/z 307.1153     0   2   4   6E+3   5.5   6   6.5  RT (min)      lcorbi_t3_pos_4_094_60    m/z 307.1687     0   1   2   3   4E+4   4.5   5  RT (min)      lcorbi_t3_pos_4_106_62    m/z 307.1755     0   1   2E+4   9   9.5  RT (min)      lcorbi_t3_pos_4_098_61    m/z 307.1767     0   2   4   6E+4   2.5   3  RT (min)      lcorbi_t3_pos_4_001_49    m/z 307.2421     0   1   2   3   4E+4   14   14.5  RT (min)       lcorbi_t3_pos_2_027_05    m/z 307.3038     0   1E+5   11   11.5   12  RT (min)       lcorbi_t3_pos_3_020_35    m/z 307.5850     0   1   2   2.5  RT (min)      lcorbi_t3_pos_4_082_59    m/z 307.9976     0   1   2   3E+4   1   1.5  RT (min)      lcorbi_t3_pos_4_075_58    m/z 308.0088     0   1   2   3   4E+3   2.5   3  RT (min)      lcorbi_t3_pos_1_002_01    m/z 308.0208     0   1   2.5   3  RT (min)      lcorbi_t3_pos_1_001_49    m/z 308.1219     0   1E+5   2.5   3  RT (min)      lcorbi_t3_pos_4_051_34    m/z 308.1258     0   1   2E+4   9   9.5  RT (min)      lcorbi_t3_pos_4_080_58    m/z 308.1305     0   1   2E+5   4.5   5  RT (min)       lcorbi_t3_pos_1_036_08    m/z 308.1331     0   1   1   1.5  RT (min)      lcorbi_t3_pos_2_023_26    m/z 308.1645     0   1E+4   7   7.5  RT (min)      lcorbi_t3_pos_3_023_05    m/z 308.2068     0   2   4   6   8E+3   4.5   5  RT (min)      lcorbi_t3_pos_4_075_58    m/z 308.2117     0   1E+4   9.5   10  RT (min)      lcorbi_t3_pos_4_100_61    m/z 308.2325     0   1   9.5   10  RT (min)      lcorbi_t3_pos_4_101_61    m/z 308.2796     0   1E+5   10.5   11  RT (min)       lcorbi_t3_pos_4_084_59    m/z 308.7625     0   2   4   6E+4   1   1.5  RT (min)       lcorbi_t3_pos_4_073_57    m/z 308.9565     0   2   4   6E+3   0.5   1  RT (min)      lcorbi_t3_pos_1_006_15    m/z 309.0263     0   2   4E+4   18   18.5  RT (min)      lcorbi_t3_pos_4_071_57    m/z 309.0644     0   1   1   1.5  RT (min)       lcorbi_t3_pos_1_032_26    m/z 309.0945     0   2   4   6E+4   1.5   2  RT (min)      lcorbi_t3_pos_1_031_40    m/z 309.1146     0   1   2.5   3  RT (min)      lcorbi_t3_pos_4_090_60    m/z 309.1171     0   2   4   6   8E+4   1   1.5  RT (min)      lcorbi_t3_pos_1_050_13    m/z 309.1293     0   1E+5   1   1.5  RT (min)       lcorbi_t3_pos_4_084_59    m/z 309.1445     0   2   4   6   8E+2   5.5   6  RT (min)      lcorbi_t3_pos_4_052_06    m/z 309.1486     0   1   2E+4   9   9.5  RT (min)      lcorbi_t3_pos_4_106_62    m/z 309.1516     0   1E+4   2.5   3  RT (min)      lcorbi_t3_pos_1_036_08    m/z 309.1598     0   1   2   3E+4   4.5   5  RT (min)       lcorbi_t3_pos_4_113_62    m/z 309.1770     0   2   4   6   8E+4   0   1  RT (min)       lcorbi_t3_pos_3_027_27    m/z 309.2037     0   1E+3   1   1.5  RT (min)      lcorbi_t3_pos_1_050_13    m/z 309.2039     0   1   2   3E+4   11   11.5  RT (min)      lcorbi_t3_pos_4_098_61    m/z 309.2042     0   1   2   3E+4   4.5   5  RT (min)       lcorbi_t3_pos_4_090_60    m/z 309.2149     0   1E+4   7.5   8  RT (min)      lcorbi_t3_pos_2_013_33    m/z 309.2271     0   1E+4   6.5   7  RT (min)      lcorbi_t3_pos_4_089_59    m/z 309.9889     0   1E+5   1   1.5  RT (min)      lcorbi_t3_pos_1_003_45    m/z 310.0869     0   1   1.5   2  RT (min)      lcorbi_t3_pos_2_037_27    m/z 310.1134     0   2   4E+4   1   1.5  RT (min)       lcorbi_t3_pos_4_086_59    m/z 310.1310     0   1   2   3   4E+4   4.5   5  RT (min)       lcorbi_t3_pos_2_024_15    m/z 310.1497     0   1   1   1.5  RT (min)       lcorbi_t3_pos_4_090_60    m/z 310.1517     0   1   5   5.5  RT (min)      lcorbi_t3_pos_1_001_49    m/z 310.1761     0   1   2E+3   1   1.5  RT (min)      lcorbi_t3_pos_1_002_01    m/z 310.1803     0   1   2   3E+3   1   1.5  RT (min)      lcorbi_t3_pos_1_001_49    m/z 310.2013     0   1   2   3E+4   7   7.5  RT (min)       lcorbi_t3_pos_1_002_01    m/z 310.2014     0   1   2   3   4E+4   10.5   11  RT (min)      lcorbi_t3_pos_1_016_05    m/z 310.2075     0   1   2   3E+5   4.5   5  RT (min)       lcorbi_t3_pos_4_090_60    m/z 310.2161     0   1   2E+4   8   8.5  RT (min)       lcorbi_t3_pos_4_053_19    m/z 310.2225     0   2   4   6E+3   6   6.5  RT (min)      lcorbi_t3_pos_1_001_49    m/z 310.2353     0   1E+5   9.5   10  RT (min)       lcorbi_t3_pos_4_101_61    m/z 310.2353     0   1   2E+4   8   8.5  RT (min)       lcorbi_t3_pos_3_052_47    m/z 310.7595     0   2   4E+4   1   1.5  RT (min)      lcorbi_t3_pos_4_021_25    m/z 310.9129     0   2   4   6E+5   1   1.5  RT (min)       lcorbi_t3_pos_1_029_29    m/z 310.9538     0   1   2E+3   1.5   2  RT (min)      lcorbi_t3_pos_1_029_29    m/z 310.9815     0   1   1.5   2  RT (min)      lcorbi_t3_pos_2_040_25    m/z 310.9992     0   1   1   1.5  RT (min)      lcorbi_t3_pos_1_029_29    m/z 311.0075     0   2   4E+4   18   18.5  RT (min)       lcorbi_t3_pos_4_080_58    m/z 311.0170     0   2   4   6E+4   0   0.5  RT (min)      lcorbi_t3_pos_1_002_01    m/z 311.0315     0   1E+4   4.5   5  RT (min)      lcorbi_t3_pos_4_081_58    m/z 311.0354     0   2   4E+4   1.5   2  RT (min)       lcorbi_t3_pos_2_040_25    m/z 311.0738     0   1E+4   2.5   3  RT (min)      lcorbi_t3_pos_4_108_62    m/z 311.1044     0   1E+5   11.5   12  RT (min)       lcorbi_t3_pos_1_029_29    m/z 311.1278     0   1E+4   8   8.5  RT (min)      lcorbi_t3_pos_2_026_44    m/z 311.1296     0   1E+5   4.5   5  RT (min)       lcorbi_t3_pos_1_036_08    m/z 311.1337     0   1E+4   4.5   5  RT (min)      lcorbi_t3_pos_4_066_57    m/z 311.1479     0   2   4E+4   5   5.5  RT (min)       lcorbi_t3_pos_4_008_30    m/z 311.1854     0   1   2   3E+4   9   9.5  RT (min)       lcorbi_t3_pos_4_066_57    m/z 311.2047     0   1E+4   7   7.5  RT (min)      lcorbi_t3_pos_4_075_58    m/z 311.2065     0   1E+4   5   5.5  RT (min)      lcorbi_t3_pos_2_021_09    m/z 311.2387     0   1   2   3   4E+4   10   10.5  RT (min)      lcorbi_t3_pos_2_044_21    m/z 311.2558     0   1E+5   11.5   12  RT (min)       lcorbi_t3_pos_4_083_59    m/z 311.2582     0   1   2E+5   14.5   15  RT (min)       lcorbi_t3_pos_1_032_26    m/z 311.2946     0   2   4   6E+4   13   13.5  RT (min)       lcorbi_t3_pos_4_073_57    m/z 311.3273     0   1   2E+4   9   9.5  RT (min)      lcorbi_t3_pos_3_055_10    m/z 311.9164     0   1   2E+4   1   1.5  RT (min)      lcorbi_t3_pos_1_029_29    m/z 311.9568     0   1E+3   1   1.5  RT (min)      lcorbi_t3_pos_2_020_22    m/z 311.9731     0   2   4E+4   18   18.5  RT (min)       lcorbi_t3_pos_4_085_59    m/z 311.9832     0   1E+5   1   1.5  RT (min)      lcorbi_t3_pos_1_034_34    m/z 312.0358     0   1E+4   1   1.5  RT (min)      lcorbi_t3_pos_2_024_15    m/z 312.0867     0   2   4   6E+3   6.5   7  RT (min)      lcorbi_t3_pos_3_025_52    m/z 312.0989     0   1   2   3   4E+4   1.5   2  RT (min)       lcorbi_t3_pos_4_093_60    m/z 312.1166     0   2   4   6   8E+3   5   5.5  RT (min)       lcorbi_t3_pos_1_012_14    m/z 312.1168     0   1   2E+4   1.5   2  RT (min)      lcorbi_t3_pos_4_032_40    m/z 312.1338     0   1   2E+5   5   5.5  RT (min)      lcorbi_t3_pos_4_110_62    m/z 312.1594     0   1   2   3E+4   9.5   10  RT (min)      lcorbi_t3_pos_4_071_57    m/z 312.1655     0   1   2   3E+4   1.5   2  RT (min)      lcorbi_t3_pos_3_016_21    m/z 312.1767     0   1   2E+4   5   5.5  RT (min)      lcorbi_t3_pos_1_012_14    m/z 312.1872     0   1E+4   5   5.5  RT (min)      lcorbi_t3_pos_2_053_28    m/z 312.2016     0   1E+4   5   5.5  RT (min)      lcorbi_t3_pos_3_048_30    m/z 312.3010     0   1   2E+4   9   9.5  RT (min)       lcorbi_t3_pos_4_113_62    m/z 312.3626     0   2   4   6   8E+4   11   11.5  RT (min)       lcorbi_t3_pos_1_004_24    m/z 312.7566     0   1E+4   1   1.5  RT (min)      lcorbi_t3_pos_1_051_09    m/z 312.8581     0   2   4   6E+4   1   1.5  RT (min)      lcorbi_t3_pos_1_036_08    m/z 312.9023     0   2   4   6E+3   1   1.5  RT (min)      lcorbi_t3_pos_4_045_08    m/z 312.9111     0   1   2   3   4E+4   1   1.5  RT (min)      lcorbi_t3_pos_1_016_05    m/z 313.0075     0   1   2   3E+3   0.5   1  RT (min)      lcorbi_t3_pos_4_083_59    m/z 313.0359     0   1   2   3   4E+4   1   1.5  RT (min)       lcorbi_t3_pos_1_032_26    m/z 313.0456     0   1   1.5   2  RT (min)      lcorbi_t3_pos_1_026_17    m/z 313.0596     0   2   4E+3   5.5   6  RT (min)      lcorbi_t3_pos_4_113_62    m/z 313.0894     0   2   4   6   8E+4   4.5   5  RT (min)       lcorbi_t3_pos_4_066_57    m/z 313.1175     0   1   2   3   4E+4   7.5   8  RT (min)       lcorbi_t3_pos_4_093_60    m/z 313.1194     0   1   2   3E+3   5   5.5  RT (min)      lcorbi_t3_pos_4_014_48    m/z 313.1435     0   1   2   3E+4   8   8.5  RT (min)       lcorbi_t3_pos_4_103_61    m/z 313.1435     0   1   2   3E+4   10.5   11  RT (min)      lcorbi_t3_pos_1_032_26    m/z 313.1499     0   1   1   1.5  RT (min)       lcorbi_t3_pos_4_084_59    m/z 313.2142     0   2   4   6   8E+4   11.5   12  RT (min)       lcorbi_t3_pos_4_083_59    m/z 313.3214     0   1   2   3E+5   9.5   10  RT (min)       lcorbi_t3_pos_4_087_59    m/z 313.3215     0   1   2E+5   13   13.5  RT (min)       lcorbi_t3_pos_2_044_21    m/z 313.9539     0   1   2   3   4E+4   1   1.5  RT (min)      lcorbi_t3_pos_2_024_15    m/z 314.0637     0   1E+4   4.5   5  RT (min)      lcorbi_t3_pos_4_090_60    m/z 314.0639     0   1   2E+3   2.5   3  RT (min)      lcorbi_t3_pos_4_013_36    m/z 314.0918     0   1   2   3E+5   2.5   3  RT (min)       lcorbi_t3_pos_4_013_36    m/z 314.0919     0   2   4E+5   1   1.5   2  RT (min)       lcorbi_t3_pos_3_029_13    m/z 314.1081     0   2   4E+3   1   1.5  RT (min)       lcorbi_t3_pos_3_055_10    m/z 314.1298     0   2   4   6E+3   9   9.5  RT (min)      lcorbi_t3_pos_4_021_25    m/z 314.1340     0   1E+5   1   1.5  RT (min)       lcorbi_t3_pos_4_090_60    m/z 314.1534     0   2   4   6E+3   6   6.5  RT (min)      lcorbi_t3_pos_4_109_62    m/z 314.1810     0   2   4   6E+3   4.5   5  RT (min)      lcorbi_t3_pos_4_076_58    m/z 314.1938     0   1   2   3E+5   1   1.5  RT (min)      lcorbi_t3_pos_4_007_46    m/z 314.2174     0   1E+4   4.5   5  RT (min)      lcorbi_t3_pos_4_106_62    m/z 314.2439     0   1   5   5.5   6  RT (min)      lcorbi_t3_pos_4_006_12    m/z 314.2480     0   2   4   6   8E+4   12   12.5  RT (min)       lcorbi_t3_pos_2_020_22    m/z 314.2772     0   2   4   6   8E+4   14.5   15  RT (min)      lcorbi_t3_pos_2_013_33    m/z 314.3248     0   2   4   6E+4   9.5   10  RT (min)      lcorbi_t3_pos_4_086_59    m/z 314.3418     0   1   2E+4   10   10.5  RT (min)      lcorbi_t3_pos_2_011_10    m/z 314.3611     0   1   2   3   4E+4   9.5   10  RT (min)      lcorbi_t3_pos_4_082_59    m/z 314.8499     0   1   2   3E+5   0   0.5  RT (min)      lcorbi_t3_pos_1_005_42    m/z 314.9500     0   2   4   6E+4   1   1.5  RT (min)       lcorbi_t3_pos_4_076_58    m/z 315.0067     0   1E+4   1   1.5  RT (min)      lcorbi_t3_pos_2_021_09    m/z 315.0931     0   1E+5   10   10.5  RT (min)       lcorbi_t3_pos_4_113_62    m/z 315.0952     0   1   1   1.5  RT (min)       lcorbi_t3_pos_2_040_25    m/z 315.0952     0   1   2.5   3  RT (min)      lcorbi_t3_pos_4_082_59    m/z 315.1089     0   1E+4   4.5   5  RT (min)      lcorbi_t3_pos_1_004_24    m/z 315.1228     0   1E+4   9   9.5  RT (min)      lcorbi_t3_pos_4_106_62    m/z 315.1288     0   1   1.5   2  RT (min)      lcorbi_t3_pos_2_040_25    m/z 315.1349     0   1E+4   5   5.5  RT (min)      lcorbi_t3_pos_4_074_58    m/z 315.1568     0   1   2E+4   9.5   10  RT (min)      lcorbi_t3_pos_4_082_59    m/z 315.1630     0   2   4   6E+4   5   5.5  RT (min)       lcorbi_t3_pos_4_113_62    m/z 315.1666     0   1E+5   1.5   2  RT (min)       lcorbi_t3_pos_1_050_13    m/z 315.1779     0   2   4   6E+4   8.5   9  RT (min)       lcorbi_t3_pos_4_086_59    m/z 315.1873     0   1   2E+3   2.5   3  RT (min)      lcorbi_t3_pos_4_094_60    m/z 315.1956     0   1   2E+5   10.5   11  RT (min)       lcorbi_t3_pos_1_053_39    m/z 315.1985     0   1   2E+5   1   1.5  RT (min)       lcorbi_t3_pos_4_007_46    m/z 315.2360     0   1   2E+4   7.5   8  RT (min)      lcorbi_t3_pos_4_080_58    m/z 315.2838     0   1   2   3   4E+4   4.5   5  RT (min)      lcorbi_t3_pos_4_089_59    m/z 315.3393     0   2   4   6E+4   12   12.5  RT (min)       lcorbi_t3_pos_4_093_60    m/z 315.9080     0   2   4   6   8E+4   1   1.5  RT (min)      lcorbi_t3_pos_1_028_32    m/z 315.9984     0   1   2   3E+4   1   1.5  RT (min)      lcorbi_t3_pos_1_036_08    m/z 316.0833     0   1   2   3   4E+5   1   1.5  RT (min)       lcorbi_t3_pos_4_076_58    m/z 316.0877     0   1E+4   2.5   3  RT (min)      lcorbi_t3_pos_4_013_36    m/z 316.1003     0   1E+3   2.5   3  RT (min)      lcorbi_t3_pos_2_005_11    m/z 316.1004     0   1   2E+4   4.5   5  RT (min)      lcorbi_t3_pos_4_082_59    m/z 316.1156     0   1E+4   5   5.5  RT (min)      lcorbi_t3_pos_3_039_03    m/z 316.1365     0   2   4E+4   9   9.5  RT (min)       lcorbi_t3_pos_4_109_62    m/z 316.1809     0   1   4.5   5  RT (min)      lcorbi_t3_pos_4_020_41    m/z 316.1884     0   1E+4   9   9.5  RT (min)      lcorbi_t3_pos_4_074_58    m/z 316.1907     0   1E+4   1   1.5  RT (min)      lcorbi_t3_pos_4_091_60    m/z 316.2564     0   1   2   3E+4   10   10.5  RT (min)      lcorbi_t3_pos_2_012_01    m/z 316.2654     0   1E+5   9   9.5  RT (min)       lcorbi_t3_pos_4_113_62    m/z 316.3211     0   1E+5   9.5   10  RT (min)       lcorbi_t3_pos_2_044_21    m/z 316.6153     0   1   2   2.5  RT (min)       lcorbi_t3_pos_4_090_60    m/z 316.8240     0   1   1   1.5  RT (min)      lcorbi_t3_pos_3_012_31    m/z 316.8695     0   1   2E+4   1   1.5  RT (min)      lcorbi_t3_pos_3_001_49    m/z 316.8789     0   2   4   6E+4   1   1.5  RT (min)      lcorbi_t3_pos_2_024_15    m/z 316.9284     0   2   4E+4   18   18.5  RT (min)       lcorbi_t3_pos_4_099_61    m/z 316.9864     0   2   4   6E+3   1   1.5  RT (min)      lcorbi_t3_pos_3_010_44    m/z 316.9998     0   1E+5   1   1.5  RT (min)      lcorbi_t3_pos_1_036_08    m/z 317.0395     0   2   4   6   8E+4   18   18.5  RT (min)       lcorbi_t3_pos_4_077_58    m/z 317.0843     0   1   1   1.5  RT (min)      lcorbi_t3_pos_4_023_16    m/z 317.0910     0   2   4   6E+3   6.5   7  RT (min)      lcorbi_t3_pos_4_104_61    m/z 317.0934     0   1   2   3   4E+3   4.5   5  RT (min)      lcorbi_t3_pos_4_066_57    m/z 317.1027     0   1   2   3E+4   2.5   3  RT (min)       lcorbi_t3_pos_1_036_08    m/z 317.1109     0   1E+5   5   5.5  RT (min)       lcorbi_t3_pos_4_104_61    m/z 317.1130     0   2   4E+4   6.5   7  RT (min)      lcorbi_t3_pos_4_005_33    m/z 317.1170     0   1E+5   1.5   2  RT (min)       lcorbi_t3_pos_4_097_60    m/z 317.1336     0   1   1   1.5   2  RT (min)       lcorbi_t3_pos_2_048_36    m/z 317.1724     0   2   4   6   8E+4   9.5   10  RT (min)       lcorbi_t3_pos_1_019_16    m/z 317.1821     0   2   4   6E+3   2.5   3  RT (min)      lcorbi_t3_pos_4_005_33    m/z 317.2021     0   1   2E+4   7   7.5  RT (min)      lcorbi_t3_pos_4_098_61    m/z 317.2153     0   1   2E+4   4.5   5  RT (min)      lcorbi_t3_pos_4_052_06    m/z 317.2185     0   1E+4   1   1.5  RT (min)      lcorbi_t3_pos_4_084_59    m/z 317.2386     0   1E+5   11.5   12  RT (min)      lcorbi_t3_pos_1_054_48    m/z 317.2435     0   1   2E+3   5.5   6  RT (min)      lcorbi_t3_pos_4_055_02    m/z 317.2516     0   1   2   3   4E+4   8   8.5  RT (min)      lcorbi_t3_pos_4_069_57    m/z 317.2688     0   2   4   6   8E+4   13   13.5  RT (min)       lcorbi_t3_pos_4_077_58    m/z 317.2799     0   1   2E+4   10.5   11  RT (min)      lcorbi_t3_pos_3_016_21    m/z 317.2847     0   1E+5   8.5   9  RT (min)       lcorbi_t3_pos_2_044_21    m/z 317.9050     0   1   2   3   4E+4   1   1.5  RT (min)      lcorbi_t3_pos_1_028_32    m/z 317.9549     0   2   4   6E+4   1   1.5  RT (min)      lcorbi_t3_pos_1_036_08    m/z 318.0175     0   2   4E+3   2.5   3  RT (min)      lcorbi_t3_pos_4_006_12    m/z 318.0594     0   1   1   1.5  RT (min)      lcorbi_t3_pos_3_027_27    m/z 318.1142     0   1E+4   5   5.5  RT (min)      lcorbi_t3_pos_4_104_61    m/z 318.1149     0   2   4E+3   1.5   2   2.5  RT (min)      lcorbi_t3_pos_1_036_08    m/z 318.1184     0   1E+5   10   10.5  RT (min)      lcorbi_t3_pos_4_080_58    m/z 318.1248     0   1   2   3E+5   1.5   2  RT (min)       lcorbi_t3_pos_1_036_08    m/z 318.1758     0   1E+4   9.5   10  RT (min)      lcorbi_t3_pos_1_019_16    m/z 318.1912     0   1   4.5   5  RT (min)      lcorbi_t3_pos_4_098_61    m/z 318.3502     0   1   10.5   11  RT (min)       lcorbi_t3_pos_1_032_26    m/z 318.7736     0   2   4   6E+4   1   1.5  RT (min)      lcorbi_t3_pos_2_028_24    m/z 318.7915     0   1E+4   1   1.5  RT (min)      lcorbi_t3_pos_1_008_36    m/z 318.8032     0   1   1   1.5  RT (min)      lcorbi_t3_pos_3_039_03    m/z 318.8211     0   1   1   1.5  RT (min)      lcorbi_t3_pos_1_023_02    m/z 318.9375     0   1E+4   1   1.5  RT (min)      lcorbi_t3_pos_4_080_58    m/z 319.0637     0   1   2E+4   1   1.5  RT (min)      lcorbi_t3_pos_4_084_59    m/z 319.1001     0   2   4   6   8E+4   2.5   3  RT (min)      lcorbi_t3_pos_3_021_39    m/z 319.1081     0   1E+5   6.5   7  RT (min)      lcorbi_t3_pos_4_098_61    m/z 319.1132     0   2   4   6E+4   10   10.5  RT (min)      lcorbi_t3_pos_4_078_58    m/z 319.1365     0   2   4   6   8E+3   4.5   5  RT (min)      lcorbi_t3_pos_4_069_57    m/z 319.1366     0   2   4E+3   2.5   3  RT (min)      lcorbi_t3_pos_1_021_18    m/z 319.1541     0   2   4   6   8E+3   8   8.5  RT (min)      lcorbi_t3_pos_4_013_36    m/z 319.1583     0   1   2   3   4E+4   4.5   5  RT (min)       lcorbi_t3_pos_4_098_61    m/z 319.1615     0   1   2.5   3  RT (min)      lcorbi_t3_pos_4_024_27    m/z 319.1822     0   2   4   6   8E+4   1   1.5  RT (min)       lcorbi_t3_pos_4_091_60    m/z 319.1905     0   1   2E+4   11   11.5  RT (min)      lcorbi_t3_pos_4_074_58    m/z 319.1946     0   1   2   3   4E+4   12.5   13  RT (min)      lcorbi_t3_pos_4_074_58    m/z 319.2018     0   1E+4   7.5   8  RT (min)      lcorbi_t3_pos_4_113_62    m/z 319.2058     0   1E+4   6   6.5  RT (min)      lcorbi_t3_pos_2_053_28    m/z 319.2105     0   1E+5   1   1.5  RT (min)       lcorbi_t3_pos_4_097_60    m/z 319.2268     0   1E+5   11   11.5   12  RT (min)       lcorbi_t3_pos_4_098_61    m/z 319.2592     0   2   4   6   8E+4   6.5   7  RT (min)       lcorbi_t3_pos_1_043_06    m/z 319.2633     0   2   4   6   8E+4   12   12.5  RT (min)       lcorbi_t3_pos_4_087_59    m/z 319.2845     0   2   4   6   8E+4   8.5   9  RT (min)      lcorbi_t3_pos_2_044_21    m/z 319.3625     0   2   4E+4   10.5   11  RT (min)       lcorbi_t3_pos_1_036_08    m/z 319.9213     0   1E+4   1   1.5  RT (min)      lcorbi_t3_pos_1_034_34    m/z 319.9496     0   1E+5   1   1.5  RT (min)      lcorbi_t3_pos_2_020_22    m/z 320.0531     0   1E+5   1   1.5  RT (min)      lcorbi_t3_pos_1_036_08    m/z 320.0789     0   2   4   6   8E+4   4.5   5  RT (min)       lcorbi_t3_pos_4_004_14    m/z 320.0913     0   2   4   6E+4   1   2  RT (min)       lcorbi_t3_pos_2_020_22    m/z 320.1097     0   1E+4   6.5   7  RT (min)      lcorbi_t3_pos_3_011_42    m/z 320.1455     0   2   4   6   8E+4   1.5   2  RT (min)      lcorbi_t3_pos_2_046_07    m/z 320.1493     0   1   2   3E+4   7   7.5  RT (min)      lcorbi_t3_pos_3_034_26    m/z 320.1494     0   2   4   6E+3   5   5.5  RT (min)      lcorbi_t3_pos_4_080_58    m/z 320.1605     0   2   4   6E+3   5   5.5  RT (min)      lcorbi_t3_pos_4_082_59    m/z 320.2139     0   1   2E+6   1   1.5  RT (min)       lcorbi_t3_pos_3_055_10    m/z 320.2302     0   2   4   6   8E+4   10.5   11  RT (min)      lcorbi_t3_pos_1_037_19    m/z 320.2585     0   1   2   3   4E+4   11.5   12  RT (min)      lcorbi_t3_pos_1_029_29    m/z 320.2626     0   1E+4   6.5   7  RT (min)      lcorbi_t3_pos_3_015_06    m/z 320.3030     0   2   4   6E+4   13   13.5  RT (min)       lcorbi_t3_pos_2_044_21    m/z 320.8004     0   1E+4   1   1.5  RT (min)      lcorbi_t3_pos_2_024_15    m/z 320.8656     0   1   2   3   4E+4   1   1.5  RT (min)      lcorbi_t3_pos_1_036_08    m/z 320.8698     0   2   4   6E+4   1   1.5  RT (min)       lcorbi_t3_pos_4_090_60    m/z 321.0525     0   2   4   6E+4   18   18.5  RT (min)       lcorbi_t3_pos_4_094_60    m/z 321.0952     0   2   4   6E+4   5   5.5  RT (min)      lcorbi_t3_pos_4_101_61    m/z 321.1034     0   1E+3   1   1.5  RT (min)      lcorbi_t3_pos_4_007_46    m/z 321.1294     0   1   2.5   3  RT (min)      lcorbi_t3_pos_1_050_13    m/z 321.1311     0   1E+4   4.5   5  RT (min)      lcorbi_t3_pos_4_113_62    m/z 321.1462     0   1E+4   10.5   11  RT (min)      lcorbi_t3_pos_4_012_37    m/z 321.1480     0   1   2   3E+3   5   5.5  RT (min)      lcorbi_t3_pos_4_102_61    m/z 321.1502     0   2   4   6   8E+4   1   1.5  RT (min)       lcorbi_t3_pos_2_037_27    m/z 321.1599     0   2   4   6E+4   4.5   5  RT (min)       lcorbi_t3_pos_4_112_62    m/z 321.2061     0   1E+4   11.5   12  RT (min)      lcorbi_t3_pos_4_082_59    m/z 321.2062     0   2   4   6   8E+4   13   13.5  RT (min)      lcorbi_t3_pos_4_077_58    m/z 321.2425     0   1   2   3E+4   9   9.5  RT (min)       lcorbi_t3_pos_4_100_61    m/z 321.2425     0   2   4   6E+4   7.5   8  RT (min)       lcorbi_t3_pos_4_105_61    m/z 321.2514     0   1   12   12.5  RT (min)       lcorbi_t3_pos_4_085_59    m/z 321.2909     0   1E+5   11.5   12  RT (min)      lcorbi_t3_pos_4_085_59    m/z 321.3265     0   1   2   3E+4   9.5   10  RT (min)      lcorbi_t3_pos_1_025_52    m/z 321.9184     0   2   4   6E+4   1   1.5  RT (min)      lcorbi_t3_pos_1_036_08    m/z 322.0024     0   1   0   0.5  RT (min)      lcorbi_t3_pos_1_001_49    m/z 322.0685     0   1E+4   4.5   5  RT (min)      lcorbi_t3_pos_4_106_62    m/z 322.0985     0   1E+5   1   1.5  RT (min)       lcorbi_t3_pos_4_086_59    m/z 322.0985     0   1   2   3   4E+3   5   5.5  RT (min)      lcorbi_t3_pos_1_028_32    m/z 322.1635     0   1E+4   4.5   5  RT (min)       lcorbi_t3_pos_4_111_62    m/z 322.1674     0   1   2   3E+4   1   1.5  RT (min)      lcorbi_t3_pos_3_055_10    m/z 322.1675     0   1E+4   7   7.5  RT (min)      lcorbi_t3_pos_4_099_61    m/z 322.1800     0   1E+4   7   7.5  RT (min)      lcorbi_t3_pos_4_030_05    m/z 322.1873     0   2   4   6   8E+4   1   2   3  RT (min)       lcorbi_t3_pos_4_091_60    m/z 322.2102     0   1E+4   7   7.5  RT (min)       lcorbi_t3_pos_4_106_62    m/z 322.2287     0   2   4   6   8E+3   4.5   5  RT (min)      lcorbi_t3_pos_4_106_62    m/z 322.2459     0   1E+4   7.5   8  RT (min)      lcorbi_t3_pos_4_101_61    m/z 322.2718     0   1   2   3   4E+4   10.5   11  RT (min)       lcorbi_t3_pos_1_032_26    m/z 322.2854     0   1   2E+4   7   7.5  RT (min)       lcorbi_t3_pos_4_082_59    m/z 322.2954     0   1   2   3E+4   11   11.5  RT (min)      lcorbi_t3_pos_4_089_59    m/z 322.3106     0   2   4   6E+4   11   11.5   12  RT (min)       lcorbi_t3_pos_4_083_59    m/z 322.8113     0   2   4   6E+4   1   1.5  RT (min)      lcorbi_t3_pos_1_020_31    m/z 322.9330     0   1   2   3E+4   12   12.5  RT (min)      lcorbi_t3_pos_3_045_43    m/z 322.9607     0   1   2   3E+3   1.5   2  RT (min)      lcorbi_t3_pos_1_023_02    m/z 322.9818     0   1E+4   1   1.5  RT (min)      lcorbi_t3_pos_2_023_26    m/z 323.0020     0   1E+4   1   1.5  RT (min)      lcorbi_t3_pos_1_032_26    m/z 323.0115     0   1   2E+5   1   1.5  RT (min)      lcorbi_t3_pos_2_020_22    m/z 323.0695     0   1   2.5   3  RT (min)      lcorbi_t3_pos_4_082_59    m/z 323.0915     0   2   4   6   8E+4   5   5.5  RT (min)       lcorbi_t3_pos_4_014_48    m/z 323.1279     0   1   2E+4   8   8.5  RT (min)       lcorbi_t3_pos_4_098_61    m/z 323.1412     0   1E+5   4.5   5  RT (min)      lcorbi_t3_pos_4_110_62    m/z 323.1531     0   1   2E+4   9   9.5  RT (min)      lcorbi_t3_pos_4_066_57    m/z 323.1537     0   2   4   6E+3   1.5   2  RT (min)      lcorbi_t3_pos_1_036_08    m/z 323.1638     0   1E+5   1   1.5  RT (min)       lcorbi_t3_pos_3_055_10    m/z 323.2044     0   2   4   6E+4   4.5   5  RT (min)      lcorbi_t3_pos_3_004_24    m/z 323.2524     0   1   2E+4   5   5.5  RT (min)      lcorbi_t3_pos_4_084_59    m/z 323.9630     0   1   2E+5   1   1.5  RT (min)      lcorbi_t3_pos_1_036_08    m/z 323.9827     0   1   2E+3   3   3.5  RT (min)      lcorbi_t3_pos_1_002_01    m/z 324.0016     0   1E+5   1   1.5  RT (min)       lcorbi_t3_pos_4_076_58    m/z 324.0960     0   1   2.5   3  RT (min)      lcorbi_t3_pos_4_091_60    m/z 324.1071     0   1E+5   1   1.5  RT (min)      lcorbi_t3_pos_4_007_46    m/z 324.1169     0   2   4   6E+4   2.5   3  RT (min)      lcorbi_t3_pos_3_021_39    m/z 324.1251     0   1   2.5   3  RT (min)       lcorbi_t3_pos_4_090_60    m/z 324.1265     0   2   4E+4   10   10.5  RT (min)       lcorbi_t3_pos_4_078_58    m/z 324.1281     0   2   4E+4   1.5   2  RT (min)       lcorbi_t3_pos_4_095_60    m/z 324.1419     0   2   4E+4   1   1.5  RT (min)       lcorbi_t3_pos_4_094_60    m/z 324.1533     0   1   2   3E+4   2.5   3  RT (min)      lcorbi_t3_pos_3_026_04    m/z 324.1584     0   2   4   6E+4   5.5   6  RT (min)      lcorbi_t3_pos_4_105_61    m/z 324.1654     0   1E+5   1   1.5  RT (min)       lcorbi_t3_pos_4_068_57    m/z 324.1739     0   2   4   6E+5   4.5   5  RT (min)      lcorbi_t3_pos_2_053_28    m/z 324.1900     0   1E+5   4.5   5  RT (min)      lcorbi_t3_pos_4_008_30    m/z 324.2170     0   1E+5   10   10.5  RT (min)       lcorbi_t3_pos_1_002_01    m/z 324.2232     0   1   4.5   5  RT (min)      lcorbi_t3_pos_4_112_62    m/z 324.2511     0   2   4   6E+4   10.5   11  RT (min)       lcorbi_t3_pos_4_111_62    m/z 324.2534     0   2   4   6   8E+3   8.5   9  RT (min)      lcorbi_t3_pos_1_019_16    m/z 324.2534     0   1   2   3   4E+4   11.5   12  RT (min)      lcorbi_t3_pos_3_023_05    m/z 324.2875     0   2   4   6E+4   10.5   11  RT (min)       lcorbi_t3_pos_3_020_35    m/z 324.3263     0   2   4   6   8E+4   13   13.5  RT (min)      lcorbi_t3_pos_2_044_21    m/z 324.3373     0   1   2   3E+4   10.5   11  RT (min)      lcorbi_t3_pos_1_036_08    m/z 324.7364     0   1   2E+4   1   1.5  RT (min)      lcorbi_t3_pos_4_006_12    m/z 324.9305     0   1   2E+6   0   0.5  RT (min)      lcorbi_t3_pos_1_005_42    m/z 325.0435     0   1E+4   1   1.5  RT (min)      lcorbi_t3_pos_4_013_36    m/z 325.1130     0   1   1   1.5  RT (min)       lcorbi_t3_pos_4_093_60    m/z 325.1131     0   1   2.5   3  RT (min)      lcorbi_t3_pos_4_092_60    m/z 325.1156     0   2   4   6E+4   5.5   6  RT (min)      lcorbi_t3_pos_2_044_21    m/z 325.1391     0   2   4   6E+4   5.5   6  RT (min)      lcorbi_t3_pos_1_037_19    m/z 325.1414     0   1E+4   7.5   8  RT (min)      lcorbi_t3_pos_4_074_58    m/z 325.1687     0   1   2   3E+4   1   1.5  RT (min)      lcorbi_t3_pos_1_020_31    m/z 325.1711     0   2   4   6E+3   6.5   7  RT (min)      lcorbi_t3_pos_4_071_57    m/z 325.1911     0   1   2   3E+4   5   5.5  RT (min)      lcorbi_t3_pos_4_008_30    m/z 325.2011     0   1E+4   9.5   10  RT (min)      lcorbi_t3_pos_4_070_57    m/z 325.2140     0   1E+5   11.5   12  RT (min)      lcorbi_t3_pos_1_029_29    m/z 325.2222     0   2   4E+3   5.5   6  RT (min)      lcorbi_t3_pos_4_066_57    m/z 325.2276     0   2   4   6E+3   6.5   7  RT (min)      lcorbi_t3_pos_4_038_38    m/z 325.2351     0   2   4E+4   11.5   12  RT (min)       lcorbi_t3_pos_4_068_57    m/z 325.3215     0   2   4E+4   10   10.5  RT (min)       lcorbi_t3_pos_4_089_59    m/z 326.0377     0   1   2   3   4E+4   4.5   5  RT (min)       lcorbi_t3_pos_4_066_57    m/z 326.0570     0   1   2   2.5  RT (min)      lcorbi_t3_pos_4_082_59    m/z 326.1083     0   1   2   3   4E+4   1.5   2  RT (min)       lcorbi_t3_pos_2_020_22    m/z 326.1326     0   1   2.5   3  RT (min)      lcorbi_t3_pos_4_092_60    m/z 326.1812     0   2   4E+3   5   5.5   6  RT (min)       lcorbi_t3_pos_1_012_14    m/z 326.1946     0   1E+4   5   5.5  RT (min)      lcorbi_t3_pos_4_111_62    m/z 326.2078     0   1   4.5   5  RT (min)      lcorbi_t3_pos_2_044_21    m/z 326.2439     0   2   4   6E+3   5.5   6  RT (min)      lcorbi_t3_pos_4_084_59    m/z 326.2693     0   1   2E+4   11   11.5  RT (min)       lcorbi_t3_pos_4_087_59    m/z 326.3055     0   2   4   6E+4   12   12.5  RT (min)      lcorbi_t3_pos_3_021_39    m/z 326.3783     0   2   4   6E+4   10.5   11  RT (min)       lcorbi_t3_pos_3_032_09    m/z 326.8054     0   1   2   3   4E+4   1   1.5  RT (min)      lcorbi_t3_pos_1_020_31    m/z 326.8232     0   1   2E+4   1   1.5  RT (min)      lcorbi_t3_pos_1_020_31    m/z 326.8870     0   1E+5   1   1.5  RT (min)       lcorbi_t3_pos_1_029_29    m/z 326.9316     0   1   2   3   4E+3   1.5   2  RT (min)      lcorbi_t3_pos_4_074_58    m/z 327.0083     0   1   2E+5   9   9.5  RT (min)       lcorbi_t3_pos_1_005_42    m/z 327.0153     0   2   4E+3   1   1.5  RT (min)      lcorbi_t3_pos_1_001_49    m/z 327.0517     0   2   4   6   8E+4   1   1.5  RT (min)       lcorbi_t3_pos_3_011_42    m/z 327.1151     0   2   4   6E+3   1   1.5  RT (min)       lcorbi_t3_pos_2_023_26    m/z 327.1264     0   1   2   3   4E+5   1   1.5  RT (min)       lcorbi_t3_pos_1_031_40    m/z 327.1312     0   1E+5   5.5   6  RT (min)       lcorbi_t3_pos_4_098_61    m/z 327.1357     0   1   2   3   4E+4   12   12.5  RT (min)       lcorbi_t3_pos_1_001_49    m/z 327.1381     0   1   1.5   2  RT (min)      lcorbi_t3_pos_1_052_38    m/z 327.1415     0   2   4   6E+3   8.5   9  RT (min)      lcorbi_t3_pos_1_001_49    m/z 327.1592     0   2   4E+3   8   8.5  RT (min)       lcorbi_t3_pos_4_008_30    m/z 327.1889     0   1E+4   5   5.5  RT (min)      lcorbi_t3_pos_4_082_59    m/z 327.2014     0   2   4   6E+4   4.5   5  RT (min)       lcorbi_t3_pos_4_112_62    m/z 327.2321     0   2   4   6E+4   15   15.5  RT (min)       lcorbi_t3_pos_4_075_58    m/z 327.2718     0   2   4   6   8E+4   12.5   13  RT (min)       lcorbi_t3_pos_4_066_57    m/z 327.3008     0   2   4   6   8E+4   11.5   12  RT (min)       lcorbi_t3_pos_4_097_60    m/z 327.3781     0   1   2   3   4E+4   11   11.5  RT (min)      lcorbi_t3_pos_2_037_27    m/z 328.0551     0   2   4   6E+3   1   1.5  RT (min)      lcorbi_t3_pos_4_018_01    m/z 328.0754     0   2   4   6   8E+2   1.5   2  RT (min)      lcorbi_t3_pos_1_031_40    m/z 328.1028     0   1   2   3E+3   2.5   3  RT (min)      lcorbi_t3_pos_1_036_08    m/z 328.1297     0   2   4E+4   1   1.5  RT (min)      lcorbi_t3_pos_1_031_40    m/z 328.1303     0   1   2E+4   2.5   3  RT (min)      lcorbi_t3_pos_4_091_60    m/z 328.1332     0   1   2E+4   9   9.5  RT (min)      lcorbi_t3_pos_4_100_61    m/z 328.1579     0   1   2E+4   8   8.5  RT (min)      lcorbi_t3_pos_4_074_58    m/z 328.1603     0   1E+5   4.5   5  RT (min)       lcorbi_t3_pos_4_090_60    m/z 328.1756     0   1   2E+4   6   6.5  RT (min)       lcorbi_t3_pos_4_078_58    m/z 328.1967     0   1E+4   4.5   5  RT (min)      lcorbi_t3_pos_4_082_59    m/z 328.2049     0   1   4.5   5  RT (min)      lcorbi_t3_pos_4_106_62    m/z 328.2331     0   1E+4   5   5.5  RT (min)      lcorbi_t3_pos_4_106_62    m/z 328.2354     0   2   4   6E+4   12   12.5  RT (min)      lcorbi_t3_pos_4_078_58    m/z 328.2484     0   1   2   3E+4   8   8.5  RT (min)       lcorbi_t3_pos_4_081_58    m/z 328.2958     0   1E+5   4.5   5  RT (min)       lcorbi_t3_pos_4_085_59    m/z 328.3212     0   2   4E+4   12.5   13  RT (min)       lcorbi_t3_pos_2_020_22    m/z 328.7564     0   2   4E+4   1   1.5  RT (min)      lcorbi_t3_pos_2_051_35    m/z 328.8202     0   1E+4   1   1.5  RT (min)      lcorbi_t3_pos_1_034_34    m/z 328.8851     0   1   2E+4   1   1.5  RT (min)      lcorbi_t3_pos_1_029_29    m/z 328.8981     0   1   2E+4   1   1.5  RT (min)       lcorbi_t3_pos_4_074_58    m/z 328.9702     0   2   4   6E+4   18   18.5  RT (min)       lcorbi_t3_pos_4_070_57    m/z 329.0037     0   1   2   3E+4   1   1.5  RT (min)       lcorbi_t3_pos_1_015_11    m/z 329.0053     0   1E+5   9   9.5  RT (min)       lcorbi_t3_pos_1_005_42    m/z 329.0101     0   1   2   3   4E+3   1   1.5  RT (min)      lcorbi_t3_pos_1_001_49    m/z 329.0847     0   1E+4   10   10.5  RT (min)      lcorbi_t3_pos_4_077_58    m/z 329.0957     0   1   2   3E+5   1.5   2  RT (min)       lcorbi_t3_pos_4_078_58    m/z 329.1125     0   2   4   6E+4   7.5   8  RT (min)       lcorbi_t3_pos_4_091_60    m/z 329.1239     0   2   4   6E+5   1   1.5  RT (min)       lcorbi_t3_pos_3_055_10    m/z 329.1596     0   1   2E+4   9   9.5  RT (min)      lcorbi_t3_pos_4_082_59    m/z 329.1637     0   1E+4   4.5   5  RT (min)      lcorbi_t3_pos_4_090_60    m/z 329.2113     0   1E+4   11   11.5  RT (min)       lcorbi_t3_pos_4_074_58    m/z 329.2436     0   1   2E+4   6.5   7  RT (min)       lcorbi_t3_pos_4_084_59    m/z 329.2477     0   2   4   6   8E+4   12   12.5  RT (min)       lcorbi_t3_pos_4_077_58    m/z 329.2589     0   1E+5   11.5   12  RT (min)       lcorbi_t3_pos_1_007_07    m/z 329.2630     0   1   2E+4   6   6.5  RT (min)      lcorbi_t3_pos_4_089_59    m/z 329.2994     0   1   2E+4   4.5   5  RT (min)      lcorbi_t3_pos_4_089_59    m/z 329.3528     0   2   4   6E+4   9.5   10  RT (min)       lcorbi_t3_pos_4_074_58    m/z 329.9237     0   2   4   6   8E+4   1   1.5  RT (min)      lcorbi_t3_pos_1_052_38    m/z 330.0599     0   1E+4   4.5   5  RT (min)      lcorbi_t3_pos_4_082_59    m/z 330.0600     0   2   4   6E+3   1.5   2  RT (min)      lcorbi_t3_pos_2_007_02    m/z 330.0732     0   1   1.5   2  RT (min)      lcorbi_t3_pos_4_042_09    m/z 330.0990     0   1   2   3   4E+3   1.5   2  RT (min)      lcorbi_t3_pos_2_042_45    m/z 330.1313     0   1E+5   1   1.5  RT (min)       lcorbi_t3_pos_4_042_09    m/z 330.1523     0   1   9.5   10  RT (min)      lcorbi_t3_pos_1_037_19    m/z 330.1701     0   1E+4   4.5   5  RT (min)      lcorbi_t3_pos_4_071_57    m/z 330.2276     0   2   4   6   8E+3   4.5   5  RT (min)      lcorbi_t3_pos_4_006_12    m/z 330.2276     0   1   2   3   4E+3   6.5   7  RT (min)      lcorbi_t3_pos_4_006_12    m/z 330.2640     0   1   2E+4   8.5   9  RT (min)      lcorbi_t3_pos_4_075_58    m/z 330.2811     0   1   2   3E+4   8   8.5  RT (min)       lcorbi_t3_pos_4_053_19    m/z 330.2834     0   2   4E+3   10.5   11  RT (min)      lcorbi_t3_pos_4_032_40    m/z 330.3220     0   1   2E+5   14.5   15  RT (min)       lcorbi_t3_pos_1_036_08    m/z 330.3368     0   1E+5   10.5   11  RT (min)      lcorbi_t3_pos_3_015_06    m/z 330.3561     0   1E+4   9.5   10  RT (min)      lcorbi_t3_pos_4_082_59    m/z 330.7535     0   2   4   6   8E+4   1   1.5  RT (min)       lcorbi_t3_pos_1_029_29    m/z 330.9032     0   1E+4   1   1.5  RT (min)      lcorbi_t3_pos_1_020_31    m/z 330.9311     0   1E+4   1   1.5  RT (min)      lcorbi_t3_pos_4_074_58    m/z 331.0023     0   2   4   6E+4   9   9.5  RT (min)      lcorbi_t3_pos_1_005_42    m/z 331.0638     0   1   2   3   4E+4   1.5   2  RT (min)      lcorbi_t3_pos_1_032_26    m/z 331.1038     0   1E+4   4.5   5   5.5  RT (min)      lcorbi_t3_pos_4_066_57    m/z 331.1158     0   2   4E+4   4.5   5  RT (min)       lcorbi_t3_pos_2_024_15    m/z 331.1266     0   2   4   6   8E+3   5.5   6  RT (min)      lcorbi_t3_pos_3_001_49    m/z 331.1346     0   1   1.5   2  RT (min)      lcorbi_t3_pos_2_020_22    m/z 331.1653     0   1   2   3E+4   11   11.5  RT (min)      lcorbi_t3_pos_4_074_58    m/z 331.2092     0   1E+4   6.5   7  RT (min)      lcorbi_t3_pos_4_083_59    m/z 331.2229     0   1   2E+4   5   5.5  RT (min)      lcorbi_t3_pos_4_091_60    m/z 331.2480     0   2   4E+4   9.5   10  RT (min)       lcorbi_t3_pos_4_078_58    m/z 331.2545     0   1   2E+4   6   6.5  RT (min)       lcorbi_t3_pos_4_084_59    m/z 331.2844     0   1   2   3E+4   12   12.5  RT (min)      lcorbi_t3_pos_3_003_33    m/z 331.2956     0   1E+4   8   8.5  RT (min)      lcorbi_t3_pos_4_090_60    m/z 331.7560     0   2   4   6E+3   6   6.5  RT (min)      lcorbi_t3_pos_2_027_05    m/z 332.0756     0   1   2E+3   1   1.5  RT (min)       lcorbi_t3_pos_4_054_18    m/z 332.1197     0   1E+5   10   10.5  RT (min)       lcorbi_t3_pos_4_113_62    m/z 332.1313     0   2   4   6E+3   1   1.5  RT (min)       lcorbi_t3_pos_1_030_21    m/z 332.1494     0   1   2E+4   9   9.5  RT (min)      lcorbi_t3_pos_1_032_26    m/z 332.1553     0   2   4E+4   1   1.5  RT (min)      lcorbi_t3_pos_1_021_18    m/z 332.2106     0   2   4   6E+4   9.5   10  RT (min)      lcorbi_t3_pos_4_101_61    m/z 332.2433     0   1   2E+4   7.5   8  RT (min)       lcorbi_t3_pos_4_066_57    m/z 332.2927     0   1   2E+5   12.5   13  RT (min)       lcorbi_t3_pos_2_044_21    m/z 332.2968     0   1   2   3E+4   8   8.5  RT (min)      lcorbi_t3_pos_2_044_21    m/z 332.3377     0   1   2E+5   14.5   15  RT (min)       lcorbi_t3_pos_4_092_60    m/z 332.7507     0   2   4   6E+4   1   1.5  RT (min)      lcorbi_t3_pos_1_029_29    m/z 332.8401     0   1E+5   1   1.5  RT (min)       lcorbi_t3_pos_1_020_31    m/z 332.9415     0   1   2E+5   18   18.5  RT (min)       lcorbi_t3_pos_4_099_61    m/z 332.9416     0   1   2   3   4E+4   10   10.5  RT (min)      lcorbi_t3_pos_4_088_59    m/z 332.9484     0   1   2   2.5  RT (min)      lcorbi_t3_pos_2_031_20    m/z 333.0718     0   2   4   6E+4   1   1.5  RT (min)      lcorbi_t3_pos_4_052_06    m/z 333.0759     0   1   2E+4   5   5.5  RT (min)       lcorbi_t3_pos_4_098_61    m/z 333.0794     0   1   2   3E+3   1   1.5  RT (min)      lcorbi_t3_pos_1_003_45    m/z 333.0849     0   2   4E+3   5   5.5  RT (min)      lcorbi_t3_pos_1_023_02    m/z 333.0904     0   1E+5   1   1.5  RT (min)       lcorbi_t3_pos_4_097_60    m/z 333.1030     0   1   2.5   3  RT (min)      lcorbi_t3_pos_4_090_60    m/z 333.1158     0   2   4   6E+4   4.5   5  RT (min)       lcorbi_t3_pos_4_066_57    m/z 333.1408     0   1E+3   1   1.5  RT (min)      lcorbi_t3_pos_3_010_44    m/z 333.1464     0   1   2E+4   9.5   10  RT (min)      lcorbi_t3_pos_3_016_21    m/z 333.1632     0   2   4E+4   9   9.5  RT (min)      lcorbi_t3_pos_1_011_44    m/z 333.1674     0   2   4   6   8E+4   6   6.5  RT (min)      lcorbi_t3_pos_4_106_62    m/z 333.1885     0   1   2   3   4E+3   5   5.5  RT (min)      lcorbi_t3_pos_4_098_61    m/z 333.1971     0   1   11.5   12  RT (min)      lcorbi_t3_pos_1_054_48    m/z 333.2579     0   2   4E+4   14   14.5  RT (min)       lcorbi_t3_pos_2_027_05    m/z 333.2749     0   1E+4   6.5   7  RT (min)      lcorbi_t3_pos_4_093_60    m/z 333.3195     0   2   4   6   8E+4   10.5   11  RT (min)       lcorbi_t3_pos_1_031_40    m/z 333.7061     0   1   2   3E+4   4.5   5  RT (min)       lcorbi_t3_pos_4_006_12    m/z 333.8791     0   1   2E+4   1   1.5  RT (min)      lcorbi_t3_pos_2_020_22    m/z 333.9653     0   1   2   3E+5   1   1.5  RT (min)      lcorbi_t3_pos_1_031_40    m/z 334.0268     0   1   2   3   4E+4   1   1.5  RT (min)      lcorbi_t3_pos_4_052_06    m/z 334.0736     0   1   1   1.5  RT (min)       lcorbi_t3_pos_3_048_30    m/z 334.0827     0   1   2   3E+3   1   1.5  RT (min)      lcorbi_t3_pos_3_020_35    m/z 334.1158     0   1   2E+5   1   1.5  RT (min)       lcorbi_t3_pos_4_084_59    m/z 334.1191     0   1   4.5   5  RT (min)      lcorbi_t3_pos_1_037_19    m/z 334.1286     0   2   4   6   8E+4   7.5   8  RT (min)       lcorbi_t3_pos_4_100_61    m/z 334.1374     0   2   4   6E+4   5   5.5  RT (min)       lcorbi_t3_pos_4_008_30    m/z 334.1598     0   1   2E+4   6   6.5  RT (min)      lcorbi_t3_pos_4_108_62    m/z 334.2078     0   1E+4   4.5   5  RT (min)      lcorbi_t3_pos_4_075_58    m/z 334.2095     0   1E+5   7   7.5  RT (min)      lcorbi_t3_pos_4_005_33    m/z 334.2460     0   1E+5   11.5   12  RT (min)       lcorbi_t3_pos_4_082_59    m/z 334.2568     0   1   2E+4   8   8.5  RT (min)      lcorbi_t3_pos_4_038_38    m/z 334.2596     0   1   2   3   4E+4   6.5   7  RT (min)       lcorbi_t3_pos_4_085_59    m/z 334.2671     0   2   4   6E+4   9.5   10  RT (min)       lcorbi_t3_pos_4_066_57    m/z 334.2760     0   1E+4   9   9.5  RT (min)      lcorbi_t3_pos_4_109_62    m/z 334.2953     0   1   2   3   4E+4   9   9.5  RT (min)      lcorbi_t3_pos_2_044_21    m/z 334.3380     0   1   2   3E+4   10   10.5  RT (min)      lcorbi_t3_pos_2_049_55    m/z 334.3541     0   1   2   3   4E+4   10   10.5  RT (min)      lcorbi_t3_pos_1_036_08    m/z 334.6901     0   1   2   3E+4   5   5.5  RT (min)       lcorbi_t3_pos_4_078_58    m/z 334.7478     0   1   2E+4   1   1.5  RT (min)      lcorbi_t3_pos_1_029_29    m/z 334.7613     0   1   2E+4   6.5   7  RT (min)      lcorbi_t3_pos_4_086_59    m/z 334.8372     0   1E+5   1   1.5  RT (min)       lcorbi_t3_pos_1_020_31    m/z 334.9151     0   1   2   3   4E+4   9   9.5  RT (min)      lcorbi_t3_pos_1_043_06    m/z 334.9151     0   1   2   3   4E+5   18   18.5  RT (min)       lcorbi_t3_pos_4_074_58    m/z 335.0116     0   2   4   6   8E+4   1   1.5  RT (min)      lcorbi_t3_pos_1_052_38    m/z 335.0276     0   1E+5   0   0.5  RT (min)      lcorbi_t3_pos_1_006_15    m/z 335.0641     0   1E+4   1   1.5  RT (min)       lcorbi_t3_pos_2_023_26    m/z 335.1303     0   1E+4   4.5   5  RT (min)      lcorbi_t3_pos_2_024_15    m/z 335.1320     0   1E+4   7.5   8  RT (min)      lcorbi_t3_pos_2_044_21    m/z 335.1684     0   2   4   6   8E+4   7   7.5  RT (min)       lcorbi_t3_pos_4_098_61    m/z 335.1736     0   2   4   6E+4   1.5   2  RT (min)      lcorbi_t3_pos_3_051_12    m/z 335.1918     0   2   4   6   8E+3   5   5.5  RT (min)      lcorbi_t3_pos_4_077_58    m/z 335.2195     0   1   2   3E+4   10.5   11  RT (min)      lcorbi_t3_pos_4_066_57    m/z 335.2218     0   1   2   3E+4   12   12.5  RT (min)       lcorbi_t3_pos_4_082_59    m/z 335.2583     0   1E+5   14   14.5  RT (min)       lcorbi_t3_pos_1_031_40    m/z 335.2794     0   2   4   6   8E+4   10.5   11  RT (min)       lcorbi_t3_pos_4_085_59    m/z 335.3185     0   1   2E+4   12   12.5  RT (min)      lcorbi_t3_pos_4_095_60    m/z 335.8954     0   1   2   3   4E+4   1   1.5  RT (min)      lcorbi_t3_pos_1_036_08    m/z 335.9165     0   2   4   6   8E+4   18   18.5  RT (min)       lcorbi_t3_pos_4_088_59    m/z 335.9235     0   1   2   3   4E+4   1   1.5  RT (min)      lcorbi_t3_pos_1_034_34    m/z 336.0157     0   1   2   3   4E+4   1   1.5  RT (min)      lcorbi_t3_pos_3_024_07    m/z 336.0270     0   1   2E+3   1   1.5  RT (min)      lcorbi_t3_pos_2_040_25    m/z 336.0716     0   1E+5   8   8.5  RT (min)       lcorbi_t3_pos_1_051_09    m/z 336.0739     0   1   2.5   3  RT (min)      lcorbi_t3_pos_4_082_59    m/z 336.0868     0   1E+4   6   6.5  RT (min)      lcorbi_t3_pos_3_034_26    m/z 336.0906     0   1   2E+4   2   2.5  RT (min)       lcorbi_t3_pos_4_097_60    m/z 336.1155     0   2   4   6E+4   1.5   2  RT (min)      lcorbi_t3_pos_4_087_59    m/z 336.1564     0   2   4   6E+4   1   1.5  RT (min)       lcorbi_t3_pos_1_036_08    m/z 336.2535     0   2   4   6   8E+3   7.5   8  RT (min)      lcorbi_t3_pos_4_068_57    m/z 336.2917     0   1E+4   11.5   12  RT (min)       lcorbi_t3_pos_4_090_60    m/z 336.2923     0   1E+4   12.5   13  RT (min)       lcorbi_t3_pos_4_094_60    m/z 336.3011     0   1E+4   7.5   8  RT (min)      lcorbi_t3_pos_4_098_61    m/z 336.8343     0   1   2   3E+4   1   1.5  RT (min)      lcorbi_t3_pos_1_007_07    m/z 336.8519     0   1   1   1.5  RT (min)       lcorbi_t3_pos_4_090_60    m/z 336.9122     0   2   4   6E+4   18   18.5  RT (min)       lcorbi_t3_pos_4_092_60    m/z 337.0148     0   1E+5   1   1.5   2  RT (min)       lcorbi_t3_pos_3_008_20    m/z 337.0620     0   1   2E+5   1   1.5  RT (min)       lcorbi_t3_pos_4_075_58    m/z 337.0751     0   1E+5   10   10.5  RT (min)       lcorbi_t3_pos_4_113_62    m/z 337.1183     0   1   2   3   4E+4   6.5   7  RT (min)      lcorbi_t3_pos_4_105_61    m/z 337.1454     0   1   2E+5   1.5   2  RT (min)       lcorbi_t3_pos_4_090_60    m/z 337.1599     0   1E+6   1   1.5  RT (min)       lcorbi_t3_pos_1_036_08    m/z 337.1720     0   1E+4   1   1.5  RT (min)       lcorbi_t3_pos_4_054_18    m/z 337.1911     0   1E+4   6.5   7  RT (min)      lcorbi_t3_pos_4_101_61    m/z 337.2739     0   1   2   3   4E+4   12   12.5  RT (min)       lcorbi_t3_pos_3_027_27    m/z 337.3103     0   1E+4   13   13.5  RT (min)       lcorbi_t3_pos_2_044_21    m/z 337.3216     0   1   2   3E+4   13   13.5  RT (min)      lcorbi_t3_pos_2_044_21    m/z 337.3297     0   1   11.5   12  RT (min)      lcorbi_t3_pos_1_032_26    m/z 337.9748     0   1   2   3E+5   0   0.5  RT (min)      lcorbi_t3_pos_1_005_42    m/z 338.0508     0   2   4E+4   1   1.5  RT (min)      lcorbi_t3_pos_4_077_58    m/z 338.0686     0   1   2   3E+4   8   8.5  RT (min)      lcorbi_t3_pos_1_051_09    m/z 338.0760     0   1   2E+5   1  RT (min)       lcorbi_t3_pos_4_087_59    m/z 338.0961     0   1   1.5   2  RT (min)      lcorbi_t3_pos_2_005_11    m/z 338.1141     0   2   4   6E+3   1.5   2  RT (min)      lcorbi_t3_pos_2_005_11    m/z 338.1186     0   1   2   3E+4   9   9.5  RT (min)      lcorbi_t3_pos_4_079_58    m/z 338.1560     0   2   4   6   8E+4   1   1.5  RT (min)       lcorbi_t3_pos_4_079_58    m/z 338.1812     0   1   2E+4   1   1.5  RT (min)       lcorbi_t3_pos_4_068_57    m/z 338.2197     0   1   2   3   4E+4   8   8.5  RT (min)      lcorbi_t3_pos_1_014_03    m/z 338.2197     0   2   4   6E+4   6   6.5  RT (min)      lcorbi_t3_pos_4_112_62    m/z 338.2475     0   1   2   3E+4   9   9.5  RT (min)       lcorbi_t3_pos_4_112_62    m/z 338.2667     0   1   2   3E+4   9.5   10  RT (min)      lcorbi_t3_pos_4_066_57    m/z 338.2774     0   1   14.5   15  RT (min)       lcorbi_t3_pos_1_045_12    m/z 338.3174     0   2   4   6   8E+4   11.5   12  RT (min)      lcorbi_t3_pos_4_084_59    m/z 338.7853     0   1   2E+4   1   1.5  RT (min)      lcorbi_t3_pos_1_034_34    m/z 338.7965     0   2   4   6E+3   1   1.5  RT (min)      lcorbi_t3_pos_1_032_26    m/z 338.8060     0   1E+5   1   1.5  RT (min)      lcorbi_t3_pos_2_020_22    m/z 338.8490     0   2   4   6   8E+3   1   1.5  RT (min)      lcorbi_t3_pos_2_023_26    m/z 338.8609     0   2   4   6   8E+4   1   1.5  RT (min)      lcorbi_t3_pos_2_024_15    m/z 338.9102     0   1E+5   18   18.5  RT (min)       lcorbi_t3_pos_4_072_57    m/z 338.9820     0   1E+5   1   1.5  RT (min)      lcorbi_t3_pos_1_036_08    m/z 339.0794     0   1   2   3E+4   1   1.5  RT (min)      lcorbi_t3_pos_4_084_59    m/z 339.0840     0   1   2E+4   8   8.5  RT (min)      lcorbi_t3_pos_3_034_26    m/z 339.0929     0   1E+4   5   5.5  RT (min)      lcorbi_t3_pos_4_101_61    m/z 339.0971     0   2   4E+4   10   10.5  RT (min)       lcorbi_t3_pos_1_029_29    m/z 339.1257     0   2   4   6   8E+3   1.5   2  RT (min)      lcorbi_t3_pos_1_033_53    m/z 339.1325     0   2   4   6   8E+4   1   1.5  RT (min)      lcorbi_t3_pos_3_041_54    m/z 339.1544     0   1E+4   9.5   10  RT (min)      lcorbi_t3_pos_2_042_45    m/z 339.1703     0   2   4   6   8E+4   1   1.5  RT (min)       lcorbi_t3_pos_4_068_57    m/z 339.1704     0   1E+5   4.5   5  RT (min)       lcorbi_t3_pos_4_112_62    m/z 339.2229     0   1E+4   6   6.5  RT (min)      lcorbi_t3_pos_4_107_62    m/z 339.2319     0   1E+4   9   9.5  RT (min)      lcorbi_t3_pos_3_005_25    m/z 339.2894     0   1   2   3E+4   9.5   10   10.5  RT (min)       lcorbi_t3_pos_2_002_06    m/z 339.3089     0   2   4E+4   10.5   11  RT (min)      lcorbi_t3_pos_4_110_62    m/z 339.3261     0   1   2E+4   13   13.5  RT (min)       lcorbi_t3_pos_4_066_57    m/z 339.3401     0   1E+5   13   13.5  RT (min)      lcorbi_t3_pos_2_002_06    m/z 339.9369     0   1E+5   1   1.5  RT (min)      lcorbi_t3_pos_1_036_08    m/z 340.0478     0   1   2E+4   1   1.5  RT (min)      lcorbi_t3_pos_4_075_58    m/z 340.0876     0   1   2E+4   4.5   5  RT (min)      lcorbi_t3_pos_4_098_61    m/z 340.1068     0   2   4   6E+4   1.5   2  RT (min)      lcorbi_t3_pos_1_036_08    m/z 340.1181     0   1E+4   6   6.5  RT (min)      lcorbi_t3_pos_4_101_61    m/z 340.1544     0   1E+5   5.5   6  RT (min)       lcorbi_t3_pos_4_100_61    m/z 340.1669     0   1   2   3E+4   9   9.5   10  RT (min)      lcorbi_t3_pos_4_048_24    m/z 340.1738     0   1   2   3E+4   4.5   5  RT (min)      lcorbi_t3_pos_4_110_62    m/z 340.1904     0   1E+5   1   1.5  RT (min)       lcorbi_t3_pos_3_055_10    m/z 340.1967     0   2   4   6E+3   6   6.5  RT (min)      lcorbi_t3_pos_4_098_61    m/z 340.2399     0   1E+5   12.5   13  RT (min)       lcorbi_t3_pos_4_072_57    m/z 340.2479     0   1   2   3E+4   5   5.5  RT (min)      lcorbi_t3_pos_3_020_35    m/z 340.2484     0   1E+4   8.5   9  RT (min)      lcorbi_t3_pos_4_073_57    m/z 340.2716     0   1   2E+4   5   5.5  RT (min)      lcorbi_t3_pos_4_034_42    m/z 340.2929     0   1E+4   12   12.5  RT (min)      lcorbi_t3_pos_4_007_46    m/z 340.3322     0   2   4   6E+4   10.5   11  RT (min)       lcorbi_t3_pos_1_036_08    m/z 340.3768     0   2   4E+4   9.5   10  RT (min)       lcorbi_t3_pos_4_066_57    m/z 340.7824     0   1   2   3E+4   1   1.5  RT (min)      lcorbi_t3_pos_1_034_34    m/z 340.7935     0   1E+4   1   1.5  RT (min)      lcorbi_t3_pos_2_023_26    m/z 340.8031     0   1E+5   1   1.5  RT (min)      lcorbi_t3_pos_1_031_40    m/z 340.9320     0   1E+5   1   1.5  RT (min)       lcorbi_t3_pos_4_077_58    m/z 341.0116     0   2   4E+3   4.5   5  RT (min)      lcorbi_t3_pos_1_002_01    m/z 341.0822     0   1E+4   4.5   5  RT (min)      lcorbi_t3_pos_4_098_61    m/z 341.0994     0   1   2   3E+3   4.5   5  RT (min)      lcorbi_t3_pos_4_098_61    m/z 341.1230     0   1   2E+4   5   5.5  RT (min)       lcorbi_t3_pos_4_014_48    m/z 341.1319     0   1E+4   1   1.5  RT (min)      lcorbi_t3_pos_2_046_07    m/z 341.1384     0   1   2   3E+4   8   8.5  RT (min)       lcorbi_t3_pos_4_099_61    m/z 341.1393     0   2   4   6   8E+3   2.5   3  RT (min)      lcorbi_t3_pos_2_020_22    m/z 341.1579     0   1   2E+4   5.5   6  RT (min)      lcorbi_t3_pos_4_102_61    m/z 341.1748     0   2   4   6   8E+4   9.5   10  RT (min)      lcorbi_t3_pos_4_080_58    m/z 341.1942     0   1E+5   8.5   9  RT (min)      lcorbi_t3_pos_4_100_61    m/z 341.2090     0   2   4E+4   10.5   11  RT (min)       lcorbi_t3_pos_1_037_19    m/z 341.2171     0   2   4   6E+3   5   5.5  RT (min)      lcorbi_t3_pos_4_090_60    m/z 341.2434     0   1   2E+3   6.5   7  RT (min)      lcorbi_t3_pos_2_051_35    m/z 341.2471     0   1E+5   11.5   12   12.5  RT (min)       lcorbi_t3_pos_4_086_59    m/z 341.2571     0   1   5   5.5  RT (min)      lcorbi_t3_pos_4_089_59    m/z 341.2857     0   2   4   6   8E+4   10   10.5  RT (min)      lcorbi_t3_pos_4_086_59    m/z 341.3052     0   1   2E+5   14.5   15  RT (min)       lcorbi_t3_pos_1_032_26    m/z 341.3519     0   2   4   6   8E+4   13   13.5  RT (min)       lcorbi_t3_pos_2_002_06    m/z 341.3528     0   2   4   6E+4   10.5   11  RT (min)       lcorbi_t3_pos_4_089_59    m/z 341.3892     0   2   4E+4   10   10.5  RT (min)       lcorbi_t3_pos_4_106_62    m/z 341.7643     0   1E+4   6   6.5  RT (min)      lcorbi_t3_pos_4_087_59    m/z 341.9331     0   2   4E+4   1   1.5  RT (min)      lcorbi_t3_pos_1_031_40    m/z 342.0327     0   1E+4   4.5   5  RT (min)      lcorbi_t3_pos_4_066_57    m/z 342.0357     0   2   4E+4   1   1.5  RT (min)      lcorbi_t3_pos_1_036_08    m/z 342.0732     0   1   2   3   4E+3   1.5   2  RT (min)      lcorbi_t3_pos_2_009_50    m/z 342.0900     0   1   2E+4   11   11.5  RT (min)      lcorbi_t3_pos_4_074_58    m/z 342.1005     0   2   4   6   8E+4   9   9.5  RT (min)       lcorbi_t3_pos_4_099_61    m/z 342.1007     0   2   4   6   8E+3   1.5   2  RT (min)      lcorbi_t3_pos_1_005_42    m/z 342.1396     0   1E+4   1   1.5  RT (min)      lcorbi_t3_pos_1_021_18    m/z 342.1562     0   1   2   3   4E+3   3   3.5  RT (min)      lcorbi_t3_pos_4_034_42    m/z 342.1760     0   2   4E+3   4.5   5  RT (min)      lcorbi_t3_pos_4_066_57    m/z 342.1783     0   1   2E+4   9.5   10  RT (min)      lcorbi_t3_pos_1_032_26    m/z 342.1912     0   1E+4   7.5   8  RT (min)      lcorbi_t3_pos_3_001_49    m/z 342.2389     0   2   4   6   8E+3   5.5   6  RT (min)      lcorbi_t3_pos_4_087_59    m/z 342.2488     0   2   4   6   8E+3   5.5   6  RT (min)      lcorbi_t3_pos_4_066_57    m/z 342.2640     0   2   4   6E+4   8.5   9  RT (min)       lcorbi_t3_pos_4_068_57    m/z 342.2659     0   1E+5   5   5.5  RT (min)      lcorbi_t3_pos_4_085_59    m/z 342.2698     0   2   4   6E+4   11.5   12  RT (min)      lcorbi_t3_pos_4_084_59    m/z 342.3368     0   2   4   6E+4   10.5   11  RT (min)      lcorbi_t3_pos_1_050_13    m/z 342.3479     0   2   4E+4   10.5   11  RT (min)      lcorbi_t3_pos_1_036_08    m/z 342.8001     0   1   2E+4   1   1.5  RT (min)      lcorbi_t3_pos_1_031_40    m/z 342.8476     0   2   4   6E+4   1   1.5  RT (min)      lcorbi_t3_pos_1_036_08    m/z 342.8610     0   1   2   3E+3   1   1.5  RT (min)      lcorbi_t3_pos_1_002_01    m/z 342.9296     0   1E+4   1   1.5  RT (min)      lcorbi_t3_pos_4_077_58    m/z 343.0222     0   2   4E+4   18   18.5  RT (min)       lcorbi_t3_pos_4_112_62    m/z 343.0257     0   1E+4   1   1.5  RT (min)      lcorbi_t3_pos_4_005_33    m/z 343.0731     0   1E+4   9   9.5  RT (min)      lcorbi_t3_pos_4_099_61    m/z 343.0928     0   2   4   6   8E+4   1   1.5  RT (min)       lcorbi_t3_pos_1_043_06    m/z 343.1038     0   1   2E+4   9   9.5  RT (min)      lcorbi_t3_pos_4_099_61    m/z 343.1114     0   2   4E+4   2.5   3  RT (min)      lcorbi_t3_pos_4_042_09    m/z 343.1299     0   1E+4   5   5.5  RT (min)      lcorbi_t3_pos_4_082_59    m/z 343.1308     0   2   4   6   8E+4   1   1.5  RT (min)       lcorbi_t3_pos_3_021_39    m/z 343.1477     0   1E+4   4.5   5  RT (min)      lcorbi_t3_pos_4_066_57    m/z 343.1478     0   1   2.5   3  RT (min)      lcorbi_t3_pos_4_082_59    m/z 343.1921     0   2   4   6E+4   1   1.5  RT (min)      lcorbi_t3_pos_2_046_07    m/z 343.2093     0   1   2   3E+4   9.5   10  RT (min)       lcorbi_t3_pos_4_088_59    m/z 343.2246     0   2   4   6E+4   11   11.5  RT (min)       lcorbi_t3_pos_1_029_29    m/z 343.2633     0   1   2   3   4E+4   12   12.5  RT (min)       lcorbi_t3_pos_4_074_58    m/z 343.2674     0   1E+4   8.5   9  RT (min)      lcorbi_t3_pos_4_068_57    m/z 343.2684     0   2   4   6   8E+3   5   5.5  RT (min)      lcorbi_t3_pos_4_088_59    m/z 343.3109     0   1E+4   10   10.5  RT (min)      lcorbi_t3_pos_4_066_57    m/z 343.3321     0   1E+5   10.5   11  RT (min)       lcorbi_t3_pos_1_028_32    m/z 344.0142     0   1   2   3   4E+4   1   1.5  RT (min)       lcorbi_t3_pos_4_081_58    m/z 344.1047     0   1   2   3E+4   10   10.5  RT (min)      lcorbi_t3_pos_4_082_59    m/z 344.1425     0   1   2   3   4E+4   1.5   2  RT (min)      lcorbi_t3_pos_4_024_27    m/z 344.1494     0   1   2E+5   1   1.5  RT (min)       lcorbi_t3_pos_2_024_15    m/z 344.1575     0   2   4   6   8E+4   10   10.5  RT (min)      lcorbi_t3_pos_4_082_59    m/z 344.1681     0   2   4E+4   12   12.5  RT (min)      lcorbi_t3_pos_3_026_04    m/z 344.2150     0   1   2   3E+4   11.5   12  RT (min)      lcorbi_t3_pos_4_090_60    m/z 344.2280     0   1E+5   4.5   5  RT (min)       lcorbi_t3_pos_4_106_62    m/z 344.2432     0   2   4   6E+3   5   5.5  RT (min)      lcorbi_t3_pos_4_081_58    m/z 344.2439     0   1E+4   6.5   7  RT (min)      lcorbi_t3_pos_4_088_59    m/z 344.2545     0   1E+4   4.5   5  RT (min)      lcorbi_t3_pos_4_088_59    m/z 344.2587     0   2   4   6   8E+4   12   12.5  RT (min)       lcorbi_t3_pos_4_076_58    m/z 344.2968     0   1   2   3   4E+4   10   10.5  RT (min)       lcorbi_t3_pos_1_034_34    m/z 344.3354     0   1   2   3E+4   10.5   11  RT (min)      lcorbi_t3_pos_4_086_59    m/z 344.8447     0   1   2E+4   1   1.5  RT (min)      lcorbi_t3_pos_3_011_42    m/z 344.8660     0   1E+5   1   1.5  RT (min)      lcorbi_t3_pos_1_020_31    m/z 344.8756     0   1   2   3   4E+4   18   18.5  RT (min)       lcorbi_t3_pos_4_088_59    m/z 344.8757     0   1   2   3E+3   1   1.5  RT (min)       lcorbi_t3_pos_4_070_57    m/z 344.9477     0   2   4   6   8E+4   18   18.5  RT (min)       lcorbi_t3_pos_4_078_58    m/z 344.9839     0   1   2   3E+4   1   1.5  RT (min)      lcorbi_t3_pos_1_044_25    m/z 345.1295     0   1E+4   2.5   3  RT (min)      lcorbi_t3_pos_4_069_57    m/z 345.1365     0   1E+4   8   8.5  RT (min)      lcorbi_t3_pos_4_073_57    m/z 345.1576     0   2   4   6   8E+4   4.5   5  RT (min)       lcorbi_t3_pos_4_014_48    m/z 345.1698     0   1   2   3E+4   8   8.5  RT (min)       lcorbi_t3_pos_4_101_61    m/z 345.1714     0   2   4   6   8E+4   10.5   11  RT (min)      lcorbi_t3_pos_1_028_32    m/z 345.2314     0   2   4   6E+3   4.5   5  RT (min)      lcorbi_t3_pos_4_014_48    m/z 345.2831     0   1   2E+4   9   9.5  RT (min)      lcorbi_t3_pos_4_080_58    m/z 345.3001     0   1   2   3E+4   12.5   13  RT (min)       lcorbi_t3_pos_4_077_58    m/z 345.3022     0   1   2   3   4E+4   9.5   10  RT (min)      lcorbi_t3_pos_4_089_59    m/z 345.9450     0   2   4E+4   1   1.5  RT (min)      lcorbi_t3_pos_1_036_08    m/z 346.0075     0   2   4   6E+4   8   8.5  RT (min)      lcorbi_t3_pos_1_051_09    m/z 346.0382     0   1   1.5   2  RT (min)      lcorbi_t3_pos_1_029_29    m/z 346.0549     0   1   2   3E+3   1   1.5  RT (min)      lcorbi_t3_pos_3_024_07    m/z 346.0549     0   2   4   6E+3   4   4.5  RT (min)      lcorbi_t3_pos_4_098_61    m/z 346.1010     0   1E+5   1   1.5  RT (min)       lcorbi_t3_pos_4_093_60    m/z 346.2953     0   1   2E+4   7.5   8  RT (min)       lcorbi_t3_pos_4_081_58    m/z 346.3727     0   2   4   6E+4   12   12.5  RT (min)       lcorbi_t3_pos_1_036_08    m/z 346.8808     0   1   2   3E+4   1   1.5  RT (min)      lcorbi_t3_pos_1_023_02    m/z 346.9087     0   1   2   3   4E+4   1   1.5  RT (min)       lcorbi_t3_pos_4_074_58    m/z 346.9528     0   2   4E+4   1   1.5  RT (min)      lcorbi_t3_pos_4_073_57    m/z 347.0127     0   2   4   6E+4   1   1.5  RT (min)      lcorbi_t3_pos_4_080_58    m/z 347.0302     0   1E+5   9   9.5  RT (min)       lcorbi_t3_pos_1_036_08    m/z 347.1107     0   1   2   3   4E+3   2  RT (min)       lcorbi_t3_pos_4_092_60    m/z 347.1315     0   1   2.5   3  RT (min)      lcorbi_t3_pos_4_093_60    m/z 347.1340     0   1   2E+4   10   10.5  RT (min)       lcorbi_t3_pos_4_081_58    m/z 347.1451     0   2   4E+4   4.5   5  RT (min)       lcorbi_t3_pos_4_066_57    m/z 347.1500     0   2   4   6   8E+3   1   1.5  RT (min)      lcorbi_t3_pos_1_003_45    m/z 347.1702     0   1   2   3E+4   9   9.5  RT (min)       lcorbi_t3_pos_4_066_57    m/z 347.2906     0   1E+4   7.5   8  RT (min)      lcorbi_t3_pos_3_016_21    m/z 347.3351     0   2   4E+4   11   11.5  RT (min)      lcorbi_t3_pos_1_028_32    m/z 348.0054     0   2   4   6E+4   8   8.5  RT (min)      lcorbi_t3_pos_1_051_09    m/z 348.0197     0   1E+4   4.5   5  RT (min)      lcorbi_t3_pos_4_074_58    m/z 348.0706     0   1E+5   1.5   2  RT (min)       lcorbi_t3_pos_1_054_48    m/z 348.0797     0   2   4   6E+5   1   1.5  RT (min)       lcorbi_t3_pos_4_006_12    m/z 348.1091     0   1E+5   1   1.5  RT (min)       lcorbi_t3_pos_4_097_60    m/z 348.1476     0   1   1   1.5  RT (min)       lcorbi_t3_pos_4_013_36    m/z 348.1483     0   2   4   6E+3   4.5   5  RT (min)      lcorbi_t3_pos_4_098_61    m/z 348.1761     0   1   2   3   4E+4   9.5   10  RT (min)      lcorbi_t3_pos_1_028_32    m/z 348.2283     0   1E+4   6   6.5  RT (min)       lcorbi_t3_pos_4_089_59    m/z 348.2616     0   1   2   3E+4   12.5   13  RT (min)      lcorbi_t3_pos_3_005_25    m/z 348.2616     0   2   4   6E+4   11   11.5  RT (min)      lcorbi_t3_pos_4_108_62    m/z 348.3476     0   1E+5   13   13.5  RT (min)       lcorbi_t3_pos_4_089_59    m/z 348.7729     0   2   4   6E+4   5   5.5  RT (min)       lcorbi_t3_pos_4_004_14    m/z 348.8141     0   1   2   3   4E+4   1   1.5  RT (min)      lcorbi_t3_pos_1_016_05    m/z 348.8254     0   2   4   6E+3   1   1.5  RT (min)      lcorbi_t3_pos_1_029_29    m/z 348.9190     0   1   2E+5   18   18.5  RT (min)      lcorbi_t3_pos_4_073_57    m/z 348.9191     0   2   4E+4   10   10.5  RT (min)      lcorbi_t3_pos_1_026_17    m/z 349.0458     0   1   1   1.5  RT (min)      lcorbi_t3_pos_4_078_58    m/z 349.0602     0   1   2   3   4E+4   10   10.5  RT (min)      lcorbi_t3_pos_4_082_59    m/z 349.0739     0   1   1.5   2  RT (min)      lcorbi_t3_pos_2_028_24    m/z 349.0897     0   1E+4   4.5   5  RT (min)      lcorbi_t3_pos_4_090_60    m/z 349.1106     0   1   2   3   4E+4   1   1.5   2  RT (min)       lcorbi_t3_pos_4_068_57    m/z 349.1834     0   1   2E+4   4.5   5  RT (min)      lcorbi_t3_pos_4_111_62    m/z 349.1840     0   1   2   3   4E+4   7   7.5  RT (min)      lcorbi_t3_pos_4_101_61    m/z 349.2020     0   1E+4   5   5.5  RT (min)       lcorbi_t3_pos_4_074_58    m/z 349.2123     0   1   2   3   4E+4   6.5   7  RT (min)       lcorbi_t3_pos_4_085_59    m/z 349.2140     0   2   4   6E+4   12   12.5  RT (min)       lcorbi_t3_pos_4_074_58    m/z 349.2308     0   1E+5   0   1  RT (min)       lcorbi_t3_pos_4_089_59    m/z 349.2467     0   1E+4   5.5   6  RT (min)       lcorbi_t3_pos_4_088_59    m/z 349.2649     0   2   4   6E+4   5.5   6  RT (min)       lcorbi_t3_pos_4_082_59    m/z 349.2747     0   1   2   3E+4   5   5.5  RT (min)       lcorbi_t3_pos_4_032_40    m/z 349.3223     0   1E+5   12   12.5  RT (min)      lcorbi_t3_pos_4_087_59    m/z 349.3579     0   1   2   3   4E+4   10.5   11  RT (min)      lcorbi_t3_pos_1_023_02    m/z 349.7037     0   2   4E+3   5   5.5  RT (min)      lcorbi_t3_pos_4_074_58    m/z 349.7484     0   2   4   6   8E+3   5.5   6  RT (min)      lcorbi_t3_pos_4_086_59    m/z 349.7666     0   1   2E+4   5.5   6  RT (min)      lcorbi_t3_pos_4_087_59    m/z 349.9392     0   2   4   6   8E+4   1   1.5  RT (min)      lcorbi_t3_pos_2_020_22    m/z 350.0849     0   1   2.5   3  RT (min)      lcorbi_t3_pos_3_029_13    m/z 350.2158     0   1E+4   6.5   7  RT (min)      lcorbi_t3_pos_4_083_59    m/z 350.2331     0   1E+4   7   7.5  RT (min)      lcorbi_t3_pos_4_005_33    m/z 350.2681     0   2   4   6E+3   5.5   6  RT (min)      lcorbi_t3_pos_4_084_59    m/z 350.2903     0   1   2   3E+4   9.5   10  RT (min)      lcorbi_t3_pos_4_098_61    m/z 350.7437     0   1   2   3   4E+4   1   1.5  RT (min)      lcorbi_t3_pos_4_068_57    m/z 350.8112     0   1   2   3   4E+4   1   1.5  RT (min)      lcorbi_t3_pos_1_029_29    m/z 351.0305     0   1E+4   2   2.5  RT (min)      lcorbi_t3_pos_1_044_25    m/z 351.0861     0   1E+5   1   1.5  RT (min)       lcorbi_t3_pos_4_084_59    m/z 351.1341     0   1   2   3   4E+4   7.5   8  RT (min)      lcorbi_t3_pos_4_099_61    m/z 351.1866     0   2   4E+4   7   7.5  RT (min)      lcorbi_t3_pos_2_044_21    m/z 351.1940     0   1   7   7.5  RT (min)      lcorbi_t3_pos_4_098_61    m/z 351.2297     0   1   2   3   4E+4   12.5   13  RT (min)       lcorbi_t3_pos_4_074_58    m/z 351.2324     0   1   2E+4   7   7.5  RT (min)      lcorbi_t3_pos_1_037_19    m/z 351.2409     0   1   2   3   4E+4   11.5   12  RT (min)       lcorbi_t3_pos_3_054_15    m/z 351.2506     0   1   6   6.5  RT (min)      lcorbi_t3_pos_4_105_61    m/z 351.2508     0   1   11.5   12  RT (min)       lcorbi_t3_pos_4_089_59    m/z 351.2985     0   2   4   6   8E+4   12.5   13  RT (min)       lcorbi_t3_pos_4_074_58    m/z 351.3799     0   2   4E+4   10.5   11  RT (min)       lcorbi_t3_pos_1_036_08    m/z 351.9752     0   1   2E+4   1.5   2  RT (min)      lcorbi_t3_pos_1_036_08    m/z 351.9896     0   1E+4   1   1.5  RT (min)      lcorbi_t3_pos_4_005_33    m/z 352.0552     0   1   2E+4   1.5   2  RT (min)      lcorbi_t3_pos_1_032_26    m/z 352.1289     0   1E+4   1.5   2  RT (min)      lcorbi_t3_pos_3_036_37    m/z 352.1343     0   1   2E+4   9.5   10  RT (min)      lcorbi_t3_pos_3_054_15    m/z 352.1657     0   1   5.5   6  RT (min)      lcorbi_t3_pos_4_104_61    m/z 352.2049     0   2   4   6E+4   7   7.5  RT (min)      lcorbi_t3_pos_3_023_05    m/z 352.2291     0   1   2E+4   9.5   10  RT (min)      lcorbi_t3_pos_4_103_61    m/z 352.2331     0   2   4   6   8E+4   4.5   5  RT (min)       lcorbi_t3_pos_4_090_60    m/z 352.2460     0   1   2E+4   11.5   12  RT (min)      lcorbi_t3_pos_1_003_45    m/z 352.2849     0   1E+4   11.5   12  RT (min)       lcorbi_t3_pos_4_074_58    m/z 352.3060     0   2   4   6E+4   10.5   11  RT (min)       lcorbi_t3_pos_1_032_26    m/z 352.3212     0   1E+5   11.5   12  RT (min)       lcorbi_t3_pos_2_002_06    m/z 352.3577     0   2   4E+4   13   14  RT (min)       lcorbi_t3_pos_2_002_06    m/z 352.7408     0   1   2   3E+4   1   1.5  RT (min)      lcorbi_t3_pos_2_028_24    m/z 352.9165     0   1   1   1.5  RT (min)      lcorbi_t3_pos_1_002_01    m/z 353.0845     0   2   4E+4   1   1.5  RT (min)      lcorbi_t3_pos_3_021_39    m/z 353.0892     0   1   2   3E+4   9   9.5  RT (min)       lcorbi_t3_pos_4_074_58    m/z 353.1249     0   1E+5   1   1.5  RT (min)       lcorbi_t3_pos_4_095_60    m/z 353.1345     0   1E+4   5   5.5  RT (min)       lcorbi_t3_pos_4_066_57    m/z 353.1864     0   1   4.5   5  RT (min)      lcorbi_t3_pos_4_094_60    m/z 353.2302     0   2   4   6E+4   10   11  RT (min)       lcorbi_t3_pos_4_076_58    m/z 353.2365     0   1   2   3E+4   4.5   5  RT (min)      lcorbi_t3_pos_4_090_60    m/z 353.2800     0   1   2E+5   9.5   10  RT (min)       lcorbi_t3_pos_1_004_24    m/z 353.3053     0   2   4E+4   14.5   15  RT (min)       lcorbi_t3_pos_4_066_57    m/z 353.3097     0   1E+5   11.5   12  RT (min)       lcorbi_t3_pos_4_094_60    m/z 353.3416     0   2   4E+4   13.5   14  RT (min)       lcorbi_t3_pos_4_113_62    m/z 353.3528     0   1   2   3E+4   10.5   11  RT (min)      lcorbi_t3_pos_4_085_59    m/z 353.9012     0   1   2   3E+4   1   1.5  RT (min)      lcorbi_t3_pos_1_020_31    m/z 354.0413     0   2   4   6E+4   1.5   2  RT (min)       lcorbi_t3_pos_1_035_28    m/z 354.0700     0   1E+4   1   1.5  RT (min)      lcorbi_t3_pos_2_046_07    m/z 354.1337     0   1E+4   5   5.5  RT (min)      lcorbi_t3_pos_4_099_61    m/z 354.1451     0   1E+4   6   6.5  RT (min)       lcorbi_t3_pos_4_102_61    m/z 354.1661     0   1   2   3   4E+4   4.5   5  RT (min)       lcorbi_t3_pos_4_088_59    m/z 354.1889     0   1   2.5   3  RT (min)      lcorbi_t3_pos_4_090_60    m/z 354.1974     0   1   4.5   5  RT (min)      lcorbi_t3_pos_4_082_59    m/z 354.2125     0   1   2   3E+4   6.5   7  RT (min)       lcorbi_t3_pos_1_051_09    m/z 354.2125     0   1E+4   10   10.5  RT (min)      lcorbi_t3_pos_2_012_01    m/z 354.7800     0   2   4   6E+4   1   1.5  RT (min)      lcorbi_t3_pos_1_036_08    m/z 354.8948     0   1   2E+5   1   1.5  RT (min)       lcorbi_t3_pos_1_020_31    m/z 355.0116     0   1   2E+4   1   1.5  RT (min)       lcorbi_t3_pos_4_006_12    m/z 355.0253     0   1   2E+5   1.5   2  RT (min)       lcorbi_t3_pos_2_040_25    m/z 355.0579     0   1   2E+5   1   1.5  RT (min)      lcorbi_t3_pos_3_024_07    m/z 355.0850     0   1E+4   8   8.5  RT (min)      lcorbi_t3_pos_4_102_61    m/z 355.1212     0   1   2   3   4E+5   1   1.5  RT (min)      lcorbi_t3_pos_1_031_40    m/z 355.1306     0   1   2   3   4E+4   7   7.5  RT (min)      lcorbi_t3_pos_4_072_57    m/z 355.1493     0   1   6   6.5  RT (min)      lcorbi_t3_pos_3_054_15    m/z 355.1600     0   1   2E+4   4.5   5  RT (min)      lcorbi_t3_pos_4_090_60    m/z 355.1695     0   1   4.5   5  RT (min)      lcorbi_t3_pos_4_090_60    m/z 355.1790     0   1   11.5   12  RT (min)      lcorbi_t3_pos_4_082_59    m/z 355.2008     0   1E+5   4.5   5  RT (min)       lcorbi_t3_pos_4_090_60    m/z 355.2099     0   1E+4   8.5   9  RT (min)      lcorbi_t3_pos_1_021_18    m/z 355.2246     0   1E+5   11.5   12  RT (min)       lcorbi_t3_pos_4_087_59    m/z 355.2328     0   2   4   6   8E+3   5   5.5  RT (min)      lcorbi_t3_pos_4_066_57    m/z 355.2341     0   1E+5   1  RT (min)       lcorbi_t3_pos_4_084_59    m/z 355.2510     0   1E+5   1   1.5  RT (min)      lcorbi_t3_pos_3_027_27    m/z 355.3039     0   1   2E+5   12   12.5  RT (min)      lcorbi_t3_pos_4_087_59    m/z 355.3209     0   2   4   6E+4   13   13.5  RT (min)       lcorbi_t3_pos_4_083_59    m/z 355.7625     0   1   2E+4   5   5.5  RT (min)       lcorbi_t3_pos_4_089_59    m/z 355.9473     0   2   4   6E+4   1   1.5  RT (min)      lcorbi_t3_pos_3_048_30    m/z 356.0613     0   1   1   1.5  RT (min)      lcorbi_t3_pos_4_092_60    m/z 356.1029     0   1E+5   1   1.5  RT (min)      lcorbi_t3_pos_3_051_12    m/z 356.1229     0   1E+4   7   7.5  RT (min)      lcorbi_t3_pos_4_068_57    m/z 356.1246     0   1   1   1.5  RT (min)      lcorbi_t3_pos_3_051_12    m/z 356.1575     0   1   2E+4   8   8.5  RT (min)      lcorbi_t3_pos_4_101_61    m/z 356.1917     0   1E+4   5   5.5  RT (min)      lcorbi_t3_pos_4_090_60    m/z 356.2303     0   2   4   6   8E+4   8   8.5  RT (min)      lcorbi_t3_pos_1_014_03    m/z 356.2546     0   2   4   6E+4   5.5   6  RT (min)       lcorbi_t3_pos_4_088_59    m/z 356.2642     0   1E+4   5   5.5  RT (min)      lcorbi_t3_pos_4_088_59    m/z 356.2797     0   2   4   6   8E+4   7.5   8  RT (min)       lcorbi_t3_pos_4_098_61    m/z 356.2797     0   1   2   3   4E+4   9   9.5  RT (min)       lcorbi_t3_pos_4_071_57    m/z 356.2798     0   2   4   6   8E+4   10.5   11  RT (min)       lcorbi_t3_pos_4_087_59    m/z 356.2878     0   1   2E+4   12   12.5  RT (min)      lcorbi_t3_pos_2_028_24    m/z 356.3272     0   2   4   6E+4   4.5   5  RT (min)       lcorbi_t3_pos_4_082_59    m/z 356.3717     0   1   2   3   4E+4   13   13.5  RT (min)      lcorbi_t3_pos_2_002_06    m/z 356.3718     0   1   2   3E+4   9.5   10  RT (min)       lcorbi_t3_pos_4_074_58    m/z 356.7465     0   2   4   6E+3   6   6.5  RT (min)      lcorbi_t3_pos_3_020_35    m/z 356.7563     0   1   2E+4   5.5   6  RT (min)      lcorbi_t3_pos_4_083_59    m/z 356.7771     0   2   4   6E+4   1   1.5  RT (min)      lcorbi_t3_pos_1_036_08    m/z 356.8487     0   1E+5   18.5   19  RT (min)       lcorbi_t3_pos_4_091_60    m/z 356.9093     0   2   4   6   8E+4   18   18.5  RT (min)       lcorbi_t3_pos_4_084_59    m/z 357.0014     0   1E+4   1   1.5  RT (min)      lcorbi_t3_pos_4_075_58    m/z 357.0083     0   1E+4   1   1.5  RT (min)      lcorbi_t3_pos_3_008_20    m/z 357.0816     0   1   2   3E+4   8   8.5  RT (min)      lcorbi_t3_pos_1_051_09    m/z 357.1008     0   1   2   3E+4   1.5   2  RT (min)       lcorbi_t3_pos_2_037_27    m/z 357.2147     0   2   4   6E+3   4.5   5  RT (min)      lcorbi_t3_pos_4_018_01    m/z 357.2335     0   1   2   3E+4   6   6.5  RT (min)      lcorbi_t3_pos_4_109_62    m/z 357.2335     0   2   4E+3   8   8.5  RT (min)      lcorbi_t3_pos_1_021_18    m/z 357.2790     0   2   4E+4   10   10.5  RT (min)       lcorbi_t3_pos_4_080_58    m/z 357.2830     0   1E+4   7.5   8  RT (min)      lcorbi_t3_pos_4_101_61    m/z 357.2862     0   2   4   6E+4   5   5.5  RT (min)       lcorbi_t3_pos_1_031_40    m/z 357.3841     0   1   2   3E+4   10   10.5  RT (min)      lcorbi_t3_pos_4_066_57    m/z 357.8720     0   1   2   3E+4   1   1.5  RT (min)      lcorbi_t3_pos_4_069_57    m/z 357.9130     0   2   4   6E+3   1   1.5  RT (min)      lcorbi_t3_pos_4_068_57    m/z 357.9921     0   1E+5   1   1.5  RT (min)      lcorbi_t3_pos_1_036_08    m/z 358.1041     0   1   2   3E+5   1.5   2  RT (min)       lcorbi_t3_pos_4_095_60    m/z 358.1754     0   1E+4   9.5   10  RT (min)      lcorbi_t3_pos_1_039_23    m/z 358.1885     0   1E+4   9   9.5   10  RT (min)      lcorbi_t3_pos_4_090_60    m/z 358.2015     0   2   4   6E+4   0   0.5  RT (min)      lcorbi_t3_pos_3_054_15    m/z 358.2015     0   1   2   3E+6   15   15.5  RT (min)       lcorbi_t3_pos_4_040_21    m/z 358.2077     0   1E+4   4.5   5  RT (min)      lcorbi_t3_pos_4_079_58    m/z 358.2139     0   1   9.5   10  RT (min)      lcorbi_t3_pos_4_090_60    m/z 358.2276     0   1   9.5   10  RT (min)      lcorbi_t3_pos_1_039_23    m/z 358.2338     0   1   2E+4   4.5   5  RT (min)       lcorbi_t3_pos_4_086_59    m/z 358.2436     0   1   12   12.5  RT (min)       lcorbi_t3_pos_1_038_30    m/z 358.2437     0   1E+4   5   5.5  RT (min)      lcorbi_t3_pos_4_106_62    m/z 358.2590     0   2   4   6   8E+3   8   8.5  RT (min)      lcorbi_t3_pos_4_075_58    m/z 358.2702     0   2   4   6   8E+4   4.5   5  RT (min)       lcorbi_t3_pos_4_088_59    m/z 358.2743     0   2   4   6   8E+4   12.5   13  RT (min)       lcorbi_t3_pos_4_067_57    m/z 358.2847     0   1   2E+4   10   10.5  RT (min)      lcorbi_t3_pos_4_078_58    m/z 358.2895     0   1E+4   5   5.5  RT (min)      lcorbi_t3_pos_1_031_40    m/z 358.2953     0   1   2E+4   9   9.5  RT (min)       lcorbi_t3_pos_4_073_57    m/z 358.3035     0   1   2   3   4E+4   12   12.5  RT (min)       lcorbi_t3_pos_3_012_31    m/z 358.3084     0   1   2E+5   13   13.5  RT (min)       lcorbi_t3_pos_2_044_21    m/z 358.3125     0   2   4   6E+4   9   9.5  RT (min)       lcorbi_t3_pos_4_111_62    m/z 358.3429     0   1E+5   10   10.5  RT (min)       lcorbi_t3_pos_3_027_27    m/z 358.7743     0   1   1   1.5  RT (min)      lcorbi_t3_pos_1_023_02    m/z 358.8216     0   1   2E+4   1   1.5  RT (min)      lcorbi_t3_pos_1_031_40    m/z 358.8429     0   1E+5   1   1.5  RT (min)       lcorbi_t3_pos_1_029_29    m/z 358.8541     0   2   4E+3   1   1.5  RT (min)      lcorbi_t3_pos_1_001_49    m/z 358.9967     0   1E+5   1   1.5  RT (min)       lcorbi_t3_pos_2_051_35    m/z 358.9997     0   2   4   6E+4   18   18.5  RT (min)       lcorbi_t3_pos_4_078_58    m/z 359.0440     0   1E+5   1   1.5  RT (min)       lcorbi_t3_pos_4_075_58    m/z 359.0971     0   1   2E+4   9   9.5  RT (min)       lcorbi_t3_pos_1_035_28    m/z 359.1002     0   1   2   3E+4   6.5   7  RT (min)      lcorbi_t3_pos_4_099_61    m/z 359.1217     0   2   4E+3   4.5   5  RT (min)      lcorbi_t3_pos_4_070_57    m/z 359.1443     0   1   2.5   3  RT (min)       lcorbi_t3_pos_4_090_60    m/z 359.1522     0   2   4   6E+4   5   5.5  RT (min)      lcorbi_t3_pos_4_048_24    m/z 359.1678     0   2   4   6E+3   6.5   7  RT (min)      lcorbi_t3_pos_4_074_58    m/z 359.1716     0   1   4.5   5  RT (min)      lcorbi_t3_pos_4_090_60    m/z 359.2049     0   1   2   3   4E+5   12   12.5  RT (min)      lcorbi_t3_pos_4_101_61    m/z 359.2049     0   2   4   6   8E+5   15   15.5  RT (min)      lcorbi_t3_pos_4_089_59    m/z 359.2164     0   1   10   10.5  RT (min)      lcorbi_t3_pos_4_091_60    m/z 359.2291     0   1E+4   5.5   6  RT (min)      lcorbi_t3_pos_4_085_59    m/z 359.2330     0   2   4   6E+3   6   6.5  RT (min)      lcorbi_t3_pos_4_089_59    m/z 359.2406     0   1E+4   9.5   10  RT (min)      lcorbi_t3_pos_4_090_60    m/z 359.2736     0   1E+4   4.5   5  RT (min)      lcorbi_t3_pos_4_085_59    m/z 359.3158     0   2   4E+4   12.5   13  RT (min)      lcorbi_t3_pos_2_012_01    m/z 359.3568     0   1   2E+5   14.5   15  RT (min)       lcorbi_t3_pos_1_036_08    m/z 359.3842     0   2   4   6   8E+4   13   13.5  RT (min)       lcorbi_t3_pos_2_037_27    m/z 359.8691     0   1   2   3E+4   1   1.5  RT (min)      lcorbi_t3_pos_4_071_57    m/z 360.0252     0   1E+4   1.5   2  RT (min)      lcorbi_t3_pos_4_079_58    m/z 360.0748     0   1   2   3E+4   1   1.5  RT (min)      lcorbi_t3_pos_1_028_32    m/z 360.1138     0   1   1.5   2  RT (min)       lcorbi_t3_pos_1_036_08    m/z 360.1234     0   2   4   6E+3   1   1.5  RT (min)       lcorbi_t3_pos_2_003_19    m/z 360.1380     0   1   2   3   4E+5   1   1.5  RT (min)      lcorbi_t3_pos_1_036_08    m/z 360.1569     0   1E+4   6   6.5  RT (min)      lcorbi_t3_pos_3_048_30    m/z 360.1807     0   1E+5   10   10.5  RT (min)      lcorbi_t3_pos_1_032_26    m/z 360.2080     0   1E+5   14.5   15  RT (min)      lcorbi_t3_pos_4_073_57    m/z 360.2244     0   1   1   1.5  RT (min)       lcorbi_t3_pos_4_012_37    m/z 360.3626     0   1E+5   10.5   11  RT (min)       lcorbi_t3_pos_1_050_13    m/z 360.8401     0   2   4   6   8E+4   1   1.5  RT (min)      lcorbi_t3_pos_1_029_29    m/z 361.0201     0   1E+5   1   1.5  RT (min)      lcorbi_t3_pos_4_080_58    m/z 361.0780     0   1   2   3   4E+5   1   1.5  RT (min)      lcorbi_t3_pos_3_010_44    m/z 361.0919     0   1   2   3   4E+4   7   7.5  RT (min)      lcorbi_t3_pos_2_022_38    m/z 361.1090     0   1   2E+4   6.5   7  RT (min)       lcorbi_t3_pos_4_005_33    m/z 361.1197     0   2   4E+5   1   1.5  RT (min)       lcorbi_t3_pos_2_037_27    m/z 361.1388     0   1   2   3E+4   5   5.5  RT (min)      lcorbi_t3_pos_4_110_62    m/z 361.1536     0   2   4E+4   1   1.5  RT (min)      lcorbi_t3_pos_4_071_57    m/z 361.1841     0   1   2   3E+4   10   10.5  RT (min)      lcorbi_t3_pos_4_086_59    m/z 361.2010     0   1E+4   7   7.5  RT (min)      lcorbi_t3_pos_4_079_58    m/z 361.2103     0   1   2E+4   9.5   10  RT (min)      lcorbi_t3_pos_4_083_59    m/z 361.2711     0   1   2   3E+5   1   1.5  RT (min)       lcorbi_t3_pos_3_055_10    m/z 361.2740     0   1   2   3E+4   14.5   15  RT (min)      lcorbi_t3_pos_4_052_06    m/z 361.3660     0   1   2E+4   10.5   11  RT (min)      lcorbi_t3_pos_4_105_61    m/z 361.3999     0   1E+5   13.5   14  RT (min)       lcorbi_t3_pos_1_003_45    m/z 362.1108     0   2   4E+4   1.5   2  RT (min)      lcorbi_t3_pos_3_048_30    m/z 362.2075     0   2   4E+3   6.5   7  RT (min)      lcorbi_t3_pos_4_066_57    m/z 362.2207     0   2   4   6E+3   6   6.5  RT (min)      lcorbi_t3_pos_4_087_59    m/z 362.2385     0   2   4   6E+4   7.5   8  RT (min)      lcorbi_t3_pos_1_028_32    m/z 362.2773     0   1   2E+4   11   11.5  RT (min)      lcorbi_t3_pos_4_039_35    m/z 362.2773     0   2   4   6E+4   12.5   13  RT (min)       lcorbi_t3_pos_4_087_59    m/z 362.2902     0   2   4   6E+4   7   7.5  RT (min)       lcorbi_t3_pos_4_081_58    m/z 362.3307     0   1E+5   13   13.5  RT (min)      lcorbi_t3_pos_2_002_06    m/z 362.3396     0   2   4   6E+4   13.5   14  RT (min)      lcorbi_t3_pos_2_044_21    m/z 362.3758     0   1   2   3   4E+5   14.5   15  RT (min)       lcorbi_t3_pos_1_036_08    m/z 362.3765     0   1E+5   10.5   11  RT (min)       lcorbi_t3_pos_4_094_60    m/z 362.7704     0   1   2E+4   5.5   6  RT (min)       lcorbi_t3_pos_4_086_59    m/z 362.8274     0   1   2   3E+4   1   1.5  RT (min)      lcorbi_t3_pos_4_021_25    m/z 362.8660     0   1   2   3   4E+4   11.5   12  RT (min)      lcorbi_t3_pos_2_017_51    m/z 362.8660     0   1E+4   1   1.5  RT (min)       lcorbi_t3_pos_4_070_57    m/z 362.8863     0   1   2   3E+3   1   1.5  RT (min)       lcorbi_t3_pos_4_067_57    m/z 362.8863     0   1   2   3   4E+4   18   18.5  RT (min)       lcorbi_t3_pos_4_108_62    m/z 362.9133     0   1E+4   1   1.5  RT (min)      lcorbi_t3_pos_1_031_40    m/z 362.9396     0   1E+4   1   1.5  RT (min)      lcorbi_t3_pos_3_012_31    m/z 362.9936     0   1   2   3   4E+3   4.5   5  RT (min)      lcorbi_t3_pos_1_029_29    m/z 363.0361     0   1   2   3E+4   1   1.5  RT (min)       lcorbi_t3_pos_4_078_58    m/z 363.1039     0   1E+4   5   5.5  RT (min)      lcorbi_t3_pos_4_104_61    m/z 363.1071     0   1E+5   1.5   2  RT (min)      lcorbi_t3_pos_3_048_30    m/z 363.1116     0   2   4E+4   10   10.5  RT (min)       lcorbi_t3_pos_4_079_58    m/z 363.1440     0   1   2E+4   8   8.5  RT (min)       lcorbi_t3_pos_4_098_61    m/z 363.1569     0   1   2E+5   12   13  RT (min)       lcorbi_t3_pos_3_005_25    m/z 363.1569     0   1   2   3   4E+5   16  RT (min)       lcorbi_t3_pos_4_102_61    m/z 363.2167     0   2   4   6E+4   7   7.5  RT (min)       lcorbi_t3_pos_4_079_58    m/z 363.2281     0   1E+4   10.5   11  RT (min)      lcorbi_t3_pos_4_099_61    m/z 363.2414     0   2   4   6   8E+3   7.5   8  RT (min)      lcorbi_t3_pos_4_104_61    m/z 363.2449     0   2   4   6E+4   5   5.5  RT (min)      lcorbi_t3_pos_4_088_59    m/z 363.2719     0   2   4   6E+3   5.5   6  RT (min)      lcorbi_t3_pos_4_088_59    m/z 363.2745     0   2   4   6   8E+4   12.5   13  RT (min)       lcorbi_t3_pos_4_074_58    m/z 363.2936     0   1E+4   7   7.5  RT (min)      lcorbi_t3_pos_4_096_60    m/z 364.0146     0   1   2E+4   4.5   5  RT (min)      lcorbi_t3_pos_4_082_59    m/z 364.0316     0   1   2E+4   1   1.5  RT (min)      lcorbi_t3_pos_2_024_15    m/z 364.2201     0   1E+4   7   7.5  RT (min)      lcorbi_t3_pos_4_077_58    m/z 364.2236     0   2   4   6E+4   11   11.5  RT (min)       lcorbi_t3_pos_4_096_60    m/z 364.3170     0   1E+4   10   10.5  RT (min)      lcorbi_t3_pos_2_030_14    m/z 364.3231     0   1   2   3E+4   12   12.5  RT (min)       lcorbi_t3_pos_1_008_36    m/z 364.3324     0   1E+4   8   8.5  RT (min)      lcorbi_t3_pos_2_035_34    m/z 364.7242     0   1   2   3E+4   1   1.5  RT (min)      lcorbi_t3_pos_2_028_24    m/z 364.8295     0   1   2E+4   1   1.5  RT (min)      lcorbi_t3_pos_2_024_15    m/z 364.9164     0   1   2   3E+4   1   1.5  RT (min)      lcorbi_t3_pos_1_028_32    m/z 364.9308     0   2   4   6E+4   1   1.5  RT (min)      lcorbi_t3_pos_4_068_57    m/z 365.0692     0   1   1   1.5  RT (min)      lcorbi_t3_pos_3_054_15    m/z 365.1361     0   1E+5   10   10.5  RT (min)       lcorbi_t3_pos_4_088_59    m/z 365.1446     0   1   10   10.5  RT (min)      lcorbi_t3_pos_4_082_59    m/z 365.1632     0   1   2   3   4E+4   9   9.5  RT (min)      lcorbi_t3_pos_4_071_57    m/z 365.1936     0   1E+5   11.5   12  RT (min)       lcorbi_t3_pos_4_082_59    m/z 365.2074     0   1   2E+4   7   7.5  RT (min)      lcorbi_t3_pos_2_029_43    m/z 365.2776     0   1E+5   9.5   10  RT (min)       lcorbi_t3_pos_4_089_59    m/z 366.1090     0   1E+5   1   1.5  RT (min)       lcorbi_t3_pos_4_071_57    m/z 366.1759     0   1   2   3E+4   1   1.5  RT (min)       lcorbi_t3_pos_4_072_57    m/z 366.2640     0   1E+4   10   10.5  RT (min)       lcorbi_t3_pos_2_030_14    m/z 366.2809     0   1E+4   9.5   10  RT (min)      lcorbi_t3_pos_4_089_59    m/z 366.2852     0   1   10   10.5  RT (min)       lcorbi_t3_pos_4_076_58    m/z 366.3368     0   2   4   6   8E+4   10.5   11  RT (min)      lcorbi_t3_pos_1_050_13    m/z 366.3488     0   2   4   6E+4   12   12.5  RT (min)      lcorbi_t3_pos_4_088_59    m/z 366.3989     0   1E+5   11   11.5  RT (min)       lcorbi_t3_pos_1_036_08    m/z 366.7212     0   2   4   6E+4   1   1.5  RT (min)      lcorbi_t3_pos_4_070_57    m/z 366.8665     0   2   4   6E+4   18   18.5  RT (min)       lcorbi_t3_pos_4_091_60    m/z 366.9134     0   1   2   3E+4   1   1.5  RT (min)      lcorbi_t3_pos_2_020_22    m/z 366.9775     0   2   4   6   8E+5   0   0.5  RT (min)      lcorbi_t3_pos_1_006_15    m/z 367.1047     0   2   4   6E+4   9.5   10  RT (min)       lcorbi_t3_pos_3_001_49    m/z 367.1212     0   2   4   6   8E+4   1   1.5  RT (min)       lcorbi_t3_pos_4_076_58    m/z 367.1276     0   1   1   1.5  RT (min)      lcorbi_t3_pos_2_028_24    m/z 367.1881     0   1   2   3E+4   9   9.5  RT (min)      lcorbi_t3_pos_4_066_57    m/z 367.2457     0   1   2E+4   10   10.5  RT (min)      lcorbi_t3_pos_3_054_15    m/z 367.2691     0   1E+4   7   7.5  RT (min)      lcorbi_t3_pos_4_082_59    m/z 367.2821     0   2   4E+4   12.5   13  RT (min)       lcorbi_t3_pos_4_073_57    m/z 367.2957     0   1   2E+4   9.5   10  RT (min)      lcorbi_t3_pos_4_101_61    m/z 367.3254     0   1E+5   12.5   13  RT (min)       lcorbi_t3_pos_4_093_60    m/z 367.3685     0   2   4   6   8E+4   10.5   11  RT (min)       lcorbi_t3_pos_4_086_59    m/z 367.7828     0   1   2E+4   1   1.5  RT (min)      lcorbi_t3_pos_4_021_25    m/z 368.0278     0   1   2E+3   2   2.5  RT (min)      lcorbi_t3_pos_4_077_58    m/z 368.1131     0   1E+4   5.5   6  RT (min)      lcorbi_t3_pos_4_100_61    m/z 368.1239     0   2   4   6   8E+4   1   1.5  RT (min)      lcorbi_t3_pos_4_045_08    m/z 368.1606     0   2   4   6   8E+4   7.5   8  RT (min)      lcorbi_t3_pos_4_098_61    m/z 368.2199     0   1   2E+4   11.5   12  RT (min)      lcorbi_t3_pos_3_042_14    m/z 368.2281     0   1E+4   7   7.5  RT (min)      lcorbi_t3_pos_4_066_57    m/z 368.2797     0   1E+5   9   9.5  RT (min)       lcorbi_t3_pos_4_075_58    m/z 368.3161     0   2   4   6   8E+4   12   12.5  RT (min)       lcorbi_t3_pos_4_086_59    m/z 368.3800     0   2   4E+4   13.5   14  RT (min)      lcorbi_t3_pos_2_044_21    m/z 368.3889     0   1E+5   14   14.5  RT (min)      lcorbi_t3_pos_3_052_47    m/z 368.4253     0   1   2   3E+5   12   12.5  RT (min)       lcorbi_t3_pos_2_046_07    m/z 368.7182     0   2   4E+4   1   1.5  RT (min)      lcorbi_t3_pos_4_070_57    m/z 368.8223     0   2   4   6E+4   11.5   12  RT (min)       lcorbi_t3_pos_4_066_57    m/z 368.8829     0   2   4   6E+4   1   1.5  RT (min)       lcorbi_t3_pos_4_067_57    m/z 368.8925     0   1   2   3   4E+3   1   1.5  RT (min)      lcorbi_t3_pos_1_023_02    m/z 369.0256     0   1E+4   1   1.5  RT (min)      lcorbi_t3_pos_2_051_35    m/z 369.0410     0   2   4   6   8E+3   1   1.5  RT (min)      lcorbi_t3_pos_3_024_07    m/z 369.1032     0   2   4   6E+3   4.5   5  RT (min)      lcorbi_t3_pos_4_103_61    m/z 369.1253     0   1E+5   11   11.5  RT (min)       lcorbi_t3_pos_4_087_59    m/z 369.1269     0   1   2E+4   4.5   5  RT (min)      lcorbi_t3_pos_4_090_60    m/z 369.1334     0   1   2   3E+4   10.5   11  RT (min)      lcorbi_t3_pos_1_049_55    m/z 369.1521     0   1E+4   9   9.5  RT (min)      lcorbi_t3_pos_4_093_60    m/z 369.1698     0   1   2E+4   9   9.5  RT (min)      lcorbi_t3_pos_1_056_56    m/z 369.2174     0   1   2   3E+4   6   6.5  RT (min)      lcorbi_t3_pos_1_025_52    m/z 369.2404     0   2   4   6   8E+4   12   12.5  RT (min)       lcorbi_t3_pos_1_032_26    m/z 369.2485     0   2   4   6E+3   5.5   6  RT (min)      lcorbi_t3_pos_4_090_60    m/z 369.2785     0   2   4   6E+4   12   12.5  RT (min)       lcorbi_t3_pos_4_087_59    m/z 369.2831     0   1   2E+4   9   9.5  RT (min)      lcorbi_t3_pos_4_080_58    m/z 369.2978     0   1E+5   12   12.5  RT (min)       lcorbi_t3_pos_4_086_59    m/z 369.3841     0   2   4   6E+4   13   13.5  RT (min)       lcorbi_t3_pos_2_002_06    m/z 369.4287     0   1E+5   12   12.5  RT (min)      lcorbi_t3_pos_2_046_07    m/z 370.0400     0   2   4E+3   5.5   6  RT (min)      lcorbi_t3_pos_2_025_52    m/z 370.0525     0   1E+4   1.5   2  RT (min)      lcorbi_t3_pos_4_013_36    m/z 370.1199     0   2   4   6E+3   6   6.5  RT (min)      lcorbi_t3_pos_4_106_62    m/z 370.1498     0   2   4   6   8E+4   11.5   12  RT (min)      lcorbi_t3_pos_4_075_58    m/z 370.2073     0   1E+4   5.5   6  RT (min)      lcorbi_t3_pos_4_106_62    m/z 370.2198     0   2   4E+4   4.5   5  RT (min)       lcorbi_t3_pos_1_036_08    m/z 370.2208     0   2   4   6   8E+3   6   6.5  RT (min)      lcorbi_t3_pos_3_025_52    m/z 370.2496     0   1   2E+4   7   7.5  RT (min)      lcorbi_t3_pos_4_104_61    m/z 370.2589     0   2   4   6E+3   9   9.5  RT (min)      lcorbi_t3_pos_3_025_52    m/z 370.2755     0   1E+5   4.5   5  RT (min)       lcorbi_t3_pos_4_090_60    m/z 370.2756     0   1   2E+4   2.5   3  RT (min)      lcorbi_t3_pos_4_052_06    m/z 370.2954     0   2   4   6   8E+4   11   11.5  RT (min)       lcorbi_t3_pos_4_071_57    m/z 370.3317     0   1E+4   9   9.5  RT (min)      lcorbi_t3_pos_3_033_53    m/z 370.3551     0   2   4   6   8E+4   12.5   13  RT (min)      lcorbi_t3_pos_4_068_57    m/z 370.3682     0   1E+5   12   12.5  RT (min)       lcorbi_t3_pos_1_003_45    m/z 370.7152     0   2   4   6   8E+3   1   1.5  RT (min)      lcorbi_t3_pos_1_029_29    m/z 370.8689     0   1E+5   1   1.5  RT (min)      lcorbi_t3_pos_1_029_29    m/z 370.9117     0   1   2   3E+3   1   1.5  RT (min)      lcorbi_t3_pos_1_034_34    m/z 371.0202     0   1E+5   1.5   2  RT (min)       lcorbi_t3_pos_2_040_25    m/z 371.0319     0   2   4E+3   1   1.5  RT (min)       lcorbi_t3_pos_3_008_20    m/z 371.0907     0   1   2E+4   9   9.5  RT (min)      lcorbi_t3_pos_4_076_58    m/z 371.1015     0   1   2E+5   11.5   12  RT (min)       lcorbi_t3_pos_4_111_62    m/z 371.1422     0   2   4   6E+4   1.5   2  RT (min)      lcorbi_t3_pos_4_047_28    m/z 371.1451     0   2   4   6E+3   2.5   3  RT (min)      lcorbi_t3_pos_4_067_57    m/z 371.1508     0   2   4   6E+4   1   1.5  RT (min)       lcorbi_t3_pos_4_095_60    m/z 371.1962     0   1E+4   10.5   11  RT (min)       lcorbi_t3_pos_1_032_26    m/z 371.2128     0   1   1   1.5  RT (min)      lcorbi_t3_pos_4_090_60    m/z 371.2152     0   1   2E+4   5   5.5  RT (min)       lcorbi_t3_pos_4_066_57    m/z 371.2197     0   1   2E+4   9.5   10  RT (min)      lcorbi_t3_pos_4_082_59    m/z 371.2218     0   1E+4   8   8.5  RT (min)      lcorbi_t3_pos_4_014_48    m/z 371.2277     0   1E+4   4.5   5  RT (min)      lcorbi_t3_pos_4_086_59    m/z 371.2649     0   1   13.5   14  RT (min)      lcorbi_t3_pos_1_004_24    m/z 371.2654     0   1   4.5   5  RT (min)       lcorbi_t3_pos_1_002_01    m/z 371.2987     0   1E+4   9.5   10  RT (min)      lcorbi_t3_pos_1_001_49    m/z 371.3269     0   1   2   3E+4   12   12.5  RT (min)       lcorbi_t3_pos_1_001_49    m/z 371.3396     0   1   11.5   12  RT (min)       lcorbi_t3_pos_1_001_49    m/z 371.3634     0   1   2   3E+4   11   11.5  RT (min)      lcorbi_t3_pos_1_054_48    m/z 371.7168     0   2   4   6E+3   5   5.5  RT (min)      lcorbi_t3_pos_4_074_58    m/z 372.0380     0   2   4E+3   5   5.5  RT (min)      lcorbi_t3_pos_2_017_51    m/z 372.1135     0   1   2   3   4E+4   7.5   8  RT (min)       lcorbi_t3_pos_4_098_61    m/z 372.1380     0   1   1.5   2  RT (min)      lcorbi_t3_pos_1_031_40    m/z 372.1587     0   1E+5   6   6.5  RT (min)       lcorbi_t3_pos_4_068_57    m/z 372.1905     0   1E+5   1   1.5  RT (min)       lcorbi_t3_pos_4_096_60    m/z 372.1994     0   1   2E+5   11   11.5  RT (min)      lcorbi_t3_pos_1_014_03    m/z 372.2170     0   1   2   3E+4   9.5   10  RT (min)      lcorbi_t3_pos_1_046_22    m/z 372.2230     0   1E+4   4.5   5  RT (min)      lcorbi_t3_pos_4_082_59    m/z 372.2437     0   1   2   3E+4   10   10.5  RT (min)       lcorbi_t3_pos_3_020_35    m/z 372.2494     0   2   4   6E+4   5   5.5  RT (min)       lcorbi_t3_pos_4_089_59    m/z 372.2746     0   1E+4   10   10.5  RT (min)      lcorbi_t3_pos_1_001_49    m/z 372.2912     0   1   1   1.5  RT (min)       lcorbi_t3_pos_1_002_01    m/z 372.3110     0   2   4   6   8E+3   10   10.5  RT (min)      lcorbi_t3_pos_1_003_45    m/z 372.3192     0   1E+5   12   12.5  RT (min)      lcorbi_t3_pos_1_025_52    m/z 372.3192     0   1E+5   15   15.5  RT (min)      lcorbi_t3_pos_1_050_13    m/z 372.3474     0   1   2   3E+5   10.5   11  RT (min)      lcorbi_t3_pos_4_050_31    m/z 372.3475     0   1   13   13.5  RT (min)      lcorbi_t3_pos_1_001_49    m/z 372.7382     0   1   2   3E+5   1   1.5  RT (min)       lcorbi_t3_pos_2_028_24    m/z 372.8834     0   1   2   3   4E+4   1   1.5  RT (min)      lcorbi_t3_pos_1_029_29    m/z 372.9087     0   1   2   3E+4   1   1.5  RT (min)      lcorbi_t3_pos_4_078_58    m/z 372.9340     0   1E+5   0   0.5  RT (min)      lcorbi_t3_pos_1_006_15    m/z 372.9781     0   2   4   6   8E+3   1   1.5  RT (min)      lcorbi_t3_pos_4_078_58    m/z 373.1057     0   1E+5   8   8.5  RT (min)       lcorbi_t3_pos_1_051_09    m/z 373.1493     0   1   2E+4   5   5.5  RT (min)      lcorbi_t3_pos_4_014_48    m/z 373.1621     0   1   6   6.5  RT (min)       lcorbi_t3_pos_2_029_43    m/z 373.1884     0   1   2E+4   7   7.5  RT (min)       lcorbi_t3_pos_4_099_61    m/z 373.1900     0   1   2   3E+4   4.5   5  RT (min)      lcorbi_t3_pos_4_090_60    m/z 373.2529     0   1E+4   5   5.5  RT (min)      lcorbi_t3_pos_3_029_13    m/z 373.3145     0   1   2   3E+4   12   12.5  RT (min)      lcorbi_t3_pos_4_071_57    m/z 373.3314     0   1   2   3   4E+4   13   13.5  RT (min)      lcorbi_t3_pos_2_020_22    m/z 373.3336     0   1E+4   10   10.5  RT (min)      lcorbi_t3_pos_1_050_13    m/z 373.3553     0   1E+5   11.5   12  RT (min)       lcorbi_t3_pos_4_095_60    m/z 373.8667     0   2   4   6E+4   1   1.5  RT (min)      lcorbi_t3_pos_1_031_40    m/z 373.9572     0   1   2   3E+4   1   1.5   2  RT (min)      lcorbi_t3_pos_1_036_08    m/z 374.0719     0   2   4   6   8E+4   6.5   7  RT (min)      lcorbi_t3_pos_4_005_33    m/z 374.1539     0   1   2  RT (min)       lcorbi_t3_pos_1_048_47    m/z 374.1558     0   1   6   6.5  RT (min)       lcorbi_t3_pos_4_110_62    m/z 374.1660     0   2   4   6   8E+4   1   1.5  RT (min)       lcorbi_t3_pos_1_048_47    m/z 374.1916     0   2   4   6   8E+3   4.5   5  RT (min)      lcorbi_t3_pos_4_093_60    m/z 374.2328     0   1   2   3   4E+4   9   9.5  RT (min)       lcorbi_t3_pos_4_091_60    m/z 374.3074     0   1   2E+4   12   12.5  RT (min)      lcorbi_t3_pos_3_003_33    m/z 374.3266     0   1   2E+3   8   8.5  RT (min)      lcorbi_t3_pos_4_050_31    m/z 374.3676     0   1E+5   12   12.5  RT (min)       lcorbi_t3_pos_4_096_60    m/z 374.8593     0   2   4   6E+4   18   18.5  RT (min)       lcorbi_t3_pos_4_098_61    m/z 375.0607     0   1E+5   8   8.5  RT (min)       lcorbi_t3_pos_3_034_26    m/z 375.0666     0   1   2   3E+4   6.5   7  RT (min)      lcorbi_t3_pos_4_005_33    m/z 375.0739     0   1   6.5   7  RT (min)      lcorbi_t3_pos_4_104_61    m/z 375.1076     0   1   2   3E+4   7.5   8  RT (min)       lcorbi_t3_pos_4_098_61    m/z 375.1300     0   1E+4   4.5   5  RT (min)      lcorbi_t3_pos_4_066_57    m/z 375.1301     0   1   2.5   3  RT (min)      lcorbi_t3_pos_4_092_60    m/z 375.1481     0   1   2E+4   10   10.5  RT (min)       lcorbi_t3_pos_4_080_58    m/z 375.1591     0   1   6   6.5  RT (min)      lcorbi_t3_pos_4_110_62    m/z 375.1708     0   2   4   6   8E+4   8   8.5  RT (min)      lcorbi_t3_pos_4_101_61    m/z 375.2048     0   1   2E+4   7   7.5  RT (min)      lcorbi_t3_pos_4_098_61    m/z 375.2241     0   1   2.5   3  RT (min)      lcorbi_t3_pos_4_082_59    m/z 375.2361     0   1E+4   9   9.5  RT (min)      lcorbi_t3_pos_4_080_58    m/z 375.2439     0   1   2E+4   8   8.5  RT (min)      lcorbi_t3_pos_4_112_62    m/z 375.2439     0   1   2E+4   6   6.5  RT (min)      lcorbi_t3_pos_4_111_62    m/z 375.2532     0   1E+4   8.5   9  RT (min)      lcorbi_t3_pos_4_014_48    m/z 375.2618     0   1   2   3E+4   5.5   6  RT (min)       lcorbi_t3_pos_4_080_58    m/z 375.2910     0   1E+5   11.5   12  RT (min)       lcorbi_t3_pos_1_054_48    m/z 375.3049     0   1   2   3   4E+5   11.5   12  RT (min)       lcorbi_t3_pos_1_054_48    m/z 375.3198     0   1   2   3E+4   11   11.5  RT (min)       lcorbi_t3_pos_3_054_15    m/z 375.3472     0   1E+5   13   13.5  RT (min)       lcorbi_t3_pos_4_083_59    m/z 375.4154     0   2   4   6   8E+4   13.5   14  RT (min)       lcorbi_t3_pos_4_096_60    m/z 375.7635     0   1E+4   5.5   6  RT (min)      lcorbi_t3_pos_4_078_58    m/z 375.8637     0   1   2   3   4E+4   1   1.5  RT (min)      lcorbi_t3_pos_1_031_40    m/z 375.9940     0   1   1   1.5  RT (min)      lcorbi_t3_pos_3_027_27    m/z 376.0640     0   1   2E+4   8   8.5  RT (min)      lcorbi_t3_pos_3_034_26    m/z 376.0851     0   1E+5   7.5   8  RT (min)      lcorbi_t3_pos_1_051_09    m/z 376.1110     0   2   4   6   8E+3   7.5   8  RT (min)      lcorbi_t3_pos_4_103_61    m/z 376.1280     0   2   4E+3   6   6.5  RT (min)      lcorbi_t3_pos_3_048_30    m/z 376.1392     0   1E+4   5.5   6  RT (min)      lcorbi_t3_pos_3_040_41    m/z 376.1732     0   1   8   8.5  RT (min)      lcorbi_t3_pos_4_085_59    m/z 376.2120     0   2   4   6   8E+5   11   12   13  RT (min)       lcorbi_t3_pos_4_075_58    m/z 376.2120     0   1E+3   0   0.5  RT (min)      lcorbi_t3_pos_4_091_60    m/z 376.2121     0   1E+6   15   16  RT (min)       lcorbi_t3_pos_4_036_11    m/z 376.2484     0   1   11.5   12  RT (min)       lcorbi_t3_pos_1_022_20    m/z 376.3000     0   2   4E+4   9.5   10  RT (min)      lcorbi_t3_pos_1_021_18    m/z 376.3127     0   1E+5   15   15.5  RT (min)       lcorbi_t3_pos_1_028_32    m/z 376.3260     0   1E+5   11   11.5  RT (min)       lcorbi_t3_pos_4_068_57    m/z 376.9934     0   1   2   3E+4   1   1.5  RT (min)      lcorbi_t3_pos_4_081_58    m/z 377.0520     0   1E+5   1   1.5  RT (min)      lcorbi_t3_pos_3_010_44    m/z 377.0577     0   1   2   3   4E+4   8   8.5  RT (min)       lcorbi_t3_pos_3_034_26    m/z 377.0884     0   1   2   3E+4   7.5   8  RT (min)      lcorbi_t3_pos_2_032_39    m/z 377.1427     0   1   2   3E+4   4.5   5  RT (min)      lcorbi_t3_pos_4_074_58    m/z 377.1459     0   1E+5   1   1.5  RT (min)      lcorbi_t3_pos_1_036_08    m/z 377.1638     0   2   4   6E+4   4.5   5  RT (min)      lcorbi_t3_pos_4_098_61    m/z 377.2088     0   1E+5   9   9.5  RT (min)      lcorbi_t3_pos_4_048_24    m/z 377.2090     0   1   2   3   4E+4   7.5   8  RT (min)       lcorbi_t3_pos_4_101_61    m/z 377.2155     0   1   2   3E+5   14   14.5  RT (min)      lcorbi_t3_pos_4_095_60    m/z 377.2272     0   1E+4   9.5   10  RT (min)      lcorbi_t3_pos_4_076_58    m/z 377.2688     0   1   13   13.5  RT (min)       lcorbi_t3_pos_4_084_59    m/z 377.3012     0   1   2   3   4E+4   13   13.5  RT (min)       lcorbi_t3_pos_1_008_36    m/z 377.3892     0   1   2E+4   11   11.5  RT (min)      lcorbi_t3_pos_2_044_21    m/z 377.3947     0   1E+5   13   13.5  RT (min)       lcorbi_t3_pos_4_095_60    m/z 377.8800     0   2   4E+4   1   1.5  RT (min)      lcorbi_t3_pos_1_036_08    m/z 378.1657     0   1E+4   4.5   5  RT (min)      lcorbi_t3_pos_4_103_61    m/z 378.1671     0   1   2   3E+4   10   10.5  RT (min)       lcorbi_t3_pos_4_079_58    m/z 378.1913     0   2   4   6   8E+4   5.5   6  RT (min)       lcorbi_t3_pos_4_106_62    m/z 378.1993     0   2   4   6E+3   7.5   8  RT (min)      lcorbi_t3_pos_3_002_16    m/z 378.2083     0   1   4.5   5  RT (min)       lcorbi_t3_pos_4_082_59    m/z 378.2122     0   1   2   3   4E+4   9   9.5  RT (min)      lcorbi_t3_pos_4_048_24    m/z 378.2124     0   2   4   6   8E+3   7.5   8  RT (min)      lcorbi_t3_pos_3_011_42    m/z 378.2182     0   2   4E+4   8   8.5  RT (min)      lcorbi_t3_pos_4_068_57    m/z 378.2184     0   2   4   6E+4   15.5   16  RT (min)      lcorbi_t3_pos_3_002_16    m/z 378.2276     0   2   4   6E+3   8   8.5  RT (min)      lcorbi_t3_pos_4_024_27    m/z 378.2406     0   1   2E+4   10   10.5  RT (min)      lcorbi_t3_pos_4_079_58    m/z 378.2661     0   2   4   6   8E+4   9   9.5  RT (min)      lcorbi_t3_pos_3_052_47    m/z 378.3005     0   1   2E+4   11.5   12  RT (min)       lcorbi_t3_pos_1_052_38    m/z 378.3239     0   1   2E+5   13   14  RT (min)       lcorbi_t3_pos_4_085_59    m/z 378.3580     0   2   4   6E+4   12.5   13  RT (min)       lcorbi_t3_pos_4_084_59    m/z 378.7098     0   1E+4   4.5   5  RT (min)      lcorbi_t3_pos_4_076_58    m/z 378.7294     0   1   2   3E+4   1   1.5  RT (min)      lcorbi_t3_pos_4_068_57    m/z 379.0139     0   1   2E+4   0   0.5  RT (min)      lcorbi_t3_pos_1_012_14    m/z 379.0814     0   2   4   6   8E+3   5   5.5  RT (min)      lcorbi_t3_pos_4_100_61    m/z 379.1212     0   1   2E+5   1   1.5  RT (min)       lcorbi_t3_pos_1_048_47    m/z 379.1308     0   1   2E+5   11   11.5  RT (min)      lcorbi_t3_pos_4_103_61    m/z 379.1309     0   1E+5   13.5   14  RT (min)      lcorbi_t3_pos_4_018_01    m/z 379.1309     0   1   2E+3   0   0.5  RT (min)      lcorbi_t3_pos_4_013_36    m/z 379.1883     0   1E+4   9.5   10  RT (min)      lcorbi_t3_pos_4_078_58    m/z 379.1947     0   1   2E+4   5.5   6  RT (min)      lcorbi_t3_pos_4_111_62    m/z 379.2036     0   2   4   6E+4   12.5   13  RT (min)       lcorbi_t3_pos_4_071_57    m/z 379.2086     0   1   7.5   8  RT (min)      lcorbi_t3_pos_4_110_62    m/z 379.2193     0   1   2E+3   4.5   5  RT (min)      lcorbi_t3_pos_4_089_59    m/z 379.2382     0   1   2E+4   12   12.5  RT (min)      lcorbi_t3_pos_2_020_22    m/z 379.2490     0   1E+4   5   5.5  RT (min)      lcorbi_t3_pos_1_001_49    m/z 379.2522     0   1   2   3   4E+4   12.5   13  RT (min)      lcorbi_t3_pos_1_029_29    m/z 379.2522     0   2   4   6E+4   10.5   11  RT (min)       lcorbi_t3_pos_4_098_61    m/z 379.2629     0   2   4   6E+4   12   12.5  RT (min)       lcorbi_t3_pos_1_032_26    m/z 379.2844     0   2   4E+5   12   12.5  RT (min)       lcorbi_t3_pos_4_070_57    m/z 379.3055     0   1   2   3E+4   10.5   11  RT (min)      lcorbi_t3_pos_4_088_59    m/z 379.3327     0   1E+5   11.5   12  RT (min)      lcorbi_t3_pos_4_083_59    m/z 379.3361     0   2   4   6E+4   15   15.5  RT (min)      lcorbi_t3_pos_3_020_35    m/z 379.8771     0   2   4E+4   1   1.5  RT (min)      lcorbi_t3_pos_1_036_08    m/z 379.9039     0   2   4   6E+4   1   1.5  RT (min)      lcorbi_t3_pos_4_077_58    m/z 380.0218     0   1   2E+5   0   0.5  RT (min)      lcorbi_t3_pos_1_005_42    m/z 380.0861     0   1   2   3E+4   1.5   2  RT (min)       lcorbi_t3_pos_4_091_60    m/z 380.1240     0   1   1.5   2  RT (min)      lcorbi_t3_pos_4_093_60    m/z 380.1488     0   2   4E+3   2.5   3  RT (min)      lcorbi_t3_pos_1_028_32    m/z 380.1494     0   2   4   6E+3   6   6.5  RT (min)      lcorbi_t3_pos_4_102_61    m/z 380.1705     0   1E+5   6.5   7  RT (min)       lcorbi_t3_pos_1_001_49    m/z 380.1706     0   2   4E+4   8   8.5  RT (min)       lcorbi_t3_pos_4_098_61    m/z 380.2561     0   1   2   3E+4   10.5   11  RT (min)       lcorbi_t3_pos_4_080_58    m/z 380.2797     0   1E+4   10   10.5  RT (min)       lcorbi_t3_pos_4_083_59    m/z 380.3209     0   2   4   6E+4   10   10.5  RT (min)       lcorbi_t3_pos_4_077_58    m/z 380.7700     0   2   4   6   8E+4   1   1.5  RT (min)       lcorbi_t3_pos_1_020_31    m/z 380.8899     0   1E+4   1   1.5  RT (min)      lcorbi_t3_pos_3_010_44    m/z 380.8986     0   2   4   6E+4   1   1.5  RT (min)      lcorbi_t3_pos_4_077_58    m/z 380.9051     0   1   2E+3   1   1.5  RT (min)      lcorbi_t3_pos_2_011_10    m/z 380.9227     0   2   4   6   8E+4   1   1.5  RT (min)      lcorbi_t3_pos_4_068_57    m/z 380.9702     0   2   4   6E+4   1   1.5  RT (min)      lcorbi_t3_pos_2_020_22    m/z 381.0241     0   1   2   3E+4   1   1.5  RT (min)      lcorbi_t3_pos_4_006_12    m/z 381.0795     0   2   4E+4   1   1.5  RT (min)       lcorbi_t3_pos_2_042_45    m/z 381.0817     0   1   2   3   4E+4   10   10.5  RT (min)       lcorbi_t3_pos_1_029_29    m/z 381.1289     0   1E+5   9   9.5  RT (min)      lcorbi_t3_pos_1_026_17    m/z 381.1674     0   1   2   3   4E+5   9   9.5  RT (min)      lcorbi_t3_pos_4_093_60    m/z 381.1675     0   1   2E+5   12   12.5   13  RT (min)       lcorbi_t3_pos_3_031_45    m/z 381.1675     0   2   4   6   8E+5   15.5   16  RT (min)       lcorbi_t3_pos_4_080_58    m/z 381.1739     0   1   2   3E+4   6.5   7  RT (min)      lcorbi_t3_pos_1_001_49    m/z 381.2021     0   1   11.5   12  RT (min)      lcorbi_t3_pos_1_022_20    m/z 381.2287     0   1   2   3E+4   1.5   2  RT (min)       lcorbi_t3_pos_3_048_30    m/z 381.2291     0   1E+5   7.5   8  RT (min)      lcorbi_t3_pos_4_106_62    m/z 381.2678     0   2   4   6E+4   13   13.5  RT (min)       lcorbi_t3_pos_1_034_34    m/z 381.3195     0   1E+5   12   12.5  RT (min)       lcorbi_t3_pos_4_085_59    m/z 381.3243     0   1E+4   10   10.5  RT (min)      lcorbi_t3_pos_3_051_12    m/z 381.3411     0   1   12.5   13  RT (min)       lcorbi_t3_pos_1_034_34    m/z 381.9217     0   2   4E+4   1   1.5  RT (min)      lcorbi_t3_pos_2_024_15    m/z 381.9863     0   1E+5   1   1.5  RT (min)       lcorbi_t3_pos_4_076_58    m/z 382.1322     0   1   2E+4   9   9.5   10  RT (min)      lcorbi_t3_pos_2_017_51    m/z 382.1377     0   1E+4   5   5.5  RT (min)      lcorbi_t3_pos_1_056_56    m/z 382.1707     0   1E+5   8   8.5  RT (min)      lcorbi_t3_pos_4_085_59    m/z 382.1710     0   1   2   3E+5   17.5   18  RT (min)      lcorbi_t3_pos_4_108_62    m/z 382.2313     0   1E+4   7.5   8  RT (min)      lcorbi_t3_pos_4_113_62    m/z 382.2378     0   1   2   3E+4   10   10.5  RT (min)      lcorbi_t3_pos_1_036_08    m/z 382.2437     0   1E+4   7.5   8  RT (min)      lcorbi_t3_pos_4_066_57    m/z 382.2801     0   2   4   6E+3   7   7.5  RT (min)      lcorbi_t3_pos_4_005_33    m/z 382.3106     0   1   2   3E+4   12   13  RT (min)       lcorbi_t3_pos_1_003_45    m/z 382.3435     0   1E+5   11   11.5  RT (min)      lcorbi_t3_pos_4_085_59    m/z 382.6951     0   1   2   3E+4   1   1.5  RT (min)      lcorbi_t3_pos_4_078_58    m/z 382.7671     0   1E+5   1   1.5  RT (min)       lcorbi_t3_pos_1_020_31    m/z 382.8400     0   1   2   3   4E+4   0.5   1  RT (min)      lcorbi_t3_pos_1_006_15    m/z 382.8659     0   2   4   6E+4   1   1.5  RT (min)      lcorbi_t3_pos_1_036_08    m/z 383.1038     0   2   4   6E+3   4.5   5  RT (min)      lcorbi_t3_pos_4_013_36    m/z 383.1081     0   1E+4   9   9.5  RT (min)      lcorbi_t3_pos_2_046_07    m/z 383.1210     0   1E+4   5   5.5  RT (min)      lcorbi_t3_pos_4_113_62    m/z 383.1409     0   2   4   6   8E+4   11.5   12  RT (min)      lcorbi_t3_pos_4_088_59    m/z 383.1467     0   1E+5   6   6.5  RT (min)       lcorbi_t3_pos_3_048_30    m/z 383.1540     0   1   2   3E+4   1.5   2  RT (min)      lcorbi_t3_pos_3_021_39    m/z 383.1564     0   1   2.5   3  RT (min)      lcorbi_t3_pos_4_082_59    m/z 383.1736     0   1E+4   8   8.5   9  RT (min)      lcorbi_t3_pos_4_071_57    m/z 383.2218     0   1   2E+4   9.5   10  RT (min)      lcorbi_t3_pos_4_066_57    m/z 383.2531     0   1E+5   1   1.5  RT (min)       lcorbi_t3_pos_3_055_10    m/z 383.2794     0   2   4   6   8E+4   10.5   11  RT (min)       lcorbi_t3_pos_4_082_59    m/z 383.2965     0   1   2E+4   12   12.5  RT (min)      lcorbi_t3_pos_2_024_15    m/z 383.3270     0   2   4   6E+4   10   10.5  RT (min)       lcorbi_t3_pos_4_088_59    m/z 383.3311     0   1E+5   14   14.5  RT (min)       lcorbi_t3_pos_1_020_31    m/z 383.3352     0   1E+5   12.5   13  RT (min)      lcorbi_t3_pos_4_086_59    m/z 383.4001     0   1   2E+5   13   13.5  RT (min)       lcorbi_t3_pos_1_001_49    m/z 383.6298     0   2   4   6   8E+3   5.5   6  RT (min)      lcorbi_t3_pos_4_084_59    m/z 383.8940     0   1E+4   1   1.5  RT (min)      lcorbi_t3_pos_1_004_24    m/z 383.9642     0   1   2   3   4E+3   5.5   6  RT (min)      lcorbi_t3_pos_3_020_35    m/z 383.9832     0   1   2E+4   1   1.5  RT (min)       lcorbi_t3_pos_1_029_29    m/z 384.0849     0   1   1   1.5  RT (min)      lcorbi_t3_pos_2_051_35    m/z 384.0978     0   1   1   1.5  RT (min)      lcorbi_t3_pos_2_051_35    m/z 384.1253     0   1E+4   5   5.5  RT (min)      lcorbi_t3_pos_4_113_62    m/z 384.1572     0   1E+4   1.5   2  RT (min)      lcorbi_t3_pos_2_037_27    m/z 384.1630     0   1   2   3E+3   5   5.5  RT (min)      lcorbi_t3_pos_4_108_62    m/z 384.1741     0   2   4   6   8E+4   6.5   7  RT (min)       lcorbi_t3_pos_4_066_57    m/z 384.2130     0   1E+4   6   6.5  RT (min)      lcorbi_t3_pos_4_097_60    m/z 384.2230     0   1E+4   6   6.5  RT (min)      lcorbi_t3_pos_4_074_58    m/z 384.2957     0   1E+4   7   7.5  RT (min)      lcorbi_t3_pos_4_082_59    m/z 384.2957     0   1   2E+4   9.5   10  RT (min)      lcorbi_t3_pos_4_082_59    m/z 384.6922     0   1   2E+4   1   1.5  RT (min)      lcorbi_t3_pos_2_051_35    m/z 384.7527     0   2   4   6E+3   4.5   5  RT (min)      lcorbi_t3_pos_4_085_59    m/z 384.7641     0   1E+5   1   1.5  RT (min)      lcorbi_t3_pos_1_020_31    m/z 384.7722     0   2   4   6   8E+3   4.5   5  RT (min)      lcorbi_t3_pos_4_013_36    m/z 384.8456     0   1E+4   1   1.5  RT (min)      lcorbi_t3_pos_3_034_26    m/z 384.8570     0   1   2E+4   1   1.5  RT (min)      lcorbi_t3_pos_3_010_44    m/z 384.8664     0   1   2E+4   1   1.5  RT (min)      lcorbi_t3_pos_1_036_08    m/z 385.0808     0   1E+4   4.5   5  RT (min)      lcorbi_t3_pos_4_103_61    m/z 385.0858     0   1   2E+3   5   5.5  RT (min)      lcorbi_t3_pos_1_012_14    m/z 385.1207     0   2   4   6E+4   1   1.5  RT (min)      lcorbi_t3_pos_4_079_58    m/z 385.1291     0   1   2   3E+4   1.5   2  RT (min)       lcorbi_t3_pos_1_032_26    m/z 385.1291     0   1   2E+4   2.5   3  RT (min)      lcorbi_t3_pos_3_011_42    m/z 385.1325     0   1   2E+4   5   5.5  RT (min)      lcorbi_t3_pos_2_029_43    m/z 385.1775     0   1   2E+4   6.5   7  RT (min)      lcorbi_t3_pos_4_073_57    m/z 385.1987     0   1   2E+4   7   7.5  RT (min)       lcorbi_t3_pos_4_081_58    m/z 385.2021     0   1   2   3   4E+4   11   11.5  RT (min)       lcorbi_t3_pos_4_066_57    m/z 385.2101     0   1   2   3E+4   10.5   11  RT (min)      lcorbi_t3_pos_4_090_60    m/z 385.2389     0   2   4E+3   4.5   5  RT (min)      lcorbi_t3_pos_2_024_15    m/z 385.2739     0   1   2   3   4E+4   11   12  RT (min)       lcorbi_t3_pos_4_084_59    m/z 385.2811     0   1   2E+4   5   5.5  RT (min)       lcorbi_t3_pos_4_083_59    m/z 385.2927     0   1E+5   11   11.5  RT (min)       lcorbi_t3_pos_4_084_59    m/z 385.3175     0   1   2E+4   5   5.5  RT (min)      lcorbi_t3_pos_1_008_36    m/z 385.3508     0   2   4   6E+4   12.5   13  RT (min)      lcorbi_t3_pos_4_090_60    m/z 386.0403     0   1   2   3   4E+4   1   1.5  RT (min)      lcorbi_t3_pos_3_010_44    m/z 386.0970     0   1   2E+4   10   10.5  RT (min)      lcorbi_t3_pos_4_080_58    m/z 386.1575     0   1E+5   11.5   12  RT (min)      lcorbi_t3_pos_4_074_58    m/z 386.2021     0   2   4   6   8E+3   7   7.5  RT (min)      lcorbi_t3_pos_4_080_58    m/z 386.2022     0   1E+4   4.5   5  RT (min)      lcorbi_t3_pos_4_074_58    m/z 386.2327     0   1E+5   9.5   10  RT (min)       lcorbi_t3_pos_1_001_49    m/z 386.2539     0   1   2E+4   8.5   9  RT (min)      lcorbi_t3_pos_1_004_24    m/z 386.2651     0   1   2E+3   5.5   6  RT (min)      lcorbi_t3_pos_1_002_01    m/z 386.2727     0   1E+5   13.5   14  RT (min)       lcorbi_t3_pos_1_011_44    m/z 386.2902     0   2   4   6E+3   9   9.5  RT (min)      lcorbi_t3_pos_1_001_49    m/z 386.3501     0   2   4   6   8E+4   14   14.5  RT (min)      lcorbi_t3_pos_1_051_09    m/z 386.7151     0   2   4   6   8E+4   1   1.5  RT (min)       lcorbi_t3_pos_2_051_35    m/z 386.7612     0   1   2   3E+4   1   1.5  RT (min)      lcorbi_t3_pos_1_020_31    m/z 386.9250     0   2   4   6E+4   18   18.5  RT (min)       lcorbi_t3_pos_4_075_58    m/z 386.9771     0   1E+5   1   1.5  RT (min)       lcorbi_t3_pos_4_084_59    m/z 387.0842     0   1E+5   7.5   8  RT (min)       lcorbi_t3_pos_4_099_61    m/z 387.1266     0   1   2   3E+4   10   10.5  RT (min)      lcorbi_t3_pos_1_029_29    m/z 387.1526     0   2   4   6E+4   5   5.5  RT (min)       lcorbi_t3_pos_4_113_62    m/z 387.1821     0   2   4   6E+4   12.5   13  RT (min)       lcorbi_t3_pos_4_075_58    m/z 387.1932     0   1E+5   9.5   10  RT (min)       lcorbi_t3_pos_4_090_60    m/z 387.2147     0   1   2E+4   10   10.5  RT (min)      lcorbi_t3_pos_4_084_59    m/z 387.2240     0   1   2   3E+3   5   5.5  RT (min)      lcorbi_t3_pos_3_029_13    m/z 387.2361     0   1   2   3   4E+4   9   9.5   10  RT (min)      lcorbi_t3_pos_1_002_01    m/z 387.2532     0   1   2E+4   8.5   9  RT (min)      lcorbi_t3_pos_2_001_49    m/z 387.3582     0   2   4   6E+4   9   9.5  RT (min)       lcorbi_t3_pos_2_010_08    m/z 387.9658     0   1   2   3   4E+3   2.5   3  RT (min)      lcorbi_t3_pos_4_006_12    m/z 388.0877     0   1   2E+4   7.5   8  RT (min)      lcorbi_t3_pos_4_101_61    m/z 388.1536     0   1   2E+4   4.5   5  RT (min)       lcorbi_t3_pos_4_070_57    m/z 388.1966     0   1   9.5   10  RT (min)      lcorbi_t3_pos_4_038_38    m/z 388.2543     0   2   4   6E+4   4.5   5  RT (min)       lcorbi_t3_pos_1_001_49    m/z 388.3231     0   1   10.5   11  RT (min)      lcorbi_t3_pos_1_001_49    m/z 388.3553     0   1E+5   13.5   14  RT (min)       lcorbi_t3_pos_2_002_06    m/z 388.3617     0   1E+4   9   9.5  RT (min)      lcorbi_t3_pos_3_015_06    m/z 388.8571     0   1E+4   1   1.5  RT (min)      lcorbi_t3_pos_4_090_60    m/z 388.9557     0   1   2E+4   1   1.5  RT (min)      lcorbi_t3_pos_4_077_58    m/z 389.1016     0   1   2E+5   1   1.5  RT (min)       lcorbi_t3_pos_4_095_60    m/z 389.1120     0   1   2   3E+5   10.5   11  RT (min)       lcorbi_t3_pos_4_113_62    m/z 389.1627     0   1E+4   8   8.5  RT (min)      lcorbi_t3_pos_4_073_57    m/z 389.1864     0   1   2   3E+4   8.5   9  RT (min)      lcorbi_t3_pos_1_031_40    m/z 389.2512     0   1   2   3E+3   7   7.5  RT (min)      lcorbi_t3_pos_1_003_45    m/z 389.2577     0   2   4   6   8E+3   4.5   5  RT (min)      lcorbi_t3_pos_1_004_24    m/z 389.2688     0   2   4   6E+4   13   13.5  RT (min)       lcorbi_t3_pos_1_029_29    m/z 389.2729     0   1   2E+4   9   9.5  RT (min)      lcorbi_t3_pos_4_074_58    m/z 389.9575     0   2   4E+4   18   18.5  RT (min)      lcorbi_t3_pos_4_079_58    m/z 390.1572     0   2   4   6   8E+3   1.5   2  RT (min)      lcorbi_t3_pos_2_026_44    m/z 390.1760     0   2   4   6E+4   7   7.5  RT (min)      lcorbi_t3_pos_4_005_33    m/z 390.3107     0   2   4E+4   12.5   13  RT (min)       lcorbi_t3_pos_4_093_60    m/z 390.3215     0   2   4   6E+4   8   8.5  RT (min)       lcorbi_t3_pos_4_081_58    m/z 390.3581     0   1E+4   13   13.5  RT (min)      lcorbi_t3_pos_3_017_51    m/z 390.3900     0   2   4   6E+4   12   12.5  RT (min)       lcorbi_t3_pos_4_093_60    m/z 390.7988     0   1E+5   1   1.5  RT (min)       lcorbi_t3_pos_1_020_31    m/z 390.8016     0   2   4E+4   6.5   7  RT (min)       lcorbi_t3_pos_4_089_59    m/z 390.8745     0   2   4   6   8E+4   1   1.5  RT (min)      lcorbi_t3_pos_2_024_15    m/z 390.9885     0   2   4   6E+4   18   18.5  RT (min)       lcorbi_t3_pos_4_082_59    m/z 391.0346     0   2   4   6   8E+4   8   8.5  RT (min)       lcorbi_t3_pos_1_051_09    m/z 391.1019     0   2   4   6E+4   1   1.5  RT (min)       lcorbi_t3_pos_4_096_60    m/z 391.1105     0   2   4   6E+4   10   10.5  RT (min)       lcorbi_t3_pos_4_113_62    m/z 391.1290     0   2   4   6E+4   8   8.5  RT (min)      lcorbi_t3_pos_4_101_61    m/z 391.1502     0   1   4.5   5  RT (min)      lcorbi_t3_pos_3_008_20    m/z 391.1598     0   1   2   3E+4   5   5.5  RT (min)       lcorbi_t3_pos_4_008_30    m/z 391.1752     0   1   2E+4   7   7.5  RT (min)      lcorbi_t3_pos_4_101_61    m/z 391.1809     0   1   2   3E+4   1   1.5  RT (min)       lcorbi_t3_pos_4_093_60    m/z 391.1865     0   1E+4   7   7.5  RT (min)      lcorbi_t3_pos_4_082_59    m/z 391.2035     0   2   4   6E+4   12   12.5  RT (min)       lcorbi_t3_pos_1_032_26    m/z 391.2196     0   1   2E+4   4.5   5  RT (min)       lcorbi_t3_pos_3_008_20    m/z 391.2282     0   1   2E+5   13.5   14  RT (min)      lcorbi_t3_pos_4_075_58    m/z 391.2493     0   1   9   9.5  RT (min)      lcorbi_t3_pos_4_081_58    m/z 391.2630     0   1   6   6.5  RT (min)      lcorbi_t3_pos_4_079_58    m/z 391.2674     0   1   8   8.5  RT (min)      lcorbi_t3_pos_4_079_58    m/z 391.2935     0   1   7   7.5  RT (min)      lcorbi_t3_pos_4_081_58    m/z 391.3033     0   1   6.5   7  RT (min)       lcorbi_t3_pos_4_081_58    m/z 391.3298     0   2   4   6   8E+4   13   13.5  RT (min)       lcorbi_t3_pos_2_040_25    m/z 391.7212     0   1E+4   4.5   5  RT (min)      lcorbi_t3_pos_4_079_58    m/z 391.7953     0   1E+4   7   7.5  RT (min)      lcorbi_t3_pos_4_087_59    m/z 391.8048     0   1E+4   6.5   7  RT (min)      lcorbi_t3_pos_4_085_59    m/z 391.9842     0   1E+4   1.5   2  RT (min)      lcorbi_t3_pos_1_003_45    m/z 391.9935     0   1E+5   1   1.5  RT (min)       lcorbi_t3_pos_4_096_60    m/z 392.1313     0   1E+4   8   8.5  RT (min)      lcorbi_t3_pos_4_100_61    m/z 392.1746     0   1E+4   9.5   10  RT (min)      lcorbi_t3_pos_2_016_23    m/z 392.2281     0   2   4   6E+3   8.5   9  RT (min)      lcorbi_t3_pos_3_014_29    m/z 392.2950     0   2   4   6   8E+4   7   7.5  RT (min)       lcorbi_t3_pos_4_107_62    m/z 392.3138     0   1   2E+4   11.5   12  RT (min)       lcorbi_t3_pos_1_038_30    m/z 392.3975     0   1   2E+5   12   12.5  RT (min)       lcorbi_t3_pos_4_091_60    m/z 392.7064     0   2   4   6E+4   1   1.5  RT (min)      lcorbi_t3_pos_1_023_02    m/z 392.9990     0   2   4   6E+4   18   18.5  RT (min)      lcorbi_t3_pos_4_086_59    m/z 393.0021     0   2   4   6E+4   1   1.5  RT (min)       lcorbi_t3_pos_4_076_58    m/z 393.0259     0   2   4E+4   1   1.5  RT (min)       lcorbi_t3_pos_4_091_60    m/z 393.0319     0   1   2   3E+4   8   8.5  RT (min)      lcorbi_t3_pos_1_051_09    m/z 393.1117     0   1   2E+4   7.5   8  RT (min)      lcorbi_t3_pos_1_051_09    m/z 393.1159     0   2   4   6   8E+3   4.5   5  RT (min)      lcorbi_t3_pos_4_090_60    m/z 393.1198     0   1   2   3   4E+4   1   1.5  RT (min)       lcorbi_t3_pos_4_091_60    m/z 393.1657     0   1E+4   6.5   7  RT (min)      lcorbi_t3_pos_4_103_61    m/z 393.1828     0   1   2   3   4E+4   9   9.5  RT (min)      lcorbi_t3_pos_1_021_18    m/z 393.1950     0   1   2   3E+4   9.5   10  RT (min)      lcorbi_t3_pos_1_019_16    m/z 393.2282     0   1E+4   5   5.5  RT (min)      lcorbi_t3_pos_4_066_57    m/z 393.2387     0   2   4   6   8E+4   7   7.5  RT (min)       lcorbi_t3_pos_4_108_62    m/z 393.2404     0   1   2   3E+4   12   12.5  RT (min)       lcorbi_t3_pos_1_050_13    m/z 393.2910     0   1   2E+5   17.5   18  RT (min)      lcorbi_t3_pos_4_100_61    m/z 393.2978     0   2   4   6   8E+4   1   1.5  RT (min)      lcorbi_t3_pos_1_002_01    m/z 393.3089     0   1E+4   10   10.5  RT (min)      lcorbi_t3_pos_1_001_49    m/z 393.3114     0   1E+5   11.5   12  RT (min)      lcorbi_t3_pos_1_023_02    m/z 393.3202     0   1   12   12.5  RT (min)      lcorbi_t3_pos_1_054_48    m/z 393.3484     0   2   4   6   8E+4   12   12.5  RT (min)       lcorbi_t3_pos_4_087_59    m/z 393.3772     0   1   2   3E+4   13   13.5  RT (min)       lcorbi_t3_pos_1_043_06    m/z 393.7299     0   2   4   6E+3   5   5.5  RT (min)      lcorbi_t3_pos_4_074_58    m/z 394.1862     0   1   7.5   8  RT (min)       lcorbi_t3_pos_4_090_60    m/z 394.1902     0   1E+5   4.5   5  RT (min)      lcorbi_t3_pos_4_098_61    m/z 394.2224     0   1   2E+5   8.5   9   9.5  RT (min)       lcorbi_t3_pos_4_077_58    m/z 394.2225     0   1   2   3E+5   0   0.5  RT (min)      lcorbi_t3_pos_2_039_30    m/z 394.2225     0   1   2   3   4E+5   11   11.5  RT (min)      lcorbi_t3_pos_4_075_58    m/z 394.2423     0   1   2   3E+4   7   7.5  RT (min)      lcorbi_t3_pos_4_107_62    m/z 394.2564     0   1E+4   10   10.5  RT (min)      lcorbi_t3_pos_3_007_28    m/z 394.2567     0   1E+4   6   6.5  RT (min)       lcorbi_t3_pos_4_112_62    m/z 394.2798     0   1   2   3   4E+4   7   7.5  RT (min)      lcorbi_t3_pos_2_029_43    m/z 394.2906     0   1   2   3   4E+3   7   7.5  RT (min)      lcorbi_t3_pos_4_113_62    m/z 394.2950     0   1E+4   9.5   10  RT (min)      lcorbi_t3_pos_2_027_05    m/z 394.3010     0   1E+4   1   1.5  RT (min)      lcorbi_t3_pos_1_002_01    m/z 394.3164     0   1E+5   11   11.5  RT (min)       lcorbi_t3_pos_4_082_59    m/z 394.3342     0   1   2   3E+4   7   7.5  RT (min)      lcorbi_t3_pos_2_022_38    m/z 394.3441     0   1   2   3E+4   7   7.5  RT (min)      lcorbi_t3_pos_2_029_43    m/z 394.4211     0   2   4   6   8E+4   13   13.5  RT (min)       lcorbi_t3_pos_4_096_60    m/z 394.7034     0   1E+4   1   1.5  RT (min)      lcorbi_t3_pos_4_016_15    m/z 394.7928     0   1E+5   1   1.5  RT (min)      lcorbi_t3_pos_1_020_31    m/z 394.8106     0   2   4   6   8E+3   1   1.5  RT (min)      lcorbi_t3_pos_3_039_03    m/z 394.8743     0   1   2E+5   1   1.5  RT (min)       lcorbi_t3_pos_3_008_20    m/z 395.0392     0   1   2   3E+4   1   1.5  RT (min)      lcorbi_t3_pos_1_031_40    m/z 395.1619     0   1   2E+4   10   10.5  RT (min)       lcorbi_t3_pos_1_024_04    m/z 395.1741     0   1   10   10.5  RT (min)       lcorbi_t3_pos_1_026_17    m/z 395.1920     0   1   2E+4   4.5   5  RT (min)      lcorbi_t3_pos_4_100_61    m/z 395.2258     0   1   2   3   4E+4   8   8.5  RT (min)       lcorbi_t3_pos_1_041_54    m/z 395.2259     0   2   4   6E+4   0   0.5  RT (min)      lcorbi_t3_pos_2_039_30    m/z 395.2405     0   1E+4   11   11.5  RT (min)      lcorbi_t3_pos_1_013_10    m/z 395.2559     0   1E+4   12   12.5  RT (min)       lcorbi_t3_pos_1_054_48    m/z 395.2640     0   1E+4   8.5   9  RT (min)      lcorbi_t3_pos_4_074_58    m/z 395.2957     0   1   2E+5   13   13.5  RT (min)       lcorbi_t3_pos_4_077_58    m/z 395.2982     0   1   7   7.5  RT (min)      lcorbi_t3_pos_4_108_62    m/z 395.3134     0   1   2   3E+4   13   13.5  RT (min)      lcorbi_t3_pos_4_085_59    m/z 395.3289     0   1   7   7.5  RT (min)      lcorbi_t3_pos_4_106_62    m/z 395.3332     0   1   7   7.5  RT (min)      lcorbi_t3_pos_4_108_62    m/z 395.3996     0   1   2E+4   10.5   11  RT (min)      lcorbi_t3_pos_4_069_57    m/z 395.8777     0   1E+4   1   1.5  RT (min)      lcorbi_t3_pos_4_077_58    m/z 396.0093     0   1   2   3E+4   1   1.5  RT (min)      lcorbi_t3_pos_4_078_58    m/z 396.1473     0   1   2   3   4E+4   4.5   5  RT (min)       lcorbi_t3_pos_4_090_60    m/z 396.1750     0   2   4E+4   9.5   10  RT (min)       lcorbi_t3_pos_1_043_06    m/z 396.2280     0   1E+4   7.5   8  RT (min)      lcorbi_t3_pos_1_021_18    m/z 396.2592     0   1E+4   8   8.5  RT (min)      lcorbi_t3_pos_4_074_58    m/z 396.2746     0   1   2   3E+4   7   7.5  RT (min)      lcorbi_t3_pos_1_004_24    m/z 396.3108     0   1   2E+4   9.5   10  RT (min)      lcorbi_t3_pos_4_076_58    m/z 396.3109     0   1   2E+4   12   12.5  RT (min)      lcorbi_t3_pos_3_048_30    m/z 396.3261     0   1   2   3E+4   7.5   8  RT (min)       lcorbi_t3_pos_2_053_28    m/z 396.3303     0   2   4   6   8E+4   9.5   10   10.5  RT (min)       lcorbi_t3_pos_4_085_59    m/z 396.3343     0   1   2   3E+5   13   13.5  RT (min)       lcorbi_t3_pos_1_054_48    m/z 396.3536     0   1   2   3E+4   14   14.5  RT (min)      lcorbi_t3_pos_4_011_22    m/z 396.3585     0   1E+4   9   9.5  RT (min)      lcorbi_t3_pos_3_042_14    m/z 396.3591     0   1E+5   11.5   12  RT (min)       lcorbi_t3_pos_4_085_59    m/z 396.7439     0   1   2   3E+4   1   1.5  RT (min)      lcorbi_t3_pos_2_051_35    m/z 396.7551     0   2   4E+3   1   1.5  RT (min)      lcorbi_t3_pos_2_023_26    m/z 396.7646     0   1E+5   1   1.5  RT (min)      lcorbi_t3_pos_2_020_22    m/z 396.8724     0   1   2   3   4E+4   1   1.5  RT (min)      lcorbi_t3_pos_4_077_58    m/z 396.8855     0   1   2E+4   0.5   1  RT (min)       lcorbi_t3_pos_4_075_58    m/z 397.0685     0   1   2E+4   6   6.5  RT (min)      lcorbi_t3_pos_3_024_07    m/z 397.0722     0   1E+5   1   1.5   2  RT (min)       lcorbi_t3_pos_2_040_25    m/z 397.1317     0   1   2E+3   1   1.5   2  RT (min)       lcorbi_t3_pos_2_051_35    m/z 397.1601     0   1E+4   9   9.5  RT (min)      lcorbi_t3_pos_4_081_58    m/z 397.2258     0   1E+4   8   8.5  RT (min)      lcorbi_t3_pos_3_049_55    m/z 397.2372     0   1   7.5   8  RT (min)      lcorbi_t3_pos_1_014_03    m/z 397.2716     0   2   4   6   8E+4   12.5   13  RT (min)       lcorbi_t3_pos_1_034_34    m/z 397.2926     0   2   4   6E+4   12   12.5  RT (min)      lcorbi_t3_pos_3_026_04    m/z 397.3103     0   1   2   3   4E+4   12.5   13  RT (min)      lcorbi_t3_pos_3_007_28    m/z 397.3295     0   2   4   6E+3   7.5   8  RT (min)      lcorbi_t3_pos_2_053_28    m/z 397.3424     0   1   2   3E+4   10   10.5  RT (min)      lcorbi_t3_pos_4_082_59    m/z 397.3426     0   1E+5   12   12.5  RT (min)       lcorbi_t3_pos_4_082_59    m/z 397.3539     0   2   4   6E+4   12   12.5  RT (min)      lcorbi_t3_pos_3_026_04    m/z 397.3677     0   1   2   3   4E+4   12.5   13  RT (min)      lcorbi_t3_pos_1_032_26    m/z 397.8731     0   1   2   3E+4   0.5   1  RT (min)      lcorbi_t3_pos_1_006_15    m/z 398.0929     0   1   2   3E+4   5   5.5  RT (min)      lcorbi_t3_pos_4_014_48    m/z 398.1365     0   2   4   6E+4   8   8.5  RT (min)      lcorbi_t3_pos_1_002_01    m/z 398.1446     0   2   4E+4   8   8.5  RT (min)      lcorbi_t3_pos_4_094_60    m/z 398.1533     0   1E+5   7   7.5  RT (min)       lcorbi_t3_pos_4_073_57    m/z 398.2385     0   1   2E+4   6.5   7  RT (min)       lcorbi_t3_pos_4_066_57    m/z 398.2667     0   1   2E+4   10.5   11  RT (min)       lcorbi_t3_pos_4_034_42    m/z 398.3061     0   1   2E+4   6   6.5  RT (min)      lcorbi_t3_pos_4_089_59    m/z 398.3073     0   1   2   3   4E+4   12   12.5  RT (min)       lcorbi_t3_pos_1_050_13    m/z 398.3265     0   2   4E+4   10   10.5  RT (min)       lcorbi_t3_pos_4_068_57    m/z 398.3823     0   2   4   6E+4   14   14.5  RT (min)      lcorbi_t3_pos_3_015_06    m/z 398.3994     0   1   2   3   4E+4   12.5   13  RT (min)       lcorbi_t3_pos_4_077_58    m/z 398.7409     0   2   4   6E+4   1   1.5  RT (min)       lcorbi_t3_pos_1_029_29    m/z 398.7522     0   1   2E+4   1   1.5  RT (min)      lcorbi_t3_pos_4_021_25    m/z 398.7616     0   1E+5   1   1.5  RT (min)      lcorbi_t3_pos_1_034_34    m/z 398.7619     0   1E+5   13.5   14  RT (min)       lcorbi_t3_pos_1_020_31    m/z 398.7809     0   2   4E+3   5.5   6  RT (min)      lcorbi_t3_pos_4_083_59    m/z 398.8906     0   1E+4   1   1.5  RT (min)      lcorbi_t3_pos_1_019_16    m/z 398.8907     0   2   4   6   8E+4   18   18.5  RT (min)       lcorbi_t3_pos_4_109_62    m/z 398.9183     0   2   4   6   8E+4   18   18.5  RT (min)      lcorbi_t3_pos_4_072_57    m/z 398.9891     0   2   4   6E+4   1   1.5  RT (min)       lcorbi_t3_pos_1_029_29    m/z 399.0874     0   1E+4   5   5.5  RT (min)      lcorbi_t3_pos_4_014_48    m/z 399.1205     0   1   2E+4   6   6.5  RT (min)       lcorbi_t3_pos_3_048_30    m/z 399.1303     0   1   2   3E+5   1   1.5  RT (min)       lcorbi_t3_pos_4_090_60    m/z 399.1334     0   1   2   3E+4   9.5   10  RT (min)      lcorbi_t3_pos_2_003_19    m/z 399.1393     0   1E+4   8   8.5  RT (min)      lcorbi_t3_pos_1_051_09    m/z 399.1567     0   1   2E+4   7   7.5  RT (min)      lcorbi_t3_pos_4_073_57    m/z 399.1760     0   1   2.5   3  RT (min)      lcorbi_t3_pos_4_090_60    m/z 399.1779     0   1E+5   0   0.5  RT (min)      lcorbi_t3_pos_2_039_30    m/z 399.1779     0   1E+6   14   16  RT (min)       lcorbi_t3_pos_4_076_58    m/z 399.1862     0   2   4   6E+3   4.5   5  RT (min)      lcorbi_t3_pos_4_090_60    m/z 399.1949     0   1   7   7.5  RT (min)      lcorbi_t3_pos_4_070_57    m/z 399.2148     0   1E+3   1   1.5  RT (min)      lcorbi_t3_pos_4_024_27    m/z 399.2351     0   2   4E+4   4.5   5  RT (min)       lcorbi_t3_pos_4_074_58    m/z 399.2419     0   2   4   6E+3   6.5   7  RT (min)      lcorbi_t3_pos_4_066_57    m/z 399.2507     0   1   2   3E+4   13   13.5  RT (min)       lcorbi_t3_pos_1_002_01    m/z 399.2532     0   2   4   6   8E+4   15   15.5  RT (min)       lcorbi_t3_pos_4_088_59    m/z 399.2590     0   1   2E+3   5   5.5  RT (min)      lcorbi_t3_pos_1_002_01    m/z 399.2637     0   1E+5   13   13.5   14  RT (min)       lcorbi_t3_pos_1_023_02    m/z 399.2719     0   2   4   6   8E+4   11   11.5  RT (min)       lcorbi_t3_pos_4_082_59    m/z 399.3082     0   1E+5   11.5   12  RT (min)       lcorbi_t3_pos_4_083_59    m/z 399.3279     0   2   4   6E+4   12.5   13  RT (min)       lcorbi_t3_pos_1_054_48    m/z 399.3766     0   2   4   6   8E+4   13   13.5  RT (min)       lcorbi_t3_pos_4_093_60    m/z 399.7696     0   2   4   6   8E+4   14   14.5  RT (min)      lcorbi_t3_pos_1_020_31    m/z 399.8902     0   1   2E+4   1   1.5  RT (min)      lcorbi_t3_pos_2_020_22    m/z 400.0860     0   1   2   3E+4   1   1.5   2  RT (min)      lcorbi_t3_pos_3_021_39    m/z 400.1388     0   2   4E+3   6   6.5  RT (min)      lcorbi_t3_pos_4_110_62    m/z 400.1465     0   1E+4   1.5   2  RT (min)      lcorbi_t3_pos_4_032_40    m/z 400.1473     0   1   2   3E+4   4.5   5  RT (min)      lcorbi_t3_pos_4_098_61    m/z 400.1690     0   1   2   3   4E+4   5.5   6  RT (min)       lcorbi_t3_pos_4_069_57    m/z 400.1813     0   1E+5   13.5   14  RT (min)      lcorbi_t3_pos_2_035_34    m/z 400.2178     0   1   2E+4   5   5.5  RT (min)      lcorbi_t3_pos_4_106_62    m/z 400.2245     0   1   2E+4   9.5   10  RT (min)      lcorbi_t3_pos_4_050_31    m/z 400.2384     0   1E+4   4.5   5  RT (min)      lcorbi_t3_pos_4_074_58    m/z 400.2541     0   1   2E+4   10.5   11  RT (min)       lcorbi_t3_pos_4_075_58    m/z 400.2881     0   2   4E+4   13.5   14  RT (min)       lcorbi_t3_pos_2_045_40    m/z 400.3059     0   1E+4   10.5   11  RT (min)       lcorbi_t3_pos_4_073_57    m/z 400.3230     0   2   4   6E+4   12   12.5  RT (min)       lcorbi_t3_pos_4_071_57    m/z 400.3786     0   1   2   3   4E+5   11   11.5  RT (min)       lcorbi_t3_pos_4_050_31    m/z 400.7381     0   1   2   3   4E+4   1   1.5  RT (min)      lcorbi_t3_pos_1_029_29    m/z 400.7587     0   1   1   1.5  RT (min)      lcorbi_t3_pos_1_020_31    m/z 400.7859     0   1E+4   6.5   7  RT (min)       lcorbi_t3_pos_4_089_59    m/z 400.8062     0   1   1   1.5  RT (min)      lcorbi_t3_pos_1_020_31    m/z 400.8275     0   1   2   3E+5   1   1.5  RT (min)       lcorbi_t3_pos_1_020_31    m/z 401.0593     0   1   2E+4   1   1.5  RT (min)      lcorbi_t3_pos_2_024_15    m/z 401.0631     0   1E+4   5   5.5  RT (min)      lcorbi_t3_pos_4_102_61    m/z 401.1143     0   2   4   6   8E+3   4.5   5  RT (min)      lcorbi_t3_pos_1_054_48    m/z 401.1246     0   1   2E+4   9.5   10  RT (min)      lcorbi_t3_pos_2_044_21    m/z 401.1280     0   1E+5   11.5   12  RT (min)       lcorbi_t3_pos_4_078_58    m/z 401.1645     0   1   2E+4   1.5   2  RT (min)      lcorbi_t3_pos_3_040_41    m/z 401.1724     0   1E+4   5.5   6  RT (min)      lcorbi_t3_pos_4_072_57    m/z 401.1725     0   2   4   6   8E+3   7   7.5  RT (min)      lcorbi_t3_pos_4_080_58    m/z 401.1840     0   1E+4   7.5   8  RT (min)      lcorbi_t3_pos_4_066_57    m/z 401.2323     0   2   4E+4   9.5   10  RT (min)      lcorbi_t3_pos_4_084_59    m/z 401.2620     0   1   2E+4   5   5.5  RT (min)      lcorbi_t3_pos_1_001_49    m/z 401.2664     0   1   12   12.5  RT (min)       lcorbi_t3_pos_4_086_59    m/z 401.2779     0   1   2   3E+4   7   7.5  RT (min)       lcorbi_t3_pos_4_005_33    m/z 401.2877     0   2   4   6E+3   6.5   7  RT (min)      lcorbi_t3_pos_4_088_59    m/z 401.3092     0   2   4   6   8E+3   10.5   11  RT (min)       lcorbi_t3_pos_4_073_57    m/z 401.3416     0   2   4E+4   12.5   13  RT (min)       lcorbi_t3_pos_2_012_01    m/z 401.3456     0   2   4   6E+4   10.5   11  RT (min)       lcorbi_t3_pos_4_073_57    m/z 401.3775     0   1   11.5   12  RT (min)       lcorbi_t3_pos_4_050_31    m/z 401.7795     0   1   2E+4   7   7.5  RT (min)      lcorbi_t3_pos_4_005_33    m/z 402.1912     0   1E+4   8.5   9  RT (min)      lcorbi_t3_pos_4_091_60    m/z 402.2082     0   1   7.5   8  RT (min)      lcorbi_t3_pos_4_083_59    m/z 402.2084     0   1   2   3   4E+5   14   14.5  RT (min)      lcorbi_t3_pos_1_036_08    m/z 402.2276     0   2   4E+4   10   10.5  RT (min)       lcorbi_t3_pos_1_039_23    m/z 402.2335     0   2   4   6   8E+3   4.5   5  RT (min)      lcorbi_t3_pos_4_082_59    m/z 402.2528     0   1   2   3E+4   10   10.5  RT (min)      lcorbi_t3_pos_3_020_35    m/z 402.2698     0   1   2E+4   5   5.5  RT (min)       lcorbi_t3_pos_4_066_57    m/z 402.3093     0   1   2E+5   12   12.5  RT (min)      lcorbi_t3_pos_2_006_47    m/z 402.3387     0   2   4   6   8E+4   12.5   13  RT (min)       lcorbi_t3_pos_1_034_34    m/z 402.3578     0   1   2   3   4E+5   10.5   11  RT (min)       lcorbi_t3_pos_4_050_31    m/z 402.3899     0   1E+5   12   12.5  RT (min)       lcorbi_t3_pos_4_090_60    m/z 402.6889     0   1   2E+4   1   1.5  RT (min)      lcorbi_t3_pos_2_040_25    m/z 402.8033     0   1   1   1.5  RT (min)      lcorbi_t3_pos_1_020_31    m/z 402.9025     0   1   2   3   4E+4   1   1.5  RT (min)       lcorbi_t3_pos_4_081_58    m/z 402.9425     0   1   2   3E+4   1   1.5  RT (min)      lcorbi_t3_pos_4_068_57    m/z 403.2304     0   1   2E+4   2.5   3  RT (min)      lcorbi_t3_pos_3_011_42    m/z 403.2627     0   1   2   3E+4   12   12.5  RT (min)      lcorbi_t3_pos_1_032_26    m/z 403.2732     0   1   2   3   4E+3   5   5.5  RT (min)      lcorbi_t3_pos_4_082_59    m/z 403.3055     0   2   4E+4   12   12.5  RT (min)       lcorbi_t3_pos_3_020_35    m/z 403.3987     0   1   2   3   4E+5   11.5   12  RT (min)       lcorbi_t3_pos_1_003_45    m/z 403.9037     0   1   2   3E+4   1   1.5  RT (min)      lcorbi_t3_pos_2_024_15    m/z 404.0031     0   1E+4   1   1.5  RT (min)      lcorbi_t3_pos_1_028_32    m/z 404.1914     0   1   2   3E+4   5   5.5  RT (min)      lcorbi_t3_pos_1_002_01    m/z 404.2140     0   2   4   6   8E+3   4.5   5  RT (min)      lcorbi_t3_pos_4_082_59    m/z 404.2492     0   2   4E+3   4.5   5  RT (min)      lcorbi_t3_pos_1_010_46    m/z 404.2643     0   1   2   3E+4   9.5   10  RT (min)      lcorbi_t3_pos_1_040_27    m/z 404.3263     0   2   4   6E+4   12.5   13  RT (min)       lcorbi_t3_pos_4_052_06    m/z 404.4056     0   1E+5   12.5   13  RT (min)       lcorbi_t3_pos_4_093_60    m/z 404.4065     0   1E+5   15   15.5  RT (min)      lcorbi_t3_pos_1_021_18    m/z 404.6861     0   2   4E+4   1   1.5  RT (min)      lcorbi_t3_pos_2_040_25    m/z 404.8216     0   2   4   6   8E+4   1   1.5  RT (min)      lcorbi_t3_pos_1_023_02    m/z 404.8393     0   1   2   3E+4   1   1.5  RT (min)      lcorbi_t3_pos_1_031_40    m/z 404.9332     0   1   2E+3   1   1.5  RT (min)      lcorbi_t3_pos_1_003_45    m/z 405.1909     0   1   8.5   9  RT (min)      lcorbi_t3_pos_1_036_08    m/z 405.2102     0   2   4   6   8E+4   8.5   9  RT (min)      lcorbi_t3_pos_1_036_08    m/z 405.2350     0   2   4   6E+4   1   1.5  RT (min)      lcorbi_t3_pos_4_052_06    m/z 405.2784     0   2   4   6E+4   12   12.5  RT (min)       lcorbi_t3_pos_1_036_08    m/z 405.3148     0   1   2   3E+4   5   5.5  RT (min)       lcorbi_t3_pos_4_083_59    m/z 405.7887     0   1E+4   5.5   6  RT (min)       lcorbi_t3_pos_4_088_59    m/z 405.8069     0   2   4E+4   5.5   6  RT (min)       lcorbi_t3_pos_4_089_59    m/z 405.8165     0   1E+4   5   5.5  RT (min)      lcorbi_t3_pos_4_084_59    m/z 405.9473     0   2   4   6E+4   1   1.5  RT (min)      lcorbi_t3_pos_1_029_29    m/z 406.1642     0   1E+4   5.5   6  RT (min)      lcorbi_t3_pos_4_068_57    m/z 406.2179     0   1   2   3E+4   12.5   13  RT (min)       lcorbi_t3_pos_4_052_06    m/z 406.2409     0   2   4E+4   11   11.5  RT (min)      lcorbi_t3_pos_4_090_60    m/z 406.2743     0   2   4   6E+3   6.5   7  RT (min)      lcorbi_t3_pos_4_110_62    m/z 406.2800     0   1   2   3   4E+4   10.5   11  RT (min)      lcorbi_t3_pos_4_066_57    m/z 406.2904     0   1E+4   5.5   6  RT (min)      lcorbi_t3_pos_4_087_59    m/z 406.3033     0   2   4   6   8E+4   13   13.5  RT (min)      lcorbi_t3_pos_4_086_59    m/z 406.3086     0   1   2E+4   5.5   6  RT (min)      lcorbi_t3_pos_4_086_59    m/z 406.3892     0   1E+5   11.5   12  RT (min)       lcorbi_t3_pos_4_084_59    m/z 406.6833     0   1   2   3   4E+4   1   1.5  RT (min)      lcorbi_t3_pos_1_029_29    m/z 406.7727     0   2   4E+4   1   1.5  RT (min)      lcorbi_t3_pos_1_029_29    m/z 406.7840     0   1   2   3   4E+3   1   1.5  RT (min)      lcorbi_t3_pos_1_031_40    m/z 406.8101     0   2   4   6E+3   5.5   6  RT (min)      lcorbi_t3_pos_4_088_59    m/z 406.8484     0   1   1   1.5  RT (min)      lcorbi_t3_pos_1_007_07    m/z 406.8679     0   1E+5   0.5   1  RT (min)      lcorbi_t3_pos_1_001_49    m/z 407.0713     0   1E+4   1   1.5  RT (min)      lcorbi_t3_pos_4_074_58    m/z 407.1273     0   1E+5   1   1.5  RT (min)       lcorbi_t3_pos_4_067_57    m/z 407.1295     0   2   4   6E+4   5   5.5  RT (min)       lcorbi_t3_pos_4_008_30    m/z 407.1637     0   1E+4   8   8.5  RT (min)      lcorbi_t3_pos_4_092_60    m/z 407.1914     0   1   2   3E+4   10   10.5  RT (min)      lcorbi_t3_pos_1_029_29    m/z 407.2941     0   1E+5   12.5   13  RT (min)       lcorbi_t3_pos_1_034_34    m/z 407.3134     0   1   2   3E+4   13   13.5  RT (min)      lcorbi_t3_pos_3_020_35    m/z 407.3327     0   1   2   3   4E+4   12.5   13  RT (min)      lcorbi_t3_pos_1_054_48    m/z 407.3674     0   1   2   3   4E+4   11   11.5  RT (min)      lcorbi_t3_pos_4_103_61    m/z 407.7925     0   1   2E+4   6   6.5  RT (min)       lcorbi_t3_pos_4_086_59    m/z 408.0717     0   1   2   3E+4   4.5   5  RT (min)       lcorbi_t3_pos_4_074_58    m/z 408.2017     0   1   2E+4   6.5   7  RT (min)       lcorbi_t3_pos_4_005_33    m/z 408.2299     0   1E+5   11.5   12  RT (min)       lcorbi_t3_pos_1_002_01    m/z 408.2898     0   1E+4   6.5   7  RT (min)       lcorbi_t3_pos_4_109_62    m/z 408.3261     0   1E+4   7   7.5  RT (min)      lcorbi_t3_pos_4_107_62    m/z 408.3320     0   2   4   6   8E+4   11.5   12  RT (min)       lcorbi_t3_pos_4_074_58    m/z 408.3403     0   1   2E+5   11   11.5  RT (min)      lcorbi_t3_pos_4_089_59    m/z 408.3493     0   2   4   6   8E+4   12   12.5  RT (min)       lcorbi_t3_pos_4_087_59    m/z 408.6805     0   1   2E+4   1   1.5  RT (min)      lcorbi_t3_pos_2_040_25    m/z 408.7023     0   1   2E+4   1   1.5  RT (min)      lcorbi_t3_pos_3_012_31    m/z 408.7464     0   2   4   6   8E+4   13   13.5  RT (min)       lcorbi_t3_pos_1_032_26    m/z 408.7698     0   2   4   6E+4   1   1.5  RT (min)      lcorbi_t3_pos_1_029_29    m/z 408.7810     0   2   4   6   8E+3   1   1.5  RT (min)      lcorbi_t3_pos_1_044_25    m/z 408.9766     0   2   4E+4   18   18.5  RT (min)      lcorbi_t3_pos_4_076_58    m/z 409.1124     0   1   2   3   4E+4   1   1.5  RT (min)      lcorbi_t3_pos_4_074_58    m/z 409.1318     0   1   2   3   4E+4   1.5   2  RT (min)      lcorbi_t3_pos_2_046_07    m/z 409.1622     0   1   2   3E+4   8.5   9  RT (min)       lcorbi_t3_pos_4_082_59    m/z 409.1751     0   1   2E+4   9.5   10  RT (min)      lcorbi_t3_pos_4_094_60    m/z 409.1898     0   2   4   6   8E+4   5   5.5  RT (min)      lcorbi_t3_pos_4_014_48    m/z 409.2716     0   1   2   3   4E+4   1   1.5  RT (min)      lcorbi_t3_pos_1_003_45    m/z 409.2720     0   1   13   13.5  RT (min)      lcorbi_t3_pos_4_080_58    m/z 409.2931     0   1   2   3E+3   6.5   7  RT (min)      lcorbi_t3_pos_1_001_49    m/z 409.3425     0   1E+5   9.5   10  RT (min)       lcorbi_t3_pos_4_101_61    m/z 409.7542     0   1E+5   13.5   14  RT (min)       lcorbi_t3_pos_1_032_26    m/z 409.9209     0   1   2E+4   1   1.5  RT (min)      lcorbi_t3_pos_4_079_58    m/z 410.0232     0   1E+4   1   1.5  RT (min)      lcorbi_t3_pos_2_024_15    m/z 410.1662     0   2   4   6E+3   2.5   3  RT (min)      lcorbi_t3_pos_4_109_62    m/z 410.1920     0   1E+4   5   5.5  RT (min)      lcorbi_t3_pos_4_113_62    m/z 410.2810     0   1E+4   6   6.5  RT (min)      lcorbi_t3_pos_4_112_62    m/z 410.3054     0   1E+5   6.5   7  RT (min)       lcorbi_t3_pos_4_014_48    m/z 410.3055     0   2   4E+4   8.5   9  RT (min)       lcorbi_t3_pos_4_106_62    m/z 410.3266     0   1   2   3E+4   10   10.5  RT (min)      lcorbi_t3_pos_4_071_57    m/z 410.3377     0   1E+4   8   8.5  RT (min)      lcorbi_t3_pos_4_082_59    m/z 410.3461     0   1   2E+4   9.5   10  RT (min)      lcorbi_t3_pos_4_104_61    m/z 410.3477     0   1   2E+4   10.5   11  RT (min)      lcorbi_t3_pos_2_020_22    m/z 410.3748     0   2   4   6   8E+4   12   12.5  RT (min)      lcorbi_t3_pos_4_084_59    m/z 410.6993     0   1   2   3   4E+4   1   1.5  RT (min)      lcorbi_t3_pos_2_028_24    m/z 410.7620     0   2   4   6   8E+4   13.5   14  RT (min)       lcorbi_t3_pos_1_034_34    m/z 410.7669     0   1   2   3   4E+4   1   1.5  RT (min)      lcorbi_t3_pos_1_029_29    m/z 410.8563     0   2   4   6E+5   1   1.5  RT (min)       lcorbi_t3_pos_1_020_31    m/z 410.9169     0   1   2E+4   1   1.5  RT (min)      lcorbi_t3_pos_4_079_58    m/z 411.0840     0   2   4   6E+4   7   7.5  RT (min)      lcorbi_t3_pos_1_052_38    m/z 411.0880     0   1   2   2.5  RT (min)      lcorbi_t3_pos_1_036_08    m/z 411.0985     0   2   4   6E+3   1   1.5  RT (min)      lcorbi_t3_pos_3_023_05    m/z 411.2507     0   1   2E+4   13   13.5  RT (min)      lcorbi_t3_pos_1_054_48    m/z 411.3088     0   1   2E+4   6.5   7  RT (min)      lcorbi_t3_pos_4_108_62    m/z 411.3089     0   1E+4   8.5   9  RT (min)      lcorbi_t3_pos_4_107_62    m/z 411.3623     0   2   4   6   8E+4   12   12.5  RT (min)       lcorbi_t3_pos_4_030_05    m/z 411.7187     0   1E+4   5.5   6  RT (min)      lcorbi_t3_pos_4_079_58    m/z 411.7695     0   2   4   6E+4   14   14.5  RT (min)      lcorbi_t3_pos_1_020_31    m/z 411.9251     0   2   4   6E+4   0.5   1  RT (min)      lcorbi_t3_pos_1_003_45    m/z 412.1085     0   1   2   3E+4   8   8.5  RT (min)      lcorbi_t3_pos_4_099_61    m/z 412.1886     0   2   4   6E+4   8.5   9  RT (min)       lcorbi_t3_pos_1_018_35    m/z 412.1887     0   1   2E+5   0   0.5  RT (min)      lcorbi_t3_pos_2_039_30    m/z 412.2096     0   2   4   6E+4   5.5   6  RT (min)       lcorbi_t3_pos_1_054_48    m/z 412.2196     0   1   5.5   6  RT (min)      lcorbi_t3_pos_1_054_48    m/z 412.2273     0   1   2E+4   12   12.5  RT (min)       lcorbi_t3_pos_4_030_05    m/z 412.2542     0   1E+4   7   7.5  RT (min)       lcorbi_t3_pos_4_106_62    m/z 412.3046     0   1E+4   5.5   6  RT (min)      lcorbi_t3_pos_4_089_59    m/z 412.3248     0   1   2E+5   13.5   14  RT (min)      lcorbi_t3_pos_4_106_62    m/z 412.7386     0   2   4   6E+4   1   1.5  RT (min)      lcorbi_t3_pos_1_036_08    m/z 412.7405     0   1E+4   4.5   5  RT (min)      lcorbi_t3_pos_4_079_58    m/z 412.7777     0   1E+5   14   14.5  RT (min)      lcorbi_t3_pos_1_044_25    m/z 412.7966     0   2   4E+4   5.5   6  RT (min)       lcorbi_t3_pos_4_088_59    m/z 412.8062     0   2   4   6E+3   5.5   6  RT (min)      lcorbi_t3_pos_4_086_59    m/z 412.8534     0   1   2   3   4E+5   1   1.5  RT (min)       lcorbi_t3_pos_1_020_31    m/z 412.8631     0   1   2   3   4E+3   1   1.5  RT (min)      lcorbi_t3_pos_4_070_57    m/z 413.1376     0   1   2E+4   9   9.5  RT (min)      lcorbi_t3_pos_1_032_26    m/z 413.1767     0   1   2E+4   8.5   9  RT (min)      lcorbi_t3_pos_2_024_15    m/z 413.1921     0   1E+4   8.5   9  RT (min)      lcorbi_t3_pos_2_024_15    m/z 413.2131     0   1E+4   5.5   6  RT (min)      lcorbi_t3_pos_1_054_48    m/z 413.2406     0   1E+5   13   13.5  RT (min)      lcorbi_t3_pos_4_080_58    m/z 413.2500     0   1   2E+5   13   13.5  RT (min)      lcorbi_t3_pos_1_029_29    m/z 413.2579     0   1   2   3   4E+3   7   7.5  RT (min)      lcorbi_t3_pos_4_019_47    m/z 413.2752     0   1E+3   5.5   6  RT (min)      lcorbi_t3_pos_4_046_44    m/z 413.2885     0   1   6   6.5  RT (min)      lcorbi_t3_pos_4_110_62    m/z 413.2982     0   1   2E+4   5.5   6  RT (min)      lcorbi_t3_pos_4_086_59    m/z 413.3239     0   1   12   12.5  RT (min)       lcorbi_t3_pos_4_075_58    m/z 413.3486     0   1   5   5.5  RT (min)      lcorbi_t3_pos_4_008_30    m/z 413.3779     0   1E+5   14.5   15  RT (min)       lcorbi_t3_pos_1_037_19    m/z 413.7902     0   2   4   6   8E+3   6   6.5  RT (min)      lcorbi_t3_pos_4_086_59    m/z 413.9058     0   1   2   3   4E+4   1   1.5  RT (min)      lcorbi_t3_pos_2_020_22    m/z 413.9395     0   1   2   3E+3   1   1.5  RT (min)      lcorbi_t3_pos_3_011_42    m/z 413.9753     0   1   2   3E+4   1   1.5  RT (min)      lcorbi_t3_pos_4_097_60    m/z 414.1468     0   2   4   6E+4   5.5   6  RT (min)      lcorbi_t3_pos_4_014_48    m/z 414.1486     0   2   4   6   8E+4   1.5   2  RT (min)      lcorbi_t3_pos_2_046_07    m/z 414.1579     0   1   2E+4   4.5   5  RT (min)       lcorbi_t3_pos_4_091_60    m/z 414.1833     0   1   2E+4   10   10.5  RT (min)      lcorbi_t3_pos_4_088_59    m/z 414.1856     0   1E+4   8.5   9  RT (min)      lcorbi_t3_pos_4_080_58    m/z 414.2123     0   1E+4   8.5   9  RT (min)      lcorbi_t3_pos_2_017_51    m/z 414.2358     0   1   2   3   4E+4   11   11.5  RT (min)      lcorbi_t3_pos_4_091_60    m/z 414.2698     0   2   4   6E+4   0   0.5  RT (min)      lcorbi_t3_pos_3_015_06    m/z 414.3004     0   2   4   6   8E+4   9   9.5  RT (min)       lcorbi_t3_pos_4_075_58    m/z 414.3085     0   1E+4   11.5   12  RT (min)       lcorbi_t3_pos_2_005_11    m/z 414.3216     0   2   4   6   8E+4   11   11.5  RT (min)       lcorbi_t3_pos_4_066_57    m/z 414.3368     0   2   4   6   8E+3   5.5   6  RT (min)      lcorbi_t3_pos_4_111_62    m/z 414.7358     0   2   4   6E+4   1   1.5  RT (min)       lcorbi_t3_pos_4_076_58    m/z 414.7380     0   1E+4   4.5   5  RT (min)      lcorbi_t3_pos_4_079_58    m/z 414.8505     0   1   2E+4   1   1.5  RT (min)      lcorbi_t3_pos_3_036_37    m/z 414.8681     0   1   2   3   4E+4   1   1.5  RT (min)      lcorbi_t3_pos_1_002_01    m/z 414.8761     0   1   2E+3   1   1.5  RT (min)      lcorbi_t3_pos_3_039_03    m/z 414.8961     0   1E+3   1   1.5  RT (min)      lcorbi_t3_pos_4_070_57    m/z 414.9402     0   1E+5   1   1.5  RT (min)       lcorbi_t3_pos_4_070_57    m/z 414.9840     0   1   2   3   4E+4   1   1.5  RT (min)      lcorbi_t3_pos_2_040_25    m/z 415.1074     0   1   2E+4   9.5   10  RT (min)      lcorbi_t3_pos_1_043_06    m/z 415.1196     0   2   4E+4   4.5   5  RT (min)      lcorbi_t3_pos_4_098_61    m/z 415.1518     0   2   4   6E+4   7.5   8  RT (min)       lcorbi_t3_pos_4_086_59    m/z 415.2396     0   1   2E+4   1   1.5  RT (min)       lcorbi_t3_pos_4_078_58    m/z 415.2409     0   2   4   6   8E+3   5   5.5  RT (min)      lcorbi_t3_pos_4_108_62    m/z 415.2480     0   2   4   6E+4   8   8.5  RT (min)       lcorbi_t3_pos_1_008_36    m/z 415.2542     0   1E+3   4.5   5  RT (min)      lcorbi_t3_pos_2_046_07    m/z 415.2844     0   1   6   6.5  RT (min)      lcorbi_t3_pos_4_084_59    m/z 415.2933     0   1   2E+5   11.5   12  RT (min)      lcorbi_t3_pos_1_051_09    m/z 415.3038     0   1   2   3E+4   9   9.5  RT (min)      lcorbi_t3_pos_4_079_58    m/z 415.3572     0   2   4   6E+4   13   13.5  RT (min)       lcorbi_t3_pos_4_078_58    m/z 415.3643     0   1E+4   6   6.5  RT (min)      lcorbi_t3_pos_4_089_59    m/z 415.3895     0   1   2   3E+4   10   10.5  RT (min)      lcorbi_t3_pos_1_020_31    m/z 415.7430     0   1   2   3   4E+3   5   5.5  RT (min)      lcorbi_t3_pos_4_098_61    m/z 416.0670     0   1E+5   1   1.5  RT (min)      lcorbi_t3_pos_4_006_12    m/z 416.1143     0   1   2E+4   4.5   5  RT (min)      lcorbi_t3_pos_4_098_61    m/z 416.1164     0   2   4   6E+3   6   6.5  RT (min)      lcorbi_t3_pos_4_109_62    m/z 416.1553     0   1E+4   7.5   8  RT (min)      lcorbi_t3_pos_4_083_59    m/z 416.2148     0   1E+5   9.5   10  RT (min)      lcorbi_t3_pos_3_052_47    m/z 416.2278     0   1   4.5   5  RT (min)      lcorbi_t3_pos_4_106_62    m/z 416.2314     0   1   1   1.5  RT (min)      lcorbi_t3_pos_3_029_13    m/z 416.2387     0   1E+4   6   6.5  RT (min)      lcorbi_t3_pos_4_108_62    m/z 416.2492     0   1E+4   5   5.5  RT (min)      lcorbi_t3_pos_4_082_59    m/z 416.2514     0   1   2E+4   8   8.5  RT (min)      lcorbi_t3_pos_1_014_03    m/z 416.2677     0   2   4   6E+4   10   10.5  RT (min)       lcorbi_t3_pos_3_021_39    m/z 416.2752     0   1E+5   13   13.5  RT (min)       lcorbi_t3_pos_4_080_58    m/z 416.2855     0   1   2   3E+4   5   5.5  RT (min)       lcorbi_t3_pos_4_066_57    m/z 416.3160     0   1E+4   8.5   9  RT (min)      lcorbi_t3_pos_4_069_57    m/z 416.3614     0   1   2   3E+4   10   10.5  RT (min)      lcorbi_t3_pos_1_012_14    m/z 416.3652     0   1E+5   14   14.5  RT (min)       lcorbi_t3_pos_4_076_58    m/z 416.3737     0   2   4   6E+4   13.5   14  RT (min)       lcorbi_t3_pos_1_004_24    m/z 416.7328     0   1   2   3E+4   1   1.5  RT (min)      lcorbi_t3_pos_2_003_19    m/z 416.8016     0   1E+5   1   1.5  RT (min)      lcorbi_t3_pos_1_029_29    m/z 416.8651     0   1E+4   1   1.5  RT (min)      lcorbi_t3_pos_1_044_25    m/z 417.1440     0   1   2E+4   8.5   9  RT (min)       lcorbi_t3_pos_1_040_27    m/z 417.1699     0   1E+4   7.5   8  RT (min)      lcorbi_t3_pos_1_004_24    m/z 417.1788     0   1E+4   7.5   8  RT (min)       lcorbi_t3_pos_4_014_48    m/z 417.2162     0   2   4E+4   4.5   5  RT (min)      lcorbi_t3_pos_4_106_62    m/z 417.2402     0   1E+5   10.5   11  RT (min)      lcorbi_t3_pos_1_050_13    m/z 417.2889     0   2   4   6   8E+3   5   5.5  RT (min)      lcorbi_t3_pos_4_098_61    m/z 417.3729     0   2   4   6   8E+4   14.5   15  RT (min)       lcorbi_t3_pos_4_081_58    m/z 417.8797     0   1E+4   1   1.5  RT (min)      lcorbi_t3_pos_1_007_07    m/z 418.2153     0   1E+4   9   9.5  RT (min)      lcorbi_t3_pos_3_033_53    m/z 418.2179     0   2   4   6E+3   4.5   5  RT (min)      lcorbi_t3_pos_4_074_58    m/z 418.2225     0   1   2E+4   8.5   9  RT (min)       lcorbi_t3_pos_4_082_59    m/z 418.2590     0   2   4   6E+5   12   13  RT (min)       lcorbi_t3_pos_4_107_62    m/z 418.2590     0   1   2   3   4E+4   0   0.5  RT (min)      lcorbi_t3_pos_3_054_15    m/z 418.2590     0   1E+6   15   16   17  RT (min)       lcorbi_t3_pos_2_033_53    m/z 418.2952     0   2   4   6   8E+4   8   8.5  RT (min)      lcorbi_t3_pos_1_001_49    m/z 418.3528     0   2   4   6E+4   9   9.5  RT (min)       lcorbi_t3_pos_4_078_58    m/z 418.7986     0   1E+5   1   1.5  RT (min)      lcorbi_t3_pos_1_029_29    m/z 418.8800     0   2   4E+4   1   1.5  RT (min)       lcorbi_t3_pos_4_081_58    m/z 418.9784     0   1   2   3E+4   1   1.5  RT (min)      lcorbi_t3_pos_4_076_58    m/z 419.1410     0   2   4   6   8E+3   8.5   9  RT (min)      lcorbi_t3_pos_3_026_04    m/z 419.1447     0   1   2E+4   10   10.5  RT (min)      lcorbi_t3_pos_2_040_25    m/z 419.2429     0   2   4   6E+4   10   10.5  RT (min)      lcorbi_t3_pos_4_102_61    m/z 419.2430     0   2   4   6E+4   0   0.5  RT (min)      lcorbi_t3_pos_2_039_30    m/z 419.2430     0   1E+5   14   14.5  RT (min)      lcorbi_t3_pos_1_054_48    m/z 419.2624     0   1   2   3   4E+5   14.5   15   15.5  RT (min)       lcorbi_t3_pos_1_025_52    m/z 419.2624     0   1   2E+5   13.5   14  RT (min)      lcorbi_t3_pos_4_098_61    m/z 419.2839     0   1   2   3E+3   2.5   3  RT (min)      lcorbi_t3_pos_1_001_49    m/z 419.2987     0   1   2   3E+4   11.5   12  RT (min)       lcorbi_t3_pos_1_001_49    m/z 419.3733     0   1   2E+4   12.5   13  RT (min)       lcorbi_t3_pos_4_097_60    m/z 419.3937     0   1E+5   13.5   14  RT (min)      lcorbi_t3_pos_2_030_14    m/z 419.8043     0   2   4E+3   6   6.5  RT (min)      lcorbi_t3_pos_4_034_42    m/z 419.8140     0   2   4   6E+3   5.5   6  RT (min)      lcorbi_t3_pos_4_088_59    m/z 420.1630     0   2   4   6   8E+4   1   1.5  RT (min)      lcorbi_t3_pos_4_042_09    m/z 420.1954     0   1   2   3   4E+3   7.5   8  RT (min)      lcorbi_t3_pos_4_093_60    m/z 420.2332     0   1   2   3E+4   13   13.5  RT (min)      lcorbi_t3_pos_1_043_06    m/z 420.2382     0   2   4   6E+4   9   9.5  RT (min)      lcorbi_t3_pos_1_027_43    m/z 420.2464     0   2   4   6   8E+4   15.5   16  RT (min)      lcorbi_t3_pos_1_038_30    m/z 420.2593     0   2   4   6E+3   7   7.5  RT (min)      lcorbi_t3_pos_1_001_49    m/z 420.2652     0   2   4   6E+4   9.5   10  RT (min)      lcorbi_t3_pos_1_013_10    m/z 420.3130     0   1E+4   10   10.5  RT (min)      lcorbi_t3_pos_1_001_49    m/z 420.3321     0   1E+4   7.5   8  RT (min)       lcorbi_t3_pos_1_001_49    m/z 420.4049     0   1E+5   11.5   12  RT (min)       lcorbi_t3_pos_4_088_59    m/z 420.7958     0   1   2   3E+4   1   1.5  RT (min)      lcorbi_t3_pos_1_016_05    m/z 420.8244     0   1   2E+3   1   1.5  RT (min)      lcorbi_t3_pos_1_016_05    m/z 420.9200     0   1E+5   0.5   1  RT (min)      lcorbi_t3_pos_1_005_42    m/z 420.9711     0   2   4   6   8E+4   1   1.5  RT (min)      lcorbi_t3_pos_1_029_29    m/z 421.0675     0   1E+4   9   9.5  RT (min)      lcorbi_t3_pos_2_029_43    m/z 421.0904     0   1E+5   10   10.5  RT (min)       lcorbi_t3_pos_1_029_29    m/z 421.1681     0   1E+4   4.5   5  RT (min)      lcorbi_t3_pos_4_082_59    m/z 421.2181     0   1   2E+4   4.5   5  RT (min)      lcorbi_t3_pos_1_004_24    m/z 421.2303     0   1   2   3E+4   13.5   14  RT (min)      lcorbi_t3_pos_1_045_12    m/z 421.2415     0   1   2E+4   9   9.5  RT (min)      lcorbi_t3_pos_4_073_57    m/z 421.2435     0   2   4   6E+4   10   10.5  RT (min)       lcorbi_t3_pos_1_029_29    m/z 421.2523     0   1   2E+4   12   12.5  RT (min)      lcorbi_t3_pos_1_051_09    m/z 421.2630     0   1E+4   11   11.5  RT (min)      lcorbi_t3_pos_1_001_49    m/z 421.3223     0   1   2E+5   13.5   14  RT (min)      lcorbi_t3_pos_4_082_59    m/z 421.3242     0   1E+4   5.5   6  RT (min)      lcorbi_t3_pos_1_001_49    m/z 421.3355     0   1E+5   12   12.5  RT (min)       lcorbi_t3_pos_1_032_26    m/z 421.3402     0   1   2   3   4E+4   11   11.5  RT (min)      lcorbi_t3_pos_3_020_35    m/z 421.3467     0   1   7.5   8  RT (min)      lcorbi_t3_pos_4_094_60    m/z 421.6587     0   2   4   6E+3   5.5   6  RT (min)      lcorbi_t3_pos_4_034_42    m/z 421.8886     0   2   4   6E+4   1   1.5  RT (min)      lcorbi_t3_pos_1_020_31    m/z 422.1001     0   2   4   6   8E+4   5   5.5  RT (min)       lcorbi_t3_pos_4_014_48    m/z 422.1230     0   2   4   6E+3   2.5   3  RT (min)      lcorbi_t3_pos_4_018_01    m/z 422.2462     0   1   2   3E+4   10   10.5  RT (min)      lcorbi_t3_pos_4_074_58    m/z 422.2538     0   1E+4   8   8.5  RT (min)      lcorbi_t3_pos_4_082_59    m/z 422.2691     0   2   4   6   8E+4   5.5   6  RT (min)       lcorbi_t3_pos_4_106_62    m/z 422.2902     0   1   2   3   4E+4   6   6.5  RT (min)       lcorbi_t3_pos_1_012_14    m/z 422.3034     0   1   2   3E+4   12.5   13  RT (min)       lcorbi_t3_pos_1_053_39    m/z 422.3266     0   1   2E+4   10   10.5  RT (min)      lcorbi_t3_pos_4_079_58    m/z 422.3560     0   1   2   3E+5   11   11.5  RT (min)       lcorbi_t3_pos_4_083_59    m/z 422.4355     0   1E+5   12.5   13  RT (min)       lcorbi_t3_pos_1_003_45    m/z 422.6827     0   1   2E+4   1   1.5  RT (min)      lcorbi_t3_pos_4_021_25    m/z 422.8822     0   1   2   3   4E+5   1   1.5  RT (min)       lcorbi_t3_pos_1_020_31    m/z 422.9435     0   2   4   6   8E+4   18   18.5  RT (min)       lcorbi_t3_pos_4_082_59    m/z 423.0456     0   2   4E+4   1   1.5  RT (min)      lcorbi_t3_pos_3_024_07    m/z 423.0641     0   1E+5   1   1.5  RT (min)       lcorbi_t3_pos_4_070_57    m/z 423.1034     0   1E+4   5   5.5  RT (min)      lcorbi_t3_pos_4_112_62    m/z 423.1377     0   1E+4   8   8.5  RT (min)      lcorbi_t3_pos_3_055_10    m/z 423.1692     0   1   2   3E+4   10   10.5  RT (min)       lcorbi_t3_pos_1_019_16    m/z 423.1867     0   1E+4   7.5   8  RT (min)      lcorbi_t3_pos_1_014_03    m/z 423.2054     0   1E+4   7.5   8  RT (min)      lcorbi_t3_pos_3_011_42    m/z 423.2144     0   1   2E+5   13.5   14   14.5  RT (min)       lcorbi_t3_pos_1_031_40    m/z 423.2208     0   2   4   6E+4   6.5   7  RT (min)      lcorbi_t3_pos_4_103_61    m/z 423.2749     0   1   2E+4   5   5.5  RT (min)      lcorbi_t3_pos_1_001_49    m/z 423.2937     0   1E+4   6   6.5  RT (min)      lcorbi_t3_pos_4_112_62    m/z 423.3583     0   2   4   6   8E+4   11   11.5  RT (min)      lcorbi_t3_pos_1_032_26    m/z 423.3697     0   1   12   12.5  RT (min)      lcorbi_t3_pos_2_037_27    m/z 423.8856     0   1   2E+4   1   1.5  RT (min)      lcorbi_t3_pos_1_020_31    m/z 424.2178     0   1E+5   15.5   16  RT (min)      lcorbi_t3_pos_4_072_57    m/z 424.2178     0   2   4   6   8E+4   13.5   14  RT (min)      lcorbi_t3_pos_4_052_06    m/z 424.2235     0   1E+4   6.5   7  RT (min)      lcorbi_t3_pos_4_099_61    m/z 424.2637     0   1   2   3E+4   12.5   13  RT (min)       lcorbi_t3_pos_2_002_06    m/z 424.2847     0   1   2   3E+4   5.5   6  RT (min)       lcorbi_t3_pos_1_008_36    m/z 424.6797     0   2   4E+4   1   1.5  RT (min)      lcorbi_t3_pos_4_068_57    m/z 425.0612     0   1   2   3E+4   1   1.5  RT (min)      lcorbi_t3_pos_4_071_57    m/z 425.1362     0   1   2   3E+4   8.5   9  RT (min)       lcorbi_t3_pos_4_052_06    m/z 425.1744     0   2   4   6E+3   7.5   8  RT (min)      lcorbi_t3_pos_4_027_07    m/z 425.1936     0   1E+4   9   9.5  RT (min)      lcorbi_t3_pos_3_024_07    m/z 425.1959     0   1   7   7.5  RT (min)      lcorbi_t3_pos_2_016_23    m/z 425.1977     0   1   11   11.5  RT (min)      lcorbi_t3_pos_4_082_59    m/z 425.2206     0   2   4   6E+4   9.5   10  RT (min)      lcorbi_t3_pos_4_081_58    m/z 425.2317     0   1   2E+4   11   11.5  RT (min)      lcorbi_t3_pos_4_098_61    m/z 425.2876     0   1E+5   12   12.5  RT (min)       lcorbi_t3_pos_1_007_07    m/z 425.2880     0   1E+4   5.5   6  RT (min)      lcorbi_t3_pos_4_014_48    m/z 425.3456     0   2   4   6   8E+4   10.5   11  RT (min)       lcorbi_t3_pos_4_074_58    m/z 425.3478     0   1   13   13.5  RT (min)      lcorbi_t3_pos_4_107_62    m/z 425.3853     0   1   2   3E+4   12.5   13  RT (min)      lcorbi_t3_pos_1_001_49    m/z 425.4468     0   2   4E+4   13   13.5  RT (min)       lcorbi_t3_pos_2_046_07    m/z 425.9005     0   1   2E+4   1   1.5  RT (min)      lcorbi_t3_pos_4_077_58    m/z 426.2699     0   1E+4   7.5   8  RT (min)       lcorbi_t3_pos_4_073_57    m/z 426.3004     0   1E+4   6   6.5  RT (min)       lcorbi_t3_pos_4_015_45    m/z 426.3086     0   1E+4   13   13.5  RT (min)      lcorbi_t3_pos_3_029_13    m/z 426.3123     0   1   6   6.5  RT (min)      lcorbi_t3_pos_4_108_62    m/z 426.3216     0   2   4   6E+4   9   10  RT (min)       lcorbi_t3_pos_4_088_59    m/z 426.3697     0   1E+5   11   11.5  RT (min)       lcorbi_t3_pos_4_084_59    m/z 426.4138     0   2   4   6   8E+4   14.5   15  RT (min)       lcorbi_t3_pos_4_038_38    m/z 426.4307     0   1   2   3   4E+4   13   13.5  RT (min)       lcorbi_t3_pos_4_071_57    m/z 426.6766     0   2   4   6E+4   1   1.5  RT (min)       lcorbi_t3_pos_4_070_57    m/z 426.8302     0   1   2E+5   1   1.5  RT (min)       lcorbi_t3_pos_1_029_29    m/z 426.8913     0   1E+4   1   1.5  RT (min)      lcorbi_t3_pos_1_029_29    m/z 426.9011     0   2   4   6E+3   1   1.5  RT (min)      lcorbi_t3_pos_4_039_35    m/z 426.9839     0   2   4E+4   1   1.5  RT (min)      lcorbi_t3_pos_1_029_29    m/z 427.0745     0   1   10   10.5  RT (min)      lcorbi_t3_pos_4_113_62    m/z 427.0760     0   1   2E+4   1   1.5  RT (min)      lcorbi_t3_pos_4_066_57    m/z 427.0789     0   1   2E+4   8   8.5  RT (min)       lcorbi_t3_pos_4_101_61    m/z 427.1085     0   1   2E+4   5   5.5  RT (min)      lcorbi_t3_pos_4_108_62    m/z 427.1602     0   1   2   3   4E+3   8   8.5  RT (min)      lcorbi_t3_pos_3_034_26    m/z 427.1728     0   2   4   6   8E+3   7.5   8  RT (min)      lcorbi_t3_pos_4_094_60    m/z 427.1923     0   1   2   3E+5   2.5   3  RT (min)       lcorbi_t3_pos_4_109_62    m/z 427.2115     0   1E+4   7   7.5  RT (min)      lcorbi_t3_pos_4_107_62    m/z 427.2803     0   1E+4   8   8.5  RT (min)      lcorbi_t3_pos_3_011_42    m/z 427.3612     0   2   4   6E+4   11   11.5  RT (min)      lcorbi_t3_pos_4_073_57    m/z 428.1958     0   1   2   3   4E+5   1   2   3  RT (min)       lcorbi_t3_pos_3_029_13    m/z 428.2044     0   2   4E+4   5.5   6  RT (min)       lcorbi_t3_pos_3_054_15    m/z 428.2229     0   1   2E+4   11   11.5  RT (min)      lcorbi_t3_pos_3_028_11    m/z 428.2491     0   1   2E+4   6   6.5  RT (min)      lcorbi_t3_pos_4_066_57    m/z 428.2795     0   2   4   6   8E+3   9.5   10  RT (min)      lcorbi_t3_pos_1_050_13    m/z 428.3249     0   1E+5   12   12.5  RT (min)      lcorbi_t3_pos_1_005_42    m/z 428.3677     0   1   2   3E+4   5   5.5  RT (min)      lcorbi_t3_pos_4_088_59    m/z 428.3817     0   1   2E+5   13.5   14  RT (min)      lcorbi_t3_pos_4_083_59    m/z 428.3929     0   2   4   6E+4   11.5   12  RT (min)       lcorbi_t3_pos_4_088_59    m/z 428.6736     0   1   2E+4   1   1.5  RT (min)      lcorbi_t3_pos_1_002_01    m/z 428.8274     0   1E+5   1   1.5  RT (min)      lcorbi_t3_pos_1_029_29    m/z 428.8700     0   2   4   6E+4   1   1.5  RT (min)      lcorbi_t3_pos_4_075_58    m/z 428.8900     0   1E+4   1   1.5  RT (min)      lcorbi_t3_pos_1_002_01    m/z 429.0654     0   1E+5   1   1.5  RT (min)      lcorbi_t3_pos_3_010_44    m/z 429.1157     0   2   4   6E+4   5   5.5  RT (min)       lcorbi_t3_pos_3_048_30    m/z 429.1734     0   1   2   3   4E+4   10   10.5  RT (min)       lcorbi_t3_pos_1_029_29    m/z 429.1975     0   1   2.5   3  RT (min)      lcorbi_t3_pos_3_029_13    m/z 429.2402     0   1   2E+5   0   0.5  RT (min)      lcorbi_t3_pos_3_054_15    m/z 429.2419     0   1E+4   4.5   5  RT (min)      lcorbi_t3_pos_4_086_59    m/z 429.2665     0   1   2   3E+5   5.5   6  RT (min)       lcorbi_t3_pos_4_090_60    m/z 429.3188     0   1E+5   11   11.5  RT (min)       lcorbi_t3_pos_4_083_59    m/z 429.3364     0   1E+5   12   12.5  RT (min)       lcorbi_t3_pos_3_029_13    m/z 429.3406     0   1E+5   11   11.5   12  RT (min)       lcorbi_t3_pos_1_020_31    m/z 429.3769     0   1   2   3E+4   11   11.5  RT (min)       lcorbi_t3_pos_4_072_57    m/z 429.7436     0   2   4   6   8E+3   4.5   5  RT (min)      lcorbi_t3_pos_3_029_13    m/z 430.1686     0   1   2   3E+3   6   6.5  RT (min)      lcorbi_t3_pos_1_008_36    m/z 430.2283     0   1E+4   5   5.5  RT (min)      lcorbi_t3_pos_4_066_57    m/z 430.2436     0   2   4E+4   5   5.5  RT (min)      lcorbi_t3_pos_4_091_60    m/z 430.2589     0   2   4   6   8E+4   13   13.5  RT (min)      lcorbi_t3_pos_2_029_43    m/z 430.2871     0   1   2E+4   7.5   8  RT (min)      lcorbi_t3_pos_2_046_07    m/z 430.3011     0   2   4E+4   5.5   6  RT (min)       lcorbi_t3_pos_4_098_61    m/z 430.3406     0   1E+5   12.5   13  RT (min)      lcorbi_t3_pos_2_006_47    m/z 430.8201     0   1E+5   13   13.5  RT (min)      lcorbi_t3_pos_1_036_08    m/z 430.8420     0   1E+4   1   1.5  RT (min)      lcorbi_t3_pos_1_029_29    m/z 430.8532     0   1E+4   1   1.5  RT (min)      lcorbi_t3_pos_4_012_37    m/z 430.8671     0   1   2   3   4E+3   1   1.5  RT (min)       lcorbi_t3_pos_2_011_10    m/z 430.8730     0   2   4   6E+4   18   18.5  RT (min)       lcorbi_t3_pos_4_073_57    m/z 430.8962     0   2   4E+4   1   1.5  RT (min)      lcorbi_t3_pos_3_012_31    m/z 430.9311     0   1   2E+4   1   1.5  RT (min)      lcorbi_t3_pos_3_010_44    m/z 431.2018     0   2   4   6E+4   6   6.5  RT (min)      lcorbi_t3_pos_4_107_62    m/z 431.2103     0   1   2E+3   6   6.5  RT (min)      lcorbi_t3_pos_4_107_62    m/z 431.2465     0   2   4   6   8E+4   13   13.5  RT (min)      lcorbi_t3_pos_4_080_58    m/z 431.2470     0   1E+4   5   5.5  RT (min)      lcorbi_t3_pos_4_052_06    m/z 431.2945     0   1   2   3   4E+5   12.5   13  RT (min)       lcorbi_t3_pos_2_054_29    m/z 431.3045     0   2   4   6E+3   5.5   6  RT (min)      lcorbi_t3_pos_4_074_58    m/z 431.3156     0   2   4   6   8E+3   10   10.5  RT (min)       lcorbi_t3_pos_1_054_48    m/z 431.3367     0   1   2   3E+4   9   9.5  RT (min)      lcorbi_t3_pos_3_011_42    m/z 431.3521     0   1   2E+5   11   11.5  RT (min)       lcorbi_t3_pos_2_005_11    m/z 431.8280     0   1E+5   13.5   14  RT (min)      lcorbi_t3_pos_1_036_08    m/z 432.1478     0   1   2   3E+5   2.5   3  RT (min)       lcorbi_t3_pos_4_004_14    m/z 432.1711     0   2   4   6   8E+4   1.5   2  RT (min)       lcorbi_t3_pos_4_068_57    m/z 432.2052     0   1   2E+4   6   6.5  RT (min)       lcorbi_t3_pos_4_111_62    m/z 432.2462     0   1   2   3E+4   10   10.5  RT (min)      lcorbi_t3_pos_3_053_48    m/z 432.2653     0   1E+4   11   11.5  RT (min)      lcorbi_t3_pos_2_005_11    m/z 432.2743     0   1   2E+4   9.5   10  RT (min)      lcorbi_t3_pos_1_046_22    m/z 432.2803     0   1   2   3   4E+4   4.5   5  RT (min)       lcorbi_t3_pos_1_002_01    m/z 432.2982     0   1E+5   13  RT (min)       lcorbi_t3_pos_4_108_62    m/z 432.3108     0   2   4E+4   9   9.5  RT (min)       lcorbi_t3_pos_4_081_58    m/z 432.3873     0   1E+5   14   14.5  RT (min)      lcorbi_t3_pos_4_078_58    m/z 432.8358     0   1E+5   13.5   14  RT (min)      lcorbi_t3_pos_1_036_08    m/z 432.8641     0   1   2   3E+4   1   1.5  RT (min)      lcorbi_t3_pos_1_002_01    m/z 433.0354     0   2   4   6E+4   18   18.5  RT (min)       lcorbi_t3_pos_4_091_60    m/z 433.1512     0   2   4E+4   2.5   3  RT (min)      lcorbi_t3_pos_4_004_14    m/z 433.2068     0   1   2E+4   9   9.5  RT (min)      lcorbi_t3_pos_4_066_57    m/z 433.2103     0   2   4   6E+4   1.5   2  RT (min)      lcorbi_t3_pos_3_051_12    m/z 433.2776     0   1   2E+4   13   13.5  RT (min)      lcorbi_t3_pos_3_042_14    m/z 433.2837     0   1E+4   4.5   5  RT (min)      lcorbi_t3_pos_1_001_49    m/z 433.3099     0   1   2E+4   13.5   14  RT (min)       lcorbi_t3_pos_4_087_59    m/z 433.3142     0   1   11.5   12  RT (min)      lcorbi_t3_pos_1_054_48    m/z 433.3143     0   1   2E+4   9   9.5  RT (min)      lcorbi_t3_pos_4_081_58    m/z 433.8223     0   2   4E+4   1   1.5  RT (min)      lcorbi_t3_pos_1_028_32    m/z 433.9216     0   2   4   6   8E+3   1   1.5  RT (min)      lcorbi_t3_pos_4_073_57    m/z 434.1529     0   1E+4   2.5   3  RT (min)      lcorbi_t3_pos_3_048_30    m/z 434.2187     0   2   4   6   8E+4   4.5   5  RT (min)       lcorbi_t3_pos_4_106_62    m/z 434.2384     0   2   4E+4   9   9.5  RT (min)       lcorbi_t3_pos_4_082_59    m/z 434.2749     0   2   4   6E+4   11   11.5  RT (min)       lcorbi_t3_pos_4_074_58    m/z 434.4205     0   1E+5   12   12.5  RT (min)       lcorbi_t3_pos_4_086_59    m/z 434.9915     0   1   2E+4   1.5   2  RT (min)      lcorbi_t3_pos_3_020_35    m/z 435.1222     0   1   2E+4   10   10.5  RT (min)      lcorbi_t3_pos_1_029_29    m/z 435.1241     0   1E+5   1   1.5  RT (min)       lcorbi_t3_pos_4_095_60    m/z 435.1917     0   1   2E+4   8   8.5  RT (min)      lcorbi_t3_pos_1_037_19    m/z 435.2418     0   2   4   6   8E+3   9   9.5  RT (min)      lcorbi_t3_pos_2_056_56    m/z 435.2549     0   1E+4   6   6.5  RT (min)      lcorbi_t3_pos_4_109_62    m/z 435.2647     0   1   2   3   4E+4   8   8.5  RT (min)      lcorbi_t3_pos_4_106_62    m/z 435.2854     0   1E+4   6   6.5  RT (min)      lcorbi_t3_pos_4_111_62    m/z 435.3331     0   1E+4   5.5   6  RT (min)      lcorbi_t3_pos_4_084_59    m/z 435.3378     0   2   4   6   8E+4   13.5   14  RT (min)      lcorbi_t3_pos_4_089_59    m/z 435.3599     0   1   2   3E+4   13.5   14  RT (min)      lcorbi_t3_pos_2_043_32    m/z 435.8386     0   1   2   3   4E+4   1   1.5  RT (min)      lcorbi_t3_pos_1_036_08    m/z 436.2177     0   1   2   3   4E+4   11   11.5  RT (min)       lcorbi_t3_pos_1_042_37    m/z 436.2483     0   2   4   6E+3   6   6.5  RT (min)      lcorbi_t3_pos_4_111_62    m/z 436.2661     0   2   4   6   8E+3   4.5   5  RT (min)      lcorbi_t3_pos_4_066_57    m/z 436.2694     0   1   2   3E+5   12   12.5  RT (min)       lcorbi_t3_pos_4_049_55    m/z 436.2695     0   1   2   3E+5   0   0.5  RT (min)      lcorbi_t3_pos_2_039_30    m/z 436.3670     0   1E+5   11   11.5   12  RT (min)       lcorbi_t3_pos_4_084_59    m/z 436.6879     0   2   4   6E+4   1   1.5  RT (min)      lcorbi_t3_pos_2_028_24    m/z 436.7677     0   1   2   3   4E+3   4.5   5  RT (min)      lcorbi_t3_pos_4_066_57    m/z 436.8590     0   2   4E+5   1   1.5  RT (min)       lcorbi_t3_pos_1_029_29    m/z 436.8702     0   2   4   6E+4   1   1.5  RT (min)       lcorbi_t3_pos_3_012_31    m/z 436.8798     0   1   2   3E+3   1   1.5  RT (min)      lcorbi_t3_pos_1_006_15    m/z 437.0554     0   1   2E+4   10   10.5  RT (min)      lcorbi_t3_pos_4_081_58    m/z 437.0643     0   1E+4   10   10.5  RT (min)      lcorbi_t3_pos_4_074_58    m/z 437.1033     0   2   4E+4   10   10.5  RT (min)       lcorbi_t3_pos_1_029_29    m/z 437.1265     0   2   4E+4   1.5   2  RT (min)       lcorbi_t3_pos_2_028_24    m/z 437.2210     0   2   4   6E+4   10   10.5  RT (min)       lcorbi_t3_pos_1_029_29    m/z 437.2363     0   1E+5   7.5   8  RT (min)      lcorbi_t3_pos_4_098_61    m/z 437.2543     0   2   4   6   8E+3   5   5.5  RT (min)      lcorbi_t3_pos_4_082_59    m/z 437.2728     0   2   4   6E+4   0   0.5  RT (min)      lcorbi_t3_pos_2_039_30    m/z 437.2729     0   1   2E+5   14   14.5  RT (min)      lcorbi_t3_pos_4_101_61    m/z 437.3092     0   1E+5   6   6.5  RT (min)      lcorbi_t3_pos_1_002_01    m/z 437.3207     0   1   2   3E+4   7   7.5  RT (min)      lcorbi_t3_pos_4_014_48    m/z 437.3738     0   2   4   6   8E+4   10   10.5  RT (min)      lcorbi_t3_pos_3_052_47    m/z 437.8356     0   2   4   6E+4   1   1.5  RT (min)      lcorbi_t3_pos_1_036_08    m/z 437.8624     0   1E+4   1   1.5  RT (min)      lcorbi_t3_pos_1_034_34    m/z 438.2270     0   1   2E+5   1.5   2  RT (min)      lcorbi_t3_pos_3_051_12    m/z 438.2391     0   1   2   3E+4   7.5   8  RT (min)      lcorbi_t3_pos_4_098_61    m/z 438.2756     0   2   4E+4   8.5   9  RT (min)      lcorbi_t3_pos_4_068_57    m/z 438.2980     0   2   4   6   8E+4   11.5   12  RT (min)      lcorbi_t3_pos_1_029_29    m/z 438.3115     0   2   4   6E+4   7   7.5  RT (min)       lcorbi_t3_pos_4_099_61    m/z 438.3484     0   1   2E+5   13   13.5  RT (min)       lcorbi_t3_pos_1_001_49    m/z 438.3790     0   1   2E+4   0   0.5  RT (min)      lcorbi_t3_pos_1_033_53    m/z 438.7286     0   2   4   6E+4   1   1.5  RT (min)      lcorbi_t3_pos_1_020_31    m/z 438.8562     0   1E+5   1   1.5  RT (min)       lcorbi_t3_pos_1_029_29    m/z 439.0193     0   1E+4   1   1.5  RT (min)      lcorbi_t3_pos_4_091_60    m/z 439.1438     0   1   2   3E+4   12.5   13  RT (min)       lcorbi_t3_pos_3_008_20    m/z 439.2415     0   1E+5   1   1.5  RT (min)       lcorbi_t3_pos_4_090_60    m/z 439.3205     0   1   2   3   4E+4   7   7.5  RT (min)      lcorbi_t3_pos_4_014_48    m/z 439.3207     0   1   2   3   4E+4   12.5   13  RT (min)      lcorbi_t3_pos_1_054_48    m/z 439.3461     0   2   4   6E+4   11   11.5  RT (min)       lcorbi_t3_pos_4_082_59    m/z 439.3572     0   1   2   3   4E+4   7.5   8  RT (min)       lcorbi_t3_pos_4_112_62    m/z 439.4198     0   2   4   6E+5   12.5   13  RT (min)       lcorbi_t3_pos_4_090_60    m/z 439.4260     0   2   4E+4   14.5   15  RT (min)      lcorbi_t3_pos_2_052_48    m/z 440.1915     0   2   4   6E+4   9.5   10  RT (min)      lcorbi_t3_pos_4_099_61    m/z 440.2854     0   1   2E+4   8   8.5  RT (min)      lcorbi_t3_pos_4_074_58    m/z 440.3136     0   1   2   3   4E+4   11.5   12  RT (min)      lcorbi_t3_pos_4_077_58    m/z 440.3218     0   1E+4   7.5   8  RT (min)      lcorbi_t3_pos_4_090_60    m/z 440.3582     0   2   4   6   8E+4   10   10.5  RT (min)       lcorbi_t3_pos_4_093_60    m/z 440.3603     0   2   4   6E+4   12   12.5  RT (min)       lcorbi_t3_pos_4_102_61    m/z 440.3605     0   1E+4   7.5   8  RT (min)      lcorbi_t3_pos_4_110_62    m/z 440.3852     0   1   2   3E+5   11   11.5  RT (min)      lcorbi_t3_pos_4_089_59    m/z 440.3958     0   1   2E+5   12   12.5  RT (min)       lcorbi_t3_pos_1_003_45    m/z 440.4100     0   2   4   6E+4   11   11.5  RT (min)      lcorbi_t3_pos_4_092_60    m/z 440.6536     0   1E+4   1   1.5  RT (min)      lcorbi_t3_pos_1_044_25    m/z 440.7256     0   1E+5   1   1.5  RT (min)       lcorbi_t3_pos_1_020_31    m/z 440.9652     0   2   4   6   8E+4   18   18.5  RT (min)      lcorbi_t3_pos_4_101_61    m/z 441.0693     0   1E+5   1  RT (min)       lcorbi_t3_pos_4_085_59    m/z 441.1110     0   1E+4   1.5   2  RT (min)      lcorbi_t3_pos_2_005_11    m/z 441.1862     0   1   2E+4   9.5   10  RT (min)      lcorbi_t3_pos_1_043_06    m/z 441.1883     0   2   4E+4   8   8.5  RT (min)      lcorbi_t3_pos_1_051_09    m/z 441.2076     0   1   2   3E+4   13   13.5  RT (min)      lcorbi_t3_pos_1_036_08    m/z 441.2096     0   1   2E+4   9.5   10  RT (min)      lcorbi_t3_pos_4_092_60    m/z 441.2249     0   1E+5   0   0.5  RT (min)      lcorbi_t3_pos_2_039_30    m/z 441.2249     0   1E+6   14   16  RT (min)       lcorbi_t3_pos_4_077_58    m/z 441.2998     0   1E+5   12   12.5  RT (min)       lcorbi_t3_pos_1_054_48    m/z 441.3000     0   2   4   6   8E+4   15   15.5  RT (min)       lcorbi_t3_pos_4_088_59    m/z 441.3435     0   2   4   6E+4   6   6.5  RT (min)       lcorbi_t3_pos_4_089_59    m/z 441.3617     0   1   2E+4   10   10.5  RT (min)      lcorbi_t3_pos_4_013_36    m/z 441.3728     0   1   2E+5   12.5   13  RT (min)       lcorbi_t3_pos_1_054_48    m/z 441.3799     0   2   4   6   8E+4   5   5.5  RT (min)       lcorbi_t3_pos_4_088_59    m/z 441.4133     0   1   2   3E+4   11   11.5  RT (min)      lcorbi_t3_pos_4_094_60    m/z 441.7879     0   2   4   6E+3   4.5   5  RT (min)      lcorbi_t3_pos_4_013_36    m/z 441.8741     0   1E+4   1   1.5  RT (min)      lcorbi_t3_pos_4_052_06    m/z 441.9354     0   1E+4   18.5   19  RT (min)      lcorbi_t3_pos_4_097_60    m/z 441.9357     0   2   4E+3   0.5   1  RT (min)      lcorbi_t3_pos_3_024_07    m/z 442.1805     0   2   4   6E+4   8   8.5  RT (min)       lcorbi_t3_pos_3_034_26    m/z 442.2283     0   2   4   6E+4   14   14.5  RT (min)      lcorbi_t3_pos_3_029_13    m/z 442.2283     0   1   2   3E+5   17.5   18  RT (min)      lcorbi_t3_pos_4_089_59    m/z 442.2283     0   1E+4   5   5.5  RT (min)      lcorbi_t3_pos_4_074_58    m/z 442.2647     0   1   2E+4   6.5   7  RT (min)       lcorbi_t3_pos_4_066_57    m/z 442.2799     0   1E+4   9.5   10  RT (min)      lcorbi_t3_pos_1_003_45    m/z 442.3011     0   2   4   6E+4   13.5   14  RT (min)       lcorbi_t3_pos_4_088_59    m/z 442.3034     0   1E+5   12   12.5  RT (min)       lcorbi_t3_pos_1_050_13    m/z 442.3274     0   1   2   3   4E+4   6.5   7  RT (min)       lcorbi_t3_pos_1_032_26    m/z 442.3375     0   1   2   3E+4   7.5   8  RT (min)       lcorbi_t3_pos_4_082_59    m/z 442.3470     0   1   2E+4   6   6.5  RT (min)      lcorbi_t3_pos_4_089_59    m/z 442.3700     0   1   12.5   13  RT (min)       lcorbi_t3_pos_1_054_48    m/z 442.3834     0   1   2E+4   5   5.5  RT (min)      lcorbi_t3_pos_4_088_59    m/z 442.4050     0   1   2E+5   6.5   7  RT (min)       lcorbi_t3_pos_4_014_48    m/z 442.4256     0   1   2   3   4E+4   12.5   13  RT (min)       lcorbi_t3_pos_3_029_13    m/z 442.6507     0   1E+4   1   1.5  RT (min)      lcorbi_t3_pos_2_004_42    m/z 442.7227     0   2   4   6   8E+4   1   1.5  RT (min)       lcorbi_t3_pos_1_020_31    m/z 442.8688     0   1E+4   1   1.5  RT (min)      lcorbi_t3_pos_4_074_58    m/z 442.8783     0   2   4E+4   18   18.5  RT (min)       lcorbi_t3_pos_4_098_61    m/z 442.8784     0   1   2E+4   1   1.5  RT (min)      lcorbi_t3_pos_4_077_58    m/z 443.0520     0   1   2   3E+4   10   10.5  RT (min)      lcorbi_t3_pos_1_029_29    m/z 443.0723     0   2   4   6E+4   10   10.5  RT (min)       lcorbi_t3_pos_1_029_29    m/z 443.0860     0   1   2   3   4E+4   5   5.5  RT (min)       lcorbi_t3_pos_4_110_62    m/z 443.1377     0   1E+4   8   8.5   9  RT (min)      lcorbi_t3_pos_4_067_57    m/z 443.1839     0   1   2E+4   8   8.5  RT (min)      lcorbi_t3_pos_3_034_26    m/z 443.1867     0   2   4   6   8E+4   11   11.5  RT (min)      lcorbi_t3_pos_4_098_61    m/z 443.1947     0   1   11   11.5  RT (min)      lcorbi_t3_pos_4_082_59    m/z 443.2309     0   1   2   3E+4   8.5   9  RT (min)      lcorbi_t3_pos_4_094_60    m/z 443.2439     0   2   4   6E+5   1.5   2  RT (min)       lcorbi_t3_pos_3_055_10    m/z 443.2681     0   2   4   6   8E+3   6.5   7  RT (min)      lcorbi_t3_pos_4_074_58    m/z 443.3245     0   1   12.5   13  RT (min)      lcorbi_t3_pos_2_005_11    m/z 443.3344     0   1   2   3E+5   11   11.5  RT (min)       lcorbi_t3_pos_4_083_59    m/z 443.3409     0   1E+4   7.5   8  RT (min)      lcorbi_t3_pos_4_082_59    m/z 443.4208     0   2   4E+4   10.5   11  RT (min)       lcorbi_t3_pos_4_090_60    m/z 444.1351     0   1E+4   4.5   5  RT (min)      lcorbi_t3_pos_4_100_61    m/z 444.1900     0   1E+4   11   11.5  RT (min)      lcorbi_t3_pos_3_045_43    m/z 444.2440     0   1   2E+4   5   5.5  RT (min)       lcorbi_t3_pos_4_074_58    m/z 444.2991     0   1   2E+4   11   11.5  RT (min)      lcorbi_t3_pos_3_021_39    m/z 444.3168     0   1E+4   6   6.5  RT (min)      lcorbi_t3_pos_4_106_62    m/z 444.3320     0   1E+5   10.5   11  RT (min)       lcorbi_t3_pos_4_082_59    m/z 444.4048     0   1   2E+5   6.5   7  RT (min)      lcorbi_t3_pos_2_044_21    m/z 444.4049     0   1   2E+5   13.5   14  RT (min)       lcorbi_t3_pos_4_088_59    m/z 444.6737     0   2   4   6E+4   1   1.5  RT (min)      lcorbi_t3_pos_1_023_02    m/z 444.7197     0   1   2   3E+4   1   1.5  RT (min)      lcorbi_t3_pos_1_020_31    m/z 444.8078     0   1   2E+5   0   0.5  RT (min)      lcorbi_t3_pos_1_007_07    m/z 444.9298     0   1E+4   1   1.5  RT (min)      lcorbi_t3_pos_3_012_31    m/z 445.0394     0   1   2   3E+4   1   1.5  RT (min)      lcorbi_t3_pos_3_010_44    m/z 445.1171     0   1E+4   6   6.5  RT (min)      lcorbi_t3_pos_3_004_24    m/z 445.1509     0   1   2   3E+4   10   10.5  RT (min)      lcorbi_t3_pos_1_029_29    m/z 445.1709     0   2   4E+3   7.5   8  RT (min)      lcorbi_t3_pos_3_036_37    m/z 445.2474     0   2   4   6E+3   5   5.5  RT (min)      lcorbi_t3_pos_4_082_59    m/z 445.2882     0   1   2   3E+4   5   5.5  RT (min)      lcorbi_t3_pos_1_001_49    m/z 445.3275     0   2   4   6   8E+4   12   12.5  RT (min)       lcorbi_t3_pos_4_071_57    m/z 445.3354     0   1   2   3   4E+4   10.5   11  RT (min)      lcorbi_t3_pos_4_089_59    m/z 446.0853     0   2   4   6   8E+4   1   1.5  RT (min)       lcorbi_t3_pos_1_043_06    m/z 446.2384     0   1   2E+4   5   5.5  RT (min)       lcorbi_t3_pos_4_111_62    m/z 446.2901     0   1E+5   9.5   10  RT (min)      lcorbi_t3_pos_1_003_45    m/z 446.2961     0   1E+4   5   5.5  RT (min)      lcorbi_t3_pos_4_090_60    m/z 446.3630     0   1E+5   13   13.5  RT (min)       lcorbi_t3_pos_3_029_13    m/z 447.1979     0   1   2   3E+3   7.5   8  RT (min)      lcorbi_t3_pos_4_049_55    m/z 447.2045     0   1E+5   5.5   6  RT (min)       lcorbi_t3_pos_4_030_05    m/z 447.2602     0   1   2   3   4E+4   4.5   5  RT (min)      lcorbi_t3_pos_4_112_62    m/z 447.3437     0   1   2   3   4E+4   6.5   7  RT (min)       lcorbi_t3_pos_4_089_59    m/z 447.8355     0   1   2E+4   7   7.5  RT (min)       lcorbi_t3_pos_4_088_59    m/z 447.8452     0   1   2E+4   6.5   7  RT (min)      lcorbi_t3_pos_4_085_59    m/z 447.9320     0   2   4   6E+4   1   1.5  RT (min)       lcorbi_t3_pos_4_077_58    m/z 448.1218     0   1E+5   2.5   3  RT (min)       lcorbi_t3_pos_4_004_14    m/z 448.2079     0   1   2   3E+4   5.5   6  RT (min)      lcorbi_t3_pos_4_019_47    m/z 448.2178     0   1   2   3E+4   7.5   8  RT (min)       lcorbi_t3_pos_4_074_58    m/z 448.2539     0   2   4E+4   4.5   5  RT (min)       lcorbi_t3_pos_4_110_62    m/z 448.2693     0   1E+4   6.5   7  RT (min)      lcorbi_t3_pos_1_032_26    m/z 448.2695     0   1E+5   16  RT (min)       lcorbi_t3_pos_4_070_57    m/z 448.3269     0   1E+5   10.5   11  RT (min)       lcorbi_t3_pos_4_090_60    m/z 448.3371     0   2   4   6   8E+3   7   7.5  RT (min)      lcorbi_t3_pos_3_020_35    m/z 448.3379     0   2   4E+4   10.5   11  RT (min)       lcorbi_t3_pos_3_020_35    m/z 448.3468     0   1E+4   6.5   7  RT (min)      lcorbi_t3_pos_4_084_59    m/z 448.3633     0   1   2   3E+4   8.5   9  RT (min)       lcorbi_t3_pos_4_093_60    m/z 448.7574     0   2   4   6   8E+4   1   1.5  RT (min)       lcorbi_t3_pos_1_020_31    m/z 448.9149     0   2   4   6E+4   0   0.5  RT (min)      lcorbi_t3_pos_1_001_49    m/z 449.0109     0   1   2E+4   1   1.5  RT (min)      lcorbi_t3_pos_3_011_42    m/z 449.1252     0   1E+4   2.5   3  RT (min)      lcorbi_t3_pos_4_004_14    m/z 449.1489     0   1   2E+4   8.5   9  RT (min)      lcorbi_t3_pos_3_024_07    m/z 449.1959     0   1E+4   8.5   9  RT (min)      lcorbi_t3_pos_3_048_30    m/z 449.2016     0   1   2   3E+4   5.5   6  RT (min)       lcorbi_t3_pos_4_030_05    m/z 449.2211     0   1E+4   7.5   8  RT (min)      lcorbi_t3_pos_4_090_60    m/z 449.2728     0   2   4   6E+4   10   10.5  RT (min)      lcorbi_t3_pos_1_012_14    m/z 449.2819     0   1   13.5   14  RT (min)       lcorbi_t3_pos_1_054_48    m/z 449.2939     0   2   4   6E+4   11   11.5  RT (min)      lcorbi_t3_pos_4_066_57    m/z 449.3011     0   1E+4   6   6.5  RT (min)      lcorbi_t3_pos_4_111_62    m/z 449.3303     0   1   2E+4   10.5   11  RT (min)      lcorbi_t3_pos_4_092_60    m/z 449.3600     0   2   4   6E+4   13   14  RT (min)       lcorbi_t3_pos_4_081_58    m/z 449.3679     0   1   2E+4   8.5   9  RT (min)      lcorbi_t3_pos_4_091_60    m/z 449.3850     0   1E+4   7   7.5  RT (min)      lcorbi_t3_pos_4_066_57    m/z 449.8893     0   2   4   6E+3   1   1.5  RT (min)      lcorbi_t3_pos_1_034_34    m/z 450.2042     0   2   4E+4   10   10.5  RT (min)       lcorbi_t3_pos_1_037_19    m/z 450.2123     0   1E+5   8   8.5  RT (min)       lcorbi_t3_pos_1_037_19    m/z 450.2334     0   1E+4   6.5   7  RT (min)      lcorbi_t3_pos_4_018_01    m/z 450.2698     0   1   2E+4   9   9.5  RT (min)       lcorbi_t3_pos_1_036_08    m/z 450.3125     0   1   2   3E+4   4.5   5  RT (min)       lcorbi_t3_pos_4_092_60    m/z 450.3191     0   1   2   3   4E+4   11.5   12  RT (min)       lcorbi_t3_pos_4_078_58    m/z 450.3214     0   2   4   6E+4   9   9.5  RT (min)       lcorbi_t3_pos_1_037_19    m/z 450.3215     0   1E+4   6   6.5  RT (min)      lcorbi_t3_pos_1_012_14    m/z 450.4154     0   2   4   6   8E+4   11   11.5  RT (min)       lcorbi_t3_pos_4_084_59    m/z 450.6649     0   2   4   6   8E+4   1   1.5  RT (min)      lcorbi_t3_pos_4_077_58    m/z 450.7545     0   1E+5   1   1.5  RT (min)       lcorbi_t3_pos_1_020_31    m/z 451.0475     0   1E+4   9   9.5  RT (min)      lcorbi_t3_pos_3_046_01    m/z 451.1018     0   2   4E+5   1   1.5  RT (min)       lcorbi_t3_pos_4_096_60    m/z 451.2076     0   1E+4   10   10.5  RT (min)      lcorbi_t3_pos_2_037_27    m/z 451.2156     0   1   2E+4   8   8.5  RT (min)      lcorbi_t3_pos_1_037_19    m/z 451.2520     0   1E+5   7.5   8  RT (min)       lcorbi_t3_pos_4_099_61    m/z 451.2884     0   2   4   6   8E+3   7.5   8  RT (min)      lcorbi_t3_pos_4_099_61    m/z 451.3185     0   1E+5   13   13.5  RT (min)      lcorbi_t3_pos_1_054_48    m/z 451.3631     0   1E+5   11   11.5  RT (min)       lcorbi_t3_pos_4_082_59    m/z 452.1916     0   1   2E+4   11   11.5  RT (min)      lcorbi_t3_pos_2_035_34    m/z 452.2012     0   1E+4   10   10.5  RT (min)      lcorbi_t3_pos_4_047_28    m/z 452.2432     0   1   2E+4   10.5   11  RT (min)      lcorbi_t3_pos_4_068_57    m/z 452.2772     0   1E+5   10.5   11  RT (min)       lcorbi_t3_pos_1_050_13    m/z 452.3007     0   1   2E+4   6   6.5  RT (min)       lcorbi_t3_pos_4_109_62    m/z 452.3218     0   1   2   3E+4   9.5   10  RT (min)      lcorbi_t3_pos_1_027_43    m/z 452.3271     0   1   2   3E+4   7   7.5  RT (min)       lcorbi_t3_pos_4_102_61    m/z 452.3372     0   2   4   6   8E+4   13   13.5  RT (min)      lcorbi_t3_pos_3_048_30    m/z 452.3628     0   1E+5   14   14.5  RT (min)       lcorbi_t3_pos_4_078_58    m/z 452.6621     0   1   2   3   4E+4   1   1.5  RT (min)      lcorbi_t3_pos_4_078_58    m/z 452.7376     0   1   2E+4   5.5   6  RT (min)       lcorbi_t3_pos_4_075_58    m/z 452.7515     0   1   2E+4   1   1.5  RT (min)      lcorbi_t3_pos_1_036_08    m/z 452.8331     0   1E+5   1   1.5  RT (min)      lcorbi_t3_pos_1_029_29    m/z 452.8442     0   1   2E+4   1   1.5  RT (min)      lcorbi_t3_pos_3_010_44    m/z 452.8540     0   2   4E+4   1   1.5  RT (min)      lcorbi_t3_pos_1_036_08    m/z 453.0808     0   2   4   6E+4   10   10.5  RT (min)       lcorbi_t3_pos_1_029_29    m/z 453.1732     0   2   4   6E+3   7.5   8  RT (min)      lcorbi_t3_pos_4_106_62    m/z 453.1909     0   2   4   6   8E+3   7   7.5  RT (min)      lcorbi_t3_pos_4_113_62    m/z 453.2391     0   1E+4   5.5   6  RT (min)      lcorbi_t3_pos_4_081_58    m/z 453.3041     0   1   6   6.5  RT (min)      lcorbi_t3_pos_4_084_59    m/z 453.3187     0   1   2E+4   9   9.5  RT (min)      lcorbi_t3_pos_1_028_32    m/z 453.3188     0   1E+5   12.5   13  RT (min)       lcorbi_t3_pos_1_007_07    m/z 453.3246     0   1E+3   5.5   6  RT (min)      lcorbi_t3_pos_4_089_59    m/z 453.3342     0   1   2E+5   13   13.5  RT (min)       lcorbi_t3_pos_1_054_48    m/z 453.3617     0   1   11.5   12  RT (min)      lcorbi_t3_pos_1_003_45    m/z 453.4780     0   1   2   3   4E+4   13.5   14  RT (min)      lcorbi_t3_pos_1_015_11    m/z 454.0277     0   2   4   6E+4   1   1.5  RT (min)      lcorbi_t3_pos_3_010_44    m/z 454.1991     0   2   4   6   8E+4   8   8.5  RT (min)       lcorbi_t3_pos_3_034_26    m/z 454.2281     0   2   4E+4   10   10.5  RT (min)      lcorbi_t3_pos_1_037_19    m/z 454.2356     0   1E+5   0   0.5  RT (min)      lcorbi_t3_pos_3_044_46    m/z 454.3739     0   1   2E+4   10.5   11  RT (min)      lcorbi_t3_pos_4_013_36    m/z 454.4009     0   1   12   12.5  RT (min)      lcorbi_t3_pos_4_090_60    m/z 454.4451     0   1E+5   14.5   15  RT (min)      lcorbi_t3_pos_4_038_38    m/z 454.7137     0   1   1   1.5  RT (min)      lcorbi_t3_pos_1_031_40    m/z 454.7233     0   2   4E+4   1   1.5  RT (min)      lcorbi_t3_pos_2_020_22    m/z 454.8304     0   1   2   3   4E+4   1   1.5  RT (min)      lcorbi_t3_pos_1_029_29    m/z 454.9691     0   2   4   6   8E+4   18   18.5  RT (min)       lcorbi_t3_pos_4_069_57    m/z 455.1581     0   1   2E+4   1.5   2  RT (min)      lcorbi_t3_pos_2_037_27    m/z 455.1678     0   1   2E+4   9   9.5  RT (min)      lcorbi_t3_pos_4_066_57    m/z 455.1692     0   1   2   3   4E+4   12.5   13  RT (min)      lcorbi_t3_pos_4_078_58    m/z 455.2026     0   1   2E+4   8   8.5  RT (min)       lcorbi_t3_pos_3_034_26    m/z 455.2964     0   2   4   6   8E+4   11   11.5  RT (min)      lcorbi_t3_pos_1_050_13    m/z 455.3134     0   1E+5   13.5   14  RT (min)       lcorbi_t3_pos_4_082_59    m/z 455.3158     0   2   4   6   8E+4   15   15.5  RT (min)       lcorbi_t3_pos_4_087_59    m/z 455.3330     0   1   2   3   4E+4   13.5   14  RT (min)      lcorbi_t3_pos_4_109_62    m/z 455.3593     0   1   11   11.5  RT (min)       lcorbi_t3_pos_1_003_45    m/z 455.4148     0   2   4   6   8E+5   12   12.5  RT (min)       lcorbi_t3_pos_4_097_60    m/z 456.1167     0   1E+4   4.5   5  RT (min)      lcorbi_t3_pos_1_032_26    m/z 456.1962     0   1   2   3E+4   8   8.5  RT (min)       lcorbi_t3_pos_3_034_26    m/z 456.2326     0   1E+4   10   10.5  RT (min)      lcorbi_t3_pos_3_056_56    m/z 456.2440     0   1E+4   5.5   6  RT (min)      lcorbi_t3_pos_4_074_58    m/z 456.2803     0   1E+4   7   7.5  RT (min)      lcorbi_t3_pos_4_082_59    m/z 456.3320     0   1   2E+4   8.5   9  RT (min)      lcorbi_t3_pos_4_077_58    m/z 456.3528     0   1E+4   5.5   6  RT (min)      lcorbi_t3_pos_4_088_59    m/z 456.4412     0   1   13   13.5  RT (min)       lcorbi_t3_pos_2_044_21    m/z 456.6997     0   1   2E+4   1   1.5  RT (min)      lcorbi_t3_pos_4_045_08    m/z 456.7107     0   1   2   3E+4   1   1.5  RT (min)      lcorbi_t3_pos_2_028_24    m/z 456.7204     0   2   4   6   8E+4   1   1.5  RT (min)      lcorbi_t3_pos_1_034_34    m/z 456.8446     0   1   2E+4   1   1.5  RT (min)      lcorbi_t3_pos_1_005_42    m/z 456.9736     0   1   2E+4   1   1.5  RT (min)      lcorbi_t3_pos_4_032_40    m/z 457.0643     0   1E+4   9   9.5  RT (min)      lcorbi_t3_pos_1_034_34    m/z 457.1120     0   1E+4   4.5   5  RT (min)      lcorbi_t3_pos_3_048_30    m/z 457.1366     0   1   2E+5   17.5   18  RT (min)       lcorbi_t3_pos_4_075_58    m/z 457.1737     0   1   2E+4   8   8.5  RT (min)      lcorbi_t3_pos_4_105_61    m/z 457.2368     0   1E+4   6   6.5  RT (min)      lcorbi_t3_pos_4_112_62    m/z 457.2468     0   1   2E+4   7.5   8  RT (min)      lcorbi_t3_pos_4_109_62    m/z 457.2717     0   1   2E+4   1   1.5  RT (min)      lcorbi_t3_pos_1_001_49    m/z 457.2717     0   1E+5   13.5   14  RT (min)       lcorbi_t3_pos_4_085_59    m/z 457.3022     0   1   5   5.5  RT (min)      lcorbi_t3_pos_1_002_01    m/z 457.3279     0   1   2   3   4E+3   6.5   7  RT (min)      lcorbi_t3_pos_1_001_49    m/z 457.3385     0   1   2E+3   5   5.5  RT (min)      lcorbi_t3_pos_1_001_49    m/z 457.3501     0   1   12   12.5  RT (min)       lcorbi_t3_pos_1_003_45    m/z 457.3678     0   1   2   3   4E+4   7.5   8  RT (min)       lcorbi_t3_pos_4_107_62    m/z 457.3678     0   1   12   12.5  RT (min)       lcorbi_t3_pos_1_003_45    m/z 457.4305     0   1   2E+5   12  RT (min)       lcorbi_t3_pos_4_097_60    m/z 457.8199     0   1   2E+4   7   7.5  RT (min)       lcorbi_t3_pos_4_084_59    m/z 457.8294     0   2   4   6E+3   6.5   7  RT (min)      lcorbi_t3_pos_4_084_59    m/z 457.9611     0   1   2E+4   1   1.5  RT (min)      lcorbi_t3_pos_1_029_29    m/z 458.2021     0   2   4E+4   8.5   9  RT (min)      lcorbi_t3_pos_4_084_59    m/z 458.2597     0   1   2E+4   5.5   6  RT (min)       lcorbi_t3_pos_4_066_57    m/z 458.2724     0   1   2.5   3  RT (min)       lcorbi_t3_pos_4_074_58    m/z 458.3215     0   1E+4   7   7.5  RT (min)      lcorbi_t3_pos_4_087_59    m/z 458.3323     0   2   4   6   8E+3   6.5   7  RT (min)      lcorbi_t3_pos_4_082_59    m/z 458.3477     0   1E+4   9   9.5  RT (min)      lcorbi_t3_pos_4_078_58    m/z 458.3478     0   2   4   6E+4   11   11.5  RT (min)       lcorbi_t3_pos_2_030_14    m/z 458.3712     0   1E+4   7.5   8  RT (min)      lcorbi_t3_pos_4_112_62    m/z 458.3712     0   1   2   3E+4   12   12.5  RT (min)      lcorbi_t3_pos_2_044_21    m/z 458.6968     0   1   2E+4   1   1.5  RT (min)      lcorbi_t3_pos_1_034_34    m/z 458.7172     0   2   4E+4   1   1.5  RT (min)      lcorbi_t3_pos_1_028_32    m/z 458.7175     0   2   4E+4   1   1.5  RT (min)      lcorbi_t3_pos_1_031_40    m/z 458.7862     0   1E+5   1   1.5  RT (min)       lcorbi_t3_pos_1_020_31    m/z 458.8615     0   2   4E+4   1   1.5  RT (min)       lcorbi_t3_pos_4_005_33    m/z 459.0372     0   1   2   3   4E+4   10   10.5  RT (min)      lcorbi_t3_pos_1_029_29    m/z 459.1347     0   1E+4   5   5.5  RT (min)       lcorbi_t3_pos_3_004_24    m/z 459.1535     0   1   2E+4   10   10.5  RT (min)      lcorbi_t3_pos_4_077_58    m/z 459.1909     0   1   2   3E+4   10   10.5  RT (min)      lcorbi_t3_pos_1_031_40    m/z 459.2478     0   1   2E+4   11   11.5  RT (min)      lcorbi_t3_pos_2_030_14    m/z 459.2633     0   2   4E+3   5.5   6  RT (min)      lcorbi_t3_pos_4_074_58    m/z 459.2760     0   1   2.5   3  RT (min)      lcorbi_t3_pos_4_074_58    m/z 459.3220     0   1   2   3E+5   13   13.5  RT (min)      lcorbi_t3_pos_1_002_01    m/z 459.3370     0   1   12   12.5  RT (min)      lcorbi_t3_pos_1_054_48    m/z 459.3512     0   1   2   3E+4   11   11.5  RT (min)      lcorbi_t3_pos_2_005_11    m/z 460.1127     0   1   2E+4   4.5   5  RT (min)       lcorbi_t3_pos_4_105_61    m/z 460.2544     0   1   5   5.5  RT (min)      lcorbi_t3_pos_4_090_60    m/z 460.3058     0   1   2   3   4E+4   13.5   14  RT (min)      lcorbi_t3_pos_1_039_23    m/z 460.3117     0   1   2E+4   5.5   6  RT (min)       lcorbi_t3_pos_4_074_58    m/z 460.3422     0   1   2E+4   8   8.5  RT (min)       lcorbi_t3_pos_4_049_55    m/z 460.7833     0   1E+5   1   1.5  RT (min)       lcorbi_t3_pos_1_020_31    m/z 460.9717     0   1   2   3E+4   0   0.5  RT (min)      lcorbi_t3_pos_1_001_49    m/z 461.1079     0   2   4   6   8E+4   7   7.5  RT (min)       lcorbi_t3_pos_4_014_48    m/z 461.1258     0   1   2   3E+4   12.5   13  RT (min)      lcorbi_t3_pos_2_040_25    m/z 461.1483     0   1E+4   7.5   8  RT (min)      lcorbi_t3_pos_4_090_60    m/z 461.2202     0   1E+5   5.5   6  RT (min)       lcorbi_t3_pos_4_030_05    m/z 461.2424     0   1   2E+4   4.5   5  RT (min)      lcorbi_t3_pos_4_082_59    m/z 461.2687     0   1   2   3E+4   8.5   9  RT (min)      lcorbi_t3_pos_4_100_61    m/z 461.2760     0   2   4E+4   4.5   5  RT (min)      lcorbi_t3_pos_4_112_62    m/z 461.2978     0   1   10   10.5  RT (min)      lcorbi_t3_pos_4_074_58    m/z 461.3150     0   1   2   3E+3   5.5   6  RT (min)      lcorbi_t3_pos_4_086_59    m/z 461.3456     0   2   4   6   8E+3   8   8.5  RT (min)      lcorbi_t3_pos_4_041_54    m/z 461.3628     0   2   4   6   8E+4   14   14.5  RT (min)       lcorbi_t3_pos_4_082_59    m/z 461.4019     0   1E+5   12.5   13  RT (min)       lcorbi_t3_pos_4_092_60    m/z 461.6981     0   2   4   6E+3   4.5   5  RT (min)      lcorbi_t3_pos_4_082_59    m/z 461.8569     0   1   2   3E+4   5   5.5  RT (min)       lcorbi_t3_pos_4_088_59    m/z 462.1112     0   1   2E+4   7   7.5  RT (min)      lcorbi_t3_pos_4_014_48    m/z 462.1972     0   1   2   3   4E+3   5.5   6  RT (min)      lcorbi_t3_pos_4_098_61    m/z 462.2236     0   1   2   3E+4   5.5   6  RT (min)      lcorbi_t3_pos_4_100_61    m/z 462.2444     0   2   4   6   8E+3   4.5   5  RT (min)      lcorbi_t3_pos_4_106_62    m/z 462.2660     0   1   2E+5   14   15   16  RT (min)       lcorbi_t3_pos_1_054_48    m/z 462.2674     0   2   4E+4   8.5   9  RT (min)      lcorbi_t3_pos_4_092_60    m/z 462.2696     0   1   2   3   4E+4   4.5   5  RT (min)      lcorbi_t3_pos_4_107_62    m/z 462.2932     0   2   4E+4   10   10.5  RT (min)       lcorbi_t3_pos_4_076_58    m/z 462.2963     0   1   2   3E+4   6.5   7  RT (min)       lcorbi_t3_pos_4_084_59    m/z 462.3063     0   2   4E+4   11   11.5  RT (min)       lcorbi_t3_pos_4_090_60    m/z 462.3308     0   1E+4   5.5   6  RT (min)      lcorbi_t3_pos_1_028_32    m/z 462.3426     0   1   2   3   4E+4   10.5   11  RT (min)      lcorbi_t3_pos_4_090_60    m/z 462.3490     0   1   2   3   4E+4   5.5   6  RT (min)       lcorbi_t3_pos_3_024_07    m/z 462.3580     0   2   4   6E+4   12.5   13  RT (min)       lcorbi_t3_pos_3_029_13    m/z 462.3585     0   1E+4   5   5.5  RT (min)      lcorbi_t3_pos_4_089_59    m/z 462.4307     0   2   4E+4   13.5   14  RT (min)       lcorbi_t3_pos_4_084_59    m/z 462.6448     0   1   2   3   4E+4   1   1.5  RT (min)      lcorbi_t3_pos_4_076_58    m/z 462.7804     0   2   4   6E+4   1   1.5  RT (min)      lcorbi_t3_pos_1_020_31    m/z 462.8325     0   2   4   6E+3   5.5   6  RT (min)      lcorbi_t3_pos_4_089_59    m/z 462.8506     0   1   2E+4   5.5   6  RT (min)      lcorbi_t3_pos_4_089_59    m/z 462.8597     0   2   4   6E+3   5   5.5  RT (min)      lcorbi_t3_pos_4_089_59    m/z 462.9669     0   1   2   3   4E+5   0   0.5  RT (min)      lcorbi_t3_pos_1_005_42    m/z 463.0266     0   1   2   3E+4   1   1.5  RT (min)      lcorbi_t3_pos_4_005_33    m/z 463.0873     0   2   4   6   8E+3   5.5   6  RT (min)      lcorbi_t3_pos_3_004_24    m/z 463.1308     0   1   2   3E+5   11.5   12  RT (min)       lcorbi_t3_pos_4_113_62    m/z 463.1754     0   1   2E+4   7.5   8  RT (min)       lcorbi_t3_pos_4_110_62    m/z 463.2173     0   1   2   3E+4   5.5   6  RT (min)       lcorbi_t3_pos_4_030_05    m/z 463.2456     0   1   2E+4   10   10.5  RT (min)      lcorbi_t3_pos_4_074_58    m/z 463.2997     0   1E+4   6.5   7  RT (min)      lcorbi_t3_pos_4_084_59    m/z 463.3521     0   2   4E+3   5.5   6  RT (min)      lcorbi_t3_pos_1_054_48    m/z 463.4260     0   2   4   6   8E+4   13.5   14  RT (min)       lcorbi_t3_pos_4_106_62    m/z 463.8652     0   1   2   3E+4   1   1.5  RT (min)      lcorbi_t3_pos_4_077_58    m/z 463.9696     0   1   2   3   4E+4   0   0.5  RT (min)      lcorbi_t3_pos_1_005_42    m/z 464.1915     0   1E+4   6.5   7  RT (min)      lcorbi_t3_pos_4_077_58    m/z 464.2126     0   1   2E+4   10.5   11  RT (min)      lcorbi_t3_pos_3_025_52    m/z 464.2642     0   1   2E+4   9   9.5  RT (min)       lcorbi_t3_pos_4_093_60    m/z 464.2853     0   2   4   6   8E+3   7   7.5  RT (min)      lcorbi_t3_pos_3_023_05    m/z 464.3005     0   1   2   3E+4   8.5   9  RT (min)      lcorbi_t3_pos_1_001_49    m/z 464.3007     0   1   2E+4   7.5   8  RT (min)      lcorbi_t3_pos_1_037_19    m/z 464.3370     0   1E+4   6   6.5  RT (min)      lcorbi_t3_pos_4_110_62    m/z 464.3582     0   2   4E+4   11.5   12  RT (min)      lcorbi_t3_pos_4_089_59    m/z 464.3676     0   1   2   3E+5   9   9.5  RT (min)      lcorbi_t3_pos_3_052_47    m/z 464.4294     0   1   2   3E+4   13.5   14  RT (min)      lcorbi_t3_pos_1_002_01    m/z 464.4310     0   1E+5   11.5   12  RT (min)       lcorbi_t3_pos_4_086_59    m/z 464.6417     0   2   4E+4   1   1.5  RT (min)      lcorbi_t3_pos_4_078_58    m/z 464.8599     0   2   4   6E+4   1   1.5  RT (min)      lcorbi_t3_pos_4_075_58    m/z 465.0542     0   1   2   3E+4   10   10.5  RT (min)      lcorbi_t3_pos_1_035_28    m/z 465.1802     0   2   4E+4   9.5   10  RT (min)      lcorbi_t3_pos_1_036_08    m/z 465.1943     0   1   2   3   4E+3   4.5   5  RT (min)      lcorbi_t3_pos_4_074_58    m/z 465.2959     0   1E+5   11.5   12  RT (min)       lcorbi_t3_pos_4_084_59    m/z 465.3787     0   1   2   3E+5   11   11.5  RT (min)       lcorbi_t3_pos_4_085_59    m/z 465.8743     0   2   4E+4   18   18.5  RT (min)      lcorbi_t3_pos_4_072_57    m/z 466.1754     0   1E+4   8.5   9  RT (min)      lcorbi_t3_pos_2_044_21    m/z 466.2588     0   1E+6   10.5   11  RT (min)       lcorbi_t3_pos_1_029_29    m/z 466.2589     0   1E+4   0   0.5  RT (min)      lcorbi_t3_pos_2_011_10    m/z 466.2589     0   2   4   6   8E+5   14   15  RT (min)       lcorbi_t3_pos_1_023_02    m/z 466.2799     0   1E+4   8   8.5  RT (min)      lcorbi_t3_pos_4_082_59    m/z 466.3163     0   1   2   3E+5   8.5   9  RT (min)       lcorbi_t3_pos_1_037_19    m/z 466.3293     0   1   12   12.5  RT (min)       lcorbi_t3_pos_4_090_60    m/z 466.3315     0   1E+4   7.5   8  RT (min)      lcorbi_t3_pos_4_048_24    m/z 466.6390     0   1   2E+4   1   1.5  RT (min)      lcorbi_t3_pos_2_040_25    m/z 466.6609     0   1   2   3   4E+3   1   1.5  RT (min)      lcorbi_t3_pos_1_029_29    m/z 466.7284     0   1   2   3   4E+4   1   1.5  RT (min)      lcorbi_t3_pos_1_029_29    m/z 467.2426     0   1E+4   9   9.5  RT (min)      lcorbi_t3_pos_2_041_54    m/z 467.2623     0   1   2E+5   14.5   15  RT (min)      lcorbi_t3_pos_4_089_59    m/z 467.2625     0   1   2E+5   14   14.5  RT (min)      lcorbi_t3_pos_4_084_59    m/z 467.2853     0   1E+4   7.5   8  RT (min)      lcorbi_t3_pos_1_028_32    m/z 467.2923     0   1   13   13.5  RT (min)       lcorbi_t3_pos_4_096_60    m/z 467.3012     0   1   2   3E+4   5   5.5  RT (min)      lcorbi_t3_pos_1_001_49    m/z 467.3103     0   1E+5   12   12.5  RT (min)       lcorbi_t3_pos_2_005_11    m/z 467.3157     0   1   15   15.5  RT (min)       lcorbi_t3_pos_4_099_61    m/z 467.3480     0   1E+4   9   9.5  RT (min)      lcorbi_t3_pos_4_079_58    m/z 467.4573     0   2   4   6   8E+4   15   15.5  RT (min)      lcorbi_t3_pos_2_052_48    m/z 468.1102     0   1   2   3E+3   4.5   5  RT (min)      lcorbi_t3_pos_4_014_48    m/z 468.2652     0   2   4   6E+4   10   10.5  RT (min)       lcorbi_t3_pos_1_023_02    m/z 468.3452     0   1   2   3   4E+4   11   11.5  RT (min)       lcorbi_t3_pos_4_079_58    m/z 468.3703     0   1   2   3   4E+4   11   11.5  RT (min)      lcorbi_t3_pos_1_032_26    m/z 468.4049     0   1   2E+4   11   11.5  RT (min)      lcorbi_t3_pos_4_088_59    m/z 468.4412     0   2   4   6   8E+4   12   12.5  RT (min)       lcorbi_t3_pos_4_092_60    m/z 468.6580     0   1   2E+4   1   1.5  RT (min)      lcorbi_t3_pos_4_070_57    m/z 468.8150     0   1   2E+5   1   1.5  RT (min)      lcorbi_t3_pos_1_020_31    m/z 468.8466     0   1E+4   5.5   6  RT (min)      lcorbi_t3_pos_4_086_59    m/z 469.0469     0   1   2E+4   1   1.5  RT (min)      lcorbi_t3_pos_4_052_06    m/z 469.1986     0   2   4E+4   10   10.5  RT (min)       lcorbi_t3_pos_4_097_60    m/z 469.2193     0   1   2E+4   9   9.5  RT (min)      lcorbi_t3_pos_4_097_60    m/z 469.3290     0   1E+5   13.5   14  RT (min)       lcorbi_t3_pos_4_103_61    m/z 469.3385     0   1   2   3E+4   6   6.5  RT (min)       lcorbi_t3_pos_4_083_59    m/z 469.3482     0   2   4E+3   5.5   6  RT (min)      lcorbi_t3_pos_4_083_59    m/z 469.3654     0   1E+5   13.5   14  RT (min)       lcorbi_t3_pos_4_073_57    m/z 469.8306     0   1E+4   6   6.5  RT (min)      lcorbi_t3_pos_4_084_59    m/z 469.8402     0   1   2E+4   6   6.5  RT (min)      lcorbi_t3_pos_4_088_59    m/z 470.0016     0   1   2E+4   1   1.5  RT (min)      lcorbi_t3_pos_4_052_06    m/z 470.2020     0   1E+4   10   10.5  RT (min)      lcorbi_t3_pos_4_043_13    m/z 470.2537     0   1E+4   9.5   10  RT (min)      lcorbi_t3_pos_1_038_30    m/z 470.2596     0   1   2   3   4E+3   5.5   6  RT (min)      lcorbi_t3_pos_4_082_59    m/z 470.2611     0   1E+4   1   1.5  RT (min)      lcorbi_t3_pos_4_087_59    m/z 470.2748     0   1   2   3E+4   10   10.5  RT (min)      lcorbi_t3_pos_1_034_34    m/z 470.2960     0   2   4   6   8E+3   7.5   8  RT (min)      lcorbi_t3_pos_4_066_57    m/z 470.3144     0   2   4E+4   10   10.5  RT (min)       lcorbi_t3_pos_4_072_57    m/z 470.3322     0   2   4   6E+3   6   6.5  RT (min)      lcorbi_t3_pos_4_089_59    m/z 470.3506     0   2   4   6E+4   10   10.5  RT (min)       lcorbi_t3_pos_4_087_59    m/z 470.3701     0   1E+5   5.5   6  RT (min)       lcorbi_t3_pos_1_008_36    m/z 470.3922     0   1   12.5   13  RT (min)      lcorbi_t3_pos_4_095_60    m/z 470.3955     0   1E+5   11   11.5  RT (min)      lcorbi_t3_pos_4_087_59    m/z 470.6972     0   1   2E+4   1   1.5  RT (min)      lcorbi_t3_pos_2_024_15    m/z 470.8120     0   1   2E+5   1   1.5  RT (min)      lcorbi_t3_pos_1_020_31    m/z 470.8899     0   1   2   3E+4   1   1.5  RT (min)       lcorbi_t3_pos_4_077_58    m/z 470.9468     0   2   4   6   8E+4   18   18.5  RT (min)       lcorbi_t3_pos_4_096_60    m/z 471.2143     0   1   2E+5   15.5   16   16.5  RT (min)       lcorbi_t3_pos_2_052_48    m/z 471.2143     0   1E+5   13.5   14  RT (min)       lcorbi_t3_pos_4_001_49    m/z 471.2143     0   1   2   3   4E+5   17.5   18  RT (min)       lcorbi_t3_pos_4_085_59    m/z 471.2489     0   1   2E+4   5   5.5  RT (min)       lcorbi_t3_pos_4_107_62    m/z 471.2717     0   1E+4   6   6.5  RT (min)      lcorbi_t3_pos_2_029_43    m/z 471.3177     0   1   2E+4   5   5.5  RT (min)       lcorbi_t3_pos_3_048_30    m/z 471.3446     0   1   2   3   4E+4   12.5   13  RT (min)       lcorbi_t3_pos_4_077_58    m/z 471.3541     0   2   4   6   8E+4   5   5.5  RT (min)       lcorbi_t3_pos_4_089_59    m/z 471.3735     0   1   2E+4   5.5   6  RT (min)      lcorbi_t3_pos_4_089_59    m/z 471.3834     0   2   4   6E+4   14.5   15  RT (min)       lcorbi_t3_pos_4_099_61    m/z 471.4521     0   1   2   3E+4   12.5   13  RT (min)       lcorbi_t3_pos_4_092_60    m/z 471.9905     0   1   2E+4   1   1.5  RT (min)      lcorbi_t3_pos_3_048_30    m/z 472.1449     0   1   2   3   4E+4   8   8.5  RT (min)      lcorbi_t3_pos_4_099_61    m/z 472.2047     0   2   4   6   8E+4   7   7.5  RT (min)      lcorbi_t3_pos_4_107_62    m/z 472.2176     0   2   4   6E+4   7.5   8  RT (min)      lcorbi_t3_pos_4_106_62    m/z 472.2752     0   1E+4   6   6.5  RT (min)      lcorbi_t3_pos_4_106_62    m/z 472.3576     0   1   2E+4   5   5.5  RT (min)      lcorbi_t3_pos_4_089_59    m/z 472.3912     0   1   2   3E+5   14.5   15  RT (min)       lcorbi_t3_pos_4_100_61    m/z 472.6943     0   2   4   6E+4   1   1.5  RT (min)      lcorbi_t3_pos_1_036_08    m/z 472.8091     0   2   4   6E+4   1   1.5  RT (min)      lcorbi_t3_pos_1_020_31    m/z 472.8184     0   1E+3   1   1.5  RT (min)      lcorbi_t3_pos_1_051_09    m/z 472.9012     0   1   2   3E+4   1   1.5  RT (min)      lcorbi_t3_pos_4_070_57    m/z 472.9202     0   2   4   6E+4   18   18.5  RT (min)       lcorbi_t3_pos_4_112_62    m/z 473.0293     0   1E+4   9   9.5  RT (min)      lcorbi_t3_pos_1_029_29    m/z 473.1394     0   1E+4   8   8.5  RT (min)      lcorbi_t3_pos_4_099_61    m/z 473.2076     0   1   2E+4   7   7.5  RT (min)      lcorbi_t3_pos_4_110_62    m/z 473.2206     0   1   2   3   4E+4   10   10.5  RT (min)      lcorbi_t3_pos_4_074_58    m/z 473.2874     0   2   4   6   8E+3   6   6.5  RT (min)      lcorbi_t3_pos_1_012_14    m/z 473.2899     0   2   4   6E+4   11   11.5  RT (min)       lcorbi_t3_pos_4_091_60    m/z 473.3476     0   1E+5   10.5   11  RT (min)       lcorbi_t3_pos_1_050_13    m/z 473.3657     0   1E+5   13   14  RT (min)       lcorbi_t3_pos_4_103_61    m/z 474.1689     0   1E+4   4.5   5  RT (min)      lcorbi_t3_pos_1_003_45    m/z 474.1769     0   1E+4   4.5   5  RT (min)      lcorbi_t3_pos_2_024_15    m/z 474.2545     0   1E+4   5   5.5  RT (min)      lcorbi_t3_pos_4_082_59    m/z 474.2672     0   1   2E+4   6.5   7  RT (min)      lcorbi_t3_pos_2_054_29    m/z 474.2850     0   2   4E+4   8.5   9  RT (min)      lcorbi_t3_pos_4_052_06    m/z 474.2932     0   1   2E+4   11   11.5  RT (min)      lcorbi_t3_pos_4_083_59    m/z 474.3273     0   1   2   3   4E+4   5.5   6  RT (min)       lcorbi_t3_pos_4_098_61    m/z 474.3577     0   1   2E+4   8   8.5  RT (min)      lcorbi_t3_pos_4_049_55    m/z 474.3943     0   2   4   6   8E+4   12   12.5  RT (min)       lcorbi_t3_pos_1_051_09    m/z 474.4570     0   1E+5   13   13.5  RT (min)       lcorbi_t3_pos_4_092_60    m/z 474.6914     0   1   2   3E+4   1   1.5  RT (min)      lcorbi_t3_pos_2_003_19    m/z 474.8333     0   2   4   6   8E+4   18   18.5  RT (min)       lcorbi_t3_pos_4_072_57    m/z 474.8337     0   2   4   6   8E+3   1   1.5  RT (min)      lcorbi_t3_pos_1_016_05    m/z 475.1123     0   1   2   3   4E+4   5   5.5  RT (min)       lcorbi_t3_pos_4_014_48    m/z 475.1309     0   1   2   3   4E+4   10   10.5  RT (min)       lcorbi_t3_pos_1_029_29    m/z 475.2369     0   1E+5   13   13.5  RT (min)       lcorbi_t3_pos_4_081_58    m/z 475.2972     0   2   4E+4   12   12.5  RT (min)       lcorbi_t3_pos_4_075_58    m/z 475.3046     0   2   4   6E+3   6   6.5  RT (min)      lcorbi_t3_pos_4_092_60    m/z 475.3783     0   1   2E+4   7.5   8  RT (min)       lcorbi_t3_pos_4_106_62    m/z 475.8544     0   2   4   6   8E+3   5.5   6  RT (min)      lcorbi_t3_pos_4_089_59    m/z 476.1611     0   1E+5   1   2  RT (min)       lcorbi_t3_pos_4_086_59    m/z 476.1875     0   1E+4   4.5   5  RT (min)      lcorbi_t3_pos_4_098_61    m/z 476.3005     0   1E+4   12   12.5  RT (min)      lcorbi_t3_pos_3_044_46    m/z 476.3065     0   1   2   3   4E+4   4.5   5  RT (min)       lcorbi_t3_pos_1_001_49    m/z 476.3221     0   1   2   3   4E+4   11   11.5  RT (min)      lcorbi_t3_pos_4_090_60    m/z 476.3289     0   2   4   6E+4   5.5   6  RT (min)      lcorbi_t3_pos_4_086_59    m/z 476.3290     0   2   4   6E+4   13.5   14  RT (min)      lcorbi_t3_pos_1_011_44    m/z 476.3817     0   2   4   6   8E+4   14   14.5  RT (min)      lcorbi_t3_pos_4_086_59    m/z 476.7573     0   2   4   6E+4   1   1.5  RT (min)      lcorbi_t3_pos_1_029_29    m/z 477.1028     0   2   4   6E+3   6   6.5  RT (min)      lcorbi_t3_pos_4_112_62    m/z 477.2805     0   2   4   6   8E+4   10.5   11  RT (min)       lcorbi_t3_pos_2_030_14    m/z 477.3100     0   1E+4   4.5   5  RT (min)      lcorbi_t3_pos_1_001_49    m/z 477.3320     0   2   4   6   8E+3   5.5   6  RT (min)      lcorbi_t3_pos_4_085_59    m/z 477.9087     0   1   2E+4   1   1.5  RT (min)      lcorbi_t3_pos_4_077_58    m/z 478.0107     0   1E+4   1   1.5  RT (min)      lcorbi_t3_pos_4_052_06    m/z 478.2435     0   2   4E+4   8   8.5  RT (min)       lcorbi_t3_pos_1_004_24    m/z 478.2646     0   2   4E+4   9   9.5  RT (min)       lcorbi_t3_pos_2_036_12    m/z 478.2799     0   1E+4   9   9.5  RT (min)      lcorbi_t3_pos_2_036_12    m/z 478.2964     0   1   2   3   4E+4   13.5   14  RT (min)       lcorbi_t3_pos_4_091_60    m/z 478.3163     0   1E+5   7.5   8  RT (min)      lcorbi_t3_pos_1_001_49    m/z 478.3164     0   1   2   3   4E+5   0   0.5  RT (min)      lcorbi_t3_pos_4_010_43    m/z 478.3292     0   1E+4   11   11.5  RT (min)       lcorbi_t3_pos_3_027_27    m/z 478.3833     0   1   2   3E+5   9   9.5  RT (min)      lcorbi_t3_pos_3_052_47    m/z 478.4467     0   1E+4   12   12.5  RT (min)       lcorbi_t3_pos_1_054_48    m/z 478.8357     0   1E+4   1   1.5  RT (min)      lcorbi_t3_pos_3_034_26    m/z 478.8437     0   1E+5   18   18.5  RT (min)       lcorbi_t3_pos_4_076_58    m/z 478.8438     0   2   4E+5   1   1.5  RT (min)       lcorbi_t3_pos_2_028_24    m/z 478.8765     0   2   4   6E+4   18   18.5  RT (min)       lcorbi_t3_pos_4_070_57    m/z 478.8816     0   1   2   3E+4   1   1.5  RT (min)      lcorbi_t3_pos_4_034_42    m/z 478.9044     0   1E+4   1   1.5  RT (min)      lcorbi_t3_pos_4_079_58    m/z 478.9298     0   1   2E+4   1   1.5  RT (min)      lcorbi_t3_pos_1_029_29    m/z 479.0005     0   2   4   6   8E+3   1   1.5  RT (min)      lcorbi_t3_pos_4_034_42    m/z 479.2681     0   1   2E+4   9   9.5  RT (min)      lcorbi_t3_pos_2_031_20    m/z 479.3198     0   1E+5   0   0.5  RT (min)      lcorbi_t3_pos_4_010_43    m/z 479.3199     0   1   2E+5   14   14.5  RT (min)      lcorbi_t3_pos_4_112_62    m/z 479.3331     0   2   4   6E+3   11   11.5  RT (min)      lcorbi_t3_pos_3_027_27    m/z 479.3474     0   1E+4   6   6.5  RT (min)      lcorbi_t3_pos_3_004_24    m/z 479.3497     0   1   12   12.5  RT (min)       lcorbi_t3_pos_1_003_45    m/z 479.3562     0   2   4   6   8E+4   6   6.5  RT (min)      lcorbi_t3_pos_4_107_62    m/z 479.3926     0   1   2   3E+5   17.5   18  RT (min)      lcorbi_t3_pos_4_075_58    m/z 479.4123     0   1E+5   13   13.5  RT (min)       lcorbi_t3_pos_4_096_60    m/z 480.1073     0   1E+5   5.5   6  RT (min)      lcorbi_t3_pos_1_008_36    m/z 480.1573     0   1E+5   11.5   12  RT (min)       lcorbi_t3_pos_4_113_62    m/z 480.2381     0   1E+4   10   10.5  RT (min)      lcorbi_t3_pos_2_011_10    m/z 480.2763     0   2   4   6   8E+3   8.5   9  RT (min)      lcorbi_t3_pos_4_044_32    m/z 480.2825     0   1   2   3E+4   8.5   9  RT (min)      lcorbi_t3_pos_4_101_61    m/z 480.2990     0   2   4E+4   11   11.5  RT (min)      lcorbi_t3_pos_2_030_14    m/z 480.3226     0   1E+5   10   10.5  RT (min)      lcorbi_t3_pos_4_074_58    m/z 480.3530     0   1   2   3E+4   12   12.5  RT (min)      lcorbi_t3_pos_3_023_05    m/z 480.3594     0   1E+4   6   6.5  RT (min)      lcorbi_t3_pos_4_106_62    m/z 480.4259     0   2   4   6   8E+4   12.5   13  RT (min)       lcorbi_t3_pos_4_086_59    m/z 480.7773     0   1E+4   4.5   5  RT (min)      lcorbi_t3_pos_4_079_58    m/z 480.8408     0   1   2E+5   1   1.5  RT (min)       lcorbi_t3_pos_1_020_31    m/z 481.1107     0   1   2   3E+4   5.5   6  RT (min)      lcorbi_t3_pos_1_008_36    m/z 481.1165     0   1   2E+4   1   1.5  RT (min)       lcorbi_t3_pos_4_082_59    m/z 481.1353     0   1   2   3E+4   10   10.5  RT (min)      lcorbi_t3_pos_1_029_29    m/z 481.1780     0   1   2E+4   1.5   2  RT (min)      lcorbi_t3_pos_1_043_06    m/z 481.2535     0   1E+5   13   13.5  RT (min)       lcorbi_t3_pos_4_074_58    m/z 481.3120     0   1   2E+4   11.5   12  RT (min)       lcorbi_t3_pos_1_040_27    m/z 481.3253     0   1E+4   10   10.5  RT (min)      lcorbi_t3_pos_2_005_11    m/z 481.3314     0   1   2   3E+4   12.5   13  RT (min)       lcorbi_t3_pos_4_087_59    m/z 481.3654     0   1E+5   14   14.5  RT (min)       lcorbi_t3_pos_4_072_57    m/z 481.4281     0   1   2E+5   13   13.5  RT (min)       lcorbi_t3_pos_1_003_45    m/z 482.2150     0   1   2   3   4E+3   6.5   7  RT (min)      lcorbi_t3_pos_4_098_61    m/z 482.4764     0   1E+5   15   15.5  RT (min)       lcorbi_t3_pos_4_038_38    m/z 482.6382     0   2   4E+4   1   1.5  RT (min)      lcorbi_t3_pos_4_068_57    m/z 482.7643     0   2   4   6E+3   5   5.5  RT (min)      lcorbi_t3_pos_4_079_58    m/z 482.7948     0   2   4   6E+3   4.5   5  RT (min)      lcorbi_t3_pos_4_082_59    m/z 482.8554     0   1   2   3E+4   1   1.5  RT (min)      lcorbi_t3_pos_1_019_16    m/z 482.9277     0   1E+5   1   1.5  RT (min)       lcorbi_t3_pos_4_067_57    m/z 483.2629     0   1E+4   7.5   8  RT (min)      lcorbi_t3_pos_1_028_32    m/z 483.2717     0   2   4   6E+4   0   0.5  RT (min)      lcorbi_t3_pos_3_054_15    m/z 483.2719     0   2   4E+5   16   17   18  RT (min)       lcorbi_t3_pos_4_092_60    m/z 483.2719     0   1E+5   13.5   14  RT (min)      lcorbi_t3_pos_4_096_60    m/z 483.2989     0   1E+5   11.5   12  RT (min)       lcorbi_t3_pos_1_054_48    m/z 483.3470     0   2   4   6E+4   12.5   13  RT (min)       lcorbi_t3_pos_4_094_60    m/z 483.3581     0   1   2   3E+4   12.5   13  RT (min)      lcorbi_t3_pos_2_035_34    m/z 483.3729     0   1E+4   11   11.5  RT (min)       lcorbi_t3_pos_3_020_35    m/z 483.4048     0   2   4   6   8E+4   13   13.5  RT (min)       lcorbi_t3_pos_3_054_15    m/z 484.1169     0   2   4   6E+3   1.5   2  RT (min)      lcorbi_t3_pos_4_086_59    m/z 484.2039     0   2   4   6E+3   2.5   3  RT (min)      lcorbi_t3_pos_1_034_34    m/z 484.2696     0   1   2E+5   0   0.5  RT (min)      lcorbi_t3_pos_2_039_30    m/z 484.2696     0   2   4E+5   14   14.5  RT (min)       lcorbi_t3_pos_3_027_27    m/z 484.3117     0   2   4   6   8E+3   8   8.5  RT (min)      lcorbi_t3_pos_4_066_57    m/z 484.3301     0   1   10.5   11  RT (min)      lcorbi_t3_pos_4_093_60    m/z 484.3494     0   1E+4   5   5.5  RT (min)      lcorbi_t3_pos_4_084_59    m/z 484.3504     0   1   2E+4   12.5   13  RT (min)      lcorbi_t3_pos_4_094_60    m/z 484.3668     0   1   11.5   12  RT (min)      lcorbi_t3_pos_4_082_59    m/z 484.3844     0   2   4   6   8E+4   10   10.5  RT (min)       lcorbi_t3_pos_4_013_36    m/z 484.4115     0   1   2   3   4E+5   11   12  RT (min)       lcorbi_t3_pos_4_085_59    m/z 484.6352     0   1   2   3   4E+4   1   1.5  RT (min)      lcorbi_t3_pos_4_073_57    m/z 484.7889     0   2   4   6E+4   1   1.5  RT (min)      lcorbi_t3_pos_2_003_19    m/z 485.1572     0   1E+4   7.5   8  RT (min)      lcorbi_t3_pos_4_113_62    m/z 485.2647     0   1   2   3E+4   5   5.5  RT (min)       lcorbi_t3_pos_4_106_62    m/z 485.2729     0   2   4E+4   0   0.5  RT (min)      lcorbi_t3_pos_2_039_30    m/z 485.2730     0   1   2E+5   14   14.5  RT (min)      lcorbi_t3_pos_4_071_57    m/z 485.2896     0   1   2   3E+4   12   12.5  RT (min)      lcorbi_t3_pos_4_098_61    m/z 485.3333     0   1   2   3   4E+4   5   5.5  RT (min)       lcorbi_t3_pos_1_054_48    m/z 485.3374     0   1   11.5   12  RT (min)      lcorbi_t3_pos_4_108_62    m/z 486.2757     0   2   4   6E+4   9   9.5  RT (min)      lcorbi_t3_pos_4_097_60    m/z 486.2909     0   1E+4   6.5   7  RT (min)      lcorbi_t3_pos_4_066_57    m/z 486.3174     0   1   5.5   6  RT (min)      lcorbi_t3_pos_4_092_60    m/z 486.3367     0   1E+4   5   5.5  RT (min)      lcorbi_t3_pos_4_082_59    m/z 486.3710     0   2   4   6E+4   13   13.5  RT (min)       lcorbi_t3_pos_4_075_58    m/z 486.4205     0   2   4   6   8E+4   12   12.5  RT (min)       lcorbi_t3_pos_1_003_45    m/z 486.6322     0   1   2E+4   1   1.5  RT (min)      lcorbi_t3_pos_4_067_57    m/z 486.6584     0   2   4   6E+4   1   1.5  RT (min)       lcorbi_t3_pos_2_028_24    m/z 486.7860     0   2   4   6   8E+4   1   1.5  RT (min)      lcorbi_t3_pos_1_029_29    m/z 486.8064     0   2   4   6E+4   18   18.5  RT (min)       lcorbi_t3_pos_4_068_57    m/z 486.8675     0   1   2   3   4E+4   1   1.5  RT (min)       lcorbi_t3_pos_4_075_58    m/z 487.3608     0   1   2E+5   11   11.5  RT (min)       lcorbi_t3_pos_4_086_59    m/z 488.1915     0   2   4   6E+4   10   10.5  RT (min)       lcorbi_t3_pos_4_092_60    m/z 488.2702     0   1   2E+4   5   5.5  RT (min)      lcorbi_t3_pos_4_082_59    m/z 488.3007     0   1   2   3E+4   8.5   9  RT (min)      lcorbi_t3_pos_1_037_19    m/z 488.3036     0   1   2E+4   5  RT (min)       lcorbi_t3_pos_4_090_60    m/z 488.3191     0   1   2E+4   4.5   5  RT (min)       lcorbi_t3_pos_4_075_58    m/z 488.3254     0   2   4   6   8E+4   11   11.5  RT (min)       lcorbi_t3_pos_4_089_59    m/z 488.3371     0   1E+4   8   8.5  RT (min)      lcorbi_t3_pos_4_067_57    m/z 488.3430     0   1   2E+4   6   6.5  RT (min)      lcorbi_t3_pos_4_106_62    m/z 488.3582     0   1E+5   10.5   11  RT (min)       lcorbi_t3_pos_4_083_59    m/z 488.3947     0   2   4   6E+4   13.5   14  RT (min)       lcorbi_t3_pos_4_087_59    m/z 488.4100     0   1E+5   14.5   15  RT (min)       lcorbi_t3_pos_4_101_61    m/z 488.4362     0   1   2   3E+5   11.5   12  RT (min)       lcorbi_t3_pos_4_096_60    m/z 488.6554     0   1E+5   1   1.5  RT (min)       lcorbi_t3_pos_2_028_24    m/z 489.1859     0   1E+4   10   10.5  RT (min)      lcorbi_t3_pos_1_042_37    m/z 489.1941     0   2   4   6   8E+3   1.5   2  RT (min)      lcorbi_t3_pos_4_013_36    m/z 489.2250     0   1E+5   13.5   14  RT (min)       lcorbi_t3_pos_4_067_57    m/z 489.2250     0   1   2   3E+5   15.5   16  RT (min)       lcorbi_t3_pos_4_101_61    m/z 489.2738     0   1   2   3   4E+3   5   5.5  RT (min)      lcorbi_t3_pos_4_090_60    m/z 489.3144     0   1   2   3E+4   5   5.5  RT (min)      lcorbi_t3_pos_1_001_49    m/z 489.3227     0   2   4   6E+3   4.5   5  RT (min)      lcorbi_t3_pos_4_078_58    m/z 489.3464     0   2   4E+3   6   6.5  RT (min)      lcorbi_t3_pos_4_066_57    m/z 489.8760     0   2   4E+4   1   1.5  RT (min)      lcorbi_t3_pos_1_020_31    m/z 490.1409     0   1E+4   5.5   6  RT (min)      lcorbi_t3_pos_1_047_41    m/z 490.2330     0   1E+4   4.5   5  RT (min)      lcorbi_t3_pos_4_106_62    m/z 490.2332     0   1   2   3   4E+3   2.5   3  RT (min)      lcorbi_t3_pos_4_032_40    m/z 490.2340     0   1   2E+4   7   7.5  RT (min)      lcorbi_t3_pos_4_110_62    m/z 490.2859     0   1E+4   4.5   5  RT (min)      lcorbi_t3_pos_4_074_58    m/z 490.2906     0   1   2E+4   10   10.5  RT (min)       lcorbi_t3_pos_2_030_14    m/z 490.3222     0   1   2   3   4E+3   5   5.5  RT (min)      lcorbi_t3_pos_4_074_58    m/z 490.3891     0   1   2E+5   13   13.5  RT (min)       lcorbi_t3_pos_1_002_01    m/z 490.4042     0   1   14.5   15  RT (min)      lcorbi_t3_pos_4_099_61    m/z 490.4467     0   1   14.5   15  RT (min)      lcorbi_t3_pos_4_099_61    m/z 490.4980     0   1E+5   12   12.5  RT (min)       lcorbi_t3_pos_4_090_60    m/z 491.0329     0   2   4   6E+4   1   1.5  RT (min)      lcorbi_t3_pos_4_005_33    m/z 491.1440     0   1   2   3E+4   10   10.5  RT (min)      lcorbi_t3_pos_1_035_28    m/z 491.2064     0   1   2E+4   9   9.5  RT (min)       lcorbi_t3_pos_4_108_62    m/z 491.2311     0   1   2   3E+4   9   9.5  RT (min)      lcorbi_t3_pos_4_102_61    m/z 491.2462     0   1   2   3   4E+4   1.5   2  RT (min)       lcorbi_t3_pos_4_084_59    m/z 491.2995     0   1   2E+4   5.5   6  RT (min)       lcorbi_t3_pos_4_086_59    m/z 491.3199     0   2   4   6E+4   11   11.5  RT (min)       lcorbi_t3_pos_4_095_60    m/z 491.3317     0   1   2E+4   1.5   2  RT (min)      lcorbi_t3_pos_4_093_60    m/z 491.3754     0   2   4   6   8E+4   12   12.5  RT (min)       lcorbi_t3_pos_4_092_60    m/z 491.4572     0   2   4   6E+4   14   14.5  RT (min)       lcorbi_t3_pos_4_110_62    m/z 492.2149     0   1   2E+5   1.5   2  RT (min)       lcorbi_t3_pos_4_097_60    m/z 492.3085     0   1   2E+5   10   10.5  RT (min)       lcorbi_t3_pos_1_050_13    m/z 492.3238     0   1   2   3E+4   12   12.5  RT (min)       lcorbi_t3_pos_4_066_57    m/z 492.3439     0   1   2E+4   11   11.5  RT (min)      lcorbi_t3_pos_4_090_60    m/z 492.3637     0   2   4E+4   10   10.5  RT (min)       lcorbi_t3_pos_4_083_59    m/z 492.4412     0   2   4E+4   14   14.5  RT (min)       lcorbi_t3_pos_4_108_62    m/z 492.8457     0   2   4   6   8E+4   1   1.5  RT (min)      lcorbi_t3_pos_1_020_31    m/z 492.9775     0   2   4E+3   0   0.5  RT (min)      lcorbi_t3_pos_4_048_24    m/z 493.1128     0   2   4E+4   7   7.5  RT (min)       lcorbi_t3_pos_4_104_61    m/z 493.3718     0   2   4   6E+4   6   6.5  RT (min)      lcorbi_t3_pos_4_112_62    m/z 493.4084     0   1E+5   17.5   18  RT (min)      lcorbi_t3_pos_4_080_58    m/z 493.8878     0   1   2E+4   1   1.5  RT (min)      lcorbi_t3_pos_4_077_58    m/z 494.1183     0   1   2.5   3  RT (min)      lcorbi_t3_pos_4_106_62    m/z 494.1867     0   1E+5   7   7.5  RT (min)      lcorbi_t3_pos_4_112_62    m/z 494.2749     0   1E+4   8.5   9  RT (min)      lcorbi_t3_pos_4_070_57    m/z 494.2902     0   2   4   6   8E+4   10   10.5  RT (min)      lcorbi_t3_pos_1_002_01    m/z 494.3265     0   2   4   6   8E+3   8   8.5  RT (min)      lcorbi_t3_pos_4_078_58    m/z 494.3748     0   1E+4   6   6.5  RT (min)      lcorbi_t3_pos_4_106_62    m/z 494.3922     0   2   4   6E+4   9   9.5  RT (min)      lcorbi_t3_pos_1_021_18    m/z 494.6466     0   1   2   3   4E+4   1   1.5  RT (min)      lcorbi_t3_pos_2_028_24    m/z 494.8177     0   1E+5   1   1.5  RT (min)      lcorbi_t3_pos_1_029_29    m/z 494.8891     0   2   4   6E+3   1   1.5  RT (min)      lcorbi_t3_pos_2_051_35    m/z 495.0111     0   1E+4   9   9.5  RT (min)      lcorbi_t3_pos_1_035_28    m/z 495.1663     0   2   4   6   8E+4   1  RT (min)       lcorbi_t3_pos_4_082_59    m/z 495.1895     0   1E+4   7   7.5  RT (min)      lcorbi_t3_pos_4_112_62    m/z 495.4048     0   1   2E+5   12   13  RT (min)       lcorbi_t3_pos_4_095_60    m/z 495.4409     0   2   4E+4   13.5   14   14.5  RT (min)       lcorbi_t3_pos_4_097_60    m/z 495.4886     0   1E+5   15   15.5  RT (min)       lcorbi_t3_pos_4_038_38    m/z 495.8336     0   1   2E+4   0   0.5  RT (min)      lcorbi_t3_pos_4_018_01    m/z 496.2753     0   1   2   3   4E+4   11   11.5  RT (min)      lcorbi_t3_pos_1_029_29    m/z 496.3262     0   1   2   3E+4   13.5   14  RT (min)      lcorbi_t3_pos_2_045_40    m/z 496.3303     0   1E+5   10.5   11  RT (min)      lcorbi_t3_pos_4_068_57    m/z 496.3399     0   1   2   3E+5   0   0.5  RT (min)      lcorbi_t3_pos_1_037_19    m/z 496.3917     0   2   4E+4   11   11.5  RT (min)      lcorbi_t3_pos_1_032_26    m/z 496.4035     0   1   2   3E+4   11   11.5  RT (min)      lcorbi_t3_pos_1_032_26    m/z 496.8149     0   1E+5   1   1.5  RT (min)      lcorbi_t3_pos_1_029_29    m/z 496.8398     0   2   4   6   8E+4   1   1.5  RT (min)       lcorbi_t3_pos_1_023_02    m/z 497.0529     0   1E+5   1   1.5  RT (min)      lcorbi_t3_pos_3_010_44    m/z 497.1128     0   1   2   3   4E+4   10   10.5  RT (min)      lcorbi_t3_pos_1_029_29    m/z 497.1704     0   1   2   3   4E+4   1.5   2  RT (min)       lcorbi_t3_pos_4_090_60    m/z 497.1978     0   2   4   6E+3   4.5   5  RT (min)      lcorbi_t3_pos_4_074_58    m/z 497.3434     0   2   4   6E+4   0   0.5  RT (min)      lcorbi_t3_pos_1_008_36    m/z 497.3879     0   1E+5   11   11.5  RT (min)       lcorbi_t3_pos_4_076_58    m/z 497.4164     0   1E+5   15   15.5  RT (min)       lcorbi_t3_pos_1_036_08    m/z 498.2487     0   2   4   6   8E+3   9   9.5  RT (min)      lcorbi_t3_pos_4_009_50    m/z 498.2698     0   1   2   3E+4   10   10.5  RT (min)      lcorbi_t3_pos_1_054_48    m/z 498.2699     0   1   2   3E+5   11.5   12  RT (min)       lcorbi_t3_pos_1_054_48    m/z 498.2932     0   1E+5   8.5   9  RT (min)      lcorbi_t3_pos_4_101_61    m/z 498.3637     0   1E+4   8   8.5  RT (min)      lcorbi_t3_pos_4_088_59    m/z 498.3650     0   1E+4   5   5.5  RT (min)      lcorbi_t3_pos_4_083_59    m/z 498.4000     0   1E+4   10.5   11  RT (min)      lcorbi_t3_pos_1_050_13    m/z 498.4013     0   1E+4   5.5   6  RT (min)      lcorbi_t3_pos_1_008_36    m/z 498.4270     0   2   4   6   8E+4   11.5   12  RT (min)      lcorbi_t3_pos_4_083_59    m/z 498.6844     0   2   4E+4   1   1.5  RT (min)      lcorbi_t3_pos_1_020_31    m/z 498.8407     0   1   2   3E+4   1   1.5  RT (min)      lcorbi_t3_pos_2_028_24    m/z 499.0737     0   1   2E+4   10   10.5  RT (min)      lcorbi_t3_pos_4_078_58    m/z 499.1673     0   1   2E+3   1.5   2  RT (min)      lcorbi_t3_pos_3_021_39    m/z 499.2749     0   2   4   6E+4   13.5   14  RT (min)      lcorbi_t3_pos_2_039_30    m/z 499.2861     0   2   4   6   8E+4   10.5   11  RT (min)      lcorbi_t3_pos_1_050_13    m/z 499.2961     0   1   2E+4   8.5   9  RT (min)      lcorbi_t3_pos_4_105_61    m/z 499.3053     0   2   4   6   8E+4   10   11  RT (min)       lcorbi_t3_pos_2_005_11    m/z 499.3055     0   1   2   3   4E+4   8   8.5  RT (min)       lcorbi_t3_pos_1_014_03    m/z 499.3419     0   1   2   3   4E+4   12.5   13  RT (min)       lcorbi_t3_pos_4_083_59    m/z 499.3491     0   2   4   6E+3   5.5   6  RT (min)      lcorbi_t3_pos_4_089_59    m/z 499.9048     0   1   2   3E+5   1   1.5  RT (min)      lcorbi_t3_pos_4_067_57    m/z 500.1681     0   1   2   3E+4   5.5   6  RT (min)      lcorbi_t3_pos_3_024_07    m/z 500.2490     0   1   2   3   4E+4   10   10.5  RT (min)      lcorbi_t3_pos_4_094_60    m/z 500.2771     0   1E+5   10.5   11  RT (min)       lcorbi_t3_pos_4_068_57    m/z 500.3067     0   2   4   6E+3   7   7.5  RT (min)      lcorbi_t3_pos_1_037_19    m/z 500.3514     0   1   11   11.5  RT (min)       lcorbi_t3_pos_3_027_27    m/z 500.3668     0   1   2E+5   11   11.5  RT (min)       lcorbi_t3_pos_3_027_27    m/z 500.3793     0   1   2   3E+4   8   8.5  RT (min)       lcorbi_t3_pos_4_074_58    m/z 500.6094     0   1   2E+4   1   1.5  RT (min)      lcorbi_t3_pos_4_078_58    m/z 500.6815     0   2   4   6E+4   1   1.5  RT (min)      lcorbi_t3_pos_1_020_31    m/z 500.7567     0   1E+4   4.5   5  RT (min)      lcorbi_t3_pos_4_074_58    m/z 500.9058     0   1E+5   1   1.5  RT (min)      lcorbi_t3_pos_4_067_57    m/z 501.2806     0   1   2   3   4E+4   10.5   11  RT (min)      lcorbi_t3_pos_4_071_57    m/z 501.3830     0   2   4   6E+4   8   8.5   9  RT (min)       lcorbi_t3_pos_4_093_60    m/z 501.8797     0   1   2   3E+4   1   1.5  RT (min)      lcorbi_t3_pos_4_072_57    m/z 501.9092     0   1   1   1.5  RT (min)      lcorbi_t3_pos_4_084_59    m/z 502.2355     0   1   2   3   4E+4   10.5   11  RT (min)       lcorbi_t3_pos_4_071_57    m/z 502.2859     0   1   2   3E+4   5.5   6  RT (min)       lcorbi_t3_pos_4_095_60    m/z 502.3165     0   1   2E+4   12   12.5  RT (min)       lcorbi_t3_pos_4_042_09    m/z 502.3291     0   1   2   3   4E+4   11   11.5  RT (min)       lcorbi_t3_pos_1_023_02    m/z 502.3586     0   2   4   6E+3   6.5   7  RT (min)      lcorbi_t3_pos_4_066_57    m/z 502.3741     0   2   4E+4   11   11.5  RT (min)       lcorbi_t3_pos_4_096_60    m/z 502.3825     0   1E+5   11.5   12  RT (min)       lcorbi_t3_pos_3_027_27    m/z 502.6323     0   1   2   3   4E+4   1   1.5  RT (min)      lcorbi_t3_pos_4_078_58    m/z 503.2894     0   2   4   6   8E+3   5.5   6  RT (min)      lcorbi_t3_pos_4_090_60    m/z 503.3771     0   1   2   3E+4   11   11.5  RT (min)      lcorbi_t3_pos_4_092_60    m/z 503.8855     0   2   4E+4   6.5   7  RT (min)       lcorbi_t3_pos_4_034_42    m/z 504.1802     0   1E+5   1   1.5  RT (min)      lcorbi_t3_pos_3_021_39    m/z 504.2038     0   2   4   6E+3   1   1.5  RT (min)      lcorbi_t3_pos_1_009_50    m/z 504.2978     0   1   2   3E+4   5.5   6  RT (min)       lcorbi_t3_pos_4_090_60    m/z 504.2991     0   2   4   6   8E+4   11   12  RT (min)       lcorbi_t3_pos_4_068_57    m/z 504.3085     0   2   4   6   8E+4   11  RT (min)       lcorbi_t3_pos_4_070_57    m/z 504.3379     0   1E+4   5.5   6  RT (min)      lcorbi_t3_pos_4_066_57    m/z 504.3429     0   1   2   3   4E+4   11.5   12  RT (min)       lcorbi_t3_pos_4_072_57    m/z 504.3868     0   1   2E+4   6.5   7  RT (min)      lcorbi_t3_pos_4_087_59    m/z 504.6294     0   2   4   6   8E+4   1   1.5  RT (min)      lcorbi_t3_pos_4_077_58    m/z 504.8465     0   1   2   3E+5   1   1.5  RT (min)       lcorbi_t3_pos_1_029_29    m/z 504.8578     0   2   4E+4   1   1.5  RT (min)       lcorbi_t3_pos_3_012_31    m/z 504.8676     0   1   2   3   4E+3   1   1.5  RT (min)      lcorbi_t3_pos_3_012_31    m/z 505.1999     0   2   4   6E+4   1   1.5  RT (min)      lcorbi_t3_pos_3_045_43    m/z 505.2686     0   1E+4   4.5   5  RT (min)      lcorbi_t3_pos_4_074_58    m/z 505.3120     0   1   11   11.5  RT (min)      lcorbi_t3_pos_2_030_14    m/z 505.3235     0   2   4E+4   10   10.5  RT (min)      lcorbi_t3_pos_4_088_59    m/z 505.3565     0   1E+4   11   11.5  RT (min)       lcorbi_t3_pos_1_003_45    m/z 505.4015     0   1E+5   11.5   12  RT (min)       lcorbi_t3_pos_4_091_60    m/z 505.4448     0   1E+5   14.5   15  RT (min)      lcorbi_t3_pos_4_099_61    m/z 505.5094     0   2   4   6   8E+4   14.5   15  RT (min)       lcorbi_t3_pos_3_016_21    m/z 505.8904     0   2   4E+4   1   1.5  RT (min)      lcorbi_t3_pos_4_076_58    m/z 506.1678     0   1E+5   8.5   9  RT (min)      lcorbi_t3_pos_4_104_61    m/z 506.2019     0   1   2   3   4E+4   9   9.5  RT (min)      lcorbi_t3_pos_4_103_61    m/z 506.2957     0   1E+4   9.5   10  RT (min)      lcorbi_t3_pos_3_006_38    m/z 506.3241     0   1   2   3E+5   11   11.5  RT (min)       lcorbi_t3_pos_1_050_13    m/z 506.3393     0   1   2E+4   13.5   14  RT (min)       lcorbi_t3_pos_1_001_49    m/z 506.3475     0   2   4   6E+4   10   10.5  RT (min)      lcorbi_t3_pos_1_002_01    m/z 506.3605     0   1E+5   11.5   12  RT (min)       lcorbi_t3_pos_4_078_58    m/z 506.3838     0   1E+4   6   6.5  RT (min)      lcorbi_t3_pos_4_113_62    m/z 506.4204     0   2   4   6E+3   4.5   5  RT (min)      lcorbi_t3_pos_1_011_44    m/z 506.4721     0   1   2   3E+4   15   15.5  RT (min)      lcorbi_t3_pos_3_030_22    m/z 506.8437     0   1E+5   1   1.5  RT (min)      lcorbi_t3_pos_4_078_58    m/z 507.0068     0   1   2E+4   1   1.5  RT (min)      lcorbi_t3_pos_3_024_07    m/z 507.1912     0   1E+4   10   10.5  RT (min)      lcorbi_t3_pos_3_002_16    m/z 507.3241     0   1E+5   11   11.5  RT (min)       lcorbi_t3_pos_1_050_13    m/z 507.4239     0   1E+5   13.5   14  RT (min)      lcorbi_t3_pos_4_082_59    m/z 507.8875     0   1   2E+4   1   1.5  RT (min)      lcorbi_t3_pos_1_029_29    m/z 508.2903     0   1   2E+4   9   9.5  RT (min)       lcorbi_t3_pos_4_067_57    m/z 508.3114     0   1E+4   7   7.5  RT (min)      lcorbi_t3_pos_4_074_58    m/z 508.5453     0   1   2   3E+5   14.5   15  RT (min)       lcorbi_t3_pos_2_044_21    m/z 508.6234     0   2   4   6E+4   1   1.5  RT (min)      lcorbi_t3_pos_4_080_58    m/z 508.7130     0   2   4   6   8E+4   1   1.5  RT (min)      lcorbi_t3_pos_1_020_31    m/z 508.8581     0   2   4   6   8E+4   1   1.5  RT (min)       lcorbi_t3_pos_4_077_58    m/z 509.1853     0   1E+4   4.5   5  RT (min)      lcorbi_t3_pos_4_098_61    m/z 509.2203     0   2   4E+4   1   1.5  RT (min)       lcorbi_t3_pos_3_055_10    m/z 509.2938     0   1E+4   9   9.5  RT (min)      lcorbi_t3_pos_4_074_58    m/z 509.3432     0   1E+5   11   11.5  RT (min)      lcorbi_t3_pos_3_027_27    m/z 509.3626     0   1E+5   15   15.5  RT (min)       lcorbi_t3_pos_1_028_32    m/z 509.5041     0   2   4   6E+4   14   14.5  RT (min)       lcorbi_t3_pos_4_106_62    m/z 509.8656     0   1E+4   1   1.5  RT (min)      lcorbi_t3_pos_4_080_58    m/z 509.8854     0   2   4   6E+4   6   6.5  RT (min)       lcorbi_t3_pos_4_034_42    m/z 510.1392     0   1   2   3   4E+4   7   7.5  RT (min)      lcorbi_t3_pos_3_034_26    m/z 510.1605     0   2   4E+4   7   7.5  RT (min)      lcorbi_t3_pos_4_107_62    m/z 510.2167     0   2   4   6   8E+4   1   1.5  RT (min)       lcorbi_t3_pos_3_055_10    m/z 510.2204     0   1   2   3E+4   8.5   9  RT (min)      lcorbi_t3_pos_2_027_05    m/z 510.2627     0   1   2   3E+4   10.5   11  RT (min)      lcorbi_t3_pos_4_074_58    m/z 510.3034     0   1E+5   5   5.5  RT (min)       lcorbi_t3_pos_4_079_58    m/z 510.3060     0   1   2   3   4E+3   8   8.5  RT (min)      lcorbi_t3_pos_1_054_48    m/z 510.3099     0   2   4   6E+4   1   1.5   2  RT (min)       lcorbi_t3_pos_4_093_60    m/z 510.3823     0   2   4E+4   11.5   12  RT (min)      lcorbi_t3_pos_1_035_28    m/z 510.3870     0   1   2   3E+4   6   6.5  RT (min)      lcorbi_t3_pos_4_034_42    m/z 510.5075     0   1   2E+4   14   14.5  RT (min)      lcorbi_t3_pos_2_055_46    m/z 510.6206     0   1   2E+4   1   1.5  RT (min)      lcorbi_t3_pos_4_077_58    m/z 510.7102     0   2   4   6E+4   1   1.5  RT (min)      lcorbi_t3_pos_2_028_24    m/z 510.8655     0   2   4   6   8E+4   18   18.5  RT (min)       lcorbi_t3_pos_4_093_60    m/z 510.8886     0   1E+4   6   6.5  RT (min)      lcorbi_t3_pos_4_045_08    m/z 511.0475     0   1E+4   10   10.5  RT (min)      lcorbi_t3_pos_4_074_58    m/z 511.1426     0   1E+4   7   7.5  RT (min)      lcorbi_t3_pos_3_011_42    m/z 511.1550     0   1E+4   7   7.5  RT (min)      lcorbi_t3_pos_4_112_62    m/z 511.2457     0   1   2E+4   11   11.5  RT (min)      lcorbi_t3_pos_4_013_36    m/z 511.3068     0   1   2   3E+4   5   5.5  RT (min)      lcorbi_t3_pos_4_080_58    m/z 511.3274     0   1   2E+4   5   5.5  RT (min)      lcorbi_t3_pos_1_001_49    m/z 511.4427     0   2   4   6E+4   12   13  RT (min)       lcorbi_t3_pos_4_095_60    m/z 511.7850     0   1   2E+4   4.5   5  RT (min)       lcorbi_t3_pos_4_082_59    m/z 512.2277     0   2   4   6E+4   10.5   11  RT (min)       lcorbi_t3_pos_2_028_24    m/z 512.2641     0   1E+4   9.5   10  RT (min)      lcorbi_t3_pos_4_082_59    m/z 512.2866     0   1E+4   4.5   5  RT (min)      lcorbi_t3_pos_4_074_58    m/z 512.3092     0   2   4   6E+3   5   5.5  RT (min)      lcorbi_t3_pos_4_080_58    m/z 512.3347     0   2   4   6E+4   8.5   9  RT (min)       lcorbi_t3_pos_4_071_57    m/z 512.3369     0   1E+4   7   7.5  RT (min)      lcorbi_t3_pos_4_086_59    m/z 512.3586     0   2   4   6E+4   14.5   15  RT (min)      lcorbi_t3_pos_4_101_61    m/z 512.3616     0   2   4   6E+4   11.5   12  RT (min)      lcorbi_t3_pos_4_077_58    m/z 512.3981     0   1   2   3E+4   12   12.5  RT (min)      lcorbi_t3_pos_4_006_12    m/z 512.4674     0   1   13   13.5  RT (min)       lcorbi_t3_pos_1_024_04    m/z 512.5039     0   2   4   6E+4   13   13.5  RT (min)      lcorbi_t3_pos_2_044_21    m/z 512.6819     0   1   2   3E+4   1   1.5  RT (min)      lcorbi_t3_pos_1_028_32    m/z 512.8754     0   1   2   3E+4   1   1.5  RT (min)      lcorbi_t3_pos_3_024_07    m/z 513.0267     0   1   2   3E+4   1   1.5  RT (min)      lcorbi_t3_pos_3_010_44    m/z 513.2458     0   2   4   6   8E+3   9   9.5  RT (min)      lcorbi_t3_pos_4_027_07    m/z 513.3380     0   1   2E+4   8.5   9  RT (min)      lcorbi_t3_pos_4_073_57    m/z 513.3381     0   1   10.5   11  RT (min)       lcorbi_t3_pos_1_054_48    m/z 513.8697     0   2   4   6E+3   6.5   7  RT (min)      lcorbi_t3_pos_4_034_42    m/z 514.3617     0   1E+4   7   7.5  RT (min)      lcorbi_t3_pos_4_084_59    m/z 514.3764     0   2   4E+4   10   10.5  RT (min)       lcorbi_t3_pos_4_086_59    m/z 514.4063     0   2   4E+4   13.5   14  RT (min)      lcorbi_t3_pos_4_111_62    m/z 514.4220     0   2   4   6   8E+4   11   11.5  RT (min)      lcorbi_t3_pos_4_084_59    m/z 514.5194     0   2   4E+4   12.5   13  RT (min)       lcorbi_t3_pos_4_066_57    m/z 514.6789     0   2   4E+4   1   1.5  RT (min)      lcorbi_t3_pos_1_032_26    m/z 514.8633     0   1E+4   7   7.5  RT (min)      lcorbi_t3_pos_4_086_59    m/z 515.0511     0   1   2   3E+4   10   10.5  RT (min)       lcorbi_t3_pos_4_079_58    m/z 515.2210     0   1E+4   9   9.5  RT (min)      lcorbi_t3_pos_4_105_61    m/z 515.3706     0   2   4   6   8E+4   9   9.5  RT (min)      lcorbi_t3_pos_1_021_18    m/z 515.4129     0   1   2E+5   14.5   15  RT (min)       lcorbi_t3_pos_1_036_08    m/z 515.8154     0   1   2   3E+4   11   11.5  RT (min)      lcorbi_t3_pos_4_078_58    m/z 516.3014     0   2   4E+3   6   6.5  RT (min)      lcorbi_t3_pos_1_037_19    m/z 516.3664     0   1   2   3   4E+4   13.5   14  RT (min)       lcorbi_t3_pos_4_100_61    m/z 516.3739     0   1   2   3E+4   9   9.5  RT (min)      lcorbi_t3_pos_2_046_07    m/z 516.3889     0   1   2   3E+5   10   10.5  RT (min)       lcorbi_t3_pos_1_050_13    m/z 516.4261     0   1E+5   14   14.5  RT (min)       lcorbi_t3_pos_4_082_59    m/z 516.4410     0   2   4   6E+4   14.5   15  RT (min)       lcorbi_t3_pos_2_016_23    m/z 516.6759     0   1   2   3   4E+4   1   1.5  RT (min)      lcorbi_t3_pos_1_034_34    m/z 516.7449     0   2   4E+4   1   1.5  RT (min)      lcorbi_t3_pos_1_020_31    m/z 516.8737     0   1E+5   1   1.5  RT (min)       lcorbi_t3_pos_4_077_58    m/z 516.9983     0   1   2E+4   1   1.5  RT (min)      lcorbi_t3_pos_3_020_35    m/z 517.1971     0   1E+5   8.5   9  RT (min)      lcorbi_t3_pos_4_104_61    m/z 517.3063     0   1E+5   11.5   12  RT (min)       lcorbi_t3_pos_4_073_57    m/z 517.3918     0   1E+5   7   7.5  RT (min)       lcorbi_t3_pos_4_014_48    m/z 517.4013     0   1   2   3E+5   10.5   11  RT (min)       lcorbi_t3_pos_1_050_13    m/z 517.4083     0   2   4   6   8E+4   13.5   14  RT (min)      lcorbi_t3_pos_1_037_19    m/z 517.4362     0   2   4   6   8E+4   14.5   15  RT (min)       lcorbi_t3_pos_4_103_61    m/z 518.2807     0   1E+4   5   5.5  RT (min)      lcorbi_t3_pos_4_090_60    m/z 518.3535     0   1   2E+4   5.5   6  RT (min)      lcorbi_t3_pos_4_098_61    m/z 518.3952     0   1   2   3   4E+4   7   7.5  RT (min)      lcorbi_t3_pos_4_014_48    m/z 518.3988     0   1E+4   5.5   6  RT (min)      lcorbi_t3_pos_4_085_59    m/z 518.4933     0   1E+5   14   14.5  RT (min)       lcorbi_t3_pos_4_089_59    m/z 518.7419     0   1E+5   1   1.5  RT (min)      lcorbi_t3_pos_1_020_31    m/z 518.8727     0   2   4   6E+3   6   6.5  RT (min)      lcorbi_t3_pos_4_083_59    m/z 518.8909     0   1   2   3E+4   5.5   6  RT (min)       lcorbi_t3_pos_3_011_42    m/z 518.9005     0   1E+4   5.5   6  RT (min)      lcorbi_t3_pos_4_084_59    m/z 518.9132     0   1   2   3   4E+4   18   18.5  RT (min)       lcorbi_t3_pos_4_071_57    m/z 519.3222     0   1   2E+4   8   8.5  RT (min)      lcorbi_t3_pos_4_109_62    m/z 519.3315     0   2   4E+4   8.5   9  RT (min)       lcorbi_t3_pos_4_106_62    m/z 519.3889     0   1   7   7.5  RT (min)       lcorbi_t3_pos_4_097_60    m/z 519.3925     0   1   2E+4   5.5   6  RT (min)      lcorbi_t3_pos_4_005_33    m/z 519.4170     0   1   2   3   4E+5   11   11.5  RT (min)       lcorbi_t3_pos_4_091_60    m/z 519.4884     0   1   2   3E+4   12.5   13  RT (min)      lcorbi_t3_pos_4_014_48    m/z 520.2591     0   1   11.5   12  RT (min)       lcorbi_t3_pos_2_030_14    m/z 520.2903     0   1   2E+4   8   8.5  RT (min)       lcorbi_t3_pos_4_104_61    m/z 520.3290     0   1   2E+4   10.5   11  RT (min)      lcorbi_t3_pos_4_069_57    m/z 520.3327     0   1   2   3E+4   5   5.5  RT (min)       lcorbi_t3_pos_1_001_49    m/z 520.3348     0   1E+4   8.5   9  RT (min)      lcorbi_t3_pos_4_111_62    m/z 520.3399     0   1   2E+5   15.5   16  RT (min)      lcorbi_t3_pos_1_029_29    m/z 520.3399     0   2   4   6   8E+4   0   0.5  RT (min)      lcorbi_t3_pos_1_017_51    m/z 520.3632     0   1E+5   7.5   8  RT (min)       lcorbi_t3_pos_4_113_62    m/z 520.3694     0   1   11   11.5  RT (min)      lcorbi_t3_pos_4_090_60    m/z 520.4634     0   1E+5   14.5   15  RT (min)      lcorbi_t3_pos_4_098_61    m/z 520.6033     0   1   2E+4   1   1.5  RT (min)      lcorbi_t3_pos_2_040_25    m/z 520.8203     0   2   4   6E+4   1   1.5  RT (min)      lcorbi_t3_pos_1_029_29    m/z 520.8317     0   1E+4   1   1.5  RT (min)      lcorbi_t3_pos_3_026_04    m/z 520.8415     0   1   2   3   4E+4   1   1.5  RT (min)      lcorbi_t3_pos_4_052_06    m/z 520.8622     0   1   2   3   4E+4   1   1.5  RT (min)      lcorbi_t3_pos_4_052_06    m/z 521.0680     0   1   2E+4   10   10.5  RT (min)      lcorbi_t3_pos_1_029_29    m/z 521.3363     0   2   4   6   8E+3   5   5.5  RT (min)      lcorbi_t3_pos_4_074_58    m/z 521.3667     0   1   2   3E+4   7.5   8  RT (min)      lcorbi_t3_pos_4_112_62    m/z 521.4395     0   1   2   3   4E+4   14   14.5  RT (min)      lcorbi_t3_pos_1_043_06    m/z 521.4565     0   1   2   3E+4   14   14.5  RT (min)       lcorbi_t3_pos_1_054_48    m/z 521.5124     0   1   2E+5   14   14.5  RT (min)      lcorbi_t3_pos_4_082_59    m/z 521.8384     0   2   4   6E+3   5   5.5  RT (min)      lcorbi_t3_pos_4_098_61    m/z 522.0151     0   2   4   6E+4   1   1.5  RT (min)      lcorbi_t3_pos_3_010_44    m/z 522.2028     0   2   4   6E+4   1   1.5  RT (min)       lcorbi_t3_pos_4_066_57    m/z 522.3556     0   2   4   6E+4   0   0.5  RT (min)      lcorbi_t3_pos_1_033_53    m/z 522.4235     0   2   4   6   8E+4   12.5   13  RT (min)      lcorbi_t3_pos_2_040_25    m/z 522.5974     0   2   4   6   8E+4   13.5   14  RT (min)       lcorbi_t3_pos_1_004_24    m/z 522.6005     0   1   2   3   4E+4   1   1.5  RT (min)      lcorbi_t3_pos_1_029_29    m/z 523.2860     0   1E+5   10.5   11  RT (min)      lcorbi_t3_pos_1_050_13    m/z 523.3391     0   1   2   3E+4   11   11.5  RT (min)       lcorbi_t3_pos_1_003_45    m/z 523.3783     0   2   4   6E+4   15   15.5  RT (min)       lcorbi_t3_pos_4_089_59    m/z 523.4358     0   2   4E+4   13   13.5  RT (min)       lcorbi_t3_pos_4_006_12    m/z 523.4722     0   2   4   6E+4   14   14.5  RT (min)       lcorbi_t3_pos_1_032_26    m/z 524.2731     0   1E+4   6   6.5  RT (min)      lcorbi_t3_pos_4_110_62    m/z 524.2983     0   1E+5   11   11.5  RT (min)       lcorbi_t3_pos_1_050_13    m/z 524.3408     0   1   11.5   12  RT (min)      lcorbi_t3_pos_4_074_58    m/z 524.3713     0   2   4   6   8E+4   0   0.5  RT (min)      lcorbi_t3_pos_2_055_46    m/z 524.4073     0   2   4   6E+4   11.5   12  RT (min)      lcorbi_t3_pos_1_029_29    m/z 524.4158     0   2   4E+4   11   12  RT (min)       lcorbi_t3_pos_1_029_29    m/z 524.4523     0   2   4E+4   12.5   13  RT (min)       lcorbi_t3_pos_4_085_59    m/z 524.5976     0   1   2E+4   1   1.5  RT (min)      lcorbi_t3_pos_4_076_58    m/z 524.8322     0   1E+4   1   1.5  RT (min)      lcorbi_t3_pos_4_077_58    m/z 525.1390     0   1   2   3   4E+4   7   7.5  RT (min)       lcorbi_t3_pos_4_101_61    m/z 525.2782     0   1   11   11.5  RT (min)      lcorbi_t3_pos_1_050_13    m/z 525.3363     0   1E+5   12.5   13  RT (min)       lcorbi_t3_pos_1_011_44    m/z 525.3508     0   1   2   3   4E+4   11.5   12  RT (min)      lcorbi_t3_pos_1_029_29    m/z 525.3977     0   2   4E+4   11.5   12  RT (min)       lcorbi_t3_pos_1_029_29    m/z 525.8805     0   1   2   3E+4   6   6.5  RT (min)       lcorbi_t3_pos_4_087_59    m/z 526.2646     0   1   2   3E+4   9.5   10  RT (min)      lcorbi_t3_pos_4_076_58    m/z 526.2798     0   1   9.5   10  RT (min)      lcorbi_t3_pos_4_078_58    m/z 526.3165     0   1   2E+5   13   14  RT (min)       lcorbi_t3_pos_1_010_46    m/z 526.3165     0   2   4   6   8E+4   0   0.5  RT (min)      lcorbi_t3_pos_4_010_43    m/z 526.3724     0   2   4   6   8E+3   6   6.5  RT (min)      lcorbi_t3_pos_4_018_01    m/z 526.3821     0   1E+4   6   6.5  RT (min)      lcorbi_t3_pos_4_086_59    m/z 526.3890     0   2   4   6E+3   4.5   5  RT (min)      lcorbi_t3_pos_4_005_33    m/z 526.3949     0   1   11   11.5  RT (min)       lcorbi_t3_pos_1_032_26    m/z 526.7737     0   2   4   6   8E+4   1   1.5  RT (min)      lcorbi_t3_pos_1_020_31    m/z 526.7834     0   1   2   3E+4   4.5   5  RT (min)       lcorbi_t3_pos_4_080_58    m/z 526.8492     0   2   4   6E+4   1   1.5  RT (min)      lcorbi_t3_pos_4_052_06    m/z 526.8741     0   1   2   3   4E+3   6   6.5  RT (min)      lcorbi_t3_pos_4_034_42    m/z 527.0243     0   1   2   3E+4   10   10.5  RT (min)      lcorbi_t3_pos_1_029_29    m/z 527.1582     0   2   4   6E+4   1   1.5  RT (min)       lcorbi_t3_pos_4_070_57    m/z 527.2786     0   1   2E+4   10   10.5  RT (min)      lcorbi_t3_pos_4_094_60    m/z 527.2846     0   1E+4   4.5   5  RT (min)      lcorbi_t3_pos_4_077_58    m/z 527.3975     0   1   2E+4   11   11.5  RT (min)       lcorbi_t3_pos_4_086_59    m/z 527.4781     0   2   4   6E+4   11   11.5  RT (min)       lcorbi_t3_pos_4_084_59    m/z 528.1497     0   1E+5   8   8.5  RT (min)      lcorbi_t3_pos_4_106_62    m/z 528.2509     0   2   4   6   8E+3   10.5   11  RT (min)      lcorbi_t3_pos_2_044_21    m/z 528.2796     0   2   4   6E+4   6   6.5  RT (min)       lcorbi_t3_pos_1_029_29    m/z 528.2989     0   1   2   3   4E+4   11   11.5  RT (min)      lcorbi_t3_pos_4_069_57    m/z 528.3227     0   1   2   3   4E+4   10.5   11  RT (min)      lcorbi_t3_pos_4_097_60    m/z 528.3826     0   2   4E+4   11.5   12  RT (min)       lcorbi_t3_pos_4_077_58    m/z 528.4048     0   1E+5   12   12.5  RT (min)       lcorbi_t3_pos_4_087_59    m/z 528.7707     0   1E+5   1   1.5  RT (min)      lcorbi_t3_pos_1_020_31    m/z 528.7811     0   1E+3   6   6.5  RT (min)      lcorbi_t3_pos_4_030_05    m/z 528.7935     0   1   2   3E+4   4.5   5  RT (min)       lcorbi_t3_pos_4_078_58    m/z 529.2826     0   1   6   6.5  RT (min)      lcorbi_t3_pos_3_023_05    m/z 529.2950     0   1E+4   4.5   5  RT (min)      lcorbi_t3_pos_4_077_58    m/z 529.3119     0   2   4   6E+4   11   11.5  RT (min)      lcorbi_t3_pos_4_067_57    m/z 529.3923     0   1   2E+5   10   10.5  RT (min)       lcorbi_t3_pos_1_050_13    m/z 530.3171     0   1E+4   6.5   7  RT (min)      lcorbi_t3_pos_4_066_57    m/z 530.3241     0   1E+5   11.5   12  RT (min)       lcorbi_t3_pos_4_069_57    m/z 530.3582     0   1   2   3   4E+4   11.5   12  RT (min)       lcorbi_t3_pos_4_075_58    m/z 530.4569     0   1   14.5   15  RT (min)      lcorbi_t3_pos_4_007_46    m/z 530.6530     0   1   2   3E+4   1   1.5  RT (min)      lcorbi_t3_pos_1_032_26    m/z 530.7880     0   1E+5   5   5.5  RT (min)       lcorbi_t3_pos_4_079_58    m/z 531.0141     0   1   2   3E+4   1   1.5  RT (min)      lcorbi_t3_pos_4_005_33    m/z 531.2895     0   2   4   6E+4   5   5.5  RT (min)      lcorbi_t3_pos_4_074_58    m/z 531.3292     0   1   2   3   4E+4   10   10.5  RT (min)       lcorbi_t3_pos_4_066_57    m/z 531.4076     0   2   4   6   8E+4   7   7.5  RT (min)      lcorbi_t3_pos_4_108_62    m/z 531.4080     0   1   2E+5   17.5   18  RT (min)      lcorbi_t3_pos_4_103_61    m/z 531.4081     0   2   4E+5   11   11.5  RT (min)      lcorbi_t3_pos_1_050_13    m/z 531.5249     0   2   4   6   8E+4   14.5   15  RT (min)       lcorbi_t3_pos_4_068_57    m/z 531.7572     0   1   2E+4   5   5.5  RT (min)      lcorbi_t3_pos_4_074_58    m/z 531.7908     0   1   2E+4   5   5.5  RT (min)      lcorbi_t3_pos_4_074_58    m/z 531.8527     0   1   2E+4   1   1.5  RT (min)      lcorbi_t3_pos_4_076_58    m/z 532.2586     0   1E+4   5   5.5  RT (min)      lcorbi_t3_pos_3_004_24    m/z 532.2963     0   1   2E+4   5   5.5  RT (min)      lcorbi_t3_pos_4_066_57    m/z 532.3373     0   1   2   3   4E+4   11.5   12  RT (min)       lcorbi_t3_pos_4_078_58    m/z 532.3515     0   1E+5   11   11.5  RT (min)       lcorbi_t3_pos_4_084_59    m/z 532.3576     0   1   14   14.5  RT (min)      lcorbi_t3_pos_4_093_60    m/z 532.3691     0   1E+4   6   6.5  RT (min)      lcorbi_t3_pos_4_066_57    m/z 532.4109     0   1   2   3   4E+4   7   7.5  RT (min)      lcorbi_t3_pos_4_014_48    m/z 532.6500     0   1   2E+4   1   1.5  RT (min)      lcorbi_t3_pos_1_028_32    m/z 532.7988     0   2   4   6E+4   18   18.5  RT (min)       lcorbi_t3_pos_4_080_58    m/z 532.8476     0   2   4E+4   1   1.5  RT (min)      lcorbi_t3_pos_4_077_58    m/z 532.8951     0   1   2   3   4E+4   1   1.5  RT (min)      lcorbi_t3_pos_4_072_57    m/z 533.2063     0   2   4   6E+4   10.5   11  RT (min)       lcorbi_t3_pos_4_071_57    m/z 533.2780     0   1   2E+4   10.5   11  RT (min)      lcorbi_t3_pos_4_091_60    m/z 533.2997     0   2   4E+3   5   5.5  RT (min)      lcorbi_t3_pos_4_082_59    m/z 533.3405     0   1E+4   5.5   6  RT (min)      lcorbi_t3_pos_1_001_49    m/z 533.3549     0   1E+4   11   11.5  RT (min)      lcorbi_t3_pos_1_050_13    m/z 533.4237     0   1   2   3   4E+5   11   11.5  RT (min)       lcorbi_t3_pos_1_050_13    m/z 533.4429     0   2   4   6   8E+4   14.5   15  RT (min)      lcorbi_t3_pos_2_024_15    m/z 533.4761     0   1E+5   14.5   15  RT (min)      lcorbi_t3_pos_4_103_61    m/z 533.5406     0   2   4   6E+4   15   15.5  RT (min)       lcorbi_t3_pos_3_038_18    m/z 534.2544     0   1   2E+4   8   8.5  RT (min)      lcorbi_t3_pos_4_090_60    m/z 534.3061     0   1E+4   8   8.5  RT (min)      lcorbi_t3_pos_4_104_61    m/z 534.3120     0   1E+4   5   5.5  RT (min)      lcorbi_t3_pos_4_075_58    m/z 534.3789     0   2   4E+4   7.5   8  RT (min)       lcorbi_t3_pos_4_106_62    m/z 534.3914     0   2   4   6   8E+4   11.5   12  RT (min)       lcorbi_t3_pos_4_075_58    m/z 534.4176     0   1   2E+4   13.5   14  RT (min)      lcorbi_t3_pos_4_106_62    m/z 534.4519     0   2   4   6E+4   13.5   14  RT (min)       lcorbi_t3_pos_3_002_16    m/z 534.5246     0   1E+5   14   14.5  RT (min)       lcorbi_t3_pos_4_082_59    m/z 535.0225     0   1   2E+4   10   10.5  RT (min)      lcorbi_t3_pos_1_029_29    m/z 535.3784     0   1E+5   15   15.5  RT (min)       lcorbi_t3_pos_4_089_59    m/z 535.3823     0   1   2E+4   7.5   8  RT (min)      lcorbi_t3_pos_4_108_62    m/z 535.4358     0   1E+5   12.5   13  RT (min)       lcorbi_t3_pos_4_075_58    m/z 535.4723     0   2   4   6   8E+4   13.5   14  RT (min)       lcorbi_t3_pos_4_106_62    m/z 535.8731     0   1   2   3E+4   1   1.5  RT (min)      lcorbi_t3_pos_4_067_57    m/z 536.2903     0   1E+5   11   11.5  RT (min)       lcorbi_t3_pos_2_030_14    m/z 536.3346     0   1   2   3   4E+4   9   9.5  RT (min)       lcorbi_t3_pos_4_006_12    m/z 536.3581     0   1   2E+4   8.5   9  RT (min)      lcorbi_t3_pos_1_008_36    m/z 536.3711     0   1E+5   11.5   12  RT (min)       lcorbi_t3_pos_4_075_58    m/z 536.3897     0   1   2   3   4E+4   10   10.5  RT (min)      lcorbi_t3_pos_4_086_59    m/z 536.4075     0   1E+5   12   12.5  RT (min)       lcorbi_t3_pos_4_076_58    m/z 536.4391     0   2   4   6E+4   12.5   13  RT (min)      lcorbi_t3_pos_4_077_58    m/z 536.4757     0   1   2   3E+4   13.5   14  RT (min)      lcorbi_t3_pos_4_072_57    m/z 536.5039     0   1   2   3   4E+5   13   13.5   14  RT (min)       lcorbi_t3_pos_4_089_59    m/z 536.5767     0   1   2E+5   14.5   15  RT (min)       lcorbi_t3_pos_2_044_21    m/z 536.8026     0   1E+5   1   1.5  RT (min)      lcorbi_t3_pos_1_020_31    m/z 537.0342     0   1   2E+4   1   1.5  RT (min)      lcorbi_t3_pos_3_010_44    m/z 537.3342     0   2   4   6E+4   13.5   14  RT (min)       lcorbi_t3_pos_4_103_61    m/z 537.3505     0   2   4   6   8E+4   13.5   14  RT (min)       lcorbi_t3_pos_1_015_11    m/z 537.3744     0   1   2E+4   11.5   12  RT (min)      lcorbi_t3_pos_3_038_18    m/z 537.4114     0   1   2   3   4E+4   12   12.5  RT (min)      lcorbi_t3_pos_4_076_58    m/z 537.5073     0   1E+5   14   14.5  RT (min)      lcorbi_t3_pos_4_083_59    m/z 537.5356     0   1   2E+5   17.5   18  RT (min)       lcorbi_t3_pos_4_076_58    m/z 537.9891     0   1   2   3E+4   1   1.5  RT (min)      lcorbi_t3_pos_3_010_44    m/z 538.1765     0   2   4   6E+4   8.5   9  RT (min)      lcorbi_t3_pos_4_101_61    m/z 538.4231     0   2   4   6   8E+4   12.5   13  RT (min)       lcorbi_t3_pos_4_081_58    m/z 538.4467     0   2   4   6   8E+4   12.5   13  RT (min)       lcorbi_t3_pos_1_029_29    m/z 538.5753     0   1E+5   14.5   15  RT (min)      lcorbi_t3_pos_2_044_21    m/z 538.7855     0   1E+4   5   5.5  RT (min)      lcorbi_t3_pos_4_075_58    m/z 538.7995     0   1E+5   1   1.5  RT (min)      lcorbi_t3_pos_1_020_31    m/z 538.8152     0   2   4   6   8E+4   18   18.5  RT (min)       lcorbi_t3_pos_4_080_58    m/z 538.8774     0   1   2E+4   1   1.5  RT (min)      lcorbi_t3_pos_4_077_58    m/z 539.0598     0   2   4   6E+4   1   1.5  RT (min)      lcorbi_t3_pos_1_029_29    m/z 539.3197     0   2   4   6E+4   11.5   12  RT (min)       lcorbi_t3_pos_4_093_60    m/z 539.3901     0   2   4E+4   12   12.5  RT (min)      lcorbi_t3_pos_1_029_29    m/z 539.4266     0   1   2E+4   12.5   13  RT (min)      lcorbi_t3_pos_4_069_57    m/z 539.9780     0   1   2E+4   1   1.5  RT (min)      lcorbi_t3_pos_4_005_33    m/z 540.3658     0   1   2E+4   9.5   10  RT (min)      lcorbi_t3_pos_4_074_58    m/z 540.3660     0   1   11   11.5  RT (min)       lcorbi_t3_pos_4_090_60    m/z 540.4108     0   2   4E+4   11   11.5  RT (min)       lcorbi_t3_pos_4_112_62    m/z 540.4484     0   2   4E+4   5.5   6  RT (min)       lcorbi_t3_pos_4_088_59    m/z 540.4623     0   1   2   3   4E+4   13   13.5  RT (min)       lcorbi_t3_pos_4_006_12    m/z 540.4989     0   1   2   3   4E+4   13.5   14  RT (min)       lcorbi_t3_pos_4_069_57    m/z 540.5258     0   1   2   3E+4   14   14.5  RT (min)      lcorbi_t3_pos_4_069_57    m/z 540.5454     0   2   4E+4   14   14.5  RT (min)       lcorbi_t3_pos_1_032_26    m/z 540.5967     0   1   2   3E+4   1   1.5  RT (min)      lcorbi_t3_pos_4_070_57    m/z 540.8891     0   1   2   3   4E+4   1   1.5  RT (min)      lcorbi_t3_pos_4_071_57    m/z 541.2606     0   1E+4   2.5   3  RT (min)      lcorbi_t3_pos_3_048_30    m/z 541.3064     0   1E+4   11.5   12  RT (min)       lcorbi_t3_pos_2_030_14    m/z 541.3135     0   1   2E+4   8.5   9  RT (min)       lcorbi_t3_pos_4_108_62    m/z 541.3250     0   2   4   6E+4   11.5   12  RT (min)       lcorbi_t3_pos_4_072_57    m/z 541.4294     0   2   4E+4   12.5   13  RT (min)      lcorbi_t3_pos_4_013_36    m/z 541.4367     0   1   2   3   4E+5   14.5   15  RT (min)      lcorbi_t3_pos_1_004_24    m/z 541.4504     0   1   2   3E+5   11.5   12  RT (min)       lcorbi_t3_pos_4_082_59    m/z 541.4519     0   1   2E+4   5.5   6  RT (min)      lcorbi_t3_pos_4_086_59    m/z 541.4616     0   1   2   3E+4   13.5   14  RT (min)       lcorbi_t3_pos_3_050_36    m/z 541.5386     0   1E+5   14   14.5  RT (min)       lcorbi_t3_pos_4_087_59    m/z 542.4047     0   1   2E+5   10.5   11  RT (min)       lcorbi_t3_pos_1_050_13    m/z 542.4205     0   2   4   6   8E+4   12   12.5  RT (min)       lcorbi_t3_pos_4_086_59    m/z 542.4734     0   2   4   6   8E+4   15   15.5  RT (min)       lcorbi_t3_pos_2_024_15    m/z 542.5937     0   1   2   3E+4   1   1.5  RT (min)      lcorbi_t3_pos_4_068_57    m/z 542.8202     0   1E+5   18   18.5  RT (min)       lcorbi_t3_pos_4_082_59    m/z 542.8230     0   1   2   3E+4   1   1.5  RT (min)      lcorbi_t3_pos_3_010_44    m/z 543.2718     0   1E+5   11.5   12  RT (min)       lcorbi_t3_pos_4_090_60    m/z 543.3407     0   1E+5   12   12.5  RT (min)      lcorbi_t3_pos_4_073_57    m/z 543.4022     0   2   4   6E+4   12.5   13  RT (min)       lcorbi_t3_pos_3_008_20    m/z 543.4774     0   2   4   6E+4   14   14.5  RT (min)      lcorbi_t3_pos_4_074_58    m/z 543.8602     0   1E+4   1   1.5  RT (min)      lcorbi_t3_pos_4_079_58    m/z 544.2751     0   1   2E+4   11.5   12  RT (min)      lcorbi_t3_pos_4_073_57    m/z 544.3150     0   1   11   11.5  RT (min)      lcorbi_t3_pos_4_071_57    m/z 544.3302     0   1   2   3   4E+4   10.5   11  RT (min)      lcorbi_t3_pos_4_066_57    m/z 544.3328     0   2   4   6E+3   7   7.5  RT (min)      lcorbi_t3_pos_4_074_58    m/z 544.3399     0   1   2   3E+5   15.5   16  RT (min)      lcorbi_t3_pos_1_014_03    m/z 544.3400     0   2   4   6   8E+4   0   0.5  RT (min)      lcorbi_t3_pos_1_008_36    m/z 544.3644     0   1E+4   11   11.5  RT (min)      lcorbi_t3_pos_2_030_14    m/z 544.4204     0   1   2E+5   11   11.5  RT (min)       lcorbi_t3_pos_1_050_13    m/z 544.4574     0   2   4   6   8E+4   14   14.5  RT (min)       lcorbi_t3_pos_4_083_59    m/z 544.5908     0   1   2   3E+4   1   1.5  RT (min)      lcorbi_t3_pos_2_028_24    m/z 544.6172     0   1   2   3   4E+4   1   1.5  RT (min)      lcorbi_t3_pos_4_067_57    m/z 544.8945     0   2   4   6   8E+4   1   1.5  RT (min)       lcorbi_t3_pos_4_077_58    m/z 545.3109     0   1   2   3E+4   8   8.5  RT (min)       lcorbi_t3_pos_4_107_62    m/z 545.3599     0   1E+5   10.5   11  RT (min)       lcorbi_t3_pos_1_050_13    m/z 545.4174     0   1E+5   10.5   11  RT (min)       lcorbi_t3_pos_1_050_13    m/z 545.8962     0   1E+4   1   1.5  RT (min)      lcorbi_t3_pos_4_079_58    m/z 545.9983     0   1E+4   1   1.5  RT (min)      lcorbi_t3_pos_3_010_44    m/z 546.2567     0   1   2E+4   11   11.5  RT (min)      lcorbi_t3_pos_4_066_57    m/z 546.3121     0   1   2E+4   5.5   6  RT (min)      lcorbi_t3_pos_4_098_61    m/z 546.3359     0   2   4   6E+4   10.5   11  RT (min)       lcorbi_t3_pos_4_073_57    m/z 546.3765     0   1   9.5   10  RT (min)      lcorbi_t3_pos_4_095_60    m/z 546.3849     0   2   4   6E+3   6.5   7  RT (min)      lcorbi_t3_pos_4_066_57    m/z 546.4001     0   1   2   3   4E+4   10.5   11  RT (min)       lcorbi_t3_pos_1_033_53    m/z 546.4154     0   1   10.5   11  RT (min)       lcorbi_t3_pos_4_086_59    m/z 546.5246     0   2   4   6E+4   14   14.5  RT (min)       lcorbi_t3_pos_4_086_59    m/z 546.6142     0   2   4   6   8E+4   1   1.5  RT (min)       lcorbi_t3_pos_1_020_31    m/z 546.8312     0   1   2E+5   1   1.5  RT (min)      lcorbi_t3_pos_1_020_31    m/z 546.8691     0   1   2   3E+4   1   1.5  RT (min)      lcorbi_t3_pos_4_005_33    m/z 546.8919     0   1E+4   1   1.5  RT (min)      lcorbi_t3_pos_4_075_58    m/z 546.9879     0   1E+4   1   1.5  RT (min)      lcorbi_t3_pos_3_011_42    m/z 547.2456     0   2   4   6E+4   10.5   11  RT (min)       lcorbi_t3_pos_4_090_60    m/z 547.3487     0   1E+5   11   11.5  RT (min)       lcorbi_t3_pos_4_073_57    m/z 547.4035     0   1E+4   10.5   11  RT (min)      lcorbi_t3_pos_1_050_13    m/z 547.4916     0   1E+5   14.5   15  RT (min)      lcorbi_t3_pos_4_104_61    m/z 547.5197     0   1   13   13.5  RT (min)      lcorbi_t3_pos_4_082_59    m/z 548.2490     0   1   2   3E+4   10.5   11  RT (min)      lcorbi_t3_pos_4_093_60    m/z 548.2748     0   1   2   3E+4   11   11.5  RT (min)      lcorbi_t3_pos_4_072_57    m/z 548.2913     0   1   2   3   4E+4   5   5.5  RT (min)       lcorbi_t3_pos_4_106_62    m/z 548.3346     0   2   4E+4   11   11.5  RT (min)      lcorbi_t3_pos_1_008_36    m/z 548.3641     0   1   2   3   4E+3   5.5   6  RT (min)      lcorbi_t3_pos_4_106_62    m/z 548.3789     0   1   2   3E+5   11   11.5  RT (min)       lcorbi_t3_pos_4_097_60    m/z 548.4030     0   1E+5   14   14.5  RT (min)       lcorbi_t3_pos_4_086_59    m/z 548.4312     0   1   2E+5   11   11.5  RT (min)       lcorbi_t3_pos_1_050_13    m/z 548.4756     0   2   4   6E+4   13.5   14   14.5  RT (min)       lcorbi_t3_pos_1_008_36    m/z 548.4948     0   1E+5   13   13.5  RT (min)      lcorbi_t3_pos_4_066_57    m/z 548.8284     0   1E+5   1   1.5  RT (min)      lcorbi_t3_pos_1_020_31    m/z 548.8693     0   1E+4   1   1.5  RT (min)      lcorbi_t3_pos_4_081_58    m/z 549.0579     0   1E+5   8.5   9  RT (min)       lcorbi_t3_pos_4_100_61    m/z 549.1803     0   1   2   3   4E+4   10.5   11  RT (min)       lcorbi_t3_pos_2_028_24    m/z 549.2948     0   1E+4   5   5.5  RT (min)      lcorbi_t3_pos_4_074_58    m/z 549.3496     0   2   4E+4   10   10.5  RT (min)      lcorbi_t3_pos_4_086_59    m/z 549.4161     0   1E+5   12   12.5  RT (min)      lcorbi_t3_pos_3_021_39    m/z 549.7882     0   1   2E+4   5   5.5  RT (min)       lcorbi_t3_pos_4_081_58    m/z 550.2897     0   1   2E+4   5   5.5  RT (min)      lcorbi_t3_pos_4_081_58    m/z 550.3349     0   2   4   6E+3   6   6.5  RT (min)      lcorbi_t3_pos_4_006_12    m/z 550.3468     0   1   2E+5   4.5   5  RT (min)      lcorbi_t3_pos_3_048_30    m/z 550.3672     0   1   2   3   4E+4   11.5   12  RT (min)       lcorbi_t3_pos_1_031_40    m/z 550.3773     0   1E+4   11.5   12  RT (min)      lcorbi_t3_pos_4_090_60    m/z 550.3946     0   2   4   6   8E+4   11.5   12  RT (min)       lcorbi_t3_pos_4_095_60    m/z 550.4171     0   1E+5   14.5   15  RT (min)       lcorbi_t3_pos_4_081_58    m/z 550.5195     0   1   2E+5   14   14.5  RT (min)       lcorbi_t3_pos_4_089_59    m/z 550.6083     0   1E+5   1   1.5  RT (min)       lcorbi_t3_pos_4_067_57    m/z 550.8429     0   1   2E+4   1   1.5  RT (min)      lcorbi_t3_pos_4_077_58    m/z 550.9152     0   1E+5   1   1.5  RT (min)       lcorbi_t3_pos_4_067_57    m/z 551.1679     0   1   9   9.5  RT (min)      lcorbi_t3_pos_4_108_62    m/z 551.5231     0   2   4   6   8E+4   14   14.5  RT (min)      lcorbi_t3_pos_4_086_59    m/z 551.6322     0   1E+5   13.5   14  RT (min)      lcorbi_t3_pos_1_055_33    m/z 551.8466     0   1E+4   1   1.5  RT (min)      lcorbi_t3_pos_4_080_58    m/z 552.2310     0   2   4   6E+4   9   9.5  RT (min)      lcorbi_t3_pos_4_107_62    m/z 552.3014     0   1E+4   8.5   9  RT (min)      lcorbi_t3_pos_3_022_02    m/z 552.3296     0   1   2E+5   9   9.5  RT (min)       lcorbi_t3_pos_1_043_06    m/z 552.4326     0   1E+5   14.5   15  RT (min)       lcorbi_t3_pos_4_078_58    m/z 552.4835     0   1   2E+4   13   13.5  RT (min)      lcorbi_t3_pos_4_086_59    m/z 552.5069     0   1   2E+4   14   14.5   15  RT (min)       lcorbi_t3_pos_2_030_14    m/z 552.5545     0   2   4   6E+4   14   14.5  RT (min)      lcorbi_t3_pos_4_077_58    m/z 552.6054     0   2   4   6E+4   1   1.5  RT (min)      lcorbi_t3_pos_4_067_57    m/z 552.8012     0   1   2   3E+4   1   1.5  RT (min)      lcorbi_t3_pos_1_044_25    m/z 553.2340     0   1E+4   9   9.5  RT (min)      lcorbi_t3_pos_4_112_62    m/z 553.4059     0   2   4   6E+4   12.5   13  RT (min)      lcorbi_t3_pos_1_029_29    m/z 553.5176     0   2   4   6E+4   14.5   15  RT (min)      lcorbi_t3_pos_4_084_59    m/z 553.5386     0   2   4   6E+4   14   14.5  RT (min)      lcorbi_t3_pos_4_089_59    m/z 554.4277     0   1   2   3   4E+4   6.5   7  RT (min)       lcorbi_t3_pos_4_088_59    m/z 554.4498     0   2   4   6E+4   12.5   13  RT (min)      lcorbi_t3_pos_4_079_58    m/z 554.4642     0   1   2   3E+4   5.5   6  RT (min)       lcorbi_t3_pos_4_086_59    m/z 554.5145     0   1   2E+5   14   14.5  RT (min)       lcorbi_t3_pos_4_083_59    m/z 554.6232     0   1E+5   14   14.5  RT (min)       lcorbi_t3_pos_4_093_60    m/z 554.7932     0   2   4   6E+4   18   18.5  RT (min)      lcorbi_t3_pos_4_085_59    m/z 554.8553     0   1   2   3   4E+4   1   1.5  RT (min)       lcorbi_t3_pos_4_052_06    m/z 555.1768     0   1   2   3   4E+5   12   12.5  RT (min)      lcorbi_t3_pos_4_113_62    m/z 555.3853     0   1   2   3E+4   13.5   14  RT (min)      lcorbi_t3_pos_4_106_62    m/z 555.3966     0   2   4   6   8E+4   13.5   14  RT (min)      lcorbi_t3_pos_4_099_61    m/z 555.4116     0   1E+4   7   7.5  RT (min)      lcorbi_t3_pos_4_084_59    m/z 555.4311     0   1E+4   6.5   7  RT (min)      lcorbi_t3_pos_4_084_59    m/z 555.4675     0   1E+4   5.5   6  RT (min)      lcorbi_t3_pos_4_088_59    m/z 555.5179     0   1E+5   14   14.5  RT (min)      lcorbi_t3_pos_1_054_48    m/z 555.5542     0   2   4   6E+4   14   14.5  RT (min)      lcorbi_t3_pos_1_032_26    m/z 556.1744     0   1   2E+5   12   12.5  RT (min)       lcorbi_t3_pos_4_113_62    m/z 556.3634     0   2   4   6E+3   6.5   7  RT (min)      lcorbi_t3_pos_4_048_24    m/z 556.4055     0   2   4   6   8E+3   8   8.5  RT (min)      lcorbi_t3_pos_4_087_59    m/z 556.4938     0   1E+5   14   14.5  RT (min)       lcorbi_t3_pos_1_008_36    m/z 556.4988     0   1   2E+5   12   12.5  RT (min)       lcorbi_t3_pos_4_090_60    m/z 557.0042     0   1E+4   10   10.5  RT (min)      lcorbi_t3_pos_3_008_20    m/z 557.3223     0   1E+4   6.5   7  RT (min)      lcorbi_t3_pos_4_048_24    m/z 557.4454     0   1   2E+5   11   11.5  RT (min)       lcorbi_t3_pos_4_084_59    m/z 557.4972     0   1   2   3   4E+4   14   14.5  RT (min)      lcorbi_t3_pos_1_036_08    m/z 557.8631     0   1E+5   1   1.5  RT (min)      lcorbi_t3_pos_4_067_57    m/z 558.4211     0   1   2E+4   8.5   9  RT (min)      lcorbi_t3_pos_4_074_58    m/z 558.5094     0   1E+5   14   14.5  RT (min)       lcorbi_t3_pos_1_032_26    m/z 558.8573     0   1   2E+5   1   1.5  RT (min)       lcorbi_t3_pos_4_070_57    m/z 559.0207     0   2   4   6E+4   1   1.5  RT (min)      lcorbi_t3_pos_3_048_30    m/z 559.2457     0   1   2E+4   11.5   12  RT (min)      lcorbi_t3_pos_4_067_57    m/z 559.2992     0   1   2   3   4E+4   11   11.5  RT (min)      lcorbi_t3_pos_4_069_57    m/z 559.3267     0   1   2E+4   8   8.5  RT (min)      lcorbi_t3_pos_1_014_03    m/z 560.2062     0   2   4   6   8E+3   6.5   7  RT (min)      lcorbi_t3_pos_4_066_57    m/z 560.3067     0   2   4   6E+3   6.5   7  RT (min)      lcorbi_t3_pos_4_077_58    m/z 560.3115     0   2   4   6E+4   11   11.5  RT (min)       lcorbi_t3_pos_4_077_58    m/z 560.3279     0   2   4   6   8E+3   6   6.5  RT (min)      lcorbi_t3_pos_4_074_58    m/z 560.3346     0   1E+5   9.5   10  RT (min)       lcorbi_t3_pos_4_076_58    m/z 560.4274     0   1E+4   6.5   7  RT (min)       lcorbi_t3_pos_4_018_01    m/z 560.4757     0   1   2E+4   13.5   14  RT (min)      lcorbi_t3_pos_3_015_06    m/z 560.9195     0   2   4   6E+3   7   7.5  RT (min)      lcorbi_t3_pos_3_024_07    m/z 560.9292     0   1   2E+4   6.5   7  RT (min)      lcorbi_t3_pos_1_032_26    m/z 561.0418     0   1   2   3   4E+4   1   1.5  RT (min)      lcorbi_t3_pos_4_078_58    m/z 561.4880     0   1E+5   14   14.5  RT (min)       lcorbi_t3_pos_4_086_59    m/z 561.5241     0   1   2   3   4E+4   14   14.5  RT (min)      lcorbi_t3_pos_1_026_17    m/z 561.8749     0   1E+5   1   1.5  RT (min)       lcorbi_t3_pos_4_067_57    m/z 562.3071     0   1E+4   5   5.5  RT (min)      lcorbi_t3_pos_4_074_58    m/z 562.3776     0   1E+5   11   11.5  RT (min)       lcorbi_t3_pos_1_050_13    m/z 562.3800     0   2   4   6E+3   5.5   6  RT (min)      lcorbi_t3_pos_4_066_57    m/z 562.4232     0   1E+5   12.5   13  RT (min)       lcorbi_t3_pos_4_079_58    m/z 562.5104     0   1   2   3E+4   13   13.5  RT (min)      lcorbi_t3_pos_1_038_30    m/z 562.5196     0   1   2E+5   14   14.5  RT (min)       lcorbi_t3_pos_4_089_59    m/z 562.5882     0   2   4   6E+4   1   1.5  RT (min)      lcorbi_t3_pos_4_078_58    m/z 562.8053     0   1E+5   1   1.5  RT (min)      lcorbi_t3_pos_1_029_29    m/z 562.8746     0   2   4E+4   1   1.5  RT (min)      lcorbi_t3_pos_4_067_57    m/z 563.1639     0   1   2E+5   9   9.5  RT (min)       lcorbi_t3_pos_4_098_61    m/z 563.3094     0   1E+5   11.5   12  RT (min)      lcorbi_t3_pos_4_072_57    m/z 563.3215     0   1   2E+4   8   8.5  RT (min)       lcorbi_t3_pos_2_053_28    m/z 563.3306     0   2   4   6E+4   11.5   12  RT (min)      lcorbi_t3_pos_1_029_29    m/z 563.4656     0   1   2E+4   14   14.5  RT (min)       lcorbi_t3_pos_4_102_61    m/z 563.5225     0   1   14   14.5  RT (min)      lcorbi_t3_pos_4_109_62    m/z 563.5513     0   1   2   3   4E+4   14   14.5  RT (min)       lcorbi_t3_pos_4_102_61    m/z 563.8803     0   1   2   3E+4   1   1.5  RT (min)      lcorbi_t3_pos_4_072_57    m/z 564.0865     0   1   2E+5   8.5   9  RT (min)       lcorbi_t3_pos_4_100_61    m/z 564.1857     0   1E+4   4.5   5  RT (min)      lcorbi_t3_pos_4_073_57    m/z 564.3030     0   1E+5   11.5   12  RT (min)       lcorbi_t3_pos_4_066_57    m/z 564.3249     0   2   4   6   8E+3   8   8.5  RT (min)      lcorbi_t3_pos_2_053_28    m/z 564.3591     0   1   2E+4   5   5.5  RT (min)       lcorbi_t3_pos_1_003_45    m/z 564.3837     0   2   4   6   8E+4   11   11.5  RT (min)       lcorbi_t3_pos_4_075_58    m/z 564.3933     0   1E+5   11.5   12  RT (min)       lcorbi_t3_pos_1_050_13    m/z 564.5853     0   2   4   6E+4   1   1.5  RT (min)      lcorbi_t3_pos_4_077_58    m/z 564.8025     0   2   4   6   8E+4   1   1.5  RT (min)      lcorbi_t3_pos_1_029_29    m/z 565.0406     0   1E+5   1   1.5  RT (min)      lcorbi_t3_pos_3_010_44    m/z 565.0879     0   1E+4   10   10.5  RT (min)      lcorbi_t3_pos_4_006_12    m/z 565.1619     0   1E+5   9   9.5  RT (min)       lcorbi_t3_pos_4_101_61    m/z 565.3066     0   2   4   6E+4   11.5   12  RT (min)       lcorbi_t3_pos_4_068_57    m/z 565.3171     0   1   2   3   4E+4   11   11.5  RT (min)       lcorbi_t3_pos_4_077_58    m/z 565.3626     0   2   4   6   8E+3   5   5.5  RT (min)      lcorbi_t3_pos_1_001_49    m/z 565.4014     0   2   4E+4   13.5   14  RT (min)       lcorbi_t3_pos_4_098_61    m/z 565.4426     0   1   2   3   4E+4   12.5   13  RT (min)       lcorbi_t3_pos_4_077_58    m/z 565.4829     0   1   2   3E+4   13   13.5  RT (min)      lcorbi_t3_pos_4_068_57    m/z 565.4926     0   1   2   3   4E+4   12.5   13  RT (min)       lcorbi_t3_pos_4_082_59    m/z 565.5103     0   1E+5   13.5   14  RT (min)      lcorbi_t3_pos_4_098_61    m/z 565.5669     0   1E+5   17.5   18  RT (min)       lcorbi_t3_pos_4_079_58    m/z 566.0844     0   1   2E+5   8.5   9  RT (min)       lcorbi_t3_pos_4_101_61    m/z 566.3089     0   1E+5   7   7.5  RT (min)       lcorbi_t3_pos_1_054_48    m/z 566.3934     0   1   2   3E+5   10.5   11  RT (min)       lcorbi_t3_pos_1_050_13    m/z 566.4416     0   1E+5   13   13.5  RT (min)       lcorbi_t3_pos_4_070_57    m/z 566.5824     0   2   4E+4   1   1.5  RT (min)      lcorbi_t3_pos_4_076_58    m/z 566.8285     0   1   2   3   4E+3   1   1.5  RT (min)      lcorbi_t3_pos_3_032_09    m/z 566.9293     0   1E+4   6.5   7  RT (min)      lcorbi_t3_pos_2_003_19    m/z 567.3253     0   1E+5   11   11.5  RT (min)       lcorbi_t3_pos_4_071_57    m/z 567.3907     0   1   2   3   4E+4   1   1.5  RT (min)       lcorbi_t3_pos_4_073_57    m/z 567.4386     0   2   4   6   8E+4   14   14.5  RT (min)       lcorbi_t3_pos_4_106_62    m/z 567.6100     0   1E+5   14.5   15  RT (min)      lcorbi_t3_pos_2_044_21    m/z 568.2350     0   1   2E+4   4.5   5  RT (min)      lcorbi_t3_pos_4_106_62    m/z 568.4091     0   1   2   3E+5   11   11.5  RT (min)       lcorbi_t3_pos_1_050_13    m/z 568.4574     0   1E+5   13   13.5  RT (min)       lcorbi_t3_pos_1_054_48    m/z 568.4702     0   1   2   3   4E+4   13   13.5  RT (min)      lcorbi_t3_pos_4_080_58    m/z 568.5574     0   1E+5   14   14.5  RT (min)      lcorbi_t3_pos_4_084_59    m/z 568.5769     0   1   2   3   4E+4   14.5   15  RT (min)       lcorbi_t3_pos_1_032_26    m/z 568.5796     0   1   2E+4   1   1.5  RT (min)      lcorbi_t3_pos_2_051_35    m/z 569.4368     0   1   2   3E+4   5.5   6  RT (min)      lcorbi_t3_pos_4_005_33    m/z 569.4543     0   2   4   6   8E+4   14   14.5  RT (min)       lcorbi_t3_pos_4_106_62    m/z 569.4679     0   1E+5   13.5   14  RT (min)      lcorbi_t3_pos_1_023_02    m/z 569.5336     0   1E+5   14   14.5  RT (min)      lcorbi_t3_pos_4_082_59    m/z 569.8654     0   1   2   3E+4   1   1.5  RT (min)       lcorbi_t3_pos_4_068_57    m/z 569.8964     0   2   4   6E+3   1   1.5  RT (min)      lcorbi_t3_pos_2_021_09    m/z 570.0367     0   2   4E+4   1   1.5  RT (min)      lcorbi_t3_pos_4_081_58    m/z 570.2417     0   2   4   6E+4   8.5   9  RT (min)      lcorbi_t3_pos_4_112_62    m/z 570.2634     0   2   4   6E+4   11   11.5  RT (min)       lcorbi_t3_pos_1_036_08    m/z 570.3356     0   2   4   6E+4   10.5   11  RT (min)      lcorbi_t3_pos_4_073_57    m/z 570.3462     0   1E+5   11   11.5  RT (min)      lcorbi_t3_pos_4_070_57    m/z 570.3673     0   2   4   6E+4   13.5   14  RT (min)      lcorbi_t3_pos_4_098_61    m/z 570.3882     0   1   2   3   4E+4   13.5   14  RT (min)      lcorbi_t3_pos_4_099_61    m/z 570.4218     0   1E+5   10.5   11  RT (min)       lcorbi_t3_pos_4_091_60    m/z 570.4730     0   2   4   6   8E+4   12.5   13  RT (min)       lcorbi_t3_pos_1_044_25    m/z 570.4959     0   1E+5   15   15.5  RT (min)       lcorbi_t3_pos_1_036_08    m/z 570.5094     0   2   4   6E+4   14   14.5  RT (min)       lcorbi_t3_pos_1_034_34    m/z 570.5145     0   1E+5   12   12.5  RT (min)       lcorbi_t3_pos_4_090_60    m/z 570.5458     0   1   2E+4   14   14.5  RT (min)      lcorbi_t3_pos_3_027_27    m/z 570.6181     0   1   2E+5   14   14.5  RT (min)       lcorbi_t3_pos_4_094_60    m/z 570.9040     0   1E+4   7   7.5  RT (min)      lcorbi_t3_pos_4_085_59    m/z 571.2444     0   1   2E+4   8.5   9  RT (min)      lcorbi_t3_pos_4_112_62    m/z 571.3971     0   1   2   3E+5   13   13.5  RT (min)       lcorbi_t3_pos_4_067_57    m/z 571.4724     0   1E+5   14   14.5  RT (min)       lcorbi_t3_pos_4_082_59    m/z 571.6304     0   2   4   6E+4   14   14.5  RT (min)       lcorbi_t3_pos_4_094_60    m/z 571.8674     0   1   2E+4   1   1.5  RT (min)      lcorbi_t3_pos_4_075_58    m/z 572.3348     0   2   4E+4   11   11.5  RT (min)      lcorbi_t3_pos_1_008_36    m/z 572.3617     0   2   4   6E+4   11   11.5  RT (min)      lcorbi_t3_pos_1_020_31    m/z 572.4004     0   1E+5   13   13.5  RT (min)      lcorbi_t3_pos_4_067_57    m/z 572.4639     0   1   2   3E+5   11   11.5  RT (min)      lcorbi_t3_pos_4_088_59    m/z 572.5252     0   2   4   6E+4   14.5   15  RT (min)       lcorbi_t3_pos_4_089_59    m/z 572.6378     0   1   2   3E+4   1   1.5  RT (min)      lcorbi_t3_pos_2_020_22    m/z 572.8341     0   1   2E+5   1   1.5  RT (min)      lcorbi_t3_pos_1_029_29    m/z 572.8456     0   2   4E+4   1   1.5  RT (min)       lcorbi_t3_pos_4_066_57    m/z 574.1986     0   1E+4   1.5   2  RT (min)      lcorbi_t3_pos_4_095_60    m/z 574.3741     0   1E+5   13.5   14  RT (min)      lcorbi_t3_pos_2_044_21    m/z 574.3776     0   2   4E+4   11.5   12  RT (min)      lcorbi_t3_pos_4_071_57    m/z 574.3964     0   1E+4   6   6.5  RT (min)      lcorbi_t3_pos_4_088_59    m/z 574.6350     0   1   2   3E+4   1   1.5  RT (min)      lcorbi_t3_pos_1_034_34    m/z 574.6494     0   1E+5   14   14.5  RT (min)       lcorbi_t3_pos_4_095_60    m/z 574.8315     0   2   4   6   8E+4   1   1.5  RT (min)      lcorbi_t3_pos_4_075_58    m/z 574.8760     0   1E+5   1   1.5  RT (min)       lcorbi_t3_pos_4_075_58    m/z 574.9411     0   2   4   6E+3   5.5   6  RT (min)      lcorbi_t3_pos_4_034_42    m/z 574.9944     0   1   2E+4   1   1.5  RT (min)      lcorbi_t3_pos_3_048_30    m/z 575.3776     0   1   2   3E+4   1   1.5  RT (min)      lcorbi_t3_pos_4_080_58    m/z 575.3803     0   1   2   3E+4   6.5   7  RT (min)       lcorbi_t3_pos_4_018_01    m/z 575.4283     0   1E+5   12.5   13  RT (min)       lcorbi_t3_pos_4_080_58    m/z 575.4332     0   1E+4   5.5   6   6.5  RT (min)       lcorbi_t3_pos_4_085_59    m/z 575.4672     0   2   4   6   8E+4   13   13.5  RT (min)      lcorbi_t3_pos_4_081_58    m/z 575.4914     0   1   2   3   4E+4   14   14.5  RT (min)      lcorbi_t3_pos_2_045_40    m/z 575.8769     0   1   2E+4   1   1.5  RT (min)      lcorbi_t3_pos_4_074_58    m/z 575.9348     0   1E+4   6   6.5  RT (min)      lcorbi_t3_pos_2_013_33    m/z 576.2807     0   2   4   6   8E+4   11   11.5  RT (min)       lcorbi_t3_pos_1_035_28    m/z 576.3228     0   1E+4   5   5.5  RT (min)      lcorbi_t3_pos_4_106_62    m/z 576.3229     0   1   2E+4   11   11.5  RT (min)      lcorbi_t3_pos_4_073_57    m/z 576.3438     0   2   4   6E+4   8.5   9  RT (min)      lcorbi_t3_pos_4_101_61    m/z 576.3777     0   2   4   6E+4   11   11.5  RT (min)       lcorbi_t3_pos_4_086_59    m/z 576.3836     0   2   4   6   8E+3   6.5   7  RT (min)      lcorbi_t3_pos_3_020_35    m/z 576.3956     0   2   4   6E+3   6   6.5  RT (min)      lcorbi_t3_pos_4_106_62    m/z 576.4180     0   1   2   3   4E+4   13.5   14  RT (min)       lcorbi_t3_pos_4_108_62    m/z 576.5264     0   1   2E+4   14.5   15  RT (min)      lcorbi_t3_pos_4_086_59    m/z 576.5352     0   1E+5   14   14.5  RT (min)      lcorbi_t3_pos_4_086_59    m/z 576.8457     0   2   4   6E+4   1   1.5  RT (min)       lcorbi_t3_pos_4_077_58    m/z 577.3261     0   1   2   3E+3   5   5.5  RT (min)      lcorbi_t3_pos_4_082_59    m/z 577.4253     0   1E+5   15   15.5  RT (min)       lcorbi_t3_pos_4_084_59    m/z 577.4692     0   1   2E+4   13.5   14  RT (min)      lcorbi_t3_pos_4_086_59    m/z 577.4828     0   1   2   3E+4   13.5   14  RT (min)      lcorbi_t3_pos_4_082_59    m/z 577.5103     0   1E+5   14.5   15  RT (min)      lcorbi_t3_pos_4_082_59    m/z 577.8519     0   1   2   3   4E+4   1   1.5  RT (min)      lcorbi_t3_pos_4_075_58    m/z 578.1385     0   2   4   6E+4   13   13.5  RT (min)       lcorbi_t3_pos_1_041_54    m/z 578.3386     0   2   4   6   8E+3   5   5.5  RT (min)      lcorbi_t3_pos_4_080_58    m/z 578.4136     0   2   4   6   8E+4   13.5   14  RT (min)       lcorbi_t3_pos_1_039_23    m/z 578.4808     0   1E+5   12   12.5  RT (min)       lcorbi_t3_pos_2_030_14    m/z 578.8528     0   2   4E+4   18   18.5  RT (min)      lcorbi_t3_pos_4_074_58    m/z 578.9858     0   1   2E+4   10   10.5  RT (min)      lcorbi_t3_pos_4_076_58    m/z 579.2928     0   2   4   6   8E+4   8.5   9  RT (min)       lcorbi_t3_pos_4_101_61    m/z 579.5350     0   1E+5   14.5   15  RT (min)       lcorbi_t3_pos_4_082_59    m/z 580.2752     0   1   2   3   4E+4   11   11.5  RT (min)      lcorbi_t3_pos_1_036_08    m/z 580.2962     0   1   2   3E+4   8.5   9  RT (min)      lcorbi_t3_pos_4_101_61    m/z 580.3462     0   1   2   3   4E+4   8   8.5  RT (min)       lcorbi_t3_pos_4_107_62    m/z 580.4338     0   2   4   6E+4   13   13.5  RT (min)       lcorbi_t3_pos_4_070_57    m/z 580.5019     0   1   2   3E+4   13.5   14  RT (min)       lcorbi_t3_pos_1_034_34    m/z 580.5593     0   1   2E+4   1   1.5  RT (min)      lcorbi_t3_pos_4_078_58    m/z 580.5859     0   1   2   3   4E+4   14.5   15  RT (min)       lcorbi_t3_pos_3_006_38    m/z 580.8630     0   1   2E+4   1   1.5  RT (min)      lcorbi_t3_pos_4_005_33    m/z 581.0145     0   1   2   3   4E+4   1   1.5  RT (min)      lcorbi_t3_pos_3_010_44    m/z 581.2958     0   1   2E+4   11   11.5  RT (min)      lcorbi_t3_pos_3_008_20    m/z 581.4973     0   2   4E+4   14   14.5  RT (min)      lcorbi_t3_pos_4_087_59    m/z 581.5142     0   1   2   3E+4   14.5   15  RT (min)      lcorbi_t3_pos_4_106_62    m/z 581.5336     0   1E+5   14   14.5  RT (min)       lcorbi_t3_pos_4_082_59    m/z 582.2959     0   2   4   6E+4   10.5   11   11.5  RT (min)       lcorbi_t3_pos_4_071_57    m/z 582.4226     0   1   2E+4   6   6.5  RT (min)       lcorbi_t3_pos_4_087_59    m/z 582.9145     0   2   4E+3   6.5   7  RT (min)      lcorbi_t3_pos_2_020_22    m/z 582.9243     0   1E+4   6   6.5  RT (min)      lcorbi_t3_pos_4_086_59    m/z 583.2550     0   2   4   6   8E+4   9   9.5  RT (min)       lcorbi_t3_pos_4_073_57    m/z 583.3242     0   1   2E+4   11   11.5  RT (min)      lcorbi_t3_pos_2_032_39    m/z 583.3645     0   1E+4   1   1.5  RT (min)      lcorbi_t3_pos_4_074_58    m/z 583.4543     0   1E+5   5.5   6  RT (min)       lcorbi_t3_pos_4_005_33    m/z 583.6016     0   1E+5   14.5   15  RT (min)       lcorbi_t3_pos_4_094_60    m/z 583.8661     0   1E+5   1   1.5  RT (min)      lcorbi_t3_pos_4_075_58    m/z 584.4019     0   1E+4   5   5.5  RT (min)      lcorbi_t3_pos_4_087_59    m/z 584.4383     0   2   4E+4   5   5.5  RT (min)       lcorbi_t3_pos_4_082_59    m/z 584.4576     0   2   4   6E+4   5.5   6  RT (min)      lcorbi_t3_pos_4_005_33    m/z 584.5159     0   1E+5   14   14.5  RT (min)      lcorbi_t3_pos_1_028_32    m/z 584.5522     0   1E+5   14   14.5  RT (min)      lcorbi_t3_pos_4_083_59    m/z 584.6611     0   1E+5   14   14.5  RT (min)       lcorbi_t3_pos_4_093_60    m/z 584.8092     0   1E+4   4.5   5  RT (min)      lcorbi_t3_pos_4_074_58    m/z 584.8620     0   1E+5   1   1.5  RT (min)      lcorbi_t3_pos_4_081_58    m/z 584.9860     0   1E+4   1   1.5  RT (min)      lcorbi_t3_pos_3_048_30    m/z 585.2708     0   1   2   3E+4   7.5   8  RT (min)       lcorbi_t3_pos_1_029_29    m/z 585.3034     0   1E+4   8   8.5  RT (min)      lcorbi_t3_pos_4_106_62    m/z 585.3106     0   1E+4   4.5   5  RT (min)      lcorbi_t3_pos_4_078_58    m/z 585.3587     0   1   2   3   4E+4   13.5   14  RT (min)      lcorbi_t3_pos_4_102_61    m/z 585.4416     0   1   2E+4   5   5.5  RT (min)      lcorbi_t3_pos_4_088_59    m/z 585.4765     0   1   2   3E+4   12.5   13  RT (min)      lcorbi_t3_pos_4_087_59    m/z 585.5649     0   1E+5   14   14.5  RT (min)       lcorbi_t3_pos_4_086_59    m/z 585.8396     0   1   2E+4   1   1.5  RT (min)      lcorbi_t3_pos_4_079_58    m/z 586.2440     0   2   4   6   8E+4   13   13.5  RT (min)       lcorbi_t3_pos_4_100_61    m/z 586.2742     0   1E+4   7.5   8  RT (min)      lcorbi_t3_pos_4_079_58    m/z 586.2968     0   1   2   3   4E+4   11   11.5  RT (min)      lcorbi_t3_pos_1_035_28    m/z 586.3094     0   2   4   6   8E+4   11   11.5  RT (min)       lcorbi_t3_pos_1_035_28    m/z 586.5316     0   1E+5   14.5   15  RT (min)      lcorbi_t3_pos_1_028_32    m/z 586.6130     0   1E+5   13.5   14  RT (min)       lcorbi_t3_pos_4_096_60    m/z 587.5037     0   1   14.5   15  RT (min)       lcorbi_t3_pos_4_106_62    m/z 588.1106     0   1E+4   2   2.5  RT (min)      lcorbi_t3_pos_3_020_35    m/z 588.1707     0   1   2E+4   1   1.5  RT (min)      lcorbi_t3_pos_4_005_33    m/z 588.3147     0   2   4E+4   11   11.5  RT (min)       lcorbi_t3_pos_4_070_57    m/z 588.4097     0   1E+5   5.5   6  RT (min)       lcorbi_t3_pos_4_005_33    m/z 588.4682     0   2   4E+4   14   14.5  RT (min)       lcorbi_t3_pos_3_031_45    m/z 588.4796     0   2   4   6   8E+4   12.5   13  RT (min)       lcorbi_t3_pos_4_077_58    m/z 588.6287     0   1E+5   14   14.5  RT (min)       lcorbi_t3_pos_4_094_60    m/z 588.8083     0   2   4   6E+4   1   1.5  RT (min)      lcorbi_t3_pos_1_029_29    m/z 588.8196     0   1E+4   1   1.5  RT (min)      lcorbi_t3_pos_3_025_52    m/z 588.8499     0   1   2   3E+4   1   1.5  RT (min)      lcorbi_t3_pos_2_012_01    m/z 589.2114     0   1E+5   10   10.5  RT (min)       lcorbi_t3_pos_1_020_31    m/z 589.4130     0   1   2   3   4E+4   5.5   6  RT (min)      lcorbi_t3_pos_4_005_33    m/z 589.4830     0   1   2   3   4E+4   12.5   13  RT (min)      lcorbi_t3_pos_4_079_58    m/z 589.5363     0   2   4   6   8E+4   14   14.5  RT (min)      lcorbi_t3_pos_1_008_36    m/z 589.5669     0   2   4   6E+4   14   14.5  RT (min)       lcorbi_t3_pos_2_037_27    m/z 589.8859     0   1E+4   1   1.5  RT (min)      lcorbi_t3_pos_4_068_57    m/z 590.0029     0   2   4E+4   1   1.5  RT (min)      lcorbi_t3_pos_4_052_06    m/z 590.2148     0   1   2   3E+4   10   10.5  RT (min)      lcorbi_t3_pos_4_021_25    m/z 590.3218     0   2   4E+4   11   11.5  RT (min)       lcorbi_t3_pos_4_068_57    m/z 590.3303     0   1   2   3E+4   11   11.5  RT (min)       lcorbi_t3_pos_4_078_58    m/z 590.3385     0   2   4   6E+3   5.5   6  RT (min)      lcorbi_t3_pos_4_066_57    m/z 590.3582     0   2   4E+4   10.5   11  RT (min)      lcorbi_t3_pos_1_032_26    m/z 590.4112     0   2   4E+3   6.5   7  RT (min)      lcorbi_t3_pos_4_091_60    m/z 590.4263     0   2   4   6E+4   10.5   11  RT (min)       lcorbi_t3_pos_4_093_60    m/z 590.4335     0   2   4   6E+4   13.5   14  RT (min)       lcorbi_t3_pos_4_100_61    m/z 590.5140     0   2   4E+4   14   14.5  RT (min)       lcorbi_t3_pos_3_054_15    m/z 590.5226     0   1E+5   14.5   15  RT (min)      lcorbi_t3_pos_4_085_59    m/z 590.5487     0   2   4   6E+4   14.5   15  RT (min)       lcorbi_t3_pos_1_034_34    m/z 590.8511     0   2   4E+4   1   1.5  RT (min)      lcorbi_t3_pos_4_081_58    m/z 591.1994     0   2   4   6   8E+4   13   13.5  RT (min)      lcorbi_t3_pos_4_111_62    m/z 591.4297     0   1   2E+4   10.5   11  RT (min)      lcorbi_t3_pos_2_045_40    m/z 591.4411     0   1E+5   15   15.5  RT (min)       lcorbi_t3_pos_4_089_59    m/z 591.4585     0   2   4   6   8E+4   13   13.5  RT (min)      lcorbi_t3_pos_4_079_58    m/z 591.5349     0   1E+5   14   14.5  RT (min)       lcorbi_t3_pos_4_084_59    m/z 591.8518     0   1E+4   1   1.5  RT (min)      lcorbi_t3_pos_4_075_58    m/z 592.3177     0   1   2   3E+4   5   5.5  RT (min)       lcorbi_t3_pos_4_066_57    m/z 592.3461     0   2   4   6   8E+4   11.5   12  RT (min)       lcorbi_t3_pos_1_029_29    m/z 592.5000     0   2   4   6E+4   14   14.5  RT (min)       lcorbi_t3_pos_4_084_59    m/z 592.5414     0   1   2E+4   13   13.5  RT (min)      lcorbi_t3_pos_1_036_08    m/z 592.8200     0   1E+4   1   1.5  RT (min)      lcorbi_t3_pos_4_075_58    m/z 593.3211     0   1E+4   5   5.5  RT (min)      lcorbi_t3_pos_4_074_58    m/z 593.3484     0   1   2E+4   11.5   12  RT (min)      lcorbi_t3_pos_3_038_18    m/z 593.3759     0   1   2   3E+4   10   10.5  RT (min)      lcorbi_t3_pos_4_086_59    m/z 593.4424     0   1E+5   12   12.5  RT (min)       lcorbi_t3_pos_3_021_39    m/z 594.1606     0   2   4E+4   9   9.5  RT (min)       lcorbi_t3_pos_4_084_59    m/z 594.4860     0   1E+5   13.5   14  RT (min)       lcorbi_t3_pos_4_069_57    m/z 594.5095     0   2   4   6   8E+4   13.5   14  RT (min)       lcorbi_t3_pos_4_082_59    m/z 594.5176     0   2   4   6E+4   14.5   15  RT (min)      lcorbi_t3_pos_2_020_22    m/z 594.5458     0   1E+5   14   14.5   15  RT (min)       lcorbi_t3_pos_4_070_57    m/z 594.5631     0   1E+5   14.5   15  RT (min)      lcorbi_t3_pos_3_027_27    m/z 594.8371     0   2   4   6   8E+4   1   1.5  RT (min)      lcorbi_t3_pos_3_010_44    m/z 595.2458     0   1   2   3E+4   11   11.5  RT (min)       lcorbi_t3_pos_4_091_60    m/z 595.2577     0   2   4   6   8E+4   10   10.5  RT (min)       lcorbi_t3_pos_4_077_58    m/z 595.2668     0   1   2   3   4E+4   8.5   9  RT (min)       lcorbi_t3_pos_4_101_61    m/z 595.4699     0   2   4   6E+4   14.5   15  RT (min)       lcorbi_t3_pos_4_102_61    m/z 595.5129     0   1   2E+4   13.5   14  RT (min)      lcorbi_t3_pos_4_032_40    m/z 595.5492     0   1   2   3   4E+4   14   14.5  RT (min)      lcorbi_t3_pos_4_086_59    m/z 595.8685     0   1E+5   1   1.5  RT (min)       lcorbi_t3_pos_4_067_57    m/z 596.2702     0   1E+4   8.5   9  RT (min)      lcorbi_t3_pos_4_103_61    m/z 596.4733     0   1   14.5   15  RT (min)      lcorbi_t3_pos_1_028_32    m/z 596.4969     0   2   4   6E+4   12.5   13  RT (min)      lcorbi_t3_pos_4_076_58    m/z 596.5098     0   1   13   13.5  RT (min)      lcorbi_t3_pos_4_078_58    m/z 596.5333     0   1   2   3E+4   14.5   15  RT (min)      lcorbi_t3_pos_2_020_22    m/z 596.5615     0   2   4   6   8E+4   15   15.5  RT (min)       lcorbi_t3_pos_4_067_57    m/z 596.8677     0   2   4   6   8E+4   1   1.5  RT (min)      lcorbi_t3_pos_4_070_57    m/z 597.3356     0   2   4   6E+3   5   5.5  RT (min)      lcorbi_t3_pos_4_076_58    m/z 597.3726     0   1   2E+4   8   8.5  RT (min)      lcorbi_t3_pos_4_107_62    m/z 597.4492     0   2   4   6   8E+4   14   14.5  RT (min)       lcorbi_t3_pos_4_106_62    m/z 597.4879     0   1E+5   14   14.5  RT (min)       lcorbi_t3_pos_1_032_26    m/z 597.5094     0   2   4   6   8E+4   13   13.5  RT (min)       lcorbi_t3_pos_4_074_58    m/z 597.6368     0   1E+5   14   14.5  RT (min)       lcorbi_t3_pos_4_093_60    m/z 597.8740     0   2   4E+4   1   1.5  RT (min)       lcorbi_t3_pos_4_068_57    m/z 598.2949     0   1   2E+4   11.5   12  RT (min)      lcorbi_t3_pos_2_051_35    m/z 598.3757     0   1E+4   1   1.5  RT (min)      lcorbi_t3_pos_4_071_57    m/z 598.3873     0   2   4   6E+4   10.5   11  RT (min)       lcorbi_t3_pos_4_092_60    m/z 598.4177     0   1   2E+4   5.5   6  RT (min)       lcorbi_t3_pos_1_054_48    m/z 598.4316     0   2   4   6   8E+4   14   14.5  RT (min)       lcorbi_t3_pos_4_074_58    m/z 598.4808     0   2   4E+4   12.5   13  RT (min)       lcorbi_t3_pos_4_098_61    m/z 598.4913     0   2   4   6E+4   14   14.5  RT (min)      lcorbi_t3_pos_4_086_59    m/z 598.6289     0   1   2   3E+4   13.5   14  RT (min)      lcorbi_t3_pos_2_029_43    m/z 598.6405     0   1   2   3E+5   14   14.5  RT (min)       lcorbi_t3_pos_4_092_60    m/z 598.8372     0   1   2   3E+5   1   1.5  RT (min)       lcorbi_t3_pos_4_075_58    m/z 599.0018     0   1   2E+4   1   1.5  RT (min)      lcorbi_t3_pos_3_003_33    m/z 599.2500     0   2   4E+3   8   8.5  RT (min)      lcorbi_t3_pos_4_107_62    m/z 599.3905     0   2   4   6   8E+3   4.5   5  RT (min)      lcorbi_t3_pos_4_091_60    m/z 599.4019     0   1E+4   6   6.5  RT (min)      lcorbi_t3_pos_4_087_59    m/z 599.4211     0   1E+4   5.5   6  RT (min)      lcorbi_t3_pos_4_084_59    m/z 599.6527     0   1E+5   14   14.5  RT (min)       lcorbi_t3_pos_4_096_60    m/z 599.8401     0   2   4E+4   1   1.5  RT (min)      lcorbi_t3_pos_4_080_58    m/z 600.2808     0   1   2E+4   11   11.5  RT (min)      lcorbi_t3_pos_2_054_29    m/z 600.3897     0   1   2   3   4E+4   13.5   14  RT (min)      lcorbi_t3_pos_2_032_39    m/z 600.4683     0   2   4   6   8E+4   14   15  RT (min)       lcorbi_t3_pos_4_098_61    m/z 600.5473     0   1E+5   14.5   15  RT (min)      lcorbi_t3_pos_4_087_59    m/z 600.5525     0   1   2E+4   1   1.5  RT (min)      lcorbi_t3_pos_4_070_57    m/z 600.7857     0   2   4   6E+4   18   18.5  RT (min)       lcorbi_t3_pos_4_075_58    m/z 600.8357     0   2   4E+4   1   1.5  RT (min)      lcorbi_t3_pos_4_075_58    m/z 601.2749     0   1E+5   10   10.5  RT (min)       lcorbi_t3_pos_4_078_58    m/z 601.3845     0   1E+5   1   1.5  RT (min)      lcorbi_t3_pos_4_067_57    m/z 601.4716     0   1E+5   10.5   11  RT (min)      lcorbi_t3_pos_4_082_59    m/z 601.4806     0   2   4   6   8E+4   14   14.5  RT (min)       lcorbi_t3_pos_4_089_59    m/z 601.6684     0   1E+5   14   14.5  RT (min)       lcorbi_t3_pos_1_036_08    m/z 601.8851     0   2   4   6E+4   1   1.5  RT (min)      lcorbi_t3_pos_4_072_57    m/z 602.2744     0   1   2   3E+4   11   11.5  RT (min)      lcorbi_t3_pos_4_080_58    m/z 602.2965     0   1   2E+4   11   11.5  RT (min)      lcorbi_t3_pos_2_051_35    m/z 602.3280     0   1   2E+4   8   8.5  RT (min)      lcorbi_t3_pos_4_106_62    m/z 602.4911     0   2   4   6E+4   15   15.5  RT (min)      lcorbi_t3_pos_3_054_15    m/z 602.5358     0   2   4E+4   14   14.5  RT (min)      lcorbi_t3_pos_4_090_60    m/z 602.5496     0   1E+4   1   1.5  RT (min)      lcorbi_t3_pos_1_020_31    m/z 603.2929     0   1E+4   10.5   11  RT (min)      lcorbi_t3_pos_4_081_58    m/z 603.4411     0   2   4   6E+4   15   15.5  RT (min)       lcorbi_t3_pos_4_088_59    m/z 603.5249     0   2   4   6   8E+4   14.5   15  RT (min)      lcorbi_t3_pos_4_087_59    m/z 603.5480     0   1   2E+5   14.5   15  RT (min)      lcorbi_t3_pos_1_032_26    m/z 603.8562     0   2   4   6E+4   1   1.5  RT (min)      lcorbi_t3_pos_4_067_57    m/z 604.2924     0   2   4   6E+4   11   11.5  RT (min)       lcorbi_t3_pos_4_068_57    m/z 604.3119     0   2   4   6E+4   11.5   12  RT (min)       lcorbi_t3_pos_1_029_29    m/z 604.3541     0   2   4   6E+3   6   6.5  RT (min)      lcorbi_t3_pos_4_066_57    m/z 604.3845     0   1   2E+5   10.5   11  RT (min)      lcorbi_t3_pos_1_001_49    m/z 604.5732     0   2   4   6E+4   1   1.5  RT (min)      lcorbi_t3_pos_4_067_57    m/z 604.6874     0   1E+5   14.5   15  RT (min)       lcorbi_t3_pos_1_036_08    m/z 604.7902     0   1E+5   1   1.5  RT (min)      lcorbi_t3_pos_2_028_24    m/z 604.8544     0   2   4E+4   1   1.5  RT (min)      lcorbi_t3_pos_4_079_58    m/z 605.2959     0   1   2E+4   11   11.5  RT (min)      lcorbi_t3_pos_4_067_57    m/z 605.4546     0   2   4E+4   14.5   15  RT (min)      lcorbi_t3_pos_4_032_40    m/z 605.4755     0   1   2   3E+4   14   14.5  RT (min)      lcorbi_t3_pos_4_084_59    m/z 605.5311     0   1   2E+5   14   14.5  RT (min)       lcorbi_t3_pos_4_083_59    m/z 605.5416     0   1E+5   14.5   15  RT (min)      lcorbi_t3_pos_4_085_59    m/z 605.8329     0   2   4   6   8E+3   6   6.5  RT (min)      lcorbi_t3_pos_4_076_58    m/z 605.8611     0   2   4E+4   1   1.5  RT (min)      lcorbi_t3_pos_4_081_58    m/z 606.3082     0   1   2   3   4E+4   11   11.5  RT (min)       lcorbi_t3_pos_4_070_57    m/z 606.3330     0   1E+5   5   5.5  RT (min)       lcorbi_t3_pos_4_078_58    m/z 606.4495     0   1   2   3E+4   13   13.5  RT (min)      lcorbi_t3_pos_4_070_57    m/z 606.5702     0   1E+5   1   1.5  RT (min)       lcorbi_t3_pos_4_067_57    m/z 606.8321     0   1E+5   5.5   6  RT (min)      lcorbi_t3_pos_4_074_58    m/z 606.8643     0   1E+4   1   1.5  RT (min)      lcorbi_t3_pos_4_080_58    m/z 607.2414     0   2   4   6E+4   10   10.5  RT (min)      lcorbi_t3_pos_1_020_31    m/z 607.2528     0   1   2   3E+4   12.5   13  RT (min)      lcorbi_t3_pos_2_053_28    m/z 607.3095     0   1   2   3   4E+4   5.5   6  RT (min)       lcorbi_t3_pos_4_098_61    m/z 607.3336     0   1   2   3E+4   5.5   6  RT (min)      lcorbi_t3_pos_4_075_58    m/z 607.3912     0   2   4   6E+4   14.5   15  RT (min)      lcorbi_t3_pos_4_082_59    m/z 607.4507     0   1   2   3   4E+4   14   14.5  RT (min)       lcorbi_t3_pos_3_043_23    m/z 607.5299     0   1E+5   13.5   14  RT (min)       lcorbi_t3_pos_4_099_61    m/z 607.5662     0   1   2   3   4E+4   15   15.5  RT (min)      lcorbi_t3_pos_4_018_01    m/z 607.8109     0   1   2E+4   5.5   6  RT (min)      lcorbi_t3_pos_4_106_62    m/z 607.8350     0   1E+4   5.5   6  RT (min)      lcorbi_t3_pos_4_075_58    m/z 607.9659     0   1   2E+4   1   1.5  RT (min)      lcorbi_t3_pos_3_048_30    m/z 608.2379     0   2   4E+4   1   1.5   2  RT (min)       lcorbi_t3_pos_4_091_60    m/z 608.3122     0   2   4   6   8E+3   5.5   6  RT (min)      lcorbi_t3_pos_4_106_62    m/z 608.3236     0   2   4   6   8E+4   11.5   12  RT (min)       lcorbi_t3_pos_1_029_29    m/z 608.3794     0   1   8   8.5  RT (min)       lcorbi_t3_pos_4_094_60    m/z 608.3854     0   1   2E+4   5   5.5  RT (min)       lcorbi_t3_pos_1_001_49    m/z 608.3910     0   1   2   3E+5   7   7.5  RT (min)       lcorbi_t3_pos_4_091_60    m/z 608.3991     0   1   2E+4   11   11.5  RT (min)       lcorbi_t3_pos_1_054_48    m/z 608.4654     0   2   4   6E+4   13   13.5  RT (min)       lcorbi_t3_pos_4_071_57    m/z 608.5161     0   1E+5   14   14.5  RT (min)      lcorbi_t3_pos_4_083_59    m/z 608.5423     0   1   2E+5   14   14.5  RT (min)       lcorbi_t3_pos_4_086_59    m/z 608.5673     0   2   4   6E+4   1   1.5  RT (min)      lcorbi_t3_pos_2_028_24    m/z 608.5965     0   2   4E+4   14   14.5  RT (min)       lcorbi_t3_pos_3_055_10    m/z 609.3072     0   2   4   6E+3   5   5.5  RT (min)      lcorbi_t3_pos_1_024_04    m/z 609.3271     0   1   2   3E+4   11.5   12  RT (min)      lcorbi_t3_pos_1_036_08    m/z 609.3716     0   2   4   6   8E+4   1   1.5  RT (min)      lcorbi_t3_pos_4_077_58    m/z 609.3829     0   1E+4   8   8.5  RT (min)      lcorbi_t3_pos_4_107_62    m/z 609.3889     0   2   4   6   8E+3   5   5.5  RT (min)      lcorbi_t3_pos_1_001_49    m/z 609.4666     0   1   2   3E+4   14   14.5  RT (min)      lcorbi_t3_pos_4_030_05    m/z 609.5459     0   1E+5   14   14.5  RT (min)       lcorbi_t3_pos_4_088_59    m/z 609.6088     0   1E+5   14   14.5  RT (min)       lcorbi_t3_pos_1_036_08    m/z 609.8708     0   2   4E+4   1   1.5  RT (min)      lcorbi_t3_pos_4_075_58    m/z 610.3098     0   1   2E+4   11   11.5  RT (min)      lcorbi_t3_pos_1_016_05    m/z 610.3717     0   1   2   3   4E+4   11   11.5  RT (min)       lcorbi_t3_pos_3_052_47    m/z 610.4316     0   2   4   6   8E+4   13   13.5  RT (min)       lcorbi_t3_pos_4_079_58    m/z 610.6247     0   1   2E+5   14.5   15  RT (min)       lcorbi_t3_pos_1_001_49    m/z 610.8111     0   1   2   3E+4   1   1.5  RT (min)      lcorbi_t3_pos_3_010_44    m/z 611.4650     0   1   2E+5   14   14.5  RT (min)       lcorbi_t3_pos_4_087_59    m/z 611.8451     0   2   4   6E+4   1   1.5  RT (min)      lcorbi_t3_pos_4_079_58    m/z 612.1639     0   1   2   3   4E+5   13.5   14  RT (min)       lcorbi_t3_pos_4_113_62    m/z 612.3256     0   1   2E+4   11   11.5  RT (min)      lcorbi_t3_pos_1_019_16    m/z 612.4683     0   2   4   6   8E+4   14   14.5  RT (min)       lcorbi_t3_pos_4_088_59    m/z 612.5201     0   1E+5   12.5   13  RT (min)       lcorbi_t3_pos_4_076_58    m/z 612.8823     0   2   4   6E+4   1   1.5  RT (min)       lcorbi_t3_pos_4_080_58    m/z 613.1599     0   1   2   3E+5   1.5   2  RT (min)       lcorbi_t3_pos_4_083_59    m/z 613.5503     0   1E+5   14.5   15  RT (min)       lcorbi_t3_pos_1_034_34    m/z 613.6596     0   1E+5   14   14.5  RT (min)       lcorbi_t3_pos_4_097_60    m/z 613.8481     0   1   2   3E+4   1   1.5  RT (min)      lcorbi_t3_pos_4_079_58    m/z 613.8844     0   1E+4   1   1.5  RT (min)      lcorbi_t3_pos_4_076_58    m/z 614.1631     0   1   2E+3   1.5   2  RT (min)      lcorbi_t3_pos_4_082_59    m/z 614.3276     0   1   2   3   4E+4   11.5   12  RT (min)      lcorbi_t3_pos_1_029_29    m/z 614.3409     0   2   4   6E+4   11.5   12  RT (min)       lcorbi_t3_pos_1_029_29    m/z 614.4476     0   1E+4   8.5   9  RT (min)      lcorbi_t3_pos_4_073_57    m/z 614.6436     0   1E+6   14   14.5  RT (min)       lcorbi_t3_pos_4_096_60    m/z 614.8112     0   2   4E+3   1   1.5  RT (min)      lcorbi_t3_pos_3_039_03    m/z 614.8189     0   1   2E+5   1   1.5  RT (min)       lcorbi_t3_pos_4_067_57    m/z 614.8802     0   1E+4   1   1.5  RT (min)      lcorbi_t3_pos_4_075_58    m/z 615.1559     0   1   1.5   2  RT (min)      lcorbi_t3_pos_4_082_59    m/z 615.2672     0   2   4   6   8E+4   13.5   14  RT (min)      lcorbi_t3_pos_4_110_62    m/z 615.3071     0   1E+4   5   5.5  RT (min)      lcorbi_t3_pos_4_066_57    m/z 615.4864     0   1   2   3   4E+4   14   14.5  RT (min)      lcorbi_t3_pos_4_086_59    m/z 615.4957     0   1E+5   12.5   13  RT (min)      lcorbi_t3_pos_1_036_08    m/z 615.5530     0   1E+5   15   15.5  RT (min)       lcorbi_t3_pos_1_036_08    m/z 615.5661     0   2   4E+4   14.5   15  RT (min)      lcorbi_t3_pos_2_027_05    m/z 615.5767     0   2   4   6   8E+4   14.5   15  RT (min)       lcorbi_t3_pos_4_089_59    m/z 615.6115     0   1E+5   14   14.5  RT (min)       lcorbi_t3_pos_1_036_08    m/z 615.6478     0   1E+5   14   14.5  RT (min)       lcorbi_t3_pos_4_097_60    m/z 615.8086     0   2   4   6E+3   5   5.5  RT (min)      lcorbi_t3_pos_4_066_57    m/z 616.1592     0   1E+4   2   2.5  RT (min)      lcorbi_t3_pos_4_083_59    m/z 616.2896     0   2   4   6   8E+4   11   11.5  RT (min)       lcorbi_t3_pos_1_029_29    m/z 616.4902     0   1   2   3E+5   11   11.5  RT (min)      lcorbi_t3_pos_4_084_59    m/z 616.5422     0   1   2E+4   14   14.5  RT (min)      lcorbi_t3_pos_1_050_13    m/z 616.8162     0   1E+5   1   1.5  RT (min)      lcorbi_t3_pos_1_020_31    m/z 616.8569     0   1   2E+5   1   1.5  RT (min)       lcorbi_t3_pos_4_075_58    m/z 616.9695     0   2   4   6   8E+3   7   7.5  RT (min)      lcorbi_t3_pos_4_034_42    m/z 617.2397     0   2   4   6   8E+4   10   10.5  RT (min)       lcorbi_t3_pos_4_075_58    m/z 617.2911     0   2   4E+4   8   8.5  RT (min)      lcorbi_t3_pos_4_107_62    m/z 617.2933     0   1   2   3E+4   11   11.5  RT (min)      lcorbi_t3_pos_1_036_08    m/z 617.3045     0   2   4   6   8E+4   9   9.5  RT (min)       lcorbi_t3_pos_4_074_58    m/z 617.3585     0   2   4E+4   1   1.5  RT (min)      lcorbi_t3_pos_4_079_58    m/z 617.4433     0   1   2   3E+4   9.5   10  RT (min)      lcorbi_t3_pos_4_099_61    m/z 617.5022     0   1E+5   14.5   15  RT (min)      lcorbi_t3_pos_4_084_59    m/z 617.8573     0   2   4E+4   1   1.5  RT (min)      lcorbi_t3_pos_4_075_58    m/z 618.2951     0   1   2E+4   8   8.5  RT (min)      lcorbi_t3_pos_4_113_62    m/z 618.3080     0   1   2   3E+4   9   9.5  RT (min)      lcorbi_t3_pos_4_074_58    m/z 618.4790     0   2   4   6   8E+4   12.5   13  RT (min)       lcorbi_t3_pos_4_077_58    m/z 618.8307     0   1   2E+4   1   1.5  RT (min)      lcorbi_t3_pos_4_079_58    m/z 618.9030     0   2   4   6   8E+4   1   1.5  RT (min)       lcorbi_t3_pos_4_070_57    m/z 619.4703     0   2   4   6E+4   14.5   15  RT (min)      lcorbi_t3_pos_1_032_26    m/z 619.8330     0   1   2   3   4E+4   1   1.5  RT (min)      lcorbi_t3_pos_4_076_58    m/z 620.3492     0   1E+4   5   5.5  RT (min)      lcorbi_t3_pos_4_090_60    m/z 620.4041     0   2   4E+4   10.5   11  RT (min)       lcorbi_t3_pos_4_086_59    m/z 620.4946     0   1E+5   13   14  RT (min)       lcorbi_t3_pos_4_074_58    m/z 620.5335     0   1E+5   14.5   15  RT (min)       lcorbi_t3_pos_1_032_26    m/z 620.5617     0   2   4   6   8E+4   15   15.5  RT (min)       lcorbi_t3_pos_4_066_57    m/z 620.5980     0   1E+5   14   14.5  RT (min)       lcorbi_t3_pos_4_071_57    m/z 620.6095     0   1   14.5   15  RT (min)      lcorbi_t3_pos_1_050_13    m/z 620.6825     0   2   4   6   8E+4   14   14.5  RT (min)       lcorbi_t3_pos_4_092_60    m/z 621.3747     0   2   4   6   8E+3   8   8.5  RT (min)      lcorbi_t3_pos_4_109_62    m/z 621.6014     0   1E+5   14   14.5  RT (min)      lcorbi_t3_pos_4_070_57    m/z 622.2321     0   1E+4   4.5   5  RT (min)      lcorbi_t3_pos_4_098_61    m/z 622.3952     0   1   2   3E+5   8   8.5  RT (min)       lcorbi_t3_pos_1_002_01    m/z 622.4083     0   2   4   6E+4   11.5   12  RT (min)       lcorbi_t3_pos_4_073_57    m/z 622.5444     0   1   2   3   4E+4   1   1.5  RT (min)      lcorbi_t3_pos_4_078_58    m/z 622.5490     0   1E+5   14.5   15  RT (min)       lcorbi_t3_pos_4_089_59    m/z 622.6137     0   1E+5   14.5   15  RT (min)       lcorbi_t3_pos_4_073_57    m/z 622.8430     0   1E+4   1   1.5  RT (min)      lcorbi_t3_pos_1_029_29    m/z 622.9700     0   1E+4   6.5   7  RT (min)      lcorbi_t3_pos_1_031_40    m/z 623.1938     0   1E+4   2   2.5  RT (min)      lcorbi_t3_pos_3_051_12    m/z 623.2569     0   2   4   6E+4   10   10.5  RT (min)       lcorbi_t3_pos_4_076_58    m/z 623.3128     0   2   4   6E+4   13.5   14  RT (min)       lcorbi_t3_pos_4_109_62    m/z 623.3986     0   1E+5   8   8.5  RT (min)      lcorbi_t3_pos_1_002_01    m/z 623.5613     0   1   2   3E+4   14.5   15  RT (min)       lcorbi_t3_pos_3_027_27    m/z 623.5947     0   1E+5   14.5   15  RT (min)       lcorbi_t3_pos_3_027_27    m/z 623.8797     0   1   2E+4   1   1.5  RT (min)      lcorbi_t3_pos_4_073_57    m/z 624.3860     0   1E+5   7   7.5  RT (min)       lcorbi_t3_pos_4_097_60    m/z 624.3941     0   2   4   6E+4   10.5   11  RT (min)       lcorbi_t3_pos_1_050_13    m/z 624.5414     0   1   2   3E+4   1   1.5  RT (min)      lcorbi_t3_pos_4_078_58    m/z 625.3465     0   1   2E+4   1   1.5  RT (min)      lcorbi_t3_pos_4_080_58    m/z 625.4806     0   1E+5   14   14.5  RT (min)       lcorbi_t3_pos_4_085_59    m/z 625.8491     0   1E+5   1   1.5  RT (min)      lcorbi_t3_pos_4_070_57    m/z 626.5358     0   2   4   6   8E+4   14   14.5  RT (min)      lcorbi_t3_pos_1_054_48    m/z 626.5630     0   1E+5   14.5   15  RT (min)      lcorbi_t3_pos_4_082_59    m/z 626.6072     0   2   4E+4   14   14.5  RT (min)       lcorbi_t3_pos_4_090_60    m/z 626.6603     0   1   2   3E+4   13.5   14  RT (min)      lcorbi_t3_pos_3_030_22    m/z 626.8453     0   1   2E+5   1   1.5  RT (min)       lcorbi_t3_pos_4_067_57    m/z 627.0084     0   1   2   3   4E+4   1   1.5  RT (min)      lcorbi_t3_pos_3_048_30    m/z 627.1756     0   2   4   6   8E+3   2.5   3  RT (min)      lcorbi_t3_pos_1_054_48    m/z 627.6471     0   1E+5   14.5   15  RT (min)       lcorbi_t3_pos_4_091_60    m/z 628.2925     0   1   2E+4   11   11.5  RT (min)      lcorbi_t3_pos_4_068_57    m/z 628.3748     0   2   4   6   8E+4   15.5   16  RT (min)       lcorbi_t3_pos_1_002_01    m/z 628.4996     0   1E+5   11.5   12  RT (min)       lcorbi_t3_pos_4_050_31    m/z 628.5323     0   1   2   3   4E+4   14.5   15  RT (min)      lcorbi_t3_pos_4_085_59    m/z 628.5516     0   1   2E+5   14.5   15  RT (min)       lcorbi_t3_pos_3_023_05    m/z 628.6228     0   1   2E+5   14   14.5  RT (min)       lcorbi_t3_pos_1_036_08    m/z 629.1956     0   1   2   3   4E+5   12.5   13  RT (min)       lcorbi_t3_pos_4_113_62    m/z 629.3583     0   2   4   6   8E+3   5   5.5  RT (min)      lcorbi_t3_pos_4_082_59    m/z 629.4546     0   1E+5   14   14.5  RT (min)       lcorbi_t3_pos_1_052_38    m/z 629.5031     0   2   4   6E+4   11.5   12  RT (min)      lcorbi_t3_pos_4_050_31    m/z 629.6926     0   2   4   6   8E+3   5   5.5  RT (min)      lcorbi_t3_pos_4_098_61    m/z 629.8625     0   1   2E+5   1   1.5  RT (min)       lcorbi_t3_pos_4_070_57    m/z 630.1936     0   1   2E+5   12.5   13  RT (min)       lcorbi_t3_pos_4_113_62    m/z 630.4579     0   2   4   6E+4   14   14.5  RT (min)      lcorbi_t3_pos_4_087_59    m/z 630.6187     0   1E+5   14.5   15  RT (min)       lcorbi_t3_pos_2_030_14    m/z 630.6386     0   1   2E+5   14.5   15  RT (min)       lcorbi_t3_pos_1_036_08    m/z 630.8619     0   2   4   6   8E+4   1   1.5  RT (min)      lcorbi_t3_pos_4_067_57    m/z 631.2207     0   1   2   2.5  RT (min)      lcorbi_t3_pos_4_090_60    m/z 631.4933     0   1   2E+4   14.5   15  RT (min)      lcorbi_t3_pos_3_008_20    m/z 631.5274     0   1E+5   14.5   15  RT (min)       lcorbi_t3_pos_4_082_59    m/z 631.6702     0   1   2   3   4E+5   14   14.5  RT (min)       lcorbi_t3_pos_4_092_60    m/z 631.8679     0   2   4   6E+4   1   1.5  RT (min)       lcorbi_t3_pos_4_073_57    m/z 631.9754     0   2   4   6E+3   6   6.5  RT (min)      lcorbi_t3_pos_4_084_59    m/z 632.2239     0   1E+5   1.5   2  RT (min)       lcorbi_t3_pos_4_097_60    m/z 632.2544     0   2   4E+4   11   11.5  RT (min)       lcorbi_t3_pos_4_081_58    m/z 632.2772     0   2   4   6   8E+4   13.5   14  RT (min)       lcorbi_t3_pos_1_018_35    m/z 632.3692     0   1E+4   1   1.5  RT (min)      lcorbi_t3_pos_4_070_57    m/z 632.4737     0   1E+5   14.5   15  RT (min)       lcorbi_t3_pos_1_028_32    m/z 632.5163     0   1E+5   14   14.5  RT (min)      lcorbi_t3_pos_4_082_59    m/z 632.6541     0   1E+5   14.5   15  RT (min)       lcorbi_t3_pos_4_094_60    m/z 632.8331     0   2   4E+4   1   1.5  RT (min)      lcorbi_t3_pos_4_080_58    m/z 633.0284     0   2   4   6   8E+4   1   1.5  RT (min)      lcorbi_t3_pos_3_010_44    m/z 633.4625     0   2   4E+4   15   15.5  RT (min)       lcorbi_t3_pos_4_013_36    m/z 634.2249     0   1   2   3E+4   2   2.5  RT (min)       lcorbi_t3_pos_4_097_60    m/z 634.4526     0   2   4E+4   10.5   11  RT (min)       lcorbi_t3_pos_1_042_37    m/z 634.4895     0   2   4E+4   14.5   15  RT (min)       lcorbi_t3_pos_4_082_59    m/z 634.6137     0   1E+5   14   14.5  RT (min)       lcorbi_t3_pos_4_069_57    m/z 634.6618     0   1E+5   14.5   15  RT (min)       lcorbi_t3_pos_4_095_60    m/z 635.1423     0   1   2E+4   1.5   2  RT (min)       lcorbi_t3_pos_4_083_59    m/z 635.2048     0   1E+5   13.5   14  RT (min)       lcorbi_t3_pos_4_099_61    m/z 635.2234     0   2   4   6E+4   1.5   2  RT (min)       lcorbi_t3_pos_4_092_60    m/z 635.3785     0   1E+5   1   1.5  RT (min)       lcorbi_t3_pos_4_070_57    m/z 635.4560     0   1E+4   10.5   11  RT (min)      lcorbi_t3_pos_4_027_07    m/z 635.4651     0   1   2   3E+5   14   14.5  RT (min)       lcorbi_t3_pos_4_088_59    m/z 635.5347     0   2   4   6E+4   14   14.5  RT (min)       lcorbi_t3_pos_1_032_26    m/z 635.5932     0   2   4   6E+4   14.5   15  RT (min)       lcorbi_t3_pos_3_055_10    m/z 635.6155     0   2   4E+4   14   14.5  RT (min)      lcorbi_t3_pos_2_024_15    m/z 636.3441     0   1E+4   5   5.5  RT (min)      lcorbi_t3_pos_4_082_59    m/z 636.3988     0   1E+5   10   10.5  RT (min)       lcorbi_t3_pos_4_086_59    m/z 636.4109     0   1E+4   8   8.5  RT (min)      lcorbi_t3_pos_1_008_36    m/z 636.4239     0   1E+5   11.5   12  RT (min)       lcorbi_t3_pos_1_054_48    m/z 636.4684     0   1E+5   14   14.5  RT (min)      lcorbi_t3_pos_4_088_59    m/z 636.8812     0   2   4   6   8E+4   1   1.5  RT (min)       lcorbi_t3_pos_4_068_57    m/z 637.2325     0   2   4   6   8E+3   5   5.5  RT (min)      lcorbi_t3_pos_1_049_55    m/z 637.3475     0   2   4   6E+3   5   5.5  RT (min)      lcorbi_t3_pos_4_066_57    m/z 637.4022     0   1   2   3   4E+4   10   10.5  RT (min)      lcorbi_t3_pos_4_085_59    m/z 637.4187     0   1E+5   9.5   10  RT (min)       lcorbi_t3_pos_4_101_61    m/z 637.4686     0   1E+5   12   12.5  RT (min)       lcorbi_t3_pos_3_021_39    m/z 637.5505     0   1   2   3   4E+4   14.5   15  RT (min)       lcorbi_t3_pos_1_036_08    m/z 637.6089     0   2   4   6   8E+4   15   15.5  RT (min)       lcorbi_t3_pos_4_093_60    m/z 637.6312     0   1   2   3E+4   14.5   15  RT (min)       lcorbi_t3_pos_1_052_38    m/z 637.8502     0   1E+5   1   1.5  RT (min)       lcorbi_t3_pos_4_067_57    m/z 638.2714     0   1E+5   11   11.5  RT (min)       lcorbi_t3_pos_1_029_29    m/z 638.3903     0   1E+4   8   8.5  RT (min)      lcorbi_t3_pos_4_113_62    m/z 638.4222     0   1   2   3E+4   9.5   10  RT (min)      lcorbi_t3_pos_4_102_61    m/z 638.8487     0   2   4E+4   1   1.5  RT (min)      lcorbi_t3_pos_4_070_57    m/z 638.9650     0   2   4   6E+3   6   6.5  RT (min)      lcorbi_t3_pos_4_083_59    m/z 639.2747     0   1   2   3E+4   11   11.5  RT (min)      lcorbi_t3_pos_1_046_22    m/z 639.3121     0   1E+5   12   12.5  RT (min)      lcorbi_t3_pos_4_086_59    m/z 639.4870     0   2   4   6   8E+4   14   14.5  RT (min)       lcorbi_t3_pos_4_083_59    m/z 639.5661     0   2   4E+4   14.5   15  RT (min)       lcorbi_t3_pos_1_036_08    m/z 639.8549     0   2   4E+4   1   1.5  RT (min)      lcorbi_t3_pos_4_080_58    m/z 640.2895     0   1   2E+4   11   11.5  RT (min)       lcorbi_t3_pos_3_012_31    m/z 640.3567     0   1E+4   1   1.5  RT (min)      lcorbi_t3_pos_4_079_58    m/z 640.5361     0   1   2   3E+4   12.5   13  RT (min)      lcorbi_t3_pos_3_047_40    m/z 640.6629     0   2   4   6E+4   14   14.5  RT (min)       lcorbi_t3_pos_4_096_60    m/z 640.8219     0   1   2E+5   1   1.5  RT (min)      lcorbi_t3_pos_4_080_58    m/z 640.8335     0   1   2   3   4E+4   1   1.5  RT (min)       lcorbi_t3_pos_4_067_57    m/z 641.6181     0   1E+5   14   14.5  RT (min)       lcorbi_t3_pos_1_050_13    m/z 641.7800     0   1   2E+4   4.5   5  RT (min)      lcorbi_t3_pos_4_082_59    m/z 642.2818     0   1E+4   4.5   5  RT (min)      lcorbi_t3_pos_4_106_62    m/z 642.3053     0   1   11   11.5  RT (min)      lcorbi_t3_pos_4_090_60    m/z 642.3182     0   1   2   3E+4   14   14.5  RT (min)      lcorbi_t3_pos_4_110_62    m/z 642.3699     0   2   4   6   8E+4   7   7.5  RT (min)      lcorbi_t3_pos_4_103_61    m/z 642.5480     0   1   2   3   4E+4   14.5   15  RT (min)      lcorbi_t3_pos_1_032_26    m/z 642.8194     0   2   4   6E+4   1   1.5  RT (min)      lcorbi_t3_pos_4_075_58    m/z 643.3462     0   1E+5   11   11.5  RT (min)      lcorbi_t3_pos_4_097_60    m/z 643.3655     0   2   4   6   8E+4   1   1.5  RT (min)      lcorbi_t3_pos_4_080_58    m/z 643.5184     0   1   2E+5   14.5   15  RT (min)      lcorbi_t3_pos_1_031_40    m/z 643.6069     0   2   4   6E+4   15   15.5  RT (min)       lcorbi_t3_pos_4_084_59    m/z 643.6433     0   1E+5   14   14.5  RT (min)       lcorbi_t3_pos_4_093_60    m/z 643.8647     0   2   4   6E+4   1   1.5  RT (min)      lcorbi_t3_pos_4_075_58    m/z 644.3026     0   2   4   6E+4   14   14.5  RT (min)       lcorbi_t3_pos_4_103_61    m/z 644.3207     0   2   4   6   8E+4   11.5   12  RT (min)       lcorbi_t3_pos_1_029_29    m/z 644.3493     0   1   2   3E+4   11   11.5  RT (min)      lcorbi_t3_pos_4_095_60    m/z 644.4945     0   1E+5   10.5   11  RT (min)       lcorbi_t3_pos_4_088_59    m/z 644.5425     0   2   4   6E+4   14   14.5  RT (min)      lcorbi_t3_pos_4_090_60    m/z 644.8335     0   2   4E+4   1   1.5  RT (min)      lcorbi_t3_pos_4_075_58    m/z 645.1320     0   1   2.5   3  RT (min)       lcorbi_t3_pos_4_082_59    m/z 645.3031     0   2   4E+4   10   10.5  RT (min)       lcorbi_t3_pos_4_073_57    m/z 645.3035     0   1E+4   6   6.5  RT (min)      lcorbi_t3_pos_1_007_07    m/z 645.3244     0   1   2E+4   11.5   12  RT (min)      lcorbi_t3_pos_4_080_58    m/z 645.3436     0   2   4   6   8E+4   4.5   5  RT (min)       lcorbi_t3_pos_4_014_48    m/z 645.4979     0   2   4   6E+4   10.5   11  RT (min)      lcorbi_t3_pos_4_050_31    m/z 645.8378     0   2   4   6   8E+4   1   1.5  RT (min)      lcorbi_t3_pos_4_075_58    m/z 646.1349     0   1   2.5   3  RT (min)      lcorbi_t3_pos_4_082_59    m/z 646.4915     0   2   4   6   8E+4   15   15.5  RT (min)      lcorbi_t3_pos_4_082_59    m/z 646.5644     0   1E+5   15   15.5  RT (min)       lcorbi_t3_pos_1_029_29    m/z 646.8351     0   2   4E+4   1   1.5  RT (min)      lcorbi_t3_pos_4_075_58    m/z 647.2427     0   1   2E+4   2.5   3  RT (min)      lcorbi_t3_pos_3_001_49    m/z 647.2660     0   1   2   3   4E+3   5   5.5  RT (min)      lcorbi_t3_pos_4_051_34    m/z 647.2983     0   2   4   6E+4   11   11.5  RT (min)       lcorbi_t3_pos_4_095_60    m/z 647.5036     0   2   4E+4   14.5   15  RT (min)       lcorbi_t3_pos_4_067_57    m/z 647.5226     0   1E+5   14.5   15  RT (min)       lcorbi_t3_pos_4_106_62    m/z 647.5499     0   2   4   6E+4   14.5   15  RT (min)      lcorbi_t3_pos_4_084_59    m/z 647.6172     0   1E+5   14   14.5  RT (min)       lcorbi_t3_pos_4_072_57    m/z 647.8421     0   1   2   3E+4   1   1.5  RT (min)      lcorbi_t3_pos_4_076_58    m/z 648.2538     0   1E+5   14   14.5  RT (min)      lcorbi_t3_pos_4_106_62    m/z 648.2662     0   1E+5   14   14.5  RT (min)      lcorbi_t3_pos_4_109_62    m/z 648.3017     0   1   2   3E+4   11   11.5  RT (min)      lcorbi_t3_pos_4_090_60    m/z 648.4757     0   2   4   6   8E+4   14.5   15  RT (min)       lcorbi_t3_pos_4_089_59    m/z 648.5160     0   2   4   6E+4   13   13.5  RT (min)      lcorbi_t3_pos_4_073_57    m/z 648.5258     0   2   4E+4   14.5   15  RT (min)      lcorbi_t3_pos_4_086_59    m/z 649.3038     0   1E+5   13.5   14  RT (min)       lcorbi_t3_pos_1_018_35    m/z 649.5833     0   1E+5   15   15.5  RT (min)      lcorbi_t3_pos_2_040_25    m/z 649.6328     0   1   2E+5   14.5   15  RT (min)      lcorbi_t3_pos_4_072_57    m/z 649.6695     0   1E+5   14.5   15  RT (min)       lcorbi_t3_pos_1_036_08    m/z 650.2233     0   1   2E+4   8.5   9  RT (min)      lcorbi_t3_pos_4_099_61    m/z 650.4266     0   1   2E+4   8   8.5  RT (min)       lcorbi_t3_pos_4_110_62    m/z 650.5360     0   1E+4   14   14.5  RT (min)      lcorbi_t3_pos_4_006_12    m/z 650.5724     0   2   4E+4   14.5   15  RT (min)      lcorbi_t3_pos_4_034_42    m/z 650.5897     0   2   4   6   8E+4   14   14.5  RT (min)       lcorbi_t3_pos_3_027_27    m/z 650.6450     0   1E+5   14.5   15  RT (min)       lcorbi_t3_pos_4_071_57    m/z 650.6866     0   1E+5   14.5   15  RT (min)       lcorbi_t3_pos_4_094_60    m/z 651.1073     0   1   2   2.5  RT (min)       lcorbi_t3_pos_4_082_59    m/z 651.3441     0   1E+5   14   14.5  RT (min)       lcorbi_t3_pos_4_107_62    m/z 651.3525     0   2   4   6E+4   1   1.5  RT (min)      lcorbi_t3_pos_4_081_58    m/z 651.4795     0   2   4   6   8E+4   13.5   14  RT (min)      lcorbi_t3_pos_4_106_62    m/z 651.4948     0   1E+5   14.5   15  RT (min)      lcorbi_t3_pos_4_087_59    m/z 651.6484     0   2   4E+4   14.5   15  RT (min)      lcorbi_t3_pos_4_070_57    m/z 651.8533     0   1E+5   1   1.5  RT (min)       lcorbi_t3_pos_4_075_58    m/z 652.1104     0   1   2   2.5  RT (min)      lcorbi_t3_pos_4_082_59    m/z 652.4118     0   1E+4   5   5.5  RT (min)      lcorbi_t3_pos_1_001_49    m/z 652.4883     0   1E+5   14.5   15  RT (min)      lcorbi_t3_pos_4_087_59    m/z 652.5880     0   1   14.5   15  RT (min)      lcorbi_t3_pos_4_096_60    m/z 652.6140     0   1E+6   14   14.5  RT (min)       lcorbi_t3_pos_1_050_13    m/z 652.7082     0   1   2E+5   14.5   15  RT (min)       lcorbi_t3_pos_2_044_21    m/z 652.8505     0   2   4   6E+4   1   1.5  RT (min)      lcorbi_t3_pos_4_075_58    m/z 653.2828     0   1   2   3E+5   13   13.5  RT (min)       lcorbi_t3_pos_4_090_60    m/z 653.4199     0   1   10   10.5  RT (min)      lcorbi_t3_pos_1_054_48    m/z 653.5327     0   1   2   3E+4   5.5   6  RT (min)      lcorbi_t3_pos_4_005_33    m/z 653.5356     0   2   4E+4   14   14.5  RT (min)      lcorbi_t3_pos_4_086_59    m/z 653.5516     0   1E+5   14.5   15  RT (min)       lcorbi_t3_pos_1_032_26    m/z 653.6261     0   1   2E+5   14   14.5  RT (min)      lcorbi_t3_pos_1_036_08    m/z 653.6522     0   1E+5   14   14.5  RT (min)       lcorbi_t3_pos_4_090_60    m/z 653.8260     0   2   4E+4   1   1.5  RT (min)      lcorbi_t3_pos_4_080_58    m/z 654.2592     0   1E+5   13.5   14  RT (min)      lcorbi_t3_pos_1_018_35    m/z 654.2862     0   2   4   6   8E+4   13   13.5  RT (min)      lcorbi_t3_pos_4_096_60    m/z 654.4580     0   2   4   6E+4   14   14.5  RT (min)      lcorbi_t3_pos_4_089_59    m/z 654.5040     0   1E+5   15   15.5  RT (min)      lcorbi_t3_pos_4_087_59    m/z 654.6035     0   1E+5   14.5   15  RT (min)       lcorbi_t3_pos_4_086_59    m/z 655.1957     0   1E+3   2   2.5  RT (min)      lcorbi_t3_pos_4_091_60    m/z 655.2096     0   1   2   3   4E+4   1.5   2   2.5  RT (min)       lcorbi_t3_pos_4_097_60    m/z 655.4315     0   1E+5   13.5   14  RT (min)       lcorbi_t3_pos_4_074_58    m/z 655.4418     0   1   2   3   4E+4   7.5   8  RT (min)       lcorbi_t3_pos_1_041_54    m/z 655.5514     0   1E+5   14.5   15  RT (min)       lcorbi_t3_pos_1_032_26    m/z 655.5704     0   1   2   3E+4   14   14.5  RT (min)      lcorbi_t3_pos_1_036_08    m/z 655.6419     0   1E+5   14.5   15  RT (min)       lcorbi_t3_pos_1_036_08    m/z 656.4452     0   1   2E+4   7.5   8  RT (min)      lcorbi_t3_pos_1_025_52    m/z 656.4737     0   1E+5   14.5   15  RT (min)      lcorbi_t3_pos_1_032_26    m/z 656.5249     0   1   2E+4   14   14.5  RT (min)       lcorbi_t3_pos_4_089_59    m/z 656.6373     0   1E+5   14   14.5  RT (min)       lcorbi_t3_pos_1_036_08    m/z 656.8075     0   2   4   6   8E+3   1   1.5  RT (min)      lcorbi_t3_pos_1_034_34    m/z 657.3611     0   1E+4   8   8.5  RT (min)      lcorbi_t3_pos_1_055_33    m/z 657.5272     0   2   4   6E+4   14   14.5   15  RT (min)       lcorbi_t3_pos_4_082_59    m/z 657.5672     0   1E+5   14.5   15  RT (min)       lcorbi_t3_pos_1_032_26    m/z 657.5863     0   1E+5   15   15.5  RT (min)      lcorbi_t3_pos_3_023_05    m/z 657.6495     0   1E+5   14   14.5  RT (min)       lcorbi_t3_pos_1_036_08    m/z 657.6577     0   2   4   6   8E+4   14.5   15  RT (min)       lcorbi_t3_pos_1_036_08    m/z 657.8736     0   1   2E+4   1   1.5  RT (min)      lcorbi_t3_pos_4_067_57    m/z 657.9908     0   1   2   3E+4   1   1.5  RT (min)      lcorbi_t3_pos_4_052_06    m/z 658.4895     0   1E+5   14.5   15  RT (min)      lcorbi_t3_pos_1_032_26    m/z 658.5540     0   1E+5   15   15.5  RT (min)       lcorbi_t3_pos_2_005_11    m/z 658.8383     0   1E+5   1   1.5  RT (min)       lcorbi_t3_pos_4_075_58    m/z 659.2961     0   1E+5   12.5   13  RT (min)      lcorbi_t3_pos_2_044_21    m/z 659.3399     0   1   2E+4   1   1.5  RT (min)      lcorbi_t3_pos_4_077_58    m/z 659.3446     0   2   4   6   8E+4   14   14.5  RT (min)      lcorbi_t3_pos_1_018_35    m/z 659.4782     0   2   4E+4   15   15.5  RT (min)      lcorbi_t3_pos_4_069_57    m/z 659.5227     0   1   2   3E+4   14   14.5  RT (min)       lcorbi_t3_pos_4_081_58    m/z 659.6652     0   1E+5   14.5   15  RT (min)       lcorbi_t3_pos_1_036_08    m/z 659.6733     0   2   4   6   8E+4   14.5   15  RT (min)       lcorbi_t3_pos_2_024_15    m/z 659.8386     0   1   2   3   4E+4   1   1.5  RT (min)      lcorbi_t3_pos_4_080_58    m/z 660.2909     0   2   4   6E+4   12   12.5  RT (min)       lcorbi_t3_pos_4_086_59    m/z 660.5050     0   1E+5   14.5   15  RT (min)      lcorbi_t3_pos_1_052_38    m/z 660.5165     0   1   2E+5   10.5   11  RT (min)      lcorbi_t3_pos_4_083_59    m/z 660.5471     0   2   4   6E+4   14.5   15  RT (min)      lcorbi_t3_pos_4_078_58    m/z 660.8079     0   2   4   6   8E+3   1   1.5  RT (min)      lcorbi_t3_pos_4_079_58    m/z 661.3118     0   1   2E+5   14   14.5  RT (min)       lcorbi_t3_pos_1_018_35    m/z 661.3565     0   2   4E+4   10   10.5  RT (min)      lcorbi_t3_pos_4_075_58    m/z 661.4937     0   2   4E+4   15   15.5  RT (min)       lcorbi_t3_pos_4_013_36    m/z 661.5176     0   2   4   6E+4   14.5   15  RT (min)       lcorbi_t3_pos_3_020_35    m/z 661.5282     0   2   4   6   8E+4   13   13.5  RT (min)       lcorbi_t3_pos_4_073_57    m/z 661.5999     0   2   4   6E+4   14.5   15  RT (min)      lcorbi_t3_pos_3_055_10    m/z 661.6809     0   1   2E+5   14.5   15  RT (min)       lcorbi_t3_pos_4_092_60    m/z 662.2716     0   1   2   3   4E+4   11   11.5  RT (min)       lcorbi_t3_pos_4_068_57    m/z 662.5632     0   1E+5   14.5   15  RT (min)      lcorbi_t3_pos_4_078_58    m/z 662.5876     0   1E+5   15   16  RT (min)       lcorbi_t3_pos_4_088_59    m/z 662.6919     0   1E+5   14.5   15  RT (min)       lcorbi_t3_pos_4_092_60    m/z 662.8249     0   1   2   3   4E+4   1   1.5  RT (min)      lcorbi_t3_pos_4_052_06    m/z 663.2725     0   2   4   6E+4   11   11.5  RT (min)       lcorbi_t3_pos_4_075_58    m/z 663.2832     0   1E+5   13.5   14  RT (min)       lcorbi_t3_pos_2_044_21    m/z 663.4870     0   2   4E+4   13.5   14  RT (min)      lcorbi_t3_pos_4_112_62    m/z 663.8563     0   1   2E+5   1   1.5  RT (min)       lcorbi_t3_pos_4_067_57    m/z 664.1169     0   2   4   6E+5   1.5   2  RT (min)       lcorbi_t3_pos_1_052_38    m/z 664.2301     0   1E+5   14   14.5  RT (min)       lcorbi_t3_pos_4_101_61    m/z 664.2756     0   1   2E+4   11   11.5  RT (min)      lcorbi_t3_pos_4_072_57    m/z 664.2872     0   1   2   3E+4   11   11.5  RT (min)       lcorbi_t3_pos_4_072_57    m/z 664.3000     0   1E+5   14   14.5  RT (min)       lcorbi_t3_pos_1_018_35    m/z 664.6278     0   1   2E+5   15   15.5  RT (min)       lcorbi_t3_pos_3_055_10    m/z 664.6997     0   1E+5   14.5   15  RT (min)       lcorbi_t3_pos_4_090_60    m/z 664.8555     0   1E+5   1   1.5  RT (min)      lcorbi_t3_pos_4_067_57    m/z 665.1201     0   1E+5   1.5   2  RT (min)      lcorbi_t3_pos_1_052_38    m/z 665.1992     0   1E+3   2.5   3  RT (min)       lcorbi_t3_pos_4_090_60    m/z 665.3086     0   1   2E+4   10   10.5  RT (min)      lcorbi_t3_pos_4_092_60    m/z 665.4758     0   1E+5   13   14  RT (min)       lcorbi_t3_pos_4_097_60    m/z 665.4952     0   2   4E+4   13.5   14  RT (min)       lcorbi_t3_pos_3_002_16    m/z 665.5914     0   1E+5   14.5   15  RT (min)       lcorbi_t3_pos_1_001_49    m/z 665.6127     0   2   4   6   8E+4   14.5   15   15.5  RT (min)       lcorbi_t3_pos_3_055_10    m/z 665.8617     0   2   4   6E+4   1   1.5  RT (min)       lcorbi_t3_pos_4_068_57    m/z 666.1221     0   1   2   3E+4   1.5   2  RT (min)      lcorbi_t3_pos_1_052_38    m/z 666.3027     0   1E+5   11.5   12  RT (min)       lcorbi_t3_pos_1_029_29    m/z 666.3631     0   1E+4   1   1.5  RT (min)      lcorbi_t3_pos_4_070_57    m/z 666.4345     0   1   2   3   4E+5   11   11.5  RT (min)       lcorbi_t3_pos_1_054_48    m/z 666.4484     0   1   2   3E+4   8   8.5  RT (min)      lcorbi_t3_pos_4_107_62    m/z 666.5154     0   1   14.5   15  RT (min)       lcorbi_t3_pos_4_082_59    m/z 666.5788     0   1   2E+5   14   14.5  RT (min)       lcorbi_t3_pos_1_036_08    m/z 666.6037     0   1   2E+5   14.5   15  RT (min)       lcorbi_t3_pos_1_002_01    m/z 666.6400     0   1   2E+4   14.5   15  RT (min)       lcorbi_t3_pos_4_055_02    m/z 666.7239     0   1E+5   14.5   15  RT (min)      lcorbi_t3_pos_2_044_21    m/z 666.9897     0   1E+4   1   1.5  RT (min)      lcorbi_t3_pos_3_048_30    m/z 667.4380     0   1E+5   11   11.5  RT (min)      lcorbi_t3_pos_1_054_48    m/z 667.5107     0   1E+5   13.5   14  RT (min)      lcorbi_t3_pos_4_098_61    m/z 667.5120     0   1E+4   6.5   7  RT (min)      lcorbi_t3_pos_4_088_59    m/z 667.5280     0   1   14.5   15  RT (min)       lcorbi_t3_pos_2_005_11    m/z 667.5912     0   2   4   6   8E+4   14.5   15  RT (min)       lcorbi_t3_pos_2_024_15    m/z 667.6432     0   1   2E+5   14.5   15  RT (min)       lcorbi_t3_pos_4_092_60    m/z 667.8274     0   1   2   3   4E+4   1   1.5  RT (min)      lcorbi_t3_pos_4_075_58    m/z 668.3732     0   1E+4   5   5.5  RT (min)       lcorbi_t3_pos_4_081_58    m/z 668.4959     0   1E+4   7   7.5  RT (min)      lcorbi_t3_pos_4_084_59    m/z 668.5638     0   2   4E+4   14.5   15  RT (min)       lcorbi_t3_pos_1_036_08    m/z 668.6555     0   2   4   6   8E+4   14.5   15  RT (min)       lcorbi_t3_pos_1_031_40    m/z 668.6943     0   1E+5   14.5   15  RT (min)      lcorbi_t3_pos_4_082_59    m/z 668.8241     0   1   2E+4   1   1.5  RT (min)      lcorbi_t3_pos_4_052_06    m/z 669.3724     0   1E+5   1   1.5  RT (min)      lcorbi_t3_pos_4_070_57    m/z 669.4472     0   2   4   6E+4   13.5   14  RT (min)       lcorbi_t3_pos_4_101_61    m/z 669.6500     0   1   2   3E+5   14   14.5  RT (min)      lcorbi_t3_pos_1_036_08    m/z 669.8731     0   2   4   6   8E+4   1   1.5  RT (min)      lcorbi_t3_pos_4_071_57    m/z 670.5642     0   1   2E+5   15   16  RT (min)       lcorbi_t3_pos_4_079_58    m/z 670.5803     0   2   4E+4   14.5   15  RT (min)       lcorbi_t3_pos_1_036_08    m/z 670.6102     0   1   2E+5   14.5   15  RT (min)       lcorbi_t3_pos_4_090_60    m/z 670.6349     0   1   2E+4   14.5   15  RT (min)      lcorbi_t3_pos_4_096_60    m/z 671.1745     0   1   2   3   4E+4   2   2.5  RT (min)       lcorbi_t3_pos_4_092_60    m/z 671.3128     0   1   2E+5   13   13.5  RT (min)      lcorbi_t3_pos_4_090_60    m/z 671.4628     0   2   4E+4   13.5   14  RT (min)       lcorbi_t3_pos_4_106_62    m/z 671.8440     0   2   4   6   8E+4   1   1.5  RT (min)      lcorbi_t3_pos_4_068_57    m/z 672.5260     0   1E+5   11.5   12  RT (min)       lcorbi_t3_pos_4_050_31    m/z 672.8425     0   1   2E+4   1   1.5  RT (min)      lcorbi_t3_pos_2_054_29    m/z 673.4550     0   2   4E+4   14.5   15  RT (min)       lcorbi_t3_pos_2_020_22    m/z 673.5293     0   1   2   3   4E+4   11.5   12  RT (min)      lcorbi_t3_pos_4_050_31    m/z 673.5929     0   2   4   6   8E+4   14   14.5  RT (min)       lcorbi_t3_pos_1_050_13    m/z 673.8486     0   2   4E+4   1   1.5  RT (min)      lcorbi_t3_pos_4_079_58    m/z 674.3502     0   1E+4   1   1.5  RT (min)      lcorbi_t3_pos_4_076_58    m/z 674.4840     0   2   4   6   8E+4   13.5   14  RT (min)       lcorbi_t3_pos_4_085_59    m/z 674.5048     0   1   2E+4   9   9.5  RT (min)      lcorbi_t3_pos_4_082_59    m/z 674.5866     0   1   15   15.5  RT (min)       lcorbi_t3_pos_4_084_59    m/z 674.8512     0   2   4   6E+3   1   1.5  RT (min)      lcorbi_t3_pos_1_007_07    m/z 675.2617     0   1   2   3E+4   12   12.5  RT (min)       lcorbi_t3_pos_1_036_08    m/z 675.4429     0   2   4E+5   11   11.5  RT (min)      lcorbi_t3_pos_1_054_48    m/z 675.5347     0   2   4E+4   13   13.5  RT (min)      lcorbi_t3_pos_4_066_57    m/z 675.9540     0   1E+4   1   1.5  RT (min)      lcorbi_t3_pos_2_022_38    m/z 676.4267     0   1E+4   8   8.5  RT (min)      lcorbi_t3_pos_4_100_61    m/z 677.3065     0   1   2E+5   12.5   13  RT (min)       lcorbi_t3_pos_2_044_21    m/z 677.3592     0   2   4   6E+4   1   1.5  RT (min)      lcorbi_t3_pos_4_075_58    m/z 677.4236     0   1   2E+4   7.5   8  RT (min)      lcorbi_t3_pos_2_056_56    m/z 677.4948     0   1   2   3E+4   13.5   14  RT (min)      lcorbi_t3_pos_4_107_62    m/z 677.5599     0   1   13.5   14  RT (min)       lcorbi_t3_pos_1_003_45    m/z 677.8586     0   2   4   6E+4   1   1.5  RT (min)      lcorbi_t3_pos_4_075_58    m/z 678.3636     0   2   4E+4   9   9.5  RT (min)       lcorbi_t3_pos_4_088_59    m/z 678.4707     0   1   2E+5   12   12.5  RT (min)       lcorbi_t3_pos_2_005_11    m/z 678.4787     0   2   4   6E+4   10.5   11  RT (min)       lcorbi_t3_pos_2_045_40    m/z 679.2675     0   1E+4   4.5   5  RT (min)      lcorbi_t3_pos_4_082_59    m/z 679.3670     0   1   2E+4   9   9.5  RT (min)      lcorbi_t3_pos_4_089_59    m/z 679.4312     0   1E+5   15   15.5  RT (min)       lcorbi_t3_pos_4_086_59    m/z 679.4611     0   1E+5   8   8.5  RT (min)      lcorbi_t3_pos_4_109_62    m/z 679.4819     0   1   2   3E+4   13.5   14  RT (min)      lcorbi_t3_pos_1_054_48    m/z 679.4821     0   1E+4   10.5   11  RT (min)      lcorbi_t3_pos_1_003_45    m/z 679.5513     0   1E+5   14.5   15  RT (min)      lcorbi_t3_pos_1_032_26    m/z 679.5673     0   2   4E+4   14.5   15  RT (min)      lcorbi_t3_pos_1_036_08    m/z 679.7073     0   1   2E+5   14.5   15  RT (min)       lcorbi_t3_pos_1_036_08    m/z 679.8317     0   2   4   6   8E+4   1   1.5  RT (min)      lcorbi_t3_pos_4_075_58    m/z 680.3785     0   2   4   6E+4   14   14.5  RT (min)       lcorbi_t3_pos_4_110_62    m/z 680.4249     0   2   4   6E+4   10   10.5  RT (min)       lcorbi_t3_pos_4_086_59    m/z 680.4501     0   1   2E+5   11.5   12  RT (min)      lcorbi_t3_pos_1_054_48    m/z 680.4641     0   1   2E+4   8   8.5  RT (min)      lcorbi_t3_pos_4_107_62    m/z 680.5827     0   1E+5   15   15.5  RT (min)       lcorbi_t3_pos_4_070_57    m/z 680.5980     0   1E+5   15   15.5  RT (min)       lcorbi_t3_pos_1_029_29    m/z 680.7244     0   1   2E+5   14.5   15  RT (min)       lcorbi_t3_pos_1_036_08    m/z 680.8699     0   1   2   3   4E+4   1   1.5  RT (min)      lcorbi_t3_pos_4_075_58    m/z 681.4282     0   1   2   3E+4   10   10.5  RT (min)      lcorbi_t3_pos_4_088_59    m/z 681.4470     0   1   2   3E+4   13.5   14  RT (min)       lcorbi_t3_pos_3_015_06    m/z 681.5165     0   2   4   6E+4   13.5   14  RT (min)       lcorbi_t3_pos_4_103_61    m/z 681.5185     0   1   2   3E+4   6   6.5  RT (min)      lcorbi_t3_pos_4_089_59    m/z 681.5670     0   1   2E+5   14.5   15  RT (min)       lcorbi_t3_pos_1_032_26    m/z 681.5789     0   1E+5   14  RT (min)       lcorbi_t3_pos_1_032_26    m/z 681.7066     0   2   4E+4   14.5   15  RT (min)       lcorbi_t3_pos_3_036_37    m/z 681.8357     0   1   2   3E+4   1   1.5  RT (min)      lcorbi_t3_pos_4_076_58    m/z 681.8720     0   1E+4   1   1.5  RT (min)      lcorbi_t3_pos_4_073_57    m/z 682.3636     0   1E+5   12   12.5  RT (min)       lcorbi_t3_pos_1_003_45    m/z 682.5022     0   2   4   6E+4   13.5   14  RT (min)       lcorbi_t3_pos_4_106_62    m/z 682.5409     0   2   4E+4   14.5   15  RT (min)       lcorbi_t3_pos_4_082_59    m/z 682.8063     0   1   2E+5   1   1.5  RT (min)      lcorbi_t3_pos_4_067_57    m/z 683.3502     0   2   4   6E+4   14   14.5  RT (min)       lcorbi_t3_pos_4_105_61    m/z 683.4899     0   2   4   6   8E+4   14.5   15  RT (min)       lcorbi_t3_pos_4_082_59    m/z 683.5035     0   2   4   6   8E+4   14.5   15  RT (min)       lcorbi_t3_pos_1_052_38    m/z 683.5441     0   2   4E+4   14.5   15  RT (min)       lcorbi_t3_pos_4_078_58    m/z 683.6016     0   1   2   3   4E+4   14.5   15  RT (min)      lcorbi_t3_pos_1_032_26    m/z 683.6170     0   1E+5   15   15.5  RT (min)      lcorbi_t3_pos_1_029_29    m/z 683.8838     0   1E+5   5.5   6  RT (min)       lcorbi_t3_pos_4_076_58    m/z 684.1841     0   1   2   3   4E+4   1.5   2  RT (min)      lcorbi_t3_pos_1_036_08    m/z 684.2189     0   1   2E+5   5.5   6  RT (min)      lcorbi_t3_pos_4_030_05    m/z 684.3205     0   1E+5   4.5   5  RT (min)       lcorbi_t3_pos_4_014_48    m/z 684.3852     0   2   4   6   8E+4   5.5   6  RT (min)      lcorbi_t3_pos_4_075_58    m/z 684.5619     0   2   4   6E+4   12   13  RT (min)       lcorbi_t3_pos_4_090_60    m/z 684.5950     0   2   4   6   8E+4   14.5   15  RT (min)       lcorbi_t3_pos_1_016_05    m/z 684.6141     0   1   2E+5   14   15  RT (min)       lcorbi_t3_pos_3_023_05    m/z 684.8446     0   1   2E+5   1   1.5  RT (min)       lcorbi_t3_pos_4_075_58    m/z 684.8866     0   1   2   3E+4   5.5   6  RT (min)      lcorbi_t3_pos_4_075_58    m/z 685.1873     0   1   2   3E+5   1.5   2  RT (min)       lcorbi_t3_pos_1_036_08    m/z 685.2128     0   1E+5   5.5   6  RT (min)      lcorbi_t3_pos_4_100_61    m/z 685.3463     0   2   4E+4   1   1.5  RT (min)      lcorbi_t3_pos_4_079_58    m/z 685.4420     0   2   4   6E+4   13   13.5  RT (min)       lcorbi_t3_pos_4_066_57    m/z 685.4691     0   1E+5   13.5   14  RT (min)      lcorbi_t3_pos_4_098_61    m/z 685.5190     0   1E+5   14.5   15  RT (min)       lcorbi_t3_pos_1_008_36    m/z 685.5594     0   2   4   6E+4   14.5   15  RT (min)       lcorbi_t3_pos_4_080_58    m/z 685.5647     0   2   4   6   8E+4   13   13.5  RT (min)       lcorbi_t3_pos_4_070_57    m/z 685.5983     0   2   4   6E+4   14.5   15  RT (min)       lcorbi_t3_pos_2_013_33    m/z 685.8448     0   2   4E+4   1   1.5  RT (min)      lcorbi_t3_pos_4_075_58    m/z 686.0987     0   2   4   6   8E+4   1.5   2  RT (min)      lcorbi_t3_pos_1_052_38    m/z 686.2160     0   2   4   6E+4   5.5   6  RT (min)       lcorbi_t3_pos_4_030_05    m/z 686.2445     0   2   4E+4   6.5   7  RT (min)       lcorbi_t3_pos_4_005_33    m/z 686.4762     0   2   4   6E+4   13   13.5  RT (min)       lcorbi_t3_pos_4_084_59    m/z 686.5573     0   1E+5   15   15.5   16  RT (min)       lcorbi_t3_pos_4_075_58    m/z 686.5875     0   2   4   6E+4   15   15.5  RT (min)      lcorbi_t3_pos_1_015_11    m/z 686.8907     0   2   4   6   8E+4   1   1.5  RT (min)       lcorbi_t3_pos_4_070_57    m/z 687.2033     0   1   2E+4   2.5   3  RT (min)      lcorbi_t3_pos_1_016_05    m/z 687.2477     0   1   2E+4   6.5   7  RT (min)      lcorbi_t3_pos_4_005_33    m/z 687.4426     0   1   2   3   4E+4   14.5   15  RT (min)      lcorbi_t3_pos_3_050_36    m/z 687.4577     0   1   2   3E+4   13.5   14  RT (min)       lcorbi_t3_pos_4_100_61    m/z 687.4965     0   1   2   3   4E+4   14.5   15  RT (min)      lcorbi_t3_pos_1_032_26    m/z 687.5349     0   1E+5   14.5   15  RT (min)       lcorbi_t3_pos_1_032_26    m/z 687.5442     0   2   4E+4   13   13.5  RT (min)       lcorbi_t3_pos_4_068_57    m/z 687.6065     0   1E+5   14.5   15  RT (min)       lcorbi_t3_pos_1_050_13    m/z 687.8199     0   1   2   3   4E+4   1   1.5  RT (min)      lcorbi_t3_pos_4_079_58    m/z 688.4644     0   1   2E+4   6.5   7  RT (min)       lcorbi_t3_pos_4_018_01    m/z 688.5207     0   1E+5   10.5   11  RT (min)       lcorbi_t3_pos_4_050_31    m/z 688.5880     0   1   14.5   15  RT (min)       lcorbi_t3_pos_1_054_48    m/z 688.6192     0   1E+5   14.5   15  RT (min)      lcorbi_t3_pos_3_027_27    m/z 688.6325     0   1   15   15.5  RT (min)       lcorbi_t3_pos_1_029_29    m/z 689.4757     0   2   4   6E+4   13.5   14  RT (min)       lcorbi_t3_pos_4_066_57    m/z 689.4954     0   1E+5   13.5   14  RT (min)       lcorbi_t3_pos_4_087_59    m/z 689.5757     0   2   4   6E+4   15   15.5  RT (min)       lcorbi_t3_pos_1_036_08    m/z 689.6562     0   1   2E+5   14   14.5  RT (min)       lcorbi_t3_pos_1_036_08    m/z 690.6098     0   1   15   15.5  RT (min)      lcorbi_t3_pos_4_066_57    m/z 690.6723     0   1E+5   14.5   15  RT (min)       lcorbi_t3_pos_4_094_60    m/z 690.8304     0   1E+4   1   1.5  RT (min)      lcorbi_t3_pos_4_081_58    m/z 691.1998     0   1   2   3E+4   6.5   7  RT (min)       lcorbi_t3_pos_4_005_33    m/z 691.4913     0   1   2E+4   13.5   14  RT (min)      lcorbi_t3_pos_1_008_36    m/z 691.6194     0   2   4   6E+4   14.5   15  RT (min)       lcorbi_t3_pos_4_095_60    m/z 691.7187     0   1E+5   14.5   15  RT (min)       lcorbi_t3_pos_4_092_60    m/z 691.8674     0   1   2E+4   1   1.5  RT (min)      lcorbi_t3_pos_4_068_57    m/z 692.2982     0   1   2   3E+5   12.5   13  RT (min)       lcorbi_t3_pos_2_044_21    m/z 692.4612     0   1   2   3E+4   11   11.5  RT (min)      lcorbi_t3_pos_4_021_25    m/z 692.4735     0   1E+4   8   8.5  RT (min)       lcorbi_t3_pos_4_106_62    m/z 692.4866     0   2   4E+4   13   13.5  RT (min)       lcorbi_t3_pos_4_098_61    m/z 692.5139     0   1E+5   13.5   14  RT (min)      lcorbi_t3_pos_4_086_59    m/z 692.5638     0   1E+5   14.5   15  RT (min)       lcorbi_t3_pos_1_031_40    m/z 693.3016     0   1E+5   12.5   13  RT (min)      lcorbi_t3_pos_2_044_21    m/z 693.3337     0   1   2E+4   1   1.5  RT (min)      lcorbi_t3_pos_4_074_58    m/z 693.5265     0   1   2   3E+5   14   14.5  RT (min)       lcorbi_t3_pos_4_089_59    m/z 693.5978     0   1E+5   14.5   15  RT (min)       lcorbi_t3_pos_1_036_08    m/z 693.6877     0   1   2E+5   14.5   15  RT (min)       lcorbi_t3_pos_4_090_60    m/z 693.8343     0   2   4E+4   1   1.5  RT (min)      lcorbi_t3_pos_4_075_58    m/z 694.4656     0   2   4   6E+4   12   12.5  RT (min)       lcorbi_t3_pos_3_029_13    m/z 694.4966     0   1E+5   14.5   15  RT (min)       lcorbi_t3_pos_4_082_59    m/z 694.5794     0   1E+5   14.5   15  RT (min)       lcorbi_t3_pos_2_020_22    m/z 694.6137     0   2   4   6   8E+4   15.5   16  RT (min)       lcorbi_t3_pos_4_075_58    m/z 694.6659     0   1   2E+5   15   15.5  RT (min)       lcorbi_t3_pos_4_093_60    m/z 694.8330     0   1E+5   1   1.5  RT (min)       lcorbi_t3_pos_4_070_57    m/z 694.9958     0   1E+4   1   1.5  RT (min)      lcorbi_t3_pos_4_005_33    m/z 695.2924     0   1E+5   5   5.5  RT (min)       lcorbi_t3_pos_4_014_48    m/z 695.4999     0   2   4E+4   14.5   15  RT (min)      lcorbi_t3_pos_4_086_59    m/z 695.6134     0   1E+5   14.5   15  RT (min)       lcorbi_t3_pos_1_036_08    m/z 695.7222     0   2   4   6   8E+4   14.5   15  RT (min)       lcorbi_t3_pos_4_066_57    m/z 696.2957     0   1   2   3E+4   5   5.5  RT (min)       lcorbi_t3_pos_4_014_48    m/z 696.4383     0   1E+4   5   5.5  RT (min)      lcorbi_t3_pos_1_001_49    m/z 696.4664     0   2   4   6E+4   13.5   14  RT (min)       lcorbi_t3_pos_4_098_61    m/z 696.5087     0   2   4   6   8E+4   13.5   14  RT (min)      lcorbi_t3_pos_4_098_61    m/z 696.5390     0   1   2   3E+4   6   6.5  RT (min)       lcorbi_t3_pos_4_088_59    m/z 696.5798     0   1   2   3E+5   15   16  RT (min)       lcorbi_t3_pos_4_069_57    m/z 696.5951     0   1E+5   14.5   15  RT (min)       lcorbi_t3_pos_2_013_33    m/z 696.6867     0   2   4   6   8E+4   15   15.5  RT (min)       lcorbi_t3_pos_1_028_32    m/z 696.8471     0   1E+5   1   1.5  RT (min)       lcorbi_t3_pos_4_075_58    m/z 697.2731     0   1E+5   13.5   14  RT (min)      lcorbi_t3_pos_1_018_35    m/z 697.2778     0   1   2   3   4E+4   4.5   5  RT (min)       lcorbi_t3_pos_4_066_57    m/z 697.2781     0   1   2E+3   2.5   3  RT (min)      lcorbi_t3_pos_4_001_49    m/z 697.3632     0   2   4E+4   12   12.5  RT (min)       lcorbi_t3_pos_1_028_32    m/z 697.4784     0   2   4   6   8E+4   14   14.5  RT (min)       lcorbi_t3_pos_4_086_59    m/z 697.5226     0   2   4   6E+3   5.5   6  RT (min)      lcorbi_t3_pos_4_088_59    m/z 697.5422     0   1E+4   6   6.5  RT (min)      lcorbi_t3_pos_4_084_59    m/z 697.6848     0   1   2E+5   15   15.5  RT (min)       lcorbi_t3_pos_4_093_60    m/z 697.8501     0   1E+5   1   1.5  RT (min)       lcorbi_t3_pos_4_067_57    m/z 698.2812     0   2   4   6E+3   4.5   5  RT (min)      lcorbi_t3_pos_4_066_57    m/z 698.2852     0   1   2E+5   12.5   13  RT (min)       lcorbi_t3_pos_2_044_21    m/z 698.4735     0   1   2   3   4E+4   13.5   14  RT (min)      lcorbi_t3_pos_4_098_61    m/z 698.5293     0   2   4   6E+4   13   13.5  RT (min)      lcorbi_t3_pos_4_072_57    m/z 698.5552     0   2   4   6   8E+4   14.5   15  RT (min)      lcorbi_t3_pos_1_028_32    m/z 698.6104     0   1E+5   14.5   15  RT (min)       lcorbi_t3_pos_4_092_60    m/z 698.8494     0   2   4   6E+4   1   1.5  RT (min)      lcorbi_t3_pos_4_068_57    m/z 699.2677     0   1   2   3E+4   14   14.5  RT (min)      lcorbi_t3_pos_3_044_46    m/z 699.4366     0   1E+5   13.5   14  RT (min)       lcorbi_t3_pos_4_098_61    m/z 699.4709     0   2   4   6E+4   14.5   15  RT (min)      lcorbi_t3_pos_1_032_26    m/z 699.6873     0   1E+5   14   15  RT (min)       lcorbi_t3_pos_1_003_45    m/z 699.8553     0   2   4E+4   1   1.5  RT (min)      lcorbi_t3_pos_4_068_57    m/z 700.3569     0   1E+4   1   1.5  RT (min)      lcorbi_t3_pos_4_071_57    m/z 700.4916     0   2   4   6   8E+4   13   13.5  RT (min)       lcorbi_t3_pos_1_032_26    m/z 700.5670     0   2   4   6E+4   15   15.5  RT (min)       lcorbi_t3_pos_1_032_26    m/z 700.5994     0   2   4   6E+4   15   15.5  RT (min)      lcorbi_t3_pos_4_083_59    m/z 700.6274     0   1E+5   15.5   16  RT (min)       lcorbi_t3_pos_4_093_60    m/z 700.6480     0   1   2E+5   14.5   15  RT (min)       lcorbi_t3_pos_4_091_60    m/z 700.8200     0   1   2   3   4E+4   1   1.5  RT (min)      lcorbi_t3_pos_4_075_58    m/z 701.0159     0   1   2   3E+4   1   1.5  RT (min)      lcorbi_t3_pos_4_052_06    m/z 701.4063     0   2   4   6   8E+4   7   7.5  RT (min)       lcorbi_t3_pos_4_102_61    m/z 701.4099     0   2   4   6E+4   14.5   15  RT (min)      lcorbi_t3_pos_4_085_59    m/z 701.4639     0   1   2   3E+4   13.5   14  RT (min)       lcorbi_t3_pos_4_071_57    m/z 701.4864     0   2   4E+4   14.5   15  RT (min)       lcorbi_t3_pos_2_020_22    m/z 701.4944     0   1   2E+4   6   6.5  RT (min)       lcorbi_t3_pos_4_088_59    m/z 701.6941     0   2   4   6   8E+4   14   14.5  RT (min)       lcorbi_t3_pos_4_092_60    m/z 701.7030     0   1   2E+5   14.5   15  RT (min)       lcorbi_t3_pos_1_001_49    m/z 702.4974     0   2   4   6   8E+3   6   6.5  RT (min)      lcorbi_t3_pos_4_084_59    m/z 702.5880     0   1   2   3E+5   15.5   16  RT (min)       lcorbi_t3_pos_4_079_58    m/z 703.2144     0   1   2   3E+5   13   13.5  RT (min)       lcorbi_t3_pos_4_113_62    m/z 703.3661     0   1E+5   1   1.5  RT (min)       lcorbi_t3_pos_4_071_57    m/z 703.4038     0   2   4   6E+4   11   11.5  RT (min)      lcorbi_t3_pos_4_097_60    m/z 703.4914     0   1   2   3E+4   13.5   14  RT (min)      lcorbi_t3_pos_4_073_57    m/z 703.5105     0   1E+5   13.5   14  RT (min)       lcorbi_t3_pos_4_083_59    m/z 703.5182     0   2   4   6E+4   14   14.5  RT (min)      lcorbi_t3_pos_4_100_61    m/z 703.5385     0   1   13.5   14  RT (min)      lcorbi_t3_pos_4_072_57    m/z 703.5398     0   1   2   3   4E+5   10.5   11  RT (min)      lcorbi_t3_pos_4_084_59    m/z 703.5429     0   2   4   6E+4   15   15.5  RT (min)       lcorbi_t3_pos_4_006_12    m/z 703.6115     0   1   13.5   14  RT (min)      lcorbi_t3_pos_4_069_57    m/z 703.6512     0   1   14.5   15  RT (min)       lcorbi_t3_pos_4_073_57    m/z 703.7098     0   2   4   6   8E+4   14   14.5  RT (min)       lcorbi_t3_pos_4_052_06    m/z 703.7186     0   1   2   3E+5   14.5   15  RT (min)       lcorbi_t3_pos_1_003_45    m/z 704.2127     0   1   2E+5   13   13.5  RT (min)       lcorbi_t3_pos_4_113_62    m/z 704.3155     0   1E+5   14.5   15  RT (min)      lcorbi_t3_pos_4_100_61    m/z 704.5980     0   1E+5   15   15.5  RT (min)       lcorbi_t3_pos_4_080_58    m/z 704.6191     0   2   4   6E+4   14.5   15  RT (min)       lcorbi_t3_pos_1_032_26    m/z 704.8690     0   2   4   6E+4   1   1.5  RT (min)       lcorbi_t3_pos_4_070_57    m/z 705.6015     0   2   4   6   8E+4   15   15.5  RT (min)       lcorbi_t3_pos_4_078_58    m/z 705.6878     0   1E+5   14.5   15  RT (min)       lcorbi_t3_pos_1_036_08    m/z 705.8377     0   2   4   6   8E+4   1   1.5  RT (min)      lcorbi_t3_pos_4_073_57    m/z 707.3093     0   1E+5   13.5   14  RT (min)      lcorbi_t3_pos_2_044_21    m/z 707.3559     0   2   4   6   8E+4   11   11.5  RT (min)       lcorbi_t3_pos_4_093_60    m/z 707.4512     0   1   2E+4   13.5   14  RT (min)      lcorbi_t3_pos_1_020_31    m/z 707.5322     0   1E+5   14   14.5  RT (min)       lcorbi_t3_pos_4_106_62    m/z 707.5878     0   2   4   6   8E+4   13.5   14  RT (min)       lcorbi_t3_pos_4_066_57    m/z 707.5971     0   1E+5   13.5   14  RT (min)       lcorbi_t3_pos_4_071_57    m/z 707.7411     0   1E+5   14.5   15  RT (min)       lcorbi_t3_pos_4_096_60    m/z 707.8425     0   2   4E+4   1   1.5  RT (min)      lcorbi_t3_pos_4_075_58    m/z 708.3593     0   1   2   3E+4   11   11.5  RT (min)       lcorbi_t3_pos_4_093_60    m/z 708.4042     0   1E+4   5.5   6  RT (min)      lcorbi_t3_pos_1_035_28    m/z 708.4099     0   2   4   6   8E+4   14.5   15  RT (min)      lcorbi_t3_pos_2_041_54    m/z 708.6125     0   1   2E+5   14.5   15  RT (min)       lcorbi_t3_pos_1_044_25    m/z 708.6204     0   1E+5   15   15.5  RT (min)      lcorbi_t3_pos_4_080_58    m/z 708.6507     0   2   4   6   8E+4   14.5   15  RT (min)      lcorbi_t3_pos_4_084_59    m/z 708.8094     0   1E+5   1   1.5  RT (min)       lcorbi_t3_pos_4_077_58    m/z 708.8212     0   2   4   6E+3   1   1.5  RT (min)      lcorbi_t3_pos_1_029_29    m/z 709.3248     0   1E+5   12.5   13  RT (min)       lcorbi_t3_pos_2_044_21    m/z 709.4784     0   1   2   3E+4   13.5   14  RT (min)      lcorbi_t3_pos_4_106_62    m/z 709.5539     0   1E+5   15   15.5  RT (min)       lcorbi_t3_pos_4_078_58    m/z 709.6156     0   2   4   6   8E+4   14.5   15  RT (min)      lcorbi_t3_pos_4_078_58    m/z 709.6522     0   2   4   6   8E+4   14.5   15  RT (min)       lcorbi_t3_pos_4_087_59    m/z 709.7567     0   1E+5   15   16  RT (min)       lcorbi_t3_pos_1_036_08    m/z 710.4761     0   1   2E+5   13   13.5  RT (min)       lcorbi_t3_pos_4_087_59    m/z 710.5565     0   1   13.5   14  RT (min)       lcorbi_t3_pos_1_003_45    m/z 710.5572     0   1   15   15.5  RT (min)      lcorbi_t3_pos_4_066_57    m/z 710.6050     0   1E+5   13.5   14  RT (min)       lcorbi_t3_pos_4_091_60    m/z 710.6296     0   1E+5   15   15.5  RT (min)       lcorbi_t3_pos_4_066_57    m/z 711.1277     0   1E+4   2   2.5  RT (min)      lcorbi_t3_pos_4_083_59    m/z 711.3531     0   2   4   6E+4   1   1.5  RT (min)      lcorbi_t3_pos_4_075_58    m/z 711.3816     0   1E+5   14.5   15  RT (min)       lcorbi_t3_pos_4_085_59    m/z 711.4577     0   1   2   3   4E+4   13.5   14  RT (min)      lcorbi_t3_pos_4_066_57    m/z 711.4794     0   2   4   6   8E+4   13   13.5  RT (min)      lcorbi_t3_pos_4_088_59    m/z 711.4941     0   1   2   3   4E+4   14   14.5  RT (min)       lcorbi_t3_pos_3_023_05    m/z 711.5417     0   1E+5   13.5   14  RT (min)       lcorbi_t3_pos_4_066_57    m/z 711.6503     0   1E+5   14.5   15  RT (min)      lcorbi_t3_pos_1_031_40    m/z 711.8525     0   2   4E+4   1   1.5  RT (min)      lcorbi_t3_pos_4_081_58    m/z 712.4050     0   1   2E+4   5   5.5  RT (min)       lcorbi_t3_pos_4_076_58    m/z 712.5450     0   1   13.5   14  RT (min)      lcorbi_t3_pos_4_082_59    m/z 712.6456     0   1   15   15.5  RT (min)       lcorbi_t3_pos_4_076_58    m/z 712.8212     0   1   2   3E+4   1   1.5  RT (min)      lcorbi_t3_pos_4_075_58    m/z 712.9067     0   1E+4   5   5.5  RT (min)      lcorbi_t3_pos_4_079_58    m/z 713.4951     0   1   2E+5   13.5   14  RT (min)       lcorbi_t3_pos_4_082_59    m/z 713.5759     0   1E+5   15   15.5  RT (min)      lcorbi_t3_pos_4_076_58    m/z 713.8255     0   2   4E+4   1   1.5  RT (min)      lcorbi_t3_pos_4_076_58    m/z 714.2757     0   2   4   6E+4   7.5   8  RT (min)       lcorbi_t3_pos_4_098_61    m/z 714.2802     0   1E+5   12.5   13  RT (min)      lcorbi_t3_pos_2_044_21    m/z 714.4005     0   1   2E+4   12   12.5  RT (min)      lcorbi_t3_pos_3_056_56    m/z 714.4977     0   2   4   6   8E+4   13.5   14  RT (min)      lcorbi_t3_pos_4_082_59    m/z 714.5522     0   1E+5   13.5   14  RT (min)       lcorbi_t3_pos_1_054_48    m/z 714.5884     0   2   4   6E+4   14.5   15  RT (min)       lcorbi_t3_pos_2_020_22    m/z 715.5551     0   1   13.5   14  RT (min)      lcorbi_t3_pos_4_072_57    m/z 715.8293     0   1   2   3E+4   1   1.5  RT (min)      lcorbi_t3_pos_4_078_58    m/z 716.3794     0   1E+4   10   10.5  RT (min)      lcorbi_t3_pos_3_007_28    m/z 716.5135     0   1   2   3E+5   13.5   14  RT (min)      lcorbi_t3_pos_4_083_59    m/z 716.5522     0   2   4   6   8E+4   11.5   12  RT (min)       lcorbi_t3_pos_4_050_31    m/z 716.5597     0   2   4   6E+4   13.5   14  RT (min)       lcorbi_t3_pos_4_082_59    m/z 716.5789     0   1E+5   13.5   14  RT (min)       lcorbi_t3_pos_4_072_57    m/z 716.6706     0   2   4E+4   15   15.5  RT (min)      lcorbi_t3_pos_1_012_14    m/z 716.6785     0   1E+5   14.5   15  RT (min)       lcorbi_t3_pos_1_036_08    m/z 717.4447     0   2   4E+4   13.5   14  RT (min)       lcorbi_t3_pos_4_035_17    m/z 717.7432     0   2   4   6   8E+4   14.5   15  RT (min)       lcorbi_t3_pos_4_090_60    m/z 718.5022     0   1   2   3E+4   13.5   14  RT (min)      lcorbi_t3_pos_4_071_57    m/z 718.5103     0   1   2   3   4E+4   13.5   14  RT (min)       lcorbi_t3_pos_4_083_59    m/z 718.6138     0   2   4   6   8E+4   15.5   16  RT (min)       lcorbi_t3_pos_4_074_58    m/z 718.6945     0   1   2E+5   14.5   15  RT (min)       lcorbi_t3_pos_1_036_08    m/z 718.8676     0   1E+4   6   6.5  RT (min)      lcorbi_t3_pos_4_074_58    m/z 719.3403     0   1   2   3   4E+4   1   1.5  RT (min)      lcorbi_t3_pos_4_079_58    m/z 719.5327     0   2   4   6E+5   14   14.5  RT (min)       lcorbi_t3_pos_4_086_59    m/z 719.5702     0   2   4   6E+4   13.5   14  RT (min)       lcorbi_t3_pos_4_066_57    m/z 719.5783     0   1E+5   14   14.5  RT (min)      lcorbi_t3_pos_1_019_16    m/z 719.5980     0   2   4   6E+4   13.5   14  RT (min)      lcorbi_t3_pos_4_070_57    m/z 719.6745     0   1E+5   14.5   15  RT (min)      lcorbi_t3_pos_1_036_08    m/z 719.8406     0   1E+5   1   1.5  RT (min)      lcorbi_t3_pos_4_075_58    m/z 720.7099     0   1   2E+5   14.5   15  RT (min)       lcorbi_t3_pos_1_036_08    m/z 720.8388     0   2   4E+4   1   1.5  RT (min)      lcorbi_t3_pos_4_075_58    m/z 721.2887     0   2   4   6E+4   13.5   14  RT (min)      lcorbi_t3_pos_2_044_21    m/z 721.3979     0   1   2E+4   5.5   6  RT (min)       lcorbi_t3_pos_4_075_58    m/z 721.4141     0   1   2   3   4E+4   10.5   11  RT (min)      lcorbi_t3_pos_4_084_59    m/z 721.4179     0   2   4E+4   13.5   14  RT (min)       lcorbi_t3_pos_4_109_62    m/z 721.5478     0   1E+5   14   14.5  RT (min)       lcorbi_t3_pos_4_085_59    m/z 721.5945     0   1E+5   14   14.5  RT (min)      lcorbi_t3_pos_4_079_58    m/z 721.6516     0   1E+4   14.5   15  RT (min)       lcorbi_t3_pos_4_052_06    m/z 721.8995     0   1E+4   5.5   6  RT (min)      lcorbi_t3_pos_4_078_58    m/z 722.4431     0   2   4E+4   13.5   14  RT (min)       lcorbi_t3_pos_4_098_61    m/z 722.4761     0   2   4E+4   13   13.5  RT (min)       lcorbi_t3_pos_4_085_59    m/z 722.5050     0   1   2   3E+4   10.5   11  RT (min)      lcorbi_t3_pos_2_045_40    m/z 722.5971     0   1   2   3   4E+4   14   14.5  RT (min)      lcorbi_t3_pos_4_081_58    m/z 722.6341     0   1E+5   14.5   15  RT (min)       lcorbi_t3_pos_2_024_15    m/z 722.6361     0   1E+5   15.5   16  RT (min)      lcorbi_t3_pos_4_079_58    m/z 722.6663     0   1E+5   14.5   15  RT (min)       lcorbi_t3_pos_4_086_59    m/z 722.6938     0   1   2E+5   14.5   15  RT (min)      lcorbi_t3_pos_2_024_15    m/z 723.0296     0   1   2   3   4E+4   6   6.5  RT (min)       lcorbi_t3_pos_4_079_58    m/z 723.3043     0   1E+5   13.5   14  RT (min)       lcorbi_t3_pos_2_044_21    m/z 723.3639     0   2   4E+4   6   6.5  RT (min)      lcorbi_t3_pos_4_079_58    m/z 723.5308     0   1E+5   14   14.5   15  RT (min)       lcorbi_t3_pos_4_100_61    m/z 723.6131     0   1E+5   14.5   15  RT (min)       lcorbi_t3_pos_1_032_26    m/z 723.6981     0   1   2   3E+4   6   6.5  RT (min)      lcorbi_t3_pos_4_076_58    m/z 724.0322     0   1E+4   6   6.5  RT (min)      lcorbi_t3_pos_4_078_58    m/z 724.0480     0   1E+4   6   6.5  RT (min)      lcorbi_t3_pos_4_075_58    m/z 724.4512     0   2   4E+4   10   11  RT (min)       lcorbi_t3_pos_2_008_16    m/z 724.5282     0   1   2E+4   14   14.5  RT (min)      lcorbi_t3_pos_4_106_62    m/z 724.5714     0   1E+5   14   14.5   15  RT (min)       lcorbi_t3_pos_4_086_59    m/z 724.6499     0   1E+5   14.5   15  RT (min)       lcorbi_t3_pos_1_036_08    m/z 724.6627     0   2   4E+4   14.5   15  RT (min)       lcorbi_t3_pos_4_079_58    m/z 724.6817     0   2   4   6E+4   15   15.5  RT (min)       lcorbi_t3_pos_4_087_59    m/z 725.3661     0   1   2   3   4E+4   10   10.5   11  RT (min)       lcorbi_t3_pos_4_078_58    m/z 725.4139     0   2   4   6E+4   12   12.5  RT (min)      lcorbi_t3_pos_4_088_59    m/z 725.5313     0   1E+5   14   14.5  RT (min)      lcorbi_t3_pos_1_008_36    m/z 725.5848     0   1E+5   15.5   16  RT (min)       lcorbi_t3_pos_4_077_58    m/z 725.6159     0   1   13.5   14  RT (min)       lcorbi_t3_pos_4_073_57    m/z 725.6296     0   1E+5   14.5   15  RT (min)       lcorbi_t3_pos_1_032_26    m/z 725.6658     0   2   4   6E+4   14.5   15  RT (min)       lcorbi_t3_pos_4_096_60    m/z 725.8612     0   2   4   6   8E+3   1   1.5  RT (min)      lcorbi_t3_pos_4_075_58    m/z 725.9362     0   1   2   3E+4   1   1.5  RT (min)      lcorbi_t3_pos_4_021_25    m/z 726.3696     0   1   2   3E+4   10   10.5  RT (min)      lcorbi_t3_pos_4_074_58    m/z 726.5441     0   1   2   3E+4   14   14.5  RT (min)       lcorbi_t3_pos_4_106_62    m/z 726.5868     0   1E+5   14   14.5   15  RT (min)       lcorbi_t3_pos_4_089_59    m/z 726.5882     0   1E+5   15.5   16  RT (min)      lcorbi_t3_pos_4_077_58    m/z 726.6281     0   1   2E+5   13.5   14  RT (min)       lcorbi_t3_pos_4_092_60    m/z 726.6421     0   2   4E+4   14.5   15  RT (min)       lcorbi_t3_pos_3_011_42    m/z 726.6656     0   1E+5   14.5   15  RT (min)       lcorbi_t3_pos_1_036_08    m/z 726.8260     0   1E+5   1   1.5  RT (min)       lcorbi_t3_pos_4_075_58    m/z 727.5462     0   1   2E+4   14   14.5  RT (min)      lcorbi_t3_pos_3_008_20    m/z 727.5616     0   1   2   3E+5   13.5   14  RT (min)      lcorbi_t3_pos_4_067_57    m/z 727.7008     0   2   4   6E+4   15   15.5  RT (min)      lcorbi_t3_pos_1_031_40    m/z 727.8261     0   1   2   3E+4   1   1.5  RT (min)      lcorbi_t3_pos_4_080_58    m/z 727.9331     0   1   2   3   4E+4   1   1.5  RT (min)      lcorbi_t3_pos_4_073_57    m/z 728.5114     0   1   14   14.5  RT (min)       lcorbi_t3_pos_4_088_59    m/z 728.5675     0   1   2E+5   13.5   14  RT (min)       lcorbi_t3_pos_4_084_59    m/z 728.6439     0   1   2E+5   14   14.5  RT (min)       lcorbi_t3_pos_4_096_60    m/z 729.4877     0   1   13.5   14  RT (min)       lcorbi_t3_pos_1_034_34    m/z 729.5628     0   2   4   6   8E+4   14   14.5  RT (min)      lcorbi_t3_pos_1_031_40    m/z 729.5789     0   1   2   3   4E+5   14.5   15  RT (min)      lcorbi_t3_pos_4_004_14    m/z 729.5819     0   2   4   6   8E+4   13.5   14  RT (min)       lcorbi_t3_pos_4_069_57    m/z 730.4528     0   2   4   6E+4   12.5   13  RT (min)       lcorbi_t3_pos_1_013_10    m/z 730.5000     0   1   2   3E+4   13   13.5   14  RT (min)       lcorbi_t3_pos_1_034_34    m/z 730.8916     0   2   4E+4   1   1.5  RT (min)      lcorbi_t3_pos_4_021_25    m/z 731.1951     0   1   2   2.5  RT (min)       lcorbi_t3_pos_4_090_60    m/z 731.4562     0   1   2   3E+4   12.5   13  RT (min)      lcorbi_t3_pos_2_012_01    m/z 731.6171     0   1E+5   15   15.5  RT (min)       lcorbi_t3_pos_1_029_29    m/z 731.6948     0   1E+5   14.5   15  RT (min)       lcorbi_t3_pos_1_036_08    m/z 731.8439     0   1E+5   1   1.5  RT (min)      lcorbi_t3_pos_4_068_57    m/z 732.4112     0   1E+4   11   11.5  RT (min)      lcorbi_t3_pos_3_053_48    m/z 732.5468     0   2   4   6   8E+4   10.5   11  RT (min)      lcorbi_t3_pos_4_050_31    m/z 732.5843     0   1E+5   14   15  RT (min)       lcorbi_t3_pos_1_003_45    m/z 732.5972     0   2   4   6   8E+4   13.5   14  RT (min)       lcorbi_t3_pos_4_071_57    m/z 732.6290     0   1   14.5   15  RT (min)       lcorbi_t3_pos_4_072_57    m/z 732.8887     0   2   4E+4   1   1.5  RT (min)       lcorbi_t3_pos_3_012_31    m/z 733.4849     0   1E+5   13   13.5  RT (min)       lcorbi_t3_pos_4_089_59    m/z 733.6140     0   1   14   14.5  RT (min)      lcorbi_t3_pos_4_083_59    m/z 733.7189     0   1E+5   14.5   15  RT (min)       lcorbi_t3_pos_4_097_60    m/z 733.8494     0   1   2   3   4E+4   1   1.5  RT (min)      lcorbi_t3_pos_4_070_57    m/z 734.3900     0   1   2E+4   9.5   10  RT (min)      lcorbi_t3_pos_4_089_59    m/z 734.6452     0   1   2E+5   15   15.5  RT (min)       lcorbi_t3_pos_1_002_01    m/z 734.7231     0   1   2E+5   14.5   15  RT (min)       lcorbi_t3_pos_4_092_60    m/z 734.8129     0   1E+5   1   1.5  RT (min)       lcorbi_t3_pos_4_075_58    m/z 734.8857     0   1   2   3   4E+4   1   1.5  RT (min)      lcorbi_t3_pos_3_012_31    m/z 735.5009     0   1   2E+5   13   13.5  RT (min)       lcorbi_t3_pos_4_087_59    m/z 735.5176     0   1   2   3E+4   13.5   14  RT (min)      lcorbi_t3_pos_4_066_57    m/z 735.7260     0   1   2E+5   14.5   15  RT (min)       lcorbi_t3_pos_1_036_08    m/z 735.8470     0   2   4   6E+4   1   1.5  RT (min)      lcorbi_t3_pos_3_012_31    m/z 736.4415     0   2   4   6E+4   14.5   15  RT (min)      lcorbi_t3_pos_3_041_54    m/z 736.5433     0   1E+5   14.5   15  RT (min)       lcorbi_t3_pos_4_082_59    m/z 736.5673     0   2   4   6E+4   13.5   14  RT (min)       lcorbi_t3_pos_1_029_29    m/z 736.6457     0   1E+5   15   15.5  RT (min)       lcorbi_t3_pos_4_066_57    m/z 736.8583     0   1   2   3E+5   1   1.5  RT (min)       lcorbi_t3_pos_4_068_57    m/z 737.3600     0   2   4   6   8E+4   1   1.5  RT (min)      lcorbi_t3_pos_4_068_57    m/z 737.5470     0   2   4   6E+4   14.5   15  RT (min)      lcorbi_t3_pos_4_082_59    m/z 737.5792     0   2   4E+4   14   14.5  RT (min)       lcorbi_t3_pos_4_078_58    m/z 737.8441     0   2   4   6E+4   1   1.5  RT (min)       lcorbi_t3_pos_4_073_57    m/z 738.3152     0   2   4   6E+4   13.5   14  RT (min)      lcorbi_t3_pos_2_044_21    m/z 738.4985     0   1E+6   13.5   14  RT (min)      lcorbi_t3_pos_4_082_59    m/z 738.6020     0   2   4   6E+4   14.5   15  RT (min)      lcorbi_t3_pos_2_024_15    m/z 738.6609     0   1E+5   15   15.5  RT (min)       lcorbi_t3_pos_4_069_57    m/z 738.6785     0   1E+5   14.5   15  RT (min)       lcorbi_t3_pos_1_032_26    m/z 738.8054     0   1   2   3E+4   1   1.5  RT (min)      lcorbi_t3_pos_4_073_57    m/z 739.4130     0   2   4   6   8E+4   14.5   15  RT (min)       lcorbi_t3_pos_4_110_62    m/z 739.4297     0   2   4E+4   12   12.5  RT (min)      lcorbi_t3_pos_4_088_59    m/z 739.4671     0   1   2   3E+4   14   14.5  RT (min)       lcorbi_t3_pos_4_084_59    m/z 739.4890     0   2   4   6E+3   13.5   14  RT (min)       lcorbi_t3_pos_1_034_34    m/z 739.5924     0   1   2E+5   14   14.5  RT (min)       lcorbi_t3_pos_4_089_59    m/z 739.6644     0   2   4   6E+4   15   15.5  RT (min)      lcorbi_t3_pos_4_067_57    m/z 739.6817     0   2   4   6E+4   14.5   15  RT (min)      lcorbi_t3_pos_1_031_40    m/z 739.8318     0   2   4   6E+4   1   1.5  RT (min)      lcorbi_t3_pos_4_067_57    m/z 739.8411     0   1   2   3   4E+4   1   1.5  RT (min)      lcorbi_t3_pos_4_067_57    m/z 740.3877     0   1E+5   13   13.5  RT (min)       lcorbi_t3_pos_4_096_60    m/z 740.4925     0   1   2   3E+4   13.5   14  RT (min)      lcorbi_t3_pos_4_083_59    m/z 740.5592     0   1   13.5   14  RT (min)       lcorbi_t3_pos_4_096_60    m/z 740.6174     0   2   4   6   8E+4   14.5   15  RT (min)       lcorbi_t3_pos_4_038_38    m/z 740.6346     0   1E+5   13.5   14  RT (min)       lcorbi_t3_pos_4_096_60    m/z 740.6430     0   2   4   6   8E+4   14   14.5  RT (min)       lcorbi_t3_pos_4_090_60    m/z 740.6766     0   2   4   6E+4   15.5   16  RT (min)       lcorbi_t3_pos_2_016_23    m/z 740.7131     0   2   4   6   8E+4   15   15.5  RT (min)       lcorbi_t3_pos_1_031_40    m/z 740.8024     0   2   4   6   8E+4   1   1.5  RT (min)       lcorbi_t3_pos_3_012_31    m/z 741.5172     0   1   2   3E+5   13.5   14  RT (min)       lcorbi_t3_pos_4_084_59    m/z 741.8369     0   1   2   3   4E+4   1   1.5  RT (min)      lcorbi_t3_pos_4_068_57    m/z 741.9580     0   1E+4   6.5   7  RT (min)       lcorbi_t3_pos_4_104_61    m/z 742.2706     0   1   2   3E+4   5   5.5  RT (min)       lcorbi_t3_pos_4_014_48    m/z 742.3525     0   1E+4   4.5   5  RT (min)      lcorbi_t3_pos_4_078_58    m/z 742.4787     0   1   2E+4   14   14.5  RT (min)       lcorbi_t3_pos_4_082_59    m/z 742.5301     0   1E+6   13.5   14  RT (min)      lcorbi_t3_pos_4_086_59    m/z 742.6507     0   1   2E+5   13.5   14  RT (min)       lcorbi_t3_pos_4_090_60    m/z 742.7864     0   2   4   6E+4   14.5   15  RT (min)       lcorbi_t3_pos_2_024_15    m/z 742.7994     0   2   4   6   8E+4   1   1.5  RT (min)       lcorbi_t3_pos_4_073_57    m/z 743.3939     0   2   4   6E+3   5   5.5  RT (min)      lcorbi_t3_pos_4_080_58    m/z 743.4350     0   1E+5   7.5   8  RT (min)       lcorbi_t3_pos_4_098_61    m/z 743.4823     0   1   2E+4   14   14.5  RT (min)      lcorbi_t3_pos_4_082_59    m/z 743.5859     0   1   2E+5   13.5   14  RT (min)       lcorbi_t3_pos_4_083_59    m/z 743.6061     0   1   13.5   14  RT (min)      lcorbi_t3_pos_1_054_48    m/z 743.7502     0   1E+5   14.5   15  RT (min)       lcorbi_t3_pos_2_024_15    m/z 743.7608     0   1   2   3E+4   1   1.5  RT (min)      lcorbi_t3_pos_4_067_57    m/z 744.4381     0   1   2   3   4E+4   7.5   8  RT (min)      lcorbi_t3_pos_4_100_61    m/z 744.5064     0   2   4   6   8E+4   13.5   14  RT (min)      lcorbi_t3_pos_4_099_61    m/z 744.5262     0   1   13.5   14  RT (min)      lcorbi_t3_pos_4_082_59    m/z 744.7965     0   2   4   6   8E+4   1   1.5  RT (min)       lcorbi_t3_pos_4_021_25    m/z 744.8455     0   1E+5   1   1.5  RT (min)       lcorbi_t3_pos_4_075_58    m/z 745.3472     0   1   2   3   4E+4   1   1.5  RT (min)      lcorbi_t3_pos_4_068_57    m/z 745.5483     0   2   4   6E+4   13.5   14  RT (min)       lcorbi_t3_pos_4_082_59    m/z 745.6228     0   1   14   14.5  RT (min)      lcorbi_t3_pos_4_082_59    m/z 745.7578     0   2   4   6   8E+4   1   1.5  RT (min)       lcorbi_t3_pos_4_070_57    m/z 745.7659     0   1E+5   14.5   15  RT (min)       lcorbi_t3_pos_2_024_15    m/z 745.8466     0   1   2E+4   1   1.5  RT (min)      lcorbi_t3_pos_4_074_58    m/z 746.5413     0   1E+5   14   14.5  RT (min)      lcorbi_t3_pos_4_106_62    m/z 746.6245     0   1   2E+4   14   14.5  RT (min)      lcorbi_t3_pos_4_082_59    m/z 746.7935     0   1   2   3   4E+4   1   1.5  RT (min)      lcorbi_t3_pos_3_012_31    m/z 747.5184     0   1E+5   13.5   14  RT (min)       lcorbi_t3_pos_4_082_59    m/z 747.5371     0   1E+4   13.5   14  RT (min)      lcorbi_t3_pos_3_021_39    m/z 747.5638     0   1   2   3   4E+5   14   14.5  RT (min)       lcorbi_t3_pos_4_089_59    m/z 747.5660     0   1   2E+5   10.5   11  RT (min)      lcorbi_t3_pos_4_082_59    m/z 747.5993     0   1   14   14.5  RT (min)      lcorbi_t3_pos_4_084_59    m/z 747.7549     0   2   4   6   8E+4   1   1.5  RT (min)       lcorbi_t3_pos_4_070_57    m/z 747.7814     0   1E+5   14.5   15  RT (min)       lcorbi_t3_pos_4_091_60    m/z 748.3249     0   2   4   6   8E+4   14.5   15  RT (min)       lcorbi_t3_pos_4_110_62    m/z 748.5040     0   1   2E+4   13   13.5  RT (min)       lcorbi_t3_pos_1_043_06    m/z 748.5277     0   1   14   14.5  RT (min)      lcorbi_t3_pos_4_106_62    m/z 748.5856     0   1E+5   14   14.5  RT (min)       lcorbi_t3_pos_4_084_59    m/z 748.6058     0   1E+5   15   15.5  RT (min)       lcorbi_t3_pos_4_089_59    m/z 748.6216     0   1E+5   14   14.5  RT (min)       lcorbi_t3_pos_4_072_57    m/z 748.6607     0   2   4   6E+4   15   15.5  RT (min)       lcorbi_t3_pos_4_006_12    m/z 748.7163     0   2   4E+4   1   1.5  RT (min)      lcorbi_t3_pos_4_073_57    m/z 748.7733     0   1E+5   14.5   15  RT (min)       lcorbi_t3_pos_1_036_08    m/z 748.8576     0   1   2   3E+4   1   1.5  RT (min)      lcorbi_t3_pos_4_077_58    m/z 749.4707     0   2   4   6E+4   13.5   14  RT (min)       lcorbi_t3_pos_4_101_61    m/z 749.5329     0   1E+5   13.5   14  RT (min)       lcorbi_t3_pos_4_082_59    m/z 749.5890     0   1   2E+4   14   14.5  RT (min)       lcorbi_t3_pos_4_098_61    m/z 749.6070     0   1E+5   13   13.5  RT (min)       lcorbi_t3_pos_1_036_08    m/z 749.7518     0   2   4   6E+4   1   1.5  RT (min)       lcorbi_t3_pos_4_067_57    m/z 750.3125     0   1E+4   6.5   7  RT (min)      lcorbi_t3_pos_1_014_03    m/z 750.4003     0   1   2E+4   11.5   12  RT (min)      lcorbi_t3_pos_2_044_21    m/z 750.4661     0   1   13.5   14  RT (min)      lcorbi_t3_pos_4_084_59    m/z 750.5432     0   1   2   3   4E+5   14   14.5  RT (min)       lcorbi_t3_pos_1_031_40    m/z 750.6198     0   1E+5   13.5   14  RT (min)       lcorbi_t3_pos_1_036_08    m/z 750.6211     0   1   15   15.5  RT (min)       lcorbi_t3_pos_4_084_59    m/z 750.6761     0   1E+5   15.5   16  RT (min)       lcorbi_t3_pos_4_075_58    m/z 750.7131     0   2   4   6E+4   1   1.5  RT (min)       lcorbi_t3_pos_4_068_57    m/z 751.5467     0   1   2E+5   14   14.5  RT (min)      lcorbi_t3_pos_1_031_40    m/z 751.7488     0   2   4   6E+4   1   1.5  RT (min)      lcorbi_t3_pos_2_028_24    m/z 752.4005     0   1   2E+4   8.5   9  RT (min)      lcorbi_t3_pos_1_054_48    m/z 752.5137     0   1E+5   13   13.5  RT (min)      lcorbi_t3_pos_1_032_26    m/z 752.5514     0   1E+5   15   15.5  RT (min)       lcorbi_t3_pos_4_074_58    m/z 752.5764     0   1   2E+4   13.5   14  RT (min)      lcorbi_t3_pos_1_036_08    m/z 752.6077     0   2   4   6E+4   13   13.5  RT (min)      lcorbi_t3_pos_2_044_21    m/z 752.7101     0   1E+5   1   1.5  RT (min)       lcorbi_t3_pos_4_068_57    m/z 752.8322     0   1E+5   1   1.5  RT (min)       lcorbi_t3_pos_4_075_58    m/z 753.5473     0   1   2   3   4E+4   13.5   14  RT (min)       lcorbi_t3_pos_1_008_36    m/z 753.5682     0   1   2   3E+5   14.5   15  RT (min)       lcorbi_t3_pos_1_036_08    m/z 753.5905     0   1   2   3   4E+4   13.5   14  RT (min)       lcorbi_t3_pos_1_036_08    m/z 753.6716     0   2   4   6E+4   1   1.5  RT (min)      lcorbi_t3_pos_2_028_24    m/z 753.6954     0   1E+5   15   15.5  RT (min)       lcorbi_t3_pos_3_008_20    m/z 754.4798     0   1E+5   14   14.5  RT (min)       lcorbi_t3_pos_4_086_59    m/z 754.4999     0   1   13.5   14  RT (min)       lcorbi_t3_pos_4_083_59    m/z 754.7072     0   2   4   6   8E+4   1   1.5  RT (min)      lcorbi_t3_pos_1_020_31    m/z 755.4234     0   2   4   6E+5   14.5   15  RT (min)      lcorbi_t3_pos_4_095_60    m/z 755.5325     0   1E+5   13   13.5  RT (min)       lcorbi_t3_pos_4_086_59    m/z 755.5784     0   2   4   6E+4   14   14.5  RT (min)      lcorbi_t3_pos_1_031_40    m/z 755.6685     0   2   4   6   8E+4   1   1.5  RT (min)       lcorbi_t3_pos_4_073_57    m/z 756.4812     0   1E+5   13   13.5   14  RT (min)       lcorbi_t3_pos_4_089_59    m/z 756.4958     0   1E+5   14   14.5  RT (min)       lcorbi_t3_pos_1_028_32    m/z 756.5130     0   1   2   3E+5   13.5   14  RT (min)      lcorbi_t3_pos_1_028_32    m/z 756.6091     0   1E+5   13.5   14  RT (min)      lcorbi_t3_pos_1_034_34    m/z 756.7044     0   1   2   3   4E+4   1   1.5  RT (min)       lcorbi_t3_pos_4_067_57    m/z 757.4846     0   2   4E+4   13   13.5  RT (min)      lcorbi_t3_pos_4_086_59    m/z 757.4996     0   2   4   6E+4   13.5   14  RT (min)       lcorbi_t3_pos_1_034_34    m/z 757.6581     0   1   2E+5   13.5   14  RT (min)      lcorbi_t3_pos_4_096_60    m/z 757.6655     0   1E+5   1   1.5  RT (min)       lcorbi_t3_pos_2_028_24    m/z 757.7101     0   1   2   3   4E+4   14   14.5  RT (min)      lcorbi_t3_pos_4_092_60    m/z 758.0298     0   1E+5   11   11.5  RT (min)       lcorbi_t3_pos_1_041_54    m/z 758.4651     0   1   2   3   4E+4   13.5   14  RT (min)      lcorbi_t3_pos_1_035_28    m/z 758.4740     0   2   4   6   8E+4   13.5   14  RT (min)       lcorbi_t3_pos_1_031_40    m/z 758.5164     0   1   13.5   14  RT (min)      lcorbi_t3_pos_4_074_58    m/z 758.5508     0   1E+5   13.5   14  RT (min)       lcorbi_t3_pos_1_031_40    m/z 758.5702     0   1E+5   0   0.5  RT (min)      lcorbi_t3_pos_3_004_24    m/z 758.6020     0   1   2E+5   14   14.5  RT (min)      lcorbi_t3_pos_1_019_16    m/z 758.6270     0   1   2   3   4E+4   1   1.5  RT (min)      lcorbi_t3_pos_4_073_57    m/z 758.6754     0   1   2   3E+5   14   14.5  RT (min)       lcorbi_t3_pos_4_093_60    m/z 759.5202     0   1   13.5   14  RT (min)      lcorbi_t3_pos_4_074_58    m/z 759.5737     0   2   4   6E+4   0   0.5  RT (min)      lcorbi_t3_pos_3_004_24    m/z 759.6054     0   1   2E+5   13.5   14  RT (min)      lcorbi_t3_pos_1_014_03    m/z 759.6119     0   1   2   3E+4   15   15.5  RT (min)      lcorbi_t3_pos_4_066_57    m/z 759.6251     0   1E+5   13.5   14  RT (min)      lcorbi_t3_pos_1_019_16    m/z 759.6626     0   1E+5   1   1.5  RT (min)       lcorbi_t3_pos_1_020_31    m/z 759.6740     0   1   2E+5   14   14.5  RT (min)       lcorbi_t3_pos_4_093_60    m/z 759.7257     0   1   2E+5   14.5   15  RT (min)       lcorbi_t3_pos_1_036_08    m/z 760.5668     0   1   2E+5   13.5   14  RT (min)       lcorbi_t3_pos_1_031_40    m/z 760.6238     0   1E+5   1   1.5  RT (min)       lcorbi_t3_pos_1_020_31    m/z 760.6604     0   1   2   3E+5   14   14.5  RT (min)      lcorbi_t3_pos_4_075_58    m/z 760.8220     0   2   4   6   8E+4   1   1.5  RT (min)      lcorbi_t3_pos_4_075_58    m/z 761.3925     0   1   2   3   4E+4   13.5   14  RT (min)      lcorbi_t3_pos_2_044_21    m/z 761.4499     0   1   14   14.5  RT (min)      lcorbi_t3_pos_1_020_31    m/z 761.4709     0   1   2E+4   13.5   14  RT (min)      lcorbi_t3_pos_1_054_48    m/z 761.5066     0   1   13   13.5  RT (min)       lcorbi_t3_pos_4_075_58    m/z 761.5352     0   2   4   6   8E+4   14   14.5  RT (min)      lcorbi_t3_pos_1_020_31    m/z 761.6292     0   1E+5   14   14.5  RT (min)      lcorbi_t3_pos_1_030_21    m/z 761.6405     0   1E+5   14   14.5  RT (min)      lcorbi_t3_pos_1_007_07    m/z 761.6447     0   1E+5   14   14.5  RT (min)       lcorbi_t3_pos_1_028_32    m/z 761.6596     0   2   4E+4   1   1.5  RT (min)       lcorbi_t3_pos_3_012_31    m/z 761.7415     0   1   2   3E+5   14.5   15  RT (min)       lcorbi_t3_pos_1_036_08    m/z 761.7969     0   1E+5   14.5   15  RT (min)       lcorbi_t3_pos_4_090_60    m/z 762.6209     0   1E+5   1   1.5  RT (min)       lcorbi_t3_pos_2_028_24    m/z 762.6608     0   2   4   6   8E+4   15   15.5  RT (min)       lcorbi_t3_pos_4_067_57    m/z 762.7572     0   1E+5   14.5   15  RT (min)       lcorbi_t3_pos_2_024_15    m/z 763.4032     0   2   4E+4   13.5   14  RT (min)       lcorbi_t3_pos_4_108_62    m/z 763.4613     0   2   4E+4   11   11.5  RT (min)      lcorbi_t3_pos_4_090_60    m/z 763.5081     0   1   2E+5   13.5   14  RT (min)       lcorbi_t3_pos_1_054_48    m/z 763.5679     0   1   14   14.5  RT (min)       lcorbi_t3_pos_1_020_31    m/z 763.5825     0   2   4   6E+4   1   1.5  RT (min)      lcorbi_t3_pos_1_020_31    m/z 763.5853     0   2   4   6E+4   13.5   14  RT (min)       lcorbi_t3_pos_1_007_07    m/z 763.6224     0   1E+5   13.5   14  RT (min)       lcorbi_t3_pos_1_036_08    m/z 763.7287     0   1   2   3E+4   1   1.5  RT (min)      lcorbi_t3_pos_2_051_35    m/z 763.7572     0   1   2   3   4E+5   14.5   15  RT (min)       lcorbi_t3_pos_1_036_08    m/z 764.4610     0   1   14   14.5  RT (min)      lcorbi_t3_pos_4_084_59    m/z 764.6074     0   2   4   6E+4   14   14.5  RT (min)       lcorbi_t3_pos_4_070_57    m/z 764.6180     0   2   4   6   8E+4   1   1.5  RT (min)      lcorbi_t3_pos_1_020_31    m/z 764.8347     0   1   2E+4   1   1.5  RT (min)      lcorbi_t3_pos_4_081_58    m/z 765.4851     0   1   13.5   14  RT (min)      lcorbi_t3_pos_4_083_59    m/z 765.5382     0   1E+5   13.5   14  RT (min)       lcorbi_t3_pos_4_089_59    m/z 765.5479     0   1   2E+5   13.5   14  RT (min)       lcorbi_t3_pos_1_031_40    m/z 765.5794     0   2   4   6   8E+4   1   1.5  RT (min)      lcorbi_t3_pos_1_020_31    m/z 765.5984     0   1   2   3   4E+4   14   14.5  RT (min)      lcorbi_t3_pos_1_035_28    m/z 765.6381     0   1E+5   13   13.5  RT (min)      lcorbi_t3_pos_4_052_06    m/z 765.6799     0   1E+5   15   15.5  RT (min)       lcorbi_t3_pos_4_066_57    m/z 765.7170     0   2   4   6E+4   14.5   15  RT (min)      lcorbi_t3_pos_4_097_60    m/z 765.7726     0   1E+5   14.5   15  RT (min)       lcorbi_t3_pos_1_036_08    m/z 766.5310     0   1   2E+4   10.5   11  RT (min)      lcorbi_t3_pos_4_027_07    m/z 766.6150     0   2   4   6E+4   1   1.5  RT (min)       lcorbi_t3_pos_1_020_31    m/z 766.6505     0   1E+5   13   13.5  RT (min)       lcorbi_t3_pos_1_036_08    m/z 766.7076     0   1E+5   15.5   16  RT (min)       lcorbi_t3_pos_4_077_58    m/z 766.7132     0   1E+5   13.5   14  RT (min)       lcorbi_t3_pos_4_092_60    m/z 767.2811     0   1E+5   5   5.5  RT (min)       lcorbi_t3_pos_4_014_48    m/z 767.4134     0   1   2   3   4E+4   11   11.5  RT (min)       lcorbi_t3_pos_4_096_60    m/z 767.4141     0   2   4   6E+4   13.5   14  RT (min)      lcorbi_t3_pos_4_113_62    m/z 767.5203     0   1   14   14.5  RT (min)       lcorbi_t3_pos_4_089_59    m/z 767.5327     0   2   4   6   8E+4   13   13.5  RT (min)       lcorbi_t3_pos_4_086_59    m/z 767.5638     0   1E+5   13.5   14  RT (min)      lcorbi_t3_pos_1_054_48    m/z 767.5764     0   1E+5   1   1.5  RT (min)       lcorbi_t3_pos_1_020_31    m/z 767.5954     0   1   2E+4   8.5   9  RT (min)      lcorbi_t3_pos_4_098_61    m/z 767.6955     0   1E+5   15.5   16  RT (min)      lcorbi_t3_pos_4_066_57    m/z 767.7110     0   2   4E+4   15   15.5  RT (min)      lcorbi_t3_pos_4_079_58    m/z 767.8172     0   1E+5   15   15.5  RT (min)       lcorbi_t3_pos_1_036_08    m/z 767.8428     0   1   2E+4   1   1.5  RT (min)      lcorbi_t3_pos_4_068_57    m/z 768.2844     0   2   4   6E+4   5   5.5  RT (min)      lcorbi_t3_pos_4_008_30    m/z 768.4108     0   1   2E+4   10.5   11  RT (min)       lcorbi_t3_pos_4_102_61    m/z 768.4796     0   1E+5   13   13.5  RT (min)       lcorbi_t3_pos_4_083_59    m/z 768.5429     0   2   4E+4   14   14.5  RT (min)      lcorbi_t3_pos_1_034_34    m/z 768.6842     0   1   2   3E+4   1   1.5  RT (min)      lcorbi_t3_pos_4_039_35    m/z 768.7009     0   2   4   6   8E+4   13.5   14  RT (min)       lcorbi_t3_pos_4_091_60    m/z 768.7078     0   1   15.5   16  RT (min)       lcorbi_t3_pos_4_092_60    m/z 768.8506     0   1E+4   5   5.5  RT (min)      lcorbi_t3_pos_4_076_58    m/z 769.3518     0   1E+4   5   5.5  RT (min)      lcorbi_t3_pos_4_078_58    m/z 769.4140     0   1E+4   10.5   11  RT (min)      lcorbi_t3_pos_3_032_09    m/z 769.4998     0   2   4   6E+4   13.5   14  RT (min)       lcorbi_t3_pos_1_054_48    m/z 769.5350     0   1   13.5   14  RT (min)       lcorbi_t3_pos_4_087_59    m/z 769.5483     0   1E+5   13.5   14  RT (min)       lcorbi_t3_pos_4_084_59    m/z 769.5736     0   2   4   6   8E+4   1   1.5  RT (min)      lcorbi_t3_pos_2_028_24    m/z 769.7054     0   1   2   3E+5   13.5   14  RT (min)       lcorbi_t3_pos_4_090_60    m/z 769.8327     0   1   2E+5   15   15.5  RT (min)       lcorbi_t3_pos_4_091_60    m/z 770.4971     0   1   2   3   4E+5   12.5   13  RT (min)       lcorbi_t3_pos_2_005_11    m/z 770.5106     0   2   4   6E+4   13.5   14  RT (min)       lcorbi_t3_pos_4_071_57    m/z 770.5350     0   2   4   6E+4   1   1.5  RT (min)      lcorbi_t3_pos_2_028_24    m/z 771.0677     0   1E+5   14   14.5  RT (min)       lcorbi_t3_pos_1_007_07    m/z 771.5159     0   1E+5   13.5   14  RT (min)       lcorbi_t3_pos_1_054_48    m/z 771.5503     0   1   13.5   14  RT (min)       lcorbi_t3_pos_4_083_59    m/z 771.5642     0   1   2   3E+5   13.5   14  RT (min)       lcorbi_t3_pos_4_084_59    m/z 771.5709     0   1   2   3   4E+4   1   1.5  RT (min)      lcorbi_t3_pos_2_028_24    m/z 771.6218     0   1E+5   13   13.5  RT (min)       lcorbi_t3_pos_4_091_60    m/z 771.6385     0   1E+5   14   14.5  RT (min)      lcorbi_t3_pos_1_016_05    m/z 771.6425     0   1   2   3E+4   1   1.5  RT (min)      lcorbi_t3_pos_4_039_35    m/z 771.8553     0   2   4   6E+4   1   1.5  RT (min)      lcorbi_t3_pos_4_068_57    m/z 772.0768     0   1   2E+5   14   14.5  RT (min)       lcorbi_t3_pos_1_020_31    m/z 772.4980     0   1   2   3   4E+4   14   14.5  RT (min)      lcorbi_t3_pos_2_015_17    m/z 772.5318     0   2   4   6   8E+4   1   1.5  RT (min)      lcorbi_t3_pos_1_020_31    m/z 772.5493     0   2   4   6E+5   13   13.5   14  RT (min)       lcorbi_t3_pos_1_054_48    m/z 772.5679     0   1   13.5   14  RT (min)      lcorbi_t3_pos_4_073_57    m/z 772.6126     0   1   14   14.5  RT (min)      lcorbi_t3_pos_4_086_59    m/z 772.6423     0   1   14   14.5  RT (min)      lcorbi_t3_pos_4_084_59    m/z 772.6768     0   2   4   6   8E+4   13.5   14  RT (min)       lcorbi_t3_pos_4_091_60    m/z 773.0794     0   1   2   3E+4   14   14.5  RT (min)       lcorbi_t3_pos_4_073_57    m/z 773.4321     0   1   2   3   4E+4   13.5   14  RT (min)      lcorbi_t3_pos_4_112_62    m/z 773.5164     0   1E+5   13.5   14  RT (min)      lcorbi_t3_pos_1_054_48    m/z 773.5526     0   1   2   3E+5   13   13.5   14  RT (min)       lcorbi_t3_pos_2_005_11    m/z 773.6141     0   1   14   14.5  RT (min)      lcorbi_t3_pos_4_082_59    m/z 773.6397     0   1   2   3E+4   1   1.5  RT (min)      lcorbi_t3_pos_3_039_03    m/z 773.6933     0   1E+5   13.5   14  RT (min)       lcorbi_t3_pos_4_092_60    m/z 774.5050     0   1E+5   13   13.5  RT (min)       lcorbi_t3_pos_4_086_59    m/z 774.5290     0   1   2   3   4E+4   1   1.5  RT (min)      lcorbi_t3_pos_2_028_24    m/z 774.5414     0   1E+5   14   14.5  RT (min)      lcorbi_t3_pos_1_036_08    m/z 775.4902     0   2   4E+4   1   1.5  RT (min)      lcorbi_t3_pos_2_028_24    m/z 775.5086     0   1E+5   13   13.5  RT (min)       lcorbi_t3_pos_4_089_59    m/z 775.5956     0   1   2   3E+5   14   14.5  RT (min)       lcorbi_t3_pos_4_083_59    m/z 775.6213     0   1E+5   13.5   14  RT (min)      lcorbi_t3_pos_4_092_60    m/z 775.6297     0   1   2   3   4E+4   14   14.5  RT (min)      lcorbi_t3_pos_1_034_34    m/z 775.6605     0   1   2   3   4E+4   14   14.5  RT (min)       lcorbi_t3_pos_2_053_28    m/z 776.2328     0   1   2E+5   13   13.5  RT (min)       lcorbi_t3_pos_4_113_62    m/z 776.3068     0   2   4   6E+4   8.5   9  RT (min)       lcorbi_t3_pos_4_111_62    m/z 776.3584     0   1E+5   14.5   15  RT (min)      lcorbi_t3_pos_4_099_61    m/z 776.4370     0   1   2E+4   10.5   11  RT (min)      lcorbi_t3_pos_4_077_58    m/z 776.4630     0   2   4   6   8E+4   13.5   14  RT (min)       lcorbi_t3_pos_4_084_59    m/z 776.5358     0   1   2E+5   13.5   14  RT (min)      lcorbi_t3_pos_4_082_59    m/z 776.5437     0   1   2   3E+5   13.5   14  RT (min)       lcorbi_t3_pos_4_083_59    m/z 776.5590     0   1E+5   14   14.5  RT (min)      lcorbi_t3_pos_1_036_08    m/z 776.5714     0   1E+5   12.5   13  RT (min)      lcorbi_t3_pos_1_029_29    m/z 776.6328     0   1   2E+5   14   14.5  RT (min)       lcorbi_t3_pos_4_090_60    m/z 776.6441     0   1   2   3E+4   14   14.5  RT (min)      lcorbi_t3_pos_2_024_15    m/z 776.6682     0   1E+5   13.5   14  RT (min)       lcorbi_t3_pos_4_091_60    m/z 777.3102     0   1   2E+4   8.5   9  RT (min)      lcorbi_t3_pos_4_113_62    m/z 777.4876     0   2   4E+4   1   1.5  RT (min)       lcorbi_t3_pos_1_020_31    m/z 777.5426     0   1E+5   13.5   14  RT (min)       lcorbi_t3_pos_4_087_59    m/z 777.5482     0   1   13.5   14  RT (min)       lcorbi_t3_pos_4_091_60    m/z 777.5624     0   1E+5   14   14.5  RT (min)      lcorbi_t3_pos_1_036_08    m/z 777.6124     0   1   14   14.5  RT (min)      lcorbi_t3_pos_4_092_60    m/z 777.6954     0   1E+5   15   15.5  RT (min)       lcorbi_t3_pos_4_079_58    m/z 777.7384     0   1   2   3E+5   14.5   15  RT (min)       lcorbi_t3_pos_1_016_05    m/z 778.0439     0   2   4   6   8E+4   13.5   14  RT (min)      lcorbi_t3_pos_1_007_07    m/z 778.4957     0   1   2   3   4E+5   13.5   14  RT (min)      lcorbi_t3_pos_1_034_34    m/z 778.5468     0   1   2E+4   8   8.5  RT (min)       lcorbi_t3_pos_4_112_62    m/z 778.5951     0   1   2   3   4E+4   1   1.5  RT (min)      lcorbi_t3_pos_4_016_15    m/z 778.6041     0   1E+5   14  RT (min)       lcorbi_t3_pos_1_034_34    m/z 778.7389     0   1   2   3E+5   15   15.5  RT (min)      lcorbi_t3_pos_1_036_08    m/z 778.8114     0   1E+5   14.5   15  RT (min)       lcorbi_t3_pos_1_036_08    m/z 778.8392     0   2   4   6   8E+4   1   1.5  RT (min)       lcorbi_t3_pos_4_075_58    m/z 779.5590     0   1   14   14.5  RT (min)      lcorbi_t3_pos_1_054_48    m/z 779.5833     0   1   2   3E+5   14   14.5  RT (min)       lcorbi_t3_pos_1_054_48    m/z 779.5939     0   2   4   6E+4   14.5   15  RT (min)       lcorbi_t3_pos_4_082_59    m/z 779.6052     0   1   14   14.5  RT (min)       lcorbi_t3_pos_1_054_48    m/z 780.0464     0   1E+5   13.5   14  RT (min)       lcorbi_t3_pos_1_008_36    m/z 780.4457     0   2   4E+4   1   1.5  RT (min)      lcorbi_t3_pos_2_028_24    m/z 780.4872     0   1   2E+5   13.5   14  RT (min)       lcorbi_t3_pos_1_028_32    m/z 780.4984     0   2   4E+4   13.5   14  RT (min)       lcorbi_t3_pos_4_085_59    m/z 780.5107     0   1   2   3E+4   13.5   14  RT (min)      lcorbi_t3_pos_4_082_59    m/z 780.5703     0   1   2   3E+5   15   15.5  RT (min)       lcorbi_t3_pos_4_044_32    m/z 780.6081     0   1   13.5   14  RT (min)      lcorbi_t3_pos_4_082_59    m/z 780.6197     0   1   2E+5   13.5   14  RT (min)      lcorbi_t3_pos_1_031_40    m/z 780.7286     0   2   4E+4   14   14.5  RT (min)       lcorbi_t3_pos_1_033_53    m/z 781.5010     0   1E+5   13.5   14  RT (min)       lcorbi_t3_pos_1_034_34    m/z 781.5536     0   1   2   3E+4   1   1.5  RT (min)      lcorbi_t3_pos_3_004_24    m/z 781.5992     0   1   2E+4   13.5   14  RT (min)      lcorbi_t3_pos_4_082_59    m/z 781.6210     0   1   14   14.5  RT (min)       lcorbi_t3_pos_4_088_59    m/z 781.6493     0   1   2   3E+4   13.5   14  RT (min)       lcorbi_t3_pos_1_034_34    m/z 782.4427     0   2   4E+4   1   1.5  RT (min)      lcorbi_t3_pos_1_020_31    m/z 782.4729     0   1   13.5   14  RT (min)      lcorbi_t3_pos_4_085_59    m/z 782.5103     0   1E+5   14   14.5  RT (min)       lcorbi_t3_pos_4_013_36    m/z 782.5702     0   1E+5   0   0.5  RT (min)      lcorbi_t3_pos_3_004_24    m/z 782.6239     0   1   2E+5   13.5   14  RT (min)      lcorbi_t3_pos_1_028_32    m/z 782.6356     0   2   4   6   8E+4   13.5   14  RT (min)      lcorbi_t3_pos_1_008_36    m/z 782.6638     0   1   2   3E+5   13.5   14  RT (min)      lcorbi_t3_pos_4_091_60    m/z 782.7236     0   2   4   6E+4   15.5   16  RT (min)      lcorbi_t3_pos_2_008_16    m/z 783.0683     0   1E+5   14   14.5  RT (min)       lcorbi_t3_pos_1_007_07    m/z 783.5299     0   1   2E+5   13.5   14  RT (min)      lcorbi_t3_pos_1_008_36    m/z 783.5541     0   2   4   6   8E+4   13   13.5  RT (min)       lcorbi_t3_pos_1_032_26    m/z 783.6103     0   1   2E+4   14   14.5  RT (min)       lcorbi_t3_pos_4_087_59    m/z 783.6845     0   2   4   6E+6   13.5   14  RT (min)      lcorbi_t3_pos_4_091_60    m/z 783.7214     0   1E+5   13.5   14  RT (min)       lcorbi_t3_pos_1_050_13    m/z 784.0768     0   1E+5   14   14.5  RT (min)       lcorbi_t3_pos_1_020_31    m/z 784.5005     0   1   2   3E+5   13.5   14  RT (min)      lcorbi_t3_pos_4_082_59    m/z 784.5285     0   1   2   3E+4   14   14.5  RT (min)      lcorbi_t3_pos_4_089_59    m/z 784.5405     0   1   14   14.5  RT (min)      lcorbi_t3_pos_4_083_59    m/z 784.6268     0   1   2   3E+5   14   14.5  RT (min)      lcorbi_t3_pos_1_020_31    m/z 784.6407     0   1   14   14.5  RT (min)      lcorbi_t3_pos_4_082_59    m/z 784.7332     0   1   2   3   4E+4   15.5   16  RT (min)      lcorbi_t3_pos_2_043_32    m/z 785.0848     0   1E+5   14   14.5  RT (min)       lcorbi_t3_pos_1_044_25    m/z 785.5323     0   1   2E+4   14   14.5  RT (min)      lcorbi_t3_pos_4_089_59    m/z 785.5443     0   1   2E+4   14   14.5  RT (min)      lcorbi_t3_pos_4_089_59    m/z 785.5691     0   1E+5   13.5   14  RT (min)       lcorbi_t3_pos_1_028_32    m/z 785.6434     0   2   4   6E+4   14   14.5  RT (min)      lcorbi_t3_pos_1_007_07    m/z 786.5048     0   1E+5   13   13.5  RT (min)      lcorbi_t3_pos_4_084_59    m/z 786.5194     0   1   2   3E+4   13.5   14  RT (min)      lcorbi_t3_pos_4_079_58    m/z 786.5271     0   1E+4   11   11.5  RT (min)      lcorbi_t3_pos_1_054_48    m/z 786.5452     0   1E+4   14   14.5  RT (min)      lcorbi_t3_pos_4_074_58    m/z 786.5572     0   1   2   3E+5   14   14.5  RT (min)      lcorbi_t3_pos_1_007_07    m/z 786.6672     0   2   4   6E+5   13   14  RT (min)       lcorbi_t3_pos_4_075_58    m/z 786.6861     0   1   14   14.5  RT (min)      lcorbi_t3_pos_4_092_60    m/z 786.7390     0   1   2   3E+5   14   14.5  RT (min)       lcorbi_t3_pos_4_091_60    m/z 786.8262     0   1E+5   1   1.5  RT (min)       lcorbi_t3_pos_4_075_58    m/z 787.5361     0   1   2E+5   14   14.5  RT (min)      lcorbi_t3_pos_4_076_58    m/z 787.5486     0   1   14   14.5  RT (min)      lcorbi_t3_pos_4_075_58    m/z 787.6236     0   1   15   15.5  RT (min)      lcorbi_t3_pos_4_071_57    m/z 787.6446     0   1   2E+5   13   14  RT (min)       lcorbi_t3_pos_1_019_16    m/z 788.4203     0   2   4   6   8E+4   15   15.5  RT (min)       lcorbi_t3_pos_4_104_61    m/z 788.4928     0   1   2   3E+4   13.5   14  RT (min)      lcorbi_t3_pos_1_008_36    m/z 788.5355     0   1   2   3   4E+4   13.5   14  RT (min)       lcorbi_t3_pos_4_072_57    m/z 788.5978     0   1   2E+5   14   14.5  RT (min)       lcorbi_t3_pos_4_084_59    m/z 788.6240     0   1E+7   14   14.5  RT (min)       lcorbi_t3_pos_4_068_57    m/z 789.4763     0   2   4E+4   11   11.5  RT (min)       lcorbi_t3_pos_4_099_61    m/z 790.0295     0   1E+5   13   13.5  RT (min)       lcorbi_t3_pos_1_008_36    m/z 790.0432     0   2   4   6E+4   13.5   14  RT (min)       lcorbi_t3_pos_1_007_07    m/z 790.3226     0   2   4   6E+4   9   9.5  RT (min)       lcorbi_t3_pos_4_108_62    m/z 790.4798     0   1   2E+4   11   11.5  RT (min)      lcorbi_t3_pos_2_005_11    m/z 790.5361     0   1   2   3E+5   13.5   14  RT (min)       lcorbi_t3_pos_4_082_59    m/z 790.5507     0   2   4   6   8E+4   13.5   14  RT (min)      lcorbi_t3_pos_1_054_48    m/z 790.5871     0   1   2E+4   14   14.5  RT (min)       lcorbi_t3_pos_1_020_31    m/z 790.5887     0   1E+5   10.5   11  RT (min)       lcorbi_t3_pos_4_084_59    m/z 790.6394     0   1E+5   13   13.5  RT (min)       lcorbi_t3_pos_1_036_08    m/z 790.6785     0   2   4E+4   14   14.5  RT (min)      lcorbi_t3_pos_4_071_57    m/z 790.7077     0   1E+5   15   15.5  RT (min)       lcorbi_t3_pos_4_079_58    m/z 790.7474     0   1   2E+5   15   15.5  RT (min)       lcorbi_t3_pos_1_036_08    m/z 791.0531     0   1   2   3   4E+4   14   14.5  RT (min)      lcorbi_t3_pos_1_019_16    m/z 791.3261     0   1   2   3E+4   9   9.5  RT (min)       lcorbi_t3_pos_4_111_62    m/z 791.5006     0   1E+5   15   15.5  RT (min)       lcorbi_t3_pos_4_109_62    m/z 791.5417     0   1   2   3E+5   13.5   14  RT (min)       lcorbi_t3_pos_4_085_59    m/z 791.5628     0   1E+5   14   14.5  RT (min)       lcorbi_t3_pos_1_054_48    m/z 791.5761     0   1E+5   14   14.5  RT (min)       lcorbi_t3_pos_1_029_29    m/z 791.6872     0   1E+5   13.5   14  RT (min)       lcorbi_t3_pos_4_092_60    m/z 791.6956     0   1E+5   15   15.5  RT (min)       lcorbi_t3_pos_4_073_57    m/z 791.7112     0   1   2   3E+4   15   15.5  RT (min)      lcorbi_t3_pos_1_036_08    m/z 792.4947     0   1   2   3E+4   13.5   14  RT (min)       lcorbi_t3_pos_1_043_06    m/z 792.5132     0   1E+5   13   14  RT (min)       lcorbi_t3_pos_4_082_59    m/z 792.5753     0   1   13.5   14  RT (min)       lcorbi_t3_pos_4_083_59    m/z 792.5961     0   1E+4   6   6.5  RT (min)      lcorbi_t3_pos_4_086_59    m/z 792.6292     0   2   4   6   8E+4   14   14.5  RT (min)       lcorbi_t3_pos_1_008_36    m/z 792.7619     0   2   4   6E+6   14.5   15  RT (min)      lcorbi_t3_pos_2_024_15    m/z 793.0531     0   1E+5   13.5   14  RT (min)       lcorbi_t3_pos_1_029_29    m/z 793.4982     0   1   2   3E+4   13.5   14  RT (min)      lcorbi_t3_pos_2_005_11    m/z 793.5447     0   1E+5   13.5   14  RT (min)       lcorbi_t3_pos_4_067_57    m/z 794.0609     0   1   2E+5   13.5   14  RT (min)       lcorbi_t3_pos_4_076_58    m/z 794.3174     0   1E+5   8.5   9  RT (min)       lcorbi_t3_pos_4_102_61    m/z 794.4472     0   1   2E+4   9.5   10  RT (min)      lcorbi_t3_pos_4_087_59    m/z 794.4798     0   1   2   3   4E+4   14   14.5  RT (min)      lcorbi_t3_pos_3_035_08    m/z 794.5098     0   1   2E+5   13.5   14  RT (min)       lcorbi_t3_pos_4_083_59    m/z 794.5490     0   1   13.5   14  RT (min)      lcorbi_t3_pos_1_008_36    m/z 794.5974     0   1E+5   14   14.5  RT (min)      lcorbi_t3_pos_1_029_29    m/z 795.0688     0   1   2E+5   14   14.5  RT (min)       lcorbi_t3_pos_1_020_31    m/z 795.3208     0   2   4E+4   8.5   9  RT (min)      lcorbi_t3_pos_4_103_61    m/z 795.5144     0   1E+5   13.5   14  RT (min)      lcorbi_t3_pos_4_082_59    m/z 795.7618     0   1E+5   15   15.5  RT (min)       lcorbi_t3_pos_1_036_08    m/z 796.0758     0   1E+5   14   14.5  RT (min)       lcorbi_t3_pos_1_020_31    m/z 796.3238     0   1   2   3E+4   8.5   9  RT (min)      lcorbi_t3_pos_4_102_61    m/z 796.5106     0   1E+5   13.5   14  RT (min)       lcorbi_t3_pos_1_054_48    m/z 796.5985     0   2   4E+4   13.5   14  RT (min)       lcorbi_t3_pos_1_036_08    m/z 796.6351     0   1   2   3E+5   13.5   14  RT (min)      lcorbi_t3_pos_1_036_08    m/z 796.6684     0   1E+5   13   13.5  RT (min)      lcorbi_t3_pos_1_036_08    m/z 796.7060     0   1E+5   15   15.5  RT (min)       lcorbi_t3_pos_4_052_06    m/z 796.7231     0   1   2   3E+4   13.5   14  RT (min)      lcorbi_t3_pos_4_091_60    m/z 796.7392     0   1E+5   15.5   16  RT (min)       lcorbi_t3_pos_4_050_31    m/z 797.0841     0   2   4   6   8E+4   14   14.5  RT (min)       lcorbi_t3_pos_1_020_31    m/z 797.5520     0   1   2   3   4E+5   12.5   13  RT (min)       lcorbi_t3_pos_1_054_48    m/z 797.6158     0   1   2   3E+5   14   14.5  RT (min)      lcorbi_t3_pos_4_074_58    m/z 797.6365     0   1   2E+5   13.5   14  RT (min)       lcorbi_t3_pos_4_091_60    m/z 797.7367     0   1   2   3   4E+4   14   14.5  RT (min)       lcorbi_t3_pos_1_025_52    m/z 797.7427     0   2   4   6E+4   15.5   16  RT (min)      lcorbi_t3_pos_4_067_57    m/z 797.7775     0   1E+5   14.5   15  RT (min)      lcorbi_t3_pos_2_024_15    m/z 798.0929     0   1E+5   14   14.5  RT (min)       lcorbi_t3_pos_1_020_31    m/z 798.5314     0   1E+5   13.5   14  RT (min)       lcorbi_t3_pos_4_079_58    m/z 798.6137     0   1   2E+5   13.5   14  RT (min)      lcorbi_t3_pos_4_091_60    m/z 798.6556     0   1   14   14.5  RT (min)      lcorbi_t3_pos_4_097_60    m/z 798.7209     0   2   4   6E+5   14.5   15  RT (min)      lcorbi_t3_pos_4_096_60    m/z 799.1004     0   2   4   6E+4   14   14.5  RT (min)      lcorbi_t3_pos_1_020_31    m/z 799.4701     0   1E+5   13.5   14  RT (min)       lcorbi_t3_pos_4_084_59    m/z 799.5451     0   1E+5   14   14.5  RT (min)       lcorbi_t3_pos_1_020_31    m/z 799.8707     0   1E+5   15.5   16  RT (min)       lcorbi_t3_pos_4_094_60    m/z 800.6074     0   1E+5   14   14.5  RT (min)      lcorbi_t3_pos_4_074_58    m/z 801.0224     0   1E+5   13   13.5  RT (min)       lcorbi_t3_pos_1_031_40    m/z 801.0594     0   1E+4   7   7.5  RT (min)      lcorbi_t3_pos_1_037_19    m/z 802.4613     0   1E+5   13   13.5  RT (min)       lcorbi_t3_pos_4_089_59    m/z 802.5875     0   2   4E+4   13   13.5  RT (min)       lcorbi_t3_pos_1_054_48    m/z 802.6377     0   1E+5   13.5   14  RT (min)      lcorbi_t3_pos_4_077_58    m/z 802.6787     0   1   2E+4   14   14.5  RT (min)       lcorbi_t3_pos_4_093_60    m/z 803.0382     0   1E+5   13.5   14  RT (min)       lcorbi_t3_pos_4_083_59    m/z 803.3926     0   2   4   6   8E+4   15   15.5  RT (min)       lcorbi_t3_pos_4_110_62    m/z 803.4121     0   2   4   6   8E+4   13.5   14  RT (min)       lcorbi_t3_pos_4_099_61    m/z 803.4347     0   1   2   3   4E+4   14   14.5  RT (min)      lcorbi_t3_pos_4_037_26    m/z 803.4648     0   1   13   13.5  RT (min)       lcorbi_t3_pos_4_074_58    m/z 803.4711     0   1   2E+4   6   6.5  RT (min)       lcorbi_t3_pos_4_113_62    m/z 803.5064     0   1E+5   7.5   8  RT (min)      lcorbi_t3_pos_4_107_62    m/z 803.5996     0   1   2   3E+5   13   13.5  RT (min)      lcorbi_t3_pos_1_029_29    m/z 803.6198     0   1   2   3E+4   6.5   7  RT (min)       lcorbi_t3_pos_4_066_57    m/z 803.6369     0   1   2   3   4E+5   13   14  RT (min)       lcorbi_t3_pos_4_091_60    m/z 803.6678     0   1   2   3E+5   13.5   14  RT (min)       lcorbi_t3_pos_4_091_60    m/z 803.6750     0   1E+5   13.5   14  RT (min)       lcorbi_t3_pos_1_036_08    m/z 803.6916     0   1E+5   14.5   15  RT (min)      lcorbi_t3_pos_4_072_57    m/z 803.7035     0   1   2   3   4E+4   6.5   7  RT (min)       lcorbi_t3_pos_4_066_57    m/z 803.7317     0   1   2   3E+5   14   14.5  RT (min)       lcorbi_t3_pos_4_090_60    m/z 803.7871     0   2   4E+4   6.5   7  RT (min)       lcorbi_t3_pos_4_074_58    m/z 803.8706     0   1   2   3   4E+4   6.5   7  RT (min)       lcorbi_t3_pos_4_066_57    m/z 803.9542     0   1   2   3   4E+4   6.5   7  RT (min)       lcorbi_t3_pos_4_082_59    m/z 804.0376     0   1   2E+4   6.5   7  RT (min)       lcorbi_t3_pos_4_099_61    m/z 804.2927     0   1   2   3E+4   4.5   5  RT (min)       lcorbi_t3_pos_1_008_36    m/z 804.4154     0   1   2   3   4E+4   13.5   14  RT (min)       lcorbi_t3_pos_4_109_62    m/z 804.5095     0   1   2E+4   7.5   8  RT (min)      lcorbi_t3_pos_4_108_62    m/z 804.5131     0   1   2   3E+4   14   14.5  RT (min)      lcorbi_t3_pos_3_052_47    m/z 804.6414     0   1   2   3E+4   13.5   14  RT (min)      lcorbi_t3_pos_4_074_58    m/z 804.6549     0   1E+5   14   14.5  RT (min)       lcorbi_t3_pos_4_096_60    m/z 804.6669     0   1E+5   13.5   14  RT (min)       lcorbi_t3_pos_4_090_60    m/z 804.7309     0   1   2E+5   14   14.5  RT (min)       lcorbi_t3_pos_4_091_60    m/z 805.0533     0   1E+5   13.5   14  RT (min)       lcorbi_t3_pos_4_075_58    m/z 805.2960     0   1E+4   4.5   5  RT (min)      lcorbi_t3_pos_1_012_14    m/z 805.4800     0   1E+5   13.5   14  RT (min)       lcorbi_t3_pos_1_054_48    m/z 805.6908     0   1E+5   13.5   14  RT (min)       lcorbi_t3_pos_4_090_60    m/z 806.0613     0   2   4   6   8E+4   13.5   14   14.5  RT (min)       lcorbi_t3_pos_4_076_58    m/z 806.4456     0   2   4   6E+4   14   14.5  RT (min)       lcorbi_t3_pos_4_098_61    m/z 806.4949     0   1E+5   13   13.5  RT (min)       lcorbi_t3_pos_4_083_59    m/z 806.7060     0   1   2   3E+5   13.5   14  RT (min)       lcorbi_t3_pos_4_091_60    m/z 806.7236     0   1E+5   15.5   16  RT (min)       lcorbi_t3_pos_2_008_16    m/z 807.0686     0   2   4   6   8E+4   14   14.5  RT (min)       lcorbi_t3_pos_4_075_58    m/z 807.4476     0   1   2   3E+4   14   14.5  RT (min)      lcorbi_t3_pos_4_035_17    m/z 807.4991     0   2   4   6   8E+4   13   13.5  RT (min)      lcorbi_t3_pos_4_084_59    m/z 807.5563     0   1E+4   13   13.5  RT (min)       lcorbi_t3_pos_1_054_48    m/z 807.7731     0   1E+5   15   15.5  RT (min)       lcorbi_t3_pos_2_024_15    m/z 808.0757     0   2   4   6   8E+4   14   14.5  RT (min)       lcorbi_t3_pos_1_019_16    m/z 808.3332     0   1   2E+4   9   9.5  RT (min)      lcorbi_t3_pos_4_112_62    m/z 808.5116     0   2   4   6   8E+4   13.5   14  RT (min)       lcorbi_t3_pos_1_054_48    m/z 808.6391     0   2   4   6E+4   14   14.5  RT (min)       lcorbi_t3_pos_1_007_07    m/z 808.7393     0   1E+5   15.5   16  RT (min)       lcorbi_t3_pos_2_008_16    m/z 809.0848     0   1E+5   14   14.5  RT (min)       lcorbi_t3_pos_4_075_58    m/z 809.7888     0   1   2   3E+4   14.5   15  RT (min)      lcorbi_t3_pos_4_095_60    m/z 810.0926     0   1   2E+5   14   14.5  RT (min)       lcorbi_t3_pos_1_035_28    m/z 810.5264     0   1   2E+5   13.5   14  RT (min)       lcorbi_t3_pos_1_054_48    m/z 810.5424     0   1   2E+4   14   14.5  RT (min)      lcorbi_t3_pos_4_078_58    m/z 810.5553     0   1   2E+5   14   14.5  RT (min)       lcorbi_t3_pos_1_008_36    m/z 810.5627     0   1   2E+5   14   14.5  RT (min)       lcorbi_t3_pos_1_007_07    m/z 810.6577     0   1   2   3   4E+5   14   14.5  RT (min)      lcorbi_t3_pos_1_023_02    m/z 810.7549     0   2   4   6E+4   15.5   16  RT (min)       lcorbi_t3_pos_2_008_16    m/z 811.0991     0   1E+5   14   14.5  RT (min)       lcorbi_t3_pos_4_073_57    m/z 811.5589     0   1E+5   14   14.5  RT (min)      lcorbi_t3_pos_1_007_07    m/z 811.6510     0   2   4   6   8E+4   14   14.5  RT (min)       lcorbi_t3_pos_1_002_01    m/z 811.6874     0   1E+5   13.5   14  RT (min)      lcorbi_t3_pos_1_036_08    m/z 812.4949     0   2   4   6   8E+4   15   15.5  RT (min)       lcorbi_t3_pos_4_066_57    m/z 812.5290     0   1E+5   14   14.5  RT (min)      lcorbi_t3_pos_4_077_58    m/z 812.5806     0   1   2E+5   13.5   14  RT (min)       lcorbi_t3_pos_4_074_58    m/z 813.4915     0   2   4   6E+4   13   13.5  RT (min)       lcorbi_t3_pos_4_096_60    m/z 813.5840     0   1E+5   13.5   14  RT (min)       lcorbi_t3_pos_4_081_58    m/z 813.6504     0   2   4   6   8E+5   13   14  RT (min)       lcorbi_t3_pos_1_043_06    m/z 813.6590     0   1E+5   15.5   16  RT (min)      lcorbi_t3_pos_4_082_59    m/z 813.7022     0   1E+5   13.5   14  RT (min)       lcorbi_t3_pos_1_036_08    m/z 814.1208     0   1E+5   13.5   14  RT (min)      lcorbi_t3_pos_1_016_05    m/z 814.5958     0   1   2E+5   13.5   14  RT (min)       lcorbi_t3_pos_4_074_58    m/z 814.6148     0   2   4E+5   14   14.5  RT (min)       lcorbi_t3_pos_4_075_58    m/z 814.6623     0   1   2E+5   13   14  RT (min)       lcorbi_t3_pos_1_003_45    m/z 815.6268     0   2   4   6   8E+5   14   14.5  RT (min)       lcorbi_t3_pos_4_074_58    m/z 815.7192     0   2   4   6   8E+4   13.5   14  RT (min)       lcorbi_t3_pos_4_092_60    m/z 816.1361     0   2   4   6E+4   13.5   14  RT (min)      lcorbi_t3_pos_1_016_05    m/z 816.5889     0   1   2E+5   14   14.5  RT (min)       lcorbi_t3_pos_4_075_58    m/z 816.6120     0   1   2E+5   13.5   14  RT (min)       lcorbi_t3_pos_4_077_58    m/z 816.6299     0   1   2E+5   14   14.5  RT (min)       lcorbi_t3_pos_1_032_26    m/z 816.6532     0   1   13.5   14  RT (min)       lcorbi_t3_pos_1_020_31    m/z 816.7416     0   1   2E+5   14   14.5  RT (min)       lcorbi_t3_pos_4_096_60    m/z 817.4984     0   1   2E+5   13   13.5   14  RT (min)      lcorbi_t3_pos_1_032_26    m/z 817.6008     0   1E+5   13.5   14  RT (min)       lcorbi_t3_pos_4_076_58    m/z 817.6154     0   1E+5   13.5   14  RT (min)       lcorbi_t3_pos_4_080_58    m/z 817.6419     0   1E+5   14   14.5  RT (min)      lcorbi_t3_pos_4_067_57    m/z 817.6815     0   1   2   3E+5   13   13.5   14  RT (min)       lcorbi_t3_pos_4_097_60    m/z 817.7455     0   1   2   3   4E+5   14   14.5  RT (min)       lcorbi_t3_pos_4_096_60    m/z 818.5008     0   1E+5   13   13.5   14  RT (min)      lcorbi_t3_pos_4_087_59    m/z 818.5264     0   1   2   3   4E+4   13   13.5  RT (min)      lcorbi_t3_pos_2_005_11    m/z 818.5683     0   2   4   6   8E+4   14   14.5  RT (min)       lcorbi_t3_pos_1_029_29    m/z 818.6712     0   1E+5   13.5   14  RT (min)       lcorbi_t3_pos_4_097_60    m/z 818.7148     0   1   2E+5   15   15.5  RT (min)      lcorbi_t3_pos_3_020_35    m/z 819.4072     0   2   4   6E+4   13.5   14  RT (min)       lcorbi_t3_pos_4_101_61    m/z 819.5320     0   1   15   15.5  RT (min)      lcorbi_t3_pos_3_027_27    m/z 820.5471     0   1E+5   13   13.5  RT (min)       lcorbi_t3_pos_4_077_58    m/z 820.5850     0   1   2   3E+5   14   14.5  RT (min)       lcorbi_t3_pos_1_029_29    m/z 820.7858     0   1   2E+5   15   15.5  RT (min)       lcorbi_t3_pos_1_001_49    m/z 821.0846     0   2   4   6   8E+4   14   14.5  RT (min)       lcorbi_t3_pos_4_074_58    m/z 821.5094     0   1E+5   13.5   14  RT (min)       lcorbi_t3_pos_4_067_57    m/z 821.5303     0   1E+5   13.5   14  RT (min)      lcorbi_t3_pos_4_082_59    m/z 821.5504     0   2   4E+4   13   13.5  RT (min)       lcorbi_t3_pos_2_040_25    m/z 821.5890     0   1E+5   14   14.5  RT (min)       lcorbi_t3_pos_1_014_03    m/z 822.0924     0   2   4   6   8E+4   14   14.5  RT (min)       lcorbi_t3_pos_4_075_58    m/z 822.4406     0   2   4E+4   13.5   14  RT (min)      lcorbi_t3_pos_4_099_61    m/z 822.5220     0   1E+5   13   13.5  RT (min)       lcorbi_t3_pos_4_077_58    m/z 822.5417     0   1E+5   14   14.5  RT (min)       lcorbi_t3_pos_4_089_59    m/z 822.5990     0   1E+5   14   14.5  RT (min)       lcorbi_t3_pos_1_029_29    m/z 822.6393     0   1E+6   13.5   14  RT (min)       lcorbi_t3_pos_4_072_57    m/z 822.7319     0   1   2E+5   13.5   14  RT (min)      lcorbi_t3_pos_1_050_13    m/z 822.7462     0   1   2   3E+5   15.5   16  RT (min)      lcorbi_t3_pos_4_066_57    m/z 823.0992     0   2   4   6E+4   14   14.5  RT (min)       lcorbi_t3_pos_1_020_31    m/z 823.4822     0   2   4   6   8E+4   7.5   8  RT (min)       lcorbi_t3_pos_4_106_62    m/z 823.5456     0   2   4   6   8E+4   14   14.5  RT (min)       lcorbi_t3_pos_4_084_59    m/z 823.6050     0   1E+5   14   14.5  RT (min)       lcorbi_t3_pos_4_076_58    m/z 823.6520     0   1E+5   13.5   14  RT (min)       lcorbi_t3_pos_4_091_60    m/z 823.6664     0   1   2E+4   14.5   15  RT (min)      lcorbi_t3_pos_4_052_06    m/z 823.6790     0   1E+5   15.5   16  RT (min)       lcorbi_t3_pos_4_066_57    m/z 823.8043     0   1   2   3E+5   14.5   15  RT (min)      lcorbi_t3_pos_1_001_49    m/z 824.4725     0   2   4   6E+4   14   14.5  RT (min)      lcorbi_t3_pos_4_098_61    m/z 824.4856     0   1   2   3E+4   7.5   8  RT (min)      lcorbi_t3_pos_4_108_62    m/z 824.5281     0   1E+5   13.5   14  RT (min)       lcorbi_t3_pos_4_066_57    m/z 824.5656     0   1E+5   13.5   14  RT (min)       lcorbi_t3_pos_4_082_59    m/z 824.6439     0   2   4   6E+5   13.5   14  RT (min)       lcorbi_t3_pos_1_003_45    m/z 824.6824     0   1   15.5   16  RT (min)      lcorbi_t3_pos_1_036_08    m/z 824.7605     0   1   2   3   4E+4   13.5   14  RT (min)      lcorbi_t3_pos_4_095_60    m/z 824.7619     0   1E+5   15.5   16  RT (min)       lcorbi_t3_pos_4_067_57    m/z 824.7704     0   1E+5   15.5   16  RT (min)       lcorbi_t3_pos_4_067_57    m/z 825.3943     0   2   4   6E+4   13.5   14  RT (min)       lcorbi_t3_pos_4_099_61    m/z 825.5610     0   1E+5   14   14.5  RT (min)      lcorbi_t3_pos_1_014_03    m/z 825.5835     0   1   2E+5   13   13.5  RT (min)       lcorbi_t3_pos_4_077_58    m/z 825.6461     0   1   2   3   4E+4   14   14.5  RT (min)      lcorbi_t3_pos_1_034_34    m/z 825.6846     0   2   4   6E+4   13   14  RT (min)       lcorbi_t3_pos_4_072_57    m/z 825.7740     0   2   4   6   8E+4   15.5   16  RT (min)       lcorbi_t3_pos_4_050_31    m/z 826.5721     0   2   4   6   8E+4   14   14.5  RT (min)       lcorbi_t3_pos_1_034_34    m/z 826.6881     0   1   2E+4   13   14  RT (min)       lcorbi_t3_pos_3_010_44    m/z 826.7765     0   1   13.5   14  RT (min)      lcorbi_t3_pos_4_091_60    m/z 827.6357     0   1E+5   14   14.5  RT (min)       lcorbi_t3_pos_4_095_60    m/z 827.6729     0   1E+5   14.5   15  RT (min)      lcorbi_t3_pos_4_070_57    m/z 827.7104     0   1E+5   15.5   16  RT (min)       lcorbi_t3_pos_4_073_57    m/z 828.4440     0   2   4   6E+4   11   11.5  RT (min)      lcorbi_t3_pos_4_074_58    m/z 828.6109     0   1   2E+5   13   13.5  RT (min)       lcorbi_t3_pos_4_078_58    m/z 828.7042     0   1   14.5   15  RT (min)      lcorbi_t3_pos_4_092_60    m/z 829.6152     0   1   2E+4   13   13.5  RT (min)       lcorbi_t3_pos_1_003_45    m/z 829.6553     0   2   4   6   8E+4   14   14.5  RT (min)       lcorbi_t3_pos_1_040_27    m/z 829.7157     0   2   4   6   8E+4   14   15  RT (min)       lcorbi_t3_pos_4_094_60    m/z 830.6657     0   1   2   3E+4   14.5   15  RT (min)       lcorbi_t3_pos_4_093_60    m/z 830.7239     0   1E+5   15.5   16  RT (min)       lcorbi_t3_pos_4_087_59    m/z 831.4239     0   2   4E+4   15   15.5  RT (min)      lcorbi_t3_pos_4_109_62    m/z 831.6650     0   1E+5   13.5   14  RT (min)       lcorbi_t3_pos_1_041_54    m/z 831.7302     0   1   2   3E+5   15   15.5  RT (min)       lcorbi_t3_pos_4_091_60    m/z 831.7499     0   1E+5   14.5   15  RT (min)      lcorbi_t3_pos_3_025_52    m/z 833.6798     0   1   2E+5   13.5   14  RT (min)       lcorbi_t3_pos_4_093_60    m/z 833.7427     0   1E+5   15.5   16  RT (min)       lcorbi_t3_pos_1_032_26    m/z 834.5865     0   1   13.5   14  RT (min)       lcorbi_t3_pos_1_054_48    m/z 834.6149     0   2   4   6E+4   10.5   11  RT (min)       lcorbi_t3_pos_4_082_59    m/z 834.6356     0   1E+5   13.5   14  RT (min)       lcorbi_t3_pos_1_036_08    m/z 835.1903     0   1   2E+5   13.5   14  RT (min)       lcorbi_t3_pos_1_036_08    m/z 835.5343     0   1E+5   13.5   14  RT (min)       lcorbi_t3_pos_4_084_59    m/z 835.6572     0   2   4   6   8E+4   14   14.5  RT (min)       lcorbi_t3_pos_4_074_58    m/z 835.7585     0   1E+5   15.5   16  RT (min)       lcorbi_t3_pos_2_008_16    m/z 836.4946     0   2   4E+4   10.5   11  RT (min)      lcorbi_t3_pos_1_054_48    m/z 836.5372     0   2   4E+4   13.5   14  RT (min)      lcorbi_t3_pos_2_005_11    m/z 836.5805     0   2   4   6   8E+4   13.5   14  RT (min)       lcorbi_t3_pos_4_074_58    m/z 836.6509     0   1E+5   13.5   14  RT (min)       lcorbi_t3_pos_4_091_60    m/z 836.7704     0   1E+5   15.5   16  RT (min)       lcorbi_t3_pos_2_008_16    m/z 836.7881     0   1E+5   15   15.5  RT (min)      lcorbi_t3_pos_1_036_08    m/z 837.3311     0   2   4   6E+4   8.5   9  RT (min)       lcorbi_t3_pos_4_112_62    m/z 837.6649     0   2   4   6E+4   14   14.5  RT (min)      lcorbi_t3_pos_4_092_60    m/z 837.6741     0   1E+5   14.5   15  RT (min)       lcorbi_t3_pos_4_068_57    m/z 837.6844     0   2   4   6E+4   14   14.5  RT (min)       lcorbi_t3_pos_2_007_02    m/z 837.7114     0   1   2   3   4E+5   14   14.5  RT (min)       lcorbi_t3_pos_4_090_60    m/z 837.7739     0   2   4   6   8E+4   15.5   16  RT (min)      lcorbi_t3_pos_2_008_16    m/z 837.9630     0   1   2   3E+4   4.5   5  RT (min)       lcorbi_t3_pos_4_106_62    m/z 838.2367     0   1   2E+5   14.5   15  RT (min)       lcorbi_t3_pos_4_105_61    m/z 838.3343     0   1   2E+4   8.5   9  RT (min)      lcorbi_t3_pos_4_109_62    m/z 838.6863     0   2   4E+4   14.5   15  RT (min)      lcorbi_t3_pos_4_069_57    m/z 839.7009     0   2   4   6   8E+4   14.5   15  RT (min)       lcorbi_t3_pos_4_068_57    m/z 840.6116     0   1E+5   13.5   14  RT (min)       lcorbi_t3_pos_4_074_58    m/z 841.6155     0   2   4   6   8E+4   13.5   14  RT (min)       lcorbi_t3_pos_4_080_58    m/z 841.6420     0   1E+5   14   14.5  RT (min)       lcorbi_t3_pos_4_068_57    m/z 841.6507     0   1E+5   14   14.5  RT (min)       lcorbi_t3_pos_4_067_57    m/z 841.7366     0   1   2E+5   13.5   14  RT (min)       lcorbi_t3_pos_4_091_60    m/z 842.5953     0   2   4   6   8E+4   13   13.5  RT (min)       lcorbi_t3_pos_4_090_60    m/z 842.6617     0   1   2E+5   13   13.5  RT (min)       lcorbi_t3_pos_1_036_08    m/z 842.6639     0   2   4   6   8E+4   14.5   15  RT (min)       lcorbi_t3_pos_4_072_57    m/z 843.4435     0   2   4   6E+4   14   14.5  RT (min)       lcorbi_t3_pos_4_101_61    m/z 843.4809     0   1E+5   15   15.5  RT (min)       lcorbi_t3_pos_4_069_57    m/z 843.4915     0   2   4   6E+4   13.5   14  RT (min)       lcorbi_t3_pos_4_070_57    m/z 843.6078     0   2   4   6   8E+4   13.5   14  RT (min)       lcorbi_t3_pos_1_003_45    m/z 843.7523     0   1   2E+5   14   14.5  RT (min)       lcorbi_t3_pos_4_094_60    m/z 844.6195     0   2   4   6   8E+4   14   14.5  RT (min)       lcorbi_t3_pos_4_067_57    m/z 845.5068     0   1E+5   13.5   14  RT (min)       lcorbi_t3_pos_4_066_57    m/z 845.7679     0   1E+5   14   14.5  RT (min)       lcorbi_t3_pos_4_093_60    m/z 846.4542     0   2   4   6   8E+4   14   14.5  RT (min)       lcorbi_t3_pos_4_099_61    m/z 846.4766     0   1   2   3   4E+4   14   14.5  RT (min)      lcorbi_t3_pos_2_047_31    m/z 846.5122     0   2   4   6   8E+4   13.5   14  RT (min)       lcorbi_t3_pos_4_083_59    m/z 846.5577     0   2   4   6   8E+4   13   13.5  RT (min)       lcorbi_t3_pos_4_074_58    m/z 847.4576     0   1   2   3   4E+4   14   14.5  RT (min)       lcorbi_t3_pos_4_098_61    m/z 847.6407     0   2   4   6   8E+4   14   14.5  RT (min)       lcorbi_t3_pos_4_067_57    m/z 847.7493     0   2   4E+5   15.5   16  RT (min)       lcorbi_t3_pos_4_084_59    m/z 848.5575     0   1E+5   14   14.5  RT (min)       lcorbi_t3_pos_1_029_29    m/z 848.6356     0   1E+5   13   13.5  RT (min)       lcorbi_t3_pos_4_091_60    m/z 848.6823     0   1E+5   15.5   16  RT (min)       lcorbi_t3_pos_3_027_27    m/z 849.5602     0   2   4   6   8E+4   14   14.5  RT (min)       lcorbi_t3_pos_1_035_28    m/z 850.3435     0   1E+5   9   9.5  RT (min)       lcorbi_t3_pos_4_111_62    m/z 850.5449     0   2   4   6   8E+4   13   13.5  RT (min)       lcorbi_t3_pos_4_087_59    m/z 850.5719     0   1E+5   14   14.5  RT (min)       lcorbi_t3_pos_4_081_58    m/z 850.5961     0   1   2E+5   13   13.5  RT (min)       lcorbi_t3_pos_4_075_58    m/z 850.6310     0   2   4E+4   14   14.5  RT (min)       lcorbi_t3_pos_1_014_03    m/z 850.6520     0   1E+5   13.5   14  RT (min)       lcorbi_t3_pos_4_090_60    m/z 850.6601     0   2   4E+5   14   14.5  RT (min)      lcorbi_t3_pos_4_067_57    m/z 851.3469     0   1E+5   9   9.5  RT (min)       lcorbi_t3_pos_4_107_62    m/z 851.5996     0   1E+5   13   13.5  RT (min)      lcorbi_t3_pos_1_029_29    m/z 851.7015     0   1   2E+5   15.5   16  RT (min)      lcorbi_t3_pos_4_089_59    m/z 851.7215     0   1   2E+5   13.5   14  RT (min)       lcorbi_t3_pos_4_092_60    m/z 851.7806     0   1   2E+5   15.5   16  RT (min)       lcorbi_t3_pos_4_073_57    m/z 852.3501     0   1   2E+4   9   9.5  RT (min)      lcorbi_t3_pos_4_113_62    m/z 852.6112     0   2   4   6E+4   13   13.5  RT (min)       lcorbi_t3_pos_1_031_40    m/z 852.6676     0   1   2   3E+5   13.5   14  RT (min)       lcorbi_t3_pos_4_092_60    m/z 853.5638     0   1   2E+5   13.5   14  RT (min)       lcorbi_t3_pos_4_082_59    m/z 853.7171     0   1E+5   15.5   16  RT (min)      lcorbi_t3_pos_4_083_59    m/z 853.7370     0   1   2E+5   13.5   14  RT (min)       lcorbi_t3_pos_4_091_60    m/z 855.5740     0   1E+5   13.5   14  RT (min)      lcorbi_t3_pos_1_054_48    m/z 855.7033     0   1E+5   14.5   15  RT (min)       lcorbi_t3_pos_4_067_57    m/z 855.7269     0   1E+5   15.5   16  RT (min)       lcorbi_t3_pos_1_032_26    m/z 856.5833     0   1E+5   14   14.5  RT (min)       lcorbi_t3_pos_1_029_29    m/z 856.6428     0   2   4   6   8E+4   14   14.5  RT (min)       lcorbi_t3_pos_4_066_57    m/z 856.7154     0   2   4   6E+4   14.5   15  RT (min)       lcorbi_t3_pos_4_068_57    m/z 857.3474     0   2   4E+4   8.5   9  RT (min)       lcorbi_t3_pos_4_108_62    m/z 857.5155     0   1E+5   13.5   14  RT (min)       lcorbi_t3_pos_4_083_59    m/z 857.5874     0   2   4   6E+4   14   14.5  RT (min)      lcorbi_t3_pos_1_048_47    m/z 858.4515     0   2   4E+4   15   15.5  RT (min)       lcorbi_t3_pos_4_070_57    m/z 858.4796     0   1E+5   13   13.5  RT (min)       lcorbi_t3_pos_4_082_59    m/z 859.6027     0   2   4   6E+4   14   14.5  RT (min)       lcorbi_t3_pos_1_019_16    m/z 859.6388     0   1E+5   13   13.5  RT (min)       lcorbi_t3_pos_1_036_08    m/z 859.7833     0   2   4   6   8E+4   14   14.5  RT (min)       lcorbi_t3_pos_4_095_60    m/z 860.7340     0   1E+5   14.5   15  RT (min)       lcorbi_t3_pos_3_008_20    m/z 861.6180     0   2   4   6   8E+4   14   14.5  RT (min)       lcorbi_t3_pos_4_073_57    m/z 861.6554     0   2   4   6   8E+4   13.5   14  RT (min)       lcorbi_t3_pos_1_036_08    m/z 861.6947     0   2   4   6   8E+4   15.5   16  RT (min)      lcorbi_t3_pos_4_082_59    m/z 862.4491     0   2   4   6E+4   13.5   14  RT (min)       lcorbi_t3_pos_4_106_62    m/z 862.6259     0   2   4   6E+4   13.5   14  RT (min)       lcorbi_t3_pos_4_070_57    m/z 862.6320     0   1E+5   14   14.5  RT (min)       lcorbi_t3_pos_4_072_57    m/z 863.5646     0   2   4   6E+4   13.5   14  RT (min)       lcorbi_t3_pos_4_072_57    m/z 863.6362     0   2   4   6   8E+4   14   14.5  RT (min)       lcorbi_t3_pos_4_070_57    m/z 863.7104     0   1E+5   15.5   16  RT (min)       lcorbi_t3_pos_4_089_59    m/z 863.7531     0   2   4   6   8E+4   15   15.5  RT (min)      lcorbi_t3_pos_1_031_40    m/z 864.5035     0   2   4   6E+4   14   14.5  RT (min)       lcorbi_t3_pos_4_087_59    m/z 864.6733     0   1   2E+5   13.5   14  RT (min)       lcorbi_t3_pos_4_091_60    m/z 864.7649     0   2   4   6   8E+4   15   15.5  RT (min)       lcorbi_t3_pos_3_020_35    m/z 864.7934     0   2   4   6E+4   15.5   16  RT (min)      lcorbi_t3_pos_4_086_59    m/z 864.8404     0   1E+5   15   15.5  RT (min)       lcorbi_t3_pos_1_036_08    m/z 865.6514     0   1E+5   14   14.5  RT (min)       lcorbi_t3_pos_4_072_57    m/z 865.6684     0   1E+5   15.5   16  RT (min)       lcorbi_t3_pos_3_020_35    m/z 865.7112     0   1   2   3   4E+5   15   15.5  RT (min)       lcorbi_t3_pos_3_027_27    m/z 865.7259     0   1E+5   15.5   16  RT (min)       lcorbi_t3_pos_4_082_59    m/z 866.6636     0   2   4   6   8E+4   14.5   15  RT (min)       lcorbi_t3_pos_4_068_57    m/z 866.7061     0   1E+5   13.5   14  RT (min)       lcorbi_t3_pos_4_090_60    m/z 866.7289     0   1E+6   15.5   16  RT (min)      lcorbi_t3_pos_4_084_59    m/z 866.8365     0   1E+5   15.5   16  RT (min)       lcorbi_t3_pos_1_036_08    m/z 867.6842     0   1E+5   15.5   16  RT (min)       lcorbi_t3_pos_1_031_40    m/z 867.8791     0   1E+5   15   15.5  RT (min)       lcorbi_t3_pos_2_024_15    m/z 868.5092     0   1E+5   13   13.5  RT (min)       lcorbi_t3_pos_4_083_59    m/z 868.6876     0   2   4   6E+4   15.5   16  RT (min)      lcorbi_t3_pos_1_028_32    m/z 869.3574     0   1   2E+5   9   9.5  RT (min)       lcorbi_t3_pos_4_107_62    m/z 869.3737     0   2   4   6   8E+4   14   14.5  RT (min)       lcorbi_t3_pos_4_110_62    m/z 869.6998     0   2   4   6   8E+4   15.5   16  RT (min)       lcorbi_t3_pos_4_066_57    m/z 869.8947     0   1   2E+5   15.5   16  RT (min)       lcorbi_t3_pos_1_036_08    m/z 870.3604     0   2   4   6E+4   9   9.5  RT (min)      lcorbi_t3_pos_4_108_62    m/z 870.5089     0   2   4   6E+4   15.5   16  RT (min)       lcorbi_t3_pos_4_107_62    m/z 870.6950     0   2   4   6   8E+4   14.5   15  RT (min)      lcorbi_t3_pos_1_031_40    m/z 871.3648     0   1E+5   8.5   9  RT (min)       lcorbi_t3_pos_1_014_03    m/z 871.9103     0   1   2E+5   15.5   16  RT (min)       lcorbi_t3_pos_4_091_60    m/z 872.3487     0   2   4   6E+4   8.5   9  RT (min)       lcorbi_t3_pos_4_108_62    m/z 872.3683     0   2   4   6E+4   8.5   9  RT (min)       lcorbi_t3_pos_1_014_03    m/z 872.6530     0   1E+5   14   14.5  RT (min)       lcorbi_t3_pos_4_068_57    m/z 872.6822     0   1E+5   15   15.5  RT (min)       lcorbi_t3_pos_4_086_59    m/z 873.3520     0   1   2   3   4E+4   8.5   9  RT (min)       lcorbi_t3_pos_4_108_62    m/z 873.3713     0   1   2   3E+4   8.5   9  RT (min)      lcorbi_t3_pos_1_014_03    m/z 873.6566     0   2   4   6E+4   14   14.5  RT (min)      lcorbi_t3_pos_4_066_57    m/z 875.6723     0   1E+5   14.5   15  RT (min)       lcorbi_t3_pos_4_069_57    m/z 876.3203     0   1E+5   8.5   9  RT (min)       lcorbi_t3_pos_4_109_62    m/z 876.5847     0   1   2   3E+4   6.5   7  RT (min)       lcorbi_t3_pos_4_102_61    m/z 876.6763     0   1   2   3E+4   6.5   7  RT (min)       lcorbi_t3_pos_4_066_57    m/z 876.7081     0   1   2   3   4E+5   15   15.5  RT (min)       lcorbi_t3_pos_1_032_26    m/z 876.7651     0   2   4   6E+4   15   15.5  RT (min)       lcorbi_t3_pos_4_075_58    m/z 876.7675     0   2   4E+4   6.5   7  RT (min)       lcorbi_t3_pos_4_106_62    m/z 876.7835     0   1   2E+5   15.5   16  RT (min)      lcorbi_t3_pos_4_085_59    m/z 876.8479     0   1E+5   15   15.5  RT (min)       lcorbi_t3_pos_1_036_08    m/z 876.8584     0   1   2   3   4E+4   6.5   7  RT (min)       lcorbi_t3_pos_4_106_62    m/z 876.9495     0   1   2   3E+4   6.5   7  RT (min)       lcorbi_t3_pos_4_106_62    m/z 877.3237     0   2   4   6E+4   8.5   9  RT (min)       lcorbi_t3_pos_4_106_62    m/z 877.5636     0   2   4   6   8E+4   13.5   14  RT (min)       lcorbi_t3_pos_4_072_57    m/z 877.6746     0   1E+5   13.5   14  RT (min)       lcorbi_t3_pos_4_094_60    m/z 878.3266     0   1   2   3E+4   8.5   9  RT (min)      lcorbi_t3_pos_4_107_62    m/z 878.5755     0   2   4   6   8E+4   13.5   14  RT (min)       lcorbi_t3_pos_4_067_57    m/z 879.8117     0   1E+5   15.5   16  RT (min)       lcorbi_t3_pos_4_067_57    m/z 880.6935     0   1   2   3   4E+5   13.5   14  RT (min)       lcorbi_t3_pos_4_091_60    m/z 881.5157     0   2   4   6E+4   13.5   14  RT (min)       lcorbi_t3_pos_4_071_57    m/z 881.5948     0   1E+5   13.5   14  RT (min)      lcorbi_t3_pos_4_078_58    m/z 881.7067     0   1E+5   14.5   15  RT (min)       lcorbi_t3_pos_4_103_61    m/z 882.7246     0   2   4   6E+4   15.5   16  RT (min)       lcorbi_t3_pos_4_071_57    m/z 882.7337     0   2   4   6E+4   15   15.5  RT (min)      lcorbi_t3_pos_3_020_35    m/z 883.7369     0   2   4E+4   15   15.5  RT (min)      lcorbi_t3_pos_3_039_03    m/z 884.4313     0   1   2   3   4E+4   13.5   14  RT (min)      lcorbi_t3_pos_4_098_61    m/z 884.7343     0   2   4   6E+4   15   15.5  RT (min)       lcorbi_t3_pos_3_024_07    m/z 885.5472     0   2   4   6   8E+4   13.5   14  RT (min)       lcorbi_t3_pos_4_077_58    m/z 885.6945     0   1E+5   15.5   16  RT (min)       lcorbi_t3_pos_4_088_59    m/z 886.3645     0   1   2   3   4E+4   8.5   9  RT (min)       lcorbi_t3_pos_4_106_62    m/z 886.4004     0   2   4   6   8E+4   14   14.5  RT (min)       lcorbi_t3_pos_4_111_62    m/z 886.4855     0   2   4   6E+4   14   14.5  RT (min)       lcorbi_t3_pos_4_084_59    m/z 886.6979     0   2   4   6E+4   15.5   16  RT (min)       lcorbi_t3_pos_4_085_59    m/z 887.3678     0   1   2E+4   8.5   9  RT (min)      lcorbi_t3_pos_4_109_62    m/z 887.3870     0   1E+5   9   9.5  RT (min)      lcorbi_t3_pos_1_014_03    m/z 887.5650     0   1E+5   13.5   14  RT (min)       lcorbi_t3_pos_1_044_25    m/z 888.3778     0   1   2   3E+4   13   13.5  RT (min)      lcorbi_t3_pos_3_048_30    m/z 888.3902     0   1   2   3E+4   9   9.5  RT (min)       lcorbi_t3_pos_4_110_62    m/z 888.5681     0   1E+5   13.5   14  RT (min)      lcorbi_t3_pos_4_073_57    m/z 888.7137     0   1E+5   15.5   16  RT (min)      lcorbi_t3_pos_4_089_59    m/z 889.3752     0   2   4   6E+4   8.5   9  RT (min)       lcorbi_t3_pos_4_111_62    m/z 890.3360     0   2   4   6E+4   9   9.5  RT (min)       lcorbi_t3_pos_1_030_21    m/z 890.3787     0   1   2   3E+4   8.5   9  RT (min)      lcorbi_t3_pos_4_108_62    m/z 890.6720     0   2   4   6   8E+4   15.5   16  RT (min)       lcorbi_t3_pos_1_008_36    m/z 891.3393     0   1   2   3   4E+4   9   9.5  RT (min)      lcorbi_t3_pos_1_030_21    m/z 891.7414     0   1   2E+5   15.5   16  RT (min)       lcorbi_t3_pos_4_084_59    m/z 892.2943     0   1   2   3   4E+4   8.5   9  RT (min)       lcorbi_t3_pos_4_107_62    m/z 892.6876     0   2   4   6   8E+4   15.5   16  RT (min)       lcorbi_t3_pos_4_005_33    m/z 893.2978     0   1   2   3E+4   8.5   9  RT (min)      lcorbi_t3_pos_4_108_62    m/z 893.5849     0   1E+5   15.5   16  RT (min)       lcorbi_t3_pos_1_017_51    m/z 893.6257     0   2   4   6E+4   14.5   15  RT (min)       lcorbi_t3_pos_4_068_57    m/z 893.6891     0   1   2E+5   13   13.5  RT (min)       lcorbi_t3_pos_4_096_60    m/z 893.7207     0   1E+5   15   15.5  RT (min)       lcorbi_t3_pos_4_081_58    m/z 893.7568     0   1E+5   15.5   16  RT (min)       lcorbi_t3_pos_4_088_59    m/z 893.7996     0   1E+5   15   15.5  RT (min)       lcorbi_t3_pos_4_086_59    m/z 893.8858     0   1E+5   15   15.5  RT (min)       lcorbi_t3_pos_1_036_08    m/z 894.3306     0   2   4   6   8E+4   8.5   9  RT (min)       lcorbi_t3_pos_4_108_62    m/z 894.7033     0   1E+5   15.5   16  RT (min)       lcorbi_t3_pos_2_020_22    m/z 894.7243     0   1E+5   15   15.5  RT (min)       lcorbi_t3_pos_4_076_58    m/z 895.3342     0   2   4E+4   8.5   9  RT (min)       lcorbi_t3_pos_4_106_62    m/z 895.7362     0   1E+5   15   15.5  RT (min)       lcorbi_t3_pos_4_078_58    m/z 896.6822     0   1E+5   15   15.5  RT (min)       lcorbi_t3_pos_4_083_59    m/z 896.7168     0   1   2   3E+5   13.5   14  RT (min)       lcorbi_t3_pos_4_091_60    m/z 896.7397     0   2   4   6   8E+4   15   15.5  RT (min)      lcorbi_t3_pos_4_080_58    m/z 897.7308     0   1E+5   15.5   16  RT (min)       lcorbi_t3_pos_4_066_57    m/z 898.4876     0   1E+5   13   13.5  RT (min)       lcorbi_t3_pos_4_083_59    m/z 898.6663     0   2   4E+4   14.5   15  RT (min)      lcorbi_t3_pos_4_073_57    m/z 898.7346     0   2   4E+4   15.5   16  RT (min)       lcorbi_t3_pos_4_071_57    m/z 898.7680     0   1E+5   15.5   16  RT (min)       lcorbi_t3_pos_4_087_59    m/z 899.6569     0   1E+5   13.5   14  RT (min)       lcorbi_t3_pos_4_092_60    m/z 900.6844     0   1E+5   14.5   15  RT (min)       lcorbi_t3_pos_3_027_27    m/z 900.7652     0   1E+5   15   15.5  RT (min)       lcorbi_t3_pos_4_078_58    m/z 900.7837     0   1E+5   15.5   16  RT (min)       lcorbi_t3_pos_4_085_59    m/z 900.8568     0   1E+5   15.5   16  RT (min)       lcorbi_t3_pos_2_024_15    m/z 900.9449     0   1   2E+5   15.5   16  RT (min)       lcorbi_t3_pos_4_097_60    m/z 901.6880     0   1   2   3E+5   14.5   15  RT (min)      lcorbi_t3_pos_1_050_13    m/z 901.7687     0   2   4   6   8E+4   15   15.5  RT (min)       lcorbi_t3_pos_3_038_18    m/z 902.3192     0   1E+5   14.5   15  RT (min)       lcorbi_t3_pos_4_105_61    m/z 902.5757     0   1E+5   13.5   14  RT (min)       lcorbi_t3_pos_4_073_57    m/z 902.6266     0   2   4   6   8E+4   13   13.5  RT (min)       lcorbi_t3_pos_4_095_60    m/z 902.6764     0   1   2E+5   13.5   14  RT (min)       lcorbi_t3_pos_1_003_45    m/z 902.8725     0   1E+5   15.5   16  RT (min)       lcorbi_t3_pos_2_024_15    m/z 903.3202     0   1E+5   14.5   15  RT (min)       lcorbi_t3_pos_4_105_61    m/z 903.3908     0   2   4E+4   8.5   9  RT (min)       lcorbi_t3_pos_4_111_62    m/z 903.5795     0   2   4   6   8E+4   13.5   14  RT (min)      lcorbi_t3_pos_4_071_57    m/z 903.7036     0   1E+5   14.5   15  RT (min)       lcorbi_t3_pos_4_066_57    m/z 904.3943     0   1   2   3E+4   8.5   9  RT (min)      lcorbi_t3_pos_1_023_02    m/z 904.8881     0   1E+5   15.5   16  RT (min)       lcorbi_t3_pos_4_096_60    m/z 905.4045     0   2   4   6E+4   13   13.5  RT (min)       lcorbi_t3_pos_1_029_29    m/z 905.7191     0   2   4   6   8E+4   14.5   15  RT (min)       lcorbi_t3_pos_4_066_57    m/z 906.3102     0   1   2   3   4E+4   9   9.5  RT (min)       lcorbi_t3_pos_4_107_62    m/z 906.5728     0   1   2   3E+4   13   13.5  RT (min)      lcorbi_t3_pos_2_017_51    m/z 906.5984     0   1E+5   13.5   14  RT (min)      lcorbi_t3_pos_4_068_57    m/z 906.7306     0   1E+5   14.5   15  RT (min)       lcorbi_t3_pos_4_073_57    m/z 907.7356     0   1E+5   15   15.5  RT (min)       lcorbi_t3_pos_4_080_58    m/z 908.3463     0   2   4E+4   8.5   9  RT (min)       lcorbi_t3_pos_1_023_02    m/z 909.3567     0   2   4   6E+4   13   13.5  RT (min)      lcorbi_t3_pos_3_020_35    m/z 909.7737     0   1E+5   15.5   16  RT (min)       lcorbi_t3_pos_4_082_59    m/z 910.5504     0   1E+5   13.5   14  RT (min)      lcorbi_t3_pos_1_023_02    m/z 910.7858     0   1E+5   15.5   16  RT (min)       lcorbi_t3_pos_4_089_59    m/z 910.9204     0   1   2E+5   15.5   16  RT (min)       lcorbi_t3_pos_1_036_08    m/z 911.7531     0   2   4   6   8E+4   14.5   15  RT (min)       lcorbi_t3_pos_1_034_34    m/z 911.7892     0   2   4   6   8E+4   15.5   16  RT (min)       lcorbi_t3_pos_4_085_59    m/z 912.6561     0   2   4E+4   13   13.5  RT (min)      lcorbi_t3_pos_4_093_60    m/z 912.8015     0   1E+5   15.5   16  RT (min)       lcorbi_t3_pos_4_088_59    m/z 912.9361     0   1E+5   15.5   16  RT (min)       lcorbi_t3_pos_4_052_06    m/z 914.6717     0   1E+5   15.5   16  RT (min)      lcorbi_t3_pos_3_020_35    m/z 914.9517     0   1E+5   15.5   16  RT (min)       lcorbi_t3_pos_4_097_60    m/z 917.6893     0   1   2E+5   15.5   16  RT (min)      lcorbi_t3_pos_4_084_59    m/z 918.6153     0   1   2   3E+4   13.5   14  RT (min)      lcorbi_t3_pos_4_029_23    m/z 918.8483     0   2   4   6   8E+4   15.5   16  RT (min)       lcorbi_t3_pos_4_067_57    m/z 919.6950     0   1E+5   13   13.5  RT (min)       lcorbi_t3_pos_3_010_44    m/z 919.7051     0   1E+5   15.5   16  RT (min)      lcorbi_t3_pos_4_083_59    m/z 919.7360     0   2   4   6   8E+4   15   15.5  RT (min)       lcorbi_t3_pos_4_081_58    m/z 919.7580     0   1E+5   15.5   16  RT (min)       lcorbi_t3_pos_3_047_40    m/z 919.8520     0   2   4E+4   15.5   16  RT (min)      lcorbi_t3_pos_4_068_57    m/z 919.8926     0   1   2E+5   15   15.5  RT (min)       lcorbi_t3_pos_2_024_15    m/z 920.8226     0   1E+5   15   15.5  RT (min)       lcorbi_t3_pos_4_082_59    m/z 920.8633     0   2   4   6E+4   15   16  RT (min)       lcorbi_t3_pos_4_052_06    m/z 921.7206     0   1E+5   15.5   16  RT (min)      lcorbi_t3_pos_4_088_59    m/z 921.7514     0   2   4   6E+4   15   15.5  RT (min)      lcorbi_t3_pos_2_020_22    m/z 922.6389     0   2   4   6   8E+4   13   13.5  RT (min)       lcorbi_t3_pos_2_053_28    m/z 922.7345     0   1E+5   15.5   16  RT (min)       lcorbi_t3_pos_4_087_59    m/z 923.7361     0   1E+5   15.5   16  RT (min)      lcorbi_t3_pos_4_088_59    m/z 924.3274     0   2   4   6E+4   13   13.5  RT (min)      lcorbi_t3_pos_3_038_18    m/z 925.7258     0   1   2   3   4E+4   15.5   16  RT (min)       lcorbi_t3_pos_4_016_15    m/z 926.6693     0   1   2E+5   14   14.5  RT (min)       lcorbi_t3_pos_1_054_48    m/z 926.7808     0   1E+5   15   15.5  RT (min)       lcorbi_t3_pos_4_080_58    m/z 926.8632     0   1E+5   15   15.5  RT (min)      lcorbi_t3_pos_2_024_15    m/z 927.7412     0   2   4   6E+4   15.5   16  RT (min)       lcorbi_t3_pos_3_038_18    m/z 927.7843     0   2   4   6   8E+4   15   15.5  RT (min)       lcorbi_t3_pos_4_075_58    m/z 927.8202     0   1E+5   15.5   16  RT (min)       lcorbi_t3_pos_1_001_49    m/z 927.8755     0   1E+5   15.5   16  RT (min)      lcorbi_t3_pos_4_052_06    m/z 929.7568     0   1   15.5   16  RT (min)       lcorbi_t3_pos_4_085_59    m/z 929.7996     0   1E+5   15   15.5  RT (min)       lcorbi_t3_pos_4_078_58    m/z 929.8914     0   1E+5   15.5   16  RT (min)       lcorbi_t3_pos_4_052_06    m/z 930.4261     0   1E+5   13   13.5  RT (min)       lcorbi_t3_pos_1_002_01    m/z 930.8114     0   1E+5   15.5   16  RT (min)       lcorbi_t3_pos_4_081_58    m/z 930.9824     0   1   2E+5   15.5   16  RT (min)       lcorbi_t3_pos_4_090_60    m/z 931.4295     0   1E+5   13   13.5  RT (min)      lcorbi_t3_pos_1_002_01    m/z 931.8152     0   2   4   6E+4   15.5   16  RT (min)       lcorbi_t3_pos_4_074_58    m/z 931.9069     0   1E+5   15.5   16  RT (min)       lcorbi_t3_pos_4_052_06    m/z 932.8642     0   1   2   3E+4   15.5   16  RT (min)       lcorbi_t3_pos_2_024_15    m/z 933.6343     0   2   4   6E+4   14.5   15  RT (min)       lcorbi_t3_pos_4_069_57    m/z 933.9225     0   1E+5   15.5   16  RT (min)       lcorbi_t3_pos_4_092_60    m/z 934.9114     0   1E+5   15   15.5  RT (min)       lcorbi_t3_pos_2_024_15    m/z 935.9237     0   1E+5   15.5   16  RT (min)      lcorbi_t3_pos_2_024_15    m/z 936.7652     0   1   2E+5   15   15.5  RT (min)       lcorbi_t3_pos_4_076_58    m/z 936.8016     0   2   4E+4   15.5   16  RT (min)       lcorbi_t3_pos_4_032_40    m/z 937.7688     0   1E+5   15   15.5  RT (min)       lcorbi_t3_pos_4_081_58    m/z 937.9394     0   1E+5   15.5   16  RT (min)       lcorbi_t3_pos_2_024_15    m/z 939.7842     0   1   2E+5   15   15.5  RT (min)       lcorbi_t3_pos_4_079_58    m/z 939.9552     0   1E+5   15.5   16  RT (min)       lcorbi_t3_pos_2_024_15    m/z 940.7959     0   1   2E+5   15   15.5  RT (min)       lcorbi_t3_pos_4_076_58    m/z 940.8329     0   1   2E+5   15.5   16  RT (min)       lcorbi_t3_pos_4_083_59    m/z 941.4436     0   1   2   3E+4   8.5   9  RT (min)       lcorbi_t3_pos_4_109_62    m/z 941.8001     0   1E+5   15   15.5  RT (min)       lcorbi_t3_pos_4_074_58    m/z 941.8366     0   1E+5   15.5   16  RT (min)       lcorbi_t3_pos_4_089_59    m/z 941.9709     0   2   4   6   8E+4   15.5   16  RT (min)       lcorbi_t3_pos_2_024_15    m/z 942.4468     0   1   2   3E+4   8.5   9  RT (min)      lcorbi_t3_pos_4_106_62    m/z 942.6360     0   2   4   6   8E+4   13   13.5  RT (min)       lcorbi_t3_pos_4_093_60    m/z 942.7549     0   1   2   3E+4   15.5   16  RT (min)       lcorbi_t3_pos_4_031_29    m/z 943.8522     0   1   2E+5   15.5   16  RT (min)       lcorbi_t3_pos_4_086_59    m/z 943.9503     0   1E+5   15   15.5  RT (min)      lcorbi_t3_pos_4_052_06    m/z 944.9899     0   1   2E+5   15.5   16  RT (min)       lcorbi_t3_pos_4_091_60    m/z 945.7740     0   2   4   6   8E+4   15.5   16  RT (min)       lcorbi_t3_pos_4_076_58    m/z 945.9081     0   1E+5   15   15.5  RT (min)      lcorbi_t3_pos_2_024_15    m/z 945.9661     0   1E+5   15   15.5  RT (min)       lcorbi_t3_pos_1_036_08    m/z 946.6388     0   1E+5   13.5   14  RT (min)       lcorbi_t3_pos_1_014_03    m/z 946.8702     0   1E+5   15.5   16  RT (min)       lcorbi_t3_pos_4_052_06    m/z 946.8795     0   1E+5   15.5   16  RT (min)       lcorbi_t3_pos_4_066_57    m/z 947.6420     0   2   4   6   8E+4   13.5   14  RT (min)       lcorbi_t3_pos_3_004_24    m/z 947.7896     0   2   4   6E+4   15.5   16  RT (min)       lcorbi_t3_pos_4_039_35    m/z 947.8834     0   2   4   6   8E+4   15.5   16  RT (min)      lcorbi_t3_pos_4_066_57    m/z 948.8014     0   2   4   6E+4   15.5   16  RT (min)       lcorbi_t3_pos_4_006_12    m/z 949.8052     0   1   2   3   4E+4   15.5   16  RT (min)      lcorbi_t3_pos_3_054_15    m/z 952.7964     0   2   4   6E+4   15.5   16  RT (min)       lcorbi_t3_pos_3_042_14    m/z 952.8329     0   2   4   6   8E+4   15.5   16  RT (min)       lcorbi_t3_pos_4_085_59    m/z 952.8704     0   1E+5   15   15.5  RT (min)      lcorbi_t3_pos_1_036_08    m/z 953.8824     0   1E+5   15.5   16  RT (min)      lcorbi_t3_pos_1_036_08    m/z 954.7013     0   1E+5   14   14.5  RT (min)       lcorbi_t3_pos_1_054_48    m/z 954.8486     0   1   2E+5   15.5   16  RT (min)       lcorbi_t3_pos_4_083_59    m/z 955.8523     0   1E+5   15.5   16  RT (min)       lcorbi_t3_pos_4_089_59    m/z 955.8983     0   1   2E+5   15.5   16  RT (min)       lcorbi_t3_pos_4_052_06    m/z 956.8643     0   1   2E+5   15.5   16  RT (min)       lcorbi_t3_pos_4_083_59    m/z 957.8679     0   1E+5   15.5   16  RT (min)       lcorbi_t3_pos_4_086_59    m/z 957.9140     0   1   2E+5   15.5   16  RT (min)       lcorbi_t3_pos_1_036_08    m/z 959.8835     0   1E+5   15.5   16  RT (min)      lcorbi_t3_pos_4_084_59    m/z 959.9296     0   1   2E+5   15.5   16  RT (min)       lcorbi_t3_pos_2_024_15    m/z 960.7654     0   1E+5   15   15.5  RT (min)       lcorbi_t3_pos_4_080_58    m/z 960.8950     0   1E+5   15.5   16  RT (min)       lcorbi_t3_pos_4_071_57    m/z 961.9452     0   1E+5   15.5   16  RT (min)       lcorbi_t3_pos_2_024_15    m/z 962.7807     0   1E+5   15   15.5  RT (min)       lcorbi_t3_pos_4_081_58    m/z 964.8248     0   1E+5   14.5   15  RT (min)      lcorbi_t3_pos_4_099_61    m/z 965.7423     0   2   4   6E+4   16   16.5  RT (min)       lcorbi_t3_pos_4_078_58    m/z 969.7737     0   2   4   6   8E+4   16.5   17  RT (min)       lcorbi_t3_pos_4_081_58    m/z 970.8597     0   1E+5   15.5   16  RT (min)      lcorbi_t3_pos_3_010_44    m/z 971.7316     0   2   4E+4   14   14.5  RT (min)      lcorbi_t3_pos_4_103_61    m/z 971.8835     0   1E+5   15.5   16  RT (min)       lcorbi_t3_pos_4_085_59    m/z 971.9725     0   1E+5   15   15.5  RT (min)      lcorbi_t3_pos_2_024_15    m/z 972.8754     0   1E+5   15.5   16  RT (min)       lcorbi_t3_pos_2_024_15    m/z 972.8928     0   1   2E+5   15.5   16  RT (min)       lcorbi_t3_pos_4_086_59    m/z 973.9884     0   1   2E+5   15   15.5  RT (min)       lcorbi_t3_pos_2_024_15    m/z 974.4797     0   2   4   6E+4   13   13.5  RT (min)      lcorbi_t3_pos_4_045_08    m/z 974.8309     0   1   2   3E+5   14   14.5  RT (min)       lcorbi_t3_pos_4_091_60    m/z 974.8911     0   1E+5   15.5   16  RT (min)       lcorbi_t3_pos_4_052_06    m/z 976.0039     0   1E+5   15   15.5  RT (min)       lcorbi_t3_pos_2_024_15    m/z 976.3382     0   1E+5   15   15.5  RT (min)       lcorbi_t3_pos_4_084_59    m/z 976.8330     0   1E+5   15.5   16  RT (min)       lcorbi_t3_pos_4_084_59    m/z 976.9068     0   1E+5   15.5   16  RT (min)       lcorbi_t3_pos_2_024_15    m/z 977.3390     0   1E+5   15   15.5  RT (min)       lcorbi_t3_pos_4_089_59    m/z 977.8365     0   2   4   6   8E+4   15.5   16  RT (min)      lcorbi_t3_pos_4_088_59    m/z 978.3374     0   1E+5   15   15.5  RT (min)       lcorbi_t3_pos_4_084_59    m/z 978.9224     0   1E+5   15.5   16  RT (min)       lcorbi_t3_pos_2_024_15    m/z 979.8522     0   1E+5   15.5   16  RT (min)      lcorbi_t3_pos_4_089_59    m/z 980.8643     0   1   2E+5   15.5   16  RT (min)       lcorbi_t3_pos_4_087_59    m/z 981.8678     0   1E+5   15.5   16  RT (min)      lcorbi_t3_pos_4_089_59    m/z 983.8835     0   1E+5   15.5   16  RT (min)      lcorbi_t3_pos_4_087_59    m/z 984.8954     0   1E+5   15.5   16  RT (min)       lcorbi_t3_pos_4_083_59    m/z 985.8991     0   1E+5   15.5   16  RT (min)      lcorbi_t3_pos_4_087_59    m/z 987.4921     0   1E+5   13   13.5  RT (min)      lcorbi_t3_pos_4_076_58    m/z 1004.8687     0   1E+5   14   14.5  RT (min)       lcorbi_t3_pos_1_036_08    m/z 1022.0248     0   1E+5   15.5   16  RT (min)      lcorbi_t3_pos_4_052_06    m/z 1023.5425     0   1E+4   6.5   7  RT (min)      lcorbi_t3_pos_1_049_55    m/z 1024.0402     0   1E+5   15.5   16  RT (min)       lcorbi_t3_pos_4_052_06    m/z 1026.0557     0   1E+5   15.5   16  RT (min)       lcorbi_t3_pos_2_024_15    m/z 1050.3568     0   1E+5   15   15.5  RT (min)       lcorbi_t3_pos_4_086_59    m/z 1051.3577     0   1E+5   15   15.5  RT (min)       lcorbi_t3_pos_4_089_59    m/z 1052.3562     0   1E+5   15   15.5  RT (min)       lcorbi_t3_pos_4_086_59    m/z 1058.8753     0   1E+5   12.5   13  RT (min)       lcorbi_t3_pos_4_066_57    m/z 1059.0005     0   1E+5   12.5   13  RT (min)       lcorbi_t3_pos_4_079_58    m/z 1059.3766     0   1E+5   12.5   13  RT (min)       lcorbi_t3_pos_4_070_57    m/z 1059.5020     0   1E+5   12.5   13  RT (min)       lcorbi_t3_pos_4_074_58    m/z 1059.6278     0   1E+5   12.5   13  RT (min)       lcorbi_t3_pos_4_087_59    m/z 1059.7527     0   2   4   6   8E+4   12.5   13  RT (min)       lcorbi_t3_pos_4_098_61    m/z 1060.2936     0   1   2E+5   15   15.5  RT (min)       lcorbi_t3_pos_4_084_59    m/z 1061.0005     0   1E+5   12.5   13  RT (min)       lcorbi_t3_pos_4_109_62    m/z 1061.1253     0   1E+5   12.5   13  RT (min)       lcorbi_t3_pos_4_078_58    m/z 1061.2506     0   1E+5   12.5   13  RT (min)       lcorbi_t3_pos_4_113_62    m/z 1061.3762     0   2   4   6   8E+4   12.5   13  RT (min)       lcorbi_t3_pos_4_081_58    m/z 1061.5013     0   1E+5   12.5   13  RT (min)       lcorbi_t3_pos_4_099_61    m/z 1094.6451     0   1E+5   11   11.5  RT (min)      lcorbi_t3_pos_1_001_49    m/z 1094.8957     0   1E+5   11   11.5  RT (min)      lcorbi_t3_pos_1_001_49    m/z 1095.1466     0   1E+5   11   11.5  RT (min)       lcorbi_t3_pos_1_001_49    m/z 1134.3129     0   1E+5   15   15.5  RT (min)       lcorbi_t3_pos_4_089_59   
