## Supplementary material for "The Everything Bagel Feature Finder: Ultra-fast automated feature finding for untargeted metabolomics": SI Data File: tool_unique_xic_st_XCMS_ST004626.html

XCMS-unique features (not detected by the other two tools) - ST004626

### XCMS-unique features (not detected by the other two tools) - ST004626

401 features · qtof · XIC window m/z ± 25 ppm (≥ 0.002 Da) · ± 0.5 min

Every panel is a feature **XCMS** detected that neither of the other two tools detected — three-way unique across EB (standard mode (min\_detection\_rate=0.1)), Masscube and XCMS, under the standard matching rule (m/z ± 25 ppm; apex within the reference RT window ± 0.2 min). It shows the MS1 extracted-ion chromatogram at the feature's m/z, drawn from the raw mzML named in `alignment_reference_file`, zoomed to ± 0.5 min around the feature's reported apex RT. **Gray solid line** = the feature's reported apex RT; **red dashed line** = XCMS's assigned MS/MS — `RTINSECONDS` in its `aligned_features.mgf`; the panel widens to keep an off-apex MS/MS on-screen. Features with no assigned MS/MS show only the gray line. y is per-cell autoscaled. Rows keep the table order (m/z ascending).

m/z 61.9876

0123E+377.5RT (min)

lcqtof\_t3\_neg\_3\_077\_58

m/z 68.9952

0123E+314.515RT (min)

lcqtof\_t3\_neg\_4\_077\_58

m/z 78.9181

01234E+31415RT (min)

lcqtof\_t3\_neg\_4\_081\_58

m/z 80.9161

0246E+314.515RT (min)

lcqtof\_t3\_neg\_4\_103\_61

m/z 99.0192

0123E+31414.5RT (min)

lcqtof\_t3\_neg\_4\_077\_58

m/z 100.9328

0246E+311.512RT (min)

lcqtof\_t3\_neg\_1\_012\_16

m/z 100.9329

02468E+31515.5RT (min)

lcqtof\_t3\_neg\_1\_031\_24

m/z 102.9562

012E+31515.5RT (min)

lcqtof\_t3\_neg\_4\_113\_62

m/z 114.9882

012E+316.517RT (min)

lcqtof\_t3\_neg\_1\_047\_22

m/z 115.0200

01234E+31.52RT (min)

lcqtof\_t3\_neg\_4\_016\_14

m/z 119.9463

012E+31515.5RT (min)

lcqtof\_t3\_neg\_4\_082\_59

m/z 128.9277

0123E+31010.5RT (min)

lcqtof\_t3\_neg\_4\_031\_42

m/z 128.9278

02468E+316.517RT (min)

lcqtof\_t3\_neg\_4\_002\_17

m/z 129.9751

012E+31515.5RT (min)

lcqtof\_t3\_neg\_4\_071\_57

m/z 129.9752

0123E+388.5RT (min)

lcqtof\_t3\_neg\_1\_025\_52

m/z 129.9752

01234E+310.511RT (min)

lcqtof\_t3\_neg\_1\_005\_18

m/z 129.9996

0123E+31313.5RT (min)

lcqtof\_t3\_neg\_4\_100\_61

m/z 130.0162

0123E+311.512RT (min)

lcqtof\_t3\_neg\_4\_015\_36

m/z 130.9655

0123E+389RT (min)

lcqtof\_t3\_neg\_4\_103\_61

m/z 130.9657

0246E+31718RT (min)

lcqtof\_t3\_neg\_3\_093\_60

m/z 130.9961

0246E+31313.5RT (min)

lcqtof\_t3\_neg\_4\_079\_58

m/z 131.0128

0246E+311.512RT (min)

lcqtof\_t3\_neg\_4\_102\_61

m/z 136.9831

0123E+31717.5RT (min)

lcqtof\_t3\_neg\_1\_040\_08

m/z 146.0114

01234E+31212.5RT (min)

lcqtof\_t3\_neg\_4\_106\_62

m/z 146.9390

01E+43.54RT (min)

lcqtof\_t3\_neg\_4\_113\_62

m/z 157.0150

02468E+31111.5RT (min)

lcqtof\_t3\_neg\_4\_108\_62

m/z 158.9764

0123E+38.59RT (min)

lcqtof\_t3\_neg\_4\_095\_60

m/z 158.9775

01234E+316.517RT (min)

lcqtof\_t3\_neg\_4\_003\_37

m/z 159.0766

01E+41414.5RT (min)

lcqtof\_t3\_neg\_4\_078\_58

m/z 159.8930

01E+31616.5RT (min)

lcqtof\_t3\_neg\_3\_093\_60

m/z 160.0254

01234E+311.512RT (min)

lcqtof\_t3\_neg\_4\_112\_62

m/z 160.8414

0123E+313.514RT (min)

lcqtof\_t3\_neg\_4\_109\_62

m/z 160.8920

012E+315.516RT (min)

lcqtof\_t3\_neg\_3\_084\_59

m/z 161.0225

0246E+31111.5RT (min)

lcqtof\_t3\_neg\_4\_075\_58

m/z 162.8386

0123E+313.514RT (min)

lcqtof\_t3\_neg\_4\_054\_44

m/z 162.8927

012E+311.512RT (min)

lcqtof\_t3\_neg\_4\_107\_62

m/z 162.8929

012E+31515.5RT (min)

lcqtof\_t3\_neg\_3\_091\_60

m/z 162.9822

01E+41717.5RT (min)

lcqtof\_t3\_neg\_4\_001\_49

m/z 168.8354

012E+31111.5RT (min)

lcqtof\_t3\_neg\_4\_087\_59

m/z 170.8324

02468E+31111.5RT (min)

lcqtof\_t3\_neg\_4\_014\_13

m/z 170.9434

0123E+31515.5RT (min)

lcqtof\_t3\_neg\_4\_047\_07

m/z 171.0515

012E+31515.5RT (min)

lcqtof\_t3\_neg\_4\_103\_61

m/z 172.8299

0246E+31111.5RT (min)

lcqtof\_t3\_neg\_4\_012\_27

m/z 176.9785

01E+416.517RT (min)

lcqtof\_t3\_neg\_4\_007\_09

m/z 176.9785

01234E+377.5RT (min)

lcqtof\_t3\_neg\_4\_008\_35

m/z 178.0175

012E+310.511RT (min)

lcqtof\_t3\_neg\_4\_111\_62

m/z 178.8143

012E+30.51RT (min)

lcqtof\_t3\_neg\_4\_081\_58

m/z 178.8144

0246E+31818.5RT (min)

lcqtof\_t3\_neg\_4\_010\_46

m/z 179.0139

012E+310.511RT (min)

lcqtof\_t3\_neg\_4\_100\_61

m/z 180.8123

01234E+317.518RT (min)

lcqtof\_t3\_neg\_4\_012\_27

m/z 180.9116

01E+316.517RT (min)

lcqtof\_t3\_neg\_4\_094\_60

m/z 180.9723

0246E+310.511RT (min)

lcqtof\_t3\_neg\_4\_100\_61

m/z 180.9723

0246E+388.5RT (min)

lcqtof\_t3\_neg\_4\_071\_57

m/z 180.9729

0123E+31818.5RT (min)

lcqtof\_t3\_neg\_4\_110\_62

m/z 182.9539

01234E+317.518RT (min)

lcqtof\_t3\_neg\_1\_021\_39

m/z 183.0113

01234E+31515.5RT (min)

lcqtof\_t3\_neg\_4\_036\_47

m/z 184.9499

0123E+317.518RT (min)

lcqtof\_t3\_neg\_1\_011\_45

m/z 187.9407

0246E+314.515RT (min)

lcqtof\_t3\_neg\_4\_011\_32

m/z 188.0341

0246E+316.517RT (min)

lcqtof\_t3\_neg\_4\_003\_37

m/z 189.0079

0123E+399.5RT (min)

lcqtof\_t3\_neg\_1\_052\_12

m/z 189.0219

01234E+31212.5RT (min)

lcqtof\_t3\_neg\_4\_107\_62

m/z 191.9457

01234E+38.59RT (min)

lcqtof\_t3\_neg\_3\_085\_59

m/z 201.0224

0123E+311.512RT (min)

lcqtof\_t3\_neg\_4\_108\_62

m/z 202.0199

01E+41111.5RT (min)

lcqtof\_t3\_neg\_4\_078\_58

m/z 203.0200

012E+41111.5RT (min)

lcqtof\_t3\_neg\_4\_076\_58

m/z 204.0171

024E+411.512RT (min)

lcqtof\_t3\_neg\_4\_112\_62

m/z 204.9723

01E+36.57RT (min)

lcqtof\_t3\_neg\_4\_098\_61

m/z 206.0174

01234E+311.512RT (min)

lcqtof\_t3\_neg\_4\_087\_59

m/z 206.9721

012E+41818.5RT (min)

lcqtof\_t3\_neg\_4\_002\_17

m/z 207.0143

0123E+311.512RT (min)

lcqtof\_t3\_neg\_4\_072\_57

m/z 208.9360

0123E+388.5RT (min)

lcqtof\_t3\_neg\_1\_035\_31

m/z 209.0286

024E+31515.5RT (min)

lcqtof\_t3\_neg\_4\_071\_57

m/z 210.9489

01234E+35.56RT (min)

lcqtof\_t3\_neg\_1\_026\_19

m/z 210.9491

0246E+31818.5RT (min)

lcqtof\_t3\_neg\_4\_001\_49

m/z 214.7824

02468E+310.511RT (min)

lcqtof\_t3\_neg\_4\_006\_16

m/z 216.0702

0123E+311.512RT (min)

lcqtof\_t3\_neg\_4\_112\_62

m/z 216.7798

024E+310.511RT (min)

lcqtof\_t3\_neg\_4\_024\_26

m/z 217.0348

01234E+31212.5RT (min)

lcqtof\_t3\_neg\_4\_099\_61

m/z 217.0667

0123E+311.512RT (min)

lcqtof\_t3\_neg\_4\_076\_58

m/z 218.7778

0123E+310.511RT (min)

lcqtof\_t3\_neg\_4\_030\_31

m/z 220.0275

012E+311.512RT (min)

lcqtof\_t3\_neg\_4\_071\_57

m/z 224.0838

024E+399.5RT (min)

lcqtof\_t3\_neg\_4\_034\_34

m/z 226.9641

012E+388.5RT (min)

lcqtof\_t3\_neg\_4\_087\_59

m/z 228.9022

012E+31616.5RT (min)

lcqtof\_t3\_neg\_4\_049\_55

m/z 228.9594

0123E+31818.5RT (min)

lcqtof\_t3\_neg\_3\_107\_62

m/z 230.9555

0123E+317.518RT (min)

lcqtof\_t3\_neg\_1\_031\_24

m/z 234.0258

01234E+312.513RT (min)

lcqtof\_t3\_neg\_4\_112\_62

m/z 235.0224

0123E+31212.5RT (min)

lcqtof\_t3\_neg\_4\_044\_48

m/z 240.9281

0123E+315.516RT (min)

lcqtof\_t3\_neg\_4\_050\_43

m/z 242.9431

0246E+31111.5RT (min)

lcqtof\_t3\_neg\_4\_105\_61

m/z 243.1090

0123E+34.55RT (min)

lcqtof\_t3\_neg\_4\_002\_17

m/z 243.1091

0123E+31515.5RT (min)

lcqtof\_t3\_neg\_4\_080\_58

m/z 246.0274

0123E+311.512RT (min)

lcqtof\_t3\_neg\_4\_106\_62

m/z 256.9545

024E+35.56RT (min)

lcqtof\_t3\_neg\_1\_026\_19

m/z 259.0364

0123E+31111.5RT (min)

lcqtof\_t3\_neg\_4\_023\_45

m/z 259.9332

012E+388.5RT (min)

lcqtof\_t3\_neg\_3\_048\_01

m/z 261.0408

012E+31212.5RT (min)

lcqtof\_t3\_neg\_4\_069\_57

m/z 262.0379

0123E+31212.5RT (min)

lcqtof\_t3\_neg\_4\_001\_49

m/z 263.0347

0123E+311.51212.5RT (min)

lcqtof\_t3\_neg\_4\_104\_61

m/z 267.0862

012E+314.515RT (min)

lcqtof\_t3\_neg\_4\_007\_09

m/z 272.8979

012E+377.5RT (min)

lcqtof\_t3\_neg\_4\_071\_57

m/z 287.0636

01234E+31515.5RT (min)

lcqtof\_t3\_neg\_4\_012\_27

m/z 293.1436

01234E+315RT (min)

lcqtof\_t3\_neg\_4\_076\_58

m/z 293.1777

0246E+315.516RT (min)

lcqtof\_t3\_neg\_4\_013\_39

m/z 295.0684

01E+311.5RT (min)

lcqtof\_t3\_neg\_4\_055\_38

m/z 296.8897

012E+315.516RT (min)

lcqtof\_t3\_neg\_4\_077\_58

m/z 298.1559

024E+31515.5RT (min)

lcqtof\_t3\_neg\_4\_029\_40

m/z 299.1504

012E+31515.5RT (min)

lcqtof\_t3\_neg\_4\_109\_62

m/z 299.9185

012E+314.515RT (min)

lcqtof\_t3\_neg\_4\_089\_59

m/z 301.0394

0123E+366.5RT (min)

lcqtof\_t3\_neg\_4\_048\_19

m/z 310.9302

024E+38.59RT (min)

lcqtof\_t3\_neg\_4\_087\_59

m/z 313.1139

0123E+31515.5RT (min)

lcqtof\_t3\_neg\_3\_086\_59

m/z 313.1671

0246E+314.515RT (min)

lcqtof\_t3\_neg\_4\_106\_62

m/z 321.1758

01234E+314.515RT (min)

lcqtof\_t3\_neg\_4\_079\_58

m/z 321.9034

01234E+37.58RT (min)

lcqtof\_t3\_neg\_1\_024\_10

m/z 323.0119

0246E+366.5RT (min)

lcqtof\_t3\_neg\_4\_007\_09

m/z 323.9088

012E+31616.5RT (min)

lcqtof\_t3\_neg\_4\_026\_30

m/z 326.1870

01E+41515.5RT (min)

lcqtof\_t3\_neg\_4\_071\_57

m/z 327.1829

01E+41515.5RT (min)

lcqtof\_t3\_neg\_4\_003\_37

m/z 331.1261

0123E+31515.5RT (min)

lcqtof\_t3\_neg\_4\_002\_17

m/z 340.2027

01E+41515.5RT (min)

lcqtof\_t3\_neg\_4\_004\_20

m/z 341.1453

012E+315.516RT (min)

lcqtof\_t3\_neg\_3\_105\_61

m/z 341.1989

01234E+31515.5RT (min)

lcqtof\_t3\_neg\_4\_113\_62

m/z 345.1029

012E+31616.5RT (min)

lcqtof\_t3\_neg\_4\_095\_60

m/z 355.1109

0123E+31515.5RT (min)

lcqtof\_t3\_neg\_4\_028\_25

m/z 355.1454

01234E+399.5RT (min)

lcqtof\_t3\_neg\_4\_005\_18

m/z 355.1992

02468E+39.510RT (min)

lcqtof\_t3\_neg\_4\_003\_37

m/z 358.2100

0123E+315.516RT (min)

lcqtof\_t3\_neg\_4\_037\_01

m/z 364.1608

012E+39.510RT (min)

lcqtof\_t3\_neg\_4\_081\_58

m/z 365.1392

0123E+31515.5RT (min)

lcqtof\_t3\_neg\_4\_099\_61

m/z 367.1009

0246E+31515.5RT (min)

lcqtof\_t3\_neg\_4\_001\_49

m/z 367.1931

012E+399.5RT (min)

lcqtof\_t3\_neg\_4\_105\_61

m/z 371.1523

012E+399.5RT (min)

lcqtof\_t3\_neg\_4\_098\_61

m/z 378.9175

0246E+38.59RT (min)

lcqtof\_t3\_neg\_4\_085\_59

m/z 379.0797

01234E+37.58RT (min)

lcqtof\_t3\_neg\_4\_048\_19

m/z 380.1582

0123E+31515.5RT (min)

lcqtof\_t3\_neg\_4\_107\_62

m/z 382.1716

012E+39.510RT (min)

lcqtof\_t3\_neg\_4\_016\_14

m/z 382.2344

0246E+31010.5RT (min)

lcqtof\_t3\_neg\_4\_019\_05

m/z 383.1417

02468E+315.516RT (min)

lcqtof\_t3\_neg\_4\_001\_49

m/z 384.1759

024E+39.510RT (min)

lcqtof\_t3\_neg\_4\_016\_14

m/z 384.9343

0246E+399.5RT (min)

lcqtof\_t3\_neg\_4\_111\_62

m/z 384.9348

0246E+388.5RT (min)

lcqtof\_t3\_neg\_4\_080\_58

m/z 384.9349

0246E+31111.5RT (min)

lcqtof\_t3\_neg\_4\_113\_62

m/z 390.8714

0123E+316.517RT (min)

lcqtof\_t3\_neg\_4\_001\_49

m/z 394.1740

012E+31515.5RT (min)

lcqtof\_t3\_neg\_4\_073\_57

m/z 394.8901

012E+366.5RT (min)

lcqtof\_t3\_neg\_4\_066\_57

m/z 395.1056

0246E+299.5RT (min)

lcqtof\_t3\_neg\_4\_005\_18

m/z 395.9076

012E+38.59RT (min)

lcqtof\_t3\_neg\_1\_018\_33

m/z 396.1457

0246E+31515.5RT (min)

lcqtof\_t3\_neg\_4\_007\_09

m/z 397.1565

0246E+31616.5RT (min)

lcqtof\_t3\_neg\_4\_003\_37

m/z 406.0041

012E+399.5RT (min)

lcqtof\_t3\_neg\_4\_005\_18

m/z 407.1868

01E+41515.5RT (min)

lcqtof\_t3\_neg\_4\_021\_21

m/z 410.1262

0123E+399.5RT (min)

lcqtof\_t3\_neg\_1\_015\_34

m/z 410.1613

0246E+315.516RT (min)

lcqtof\_t3\_neg\_4\_005\_18

m/z 413.0078

02468E+39.510RT (min)

lcqtof\_t3\_neg\_4\_016\_14

m/z 413.2354

01234E+311.512RT (min)

lcqtof\_t3\_neg\_4\_003\_37

m/z 417.2104

0123E+31515.5RT (min)

lcqtof\_t3\_neg\_4\_110\_62

m/z 424.1770

01234E+315.516RT (min)

lcqtof\_t3\_neg\_4\_007\_09

m/z 437.2543

0246E+311.512RT (min)

lcqtof\_t3\_neg\_4\_016\_14

m/z 439.2694

0246E+31212.5RT (min)

lcqtof\_t3\_neg\_4\_037\_01

m/z 440.2524

01E+411.512RT (min)

lcqtof\_t3\_neg\_4\_016\_14

m/z 441.2849

012E+312.513RT (min)

lcqtof\_t3\_neg\_4\_026\_30

m/z 442.2558

0246E+29.510RT (min)

lcqtof\_t3\_neg\_4\_043\_11

m/z 445.1206

012E+316.517RT (min)

lcqtof\_t3\_neg\_4\_053\_12

m/z 452.9221

0246E+38.59RT (min)

lcqtof\_t3\_neg\_4\_092\_60

m/z 453.1615

0246E+39.510RT (min)

lcqtof\_t3\_neg\_4\_016\_14

m/z 455.4150

024E+31212.5RT (min)

lcqtof\_t3\_neg\_4\_040\_24

m/z 456.9580

0123E+39.510RT (min)

lcqtof\_t3\_neg\_4\_003\_37

m/z 461.1587

0246E+315.516RT (min)

lcqtof\_t3\_neg\_4\_022\_28

m/z 463.2491

0246E+311.512RT (min)

lcqtof\_t3\_neg\_4\_016\_14

m/z 464.0765

024E+39.510RT (min)

lcqtof\_t3\_neg\_4\_016\_14

m/z 469.3639

0123E+310.511RT (min)

lcqtof\_t3\_neg\_4\_018\_23

m/z 475.1746

0123E+31515.5RT (min)

lcqtof\_t3\_neg\_4\_001\_49

m/z 479.0821

012E+399.5RT (min)

lcqtof\_t3\_neg\_4\_099\_61

m/z 482.1111

024E+39.510RT (min)

lcqtof\_t3\_neg\_4\_075\_58

m/z 483.2781

01234E+31212.5RT (min)

lcqtof\_t3\_neg\_4\_016\_14

m/z 486.2823

01234E+39.510RT (min)

lcqtof\_t3\_neg\_4\_018\_23

m/z 489.1005

01234E+39.510RT (min)

lcqtof\_t3\_neg\_4\_002\_17

m/z 492.9310

0246E+38.59RT (min)

lcqtof\_t3\_neg\_4\_004\_20

m/z 497.1665

02468E+31515.5RT (min)

lcqtof\_t3\_neg\_4\_034\_34

m/z 503.3316

024E+31212.5RT (min)

lcqtof\_t3\_neg\_4\_040\_24

m/z 514.8923

024E+378RT (min)

lcqtof\_t3\_neg\_4\_088\_59

m/z 515.4099

024E+312.513RT (min)

lcqtof\_t3\_neg\_4\_016\_14

m/z 520.1513

01234E+31515.5RT (min)

lcqtof\_t3\_neg\_4\_014\_13

m/z 520.9095

0246E+388.5RT (min)

lcqtof\_t3\_neg\_4\_111\_62

m/z 523.0387

01234E+377.5RT (min)

lcqtof\_t3\_neg\_4\_041\_54

m/z 525.0744

024E+38.59RT (min)

lcqtof\_t3\_neg\_4\_004\_20

m/z 527.2181

02468E+377.5RT (min)

lcqtof\_t3\_neg\_4\_016\_14

m/z 529.1465

01234E+315.516RT (min)

lcqtof\_t3\_neg\_4\_031\_42

m/z 535.5455

01234E+314.515RT (min)

lcqtof\_t3\_neg\_4\_006\_16

m/z 538.4190

024E+311.512RT (min)

lcqtof\_t3\_neg\_4\_016\_14

m/z 539.0659

0123E+39.510RT (min)

lcqtof\_t3\_neg\_4\_008\_35

m/z 543.2534

02468E+29.510RT (min)

lcqtof\_t3\_neg\_4\_040\_24

m/z 549.5607

01234E+314.515RT (min)

lcqtof\_t3\_neg\_4\_050\_43

m/z 550.0973

0123E+39.510RT (min)

lcqtof\_t3\_neg\_4\_046\_33

m/z 556.3260

0123E+312.513RT (min)

lcqtof\_t3\_neg\_4\_004\_20

m/z 556.3373

024E+31313.5RT (min)

lcqtof\_t3\_neg\_4\_018\_23

m/z 571.1396

01E+377.5RT (min)

lcqtof\_t3\_neg\_4\_103\_61

m/z 573.3315

02468E+39.510RT (min)

lcqtof\_t3\_neg\_4\_003\_37

m/z 577.4228

0246E+312.513RT (min)

lcqtof\_t3\_neg\_4\_016\_14

m/z 581.4994

012E+31313.5RT (min)

lcqtof\_t3\_neg\_4\_018\_23

m/z 586.2375

02468E+31010.5RT (min)

lcqtof\_t3\_neg\_4\_019\_05

m/z 588.8969

024E+388.5RT (min)

lcqtof\_t3\_neg\_4\_110\_62

m/z 590.4802

01234E+31313.5RT (min)

lcqtof\_t3\_neg\_1\_019\_23

m/z 591.1335

01234E+38.59RT (min)

lcqtof\_t3\_neg\_4\_004\_20

m/z 613.1436

024E+333.5RT (min)

lcqtof\_t3\_neg\_4\_087\_59

m/z 614.5634

0123E+313.514RT (min)

lcqtof\_t3\_neg\_4\_003\_37

m/z 616.3732

0123E+31313.5RT (min)

lcqtof\_t3\_neg\_4\_014\_13

m/z 620.5743

0123E+31313.5RT (min)

lcqtof\_t3\_neg\_4\_040\_24

m/z 622.5893

024E+31313.5RT (min)

lcqtof\_t3\_neg\_4\_040\_24

m/z 623.3093

0123E+312.513RT (min)

lcqtof\_t3\_neg\_4\_005\_18

m/z 627.3234

0123E+31111.5RT (min)

lcqtof\_t3\_neg\_4\_014\_13

m/z 630.5130

0123E+31313.5RT (min)

lcqtof\_t3\_neg\_1\_019\_23

m/z 636.2382

0246E+39.510RT (min)

lcqtof\_t3\_neg\_4\_016\_14

m/z 642.6459

0246E+313.514RT (min)

lcqtof\_t3\_neg\_4\_040\_24

m/z 643.4336

0246E+31212.5RT (min)

lcqtof\_t3\_neg\_4\_003\_37

m/z 644.6484

01234E+313.514RT (min)

lcqtof\_t3\_neg\_4\_011\_32

m/z 645.5361

024E+313.514RT (min)

lcqtof\_t3\_neg\_2\_030\_23

m/z 645.6228

024E+31313.5RT (min)

lcqtof\_t3\_neg\_4\_040\_24

m/z 649.6091

0246E+31313.5RT (min)

lcqtof\_t3\_neg\_1\_031\_24

m/z 650.1600

01E+377.5RT (min)

lcqtof\_t3\_neg\_4\_104\_61

m/z 650.5164

01234E+313.514RT (min)

lcqtof\_t3\_neg\_4\_087\_59

m/z 657.2391

012E+39.510RT (min)

lcqtof\_t3\_neg\_4\_016\_14

m/z 671.8673

01E+41313.5RT (min)

lcqtof\_t3\_neg\_4\_034\_34

m/z 672.3562

0246E+312.513RT (min)

lcqtof\_t3\_neg\_4\_055\_38

m/z 673.4798

0123E+316.517RT (min)

lcqtof\_t3\_neg\_4\_089\_59

m/z 677.2381

0246E+31515.5RT (min)

lcqtof\_t3\_neg\_4\_103\_61

m/z 677.2383

024E+388.5RT (min)

lcqtof\_t3\_neg\_4\_089\_59

m/z 678.3127

01234E+310.511RT (min)

lcqtof\_t3\_neg\_4\_055\_38

m/z 679.2884

024E+39.510RT (min)

lcqtof\_t3\_neg\_4\_002\_17

m/z 682.2994

024E+311.512RT (min)

lcqtof\_t3\_neg\_4\_032\_04

m/z 685.8699

0246E+31313.5RT (min)

lcqtof\_t3\_neg\_4\_055\_38

m/z 687.3365

0246E+311.512RT (min)

lcqtof\_t3\_neg\_4\_019\_05

m/z 691.5540

012E+315.516RT (min)

lcqtof\_t3\_neg\_3\_101\_61

m/z 693.5157

024E+313.514RT (min)

lcqtof\_t3\_neg\_4\_003\_37

m/z 694.9601

01234E+31313.5RT (min)

lcqtof\_t3\_neg\_4\_011\_32

m/z 696.4855

012E+313.514RT (min)

lcqtof\_t3\_neg\_4\_085\_59

m/z 698.6278

01234E+313.514RT (min)

lcqtof\_t3\_neg\_4\_003\_37

m/z 699.6156

01234E+315.516RT (min)

lcqtof\_t3\_neg\_1\_012\_16

m/z 701.5889

0246E+31516RT (min)

lcqtof\_t3\_neg\_1\_029\_44

m/z 701.5901

0246E+31515.5RT (min)

lcqtof\_t3\_neg\_2\_037\_42

m/z 705.6803

012E+314.515RT (min)

lcqtof\_t3\_neg\_4\_020\_22

m/z 707.4635

01234E+312.513RT (min)

lcqtof\_t3\_neg\_4\_018\_23

m/z 708.5898

024E+316.517RT (min)

lcqtof\_t3\_neg\_4\_095\_60

m/z 711.6100

0123E+31313.514RT (min)

lcqtof\_t3\_neg\_4\_096\_60

m/z 715.2494

01E+34.55RT (min)

lcqtof\_t3\_neg\_4\_088\_59

m/z 716.4838

024E+21313.5RT (min)

lcqtof\_t3\_neg\_4\_073\_57

m/z 716.6146

024E+315.516RT (min)

lcqtof\_t3\_neg\_3\_112\_62

m/z 716.9743

024E+31313.5RT (min)

lcqtof\_t3\_neg\_4\_011\_32

m/z 717.6092

024E+315.516RT (min)

lcqtof\_t3\_neg\_3\_105\_61

m/z 719.2074

02468E+39.510RT (min)

lcqtof\_t3\_neg\_4\_016\_14

m/z 721.5848

02468E+315.516RT (min)

lcqtof\_t3\_neg\_1\_022\_28

m/z 722.5861

0123E+315.516RT (min)

lcqtof\_t3\_neg\_2\_051\_14

m/z 723.5171

01234E+212.513RT (min)

lcqtof\_t3\_neg\_2\_019\_32

m/z 724.9847

0246E+31414.5RT (min)

lcqtof\_t3\_neg\_4\_003\_37

m/z 726.6415

0246E+315.516RT (min)

lcqtof\_t3\_neg\_3\_113\_62

m/z 728.4592

024E+31212.5RT (min)

lcqtof\_t3\_neg\_4\_003\_37

m/z 728.5459

01E+413.514RT (min)

lcqtof\_t3\_neg\_4\_015\_36

m/z 731.5778

0113.514RT (min)

lcqtof\_t3\_neg\_2\_050\_01

m/z 733.5634

0123E+315.516RT (min)

lcqtof\_t3\_neg\_1\_029\_44

m/z 735.4341

0246E+312.513RT (min)

lcqtof\_t3\_neg\_4\_030\_31

m/z 735.5795

01234E+41515.516RT (min)

lcqtof\_t3\_neg\_3\_069\_57

m/z 736.2531

01E+39.510RT (min)

lcqtof\_t3\_neg\_4\_005\_18

m/z 737.5793

01E+41516RT (min)

lcqtof\_t3\_neg\_3\_076\_58

m/z 738.5803

024E+315.516RT (min)

lcqtof\_t3\_neg\_3\_113\_62

m/z 741.2200

0123E+39.510RT (min)

lcqtof\_t3\_neg\_1\_036\_14

m/z 742.6311

02468E+315.516RT (min)

lcqtof\_t3\_neg\_1\_025\_52

m/z 747.2339

0123E+39.510RT (min)

lcqtof\_t3\_neg\_1\_036\_14

m/z 749.4945

0246E+312.513RT (min)

lcqtof\_t3\_neg\_4\_034\_34

m/z 755.5909

024E+31313.5RT (min)

lcqtof\_t3\_neg\_4\_037\_01

m/z 759.2712

0123E+312.513RT (min)

lcqtof\_t3\_neg\_4\_084\_59

m/z 759.5611

024E+31515.516RT (min)

lcqtof\_t3\_neg\_3\_091\_60

m/z 760.5743

01234E+315.516RT (min)

lcqtof\_t3\_neg\_3\_086\_59

m/z 761.5660

02468E+315.516RT (min)

lcqtof\_t3\_neg\_3\_090\_60

m/z 764.5378

0246E+315.516RT (min)

lcqtof\_t3\_neg\_2\_007\_11

m/z 764.6050

024E+315.516RT (min)

lcqtof\_t3\_neg\_3\_086\_59

m/z 766.5362

01234E+315.516RT (min)

lcqtof\_t3\_neg\_3\_106\_62

m/z 777.5788

01E+41414.5RT (min)

lcqtof\_t3\_neg\_4\_016\_14

m/z 780.3715

012E+313.514RT (min)

lcqtof\_t3\_neg\_4\_083\_59

m/z 780.4783

01234E+316.517RT (min)

lcqtof\_t3\_neg\_4\_089\_59

m/z 780.5324

01E+415.516RT (min)

lcqtof\_t3\_neg\_2\_015\_33

m/z 781.5364

01E+413.514RT (min)

lcqtof\_t3\_neg\_1\_019\_23

m/z 782.5312

01E+415.516RT (min)

lcqtof\_t3\_neg\_2\_043\_31

m/z 782.6014

01E+413.514RT (min)

lcqtof\_t3\_neg\_4\_016\_14

m/z 783.5344

0123E+315.516RT (min)

lcqtof\_t3\_neg\_2\_002\_35

m/z 785.7736

02468E+313.514RT (min)

lcqtof\_t3\_neg\_4\_011\_32

m/z 793.5201

024E+313.514RT (min)

lcqtof\_t3\_neg\_4\_030\_31

m/z 793.6285

0123E+315.516RT (min)

lcqtof\_t3\_neg\_3\_076\_58

m/z 797.4602

01234E+312.513RT (min)

lcqtof\_t3\_neg\_1\_032\_36

m/z 798.6056

012E+315.516RT (min)

lcqtof\_t3\_neg\_3\_094\_60

m/z 807.5454

02468E+313.514RT (min)

lcqtof\_t3\_neg\_4\_030\_31

m/z 808.6217

01E+413.514RT (min)

lcqtof\_t3\_neg\_4\_037\_01

m/z 809.5723

0246E+312.513RT (min)

lcqtof\_t3\_neg\_4\_034\_34

m/z 812.6738

024E+313.514RT (min)

lcqtof\_t3\_neg\_4\_016\_14

m/z 813.2811

0246E+310.511RT (min)

lcqtof\_t3\_neg\_4\_076\_58

m/z 814.6002

02468E+315.516RT (min)

lcqtof\_t3\_neg\_2\_027\_15

m/z 816.5135

01E+412.513RT (min)

lcqtof\_t3\_neg\_4\_034\_34

m/z 818.6589

024E+31212.5RT (min)

lcqtof\_t3\_neg\_4\_092\_60

m/z 819.5022

0246E+31212.5RT (min)

lcqtof\_t3\_neg\_4\_034\_34

m/z 821.5153

024E+312.513RT (min)

lcqtof\_t3\_neg\_1\_032\_36

m/z 822.6698

02468E+312.513RT (min)

lcqtof\_t3\_neg\_4\_034\_34

m/z 826.6562

024E+316.517RT (min)

lcqtof\_t3\_neg\_4\_097\_60

m/z 830.6859

0246E+316.517RT (min)

lcqtof\_t3\_neg\_4\_097\_60

m/z 832.6365

024E+31212.5RT (min)

lcqtof\_t3\_neg\_4\_011\_32

m/z 836.6498

012E+213.514RT (min)

lcqtof\_t3\_neg\_1\_031\_24

m/z 838.4358

024E+31212.5RT (min)

lcqtof\_t3\_neg\_4\_086\_59

m/z 839.6186

0123E+315.516RT (min)

lcqtof\_t3\_neg\_2\_011\_25

m/z 842.6151

01E+412.513RT (min)

lcqtof\_t3\_neg\_4\_016\_14

m/z 843.5920

0246E+315.516RT (min)

lcqtof\_t3\_neg\_1\_034\_30

m/z 844.4347

0246E+31212.5RT (min)

lcqtof\_t3\_neg\_4\_034\_34

m/z 844.5945

01234E+315.516RT (min)

lcqtof\_t3\_neg\_1\_006\_05

m/z 847.7226

0246E+31313.5RT (min)

lcqtof\_t3\_neg\_2\_028\_24

m/z 850.2318

0123E+39.510RT (min)

lcqtof\_t3\_neg\_4\_076\_58

m/z 853.5034

02468E+31313.5RT (min)

lcqtof\_t3\_neg\_4\_089\_59

m/z 859.5865

01E+415.516RT (min)

lcqtof\_t3\_neg\_1\_026\_19

m/z 860.5895

01E+415.516RT (min)

lcqtof\_t3\_neg\_1\_026\_19

m/z 861.5917

0123E+315.516RT (min)

lcqtof\_t3\_neg\_1\_015\_34

m/z 862.4253

0246E+312.513RT (min)

lcqtof\_t3\_neg\_4\_088\_59

m/z 862.4866

02468E+31212.5RT (min)

lcqtof\_t3\_neg\_4\_034\_34

m/z 866.5916

01E+21313.5RT (min)

lcqtof\_t3\_neg\_1\_003\_07

m/z 885.7268

0246E+31313.5RT (min)

lcqtof\_t3\_neg\_4\_040\_24

m/z 887.6482

02468E+31212.5RT (min)

lcqtof\_t3\_neg\_4\_040\_24

m/z 888.7116

0246E+31313.5RT (min)

lcqtof\_t3\_neg\_1\_031\_24

m/z 889.7575

0246E+313.514RT (min)

lcqtof\_t3\_neg\_4\_040\_24

m/z 895.6952

0246E+31313.5RT (min)

lcqtof\_t3\_neg\_4\_095\_60

m/z 899.4256

01234E+312.513RT (min)

lcqtof\_t3\_neg\_4\_018\_23

m/z 900.4957

01234E+31212.5RT (min)

lcqtof\_t3\_neg\_4\_055\_38

m/z 901.6733

0246E+313.514RT (min)

lcqtof\_t3\_neg\_4\_002\_17

m/z 909.6638

01E+41313.5RT (min)

lcqtof\_t3\_neg\_1\_001\_49

m/z 912.4399

02468E+39.510RT (min)

lcqtof\_t3\_neg\_4\_016\_14

m/z 917.4589

02468E+312.513RT (min)

lcqtof\_t3\_neg\_4\_035\_08

m/z 925.2853

012E+31111.5RT (min)

lcqtof\_t3\_neg\_1\_010\_27

m/z 928.6290

02468E+313.514RT (min)

lcqtof\_t3\_neg\_4\_037\_01

m/z 931.5388

0123E+312.513RT (min)

lcqtof\_t3\_neg\_4\_072\_57

m/z 932.6011

0246E+31212.5RT (min)

lcqtof\_t3\_neg\_4\_091\_60

m/z 932.6991

0246E+313.514RT (min)

lcqtof\_t3\_neg\_4\_040\_24

m/z 933.6949

0246E+31111.5RT (min)

lcqtof\_t3\_neg\_4\_005\_18

m/z 936.6409

0123E+313.514RT (min)

lcqtof\_t3\_neg\_4\_094\_60

m/z 936.6903

0246E+313.514RT (min)

lcqtof\_t3\_neg\_4\_018\_23

m/z 937.6486

0123E+313.514RT (min)

lcqtof\_t3\_neg\_4\_006\_16

m/z 940.6761

0246E+31313.5RT (min)

lcqtof\_t3\_neg\_4\_094\_60

m/z 941.6527

011313.5RT (min)

lcqtof\_t3\_neg\_4\_040\_24

m/z 944.4884

02468E+312.513RT (min)

lcqtof\_t3\_neg\_4\_034\_34

m/z 947.4079

0246E+39.510RT (min)

lcqtof\_t3\_neg\_4\_016\_14

m/z 947.4595

0246E+31010.5RT (min)

lcqtof\_t3\_neg\_4\_102\_61

m/z 947.6940

024E+31414.5RT (min)

lcqtof\_t3\_neg\_4\_037\_01

m/z 948.4735

024E+312.513RT (min)

lcqtof\_t3\_neg\_4\_018\_23

m/z 952.4859

01E+312.513RT (min)

lcqtof\_t3\_neg\_4\_068\_57

m/z 960.5480

01E+313.514RT (min)

lcqtof\_t3\_neg\_4\_003\_37

m/z 961.5034

0246E+312.513RT (min)

lcqtof\_t3\_neg\_4\_003\_37

m/z 969.3259

02468E+388.5RT (min)

lcqtof\_t3\_neg\_4\_004\_20

m/z 970.3055

0246E+388.5RT (min)

lcqtof\_t3\_neg\_4\_004\_20

m/z 973.6290

012E+313.514RT (min)

lcqtof\_t3\_neg\_4\_003\_37

m/z 977.5308

0123E+31313.5RT (min)

lcqtof\_t3\_neg\_4\_066\_57

m/z 980.5319

0246E+312.513RT (min)

lcqtof\_t3\_neg\_4\_003\_37

m/z 980.7157

024E+313.514RT (min)

lcqtof\_t3\_neg\_4\_018\_23

m/z 981.7191

01234E+213.514RT (min)

lcqtof\_t3\_neg\_4\_091\_60

m/z 982.2885

012E+388.5RT (min)

lcqtof\_t3\_neg\_4\_108\_62

m/z 983.3017

012E+388.5RT (min)

lcqtof\_t3\_neg\_4\_004\_20

m/z 983.4343

024E+312.513RT (min)

lcqtof\_t3\_neg\_4\_084\_59

m/z 984.7382

012E+313.514RT (min)

lcqtof\_t3\_neg\_4\_040\_24

m/z 986.6350

012E+313.514RT (min)

lcqtof\_t3\_neg\_4\_046\_33

m/z 1000.6655

024E+31313.5RT (min)

lcqtof\_t3\_neg\_4\_015\_36

m/z 1004.5352

0246E+312.513RT (min)

lcqtof\_t3\_neg\_4\_003\_37

m/z 1008.6141

0123E+313.514RT (min)

lcqtof\_t3\_neg\_1\_008\_02

m/z 1012.7304

0246E+313.514RT (min)

lcqtof\_t3\_neg\_4\_091\_60

m/z 1018.8400

0246E+315.516RT (min)

lcqtof\_t3\_neg\_1\_005\_18

m/z 1019.8380

024E+315.516RT (min)

lcqtof\_t3\_neg\_1\_017\_51

m/z 1024.3595

0123E+39.510RT (min)

lcqtof\_t3\_neg\_1\_036\_14

m/z 1026.8537

024E+31313.5RT (min)

lcqtof\_t3\_neg\_4\_011\_32

m/z 1033.6064

012E+313.514RT (min)

lcqtof\_t3\_neg\_4\_074\_58

m/z 1042.6138

024E+312.513RT (min)

lcqtof\_t3\_neg\_4\_011\_32

m/z 1044.8554

02468E+315.516RT (min)

lcqtof\_t3\_neg\_1\_047\_22

m/z 1050.4183

024E+312.513RT (min)

lcqtof\_t3\_neg\_4\_082\_59

m/z 1050.7742

0246E+31212.5RT (min)

lcqtof\_t3\_neg\_4\_040\_24

m/z 1054.5223

01234E+31313.5RT (min)

lcqtof\_t3\_neg\_4\_032\_04

m/z 1061.7870

01234E+315.516RT (min)

lcqtof\_t3\_neg\_2\_011\_25

m/z 1063.7868

024E+315.516RT (min)

lcqtof\_t3\_neg\_2\_025\_52

m/z 1069.6357

024E+312.513RT (min)

lcqtof\_t3\_neg\_4\_047\_07

m/z 1069.7290

024E+313.514RT (min)

lcqtof\_t3\_neg\_4\_003\_37

m/z 1069.8388

0246E+313.514RT (min)

lcqtof\_t3\_neg\_4\_040\_24

m/z 1073.6942

0246E+31313.5RT (min)

lcqtof\_t3\_neg\_4\_018\_23

m/z 1076.3616

012E+39.510RT (min)

lcqtof\_t3\_neg\_4\_081\_58

m/z 1085.7249

0246E+313.514RT (min)

lcqtof\_t3\_neg\_4\_008\_35

m/z 1087.5001

0123E+312.513RT (min)

lcqtof\_t3\_neg\_4\_066\_57

m/z 1088.7153

0246E+316.517RT (min)

lcqtof\_t3\_neg\_4\_055\_38

m/z 1095.6237

024E+310.511RT (min)

lcqtof\_t3\_neg\_4\_037\_01

m/z 1096.7290

0246E+31414.5RT (min)

lcqtof\_t3\_neg\_4\_003\_37

m/z 1099.7491

0246E+313.514RT (min)

lcqtof\_t3\_neg\_4\_018\_23

m/z 1106.8808

0123E+31313.5RT (min)

lcqtof\_t3\_neg\_2\_016\_05

m/z 1110.6138

012E+310.511RT (min)

lcqtof\_t3\_neg\_1\_036\_14

m/z 1110.7437

02468E+31414.5RT (min)

lcqtof\_t3\_neg\_4\_003\_37

m/z 1110.8483

024E+31313.5RT (min)

lcqtof\_t3\_neg\_4\_011\_32

m/z 1125.8707

02468E+31313.5RT (min)

lcqtof\_t3\_neg\_4\_040\_24

m/z 1136.8931

0123E+21414.5RT (min)

lcqtof\_t3\_neg\_1\_010\_27

m/z 1137.8591

0246E+313.514RT (min)

lcqtof\_t3\_neg\_4\_011\_32

m/z 1138.6245

0246E+312.513RT (min)

lcqtof\_t3\_neg\_4\_035\_08

m/z 1140.7351

012E+413.514RT (min)

lcqtof\_t3\_neg\_4\_018\_23

m/z 1143.3433

0123E+39.510RT (min)

lcqtof\_t3\_neg\_4\_081\_58

m/z 1149.3597

01234E+39.510RT (min)

lcqtof\_t3\_neg\_4\_077\_58

m/z 1154.8743

0246E+41313.5RT (min)

lcqtof\_t3\_neg\_4\_040\_24
