## Supplementary material for "The Everything Bagel Feature Finder: Ultra-fast automated feature finding for untargeted metabolomics": SI Data File: tool_unique_xic_st_XCMS_ST004650.html

XCMS-unique features (not detected by the other two tools) - ST004650

### XCMS-unique features (not detected by the other two tools) - ST004650

841 features · qtof · XIC window m/z ± 25 ppm (≥ 0.002 Da) · ± 0.5 min

Every panel is a feature **XCMS** detected that neither of the other two tools detected — three-way unique across EB (standard mode (min\_detection\_rate=0.1)), Masscube and XCMS, under the standard matching rule (m/z ± 25 ppm; apex within the reference RT window ± 0.2 min). It shows the MS1 extracted-ion chromatogram at the feature's m/z, drawn from the raw mzML named in `alignment_reference_file`, zoomed to ± 0.5 min around the feature's reported apex RT. **Gray solid line** = the feature's reported apex RT; **red dashed line** = XCMS's assigned MS/MS — `RTINSECONDS` in its `aligned_features.mgf`; the panel widens to keep an off-apex MS/MS on-screen. Features with no assigned MS/MS show only the gray line. y is per-cell autoscaled. Rows keep the table order (m/z ascending).

m/z 57.0704

012E+316.517RT (min)

lcqtof\_hilic\_pos\_4\_091\_60

m/z 87.0454

0123E+344.5RT (min)

lcqtof\_hilic\_pos\_4\_026\_01

m/z 92.0379

0246E+300.5RT (min)

lcqtof\_hilic\_pos\_4\_111\_62

m/z 92.0380

012E+318.519RT (min)

lcqtof\_hilic\_pos\_4\_073\_57

m/z 92.0380

024E+322.5RT (min)

lcqtof\_hilic\_pos\_4\_088\_59

m/z 99.0450

01E+30.51RT (min)

lcqtof\_hilic\_pos\_1\_039\_06

m/z 99.0579

024E+233.5RT (min)

lcqtof\_hilic\_pos\_4\_004\_13

m/z 100.0768

01E+41.52RT (min)

lcqtof\_hilic\_pos\_4\_080\_58

m/z 100.0769

0246E+318.519RT (min)

lcqtof\_hilic\_pos\_4\_083\_59

m/z 100.0864

012E+36.57RT (min)

lcqtof\_hilic\_pos\_2\_041\_54

m/z 101.0644

0123E+34.55RT (min)

lcqtof\_hilic\_pos\_4\_036\_24

m/z 101.1086

02468E+333.5RT (min)

lcqtof\_hilic\_pos\_4\_036\_24

m/z 103.9566

0123E+48.59RT (min)

lcqtof\_hilic\_pos\_4\_101\_61

m/z 103.9567

024E+42.53RT (min)

lcqtof\_hilic\_pos\_4\_035\_41

m/z 104.0619

01234E+211.5RT (min)

lcqtof\_hilic\_pos\_1\_026\_20

m/z 105.0707

0123E+23.54RT (min)

lcqtof\_hilic\_pos\_3\_023\_01

m/z 105.9549

01E+48.59RT (min)

lcqtof\_hilic\_pos\_4\_068\_57

m/z 105.9552

0123E+318.519RT (min)

lcqtof\_hilic\_pos\_3\_113\_62

m/z 107.0616

012E+31919.5RT (min)

lcqtof\_hilic\_pos\_4\_081\_58

m/z 107.0667

01E+23.54RT (min)

lcqtof\_hilic\_pos\_4\_076\_58

m/z 109.0297

0246E+21.52RT (min)

lcqtof\_hilic\_pos\_4\_023\_19

m/z 111.0460

012E+322.5RT (min)

lcqtof\_hilic\_pos\_4\_088\_59

m/z 111.0926

01234E+33.54RT (min)

lcqtof\_hilic\_pos\_1\_003\_44

m/z 113.0793

02468E+11.52RT (min)

lcqtof\_hilic\_pos\_4\_111\_62

m/z 113.0817

02468E+166.5RT (min)

lcqtof\_hilic\_pos\_4\_074\_58

m/z 114.1013

0246E+377.5RT (min)

lcqtof\_hilic\_pos\_4\_013\_36

m/z 114.1368

0123E+31818.5RT (min)

lcqtof\_hilic\_pos\_4\_098\_61

m/z 115.1240

0123E+31919.5RT (min)

lcqtof\_hilic\_pos\_4\_013\_36

m/z 115.1240

012E+31.52RT (min)

lcqtof\_hilic\_pos\_4\_009\_50

m/z 118.0490

02468E+255.5RT (min)

lcqtof\_hilic\_pos\_1\_005\_01

m/z 119.0732

0246E+45.56RT (min)

lcqtof\_hilic\_pos\_4\_036\_24

m/z 119.0902

0246E+311.512RT (min)

lcqtof\_hilic\_pos\_3\_038\_03

m/z 124.0462

012E+31.52RT (min)

lcqtof\_hilic\_pos\_4\_027\_05

m/z 124.0549

024E+21.52RT (min)

lcqtof\_hilic\_pos\_4\_015\_42

m/z 124.1144

01E+32.533.5RT (min)

lcqtof\_hilic\_pos\_4\_003\_21

m/z 125.0732

02468E+42.53RT (min)

lcqtof\_hilic\_pos\_4\_016\_09

m/z 125.0960

01E+44.55RT (min)

lcqtof\_hilic\_pos\_4\_106\_62

m/z 125.9874

0246E+39.510RT (min)

lcqtof\_hilic\_pos\_4\_077\_58

m/z 126.0736

0246E+318.519RT (min)

lcqtof\_hilic\_pos\_4\_075\_58

m/z 126.0740

0246E+300.5RT (min)

lcqtof\_hilic\_pos\_4\_043\_39

m/z 126.0741

0246E+32.53RT (min)

lcqtof\_hilic\_pos\_4\_004\_13

m/z 126.9903

012E+38.59RT (min)

lcqtof\_hilic\_pos\_4\_079\_58

m/z 127.0077

01234E+333.5RT (min)

lcqtof\_hilic\_pos\_4\_096\_60

m/z 127.0735

012E+488.5RT (min)

lcqtof\_hilic\_pos\_4\_047\_40

m/z 127.0870

01234E+322.5RT (min)

lcqtof\_hilic\_pos\_3\_001\_49

m/z 127.0870

01234E+318.519RT (min)

lcqtof\_hilic\_pos\_3\_105\_61

m/z 130.0616

012E+31919.5RT (min)

lcqtof\_hilic\_pos\_4\_090\_60

m/z 130.0868

02468E+11.52RT (min)

lcqtof\_hilic\_pos\_4\_005\_11

m/z 131.1072

01E+30.51RT (min)

lcqtof\_hilic\_pos\_4\_048\_04

m/z 132.0777

01E+33.54RT (min)

lcqtof\_hilic\_pos\_4\_040\_06

m/z 133.0580

01E+455.5RT (min)

lcqtof\_hilic\_pos\_4\_023\_19

m/z 133.0631

01234E+366.5RT (min)

lcqtof\_hilic\_pos\_3\_083\_59

m/z 134.0615

024E+31819RT (min)

lcqtof\_hilic\_pos\_4\_089\_59

m/z 135.0564

0246E+31.52RT (min)

lcqtof\_hilic\_pos\_4\_050\_16

m/z 136.0626

012E+41818.5RT (min)

lcqtof\_hilic\_pos\_4\_037\_35

m/z 136.0629

0123E+36.57RT (min)

lcqtof\_hilic\_pos\_4\_008\_25

m/z 137.1077

0246E+30.51RT (min)

lcqtof\_hilic\_pos\_4\_030\_03

m/z 138.1037

01234E+311.52RT (min)

lcqtof\_hilic\_pos\_4\_112\_62

m/z 138.1047

012E+319.520RT (min)

lcqtof\_hilic\_pos\_4\_080\_58

m/z 140.0696

012E+31010.5RT (min)

lcqtof\_hilic\_pos\_4\_038\_38

m/z 141.9596

024E+31214RT (min)

lcqtof\_hilic\_pos\_1\_039\_06

m/z 142.1240

012E+31818.5RT (min)

lcqtof\_hilic\_pos\_3\_111\_62

m/z 143.0753

0123E+20.51RT (min)

lcqtof\_hilic\_pos\_4\_113\_62

m/z 144.9830

01234E+48.59RT (min)

lcqtof\_hilic\_pos\_4\_041\_54

m/z 146.9812

012E+48.59RT (min)

lcqtof\_hilic\_pos\_4\_073\_57

m/z 146.9813

0123E+42.53RT (min)

lcqtof\_hilic\_pos\_4\_048\_04

m/z 146.9814

01E+418.519RT (min)

lcqtof\_hilic\_pos\_4\_037\_35

m/z 148.0805

02468E+311.5RT (min)

lcqtof\_hilic\_pos\_4\_103\_61

m/z 148.0876

024E+318.519RT (min)

lcqtof\_hilic\_pos\_4\_089\_59

m/z 149.0815

01E+418.519RT (min)

lcqtof\_hilic\_pos\_4\_012\_02

m/z 150.0272

01E+418.519RT (min)

lcqtof\_hilic\_pos\_4\_083\_59

m/z 150.0273

01E+41616.5RT (min)

lcqtof\_hilic\_pos\_4\_008\_25

m/z 150.0668

012E+418.519RT (min)

lcqtof\_hilic\_pos\_3\_094\_60

m/z 150.0862

01E+34.55RT (min)

lcqtof\_hilic\_pos\_4\_031\_33

m/z 151.0705

01E+318.519RT (min)

lcqtof\_hilic\_pos\_4\_108\_62

m/z 153.0653

012E+366.5RT (min)

lcqtof\_hilic\_pos\_4\_036\_24

m/z 153.1122

024E+24.55RT (min)

lcqtof\_hilic\_pos\_3\_034\_24

m/z 155.0677

02468E+188.5RT (min)

lcqtof\_hilic\_pos\_4\_112\_62

m/z 155.0738

02468E+311.5RT (min)

lcqtof\_hilic\_pos\_4\_113\_62

m/z 157.1335

01E+333.5RT (min)

lcqtof\_hilic\_pos\_4\_054\_18

m/z 158.1548

01E+42.53RT (min)

lcqtof\_hilic\_pos\_4\_001\_49

m/z 159.0672

01E+311.5RT (min)

lcqtof\_hilic\_pos\_4\_050\_16

m/z 159.1138

024E+36.57RT (min)

lcqtof\_hilic\_pos\_3\_111\_62

m/z 160.0629

0123E+22.53RT (min)

lcqtof\_hilic\_pos\_4\_002\_23

m/z 160.1099

01E+33.54RT (min)

lcqtof\_hilic\_pos\_4\_006\_29

m/z 163.0764

0123E+31818.5RT (min)

lcqtof\_hilic\_pos\_4\_075\_58

m/z 163.0835

01E+511.5RT (min)

lcqtof\_hilic\_pos\_2\_002\_32

m/z 164.1010

012E+344.5RT (min)

lcqtof\_hilic\_pos\_4\_090\_60

m/z 165.1194

0246E+40.51RT (min)

lcqtof\_hilic\_pos\_4\_015\_42

m/z 167.0998

02468E+318.519RT (min)

lcqtof\_hilic\_pos\_4\_076\_58

m/z 168.0374

0123E+31818.5RT (min)

lcqtof\_hilic\_pos\_4\_113\_62

m/z 168.0471

01E+40.51RT (min)

lcqtof\_hilic\_pos\_2\_001\_49

m/z 168.0681

0123E+316.517RT (min)

lcqtof\_hilic\_pos\_4\_039\_15

m/z 168.1400

0122.5RT (min)

lcqtof\_hilic\_pos\_3\_105\_61

m/z 169.1175

0123E+30.51RT (min)

lcqtof\_hilic\_pos\_2\_023\_22

m/z 169.1177

012E+318.519RT (min)

lcqtof\_hilic\_pos\_3\_097\_60

m/z 170.1546

0123E+333.5RT (min)

lcqtof\_hilic\_pos\_4\_006\_29

m/z 171.1487

0246E+318.519RT (min)

lcqtof\_hilic\_pos\_3\_106\_62

m/z 172.0616

01E+34.55RT (min)

lcqtof\_hilic\_pos\_4\_050\_16

m/z 173.1299

01234E+318.519RT (min)

lcqtof\_hilic\_pos\_4\_089\_59

m/z 177.0528

01E+20.51RT (min)

lcqtof\_hilic\_pos\_4\_082\_59

m/z 178.1234

01234E+31111.5RT (min)

lcqtof\_hilic\_pos\_4\_053\_30

m/z 178.1234

012E+31414.5RT (min)

lcqtof\_hilic\_pos\_3\_039\_22

m/z 180.0769

0123E+318.519RT (min)

lcqtof\_hilic\_pos\_4\_107\_62

m/z 181.0518

01E+311.5RT (min)

lcqtof\_hilic\_pos\_4\_083\_59

m/z 181.1333

01E+211.5RT (min)

lcqtof\_hilic\_pos\_4\_111\_62

m/z 182.1915

012E+31.52RT (min)

lcqtof\_hilic\_pos\_1\_018\_08

m/z 182.1917

012E+31919.5RT (min)

lcqtof\_hilic\_pos\_2\_027\_18

m/z 183.0497

02468E+399.5RT (min)

lcqtof\_hilic\_pos\_4\_022\_10

m/z 183.0573

024E+344.5RT (min)

lcqtof\_hilic\_pos\_4\_113\_62

m/z 183.1946

0246E+32.53RT (min)

lcqtof\_hilic\_pos\_4\_040\_06

m/z 185.0861

012E+344.5RT (min)

lcqtof\_hilic\_pos\_4\_015\_42

m/z 185.1601

02468E+233.5RT (min)

lcqtof\_hilic\_pos\_4\_107\_62

m/z 186.0748

02468E+311.5RT (min)

lcqtof\_hilic\_pos\_4\_036\_24

m/z 186.1246

01E+488.5RT (min)

lcqtof\_hilic\_pos\_3\_015\_10

m/z 187.0545

01234E+444.5RT (min)

lcqtof\_hilic\_pos\_4\_082\_59

m/z 189.1140

0123E+333.5RT (min)

lcqtof\_hilic\_pos\_4\_091\_60

m/z 189.1144

0123E+318.519RT (min)

lcqtof\_hilic\_pos\_4\_106\_62

m/z 189.1216

01E+244.5RT (min)

lcqtof\_hilic\_pos\_4\_015\_42

m/z 190.0988

024E+318.519RT (min)

lcqtof\_hilic\_pos\_4\_098\_61

m/z 190.1201

01234E+43.54RT (min)

lcqtof\_hilic\_pos\_4\_050\_16

m/z 193.0644

01234E+311.5RT (min)

lcqtof\_hilic\_pos\_4\_109\_62

m/z 193.0753

0123E+11.52RT (min)

lcqtof\_hilic\_pos\_4\_105\_61

m/z 194.1182

0123E+31111.5RT (min)

lcqtof\_hilic\_pos\_4\_002\_23

m/z 194.1185

0123E+31414.5RT (min)

lcqtof\_hilic\_pos\_4\_006\_29

m/z 196.1708

01234E+322.5RT (min)

lcqtof\_hilic\_pos\_4\_036\_24

m/z 197.0614

01E+30.51RT (min)

lcqtof\_hilic\_pos\_4\_010\_37

m/z 197.0659

012E+31010.5RT (min)

lcqtof\_hilic\_pos\_4\_092\_60

m/z 198.1866

012E+31818.5RT (min)

lcqtof\_hilic\_pos\_3\_108\_62

m/z 198.9545

012E+31313.5RT (min)

lcqtof\_hilic\_pos\_2\_049\_55

m/z 198.9551

012E+31313.5RT (min)

lcqtof\_hilic\_pos\_3\_043\_46

m/z 199.1787

01234E+30.51RT (min)

lcqtof\_hilic\_pos\_4\_043\_39

m/z 199.1804

012E+31818.5RT (min)

lcqtof\_hilic\_pos\_3\_093\_60

m/z 200.2380

0123E+31919.5RT (min)

lcqtof\_hilic\_pos\_4\_079\_58

m/z 201.0877

01234E+34.55RT (min)

lcqtof\_hilic\_pos\_4\_026\_01

m/z 202.0579

0246E+30.51RT (min)

lcqtof\_hilic\_pos\_4\_050\_16

m/z 202.0778

0246E+30.51RT (min)

lcqtof\_hilic\_pos\_4\_053\_30

m/z 203.0822

01E+411.5RT (min)

lcqtof\_hilic\_pos\_4\_003\_21

m/z 204.1247

012E+21.52RT (min)

lcqtof\_hilic\_pos\_4\_099\_61

m/z 205.0952

02468E+311.5RT (min)

lcqtof\_hilic\_pos\_4\_053\_30

m/z 206.1084

01E+333.5RT (min)

lcqtof\_hilic\_pos\_4\_002\_23

m/z 206.1400

01234E+444.5RT (min)

lcqtof\_hilic\_pos\_4\_001\_49

m/z 207.1324

01E+50.51RT (min)

lcqtof\_hilic\_pos\_4\_055\_43

m/z 209.1297

01E+355.5RT (min)

lcqtof\_hilic\_pos\_4\_036\_24

m/z 209.1546

01E+317.518RT (min)

lcqtof\_hilic\_pos\_3\_096\_60

m/z 212.1660

012E+318.519RT (min)

lcqtof\_hilic\_pos\_4\_097\_60

m/z 213.1145

01E+42.53RT (min)

lcqtof\_hilic\_pos\_4\_007\_45

m/z 213.1755

01E+40.51RT (min)

lcqtof\_hilic\_pos\_4\_048\_04

m/z 213.1963

0123E+311.5RT (min)

lcqtof\_hilic\_pos\_4\_011\_48

m/z 214.2536

01234E+31919.5RT (min)

lcqtof\_hilic\_pos\_4\_105\_61

m/z 215.0663

01E+555.5RT (min)

lcqtof\_hilic\_pos\_3\_091\_60

m/z 215.1380

02468E+30.51RT (min)

lcqtof\_hilic\_pos\_4\_023\_19

m/z 218.1091

0246E+211.5RT (min)

lcqtof\_hilic\_pos\_4\_054\_18

m/z 218.9410

0123E+31313.5RT (min)

lcqtof\_hilic\_pos\_1\_008\_11

m/z 219.0887

0246E+311.5RT (min)

lcqtof\_hilic\_pos\_4\_050\_16

m/z 219.1750

024E+31818.5RT (min)

lcqtof\_hilic\_pos\_4\_088\_59

m/z 220.9355

012E+39.510RT (min)

lcqtof\_hilic\_pos\_4\_002\_23

m/z 220.9364

0123E+313.514RT (min)

lcqtof\_hilic\_pos\_3\_113\_62

m/z 221.0362

01E+33.54RT (min)

lcqtof\_hilic\_pos\_4\_096\_60

m/z 221.1335

01234E+333.5RT (min)

lcqtof\_hilic\_pos\_4\_023\_19

m/z 222.1275

01E+466.5RT (min)

lcqtof\_hilic\_pos\_1\_001\_49

m/z 223.1115

01E+311.5RT (min)

lcqtof\_hilic\_pos\_4\_095\_60

m/z 225.1327

0246E+21.52RT (min)

lcqtof\_hilic\_pos\_4\_085\_59

m/z 225.1349

024E+310.511RT (min)

lcqtof\_hilic\_pos\_4\_009\_50

m/z 225.1354

01234E+310.511RT (min)

lcqtof\_hilic\_pos\_4\_002\_23

m/z 226.1085

01E+32.53RT (min)

lcqtof\_hilic\_pos\_3\_094\_60

m/z 226.1086

01E+318.519RT (min)

lcqtof\_hilic\_pos\_3\_105\_61

m/z 226.1812

0246E+318.519RT (min)

lcqtof\_hilic\_pos\_3\_007\_31

m/z 226.1913

02468E+33.54RT (min)

lcqtof\_hilic\_pos\_4\_010\_37

m/z 228.2325

01234E+31818.5RT (min)

lcqtof\_hilic\_pos\_4\_087\_59

m/z 228.2693

012E+31919.5RT (min)

lcqtof\_hilic\_pos\_4\_029\_44

m/z 229.0956

0246E+333.5RT (min)

lcqtof\_hilic\_pos\_4\_031\_33

m/z 231.1274

01234E+422.5RT (min)

lcqtof\_hilic\_pos\_4\_106\_62

m/z 231.1345

0123E+40.51RT (min)

lcqtof\_hilic\_pos\_4\_101\_61

m/z 231.1386

0246E+13.54RT (min)

lcqtof\_hilic\_pos\_4\_103\_61

m/z 231.1401

01E+25.56RT (min)

lcqtof\_hilic\_pos\_4\_109\_62

m/z 231.2519

012E+31919.5RT (min)

lcqtof\_hilic\_pos\_4\_091\_60

m/z 232.1203

01E+418.519RT (min)

lcqtof\_hilic\_pos\_1\_030\_33

m/z 232.1204

012E+422.5RT (min)

lcqtof\_hilic\_pos\_1\_026\_20

m/z 232.1278

02468E+30.51RT (min)

lcqtof\_hilic\_pos\_4\_031\_33

m/z 233.1219

01E+322.5RT (min)

lcqtof\_hilic\_pos\_4\_100\_61

m/z 233.1226

012E+318.519RT (min)

lcqtof\_hilic\_pos\_2\_014\_25

m/z 233.1285

012E+333.5RT (min)

lcqtof\_hilic\_pos\_2\_001\_49

m/z 234.0978

012E+31818.5RT (min)

lcqtof\_hilic\_pos\_4\_088\_59

m/z 234.0980

0246E+30.51RT (min)

lcqtof\_hilic\_pos\_3\_098\_61

m/z 234.1285

01E+41.52RT (min)

lcqtof\_hilic\_pos\_4\_054\_18

m/z 235.1310

012E+318.519RT (min)

lcqtof\_hilic\_pos\_4\_100\_61

m/z 238.0817

012E+244.5RT (min)

lcqtof\_hilic\_pos\_4\_095\_60

m/z 238.1572

0123E+411.5RT (min)

lcqtof\_hilic\_pos\_4\_044\_32

m/z 240.1090

01E+36.57RT (min)

lcqtof\_hilic\_pos\_4\_036\_24

m/z 241.1815

0246E+40.51RT (min)

lcqtof\_hilic\_pos\_4\_012\_02

m/z 242.2845

0123E+318.519RT (min)

lcqtof\_hilic\_pos\_4\_099\_61

m/z 242.2852

0123E+31919.5RT (min)

lcqtof\_hilic\_pos\_4\_108\_62

m/z 244.1156

01234E+333.5RT (min)

lcqtof\_hilic\_pos\_4\_004\_13

m/z 244.1207

0123E+316.517RT (min)

lcqtof\_hilic\_pos\_4\_109\_62

m/z 244.1919

012E+31818.5RT (min)

lcqtof\_hilic\_pos\_3\_091\_60

m/z 245.1042

01E+344.5RT (min)

lcqtof\_hilic\_pos\_4\_018\_34

m/z 245.1238

02468E+333.5RT (min)

lcqtof\_hilic\_pos\_4\_036\_24

m/z 246.2436

0123E+377.5RT (min)

lcqtof\_hilic\_pos\_4\_056\_56

m/z 246.9358

01E+41313.5RT (min)

lcqtof\_hilic\_pos\_1\_019\_19

m/z 247.1483

0144.5RT (min)

lcqtof\_hilic\_pos\_4\_003\_21

m/z 248.0940

01E+26.57RT (min)

lcqtof\_hilic\_pos\_4\_036\_24

m/z 248.1701

02468E+399.5RT (min)

lcqtof\_hilic\_pos\_4\_051\_17

m/z 248.1703

02468E+311.512RT (min)

lcqtof\_hilic\_pos\_4\_095\_60

m/z 250.1276

01234E+318.519RT (min)

lcqtof\_hilic\_pos\_4\_108\_62

m/z 251.1055

012E+20.51RT (min)

lcqtof\_hilic\_pos\_4\_111\_62

m/z 251.1284

012E+311.5RT (min)

lcqtof\_hilic\_pos\_4\_095\_60

m/z 251.1370

01E+355.5RT (min)

lcqtof\_hilic\_pos\_3\_045\_32

m/z 253.0774

012E+211.5RT (min)

lcqtof\_hilic\_pos\_4\_084\_59

m/z 253.1919

01234E+333.5RT (min)

lcqtof\_hilic\_pos\_4\_004\_13

m/z 254.1399

0123E+31818.5RT (min)

lcqtof\_hilic\_pos\_4\_095\_60

m/z 254.1402

0246E+322.5RT (min)

lcqtof\_hilic\_pos\_4\_005\_11

m/z 254.1404

01234E+318.519RT (min)

lcqtof\_hilic\_pos\_4\_106\_62

m/z 255.2407

012E+50.51RT (min)

lcqtof\_hilic\_pos\_4\_111\_62

m/z 256.2640

01E+418.519RT (min)

lcqtof\_hilic\_pos\_4\_098\_61

m/z 257.0181

0123E+34.55RT (min)

lcqtof\_hilic\_pos\_4\_008\_25

m/z 257.1252

012E+377.5RT (min)

lcqtof\_hilic\_pos\_1\_016\_14

m/z 257.2673

012E+31818.5RT (min)

lcqtof\_hilic\_pos\_4\_074\_58

m/z 258.0508

024E+10.51RT (min)

lcqtof\_hilic\_pos\_4\_079\_58

m/z 258.2794

01E+41919.5RT (min)

lcqtof\_hilic\_pos\_4\_082\_59

m/z 259.2831

012E+31919.5RT (min)

lcqtof\_hilic\_pos\_4\_072\_57

m/z 259.2832

01234E+322.5RT (min)

lcqtof\_hilic\_pos\_4\_103\_61

m/z 259.9674

012E+312.513RT (min)

lcqtof\_hilic\_pos\_1\_034\_15

m/z 260.9376

024E+30.51RT (min)

lcqtof\_hilic\_pos\_4\_077\_58

m/z 261.0370

0123E+37.58RT (min)

lcqtof\_hilic\_pos\_4\_010\_37

m/z 261.0780

0123E+222.5RT (min)

lcqtof\_hilic\_pos\_4\_106\_62

m/z 261.1608

0133.5RT (min)

lcqtof\_hilic\_pos\_4\_094\_60

m/z 262.1125

01234E+366.5RT (min)

lcqtof\_hilic\_pos\_1\_031\_32

m/z 262.1526

02468E+318.519RT (min)

lcqtof\_hilic\_pos\_4\_106\_62

m/z 262.1526

02468E+31616.5RT (min)

lcqtof\_hilic\_pos\_4\_009\_50

m/z 263.0818

0123E+31818.5RT (min)

lcqtof\_hilic\_pos\_1\_034\_15

m/z 263.0867

012E+36.57RT (min)

lcqtof\_hilic\_pos\_4\_001\_49

m/z 263.0906

01E+33.54RT (min)

lcqtof\_hilic\_pos\_4\_001\_49

m/z 265.0050

01234E+318.519RT (min)

lcqtof\_hilic\_pos\_4\_106\_62

m/z 265.1248

024E+21.52RT (min)

lcqtof\_hilic\_pos\_2\_002\_32

m/z 266.1554

01E+30.51RT (min)

lcqtof\_hilic\_pos\_4\_048\_04

m/z 267.0819

012E+31.52RT (min)

lcqtof\_hilic\_pos\_1\_052\_21

m/z 268.0550

012E+466.5RT (min)

lcqtof\_hilic\_pos\_4\_039\_15

m/z 268.0552

012E+48.59RT (min)

lcqtof\_hilic\_pos\_4\_015\_42

m/z 268.2640

012E+318.519RT (min)

lcqtof\_hilic\_pos\_4\_028\_31

m/z 268.2641

0123E+31616.5RT (min)

lcqtof\_hilic\_pos\_4\_004\_13

m/z 269.0578

024E+35.56RT (min)

lcqtof\_hilic\_pos\_4\_004\_13

m/z 269.2585

012E+318.519RT (min)

lcqtof\_hilic\_pos\_4\_108\_62

m/z 270.1329

012E+344.5RT (min)

lcqtof\_hilic\_pos\_1\_002\_38

m/z 270.2798

0123E+31616.5RT (min)

lcqtof\_hilic\_pos\_4\_084\_59

m/z 271.1666

024E+31818.5RT (min)

lcqtof\_hilic\_pos\_4\_069\_57

m/z 272.1297

0246E+34.55RT (min)

lcqtof\_hilic\_pos\_4\_021\_46

m/z 272.1308

0111.5RT (min)

lcqtof\_hilic\_pos\_4\_086\_59

m/z 273.1499

0246E+322.5RT (min)

lcqtof\_hilic\_pos\_4\_053\_30

m/z 273.1500

0246E+318.519RT (min)

lcqtof\_hilic\_pos\_4\_066\_57

m/z 274.0022

0123E+35.56RT (min)

lcqtof\_hilic\_pos\_4\_016\_09

m/z 274.2749

01E+477.5RT (min)

lcqtof\_hilic\_pos\_4\_093\_60

m/z 274.9511

0123E+38.59RT (min)

lcqtof\_hilic\_pos\_4\_006\_29

m/z 275.0009

01234E+30.51RT (min)

lcqtof\_hilic\_pos\_4\_075\_58

m/z 275.1048

024E+344.5RT (min)

lcqtof\_hilic\_pos\_1\_012\_10

m/z 275.1199

0246E+25.56RT (min)

lcqtof\_hilic\_pos\_3\_092\_60

m/z 275.2411

012E+30.51RT (min)

lcqtof\_hilic\_pos\_1\_018\_08

m/z 275.2781

01234E+37.58RT (min)

lcqtof\_hilic\_pos\_4\_040\_06

m/z 276.2331

02468E+30.51RT (min)

lcqtof\_hilic\_pos\_4\_035\_41

m/z 279.0939

0123E+45.56RT (min)

lcqtof\_hilic\_pos\_4\_084\_59

m/z 280.0973

02468E+311.5RT (min)

lcqtof\_hilic\_pos\_4\_111\_62

m/z 280.0976

02468E+318.519RT (min)

lcqtof\_hilic\_pos\_4\_090\_60

m/z 280.1098

0111.5RT (min)

lcqtof\_hilic\_pos\_4\_098\_61

m/z 280.1630

012E+416.517RT (min)

lcqtof\_hilic\_pos\_4\_008\_25

m/z 281.1646

0123E+31818.5RT (min)

lcqtof\_hilic\_pos\_4\_080\_58

m/z 281.1657

012E+317.518RT (min)

lcqtof\_hilic\_pos\_4\_098\_61

m/z 281.1982

0123E+455.5RT (min)

lcqtof\_hilic\_pos\_4\_056\_56

m/z 282.1207

01E+322.5RT (min)

lcqtof\_hilic\_pos\_4\_012\_02

m/z 282.1700

01234E+377.5RT (min)

lcqtof\_hilic\_pos\_4\_036\_24

m/z 282.1739

01234E+34.55RT (min)

lcqtof\_hilic\_pos\_1\_020\_39

m/z 282.2425

01E+40.51RT (min)

lcqtof\_hilic\_pos\_1\_005\_01

m/z 283.0661

02468E+31818.5RT (min)

lcqtof\_hilic\_pos\_4\_017\_51

m/z 283.0987

01E+311.5RT (min)

lcqtof\_hilic\_pos\_4\_005\_11

m/z 283.1274

01E+31.52RT (min)

lcqtof\_hilic\_pos\_4\_005\_11

m/z 283.2732

012E+318.519RT (min)

lcqtof\_hilic\_pos\_4\_089\_59

m/z 284.0878

012E+34.55RT (min)

lcqtof\_hilic\_pos\_4\_002\_23

m/z 284.1443

01234E+36.57RT (min)

lcqtof\_hilic\_pos\_4\_036\_24

m/z 284.2943

024E+318.519RT (min)

lcqtof\_hilic\_pos\_4\_077\_58

m/z 284.3299

01E+418.519RT (min)

lcqtof\_hilic\_pos\_4\_028\_31

m/z 284.8893

012E+36.57RT (min)

lcqtof\_hilic\_pos\_2\_047\_07

m/z 285.0314

0123E+34.55RT (min)

lcqtof\_hilic\_pos\_4\_008\_25

m/z 285.3353

012E+422.5RT (min)

lcqtof\_hilic\_pos\_4\_112\_62

m/z 285.3356

024E+318.519RT (min)

lcqtof\_hilic\_pos\_4\_108\_62

m/z 286.3114

012E+31919.5RT (min)

lcqtof\_hilic\_pos\_4\_085\_59

m/z 286.3115

01234E+322.5RT (min)

lcqtof\_hilic\_pos\_4\_008\_25

m/z 287.1261

02468E+31.52RT (min)

lcqtof\_hilic\_pos\_4\_094\_60

m/z 287.1459

01E+32.53RT (min)

lcqtof\_hilic\_pos\_4\_004\_13

m/z 287.1558

01234E+21.52RT (min)

lcqtof\_hilic\_pos\_2\_004\_13

m/z 287.9626

012E+312.513RT (min)

lcqtof\_hilic\_pos\_1\_026\_20

m/z 288.9230

012E+31010.5RT (min)

lcqtof\_hilic\_pos\_4\_092\_60

m/z 291.1628

0123E+31818.5RT (min)

lcqtof\_hilic\_pos\_4\_092\_60

m/z 293.1927

012E+344.5RT (min)

lcqtof\_hilic\_pos\_4\_075\_58

m/z 293.2238

024E+311.5RT (min)

lcqtof\_hilic\_pos\_4\_052\_12

m/z 295.1515

01E+37.58RT (min)

lcqtof\_hilic\_pos\_4\_027\_05

m/z 296.1070

0246E+277.5RT (min)

lcqtof\_hilic\_pos\_4\_023\_19

m/z 296.1539

0123E+34.55RT (min)

lcqtof\_hilic\_pos\_3\_028\_34

m/z 296.2103

01E+40.51RT (min)

lcqtof\_hilic\_pos\_4\_030\_03

m/z 297.1315

01234E+411.5RT (min)

lcqtof\_hilic\_pos\_4\_050\_16

m/z 297.1723

024E+25.56RT (min)

lcqtof\_hilic\_pos\_4\_036\_24

m/z 297.2907

02468E+318.519RT (min)

lcqtof\_hilic\_pos\_4\_021\_46

m/z 298.3124

0246E+32.53RT (min)

lcqtof\_hilic\_pos\_4\_013\_36

m/z 299.1878

01234E+311.5RT (min)

lcqtof\_hilic\_pos\_4\_100\_61

m/z 300.2028

0123E+31919.5RT (min)

lcqtof\_hilic\_pos\_4\_084\_59

m/z 302.3057

0123E+37.58RT (min)

lcqtof\_hilic\_pos\_4\_029\_44

m/z 304.3019

0123E+318.519RT (min)

lcqtof\_hilic\_pos\_4\_074\_58

m/z 307.2043

01234E+333.5RT (min)

lcqtof\_hilic\_pos\_4\_004\_13

m/z 307.2277

01234E+20.51RT (min)

lcqtof\_hilic\_pos\_4\_023\_19

m/z 310.3120

0123E+32.53RT (min)

lcqtof\_hilic\_pos\_3\_012\_12

m/z 311.1348

01E+366.5RT (min)

lcqtof\_hilic\_pos\_4\_050\_16

m/z 311.1603

0246E+344.5RT (min)

lcqtof\_hilic\_pos\_4\_008\_25

m/z 311.2326

02468E+311.5RT (min)

lcqtof\_hilic\_pos\_4\_023\_19

m/z 312.1083

01E+33.54RT (min)

lcqtof\_hilic\_pos\_4\_072\_57

m/z 312.3132

01234E+32.53RT (min)

lcqtof\_hilic\_pos\_4\_022\_10

m/z 313.1762

024E+318.519RT (min)

lcqtof\_hilic\_pos\_4\_105\_61

m/z 313.1765

024E+322.5RT (min)

lcqtof\_hilic\_pos\_4\_049\_55

m/z 316.1007

01234E+366.5RT (min)

lcqtof\_hilic\_pos\_3\_043\_46

m/z 318.1480

01E+244.5RT (min)

lcqtof\_hilic\_pos\_4\_024\_26

m/z 319.2235

01E+333.5RT (min)

lcqtof\_hilic\_pos\_4\_005\_11

m/z 321.1176

01234E+255.5RT (min)

lcqtof\_hilic\_pos\_4\_002\_23

m/z 321.3149

024E+31818.5RT (min)

lcqtof\_hilic\_pos\_4\_015\_42

m/z 321.3158

01234E+32.53RT (min)

lcqtof\_hilic\_pos\_4\_030\_03

m/z 321.3163

024E+31818.5RT (min)

lcqtof\_hilic\_pos\_4\_003\_21

m/z 321.9551

0246E+30.51RT (min)

lcqtof\_hilic\_pos\_4\_077\_58

m/z 322.3157

0123E+31818.5RT (min)

lcqtof\_hilic\_pos\_4\_007\_45

m/z 324.1168

01E+433.5RT (min)

lcqtof\_hilic\_pos\_2\_027\_18

m/z 325.1016

0246E+244.5RT (min)

lcqtof\_hilic\_pos\_4\_097\_60

m/z 326.1331

0246E+35.56RT (min)

lcqtof\_hilic\_pos\_4\_044\_32

m/z 326.1652

0246E+24.55RT (min)

lcqtof\_hilic\_pos\_4\_103\_61

m/z 326.3258

0123E+31919.5RT (min)

lcqtof\_hilic\_pos\_4\_006\_29

m/z 326.3261

024E+32.53RT (min)

lcqtof\_hilic\_pos\_4\_019\_08

m/z 327.0799

012E+316.517RT (min)

lcqtof\_hilic\_pos\_4\_084\_59

m/z 327.0817

012E+31919.5RT (min)

lcqtof\_hilic\_pos\_4\_085\_59

m/z 327.2032

0123E+31919.5RT (min)

lcqtof\_hilic\_pos\_4\_090\_60

m/z 328.7403

012E+511.5RT (min)

lcqtof\_hilic\_pos\_4\_099\_61

m/z 329.0970

0246E+36.57RT (min)

lcqtof\_hilic\_pos\_3\_079\_58

m/z 330.2044

0246E+318.519RT (min)

lcqtof\_hilic\_pos\_4\_108\_62

m/z 330.2047

0246E+31.52RT (min)

lcqtof\_hilic\_pos\_4\_047\_40

m/z 330.2651

0123E+388.5RT (min)

lcqtof\_hilic\_pos\_1\_002\_38

m/z 332.1229

01E+377.5RT (min)

lcqtof\_hilic\_pos\_3\_034\_24

m/z 332.2091

01234E+22.53RT (min)

lcqtof\_hilic\_pos\_4\_050\_16

m/z 332.3315

012E+318.519RT (min)

lcqtof\_hilic\_pos\_4\_089\_59

m/z 333.1443

014.55RT (min)

lcqtof\_hilic\_pos\_4\_112\_62

m/z 335.0969

01234E+35.56RT (min)

lcqtof\_hilic\_pos\_3\_067\_57

m/z 335.2967

012E+31818.5RT (min)

lcqtof\_hilic\_pos\_4\_113\_62

m/z 336.1946

012E+344.5RT (min)

lcqtof\_hilic\_pos\_2\_002\_32

m/z 336.2746

01234E+333.5RT (min)

lcqtof\_hilic\_pos\_4\_008\_25

m/z 336.3250

012E+31818.5RT (min)

lcqtof\_hilic\_pos\_4\_082\_59

m/z 337.1133

0123E+317.518RT (min)

lcqtof\_hilic\_pos\_4\_027\_05

m/z 337.3228

012E+31818.5RT (min)

lcqtof\_hilic\_pos\_4\_099\_61

m/z 339.1018

01234E+24.55RT (min)

lcqtof\_hilic\_pos\_4\_036\_24

m/z 339.1595

0123E+377.5RT (min)

lcqtof\_hilic\_pos\_4\_026\_01

m/z 340.3521

01234E+318.519RT (min)

lcqtof\_hilic\_pos\_4\_074\_58

m/z 341.0716

01E+31818.5RT (min)

lcqtof\_hilic\_pos\_4\_090\_60

m/z 341.1418

0123E+35.56RT (min)

lcqtof\_hilic\_pos\_4\_088\_59

m/z 341.1821

024E+322.5RT (min)

lcqtof\_hilic\_pos\_4\_017\_51

m/z 342.2771

024E+21.52RT (min)

lcqtof\_hilic\_pos\_4\_008\_25

m/z 343.1605

0123E+366.5RT (min)

lcqtof\_hilic\_pos\_4\_010\_37

m/z 344.2279

0123E+31919.5RT (min)

lcqtof\_hilic\_pos\_4\_081\_58

m/z 346.1849

0123E+344.5RT (min)

lcqtof\_hilic\_pos\_4\_017\_51

m/z 347.1512

024E+311.5RT (min)

lcqtof\_hilic\_pos\_4\_033\_53

m/z 348.3261

01E+318.519RT (min)

lcqtof\_hilic\_pos\_4\_097\_60

m/z 349.1771

0246E+333.5RT (min)

lcqtof\_hilic\_pos\_4\_027\_05

m/z 352.3284

01E+416.517RT (min)

lcqtof\_hilic\_pos\_4\_001\_49

m/z 353.0565

0246E+35.56RT (min)

lcqtof\_hilic\_pos\_4\_033\_53

m/z 353.1065

01234E+266.5RT (min)

lcqtof\_hilic\_pos\_4\_044\_32

m/z 353.2210

0123E+344.5RT (min)

lcqtof\_hilic\_pos\_3\_040\_45

m/z 354.1055

01E+311.5RT (min)

lcqtof\_hilic\_pos\_3\_105\_61

m/z 354.2043

024E+31818.5RT (min)

lcqtof\_hilic\_pos\_4\_095\_60

m/z 354.2053

024E+31919.5RT (min)

lcqtof\_hilic\_pos\_4\_011\_48

m/z 354.2055

02468E+322.5RT (min)

lcqtof\_hilic\_pos\_4\_010\_37

m/z 354.3456

012E+318.519RT (min)

lcqtof\_hilic\_pos\_4\_088\_59

m/z 354.3464

0246E+400.5RT (min)

lcqtof\_hilic\_pos\_2\_004\_13

m/z 355.0720

01E+31515.5RT (min)

lcqtof\_hilic\_pos\_1\_006\_04

m/z 355.1948

0123E+31919.5RT (min)

lcqtof\_hilic\_pos\_4\_088\_59

m/z 356.3284

01E+31515.5RT (min)

lcqtof\_hilic\_pos\_4\_081\_58

m/z 358.1207

01E+366.5RT (min)

lcqtof\_hilic\_pos\_1\_053\_46

m/z 359.1627

012E+266.5RT (min)

lcqtof\_hilic\_pos\_4\_004\_13

m/z 359.2191

0123E+32.53RT (min)

lcqtof\_hilic\_pos\_4\_089\_59

m/z 362.2538

012E+322.5RT (min)

lcqtof\_hilic\_pos\_1\_004\_34

m/z 363.2641

01234E+311.5RT (min)

lcqtof\_hilic\_pos\_4\_010\_37

m/z 364.3285

012E+31515.5RT (min)

lcqtof\_hilic\_pos\_4\_110\_62

m/z 364.9318

0123E+388.5RT (min)

lcqtof\_hilic\_pos\_4\_042\_14

m/z 365.1576

012E+34.55RT (min)

lcqtof\_hilic\_pos\_4\_027\_05

m/z 365.2309

01E+30.51RT (min)

lcqtof\_hilic\_pos\_4\_078\_58

m/z 366.0743

0246E+15.56RT (min)

lcqtof\_hilic\_pos\_3\_070\_57

m/z 367.2036

0246E+322.5RT (min)

lcqtof\_hilic\_pos\_4\_104\_61

m/z 368.4246

0123E+318.519RT (min)

lcqtof\_hilic\_pos\_4\_075\_58

m/z 369.3013

0123E+315.516RT (min)

lcqtof\_hilic\_pos\_1\_018\_08

m/z 373.0577

01234E+266.5RT (min)

lcqtof\_hilic\_pos\_4\_033\_53

m/z 373.2697

012E+32.53RT (min)

lcqtof\_hilic\_pos\_4\_054\_18

m/z 376.2645

024E+34.55RT (min)

lcqtof\_hilic\_pos\_4\_042\_14

m/z 376.8945

012E+31313.5RT (min)

lcqtof\_hilic\_pos\_2\_025\_52

m/z 377.3276

012E+31919.5RT (min)

lcqtof\_hilic\_pos\_4\_016\_09

m/z 378.2357

01E+377.5RT (min)

lcqtof\_hilic\_pos\_3\_014\_48

m/z 379.1399

024E+32.53RT (min)

lcqtof\_hilic\_pos\_4\_045\_20

m/z 381.3821

012E+31919.5RT (min)

lcqtof\_hilic\_pos\_4\_083\_59

m/z 382.2439

01E+31818.5RT (min)

lcqtof\_hilic\_pos\_4\_081\_58

m/z 382.3339

024E+31515.5RT (min)

lcqtof\_hilic\_pos\_4\_002\_23

m/z 383.2342

01234E+32.53RT (min)

lcqtof\_hilic\_pos\_4\_001\_49

m/z 383.2649

0246E+23.54RT (min)

lcqtof\_hilic\_pos\_1\_018\_08

m/z 383.3197

012E+4161718RT (min)

lcqtof\_hilic\_pos\_3\_105\_61

m/z 385.3502

0123E+31818.5RT (min)

lcqtof\_hilic\_pos\_4\_092\_60

m/z 386.1680

0123E+316.517RT (min)

lcqtof\_hilic\_pos\_4\_007\_45

m/z 386.3266

0123E+222.5RT (min)

lcqtof\_hilic\_pos\_4\_052\_12

m/z 391.1618

02468E+233.5RT (min)

lcqtof\_hilic\_pos\_4\_036\_24

m/z 391.2199

01234E+377.5RT (min)

lcqtof\_hilic\_pos\_3\_080\_58

m/z 391.2965

024E+611.5RT (min)

lcqtof\_hilic\_pos\_2\_002\_32

m/z 391.4850

02468E+31818.5RT (min)

lcqtof\_hilic\_pos\_4\_001\_49

m/z 392.1061

01234E+366.5RT (min)

lcqtof\_hilic\_pos\_4\_094\_60

m/z 392.3632

01E+41919.5RT (min)

lcqtof\_hilic\_pos\_4\_009\_50

m/z 393.2040

012E+333.5RT (min)

lcqtof\_hilic\_pos\_4\_023\_19

m/z 393.2984

024E+366.5RT (min)

lcqtof\_hilic\_pos\_1\_016\_14

m/z 393.3077

01E+23.54RT (min)

lcqtof\_hilic\_pos\_4\_087\_59

m/z 393.3661

0246E+31919.5RT (min)

lcqtof\_hilic\_pos\_4\_002\_23

m/z 395.3667

01E+333.5RT (min)

lcqtof\_hilic\_pos\_3\_100\_61

m/z 396.4038

01E+31515.5RT (min)

lcqtof\_hilic\_pos\_4\_082\_59

m/z 398.3272

012E+31818.5RT (min)

lcqtof\_hilic\_pos\_4\_099\_61

m/z 400.0215

0246E+311.5RT (min)

lcqtof\_hilic\_pos\_1\_033\_53

m/z 400.2549

024E+31818.5RT (min)

lcqtof\_hilic\_pos\_3\_094\_60

m/z 400.2550

024E+322.5RT (min)

lcqtof\_hilic\_pos\_1\_009\_50

m/z 401.3004

024E+22.53RT (min)

lcqtof\_hilic\_pos\_4\_026\_01

m/z 402.1061

02468E+377.5RT (min)

lcqtof\_hilic\_pos\_4\_007\_45

m/z 402.2472

01E+46.57RT (min)

lcqtof\_hilic\_pos\_4\_002\_23

m/z 404.3018

012E+318.519RT (min)

lcqtof\_hilic\_pos\_3\_110\_62

m/z 407.3331

0246E+35.56RT (min)

lcqtof\_hilic\_pos\_4\_001\_49

m/z 408.3103

012E+35.56RT (min)

lcqtof\_hilic\_pos\_3\_050\_21

m/z 409.3328

01234E+32.53RT (min)

lcqtof\_hilic\_pos\_4\_023\_19

m/z 411.3775

024E+310.511RT (min)

lcqtof\_hilic\_pos\_4\_024\_26

m/z 413.1915

012E+31515.5RT (min)

lcqtof\_hilic\_pos\_4\_031\_33

m/z 415.2724

02468E+388.5RT (min)

lcqtof\_hilic\_pos\_4\_054\_18

m/z 416.2148

0246E+31818.5RT (min)

lcqtof\_hilic\_pos\_4\_029\_44

m/z 416.2158

0246E+31818.5RT (min)

lcqtof\_hilic\_pos\_4\_079\_58

m/z 416.2162

0246E+322.5RT (min)

lcqtof\_hilic\_pos\_4\_040\_06

m/z 418.3293

012E+35.56RT (min)

lcqtof\_hilic\_pos\_4\_097\_60

m/z 419.1822

024E+333.5RT (min)

lcqtof\_hilic\_pos\_4\_046\_07

m/z 420.3567

012E+300.5RT (min)

lcqtof\_hilic\_pos\_4\_084\_59

m/z 421.2341

012E+355.5RT (min)

lcqtof\_hilic\_pos\_1\_015\_45

m/z 422.1552

0123E+344.5RT (min)

lcqtof\_hilic\_pos\_4\_033\_53

m/z 423.1622

012E+34.55RT (min)

lcqtof\_hilic\_pos\_4\_036\_24

m/z 423.3788

02468E+31919.5RT (min)

lcqtof\_hilic\_pos\_4\_015\_42

m/z 425.4465

0246E+322.5RT (min)

lcqtof\_hilic\_pos\_2\_031\_24

m/z 428.2478

012E+31515.5RT (min)

lcqtof\_hilic\_pos\_4\_010\_37

m/z 428.2855

012E+36.57RT (min)

lcqtof\_hilic\_pos\_4\_087\_59

m/z 428.3369

01E+41818.5RT (min)

lcqtof\_hilic\_pos\_4\_009\_50

m/z 429.3415

012E+31515.5RT (min)

lcqtof\_hilic\_pos\_4\_101\_61

m/z 432.3535

01E+31515.5RT (min)

lcqtof\_hilic\_pos\_4\_096\_60

m/z 432.3606

01E+23.54RT (min)

lcqtof\_hilic\_pos\_4\_023\_19

m/z 432.7973

0123E+33.54RT (min)

lcqtof\_hilic\_pos\_4\_026\_01

m/z 434.3042

0123E+34.55RT (min)

lcqtof\_hilic\_pos\_4\_008\_25

m/z 435.3225

02468E+377.5RT (min)

lcqtof\_hilic\_pos\_4\_046\_07

m/z 439.0198

01234E+244.5RT (min)

lcqtof\_hilic\_pos\_4\_024\_26

m/z 439.2939

01234E+322.5RT (min)

lcqtof\_hilic\_pos\_4\_002\_23

m/z 439.3803

024E+4161718RT (min)

lcqtof\_hilic\_pos\_3\_102\_61

m/z 440.2869

0123E+31818.5RT (min)

lcqtof\_hilic\_pos\_4\_108\_62

m/z 440.2870

02468E+322.5RT (min)

lcqtof\_hilic\_pos\_4\_003\_21

m/z 440.3354

024E+31515.5RT (min)

lcqtof\_hilic\_pos\_4\_003\_21

m/z 441.3089

01E+344.5RT (min)

lcqtof\_hilic\_pos\_4\_016\_09

m/z 441.4345

0246E+20.51RT (min)

lcqtof\_hilic\_pos\_4\_028\_31

m/z 442.2560

02468E+23.54RT (min)

lcqtof\_hilic\_pos\_3\_005\_13

m/z 442.2970

0246E+33.54RT (min)

lcqtof\_hilic\_pos\_4\_108\_62

m/z 443.1464

01E+244.5RT (min)

lcqtof\_hilic\_pos\_4\_092\_60

m/z 448.1370

012E+24.55RT (min)

lcqtof\_hilic\_pos\_4\_045\_20

m/z 448.2193

012E+27.58RT (min)

lcqtof\_hilic\_pos\_4\_036\_24

m/z 448.3302

012E+31818.5RT (min)

lcqtof\_hilic\_pos\_3\_091\_60

m/z 448.9751

012E+388.5RT (min)

lcqtof\_hilic\_pos\_3\_002\_23

m/z 449.3442

024E+32.53RT (min)

lcqtof\_hilic\_pos\_4\_026\_01

m/z 450.0199

0246E+37.58RT (min)

lcqtof\_hilic\_pos\_3\_011\_37

m/z 450.4008

012E+31515.5RT (min)

lcqtof\_hilic\_pos\_4\_003\_21

m/z 451.4598

012E+31515.5RT (min)

lcqtof\_hilic\_pos\_4\_001\_49

m/z 452.3674

024E+33.54RT (min)

lcqtof\_hilic\_pos\_3\_083\_59

m/z 452.7380

01234E+35.56RT (min)

lcqtof\_hilic\_pos\_4\_081\_58

m/z 455.3341

024E+344.5RT (min)

lcqtof\_hilic\_pos\_4\_002\_23

m/z 455.3838

012E+311.5RT (min)

lcqtof\_hilic\_pos\_4\_036\_24

m/z 455.3946

01E+32.53RT (min)

lcqtof\_hilic\_pos\_3\_067\_57

m/z 457.1633

024E+34.55RT (min)

lcqtof\_hilic\_pos\_1\_039\_06

m/z 458.0052

024E+288.5RT (min)

lcqtof\_hilic\_pos\_4\_081\_58

m/z 462.3204

012E+31.52RT (min)

lcqtof\_hilic\_pos\_4\_088\_59

m/z 462.3284

0246E+21515.5RT (min)

lcqtof\_hilic\_pos\_4\_073\_57

m/z 464.3335

01234E+30.51RT (min)

lcqtof\_hilic\_pos\_4\_013\_36

m/z 465.3088

01E+31515.5RT (min)

lcqtof\_hilic\_pos\_4\_099\_61

m/z 465.4597

01E+315.516RT (min)

lcqtof\_hilic\_pos\_1\_037\_36

m/z 466.2946

012E+31515.5RT (min)

lcqtof\_hilic\_pos\_4\_109\_62

m/z 467.1166

01E+44.55RT (min)

lcqtof\_hilic\_pos\_4\_012\_02

m/z 467.2082

0246E+30.51RT (min)

lcqtof\_hilic\_pos\_4\_109\_62

m/z 467.4108

01234E+4161718RT (min)

lcqtof\_hilic\_pos\_3\_112\_62

m/z 468.3843

02468E+318.519RT (min)

lcqtof\_hilic\_pos\_4\_016\_09

m/z 468.3863

024E+31818.5RT (min)

lcqtof\_hilic\_pos\_4\_098\_61

m/z 468.3974

0123E+31818.5RT (min)

lcqtof\_hilic\_pos\_3\_106\_62

m/z 468.4113

01E+344.5RT (min)

lcqtof\_hilic\_pos\_4\_074\_58

m/z 469.1994

0123E+31919.5RT (min)

lcqtof\_hilic\_pos\_4\_009\_50

m/z 469.2007

01234E+319.520RT (min)

lcqtof\_hilic\_pos\_4\_015\_42

m/z 469.2016

012E+322.5RT (min)

lcqtof\_hilic\_pos\_4\_028\_31

m/z 469.3841

0123E+31818.5RT (min)

lcqtof\_hilic\_pos\_4\_099\_61

m/z 470.3333

012E+31212.5RT (min)

lcqtof\_hilic\_pos\_4\_050\_16

m/z 472.3270

01234E+31818.5RT (min)

lcqtof\_hilic\_pos\_4\_095\_60

m/z 474.1705

024E+366.5RT (min)

lcqtof\_hilic\_pos\_4\_036\_24

m/z 475.2213

01E+35.56RT (min)

lcqtof\_hilic\_pos\_1\_027\_05

m/z 475.3242

0246E+31.52RT (min)

lcqtof\_hilic\_pos\_4\_002\_23

m/z 475.3958

01E+31515.5RT (min)

lcqtof\_hilic\_pos\_4\_095\_60

m/z 476.3300

02468E+333.5RT (min)

lcqtof\_hilic\_pos\_4\_085\_59

m/z 476.8108

012E+36.57RT (min)

lcqtof\_hilic\_pos\_4\_040\_06

m/z 476.8466

0123E+344.5RT (min)

lcqtof\_hilic\_pos\_4\_084\_59

m/z 481.2549

01E+355.5RT (min)

lcqtof\_hilic\_pos\_4\_036\_24

m/z 481.3254

01E+255.5RT (min)

lcqtof\_hilic\_pos\_4\_079\_58

m/z 482.4066

02468E+611.5RT (min)

lcqtof\_hilic\_pos\_4\_088\_59

m/z 483.3450

0123E+31515.5RT (min)

lcqtof\_hilic\_pos\_4\_110\_62

m/z 484.4350

02468E+22.53RT (min)

lcqtof\_hilic\_pos\_2\_001\_49

m/z 486.1352

0246E+366.5RT (min)

lcqtof\_hilic\_pos\_4\_040\_06

m/z 487.0784

01E+233.5RT (min)

lcqtof\_hilic\_pos\_3\_019\_16

m/z 487.4019

012E+31515.5RT (min)

lcqtof\_hilic\_pos\_4\_018\_34

m/z 489.1130

01234E+37.58RT (min)

lcqtof\_hilic\_pos\_4\_010\_37

m/z 490.3621

012E+318.519RT (min)

lcqtof\_hilic\_pos\_4\_074\_58

m/z 490.3629

012E+30.51RT (min)

lcqtof\_hilic\_pos\_4\_098\_61

m/z 490.4261

01E+31515.5RT (min)

lcqtof\_hilic\_pos\_4\_003\_21

m/z 491.9699

012E+37.58RT (min)

lcqtof\_hilic\_pos\_3\_010\_02

m/z 493.4943

01E+315.516RT (min)

lcqtof\_hilic\_pos\_3\_093\_60

m/z 496.4188

01234E+318.519RT (min)

lcqtof\_hilic\_pos\_4\_035\_41

m/z 496.4269

01234E+31818.5RT (min)

lcqtof\_hilic\_pos\_4\_107\_62

m/z 498.3277

01E+41818.5RT (min)

lcqtof\_hilic\_pos\_4\_089\_59

m/z 499.3313

0123E+31818.5RT (min)

lcqtof\_hilic\_pos\_4\_087\_59

m/z 499.3316

0246E+322.5RT (min)

lcqtof\_hilic\_pos\_4\_012\_02

m/z 503.3672

0123E+31515.5RT (min)

lcqtof\_hilic\_pos\_4\_001\_49

m/z 504.0702

01234E+366.5RT (min)

lcqtof\_hilic\_pos\_4\_026\_01

m/z 506.3792

01E+33.54RT (min)

lcqtof\_hilic\_pos\_1\_005\_01

m/z 506.8122

01E+31515.5RT (min)

lcqtof\_hilic\_pos\_4\_018\_34

m/z 510.2832

012E+24.55RT (min)

lcqtof\_hilic\_pos\_3\_024\_18

m/z 515.0641

024E+366.5RT (min)

lcqtof\_hilic\_pos\_4\_001\_49

m/z 517.1749

012E+355.5RT (min)

lcqtof\_hilic\_pos\_1\_005\_01

m/z 517.4589

01E+315.516RT (min)

lcqtof\_hilic\_pos\_4\_082\_59

m/z 519.9460

0246E+10.51RT (min)

lcqtof\_hilic\_pos\_2\_022\_09

m/z 521.4415

01234E+22.53RT (min)

lcqtof\_hilic\_pos\_2\_002\_32

m/z 521.5216

012E+315.516RT (min)

lcqtof\_hilic\_pos\_1\_018\_08

m/z 522.2246

01234E+35.56RT (min)

lcqtof\_hilic\_pos\_4\_004\_13

m/z 525.3883

02468E+34.55RT (min)

lcqtof\_hilic\_pos\_2\_038\_33

m/z 525.3986

02468E+34.55RT (min)

lcqtof\_hilic\_pos\_1\_027\_05

m/z 526.1429

0123E+377.5RT (min)

lcqtof\_hilic\_pos\_4\_044\_32

m/z 526.4096

012E+31515.5RT (min)

lcqtof\_hilic\_pos\_4\_001\_49

m/z 526.4325

01234E+318.519RT (min)

lcqtof\_hilic\_pos\_4\_097\_60

m/z 527.3335

012E+318.519RT (min)

lcqtof\_hilic\_pos\_4\_020\_27

m/z 527.3337

0123E+32.53RT (min)

lcqtof\_hilic\_pos\_4\_003\_21

m/z 530.2362

0123E+34.55RT (min)

lcqtof\_hilic\_pos\_4\_031\_33

m/z 531.2753

0123E+31515.5RT (min)

lcqtof\_hilic\_pos\_4\_073\_57

m/z 531.2900

02468E+36.57RT (min)

lcqtof\_hilic\_pos\_3\_074\_58

m/z 532.0886

0123E+366.5RT (min)

lcqtof\_hilic\_pos\_4\_082\_59

m/z 535.4213

0123E+31515.5RT (min)

lcqtof\_hilic\_pos\_1\_018\_08

m/z 535.4298

0123E+31515.5RT (min)

lcqtof\_hilic\_pos\_1\_018\_08

m/z 535.4918

024E+32.53RT (min)

lcqtof\_hilic\_pos\_4\_002\_23

m/z 537.4154

02468E+311.5RT (min)

lcqtof\_hilic\_pos\_4\_042\_14

m/z 542.0442

01234E+366.5RT (min)

lcqtof\_hilic\_pos\_1\_031\_32

m/z 542.2541

01234E+244.5RT (min)

lcqtof\_hilic\_pos\_1\_005\_01

m/z 543.0681

0246E+36.57RT (min)

lcqtof\_hilic\_pos\_4\_053\_30

m/z 546.4360

0123E+31515.5RT (min)

lcqtof\_hilic\_pos\_4\_106\_62

m/z 547.2438

012E+35.56RT (min)

lcqtof\_hilic\_pos\_4\_004\_13

m/z 548.1242

012E+377.5RT (min)

lcqtof\_hilic\_pos\_4\_033\_53

m/z 548.3936

0123E+411.5RT (min)

lcqtof\_hilic\_pos\_3\_041\_54

m/z 548.5045

01E+315.516RT (min)

lcqtof\_hilic\_pos\_3\_092\_60

m/z 549.3452

01234E+21515.5RT (min)

lcqtof\_hilic\_pos\_1\_018\_08

m/z 550.4910

01234E+30.51RT (min)

lcqtof\_hilic\_pos\_4\_018\_34

m/z 550.5170

024E+322.5RT (min)

lcqtof\_hilic\_pos\_4\_002\_23

m/z 550.6292

0123E+31818.5RT (min)

lcqtof\_hilic\_pos\_4\_005\_11

m/z 551.4508

0123E+31515.5RT (min)

lcqtof\_hilic\_pos\_4\_105\_61

m/z 551.5455

0123E+215.516RT (min)

lcqtof\_hilic\_pos\_3\_071\_57

m/z 552.3298

02468E+244.5RT (min)

lcqtof\_hilic\_pos\_4\_016\_09

m/z 555.4220

012E+315.516RT (min)

lcqtof\_hilic\_pos\_3\_074\_58

m/z 556.3692

024E+41818.5RT (min)

lcqtof\_hilic\_pos\_4\_015\_42

m/z 556.3693

0123E+42.53RT (min)

lcqtof\_hilic\_pos\_4\_039\_15

m/z 557.3731

0246E+31818.5RT (min)

lcqtof\_hilic\_pos\_4\_108\_62

m/z 557.3732

01E+42.53RT (min)

lcqtof\_hilic\_pos\_4\_025\_52

m/z 557.3732

01E+40.51RT (min)

lcqtof\_hilic\_pos\_4\_014\_22

m/z 558.3781

0123E+31818.5RT (min)

lcqtof\_hilic\_pos\_4\_014\_22

m/z 558.3783

01234E+322.5RT (min)

lcqtof\_hilic\_pos\_4\_006\_29

m/z 563.2171

01234E+31010.5RT (min)

lcqtof\_hilic\_pos\_4\_105\_61

m/z 563.4419

0246E+33.54RT (min)

lcqtof\_hilic\_pos\_4\_003\_21

m/z 564.3655

012E+31515.5RT (min)

lcqtof\_hilic\_pos\_4\_005\_11

m/z 566.0356

0246E+36.57RT (min)

lcqtof\_hilic\_pos\_4\_090\_60

m/z 566.5302

024E+322.5RT (min)

lcqtof\_hilic\_pos\_4\_002\_23

m/z 570.4348

0123E+31515.5RT (min)

lcqtof\_hilic\_pos\_4\_006\_29

m/z 570.4594

0123E+318.519RT (min)

lcqtof\_hilic\_pos\_4\_067\_57

m/z 571.0953

012E+25.56RT (min)

lcqtof\_hilic\_pos\_4\_036\_24

m/z 571.3591

0123E+318.519RT (min)

lcqtof\_hilic\_pos\_4\_019\_08

m/z 571.3605

0123E+318.519RT (min)

lcqtof\_hilic\_pos\_4\_020\_27

m/z 571.3606

01E+222.5RT (min)

lcqtof\_hilic\_pos\_2\_038\_33

m/z 572.4877

0123E+318.519RT (min)

lcqtof\_hilic\_pos\_3\_096\_60

m/z 575.4764

012E+315.516RT (min)

lcqtof\_hilic\_pos\_3\_082\_59

m/z 575.5054

0246E+316.517RT (min)

lcqtof\_hilic\_pos\_1\_019\_19

m/z 579.5104

0246E+31515.5RT (min)

lcqtof\_hilic\_pos\_4\_001\_49

m/z 583.4002

0123E+31515.5RT (min)

lcqtof\_hilic\_pos\_4\_001\_49

m/z 584.4718

01E+318.519RT (min)

lcqtof\_hilic\_pos\_4\_103\_61

m/z 586.2948

012E+43.54RT (min)

lcqtof\_hilic\_pos\_4\_004\_13

m/z 590.4864

012E+315.516RT (min)

lcqtof\_hilic\_pos\_4\_104\_61

m/z 592.3752

01E+311.5RT (min)

lcqtof\_hilic\_pos\_4\_042\_14

m/z 592.8038

0246E+28.59RT (min)

lcqtof\_hilic\_pos\_1\_003\_44

m/z 595.4197

012E+31515.5RT (min)

lcqtof\_hilic\_pos\_4\_099\_61

m/z 599.4208

0123E+31515.5RT (min)

lcqtof\_hilic\_pos\_4\_002\_23

m/z 599.5040

012E+51617RT (min)

lcqtof\_hilic\_pos\_1\_024\_25

m/z 599.5055

0246E+317.518RT (min)

lcqtof\_hilic\_pos\_1\_021\_09

m/z 600.4705

0118.519RT (min)

lcqtof\_hilic\_pos\_1\_018\_08

m/z 602.3563

01E+444.5RT (min)

lcqtof\_hilic\_pos\_1\_001\_49

m/z 602.5645

012E+315.516RT (min)

lcqtof\_hilic\_pos\_4\_069\_57

m/z 603.4918

0123E+31515.5RT (min)

lcqtof\_hilic\_pos\_4\_013\_36

m/z 604.4043

0123E+31818.5RT (min)

lcqtof\_hilic\_pos\_4\_093\_60

m/z 604.5404

012E+31717.5RT (min)

lcqtof\_hilic\_pos\_1\_050\_22

m/z 605.4502

02468E+355.5RT (min)

lcqtof\_hilic\_pos\_1\_024\_25

m/z 606.2740

0246E+35.56RT (min)

lcqtof\_hilic\_pos\_4\_004\_13

m/z 606.3853

0123E+344.5RT (min)

lcqtof\_hilic\_pos\_4\_036\_24

m/z 607.4595

0123E+31515.5RT (min)

lcqtof\_hilic\_pos\_4\_110\_62

m/z 607.9102

012E+36.57RT (min)

lcqtof\_hilic\_pos\_4\_039\_15

m/z 610.4339

012E+31515.5RT (min)

lcqtof\_hilic\_pos\_4\_103\_61

m/z 611.1291

0123E+377.5RT (min)

lcqtof\_hilic\_pos\_4\_084\_59

m/z 612.5552

0123E+315.516RT (min)

lcqtof\_hilic\_pos\_1\_018\_08

m/z 614.2686

01234E+32.53RT (min)

lcqtof\_hilic\_pos\_4\_046\_07

m/z 614.2687

024E+31919.5RT (min)

lcqtof\_hilic\_pos\_4\_015\_42

m/z 614.4113

0246E+42.53RT (min)

lcqtof\_hilic\_pos\_4\_003\_21

m/z 614.4113

0246E+40.51RT (min)

lcqtof\_hilic\_pos\_4\_002\_23

m/z 615.4112

012E+41818.5RT (min)

lcqtof\_hilic\_pos\_4\_020\_27

m/z 616.4136

0246E+31818.5RT (min)

lcqtof\_hilic\_pos\_4\_004\_13

m/z 622.3654

01E+31.52RT (min)

lcqtof\_hilic\_pos\_4\_026\_01

m/z 625.2677

0246E+31919.5RT (min)

lcqtof\_hilic\_pos\_4\_018\_34

m/z 626.2574

012E+35.56RT (min)

lcqtof\_hilic\_pos\_4\_005\_11

m/z 626.2677

012E+31919.5RT (min)

lcqtof\_hilic\_pos\_4\_026\_01

m/z 626.5643

01E+315.516RT (min)

lcqtof\_hilic\_pos\_3\_107\_62

m/z 629.5491

01E+40.51RT (min)

lcqtof\_hilic\_pos\_1\_019\_19

m/z 629.5551

024E+31818.5RT (min)

lcqtof\_hilic\_pos\_3\_090\_60

m/z 629.5841

02468E+115.516RT (min)

lcqtof\_hilic\_pos\_1\_018\_08

m/z 629.8924

012E+36.57RT (min)

lcqtof\_hilic\_pos\_4\_074\_58

m/z 632.0767

0123E+377.5RT (min)

lcqtof\_hilic\_pos\_4\_028\_31

m/z 632.9733

012E+355.5RT (min)

lcqtof\_hilic\_pos\_3\_076\_58

m/z 634.6332

01E+31515.5RT (min)

lcqtof\_hilic\_pos\_4\_016\_09

m/z 635.2486

01E+49.510RT (min)

lcqtof\_hilic\_pos\_4\_095\_60

m/z 635.4606

0123E+31515.5RT (min)

lcqtof\_hilic\_pos\_4\_112\_62

m/z 636.2510

01234E+39.510RT (min)

lcqtof\_hilic\_pos\_4\_024\_26

m/z 643.6042

0246E+215.516RT (min)

lcqtof\_hilic\_pos\_4\_002\_23

m/z 646.5193

01234E+31818.5RT (min)

lcqtof\_hilic\_pos\_4\_051\_17

m/z 647.5456

01234E+315.516RT (min)

lcqtof\_hilic\_pos\_4\_102\_61

m/z 648.4894

024E+315.516RT (min)

lcqtof\_hilic\_pos\_4\_102\_61

m/z 649.6367

01E+511.5RT (min)

lcqtof\_hilic\_pos\_4\_027\_05

m/z 652.2720

0123E+366.5RT (min)

lcqtof\_hilic\_pos\_4\_028\_31

m/z 653.3846

02468E+344.5RT (min)

lcqtof\_hilic\_pos\_4\_087\_59

m/z 654.0585

012E+377.5RT (min)

lcqtof\_hilic\_pos\_4\_040\_06

m/z 654.5688

01234E+318.519RT (min)

lcqtof\_hilic\_pos\_1\_021\_09

m/z 655.5421

012E+31515.5RT (min)

lcqtof\_hilic\_pos\_4\_010\_37

m/z 657.5865

0246E+31818.5RT (min)

lcqtof\_hilic\_pos\_3\_099\_61

m/z 659.4109

0246E+32.53RT (min)

lcqtof\_hilic\_pos\_4\_016\_09

m/z 659.4114

01234E+31919.5RT (min)

lcqtof\_hilic\_pos\_4\_015\_42

m/z 660.4138

02468E+32.53RT (min)

lcqtof\_hilic\_pos\_4\_013\_36

m/z 660.4181

02468E+32.53RT (min)

lcqtof\_hilic\_pos\_4\_085\_59

m/z 664.4606

012E+318.519RT (min)

lcqtof\_hilic\_pos\_3\_105\_61

m/z 664.6247

012E+315.516RT (min)

lcqtof\_hilic\_pos\_4\_029\_44

m/z 668.6455

0246E+215.516RT (min)

lcqtof\_hilic\_pos\_1\_018\_08

m/z 671.2721

0123E+31919.5RT (min)

lcqtof\_hilic\_pos\_4\_021\_46

m/z 671.4747

012E+31515.5RT (min)

lcqtof\_hilic\_pos\_4\_015\_42

m/z 672.4524

012E+42.53RT (min)

lcqtof\_hilic\_pos\_4\_041\_54

m/z 672.4530

024E+41818.5RT (min)

lcqtof\_hilic\_pos\_4\_001\_49

m/z 673.4564

01E+42.53RT (min)

lcqtof\_hilic\_pos\_4\_008\_25

m/z 673.4566

01E+41818.5RT (min)

lcqtof\_hilic\_pos\_4\_012\_02

m/z 673.4568

01E+40.51RT (min)

lcqtof\_hilic\_pos\_4\_030\_03

m/z 674.4631

01234E+31818.5RT (min)

lcqtof\_hilic\_pos\_4\_013\_36

m/z 675.4992

01E+315.516RT (min)

lcqtof\_hilic\_pos\_4\_106\_62

m/z 679.4292

012E+315.516RT (min)

lcqtof\_hilic\_pos\_4\_077\_58

m/z 679.6043

012E+315.516RT (min)

lcqtof\_hilic\_pos\_4\_001\_49

m/z 682.5995

01234E+31818.5RT (min)

lcqtof\_hilic\_pos\_3\_101\_61

m/z 684.6131

01E+41818.5RT (min)

lcqtof\_hilic\_pos\_3\_104\_61

m/z 685.6175

024E+31818.5RT (min)

lcqtof\_hilic\_pos\_3\_109\_62

m/z 686.5579

0123E+31818.5RT (min)

lcqtof\_hilic\_pos\_4\_073\_57

m/z 687.2922

01234E+32.53RT (min)

lcqtof\_hilic\_pos\_4\_011\_48

m/z 691.6590

0123E+315.516RT (min)

lcqtof\_hilic\_pos\_4\_107\_62

m/z 693.5552

01234E+233.5RT (min)

lcqtof\_hilic\_pos\_4\_094\_60

m/z 693.5775

0246E+315.516RT (min)

lcqtof\_hilic\_pos\_4\_002\_23

m/z 694.3870

0123E+34.55RT (min)

lcqtof\_hilic\_pos\_4\_006\_29

m/z 694.8747

01234E+36.57RT (min)

lcqtof\_hilic\_pos\_3\_075\_58

m/z 698.5232

012E+31515.5RT (min)

lcqtof\_hilic\_pos\_4\_094\_60

m/z 698.6596

0123E+31616.5RT (min)

lcqtof\_hilic\_pos\_4\_085\_59

m/z 699.5010

01E+31515.5RT (min)

lcqtof\_hilic\_pos\_4\_104\_61

m/z 700.5294

024E+233.5RT (min)

lcqtof\_hilic\_pos\_4\_002\_23

m/z 702.4453

012E+344.5RT (min)

lcqtof\_hilic\_pos\_4\_002\_23

m/z 703.4711

01234E+33.54RT (min)

lcqtof\_hilic\_pos\_3\_072\_57

m/z 703.6305

012E+31515.5RT (min)

lcqtof\_hilic\_pos\_4\_106\_62

m/z 704.4387

0123E+31919.5RT (min)

lcqtof\_hilic\_pos\_4\_006\_29

m/z 704.4622

0123E+344.5RT (min)

lcqtof\_hilic\_pos\_4\_004\_13

m/z 705.4908

01234E+43.54RT (min)

lcqtof\_hilic\_pos\_4\_026\_01

m/z 705.5023

02468E+317.518RT (min)

lcqtof\_hilic\_pos\_4\_113\_62

m/z 705.5056

01E+51618RT (min)

lcqtof\_hilic\_pos\_4\_039\_15

m/z 707.2707

01E+48.59RT (min)

lcqtof\_hilic\_pos\_4\_010\_37

m/z 709.2744

012E+35.56RT (min)

lcqtof\_hilic\_pos\_4\_104\_61

m/z 709.5249

012E+315.516RT (min)

lcqtof\_hilic\_pos\_4\_086\_59

m/z 709.6056

012E+133.5RT (min)

lcqtof\_hilic\_pos\_4\_095\_60

m/z 710.6301

0123E+31818.5RT (min)

lcqtof\_hilic\_pos\_3\_106\_62

m/z 711.3808

01E+315.516RT (min)

lcqtof\_hilic\_pos\_4\_006\_29

m/z 712.6447

012E+41919.5RT (min)

lcqtof\_hilic\_pos\_1\_018\_08

m/z 713.6504

02468E+318.519RT (min)

lcqtof\_hilic\_pos\_1\_024\_25

m/z 714.5897

01234E+31818.5RT (min)

lcqtof\_hilic\_pos\_4\_088\_59

m/z 714.5997

01234E+344.5RT (min)

lcqtof\_hilic\_pos\_4\_085\_59

m/z 722.4958

012E+315.516RT (min)

lcqtof\_hilic\_pos\_4\_081\_58

m/z 723.5988

01E+31515.5RT (min)

lcqtof\_hilic\_pos\_4\_097\_60

m/z 724.0313

0246E+26.57RT (min)

lcqtof\_hilic\_pos\_4\_080\_58

m/z 726.2461

0123E+34.55RT (min)

lcqtof\_hilic\_pos\_4\_008\_25

m/z 730.4953

012E+41818.5RT (min)

lcqtof\_hilic\_pos\_4\_017\_51

m/z 731.4987

01E+41818.5RT (min)

lcqtof\_hilic\_pos\_4\_023\_19

m/z 735.4507

024E+355.5RT (min)

lcqtof\_hilic\_pos\_4\_006\_29

m/z 737.3078

01234E+366.5RT (min)

lcqtof\_hilic\_pos\_4\_072\_57

m/z 738.6392

024E+415.516RT (min)

lcqtof\_hilic\_pos\_3\_023\_01

m/z 740.6759

024E+31818.5RT (min)

lcqtof\_hilic\_pos\_3\_101\_61

m/z 741.6801

012E+316.517RT (min)

lcqtof\_hilic\_pos\_3\_104\_61

m/z 741.9923

012E+37.58RT (min)

lcqtof\_hilic\_pos\_3\_026\_15

m/z 742.4981

012E+315.516RT (min)

lcqtof\_hilic\_pos\_4\_003\_21

m/z 742.5774

01E+44.55RT (min)

lcqtof\_hilic\_pos\_4\_013\_36

m/z 745.6793

012E+31515.5RT (min)

lcqtof\_hilic\_pos\_4\_019\_08

m/z 747.4642

01234E+37.58RT (min)

lcqtof\_hilic\_pos\_4\_043\_39

m/z 747.6611

0123E+31515.5RT (min)

lcqtof\_hilic\_pos\_4\_003\_21

m/z 751.4093

0123E+411.5RT (min)

lcqtof\_hilic\_pos\_4\_046\_07

m/z 752.7495

012E+215.516RT (min)

lcqtof\_hilic\_pos\_1\_018\_08

m/z 755.5767

01E+333.5RT (min)

lcqtof\_hilic\_pos\_1\_015\_45

m/z 760.4086

012E+344.5RT (min)

lcqtof\_hilic\_pos\_4\_084\_59

m/z 760.6298

01234E+315.516RT (min)

lcqtof\_hilic\_pos\_4\_053\_30

m/z 764.5953

012E+315.516RT (min)

lcqtof\_hilic\_pos\_4\_071\_57

m/z 764.9733

012E+37.58RT (min)

lcqtof\_hilic\_pos\_3\_077\_58

m/z 765.6653

0246E+333.5RT (min)

lcqtof\_hilic\_pos\_4\_070\_57

m/z 766.3048

012E+35.56RT (min)

lcqtof\_hilic\_pos\_4\_001\_49

m/z 768.4943

01234E+233.5RT (min)

lcqtof\_hilic\_pos\_4\_016\_09

m/z 769.5023

012.53RT (min)

lcqtof\_hilic\_pos\_3\_002\_23

m/z 771.3364

0246E+366.5RT (min)

lcqtof\_hilic\_pos\_4\_002\_23

m/z 774.6265

0123E+315.516RT (min)

lcqtof\_hilic\_pos\_4\_090\_60

m/z 775.3398

0123E+35.56RT (min)

lcqtof\_hilic\_pos\_4\_004\_13

m/z 777.5845

024E+315.516RT (min)

lcqtof\_hilic\_pos\_4\_106\_62

m/z 778.4313

0123E+333.5RT (min)

lcqtof\_hilic\_pos\_4\_084\_59

m/z 780.4411

012E+311.5RT (min)

lcqtof\_hilic\_pos\_3\_002\_23

m/z 780.4450

01E+333.5RT (min)

lcqtof\_hilic\_pos\_2\_032\_35

m/z 785.5565

012E+33.54RT (min)

lcqtof\_hilic\_pos\_2\_002\_32

m/z 786.4995

012E+333.5RT (min)

lcqtof\_hilic\_pos\_3\_079\_58

m/z 788.5387

0246E+31818.5RT (min)

lcqtof\_hilic\_pos\_4\_093\_60

m/z 789.5427

0246E+31818.5RT (min)

lcqtof\_hilic\_pos\_4\_009\_50

m/z 789.5484

0123E+31818.5RT (min)

lcqtof\_hilic\_pos\_4\_079\_58

m/z 790.5902

01E+31515.5RT (min)

lcqtof\_hilic\_pos\_4\_026\_01

m/z 792.5569

02468E+133.5RT (min)

lcqtof\_hilic\_pos\_4\_036\_24

m/z 794.4909

01E+333.5RT (min)

lcqtof\_hilic\_pos\_3\_082\_59

m/z 794.7236

01234E+31919.5RT (min)

lcqtof\_hilic\_pos\_3\_089\_59

m/z 800.3888

0246E+33.54RT (min)

lcqtof\_hilic\_pos\_4\_091\_60

m/z 800.7760

01234E+344.5RT (min)

lcqtof\_hilic\_pos\_4\_036\_24

m/z 804.0533

0123E+31515.5RT (min)

lcqtof\_hilic\_pos\_4\_111\_62

m/z 804.5467

02468E+388.5RT (min)

lcqtof\_hilic\_pos\_4\_040\_06

m/z 806.6104

012E+315.516RT (min)

lcqtof\_hilic\_pos\_4\_077\_58

m/z 806.7073

01E+333.5RT (min)

lcqtof\_hilic\_pos\_3\_090\_60

m/z 808.2909

02468E+366.5RT (min)

lcqtof\_hilic\_pos\_4\_004\_13

m/z 808.6039

012E+31515.5RT (min)

lcqtof\_hilic\_pos\_4\_004\_13

m/z 810.4884

012E+311.5RT (min)

lcqtof\_hilic\_pos\_4\_079\_58

m/z 811.5476

01234E+33.54RT (min)

lcqtof\_hilic\_pos\_4\_044\_32

m/z 811.5555

024E+33.54RT (min)

lcqtof\_hilic\_pos\_4\_010\_37

m/z 812.4654

012E+43.54RT (min)

lcqtof\_hilic\_pos\_4\_031\_33

m/z 813.5595

0246E+33.54RT (min)

lcqtof\_hilic\_pos\_4\_010\_37

m/z 818.5692

012E+31515.5RT (min)

lcqtof\_hilic\_pos\_4\_103\_61

m/z 819.5159

0123E+34.55RT (min)

lcqtof\_hilic\_pos\_4\_089\_59

m/z 820.4522

01234E+344.5RT (min)

lcqtof\_hilic\_pos\_3\_083\_59

m/z 820.7139

0246E+315.516RT (min)

lcqtof\_hilic\_pos\_3\_044\_04

m/z 821.5163

012E+31515.5RT (min)

lcqtof\_hilic\_pos\_4\_092\_60

m/z 822.6656

0123E+31515.5RT (min)

lcqtof\_hilic\_pos\_4\_036\_24

m/z 822.7558

01E+41919.5RT (min)

lcqtof\_hilic\_pos\_1\_020\_39

m/z 824.8351

01E+411.5RT (min)

lcqtof\_hilic\_pos\_4\_093\_60

m/z 825.6513

0123E+31515.5RT (min)

lcqtof\_hilic\_pos\_4\_001\_49

m/z 825.7732

02468E+31717.5RT (min)

lcqtof\_hilic\_pos\_1\_019\_19

m/z 826.6760

0122.5RT (min)

lcqtof\_hilic\_pos\_4\_008\_25

m/z 828.6764

024E+53.54RT (min)

lcqtof\_hilic\_pos\_4\_095\_60

m/z 833.5831

012E+31515.5RT (min)

lcqtof\_hilic\_pos\_4\_099\_61

m/z 834.6000

0123E+422.5RT (min)

lcqtof\_hilic\_pos\_4\_071\_57

m/z 835.5149

0123E+31515.5RT (min)

lcqtof\_hilic\_pos\_4\_022\_10

m/z 835.8836

024E+30.51RT (min)

lcqtof\_hilic\_pos\_4\_074\_58

m/z 836.6144

012E+42.53RT (min)

lcqtof\_hilic\_pos\_4\_071\_57

m/z 836.7446

0246E+315.516RT (min)

lcqtof\_hilic\_pos\_3\_076\_58

m/z 837.3171

0246E+366.5RT (min)

lcqtof\_hilic\_pos\_4\_019\_08

m/z 844.6593

0123E+315.516RT (min)

lcqtof\_hilic\_pos\_4\_076\_58

m/z 845.5896

012E+31515.5RT (min)

lcqtof\_hilic\_pos\_4\_094\_60

m/z 846.5786

0246E+322.5RT (min)

lcqtof\_hilic\_pos\_4\_011\_48

m/z 846.5788

0123E+31818.5RT (min)

lcqtof\_hilic\_pos\_4\_095\_60

m/z 847.5834

01E+31818.5RT (min)

lcqtof\_hilic\_pos\_4\_090\_60

m/z 848.7717

024E+31818.5RT (min)

lcqtof\_hilic\_pos\_3\_110\_62

m/z 850.7863

0246E+31818.5RT (min)

lcqtof\_hilic\_pos\_3\_105\_61

m/z 851.7912

0123E+31818.5RT (min)

lcqtof\_hilic\_pos\_3\_112\_62

m/z 852.6797

0123E+315.516RT (min)

lcqtof\_hilic\_pos\_4\_001\_49

m/z 856.7211

012E+318.519RT (min)

lcqtof\_hilic\_pos\_4\_010\_37

m/z 860.5839

02468E+344.5RT (min)

lcqtof\_hilic\_pos\_4\_036\_24

m/z 870.7393

0123E+500.5RT (min)

lcqtof\_hilic\_pos\_3\_053\_36

m/z 872.6662

01E+31616.5RT (min)

lcqtof\_hilic\_pos\_1\_035\_18

m/z 872.7020

01234E+315.516RT (min)

lcqtof\_hilic\_pos\_4\_101\_61

m/z 872.7695

02468E+31818.5RT (min)

lcqtof\_hilic\_pos\_3\_110\_62

m/z 873.7738

0123E+31818.5RT (min)

lcqtof\_hilic\_pos\_3\_110\_62

m/z 873.7755

0123E+318.519RT (min)

lcqtof\_hilic\_pos\_2\_022\_09

m/z 874.7852

01E+41818.5RT (min)

lcqtof\_hilic\_pos\_3\_110\_62

m/z 875.7906

024E+31818.5RT (min)

lcqtof\_hilic\_pos\_3\_113\_62

m/z 876.8008

01E+41818.5RT (min)

lcqtof\_hilic\_pos\_3\_110\_62

m/z 877.8061

0246E+31818.5RT (min)

lcqtof\_hilic\_pos\_3\_112\_62

m/z 878.8147

01E+418.519RT (min)

lcqtof\_hilic\_pos\_1\_019\_19

m/z 878.8163

01E+418.519RT (min)

lcqtof\_hilic\_pos\_1\_019\_19

m/z 880.5378

0123E+31515.5RT (min)

lcqtof\_hilic\_pos\_4\_002\_23

m/z 884.7258

02468E+315.516RT (min)

lcqtof\_hilic\_pos\_4\_050\_16

m/z 885.3672

0123E+318.519RT (min)

lcqtof\_hilic\_pos\_4\_014\_22

m/z 887.3434

0123E+35.56RT (min)

lcqtof\_hilic\_pos\_4\_031\_33

m/z 891.7159

0123E+315.516RT (min)

lcqtof\_hilic\_pos\_4\_013\_36

m/z 894.7535

0123E+31818.5RT (min)

lcqtof\_hilic\_pos\_1\_050\_22

m/z 895.7274

01E+41616.5RT (min)

lcqtof\_hilic\_pos\_1\_019\_19

m/z 895.8213

01E+21616.5RT (min)

lcqtof\_hilic\_pos\_1\_020\_39

m/z 896.7691

01E+41818.5RT (min)

lcqtof\_hilic\_pos\_3\_109\_62

m/z 897.7735

0246E+317.518RT (min)

lcqtof\_hilic\_pos\_3\_112\_62

m/z 897.7754

012E+418.519RT (min)

lcqtof\_hilic\_pos\_1\_018\_08

m/z 898.6711

0123E+31616.5RT (min)

lcqtof\_hilic\_pos\_1\_023\_27

m/z 898.7853

01234E+418.519RT (min)

lcqtof\_hilic\_pos\_1\_023\_27

m/z 899.7885

02468E+31818.5RT (min)

lcqtof\_hilic\_pos\_3\_110\_62

m/z 900.6799

01234E+31616.5RT (min)

lcqtof\_hilic\_pos\_1\_019\_19

m/z 900.8007

01E+418.519RT (min)

lcqtof\_hilic\_pos\_3\_088\_59

m/z 901.8059

02468E+318.519RT (min)

lcqtof\_hilic\_pos\_3\_086\_59

m/z 903.8206

01E+41818.5RT (min)

lcqtof\_hilic\_pos\_3\_113\_62

m/z 904.8300

02468E+31818.5RT (min)

lcqtof\_hilic\_pos\_3\_002\_23

m/z 910.7212

012E+315.516RT (min)

lcqtof\_hilic\_pos\_4\_003\_21

m/z 910.7897

012E+31616.5RT (min)

lcqtof\_hilic\_pos\_4\_104\_61

m/z 922.7364

01E+31616.5RT (min)

lcqtof\_hilic\_pos\_3\_110\_62

m/z 926.4512

0246E+35.56RT (min)

lcqtof\_hilic\_pos\_3\_077\_58

m/z 936.8965

0123E+31717.5RT (min)

lcqtof\_hilic\_pos\_1\_021\_09

m/z 943.7703

012E+31616.5RT (min)

lcqtof\_hilic\_pos\_4\_004\_13

m/z 944.7610

01234E+30.51RT (min)

lcqtof\_hilic\_pos\_4\_069\_57

m/z 962.9113

0246E+31717.5RT (min)

lcqtof\_hilic\_pos\_1\_048\_26

m/z 963.9162

0123E+31717.5RT (min)

lcqtof\_hilic\_pos\_1\_036\_13

m/z 975.8369

0123E+31616.5RT (min)

lcqtof\_hilic\_pos\_4\_002\_23

m/z 977.6947

024E+315.516RT (min)

lcqtof\_hilic\_pos\_4\_106\_62

m/z 979.7200

012E+315.516RT (min)

lcqtof\_hilic\_pos\_4\_077\_58

m/z 1008.7232

01E+31515.5RT (min)

lcqtof\_hilic\_pos\_4\_075\_58

m/z 1010.7504

01E+31515.5RT (min)

lcqtof\_hilic\_pos\_4\_001\_49

m/z 1015.5461

01234E+344.5RT (min)

lcqtof\_hilic\_pos\_4\_050\_16

m/z 1015.6995

01E+31515.5RT (min)

lcqtof\_hilic\_pos\_4\_082\_59

m/z 1016.5653

01234E+344.5RT (min)

lcqtof\_hilic\_pos\_1\_005\_01

m/z 1019.6900

01E+315.516RT (min)

lcqtof\_hilic\_pos\_4\_091\_60

m/z 1021.7053

012E+31515.5RT (min)

lcqtof\_hilic\_pos\_4\_003\_21

m/z 1042.7592

01E+31515.5RT (min)

lcqtof\_hilic\_pos\_4\_098\_61

m/z 1056.7382

01E+31515.5RT (min)

lcqtof\_hilic\_pos\_4\_109\_62

m/z 1056.7542

01234E+32.53RT (min)

lcqtof\_hilic\_pos\_4\_006\_29

m/z 1060.6903

01E+31515.5RT (min)

lcqtof\_hilic\_pos\_4\_104\_61

m/z 1092.0005

012E+37.58RT (min)

lcqtof\_hilic\_pos\_3\_076\_58

m/z 1098.5456

02468E+35.56RT (min)

lcqtof\_hilic\_pos\_4\_005\_11

m/z 1099.0754

01234E+366.5RT (min)

lcqtof\_hilic\_pos\_4\_012\_02

m/z 1144.0689

01234E+34.55RT (min)

lcqtof\_hilic\_pos\_3\_083\_59

m/z 1144.2691

0123E+34.55RT (min)

lcqtof\_hilic\_pos\_4\_086\_59

m/z 1144.4691

024E+34.55RT (min)

lcqtof\_hilic\_pos\_3\_088\_59

m/z 1199.6545

0123E+34.55RT (min)

lcqtof\_hilic\_pos\_3\_095\_60

m/z 1199.8551

01234E+34.55RT (min)

lcqtof\_hilic\_pos\_3\_091\_60
