## Supplementary material for "The Everything Bagel Feature Finder: Ultra-fast automated feature finding for untargeted metabolomics": SI Data File: tool_unique_xic_st_XCMS_ST004651.html

XCMS-unique features (not detected by the other two tools) - ST004651

### XCMS-unique features (not detected by the other two tools) - ST004651

500 features · qtof · XIC window m/z ± 25 ppm (≥ 0.002 Da) · ± 0.5 min

Every panel is a feature **XCMS** detected that neither of the other two tools detected — three-way unique across EB (standard mode (min\_detection\_rate=0.1)), Masscube and XCMS, under the standard matching rule (m/z ± 25 ppm; apex within the reference RT window ± 0.2 min). It shows the MS1 extracted-ion chromatogram at the feature's m/z, drawn from the raw mzML named in `alignment_reference_file`, zoomed to ± 0.5 min around the feature's reported apex RT. **Gray solid line** = the feature's reported apex RT; **red dashed line** = XCMS's assigned MS/MS — `RTINSECONDS` in its `aligned_features.mgf`; the panel widens to keep an off-apex MS/MS on-screen. Features with no assigned MS/MS show only the gray line. y is per-cell autoscaled. Rows keep the table order (m/z ascending).

m/z 59.0135

012E+35.56RT (min)

lcqtof\_hilic\_neg\_4\_011\_35

m/z 61.9877

0123E+31818.5RT (min)

lcqtof\_hilic\_neg\_4\_075\_58

m/z 68.9954

01E+411.5RT (min)

lcqtof\_hilic\_neg\_4\_050\_02

m/z 78.9584

012E+31313.5RT (min)

lcqtof\_hilic\_neg\_4\_056\_56

m/z 80.9645

02468E+311.5RT (min)

lcqtof\_hilic\_neg\_4\_047\_06

m/z 83.0313

0246E+311.5RT (min)

lcqtof\_hilic\_neg\_4\_104\_61

m/z 84.9380

024E+319.520RT (min)

lcqtof\_hilic\_neg\_1\_033\_53

m/z 84.9380

024E+35.56RT (min)

lcqtof\_hilic\_neg\_3\_017\_51

m/z 87.0269

024E+31.52RT (min)

lcqtof\_hilic\_neg\_4\_055\_32

m/z 89.0237

0123E+31919.5RT (min)

lcqtof\_hilic\_neg\_4\_092\_60

m/z 96.9595

0123E+318.519RT (min)

lcqtof\_hilic\_neg\_4\_083\_59

m/z 98.9387

012E+311.5RT (min)

lcqtof\_hilic\_neg\_2\_002\_36

m/z 99.9250

0246E+40.51RT (min)

lcqtof\_hilic\_neg\_4\_050\_02

m/z 99.9250

01234E+433.5RT (min)

lcqtof\_hilic\_neg\_4\_050\_02

m/z 101.9336

024E+319.520RT (min)

lcqtof\_hilic\_neg\_1\_054\_01

m/z 101.9339

0246E+311.5RT (min)

lcqtof\_hilic\_neg\_3\_111\_62

m/z 102.9326

012E+31.52RT (min)

lcqtof\_hilic\_neg\_2\_014\_08

m/z 112.9850

024E+31818.5RT (min)

lcqtof\_hilic\_neg\_4\_056\_56

m/z 113.0602

012E+31.52RT (min)

lcqtof\_hilic\_neg\_4\_089\_59

m/z 114.9485

0123E+411.5RT (min)

lcqtof\_hilic\_neg\_3\_109\_62

m/z 119.0806

0123E+31.52RT (min)

lcqtof\_hilic\_neg\_4\_090\_60

m/z 125.8991

01234E+33.54RT (min)

lcqtof\_hilic\_neg\_4\_016\_37

m/z 128.0347

0123E+31919.5RT (min)

lcqtof\_hilic\_neg\_4\_093\_60

m/z 128.0348

02468E+300.5RT (min)

lcqtof\_hilic\_neg\_4\_006\_30

m/z 129.9315

0123E+319.520RT (min)

lcqtof\_hilic\_neg\_3\_066\_57

m/z 129.9321

01234E+333.5RT (min)

lcqtof\_hilic\_neg\_3\_068\_57

m/z 129.9750

012E+39.510RT (min)

lcqtof\_hilic\_neg\_4\_099\_61

m/z 130.9655

02468E+39.510RT (min)

lcqtof\_hilic\_neg\_4\_050\_02

m/z 132.0204

01E+311.5RT (min)

lcqtof\_hilic\_neg\_4\_008\_24

m/z 132.8669

01E+41212.5RT (min)

lcqtof\_hilic\_neg\_4\_094\_60

m/z 132.8672

0246E+318.519RT (min)

lcqtof\_hilic\_neg\_2\_040\_20

m/z 134.8645

012E+411.512RT (min)

lcqtof\_hilic\_neg\_4\_082\_59

m/z 134.8647

0246E+318.519RT (min)

lcqtof\_hilic\_neg\_4\_003\_16

m/z 135.9704

0246E+317.518RT (min)

lcqtof\_hilic\_neg\_4\_084\_59

m/z 136.8620

01E+412.513RT (min)

lcqtof\_hilic\_neg\_4\_006\_30

m/z 136.8625

01E+31818.5RT (min)

lcqtof\_hilic\_neg\_3\_050\_41

m/z 138.0191

012E+31818.5RT (min)

lcqtof\_hilic\_neg\_4\_099\_61

m/z 144.0524

024E+333.5RT (min)

lcqtof\_hilic\_neg\_4\_084\_59

m/z 144.9230

01E+319.520RT (min)

lcqtof\_hilic\_neg\_1\_051\_35

m/z 144.9230

012E+31.52RT (min)

lcqtof\_hilic\_neg\_2\_050\_19

m/z 145.9304

012E+319.520RT (min)

lcqtof\_hilic\_neg\_3\_046\_03

m/z 145.9304

0123E+333.5RT (min)

lcqtof\_hilic\_neg\_4\_050\_02

m/z 146.0815

024E+322.5RT (min)

lcqtof\_hilic\_neg\_4\_048\_23

m/z 146.9370

012E+414.515RT (min)

lcqtof\_hilic\_neg\_4\_107\_62

m/z 146.9380

012E+41111.5RT (min)

lcqtof\_hilic\_neg\_4\_104\_61

m/z 146.9649

0246E+39.510RT (min)

lcqtof\_hilic\_neg\_3\_066\_57

m/z 146.9650

0246E+31313.5RT (min)

lcqtof\_hilic\_neg\_1\_050\_12

m/z 146.9651

0246E+31818.5RT (min)

lcqtof\_hilic\_neg\_4\_101\_61

m/z 149.0715

02468E+311.5RT (min)

lcqtof\_hilic\_neg\_4\_055\_32

m/z 154.0503

02468E+322.5RT (min)

lcqtof\_hilic\_neg\_4\_052\_13

m/z 155.9507

0123E+25.56RT (min)

lcqtof\_hilic\_neg\_4\_011\_35

m/z 157.0244

0123E+255.5RT (min)

lcqtof\_hilic\_neg\_4\_047\_06

m/z 157.0502

0123E+311.5RT (min)

lcqtof\_hilic\_neg\_4\_085\_59

m/z 157.9306

012E+32.53RT (min)

lcqtof\_hilic\_neg\_3\_113\_62

m/z 159.8972

01E+40.51RT (min)

lcqtof\_hilic\_neg\_4\_050\_02

m/z 160.0342

01234E+35.56RT (min)

lcqtof\_hilic\_neg\_4\_013\_20

m/z 160.9352

01234E+31919.5RT (min)

lcqtof\_hilic\_neg\_1\_022\_36

m/z 162.9686

0246E+34.55RT (min)

lcqtof\_hilic\_neg\_3\_066\_57

m/z 164.8357

0123E+399.5RT (min)

lcqtof\_hilic\_neg\_2\_031\_01

m/z 165.0309

0123E+30.51RT (min)

lcqtof\_hilic\_neg\_4\_026\_04

m/z 167.8362

01234E+318.519RT (min)

lcqtof\_hilic\_neg\_1\_033\_53

m/z 168.8864

01234E+311.5RT (min)

lcqtof\_hilic\_neg\_4\_036\_11

m/z 168.9885

01234E+318.519RT (min)

lcqtof\_hilic\_neg\_4\_084\_59

m/z 169.0244

01E+322.5RT (min)

lcqtof\_hilic\_neg\_2\_021\_17

m/z 169.0246

0123E+31919.5RT (min)

lcqtof\_hilic\_neg\_4\_010\_28

m/z 170.0174

012E+311.5RT (min)

lcqtof\_hilic\_neg\_4\_032\_01

m/z 170.0725

01E+411.5RT (min)

lcqtof\_hilic\_neg\_4\_052\_13

m/z 170.9779

024E+31.52RT (min)

lcqtof\_hilic\_neg\_4\_004\_47

m/z 171.0484

01234E+333.5RT (min)

lcqtof\_hilic\_neg\_4\_087\_59

m/z 171.8310

01E+318.519RT (min)

lcqtof\_hilic\_neg\_3\_078\_58

m/z 171.9461

012E+300.5RT (min)

lcqtof\_hilic\_neg\_3\_024\_35

m/z 171.9463

012E+32.53RT (min)

lcqtof\_hilic\_neg\_1\_029\_06

m/z 171.9463

012E+31919.5RT (min)

lcqtof\_hilic\_neg\_3\_072\_57

m/z 172.0411

012E+433.5RT (min)

lcqtof\_hilic\_neg\_4\_088\_59

m/z 173.0558

0246E+344.5RT (min)

lcqtof\_hilic\_neg\_4\_048\_23

m/z 173.8286

01E+33.54RT (min)

lcqtof\_hilic\_neg\_1\_043\_41

m/z 174.0015

0246E+266.5RT (min)

lcqtof\_hilic\_neg\_4\_040\_05

m/z 174.0768

02468E+322.5RT (min)

lcqtof\_hilic\_neg\_4\_048\_23

m/z 176.8164

01E+41313.5RT (min)

lcqtof\_hilic\_neg\_4\_016\_37

m/z 176.8165

01E+499.5RT (min)

lcqtof\_hilic\_neg\_4\_013\_20

m/z 176.8169

02468E+30.51RT (min)

lcqtof\_hilic\_neg\_4\_036\_11

m/z 177.0654

01E+31.52RT (min)

lcqtof\_hilic\_neg\_4\_055\_32

m/z 177.8911

017.58RT (min)

lcqtof\_hilic\_neg\_1\_008\_13

m/z 178.8143

01E+499.5RT (min)

lcqtof\_hilic\_neg\_4\_052\_13

m/z 178.8146

012E+40.51RT (min)

lcqtof\_hilic\_neg\_4\_015\_09

m/z 180.0614

0246E+33.54RT (min)

lcqtof\_hilic\_neg\_4\_096\_60

m/z 180.8121

02468E+31313.5RT (min)

lcqtof\_hilic\_neg\_4\_079\_58

m/z 180.8121

02468E+399.5RT (min)

lcqtof\_hilic\_neg\_4\_005\_29

m/z 180.9227

012E+411.512RT (min)

lcqtof\_hilic\_neg\_4\_106\_62

m/z 180.9729

012E+313.514RT (min)

lcqtof\_hilic\_neg\_4\_080\_58

m/z 181.0245

0123E+31919.5RT (min)

lcqtof\_hilic\_neg\_4\_002\_25

m/z 181.0606

012E+355.5RT (min)

lcqtof\_hilic\_neg\_4\_054\_48

m/z 181.8371

012E+344.5RT (min)

lcqtof\_hilic\_neg\_4\_009\_50

m/z 182.8099

01E+31313.5RT (min)

lcqtof\_hilic\_neg\_4\_081\_58

m/z 182.9204

01E+411.512RT (min)

lcqtof\_hilic\_neg\_4\_067\_57

m/z 182.9204

01E+46.57RT (min)

lcqtof\_hilic\_neg\_4\_112\_62

m/z 182.9539

012E+313.514RT (min)

lcqtof\_hilic\_neg\_4\_056\_56

m/z 182.9877

012E+31818.5RT (min)

lcqtof\_hilic\_neg\_4\_076\_58

m/z 183.0109

024E+31818.5RT (min)

lcqtof\_hilic\_neg\_4\_074\_58

m/z 184.9180

01234E+31212.5RT (min)

lcqtof\_hilic\_neg\_4\_082\_59

m/z 187.0972

012E+318.519RT (min)

lcqtof\_hilic\_neg\_4\_112\_62

m/z 187.8611

01E+31010.5RT (min)

lcqtof\_hilic\_neg\_3\_102\_61

m/z 187.9412

024E+32.53RT (min)

lcqtof\_hilic\_neg\_1\_054\_01

m/z 187.9413

01E+318.519RT (min)

lcqtof\_hilic\_neg\_3\_102\_61

m/z 187.9646

012E+266.5RT (min)

lcqtof\_hilic\_neg\_2\_052\_24

m/z 188.0337

012E+31.52RT (min)

lcqtof\_hilic\_neg\_4\_048\_23

m/z 190.0041

0246E+31818.5RT (min)

lcqtof\_hilic\_neg\_4\_050\_02

m/z 190.0041

01E+41.52RT (min)

lcqtof\_hilic\_neg\_3\_066\_57

m/z 190.0513

02468E+366.5RT (min)

lcqtof\_hilic\_neg\_4\_096\_60

m/z 190.9279

012E+310.511RT (min)

lcqtof\_hilic\_neg\_3\_107\_62

m/z 191.0000

0123E+411.5RT (min)

lcqtof\_hilic\_neg\_4\_050\_02

m/z 191.0003

0246E+41818.5RT (min)

lcqtof\_hilic\_neg\_3\_067\_57

m/z 193.0624

024E+222.5RT (min)

lcqtof\_hilic\_neg\_4\_052\_13

m/z 193.8973

012E+37.58RT (min)

lcqtof\_hilic\_neg\_4\_017\_51

m/z 194.8920

01E+37.58RT (min)

lcqtof\_hilic\_neg\_1\_005\_32

m/z 194.9960

0246E+333.5RT (min)

lcqtof\_hilic\_neg\_4\_037\_39

m/z 195.0100

01E+30.51RT (min)

lcqtof\_hilic\_neg\_4\_050\_02

m/z 195.9757

02468E+24.55RT (min)

lcqtof\_hilic\_neg\_3\_066\_57

m/z 197.0235

012E+318.519RT (min)

lcqtof\_hilic\_neg\_4\_079\_58

m/z 197.9917

012E+31818.5RT (min)

lcqtof\_hilic\_neg\_4\_085\_59

m/z 197.9919

01E+300.5RT (min)

lcqtof\_hilic\_neg\_4\_088\_59

m/z 199.0978

0123E+311.5RT (min)

lcqtof\_hilic\_neg\_4\_034\_31

m/z 199.8506

024E+211.5RT (min)

lcqtof\_hilic\_neg\_4\_046\_41

m/z 200.0481

0123E+322.5RT (min)

lcqtof\_hilic\_neg\_4\_031\_38

m/z 200.8581

0123E+31919.5RT (min)

lcqtof\_hilic\_neg\_2\_024\_10

m/z 200.9859

012E+31919.5RT (min)

lcqtof\_hilic\_neg\_4\_111\_62

m/z 203.0202

01E+433.5RT (min)

lcqtof\_hilic\_neg\_4\_045\_26

m/z 206.9721

0246E+312.513RT (min)

lcqtof\_hilic\_neg\_3\_067\_57

m/z 208.0461

02468E+34.55RT (min)

lcqtof\_hilic\_neg\_4\_016\_37

m/z 208.9759

012E+30.51RT (min)

lcqtof\_hilic\_neg\_4\_017\_51

m/z 210.9491

0246E+31313.514RT (min)

lcqtof\_hilic\_neg\_3\_066\_57

m/z 212.9105

0123E+319.520RT (min)

lcqtof\_hilic\_neg\_3\_068\_57

m/z 214.8598

0111.5RT (min)

lcqtof\_hilic\_neg\_4\_112\_62

m/z 214.9459

012E+21.52RT (min)

lcqtof\_hilic\_neg\_3\_066\_57

m/z 215.9666

0123E+244.5RT (min)

lcqtof\_hilic\_neg\_4\_091\_60

m/z 218.9056

0123E+37.58RT (min)

lcqtof\_hilic\_neg\_4\_002\_25

m/z 219.0525

01E+21.52RT (min)

lcqtof\_hilic\_neg\_4\_013\_20

m/z 221.7999

0246E+33.54RT (min)

lcqtof\_hilic\_neg\_4\_020\_42

m/z 222.7644

012E+313.514RT (min)

lcqtof\_hilic\_neg\_4\_056\_56

m/z 223.8394

01E+388.5RT (min)

lcqtof\_hilic\_neg\_1\_024\_21

m/z 223.9780

01E+31818.5RT (min)

lcqtof\_hilic\_neg\_4\_087\_59

m/z 224.7623

01E+313.514RT (min)

lcqtof\_hilic\_neg\_4\_104\_61

m/z 224.8725

0123E+312.513RT (min)

lcqtof\_hilic\_neg\_4\_076\_58

m/z 225.5696

0246E+333.5RT (min)

lcqtof\_hilic\_neg\_4\_050\_02

m/z 226.0830

02468E+344.5RT (min)

lcqtof\_hilic\_neg\_4\_056\_56

m/z 226.8705

01234E+312.513RT (min)

lcqtof\_hilic\_neg\_4\_113\_62

m/z 227.0670

024E+32.53RT (min)

lcqtof\_hilic\_neg\_4\_016\_37

m/z 228.1093

0123E+34.55RT (min)

lcqtof\_hilic\_neg\_4\_049\_55

m/z 228.8687

01E+312.513RT (min)

lcqtof\_hilic\_neg\_4\_070\_57

m/z 228.9771

0123E+31111.5RT (min)

lcqtof\_hilic\_neg\_4\_107\_62

m/z 230.9557

01E+313.514RT (min)

lcqtof\_hilic\_neg\_4\_067\_57

m/z 231.9319

01E+31919.5RT (min)

lcqtof\_hilic\_neg\_2\_028\_05

m/z 232.0074

01E+411.5RT (min)

lcqtof\_hilic\_neg\_4\_011\_35

m/z 235.1350

012E+30.51RT (min)

lcqtof\_hilic\_neg\_4\_094\_60

m/z 236.0560

01E+311.5RT (min)

lcqtof\_hilic\_neg\_4\_035\_33

m/z 236.7587

0246E+30.51RT (min)

lcqtof\_hilic\_neg\_4\_015\_09

m/z 237.0490

01234E+366.5RT (min)

lcqtof\_hilic\_neg\_4\_053\_14

m/z 239.7603

0123E+32.53RT (min)

lcqtof\_hilic\_neg\_3\_066\_57

m/z 240.0513

02468E+30.51RT (min)

lcqtof\_hilic\_neg\_4\_002\_25

m/z 241.9759

0133.5RT (min)

lcqtof\_hilic\_neg\_4\_050\_02

m/z 242.1760

012E+31818.5RT (min)

lcqtof\_hilic\_neg\_4\_098\_61

m/z 242.9428

0123E+39.510RT (min)

lcqtof\_hilic\_neg\_4\_050\_02

m/z 243.0179

0246E+344.5RT (min)

lcqtof\_hilic\_neg\_4\_052\_13

m/z 243.0553

0246E+311.5RT (min)

lcqtof\_hilic\_neg\_4\_028\_36

m/z 245.0625

024E+311.5RT (min)

lcqtof\_hilic\_neg\_4\_013\_20

m/z 246.0280

012E+26.57RT (min)

lcqtof\_hilic\_neg\_4\_053\_14

m/z 246.9375

01234E+31.52RT (min)

lcqtof\_hilic\_neg\_4\_023\_15

m/z 248.5806

012E+33.54RT (min)

lcqtof\_hilic\_neg\_4\_050\_02

m/z 248.9726

012E+416.517RT (min)

lcqtof\_hilic\_neg\_4\_050\_02

m/z 249.0786

024E+344.5RT (min)

lcqtof\_hilic\_neg\_4\_083\_59

m/z 251.0954

01234E+318.519RT (min)

lcqtof\_hilic\_neg\_4\_105\_61

m/z 253.0495

024E+31.52RT (min)

lcqtof\_hilic\_neg\_4\_016\_37

m/z 253.1078

01234E+411.5RT (min)

lcqtof\_hilic\_neg\_4\_052\_13

m/z 253.7939

012E+30.51RT (min)

lcqtof\_hilic\_neg\_1\_026\_10

m/z 257.0082

01E+37.58RT (min)

lcqtof\_hilic\_neg\_4\_034\_31

m/z 258.0671

0246E+222.5RT (min)

lcqtof\_hilic\_neg\_3\_054\_37

m/z 258.1087

024E+24.55RT (min)

lcqtof\_hilic\_neg\_4\_018\_27

m/z 258.9457

0246E+32.53RT (min)

lcqtof\_hilic\_neg\_4\_034\_31

m/z 259.1297

024E+32.53RT (min)

lcqtof\_hilic\_neg\_4\_048\_23

m/z 260.0162

0246E+34.55RT (min)

lcqtof\_hilic\_neg\_4\_050\_02

m/z 260.9436

0246E+32.53RT (min)

lcqtof\_hilic\_neg\_4\_034\_31

m/z 260.9699

024E+222.5RT (min)

lcqtof\_hilic\_neg\_4\_038\_10

m/z 265.0733

01E+31.52RT (min)

lcqtof\_hilic\_neg\_4\_098\_61

m/z 265.0826

012E+34.55RT (min)

lcqtof\_hilic\_neg\_4\_096\_60

m/z 265.1440

01E+41010.5RT (min)

lcqtof\_hilic\_neg\_4\_099\_61

m/z 266.1507

01234E+31919.5RT (min)

lcqtof\_hilic\_neg\_4\_074\_58

m/z 266.1509

01234E+31717.5RT (min)

lcqtof\_hilic\_neg\_4\_083\_59

m/z 267.0290

01234E+31.52RT (min)

lcqtof\_hilic\_neg\_4\_039\_18

m/z 267.0721

0123E+311.5RT (min)

lcqtof\_hilic\_neg\_4\_094\_60

m/z 267.1460

012E+318.519RT (min)

lcqtof\_hilic\_neg\_4\_084\_59

m/z 267.1566

01234E+40.51RT (min)

lcqtof\_hilic\_neg\_3\_035\_46

m/z 268.9631

012E+333.5RT (min)

lcqtof\_hilic\_neg\_4\_084\_59

m/z 270.8285

0123E+37.58RT (min)

lcqtof\_hilic\_neg\_4\_052\_13

m/z 271.0813

024E+377.5RT (min)

lcqtof\_hilic\_neg\_4\_084\_59

m/z 271.1286

012E+41.52RT (min)

lcqtof\_hilic\_neg\_4\_046\_41

m/z 277.0602

01234E+222.5RT (min)

lcqtof\_hilic\_neg\_4\_050\_02

m/z 280.0618

02468E+22.53RT (min)

lcqtof\_hilic\_neg\_4\_055\_32

m/z 280.1663

01E+318.519RT (min)

lcqtof\_hilic\_neg\_4\_067\_57

m/z 281.0473

012E+33.54RT (min)

lcqtof\_hilic\_neg\_4\_078\_58

m/z 282.3676

02468E+40.51RT (min)

lcqtof\_hilic\_neg\_4\_050\_02

m/z 283.0943

024E+344.5RT (min)

lcqtof\_hilic\_neg\_4\_085\_59

m/z 284.0740

0246E+311.5RT (min)

lcqtof\_hilic\_neg\_4\_102\_61

m/z 284.9779

012E+30.51RT (min)

lcqtof\_hilic\_neg\_4\_038\_10

m/z 285.0960

024E+21.52RT (min)

lcqtof\_hilic\_neg\_4\_053\_14

m/z 286.1051

02468E+311.5RT (min)

lcqtof\_hilic\_neg\_4\_005\_29

m/z 291.0670

0123E+244.5RT (min)

lcqtof\_hilic\_neg\_4\_035\_33

m/z 293.1425

02468E+318.519RT (min)

lcqtof\_hilic\_neg\_4\_091\_60

m/z 293.9971

012E+36.57RT (min)

lcqtof\_hilic\_neg\_4\_053\_14

m/z 294.1412

0246E+277.5RT (min)

lcqtof\_hilic\_neg\_4\_034\_31

m/z 294.1817

012E+31919.5RT (min)

lcqtof\_hilic\_neg\_4\_001\_49

m/z 294.8309

012E+311.5RT (min)

lcqtof\_hilic\_neg\_1\_020\_09

m/z 295.0114

01234E+222.5RT (min)

lcqtof\_hilic\_neg\_4\_050\_02

m/z 295.1381

012E+344.5RT (min)

lcqtof\_hilic\_neg\_4\_087\_59

m/z 296.0934

0246E+35.56RT (min)

lcqtof\_hilic\_neg\_4\_052\_13

m/z 298.0913

0246E+233.5RT (min)

lcqtof\_hilic\_neg\_4\_047\_06

m/z 298.0996

0246E+25.56RT (min)

lcqtof\_hilic\_neg\_4\_087\_59

m/z 298.1559

024E+318.519RT (min)

lcqtof\_hilic\_neg\_4\_101\_61

m/z 299.0970

012E+355.5RT (min)

lcqtof\_hilic\_neg\_4\_052\_13

m/z 299.1366

0246E+35.56RT (min)

lcqtof\_hilic\_neg\_4\_040\_05

m/z 299.1452

0123E+318.519RT (min)

lcqtof\_hilic\_neg\_4\_001\_49

m/z 300.0926

01E+444.5RT (min)

lcqtof\_hilic\_neg\_4\_033\_53

m/z 300.9637

0123E+322.5RT (min)

lcqtof\_hilic\_neg\_4\_013\_20

m/z 305.1864

012E+317.518RT (min)

lcqtof\_hilic\_neg\_4\_083\_59

m/z 306.0998

0246E+35.56RT (min)

lcqtof\_hilic\_neg\_4\_050\_02

m/z 306.9367

01E+313.514RT (min)

lcqtof\_hilic\_neg\_4\_046\_41

m/z 307.1964

0123E+322.5RT (min)

lcqtof\_hilic\_neg\_4\_052\_13

m/z 308.1612

012E+318.519RT (min)

lcqtof\_hilic\_neg\_4\_102\_61

m/z 309.0770

024E+266.5RT (min)

lcqtof\_hilic\_neg\_4\_008\_24

m/z 309.0992

012E+31.52RT (min)

lcqtof\_hilic\_neg\_4\_003\_16

m/z 310.1313

01E+30.51RT (min)

lcqtof\_hilic\_neg\_4\_052\_13

m/z 310.9301

012E+399.5RT (min)

lcqtof\_hilic\_neg\_4\_040\_05

m/z 311.0217

01234E+244.5RT (min)

lcqtof\_hilic\_neg\_4\_039\_18

m/z 311.1680

01E+466.5RT (min)

lcqtof\_hilic\_neg\_4\_034\_31

m/z 312.0839

0123E+33.54RT (min)

lcqtof\_hilic\_neg\_4\_013\_20

m/z 312.1714

012E+418.519RT (min)

lcqtof\_hilic\_neg\_4\_113\_62

m/z 313.1154

01234E+318.519RT (min)

lcqtof\_hilic\_neg\_4\_003\_16

m/z 313.1671

0246E+318.519RT (min)

lcqtof\_hilic\_neg\_4\_076\_58

m/z 313.9081

01E+34.55RT (min)

lcqtof\_hilic\_neg\_4\_050\_02

m/z 315.1805

0123E+31818.5RT (min)

lcqtof\_hilic\_neg\_4\_102\_61

m/z 316.0439

0246E+366.5RT (min)

lcqtof\_hilic\_neg\_4\_081\_58

m/z 316.7786

024E+311.5RT (min)

lcqtof\_hilic\_neg\_1\_021\_42

m/z 316.8899

01234E+344.5RT (min)

lcqtof\_hilic\_neg\_1\_043\_41

m/z 318.1033

01E+55.56RT (min)

lcqtof\_hilic\_neg\_1\_007\_05

m/z 320.0961

01E+55.56RT (min)

lcqtof\_hilic\_neg\_4\_047\_06

m/z 321.0624

02468E+222.5RT (min)

lcqtof\_hilic\_neg\_1\_031\_16

m/z 321.1735

012E+31819RT (min)

lcqtof\_hilic\_neg\_4\_001\_49

m/z 321.2104

0123E+31919.5RT (min)

lcqtof\_hilic\_neg\_4\_085\_59

m/z 322.0132

01234E+366.5RT (min)

lcqtof\_hilic\_neg\_4\_052\_13

m/z 323.1756

0123E+318.519RT (min)

lcqtof\_hilic\_neg\_4\_096\_60

m/z 324.1411

01E+25.56RT (min)

lcqtof\_hilic\_neg\_4\_087\_59

m/z 326.1869

0118.519RT (min)

lcqtof\_hilic\_neg\_4\_001\_49

m/z 327.0629

024E+34.55RT (min)

lcqtof\_hilic\_neg\_3\_073\_57

m/z 327.1286

0123E+318.519RT (min)

lcqtof\_hilic\_neg\_4\_056\_56

m/z 327.1828

02468E+31818.5RT (min)

lcqtof\_hilic\_neg\_4\_108\_62

m/z 328.0799

0123E+33.54RT (min)

lcqtof\_hilic\_neg\_4\_010\_28

m/z 328.8626

01234E+37.58RT (min)

lcqtof\_hilic\_neg\_4\_112\_62

m/z 329.1893

024E+20.51RT (min)

lcqtof\_hilic\_neg\_2\_051\_14

m/z 329.2170

01234E+20.51RT (min)

lcqtof\_hilic\_neg\_4\_050\_02

m/z 331.0389

0246E+31.52RT (min)

lcqtof\_hilic\_neg\_4\_018\_27

m/z 335.0533

01234E+32.53RT (min)

lcqtof\_hilic\_neg\_2\_020\_23

m/z 335.1892

01234E+31818.5RT (min)

lcqtof\_hilic\_neg\_4\_090\_60

m/z 335.1902

01234E+31818.5RT (min)

lcqtof\_hilic\_neg\_4\_086\_59

m/z 336.1559

0246E+31.52RT (min)

lcqtof\_hilic\_neg\_4\_004\_47

m/z 337.0987

0123E+211.5RT (min)

lcqtof\_hilic\_neg\_4\_089\_59

m/z 338.8549

012E+37.58RT (min)

lcqtof\_hilic\_neg\_4\_002\_25

m/z 340.1289

0246E+311.5RT (min)

lcqtof\_hilic\_neg\_4\_008\_24

m/z 340.2027

01E+418.519RT (min)

lcqtof\_hilic\_neg\_4\_079\_58

m/z 341.0857

01E+36.57RT (min)

lcqtof\_hilic\_neg\_4\_040\_05

m/z 341.1975

024E+318.519RT (min)

lcqtof\_hilic\_neg\_4\_112\_62

m/z 342.2277

024E+20.51RT (min)

lcqtof\_hilic\_neg\_4\_002\_25

m/z 343.1204

01E+300.5RT (min)

lcqtof\_hilic\_neg\_4\_071\_57

m/z 343.1211

012E+318.519RT (min)

lcqtof\_hilic\_neg\_4\_068\_57

m/z 343.1228

0246E+366.5RT (min)

lcqtof\_hilic\_neg\_4\_054\_48

m/z 343.2120

012E+31818.5RT (min)

lcqtof\_hilic\_neg\_4\_077\_58

m/z 344.0565

0246E+24.55RT (min)

lcqtof\_hilic\_neg\_4\_053\_14

m/z 345.1281

02468E+31.52RT (min)

lcqtof\_hilic\_neg\_4\_079\_58

m/z 346.8420

012E+36.57RT (min)

lcqtof\_hilic\_neg\_4\_035\_33

m/z 350.1250

0123E+31717.5RT (min)

lcqtof\_hilic\_neg\_4\_113\_62

m/z 353.0353

02468E+366.5RT (min)

lcqtof\_hilic\_neg\_4\_048\_23

m/z 353.2082

0123E+31818.5RT (min)

lcqtof\_hilic\_neg\_4\_076\_58

m/z 354.0606

0246E+377.5RT (min)

lcqtof\_hilic\_neg\_4\_012\_07

m/z 355.1934

01E+411.5RT (min)

lcqtof\_hilic\_neg\_4\_050\_02

m/z 355.2251

0123E+20.51RT (min)

lcqtof\_hilic\_neg\_2\_002\_36

m/z 357.1089

02468E+35.56RT (min)

lcqtof\_hilic\_neg\_4\_032\_01

m/z 358.0142

01234E+30.51RT (min)

lcqtof\_hilic\_neg\_4\_017\_51

m/z 358.2098

0246E+31818.5RT (min)

lcqtof\_hilic\_neg\_4\_001\_49

m/z 358.2103

0246E+318.519RT (min)

lcqtof\_hilic\_neg\_4\_037\_39

m/z 359.2169

012E+317.518RT (min)

lcqtof\_hilic\_neg\_4\_003\_16

m/z 360.0482

01234E+166.5RT (min)

lcqtof\_hilic\_neg\_4\_095\_60

m/z 361.0071

01E+34.55RT (min)

lcqtof\_hilic\_neg\_2\_004\_26

m/z 364.0926

01234E+233.5RT (min)

lcqtof\_hilic\_neg\_4\_003\_16

m/z 366.1396

0246E+35.56RT (min)

lcqtof\_hilic\_neg\_4\_014\_46

m/z 367.2261

01E+318.519RT (min)

lcqtof\_hilic\_neg\_4\_084\_59

m/z 368.1057

0246E+333.5RT (min)

lcqtof\_hilic\_neg\_1\_049\_55

m/z 369.2186

01E+40.51RT (min)

lcqtof\_hilic\_neg\_4\_005\_29

m/z 371.2437

012E+31818.5RT (min)

lcqtof\_hilic\_neg\_4\_001\_49

m/z 375.2385

012E+311.5RT (min)

lcqtof\_hilic\_neg\_4\_005\_29

m/z 377.2363

01E+318.519RT (min)

lcqtof\_hilic\_neg\_4\_066\_57

m/z 378.9173

012E+39.510RT (min)

lcqtof\_hilic\_neg\_4\_093\_60

m/z 379.1558

01234E+31616.5RT (min)

lcqtof\_hilic\_neg\_4\_050\_02

m/z 379.6001

0123E+32.53RT (min)

lcqtof\_hilic\_neg\_4\_052\_13

m/z 382.1309

01234E+317.518RT (min)

lcqtof\_hilic\_neg\_4\_111\_62

m/z 383.1211

01E+41.52RT (min)

lcqtof\_hilic\_neg\_4\_050\_02

m/z 383.1413

012E+31717.5RT (min)

lcqtof\_hilic\_neg\_4\_099\_61

m/z 384.0125

0123E+43.54RT (min)

lcqtof\_hilic\_neg\_4\_050\_02

m/z 384.0909

024E+31.52RT (min)

lcqtof\_hilic\_neg\_4\_005\_29

m/z 384.1492

01E+318.519RT (min)

lcqtof\_hilic\_neg\_4\_085\_59

m/z 384.1504

0123E+31818.5RT (min)

lcqtof\_hilic\_neg\_4\_017\_51

m/z 384.9836

02468E+37.58RT (min)

lcqtof\_hilic\_neg\_4\_016\_37

m/z 388.1720

02468E+33.54RT (min)

lcqtof\_hilic\_neg\_4\_048\_23

m/z 389.1962

01E+41818.5RT (min)

lcqtof\_hilic\_neg\_4\_099\_61

m/z 390.1998

0123E+31818.5RT (min)

lcqtof\_hilic\_neg\_4\_099\_61

m/z 391.2258

012E+311.5RT (min)

lcqtof\_hilic\_neg\_4\_069\_57

m/z 392.1355

012E+31717.5RT (min)

lcqtof\_hilic\_neg\_4\_101\_61

m/z 393.1708

0123E+31616.5RT (min)

lcqtof\_hilic\_neg\_4\_069\_57

m/z 397.1301

01E+30.51RT (min)

lcqtof\_hilic\_neg\_4\_023\_15

m/z 397.1527

012E+317.518RT (min)

lcqtof\_hilic\_neg\_4\_093\_60

m/z 397.1536

0123E+30.51RT (min)

lcqtof\_hilic\_neg\_4\_016\_37

m/z 397.1555

012E+31616.5RT (min)

lcqtof\_hilic\_neg\_1\_054\_01

m/z 397.3318

012E+316.517RT (min)

lcqtof\_hilic\_neg\_4\_077\_58

m/z 397.3320

012E+318.519RT (min)

lcqtof\_hilic\_neg\_4\_084\_59

m/z 402.1336

01E+455.5RT (min)

lcqtof\_hilic\_neg\_4\_050\_02

m/z 403.1862

0246E+244.5RT (min)

lcqtof\_hilic\_neg\_4\_099\_61

m/z 411.1641

0123E+317.518RT (min)

lcqtof\_hilic\_neg\_4\_106\_62

m/z 411.3471

012E+316.517RT (min)

lcqtof\_hilic\_neg\_4\_024\_40

m/z 412.0234

0123E+311.5RT (min)

lcqtof\_hilic\_neg\_4\_037\_39

m/z 417.2110

01234E+31818.5RT (min)

lcqtof\_hilic\_neg\_4\_112\_62

m/z 424.1772

0246E+31819RT (min)

lcqtof\_hilic\_neg\_4\_083\_59

m/z 425.1105

012E+31919.5RT (min)

lcqtof\_hilic\_neg\_3\_109\_62

m/z 425.3631

012E+316.517RT (min)

lcqtof\_hilic\_neg\_4\_082\_59

m/z 431.2259

01E+31818.5RT (min)

lcqtof\_hilic\_neg\_4\_079\_58

m/z 432.1445

024E+344.5RT (min)

lcqtof\_hilic\_neg\_4\_048\_23

m/z 433.2804

0123E+32.53RT (min)

lcqtof\_hilic\_neg\_4\_032\_01

m/z 434.1412

024E+344.5RT (min)

lcqtof\_hilic\_neg\_4\_008\_24

m/z 435.0053

0246E+30.51RT (min)

lcqtof\_hilic\_neg\_4\_097\_60

m/z 436.1700

0246E+333.5RT (min)

lcqtof\_hilic\_neg\_4\_013\_20

m/z 439.3239

0246E+30.51RT (min)

lcqtof\_hilic\_neg\_4\_048\_23

m/z 442.3379

01234E+411.5RT (min)

lcqtof\_hilic\_neg\_3\_003\_20

m/z 444.0905

0246E+344.5RT (min)

lcqtof\_hilic\_neg\_4\_048\_23

m/z 444.1415

0123E+34.55RT (min)

lcqtof\_hilic\_neg\_4\_108\_62

m/z 447.1476

012E+31818.5RT (min)

lcqtof\_hilic\_neg\_4\_113\_62

m/z 449.1281

024E+311.5RT (min)

lcqtof\_hilic\_neg\_3\_098\_61

m/z 451.1087

024E+322.5RT (min)

lcqtof\_hilic\_neg\_4\_023\_15

m/z 456.0684

0246E+35.56RT (min)

lcqtof\_hilic\_neg\_1\_022\_36

m/z 457.3161

012E+31818.5RT (min)

lcqtof\_hilic\_neg\_4\_101\_61

m/z 458.8050

0123E+37.58RT (min)

lcqtof\_hilic\_neg\_4\_002\_25

m/z 459.2938

0246E+318.519RT (min)

lcqtof\_hilic\_neg\_4\_003\_16

m/z 469.1186

012E+255.5RT (min)

lcqtof\_hilic\_neg\_4\_096\_60

m/z 472.2452

0123E+31.52RT (min)

lcqtof\_hilic\_neg\_4\_053\_14

m/z 477.0829

01E+466.5RT (min)

lcqtof\_hilic\_neg\_4\_013\_20

m/z 482.2952

0123E+23.54RT (min)

lcqtof\_hilic\_neg\_4\_039\_18

m/z 485.3308

01E+311.5RT (min)

lcqtof\_hilic\_neg\_1\_054\_01

m/z 491.3357

01E+411.5RT (min)

lcqtof\_hilic\_neg\_4\_032\_01

m/z 492.2041

0246E+333.5RT (min)

lcqtof\_hilic\_neg\_1\_027\_27

m/z 495.0772

01E+46.57RT (min)

lcqtof\_hilic\_neg\_4\_048\_23

m/z 498.1470

0246E+222.5RT (min)

lcqtof\_hilic\_neg\_4\_039\_18

m/z 504.0512

0246E+311.5RT (min)

lcqtof\_hilic\_neg\_4\_027\_19

m/z 507.3163

01234E+31.52RT (min)

lcqtof\_hilic\_neg\_4\_070\_57

m/z 508.0320

0246E+30.51RT (min)

lcqtof\_hilic\_neg\_4\_040\_05

m/z 510.9216

01234E+37.58RT (min)

lcqtof\_hilic\_neg\_4\_018\_27

m/z 521.1629

012E+30.51RT (min)

lcqtof\_hilic\_neg\_4\_077\_58

m/z 528.0705

024E+30.51RT (min)

lcqtof\_hilic\_neg\_4\_046\_41

m/z 529.0033

012E+36.57RT (min)

lcqtof\_hilic\_neg\_4\_040\_05

m/z 529.3600

012E+31515.5RT (min)

lcqtof\_hilic\_neg\_4\_050\_02

m/z 531.3174

01E+311.5RT (min)

lcqtof\_hilic\_neg\_4\_032\_01

m/z 535.2551

01234E+322.5RT (min)

lcqtof\_hilic\_neg\_4\_018\_27

m/z 541.1807

012E+34.55RT (min)

lcqtof\_hilic\_neg\_1\_006\_22

m/z 552.2437

01E+30.51RT (min)

lcqtof\_hilic\_neg\_4\_050\_02

m/z 553.0367

012E+30.51RT (min)

lcqtof\_hilic\_neg\_4\_070\_57

m/z 553.3971

01234E+32.53RT (min)

lcqtof\_hilic\_neg\_4\_016\_37

m/z 555.3190

0246E+211.5RT (min)

lcqtof\_hilic\_neg\_4\_056\_56

m/z 557.3340

012E+31819RT (min)

lcqtof\_hilic\_neg\_4\_079\_58

m/z 557.3356

01E+319RT (min)

lcqtof\_hilic\_neg\_4\_102\_61

m/z 564.1963

0123E+344.5RT (min)

lcqtof\_hilic\_neg\_4\_047\_06

m/z 565.1920

01E+344.5RT (min)

lcqtof\_hilic\_neg\_4\_040\_05

m/z 571.3382

0123E+317.518RT (min)

lcqtof\_hilic\_neg\_1\_009\_50

m/z 571.3480

012E+3181920RT (min)

lcqtof\_hilic\_neg\_4\_111\_62

m/z 573.4262

012E+316.517RT (min)

lcqtof\_hilic\_neg\_4\_104\_61

m/z 584.1314

01234E+399.5RT (min)

lcqtof\_hilic\_neg\_4\_016\_37

m/z 585.3104

02468E+33.54RT (min)

lcqtof\_hilic\_neg\_4\_052\_13

m/z 585.3393

02468E+30.51RT (min)

lcqtof\_hilic\_neg\_1\_040\_20

m/z 585.3584

01E+40.51RT (min)

lcqtof\_hilic\_neg\_2\_042\_27

m/z 585.3586

01234E+31818.519RT (min)

lcqtof\_hilic\_neg\_4\_099\_61

m/z 586.3512

0123E+33.54RT (min)

lcqtof\_hilic\_neg\_4\_069\_57

m/z 588.2349

024E+33.54RT (min)

lcqtof\_hilic\_neg\_1\_035\_23

m/z 592.8068

024E+377.5RT (min)

lcqtof\_hilic\_neg\_4\_047\_06

m/z 598.2091

024E+344.5RT (min)

lcqtof\_hilic\_neg\_3\_045\_18

m/z 599.3803

012E+31819RT (min)

lcqtof\_hilic\_neg\_4\_084\_59

m/z 601.3230

012E+318.519RT (min)

lcqtof\_hilic\_neg\_1\_056\_56

m/z 602.3278

024E+344.5RT (min)

lcqtof\_hilic\_neg\_4\_008\_24

m/z 606.3769

01E+42.53RT (min)

lcqtof\_hilic\_neg\_4\_048\_23

m/z 607.4593

024E+311.5RT (min)

lcqtof\_hilic\_neg\_4\_075\_58

m/z 608.3879

0246E+322.5RT (min)

lcqtof\_hilic\_neg\_4\_032\_01

m/z 612.4944

01E+30.51RT (min)

lcqtof\_hilic\_neg\_1\_035\_23

m/z 613.3951

0123E+31819RT (min)

lcqtof\_hilic\_neg\_4\_001\_49

m/z 614.3595

01234E+355.5RT (min)

lcqtof\_hilic\_neg\_4\_034\_31

m/z 616.3484

024E+333.5RT (min)

lcqtof\_hilic\_neg\_4\_052\_13

m/z 626.3186

024E+33.54RT (min)

lcqtof\_hilic\_neg\_4\_026\_04

m/z 627.4305

012E+30.51RT (min)

lcqtof\_hilic\_neg\_4\_013\_20

m/z 630.0603

024E+37.58RT (min)

lcqtof\_hilic\_neg\_4\_031\_38

m/z 634.1556

02468E+35.56RT (min)

lcqtof\_hilic\_neg\_4\_048\_23

m/z 635.3800

024E+32.53RT (min)

lcqtof\_hilic\_neg\_4\_053\_14

m/z 644.0294

0246E+37.58RT (min)

lcqtof\_hilic\_neg\_4\_028\_36

m/z 648.2783

01E+411.5RT (min)

lcqtof\_hilic\_neg\_4\_006\_30

m/z 652.4231

0123E+322.5RT (min)

lcqtof\_hilic\_neg\_4\_011\_35

m/z 653.4614

0123E+33.54RT (min)

lcqtof\_hilic\_neg\_4\_053\_14

m/z 653.4884

01E+41617RT (min)

lcqtof\_hilic\_neg\_4\_069\_57

m/z 655.4347

024E+333.5RT (min)

lcqtof\_hilic\_neg\_4\_011\_35

m/z 655.4452

012E+344.5RT (min)

lcqtof\_hilic\_neg\_4\_047\_06

m/z 659.3431

012E+31718RT (min)

lcqtof\_hilic\_neg\_4\_044\_08

m/z 664.6753

01E+388.5RT (min)

lcqtof\_hilic\_neg\_4\_017\_51

m/z 665.1199

024E+377.5RT (min)

lcqtof\_hilic\_neg\_4\_016\_37

m/z 665.3538

01234E+333.5RT (min)

lcqtof\_hilic\_neg\_4\_039\_18

m/z 667.5143

012E+31818.5RT (min)

lcqtof\_hilic\_neg\_4\_001\_49

m/z 668.8034

024E+36.57RT (min)

lcqtof\_hilic\_neg\_4\_012\_07

m/z 669.4585

0123E+31819RT (min)

lcqtof\_hilic\_neg\_4\_033\_53

m/z 671.4646

0246E+533.54RT (min)

lcqtof\_hilic\_neg\_4\_034\_31

m/z 676.1653

012E+388.5RT (min)

lcqtof\_hilic\_neg\_4\_047\_06

m/z 683.0576

0246E+377.5RT (min)

lcqtof\_hilic\_neg\_3\_079\_58

m/z 693.9977

0123E+34.55RT (min)

lcqtof\_hilic\_neg\_4\_032\_01

m/z 695.5457

0246E+41616.5RT (min)

lcqtof\_hilic\_neg\_4\_047\_06

m/z 695.5461

01234E+31920RT (min)

lcqtof\_hilic\_neg\_4\_003\_16

m/z 695.5756

024E+333.5RT (min)

lcqtof\_hilic\_neg\_4\_093\_60

m/z 696.7065

01234E+37.58RT (min)

lcqtof\_hilic\_neg\_4\_001\_49

m/z 699.4837

01E+533.5RT (min)

lcqtof\_hilic\_neg\_4\_084\_59

m/z 702.5072

02468E+42.533.5RT (min)

lcqtof\_hilic\_neg\_1\_051\_35

m/z 706.4128

01E+40.51RT (min)

lcqtof\_hilic\_neg\_4\_085\_59

m/z 712.7595

024E+377.5RT (min)

lcqtof\_hilic\_neg\_4\_012\_07

m/z 713.5205

012E+315.516RT (min)

lcqtof\_hilic\_neg\_4\_020\_42

m/z 720.0194

0123E+366.5RT (min)

lcqtof\_hilic\_neg\_4\_034\_31

m/z 725.0398

01E+26.57RT (min)

lcqtof\_hilic\_neg\_4\_053\_14

m/z 727.5183

0246E+20.51RT (min)

lcqtof\_hilic\_neg\_4\_038\_10

m/z 741.9523

01E+40.51RT (min)

lcqtof\_hilic\_neg\_4\_004\_47

m/z 742.9509

02468E+30.51RT (min)

lcqtof\_hilic\_neg\_4\_004\_47

m/z 748.9981

01E+40.51RT (min)

lcqtof\_hilic\_neg\_4\_077\_58

m/z 753.6105

012E+315.516RT (min)

lcqtof\_hilic\_neg\_4\_099\_61

m/z 757.6355

01E+315.516RT (min)

lcqtof\_hilic\_neg\_4\_109\_62

m/z 759.5517

012E+315.516RT (min)

lcqtof\_hilic\_neg\_1\_005\_32

m/z 760.4819

01234E+322.5RT (min)

lcqtof\_hilic\_neg\_4\_053\_14

m/z 768.6721

01E+315.516RT (min)

lcqtof\_hilic\_neg\_4\_111\_62

m/z 775.0889

02468E+288.5RT (min)

lcqtof\_hilic\_neg\_4\_016\_37

m/z 797.0476

0246E+388.5RT (min)

lcqtof\_hilic\_neg\_4\_083\_59

m/z 799.0314

0123E+355.5RT (min)

lcqtof\_hilic\_neg\_4\_039\_18

m/z 801.5010

0123E+411.5RT (min)

lcqtof\_hilic\_neg\_1\_032\_18

m/z 801.5543

0123E+42.53RT (min)

lcqtof\_hilic\_neg\_4\_022\_34

m/z 801.8471

01234E+36.57RT (min)

lcqtof\_hilic\_neg\_4\_034\_31

m/z 803.4897

01E+315.516RT (min)

lcqtof\_hilic\_neg\_4\_024\_40

m/z 806.5398

02468E+22.53RT (min)

lcqtof\_hilic\_neg\_4\_005\_29

m/z 808.5366

01E+43.54RT (min)

lcqtof\_hilic\_neg\_4\_053\_14

m/z 810.4499

0246E+344.5RT (min)

lcqtof\_hilic\_neg\_4\_086\_59

m/z 812.4656

024E+33.54RT (min)

lcqtof\_hilic\_neg\_4\_087\_59

m/z 816.4175

0246E+333.5RT (min)

lcqtof\_hilic\_neg\_1\_014\_31

m/z 816.9616

012E+37.58RT (min)

lcqtof\_hilic\_neg\_2\_016\_33

m/z 817.1140

0246E+388.5RT (min)

lcqtof\_hilic\_neg\_4\_090\_60

m/z 817.6162

02468E+133.5RT (min)

lcqtof\_hilic\_neg\_3\_086\_59

m/z 825.1464

024E+20.51RT (min)

lcqtof\_hilic\_neg\_4\_077\_58

m/z 834.6148

01E+315.516RT (min)

lcqtof\_hilic\_neg\_4\_005\_29

m/z 848.4465

0246E+311.5RT (min)

lcqtof\_hilic\_neg\_1\_035\_23

m/z 851.2525

01E+34.55RT (min)

lcqtof\_hilic\_neg\_4\_071\_57

m/z 852.5604

0246E+34.55RT (min)

lcqtof\_hilic\_neg\_4\_016\_37

m/z 853.6043

01234E+311.5RT (min)

lcqtof\_hilic\_neg\_4\_016\_37

m/z 856.4492

02468E+333.5RT (min)

lcqtof\_hilic\_neg\_4\_011\_35

m/z 862.5759

01E+43.54RT (min)

lcqtof\_hilic\_neg\_4\_008\_24

m/z 868.5793

01E+411.5RT (min)

lcqtof\_hilic\_neg\_4\_053\_14

m/z 871.5688

01234E+344.5RT (min)

lcqtof\_hilic\_neg\_4\_052\_13

m/z 873.3374

0123E+34.55RT (min)

lcqtof\_hilic\_neg\_4\_053\_14

m/z 874.5769

01E+33.54RT (min)

lcqtof\_hilic\_neg\_4\_008\_24

m/z 882.5549

0246E+211.5RT (min)

lcqtof\_hilic\_neg\_4\_084\_59

m/z 885.4807

01234E+333.5RT (min)

lcqtof\_hilic\_neg\_4\_032\_01

m/z 897.6199

024E+333.5RT (min)

lcqtof\_hilic\_neg\_4\_053\_14

m/z 898.7196

02468E+322.5RT (min)

lcqtof\_hilic\_neg\_4\_004\_47

m/z 906.9295

02468E+30.51RT (min)

lcqtof\_hilic\_neg\_4\_075\_58

m/z 910.5201

012E+244.5RT (min)

lcqtof\_hilic\_neg\_1\_012\_04

m/z 910.7249

024E+322.5RT (min)

lcqtof\_hilic\_neg\_1\_005\_32

m/z 915.5231

0123E+333.5RT (min)

lcqtof\_hilic\_neg\_4\_032\_01

m/z 916.5309

01234E+333.5RT (min)

lcqtof\_hilic\_neg\_4\_032\_01

m/z 934.6238

01E+411.5RT (min)

lcqtof\_hilic\_neg\_4\_052\_13

m/z 934.6662

0123E+311.5RT (min)

lcqtof\_hilic\_neg\_2\_001\_49

m/z 938.6955

0123E+34.55RT (min)

lcqtof\_hilic\_neg\_4\_089\_59

m/z 942.6195

0246E+233.5RT (min)

lcqtof\_hilic\_neg\_4\_008\_24

m/z 945.0058

012E+34.55RT (min)

lcqtof\_hilic\_neg\_4\_023\_15

m/z 946.5140

01E+311.5RT (min)

lcqtof\_hilic\_neg\_4\_048\_23

m/z 951.7499

012E+43.54RT (min)

lcqtof\_hilic\_neg\_4\_008\_24

m/z 957.6155

01234E+322.5RT (min)

lcqtof\_hilic\_neg\_4\_048\_23

m/z 967.3523

0123E+344.5RT (min)

lcqtof\_hilic\_neg\_1\_040\_20

m/z 968.6168

01E+422.5RT (min)

lcqtof\_hilic\_neg\_4\_034\_31

m/z 978.6773

0123E+34.55RT (min)

lcqtof\_hilic\_neg\_4\_029\_17

m/z 994.9173

012E+355.5RT (min)

lcqtof\_hilic\_neg\_4\_026\_04

m/z 1010.5596

0246E+31.52RT (min)

lcqtof\_hilic\_neg\_4\_032\_01

m/z 1012.7558

0123E+34.55RT (min)

lcqtof\_hilic\_neg\_4\_029\_17

m/z 1018.6457

01E+31.52RT (min)

lcqtof\_hilic\_neg\_4\_032\_01

m/z 1021.6699

01E+32.53RT (min)

lcqtof\_hilic\_neg\_4\_053\_14

m/z 1052.7436

0246E+322.5RT (min)

lcqtof\_hilic\_neg\_4\_016\_37

m/z 1072.0308

024E+366.5RT (min)

lcqtof\_hilic\_neg\_4\_034\_31

m/z 1080.0073

012E+34.55RT (min)

lcqtof\_hilic\_neg\_4\_029\_17

m/z 1082.7853

0246E+322.5RT (min)

lcqtof\_hilic\_neg\_4\_016\_37

m/z 1092.9836

01E+37.58RT (min)

lcqtof\_hilic\_neg\_3\_040\_14

m/z 1094.8849

024E+322.5RT (min)

lcqtof\_hilic\_neg\_4\_008\_24

m/z 1122.2173

0123E+30.51RT (min)

lcqtof\_hilic\_neg\_4\_026\_04

m/z 1132.9520

0123E+37.58RT (min)

lcqtof\_hilic\_neg\_3\_029\_29
