## Supplementary material for "The Everything Bagel Feature Finder: Ultra-fast automated feature finding for untargeted metabolomics": SI Data File: yeast_native_unique_xic_EB.html

EB tool-unique native yeast analytes

### EB-unique native yeast analytes (not detected by the other two tools)

123 analytes across 7 datasets · Orbitrap 10 ppm / Q-ToF 25 ppm · ± 0.5 min

Every panel shows a XIC of a reference yeast analyte that EB detected as a reference yeast analyte but XCMS or Masscube did not. For each analyte we show one representative feature, the matched feature with the highest abundance in the yeast 12C 1000 µL samples. If more than one feature is matched, the caption reads "1 of N". The XIC is the MS1 extracted-ion chromatogram at that feature's m/z ± the instrument ppm (Orbitrap 10 / Q-ToF 25, shown per section), drawn from the raw mzML of the yeast 12C 1000 µL sample in which the feature is most abundant, zoomed to ± 0.5 min around the analyte's consensus RT. A flat/empty trace means the analyte has no real signal in the sample group, i.e. it is a gap-fill artifact. **Gray solid line** = the feature's per-file apex in the file (`RT_apex`) when reported, else the consensus RT. **Red dashed line** = the MS/MS EB assigned to the feature (`source_scan`, a 1-based spectrum index, in `source_file`). The y axis is per-cell autoscaled. Panels in m/z order.

#### ST004500 · Orbitrap · 19 analytes · m/z ±10 ppm

m/z 115.0038

ST004500\_NATIVE\_00002 · id 2118 · area 1.35e+05

01E+511.5RT (min)

lcorbi\_t3\_neg\_1\_055\_05

m/z 133.0335

ST004500\_NATIVE\_00006 · id 2319 · area 1.44e+05

01E+511.5RT (min)

lcorbi\_t3\_neg\_1\_055\_05

m/z 134.0175

ST004500\_NATIVE\_00007 · id 3663 · area 6.77e+04

0246E+411.5RT (min)

lcorbi\_t3\_neg\_1\_055\_05

m/z 142.0513

ST004500\_NATIVE\_00008 · id 691 · area 3.11e+05 · 1 of 2

0123E+52.53RT (min)

lcorbi\_t3\_neg\_4\_016\_05

m/z 147.0301

ST004500\_NATIVE\_00009 · id 1695 · area 7.57e+04

0246E+422.5RT (min)

lcorbi\_t3\_neg\_1\_055\_05

m/z 221.0601

ST004500\_NATIVE\_00014 · id 2302 · area 1.53e+05

01E+511.5RT (min)

lcorbi\_t3\_neg\_1\_055\_05

m/z 251.1035

ST004500\_NATIVE\_00019 · id 2223 · area 1.06e+05

01E+566.5RT (min)

lcorbi\_t3\_neg\_1\_055\_05

m/z 305.0639

ST004500\_NATIVE\_00031 · id 3722 · area 4.52e+04

01234E+411.5RT (min)

lcorbi\_t3\_neg\_1\_055\_05

m/z 363.2941

ST004500\_NATIVE\_00037 · id 7652 · area 2.21e+04

012E+410.511RT (min)

lcorbi\_t3\_neg\_1\_055\_05

m/z 596.4902

ST004500\_NATIVE\_00050 · id 4652 · area 6.34e+04

0246E+41414.5RT (min)

lcorbi\_t3\_neg\_1\_055\_05

m/z 610.5054

ST004500\_NATIVE\_00056 · id 6100 · area 4.3e+04

01234E+41414.5RT (min)

lcorbi\_t3\_neg\_1\_055\_05

m/z 612.2529

ST004500\_NATIVE\_00058 · id 5372 · area 6.93e+04 · 1 of 2

0246E+41111.5RT (min)

lcorbi\_t3\_neg\_3\_040\_05

m/z 665.4188

ST004500\_NATIVE\_00068 · id 6722 · area 2.67e+04

012E+41313.5RT (min)

lcorbi\_t3\_neg\_1\_055\_05

m/z 709.4647

ST004500\_NATIVE\_00097 · id 4141 · area 6.1e+04

0246E+413.514RT (min)

lcorbi\_t3\_neg\_1\_055\_05

m/z 768.4824

ST004500\_NATIVE\_00124 · id 1972 · area 1.51e+05

01E+513.514RT (min)

lcorbi\_t3\_neg\_1\_055\_05

m/z 769.4855

ST004500\_NATIVE\_00125 · id 3911 · area 6.67e+04

0246E+413.514RT (min)

lcorbi\_t3\_neg\_1\_055\_05

m/z 773.4996

ST004500\_NATIVE\_00127 · id 2357 · area 7.72e+04

0246E+414.515RT (min)

lcorbi\_t3\_neg\_1\_055\_05

m/z 807.5027

ST004500\_NATIVE\_00135 · id 2112 · area 5.29e+04

024E+413.514RT (min)

lcorbi\_t3\_neg\_1\_055\_05

m/z 820.5346

ST004500\_NATIVE\_00138 · id 5594 · area 2.58e+04

012E+413.514RT (min)

lcorbi\_t3\_neg\_1\_055\_05

#### ST004501 · Orbitrap · 21 analytes · m/z ±10 ppm

m/z 84.0443

ST004501\_NATIVE\_00001 · id 2087 · area 3.49e+04

0123E+433.5RT (min)

lcorbi\_hilic\_pos\_1\_050\_05

m/z 97.0760

ST004501\_NATIVE\_00003 · id 683 · area 8.6e+04

02468E+43.54RT (min)

lcorbi\_hilic\_pos\_1\_050\_05

m/z 124.0586

ST004501\_NATIVE\_00007 · id 814 · area 1.31e+05

01E+511.5RT (min)

lcorbi\_hilic\_pos\_1\_050\_05

m/z 133.0609

ST004501\_NATIVE\_00011 · id 1224 · area 5.12e+04

024E+46.57RT (min)

lcorbi\_hilic\_pos\_1\_050\_05

m/z 148.0798

ST004501\_NATIVE\_00016 · id 1326 · area 4.37e+04

01234E+46.57RT (min)

lcorbi\_hilic\_pos\_2\_030\_05

m/z 158.0812

ST004501\_NATIVE\_00018 · id 1261 · area 5.26e+04

024E+41.52RT (min)

lcorbi\_hilic\_pos\_1\_050\_05

m/z 162.0760

ST004501\_NATIVE\_00021 · id 1041 · area 4.74e+04

01234E+455.5RT (min)

lcorbi\_hilic\_pos\_1\_050\_05

m/z 162.0761

ST004501\_NATIVE\_00022 · id 826 · area 1.06e+05

01E+566.5RT (min)

lcorbi\_hilic\_pos\_2\_030\_05

m/z 181.0584

ST004501\_NATIVE\_00026 · id 1264 · area 6.8e+04

0246E+45.56RT (min)

lcorbi\_hilic\_pos\_1\_050\_05

m/z 187.1077

ST004501\_NATIVE\_00028 · id 1211 · area 6.22e+04

0246E+45.56RT (min)

lcorbi\_hilic\_pos\_1\_050\_05

m/z 191.0486

ST004501\_NATIVE\_00029 · id 2055 · area 2.88e+04

012E+41.52RT (min)

lcorbi\_hilic\_pos\_1\_050\_05

m/z 286.3102

ST004501\_NATIVE\_00053 · id 2276 · area 1.78e+04

01E+433.5RT (min)

lcorbi\_hilic\_pos\_1\_050\_05

m/z 299.1000

ST004501\_NATIVE\_00059 · id 887 · area 6.26e+04

0246E+422.5RT (min)

lcorbi\_hilic\_pos\_3\_027\_05

m/z 300.0927

ST004501\_NATIVE\_00060 · id 1762 · area 2.36e+04

012E+422.5RT (min)

lcorbi\_hilic\_pos\_4\_007\_05

m/z 300.2896

ST004501\_NATIVE\_00061 · id 1495 · area 3.73e+04

0123E+433.5RT (min)

lcorbi\_hilic\_pos\_4\_007\_05

m/z 345.3192

ST004501\_NATIVE\_00065 · id 1301 · area 5.28e+04

024E+41.52RT (min)

lcorbi\_hilic\_pos\_1\_050\_05

m/z 392.3459

ST004501\_NATIVE\_00080 · id 696 · area 6.21e+04

0246E+40.51RT (min)

lcorbi\_hilic\_pos\_1\_050\_05

m/z 393.3156

ST004501\_NATIVE\_00081 · id 1606 · area 5.09e+04

024E+40.51RT (min)

lcorbi\_hilic\_pos\_2\_030\_05

m/z 408.3414

ST004501\_NATIVE\_00083 · id 1555 · area 2.73e+04

012E+40.51RT (min)

lcorbi\_hilic\_pos\_4\_007\_05

m/z 429.3191

ST004501\_NATIVE\_00090 · id 2237 · area 3.24e+04

0123E+40.51RT (min)

lcorbi\_hilic\_pos\_1\_050\_05

m/z 441.3006

ST004501\_NATIVE\_00091 · id 1368 · area 3.85e+04

0123E+40.51RT (min)

lcorbi\_hilic\_pos\_2\_030\_05

#### ST004502 · Orbitrap · 10 analytes · m/z ±10 ppm

m/z 109.0408

ST004502\_NATIVE\_00003 · id 1460 · area 1.75e+05

01E+555.5RT (min)

lcorbi\_hilic\_neg\_1\_047\_05

m/z 129.0193

ST004502\_NATIVE\_00005 · id 772 · area 1.53e+05

01E+51.52RT (min)

lcorbi\_hilic\_neg\_1\_047\_05

m/z 130.0392

ST004502\_NATIVE\_00006 · id 2729 · area 4.84e+04

01234E+433.5RT (min)

lcorbi\_hilic\_neg\_1\_047\_05

m/z 145.0506

ST004502\_NATIVE\_00008 · id 1657 · area 1.26e+05

01E+511.5RT (min)

lcorbi\_hilic\_neg\_1\_047\_05

m/z 189.0598

ST004502\_NATIVE\_00015 · id 1400 · area 1.7e+05

01E+533.5RT (min)

lcorbi\_hilic\_neg\_1\_047\_05

m/z 243.0623

ST004502\_NATIVE\_00019 · id 1218 · area 1.66e+05

01E+54.55RT (min)

lcorbi\_hilic\_neg\_1\_047\_05

m/z 269.0779

ST004502\_NATIVE\_00024 · id 1268 · area 1.35e+05 · 1 of 2

01E+511.5RT (min)

lcorbi\_hilic\_neg\_1\_047\_05

m/z 271.0813

ST004502\_NATIVE\_00025 · id 1593 · area 9.03e+04

02468E+477.5RT (min)

lcorbi\_hilic\_neg\_1\_047\_05

m/z 287.0520

ST004502\_NATIVE\_00028 · id 1341 · area 1.77e+05

01E+55.56RT (min)

lcorbi\_hilic\_neg\_1\_047\_05

m/z 341.1088

ST004502\_NATIVE\_00037 · id 1313 · area 1.32e+05

01E+577.5RT (min)

lcorbi\_hilic\_neg\_1\_047\_05

#### ST004503 · Q-ToF · 12 analytes · m/z ±25 ppm

m/z 130.0529

ST004503\_NATIVE\_00007 · id 14423 · area 5.67e+03

024E+344.5RT (min)

lcqtof\_t3\_pos\_4\_003\_05

m/z 143.0826

ST004503\_NATIVE\_00011 · id 11496 · area 5.38e+03

024E+311.5RT (min)

lcqtof\_t3\_pos\_2\_023\_05

m/z 258.1794

ST004503\_NATIVE\_00031 · id 11888 · area 5.36e+03

024E+344.5RT (min)

lcqtof\_t3\_pos\_4\_003\_05

m/z 261.0926

ST004503\_NATIVE\_00036 · id 7583 · area 2.98e+04

012E+444.5RT (min)

lcqtof\_t3\_pos\_4\_003\_05

m/z 291.1438

ST004503\_NATIVE\_00043 · id 11859 · area 5.98e+03

0246E+311.5RT (min)

lcqtof\_t3\_pos\_4\_003\_05

m/z 301.1717

ST004503\_NATIVE\_00048 · id 3809 · area 5.52e+04

024E+411.5RT (min)

lcqtof\_t3\_pos\_4\_003\_05

m/z 428.2803

ST004503\_NATIVE\_00087 · id 11213 · area 6.35e+03

0246E+344.5RT (min)

lcqtof\_t3\_pos\_4\_003\_05

m/z 438.1504

ST004503\_NATIVE\_00089 · id 12308 · area 5.36e+03

024E+311.5RT (min)

lcqtof\_t3\_pos\_4\_003\_05

m/z 590.5233

ST004503\_NATIVE\_00124 · id 6688 · area 1.8e+04

01E+413.514RT (min)

lcqtof\_t3\_pos\_4\_003\_05

m/z 598.2980

ST004503\_NATIVE\_00130 · id 11755 · area 8.57e+03

02468E+36.57RT (min)

lcqtof\_t3\_pos\_4\_003\_05

m/z 900.7124

ST004503\_NATIVE\_00244 · id 1822 · area 1.34e+05

01E+514.515RT (min)

lcqtof\_t3\_pos\_4\_003\_05

m/z 990.8117

ST004503\_NATIVE\_00262 · id 10291 · area 8.2e+03

02468E+314.515RT (min)

lcqtof\_t3\_pos\_1\_004\_05

#### ST004626 · Q-ToF · 7 analytes · m/z ±25 ppm

m/z 122.0604

ST004626\_NATIVE\_00001 · id 5944 · area 3.04e+03

0123E+344.5RT (min)

lcqtof\_t3\_neg\_3\_037\_05

m/z 155.0642

ST004626\_NATIVE\_00005 · id 5412 · area 6.56e+03

0246E+311.5RT (min)

lcqtof\_t3\_neg\_1\_006\_05

m/z 270.0801

ST004626\_NATIVE\_00016 · id 5907 · area 5.68e+03

024E+344.5RT (min)

lcqtof\_t3\_neg\_3\_037\_05

m/z 272.0883

ST004626\_NATIVE\_00017 · id 6189 · area 4.97e+03

01234E+311.5RT (min)

lcqtof\_t3\_neg\_4\_019\_05

m/z 472.1870

ST004626\_NATIVE\_00023 · id 1246 · area 2.58e+04

012E+411.5RT (min)

lcqtof\_t3\_neg\_1\_006\_05

m/z 614.1431

ST004626\_NATIVE\_00036 · id 6176 · area 2.86e+03

012E+311.5RT (min)

lcqtof\_t3\_neg\_4\_019\_05

m/z 808.1193

ST004626\_NATIVE\_00066 · id 4556 · area 2.43e+03

012E+344.5RT (min)

lcqtof\_t3\_neg\_1\_006\_05

#### ST004650 · Q-ToF · 33 analytes · m/z ±25 ppm

m/z 116.0650

ST004650\_NATIVE\_00004 · id 3184 · area 6.92e+04

0246E+444.5RT (min)

lcqtof\_hilic\_pos\_4\_027\_05

m/z 168.1144

ST004650\_NATIVE\_00011 · id 14994 · area 5.92e+03

024E+35.56RT (min)

lcqtof\_hilic\_pos\_4\_027\_05

m/z 195.0057

ST004650\_NATIVE\_00017 · id 19291 · area 3.57e+03

01234E+36.57RT (min)

lcqtof\_hilic\_pos\_4\_027\_05

m/z 217.1309

ST004650\_NATIVE\_00021 · id 14005 · area 4.89e+03

01234E+35.56RT (min)

lcqtof\_hilic\_pos\_4\_027\_05

m/z 222.0139

ST004650\_NATIVE\_00022 · id 18599 · area 2.84e+03

012E+355.5RT (min)

lcqtof\_hilic\_pos\_4\_027\_05

m/z 229.1557

ST004650\_NATIVE\_00025 · id 12925 · area 7.52e+03

0246E+34.55RT (min)

lcqtof\_hilic\_pos\_4\_027\_05

m/z 230.0796

ST004650\_NATIVE\_00026 · id 13726 · area 5.2e+03

024E+366.5RT (min)

lcqtof\_hilic\_pos\_1\_027\_05

m/z 244.0796

ST004650\_NATIVE\_00031 · id 22155 · area 1.7e+03

01E+35.56RT (min)

lcqtof\_hilic\_pos\_2\_048\_05

m/z 260.0537

ST004650\_NATIVE\_00036 · id 6701 · area 3.78e+03

0123E+34.55RT (min)

lcqtof\_hilic\_pos\_1\_027\_05

m/z 269.1632

ST004650\_NATIVE\_00039 · id 5490 · area 2.62e+04

012E+46.57RT (min)

lcqtof\_hilic\_pos\_4\_027\_05

m/z 307.5857

ST004650\_NATIVE\_00051 · id 15029 · area 4.72e+03

01234E+388.5RT (min)

lcqtof\_hilic\_pos\_2\_048\_05

m/z 332.3329

ST004650\_NATIVE\_00060 · id 5351 · area 4.16e+04

01234E+40.51RT (min)

lcqtof\_hilic\_pos\_4\_027\_05

m/z 334.2320

ST004650\_NATIVE\_00061 · id 13787 · area 2.89e+03

02468E+20.51RT (min)

lcqtof\_hilic\_pos\_1\_027\_05

m/z 345.3167

ST004650\_NATIVE\_00063 · id 3256 · area 7.31e+04

0246E+41.52RT (min)

lcqtof\_hilic\_pos\_1\_027\_05

m/z 397.3378

ST004650\_NATIVE\_00078 · id 12510 · area 1.04e+04

01E+40.51RT (min)

lcqtof\_hilic\_pos\_1\_027\_05

m/z 416.3749

ST004650\_NATIVE\_00084 · id 14070 · area 5.83e+03

02468E+311.5RT (min)

lcqtof\_hilic\_pos\_4\_027\_05

m/z 515.4131

ST004650\_NATIVE\_00118 · id 5180 · area 2.77e+04

012E+40.51RT (min)

lcqtof\_hilic\_pos\_4\_027\_05

m/z 518.4940

ST004650\_NATIVE\_00120 · id 2965 · area 5.77e+04

024E+40.51RT (min)

lcqtof\_hilic\_pos\_4\_027\_05

m/z 523.5217

ST004650\_NATIVE\_00123 · id 11752 · area 6.85e+03 · 1 of 2

0246E+30.51RT (min)

lcqtof\_hilic\_pos\_4\_027\_05

m/z 525.3147

ST004650\_NATIVE\_00125 · id 17129 · area 5.04e+03

024E+36.57RT (min)

lcqtof\_hilic\_pos\_4\_027\_05

m/z 538.1296

ST004650\_NATIVE\_00133 · id 13478 · area 5.5e+03

024E+388.5RT (min)

lcqtof\_hilic\_pos\_2\_048\_05

m/z 551.3895

ST004650\_NATIVE\_00141 · id 16607 · area 5.06e+03

0246E+311.5RT (min)

lcqtof\_hilic\_pos\_4\_027\_05

m/z 561.4310

ST004650\_NATIVE\_00147 · id 18459 · area 2.78e+03

012E+344.5RT (min)

lcqtof\_hilic\_pos\_1\_027\_05

m/z 573.4693

ST004650\_NATIVE\_00155 · id 2673 · area 4.69e+04

01234E+40.51RT (min)

lcqtof\_hilic\_pos\_4\_027\_05

m/z 613.4388

ST004650\_NATIVE\_00167 · id 23216 · area 974

02468E+23.54RT (min)

lcqtof\_hilic\_pos\_2\_048\_05

m/z 622.3170

ST004650\_NATIVE\_00173 · id 21271 · area 2.89e+03

012E+34.55RT (min)

lcqtof\_hilic\_pos\_1\_027\_05

m/z 645.1337

ST004650\_NATIVE\_00178 · id 11524 · area 6.27e+03

0246E+388.5RT (min)

lcqtof\_hilic\_pos\_2\_048\_05

m/z 667.1133

ST004650\_NATIVE\_00182 · id 13467 · area 3.55e+03

0123E+388.5RT (min)

lcqtof\_hilic\_pos\_2\_048\_05

m/z 671.1415

ST004650\_NATIVE\_00184 · id 21606 · area 1.75e+03

01E+377.5RT (min)

lcqtof\_hilic\_pos\_4\_027\_05

m/z 774.5941

ST004650\_NATIVE\_00217 · id 7857 · area 1.13e+04

01E+422.5RT (min)

lcqtof\_hilic\_pos\_4\_027\_05

m/z 832.5932

ST004650\_NATIVE\_00237 · id 18475 · area 1.96e+03

01E+34.55RT (min)

lcqtof\_hilic\_pos\_4\_027\_05

m/z 1012.7195

ST004650\_NATIVE\_00250 · id 19733 · area 3.43e+03

0123E+30.51RT (min)

lcqtof\_hilic\_pos\_4\_027\_05

m/z 1183.7148

ST004650\_NATIVE\_00251 · id 22297 · area 2.26e+03

012E+33.54RT (min)

lcqtof\_hilic\_pos\_4\_027\_05

#### ST004651 · Q-ToF · 21 analytes · m/z ±25 ppm

m/z 142.0504

ST004651\_NATIVE\_00001 · id 11535 · area 5.4e+03

024E+366.5RT (min)

lcqtof\_hilic\_neg\_2\_028\_05

m/z 280.0757

ST004651\_NATIVE\_00020 · id 14565 · area 3.85e+03

0123E+35.56RT (min)

lcqtof\_hilic\_neg\_4\_040\_05

m/z 281.0431

ST004651\_NATIVE\_00021 · id 16856 · area 2.12e+03

012E+34.55RT (min)

lcqtof\_hilic\_neg\_1\_007\_05

m/z 356.3251

ST004651\_NATIVE\_00032 · id 9815 · area 7.68e+03

0246E+30.51RT (min)

lcqtof\_hilic\_neg\_2\_028\_05

m/z 381.1407

ST004651\_NATIVE\_00034 · id 6801 · area 1.73e+04

01E+466.5RT (min)

lcqtof\_hilic\_neg\_4\_040\_05

m/z 389.3096

ST004651\_NATIVE\_00036 · id 15298 · area 3.23e+03

0123E+311.5RT (min)

lcqtof\_hilic\_neg\_1\_007\_05

m/z 566.5493

ST004651\_NATIVE\_00060 · id 14481 · area 6.3e+03

0246E+30.51RT (min)

lcqtof\_hilic\_neg\_4\_040\_05

m/z 646.3924

ST004651\_NATIVE\_00077 · id 5895 · area 2.63e+04

012E+40.51RT (min)

lcqtof\_hilic\_neg\_4\_040\_05

m/z 669.1215

ST004651\_NATIVE\_00082 · id 19003 · area 1.47e+03

01E+377.5RT (min)

lcqtof\_hilic\_neg\_3\_048\_05

m/z 699.5990

ST004651\_NATIVE\_00090 · id 15189 · area 7.01e+03

0246E+30.51RT (min)

lcqtof\_hilic\_neg\_2\_028\_05

m/z 740.4332

ST004651\_NATIVE\_00095 · id 17090 · area 3.91e+03

0123E+344.5RT (min)

lcqtof\_hilic\_neg\_4\_040\_05

m/z 819.7090

ST004651\_NATIVE\_00104 · id 17410 · area 3.72e+03

0123E+30.51RT (min)

lcqtof\_hilic\_neg\_4\_040\_05

m/z 887.6078

ST004651\_NATIVE\_00110 · id 16326 · area 4.69e+03

01234E+34.55RT (min)

lcqtof\_hilic\_neg\_4\_040\_05

m/z 968.5723

ST004651\_NATIVE\_00125 · id 7655 · area 2.34e+04

012E+44.55RT (min)

lcqtof\_hilic\_neg\_4\_040\_05

m/z 972.5699

ST004651\_NATIVE\_00126 · id 20006 · area 4.13e+03 · 1 of 2

0246E+34.55RT (min)

lcqtof\_hilic\_neg\_4\_040\_05

m/z 1006.8380

ST004651\_NATIVE\_00130 · id 7490 · area 2.53e+04

012E+40.51RT (min)

lcqtof\_hilic\_neg\_4\_040\_05

m/z 1007.7317

ST004651\_NATIVE\_00131 · id 15127 · area 5.21e+03 · 1 of 2

024E+311.5RT (min)

lcqtof\_hilic\_neg\_2\_028\_05

m/z 1010.5503

ST004651\_NATIVE\_00132 · id 16557 · area 5.2e+03

024E+34.55RT (min)

lcqtof\_hilic\_neg\_4\_040\_05

m/z 1056.5280

ST004651\_NATIVE\_00133 · id 15283 · area 6.89e+03 · 1 of 2

0246E+34.55RT (min)

lcqtof\_hilic\_neg\_4\_040\_05

m/z 1069.7573

ST004651\_NATIVE\_00134 · id 19666 · area 2.5e+03

012E+355.5RT (min)

lcqtof\_hilic\_neg\_4\_040\_05

m/z 1173.0477

ST004651\_NATIVE\_00138 · id 15821 · area 2.74e+03

012E+30.51RT (min)

lcqtof\_hilic\_neg\_4\_040\_05
