## Supplementary material for "The Everything Bagel Feature Finder: Ultra-fast automated feature finding for untargeted metabolomics": SI Data File: yeast_native_unique_xic_Masscube.html

Masscube tool-unique native yeast analytes

### Masscube-unique native yeast analytes (not detected by the other two tools)

78 analytes across 4 datasets · Orbitrap 10 ppm / Q-ToF 25 ppm · ± 0.5 min

Every panel shows a XIC of a reference yeast analyte that Masscube detected as a reference yeast analyte but EB or XCMS did not. For each analyte we show one representative feature, the matched feature with the highest abundance in the yeast 12C 1000 µL samples. If more than one feature is matched, the caption reads "1 of N". The XIC is the MS1 extracted-ion chromatogram at that feature's m/z ± the instrument ppm (Orbitrap 10 / Q-ToF 25, shown per section), drawn from the raw mzML of the yeast 12C 1000 µL sample in which the feature is most abundant, zoomed to ± 0.5 min around the analyte's consensus RT. A flat/empty trace means the analyte has no real signal in the sample group, i.e. it is a gap-fill artifact. **Gray solid line** = the reported consensus RT. Masscube carries no MS/MS in the benchmark aligned table, so panels show only the apex line. The y axis is per-cell autoscaled. Panels in m/z order.

#### ST004503 · Q-ToF · 42 analytes · m/z ±25 ppm

m/z 184.0739

ST004503\_NATIVE\_00020 · id 14843 · area 2.69e+04 · 1 of 4

012E+412.513RT (min)

lcqtof\_t3\_pos\_4\_003\_05

m/z 266.1248

ST004503\_NATIVE\_00037 · id 33147 · area 2.16e+04

012E+311.5RT (min)

lcqtof\_t3\_pos\_1\_004\_05

m/z 284.2960

ST004503\_NATIVE\_00041 · id 9434 · area 2.3e+04 · 1 of 4

012E+41010.5RT (min)

lcqtof\_t3\_pos\_2\_023\_05

m/z 300.2907

ST004503\_NATIVE\_00047 · id 17172 · area 1.6e+04 · 1 of 5

012E+41010.5RT (min)

lcqtof\_t3\_pos\_2\_023\_05

m/z 302.3061

ST004503\_NATIVE\_00049 · id 30092 · area 1.84e+04 · 1 of 5

012E+41010.5RT (min)

lcqtof\_t3\_pos\_2\_023\_05

m/z 320.0792

ST004503\_NATIVE\_00057 · id 5480 · area 6.78e+04

0246E+44.55RT (min)

lcqtof\_t3\_pos\_4\_003\_05

m/z 344.3168

ST004503\_NATIVE\_00067 · id 16994 · area 6.14e+04 · 1 of 2

0246E+41010.5RT (min)

lcqtof\_t3\_pos\_2\_023\_05

m/z 347.3160

ST004503\_NATIVE\_00070 · id 11588 · area 6.15e+04

0246E+411.512RT (min)

lcqtof\_t3\_pos\_4\_003\_05

m/z 350.3265

ST004503\_NATIVE\_00071 · id 23826 · area 3.87e+04 · 1 of 2

024E+410.511RT (min)

lcqtof\_t3\_pos\_4\_003\_05

m/z 355.2813

ST004503\_NATIVE\_00073 · id 19467 · area 1.88e+04 · 1 of 6

012E+41111.5RT (min)

lcqtof\_t3\_pos\_4\_003\_05

m/z 355.3027

ST004503\_NATIVE\_00074 · id 24316 · area 2.71e+04 · 1 of 3

012E+41111.5RT (min)

lcqtof\_t3\_pos\_4\_003\_05

m/z 364.3426

ST004503\_NATIVE\_00075 · id 4610 · area 2.41e+04

012E+41111.5RT (min)

lcqtof\_t3\_pos\_1\_004\_05

m/z 378.3302

ST004503\_NATIVE\_00077 · id 5873 · area 5.83e+04 · 1 of 3

024E+410.511RT (min)

lcqtof\_t3\_pos\_4\_003\_05

m/z 380.3378

ST004503\_NATIVE\_00078 · id 17589 · area 3.61e+04

0123E+410.511RT (min)

lcqtof\_t3\_pos\_1\_004\_05

m/z 381.3522

ST004503\_NATIVE\_00080 · id 14391 · area 2.5e+04 · 1 of 2

012E+41515.5RT (min)

lcqtof\_t3\_pos\_4\_003\_05

m/z 395.3335

ST004503\_NATIVE\_00082 · id 27500 · area 2.23e+04 · 1 of 5

012E+41212.5RT (min)

lcqtof\_t3\_pos\_4\_003\_05

m/z 397.3479

ST004503\_NATIVE\_00083 · id 22576 · area 2.69e+04

012E+41313.5RT (min)

lcqtof\_t3\_pos\_2\_023\_05

m/z 512.4161

ST004503\_NATIVE\_00097 · id 53435 · area 2.04e+04 · 1 of 2

0123E+49.510RT (min)

lcqtof\_t3\_pos\_4\_003\_05

m/z 547.4794

ST004503\_NATIVE\_00099 · id 23173 · area 1.11e+04 · 1 of 2

01E+41212.5RT (min)

lcqtof\_t3\_pos\_4\_003\_05

m/z 573.4868

ST004503\_NATIVE\_00106 · id 14007 · area 5.08e+04 · 1 of 6

01E+512.513RT (min)

lcqtof\_t3\_pos\_4\_003\_05

m/z 574.4912

ST004503\_NATIVE\_00108 · id 16717 · area 1.75e+04

01E+414.515RT (min)

lcqtof\_t3\_pos\_4\_003\_05

m/z 574.4920

ST004503\_NATIVE\_00109 · id 43357 · area 3.64e+04 · 1 of 2

0123E+413.514RT (min)

lcqtof\_t3\_pos\_4\_003\_05

m/z 576.5064

ST004503\_NATIVE\_00112 · id 3572 · area 1.32e+05

01E+513.514RT (min)

lcqtof\_t3\_pos\_3\_037\_05

m/z 589.4829

ST004503\_NATIVE\_00121 · id 31747 · area 1.38e+04 · 1 of 3

01E+41313.5RT (min)

lcqtof\_t3\_pos\_4\_003\_05

m/z 591.5048

ST004503\_NATIVE\_00125 · id 47960 · area 3.04e+04 · 1 of 3

0123E+413.514RT (min)

lcqtof\_t3\_pos\_4\_003\_05

m/z 597.4872

ST004503\_NATIVE\_00128 · id 15307 · area 1.61e+04 · 1 of 2

01234E+41414.5RT (min)

lcqtof\_t3\_pos\_4\_003\_05

m/z 598.4908

ST004503\_NATIVE\_00131 · id 16907 · area 2.31e+04

012E+413.514RT (min)

lcqtof\_t3\_pos\_3\_037\_05

m/z 631.5126

ST004503\_NATIVE\_00146 · id 21548 · area 3.04e+04

0123E+413.514RT (min)

lcqtof\_t3\_pos\_3\_037\_05

m/z 674.5727

ST004503\_NATIVE\_00158 · id 32038 · area 1.14e+04

012E+413.514RT (min)

lcqtof\_t3\_pos\_2\_023\_05

m/z 702.5350

ST004503\_NATIVE\_00166 · id 2235 · area 5.61e+04 · 1 of 2

024E+41010.5RT (min)

lcqtof\_t3\_pos\_4\_003\_05

m/z 727.5108

ST004503\_NATIVE\_00169 · id 18930 · area 2.41e+04

012E+41212.5RT (min)

lcqtof\_t3\_pos\_4\_003\_05

m/z 730.5270

ST004503\_NATIVE\_00170 · id 24143 · area 1.12e+05 · 1 of 2

0123E+51313.5RT (min)

lcqtof\_t3\_pos\_2\_023\_05

m/z 738.5078

ST004503\_NATIVE\_00173 · id 30004 · area 2e+04 · 1 of 3

01234E+51212.5RT (min)

lcqtof\_t3\_pos\_4\_003\_05

m/z 765.5259

ST004503\_NATIVE\_00181 · id 11188 · area 2.09e+05 · 1 of 3

012E+512.513RT (min)

lcqtof\_t3\_pos\_2\_023\_05

m/z 794.5326

ST004503\_NATIVE\_00192 · id 20191 · area 1.96e+04 · 1 of 5

01E+411.512RT (min)

lcqtof\_t3\_pos\_4\_003\_05

m/z 801.6166

ST004503\_NATIVE\_00194 · id 13969 · area 1.11e+04

01E+413.514RT (min)

lcqtof\_t3\_pos\_4\_003\_05

m/z 840.7091

ST004503\_NATIVE\_00195 · id 11572 · area 2.19e+04

012E+414.515RT (min)

lcqtof\_t3\_pos\_2\_023\_05

m/z 883.6754

ST004503\_NATIVE\_00223 · id 31643 · area 9.92e+03

02468E+314.515RT (min)

lcqtof\_t3\_pos\_2\_023\_05

m/z 898.9272

ST004503\_NATIVE\_00242 · id 6441 · area 4.88e+04

01234E+414.515RT (min)

lcqtof\_t3\_pos\_4\_003\_05

m/z 901.9465

ST004503\_NATIVE\_00247 · id 4545 · area 8.26e+04

02468E+41515.5RT (min)

lcqtof\_t3\_pos\_4\_003\_05

m/z 913.7668

ST004503\_NATIVE\_00250 · id 14066 · area 1.39e+04

01E+41414.5RT (min)

lcqtof\_t3\_pos\_4\_003\_05

m/z 915.7813

ST004503\_NATIVE\_00253 · id 9368 · area 1.99e+04

01E+41414.5RT (min)

lcqtof\_t3\_pos\_3\_037\_05

#### ST004626 · Q-ToF · 1 analyte · m/z ±25 ppm

m/z 795.6118

ST004626\_NATIVE\_00065 · id 3537 · area 8.37e+04

02468E+412.513RT (min)

lcqtof\_t3\_neg\_4\_019\_05

#### ST004650 · Q-ToF · 28 analytes · m/z ±25 ppm

m/z 142.0983

ST004650\_NATIVE\_00007 · id 3686 · area 1.32e+04 · 1 of 6

01E+577.5RT (min)

lcqtof\_hilic\_pos\_3\_047\_05

m/z 364.3071

ST004650\_NATIVE\_00068 · id 22468 · area 3.36e+05 · 1 of 2

0123E+50.51RT (min)

lcqtof\_hilic\_pos\_4\_027\_05

m/z 394.3501

ST004650\_NATIVE\_00077 · id 15061 · area 9.65e+04

01E+60.51RT (min)

lcqtof\_hilic\_pos\_4\_027\_05

m/z 403.3030

ST004650\_NATIVE\_00081 · id 12845 · area 5.06e+04

024E+40.51RT (min)

lcqtof\_hilic\_pos\_4\_027\_05

m/z 428.3796

ST004650\_NATIVE\_00090 · id 16975 · area 9.61e+03

02468E+30.51RT (min)

lcqtof\_hilic\_pos\_2\_048\_05

m/z 510.3873

ST004650\_NATIVE\_00114 · id 13163 · area 2.96e+04

012E+444.5RT (min)

lcqtof\_hilic\_pos\_4\_027\_05

m/z 539.4169

ST004650\_NATIVE\_00134 · id 24205 · area 1.48e+04

01E+40.51RT (min)

lcqtof\_hilic\_pos\_4\_027\_05

m/z 558.4420

ST004650\_NATIVE\_00145 · id 15096 · area 4.04e+04

01234E+40.51RT (min)

lcqtof\_hilic\_pos\_4\_027\_05

m/z 575.5038

ST004650\_NATIVE\_00156 · id 4812 · area 7.63e+04 · 1 of 4

02468E+422.5RT (min)

lcqtof\_hilic\_pos\_4\_027\_05

m/z 597.4880

ST004650\_NATIVE\_00163 · id 9468 · area 3.91e+04 · 1 of 5

0123E+422.5RT (min)

lcqtof\_hilic\_pos\_4\_027\_05

m/z 689.5202

ST004650\_NATIVE\_00188 · id 13397 · area 4.66e+04 · 1 of 3

01234E+40.51RT (min)

lcqtof\_hilic\_pos\_4\_027\_05

m/z 718.5363

ST004650\_NATIVE\_00194 · id 13926 · area 3e+04 · 1 of 7

0123E+422.5RT (min)

lcqtof\_hilic\_pos\_4\_027\_05

m/z 740.5190

ST004650\_NATIVE\_00199 · id 13956 · area 2.83e+04 · 1 of 4

01234E+42.53RT (min)

lcqtof\_hilic\_pos\_4\_027\_05

m/z 740.5205

ST004650\_NATIVE\_00200 · id 7732 · area 3.96e+04 · 1 of 5

0123E+422.5RT (min)

lcqtof\_hilic\_pos\_4\_027\_05

m/z 743.5411

ST004650\_NATIVE\_00202 · id 5529 · area 7.38e+04 · 1 of 4

0246E+422.5RT (min)

lcqtof\_hilic\_pos\_4\_027\_05

m/z 754.5339

ST004650\_NATIVE\_00206 · id 9747 · area 3.07e+04 · 1 of 5

0123E+422.5RT (min)

lcqtof\_hilic\_pos\_1\_027\_05

m/z 756.5533

ST004650\_NATIVE\_00207 · id 4854 · area 1.7e+05 · 1 of 7

012E+52.53RT (min)

lcqtof\_hilic\_pos\_4\_027\_05

m/z 766.5478

ST004650\_NATIVE\_00211 · id 17299 · area 2.56e+04 · 1 of 6

012E+42.53RT (min)

lcqtof\_hilic\_pos\_4\_027\_05

m/z 778.5351

ST004650\_NATIVE\_00219 · id 5294 · area 1.55e+05 · 1 of 5

012E+522.5RT (min)

lcqtof\_hilic\_pos\_4\_027\_05

m/z 779.5341

ST004650\_NATIVE\_00220 · id 11722 · area 3.74e+04 · 1 of 3

01234E+422.5RT (min)

lcqtof\_hilic\_pos\_1\_027\_05

m/z 780.5524

ST004650\_NATIVE\_00221 · id 4518 · area 1.73e+05 · 1 of 13

01E+522.5RT (min)

lcqtof\_hilic\_pos\_1\_027\_05

m/z 781.2100

ST004650\_NATIVE\_00223 · id 8018 · area 2.01e+04

012E+40.51RT (min)

lcqtof\_hilic\_pos\_4\_027\_05

m/z 781.5563

ST004650\_NATIVE\_00224 · id 4181 · area 5.9e+04 · 1 of 2

02468E+41.52RT (min)

lcqtof\_hilic\_pos\_1\_027\_05

m/z 781.5564

ST004650\_NATIVE\_00225 · id 5939 · area 1.56e+05 · 1 of 3

01E+52.53RT (min)

lcqtof\_hilic\_pos\_4\_027\_05

m/z 782.5682

ST004650\_NATIVE\_00226 · id 1171 · area 1.48e+05 · 1 of 2

01E+522.5RT (min)

lcqtof\_hilic\_pos\_1\_027\_05

m/z 786.3844

ST004650\_NATIVE\_00229 · id 8637 · area 2.18e+04 · 1 of 2

012E+433.5RT (min)

lcqtof\_hilic\_pos\_4\_027\_05

m/z 821.6004

ST004650\_NATIVE\_00235 · id 26898 · area 1.67e+04

01E+40.51RT (min)

lcqtof\_hilic\_pos\_4\_027\_05

m/z 864.6240

ST004650\_NATIVE\_00238 · id 22781 · area 1.85e+04

01E+40.51RT (min)

lcqtof\_hilic\_pos\_4\_027\_05

#### ST004651 · Q-ToF · 7 analytes · m/z ±25 ppm

m/z 346.1438

ST004651\_NATIVE\_00030 · id 3467 · area 5.91e+04

024E+40.51RT (min)

lcqtof\_hilic\_neg\_2\_028\_05

m/z 558.3388

ST004651\_NATIVE\_00057 · id 4404 · area 5.71e+04

024E+40.51RT (min)

lcqtof\_hilic\_neg\_4\_040\_05

m/z 610.5008

ST004651\_NATIVE\_00069 · id 19114 · area 3.46e+04

0123E+40.51RT (min)

lcqtof\_hilic\_neg\_4\_040\_05

m/z 635.2906

ST004651\_NATIVE\_00073 · id 22566 · area 1.53e+04

01E+40.51RT (min)

lcqtof\_hilic\_neg\_4\_040\_05

m/z 674.4839

ST004651\_NATIVE\_00084 · id 5436 · area 2.64e+04

01234E+433.5RT (min)

lcqtof\_hilic\_neg\_4\_040\_05

m/z 822.5318

ST004651\_NATIVE\_00106 · id 12624 · area 2.16e+04 · 1 of 5

0246E+42.53RT (min)

lcqtof\_hilic\_neg\_2\_028\_05

m/z 877.5848

ST004651\_NATIVE\_00108 · id 18832 · area 1.92e+04 · 1 of 2

01E+43.54RT (min)

lcqtof\_hilic\_neg\_4\_040\_05
