## Supplementary material for "The Everything Bagel Feature Finder: Ultra-fast automated feature finding for untargeted metabolomics": SI Data File: yeast_native_unique_xic_XCMS.html

XCMS tool-unique native yeast analytes

### XCMS-unique native yeast analytes (not detected by the other two tools)

190 analytes across 7 datasets · Orbitrap 10 ppm / Q-ToF 25 ppm · ± 0.5 min

Every panel shows a XIC of a reference yeast analyte that XCMS detected as a reference yeast analyte but EB or Masscube did not. For each analyte we show one representative feature, the matched feature with the highest abundance in the yeast 12C 1000 µL samples. If more than one feature is matched, the caption reads "1 of N". The XIC is the MS1 extracted-ion chromatogram at that feature's m/z ± the instrument ppm (Orbitrap 10 / Q-ToF 25, shown per section), drawn from the raw mzML of the yeast 12C 1000 µL sample in which the feature is most abundant, zoomed to ± 0.5 min around the analyte's consensus RT. A flat/empty trace means the analyte has no real signal in the sample group, i.e. it is a gap-fill artifact. **Gray solid line** = the reported consensus RT. XCMS carries no MS/MS in the benchmark aligned table, so panels show only the apex line. The y axis is per-cell autoscaled. Panels in m/z order.

#### ST004500 · Orbitrap · 69 analytes · m/z ±10 ppm

m/z 189.0598

ST004500\_NATIVE\_00012 · id 1139 · area 4.52e+04

02468E+311.5RT (min)

lcorbi\_t3\_neg\_1\_055\_05

m/z 208.9621

ST004500\_NATIVE\_00013 · id 1594 · area 2.62e+05

0246E+411.5RT (min)

lcorbi\_t3\_neg\_1\_055\_05

m/z 228.9865

ST004500\_NATIVE\_00015 · id 2045 · area 2.46e+04

0111.5RT (min)

lcorbi\_t3\_neg\_1\_055\_05

m/z 238.0912

ST004500\_NATIVE\_00016 · id 2224 · area 2.37e+04

0246E+355.5RT (min)

lcorbi\_t3\_neg\_1\_055\_05

m/z 244.0225

ST004500\_NATIVE\_00017 · id 2312 · area 8.23e+05

012E+511.5RT (min)

lcorbi\_t3\_neg\_1\_055\_05

m/z 259.0587

ST004500\_NATIVE\_00022 · id 2588 · area 1.6e+05

01E+311.5RT (min)

lcorbi\_t3\_neg\_1\_055\_05

m/z 267.0718

ST004500\_NATIVE\_00023 · id 2736 · area 6.52e+04

01E+411.5RT (min)

lcorbi\_t3\_neg\_1\_055\_05

m/z 273.0854

ST004500\_NATIVE\_00024 · id 2827 · area 6.44e+04

01E+466.5RT (min)

lcorbi\_t3\_neg\_4\_016\_05

m/z 274.1043

ST004500\_NATIVE\_00025 · id 2847 · area 5.88e+04

01E+411.5RT (min)

lcorbi\_t3\_neg\_1\_055\_05

m/z 275.1245

ST004500\_NATIVE\_00026 · id 2865 · area 2.43e+05

0246E+411.5RT (min)

lcorbi\_t3\_neg\_1\_055\_05

m/z 289.0681

ST004500\_NATIVE\_00028 · id 3120 · area 1.26e+05

01E+311.5RT (min)

lcorbi\_t3\_neg\_1\_055\_05

m/z 289.1232

ST004500\_NATIVE\_00029 · id 3126 · area 8.09e+04

01E+422.5RT (min)

lcorbi\_t3\_neg\_1\_055\_05

m/z 334.9112

ST004500\_NATIVE\_00032 · id 4065 · area 1.11e+05

02468E+211.5RT (min)

lcorbi\_t3\_neg\_1\_055\_05

m/z 343.2850

ST004500\_NATIVE\_00033 · id 4265 · area 4.74e+04

02468E+31111.5RT (min)

lcorbi\_t3\_neg\_1\_055\_05

m/z 357.2138

ST004500\_NATIVE\_00035 · id 4623 · area 1.39e+05

0123E+45.56RT (min)

lcorbi\_t3\_neg\_1\_055\_05

m/z 367.2460

ST004500\_NATIVE\_00038 · id 4826 · area 1.75e+05

0123E+412.513RT (min)

lcorbi\_t3\_neg\_1\_055\_05

m/z 387.0098

ST004500\_NATIVE\_00039 · id 5307 · area 7.54e+04

01E+411.5RT (min)

lcorbi\_t3\_neg\_1\_055\_05

m/z 425.2575

ST004500\_NATIVE\_00040 · id 6288 · area 1.06e+06

01E+51111.5RT (min)

lcorbi\_t3\_neg\_1\_055\_05

m/z 433.1724

ST004500\_NATIVE\_00042 · id 6526 · area 1.3e+04

0123E+355.5RT (min)

lcorbi\_t3\_neg\_1\_055\_05

m/z 439.0852

ST004500\_NATIVE\_00043 · id 6672 · area 1.34e+05

012E+411.5RT (min)

lcorbi\_t3\_neg\_1\_055\_05

m/z 441.3371

ST004500\_NATIVE\_00044 · id 6752 · area 9.9e+04

01E+413.514RT (min)

lcorbi\_t3\_neg\_1\_055\_05

m/z 534.2043

ST004500\_NATIVE\_00045 · id 9302 · area 8.88e+03

01E+31010.5RT (min)

lcorbi\_t3\_neg\_1\_055\_05

m/z 597.4935

ST004500\_NATIVE\_00051 · id 10837 · area 1.02e+05

012E+41414.5RT (min)

lcorbi\_t3\_neg\_1\_055\_05

m/z 609.5297

ST004500\_NATIVE\_00055 · id 11127 · area 7.99e+04

01E+414.515RT (min)

lcorbi\_t3\_neg\_1\_055\_05

m/z 612.1477

ST004500\_NATIVE\_00057 · id 11190 · area 7.69e+05

01E+522.5RT (min)

lcorbi\_t3\_neg\_1\_055\_05

m/z 613.3900

ST004500\_NATIVE\_00060 · id 11236 · area 4.32e+04

01E+414.515RT (min)

lcorbi\_t3\_neg\_1\_055\_05

m/z 620.3009

ST004500\_NATIVE\_00061 · id 11402 · area 4.77e+04

0246E+31010.5RT (min)

lcorbi\_t3\_neg\_1\_055\_05

m/z 640.5523

ST004500\_NATIVE\_00063 · id 11848 · area 1.15e+05

012E+414.515RT (min)

lcorbi\_t3\_neg\_1\_055\_05

m/z 645.4676

ST004500\_NATIVE\_00064 · id 11964 · area 1.91e+04

024E+314.515RT (min)

lcorbi\_t3\_neg\_1\_055\_05

m/z 653.4809

ST004500\_NATIVE\_00065 · id 12150 · area 1.36e+05

012E+414.515RT (min)

lcorbi\_t3\_neg\_1\_055\_05

m/z 660.5576

ST004500\_NATIVE\_00066 · id 12319 · area 9.28e+04

01E+414.515RT (min)

lcorbi\_t3\_neg\_1\_055\_05

m/z 660.6421

ST004500\_NATIVE\_00067 · id 12320 · area 2.57e+06

024E+51414.5RT (min)

lcorbi\_t3\_neg\_1\_055\_05

m/z 667.4965

ST004500\_NATIVE\_00069 · id 12506 · area 5.95e+04

01E+414.515RT (min)

lcorbi\_t3\_neg\_1\_055\_05

m/z 667.5521

ST004500\_NATIVE\_00070 · id 12510 · area 7.67e+04

0123E+314.515RT (min)

lcorbi\_t3\_neg\_4\_016\_05

m/z 670.4869

ST004500\_NATIVE\_00071 · id 12566 · area 5.01e+04

01E+414.515RT (min)

lcorbi\_t3\_neg\_1\_055\_05

m/z 672.2707

ST004500\_NATIVE\_00074 · id 12599 · area 7.16e+04

01E+412.513RT (min)

lcorbi\_t3\_neg\_1\_055\_05

m/z 672.4864

ST004500\_NATIVE\_00075 · id 12608 · area 9.8e+04

012E+414.515RT (min)

lcorbi\_t3\_neg\_1\_055\_05

m/z 673.5050

ST004500\_NATIVE\_00076 · id 12633 · area 8.53e+04

01E+414.515RT (min)

lcorbi\_t3\_neg\_1\_055\_05

m/z 678.4842

ST004500\_NATIVE\_00077 · id 12738 · area 4.52e+04

02468E+314.515RT (min)

lcorbi\_t3\_neg\_1\_055\_05

m/z 680.5843

ST004500\_NATIVE\_00080 · id 12800 · area 3.99e+04

0246E+314.515RT (min)

lcorbi\_t3\_neg\_1\_055\_05

m/z 696.5312

ST004500\_NATIVE\_00086 · id 13167 · area 6.72e+04

01E+414.515RT (min)

lcorbi\_t3\_neg\_1\_055\_05

m/z 697.5438

ST004500\_NATIVE\_00087 · id 13193 · area 3.99e+04

02468E+31515.5RT (min)

lcorbi\_t3\_neg\_1\_055\_05

m/z 698.5020

ST004500\_NATIVE\_00088 · id 13210 · area 9.27e+04

01E+414.515RT (min)

lcorbi\_t3\_neg\_1\_055\_05

m/z 700.5178

ST004500\_NATIVE\_00089 · id 13250 · area 8.11e+04

01E+414.515RT (min)

lcorbi\_t3\_neg\_1\_055\_05

m/z 702.6173

ST004500\_NATIVE\_00090 · id 13301 · area 1.57e+04

0123E+31515.5RT (min)

lcorbi\_t3\_neg\_1\_055\_05

m/z 705.4505

ST004500\_NATIVE\_00093 · id 13360 · area 9.55e+04

01E+413.514RT (min)

lcorbi\_t3\_neg\_1\_055\_05

m/z 710.5469

ST004500\_NATIVE\_00098 · id 13488 · area 5.48e+05

01E+514.515RT (min)

lcorbi\_t3\_neg\_1\_055\_05

m/z 711.5221

ST004500\_NATIVE\_00099 · id 13511 · area 8.8e+04

01E+314.515RT (min)

lcorbi\_t3\_neg\_1\_055\_05

m/z 716.3637

ST004500\_NATIVE\_00102 · id 13609 · area 1.03e+05

01E+41111.5RT (min)

lcorbi\_t3\_neg\_3\_040\_05

m/z 720.4585

ST004500\_NATIVE\_00103 · id 13714 · area 3.02e+04

0246E+314.515RT (min)

lcorbi\_t3\_neg\_1\_055\_05

m/z 727.5357

ST004500\_NATIVE\_00104 · id 13889 · area 2.06e+04

0246E+314.515RT (min)

lcorbi\_t3\_neg\_1\_055\_05

m/z 732.6545

ST004500\_NATIVE\_00106 · id 14008 · area 4.09e+04

01E+414.515RT (min)

lcorbi\_t3\_neg\_1\_055\_05

m/z 733.4756

ST004500\_NATIVE\_00107 · id 14018 · area 3.84e+04

02468E+31313.5RT (min)

lcorbi\_t3\_neg\_1\_055\_05

m/z 736.6295

ST004500\_NATIVE\_00108 · id 14097 · area 3.24e+05

0246E+414.515RT (min)

lcorbi\_t3\_neg\_1\_055\_05

m/z 737.6452

ST004500\_NATIVE\_00109 · id 14123 · area 1.26e+05

012E+414.515RT (min)

lcorbi\_t3\_neg\_1\_055\_05

m/z 738.6453

ST004500\_NATIVE\_00110 · id 14149 · area 3.45e+05

0246E+414.515RT (min)

lcorbi\_t3\_neg\_1\_055\_05

m/z 750.5025

ST004500\_NATIVE\_00115 · id 14418 · area 1.25e+05

012E+414.515RT (min)

lcorbi\_t3\_neg\_1\_055\_05

m/z 750.6444

ST004500\_NATIVE\_00116 · id 14427 · area 3.41e+04

0246E+31515.5RT (min)

lcorbi\_t3\_neg\_1\_055\_05

m/z 750.6894

ST004500\_NATIVE\_00117 · id 14429 · area 5.4e+05

01E+51515.5RT (min)

lcorbi\_t3\_neg\_1\_055\_05

m/z 751.5540

ST004500\_NATIVE\_00118 · id 14445 · area 1.97e+04

0123E+314.515RT (min)

lcorbi\_t3\_neg\_1\_055\_05

m/z 753.3439

ST004500\_NATIVE\_00120 · id 14503 · area 8.49e+03

012E+314.515RT (min)

lcorbi\_t3\_neg\_4\_016\_05

m/z 765.5309

ST004500\_NATIVE\_00123 · id 14818 · area 1.04e+04

012E+31515.5RT (min)

lcorbi\_t3\_neg\_1\_055\_05

m/z 779.5328

ST004500\_NATIVE\_00130 · id 15161 · area 3.64e+04

0246E+314.515RT (min)

lcorbi\_t3\_neg\_1\_055\_05

m/z 796.5040

ST004500\_NATIVE\_00134 · id 15588 · area 9.73e+04

012E+41313.5RT (min)

lcorbi\_t3\_neg\_1\_055\_05

m/z 825.5679

ST004500\_NATIVE\_00139 · id 16300 · area 1.01e+06

01234E+313.514RT (min)

lcorbi\_t3\_neg\_1\_055\_05

m/z 838.5222

ST004500\_NATIVE\_00141 · id 16613 · area 2.25e+05

0246E+313.514RT (min)

lcorbi\_t3\_neg\_1\_055\_05

m/z 872.6662

ST004500\_NATIVE\_00144 · id 17375 · area 8.88e+04

01E+414.515RT (min)

lcorbi\_t3\_neg\_1\_055\_05

m/z 942.6546

ST004500\_NATIVE\_00145 · id 18560 · area 2.19e+04

0123E+214.515RT (min)

lcorbi\_t3\_neg\_1\_055\_05

m/z 985.7478

ST004500\_NATIVE\_00146 · id 19027 · area 5.96e+04

01E+414.515RT (min)

lcorbi\_t3\_neg\_4\_016\_05

#### ST004501 · Orbitrap · 35 analytes · m/z ±10 ppm

m/z 98.0600

ST004501\_NATIVE\_00004 · id 286 · area 7.4e+04

01E+433.5RT (min)

lcorbi\_hilic\_pos\_2\_030\_05

m/z 131.0531

ST004501\_NATIVE\_00009 · id 669 · area 2.33e+04

01E+433.5RT (min)

lcorbi\_hilic\_pos\_2\_030\_05

m/z 147.1129

ST004501\_NATIVE\_00015 · id 936 · area 2.62e+05

01E+57.58RT (min)

lcorbi\_hilic\_pos\_1\_050\_05

m/z 159.0764

ST004501\_NATIVE\_00019 · id 1113 · area 2.94e+05

01E+56.57RT (min)

lcorbi\_hilic\_pos\_1\_050\_05

m/z 160.0798

ST004501\_NATIVE\_00020 · id 1131 · area 1.29e+04

01234E+36.57RT (min)

lcorbi\_hilic\_pos\_1\_050\_05

m/z 197.0921

ST004501\_NATIVE\_00030 · id 1808 · area 8.63e+04

012E+41.52RT (min)

lcorbi\_hilic\_pos\_1\_050\_05

m/z 198.0849

ST004501\_NATIVE\_00031 · id 1829 · area 3.34e+05

01E+56.57RT (min)

lcorbi\_hilic\_pos\_1\_050\_05

m/z 205.0819

ST004501\_NATIVE\_00032 · id 1950 · area 6.66e+03

012E+377.5RT (min)

lcorbi\_hilic\_pos\_2\_030\_05

m/z 208.0798

ST004501\_NATIVE\_00035 · id 2002 · area 2.7e+04

01E+41.52RT (min)

lcorbi\_hilic\_pos\_1\_050\_05

m/z 211.1077

ST004501\_NATIVE\_00036 · id 2047 · area 3.18e+04

01E+444.5RT (min)

lcorbi\_hilic\_pos\_1\_050\_05

m/z 225.0159

ST004501\_NATIVE\_00038 · id 2269 · area 2.16e+04

01234E+37.58RT (min)

lcorbi\_hilic\_pos\_1\_050\_05

m/z 227.0639

ST004501\_NATIVE\_00040 · id 2309 · area 4.16e+03

01E+377.5RT (min)

lcorbi\_hilic\_pos\_1\_050\_05

m/z 247.0578

ST004501\_NATIVE\_00042 · id 2680 · area 3.03e+04

01E+466.5RT (min)

lcorbi\_hilic\_pos\_4\_007\_05

m/z 254.1216

ST004501\_NATIVE\_00044 · id 2800 · area 1.44e+05

0246E+411.5RT (min)

lcorbi\_hilic\_pos\_1\_050\_05

m/z 254.1612

ST004501\_NATIVE\_00045 · id 2804 · area 1.7e+04

0246E+355.5RT (min)

lcorbi\_hilic\_pos\_1\_050\_05

m/z 255.0976

ST004501\_NATIVE\_00046 · id 2818 · area 3.16e+04

02468E+35.56RT (min)

lcorbi\_hilic\_pos\_1\_050\_05

m/z 265.1112

ST004501\_NATIVE\_00047 · id 3015 · area 5.15e+04

01E+444.5RT (min)

lcorbi\_hilic\_pos\_2\_030\_05

m/z 269.1132

ST004501\_NATIVE\_00048 · id 3085 · area 9.26e+04

012E+42.53RT (min)

lcorbi\_hilic\_pos\_1\_050\_05

m/z 271.0900

ST004501\_NATIVE\_00049 · id 3120 · area 3.68e+03

012E+34.55RT (min)

lcorbi\_hilic\_pos\_1\_050\_05

m/z 291.1300

ST004501\_NATIVE\_00055 · id 3470 · area 5.59e+04

01E+47.58RT (min)

lcorbi\_hilic\_pos\_1\_050\_05

m/z 291.1381

ST004501\_NATIVE\_00056 · id 3471 · area 2.02e+04

02468E+36.57RT (min)

lcorbi\_hilic\_pos\_1\_050\_05

m/z 298.0879

ST004501\_NATIVE\_00057 · id 3589 · area 2.23e+06

01234E+322.5RT (min)

lcorbi\_hilic\_pos\_3\_027\_05

m/z 301.2932

ST004501\_NATIVE\_00062 · id 3661 · area 1.3e+04

02468E+311.5RT (min)

lcorbi\_hilic\_pos\_4\_007\_05

m/z 307.3109

ST004501\_NATIVE\_00063 · id 3774 · area 2.18e+04

0246E+322.5RT (min)

lcorbi\_hilic\_pos\_3\_027\_05

m/z 346.3226

ST004501\_NATIVE\_00067 · id 4417 · area 2.18e+04

01E+411.5RT (min)

lcorbi\_hilic\_pos\_2\_030\_05

m/z 354.1662

ST004501\_NATIVE\_00070 · id 4520 · area 1.37e+04

0246E+35.56RT (min)

lcorbi\_hilic\_pos\_1\_050\_05

m/z 354.3005

ST004501\_NATIVE\_00071 · id 4523 · area 4.39e+05

012E+50.51RT (min)

lcorbi\_hilic\_pos\_1\_050\_05

m/z 355.3038

ST004501\_NATIVE\_00072 · id 4539 · area 9.38e+04

024E+40.51RT (min)

lcorbi\_hilic\_pos\_1\_050\_05

m/z 370.0527

ST004501\_NATIVE\_00075 · id 4716 · area 4.14e+04

01E+477.5RT (min)

lcorbi\_hilic\_pos\_1\_050\_05

m/z 410.3182

ST004501\_NATIVE\_00085 · id 5104 · area 1.42e+04

01234E+30.51RT (min)

lcorbi\_hilic\_pos\_1\_050\_05

m/z 426.3089

ST004501\_NATIVE\_00088 · id 5210 · area 4.88e+03

012E+30.51RT (min)

lcorbi\_hilic\_pos\_3\_027\_05

m/z 429.2425

ST004501\_NATIVE\_00089 · id 5236 · area 6.26e+03

012E+36.57RT (min)

lcorbi\_hilic\_pos\_2\_030\_05

m/z 442.3036

ST004501\_NATIVE\_00092 · id 5309 · area 2.19e+04

01E+40.51RT (min)

lcorbi\_hilic\_pos\_2\_030\_05

m/z 463.3431

ST004501\_NATIVE\_00093 · id 5401 · area 2.06e+04

01E+40.51RT (min)

lcorbi\_hilic\_pos\_1\_050\_05

m/z 580.5308

ST004501\_NATIVE\_00094 · id 5736 · area 1.31e+04

02468E+30.51RT (min)

lcorbi\_hilic\_pos\_2\_030\_05

#### ST004502 · Orbitrap · 31 analytes · m/z ±10 ppm

m/z 84.0455

ST004502\_NATIVE\_00001 · id 230 · area 1.95e+04

02468E+36.57RT (min)

lcorbi\_hilic\_neg\_1\_047\_05

m/z 101.0720

ST004502\_NATIVE\_00002 · id 445 · area 2.13e+04

02468E+36.57RT (min)

lcorbi\_hilic\_neg\_1\_047\_05

m/z 122.0610

ST004502\_NATIVE\_00004 · id 719 · area 4.15e+04

01E+43.54RT (min)

lcorbi\_hilic\_neg\_1\_047\_05

m/z 143.0543

ST004502\_NATIVE\_00007 · id 1146 · area 1.1e+06

01E+533.5RT (min)

lcorbi\_hilic\_neg\_1\_047\_05

m/z 163.1239

ST004502\_NATIVE\_00011 · id 1646 · area 1.49e+04

0246E+311.5RT (min)

lcorbi\_hilic\_neg\_1\_047\_05

m/z 174.1077

ST004502\_NATIVE\_00012 · id 1923 · area 4.55e+05

01E+577.5RT (min)

lcorbi\_hilic\_neg\_1\_047\_05

m/z 178.0564

ST004502\_NATIVE\_00013 · id 2046 · area 3.84e+04

01E+466.5RT (min)

lcorbi\_hilic\_neg\_3\_042\_05

m/z 185.0930

ST004502\_NATIVE\_00014 · id 2271 · area 7.63e+04

0123E+455.5RT (min)

lcorbi\_hilic\_neg\_1\_047\_05

m/z 224.0765

ST004502\_NATIVE\_00016 · id 3523 · area 1.24e+05

01E+41.52RT (min)

lcorbi\_hilic\_neg\_4\_030\_05

m/z 232.0937

ST004502\_NATIVE\_00017 · id 3782 · area 9.1e+03

012E+36.57RT (min)

lcorbi\_hilic\_neg\_4\_030\_05

m/z 238.0912

ST004502\_NATIVE\_00018 · id 3981 · area 4.65e+04

012E+411.5RT (min)

lcorbi\_hilic\_neg\_1\_047\_05

m/z 276.1279

ST004502\_NATIVE\_00027 · id 5227 · area 7.53e+04

012E+47.58RT (min)

lcorbi\_hilic\_neg\_1\_047\_05

m/z 289.1154

ST004502\_NATIVE\_00030 · id 5653 · area 4.66e+04

01E+47.58RT (min)

lcorbi\_hilic\_neg\_1\_047\_05

m/z 289.1235

ST004502\_NATIVE\_00031 · id 5655 · area 9.66e+04

0123E+46.57RT (min)

lcorbi\_hilic\_neg\_1\_047\_05

m/z 289.1400

ST004502\_NATIVE\_00032 · id 5656 · area 1.87e+04

0246E+37.58RT (min)

lcorbi\_hilic\_neg\_1\_047\_05

m/z 291.1310

ST004502\_NATIVE\_00033 · id 5717 · area 4.05e+04

0123E+46.57RT (min)

lcorbi\_hilic\_neg\_1\_047\_05

m/z 319.0801

ST004502\_NATIVE\_00035 · id 6539 · area 5.27e+04

012E+46.57RT (min)

lcorbi\_hilic\_neg\_4\_030\_05

m/z 338.1355

ST004502\_NATIVE\_00036 · id 7074 · area 3.78e+04

01E+45.56RT (min)

lcorbi\_hilic\_neg\_1\_047\_05

m/z 357.2141

ST004502\_NATIVE\_00039 · id 7614 · area 3.28e+04

01E+44.55RT (min)

lcorbi\_hilic\_neg\_1\_047\_05

m/z 374.1332

ST004502\_NATIVE\_00040 · id 7989 · area 1.11e+04

0123E+35.56RT (min)

lcorbi\_hilic\_neg\_3\_042\_05

m/z 377.0410

ST004502\_NATIVE\_00041 · id 8053 · area 1.91e+06

01E+366.5RT (min)

lcorbi\_hilic\_neg\_1\_047\_05

m/z 387.0766

ST004502\_NATIVE\_00044 · id 8311 · area 1.26e+05

01E+466.5RT (min)

lcorbi\_hilic\_neg\_1\_047\_05

m/z 396.1154

ST004502\_NATIVE\_00045 · id 8507 · area 6.99e+03

01234E+35.56RT (min)

lcorbi\_hilic\_neg\_1\_047\_05

m/z 427.2635

ST004502\_NATIVE\_00046 · id 9065 · area 1.66e+04

01E+40.51RT (min)

lcorbi\_hilic\_neg\_1\_047\_05

m/z 582.5113

ST004502\_NATIVE\_00047 · id 10933 · area 1.52e+05

01E+50.51RT (min)

lcorbi\_hilic\_neg\_2\_021\_05

m/z 585.5304

ST004502\_NATIVE\_00048 · id 10955 · area 1.34e+04

0246E+30.51RT (min)

lcorbi\_hilic\_neg\_1\_047\_05

m/z 606.0765

ST004502\_NATIVE\_00049 · id 11096 · area 7.74e+04

01E+488.5RT (min)

lcorbi\_hilic\_neg\_3\_042\_05

m/z 610.5426

ST004502\_NATIVE\_00050 · id 11132 · area 9.68e+04

01234E+40.51RT (min)

lcorbi\_hilic\_neg\_1\_047\_05

m/z 611.5461

ST004502\_NATIVE\_00051 · id 11140 · area 3.85e+04

012E+40.51RT (min)

lcorbi\_hilic\_neg\_2\_021\_05

m/z 614.4929

ST004502\_NATIVE\_00052 · id 11154 · area 2.3e+04

01E+40.51RT (min)

lcorbi\_hilic\_neg\_2\_021\_05

m/z 669.4508

ST004502\_NATIVE\_00053 · id 11304 · area 4.62e+04

012E+40.51RT (min)

lcorbi\_hilic\_neg\_1\_047\_05

#### ST004503 · Q-ToF · 14 analytes · m/z ±25 ppm

m/z 184.0634

ST004503\_NATIVE\_00019 · id 838 · area 2.68e+04

0246E+311.5RT (min)

lcqtof\_t3\_pos\_4\_003\_05

m/z 258.1108

ST004503\_NATIVE\_00030 · id 1834 · area 5.75e+06

01E+611.5RT (min)

lcqtof\_t3\_pos\_2\_023\_05

m/z 260.1284

ST004503\_NATIVE\_00035 · id 1868 · area 7.99e+04

012E+411.5RT (min)

lcqtof\_t3\_pos\_4\_003\_05

m/z 432.0268

ST004503\_NATIVE\_00088 · id 4636 · area 2.08e+04

01234E+344.5RT (min)

lcqtof\_t3\_pos\_4\_003\_05

m/z 441.3075

ST004503\_NATIVE\_00090 · id 4764 · area 1.79e+04

012E+355.5RT (min)

lcqtof\_t3\_pos\_4\_003\_05

m/z 577.4801

ST004503\_NATIVE\_00113 · id 6606 · area 1.17e+06

0123E+313.514RT (min)

lcqtof\_t3\_pos\_4\_003\_05

m/z 577.4970

ST004503\_NATIVE\_00114 · id 6608 · area 9.6e+05

0123E+413.514RT (min)

lcqtof\_t3\_pos\_4\_003\_05

m/z 578.4822

ST004503\_NATIVE\_00116 · id 6623 · area 4.31e+05

01E+313.514RT (min)

lcqtof\_t3\_pos\_4\_003\_05

m/z 676.6099

ST004503\_NATIVE\_00160 · id 7967 · area 2.35e+06

0123E+513.514RT (min)

lcqtof\_t3\_pos\_2\_023\_05

m/z 678.6260

ST004503\_NATIVE\_00161 · id 7997 · area 3.36e+06

01234E+513.514RT (min)

lcqtof\_t3\_pos\_2\_023\_05

m/z 734.7374

ST004503\_NATIVE\_00172 · id 8931 · area 1.58e+05

012E+41414.5RT (min)

lcqtof\_t3\_pos\_4\_003\_05

m/z 759.4994

ST004503\_NATIVE\_00179 · id 9302 · area 8.9e+05

02468E+412.513RT (min)

lcqtof\_t3\_pos\_4\_003\_05

m/z 872.7450

ST004503\_NATIVE\_00215 · id 11159 · area 2.99e+06

01E+514.515RT (min)

lcqtof\_t3\_pos\_4\_003\_05

m/z 883.7416

ST004503\_NATIVE\_00224 · id 11279 · area 2e+05

02468E+414.515RT (min)

lcqtof\_t3\_pos\_4\_003\_05

#### ST004626 · Q-ToF · 5 analytes · m/z ±25 ppm

m/z 578.5144

ST004626\_NATIVE\_00024 · id 2445 · area 1.98e+04

0123E+31313.5RT (min)

lcqtof\_t3\_neg\_1\_006\_05

m/z 583.5135

ST004626\_NATIVE\_00027 · id 2478 · area 6.94e+04

02468E+31313.5RT (min)

lcqtof\_t3\_neg\_4\_019\_05

m/z 615.4948

ST004626\_NATIVE\_00037 · id 2692 · area 4.12e+04

01234E+31313.5RT (min)

lcqtof\_t3\_neg\_2\_016\_05

m/z 629.5104

ST004626\_NATIVE\_00044 · id 2771 · area 3.55e+04

01234E+31313.5RT (min)

lcqtof\_t3\_neg\_1\_006\_05

m/z 756.5899

ST004626\_NATIVE\_00053 · id 3779 · area 3.46e+04

0246E+213.514RT (min)

lcqtof\_t3\_neg\_4\_019\_05

#### ST004650 · Q-ToF · 18 analytes · m/z ±25 ppm

m/z 187.1093

ST004650\_NATIVE\_00016 · id 1326 · area 1.74e+05

012E+24.55RT (min)

lcqtof\_hilic\_pos\_4\_027\_05

m/z 379.2854

ST004650\_NATIVE\_00071 · id 5550 · area 2.99e+05

012E+30.51RT (min)

lcqtof\_hilic\_pos\_4\_027\_05

m/z 467.1166

ST004650\_NATIVE\_00102 · id 7307 · area 2.2e+03

02468E+24.55RT (min)

lcqtof\_hilic\_pos\_3\_047\_05

m/z 486.2767

ST004650\_NATIVE\_00108 · id 7617 · area 2.81e+04

02468E+23.54RT (min)

lcqtof\_hilic\_pos\_4\_027\_05

m/z 534.5234

ST004650\_NATIVE\_00131 · id 8573 · area 1.22e+05 · 1 of 2

0246E+40.51RT (min)

lcqtof\_hilic\_pos\_4\_027\_05

m/z 541.2531

ST004650\_NATIVE\_00136 · id 8707 · area 3.42e+04

01E+344.5RT (min)

lcqtof\_hilic\_pos\_1\_027\_05

m/z 546.2985

ST004650\_NATIVE\_00137 · id 8796 · area 7.97e+03

01234E+33.54RT (min)

lcqtof\_hilic\_pos\_1\_027\_05

m/z 550.5488

ST004650\_NATIVE\_00140 · id 8901 · area 7.91e+04

01234E+40.51RT (min)

lcqtof\_hilic\_pos\_4\_027\_05

m/z 562.4914

ST004650\_NATIVE\_00148 · id 9106 · area 3.13e+05

0246E+20.51RT (min)

lcqtof\_hilic\_pos\_4\_027\_05

m/z 585.3858

ST004650\_NATIVE\_00158 · id 9587 · area 1.12e+04

01234E+33.54RT (min)

lcqtof\_hilic\_pos\_4\_027\_05

m/z 588.1715

ST004650\_NATIVE\_00159 · id 9640 · area 1.21e+04

0123E+37.58RT (min)

lcqtof\_hilic\_pos\_4\_027\_05

m/z 616.4611

ST004650\_NATIVE\_00171 · id 10200 · area 1.73e+05

01E+40.51RT (min)

lcqtof\_hilic\_pos\_1\_027\_05

m/z 671.4290

ST004650\_NATIVE\_00185 · id 11148 · area 9.32e+03

024E+30.51RT (min)

lcqtof\_hilic\_pos\_4\_027\_05

m/z 685.5096

ST004650\_NATIVE\_00187 · id 11405 · area 1.29e+04

02468E+30.51RT (min)

lcqtof\_hilic\_pos\_4\_027\_05

m/z 703.5366

ST004650\_NATIVE\_00189 · id 11739 · area 3.82e+05

012E+50.51RT (min)

lcqtof\_hilic\_pos\_4\_027\_05

m/z 704.5164

ST004650\_NATIVE\_00190 · id 11772 · area 9.34e+04

012E+40.51RT (min)

lcqtof\_hilic\_pos\_4\_027\_05

m/z 790.6901

ST004650\_NATIVE\_00231 · id 13438 · area 3.94e+04

012E+40.51RT (min)

lcqtof\_hilic\_pos\_4\_027\_05

m/z 998.7149

ST004650\_NATIVE\_00249 · id 16007 · area 4.2e+03

01234E+233.5RT (min)

lcqtof\_hilic\_pos\_4\_027\_05

#### ST004651 · Q-ToF · 18 analytes · m/z ±25 ppm

m/z 216.0369

ST004651\_NATIVE\_00014 · id 1305 · area 2.92e+04

012E+322.5RT (min)

lcqtof\_hilic\_neg\_2\_028\_05

m/z 413.2427

ST004651\_NATIVE\_00038 · id 4554 · area 2.37e+04

01E+30.51RT (min)

lcqtof\_hilic\_neg\_4\_040\_05

m/z 489.3031

ST004651\_NATIVE\_00047 · id 5700 · area 7.38e+05

01E+533.5RT (min)

lcqtof\_hilic\_neg\_4\_040\_05

m/z 564.4783

ST004651\_NATIVE\_00058 · id 6752 · area 1.32e+04

024E+20.51RT (min)

lcqtof\_hilic\_neg\_4\_040\_05

m/z 565.3214

ST004651\_NATIVE\_00059 · id 6770 · area 1.71e+05

0246E+433.5RT (min)

lcqtof\_hilic\_neg\_1\_007\_05

m/z 568.4572

ST004651\_NATIVE\_00061 · id 6822 · area 1.19e+04

02468E+20.51RT (min)

lcqtof\_hilic\_neg\_4\_040\_05

m/z 585.4489

ST004651\_NATIVE\_00064 · id 7082 · area 8.83e+04

012E+30.51RT (min)

lcqtof\_hilic\_neg\_4\_040\_05

m/z 592.1827

ST004651\_NATIVE\_00066 · id 7172 · area 5.11e+03

012E+34.55RT (min)

lcqtof\_hilic\_neg\_2\_028\_05

m/z 612.4944

ST004651\_NATIVE\_00070 · id 7491 · area 1.39e+05

01E+30.51RT (min)

lcqtof\_hilic\_neg\_2\_028\_05

m/z 655.5299

ST004651\_NATIVE\_00079 · id 8141 · area 3.05e+04

01E+40.51RT (min)

lcqtof\_hilic\_neg\_4\_040\_05

m/z 872.6949

ST004651\_NATIVE\_00107 · id 11530 · area 777

01E+30.51RT (min)

lcqtof\_hilic\_neg\_1\_007\_05

m/z 879.5853

ST004651\_NATIVE\_00109 · id 11638 · area 7.22e+03

012E+33.54RT (min)

lcqtof\_hilic\_neg\_4\_040\_05

m/z 895.8088

ST004651\_NATIVE\_00112 · id 11869 · area 1.52e+04

02468E+30.51RT (min)

lcqtof\_hilic\_neg\_4\_040\_05

m/z 911.4693

ST004651\_NATIVE\_00117 · id 12101 · area 5.08e+03

012E+355.5RT (min)

lcqtof\_hilic\_neg\_4\_040\_05

m/z 920.6756

ST004651\_NATIVE\_00121 · id 12228 · area 5.19e+04

012E+30.51RT (min)

lcqtof\_hilic\_neg\_1\_007\_05

m/z 934.8209

ST004651\_NATIVE\_00122 · id 12415 · area 1.4e+04 · 1 of 2

01E+30.51RT (min)

lcqtof\_hilic\_neg\_4\_040\_05

m/z 984.7356

ST004651\_NATIVE\_00128 · id 12900 · area 2.82e+03

012E+311.5RT (min)

lcqtof\_hilic\_neg\_2\_028\_05

m/z 990.9405

ST004651\_NATIVE\_00129 · id 12945 · area 4.05e+03

0246E+20.51RT (min)

lcqtof\_hilic\_neg\_4\_040\_05
